# Targeted Transcriptome Analysis using Synthetic Long Read Sequencing Uncovers Isoform Reprograming in the Progression of Colon Cancer

**DOI:** 10.1101/2020.08.07.240069

**Authors:** Silvia Liu, Indira Wu, Yan-Ping Yu, Michael Balamotis, Baoguo Ren, Tuval Ben Yehezkel, Jian-Hua Luo

**Affiliations:** Department of Pathology, 5941 Optical Court, San Jose, CA 95138; High Throughput Genome Center, 5941 Optical Court, San Jose, CA 95138; Pittsburgh Liver Research Center, 5941 Optical Court, San Jose, CA 95138; University of Pittsburgh School of Medicine, 3550 Terrace Street, Pittsburgh, PA 15261, Loop Genomics, Inc., 5941 Optical Court, San Jose, CA 95138

## Abstract

Diversity in human gene expression stems, to a large extent, from splicing exons into multiple mRNA isoforms. Characterization of isoforms requires accurate long-read sequencing. However, read lengths, high error rates, low throughput and large input requirements are some of the challenges that remain to be addressed in sequencing technologies.

In this study, we used a barcoding-based synthetic long read (SLR) isoform sequencing approach, LoopSeq, to generate sequencing reads sufficiently long and accurate to identify isoforms using standard short read Illumina sequencers. The method identifies isoforms from control RNA samples with 99.4% accuracy and a 0.01% per-base error rate, exceeding the accuracy reported for other long-read sequencing technologies.

Applied to targeted transcriptome sequencing of over 10,000 genes from colon cancers and their metastatic counterparts, LoopSeq revealed large scale isoform redistributions from benign colon mucosa to primary colon cancer and metastatic cancer and identified several novel gene fusion isoforms in the colon cancer samples. Strikingly, our data showed that most single nucleotide variants (SNV’s) occurred dominantly in specific isoforms and that some SNVs underwent isoform switching in cancer progression.

The ability to use short read sequencers to generate accurate long-read isoform information as the raw unit of transcriptional information holds promise as a new and widely accessible approach in RNA isoform analyses.

## Introduction

The development of massively parallel short read sequencing and their overlapping alignment in the last 20 years has made it possible to decipher the genome sequences of numerous organisms^1-3^. It has also enhanced the quantification of gene expression by allowing the sequencing of millions of transcripts at an affordable price point. For mammalian cells, the transcription of a gene involves an alternative splicing process that selectively utilizes specific exons while removing other exons and introns from the final transcript^4^. This generates many different transcripts (isoforms) with altered amino acid sequences from the same gene and dramatically increases the diversity of the gene products^5^. However, given the high sequence homology between the different isoforms from the same gene, the characterization and quantification of isoforms using short-read sequencing is challenging as the span of the short reads typically renders isoform mapping and identification ambiguous. Similarly, determining the exact exon composition of a single mRNA molecule is difficult without longer sequencing lengths or the ability to trace the short reads to the originating individual transcripts.

To overcome the read-length limitation of short read sequencers, various approaches of synthetic long read (SLR) sequencing have been developed and applied to various difficult-to-sequence applications, including transcriptome sequencing^6^. SLR sequencing methods rely on a molecule-specific barcode sequence to be assigned to individual transcripts during library preparation that identifies the molecular origin of the short reads. Subsequently, synthetic long reads are reconstructed from clusters of short reads that share the same barcode. One approach of SLR involves physically partitioning DNA molecules into 384-well plates or microfluidic droplets, typically in the order of hundreds of molecules per partition, and the DNA molecules within each partition are amplified, fragmented, and barcoded for a highly-parallel library preparation^7-9^. While this approach demonstrates the power of SLR by resolving highly repetitive or difficult to sequence regions, the process is labor-intensive and cannot be used on samples with high sequence homology between RNA molecules, such as mRNA isoforms. Here we present the application of synthetic long reads to isoform sequencing using LoopSeq, a SLR technology that builds upon previously developed SLR approaches^10,11^ and leverages Illumina short read sequencing platforms to generate accurate long reads from mRNA. We validated that LoopSeq is highly accurate by identifying known isoforms in control samples and demonstrated its utility in discovering novel fusion gene isoforms, in quantifying isoform distributions, and in discovering novel mutation isoform expression patterns in clinical samples.

## Results

### LoopSeq accuracy in sequencing full-length transcripts

LoopSeq employs a SLR sample preparation method that enables long read assembly, building upon previously published synthetic long read sequencing methodologies^10,11^. As shown in Figure 1A, it first assigns a unique molecular identifier (UMI) to each first-strand cDNA molecule during reverse transcription. Following the barcoding step, an optional probe capture-based target enrichment step is applied to select for full-length cDNA molecules of interest. Following capture and PCR amplification of the barcoded cDNA, UMIs are randomly transposed to various internal positions of the molecules, and the sequence immediately adjacent to the UMI insertion site is converted into an Illumina short read that contains both the UMI and the adjacent sequence. After short read sequencing, short reads tagged with identical sample indices and UMI’s are binned and used for de novo synthetic long read assembly which generates a single long read for each barcoded cDNA molecule.

**Figure 1.**
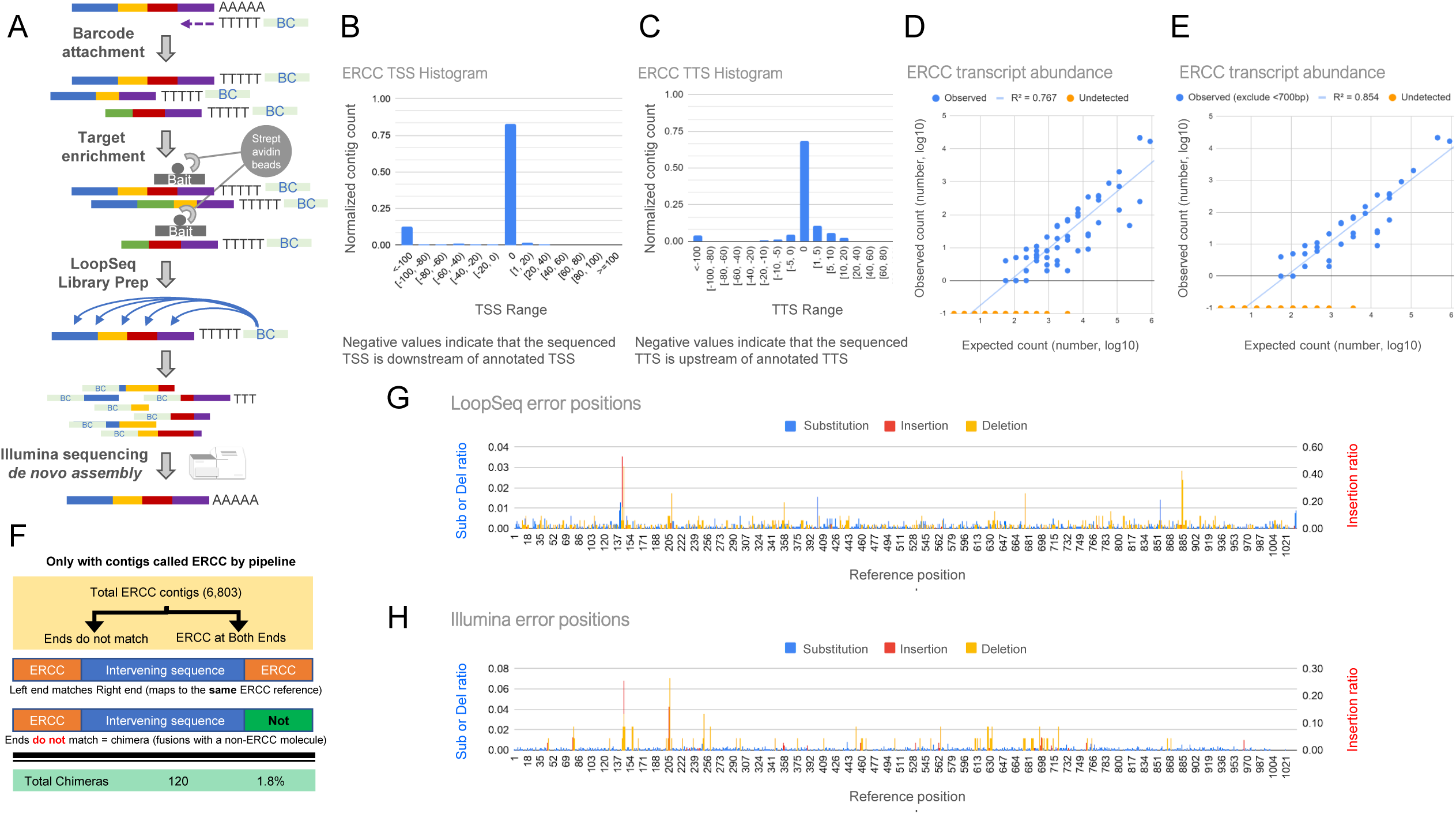
Schematics of LoopSeq library preparation and validation data. (A) Overview of the library preparation for isoform sequencing using LoopSeq, including an optional target enrichment step to focus the sequencing depth on genes or isoforms of interest. (B) The transcription start site (TSS) of reconstructed ERCC contigs as compared to the reference annotation. (C) The transcription termination site (TTS) of reconstructed ERCC contigs as compared to the reference annotation. (D) Comparison of ERCC transcript counts between the observed abundance as determined by reconstructed contigs and the expected abundance given the input into the library prep. (E) Comparison of ERCC transcript counts between the observed and the expected abundance, excluding references <700bp. (F) Overview of the chimera contig detection. (G) Demonstration of the positional bias of LoopSeq errors along an ERCC reference and (H) of the positional bias of Illumina short read errors along the same ERCC reference. The ratio of substitution (left axis), deletion (left axis), and insertion errors (right axis) against the position on ERCC-0002 reference are shown. The plotted values are the ratio of each error at a given reference position normalized by the overall error rate of a given error type. The Illumina short reads used for error analysis are obtained from previously published data (Oikonomopoulos 2016).

To demonstrate that LoopSeq SLRs cover the full-length of cDNA molecules, we prepared and sequenced 7,481 synthetic RNA control ERCC cDNA molecules alongside 27,426 mRNA molecules from Hela total RNA. After SLR reconstruction for each uniquely tagged cDNA molecule, the transcription start site (TSS) and transcription termination site (TTS) of each ERCC transcript identified as full-length were compared to the reference sequences. Histograms of the TSS and TTS differences are shown in Figure 1B and 1C. For TSS, 82.6% of the full-length contigs correctly identified the start site of the cDNA, and 12.6% of the full-length contigs report a TSS that is downstream of the annotated TSS. Note that the designation of full-length contigs is that the reconstructed sequence reaches the expected adaptor sequence at both the 5’ and the 3’ of the cDNA molecule. If a cDNA molecule is prematurely terminated during reverse transcription, either due to degraded RNA or non-specific terminal transferase activity of the reverse transcriptase, the reconstructed molecule would include the 5’ adaptor sequence, correctly identifying that the full-length cDNA molecule is reconstructed, while having a TSS that is downstream of the annotated TSS. For TTS, 68.7% of the full-length contigs correctly identified the termination site of the cDNA, while 15.6% of the full-length contigs had a TTS that was within 5 nucleotides from the annotated TTS.

Next we analyzed the fidelity of the reconstructed sequences, both by examining the rate of chimeric sequence formation resulting in large sections of long-reads being incorrect, and the rate of substitution and indels, i.e. single-nucleotide edits. Known sources of chimeric sequence formation in mRNA sequencing are cDNA synthesis and PCR amplification. Reverse transcriptase with template switching activity has been reported to jump within or between different templates without terminating DNA synthesis activity, resulting in chimeric cDNA formation. Over-amplification during PCR also leads to formation of chimeric molecules. While care is taken to not over-amplify cDNA molecules during PCR, chimeric molecules can sometimes be made during cDNA amplification. LoopSeq employs consensus sequence correction to remove chimeric sequences that are introduced during PCR, but it does not completely eliminate it. If a chimeric molecule is to form, we presume the chimeric junction is likely to occur once in the middle of the molecules, and one of the ends of the molecules would not map to the expected reference. To measure the rate of chimera formation, we examined 6,803 reconstructed full-length ERCC contigs and separately mapped the ends to the reference database. As illustrated in Figure 1F, 120 contigs were found to have ends that do not map to ERCC, which indicates a chimera rate of 1.8%. Most of the ends that did not map to ERCC mapped to Hela mRNA molecules, though some sequences were found to map to E. coli plasmid, most likely the plasmid from which the ERCC mRNA was transcribed

To examine LoopSeq’s error rate, we compared LoopSeq’s contigs to the expected ERCC sequences. A variant table of single-nucleotide edits, either from substitution, insertion, or deletion were constructed by comparing the ERCC full-length contig sequences to the reference sequence of ERCC-00002. As shown in Figure 1G, plotting the frequency of single-nucleotide edits against the reference shows that while the positions of substitutions are mostly random, the positions of insertions and deletions are concentrated at specific locations. Specifically, 53% of insertion errors were found near position 142, at a homopolymer region of seven As.

Substitution errors were also concentrated near homopolymer regions. for example positions 142 (seven As), 203 (five As), 672 (four As), and 882 (four Ts). To examine whether substitution errors originated from LoopSeq sequencing, we performed the same variant sequence analysis on a previously published ERCC Illumina short read dataset. As shown in Figure 1H, the published ERCC Illumina short read dataset shares many of the same positions prone to have high insertion and deletion rates, including positions 142 and 203, but not all the positions. This suggests some of the indel errors from LoopSeq are due to errors introduced during synthetic RNA synthesis or the use of template-switching reverse transcriptase, as the errors are shared across two different methods.

Next, we performed a comparative error rate between LoopSeq, PacBio CCS reads, Oxford Nanopore 2D reads and Illumina short read sequencing. Table 1 summarizes the error rates across the different sequencing methods. When sequencing synthetic RNA, LoopSeq exhibits at least two orders of magnitude lower error rate compared to PacBio CCS reads or Oxford Nanopore 2D reads, and an order of magnitude lower error rates compared to Illumina short read sequencing. As suggested by the non-random nature of the indel errors observed in the LoopSeq RNA sequencing data, the errors are not characteristic to the technology but rather to how the RNA is converted to cDNA using template switching. When sequencing DNA templates directly without reverse transcription^12^ LoopSeq’s indel errors are reduced to a negligible rate, roughly 1 in 220,000 bases assembled.

**Table 1.** Comparison of error rates across different sequencing platforms. Error rates of PacBio CCS, Oxford Nanopore ONT-2D are obtained from previously published data^53^. Error rate of Illumina short reads are computed from previously published data^54^.

| Error type | PacBio-CCS RNA | ONT-2D RNA | Illumina RNA | LoopSeq RNA | LoopSeq DNA |
| --- | --- | --- | --- | --- | --- |
| Match | 9.83E-01 | 8.66E-01 | 9.95E-01 | 9.992E-01 | 9.998E-01 |
| Substitution | 1.3E-02 | 5.50E-02 | 4.15E-03 | 3.475E-04 | 1.476E-04 |
| Insertion | 8.7E-04 | 3.12E-02 | 3.84E-04 | 2.060E-04 | 2.148E-06 |
| Deletion | 3.4E-03 | 4.79E-02 | 4.80E-04 | 2.691E-04 | 2.387E-06 |
| **Sum error** | **1.72E-02** | **1.34E-01** | **5.018E-03** | **8.226E-04** | **1.521E-04** |

Lastly, to demonstrate the accuracy in transcript quantification using LoopSeq, we prepared and sequenced a separate library of 66,308 synthetic RNA control ERCC cDNA molecules, and the observed abundances of ERCC molecules were compared with the expected abundance. As shown in Figure 1D, the agreement between the observed abundance and the expected abundance is moderate. Similar to what has been reported previously with other long read technologies such as PacBio sequencing, there is an intrinsic bias against short transcripts that are at the lower length limit that LoopSeq can prepare and sequence. As shown in Figure 1E, only considering transcripts that are expected to be at least 700bp in length increases the agreement between the observed and expected abundances.

### Quantification of gene and isoform expressions in human colon cancer

To examine the utility of LoopSeq in quantifying the expressions of genes and isoforms in human cancers, we prepared and sequenced three pairs of primary colon cancers and their matched metastases in the lymph nodes. To enrich for cancer-specific gene fusions and isoforms, a panel of 2149 probe capture oligos representing 2193 genes was designed (Supplementary Table S1). These oligos represent the split regions of the most frequent cancer-related gene fusions found in the TCGA databases. Using the fusion junction probe capture oligos, we obtained transcripts of 12127 cancer-related genes from the cancer samples (Supplementary Table S2). As shown in Figure 2A, we found 2682 genes that were differentially expressed between the metastasis and primary cancer samples or between the tumor samples and their corresponding benign colon tissue adjacent to cancer (Supplementary Table S3). When hierarchical clustering analyses were performed, differentially expressed genes (DEGs) showed proper segregation between primary cancer/metastases and normal colon samples (Figure 2A and Supplementary Table S3). However, the separation between primary cancer and metastasis samples is inadequate, and we were unable to differentiate between primary cancer and metastasis samples with gene expression patterns alone. When leveraging the isoform mapping from reconstructed long reads, we found 5941 isoforms differential expressions, across 4643 unique genes among these samples (Supplementary Table S4). Unlike DEGs, differentially expressed isoforms (DEIs) showed excellent segregation of all three groups of tissues (Figure 2A and Supplementary Table S4), demonstrating the power of performing differential expression analysis using isoform expression data over gene expression data.

**Figure 2.**
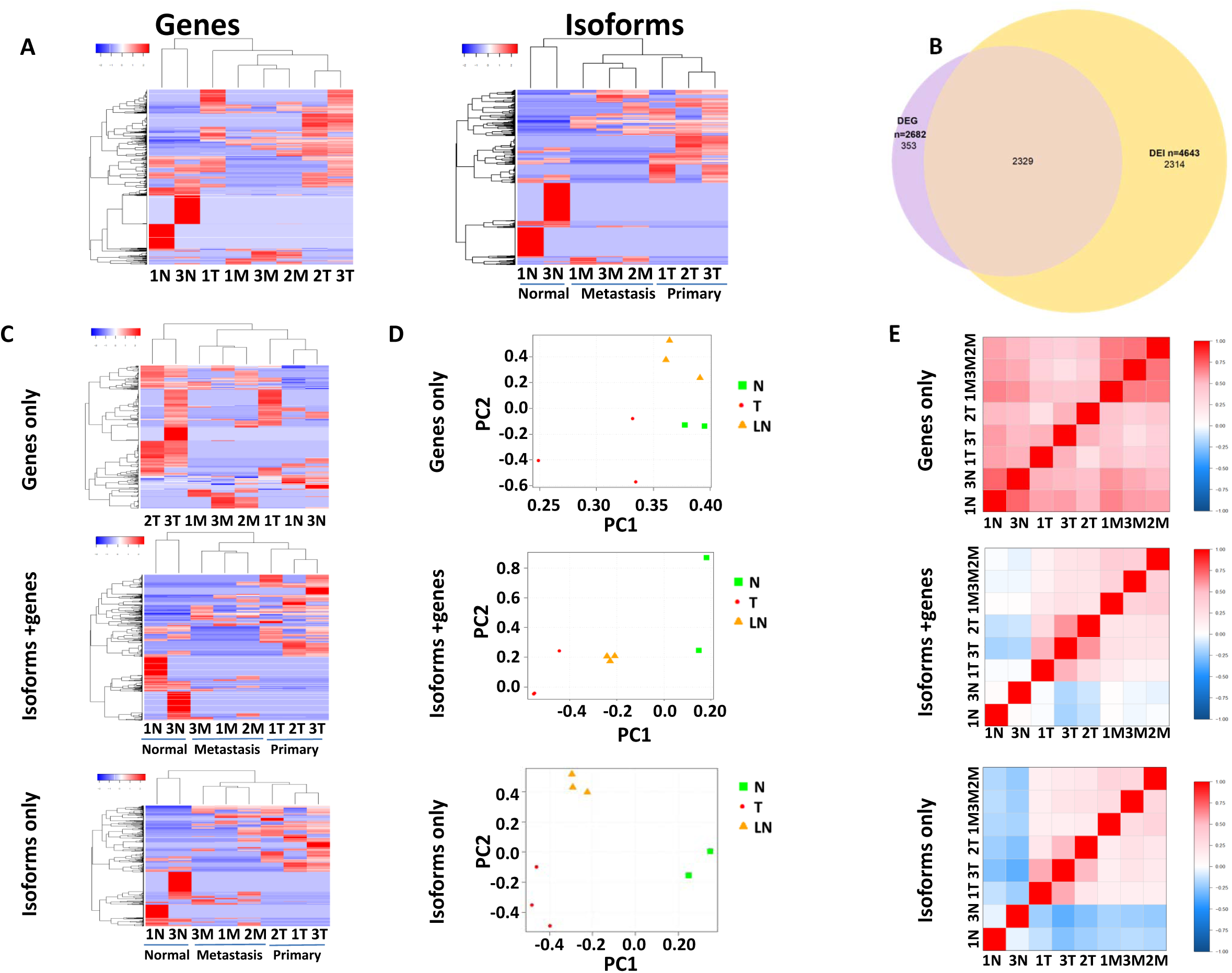
Quantification of isoform and gene expression to distinguish benign colon, primary colon cancer and metastatic colon cancers. (A) Hierarchical clustering of benign colon samples adjacent to cancer (1N and 3N), primary colon cancer samples (1-3T) and metastatic colon cancer samples (1-3M) based on differentially expressed genes (left) or isoforms (right). (B) Venn diagram of overlapping differentially expressed genes and isoforms in colon cancers, metastases and benign colon tissues adjacent to cancer. (C) Hierarchical clustering of colon samples based on differential expressed genes but not isoforms (top), or differential expressed genes accompanied with concomitant isoform differential expression (middle), or different isoform expressions without alteration of gene expression (bottom). (D) Principal component analyses of benign colon tissues adjacent to cancer, primary colon cancers and metastatic colon cancers based on differential gene expression without isoform expression alteration (top), or differential gene expression with concomitant isoform alteration (middle), or differential isoform expression without the alteration of gene expression (bottom). (E) Pearson’s correlation of benign colon tissues adjacent to cancer, primary colon cancers and metastatic colon cancers based on differential gene expression without isoform expression alteration (top), or differential gene expression with concomitant isoform alteration (middle), or differential isoform expression without the alteration of gene expression (bottom).

To further elucidate the significance of gene versus isoform expression patterns, next we examined DEGs with no isoform expression change and DEIs with no gene expression changes. While only 13.3% (353 of 2682) DEGs have no change in isoform distribution (Figure 2B), nearly half (49.9% or 2316/4643) of DEIs belong to genes with no gene expression changes (Supplementary Table S5). To investigate the impact of DEG with no change in isoform distribution on tissue differentiation, a hierarchical clustering analysis was performed on the primary colon cancer, the lymph node metastasis and their matched benign colon tissues adjacent to cancer using 353 DEGs that have no isoform alterations. The results indicated that these DEGs are not able to segregate the normal samples from the cancer samples (Figure 2C and Supplementary Table S3 and S5). Both principal component and Pearson’s correlation analyses confirmed the inadequacy of tissue segregation based on these genes (Figure 2D and 2E). In contrast, clustering analysis using DEGs with isoform redistribution produced a far better segregation between the benign colon and cancer samples (Figure 2C-E, mid panel and Supplementary Table S3 - S5). Interestingly, isoform redistribution without a change in gene expression produced the best tissue-differentiation results: all benign colon, primary cancer and metastases samples were segregated into different groups (Figure 2c, lower panel and Supplementary Table S4 and S5). These results indicate that DEI analysis produced robust sample segregation not obtainable by DEG analysis. These DEIs, which might have previously been inaccessible and were hidden within comparable gene expression levels, represent a new dimension in differential expression analysis.

Genes involved in cancer metastasis such as CD44^13^ are among DEIs that are not accompanied with a change in gene expression (Supplementary Figure S1). There were 26 different isoforms of CD44 detected in the colon cancer tissues, with the protein length ranging from 139 aa to 743 aa. Isoform analysis indicates that two isoforms (XM_005253231.3 and XM_011520484.2) emerged in the colon cancer and cancer metastasis samples but were absent in the benign colon tissues. Another gene of interest is ATP1A1, a Na^+^/K^+^ ATPase that is a subunit of Na^+^/K^+^ pump essential for maintaining ionic homeostasis for a cell^14^. ATP1A1 produced two additional isoforms (NM_000701.7 and NM_001160233.1) in the primary colon cancer samples and their corresponding metastasis, but not in the benign colon tissues (Supplementary Figure S2). The distribution of ATP1A1 isoforms were validated by Taqman qRT-PCR (Supplementary Figure S3).

### Expression pattern analyses of isoform-specific single nucleotide variants

Mutation is the hallmark of human malignancies. However, little is known about isoform-specific mutations due to the difficulty of identifying mutations and isoforms simultaneously. Taking the advantage of the read-length and the accuracy of LoopSeq synthetic long reads, we examined the single nucleotide variants (SNVs) in assembled contigs in the context of isoforms. A total of 4042 SNVs was identified in 6 cancer samples using LoopSeq that were cross-validated by standard short read whole exome sequencing (Supplementary Table S6). These SNVs were distributed among 1340 genes and 8712 isoforms. Interestingly, many SNVs were found in specific isoforms for a given gene. Of the 1509 SNVs found with at least 2 isoforms and 5 assembled contigs, 1297 SNVs were not distributed evenly among the isoforms of these genes but were predominantly found in specific isoforms (Supplementary Table S7). While the majority of the SNV isoform distribution is comparable to the wild-type isoform distribution, the isoform expression patterns of 113 SNVs were significantly different from their wild type counterparts (Supplementary Table S8), suggesting the alterations of splicing patterns for the variants. To validate the SNV isoform expression patterns observed in the long read data, isoforms of two genes were selected for targeted and isoform-specific short-read sequencing, made possible by the close proximity of the mutation and the alternative-splicing junction. For FAM104A, a protein involved in centriole biogenesis^15^, the SNV isoform expression pattern is comparable to the wild-type counterpart of NM_001098832, NM_032837 and NM_001289410, and the short read data was largely consistent with the LoopSeq data (Supplementary Table S9). For PABPC1, a poly-A binding protein^16^, the SNV isoform distribution of NM_002568 and XM_005250861 was different from the wild-type distribution, and again similar observation was made with the short read data (Supplementary Table S9).

To identify SNVs that change their expression in a given isoform during the progression of colon cancers, we screened for isoforms that uniformly have high SNV rate (>=0.5) or low SNV rate (<=0.5) across all metastatic colon cancer samples versus matched primary cancer samples. SNV rate was computed by normalizing the SNV counts with the total transcript counts of an isoform. Twenty-three SNV containing isoforms were identified that match the search criteria (Supplementary Table S10). The hierarchical clustering analysis based on the SNV rates of 23 SNV-containing isoforms confirmed that these isoforms produced a complete separation between cancer and metastatic samples (Figure 3A and Supplementary Table S10). Similar results were obtained by the principal component and Pearson’s correlation analyses (Figure 3B and 3C). In contrast, hierarchical clustering analysis based on the SNV rates of all 8712 isoforms failed to yield appreciable separation of metastatic and primary cancer samples (Supplementary Figure S4 and Supplementary Table S6). The ingenuity pathway analysis indicates that many of the 23 SNV-containing isoforms belong to genes involved in DNA repairing signaling and antigen-presentation signaling (Figure 3D).

**Figure 3.**
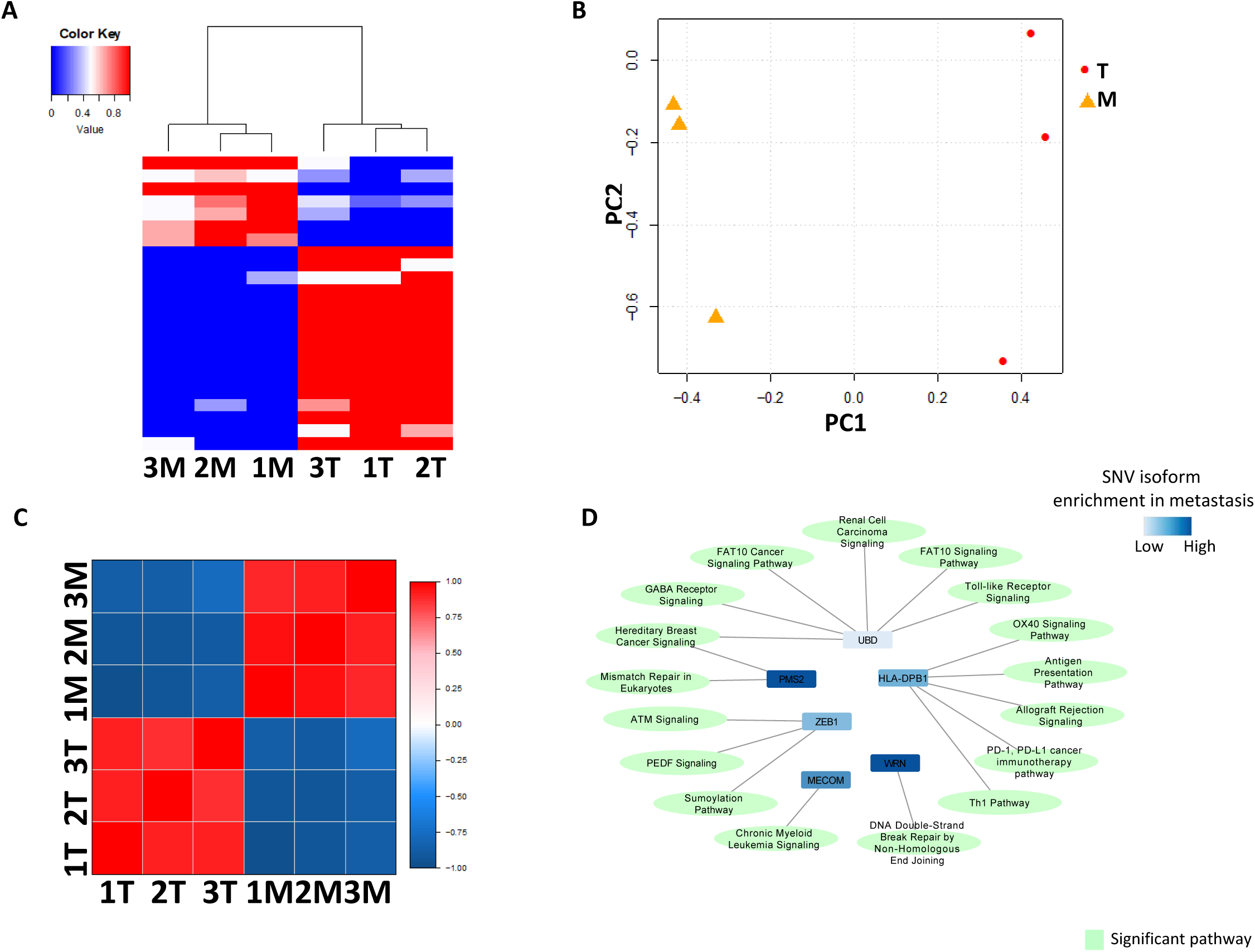
Isoform switches of single nucleotide variant between primary colon cancers and metastatic colon cancers. (A) Hierarchical clustering between primary colon cancers and metastatic colon cancers based on the quantities of non-synonymous single nucleotide variants of 23 isoforms in each sample. (B) Principal component analyses of primary colon cancers and metastatic colon cancers based on the quantities of non-synonymous single nucleotide variants of (A). (C) Pearson’s correlation of primary colon cancers and metastatic colon cancers based on the quantities of non-synonymous single nucleotide variants of (A). (D) Pathway analysis of 23 single nucleotide variant isoforms showed enrichment in genes involved in HLA/CD74 antigen presentation pathways.

To study the potential pathological SNVs in the development of colon cancer, we cross-referenced the 4042 validated SNVs against the database of Catalogue of Somatic Mutations in Cancers (COSMIC). Four hundred and one SNVs were identical to the mutations in the colon cancer database of COSMIC, suggesting many SNVs we discovered in our dataset may be involved in the pathological progress of colon cancer (Supplementary Table S11). One hundred ninety of these potential pathological SNVs were present predominantly in some specific isoforms of the residing genes. Twelve SNVs displayed different isoform expression patterns in comparison with their wild type patterns. Two SNV examples we investigated are BRAF and KRAS. The V600E variant^17^ of BRAF was detected in sample 3T (primary tumor sample) and 3M (metastasis tumor sample). The predominant isoform for both wild-type and the V600E variant was NM_04333 in both samples (p<0.05). However, new isoforms for BRAF (NM_001354609, XM_017012558 and XR_001744857) emerged in both primary cancer and the corresponding metastasis and all contained the V600E variant (Supplementary Figure S5). In contrast, wild-type isoform distribution of KRAS proto-oncogene^18^ is distinct between sample 2T (primary tumor sample) and 2M (metastasis tumor sample). Similarly, the G12V variant also had different isoform distributions. This suggests that KRAS may have undergone a change in isoform distribution when the colon cancer evolved from its primary site (2T, NM_033360) to the lymph node site (2M, NM_004985 and XM_011520653)(Supplementary Figure S6).

### Discovery of fusion gene isoforms

Discovery of fusion isoforms remains difficult due to the requirement that the fusion junction needs to be sequenced alongside exons that can differentiate one isoform from another. To investigate the utility of LoopSeq in identifying fusion gene isoforms, we used SQANTI, a bioinformatic pipeline for classifying long reads by splice-junctions, to detect fusion transcripts in the synthetic long read data. We applied two additional filtering criteria on the fusion isoform candidates: (i) the fusion gene partners are of trans direction or cis direction separated by >40 kb with at least one gene in between, and (ii) the fusion junction point is derived from the exon junctures between the fusion partners (Supplementary Figure S7). Among 6 samples of colon cancers, 4 novel fusion isoforms were found to meet these criteria. These fusion junctions were validated using Taqman qRT-PCR and Sanger’s sequencing (Figure 4 and Supplemental Figure 8).

**Figure 4.**
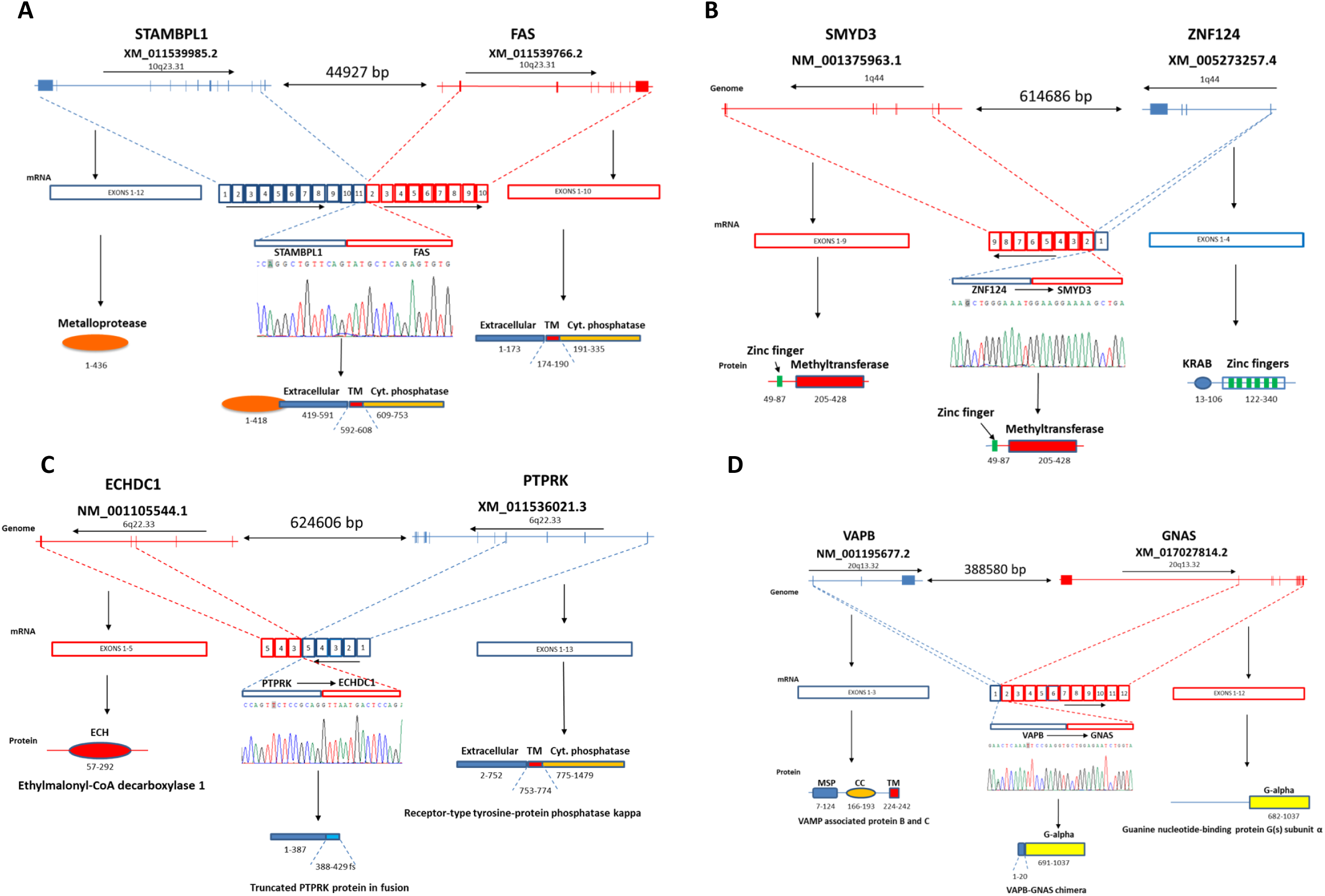
Fusion gene isoforms identified in colon cancers through LOOP sequencing. (A) STAMBPL1-FAS fusion. Top: diagram of mini genomes of STAMBPL1 and FAS. Direction of transcription and distance between the 2 genes are indicated. Mid: mRNA represented by exons from each gene. Bottom: diagram of functional protein domains of STAMBPL1 and FAS. (B) ZNF124-SMYD3 fusion. Top: diagram of minigenomes of ZNF124 and SMYD3. Direction of transcription and distance between the 2 genes are indicated. Mid: mRNA represented by exons from each gene. Bottom: diagram of functional protein domains of ZNF124 and SMYD3. (C) PTPRK-ECHCD1 fusion. Top: diagram of minigenomes of PTPRK and ECHCD1. Direction of transcription and distance between the 2 genes are indicated. Mid: mRNA represented by exons from each gene. Bottom: diagram of functional protein domains of PTPRK and ECHCD1. (D) VAPB-GNAS fusion. Top: diagram of minigenomes of VAPB and GNAS. Direction of transcription and distance between the 2 genes are indicated. Mid: mRNA represented by exons from each gene. Bottom: diagram of functional protein domains of VAPB and GNAS.

STAMBPL1-FAS fusion isoform was identified in sample 1M (metastasis cancer) and 2M (the metastasis cancer): XM_011539985 from STAMBPL1 and XM_011539766.2 from FAS (Figure 4A). However, subsequent analysis using Taqman qPCR showed that the STAMBPL1-FAS fusion isoform can be found in all 6 cancer samples (Supplemental Figure S8), implying a wider distribution of this gene fusion in colon cancers. STAMBPL1 is a deubiquitinase involving NF-kB activation and the inhibition of apoptosis^19^. FAS, on the other hand, is a cell surface death receptor^20^. The FAS isoform (XM_011539766.2) in the fusion protein contains the transmembrane domain and death domain in the cytoplasmic portion of the protein. The fusion isoform is a chimera of largely the intact metalloprotease domain from STAMBPL1 fused with the extracellular domain of FAS. Since FAS is a transmembrane protein while STAMBPL1 is an endosomal one, it is unclear where the ultimate subcellular localization of the chimera protein is or whether the activity of FAS is neutralized by STAMBPL1. In contrast, both the PTPRK-ECHDC1 gene fusion isoform and the ZNF124-SMYD3 gene fusion isoform introduced a frame-shift from the fusion event (Figure 4B and C, Supplementary Figure S8). As a result, only truncated PTPRK and ZNF124 proteins are produced. The tail gene expressions are eliminated. GNAS is a component of the guanine nucleotide binding protein^21^, and is frequently mutated in colon and pancreatic cancers^22^. In our analysis, GNAS forms a fusion gene with VAPB^23^ (Figure 4D and Supplementary Figure S8). The fusion generated a chimeric protein resulting in a loss of the regulatory domain in the N-terminus of GNAS while leaving the G-binding domain intact. The chimera protein may have a new function in the cancer cells.

## Discussion

The survival of eukaryotic cells requires rapid adaptability to the constant changes of their environment. By utilizing different combinations of exons of a gene, eukaryotic cells generate an array of different isoforms and proteins from a single gene, which allow them to cope with the challenge of the environment^24^. Even though alternative splicing of mRNA has long been known and is robust in most genes, little is known about the isoform distribution in cells. Isoform quantification has been hampered by the complexity of alternative splicing and the lack of adequate tools to identify the different isoforms. In previous studies, long read sequencing technologies from Pacific Biosciences (PacBio) or Oxford Nanopore (ONT) were used to sequence full-length transcripts, either on cDNA synthesized from RNA or directly on RNA molecules^25^. However, the single long-molecule readout offered by PacBio SMRTseq must be accompanied by short read RNAseq for error correction, effectively requiring independent rounds of sequencing per sample. Without short reads for correction, downstream informatics processing is needed, with varying levels of error reduction^12,26^. ONT has native RNA read ability, skipping the need for cDNA library construction, but is hampered by low throughput, high error rate, and incomplete read lengths^27,28^. The relatively high error rates of both long-read sequencing methods hamper their usefulness in accurate isoform mapping and quantification. In comparison, LoopSeq combines the low error rate of Illumina short reads with long-molecule data that facilitates transcriptome profiling and isoform discovery. Short read densities that cover each contig/UMI allow for error correction by base-pair consensus. Additionally, UMI tagging enables accurate assessment of relative transcript abundance. Our results demonstrate that isoform characterization with LoopSeq enables obtaining detailed granularity in isoform expression regulation, isoform-specific mutation expression and fusion gene isoform expression that were previously inaccessible.

Comprehensive quantification of isoform-specific mutations were seldom performed, even though mutations at the genome levels have been extensively studied in human cancers. Our analysis showed that most mutations are not evenly distributed among the isoforms, but rather are dominantly present in some isoforms. A significant number of mutations undergo a change in isoform distribution as the cancer evolves. These changes in the expression of dominant mutation isoform may bear important clinical significance, as it may alter signaling pathways and adapt cancer cells to their new environment. Lastly, drug targeting design relies on accurate assessment of the physical interaction between the drug and the structure of the protein target. Subtle variations in the amino acid sequences among different protein isoforms can have a profound impact on the interaction between the drug and its targets. Isoform switching can also impact the signaling mechanism of cancer driver proteins, leading to resistance to cancer treatment^29,30^. The ability to accurately characterize and quantify isoforms of target proteins will undoubtedly provide new insight in cancer drug design and shed light into the mechanism of cancer drug resistance.

## Acknowledgement

The work was supported in part by grants from National Cancer Institute (1R56CA229262-01 to JHL), Department of Defense (W81XWH-16-1-0541 to JHL) and University of Pittsburgh Medical Center (Endowed Chair of Molecular Carcinogenesis).

## Methods

### Samples collection for internal quality control

ERCC synthetic RNA and Hela Total RNA were obtained from ThermoFisher Scientific, Inc., and used as internal quality control to demonstrate the error rate, chimera rate, and transcript quantification using LoopSeq.

### Colon cancer sample collection and RNA extraction

Frozen tissue samples were collected from 3 colon cancer patients, including benign colon tissue adjacent to cancer samples, primary colon cancer samples, and lymph node metastasis samples. The procedure of obtaining the tissues was approved by the institutional review board of University of Pittsburgh. Total RNA was extracted using TRIzol (Invitrogen, CA) and RNeasy column methods. The methods were described in several previous publications^31-39^.

### Target sequence selection for Loop Sequencing

Fusion transcript candidates were selected from University of Pittsburgh Medical Center (UPMC) cohort and TCGA database. For the UPMC cohort, 14 novel fusion transcripts were detected by our previous study^36,38^. For the TCGA panel, a list of 17754 fusion transcripts were downloaded from Tumor fusion gene data portal (https://www.tumorfusions.org/). Based on the Cancer gene census (https://cancer.sanger.ac.uk/census), 315 oncogenes and 315 tumor suppressor genes (TSGs) were defined. Eventually, 2135 fusions were selected by satisfying one of the following criteria: (1) Fusions can be detected in more than one sample; (2) Two genes involved in one fusion transcripts are either oncogenes or TSGs; (3) Only one fusion gene is a oncogene or TSG, and the fusion event is either in-frame or out-of-frame. To sum up the two cohorts, 2149 fusion transcripts in total were selected (Supplementary Table S1: Target sequence selection), which involved 2193 unique genes. The 100bp sequences surrounding the junction point (50bp upstream to the junction and 50bp downstream to the junction) were extracted and used for enrichment. The oligonucleotides were provided by Twist Bioscience, Inc., CA.

### LoopSeq sequencing library preparation

LoopSeq sequencing libraries were prepared from ERCC synthetic RNA, a blend of ERCC synthetic RNA and Hela Total RNA, or Total RNA extracted from tissue samples using LoopSeq Transcriptome Kit according to manufacturer’s protocol except when specified. Two hundred nanograms of total RNA extracted from cancer patient tissues was reverse transcribed and barcoded using LoopSeq Transcriptome Kit, and an enrichment step using Twist Biosciences probes targeting 2149 fusion junction sequences (selected in previous step) was applied prior to Barcode Amplification in LoopSeq library prep. Specifically, the barcoding reaction was purified using 0.6X SPRISelect ratio and then eluted into 20 μL of Hybridization Mix per manufacturer’s pre-capture concentration protocol. The hybridization reactions contain 20 μL of Hybridization Mix, 5 μL of Blocker Solution, 9 μL of LG Adaptor Blocker (Loop Genomics), 4 μL of Twist Custom Enrichment Panel, 2 μL of Buffer EB (Qiagen), and 30 μL of Hybridization Enhancer. All probe hybridization and washes were conducted per manufacturer’s protocol. All target enrichment reagents are from Twist Biosciences except when specified. The captured barcoded cDNA was amplified following the LoopSeq Transcriptome protocol starting from Barcode Amplification.

### Probe hybridization analysis

Selected bases on-target rate is computed by aligning the contigs to GRCh37 reference genome using minimap2 (Li 2018) and comparing the alignment against the probe BED file using Picard. As shown in supplementary table S12, the on-target reads of the transcriptome sequencing ranged from 83% to 92% in long-read level, and 93-96% in base level per sample, indicating a high level of enrichment by the probes. In addition, two known fusion genes (AGRN-NOC4L and CTNND1-TMX2) in the cancer sample (3T) were enriched and detected in the dataset, suggesting that the probe design for the fusion transcript is adequate.

### Isoform expression analysis

Transcriptome samples of 8 samples across 3 colon cancer patients were measured by LoopSeq technology, including 2 benign colon tissue samples (1N and 3N), 3 tumor samples (1T, 2T and 3T) and 3 metastasis samples (1M, 2M and 3M). LoopSeq long-molecules were analyzed by tool SQANTI for transcriptome isoform identification and quantification^40^. Isoforms were quantified by read count across all the eight samples. Differential expression analysis was performed by R tool ‘DEseq2’^41^ to compare normal samples versus tumor samples, and tumor samples versus metastasis samples respectively. Top differentially expressed isoforms (DEIs) were selected by false discovery rate FDR=5% and absolute fold change greater than 2-fold. Then hierarchical clustering^42^ was applied on these 8 samples based on the DEIs pooling from the two comparisons. Besides isoform level analysis, similar analysis was performed at gene level to detect differentially expressed genes (DEGs) and cluster samples based on these DEGs. To compare isoform-level and gene-level analysis, these DEIs/DEGs were categorized into three groups: DEG only, DEG/DEI intersect, and DEI only. Hierarchical clustering, principal component analysis (PCA) and Pearson correlation analysis was performed based on genes/isoforms within each category.

### SNV isoform analysis

Loop-seq long-reads were aligned to the human reference genome hg38 by Minimap2 aligner^43^. For each sample, mutations/SNPs were called and quantified by SAMtools mpileup function^44,45^. These mutations/SNVs were then annotated by ANNOVAR tool, dbSNP and Cosmic database to identify known SNPs and somatic mutations in human cancer^46-48^. Those detected SNVs were further filtered by the following criteria: (1) validated by WES (see next section) and (2) either stop-gain or non-synonymous mutations/SNVs. For a given SNV position of interest, reads with reference base (wild type) and alteration base were able to be identified and annotated at isoform level by SQANTI^40^.

Several statistical tests were applied to the SNV isoform data. (1) When defining the unevenly distributed SNV isoforms per gene, only SNVs involved with more than one isoforms and covered by at least 5 contigs were analyzed by Chi-squared test. p-values were adjusted by Benjamini-Hochberg (BH) method and FDR=5% were applied to define significant unevenly distributed SNV isoforms. (2) When detecting differentially expressed SNV isoforms between reference (wild-type) and altered alleles, Fisher’s exact tests were applied to test the long-molecule read count of reference/altered alleles across multiple SNV isoforms. BH adjustment and FDR=5% were applied. (3) When comparing tumor samples and metastasis samples, SNV rate was defined as SNV count divided by total count (SNV count + wild type count). Based on this, SNV isoforms with SNV rate low in tumor samples and high in metastasis samples were detected. Similar or the other way around. Hierarchical clustering, PCA and Pearson correlation analysis were applied on these selected switching isoforms. We further applied these top switching isoforms for IPA pathway analysis (QIAGEN Inc., https://www.qiagenbioinformatics.com/products/ingenuitypathway-analysis). Significant pathways were visualized by network plot drawn by Cytoscape^49^.

### Single nucleotide variant calling from WES as validation

Whole Exome Sequencing (WES) was performed on the same three individuals for mutation validation at DNA level. Illumina TruSeq Exome kit was used to prepare the exome DNA libraries of 2T, 2M, 1T, 1M, 3T and 3M. The genome DNA was sheared to 150 bp using Covaris sonicator. The fragmented DNA was end-repaired, polyadenylated and ligated with Illumina adaptors. The adaptor ligated DNA will be amplified by PCR for 8 cycles in the following condition: 98°C for 20 seconds, 60°C for 15 seconds and 72°C for 30 seconds. The amplified libraries were then pooled and bound with Coding Exome oligos. The hybrids were then captured by Streptavidin Magnetic Beads provided by Illumina, inc. The beads were then washed. The captured libraries were eluted. The capture procedure was repeated once. The eluted libraries were amplified for 8 cycles in the same condition as above. The libraries were cleaned up and assessed for quantity and quality based on Agilent’s Bioanalyzer 2000 and Qbit. The libraries were sequenced on a Illumina NextSeq Dx550 sequencer.

For raw sequencing reads pre-processing, quality control was applied by tool FastQC (https://qubeshub.org/resources/fastqc). Low-quality reads and adapter sequences were trimmed by tool Trimmomatic^50^. Surviving reads were then mapped to human reference genome hg38 by Burrows-Wheeler Aligner^51^. Aligned reads were sorted and marked duplicates by tool Picard (http://broadinstitute.github.io/picard). Mutation/SNP calling on individual samples were performed by SAMtools mpileup function^44,45^.

### Amplicon sequencing validation of mutation isoform expression

Transcriptome sequencing on the same samples as loop-seq were performed for mutation isoform validation. Amplicon sequencing was specifically targeted on two candidate genes: FAM104A and PABPC1 using primer sets ACAACCCCCTCTGTTCCCTCT/ATGGTCTGGCTCAAGCTGCCT for FAM104A and AGCAAATGTTGGGTGAACGGC/TTCTTCGGTGAAGCACAAGTTTC.

For bioinformatics analysis, raw sequencing reads first went through the pipeline of quality control by FastQC and then low-quality reads and adapter sequences were filtered out by tool Trimmomatic^50^. Surviving reads were then aligned to human reference genome hg38 by HISAT2 aligner^52^. Mutation calling was performed by SAMtools mpileup function^44,45^ and isoform identifications were supported by the reads exactly splitting across more than one exon and their counterpart paired-ends spanning.

### Taqman qRT-PCR to validate isoform expressions

Two micrograms of RNA were used to synthesize first-strand cDNA with random hexamer primers and Superscript II™ (Invitrogen, CA). For NM_000701.8//XR_002956654.1 of ATP1A1 detection, one microliter of each cDNA sample was used for TaqMan PCR with 50 heating cycles at 94°C for 30 seconds, 61°C for 30 seconds, and 72°C for 30 seconds using the primers and probes listed in Supplementary Table S12 (Primers and probes design). At least one negative control and a synthetic positive control were included in each reaction batch. The PCR products were gel purified, and Sanger sequencing was performed on the positive samples. The procedure of fusion gene validation also followed the similar process except using the primers and probes listed in Supplementary Table S13 (Primers and probes design).

### Fusion Transcript detection

Fusion transcripts were detected from the LoopSeq long-molecules in two methods. For the first method, 2149 targeted fusion junctions (Supplementary Table S1) were specifically checked. Text search based on the 15bp ahead of and 15bp right after the junction point were applied to the long-read molecules. For the second method, SQANTI tool^40^ were applied to for novel fusion isoforms detection.

### Data Availability

Data for LoopSeq quality control, LoopSeq colon cancer samples and RNA amplicon sequencing were submitted to the GEO database, which can be accessed by GSE155375. Raw whole exome sequencing data for colon cancer samples were submitted to the SRA database with Bioproject accession number PRJNA648918.

