## Supplementary material for "Targeted Transcriptome Analysis using Synthetic Long Read Sequencing Uncovers Isoform Reprograming in the Progression of Colon Cancer": legends

**Supplementary Tables**

Supplementary Table S1: Target sequence selection.

Supplementary Table S2: Gene short-read count.

Supplementary Table S3: Differentially expressed genes.

Supplementary Table S4: Differentially expressed isoforms.

Supplementary Table S5: Differentially expressed genes and isoforms.

Supplementary Table S6: Total SNVs identified in colon cancer samples.

Supplementary Table S7: Unevenly distributed SNV isoforms within a gene.

Supplementary Table S8: Differential expression of SNV isoforms compared with wild type counterparts.

Supplementary Table S9: SNV isoform validation via targeted RNA sequencing. S10A: Gene FAM104A. S10B: Gene PABPC1.

Supplementary Table S10: Switching SNV isoforms between tumor and metastasis.

Supplementary Table S11: Colon cancer related mutations/SNVs.

Supplementary Table S12: LoopSeq targeted sequencing on-target rate

Supplementary Table S13: Primers and probes design.

**Supplementary Figure S1. Isoform expression of CD44.** Isoform distribution of CD44 is normalized to FPKM scale. Arrows indicate isoforms that are overexpressed in cancers and metastases.

**Supplementary Figure S2. Isoform expression of ATP1A1.** Isoform distribution of ATP1A1 is normalized to FPKM scale. Arrows indicate isoforms that are overexpressed in cancers.

**Supplementary Figure S3. Taqman qRT-PCR to quantify isoforms of ATP1A1.** Five sets of primers and probes for the indicated isoforms were designed. Taqman qRT-PCRs were performed triplicate to quantify the isoform expression. The results were normalized to the mRNA of β-actin.

**Supplementary Figure S4. Non-switching SNV isoforms do not segregate primary cancer samples and metastasis samples.** Hierarchical clustering between primary colon cancers and metastatic colon cancers based on the quantities of total (top) or non-switching (bottom) non-synonymous single nucleotide variants of all isoforms in each sample. (B) Principal component analyses of primary colon cancers and metastatic colon cancers based on the quantities of total (top) or non-switching (bottom) non-synonymous single nucleotide variants of (A). (C) Pearson's correlation of primary colon cancers and metastatic colon cancers based on the quantities of total (top) or non-switching (bottom) non-synonymous single nucleotide variants of (A).

**Supplementary Figure S5. Isoform distribution of V600E of BRAF in colon cancer.**

**Supplementary Figure S6. Isoform distribution of G12V of KRAS in colon cancer.**

**Supplementary Figure S7. Schematic diagram of fusion gene screening criteria.**

**Supplementary Figure S8. Validation and screening analyses of STAMBPL1-FAS, ZNF124-SMYD3, PTPRK-ECHDC1 and VAPB-GNAS.** Taqman qRT-PCRs were performed on benign colon tissues adjacent cancer, colon cancer and lymph node metastasis samples using the primers and probes described in the methods. Taqman qRT-PCRs on β-actin are the controls. The positions of primers and probes are indicated.

#### Supplemental figure 1

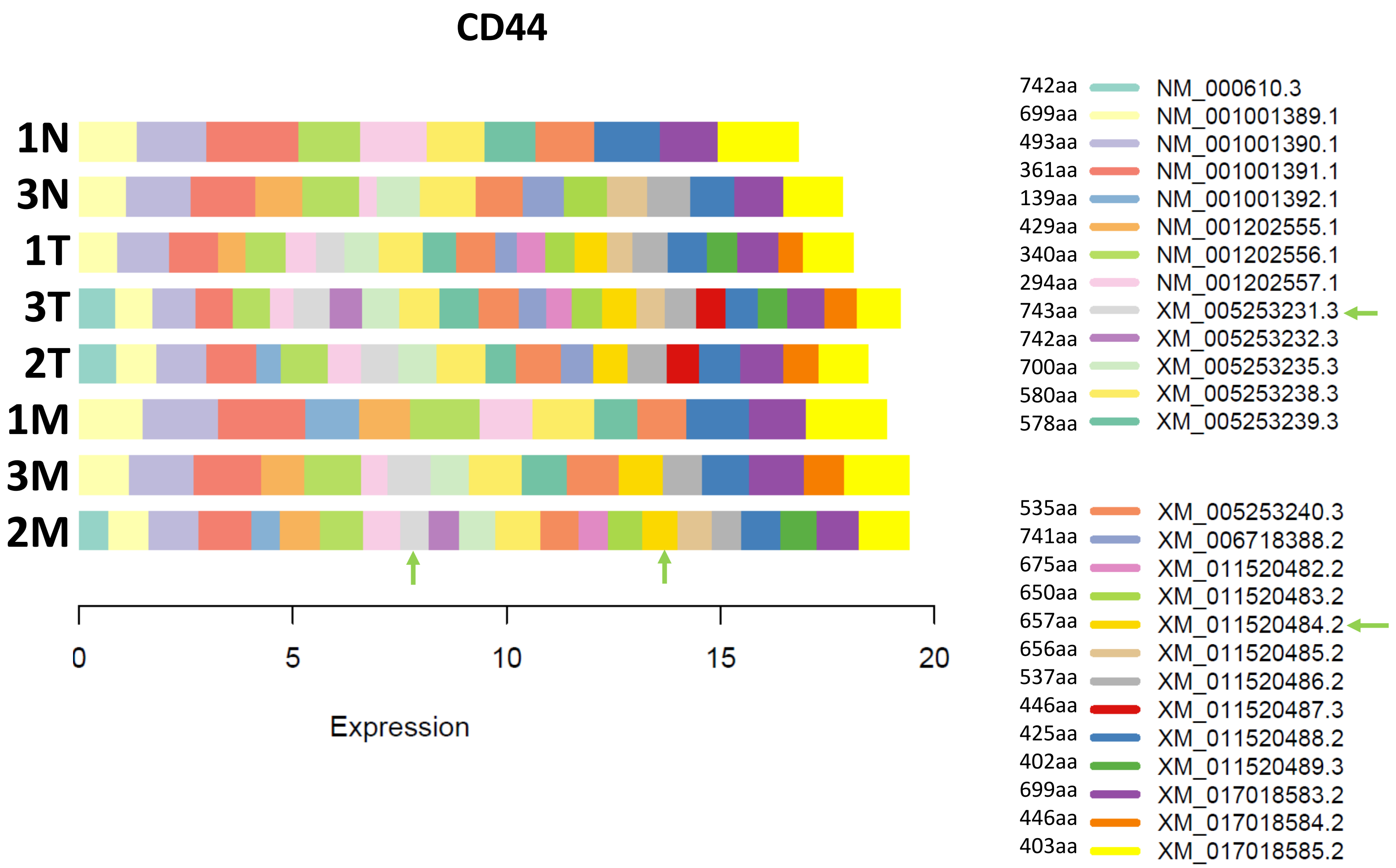

**Supplemental figure 2**

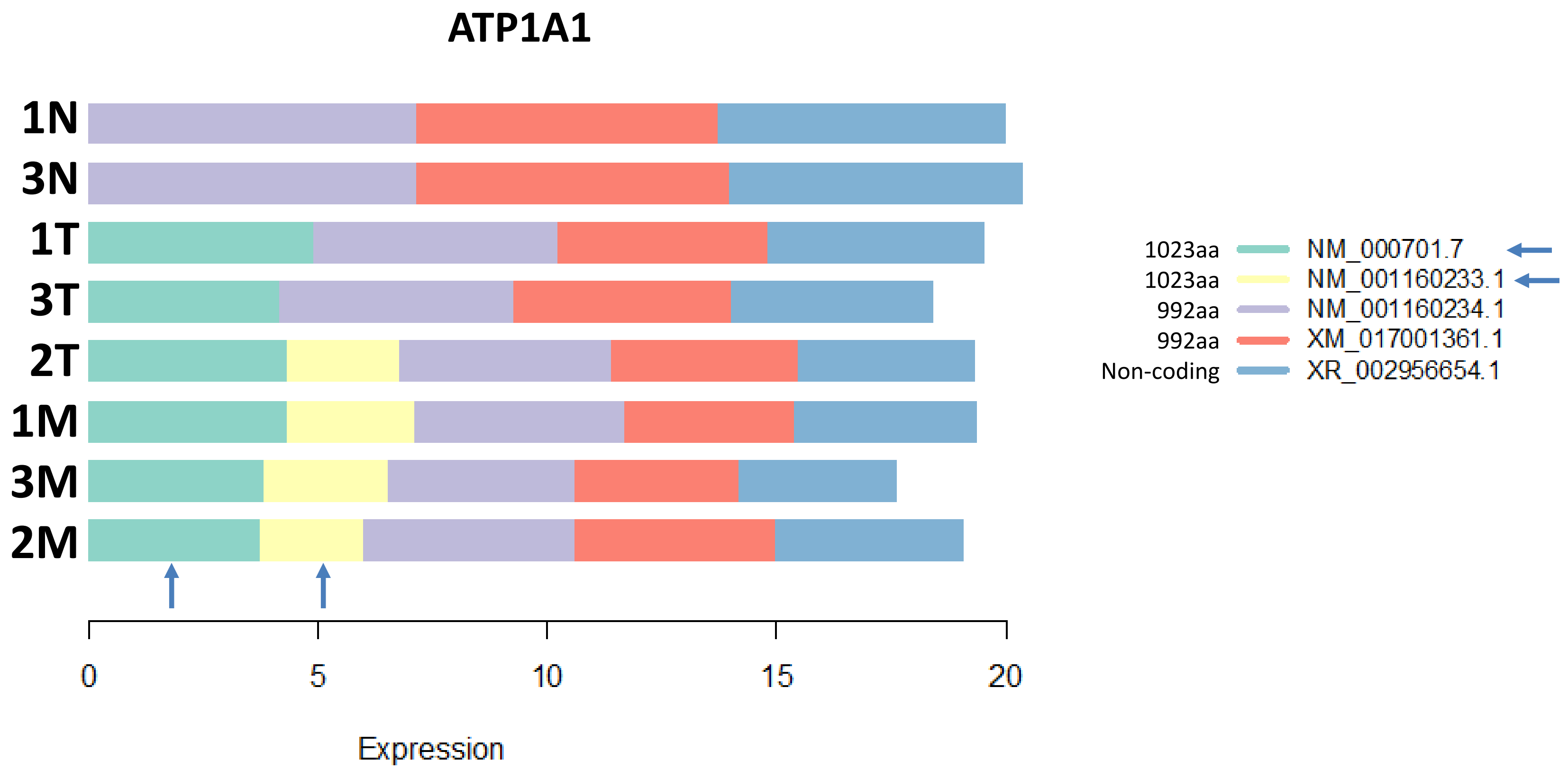

### Supplemental figure 3

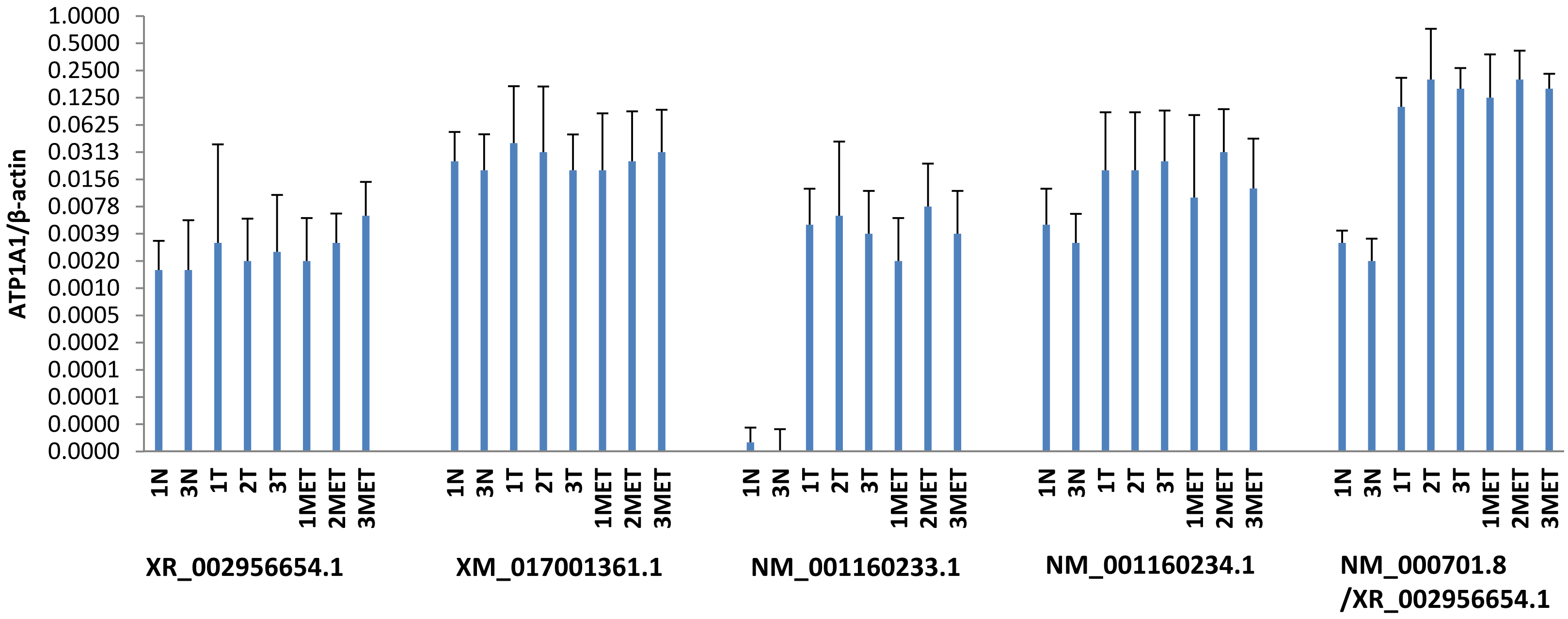

### Supplemental figure 4

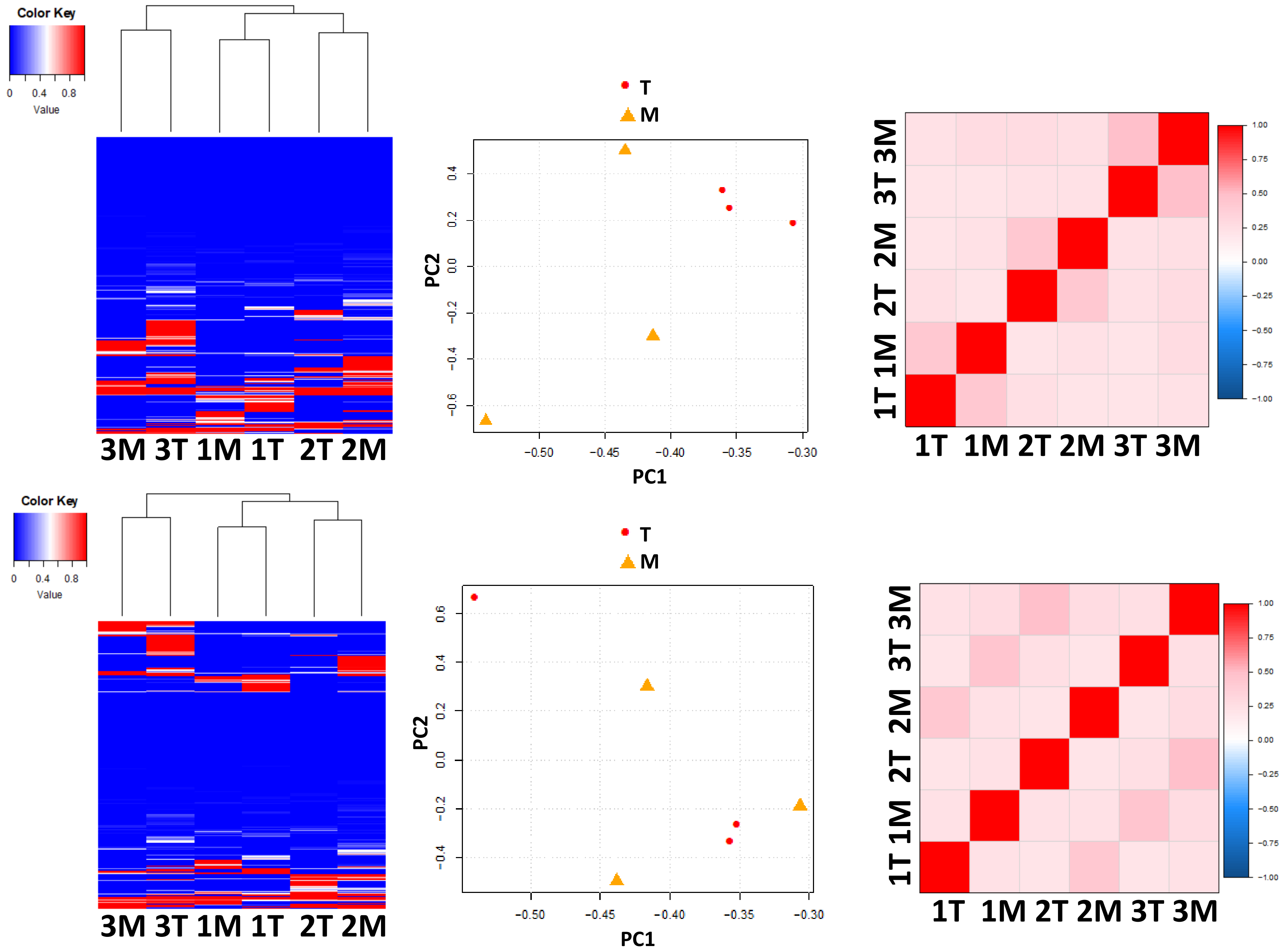

**Supplemental figure 5**

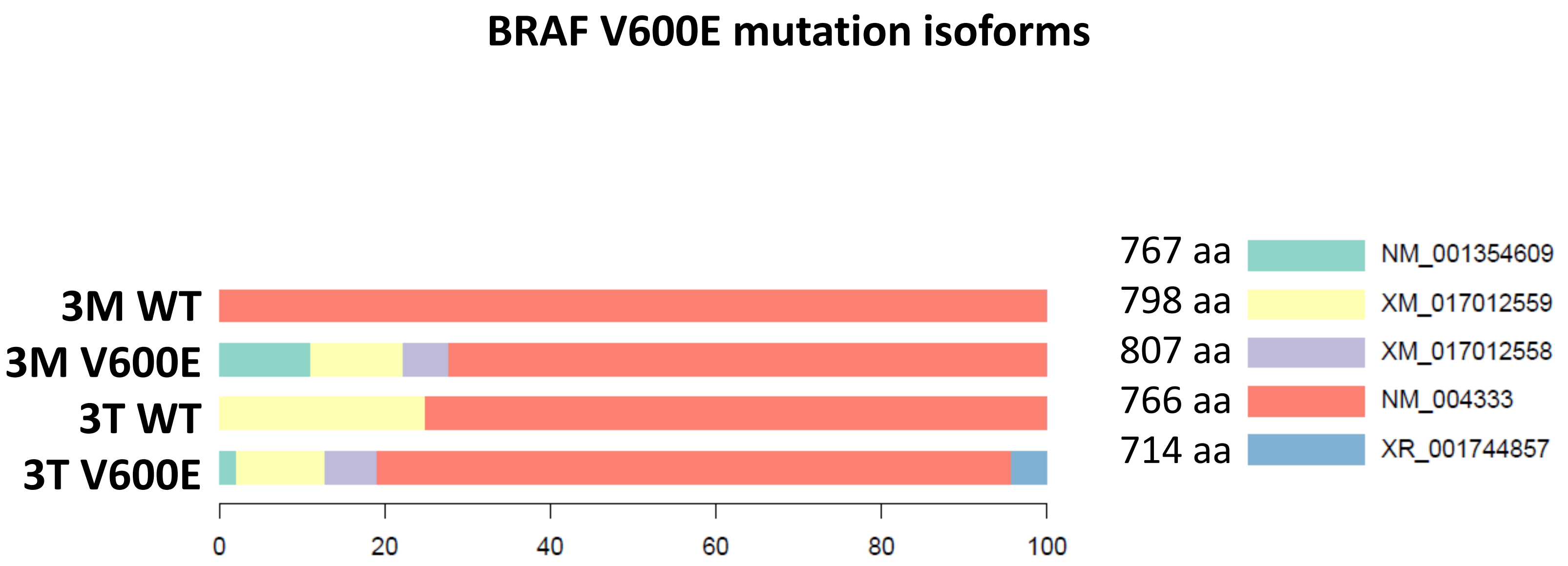

Supplemental figure 6

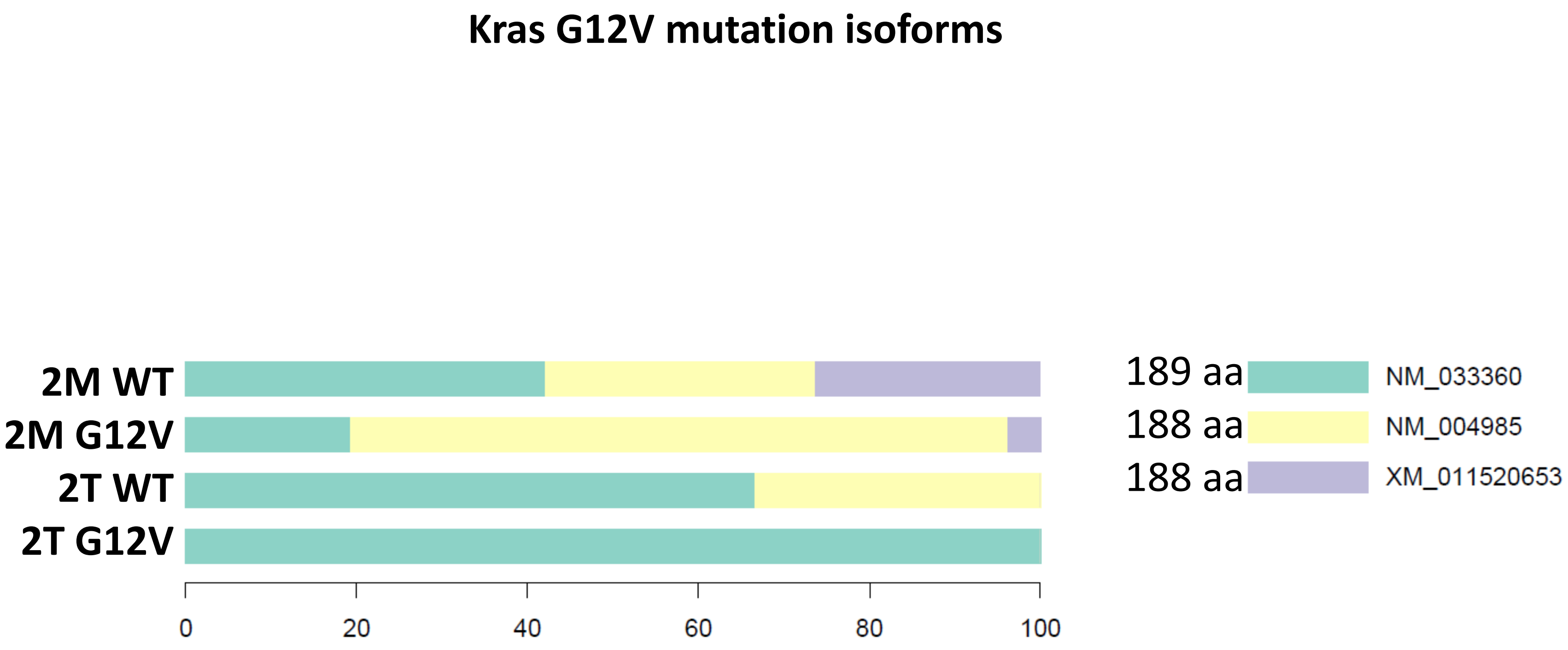

**Supplemental figure 7**

### Detection of fusion transcripts

#### Criteria for filtering chromosome rearrangement based fusion transcripts after SQANTI mapping

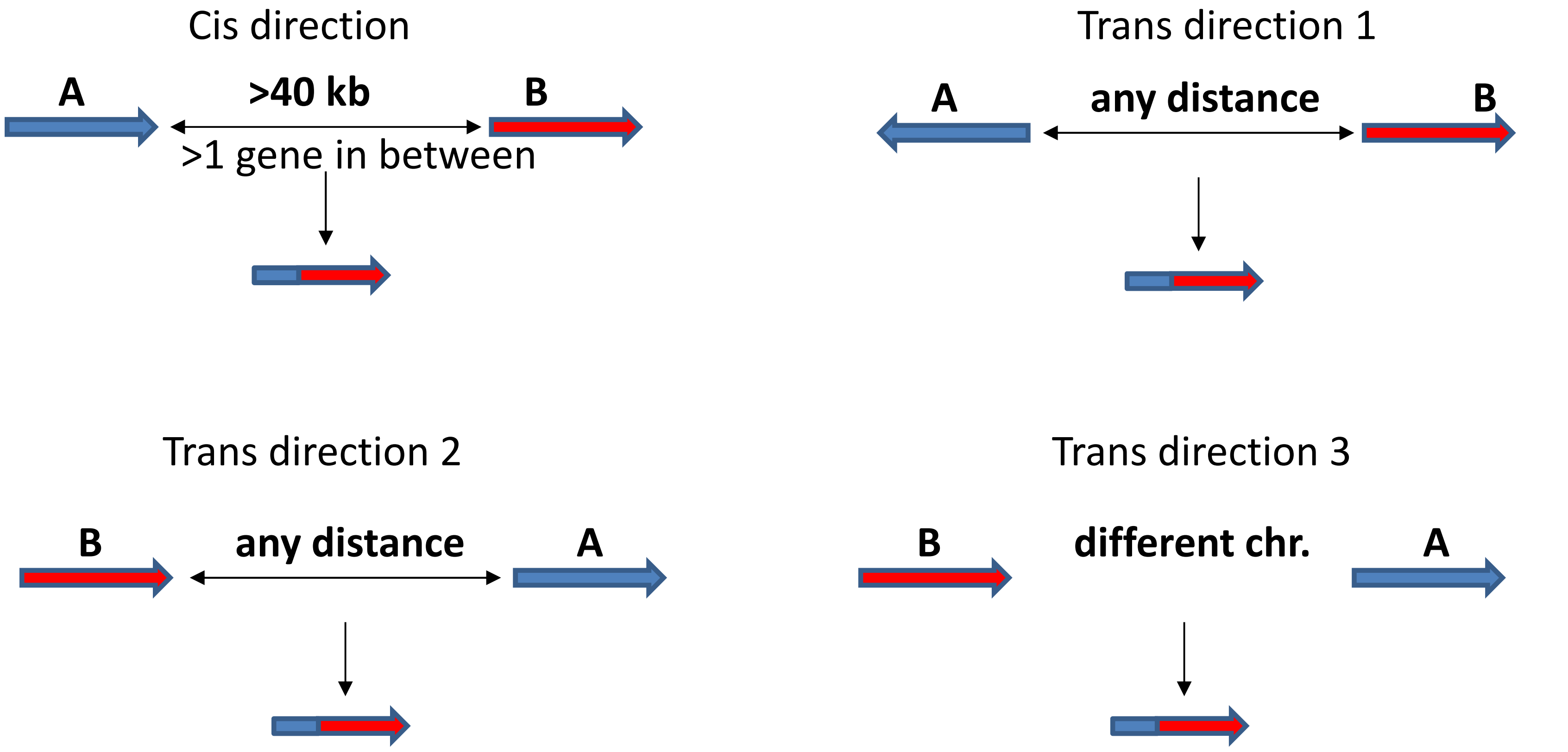

### Supplemental figure 8

#### STAMBPL1-FAS

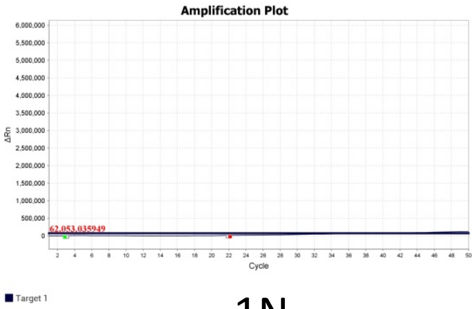

1N

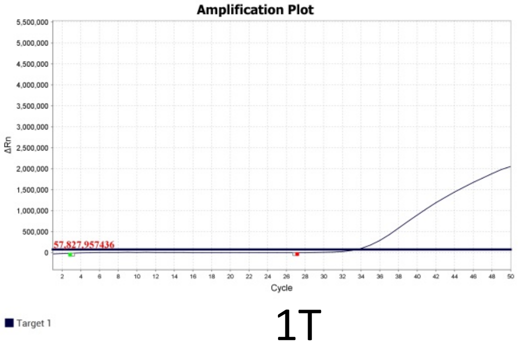

1T

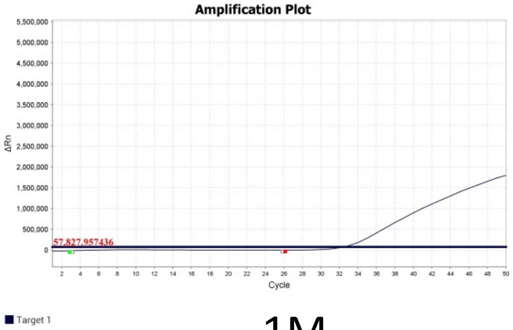

1M

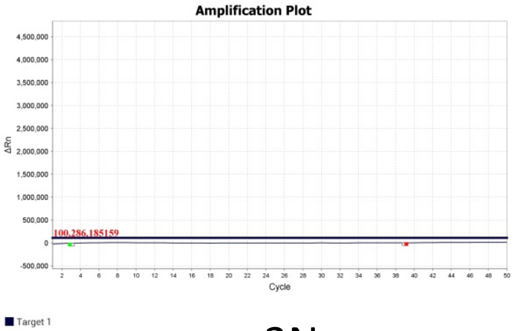

3N

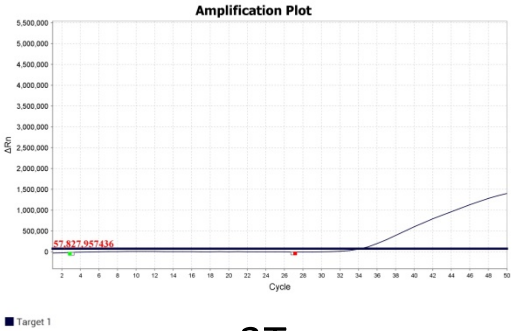

3T

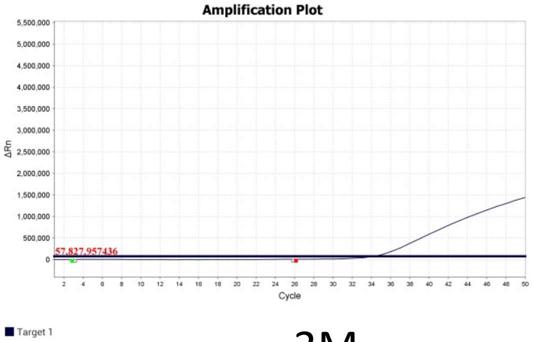

3M

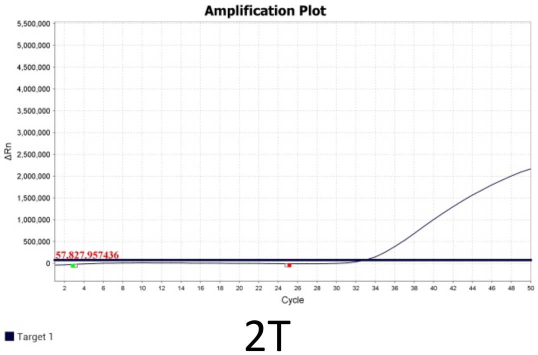

2T

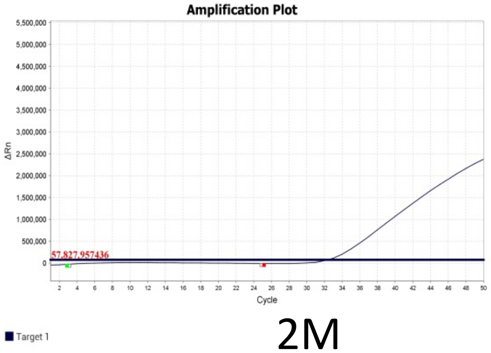

2M

### Supplemental figure 8

#### ZNF124-SMYD3

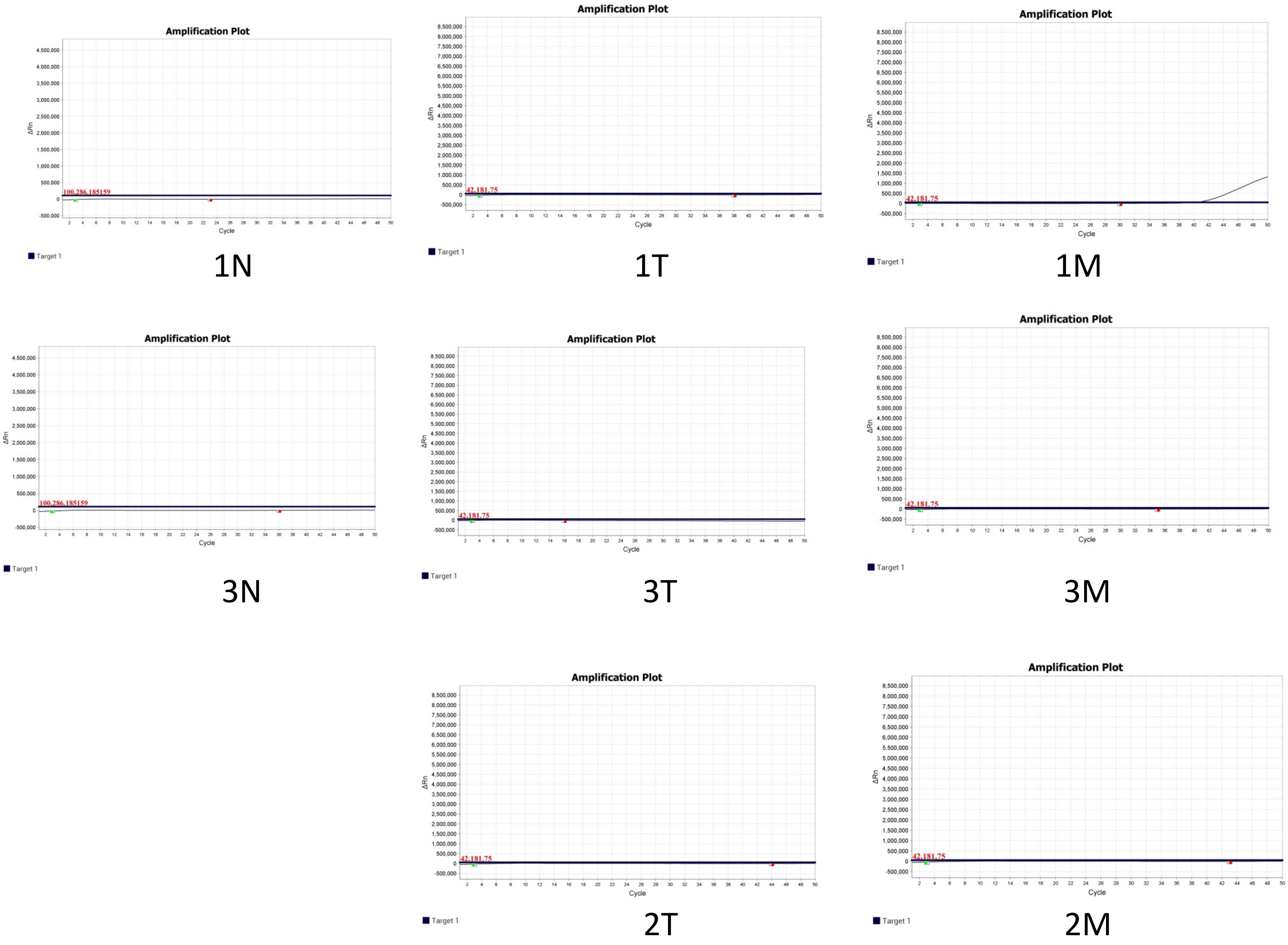

### Supplemental figure 8

#### PTPRK-ECHDC1

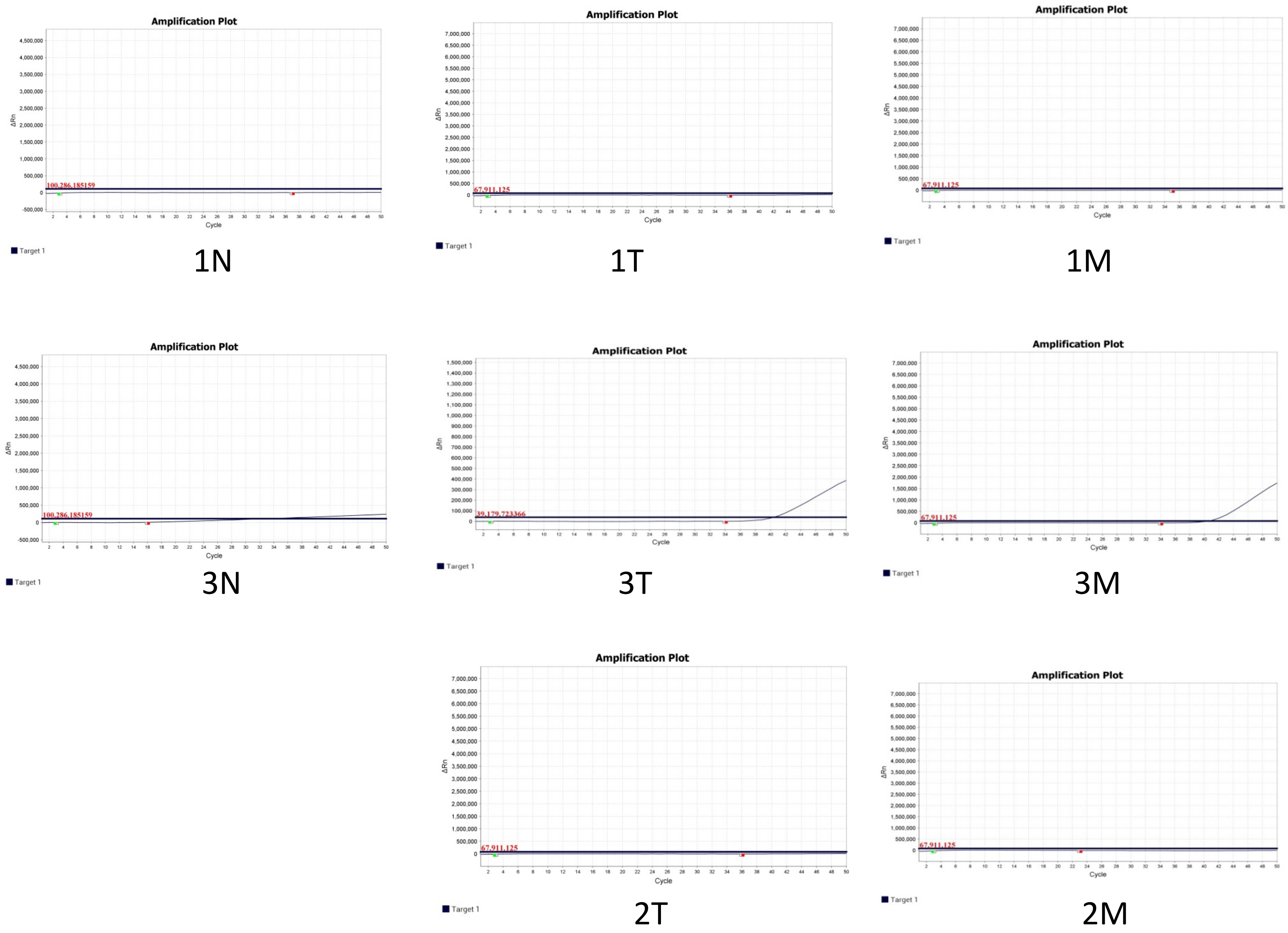

### Supplemental figure 8

#### VAPB-GNAS

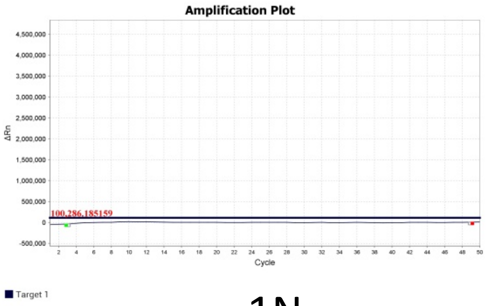

1N

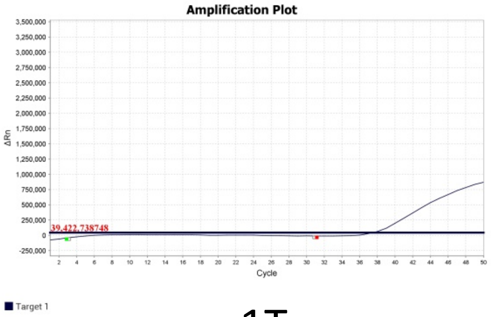

1T

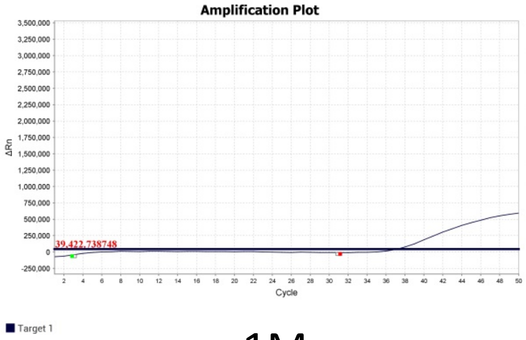

1M

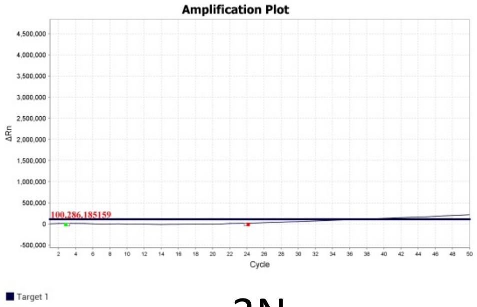

3N

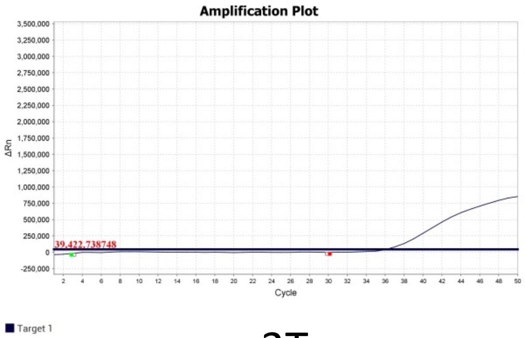

3T

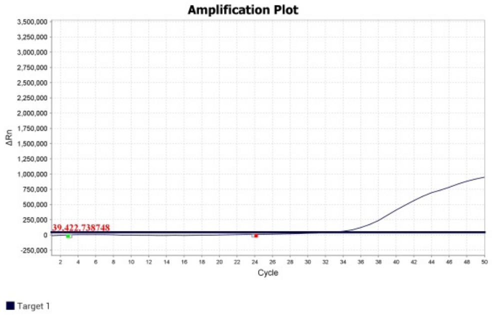

3M

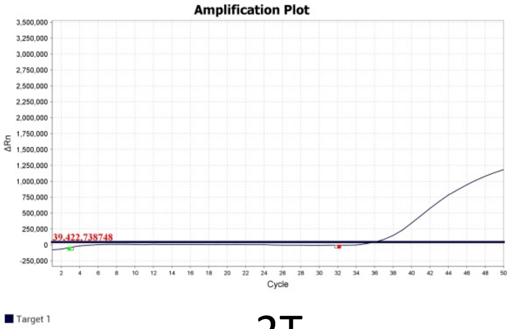

2T

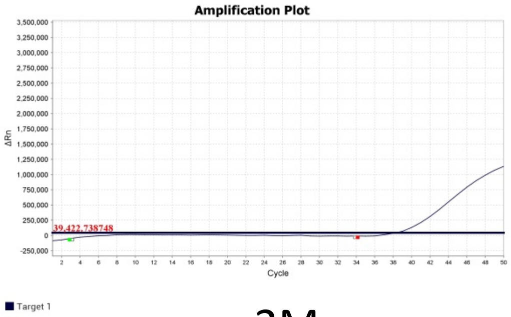

2M

### Supplemental figure 8

#### β-ACTIN

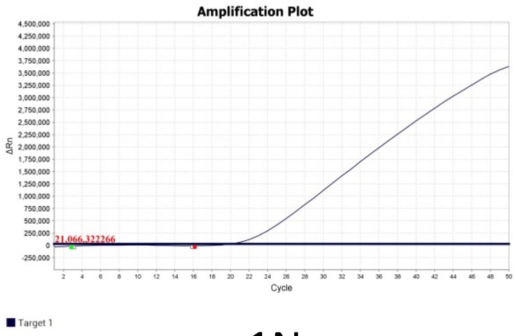

1N

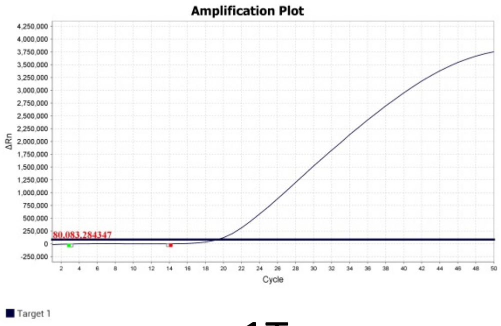

1T

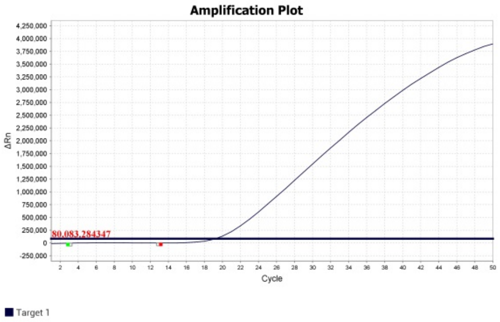

1M

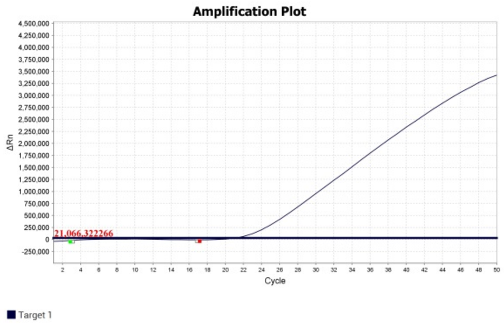

3N

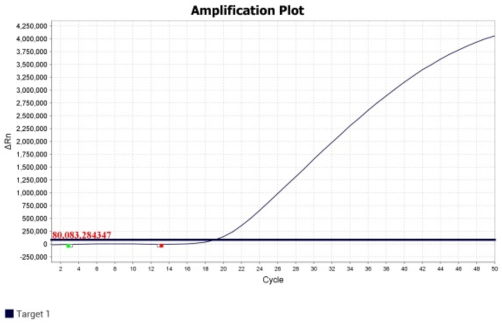

3T

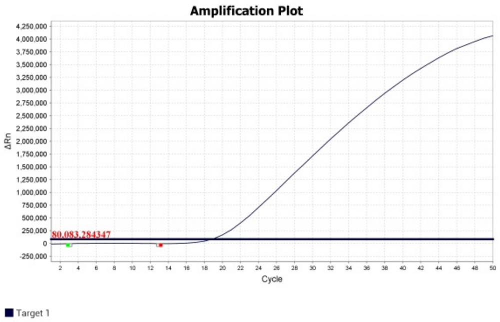

3M

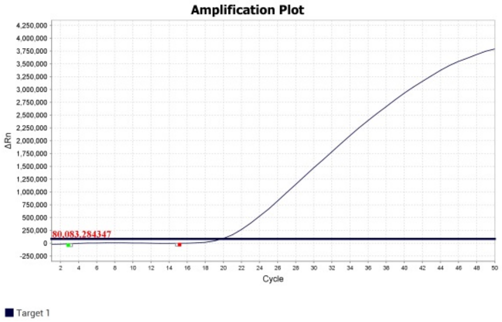

2T

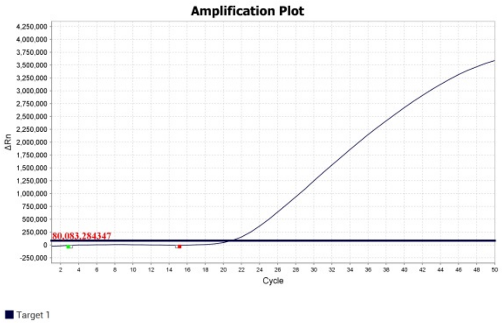

2M

Supplementary Table S1: Target sequence selection

| Gene_A | Gene_B | A_chr | B_chr | A_strand | B_strand | Junction_A | Junction_B | A_pos | B_pos |
|---|---|---|---|---|---|---|---|---|---|
| MAN2A1 | FER | 5 | 5 | 1 | 1 | 109153139 | 108380381 | MAN2A1::chr5:109153089-109153139(+) | FER::chr5:108380380-108380430(+) |
| TMEM135 | CCDC67 | 11 | 11 | 1 | 1 | 87030419 | 93127625 | TMEM135::chr11:87030369-87030419(+) | CCDC67::chr11:93127624-93127674(+) |
| LRRC59 | FLJ60017 | 17 | 11 | -1 | 1 | 48469759 | 63129852 | LRRC59::chr17:48469758-48469808(-) | FLJ60017::chr11:62987386-63137190(+) |
| TRMT11 | GRIK2 | 6 | 6 | 1 | 1 | 126307768 | 102069824 | TRMT11::chr6:126307718-126307768(+) | GRIK2::chr6:102069823-102069873(+) |
| MTOR | TP53BP1 | 1 | 15 | -1 | -1 | 11290982 | 43773220 | MTOR::chr1:11290981-11291031(-) | TP53BP1::chr15:43773170-43773220(-) |
| CCNH | C5ORF30 | 5 | 5 | -1 | 1 | 86697519 | 102601609 | CCNH::chr5:86697518-86697568(-) | C5ORF30::chr5:102601608-102601658(+) |
| SLC45A2 | AMACR | 5 | 5 | -1 | -1 | 33982341 | 34006004 | SLC45A2::chr5:33982340-33982390(-) | AMACR::chr5:34005954-34006004(-) |
| KDM4B | AC011523.2 | 19 | 19 | 1 | 1 | 5047680 | 51354167 | KDM4B::chr19:5047630-5047680(+) | AC011523.2::chr19:51354166-51354216(+) |
| PTEN | NOLC1 | 10 | 10 | 1 | 1 | 89720875 | 103916776 | PTEN::chr10:89720825-89720875(+) | NOLC1::chr10:103916775-103916825(+) |
| DOCK7 | OLR1 | 1 | 12 | -1 | -1 | 62941378 | 10324689 | DOCK7::chr1:62941377-62941427(-) | OLR1::chr12:10324639-10324689(-) |
| PCMTD1 | SNTG1 | 8 | 8 | -1 | 1 | 52758221 | 51085121 | PCMTD1::chr8:52758220-52758270(-) | SNTG1::chr8:51085120-51085170(+) |
| ACPP | SEC13 | 3 | 3 | 1 | -1 | 132047206 | 10347376 | ACPP::chr3:132047156-132047206(+) | SEC13::chr3:10347326-10347376(-) |
| ZMPSTE24 | ZMYM4 | 1 | 1 | 1 | 1 | 40747199 | 35879573 | ZMPSTE24::chr1:40747149-40747199(+) | ZMYM4::chr1:35879572-35879622(+) |
| CLTC | ETV1 | 17 | 7 | 1 | -1 | 57759200 | 13978871 | CLTC::chr17:57759150-57759200(+) | ETV1::chr7:13978821-13978871(-) |
| AAGAB | SMAD3 | 15 | 15 | -1 | 1 | 67500900 | 67457233 | AAGAB::chr15:67500899-67500949(-) | SMAD3::chr15:67457232-67457282(+) |
| ABCB8 | NOS3 | 7 | 7 | 1 | 1 | 150725697 | 150706018 | ABCB8::chr7:150725647-150725697(+) | NOS3::chr7:150706017-150706067(+) |
| ABHD6 | FLNB | 3 | 3 | 1 | 1 | 58242432 | 58104599 | ABHD6::chr3:58242382-58242432(+) | FLNB::chr3:58104598-58104648(+) |
| ABI1 | PTP4A3 | 10 | 8 | -1 | 1 | 27112067 | 142431488 | ABI1::chr10:27112066-27112116(-) | PTP4A3::chr8:142431487-142431537(+) |
| ABL1 | BCR | 9 | 22 | 1 | 1 | 133589842 | 23634728 | ABL1::chr9:133589792-133589842(+) | BCR::chr22:23634727-23634777(+) |
| ABL1 | FUBP3 | 9 | 9 | 1 | 1 | 133589842 | 133493184 | ABL1::chr9:133589792-133589842(+) | FUBP3::chr9:133493183-133493233(+) |
| ACACB | UNG | 12 | 12 | 1 | 1 | 109665294 | 109547634 | ACACB::chr12:109665244-109665294(+) | UNG::chr12:109547633-109547683(+) |
| ACOT9 | APOO | X | X | -1 | -1 | 23739998 | 23899069 | ACOT9::chrX:23739997-23740047(-) | APOO::chrX:23899019-23899069(-) |
| ACPP | ZNF462 | 3 | 9 | 1 | 1 | 132036420 | 109685635 | ACPP::chr3:132036370-132036420(+) | ZNF462::chr9:109685634-109685684(+) |
| ACSL3 | SGPP2 | 2 | 2 | 1 | 1 | 223725976 | 223386486 | ACSL3::chr2:223725926-223725976(+) | SGPP2::chr2:223386485-223386535(+) |
| ACTB | CELA3A | 7 | 1 | -1 | 1 | 5567255 | 22328149 | ACTB::chr7:5567254-5567304(-) | CELA3A::chr1:22328148-22328198(+) |
| ACTB | GNB2L1 | 7 | 5 | -1 | -1 | 5567255 | 180670914 | ACTB::chr7:5567254-5567304(-) | GNB2L1::chr5:180670864-180670914(-) |
| ACTB | TM4SF4 | 7 | 3 | -1 | 1 | 5567255 | 149192486 | ACTB::chr7:5567254-5567304(-) | TM4SF4::chr3:149192485-149192535(+) |
| ACTB | YWHAE | 7 | 17 | -1 | -1 | 5567255 | 1303494 | ACTB::chr7:5567254-5567304(-) | YWHAE::chr17:1303444-1303494(-) |
| ACTG1 | ANKS1A | 17 | 6 | -1 | 1 | 79476999 | 34977682 | ACTG1::chr17:79476998-79477048(-) | ANKS1A::chr6:34977681-34977731(+) |
| ACTG1 | CLCN3 | 17 | 4 | -1 | 1 | 79476999 | 170623929 | ACTG1::chr17:79476998-79477048(-) | CLCN3::chr4:170623928-170623978(+) |
| ACTG1 | STAT6 | 17 | 12 | -1 | -1 | 79476999 | 57496773 | ACTG1::chr17:79476998-79477048(-) | STAT6::chr12:57496723-57496773(-) |
| ACTN4 | EIF3K | 19 | 19 | 1 | 1 | 39138547 | 39123070 | ACTN4::chr19:39138497-39138547(+) | EIF3K::chr19:39123069-39123119(+) |
| ADAM15 | DCST1 | 1 | 1 | 1 | 1 | 155025253 | 155006448 | ADAM15::chr1:155025203-155025253(+) | DCST1::chr1:155006447-155006497(+) |
| ADD1 | FAM193A | 4 | 4 | 1 | 1 | 2845751 | 2733502 | ADD1::chr4:2845701-2845751(+) | FAM193A::chr4:2733501-2733551(+) |
| AFAP1 | AKR1C1 | 4 | 10 | -1 | 1 | 7774547 | 4975429 | AFAP1::chr4:7774546-7774596(-) | AKR1C1::chr10:4975428-4975478(+) |
| AFF1 | PTPN13 | 4 | 4 | 1 | 1 | 87869723 | 87556405 | AFF1::chr4:87869673-87869723(+) | PTPN13::chr4:87556404-87556454(+) |
| AGBL4 | FAF1 | 1 | 1 | -1 | -1 | 50489435 | 51121200 | AGBL4::chr1:50489434-50489484(-) | FAF1::chr1:51121150-51121200(-) |
| AGK | BRAF | 7 | 7 | 1 | -1 | 141255367 | 140494267 | AGK::chr7:141255317-141255367(+) | BRAF::chr7:140494217-140494267(-) |
| AGPAT3 | PDXK | 21 | 21 | 1 | 1 | 45323900 | 45153950 | AGPAT3::chr21:45323850-45323900(+) | PDXK::chr21:45153949-45153999(+) |
| AGRN | NOC4L | 1 | 12 | 1 | 1 | 970704 | 132635526 | AGRN::chr1:970654-970704(+) | NOC4L::chr12:132635525-132635575(+) |
| ALB | EXOSC9 | 4 | 4 | 1 | 1 | 74270123 | 122737927 | ALB::chr4:74270073-74270123(+) | EXOSC9::chr4:122737926-122737976(+) |

| | | | | | | | | | |
|---|---|---|---|---|---|---|---|---|---|
| ALDOA | NDUFV1 | 16 | 11 | 1 | 1 | 30081734 | 67375697 | ALDOA::chr16:30081684-30081734(+) | NDUFV1::chr11:67375696-67375746(+) |
| ALDOA | SIAE | 16 | 11 | 1 | -1 | 30081734 | 124543777 | ALDOA::chr16:30081684-30081734(+) | SIAE::chr11:124543727-124543777(-) |
| AMBRA1 | HARBI1 | 11 | 11 | -1 | -1 | 46615378 | 46625459 | AMBRA1::chr11:46615377-46615427(-) | HARBI1::chr11:46625409-46625459(-) |
| AMBRA1 | HARBI1 | 11 | 11 | -1 | -1 | 46615378 | 46637931 | AMBRA1::chr11:46615377-46615427(-) | HARBI1::chr11:46637881-46637931(-) |
| AMOTL1 | CCDC88B | 11 | 11 | 1 | 1 | 94587247 | 64118948 | AMOTL1::chr11:94587197-94587247(+) | CCDC88B::chr11:64118947-64118997(+) |
| ANKRD11 | CHMP1A | 16 | 16 | -1 | -1 | 89484692 | 89720331 | ANKRD11::chr16:89484691-89484741(-) | CHMP1A::chr16:89720281-89720331(-) |
| AP2M1 | SEC63 | 3 | 6 | 1 | -1 | 183901878 | 108214782 | AP2M1::chr3:183901828-183901878(+) | SEC63::chr6:108214732-108214782(-) |
| APIP | S100A13 | 11 | 1 | -1 | -1 | 34937775 | 153599009 | APIP::chr11:34937774-34937824(-) | S100A13::chr1:153598959-153599009(-) |
| APLP2 | C7 | 11 | 5 | 1 | 1 | 130014699 | 40954982 | APLP2::chr11:130014649-130014699(+) | C7::chr5:40954981-40955031(+) |
| APLP2 | EIF2AK2 | 11 | 2 | 1 | -1 | 130014699 | 37342384 | APLP2::chr11:130014649-130014699(+) | EIF2AK2::chr2:37342334-37342384(-) |
| APOA2 | ITPR2 | 1 | 12 | -1 | -1 | 161192082 | 26775338 | APOA2::chr1:161192081-161192131(-) | ITPR2::chr12:26775288-26775338(-) |
| APOE | CCT5 | 19 | 5 | 1 | 1 | 45412650 | 10250033 | APOE::chr19:45412600-45412650(+) | CCT5::chr5:10250032-10250082(+) |
| ARHGDIA | SQSTM1 | 17 | 5 | -1 | 1 | 79825598 | 179250858 | ARHGDIA::chr17:79825597-79825647(-) | SQSTM1::chr5:179250857-179250907(+) |
| ARHGEF1 | SNRNP70 | 19 | 19 | 1 | 1 | 42399559 | 49604647 | ARHGEF1::chr19:42399509-42399559(+) | SNRNP70::chr19:49604646-49604696(+) |
| ARHGEF1 | UCKL1 | 19 | 20 | 1 | -1 | 42400555 | 62572561 | ARHGEF1::chr19:42400505-42400555(+) | UCKL1::chr20:62572511-62572561(-) |
| ARID1A | RNF31 | 1 | 14 | 1 | 1 | 27094490 | 24624366 | ARID1A::chr1:27094440-27094490(+) | RNF31::chr14:24624365-24624415(+) |
| ARL8B | ITPR1 | 3 | 3 | 1 | 1 | 5164273 | 4808214 | ARL8B::chr3:5164223-5164273(+) | ITPR1::chr3:4808213-4808263(+) |
| ARRB1 | GDPD5 | 11 | 11 | -1 | -1 | 75062632 | 75201365 | ARRB1::chr11:75062631-75062681(-) | GDPD5::chr11:75201315-75201365(-) |
| ASH2L | C20orf72 | 8 | 20 | 1 | 1 | 37971805 | 17968809 | ASH2L::chr8:37971755-37971805(+) | C20orf72::chr20:17968808-17968858(+) |
| ASXL1 | TM9SF4 | 20 | 20 | 1 | 1 | 30956926 | 30720816 | ASXL1::chr20:30956876-30956926(+) | TM9SF4::chr20:30720815-30720865(+) |
| ATF6 | ODZ3 | 1 | 4 | 1 | 1 | 161762117 | 183522077 | ATF6::chr1:161762067-161762117(+) | ODZ3::chr4:183522076-183522126(+) |
| ATL2 | HNRPLL | 2 | 2 | -1 | -1 | 38604285 | 38800569 | ATL2::chr2:38604284-38604334(-) | HNRPLL::chr2:38800519-38800569(-) |
| ATP10D | FAM133A | 4 | X | 1 | 1 | 47538955 | 92953396 | ATP10D::chr4:47538905-47538955(+) | FAM133A::chrX:92953395-92953445(+) |
| ATP11A | C13orf35 | 13 | 13 | 1 | 1 | 113439571 | 113337588 | ATP11A::chr13:113439521-113439571(+) | C13orf35::chr13:113337587-113337637(+) |
| ATP1A1 | DDX21 | 1 | 10 | 1 | 1 | 116947394 | 70723858 | ATP1A1::chr1:116947344-116947394(+) | DDX21::chr10:70723857-70723907(+) |
| ATRN | UQCRB | 20 | 8 | 1 | -1 | 3541552 | 97245455 | ATRN::chr20:3541502-3541552(+) | UQCRB::chr8:97245405-97245455(-) |
| B4GALT1 | BAG1 | 9 | 9 | -1 | -1 | 33166756 | 33262828 | B4GALT1::chr9:33166755-33166805(-) | BAG1::chr9:33262778-33262828(-) |
| BCOR | L3MBTL2 | X | 22 | -1 | 1 | 39931602 | 41605700 | BCOR::chrX:39931601-39931651(-) | L3MBTL2::chr22:41605699-41605749(+) |
| BCR | ABL1 | 22 | 9 | 1 | 1 | 23632600 | 133729451 | BCR::chr22:23632550-23632600(+) | ABL1::chr9:133729450-133729500(+) |
| BGN | CLCN3 | X | 4 | 1 | 1 | 152775004 | 170623929 | BGN::chrX:152774954-152775004(+) | CLCN3::chr4:170623928-170623978(+) |
| BGN | TTN | X | 2 | 1 | -1 | 152774998 | 179511286 | BGN::chrX:152774948-152774998(+) | TTN::chr2:179511236-179511286(-) |
| BPHL | C6orf125 | 6 | 6 | 1 | -1 | 3119081 | 33665527 | BPHL::chr6:3119031-3119081(+) | C6orf125::chr6:33665477-33665527(-) |
| BRAF | AGK | 7 | 7 | -1 | 1 | 140500162 | 141292946 | BRAF::chr7:140500161-140500211(-) | AGK::chr7:141292945-141292995(+) |
| BRAF | MRPS33 | 7 | 7 | -1 | -1 | 140624366 | 140710460 | BRAF::chr7:140624365-140624415(-) | MRPS33::chr7:140710410-140710460(-) |
| C10orf11 | ADK | 10 | 10 | 1 | 1 | 77198252 | 76285014 | C10orf11::chr10:77198202-77198252(+) | ADK::chr10:76285013-76285063(+) |
| C10orf26 | SUFU | 10 | 10 | 1 | 1 | 104503900 | 104352339 | C10orf26::chr10:104503850-104503900(+) | SUFU::chr10:104352338-104352388(+) |
| C10orf32 | C10orf26 | 10 | 10 | 1 | 1 | 104622689 | 104569650 | C10orf32::chr10:104622639-104622689(+) | C10orf26::chr10:104569649-104569699(+) |
| C10orf68 | CCDC7 | 10 | 10 | 1 | 1 | 32873232 | 32832228 | C10orf68::chr10:32873182-32873232(+) | CCDC7::chr10:32832227-32832277(+) |
| C10orf68 | CCDC7 | 10 | 10 | 1 | 1 | 32873232 | 32854486 | C10orf68::chr10:32873182-32873232(+) | CCDC7::chr10:32854485-32854535(+) |
| C19orf2 | PLEKHF1 | 19 | 19 | 1 | 1 | 30477324 | 30164731 | C19orf2::chr19:30477274-30477324(+) | PLEKHF1::chr19:30164730-30164780(+) |
| C5orf45 | APOA2 | 5 | 1 | -1 | -1 | 179263802 | 161193418 | C5orf45::chr5:179263801-179263851(-) | APOA2::chr1:161193368-161193418(-) |
| C7orf50 | MAD1L1 | 7 | 7 | -1 | -1 | 1166893 | 1855864 | C7orf50::chr7:1166892-1166942(-) | MAD1L1::chr7:1855814-1855864(-) |
| CALD1 | CGN | 7 | 1 | 1 | 1 | 134576424 | 151490982 | CALD1::chr7:134576374-134576424(+) | CGN::chr1:151490981-151491031(+) |

| | | | | | | | | | |
|---|---|---|---|---|---|---|---|---|---|
| CASP8 | CFLAR | 2 | 2 | 1 | 1 | 202098835 | 201994452 | CASP8::chr2:202098785-202098835(+) | CFLAR::chr2:201994451-201994501(+) |
| CATSPERB | TC2N | 14 | 14 | -1 | -1 | 92246920 | 92280169 | CATSPERB::chr14:92246919-92246969(-) | TC2N::chr14:92280119-92280169(-) |
| CBFB | MYH11 | 16 | 16 | 1 | -1 | 67116211 | 15814908 | CBFB::chr16:67116161-67116211(+) | MYH11::chr16:15814858-15814908(-) |
| CCDC6 | ANK3 | 10 | 10 | -1 | -1 | 61665880 | 61789584 | CCDC6::chr10:61665879-61665929(-) | ANK3::chr10:61789534-61789584(-) |
| CCDC6 | RET | 10 | 10 | -1 | 1 | 61665880 | 43612032 | CCDC6::chr10:61665879-61665929(-) | RET::chr10:43612031-43612081(+) |
| CCND3 | TRERF1 | 6 | 6 | -1 | -1 | 42016239 | 42268487 | CCND3::chr6:42016238-42016288(-) | TRERF1::chr6:42268437-42268487(-) |
| CD74 | ROS1 | 5 | 6 | -1 | -1 | 149784243 | 117645578 | CD74::chr5:149784242-149784292(-) | ROS1::chr6:117645528-117645578(-) |
| CD74 | SUPT16H | 5 | 14 | -1 | -1 | 149781200 | 21829491 | CD74::chr5:149781199-149781249(-) | SUPT16H::chr14:21829441-21829491(-) |
| CDC42EP4 | SDK2 | 17 | 17 | -1 | -1 | 71308032 | 71348776 | CDC42EP4::chr17:71308031-71308081(-) | SDK2::chr17:71348726-71348776(-) |
| CELF1 | BBOX1 | 11 | 11 | -1 | 1 | 47586826 | 27141196 | CELF1::chr11:47586825-47586875(-) | BBOX1::chr11:27141195-27141245(+) |
| CEP70 | TNFSF10 | 3 | 3 | -1 | -1 | 138216873 | 172227111 | CEP70::chr3:138216872-138216922(-) | TNFSF10::chr3:172227061-172227111(-) |
| CFB | SERPINA1 | 6 | 14 | 1 | -1 | 31919830 | 94857030 | CFB::chr6:31919780-31919830(+) | SERPINA1::chr14:94856980-94857030(-) |
| CHD7 | TOX | 8 | 8 | 1 | -1 | 61655656 | 59872567 | CHD7::chr8:61655606-61655656(+) | TOX::chr8:59872517-59872567(-) |
| CHGB | IGF2 | 20 | 11 | 1 | -1 | 5903719 | 2154895 | CHGB::chr20:5903669-5903719(+) | IGF2::chr11:2154845-2154895(-) |
| CKAP5 | TMEM135 | 11 | 11 | -1 | 1 | 46797805 | 86778736 | CKAP5::chr11:46797804-46797854(-) | TMEM135::chr11:86778735-86778785(+) |
| CLCF1 | PPP1CA | 11 | 11 | -1 | -1 | 67140995 | 67168670 | CLCF1::chr11:67140994-67141044(-) | PPP1CA::chr11:67168620-67168670(-) |
| CLNK | WDR70 | 4 | 5 | -1 | 1 | 10542118 | 37438024 | CLNK::chr4:10542117-10542167(-) | WDR70::chr5:37438023-37438073(+) |
| CNN3 | ALG14 | 1 | 1 | -1 | -1 | 95392394 | 95448862 | CNN3::chr1:95392393-95392443(-) | ALG14::chr1:95448812-95448862(-) |
| COL16A1 | CPSF6 | 1 | 12 | -1 | 1 | 32119200 | 69663292 | COL16A1::chr1:32119199-32119249(-) | CPSF6::chr12:69663291-69663341(+) |
| COL3A1 | PLAT | 2 | 8 | 1 | -1 | 189877472 | 42051245 | COL3A1::chr2:189877422-189877472(+) | PLAT::chr8:42051195-42051245(-) |
| COMMD10 | AP3S1 | 5 | 5 | 1 | 1 | 115428397 | 115230784 | COMMD10::chr5:115428347-115428397(+) | AP3S1::chr5:115230783-115230833(+) |
| COQ10A | OBFC2B | 12 | 12 | 1 | 1 | 56663345 | 56617368 | COQ10A::chr12:56663295-56663345(+) | OBFC2B::chr12:56617367-56617417(+) |
| CPM | GOLT1B | 12 | 12 | -1 | 1 | 69326458 | 21659819 | CPM::chr12:69326457-69326507(-) | GOLT1B::chr12:21659818-21659868(+) |
| CPSF6 | CCNL2 | 12 | 1 | 1 | -1 | 69656342 | 1326245 | CPSF6::chr12:69656292-69656342(+) | CCNL2::chr1:1326195-1326245(-) |
| CPSF6 | COL16A1 | 12 | 1 | 1 | -1 | 69656342 | 32118454 | CPSF6::chr12:69656292-69656342(+) | COL16A1::chr1:32118404-32118454(-) |
| CPT1A | ANPEP | 11 | 15 | -1 | -1 | 68609243 | 90328732 | CPT1A::chr11:68609242-68609292(-) | ANPEP::chr15:90328682-90328732(-) |
| CRADD | MRPL42 | 12 | 12 | 1 | 1 | 94072848 | 93894952 | CRADD::chr12:94072798-94072848(+) | MRPL42::chr12:93894951-93895001(+) |
| CRTC1 | KLHL26 | 19 | 19 | 1 | 1 | 18794638 | 18775071 | CRTC1::chr19:18794588-18794638(+) | KLHL26::chr19:18775070-18775120(+) |
| CTNND1 | TMX2 | 11 | 11 | 1 | 1 | 57529518 | 57505080 | CTNND1::chr11:57529468-57529518(+) | TMX2::chr11:57505079-57505129(+) |
| CTSB | EIF4A2 | 8 | 3 | -1 | 1 | 11701564 | 186501336 | CTSB::chr8:11701563-11701613(-) | EIF4A2::chr3:186501335-186501385(+) |
| CTTN | FOLR1 | 11 | 11 | 1 | 1 | 70244720 | 71903210 | CTTN::chr11:70244670-70244720(+) | FOLR1::chr11:71903209-71903259(+) |
| CWC25 | B4GALNT2 | 17 | 17 | -1 | 1 | 36977154 | 47218609 | CWC25::chr17:36977153-36977203(-) | B4GALNT2::chr17:47218608-47218658(+) |
| CYP2C18 | HELLS | 10 | 10 | 1 | 1 | 96466717 | 96331145 | CYP2C18::chr10:96466667-96466717(+) | HELLS::chr10:96331144-96331194(+) |
| DDX23 | RND1 | 12 | 12 | -1 | -1 | 49245867 | 49255901 | DDX23::chr12:49245866-49245916(-) | RND1::chr12:49255851-49255901(-) |
| DGKH | MED13L | 13 | 12 | 1 | -1 | 42795530 | 116460406 | DGKH::chr13:42795480-42795530(+) | MED13L::chr12:116460356-116460406(-) |
| DHCR24 | EPHX1 | 1 | 1 | -1 | 1 | 55315306 | 225997794 | DHCR24::chr1:55315305-55315355(-) | EPHX1::chr1:225997793-225997843(+) |
| DHCR24 | SFTPC | 1 | 8 | -1 | 1 | 55317396 | 22019313 | DHCR24::chr1:55317395-55317445(-) | SFTPC::chr8:22019312-22019362(+) |
| DHCR24 | THBS1 | 1 | 15 | -1 | 1 | 55315306 | 39873127 | DHCR24::chr1:55315305-55315355(-) | THBS1::chr15:39873126-39873176(+) |
| DLG3 | UNC5D | X | 8 | 1 | 1 | 69699114 | 35606045 | DLG3::chrX:69699064-69699114(+) | UNC5D::chr8:35606044-35606094(+) |
| DLK1 | STAT6 | 14 | 12 | 1 | -1 | 101201464 | 57496773 | DLK1::chr14:101201414-101201464(+) | STAT6::chr12:57496723-57496773(-) |
| DMTF1 | SEMA3D | 7 | 7 | 1 | -1 | 86800405 | 84702397 | DMTF1::chr7:86800355-86800405(+) | SEMA3D::chr7:84702347-84702397(-) |
| DNAJC3 | CLDN10 | 13 | 13 | 1 | 1 | 96329591 | 96212386 | DNAJC3::chr13:96329541-96329591(+) | CLDN10::chr13:96212385-96212435(+) |
| DNM2 | RELB | 19 | 19 | 1 | 1 | 10829079 | 45537502 | DNM2::chr19:10829029-10829079(+) | RELB::chr19:45537501-45537551(+) |

| | | | | | | | | | |
|---|---|---|---|---|---|---|---|---|---|
| DTL | INTS7 | 1 | 1 | 1 | -1 | 212220759 | 212194554 | DTL::chr1:212220709-212220759(+) | INTS7::chr1:212194504-212194554(-) |
| EDEM2 | UQCC | 20 | 20 | -1 | -1 | 33859808 | 33935075 | EDEM2::chr20:33859807-33859857(-) | UQCC::chr20:33935025-33935075(-) |
| EEF2 | ANKS1A | 19 | 6 | -1 | 1 | 3976055 | 34977682 | EEF2::chr19:3976054-3976104(-) | ANKS1A::chr6:34977681-34977731(+) |
| EEF2 | ARFGEF2 | 19 | 20 | -1 | 1 | 3976055 | 47615777 | EEF2::chr19:3976054-3976104(-) | ARFGEF2::chr20:47615776-47615826(+) |
| EEF2 | CLCN3 | 19 | 4 | -1 | 1 | 3976055 | 170623929 | EEF2::chr19:3976054-3976104(-) | CLCN3::chr4:170623928-170623978(+) |
| EEF2 | DDX27 | 19 | 20 | -1 | 1 | 3976055 | 47849303 | EEF2::chr19:3976054-3976104(-) | DDX27::chr20:47849302-47849352(+) |
| EEF2 | EIF2AK2 | 19 | 2 | -1 | -1 | 3976055 | 37342384 | EEF2::chr19:3976054-3976104(-) | EIF2AK2::chr2:37342334-37342384(-) |
| EEF2 | SEC63 | 19 | 6 | -1 | -1 | 3976055 | 108214782 | EEF2::chr19:3976054-3976104(-) | SEC63::chr6:108214732-108214782(-) |
| EEF2 | THSD4 | 19 | 15 | -1 | 1 | 3976055 | 72072881 | EEF2::chr19:3976054-3976104(-) | THSD4::chr15:72072880-72072930(+) |
| EEF2 | TYK2 | 19 | 19 | -1 | -1 | 3976055 | 10491352 | EEF2::chr19:3976054-3976104(-) | TYK2::chr19:10491302-10491352(-) |
| EGFR | 14-Sep | 7 | 7 | 1 | -1 | 55268106 | 55863785 | EGFR::chr7:55268056-55268106(+) | Sep-14::chr7:55863735-55863785(-) |
| EIF2AK1 | GCK | 7 | 7 | -1 | -1 | 6094177 | 44189663 | EIF2AK1::chr7:6094176-6094226(-) | GCK::chr7:44189613-44189663(-) |
| EIF4G3 | ECE1 | 1 | 1 | -1 | -1 | 21377359 | 21616649 | EIF4G3::chr1:21377358-21377408(-) | ECE1::chr1:21616599-21616649(-) |
| EML4 | ALK | 2 | 2 | 1 | -1 | 42492091 | 29446394 | EML4::chr2:42492041-42492091(+) | ALK::chr2:29446344-29446394(-) |
| ENO1 | SNW1 | 1 | 14 | -1 | -1 | 8921061 | 78189203 | ENO1::chr1:8921060-8921110(-) | SNW1::chr14:78189153-78189203(-) |
| EP300 | BCOR | 22 | X | 1 | -1 | 41572532 | 39930412 | EP300::chr22:41572482-41572532(+) | BCOR::chrX:39930362-39930412(-) |
| ERBB2 | PPP1R1B | 17 | 17 | 1 | 1 | 37883800 | 37791860 | ERBB2::chr17:37883750-37883800(+) | PPP1R1B::chr17:37791859-37791909(+) |
| ERGIC1 | RANBP17 | 5 | 5 | 1 | 1 | 172341841 | 170720887 | ERGIC1::chr5:172341791-172341841(+) | RANBP17::chr5:170720886-170720936(+) |
| ESR1 | C6orf97 | 6 | 6 | 1 | 1 | 152023140 | 151936578 | ESR1::chr6:152023090-152023140(+) | C6orf97::chr6:151936577-151936627(+) |
| ESR1 | C6orf97 | 6 | 6 | 1 | 1 | 152023140 | 151917470 | ESR1::chr6:152023090-152023140(+) | C6orf97::chr6:151917469-151917519(+) |
| ESR1 | C6orf97 | 6 | 6 | 1 | 1 | 152023140 | 151894309 | ESR1::chr6:152023090-152023140(+) | C6orf97::chr6:151894308-151894358(+) |
| ESR1 | C6orf97 | 6 | 6 | 1 | 1 | 152023140 | 151857453 | ESR1::chr6:152023090-152023140(+) | C6orf97::chr6:151857452-151857502(+) |
| ETV6 | NTRK3 | 12 | 15 | 1 | -1 | 12022903 | 88483984 | ETV6::chr12:12022853-12022903(+) | NTRK3::chr15:88483934-88483984(-) |
| ETV6 | NTRK3 | 12 | 15 | 1 | -1 | 12006495 | 88576276 | ETV6::chr12:12006445-12006495(+) | NTRK3::chr15:88576226-88576276(-) |
| EXT1 | SAMD12 | 8 | 8 | -1 | -1 | 119122324 | 119209971 | EXT1::chr8:119122323-119122373(-) | SAMD12::chr8:119209921-119209971(-) |
| EYA3 | GUCA1C | 1 | 3 | -1 | -1 | 28415035 | 108639432 | EYA3::chr1:28415034-28415084(-) | GUCA1C::chr3:108639382-108639432(-) |
| EYS | KIAA1543 | 6 | 19 | -1 | 1 | 65300116 | 7670112 | EYS::chr6:65300115-65300165(-) | KIAA1543::chr19:7670111-7670161(+) |
| EZR | ROS1 | 6 | 6 | -1 | -1 | 159191796 | 117645578 | EZR::chr6:159191795-159191845(-) | ROS1::chr6:117645528-117645578(-) |
| FAM104A | CDC42EP4 | 17 | 17 | -1 | -1 | 71228225 | 71282751 | FAM104A::chr17:71228224-71228274(-) | CDC42EP4::chr17:71282701-71282751(-) |
| FAM126A | TRA2A | 7 | 7 | -1 | -1 | 23023563 | 23547154 | FAM126A::chr7:23023562-23023612(-) | TRA2A::chr7:23547104-23547154(-) |
| FAM168A | PXDNL | 11 | 8 | -1 | -1 | 73308968 | 52339318 | FAM168A::chr11:73308967-73309017(-) | PXDNL::chr8:52339268-52339318(-) |
| FAM190A | MMRN1 | 4 | 4 | 1 | 1 | 91048982 | 90815875 | FAM190A::chr4:91048932-91048982(+) | MMRN1::chr4:90815874-90815924(+) |
| FAM193A | PPP2R2C | 4 | 4 | 1 | -1 | 2632896 | 6335458 | FAM193A::chr4:2632846-2632896(+) | PPP2R2C::chr4:6335408-6335458(-) |
| FBXL20 | GRB7 | 17 | 17 | -1 | 1 | 37557614 | 37898505 | FBXL20::chr17:37557613-37557663(-) | GRB7::chr17:37898504-37898554(+) |
| FBXL20 | GSDMB | 17 | 17 | -1 | -1 | 37557614 | 38068750 | FBXL20::chr17:37557613-37557663(-) | GSDMB::chr17:38068700-38068750(-) |
| FCHSD2 | ATP6V1H | 11 | 8 | -1 | -1 | 72851076 | 54682303 | FCHSD2::chr11:72851075-72851125(-) | ATP6V1H::chr8:54682253-54682303(-) |
| FDFT1 | INTS9 | 8 | 8 | 1 | -1 | 11660440 | 28707790 | FDFT1::chr8:11660390-11660440(+) | INTS9::chr8:28707740-28707790(-) |
| FGF12 | LAMP3 | 3 | 3 | -1 | -1 | 192444970 | 182872179 | FGF12::chr3:192444969-192445019(-) | LAMP3::chr3:182872129-182872179(-) |
| FGFR2 | BICC1 | 10 | 10 | -1 | 1 | 123243212 | 60461834 | FGFR2::chr10:123243211-123243261(-) | BICC1::chr10:60461833-60461883(+) |
| FGFR2 | KIAA1598 | 10 | 10 | -1 | -1 | 123243212 | 118708508 | FGFR2::chr10:123243211-123243261(-) | KIAA1598::chr10:118708458-118708508(-) |
| FGFR3 | TACC3 | 4 | 4 | 1 | 1 | 1808661 | 1741429 | FGFR3::chr4:1808611-1808661(+) | TACC3::chr4:1741428-1741478(+) |
| FGFR3 | TACC3 | 4 | 4 | 1 | 1 | 1808661 | 1737458 | FGFR3::chr4:1808611-1808661(+) | TACC3::chr4:1737457-1737507(+) |
| FKBP8 | SLC1A6 | 19 | 19 | -1 | -1 | 18644080 | 15061202 | FKBP8::chr19:18644079-18644129(-) | SLC1A6::chr19:15061152-15061202(-) |

| | | | | | | | | | |
|---|---|---|---|---|---|---|---|---|---|
| FOXP1 | EIF4E3 | 3 | 3 | -1 | -1 | 71542577 | 71777774 | FOXP1::chr3:71542576-71542626(-) | EIF4E3::chr3:71777724-71777774(-) |
| FZR1 | TBC1D17 | 19 | 19 | 1 | 1 | 3532648 | 50390733 | FZR1::chr19:3532598-3532648(+) | TBC1D17::chr19:50390732-50390782(+) |
| GAPDH | APOA2 | 12 | 1 | 1 | -1 | 6647481 | 161193418 | GAPDH::chr12:6647431-6647481(+) | APOA2::chr1:161193368-161193418(-) |
| GAS7 | MYH1 | 17 | 17 | -1 | -1 | 10101525 | 10415863 | GAS7::chr17:10101524-10101574(-) | MYH1::chr17:10415813-10415863(-) |
| GATAD2B | KCNN3 | 1 | 1 | -1 | -1 | 153895209 | 154744869 | GATAD2B::chr1:153895208-153895258(-) | KCNN3::chr1:154744819-154744869(-) |
| GLB1 | INSC | 3 | 11 | -1 | 1 | 33109722 | 15260465 | GLB1::chr3:33109721-33109771(-) | INSC::chr11:15260464-15260514(+) |
| GLI3 | C7orf23 | 7 | 7 | -1 | -1 | 42063067 | 86828373 | GLI3::chr7:42063066-42063116(-) | C7orf23::chr7:86828323-86828373(-) |
| GMCL1 | ANXA4 | 2 | 2 | 1 | 1 | 70057284 | 70008652 | GMCL1::chr2:70057234-70057284(+) | ANXA4::chr2:70008651-70008701(+) |
| GNAS | CLIP1 | 20 | 12 | 1 | -1 | 57486241 | 122819252 | GNAS::chr20:57486191-57486241(+) | CLIP1::chr12:122819202-122819252(-) |
| GNAS | FOXP1 | 20 | 3 | 1 | -1 | 57486241 | 71007472 | GNAS::chr20:57486191-57486241(+) | FOXP1::chr3:71007422-71007472(-) |
| GNAS | HEG1 | 20 | 3 | 1 | -1 | 57486241 | 124692749 | GNAS::chr20:57486191-57486241(+) | HEG1::chr3:124692699-124692749(-) |
| GNAS | SIAE | 20 | 11 | 1 | -1 | 57486241 | 124543777 | GNAS::chr20:57486191-57486241(+) | SIAE::chr11:124543727-124543777(-) |
| GNAS | SUPT16H | 20 | 14 | 1 | -1 | 57486241 | 21829491 | GNAS::chr20:57486191-57486241(+) | SUPT16H::chr14:21829441-21829491(-) |
| GNAS | TSPAN7 | 20 | X | 1 | 1 | 57486241 | 38505515 | GNAS::chr20:57486191-57486241(+) | TSPAN7::chrX:38505514-38505564(+) |
| GNB2L1 | APOB | 5 | 2 | -1 | -1 | 180663934 | 21250937 | GNB2L1::chr5:180663933-180663983(-) | APOB::chr2:21250887-21250937(-) |
| GNB2L1 | DECR1 | 5 | 8 | -1 | 1 | 180663934 | 91017676 | GNB2L1::chr5:180663933-180663983(-) | DECR1::chr8:91017675-91017725(+) |
| GNG12 | DIRAS3 | 1 | 1 | -1 | -1 | 68299027 | 68513045 | GNG12::chr1:68299026-68299076(-) | DIRAS3::chr1:68512995-68513045(-) |
| GNG12 | PLXNA2 | 1 | 1 | -1 | -1 | 68299027 | 208257925 | GNG12::chr1:68299026-68299076(-) | PLXNA2::chr1:208257875-208257925(-) |
| GPC6 | GPC5 | 13 | 13 | 1 | 1 | 93879869 | 93518535 | GPC6::chr13:93879819-93879869(+) | GPC5::chr13:93518534-93518584(+) |
| GPR35 | ANKMY1 | 2 | 2 | 1 | -1 | 241558448 | 241468926 | GPR35::chr2:241558398-241558448(+) | ANKMY1::chr2:241468876-241468926(-) |
| GPX3 | CHD4 | 5 | 12 | 1 | -1 | 150408515 | 6710696 | GPX3::chr5:150408465-150408515(+) | CHD4::chr12:6710646-6710696(-) |
| GPX3 | SEC63 | 5 | 6 | 1 | -1 | 150408548 | 108214782 | GPX3::chr5:150408498-150408548(+) | SEC63::chr6:108214732-108214782(-) |
| H2AFY | CNOT6 | 5 | 5 | -1 | 1 | 134688636 | 179991489 | H2AFY::chr5:134688635-134688685(-) | CNOT6::chr5:179991488-179991538(+) |
| HARBI1 | TNS3 | 11 | 7 | -1 | -1 | 46637118 | 47344601 | HARBI1::chr11:46637117-46637167(-) | TNS3::chr7:47344551-47344601(-) |
| HERPUD2 | AOAH | 7 | 7 | -1 | -1 | 35733794 | 36580097 | HERPUD2::chr7:35733793-35733843(-) | AOAH::chr7:36580047-36580097(-) |
| HEYL | COL9A2 | 1 | 1 | -1 | -1 | 40105218 | 40777387 | HEYL::chr1:40105217-40105267(-) | COL9A2::chr1:40777337-40777387(-) |
| HHIPL1 | CCNK | 14 | 14 | 1 | 1 | 100111799 | 99973413 | HHIPL1::chr14:100111749-100111799(+) | CCNK::chr14:99973412-99973462(+) |
| HLA-B | CCT5 | 6 | 5 | -1 | 1 | 31321649 | 10250033 | HLA-B::chr6:31321648-31321698(-) | CCT5::chr5:10250032-10250082(+) |
| HTATSF1 | BRS3 | X | X | 1 | 1 | 135586622 | 135572292 | HTATSF1::chrX:135586572-135586622(+) | BRS3::chrX:135572291-135572341(+) |
| IFT80 | RARRES1 | 3 | 3 | -1 | -1 | 160093600 | 158422716 | IFT80::chr3:160093599-160093649(-) | RARRES1::chr3:158422666-158422716(-) |
| IGF2 | CCT5 | 11 | 5 | -1 | 1 | 2150351 | 10250033 | IGF2::chr11:2150350-2150400(-) | CCT5::chr5:10250032-10250082(+) |
| IL27RA | RLN3 | 19 | 19 | 1 | 1 | 14153601 | 14141522 | IL27RA::chr19:14153551-14153601(+) | RLN3::chr19:14141521-14141571(+) |
| IL27RA | RLN3 | 19 | 19 | 1 | 1 | 14160126 | 14141522 | IL27RA::chr19:14160076-14160126(+) | RLN3::chr19:14141521-14141571(+) |
| IPO7 | TMEM41B | 11 | 11 | 1 | -1 | 9406394 | 9321248 | IPO7::chr11:9406344-9406394(+) | TMEM41B::chr11:9321198-9321248(-) |
| IQGAP1 | SEMA4B | 15 | 15 | 1 | 1 | 90934105 | 90760671 | IQGAP1::chr15:90934055-90934105(+) | SEMA4B::chr15:90760670-90760720(+) |
| IQGAP1 | ZNF774 | 15 | 15 | 1 | 1 | 90934105 | 90897874 | IQGAP1::chr15:90934055-90934105(+) | ZNF774::chr15:90897873-90897923(+) |
| ITGB6 | RBMS1 | 2 | 2 | -1 | -1 | 161051880 | 161174738 | ITGB6::chr2:161051879-161051929(-) | RBMS1::chr2:161174688-161174738(-) |
| ITM2B | RPL30 | 13 | 8 | 1 | -1 | 48835524 | 99058697 | ITM2B::chr13:48835474-48835524(+) | RPL30::chr8:99058647-99058697(-) |
| ITPKC | LTBP4 | 19 | 19 | 1 | 1 | 41231344 | 41110941 | ITPKC::chr19:41231294-41231344(+) | LTBP4::chr19:41110940-41110990(+) |
| ITPKC | SPTBN4 | 19 | 19 | 1 | 1 | 41231344 | 41071329 | ITPKC::chr19:41231294-41231344(+) | SPTBN4::chr19:41071328-41071378(+) |
| KDM2A | RHOD | 11 | 11 | 1 | 1 | 66888829 | 66833367 | KDM2A::chr11:66888779-66888829(+) | RHOD::chr11:66833366-66833416(+) |
| KIAA0391 | FAM177A1 | 14 | 14 | 1 | 1 | 35649983 | 35546356 | KIAA0391::chr14:35649933-35649983(+) | FAM177A1::chr14:35546355-35546405(+) |
| KIAA1109 | CCNA2 | 4 | 4 | 1 | -1 | 123075267 | 122742246 | KIAA1109::chr4:123075217-123075267(+) | CCNA2::chr4:122742196-122742246(-) |

| | | | | | | | | | |
|---|---|---|---|---|---|---|---|---|---|
| KIAA1429 | SYBU | 8 | 8 | -1 | -1 | 95547067 | 110631268 | KIAA1429::chr8:95547066-95547116(-) | SYBU::chr8:110631218-110631268(-) |
| KIF26B | SMYD3 | 1 | 1 | 1 | -1 | 245319985 | 246093239 | KIF26B::chr1:245319935-245319985(+) | SMYD3::chr1:246093189-246093239(-) |
| KIF26B | SMYD3 | 1 | 1 | 1 | -1 | 245319985 | 246027188 | KIF26B::chr1:245319935-245319985(+) | SMYD3::chr1:246027138-246027188(-) |
| KLF5 | PIBF1 | 13 | 13 | 1 | 1 | 73651676 | 73410241 | KLF5::chr13:73651626-73651676(+) | PIBF1::chr13:73410240-73410290(+) |
| KLK2 | FGFR2 | 19 | 10 | 1 | -1 | 51376775 | 123310973 | KLK2::chr19:51376725-51376775(+) | FGFR2::chr10:123310923-123310973(-) |
| KLK2 | NDUFV1 | 19 | 11 | 1 | 1 | 51383822 | 67375697 | KLK2::chr19:51383772-51383822(+) | NDUFV1::chr11:67375696-67375746(+) |
| KRT14 | SUPT16H | 17 | 14 | -1 | -1 | 39738531 | 21829491 | KRT14::chr17:39738530-39738580(-) | SUPT16H::chr14:21829441-21829491(-) |
| KRT19 | CTNND1 | 17 | 11 | -1 | 1 | 39679869 | 57574387 | KRT19::chr17:39679868-39679918(-) | CTNND1::chr11:57574386-57574436(+) |
| LAMA5 | CLCN7 | 20 | 16 | -1 | -1 | 60900372 | 1509188 | LAMA5::chr20:60900371-60900421(-) | CLCN7::chr16:1509138-1509188(-) |
| LATS2 | MYLK | 13 | 3 | -1 | -1 | 21635485 | 123386034 | LATS2::chr13:21635484-21635534(-) | MYLK::chr3:123385984-123386034(-) |
| LBP | BPI | 20 | 20 | 1 | 1 | 36983809 | 36952272 | LBP::chr20:36983759-36983809(+) | BPI::chr20:36952271-36952321(+) |
| LCN2 | ZNFX1 | 9 | 20 | 1 | -1 | 130915734 | 47863418 | LCN2::chr9:130915684-130915734(+) | ZNFX1::chr20:47863368-47863418(-) |
| LIMCH1 | UCHL1 | 4 | 4 | 1 | 1 | 41526495 | 41258610 | LIMCH1::chr4:41526445-41526495(+) | UCHL1::chr4:41258609-41258659(+) |
| LRP12 | LAPTM4B | 8 | 8 | -1 | 1 | 105601047 | 98863625 | LRP12::chr8:105601046-105601096(-) | LAPTM4B::chr8:98863624-98863674(+) |
| LRRC1 | SH3GL1 | 6 | 19 | 1 | -1 | 53707025 | 4366991 | LRRC1::chr6:53706975-53707025(+) | SH3GL1::chr19:4366941-4366991(-) |
| LTBP1 | BIRC6 | 2 | 2 | 1 | 1 | 33246273 | 32631567 | LTBP1::chr2:33246223-33246273(+) | BIRC6::chr2:32631566-32631616(+) |
| LTF | CFB | 3 | 6 | -1 | 1 | 46477510 | 31913427 | LTF::chr3:46477509-46477559(-) | CFB::chr6:31913426-31913476(+) |
| LUC7L2 | ETV1 | 7 | 7 | 1 | -1 | 139045068 | 13978871 | LUC7L2::chr7:139045018-139045068(+) | ETV1::chr7:13978821-13978871(-) |
| LYZ | ANXA1 | 12 | 9 | 1 | 1 | 69748014 | 75772570 | LYZ::chr12:69747964-69748014(+) | ANXA1::chr9:75772569-75772619(+) |
| LYZ | DDX21 | 12 | 10 | 1 | 1 | 69748014 | 70723858 | LYZ::chr12:69747964-69748014(+) | DDX21::chr10:70723857-70723907(+) |
| MAMSTR | RASIP1 | 19 | 19 | -1 | -1 | 49222231 | 49230687 | MAMSTR::chr19:49222230-49222280(-) | RASIP1::chr19:49230637-49230687(-) |
| MAP4 | SPINK8 | 3 | 3 | -1 | -1 | 48130263 | 48351436 | MAP4::chr3:48130262-48130312(-) | SPINK8::chr3:48351386-48351436(-) |
| MAP4 | SPINK8 | 3 | 3 | -1 | -1 | 48019355 | 48351436 | MAP4::chr3:48019354-48019404(-) | SPINK8::chr3:48351386-48351436(-) |
| MAPK10 | ARHGAP24 | 4 | 4 | -1 | 1 | 87275683 | 86893189 | MAPK10::chr4:87275682-87275732(-) | ARHGAP24::chr4:86893188-86893238(+) |
| MAPKAPK3 | HEMK1 | 3 | 3 | 1 | 1 | 50655215 | 50609141 | MAPKAPK3::chr3:50655165-50655215(+) | HEMK1::chr3:50609140-50609190(+) |
| MARK2 | C11orf84 | 11 | 11 | 1 | 1 | 63607032 | 63585277 | MARK2::chr11:63606982-63607032(+) | C11orf84::chr11:63585276-63585326(+) |
| MAST2 | NASP | 1 | 1 | 1 | 1 | 46348067 | 46056895 | MAST2::chr1:46348017-46348067(+) | NASP::chr1:46056894-46056944(+) |
| MCFD2 | TTC7A | 2 | 2 | -1 | 1 | 47142862 | 47177502 | MCFD2::chr2:47142861-47142911(-) | TTC7A::chr2:47177501-47177551(+) |
| MED1 | KRT222 | 17 | 17 | -1 | -1 | 37604051 | 38818296 | MED1::chr17:37604050-37604100(-) | KRT222::chr17:38818246-38818296(-) |
| MGAT4A | C2orf55 | 2 | 2 | -1 | -1 | 99342702 | 99463273 | MGAT4A::chr2:99342701-99342751(-) | C2orf55::chr2:99463223-99463273(-) |
| MKL1 | KIAA0146 | 22 | 8 | -1 | 1 | 41032482 | 48586363 | MKL1::chr22:41032481-41032531(-) | KIAA0146::chr8:48586362-48586412(+) |
| MKRN2 | PPARG | 3 | 3 | 1 | 1 | 12598675 | 12353879 | MKRN2::chr3:12598625-12598675(+) | PPARG::chr3:12353878-12353928(+) |
| MLL5 | LHFPL3 | 7 | 7 | 1 | 1 | 104654982 | 104546634 | MLL5::chr7:104654932-104654982(+) | LHFPL3::chr7:104546633-104546683(+) |
| MLL5 | LHFPL3 | 7 | 7 | 1 | 1 | 104707254 | 104546634 | MLL5::chr7:104707204-104707254(+) | LHFPL3::chr7:104546633-104546683(+) |
| MLL5 | LHFPL3 | 7 | 7 | 1 | 1 | 104681470 | 104546634 | MLL5::chr7:104681420-104681470(+) | LHFPL3::chr7:104546633-104546683(+) |
| MLLT10 | PICALM | 10 | 11 | 1 | -1 | 21906136 | 85670103 | MLLT10::chr10:21906086-21906136(+) | PICALM::chr11:85670053-85670103(-) |
| MLLT10 | WAC | 10 | 10 | 1 | 1 | 21906136 | 28824491 | MLLT10::chr10:21906086-21906136(+) | WAC::chr10:28824490-28824540(+) |
| MPP2 | PPY | 17 | 17 | -1 | -1 | 41975630 | 42019022 | MPP2::chr17:41975629-41975679(-) | PPY::chr17:42018972-42019022(-) |
| MPP5 | GPHN | 14 | 14 | 1 | 1 | 67746254 | 67635650 | MPP5::chr14:67746204-67746254(+) | GPHN::chr14:67635649-67635699(+) |
| MPPED2 | AMOTL1 | 11 | 11 | -1 | 1 | 30601793 | 94554696 | MPPED2::chr11:30601792-30601842(-) | AMOTL1::chr11:94554695-94554745(+) |
| MRPL48 | DTX4 | 11 | 11 | 1 | 1 | 73536841 | 58972149 | MRPL48::chr11:73536791-73536841(+) | DTX4::chr11:58972148-58972198(+) |
| MRPS23 | FAM117A | 17 | 17 | -1 | -1 | 55926601 | 47810082 | MRPS23::chr17:55926600-55926650(-) | FAM117A::chr17:47810032-47810082(-) |
| MSMB | KLK3 | 10 | 19 | 1 | 1 | 51562517 | 51361285 | MSMB::chr10:51562467-51562517(+) | KLK3::chr19:51361284-51361334(+) |

| | | | | | | | | | |
|---|---|---|---|---|---|---|---|---|---|
| MTHFS | BCL2A1 | 15 | 15 | -1 | -1 | 80181435 | 80253516 | MTHFS::chr15:80181434-80181484(-) | BCL2A1::chr15:80253466-80253516(-) |
| MVP | KIF22 | 16 | 16 | 1 | 1 | 29851780 | 29814770 | MVP::chr16:29851730-29851780(+) | KIF22::chr16:29814769-29814819(+) |
| MYH9 | TXN2 | 22 | 22 | -1 | -1 | 36783852 | 36863964 | MYH9::chr22:36783851-36783901(-) | TXN2::chr22:36863914-36863964(-) |
| MYO1D | ACCN1 | 17 | 17 | -1 | -1 | 31203796 | 31355410 | MYO1D::chr17:31203795-31203845(-) | ACCN1::chr17:31355360-31355410(-) |
| MYO1D | TMIGD1 | 17 | 17 | -1 | -1 | 31203796 | 28656547 | MYO1D::chr17:31203795-31203845(-) | TMIGD1::chr17:28656497-28656547(-) |
| MYO6 | SENP6 | 6 | 6 | 1 | 1 | 76459140 | 76405520 | MYO6::chr6:76459090-76459140(+) | SENP6::chr6:76405519-76405569(+) |
| NBAS | ALB | 2 | 4 | -1 | 1 | 15307033 | 74270025 | NBAS::chr2:15307032-15307082(-) | ALB::chr4:74270024-74270074(+) |
| NCKAP5 | DYNC2H1 | 2 | 11 | -1 | 1 | 133971288 | 103306671 | NCKAP5::chr2:133971287-133971337(-) | DYNC2H1::chr11:103306670-103306720(+) |
| NCL | RBM25 | 2 | 14 | -1 | 1 | 232319462 | 73572567 | NCL::chr2:232319461-232319511(-) | RBM25::chr14:73572566-73572616(+) |
| NCOA4 | RET | 10 | 10 | 1 | 1 | 51582939 | 43612032 | NCOA4::chr10:51582889-51582939(+) | RET::chr10:43612031-43612081(+) |
| NCOA6 | GSS | 20 | 20 | -1 | -1 | 33413275 | 33539663 | NCOA6::chr20:33413274-33413324(-) | GSS::chr20:33539613-33539663(-) |
| NCOR2 | SCARB1 | 12 | 12 | -1 | -1 | 125051854 | 125302253 | NCOR2::chr12:125051853-125051903(-) | SCARB1::chr12:125302203-125302253(-) |
| NCOR2 | SCARB1 | 12 | 12 | -1 | -1 | 124904503 | 125302253 | NCOR2::chr12:124904502-124904552(-) | SCARB1::chr12:125302203-125302253(-) |
| NCOR2 | SCARB1 | 12 | 12 | -1 | -1 | 125020111 | 125302253 | NCOR2::chr12:125020110-125020160(-) | SCARB1::chr12:125302203-125302253(-) |
| NDUFA10 | ANXA4 | 2 | 2 | -1 | 1 | 240929491 | 70008652 | NDUFA10::chr2:240929490-240929540(-) | ANXA4::chr2:70008651-70008701(+) |
| NDUFB9 | TDRKH | 8 | 1 | 1 | -1 | 125551528 | 151755525 | NDUFB9::chr8:125551478-125551528(+) | TDRKH::chr1:151755475-151755525(-) |
| NEBL | C10orf140 | 10 | 10 | -1 | -1 | 21461312 | 21812800 | NEBL::chr10:21461311-21461361(-) | C10orf140::chr10:21812750-21812800(-) |
| NFASC | PRELP | 1 | 1 | 1 | 1 | 204797910 | 203452297 | NFASC::chr1:204797860-204797910(+) | PRELP::chr1:203452296-203452346(+) |
| NMD3 | USP40 | 3 | 2 | 1 | -1 | 160953000 | 234407258 | NMD3::chr3:160952950-160953000(+) | USP40::chr2:234407208-234407258(-) |
| NOS1AP | ATF6 | 1 | 1 | 1 | 1 | 162040072 | 161928236 | NOS1AP::chr1:162040022-162040072(+) | ATF6::chr1:161928235-161928285(+) |
| NXN | ABR | 17 | 17 | -1 | -1 | 882559 | 916404 | NXN::chr17:882558-882608(-) | ABR::chr17:916354-916404(-) |
| P4HB | NDUFV1 | 17 | 11 | -1 | 1 | 79801037 | 67375697 | P4HB::chr17:79801036-79801086(-) | NDUFV1::chr11:67375696-67375746(+) |
| P4HB | NHP2L1 | 17 | 22 | -1 | -1 | 79801042 | 42071199 | P4HB::chr17:79801041-79801091(-) | NHP2L1::chr22:42071149-42071199(-) |
| PACSIN2 | TTLL1 | 22 | 22 | -1 | -1 | 43411027 | 43447893 | PACSIN2::chr22:43411026-43411076(-) | TTLL1::chr22:43447843-43447893(-) |
| PARD3 | LYAR | 10 | 4 | -1 | -1 | 35103804 | 4276496 | PARD3::chr10:35103803-35103853(-) | LYAR::chr4:4276446-4276496(-) |
| PAX8 | PPARG | 2 | 3 | -1 | 1 | 113992971 | 12421203 | PAX8::chr2:113992970-113993020(-) | PPARG::chr3:12421202-12421252(+) |
| PCMTD1 | ST18 | 8 | 8 | -1 | -1 | 52811490 | 53159224 | PCMTD1::chr8:52811489-52811539(-) | ST18::chr8:53159174-53159224(-) |
| PCYT1A | TCTEX1D2 | 3 | 3 | -1 | -1 | 196014486 | 196022940 | PCYT1A::chr3:196014485-196014535(-) | TCTEX1D2::chr3:196022890-196022940(-) |
| PDE4D | DEPDC1B | 5 | 5 | -1 | -1 | 59783769 | 59943360 | PDE4D::chr5:59783768-59783818(-) | DEPDC1B::chr5:59943310-59943360(-) |
| PDE4D | DEPDC1B | 5 | 5 | -1 | -1 | 59481384 | 59983054 | PDE4D::chr5:59481383-59481433(-) | DEPDC1B::chr5:59983004-59983054(-) |
| PGD | CAMTA1 | 1 | 1 | 1 | 1 | 10459761 | 7309551 | PGD::chr1:10459711-10459761(+) | CAMTA1::chr1:7309550-7309600(+) |
| PHACTR2 | AIG1 | 6 | 6 | 1 | 1 | 143858344 | 143605247 | PHACTR2::chr6:143858294-143858344(+) | AIG1::chr6:143605246-143605296(+) |
| PHACTR4 | SESN2 | 1 | 1 | 1 | 1 | 28696336 | 28595694 | PHACTR4::chr1:28696286-28696336(+) | SESN2::chr1:28595693-28595743(+) |
| PHF21A | ARHGAP1 | 11 | 11 | -1 | -1 | 46142486 | 46717706 | PHF21A::chr11:46142485-46142535(-) | ARHGAP1::chr11:46717656-46717706(-) |
| PHF21A | GLIS2 | 11 | 16 | -1 | 1 | 45967391 | 4386726 | PHF21A::chr11:45967390-45967440(-) | GLIS2::chr16:4386725-4386775(+) |
| PICALM | MLLT10 | 11 | 10 | -1 | 1 | 85685751 | 21940602 | PICALM::chr11:85685750-85685800(-) | MLLT10::chr10:21940601-21940651(+) |
| PKM2 | FOXP1 | 15 | 3 | -1 | -1 | 72491373 | 71007472 | PKM2::chr15:72491372-72491422(-) | FOXP1::chr3:71007422-71007472(-) |
| PML | RARA | 15 | 17 | 1 | 1 | 74325755 | 38504568 | PML::chr15:74325705-74325755(+) | RARA::chr17:38504567-38504617(+) |
| PML | RARA | 15 | 17 | 1 | 1 | 74315749 | 38504568 | PML::chr15:74315699-74315749(+) | RARA::chr17:38504567-38504617(+) |
| POLR1A | REEP1 | 2 | 2 | -1 | -1 | 86292397 | 86509365 | POLR1A::chr2:86292396-86292446(-) | REEP1::chr2:86509315-86509365(-) |
| POLR2B | FIP1L1 | 4 | 4 | 1 | 1 | 57845170 | 54308820 | POLR2B::chr4:57845120-57845170(+) | FIP1L1::chr4:54308819-54308869(+) |
| PPAP2A | SLC38A9 | 5 | 5 | -1 | -1 | 54830400 | 54993820 | PPAP2A::chr5:54830399-54830449(-) | SLC38A9::chr5:54993770-54993820(-) |
| PPARG | PAX8 | 3 | 2 | 1 | -1 | 12353952 | 113984833 | PPARG::chr3:12353902-12353952(+) | PAX8::chr2:113984783-113984833(-) |

| | | | | | | | | | |
|---|---|---|---|---|---|---|---|---|---|
| PPP1R13B | C14orf2 | 14 | 14 | -1 | -1 | 104313636 | 104381526 | PPP1R13B::chr14:104313635-104313685(-) | C14orf2::chr14:104381476-104381526(-) |
| PRCC | TFE3 | 1 | X | 1 | -1 | 156738031 | 48895967 | PRCC::chr1:156737981-156738031(+) | TFE3::chrX:48895917-48895967(-) |
| PRR12 | PRRG2 | 19 | 19 | 1 | 1 | 50124879 | 50091754 | PRR12::chr19:50124829-50124879(+) | PRRG2::chr19:50091753-50091803(+) |
| PTGES3 | NACA | 12 | 12 | -1 | -1 | 57081782 | 57108471 | PTGES3::chr12:57081781-57081831(-) | NACA::chr12:57108421-57108471(-) |
| PTPRG | FHIT | 3 | 3 | 1 | -1 | 61734656 | 60522712 | PTPRG::chr3:61734606-61734656(+) | FHIT::chr3:60522662-60522712(-) |
| PTPRK | RSPO3 | 6 | 6 | -1 | 1 | 128841404 | 127469793 | PTPRK::chr6:128841403-128841453(-) | RSPO3::chr6:127469792-127469842(+) |
| PTPRK | RSPO3 | 6 | 6 | -1 | 1 | 128505577 | 127469793 | PTPRK::chr6:128505576-128505626(-) | RSPO3::chr6:127469792-127469842(+) |
| PTPRZ1 | MET | 7 | 7 | 1 | 1 | 121513611 | 116339125 | PTPRZ1::chr7:121513561-121513611(+) | MET::chr7:116339124-116339174(+) |
| PXDNL | USP8 | 8 | 15 | -1 | 1 | 52252184 | 50769473 | PXDNL::chr8:52252183-52252233(-) | USP8::chr15:50769472-50769522(+) |
| PXN | PLA2G1B | 12 | 12 | -1 | -1 | 120703420 | 120763823 | PXN::chr12:120703419-120703469(-) | PLA2G1B::chr12:120763773-120763823(-) |
| QKI | PACRG | 6 | 6 | 1 | 1 | 163836367 | 163483182 | QKI::chr6:163836317-163836367(+) | PACRG::chr6:163483181-163483231(+) |
| QKI | PACRG | 6 | 6 | 1 | 1 | 163899928 | 163483182 | QKI::chr6:163899878-163899928(+) | PACRG::chr6:163483181-163483231(+) |
| RABEP1 | RNF167 | 17 | 17 | 1 | 1 | 5235447 | 4843782 | RABEP1::chr17:5235397-5235447(+) | RNF167::chr17:4843781-4843831(+) |
| RAF1 | TMEM40 | 3 | 3 | -1 | -1 | 12632297 | 12791339 | RAF1::chr3:12632296-12632346(-) | TMEM40::chr3:12791289-12791339(-) |
| RARA | PML | 17 | 15 | 1 | 1 | 38487648 | 74317198 | RARA::chr17:38487598-38487648(+) | PML::chr15:74317197-74317247(+) |
| RARA | PML | 17 | 15 | 1 | 1 | 38487648 | 74326819 | RARA::chr17:38487598-38487648(+) | PML::chr15:74326818-74326868(+) |
| RASA2 | ACPL2 | 3 | 3 | 1 | 1 | 141206058 | 140997184 | RASA2::chr3:141206008-141206058(+) | ACPL2::chr3:140997183-140997233(+) |
| RASA2 | ZBTB38 | 3 | 3 | 1 | 1 | 141206058 | 141161231 | RASA2::chr3:141206008-141206058(+) | ZBTB38::chr3:141161230-141161280(+) |
| RB1 | MED4 | 13 | 13 | 1 | -1 | 48916850 | 48664554 | RB1::chr13:48916800-48916850(+) | MED4::chr13:48664504-48664554(-) |
| RBM47 | APBB2 | 4 | 4 | -1 | -1 | 40631413 | 40832594 | RBM47::chr4:40631412-40631462(-) | APBB2::chr4:40832544-40832594(-) |
| RCC1 | SERINC2 | 1 | 1 | 1 | 1 | 28834672 | 31905814 | RCC1::chr1:28834622-28834672(+) | SERINC2::chr1:31905813-31905863(+) |
| REV3L | FYN | 6 | 6 | -1 | -1 | 111803953 | 112041265 | REV3L::chr6:111803952-111804002(-) | FYN::chr6:112041215-112041265(-) |
| RIMS2 | BAALC | 8 | 8 | 1 | 1 | 104513290 | 104225147 | RIMS2::chr8:104513240-104513290(+) | BAALC::chr8:104225146-104225196(+) |
| RIOK1 | EYS | 6 | 6 | 1 | -1 | 7414623 | 66006012 | RIOK1::chr6:7414573-7414623(+) | EYS::chr6:66005962-66006012(-) |
| RNASE10 | CD38 | 14 | 4 | 1 | 1 | 20974178 | 15818134 | RNASE10::chr14:20974128-20974178(+) | CD38::chr4:15818133-15818183(+) |
| RNF213 | SLC26A11 | 17 | 17 | 1 | 1 | 78324196 | 78225128 | RNF213::chr17:78324146-78324196(+) | SLC26A11::chr17:78225127-78225177(+) |
| RPL18 | SLC38A5 | 19 | X | -1 | -1 | 49118590 | 48327824 | RPL18::chr19:49118589-49118639(-) | SLC38A5::chrX:48327774-48327824(-) |
| RPS6 | GNB2L1 | 9 | 5 | -1 | -1 | 19376283 | 180670914 | RPS6::chr9:19376282-19376332(-) | GNB2L1::chr5:180670864-180670914(-) |
| RRBP1 | TASP1 | 20 | 20 | -1 | -1 | 17614105 | 13398168 | RRBP1::chr20:17614104-17614154(-) | TASP1::chr20:13398118-13398168(-) |
| RUNX1 | RUNX1T1 | 21 | 8 | -1 | -1 | 36231771 | 93029591 | RUNX1::chr21:36231770-36231820(-) | RUNX1T1::chr8:93029541-93029591(-) |
| RYBP | FHIT | 3 | 3 | -1 | -1 | 72495647 | 59738047 | RYBP::chr3:72495646-72495696(-) | FHIT::chr3:59737997-59738047(-) |
| S100A2 | RPL30 | 1 | 8 | -1 | -1 | 153533816 | 99058697 | S100A2::chr1:153533815-153533865(-) | RPL30::chr8:99058647-99058697(-) |
| S100A6 | PLEKHB2 | 1 | 2 | -1 | 1 | 153507075 | 131876464 | S100A6::chr1:153507074-153507124(-) | PLEKHB2::chr2:131876463-131876513(+) |
| SARNP | MMP19 | 12 | 12 | -1 | -1 | 56211450 | 56236222 | SARNP::chr12:56211449-56211499(-) | MMP19::chr12:56236172-56236222(-) |
| SBNO2 | C19orf20 | 19 | 19 | -1 | 1 | 1149368 | 518889 | SBNO2::chr19:1149367-1149417(-) | C19orf20::chr19:518888-518938(+) |
| SBNO2 | CLEC4M | 19 | 19 | -1 | 1 | 1147308 | 7830524 | SBNO2::chr19:1147307-1147357(-) | CLEC4M::chr19:7830523-7830573(+) |
| SEC23B | CSRP2BP | 20 | 20 | 1 | 1 | 18496380 | 18162354 | SEC23B::chr20:18496330-18496380(+) | CSRP2BP::chr20:18162353-18162403(+) |
| SERPINA1 | CCT5 | 14 | 5 | -1 | 1 | 94843084 | 10250033 | SERPINA1::chr14:94843083-94843133(-) | CCT5::chr5:10250032-10250082(+) |
| SERPINA1 | CD46 | 14 | 1 | -1 | 1 | 94844713 | 207940358 | SERPINA1::chr14:94844712-94844762(-) | CD46::chr1:207940357-207940407(+) |
| SERPINA1 | CD63 | 14 | 12 | -1 | -1 | 94844713 | 56123444 | SERPINA1::chr14:94844712-94844762(-) | CD63::chr12:56123394-56123444(-) |
| SERPINA1 | CPD | 14 | 17 | -1 | 1 | 94844713 | 28783274 | SERPINA1::chr14:94844712-94844762(-) | CPD::chr17:28783273-28783323(+) |
| SETD2 | KIF9 | 3 | 3 | -1 | -1 | 47205344 | 47289576 | SETD2::chr3:47205343-47205393(-) | KIF9::chr3:47289526-47289576(-) |
| SF3B3 | DDX19B | 16 | 16 | 1 | 1 | 70573110 | 70346512 | SF3B3::chr16:70573060-70573110(+) | DDX19B::chr16:70346511-70346561(+) |

| | | | | | | | | | |
|---|---|---|---|---|---|---|---|---|---|
| SFN | ELK3 | 1 | 12 | 1 | 1 | 27190947 | 96588323 | SFN::chr1:27190897-27190947(+) | ELK3::chr12:96588322-96588372(+) |
| SFPQ | TFE3 | 1 | X | -1 | -1 | 35654604 | 48898095 | SFPQ::chr1:35654603-35654653(-) | TFE3::chrX:48898045-48898095(-) |
| SFPQ | TFE3 | 1 | X | -1 | -1 | 35652602 | 48891766 | SFPQ::chr1:35652601-35652651(-) | TFE3::chrX:48891716-48891766(-) |
| SH3PXD2A | OBFC1 | 10 | 10 | -1 | -1 | 105614953 | 105670380 | SH3PXD2A::chr10:105614952-105615002(-) | OBFC1::chr10:105670330-105670380(-) |
| SLAMF1 | CD1A | 1 | 1 | -1 | 1 | 160616660 | 158224874 | SLAMF1::chr1:160616659-160616709(-) | CD1A::chr1:158224873-158224923(+) |
| SLC12A7 | TERT | 5 | 5 | -1 | -1 | 1111983 | 1282739 | SLC12A7::chr5:1111982-1112032(-) | TERT::chr5:1282689-1282739(-) |
| SLC1A5 | AP2A1 | 19 | 19 | -1 | 1 | 47285640 | 50285014 | SLC1A5::chr19:47285639-47285689(-) | AP2A1::chr19:50285013-50285063(+) |
| SLC22A23 | C6orf145 | 6 | 6 | -1 | -1 | 3410422 | 3738382 | SLC22A23::chr6:3410421-3410471(-) | C6orf145::chr6:3738332-3738382(-) |
| SLC25A32 | SMYD3 | 8 | 1 | -1 | -1 | 104419862 | 246518396 | SLC25A32::chr8:104419861-104419911(-) | SMYD3::chr1:246518346-246518396(-) |
| SLC25A44 | SEMA4A | 1 | 1 | 1 | 1 | 156164038 | 156142617 | SLC25A44::chr1:156163988-156164038(+) | SEMA4A::chr1:156142616-156142666(+) |
| SLC34A2 | CLDN7 | 4 | 17 | 1 | -1 | 25680370 | 7166264 | SLC34A2::chr4:25680320-25680370(+) | CLDN7::chr17:7166214-7166264(-) |
| SLC39A3 | SGTA | 19 | 19 | -1 | -1 | 2737046 | 2765283 | SLC39A3::chr19:2737045-2737095(-) | SGTA::chr19:2765233-2765283(-) |
| SLC45A3 | ERG | 1 | 21 | -1 | -1 | 205649522 | 39817544 | SLC45A3::chr1:205649521-205649571(-) | ERG::chr21:39817494-39817544(-) |
| SLC45A3 | ETV1 | 1 | 7 | -1 | -1 | 205649522 | 13978871 | SLC45A3::chr1:205649521-205649571(-) | ETV1::chr7:13978821-13978871(-) |
| SLC45A3 | STAT6 | 1 | 12 | -1 | -1 | 205626979 | 57496773 | SLC45A3::chr1:205626978-205627028(-) | STAT6::chr12:57496723-57496773(-) |
| SMAD3 | AAGAB | 15 | 15 | 1 | -1 | 67358698 | 67495236 | SMAD3::chr15:67358648-67358698(+) | AAGAB::chr15:67495186-67495236(-) |
| SMAD3 | SMAD6 | 15 | 15 | 1 | 1 | 67358698 | 67073335 | SMAD3::chr15:67358648-67358698(+) | SMAD6::chr15:67073334-67073384(+) |
| SMEK1 | FBLN5 | 14 | 14 | -1 | -1 | 91975850 | 92361416 | SMEK1::chr14:91975849-91975899(-) | FBLN5::chr14:92361366-92361416(-) |
| SMURF1 | GJC3 | 7 | 7 | -1 | -1 | 98741349 | 99521226 | SMURF1::chr7:98741348-98741398(-) | GJC3::chr7:99521176-99521226(-) |
| SMURF1 | TRIM4 | 7 | 7 | -1 | -1 | 98741349 | 99507283 | SMURF1::chr7:98741348-98741398(-) | TRIM4::chr7:99507233-99507283(-) |
| SMYD3 | ZNF695 | 1 | 1 | -1 | -1 | 246670356 | 247131094 | SMYD3::chr1:246670355-246670405(-) | ZNF695::chr1:247131044-247131094(-) |
| SNAI1 | PFDN4 | 20 | 20 | 1 | 1 | 48600888 | 52835558 | SNAI1::chr20:48600838-48600888(+) | PFDN4::chr20:52835557-52835607(+) |
| SND1 | BRAF | 7 | 7 | 1 | -1 | 127361454 | 140487384 | SND1::chr7:127361404-127361454(+) | BRAF::chr7:140487334-140487384(-) |
| SNX30 | KIAA1958 | 9 | 9 | 1 | 1 | 115580095 | 115380151 | SNX30::chr9:115580045-115580095(+) | KIAA1958::chr9:115380150-115380200(+) |
| SOS1 | CDKL4 | 2 | 2 | -1 | -1 | 39347477 | 39417643 | SOS1::chr2:39347476-39347526(-) | CDKL4::chr2:39417593-39417643(-) |
| SOX6 | PTPRJ | 11 | 11 | -1 | 1 | 16362558 | 48134299 | SOX6::chr11:16362557-16362607(-) | PTPRJ::chr11:48134298-48134348(+) |
| SPAG7 | CAMTA2 | 17 | 17 | -1 | -1 | 4871015 | 4886338 | SPAG7::chr17:4871014-4871064(-) | CAMTA2::chr17:4886288-4886338(-) |
| SPAG9 | MBTD1 | 17 | 17 | -1 | -1 | 49156945 | 49258055 | SPAG9::chr17:49156944-49156994(-) | MBTD1::chr17:49258005-49258055(-) |
| SPSB1 | H6PD | 1 | 1 | 1 | 1 | 9353130 | 9322118 | SPSB1::chr1:9353080-9353130(+) | H6PD::chr1:9322117-9322167(+) |
| SRGAP3 | LHFPL4 | 3 | 3 | -1 | -1 | 9290569 | 9547887 | SRGAP3::chr3:9290568-9290618(-) | LHFPL4::chr3:9547837-9547887(-) |
| SRPK2 | PUS7 | 7 | 7 | -1 | -1 | 105029095 | 105148991 | SRPK2::chr7:105029094-105029144(-) | PUS7::chr7:105148941-105148991(-) |
| SS18 | SSX1 | 18 | X | -1 | 1 | 23612363 | 48123217 | SS18::chr18:23612362-23612412(-) | SSX1::chrX:48123216-48123266(+) |
| ST7 | CAPZA2 | 7 | 7 | 1 | 1 | 116593745 | 116556114 | ST7::chr7:116593695-116593745(+) | CAPZA2::chr7:116556113-116556163(+) |
| ST7 | MET | 7 | 7 | 1 | 1 | 116593745 | 116339125 | ST7::chr7:116593695-116593745(+) | MET::chr7:116339124-116339174(+) |
| ST7OT4 | CAPZA2 | 7 | 7 | 1 | 1 | 116595207 | 116556114 | ST7OT4::chr7:116595157-116595207(+) | CAPZA2::chr7:116556113-116556163(+) |
| STK24 | SLC15A1 | 13 | 13 | -1 | -1 | 99228999 | 99379113 | STK24::chr13:99228998-99229048(-) | SLC15A1::chr13:99379063-99379113(-) |
| STK35 | GP5 | 20 | 3 | 1 | -1 | 2098061 | 194119013 | STK35::chr20:2098011-2098061(+) | GP5::chr3:194118963-194119013(-) |
| STRBP | DENND1A | 9 | 9 | -1 | -1 | 126030715 | 126165769 | STRBP::chr9:126030714-126030764(-) | DENND1A::chr9:126165719-126165769(-) |
| STRN | ALK | 2 | 2 | -1 | -1 | 37143221 | 29446394 | STRN::chr2:37143220-37143270(-) | ALK::chr2:29446344-29446394(-) |
| STX5 | WDR74 | 11 | 11 | -1 | -1 | 62599426 | 62607131 | STX5::chr11:62599425-62599475(-) | WDR74::chr11:62607081-62607131(-) |
| SUFU | GBF1 | 10 | 10 | 1 | 1 | 104309863 | 104111013 | SUFU::chr10:104309813-104309863(+) | GBF1::chr10:104111012-104111062(+) |
| TACC2 | PPAPDC1A | 10 | 10 | 1 | 1 | 123903221 | 122263330 | TACC2::chr10:123903171-123903221(+) | PPAPDC1A::chr10:122263329-122263379(+) |
| TAF4 | TSHZ2 | 20 | 20 | -1 | 1 | 60572606 | 52103683 | TAF4::chr20:60572605-60572655(-) | TSHZ2::chr20:52103682-52103732(+) |

| | | | | | | | | | |
|---|---|---|---|---|---|---|---|---|---|
| TAF8 | COL5A3 | 6 | 19 | 1 | -1 | 42044977 | 10116907 | TAF8::chr6:42044927-42044977(+) | COL5A3::chr19:10116857-10116907(-) |
| TANC2 | CRB3 | 17 | 19 | 1 | 1 | 61345234 | 6465556 | TANC2::chr17:61345184-61345234(+) | CRB3::chr19:6465555-6465605(+) |
| TBL1XR1 | PIK3CA | 3 | 3 | -1 | 1 | 176914909 | 178916538 | TBL1XR1::chr3:176914908-176914958(-) | PIK3CA::chr3:178916537-178916587(+) |
| TCF7L2 | VTI1A | 10 | 10 | 1 | 1 | 114799885 | 114286846 | TCF7L2::chr10:114799835-114799885(+) | VTI1A::chr10:114286845-114286895(+) |
| TEAD1 | PARVA | 11 | 11 | 1 | 1 | 12697108 | 12495292 | TEAD1::chr11:12697058-12697108(+) | PARVA::chr11:12495291-12495341(+) |
| THOC1 | GLI3 | 18 | 7 | -1 | -1 | 223440 | 42262894 | THOC1::chr18:223439-223489(-) | GLI3::chr7:42262844-42262894(-) |
| TJP2 | FXN | 9 | 9 | 1 | 1 | 71766687 | 71661301 | TJP2::chr9:71766637-71766687(+) | FXN::chr9:71661300-71661350(+) |
| TMBIM6 | CLCN3 | 12 | 4 | 1 | 1 | 50158717 | 170623929 | TMBIM6::chr12:50158667-50158717(+) | CLCN3::chr4:170623928-170623978(+) |
| TMEM135 | GPD1L | 11 | 3 | 1 | 1 | 86749228 | 32169568 | TMEM135::chr11:86749178-86749228(+) | GPD1L::chr3:32169567-32169617(+) |
| TMEM87B | MERTK | 2 | 2 | 1 | 1 | 112813333 | 112751828 | TMEM87B::chr2:112813283-112813333(+) | MERTK::chr2:112751827-112751877(+) |
| TMPRSS2 | ERG | 21 | 21 | -1 | -1 | 42866283 | 39817544 | TMPRSS2::chr21:42866282-42866332(-) | ERG::chr21:39817494-39817544(-) |
| TMPRSS2 | ERG | 21 | 21 | -1 | -1 | 42880008 | 39817544 | TMPRSS2::chr21:42880007-42880057(-) | ERG::chr21:39817494-39817544(-) |
| TMPRSS2 | ERG | 21 | 21 | -1 | -1 | 42870046 | 39817544 | TMPRSS2::chr21:42870045-42870095(-) | ERG::chr21:39817494-39817544(-) |
| TMPRSS2 | ERG | 21 | 21 | -1 | -1 | 42880008 | 39956869 | TMPRSS2::chr21:42880007-42880057(-) | ERG::chr21:39956819-39956869(-) |
| TMPRSS2 | ERG | 21 | 21 | -1 | -1 | 42860321 | 39817544 | TMPRSS2::chr21:42860320-42860370(-) | ERG::chr21:39817494-39817544(-) |
| TMPRSS2 | ERG | 21 | 21 | -1 | -1 | 42878372 | 39817544 | TMPRSS2::chr21:42878371-42878421(-) | ERG::chr21:39817494-39817544(-) |
| TMPRSS2 | ERG | 21 | 21 | -1 | -1 | 42879877 | 39817544 | TMPRSS2::chr21:42879876-42879926(-) | ERG::chr21:39817494-39817544(-) |
| TMPRSS2 | ERG | 21 | 21 | -1 | -1 | 42880008 | 39795483 | TMPRSS2::chr21:42880007-42880057(-) | ERG::chr21:39795433-39795483(-) |
| TMPRSS2 | ERG | 21 | 21 | -1 | -1 | 42879877 | 39956869 | TMPRSS2::chr21:42879876-42879926(-) | ERG::chr21:39956819-39956869(-) |
| TMPRSS2 | ERG | 21 | 21 | -1 | -1 | 42870046 | 39956869 | TMPRSS2::chr21:42870045-42870095(-) | ERG::chr21:39956819-39956869(-) |
| TMPRSS2 | ETV1 | 21 | 7 | -1 | -1 | 42879877 | 13978871 | TMPRSS2::chr21:42879876-42879926(-) | ETV1::chr7:13978821-13978871(-) |
| TMPRSS2 | ETV4 | 21 | 17 | -1 | -1 | 42870046 | 41623036 | TMPRSS2::chr21:42870045-42870095(-) | ETV4::chr17:41622986-41623036(-) |
| TMPRSS2 | ETV4 | 21 | 17 | -1 | -1 | 42880008 | 41622735 | TMPRSS2::chr21:42880007-42880057(-) | ETV4::chr17:41622685-41622735(-) |
| TNRC18 | RNF216 | 7 | 7 | -1 | -1 | 5460691 | 5681007 | TNRC18::chr7:5460690-5460740(-) | RNF216::chr7:5680957-5681007(-) |
| TNRC6B | FAM83F | 22 | 22 | 1 | 1 | 40441002 | 40415172 | TNRC6B::chr22:40440952-40441002(+) | FAM83F::chr22:40415171-40415221(+) |
| TOP3A | TEKT1 | 17 | 17 | -1 | -1 | 18217913 | 6719281 | TOP3A::chr17:18217912-18217962(-) | TEKT1::chr17:6719231-6719281(-) |
| TOPBP1 | SPDEF | 3 | 6 | -1 | -1 | 133341935 | 34512261 | TOPBP1::chr3:133341934-133341984(-) | SPDEF::chr6:34512211-34512261(-) |
| TPST2 | CRYBB1 | 22 | 22 | -1 | -1 | 26932254 | 26997985 | TPST2::chr22:26932253-26932303(-) | CRYBB1::chr22:26997935-26997985(-) |
| TPT1 | ERBB3 | 13 | 12 | -1 | 1 | 45911313 | 56492035 | TPT1::chr13:45911312-45911362(-) | ERBB3::chr12:56492034-56492084(+) |
| TPT1 | NCALD | 13 | 8 | -1 | -1 | 45911313 | 102813898 | TPT1::chr13:45911312-45911362(-) | NCALD::chr8:102813848-102813898(-) |
| TPX2 | DEFB123 | 20 | 20 | 1 | 1 | 30330450 | 30037832 | TPX2::chr20:30330400-30330450(+) | DEFB123::chr20:30037831-30037881(+) |
| TRAPPC2 | ATP2B3 | X | X | -1 | 1 | 13737990 | 152845436 | TRAPPC2::chrX:13737989-13738039(-) | ATP2B3::chrX:152845435-152845485(+) |
| TRHDE | NUP107 | 12 | 12 | 1 | 1 | 72962436 | 69128488 | TRHDE::chr12:72962386-72962436(+) | NUP107::chr12:69128487-69128537(+) |
| TRIM33 | CSDE1 | 1 | 1 | -1 | -1 | 114945382 | 115282511 | TRIM33::chr1:114945381-114945431(-) | CSDE1::chr1:115282461-115282511(-) |
| TRIO | CLNK | 5 | 4 | 1 | -1 | 14401173 | 10586579 | TRIO::chr5:14401123-14401173(+) | CLNK::chr4:10586529-10586579(-) |
| TRIO | TERT | 5 | 5 | 1 | -1 | 14420130 | 1282739 | TRIO::chr5:14420080-14420130(+) | TERT::chr5:1282689-1282739(-) |
| TRIP12 | SLC16A14 | 2 | 2 | -1 | -1 | 230723488 | 230902247 | TRIP12::chr2:230723487-230723537(-) | SLC16A14::chr2:230902197-230902247(-) |
| TRMT11 | HINT3 | 6 | 6 | 1 | 1 | 126307768 | 126288033 | TRMT11::chr6:126307718-126307768(+) | HINT3::chr6:126288032-126288082(+) |
| TRPM4 | PPFIA3 | 19 | 19 | 1 | 1 | 49675365 | 49636258 | TRPM4::chr19:49675315-49675365(+) | PPFIA3::chr19:49636257-49636307(+) |
| TTYH3 | MAD1L1 | 7 | 7 | 1 | -1 | 2671912 | 1938026 | TTYH3::chr7:2671862-2671912(+) | MAD1L1::chr7:1937976-1938026(-) |
| TTYH3 | NOC4L | 7 | 12 | 1 | 1 | 2671912 | 132635526 | TTYH3::chr7:2671862-2671912(+) | NOC4L::chr12:132635525-132635575(+) |
| UBA3 | FRMD4B | 3 | 3 | -1 | -1 | 69120686 | 69225807 | UBA3::chr3:69120685-69120735(-) | FRMD4B::chr3:69225757-69225807(-) |
| UBAC1 | SHANK2 | 9 | 11 | -1 | -1 | 138852871 | 70935842 | UBAC1::chr9:138852870-138852920(-) | SHANK2::chr11:70935792-70935842(-) |

| | | | | | | | | | |
|---|---|---|---|---|---|---|---|---|---|
| UBTF | MAML3 | 17 | 4 | -1 | -1 | 42287492 | 140812121 | UBTF::chr17:42287491-42287541(-) | MAML3::chr4:140812071-140812121(-) |
| UBTF | MAML3 | 17 | 4 | -1 | -1 | 42286720 | 140812121 | UBTF::chr17:42286719-42286769(-) | MAML3::chr4:140812071-140812121(-) |
| UNK | KIAA0195 | 17 | 17 | 1 | 1 | 73781065 | 73467881 | UNK::chr17:73781015-73781065(+) | KIAA0195::chr17:73467880-73467930(+) |
| UNK | ZNF266 | 17 | 19 | 1 | -1 | 73781065 | 9530894 | UNK::chr17:73781015-73781065(+) | ZNF266::chr19:9530844-9530894(-) |
| UPF3A | CDC16 | 13 | 13 | 1 | 1 | 115052104 | 115027362 | UPF3A::chr13:115052054-115052104(+) | CDC16::chr13:115027361-115027411(+) |
| USP22 | MYH10 | 17 | 17 | -1 | -1 | 20919065 | 8508300 | USP22::chr17:20919064-20919114(-) | MYH10::chr17:8508250-8508300(-) |
| USP34 | C2orf74 | 2 | 2 | -1 | 1 | 61417406 | 61391489 | USP34::chr2:61417405-61417455(-) | C2orf74::chr2:61391488-61391538(+) |
| USP53 | MYOZ2 | 4 | 4 | 1 | 1 | 120138828 | 120107121 | USP53::chr4:120138778-120138828(+) | MYOZ2::chr4:120107120-120107170(+) |
| VEGFA | STK38 | 6 | 6 | 1 | -1 | 43745402 | 36467767 | VEGFA::chr6:43745352-43745402(+) | STK38::chr6:36467717-36467767(-) |
| VGF | SIAE | 7 | 11 | -1 | -1 | 100805797 | 124543777 | VGF::chr7:100805796-100805846(-) | SIAE::chr11:124543727-124543777(-) |
| VIM | LAMA4 | 10 | 6 | 1 | -1 | 17279584 | 112507969 | VIM::chr10:17279534-17279584(+) | LAMA4::chr6:112507919-112507969(-) |
| VPS52 | ZNF358 | 6 | 19 | -1 | 1 | 33231261 | 7584091 | VPS52::chr6:33231260-33231310(-) | ZNF358::chr19:7584090-7584140(+) |
| VPS53 | GLOD4 | 17 | 17 | -1 | -1 | 613773 | 681984 | VPS53::chr17:613772-613822(-) | GLOD4::chr17:681934-681984(-) |
| VSIG1 | MID2 | X | X | 1 | 1 | 107288409 | 107169324 | VSIG1::chrX:107288359-107288409(+) | MID2::chrX:107169323-107169373(+) |
| WASF2 | AHDC1 | 1 | 1 | -1 | -1 | 27816497 | 27879696 | WASF2::chr1:27816496-27816546(-) | AHDC1::chr1:27879646-27879696(-) |
| WASF2 | AHDC1 | 1 | 1 | -1 | -1 | 27816497 | 27885394 | WASF2::chr1:27816496-27816546(-) | AHDC1::chr1:27885344-27885394(-) |
| WASF2 | FGR | 1 | 1 | -1 | -1 | 27816497 | 27951662 | WASF2::chr1:27816496-27816546(-) | FGR::chr1:27951612-27951662(-) |
| WHSC1 | CPLX1 | 4 | 4 | 1 | -1 | 1920350 | 786396 | WHSC1::chr4:1920300-1920350(+) | CPLX1::chr4:786346-786396(-) |
| WHSC1L1 | CSMD1 | 8 | 8 | -1 | -1 | 38172165 | 2796266 | WHSC1L1::chr8:38172164-38172214(-) | CSMD1::chr8:2796216-2796266(-) |
| WHSC1L1 | FGFR1 | 8 | 8 | -1 | -1 | 38239317 | 38315052 | WHSC1L1::chr8:38239316-38239366(-) | FGFR1::chr8:38315002-38315052(-) |
| WSCD1 | NLRP1 | 17 | 17 | 1 | -1 | 5998543 | 5405192 | WSCD1::chr17:5998493-5998543(+) | NLRP1::chr17:5405142-5405192(-) |
| YEATS4 | CPSF6 | 12 | 12 | 1 | 1 | 69764755 | 69663292 | YEATS4::chr12:69764705-69764755(+) | CPSF6::chr12:69663291-69663341(+) |
| YEATS4 | LYZ | 12 | 12 | 1 | 1 | 69764755 | 69743888 | YEATS4::chr12:69764705-69764755(+) | LYZ::chr12:69743887-69743937(+) |
| ZBTB38 | ACPL2 | 3 | 3 | 1 | 1 | 141122873 | 140978965 | ZBTB38::chr3:141122823-141122873(+) | ACPL2::chr3:140978964-140979014(+) |
| ZC3H18 | CHMP1A | 16 | 16 | 1 | -1 | 88636974 | 89713739 | ZC3H18::chr16:88636924-88636974(+) | CHMP1A::chr16:89713689-89713739(-) |
| ZCCHC8 | RSRC2 | 12 | 12 | -1 | -1 | 122983374 | 122990253 | ZCCHC8::chr12:122983373-122983423(-) | RSRC2::chr12:122990203-122990253(-) |
| ZDHHC20 | EFHA1 | 13 | 13 | -1 | -1 | 21987791 | 22084240 | ZDHHC20::chr13:21987790-21987840(-) | EFHA1::chr13:22084190-22084240(-) |
| ZFP91 | RAB6A | 11 | 11 | 1 | -1 | 58352357 | 73390765 | ZFP91::chr11:58352307-58352357(+) | RAB6A::chr11:73390715-73390765(-) |
| ZMYND11 | SORBS2 | 10 | 4 | 1 | -1 | 293406 | 186515089 | ZMYND11::chr10:293356-293406(+) | SORBS2::chr4:186515039-186515089(-) |
| ZNF189 | BAAT | 9 | 9 | 1 | -1 | 104162292 | 104133745 | ZNF189::chr9:104162242-104162292(+) | BAAT::chr9:104133695-104133745(-) |
| ZNF236 | QRSL1 | 18 | 6 | 1 | 1 | 74563895 | 107088224 | ZNF236::chr18:74563845-74563895(+) | QRSL1::chr6:107088223-107088273(+) |
| ZNF253 | HSD17B14 | 19 | 19 | 1 | -1 | 19976808 | 49337615 | ZNF253::chr19:19976758-19976808(+) | HSD17B14::chr19:49337565-49337615(-) |
| ZNF292 | GRM4 | 6 | 6 | 1 | -1 | 87865477 | 34101636 | ZNF292::chr6:87865427-87865477(+) | GRM4::chr6:34101586-34101636(-) |
| ZNF91 | CCDC88A | 19 | 2 | -1 | -1 | 23556544 | 55523659 | ZNF91::chr19:23556543-23556593(-) | CCDC88A::chr2:55523609-55523659(-) |
| AR | NR4A3 | X | 9 | 1 | 1 | 66863249 | 102587941 | AR::chrX:66863199-66863249(+) | NR4A3::chr9:102587940-102587990(+) |
| ATP1A1 | NOTCH2 | 1 | 1 | 1 | -1 | 116926746 | 120465401 | ATP1A1::chr1:116926696-116926746(+) | NOTCH2::chr1:120465351-120465401(-) |
| BRAF | CUX1 | 7 | 7 | -1 | 1 | 140500162 | 101801840 | BRAF::chr7:140500161-140500211(-) | CUX1::chr7:101801839-101801889(+) |
| BRAF | SND1 | 7 | 7 | -1 | 1 | 140487348 | 127724776 | BRAF::chr7:140487347-140487397(-) | SND1::chr7:127724775-127724825(+) |
| BRAF | TRIM24 | 7 | 7 | -1 | 1 | 140624366 | 138258252 | BRAF::chr7:140624365-140624415(-) | TRIM24::chr7:138258251-138258301(+) |
| CD74 | ERBB3 | 5 | 12 | -1 | 1 | 149781200 | 56492035 | CD74::chr5:149781199-149781249(-) | ERBB3::chr12:56492034-56492084(+) |
| CUX1 | BRAF | 7 | 7 | 1 | -1 | 101758553 | 140494267 | CUX1::chr7:101758503-101758553(+) | BRAF::chr7:140494217-140494267(-) |
| DDIT3 | GLI1 | 12 | 12 | -1 | 1 | 57914201 | 57857448 | DDIT3::chr12:57914200-57914250(-) | GLI1::chr12:57857447-57857497(+) |
| DDX6 | ALK | 11 | 2 | -1 | -1 | 118656761 | 29606725 | DDX6::chr11:118656760-118656810(-) | ALK::chr2:29606675-29606725(-) |

| | | | | | | | | | |
|---|---|---|---|---|---|---|---|---|---|
| DDX6 | FOXR1 | 11 | 11 | -1 | 1 | 118651830 | 118849492 | DDX6::chr11:118651829-118651879(-) | FOXR1::chr11:118849491-118849541(+) |
| ERBB3 | FOXA1 | 12 | 14 | 1 | -1 | 56497283 | 38069245 | ERBB3::chr12:56497233-56497283(+) | FOXA1::chr14:38069195-38069245(-) |
| GNAS | BIRC6 | 20 | 2 | 1 | 1 | 57486247 | 32820101 | GNAS::chr20:57486197-57486247(+) | BIRC6::chr2:32820100-32820150(+) |
| GNAS | BIRC6 | 20 | 2 | 1 | 1 | 57486241 | 32820101 | GNAS::chr20:57486191-57486241(+) | BIRC6::chr2:32820100-32820150(+) |
| MAP2K1 | CUX1 | 15 | 7 | 1 | 1 | 66679765 | 101870647 | MAP2K1::chr15:66679715-66679765(+) | CUX1::chr7:101870646-101870696(+) |
| MAP2K4 | CD79B | 17 | 17 | 1 | -1 | 11958308 | 62008745 | MAP2K4::chr17:11958258-11958308(+) | CD79B::chr17:62008695-62008745(-) |
| MAPK1 | NOTCH1 | 22 | 9 | -1 | -1 | 22153301 | 139414017 | MAPK1::chr22:22153300-22153350(-) | NOTCH1::chr9:139413967-139414017(-) |
| MDM2 | EGFR | 12 | 7 | 1 | 1 | 69203072 | 55231426 | MDM2::chr12:69203022-69203072(+) | EGFR::chr7:55231425-55231475(+) |
| MITF | FOXP1 | 3 | 3 | 1 | -1 | 70017484 | 71007472 | MITF::chr3:70017434-70017484(+) | FOXP1::chr3:71007422-71007472(-) |
| MLLT4 | SYK | 6 | 9 | 1 | 1 | 168227917 | 93606140 | MLLT4::chr6:168227867-168227917(+) | SYK::chr9:93606139-93606189(+) |
| NOTCH1 | MAPK1 | 9 | 22 | -1 | -1 | 139405604 | 22143097 | NOTCH1::chr9:139405603-139405653(-) | MAPK1::chr22:22143047-22143097(-) |
| PDGFRA | USP8 | 4 | 15 | 1 | 1 | 55156721 | 50751197 | PDGFRA::chr4:55156671-55156721(+) | USP8::chr15:50751196-50751246(+) |
| PRKAR1A | FOXP1 | 17 | 3 | 1 | -1 | 66528899 | 71007472 | PRKAR1A::chr17:66528849-66528899(+) | FOXP1::chr3:71007422-71007472(-) |
| RARA | STAT3 | 17 | 17 | 1 | -1 | 38465538 | 40500557 | RARA::chr17:38465488-38465538(+) | STAT3::chr17:40500507-40500557(-) |
| RUNX1T1 | RUNX1 | 8 | 21 | -1 | -1 | 93074774 | 36231875 | RUNX1T1::chr8:93074773-93074823(-) | RUNX1::chr21:36231825-36231875(-) |
| SND1 | BRAF | 7 | 7 | 1 | -1 | 127721553 | 140482957 | SND1::chr7:127721503-127721553(+) | BRAF::chr7:140482907-140482957(-) |
| STAT3 | ETV4 | 17 | 17 | -1 | -1 | 40468807 | 41611353 | STAT3::chr17:40468806-40468856(-) | ETV4::chr17:41611303-41611353(-) |
| STAT5B | STAT3 | 17 | 17 | -1 | -1 | 40354358 | 40477079 | STAT5B::chr17:40354357-40354407(-) | STAT3::chr17:40477029-40477079(-) |
| TBL1XR1 | RET | 3 | 10 | -1 | 1 | 176765088 | 43612032 | TBL1XR1::chr3:176765087-176765137(-) | RET::chr10:43612031-43612081(+) |
| TBL1XR1 | STAT3 | 3 | 17 | -1 | -1 | 176914909 | 40500557 | TBL1XR1::chr3:176914908-176914958(-) | STAT3::chr17:40500507-40500557(-) |
| TRIM24 | BRAF | 7 | 7 | 1 | -1 | 138239609 | 140550012 | TRIM24::chr7:138239559-138239609(+) | BRAF::chr7:140549962-140550012(-) |
| TRIM24 | BRAF | 7 | 7 | 1 | -1 | 138255748 | 140550012 | TRIM24::chr7:138255698-138255748(+) | BRAF::chr7:140549962-140550012(-) |
| TRIM27 | RET | 6 | 10 | -1 | 1 | 28887789 | 43612032 | TRIM27::chr6:28887788-28887838(-) | RET::chr10:43612031-43612081(+) |
| WHSC1 | QKI | 4 | 6 | 1 | 1 | 1959763 | 163876311 | WHSC1::chr4:1959713-1959763(+) | QKI::chr6:163876310-163876360(+) |
| ACVR2A | CASP8 | 2 | 2 | 1 | 1 | 148674995 | 202131184 | ACVR2A::chr2:148674945-148674995(+) | CASP8::chr2:202131183-202131233(+) |
| ARID2 | PTPRB | 12 | 12 | 1 | -1 | 46125097 | 70933802 | ARID2::chr12:46125047-46125097(+) | PTPRB::chr12:70933752-70933802(-) |
| BAP1 | PBRM1 | 3 | 3 | -1 | -1 | 52438469 | 52663051 | BAP1::chr3:52438468-52438518(-) | PBRM1::chr3:52663001-52663051(-) |
| BAZ1A | NKX2-1 | 14 | 14 | -1 | -1 | 35331250 | 36988575 | BAZ1A::chr14:35331249-35331299(-) | NKX2-1::chr14:36988525-36988575(-) |
| BRCA1 | NF1 | 17 | 17 | -1 | 1 | 41199660 | 29654517 | BRCA1::chr17:41199659-41199709(-) | NF1::chr17:29654516-29654566(+) |
| NF1 | BRCA1 | 17 | 17 | 1 | -1 | 29653270 | 41215968 | NF1::chr17:29653220-29653270(+) | BRCA1::chr17:41215918-41215968(-) |
| NF1 | CLTC | 17 | 17 | 1 | 1 | 29563039 | 57721637 | NF1::chr17:29562989-29563039(+) | CLTC::chr17:57721636-57721686(+) |
| NF1 | EZH2 | 17 | 7 | 1 | -1 | 29592357 | 148544397 | NF1::chr17:29592307-29592357(+) | EZH2::chr7:148544347-148544397(-) |
| NF1 | MAP2K4 | 17 | 17 | 1 | 1 | 29509683 | 12028611 | NF1::chr17:29509633-29509683(+) | MAP2K4::chr17:12028610-12028660(+) |
| SMAD4 | NRG1 | 18 | 8 | 1 | 1 | 48556993 | 32585467 | SMAD4::chr18:48556943-48556993(+) | NRG1::chr8:32585466-32585516(+) |
| SMARCA4 | DNM2 | 19 | 19 | 1 | 1 | 11071850 | 10916592 | SMARCA4::chr19:11071800-11071850(+) | DNM2::chr19:10916591-10916641(+) |
| SMARCA4 | DNM2 | 19 | 19 | 1 | 1 | 11170863 | 10939712 | SMARCA4::chr19:11170813-11170863(+) | DNM2::chr19:10939711-10939761(+) |
| SUZ12 | NF1 | 17 | 17 | 1 | 1 | 30303633 | 29576002 | SUZ12::chr17:30303583-30303633(+) | NF1::chr17:29576001-29576051(+) |
| TGFBR2 | XPC | 3 | 3 | 1 | -1 | 30648469 | 14214562 | TGFBR2::chr3:30648419-30648469(+) | XPC::chr3:14214512-14214562(-) |
| TPM3 | NTRK1 | 1 | 1 | -1 | 1 | 154142876 | 156845312 | TPM3::chr1:154142875-154142925(-) | NTRK1::chr1:156845311-156845361(+) |
| TPM3 | NTRK1 | 1 | 1 | -1 | 1 | 154142876 | 156844363 | TPM3::chr1:154142875-154142925(-) | NTRK1::chr1:156844362-156844412(+) |
| WRN | NRG1 | 8 | 8 | 1 | 1 | 31024746 | 32585467 | WRN::chr8:31024696-31024746(+) | NRG1::chr8:32585466-32585516(+) |
| ACVR2A | AKT3 | 2 | 1 | 1 | -1 | 148602776 | 243828185 | ACVR2A::chr2:148602726-148602776(+) | AKT3::chr1:243828135-243828185(-) |
| ATRX | BCL2 | X | 18 | -1 | -1 | 77041468 | 60795992 | ATRX::chrX:77041467-77041517(-) | BCL2::chr18:60795942-60795992(-) |

| | | | | | | | | | |
|---|---|---|---|---|---|---|---|---|---|
| BRAF | CIITA | 7 | 16 | -1 | 1 | 140487348 | 10995371 | BRAF::chr7:140487347-140487397(-) | CIITA::chr16:10995370-10995420(+) |
| CCDC6 | RET | 10 | 10 | -1 | 1 | 61554231 | 43612032 | CCDC6::chr10:61554230-61554280(-) | RET::chr10:43612031-43612081(+) |
| CDK12 | TAF15 | 17 | 17 | 1 | 1 | 37682569 | 34169371 | CDK12::chr17:37682519-37682569(+) | TAF15::chr17:34169370-34169420(+) |
| CIITA | PRDM16 | 16 | 1 | 1 | 1 | 10971239 | 3301716 | CIITA::chr16:10971189-10971239(+) | PRDM16::chr1:3301715-3301765(+) |
| CLTC | ETV4 | 17 | 17 | 1 | -1 | 57744338 | 41613847 | CLTC::chr17:57744288-57744338(+) | ETV4::chr17:41613797-41613847(-) |
| CLTC | ROS1 | 17 | 6 | 1 | -1 | 57768072 | 117642557 | CLTC::chr17:57768022-57768072(+) | ROS1::chr6:117642507-117642557(-) |
| CTNNB1 | IGF2BP2 | 3 | 3 | 1 | -1 | 41241161 | 185416135 | CTNNB1::chr3:41241111-41241161(+) | IGF2BP2::chr3:185416085-185416135(-) |
| ERBB2 | CDK12 | 17 | 17 | 1 | 1 | 37868701 | 37680927 | ERBB2::chr17:37868651-37868701(+) | CDK12::chr17:37680926-37680976(+) |
| ERG | ZNRF3 | 21 | 22 | -1 | 1 | 39947586 | 29383064 | ERG::chr21:39947585-39947635(-) | ZNRF3::chr22:29383063-29383113(+) |
| FANCC | PSIP1 | 9 | 9 | -1 | -1 | 97912205 | 15490122 | FANCC::chr9:97912204-97912254(-) | PSIP1::chr9:15490072-15490122(-) |
| FAS | CTNND2 | 10 | 5 | 1 | -1 | 90762951 | 11023091 | FAS::chr10:90762901-90762951(+) | CTNND2::chr5:11023041-11023091(-) |
| FAT1 | NTRK3 | 4 | 15 | -1 | -1 | 187627717 | 88680792 | FAT1::chr4:187627716-187627766(-) | NTRK3::chr15:88680742-88680792(-) |
| FAT1 | STAT6 | 4 | 12 | -1 | -1 | 187508949 | 57496773 | FAT1::chr4:187508948-187508998(-) | STAT6::chr12:57496723-57496773(-) |
| FGFR2 | CCDC6 | 10 | 10 | -1 | -1 | 123243212 | 61612460 | FGFR2::chr10:123243211-123243261(-) | CCDC6::chr10:61612410-61612460(-) |
| FOXP1 | FHIT | 3 | 3 | -1 | -1 | 71348971 | 59908140 | FOXP1::chr3:71348970-71349020(-) | FHIT::chr3:59908090-59908140(-) |
| FUS | DDIT3 | 16 | 12 | 1 | -1 | 31199678 | 57911536 | FUS::chr16:31199628-31199678(+) | DDIT3::chr12:57911486-57911536(-) |
| MYH9 | BIRC6 | 22 | 2 | -1 | 1 | 36677327 | 32820101 | MYH9::chr22:36677326-36677376(-) | BIRC6::chr2:32820100-32820150(+) |
| MYH9 | CD74 | 22 | 5 | -1 | -1 | 36685235 | 149792314 | MYH9::chr22:36685234-36685284(-) | CD74::chr5:149792264-149792314(-) |
| NAB2 | STAT6 | 12 | 12 | 1 | -1 | 57487381 | 57493223 | NAB2::chr12:57487331-57487381(+) | STAT6::chr12:57493173-57493223(-) |
| NCOA2 | LEPROTL1 | 8 | 8 | -1 | 1 | 71315935 | 29959414 | NCOA2::chr8:71315934-71315984(-) | LEPROTL1::chr8:29959413-29959463(+) |
| NDRG1 | MYC | 8 | 8 | -1 | 1 | 134292475 | 128750497 | NDRG1::chr8:134292474-134292524(-) | MYC::chr8:128750496-128750546(+) |
| NTRK3 | ETV6 | 15 | 12 | -1 | 1 | 88669502 | 12022358 | NTRK3::chr15:88669501-88669551(-) | ETV6::chr12:12022357-12022407(+) |
| RAF1 | XPC | 3 | 3 | -1 | -1 | 12705312 | 14189501 | RAF1::chr3:12705311-12705361(-) | XPC::chr3:14189451-14189501(-) |
| RARA | CDK12 | 17 | 17 | 1 | 1 | 38504716 | 37686857 | RARA::chr17:38504666-38504716(+) | CDK12::chr17:37686856-37686906(+) |
| RARA | PML | 17 | 15 | 1 | 1 | 38487648 | 74325497 | RARA::chr17:38487598-38487648(+) | PML::chr15:74325496-74325546(+) |
| RARA | PML | 17 | 15 | 1 | 1 | 38499119 | 74326819 | RARA::chr17:38499069-38499119(+) | PML::chr15:74326818-74326868(+) |
| RBM10 | TFE3 | X | X | 1 | -1 | 47041725 | 48895639 | RBM10::chrX:47041675-47041725(+) | TFE3::chrX:48895589-48895639(-) |
| RET | CCDC6 | 10 | 10 | 1 | -1 | 43610184 | 61612460 | RET::chr10:43610134-43610184(+) | CCDC6::chr10:61612410-61612460(-) |
| RET | NCOA4 | 10 | 10 | 1 | 1 | 43610184 | 51584616 | RET::chr10:43610134-43610184(+) | NCOA4::chr10:51584615-51584665(+) |
| RNF43 | RARA | 17 | 17 | -1 | 1 | 56492687 | 38487109 | RNF43::chr17:56492686-56492736(-) | RARA::chr17:38487108-38487158(+) |
| RPL22 | PRDM16 | 1 | 1 | -1 | 1 | 6257712 | 3342145 | RPL22::chr1:6257711-6257761(-) | PRDM16::chr1:3342144-3342194(+) |
| SMARCE1 | RARA | 17 | 17 | -1 | 1 | 38792183 | 38511515 | SMARCE1::chr17:38792182-38792232(-) | RARA::chr17:38511514-38511564(+) |
| SPOP | PRDM16 | 17 | 1 | -1 | 1 | 47755295 | 3301716 | SPOP::chr17:47755294-47755344(-) | PRDM16::chr1:3301715-3301765(+) |
| SPOP | RARA | 17 | 17 | -1 | 1 | 47755295 | 38487109 | SPOP::chr17:47755294-47755344(-) | RARA::chr17:38487108-38487158(+) |
| TFE3 | SFPQ | X | 1 | -1 | -1 | 48900637 | 35653691 | TFE3::chrX:48900636-48900686(-) | SFPQ::chr1:35653641-35653691(-) |
| TRIM33 | RET | 1 | 10 | -1 | 1 | 114964058 | 43612032 | TRIM33::chr1:114964057-114964107(-) | RET::chr10:43612031-43612081(+) |
| WWTR1 | ATR | 3 | 3 | -1 | -1 | 149374663 | 142168444 | WWTR1::chr3:149374662-149374712(-) | ATR::chr3:142168394-142168444(-) |
| ABI1 | CUBN | 10 | 10 | -1 | -1 | 27149676 | 17032514 | ABI1::chr10:27149675-27149725(-) | CUBN::chr10:17032464-17032514(-) |
| ACVR2A | KANK1 | 2 | 9 | 1 | 1 | 148672903 | 676890 | ACVR2A::chr2:148672853-148672903(+) | KANK1::chr9:676889-676939(+) |
| ACVR2A | SUMO1 | 2 | 2 | 1 | -1 | 148680680 | 203072044 | ACVR2A::chr2:148680630-148680680(+) | SUMO1::chr2:203071994-203072044(-) |
| ADAMTS16 | TERT | 5 | 5 | 1 | -1 | 5191903 | 1282739 | ADAMTS16::chr5:5191853-5191903(+) | TERT::chr5:1282689-1282739(-) |
| ADRBK2 | EP300 | 22 | 22 | 1 | 1 | 25961120 | 41550999 | ADRBK2::chr22:25961070-25961120(+) | EP300::chr22:41550998-41551048(+) |
| ADRBK2 | ZNRF3 | 22 | 22 | 1 | 1 | 25961120 | 29383064 | ADRBK2::chr22:25961070-25961120(+) | ZNRF3::chr22:29383063-29383113(+) |

| | | | | | | | | | |
|---|---|---|---|---|---|---|---|---|---|
| AFF1 | PTPN13 | 4 | 4 | 1 | 1 | 87928576 | 87718027 | AFF1::chr4:87928526-87928576(+) | PTPN13::chr4:87718026-87718076(+) |
| AFF1 | PTPN13 | 4 | 4 | 1 | 1 | 87869723 | 87718027 | AFF1::chr4:87869673-87869723(+) | PTPN13::chr4:87718026-87718076(+) |
| AFF1 | PTPN13 | 4 | 4 | 1 | 1 | 87869723 | 87637683 | AFF1::chr4:87869673-87869723(+) | PTPN13::chr4:87637682-87637732(+) |
| AGAP1 | ERBB4 | 2 | 2 | 1 | -1 | 236715975 | 212289026 | AGAP1::chr2:236715925-236715975(+) | ERBB4::chr2:212288976-212289026(-) |
| AKAP13 | CHD2 | 15 | 15 | 1 | 1 | 86077111 | 93555575 | AKAP13::chr15:86077061-86077111(+) | CHD2::chr15:93555574-93555624(+) |
| AKAP9 | RUNX1 | 7 | 21 | 1 | -1 | 91699496 | 36171759 | AKAP9::chr7:91699446-91699496(+) | RUNX1::chr21:36171709-36171759(-) |
| APC | ALDH1L1 | 5 | 3 | 1 | -1 | 112043579 | 125879845 | APC::chr5:112043529-112043579(+) | ALDH1L1::chr3:125879795-125879845(-) |
| APC | C5orf13 | 5 | 5 | 1 | -1 | 112043579 | 111071204 | APC::chr5:112043529-112043579(+) | C5orf13::chr5:111071154-111071204(-) |
| APC | REEP2 | 5 | 5 | 1 | 1 | 112137080 | 137781372 | APC::chr5:112137030-112137080(+) | REEP2::chr5:137781371-137781421(+) |
| APC | SLC27A6 | 5 | 5 | 1 | 1 | 112043579 | 128364038 | APC::chr5:112043529-112043579(+) | SLC27A6::chr5:128364037-128364087(+) |
| APPBP2 | SETD2 | 17 | 3 | -1 | -1 | 58571827 | 47061330 | APPBP2::chr17:58571826-58571876(-) | SETD2::chr3:47061280-47061330(-) |
| ARHGEF10 | C6orf103 | 8 | 6 | 1 | 1 | 1791602 | 147022077 | ARHGEF10::chr8:1791552-1791602(+) | C6orf103::chr6:147022076-147022126(+) |
| ARHGEF10 | RALYL | 8 | 8 | 1 | 1 | 1857636 | 85686814 | ARHGEF10::chr8:1857586-1857636(+) | RALYL::chr8:85686813-85686863(+) |
| ARHGEF10L | CSDE1 | 1 | 1 | 1 | -1 | 18014242 | 115292828 | ARHGEF10L::chr1:18014192-18014242(+) | CSDE1::chr1:115292778-115292828(-) |
| ARHGEF10L | EYA2 | 1 | 20 | 1 | 1 | 17914974 | 45618640 | ARHGEF10L::chr1:17914924-17914974(+) | EYA2::chr20:45618639-45618689(+) |
| ARHGEF10L | PADI4 | 1 | 1 | 1 | 1 | 17866445 | 17681028 | ARHGEF10L::chr1:17866395-17866445(+) | PADI4::chr1:17681027-17681077(+) |
| ARHGEF10L | PADI4 | 1 | 1 | 1 | 1 | 17991090 | 17657464 | ARHGEF10L::chr1:17991040-17991090(+) | PADI4::chr1:17657463-17657513(+) |
| ARHGEF10L | PADI4 | 1 | 1 | 1 | 1 | 17983203 | 17657464 | ARHGEF10L::chr1:17983153-17983203(+) | PADI4::chr1:17657463-17657513(+) |
| ARHGEF10L | PADI4 | 1 | 1 | 1 | 1 | 18021775 | 17668438 | ARHGEF10L::chr1:18021725-18021775(+) | PADI4::chr1:17668437-17668487(+) |
| ARHGEF10L | PADI4 | 1 | 1 | 1 | 1 | 17965184 | 17664533 | ARHGEF10L::chr1:17965134-17965184(+) | PADI4::chr1:17664532-17664582(+) |
| ARHGEF10L | PAK4 | 1 | 19 | 1 | 1 | 17991090 | 39660172 | ARHGEF10L::chr1:17991040-17991090(+) | PAK4::chr19:39660171-39660221(+) |
| ARHGEF12 | CEP164 | 11 | 11 | 1 | 1 | 120207984 | 117209282 | ARHGEF12::chr11:120207934-120207984(+) | CEP164::chr11:117209281-117209331(+) |
| ARHGEF12 | CNTN5 | 11 | 11 | 1 | 1 | 120280159 | 99786786 | ARHGEF12::chr11:120280109-120280159(+) | CNTN5::chr11:99786785-99786835(+) |
| ARHGEF12 | OAF | 11 | 11 | 1 | 1 | 120207984 | 120096370 | ARHGEF12::chr11:120207934-120207984(+) | OAF::chr11:120096369-120096419(+) |
| ARHGEF12 | POU2F3 | 11 | 11 | 1 | 1 | 120207984 | 120139919 | ARHGEF12::chr11:120207934-120207984(+) | POU2F3::chr11:120139918-120139968(+) |
| ARHGEF12 | POU2F3 | 11 | 11 | 1 | 1 | 120207984 | 120186070 | ARHGEF12::chr11:120207934-120207984(+) | POU2F3::chr11:120186069-120186119(+) |
| ARHGEF12 | SIK3 | 11 | 11 | 1 | -1 | 120207984 | 116827780 | ARHGEF12::chr11:120207934-120207984(+) | SIK3::chr11:116827730-116827780(-) |
| ARID1A | BEND5 | 1 | 1 | 1 | -1 | 27024031 | 49202124 | ARID1A::chr1:27023981-27024031(+) | BEND5::chr1:49202074-49202124(-) |
| ARID1A | DHDDS | 1 | 1 | 1 | 1 | 27059283 | 26764659 | ARID1A::chr1:27059233-27059283(+) | DHDDS::chr1:26764658-26764708(+) |
| ARID1A | NIPBL | 1 | 5 | 1 | 1 | 27102198 | 36995724 | ARID1A::chr1:27102148-27102198(+) | NIPBL::chr5:36995723-36995773(+) |
| ARID1A | PRKCZ | 1 | 1 | 1 | 1 | 27088810 | 2066701 | ARID1A::chr1:27088760-27088810(+) | PRKCZ::chr1:2066700-2066750(+) |
| ARID1A | RPS6KA1 | 1 | 1 | 1 | 1 | 27102198 | 26897940 | ARID1A::chr1:27102148-27102198(+) | RPS6KA1::chr1:26897939-26897989(+) |
| ARID1A | SEC11A | 1 | 15 | 1 | -1 | 27094490 | 85234875 | ARID1A::chr1:27094440-27094490(+) | SEC11A::chr15:85234825-85234875(-) |
| ARID1A | THSD7B | 1 | 2 | 1 | 1 | 27059283 | 137639714 | ARID1A::chr1:27059233-27059283(+) | THSD7B::chr2:137639713-137639763(+) |
| ARID1A | UBE2U | 1 | 1 | 1 | 1 | 27059283 | 64686542 | ARID1A::chr1:27059233-27059283(+) | UBE2U::chr1:64686541-64686591(+) |
| ARID1A | YTHDF2 | 1 | 1 | 1 | 1 | 27059283 | 29064170 | ARID1A::chr1:27059233-27059283(+) | YTHDF2::chr1:29064169-29064219(+) |
| ARID1B | C6orf35 | 6 | 6 | 1 | -1 | 157358010 | 157714031 | ARID1B::chr6:157357960-157358010(+) | C6orf35::chr6:157713981-157714031(-) |
| ARID1B | ENPP4 | 6 | 6 | 1 | 1 | 157406039 | 46107288 | ARID1B::chr6:157405989-157406039(+) | ENPP4::chr6:46107287-46107337(+) |
| ARID1B | EZR | 6 | 6 | 1 | -1 | 157256710 | 159210403 | ARID1B::chr6:157256660-157256710(+) | EZR::chr6:159210353-159210403(-) |
| ARID1B | PARK2 | 6 | 6 | 1 | -1 | 157150555 | 162864505 | ARID1B::chr6:157150505-157150555(+) | PARK2::chr6:162864455-162864505(-) |
| ARID1B | PREP | 6 | 6 | 1 | -1 | 157150555 | 105845802 | ARID1B::chr6:157150505-157150555(+) | PREP::chr6:105845752-105845802(-) |
| ARID1B | SAMD5 | 6 | 6 | 1 | 1 | 157150555 | 147885530 | ARID1B::chr6:157150505-157150555(+) | SAMD5::chr6:147885529-147885579(+) |
| ARID1B | SASH1 | 6 | 6 | 1 | 1 | 157150555 | 148711270 | ARID1B::chr6:157150505-157150555(+) | SASH1::chr6:148711269-148711319(+) |

| | | | | | | | | | |
|---|---|---|---|---|---|---|---|---|---|
| ARID1B | SMOC2 | 6 | 6 | 1 | 1 | 157150555 | 169051361 | ARID1B::chr6:157150505-157150555(+) | SMOC2::chr6:169051360-169051410(+) |
| ARID1B | SNX9 | 6 | 6 | 1 | 1 | 157150555 | 158288579 | ARID1B::chr6:157150505-157150555(+) | SNX9::chr6:158288578-158288628(+) |
| ARID1B | SYNE1 | 6 | 6 | 1 | -1 | 157406039 | 152464900 | ARID1B::chr6:157405989-157406039(+) | SYNE1::chr6:152464850-152464900(-) |
| ARID1B | TIAM2 | 6 | 6 | 1 | 1 | 157100605 | 155411423 | ARID1B::chr6:157100555-157100605(+) | TIAM2::chr6:155411422-155411472(+) |
| ARID1B | TIAM2 | 6 | 6 | 1 | 1 | 157470085 | 155265250 | ARID1B::chr6:157470035-157470085(+) | TIAM2::chr6:155265249-155265299(+) |
| ARID2 | C12orf56 | 12 | 12 | 1 | -1 | 46125097 | 64746836 | ARID2::chr12:46125047-46125097(+) | C12orf56::chr12:64746786-64746836(-) |
| ARID2 | SPATS1 | 12 | 6 | 1 | 1 | 46287504 | 44336116 | ARID2::chr12:46287454-46287504(+) | SPATS1::chr6:44336115-44336165(+) |
| ARID2 | SPSB2 | 12 | 12 | 1 | -1 | 46205334 | 6991451 | ARID2::chr12:46205284-46205334(+) | SPSB2::chr12:6991401-6991451(-) |
| ARID2 | TMPRSS2 | 12 | 21 | 1 | -1 | 46233279 | 42902145 | ARID2::chr12:46233229-46233279(+) | TMPRSS2::chr21:42902095-42902145(-) |
| ARNT | F5 | 1 | 1 | -1 | -1 | 150825238 | 169494146 | ARNT::chr1:150825237-150825287(-) | F5::chr1:169494096-169494146(-) |
| AS3MT | SUFU | 10 | 10 | 1 | 1 | 104650435 | 104356897 | AS3MT::chr10:104650385-104650435(+) | SUFU::chr10:104356896-104356946(+) |
| ASXL1 | C20orf160 | 20 | 20 | 1 | 1 | 30956926 | 30613659 | ASXL1::chr20:30956876-30956926(+) | C20orf160::chr20:30613658-30613708(+) |
| ASXL1 | DEFB118 | 20 | 20 | 1 | 1 | 30954269 | 29960660 | ASXL1::chr20:30954219-30954269(+) | DEFB118::chr20:29960659-29960709(+) |
| ASXL1 | PDRG1 | 20 | 20 | 1 | -1 | 30959677 | 30536685 | ASXL1::chr20:30959627-30959677(+) | PDRG1::chr20:30536635-30536685(-) |
| ASXL1 | TPX2 | 20 | 20 | 1 | 1 | 30956926 | 30359363 | ASXL1::chr20:30956876-30956926(+) | TPX2::chr20:30359362-30359412(+) |
| ASXL1 | XKR7 | 20 | 20 | 1 | 1 | 30956926 | 30582871 | ASXL1::chr20:30956876-30956926(+) | XKR7::chr20:30582870-30582920(+) |
| ASXL2 | C2orf44 | 2 | 2 | -1 | -1 | 26058407 | 24262382 | ASXL2::chr2:26058406-26058456(-) | C2orf44::chr2:24262332-24262382(-) |
| ASXL2 | DENND2A | 2 | 7 | -1 | -1 | 26101035 | 140302342 | ASXL2::chr2:26101034-26101084(-) | DENND2A::chr7:140302292-140302342(-) |
| ASXL2 | FKBP8 | 2 | 19 | -1 | -1 | 26058407 | 18652805 | ASXL2::chr2:26058406-26058456(-) | FKBP8::chr19:18652755-18652805(-) |
| ASXL2 | HSPH1 | 2 | 13 | -1 | -1 | 26101035 | 31733000 | ASXL2::chr2:26101034-26101084(-) | HSPH1::chr13:31732950-31733000(-) |
| ASXL2 | KIF3C | 2 | 2 | -1 | -1 | 26101035 | 26179313 | ASXL2::chr2:26101034-26101084(-) | KIF3C::chr2:26179263-26179313(-) |
| ASXL2 | MINPP1 | 2 | 10 | -1 | 1 | 26068350 | 89272881 | ASXL2::chr2:26068349-26068399(-) | MINPP1::chr10:89272880-89272930(+) |
| ASXL2 | MORC3 | 2 | 21 | -1 | 1 | 26101035 | 37744672 | ASXL2::chr2:26101034-26101084(-) | MORC3::chr21:37744671-37744721(+) |
| ASXL2 | PPWD1 | 2 | 5 | -1 | 1 | 26029098 | 64863340 | ASXL2::chr2:26029097-26029147(-) | PPWD1::chr5:64863339-64863389(+) |
| ASXL2 | SRBD1 | 2 | 2 | -1 | -1 | 26101035 | 45715470 | ASXL2::chr2:26101034-26101084(-) | SRBD1::chr2:45715420-45715470(-) |
| ATAD2 | ERBB4 | 8 | 2 | -1 | -1 | 124381298 | 212530202 | ATAD2::chr8:124381297-124381347(-) | ERBB4::chr2:212530152-212530202(-) |
| ATG7 | PPARG | 3 | 3 | 1 | 1 | 11468400 | 12475397 | ATG7::chr3:11468350-11468400(+) | PPARG::chr3:12475396-12475446(+) |
| ATM | CARD18 | 11 | 11 | 1 | -1 | 108218092 | 105009805 | ATM::chr11:108218042-108218092(+) | CARD18::chr11:105009755-105009805(-) |
| ATP1A1 | FN1 | 1 | 2 | 1 | -1 | 116947396 | 216274837 | ATP1A1::chr1:116947346-116947396(+) | FN1::chr2:216274787-216274837(-) |
| ATP1A1 | IGSF3 | 1 | 1 | 1 | -1 | 116946597 | 117122499 | ATP1A1::chr1:116946547-116946597(+) | IGSF3::chr1:117122449-117122499(-) |
| ATP1A1 | RABEP1 | 1 | 17 | 1 | 1 | 116947394 | 5283828 | ATP1A1::chr1:116947344-116947394(+) | RABEP1::chr17:5283827-5283877(+) |
| ATP1A1 | VANGL1 | 1 | 1 | 1 | 1 | 116916145 | 116206282 | ATP1A1::chr1:116916095-116916145(+) | VANGL1::chr1:116206281-116206331(+) |
| ATR | DBR1 | 3 | 3 | -1 | -1 | 142297488 | 137892468 | ATR::chr3:142297487-142297537(-) | DBR1::chr3:137892418-137892468(-) |
| ATR | R3HDM2 | 3 | 12 | -1 | -1 | 142284963 | 57650294 | ATR::chr3:142284962-142285012(-) | R3HDM2::chr12:57650244-57650294(-) |
| ATRX | ABCB7 | X | X | -1 | -1 | 77041468 | 74334669 | ATRX::chrX:77041467-77041517(-) | ABCB7::chrX:74334619-74334669(-) |
| ATRX | ARHGEF9 | X | X | -1 | -1 | 77041468 | 62875617 | ATRX::chrX:77041467-77041517(-) | ARHGEF9::chrX:62875567-62875617(-) |
| ATRX | EFCAB4B | X | 12 | -1 | -1 | 77041468 | 3822483 | ATRX::chrX:77041467-77041517(-) | EFCAB4B::chr12:3822433-3822483(-) |
| ATRX | LPAR4 | X | X | -1 | 1 | 77041468 | 78010288 | ATRX::chrX:77041467-77041517(-) | LPAR4::chrX:78010287-78010337(+) |
| ATRX | MAGT1 | X | X | -1 | -1 | 76972608 | 77112994 | ATRX::chrX:76972607-76972657(-) | MAGT1::chrX:77112944-77112994(-) |
| ATRX | SNTB2 | X | 16 | -1 | 1 | 76953071 | 69304046 | ATRX::chrX:76953070-76953120(-) | SNTB2::chr16:69304045-69304095(+) |
| ATRX | ZNF304 | X | 19 | -1 | 1 | 77041468 | 57865093 | ATRX::chrX:77041467-77041517(-) | ZNF304::chr19:57865092-57865142(+) |
| AXIN1 | BBX | 16 | 3 | -1 | 1 | 396148 | 107466812 | AXIN1::chr16:396147-396197(-) | BBX::chr3:107466811-107466861(+) |
| AXIN1 | IFT140 | 16 | 16 | -1 | -1 | 354304 | 1576797 | AXIN1::chr16:354303-354353(-) | IFT140::chr16:1576747-1576797(-) |

| | | | | | | | | | |
|---|---|---|---|---|---|---|---|---|---|
| AXIN1 | TMEM207 | 16 | 3 | -1 | -1 | 354304 | 190165616 | AXIN1::chr16:354303-354353(-) | TMEM207::chr3:190165566-190165616(-) |
| AXIN1 | TMEM8A | 16 | 16 | -1 | -1 | 364543 | 427848 | AXIN1::chr16:364542-364592(-) | TMEM8A::chr16:427798-427848(-) |
| AXIN2 | EIF2AK2 | 17 | 2 | -1 | -1 | 63524685 | 37342384 | AXIN2::chr17:63524684-63524734(-) | EIF2AK2::chr2:37342334-37342384(-) |
| AXIN2 | VWA3B | 17 | 2 | -1 | 1 | 63553924 | 98872587 | AXIN2::chr17:63553923-63553973(-) | VWA3B::chr2:98872586-98872636(+) |
| B2M | CCT5 | 15 | 5 | 1 | 1 | 45010359 | 10250033 | B2M::chr15:45010309-45010359(+) | CCT5::chr5:10250032-10250082(+) |
| B2M | PPP4R1 | 15 | 18 | 1 | -1 | 45003811 | 9562073 | B2M::chr15:45003761-45003811(+) | PPP4R1::chr18:9562023-9562073(-) |
| BAP1 | DNAH1 | 3 | 3 | -1 | 1 | 52439781 | 52378506 | BAP1::chr3:52439780-52439830(-) | DNAH1::chr3:52378505-52378555(+) |
| BAP1 | PLXNB1 | 3 | 3 | -1 | -1 | 52437154 | 48455480 | BAP1::chr3:52437153-52437203(-) | PLXNB1::chr3:48455430-48455480(-) |
| BAP1 | RILP | 3 | 17 | -1 | -1 | 52443570 | 1551789 | BAP1::chr3:52443569-52443619(-) | RILP::chr17:1551739-1551789(-) |
| BAP1 | TCEA3 | 3 | 1 | -1 | -1 | 52440269 | 23745632 | BAP1::chr3:52440268-52440318(-) | TCEA3::chr1:23745582-23745632(-) |
| BARD1 | C6orf222 | 2 | 6 | -1 | -1 | 215645284 | 36291967 | BARD1::chr2:215645283-215645333(-) | C6orf222::chr6:36291917-36291967(-) |
| BAX | VAV2 | 19 | 9 | 1 | -1 | 49464171 | 136629231 | BAX::chr19:49464121-49464171(+) | VAV2::chr9:136629181-136629231(-) |
| BAZ1A | RNF115 | 14 | 1 | -1 | 1 | 35343698 | 145646115 | BAZ1A::chr14:35343697-35343747(-) | RNF115::chr1:145646114-145646164(+) |
| BCL10 | DPYD | 1 | 1 | -1 | -1 | 85741979 | 98015300 | BCL10::chr1:85741978-85742028(-) | DPYD::chr1:98015250-98015300(-) |
| BCL11B | MIPOL1 | 14 | 14 | -1 | 1 | 99723808 | 37688758 | BCL11B::chr14:99723807-99723857(-) | MIPOL1::chr14:37688757-37688807(+) |
| BCL9L | DGKI | 11 | 7 | -1 | -1 | 118795947 | 137374748 | BCL9L::chr11:118795946-118795996(-) | DGKI::chr7:137374698-137374748(-) |
| BCOR | PHEX | X | X | -1 | 1 | 40036260 | 22231021 | BCOR::chrX:40036259-40036309(-) | PHEX::chrX:22231020-22231070(+) |
| BCORL1 | UTP14A | X | X | 1 | 1 | 129173257 | 129041343 | BCORL1::chrX:129173207-129173257(+) | UTP14A::chrX:129041342-129041392(+) |
| BIRC3 | CCT5 | 11 | 5 | 1 | 1 | 102208465 | 10250033 | BIRC3::chr11:102208415-102208465(+) | CCT5::chr5:10250032-10250082(+) |
| BLM | ABHD2 | 15 | 15 | 1 | 1 | 91260671 | 89694908 | BLM::chr15:91260621-91260671(+) | ABHD2::chr15:89694907-89694957(+) |
| BMPR1A | C10orf11 | 10 | 10 | 1 | 1 | 88659883 | 77818424 | BMPR1A::chr10:88659833-88659883(+) | C10orf11::chr10:77818423-77818473(+) |
| BMPR1A | ENTPD1 | 10 | 10 | 1 | 1 | 88659883 | 97604233 | BMPR1A::chr10:88659833-88659883(+) | ENTPD1::chr10:97604232-97604282(+) |
| BPTF | RAD17 | 17 | 5 | 1 | 1 | 65871860 | 68692224 | BPTF::chr17:65871810-65871860(+) | RAD17::chr5:68692223-68692273(+) |
| BRIP1 | CUEDC1 | 17 | 17 | -1 | -1 | 59853762 | 55963240 | BRIP1::chr17:59853761-59853811(-) | CUEDC1::chr17:55963190-55963240(-) |
| BUB1B | EIF2AK4 | 15 | 15 | 1 | 1 | 40468874 | 40301815 | BUB1B::chr15:40468824-40468874(+) | EIF2AK4::chr15:40301814-40301864(+) |
| BUB1B | EIF2AK4 | 15 | 15 | 1 | 1 | 40510763 | 40278006 | BUB1B::chr15:40510713-40510763(+) | EIF2AK4::chr15:40278005-40278055(+) |
| BUB1B | PCM1 | 15 | 8 | 1 | 1 | 40462882 | 17842956 | BUB1B::chr15:40462832-40462882(+) | PCM1::chr8:17842955-17843005(+) |
| C10orf26 | SUFU | 10 | 10 | 1 | 1 | 104536174 | 104352339 | C10orf26::chr10:104536124-104536174(+) | SUFU::chr10:104352338-104352388(+) |
| C10orf26 | SUFU | 10 | 10 | 1 | 1 | 104503900 | 104375025 | C10orf26::chr10:104503850-104503900(+) | SUFU::chr10:104375024-104375074(+) |
| C11orf84 | BUB1B | 11 | 15 | 1 | 1 | 63581285 | 40468678 | C11orf84::chr11:63581235-63581285(+) | BUB1B::chr15:40468677-40468727(+) |
| C2CD2L | DDX10 | 11 | 11 | 1 | 1 | 118985075 | 108709173 | C2CD2L::chr11:118985025-118985075(+) | DDX10::chr11:108709172-108709222(+) |
| C3orf67 | FHIT | 3 | 3 | -1 | -1 | 58899434 | 59738047 | C3orf67::chr3:58899433-58899483(-) | FHIT::chr3:59737997-59738047(-) |
| CABIN1 | GRIN2A | 22 | 16 | 1 | -1 | 24515665 | 9984957 | CABIN1::chr22:24515615-24515665(+) | GRIN2A::chr16:9984907-9984957(-) |
| CAMSAP1L1 | CARS | 1 | 11 | 1 | -1 | 200824017 | 3022478 | CAMSAP1L1::chr1:200823967-200824017(+) | CARS::chr11:3022428-3022478(-) |
| CAMTA1 | DHRS3 | 1 | 1 | 1 | -1 | 6845635 | 12640694 | CAMTA1::chr1:6845585-6845635(+) | DHRS3::chr1:12640644-12640694(-) |
| CAMTA1 | THAP3 | 1 | 1 | 1 | 1 | 6845635 | 6688559 | CAMTA1::chr1:6845585-6845635(+) | THAP3::chr1:6688558-6688608(+) |
| CARS | OSBPL5 | 11 | 11 | -1 | -1 | 3059280 | 3115564 | CARS::chr11:3059279-3059329(-) | OSBPL5::chr11:3115514-3115564(-) |
| CARS | TSHZ3 | 11 | 19 | -1 | -1 | 3078573 | 31770658 | CARS::chr11:3078572-3078622(-) | TSHZ3::chr19:31770608-31770658(-) |
| CASC3 | CDK12 | 17 | 17 | 1 | 1 | 38297860 | 37686884 | CASC3::chr17:38297810-38297860(+) | CDK12::chr17:37686883-37686933(+) |
| CASC5 | LIPC | 15 | 15 | 1 | 1 | 40886590 | 58855704 | CASC5::chr15:40886540-40886590(+) | LIPC::chr15:58855703-58855753(+) |
| CASP8 | NRP2 | 2 | 2 | 1 | 1 | 202098835 | 206547815 | CASP8::chr2:202098785-202098835(+) | NRP2::chr2:206547814-206547864(+) |
| CASP8 | UBR3 | 2 | 2 | 1 | 1 | 202131514 | 170789422 | CASP8::chr2:202131464-202131514(+) | UBR3::chr2:170789421-170789471(+) |
| CBFA2T3 | ACSF3 | 16 | 16 | -1 | 1 | 89007428 | 89169012 | CBFA2T3::chr16:89007427-89007477(-) | ACSF3::chr16:89169011-89169061(+) |

| | | | | | | | | | |
|---|---|---|---|---|---|---|---|---|---|
| CBFB | ATP13A3 | 16 | 3 | 1 | -1 | 67070658 | 194182965 | CBFB::chr16:67070608-67070658(+) | ATP13A3::chr3:194182915-194182965(-) |
| CBFB | CA7 | 16 | 16 | 1 | 1 | 67100701 | 66880933 | CBFB::chr16:67100651-67100701(+) | CA7::chr16:66880932-66880982(+) |
| CBFB | COG4 | 16 | 16 | 1 | -1 | 67100701 | 70553634 | CBFB::chr16:67100651-67100701(+) | COG4::chr16:70553584-70553634(-) |
| CBFB | NAE1 | 16 | 16 | 1 | -1 | 67070658 | 66857503 | CBFB::chr16:67070608-67070658(+) | NAE1::chr16:66857453-66857503(-) |
| CBL | CXCR5 | 11 | 11 | 1 | 1 | 119103405 | 118764305 | CBL::chr11:119103355-119103405(+) | CXCR5::chr11:118764304-118764354(+) |
| CBL | PDZD3 | 11 | 11 | 1 | 1 | 119077322 | 119059349 | CBL::chr11:119077272-119077322(+) | PDZD3::chr11:119059348-119059398(+) |
| CBL | PHLDB1 | 11 | 11 | 1 | 1 | 119156276 | 118514518 | CBL::chr11:119156226-119156276(+) | PHLDB1::chr11:118514517-118514567(+) |
| CBL | PHLDB1 | 11 | 11 | 1 | 1 | 119077322 | 118484531 | CBL::chr11:119077272-119077322(+) | PHLDB1::chr11:118484530-118484580(+) |
| CBL | SC5DL | 11 | 11 | 1 | 1 | 119103405 | 121177095 | CBL::chr11:119103355-119103405(+) | SC5DL::chr11:121177094-121177144(+) |
| CBL | TRAPPC4 | 11 | 11 | 1 | 1 | 119158656 | 118894031 | CBL::chr11:119158606-119158656(+) | TRAPPC4::chr11:118894030-118894080(+) |
| CBL | VWA5A | 11 | 11 | 1 | 1 | 119077322 | 123988204 | CBL::chr11:119077272-119077322(+) | VWA5A::chr11:123988203-123988253(+) |
| CBLB | IL17RE | 3 | 3 | -1 | 1 | 105572258 | 9944574 | CBLB::chr3:105572257-105572307(-) | IL17RE::chr3:9944573-9944623(+) |
| CBLB | KIAA1524 | 3 | 3 | -1 | -1 | 105572258 | 108304059 | CBLB::chr3:105572257-105572307(-) | KIAA1524::chr3:108304009-108304059(-) |
| CBLC | PVR | 19 | 19 | 1 | 1 | 45303707 | 45164559 | CBLC::chr19:45303657-45303707(+) | PVR::chr19:45164558-45164608(+) |
| CCAR1 | TET1 | 10 | 10 | 1 | 1 | 70520949 | 70432652 | CCAR1::chr10:70520899-70520949(+) | TET1::chr10:70432651-70432701(+) |
| CCAR1 | TET1 | 10 | 10 | 1 | 1 | 70517134 | 70432652 | CCAR1::chr10:70517084-70517134(+) | TET1::chr10:70432651-70432701(+) |
| CCDC6 | ANK3 | 10 | 10 | -1 | -1 | 61665880 | 61827767 | CCDC6::chr10:61665879-61665929(-) | ANK3::chr10:61827717-61827767(-) |
| CCDC6 | ANK3 | 10 | 10 | -1 | -1 | 61665880 | 61802514 | CCDC6::chr10:61665879-61665929(-) | ANK3::chr10:61802464-61802514(-) |
| CCDC6 | ANK3 | 10 | 10 | -1 | -1 | 61665880 | 61802517 | CCDC6::chr10:61665879-61665929(-) | ANK3::chr10:61802467-61802517(-) |
| CCDC6 | ANK3 | 10 | 10 | -1 | -1 | 61665880 | 61842495 | CCDC6::chr10:61665879-61665929(-) | ANK3::chr10:61842445-61842495(-) |
| CCDC6 | ANK3 | 10 | 10 | -1 | -1 | 61564178 | 61848116 | CCDC6::chr10:61564177-61564227(-) | ANK3::chr10:61848066-61848116(-) |
| CCDC6 | ANK3 | 10 | 10 | -1 | -1 | 61574410 | 61789584 | CCDC6::chr10:61574409-61574459(-) | ANK3::chr10:61789534-61789584(-) |
| CCDC6 | LIPI | 10 | 21 | -1 | -1 | 61612311 | 15481401 | CCDC6::chr10:61612310-61612360(-) | LIPI::chr21:15481351-15481401(-) |
| CCDC6 | UBE2D1 | 10 | 10 | -1 | 1 | 61665880 | 60121098 | CCDC6::chr10:61665879-61665929(-) | UBE2D1::chr10:60121097-60121147(+) |
| CCNC | MICAL1 | 6 | 6 | -1 | -1 | 100016372 | 109767975 | CCNC::chr6:100016371-100016421(-) | MICAL1::chr6:109767925-109767975(-) |
| CD274 | IGFBPL1 | 9 | 9 | 1 | -1 | 5465606 | 38409214 | CD274::chr9:5465556-5465606(+) | IGFBPL1::chr9:38409164-38409214(-) |
| CD44 | EXT2 | 11 | 11 | 1 | 1 | 35160917 | 44265699 | CD44::chr11:35160867-35160917(+) | EXT2::chr11:44265698-44265748(+) |
| CDC73 | TROVE2 | 1 | 1 | 1 | 1 | 193121574 | 193060781 | CDC73::chr1:193121524-193121574(+) | TROVE2::chr1:193060780-193060830(+) |
| CDH1 | CA1 | 16 | 8 | 1 | -1 | 68772314 | 86245849 | CDH1::chr16:68772264-68772314(+) | CA1::chr8:86245799-86245849(-) |
| CDH1 | CCDC132 | 16 | 7 | 1 | 1 | 68857529 | 92952923 | CDH1::chr16:68857479-68857529(+) | CCDC132::chr7:92952922-92952972(+) |
| CDH1 | CDKL3 | 16 | 5 | 1 | -1 | 68856128 | 133644157 | CDH1::chr16:68856078-68856128(+) | CDKL3::chr5:133644107-133644157(-) |
| CDH1 | DEXI | 16 | 16 | 1 | -1 | 68842751 | 11023416 | CDH1::chr16:68842701-68842751(+) | DEXI::chr16:11023366-11023416(-) |
| CDH1 | NPFFR2 | 16 | 4 | 1 | 1 | 68772314 | 73003757 | CDH1::chr16:68772264-68772314(+) | NPFFR2::chr4:73003756-73003806(+) |
| CDH1 | TRHDE | 16 | 12 | 1 | 1 | 68835796 | 72771775 | CDH1::chr16:68835746-68835796(+) | TRHDE::chr12:72771774-72771824(+) |
| CDH11 | KIFC3 | 16 | 16 | -1 | -1 | 64982512 | 57795448 | CDH11::chr16:64982511-64982561(-) | KIFC3::chr16:57795398-57795448(-) |
| CDH11 | PRC1 | 16 | 15 | -1 | -1 | 65005482 | 91512853 | CDH11::chr16:65005481-65005531(-) | PRC1::chr15:91512803-91512853(-) |
| CDK12 | CACNB1 | 17 | 17 | 1 | -1 | 37646986 | 37347846 | CDK12::chr17:37646936-37646986(+) | CACNB1::chr17:37347796-37347846(-) |
| CDK12 | FAM20A | 17 | 17 | 1 | -1 | 37628016 | 66551884 | CDK12::chr17:37627966-37628016(+) | FAM20A::chr17:66551834-66551884(-) |
| CDK12 | FBXL20 | 17 | 17 | 1 | -1 | 37628016 | 37439108 | CDK12::chr17:37627966-37628016(+) | FBXL20::chr17:37439058-37439108(-) |
| CDK12 | IKZF3 | 17 | 17 | 1 | -1 | 37657692 | 37988404 | CDK12::chr17:37657642-37657692(+) | IKZF3::chr17:37988354-37988404(-) |
| CDK12 | MED1 | 17 | 17 | 1 | -1 | 37619370 | 37571592 | CDK12::chr17:37619320-37619370(+) | MED1::chr17:37571542-37571592(-) |
| CDK12 | PLXDC1 | 17 | 17 | 1 | -1 | 37619370 | 37265644 | CDK12::chr17:37619320-37619370(+) | PLXDC1::chr17:37265594-37265644(-) |
| CDK12 | SMG6 | 17 | 17 | 1 | -1 | 37657692 | 1985250 | CDK12::chr17:37657642-37657692(+) | SMG6::chr17:1985200-1985250(-) |

| | | | | | | | | | |
|---|---|---|---|---|---|---|---|---|---|
| CDK5RAP3 | RNF43 | 17 | 17 | 1 | -1 | 46051814 | 56448394 | CDK5RAP3::chr17:46051764-46051814(+) | RNF43::chr17:56448344-56448394(-) |
| CDKN2A | CAPZB | 9 | 1 | -1 | -1 | 21974677 | 19671746 | CDKN2A::chr9:21974676-21974726(-) | CAPZB::chr1:19671696-19671746(-) |
| CDKN2A | EXD2 | 9 | 14 | -1 | 1 | 21974677 | 69695533 | CDKN2A::chr9:21974676-21974726(-) | EXD2::chr14:69695532-69695582(+) |
| CDKN2A | FASTKD3 | 9 | 5 | -1 | -1 | 21994138 | 7866096 | CDKN2A::chr9:21994137-21994187(-) | FASTKD3::chr5:7866046-7866096(-) |
| CDKN2A | KIAA1797 | 9 | 9 | -1 | 1 | 21994138 | 20881870 | CDKN2A::chr9:21994137-21994187(-) | KIAA1797::chr9:20881869-20881919(+) |
| CDKN2A | MGA | 9 | 15 | -1 | 1 | 21970901 | 42019378 | CDKN2A::chr9:21970900-21970950(-) | MGA::chr15:42019377-42019427(+) |
| CDKN2A | MTAP | 9 | 9 | -1 | 1 | 21974677 | 21815432 | CDKN2A::chr9:21974676-21974726(-) | MTAP::chr9:21815431-21815481(+) |
| CDKN2A | MYH10 | 9 | 17 | -1 | -1 | 21994138 | 8508300 | CDKN2A::chr9:21994137-21994187(-) | MYH10::chr17:8508250-8508300(-) |
| CDKN2A | SOBP | 9 | 6 | -1 | 1 | 21994138 | 107908284 | CDKN2A::chr9:21994137-21994187(-) | SOBP::chr6:107908283-107908333(+) |
| CDKN2A | SRGAP2 | 9 | 1 | -1 | 1 | 21994138 | 206634382 | CDKN2A::chr9:21994137-21994187(-) | SRGAP2::chr1:206634381-206634431(+) |
| CHD2 | ACSBG1 | 15 | 15 | 1 | -1 | 93444529 | 78466181 | CHD2::chr15:93444479-93444529(+) | ACSBG1::chr15:78466131-78466181(-) |
| CHD2 | DYX1C1 | 15 | 15 | 1 | -1 | 93444529 | 55677912 | CHD2::chr15:93444479-93444529(+) | DYX1C1::chr15:55677862-55677912(-) |
| CHD2 | HOMER2 | 15 | 15 | 1 | -1 | 93496803 | 83533011 | CHD2::chr15:93496753-93496803(+) | HOMER2::chr15:83532961-83533011(-) |
| CHD2 | ZNF880 | 15 | 19 | 1 | 1 | 93444529 | 52887102 | CHD2::chr15:93444479-93444529(+) | ZNF880::chr19:52887101-52887151(+) |
| CHEK2 | C22orf31 | 22 | 22 | -1 | -1 | 29130391 | 29456831 | CHEK2::chr22:29130390-29130440(-) | C22orf31::chr22:29456781-29456831(-) |
| CHMP1A | FANCA | 16 | 16 | -1 | -1 | 89720312 | 89833645 | CHMP1A::chr16:89720311-89720361(-) | FANCA::chr16:89833595-89833645(-) |
| CHMP1A | FANCA | 16 | 16 | -1 | -1 | 89723990 | 89851372 | CHMP1A::chr16:89723989-89724039(-) | FANCA::chr16:89851322-89851372(-) |
| CIC | DLK1 | 19 | 14 | 1 | 1 | 42798456 | 101193333 | CIC::chr19:42798406-42798456(+) | DLK1::chr14:101193332-101193382(+) |
| CIC | KLK11 | 19 | 19 | 1 | -1 | 42793558 | 51528922 | CIC::chr19:42793508-42793558(+) | KLK11::chr19:51528872-51528922(-) |
| CIC | LMNA | 19 | 1 | 1 | 1 | 42798456 | 156084592 | CIC::chr19:42798406-42798456(+) | LMNA::chr1:156084591-156084641(+) |
| CLASP1 | FBXO11 | 2 | 2 | -1 | -1 | 122363277 | 48047595 | CLASP1::chr2:122363276-122363326(-) | FBXO11::chr2:48047545-48047595(-) |
| CLEC16A | GRIN2A | 16 | 16 | 1 | -1 | 11154879 | 10032408 | CLEC16A::chr16:11154829-11154879(+) | GRIN2A::chr16:10032358-10032408(-) |
| CLTC | DNAH17 | 17 | 17 | 1 | -1 | 57697534 | 76511104 | CLTC::chr17:57697484-57697534(+) | DNAH17::chr17:76511054-76511104(-) |
| CLTC | FAM129C | 17 | 19 | 1 | 1 | 57697534 | 17648185 | CLTC::chr17:57697484-57697534(+) | FAM129C::chr19:17648184-17648234(+) |
| CLTC | PTRH2 | 17 | 17 | 1 | -1 | 57697534 | 57775339 | CLTC::chr17:57697484-57697534(+) | PTRH2::chr17:57775289-57775339(-) |
| CLTC | USP32 | 17 | 17 | 1 | -1 | 57697534 | 58422969 | CLTC::chr17:57697484-57697534(+) | USP32::chr17:58422919-58422969(-) |
| CNBP | ALDH1L1 | 3 | 3 | -1 | -1 | 128902619 | 125831723 | CNBP::chr3:128902618-128902668(-) | ALDH1L1::chr3:125831673-125831723(-) |
| CNTNAP2 | GSTK1 | 7 | 7 | 1 | 1 | 146536996 | 142960241 | CNTNAP2::chr7:146536946-146536996(+) | GSTK1::chr7:142960240-142960290(+) |
| COG6 | GPC5 | 13 | 13 | 1 | 1 | 40301685 | 93518535 | COG6::chr13:40301635-40301685(+) | GPC5::chr13:93518534-93518584(+) |
| CPSF6 | PTPRB | 12 | 12 | 1 | -1 | 69633486 | 70975082 | CPSF6::chr12:69633436-69633486(+) | PTPRB::chr12:70975032-70975082(-) |
| CREBBP | ADCY9 | 16 | 16 | -1 | -1 | 3929833 | 4016967 | CREBBP::chr16:3929832-3929882(-) | ADCY9::chr16:4016917-4016967(-) |
| CREBBP | ATG10 | 16 | 5 | -1 | 1 | 3900298 | 81354314 | CREBBP::chr16:3900297-3900347(-) | ATG10::chr5:81354313-81354363(+) |
| CREBBP | C16orf59 | 16 | 16 | -1 | 1 | 3788560 | 2514040 | CREBBP::chr16:3788559-3788609(-) | C16orf59::chr16:2514039-2514089(+) |
| CREBBP | CASKIN1 | 16 | 16 | -1 | -1 | 3794895 | 2240359 | CREBBP::chr16:3794894-3794944(-) | CASKIN1::chr16:2240309-2240359(-) |
| CREBBP | GLYR1 | 16 | 16 | -1 | -1 | 3900298 | 4882913 | CREBBP::chr16:3900297-3900347(-) | GLYR1::chr16:4882863-4882913(-) |
| CREBBP | HCN1 | 16 | 5 | -1 | -1 | 3900298 | 45645710 | CREBBP::chr16:3900297-3900347(-) | HCN1::chr5:45645660-45645710(-) |
| CREBBP | IFT140 | 16 | 16 | -1 | -1 | 3860604 | 1642589 | CREBBP::chr16:3860603-3860653(-) | IFT140::chr16:1642539-1642589(-) |
| CREBBP | LUC7L | 16 | 16 | -1 | -1 | 3828012 | 243014 | CREBBP::chr16:3828011-3828061(-) | LUC7L::chr16:242964-243014(-) |
| CREBBP | NUBP1 | 16 | 16 | -1 | 1 | 3819175 | 10846467 | CREBBP::chr16:3819174-3819224(-) | NUBP1::chr16:10846466-10846516(+) |
| CREBBP | PRSS33 | 16 | 16 | -1 | -1 | 3929833 | 2836122 | CREBBP::chr16:3929832-3929882(-) | PRSS33::chr16:2836072-2836122(-) |
| CREBBP | RANBP3 | 16 | 19 | -1 | -1 | 3900298 | 5941846 | CREBBP::chr16:3900297-3900347(-) | RANBP3::chr19:5941796-5941846(-) |
| CREBBP | SRRM2 | 16 | 16 | -1 | 1 | 3929833 | 2806335 | CREBBP::chr16:3929832-3929882(-) | SRRM2::chr16:2806334-2806384(+) |
| CREBBP | WRAP53 | 16 | 17 | -1 | 1 | 3860604 | 7591966 | CREBBP::chr16:3860603-3860653(-) | WRAP53::chr17:7591965-7592015(+) |

| | | | | | | | | | |
|---|---|---|---|---|---|---|---|---|---|
| CSMD3 | ADCY8 | 8 | 8 | -1 | -1 | 114290821 | 131964244 | CSMD3::chr8:114290820-114290870(-) | ADCY8::chr8:131964194-131964244(-) |
| CSMD3 | RIMS2 | 8 | 8 | -1 | 1 | 113599295 | 104955001 | CSMD3::chr8:113599294-113599344(-) | RIMS2::chr8:104955000-104955050(+) |
| CTCF | CES2 | 16 | 16 | 1 | 1 | 67605167 | 66971940 | CTCF::chr16:67605117-67605167(+) | CES2::chr16:66971939-66971989(+) |
| CTCF | E2F4 | 16 | 16 | 1 | 1 | 67605167 | 67228589 | CTCF::chr16:67605117-67605167(+) | E2F4::chr16:67228588-67228638(+) |
| CTCF | GFOD2 | 16 | 16 | 1 | -1 | 67663436 | 67719705 | CTCF::chr16:67663386-67663436(+) | GFOD2::chr16:67719655-67719705(-) |
| CTCF | HSD11B2 | 16 | 16 | 1 | 1 | 67605167 | 67469531 | CTCF::chr16:67605117-67605167(+) | HSD11B2::chr16:67469530-67469580(+) |
| CTDSPL | WRN | 3 | 8 | 1 | 1 | 37903769 | 30954284 | CTDSPL::chr3:37903719-37903769(+) | WRN::chr8:30954283-30954333(+) |
| CUL3 | IRS1 | 2 | 2 | -1 | -1 | 225422376 | 227600966 | CUL3::chr2:225422375-225422425(-) | IRS1::chr2:227600916-227600966(-) |
| CUL3 | SPAG16 | 2 | 2 | -1 | 1 | 225360549 | 214174783 | CUL3::chr2:225360548-225360598(-) | SPAG16::chr2:214174782-214174832(+) |
| CUX1 | AHCYL2 | 7 | 7 | 1 | 1 | 101459373 | 129019482 | CUX1::chr7:101459323-101459373(+) | AHCYL2::chr7:129019481-129019531(+) |
| CUX1 | CDK14 | 7 | 7 | 1 | 1 | 101758553 | 90376996 | CUX1::chr7:101758503-101758553(+) | CDK14::chr7:90376995-90377045(+) |
| CUX1 | CLCN3 | 7 | 4 | 1 | 1 | 101927249 | 170623929 | CUX1::chr7:101927199-101927249(+) | CLCN3::chr4:170623928-170623978(+) |
| CUX1 | EHBP1L1 | 7 | 11 | 1 | 1 | 101459373 | 65351712 | CUX1::chr7:101459323-101459373(+) | EHBP1L1::chr11:65351711-65351761(+) |
| CUX1 | EMID2 | 7 | 7 | 1 | 1 | 101559505 | 101176363 | CUX1::chr7:101559455-101559505(+) | EMID2::chr7:101176362-101176412(+) |
| CUX1 | POP7 | 7 | 7 | 1 | 1 | 101801888 | 100304444 | CUX1::chr7:101801838-101801888(+) | POP7::chr7:100304443-100304493(+) |
| CUX1 | THSD7A | 7 | 7 | 1 | -1 | 101713697 | 11676588 | CUX1::chr7:101713647-101713697(+) | THSD7A::chr7:11676538-11676588(-) |
| CUX1 | YEATS2 | 7 | 3 | 1 | 1 | 101459373 | 183490093 | CUX1::chr7:101459323-101459373(+) | YEATS2::chr3:183490092-183490142(+) |
| CYLD | FAM49B | 16 | 8 | 1 | -1 | 50776752 | 130983241 | CYLD::chr16:50776702-50776752(+) | FAM49B::chr8:130983191-130983241(-) |
| D2HGDH | CUL3 | 2 | 2 | 1 | -1 | 242674931 | 225376299 | D2HGDH::chr2:242674881-242674931(+) | CUL3::chr2:225376249-225376299(-) |
| DCAF10 | PAX5 | 9 | 9 | 1 | -1 | 37801402 | 37020798 | DCAF10::chr9:37801352-37801402(+) | PAX5::chr9:37020748-37020798(-) |
| DCUN1D4 | IKZF1 | 4 | 7 | 1 | 1 | 52758017 | 50444231 | DCUN1D4::chr4:52757967-52758017(+) | IKZF1::chr7:50444230-50444280(+) |
| DCUN1D4 | PTPRT | 4 | 20 | 1 | -1 | 52777340 | 41101202 | DCUN1D4::chr4:52777290-52777340(+) | PTPRT::chr20:41101152-41101202(-) |
| DDX10 | ARNTL | 11 | 11 | 1 | 1 | 108594189 | 13387047 | DDX10::chr11:108594139-108594189(+) | ARNTL::chr11:13387046-13387096(+) |
| DGCR2 | CLTCL1 | 22 | 22 | -1 | -1 | 19076881 | 19178947 | DGCR2::chr22:19076880-19076930(-) | CLTCL1::chr22:19178897-19178947(-) |
| DICER1 | CLMN | 14 | 14 | -1 | -1 | 95623567 | 95663031 | DICER1::chr14:95623566-95623616(-) | CLMN::chr14:95662981-95663031(-) |
| DICER1 | PAPOLA | 14 | 14 | -1 | 1 | 95569683 | 97008580 | DICER1::chr14:95569682-95569732(-) | PAPOLA::chr14:97008579-97008629(+) |
| DICER1 | SAMD4A | 14 | 14 | -1 | 1 | 95596395 | 55034214 | DICER1::chr14:95596394-95596444(-) | SAMD4A::chr14:55034213-55034263(+) |
| DIP2C | LARP4B | 10 | 10 | -1 | -1 | 735434 | 910210 | DIP2C::chr10:735433-735483(-) | LARP4B::chr10:910160-910210(-) |
| DNM2 | ANGPTL4 | 19 | 19 | 1 | 1 | 10829079 | 8438589 | DNM2::chr19:10829029-10829079(+) | ANGPTL4::chr19:8438588-8438638(+) |
| DNM2 | ARHGAP25 | 19 | 2 | 1 | 1 | 10829079 | 69002353 | DNM2::chr19:10829029-10829079(+) | ARHGAP25::chr2:69002352-69002402(+) |
| DNM2 | ARHGEF7 | 19 | 13 | 1 | 1 | 10930765 | 111953095 | DNM2::chr19:10930715-10930765(+) | ARHGEF7::chr13:111953094-111953144(+) |
| DNM2 | ATG4D | 19 | 19 | 1 | 1 | 10897382 | 10662573 | DNM2::chr19:10897332-10897382(+) | ATG4D::chr19:10662572-10662622(+) |
| DNM2 | C19orf66 | 19 | 19 | 1 | 1 | 10829079 | 10197605 | DNM2::chr19:10829029-10829079(+) | C19orf66::chr19:10197604-10197654(+) |
| DNM2 | DAND5 | 19 | 19 | 1 | 1 | 10870487 | 13084203 | DNM2::chr19:10870437-10870487(+) | DAND5::chr19:13084202-13084252(+) |
| DNM2 | FDX1L | 19 | 19 | 1 | -1 | 10829079 | 10419226 | DNM2::chr19:10829029-10829079(+) | FDX1L::chr19:10419176-10419226(-) |
| DNM2 | ILF3 | 19 | 19 | 1 | 1 | 10829079 | 10787839 | DNM2::chr19:10829029-10829079(+) | ILF3::chr19:10787838-10787888(+) |
| DNM2 | MRPL4 | 19 | 19 | 1 | 1 | 10829079 | 10363073 | DNM2::chr19:10829029-10829079(+) | MRPL4::chr19:10363072-10363122(+) |
| DNM2 | NFIX | 19 | 19 | 1 | 1 | 10870487 | 13135835 | DNM2::chr19:10870437-10870487(+) | NFIX::chr19:13135834-13135884(+) |
| DNM2 | PDE4A | 19 | 19 | 1 | 1 | 10829079 | 10556894 | DNM2::chr19:10829029-10829079(+) | PDE4A::chr19:10556893-10556943(+) |
| DNM2 | PTH2R | 19 | 2 | 1 | 1 | 10829079 | 209292926 | DNM2::chr19:10829029-10829079(+) | PTH2R::chr2:209292925-209292975(+) |
| DNM2 | QTRT1 | 19 | 19 | 1 | 1 | 10829079 | 10812626 | DNM2::chr19:10829029-10829079(+) | QTRT1::chr19:10812625-10812675(+) |
| DNM2 | QTRT1 | 19 | 19 | 1 | 1 | 10829079 | 10817978 | DNM2::chr19:10829029-10829079(+) | QTRT1::chr19:10817977-10818027(+) |
| DNM2 | RFX1 | 19 | 19 | 1 | -1 | 10870487 | 14088898 | DNM2::chr19:10870437-10870487(+) | RFX1::chr19:14088848-14088898(-) |

| | | | | | | | | | |
|---|---|---|---|---|---|---|---|---|---|
| DNM2 | RFX2 | 19 | 19 | 1 | -1 | 10916643 | 5997224 | DNM2::chr19:10916593-10916643(+) | RFX2::chr19:5997174-5997224(-) |
| DNM2 | SLC44A2 | 19 | 19 | 1 | 1 | 10893796 | 10745432 | DNM2::chr19:10893746-10893796(+) | SLC44A2::chr19:10745431-10745481(+) |
| DNM2 | SLC44A2 | 19 | 19 | 1 | 1 | 10870487 | 10736929 | DNM2::chr19:10870437-10870487(+) | SLC44A2::chr19:10736928-10736978(+) |
| DNM2 | SLC44A2 | 19 | 19 | 1 | 1 | 10930765 | 10736929 | DNM2::chr19:10930715-10930765(+) | SLC44A2::chr19:10736928-10736978(+) |
| DNM2 | SLC44A2 | 19 | 19 | 1 | 1 | 10886582 | 10736929 | DNM2::chr19:10886532-10886582(+) | SLC44A2::chr19:10736928-10736978(+) |
| DNM2 | SLC47A1 | 19 | 17 | 1 | 1 | 10942573 | 19482057 | DNM2::chr19:10942523-10942573(+) | SLC47A1::chr17:19482056-19482106(+) |
| DNM2 | TGIF1 | 19 | 18 | 1 | 1 | 10870487 | 3418141 | DNM2::chr19:10870437-10870487(+) | TGIF1::chr18:3418140-3418190(+) |
| DNM2 | ZGLP1 | 19 | 19 | 1 | -1 | 10942579 | 10420556 | DNM2::chr19:10942529-10942579(+) | ZGLP1::chr19:10420506-10420556(-) |
| DNMT3A | DTNB | 2 | 2 | -1 | -1 | 25564695 | 25602192 | DNMT3A::chr2:25564694-25564744(-) | DTNB::chr2:25602142-25602192(-) |
| DNMT3A | GTF3C2 | 2 | 2 | -1 | -1 | 25458576 | 27566445 | DNMT3A::chr2:25458575-25458625(-) | GTF3C2::chr2:27566395-27566445(-) |
| DNMT3A | HADHA | 2 | 2 | -1 | -1 | 25475063 | 26438044 | DNMT3A::chr2:25475062-25475112(-) | HADHA::chr2:26437994-26438044(-) |
| DOCK1 | MGMT | 10 | 10 | 1 | 1 | 128769049 | 131506159 | DOCK1::chr10:128768999-128769049(+) | MGMT::chr10:131506158-131506208(+) |
| DTNB | ASXL2 | 2 | 2 | -1 | -1 | 25818955 | 25973282 | DTNB::chr2:25818954-25819004(-) | ASXL2::chr2:25973232-25973282(-) |
| DTNB | ASXL2 | 2 | 2 | -1 | -1 | 25754342 | 25994409 | DTNB::chr2:25754341-25754391(-) | ASXL2::chr2:25994359-25994409(-) |
| DYRK1A | RUNX1 | 21 | 21 | 1 | -1 | 38868560 | 36265260 | DYRK1A::chr21:38868510-38868560(+) | RUNX1::chr21:36265210-36265260(-) |
| EIF3H | RAD21 | 8 | 8 | -1 | -1 | 117738255 | 117870697 | EIF3H::chr8:117738254-117738304(-) | RAD21::chr8:117870647-117870697(-) |
| ELF3 | DDX21 | 1 | 10 | 1 | 1 | 201985131 | 70723858 | ELF3::chr1:201985081-201985131(+) | DDX21::chr10:70723857-70723907(+) |
| ELL | ARL4D | 19 | 17 | -1 | 1 | 18632731 | 41477029 | ELL::chr19:18632730-18632780(-) | ARL4D::chr17:41477028-41477078(+) |
| ELL | C19orf42 | 19 | 19 | -1 | -1 | 18632731 | 16758072 | ELL::chr19:18632730-18632780(-) | C19orf42::chr19:16758022-16758072(-) |
| ELL | CISD3 | 19 | 17 | -1 | 1 | 18576222 | 36887013 | ELL::chr19:18576221-18576271(-) | CISD3::chr17:36887012-36887062(+) |
| ELL | CRLF1 | 19 | 19 | -1 | -1 | 18632731 | 18710656 | ELL::chr19:18632730-18632780(-) | CRLF1::chr19:18710606-18710656(-) |
| ELL | FAM125A | 19 | 19 | -1 | 1 | 18632731 | 17516886 | ELL::chr19:18632730-18632780(-) | FAM125A::chr19:17516885-17516935(+) |
| ELL | FKBP8 | 19 | 19 | -1 | -1 | 18632731 | 18652805 | ELL::chr19:18632730-18632780(-) | FKBP8::chr19:18652755-18652805(-) |
| ELL | FKBP8 | 19 | 19 | -1 | -1 | 18583645 | 18644157 | ELL::chr19:18583644-18583694(-) | FKBP8::chr19:18644107-18644157(-) |
| ELL | INSR | 19 | 19 | -1 | -1 | 18632731 | 7267907 | ELL::chr19:18632730-18632780(-) | INSR::chr19:7267857-7267907(-) |
| ELL | LDLRAD1 | 19 | 1 | -1 | -1 | 18572388 | 54476083 | ELL::chr19:18572387-18572437(-) | LDLRAD1::chr1:54476033-54476083(-) |
| ELL | MKNK2 | 19 | 19 | -1 | -1 | 18569015 | 2046690 | ELL::chr19:18569014-18569064(-) | MKNK2::chr19:2046640-2046690(-) |
| ELL | RABGGTA | 19 | 14 | -1 | -1 | 18557558 | 24740375 | ELL::chr19:18557557-18557607(-) | RABGGTA::chr14:24740325-24740375(-) |
| EP300 | FAM55B | 22 | 11 | 1 | 1 | 41546202 | 114568767 | EP300::chr22:41546152-41546202(+) | FAM55B::chr11:114568766-114568816(+) |
| EP300 | FBLN1 | 22 | 22 | 1 | 1 | 41551117 | 45996187 | EP300::chr22:41551067-41551117(+) | FBLN1::chr22:45996186-45996236(+) |
| EP300 | HAUS8 | 22 | 19 | 1 | -1 | 41513825 | 17166812 | EP300::chr22:41513775-41513825(+) | HAUS8::chr19:17166762-17166812(-) |
| EP300 | MCHR1 | 22 | 22 | 1 | 1 | 41489102 | 41076953 | EP300::chr22:41489052-41489102(+) | MCHR1::chr22:41076952-41077002(+) |
| EP300 | PAICS | 22 | 4 | 1 | 1 | 41513825 | 57325538 | EP300::chr22:41513775-41513825(+) | PAICS::chr4:57325537-57325587(+) |
| EP300 | RBX1 | 22 | 22 | 1 | 1 | 41551117 | 41363803 | EP300::chr22:41551067-41551117(+) | RBX1::chr22:41363802-41363852(+) |
| EP300 | TNRC6B | 22 | 22 | 1 | 1 | 41489102 | 40642019 | EP300::chr22:41489052-41489102(+) | TNRC6B::chr22:40642018-40642068(+) |
| EP300 | TNRC6B | 22 | 22 | 1 | 1 | 41546202 | 40681649 | EP300::chr22:41546152-41546202(+) | TNRC6B::chr22:40681648-40681698(+) |
| EP300 | TTC28 | 22 | 22 | 1 | -1 | 41523752 | 28702631 | EP300::chr22:41523702-41523752(+) | TTC28::chr22:28702581-28702631(-) |
| EP300 | XPNPEP3 | 22 | 22 | 1 | 1 | 41551117 | 41363803 | EP300::chr22:41551067-41551117(+) | XPNPEP3::chr22:41363802-41363852(+) |
| EPAS1 | SYPL2 | 2 | 1 | 1 | 1 | 46525076 | 110009681 | EPAS1::chr2:46525026-46525076(+) | SYPL2::chr1:110009680-110009730(+) |
| EPB41L2 | PTPRK | 6 | 6 | -1 | -1 | 131276245 | 128330413 | EPB41L2::chr6:131276244-131276294(-) | PTPRK::chr6:128330363-128330413(-) |
| EPS15 | C1orf159 | 1 | 1 | -1 | -1 | 51929345 | 1027483 | EPS15::chr1:51929344-51929394(-) | C1orf159::chr1:1027433-1027483(-) |
| EPS15 | CCDC18 | 1 | 1 | -1 | 1 | 51934145 | 93676359 | EPS15::chr1:51934144-51934194(-) | CCDC18::chr1:93676358-93676408(+) |
| EPS15 | INADL | 1 | 1 | -1 | 1 | 51984871 | 62503649 | EPS15::chr1:51984870-51984920(-) | INADL::chr1:62503648-62503698(+) |

| | | | | | | | | | |
|---|---|---|---|---|---|---|---|---|---|
| EPS15 | OSBPL9 | 1 | 1 | -1 | 1 | 51946945 | 52246836 | EPS15::chr1:51946944-51946994(-) | OSBPL9::chr1:52246835-52246885(+) |
| ERC1 | GPC5 | 12 | 13 | 1 | 1 | 1250953 | 93518535 | ERC1::chr12:1250903-1250953(+) | GPC5::chr13:93518534-93518584(+) |
| ERCC2 | C12orf40 | 19 | 12 | -1 | 1 | 45872188 | 40110558 | ERCC2::chr19:45872187-45872237(-) | C12orf40::chr12:40110557-40110607(+) |
| ERCC2 | MYH14 | 19 | 19 | -1 | 1 | 45864782 | 50810306 | ERCC2::chr19:45864781-45864831(-) | MYH14::chr19:50810305-50810355(+) |
| ERCC2 | NOVA2 | 19 | 19 | -1 | -1 | 45871888 | 46444203 | ERCC2::chr19:45871887-45871937(-) | NOVA2::chr19:46444153-46444203(-) |
| ERCC3 | MAP3K2 | 2 | 2 | -1 | -1 | 128038023 | 128072459 | ERCC3::chr2:128038022-128038072(-) | MAP3K2::chr2:128072409-128072459(-) |
| ESR1 | ASPH | 6 | 8 | 1 | -1 | 152201906 | 62416068 | ESR1::chr6:152201856-152201906(+) | ASPH::chr8:62416018-62416068(-) |
| ESR1 | BNC2 | 6 | 9 | 1 | -1 | 152163922 | 16738483 | ESR1::chr6:152163872-152163922(+) | BNC2::chr9:16738433-16738483(-) |
| ESR1 | C6orf97 | 6 | 6 | 1 | 1 | 152023140 | 151859180 | ESR1::chr6:152023090-152023140(+) | C6orf97::chr6:151859179-151859229(+) |
| ESR1 | C6orf97 | 6 | 6 | 1 | 1 | 152023140 | 151869439 | ESR1::chr6:152023090-152023140(+) | C6orf97::chr6:151869438-151869488(+) |
| ESR1 | C6orf97 | 6 | 6 | 1 | 1 | 152129499 | 151857453 | ESR1::chr6:152129449-152129499(+) | C6orf97::chr6:151857452-151857502(+) |
| ESR1 | C6orf97 | 6 | 6 | 1 | 1 | 152023140 | 151865707 | ESR1::chr6:152023090-152023140(+) | C6orf97::chr6:151865706-151865756(+) |
| ESR1 | C6orf97 | 6 | 6 | 1 | 1 | 152023140 | 151907024 | ESR1::chr6:152023090-152023140(+) | C6orf97::chr6:151907023-151907073(+) |
| ESR1 | C6orf97 | 6 | 6 | 1 | 1 | 152011799 | 151936578 | ESR1::chr6:152011749-152011799(+) | C6orf97::chr6:151936577-151936627(+) |
| ESR1 | C6orf97 | 6 | 6 | 1 | 1 | 152201906 | 151857453 | ESR1::chr6:152201856-152201906(+) | C6orf97::chr6:151857452-151857502(+) |
| ESR1 | MTHFD1L | 6 | 6 | 1 | 1 | 152265643 | 151330955 | ESR1::chr6:152265593-152265643(+) | MTHFD1L::chr6:151330954-151331004(+) |
| ESR1 | PDE7B | 6 | 6 | 1 | 1 | 152129499 | 136429869 | ESR1::chr6:152129449-152129499(+) | PDE7B::chr6:136429868-136429918(+) |
| ESR1 | SIM1 | 6 | 6 | 1 | -1 | 152023140 | 100841765 | ESR1::chr6:152023090-152023140(+) | SIM1::chr6:100841715-100841765(-) |
| ESR1 | SYNE1 | 6 | 6 | 1 | -1 | 152265643 | 152694354 | ESR1::chr6:152265593-152265643(+) | SYNE1::chr6:152694304-152694354(-) |
| ESR1 | TFB1M | 6 | 6 | 1 | -1 | 152265643 | 155581534 | ESR1::chr6:152265593-152265643(+) | TFB1M::chr6:155581484-155581534(-) |
| ETNK1 | CMAS | 12 | 12 | 1 | 1 | 22812088 | 22208083 | ETNK1::chr12:22812038-22812088(+) | CMAS::chr12:22208082-22208132(+) |
| ETNK1 | GLT1D1 | 12 | 12 | 1 | 1 | 22778520 | 129429871 | ETNK1::chr12:22778470-22778520(+) | GLT1D1::chr12:129429870-129429920(+) |
| ETNK1 | ITPR2 | 12 | 12 | 1 | -1 | 22826594 | 26634174 | ETNK1::chr12:22826544-22826594(+) | ITPR2::chr12:26634124-26634174(-) |
| ETNK1 | MYO9B | 12 | 19 | 1 | 1 | 22778520 | 17298744 | ETNK1::chr12:22778470-22778520(+) | MYO9B::chr19:17298743-17298793(+) |
| ETNK1 | RCAN2 | 12 | 6 | 1 | -1 | 22778520 | 46216633 | ETNK1::chr12:22778470-22778520(+) | RCAN2::chr6:46216583-46216633(-) |
| ETV6 | APOLD1 | 12 | 12 | 1 | 1 | 11803094 | 12939843 | ETV6::chr12:11803044-11803094(+) | APOLD1::chr12:12939842-12939892(+) |
| ETV6 | CIDEA | 12 | 18 | 1 | 1 | 11803094 | 12274092 | ETV6::chr12:11803044-11803094(+) | CIDEA::chr18:12274091-12274141(+) |
| ETV6 | DUSP16 | 12 | 12 | 1 | -1 | 11905513 | 12672934 | ETV6::chr12:11905463-11905513(+) | DUSP16::chr12:12672884-12672934(-) |
| ETV6 | EPB41L2 | 12 | 6 | 1 | -1 | 11803094 | 131222320 | ETV6::chr12:11803044-11803094(+) | EPB41L2::chr6:131222270-131222320(-) |
| ETV6 | ITLN1 | 12 | 1 | 1 | -1 | 11803094 | 160854673 | ETV6::chr12:11803044-11803094(+) | ITLN1::chr1:160854623-160854673(-) |
| ETV6 | PDE6H | 12 | 12 | 1 | 1 | 11803094 | 15130906 | ETV6::chr12:11803044-11803094(+) | PDE6H::chr12:15130905-15130955(+) |
| ETV6 | PLEKHG6 | 12 | 12 | 1 | 1 | 11803094 | 6424171 | ETV6::chr12:11803044-11803094(+) | PLEKHG6::chr12:6424170-6424220(+) |
| EXT1 | FAM155A | 8 | 13 | -1 | -1 | 119122324 | 107823134 | EXT1::chr8:119122323-119122373(-) | FAM155A::chr13:107823084-107823134(-) |
| EXT1 | OC90 | 8 | 8 | -1 | -1 | 118830674 | 133045411 | EXT1::chr8:118830673-118830723(-) | OC90::chr8:133045361-133045411(-) |
| EXT1 | RSF1 | 8 | 11 | -1 | -1 | 118825111 | 77451981 | EXT1::chr8:118825110-118825160(-) | RSF1::chr11:77451931-77451981(-) |
| EXT1 | SAMD12 | 8 | 8 | -1 | -1 | 119122324 | 119391939 | EXT1::chr8:119122323-119122373(-) | SAMD12::chr8:119391889-119391939(-) |
| EXT1 | TSNARE1 | 8 | 8 | -1 | -1 | 119122324 | 143293674 | EXT1::chr8:119122323-119122373(-) | TSNARE1::chr8:143293624-143293674(-) |
| EXT1 | WDYHV1 | 8 | 8 | -1 | 1 | 119122324 | 124440164 | EXT1::chr8:119122323-119122373(-) | WDYHV1::chr8:124440163-124440213(+) |
| EXT2 | ALKBH3 | 11 | 11 | 1 | 1 | 44135851 | 43940588 | EXT2::chr11:44135801-44135851(+) | ALKBH3::chr11:43940587-43940637(+) |
| EXT2 | ALKBH3 | 11 | 11 | 1 | 1 | 44219568 | 43940588 | EXT2::chr11:44219518-44219568(+) | ALKBH3::chr11:43940587-43940637(+) |
| EXT2 | ALX4 | 11 | 11 | 1 | -1 | 44257925 | 44289172 | EXT2::chr11:44257875-44257925(+) | ALX4::chr11:44289122-44289172(-) |
| EXT2 | PRMT10 | 11 | 4 | 1 | -1 | 44151688 | 148601622 | EXT2::chr11:44151638-44151688(+) | PRMT10::chr4:148601572-148601622(-) |
| EZH2 | AUTS2 | 7 | 7 | -1 | 1 | 148543562 | 70227856 | EZH2::chr7:148543561-148543611(-) | AUTS2::chr7:70227855-70227905(+) |

| | | | | | | | | | |
|---|---|---|---|---|---|---|---|---|---|
| EZH2 | TMEM176B | 7 | 7 | -1 | -1 | 148581256 | 150493662 | EZH2::chr7:148581255-148581305(-) | TMEM176B::chr7:150493612-150493662(-) |
| FAM134B | TERT | 5 | 5 | -1 | -1 | 16508687 | 1282739 | FAM134B::chr5:16508686-16508736(-) | TERT::chr5:1282689-1282739(-) |
| FANCA | CACNA2D4 | 16 | 12 | -1 | -1 | 89818546 | 1955855 | FANCA::chr16:89818545-89818595(-) | CACNA2D4::chr12:1955805-1955855(-) |
| FANCA | SPIRE2 | 16 | 16 | -1 | 1 | 89836245 | 89911730 | FANCA::chr16:89836244-89836294(-) | SPIRE2::chr16:89911729-89911779(+) |
| FANCC | C9orf3 | 9 | 9 | -1 | 1 | 98002931 | 97822976 | FANCC::chr9:98002930-98002980(-) | C9orf3::chr9:97822975-97823025(+) |
| FANCC | DDX31 | 9 | 9 | -1 | -1 | 98079808 | 135538082 | FANCC::chr9:98079807-98079857(-) | DDX31::chr9:135538032-135538082(-) |
| FANCC | ROR2 | 9 | 9 | -1 | -1 | 98002931 | 94538100 | FANCC::chr9:98002930-98002980(-) | ROR2::chr9:94538050-94538100(-) |
| FANCD2 | MTMR14 | 3 | 3 | 1 | 1 | 10094181 | 9703951 | FANCD2::chr3:10094131-10094181(+) | MTMR14::chr3:9703950-9704000(+) |
| FANCD2 | MTMR14 | 3 | 3 | 1 | 1 | 10074656 | 9719001 | FANCD2::chr3:10074606-10074656(+) | MTMR14::chr3:9719000-9719050(+) |
| FAT1 | ACSL1 | 4 | 4 | -1 | -1 | 187627717 | 185724700 | FAT1::chr4:187627716-187627766(-) | ACSL1::chr4:185724650-185724700(-) |
| FAT1 | IRF2 | 4 | 4 | -1 | -1 | 187530337 | 185350224 | FAT1::chr4:187530336-187530386(-) | IRF2::chr4:185350174-185350224(-) |
| FAT1 | KIAA1430 | 4 | 4 | -1 | -1 | 187554838 | 186085333 | FAT1::chr4:187554837-187554887(-) | KIAA1430::chr4:186085283-186085333(-) |
| FAT1 | NPY1R | 4 | 4 | -1 | -1 | 187647508 | 164247857 | FAT1::chr4:187647507-187647557(-) | NPY1R::chr4:164247807-164247857(-) |
| FBXL20 | NOTCH2 | 17 | 1 | -1 | -1 | 37557614 | 120465401 | FBXL20::chr17:37557613-37557663(-) | NOTCH2::chr1:120465351-120465401(-) |
| FBXO11 | INTS9 | 2 | 8 | -1 | -1 | 48132628 | 28671129 | FBXO11::chr2:48132627-48132677(-) | INTS9::chr8:28671079-28671129(-) |
| FBXW7 | ARHGAP10 | 4 | 4 | -1 | 1 | 153456093 | 148993144 | FBXW7::chr4:153456092-153456142(-) | ARHGAP10::chr4:148993143-148993193(+) |
| FBXW7 | INPP4B | 4 | 4 | -1 | -1 | 153249360 | 143383879 | FBXW7::chr4:153249359-153249409(-) | INPP4B::chr4:143383829-143383879(-) |
| FBXW7 | SH3D19 | 4 | 4 | -1 | -1 | 153249360 | 152147238 | FBXW7::chr4:153249359-153249409(-) | SH3D19::chr4:152147188-152147238(-) |
| FBXW7 | TMEM126B | 4 | 11 | -1 | 1 | 153456093 | 85345130 | FBXW7::chr4:153456092-153456142(-) | TMEM126B::chr11:85345129-85345179(+) |
| FBXW7 | TMEM154 | 4 | 4 | -1 | -1 | 153456093 | 153574079 | FBXW7::chr4:153456092-153456142(-) | TMEM154::chr4:153574029-153574079(-) |
| FBXW7 | TMEM154 | 4 | 4 | -1 | -1 | 153456093 | 153549713 | FBXW7::chr4:153456092-153456142(-) | TMEM154::chr4:153549663-153549713(-) |
| FEN1 | GDPD4 | 11 | 11 | 1 | -1 | 61560500 | 76940277 | FEN1::chr11:61560450-61560500(+) | GDPD4::chr11:76940227-76940277(-) |
| FHIT | CADPS | 3 | 3 | -1 | -1 | 61027719 | 62484943 | FHIT::chr3:61027718-61027768(-) | CADPS::chr3:62484893-62484943(-) |
| FHIT | CAMK4 | 3 | 5 | -1 | 1 | 61027719 | 110679722 | FHIT::chr3:61027718-61027768(-) | CAMK4::chr5:110679721-110679771(+) |
| FHIT | FAM19A4 | 3 | 3 | -1 | -1 | 60807613 | 68934461 | FHIT::chr3:60807612-60807662(-) | FAM19A4::chr3:68934411-68934461(-) |
| FHIT | KHDRBS3 | 3 | 8 | -1 | 1 | 61027719 | 136554897 | FHIT::chr3:61027718-61027768(-) | KHDRBS3::chr8:136554896-136554946(+) |
| FHIT | PTPRG | 3 | 3 | -1 | 1 | 61236975 | 61734552 | FHIT::chr3:61236974-61237024(-) | PTPRG::chr3:61734551-61734601(+) |
| FHIT | STT3B | 3 | 3 | -1 | 1 | 60807613 | 31617888 | FHIT::chr3:60807612-60807662(-) | STT3B::chr3:31617887-31617937(+) |
| FHIT | SYNPR | 3 | 3 | -1 | 1 | 61186291 | 63466508 | FHIT::chr3:61186290-61186340(-) | SYNPR::chr3:63466507-63466557(+) |
| FLCN | COPS3 | 17 | 17 | -1 | -1 | 17140226 | 17152353 | FLCN::chr17:17140225-17140275(-) | COPS3::chr17:17152303-17152353(-) |
| FOXO1 | ELF1 | 13 | 13 | -1 | -1 | 41239720 | 41533152 | FOXO1::chr13:41239719-41239769(-) | ELF1::chr13:41533102-41533152(-) |
| FOXO1 | MRPS31 | 13 | 13 | -1 | -1 | 41239720 | 41303737 | FOXO1::chr13:41239719-41239769(-) | MRPS31::chr13:41303687-41303737(-) |
| FOXO1 | WDFY2 | 13 | 13 | -1 | 1 | 41239720 | 52234732 | FOXO1::chr13:41239719-41239769(-) | WDFY2::chr13:52234731-52234781(+) |
| FOXO3 | CHD7 | 6 | 8 | 1 | 1 | 108883032 | 61653818 | FOXO3::chr6:108882982-108883032(+) | CHD7::chr8:61653817-61653867(+) |
| FOXO3 | LACE1 | 6 | 6 | 1 | 1 | 108883032 | 108840900 | FOXO3::chr6:108882982-108883032(+) | LACE1::chr6:108840899-108840949(+) |
| FRAS1 | MGMT | 4 | 10 | 1 | 1 | 79343154 | 131334505 | FRAS1::chr4:79343104-79343154(+) | MGMT::chr10:131334504-131334554(+) |
| FUS | PRSS36 | 16 | 16 | 1 | -1 | 31195717 | 31161052 | FUS::chr16:31195667-31195717(+) | PRSS36::chr16:31161002-31161052(-) |
| GAB2 | PTPRT | 11 | 20 | -1 | -1 | 78128692 | 40828028 | GAB2::chr11:78128691-78128741(-) | PTPRT::chr20:40827978-40828028(-) |
| GALNTL4 | PRDM2 | 11 | 1 | -1 | 1 | 11642906 | 14142922 | GALNTL4::chr11:11642905-11642955(-) | PRDM2::chr1:14142921-14142971(+) |
| GAS6 | GPC5 | 13 | 13 | -1 | 1 | 114566548 | 93518535 | GAS6::chr13:114566547-114566597(-) | GPC5::chr13:93518534-93518584(+) |
| GATA3 | TAF3 | 10 | 10 | 1 | 1 | 8111561 | 8055694 | GATA3::chr10:8111511-8111561(+) | TAF3::chr10:8055693-8055743(+) |
| GLIS3 | TERT | 9 | 5 | -1 | -1 | 4117768 | 1282739 | GLIS3::chr9:4117767-4117817(-) | TERT::chr5:1282689-1282739(-) |
| GNL3 | BAP1 | 3 | 3 | 1 | -1 | 52725645 | 52442093 | GNL3::chr3:52725595-52725645(+) | BAP1::chr3:52442043-52442093(-) |

| | | | | | | | | | |
|---|---|---|---|---|---|---|---|---|---|
| GOLGA4 | MLH1 | 3 | 3 | 1 | 1 | 37360685 | 37061801 | GOLGA4::chr3:37360635-37360685(+) | MLH1::chr3:37061800-37061850(+) |
| GOLPH3L | ARNT | 1 | 1 | -1 | -1 | 150667132 | 150790506 | GOLPH3L::chr1:150667131-150667181(-) | ARNT::chr1:150790456-150790506(-) |
| GPC6 | GPC5 | 13 | 13 | 1 | 1 | 94482798 | 93518535 | GPC6::chr13:94482748-94482798(+) | GPC5::chr13:93518534-93518584(+) |
| GPC6 | GPC5 | 13 | 13 | 1 | 1 | 93879869 | 92797083 | GPC6::chr13:93879819-93879869(+) | GPC5::chr13:92797082-92797132(+) |
| GPR98 | ATR | 5 | 3 | 1 | -1 | 89924649 | 142286996 | GPR98::chr5:89924599-89924649(+) | ATR::chr3:142286946-142286996(-) |
| GRB7 | CDK12 | 17 | 17 | 1 | 1 | 37902455 | 37680927 | GRB7::chr17:37902405-37902455(+) | CDK12::chr17:37680926-37680976(+) |
| GRIA1 | RAD17 | 5 | 5 | 1 | 1 | 153085627 | 68677728 | GRIA1::chr5:153085577-153085627(+) | RAD17::chr5:68677727-68677777(+) |
| GRIN2A | MTHFD1L | 16 | 6 | -1 | 1 | 10031816 | 151413603 | GRIN2A::chr16:10031815-10031865(-) | MTHFD1L::chr6:151413602-151413652(+) |
| GRIN2A | RASSF8 | 16 | 12 | -1 | 1 | 10273855 | 26208212 | GRIN2A::chr16:10273854-10273904(-) | RASSF8::chr12:26208211-26208261(+) |
| GTF2H3 | NCOR2 | 12 | 12 | 1 | -1 | 124130108 | 124862930 | GTF2H3::chr12:124130058-124130108(+) | NCOR2::chr12:124862880-124862930(-) |
| HERC4 | CCDC6 | 10 | 10 | -1 | -1 | 69692354 | 61554355 | HERC4::chr10:69692353-69692403(-) | CCDC6::chr10:61554305-61554355(-) |
| HIRA | ZBTB16 | 22 | 11 | -1 | 1 | 19384310 | 114027059 | HIRA::chr22:19384309-19384359(-) | ZBTB16::chr11:114027058-114027108(+) |
| IGF2BP2 | IFITM2 | 3 | 11 | -1 | 1 | 185540941 | 308231 | IGF2BP2::chr3:185540940-185540990(-) | IFITM2::chr11:308230-308280(+) |
| IGF2R | ACVR2A | 6 | 2 | 1 | 1 | 160390427 | 148672760 | IGF2R::chr6:160390377-160390427(+) | ACVR2A::chr2:148672759-148672809(+) |
| IGF2R | QKI | 6 | 6 | 1 | 1 | 160511135 | 163956014 | IGF2R::chr6:160511085-160511135(+) | QKI::chr6:163956013-163956063(+) |
| IKZF1 | LRBA | 7 | 4 | 1 | -1 | 50367353 | 151207186 | IKZF1::chr7:50367303-50367353(+) | LRBA::chr4:151207136-151207186(-) |
| ITPR2 | RAD17 | 12 | 5 | -1 | 1 | 26553061 | 68706320 | ITPR2::chr12:26553060-26553110(-) | RAD17::chr5:68706319-68706369(+) |
| JAK1 | HSD11B1 | 1 | 1 | -1 | 1 | 65432016 | 209878262 | JAK1::chr1:65432015-65432065(-) | HSD11B1::chr1:209878261-209878311(+) |
| KAT2A | STAT5B | 17 | 17 | -1 | -1 | 40269053 | 40354826 | KAT2A::chr17:40269052-40269102(-) | STAT5B::chr17:40354776-40354826(-) |
| KATNA1 | LATS1 | 6 | 6 | -1 | -1 | 149944239 | 150005728 | KATNA1::chr6:149944238-149944288(-) | LATS1::chr6:150005678-150005728(-) |
| KDM5C | HUWE1 | X | X | -1 | -1 | 53245256 | 53707139 | KDM5C::chrX:53245255-53245305(-) | HUWE1::chrX:53707089-53707139(-) |
| KDM5C | MAGT1 | X | X | -1 | -1 | 53222619 | 77086392 | KDM5C::chrX:53222618-53222668(-) | MAGT1::chrX:77086342-77086392(-) |
| KDM5C | PPP4R1 | X | 18 | -1 | -1 | 53240679 | 9550396 | KDM5C::chrX:53240678-53240728(-) | PPP4R1::chr18:9550346-9550396(-) |
| KDM6A | ARHGAP6 | X | X | 1 | -1 | 44918711 | 11272827 | KDM6A::chrX:44918661-44918711(+) | ARHGAP6::chrX:11272777-11272827(-) |
| KDM6A | DCAF7 | X | 17 | 1 | 1 | 44733233 | 61655831 | KDM6A::chrX:44733183-44733233(+) | DCAF7::chr17:61655830-61655880(+) |
| KDM6A | DLL1 | X | 6 | 1 | -1 | 44833960 | 170595386 | KDM6A::chrX:44833910-44833960(+) | DLL1::chr6:170595336-170595386(-) |
| KDM6A | DMD | X | X | 1 | -1 | 44833960 | 31497220 | KDM6A::chrX:44833910-44833960(+) | DMD::chrX:31497170-31497220(-) |
| KDM6A | FUNDC1 | X | X | 1 | -1 | 44833960 | 44401347 | KDM6A::chrX:44833910-44833960(+) | FUNDC1::chrX:44401297-44401347(-) |
| KDM6A | KAL1 | X | X | 1 | -1 | 44945224 | 8591711 | KDM6A::chrX:44945174-44945224(+) | KAL1::chrX:8591661-8591711(-) |
| KDM6A | PDHX | X | 11 | 1 | 1 | 44950109 | 34952951 | KDM6A::chrX:44950059-44950109(+) | PDHX::chr11:34952950-34953000(+) |
| KDM6A | RPS6KA3 | X | X | 1 | -1 | 44950109 | 20252932 | KDM6A::chrX:44950059-44950109(+) | RPS6KA3::chrX:20252882-20252932(-) |
| KDM6A | ZNF674 | X | X | 1 | -1 | 44733233 | 46360770 | KDM6A::chrX:44733183-44733233(+) | ZNF674::chrX:46360720-46360770(-) |
| KIAA0146 | CSMD3 | 8 | 8 | 1 | -1 | 48511758 | 113569183 | KIAA0146::chr8:48511708-48511758(+) | CSMD3::chr8:113569133-113569183(-) |
| LAMA5 | NCOR2 | 20 | 12 | -1 | -1 | 60926760 | 124841352 | LAMA5::chr20:60926759-60926809(-) | NCOR2::chr12:124841302-124841352(-) |
| LARP4B | C12orf69 | 10 | 12 | -1 | -1 | 882343 | 14959630 | LARP4B::chr10:882342-882392(-) | C12orf69::chr12:14959580-14959630(-) |
| LARP4B | PAMR1 | 10 | 11 | -1 | -1 | 909683 | 35515820 | LARP4B::chr10:909682-909732(-) | PAMR1::chr11:35515770-35515820(-) |
| LARP4B | PFKFB3 | 10 | 10 | -1 | 1 | 909683 | 6255586 | LARP4B::chr10:909682-909732(-) | PFKFB3::chr10:6255585-6255635(+) |
| LATS1 | LACE1 | 6 | 6 | -1 | 1 | 150038985 | 108723200 | LATS1::chr6:150038984-150039034(-) | LACE1::chr6:108723199-108723249(+) |
| LDLR | DNM2 | 19 | 19 | 1 | 1 | 11200291 | 10935733 | LDLR::chr19:11200241-11200291(+) | DNM2::chr19:10935732-10935782(+) |
| LDLR | DNM2 | 19 | 19 | 1 | 1 | 11231198 | 10922940 | LDLR::chr19:11231148-11231198(+) | DNM2::chr19:10922939-10922989(+) |
| LDLR | SMARCA4 | 19 | 19 | 1 | 1 | 11234020 | 11172460 | LDLR::chr19:11233970-11234020(+) | SMARCA4::chr19:11172459-11172509(+) |
| LRIG3 | ATF7 | 12 | 12 | -1 | -1 | 59307763 | 53994805 | LRIG3::chr12:59307762-59307812(-) | ATF7::chr12:53994755-53994805(-) |
| LRIG3 | PIP4K2C | 12 | 12 | -1 | 1 | 59307763 | 57995709 | LRIG3::chr12:59307762-59307812(-) | PIP4K2C::chr12:57995708-57995758(+) |

| | | | | | | | | | |
|---|---|---|---|---|---|---|---|---|---|
| LRIG3 | SLC16A7 | 12 | 12 | -1 | 1 | 59267837 | 60165000 | LRIG3::chr12:59267836-59267886(-) | SLC16A7::chr12:60164999-60165049(+) |
| LRIG3 | XRCC6BP1 | 12 | 12 | -1 | 1 | 59307763 | 58339411 | LRIG3::chr12:59307762-59307812(-) | XRCC6BP1::chr12:58339410-58339460(+) |
| LRP1B | ADAP1 | 2 | 7 | -1 | -1 | 142237965 | 975141 | LRP1B::chr2:142237964-142238014(-) | ADAP1::chr7:975091-975141(-) |
| LRP1B | HECW2 | 2 | 2 | -1 | -1 | 142567848 | 197208488 | LRP1B::chr2:142567847-142567897(-) | HECW2::chr2:197208438-197208488(-) |
| LRRC6 | EXT1 | 8 | 8 | -1 | -1 | 133687730 | 118849440 | LRRC6::chr8:133687729-133687779(-) | EXT1::chr8:118849390-118849440(-) |
| LUC7L | AXIN1 | 16 | 16 | -1 | -1 | 249061 | 364683 | LUC7L::chr16:249060-249110(-) | AXIN1::chr16:364633-364683(-) |
| LUC7L | AXIN1 | 16 | 16 | -1 | -1 | 277241 | 364683 | LUC7L::chr16:277240-277290(-) | AXIN1::chr16:364633-364683(-) |
| LUC7L | AXIN1 | 16 | 16 | -1 | -1 | 270648 | 364683 | LUC7L::chr16:270647-270697(-) | AXIN1::chr16:364633-364683(-) |
| MAP2K4 | DNAH9 | 17 | 17 | 1 | 1 | 12032604 | 11865184 | MAP2K4::chr17:12032554-12032604(+) | DNAH9::chr17:11865183-11865233(+) |
| MAP2K4 | PSMD12 | 17 | 17 | 1 | -1 | 11924318 | 65353694 | MAP2K4::chr17:11924268-11924318(+) | PSMD12::chr17:65353644-65353694(-) |
| MAP3K1 | OR4N2 | 5 | 14 | 1 | 1 | 56155742 | 20201528 | MAP3K1::chr5:56155692-56155742(+) | OR4N2::chr14:20201527-20201577(+) |
| MAP3K1 | PARN | 5 | 16 | 1 | -1 | 56111882 | 14540938 | MAP3K1::chr5:56111832-56111882(+) | PARN::chr16:14540888-14540938(-) |
| MAP3K13 | CD37 | 3 | 19 | 1 | 1 | 185003431 | 49840166 | MAP3K13::chr3:185003381-185003431(+) | CD37::chr19:49840165-49840215(+) |
| MAP3K13 | FETUB | 3 | 3 | 1 | 1 | 185003431 | 186369007 | MAP3K13::chr3:185003381-185003431(+) | FETUB::chr3:186369006-186369056(+) |
| MAP3K13 | NSUN2 | 3 | 5 | 1 | -1 | 185046750 | 6625782 | MAP3K13::chr3:185046700-185046750(+) | NSUN2::chr5:6625732-6625782(-) |
| MAP3K13 | P2RX5 | 3 | 17 | 1 | -1 | 185081156 | 3577171 | MAP3K13::chr3:185081106-185081156(+) | P2RX5::chr17:3577121-3577171(-) |
| MAP3K13 | UTS2D | 3 | 3 | 1 | -1 | 185003431 | 191046498 | MAP3K13::chr3:185003381-185003431(+) | UTS2D::chr3:191046448-191046498(-) |
| MAP3K13 | VPS8 | 3 | 3 | 1 | 1 | 185081156 | 184769664 | MAP3K13::chr3:185081106-185081156(+) | VPS8::chr3:184769663-184769713(+) |
| MAX | ACTN1 | 14 | 14 | -1 | -1 | 65560426 | 69392389 | MAX::chr14:65560425-65560475(-) | ACTN1::chr14:69392339-69392389(-) |
| MAX | RUNX2 | 14 | 6 | -1 | 1 | 65560426 | 45405684 | MAX::chr14:65560425-65560475(-) | RUNX2::chr6:45405683-45405733(+) |
| MAX | TEX10 | 14 | 9 | -1 | -1 | 65560426 | 103111654 | MAX::chr14:65560425-65560475(-) | TEX10::chr9:103111604-103111654(-) |
| MCTP1 | CBLB | 5 | 3 | -1 | -1 | 94204038 | 105495386 | MCTP1::chr5:94204037-94204087(-) | CBLB::chr3:105495336-105495386(-) |
| MED1 | CDK12 | 17 | 17 | -1 | 1 | 37607291 | 37627132 | MED1::chr17:37607290-37607340(-) | CDK12::chr17:37627131-37627181(+) |
| MEN1 | CDC42BPG | 11 | 11 | -1 | -1 | 64572506 | 64608120 | MEN1::chr11:64572505-64572555(-) | CDC42BPG::chr11:64608070-64608120(-) |
| MEX3D | TCF3 | 19 | 19 | -1 | -1 | 1567463 | 1646426 | MEX3D::chr19:1567462-1567512(-) | TCF3::chr19:1646376-1646426(-) |
| MGMT | DOCK1 | 10 | 10 | 1 | 1 | 131334641 | 129237291 | MGMT::chr10:131334591-131334641(+) | DOCK1::chr10:129237290-129237340(+) |
| MINPP1 | ASXL2 | 10 | 2 | 1 | -1 | 89268290 | 26068432 | MINPP1::chr10:89268240-89268290(+) | ASXL2::chr2:26068382-26068432(-) |
| MKL1 | APOL4 | 22 | 22 | -1 | -1 | 41032482 | 36598101 | MKL1::chr22:41032481-41032531(-) | APOL4::chr22:36598051-36598101(-) |
| MKL1 | ARAP1 | 22 | 11 | -1 | -1 | 41032482 | 72414124 | MKL1::chr22:41032481-41032531(-) | ARAP1::chr11:72414074-72414124(-) |
| MKL1 | CSNK1E | 22 | 22 | -1 | -1 | 40948110 | 38710174 | MKL1::chr22:40948109-40948159(-) | CSNK1E::chr22:38710124-38710174(-) |
| MKL1 | INPP5D | 22 | 2 | -1 | 1 | 40948110 | 234077951 | MKL1::chr22:40948109-40948159(-) | INPP5D::chr2:234077950-234078000(+) |
| MKL1 | NTNG1 | 22 | 1 | -1 | 1 | 41032482 | 107937776 | MKL1::chr22:41032481-41032531(-) | NTNG1::chr1:107937775-107937825(+) |
| MKL1 | SCGB1C1 | 22 | 11 | -1 | 1 | 40859225 | 194418 | MKL1::chr22:40859224-40859274(-) | SCGB1C1::chr11:194417-194467(+) |
| MKL1 | SERPIND1 | 22 | 22 | -1 | 1 | 40859225 | 21133585 | MKL1::chr22:40859224-40859274(-) | SERPIND1::chr22:21133584-21133634(+) |
| MKL1 | SLC25A17 | 22 | 22 | -1 | -1 | 40990678 | 41195087 | MKL1::chr22:40990677-40990727(-) | SLC25A17::chr22:41195037-41195087(-) |
| MKL1 | SLC25A17 | 22 | 22 | -1 | -1 | 40948110 | 41175129 | MKL1::chr22:40948109-40948159(-) | SLC25A17::chr22:41175079-41175129(-) |
| MKL1 | SLC25A17 | 22 | 22 | -1 | -1 | 40859225 | 41175129 | MKL1::chr22:40859224-40859274(-) | SLC25A17::chr22:41175079-41175129(-) |
| MKL1 | SLC25A17 | 22 | 22 | -1 | -1 | 41032482 | 41175129 | MKL1::chr22:41032481-41032531(-) | SLC25A17::chr22:41175079-41175129(-) |
| MKL1 | TMEM218 | 22 | 11 | -1 | -1 | 40819540 | 124972705 | MKL1::chr22:40819539-40819589(-) | TMEM218::chr11:124972655-124972705(-) |
| MKL1 | WFIKKN2 | 22 | 17 | -1 | 1 | 40990678 | 48916860 | MKL1::chr22:40990677-40990727(-) | WFIKKN2::chr17:48916859-48916909(+) |
| MLF1 | RSRC1 | 3 | 3 | 1 | 1 | 158310370 | 158261124 | MLF1::chr3:158310320-158310370(+) | RSRC1::chr3:158261123-158261173(+) |
| MLL3 | CNTNAP2 | 7 | 7 | -1 | 1 | 152027686 | 146740999 | MLL3::chr7:152027685-152027735(-) | CNTNAP2::chr7:146740998-146741048(+) |
| MLL3 | CNTNAP2 | 7 | 7 | -1 | 1 | 152132711 | 146471363 | MLL3::chr7:152132710-152132760(-) | CNTNAP2::chr7:146471362-146471412(+) |

| | | | | | | | | | |
|---|---|---|---|---|---|---|---|---|---|
| MOBKL1B | GRIN2A | 2 | 16 | -1 | -1 | 74405788 | 9928087 | MOBKL1B::chr2:74405787-74405837(-) | GRIN2A::chr16:9928037-9928087(-) |
| MOBKL2A | STK11 | 19 | 19 | -1 | 1 | 2076810 | 1218416 | MOBKL2A::chr19:2076809-2076859(-) | STK11::chr19:1218415-1218465(+) |
| MSH2 | CCDC57 | 2 | 17 | 1 | -1 | 47637511 | 80121233 | MSH2::chr2:47637461-47637511(+) | CCDC57::chr17:80121183-80121233(-) |
| MSH2 | EPCAM | 2 | 2 | 1 | 1 | 47643568 | 47612305 | MSH2::chr2:47643518-47643568(+) | EPCAM::chr2:47612304-47612354(+) |
| MSH2 | EPCAM | 2 | 2 | 1 | 1 | 47630541 | 47612305 | MSH2::chr2:47630491-47630541(+) | EPCAM::chr2:47612304-47612354(+) |
| MSH2 | SLC3A1 | 2 | 2 | 1 | 1 | 47637511 | 44539725 | MSH2::chr2:47637461-47637511(+) | SLC3A1::chr2:44539724-44539774(+) |
| MSH2 | SLCO6A1 | 2 | 5 | 1 | -1 | 47672796 | 101709198 | MSH2::chr2:47672746-47672796(+) | SLCO6A1::chr5:101709148-101709198(-) |
| MSH2 | SPTLC1 | 2 | 9 | 1 | -1 | 47672796 | 94800647 | MSH2::chr2:47672746-47672796(+) | SPTLC1::chr9:94800597-94800647(-) |
| MSH2 | TTC7A | 2 | 2 | 1 | 1 | 47630541 | 47177502 | MSH2::chr2:47630491-47630541(+) | TTC7A::chr2:47177501-47177551(+) |
| MSL2 | STAG1 | 3 | 3 | -1 | -1 | 135913814 | 136068205 | MSL2::chr3:135913813-135913863(-) | STAG1::chr3:136068155-136068205(-) |
| MTAP | CDKN2A | 9 | 9 | 1 | -1 | 21815518 | 21968241 | MTAP::chr9:21815468-21815518(+) | CDKN2A::chr9:21968191-21968241(-) |
| MTAP | CDKN2A | 9 | 9 | 1 | -1 | 21818201 | 21971207 | MTAP::chr9:21818151-21818201(+) | CDKN2A::chr9:21971157-21971207(-) |
| MTF2 | RPL5 | 1 | 1 | 1 | 1 | 93581175 | 93306108 | MTF2::chr1:93581125-93581175(+) | RPL5::chr1:93306107-93306157(+) |
| MTMR12 | TERT | 5 | 5 | -1 | -1 | 32263219 | 1294781 | MTMR12::chr5:32263218-32263268(-) | TERT::chr5:1294731-1294781(-) |
| MYH14 | FBLN2 | 19 | 3 | 1 | 1 | 50805126 | 13649562 | MYH14::chr19:50805076-50805126(+) | FBLN2::chr3:13649561-13649611(+) |
| MYH9 | CACNA1C | 22 | 12 | -1 | 1 | 36783852 | 2224390 | MYH9::chr22:36783851-36783901(-) | CACNA1C::chr12:2224389-2224439(+) |
| MYH9 | CACNG2 | 22 | 22 | -1 | -1 | 36705327 | 36983595 | MYH9::chr22:36705326-36705376(-) | CACNG2::chr22:36983545-36983595(-) |
| MYH9 | CACNG2 | 22 | 22 | -1 | -1 | 36783852 | 36983595 | MYH9::chr22:36783851-36783901(-) | CACNG2::chr22:36983545-36983595(-) |
| MYH9 | CACNG2 | 22 | 22 | -1 | -1 | 36722613 | 36962540 | MYH9::chr22:36722612-36722662(-) | CACNG2::chr22:36962490-36962540(-) |
| MYH9 | CACNG2 | 22 | 22 | -1 | -1 | 36783852 | 36962540 | MYH9::chr22:36783851-36783901(-) | CACNG2::chr22:36962490-36962540(-) |
| MYH9 | DDX21 | 22 | 10 | -1 | 1 | 36677327 | 70723858 | MYH9::chr22:36677326-36677376(-) | DDX21::chr10:70723857-70723907(+) |
| MYH9 | DNASE1 | 22 | 16 | -1 | 1 | 36697580 | 3704962 | MYH9::chr22:36697579-36697629(-) | DNASE1::chr16:3704961-3705011(+) |
| MYH9 | HSP90AA1 | 22 | 14 | -1 | -1 | 36783852 | 102550901 | MYH9::chr22:36783851-36783901(-) | HSP90AA1::chr14:102550851-102550901(-) |
| MYH9 | KLK3 | 22 | 19 | -1 | 1 | 36685235 | 51358191 | MYH9::chr22:36685234-36685284(-) | KLK3::chr19:51358190-51358240(+) |
| MYH9 | ORAOV1 | 22 | 11 | -1 | -1 | 36744949 | 69488089 | MYH9::chr22:36744948-36744998(-) | ORAOV1::chr11:69488039-69488089(-) |
| MYH9 | PLA2G6 | 22 | 22 | -1 | -1 | 36783852 | 38565478 | MYH9::chr22:36783851-36783901(-) | PLA2G6::chr22:38565428-38565478(-) |
| MYH9 | PLAT | 22 | 8 | -1 | -1 | 36677327 | 42051245 | MYH9::chr22:36677326-36677376(-) | PLAT::chr8:42051195-42051245(-) |
| MYH9 | PVALB | 22 | 22 | -1 | -1 | 36783852 | 37213056 | MYH9::chr22:36783851-36783901(-) | PVALB::chr22:37213006-37213056(-) |
| MYH9 | TXN2 | 22 | 22 | -1 | -1 | 36783852 | 36876884 | MYH9::chr22:36783851-36783901(-) | TXN2::chr22:36876834-36876884(-) |
| MYH9 | TXN2 | 22 | 22 | -1 | -1 | 36718474 | 36876884 | MYH9::chr22:36718473-36718523(-) | TXN2::chr22:36876834-36876884(-) |
| MYH9 | TXN2 | 22 | 22 | -1 | -1 | 36705327 | 36863964 | MYH9::chr22:36705326-36705376(-) | TXN2::chr22:36863914-36863964(-) |
| MYRIP | MLH1 | 3 | 3 | 1 | 1 | 39942417 | 37089010 | MYRIP::chr3:39942367-39942417(+) | MLH1::chr3:37089009-37089059(+) |
| NAP1L4 | CARS | 11 | 11 | -1 | -1 | 2999514 | 3023830 | NAP1L4::chr11:2999513-2999563(-) | CARS::chr11:3023780-3023830(-) |
| NBEA | BRCA2 | 13 | 13 | 1 | 1 | 35770439 | 32899213 | NBEA::chr13:35770389-35770439(+) | BRCA2::chr13:32899212-32899262(+) |
| NBN | RIPK2 | 8 | 8 | -1 | 1 | 90976638 | 90796278 | NBN::chr8:90976637-90976687(-) | RIPK2::chr8:90796277-90796327(+) |
| NBN | SLC26A7 | 8 | 8 | -1 | 1 | 90994950 | 92406025 | NBN::chr8:90994949-90994999(-) | SLC26A7::chr8:92406024-92406074(+) |
| NCKAP5 | ERBB4 | 2 | 2 | -1 | -1 | 133887550 | 212812341 | NCKAP5::chr2:133887549-133887599(-) | ERBB4::chr2:212812291-212812341(-) |
| NCOA4 | C10orf25 | 10 | 10 | 1 | -1 | 51565296 | 45495864 | NCOA4::chr10:51565246-51565296(+) | C10orf25::chr10:45495814-45495864(-) |
| NCOA4 | MSMB | 10 | 10 | 1 | 1 | 51585599 | 51555731 | NCOA4::chr10:51585549-51585599(+) | MSMB::chr10:51555730-51555780(+) |
| NCOA4 | MSMB | 10 | 10 | 1 | 1 | 51581378 | 51555731 | NCOA4::chr10:51581328-51581378(+) | MSMB::chr10:51555730-51555780(+) |
| NCOA4 | MSMB | 10 | 10 | 1 | 1 | 51565296 | 51555731 | NCOA4::chr10:51565246-51565296(+) | MSMB::chr10:51555730-51555780(+) |
| NCOA4 | OR13A1 | 10 | 10 | 1 | -1 | 51565296 | 45803256 | NCOA4::chr10:51565246-51565296(+) | OR13A1::chr10:45803206-45803256(-) |
| NCOA4 | RSU1 | 10 | 10 | 1 | -1 | 51565296 | 16635495 | NCOA4::chr10:51565246-51565296(+) | RSU1::chr10:16635445-16635495(-) |

| | | | | | | | | | |
|---|---|---|---|---|---|---|---|---|---|
| NCOR1 | ACACA | 17 | 17 | -1 | -1 | 15960828 | 35627752 | NCOR1::chr17:15960827-15960877(-) | ACACA::chr17:35627702-35627752(-) |
| NCOR1 | DPM1 | 17 | 20 | -1 | -1 | 16055260 | 49571822 | NCOR1::chr17:16055259-16055309(-) | DPM1::chr20:49571772-49571822(-) |
| NCOR1 | FBXO3 | 17 | 11 | -1 | -1 | 15960828 | 33792376 | NCOR1::chr17:15960827-15960877(-) | FBXO3::chr11:33792326-33792376(-) |
| NCOR1 | HIC1 | 17 | 17 | -1 | 1 | 15973479 | 1959965 | NCOR1::chr17:15973478-15973528(-) | HIC1::chr17:1959964-1960014(+) |
| NCOR1 | PIGL | 17 | 17 | -1 | 1 | 16062074 | 16203202 | NCOR1::chr17:16062073-16062123(-) | PIGL::chr17:16203201-16203251(+) |
| NCOR1 | PRR11 | 17 | 17 | -1 | 1 | 16089868 | 57247109 | NCOR1::chr17:16089867-16089917(-) | PRR11::chr17:57247108-57247158(+) |
| NCOR1 | RHOU | 17 | 1 | -1 | 1 | 16097776 | 228873420 | NCOR1::chr17:16097775-16097825(-) | RHOU::chr1:228873419-228873469(+) |
| NCOR1 | TFDP2 | 17 | 3 | -1 | -1 | 15960828 | 141693033 | NCOR1::chr17:15960827-15960877(-) | TFDP2::chr3:141692983-141693033(-) |
| NCOR2 | DDX27 | 12 | 20 | -1 | 1 | 124808962 | 47849303 | NCOR2::chr12:124808961-124809011(-) | DDX27::chr20:47849302-47849352(+) |
| NCOR2 | DHX37 | 12 | 12 | -1 | -1 | 124950719 | 125450358 | NCOR2::chr12:124950718-124950768(-) | DHX37::chr12:125450308-125450358(-) |
| NCOR2 | DHX37 | 12 | 12 | -1 | -1 | 124970987 | 125470811 | NCOR2::chr12:124970986-124971036(-) | DHX37::chr12:125470761-125470811(-) |
| NCOR2 | DHX37 | 12 | 12 | -1 | -1 | 124846672 | 125435350 | NCOR2::chr12:124846671-124846721(-) | DHX37::chr12:125435300-125435350(-) |
| NCOR2 | FMR1NB | 12 | X | -1 | 1 | 124882665 | 147084721 | NCOR2::chr12:124882664-124882714(-) | FMR1NB::chrX:147084720-147084770(+) |
| NCOR2 | GATA4 | 12 | 8 | -1 | 1 | 124862783 | 11606428 | NCOR2::chr12:124862782-124862832(-) | GATA4::chr8:11606427-11606477(+) |
| NCOR2 | ITGB5 | 12 | 3 | -1 | -1 | 125002686 | 124528003 | NCOR2::chr12:125002685-125002735(-) | ITGB5::chr3:124527953-124528003(-) |
| NCOR2 | MORN3 | 12 | 12 | -1 | -1 | 124979693 | 122097254 | NCOR2::chr12:124979692-124979742(-) | MORN3::chr12:122097204-122097254(-) |
| NCOR2 | NUP37 | 12 | 12 | -1 | -1 | 124904503 | 102512361 | NCOR2::chr12:124904502-124904552(-) | NUP37::chr12:102512311-102512361(-) |
| NCOR2 | PGC | 12 | 6 | -1 | -1 | 124809926 | 41715135 | NCOR2::chr12:124809925-124809975(-) | PGC::chr6:41715085-41715135(-) |
| NCOR2 | PIP4K2B | 12 | 17 | -1 | -1 | 125051854 | 36943173 | NCOR2::chr12:125051853-125051903(-) | PIP4K2B::chr17:36943123-36943173(-) |
| NCOR2 | S100A4 | 12 | 1 | -1 | -1 | 124831139 | 153518278 | NCOR2::chr12:124831138-124831188(-) | S100A4::chr1:153518228-153518278(-) |
| NCOR2 | SCARB1 | 12 | 12 | -1 | -1 | 125020111 | 125271049 | NCOR2::chr12:125020110-125020160(-) | SCARB1::chr12:125270999-125271049(-) |
| NCOR2 | SCARB1 | 12 | 12 | -1 | -1 | 125020111 | 125284788 | NCOR2::chr12:125020110-125020160(-) | SCARB1::chr12:125284738-125284788(-) |
| NCOR2 | SCARB1 | 12 | 12 | -1 | -1 | 125051854 | 125279814 | NCOR2::chr12:125051853-125051903(-) | SCARB1::chr12:125279764-125279814(-) |
| NCOR2 | SCARB1 | 12 | 12 | -1 | -1 | 124882665 | 125263132 | NCOR2::chr12:124882664-124882714(-) | SCARB1::chr12:125263082-125263132(-) |
| NCOR2 | SCARB1 | 12 | 12 | -1 | -1 | 124979693 | 125302253 | NCOR2::chr12:124979692-124979742(-) | SCARB1::chr12:125302203-125302253(-) |
| NCOR2 | TMEM132D | 12 | 12 | -1 | -1 | 124885047 | 130015750 | NCOR2::chr12:124885046-124885096(-) | TMEM132D::chr12:130015700-130015750(-) |
| NDRG1 | CLCN3 | 8 | 4 | -1 | 1 | 134249419 | 170623929 | NDRG1::chr8:134249418-134249468(-) | CLCN3::chr4:170623928-170623978(+) |
| NDRG1 | DACT3 | 8 | 19 | -1 | -1 | 134292475 | 47156332 | NDRG1::chr8:134292474-134292524(-) | DACT3::chr19:47156282-47156332(-) |
| NDRG1 | LARP1 | 8 | 5 | -1 | 1 | 134249419 | 154188019 | NDRG1::chr8:134249418-134249468(-) | LARP1::chr5:154188018-154188068(+) |
| NDRG1 | PLEKHA5 | 8 | 12 | -1 | 1 | 134249419 | 19441012 | NDRG1::chr8:134249418-134249468(-) | PLEKHA5::chr12:19441011-19441061(+) |
| NDRG1 | RABEP1 | 8 | 17 | -1 | 1 | 134249419 | 5283828 | NDRG1::chr8:134249418-134249468(-) | RABEP1::chr17:5283827-5283877(+) |
| NDRG1 | ST3GAL1 | 8 | 8 | -1 | -1 | 134296492 | 134488843 | NDRG1::chr8:134296491-134296541(-) | ST3GAL1::chr8:134488793-134488843(-) |
| NDUFB9 | CSMD3 | 8 | 8 | 1 | -1 | 125555520 | 114327022 | NDUFB9::chr8:125555470-125555520(+) | CSMD3::chr8:114326972-114327022(-) |
| NEBL | FBXO11 | 10 | 2 | -1 | -1 | 21169723 | 48066908 | NEBL::chr10:21169722-21169772(-) | FBXO11::chr2:48066858-48066908(-) |
| NF1 | ACACA | 17 | 17 | 1 | -1 | 29422387 | 35518964 | NF1::chr17:29422337-29422387(+) | ACACA::chr17:35518914-35518964(-) |
| NF1 | ACCN1 | 17 | 17 | 1 | -1 | 29560231 | 31439085 | NF1::chr17:29560181-29560231(+) | ACCN1::chr17:31439035-31439085(-) |
| NF1 | ADAP2 | 17 | 17 | 1 | 1 | 29422387 | 29271925 | NF1::chr17:29422337-29422387(+) | ADAP2::chr17:29271924-29271974(+) |
| NF1 | AGTR1 | 17 | 3 | 1 | 1 | 29662049 | 148425731 | NF1::chr17:29661999-29662049(+) | AGTR1::chr3:148425730-148425780(+) |
| NF1 | ANKFN1 | 17 | 17 | 1 | 1 | 29592357 | 54520204 | NF1::chr17:29592307-29592357(+) | ANKFN1::chr17:54520203-54520253(+) |
| NF1 | ATAD5 | 17 | 17 | 1 | 1 | 29422387 | 29203392 | NF1::chr17:29422337-29422387(+) | ATAD5::chr17:29203391-29203441(+) |
| NF1 | ATAD5 | 17 | 17 | 1 | 1 | 29533389 | 29161166 | NF1::chr17:29533339-29533389(+) | ATAD5::chr17:29161165-29161215(+) |
| NF1 | ATAD5 | 17 | 17 | 1 | 1 | 29497015 | 29192722 | NF1::chr17:29496965-29497015(+) | ATAD5::chr17:29192721-29192771(+) |
| NF1 | CACNA1G | 17 | 17 | 1 | 1 | 29592357 | 48676789 | NF1::chr17:29592307-29592357(+) | CACNA1G::chr17:48676788-48676838(+) |

| | | | | | | | | | |
|---|---|---|---|---|---|---|---|---|---|
| NF1 | GFPT1 | 17 | 2 | 1 | -1 | 29664600 | 69556930 | NF1::chr17:29664550-29664600(+) | GFPT1::chr2:69556880-69556930(-) |
| NF1 | IKZF3 | 17 | 17 | 1 | -1 | 29563039 | 37988404 | NF1::chr17:29562989-29563039(+) | IKZF3::chr17:37988354-37988404(-) |
| NF1 | PDS5B | 17 | 13 | 1 | 1 | 29422387 | 33281071 | NF1::chr17:29422337-29422387(+) | PDS5B::chr13:33281070-33281120(+) |
| NF1 | PSMD11 | 17 | 17 | 1 | 1 | 29497015 | 30773963 | NF1::chr17:29496965-29497015(+) | PSMD11::chr17:30773962-30774012(+) |
| NF1 | RNF135 | 17 | 17 | 1 | 1 | 29422387 | 29324260 | NF1::chr17:29422337-29422387(+) | RNF135::chr17:29324259-29324309(+) |
| NF1 | RNF135 | 17 | 17 | 1 | 1 | 29497015 | 29324260 | NF1::chr17:29496965-29497015(+) | RNF135::chr17:29324259-29324309(+) |
| NF1 | RPTOR | 17 | 17 | 1 | 1 | 29422387 | 78599491 | NF1::chr17:29422337-29422387(+) | RPTOR::chr17:78599490-78599540(+) |
| NF1 | SPACA3 | 17 | 17 | 1 | 1 | 29497015 | 31323861 | NF1::chr17:29496965-29497015(+) | SPACA3::chr17:31323860-31323910(+) |
| NF1 | TPMT | 17 | 6 | 1 | -1 | 29422387 | 18149402 | NF1::chr17:29422337-29422387(+) | TPMT::chr6:18149352-18149402(-) |
| NF1 | TRMT61B | 17 | 2 | 1 | -1 | 29497015 | 29084174 | NF1::chr17:29496965-29497015(+) | TRMT61B::chr2:29084124-29084174(-) |
| NF1 | YPEL5 | 17 | 2 | 1 | 1 | 29528503 | 30379494 | NF1::chr17:29528453-29528503(+) | YPEL5::chr2:30379493-30379543(+) |
| NF2 | EIF3D | 22 | 22 | 1 | -1 | 30000101 | 36908649 | NF2::chr22:30000051-30000101(+) | EIF3D::chr22:36908599-36908649(-) |
| NF2 | GRAP2 | 22 | 22 | 1 | 1 | 30035201 | 40351823 | NF2::chr22:30035151-30035201(+) | GRAP2::chr22:40351822-40351872(+) |
| NF2 | PIAS1 | 22 | 15 | 1 | 1 | 30000101 | 68378644 | NF2::chr22:30000051-30000101(+) | PIAS1::chr15:68378643-68378693(+) |
| NFE2L2 | AGPS | 2 | 2 | -1 | 1 | 178128165 | 178386057 | NFE2L2::chr2:178128164-178128214(-) | AGPS::chr2:178386056-178386106(+) |
| NFE2L2 | GAPVD1 | 2 | 9 | -1 | 1 | 178097120 | 128083712 | NFE2L2::chr2:178097119-178097169(-) | GAPVD1::chr9:128083711-128083761(+) |
| NFE2L2 | PDE11A | 2 | 2 | -1 | -1 | 178097120 | 178634101 | NFE2L2::chr2:178097119-178097169(-) | PDE11A::chr2:178634051-178634101(-) |
| NFE2L2 | RFTN2 | 2 | 2 | -1 | -1 | 178129260 | 198437004 | NFE2L2::chr2:178129259-178129309(-) | RFTN2::chr2:198436954-198437004(-) |
| NFE2L2 | TMEM194B | 2 | 2 | -1 | -1 | 178129260 | 191390104 | NFE2L2::chr2:178129259-178129309(-) | TMEM194B::chr2:191390054-191390104(-) |
| NFE2L2 | ZC3H13 | 2 | 13 | -1 | -1 | 178129260 | 46554139 | NFE2L2::chr2:178129259-178129309(-) | ZC3H13::chr13:46554089-46554139(-) |
| NFKB2 | GBF1 | 10 | 10 | 1 | 1 | 104156246 | 104128496 | NFKB2::chr10:104156196-104156246(+) | GBF1::chr10:104128495-104128545(+) |
| NFKBIE | FBXO9 | 6 | 6 | -1 | 1 | 44232719 | 52935855 | NFKBIE::chr6:44232718-44232768(-) | FBXO9::chr6:52935854-52935904(+) |
| NFKBIE | SLC29A1 | 6 | 6 | -1 | 1 | 44227780 | 44194997 | NFKBIE::chr6:44227779-44227829(-) | SLC29A1::chr6:44194996-44195046(+) |
| NHSL1 | PRDM1 | 6 | 6 | -1 | 1 | 138817356 | 106536076 | NHSL1::chr6:138817355-138817405(-) | PRDM1::chr6:106536075-106536125(+) |
| NKX2-1 | SLC25A21 | 14 | 14 | -1 | -1 | 36988190 | 37153085 | NKX2-1::chr14:36988189-36988239(-) | SLC25A21::chr14:37153035-37153085(-) |
| NLK | NF1 | 17 | 17 | 1 | 1 | 26370357 | 29687505 | NLK::chr17:26370307-26370357(+) | NF1::chr17:29687504-29687554(+) |
| NOTCH1 | HNF1B | 9 | 17 | -1 | -1 | 139397634 | 36093736 | NOTCH1::chr9:139397633-139397683(-) | HNF1B::chr17:36093686-36093736(-) |
| NOTCH1 | LCN10 | 9 | 9 | -1 | -1 | 139438476 | 139634500 | NOTCH1::chr9:139438475-139438525(-) | LCN10::chr9:139634450-139634500(-) |
| NOTCH1 | UHRF1BP1L | 9 | 12 | -1 | -1 | 139408962 | 100463947 | NOTCH1::chr9:139408961-139409011(-) | UHRF1BP1L::chr12:100463897-100463947(-) |
| NOTCH2 | CWH43 | 1 | 4 | -1 | 1 | 120506197 | 49032842 | NOTCH2::chr1:120506196-120506246(-) | CWH43::chr4:49032841-49032891(+) |
| NOTCH2 | MNDA | 1 | 1 | -1 | 1 | 120478095 | 158815377 | NOTCH2::chr1:120478094-120478144(-) | MNDA::chr1:158815376-158815426(+) |
| NOTCH2 | PSMA5 | 1 | 1 | -1 | -1 | 120611948 | 109964548 | NOTCH2::chr1:120611947-120611997(-) | PSMA5::chr1:109964498-109964548(-) |
| NOTCH2 | SPAG17 | 1 | 1 | -1 | -1 | 120611948 | 118524024 | NOTCH2::chr1:120611947-120611997(-) | SPAG17::chr1:118523974-118524024(-) |
| NRG1 | KRT78 | 8 | 12 | 1 | -1 | 32406344 | 53233768 | NRG1::chr8:32406294-32406344(+) | KRT78::chr12:53233718-53233768(-) |
| NRG1 | PMEPA1 | 8 | 20 | 1 | -1 | 31498245 | 56234753 | NRG1::chr8:31498195-31498245(+) | PMEPA1::chr20:56234703-56234753(-) |
| NRG1 | PROSC | 8 | 8 | 1 | 1 | 32406344 | 37630273 | NRG1::chr8:32406294-32406344(+) | PROSC::chr8:37630272-37630322(+) |
| NRG1 | STMN2 | 8 | 8 | 1 | 1 | 32463201 | 80549037 | NRG1::chr8:32463151-32463201(+) | STMN2::chr8:80549036-80549086(+) |
| NRP1 | PTEN | 10 | 10 | -1 | 1 | 33619636 | 89717610 | NRP1::chr10:33619635-33619685(-) | PTEN::chr10:89717609-89717659(+) |
| NSMCE2 | EXT1 | 8 | 8 | 1 | -1 | 126163519 | 118817132 | NSMCE2::chr8:126163469-126163519(+) | EXT1::chr8:118817082-118817132(-) |
| NTHL1 | KLRC3 | 16 | 12 | -1 | -1 | 2097710 | 10565215 | NTHL1::chr16:2097709-2097759(-) | KLRC3::chr12:10565165-10565215(-) |
| NTRK1 | DYNC2H1 | 1 | 11 | 1 | 1 | 156849949 | 103306671 | NTRK1::chr1:156849899-156849949(+) | DYNC2H1::chr11:103306670-103306720(+) |
| NUMB | DNM2 | 14 | 19 | -1 | 1 | 73822334 | 10893636 | NUMB::chr14:73822333-73822383(-) | DNM2::chr19:10893635-10893685(+) |
| NUP210 | FBLN2 | 3 | 3 | -1 | 1 | 13393379 | 13679062 | NUP210::chr3:13393378-13393428(-) | FBLN2::chr3:13679061-13679111(+) |

| | | | | | | | | | |
|---|---|---|---|---|---|---|---|---|---|
| OXR1 | CSMD3 | 8 | 8 | 1 | -1 | 107726213 | 114327022 | OXR1::chr8:107726163-107726213(+) | CSMD3::chr8:114326972-114327022(-) |
| PABPC1 | VPS13B | 8 | 8 | -1 | 1 | 101716525 | 100779002 | PABPC1::chr8:101716524-101716574(-) | VPS13B::chr8:100779001-100779051(+) |
| PAX5 | TMEM9B | 9 | 11 | -1 | -1 | 37033983 | 8977845 | PAX5::chr9:37033982-37034032(-) | TMEM9B::chr11:8977795-8977845(-) |
| PBRM1 | CACNA2D3 | 3 | 3 | -1 | 1 | 52637537 | 54420742 | PBRM1::chr3:52637536-52637586(-) | CACNA2D3::chr3:54420741-54420791(+) |
| PBRM1 | FBXW12 | 3 | 3 | -1 | 1 | 52595783 | 48416844 | PBRM1::chr3:52595782-52595832(-) | FBXW12::chr3:48416843-48416893(+) |
| PBRM1 | GLT8D1 | 3 | 3 | -1 | -1 | 52712516 | 52734512 | PBRM1::chr3:52712515-52712565(-) | GLT8D1::chr3:52734462-52734512(-) |
| PBRM1 | IGSF11 | 3 | 3 | -1 | -1 | 52712516 | 118647563 | PBRM1::chr3:52712515-52712565(-) | IGSF11::chr3:118647513-118647563(-) |
| PBRM1 | KYNU | 3 | 2 | -1 | 1 | 52692215 | 143742653 | PBRM1::chr3:52692214-52692264(-) | KYNU::chr2:143742652-143742702(+) |
| PBRM1 | NEK4 | 3 | 3 | -1 | -1 | 52637537 | 52773650 | PBRM1::chr3:52637536-52637586(-) | NEK4::chr3:52773600-52773650(-) |
| PDCD6 | TERT | 5 | 5 | 1 | -1 | 271936 | 1294781 | PDCD6::chr5:271886-271936(+) | TERT::chr5:1294731-1294781(-) |
| PDE12 | ROBO2 | 3 | 3 | 1 | 1 | 57545115 | 77671378 | PDE12::chr3:57545065-57545115(+) | ROBO2::chr3:77671377-77671427(+) |
| PDHX | KDM6A | 11 | X | 1 | 1 | 34938362 | 44966655 | PDHX::chr11:34938312-34938362(+) | KDM6A::chrX:44966654-44966704(+) |
| PER1 | ENOPH1 | 17 | 4 | -1 | 1 | 8045967 | 83369073 | PER1::chr17:8045966-8046016(-) | ENOPH1::chr4:83369072-83369122(+) |
| PI4KA | CLTCL1 | 22 | 22 | -1 | -1 | 21188850 | 19230459 | PI4KA::chr22:21188849-21188899(-) | CLTCL1::chr22:19230409-19230459(-) |
| PIK3R1 | MRPL42 | 5 | 12 | 1 | 1 | 67522837 | 93862965 | PIK3R1::chr5:67522787-67522837(+) | MRPL42::chr12:93862964-93863014(+) |
| PIK3R1 | NDUFB7 | 5 | 19 | 1 | -1 | 67576834 | 14677745 | PIK3R1::chr5:67576784-67576834(+) | NDUFB7::chr19:14677695-14677745(-) |
| PML | POLN | 15 | 4 | 1 | -1 | 74325755 | 2097660 | PML::chr15:74325705-74325755(+) | POLN::chr4:2097610-2097660(-) |
| PMS2 | AIMP2 | 7 | 7 | -1 | 1 | 6048628 | 6054777 | PMS2::chr7:6048627-6048677(-) | AIMP2::chr7:6054776-6054826(+) |
| PMS2 | DNAH11 | 7 | 7 | -1 | 1 | 6048628 | 21813384 | PMS2::chr7:6048627-6048677(-) | DNAH11::chr7:21813383-21813433(+) |
| PMS2 | USP42 | 7 | 7 | -1 | 1 | 6048628 | 6150756 | PMS2::chr7:6048627-6048677(-) | USP42::chr7:6150755-6150805(+) |
| POLE | ANKS1B | 12 | 12 | -1 | -1 | 133263840 | 99225914 | POLE::chr12:133263839-133263889(-) | ANKS1B::chr12:99225864-99225914(-) |
| POLG | CGNL1 | 15 | 15 | -1 | 1 | 89866635 | 57808978 | POLG::chr15:89866634-89866684(-) | CGNL1::chr15:57808977-57809027(+) |
| POLR1A | SETD2 | 2 | 3 | -1 | -1 | 86292397 | 47079267 | POLR1A::chr2:86292396-86292446(-) | SETD2::chr3:47079217-47079267(-) |
| POT1 | RTKN2 | 7 | 10 | -1 | -1 | 124568869 | 63959620 | POT1::chr7:124568868-124568918(-) | RTKN2::chr10:63959570-63959620(-) |
| PPARG | SYN2 | 3 | 3 | 1 | 1 | 12330633 | 12187179 | PPARG::chr3:12330583-12330633(+) | SYN2::chr3:12187178-12187228(+) |
| PPARG | SYN2 | 3 | 3 | 1 | 1 | 12330633 | 12228869 | PPARG::chr3:12330583-12330633(+) | SYN2::chr3:12228868-12228918(+) |
| PPFIA1 | CBLB | 11 | 3 | 1 | -1 | 70208594 | 105459475 | PPFIA1::chr11:70208544-70208594(+) | CBLB::chr3:105459425-105459475(-) |
| PPM1H | WIF1 | 12 | 12 | -1 | -1 | 63131282 | 65462684 | PPM1H::chr12:63131281-63131331(-) | WIF1::chr12:65462634-65462684(-) |
| PPP2R1A | CENPP | 19 | 9 | 1 | 1 | 52693427 | 95373596 | PPP2R1A::chr19:52693377-52693427(+) | CENPP::chr9:95373595-95373645(+) |
| PPP6C | C11orf84 | 9 | 11 | -1 | 1 | 127933364 | 63594400 | PPP6C::chr9:127933363-127933413(-) | C11orf84::chr11:63594399-63594449(+) |
| PRDM1 | SLC22A16 | 6 | 6 | 1 | -1 | 106554374 | 110763978 | PRDM1::chr6:106554324-106554374(+) | SLC22A16::chr6:110763928-110763978(-) |
| PREP | PTPRK | 6 | 6 | -1 | -1 | 105729621 | 128643455 | PREP::chr6:105729620-105729670(-) | PTPRK::chr6:128643405-128643455(-) |
| PRKAR1A | PITPNC1 | 17 | 17 | 1 | 1 | 66511717 | 65528918 | PRKAR1A::chr17:66511667-66511717(+) | PITPNC1::chr17:65528917-65528967(+) |
| PRKCA | NF1 | 17 | 17 | 1 | 1 | 64302245 | 29652838 | PRKCA::chr17:64302195-64302245(+) | NF1::chr17:29652837-29652887(+) |
| PTCH1 | IPPK | 9 | 9 | -1 | -1 | 98268689 | 95381847 | PTCH1::chr9:98268688-98268738(-) | IPPK::chr9:95381797-95381847(-) |
| PTEN | BTRC | 10 | 10 | 1 | 1 | 89693008 | 103290994 | PTEN::chr10:89692958-89693008(+) | BTRC::chr10:103290993-103291043(+) |
| PTEN | C1orf226 | 10 | 1 | 1 | 1 | 89653866 | 162352972 | PTEN::chr10:89653816-89653866(+) | C1orf226::chr1:162352971-162353021(+) |
| PTEN | CTNNA3 | 10 | 10 | 1 | -1 | 89690846 | 68979628 | PTEN::chr10:89690796-89690846(+) | CTNNA3::chr10:68979578-68979628(-) |
| PTEN | GRID1 | 10 | 10 | 1 | -1 | 89690846 | 87489371 | PTEN::chr10:89690796-89690846(+) | GRID1::chr10:87489321-87489371(-) |
| PTEN | KIF20B | 10 | 10 | 1 | 1 | 89653866 | 91497144 | PTEN::chr10:89653816-89653866(+) | KIF20B::chr10:91497143-91497193(+) |
| PTEN | NRP1 | 10 | 10 | 1 | -1 | 89693008 | 33559784 | PTEN::chr10:89692958-89693008(+) | NRP1::chr10:33559734-33559784(-) |
| PTEN | PYGB | 10 | 20 | 1 | 1 | 89693008 | 25249765 | PTEN::chr10:89692958-89693008(+) | PYGB::chr20:25249764-25249814(+) |
| PTEN | RNLS | 10 | 10 | 1 | -1 | 89693008 | 90045263 | PTEN::chr10:89692958-89693008(+) | RNLS::chr10:90045213-90045263(-) |

| | | | | | | | | | |
|---|---|---|---|---|---|---|---|---|---|
| PTPN13 | MAPK10 | 4 | 4 | 1 | -1 | 87515942 | 87115560 | PTPN13::chr4:87515892-87515942(+) | MAPK10::chr4:87115510-87115560(-) |
| PTPN6 | LEPREL2 | 12 | 12 | 1 | 1 | 7066948 | 6948164 | PTPN6::chr12:7066898-7066948(+) | LEPREL2::chr12:6948163-6948213(+) |
| PTPN6 | LRRC23 | 12 | 12 | 1 | 1 | 7061340 | 7014749 | PTPN6::chr12:7061290-7061340(+) | LRRC23::chr12:7014748-7014798(+) |
| PTPRD | CYGB | 9 | 17 | -1 | -1 | 9938507 | 74527224 | PTPRD::chr9:9938506-9938556(-) | CYGB::chr17:74527174-74527224(-) |
| PTPRD | IDO1 | 9 | 8 | -1 | 1 | 8733780 | 39771408 | PTPRD::chr9:8733779-8733829(-) | IDO1::chr8:39771407-39771457(+) |
| PTPRD | RAC2 | 9 | 22 | -1 | -1 | 10612398 | 37637698 | PTPRD::chr9:10612397-10612447(-) | RAC2::chr22:37637648-37637698(-) |
| PTPRD | TRHDE | 9 | 12 | -1 | 1 | 8314247 | 72666463 | PTPRD::chr9:8314246-8314296(-) | TRHDE::chr12:72666462-72666512(+) |
| PTPRK | ARHGAP18 | 6 | 6 | -1 | -1 | 128505577 | 129939997 | PTPRK::chr6:128505576-128505626(-) | ARHGAP18::chr6:129939947-129939997(-) |
| PTPRK | BCKDHB | 6 | 6 | -1 | 1 | 128841404 | 80837264 | PTPRK::chr6:128841403-128841453(-) | BCKDHB::chr6:80837263-80837313(+) |
| PTPRK | ECHDC1 | 6 | 6 | -1 | -1 | 128841404 | 127652175 | PTPRK::chr6:128841403-128841453(-) | ECHDC1::chr6:127652125-127652175(-) |
| PTPRK | ECHDC1 | 6 | 6 | -1 | -1 | 128561180 | 127611422 | PTPRK::chr6:128561179-128561229(-) | ECHDC1::chr6:127611372-127611422(-) |
| PTPRK | EPB41L2 | 6 | 6 | -1 | -1 | 128841404 | 131184858 | PTPRK::chr6:128841403-128841453(-) | EPB41L2::chr6:131184808-131184858(-) |
| PTPRK | EYA4 | 6 | 6 | -1 | 1 | 128505577 | 133782252 | PTPRK::chr6:128505576-128505626(-) | EYA4::chr6:133782251-133782301(+) |
| PTPRK | EYS | 6 | 6 | -1 | -1 | 128841404 | 65303202 | PTPRK::chr6:128841403-128841453(-) | EYS::chr6:65303152-65303202(-) |
| PTPRK | FBXO5 | 6 | 6 | -1 | -1 | 128718711 | 153296756 | PTPRK::chr6:128718710-128718760(-) | FBXO5::chr6:153296706-153296756(-) |
| PTPRK | IL20RA | 6 | 6 | -1 | -1 | 128841404 | 137338240 | PTPRK::chr6:128841403-128841453(-) | IL20RA::chr6:137338190-137338240(-) |
| PTPRK | OXR1 | 6 | 8 | -1 | 1 | 128841404 | 107531171 | PTPRK::chr6:128841403-128841453(-) | OXR1::chr8:107531170-107531220(+) |
| PTPRK | PEX5L | 6 | 3 | -1 | -1 | 128505577 | 179605572 | PTPRK::chr6:128505576-128505626(-) | PEX5L::chr3:179605522-179605572(-) |
| PTPRK | POPDC3 | 6 | 6 | -1 | -1 | 128718711 | 105610035 | PTPRK::chr6:128718710-128718760(-) | POPDC3::chr6:105609985-105610035(-) |
| PTPRK | SAMD3 | 6 | 6 | -1 | -1 | 128718711 | 130505768 | PTPRK::chr6:128718710-128718760(-) | SAMD3::chr6:130505718-130505768(-) |
| PTPRK | SMNDC1 | 6 | 10 | -1 | -1 | 128561180 | 112058548 | PTPRK::chr6:128561179-128561229(-) | SMNDC1::chr10:112058498-112058548(-) |
| PTPRK | SPARCL1 | 6 | 4 | -1 | -1 | 128505577 | 88420737 | PTPRK::chr6:128505576-128505626(-) | SPARCL1::chr4:88420687-88420737(-) |
| PTPRT | EXD2 | 20 | 14 | -1 | 1 | 41385102 | 69695533 | PTPRT::chr20:41385101-41385151(-) | EXD2::chr14:69695532-69695582(+) |
| PTPRT | NCOA3 | 20 | 20 | -1 | 1 | 40944363 | 46211927 | PTPRT::chr20:40944362-40944412(-) | NCOA3::chr20:46211926-46211976(+) |
| QKI | ATG5 | 6 | 6 | 1 | -1 | 163836367 | 106764141 | QKI::chr6:163836317-163836367(+) | ATG5::chr6:106764091-106764141(-) |
| QKI | PACRG | 6 | 6 | 1 | 1 | 163899928 | 163733811 | QKI::chr6:163899878-163899928(+) | PACRG::chr6:163733810-163733860(+) |
| QKI | ZNF214 | 6 | 11 | 1 | -1 | 163836367 | 7024086 | QKI::chr6:163836317-163836367(+) | ZNF214::chr11:7024036-7024086(-) |
| RAB3GAP1 | LRP1B | 2 | 2 | 1 | -1 | 135851263 | 141739845 | RAB3GAP1::chr2:135851213-135851263(+) | LRP1B::chr2:141739795-141739845(-) |
| RAB3IL1 | NRG1 | 11 | 8 | -1 | 1 | 61669914 | 32585467 | RAB3IL1::chr11:61669913-61669963(-) | NRG1::chr8:32585466-32585516(+) |
| RAD17 | CNTN5 | 5 | 11 | 1 | 1 | 68677884 | 100226848 | RAD17::chr5:68677834-68677884(+) | CNTN5::chr11:100226847-100226897(+) |
| RAD17 | SEC24A | 5 | 5 | 1 | 1 | 68670538 | 134050714 | RAD17::chr5:68670488-68670538(+) | SEC24A::chr5:134050713-134050763(+) |
| RANBP2 | BMP5 | 2 | 6 | 1 | -1 | 109371715 | 55659225 | RANBP2::chr2:109371665-109371715(+) | BMP5::chr6:55659175-55659225(-) |
| RB1 | CLEC2B | 13 | 12 | 1 | -1 | 48881542 | 10015172 | RB1::chr13:48881492-48881542(+) | CLEC2B::chr12:10015122-10015172(-) |
| RB1 | CNIH3 | 13 | 1 | 1 | 1 | 48881542 | 224868660 | RB1::chr13:48881492-48881542(+) | CNIH3::chr1:224868659-224868709(+) |
| RB1 | DIAPH3 | 13 | 13 | 1 | -1 | 48881542 | 60707123 | RB1::chr13:48881492-48881542(+) | DIAPH3::chr13:60707073-60707123(-) |
| RB1 | ESR2 | 13 | 14 | 1 | -1 | 48955579 | 64749793 | RB1::chr13:48955529-48955579(+) | ESR2::chr14:64749743-64749793(-) |
| RB1 | FRYL | 13 | 4 | 1 | -1 | 49039504 | 48636507 | RB1::chr13:49039454-49039504(+) | FRYL::chr4:48636457-48636507(-) |
| RB1 | GADD45GIP1 | 13 | 19 | 1 | -1 | 48955579 | 13065340 | RB1::chr13:48955529-48955579(+) | GADD45GIP1::chr19:13065290-13065340(-) |
| RB1 | HTR2C | 13 | X | 1 | 1 | 48881542 | 114141152 | RB1::chr13:48881492-48881542(+) | HTR2C::chrX:114141151-114141201(+) |
| RB1 | KCNRG | 13 | 13 | 1 | 1 | 48939107 | 50594350 | RB1::chr13:48939057-48939107(+) | KCNRG::chr13:50594349-50594399(+) |
| RB1 | WDFY2 | 13 | 13 | 1 | 1 | 48881542 | 52325446 | RB1::chr13:48881492-48881542(+) | WDFY2::chr13:52325445-52325495(+) |
| RBM10 | CMAS | X | 12 | 1 | 1 | 47006897 | 22208083 | RBM10::chrX:47006847-47006897(+) | CMAS::chr12:22208082-22208132(+) |
| RBM10 | PHF16 | X | X | 1 | 1 | 47028897 | 46857522 | RBM10::chrX:47028847-47028897(+) | PHF16::chrX:46857521-46857571(+) |

| | | | | | | | | | |
|---|---|---|---|---|---|---|---|---|---|
| RCAN1 | RUNX1 | 21 | 21 | -1 | -1 | 35987059 | 36171759 | RCAN1::chr21:35987058-35987108(-) | RUNX1::chr21:36171709-36171759(-) |
| RFWD3 | STK31 | 16 | 7 | -1 | 1 | 74694830 | 23757100 | RFWD3::chr16:74694829-74694879(-) | STK31::chr7:23757099-23757149(+) |
| RHOA | C3orf43 | 3 | 3 | -1 | -1 | 49449253 | 196236540 | RHOA::chr3:49449252-49449302(-) | C3orf43::chr3:196236490-196236540(-) |
| RHOA | NICN1 | 3 | 3 | -1 | -1 | 49449253 | 49463861 | RHOA::chr3:49449252-49449302(-) | NICN1::chr3:49463811-49463861(-) |
| RHOA | TMEM111 | 3 | 3 | -1 | -1 | 49405861 | 10011485 | RHOA::chr3:49405860-49405910(-) | TMEM111::chr3:10011435-10011485(-) |
| RNF217 | PRDM1 | 6 | 6 | 1 | 1 | 125284572 | 106552700 | RNF217::chr6:125284522-125284572(+) | PRDM1::chr6:106552699-106552749(+) |
| RNF217 | PTPRK | 6 | 6 | 1 | -1 | 125284572 | 128718833 | RNF217::chr6:125284522-125284572(+) | PTPRK::chr6:128718783-128718833(-) |
| RNF43 | HSF5 | 17 | 17 | -1 | -1 | 56492687 | 56499791 | RNF43::chr17:56492686-56492736(-) | HSF5::chr17:56499741-56499791(-) |
| RNF43 | LIMCH1 | 17 | 4 | -1 | 1 | 56492687 | 41496553 | RNF43::chr17:56492686-56492736(-) | LIMCH1::chr4:41496552-41496602(+) |
| ROBO2 | ERLIN2 | 3 | 8 | 1 | 1 | 77147491 | 37607077 | ROBO2::chr3:77147441-77147491(+) | ERLIN2::chr8:37607076-37607126(+) |
| ROBO2 | TNNT2 | 3 | 1 | 1 | -1 | 77089997 | 201328791 | ROBO2::chr3:77089947-77089997(+) | TNNT2::chr1:201328741-201328791(-) |
| RPL22 | ACOT7 | 1 | 1 | -1 | -1 | 6252990 | 6324755 | RPL22::chr1:6252989-6253039(-) | ACOT7::chr1:6324705-6324755(-) |
| RPS6KA3 | KDM6A | X | X | -1 | 1 | 20284682 | 44966655 | RPS6KA3::chrX:20284681-20284731(-) | KDM6A::chrX:44966654-44966704(+) |
| RPTOR | GPC3 | 17 | X | 1 | -1 | 78519591 | 133087238 | RPTOR::chr17:78519541-78519591(+) | GPC3::chrX:133087188-133087238(-) |
| RSU1 | LARP4B | 10 | 10 | -1 | -1 | 16858972 | 863829 | RSU1::chr10:16858971-16859021(-) | LARP4B::chr10:863779-863829(-) |
| RUNX1 | CEP76 | 21 | 18 | -1 | -1 | 36206707 | 12686449 | RUNX1::chr21:36206706-36206756(-) | CEP76::chr18:12686399-12686449(-) |
| RUNX1 | DTD1 | 21 | 20 | -1 | 1 | 36252854 | 18724744 | RUNX1::chr21:36252853-36252903(-) | DTD1::chr20:18724743-18724793(+) |
| RUNX1 | IFNAR1 | 21 | 21 | -1 | 1 | 36206707 | 34725064 | RUNX1::chr21:36206706-36206756(-) | IFNAR1::chr21:34725063-34725113(+) |
| RUNX1 | PROS1 | 21 | 3 | -1 | -1 | 36206707 | 93646251 | RUNX1::chr21:36206706-36206756(-) | PROS1::chr3:93646201-93646251(-) |
| RUNX1 | ZMAT4 | 21 | 8 | -1 | -1 | 36259140 | 40532450 | RUNX1::chr21:36259139-36259189(-) | ZMAT4::chr8:40532400-40532450(-) |
| RUNX1T1 | E2F5 | 8 | 8 | -1 | 1 | 93115085 | 86114382 | RUNX1T1::chr8:93115084-93115134(-) | E2F5::chr8:86114381-86114431(+) |
| RUNX1T1 | FAM164A | 8 | 8 | -1 | 1 | 93088193 | 79590798 | RUNX1T1::chr8:93088192-93088242(-) | FAM164A::chr8:79590797-79590847(+) |
| RUNX2 | MAX | 6 | 14 | 1 | -1 | 45399756 | 65544754 | RUNX2::chr6:45399706-45399756(+) | MAX::chr14:65544704-65544754(-) |
| RYR3 | BUB1B | 15 | 15 | 1 | 1 | 33765739 | 40457254 | RYR3::chr15:33765689-33765739(+) | BUB1B::chr15:40457253-40457303(+) |
| SDHAF2 | RIN1 | 11 | 11 | 1 | -1 | 61205585 | 66101129 | SDHAF2::chr11:61205535-61205585(+) | RIN1::chr11:66101079-66101129(-) |
| SDHAF2 | TPH1 | 11 | 11 | 1 | -1 | 61205585 | 18062335 | SDHAF2::chr11:61205535-61205585(+) | TPH1::chr11:18062285-18062335(-) |
| SDHB | PLN | 1 | 6 | -1 | 1 | 17380443 | 118879988 | SDHB::chr1:17380442-17380492(-) | PLN::chr6:118879987-118880037(+) |
| SEC16A | NOTCH1 | 9 | 9 | -1 | -1 | 139357339 | 139438554 | SEC16A::chr9:139357338-139357388(-) | NOTCH1::chr9:139438504-139438554(-) |
| SEC16A | NOTCH1 | 9 | 9 | -1 | -1 | 139353586 | 139418431 | SEC16A::chr9:139353585-139353635(-) | NOTCH1::chr9:139418381-139418431(-) |
| SETD2 | CCRL2 | 3 | 3 | -1 | 1 | 47139445 | 46449559 | SETD2::chr3:47139444-47139494(-) | CCRL2::chr3:46449558-46449608(+) |
| SETD2 | MACROD2 | 3 | 20 | -1 | 1 | 47125210 | 14066267 | SETD2::chr3:47125209-47125259(-) | MACROD2::chr20:14066266-14066316(+) |
| SETD2 | NISCH | 3 | 3 | -1 | 1 | 47125210 | 52504875 | SETD2::chr3:47125209-47125259(-) | NISCH::chr3:52504874-52504924(+) |
| SETD2 | PALMD | 3 | 1 | -1 | 1 | 47205344 | 100159575 | SETD2::chr3:47205343-47205393(-) | PALMD::chr1:100159574-100159624(+) |
| SETD2 | QRICH1 | 3 | 3 | -1 | -1 | 47205344 | 49070661 | SETD2::chr3:47205343-47205393(-) | QRICH1::chr3:49070611-49070661(-) |
| SETD2 | RBM6 | 3 | 3 | -1 | 1 | 47205344 | 50112634 | SETD2::chr3:47205343-47205393(-) | RBM6::chr3:50112633-50112683(+) |
| SETD2 | SCAP | 3 | 3 | -1 | -1 | 47139445 | 47484581 | SETD2::chr3:47139444-47139494(-) | SCAP::chr3:47484531-47484581(-) |
| SETD2 | SMARCC1 | 3 | 3 | -1 | -1 | 47205344 | 47632327 | SETD2::chr3:47205343-47205393(-) | SMARCC1::chr3:47632277-47632327(-) |
| SETD2 | SPATA12 | 3 | 3 | -1 | 1 | 47155366 | 57107394 | SETD2::chr3:47155365-47155415(-) | SPATA12::chr3:57107393-57107443(+) |
| SH2B3 | ATXN2 | 12 | 12 | 1 | -1 | 111844081 | 111993723 | SH2B3::chr12:111844031-111844081(+) | ATXN2::chr12:111993673-111993723(-) |
| SH2B3 | CUX2 | 12 | 12 | 1 | 1 | 111844081 | 111757816 | SH2B3::chr12:111844031-111844081(+) | CUX2::chr12:111757815-111757865(+) |
| SHB | CDKN2A | 9 | 9 | -1 | -1 | 38067926 | 21968241 | SHB::chr9:38067925-38067975(-) | CDKN2A::chr9:21968191-21968241(-) |
| SHB | GRIN2A | 9 | 16 | -1 | -1 | 37955880 | 9943818 | SHB::chr9:37955879-37955929(-) | GRIN2A::chr16:9943768-9943818(-) |
| SHC2 | PTEN | 19 | 10 | -1 | 1 | 438718 | 89685270 | SHC2::chr19:438717-438767(-) | PTEN::chr10:89685269-89685319(+) |

| | | | | | | | | | |
|---|---|---|---|---|---|---|---|---|---|
| SLC12A7 | TERT | 5 | 5 | -1 | -1 | 1060459 | 1282739 | SLC12A7::chr5:1060458-1060508(-) | TERT::chr5:1282689-1282739(-) |
| SLC22A23 | CAMTA1 | 6 | 1 | -1 | 1 | 3410422 | 6880241 | SLC22A23::chr6:3410421-3410471(-) | CAMTA1::chr1:6880240-6880290(+) |
| SLC25A14 | BCORL1 | X | X | 1 | 1 | 129484705 | 129184692 | SLC25A14::chrX:129484655-129484705(+) | BCORL1::chrX:129184691-129184741(+) |
| SLC2A13 | TERT | 12 | 5 | -1 | -1 | 40345059 | 1282739 | SLC2A13::chr12:40345058-40345108(-) | TERT::chr5:1282689-1282739(-) |
| SLC34A2 | EEA1 | 4 | 12 | 1 | -1 | 25680370 | 93229203 | SLC34A2::chr4:25680320-25680370(+) | EEA1::chr12:93229153-93229203(-) |
| SLC34A2 | NDUFV1 | 4 | 11 | 1 | 1 | 25680370 | 67375697 | SLC34A2::chr4:25680320-25680370(+) | NDUFV1::chr11:67375696-67375746(+) |
| SMAD2 | C10orf90 | 18 | 10 | -1 | -1 | 45394694 | 128202508 | SMAD2::chr18:45394693-45394743(-) | C10orf90::chr10:128202458-128202508(-) |
| SMAD2 | CABLES1 | 18 | 18 | -1 | 1 | 45456732 | 20768802 | SMAD2::chr18:45456731-45456781(-) | CABLES1::chr18:20768801-20768851(+) |
| SMAD2 | CD226 | 18 | 18 | -1 | -1 | 45456732 | 67563281 | SMAD2::chr18:45456731-45456781(-) | CD226::chr18:67563231-67563281(-) |
| SMAD2 | ZBTB7C | 18 | 18 | -1 | -1 | 45456732 | 45864769 | SMAD2::chr18:45456731-45456781(-) | ZBTB7C::chr18:45864719-45864769(-) |
| SMAD2 | ZBTB7C | 18 | 18 | -1 | -1 | 45456732 | 45567494 | SMAD2::chr18:45456731-45456781(-) | ZBTB7C::chr18:45567444-45567494(-) |
| SMAD3 | AAGAB | 15 | 15 | 1 | -1 | 67430438 | 67529158 | SMAD3::chr15:67430388-67430438(+) | AAGAB::chr15:67529108-67529158(-) |
| SMAD3 | C15orf44 | 15 | 15 | 1 | -1 | 67358698 | 65877226 | SMAD3::chr15:67358648-67358698(+) | C15orf44::chr15:65877176-65877226(-) |
| SMAD3 | DLGAP4 | 15 | 20 | 1 | 1 | 67358698 | 35125108 | SMAD3::chr15:67358648-67358698(+) | DLGAP4::chr20:35125107-35125157(+) |
| SMAD3 | IL17RA | 15 | 22 | 1 | 1 | 67358698 | 17577952 | SMAD3::chr15:67358648-67358698(+) | IL17RA::chr22:17577951-17578001(+) |
| SMAD3 | SENP8 | 15 | 15 | 1 | 1 | 67358698 | 72431918 | SMAD3::chr15:67358648-67358698(+) | SENP8::chr15:72431917-72431967(+) |
| SMAD4 | SKAP1 | 18 | 17 | 1 | -1 | 48593557 | 46474147 | SMAD4::chr18:48593507-48593557(+) | SKAP1::chr17:46474097-46474147(-) |
| SMARCA4 | C19orf38 | 19 | 19 | 1 | 1 | 11101999 | 10979894 | SMARCA4::chr19:11101949-11101999(+) | C19orf38::chr19:10979893-10979943(+) |
| SMARCA4 | C19orf38 | 19 | 19 | 1 | 1 | 11098600 | 10979894 | SMARCA4::chr19:11098550-11098600(+) | C19orf38::chr19:10979893-10979943(+) |
| SMARCA4 | CARM1 | 19 | 19 | 1 | 1 | 11100119 | 11022860 | SMARCA4::chr19:11100069-11100119(+) | CARM1::chr19:11022859-11022909(+) |
| SMARCA4 | DOCK6 | 19 | 19 | 1 | -1 | 11130377 | 11311728 | SMARCA4::chr19:11130327-11130377(+) | DOCK6::chr19:11311678-11311728(-) |
| SMARCA4 | KDM4B | 19 | 19 | 1 | 1 | 11100119 | 5071021 | SMARCA4::chr19:11100069-11100119(+) | KDM4B::chr19:5071020-5071070(+) |
| SMARCA4 | RNF216 | 19 | 7 | 1 | -1 | 11152236 | 5800769 | SMARCA4::chr19:11152186-11152236(+) | RNF216::chr7:5800719-5800769(-) |
| SMARCA4 | SLC44A2 | 19 | 19 | 1 | 1 | 11107220 | 10745432 | SMARCA4::chr19:11107170-11107220(+) | SLC44A2::chr19:10745431-10745481(+) |
| SMARCA4 | SLC44A2 | 19 | 19 | 1 | 1 | 11114073 | 10736929 | SMARCA4::chr19:11114023-11114073(+) | SLC44A2::chr19:10736928-10736978(+) |
| SMARCB1 | HRASLS2 | 22 | 11 | 1 | -1 | 24159123 | 63320537 | SMARCB1::chr22:24159073-24159123(+) | HRASLS2::chr11:63320487-63320537(-) |
| SMARCB1 | SNW1 | 22 | 14 | 1 | -1 | 24129449 | 78201355 | SMARCB1::chr22:24129399-24129449(+) | SNW1::chr14:78201305-78201355(-) |
| SPEN | CTRC | 1 | 1 | 1 | 1 | 16199631 | 15768943 | SPEN::chr1:16199581-16199631(+) | CTRC::chr1:15768942-15768992(+) |
| SPEN | EFHD2 | 1 | 1 | 1 | 1 | 16174645 | 15752367 | SPEN::chr1:16174595-16174645(+) | EFHD2::chr1:15752366-15752416(+) |
| SPEN | FBLIM1 | 1 | 1 | 1 | 1 | 16203173 | 16101113 | SPEN::chr1:16203123-16203173(+) | FBLIM1::chr1:16101112-16101162(+) |
| SPEN | FBLIM1 | 1 | 1 | 1 | 1 | 16203173 | 16090991 | SPEN::chr1:16203123-16203173(+) | FBLIM1::chr1:16090990-16091040(+) |
| SPEN | FBLIM1 | 1 | 1 | 1 | 1 | 16174645 | 16096904 | SPEN::chr1:16174595-16174645(+) | FBLIM1::chr1:16096903-16096953(+) |
| SPEN | FHAD1 | 1 | 1 | 1 | 1 | 16174645 | 15668232 | SPEN::chr1:16174595-16174645(+) | FHAD1::chr1:15668231-15668281(+) |
| SPEN | PLEKHM2 | 1 | 1 | 1 | 1 | 16203173 | 16060292 | SPEN::chr1:16203123-16203173(+) | PLEKHM2::chr1:16060291-16060341(+) |
| SPEN | PLEKHM2 | 1 | 1 | 1 | 1 | 16203173 | 16042732 | SPEN::chr1:16203123-16203173(+) | PLEKHM2::chr1:16042731-16042781(+) |
| SPEN | TMEM51 | 1 | 1 | 1 | 1 | 16203173 | 15536986 | SPEN::chr1:16203123-16203173(+) | TMEM51::chr1:15536985-15537035(+) |
| SPEN | TNFRSF1B | 1 | 1 | 1 | 1 | 16242774 | 12262024 | SPEN::chr1:16242724-16242774(+) | TNFRSF1B::chr1:12262023-12262073(+) |
| SPOCK1 | ARHGAP26 | 5 | 5 | -1 | 1 | 136834062 | 142500553 | SPOCK1::chr5:136834061-136834111(-) | ARHGAP26::chr5:142500552-142500602(+) |
| SPOP | AARSD1 | 17 | 17 | -1 | -1 | 47688642 | 41116254 | SPOP::chr17:47688641-47688691(-) | AARSD1::chr17:41116204-41116254(-) |
| SPOP | ARL5C | 17 | 17 | -1 | -1 | 47755295 | 37322870 | SPOP::chr17:47755294-47755344(-) | ARL5C::chr17:37322820-37322870(-) |
| SPOP | C17orf57 | 17 | 17 | -1 | 1 | 47755295 | 45351785 | SPOP::chr17:47755294-47755344(-) | C17orf57::chr17:45351784-45351834(+) |
| SPOP | CA10 | 17 | 17 | -1 | -1 | 47755295 | 50008492 | SPOP::chr17:47755294-47755344(-) | CA10::chr17:50008442-50008492(-) |
| SPOP | EPHX2 | 17 | 8 | -1 | 1 | 47755295 | 27358443 | SPOP::chr17:47755294-47755344(-) | EPHX2::chr8:27358442-27358492(+) |

| | | | | | | | | | |
|---|---|---|---|---|---|---|---|---|---|
| SPOP | FAM117A | 17 | 17 | -1 | -1 | 47745389 | 47810082 | SPOP::chr17:47745388-47745438(-) | FAM117A::chr17:47810032-47810082(-) |
| SPOP | FAM117A | 17 | 17 | -1 | -1 | 47755295 | 47799956 | SPOP::chr17:47755294-47755344(-) | FAM117A::chr17:47799906-47799956(-) |
| SPOP | GPATCH1 | 17 | 19 | -1 | 1 | 47755295 | 33609916 | SPOP::chr17:47755294-47755344(-) | GPATCH1::chr19:33609915-33609965(+) |
| SPOP | NDUFS5 | 17 | 1 | -1 | 1 | 47755295 | 39494395 | SPOP::chr17:47755294-47755344(-) | NDUFS5::chr1:39494394-39494444(+) |
| SPOP | SLC35B1 | 17 | 17 | -1 | -1 | 47755295 | 47783671 | SPOP::chr17:47755294-47755344(-) | SLC35B1::chr17:47783621-47783671(-) |
| SPOP | SLC35B1 | 17 | 17 | -1 | -1 | 47684612 | 47781588 | SPOP::chr17:47684611-47684661(-) | SLC35B1::chr17:47781538-47781588(-) |
| SPOP | TAC4 | 17 | 17 | -1 | -1 | 47755295 | 47921519 | SPOP::chr17:47755294-47755344(-) | TAC4::chr17:47921469-47921519(-) |
| SPOP | XYLT2 | 17 | 17 | -1 | 1 | 47755295 | 48430991 | SPOP::chr17:47755294-47755344(-) | XYLT2::chr17:48430990-48431040(+) |
| ST6GALNAC2 | BRIP1 | 17 | 17 | -1 | -1 | 74574838 | 59820495 | ST6GALNAC2::chr17:74574837-74574887(-) | BRIP1::chr17:59820445-59820495(-) |
| STAG1 | ANKS1B | 3 | 12 | -1 | -1 | 136183723 | 99548173 | STAG1::chr3:136183722-136183772(-) | ANKS1B::chr12:99548123-99548173(-) |
| STAG1 | CCDC77 | 3 | 12 | -1 | 1 | 136471037 | 514677 | STAG1::chr3:136471036-136471086(-) | CCDC77::chr12:514676-514726(+) |
| STAG1 | FAM168A | 3 | 11 | -1 | -1 | 136471037 | 73179537 | STAG1::chr3:136471036-136471086(-) | FAM168A::chr11:73179487-73179537(-) |
| STAG1 | NCK1 | 3 | 3 | -1 | 1 | 136221470 | 136646826 | STAG1::chr3:136221469-136221519(-) | NCK1::chr3:136646825-136646875(+) |
| STAG1 | P2RY13 | 3 | 3 | -1 | -1 | 136471037 | 151046795 | STAG1::chr3:136471036-136471086(-) | P2RY13::chr3:151046745-151046795(-) |
| STAG1 | SPSB4 | 3 | 3 | -1 | 1 | 136117591 | 140865984 | STAG1::chr3:136117590-136117640(-) | SPSB4::chr3:140865983-140866033(+) |
| STAG2 | AMMECR1 | X | X | 1 | -1 | 123094716 | 109507827 | STAG2::chrX:123094666-123094716(+) | AMMECR1::chrX:109507777-109507827(-) |
| STAG2 | C16orf45 | X | 16 | 1 | 1 | 123220620 | 15609162 | STAG2::chrX:123220570-123220620(+) | C16orf45::chr16:15609161-15609211(+) |
| STAG2 | FAM49B | X | 8 | 1 | -1 | 123164975 | 130916831 | STAG2::chrX:123164925-123164975(+) | FAM49B::chr8:130916781-130916831(-) |
| STAG2 | GRIA3 | X | X | 1 | 1 | 123164975 | 122459877 | STAG2::chrX:123164925-123164975(+) | GRIA3::chrX:122459876-122459926(+) |
| STAG2 | RGAG1 | X | X | 1 | 1 | 123094716 | 109693795 | STAG2::chrX:123094666-123094716(+) | RGAG1::chrX:109693794-109693844(+) |
| STAT5B | ODZ4 | 17 | 11 | -1 | -1 | 40368032 | 78926954 | STAT5B::chr17:40368031-40368081(-) | ODZ4::chr11:78926904-78926954(-) |
| STAT5B | PFAS | 17 | 17 | -1 | 1 | 40362189 | 8166238 | STAT5B::chr17:40362188-40362238(-) | PFAS::chr17:8166237-8166287(+) |
| STK11 | C19orf26 | 19 | 19 | 1 | -1 | 1207202 | 1231274 | STK11::chr19:1207152-1207202(+) | C19orf26::chr19:1231224-1231274(-) |
| STK11 | HMHA1 | 19 | 19 | 1 | 1 | 1222005 | 1080254 | STK11::chr19:1221955-1222005(+) | HMHA1::chr19:1080253-1080303(+) |
| STK11 | SRFBP1 | 19 | 5 | 1 | 1 | 1207202 | 121354955 | STK11::chr19:1207152-1207202(+) | SRFBP1::chr5:121354954-121355004(+) |
| SUFU | ADK | 10 | 10 | 1 | 1 | 104309863 | 76468079 | SUFU::chr10:104309813-104309863(+) | ADK::chr10:76468078-76468128(+) |
| SUFU | CYP26A1 | 10 | 10 | 1 | 1 | 104377185 | 94835583 | SUFU::chr10:104377135-104377185(+) | CYP26A1::chr10:94835582-94835632(+) |
| SUFU | KCNH2 | 10 | 7 | 1 | -1 | 104309863 | 150672029 | SUFU::chr10:104309813-104309863(+) | KCNH2::chr7:150671979-150672029(-) |
| SUFU | ZCCHC24 | 10 | 10 | 1 | -1 | 104309863 | 81154196 | SUFU::chr10:104309813-104309863(+) | ZCCHC24::chr10:81154146-81154196(-) |
| SUZ12 | ADAP2 | 17 | 17 | 1 | 1 | 30267505 | 29271925 | SUZ12::chr17:30267455-30267505(+) | ADAP2::chr17:29271924-29271974(+) |
| SUZ12 | VANGL1 | 17 | 1 | 1 | 1 | 30323896 | 116202262 | SUZ12::chr17:30323846-30323896(+) | VANGL1::chr1:116202261-116202311(+) |
| SYNE1 | ESR1 | 6 | 6 | -1 | 1 | 152565672 | 152415520 | SYNE1::chr6:152565671-152565721(-) | ESR1::chr6:152415519-152415569(+) |
| SYT7 | SDHAF2 | 11 | 11 | -1 | 1 | 61290547 | 61213413 | SYT7::chr11:61290546-61290596(-) | SDHAF2::chr11:61213412-61213462(+) |
| TAOK1 | NF1 | 17 | 17 | 1 | 1 | 27829741 | 29652838 | TAOK1::chr17:27829691-27829741(+) | NF1::chr17:29652837-29652887(+) |
| TBL1XR1 | ADAM33 | 3 | 20 | -1 | -1 | 176914909 | 3657790 | TBL1XR1::chr3:176914908-176914958(-) | ADAM33::chr20:3657740-3657790(-) |
| TBL1XR1 | ATG7 | 3 | 3 | -1 | 1 | 176914909 | 11596285 | TBL1XR1::chr3:176914908-176914958(-) | ATG7::chr3:11596284-11596334(+) |
| TBL1XR1 | CACNA1C | 3 | 12 | -1 | 1 | 176816254 | 2721069 | TBL1XR1::chr3:176816253-176816303(-) | CACNA1C::chr12:2721068-2721118(+) |
| TBL1XR1 | DCUN1D1 | 3 | 3 | -1 | -1 | 176914909 | 182683541 | TBL1XR1::chr3:176914908-176914958(-) | DCUN1D1::chr3:182683491-182683541(-) |
| TBL1XR1 | MAST3 | 3 | 19 | -1 | 1 | 176914909 | 18245389 | TBL1XR1::chr3:176914908-176914958(-) | MAST3::chr19:18245388-18245438(+) |
| TBL1XR1 | MCF2L2 | 3 | 3 | -1 | -1 | 176914909 | 183107567 | TBL1XR1::chr3:176914908-176914958(-) | MCF2L2::chr3:183107517-183107567(-) |
| TBL1XR1 | MIPEP | 3 | 13 | -1 | -1 | 176914909 | 24330757 | TBL1XR1::chr3:176914908-176914958(-) | MIPEP::chr13:24330707-24330757(-) |
| TBL1XR1 | ROCK1 | 3 | 18 | -1 | -1 | 176914909 | 18619524 | TBL1XR1::chr3:176914908-176914958(-) | ROCK1::chr18:18619474-18619524(-) |
| TBL1XR1 | TP73 | 3 | 1 | -1 | 1 | 176771561 | 3624113 | TBL1XR1::chr3:176771560-176771610(-) | TP73::chr1:3624112-3624162(+) |

| | | | | | | | | | |
|---|---|---|---|---|---|---|---|---|---|
| TBL1XR1 | USP13 | 3 | 3 | -1 | 1 | 176763917 | 179478900 | TBL1XR1::chr3:176763916-176763966(-) | USP13::chr3:179478899-179478949(+) |
| TCF20 | MKL1 | 22 | 22 | -1 | -1 | 42605657 | 40831558 | TCF20::chr22:42605656-42605706(-) | MKL1::chr22:40831508-40831558(-) |
| TCF3 | AP3D1 | 19 | 19 | -1 | -1 | 1621837 | 2111827 | TCF3::chr19:1621836-1621886(-) | AP3D1::chr19:2111777-2111827(-) |
| TCF3 | APC2 | 19 | 19 | -1 | 1 | 1619110 | 1452983 | TCF3::chr19:1619109-1619159(-) | APC2::chr19:1452982-1453032(+) |
| TCF3 | DNAJC11 | 19 | 1 | -1 | -1 | 1619779 | 6714099 | TCF3::chr19:1619778-1619828(-) | DNAJC11::chr1:6714049-6714099(-) |
| TET1 | EFHC2 | 10 | X | 1 | -1 | 70442730 | 44172002 | TET1::chr10:70442680-70442730(+) | EFHC2::chrX:44171952-44172002(-) |
| TET1 | TMEM222 | 10 | 1 | 1 | 1 | 70411693 | 27660449 | TET1::chr10:70411643-70411693(+) | TMEM222::chr1:27660448-27660498(+) |
| TET2 | ADH6 | 4 | 4 | 1 | -1 | 106111662 | 100137419 | TET2::chr4:106111612-106111662(+) | ADH6::chr4:100137369-100137419(-) |
| TET2 | KLK3 | 4 | 19 | 1 | 1 | 106200973 | 51358191 | TET2::chr4:106200923-106200973(+) | KLK3::chr19:51358190-51358240(+) |
| TET2 | LARP7 | 4 | 4 | 1 | 1 | 106068136 | 113565824 | TET2::chr4:106068086-106068136(+) | LARP7::chr4:113565823-113565873(+) |
| TFR2 | NCOR2 | 7 | 12 | -1 | -1 | 100224386 | 124848345 | TFR2::chr7:100224385-100224435(-) | NCOR2::chr12:124848295-124848345(-) |
| TGFBR2 | NR2C2 | 3 | 3 | 1 | 1 | 30648469 | 15055096 | TGFBR2::chr3:30648419-30648469(+) | NR2C2::chr3:15055095-15055145(+) |
| THAP7 | NRG1 | 22 | 8 | -1 | 1 | 21354936 | 32453346 | THAP7::chr22:21354935-21354985(-) | NRG1::chr8:32453345-32453395(+) |
| THBS1 | NRG1 | 15 | 8 | 1 | 1 | 39876623 | 32585467 | THBS1::chr15:39876573-39876623(+) | NRG1::chr8:32585466-32585516(+) |
| THRA | CDK12 | 17 | 17 | 1 | 1 | 38245586 | 37686884 | THRA::chr17:38245536-38245586(+) | CDK12::chr17:37686883-37686933(+) |
| TMCO7 | CDH1 | 16 | 16 | 1 | 1 | 68877614 | 68772200 | TMCO7::chr16:68877564-68877614(+) | CDH1::chr16:68772199-68772249(+) |
| TMEM65 | EXT1 | 8 | 8 | -1 | -1 | 125384095 | 118849440 | TMEM65::chr8:125384094-125384144(-) | EXT1::chr8:118849390-118849440(-) |
| TNFAIP3 | IL26 | 6 | 12 | 1 | -1 | 138192659 | 68595863 | TNFAIP3::chr6:138192609-138192659(+) | IL26::chr12:68595813-68595863(-) |
| TNFRSF14 | TXLNA | 1 | 1 | 1 | 1 | 2489907 | 32659666 | TNFRSF14::chr1:2489857-2489907(+) | TXLNA::chr1:32659665-32659715(+) |
| TNRC6B | EP300 | 22 | 22 | 1 | 1 | 40552186 | 41572251 | TNRC6B::chr22:40552136-40552186(+) | EP300::chr22:41572250-41572300(+) |
| TNRC6B | STK11 | 22 | 19 | 1 | 1 | 40697337 | 1222984 | TNRC6B::chr22:40697287-40697337(+) | STK11::chr19:1222983-1223033(+) |
| TP53 | C17orf39 | 17 | 17 | -1 | 1 | 7590695 | 17948461 | TP53::chr17:7590694-7590744(-) | C17orf39::chr17:17948460-17948510(+) |
| TP53 | CHD3 | 17 | 17 | -1 | 1 | 7590695 | 7798657 | TP53::chr17:7590694-7590744(-) | CHD3::chr17:7798656-7798706(+) |
| TP53 | ECSIT | 17 | 19 | -1 | -1 | 7576853 | 11630008 | TP53::chr17:7576852-7576902(-) | ECSIT::chr19:11629958-11630008(-) |
| TP53 | FARP2 | 17 | 2 | -1 | 1 | 7590695 | 242312499 | TP53::chr17:7590694-7590744(-) | FARP2::chr2:242312498-242312548(+) |
| TP53 | KIAA0753 | 17 | 17 | -1 | -1 | 7590695 | 6498373 | TP53::chr17:7590694-7590744(-) | KIAA0753::chr17:6498323-6498373(-) |
| TP63 | LEPREL1 | 3 | 3 | 1 | -1 | 189526315 | 189713231 | TP63::chr3:189526265-189526315(+) | LEPREL1::chr3:189713181-189713231(-) |
| TP63 | TPRG1 | 3 | 3 | 1 | 1 | 189349366 | 189028175 | TP63::chr3:189349316-189349366(+) | TPRG1::chr3:189028174-189028224(+) |
| TRAF7 | LRRC1 | 16 | 6 | 1 | 1 | 2215937 | 53743791 | TRAF7::chr16:2215887-2215937(+) | LRRC1::chr6:53743790-53743840(+) |
| TRAF7 | MAPK8IP3 | 16 | 16 | 1 | 1 | 2214002 | 1816905 | TRAF7::chr16:2213952-2214002(+) | MAPK8IP3::chr16:1816904-1816954(+) |
| TRAF7 | WDR24 | 16 | 16 | 1 | -1 | 2205859 | 737739 | TRAF7::chr16:2205809-2205859(+) | WDR24::chr16:737689-737739(-) |
| TRAP1 | CREBBP | 16 | 16 | -1 | -1 | 3740828 | 3799684 | TRAP1::chr16:3740827-3740877(-) | CREBBP::chr16:3799634-3799684(-) |
| TRIM24 | NTRK2 | 7 | 9 | 1 | 1 | 138258387 | 87482158 | TRIM24::chr7:138258337-138258387(+) | NTRK2::chr9:87482157-87482207(+) |
| TRIM24 | SVOPL | 7 | 7 | 1 | -1 | 138258387 | 138281275 | TRIM24::chr7:138258337-138258387(+) | SVOPL::chr7:138281225-138281275(-) |
| TRIM24 | ULK4 | 7 | 3 | 1 | -1 | 138189153 | 41497086 | TRIM24::chr7:138189103-138189153(+) | ULK4::chr3:41497036-41497086(-) |
| TRIM33 | BCAR3 | 1 | 1 | -1 | -1 | 115053172 | 94171698 | TRIM33::chr1:115053171-115053221(-) | BCAR3::chr1:94171648-94171698(-) |
| TRIM33 | COL24A1 | 1 | 1 | -1 | -1 | 115005726 | 86334309 | TRIM33::chr1:115005725-115005775(-) | COL24A1::chr1:86334259-86334309(-) |
| TRIM33 | CSMD2 | 1 | 1 | -1 | -1 | 114948032 | 34128692 | TRIM33::chr1:114948031-114948081(-) | CSMD2::chr1:34128642-34128692(-) |
| TRIM33 | SHC1 | 1 | 1 | -1 | -1 | 114963002 | 154942675 | TRIM33::chr1:114963001-114963051(-) | SHC1::chr1:154942625-154942675(-) |
| TRIO | TERT | 5 | 5 | 1 | -1 | 14406781 | 1282739 | TRIO::chr5:14406731-14406781(+) | TERT::chr5:1282689-1282739(-) |
| TSC1 | CEL | 9 | 9 | -1 | 1 | 135796750 | 135945848 | TSC1::chr9:135796749-135796799(-) | CEL::chr9:135945847-135945897(+) |
| TSC1 | DOLPP1 | 9 | 9 | -1 | 1 | 135796750 | 131848988 | TSC1::chr9:135796749-135796799(-) | DOLPP1::chr9:131848987-131849037(+) |
| TSC1 | SARDH | 9 | 9 | -1 | -1 | 135796750 | 136578246 | TSC1::chr9:135796749-135796799(-) | SARDH::chr9:136578196-136578246(-) |

| | | | | | | | | | |
|---|---|---|---|---|---|---|---|---|---|
| TSC2 | MAPK8IP3 | 16 | 16 | 1 | 1 | 2136872 | 1809959 | TSC2::chr16:2136822-2136872(+) | MAPK8IP3::chr16:1809958-1810008(+) |
| TTC28 | EP300 | 22 | 22 | -1 | 1 | 29025540 | 41531817 | TTC28::chr22:29025539-29025589(-) | EP300::chr22:41531816-41531866(+) |
| TTC7B | RUNX1 | 14 | 21 | -1 | -1 | 91110392 | 36265260 | TTC7B::chr14:91110391-91110441(-) | RUNX1::chr21:36265210-36265260(-) |
| TUBD1 | CDK12 | 17 | 17 | -1 | 1 | 57943970 | 37667782 | TUBD1::chr17:57943969-57944019(-) | CDK12::chr17:37667781-37667831(+) |
| TULP4 | ESR1 | 6 | 6 | 1 | 1 | 158735300 | 152201790 | TULP4::chr6:158735250-158735300(+) | ESR1::chr6:152201789-152201839(+) |
| TXNL1 | LRP1B | 18 | 2 | -1 | -1 | 54305574 | 141299540 | TXNL1::chr18:54305573-54305623(-) | LRP1B::chr2:141299490-141299540(-) |
| UACA | PTPN6 | 15 | 12 | -1 | 1 | 70987356 | 7069090 | UACA::chr15:70987355-70987405(-) | PTPN6::chr12:7069089-7069139(+) |
| UGGT2 | GPC5 | 13 | 13 | -1 | 1 | 96529533 | 93518535 | UGGT2::chr13:96529532-96529582(-) | GPC5::chr13:93518534-93518584(+) |
| UNC13B | PTPRD | 9 | 9 | 1 | -1 | 35386290 | 8636844 | UNC13B::chr9:35386240-35386290(+) | PTPRD::chr9:8636794-8636844(-) |
| UNC45A | BLM | 15 | 15 | 1 | 1 | 91483703 | 91358332 | UNC45A::chr15:91483653-91483703(+) | BLM::chr15:91358331-91358381(+) |
| USP4 | RHOA | 3 | 3 | -1 | -1 | 49377357 | 49405981 | USP4::chr3:49377356-49377406(-) | RHOA::chr3:49405931-49405981(-) |
| USP9X | KDM6A | X | X | 1 | 1 | 41031211 | 44870206 | USP9X::chrX:41031161-41031211(+) | KDM6A::chrX:44870205-44870255(+) |
| VHL | C3orf24 | 3 | 3 | 1 | -1 | 10183871 | 10123215 | VHL::chr3:10183821-10183871(+) | C3orf24::chr3:10123165-10123215(-) |
| VHL | KIAA1430 | 3 | 4 | 1 | -1 | 10188320 | 186128945 | VHL::chr3:10188270-10188320(+) | KIAA1430::chr4:186128895-186128945(-) |
| VNN3 | PTPRK | 6 | 6 | -1 | -1 | 133055392 | 128563747 | VNN3::chr6:133055391-133055441(-) | PTPRK::chr6:128563697-128563747(-) |
| VPS13B | CCDC6 | 8 | 10 | 1 | -1 | 100287482 | 61592411 | VPS13B::chr8:100287432-100287482(+) | CCDC6::chr10:61592361-61592411(-) |
| WDFY2 | FOXO1 | 13 | 13 | 1 | -1 | 52158960 | 41134997 | WDFY2::chr13:52158910-52158960(+) | FOXO1::chr13:41134947-41134997(-) |
| WDR59 | CDH11 | 16 | 16 | -1 | -1 | 75018862 | 64982690 | WDR59::chr16:75018861-75018911(-) | CDH11::chr16:64982640-64982690(-) |
| WLS | TBX3 | 1 | 12 | -1 | -1 | 68603463 | 115110107 | WLS::chr1:68603462-68603512(-) | TBX3::chr12:115110057-115110107(-) |
| WNK2 | C9orf89 | 9 | 9 | 1 | 1 | 96052329 | 95869956 | WNK2::chr9:96052279-96052329(+) | C9orf89::chr9:95869955-95870005(+) |
| WNK2 | CENPP | 9 | 9 | 1 | 1 | 96070866 | 95373596 | WNK2::chr9:96070816-96070866(+) | CENPP::chr9:95373595-95373645(+) |
| WNK2 | SECISBP2 | 9 | 9 | 1 | 1 | 96026315 | 91972326 | WNK2::chr9:96026265-96026315(+) | SECISBP2::chr9:91972325-91972375(+) |
| WNK2 | SUSD3 | 9 | 9 | 1 | 1 | 96031028 | 95838066 | WNK2::chr9:96030978-96031028(+) | SUSD3::chr9:95838065-95838115(+) |
| WRN | LETM2 | 8 | 8 | 1 | 1 | 30891489 | 38245468 | WRN::chr8:30891439-30891489(+) | LETM2::chr8:38245467-38245517(+) |
| XPA | CORO2A | 9 | 9 | -1 | -1 | 100449378 | 100919942 | XPA::chr9:100449377-100449427(-) | CORO2A::chr9:100919892-100919942(-) |
| XPC | DAG1 | 3 | 3 | -1 | 1 | 14219966 | 49547852 | XPC::chr3:14219965-14220015(-) | DAG1::chr3:49547851-49547901(+) |
| YWHAE | GCGR | 17 | 17 | -1 | 1 | 1303341 | 79766718 | YWHAE::chr17:1303340-1303390(-) | GCGR::chr17:79766717-79766767(+) |
| YWHAE | INPP5K | 17 | 17 | -1 | -1 | 1303341 | 1417273 | YWHAE::chr17:1303340-1303390(-) | INPP5K::chr17:1417223-1417273(-) |
| YWHAE | PITPNA | 17 | 17 | -1 | -1 | 1303341 | 1424939 | YWHAE::chr17:1303340-1303390(-) | PITPNA::chr17:1424889-1424939(-) |
| YWHAE | SLC43A2 | 17 | 17 | -1 | -1 | 1257505 | 1489345 | YWHAE::chr17:1257504-1257554(-) | SLC43A2::chr17:1489295-1489345(-) |
| ZDHHC14 | ARID1B | 6 | 6 | 1 | 1 | 157803298 | 157431606 | ZDHHC14::chr6:157803248-157803298(+) | ARID1B::chr6:157431605-157431655(+) |
| ZFHX3 | ALS2CR11 | 16 | 2 | -1 | -1 | 72984368 | 202440049 | ZFHX3::chr16:72984367-72984417(-) | ALS2CR11::chr2:202439999-202440049(-) |
| ZFHX3 | ARL15 | 16 | 5 | -1 | -1 | 72816784 | 53183213 | ZFHX3::chr16:72816783-72816833(-) | ARL15::chr5:53183163-53183213(-) |
| ZFHX3 | ENY2 | 16 | 8 | -1 | 1 | 72845476 | 110355634 | ZFHX3::chr16:72845475-72845525(-) | ENY2::chr8:110355633-110355683(+) |
| ZFHX3 | WWP2 | 16 | 16 | -1 | 1 | 73081651 | 69874029 | ZFHX3::chr16:73081650-73081700(-) | WWP2::chr16:69874028-69874078(+) |
| ZFHX3 | ZMAT4 | 16 | 8 | -1 | -1 | 72845476 | 40438780 | ZFHX3::chr16:72845475-72845525(-) | ZMAT4::chr8:40438730-40438780(-) |
| ZFHX3 | ZNF821 | 16 | 16 | -1 | -1 | 72923630 | 71901940 | ZFHX3::chr16:72923629-72923679(-) | ZNF821::chr16:71901890-71901940(-) |
| ZMAT3 | TP63 | 3 | 3 | -1 | 1 | 178785271 | 189455529 | ZMAT3::chr3:178785270-178785320(-) | TP63::chr3:189455528-189455578(+) |
| ZMYND8 | RNF43 | 20 | 17 | -1 | -1 | 45976600 | 56448394 | ZMYND8::chr20:45976599-45976649(-) | RNF43::chr17:56448344-56448394(-) |
| ZNF253 | BAZ1A | 19 | 14 | 1 | -1 | 19976808 | 35280242 | ZNF253::chr19:19976758-19976808(+) | BAZ1A::chr14:35280192-35280242(-) |
| ZNF331 | NLRP11 | 19 | 19 | 1 | -1 | 54074984 | 56313105 | ZNF331::chr19:54074934-54074984(+) | NLRP11::chr19:56313055-56313105(-) |
| ZNF331 | NLRP4 | 19 | 19 | 1 | 1 | 54025938 | 56363382 | ZNF331::chr19:54025888-54025938(+) | NLRP4::chr19:56363381-56363431(+) |
| ZNF496 | WIF1 | 1 | 12 | -1 | -1 | 247491985 | 65471634 | ZNF496::chr1:247491984-247492034(-) | WIF1::chr12:65471584-65471634(-) |

| | | | | | | | | | |
|---|---|---|---|---|---|---|---|---|---|
| ZNF709 | ELL | 19 | 19 | -1 | -1 | 12595469 | 18557624 | ZNF709::chr19:12595468-12595518(-) | ELL::chr19:18557574-18557624(-) |
| ZNRF3 | CRYBB2 | 22 | 22 | 1 | 1 | 29280054 | 25617371 | ZNRF3::chr22:29280004-29280054(+) | CRYBB2::chr22:25617370-25617420(+) |
| ZNRF3 | FAM19A5 | 22 | 22 | 1 | 1 | 29383189 | 49042409 | ZNRF3::chr22:29383139-29383189(+) | FAM19A5::chr22:49042408-49042458(+) |
| AARSD1 | CCR7 | 17 | 17 | -1 | -1 | 41113209 | 38715194 | AARSD1::chr17:41113208-41113258(-) | CCR7::chr17:38715144-38715194(-) |
| ABL1 | ADAMTSL2 | 9 | 9 | 1 | 1 | 133589842 | 136405720 | ABL1::chr9:133589792-133589842(+) | ADAMTSL2::chr9:136405719-136405769(+) |
| ABL1 | BCR | 9 | 22 | 1 | 1 | 133589842 | 23595986 | ABL1::chr9:133589792-133589842(+) | BCR::chr22:23595985-23596035(+) |
| ABL1 | CCBL1 | 9 | 9 | 1 | -1 | 133589842 | 131607690 | ABL1::chr9:133589792-133589842(+) | CCBL1::chr9:131607640-131607690(-) |
| ABL1 | DDX27 | 9 | 20 | 1 | 1 | 133763062 | 47849303 | ABL1::chr9:133763012-133763062(+) | DDX27::chr20:47849302-47849352(+) |
| ABL1 | EXOSC2 | 9 | 9 | 1 | 1 | 133589842 | 133570879 | ABL1::chr9:133589792-133589842(+) | EXOSC2::chr9:133570878-133570928(+) |
| ABL1 | PRDM12 | 9 | 9 | 1 | 1 | 133589842 | 133553916 | ABL1::chr9:133589792-133589842(+) | PRDM12::chr9:133553915-133553965(+) |
| ABL1 | PTPRN2 | 9 | 7 | 1 | -1 | 133589842 | 157475629 | ABL1::chr9:133589792-133589842(+) | PTPRN2::chr7:157475579-157475629(-) |
| ABL1 | SLC38A10 | 9 | 17 | 1 | -1 | 133730483 | 79246427 | ABL1::chr9:133730433-133730483(+) | SLC38A10::chr17:79246377-79246427(-) |
| ABL2 | XRRA1 | 1 | 11 | -1 | -1 | 179198376 | 74559507 | ABL2::chr1:179198375-179198425(-) | XRRA1::chr11:74559457-74559507(-) |
| ACLY | TAF15 | 17 | 17 | -1 | 1 | 40054002 | 34163136 | ACLY::chr17:40054001-40054051(-) | TAF15::chr17:34163135-34163185(+) |
| ACVR1 | CCDC148 | 2 | 2 | -1 | -1 | 158674897 | 159035508 | ACVR1::chr2:158674896-158674946(-) | CCDC148::chr2:159035458-159035508(-) |
| ACVR1 | CCDC148 | 2 | 2 | -1 | -1 | 158655939 | 159145348 | ACVR1::chr2:158655938-158655988(-) | CCDC148::chr2:159145298-159145348(-) |
| ACVR1 | NFATC1 | 2 | 18 | -1 | 1 | 158674897 | 77246248 | ACVR1::chr2:158674896-158674946(-) | NFATC1::chr18:77246247-77246297(+) |
| AFF3 | ASZ1 | 2 | 7 | -1 | -1 | 100453939 | 117066989 | AFF3::chr2:100453938-100453988(-) | ASZ1::chr7:117066939-117066989(-) |
| AFF3 | HDAC5 | 2 | 17 | -1 | -1 | 100623094 | 42171202 | AFF3::chr2:100623093-100623143(-) | HDAC5::chr17:42171152-42171202(-) |
| AFF3 | NPAS2 | 2 | 2 | -1 | 1 | 100623094 | 101604541 | AFF3::chr2:100623093-100623143(-) | NPAS2::chr2:101604540-101604590(+) |
| AFF3 | TMEM182 | 2 | 2 | -1 | 1 | 100209657 | 103379046 | AFF3::chr2:100209656-100209706(-) | TMEM182::chr2:103379045-103379095(+) |
| AFF4 | FSTL4 | 5 | 5 | -1 | -1 | 132227856 | 132561511 | AFF4::chr5:132227855-132227905(-) | FSTL4::chr5:132561461-132561511(-) |
| AFF4 | SNTG2 | 5 | 2 | -1 | 1 | 132227856 | 1204789 | AFF4::chr5:132227855-132227905(-) | SNTG2::chr2:1204788-1204838(+) |
| AFF4 | SPATA21 | 5 | 1 | -1 | -1 | 132298951 | 16757917 | AFF4::chr5:132298950-132299000(-) | SPATA21::chr1:16757867-16757917(-) |
| AHCYL2 | SND1 | 7 | 7 | 1 | 1 | 128865280 | 127714554 | AHCYL2::chr7:128865230-128865280(+) | SND1::chr7:127714553-127714603(+) |
| AHI1 | MYB | 6 | 6 | -1 | 1 | 135679270 | 135539002 | AHI1::chr6:135679269-135679319(-) | MYB::chr6:135539001-135539051(+) |
| AKAP13 | NTRK3 | 15 | 15 | 1 | -1 | 86213061 | 88576276 | AKAP13::chr15:86213011-86213061(+) | NTRK3::chr15:88576226-88576276(-) |
| AKAP13 | RET | 15 | 10 | 1 | 1 | 86284726 | 43612032 | AKAP13::chr15:86284676-86284726(+) | RET::chr10:43612031-43612081(+) |
| AKT1 | ANKS1A | 14 | 6 | -1 | 1 | 105235689 | 34977682 | AKT1::chr14:105235688-105235738(-) | ANKS1A::chr6:34977681-34977731(+) |
| AKT2 | ELK3 | 19 | 12 | -1 | 1 | 40736224 | 96588323 | AKT2::chr19:40736223-40736273(-) | ELK3::chr12:96588322-96588372(+) |
| AKT2 | SRRM5 | 19 | 19 | -1 | 1 | 40771129 | 44101246 | AKT2::chr19:40771128-40771178(-) | SRRM5::chr19:44101245-44101295(+) |
| AKT2 | UPF1 | 19 | 19 | -1 | 1 | 40791088 | 18956789 | AKT2::chr19:40791087-40791137(-) | UPF1::chr19:18956788-18956838(+) |
| AKT2 | YIF1B | 19 | 19 | -1 | -1 | 40791088 | 38800283 | AKT2::chr19:40791087-40791137(-) | YIF1B::chr19:38800233-38800283(-) |
| AKT3 | ADSS | 1 | 1 | -1 | -1 | 244006427 | 244601070 | AKT3::chr1:244006426-244006476(-) | ADSS::chr1:244601020-244601070(-) |
| AKT3 | C1orf150 | 1 | 1 | -1 | 1 | 244006427 | 247690242 | AKT3::chr1:244006426-244006476(-) | C1orf150::chr1:247690241-247690291(+) |
| AKT3 | HEATR1 | 1 | 1 | -1 | -1 | 243800913 | 236714290 | AKT3::chr1:243800912-243800962(-) | HEATR1::chr1:236714240-236714290(-) |
| AKT3 | PPP2R2A | 1 | 8 | -1 | 1 | 244006427 | 26211984 | AKT3::chr1:244006426-244006476(-) | PPP2R2A::chr8:26211983-26212033(+) |
| AKT3 | PTPRR | 1 | 12 | -1 | -1 | 243776973 | 71078044 | AKT3::chr1:243776972-243777022(-) | PTPRR::chr12:71077994-71078044(-) |
| AKT3 | RGS7 | 1 | 1 | -1 | -1 | 244006427 | 240990472 | AKT3::chr1:244006426-244006476(-) | RGS7::chr1:240990422-240990472(-) |
| ALG14 | JAK1 | 1 | 1 | -1 | -1 | 95492685 | 65313358 | ALG14::chr1:95492684-95492734(-) | JAK1::chr1:65313308-65313358(-) |
| ALK | GALNT14 | 2 | 2 | -1 | -1 | 29519754 | 31152353 | ALK::chr2:29519753-29519803(-) | GALNT14::chr2:31152303-31152353(-) |
| ALK | SCEL | 2 | 13 | -1 | 1 | 29754781 | 78202082 | ALK::chr2:29754780-29754830(-) | SCEL::chr13:78202081-78202131(+) |
| ALK | STK39 | 2 | 2 | -1 | -1 | 29940444 | 169038600 | ALK::chr2:29940443-29940493(-) | STK39::chr2:169038550-169038600(-) |

| | | | | | | | | | |
|---|---|---|---|---|---|---|---|---|---|
| ARHGAP5 | C18orf10 | 14 | 18 | 1 | -1 | 32546641 | 34387897 | ARHGAP5::chr14:32546591-32546641(+) | C18orf10::chr18:34387847-34387897(-) |
| ARHGAP5 | L2HGDH | 14 | 14 | 1 | -1 | 32563592 | 50760964 | ARHGAP5::chr14:32563542-32563592(+) | L2HGDH::chr14:50760914-50760964(-) |
| ARHGAP5 | LSAMP | 14 | 3 | 1 | -1 | 32546641 | 116163802 | ARHGAP5::chr14:32546591-32546641(+) | LSAMP::chr3:116163752-116163802(-) |
| ARHGAP5 | NPAS3 | 14 | 14 | 1 | 1 | 32563592 | 33836392 | ARHGAP5::chr14:32563542-32563592(+) | NPAS3::chr14:33836391-33836441(+) |
| ARHGAP5 | NUBPL | 14 | 14 | 1 | 1 | 32586493 | 32142561 | ARHGAP5::chr14:32586443-32586493(+) | NUBPL::chr14:32142560-32142610(+) |
| ATF1 | GRIP1 | 12 | 12 | 1 | -1 | 51174021 | 66990707 | ATF1::chr12:51173971-51174021(+) | GRIP1::chr12:66990657-66990707(-) |
| ATG7 | BRAF | 3 | 7 | 1 | -1 | 11468400 | 140487384 | ATG7::chr3:11468350-11468400(+) | BRAF::chr7:140487334-140487384(-) |
| B4GALT5 | GNAS | 20 | 20 | -1 | 1 | 48330113 | 57470667 | B4GALT5::chr20:48330112-48330162(-) | GNAS::chr20:57470666-57470716(+) |
| BAG4 | FGFR1 | 8 | 8 | 1 | -1 | 38050313 | 38283763 | BAG4::chr8:38050263-38050313(+) | FGFR1::chr8:38283713-38283763(-) |
| BCL11A | PTPLA | 2 | 10 | -1 | -1 | 60773106 | 17646046 | BCL11A::chr2:60773105-60773155(-) | PTPLA::chr10:17645996-17646046(-) |
| BCL2L11 | BRAF | 2 | 7 | 1 | -1 | 111881716 | 140482957 | BCL2L11::chr2:111881666-111881716(+) | BRAF::chr7:140482907-140482957(-) |
| BCL2L12 | FLT3LG | 19 | 19 | 1 | 1 | 50173745 | 49989272 | BCL2L12::chr19:50173695-50173745(+) | FLT3LG::chr19:49989271-49989321(+) |
| BCL3 | ATP9B | 19 | 18 | 1 | 1 | 45251385 | 77013381 | BCL3::chr19:45251335-45251385(+) | ATP9B::chr18:77013380-77013430(+) |
| BCL9 | BEST3 | 1 | 12 | 1 | -1 | 147013444 | 70049593 | BCL9::chr1:147013394-147013444(+) | BEST3::chr12:70049543-70049593(-) |
| BCL9 | C1orf21 | 1 | 1 | 1 | 1 | 147087702 | 184588652 | BCL9::chr1:147087652-147087702(+) | C1orf21::chr1:184588651-184588701(+) |
| BCL9 | GOLPH3L | 1 | 1 | 1 | -1 | 147013444 | 150636239 | BCL9::chr1:147013394-147013444(+) | GOLPH3L::chr1:150636189-150636239(-) |
| BCR | ABL1 | 22 | 9 | 1 | 1 | 23524426 | 133729451 | BCR::chr22:23524376-23524426(+) | ABL1::chr9:133729450-133729500(+) |
| BICC1 | FGFR2 | 10 | 10 | 1 | -1 | 60563002 | 123239535 | BICC1::chr10:60562952-60563002(+) | FGFR2::chr10:123239485-123239535(-) |
| BIRC6 | ABCC11 | 2 | 16 | 1 | -1 | 32738221 | 48211024 | BIRC6::chr2:32738171-32738221(+) | ABCC11::chr16:48210974-48211024(-) |
| BIRC6 | EML4 | 2 | 2 | 1 | 1 | 32774548 | 42515367 | BIRC6::chr2:32774498-32774548(+) | EML4::chr2:42515366-42515416(+) |
| BIRC6 | EML6 | 2 | 2 | 1 | 1 | 32679010 | 55071184 | BIRC6::chr2:32678960-32679010(+) | EML6::chr2:55071183-55071233(+) |
| BIRC6 | FAM82A1 | 2 | 2 | 1 | 1 | 32823035 | 38294125 | BIRC6::chr2:32822985-32823035(+) | FAM82A1::chr2:38294124-38294174(+) |
| BIRC6 | MATN3 | 2 | 2 | 1 | -1 | 32641231 | 20206071 | BIRC6::chr2:32641181-32641231(+) | MATN3::chr2:20206021-20206071(-) |
| BIRC6 | SLC30A6 | 2 | 2 | 1 | 1 | 32582554 | 32396356 | BIRC6::chr2:32582504-32582554(+) | SLC30A6::chr2:32396355-32396405(+) |
| BIRC6 | SLC30A6 | 2 | 2 | 1 | 1 | 32774548 | 32409342 | BIRC6::chr2:32774498-32774548(+) | SLC30A6::chr2:32409341-32409391(+) |
| BIRC6 | SPAST | 2 | 2 | 1 | 1 | 32631625 | 32339707 | BIRC6::chr2:32631575-32631625(+) | SPAST::chr2:32339706-32339756(+) |
| BIRC6 | SPAST | 2 | 2 | 1 | 1 | 32658872 | 32323865 | BIRC6::chr2:32658822-32658872(+) | SPAST::chr2:32323864-32323914(+) |
| BIRC6 | SPAST | 2 | 2 | 1 | 1 | 32774548 | 32372287 | BIRC6::chr2:32774498-32774548(+) | SPAST::chr2:32372286-32372336(+) |
| BIRC6 | YIPF4 | 2 | 2 | 1 | 1 | 32582554 | 32515552 | BIRC6::chr2:32582504-32582554(+) | YIPF4::chr2:32515551-32515601(+) |
| BIRC6 | YIPF4 | 2 | 2 | 1 | 1 | 32664754 | 32515552 | BIRC6::chr2:32664704-32664754(+) | YIPF4::chr2:32515551-32515601(+) |
| BRAF | AP3B1 | 7 | 5 | -1 | -1 | 140494108 | 77335098 | BRAF::chr7:140494107-140494157(-) | AP3B1::chr5:77335048-77335098(-) |
| BRAF | ATG7 | 7 | 3 | -1 | 1 | 140507760 | 11596285 | BRAF::chr7:140507759-140507809(-) | ATG7::chr3:11596284-11596334(+) |
| BRAF | C7orf10 | 7 | 7 | -1 | 1 | 140624366 | 40789033 | BRAF::chr7:140624365-140624415(-) | C7orf10::chr7:40789032-40789082(+) |
| BRAF | CUL1 | 7 | 7 | -1 | 1 | 140549911 | 148427054 | BRAF::chr7:140549910-140549960(-) | CUL1::chr7:148427053-148427103(+) |
| BRAF | FAM114A2 | 7 | 5 | -1 | -1 | 140482821 | 153382529 | BRAF::chr7:140482820-140482870(-) | FAM114A2::chr5:153382479-153382529(-) |
| BRAF | HIBADH | 7 | 7 | -1 | -1 | 140500162 | 27582719 | BRAF::chr7:140500161-140500211(-) | HIBADH::chr7:27582669-27582719(-) |
| BRAF | MACF1 | 7 | 1 | -1 | 1 | 140494108 | 39898207 | BRAF::chr7:140494107-140494157(-) | MACF1::chr1:39898206-39898256(+) |
| BRAF | SLC26A4 | 7 | 7 | -1 | 1 | 140534409 | 107323647 | BRAF::chr7:140534408-140534458(-) | SLC26A4::chr7:107323646-107323696(+) |
| BRD4 | AKAP8L | 19 | 19 | -1 | -1 | 15443101 | 15491423 | BRD4::chr19:15443100-15443150(-) | AKAP8L::chr19:15491373-15491423(-) |
| BRD4 | AKAP8L | 19 | 19 | -1 | -1 | 15443101 | 15514853 | BRD4::chr19:15443100-15443150(-) | AKAP8L::chr19:15514803-15514853(-) |
| BRD4 | CSN2 | 19 | 4 | -1 | -1 | 15443101 | 70826725 | BRD4::chr19:15443100-15443150(-) | CSN2::chr4:70826675-70826725(-) |
| BRD4 | ZC3H15 | 19 | 2 | -1 | 1 | 15443101 | 187367223 | BRD4::chr19:15443100-15443150(-) | ZC3H15::chr2:187367222-187367272(+) |
| BRD9 | TERT | 5 | 5 | -1 | -1 | 892721 | 1282739 | BRD9::chr5:892720-892770(-) | TERT::chr5:1282689-1282739(-) |

| | | | | | | | | | |
|---|---|---|---|---|---|---|---|---|---|
| BRD9 | TERT | 5 | 5 | -1 | -1 | 891270 | 1282739 | BRD9::chr5:891269-891319(-) | TERT::chr5:1282689-1282739(-) |
| BSPRY | NPM1 | 9 | 5 | 1 | 1 | 116123017 | 170837531 | BSPRY::chr9:116122967-116123017(+) | NPM1::chr5:170837530-170837580(+) |
| C11orf49 | NUP98 | 11 | 11 | 1 | -1 | 47182741 | 3735225 | C11orf49::chr11:47182691-47182741(+) | NUP98::chr11:3735175-3735225(-) |
| C12orf49 | MDM2 | 12 | 12 | -1 | 1 | 117175595 | 69229609 | C12orf49::chr12:117175594-117175644(-) | MDM2::chr12:69229608-69229658(+) |
| C3orf21 | ETV5 | 3 | 3 | -1 | -1 | 194877178 | 185783861 | C3orf21::chr3:194877177-194877227(-) | ETV5::chr3:185783811-185783861(-) |
| C6orf204 | ROS1 | 6 | 6 | -1 | -1 | 118802942 | 117641193 | C6orf204::chr6:118802941-118802991(-) | ROS1::chr6:117641143-117641193(-) |
| C8orf34 | MET | 8 | 7 | 1 | 1 | 69351881 | 116414935 | C8orf34::chr8:69351831-69351881(+) | MET::chr7:116414934-116414984(+) |
| CACNA1D | GNL3 | 3 | 3 | 1 | 1 | 53804579 | 52724608 | CACNA1D::chr3:53804529-53804579(+) | GNL3::chr3:52724607-52724657(+) |
| CACNA1D | PRKCD | 3 | 3 | 1 | 1 | 53535747 | 53217468 | CACNA1D::chr3:53535697-53535747(+) | PRKCD::chr3:53217467-53217517(+) |
| CALR | BRD2 | 19 | 6 | 1 | 1 | 13055304 | 32944339 | CALR::chr19:13055254-13055304(+) | BRD2::chr6:32944338-32944388(+) |
| CALR | PLEKHB2 | 19 | 2 | 1 | 1 | 13055304 | 131876464 | CALR::chr19:13055254-13055304(+) | PLEKHB2::chr2:131876463-131876513(+) |
| CALR | TNNT1 | 19 | 19 | 1 | -1 | 13055304 | 55646910 | CALR::chr19:13055254-13055304(+) | TNNT1::chr19:55646860-55646910(-) |
| CAPZA2 | MET | 7 | 7 | 1 | 1 | 116528244 | 116414935 | CAPZA2::chr7:116528194-116528244(+) | MET::chr7:116414934-116414984(+) |
| CAPZA2 | MET | 7 | 7 | 1 | 1 | 116550365 | 116409699 | CAPZA2::chr7:116550315-116550365(+) | MET::chr7:116409698-116409748(+) |
| CARD11 | ZNRD1 | 7 | 6 | -1 | 1 | 3083301 | 30029292 | CARD11::chr7:3083300-3083350(-) | ZNRD1::chr6:30029291-30029341(+) |
| CCDC127 | TERT | 5 | 5 | -1 | -1 | 216844 | 1282739 | CCDC127::chr5:216843-216893(-) | TERT::chr5:1282689-1282739(-) |
| CCND3 | C6orf89 | 6 | 6 | -1 | 1 | 42016239 | 36846784 | CCND3::chr6:42016238-42016288(-) | C6orf89::chr6:36846783-36846833(+) |
| CCND3 | EHMT2 | 6 | 6 | -1 | -1 | 42016239 | 31848625 | CCND3::chr6:42016238-42016288(-) | EHMT2::chr6:31848575-31848625(-) |
| CCND3 | EXOC4 | 6 | 7 | -1 | 1 | 42016239 | 133682245 | CCND3::chr6:42016238-42016288(-) | EXOC4::chr7:133682244-133682294(+) |
| CCND3 | GUCA1B | 6 | 6 | -1 | -1 | 42016239 | 42156469 | CCND3::chr6:42016238-42016288(-) | GUCA1B::chr6:42156419-42156469(-) |
| CCND3 | LUZP1 | 6 | 1 | -1 | -1 | 42016239 | 23415546 | CCND3::chr6:42016238-42016288(-) | LUZP1::chr1:23415496-23415546(-) |
| CCND3 | TRERF1 | 6 | 6 | -1 | -1 | 42016239 | 42330817 | CCND3::chr6:42016238-42016288(-) | TRERF1::chr6:42330767-42330817(-) |
| CCND3 | TRERF1 | 6 | 6 | -1 | -1 | 42016239 | 42222664 | CCND3::chr6:42016238-42016288(-) | TRERF1::chr6:42222614-42222664(-) |
| CCND3 | VPS52 | 6 | 6 | -1 | -1 | 42016239 | 33237596 | CCND3::chr6:42016238-42016288(-) | VPS52::chr6:33237546-33237596(-) |
| CCNY | BRAF | 10 | 7 | 1 | -1 | 35626135 | 140482957 | CCNY::chr10:35626085-35626135(+) | BRAF::chr7:140482907-140482957(-) |
| CCT4 | BIRC6 | 2 | 2 | -1 | 1 | 62107421 | 32756422 | CCT4::chr2:62107420-62107470(-) | BIRC6::chr2:32756421-32756471(+) |
| CD74 | CD40 | 5 | 20 | -1 | 1 | 149781200 | 44754189 | CD74::chr5:149781199-149781249(-) | CD40::chr20:44754188-44754238(+) |
| CD74 | CXCR5 | 5 | 11 | -1 | 1 | 149781200 | 118754475 | CD74::chr5:149781199-149781249(-) | CXCR5::chr11:118754474-118754524(+) |
| CD74 | DDX27 | 5 | 20 | -1 | 1 | 149781200 | 47849303 | CD74::chr5:149781199-149781249(-) | DDX27::chr20:47849302-47849352(+) |
| CD74 | IQCE | 5 | 7 | -1 | 1 | 149781200 | 2641075 | CD74::chr5:149781199-149781249(-) | IQCE::chr7:2641074-2641124(+) |
| CD74 | NDUFV1 | 5 | 11 | -1 | 1 | 149781200 | 67375697 | CD74::chr5:149781199-149781249(-) | NDUFV1::chr11:67375696-67375746(+) |
| CD74 | THSD4 | 5 | 15 | -1 | 1 | 149781200 | 72072881 | CD74::chr5:149781199-149781249(-) | THSD4::chr15:72072880-72072930(+) |
| CD74 | UTRN | 5 | 6 | -1 | 1 | 149781200 | 144665237 | CD74::chr5:149781199-149781249(-) | UTRN::chr6:144665236-144665286(+) |
| CDC27 | BRAF | 17 | 7 | -1 | -1 | 45206759 | 140487384 | CDC27::chr17:45206758-45206808(-) | BRAF::chr7:140487334-140487384(-) |
| CDH17 | RAD54B | 8 | 8 | -1 | -1 | 95160972 | 95423543 | CDH17::chr8:95160971-95161021(-) | RAD54B::chr8:95423493-95423543(-) |
| CDH18 | TERT | 5 | 5 | -1 | -1 | 19612543 | 1282739 | CDH18::chr5:19612542-19612592(-) | TERT::chr5:1282689-1282739(-) |
| CDK6 | ANKIB1 | 7 | 7 | -1 | 1 | 92404010 | 91924203 | CDK6::chr7:92404009-92404059(-) | ANKIB1::chr7:91924202-91924252(+) |
| CDK6 | ANKIB1 | 7 | 7 | -1 | 1 | 92462405 | 91936673 | CDK6::chr7:92462404-92462454(-) | ANKIB1::chr7:91936672-91936722(+) |
| CDK6 | HEPACAM2 | 7 | 7 | -1 | -1 | 92404010 | 92821676 | CDK6::chr7:92404009-92404059(-) | HEPACAM2::chr7:92821626-92821676(-) |
| CDK6 | STXBP6 | 7 | 14 | -1 | -1 | 92462405 | 25281938 | CDK6::chr7:92462404-92462454(-) | STXBP6::chr14:25281888-25281938(-) |
| CEP170 | AKT3 | 1 | 1 | -1 | -1 | 243349081 | 243736350 | CEP170::chr1:243349080-243349130(-) | AKT3::chr1:243736300-243736350(-) |
| CHD4 | CLEC9A | 12 | 12 | -1 | 1 | 6686951 | 10194065 | CHD4::chr12:6686950-6687000(-) | CLEC9A::chr12:10194064-10194114(+) |
| CHD4 | CPM | 12 | 12 | -1 | -1 | 6700632 | 69265736 | CHD4::chr12:6700631-6700681(-) | CPM::chr12:69265686-69265736(-) |

| | | | | | | | | | |
|---|---|---|---|---|---|---|---|---|---|
| CHD4 | DTX1 | 12 | 12 | -1 | 1 | 6690210 | 113515229 | CHD4::chr12:6690209-6690259(-) | DTX1::chr12:113515228-113515278(+) |
| CHD7 | TCF7L2 | 8 | 10 | 1 | 1 | 61655656 | 114900943 | CHD7::chr8:61655606-61655656(+) | TCF7L2::chr10:114900942-114900992(+) |
| CHST11 | ATP1B4 | 12 | X | 1 | 1 | 104995769 | 119500380 | CHST11::chr12:104995719-104995769(+) | ATP1B4::chrX:119500379-119500429(+) |
| CHST11 | HCFC2 | 12 | 12 | 1 | 1 | 104851292 | 104459945 | CHST11::chr12:104851242-104851292(+) | HCFC2::chr12:104459944-104459994(+) |
| CHST11 | PTPRR | 12 | 12 | 1 | -1 | 104995769 | 71158558 | CHST11::chr12:104995719-104995769(+) | PTPRR::chr12:71158508-71158558(-) |
| CHST11 | RFX4 | 12 | 12 | 1 | 1 | 104995769 | 107075771 | CHST11::chr12:104995719-104995769(+) | RFX4::chr12:107075770-107075820(+) |
| CHST11 | TXNRD1 | 12 | 12 | 1 | 1 | 104995769 | 104732917 | CHST11::chr12:104995719-104995769(+) | TXNRD1::chr12:104732916-104732966(+) |
| CLCN6 | RAF1 | 1 | 3 | 1 | -1 | 11867247 | 12641914 | CLCN6::chr1:11867197-11867247(+) | RAF1::chr3:12641864-12641914(-) |
| CLPTM1 | CBLC | 19 | 19 | 1 | 1 | 45465332 | 45293261 | CLPTM1::chr19:45465282-45465332(+) | CBLC::chr19:45293260-45293310(+) |
| CREB1 | SUCNR1 | 2 | 3 | 1 | 1 | 208425884 | 151597640 | CREB1::chr2:208425834-208425884(+) | SUCNR1::chr3:151597639-151597689(+) |
| CREB3L2 | CPA2 | 7 | 7 | -1 | 1 | 137686350 | 129921944 | CREB3L2::chr7:137686349-137686399(-) | CPA2::chr7:129921943-129921993(+) |
| CREB3L2 | SHANK2 | 7 | 11 | -1 | -1 | 137686350 | 70349036 | CREB3L2::chr7:137686349-137686399(-) | SHANK2::chr11:70348986-70349036(-) |
| CRTC1 | KCNN1 | 19 | 19 | 1 | 1 | 18794638 | 18084617 | CRTC1::chr19:18794588-18794638(+) | KCNN1::chr19:18084616-18084666(+) |
| CRTC1 | LYPD4 | 19 | 19 | 1 | -1 | 18860685 | 42343098 | CRTC1::chr19:18860635-18860685(+) | LYPD4::chr19:42343048-42343098(-) |
| CTNNA2 | REG3G | 2 | 2 | 1 | 1 | 79878784 | 79253109 | CTNNA2::chr2:79878734-79878784(+) | REG3G::chr2:79253108-79253158(+) |
| CTNNB1 | DOCK3 | 3 | 3 | 1 | 1 | 41280845 | 50927457 | CTNNB1::chr3:41280795-41280845(+) | DOCK3::chr3:50927456-50927506(+) |
| CTNNB1 | IQCE | 3 | 7 | 1 | 1 | 41281934 | 2641075 | CTNNB1::chr3:41281884-41281934(+) | IQCE::chr7:2641074-2641124(+) |
| CTNNB1 | SLC38A5 | 3 | X | 1 | -1 | 41281934 | 48327824 | CTNNB1::chr3:41281884-41281934(+) | SLC38A5::chrX:48327774-48327824(-) |
| CTNNB1 | TCOF1 | 3 | 5 | 1 | 1 | 41281934 | 149777917 | CTNNB1::chr3:41281884-41281934(+) | TCOF1::chr5:149777916-149777966(+) |
| CTNNB1 | THRB | 3 | 3 | 1 | -1 | 41241161 | 24270492 | CTNNB1::chr3:41241111-41241161(+) | THRB::chr3:24270442-24270492(-) |
| CTNND2 | DAP | 5 | 5 | -1 | -1 | 11565056 | 10683683 | CTNND2::chr5:11565055-11565105(-) | DAP::chr5:10683633-10683683(-) |
| CTNND2 | NRBP1 | 5 | 2 | -1 | 1 | 11732248 | 27656540 | CTNND2::chr5:11732247-11732297(-) | NRBP1::chr2:27656539-27656589(+) |
| CTNND2 | P4HA2 | 5 | 5 | -1 | -1 | 11903929 | 131554337 | CTNND2::chr5:11903928-11903978(-) | P4HA2::chr5:131554287-131554337(-) |
| CTNND2 | POLR1A | 5 | 2 | -1 | -1 | 11732248 | 86257519 | CTNND2::chr5:11732247-11732297(-) | POLR1A::chr2:86257469-86257519(-) |
| CTNND2 | PPARGC1B | 5 | 5 | -1 | 1 | 11082808 | 149199996 | CTNND2::chr5:11082807-11082857(-) | PPARGC1B::chr5:149199995-149200045(+) |
| CXCR4 | CFLAR | 2 | 2 | -1 | 1 | 136875616 | 201994452 | CXCR4::chr2:136875615-136875665(-) | CFLAR::chr2:201994451-201994501(+) |
| DDR2 | CHD1L | 1 | 1 | 1 | 1 | 162625164 | 146724278 | DDR2::chr1:162625114-162625164(+) | CHD1L::chr1:146724277-146724327(+) |
| DDR2 | FMO4 | 1 | 1 | 1 | 1 | 162625164 | 171293277 | DDR2::chr1:162625114-162625164(+) | FMO4::chr1:171293276-171293326(+) |
| DDX5 | PFKP | 17 | 10 | -1 | 1 | 62495741 | 3177928 | DDX5::chr17:62495740-62495790(-) | PFKP::chr10:3177927-3177977(+) |
| DDX52 | STAT5B | 17 | 17 | -1 | -1 | 35988605 | 40354826 | DDX52::chr17:35988604-35988654(-) | STAT5B::chr17:40354776-40354826(-) |
| DDX6 | TSPAN18 | 11 | 11 | -1 | 1 | 118627880 | 44927958 | DDX6::chr11:118627879-118627929(-) | TSPAN18::chr11:44927957-44928007(+) |
| DEK | KIF25 | 6 | 6 | -1 | 1 | 18264074 | 168431453 | DEK::chr6:18264073-18264123(-) | KIF25::chr6:168431452-168431502(+) |
| DEK | NUP210L | 6 | 1 | -1 | -1 | 18249882 | 154000073 | DEK::chr6:18249881-18249931(-) | NUP210L::chr1:154000023-154000073(-) |
| DGCR8 | COMT | 22 | 22 | 1 | 1 | 20067906 | 19950050 | DGCR8::chr22:20067856-20067906(+) | COMT::chr22:19950049-19950099(+) |
| DGCR8 | CRKL | 22 | 22 | 1 | 1 | 20096526 | 21288067 | DGCR8::chr22:20096476-20096526(+) | CRKL::chr22:21288066-21288116(+) |
| DGCR8 | DGCR14 | 22 | 22 | 1 | -1 | 20082318 | 19127242 | DGCR8::chr22:20082268-20082318(+) | DGCR14::chr22:19127192-19127242(-) |
| DGCR8 | ITGA2 | 22 | 5 | 1 | 1 | 20080430 | 52385776 | DGCR8::chr22:20080380-20080430(+) | ITGA2::chr5:52385775-52385825(+) |
| DGCR8 | SGSM1 | 22 | 22 | 1 | 1 | 20067906 | 25240864 | DGCR8::chr22:20067856-20067906(+) | SGSM1::chr22:25240863-25240913(+) |
| DHX15 | ETV1 | 4 | 7 | -1 | -1 | 24585946 | 13975521 | DHX15::chr4:24585945-24585995(-) | ETV1::chr7:13975471-13975521(-) |
| DHX8 | MSI2 | 17 | 17 | 1 | 1 | 41567831 | 55674229 | DHX8::chr17:41567781-41567831(+) | MSI2::chr17:55674228-55674278(+) |
| DLG5 | RET | 10 | 10 | -1 | 1 | 79588640 | 43612032 | DLG5::chr10:79588639-79588689(-) | RET::chr10:43612031-43612081(+) |
| DNAH1 | CACNA1D | 3 | 3 | 1 | 1 | 52418047 | 53834276 | DNAH1::chr3:52417997-52418047(+) | CACNA1D::chr3:53834275-53834325(+) |
| DSTYK | BRAF | 1 | 7 | -1 | -1 | 205180399 | 140482957 | DSTYK::chr1:205180398-205180448(-) | BRAF::chr7:140482907-140482957(-) |

| | | | | | | | | | |
|---|---|---|---|---|---|---|---|---|---|
| DTNA | ZNF521 | 18 | 18 | 1 | -1 | 32346005 | 22902151 | DTNA::chr18:32345955-32346005(+) | ZNF521::chr18:22902101-22902151(-) |
| DVL2 | TFE3 | 17 | X | -1 | -1 | 7133127 | 48896935 | DVL2::chr17:7133126-7133176(-) | TFE3::chrX:48896885-48896935(-) |
| EGFR | ACADM | 7 | 1 | 1 | 1 | 55087058 | 76198329 | EGFR::chr7:55087008-55087058(+) | ACADM::chr1:76198328-76198378(+) |
| EGFR | DYM | 7 | 18 | 1 | -1 | 55087058 | 46918015 | EGFR::chr7:55087008-55087058(+) | DYM::chr18:46917965-46918015(-) |
| EGFR | ERP44 | 7 | 9 | 1 | -1 | 55087058 | 102784508 | EGFR::chr7:55087008-55087058(+) | ERP44::chr9:102784458-102784508(-) |
| EGFR | GNS | 7 | 12 | 1 | -1 | 55240817 | 65110599 | EGFR::chr7:55240767-55240817(+) | GNS::chr12:65110549-65110599(-) |
| EGFR | GRB10 | 7 | 7 | 1 | -1 | 55087058 | 50742355 | EGFR::chr7:55087008-55087058(+) | GRB10::chr7:50742305-50742355(-) |
| EGFR | LYST | 7 | 1 | 1 | -1 | 55087058 | 235996967 | EGFR::chr7:55087008-55087058(+) | LYST::chr1:235996917-235996967(-) |
| EGFR | PPM1H | 7 | 12 | 1 | -1 | 55087058 | 63195940 | EGFR::chr7:55087008-55087058(+) | PPM1H::chr12:63195890-63195940(-) |
| EGFR | 14-Sep | 7 | 7 | 1 | -1 | 55087058 | 55874951 | EGFR::chr7:55087008-55087058(+) | Sep-14::chr7:55874901-55874951(-) |
| EGFR | TEAD2 | 7 | 19 | 1 | -1 | 55087058 | 49862756 | EGFR::chr7:55087008-55087058(+) | TEAD2::chr19:49862706-49862756(-) |
| ELK4 | SLC26A9 | 1 | 1 | -1 | -1 | 205592804 | 205904966 | ELK4::chr1:205592803-205592853(-) | SLC26A9::chr1:205904916-205904966(-) |
| ELK4 | VCL | 1 | 10 | -1 | 1 | 205592804 | 75855414 | ELK4::chr1:205592803-205592853(-) | VCL::chr10:75855413-75855463(+) |
| ELK4 | ZNF350 | 1 | 19 | -1 | -1 | 205588085 | 52477784 | ELK4::chr1:205588084-205588134(-) | ZNF350::chr19:52477734-52477784(-) |
| EML4 | ALK | 2 | 2 | 1 | -1 | 42472827 | 29446394 | EML4::chr2:42472777-42472827(+) | ALK::chr2:29446344-29446394(-) |
| EML4 | ALK | 2 | 2 | 1 | -1 | 42491871 | 29446394 | EML4::chr2:42491821-42491871(+) | ALK::chr2:29446344-29446394(-) |
| EML4 | ALK | 2 | 2 | 1 | -1 | 42522656 | 29446394 | EML4::chr2:42522606-42522656(+) | ALK::chr2:29446344-29446394(-) |
| EML4 | ALK | 2 | 2 | 1 | -1 | 42552694 | 29446394 | EML4::chr2:42552644-42552694(+) | ALK::chr2:29446344-29446394(-) |
| EML4 | ALK | 2 | 2 | 1 | -1 | 42492091 | 29449940 | EML4::chr2:42492041-42492091(+) | ALK::chr2:29449890-29449940(-) |
| EML4 | NTRK3 | 2 | 15 | 1 | -1 | 42491871 | 88576276 | EML4::chr2:42491821-42491871(+) | NTRK3::chr15:88576226-88576276(-) |
| EPHB2 | NTRK1 | 1 | 1 | 1 | 1 | 23111569 | 156843425 | EPHB2::chr1:23111519-23111569(+) | NTRK1::chr1:156843424-156843474(+) |
| ERBB2 | CLDN7 | 17 | 17 | 1 | -1 | 37884915 | 7166264 | ERBB2::chr17:37884865-37884915(+) | CLDN7::chr17:7166214-7166264(-) |
| ERBB2 | DNAJC7 | 17 | 17 | 1 | -1 | 37844531 | 40134419 | ERBB2::chr17:37844481-37844531(+) | DNAJC7::chr17:40134369-40134419(-) |
| ERBB2 | IKZF3 | 17 | 17 | 1 | -1 | 37872858 | 37985741 | ERBB2::chr17:37872808-37872858(+) | IKZF3::chr17:37985691-37985741(-) |
| ERBB2 | NEUROD2 | 17 | 17 | 1 | -1 | 37856564 | 37762857 | ERBB2::chr17:37856514-37856564(+) | NEUROD2::chr17:37762807-37762857(-) |
| ERBB2 | PPP1R1B | 17 | 17 | 1 | 1 | 37866734 | 37790136 | ERBB2::chr17:37866684-37866734(+) | PPP1R1B::chr17:37790135-37790185(+) |
| ERBB2 | PSMB3 | 17 | 17 | 1 | 1 | 37876087 | 36916684 | ERBB2::chr17:37876037-37876087(+) | PSMB3::chr17:36916683-36916733(+) |
| ERBB2 | SLC29A3 | 17 | 10 | 1 | 1 | 37883256 | 73121711 | ERBB2::chr17:37883206-37883256(+) | SLC29A3::chr10:73121710-73121760(+) |
| ERBB2 | SLC39A11 | 17 | 17 | 1 | -1 | 37871789 | 71084914 | ERBB2::chr17:37871739-37871789(+) | SLC39A11::chr17:71084864-71084914(-) |
| ERBB2 | SUPT16H | 17 | 14 | 1 | -1 | 37884915 | 21829491 | ERBB2::chr17:37884865-37884915(+) | SUPT16H::chr14:21829441-21829491(-) |
| ERBB2 | TCAP | 17 | 17 | 1 | 1 | 37872858 | 37821969 | ERBB2::chr17:37872808-37872858(+) | TCAP::chr17:37821968-37822018(+) |
| ERBB3 | COX4I2 | 12 | 20 | 1 | 1 | 56474166 | 30231207 | ERBB3::chr12:56474116-56474166(+) | COX4I2::chr20:30231206-30231256(+) |
| ERBB3 | RPL30 | 12 | 8 | 1 | -1 | 56497283 | 99058697 | ERBB3::chr12:56497233-56497283(+) | RPL30::chr8:99058647-99058697(-) |
| ERBB3 | SUOX | 12 | 12 | 1 | 1 | 56494029 | 56395996 | ERBB3::chr12:56493979-56494029(+) | SUOX::chr12:56395995-56396045(+) |
| ERC1 | RET | 12 | 10 | 1 | 1 | 1250953 | 43612032 | ERC1::chr12:1250903-1250953(+) | RET::chr10:43612031-43612081(+) |
| ERC1 | RET | 12 | 10 | 1 | 1 | 1346070 | 43612032 | ERC1::chr12:1346020-1346070(+) | RET::chr10:43612031-43612081(+) |
| ERC1 | RET | 12 | 10 | 1 | 1 | 1553916 | 43612032 | ERC1::chr12:1553866-1553916(+) | RET::chr10:43612031-43612081(+) |
| ERG | ABCC4 | 21 | 13 | -1 | -1 | 39947586 | 95900007 | ERG::chr21:39947585-39947635(-) | ABCC4::chr13:95899957-95900007(-) |
| ERG | KCNJ6 | 21 | 21 | -1 | -1 | 39947586 | 38997786 | ERG::chr21:39947585-39947635(-) | KCNJ6::chr21:38997736-38997786(-) |
| ERG | PLEKHH1 | 21 | 14 | -1 | 1 | 39947586 | 68022542 | ERG::chr21:39947585-39947635(-) | PLEKHH1::chr14:68022541-68022591(+) |
| ERG | SON | 21 | 21 | -1 | 1 | 39947586 | 34918519 | ERG::chr21:39947585-39947635(-) | SON::chr21:34918518-34918568(+) |
| ERG | TAF8 | 21 | 6 | -1 | 1 | 39947586 | 42019095 | ERG::chr21:39947585-39947635(-) | TAF8::chr6:42019094-42019144(+) |
| ERG | TMPRSS2 | 21 | 21 | -1 | -1 | 39947586 | 42870116 | ERG::chr21:39947585-39947635(-) | TMPRSS2::chr21:42870066-42870116(-) |

| | | | | | | | | | |
|---|---|---|---|---|---|---|---|---|---|
| ERG | TMPRSS2 | 21 | 21 | -1 | -1 | 39947586 | 42852529 | ERG::chr21:39947585-39947635(-) | TMPRSS2::chr21:42852479-42852529(-) |
| ETV1 | BTBD9 | 7 | 6 | -1 | -1 | 13971127 | 38224292 | ETV1::chr7:13971126-13971176(-) | BTBD9::chr6:38224242-38224292(-) |
| ETV1 | CACNA2D1 | 7 | 7 | -1 | -1 | 14017052 | 81695840 | ETV1::chr7:14017051-14017101(-) | CACNA2D1::chr7:81695790-81695840(-) |
| ETV4 | CCL16 | 17 | 17 | -1 | -1 | 41610042 | 34305299 | ETV4::chr17:41610041-41610091(-) | CCL16::chr17:34305249-34305299(-) |
| ETV4 | TCAP | 17 | 17 | -1 | 1 | 41610042 | 37821969 | ETV4::chr17:41610041-41610091(-) | TCAP::chr17:37821968-37822018(+) |
| ETV5 | DGKG | 3 | 3 | -1 | -1 | 185797606 | 185929674 | ETV5::chr3:185797605-185797655(-) | DGKG::chr3:185929624-185929674(-) |
| EWSR1 | ABR | 22 | 17 | 1 | -1 | 29669853 | 1028702 | EWSR1::chr22:29669803-29669853(+) | ABR::chr17:1028652-1028702(-) |
| EXOSC10 | MTOR | 1 | 1 | -1 | -1 | 11139768 | 11190834 | EXOSC10::chr1:11139767-11139817(-) | MTOR::chr1:11190784-11190834(-) |
| FAM114A2 | BRAF | 5 | 7 | -1 | -1 | 153390801 | 140481493 | FAM114A2::chr5:153390800-153390850(-) | BRAF::chr7:140481443-140481493(-) |
| FARSA | LYL1 | 19 | 19 | -1 | -1 | 13039156 | 13210548 | FARSA::chr19:13039155-13039205(-) | LYL1::chr19:13210498-13210548(-) |
| FGFR1 | ADAM18 | 8 | 8 | -1 | 1 | 38314874 | 39463826 | FGFR1::chr8:38314873-38314923(-) | ADAM18::chr8:39463825-39463875(+) |
| FGFR1 | SLC20A2 | 8 | 8 | -1 | -1 | 38277057 | 42330172 | FGFR1::chr8:38277056-38277106(-) | SLC20A2::chr8:42330122-42330172(-) |
| FGFR2 | ATE1 | 10 | 10 | -1 | -1 | 123298106 | 123683844 | FGFR2::chr10:123298105-123298155(-) | ATE1::chr10:123683794-123683844(-) |
| FGFR2 | BICC1 | 10 | 10 | -1 | 1 | 123243212 | 60566344 | FGFR2::chr10:123243211-123243261(-) | BICC1::chr10:60566343-60566393(+) |
| FGFR2 | C10orf118 | 10 | 10 | -1 | -1 | 123243212 | 115910979 | FGFR2::chr10:123243211-123243261(-) | C10orf118::chr10:115910929-115910979(-) |
| FGFR2 | ENPP2 | 10 | 8 | -1 | -1 | 123274637 | 120650767 | FGFR2::chr10:123274636-123274686(-) | ENPP2::chr8:120650717-120650767(-) |
| FGFR2 | KIAA1967 | 10 | 8 | -1 | 1 | 123243212 | 22464120 | FGFR2::chr10:123243211-123243261(-) | KIAA1967::chr8:22464119-22464169(+) |
| FGFR2 | OFD1 | 10 | X | -1 | 1 | 123243212 | 13754597 | FGFR2::chr10:123243211-123243261(-) | OFD1::chrX:13754596-13754646(+) |
| FGFR2 | TACC2 | 10 | 10 | -1 | 1 | 123243212 | 123996910 | FGFR2::chr10:123243211-123243261(-) | TACC2::chr10:123996909-123996959(+) |
| FGFR2 | USP10 | 10 | 16 | -1 | 1 | 123353223 | 84792322 | FGFR2::chr10:123353222-123353272(-) | USP10::chr16:84792321-84792371(+) |
| FGFR3 | AES | 4 | 19 | 1 | -1 | 1808661 | 3061255 | FGFR3::chr4:1808611-1808661(+) | AES::chr19:3061205-3061255(-) |
| FGFR3 | ELAVL3 | 4 | 19 | 1 | -1 | 1808661 | 11577642 | FGFR3::chr4:1808611-1808661(+) | ELAVL3::chr19:11577592-11577642(-) |
| FGFR3 | FBXO28 | 4 | 1 | 1 | 1 | 1808661 | 224340844 | FGFR3::chr4:1808611-1808661(+) | FBXO28::chr1:224340843-224340893(+) |
| FGFR3 | TACC3 | 4 | 4 | 1 | 1 | 1808661 | 1737000 | FGFR3::chr4:1808611-1808661(+) | TACC3::chr4:1736999-1737049(+) |
| FGFR3 | TACC3 | 4 | 4 | 1 | 1 | 1803470 | 1746245 | FGFR3::chr4:1803420-1803470(+) | TACC3::chr4:1746244-1746294(+) |
| FGFR3 | TACC3 | 4 | 4 | 1 | 1 | 1808661 | 1746245 | FGFR3::chr4:1808611-1808661(+) | TACC3::chr4:1746244-1746294(+) |
| FGFR3 | TACC3 | 4 | 4 | 1 | 1 | 1808661 | 1739325 | FGFR3::chr4:1808611-1808661(+) | TACC3::chr4:1739324-1739374(+) |
| FGFR3 | TACC3 | 4 | 4 | 1 | 1 | 1801539 | 1746245 | FGFR3::chr4:1801489-1801539(+) | TACC3::chr4:1746244-1746294(+) |
| FGFR3 | TACC3 | 4 | 4 | 1 | 1 | 1808661 | 1732899 | FGFR3::chr4:1808611-1808661(+) | TACC3::chr4:1732898-1732948(+) |
| FGFR3 | TACC3 | 4 | 4 | 1 | 1 | 1795770 | 1746245 | FGFR3::chr4:1795720-1795770(+) | TACC3::chr4:1746244-1746294(+) |
| FGFR3 | TPRG1 | 4 | 3 | 1 | 1 | 1795770 | 189028175 | FGFR3::chr4:1795720-1795770(+) | TPRG1::chr3:189028174-189028224(+) |
| FGGY | CDH17 | 1 | 8 | 1 | -1 | 59978111 | 95174390 | FGGY::chr1:59978061-59978111(+) | CDH17::chr8:95174340-95174390(-) |
| FKBP15 | RET | 9 | 10 | -1 | 1 | 115932802 | 43612032 | FKBP15::chr9:115932801-115932851(-) | RET::chr10:43612031-43612081(+) |
| FOXP1 | ACTR3B | 3 | 7 | -1 | 1 | 71408301 | 152549211 | FOXP1::chr3:71408300-71408350(-) | ACTR3B::chr7:152549210-152549260(+) |
| FOXP1 | ARMC8 | 3 | 3 | -1 | 1 | 71247353 | 137928659 | FOXP1::chr3:71247352-71247402(-) | ARMC8::chr3:137928658-137928708(+) |
| FOXP1 | C5orf51 | 3 | 5 | -1 | 1 | 71348971 | 41907838 | FOXP1::chr3:71348970-71349020(-) | C5orf51::chr5:41907837-41907887(+) |
| FOXP1 | COQ10B | 3 | 2 | -1 | 1 | 71090479 | 198334794 | FOXP1::chr3:71090478-71090528(-) | COQ10B::chr2:198334793-198334843(+) |
| FOXP1 | EIF4E3 | 3 | 3 | -1 | -1 | 71542577 | 71759635 | FOXP1::chr3:71542576-71542626(-) | EIF4E3::chr3:71759585-71759635(-) |
| FOXP1 | HRH1 | 3 | 3 | -1 | 1 | 71247353 | 11300689 | FOXP1::chr3:71247352-71247402(-) | HRH1::chr3:11300688-11300738(+) |
| FOXP1 | PXDNL | 3 | 8 | -1 | -1 | 71542577 | 52412330 | FOXP1::chr3:71542576-71542626(-) | PXDNL::chr8:52412280-52412330(-) |
| FOXP1 | RGAG4 | 3 | X | -1 | -1 | 71542577 | 71347560 | FOXP1::chr3:71542576-71542626(-) | RGAG4::chrX:71347510-71347560(-) |
| FOXP1 | RUFY1 | 3 | 5 | -1 | 1 | 71348971 | 179036377 | FOXP1::chr3:71348970-71349020(-) | RUFY1::chr5:179036376-179036426(+) |
| FOXP1 | SH3RF2 | 3 | 5 | -1 | 1 | 71090479 | 145379621 | FOXP1::chr3:71090478-71090528(-) | SH3RF2::chr5:145379620-145379670(+) |

| | | | | | | | | | |
|---|---|---|---|---|---|---|---|---|---|
| FOXP1 | UBA3 | 3 | 3 | -1 | -1 | 71542577 | 69112654 | FOXP1::chr3:71542576-71542626(-) | UBA3::chr3:69112604-69112654(-) |
| FOXP1 | UGGT1 | 3 | 2 | -1 | 1 | 71037145 | 128941249 | FOXP1::chr3:71037144-71037194(-) | UGGT1::chr2:128941248-128941298(+) |
| FUBP1 | DNAJB4 | 1 | 1 | -1 | 1 | 78444569 | 78478735 | FUBP1::chr1:78444568-78444618(-) | DNAJB4::chr1:78478734-78478784(+) |
| FUBP1 | LRRC7 | 1 | 1 | -1 | 1 | 78432378 | 70144064 | FUBP1::chr1:78432377-78432427(-) | LRRC7::chr1:70144063-70144113(+) |
| FUBP1 | SLC44A5 | 1 | 1 | -1 | -1 | 78444569 | 75766370 | FUBP1::chr1:78444568-78444618(-) | SLC44A5::chr1:75766320-75766370(-) |
| FXR2 | TP53 | 17 | 17 | -1 | -1 | 7504727 | 7577608 | FXR2::chr17:7504726-7504776(-) | TP53::chr17:7577558-7577608(-) |
| GAS2L3 | SND1 | 12 | 7 | 1 | 1 | 100995531 | 127334882 | GAS2L3::chr12:100995481-100995531(+) | SND1::chr7:127334881-127334931(+) |
| GBF1 | NT5C2 | 10 | 10 | 1 | -1 | 104019873 | 104866463 | GBF1::chr10:104019823-104019873(+) | NT5C2::chr10:104866413-104866463(-) |
| GLIS3 | CTNNA2 | 9 | 2 | -1 | 1 | 4117768 | 80874710 | GLIS3::chr9:4117767-4117817(-) | CTNNA2::chr2:80874709-80874759(+) |
| GMDS | CCND3 | 6 | 6 | -1 | -1 | 2245555 | 41905132 | GMDS::chr6:2245554-2245604(-) | CCND3::chr6:41905082-41905132(-) |
| GNA11 | TMCO3 | 19 | 13 | 1 | 1 | 3094785 | 114174931 | GNA11::chr19:3094735-3094785(+) | TMCO3::chr13:114174930-114174980(+) |
| GNAQ | C9orf64 | 9 | 9 | -1 | -1 | 80409379 | 86559904 | GNAQ::chr9:80409378-80409428(-) | C9orf64::chr9:86559854-86559904(-) |
| GNAS | AP1G2 | 20 | 14 | 1 | -1 | 57486247 | 24037279 | GNAS::chr20:57486197-57486247(+) | AP1G2::chr14:24037229-24037279(-) |
| GNAS | ARFGEF2 | 20 | 20 | 1 | 1 | 57486241 | 47615777 | GNAS::chr20:57486191-57486241(+) | ARFGEF2::chr20:47615776-47615826(+) |
| GNAS | CLCN3 | 20 | 4 | 1 | 1 | 57486241 | 170623929 | GNAS::chr20:57486191-57486241(+) | CLCN3::chr4:170623928-170623978(+) |
| GNAS | EEA1 | 20 | 12 | 1 | -1 | 57486241 | 93229203 | GNAS::chr20:57486191-57486241(+) | EEA1::chr12:93229153-93229203(-) |
| GNAS | EIF2AK2 | 20 | 2 | 1 | -1 | 57486241 | 37342384 | GNAS::chr20:57486191-57486241(+) | EIF2AK2::chr2:37342334-37342384(-) |
| GNAS | ELK3 | 20 | 12 | 1 | 1 | 57486247 | 96588323 | GNAS::chr20:57486197-57486247(+) | ELK3::chr12:96588322-96588372(+) |
| GNAS | FN1 | 20 | 2 | 1 | -1 | 57486247 | 216274837 | GNAS::chr20:57486197-57486247(+) | FN1::chr2:216274787-216274837(-) |
| GNAS | ITGAV | 20 | 2 | 1 | 1 | 57486241 | 187540547 | GNAS::chr20:57486191-57486241(+) | ITGAV::chr2:187540546-187540596(+) |
| GNAS | PLXNC1 | 20 | 12 | 1 | 1 | 57486241 | 94642003 | GNAS::chr20:57486191-57486241(+) | PLXNC1::chr12:94642002-94642052(+) |
| GNAS | PSAP | 20 | 10 | 1 | -1 | 57486241 | 73579657 | GNAS::chr20:57486191-57486241(+) | PSAP::chr10:73579607-73579657(-) |
| GNAS | SEC63 | 20 | 6 | 1 | -1 | 57486241 | 108214782 | GNAS::chr20:57486191-57486241(+) | SEC63::chr6:108214732-108214782(-) |
| GNAS | SMAP2 | 20 | 1 | 1 | 1 | 57486241 | 40878688 | GNAS::chr20:57486191-57486241(+) | SMAP2::chr1:40878687-40878737(+) |
| GNAS | SNW1 | 20 | 14 | 1 | -1 | 57486241 | 78189203 | GNAS::chr20:57486191-57486241(+) | SNW1::chr14:78189153-78189203(-) |
| GNAS | THSD4 | 20 | 15 | 1 | 1 | 57486241 | 72072881 | GNAS::chr20:57486191-57486241(+) | THSD4::chr15:72072880-72072930(+) |
| GNAS | ZNFX1 | 20 | 20 | 1 | -1 | 57486241 | 47863418 | GNAS::chr20:57486191-57486241(+) | ZNFX1::chr20:47863368-47863418(-) |
| GNAS | ZNFX1 | 20 | 20 | 1 | -1 | 57486247 | 47863418 | GNAS::chr20:57486197-57486247(+) | ZNFX1::chr20:47863368-47863418(-) |
| GOLIM4 | MECOM | 3 | 3 | -1 | -1 | 167742314 | 168861620 | GOLIM4::chr3:167742313-167742363(-) | MECOM::chr3:168861570-168861620(-) |
| GPC4 | GPC3 | X | X | -1 | -1 | 132548834 | 132670321 | GPC4::chrX:132548833-132548883(-) | GPC3::chrX:132670271-132670321(-) |
| GTF2IRD1 | ALK | 7 | 2 | 1 | -1 | 73935627 | 29446394 | GTF2IRD1::chr7:73935577-73935627(+) | ALK::chr2:29446344-29446394(-) |
| H3F3B | CCT5 | 17 | 5 | -1 | 1 | 73772517 | 10250033 | H3F3B::chr17:73772516-73772566(-) | CCT5::chr5:10250032-10250082(+) |
| HGD | CACNA1D | 3 | 3 | -1 | 1 | 120389274 | 53777038 | HGD::chr3:120389273-120389323(-) | CACNA1D::chr3:53777037-53777087(+) |
| HIBCH | CTNNA2 | 2 | 2 | -1 | 1 | 191175480 | 80620336 | HIBCH::chr2:191175479-191175529(-) | CTNNA2::chr2:80620335-80620385(+) |
| HIF1A | C14orf37 | 14 | 14 | 1 | -1 | 62162557 | 58606132 | HIF1A::chr14:62162507-62162557(+) | C14orf37::chr14:58606082-58606132(-) |
| HIF1A | PRKCH | 14 | 14 | 1 | 1 | 62162557 | 61909829 | HIF1A::chr14:62162507-62162557(+) | PRKCH::chr14:61909828-61909878(+) |
| HIF1A | PRKCH | 14 | 14 | 1 | 1 | 62207906 | 61995793 | HIF1A::chr14:62207856-62207906(+) | PRKCH::chr14:61995792-61995842(+) |
| HIF1A | PRKCH | 14 | 14 | 1 | 1 | 62162557 | 61995793 | HIF1A::chr14:62162507-62162557(+) | PRKCH::chr14:61995792-61995842(+) |
| HIF1A | PRKCH | 14 | 14 | 1 | 1 | 62162557 | 61857943 | HIF1A::chr14:62162507-62162557(+) | PRKCH::chr14:61857942-61857992(+) |
| HIF1A | VRK1 | 14 | 14 | 1 | 1 | 62162557 | 97299804 | HIF1A::chr14:62162507-62162557(+) | VRK1::chr14:97299803-97299853(+) |
| HLTF | WWTR1 | 3 | 3 | -1 | -1 | 148777505 | 149260324 | HLTF::chr3:148777504-148777554(-) | WWTR1::chr3:149260274-149260324(-) |
| HMGA1 | EEA1 | 6 | 12 | 1 | -1 | 34214008 | 93229203 | HMGA1::chr6:34213958-34214008(+) | EEA1::chr12:93229153-93229203(-) |
| HMGA2 | ARFGAP2 | 12 | 11 | 1 | -1 | 66219161 | 47193884 | HMGA2::chr12:66219111-66219161(+) | ARFGAP2::chr11:47193834-47193884(-) |

| | | | | | | | | | |
|---|---|---|---|---|---|---|---|---|---|
| HMGA2 | C10orf11 | 12 | 10 | 1 | 1 | 66232349 | 78084159 | HMGA2::chr12:66232299-66232349(+) | C10orf11::chr10:78084158-78084208(+) |
| HMGA2 | C1orf87 | 12 | 1 | 1 | -1 | 66232349 | 60463488 | HMGA2::chr12:66232299-66232349(+) | C1orf87::chr1:60463438-60463488(-) |
| HMGA2 | DAB1 | 12 | 1 | 1 | -1 | 66232349 | 57463801 | HMGA2::chr12:66232299-66232349(+) | DAB1::chr1:57463751-57463801(-) |
| HMGA2 | GABRA3 | 12 | X | 1 | -1 | 66232349 | 151393317 | HMGA2::chr12:66232299-66232349(+) | GABRA3::chrX:151393267-151393317(-) |
| HMGA2 | KALRN | 12 | 3 | 1 | 1 | 66345195 | 124281690 | HMGA2::chr12:66345145-66345195(+) | KALRN::chr3:124281689-124281739(+) |
| HMGA2 | NELL2 | 12 | 12 | 1 | -1 | 66232349 | 45097637 | HMGA2::chr12:66232299-66232349(+) | NELL2::chr12:45097587-45097637(-) |
| HMGA2 | NR2C1 | 12 | 12 | 1 | -1 | 66232349 | 95461167 | HMGA2::chr12:66232299-66232349(+) | NR2C1::chr12:95461117-95461167(-) |
| HMGA2 | NUP107 | 12 | 12 | 1 | 1 | 66232349 | 69109407 | HMGA2::chr12:66232299-66232349(+) | NUP107::chr12:69109406-69109456(+) |
| HMGA2 | PCBP2 | 12 | 12 | 1 | 1 | 66232349 | 53856277 | HMGA2::chr12:66232299-66232349(+) | PCBP2::chr12:53856276-53856326(+) |
| HMGA2 | PPFIA2 | 12 | 12 | 1 | -1 | 66232349 | 81851645 | HMGA2::chr12:66232299-66232349(+) | PPFIA2::chr12:81851595-81851645(-) |
| HMGA2 | SARNP | 12 | 12 | 1 | -1 | 66345195 | 56146624 | HMGA2::chr12:66345145-66345195(+) | SARNP::chr12:56146574-56146624(-) |
| HNF1B | NOTCH1 | 17 | 9 | -1 | -1 | 36099431 | 139396940 | HNF1B::chr17:36099430-36099480(-) | NOTCH1::chr9:139396890-139396940(-) |
| HNRNPA2B1 | CANT1 | 7 | 17 | -1 | -1 | 26236208 | 76994045 | HNRNPA2B1::chr7:26236207-26236257(-) | CANT1::chr17:76993995-76994045(-) |
| HNRNPA2B1 | CLASP2 | 7 | 3 | -1 | -1 | 26231467 | 33602368 | HNRNPA2B1::chr7:26231466-26231516(-) | CLASP2::chr3:33602318-33602368(-) |
| HNRNPA2B1 | DSP | 7 | 6 | -1 | 1 | 26231467 | 7575528 | HNRNPA2B1::chr7:26231466-26231516(-) | DSP::chr6:7575527-7575577(+) |
| HNRNPA2B1 | RABEP1 | 7 | 17 | -1 | 1 | 26231467 | 5283828 | HNRNPA2B1::chr7:26231466-26231516(-) | RABEP1::chr17:5283827-5283877(+) |
| HNRNPA2B1 | SIAE | 7 | 11 | -1 | -1 | 26231467 | 124543777 | HNRNPA2B1::chr7:26231466-26231516(-) | SIAE::chr11:124543727-124543777(-) |
| HNRNPA2B1 | SIAE | 7 | 11 | -1 | -1 | 26231464 | 124543777 | HNRNPA2B1::chr7:26231463-26231513(-) | SIAE::chr11:124543727-124543777(-) |
| HNRNPH1 | ERG | 5 | 21 | -1 | -1 | 179047893 | 39755845 | HNRNPH1::chr5:179047892-179047942(-) | ERG::chr21:39755795-39755845(-) |
| HOOK3 | IKBKB | 8 | 8 | 1 | 1 | 42798588 | 42162705 | HOOK3::chr8:42798538-42798588(+) | IKBKB::chr8:42162704-42162754(+) |
| HSD17B12 | WT1 | 11 | 11 | 1 | -1 | 43702537 | 32450165 | HSD17B12::chr11:43702487-43702537(+) | WT1::chr11:32450115-32450165(-) |
| IKBKB | ANK1 | 8 | 8 | 1 | -1 | 42129723 | 41615655 | IKBKB::chr8:42129673-42129723(+) | ANK1::chr8:41615605-41615655(-) |
| IKBKB | ANK1 | 8 | 8 | 1 | -1 | 42129723 | 41566493 | IKBKB::chr8:42129673-42129723(+) | ANK1::chr8:41566443-41566493(-) |
| IKBKB | ANK1 | 8 | 8 | 1 | -1 | 42129723 | 41591587 | IKBKB::chr8:42129673-42129723(+) | ANK1::chr8:41591537-41591587(-) |
| IKBKB | FADD | 8 | 11 | 1 | 1 | 42166543 | 70052239 | IKBKB::chr8:42166493-42166543(+) | FADD::chr11:70052238-70052288(+) |
| IKBKB | LETM2 | 8 | 8 | 1 | 1 | 42129723 | 38257790 | IKBKB::chr8:42129673-42129723(+) | LETM2::chr8:38257789-38257839(+) |
| IKBKB | UNC5D | 8 | 8 | 1 | 1 | 42129723 | 35406810 | IKBKB::chr8:42129673-42129723(+) | UNC5D::chr8:35406809-35406859(+) |
| IL6ST | ANKRD55 | 5 | 5 | -1 | -1 | 55272043 | 55412609 | IL6ST::chr5:55272042-55272092(-) | ANKRD55::chr5:55412559-55412609(-) |
| IL6ST | DDX4 | 5 | 5 | -1 | 1 | 55272043 | 55034778 | IL6ST::chr5:55272042-55272092(-) | DDX4::chr5:55034777-55034827(+) |
| IL6ST | LAMC2 | 5 | 1 | -1 | 1 | 55247292 | 183177016 | IL6ST::chr5:55247291-55247341(-) | LAMC2::chr1:183177015-183177065(+) |
| IQCH | CTNND2 | 15 | 5 | 1 | -1 | 67553732 | 11199773 | IQCH::chr15:67553682-67553732(+) | CTNND2::chr5:11199723-11199773(-) |
| IRF2BP2 | NTRK1 | 1 | 1 | -1 | 1 | 234744241 | 156844363 | IRF2BP2::chr1:234744240-234744290(-) | NTRK1::chr1:156844362-156844412(+) |
| JAK2 | CSTF3 | 9 | 11 | 1 | -1 | 4986022 | 33108687 | JAK2::chr9:4985972-4986022(+) | CSTF3::chr11:33108637-33108687(-) |
| JAK2 | DOCK8 | 9 | 9 | 1 | 1 | 5055788 | 286461 | JAK2::chr9:5055738-5055788(+) | DOCK8::chr9:286460-286510(+) |
| JAK2 | GLDC | 9 | 9 | 1 | -1 | 5029906 | 6565429 | JAK2::chr9:5029856-5029906(+) | GLDC::chr9:6565379-6565429(-) |
| JAK2 | PKNOX2 | 9 | 11 | 1 | 1 | 4986022 | 125201715 | JAK2::chr9:4985972-4986022(+) | PKNOX2::chr11:125201714-125201764(+) |
| JAK2 | RCL1 | 9 | 9 | 1 | 1 | 5090911 | 4834141 | JAK2::chr9:5090861-5090911(+) | RCL1::chr9:4834140-4834190(+) |
| JHDM1D | BRAF | 7 | 7 | -1 | -1 | 139810895 | 140481493 | JHDM1D::chr7:139810894-139810944(-) | BRAF::chr7:140481443-140481493(-) |
| KCNQ5 | ALK | 6 | 2 | 1 | -1 | 73332315 | 29498362 | KCNQ5::chr6:73332265-73332315(+) | ALK::chr2:29498312-29498362(-) |
| KCTD15 | MSI2 | 19 | 17 | 1 | 1 | 34302457 | 55693331 | KCTD15::chr19:34302407-34302457(+) | MSI2::chr17:55693330-55693380(+) |
| KDM2A | DDR2 | 11 | 1 | 1 | 1 | 67007840 | 162749902 | KDM2A::chr11:67007790-67007840(+) | DDR2::chr1:162749901-162749951(+) |
| KDM4A | MPL | 1 | 1 | 1 | 1 | 44169783 | 43805635 | KDM4A::chr1:44169733-44169783(+) | MPL::chr1:43805634-43805684(+) |
| KDM5A | ANO2 | 12 | 12 | -1 | -1 | 427272 | 5708795 | KDM5A::chr12:427271-427321(-) | ANO2::chr12:5708745-5708795(-) |

| | | | | | | | | | |
|---|---|---|---|---|---|---|---|---|---|
| KDM5A | IQSEC3 | 12 | 12 | -1 | 1 | 442653 | 266191 | KDM5A::chr12:442652-442702(-) | IQSEC3::chr12:266190-266240(+) |
| KDM5A | NINJ2 | 12 | 12 | -1 | -1 | 404739 | 675344 | KDM5A::chr12:404738-404788(-) | NINJ2::chr12:675294-675344(-) |
| KDM5A | NINJ2 | 12 | 12 | -1 | -1 | 430161 | 675344 | KDM5A::chr12:430160-430210(-) | NINJ2::chr12:675294-675344(-) |
| KDM5A | RAE1 | 12 | 20 | -1 | 1 | 459787 | 55940412 | KDM5A::chr12:459786-459836(-) | RAE1::chr20:55940411-55940461(+) |
| KIAA0947 | TERT | 5 | 5 | 1 | -1 | 5448010 | 1282739 | KIAA0947::chr5:5447960-5448010(+) | TERT::chr5:1282689-1282739(-) |
| KIAA1549 | BRAF | 7 | 7 | -1 | -1 | 138545885 | 140487384 | KIAA1549::chr7:138545884-138545934(-) | BRAF::chr7:140487334-140487384(-) |
| KIF5B | MET | 10 | 7 | -1 | 1 | 32306071 | 116411903 | KIF5B::chr10:32306070-32306120(-) | MET::chr7:116411902-116411952(+) |
| KIT | DNAJB14 | 4 | 4 | 1 | -1 | 55598164 | 100851778 | KIT::chr4:55598114-55598164(+) | DNAJB14::chr4:100851728-100851778(-) |
| KIT | FAR1 | 4 | 11 | 1 | 1 | 55606881 | 13744912 | KIT::chr4:55606831-55606881(+) | FAR1::chr11:13744911-13744961(+) |
| KIT | HOPX | 4 | 4 | 1 | -1 | 55524248 | 57522178 | KIT::chr4:55524198-55524248(+) | HOPX::chr4:57522128-57522178(-) |
| KIT | SLC4A4 | 4 | 4 | 1 | 1 | 55599358 | 72102293 | KIT::chr4:55599308-55599358(+) | SLC4A4::chr4:72102292-72102342(+) |
| KIT | UGT2B7 | 4 | 4 | 1 | 1 | 55524248 | 69955190 | KIT::chr4:55524198-55524248(+) | UGT2B7::chr4:69955189-69955239(+) |
| KRAS | IFLTD1 | 12 | 12 | -1 | -1 | 25398208 | 25656734 | KRAS::chr12:25398207-25398257(-) | IFLTD1::chr12:25656684-25656734(-) |
| KRAS | RERGL | 12 | 12 | -1 | -1 | 25398208 | 18241890 | KRAS::chr12:25398207-25398257(-) | RERGL::chr12:18241840-18241890(-) |
| LAMA3 | ZNF521 | 18 | 18 | 1 | -1 | 21529888 | 22807661 | LAMA3::chr18:21529838-21529888(+) | ZNF521::chr18:22807611-22807661(-) |
| LARS2 | TBL1XR1 | 3 | 3 | 1 | -1 | 45461221 | 176769514 | LARS2::chr3:45461171-45461221(+) | TBL1XR1::chr3:176769464-176769514(-) |
| LDHB | CTNNA2 | 12 | 2 | -1 | 1 | 21790005 | 80620336 | LDHB::chr12:21790004-21790054(-) | CTNNA2::chr2:80620335-80620385(+) |
| LEPREL1 | TP63 | 3 | 3 | -1 | 1 | 189838041 | 189526061 | LEPREL1::chr3:189838040-189838090(-) | TP63::chr3:189526060-189526110(+) |
| LMBRD2 | CTNND2 | 5 | 5 | -1 | -1 | 36108636 | 11365002 | LMBRD2::chr5:36108635-36108685(-) | CTNND2::chr5:11364952-11365002(-) |
| LNPEP | FLT4 | 5 | 5 | 1 | -1 | 96271878 | 180048904 | LNPEP::chr5:96271828-96271878(+) | FLT4::chr5:180048854-180048904(-) |
| LPP | CASR | 3 | 3 | 1 | 1 | 188202492 | 121980375 | LPP::chr3:188202442-188202492(+) | CASR::chr3:121980374-121980424(+) |
| LPP | CLPTM1L | 3 | 5 | 1 | -1 | 188478070 | 1338097 | LPP::chr3:188478020-188478070(+) | CLPTM1L::chr5:1338047-1338097(-) |
| LPP | CLSTN2 | 3 | 3 | 1 | 1 | 187943315 | 140251166 | LPP::chr3:187943265-187943315(+) | CLSTN2::chr3:140251165-140251215(+) |
| LPP | F2 | 3 | 11 | 1 | 1 | 188059507 | 46741252 | LPP::chr3:188059457-188059507(+) | F2::chr11:46741251-46741301(+) |
| LPP | FAM108C1 | 3 | 15 | 1 | 1 | 187943315 | 81046492 | LPP::chr3:187943265-187943315(+) | FAM108C1::chr15:81046491-81046541(+) |
| LPP | GOLIM4 | 3 | 3 | 1 | -1 | 187943315 | 167766153 | LPP::chr3:187943265-187943315(+) | GOLIM4::chr3:167766103-167766153(-) |
| LPP | HRASLS | 3 | 3 | 1 | 1 | 188202492 | 192973440 | LPP::chr3:188202442-188202492(+) | HRASLS::chr3:192973439-192973489(+) |
| LPP | KCNIP4 | 3 | 4 | 1 | -1 | 187943315 | 20852290 | LPP::chr3:187943265-187943315(+) | KCNIP4::chr4:20852240-20852290(-) |
| LPP | LEPREL1 | 3 | 3 | 1 | -1 | 188327632 | 189713231 | LPP::chr3:188327582-188327632(+) | LEPREL1::chr3:189713181-189713231(-) |
| LPP | LIPJ | 3 | 10 | 1 | 1 | 188426181 | 90347023 | LPP::chr3:188426131-188426181(+) | LIPJ::chr10:90347022-90347072(+) |
| LPP | ST6GAL1 | 3 | 3 | 1 | 1 | 187871809 | 186790637 | LPP::chr3:187871759-187871809(+) | ST6GAL1::chr3:186790636-186790686(+) |
| LPP | ZNRF1 | 3 | 16 | 1 | 1 | 187943315 | 75127470 | LPP::chr3:187943265-187943315(+) | ZNRF1::chr16:75127469-75127519(+) |
| LTBP1 | BIRC6 | 2 | 2 | 1 | 1 | 33447218 | 32770710 | LTBP1::chr2:33447168-33447218(+) | BIRC6::chr2:32770709-32770759(+) |
| LYN | NTRK3 | 8 | 15 | 1 | -1 | 56866543 | 88576276 | LYN::chr8:56866493-56866543(+) | NTRK3::chr15:88576226-88576276(-) |
| MACF1 | BRAF | 1 | 7 | 1 | -1 | 39896580 | 140487384 | MACF1::chr1:39896530-39896580(+) | BRAF::chr7:140487334-140487384(-) |
| MADD | DDB2 | 11 | 11 | 1 | 1 | 47317606 | 47254365 | MADD::chr11:47317556-47317606(+) | DDB2::chr11:47254364-47254414(+) |
| MALT1 | NOL4 | 18 | 18 | 1 | -1 | 56367823 | 31432999 | MALT1::chr18:56367773-56367823(+) | NOL4::chr18:31432949-31432999(-) |
| MAML2 | ANKMY2 | 11 | 7 | -1 | -1 | 96074547 | 16666803 | MAML2::chr11:96074546-96074596(-) | ANKMY2::chr7:16666753-16666803(-) |
| MAML2 | BCAS4 | 11 | 20 | -1 | 1 | 95825056 | 49493023 | MAML2::chr11:95825055-95825105(-) | BCAS4::chr20:49493022-49493072(+) |
| MAML2 | CNTN5 | 11 | 11 | -1 | 1 | 95825056 | 99715584 | MAML2::chr11:95825055-95825105(-) | CNTN5::chr11:99715583-99715633(+) |
| MAML2 | YAP1 | 11 | 11 | -1 | 1 | 96074547 | 102094353 | MAML2::chr11:96074546-96074596(-) | YAP1::chr11:102094352-102094402(+) |
| MANBAL | UBR5 | 20 | 8 | 1 | -1 | 35929816 | 103341627 | MANBAL::chr20:35929766-35929816(+) | UBR5::chr8:103341577-103341627(-) |
| MAP1S | CRTC1 | 19 | 19 | 1 | 1 | 17844237 | 18853720 | MAP1S::chr19:17844187-17844237(+) | CRTC1::chr19:18853719-18853769(+) |

| | | | | | | | | | |
|---|---|---|---|---|---|---|---|---|---|
| MAP2K2 | PSMD9 | 19 | 12 | -1 | 1 | 4110507 | 122353762 | MAP2K2::chr19:4110506-4110556(-) | PSMD9::chr12:122353761-122353811(+) |
| MAP2K2 | RP11-87C12.2 | 19 | 12 | -1 | 1 | 4110507 | 122353762 | MAP2K2::chr19:4110506-4110556(-) | RP11-87C12.2::chr12:122353761-122353811(+) |
| MAPK1 | ARHGEF7 | 22 | 13 | -1 | 1 | 22221612 | 111806252 | MAPK1::chr22:22221611-22221661(-) | ARHGEF7::chr13:111806251-111806301(+) |
| MAPK1 | KCNJ4 | 22 | 22 | -1 | -1 | 22160139 | 38824176 | MAPK1::chr22:22160138-22160188(-) | KCNJ4::chr22:38824126-38824176(-) |
| MAPK1 | PRODH | 22 | 22 | -1 | -1 | 22221612 | 18918711 | MAPK1::chr22:22221611-22221661(-) | PRODH::chr22:18918661-18918711(-) |
| MAPK1 | TOP3B | 22 | 22 | -1 | -1 | 22221612 | 22313604 | MAPK1::chr22:22221611-22221661(-) | TOP3B::chr22:22313554-22313604(-) |
| MDM2 | DIP2B | 12 | 12 | 1 | 1 | 69207408 | 51053977 | MDM2::chr12:69207358-69207408(+) | DIP2B::chr12:51053976-51054026(+) |
| MDM2 | FGF18 | 12 | 5 | 1 | 1 | 69214154 | 170883543 | MDM2::chr12:69214104-69214154(+) | FGF18::chr5:170883542-170883592(+) |
| MDM2 | NUP107 | 12 | 12 | 1 | 1 | 69229764 | 69113121 | MDM2::chr12:69229714-69229764(+) | NUP107::chr12:69113120-69113170(+) |
| MDM2 | NUP107 | 12 | 12 | 1 | 1 | 69229764 | 69135593 | MDM2::chr12:69229714-69229764(+) | NUP107::chr12:69135592-69135642(+) |
| MDM2 | NUP107 | 12 | 12 | 1 | 1 | 69202271 | 69082742 | MDM2::chr12:69202221-69202271(+) | NUP107::chr12:69082741-69082791(+) |
| MDM2 | PTPRR | 12 | 12 | 1 | -1 | 69207408 | 71078563 | MDM2::chr12:69207358-69207408(+) | PTPRR::chr12:71078513-71078563(-) |
| MDM2 | SLAMF9 | 12 | 1 | 1 | -1 | 69230529 | 159923443 | MDM2::chr12:69230479-69230529(+) | SLAMF9::chr1:159923393-159923443(-) |
| MECOM | ANKRD10 | 3 | 13 | -1 | -1 | 168849217 | 111536140 | MECOM::chr3:168849216-168849266(-) | ANKRD10::chr13:111536090-111536140(-) |
| MECOM | ARMC8 | 3 | 3 | -1 | 1 | 169098975 | 138007890 | MECOM::chr3:169098974-169099024(-) | ARMC8::chr3:138007889-138007939(+) |
| MECOM | CD86 | 3 | 3 | -1 | 1 | 169381124 | 121810448 | MECOM::chr3:169381123-169381173(-) | CD86::chr3:121810447-121810497(+) |
| MECOM | FRMD5 | 3 | 15 | -1 | -1 | 168810746 | 44180464 | MECOM::chr3:168810745-168810795(-) | FRMD5::chr15:44180414-44180464(-) |
| MECOM | LMAN2L | 3 | 2 | -1 | -1 | 169381124 | 97400263 | MECOM::chr3:169381123-169381173(-) | LMAN2L::chr2:97400213-97400263(-) |
| MECOM | LRRC34 | 3 | 3 | -1 | -1 | 169381124 | 169526533 | MECOM::chr3:169381123-169381173(-) | LRRC34::chr3:169526483-169526533(-) |
| MECOM | LRRC34 | 3 | 3 | -1 | -1 | 169098975 | 169521969 | MECOM::chr3:169098974-169099024(-) | LRRC34::chr3:169521919-169521969(-) |
| MECOM | NCEH1 | 3 | 3 | -1 | -1 | 169381124 | 172363482 | MECOM::chr3:169381123-169381173(-) | NCEH1::chr3:172363432-172363482(-) |
| MECOM | SLC7A14 | 3 | 3 | -1 | -1 | 169381124 | 170219134 | MECOM::chr3:169381123-169381173(-) | SLC7A14::chr3:170219084-170219134(-) |
| MED15 | TFE3 | 22 | X | 1 | -1 | 20929519 | 48891766 | MED15::chr22:20929469-20929519(+) | TFE3::chrX:48891716-48891766(-) |
| MET | C8orf34 | 7 | 8 | 1 | 1 | 116412043 | 69358564 | MET::chr7:116411993-116412043(+) | C8orf34::chr8:69358563-69358613(+) |
| MET | CAV1 | 7 | 7 | 1 | 1 | 116381079 | 116199000 | MET::chr7:116381029-116381079(+) | CAV1::chr7:116198999-116199049(+) |
| MET | CAV1 | 7 | 7 | 1 | 1 | 116312631 | 116199000 | MET::chr7:116312581-116312631(+) | CAV1::chr7:116198999-116199049(+) |
| MET | CAV1 | 7 | 7 | 1 | 1 | 116340338 | 116199000 | MET::chr7:116340288-116340338(+) | CAV1::chr7:116198999-116199049(+) |
| MET | DYNC1I1 | 7 | 7 | 1 | 1 | 116371913 | 95434033 | MET::chr7:116371863-116371913(+) | DYNC1I1::chr7:95434032-95434082(+) |
| MET | TFG | 7 | 3 | 1 | 1 | 116412043 | 100455420 | MET::chr7:116411993-116412043(+) | TFG::chr3:100455419-100455469(+) |
| MET | WNT2 | 7 | 7 | 1 | -1 | 116417523 | 116937930 | MET::chr7:116417473-116417523(+) | WNT2::chr7:116937880-116937930(-) |
| MITF | MTERFD1 | 3 | 8 | 1 | -1 | 69788852 | 97270928 | MITF::chr3:69788802-69788852(+) | MTERFD1::chr8:97270878-97270928(-) |
| MITF | PRICKLE2 | 3 | 3 | 1 | -1 | 69788852 | 64085601 | MITF::chr3:69788802-69788852(+) | PRICKLE2::chr3:64085551-64085601(-) |
| MKL2 | FSTL3 | 16 | 19 | 1 | 1 | 14234617 | 681650 | MKL2::chr16:14234567-14234617(+) | FSTL3::chr19:681649-681699(+) |
| MKRN1 | BRAF | 7 | 7 | -1 | -1 | 140158807 | 140487384 | MKRN1::chr7:140158806-140158856(-) | BRAF::chr7:140487334-140487384(-) |
| MLL | MLLT10 | 11 | 10 | 1 | 1 | 118355029 | 21959378 | MLL::chr11:118354979-118355029(+) | MLLT10::chr10:21959377-21959427(+) |
| MLL | MLLT10 | 11 | 10 | 1 | 1 | 118353210 | 21875223 | MLL::chr11:118353160-118353210(+) | MLLT10::chr10:21875222-21875272(+) |
| MLL | MLLT10 | 11 | 10 | 1 | 1 | 118355690 | 21940602 | MLL::chr11:118355640-118355690(+) | MLLT10::chr10:21940601-21940651(+) |
| MLL | MLLT10 | 11 | 10 | 1 | 1 | 118350953 | 22002701 | MLL::chr11:118350903-118350953(+) | MLLT10::chr10:22002700-22002750(+) |
| MLL | MLLT4 | 11 | 6 | 1 | 1 | 118355029 | 168265231 | MLL::chr11:118354979-118355029(+) | MLLT4::chr6:168265230-168265280(+) |
| MLL | MLLT4 | 11 | 6 | 1 | 1 | 118353210 | 168265231 | MLL::chr11:118353160-118353210(+) | MLLT4::chr6:168265230-168265280(+) |
| MLLT10 | ASCC1 | 10 | 10 | 1 | -1 | 21906136 | 73892939 | MLLT10::chr10:21906086-21906136(+) | ASCC1::chr10:73892889-73892939(-) |
| MLLT10 | C10orf140 | 10 | 10 | 1 | -1 | 21906136 | 21812800 | MLLT10::chr10:21906086-21906136(+) | C10orf140::chr10:21812750-21812800(-) |
| MLLT10 | KCNJ6 | 10 | 21 | 1 | -1 | 21827841 | 39087434 | MLLT10::chr10:21827791-21827841(+) | KCNJ6::chr21:39087384-39087434(-) |

| | | | | | | | | | |
|---|---|---|---|---|---|---|---|---|---|
| MLLT10 | MINPP1 | 10 | 10 | 1 | 1 | 21823733 | 89311839 | MLLT10::chr10:21823683-21823733(+) | MINPP1::chr10:89311838-89311888(+) |
| MLLT10 | SLC16A1 | 10 | 1 | 1 | -1 | 21823733 | 113471974 | MLLT10::chr10:21823683-21823733(+) | SLC16A1::chr1:113471924-113471974(-) |
| MLLT10 | SNTG1 | 10 | 8 | 1 | 1 | 22024164 | 51415338 | MLLT10::chr10:22024114-22024164(+) | SNTG1::chr8:51415337-51415387(+) |
| MLLT4 | EXOC6B | 6 | 2 | 1 | -1 | 168303066 | 72802797 | MLLT4::chr6:168303016-168303066(+) | EXOC6B::chr2:72802747-72802797(-) |
| MLLT4 | GMDS | 6 | 6 | 1 | -1 | 168227917 | 1742820 | MLLT4::chr6:168227867-168227917(+) | GMDS::chr6:1742770-1742820(-) |
| MLLT4 | HMGCLL1 | 6 | 6 | 1 | -1 | 168271178 | 55406938 | MLLT4::chr6:168271128-168271178(+) | HMGCLL1::chr6:55406888-55406938(-) |
| MLLT4 | KCNAB1 | 6 | 3 | 1 | 1 | 168227917 | 156232153 | MLLT4::chr6:168227867-168227917(+) | KCNAB1::chr3:156232152-156232202(+) |
| MLLT4 | MLL | 6 | 11 | 1 | 1 | 168227917 | 118352430 | MLLT4::chr6:168227867-168227917(+) | MLL::chr11:118352429-118352479(+) |
| MLLT4 | SASH1 | 6 | 6 | 1 | 1 | 168281197 | 148711270 | MLLT4::chr6:168281147-168281197(+) | SASH1::chr6:148711269-148711319(+) |
| MLLT4 | TTLL2 | 6 | 6 | 1 | 1 | 168227917 | 167752135 | MLLT4::chr6:168227867-168227917(+) | TTLL2::chr6:167752134-167752184(+) |
| MLLT4 | UNC93A | 6 | 6 | 1 | 1 | 168227917 | 167717407 | MLLT4::chr6:168227867-168227917(+) | UNC93A::chr6:167717406-167717456(+) |
| MNAT1 | SIX1 | 14 | 14 | 1 | -1 | 61201669 | 61113295 | MNAT1::chr14:61201619-61201669(+) | SIX1::chr14:61113245-61113295(-) |
| MRPL35 | CTNNA2 | 2 | 2 | 1 | 1 | 86434450 | 80772107 | MRPL35::chr2:86434400-86434450(+) | CTNNA2::chr2:80772106-80772156(+) |
| MSI2 | AKAP1 | 17 | 17 | 1 | 1 | 55607085 | 55182802 | MSI2::chr17:55607035-55607085(+) | AKAP1::chr17:55182801-55182851(+) |
| MSI2 | BCAS3 | 17 | 17 | 1 | 1 | 55729522 | 58885365 | MSI2::chr17:55729472-55729522(+) | BCAS3::chr17:58885364-58885414(+) |
| MSI2 | BRD7 | 17 | 16 | 1 | -1 | 55607085 | 50355949 | MSI2::chr17:55607035-55607085(+) | BRD7::chr16:50355899-50355949(-) |
| MSI2 | C18orf34 | 17 | 18 | 1 | -1 | 55339553 | 30977172 | MSI2::chr17:55339503-55339553(+) | C18orf34::chr18:30977122-30977172(-) |
| MSI2 | CA10 | 17 | 17 | 1 | -1 | 55339553 | 50008492 | MSI2::chr17:55339503-55339553(+) | CA10::chr17:50008442-50008492(-) |
| MSI2 | CA10 | 17 | 17 | 1 | -1 | 55607085 | 50149753 | MSI2::chr17:55607035-55607085(+) | CA10::chr17:50149703-50149753(-) |
| MSI2 | CISD3 | 17 | 17 | 1 | 1 | 55478832 | 36887013 | MSI2::chr17:55478782-55478832(+) | CISD3::chr17:36887012-36887062(+) |
| MSI2 | EFCAB5 | 17 | 17 | 1 | 1 | 55607085 | 28378136 | MSI2::chr17:55607035-55607085(+) | EFCAB5::chr17:28378135-28378185(+) |
| MSI2 | HNF1B | 17 | 17 | 1 | -1 | 55478832 | 36070671 | MSI2::chr17:55478782-55478832(+) | HNF1B::chr17:36070621-36070671(-) |
| MSI2 | ICAM2 | 17 | 17 | 1 | -1 | 55704664 | 62081324 | MSI2::chr17:55704614-55704664(+) | ICAM2::chr17:62081274-62081324(-) |
| MSI2 | RAD51C | 17 | 17 | 1 | 1 | 55339553 | 56772292 | MSI2::chr17:55339503-55339553(+) | RAD51C::chr17:56772291-56772341(+) |
| MSI2 | SMYD3 | 17 | 1 | 1 | -1 | 55607085 | 246518396 | MSI2::chr17:55607035-55607085(+) | SMYD3::chr1:246518346-246518396(-) |
| MSI2 | SYNPO2 | 17 | 4 | 1 | 1 | 55478832 | 119944585 | MSI2::chr17:55478782-55478832(+) | SYNPO2::chr4:119944584-119944634(+) |
| MSI2 | TEX2 | 17 | 17 | 1 | -1 | 55607085 | 62272455 | MSI2::chr17:55607035-55607085(+) | TEX2::chr17:62272405-62272455(-) |
| MSI2 | TOB1 | 17 | 17 | 1 | -1 | 55704664 | 48944963 | MSI2::chr17:55704614-55704664(+) | TOB1::chr17:48944913-48944963(-) |
| MSI2 | UTP18 | 17 | 17 | 1 | 1 | 55478832 | 49374326 | MSI2::chr17:55478782-55478832(+) | UTP18::chr17:49374325-49374375(+) |
| MSI2 | YPEL2 | 17 | 17 | 1 | 1 | 55339553 | 57430576 | MSI2::chr17:55339503-55339553(+) | YPEL2::chr17:57430575-57430625(+) |
| MSI2 | YPEL2 | 17 | 17 | 1 | 1 | 55607085 | 57430576 | MSI2::chr17:55607035-55607085(+) | YPEL2::chr17:57430575-57430625(+) |
| MTMR2 | MAML2 | 11 | 11 | -1 | -1 | 95621320 | 95826681 | MTMR2::chr11:95621319-95621369(-) | MAML2::chr11:95826631-95826681(-) |
| MTOR | CR1L | 1 | 1 | -1 | 1 | 11288725 | 207857217 | MTOR::chr1:11288724-11288774(-) | CR1L::chr1:207857216-207857266(+) |
| MTOR | GARNL3 | 1 | 9 | -1 | 1 | 11217209 | 130151183 | MTOR::chr1:11217208-11217258(-) | GARNL3::chr9:130151182-130151232(+) |
| MTOR | MAD2L2 | 1 | 1 | -1 | -1 | 11227499 | 11740670 | MTOR::chr1:11227498-11227548(-) | MAD2L2::chr1:11740620-11740670(-) |
| MUC4 | PCYT1A | 3 | 3 | -1 | -1 | 195538607 | 195941244 | MUC4::chr3:195538606-195538656(-) | PCYT1A::chr3:195941194-195941244(-) |
| MUC4 | TNK2 | 3 | 3 | -1 | -1 | 195495893 | 195610180 | MUC4::chr3:195495892-195495942(-) | TNK2::chr3:195610130-195610180(-) |
| MYB | NFIB | 6 | 9 | 1 | -1 | 135521553 | 14307519 | MYB::chr6:135521503-135521553(+) | NFIB::chr9:14307469-14307519(-) |
| NCALD | UBR5 | 8 | 8 | -1 | -1 | 102731480 | 103266742 | NCALD::chr8:102731479-102731529(-) | UBR5::chr8:103266692-103266742(-) |
| NCOA2 | ARFGEF1 | 8 | 8 | -1 | -1 | 71315935 | 68214701 | NCOA2::chr8:71315934-71315984(-) | ARFGEF1::chr8:68214651-68214701(-) |
| NCOA2 | EIF4EBP1 | 8 | 8 | -1 | 1 | 71315935 | 37914599 | NCOA2::chr8:71315934-71315984(-) | EIF4EBP1::chr8:37914598-37914648(+) |
| NCOA2 | FAM150A | 8 | 8 | -1 | -1 | 71208979 | 53455005 | NCOA2::chr8:71208978-71209028(-) | FAM150A::chr8:53454955-53455005(-) |
| NCOA2 | NCALD | 8 | 8 | -1 | -1 | 71126138 | 102731876 | NCOA2::chr8:71126137-71126187(-) | NCALD::chr8:102731826-102731876(-) |

| | | | | | | | | | |
|---|---|---|---|---|---|---|---|---|---|
| NCOA2 | SGSM3 | 8 | 22 | -1 | 1 | 71315935 | 40796700 | NCOA2::chr8:71315934-71315984(-) | SGSM3::chr22:40796699-40796749(+) |
| NCOA2 | ST18 | 8 | 8 | -1 | -1 | 71208979 | 53159224 | NCOA2::chr8:71208978-71209028(-) | ST18::chr8:53159174-53159224(-) |
| NCOA2 | XKR9 | 8 | 8 | -1 | 1 | 71033537 | 71587053 | NCOA2::chr8:71033536-71033586(-) | XKR9::chr8:71587052-71587102(+) |
| NFASC | NTRK1 | 1 | 1 | 1 | 1 | 204951148 | 156844363 | NFASC::chr1:204951098-204951148(+) | NTRK1::chr1:156844362-156844412(+) |
| NFATC2 | DOK5 | 20 | 20 | -1 | 1 | 50133323 | 53171472 | NFATC2::chr20:50133322-50133372(-) | DOK5::chr20:53171471-53171521(+) |
| NFATC2 | PHACTR3 | 20 | 20 | -1 | 1 | 50048604 | 58420244 | NFATC2::chr20:50048603-50048653(-) | PHACTR3::chr20:58420243-58420293(+) |
| NOL11 | CARD11 | 17 | 7 | 1 | -1 | 65733261 | 2979562 | NOL11::chr17:65733211-65733261(+) | CARD11::chr7:2979512-2979562(-) |
| NOTCH3 | BRD4 | 19 | 19 | -1 | -1 | 15299800 | 15360101 | NOTCH3::chr19:15299799-15299849(-) | BRD4::chr19:15360051-15360101(-) |
| NRAS | CLDN14 | 1 | 21 | -1 | -1 | 115258671 | 37834074 | NRAS::chr1:115258670-115258720(-) | CLDN14::chr21:37834024-37834074(-) |
| NSD1 | NUP98 | 5 | 11 | 1 | -1 | 176639196 | 3756554 | NSD1::chr5:176639146-176639196(+) | NUP98::chr11:3756504-3756554(-) |
| NT5C2 | C10orf120 | 10 | 10 | -1 | -1 | 104934615 | 124458928 | NT5C2::chr10:104934614-104934664(-) | C10orf120::chr10:124458878-124458928(-) |
| NT5C2 | DLEU1 | 10 | 13 | -1 | 1 | 104899163 | 50678840 | NT5C2::chr10:104899162-104899212(-) | DLEU1::chr13:50678839-50678889(+) |
| NT5C2 | LRIT1 | 10 | 10 | -1 | -1 | 104899163 | 85994134 | NT5C2::chr10:104899162-104899212(-) | LRIT1::chr10:85994084-85994134(-) |
| NTM | CTNNA2 | 11 | 2 | 1 | 1 | 132082041 | 80085139 | NTM::chr11:132081991-132082041(+) | CTNNA2::chr2:80085138-80085188(+) |
| NTRK3 | RBPMS | 15 | 8 | -1 | 1 | 88669502 | 30402011 | NTRK3::chr15:88669501-88669551(-) | RBPMS::chr8:30402010-30402060(+) |
| NUP214 | ABL1 | 9 | 9 | 1 | 1 | 134027281 | 133747516 | NUP214::chr9:134027231-134027281(+) | ABL1::chr9:133747515-133747565(+) |
| NUP98 | DNHD1 | 11 | 11 | -1 | 1 | 3774546 | 6584212 | NUP98::chr11:3774545-3774595(-) | DNHD1::chr11:6584211-6584261(+) |
| NUP98 | MLXIP | 11 | 12 | -1 | 1 | 3789811 | 122611803 | NUP98::chr11:3789810-3789860(-) | MLXIP::chr12:122611802-122611852(+) |
| NUP98 | NSD1 | 11 | 5 | -1 | 1 | 3774546 | 176662822 | NUP98::chr11:3774545-3774595(-) | NSD1::chr5:176662821-176662871(+) |
| NUP98 | RHOG | 11 | 11 | -1 | -1 | 3797112 | 3849436 | NUP98::chr11:3797111-3797161(-) | RHOG::chr11:3849386-3849436(-) |
| NUP98 | TMEM117 | 11 | 12 | -1 | 1 | 3774546 | 44770378 | NUP98::chr11:3774545-3774595(-) | TMEM117::chr12:44770377-44770427(+) |
| OCIAD1 | KIT | 4 | 4 | 1 | 1 | 48859382 | 55589750 | OCIAD1::chr4:48859332-48859382(+) | KIT::chr4:55589749-55589799(+) |
| OLFM2 | CRTC1 | 19 | 19 | -1 | 1 | 10046980 | 18853720 | OLFM2::chr19:10046979-10047029(-) | CRTC1::chr19:18853719-18853769(+) |
| OXR1 | MET | 8 | 7 | 1 | 1 | 107726213 | 116411552 | OXR1::chr8:107726163-107726213(+) | MET::chr7:116411551-116411601(+) |
| PAPD7 | RAF1 | 5 | 3 | 1 | -1 | 6751310 | 12641307 | PAPD7::chr5:6751260-6751310(+) | RAF1::chr3:12641257-12641307(-) |
| PARD3 | ZNF521 | 10 | 18 | -1 | -1 | 34805907 | 22672045 | PARD3::chr10:34805906-34805956(-) | ZNF521::chr18:22671995-22672045(-) |
| PAX8 | NFE2L2 | 2 | 2 | -1 | -1 | 113999128 | 178097311 | PAX8::chr2:113999127-113999177(-) | NFE2L2::chr2:178097261-178097311(-) |
| PBX1 | COPG2 | 1 | 7 | 1 | -1 | 164532548 | 130148500 | PBX1::chr1:164532498-164532548(+) | COPG2::chr7:130148450-130148500(-) |
| PBX1 | CTNNA3 | 1 | 10 | 1 | -1 | 164781386 | 68535282 | PBX1::chr1:164781336-164781386(+) | CTNNA3::chr10:68535232-68535282(-) |
| PBX1 | DHX40 | 1 | 17 | 1 | 1 | 164532548 | 57679883 | PBX1::chr1:164532498-164532548(+) | DHX40::chr17:57679882-57679932(+) |
| PBX1 | DISC1 | 1 | 1 | 1 | 1 | 164532548 | 232094574 | PBX1::chr1:164532498-164532548(+) | DISC1::chr1:232094573-232094623(+) |
| PBX1 | ILDR2 | 1 | 1 | 1 | -1 | 164532548 | 166927338 | PBX1::chr1:164532498-164532548(+) | ILDR2::chr1:166927288-166927338(-) |
| PBX1 | KIAA1045 | 1 | 9 | 1 | 1 | 164532548 | 34971292 | PBX1::chr1:164532498-164532548(+) | KIAA1045::chr9:34971291-34971341(+) |
| PBX1 | LRRC52 | 1 | 1 | 1 | 1 | 164532548 | 165532742 | PBX1::chr1:164532498-164532548(+) | LRRC52::chr1:165532741-165532791(+) |
| PBX1 | RFWD2 | 1 | 1 | 1 | -1 | 164532548 | 175958615 | PBX1::chr1:164532498-164532548(+) | RFWD2::chr1:175958565-175958615(-) |
| PBX1 | RHBG | 1 | 1 | 1 | 1 | 164781386 | 156354318 | PBX1::chr1:164781336-164781386(+) | RHBG::chr1:156354317-156354367(+) |
| PCDHGC3 | QKI | 5 | 6 | 1 | 1 | 140858113 | 163956014 | PCDHGC3::chr5:140858063-140858113(+) | QKI::chr6:163956013-163956063(+) |
| PDCD6 | TERT | 5 | 5 | 1 | -1 | 272887 | 1282739 | PDCD6::chr5:272837-272887(+) | TERT::chr5:1282689-1282739(-) |
| PDCD7 | TAF15 | 15 | 17 | -1 | 1 | 65421370 | 34144720 | PDCD7::chr15:65421369-65421419(-) | TAF15::chr17:34144719-34144769(+) |
| PDGFB | DGCR2 | 22 | 22 | -1 | -1 | 39626089 | 19077003 | PDGFB::chr22:39626088-39626138(-) | DGCR2::chr22:19076953-19077003(-) |
| PDGFRB | FAR1 | 5 | 11 | -1 | 1 | 149493405 | 13744912 | PDGFRB::chr5:149493404-149493454(-) | FAR1::chr11:13744911-13744961(+) |
| PEX1 | CDK6 | 7 | 7 | -1 | -1 | 92157621 | 92355107 | PEX1::chr7:92157620-92157670(-) | CDK6::chr7:92355057-92355107(-) |
| PGAP3 | ERBB2 | 17 | 17 | -1 | 1 | 37840850 | 37880979 | PGAP3::chr17:37840849-37840899(-) | ERBB2::chr17:37880978-37881028(+) |

| | | | | | | | | | |
|---|---|---|---|---|---|---|---|---|---|
| PHACTR1 | BCL11A | 6 | 2 | 1 | -1 | 12750022 | 60679801 | PHACTR1::chr6:12749972-12750022(+) | BCL11A::chr2:60679751-60679801(-) |
| PIK3CB | BFSP2 | 3 | 3 | -1 | 1 | 138452203 | 133185672 | PIK3CB::chr3:138452202-138452252(-) | BFSP2::chr3:133185671-133185721(+) |
| PIK3CB | COPB2 | 3 | 3 | -1 | -1 | 138478015 | 139102277 | PIK3CB::chr3:138478014-138478064(-) | COPB2::chr3:139102227-139102277(-) |
| PIK3CB | ESYT3 | 3 | 3 | -1 | 1 | 138400809 | 138176410 | PIK3CB::chr3:138400808-138400858(-) | ESYT3::chr3:138176409-138176459(+) |
| PIK3CB | GRM7 | 3 | 3 | -1 | 1 | 138474596 | 7620109 | PIK3CB::chr3:138474595-138474645(-) | GRM7::chr3:7620108-7620158(+) |
| PIK3CB | HHLA2 | 3 | 3 | -1 | 1 | 138478015 | 108070635 | PIK3CB::chr3:138478014-138478064(-) | HHLA2::chr3:108070634-108070684(+) |
| PIK3CB | KCNAB1 | 3 | 3 | -1 | 1 | 138413628 | 156175242 | PIK3CB::chr3:138413627-138413677(-) | KCNAB1::chr3:156175241-156175291(+) |
| PIK3CB | PCCB | 3 | 3 | -1 | 1 | 138515305 | 136002679 | PIK3CB::chr3:138515304-138515354(-) | PCCB::chr3:136002678-136002728(+) |
| PIK3CB | RASA2 | 3 | 3 | -1 | 1 | 138383878 | 141235170 | PIK3CB::chr3:138383877-138383927(-) | RASA2::chr3:141235169-141235219(+) |
| PIK3CB | SEMA5B | 3 | 3 | -1 | -1 | 138452203 | 122667556 | PIK3CB::chr3:138452202-138452252(-) | SEMA5B::chr3:122667506-122667556(-) |
| PLCG1 | EYA2 | 20 | 20 | 1 | 1 | 39798909 | 45797787 | PLCG1::chr20:39798859-39798909(+) | EYA2::chr20:45797786-45797836(+) |
| PLCG1 | TAF4 | 20 | 20 | 1 | -1 | 39766498 | 60584230 | PLCG1::chr20:39766448-39766498(+) | TAF4::chr20:60584180-60584230(-) |
| PMEPA1 | ETV4 | 20 | 17 | -1 | -1 | 56284530 | 41613847 | PMEPA1::chr20:56284529-56284579(-) | ETV4::chr17:41613797-41613847(-) |
| POLB | IKBKB | 8 | 8 | 1 | 1 | 42202537 | 42166419 | POLB::chr8:42202487-42202537(+) | IKBKB::chr8:42166418-42166468(+) |
| POLR1A | CTNNA2 | 2 | 2 | -1 | 1 | 86265823 | 80831199 | POLR1A::chr2:86265822-86265872(-) | CTNNA2::chr2:80831198-80831248(+) |
| POLR3H | CTNND2 | 22 | 5 | -1 | -1 | 41936703 | 11732384 | POLR3H::chr22:41936702-41936752(-) | CTNND2::chr5:11732334-11732384(-) |
| POU2AF1 | BCO2 | 11 | 11 | -1 | 1 | 111228170 | 112084447 | POU2AF1::chr11:111228169-111228219(-) | BCO2::chr11:112084446-112084496(+) |
| POU2AF1 | BTG4 | 11 | 11 | -1 | -1 | 111249887 | 111369527 | POU2AF1::chr11:111249886-111249936(-) | BTG4::chr11:111369477-111369527(-) |
| PPM1D | C17orf64 | 17 | 17 | 1 | 1 | 58678247 | 58506723 | PPM1D::chr17:58678197-58678247(+) | C17orf64::chr17:58506722-58506772(+) |
| PPM1D | NUP85 | 17 | 17 | 1 | 1 | 58725443 | 73221198 | PPM1D::chr17:58725393-58725443(+) | NUP85::chr17:73221197-73221247(+) |
| PRDM16 | NARS2 | 1 | 11 | 1 | -1 | 2985860 | 78282489 | PRDM16::chr1:2985810-2985860(+) | NARS2::chr11:78282439-78282489(-) |
| PRR11 | SUZ12 | 17 | 17 | 1 | 1 | 57275150 | 30315339 | PRR11::chr17:57275100-57275150(+) | SUZ12::chr17:30315338-30315388(+) |
| PSIP1 | TBL1X | 9 | X | -1 | 1 | 15486004 | 9621585 | PSIP1::chr9:15486003-15486053(-) | TBL1X::chrX:9621584-9621634(+) |
| PTPN11 | CUX2 | 12 | 12 | 1 | 1 | 112910844 | 111779584 | PTPN11::chr12:112910794-112910844(+) | CUX2::chr12:111779583-111779633(+) |
| PTPN11 | EPYC | 12 | 12 | 1 | -1 | 112893867 | 91372039 | PTPN11::chr12:112893817-112893867(+) | EPYC::chr12:91371989-91372039(-) |
| PTPN11 | PLAGL1 | 12 | 6 | 1 | -1 | 112856929 | 144290115 | PTPN11::chr12:112856879-112856929(+) | PLAGL1::chr6:144290065-144290115(-) |
| RAF1 | C9orf153 | 3 | 9 | -1 | -1 | 12632297 | 88844544 | RAF1::chr3:12632296-12632346(-) | C9orf153::chr9:88844494-88844544(-) |
| RAF1 | GXYLT2 | 3 | 3 | -1 | 1 | 12705312 | 72957518 | RAF1::chr3:12705311-12705361(-) | GXYLT2::chr3:72957517-72957567(+) |
| RAF1 | IQSEC1 | 3 | 3 | -1 | -1 | 12705312 | 12983365 | RAF1::chr3:12705311-12705361(-) | IQSEC1::chr3:12983315-12983365(-) |
| RAF1 | NXPH1 | 3 | 7 | -1 | 1 | 12705312 | 8790638 | RAF1::chr3:12705311-12705361(-) | NXPH1::chr7:8790637-8790687(+) |
| RAF1 | PHC3 | 3 | 3 | -1 | -1 | 12641190 | 169815172 | RAF1::chr3:12641189-12641239(-) | PHC3::chr3:169815122-169815172(-) |
| RAF1 | TMEM40 | 3 | 3 | -1 | -1 | 12705312 | 12784006 | RAF1::chr3:12705311-12705361(-) | TMEM40::chr3:12783956-12784006(-) |
| RAF1 | TRAK1 | 3 | 3 | -1 | 1 | 12645635 | 42236296 | RAF1::chr3:12645634-12645684(-) | TRAK1::chr3:42236295-42236345(+) |
| RAP1GDS1 | ASTN2 | 4 | 9 | 1 | -1 | 99342545 | 119414072 | RAP1GDS1::chr4:99342495-99342545(+) | ASTN2::chr9:119414022-119414072(-) |
| RAP1GDS1 | GSTCD | 4 | 4 | 1 | 1 | 99273752 | 106766598 | RAP1GDS1::chr4:99273702-99273752(+) | GSTCD::chr4:106766597-106766647(+) |
| RAP1GDS1 | SCD5 | 4 | 4 | 1 | -1 | 99325753 | 83557976 | RAP1GDS1::chr4:99325703-99325753(+) | SCD5::chr4:83557926-83557976(-) |
| RARA | C17orf108 | 17 | 17 | 1 | -1 | 38487648 | 26209738 | RARA::chr17:38487598-38487648(+) | C17orf108::chr17:26209688-26209738(-) |
| RARA | CA10 | 17 | 17 | 1 | -1 | 38474700 | 50149753 | RARA::chr17:38474650-38474700(+) | CA10::chr17:50149703-50149753(-) |
| RARA | CDC6 | 17 | 17 | 1 | 1 | 38465538 | 38457080 | RARA::chr17:38465488-38465538(+) | CDC6::chr17:38457079-38457129(+) |
| RARA | HOXB3 | 17 | 17 | 1 | -1 | 38508759 | 46632980 | RARA::chr17:38508709-38508759(+) | HOXB3::chr17:46632930-46632980(-) |
| RARA | LTBP1 | 17 | 2 | 1 | 1 | 38474700 | 33447147 | RARA::chr17:38474650-38474700(+) | LTBP1::chr2:33447146-33447196(+) |
| RARA | MSL1 | 17 | 17 | 1 | 1 | 38487648 | 38282436 | RARA::chr17:38487598-38487648(+) | MSL1::chr17:38282435-38282485(+) |
| RARA | PCTP | 17 | 17 | 1 | 1 | 38474700 | 53848467 | RARA::chr17:38474650-38474700(+) | PCTP::chr17:53848466-53848516(+) |

| | | | | | | | | | |
|---|---|---|---|---|---|---|---|---|---|
| RARA | PGAP3 | 17 | 17 | 1 | -1 | 38474700 | 37830932 | RARA::chr17:38474650-38474700(+) | PGAP3::chr17:37830882-37830932(-) |
| RARA | PRR11 | 17 | 17 | 1 | 1 | 38474700 | 57247109 | RARA::chr17:38474650-38474700(+) | PRR11::chr17:57247108-57247158(+) |
| RARA | PSMD3 | 17 | 17 | 1 | 1 | 38487648 | 38151207 | RARA::chr17:38487598-38487648(+) | PSMD3::chr17:38151206-38151256(+) |
| RARA | SKAP1 | 17 | 17 | 1 | -1 | 38487648 | 46214699 | RARA::chr17:38487598-38487648(+) | SKAP1::chr17:46214649-46214699(-) |
| RARA | SLC9A3R1 | 17 | 17 | 1 | 1 | 38508759 | 72758151 | RARA::chr17:38508709-38508759(+) | SLC9A3R1::chr17:72758150-72758200(+) |
| RARA | STAC2 | 17 | 17 | 1 | -1 | 38474700 | 37374426 | RARA::chr17:38474650-38474700(+) | STAC2::chr17:37374376-37374426(-) |
| RBPMS | NTRK3 | 8 | 15 | 1 | -1 | 30361953 | 88576276 | RBPMS::chr8:30361903-30361953(+) | NTRK3::chr15:88576226-88576276(-) |
| RCAN1 | RUNX1 | 21 | 21 | -1 | -1 | 35987059 | 36253010 | RCAN1::chr21:35987058-35987108(-) | RUNX1::chr21:36252960-36253010(-) |
| RCOR1 | KIT | 14 | 4 | 1 | 1 | 103148315 | 55561678 | RCOR1::chr14:103148265-103148315(+) | KIT::chr4:55561677-55561727(+) |
| REL | MAN1A2 | 2 | 1 | 1 | 1 | 61147777 | 118035769 | REL::chr2:61147727-61147777(+) | MAN1A2::chr1:118035768-118035818(+) |
| REL | MVP | 2 | 16 | 1 | 1 | 61144152 | 29847025 | REL::chr2:61144102-61144152(+) | MVP::chr16:29847024-29847074(+) |
| RET | ANK3 | 10 | 10 | 1 | -1 | 43604678 | 61789584 | RET::chr10:43604628-43604678(+) | ANK3::chr10:61789534-61789584(-) |
| REV1 | AFF3 | 2 | 2 | -1 | -1 | 100065798 | 100185377 | REV1::chr2:100065797-100065847(-) | AFF3::chr2:100185327-100185377(-) |
| SASH1 | MLLT4 | 6 | 6 | 1 | 1 | 148841029 | 168343809 | SASH1::chr6:148840979-148841029(+) | MLLT4::chr6:168343808-168343858(+) |
| SBF2 | MDM2 | 11 | 12 | -1 | 1 | 9983504 | 69210592 | SBF2::chr11:9983503-9983553(-) | MDM2::chr12:69210591-69210641(+) |
| SCN8A | ATF1 | 12 | 12 | 1 | 1 | 52184286 | 51189692 | SCN8A::chr12:52184236-52184286(+) | ATF1::chr12:51189691-51189741(+) |
| SETBP1 | ARNTL | 18 | 11 | 1 | 1 | 42449248 | 13407236 | SETBP1::chr18:42449198-42449248(+) | ARNTL::chr11:13407235-13407285(+) |
| SETBP1 | NFYC | 18 | 1 | 1 | 1 | 42449248 | 41204508 | SETBP1::chr18:42449198-42449248(+) | NFYC::chr1:41204507-41204557(+) |
| SETDB1 | PKP4 | 1 | 2 | 1 | 1 | 150915164 | 159477502 | SETDB1::chr1:150915114-150915164(+) | PKP4::chr2:159477501-159477551(+) |
| SETDB1 | PLEKHO1 | 1 | 1 | 1 | 1 | 150902594 | 150128260 | SETDB1::chr1:150902544-150902594(+) | PLEKHO1::chr1:150128259-150128309(+) |
| SF3B1 | HSPD1 | 2 | 2 | -1 | -1 | 198274494 | 198355020 | SF3B1::chr2:198274493-198274543(-) | HSPD1::chr2:198354970-198355020(-) |
| SH3GL1 | DPP9 | 19 | 19 | -1 | -1 | 4400321 | 4679958 | SH3GL1::chr19:4400320-4400370(-) | DPP9::chr19:4679908-4679958(-) |
| SH3GL1 | FN1 | 19 | 2 | -1 | -1 | 4361313 | 216300790 | SH3GL1::chr19:4361312-4361362(-) | FN1::chr2:216300740-216300790(-) |
| SH3GL1 | MUM1 | 19 | 19 | -1 | 1 | 4400321 | 1356323 | SH3GL1::chr19:4400320-4400370(-) | MUM1::chr19:1356322-1356372(+) |
| SH3GL1 | TAX1BP1 | 19 | 7 | -1 | 1 | 4400321 | 27867357 | SH3GL1::chr19:4400320-4400370(-) | TAX1BP1::chr7:27867356-27867406(+) |
| SHB | PAX5 | 9 | 9 | -1 | -1 | 38016008 | 37020798 | SHB::chr9:38016007-38016057(-) | PAX5::chr9:37020748-37020798(-) |
| SKI | CA6 | 1 | 1 | 1 | 1 | 2161174 | 9027718 | SKI::chr1:2161124-2161174(+) | CA6::chr1:9027717-9027767(+) |
| SKI | CAPZB | 1 | 1 | 1 | -1 | 2161174 | 19746244 | SKI::chr1:2161124-2161174(+) | CAPZB::chr1:19746194-19746244(-) |
| SKI | PRKCZ | 1 | 1 | 1 | 1 | 2161174 | 2087434 | SKI::chr1:2161124-2161174(+) | PRKCZ::chr1:2087433-2087483(+) |
| SKI | SSU72 | 1 | 1 | 1 | -1 | 2161174 | 1500296 | SKI::chr1:2161124-2161174(+) | SSU72::chr1:1500246-1500296(-) |
| SLC12A7 | TERT | 5 | 5 | -1 | -1 | 1074682 | 1266650 | SLC12A7::chr5:1074681-1074731(-) | TERT::chr5:1266600-1266650(-) |
| SLC20A2 | HIP1 | 8 | 7 | -1 | -1 | 42286276 | 75172293 | SLC20A2::chr8:42286275-42286325(-) | HIP1::chr7:75172243-75172293(-) |
| SMEK2 | ALK | 2 | 2 | -1 | -1 | 55804451 | 29940563 | SMEK2::chr2:55804450-55804500(-) | ALK::chr2:29940513-29940563(-) |
| SMO | TSPAN33 | 7 | 7 | 1 | 1 | 128829323 | 128801520 | SMO::chr7:128829273-128829323(+) | TSPAN33::chr7:128801519-128801569(+) |
| SND1 | AKAP9 | 7 | 7 | 1 | 1 | 127447627 | 91734992 | SND1::chr7:127447577-127447627(+) | AKAP9::chr7:91734991-91735041(+) |
| SND1 | GRM8 | 7 | 7 | 1 | -1 | 127361454 | 126544737 | SND1::chr7:127361404-127361454(+) | GRM8::chr7:126544687-126544737(-) |
| SND1 | IMMP2L | 7 | 7 | 1 | -1 | 127569382 | 110303777 | SND1::chr7:127569332-127569382(+) | IMMP2L::chr7:110303727-110303777(-) |
| SNX29 | TNFRSF17 | 16 | 16 | 1 | 1 | 12172772 | 12060052 | SNX29::chr16:12172722-12172772(+) | TNFRSF17::chr16:12060051-12060101(+) |
| SNX29 | TNFRSF17 | 16 | 16 | 1 | 1 | 12155503 | 12060052 | SNX29::chr16:12155453-12155503(+) | TNFRSF17::chr16:12060051-12060101(+) |
| SQSTM1 | NTRK1 | 5 | 1 | 1 | 1 | 179252226 | 156844363 | SQSTM1::chr5:179252176-179252226(+) | NTRK1::chr1:156844362-156844412(+) |
| SSBP2 | NTRK1 | 5 | 1 | -1 | 1 | 80742687 | 156845312 | SSBP2::chr5:80742686-80742736(-) | NTRK1::chr1:156845311-156845361(+) |
| ST7 | MET | 7 | 7 | 1 | 1 | 116830990 | 116371722 | ST7::chr7:116830940-116830990(+) | MET::chr7:116371721-116371771(+) |
| STAT3 | NOS2 | 17 | 17 | -1 | -1 | 40540297 | 26116714 | STAT3::chr17:40540296-40540346(-) | NOS2::chr17:26116664-26116714(-) |

| | | | | | | | | | |
|---|---|---|---|---|---|---|---|---|---|
| STX17 | NR4A3 | 9 | 9 | 1 | 1 | 102691125 | 102606931 | STX17::chr9:102691075-102691125(+) | NR4A3::chr9:102606930-102606980(+) |
| SYK | AUH | 9 | 9 | 1 | -1 | 93564175 | 94060358 | SYK::chr9:93564125-93564175(+) | AUH::chr9:94060308-94060358(-) |
| TAF15 | AP2B1 | 17 | 17 | 1 | 1 | 34151202 | 33966614 | TAF15::chr17:34151152-34151202(+) | AP2B1::chr17:33966613-33966663(+) |
| TAX1BP1 | BRAF | 7 | 7 | 1 | -1 | 27827222 | 140481493 | TAX1BP1::chr7:27827172-27827222(+) | BRAF::chr7:140481443-140481493(-) |
| TCF7L2 | CHD7 | 10 | 8 | 1 | 1 | 114799885 | 61693559 | TCF7L2::chr10:114799835-114799885(+) | CHD7::chr8:61693558-61693608(+) |
| TCF7L2 | TACC2 | 10 | 10 | 1 | 1 | 114799885 | 123954555 | TCF7L2::chr10:114799835-114799885(+) | TACC2::chr10:123954554-123954604(+) |
| TCF7L2 | VTI1A | 10 | 10 | 1 | 1 | 114724383 | 114575049 | TCF7L2::chr10:114724333-114724383(+) | VTI1A::chr10:114575048-114575098(+) |
| TCF7L2 | VTI1A | 10 | 10 | 1 | 1 | 114905856 | 114575049 | TCF7L2::chr10:114905806-114905856(+) | VTI1A::chr10:114575048-114575098(+) |
| TCF7L2 | VTI1A | 10 | 10 | 1 | 1 | 114799885 | 114575049 | TCF7L2::chr10:114799835-114799885(+) | VTI1A::chr10:114575048-114575098(+) |
| TCF7L2 | VTI1A | 10 | 10 | 1 | 1 | 114799885 | 114427977 | TCF7L2::chr10:114799835-114799885(+) | VTI1A::chr10:114427976-114428026(+) |
| TEC | NCAM2 | 4 | 21 | -1 | 1 | 48271769 | 22746183 | TEC::chr4:48271768-48271818(-) | NCAM2::chr21:22746182-22746232(+) |
| TEC | PASD1 | 4 | X | -1 | 1 | 48271769 | 150789402 | TEC::chr4:48271768-48271818(-) | PASD1::chrX:150789401-150789451(+) |
| TFE3 | DVL2 | X | 17 | -1 | -1 | 48897982 | 7132766 | TFE3::chrX:48897981-48898031(-) | DVL2::chr17:7132716-7132766(-) |
| TFE3 | MED15 | X | 22 | -1 | 1 | 48895535 | 20936898 | TFE3::chrX:48895534-48895584(-) | MED15::chr22:20936897-20936947(+) |
| TFE3 | PRCC | X | 1 | -1 | 1 | 48896632 | 156752074 | TFE3::chrX:48896631-48896681(-) | PRCC::chr1:156752073-156752123(+) |
| TFE3 | PRCC | X | 1 | -1 | 1 | 48896632 | 156756400 | TFE3::chrX:48896631-48896681(-) | PRCC::chr1:156756399-156756449(+) |
| TFEB | COL21A1 | 6 | 6 | -1 | -1 | 41703088 | 56047454 | TFEB::chr6:41703087-41703137(-) | COL21A1::chr6:56047404-56047454(-) |
| TFEB | EPS15L1 | 6 | 19 | -1 | -1 | 41703884 | 16488065 | TFEB::chr6:41703883-41703933(-) | EPS15L1::chr19:16488015-16488065(-) |
| TFEB | MDFI | 6 | 6 | -1 | 1 | 41703088 | 41617357 | TFEB::chr6:41703087-41703137(-) | MDFI::chr6:41617356-41617406(+) |
| TFEB | PGC | 6 | 6 | -1 | -1 | 41703088 | 41711127 | TFEB::chr6:41703087-41703137(-) | PGC::chr6:41711077-41711127(-) |
| TFEB | PGC | 6 | 6 | -1 | -1 | 41703088 | 41712546 | TFEB::chr6:41703087-41703137(-) | PGC::chr6:41712496-41712546(-) |
| TFEB | SUPT3H | 6 | 6 | -1 | -1 | 41657469 | 44797594 | TFEB::chr6:41657468-41657518(-) | SUPT3H::chr6:44797544-44797594(-) |
| TFG | MET | 3 | 7 | 1 | 1 | 100451516 | 116414935 | TFG::chr3:100451466-100451516(+) | MET::chr7:116414934-116414984(+) |
| TFG | NTRK1 | 3 | 1 | 1 | 1 | 100455560 | 156844363 | TFG::chr3:100455510-100455560(+) | NTRK1::chr1:156844362-156844412(+) |
| TMEM111 | RHOA | 3 | 3 | -1 | -1 | 10012266 | 49400059 | TMEM111::chr3:10012265-10012315(-) | RHOA::chr3:49400009-49400059(-) |
| TMEM161B | CTNND2 | 5 | 5 | -1 | -1 | 87536630 | 11117679 | TMEM161B::chr5:87536629-87536679(-) | CTNND2::chr5:11117629-11117679(-) |
| TMEM9B | PSIP1 | 11 | 9 | -1 | -1 | 8983639 | 15490122 | TMEM9B::chr11:8983638-8983688(-) | PSIP1::chr9:15490072-15490122(-) |
| TMPRSS2 | ETV4 | 21 | 17 | -1 | -1 | 42866283 | 41607549 | TMPRSS2::chr21:42866282-42866332(-) | ETV4::chr17:41607499-41607549(-) |
| TNC | CDH3 | 9 | 16 | -1 | 1 | 117782839 | 68678151 | TNC::chr9:117782838-117782888(-) | CDH3::chr16:68678150-68678200(+) |
| TNFRSF1A | CHD4 | 12 | 12 | -1 | -1 | 6450942 | 6682435 | TNFRSF1A::chr12:6450941-6450991(-) | CHD4::chr12:6682385-6682435(-) |
| TNS4 | CCR7 | 17 | 17 | -1 | -1 | 38640736 | 38715194 | TNS4::chr17:38640735-38640785(-) | CCR7::chr17:38715144-38715194(-) |
| TPM1 | ALK | 15 | 2 | 1 | -1 | 63354844 | 29446394 | TPM1::chr15:63354794-63354844(+) | ALK::chr2:29446344-29446394(-) |
| TPRA1 | LPP | 3 | 3 | -1 | 1 | 127298583 | 188583988 | TPRA1::chr3:127298582-127298632(-) | LPP::chr3:188583987-188584037(+) |
| TRAK1 | RAF1 | 3 | 3 | 1 | -1 | 42235390 | 12641914 | TRAK1::chr3:42235340-42235390(+) | RAF1::chr3:12641864-12641914(-) |
| TRIM27 | PTPRR | 6 | 12 | -1 | -1 | 28879473 | 71050597 | TRIM27::chr6:28879472-28879522(-) | PTPRR::chr12:71050547-71050597(-) |
| TRRAP | ANKIB1 | 7 | 7 | 1 | 1 | 98582667 | 91924203 | TRRAP::chr7:98582617-98582667(+) | ANKIB1::chr7:91924202-91924252(+) |
| TRRAP | PRKAR2B | 7 | 7 | 1 | 1 | 98515302 | 106762397 | TRRAP::chr7:98515252-98515302(+) | PRKAR2B::chr7:106762396-106762446(+) |
| TSFM | HMGA2 | 12 | 12 | 1 | 1 | 58180074 | 66345163 | TSFM::chr12:58180024-58180074(+) | HMGA2::chr12:66345162-66345212(+) |
| TTC13 | JAK2 | 1 | 9 | -1 | 1 | 231075132 | 5064883 | TTC13::chr1:231075131-231075181(-) | JAK2::chr9:5064882-5064932(+) |
| TTC27 | BIRC6 | 2 | 2 | 1 | 1 | 32961883 | 32828073 | TTC27::chr2:32961833-32961883(+) | BIRC6::chr2:32828072-32828122(+) |
| UBE2D2 | SND1 | 5 | 7 | 1 | 1 | 138941400 | 127714554 | UBE2D2::chr5:138941350-138941400(+) | SND1::chr7:127714553-127714603(+) |
| UBR5 | ODF1 | 8 | 8 | -1 | 1 | 103424401 | 103572680 | UBR5::chr8:103424400-103424450(-) | ODF1::chr8:103572679-103572729(+) |
| UBR5 | RIMS4 | 8 | 20 | -1 | -1 | 103305833 | 43400057 | UBR5::chr8:103305832-103305882(-) | RIMS4::chr20:43400007-43400057(-) |

| | | | | | | | | | |
|---|---|---|---|---|---|---|---|---|---|
| UBR5 | TRAF3 | 8 | 14 | -1 | 1 | 103269860 | 103352526 | UBR5::chr8:103269859-103269909(-) | TRAF3::chr14:103352525-103352575(+) |
| UCK2 | PBX1 | 1 | 1 | 1 | 1 | 165875206 | 164761731 | UCK2::chr1:165875156-165875206(+) | PBX1::chr1:164761730-164761780(+) |
| UCK2 | PBX1 | 1 | 1 | 1 | 1 | 165860559 | 164532475 | UCK2::chr1:165860509-165860559(+) | PBX1::chr1:164532474-164532524(+) |
| UPF1 | CRTC1 | 19 | 19 | 1 | 1 | 18963880 | 18864310 | UPF1::chr19:18963830-18963880(+) | CRTC1::chr19:18864309-18864359(+) |
| USP13 | PIK3CA | 3 | 3 | 1 | 1 | 179418917 | 178941869 | USP13::chr3:179418867-179418917(+) | PIK3CA::chr3:178941868-178941918(+) |
| USP8 | ADAMTSL3 | 15 | 15 | 1 | 1 | 50741682 | 84700087 | USP8::chr15:50741632-50741682(+) | ADAMTSL3::chr15:84700086-84700136(+) |
| USP8 | BDP1 | 15 | 5 | 1 | 1 | 50769696 | 70786811 | USP8::chr15:50769646-50769696(+) | BDP1::chr5:70786810-70786860(+) |
| USP8 | TSPAN3 | 15 | 15 | 1 | -1 | 50763992 | 77348597 | USP8::chr15:50763942-50763992(+) | TSPAN3::chr15:77348547-77348597(-) |
| UTRN | HMGA2 | 6 | 12 | 1 | 1 | 145021379 | 66232299 | UTRN::chr6:145021329-145021379(+) | HMGA2::chr12:66232298-66232348(+) |
| VAV2 | BRD3 | 9 | 9 | -1 | -1 | 136857197 | 136910543 | VAV2::chr9:136857196-136857246(-) | BRD3::chr9:136910493-136910543(-) |
| VCL | FGFR2 | 10 | 10 | 1 | -1 | 75758133 | 123239535 | VCL::chr10:75758083-75758133(+) | FGFR2::chr10:123239485-123239535(-) |
| VPS18 | NTRK3 | 15 | 15 | 1 | -1 | 41193212 | 88423659 | VPS18::chr15:41193162-41193212(+) | NTRK3::chr15:88423609-88423659(-) |
| WHSC1 | FAM53A | 4 | 4 | 1 | -1 | 1957915 | 1619838 | WHSC1::chr4:1957865-1957915(+) | FAM53A::chr4:1619788-1619838(-) |
| WHSC1 | GSDMB | 4 | 17 | 1 | -1 | 1957915 | 38068750 | WHSC1::chr4:1957865-1957915(+) | GSDMB::chr17:38068700-38068750(-) |
| WHSC1 | NUP210L | 4 | 1 | 1 | -1 | 1902978 | 153984846 | WHSC1::chr4:1902928-1902978(+) | NUP210L::chr1:153984796-153984846(-) |
| WHSC1 | PHTF2 | 4 | 7 | 1 | 1 | 1902978 | 77522925 | WHSC1::chr4:1902928-1902978(+) | PHTF2::chr7:77522924-77522974(+) |
| WHSC1 | PIBF1 | 4 | 13 | 1 | 1 | 1873269 | 73539412 | WHSC1::chr4:1873219-1873269(+) | PIBF1::chr13:73539411-73539461(+) |
| WHSC1L1 | ADAM18 | 8 | 8 | -1 | 1 | 38137058 | 39587417 | WHSC1L1::chr8:38137057-38137107(-) | ADAM18::chr8:39587416-39587466(+) |
| WHSC1L1 | C15orf55 | 8 | 15 | -1 | 1 | 38184248 | 34640170 | WHSC1L1::chr8:38184247-38184297(-) | C15orf55::chr15:34640169-34640219(+) |
| WHSC1L1 | DDHD2 | 8 | 8 | -1 | 1 | 38178590 | 38095607 | WHSC1L1::chr8:38178589-38178639(-) | DDHD2::chr8:38095606-38095656(+) |
| WHSC1L1 | DDHD2 | 8 | 8 | -1 | 1 | 38172165 | 38090505 | WHSC1L1::chr8:38172164-38172214(-) | DDHD2::chr8:38090504-38090554(+) |
| WHSC1L1 | DUSP26 | 8 | 8 | -1 | -1 | 38162105 | 33455109 | WHSC1L1::chr8:38162104-38162154(-) | DUSP26::chr8:33455059-33455109(-) |
| WHSC1L1 | GAB2 | 8 | 11 | -1 | -1 | 38239317 | 77991947 | WHSC1L1::chr8:38239316-38239366(-) | GAB2::chr11:77991897-77991947(-) |
| WHSC1L1 | LETM2 | 8 | 8 | -1 | 1 | 38239317 | 38245468 | WHSC1L1::chr8:38239316-38239366(-) | LETM2::chr8:38245467-38245517(+) |
| WHSC1L1 | MUCL1 | 8 | 12 | -1 | 1 | 38239317 | 55248900 | WHSC1L1::chr8:38239316-38239366(-) | MUCL1::chr12:55248899-55248949(+) |
| WHSC1L1 | NEGR1 | 8 | 1 | -1 | -1 | 38239317 | 72241980 | WHSC1L1::chr8:38239316-38239366(-) | NEGR1::chr1:72241930-72241980(-) |
| WHSC1L1 | PLEKHA2 | 8 | 8 | -1 | 1 | 38146005 | 38826939 | WHSC1L1::chr8:38146004-38146054(-) | PLEKHA2::chr8:38826938-38826988(+) |
| WHSC1L1 | PSD3 | 8 | 8 | -1 | -1 | 38146005 | 18513195 | WHSC1L1::chr8:38146004-38146054(-) | PSD3::chr8:18513145-18513195(-) |
| WHSC1L1 | PURG | 8 | 8 | -1 | -1 | 38239317 | 30854213 | WHSC1L1::chr8:38239316-38239366(-) | PURG::chr8:30854163-30854213(-) |
| WHSC1L1 | SNX20 | 8 | 16 | -1 | -1 | 38239317 | 50711446 | WHSC1L1::chr8:38239316-38239366(-) | SNX20::chr16:50711396-50711446(-) |
| WHSC1L1 | TACC1 | 8 | 8 | -1 | 1 | 38176413 | 38705473 | WHSC1L1::chr8:38176412-38176462(-) | TACC1::chr8:38705472-38705522(+) |
| WHSC1L1 | ZMAT4 | 8 | 8 | -1 | -1 | 38172165 | 40389757 | WHSC1L1::chr8:38172164-38172214(-) | ZMAT4::chr8:40389707-40389757(-) |
| WHSC1L1 | ZNF721 | 8 | 4 | -1 | -1 | 38239317 | 438221 | WHSC1L1::chr8:38239316-38239366(-) | ZNF721::chr4:438171-438221(-) |
| WRB | ERG | 21 | 21 | 1 | -1 | 40752412 | 39817544 | WRB::chr21:40752362-40752412(+) | ERG::chr21:39817494-39817544(-) |
| WWTR1 | ESRRG | 3 | 1 | -1 | -1 | 149290651 | 216741440 | WWTR1::chr3:149290650-149290700(-) | ESRRG::chr1:216741390-216741440(-) |
| XPO1 | C2orf74 | 2 | 2 | -1 | 1 | 61749746 | 61389977 | XPO1::chr2:61749745-61749795(-) | C2orf74::chr2:61389976-61390026(+) |
| XPO1 | HORMAD1 | 2 | 1 | -1 | -1 | 61725808 | 150692014 | XPO1::chr2:61725807-61725857(-) | HORMAD1::chr1:150691964-150692014(-) |
| YAP1 | MAML2 | 11 | 11 | 1 | -1 | 102076805 | 95826681 | YAP1::chr11:102076755-102076805(+) | MAML2::chr11:95826631-95826681(-) |
| ZC3HAV1 | BRAF | 7 | 7 | -1 | -1 | 138758602 | 140481493 | ZC3HAV1::chr7:138758601-138758651(-) | BRAF::chr7:140481443-140481493(-) |
| ZDHHC6 | TCF7L2 | 10 | 10 | -1 | 1 | 114194055 | 114900943 | ZDHHC6::chr10:114194054-114194104(-) | TCF7L2::chr10:114900942-114900992(+) |
| ZEB1 | SYT1 | 10 | 12 | 1 | 1 | 31608221 | 79258778 | ZEB1::chr10:31608171-31608221(+) | SYT1::chr12:79258777-79258827(+) |
| ZEB1 | ZNF438 | 10 | 10 | 1 | -1 | 31608221 | 31165994 | ZEB1::chr10:31608171-31608221(+) | ZNF438::chr10:31165944-31165994(-) |
| ZFR | SND1 | 5 | 7 | -1 | 1 | 32444335 | 127527955 | ZFR::chr5:32444334-32444384(-) | SND1::chr7:127527954-127528004(+) |

| | | | | | | | | | |
|---|---|---|---|---|---|---|---|---|---|
| ZNF461 | AKT1 | 19 | 14 | -1 | -1 | 37147350 | 105246553 | ZNF461::chr19:37147349-37147399(-) | AKT1::chr14:105246503-105246553(-) |

| fusionSeq | source |
| --- | --- |
| TCAGCGCAGTTTGGGATACAGCAAATACTATTTCAGAAACAGCCTATGAG*GGAAATTTTGGTGAAGTATATAAGGGCACATTAAAGGATAAAACTTCTGT | UPMC |
| ACTTTGAGACCATCTTACTGGAAGTTCCTTTTAAGACTCACCAAGGGCAA*ATAAGAAGCCAACTCCAACAGGTGGAAGAGTACCATAACTCTGAGCAGGA | UPMC |
| TGGCAGGTGACTGCTTGGATGAGAAGCAGTGTAAGCAGTGTGCAAACAAG*GTGACTGGAAGCACCTGCTCAATGGCTGCTCATCAACAACAAACCAGAGG | UPMC |
| ACAGGTATCTGCTCCTCATGGCGCAGGAGCATCTGGAGTTCCGCCTGCCG*GTGGTATTTTTGAATATGTGGAATCTGGCCCAATGGGAGCTGAGGAACTT | UPMC |
| GGATGCCTCTGCTGTCAGCCTGTCAGAATCCAAGTCAAGTCAGGATTCCT*TGTTCTGGGAATGTCAGTGGAATCTGCTCCTGCTGTGGAGGAAGAGAAGG | UPMC |
| GCTGAAAGAGAACAGAACTTGCCTGTCACAGTTACTAGATATAATGAAAA*TACCTGGAGTAGAACAGAAAAATTATTATGTCTGTGTTCCCTTGGGACTC | UPMC |
| TCCCATCAGGACAAGGAGAAGGGCCTCCACTACCATGCCCTCTTCACAGG*TGTCATGGAGAAACTCCAGCTGGGCCCAGAGATTCTGCAGCGGGAAAATC | UPMC |
| ATGGACCTGTACAGCATCAACTACCTGCACTTTGGGGAGCCTAAGTCCTG*GACAGTAAGCAAGCCTGGATCTGAGAGAGATATCATCTTGCAAGGATGCC | UPMC |
| ACAAAGCAAATAAAGACAAAGCCAACCGATACTTTTCTCCAAATTTTAAG*ACACAGCAGGATGCCAATGCCTCTTCCCTCTTAGACATCTATAGCTTCTG | UPMC |
| ATGCCTTTCCTTATATTAAAACAAGGGTCAATGTCACTCATAAAGAAGAG*CTTGAATTTGGAAATGACTTTTGATGACCTAAAGATCCAGACTGTGAAGG | UPMC |
| GCCAAGGAAAAACTGGAGAGCTTCATCAAAAATAGTGATAGCTTTGATAA*ACTGCTCTCCAGAATGTTGAGATTGCCCGAGAAGTGACCCCAGCAAAAGA | UPMC |
| ACCCCATAAAGGAATCCTCATGGCCACAAGGATTTGGCCAACTCACCCAG*ATTGGCTGCAATGCCGTCAGCTGGGCCCCTGCTGTTGTACCTGGAAGCCT | UPMC |
| CCCGCAATGAGGAAGAAGGGAACAGTGAAGAAATAAAAGCTAAAGTTAAA*TACCTGTTTGAAAATGGTAGAATAGATAACATTTTTACTGAGCCCTATTC | UPMC |
| AGATGATCCTTCCTCCTACATGGAAGTTGTTCAGGCTGCCAATACTAGTG*TGGCTTTTCATGGCCTGCCACTGAAAATCAAGAAAGAACCCCACAGTCCA | UPMC |
| TCGGGGTGGTGCATCTAACACAACAGATGCCCAGGTTGATAGCATTGTGG*GTCCCTGGATGGCCGGTTGCAGGTGTCCCATCGGAAGGGGCTCCCTCATG | TCGA |
| CCAGGCAGGCTGCTACCGCCCCTCCGCTTCCAGACATTCTCAGCTGTCAG*GCCGCCTGTGAGACCTTCTGTGTGGGAGAGGATGCCAAGGCCGCCGCCCG | TCGA |
| GTGGCTTCATTTCTTCTGTGGCCTTCAGCACTGATAAGAATCTATTATTG*GGCAACATGCAGGTTCTGGTGACTTACGGTGGCGATCCCATCCCTAAAAG | TCGA |
| AAGCCTCTCAGCTTCGGAGAATGGAGTCTTCCATCAATCATATCTCACAG*ATGCTGTGTGCTGTGGACCCACCTGGGGTTCATGGAGTGGGCCACGGGGC | TCGA |
| GGAATCATCGAGGCATGGGGGTCCACACTGCAATGTTTTTGTGGAACATG*ATCTGTACTGCACCCTGGAGGTGGATTCCTTTGGGTATTTTGTGAATAAA | TCGA |
| GGAATCATCGAGGCATGGGGGTCCACACTGCAATGTTTTTGTGGAACATG*GAGCGGACAGGGGTGAAGATGGTCATGATCCAGGATGGCCCATTGCCCAC | TCGA |
| ACCTGCGTGGTAGAATTCCAGTTCATGCTGCCGTCCTCCCACCCAAACCG*AAGCGGCACCATGTACTACAGACGGCTCATCCCTCCCCTTTGTCAGTGTA | TCGA |
| TTAGCTGGGTCGGGAAGACATCCATGGAAGTGAAGATGCAAATGTTCCAG*GTAATTCAGAGGTCCGTGGGGCCAGCCAGCCTGAGCTTGCTCACCTTCAA | TCGA |
| TCTGGCTAGACCGAAGTGTACTAGCCAAGGAGTTGAAGTTTGTGACTTTG*GTTCCTAATGTGAGAGGCTAGACCCAGATCATGGAGGTGCTTCAGTGTGA | TCGA |
| CCAGGCCTGCGCGGGCCCGAGGCCGGAGGAACCCGGACTCCGGCGTAGCG*TTGGTGATGTATATTGGCCAAGTGGCCAAGGATGTCTTGAAGTGGCCCCG | TCGA |
| CATGGCtttatttgttttttttgttttgttttggtttttttttttttttt*TCCTATCATCACAAAACTCATGATGCTCCGGCTGCTCAGTTCCCTCCTCC | TCGA |
| CATGGCtttatttgttttttttgttttgttttggtttttttttttttttt*TTTTCTCTCTCTCTTTCACTGCAAGGCGGCGGCAGGAGAGGTTGTGGTGC | TCGA |
| CATGGCtttatttgttttttttgttttgttttggtttttttttttttttt*TTTTGAGAAGCTGAAGCAACTCCAAGGACACAGTTCACAGAAATTTGGTT | TCGA |
| CATGGCtttatttgttttttttgttttgttttggtttttttttttttttt*GCTGCCCGGACGCGGAGCGAGAGGCTGAGAGAGTCGGAGACACTATCCGC | TCGA |
| CCCCAGCAACACGTCATTGTGTAAGGAAAAATAAAAGTGCTGCCGTAACC*aaaaaaaaaaaaaaaaaaaaaaaaaaaaaaGTCCATTGCTGAGAGACGAG | TCGA |
| CCCCAGCAACACGTCATTGTGTAAGGAAAAATAAAAGTGCTGCCGTAACC*aaaaaaaaaaaaaaaaaaaagaaagccaggtgtggtggtgcacgcctgta | TCGA |
| CCCCAGCAACACGTCATTGTGTAAGGAAAAATAAAAGTGCTGCCGTAACC*aaaaaaaaaaaaaaaaaaaaaaaaaggaaagaaagaaagaaaagaaaCCC | TCGA |
| GGGACCGGGACCTGCTGCTGGACCCGGCCTGGGAGAAGCAGCAGCGCAAG*CAAGCCCTGGATGAAAACATGGACCTCTTGGAAGGTATAACTGGCTTTGA | TCGA |
| CCCAGGTCCTTCAGGACGATCTCCCAATTAGCCTCAAAAAGGTGCTTCAG*AACCTTGTGTCCAAGGAGGTCCTTCAGGAGACCTGGGATGAGTGGTGCTT | TCGA |
| CCGCACTGGGACCCCGGAATCCCGCGCGCTGCCCACGATTCGCTTCTGAG*GTTCTGCTTGGATTCTGCTAGACAGACCCGACAAAGACTGTCTATCAACT | TCGA |
| TCACCCTGGGGCTGGCCATCGAGCCCAAGTCAGGGACATCGAGTCCACAG*AACCTTGGCCATTCTGATGTTTAACCAAGAGTTGAGCTGAGTGGCTCTGA | TCGA |
| TGACTGGAAACAATGGCATTTACAGAAAGAGTCAACAGCAGTGGCAACAG*GTAATATGCACGTGTCACTAGCTGAGGCCCTGGAGGTTCGGGGTGGACCA | TCGA |
| CTGAGGGACAACAAAAATGGCGGAGGGGAGCCAGTCGGCGCCTGAGGCAG*GTAAAGAGAAATGTGTATGACCTTACAAGTATCCCCGTTCGCCACCAATT | TCGA |
| GGGCTCTGCCTGCTGACCTGGGGAGGCCATTGGCTCTATGGAAAACACTG*GCCCCAAATTCTCACCAGTCCGTCTCCTTCAAAATCCATTCCAATTCCAC | TCGA |
| GAAGCCCTGAGGGCAGCTGTTCCCACTGGCTCTGCTGACCTTGTGCCTTG*GTTTTGGGATTTGAGATTGACGCGGTGAACTCTGTCCAGTTTTCAAACCA | TCGA |
| AGATAAACCCGGGACCCACTTCACTCCAGTGCCTCCGACGCCTCCTGATG*GGGGGGCCCTCAGCCTCTTGGCCTTGAACGGGCTGTTCATCTTGATTCAC | TCGA |
| TTTTCTCTTTAGCTCGGCTTATTCCAGGGGTGTGTTTCGTCGAGATGCAC*AACACAGACCACCAGTGCAAAACAAGAAAAAGCACCAAGTAAAAAGCCAG | TCGA |

| | |
|---|---|
| GTCCATCACCCTTTCCGGCACACTGCCAAATAAACAGCTATTTAAGGGGG*aaaaaaaaaaaaTGCTTACTAAGTGGCAGTGGTATCTATGATGCTATAG | TCGA |
| GTCCATCACCCTTTCCGGCACACTGCCAAATAAACAGCTATTTAAGGGGG*AAAAAAAAAAAAAAACTACGGCGGCCGAGAAGGGGCGAGTACAGCCCAGT | TCGA |
| GGCTCAGCCCGTTCGCTGGACTGCAAACCGAGCTCCCGTCTGCCTTGCAG*GTGACAGTTCCTTCTTTCTTCGAACCTGGCTCATGACCCCACTTCACATT | TCGA |
| GGCTCAGCCCGTTCGCTGGACTGCAAACCGAGCTCCCGTCTGCCTTGCAG*AGAGAAGATTATAAATGGCAGAGCCATTTAACTGTGGTTAGCTGTTGGAT | TCGA |
| GACTGCGGACTTGGCTGGAGAGAGAGCTGGATGCACTGAGAACCCAGCAG*AATGCGATGCTGGTGGCAGAGAAGGCAGCTTTGCAGGGGCAGCTGCAGCA | TCGA |
| TGGATTTAAAAGACAGTCCTCGTCAGCACTGACTTTCAGCTATGGAATCG*ATACCCTGTTCCAGTTGAAGGTATGTACCGAATGGCCGTTCACCCCAGCC | TCGA |
| TACACGTGTGACTTCGTCCAGTTACAAACCCAATAAACTCTGTAGAGTGG*AAAAAAAAAGAAACCTTTAAAAAAAAAACCTACACCTGTGCTATTACCAC | TCGA |
| GCTGTGATGCTCGGGAGGGAGACTGTTGTTCCCGGAGATGCGGCGCGCAG*GTCAGCCCTGACAAAGGTCAGCTAGCCCCTTGAGGACATCAGCTTTGGCC | TCGA |
| CTGTGGAGAATGTACACAGTTTAAACACATCAATAAATACTTTAACTTCC*aaaaaaaaaaaaaaaaaaaaaaaaaagaaaaCCACCAAAAATAAAAtaaa | TCGA |
| CTGTGGAGAATGTACACAGTTTAAACACATCAATAAATACTTTAACTTCC*aaaaaaaaaaaaaaaaaaaaaaaaaaaactagccgggcatggtggcgggtg | TCGA |
| AGCTCCTGTCCCTACCCACTCTTTGCTACAATAAATGCTGAATGAATCCA*AAAAGAAAAAAATCCAGTTCAACTTGACGATGAAGGAGGCAGGACGTTTT | TCGA |
| TCCTCCTGGGGTGGACCCTAGTTTAATAAAGATTCACCAAGTTTCACGCA*aaaaaaaaaaaaaaCCGGAAATGGGTCCTACCATCTTCTCGGAGCCGGAG | TCGA |
| GGCGCCTGTCCCTGCTTCTTTTCTGACCGGGAAATAAATGCCCCTGAAGG*AGAAAAAAGAGTGCCGGCGGGACCACCGCCCACCGTGTGCTCAGGAGGCG | TCGA |
| TGGAGTCTCTCTGCACCCTCTGTCCCTGGACAGCCCAGACCGGGAACCAG*ATACACATGGTCTACAGTAAGCGGTCAGGAAAGCCCCGTGGCTATGCCTT | TCGA |
| GAGTCCTTGGCGCCCCCAGAGAGTACCGACGAGGGGGCCGAAACCGAGAG*GGCTGCGCTGGCCTCGGCACACCAGTGCCACCCGCTGCCCCGGACGCTGA | TCGA |
| ACCTCTATCGCCTCTATGTGTCTGTGAAGGAGATTGGTGGATTGACTCAG*ATGCAGGCCCTGACTTCCTGTGAGTGCACCATCTGTCCTGACTGCTTCCG | TCGA |
| TCGTGGGGCTGCAGTACTCGGGCAAGACCACCTTCGTCAATGTCATCGCG*CACTCCTTTTTCTGTCGCTTGACAGAAGATAAGAAGTCAGAGAAATTCTT | TCGA |
| CCGCGGACCTCCCTGCGACCGTCGCGGACCATGGGCGACAAAGGGACCCG*CTCTCCCAGTGAAGTGATATCACTGCCTCTGTGGACAAGACACCTCCAGG | TCGA |
| GACAGAGACCTGGGAAAATGACTTGGCCAAATAACATTGTTAAAACAATG*GGGCAAGCTCTGTGTGATTGATTGGAAGACATCAGAGAAACCAAAGCCTT | TCGA |
| AGGGGTTGTTTTATAAACTGCCTGGCCGAATCAGCCTTTTCACGCTCAAG*GATTGGTTGCCGTGGTCTTTACTGCTTTTCTCCCTGATGTGTGAAACAAG | TCGA |
| ATTGGCAAAGCAGCAACCAATTATCAGTTTACAACCTGCACCCACTAAAG*AGCAACCTGCAAGCAATCAAGGCCAGTCTACCCTGCAGCCCTTGCCGCCT | TCGA |
| CAGCGACCCAAGCGCCGCGGTTAACCACGTCTCGTCCACGACCTCCCTAG*GATCCCATGGTCCATTATTGCCTTTACCAAGTCGTTACAGAATGGGCTCT | TCGA |
| TTTTCGAAGATGTAGTGTGGCAGGATTTGATTACTGCCATGAAGAAAATG*tggagaacctgaaaaggaacctggaaagaagcagctggaaaaggtgagag | TCGA |
| AGGCCTACATCCCACAGAGATACCCAGACAACAGGATCGTCTCGTCCAAG*ATACCCTATCATCTTTACTCTGTTGGAGAGGAGCTATAATGGTCTATATT | TCGA |
| AATACATTTTATATCTGGATTTTTACAAATAAAGATGGCTATTATAATGG*aaaaaaaaaaaaaaaaaaagaaaaGCATGAATGGAAAGAAGGGCAAGTCA | TCGA |
| TCACTGCCCTCTCTATGGATATATAAGCAATGTGCAGGAATATGATTTGG*TTTCAGCATCAGGCAAGTGGCTGGATGGTATTCGAAAATGGTATTACAAT | TCGA |
| CACCACCGCACTGTCGCTGCCCGCCTGCCCTGAGGAGTCCCCGCTGCTTG*GCAATGAGAAGCACGACCTTCATGTTACCTCCCAGCAGGGCAGCAGTGAA | TCGA |
| AACTGTATGCAGATTCCAGTCAGCTCAGCCGGGAGCAACGGGCATTGCAG*GAGACCCCATCTTCAGAACCAATGGAGGAAGAGGAAGATGACGACTTGGA | TCGA |
| GTTTCTGAATGTCATCGTCCACTCAGCCACTGGATTTAAGCAGAGTTCAA*AAGCCCTTCAGCGGCCAGTAGCATCTGACTTTGAGCCTCAGGGTCTGAGT | TCGA |
| CTTCTGAGGTCGGTGGCTGTCTTTCCATTAAAGAAACACCGTGCAACGTG*aaaaaaaaaaaaaaaaaaaagaaagccaggtgtggtggtgcacgcctgta | TCGA |
| GGGCTGCTTCTGAGGTCGGTGGCTGTCTTTCCATTAAAGAAACACCGTGC*AACGTGAAATTGAAAAGTATATTAAACCTGAAGAGCCCGAACCTGAACCA | TCGA |
| CTGAAGCCCGGGATCCACGTCCCACGGGCCGGACCCGCGGCCGCGTTCGG*ACACCTTGGAAGAGCTTAAGGAAATAGATAAAGGCATGTGGAAGAAACTG | TCGA |
| GCCCTAACATCTGGATCATCCCCTTCCGCACCCGCCTCGGACTCTATTGG*TGATAACCTCCTAAGGAGAGCAGCCTGTCAAGAAGCTCAGGTAATACTAC | TCGA |
| gcgccggcgccgcggccTCTTCGGCTGCGGACCCTGCCATTCCGGAGGAG*GAACTCTAACTCCTTGCCACTCAAGAAATGTCCTCCCTTTCAGAATATGC | TCGA |
| GACTGTGGACATTTCGCAAAGAGGCTTGATCTGAGCTTTAACCTTCTGAG*GGCTTTTTTCTTACAGTTTCCCCAGAGTCAGTATTAAAGGTGGCTCACCA | TCGA |
| TCCTCCAGGCGCTGCCCAGCCCCTTGTCAGCCAGGGCTGAACCCCCGCAG*AGAACACCTTCTGCAGTGGGGACCATGTGTCCTGGCACAGCCCTTTGGAT | TCGA |
| TCATCGGATCTACAGGACCAGTTGAATCATCTGTTGAAATAGAATGACAT*GGTTCTGGCTGGTGTGGACCATCATCATCATCCTGAGCTGCTGCTGTGTT | TCGA |
| ACAAAAGATGACTGAAGAACAAACTTACCAAGCAGCAGAAAAATCTCAAG*AAAACTAAGCCTACAAATAATCGAACAAAGAAAGCTGTGAAAACAGTGAA | TCGA |
| ACAAAAGATGACTGAAGAACAAACTTACCAAGCAGCAGAAAAATCTCAAG*AAAGTAGCACGTCTGGAAATTGAGAACAAAGTCCTTCAGGAGCAATTGAA | TCGA |
| AAAGTGCTCAGCAAAGCAGGCTGTAGGTTTAGTTGAGCACCGGAAAGAAC*CCGCCAGCTGGAGACGATGGTGGACCACTTGGCCAACACGGAGATCAACA | TCGA |
| GCTGCAGTGAGCCTTGGGTCTCGCAGGCCCAGCCCTGCTGCAAGGAGCCA*AGGCACAGACACCAAGGACAGAGACGCTGGCTAGGTAAGATAAGGAGGCA | TCGA |
| CACTGCTGGGCCCTGAGGCTGCACCAAGTGGCGAAGGAGCCGGCTCCAAG*GCCACCAGCCCCTCGGGTTCCAAGATGCAGCTACTGGAGACAGAGTTCTC | TCGA |
| CTGGGTGGATCCGGATCGCATGGAAGACGCAGCCTGGCCGCGCTCTCCCA*GACTCCTCCTATTTATGGAGCAGGCACCCAACATGGCTGAGCCCCGGGGC | TCGA |

| | |
|---|---|
| TTGGCCGCCTGAGCCCTTGAGTTGGTCACTTGAACCTTGGGAATATTGAG*AACTCCCCCACTGGAAAGGATTCTGAAAGAAATGAAGTCAGCCCTCAGAA | TCGA |
| CTGATAGCTGATAGCTTTTACAGTGCTCATTCCTAACAGTGGATTTACTT*ATTAATATTAAGATTGGAAGTTTGTGTCTTTTGCTGGATATTGGAAATTG | TCGA |
| GGACACGCGAATTTGAAGATAGAGACAGGTCTCATCGGGAGGAAATGGAG*GTCCATGAGCTGGAGAAGTCCAAGCGGGCCCTGGAGACCCAGATGGAGGA | TCGA |
| AGGCACTGCAGGAGGAGAACCGCGACCTGCGCAAAGCCAGCGTGACCATC*GATCATAAGTTTTTACTGCCAGTATTGAGAAATTCGTGGAAGAAATGTCA | TCGA |
| AGGCACTGCAGGAGGAGAACCGCGACCTGCGCAAAGCCAGCGTGACCATC*GAGGATCCAAAGTGGGAATTCCCTCGGAAGAACTTGGTTCTTGGAAAAAC | TCGA |
| GAGCGCCTGCGACCCCAGTCGGTGCTCCGGGGAGGTCACCTGACGAGGAG*GACCAGGCCTCCTCTTCCCACCTGTCCAGCAGCATGAAAGCAGCATGATT | TCGA |
| AATGAGCAGGCACTCCTTGGAGCAAAAGCCCACTGACGCTCCACCGAAAG*ATGATTTTTGGATACCAGAAACAAGTTTCATACTTACTATTATAGTTGGA | TCGA |
| AGCTTTCAGGGCCATGGTTCACATTAGAATAAAAGGTAGTAATTAGAACA*aaaaaaaaaaaaaaaaaaaaaGCAAAGTAGAGATTAAGCTGCTTAGAAGA | TCGA |
| ACGAAGGGGACAGCGCAGCCGGCCCAGAGCTCGGGTCTCCGGGGACCGAG*ACGAGGGACTATGGGACATCCTCATCAAAGACATCCCCAAGGAGGTGAGC | TCGA |
| CCTGTGGCCGGTGACCTCCCTCCTCCGAGGGAGGGGGGTTGTGACCCCAG*GTTCAGCTTCTTCACTGCATAAAGCAAACAGTCACAGGGGGTGATTCAGA | TCGA |
| ATTTTTCCCAGCATTTCAGGCATTTACTAATGATCTACTTGAAATCTTAG*AAAAGCAACAAAATATTTCTCCCCTAGTGAGAGAAAGAGGTCCTCAGAGA | TCGA |
| CAAGATGAGGATTTGGGTTTTCTATAAGGGGTTTCCTGCTGGACAGGGGC*GTGGGCAGGAACTGGGCACTGTGCCCAGGGCATGCACTGCCTCCACGCAG | TCGA |
| AGCTCCACCCATCACCCCAGAACACCCCGCAGAAAGTGCCTGTGCATCAG*TTTGACGGTGAGAACATGTATATGAGCATGACAGAGCCGAGCCAGGACTA | TCGA |
| TCTCAAGGAGGGAGTCTTCCCTCTGAGGAAAAGGGACACCCCCAGGAGGA*GCAGGCCCGCAAGCCGTGTGAGCCGTCGCAGCCGTGGCATCGTTGAGGAG | TCGA |
| AAGATCTTTCTGAAGAGCTCAAAAAGGAAAATCCTTTCTTGAGGCAAGAG*ATTGCAGCAATTCTCCGGAAACGGAAATTAGACTATTATTTACACAAACT | TCGA |
| AGGAGCCGCACCCGGCCGGCCCGGCCCCAGCCCCATGGACCTCCGAGCAG*TGCAGGGCTCGCGGCCTGGCAAGAATGTACAGCTGACAGAGAACGAGATC | TCGA |
| CCACCTTTATCTCAGAGACACACCTTTCCAGAAGTCCAGAGAATGCCCAG*AATCCTGTTCACAAGATTCCTGACTCGCATGAGATAACGCTGAAGCATGG | TCGA |
| ACTTCAACAAGGGCCCTTCCTATGGGCTCTCGGCCGAAGTCAAGAACAAG*GTGTTGTGTAACGGACCAGGAACATGTGTTCCTATCTGTGTATCTGCCCT | TCGA |
| GGACATTGGTATTGGCATTGCAGGAGAAAATGGTCTTCCCGGCCCCCCAG*CTGAAGGAAGAGGATCACCTTCCAAGACAAAACAGTCTTCATGGGGGAAA | TCGA |
| ATCTTTTTTTTCCTTACAGACACCCATAATAAAATATCATATTAAAATTC*aaaaaaaaaaaaggacatgctacagcatggatgcaccttgaagacatttg | TCGA |
| GTCAAGAAACAGTTGAAAAGTTCCGGCAGAGAATTCTGGCTCCCTGTAAG*GTATTTGTGGAAACATTAGACAAATGTTTTGAAAATGTCTGTGAGCTGGA | TCGA |
| TCAGCCCTGGTATTCCTGCCTATCCTCGAACCTGCACTGTGGACTTTTCG*GTTCCATTTCTTCCAAGAAGGGTGAGCCTGCCTTTTGTCTCTGGTGTAAA | TCGA |
| AAACTACAGTTCTGTCACTCACTTACACAGTATTGGGAAATCTGTGAAAG*AAATTGGAATGGGATTAACAGGATTTGGAGTGTTTTTCCTGTTCTTTGGA | TCGA |
| accgtgaccgagagcgtgaccgagagcgCGAATATCGTCATCGTTAGAAG*GAATTACATGAACGACAGCCTTCGCACCGACGTCTTCGTGCGGTTCCAGC | TCGA |
| accgtgaccgagagcgtgaccgagagcgCGAATATCGTCATCGTTAGAAG*GTCCTCAAGGTCCTCCAGGCTATGGCAAGATGGGTGCAACAGGACCAATG | TCGA |
| gccgccgccgccgctgccgctgccgctgccgcACCTCCGTAGCTGACTCG*CTGGAGCAGTTCAAGAAGGACAACGAGGAAACAGGCTTCGGCTCAGGCAC | TCGA |
| GGAGAAGCTGAAGAAGGCAAGGGAAGAGGCCATGACCGACCTGCCTGCAG*GTATCACAGATGTCGTAAGAATCTGAATCCTCCAAAAGACAGATGATGCG | TCGA |
| CGGCCTTCGAGGAGGTCATGAAGGACCTGAGCCTGACGCGGGCCGCGCGG*CACGGCCGACAAGAACGGGGCCCTCAAGTGCACCTTCTCGGCACCCAGCC | TCGA |
| GCCTTGGGTTTCTTTCTTAAAGGACTGATTTTTAGAACTCCACATTTGAG*AGAGAAGTGGAGATCCTGATGTTTCTCAGTGCCATTGTGATGATGAAGAA | TCGA |
| GTGCCTGGTAAGCTTGCCCTTGCTTAAGAACTCAAAACATAGCTGTGCTC*TATAAAAAACGGGTGGTTGGGCGCCGCTGTCTTTTCAGTCGGGCGCTGAG | TCGA |
| GGCGCGGCGGAATCCAGGGCCGACCCGGGCCGGACCGACCCCAGGCGGCG*GGACAGACATGGCTCAGCGGATGACAACACAGCTGCTGCTCCTTCTAGTG | TCGA |
| AGAGCCCGGGAAGAGATGCAGCGCTATGCGGAGGATGTTGGGGCCGTCAA*CTCGAGATTTCTGTGGCTCCTCAAGATATTGGTCATAATCCTGGTACTTG | TCGA |
| TGAACAGTGCTCGGGACTTTATTGATTGTTTCCTGATCAAAATGGAACAG*GAAATTTTGTCTGTGGCTAAAAAAAATAAAAAGGAGAATGAGGTAAGAAA | TCGA |
| CGACGGCTCCGCGACGTTGAGGCCGCGTTGGGCGGTTCAGACTCAGGGTG*GATCTCCCTACTACGATAATGTCCGTCCACTCTGCTACAGCGACTCGGAT | TCGA |
| ATGACATCAGAGGGGCTGAACTTTTGCATCTGGAAAGGCGAGATCTTAAG*TGAGCATTTGTCCTGTGCTTTCACATTCTTTCTGCATGGAGAAAGTAATG | TCGA |
| CCCCATGGCTGTGGGGCCGCTGCTTCTGTTTAAATAAAAGTGGCCTGGAA*GCTGAAAACTAGCCGAGGAGAGCCAGGGAGCCGGAGAGATCGCGCGCCTG | TCGA |
| TGGAATGATGAAGGGGATACGCTGGAGTTGCCCTGCCATCGCCTCCATCA*GGAGAGGAGAGCATAGCACCTGCAGCAAGATGGATGTGGGCAGCAAAGAG | TCGA |
| CCCCATGGCTGTGGGGCCGCTGCTTCTGTTTAAATAAAAGTGGCCTGGAA*GCTGGAAAGTTGCGCGCCAGGCAGCGGGGGGCGGAGAGAGGAGCCCAGAC | TCGA |
| TCTGGGTCTGGGTCCCTCCGAACAAGTGAAAAGAGGTCCTTGTATGTCAG*CCTCCAGTCAGATGGCTCTGAGGTGCTCCTGAGTCCTGAAGTCACCTGTG | TCGA |
| CTCTTAATGCATGATACAGAATAATAATAAGAATTTCATCTTTAAATGAG*aaaaaaaaaaaaaaaaaaaaaaaaaggaaagaaagaaagaaaagaaaCCC | TCGA |
| CAGAAGTAGCAGATGATGAGGTTACTGAGGGGACTGTGACACAGATACAG*ACAGAATGTGCAAATTTCATCAGAGTACTTCAGCCCTATAACAAAACTCA | TCGA |
| GGGCTCGGTATTCCCCTTCCTGCTAGTCCTGGTGGATCTGCAGTACGAAG*GTTATATACAGGCATGTAGAGGACTTATGATCGCTGCTGTCAGCCTGGGC | TCGA |
| GGCGGCCAGAGCGCCGGCAAGAGCTCGGTGCTGGAGAACTTCGTGGGCCG*ACAGCTACGGCGTGGACAAGAAGCGGAAACGGGGGATGCCCGACGTCCTT | TCGA |

| | |
|---|---|
| AGGTCATCAATGCAGCCTCAAGTCAGTTGCCTTTTCTAAGTTTGAGAAAG*GCCTAAGATCTGGCAAACTTGGTGAACAGTGTGAAGCAGTTGTTCGCTTT | TCGA |
| TCTCTGCCGTGAATCATTCCTGCAGATGAAAAAGTGTGACTTGGCAGCTG*GATGTGTCTAGTCCGAATGAAGCAGGAAGGCCGGAGTGGGAAGTACATGT | TCGA |
| ACCTCCAGCTGTACTAGTGCCATTGGAATAATAAATTTGATAAGGTGGTG*aaaaaaaaaaaaaaaaaaaaaaaaaaaaaaGTCCATTGCTGAGAGACGAG | TCGA |
| ACCTCCAGCTGTACTAGTGCCATTGGAATAATAAATTTGATAAGGTGGTG*aaaaaaaaaTTAATAATTGTTTTTAAATCAGTAACAAGCCAAAGACCCAT | TCGA |
| ACCTCCAGCTGTACTAGTGCCATTGGAATAATAAATTTGATAAGGTGGTG*aaaaaaaaaaaaaaaaaaaagaaagccaggtgtggtggtgcacgcctgta | TCGA |
| ACCTCCAGCTGTACTAGTGCCATTGGAATAATAAATTTGATAAGGTGGTG*aaaaaaaaaaaaaaaaaaaaaaattaagtgggtgtggtggtgcatgcc | TCGA |
| ACCTCCAGCTGTACTAGTGCCATTGGAATAATAAATTTGATAAGGTGGTG*aaaaaaaaaaaaaaaaaaaaaaaaaaactagccgggcatggtggcgggtg | TCGA |
| ACCTCCAGCTGTACTAGTGCCATTGGAATAATAAATTTGATAAGGTGGTG*AAAAAAAAAGAAACCTTTAAAAAAAAACCTACACCTGTGCTATTACCAC | TCGA |
| ACCTCCAGCTGTACTAGTGCCATTGGAATAATAAATTTGATAAGGTGGTG*AAAAAAAAAAGCCATTTAAAGCCAGCCACTAGAGGGAGTCAGTTCAGTTC | TCGA |
| ACCTCCAGCTGTACTAGTGCCATTGGAATAATAAATTTGATAAGGTGGTG*AAAAAAAAAAAAATCCAATGGTGACGAGCAGGGAGAACAGAGCAGCTGCC | TCGA |
| TCGAATTCTCCAAAATGGCCCGAGACCCCCAGCGCTACCTTGTCATTCAG*CTGCAGGACAAGTTCGAGCATCTTAAAATGATTCAACAGGAGGAGATAAG | TCGA |
| GAGCCACGTGCATGAACCAAACCCACTTCGTTCAAGACAGGTGTTTAAGC*GGCATCCTTCTCAACTGGACCAAGGGCTTCAAGGCCTCAGGAGCAGAAGG | TCGA |
| ATCTTGCGGCAACGCCCCCGGGACATGCATCGTCCCCTGGACTCTCTCAA*ATGTCGACGTACAAGCGGGCCACGCTGGACGAGGAGGACCTGGTGGACTC | TCGA |
| ATTTTTGTCTCAACCAGCAAAAATGTCAACTCGCGAAAAAAACAGCCAAG*TGTACCGCCGGAAGCACCAGGAGCTGCAAGCCATGCAGATGGAGCTGCAG | TCGA |
| TCCCCGGTGGTTTTGTGCTCAAAATAAAAAGCCTCAGTGACCCATGAGAA*aaaaaaaaaaaaaaaaaaaaaaGATATGTACCAATCATGAAGAGTGGCCT | TCGA |
| ATGAATGCAAGCACCATGTGGAGACACGCTGGCACTGTACTGTCTGTGAG*CGTGCAATGATGCGCTTCTCAGAGTTGGAGATGAAAGAAAGAGAAGGTGG | TCGA |
| CTCTGAGACTGATGGCTACGTTGCCCCCCTGACCTGCAGCCCCCAGCCTG*CTGGGCAAAAGACAACCTGTGGCCAGGGTCTGGAAGGGCCCTGGGAGCGC | TCGA |
| TGAACAATGGGGCAGGCTGCCGCTTCGAGGGGCAGTTCAGCATCAACAAG*ATGATGTCTGTCCTCATGAACACCATTGTCTTTGAAGACTGTCGGAACCA | TCGA |
| TTCTTGAAAAAGGATGTAGATTTTAATCTGAACTTTGAACCATCACTGAG*ATTAAAACTTTGGAACAGACTAAAGCCATTGAAGATCTAAACAAATCCAG | TCGA |
| TTCTTGAAAAAGGATGTAGATTTTAATCTGAACTTTGAACCATCACTGAG*CTAAAGACACAGAAAGAGAGACTGGAGAGCAAAGAATTACACATGAGCCT | TCGA |
| TTCTTGAAAAAGGATGTAGATTTTAATCTGAACTTTGAACCATCACTGAG*GACCTGCTCAGTGCTGTAGAAGCAAAAGAAGCTCTTGAAAGGGAAGTTAA | TCGA |
| TTCTTGAAAAAGGATGTAGATTTTAATCTGAACTTTGAACCATCACTGAG*GAAACTTACGATCATCTTTCGGAAGTCCCGGTCACGCGGGAGCAGTTAAA | TCGA |
| CTCTGTCTCCCCGCCTGAAGAGCACGCCATGCCCATTGGGAGAATAGCAG*ATGTGCAGCACATTAAGAGGAGAGACATCGTGCTGAAGCGAGAACTGGGT | TCGA |
| CTCTATACACACACAGCCGGAGGTCATACTGCATCAGAACCATGAAGAAG*GTCCCGTGGCTGTCATCAGTGGTGAGGAGGACTCAGCCAGCCCACTGCAC | TCGA |
| CAAAAGCACAAGGATTCTCGCTGTGACAGAGACAACACCGAGTATGAGAA*CATCATCTGAGGGTTCTCCATAAACACGGTTAGCCTTCCAAGCTGCTGTC | TCGA |
| GGGTCTGGACTGATCTCCATGTCCTGTTGTGGGGCTTTTACAGCCTTTGG*GATGGATTTGTTGACTTTTTGGAGTTTATTGCTGCTGTAAATCTAATCAT | TCGA |
| TTTGTCTTCACTGGAATCCATTCTGGCACCTCAACGGCTGATGATTTCTG*AGCACGTGCCCCCGGAGCTGTGGGAGCCCTTCTATACCGACCAGTACGCG | TCGA |
| GATGCTGCGGCTGCAGGACTATGAGGAGAAGACAAAGAAGGCAGAGAGAG*ATGATTTTTGGATACCAGAAACAAGTTTCATACTTACTATTATAGTTGGA | TCGA |
| CCGCGTAGCTTCCATCCGCCAGCTGCCATGAGCGAGCGCCTCCGTCCCAG*CCTTATGATCTCCTCATTGCGTCCCCCTCTGCCCACTGGACTTGGACTTC | TCGA |
| GTTCTTTAGTTTCATCTCTCTATAAAGTTATCCAGGAGCCACAAAGTGAG*GCTATGGAAAGGGCAAATGGAATGGAGCTGGATGGTAGAAGAATTCGGGT | TCGA |
| TCGCCGCCTCAGCCGCCGCCGCCGCCGTTTTCGCCTGCAGCTGCTCACCG*CTCTTGCAGTGTTTACTCAACTTCCTCAGGATACAAGTGTCGAGGTTGGA | TCGA |
| AATCCCTGAGTGCCCGGGAGGAGCCTTAACAAGCGCACGGAGCCCTCAAG*CTTTGTCAGAGACCCTTCAGCACTTCCATAGAGCCATCCCACTAGAGGTC | TCGA |
| GCGCCCTCTGCCAGGCCGCACGCTCCATCAGCACCTTCCTTGGCACTCTG*TCTTTGAAGAGCCTGAGGACCCCAGTAACCGCTCATTCTTCTCGGAAATC | TCGA |
| ccggccccATGAGGAGGGACGTGAACGGAGTGACCAAGAGCAGGTTTGAG*GACAAGGGCACACAACTGGTTCCGTTAAGCCCCTCTCTTGCTCAGACGCC | TCGA |
| ccggccccATGAGGAGGGACGTGAACGGAGTGACCAAGAGCAGGTTTGAG*GTCAAAAGGCTGAGTTTCAAATTCTGGATAATGTAGACTCAACGGGAGAG | TCGA |
| AAACTTCAAGCCAAACATCAAGCAGAATGTGATTTGCTTGAAGATATGAG*TTCATTTGATGAATACAGTTCAGAACTTAAATCTGGAAGGTTGGAATGGA | TCGA |
| TGCGCTTCCGGATCGGGGGCAAGCGGAAGGTGATGCCCAAGATGGACCAG*TCCCAGGCTGTCCAATCTTCCTGGCTGGTCCCTGAAGGATGGAAATGCTT | TCGA |
| CTTCCATTGAACTTCCCAGCACTCGAAAGGGACCGAAATGGAGAGCAAAG*TAATTTTGCACGATGGCAGTCAAATGAGAGCAAAAGCATTTCCAGAAACC | TCGA |
| AGCAGTTGGTAGAAGACTTGGATCGAATTCTCACTCTCACAACCAATGAG*ATCATGGAGGAAACAAATACGCAGATTGCTTGGCCATCAAAACTGAAGAT | TCGA |
| AGCAGTTGGTAGAAGACTTGGATCGAATTCTCACTCTCACAACCAATGAG*GTAAATAAAGTTAAACAAGAAAAGACTGTTTTAAATTCAGAAGTTCTTGA | TCGA |
| AGCAGCTGGTGGAGGACCTGGACCGTGTCCTTACCGTGACGTCCACCGAC*GTAAAGGCGACACAGGAGGAGAACCGGGAGCTGAGGAGCAGGTGTGAGGA | TCGA |
| AGCAGCTGGTGGAGGACCTGGACCGTGTCCTTACCGTGACGTCCACCGAC*TTTAAGGAGTCGGCCTTGAGGAAGCAGTCCTTATACCTCAAGTTCGACCC | TCGA |
| AGGCCATCCCCATCCTGAGGGCAGCCCTGAAGCTGGAACCTTCCAACAAG*TGACCGGCTTCGCACAATGACCAACGTACTGGGGGACTCAATTGGAGCGG | TCGA |

| | |
|---|---|
| ATTTGCTGTCAGCCATGAACGGATGGATGTGATGCCTGCTAGCCAAAAGG*GGGCTGTGGGGTCACCTTCTCTTCCTCTCTTCTGCCACCCCAAGTCGTCT | TCGA |
| CGGGCTCCCAAGTGTGCAATCTGGCCTGGTCCAAGCACGCCAACGAGCTG*ATTCCCAGGACTCCGGCTCTCTCTGCTTCTGTTTCCGGTGGCTGCTCATC | TCGA |
| GAATCTCCCCTCCTCACAGTTGCCATGTAGACCCCTTGAAGAGGGGAGGG*AGGCACAGACACCAAGGACAGAGACGCTGGCTAGGTAAGATAAGGAGGCA | TCGA |
| AGGACGGGCTCCGTGGCTGGTTCCCGGCGAGCTACGTGCAGTTGCTGGAG*AGTGCCATTGAAATTCTGGGCTTTACTTCAGATGAAAGAGTGTCCATCTA | TCGA |
| GCCGCCGCCTGAGGAGGCGAGCTAGCCGGGAGTTACACCGCCACCGCCAG*CTCTTCGTGATCGACAATGGCGCGGATGACTGGCGGATAGCCATGACCTA | TCGA |
| TGCTTGGCTGCTAGAGAAAGAGTCTATTCTTCTCCGCTCCTCCGACCCAG*GGGTCCAGCTTATCATGGGCATGCTGTCTGAAAAACCAAGGTCTGGGACT | TCGA |
| ACTGGGAAGGCTGCGCGAGGGAGTTCGACACCCAAGAGCAGCTTGTGCAC*GATCGAATCATCAATTTAGTTGTTGGCAGCTTAACATCCTTATTGATTCT | TCGA |
| GATGAGGAGGAGGGGGACGAGCAGCAGCGGCTCCTCAACACCCCTCGAAG*AAGAACTTCTGCTTGGGTGGCTGAACTCTGATCTTGACCTAGAGTCATGG | TCGA |
| ATTGTGTTGTGCAGCattaaaaaaaatcaaaataaaaattaaatGTGAGC*AAAGAAAAGTTTGCTGAAGCTTCAGAGGAGGCAGTCTCTGTTCAGAGAAG | TCGA |
| ATTGTGTTGTGCAGCattaaaaaaaatcaaaataaaaattaaatGTGAGC*AAAAAAAAAAGAGAGACAAACTCTAGGAACAATGCCGATTCATTCCACGC | TCGA |
| ATTGTGTTGTGCAGCattaaaaaaaatcaaaataaaaattaaatGTGAGC*AAAGAATAAAAATGACATAAGCAAACTCATCTTCAAAAGTGGAGATTTCC | TCGA |
| ATTGTGTTGTGCAGCattaaaaaaaatcaaaataaaaattaaatGTGAGC*AAAAAAAAAAAAAAACTACGGCGGCCGAGAAGGGGCGAGTACAGCCCAGT | TCGA |
| ATTGTGTTGTGCAGCattaaaaaaaatcaaaataaaaattaaatGTGAGC*aaaaaaaaaaaaaaaaaaaaGCAAAGTAGAGATTAAGCTGCTTAGAAGA | TCGA |
| ATTGTGTTGTGCAGCattaaaaaaaatcaaaataaaaattaaatGTGAGC*AAAGAAAAGGCTGCAGGATTTTGTGGCAGACAGGATGTGGCTATGGCTTT | TCGA |
| ACGCTAGAAGTTTATGGCAGAGCTTTACAAATAAAAAAAAAACTGGCTTT*TCTGAAAAAGTTAGTGAAAGAAGCTCTGAAAGAATCTCAACTTCCAACTG | TCGA |
| ACGCTAGAAGTTTATGGCAGAGCTTTACAAATAAAAAAAAAACTGGCTTT*TCTGAAAAATGAGTAAAGATGGAAGCCAAAGTATCACTTTTATTGTGGTT | TCGA |
| TTAGAGACCGAGTCCCGGCGGCGACGGCGGGGCAGCGCACCGGCAGGCGG*GGGTCTCCTGCTGCCAGCTAAGTGTGGGAGAACTTGTGCACGTATCTCCC | TCGA |
| TTAGAGACCGAGTCCCGGCGGCGACGGCGGGGCAGCGCACCGGCAGGCGG*GACTGTCCCCAGCTGGTGCCCACAGAGGAGATCTTGATTCCAGTCGGGGA | TCGA |
| CGGTGCCAAGGGATTCAGCCTGGCGGACATCCCCTACCAGGAGATCGCAG*GGATGCCAGATGATATGAACTTCAGTGATGTAAAGCAAATCCATCAAACA | TCGA |
| ggcagtgaagagctgctgaagtacatgcttcatagtccttgcgtctctct*GGGCTATACAAAGCGGACCAGCGGTTTGGGCCAGGTGTCGAGACCTACCC | TCGA |
| CCTTCGGAGGACGTGCCCTCACCCCTCACTGGTCCACTGGCTTGAGACTC*ACCCCTCATTGCTGCCAAAAATCCCAAGATTGCTGTCTCCAAGATGATGA | TCGA |
| CCACTGGCTTGAGACTCACCCCGTCTGCCCAGTAAAAGCCTTTCTGCAGC*AAAAAAAAAGAAACCTTTAAAAAAAAAACCTACACCTGTGCTATTACCAC | TCGA |
| GGCCATAATCAATCCTACCAATGCTGACATTGACCTTAAAGATGACCTAG*GAAATCCCCTTACCCAGGATATATTGAACCTTTATCAGGAACCAGATGGA | TCGA |
| CCTCAGTAGTCAGTTTGAAGCGGGTATGCACAAAGATAGCTGGCTTCTGG*AACCACGAAGCTGCCCTGAGACGCTCACTCACGCTGTGGGGATGTCAGAG | TCGA |
| CCGTGGGGAAACTAAAAACGCATCTATCTAACGTTTACCCTAGCAAACCA*GAAGAGTGACCCAGTCCCAGCCATGACCACTCCTGAGAAACTCTACTCCA | TCGA |
| GACGGGGAGTCCGACGGACCCATCGACGTGGGCCAAGAGGGCCAGCTGAG*GGGGACCCTGGACCAGATGGACCATCGGGGCCCCCAGGACCCCCTGGGAA | TCGA |
| CCGCCGAGTGGGCCGCGTGCGCCGGCTACGCGAGGGACCTGCTGTGCCAG*AACCACCACCACCTAAAATCCCCAAAATTGAGACCACTCATCCACCGTTG | TCGA |
| CAATTCCTGGAATTTGAGAGAGCAAATAAAGACCTGAGAACCTTCCAGAA*aaaaaaaaaaaaaaCCGGAAATGGGTCCTACCATCTTCTCGGAGCCGGAG | TCGA |
| ATGAGCGAGTTGTCATCATCAAGAATATGTTTCATCCTATGGATTTTGAG*ATACAAGGCAGTTGTGAAGCCACTTGAGCGACAGCCCTCCAATGCCATCC | TCGA |
| AAAAATTTGGTCAAAGACTGGGATGCTTAGATCAACTTTAGCTCAGCAAG*ATAATCATGGACATATTGATCCCTCTCTGAGACTCATCTGGGATTTGGCT | TCGA |
| TTTGTCTTTAACCTTGTAACGCTTGCAATCCCAATAAAGCATTAAAAGTC*aaaaaaaaaaaaaaCCGGAAATGGGTCCTACCATCTTCTCGGAGCCGGAG | TCGA |
| GGAACCTCTGTGGGACGCCTGGAGGAGAGGAACCTTTGCTTCTATGGAAG*GAGATACCTTCCCGGATGCAGATGCTGATGAAGACAGTCTGGCAGGCGAG | TCGA |
| CTACACCTTGTGTGCACAGAGTGGAACCAGCCCCTCCGTCTGCATGAATG*GAGATACCTTCCCGGATGCAGATGCTGATGAAGACAGTCTGGCAGGCGAG | TCGA |
| GCACTATGGACCCAGCCCTGCGTGAGGCCGCGGAGCGCCAGCTCAATGAA*AAAAATCCTGGGTAGAAGCTGGATCAGCAAGAATGTCACTCCTTATATTG | TCGA |
| CGTCAGAACGTGGCTTATGAGTACCTTTGTCATTTGGAAGAAGCGAAGAG*GCTCTGAAGAGCGGCCATTCCTCAGATTCGAAGCTGAACACATCTCCAAC | TCGA |
| CGTCAGAACGTGGCTTATGAGTACCTTTGTCATTTGGAAGAAGCGAAGAG*GAACTGATGACGCTTGATAATGTGGCTGGGGACTTCAGGGAAGAGTGGGT | TCGA |
| GTATCCCCTTTTGTGAAAACAACACCAGAAGAAATTGCCAACCCTTGCAG*ATATGGGAAAATAGTCTCCACAAAGGCAATTTTGGATAAGACAACGAACA | TCGA |
| CTTTACTATCTTTTCATCTCATTAATTCAATTAAAACCATTACCTTAAAA*TTTTAAAAAATTAGACCTGTTTATCGCGCTCTCAGTGCAAACTCGGCTCA | TCGA |
| GGTCTCCTTCCGAAAACACTACCCTTGGGTCCAGCTTTCTGGACATGCTG*TCCTGTGTCCCTTGATCTGTCACAATGGCGGTGTGTGCGTGAAGCCTGAC | TCGA |
| GGTCTCCTTCCGAAAACACTACCCTTGGGTCCAGCTTTCTGGACATGCTG*GGACGTGTCATCAGTGGAGGTGCTCATGAACTACCACCAGGGCCTGAAGA | TCGA |
| AGTGAGAGATGGAACCCGAAGAAGAAAGGATTCGTTACAGCCAGAGATTG*AGCTACACCCCCACGGTGTTTGAGCGGTACATGGTCAACCTGCAAGTGAA | TCGA |
| ATGGTCTCAATGTTGCCAAAATGTTTCCTAAAGTTCGTGAATCTCAACTT*ACAAAACTTACCTGGGGTCCCTACTTATGGTTTTACATGCTTCGGGCTGC | TCGA |
| GAAAAGTAGAAGGAAAAACAGAACGTTGCAAACTCTGGAAAAAAACTTGG*AGTCACCACATACTATGGACATGTCAATTATATTAGAAGATGAAAAGCCA | TCGA |

| | |
|---|---|
| gccaccaccTCCCCAGCCACAACCAAGTTTGAAAAGGAATCCAAAACATG*ATGACACAGGCTGTCCTAGCAGCCAGTCAGTGTCTCCTGTGAAGACACCC | TCGA |
| TCAAGAGGCAGGCCCTGAGGTTGCTCCTCCCGGGGCCCTTCCCGGGCAAG*GTGATCTGCAACTCTTTCACCATCTGTAATGCGGAGATGCAGGAAGTTGG | TCGA |
| TCAAGAGGCAGGCCCTGAGGTTGCTCCTCCCGGGGCCCTTCCCGGGCAAG*GATGCTGATATGCTAACTGGTGATGAGCAAGTATGGAAGGAAGTTCAAGA | TCGA |
| TATATATTAACGACAATTTTTTTTTTGGAAAATAAAAAGTGCCTAAAAGA*aaaaaaaagaaaaaagaaaTATCATTTTTATTCATGCAGTTACTTTTAGA | TCGA |
| CAGCATGTGGGACCTGGTTCTCTCCATCGCCTTGTCTGTGGGGTGCACTG*GAGCACCATACTGGACCAACACAGAAAAGATGGAAAAGCGGCTCCATGCT | TCGA |
| gttacagatactggggcagcaaataaaactgaatcttgttttcagacctt*aaaaaaaaaaaaaTGCTTACTAAGTGGCAGTGGTATCTATGATGCTATAG | TCGA |
| CCTCACCCCCTCCTGGCAATCAATACAGCTTCATTATCTGAGTTGCATAA*aaaaaaaaaaaaaaaaaaaaGCAAAGTAGAGATTAAGCTGCTTAGAAGA | TCGA |
| ACCCCCGGCTCTTCTCCTGACCTGCCAATAAAAATTTATGGTCCAAGGGA*GGAAAAAACCTATAGAGGATCCAGCAAACGATACAGTGGATTTCCCTAAA | TCGA |
| ATGTGGAGCTGTGCCTGTGCCCCGCCAGCTACCGGGGGGACTCATGCCAG*CCGGTGGCTGCTGGCAGCGGAATCCCCCAGATCAAGTGCTTCCTCAACGG | TCGA |
| GGGAGCCGCCGGCCAGGACGCCCCCGAGGGTGTAGACCGCGCCCCTGGAG*GTGATGCGACTGTGAAAAAGAAACCTGCCCCCAAGACACCTCCGAAGGCA | TCGA |
| TCTTCCACAACCAGATTGAGTCCAAGTTCCAGAAAGTACTGGAGAGCAGG*GGGGAGTTTTACAGTGAGAACCACCACAATCCACCTCCCTTTGCTCCACC | TCGA |
| CGCTGATGGAGCCCCACCTTGTCTGCTAAATAAACATGTGCCCTCAGGCC*aaaaaaaaaTCATTCTTTTTAGTCTTAGCACCTACTTAAGGATCCACTTT | TCGA |
| CAGGACTTGTTAAAAAGATCAATAGATTGCCTACCCCCATTGCAGGACTG*AGACTCGAGCTTTAGAGTAATTGGGATGGTGAAAGGATGGGTTTCCAGAA | TCGA |
| TCCGCGGTGGAGGTCTGCGTTGCTCTTGCTTTTCCTCGCTGGGGTGTACG*GTGCTGCTACCCCCGTATGATGATGCCACTGTGAATGGTGCTGCCAAGGA | TCGA |
| AGAAATAGCAAACTTCATGCAGCTGGTGGAACTAGATGTGTCTCGAAATG*CTGGTCAGTGAGAAGGTCGGAGGGGCCGAGGGGACCAAGCTGGATGATGA | TCGA |
| ACCCTCAAGCCGAAGCCTTCAGTGGGACTCCCCCAGCAGATACATTCTCA*TGATGATTTACTGGAGGATTCAGACAGTGAAGAGCATTCCAGATCAGATT | TCGA |
| ACAATTCCCTGCTGTCGTCTTAGCAAGAAGTAAAATGAGAAATTTTGTTG*AATAAAAATCAATGGTTTCCACATCTCTCAGTGCCTCTATCTGGAGGCCA | TCGA |
| GGCCCAGATGCGCGCGATGCTGGACCAGTTGATGGGCACCTCCCGGGACG*TGGCTTTTCATGGCCTGCCACTGAAAATCAAGAAAGAACCCCACAGTCCA | TCGA |
| ATGCATGATGTAAAAAATACAAACATTCTAATTAAAGGCTTTGCAACACA*aaaaaaaaaaaaaaaaaaaaaaGCAATCATTAAGCTTTTCATCTAgccat | TCGA |
| ATGCATGATGTAAAAAATACAAACATTCTAATTAAAGGCTTTGCAACACA*aaaaaaaaaaaaaaaaaaagaaaaGCATGAATGGAAAGAAGGGCAAGTCA | TCGA |
| GGCGGCTTCCTCCCAGCGTTCCCAAATCATTCGCTCCAAGTTCCGATCTG*GAAAAGATTAAGGAAATTGGAGACCGTCAGCCAGAAAAGTAAGAGAAATT | TCGA |
| CGGCGCTCTCCTGGCTCCCTTCTGCCCCAGCTCCGTCTCGGCGGCGGCGG*ATTTGAAGGGCTTAACATAACTAAACTGTATGATGGACAATGTGTAAGTA | TCGA |
| AACACTCCTAGCCAATGGTGGTCATGGAGTAGAAGGGAGCGATACTACAG*ATTTGAAGGGCTTAACATAACTAAACTGTATGATGGACAATGTGTAAGTA | TCGA |
| ACTGTTGAATTAGACCCGTTTCCTATAGATGAGAAACCATACAAGCTGTG*CAACACAGATGTACACACGGTGGCATCACTTCTTAAGCTGTACCTCCGAG | TCGA |
| AAGTGCTGGAGTGCTTCCATCGGCGCACTGGACAGAAGTGTGCCCTGAAG*TTTCAGAGCCTGAGGCCGGCACTTTGGACCCAGCCCTTGACCTCTCAGCA | TCGA |
| CCAGCGCTCGGACCCCCCTACCCACGCTGAACGAGAGGGACACGGAGCAG*TGACCCAACAGGAGAAGACCCCACCGCCTAGACCCAGCCCGCTAGAGGCA | TCGA |
| ATATGTTTTGTTTTTCAGAAGGAGTTGAGCCTTCCAAGAAGAGGCAGCTT*AATTGAAGATGTTCCTGCTCCTTCTACATCTGCAGATAAAGTGGAGAGGT | TCGA |
| TGGTGAGGCTCACGTTGGAGGGCTTCGCGTCTGCTTCGGAGACCGTAAGG*ATGACTTTGGGAAATTGCTGCTGGCTGAGGCCCTCCTGGAGCAGTGTTTG | TCGA |
| ACCAAAATAGACCCTGGAGTGAAACCATTAAGCTTGTGCGTCAAGTCATG*CTAGAAGAAGACATAAGCAAAAAAATGGACAAAGATGAAGAGGCTTTGAA | TCGA |
| CACTTCCTTCCTTACTTTGTCTTGGTATACTACATGGCAAAATGGGAAAG*CGAGCTTGACATGATTTCCACAAGGGTGATGGACATTAAGCTTCGGGAGG | TCGA |
| GAGCCGGAGTGGGGCGAGCCGCGGAACCCGGCCGGGAGCCGCGCGAGGCG*AGCCTGCCTTCTGGTACAAGATGCCTGTGGAATGTTCGGTGAAGTGCACT | TCGA |
| CGACGGTCCCTCAGCCCAGCCACCATGAGCACCAAGCAGATCACTTGCAG*ATTTGAAAGAAGCCAACACTAAACCACAAATATACAACAAGGCCATTTTC | TCGA |
| GGGGTCGAGCACATAACGGGGTTCGGGTGTCTCGTGTGTGAACATCACAG*TTCTGCTAAGCCAATATTCTCTAGAATGAGCACAAAACAAATCGAATAAC | TCGA |
| GATAGGCAGCATATTCCTGATACATATCTATGTGAACGTTGTCAGCCTAG*TTCTGCTAAGCCAATATTCTCTAGAATGAGCACAAAACAAATCGAATAAC | TCGA |
| GGGGTTGATACAGCAGAGACGTCATACTTGGAAATGGCTGCAGGTTCAGA*TTCTGCTAAGCCAATATTCTCTAGAATGAGCACAAAACAAATCGAATAAC | TCGA |
| GTGATTCTTCCTCTCACTCTCAGGATAAACATCATGAGAAAGAGAAAAAA*ATACAGTTTATGTAACTTGATGGAAGAAAATGGAATTACTCCAAAAAGAC | TCGA |
| GTGATTCTTCCTCTCACTCTCAGGATAAACATCATGAGAAAGAGAAAAAA*GCACTTAAGTATTCATCGAAGAGTCACCCCAGTAGCGGTGATCACAGACA | TCGA |
| TCTTCCTTCGAGGCATTATGGAAAGTCCCATAGTAAGATCCCTGGCCAAG*ATGGCTGCCGCACGCCTCTGCCTCTCCCTGCTGCTCCTGTCCACCTGCGT | TCGA |
| AGAGGAAGGAATTGTCTTAGAAAGTCCTCATTATGCTGTGAAAATATTAG*GGAATCCTGTATCGGCTGTGGTCACCTGCAATCTCTTTGTTGTGCCTGCA | TCGA |
| ACGCACTACAACATCAACCAGAGCAGATTCCAGCCTCCACATGTACATAT*CCGCCCATGCCAACTTCCGTTCCCATCAACCATGCAGCAGCACAGCCCCA | TCGA |
| AGCCCACCCACGGCATTGGAAAGTACAAGCACTTAATTAAAGCAGAAGAG*GTTCTGAAGCTGCTGCTCGTGGCCTGGGATCGCCGCCTCATTTTTGCCAT | TCGA |
| GCCAAAGCTCCCATCCAAGACATCTGGTACCACGAGGATCGGATTAGAGC*CCAGCGTCCCATGCTCGGTGGCCCCAGAAAAGTCAGTGTGTAGGCCTCAG | TCGA |
| ATTGTGTAGCCATGcctatcagtaaaaagatttttgagcaaacacttgaa*CAAAAGCGTGATCTTGCTGGGTCGGCACAGCCTGTTTCATCCTGAAGACA | TCGA |

| | |
|---|---|
| TATCCCTCAGCCTGGTGAGGGTGATGTTCGGGAGGAGGCCTTGTCCACAG*GAAAATGGCTTTGTAAAGAAGTTTGAACCTAAATCTGGCTGGATGACTTT | TCGA |
| CTCTAGACGAGAACGAGGGCATCTATGTGCAGGATGTCAAGACCGGAAAG*AGGCTTAAGACGAAGCAAAAAGAACTGGAGGCCAAGATGTTGGCCCAGAA | TCGA |
| AGTCCTCTCGGGAGAAGCAGCGCCTGTTCCCGGGGCAGATCCAGGTTCAG*GTGTCAGCGGTGCCCACTGTGCTGGCCATGAAGAATGGGGACGTGGTGGA | TCGA |
| CTGATGGACACCGTCTCCATGCCCGAGTTCATGGCCAACCTCAGGCTCAG*CTCACATACCTGCCCCCACCGTGGGGTGAGTGCCGATCCTCAGAGATGGG | TCGA |
| CTGATGGACACCGTCTCCATGCCCGAGTTCATGGCCAACCTCAGGCTCAG*GTTCTGTTTTAACTGTGAATGGTAAAACTGAGAACTATATCCTGGATACT | TCGA |
| ACCCGTACAGGTAGCCCCGCCGCCGCGCACCTGCCTTCGCTCCCGCACCG*ATACTTGGTGCTTGAAAAACTGAAGAAGGAAGACGCTGACCGAATTCATA | TCGA |
| TCCATGGTTCCAACAAATTGCAAATAAAACTGTATGGAAACGATGAGCAA*CCCCACACGCCTTTGGCACAATGAAGTGGGTAACCTTTATTTCCCTTCTT | TCGA |
| TGGCCCGACTACAGCGAGAAGTTGCCCAAAGAACAAGTGAGGGGGCCATG*TGTCCTCTCGCATGGCAGAGCAAGTGGGAAGGCCCAGAAGATCCCTTACA | TCGA |
| GGTTTTTGCTTCAGCGGCTTTAGGACAAATTAAAAGTCAACTCTGGTGCC*AGAAAAAAGCAGAGATCGTGAAAGGGAACgagagcgggaaagagagagag | TCGA |
| CCGACCCCCAGGACTGGCTTACCCAAAAGCAGACCTTGGAGAACAGTCAG*GAGGATCCAAAGTGGGAATTCCCTCGGAAGAACTTGGTTCTTGGAAAAAC | TCGA |
| CCCGGGCCGGCGCTCCAAggcccgcccgggagggccggggccgcgcTCAG*GTGTTGGGATGGCCACCAACTGGGGGAGCCTCTTGCAGGATAAACAGCAG | TCGA |
| CGGCCCGAGCGCGCCCCAAGCCCGGGCGCCACCGCTGCCACCTCCGCGAG*AACGTGCGCATCGACCCCAGTAGCCTGTCCTTCAACATGTGGAAGGAGAT | TCGA |
| ACTATAAGAGCCTGGTGAGACGGAGCTATCGGCGCCGCGGCAAGAGCCAG*AACGTGCGCATCGACCCCAGTAGCCTGTCCTTCAACATGTGGAAGGAGAT | TCGA |
| CAGGTCTCCCTGAGTCTTTGAGGACACAGCCTCGCTGGAGGCAGTTTCTG*AACGTGCGCATCGACCCCAGTAGCCTGTCCTTCAACATGTGGAAGGAGAT | TCGA |
| TCACCATTGGAGCTCATCAGACTGACCGTGTCTTACATCAGTTCAGAGAG*AAGAACTTCTGCTTGGGTGGCTGAACTCTGATCTTGACCTAGAGTCATGG | TCGA |
| CGGCTTTATAAGCGGGCGCTACGCCACCTCGAGTCGTGGTGCGTCCAGAG*GATATAAGTCAAAGAGTACAGCAGAAAATGTCTACTGAACGGACTTCTTG | TCGA |
| CTCAACATGAACAACTACAAAGGCTATGAAAAGAAGCCCTATTGTAATGC*GTTCCCAGTAGCTGCAGCAGTGAAAGACAGTGATTGGCTCCAGTGCTCCC | TCGA |
| GCAGCGGACAGCTCGGACAGCGCCCAGGGCCGGAGCCCGAGCCCTTGGAG*GTGCATCACCTGGATCATGAGGTCACCCCTCTGCTGGCTCCTCCCACTTC | TCGA |
| TCTGAAATATGGAATGCATCAGAATACACTTCGTATCAAAGAGATTCATG*GAGATGCCTGGCATTTACGAAAAATGGATTGGTGCTATGAAGCTGGAGAG | TCGA |
| CCTTGCACAACGAGGACGCCTTCCAGCACGGCATCTGCTTTGAGGCCAAG*AGAATGTGATCAATGGGCAGGACTACGAAGTGATGATGCAGATTGACTGT | TCGA |
| TGCGGGACATGCCGTGGCTGGCGCTGCCCTACAAGGAGAAGCACAGGAAG*CTGGACCCACAAACCGTGGAGACCAAGAACTGGCACACGGACGTGATTGA | TCGA |
| TCACAATCGAATTGAACACATTGGCCAAATAAAGTTGAAATTTTACCACC*aaaaaaaaaaaaTGCTTACTAAGTGGCAGTGGTATCTATGATGCTATAG | TCGA |
| CTATTTCACAATCGAATTGAACACATTGGCCAAATAAAGTTGAAATTTTA*CCACCAAAACCCTCAACAGGGGCATCTCTGAGTTCATCGTGATGGCTGCA | TCGA |
| GGTCTGGAGACGCCGTGGCAGCCTGAACGGAGTGTGCGACGGATTGGGAG*CCGGTGATGAACAATGACAAGCACTGCTTTGAATGCTATGGCTACGACAT | TCGA |
| TCAGCCTCATCCAGCAGGCGGTGACCCGCTACCGGAAGGCCATCGCCAAG*GAACCAGTCAATAAGGAACAGGATCAACGGCCACTCCACCCAGTGGCAAA | TCGA |
| CCATGCTGCCTCCGTGTACGGGCAGTTCACGGGCCAGGCCCTCCTCTCAG*AAATGACCATGGTTGACACAGAGATGCCATTCTGGCCCACCAACTTTGGG | TCGA |
| ACCCCGCCGAGTCCTAGCGCCTGGCCTGCGCGCCGCTGCCCGCGCCACAG*AGCCCCCAATTTTGTGAGTGTCCCTGAAGCATTGTGACAATTCCATTTTA | TCGA |
| CGCCGGCCAGAAATCCGACCGGACCGGGCTCGGGGGAGCGTGAGTTGCAG*CATGGCTTCTCGCTGTTTCTGGGATGCTGACACTGACAACTATACTCATG | TCGA |
| GCCTTGGATGTTTCTGTCCTAAATAGCTGACATTGCATCTTCAAGACTGT*ATTTCCTCCTTCTTCACCCCTGAAACCATATCCAAAGAAGCCCCCAAACC | TCGA |
| ACACCACGATGAAAAGAAATACCTGTGATTTGCTTTCTCGGAGCAAAAGT*TGGAATGAGACCGTGGAGCTTTTTCGTGCTAAGATGCCGTTACGGAAACA | TCGA |
| CCGTCATGGGCCAGAACTTAATTCTGAACATGAATGACCACGGCTTTGTG*ACCACAGAATGGCTCAATGATACTCTACAACAGGAAGAAAGTGAAATACA | TCGA |
| GTCGTCCTCGAGCAGGGGCCGCCCGCTCCGGGTCCACATCTCCGGCTCAG*CACACGACGGTTCTGCCCTTTATATTAATCGAGATGAGGACATCGCACCA | TCGA |
| TTCCGGGAGAGGCTGCTGTGGAGGCTGAGGAGGCGGCGGCGGAGATCTGG*GAGCAGAGGGAGAGCCGGGCTCGGCGAGGCCCTCGAGGGCCCAGCGCCTT | TCGA |
| GAGTTCCTGCCTCCAGCTGCCAGGGGGGACAGCCAGCCAGCAGCAGGAGG*GTTTCTCTTAAGGTTCTGCAGGGCAAGGCTGTCTGGGACAGGCTTGGCCA | TCGA |
| TCCCTGCACCTGTTCAGCCTGTGTCCCTGCCCAGCCCCACCTCCACAGAC*GTGAGAAGCCCTACGTCTGCCCCTACGAGGGCTGCAACAAGCGCTATTCC | TCGA |
| AGCCTGTCATGAGACCTCCAAACCCCTTTGGCCCTGTATCAGGAGCACAG*AAATATAAAGAGAAGGACAAACACAAACAGAAACACAAGAAGCAGCCAGA | TCGA |
| GGCCTGTTGCTATAGAGCCTACCTGTATGTCAATAAACAACAGCTGAAGC*AAAAAAAAAAGAGAGACAAACTCTAGGAACAATGCCGATTCATTCCACGC | TCGA |
| GGTCTTCCTGCCCAACAGCAACCACGTGGCCAGTGGCGCCGGGGAGGCAG*CCATTGAGACCCAGAGCAGCAGTTCTGAAGAGATAGTGCCCAGCCCTCCC | TCGA |
| GTTCAAGGTGCGCCTGCAGGACCTCAGCTCTTGCATCACCCAGGGGAAAG*CCATTGAGACCCAGAGCAGCAGTTCTGAAGAGATAGTGCCCAGCCCTCCC | TCGA |
| TCCTTTCTCCTTCCATCCTGAAGCCCTTTCCGCTGTGGACAGGAAAACAG*GCTTATATTTGGCACCCTTTACCCTGCGTATTATTCCTACAAGGCTGTGA | TCGA |
| AGAGTTAGGAGCTCGAGAACCGTTTGGCAATATGTACGACGCGGATGAGG*GTTTTCCTCCTCCACCAGGCGCTCCACCTCCATCTCTTATACCAACAATA | TCGA |
| CAAGACGCGGCTGCCGTACGTGGCCCTCGATGTGCTCTGCGTGTTGCTGG*GACTAACTTCTTCAGTGCTTAGAGTGTGAGAAAAATGGCAAATATGAATA | TCGA |
| CACAAATATACAACAAGGCCATTTTCTCAAACGAGAGTCAGCCTTTAACG*GGCGAGAGATGGTGGGGCCCACGCTGCCCGGATACCCACCCCACATCCCC | TCGA |

| | |
|---|---|
| CTGTCAGGCGGCGGCCGAGGGCGTCGCGGACTCTCCCCGCGATGATGCCG*ATGCTTCAAAGTATTATTAAAAACATATGGATCCCCATGAAGCCCTACTA | TCGA |
| AGAGGAAAGAGCCCGTGAAGATCGCGGCGCCGGAGTTGCATAAGGGAGAT*GTGCAGACCCATCTGGAGAACCCAACGCGCTACCACCTGCAGCAGGCGCG | TCGA |
| AGCACAAGAAGAAAGTCCTGAAGCGGCTGTCGCTAAGCCCAGCCCTGCAG*GGCGTGGACGAGTGGATGTGGCCAGCCTGGCTGTGGGGCTGACAGGTGGC | TCGA |
| CCAGAGAGGCCCGCCCACCAGTTCGCCCGTCCCCCTGCCCCGTTCACAAT*GGTCTGGAACAGAATCTGACAGTGATGAATCAGTACCAGAGCTTGAAGAA | TCGA |
| GCAGATCCGCAGGCGCAAGGCTTCAGGCGACCCGTACTGGGCCTACTCTG*ACTTCAACTGTGAGGACATGTCGTTCAGATTTGGCCAACATCTCATCAAG | TCGA |
| CCTCTCTCCTTGGCCTCTCCTGGGATCGGCCCAAGGCCAGTTCTCCGCAG*TGCATCCTAACGTTAGTCAAGGCTGCCAAGGAGGCTGTGCAACATGCTCA | TCGA |
| TGGAACGGGGCTCCCAGGACCTCCACTAATCACCAGAACAAAATGTGCAG*TGCATCCTAACGTTAGTCAAGGCTGCCAAGGAGGCTGTGCAACATGCTCA | TCGA |
| CCTAAAGCGTTTCCTCGCTTGCATTCAGCTCCTCTGTGTTTGCCGCCTGG*ATAAACCTCTCATAATGAAGGCCCCCGCTGTGCTTGCACCTGGCATCCTC | TCGA |
| GCACGTTTGCAGCGGAAATTCAGGAAACCATCACAGCACTCAGAGAGCAG*TGGATTTTACTTATCCCTCATTGGAAGAATCAATTCCTTCTAAACCTGCT | TCGA |
| GCTAGCGCGACCCTGAGCCGGCGCCCGTGGTCCGGCCATGGACGACCTCG*TGGCCGCCGCCGACAGCGGCATCAGCCCTCGGGCCGTGTGGCAGTTCCGC | TCGA |
| GCCCAACTTCTGCGGGATCTTCAACCACCTCGAGCGGCTGCTGGACGAAG*GTAGAAATTGAGAAGCTGGATTACCATCATTATCTGCCTCTGTTTTTTGA | TCGA |
| GATGTAAAATCATGGTCCGAGGCAAAGGCTCAATGAGGGATAAAAAAAAG*GTAGAAATTGAGAAGCTGGATTACCATCATTATCTGCCTCTGTTTTTTGA | TCGA |
| GAAAAGACAGTGGAGAGAAGAAGTTGCTTCACTTCAGGCTGTTATGAAAG*GACCGGAAGTCCTTCATCTCAAGCATCCAATGCTGAAAGCGGCCTGATTT | TCGA |
| CAGATGTTCCAGCTAATTGACATTGCCCGGCAGACGGCTCAGGGAATGGA*GAAAAGCCATGGAGACTTCAGCATCCTCCTCCCAGCCTCAGGACAACAGT | TCGA |
| TCTCCAGCACCAGCTTCCAGTTAGTGGATATAGCACACCATCCCCAGCCA*ATGCAGCTGTATCCAAGAAAGCCAGCCCAGAGGCTGCCAGCACTCCCAGG | TCGA |
| TCTCCAGCACCAGCTTCCAGTTAGTGGATATAGCACACCATCCCCAGCCA*AGGAACGCGTTGTGGTGATCAGCAGCTCGGAAGACTCAGATGCCGAAAAC | TCGA |
| CCAGGACAGTCGCGAGGTTCGAGTGTTGCAGAGCCTGCGGGGCAAGATCT*TCCACCTGATCCCGGTGTCGACTCCTAAGAATGGAATGAGTAGCAAGAGT | TCGA |
| CCAGGACAGTCGCGAGGTTCGAGTGTTGCAGAGCCTGCGGGGCAAGATCT*ATGACAGTCATGTCCCTTTCCAGGGACCTCAAGGACGACTTTCACAGTGA | TCGA |
| GATGAGATGTCGTTCACTTTTACTGAGCTACAGAAAAACATAGAAATCAG*GGAACTTATAGAAATGCTGGCAATTTCAAGAAACCAAAAGTTGTTACAGG | TCGA |
| AATGGTGTTAATTTCTGAAGTGGGCGGCTCCAAAAACTTAACAAGGAAAG*GGATTTTGCTTATGTAGCAAGAGATAAAGATACAAGAATTTTGAAATGTC | TCGA |
| AATATCTTGTTTTGCAGGATTTGTTAAGGATTCCAAGTAACTCTTATTTG*TCTGCGCTCCTCAGACCACCGGCAGGTGAACAGCCTGATGCAGACCGAGG | TCGA |
| CCCTGTCAAGAAGGTGCCGGTGGTGCGAGTCTTCGGAGCGACCCCGGCAG*GAATTTAGATAATGGGCTGTGTGCAATGTAAGGATAAAGAAGCAACAAAA | TCGA |
| GTCATGGATAGGCAGAAGAAAGAAGAGGAGAAGGAGCAGTCCGTGCTCAA*GCATGCTGGAAGATGGACTGCCCTCCAATGGTGTGCCCCGATCTACAGCC | TCGA |
| ACCATGCCCGCCCCAAGAAACACACCACGGACCCTGACATTGATAAAAAA*ATGCAGCTGTTCTCTTAGTTACATTGGCAGATTGTGTGTTGTCAATGTTG | TCGA |
| tgtctgggatgtgaggagcgcctctgctgggccgcaaccctgtctgggag*ACATGTAGACTGCCAAAGTGTATGGGATGCTTTCAAGGGTGCATTTATTT | TCGA |
| ACAGATTTTGAAGATGGAATCCGTAGCGCCCAGCTCATTGCCTCAGCTAA*GTCCCCGTTCTCCGTGTCCTGCTGTCCGCTGACCTGAAGGGGTTCCAGTA | TCGA |
| ACAAAAACTAACCCTGGATCCTACTCTCTTATTAAAAAGATTTTTGCTGA*GAAAAAAAAAAAAGTACCGCTCCAGAGCAGGAGCCTAGGCAGCCGAGAGG | TCGA |
| CTGCGAGCTTCTACTTCTAAGTCTGAATCCAGTCAGAAATAAGATTTTTT*TTTTCTCTCTCTCTTTCACTGCAAGGCGGCGGCAGGAGAGGTTGTGGTGC | TCGA |
| CTCTGGAAGCCAAGGCAGCTGCCTTCGAGAAGCAGGTCCTGCAGCTGCAG*TGGAATTTCTGTGGAGCCACACGACGGAGAGCATGTGTGTCGGATATATG | TCGA |
| CTACCACAGAGCCATCAAAATCACAGTGGATGGGCCCCGAGAACCTCGAA*ATCGTACTGAGAAGCACTCCACAATGCCAGACTCACCTGTGGATGTGAAG | TCGA |
| GAAGCCTTTAAATGCAGCATCTGCGATGTGAGGAAAGGCACCTCCACCAG*CTCCAGAAACATGACAAGGAGGACTTTCCTGCCTCTTGGAGATCAGAGGA | TCGA |
| GATGCCTTTGAGTTTTGTATTCAATAAACTTTTTTTGTCTGTTGATAATA*TTTTAAAAAATTAGACCTGTTTATCGCGCTCTCAGTGCAAACTCGGCTCA | TCGA |
| CAAATCCAGTGGTGGGTAATTGTACAATAAATTTTTTTTGGTCAAATTTA*aaaaaaaaGCTAAAGCAGAAAACAAAAGTGGCCCATGGTCTCAGCTTCCA | TCGA |
| TGAGGGGTAACAAGATGGCGACCGAGACGGTGGAGCTCCATAAGCTAAAG*GACTACCTGTCACAATATGGGTACCTACAGAAGCCTCTAGAAGGATCTAA | TCGA |
| TGGAACACCTTCTCGCTGCCGCCATACCCTGCCTTCTCCAGCGACAGCCG*GGCCGCCTTCAACAACAACGTGAGCGTGGCCTACGAGTGCCTGAGCGCCG | TCGA |
| CCGTGGCCACCGCCTCCAGCTTGCCACCAAAGACCTGCGACTTTGCTCAG*TGTCCAAGGTCCCCAGCTCCCTAAGTCAGGAACAATCCGAGCAAGACGCA | TCGA |
| AACCTGCCGAATTGATGCCCCAGTTTTCTACAATTGAGTACGTGATACAG*ACTTCCTTGCCGTCCAGGAAGGGATTTCGACACCAGACCACCAAGTTTTT | TCGA |
| aatgtttatacccattagaacagAGAATAaatagaactacatttcttgca*aaaaaaaaaaaaaaCCGGAAATGGGTCCTACCATCTTCTCGGAGCCGGAG | TCGA |
| CCCCTCCATCCCTGGCCCCCTCCCTGGATGACATTAAAGAAGGGTTGAGC*TGGTCAAATGTCGATTTCCAGTAGTCGAAAATGGAAAACAGATATCAGGA | TCGA |
| CCCCTCCATCCCTGGCCCCCTCCCTGGATGACATTAAAGAAGGGTTGAGC*TGGAAAATGGAGAAGCTGTGCTAATAGAGGGGGGCCAGAAATCCCCACTC | TCGA |
| CCCCTCCATCCCTGGCCCCCTCCCTGGATGACATTAAAGAAGGGTTGAGC*TGGAAAATAAAGAGACTCTGAAGCATTTGGCATCTCTTTATGCAAATAAT | TCGA |
| CCGCCTCCGAAGATGGGGGATTTCTACGACCCGGAGCACCCGACCCCTGA*AGAATGGTCAAGAACCTGGAGAAGGAACTAGCACTACTCAAGCAGGAGCT | TCGA |
| TGTGTGCTTAAAACAGGGTTCCTTTTTGTAGCATCAGAATTTGGAAACCA*TTGAGCAACTTGCATCTTAAGGAAGAGAAAATCAAACCAGATACCAATGG | TCGA |

| | |
|---|---|
| CTGCTGGGTGTGACCATGTTTCCTCTCAATAAAGTTCCCCTGTGACACTC*AAAAAAAAAGAGGAGCGAGAGAAAGAAAAAAAGGGGGAAAAATCAGGATC | TCGA |
| GAAGAACTTCACAATCAAGAAATGCAGAAACGTAAAGAAATGCAATTGAG*CCTGAACTCTTTGCTTCCGGAATCCGGGATTGTTGCTGACATAGAATTAG | TCGA |
| CTGGCGTTCCACCAGCAACCATGAGTGGTTCCATGATGGGAAGTGACATG*CTGCCTGTGTCAGGGAATCTGCTTGATGTGTACAGTAGTCAAGGCGTGGC | TCGA |
| ATGCCACCGTGGTGGACGTGGAGAAGCGGAGGAACCCCTCCAAGCACTAC*GTGTATTTTTGTACAATGGACATCCAATAAAACAGGTAGATGTCTTGGGA | TCGA |
| GACCTTCGTGCTGTTTCTCTCCCTGGCTTTTGGGGCAAGCTACGGAACAG*GGCTCAAGGAGCCTCTCTCCTTCCATGTCACCTGGATCGCATCCTTTTAC | TCGA |
| GGAGGCTCCGGGCACCCCCGAGGGCCCCGAGCCCGAGCGCCCCAGCCCGG*GGGTTGGCTGTGTTCCGGCCGCAGAGCACCGTCTGCGTGAGGAGATCCTG | TCGA |
| TTCAACTCCTTCAATGAGGCCACCATGGTTCTGGTCTCCTGGATCATGTG*GTAAGAGCAAAGAGGCGGAAATTAAGAGAATCAACAAGGAACTGGCCAAC | TCGA |
| CTTTGAAGGATTTTGCCTGGCTGGAATCATTCTCACCTTGTATGCTTTAC*GACTGGTTGCCATAAAGGAAGCCCACGACATAGAGACCAGGCTTAATGAG | TCGA |
| ACCCCAAATATATGGGGTGCAGGTTTATCCTGGGGACTCTACTTTTTCTT*GAAGGAAAAGCTGATGCGATGCTCTCAGTGCCGCGTCGCCAAATACTGTA | TCGA |
| CGGGCTTGGAGTTCGCCTCAGACGCGGTGGAGCCGCCGGAGTCTGAGAAG*TGCTCAGTGGGCCCCTCCTCTGATAAGGCCCTGACCTTCATGAAGGACCA | TCGA |
| ttttttttaCTGGTTATGGGAAGGGAGAAATAAAATCATCAAACCCAAAA*GGAAAAAAAAGGAGAGCAGCGGGTGGGGGCGAAAGCAGGGCCGAGGAGAG | TCGA |
| TGTTTCTGGCCACGTGCTTCAACGCTCTGCTGCCCGCTGTGAGGGAAAAG*GAAACGAGCAGATGAAAGTGGAAAACTTTGAAGCTGCCGTGCATTTCTAC | TCGA |
| AACCAGCCTGCACGCGCTGGCTCCGGGTGACAGCCGCGCGCCTCGGCCAG*GAAGCCTTATCAGTTGTGAGTGAGGACCAGTCGTTGTTTGAGTGTGCCTA | TCGA |
| AACCAGCCTGCACGCGCTGGCTCCGGGTGACAGCCGCGCGCCTCGGCCAG*TGGCTTTTCATGGCCTGCCACTGAAAATCAAGAAAGAACCCCACAGTCCA | TCGA |
| CAGAGTATAATGTTTATGGTGACAAAATTAAAGGCTTTCTTATATGTTTA*aaaaaaaaaaaaaaaaaaaaaaaaaggaaagaaagaaagaaaagaaaCCC | TCGA |
| AAGGCCATCACCACGCAGAACGTCAACACCAAGTGCATCACCATCCCCAG*GTGGCCAAAGCATTCTGGATGGCAATCGGGGGAGACAGAGATGAAATTGA | TCGA |
| AAGGCCATCACCACGCAGAACGTCAACACCAAGTGCATCACCATCCCCAG*ACGCCAGCATGTCTCCGGACGCCACCAAGCCGAGCCACTGGTGCAGCGTG | TCGA |
| CGTGGAGCGGCTGAAGGGCATGTCCCTGCTTGTCAGGGCTGAGAGCGACG*ATGTGGACGAGTGTGCAACAGATTCCCACCAGTGCAACCCCACCCAGATC | TCGA |
| GAACCCCGGGACACGCAGGAACGGCTCCAGCATCAAGATCCGTCTGACAG*ttccaggaagaagcttagcccaggaaaaacaaagaccagttggacccaga | TCGA |
| GAACCCCGGGACACGCAGGAACGGCTCCAGCATCAAGATCCGTCTGACAG*GAGAAACTTCTTAAGTCTCAGCGTAATCTCGTGGCCAAGATGAAGAAAGT | TCGA |
| ACGGTGTGCAAGGGGAGTCGTGGCGTCGTCTGCGACCGCTGCCTTCTCGG*ccatgtgaagatgtgcttgcttcccctttgccttctgccatgattctaag | TCGA |
| GGCCTTCTCTAGGCCCTGGCTGCTACAAGGCCATGTCCGGACCCACACTG*AAAAATTTGCAAGAAGAAATTGACGCCTTAGAATCCAGAGTGGAATCAAT | TCGA |
| TTCACCTGTCCAGCATCCGACCACCGAGGCTGGAGGGGGAGAACACCCAG*GACTTGATTAGAGACCAAGGATTTCGTGGTGATGGAGGTAAGTAGTGATT | TCGA |
| ACTGGTTGAGGAGCAAACTGGAAGAATCCCAGCCCACTCATCTCATTCCC*ACCAGGATGAAAGAGCAGCTGAGCTCAGCAGGGAGCAGAACGAGAAAACC | TCGA |
| AAGAGAACGCGCCCAAGTGGCGGGGACTACTGGTGCCTGCGCTGAAAAAG*TTCCAGGAGATGCCTACACCGATTATGTAGCTACGAGATGGTACCGAGCT | TCGA |
| TTCAACAAGATGCTGACTGGGACAGCGTTCTGTCATCTCAGCAAAGAATG*CCCCTAGTCCAATTCCTGACCCTTCAGTAGCAACTGTTGCCACAGGGGAA | TCGA |
| GGAGAAGCCACCCAGCCTCGGTGACCAGGAGACTCGGCGCAAGGCCCGAG*GAGATTGCATCCTACCTGATCACCTTTGAGAAGCATGATGAGTGGCTGTC | TCGA |
| ATCTCAAACAAGACAACTTGAGCTGAAAGCGAAAAACTATGCTGACCAGA*TGACAACACTGCAGCTGAAGGAGGAGTTGCTGGATGGAGAGGATTATAAT | TCGA |
| gAGCCTCCTCGGCCTTGGAGAGCAgcggcggcggcggcaccccgggcgcg*GCCGCACCAGCTTCTATGAGGAGTACGGTGTCATTCGCGACGTCCTCCAG | TCGA |
| GTTCAAGAAAGACAAAGAGATCATTGCTGAATATGAAGCCCAAATAAAAG*CTCTGTGCCTCGTCCTGGGCTGCATGATCTTTCCTGATGGCTGGGATGCC | TCGA |
| GCCATTCAGGCCCGAAAGCGGAGGCCGAAAAGAGAGAAACATCCGAAAAA*GTTTTCTTAAATGTTTTCTTGGAGCCTTAAAGATGGAGATGACAGAAATG | TCGA |
| AGCCTGGACCACCACAGCCACCCCAGCAGAGGCCTTATGGATATGACCAG*ATCATGCCCAAGAAGCCAGCAGAGGACGAAAATGATTCGAAGGGAGTGTC | TCGA |
| GGTCTACATCCTGCGGGTGCCTTTGAAAATCAACGACAACTTGAGCACAG*AATGAAGTGCAAACAGCAAAAGAATTTATAAAGATTGTAGAAGCTGCAGA | TCGA |
| GGTCTACATCCTGCGGGTGCCTTTGAAAATCAACGACAACTTGAGCACAG*ATAAACCTCTCATAATGAAGGCCCCCGCTGTGCTTGCACCTGGCATCCTC | TCGA |
| TCTTTTTGGATGGCTGCTTGTCGCAATATTGTCTTCTTCGCTTTAGACAA*AATGAAGTGCAAACAGCAAAAGAATTTATAAAGATTGTAGAAGCTGCAGA | TCGA |
| TGAGCGCCATGGCTCACTCCCCGGTGCAGTCGGGCCTGCCCGGCATGCAG*GAATGTCCAAATCACACGTGAGTACCCTTTTTCCTTCTCTACATATCTTC | TCGA |
| ATGTGCTGCTTAAAATTCAGGGCTAAGCATTTTGGGTGATTTTAAACTAG*ACATGCTGAGGGGGACTCTACTGTGCGCGGTGCTCGGGCTTCTGCGCGCC | TCGA |
| gcggcggagcgggagcgggtaccgggcgcgggcagagcgcggcggcgcgg*GTGCGCCCACCTCGTCCACATGTTGTTAAGAGACCAAAGAGCAACATCGC | TCGA |
| CGGGACAGAATTGAATCAGGGAGATATGAAGCCTCCAAGCTATGATTCTG*TGTACCGCCGGAAGCACCAGGAGCTGCAAGCCATGCAGATGGAGCTGCAG | TCGA |
| GACGGCCGGTGAGATTGGAAGTGGCGGCGGCGGCAGGCGGCAGAGGGGAG*CAACTACACTGCGCTGCATCGGACTCGACGCCCGCTGGTGACGCACACGC | TCGA |
| ATGGCCCGCAGAGTTAATGCAGGGCTTGGCACGATACGTGTTCCAGTCAG*ATCCCACCCATGAGGGCACAGCAGAGGGCATGGAGAACATGGCAGATGCT | TCGA |
| GTAGAAGCTTCCACTCCCTCCTGCCCAGATCCGGCCAAGGACCTCAGCAG*TTTTACAGAGTTTTTGGATCCGTTCCAGAGAGTCATCCAGCCAGAAGAGA | TCGA |
| CCAGGAAAAAGAGGAAAGTGGACTGTCCGGGGCCGGGCTCAGGAGCAGAG*GACGAATGCCTTAGTTTCCACTTTCCAGCCTGGATCCCCTCACACTGAAC | TCGA |

| | |
|---|---|
| GATAACCCTTATCTGCGGCCGGTGAAGAAGCCCAAGATCCGCAGGAAGAA*ATCCTGTGGATGTCCTGAAGGCCCTGGGTGTGCAGGGAGGCCAGGCTGGG | TCGA |
| AATGCCAAGACAGAGCATGGGTGGTGCTCGCACACAGCAGGATTTAAGAG*TACAGACCACTTCTGCAAATGAGAATAGCACTGTTTTGCCTTCATCCACC | TCGA |
| TGCCTCCATTCCCGGGAGGGGGGAGCGGCGTTGGAGGCCACCGTTTCCAG*GTTTCTGCTTTGGGACAACCATACATCTAATTCCTTAAAGTAGTTTTATA | TCGA |
| GTAGACAAGCCCTCAAGGATGCCCGGTCCCCATCACCGGCACACATTGTC*AAAAGGTCACGGATCGCCTACAGTGACGAAGTACGGAATGAGCTCCTGGG | TCGA |
| CCACCAAACCCAGGAGGAGACTCTCCCTGGAAAACTTCCCTTCCCTTTCG*TGTCCGAGCTGCAGGAGGAGGGAATGAACGCCATCAACCTGCCCCTCAGC | TCGA |
| CTTTGGAATCTTTGCCCTGATAATATGGAAGCCTGTAAATCAGAGACAAG*GGCATTTTTGGTCGAAGAGAGCTGAAGTAATGAGAAGACATCATGGAGGC | TCGA |
| AGTACTCCAGGAAGCACAGAGCTGTACCAGTACACGGCCCACGTTGCCAG*AGTTCGAACCAACGTGGCCTCAACCAGATTTGGAATGTCAAAAAGCAGAG | TCGA |
| AATCTGCATAGTCTTAATTTGTAAAAAATAAAGAAAATTCCTTAACCTTT*aaaaaaaaaaaaaaaaaaaagaaagccaggtgtggtggtgcacgcctgta | TCGA |
| CCGGGGGCGCCCTGGAGGAGTCCCTGAAGATCTATGCTCCTCTGTACTTG*GGGTTCAGCTGTTGCAAAAATAATTGGTAATAATGTCAAGAAACTTCAGA | TCGA |
| CTGTGCGCGCGGTCCCTGAGCTCGGGCTCTGGTTAGAGACAGTCAACGAC*AAAGAGCTCTTGGAGGAAGTTGGCCAGAATGGCAGCCGAGCTCGGATCTC | TCGA |
| CGTCGTCTGCACGCAGCCCAAATCCCCATCCGGGACAGTGTGCACCTCAA*GAAGCCTTATCAGTTGTGAGTGAGGACCAGTCGTTGTTTGAGTGTGCCTA | TCGA |
| aggcggaggcggagggcgaggggcggggAGCGCCGCCTGGAGCGCGGCAG*GAAGCCTTATCAGTTGTGAGTGAGGACCAGTCGTTGTTTGAGTGTGCCTA | TCGA |
| ACCTATCATTACTCGATGCTGTTGATAACAGCAAGATGGCTTTGAACTCA*GAAGCCTTATCAGTTGTGAGTGAGGACCAGTCGTTGTTTGAGTGTGCCTA | TCGA |
| aggcggaggcggagggcgaggggcggggAGCGCCGCCTGGAGCGCGGCAG*GTTATTCCAGGATCTTTGGAGACCCGAGGAAAGCCGTGTTGACCAAAAGC | TCGA |
| GTGTGATGGCGTGTCACACTGCCCCGGCGGGGAGGACGAGAATCGGTGTG*GAAGCCTTATCAGTTGTGAGTGAGGACCAGTCGTTGTTTGAGTGTGCCTA | TCGA |
| CTTTTGAAGAAGTAGGACCTGCCCTGCTCCGCTACAGAACTACTGGACAG*GAAGCCTTATCAGTTGTGAGTGAGGACCAGTCGTTGTTTGAGTGTGCCTA | TCGA |
| CCCTGCCCCGCCCGGAGGTGAAAGCGGGTGTGAGGAGCGCGGCGCGGCAG*GAAGCCTTATCAGTTGTGAGTGAGGACCAGTCGTTGTTTGAGTGTGCCTA | TCGA |
| aggcggaggcggagggcgaggggcggggAGCGCCGCCTGGAGCGCGGCAG*GAACTCTCCTGATGAATGCAGTGTGGCCAAAGGCGGGAAGATGGTGGGCA | TCGA |
| CCCTGCCCCGCCCGGAGGTGAAAGCGGGTGTGAGGAGCGCGGCGCGGCAG*GTTATTCCAGGATCTTTGGAGACCCGAGGAAAGCCGTGTTGACCAAAAGC | TCGA |
| ACCTATCATTACTCGATGCTGTTGATAACAGCAAGATGGCTTTGAACTCA*GTTATTCCAGGATCTTTGGAGACCCGAGGAAAGCCGTGTTGACCAAAAGC | TCGA |
| CCCTGCCCCGCCCGGAGGTGAAAGCGGGTGTGAGGAGCGCGGCGCGGCAG*TGGCTTTTCATGGCCTGCCACTGAAAATCAAGAAAGAACCCCACAGTCCA | TCGA |
| ACCTATCATTACTCGATGCTGTTGATAACAGCAAGATGGCTTTGAACTCA*GTCTCGGCCCCCGCTTGGGGCCCCGGCCGTGCGGCCGGAGGGAGCGGCCG | TCGA |
| aggcggaggcggagggcgaggggcggggAGCGCCGCCTGGAGCGCGGCAG*AAATCGCCCGGAAATGGGAGCTTGCGCGAAGCGCTGATCGGCCCGCTGGG | TCGA |
| GCTGCCGCCCGGGAAGTACATGGCCGGCCTGAATCTCCATCCGCACCCGG*TGAAGAAAAAATGACTGCTGCCCGCATTAGAAAATGCCACAAGTGTGGGA | TCGA |
| CCTCGCCTCACAGCGAAACGCCGCGCACTCGGAGCCGGCACGGCTCGGCG*GTCATTGCTGTGGTCATGGACCTCTTCACTGATGGTGATATCTTTCAAGA | TCGA |
| ACGCGGCCAAGGGGATCGCCGACCTGCTGTCAAACGGTCGCATGAGGCGG*GGAGAAGCGCATTGGCATTGACCTGGTGCACGACACAGTGGAGCATGAGC | TCGA |
| TAAAGAGTCAGCACCATCAAATGGAAGTGGAAAGAATGACTCTAAAGGAG*ACACAGCCGCCAGCCCAAACAGCAGCGGCATGGGCAGCGCCAGCCCGGGT | TCGA |
| CCCCCAACTATGGCAACCCTGACCCCTTCGTCATCAACAACACACAGCGG*GATGCCCAGGAGCACAAAATCTCCCTGTTTGAAGGGGCCAACTTCAAGGG | TCGA |
| AAGAAAAACATGTCATGTAGGTTGTCTAAAAATAAAATGCATTTAAACTC*aaaaaaaaaaaattagccaggcgtggcggcatgcatctgtactcccagct | TCGA |
| AAGAAAAACATGTCATGTAGGTTGTCTAAAAATAAAATGCATTTAAACTC*aaaaaaaaaaaaaaaaaaaaaaaaaaGTAAGTGTGCTTGTAAATCATAG | TCGA |
| TTGCAACTTTGAGTCAGACCTGTAGGCCTGATAGACTGATTAAACCACAG*GTGGCACCCAAAGATGCTGGAATCTTTATGGCAAATGCCGTTACAGATGC | TCGA |
| ATCCAGTTTTTGAAATGGAGTTTTTGCCAGCTGGGAAGGCAGAATCCAAA*ATCCGGGTGGTGAAAGCGTTCCGTAGCTCGCTCTATGAAGGCCTGGAGAA | TCGA |
| TAGAAGCCATGTGTCTTCAGAAGCAATTATTTGGGTGTCTAACAAATCAG*GAAGCCCATGAAACCTTTAATGAGTGGTTTAAGCATATGAATTCAGTTCC | TCGA |
| ATAGTAAGAAGGGGAAAACTGCGCAGGGGTTAAGCCCCGTGGACCAAAGG*ATGAGCTTTGATCCAAACCTTCTCCACAACAATGGACATAATGGGTACCC | TCGA |
| CACTGTGGGTGGGGAGAACACCAACTTCAGATAATAAAATTGTCCTTAAG*GTCATGGCCTCGCATCAATAGTGCCACAGGTGAGTCGGGGCTCTCTGTCA | TCGA |
| CGCCCACTCCAGAAGTAGCATGGAAATGGAGGGCATCTTCAACCACAAAG*GGGTTGGCTGTGTTCCGGCCGCAGAGCACCGTCTGCGTGAGGAGATCCTG | TCGA |
| GTCAGAGACATCAAAACCTGGACCTTCTGGATTACAGGCCAAATTAGCAA*GGTGGATCTATGACATCACGCAAAAATATGATTTTTCCTTCTACATATGT | TCGA |
| ACAGGTATCTGCTCCTCATGGCGCAGGAGCATCTGGAGTTCCGCCTGCCG*AATGAGGACCTAATTTGCTTCAAAGATATCAAACCAGCAGCAACTCATCA | TCGA |
| GGATGGGTCTGAGGAATTCGAGACCATAGTTTTGAAGGCCCTTGTGAAGG*GCGCTGGCGCAGCGGGAAGATATGGAGGAGCGGATTACAACACTGGAGAA | TCGA |
| GGGAGGCCACTAGCAGCCAGTTCCGGCCCGAGGACACCGACTACCAGCAG*AGCTGAAGAAGCAGGTGGAGAGTGCCGAGCTGAAGAACCAGCGGCTCAAG | TCGA |
| GGGAGGCCACTAGCAGCCAGTTCCGGCCCGAGGACACCGACTACCAGCAG*GGGGGGCCCTCAGCCTCTTGGCCTTGAACGGGCTGTTCATCTTGATTCAC | TCGA |
| GATATGGACACTATAGATGTTTCCAATCTAAATAGGCAGTTTTTATTTAG*TGTCTTCTACAAATGCTTCTGGGAACTGGAGGACCCAGTTAACCATAGGG | TCGA |
| CCACCGAGGACACCTCGGTGGAGAAGCTCAAGGAGCGCTGCCTCAAGCAC*GTCTGGAAATCTCTCCAGGCACCCTGCTTTGGCCCTCTAGAATACAGACA | TCGA |

| | |
|---|---|
| AGAAACTGATGTGGATTAAGAAGGCAGCCGAAGACCAAAAGCGATATGAG*CTACAAGAGACTGTGAAAAGGAAGTTGGAAGGAGCTCGATCACCACTTAA | TCGA |
| CCGAGGAGCAGCAAAAGCAGTACAAGGTGCACCTGGACCTCTGGGTTAAG*CTACAAGAGACTGTGAAAAGGAAGTTGGAAGGAGCTCGATCACCACTTAA | TCGA |
| GCTACCGCTCAGGTGCTGCAGGCACAGCCCGAGAAACCGCAGCACTACAC*ATGTTGTGACTGTGACAGACTCACTGGGGTTTGTACATGCTGGGGAGGAG | TCGA |
| GCTACCGCTCAGGTGCTGCAGGCACAGCCCGAGAAACCGCAGCACTACAC*GATACCGTTAACCCTGGATAAAGAGAATGGAGGTTACCTCTGTCCGTGTA | TCGA |
| CCCTGAGACTCTGCTGGGGGAGATGGAGGCGAAGACAAGAGAGCTCATTG*GTAACAGTTGACAAATGGGAACCTTTGTTGAACAACTTGGGGCATGTCTG | TCGA |
| CCACGAGTTCCTCATCGCGGCCCTGGACGTGCTCCACCGACACTGCAAAG*ACTTATTCTGGACTCTTCTGTGTAGTTATAAACCCTTACAAGAATCTTCC | TCGA |
| ATTTCTCCAACCGAGTTGAAATTTCCAAAGCAAGTGCTTCTTTAAATGGG*GATGATGGTTTGCAGAAAATACACACATCTGTCACTAGAACTCCTTCAGT | TCGA |
| tatatttagaaggagaaaagtgccaaggaggaatgaaaaagtagaacaag*GGTTGCCACACCATTTGGAGGTTTTGAAAAAGCATCAAGAATGGTTAAAT | TCGA |
| ACGAGGGCCTGGAGTGTGTGCCCACTGAGGAGTCCAACATCACCATGCAG*GCCTTCTCCACAGTAGGCACTCCTGACTACATTGCTCCTGAGGTGTTCAT | TCGA |
| TACCCCTCACGGAGGAAATAGTTTTGCTAAGAAATAAAAGTGACTATTTT*AAAAAAAAAAAAAAACTACGGCGGCCGAGAAGGGGCGAGTACAGCCCAGT | TCGA |
| TTTTCCAGCAAGTATCCAACCAACTTGGTTCTGCTTCAATAAATCTTTGG*aaaaacaataaaaaaataatgcaacaattaaaaaagaaaaatacagtgtaa | TCGA |
| ACAGCAAAGAGGTTGAGAGCTTCCAGCAGCTGCTCAATGCTCGGACACAG*GTCTTGCCCCAGAAGCTGCGGGCACATCCACGCCTGAAATGCGGCGCTCA | TCGA |
| CAGATTTCAATGCTGTTGAGTATATCAATACCCTGTTCCCAACCGAGCAA*GTTCTGCGGCATGAGGAATTTGAAGAAGGCTGCAAAGCTGCCTGTAATGG | TCGA |
| AATGGTGTTCGCATTTTGGAAGGTCTTTCTGATCCTAAGCTGCCTTGCAG*GCCAACCCTTTAAATTGGATCCCAAAATGACTCACAAGAAGTTGAAGATC | TCGA |
| GGGCGGCCGGCCGACCCTTCCTGTCGCTGACGGCGACTGCGGGAGGCCAG*GCGCTGTCCACGCCCCGAGGCCTGGGCTCAGGCCGAAGTTGCTCCACAGC | TCGA |
| GGGCGGCCGGCCGACCCTTCCTGTCGCTGACGGCGACTGCGGGAGGCCAG*ATGGGAGCTTGCCTGATTGTCAACCCATCGGCCTGGACCCCAGAGCCCAC | TCGA |
| GGGCGGCCGGCCGACCCTTCCTGTCGCTGACGGCGACTGCGGGAGGCCAG*GTCTCTGACCCCTCCCAAGGATCATGCCGCAGCCCCACTGACCCAGGAGT | TCGA |
| GAGATGCAGCATCCCAGTTTTTGGTCTTCTGTCAAAAACACAGGGATGAG*GGGCCACCAAGGACATGGGGAAGATGCTGGGGGGTGACGAGGAGAAGGAC | TCGA |
| ATTGGCATCACTTCCTGATAGCAAGTTCATCTGCATGGAATGTAAAACTG*AACGAGACCAAAAGTTCAATACAATGGCTATGCTGGGCATGAAAACAGCA | TCGA |
| CCTGTGATCGCACTGACACGGCCGGGGGGTTAGAATGGAACAAACTGAAG*GGTCAGTTTGAAAAGGAGGATCGAGCTCACTGTGGAGTATCCATGGAGAT | TCGA |
| TACAGGCCAATGTGACCGTGGGGACTTGCTCGGGCTTTTGTTCCCAGAAA*GAAGGAACACCAGCCAGCCGTGGAACCTCAGGTGCAACAGAGACGCCAGG | TCGA |
| GACAACATCTCGTCAGCTAACATTAGGAGCCTATAAGCATGAAACAGAAT*CTGAAGGAAGAGGATCACCTTCCAAGACAAAACAGTCTTCATGGGGGAAA | TCGA |
| GACAACATCTCGTCAGCTAACATTAGGAGCCTATAAGCATGAAACAGAAT*GGATGTGTTTGGCCAAATGGGAGAGTGGTTACAACACACGAGCTACAAAC | TCGA |
| TCCAAAGCACAGGAATCTCACTCTGTTAAAGCTGGTCTGTTCTAACTGAG*GACATGTTCCCGATTTGAGGTGAAACCATGAAGAGAAAATAGAATACTTA | TCGA |
| CCGCTCCGACTCCAGGGGAGCCGTGGCCTCCTCTCCGCCCTAGCGCTGAG*GTGACCAAGAATATGGCCCAGGTGACCAAAGCCCTGGACAAGGCCCTGAG | TCGA |
| TAATAACATCAAGAACTTAAACGAAAATTGAACATTCTGACTCGACCGAG*AGTGAAGATGAAGCTGGATGTAGCTCAGTTGATGAAGAAAGTTACAAGAC | TCGA |
| GAGAAGAGCAGCAAGAGCTTTACCTATCTATACCACATCAGCTTCAAAAA*GAAGCAATAGTGAAAGAACCTGAAATTAACACAACTCTTCAGATGCGTTT | TCGA |
| TTTGATTCTTATTTCTTTCAGATGCATAGAAAAAGTAACAACTGATAAAG*CTCTTTCGACGTGTAGCAGCAGCTTTGCCGGGAATGGAAAGCACACAGGA | TCGA |
| CAAGAGCAAAGAAAGGACGACGTAATCAAAGTGTGGAGCCCAAAAAGGAA*CCACAGCGTCCTGTGTTTACTCATGAAAATATTCAAGGTGGGGGGGAACC | TCGA |
| CCTGTATAAAGATGTCATGATGGAGAATTATGGAAACCTGGTCTCACTGG*GTTTTGAGGTTTTGCTGAAGATATTCCTGAAGAATCATCCCAGGTGCCAC | TCGA |
| CCAGAGTGGCTAGTCTCAAAGCGCATATTATGCTACATGAAAAGGAAGAG*GTTTCTGCGGCACTGAAACAAGGCCAAATTACACCAACAGAGCTCTGTCA | TCGA |
| TATTGGGAGAGCCACAGCTAAACCCCGGGACCCCTGGAAGCCTAGAAATG*agtctgggggccgggcccTGGAGCAGGAGCTCCCTGGAGCTGTCTTTATC | TCGA |
| GGGTACCGGCCGTGGAAGCGGCCACCGACTACTGTCAGCAGCTGTGCCAG*GTTGGTAGGAGGAGAGGATTGGAGCTGTTTTCTCCTTGATGCCAAGATAC | TCGA |
| AAAAGAGCCCTGGAATATGAAGCAACATGAGATGGTGGATGAACCCACAG*CAGGTGCAGTTAATAACCAAAGCAGGCCACAAAGCCACAGCAGTGGAGAA | TCGA |
| AGCTCTCACATGTGGAAGCTGCAAGGTCTTCTTCAAAAGAGCCGCTGAAG*AGCCCACTGCGGAAGAGGGCAGCCCGGCAAGCCCGGGCCCTGAGCCTGGA | TCGA |
| AGGGCAAAAAAGACAGGGACATGGATGAACTGAAGAAAGAAGTTTCTATG*CTCTAAAGTCTTTCTGGAAATTGACAACCGCCAGTGTGTTCAAGACTCAG | TCGA |
| GCCCTAACATCTGGATCATCCCCTTCCGCACCCGCCTCGGACTCTATTGG*GGCCGACGAGATTGAAATGATCATGACGGACCTTGAAAGGGCAAACCAGG | TCGA |
| TATTTATTTGTAGGACTTGATTAGAGACCAAGGATTTCGTGGTGATGGAG*GCACCCAGTTGGAGAAGCTGATGGAGAACATGCGCAATGACATTGCCAGT | TCGA |
| gcgccggcgccgcggccTCTTCGGCTGCGGACCCTGCCATTCCGGAGGAG*ATTGACTGTTCAAGTACTATTATGCTGGACAATATTGTGAGGAAAGATAC | TCGA |
| AGCTTTCAGGGCCATGGTTCACATTAGAATAAAAGGTAGTAATTAGAACA*aaaaaaaaaaaaattagccaggcgtggcggcatgcatctgtactcccagct | TCGA |
| ACAGAATTATTTGACCTGAAAACCAAATACGATGAAGAAACTACTGCAAA*GCCCCAAATTCTCACCAGTCCGTCTCCTTCAAAATCCATTCCAATTCCAC | TCGA |
| GCGGGTGGCAGCGACAGAGCCAAAATCAGAGCTGGAACCTGAGGAGAGAG*TGTCCCCACACCCTCCTCTGAGACGCCATGTTCAACTCGATGACCCCACC | TCGA |
| ATCAATAATGGCACTCAGCAGCAAGCACAGAGTATGACCACCACTATTAA*TTGGACAGTGCTCCAGGGAAGAATCGGGCGTCCAGACAACCCATTTCGAG | TCGA |

| | |
|---|---|
| GGAAAAAGACTTTAAAACTCCCTCCAAAGGATCTAAGAATCAAAACTTCG*TTGCCAGGTATAAACTCCGAATTGTTAAGCCACCAAAATTACCCCTAGAG | TCGA |
| AGTTTTTTGTTTTGTTTTTATACGTGTCTGAATAAAAATGCCAAAGTTTT*AAAGCCCGAGGATCCTTTCAGCAGCACAGAGCAGAGACCGCGCTCCCCAG | TCGA |
| TTGTGCAGCattaaaaaaaatcaaaataaaaattaaatGTGAGCAAAGAA*AAAAAAAACTGGGTGAATACTCCAAGAAGGCGGCTATGAAACCCAAACCT | TCGA |
| ATTGTGTTGTGCAGCattaaaaaaaatcaaaataaaaattaaatGTGAGC*AAAAAAAACTGGGTGAATACTCCAAGAAGGCGGCTATGAAACCCAAACCT | TCGA |
| CTGAACCCGGCCCCCGACGGCTCTGCAGTTAACGGGACCAGCTCTGCGGA*AAAGCACTCCAAAGACCTCCGCCAGCTGCAGCCCTGCCCCTGAGTCCCCG | TCGA |
| GCAAGGTTTACTCTGAATCCCAATCCTACAGGAGTTCAAAACCCACACAT*CTGAGCCAGTACCAGCAGCCAGATCGGAGGACCGGTACCGGAATCCCAAA | TCGA |
| ATGTGGCCACACGTTGGTACAGGGCTCCAGAAATTATGTTGAATTCCAAG*GCTTCACCGGCCAGAACTGTGAGGAAAATATCGACGATTGTCCAGGAAAC | TCGA |
| ATGGTGCTGTAACCACCTCACAGATTCCAGCTTCGGAACAAGAGACCCTG*TGAGCCAAGGGAGTTTGTGGAGAACTCTGAGTGCATACAGTGCCACCCAG | TCGA |
| AAATAATCGACTTGTTCTTGATAGCCTCATTAAAGCATTTGGTTTTTCAC*AAAAAAAAAAGAGAGACAAACTCTAGGAACAATGCCGATTCATTCCACGC | TCGA |
| ACTGGAACGCCAACCGGCTGGACCTGTTCGAGATCAGCCAGCCGACCGAG*GTGGACACCTGCGCAGGTGTGTGCCCTCCGGCCCCTGAAGCATGGCCAGC | TCGA |
| ATCCGAGGACTATGAGAGCTTCTCCTGTGTCTGCCCCACGGGCTGGCAAG*GGCTACACCAAGTCCATTGATATTTGGTCTGTAGGCTGCATTCTGGCAGA | TCGA |
| ATTGACCCTGTCCCTGAGGAGGAGGACCTGGGCAAGAGGAACAGACACAG*ATATGAAGAAGCTGAAGTCCGGAAAAAACTTGAGGAAAAAGACAGGCAGG | TCGA |
| TATAGATTGTCTTAATGGTAGGTCAAGTAATAAAAAGAGATGAAATAATT*AAAAAAAAAAGAGAGACAAACTCTAGGAACAATGCCGATTCATTCCACGC | TCGA |
| CGGTTCTCCTGGGAACCCCATCGGCCCCCTGCCAGCACACACCTGAGCAG*TTGGGACCCCTGATTTTAGCAGGATGGCCCAATGGAATCAGCTACAGCAG | TCGA |
| TTTAGGAGACAGCCACTTTCTGTGTGGTACGCTGGATTCAAGGATGCCTG*GGAAAAGCTTCACTCTGACCATCACTGTCTTCACAAACCCACCGCAAGTC | TCGA |
| GACTGAGATCACTGATGACCTGCACTTCTACGTGCAGGATGTGGAGACCG*GATCAACCACAGGTTTGTCTGCTACCCCCCCTGCCTCATTACCTGGCTCA | TCGA |
| GTTTGGAAATAATGGTGAAGGTGCTGAACCCTCAGCAGGAGGGCAGTTTG*TAGCTTTCCACAGCCCCACCACCAGGATCAAGAAGGAGCCCCAGAGTCCC | TCGA |
| CCCTCCCCAGCTGTGTGTCCCCAGGCTCACTATAACATGTACCCACAGAA*ACCCACTCCTTGCCAGTTGTGGTGATCTCCAACATCTGTCAGATGCCAAA | TCGA |
| TAAAATGGAATAAGAAAGGAAATTTCATCCTAAGTGCTGGAGTAGACAAG*GAGGATCCAAAGTGGGAATTCCCTCGGAAGAACTTGGTTCTTGGAAAAAC | TCGA |
| TGCCTCCATTCCCGGGAGGGGGGAGCGGCGTTGGAGGCCACCGTTTCCAG*TTGGGACCCCTGATTTTAGCAGGATGGCCCAATGGAATCAGCTACAGCAG | TCGA |
| CAACACAGATCAGCCTAGCTCAATTACGGCTCCAGCATATGCAGCAACAG*GTGTGGAATATCAAACAAATGATTAAGTTGACACAGGAACATATAGAGGC | TCGA |
| ATTTGAGCTCACCAGTGGGAGGGTCTTATAATCTTCCCTCTCTTCCGGAT*GTGTGGAATATCAAACAAATGATTAAGTTGACACAGGAACATATAGAGGC | TCGA |
| TCGCTCAGCTAGAAGAGAAGCAGCAGCAGCCCACCAGGGAGCTCCTGCAG*GAGGATCCAAAGTGGGAATTCCCTCGGAAGAACTTGGTTCTTGGAAAAAC | TCGA |
| GAGAAACACAGGAGAGCGAGCGCAAGCCCCCACCATACAAGCACATCAAG*AAATTAGCAGAGTACGGAAAGACATGTACAATGACACATTAAATGGCAGT | TCGA |
| GCACCAGTGTTGATGTGGATCTTTGGCTGATCACAGCATTTCATGAAAAG*ATTATATTCTCCTGCCTTTTAAAAAGATGGACTTCAGCAGAAATCTTTAT | TCGA |
| CCCAGAAGTTGTTCTAACGCTGCCTTTGCTTTAAAACAGTATTACTTGCG*CCATGGTCGAGAAAGACCCTCTGCCCGTCTGAGCATTCGTAGGGATCGAC | TCGA |
| tatgaggatgacgaggaggatgacgTGCAGAACACCAACTCTGCCCTTAG*AACAAAACTGAAGAATCAAGAATATGAAACTTTAGATCATTTGGAGTGTG | TCGA |
| CGATATTTTGTCGAAGAAACTGTGGAAGTCATTAGGAACAATGGTGCAAG*CCGCCGCCGAATCATGTCGATGAGTCCAAAGCACACGACTCCGTTCTCAG | TCGA |
| AATTGTGGTTGTGCAGCCAGATGCCTGGACAGAGGACAATGGCTTCCATG*GTTGGTTCTACTGCTGTCCAAGTAACTTCAGCAGAGCGAACAAAAGTCCT | TCGA |
| TCCACAATGCTCTCAAGCTAGCTCACAAAGACACCAAAGTTTCTATTAAA*ATGCTGAGTTTGTGTGTGAACGGACACTGAAATATTTTCTAGGAATTGCG | TCGA |
| ATCACATCCTCTGATTGGCAACATGTTAGCTTTGAAGTGGATCCTACCAG*ctccagaacctgggtatcaacccagcaaacATTGGCTTCAGTACCCTGAC | TCGA |
| GCTGGGACTTCCAAAGCTGGGAATCCTATTTTTTATTATGTTGCACGGAG*AATAATCATGGGCCAGACTGGGAAGAAATCTGAGAAGGGACCAGTTTGTT | TCGA |
| TAATCCAGGATATATCCAAAGACGTGGTTGATGAAAACAACATGAATAAG*CCTGAAAGAATAGACCCAAGCGCATCACGACAAGGATATGATGTCCGCTC | TCGA |
| GGACCAGCAGCGCGGGAGAGCGGACTCCCCTCGCCACCGCCCGAGCCCAG*CTACATCTACATCCACCACTGGGACAAGCCATCTTGTAAAATGTGCGGAG | TCGA |
| gcggaggggcgggggaggcgccgggAAGTCGACGGCGCCGGCGGCTCCTG*TGCCCAGCAGAGGAGCACGCAGCTGAACAAGAAGAGAGCCATCCCCAATC | TCGA |
| GAGCCAAGCCGGTCGTGAGTGACGATGACAGTGAGGAGGAACAAGAGGAG*ACGAAGGCCTTCATCCACCACGAGCTGCTGGCCTACCTATACTCCTCGGC | TCGA |
| GATGGGGAAAAGACATTTGTTGCACAAATGACAGTATTTGATAAAAACAG*GTTAGAACCATCAGAGAGCCTTGAGGAAAACCAGCGGAACCTCCTTCAGA | TCGA |
| CCTGTGGACGCGTATCGCCAGCACGATCCCACCGCACGTTCAGAAGTCGG*ATGCCTTTGAAGATGAGAAACCCCCAAAGAAGAGCCTTCTCTCCAAAGTT | TCGA |
| TGCTGAGAGATCGGTAGCCAAGCTGGAAAAGACAATTGATGACCTGGAAG*GCCCGGCTGTGCTGGCTCCAGAGGATGGGCTGGCCATGTCCCTGCATTTC | TCGA |
| TGCTGAGAGATCGGTAGCCAAGCTGGAAAAGACAATTGATGACCTGGAAG*ACACTAACAGCACATCTGGAGACCCGGTGGAGAAGAAGGACGAAACACCT | TCGA |
| TCTGTTCAAGTTCTAAGAGAAGCAAGGAAGAAGTAGGCATCAATACTGAG*CTACATCTACATCCACCACTGGGACAAGCCATCTTGTAAAATGTGCGGAG | TCGA |
| AGCTGCTGCAAAGTTGGCGTTTGCCGTCTTTCTTATCTCCTGTTCTTCAG*AATGCCAGTTAATGAAAACAGAACGACCAAAGCCAAACACATTTATAATC | TCGA |
| CAGAAGCCGACAAGGCGTTCAAGCGAAAACATGACCGCTGAGCCCATGAG*GATGCCTTTGTGGAACTGTACGGCCCCAGCATGCGGCCTCTGTTTGATTT | TCGA |

| | |
|---|---|
| TATTTATTTGTAGGACTTGATTAGAGACCAAGGATTTCGTGGTGATGGAG*CCTTCCCAGAGGAGCTTCCGGCAGACCTGAAGCACTGGAAGCCAGGTGTG | TCGA |
| ATTCCCCGGGTCTTCACGTGCAGCACATGGGAACATCCCATGGTATCACA*GAGGATCCAAAGTGGGAATTCCCTCGGAAGAACTTGGTTCTTGGAAAAAC | TCGA |
| GCCACAGGGGCCCCGAAGAACTCCCACAATGCCACAGGAGGAGGCAGCAG*gaaaagaATTCCATGGCAACATCATTAAAGTGTCCTTTGCCACTAGAAGA | TCGA |
| GCGTTGCCTGGCTCCACGCCCTGCTGGGTCCTACCTGTCAGAGCCCCAAG*ATCTCCGAAGACCTGGGCAGTGAGAAGTTCTGCGTGGATGCAAATCAGGC | TCGA |
| GTCAACTCAGTCTCTGATTGAACTTTTTGAATCTTTCAAGAGTTTTGAAG*CTCAGGTACCAGACAGTGATGAGCAGTTTGTTCCTGATTTCCATTCAGAA | TCGA |
| ACTGAGAAAAGAAGAAGAACAAGCTACAGAGACACAACCCATTGTTTATG*TCTGGCATAGAAGATTAAAGAATCAAAAAAGTGCCAAGGAAGGGGTGACA | TCGA |
| AGGTCGAGGACGGTCGGACTCCCGCGGCGGGAGGAGCCTGTTCCCCTGAG*GAGCAGGAAAATTCAGATTCGAAACATCCCTCCTCACCTGCAGTGGGAGG | TCGA |
| TGCAAGAAGATCTTTGGGAGCCTGGCATTTCTGCCGGAGAGCTTTGATGG*CCTCCCCCACTGGCAGGATTGCCATGAGTTGTGGAGTAAGAAACGGCGAC | TCGA |
| TACTGAAGGACATGATTCAGACTGTCCCGGACCCAGCAGCTCATATCAAG*ATGCACCCACTGGGCCTATGTAATAACAATGACGAAGAGGACTTGTATGA | TCGA |
| GAAATCCTCCAGCCAGAGTTCTTTGAGGCTGTAAACGAGGCCATTTTGCT*TGCTTTTTTAGGACCAAAGGATATATTTCCTTACTCAGAAAATAAGGAAA | TCGA |
| GGAAGGCCTGCATCATGATGGCCAATTCTGCCATAAGCCCTGTCCTCCAG*GCAAATACGCCATGCGAGACCTAGTCCACAGGCTTCCAGGAGGGAACAAC | TCGA |
| CATTAGAGATGGCTCTGGCGTTGGTGTTTTCAAAATAGGTGAAGAGACAG*GCAGTTGGAGCAGAACTTTTTCAACTGCAGCTGTGACATCCGCTGGATGC | TCGA |
| AAACTACCATTAGCCAGTCTTTCTTACTGACAATAAATTATTAATAAAAT*aaaaaaaaaaaaaaaaaaaaaaaaaggaaagaaagaaagaaaagaaaCCC | TCGA |
| AGCAGTTGGTAGAAGACTTGGATCGAATTCTCACTCTCACAACCAATGAG*CAAGCCAGGGCTGAGCAGGAAGAAGAATTCATTAGTAACACTTTATTCAA | TCGA |
| ACCTTTTGAGGTGACTATAACTGAAGATTGCTTTACAGAAGCCAAAAAAG*CACGTTCACGTCCATGTTCTTCCCAGGAAGGCTGGAGACTTTCACAGGAA | TCGA |
| tttCCATGTCACTAAAGGCCCTCGGGACCAAGGATCACGTCATGACTCCG*TGTTCAAGAAGGAAGTGTATCTTCATACATCACCACACCTGAAAGCAGGT | TCGA |
| GGTTTACAGATGTTTTCCACTGAGCATCACAATAAAGAGAACCATGTGCT*AAAAAAAACTGGGTGAATACTCCAAGAAGGCGGCTATGAAACCCAAACCT | TCGA |
| GGCGGAGGCCCGAGAGAAGGAGACCAAGGCTCTGTCGCTGGCCCGGGCCC*AGATGCACAGGAGGAGAAGCAGGAGCTGTCGGGAAGATCAGAAGCCAGTC | TCGA |
| CCGCGTGGGCCGCCTCAGCCCCTGTGTGCCTGCGAAGCCACCTCTCGCAG*GCTCTCCACAGATAGAGAACATCCAGCCATTCTCTGCCAAAGACCTGTCC | TCGA |
| GGTCAGCGCCGACGGCAGCCGGCACCTGACGGCGTGACCGACCCGAGCCG*CTTTGATTAGTTTGTCCTTTGGAGGAGCAATCGGACTGATGTTTTTGATG | TCGA |
| TTGTCTTTTCACAGACCATCACCGGCCTCCTGCAAGAGTTTGATGTCCAG*CCTCCCGCGACGATGCCCCTCAACGTTAGCTTCACCAACAGGAACTATGA | TCGA |
| CTTCGTCATGATCAACAAATATGGTCGACGGTCCAAATTTGGAATGAAGG*ATAACTGTGTCCAGAGGACCCCCAGGCCATCCGTGGATAATGTGCACCAT | TCGA |
| GAGAAGCTGCCGCCGAACGACAGGACGTTGGGGCGGCCTGGCTCCCTCAG*GACTGATGGATACATCGTCTGCGAGGAATTCAAAGACGTGCTCCTGACTG | TCGA |
| ACAAGTCCTCAGGCTACCACTATGGGGTCAGCGCCTGTGAGGGCTGCAAG*CATGTCCTCCTCACATTCTTCCACCAGAGAAGAGGCCCCCTGAGCCCCCC | TCGA |
| TCTCCAGCACCAGCTTCCAGTTAGTGGATATAGCACACCATCCCCAGCCA*GATGTCTCCAATACAACGACAGCCCAGAAGAGGAAGTGCAGCCAGACCCA | TCGA |
| TCCGTACTCCACCCCGCTCCGGACTCCGCTTTGGAATGGCTCAAACCACT*AGGAACGCGTTGTGGTGATCAGCAGCTCGGAAGACTCAGATGCCGAAAAC | TCGA |
| AGGAGGGCAAAGAGAAGAAGGAGAAGCACAAGACCAAGACAGCTCAACAG*ATTGATGATGTCATTGATGAGATCATCAGCCTGGAGTCCAGTTACAATGA | TCGA |
| CGCTGGACTCCATGGAGAACCAGGTCTCCGTGGATGCCTTCAAGATCCTG*CAAGCCAGGGCTGAGCAGGAAGAAGAATTCATTAGTAACACTTTATTCAA | TCGA |
| CGCTGGACTCCATGGAGAACCAGGTCTCCGTGGATGCCTTCAAGATCCTG*ACTTCTTCCAGAGCCTGCAATTTCTTCAATAATGTCGGGGGAAACCTAAA | TCGA |
| TGTTTGCTGGTGTTGCTGAAATAACTCCAGCAGAAGGAAAATTAATGCAG*CATCACAGGACATGGCCCCCTCAGCCACCTAGCTGGGGCCCATCTAGGAG | TCGA |
| TTGATTGCACCCACCCTGTAGAAGATGGAATCATGGATGCTGCCAATTTT*GTGTAAGTACTGCGACCGCTCCTTCAGCATCTCTTCGAACCTCCAGCGGC | TCGA |
| GGAGAAAGGAGAACCGTACATGAGCATTCAGCCTGCTGAAGATCCAGATG*ACCGCCAGGACCTGGAGCAGCCGGACCGGGTGGACATGCTGCAGGAGCCG | TCGA |
| GAGGCAAGCGCAGGGGGACGGGGACCAGCAGCTGTCGCCGCCGCTCTCAG*ATCTCCGAAGACCTGGGCAGTGAGAAGTTCTGCGTGGATGCAAATCAGGC | TCGA |
| GAGGCAAGCGCAGGGGGACGGGGACCAGCAGCTGTCGCCGCCGCTCTCAG*CATCACAGGACATGGCCCCCTCAGCCACCTAGCTGGGGCCCATCTAGGAG | TCGA |
| TTCGTGCTGTTGGAGGAGCGCAGGCCGGCCGACTCGGCTCAGCTGCTCAG*GCAAGAGGAGGAACGACGTAGAAGAGAGGAAGAGATGATGATTCGTCAAC | TCGA |
| TTACCAATCCAGAAAACCTTCCATCGCTGCCAGATATTCCACCCATACAG*GAGGATCCAAAGTGGGAATTCCCTCGGAAGAACTTGGTTCTTGGAAAAAC | TCGA |
| CCGGGCTGGGAGATGACCTTCACGGCCACTGGCCAGAGGTACTTCCTCAA*GCCAAGACCCATGTTCTTGACATTGAGCAGCGACTACAAGGTGTAATCAA | TCGA |
| ACCAGAACCTGACTCGGGTGGCAGACTACTGTGAAAACAACTACATACAG*GTTGTGGTGGAGAGCTGTCTGGGGCCACAGGCTCCTTCAGCAGCCCCGGG | TCGA |
| AAGCCCAGTTGCTTAACGAATATGTGGCTGTCAAAATATTTCCAATACAG*GTTGAATGCCTTTGAGAACTTGATGCATAAAATTTGCATGACTCCTCACT | TCGA |
| CATGGGATTAGTCCTATGGGAACTGGCTTCTCGCTGTACTGCTGCAGATG*CTGGGAATGGAGGAAGAAGATGTGATTGAAGTTTATCAGGAACAAACGGG | TCGA |
| GATACAGGTCTTGGACTGGCCTTCACCATTGCCCATGAGTCTGGACACAA*GGGTTGGCTGTGTTCCGGCCGCAGAGCACCGTCTGCGTGAGGAGATCCTG | TCGA |
| GCGACCCCGGCCGCCCGCGCCAGCAAGAGGATCGTCCTGCCGGAGCCCAG*TTTTCAAACCAGAAGAACTACGACAGGCACTGATGCCAACTTTGGAGGCA | TCGA |
| GCGACCCCGGCCGCCCGCGCCAGCAAGAGGATCGTCCTGCCGGAGCCCAG*ATGCACCCACTGGGCCTATGTAATAACAATGACGAAGAGGACTTGTATGA | TCGA |

| | |
|---|---|
| GGACGGAAGACCCCTGGCTCTATAAGCTGAATTATGGCAGCCCAGTCAAG*GCCACCAAAATGAATGGCTGTGAAGAATATTGTGAAGAAAAAGTAAAAAG | TCGA |
| TGACTGGAAACAATGGCATTTACAGAAAGAGTCAACAGCAGTGGCAACAG*GCCACCAAAATGAATGGCTGTGAAGAATATTGTGAAGAAAAAGTAAAAAG | TCGA |
| TGACTGGAAACAATGGCATTTACAGAAAGAGTCAACAGCAGTGGCAACAG*AACCAGTTCGAAGATACAAAACTTATCATGGTGATGTCTTTAGTACCTCC | TCGA |
| AGAGTCGTGCGGACAGCATTGGGAGCGGCCGAGCCATCCCAATTAAACAG*GAGTTACTATATGGGAACTGATGACCTTTGGAGGAAAACCCTATGATGGA | TCGA |
| GAAGCTGCCCACGGAGTGGAATGTATTGGGGACAGATCAGAGTTTGCATG*GAACCTATGGATTTTTGTTTCCAAGTTTACAGAATTTGATGCTCGAAAAC | TCGA |
| AAATAGAGAAACTGGAGTTCAGAGTAAGAGAACTGGAGCAGGCGCTTCTT*ATACAAGGCAGATCCAACCATCCCCACCGTGGTCCTACGATCAGTCCTAC | TCGA |
| AGAGCCCGGGCGGCGCTCGTACTTCTGGCCACTGGGCGAGCGTCTGGCAG*GTCCTTCCAACCCTCCTGCTACCATGAAGATTGCAGTGATTGGACAGAGC | TCGA |
| AGAGCCCGGGCGGCGCTCGTACTTCTGGCCACTGGGCGAGCGTCTGGCAG*GTTTATTACCCAGAACTCTTTGTCTGGGTCAGTCAAGAACCATTTCCAAA | TCGA |
| AAGGTCAAGGAGTGGGAGAAATCAACATGGCAACTTCTGGTAATGGTCAG*CCACTGGCTTCCAAGACACTGAAGACCCGGCCCAAGAAGAAGACCTCTGG | TCGA |
| AGAGCCCGGGCGGCGCTCGTACTTCTGGCCACTGGGCGAGCGTCTGGCAG*ATGGAAAGGAGAAAATGTCGCAACCACTGAGGTTGCTGATGTTATTGGAA | TCGA |
| ATCAGATGCTGCAGTAAAGGAAAAAGCCATTCAGGTTGGCTTTGTTTTAG*GCCTCGAAAAAGCCCAAGACAGCAGAAGCAGACACCTCCAGTGAACTAGC | TCGA |
| CTGGAGCTGCAGCATGGACCAGCGAGAGCCCCTGCCTCCCGCTCCTGCAG*GCATTTTGTGCGCTCCTTAATTAAGAAAGGAATACCTCCAGGATCTGATT | TCGA |
| TGCCGTTCACCCGCCGGAGAGCCTGGCCGTGGTTGCTAACGCGAAACCAA*ATATCAACATGGCAGGAGAGCCCAAACCATACAGACCAAAACCTGGAAAC | TCGA |
| GCATCTGCAAGAGATCAACATCGCCACCAGGACCACCTTCCTCCTGCCAG*TATGAAGTGTAACAGAACAGACTTTACCACCTGAAACTGCTGCTTCAAGT | TCGA |
| TGCTCTTCTTTTGCAGACCCAGACCCAGCAGCTGCTCCACCCGGCACAGG*GTACAAGGAAATGGTAGAACTAGTGATCTCACCCAGCCTCACTGTAAACA | TCGA |
| CGCCTTTGTGAGCGGCGCGGACGACAAAGGCGCGGGCCCGGGCAGCCGAG*GGTCCAGATTTTGGCTATGTAACTCGAGGGCCCCAAACAGGGGGTATCAG | TCGA |
| GCTGTGGGCCCTGGGTCACTGTCCTGGAAGCCACCACCCTGCAGCCTCAG*CTCTGCCCCTGAGGACTGCACGTCCTTCAGCATCAACGCCTCCCCAGGGG | TCGA |
| GCTCGCTGGCCTGCAGGATGGGACCCTTGCTGCTTACCCTCGGACCAGCG*CTCTGCCCCTGAGGACTGCACGTCCTTCAGCATCAACGCCTCCCCAGGGG | TCGA |
| CATCGTCCTGCTGCCCGTGCCTCGGCTGGAAGGCATCCCCAAGATCACAG*GGGGCAAACTGTCCTCCAAGTGCAGCGTAGTCTTGGGTCCCAAGTGGCCC | TCGA |
| TCGGCCAACAAGTGCCGTCTCAGGCTCCTGCTTCCTGGGAAACCCGACAA*AGGACCTGGACCTGGGGCCCTTGTGGACAGGGTGCCATCCTGCTGGTGAA | TCGA |
| GCTGTGGGCCCTGGGTCACTGTCCTGGAAGCCACCACCCTGCAGCCTCAG*GCCGCACCGAGTCCCCGGCACCATGTTTGGGAAGAGGAAGAAGCGGGTGG | TCGA |
| CTCTCGCCAAGGGCCCCAATGAGTGGCACACAGTCTACTATCACCGACAG*GTGTTTGAGCCCAGATGAGTCATGGCTGGACGACCCCTCCGCATAGGAGA | TCGA |
| CTCCAAGAAGACAAAGTCTAGTTCAGAGGAGAGTAGATCCGAGATATATG*ATCTGGGAAATTTTAGTGGCCGGACAAGAAGTGCAGTCTCTGTGAGGGAA | TCGA |
| CTCTCGCCAAGGGCCCCAATGAGTGGCACACAGTCTACTATCACCGACAG*GATGTGAAGGTCTTCCGGGCCCTGATCCTGGGGGAGCTGGAGAAGGGGCA | TCGA |
| CTCTCGCCAAGGGCCCCAATGAGTGGCACACAGTCTACTATCACCGACAG*GAAATGATCGAAATGGCCTAGATTTCAACAGGCAGGTAAGAACTTGGGTC | TCGA |
| CTCTCGCCAAGGGCCCCAATGAGTGGCACACAGTCTACTATCACCGACAG*GTATCTCCCTCAGGGTCTCTGGGCCCCCTCTCTGTCCCTCCTGTCCACAG | TCGA |
| CTCTCGCCAAGGGCCCCAATGAGTGGCACACAGTCTACTATCACCGACAG*GTTGCTATCAAGATCATAGATAAGACCCAGCTGGATGAAGAAAACTTGAA | TCGA |
| GCCCCGGGAGCAGCGGCGGCGGGGGGCAGCCGCTCGCCCGGACCCCTCAG*GTCCATCTGGGAAGCGGGATTTGGGTTGATGAGGAGAAATGGCACCAGCT | TCGA |
| GGCAAGAAGATATGAACCTGAGCCTTCAGTCAAGACCCTCCAGCTTGCCT*GCAGGCCCAATGCCGAAACACATTGCATTCATAATGGACGGGAACCGTCG | TCGA |
| ACATCCTGCTGTATGATGACAACAGCATCATGACCTTCAACCTCAGTCAG*GTAGTATAGATCAATCAGTGTTAAAAGAATTACCCCCTGAACTCCTGGCA | TCGA |
| GAACTCCATGGGGAGCTATGGTCCCCAGGGGGGTCAGTATGGCCCACAAG*AATCTATCTACCGCCGGGGAGCCAGAAGATGGAGGAAGCTGTACCGTGCC | TCGA |
| ACATCCTGCTGTATGATGACAACAGCATCATGACCTTCAACCTCAGTCAG*GTTGTGCACAGGGACCTGAAGCCCAGCAACATCCTGTATGTGGACGAGTC | TCGA |
| ACCTCTATCGCCTCTATGTGTCTGTGAAGGAGATTGGTGGATTGACTCAG*CTCTATTATCAAGTCCTAAATTTTGGAATGATTGTCTCATCGGCACTAAT | TCGA |
| GGCAAGAAGATATGAACCTGAGCCTTCAGTCAAGACCCTCCAGCTTGCCT*GAATAGGCAAGTCAAGAGGCTGAAAAATCTGAAGCATGTTTCCAAAGAGC | TCGA |
| GGCAAGAAGATATGAACCTGAGCCTTCAGTCAAGACCCTCCAGCTTGCCT*TGAAAGATGACAGCCAGGAGTTACCTAAAGACCCACGTAAATGTATCAGG | TCGA |
| GGCAAGAAGATATGAACCTGAGCCTTCAGTCAAGACCCTCCAGCTTGCCT*AGACCAAAAGGTCAAGGAAACAAAGGTAAGTCCCGCTCCGCCGGTGGCCC | TCGA |
| GATTTCAGGTTGAAGTCTTGGCCTCGGAGGATGCAGCCTTTGGACTCAAG*ATGAACGACTTTCGAAGTAAAATGCAATCAATATTTCCAACAATTCCCAA | TCGA |
| AGGAAGCAACCAGTCTCGATCTGGCCCAATCTCTCCTGCAAGTATCCCAG*GAACCCTGATTGCTGTCCTTCAACGTGTTCATTATGAAGTTATTAGTAAT | TCGA |
| ACCATGAAGACTTGAACTTAATACAGCAAGAAAGACCATCAAGTTTACCA*ATCAATGTCCGAGTTACCACCATGGATGCAGAGCTGGAGTTTGCAATCCA | TCGA |
| AGGGTCGGACTCCCGGGGCCATGGCCGGAATGCAGTACCCTCAGCAGCAG*TGTTTGTCAGGTTCAACTCCAGCCATGGTTTCCCAGTGGAGGTCGATTCT | TCGA |
| AGGGTCGGACTCCCGGGGCCATGGCCGGAATGCAGTACCCTCAGCAGCAG*GTACAGGATTATCATGGTCATAAAATTTGTGACCCTTACGCCTGGCTTGA | TCGA |
| AGGGTCGGACTCCCGGGGCCATGGCCGGAATGCAGTACCCTCAGCAGCAG*gtcccaatggctggcatcctagagtacttaatgaattggccgaagtcatc | TCGA |
| AGGGTCGGACTCCCGGGGCCATGGCCGGAATGCAGTACCCTCAGCAGCAG*GACGGTTCACTGGGAAACATCGATGACCTGGCGCAGCAGTATGCAGATTA | TCGA |

| | |
|---|---|
| AGGGTCGGACTCCCGGGGCCATGGCCGGAATGCAGTACCCTCAGCAGCAG*GTTGTCCGGGTGCCAAAAAGCATGAGTTTCTGACCAGCGTTCTGGACGCG | TCGA |
| AGGGTCGGACTCCCGGGGCCATGGCCGGAATGCAGTACCCTCAGCAGCAG*GCTCGGGTTATGTATGATTTTGCTGCTGAACCTGGAAATAATGAACTGAC | TCGA |
| AGGAAGCAACCAGTCTCGATCTGGCCCAATCTCTCCTGCAAGTATCCCAG*GGGACCACAGTGCCCTAGAGTCACAGATCCGACAACTGGGCAAAGCCCTG | TCGA |
| ACCCGGCGATGAGCCCCGGCACCCCCGGACCGACCATGGGCAGATCCCAG*GGCTCTGTGTATGAATGACAAGGATACCTTCAGCCAGCTCATTCTGGATG | TCGA |
| GAGGCAGCCGCAGCAGTGATGCAGGCTGCTGCGAACTCAGCACAAAGCAG*GACAGAATGACTTGGAAGATAACGGAGCTTCGAAGCTTCTAGCAGAATAT | TCGA |
| CCCAGAAGTTGTTCTAACGCTGCCTTTGCTTTAAAACAGTATTACTTGCG*ATTGATGATTACCCGGAATTTTTGAGTTCGCCAGATAGAGAAATCAGCCA | TCGA |
| GCTGCCTTAATATTAAAAAATATTGGTAAATATTCAGAATGTGGTCGCAG*ATATTGATCCCAGGAATGGAATCCCAAAGTTAACTCCAGGCGACAATCCA | TCGA |
| TATTGGTGCAATTCCATCTTCCTACAATTACCAGCAACACAGTGTGTCGG*agacaaggtttccctctgtaacccaggctggagtgcagtggaacttctga | TCGA |
| ACAAGCTATAGAGCAAGTCCAAACCCAGACTCATGTAGCATCTGCCCCAG*gcagactggagctggctgtggattttatttctcaggagaaggaggtggga | TCGA |
| CTCAGGCTGGATTTTGATGATGATGGAGAAGGGAACAGTAAATTTTTGAG*ACTGTAGGATGCCAATGGGACTAAGCACTGGTATCATATCTGATTCACAG | TCGA |
| TTGGAGAGAAGTTGCCAACATCTGGAGGCTGTTCTGCTTTGGAGTTAAAG*GAGAGAGTTGACAAAGGCATCGAGACAGATGGCTCCAACCTGAGTGGTGT | TCGA |
| AGGGGTTGTTTTATAAACTGCCTGGCCGAATCAGCCTTTTCACGCTCAAG*GTGAGAGCTGCCACACAGATGGGACGTATGCCTATGATGCCGACTTCAGC | TCGA |
| CCAAAACAGATTCTGCAGGTCATAGAGGCAGAAGGACTAAAGGAAATGAG*CCTATAGTGGTGAAAAAAAATGCTGGAACAGATCAGGGCACTGCAGGAAA | TCGA |
| tcaagtgatcttcccaccacagcttcacaaagcactgggatggattacag*AAGATGTGATGGTTTGCTTCGGGAACATGTTTATCAAGATGCCTCACCCT | TCGA |
| AGGGGTTGTTTTATAAACTGCCTGGCCGAATCAGCCTTTTCACGCTCAAG*TTCTAACAACAAAAAGAAGCCAGAGGAAGAAGGCAGTGCTCATCAAGATA | TCGA |
| AGGGGTTGTTTTATAAACTGCCTGGCCGAATCAGCCTTTTCACGCTCAAG*GTACCTGCGCGCCCTGTACCTGGGGCTGCAGAGCCGCTGGCGCGGGGAGC | TCGA |
| CTTTTTTTTTCCTTCTTTCATTTTCCTGCACTGCATTCCAACTCCAGAAG*GTAACCTTTATCAAAAGGATGGAGTTGGGAAAAGGAAAACTACTCAGGAC | TCGA |
| AAAAGGGACGTAGGAAGAAGGGCAGGACCTGGGCGGAGGCCGCCAAGACG*GGTCAAGGTGGAAGGTCCAAGGACTGAGAAAGACCCTCGAAGGCATCCAC | TCGA |
| CTTTTTTTTTCCTTCTTTCATTTTCCTGCACTGCATTCCAACTCCAGAAG*GGCCAATTCCTGTCCCCCCAGCAGCATGGCATCGTGTGCTGAACCCTCTG | TCGA |
| AAAAGGGACGTAGGAAGAAGGGCAGGACCTGGGCGGAGGCCGCCAAGACG*GTCAGTCATATCATTTGGATCAAAAAATAGAACAATCGGAGTTGCAGCCA | TCGA |
| AAAAGGGACGTAGGAAGAAGGGCAGGACCTGGGCGGAGGCCGCCAAGACG*GCCATGGAGAGCAAGCTCCTCATCGGGGGCAGGAACATCATGGATCACAC | TCGA |
| CATAAAGAAATTCTTCAAGTTATCCAGAGAGAAGGACTAAAAGAAATCAG*ATTTAATTCAAGTAGCCTTTTTCACCTGTTCATTTGACCTGGCAATTAAA | TCGA |
| AAAAGGGACGTAGGAAGAAGGGCAGGACCTGGGCGGAGGCCGCCAAGACG*GTTGAATTGCTGGAAATGGAAAAGTCACAAATCCGTTCACAGTGTGAAGA | TCGA |
| AGGGCATCTTCTATAAGGTTCCAGGTAGAATGGGAGTATATACTTTGAAG*TCTTAGAGTTTGAAAGAGTCTATCTTGATAATCTCCCCAGTGCATCCATG | TCGA |
| AAAAGGGACGTAGGAAGAAGGGCAGGACCTGGGCGGAGGCCGCCAAGACG*TATCGTCAGTGAAGCAGGAGCATCAATCTACAGTGTCAGCCCTGAAGCTA | TCGA |
| CGATTATCTTCCGCAGGACCAAGAAGTCCTTACTGTAAACGAATGAACAG*GGTCCTGACAACTGTACAAAGTGCTCTCATTTTAAAGATGGCCCAAACTG | TCGA |
| AAGACTTGACTGGTCTTACATTGCTGCATCAAGAAACCCAAGCTGCTGAG*ACCGCCCAGGTTTGCTGAATGTGAAGCCCATTGAAGACATTCAAGACAAC | TCGA |
| GAATATCTTGATAAATGAGCAGTCAGCAGAACTTGTACATATAGATCTAG*ACCAACTCTTGCGTAAAAAGAGAAGAATTTTTATCCATTCAGTGGGTGCA | TCGA |
| TACATTTTATATCTGGATTTTTACAAATAAAGATGGCTATTATAATGGAA*AAAAATTCTGTAGGCCGTTGGAAGGAAGCTACCATACCAGGCCACTTAAA | TCGA |
| CTTCGTATATGACGAAGTCAGAAAACTCATCATCAGGCGACGCCCTGGCG*ATGCTTCCCTGCAGGTGGACACAGTGGTCCCCAATGCCACGGTCTCTGAG | TCGA |
| AATACATTTTATATCTGGATTTTTACAAATAAAGATGGCTATTATAATGG*AAAAAAAAATATTGGTGCTTTGAAGTGTCACCTTTTAAAAAGTTGGCAAA | TCGA |
| GCGACAGGACCCGGCGCCGGGCACTGAGCACCGCCACCATGGGGAAGGGG*GATGACAACTGGGGAGAGACCACCACGGCCATCACAGGCACCTCGGAGCA | TCGA |
| CATGGCCTGGAGCTGGCTTCCATGATCCCCGCCCTGCGGGAGCTGGGCAG*GTATTACTCTGGAGAGAAAAAGGCTCCAGTTCTCACGCTCTTCATTGGGG | TCGA |
| TATGTTTGTAAATGTGAGTGGAAGCCATGAGGCCAAAGGCAGTTGTATTG*GTGATGGGCGCTACTCCCTTTTGGGCCAGCCATTACAGTACAATCTGTCC | TCGA |
| CAGAAGCCGACAAGGCGTTCAAGCGAAAACATGACCGCTGAGCCCATGAG*CAGATTCCAGAGTCATTAAAAAGTATCACATGGCAGAGATTGGGAAAAGG | TCGA |
| CAGAAGCCGACAAGGCGTTCAAGCGAAAACATGACCGCTGAGCCCATGAG*GACCTAATCCGGAGAGACATCCTGTACTACAAAGGCCGCATTGACATGGA | TCGA |
| CAGAAGCCGACAAGGCGTTCAAGCGAAAACATGACCGCTGAGCCCATGAG*GTGGAATTCAAGTCACCAGTGGCCAAAGTACCCTCCAGGTGTGAAGAGTT | TCGA |
| CAGAAGCCGACAAGGCGTTCAAGCGAAAACATGACCGCTGAGCCCATGAG*GAGGAAAATATTTCCTACCGGTCCATAGTGTCAGAGTGGTGAACCCCTGC | TCGA |
| ATCTGAAGAAACAAGTTCTCCTCCACGACTTGCAATGAATCAAAACACAG*CTAAACATGAATTCAGCTCCAACTTTCATCAACTTTCCTGCAAAAGGGAA | TCGA |
| ACTAGCTCTTCAGAAAAATCCAAGTCTTCAGGATCGTCACGATCAAAGAG*GCAAAACTAGATGGTGGAAGACAGCAATGGAGACCTGTCCTCATGGCTGT | TCGA |
| CAGAAGCCGACAAGGCGTTCAAGCGAAAACATGACCGCTGAGCCCATGAG*AGTTGTGTGACCTTCGAGGATGTGTTCGTGTACTTCTCTCGGGAGGAGTG | TCGA |
| GCCCCGAGGGTCTCCTCCAGTAGACGGTACAGCGAAGGCAGAGAGTTCAG*ATATCTTCAAGTACGTCCCACTCTGATGCTTCTACAAAGCAGTGTCAAAC | TCGA |
| CGCGGGAGGCCGAGGAGAAGCTGGAGGAGCGGCTGAAGCGCGTGCGCATG*TGAGGCCGTCTGGGAGAACATGGCGCGCATGTGCGTGAAGACCCAGCGGC | TCGA |

| | |
|---|---|
| CGCGGGAGGCCGAGGAGAAGCTGGAGGAGCGGCTGAAGCGCGTGCGCATG*TTGGTGCTCTCGGACCTACCATGCGAAGAAGATGAAATGTAAGTAAACAG | TCGA |
| CAGAGCCTGTCCAGCGATGCAGACACCCTGTCCCTCACGGACAGCAGCGT*AGGTGGGGCTGGTGTCCGAGCACTTCTCGCAGGCCCCGCAGAGGCTGTCC | TCGA |
| TAATTATTGTAATTAAGAGATGTAGCCtttattaaaattttatatttttC*aaaaaaaaaaaaaaaaaaaaaaaaaaactagccgggcatggtggcgggtg | TCGA |
| AGGGCCACGGCGAGTGTGAGGTCCACGGAAACTGTTGACAGTGGATACAG*GCGCTTGAATAAAATTGTTTGGCGAGCATTATCTCAAGAGGAAAAAGAAA | TCGA |
| TTGTTCCACAAGTTAAATAAATCATAAAACTTGATGTGTTATCTCTTATA*aaaaaaaaaaaaaaCCGGAAATGGGTCCTACCATCTTCTCGGAGCCGGAG | TCGA |
| CTTAGCTGTGCTCGCGCTACTCTCTCTTTCTGGCCTGGAGGCTATCCAGC*AATATGGACTCCACTCTTCACTATATTCACAGCGATTCAGACTTGAGCAA | TCGA |
| GGTGCTGGAAGCAAACAGGGCCCCTGCAGCCTCTGAGGGCAACCACACAG*GGTTCTAGAGCACCTCAGCAGTCTTGCCAGAGAAGTGAGCCTGGACTATG | TCGA |
| GGATGGTGAGGCCTGGCGAGCCCTTGAGTGGGGAGAAATACTCACCCAAG*GGGGAGAACCTGGACCTTGCAATGTCCAAGGAGGAGGTGGTGGCTATGAT | TCGA |
| AAGGGGGTGCAAGTGGAGGAGATCTACGACCTTCAGAGCAAATGTCAGGG*GCGGAGGCCGCGCAGCAGCTCGGGCGCCCCTCGGAGGCAGGGCAGTGCCG | TCGA |
| TGCATGTGCTGAAGGTGAACCGTCAGACAGTACTAGAGGCTCTGCAGCAG*GAAGGGGCCCTGGACCTTCTGAAGAAGCTGCACAGCTGCCAGATGTCCAT | TCGA |
| TGAAAAGAAATCATAGAGGAGAGACTTTGCTCCATATTGCTTCTATTAAG*AAGAATTCATTCAGAAGATCATTTCCATGCTCCAAGATGCAGAAGAACAG | TCGA |
| ACTTCCTCCGGGAGCGGCTGTTGGGCTGGATCCAAGACCAGGGTGGTTGG*ATTGGCTGGTTTCCTTCAACGTACGTAGAAGAGGAGGGCATCCAGTGACG | TCGA |
| GAGGAAGTTTTCTACTGTAAAGTCACCAACGAGATCTTCCGCCACTACGA*GAATATATATGTCCCAGATGTGAATCAGGCTTTATTGAAGAAGTGACAGA | TCGA |
| CCACCGCACCGTCCCTCACCGAGGAGGACCTCACTGAAGTGAAGAAGGAC*TAAAAGAAGCCTTGAGCCCTATAAAATTTAACAGATGGGGTCTCCCAGAA | TCGA |
| CCTGCTCTCACCCACGAAAGGCATCTGTCCCAAGCAGGAGAACATTGCAG*gaacgaggtctcactatattgcccagactggtctcgaactcctgggctca | TCGA |
| TCATGGCGTCCAGCTCCGGCCTGGCGTGAGGACCCCCAATCCGCGTGAG*GAAAGCAATCTCCCGGGCAGGCCTCCAGCATCTGGCTCCTGCACATCCCC | TCGA |
| TTTGAACGGAATGCTGGAGTCAGTGGAAGACGGAGCTGGTCGGCCCGAAG*GATTTCCAGCAGGAGAGCTCCAGAAGCCTTTCTTTTGGGGAACAGAATAT | TCGA |
| GACAGCCATGAAGCTGGCCAGCAGCGACACCATGAAGCGCTTTCTCAGTG*CCTTCTGGCTTTGAGCCAACAGGAAGAACTAGCGGATTTGCCAAAAgact | TCGA |
| TAAAGAATTCTGTGAGAAAAAATTTAATAAAGCAACAAAAATTACTCTTA*aaaaaaaaaaaaaaCCGGAAATGGGTCCTACCATCTTCTCGGAGCCGGAG | TCGA |
| GCGCGGGAAGTTTGGATCCTGGTTCCGTCCGCTAGGAGTCTGCGTGCGAG*ATACATTCCACCGTTGATCTGGGGGAAAAGTGGACACATCCAGACAGCCT | TCGA |
| GCTGTCTGCATAATTGCTATGATCATCTTCTCCAGCTGCTTTTGTTACAA*ATGCTTTGTTCTGTACAAGCTGCCCAACTTGAAATTTCTGGATGCCCAGA | TCGA |
| GCTGTCTGCATAATTGCTATGATCATCTTCTCCAGCTGCTTTTGTTACAA*GATGGAAAGTGAAGAGTTGGCAGACAGGGTTCTGGATGTGGTGGAGAGGA | TCGA |
| CAACATGGCAGAGAAGAAGGTGGCATCTGAGCTCCCCCAGGATGTGCCAG*ACACGCTCTTTACTCAGGGAATATAGCACATCTATAGCTACGAGAGGTGT | TCGA |
| TGTGCCAGACTGTGAGCCAAGGAATTTTGTGTTTCTTGCCATCTTACAAG*GTGGGACCATCCGTGAAACCGGTGCCAGGATGCACCGGGGGTGTGGATGA | TCGA |
| GGGATTCAACAGAAGGCTGAACCACTAGAAAGACTACAGTCCCAGCACCG*GAAAGAAATTACAGTATTTATTTGAACCATACCATGTTATTGAAAGCAAT | TCGA |
| CAGGTCTTCTGGGATGGGTCCTTCTGGAAACTTAGCCAAAATATTTCTGA*GATGAAGATTGCAATGAAAAGAATGGCTGCCATGAAAGTGAGCCATCAGT | TCGA |
| AAAAACGATATTATAGTGATCCTCGATTTCTCAATCTCTGGCTTAAATTA*AAACTGGGAGTGATTTTTCCATGTTTGAAGCTTTGCGAGATACTATTTAT | TCGA |
| AGTGTGGACGATGCTCTTGCAGGATGCCTTTCCTTTTGGGTCTTAGACAG*AGAACACCTTCTGCAGTGGGGACCATGTGTCCTGGCACAGCCCTTTGGAT | TCGA |
| TCCTCCAGGCGCTGCCCAGCCCCTTGTCAGCCAGGGCTGAACCCCCGCAG*GAGCCGCAAAGACAGCCTGGAAAGTGACAGCTCCACGGCCATCATTCCCC | TCGA |
| GGAGGCCATGGTGGTGGCCGTAATTCCGCGGCCCGAGCCGATGCTCAGAG*GGGCGTTTATGCAATGAGCCTTTGGATATGTACAGTTACTTGCACAACCA | TCGA |
| AACGGGGTCCACTGGTGCCCTGGAGACCCGCAGCCTCAAGGATCACAAAG*AAGAAAGAACCTTCTAAATCCAGCATCAAGAAAAAAATGACCAAAGTTGC | TCGA |
| GAAGATCTTCACCTTAAAATCAAAGCCACAAGACACTGCTGATAAGGATG*CTCCAGAAACATGACAAGGAGGACTTTCCTGCCTCTTGGAGATCAGAGGA | TCGA |
| AGAGTCTCTACCGTCTGGCCTTCCTCTACACCTACAGCAAGACCCACCGG*ATTTATGGTCAATGTTACATGGGATGGCAAAGACTTATCCTTCACTGAGG | TCGA |
| CTGAACACGGGTGATAACGAGAGTGTACATTCAGGCAAGAGGACGCCAAG*GGTCTGCCCACACATGACATGGAGGGCAAAGAGCTCAGCAAAGGGCAAGC | TCGA |
| GGAGGATGTGGCGCGCGGAGGGGAAATGGCTGCCGAAAACAAGCCGGAAG*ATTGTTCTCTGGGGCCGGACTGAGAAATGCCTGAAGGAGACGACGGAGGA | TCGA |
| GGAGGATGTGGCGCGCGGAGGGGAAATGGCTGCCGAAAACAAGCCGGAAG*GTTTCCGTTCAGCCGCCCGGAGCTGCTGAAGGAATGGGTGCTGAACATCG | TCGA |
| AAGCTGTGCAGTCCAGACTCACGGGAGAGGAAGTCAACAGCTGTGTGGAG*GAATCCTGTATGGCACGATGACCCTGGAGCTGGGTGGGAAGGTCACCATC | TCGA |
| AGCTGCGGGTCCGGGATTCCCAGCCATGGCAGATTCCTCCGGGCAGCAGG*CCTATGTTTCCGAAGAGTTAAAGGCTGCTGCCCTGGTGGACGAAGGTTTA | TCGA |
| GTTTTATTTCAGCTGTTCTCTCGGATTATGAAAGTGCAGAAGACTCGGAA*AGAAGAGGCCCCCTGAGCCCCCCGGACCTCCACCGCCGCCACCTCCACCC | TCGA |
| GACCCCAAACAAGTCTGCGCAAAATTTGTCGAGGAGGTTTGCCGCGGCAG*GGGCAAAGGAATTGCTAGTAATAAAACGTATTCCTTTCTTATCACGCTGG | TCGA |
| TTGGCCGCCTGAGCCCTTGAGTTGGTCACTTGAACCTTGGGAATATTGAG*GAGCTACTCTCCTCCTGGTGAGGTGGAAATTCCAGCAAGAATAGAGGTGA | TCGA |
| ATGGAAAGGGAACTTCAGACACCAGGCAGGGCTCAAATTTCTGCCTACAG*CTGAAGTCTGGGATCAAGAGTTTGACCCCGTCATGGTCATTCTTCGAACA | TCGA |
| CGCGAGTGTGCGCCGTCCGCCGCCCGCCGGATCCCCCGGACCCCCCGACC*GTGACCGGGCTGGTCCACAAGTGGGCATGGACCAAAGACGACGTGATCCT | TCGA |

| | |
|---|---|
| GACAAACACCTAGCCGAGAGTATGTCGACTTAGAAAGAGAAGCAGGCAAG*TGATCTTCCATTGTTTGAAGCACTGGACCTTTAATCCACTGTAGGTATGG | TCGA |
| AAAGACTGGATGGTATGGGCTGTCTGGAGTTTGATGAGGAGCGAGCCCAG*GCCCCTCGCATTGGCACAAGCTGTATCCCATTGCCCAGGGAGATCGCCAA | TCGA |
| AAAGACTGGATGGTATGGGCTGTCTGGAGTTTGATGAGGAGCGAGCCCAG*AAAGTGGTGGAGAGAGAGCTGGATGCTCTTTTGGAACAGCAAAACACCAT | TCGA |
| GACAAACACCTAGCCGAGAGTATGTCGACTTAGAAAGAGAAGCAGGCAAG*AACCGAGCTGAAGCTGCCATGGAATTCTTACAAGAATTAAATAGCGATGT | TCGA |
| CTCTTCAAGGAGGGAAAAGAAAGAATGTATGAGGAGAATTCTCAGCCTAG*TTCTGGGAACTGGACAGATTGGACAACTATAACGACACCTCCCTGGTGGA | TCGA |
| GGACGGTGGACAAGAAGATGGTGGAGAAGTGCTGGAAGCTCATGGACAAG*GTTCGCCTGTCCCCACTCCTCTTCTTGGAGAACACAGAGGCTCCCGCCTC | TCGA |
| GACCAGATGTGCCTAGGCTCGGAAGCACGTTCAGTCTGGATACCTCCATG*GTTTACCGCTCCAAGATGGATGGCGAGGCCACCAGCCCCCTTCCCCGGAC | TCGA |
| GGACGGTGGACAAGAAGATGGTGGAGAAGTGCTGGAAGCTCATGGACAAG*GAGCTCTGGAGCCTTAGGACCATGGACGCTCTCAATAGGAACCAAATAGG | TCGA |
| CTCTTCAAGGAGGGAAAAGAAAGAATGTATGAGGAGAATTCTCAGCCTAG*GATTGTTTGAACTTGTCGTTAGTATAATATCTTTCCTCTTTTTCACTGAC | TCGA |
| AAAATCAAACCTTCCTCATCTGCCAATGCCATTTATTCTCTGGCTGCCAG*GTGTGAGCTCTTTGACCAGAACCTGAAGCTAGCTCTGGAGGTGGCAGAGA | TCGA |
| GGACGGTGGACAAGAAGATGGTGGAGAAGTGCTGGAAGCTCATGGACAAG*AAATCTTGCATCACCATGGTGCACTTCTGTGGCCTACTCACCCTCCACCG | TCGA |
| CTCTTTAAAGAAGGCAAGGAGAGAATGTATGAAGAACAGTCACAGGACAG*GCCATGCAGCCATGTTCCGGGAGCCCTAATTGCACAGAAGCCCATGGGGA | TCGA |
| CTCTTTAAAGAAGGCAAGGAGAGAATGTATGAAGAACAGTCACAGGACAG*GTAATTTCTGGACAGAAACTCACACGACTATTTACATCAAATCAGATATT | TCGA |
| CCCTCCAGCTGCGCTGGGACCCCAGGACCCTGCCCCGGCCTGAAGGCCAG*GTACAGAGCATGCCAGCGCCTCAGCTAATGGGGTAAGTGCAGTGTTGGAG | TCGA |
| AAAAAGAAGAGGATGAGGATGATGATAGGAAATCTGAAGACGATAAAGAG*TCGTACCTGTACATGTCAAGGAATTGATCCAGAGACTTGTGGAGCTTCAT | TCGA |
| AAATTTCTACACCTACCCATTGGTCTAAACTTGATCCAAAGACAATGAAG*TCGTACCTGTACATGTCAAGGAATTGATCCAGAGACTTGTGGAGCTTCAT | TCGA |
| AGGCACTGCAGGAGGAGAACCGCGACCTGCGCAAAGCCAGCGTGACCATC*GTCCACAGAGTCCATGTGAACGGACAGATATCAGGATGGCAATAGTAGCC | TCGA |
| AGGCACTGCAGGAGGAGAACCGCGACCTGCGCAAAGCCAGCGTGACCATC*GGAGAAGGTTTTAAGGTGAAAACGAAGAAAGAAATCCGGCATGTGGAAAA | TCGA |
| AGGCACTGCAGGAGGAGAACCGCGACCTGCGCAAAGCCAGCGTGACCATC*CAGGGAGAAGGTTTTAAGGTGAAAACGAAGAAAGAAATCCGGCATGTGGA | TCGA |
| AGGCACTGCAGGAGGAGAACCGCGACCTGCGCAAAGCCAGCGTGACCATC*ATTAGAGACACCAGCCAAGAGCCCTGTGGTCGTCTGTCTTTTCTGAAAGA | TCGA |
| CCCGATCCCTTACACACCTTCTCCGAGTTCAAGCAGGCCTATATCACCTG*AACTTGATAGCCCAGAAGAGTTAGGGAAAAAGCGTATCTGCAGGATTATC | TCGA |
| CTAGTTAATCGCCTCTGGAAAAGGATGGATAAGCTTGAAGCTGAAAAGCG*GATCATAAGTTTTTACTGCCAGTATTGAGAAATTCGTGGAAGAAATGTCA | TCGA |
| AGAAAGAAGAAGAATTCCTCACTAATGAGCTCTCCAGAAAATTGATGCAG*ACCACCACTTTGCAGGTATAATATTGTACTTAAAGACAGAGAGGAAGTGT | TCGA |
| AGGCACTGCAGGAGGAGAACCGCGACCTGCGCAAAGCCAGCGTGACCATC*GAATTGAGTGATCTACAGCGCGATCCACCTGCTCACTGTTCAGCTGGACC | TCGA |
| CAGGGCAAGCTGTGTTCCATGGCAGGGAACTTTTGGCAGAGCTCCCACTA*GACATTTCTACTGCCTCCAGCACCTGCCCCAGACAGACCACAAAGCGGAA | TCGA |
| CTTATTATGCCTTGGTGTAGCACTGACATTCATCTTCCGTTTAAGAAAAG*GGTCTTAGAAACATTGATCATGGGATGATGGAAAAGTCAAATAACGGATC | TCGA |
| GCACGCAGCCTGGGGACTCTGCCTCGTGCCGCTGAGCCTGGCGCAGATCG*GTCAGAGTGCATCAACAAGTTTGCTTCAGTCTTCGGGACCATGCCTCTCA | TCGA |
| AAATTGACACTATGGGAACCTACCATGGTATGACACTGAAATCTGTAACG*CCTTGCAAAATACCCTACTAAATAAATCATTTTAGACATGGAGTGCAGGT | TCGA |
| GTTCACGGTGCCCCGGCGCCACCTGGAGAGAGGCCGCGTCCTGGGCAGAG*CTTCACGTAGCTCACTGGAATTCTGCAAAGTACTCCAGCCTTGCTGAAGC | TCGA |
| GCAAATTCCTGCCATTCTGGGGATTCTTGGAGGAATTCTTGCTTTGCTAA*GAAAATATATGCAGATGATGAACATTCTTAAGCCAATTGCCTTTGATGTT | TCGA |
| ACTAACACACGGGGCGAGTGCCAACTGGACCATTCAGTACAACGACCCAA*GAAATAGAAAAAGAGAAAAAGCCCAAGGAGATCAAAGTCAGAGTTATTAA | TCGA |
| GGCTGAAGGTGACAGAGCCTCTGGATAGAGAACGCATTGCCACATACACT*ACCTGCTCCCCTCAAAAGCTTCGTGCCAACAGAGAGGTTCCTGTTTGAAC | TCGA |
| GTTCACGGTGCCCCGGCGCCACCTGGAGAGAGGCCGCGTCCTGGGCAGAG*GATGGCCATTTGGAAACACGATGTGCAAGATCAGTGGATTGGTCCAGGGA | TCGA |
| AAGTCACGCTGAATACAGTGGGGCACCACCACCGCCCCCCGCCCCATCAG*GTACGATTATATGCAAGACCTGATGCTATCAGAAGAGGATCCGGGGACTA | TCGA |
| AACTTCCTTTCCTCTATGTGGTGGCTACAGAGAGTCCAACCACACTTACG*GGACCTGCTAGGGAAAGAGCCTCAGGAAAAACTGGAGATCCGGCTGTGCC | TCGA |
| ACCCCCTGAAATCATTCACAATCCAAATTTCACAGTCAGAGACAACCGAG*CCAGTCGCTGCTTCCACCTGTTCAGGGAAGAAAACACCCCGTACTGGCAG | TCGA |
| GAACCAAAGGCAATCACACCACCTCAGCAACCATATAAAAAGAGACCAAA*GGCTCAGCGGAGTCCTACACCAGCCGTCCATCAGACTCTGATGTATCTCT | TCGA |
| CCTCCTTTGCCCCTCCCACCCTTATTACCTGGAGATGATGACATGGATAG*GCGACACAAGATGTACAGAGAGCAGATGAACCTTACCTCCCTGGACCCCC | TCGA |
| CCTCCTTTGCCCCTCCCACCCTTATTACCTGGAGATGATGACATGGATAG*TGAGGGATGTCCACTGTTGGAGCAGTTGAACATTTCCTGGTGTGACCAAG | TCGA |
| AATTTCCTGCATCGGGATATTAAGTGTTCTAACATTTTGCTGAATAACAG*ATATACAAACAAATGCGGAACTGAAAAGCACTCAGGAGCAGTCTGTGCCC | TCGA |
| AGCAGCCCTTTCCTGAGCAAGCGGTCTCTGAGTCGGAGTCCACTCCCCAG*ATTCTCCTGGGCTTCTCCAATTTGAAGTGTGTCCTCTCTCAGAGTCTCGT | TCGA |
| AGCAGCCCTTTCCTGAGCAAGCGGTCTCTGAGTCGGAGTCCACTCCCCAG*GAGGACAACCACAGCTATTATGTGTCCCGTCTCTATGGCCCCAGCGAGCC | TCGA |
| AATTTCCTGCATCGGGATATTAAGTGTTCTAACATTTTGCTGAATAACAG*GAGGAGGATGTGGTGATTGAAGACTTTGAGGAAGATTCAGAGGCTGAAGG | TCGA |

| | |
|---|---|
| GGATTGGCAGGAGATTATAGCTCTGTATGAGAAGGACAACACCTACTTAG*TCCCACCCGCTGTACCTGTGCAATGCCAGTGATGACGACAATCTGGAGCC | TCGA |
| CGGGGGCGCTGCCCAACGCACCGAATAGTTACGGTCGGAGGCCGATCCAG*ATGGAGAAGGATGAAACTGTGAGTGACTGCTCCCCACACATAGCCAACAT | TCGA |
| CGGGGGCGCTGCCCAACGCACCGAATAGTTACGGTCGGAGGCCGATCCAG*GTAAATTTGGAAGGCAAAGCCAGCCCTCTGTCACTTCTACAAATGGCCTC | TCGA |
| GCTACTGAGGAGCCAGCGTCTAGGGCAGCAGCCGCTTCCTAGAAGACCAG*GTAAAGAATCTCATTTGGACACATTGAGTCGGGCACAACTGACCCAACTT | TCGA |
| GCTACTGAGGAGCCAGCGTCTAGGGCAGCAGCCGCTTCCTAGAAGACCAG*CTTTGGAGGAATTTTTTACATCACTTGTGAAGCAAGAAATGGTGAATATG | TCGA |
| CACCAGAGGCAGTAACCATGCCCGCATAGATGCCGCGGAAGGTCCCTCAG*CTATATGTGAGACAGAGCCTGAACAGCCTGTTCGACATTACCCATTATGG | TCGA |
| CGGGGGCGCTGCCCAACGCACCGAATAGTTACGGTCGGAGGCCGATCCAG*ATTGGAATAATTGGTGGAACAGGCCTGGATGATCCAGAAATTTTAGAAGG | TCGA |
| GCTACTGAGGAGCCAGCGTCTAGGGCAGCAGCCGCTTCCTAGAAGACCAG*ACTTATTCTGGACTCTTCTGTGTAGTTATAAACCCTTACAAGAATCTTCC | TCGA |
| GCTACTGAGGAGCCAGCGTCTAGGGCAGCAGCCGCTTCCTAGAAGACCAG*GCTAGAGATGAAGATGGACATGCTGAAAATTTTCCCCAGCAGCACTATGC | TCGA |
| GCTACTGAGGAGCCAGCGTCTAGGGCAGCAGCCGCTTCCTAGAAGACCAG*GATATTGAGGCAACAATGAACTCGGCCCTGAATGAGCTACGGGAACTAGA | TCGA |
| AAGGACAAAAGCCAAGAGGAGGACAGTTCGCTACACAGCAATGCATCGAG*TGCACTCTGGACCCAGACACCTCTGACCAGACTGATAATCTGACTGAACA | TCGA |
| AAGGACAAAAGCCAAGAGGAGGACAGTTCGCTACACAGCAATGCATCGAG*GGGTCAGATATAGAAATGTTGAATTCTGTGACCCCCACTGACAGCTGTGA | TCGA |
| CAGAGATTAACGTAGTGGTTTACATAGGTGACCTGATGAGCAGAAATACG*TTTGCAGAGAAATTCCAGGAGGTGAAAGAAGCTGCCAAGATAGCCAAAGA | TCGA |
| AAGGACAAAAGCCAAGAGGAGGACAGTTCGCTACACAGCAATGCATCGAG*AGAGCAGCTCTAAATTGGGAAGCAATGCCGGAAACAAGTCTCTTAAAAAT | TCGA |
| TGCCCCCTGGGCTCGATTATGGGCCCTTCAGGATGGATTTGCCAATCTTG*CACCCAATCAATGTGAGACGAGACCCCAGCATCCCTATCTATGGACTCCG | TCGA |
| TCAAACCATTAATTCCCTCTTTCTTTGCAGATACCCTGTTCCAGTTGAAG*ATTTTGTACAGCAGCAATTTCTTACTCTCTCTGCAAGTTTTCTTCCCAGT | TCGA |
| CGTCCCTTCTCCCTCTCCGAGTGCTGCTCCTGTCATTGTGGCCATGGACG*GCCTCCTTTGGGAGCACACGAGGCTACCATGGCTGCAGCAAGAAGGCCCT | TCGA |
| CCCCAAGAGTGCCAAGTGCGAGGGGGACATCTTCACCTTTGACCGTACAG*AAGCCCAGTGCGGCGCCAGGAGCCGGACCCGCGCCCGCACCGCTCCCGGG | TCGA |
| TGAGGAGCGCATGGTCATCTGTGAGGAGGAAGGGGATGATGATGTCATTG*AGGAACCTGGGGCCCGCTCCTCCCCCCTCCAGGCCATGAGGATTCTGCAG | TCGA |
| CCCCAAGAGTGCCAAGTGCGAGGGGGACATCTTCACCTTTGACCGTACAG*AAGCCGAGAGCCAGCCGGCCGGCGCACTCCGACTCCGAGCAGTCTCTGTC | TCGA |
| AACTTGTGGATGGACTTGCTACCTCTTGGGTGAACTCTAGCAATTACAAG*GCAATGCATTAGCAGGAATTCAAATTAGGACAAACAGTTGTCCAATTGTT | TCGA |
| GCTGACTCGGGAGGAGGACCTGATCAAGACTGATGATGTCCTGGATCTGA*GATCCGACGTCTACCTTCTTCCAGTTTGGAGCGTCCATCCAGCAGCAAGC | TCGA |
| ACAGCGCCATGGCCCAGATTCTGCCAATTCGTTTTCAGGAGCATCTCCAG*GTAAATACCAGCATCGAGGACTGGAAGACCACCAAGTGGAAAGATATCAA | TCGA |
| ACAGCGCCATGGCCCAGATTCTGCCAATTCGTTTTCAGGAGCATCTCCAG*GAGACCATACTCAGGAAGAGCCTGACTCCCTCTTGGAAGTGCCTGTGAGC | TCGA |
| ACAGCGCCATGGCCCAGATTCTGCCAATTCGTTTTCAGGAGCATCTCCAG*ATGCCCTCCAAATCCTTGGTTATGGAATATTTGGCTCATCCCAGTACACT | TCGA |
| ACAGCGCCATGGCCCAGATTCTGCCAATTCGTTTTCAGGAGCATCTCCAG*TTACAGATGTAGAGCTAAAACGACTGAAGGATGCTTTCAAGAGGACCTGT | TCGA |
| CAGGCCTGCGTCCGAGTCTCCGCCGCTGCGGGCCCGCTCCGACGCGGAAG*GGGATGAGTTCTGTTTTCTTCAACAAAGGAGAGAATTGCATTGCAGCAGG | TCGA |
| GCGACACAGGGAGAAACTGGAAACCCTATCATCAAGATGGGAATATCTGG*ctcgtcccatgctcaggaatgaccaagggcagtttggaggttaaaggcac | TCGA |
| GACAACCTATTGATACCACAGCTGAACTTTCTTCTAAGTGCCACAGTGAA*GGATGCCAGATGATATGAACTTCAGTGATGTAAAGCAAATCCATCAAACA | TCGA |
| GCGTGGACCACATAGACATTTACGCGGATGTCGGCGAAGAGTTCAACCAG*TGCCTGCCCAAGTGACTGACTTGCATGTGGCCAACCAAGGAATGACCAGT | TCGA |
| CAAAAGAGCCAAACTCAGCTCGCCCGGTTTCTCGGCGAATGACAGCACAG*TTCCGAGAGGAACCCGTGCAATAGTTCGGTGCCGCGTGACCTCCGGCGGC | TCGA |
| TGACTGGTCACGCGGGACTGAACACCGCACAGGCAGGAGGCATGGCCAAG*GACTGTTCTGATGGCTACATGTGCAAAATACACTTTCAAATTAAGAATGG | TCGA |
| GTTTACCATGAGATCCTTATTGGATATTTAGAGTATGTGAAGAAATTAGG*CGGTGGCGGAACAGCCACCAAGACCATGTCCTGTGGGGAGGCCCCCCGGA | TCGA |
| CTTGAAGAAAACTGGCAGACCTCGAAAAGAAAACAAATTCAGTGCTAAGA*AGCTCCTGGGTTCCACCAAGAAGATCAATGTCAACTTCCAGGACCCGGAT | TCGA |
| TGACTGGTCACGCGGGACTGAACACCGCACAGGCAGGAGGCATGGCCAAG*TGCCTGGATCAAAGTGGAACAGCTGAAGCCATATCATGCTCATAAAGAGG | TCGA |
| TGACTGGTCACGCGGGACTGAACACCGCACAGGCAGGAGGCATGGCCAAG*GTTTTACTGGGATTTAATAATGCTTATAATGATGGTTGGAAATCTAGTCA | TCGA |
| CCTTCCCTACAGATATCAAGAATACTTCAGTCACCAACGTGCCAAATATG*CTTGGTGTCTTGCTCTTGTGGAGGTTGGACCAAAGGGGCCGAGTGCAAGG | TCGA |
| GGGGGCTCAGCCCCCTGTGATTCCACAGGCACAACCTGTGAGACCTCCAA*AAAACTGTCGCTGAAAAGCAGGAGAAGAGAAATCAGGATCGCTTGAGGAG | TCGA |
| AGGACACTGAGCCCGATCCTGGTGAATCCAAAGGGGAGCCCAGGTCTGAG*ATTGCTCTTCTAGACATCGATATATGTGGGCCATCGATTCCCAAGATAAT | TCGA |
| CAAAAGAGCCAAACTCAGCTCGCCCGGTTTCTCGGCGAATGACAGCACAG*GAGCTGCTGGGACTCAGGGAAGGAAGTCTGCAGGTGAGGAAGCCTCGGAG | TCGA |
| TGACTGGTCACGCGGGACTGAACACCGCACAGGCAGGAGGCATGGCCAAG*AGGTCAGCTGGCGGCTCCAGTCCTGAAGGCGGAGAAGGTAAGGAGCTCTC | TCGA |
| CAAAAGAGCCAAACTCAGCTCGCCCGGTTTCTCGGCGAATGACAGCACAG*GAGCGGTGGTGCCCCCCCCGGGCACGGGGCCATGTACAACGGGATCGGGC | TCGA |
| CCTTCCCTACAGATATCAAGAATACTTCAGTCACCAACGTGCCAAATATG*TATGAAGACTTTGGAGACTCAACCGTTAGCTCCGGACTGCTGTCCTTCAG | TCGA |

| | |
|---|---|
| GACCAGTGATTTTGCAGTTAGTGCTCATGGATTTAAGGTATATTACGAAG*GAGCGGCTCGTGCTTTCTGTGCTCCCCCGGTTTGTTGTCCTGGAAATGAT | TCGA |
| TCAATATTTCAGCCAGAACATTTCATTTAGCACAAGGAGATGTTCTTAAG*ATTTTAATTGAATTAGAAACAGCATTATTAGATGATGAGCCACATTGGTA | TCGA |
| GAGCTGGGTTCTATTTTCCCTCCTCAAACTGACTTTGCAGCCACGGAGAG*GCCAGGACTCAGCCAGTCCCATCCGGACCACACACACGGGGCAGGTGCTG | TCGA |
| GAGCTGGGTTCTATTTTCCCTCCTCAAACTGACTTTGCAGCCACGGAGAG*GGTCTCAATGGGCAGAAGAAGTACCAGATTCACCTGAAGAGTGTGAGTGG | TCGA |
| TGGAAGAAAAAGAAAGATGCGCTCTAAGAAAGAAGATTCCTCTGACAGTG*GAGCCATGTCTGACCAGGACGTTTGGAAGATCATATCCATGCCAGAGGCT | TCGA |
| GAGCTGGGTTCTATTTTCCCTCCTCAAACTGACTTTGCAGCCACGGAGAG*GCTGTGACTCTGGTTTTGGCAAGGAGACGGCCAAGAAACTGGACTCCATG | TCGA |
| GACCAACCCCAAGGAGGACGAGGGCCGGTTGCCGGGCGCGGGCGAGAAAG*AGGTAAATACCGGATTGTATACGTAACTCCAGAATACTGTTCAGGTAACA | TCGA |
| AGCTCTACACTGGACTAAGAGAAGTTGTTACCGAACATCTCATAAATAAG*AATGAAGACCTAAATGACCTCAGCAAATCCTCTTCTAACTCATGGGTACC | TCGA |
| AGATGACCATATTAATGCTCTTTAATAATAGAGAAAAATACACATTTGAG*GAACGGAAAACAGTTCTTCCTTCAAAGCATGCAGTACCTGAAGTAATAGA | TCGA |
| TGGGATCGATGTTTCAATATTGGAAGCGCTTTGATTTACAGCAGCTGCAG*ATCCAGTTTGCTGACCAGAAGCAAGAATTCAACAAACGTCCCACCAAAAT | TCGA |
| ACAGAATTATTTGACCTGAAAACCAAATACGATGAAGAAACTACTGCAAA*CCAACAAGTCCCAAATTTGGAAAAGCTGACTCATATGAAAAGCTGGAAAA | TCGA |
| GTAGTTGCATATTTAGCTTTGCACATTTGAAATAAACCACGGTTGCAGCC*aaaaaaaaaaaaaaaaaaaaagaaagccaggtgtggtggtgcacgcctgta | TCGA |
| TGGGATCGATGTTTCAATATTGGAAGCGCTTTGATTTACAGCAGCTGCAG*GCACCACCTGCCCTGGTCAGCTCCAGCCAGTCCCTGCTGGAGTGGTGCCA | TCGA |
| GAAAGCGGCTTATCGAACAGAGCCGGGAGTTCAAGAAGAACACTCCAGAG*AATGCATGAACTGCACCCGGCTCAGTGACATGAGTGAGCGACTGACCACA | TCGA |
| GGGCCGACGAGATTGAAATGATCATGACGGACCTTGAAAGGGCAAACCAG*GGCACACAGCATGGCAGAAAACCGAGAGCCCCGCGGTGCTGTGGAGGCTG | TCGA |
| AGCTGAAGCAGCTTTCTTGAATGTCTACAAAAGATTGATTGACGTCCCAG*GTCCATGGGGCCGATGTATGGGAGATGAATGTGGTCCCGGAGGCATCCAA | TCGA |
| TGGGATCGATGTTTCAATATTGGAAGCGCTTTGATTTACAGCAGCTGCAG*GTTGTCGGGGTACCAGTTGGGTCTGCTTTACCTTCAACAGTGAAGCAGGC | TCGA |
| ACACAGCCACCCGGAGTTCCTTAGTTGAAAGGTGCGCCCTGCTGTGACAG*gagaagaaactgaggcctggaatttgattaactcattcaaggttacccag | TCGA |
| CCCGGAAGCGCTGCAGGCTCCCAACGTGGACTGGTTGCGGACGCTGCGAG*ATGGAAAGCCAGAAATTTTTAGCAGAAAATAGTGCTTCAGTATATATAAA | TCGA |
| AGCACCCGCACCCACGGCGCCGTCTTCAACCTCGAGTACTCGCCCGACGG*GACATGGAGGAGTGAATCAGCTTGGGGGGGTTTTTGTGAATGGACGGCCA | TCGA |
| ATGGGTTATTTTACTCTACAGGAGTGGTTAAAAGGAATGACTTCTCTCCA*CCAGTAATGTTAAAGTAGAGACTCAGAGTGATGAAGAGAATGGGCGTGCC | TCGA |
| TATTAGGAAAAATCTGGCCCCTTTTTCCAGTTTTTCACCAATTCTTAGAG*ATCCGGTACATGGCCCACAGAACGTGGAAATCGTAGACATCAGAGCCCGG | TCGA |
| AGCGGCATAATGTGTTTGGATTGGACCTTAAAGACGAGAAAACATTACAG*GCAGCAGATGGATTTTTGTTTGTCGTAGGATGTGACCGAGGGAAGATACT | TCGA |
| TGACGGCTGGGCGACTTGCGAGGATGAGAGCGACGAAGCCAACTGTCCAG*GTTGCCAACGTCGAGCTCTGTTACAGAGCCCTGCAGTTCTATTTGGATTA | TCGA |
| CGGAGCGGGGTCCGATGGTTCTCGAAGGCCCGCGGCGCCCCGTGCTGCAG*TCCCAGGAATCCCCAAACCTGGAAAACATAGCAAACCCCCTAGAAGAAAA | TCGA |
| CTTACCCTGATGCGCATGAGGGCCGCCTTTCATATATGAGAAGCAAAAAG*GCATTATATTGTACTTCTAGCAAGTGCACCAACAGAAAAACAACGCCTGG | TCGA |
| ATGAGTGTCATCTTGCAATCCTAGACCACCCCTATCGAGAAATTATGAAG*TGGTGATCGCCGCTCGGGAATGCGGGCGTGAAGGGTCCGAGTTGCCGCCC | TCGA |
| CCTGGAGGTGCGGGCGCGCCTGGCCGAGCTGGAGCTGGAGCTGTCGGAAG*GTACCAGCAACTAAGGTTTCAGAGCTGAACCCTAATGCAGAAGTGTGGGG | TCGA |
| GGCGGCCAGAGCGCCGGCAAGAGCTCGGTGCTGGAGAACTTCGTGGGCCG*GAGGCTGGTGGTTTGGCACCTGCAGCCATTCCAACCTCAACGGCCAGTAC | TCGA |
| GGCGGCCAGAGCGCCGGCAAGAGCTCGGTGCTGGAGAACTTCGTGGGCCG*CTCGGTCAAGGAGTGTGATGACTGGCGAGCAGATGGCTGCCTTCCATCCA | TCGA |
| GGCTTCATGTCCAACAAGCACGTCTTCGCCATCTTCAACACGGAGCAGAG*GTTCACGCAAAGAATCTGCTCCACAAGTTTTGCTTCCAGAAGAAGAGAAA | TCGA |
| AAGAACTTTCGGCCCGACGACCCCACCCGCAAAACCAAAGCCCTGCTGCA*GAACTCCTGCGTTGCGAGCTGTGCCTGGGCATCATGGGTGGGAAACCGCG | TCGA |
| GGCGGCCAGAGCGCCGGCAAGAGCTCGGTGCTGGAGAACTTCGTGGGCCG*CTGGAGAAGAGCGTCCGGCGCCTCCGGGAGAAGTTTCATGGGAAGGTATC | TCGA |
| AATCGTCACCCGGCGGCCTCTCATTCTGCAGCTCATCTTCTCAAAAACAG*GTGTTCTCCCGGCCCGGCTGCTCAGCCATACGCCTCCGAAATCATCTGTG | TCGA |
| GGCGGCCAGAGCGCCGGCAAGAGCTCGGTGCTGGAGAACTTCGTGGGCCG*GTCCCTCTGGCCTGCATGCCAGGAGTCGGTCACCGCCCTGTGCTTCCTCC | TCGA |
| GGCGGCCAGAGCGCCGGCAAGAGCTCGGTGCTGGAGAACTTCGTGGGCCG*GCTGTAACAGAAGACAAGTACGAAATACTGCAATCTGTCGACGATGCTGC | TCGA |
| GGCGGCCAGAGCGCCGGCAAGAGCTCGGTGCTGGAGAACTTCGTGGGCCG*GGCCTGAGTTCCCTGGCGGAAGAGGCAGCGCGTGCGACCGAGAACCCGGA | TCGA |
| AATCGTCACCCGGCGGCCTCTCATTCTGCAGCTCATCTTCTCAAAAACAG*GATGAGTTCCACCCGTTCATCGAGGCACTGCTGCCTCACGTCCGCGCTTT | TCGA |
| GGCGGCCAGAGCGCCGGCAAGAGCTCGGTGCTGGAGAACTTCGTGGGCCG*CTTCGAGGCAGAGAATGGGCCGACACCATCTCCTGGCCGCAGCCCCCTGG | TCGA |
| GGCGGCCAGAGCGCCGGCAAGAGCTCGGTGCTGGAGAACTTCGTGGGCCG*CTGGATTCTGATGGCACCATTACTATAGAGGAGCAGATTGTCCTTGTGCT | TCGA |
| GGCGGCCAGAGCGCCGGCAAGAGCTCGGTGCTGGAGAACTTCGTGGGCCG*GGACCCGAGCTGATCCAGAAAGCCAACGGTCTCCACGGCTTCATGAATTG | TCGA |
| GGCGGCCAGAGCGCCGGCAAGAGCTCGGTGCTGGAGAACTTCGTGGGCCG*GCTCGGACATCATCATGCAGCTGGACGACGTGGTTAGCAGTACTGTGACT | TCGA |
| AATCGTCACCCGGCGGCCTCTCATTCTGCAGCTCATCTTCTCAAAAACAG*GTGCAGCAGCTCCAGCAGGTGCCCGTCCCACACGTGTACTCCAGCCAGGT | TCGA |

| | |
|---|---|
| CCCAGCAGAGGAGCACGCAGCTGAACAAGAAGAGAGCCATCCCCAATCAG*CTCCATGGTGATCCGGGACCTGACCCTGCGCAGCGCTGCCAGCTTCGGCT | TCGA |
| ACATGGCCGACCGCATGGGCACGCCACATCTGCAGAAGACGCTGAATCAG*GAATATTTCACTGCTACATGGAGTACTCCCGACTGCGTGGTGAGGCCGGC | TCGA |
| AATCGTCACCCGGCGGCCTCTCATTCTGCAGCTCATCTTCTCAAAAACAG*GAACGCCACAGAAGTATGATCCCACTTTCAAAGGACCCATTTACAATAGG | TCGA |
| GGCTTCATGTCCAACAAGCACGTCTTCGCCATCTTCAACACGGAGCAGAG*GAACGCCACAGAAGTATGATCCCACTTTCAAAGGACCCATTTACAATAGG | TCGA |
| CCTGGCCAACTCCGACGCCCTCAAGCTGGCCAAGGAAGTCGATCCCCAAG*GAACGCCACAGAAGTATGATCCCACTTTCAAAGGACCCATTTACAATAGG | TCGA |
| TGCCTGCACTCTGTATTCTATTAATAAACTAAAATAAAGGGAAGACGCTG*ctggtgaaactgcgtgctgtaagctgggaccagctttgtccataactgct | TCGA |
| AATCGTCACCCGGCGGCCTCTCATTCTGCAGCTCATCTTCTCAAAAACAG*GTTGGTGTCTTTTCCACGTGGTAGAagctggctaacctccacttccacat | TCGA |
| CACTCTGTATTCTATTAATAAACTAAAATAAAGGGAAGACGCTGCTGGTG*aaacaaaaacaaaCCACACCTGCATCTGCTGTGCTGGGCCCCCGGTTCTC | TCGA |
| CGCGGCACCAGGGCGCGCAGCCGGGCCGGCCCGACCCCACCGGCCATACG*GTTAGCAGAGGAGCCGGACACAGAGGAAGCTCAGGTGAGTACAAAGAGTC | TCGA |
| GTCTTCATGAATGAGAAAGAGGACATCTTATGGTGCACTGAAATGGAAAG*GGGCAGCCATTTTGGGGGGTGCTGATGGATACCTGCGGGGTCGGCTATGT | TCGA |
| GCGGCTCACACCTGAGCGCGACTGCAGAGGGGCTGCACCTGGCCTTATGG*GACCAGGACTAAAACCTCCAGAGGAACGGACAATAGAATACCTAGAAGAA | TCGA |
| ACTTTCTTTACAGATCGGAGACACTGTGCACATCTTAGAAACATATGAAG*TGCCGTGGAGGTCCCAGCCCCCGCTGCGGTTCTCGGAGGTCCGGAGCCCC | TCGA |
| CATCTTTTGGTTACACAGAGCACTCAGTCCGCACCTGTTTTCCACAGCAG*TTCTGGCCTGAGCCTTGAAGATTCTAAGAAATTGACAGCTTCTCCCAGTG | TCGA |
| CCTGTTTTTCCTGAGCAACCAGAGAAACCACTTGACCTTGCACATATAGT*TATCGTCGTCCTCCCCGCAGTCAGGCTGCCCATCACCCACCATTCCAGCA | TCGA |
| AGTTGCCAGATGGCACTTGGAACTTAAAGAAGACCAAAGATGGAAAACGG*AATGCATACTTGGAATGAATCCTTCTAGAGACGTCCACGGTATGTTAAAT | TCGA |
| CTGCTTTCCTTTCCCTCAGCACACAGAGGATGATGCTGACTTTGATGAAG*TGACATCGATGTGGCCCAGCAGTTCAGCTTGAATCAGAGTAGAGTGGAAG | TCGA |
| CTGTGAATGGAGGCAGAGACCTCCAATAAAGTGCCTTCTGGGCTTTTTCT*aaaaaaaaaaaaaaaaaaagaaaaGCATGAATGGAAAGAAGGGCAAGTCA | TCGA |
| AGCTCACCGACAGTGCCCTGAGGGCCTTCGAGAGCTACCGCGCCAGACAG*ATAACCCAGCTGTGCTCCCTGGAACCTTCAATTTCAAGGCCTCCCTGCCT | TCGA |
| AGCTCACCGACAGTGCCCTGAGGGCCTTCGAGAGCTACCGCGCCAGACAG*GCTGTTCGGCTCTTGAATCCCAGCGATGAAACCAGGAACTCACTTTCCCG | TCGA |
| GACGCGGAGCCGAAGTGCCATTGTCATCAAGGCTGGAGGCCGCTACCTGG*GACCTGAACCCGCGGCGGGACATCTCCTCCTGGCTGGTGAGTCCCCCCAT | TCGA |
| AGCTCACCGACAGTGCCCTGAGGGCCTTCGAGAGCTACCGCGCCAGACAG*ACACAGCTGTGATCAGTCCCCAGGATCCCACGCTTCTCATCGGCTCCTCC | TCGA |
| AGCTCACCGACAGTGCCCTGAGGGCCTTCGAGAGCTACCGCGCCAGACAG*GTGTCTCTGTGGTGACGAGCAGGGCTTCATCCAGTGCCTCTGTCCCCACC | TCGA |
| AGCTCACCGACAGTGCCCTGAGGGCCTTCGAGAGCTACCGCGCCAGACAG*GGCCAATTCCTGTCCCCCCAGCAGCATGGCATCGTGTGCTGAACCCTCTG | TCGA |
| AGGATTCTGTTTCACTGAGGCCATCTATCCGATTTCAAGGAAGCCAAGGG*GTGCTGGCCCAGCAGGGGGAGTACAGTGAGGCCATCCCCATCCTGAGGGC | TCGA |
| AGCTCACCGACAGTGCCCTGAGGGCCTTCGAGAGCTACCGCGCCAGACAG*TGTGTCCCGGCATGGATATCCGGAACAACCTCACTAGGTTGCATGAGCTG | TCGA |
| ACGGCCTGACGCAGGCGGACAAGGACGCGCTGGATGGCCTCCTCCAGCAG*GAGATGTGCCCCAGAGCCTCCCCCACTTCCTTGTGGCCCACTGTGGAGAC | TCGA |
| CCTGGCTACTCGGAGGGGGACCAGCAGCTGCTGAAGCGGGTGCTCGTCCG*GGGCAGAACCCCTTCGAGCTGGCCTTCTCCCTAGACCAGCCCGACCACGG | TCGA |
| AGCGTTTCCAGTGTTCCCACGTCCACGTCGGAGACGCCTGACTACTTGCT*ACTTCAAGGGCTACCACTGGACCCTTCCCCTGTCTTGAACCCTGAGCCGG | TCGA |
| CTGTTCCTACCCCAACAGCACCGCTGCTTCCTCCGCAGCCTGCAACTCCA*TTCTCGTTCAACTTGGAAAACCATATTATCCTGAACCAAGGGAACATCTT | TCGA |
| AATCCCTTCCCTTTCGTCAACCTGTGGACCCTCAGCTTTTAGGAATCCCT*GTGTCGTGCGCCAGGTGCGGCCCATCGTGGGCCCATTTCATGCCGTCCTG | TCGA |
| CTCCCCAGATGGGAGGACAAACAGGATTGAGAGGCCCCCAGCCTCTTAAG*ATCGAGATGCTCAGCCCCTTCGAGGCAGTGGCCACACGCTTCAAGGAGCA | TCGA |
| GCCTAAACTCTCATCTCCGGCCCTCTCGGCGTCCGCCAGCGATGGCACAG*GATCACCTCCTCGCACGGGGAGCATCTCCTACATCAACATCATCATGCCT | TCGA |
| CTCCCCAGATGGGAGGACAAACAGGATTGAGAGGCCCCCAGCCTCTTAAG*GTCTTGGCTGTTCAACCGTACTTTCTCCAGAAGGATCAGCTCAATTTGCT | TCGA |
| AATCCCTTCCCTTTCGTCAACCTGTGGACCCTCAGCTTTTAGGAATCCCT*CATGCTTTTCACTTCCACTGCATCTCTCGCTGGCTCAAAACACGACAGGT | TCGA |
| GCCTAAACTCTCATCTCCGGCCCTCTCGGCGTCCGCCAGCGATGGCACAG*agagaaggagcaagaaagggaagaacagttaatggaagacaagaaaagga | TCGA |
| CTGTTCCTACCCCAACAGCACCGCTGCTTCCTCCGCAGCCTGCAACTCCA*GGCGGTAGTCATGGTTTGTTTGGAAACAGCACAGCACAATCGAGAGGTCT | TCGA |
| GAATGTCCTAAACCACATGACACACTGCCAGTCAGGCAAGTCTTGCCAAG*GCATACTTCCGACAGGGTGTTGCCCTCCAGTACCTTGGACGTCATGCCGA | TCGA |
| AATCCCTTCCCTTTCGTCAACCTGTGGACCCTCAGCTTTTAGGAATCCCT*CATGCTTTTCACTTCCACTGCATCTCTCGCTGGCTCAAAACACGACAGGT | TCGA |
| ACGTCTCAAAGGGCCACAGCGACAATGACAGCTGACAAGGAGAAGAAAAG*GTGGACCGCCTACTCGTGGGGCTGCGCTGGCGGCGGCTGGAGGAGCCGCT | TCGA |
| AGTGTAAAGTGACCCTCTTAGATGGCACCGAATACAGCTGTGACCTGGAG*CAGCAGCAACAGAAGAACCAGAAGTGATCCCAGATCCCGCCAAGCAGACA | TCGA |
| TGAAACCAGTGTTGCTCAACTCTAAGTTACCTGTGGATATCCTTGGAAGA*GACTGGGCCGGGGTCTGCAGGGCTCAGCTGAGCCCATGAGCTCCCAGAGC | TCGA |
| TGGAAGTTTCACTAAGTAGTTTGAACCTGGCTGTTCCTCCACCAAGATTT*TTAAAAAGAGAATTATTTGGCTTTAAATCATATCTTTCTAAATACCAGAT | TCGA |
| TGCATGATGGAAACACCATGGCTGCGGCGGCCCAGCTCTCTCTGACACAG*AAACGAGGATGCAGTCAATCAGATGGCCGTTACTCCCTTTCCAGTGCCAT | TCGA |

| | |
|---|---|
| TATTTCAGTTATCAAGTGGGAATCCTGTATATGAAAAATACTATAGACAG*CATTAGTGACCAGAAGGATCCCAAGGATCGAATGGTTCAGGTTGTGAAAT | TCGA |
| AGGACATGTTGGATGTGAAGGAGCGGAAGGTTAATGTTCTTCAGAAGAAG*GGATGCCAGATGATATGAACTTCAGTGATGTAAAGCAAATCCATCAAACA | TCGA |
| AGGTGACCAAACTCATCTACTGCTCAAGAACTGTGCCAGAGATTGAGAAG*CTCTTGATTCTGCACAGAGTAGTCGGTCTACTAGTTACTCTCCAAGACCA | TCGA |
| GCTCACCCTCCTTGCTAACTTTGCCACCCTTGTCAGCACCTACGCCAAAG*AGAGCGCATCCTCTCTGGAAAGCTGGTGCGCAGAGCTGAGAAGCGGCTTA | TCGA |
| TTCTGGGACTGGCTCTGAGCTCCCGCAAAAACTTGTGTATTCACCCTGAG*GCCAAGCTGATCGTCCCCAACAGCACGGCGGGCCTGATCATCGGCAAGGG | TCGA |
| GGATGGAGCTGCAGAATAATGGCTACATCGCCAAAGTCCAGTGTGCTGAG*GGTTCAATTAAGGACCAATTAAAAGCATATGGCGCTCTTACTGAGAATGT | TCGA |
| CCAGGCCTGCCGGCTCCGTAAATGCTACGAAGTGGGAATGATGAAAGGTG*GACCTGGGAGGAAGGCAAGGTGCTCATCTTTGATGACTCCTTTGAGCACG | TCGA |
| AGTCTGGTCCTGTGAGGGCTGCAAGGCCTTCTTCAAGAGAAGTATTCAAG*GCACACCTTGGGCCCACCCCACCTCCACATAGCCTTAATTACAAATCAGA | TCGA |
| TTCTTGAAAAAGGATGTAGATTTTAATCTGAACTTTGAACCATCACTGAG*CTTCAAGACCTCCGATCCAAGATGCTTTCTAAAGAAGTCTCCTGTCAAGA | TCGA |
| TTCTTGAAAAAGGATGTAGATTTTAATCTGAACTTTGAACCATCACTGAG*CTTAGAGACCTGCGCAAAGAAAATGAATTCGTGAAAGGACAAATTGTTAT | TCGA |
| GAGCCCAGCGGCTACACGGTGCGCGAGGCCGGCCCGCCGGCATTCTACAG*GAAACTTACGATCATCTTTCGGAAGTCCCGGTCACGCGGGAGCAGTTAAA | TCGA |
| TTCTTGAAAAAGGATGTAGATTTTAATCTGAACTTTGAACCATCACTGAG*AAAGTGTTCAAAAGAAAATGAGGAGAATAAGAAACAAGTTTCAAAGAATT | TCGA |
| TTCTTGAAAAAGGATGTAGATTTTAATCTGAACTTTGAACCATCACTGAG*ATGGTCTCCCAGCTTGAAGCCCAAATATCTGAGCTTGTTGAACAGTTGGG | TCGA |
| CCATTCTATAGCATAAGAAGACAGTCTCTGAGTGATAATCTTCTCTTCAA*ATTAAAACTTTGGAACAGACTAAAGCCATTGAAGATCTAAACAAATCCAG | TCGA |
| CCAGGCCTGCCGGCTCCGTAAATGCTACGAAGTGGGAATGATGAAAGGTG*GAAACTTACGATCATCTTTCGGAAGTCCCGGTCACGCGGGAGCAGTTAAA | TCGA |
| GGCAGACAGGGAGCTGGTTCACATGATCAACTGGGCGAAGAGGGTGCCAG*TGACCGAAGCTGGCTTTGGTGCTGACATCGGAATGGAGAAATTCTTCAAC | TCGA |
| GAGCCCAGCGGCTACACGGTGCGCGAGGCCGGCCCGCCGGCATTCTACAG*GAGATATACGACTAAGGGGTCAGACGGGGGTTCGTGCTGAACGCCGTGGC | TCGA |
| TTCTTGAAAAAGGATGTAGATTTTAATCTGAACTTTGAACCATCACTGAG*TATTCGGGATTTCACACAGAAAGATCGGAATCTGATCATGACAGCCAGTG | TCGA |
| GGCAGACAGGGAGCTGGTTCACATGATCAACTGGGCGAAGAGGGTGCCAG*GAGTTTTTGTCAGAAAGTGAAAATGGACAGCACAAGCTAAACATGATGCT | TCGA |
| GGCAGACAGGGAGCTGGTTCACATGATCAACTGGGCGAAGAGGGTGCCAG*GTGGACGTGGGCGTGGTGCACTTCACTCCCTTGATACAGCCCAAGATAGA | TCGA |
| TATTTCTCTCTCATTCCCACAGGATTTGCAGATGAAGACATTAATAAAAG*TGTATGGGTTTCGACAGACCATGATGAAATTGAGAATGTGGCCAAACAAT | TCGA |
| TCCTGCAACACCTGCGGCCTCACTGGGACCCCCAGGAGGTGACCCTGCAG*GTCGATCCAGTGTTTACAAGGGAAGTGAAAGCCAAAGTGAAAAGGTAAGA | TCGA |
| CTGAAAAGGAGGTAGAAATACTCTTCATTCAAGTCAATCAGTTTGCATTG*ACCTGTATCGCTACACATGAGTCTAATGGGATTGATATCATCATTGCTTT | TCGA |
| TCCTGCAACACCTGCGGCCTCACTGGGACCCCCAGGAGGTGACCCTGCAG*AAAGAGCTGTGCTTTGACGACGAGCTGGTCCTGCAGCAGCTGCGCTACAC | TCGA |
| TCCTGCAACACCTGCGGCCTCACTGGGACCCCCAGGAGGTGACCCTGCAG*GAAAAATTTGAGGGACTGTTTCGGACTTATGATGACTGTGTGACGTTCCA | TCGA |
| CTCTCTCGCTGTGAGACATGTCTGAGACTCCTGCTCAGTGTAGCATTAAG*GGAATGGAGAGGCCGGCGGCCCGGGAGCCGCATGGGCCCGACGCGCTGCG | TCGA |
| CTCTCTCGCTGTGAGACATGTCTGAGACTCCTGCTCAGTGTAGCATTAAG*GGCAGCCAGCACGTCCCCACTTGCTCGCCGCCGAAGAGGTCGGGAATAGC | TCGA |
| TCGAGCGCTCAGGATGGAGGAAGACTCGATCCGCCTGCCTGCGCACCTGC*GTTGACATTGATTGCAGTCAGAAGGTTGTAGTTTACGATCAAAGCTCCCA | TCGA |
| CTCTCTCGCTGTGAGACATGTCTGAGACTCCTGCTCAGTGTAGCATTAAG*ATACTTCTTGTGCCTTCAGCTCCGGCAGGACATTGCCTCTGGCCGCCTGC | TCGA |
| CTCTCTCGCTGTGAGACATGTCTGAGACTCCTGCTCAGTGTAGCATTAAG*ATTACAATGAACCAACTCAGCTTCCTGCTGTTTCTCATAGCGACCACCAG | TCGA |
| CTCTCTCGCTGTGAGACATGTCTGAGACTCCTGCTCAGTGTAGCATTAAG*GAGGCTGAAAGGGAAACATCAGCCGCCCGGGGGGAGTTAAAATGAGTGAC | TCGA |
| CTCTCTCGCTGTGAGACATGTCTGAGACTCCTGCTCAGTGTAGCATTAAG*GAACTCACCAAGGCCCATGAGCTGGAGGTGAGGCTGCACACTTTCAGCAT | TCGA |
| CAAAAGCACAAGGATTCTCGCTGTGACAGAGACAACACCGAGTATGAGAA*GGCTTTATGAAACCTTTCTAACCAATGATGAACCAGAATGCTGTGACGTC | TCGA |
| GCTGGCCTGCCACTGCTGTGCCTGTCGTCGTCATTGAAGGAGAGAGCAAG*GATCTGCAGAGATAGTTGCAACAAGGGTTACAGCTAAAATTGTAACCCTT | TCGA |
| CAGACGCCGTGCTCAGCCTTGACGAGGACACGGTGCTTTCAACAACAGAG*TACCTCTGTGAGTGTCAGTTTGATGACAATCTCAAATTCAAGAATATTAT | TCGA |
| CAAAAGCACAAGGATTCTCGCTGTGACAGAGACAACACCGAGTATGAGAA*GGCGAGCCCTGCTGAGACTTACTGACAAAAAGCTCGAGCGAATGGGGATT | TCGA |
| CAAAAGCACAAGGATTCTCGCTGTGACAGAGACAACACCGAGTATGAGAA*CATCGGCCGCTGAACCGTCAGCTGGCTGTGGTGTGGGCTCTCCAGCCCCG | TCGA |
| CAAAAGCACAAGGATTCTCGCTGTGACAGAGACAACACCGAGTATGAGAA*TGAAGAAAATATTTGGAAGCTCTGTGAATACATCAAAAACCATGACCAGT | TCGA |
| GTCTATAGTCCACTGTCAGCTGAGGTGGATCTTCCAGAGAAAGGACCAGG*GAAGAGAATGGAGACTACACATATGTGGAAAGAGTGAAGATACCCTTGGA | TCGA |
| ATCCAAACTACTTGTCGTCTGGAATAATCAGAATAAAAACCCTCCAGAAG*GAAGAGAATGGAGACTACACATATGTGGAAAGAGTGAAGATACCCTTGGA | TCGA |
| TGCACAGCCATAGATGGGCTTTCACTAGACCAAACACACATGGTGGAGAG*GTCTGGTTCCAGAACCGAAGGGCCAAGTGGAGGAAGCGGGAGCGTTTTGG | TCGA |
| GTATTTTGCAGAGCATCCCCCAAAGACAGATTGAAGAAATGCAGAGACAG*GAAACTTTTCAGTACACACTTTTCAGATGGGCTGAAGAGCTTGATGCTCT | TCGA |
| AGCCTGTGCACATCCTGACTTCTGTGAGCTCATTGCGCGGGACTAGGGAG*ATCCGGAGTTAGGTGTTGGCACGCTACCAGAACATGACAGCCAGGATGCA | TCGA |

| | |
|---|---|
| cgcgcgggggcgacgCGCGGGAACAACGCGAGTCGGCGCGCGGGACGAAG*GCAGGATGACGCAAAACACGGTGATTGTGAATGGAGTTGCTATGGCCTCT | TCGA |
| TTTTAAAGGGCAGAAATGCCTGAAGGTGAAGACTTTGGACCAGGCAAAAG*GGGTTGGCTGTGTTCCGGCCGCAGAGCACCGTCTGCGTGAGGAGATCCTG | TCGA |
| TGGTCTTTGGTGGCCGCACAGGAAATGAGGATATTATTTCCAGATTGCAG*GGAGTCTGAACACGTGGTGGACATGGCCTTCCTGGGCACCCGGGCTGGCC | TCGA |
| GACAGCCTCCTGACCTGTAGGACGAGGGATTCCTTGTTCTTCTGCCTGAA*AACCTGCAACCATGGTCGTGCCACTAGCCAGCTCGGAAGCCCAGGTACTT | TCGA |
| ACAAAGAACCACAGAATTCTGGACAATCAAAACTTAACTCCTGGATACAG*CTTCTTCCAGACCAGCTGGTCTTGCTTCTGGAGCATCTCTTGGAGCAGAA | TCGA |
| CTTTCCGCCACTGCCCAAACTGCTGAAGCAGCTCCCGCGAGGACCACCCG*CAAGCAAAGGCTACGAAAAGAAAATACCAAGCGTCCAGTGAGGCTCCCCC | TCGA |
| ACAAAGAACCACAGAATTCTGGACAATCAAAACTTAACTCCTGGATACAG*GTGAAGTGGAGGTTCTGGATCCGAACGACCCTTTAGGACCCCTTGATGGG | TCGA |
| TCAGCACACTGGCATTTAGCAAACAGAATGAAGCCAGCAGCCACATCCAG*GTTTGAGAGTACCGTACAGGTGAGCAAGTTGCAAGACCTCATCCACCGCA | TCGA |
| TAAGATATCAGGAATTATTCTTAAAACGGGAGAGAGTCAGAATCAACTAG*GCTGTGAATTTTTCAAGGAATATAAAGATCGGGATTACATGGCAGAAGGG | TCGA |
| CATTAGAGATGGCTCTGGCGTTGGTGTTTTCAAAATAGGTGAAGAGACAG*AATTCAGCTTAGAGAACTATCAACACAGGACAATGCAAGCCCATGAGCTG | TCGA |
| ACCCCGTCAGGGGAGAAGTCAAAGTGACCAAACTTCTCGACCGAGAAACG*GGCACCATGCCGGTGGAAAGGATGCGCATGCGCCCGTGGCTGGAGGAGCA | TCGA |
| ACAACCTCACAGTCGAGGCTACAGATGGAACCACCACTATCCTCACTCAG*GCTTTATTGAAAAGGCTTGAGGCCGTGAAACCAACAGTTGGTATGAAACG | TCGA |
| CGGAGCTGCCGAGGACTTTAGAGCCCAGGGCCGAGGACTCGGCTAGACTG*ATTGTTCAGTTCAAGGGAATGAAGAATTCAGAATAATTTTGGTAAATGGA | TCGA |
| ccggccccATGAGGAGGGACGTGAACGGAGTGACCAAGAGCAGGTTTGAG*CTCTAAAGTCTTTCTGGAAATTGACAACCGCCAGTGTGTTCAAGACTCAG | TCGA |
| accgccgcctccgccgctgccTCAGGAGCGGAACAACGTCGGCGAGCGGG*GCTGTTACCTTCTCCTCTCAAGGATGCAGTGGAAGTCTCAACCTGGAGAA | TCGA |
| AGGAAGGGAACCAACCCACTTTCTCGGCGCCGCGGCTCTTTTCTAAAAGT*TACAAACCTCCAGGGAACCTGGCTGGCTAGAAGGGACTCTGAACGGCAAG | TCGA |
| TTATATGGGCATACTTCCACTGTGCGTTGTATGCATCTTCATGAAAAAAG*AGCCTTGAAACCAGCGAAACAATAATCAAAGAGCAGTTCTGTCAAAGAAA | TCGA |
| TTATATGGGCATACTTCCACTGTGCGTTGTATGCATCTTCATGAAAAAAG*ATCGCAACGAACGAAATAAACCAGAACATCGTTCTTCAAGGTATGAATTT | TCGA |
| AGGAAGGGAACCAACCCACTTTCTCGGCGCCGCGGCTCTTTTCTAAAAGT*GACACAAAATATATATCAAATGGCGACATTTGGAACAACAGCTGGTTTCT | TCGA |
| AGGAAGGGAACCAACCCACTTTCTCGGCGCCGCGGCTCTTTTCTAAAAGT*GTAATTATGAGGAATTAGAAAACTCAGGAGATACAACTGTGGAATCTGAA | TCGA |
| AGGAAGGGAACCAACCCACTTTCTCGGCGCCGCGGCTCTTTTCTAAAAGT*TGACAGTGAATCCTAAACCTGAATGGCGCTCATGTTTTCCAAGAGAAGCA | TCGA |
| CAACCCCGAACCAAGCTTTAGCCGCCGAGGCCGCGTGTCCCAAAGGCCAG*ACACCAAAGTTCTATGTGTTCATGTGGCTCCTTGCAGATATCATTTCTGT | TCGA |
| cgtagtgctatctacatccagacggtggaagggagagaaagagaaagaAG*AAAATGTAGGCCGGTTAATCACTCCTGCCAAAAAGCTTGAAGATACAATA | TCGA |
| cgtagtgctatctacatccagacggtggaagggagagaaagagaaagaAG*GGGTGCTACATCCATTGTGTACAGATGCAAACAGAAGGGGACCCAGAAGC | TCGA |
| CTGTGCATCCTGCCACTGAGGACTCCGAAGAGGTAGCAGTCTTCTGAAAG*GTCTTCTGAGCTACGATAATTTTTTGGAACGGCAGAAATGATTGGTTCTA | TCGA |
| cgtagtgctatctacatccagacggtggaagggagagaaagagaaagaAG*TTCAACTTTGTGGGGAAACTTTTGGGTCCACGTGGCAATTCTCTGAAGCG | TCGA |
| CCGCGCTCCCTCCCTCTGCCTTTCATTCCCAGCTGTCAACATCCTGGAAG*CGTTGACAGAAGGCTACGTTGGGGCCCTGCACGAGAATAGACACGGCAGC | TCGA |
| CTGTGCATCCTGCCACTGAGGACTCCGAAGAGGTAGCAGTCTTCTGAAAG*GTTTAACTATAGATCAACACATCATCTTGCATCTCATGGGTTCTATGAAT | TCGA |
| GCTTTGAAGCTCAGGAAAGAAGAGAAATCCACTGAGAACAGTCTGTAAAG*CTTTTTGCAATCTTTGCATTTGCAACATGCGGTGGCTATTCTGGAGGCCT | TCGA |
| ACCCCGGCTGCCGAGTCCAGGTGTCCCGCGGGCCTCGATTTGGGGAGCAG*ATAGAAGATGGTGAGATTTTTGCAAGTATTAACCAGAAGGACGGTATGGT | TCGA |
| TGCCCTACTTCAAGGATAAGGGTGACAGCAACAGCTCGGCGGGCTGGAAG*CTTGGTGATCCAGCTATTTTTCCTGCCGTAATTGTGGAACATGTTCCTGG | TCGA |
| TGCCCTACTTCAAGGATAAGGGTGACAGCAACAGCTCGGCGGGCTGGAAG*GTTTTGATGATGATGGTTCAGAATTTCATGAACATATATTTCTGGAGAAA | TCGA |
| TGCCCTACTTCAAGGATAAGGGTGACAGCAACAGCTCGGCGGGCTGGAAG*GACAGTTCGTGTTTGGTTAAAGAGAGACAGTGGACAGTATTGGCCAAGCG | TCGA |
| TGCCCTACTTCAAGGATAAGGGCGACAGCAACAGCTCTGCCGGCTGGAAG*GACCTATATCCAGACTTTGCCTGACACTGCAGGGTCCAAGAGAATTAAAG | TCGA |
| TGCCCTACTTCAAGGATAAGGGCGACAGCAACAGCTCTGCCGGCTGGAAG*GTGCGTATAATTTGCTCTGCGTCGACTCCTATATCAAGCTTATTTTTGCA | TCGA |
| TCCCCCGGCAGCACCCCACCTCCAGGAGCTCATGGCCTTCTCGTTCGCTG*GTACTTGGAAAAATGGACAAGGATTGTGAAATGAAACGCACCACACTGGA | TCGA |
| GAACCAGTACAACAGCAGCAgtggtggtggaggtggaggtggaggtggag*TCATCAGTCCCATCCCAGGAGCCTTCCAGGACTCAGGTAAGGTAGCCGGA | TCGA |
| GCACCGGCTGGCTGAGGAAATCGCCTCCCGAGAAGAAGTTGAGGCGCTAT*GAAGCTGGCCAAGAAGCAGAAGGAGACCCAGAGTGGAGCCCAGAGGGAGA | TCGA |
| TATTGAGCGGCTGGACCACCTGGAGAATGTCATCAAGCAGCACATTCAAG*CTACAGCCTCCGCTTGGCGTCCCGATGCTCTCCACCAGCGGCCCCGTACA | TCGA |
| TGTGCAGCCGCGAGGAGGCGCGGGAGGTGTTCGAGAACGACCCCGAGACG*GGATGCCAGATGATATGAACTTCAGTGATGTAAAGCAAATCCATCAAACA | TCGA |
| ATGGGGACCCTGTCTGCAATGCCTGTGGGCTCTACTACAAGCTTCACAAT*ATCCGAGATGAGTGGGGCAATCAGATCTGGATCTGCCCTGGGTGTAACAA | TCGA |
| TGCTGATCCACATGAGAGTCCACTCTGGGGAGAAGCCCAACAAGTGTACG*GGGTTGGCTGTGTTCCGGCCGCAGAGCACCGTCTGCGTGAGGAGATCCTG | TCGA |
| TTAAAACAAGAACAGATGTGTAATGTTGGTGTATCCATGGGGCTTACAAG*CTGATACCCAACTCTTGTGCAACTCATGCCTTGCTGAGCGTGCTCCTGAA | TCGA |

| | |
|---|---|
| AGCTTCAGACCCGAGAAAGGGAATTTCAGGAACAAATGAAAGTAGCTCTT*TTTAGAAATCAGTCCCCAGAATGTGGATGTTAATGTGCACCCCACAAAGC | TCGA |
| GCCTTACTCTTATGGAAGAAGTATTGCTTCTGGGACTAAAAGATAAAGAG*GAACTCTAGCCAAGAACCACGGCCTACACTCTCCAACACAATCCAGAGGC | TCGA |
| TTGTCCAGGGGCTGACTGTGGGCAGAGAAGTTGCAAACCGAGTTTCCAAG*GGATGCCAGATGATATGAACTTCAGTGATGTAAAGCAAATCCATCAAACA | TCGA |
| CGGTGCCAAGGGATTCAGCCTGGCGGACATCCCCTACCAGGAGATCGCAG*TTGTTACAGGGTAGATCACCCAAACCTGACAAGTGGGAACTTCTTCAGCT | TCGA |
| TTCTGCCTCATACAATACGAGGAGGTGCAGAAGTGAGCGAGCCAGCGGAG*TGCCACACCAGAGGAATATAATACAGTTGTACAGAAGCCAAGACAAATTC | TCGA |
| TCCTCTCTTTGTGCCACCTGCAGAAAGTGAAGCATTATCTCATCCTGCCG*CCTCCCCCACTGGCAGGATTGCCATGAGTTGTGGAGTAAGAAACGGCGAC | TCGA |
| TGGTTCTCCCTGGGAGCCTTCATGCAGCAAGGATGTGACATTTCTCCCAG*GGTGGATCTATTTTATTAATAACAGGTCCTCCTGGATGTGGAAAGACAAC | TCGA |
| TGCTACGGGCAGATGGAGAGGCCAGAGGTCCCGATGCACACCTTGCACCC*ATGAGCACCATGCCAGGACTGCCCACCCGGCCCTGCTTTTATGACATAGA | TCGA |
| TCCCCATCTTGGGCATTCATGGGGGCGCATCTATGATCATGGCTGACAAG*CTAGAGACATGACCTAACACCCTGATGACCACTCTCAGGGACCTTGAGTG | TCGA |
| ATGAATCGTTCCAACAAACTTGCTGTGATAGCAAGTCACATTCAAGAAAG*GAGAAGGAGAGGAACGCGCGGAGGAAGAAGAAGAAAGCGCCGGCGGCGGC | TCGA |
| AAGCAATGGTCATTGGAAATACAAATTATGATTGGAAGGAACTGGAAAAG*GTCTATCATATGCAAGTCACACGGTTGGTTTCACGCCACCAACTTCACTG | TCGA |
| GGACGCTGGACTGGCAGTTGGAGACCAGCATCACCAAGCCTTTTGATGAG*GAAGCTGCACAGTGGGATGAAGACGTACGGGTGCGAGCTCTGCGGGAAGC | TCGA |
| TTGCATGGGAAAATCATGGAAGTTGATTACTCAGTCTCTAAAAAGCTAAG*CGGCCAGCCTCCCAACTACGAGATGCTCAAGGAGGAGCAGGAAGTGGCTA | TCGA |
| GCCCCGGGGTCCACGCAGGCCCAGGCCGCCCCGTTCCCCGAGCTGTGCAG*GACCCAGGACCACCCCCACCTTCTCCATTACTAGGTTTGAAACCACTGCA | TCGA |
| CTTCAGTCGGAAAGTTGGAACCTCTGACAAGACCAAGTACTACCTTCAAG*GAGGAGCAAAATAGAGGCAAGCCCAATTGGGAGCATCTAAATGAAGATTT | TCGA |
| CTCCACCACCTCGGGAGGACAGCAAAGCTCCAAGAGTGACAGAGTCGTGG*GTGAGACTGCTGCTCCTCGGGCCATTTTGACCGGCCATGACTATGAGGTC | TCGA |
| TCAGGAATGGCGGTGGTGTGGGGGATGTGCTAAGAAGGCCATCGAAAGAT*CTCAGATTTCTTTTATCCAAGATATTGGAAGGCTCCCTCTGAAGCGACAC | TCGA |
| CAGCCAGCGCAGCGACGGGGCGCTGCCTGGCCCAGGCGCACACGGAAGTG*GAGTCTTCAGGCCAGCTCCCTGTCGGATGGCTTTTATGAAAAAATATCTC | TCGA |
| GCTTCACGGAGATTGTCTTCTGTGCTGTCACCTCGAATGAGCAGGTCAAG*CTAAAGCTGTTGATGGATACGTGAAGCCACAGATCAAGCAAGTGGTCCCT | TCGA |
| CTGTTCGTTGTCGTGAAAAGAAAGAACAGAATAAAGGAAGAGAGGAAAAG*GGAATGTGCAGCAATCAGTTAACCGCAAACAGAGCTGGAAAGGTTCTAAA | TCGA |
| GATGATGGGCCTGGGCCTCATGGCCAAAGACAAGACTCTGCGGAAGAAAG*GTCCTCCAGCAGCCTGACCTGAGTGGGTTAGTGATCCAGAGAAACCAGCA | TCGA |
| TGCAGGCTGGACAGCCCCCAGACCTGGAGAGGATCCGCACACTTCTGGAG*TAATGTGTGTGGCTGGTATTGGACTTGTTGTATTATTCTTCAGTTGGATG | TCGA |
| GCGGTTTCCCTGTCAGTGACAGTAAACGGCACCTAACCCCCGAAGAGGAG*ACAGCTGATTTTACTTCTAGAGTTATATAGTCCCAGAGATGTTTATGACT | TCGA |
| AAAGTTGTAGTAATACCTCTGCACTTGCAGCACGAATTAAGTATTTACAG*GGTGATTTCACCTGGAACAGCATGTCAGGCCGCAGTGTACGGCTGAGGTC | TCGA |
| TCGTGAGATTTCATGAAGATGGCGCCAGGACGAAGGCCCTACTGGGCAAG*GTTCAGCTTGTTGGTTTAGATGAGGAGAGTTCAGAGTTTATTTGCAGAAA | TCGA |
| CATTATCTGCATACCAGAGGTACTACAGTTTACAGTCTGACTACTGGAAG*CGATCTGCCTGCCTGGATGTGATGAGCAGCATGGATTTTGTGACAAACCA | TCGA |
| CATTATCTGCATACCAGAGGTACTACAGTTTACAGTCTGACTACTGGAAG*GCCTTCAAGAGGGAATTGAAAACTAAAGAACCTGTAATCATGAGTACTCT | TCGA |
| CATTATCTGCATACCAGAGGTACTACAGTTTACAGTCTGACTACTGGAAG*ACTATGAAAGTGATGACGACTCTTATGAAGTGTTGGATTTAACTGAGTAT | TCGA |
| TTTGTTGTTCCTGAAGGTTACTGGGGTGTTCTGAATGACTTCTGTGAAAA*TGCCTGGAGCCCTGCAAGGAATCAGGGGACCTGAGGAAACACCAGTGCCA | TCGA |
| ATGGCACGAAATATCAAGGTCTCAGATCCAAAGCTTTTTGAAATGATTAA*GTGATCCCATTAAGATACTAATGCCATCACTGTCTCCTACAATGGAAGAA | TCGA |
| ATGGCACGAAATATCAAGGTCTCAGATCCAAAGCTTTTTGAAATGATTAA*AATGGACAGCAAATTATGGATGAACCTATGGGAGAGGAGGAGATTAACCC | TCGA |
| TCGTGAGATTTCATGAAGATGGCGCCAGGACGAAGGCCCTACTGGGCAAG*AAGTCTGGGAAGTTGACGAGCAGATAGATCACTACAAGGAAAGCCAAGAC | TCGA |
| AGGGGTCAGCAGGGGGCCAGCTCAGGACACACAGACCCAGCTGGAACTCG*GTTTTGAACTCTCACACTGTGAAGATCCTGGCATTCCACAATTTGGATAC | TCGA |
| CACGGCCACGCGGATGCCTGCGATGCCAAAGACCCCACGGACCCGTTCAG*CAGAGAAGCCTGTGTTCTTCCCAGCCTTCGCAGCCGAGGCCCAGAAGCTG | TCGA |
| ATCGCTGCATAGTAATATTGCGTGAAATATCTGAATCTACCCCCGTGGAA*CGATCATATTGGAAATATGGCCCAAAGTGACTTCCTTTACCCAGAGAACC | TCGA |
| CGCTCTGGGTCCCTCAGAATATGACTCTCTGCCTGAAAATAGCGAGACAG*AGTACACAGTCATTAATGAAGCCTGCCCTGGAGCAGAGTGGAATATCATG | TCGA |
| CGCTCTGGGTCCCTCAGAATATGACTCTCTGCCTGAAAATAGCGAGACAG*CCTGTGGGCCAAAGCTGACCAACTCCCCCACCGTCATCGTCATGGTGGGC | TCGA |
| CCGCCCTCAGGCCCGTGGCCGCTGTCCAGGAGCTCTGCTCTCCCCTCCAG*ATCTCTATAAAAATGGACTCCAAAGAGCTAACTTTGTACCATTCATAGCA | TCGA |
| GAAATTGCGCTGGACCGTCGCCTTGCTCCTCGCCGCGGCGGGGACTGCAG*GCAGAAAACGAGGATGGGGCCCAGGAGAACACCTTCTCCATGGACCCCCA | TCGA |
| CTGCCCGGACGGCATGCTGCTGGCCAGGGACATGAGGAGCTGCCTCACAG*GTGATCCGCAGGGGCTGGCTGACCATCAACAACATCAGCCTGATGAAAGG | TCGA |
| GGCCACCCCTGGGCTCACCACGGTGGAGATAGTGACAATGTCTCACCAAG*GACCGCTCAGGAAGTGGCAGCGAAGAAGACTGAGCCCCGACATTCCAGTC | TCGA |
| ACCATTCCAAATCTGGGACCAGTCTCGGCAAATATTACACTTCTCTCCTT*CTCCTCTCTGCTATATGAAATATGGGAGACGACAGACCGTTTGTGTGCAA | TCGA |
| ACCATTCCAAATCTGGGACCAGTCTCGGCAAATATTACACTTCTCTCCTT*GAGCTAGGGGAAGGGGGTTGTTTGCCTTCTTCAGGACCTGACTGGACAGA | TCGA |

| | |
|---|---|
| TAGTGCAAATCCAGAGCCAGCGTCGGTTGCCTCGAGTAATTCTTTCATGG*GTCCTGTAAGTAGTGTTTTGGTGAATAAATACGGCAGCCGGCCGGTGGTG | TCGA |
| ACCATTCCAAATCTGGGACCAGTCTCGGCAAATATTACACTTCTCTCCTT*ATCCATATGTCAAACTTCTGCTTGATGCTATGAAACACTCAGGTTGGTAA | TCGA |
| TGGTGTCTTGGACTGCCCAGATGGGTATGACGAAGGAGTACATTGTCAGG*ATCCCGACTGGGCCTCCTACACTCTGGGCGTCTTCATCTGCCTGAGCTGC | TCGA |
| TGATGGGGACCCTGACTGCCCTGATGATTCAGACGAGTCTTTAGATACCT*ATGAGAATTCTCCAGCCAACTTCTGGGATTCTAAGAACAGGGGTGTGACT | TCGA |
| CCGGGTCCGGCTTGCGGGGTCAGCGAACTGGAGAGGCGCCATGGGCTGGA*GTATGATTATCGGGAAATGCTGCACAATGCCACTTTCTGTCTGGTTCCTC | TCGA |
| AGTTACACTTGGGGTTCATTCAGATCCGAGAGAAGCTTGATCAGTTGAGG*GTATGGATCCTGGCGGGAGCCAGTCAACCCCTATTATGTCAATGCCGGCT | TCGA |
| GCAAGAGTCACCTTCTGGACTGCTGCCCCCATGACATCCTGGCTGGGACG*GTATGGATCCTGGCGGGAGCCAGTCAACCCCTATTATGTCAATGCCGGCT | TCGA |
| CAGATTATGAGATTGCAAGTAAAGAAAGAGACCTGTTTTTTGAATTAGAT*GTATGGATCCTGGCGGGAGCCAGTCAACCCCTATTATGTCAATGCCGGCT | TCGA |
| TCTGAGGAAAGGGAATTCTCCCCGAGTTTCATCAACTTTGTCAACTTGTG*GGGGAGCTGACTATGACCAGCCACATGGAGAACTTACAGAATGCCCTGTA | TCGA |
| CCCCGTAGGGTCCCCGGCGCCAGGCCACCCGGCCGTCAGCAGCATGCAGG*GAAGGAAGACTTCAAGAAGTCATTGAAACCCTTCTCTCTCTGGAAAAGCA | TCGA |
| TGAAATCAGAATCTCCAGGAGTAAGGAGAAAAAGAGTTTCCCCAGTGCCT*TCCTGAAGATTTTCCTCAGAGAATCTTTCATCTCTTTGCTCCTGAGACTA | TCGA |
| GAGAAGCGGGCGCCCGCCGCCGAGCCGTCTCCTGCAGCGGCCCCCGCCGG*TTTTACAGCTCCCAGCACAGTAGGAAAGAGAAATTTGAGTCCTAGTCAAG | TCGA |
| ACATGAGTCAATATGTATTGTGGCATTCAAATCCTAGCACTCTGGGAGAG*GCTTGGCCTTCGTGCCTCTGCAGATCTGGTCCAAAGTCCTGGCCATCTCA | TCGA |
| ACATGAGTCAATATGTATTGTGGCATTCAAATCCTAGCACTCTGGGAGAG*CCTGTTGGTCTTTGCAAAGGTTCTCTGACTCGAACACACTGGGAAAAGTT | TCGA |
| ACTGGCGGCTGCTGGTAGTTCCATGCTGTCTGTTCAGACTTCAGCTTGGT*GTATCCTGAAGAACTTGCCTGGCACACAAATTTAAGTCGAAAAATCTTGA | TCGA |
| CTTTTCAAAGCAAGAAAATGGAACAGCATGTGTAGGAATTCTTCGTTGTT*gaGCACGTGAATTGCCTCTGCTTACGTTCAGGCCCTGTCCTAAACCCAGC | TCGA |
| ACATGAGTCAATATGTATTGTGGCATTCAAATCCTAGCACTCTGGGAGAG*ACAACCTCTCCTGTTTCATCTCCTGGCAACAGAATGCATCCTCCTTTTGG | TCGA |
| CTTTTCAAAGCAAGAAAATGGAACAGCATGTGTAGGAATTCTTCGTTGTT*CTGGCTCTCCTCACGGAACTCTCCCAGAATCGCAGCAGCGAGAGCTATAG | TCGA |
| ACAGCTTTCACAGTTTGCGGGACTCAGTCCCATCACTCCAAGGAGAGAAG*ACATTCACGGCATGGTGTAACTCCCACCTCCGGAAGGCGGGGACACAGAT | TCGA |
| ACAGCTTTCACAGTTTGCGGGACTCAGTCCCATCACTCCAAGGAGAGAAG*GCAAGAGTTTCACCTTGACCATAACCGTCTTCACAAATCCTCCCCAAGTA | TCGA |
| ACAGCTTTCACAGTTTGCGGGACTCAGTCCCATCACTCCAAGGAGAGAAG*TAGTCGAGAATGACTAAAAAAAGAAAACGCCAACATGATTTTCAAAAAGT | TCGA |
| ATAGTTGCTTTGATTGGGATTCACCCCCAAGGAGTCTCGCTGCTTTTGTG*ACGAAATCTCACAAAACTGTCCCTTATCTTCAGTCACATGCTGGCAGAAA | TCGA |
| GATCTAGCTTCTCCTCGCCTTCAGGATGAAAGCTCAGGGGGAAACCGAGG*TAGGAAATCCATGAAGTCCAGAAGTAGAAGTCCTGCATATTCAAGACATT | TCGA |
| TGGGCTGGGCCACCTTTCTTGTGCAGTCCCTAGGCCGTTTTGAGGGACAG*ACAGCCTGTTTCCGGGAGGAGCGGGATGTGCTCGTGAAAGGGGACAGCCG | TCGA |
| CGTCCCGGTGCCCAGCTCCGAGCACGTCGCCGAGATCGTGGGTCGCCAGG*ATGTTCCCGCTGCCTGTCACCAACGGGAAGGGCCGGCCCGCCTCCCTGGC | TCGA |
| CAGGGTCTGCACGAAATAAAGCTCCTGGGCAAGGGGACGTCTGCAGCTGA*GTTTGCCCTGGAGCCTCTCCTGCCAAAGAAAATGCACTCCAGGTCCCAGG | TCGA |
| AAAAAAAGTTGCAGCTACTTTGCAAGTGCCAGTAAATGATTTAAATGCAG*GTCTTAGAAAAATACCCCAATACACCCATGAGTCATAAAGAAATTCTTCA | TCGA |
| GAGCCGGAGTGGGGCGAGCCGCGGAACCCGGCCGGGAGCCGCGCGAGGCG*GAGGAGGCCCCGCAGCGACATGGAGGGAGCTGCTTTGCTGAAAATCTTTG | TCGA |
| GAGCCGGAGTGGGGCGAGCCGCGGAACCCGGCCGGGAGCCGCGCGAGGCG*GCCCTGTGTGCTGCAGTCACCACCACAGACCTGGCTGAGACCCAGGCGCT | TCGA |
| CCACCCTGGCAGGAATCCCAATTTGCCTCCACTTAGTGAGCGGAAGAATG*CAAGAGTGAGCCATGGAGCTACGTGTGGGGAACAAGTACCGCCTGGGACG | TCGA |
| CCACCCTGGCAGGAATCCCAATTTGCCTCCACTTAGTGAGCGGAAGAATG*GGAACAAGGGAGCCGTGGGGGTGTCGTTCATGTTCAATGGAACCTCCTTA | TCGA |
| GAGCCGGAGTGGGGCGAGCCGCGGAACCCGGCCGGGAGCCGCGCGAGGCG*GTGCAAGTGTAATCTCCATGCCACTGTATGTGTGTATGACAACAGCAAAT | TCGA |
| CCAGCAGCGCCGGACCCGGGAAGAACTGGTGAGCCAAGGGATCATGCCGC*GTGCAAGTGCTGGGCAGTCAGGACGGTGCCTAAGTGGACCTCAGACATGG | TCGA |
| CCAGCAGCGCCGGACCCGGGAAGAACTGGTGAGCCAAGGGATCATGCCGC*CTTTAGCTCCGCCAAAATGAAACACTCATTAAACGCACTTCTCATTTTCC | TCGA |
| ATGGTTGGCATGGAATTGAATTTCATCTGTCTGTGGGAATTGTAAGCAAG*GGAAGCGTGACAGCAATGACAGTGTTTTTTCCCCTGGATACAGCTAGACT | TCGA |
| CCACCCTGGCAGGAATCCCAATTTGCCTCCACTTAGTGAGCGGAAGAATG*GAGTGGTTAATGTGTTGCTAACAACTCCACTCTGGGTGGTAAACACCAGA | TCGA |
| CCAGCAGCGCCGGACCCGGGAAGAACTGGTGAGCCAAGGGATCATGCCGC*GAGTGGTTAATGTGTTGCTAACAACTCCACTCTGGGTGGTAAACACCAGA | TCGA |
| GAGCCGGAGTGGGGCGAGCCGCGGAACCCGGCCGGGAGCCGCGCGAGGCG*GAGTGGTTAATGTGTTGCTAACAACTCCACTCTGGGTGGTAAACACCAGA | TCGA |
| CCAATGGAACCACTATCCCCACTGCCAAGTCCACCCCCACACTCATTAAG*GACTGGGAGTTTTCATCCTCTTGAATAAGAACTCGACAACAGAGTGGGAA | TCGA |
| ATGGTTGGCATGGAATTGAATTTCATCTGTCTGTGGGAATTGTAAGCAAG*GAGTGTGAGACCTATGAGAAGTGCTGCCCCAACGTATGTGGGACCAAGAG | TCGA |
| GGAGAGCTCATAATCGTAGAGGACATAATGATGGTGAAGATTCTTTGACT*TCAGTGGAACCTAGTGAAGTGAAACAAGCAACTTCAACATCAGGACCAGC | TCGA |
| GAGGAATCGGCAAACTCCCTGGTCTCTGTTGGTGTAGAAGCCAAAATCAG*GCATTTCCCGGAAACATTAACTCTGACGGTGTGGTCCGGCACGAATTACA | TCGA |
| GGCCGGCCTCGCAAAGATGGCGCTTCCCCTTTCCAGAGAGCCAGAAAGAA*AAAAATGTGATGAGCCACTTGTCTCTGGACTCCCCATGTGGCTTTCAGC | TCGA |

| | |
|---|---|
| GCCAGGGGCCCCTCGCCTCCCCTCTGAGGACCGAAGATGAGCTTCCTCTT*AACCATTCAGCGCCTCTGTCTGGGTGATGATGTTTGTGATGCTGCTCATT | TCGA |
| ATTTCGTCAAGGAGTTCGGCCTCATCGACACCAAGGAGCTGGAGCCACTG*GGAAATTCAACTACTGAGGAGGTTACGGCACAAAAATGTCATCCAGCTGG | TCGA |
| GTGTTTTACCCGGAGGAGAGACTGCTGGAGACATGGGGAAACTGAGACAG*GAGGGTCTGCCCAAGCTGGTCAAGCCGGAGGGTCTGTCCAAGCTGGTCAA | TCGA |
| AACCAGTGGATTAAGCAGCCTCTCATGGATAAGAACAGAATAGAGGAGAG*GTTATTTCCAGAAAGAAGAGAATGGCAAAGTATGAGAAGGCTGAGGTAAA | TCGA |
| CCGGGAGGTGTTCAAGACCCAGGGGGTGATCAAGTACATGGGGCCGGCAG*GTTATTTCCAGAAAGAAGAGAATGGCAAAGTATGAGAAGGCTGAGGTAAA | TCGA |
| GTGTTTTACCCGGAGGAGAGACTGCTGGAGACATGGGGAAACTGAGACAG*ATTGGTGGACCAGACAGTTCACGGCTGACTTCGCGTTTGGGGAATCAGTA | TCGA |
| ACTTCTCCAAGTTTCAGGAAATGATAGAAACAACTTTAGATATGGATCAG*GTTTTCTTTGCAAACTATGCACTATCATCTTCACTACTATTGCATTTTTC | TCGA |
| ACTTCTCCAAGTTTCAGGAAATGATAGAAACAACTTTAGATATGGATCAG*CATTTCTGGATTAAAAGTGGTGGGGGAGTCCCTTTCTCCAGCCTTTCACC | TCGA |
| CCGGGAGGTGTTCAAGACCCAGGGGGTGATCAAGTACATGGGGCCGGCAG*ATGACTTTGGGAAATTGCTGCTGGCTGAGGCCCTCCTGGAGCAGTGTTTG | TCGA |
| TAACAGGCTCTTGCCTTACTTCCGACAGTCCCTTTCGTGCTGTGTTTGCG*TCATTCATACCTAGAGAAATTCCTTACCGAGCAGATGATGGAAAGGAGGG | TCGA |
| AAATTTTAGAAGGAAGAACTGAAAAATATGTGGATACTCCATTTGGCAAG*ACATCCCCGATTGAAAGAACCAGAGAGGCTCTGAGAAACCTCGGGAAACT | TCGA |
| GAGGAGATTCAGCCCGGCGATATTGTCATTATTGATCAGTTCATTGACAG*GTCATGATGATGGGCAGCGCCCGAGTGGCGGAGCTGCTGCTGCTCCACGG | TCGA |
| CATAAAACCAATGTCCAGCAGTGTTACTGCTATTGTGGAGGCCCTGGAGA*ATGGAGGAGATGTATAAGAAAGCTCATGCTGCTATACGAGAGAATCCAGT | TCGA |
| ATGAAGTACAAAGCAGTGAGTGTCAACGAAGGCTATAAAGTCTGTGAGAG*GTGTCCTGCCTGAAGGAGCTGGTGGCCCGAGTGCTGCAGAGGCTGTGCGA | TCGA |
| CTTGAGTCTAAGTTGGCCCAGGCTGAGGAGCAGCTAGAGCAAGAGACCAG*GCTCCACCAAGGACCTGATCGAGACTTGCTGCGCAGCCGGACAGCAGTGG | TCGA |
| AGTCCTCTCGGGAGAAGCAGCGCCTGTTCCCGGGGCAGATCCAGGTTCAG*GTTCCAACTATGGGAGCCCACGCCCCGCCCATGCCAACATGAATGCCAAT | TCGA |
| ACTGCTCCACCAGTCCTCTGACAAGTTTGTCTCGGAGCTGTGGAAGGATG*GGAATTTCAAAGGTCTGTGCAAGCAAATTGATCACTTCCCAGAGGATGCA | TCGA |
| AGTCCTCTCGGGAGAAGCAGCGCCTGTTCCCGGGGCAGATCCAGGTTCAG*GGAATTTCAAAGGTCTGTGCAAGCAAATTGATCACTTCCCAGAGGATGCA | TCGA |
| TCCAGTATCTGGCGTACGTGGCGTCCTCGCACAAGAGCAAGAAGGACCAG*GGGCCGTGAGGGCCTCCAGCATTTTCCCAATCCTGAGTGTGATTCTGCTT | TCGA |
| AGTCCTCTCGGGAGAAGCAGCGCCTGTTCCCGGGGCAGATCCAGGTTCAG*GGGCCGTGAGGGCCTCCAGCATTTTCCCAATCCTGAGTGTGATTCTGCTT | TCGA |
| GGTTTACAGATGTTTTCCACTGAGCATCACAATAAAGAGAACCATGTGCT*aaaaaaaaaaaaaaaaaaagaaaaGCATGAATGGAAAGAAGGGCAAGTCA | TCGA |
| AGCTGGCTGCGGAGAACAGGCTCACGGAGATGGAGACGCTGCAGTCTCAG*CAAAAGGAGAAAATTGTCATCAAAGGATATTCCAGATTCTTGACAGCATT | TCGA |
| AGTCCTCTCGGGAGAAGCAGCGCCTGTTCCCGGGGCAGATCCAGGTTCAG*CATTTTTCAGTTGAAGGACAGTTGGAATTCAGAGCCCTTCTATTTGTCCC | TCGA |
| GGCGGAGGCCCGAGAGAAGGAGACCAAGGCTCTGTCGCTGGCCCGGGCCC*CACCCGGAGAGCTGTGTCACCATGTGGGTCCCGGTTGTCTTCCTCACCCT | TCGA |
| CCTCGGTGCTGCACAACCTCAAGGAGCGTTACTACTCAGGGCTCATCTAC*GTTTCATGGGGAAGGGTATCGGGAAGGCTATGAAGAAGGCAGTAGTTTGG | TCGA |
| AGTCCTCTCGGGAGAAGCAGCGCCTGTTCCCGGGGCAGATCCAGGTTCAG*ACAGAGGGGGAAGACGGTGGGGCCTCCCCACCTGCCCCGCAGAAGATGCA | TCGA |
| GGTTTACAGATGTTTTCCACTGAGCATCACAATAAAGAGAACCATGTGCT*aaaaaaaaaaaaaggacatgctacagcatggatgcaccttgaagacatttg | TCGA |
| AGTCCTCTCGGGAGAAGCAGCGCCTGTTCCCGGGGCAGATCCAGGTTCAG*TTGCAGGATGTCGATGACAGACTTGCTGAACGCTGAGGACATCAAGAAGG | TCGA |
| AGTCCTCTCGGGAGAAGCAGCGCCTGTTCCCGGGGCAGATCCAGGTTCAG*ATGGCTCAGCGACTTCTTCTGAGGAGGTTCCTGGCCTCTGTCATCTCCAG | TCGA |
| AGGCCTTCGGGAACGCCAAGACCGTGAAGAATGACAACTCCTCCCGCTTC*ATGGCTCAGCGACTTCTTCTGAGGAGGTTCCTGGCCTCTGTCATCTCCAG | TCGA |
| ACTGCTCCACCAGTCCTCTGACAAGTTTGTCTCGGAGCTGTGGAAGGATG*GTGTCAGCGGTGCCCACTGTGCTGGCCATGAAGAATGGGGACGTGGTGGA | TCGA |
| GTGGTTCAAAGAGACTTCAATCTTCGCAAAAAAGAAGAAGAACGACTAAG*GAGCCAGCACCGCTCTTTGACCTTGCCATGCTTGCCTTAGATAGTCCAGA | TCGA |
| TGGGGTTCCTTCAGATTCCGTGGAAGCTGCTAAAAATGCAAGTAACACAG*GTTGAAGAGGAGAAGAGGAAGAAGAAAGAGGAGGCGGCCCGGAGGAAACA | TCGA |
| AAGAGAGAAACACAAGCTATTCTTCCTATGCAGTTTCATTCCTTTGACAG*GAAGGAATGTTCCCAATAGTAGACATAAAAGTCTTCGCACAGTGAAAACT | TCGA |
| TTTCATGACTACAAAGAATTACTGTGATCCTCAGGGCCATCCCAGTACAG*TTACAGAGTGTTTCAAGTGCCATTCACCTATGTGACAAGAAGAAAATGGA | TCGA |
| TCAGCCGAAATCATGCTGTGTTAACTGCTAACTTTTCTGTAACCAACCTG*GTTTACATGGACTGTAAAGGCAGGAGTGTGGATGTATTGTTAGCCCATTG | TCGA |
| TCTGAACGCAACAGAATTCAGATGCGTAGCTTGCAGCAGCAGTTCTCCAG*TCTGTTCGAGAAGTCACAGGCTACGTGTTAGTGGCTCTTAATCAGTTTCG | TCGA |
| TCAGTCCACCCAAGGTCTCCTCGGATCGCCTGGAGAGGCACTCGGACCTG*GCTCCTCCTCAGCCATCCTGGCCGTgctcagtcatctgaccagctgacct | TCGA |
| GCCAGTTATCTTCTGGAGAAGACAAGTGGCTGCTTCGAAAGAAGGCCCAG*AATGTTCTCCTGGGCAGCGTTGTGATCTTTGCCACCTTCGTGACTTTATG | TCGA |
| CAATTACTCTGCGCCAGACCATCACCACATTTGGGTCTCTCAAAACCATT*AATGTTCTCCTGGGCAGCGTTGTGATCTTTGCCACCTTCGTGACTTTATG | TCGA |
| TCAGTCCACCCAAGGTCTCCTCGGATCGCCTGGAGAGGCACTCGGACCTG*AATGTTCTCCTGGGCAGCGTTGTGATCTTTGCCACCTTCGTGACTTTATG | TCGA |
| TCAGTCCACCCAAGGTCTCCTCGGATCGCCTGGAGAGGCACTCGGACCTG*AAGCTTGAGTGCCCAAGGGTTTGATAAATGTGGAAAATAACCGCTGTTCG | TCGA |
| TCAGTCCACCCAAGGTCTCCTCGGATCGCCTGGAGAGGCACTCGGACCTG*CCTCTACGGCAGACACATGCAGGCCAACCCAGAACCACCGAAGAAGAATA | TCGA |

| | |
|---|---|
| CCAGGTTCAAGGGTCTCTCCAGAAAATCTTGTGGACAAATCCAGGGGAAG*GCAGCTGAGGAAGTTGGATATCCAGTAATGATCAAGGCCTCAGAGGGAGG | TCGA |
| CTGTATAACCAGCCATCAGATACCAAGGTGTACCATGAGAACATCAAGAC*TGGAATCAACTATGAAATTATAATCATAGATGATGGAAGCCCAGATGGAA | TCGA |
| CCAGGTTCAAGGGTCTCTCCAGAAAATCTTGTGGACAAATCCAGGGGAAG*CTGTTGTTATGTCAGTCGAAGACTTAGCCAGCTATCAAGTCATGATCCGC | TCGA |
| TCAAAACACCATGTCCAGAGGCTCACCCATGATGAACAGAACTTCTGATG*GAGAGTGTGCTGGGCAGACGATGCTGGACACGATGGAGGCGCCCGGCCAC | TCGA |
| TGGAGCAGAAACACCGCAGTATTGTCCAAATTATTTATGATGAGAATCGG*GGATTTCCCAGATGACCCAGGCATGCAGTGGGACACAGAGCACGTGGCCA | TCGA |
| CTTCGAAGGCGACCTTCCTTGCTTTCAGAATTTCACCCAGGTTCTGACAG*AAATCATGCCCAAGTTCAAACAACGAAGACGAAAGCTAAAAGCCAAAGCC | TCGA |
| ATCCTCCTCACTCTGTCCAGTATACATTTCCCAACACCCGCCACCAGCAG*CGGTGGTGTCTGTGGATGGGCGGCCCGTGAGACTCCAACTCTGTGACACT | TCGA |
| CCAGGTTCAAGGGTCTCTCCAGAAAATCTTGTGGACAAATCCAGGGGAAG*GCTTATGATCAGAAGAACATTAGGCGAAGAGTTTATGATGCTTTAAATGT | TCGA |
| TATTCTGTCATTTACACACGTCGTTCTAATTAAAAAGCGAATTATACTCC*aaaaaaaaaaaaaaaaaaaaaaaaaattaagtgggtgtggtggtgcatgcc | TCGA |
| TCGAGTCGAAGCACCGCAGCCTGGTGCAGATCATCTACGACGAGAACCGG*GTGCTGCCCCAGATCAACTTGGATCATTACTCGGTGTTACCGGCAGGCGA | TCGA |
| CAGCGGCGGAGGCCCTCCCTGCTGTCTGAGTTCCAGCCCGGGAATGAACG*ACAAGGACACGTTGAAGGGAGTTGATGCAAGCAACGCGCTCGTTCTACCG | TCGA |
| CAGCCCCCGGGGCAAGAGCAGGAGCCCGGCACCCCCCGCCGACAAGGAGG*ACCCCTCTCCTCGACGACCCTGTCTTCATCCACCCCAGCTCCGTCCTTTT | TCGA |
| GAATGAGAGTTCTCGCTGGACAGAAGAAGAAATGGAAACAGCCAAGAAAG*GGTCCTCATATTTTGTGCTTGCAAATGGACATATCCTGCCCAACAGTGAA | TCGA |
| GGCGTCGGGCGTGAGCGGAAATGAGGAGGAGATGGTGGAGGAGGCTGAAG*ATATGTTTGACGACTTCTCAGAAGGCAGAGAGTGTGTCAACTGTGGGGCT | TCGA |
| CTTCAAGTTGTGTTTGGAGCTGACGTGAGCGAGGTTGGTCATGTGCTCGG*AGTATCCGGTCTAAAGTGGAGTTGTCAGTCTGGGATCAGCCTGAGGATCT | TCGA |
| ACCCGCCCCACAGCCTTTCCTACCCAGTGCAGATCGCCCGGACGCACACG*GGAAAGGAACACAGGTCTGGAAGAAGAAAGGAGCCATCTATGAGGGGGAC | TCGA |
| ACTATAAGAGCCTGGTGAGACGGAGCTATCGGCGCCGCGGCAAGAGCCAG*TTTCGTGACCTTGTCCAGTAGAAGGCTATTTAATTTTCACAACTGCTTGA | TCGA |
| GTACGAGACACTCTCCGACAGCGAGTGACTCAGAACAGGGCGGGGGGGGG*ACTCCACTGCGACTGCAGAACTCAGAGCTGCTCTTCCTCTGTGGCCAGTT | TCGA |
| CGGCCCGAGCGCGCCCCAAGCCCGGGCGCCACCGCTGCCACCTCCGCGAG*ATCAATGAGCTGAGCAATGTTCCTGTTCCTGTCATGCTAATGCCAGATGA | TCGA |
| GATCCCGCGCGAGGAGCTGCGGCACACGCCCGAGCTGCCCCTGGCCCCGC*TTCCCCTCTCTACAACCCTCTCTCCTCAGCGCTTCTTCTTTCTTGGTTTG | TCGA |
| CAGGTCTCCCTGAGTCTTTGAGGACACAGCCTCGCTGGAGGCAGTTTCTG*AGCGGGGCCATGGAGGGGGAGACTCTTCACACATTCTACACTCAGCTGGT | TCGA |
| CAGGTCTCCCTGAGTCTTTGAGGACACAGCCTCGCTGGAGGCAGTTTCTG*GTGCCCCCTTGTTTCTCTCCCATCCTCACTTCCTCAACGCTGACCCGGTT | TCGA |
| CGGCCCGAGCGCGCCCCAAGCCCGGGCGCCACCGCTGCCACCTCCGCGAG*GTCACGGGAATCCCCATGAACTGCTCTGTGAAACTGCAGCTGAGCCTCTA | TCGA |
| GAATGAGAGTTCTCGCTGGACAGAAGAAGAAATGGAAACAGCCAAGAAAG*GGTCCTGAGGACACCGTGAGCCAGCCAGGCCTGGCCGCTGGGCCTGACCG | TCGA |
| ACCCGCCCCACAGCCTTTCCTACCCAGTGCAGATCGCCCGGACGCACACG*AACGTGCGCATCGACCCCAGTAGCCTGTCCTTCAACATGTGGAAGGAGAT | TCGA |
| TGAGGCCAACAGCGAGGAGGCCATCACCCCCCAGCAGAGCGCCGAGCTGG*GGCAAAGGTGAAGAAAGGCGTGAACATCATCGGCGTGCGAGCCAGCAGCC | TCGA |
| TTAATGAGATATTTGTATTTATTTCCAGACCAATAAATTTGTAACTTTGC*aaaaaaaaaaaaaaaaaaaagaaagccaggtgtggtggtgcacgcctgta | TCGA |
| TTGTCTTTTCACAGACCATCACCGGCCTCCTGCAAGAGTTTGATGTCCAG*GAGGCCCTGCCTGGTCTCGTCTGGGACCTGGGACAGCAGCTGGGAGACCT | TCGA |
| TTAATGAGATATTTGTATTTATTTCCAGACCAATAAATTTGTAACTTTGC*AGCGGAAACGCTTGGGCATTGGCCAGTCTCAGGAGATGAACACACTCTTC | TCGA |
| TTAATGAGATATTTGTATTTATTTCCAGACCAATAAATTTGTAACTTTGC*aaaaaaaaaaaaaaaaaaaagaaaCTTTATTGTATGAACTAGGTGGGTTT | TCGA |
| TTAATGAGATATTTGTATTTATTTCCAGACCAATAAATTTGTAACTTTGC*AAAAAAAAATATTGGTGCTTTGAAGTGTCACCTTTTAAAAAGTTGGCAAA | TCGA |
| TGCAGGATGTAGACCTCGCTGAGGTGAAGCCTTTGGTGGAGAAAGGGGAG*ATGACAGACAACCTTCTGTGCCTTCATCAAGCTGGTTGTGTACCCACCAT | TCGA |
| TCCCTGACTCTCCTGGGGGCACCTCCTATGAGAGATACGATTGCTACAAG*GATTTATTTATACATGTGGTGGAACTTTAAAAGGACTTAATGGCACTATA | TCGA |
| AGCCCCCTGAGGTTAAACATGCCATGGAGGTCAATAAACACCAGAGTAAT*ATGATGATGTGCCTGCAGATATGGTTGCAGAAGAATCAGGTCCTGGTGCA | TCGA |
| ACAGGCCGGTGGAATGGGTCCAGGCCGTGGTCAGCCGCTTCGACGAGCAG*ATTGGCATGGTAGCTTGGAAAATGACCTTTAAAAGTCCTGAATATCCAGA | TCGA |
| AACTCCCTATAGCGATGGCTCTGGCCAATGTGGTTCCTTGTTCTCAGTGG*GTGTTTACAAAATATGGGAAGTGTTACATGTTTAACTCAGGCGAGGATGG | TCGA |
| ACAGGCCGGTGGAATGGGTCCAGGCCGTGGTCAGCCGCTTCGACGAGCAG*GGTAAAAGCCCCAAAGCTGTCATCAGCATTAAGGACTTGAATGCCACCTT | TCGA |
| AACAGATGTACCCATCTATTCAAGCAAAAATATGGGGAAGCCTTGGGCAG*GTTTGATATTTGACAAATTGATCTAAAATGGCTGGGTTTTTATCTGAATA | TCGA |
| GCTGGGACTTCCAAAGCTGGGAATCCTATTTTTTATTATGTTGCACGGAG*GGCCAACAGTATTTTGTTCAAGTCTCGGCTTACAATATGAAAGGATGGGG | TCGA |
| ACAGGCCGGTGGAATGGGTCCAGGCCGTGGTCAGCCGCTTCGACGAGCAG*AAAAAATAAGCTCCCCTAAGAAAGTTGTTACATCACCAAGAAAAGTTCCT | TCGA |
| ATAAAGCAGTGCAAGGTTGTGGAGCACACCCAGCAATACGAATGGCACCG*CCATGCAAAAAGCGAAAGAAAGATGATGACACATCTACCTGCAAAACAAT | TCGA |
| GTTACAGTATATCAATGTGGATTGTGCAAAATTAAAACGACTCCTGAAGG*ATTCTGGAACTGAAGACATGCTTTGGACAGAAAAGTATCAACCTCAGACT | TCGA |
| GCTGGGACTTCCAAAGCTGGGAATCCTATTTTTTATTATGTTGCACGGAG*CCCAGCGCCCGCAGCTCTCCGCACAGCCCCTGGAGCGCTGCAAGCAGCTG | TCGA |

| | |
|---|---|
| CTTTGACATCCTTGGAAACAGTCACAGAAGCTTTGTTGGAGATCATGGAG*GCTTATACCAGCCAGTTTGTATCCCTTGTGATGTTTGCCCTTATGATGTG | TCGA |
| ATCACATCCTCTGATTGGCAACATGTTAGCTTTGAAGTGGATCCTACCAG*ATATACAAACAAATGCGGAACTGAAAAGCACTCAGGAGCAGTCTGTGCCC | TCGA |
| ACAGGCCGGTGGAATGGGTCCAGGCCGTGGTCAGCCGCTTCGACGAGCAG*TGCTCTTATTAAACAAGTGAACAAATCAATAGATGGAACAGCAGATGATG | TCGA |
| GTTACAGTATATCAATGTGGATTGTGCAAAATTAAAACGACTCCTGAAGG*TGAAGCGTGACATTCAGGAAAACGATGAAGAGGCAGTGCAAGTCAAAGAG | TCGA |
| ACAGGCCGGTGGAATGGGTCCAGGCCGTGGTCAGCCGCTTCGACGAGCAG*GAGAACTCCTGGAAGCCCCGTCTTCCTCCTCATGCCCATTGCCTGACCAG | TCGA |
| GTTACAGTATATCAATGTGGATTGTGCAAAATTAAAACGACTCCTGAAGG*GAGAACTCCTGGAAGCCCCGTCTTCCTCCTCATGCCCATTGCCTGACCAG | TCGA |
| ACAGGCCGGTGGAATGGGTCCAGGCCGTGGTCAGCCGCTTCGACGAGCAG*ATGAAGACAGTCAGTGTTGCCTTAGTTTTGTGCCTGAATGTTGGTGTGGA | TCGA |
| GTTACAGTATATCAATGTGGATTGTGCAAAATTAAAACGACTCCTGAAGG*GGGTCTGCCTTGCTTATTTCACAAGCGGTTTCAACGCAGCTGCTTTGGAC | TCGA |
| ACAGGCCGGTGGAATGGGTCCAGGCCGTGGTCAGCCGCTTCGACGAGCAG*GCACGGAAGACATATGCTTGTGAGACAAAGGTGTCTCTGAAACTATGGAT | TCGA |
| GTTACAGTATATCAATGTGGATTGTGCAAAATTAAAACGACTCCTGAAGG*TTGGATCACAAGGACGAGTCATAAGTTTTGAGGTACGAAAAGACCACCAT | TCGA |
| CTACATTTCACTATGTGCTGGTAAATTCACTCCATCGAATCATCACCAAT*GTTTTTAGAACTTCAGCCATAAAAATGGGCAGAATTTTCCTTGATCATAT | TCGA |
| TCACCGTGAGGATCGTCACCATGGACGCCGAGATGGAGTTCAATTGCGAG*CACTGTAATGGCGTTGACTGGCGTCAGAAGCTGGACTCTCAGCGAGGGGC | TCGA |
| CTGAAGAGGAGCTGGTTCAGGAGATCACACAACATTTATTCTTCTTACAG*ATTTTAAGTAACCAAGAGGAGTGGTTTAAGGCGGAGCTTGGGAGCCAGGA | TCGA |
| TCACCGTGAGGATCGTCACCATGGACGCCGAGATGGAGTTCAATTGCGAG*CAAATGGTTATGAGCCTTAGAGTTTCTGAACTCCAAGTACTGTTGGGCTA | TCGA |
| AGCGGCTTTGTCTTTGGATTTAGCGTTTCAGAATTGCAATTCCAAAATGT*GGGAATTAGTGACCCACTGACCGTATTTGAACAAACTGAGGTAATTTTGC | TCGA |
| AGGACATTGAGCAAGTTTGGGAGGAGCTATTATCCATTCCTGAGTTACAG*GTCCTAAACATGCAGCTTTCGGATGGAGGACAAGGAGATGTCCCTGTTGA | TCGA |
| AGGACATTGAGCAAGTTTGGGAGGAGCTATTATCCATTCCTGAGTTACAG*GTGCTATCATACCATGCAACATGTTCAAAAGCTGAAGTTGACAAGTTTAA | TCGA |
| TCATCATGATGGACTTGGAGCTGCCGCCGCCGGGACTCCCGTCCCAGCAG*AAGAAAGCCAGCCGCCACATAAAAGGTGAAGACAAGAATAAAGCCACTAG | TCGA |
| TCATCATGATGGACTTGGAGCTGCCGCCGCCGGGACTCCCGTCCCAGCAG*TTCGTAGGTGTAAAGCTTTGAAGGAAAAAGATTTAATTAGAACGTCTGAG | TCGA |
| TCATCATGATGGACTTGGAGCTGCCGCCGCCGGGACTCCCGTCCCAGCAG*GAAGTCGAGGCTCAAGAGGTTCTCAAATTGATAGTCACAGTAGTAATAGC | TCGA |
| TGTCCCCAGCTGATGGCCCCTACCTGGTGATCGTGGAACAGCCTAAGCAG*GAAGTGCGCCATGATCTCCGCCCACTATGGCCTCAGCGATGTGTTTGACA | TCGA |
| AGCCCTCAGCAGCTGGAAGCACTCACTTACATCTCCGAGGACGGAGACAC*GCAGAAGCTGAGGAAGATTGTCATTCTGATACTGTCAGAGCAGATGATGA | TCGA |
| CCCTGCTGCGGAATGTGGAGGATGAGACGCCCCAGGACCTGACTGAGGAA*CAGGCCCCTGAGGGAGGGAGCTGTCAGCCAGGGAAAACCGAGAACACCAT | TCGA |
| TGAGGACCTGCACCGCCAAGCCAAGCTCAACCTCAAATCAGTACTGAGGG*GCTGCCCCCAAGTGTAACTCCAGCACTGTGAGGTTTCAGGGATTGGCAGA | TCGA |
| CTCTGGCCCCGGATGGTACGGCGCCAACCCAGACCCGCGCTTCCCCGCCA*GATTTTAGCTTTGTACAAAGGCCTGCTTCCCAAGATTATGAGACTTGGAC | TCGA |
| GATATTGAGCCGGATAGACCTATTGGATATGGAGCCTTTGGTGTTGTCTG*CAAACACAAATTCCAGACTATGCTGAGCTTATTGTTAAGTTTCTTGATGC | TCGA |
| CGCCTCGCTGGGCAGCCTCAACATCCCCTACAAGATCGAGGCCGTGCAGA*AGTTCAGTCAACAGAGCCATGGGCCTGGGCAGTCCGATGATGCCTGCTCT | TCGA |
| TGCCTGAATGGCGGGAAGTGTGAAGCGGCCAATGGCACGGAGGCCTGCGT*ATAGGCAGAATGTTTCGAGCTTCCAGAGTCTGAAGGAATTCATGGACGCT | TCGA |
| TGCAACAGCAACCCCTGCGTCCACGGGGCCTGCCGGGACAGCCTCAACGG*CTGCCTTCCTAAGcagatgagaaaactcaagatagatgaaggaagacatc | TCGA |
| CATTGACCTGGTCAATGGCTACCAGTGCAACTGCCAGCCAGGCACGTCAG*GTGCAGATTTCATAACAATTTTGGAGAGTGATGCTTCTAAGCCCTATATG | TCGA |
| TATTGACCTTGTGAACCATTTCAAGTGCTCTTGCCCACCAGGCACTCGGG*AATCAGGAAACCCAGGCCCAACGGCAGGTGGATGCAAGAAGAAATGTTCC | TCGA |
| GCTGTGGGCGCTGCTGGCGCTCTGGCTGTGCTGCGCGGCCCCCGCGCATG*GGGCGTGAATACTTTTTCTCCCGAAGGAAGATTATTTCAAGTGGAATATG | TCGA |
| GCTGTGGGCGCTGCTGGCGCTCTGGCTGTGCTGCGCGGCCCCCGCGCATG*ATTTCAAGCCACATAAGGTTTCAGAACAGAAATCCTCAAGTGTGCCTAGT | TCGA |
| GCGAGGCTCCGGCAAGAAGCCGGAGTCCGCGGCGGGCAGCCAGAGCCCAG*AACGCCAGCCTGCAGGCCGCCATCACTGATGCTGAGCAGCGTGGGGAGCT | TCGA |
| GGGCCGGAACCTCAAGAAGGAGGTCAGCCGGGTGCTGTGCAAGCGGTGCG*CGGAGCTGGAGTTTGTTCAGATCATCATCATCGtggtggtgatgatggtg | TCGA |
| GCGAGGCTCCGGCAAGAAGCCGGAGTCCGCGGCGGGCAGCCAGAGCCCAG*CTGTCCCCAATCTCTTCATGCTGGAAACAGTGGATTCTGTGAAGTTGGCA | TCGA |
| ATTAGGAAATGACAGTGCCTCTGCCAATATCACCATCGTGGAATCAAACG*CCTACAAGGAAAAAATGAAGGAGCTGTCCATGCTGTCACTGATCTGCTCT | TCGA |
| ATTATGATCAACTTCAACCCTCACTTCGATTTGGAGGACAGAGACTGCAA*ATCCTCAGTTTGTGGTCTGCCAGCTAAAGGTGAAGATATATTCCTCCAAT | TCGA |
| ATCGGCAACTAAACCATTATGTAAAGGCTGTTCAATCTACAATAAATCAT*ATATTATCACTACCTATACTCCCATTACCTGCCAGCCAGCCTGAAGAACA | TCGA |
| GGGGTGTAGGGAGGAGCCCGGGCCTCTCCGGAGAAGAGAGGCTGCAGCAG*agacggggtttcatcatgttgaccaggctggtcttgaactcctgagctca | TCGA |
| AGATCTTCACCTACGGCAAGCAGCCCTGGTACCAGCTCTCCAACACGGAG*TGTCCTCTCGCATGGCAGAGCAAGTGGGAAGGCCCAGAAGATCCCTTACA | TCGA |
| AGACAGATGAAGAAGGCGTTCGCACCGGAAAATGTAGCTTCCCGGTTAAG*GCTACATTGGCGTGGTGAACCGCAGCCAGAAGGATATTGAGGGCAAGAAG | TCGA |
| CCGCAGATGTTGTCAAGGTGGCCTACCAGGAGGCCAGGGGTGTCGCCATG*GAAGTGCGAGCGCACCACGTGCCATGACTTCCTGGAGTGCCAGAACTCGC | TCGA |

| | |
|---|---|
| CTGAAACAATTGAGGATTCTAGTAATCAAGCAGCAGCCAGAGAATGGGAG*GATTTATTTATACATGTGGTGGAACTTTAAAAGGACTTAATGGCACTATA | TCGA |
| GGCTGCCCAGAAAGCAGTTAACAGTGCCACCGGTGTTCCAACTGTTTAAA*GTTCCTTGTAGCTTGGAATACTGGGATGAACTCCAGAAGGTTTTTGTTGC | TCGA |
| CAAAATGGATTTAGAGAAAAATTATCCGACTCCTCGGACCAGCAGGACAG*TGATTGCCTTCATGTTGTGGAGCCCATGCCTGTGCGGGGGCCTGATGTAG | TCGA |
| GCCAAAAGAAATAGAGGAAGATAAACTAAAACGAGAAGAAGAAAAAAGAG*CGTCTGGTGGAGGCTGCAGAAGAAGCACACCTGAAACATGAATTTGATGC | TCGA |
| GGGGACAGAATGGAGAAATCTTGAGACAGCCAAGAAAGCAGAATATGAAG*CATTTGAGACGGAGTTGGCTTATCTCTCAGGAAATAGACTTACAGTGGAT | TCGA |
| GAACAGGGCAGACTTCTCTGTGAGCTCTTCATTAGGGCACCAAAGCGAAG*GGTTAAAAACTAATATTTATATGACAGAAGAAAAAGATGTCATTCCGTAA | TCGA |
| GAACAGGGCAGACTTCTCTGTGAGCTCTTCATTAGGGCACCAAAGCGAAG*GTCATCCTGTATCAGGGTGGACAGATGTTTGATGGTGCCCCCCGGTTCCA | TCGA |
| CATCAGGACGTCTCATTAGCGAACTTTTTCAGAAACTGCCTTCTAAAGTG*GGTTGTTATGTTGGCTTTGATCTAGCACATGCAGTTGGAAATGTTGAACT | TCGA |
| GCCAAAAGAAATAGAGGAAGATAAACTAAAACGAGAAGAAGAAAAAAGAG*GAAAGGTCAGACAAATGAAATTAATGCCTTGGTACAATTGATGACTCAGA | TCGA |
| GCAGGCGCGGCGCTGCCGGACCAGAGCTTCCTGTGGAACGTTTTCCAGAG*GTGTCCTGCCTGAAGGAGCTGGTGGCCCGAGTGCTGCAGAGGCTGTGCGA | TCGA |
| TTCTAAAAGGATATGTGTTGCTAATACCCATCTTTACTGGCATCCTAAAG*GCACATTCAAAGCAATAATCAACCTCCACAGCCTCCAGTTCCACCGTTAG | TCGA |
| AAGCCGCGGAGCTAATTGGAGATGGTTTCACAGCACGCAGTGGCTTCGGG*GTATTGTCTTCTAAGAACTCTGAAGCAATGTCAGACATTGAGGGAAGCTC | TCGA |
| GTCTGGTTCAGGCTCCCATGAAGGGGGCAGCACCTCAGCCAGCATCACTC*GACATTTTATTTCCTTACATCGAAGAAAATGTTAAAGAGTATCTGCAGAC | TCGA |
| TCTTCCAAAAGTGTATTGGGTAGAAGAAAGCACAGCTCGGAAAGGCAGAG*CAAAACCGTGTGGTTGGAGCAATGCAGCTCTACTCTGTGGATAGGAAGGT | TCGA |
| TCCTGTTGCACCAGGTTCTTCGAAAACTGAAGCAGATGTTGAACAACAAG*CAGGAGTAGAAATTGGTATGCTTAGAAGCAGATTCTAAAAGCAGTTTCTC | TCGA |
| TATAGAAATTTTAATCTCAACTGAATGGAATGAACGACAGCCTGCACCAG*AGATGGTGGCCACACAGCAGGAGATGATGGACGCGCAGCTGAGGCTCCAG | TCGA |
| GGTCTTCCTGCCCAACAGCAACCACGTGGCCAGTGGCGCCGGGGAGGCAG*GAAGGATGTGCCCGTGGAACAGGTGACACACGCAGACAGAGAGCAAACCA | TCGA |
| CCCCGAGGCGGATCGGGTGTTGCATCCATGGAGCGAGCTGAGAGCTCGAG*GAAGAGTCTAACCTGTCTCTGCAAGCTCTTGAGTCCCGCCAAGATGATAT | TCGA |
| CCCCGAGGCGGATCGGGTGTTGCATCCATGGAGCGAGCTGAGAGCTCGAG*CCAGTGCACAAAGACTCTATTAGCCTTTTCATGGCACATGTTCACACCAC | TCGA |
| CCCCGAGGCGGATCGGGTGTTGCATCCATGGAGCGAGCTGAGAGCTCGAG*AGTTGAACAATGACCATAGTTGACAAAGCTTCTGAATCTTCAGACCCATC | TCGA |
| AgcggcgggcggcggcgcgcggACCCAGGCGCGGATGGCGAGGCCAGCAG*GTTTTAAAAATCAATTTGATTGGCCACAGGAAACGTATTTTGGCATCTCT | TCGA |
| ACAACGACGTGGACATCCCTGGCTGCTGGTTTTTCAAGCTGCCTCACAAG*AATGTCGAGGTCTTGGCGAGCAGGAGCAACACTTCAGAGCAAGACCAGGC | TCGA |
| TCCTTTCTCCTTCCATCCTGAAGCCCTTTCCGCTGTGGACAGGAAAACAG*GCAGACTCAGTGGGATCCTCCTACTTGGGAAAGCCCAGGAGATGATGCCA | TCGA |
| ATTTCGGATCCCTACAGCCAGGTTACAATTATAGCAAGAAATATATTCAG*GCCAATGGAAGCACTGTTGTGAAGAACTTATGAAAATTGAGATTATGTCA | TCGA |
| GGTGACCAGAAGCCTGCATTTCTGCATTCTGCTTAATTCCCTTTCCTTAG*GTCCTTCCGGCCAGACTTCGTGCTCATCCGGCAGCATGCATTTGGCATGG | TCGA |
| GGTGACCAGAAGCCTGCATTTCTGCATTCTGCTTAATTCCCTTTCCTTAG*GGGGCCCTGGGCAACCCCAAGGAATGCAGCCCCCAGGCAAGGTGCTGCCT | TCGA |
| AGGAGATCATGTCGCTGACCAGCCCGTCTGCCCCGCCCACATCTAGAACG*CTATATTTTCCGGTTAAGTTGCACTCGATTGGGACAGTGGGCCATTGGCT | TCGA |
| TGTCTTCAGAATTTACCCCCGAGACGGAGCGCCAGCGACTTCAGTACCTG*TTGTTCAAGTTGGTTTCCCATGTCTTGGAAAACAGGATGGGGTGGCAGCA | TCGA |
| TGTACCCCATCGCGGTGCTCATAGACGAACTCCGCAATGAGGACGTTCAG*GAAAAGTACCCAGATGCCGTGTACCTCTCGGAGGGGCCCTCCTCCTGCTC | TCGA |
| TTCAGCCAGTATCAACACCAGTAACAGTGTGTGGAGATATCCATGGACAG*CTCCCCCTCCGGGGCTCCGCGGGACACTGGATCTCCAGGTTATCCGCGTG | TCGA |
| TGCAGAAACACTACCTGGTACACACGGGAGAAAAGCCACATGAATGCCAG*GTTGCAAGTGGCTATCTTGTGGTGGGGTTTGTCTATGTGATGGAATTCAT | TCGA |
| ACTGATTATGGGTGCTCGGACAGCAAACAACACTTTGAATGGCTTGTCAA*GTTCCTATATGATAGTGGACTCTTCAGATCACGACCCTGGAGAAAAAGCC | TCGA |
| CTGAGAGACCCATGGCATTCCTCAGGGAATACTTTGAGAGGTTGGAGAAG*TACAAAATTGGACAGCTGTACATGATCAGCAAACACAGCCATGAACAGAG | TCGA |
| CTTGTTTTTGTTTTTCAGGGGGTTTGGGAAACAAGGCTTCCAGTGCCAAG*GTTCAAAACTGGTCAAATCAATGGTGATTTGCTGATATACCATGTCTTAC | TCGA |
| ATTAAAAGCAGCGAACCTCGAGACCAACGTGGAGGAGCTGTGGGTGGAAG*GTGCAGCAGTACCGCGTCGCCATGACTGCCAAGGACTGCTCCATCATGAT | TCGA |
| CACAAGAGGCCCTAGATTTCTATGGGGAAGTAAGGACCAGAGACAAAAAG*GGGACAGTATTTATTCAAAAACAAACCTCCTGACGGGAATGCTCCTCCCA | TCGA |
| GCAGAAAGACTTGAAGGCGTATACAGGAACAATATTGATGATGTAGTAAG*GTCAGTCCTGCAAGAAACATTTCCTCGGCTGGATCCTCCACCTCCCATAA | TCGA |
| TTTTAGTTGTGCTGAAAGACATTATGACACCGCCAAATTTAATTGCAGAG*GACTTAAAATCTCCAAATCAGAGAGATGAAATTGCAGGAGCCCGAGCTTC | TCGA |
| TTTTAGTTGTGCTGAAAGACATTATGACACCGCCAAATTTAATTGCAGAG*TTGGCGACATGGGACTCAGAGAAGGGCTTGAATGGCAGCTTGCAAGAGAG | TCGA |
| GCAGAAAGACTTGAAGGCGTATACAGGAACAATATTGATGATGTAGTAAG*GGTCTATCCATGTTAGTTCAGCTATCACTGAAGACCAAAAGAAAAGTGAA | TCGA |
| CACAAGAGGCCCTAGATTTCTATGGGGAAGTAAGGACCAGAGACAAAAAG*GTATGACTACGTGGAAGTCTTCGATGGAGAAAATGAAAATGGACATTTTA | TCGA |
| CACAAGAGGCCCTAGATTTCTATGGGGAAGTAAGGACCAGAGACAAAAAG*TTGGGGTTAGACTTGGAGGAACTCGAGGAGATAGAAGAAGATGCTGGCCT | TCGA |
| CACAAGAGGCCCTAGATTTCTATGGGGAAGTAAGGACCAGAGACAAAAAG*GTTACAAATGCTGCTGCCAACTGTCCTGGCCAAATGACTCTGCATCACAA | TCGA |

| | |
|---|---|
| CAGCCCTCGCCCGGCGCTCCAGTAGCAGGACCCGGTCTCGGGACCAGCCG*GTATTTATGAGCCTCCATTTCTTATACTACTGCAGTGAACCAACATTGGA | TCGA |
| ACACAACCGAATACAAACTCCGTACCTTACAGGTCTCCCCGCTGGACAAT*GCTCGGGTGCGTCCTCGCTGTGGGCGCCTTGTGGCCTTCAGCTCCGGTGT | TCGA |
| ATCATCCACCTCAAGTACCCGCTGAACTGCTCCGATCCCACTAGTGAGAG*GAGGAGGACTGAGCTTATCTGACTCCAGAGCTTTCAGGAGGGAAGAAAGA | TCGA |
| CTGACTGACAGCATGTTGGCACCGAGGGAAGCGATGGGCTTCGGCTCCAG*ATCCTCTCTGGGGTCATTCTGGAGGTGGTCGCCGAGGAATTTGCCAGTGA | TCGA |
| CAGGCTGCTGCTGCTGCTCCTCACTTTCTTCCTCCGCACGGATGCTGAGA*ACACTGAGGGGCACCAGAGGAGCAGACTACAAGAATGGCACACGCTATGG | TCGA |
| AAAGAGAAGACTCCTCAGGGTTTCCAAGGAGGACTGGAGCGGCTGCTTTA*GGCCGTGGGCAAGACCTGCCTTCTCATCAGCTACACCACCAACGCCTTTC | TCGA |
| CAAGTTCCATTTGTTACAAATGAATTCTAATAAAAATGTCAGTGTTATGC*aagaaaaagaagaagaaaaagaagaggaagaagaagaaggaggaggagga | TCGA |
| TGGAACGGGGCTCCCAGGACCTCCACTAATCACCAGAACAAAATGTGCAG*AGTTTCAGATTGCCAAAAGACAAAACGGGTACCACAAGGATTGGTGACCT | TCGA |
| CCTCTCTCCTTGGCCTCTCCTGGGATCGGCCCAAGGCCAGTTCTCCGCAG*GGCAAACTCAGAAAATGAATCTTTTCCAGTCTGTAACAAGTGCCTTGGAT | TCGA |
| CCTCTCTCCTTGGCCTCTCCTGGGATCGGCCCAAGGCCAGTTCTCCGCAG*AAATGGCGAAAAGTCTTTTGAAGACAGCCTCTCTGTCTGGAAGGACAAAA | TCGA |
| AGTGCATTGCCACAGGGAGAGATGCTGTGCATAACAAGTTATGGCTCCAG*GTTAATGACTCCAGAGAGTAAGATCAGATTCGTCCACAAAGAGATGGGCA | TCGA |
| CCTCTCTCCTTGGCCTCTCCTGGGATCGGCCCAAGGCCAGTTCTCCGCAG*ACTGTAAAAGGTGGAATTTCTGAAACAAGAATTGAGAAACGCATTGTGAT | TCGA |
| TGGAACGGGGCTCCCAGGACCTCCACTAATCACCAGAACAAAATGTGCAG*TAATTACAAGTAGTGGCTACAGCCCCAGATCAGCACATCAGTATTCCCCA | TCGA |
| CCTCTCTCCTTGGCCTCTCCTGGGATCGGCCCAAGGCCAGTTCTCCGCAG*ACCTGCTCCATTGGGCTTCTTTGTGGTGATGAAATAAGGAGAATTACCTG | TCGA |
| GCATGTTAGTGCTCAAGAGCCTCATTATCTACCACCCGAGATGCCCCAAG*GTTGTAAAGAAGAAAGTTCTACCCTTTCTGTCAAAATGAAGTGTGATTTT | TCGA |
| CCTCTCTCCTTGGCCTCTCCTGGGATCGGCCCAAGGCCAGTTCTCCGCAG*TTCCCTGTGTCTCTGGTGGTTTGCCTAAACCTGCAAACATCACCTTCTTA | TCGA |
| CCTCTCTCCTTGGCCTCTCCTGGGATCGGCCCAAGGCCAGTTCTCCGCAG*GCTGAAGAAAAAGTCCCAGTCGGTGGATATTAATGCTCCAGGGTTCAACC | TCGA |
| TGGAACGGGGCTCCCAGGACCTCCACTAATCACCAGAACAAAATGTGCAG*CTGGTGAATGAGCAACAAGAAAGCAGACCCCTCCTGAGTCCCTCCATCGA | TCGA |
| GCATGTTAGTGCTCAAGAGCCTCATTATCTACCACCCGAGATGCCCCAAG*GAAATACCAAAATATTTGCAGACTGGATCCAAATCAGGAGCCCAGATGAA | TCGA |
| GCATGTTAGTGCTCAAGAGCCTCATTATCTACCACCCGAGATGCCCCAAG*AAGAAATGTGAAACAAATTCTAGCAAGAAGCAAAGCATTACAGTGGACGA | TCGA |
| AGTGCATTGCCACAGGGAGAGATGCTGTGCATAACAAGTTATGGCTCCAG*GAAGTTATAGAACTAACCAAAGACCTTCTGTCAACTCAACCTTCTGAGAC | TCGA |
| TGGAACGGGGCTCCCAGGACCTCCACTAATCACCAGAACAAAATGTGCAG*ATCTGGAAAGCATGAAGACTGGGCTTTTTTTCCTATGTCTCTTGGGAACT | TCGA |
| CTCTGATGGTGGGTCTGGTGTGTCCAACTACGCGGAGCTGATCGTGAAAG*GTAAATTTGGAAGGCAAAGCCAGCCCTCTGTCACTTCTACAAATGGCCTC | TCGA |
| CCCTGAAAAGCTACAGCATCTACTTCCAGGCACTCAGCAAAGCCAATGGA*GATCAAAATACTTGCTGGATGGTGGACTCAGAGACCAATAAAAATAAACT | TCGA |
| GCCCAACTTCTGCGGGATCTTCAACCACCTCGAGCGGCTGCTGGACGAAG*TTTGGCTTTGGTTGAAATAAGAATTTAGCCTGTATGTACTGCTTTAACTC | TCGA |
| GATGTAAAATCATGGTCCGAGGCAAAGGCTCAATGAGGGATAAAAAAAAG*ggtcttactctttgcccaggctggagtgcagtggcgcgatcatggctcgt | TCGA |
| GCCCAACTTCTGCGGGATCTTCAACCACCTCGAGCGGCTGCTGGACGAAG*AAAGCCTGATCTTTGACCAGATGGCAGTAACATTTGAAGATGTGACTATT | TCGA |
| CTGGGTATGAATAATGACTTTCCTCCAAGAGCACATTGCCTGGTAAGATG*CCAGAACATGCCAGGTAGACCAGTTTTCTTGCGGAAATGGGCGTTGCATT | TCGA |
| TCGGGGACTCCAAAAGCCATTACTACATCTCGCCATCTTCCCGGGCCAGG*CTACATCTACATCCACCACTGGGACAAGCCATCTTGTAAAATGTGCGGAG | TCGA |
| AGTTTTACCAGACTTCCAAAAAGATGATTTCAAGGGGATGTTTAATACTG*GTGGAAAAATCACAAGTGCACAGTCGACCCTTCACTCTCTCTCCACATCT | TCGA |
| AAGTCGAAACCTGGTTAAAAGCTCAAGTTTTAGAAAGGCAACCAAAACAG*AAATCATTCCAGACTGGGACAAATGCACGTCTAGATGAACGCATTTTTGC | TCGA |
| ATCAGAATTCTTTACTGAAAATGATTTGCCAACAAGTAGAGGCCATTAAG*GGATGCAGATCTGTTCTTGTTAGACACAAGAAAGGCCCAAGCTTTAGATG | TCGA |
| AGAGAGCTTGGTTAACTTGGGAGAAAGTTTCATCTGTGGATGGAGTATTG*GTATGATGACCAAACATAAAAAGTGTTTTATAATTGTTGGTGTTTTAATA | TCGA |
| AGAGAGCTTGGTTAACTTGGGAGAAAGTTTCATCTGTGGATGGAGTATTG*ATAATTGCCTTTGATGAGTTAAGGACAGATTTTAAGAGCCCCATAGACCA | TCGA |
| AGAGAGCTTGGTTAACTTGGGAGAAAGTTTCATCTGTGGATGGAGTATTG*CATTTAAATATTAGGACACTGACGGATGATATGGTAAGATTATCCTCAAC | TCGA |
| ATTTAGAACGATGTGAACATCGAATCATGGAATCCCTTGCATGGCTCTCA*CCATTATACTTGCCCACGAATCTTTGAGAACATTATAATGACCTTTGTGC | TCGA |
| AAAATTTCAGAAGGTCTGCCAACACCAACAAAAATGACTCCAAGATCAAG*CTTCAGCGCCAGTCAGCAAAGTAAATAAATACTGTGCTTCTTCCAACTTT | TCGA |
| ATTTAGAACGATGTGAACATCGAATCATGGAATCCCTTGCATGGCTCTCA*GGAGCAGCACATCGCAGAGTGCATGGCCAAGATGCCACAGATGATTGTGA | TCGA |
| AGAGAGCTTGGTTAACTTGGGAGAAAGTTTCATCTGTGGATGGAGTATTG*GTGTATCAGTTCCTATCCCTGTGATTGGACTGAGGGACGAAGAAAAGGTG | TCGA |
| TACCTTTTATGAATTCTCTTGGACTTGTAACATCTAATGGACTTCCAGAG*GTATGTTTCTATAAAACCTGATAACCGAAAATTGGCCAACGGAACAAATG | TCGA |
| AGAGAGCTTGGTTAACTTGGGAGAAAGTTTCATCTGTGGATGGAGTATTG*CGACAGAGTCCAGGCCCTCTCCTATGCACAGCACACGCGACAATTGATCT | TCGA |
| GGCCCCAGCAGCCCCCGAAGGCCCTATCAGGACATGGAGTATGAAAGACG*TGTATGGGTTTCGACAGACCATGATGAAATTGAGAATGTGGCCAAACAAT | TCGA |
| AGGAGGGCAAGCATGACTATGACGACTCATCTGAGGAGCAGAGTGCGGAG*GTATTCCGGAAGGACCTCATCAGTGCCATGAAACTTCCAGATTCTCACCA | TCGA |

| | |
|---|---|
| CCATCGCCTGTCACCTGGACCCGCGCGTGTTCGTGGACGGCCTGTGCCGG*ATACAAGGCAGATCCAACCATCCCCACCGTGGTCCTACGATCAGTCCTAC | TCGA |
| GCTTCACGGAGGGCAAGAGCCGGAGGGTCTCAGAGGACAGACAGTGCCAG*AGTATCAATAGAAATAAGGATATCATGAAGATTGGTTGCTCACTGTCTGA | TCGA |
| GGACCGGCGTCCCCGGCGCGAAGAGGCTGGACTCGGATTCGTTGCCTGAG*AGTAGACCACAAACTCCAAGCGTTAGAAACACAGTTCAAAGAACTAGACT | TCGA |
| GGACCGGCGTCCCCGGCGCGAAGAGGCTGGACTCGGATTCGTTGCCTGAG*TTGCAGGAAATCACGTTTAAGAATTACTACACAGCTTTTTTGAGCATCCG | TCGA |
| TACCGATGTTATACTGATGTGTTTTTCCATCGACAGCCCTGATAGTTTAG*CCGCTGACCAATCACGAATGATGCAGGAGCAGATGACGGGAGCAGCCATG | TCGA |
| GCAAGAAGGCCGTGTGCGAGGAGTGCCTCAAAGTCTACCTGAGCGCCCAG*CACAAACACAGAGCAGTCTAAAGCAACCGAGCACTGAGAAAAATGAACTC | TCGA |
| GCAAGAAGGCCGTGTGCGAGGAGTGCCTCAAAGTCTACCTGAGCGCCCAG*GTGGCTGTACTTTTGATGATGGTCCAGGGGCCTGTGATTACCACCAGGAT | TCGA |
| TGTTTGCTGGTGTTGCTGAAATAACTCCAGCAGAAGGAAAATTAATGCAG*ATCTTCATCTGCTGGTGGACGTGGCCTGCAAGCAGGAGCGCTTTCCAAAG | TCGA |
| TGTTTGCTGGTGTTGCTGAAATAACTCCAGCAGAAGGAAAATTAATGCAG*CAAGTAACTGGCAGAAGTTTTGGTGATAAAGATTTTCGGACAGGTTTAGA | TCGA |
| GAACTATCTTGGTGAAGCAGTGAGTCGAAATGCGTCTCTGGAAGTGGCAT*ATCAGATTGATGAAAATCTCAAACTGGCTTTGCAACAGGACCTGACCTCC | TCGA |
| GATGTTTACACAACTACTGCTCTGTGGATTTTTATATGTTCGGGTTGATG*ATCAATGTTCTCCGAAACAGGATCAACGATAACCAGAAAGTGTAAGTGTC | TCGA |
| AGGAGCAAGAGCAAGATCACCGTGACATCCGAGGTGCCTTTCTCCAAAAG*CCCGAGACCGAGGACGAGAAGAAGCGCTTTGAGGAAGGCAAAGGGCGGTA | TCGA |
| CGGACCCGTGGCAGAAGATGGCTGTGGAGAGCCCGTCCGACAGCGCTGAG*GTATTGTCTTCTAAGAACTCTGAAGCAATGTCAGACATTGAGGGAAGCTC | TCGA |
| AAATTGAAGGCTCCAAATCCTTAGCTCAGAGCTGGAGGATGAAGGATCGG*GATCAGATTTGCAAGTATGTCTCCCTAAGGGCCCAACATGCTGCTCAAGA | TCGA |
| GGACATGAGTGACCGGGGCATCTCCAACATGCTGGATGTCAACGGCCTCT*AGCAGCCAGACACAGTCTCCAACGCCACCAAAGCCTCCGTCGCCAAGCTT | TCGA |
| CCTGAACCACTCCACTGCCTTTAACCCTCAGCCTCAGAGTCAGATGCAGG*GATGAAAATGGGATAAATAGACCAGTCTGTTCCTATGTTAAACCACTTCG | TCGA |
| CCAGGTTGCAAGATTTAATGACCTCAGGTTTGTCGGTCGAAGTGGAAGAG*CTGTCAAAGCTCGAAAAACAGCAGCAGAGGAAAGAAAAGACCAGAGCTAA | TCGA |
| CCTGAACCACTCCACTGCCTTTAACCCTCAGCCTCAGAGTCAGATGCAGG*AGAAAAATTATCGAGAAAAAAACTGATGTTACAGTTCCTAATTTGAAACC | TCGA |
| CCTGAACCACTCCACTGCCTTTAACCCTCAGCCTCAGAGTCAGATGCAGG*TTTTGTCAAAGCAACAGGCTTCACAAGTCCTGGTTAGGAAGCGTCGTGCA | TCGA |
| TGCTGCCTACGCACTGGCGCTGCAACAAGACCCTGCCCATCGCTTTCAAG*CAACACCCCTGAGCCCACTTAAGCCCCCACGGATGGACACTGCTCCGGTG | TCGA |
| TGCAGCTCGGAGCCTGTGGCGGATGGTGGGACTCTCCGGCCCTGTGCAGT*GCTGCTGATACTTTGGCTGTGAGGCAAAAAAGGAGAATTTATGATATCAC | TCGA |
| TTTAGTAATAGGTGACTGCCGGAAAAAAGGGAACTTTGAATATTGTCAAG*AAAAAACATGGACCCATTTGCCAGAAGACTGCAACTAAAAAACGGAAGAC | TCGA |
| CCAAGTAGCAAGGTTCAACGATCTGAGATTTGTGGGCCGGAGTGGACGAG*GCATCCCGGGCCCAAATCCTAGACAAAGCCACAGAATATATCCAGTATAT | TCGA |
| AGGGACTTGGGAATCGCCTGTGCTTCTTGGAACCCACTTCAGAAGCCAAG*TGAAGCCATGTCCCTGGAGGGAGATGAATGGGAACTGAGTAAAGAAAATG | TCGA |
| CCTGATTAACGAGCCTAGTAATGACTGGGATATTTACTACTGGGCCACAG*GGGAGGGCGCGGGTGCCGACGAGTTCCTGCCTCTGCTGAGCCTCGTCTTG | TCGA |
| CCTGATTAACGAGCCTAGTAATGACTGGGATATTTACTACTGGGCCACAG*TTTTAGAGAATTACTCCAAATTCATCATGATTGAAGACAATAAGGAGAAC | TCGA |
| CCTTGAGGCGCCGGTTGCCGGCCACAACCCTTGGCGGAGCCTGCCTGCAG*GCTACCTAAAAGAAGACAGTTATCTCATATTTGGCTGCCAGCTTTTTATC | TCGA |
| AGAGAGAGACCGAGAGGTTCAGGGAGCTGTTGCTGTATGGCCGTAAGAAG*GCCCGCGATGCTCCCAGCCCGGTGAGACCTGCCTGAATGGCGGGAAGTGT | TCGA |
| AGTACGCCCAGTCCCTGGGTGCCGAGACCTGCCCCCTGCCTAGTTTCCAG*CTGTGGCGGGGCCTTCGTGGGCCCGCGATGCCAGGACCCCAACCCGTGCC | TCGA |
| GAGCAGCAGGAGGGAAAATGAAGAAGGAACGATCTCGTAAGAAGGATTCA*GGCAGTCTGAAGATGGCCAATTACACGCTGGCACCAGAGGATGAATATGA | TCGA |
| ATATAAGTGATTTGGCCACCAAACTCCTGGACAGTTGGAAAGACCTAAAG*AAGAAAATACTCAGGAAACATCCCAGGTGAAGAAAAGTTTGACTGAAAAA | TCGA |
| ATATAAGTGATTTGGCCACCAAACTCCTGGACAGTTGGAAAGACCTAAAG*GAGATAAATGGCATCACCGCGGCACTGGCTGAAGAGCTCTTTGAGAAAGG | TCGA |
| CCGCCTCCGAAGATGGGGGATTTCTACGACCCGGAGCACCCGACCCCTGA*CTTTAAGGATGAGAATGGCAAAGCTGGGAAAAAAGGTGATCTAAGAGTTG | TCGA |
| CCGCCTCCGAAGATGGGGGATTTCTACGACCCGGAGCACCCGACCCCTGA*GCTATGTCTTGCCCAGCCACGTGACTGAGGAGATGCTATGGGAGTGCAAG | TCGA |
| CCGCCTCCGAAGATGGGGGATTTCTACGACCCGGAGCACCCGACCCCTGA*TGATCGTAAACTTGTTGATAAAGAAGATATCGACACTAGCAGCAAAGGAG | TCGA |
| GAGCAGCAGGAGGGAAAATGAAGAAGGAACGATCTCGTAAGAAGGATTCA*GTACCTGCACATGTTGTTCTTTGTCAGTGCTGTCAAGTGTGTGCCAGGGT | TCGA |
| CCGCCTCCGAAGATGGGGGATTTCTACGACCCGGAGCACCCGACCCCTGA*GTCAGATACCAGGCCCAGGTTCCATGATGCCCGGGCAGCACATGCCAGGC | TCGA |
| CTCACAGAAAAGAAAGGCTGGGGCTTGAGAGCTGCCAAAGACCTTCCTTC*CCAAAAGGGAAGGAAAGAGAGAGAAAGCCACTGGAGTTGGGCAGCGTGGG | TCGA |
| GGGGCGCGCGACGCCATGGGCCGGCCGGGCCCAGAGCTCCTGTCTCTCAG*AGGTCGAAACAGTAACAAAGGACTGCCTCAGTCTACGGTGAGTAACTTTA | TCGA |
| GGGGCGCGCGACGCCATGGGCCGGCCGGGCCCAGAGCTCCTGTCTCTCAG*GCGAGCCCAAGACCTCGGTGGCCCCGCTGAGCATCGCCAACGGCACGACC | TCGA |
| AGTGCGCCGCCTCAGCCGCGGAGGAGAGCGGGGCCGGCAAGAAGGACAAG*ACATCCCCGATTGAAAGAACCAGAGAGGCTCTGAGAAACCTCGGGAAACT | TCGA |
| GGGATCCTAGGAGAAAGGGTGGATCCTGCCGTCCCCCTGGAGAAGCAAAT*GTGGGCAAGTGGGAGAACCATACGCTGAGCCTGAGGCACGCCGTGTGGCC | TCGA |
| TCCACATCTCCACTGATGGCCTCAGCCTCTCCGTGCCTGCCACGCGCCAG*GTTTTTGGATTCAAAGCATAAAAACCATTACAAGATATACAATCTGTAAG | TCGA |

| | |
|---|---|
| CGCAGATGCTGAAGCAGATGCAGCTGTCCAAGAACGAGCAGGAGCGAGAG*GGGTTGGCTGTGTTCCGGCCGCAGAGCACCGTCTGCGTGAGGAGATCCTG | TCGA |
| CTTTGAAGGATTTTGCCTGGCTGGAATCATTCTCACCTTGTATGCTTTAC*agcgtttcccaaagtgtattctgcggaacTAGCACCTACTGTGTTCTCAA | TCGA |
| TGTCAGGAGTGATATCTTCCACTATAGCCAATCCCACCGATGTTCTAAAG*ATCACCTCTCGGATCTTCAGGGCCGGGCAGAGGGTGATCCCGGTGTATCC | TCGA |
| GGTTGTGGCCTACAAATGTTCCAGCAGCTCTCAGGCATTAACACCATCAT*GGGTTGGCTGTGTTCCGGCCGCAGAGCACCGTCTGCGTGAGGAGATCCTG | TCGA |
| ttttttttaCTGGTTATGGGAAGGGAGAAATAAAATCATCAAACCCAAAA*aaaaaaaaaaaaaaaaaaaaaaaaaaaGGAGTGAAACTATCTCAAGGGTG | TCGA |
| ttttttttaCTGGTTATGGGAAGGGAGAAATAAAATCATCAAACCCAAAA*aaaaaaaaaaaaaTGCTTACTAAGTGGCAGTGGTATCTATGATGCTATAG | TCGA |
| AAACACTAACTTCCCAGCAGGAATTGAGCCACAGAGTAATTATATTCCAG*GATTACGGGACAGCTACCACAGTAGAAGAGATCAAATTGCCCTTAAAAAT | TCGA |
| GTCCCGGACCCCCCGGATCTGACGGGGCGGGACCTGGCGCCACCTTGCAG*ACGGCGCCTCATCTCCCAGAGATCTTCCTTGGAGACCCTGGAAGATATTG | TCGA |
| GTCCCGGACCCCCCGGATCTGACGGGGCGGGACCTGGCGCCACCTTGCAG*ATAGTTTTGAGGCAGCTGTGCCATCAAATAGCCACATTGTTTCGGAACCT | TCGA |
| GTCCCGGACCCCCCGGATCTGACGGGGCGGGACCTGGCGCCACCTTGCAG*ATCCCCTCTGAGCAGCTGGGAGCTCAGAAGCTCTGTCCAACCCTGAGGAG | TCGA |
| GTCCCGGACCCCCCGGATCTGACGGGGCGGGACCTGGCGCCACCTTGCAG*GGCTCTGGCTGAGAACATGGCCAATGACATTGATGAGCTCATTGGCATTC | TCGA |
| ACGTGGAAAGGCGCAGCTCTGGTACACCGGAAAGCATGGTGGATGGGGAG*ATATCCTTGGAACAGAAGATCTTATTGTGGAAGTGACTTCCAATGATGCT | TCGA |
| AAGGCCATCACCACGCAGAACGTCAACACCAAGTGCATCACCATCCCCAG*ATGCTAAAGAAAACCCTTATGGCGAGGATGACAATAAGAGTCCATTCCCC | TCGA |
| AAGGCCATCACCACGCAGAACGTCAACACCAAGTGCATCACCATCCCCAG*GTTCATCATGCCTAGTGGCGTATAAGAAGACCCCGCCACCGGTCCCTCCA | TCGA |
| AAGGCCATCACCACGCAGAACGTCAACACCAAGTGCATCACCATCCCCAG*GGGCTAAACTGCACGGTCAAGAATAGTAAGTCATCTTTTTCTGTTCTTCT | TCGA |
| AAGGCCATCACCACGCAGAACGTCAACACCAAGTGCATCACCATCCCCAG*CTCTTGTTCAGCTTCTGGAATTTCTGAGCAGCCCTCGTCAGTACAAGATG | TCGA |
| GTGCACCTGGAGATGCTGTTCATAAGATCTACCCAAGTGCATATATAAAG*ATGCTGAAGAGTTTCTGGCAGAAGGTTTGCGGAATGAGAACCTCAGCGCT | TCGA |
| AGAAGCAGCAGAAGATCGAGCAGGAGCGCAAGCGCCGGCAGAAGCACCAG*AAAACGATGCCAGAAGAAGACCCGGCCACCTTGGATGATCACTCAGGCAC | TCGA |
| AAGCCGCGGGGCCTCGACCCTGTGGAGATCCTGCAGGAGCGCGAGTACAG*AAAACGATGCCAGAAGAAGACCCGGCCACCTTGGATGATCACTCAGGCAC | TCGA |
| AAGCGACCATTGAGCTCAAGGCCCTCAGGCTGCTGAACTTCCAGAGGCAG*ATCGTTCTTGATGTTGGCTGTGGCTCTGGGATCCTGTCGTTTTTTGCCGC | TCGA |
| TGCTGACGACGTACGAGTACATCATCAAAGACAAGCACATCCTCGCCAAG*ACGGTGCTGACGCCAGTGGAGGTGGCCATCGAGGACATGCAGAAGAAGAC | TCGA |
| AAGCGACCATTGAGCTCAAGGCCCTCAGGCTGCTGAACTTCCAGAGGCAG*GTACGCCATCCCACCAGAGCACGGCAAGCGCCTGGAGCGGCTGGCCATCG | TCGA |
| ACCAAGAAGATGAAGAAGATTGTGGATGCCGTGATCAAGTACAAGGACAG*TTCTTTTGAAACAGATGGTCACCATGTTTAGATATTAGCAGTCCCATATA | TCGA |
| AGGCAGAAAATGCAGAAGGACAGACGCCTGCCATTGGGCCGGATGGCGAG*GAATATTTCACTGCTACATGGAGTACTCCCGACTGCGTGGTGAGGCCGGC | TCGA |
| TAGCTCCGAGGTCTGATAGTGAAGAAAGTGGCTCAGAAGAAGAGGAAGAG*GAACGCCACAGAAGTATGATCCCACTTTCAAAGGACCCATTTACAATAGG | TCGA |
| CCACGGACAGCATCCTGGAGGACCAGTCAGACCAGCGCGTCATCATCAAG*GTCACTGGTGCAGTCACGACAGTAGGTGTGGCAGCAGGCCTGCTGGCTGC | TCGA |
| TGAAGTTCCAGCTGGAGGACGACGGCGAGTTCTACATGATCGGCTCCGAG*ATGACTGTAAAGGAACAACAAGAGTGGAAGATTCCTCCTTGTATTTCTAA | TCGA |
| CCACCGTCACTTCATGCACGAGAAGGACGTTATGAGCGGAGACTTGATGG*CAACACCCGGACCTACCGTGTGGCCGTGGGAAAGAACAACCTGGAGGTGG | TCGA |
| AACTTACCCGAGAACGTGCGGGAAGAGAAGATCATCGAGCATTTCAAACG*GTATGATGCCGGGCGGGACGGCTTCATCGACCTGATGGAGCTAAAACTCA | TCGA |
| TCCTCCAGTAGTGATTCAATCAGCAGCAGCAGTAGTACCAGCAGTGACAG*GACACACTGGAGAGGTGCGGCAAGTGTGGCGAGGTGGTCCGGGACCACAT | TCGA |
| TCCTCCAGTAGTGATTCAATCAGCAGCAGCAGTAGTACCAGCAGTGACAG*AGCTGGCCTGAACCTCCAGGACTTCAGCCCTTCTAGGAGAATCTGATCCC | TCGA |
| AACTTACCCGAGAACGTGCGGGAAGAGAAGATCATCGAGCATTTCAAACG*ACATCTGTGCCTTCTGCCACAAGACCGTGTCCCCCCGAGAGCTGGCTGTG | TCGA |
| AACTTACCCGAGAACGTGCGGGAAGAGAAGATCATCGAGCATTTCAAACG*ATCAAGTTCAACAGTTCTCAGGAAACTCAGCAGTCTTTACTGCAGGAAAA | TCGA |
| TCCTCCAGTAGTGATTCAATCAGCAGCAGCAGTAGTACCAGCAGTGACAG*GTGGACCTCCCCCACACGGCGATCCAGGAAGCCTCCAACAAGAAGAAATT | TCGA |
| TCCTCCAGTAGTGATTCAATCAGCAGCAGCAGTAGTACCAGCAGTGACAG*TTGCAGAGCTATTTTGCTGCATGTGAGGATGAGATCCCTGCCATCCGGAA | TCGA |
| TCCTCCAGTAGTGATTCAATCAGCAGCAGCAGTAGTACCAGCAGTGACAG*GTGGATCTTTAACTCAAGACTAGCATGAAGAGTTGCCTTCTGGCCTGCCC | TCGA |
| CTACTTACCATGACCTTCGCAACATCTTCCAGCGCTTTGGAGAAATTGTG*CCTCACTTGCCTGCCGATAAGGCCCGGGGTACACAGGGCCCCGAGCAGCA | TCGA |
| GCACCGTCTCCCAGTACGACCGGGACAAGTACTGGAACCGCTTTCGAGAC*TGTTGCTAACAACCACAAGCAGAATTTGATGACGGTGGCAAACCTTGGTG | TCGA |
| CTTGTGTGTTGCCGGCCAGGAATTCCAGGCTCACAAGGCTATCTTAGCAG*TTCACCACCACCGTGGTCTCCTGCTGTCCCGCGGAGCTGCAGACTGAAGG | TCGA |
| GAGGCAAGCGCAGGGGGACGGGGACCAGCAGCTGTCGCCGCCGCTCTCAG*CTTAAGCAAAGAAAGAAATTCACAAGCAACAGCAGGAAACTCCACCTTCC | TCGA |
| GAGGCAAGCGCAGGGGGACGGGGACCAGCAGCTGTCGCCGCCGCTCTCAG*GGCCCAACATCTGTACCACGCGAGGTGTGAGCTCCTGCCAGCAGTGCCTG | TCGA |
| GAGGCAAGCGCAGGGGGACGGGGACCAGCAGCTGTCGCCGCCGCTCTCAG*TTCCTTCTTTCTGGGGATTGGTGAACTCAGCTTGGAATCTTTGCTCTGTG | TCGA |
| GAGGCAAGCGCAGGGGGACGGGGACCAGCAGCTGTCGCCGCCGCTCTCAG*AGGACTTCTGAATGATGCTTTCCAGAAAGGGGGACCAGAGGGTGCCACTA | TCGA |

| | |
|---|---|
| tctgggaaccacccttctactttctgtctctaggaatttcactactctag*CCAGCGTCCCATGCTCGGTGGCCCCAGAAAAGTCAGTGTGTAGGCCTCAG | TCGA |
| GAGGCAAGCGCAGGGGGACGGGGACCAGCAGCTGTCGCCGCCGCTCTCAG*ACGCCCCTGTCCTGGCAAGAGCTAGAAGGTGAGCGTGCCAGTTCCTGTGC | TCGA |
| GAGGCAAGCGCAGGGGGACGGGGACCAGCAGCTGTCGCCGCCGCTCTCAG*CTCTTGTGCCAGCACCCCAGGAGCCGCCACCTTCCTTCCCGATACAAAAG | TCGA |
| GAGGCAAGCGCAGGGGGACGGGGACCAGCAGCTGTCGCCGCCGCTCTCAG*CCATGCCTTTCTTGGACATCCAGAAAAGGTTCGGCCTTAACATAGATCGA | TCGA |
| GAGGCAAGCGCAGGGGGACGGGGACCAGCAGCTGTCGCCGCCGCTCTCAG*GGTGGATCGTACCCGGAGCTGGCTCTATGCTGCCTGTTCTATCTCCTATC | TCGA |
| CTCCAAACCTCGACAAAATGGCTGATGATTTGCTGGCAGCTGCTGACAAG*CTATTATCGCTGACCCTGGATGGACTGACTGGTGTTTCCCAGGACCACAT | TCGA |
| GAGGCAAGCGCAGGGGGACGGGGACCAGCAGCTGTCGCCGCCGCTCTCAG*GAAGGCGCTGGCCCCAGCATTCAGCTCCAGCTGCAGGAGGTGAAGACGGG | TCGA |
| GAGGCAAGCGCAGGGGGACGGGGACCAGCAGCTGTCGCCGCCGCTCTCAG*AAAGGAAGGCAGAGGAAGCCACGGCCACTGGACCCTGGCGAGGGTTCCAA | TCGA |
| CATTTGAAGCATTCTTTCAATCCAAGGCATCGAATTCTTGGACAGGAAAG*ATGGAGCTCTCCTGGTAGCAGTTTGTCGTGGTAAAGTGAGTGAGGGTCTG | TCGA |
| TCGGCACATCGCCCTGTTGCTGTGGCAGCTGGAGAGTTCCTTCACAAAAA*GACCCAGATGCCCTGTCCAAACAGTGGGACAATGGTTGGAAAGCATTGGG | TCGA |
| TTGGGGGGAAACTAGGTCGATGGTCGGCGGCGCCCGGATGGGCAGCTGAG*GTTATTTGGACAAGAGGCACATTGACCCAATTTGGAGACGTATTTTGGAA | TCGA |
| TTGGGGGGAAACTAGGTCGATGGTCGGCGGCGCCCGGATGGGCAGCTGAG*GATCCTCAGTCTTCCACAATGAACCCTGTTTACAGCCCCGTGCAGCCTGG | TCGA |
| GGAAGAGAGCCAATGAAAGGTTGGAGTTACTACTTCAGAAACGCAAAGAG*TGCTGAAGCTGCTGAAAGATGGCAGAAGAAGTGGTGGTAGTAGCCAAATT | TCGA |
| TTGGGGGGAAACTAGGTCGATGGTCGGCGGCGCCCGGATGGGCAGCTGAG*GTGACACTGGAAGCAATGAACACCACAGTGATGCAAGGCTTCAACAGATC | TCGA |
| ATTCGATTCTTTGGCAGTTGGTGAAAATTACTGATGGCTCTCCTTCCAAA*CCGAGCCCCTGCCACTGATGGACCTGTGCCGGAGATCCATCCGCTCGGCC | TCGA |
| GGGGGAGGCGAACAGTCTCGTTGCTGCCTCTGTGTACGCTGAGGGGGGAG*CCCACTGTTTGTAACATGGAAGATTGGTCGAGACAAAAGATTACGTGGAT | TCGA |
| ACCATCTACAGGAAAACGGAAAGTGGTTGAGGGCATGCAGCTTTCACTCA*ATCAGCTGGACATCATCTCCATGGCGGAGACAACCATGATGCCAGAGGAG | TCGA |
| AAAACATGATGTTGTTTGAAGTTGTTAAAATGGGCAAGAGTGCTATGCAG*gtaattcatttgcacacctgttagcaagaaacagaagttgaaggactgga | TCGA |
| AAAACATGATGTTGTTTGAAGTTGTTAAAATGGGCAAGAGTGCTATGCAG*GATTTTCCATCCTCCAAGCGATTATGGAAGCAGCAGTGCAAAACAACTGG | TCGA |
| GGGGGAGGCGAACAGTCTCGTTGCTGCCTCTGTGTACGCTGAGGGGGGAG*ATTTTCTTGCCTCCTGGGCTCCCTCCACACTATCCTGTGGCTGTGGCAGA | TCGA |
| ACAACAATGCGACGGCCACTGTTCTCTGGGACAATGCTTTTGCAGAGCCT*AAAAGATCCCGCACCATCCTCCAGGATCCAATGGCCTTGGAGAGAGGGCT | TCGA |
| CAGTGACTCAGAAATTGGCGGCATCACCATTGCTTGGAAGTTTGATTCTC*GCATGGAAGTTGTAAAGGTTGGAGGTCCCGTCTACAGGATTGGAGTTGGA | TCGA |
| AAGAAGAAGAAGTTGCGAAGGATCCCCAACGGGGAGGCCAACGTGAAGAA*GTTCCCCCAAGCGGCACCACTTCCAGCGGCAGCGGGCAGCCAGTGAGAGC | TCGA |
| GAGTACGAACCGGCCAAGAGGTTCTCCATCCGGCAGATCCGGCAGCACAG*GTCCCCCGTCATGCGTGCCCGGAAGAGCAGCTTCAACGTGAGTGATGTGG | TCGA |
| AAGAAGAAGAAGTTGCGAAGGATCCCCAACGGGGAGGCCAACGTGAAGAA*CCAGATTCTACTGCAACTGAAAGAGCAATTGCCAGACTAGCAGTACATCC | TCGA |
| ATGGCCCGCAGAGTTAATGCAGGGCTTGGCACGATACGTGTTCCAGTCAG*GTTTTCTGTCTCAACTGGTCTCTGACAAGCCTCTGACTGAATGTATCCGT | TCGA |
| CCTTTGCCACTGAGGAGCATCCTTACGCGGCTCATGGACCCTGGTTACAA*GCACTAAAGCAATCTTCAACCGAACTCCTCTTTGGAGGACACGAAACCAC | TCGA |
| ATGGCCCGCAGAGTTAATGCAGGGCTTGGCACGATACGTGTTCCAGTCAG*GCCGTAAGTTCATCATCGCCAACGCTCGGGTGGAGAACTGCGCCGTCATC | TCGA |
| ATGGCCCGCAGAGTTAATGCAGGGCTTGGCACGATACGTGTTCCAGTCAG*GCACGCCCCAAAGGCGAGGGCCTGACTCCATACCAGGGCAAAAAGCGCTG | TCGA |
| AGAACTCTTACTTACATGTCTCATCGAAACTCCAGAACAAACATCAAAAG*GGTAAAAGCCCCAAAGCTGTCATCAGCATTAAGGACTTGAATGCCACCTT | TCGA |
| GGAGAGAAAGAAGTGATGAAACTCTGGAATCTCCATGTCATGAAGCATGG*GGAAAGAACTAGAGAGAGACACAAGTCACCCCGGAATAAAGACGGCAGAG | TCGA |
| ACAAGCTACAGGTCGAGCAGCCGTCCATGCAAGAACTCTCCAAGCTCCAG*GAGTGTACACATTTCTGTCCAGCACCCTGAAGTCTCTGGAAGAGAAGGAC | TCGA |
| TCACTGTCATGGACAAGGACAAGCTCAGCCGCAATGACGTCATCGGCAAG*AAGCTAAACCAGCCCCAGAAATATTTGAAAATGAAGTCATGGCCCTGCTG | TCGA |
| CACACTATCGTAATCGAGAACACTTTGCTACTATACGGACAGCATCACTG*GTTCAAAACTGGTCAAATCAATGGTGATTTGCTGATATACCATGTCTTAC | TCGA |
| TGCCTCCATTCCCGGGAGGGGGGAGCGGCGTTGGAGGCCACCGTTTCCAG*GTCTCGAAGCCAGACATGGGGCTGGTGGCCCTGGAGGCTGAAGGCCAGGA | TCGA |
| TGCCTCCATTCCCGGGAGGGGGGAGCGGCGTTGGAGGCCACCGTTTCCAG*ATCTGGGACATGAGCGATGATGAGACCATCTGAGATGGCCCCGCTGTGGG | TCGA |
| CGGTGACCGGATATTCAGTTGCACATCCCCACATCAATGCACTGCCAATG*CACTACGGCCAGAGCTGCCTGTTCAAAATCGCCATGAACATCCTCAACAT | TCGA |
| TGCCTCCATTCCCGGGAGGGGGGAGCGGCGTTGGAGGCCACCGTTTCCAG*AACAAGTTGAAATCATCGCAGAAGGATAAAGTTCGTCAGTTTATGATCTT | TCGA |
| TGCCTCCATTCCCGGGAGGGGGGAGCGGCGTTGGAGGCCACCGTTTCCAG*TCTGCTCATCACCTCGCTTGGCCACATCAAGCTCACGGACTTCGGCCTGT | TCGA |
| TGCCTCCATTCCCGGGAGGGGGGAGCGGCGTTGGAGGCCACCGTTTCCAG*ATGAAATTATGCAGCAGGAAGTCAGACCCCTGATGGCGGTGGAGATAATA | TCGA |
| TGCCTCCATTCCCGGGAGGGGGGAGCGGCGTTGGAGGCCACCGTTTCCAG*GGCTGCCGGGGAGCGCTATCGCAGGGAGATGCTGGCCCACGGTGGAGGCA | TCGA |
| TGCCTCCATTCCCGGGAGGGGGGAGCGGCGTTGGAGGCCACCGTTTCCAG*GGAAGTGAGGTTAGGGCGAAATGGTGTAGAAGAAATCAAACGACATCTCT | TCGA |
| TCATCCAGAAAGGTCTACAGTATGTAGAAGCAGAAGTTAGTATTAATGAG*GCCCAGTTCAATCTGCTGAGCAGCACCATGGACCAGATGAGCAGCCGCGC | TCGA |

| | |
|---|---|
| TTGGGACGCACATACTGGTGAAGCCAAGCAACAGTTTCCTTTTCATTCAG*CATCAGACATCGATGAGTCATCAGTGATGCAGCTGGCCGAGATGGGTTTC | TCGA |
| GTGGCAGGCTCTATGGCCTGCAGGAAGCGCTGGAAATAGCCAGAGAGATG*CTTTGAAAAGTCCAGCCGCATTTCATGAGCAGAGAAGGAGCTTGGAGCGG | TCGA |
| CGGCGTCTCCAGCCACACGCCGCCTGTCAGCGGGGCCGACAGCCTCCTGG*AAATCTCCCAAGCCTaagaagaagaagcacaggaaggagaaggaggagcg | TCGA |
| CCAGCCAGGCACCCTCCCTGACCTGTCTCGGCCTCCCGACTCCTACAGTG*ACGCTGCAGGAGCTGAAGATGGCGAGCTCCGTGGCGCCCTACGAGCAGCT | TCGA |
| GAGCCCCCGGTGCCTTATCGCCCAGCTACGACGGGGGTCTCCACGGCCTG*GGAACGATCAGCGTTGGAGTAGATGCCACCGACCTTTTTGATCGCTATGA | TCGA |
| GTGCTCATCCCCACAGGGACATTCACAACATGAATAATGGAAGCACTGTG*GTGGGAAAGGAGAAGTTTCACAAATCCCAACATTGGGGCTTTTGCAACAA | TCGA |
| CTTGGGGCAGGACCAAGTGTTGCTGCTGTCAGGGAAATCATGGAGAATAG*GTACTGGAAGTTGGACCCTGCTCAGGTCTATGCTAGCGGGCCCAACGCAT | TCGA |
| GCAGCCAATAGGACATGATCCAGGAAGAGCAGTAAGGGACTGAGCTGCTG*GTCATCAGATGCAAAGCAGCCATACTCTGGAAGCCTGGTGCACCATTTTC | TCGA |
| tttttaataaaatttatatttatttaatGCACTCTAAGTGTTGTCTTCCT*CACCCGGAGAGCTGTGTCACCATGTGGGTCCCGGTTGTCTTCCTCACCCT | TCGA |
| GACCCGACTGCAACTGCTGGATTGCTGCAAGGCTGAGGGACGAGAACGAG*GAATGGAAACTGAAAGTGGAAATCAGGAAAAGGTAATGGAAGAAGAAAGC | TCGA |
| AGGAGAGAGACGAGCGACTGACACGCATGTACAACGTGCGCATAATGCGG*GCTGCTGTCCCCAAGGCCCAGCCTCCTCACCCCGACTGGCGACCCCCGGG | TCGA |
| CCTGTGGACGCGTATCGCCAGCACGATCCCACCGCACGTTCAGAAGTCGG*ATTGTCACAGACCAGCAGACAGGACAGAAAATCCAGATAGTCACCGCAGT | TCGA |
| TACCCACCTGGCCCCGCTGAAGTCAGCCGGCTCAGACGATGCAGGAAGCG*CCTTGCCTCCCCGATTGAAAGAGATGAAAAGCCAGGAATCGGCTGCAGGT | TCGA |
| GAAATAACGAGGAATGGACTGTTGATAGCTGCACTGAGTGTCACTGTCAG*CTACATCTACATCCACCACTGGGACAAGCCATCTTGTAAAATGTGCGGAG | TCGA |
| ACCACCGCAAACACAACATTCCGCACTTCTGGCCCAAGCTGCTGATGAAG*AGAAGAGGCCCCCTGAGCCCCCCGGACCTCCACCGCCGCCACCTCCACCC | TCGA |
| CGGTCTGGATCGGATTTTGGAGGCATTGAAGCTGCTGCTGAGCCCGGGAG*GTCTCCTCTTGGCTCTGCCAGGAGCCGGAGCCCTGCCACCCTGGCTTTGA | TCGA |
| CTGCCTGCTCAAAGAGCTGCACCGCTTCGAGTCTATTGCCATTGCCCAAG*GTATGATTATCGGGAAATGCTGCACAATGCCACTTTCTGTCTGGTTCCTC | TCGA |
| ACTCAACTGGTGTCGAGAAGTCCGGAAGCTTGTGGCGCTGAAAACGAACG*ATTTCCTGTGCTTCATCAGCTAGAGAGATGAAATCCATTACCAGGATGAA | TCGA |
| CCACCTCAATGGCCTAAGCAAGTGTCTGCAGTGCCAAATGTGTGACCCAG*ATGACTAAGAAGATCAAGAAGCTGGAGAAAGAAACCACCATGTACCGGTC | TCGA |
| GTAATGGAAGGGCATGGCAAGACTCCACCTCCTGGTGAAGAAAGCAAACA*GTCTTCTTTGTGATCCGCCTCATTGCTGGCCCTGCTGCCAACTCCCTGCC | TCGA |
| GGGGCTCCCAGACCTTCAAACCAAAGGGCCAATACCTGGATATGGTTCTG*CTGGTTCCGGAAGAAACATCCTCCGGCTGAAGCACCAGTGCCCATCCCAC | TCGA |
| TCGGGCTGGGAGCGTGCTTTCCACGACGGTGACACGCTTCCCTGGATTGG*CACGTGGACACGGGGAACTCTTACCTTTGTGGGTACTTGAAGATTAAAGG | TCGA |
| TCGGGCTGGGAGCGTGCTTTCCACGACGGTGACACGCTTCCCTGGATTGG*TGCCCCGTGCTGAAGGGTCGAGTGCAGAAGATCCTACATTGGCGGTGGGG | TCGA |
| CTCCCCAGCCAAAGAAGAAACCACTGGATGGAGAATATTTCACCCTTCAG*GTGATTGTCTGACAAGCAGAGGCATGAGCTGGGTCCAGGCCACCCTACTG | TCGA |
| TCGGGCTGGGAGCGTGCTTTCCACGACGGTGACACGCTTCCCTGGATTGG*TGTTTTCTTCACTCATGGTGAAGAATGGGGGAGATAGAAGGAACATACAG | TCGA |
| TCGGGCTGGGAGCGTGCTTTCCACGACGGTGACACGCTTCCCTGGATTGG*AAATACCAGGAGTCTGTTCGTCAAAGATATAATAAAATCGCATATGCTGA | TCGA |
| TCGACGTGTCCTTCCAGCAGTCGAGCACCGCCAAGTCGGCCACCTGGACG*CTTAACCAGCTCGAAAAAGCAGTGGAAGCAGCTCACACATTTTTCGTGGC | TCGA |
| ACTTCACGGTGTGCCACCCTACAGTACTGCCCTGACCCTTACATCCAGCG*GAGACAAGGAGAAGGCCTTAGGATCTACTGGGGGAGTCCGGAGGAGCAGT | TCGA |
| GGAAACGACCTTCGGACCCGCCTTTTCAGCCGTCACCACCATCACAAAAG*AGATTCCTGAAATTCCAGAAAGCATTTCATTCTGTAAAGCACTGCAGGTA | TCGA |
| GCTTCTCCGGGGGGCCCAGCAATCTTCCCACCCCAGACGTCACCACAGGG*GCATGTCAAAGGCCGTGTGCTGGTGGCTCTGGCGGACGGGACCCTGGCCA | TCGA |
| CATCATGAAGCGGGCTggcggcgctgccgctcccgggcggccgcgggcgg*GCCAGTCGGAGAGCGTGCGGGACGTGCAGTTCAGTATCCGGGACTACTTC | TCGA |
| CAAGGAGGAACCCCTGCACTCGATTATCAGCAGCACAGAGAGCGTGCAGG*GACAATTTCAAAGGATCAGTTTGAAAAGAAGAAAAATGATACCTTAGACC | TCGA |
| CAGAACGGTCCAGTCACCAAATTCATCAGTGCCATCTCCAGGCCTTGCAG*GCCCAGCCTCCGTTATCAGCAATGATGATGACTCTGCCAGCCCACTCCAT | TCGA |
| CAGAACGGTCCAGTCACCAAATTCATCAGTGCCATCTCCAGGCCTTGCAG*GTTCTTATGAGTGCATCAATACTGGGGGCCCTGTGTCTCTTCTCATCTGT | TCGA |
| TGAAGGACACTACTGAGGTTCCCAGCAGTACAGTAGAAAAGTCAAATCAG*GCCCAGAAGTCTGGCTCAGGAGAGGACCCTCAGGCTGCAGAAGACCTGCT | TCGA |
| CCAGCTCAGCGTGCCCATCCCGGGGGGCAGCAACGGCGACATCCAGCAAG*GAGCTGCTTCATCTGACCCAAGCTGGCCTGGAGAAAGGAGGAAGAGGCAA | TCGA |
| GATAGATTGACATGTAGAGACTGTCAGCTATTGGAACACAAAGAACATAG*GGAGTACCAGGAGGAAGGGGACTTCCTGGAGAGGATGGAGAAAAAGGAGA | TCGA |
| AAGGTCTTTCATCTAACTTGTCATGTTCCAACACTACTTAGCTTTCCAAG*GCTACACTTCCTGGTGTTTGACACAGAGGAGGTTCACGACGTGCTGCGCA | TCGA |
| ACTCCTGGTATGAGTTTGGAGCCAGAGAGAAGACCCAGGACCAAAATGTG*GACATGAACAAGCTGAGTGGAGGCGGCGGGCGCAGGACTCGGGTGGAAGG | TCGA |
| CGATTTCCATCGCCTCACGGACGTCTCAGAACACGCTGGACAGCGATAAG*GGGTTGGCTGTGTTCCGGCCGCAGAGCACCGTCTGCGTGAGGAGATCCTG | TCGA |
| CATCCGGAATTAGTGACTGGATCCAAGGACCATGAACTGGACCCTCGAAG*GAGTGCCAAGACCTACGCCTACCTGTTTTCCCATCCCTCTCGGATGCCCG | TCGA |
| CATCCGGAATTAGTGACTGGATCCAAGGACCATGAACTGGACCCTCGAAG*GCCTGTCTCCGAGTTCTTCCTAATCCGAGACACAAGCCTCATTCCCAACG | TCGA |
| CATCCGGAATTAGTGACTGGATCCAAGGACCATGAACTGGACCCTCGAAG*AATCCTTCACGCCCGACCACAAGCCCCTGATGGGGGAGGCACCTGAGCTC | TCGA |

| | |
|---|---|
| CCATTGTCTACAATGACTCCGGTGAGGACTTCAAGCTTGGCACCATCAAG*GAATGGGCAAAGAAGTGGGGAATCTGCTACTGGAAAACTCACAGCTTCTG | TCGA |
| CACTGGATGATGCAATCAAAGCTCGACTTCTCAATCCCAAGTGGCCAAAG*GTGCTAGTCCTATGGGAGTAAATGGAGGTGTAGGAGTTCAAACGCCGAGT | TCGA |
| GCTCTGAACATCATCGACATGGCCCTGAGTGAATACCCAGAAAATTTCAT*AATGCATACTTGGAATGAATCCTTCTAGAGACGTCCACGGTATGTTAAAT | TCGA |
| TGCTAACTTGGTCATTCTTCGTGGGAAAGATGTGCAAAGTGCAGATGTGG*TCGCCCTTACACAAACAAAGTCATTACTTTGTGGTACCGACCTCCAGAAC | TCGA |
| GGAGTACTCCACAGAGGATAAATTTCAACCTCCGGGGCCACAATAGCGAG*GACATAACGACTATATGTGTCCAGCCACCAACCAGTGCACCATTGATAAA | TCGA |
| CTGAGCGGCGCGGGCTCCAGACTCGCCGTGGTCAAGTTCACCATGAGAGG*GTGATAGTTAAATCTGGGCCAGGGACTTTCCTCAGTTTGGCTGTTTATGA | TCGA |
| GAATGCCATCCTTATACATGGAGTTGATATTACAACCAGTGACACTGCAG*CGCCGGCATCGGCCGCACAGGCACCATCATTGTCATCGACATGCTCATGG | TCGA |
| ATACTGAGGTTACACCAAGGAAAATCTGAAGATATTTATCAAATAGTTGG*GGATGCCAGATGATATGAACTTCAGTGATGTAAAGCAAATCCATCAAACA | TCGA |
| AATATATCTTCAACAACTGCCACGACTTATACAGCCGCCAGTACCAGCTG*CACCTCCAAGGTTTACACGAACACCCGTTGATCAGACAGGGGTCTCTGGC | TCGA |
| TGGCGGCTCTGCGTACGCTGGTTGGCATTTGCTCTGAGCATCAGTCACGG*GGGGTCTGCCACATGTAGAAAGATATCTTCCAAAACGAAATCCTCCAGCA | TCGA |
| TCCGAGCTTGGACCCTTAATGAGGACCACACTCCAACGCGGGGCGCAGTG*GTAGAGTTGGCTTTGTGGGACACAGCTGGGCAGGAAGATTATGATCGCCT | TCGA |
| GCCACCCCTTAGAGATGGAGCAAGAGTACTTATGAAACTTATGCCGCCAG*AATGCTGCCTTTTTATATGGTCTTGGTTTGGTCTACTTCCATTATAATGC | TCGA |
| CTACCCAACGCTGCCGCCTGGCACGGGCCGCCGCATCCACAGCTACCGAG*TGCCTTGGTGACTGTCAAATAAAAGCAAGTATATACTCAGTTCTCTCCAC | TCGA |
| ATCTCTCAATGTTGACGGACAGCCTATTTTTGCCAATATCACACTGCCAG*caaccccgatggataccattatctccatctacacagcggggaactgaagc | TCGA |
| GATATACCAGACCCTGGAGTGAACTGGATTCCATGTAGAGACCCCTGGAG*GTAATATTTGAAGCTGAAGTCTCAGGAGGGAGAAGTGGTTATATTGCCAT | TCGA |
| GGCCTTCACAATCCAAGTTCCACAATATATTGACTACTGCCACAATTCCG*TTGCAGCATGAGAAAGCCGAACTAGAACAGCATCTTGAACAAGAGCAGGA | TCGA |
| GCCGTGATCGTGCCCAAAGAGGAGGGCGTCATCAGCGTCTCCGAGGACAG*AATTCAATTCGTCATAATCTGTCCCTACACAGCAAGTTCATTCGTGTGCA | TCGA |
| CGGCGCGATGGAGCAGCGAAAACGTGGTTGTAGAGTTCCGTGACTCCCAG*GTTGCCCAAGCTTAATGGAACCCCCCTCTCCCAGGGAAGACATGAGATTG | TCGA |
| GTTCTTGTATGCACCATCCCATAAAAACTATGGAGAAGACCAGTCCAATG*GGCCTGGCCATGTCCCCTTTCGGAAGCCTGTTCCCTTACCCCTACACGTA | TCGA |
| CCCCACTCGAGGGGCCGTGATGGAGCAGGGCACGTCCTCGTCAATGACAG*ATCAGACCTATTGTGACCGCCTGGTGCAGGACACGCCTTTCCTGACAGGC | TCGA |
| AAGACATGGAGGCCCAGGCAGGCTGGGCTGCCCCTGGCGAGGCGCGGGCT*GAAAAGTACCCAGATGCCGTGTACCTCTCGGAGGGGCCCTCCTCCTGCTC | TCGA |
| TCAAGTTCGACTTGGACGGGGACGCACCCGATGAAATTGCCACGTATATG*GTGGGCTGGATGACACTTTGCACACAATTATTGATTATGCCTGTGAGCAG | TCGA |
| ATCTTCGCCCCCACTTCCTCTAAGCTCCCTGCCGCCAGAAGCCAGCCAAG*GCACGTGCGCTAAGCTGCGGCTACCCCCGCAAGCAACCTTCCAAGTCCTT | TCGA |
| CTCCACAGTCATCCGGCTGAAGAAGACCTGTTGGACTGGATCTTCTCGGG*GTCCCTATGTTAACTACTTGAGACAAAAATAAATATGGCCTTCTACAGTT | TCGA |
| ATCCACATCATTCACAATGGGGTGATATGAAACTCTACTTAAAGTTACAG*ATGTCATGGCACCCCCAGTACCGGAGCTCCAAGTTCCGTCATGTCTTTGG | TCGA |
| CCAGAAATCCAAGGCCAAGAGCAAGGCCCGGCGTGAGGAGGAGGAGGAGG*GCTCTGTGTGCTCCGGGATGGAGCAGGTGTGCAGAGGGTGAGAACCCAGC | TCGA |
| GGATCTGGTGTACCAGGCGAAGCTGGCCGAGCAGGCTGAGCGATACGACG*GACCCTGAGGCTCAGAGGGGCAGCTTCAGGGGAGGACACCCCACTGGCCA | TCGA |
| GGATCTGGTGTACCAGGCGAAGCTGGCCGAGCAGGCTGAGCGATACGACG*CATACACGTCGTGACTTGGAACGTGGCTTCGGCAGCGCCCCCTCTAGATC | TCGA |
| GGATCTGGTGTACCAGGCGAAGCTGGCCGAGCAGGCTGAGCGATACGACG*ATGAGACAAAAGGACCCAGTGAAAGGAATGACAGCAGATGACTAAAGCCG | TCGA |
| GTTGTTACGTGATAATCTGACACTATGGACTTCAGACATGCAGGGTGACG*TTGGCCTCTACACCTCCATCTTCGGCGTGCTCCAGCTGCTGTGCCTGCTG | TCGA |
| CTGACGCTCGTCCTCATCCTGGTCACTAGCGGACTCTTCTTCGCCTTCGA*GTAGTCAGATGCCTCCGCAGCCACCCGGGAGCCAGTCAGAATCCAGTTCC | TCGA |
| GGCTGCACACGGTCAACTCCAGGCACGAGGCCAGCCTGAAGTTGTACAAG*AAAGGGTGCTTCATTGAAGAGGTCCAAGTGTTGCATGGAAACATAGTAAG | TCGA |
| AGTATGTTTCCTGCTCATTGTGATACATTAAAAAAAAAATATGAGCAAAA*AAAATTCAGGACTGAAAAAGAAAAAAAAAGGTAGAAGATTTTACGCATAT | TCGA |
| CCCACCTCCACAGCGTGGCACCTGACTGCGTGGAGAAGCTCATTATGACG*CCCTGGTACCTGACAGTGTAAAGAAGGAGCTCCTACAAAGAATAAGAACA | TCGA |
| CCTGCAAGTGCCGCCTCCCTCCGGAGGTGCGCGGAGGCTTGGTGTGCCAG*TGGAGAACATGCAGCTGACCCTGAACCTGCAGACGGAGAACAAAGGCAGC | TCGA |
| CCCACCTCCACAGCGTGGCACCTGACTGCGTGGAGAAGCTCATTATGACG*GTCTGAGGCGCAATTACAGATGTACCATCTGCAGTGTCTCCCTAAACTCA | TCGA |
| GGACCTGGGGCAGATCTTCACCATCCGCAGGTGCCCCTCCACGGACCCAG*GGGATCAAGTATTTGCTGGGCTAGAAGAGCAAGCCCGCCAGGCGATGATG | TCGA |
| TCTGCAATGTCACCTTGAACTCTGCACAGCAAGCCCAGGCTCATTATCAG*TTTCGTAGAAACCCCAGCTCATTTCTCTTGGAAAGAAAGTTATTACCGAT | TCGA |
| AACAGAACAGGAGGTGGTAGAGGGCATGGATATCTCTACTCGCTCCAAAG*TCCCACCCGCTGTACCTGTGCAATGCCAGTGATGACGACAATCTGGAGCC | TCGA |
| TATTGGGAGAGCCACAGCTAAACCCCGGGACCCCTGGAAGCCTAGAAATG*GAAGAAAAAAGATGCAATTGATCCCTTACTATTCAAGTATAAAGTGCAAC | TCGA |
| CCTGTACTGGGACGTGATGCTGGAGAACTACAGTAACTTGGTCTCACTGG*GTTGTCCTATGTCTCGACTGCTTCTGGTTTTGAAGACTTACTCAAGGCTT | TCGA |
| ATCCCGGAGGAGGAAGAAGAAGTGTTATATGTCCTAAAGTAAGGTTTCAG*GTTTTATTTATTTATTGTTCCTGGTCACTGTCTCTTTGAGGATTGGTATC | TCGA |
| GTGCCTGGGGCTCCCAAGCAGACCACCAAGCCAGCTCAGCGGGGACCCAG*GCAGAATACTTCTATGAATTCCTGTCCTTGCGCTCCCTGGATAAAGGCAT | TCGA |

| | |
|---|---|
| CCGCGGGAGAGGTAAACAGAACACCAGGACATCCCAGAGTCTGGGAAATG*GTTTAAACGGAACCTGCAGCGTTTCCAGTGTTCCCACGTCCACGTCGGAG | TCGA |
| GCTTCTCGCGGGCCGGGGCCACGCTCAGCGCCGAGGGCGAGATCGTGCAG*GTCATTCCTGCACAGGACAGTCCACCATGGCCTCAGATCACCAGACCCAG | TCGA |
| TGGAACAGCCAGAATTGGACCCGAAACCATGCCTCACTGTCCTAGGCAAG*GTCAGCTGGCCGCCGGCACCTGTGAGATTGTGACCTTGGACCGGGACAGC | TCGA |
| GGTCCGGGTAGATTGGGAGCGGAGGTTTGACCACATGCAGCAGCATTCAG*GGAAACCAATGAAAAGCGTGCTGGTGGTGGCTCTCCTTGTCATTTTCCAG | TCGA |
| GGAATCATCGAGGCATGGGGGTCCACACTGCAATGTTTTTGTGGAACATG*ATGACTATGTCCACATCTCCAGCAAACCGTGTGACCTGCACTGTACCACC | TCGA |
| GGAATCATCGAGGCATGGGGGTCCACACTGCAATGTTTTTGTGGAACATG*ATGGCTCGTTCGGAACACCACCTGGATACGGCTGCGCTGCAGACCGGGCA | TCGA |
| GGAATCATCGAGGCATGGGGGTCCACACTGCAATGTTTTTGTGGAACATG*CTCGCCATGGCCAAACAGCTGCAGGCCCGAAGGCTAGACGGGATCGACTA | TCGA |
| AAATCTGTCCTCTGTAGTATTTTTTAAATAAATCAGTGTTTACATTAGAA*aaaaaaaaaaaaaaaaaaaaaaaaattaagtgggtgtggtggtgcatgcc | TCGA |
| GGAATCATCGAGGCATGGGGGTCCACACTGCAATGTTTTTGTGGAACATG*GGGCCATGGAACGTATATGGGAGAAGAGAAGCTCATTGCATCTGTTGCTG | TCGA |
| GGAATCATCGAGGCATGGGGGTCCACACTGCAATGTTTTTGTGGAACATG*ATGATCCCACCTGACCAGGAACTGCTGGTGTGGTACGGAAACTCACACAA | TCGA |
| GGAATCATCGAGGCATGGGGGTCCACACTGCAATGTTTTTGTGGAACATG*AAAAGCAAACTCAAGTTCCTGCCTCCTCAGGCGGAGCAAGAAGACTCCAC | TCGA |
| ACGAAGGGAGGGTGTACCATTACAGGATCAACACTGCTTCTGATGGCAAG*CAAAAAGATGGCACCTTTGCAGCAGGGGGCTACATGCCCCCTCTCCGGTT | TCGA |
| CCCGGCGGGGCGCACCACAGAGACCGGCTTCAATATCTTCACCCAGCATG*GTGAAAAGCGAAATCCCAAAGGTGCCGAAGCAGCCTCTGGTGCTCCATCA | TCGA |
| GGCCGATGAGGTGGCGCCTGCAAAGAAGGCCAAGCCTGCCATGCCACAAG*ATTCAGAATCTGATAATTCAGATAACAACACAATCTTTGTGCAAGGACTT | TCGA |
| CACGCGGCTTGAAGGACTGTGGGCAGATGTGACCAAGAGCCTGCATTAAG*GGTTAAATATAGACAAGAATTATTGGAAAGGCGTTTGATGGAGAAAAAGG | TCGA |
| GATTCTTCCTGTGCTTATCATGATTGCTCTCCCCTCCCCTAGTATGGAAG*tctcaggtatgtctttatcagcagcatgaaaatggactaatacagtaaat | TCGA |
| CACGCGGCTTGAAGGACTGTGGGCAGATGTGACCAAGAGCCTGCATTAAG*TTCCAATTATAAAAACAGAACCCACTGATGATTATGAGCCTGCTCCAACC | TCGA |
| AGGAGAGTAGATCTGGAGAAACCAACAGCTGTGTTGAAGAAATAATCCGG*AAATTGAAAAGGCTATTACCCATTGAAGAAAAGAAAGAAAAATTTAAGAA | TCGA |
| CCAGAGCCAAGGCCAAGCTCTCCAAGTTCAGCATCCCCAAGCAGGGGGAG*TGGAGGTGAAGCCGGTGCTGCCAAGAGCCATGCCCAGTTCCATGGGGGGT | TCGA |
| CCAGAGCCAAGGCCAAGCTCTCCAAGTTCAGCATCCCCAAGCAGGGGGAG*ATGTTCCTGCAGCAGCCAGCTGTATCCCTGAGCTTCAGCAGCACCCAGCG | TCGA |
| CGGACACACCTGCAGAAAAGGCTTTGCCAAAATCCAAGAGGAAACGCAAG*ATAGAGAACGTCACTTTTCACCATGAAGGGTTCTTCTGGAGGTGTTGGTT | TCGA |
| AGCAGAAGAAGACCGAAGGGAAGACTTCCAGTAGCTCCAAGGAGGTTAAG*TGCCGCCAGTCATCCGTGTCTATCCAGAGAGCCAGGCACAGGAGCCTGGA | TCGA |
| AGCAGAAGAAGACCGAAGGGAAGACTTCCAGTAGCTCCAAGGAGGTTAAG*GCCCCATCGTCACCTTCCTCGCCCATAGCTAAGGACCCGAGGTATGAGAA | TCGA |
| CACTGCCGCAGCCAGGAGATGGTTCGGGCCTAGCGGAGCCGGGACTGGAG*GAGCACACTAGGTGATGCCCCTCACTTGGCACCCGTGTTGGCGCTGGCAT | TCGA |
| TGGTCAGCCCCGACGGCACCGTCACCGAGGCGCCGCGCACAGTCAAGAAG*GTGGAGGTGGAGGTGGAGAGCATGGACAAGGCCGGCAACTTTATCGGCTG | TCGA |
| TCAGCATAGAAAGAAAGCCTTGTAACCATCAGGTAGATACAGCACCAACG*CCTTGTAGCAGTACCTGGGAACCTGCATCCTGTGGAAAGATGGAGGAGCA | TCGA |
| ATTGACAAAATCCATCTCATTAATGACAATCAGCCATCCTGGATTGGACA*GTCCCGTGGCTGTCATCAGTGGTGAGGAGGACTCAGCCAGCCCACTGCAC | TCGA |
| GGGAGGCAGAGCGGCTCAGCCAGCGGCAGACAGAACGGGACCTGTGTCAG*GAGGATCCAAAGTGGGAATTCCCTCGGAAGAACTTGGTTCTTGGAAAAAC | TCGA |
| AATATAATATGGAACCTTCCCTCCAAATTCTTCAATAAAAGTTGCTTTTC*aaaaaaaaaaaaaaaaaaaaaaaaaaaaaGTCCATTGCTGAGAGACGAG | TCGA |
| TTCGTTTGGTGCTACCAGCAGTCCTGTAATAAACTAGCTATCCATCTGTA*AAAAAAAAAGAGGAGCGAGAGAAAGAAAAAAAGGGGGAAAAATCAGGATC | TCGA |
| CACCATGAATGAGGTGTCTGTCATCAAAGAAGGCTGGCTCCACAAGCGTG*AAGGGTCCCAGCACCATGGAGGACCCGAACCCTGAAGAGAACATGAAGCA | TCGA |
| TGGGCGGGGGTAGAGCCCCGCCGGAGAGGCTGGGCGGCTGCCGGTGACAG*GTTGGGCCCGAAGGCATCCTGCAGAACGGGGCTGTGGACGACAGTGTAGC | TCGA |
| TGGGCGGGGGTAGAGCCCCGCCGGAGAGGCTGGGCGGCTGCCGGTGACAG*CCTCGAAGCGGAGGATCCCTGTGTCCCAGCCGGGCATGGCCGACCCCCAC | TCGA |
| CATCATGAGCGATGTTACCATTGTGAAAGAAGGTTGGGTTCAGAAGAGGG*GGAGGAAATAATGCTGGCCATACAGTTGTTGTGGATTCTGTGGAATATGA | TCGA |
| CATCATGAGCGATGTTACCATTGTGAAAGAAGGTTGGGTTCAGAAGAGGG*GATTTACTGCCTCAAGATGGTTATGTTGGAGGATTCCAATAGTTCTACTG | TCGA |
| GAAAATACTATGCTATGAAGATTCTGAAGAAAGAAGTCATTATTGCAAAG*ATGAATGTGAAGAAGTAGAACATCAGTGCCAAAAGACTATTCAGCAACTG | TCGA |
| CATCATGAGCGATGTTACCATTGTGAAAGAAGGTTGGGTTCAGAAGAGGG*AACAAAATCCAGTCTCATAGCAGAGGAGAATATAATGTTTACAGCACCTT | TCGA |
| AGACAAAAGACCGTTTGTGTTTTGTGATGGAATATGTTAATGGGGGCGAG*GGCTACAGTGGCAAGGAGAAAGCCTTCATTGCCACGCAGGGCCCCATGAT | TCGA |
| CATCATGAGCGATGTTACCATTGTGAAAGAAGGTTGGGTTCAGAAGAGGG*CCTGGATGTGTAAATACAACTGAAGTGGACATTAAGAAGTCATCCAGAAT | TCGA |
| ACTCCATGTGGCTCTCCTTTCCCCTAATTCACAGGGTGAAGCCAGATTTG*GGCGAGCACCTTGGGAGAGGCACGAGAACACACATCTATTCTGGGACCCT | TCGA |
| TTGAGCCTGTGGCAGTGGATGGTGTTGCCTCTCCTCGATGTGTCTGACAG*GAACACCAAGCGGACAGCTGAAGTGTGGATGGATGAATACAAGCAATACT | TCGA |
| AACGAGGCTGCAAGAGAGATCCTCCTGATGCCCACTCCAGGGAAGCATGG*AGACCAGAACCTGGAAAATTTAATTGAAGTAAATTCTCATGTGTCTGAAA | TCGA |
| CTGGATAATGAAAGACTCCTTCCCTTTCCTGTCTCATCGCAGCCGATATG*GCAGTGGAGCTACTGCTGTGGTTCAGGCAGCCCTATGCAAACCCAGGCAA | TCGA |

| | |
|---|---|
| gccggggccgctgccgTTGAGGAGGAGACGGAGGAGACCGACGTTGTTAG*AAGAATAACTGTGTGATGCCTGAAGATGTGAAGAACTTTTACCTGATGAC | TCGA |
| ATGATAAGAAGATGAAGAAGAAAACCCACAAAGTGAAAGAAGATAAAAAG*CTTATAGTAGCTGTTGAACAAGAAGAAATTCCCAGACTTCAGGCCCTATA | TCGA |
| gccggggccgctgccgTTGAGGAGGAGACGGAGGAGACCGACGTTGTTAG*GACTGCCTGTTCGCAGCGTGGATTTTAACCGAGGCACGGACAACATCACC | TCGA |
| ATGATAAGAAGATGAAGAAGAAAACCCACAAAGTGAAAGAAGATAAAAAG*TTATAGGTGCACAGCGAAGGAGAAGCCCCAGTGCACTAGCCATTGAAGTA | TCGA |
| GCCCATACCACTATTTGTTGAGAAATGTGTGGAATTTATTGAAGATACAG*GCAACCTAATGAGGCCTCTCTTGAATTATGGTATTGCTTGGTGAGCATAT | TCGA |
| AACCTGGTTCAGCAGTTCAGGGAGCTCACATTTCTCATATTGCTCAACAG*ATGAGAGTCCCTACACTAAATCCGCCAGCCAGACAAAGCCGCCTGATGGA | TCGA |
| AAGACTTGACTGGTCTTACATTGCTGCATCAAGAAACCCAAGCTGCTGAG*GACTTGATTAGAGACCAAGGATTTCGTGGTGATGGAGGTAAGTAGTGATT | TCGA |
| TTCTCTCTCGTCCTCGCTGCTGTACTTCGTCTATGTGGCGCCCGGCATAG*GTGCTGGAGAATCTGGTAAAAGCACCATTGTGAAGCAGATGAGGATCCTG | TCGA |
| GGGCTCCTTACCCAAGTACATATCCTGTAAGACCAGAATTGCAAGGCCAG*GTCCGTTATGCCACCTGGAGCATCATAATGGACTCTGTGGTGCCCTCTGA | TCGA |
| TTTATCAACGTCATCTAGAGGAATTTGCCCCAAACAGGAACACATAGCAG*GTGGTTGGTTCTAGCTATTGCCATGGTACGTTTTTATATGGAAAAAGGAA | TCGA |
| AACCCCAAGTCCTCCTTGCCAGGCCTTCAACCACTATCTCAGTGCAATGG*GATCAACCACAGGTTTGTCTGCTACCCCCCCTGCCTCATTACCTGGCTCA | TCGA |
| TGGAGCACGTGCACAGCTTCACGCCCTGGATCCAGGCCCACGGGGGCTGG*gggaggatactgaggcacacagaggggagtcaccagccagaggatgcata | TCGA |
| ggaggaaacggctcagagaggggaagtgtttggccaaggtcacccagcga*GTTGGTTTGTTGGACCTTGAACTCAATCGGCTGACGAAAGCGCTATTTTT | TCGA |
| CCCCCCTCAGTTTCTGAAATGATTCTCCAGAATTTCTCCTCATAAAAAAG*GCTGTCTGGGTCCGACTTTCCTGACGAGGAGTGGCTGTGGGATTATGAGA | TCGA |
| ACATCCAGAACATTTCTAACAACAAGACAGAGAGAAGCACAGCGCCTCTG*GGTCGGGATTACTGTTCGGAAGAAGAGGATATCACATAGCACCAATTTTA | TCGA |
| CCCCCCTCAGTTTCTGAAATGATTCTCCAGAATTTCTCCTCATAAAAAAG*GGGTACACATCTTTCTGGAATGACTGCATATCATCAGGCCTGCGAGGGGG | TCGA |
| CGGGCAGATCTGGCCCAACGATGGCGAGGGCGCCTTCCATGGAGACGCAG*AAGCCCTTCAGCGGCCAGTAGCATCTGACTTTGAGCCTCAGGGTCTGAGT | TCGA |
| ACTCCGAAAGGGCCCACCTTGCTCCACGGTCATCATATGTCAACATGCAG*GAATACTTGGACCTCAGCCAACCTCTCGAACAGTATTCACCTAGTTACCC | TCGA |
| CTGCTTGTAGAATATGACTTACCAGCACTCCTGGACCAAGAGCTCTTTGA*ATGTGTGTCTCGGAAGCTCCTTTACACATGGAAGGCACAAGTTGTCCCCA | TCGA |
| TCAGTCCTGCCTCATCCCAGCCATGTCATCTTATCTACGAAATGATTCAG*GATTCAGGTGTTCATTTATGTATTATTGATGACTCCAATGAGCATATGCT | TCGA |
| TACTTGTGTGTCAACTAGTGATGGAACCAGAATAGAAAGGGATGATGCAA*GCCTCTCTATCCGGAGCGTGTGCTGGAAAGCAGACCGCCTTCTAGCAGGG | TCGA |
| CCTCTAGTGTGTTTGTACGCTGTGATGAGGAGCGACTTGATATCATGAAG*GAGCTGTAGTCTGGAAGATAAAGAGCCTTTGGTACCGCAAGCCCTGTCAT | TCGA |
| TTCTGTGATTTACTGTTCTGGCACAGACAGGCTGTGTGCATGCACCAAAG*GTGTTTGCAAGAGCAGACCCTTGGACCTGGTGTTTATCATTGATAGTTCT | TCGA |
| AGTCATCGACGGCACCTCGGGGGCCACACTGCAGGCCTCCGCGCTCAGTG*GGGACAATTCATCTCTTTCGAAAACCACAAAGATCCTTTTTTGGCAAGTT | TCGA |
| TCAGTCCTGCCTCATCCCAGCCATGTCATCTTATCTACGAAATGATTCAG*TTTAATGACATGTTTAATAAGTTACTGGGTAACATTGAGGAAACCTAGCC | TCGA |
| ACTGGAGGATTCAGACAGTGAAGAGCATTCCAGATCAGATTCTGTGACAG*AAAGTGGAGCTGTTCCAAAAAGAAAAGACCCCTTAACACACACTAGTAAT | TCGA |
| ACAAGTGACACTGCTGAAAAATAAAGCTCCAGGATTAGGGAAAGTCAATG*AGAAGATGCAACCAGTTTTGCCATTTTCCAAGTCACAAACGGACGTCTAT | TCGA |
| TCAGTCCTGCCTCATCCCAGCCATGTCATCTTATCTACGAAATGATTCAG*AACTAAAACCAGAACAGGTGAAGAATATGTCTGCCAGTGAGGTATAGTAT | TCGA |
| AGTCATCGACGGCACCTCGGGGGCCACACTGCAGGCCTCCGCGCTCAGTG*ATCTCAGTGGTTCAATAGCATCCCCAGATGTCAAATTAAATCTTGGTGGA | TCGA |
| ACAAGGGTTATTCACTTGCTTCACTTTTGGCTAAAGTTGCAGCAGGCAAG*ATCTCAGTGGTTCAATAGCATCCCCAGATGTCAAATTAAATCTTGGTGGA | TCGA |
| CATCAGCTCCCAATGTGCATATAAACACAATAGAACCTGTCAATATTGAT*GTCAGTACTCCTGCATTTGTACCAACGAAAACTCACGTGCTGCTTCATCG | TCGA |
| AATTGCATGTGGAAGTGTTGGAGAATGTTCCACTTACAACACACAACTTT*ATCTGGGACATGAGCGATGATGAGACCATCTGAGATGGCCCCGCTGTGGG | TCGA |
| gcgccggcgccgcggccTCTTCGGCTGCGGACCCTGCCATTCCGGAGGAG*GTATTACACAATGGCCTCGTTATGGAGATGGAGCATCCAACTGTGGGGAA | TCGA |
| TATTGGACAAATTTGGTGGGGAGCATAATCCACCATCAATATATCTGGAG*GTTTGCCTGCAATGAGATTTCATTCTCTACATTTAAAGGACATCCTTTCT | TCGA |
| CTCAGCGAGAAAGGAAGTCATCTTCATCCTCAGAAGACAGGAATCGAATG*GCAAGAAATACCGCCCACGAATGGATCAGGAAGTCTCTGACCAAGCCATT | TCGA |
| GCCCTAACATCTGGATCATCCCCTTCCGCACCCGCCTCGGACTCTATTGG*GTGTAGGAGCTGCACGATCTGGGAACCTCACGTTTATGGTGGGAGGAGTT | TCGA |
| CATCAGCTCCCAATGTGCATATAAACACAATAGAACCTGTCAATATTGAT*TATGAGCAAGCTGCCGATGCAGAACTAGCTTGGGTTGCTGAAACAAAACG | TCGA |
| CACAAAAACCTATCGTTAGAGTCTTCCTGCCCAACAAACAGAGGACAGTG*ACGCTGGTTGAGATTTTTCAAAATATTGGTGATACCAATCTTGCTGATTT | TCGA |
| cggcagcgggcggacggccggcgggggcggcgtgcggcCTAGCGTCTCAG*CTCATGATGGAGCAGTCCAAGAAGTCCTCCCTCATGGTGGCCCGCAGTAT | TCGA |
| cggcagcgggcggacggccggcgggggcggcgtgcggcCTAGCGTCTCAG*GATATGGAACTTGGAACTCTGGGACAAATAGAGGCAAGTGTCATTCCTGT | TCGA |
| cggcagcgggcggacggccggcgggggcggcgtgcggcCTAGCGTCTCAG*GACTTAGTAGCCATGAAGGTCCTCATCCTCGCCTGCCTGGTGGCTCTTGC | TCGA |
| cggcagcgggcggacggccggcgggggcggcgtgcggcCTAGCGTCTCAG*ATACTATGGATAATTGGGATGAGAAAAAGCTGGAAGAAGTAGTGAACAAG | TCGA |
| GGGCAAGAAGCACAAGAAGCACAAGGCCGAGTGGCGCTCGTCCTACGAGG*GGGTTGGCTGTGTTCCGGCCGCAGAGCACCGTCTGCGTGAGGAGATCCTG | TCGA |

| | |
|---|---|
| GGTGGAGCCGCCCCCAGATCGGCCAGTCCGAGCGTGCCGGACACAGCCAG*GGGTTGGCTGTGTTCCGGCCGCAGAGCACCGTCTGCGTGAGGAGATCCTG | TCGA |
| GCACTGGCCTGGTGGGCATGCTTACTCACCTGGATGACCTCCAGCTGATT*GCTATTCAAGATCTCTGGCAGTGGAGGAAGTCTCTTTAAGAAAATAGTTT | TCGA |
| AAGGGGGATCTGATGCACGACCCAGCAATGGATGAAGAGCTGGAACGGCT*GAAGGCTGAAGTTACATTGGATGGAGTTTGGCCAACAGATAAAACATCTC | TCGA |
| TGGTCTTCGGGCTGTCGCTCGTCTACTTCCTCAGCAGCACCTTCAAGCAG*GATCTTGATGCTGGTGTAAGTGAACATTCAGGTGATTGGTTGGATCAGGA | TCGA |
| CTGCCAGGCGCCATCATCGGCATAGCCCGGGAGATGCAGCCAGTTTACAG*ATTTCAGAGACAACTGTCTGAACCCTGCCACCCCTTCCCTCCTCAGCCAG | TCGA |
| TTAATATGCCAGAAAAAGAAAGAAAAGGAGTTAGTAACTACCGTTCAGAG*TACTCTTCCAACCCAAGAGGAGATTGAAAATCTTCCTGCCTTCCCTCGGG | TCGA |
| AAGAACTTTGTCTGGATCTGCAGCTCTATGGGCAGAAAGTGAAAAATCAG*ATCAGTTTCCTAATTCATCTCAGAACGGTTCATGCCGACAAGTGCAGTAT | TCGA |
| GGGGGGAAGGCATCCGGACATTGCTGTGGACTTTTATTAAGTCCTTTCAG*ATCTGGTACCAAAGGAGAATTTGGAGAGCTGGCTAAATTATTTGAAGAAA | TCGA |
| TTTACATCCCATTCCCTGAAGATGATTCTAATTCAACAAATCATAACTTG*TTCCAGAAAGAACGCTTCAACATCGACATGCCGCACCGCTTCAAGGTTCA | TCGA |
| AATGTAAGAACTACAAACAAAATTTCTATTAAATTAAATTTTGTGTCTCC*AAAAAAGGCGTAAAGCGGAAAGCAGATACTACCACCCCTACACCTACAGC | TCGA |
| AATGTAAGAACTACAAACAAAATTTCTATTAAATTAAATTTTGTGTCTCC*aaaaaaaaGCTAAAGCAGAAAACAAAAGTGGCCCATGGTCTCAGCTTCCA | TCGA |
| AATGTAAGAACTACAAACAAAATTTCTATTAAATTAAATTTTGTGTCTCC*aaaaaaaaaaaaaaaaaaggctgaggtgagaggatcactagagcctgagag | TCGA |
| AAAATTCATCATTCATGCCCCTCCTGGAGAATTTAATGAGGTTTTCAATG*ATCAGTTTCCTAATTCATCTCAGAACGGTTCATGCCGACAAGTGCAGTAT | TCGA |
| TTACAATCACTCCTTCAACCACTCAAGTGGTTGGCATCTTGAAAATTCAG*GGAAATGATATTGACCCTGAAGCAGTTAAAGGTGAAGTGTTAAAAGTTGG | TCGA |
| CGTGGAGGAGGCGCCCCGCGCAGCTGGCTGGCGGAGCATGAGCGCCCCAG*ACCCGACCGCATGTCTGTCATGGACCTCGCCAATACTTGCTCCAGCTTTC | TCGA |
| CAGGTGGAATTATGCCCTGTTGGTTCCAATGCTGGGATTGGCTGCTTTTC*GGGTTGGCTGTGTTCCGGCCGCAGAGCACCGTCTGCGTGAGGAGATCCTG | TCGA |
| GAGCGCCTGCGACCCCAGTCGGTGCTCCGGGGAGGTCACCTGACGAGGAG*atggtgtcaggaatgggatctaaagtggagcttctagcgacccccaggag | TCGA |
| GAGCGCCTGCGACCCCAGTCGGTGCTCCGGGGAGGTCACCTGACGAGGAG*GATGGAGAGGTGTACTGCATAGATGCCCGTTACTATGGCAACATCAGCCG | TCGA |
| GAGCGCCTGCGACCCCAGTCGGTGCTCCGGGGAGGTCACCTGACGAGGAG*TGCTTTCTCCTGCTCAAGACAGCCACACGAACACGGATCTCCCCCCAGTG | TCGA |
| GAGCGCCTGCGACCCCAGTCGGTGCTCCGGGGAGGTCACCTGACGAGGAG*GACAACACCATCGACTTCCTGGAGTACGTGGCAGCTCTGAATCTCGTGCT | TCGA |
| GAGCGCCTGCGACCCCAGTCGGTGCTCCGGGGAGGTCACCTGACGAGGAG*GAGGAAGAAGGGGAAGACTGTACACTCAGTGTCTACAGGCAACTGCACAA | TCGA |
| GAGCGCCTGCGACCCCAGTCGGTGCTCCGGGGAGGTCACCTGACGAGGAG*GCCCTGGCTGTGGAGAATGCTGAAGGAAGAAGACGCAGAAGCAGGACGAC | TCGA |
| GAGCGCCTGCGACCCCAGTCGGTGCTCCGGGGAGGTCACCTGACGAGGAG*GTTGCTCTGGAAATGCTGCTACTGCGGAAGCCTGTCAGGTTAAAATGTCA | TCGA |
| GAGCGCCTGCGACCCCAGTCGGTGCTCCGGGGAGGTCACCTGACGAGGAG*GGTGTAGATCTCCGTCACTATTCAAAGCAAGTTGAGCTGGAGCTACAGCA | TCGA |
| GGACACGGGCTGCAACCTGCAGCACATCAGCGACCGGGAGAACATAGACG*GATCAACCACAGGTTTGTCTGCTACCCCCCCTGCCTCATTACCTGGCTCA | TCGA |
| CATCATTGCTGGCTCCCTCTTAGATTCTTGTACCAAGCTTCTTCAGAAAG*ATCATGGAGACTTACTTGCTAGCTGTCCAGAAGATGAGGCTCTCACTCCA | TCGA |
| AGCTTTCAGGGCCATGGTTCACATTAGAATAAAAGGTAGTAATTAGAACA*aaaaaaaaaaaaaaaaaaaaaaGACTAAAGTACATGGTTTCTTCAAAGCT | TCGA |
| AGCTTTCAGGGCCATGGTTCACATTAGAATAAAAGGTAGTAATTAGAACA*AAAAAAAAAAAGTGATGAGTTGTGAGGCAGGTCGCGGCCCTACTGCCTCA | TCGA |
| AGCTTTCAGGGCCATGGTTCACATTAGAATAAAAGGTAGTAATTAGAACA*aaaaaaaaaaaaaaaaaaaaaaaaaattaagtgggtgtggtggtgcatgcc | TCGA |
| AGCTTTCAGGGCCATGGTTCACATTAGAATAAAAGGTAGTAATTAGAACA*AAAAAAAAGGCTGTTCTGGATGAGGTAAGCGAGGTTTATTTCTTGTTTCA | TCGA |
| AGCTTTCAGGGCCATGGTTCACATTAGAATAAAAGGTAGTAATTAGAACA*aaaaaaaaaaaaaTGCTTACTAAGTGGCAGTGGTATCTATGATGCTATAG | TCGA |
| AGCTTTCAGGGCCATGGTTCACATTAGAATAAAAGGTAGTAATTAGAACA*AAAAAAAAAAGCCATTTAAAGCCAGCCACTAGAGGGAGTCAGTTCAGTTC | TCGA |
| AGCTTTCAGGGCCATGGTTCACATTAGAATAAAAGGTAGTAATTAGAACA*aaaaaaaaaaCCCAAAATAACACAGGACATCCCAGTGTGCAGTTCGAAGG | TCGA |
| TATGCAAATTTCACAGAGCCTCAGTTTTATTTGCAAATGAAAAATATAAG*GACTTGATTAGAGACCAAGGATTTCGTGGTGATGGAGGTAAGTAGTGATT | TCGA |
| CAGAGAAGCCGGAAGTCCATATCGGGTACAAGTGGTGGCCACAGAAGTAG*TTCATTCGGCTCCTAAAGAAGTAGCAGTGTCCAAGGAACAAGAAGAGAAA | TCGA |
| CAACATCACCTTAACCGATGTCAATGACAACCCACCACGCTTTCCTCAAA*GGGTTGGCTGTGTTCCGGCCGCAGAGCACCGTCTGCGTGAGGAGATCCTG | TCGA |
| CTTACTTGGATAAAGTTCCAGAGCCTGGAGTGCCCACTGAAACCATAAAG*AATGTGAAAGATGTTGCAGAAGTGTTCCAGAAGTGGCTGAAGATAGAAGG | TCGA |
| GTGGCGGTGCTGAGGCACCTGGAGACCTTCGAGCACCCCAACGTGGTCAG*GACTTTTCTTGGTAGAGATGGAAATCCAAATAAACGGAATGTGCACAATG | TCGA |
| CTTACTTGGATAAAGTTCCAGAGCCTGGAGTGCCCACTGAAACCATAAAG*ATCCCAAGCAGGTCTGTTCCAGCCTCTGATTGTGTATCGGGGCAAGATTT | TCGA |
| GTGGCGGTGCTGAGGCACCTGGAGACCTTCGAGCACCCCAACGTGGTCAG*CTTGCCATGAAGCACAAATGTTGAGAAACTGCCTATCCTGGTGACTCTTC | TCGA |
| TAGAGAATCCCAACAGTGAGGAAGTGGAAGCAAGAAAAATGATTGACAAG*CTGTTTTTCCATTTGTCGAGAGAGCGGGTGTTCTCTGAGGACCGCACACG | TCGA |
| GTGGCAATTTCTTAGAGATCAAGAATAAATTTCTAGCTCGAAGGTTTAAG*GTGACTATAAACGCAGCCCCTGTGATGCCAACTGGACATATTATGGAGAT | TCGA |
| TGGTGGAATCACTGGGAACATGCGGCAAGAGGCCATTGACCGCTTCAATG*ACTGTTGGGCGGGAGCTGCTGCTCCATCTGATTGACTATCTCGTAACCAG | TCGA |

| | |
|---|---|
| GGCAGAGGTGAAGGAGAGAACAGAGGAACCTATGGAGACAGAGCCCAAAG*GCACCATGCGGCCCGTGCGGCGCAACTTCTACGACCCGTCGTCGGCGCCG | TCGA |
| AGCTCCACCCATCACCCCAGAACACCCCGCAGAAAGTGCCTGTGCATCAG*TCTAACAAAGTGCCAGTGGTGCAGCACCCTCACCATGTCCACCCCCTCAC | TCGA |
| GCCGGAAGGGGTCCCGAAGCCCCCTGCAGGAACTCTACAACCCAATCCAG*GATGATCCGGATGAAGCGAACCAGAACTACTTAGCAGATGAAGAGGAGGA | TCGA |
| GCTGGCCACTTGCTTGGGATCCTTTATCCTGGTCATCTTCTATTTCCAAA*CTACGAATCAGTGGTTTCTGCCAGCTGTTAGAGGAGATATCCCTCCAGGC | TCGA |
| GCCGGAAGGGGTCCCGAAGCCCCCTGCAGGAACTCTACAACCCAATCCAG*ACACTGCAAATGGATGTAAACAAGCTGAACATAACCTTGCTTCGGATCTT | TCGA |
| GCCGGAAGGGGTCCCGAAGCCCCCTGCAGGAACTCTACAACCCAATCCAG*ATCATAAGGCAGCAGTTTCCTCAGTTAACCACCAGAAGACTCGGGACCCG | TCGA |
| GCCGGAAGGGGTCCCGAAGCCCCCTGCAGGAACTCTACAACCCAATCCAG*GAACGTGTTGTGGGCTTTCACGTACTGGGTCCAAATGCTGGAGAAGTTAC | TCGA |
| GAGAAACACAGGAGGAGGAGGATGAGATTCTTCCAAGGAAAGACTATGAG*GATGCAATTCGAAGTCACAGCGAATCAGGTACTTTTCCATAGTCATTTAG | TCGA |
| GCCCAGCCGGCACCCAATGCCTGGCAGGTCATCAAAGGTGTGCTGTTTAG*CTACCTCTACCCAGATGGAAAGACCCACAACCCAGACCTGACTGAGCTCG | TCGA |
| CTTCACAGTCTTCCTGTAAGGACTTAAAAAGACTTTTCTCCGGAACACAG*GTTATGGTTTAACTCAGCAGAATTTGTTGAACAACTACGACATGCTGGGG | TCGA |
| TGAGCGAGCTGTCAGAGCCCGGGGACGGCGAGGCCCTCATGTACCACACG*AGTGAAGTGGCCCAAAAGGCTGCCCAATCTCTGAGAAGCCTGCATGGCAC | TCGA |
| TGAGCGAGCTGTCAGAGCCCGGGGACGGCGAGGCCCTCATGTACCACACG*GTTAACAATGAGAATGTTGTCAAAGTCGGCCACAGGCAGGTGGTGAACAT | TCGA |
| CGGCCTTCGAGGAGGTCATGAAGGACCTGAGCCTGACGCGGGCCGCGCGG*GTCAGTGCAGGAGCCCAGCCGCTGAGCCATGCCGGGCCCCGGGCGGCCTG | TCGA |
| GCAGCCAGAATCCTTTAGCAGTGGGTCCCAGGACGTGCACCAGAAAAGAG*GGGCTGGAGCTCTTTTGTGCTATGAAGCAACAGCCTCAAGATTCAGAGCT | TCGA |
| AAATCCGGACGCTAACAGTGGAAAGGCTGTTGGAGCCACTTGTTACACAG*TCCTAGGGGACTACAGAAGGAAAAAGACAAGAGGCAGTAGGATATCTGTG | TCGA |
| ACAGCAATCAGCTGGCCTGGTTTGATACTGACCTGTAAATCATCCTTTAG*GGGATCTTTCCTGCAAATTACATTCACTTGAAAAAGGCAATTGTCAGTAA | TCGA |
| AGAAAATGCGGTTATAAAAAATGGTTCAGAATTAAACTTTTAATTCATTC*AAAAAAAAGGCTGTTCTGGATGAGGTAAGCGAGGTTTATTTCTTGTTTCA | TCGA |
| AGAAAATGCGGTTATAAAAAATGGTTCAGAATTAAACTTTTAATTCATTC*GAAAAAAAAAAAAGTACCGCTCCAGAGCAGGAGCCTAGGCAGCCGAGAGG | TCGA |
| AGAAAATGCGGTTATAAAAAATGGTTCAGAATTAAACTTTTAATTCATTC*GaaaaaaagacaaagaaaaaaaagaaaagaagaagaaagcaaaaaaGGCC | TCGA |
| AGGTCGAGGACGGTCGGACTCCCGCGGCGGGAGGAGCCTGTTCCCCTGAG*GAGATGTCAGTGAAAAAAGGGATCCAGAATGATTACTAACCTATGACTCC | TCGA |
| CTGGAGCGATGCAAGCTCGGATCCGAGACTGGCAGCATGAGCAGCATGAG*TCCACCTAAACCCACTGTGTTCATCTCTGGGGTCATCGCCCGGGTAAGTC | TCGA |
| GCTCTGAAACAGAAACCACCTCTGCCATCCTCGCCTCAGTCAAAGAACAG*GTGAATCAACGGAATGTACCAGGTATTGACAGTGCATACCTGGCCATGGA | TCGA |
| CGGCGGAGGGTGCATGTTTGCGAGGAAGCCGCCGGGCGCCGCGCCTTTGG*ACACTTCCCTCTGTGACCATGAAACTCTGGGTGTCTGCATTGCTGATGGC | TCGA |
| GCTCTGAAACAGAAACCACCTCTGCCATCCTCGCCTCAGTCAAAGAACAG*GTGACAGTGAAGCTCCCTCTGATGAAGATCAACTTTGACATGAGCTCCCT | TCGA |
| CACTGCGCTGCGGAACATGGCCTTGGACGTCAGAAATAAGGAGCTCATCG*GGTGGAGGGTCCGGGGAGGAGCAACTCTATGCTGACTTTCCAGAACTTGA | TCGA |
| CAGTAGCCACCGCATCTGGAGAACCAGCGGTTACCATGGAGGGGATCAGT*AACTCCCCCACTGGAAAGGATTCTGAAAGAAATGAAGTCAGCCCTCAGAA | TCGA |
| AGCTGTTTTTGAGGATCCTGCTCCACAGAGAATGCTCTGCACCCGTTGAT*GGATTCACCTACGCTCTTACCAGCTGGAGGGAGTAAACTGGCTCGCCCAG | TCGA |
| AGCTGTTTTTGAGGATCCTGCTCCACAGAGAATGCTCTGCACCCGTTGAT*ACCACTGTGTGCAGCATAACGAAGCGTCCAGACTTCTCCGAAACTGGTCA | TCGA |
| CAATAAATGAAGGCAATGCATGGCCTTCCAATAAAAAATATTTGAAGACT*GAAAAAAATTTACCACCGATGATTCCATTTGTGTGCTGGGAATAAGCAAA | TCGA |
| AGCAGCAGTGGCAGCCAAGAAATTTGGACCTAAATCATCTAAAAAGTTTG*CTAAAGCTGTTGATGGATACGTGAAGCCACAGATCAAGCAAGTGGTCCCT | TCGA |
| TTTCTCAACTGGGTTATTCTTGCTTCTATATTCATGCTAAAATGAGGCAG*GTGGAGCACCATGGAAGGCGACTGTCTGAGCTGCATGAAGTATCTGATGT | TCGA |
| cgaggaggaagaggacgaggacgacgaggaggaggaggaggaggaAAAAG*GGTTTATGGTCCAGCAGAGTCTCAGAGCGCGGTCTTTGGAGATGTGTGCC | TCGA |
| AAAACAAGGAAGAGTCTTCAGATGATGAAGATAAAGAAAGTGAAGAGGAG*GTGTTTGAAAAACTCCAACTCTTCTATCCAGAGTGCCAACCAGAGCAGAT | TCGA |
| CCGGCTGTGGTTTGGCTGCGGGCGGCTTGGGCAGCCCGCGGGCGCCTCAG*ATGCCGGAGGCCCCGCCTctgctgttggcagctgtgttgctgggcctggt | TCGA |
| ACATCCTCAGCAAGCTCCAAGAGGAGATGAAGAGGCTAGCTGAGGAAAGG*GTATCCAAGCCCACCAATGGGATCTGTCTCAGCACCCAACCTGCCTACAG | TCGA |
| TTGTTAAACAGACCTCTGAAGAGAAGCCCAAAGACAGTGAAGAACTCGAG*AGGCAGAAAGATAATCTCGAACTCCCGTCAGCAGAGCACCAGGCCATCGA | TCGA |
| TTACGGATCTGGAACTGCAAGCAGCAAAAAACTTGCGAAGAATAAAGCTG*ATTCCCCTGATGATAATGAAACCTGATGAGAAAGCCGAAGTACCAACAGG | TCGA |
| CCGGCTGTGGTTTGGCTGCGGGCGGCTTGGGCAGCCCGCGGGCGCCTCAG*GTGAAGCAGATCATGGAGGAGGCTGTGACACGCAAGTTTGTCCACGAAGA | TCGA |
| GACCTAGGGGAGGATTACCCCTCTGGCAAGAAGCGTGCGGGGACCGATGG*TGCCTATGATCAGAAGCCACAAGTGGGAATGAGGCCCTCCAACCCCCCCA | TCGA |
| GCGTCTCATACAAACCATGCGGCCTCCAGCGAAGCCTTCCACTAGCAAAG*GGTTTGGCTTTGTCACTTTTGAGAATGAAGATGTTGTGGAGAAAGTCTGT | TCGA |
| GCCCTGCCGCTAAAGAAGGGTCCCTTGCTGTGGGAGACAGGATCGTTGCG*GAGGATCCAAAGTGGGAATTCCCTCGGAAGAACTTGGTTCTTGGAAAAAC | TCGA |
| TCAATGAGATCCCAGAACTGGAATCCTCCCAGGAAGAAATCCAAGGACTG*GCGGGATTAAGGACACTGCATGACATTGGGCCAGAAATCCGGCGTGCTAT | TCGA |
| ggccgagcgcggcccTGCAGGCGATGTCGCCGAAACCGGGCTGCAGGCGG*GATCAACCACAGGTTTGTCTGCTACCCCCCCTGCCTCATTACCTGGCTCA | TCGA |

| | |
|---|---|
| CACCTACAGAACAGCATGCAAGCTTAGGTTTGTTCAGAAGAAATGCAATT*ACCCCAACTGTAAACTTGAAGACAAGACTGAAGATGGAGAGGCACTAGAT | TCGA |
| CTGGAGAGTACCAGCCTGGGGGACTCGGACGAGGAGGACACCATGAGCAG*CCTCCCAATATCACTGCAGGCCACACCAGCCACCCCAGCTACACTCTCTG | TCGA |
| GCTGCTGGCTGCGCTCTGCCCGGCGAGTCGGGCTCTGGAGGAAAAGAAAG*AGTTCACCGAACAGCAGAAAGAATTTCAAGCTACTGCTCGTAAATTTGCC | TCGA |
| GCTGCTGGCTGCGCTCTGCCCGGCGAGTCGGGCTCTGGAGGAAAAGAAAG*TAGTGAGTTGAAACTCTTGGAGGAAGCAACCATTTCAGTCTGCAGGTCAT | TCGA |
| GCTGCTGGCTGCGCTCTGCCCGGCGAGTCGGGCTCTGGAGGAAAAGAAAG*CTGACATAGCCCAGAGATACAGGATAAGCAAATACCCAACCCTCAAATTG | TCGA |
| GCCACATCGTTCGGAAGCGCACGCTGCGGAGGCTGCTGCAGGAGAGGGAG*ATACAGGTTTGACCCCCGTCTCATGTTCAGCAATCGCGGCAGTGTCAGGA | TCGA |
| GCTGCTGGCTGCGCTCTGCCCGGCGAGTCGGGCTCTGGAGGAAAAGAAAG*AGGATGATGTGGACCTGGAAGCCCTGGTGAACGATATGAATGCATCCCTG | TCGA |
| GCTGCTGGCTGCGCTCTGCCCGGCGAGTCGGGCTCTGGAGGAAAAGAAAG*GTGAAGCTTTGTTCTAAGAATATTTGTTTCATCTAGTTTATGAGTCCAAA | TCGA |
| GCTGCTGGCTGCGCTCTGCCCGGCGAGTCGGGCTCTGGAGGAAAAGAAAG*GAATCCGAGGGTGTTTCCTGCCACTATTGGTCGCTGTTTGACGGGCACGC | TCGA |
| GCTGCTGGCTGCGCTCTGCCCGGCGAGTCGGGCTCTGGAGGAAAAGAAAG*TGGAAAATGAAAATCACTGTGACTTCGTTAAGCTCCGAGATATGCTTCTT | TCGA |
| GCTGCTGGCTGCGCTCTGCCCGGCGAGTCGGGCTCTGGAGGAAAAGAAAG*GTCGGAATGAACTGATCGCCCGCTACATCAAGCTGAGAACGGGGAAGACC | TCGA |
| ACATGAATTATGACAAACTCAGCCGAGCCCTCAGATACTATTATGTAAAG*CTTGTAAATGCCCCAGATATGAGCCAGCCCAGGCCCCGCTACGTGGTAGA | TCGA |
| ACATGAATTATGACAAACTCAGCCGAGCCCTCAGATACTATTATGTAAAG*GTCAGGCTGCCATCCGGGGGCTTGTGGCCGAAGGGCATCGTCTGGCTAAT | TCGA |
| TTGCTCCCCTAAGTCCAGCCAGACTGCAAGGTGCTAACACACTTTTCCAG*GTTCTGACTCCTCAGGAGCAAAAAACATAACCTGAAGAGGGAGGAAGTGG | TCGA |
| AATCTCTGAAGATCATGTGGCCTCAGTGAAAAAATCAGTCTCAAGTAAAG*TGTACCGCCGGAAGCACCAGGAGCTGCAAGCCATGCAGATGGAGCTGCAG | TCGA |
| GTGTTTCTTAAATTTCTTTTCTAGGCATAAAGATGTCATCATCAACCAAG*TGTACCGCCGGAAGCACCAGGAGCTGCAAGCCATGCAGATGGAGCTGCAG | TCGA |
| TATATGGAGCAAAACTACTGTAGAGCCCACACCTGGGAAAGGACCTAAAG*TGTACCGCCGGAAGCACCAGGAGCTGCAAGCCATGCAGATGGAGCTGCAG | TCGA |
| AGACAACAAGTATATAATGTCTAACTCGGGAGACTATGAAATATTGTACT*TGTACCGCCGGAAGCACCAGGAGCTGCAAGCCATGCAGATGGAGCTGCAG | TCGA |
| ATTTTTGTCTCAACCAGCAAAAATGTCAACTCGCGAAAAAAACAGCCAAG*TGATGGAAGGCCACGGGGAAGTGAATATTAAGCATTATCTAAACTGCAGT | TCGA |
| GTGTTTCTTAAATTTCTTTTCTAGGCATAAAGATGTCATCATCAACCAAG*GTCCCGTGGCTGTCATCAGTGGTGAGGAGGACTCAGCCAGCCCACTGCAC | TCGA |
| CATGTGCAAAGCAGGCTTCGAGGCCGTTGAGAATGGCACCGTCTGCCGAG*TCCCGGCCAGTGTGCAGCTGCACACGGCGGTGGAGATGCACCACTGGTGC | TCGA |
| TTCTCCACACCCACTTTGTCCATTTGCAAATATATTTTGGAAAACAGCTA*GGAAAAAAAAGGAGAGCAGCGGGTGGGGGCGAAAGCAGGGCCGAGGAGAG | TCGA |
| CCTGATGGGTTAATGAGCAAACTGAAGTGTTTTCCATGATCTTTTTTGAG*AACACAAACAGCTCCTAAAAAATGCGCAGCTGGAACTGAAGAAGAGTAAG | TCGA |
| GCCACCCTGAGTGTCAGCCCCAGAATGGCTCAGTGACCTGTTTTGGACCG*AAAGTGCAGCGGTTTTGAATGACTACAGTTTAACCAAATCTCATGAAATG | TCGA |
| CTGGGGGCTCCTCCTCGCCCTCTTGCCCCCCGGAGCCGCGAGCACCCAAG*GCACCATGCTGACCCGCCTGTTCAGCGAGCCCGGCCTTCTCTCGGACGTG | TCGA |
| TCCCGAGGGCCGGTATACATTCGGCGCCAGCTGTGTGACTGCCTGTCCCT*CTGTGCAGCGCATTGCTGAGTCTCACCTGCAGTCTATCAGCAATTTGAAT | TCGA |
| AGCTGTGTGGACCTGGATGACAAGGGCTGCCCCGCCGAGCAGAGAGCCAG*GTTTGGCCCTTACTACACTGAGCCAGTCATTGCCGGGTTGGACCCGAAGA | TCGA |
| CGGGCGCTGGGGGCATGGTCCACCACAGGCACCGCAGCTCATCTACCAGG*GTACTACATGAGGCCTGTTCTTGCGGCCCATGTGTTTTCTGGTGAAGAGG | TCGA |
| CTCAGCGTCTTCCAGAACCTGCAAGTAATCCGGGGACGAATTCTGCACAA*CTGTATCCAGCATGCTCCAAGGCCACAGCTCTGTGTTCCAGGCCTTGCTG | TCGA |
| TTCTCCACACCCACTTTGTCCATTTGCAAATATATTTTGGAAAACAGCTA*aaaaaaaaaaaaaaaaaaaaGCAAAGTAGAGATTAAGCTGCTTAGAAGA | TCGA |
| GCCACCCTGAGTGTCAGCCCCAGAATGGCTCAGTGACCTGTTTTGGACCG*CTGCTCCCTGCATGAGGAGGACACCCAGAGACATGAGACCTACCACCAGC | TCGA |
| CTTGCTTTTCAGCCTGGCCCGGGGCTCCGAGGTGGGCAACTCTCAGGCAG*TGTACCGGCTCCAGTTCAATGAGACCTTTGCGGAGATGAACCGTCGCTCC | TCGA |
| AGTTTTTTGTTTTGTTTTTATACGTGTCTGAATAAAAATGCCAAAGTTTT*TTTTAAAAAATTAGACCTGTTTATCGCGCTCTCAGTGCAAACTCGGCTCA | TCGA |
| CATCTGGATACATGCCCATGAACCAGGGTAATCTTGGGGAGTCTTGCCAG*GTCTGCTACAATGCTGCTGCTGCACAGAGCTGTGGTCCTCAGGCTCCAAC | TCGA |
| AGGACATGTTGGATGTGAAGGAGCGGAAGGTTAATGTTCTTCAGAAGAAG*GAGGATCCAAAGTGGGAATTCCCTCGGAAGAACTTGGTTCTTGGAAAAAC | TCGA |
| AGGCGCGACGACGGGAGGACAATCTCAACGACAGCTCTCAGCAGCTACAG*GAGGATCCAAAGTGGGAATTCCCTCGGAAGAACTTGGTTCTTGGAAAAAC | TCGA |
| ATGACCAGGCGGCTTGGGAGAATGAGCTGCAGAAGATGACCCGGGGGCAG*GAGGATCCAAAGTGGGAATTCCCTCGGAAGAACTTGGTTCTTGGAAAAAC | TCGA |
| TACTGAAGGACATGATTCAGACTGTCCCGGACCCAGCAGCTCATATCAAG*GTGGCTCAATCCCTTGTTTAAAATTGGCCATAAACGGAGATTAGAGGAAG | TCGA |
| TACTGAAGGACATGATTCAGACTGTCCCGGACCCAGCAGCTCATATCAAG*GGATGACATGCCAAGCTCGAAGCTCCTACATCACCAGTGAGATCCTGTGG | TCGA |
| TACTGAAGGACATGATTCAGACTGTCCCGGACCCAGCAGCTCATATCAAG*ATGCAGGAGCTGGAGCAGAGGCTGCTGGAGGCAGAGCAGAGAGCAGAGAA | TCGA |
| TACTGAAGGACATGATTCAGACTGTCCCGGACCCAGCAGCTCATATCAAG*TGGAAGGAATGAAGGCCAGCTGAATGGTGAAACAAATACACCCATTGAAG | TCGA |
| TACTGAAGGACATGATTCAGACTGTCCCGGACCCAGCAGCTCATATCAAG*AGATCGGGAAGTAAACAGTCCACTAACCCTGCCGATAACTATCATCTGGC | TCGA |
| TACTGAAGGACATGATTCAGACTGTCCCGGACCCAGCAGCTCATATCAAG*GTCATATTGAACATTCCAGATACCTATCATTACTCGATGCTGTTGATAAC | TCGA |

| | |
|---|---|
| TACTGAAGGACATGATTCAGACTGTCCCGGACCCAGCAGCTCATATCAAG*TTCGCCTCTACGGACCAAACTTCATCCTTCAGGTGTACTCATCTCAGAGG | TCGA |
| CCCTCCTCTGATGATTAAACAGGAACCCAGAGATTTTGCATATGACTCAG*GTTACTACTTTGGGATTGTGATGATCGAAGCTATAGCTACTACGTTGAGG | TCGA |
| GGTACCTGACAATGATGAGCAGTTTGTACCAGACTATCAGGCTGAAAGTT*CTTCACCATGGGTTGATAATAGTAGAACTCCAAATAAGATTGACCTTTAT | TCGA |
| GGCGGGGGTGGTGATCAAACAGGAACAGACGGACTTCGCCTACGACTCAG*AAGTTCCTGAGTGGGTGAACACCCCATCCACCTGCTGCCTGAAGTATTAT | TCGA |
| GGCGGGGGTGGTGATCAAACAGGAACAGACGGACTTCGCCTACGACTCAG*CTGCTCCCTGCATGAGGAGGACACCCAGAGACATGAGACCTACCACCAGC | TCGA |
| CTGCAGATGCCAAAGATGATGCCTGAAAACCAGTATCCATCAGAACAGAG*GATGAAGAACAAGCTGTGGTACTTTGAATTTGGCACCTCGGAGACTTTTG | TCGA |
| AACCTATGGGCAGACCGCCTATGCAACTTCTTATGGACAGCCTCCCACTG*ACTTCAGCTACGGGACGGACGAGTACGACGGAGAGGGGAATGAGGAGCAG | TCGA |
| ACGAAATGCACCTTTTAATCCAGCAGGCCCGAGAGATGCCCCTGCTCAAG*GCCTGGCATGCGTGGGCAGTGATGAACTTCGAAGCTGTGCTACACTACAA | TCGA |
| AGCTGTTTTCCCAGCTGCACGTTTCCTCCAAACCAGAGAAACTTGCCAGG*AAAACACTTGGTAGACGGGACTCGAGTGATGATTGGGAGATTCCTGATGG | TCGA |
| GCCCCAGCAGCACCCAGCCCGTGACCAGCACGACACCTTCTTCCTTCGAG*GGCACCAGCCCCAGAAGGTGGCCCGGCGCGTGTTCACCAACAGCCGGGAG | TCGA |
| CACAGCCACACTCTGCACCGCTAGGCCGTCCCCGACCTTGCCTGAACAAG*ATGATTTACATCATTACAATTGATGGACAACCTTACACTCTACATCTCGG | TCGA |
| GCCAGATGGCTGTGCACAAGCTGGCCAAGAGCATCCCTCTGCGCAGACAG*GAGGAGACATCACAGCTTTCCCAGATCGGGAGGAAAAATATGGAATGTGT | TCGA |
| GGAGAATGAATACGGGTCCATCAATCACACGTACCACCTGGATGTTGTGG*GCATGTGGGCACATTCCATGACAGTACAGGATTATCAGGATCTCATAGAC | TCGA |
| AGCAGTTGGTAGAAGACTTGGATCGAATTCTCACTCTCACAACCAATGAG*GCATTTGACTATGAACAGAAGAAGCTATTAGCCACCAAAGGTATGTAATA | TCGA |
| AGCAGTTGGTAGAAGACTTGGATCGAATTCTCACTCTCACAACCAATGAG*TTAGAATCAAGAATAGAAGAACTTAATAAAGAAGTTAAAGCTTCCAGAGA | TCGA |
| GCCAGCCGGCTGTGCACAAGCTGACCAAACGTATCCCCCTGCGGAGACAG*ATAATATCCCTGTTCACTTTTGCCGTTGGAGTCAATATCTGCTTAGGATT | TCGA |
| AGCAGTTGGTAGAAGACTTGGATCGAATTCTCACTCTCACAACCAATGAG*GGTGGGGAGAAACAGCGGGTCTTCACTGGTATTGTTACCAGCTTGCATGA | TCGA |
| AGCAGTTGGTAGAAGACTTGGATCGAATTCTCACTCTCACAACCAATGAG*ACACAACTTCGAAACCAGCTAATTCATGAGTTGATGCACCCTGTATTGAG | TCGA |
| AGCAGTTGGTAGAAGACTTGGATCGAATTCTCACTCTCACAACCAATGAG*AGAGAGGCTGCTCACCCAACAGACGTCTCCATCTCCAAAACAGCCTTGTA | TCGA |
| GGCCCGGCCCTCCTTCAGTTTAGTTGAGGATACCACATTAGAGCCAGAAG*AGTTGCTGGAGAATGTAACCCTAATCCATAAACCAGTGTCGTTGCAACCC | TCGA |
| AGCAGCTGGTGGAGGACCTGGACCGTGTCCTTACCGTGACGTCCACCGAC*GGCTCCTCGCACCTACCCCAGCAACTCAAATTCACCACCTCGGACTCCTG | TCGA |
| AGCAGCTGGTGGAGGACCTGGACCGTGTCCTTACCGTGACGTCCACCGAC*CAGATACTGGGGGCCATGGAGTCTCAGGTgggggggggcccggccggccc | TCGA |
| AGCAGCTGGTGGAGGACCTGGACCGTGTCCTTACCGTGACGTCCACCGAC*GTGATTGATGAGATTTATCGTGTGTTGAGATATGTCAATTCTACCAGAGC | TCGA |
| AGCAGCTGGTGGAGGACCTGGACCGTGTCCTTACCGTGACGTCCACCGAC*TTCTAGGCACGGGCGCGGAGGTGGATTACCTGGAGCAGTTTGGAACTTCC | TCGA |
| GGAGAACAAGTTTGGCAGCATCCGGCAGACGTACACGCTGGACGTGCTGG*AACGAAGAGTCACTGAAGAAGTGCGTGGAGGATTACCTGGCAAGGATCAC | TCGA |
| AGCAGCTGGTGGAGGACCTGGACCGTGTCCTTACCGTGACGTCCACCGAC*AACGAAGAGTCACTGAAGAAGTGCGTGGAGGATTACCTGGCAAGGATCAC | TCGA |
| AGCAGCTGGTGGAGGACCTGGACCGTGTCCTTACCGTGACGTCCACCGAC*GTGCCAGGCCCACCCCCAGGTGTTCCCGCGCCTGGGGGCCCACCCCTGTC | TCGA |
| AGATGACGAAGACGGGGAGGACGAGGCTGAGGACACAGGTGTGGACACAG*AACGAAGAGTCACTGAAGAAGTGCGTGGAGGATTACCTGGCAAGGATCAC | TCGA |
| AGCAGCTGGTGGAGGACCTGGACCGTGTCCTTACCGTGACGTCCACCGAC*GAGAGAGCCTTGAACTCTGCCAGCACCTCGCTTCCCACAAGCTGTCCAGG | TCGA |
| CTCCTCGGAGTCCTTGGGGACGGAGCAGCGCGTCGTGGGGCGAGCGGCAG*AACGAAGAGTCACTGAAGAAGTGCGTGGAGGATTACCTGGCAAGGATCAC | TCGA |
| CTCCTCGGAGTCCTTGGGGACGGAGCAGCGCGTCGTGGGGCGAGCGGCAG*GAGACAAGGAGAAGGCCTTAGGATCTACTGGGGGAGTCCGGAGGAGCAGT | TCGA |
| TGGGATTGCGGTCGCAGCTTCACTCATTGATGCCCATGCAGGAGGACTAG*ATTTCAAGACCCTTTGTTTTGTGCAAATCAACGTTATTGATATCAATGAT | TCGA |
| AATCTTACAATGGCAGGACCATTCTGGGAACCATCATGAATACGATCAAG*GAGGATCCAAAGTGGGAATTCCCTCGGAAGAACTTGGTTCTTGGAAAAAC | TCGA |
| CCCAGTGACCCCAGGCTTTTTATGGCTGTGAGACACGTTAAAATTTCAGG*AATGTCGTACTCTCAGGAGGCTCCACCATGTTCAGGGATTTCGGACGCCG | TCGA |
| TGGACATCGGGGCAGCTGACCTCGCCCACGCCCAGCAGCAGCAGCAACAG*GAAGTAACAGCTAGCAGTCGCCACTATGTTGACAGGCTATTTGACCCTGA | TCGA |
| ACCTTTTGAGGTGACTATAACTGAAGATTGCTTTACAGAAGCCAAAAAAG*ATCACTTTTTAATTCGGGCCTCTGCAGCATTAGAAAAATTGAAACTCCTG | TCGA |
| CATGCCTCTACCAATGGACAGCTCTCAGTCCACACTCCCAAAAGGGAAAG*GCATCTTGTACTGATGGGAGACTTTTCAATCATTTGGAGACTATTTGGCG | TCGA |
| ATTTGCTGTCAGCCATGAACGGATGGATGTGATGCCTGCTAGCCAAAAGG*ATCCCTCCCTGGTGCCACAGCAGCTGAGTGCGCATCAAATCTGAAGAAAA | TCGA |
| TGGACATCGGGGCAGCTGACCTCGCCCACGCCCAGCAGCAGCAGCAACAG*GGAGTGAGCCATAACTGGTGGCTGCTCTTGCGCCAATGAGCCTCCCCAAT | TCGA |
| ATTTGCTGTCAGCCATGAACGGATGGATGTGATGCCTGCTAGCCAAAAGG*GTATATTCATTTCAACCAACTAGAAATGCTACAGCCAGAGACCTTTGGAG | TCGA |
| ATTTGCTGTCAGCCATGAACGGATGGATGTGATGCCTGCTAGCCAAAAGG*ctgattttatctgatttcctcaagagtgcattccagaaggggaCCTGCTA | TCGA |
| ACCTTTTGAGGTGACTATAACTGAAGATTGCTTTACAGAAGCCAAAAAAG*CACCACTGCCGGAACTGTGGCCACATCTTCTGCAACACCTGCTCCAGCAA | TCGA |
| CATGCCTCTACCAATGGACAGCTCTCAGTCCACACTCCCAAAAGGGAAAG*GTGCCTCGAGCAAAGGCCTTATGCAACTACAGAGGGCAGAATCCCGGTGA | TCGA |

| | |
|---|---|
| ATTTGCTGTCAGCCATGAACGGATGGATGTGATGCCTGCTAGCCAAAAGG*AATTTCATATTATTGTATGTGGACTGGACTCTATCATCGCCAGAAGATGG | TCGA |
| TGACCCACCTGCATGTGAAGTCTACAGAACCCAAAGCCGCCCCTCAGCCC*TGCACTATATGTTGTGGATCTGAAGAAGTTTAGGAAAATAGCTGCTGGTG | TCGA |
| gtggAGGAGTTAACGACGCTTTCAAAGATGCACTGCAGAGAGCCCGGCAG*GGTTGAAAGGAGGAGCAGGAGGTACTGATGGACAAGGAGGTACCTTCCGG | TCGA |
| GGCCTTCCAGAAAGGTCCTGTATGTTAACTGGAACACCTGAATCTGTCCA*GATGGAGAACCTAATAAGGGGAAGGAATCCCCCGCAATACCAGAGAAGTC | TCGA |
| gtggAGGAGTTAACGACGCTTTCAAAGATGCACTGCAGAGAGCCCGGCAG*GAGTTGTACAGATGTTCTGTGCTGTATGATCTTCCTACTGTGTATTATTG | TCGA |
| CCAAACTGCTACTTATGTCCCGCAATGAAGAAGCTACCAAGCACCTAGAG*GTTGGCTCTGACTGTACCACCATCCACTACAACTACATGTGTAACAGTTC | TCGA |
| TTGATGTTCTTCAAAACATGGTGAAAACATGCAACTCTGAAGAATCAGGG*CCCTGGGCATTTCCAGCTCGAGAGTTCCTTCGAAAGAAGCTGATTGGGAA | TCGA |
| TCTGCTGCATAGTTTCGGTCATCTAAAGGAGGTTTTAAACAGTATAACAG*TGTACAAGTCCCCAGAGTATGAGTCCCTTGGTTTTGAGCTTACTGTGGAG | TCGA |
| TGCTGATCCACATGAGAGTCCACTCTGGGGAGAAGCCCAACAAGTGTACG*CTGGACAGTGCCACATCGCTTATCCAGGCAGCTAAAAACCTGATGAATGC | TCGA |
| AGATGGGCAAGCCCAGGAACGTGGCGCTCATCACCGGTATCACAGGCCAG*GACTGGGAGGTGCTGGTCCTAGGGAAGCTCAAGTGGGACCTGGCTGCTGT | TCGA |
| GCGGCGGGACAAGCGCGACGCCCGGCGCGAGCTCAAGCTGCTGCTGCTCG*GCGACATTGACTACAGCACCGTGCTCCTCGGCATGCTGGTGACGCAGGAC | TCGA |
| TAGTAGCGCTTAGTGAATATGATCAAGTTCTCGTGGAGTCAGACAATGAG*GGGAAAAGAGTTTCTTTTTACAAACGAGCCCAAATCCTTGTAGCAGATAC | TCGA |
| TTGTGCAGCattaaaaaaaaatcaaaataaaaattaaatGTGAGCAAAGAA*aaaaaaaatatattaataaaaaaGTTCAAAAAAGGGGGGGGATTGTCATC | TCGA |
| ATTGTGTTGTGCAGCattaaaaaaaaatcaaaataaaaattaaatGTGAGC*aaaaaaaaaTTAATAATTGTTTTTAAATCAGTAACAAGCCAAAGACCCAT | TCGA |
| ATTGTGTTGTGCAGCattaaaaaaaaatcaaaataaaaattaaatGTGAGC*aaaaaaaaaaaaaaaaaaaagaaagccaggtgtggtggtgcacgcctgta | TCGA |
| ATTGTGTTGTGCAGCattaaaaaaaaatcaaaataaaaattaaatGTGAGC*aaaaaaaaaaaaaaaaaaaaaaaaaaaaGGAGTGAAACTATCTCAAGGGTG | TCGA |
| ATTGTGTTGTGCAGCattaaaaaaaaatcaaaataaaaattaaatGTGAGC*aaaaaaaaaaaaaaaaaaaaaaaaaaaactagccgggcatggtggcgggtg | TCGA |
| TTGTGCAGCattaaaaaaaaatcaaaataaaaattaaatGTGAGCAAAGAA*AAAAAAAAAGAGGAGCGAGAGAAAGAAAAAAAGGGGGAAAAATCAGGATC | TCGA |
| TTGTGCAGCattaaaaaaaaatcaaaataaaaattaaatGTGAGCAAAGAA*AAAAATTCTGTAGGCCGTTGGAAGGAAGCTACCATACCAGGCCACTTAAA | TCGA |
| ATTGTGTTGTGCAGCattaaaaaaaaatcaaaataaaaattaaatGTGAGC*AAAGAAAATCAGAATCATTCCTATTCTCTGAAGTCGTCTGCTTCATTTAA | TCGA |
| ATTGTGTTGTGCAGCattaaaaaaaaatcaaaataaaaattaaatGTGAGC*AAAGAAAATGACAACTTCAACATTTCCAAAAAAGACATTGAAATTACTCT | TCGA |
| ATTGTGTTGTGCAGCattaaaaaaaaatcaaaataaaaattaaatGTGAGC*AAAGAAATACTCGACGCTTTTGACAAAATGTGCTCGAAGCTGCCGAAGTC | TCGA |
| ATTGTGTTGTGCAGCattaaaaaaaaatcaaaataaaaattaaatGTGAGC*AAAAAAAAAGAAACCTTTAAAAAAAAAACCTACACCTGTGCTATTACCAC | TCGA |
| ATTGTGTTGTGCAGCattaaaaaaaaatcaaaataaaaattaaatGTGAGC*AAAGAAAAAGATGACAAGTGGAAAAGAGGGAGCGAACCAGTTCCAGAAAA | TCGA |
| ATTGTGTTGTGCAGCattaaaaaaaaatcaaaataaaaattaaatGTGAGC*aaaaaaaaaaaaaaaaaaaaaaGATATGTACCAATCATGAAGAGTGGCCT | TCGA |
| ATTGTGTTGTGCAGCattaaaaaaaaatcaaaataaaaattaaatGTGAGC*AAAAAAAAAAGCCATTTAAAGCCAGCCACTAGAGGGAGTCAGTTCAGTTC | TCGA |
| ATTGTGTTGTGCAGCattaaaaaaaaatcaaaataaaaattaaatGTGAGC*aaaaaaaaaTCATTCTTTTTAGTCTTAGCACCTACTTAAGGATCCACTTT | TCGA |
| TTGTGCAGCattaaaaaaaaatcaaaataaaaattaaatGTGAGCAAAGAA*aaaaaaaaaTCATTCTTTTTAGTCTTAGCACCTACTTAAGGATCCACTTT | TCGA |
| AGCAGGAGGACAATGTTGATGAACAGTACCAGGAAGAGGCAGAAGAGGAG*ATCTTAGACGAATTTTACAATGTGAAGTTCTGCATAGATGCCAGTCAACC | TCGA |
| CGTGTCCAAAGGCTTCAACAAGAACGATGCCCCCCTCCACGAGATCAACG*AACTGGCCTATGATCTGGATGTGGATGATGCGCCTGGAAACAGTCAGCAG | TCGA |
| CGTCCATGCCTCCAAGCGCATTCTCTTCTCCATCGTCCATGACAAGTCAG*TGTACCGCCGGAAGCACCAGGAGCTGCAAGCCATGCAGATGGAGCTGCAG | TCGA |
| AGGTATCCTATGCCCTTGATGGATGAGACTAAAATAAAATTTTTGAAGTT*aaaaaaaaaaaaaaCCGGAAATGGGTCCTACCATCTTCTCGGAGCCGGAG | TCGA |
| AAGGCCAAGTCACTCACAACTGGGATGAAGTTGATCCTGATCCTAACCAG*ATGACAACTTTTGGAGCTTTCCTCCACAAAGGGGCCTTCTGCAGGAACTA | TCGA |
| GTTTCAGGTTTAATGCATTCAAAAGGACTAATACCATACTGCACCATTTG*ACTGGAAGGAAAGAAAAAGGAGATCCTCTCAACATTGCGATTGATAAGAT | TCGA |
| GGGGACCGATTCACCATGGAGGGCGCCGGCGGCGCGAACGACAAGAAAAA*GATTCAATTACTGGACTTGTCAACTCTGCCAGTGTACGTGCCATTTCTCT | TCGA |
| GGGGACCGATTCACCATGGAGGGCGCCGGCGGCGCGAACGACAAGAAAAA*CTACTCTCCAGAGAGACCGGATCTTCAAACATTTTACCAGGAAGCGCCAA | TCGA |
| AAATCTCATCCAAGAAGCCCTAACGTGTTATCTGTCGCTTTGAGTCAAAG*AGATCTGAAACTGGACAATGTCCTGTTGGACCACGAGGGTCACTGTAAAC | TCGA |
| GGGGACCGATTCACCATGGAGGGCGCCGGCGGCGCGAACGACAAGAAAAA*AGATCTGAAACTGGACAATGTCCTGTTGGACCACGAGGGTCACTGTAAAC | TCGA |
| GGGGACCGATTCACCATGGAGGGCGCCGGCGGCGCGAACGACAAGAAAAA*GTGGATCTCGAGCCAGAGGGGAAAGTATTTGTGGTAATAACCCTTACCGG | TCGA |
| GGGGACCGATTCACCATGGAGGGCGCCGGCGGCGCGAACGACAAGAAAAA*TGAAAATGCCTCGTGTAAAAGCAGCTCAAGCTGGAAGACAGAGCTCTGCA | TCGA |
| TGCATTAACTTCATCTGTTCCTACAACAAAAAAGAAAATGTTGAAAAAGG*TGATGAATCACCAACACCAGCAGCAGATGGCCCCCAGTACCCTGAGCCAG | TCGA |
| CTGGTTTCCTATTTGCAGTTACTTGAATAAAAAAAATATCCTTTTCTGGA*aaaaaaaaaaaaaaaaaaaaaaaaaaGGAGTGAAACTATCTCAAGGGTG | TCGA |
| CCCCAGCGCCTCAGAAGAGAGGACGCGGCCGCCCCAGGAAGCAGCAGCAA*AACTGAAAAGCTCCATCATTGGCAAGAAGAAGCCAGCAGCAGCTAAGAAA | TCGA |

| | |
|---|---|
| AAGCAGAAGCCACTGGAGAAAAACGGCCAAGAGGCAGACCTAGGAAATGG*GCTTCTAGTGAGGACGTTGCCAGCTCCCCGGAGCGCCACTACACGCCCTT | TCGA |
| AAGCAGAAGCCACTGGAGAAAAACGGCCAAGAGGCAGACCTAGGAAATGG*AAAACTCCAGAAGAGGAGCTGCAGCCAGAAAGCTCTCCTGCTGAAACTTC | TCGA |
| AAGCAGAAGCCACTGGAGAAAAACGGCCAAGAGGCAGACCTAGGAAATGG*GAGCCAGAGCCTCTGTACGCGCAGATCAACAGACCTAAGAAATAGCATCG | TCGA |
| AAGCAGAAGCCACTGGAGAAAAACGGCCAAGAGGCAGACCTAGGAAATGG*GTTAACAATTCATGCTGAGTGTCCCATGCATTTGGAAGATTTTCCCATGG | TCGA |
| TTTCTTTTCCTCCTTAGCCACAACAAGTTGTTCAGAAGAAGCCTGCTCAG*CTCTCTGGTGGATGTGAGCTGACAGTGGTCCTCCAGGACTTCAGTGCGGG | TCGA |
| AAGCAGAAGCCACTGGAGAAAAACGGCCAAGAGGCAGACCTAGGAAATGG*GTTATGACTTTTGTTCTGAAAGGCATAACTGCATGGAGAATTCCATCTGC | TCGA |
| AAGCAGAAGCCACTGGAGAAAAACGGCCAAGAGGCAGACCTAGGAAATGG*GTAGATCATGGCAACCATAGAAGAAATTGCACATCAAATTATTGAACAAC | TCGA |
| AAGCAGAAGCCACTGGAGAAAAACGGCCAAGAGGCAGACCTAGGAAATGG*GACCCTGATGCTCCCATAAGACAGAAAATGCCCCTTGATGATCTGGATAG | TCGA |
| AAGCAGAAGCCACTGGAGAAAAACGGCCAAGAGGCAGACCTAGGAAATGG*TCCCCCCCGAAGGGCGTGACCATCCCGTACCGGCCCAAGCCGTCCAGCTC | TCGA |
| AAGCAGAAGCCACTGGAGAAAAACGGCCAAGAGGCAGACCTAGGAAATGG*GAATTTGCTGCACTGACAAAAGAATTAAATGCCTGCAGGGAACAACTTCT | TCGA |
| TTTCTTTTCCTCCTTAGCCACAACAAGTTGTTCAGAAGAAGCCTGCTCAG*GTTCAAGTGCTGGATTGTGTCATGTGACCATCCCAAAACTCAGAGCACCC | TCGA |
| TGCCGCTCTGTACACCTGGTACGTCAGAAAGCAACGAGAGATCCTCCGAC*GTGAGACCGTGGAGCCGCCCCCGCCGGCGCAGCTGCACTTCATGTACGTG | TCGA |
| ATGGTCATAATGCAGAAGTAAGAAAGGCTTTGTCTAGACAAGAAATGCAG*AGGGACCACAGAGCACCAGCCTGCATGGAACTTCCTTCCTCACTCAGCTT | TCGA |
| GAGTTAAGAAGCTGTTGTACATTTATGATTTAATAAAATAATTCTAAAGG*AAAAAAAATGGGTGCTGATTTGCTTGGATCTGTTCAGGCAAAAGTTCAGA | TCGA |
| GAGTTAAGAAGCTGTTGTACATTTATGATTTAATAAAATAATTCTAAAGG*AAAATAAAAAATGACTTGAACTTGAAGAAGTCGTTGTTGGCCACTATGAA | TCGA |
| GAGTTAAGAAGCTGTTGTACATTTATGATTTAATAAAATAATTCTAAAGG*AAAAAAAAATATTGGTGCTTTGAAGTGTCACCTTTTAAAAAGTTGGCAAA | TCGA |
| GAGTTAAGAAGCTGTTGTACATTTATGATTTAATAAAATAATTCTAAAGG*AAAAAAAAAAAAAAACTACGGCGGCCGAGAAGGGGCGAGTACAGCCCAGT | TCGA |
| TTAAGAAGCTGTTGTACATTTATGATTTAATAAAATAATTCTAAAGGAAA*AAAAAAAAAAAAAAACTACGGCGGCCGAGAAGGGGCGAGTACAGCCCAGT | TCGA |
| AGGACTTCCCTTTGGATGTAGCAAGGAAGAAATTGTTCAGTTCTTCTCAG*GCAGTGGCCAGATCCAGCTTTGGCAGTTCCTCCTGGAGCTCCTGTCGGAC | TCGA |
| GATGCTTCAGCTCATCTTAGGCTGTGCTGTGAACTGTGAACAGAAGCAAG*CCTCTGCGCTTAGATACCTTCATGAAAACAGAATCATCCATCGGGATCTA | TCGA |
| GGGAGTGGGGAATGAGGCGGGGGTCGGCCCGGGGCTCGGAGAATGGGCAG*GTTACAGCACGGTCACCTTCGACGGGACGCCCAGCTACGGTCACACGCCC | TCGA |
| AGCGCCTTGGGACAGGGGGATTTGGAAATGTCATCCGATGGCACAATCAG*GCCGATGCTGCTACCAGCTTTCTGAGAGCAGCAAGATCAGGTAACTTGGA | TCGA |
| AGCGCCTTGGGACAGGGGGATTTGGAAATGTCATCCGATGGCACAATCAG*AATGGCTACACCCCTTTGCACATCGCTGCCAAGCAGAACCAGGTGGAGGT | TCGA |
| AGCGCCTTGGGACAGGGGGATTTGGAAATGTCATCCGATGGCACAATCAG*AATGGGTTGAATGGCTTGCATCTGGCTTCTAAGGAAGGCCATGTGAAAAT | TCGA |
| TGCATCACGGGCTTCCGGCCCTTCCTCCCCAACTGGCAGCCCGTGCAGTG*ACCTGTGTGCAGCATTTAACGTCATATGTGATAATGTGGGGAAAGATTGG | TCGA |
| AGCGCCTTGGGACAGGGGGATTTGGAAATGTCATCCGATGGCACAATCAG*GAAGAAAAACAGAAAAAGAAAATGGCTGTAAAGTTGGAACTAGCAAAATT | TCGA |
| AGCGCCTTGGGACAGGGGGATTTGGAAATGTCATCCGATGGCACAATCAG*GAACTGACAATGGCGAAGCCCTTCCCGAATCCATCCCATCAGCTCCTGGG | TCGA |
| GACTTGGCTAGTGCAAGCCTTGTTTATTTTCCTCACCACTGAATCTACAG*GACACCTCTGCACTGGGCTGCAGCTGCAGGGAAGGCCGAATGTGTCCAGT | TCGA |
| GACTTGGCTAGTGCAAGCCTTGTTTATTTTCCTCACCACTGAATCTACAG*AACTTGAAGCCACCATGGGAGATGAAGATTGGGAAGCAGAAATCAACCCT | TCGA |
| TGAAGGTGGGAAGGATGGTCCAGAATTCACTTTTACTACCCCAAAGTTTG*TCTGTGATTGCAATGGGAAGTCCAGGCAGTGTATCTTTGATCGGGAACTT | TCGA |
| ACATTCAGAGTCTTGAAACAGCAATCAAAAGGACTGAAGTGGGGTTAAGA*ATAAGGAGACAAGGAGGCATCCAGCTCCTGGTGGACCTGTTGGATCATCG | TCGA |
| CTCGGGGCCCTTCGAGAGCAAGTTTAAGAAGGAGCCGGCCCTGACTGCAG*ACACTAACAGCACATCTGGAGACCCGGTGGAGAAGAAGGACGAAACACCT | TCGA |
| ACTGTTTCTCTTCTGCAGAAAAAGAGGCTCTTCCTCCTCCTCCCGCGACG*GATGTCTCCCGTGCTAAGCTAGCAGCTATAATTCCGGACCCAGTTGTAGC | TCGA |
| CAAATGAAAGCCGAGTTGTAACTATCCATAAGCAAGATGGTAAAAATCTG*CCTCAGTTTTATGACCCTGTGGAGCCAGTGGACTTTGAAGGACTTCTGAT | TCGA |
| GTCTTCCATATAGATGAGTCAACCAGGCATAATGTACTCTACAGAATAAG*CGGAGCCCAGGGAGAATATGCTGGACTGGCCACTATCCGAGCCTACTTAA | TCGA |
| ACTGTTTCTCTTCTGCAGAAAAAGAGGCTCTTCCTCCTCCTCCCGCGACG*GTGCTTTGGCTCTAGCAGACAGGAAGAAATTCTGGCCCAGCTGGAAGTAG | TCGA |
| GACAAAGAATACTATAAAGTAAAAGAACCTGGTGAAAGTCCCATATTCTG*ATTGTGCGACGGGGAATGCCTCCCGGAGGAGGAGGCGAAGTGGTTTTCTC | TCGA |
| GACCTGGACACCTTATTAAAGAACTTTCTAAAGTAATTCGAGCAATAGAG*AAAACACTTGGTAGACGGGACTCGAGTGATGATTGGGAGATTCCTGATGG | TCGA |
| AACGTGCTGGAGAGACCCCGCGGCTGGGCGTTCATCTACCACGCTTTCGT*GTTCTGGCTGCAGATGGTCGCATGGTGGGGACAAGGATCCAGAGCCATCG | TCGA |
| ACGTCATCCGCTTCCCGCTCAATGGCTACTGCCGGCTCAACTCGGTACAG*GTAGAATGTAAGAAAGCTCAGCCGAAAGAAGTCATGTTCCCACCTGGGAC | TCGA |
| GGAAACTTCACCAAGAGCCTCGGCAGCTCCAGAGCGACGGGAAACGAAAG*ACTTACCTCCCTCAACCAGCCATTTGTCCTGACTCTGTGTATAAGCTGAT | TCGA |
| ATGTATACACACTGGATGAAGAGCTTCCCAAGAGAGTCAAATCTAGACTG*GCTTCAGCTCTGACAGCAGAGGGTGGAAGCTGCCTCATCTCAGGACTCCA | TCGA |
| ACAGTCTCTGAACGATGGGAAGAAAAGGCTAAGGTCTGCCTACAGGCAAG*GAAGCTAAAGAAACTATTTCGAAAGCTGAAAGATGAGACCGAAGCTGGAG | TCGA |

| | |
|---|---|
| AGTTAGTTACCATCATGAACCCCAACGTGCTAATGGAGCATGGTGTGCCT*CTGCGTGGTGGACGAGATGGACTTCTCCAGCATGGAGCTGGACGAGGCCC | TCGA |
| TGCAGGCCACACACCCACCCTCTGAAGACAGATTCTTGCATATCATGGAG*CCTGGAAGCTCCGACCCCAGGAGCCAGCCCATCAACCTGAACCATTACGC | TCGA |
| AACAACTTTTTAGTCTTCCGTGTGTCATCAGCCAAGCTCGGCAAGTAAAG*CCTGGAAGCTCCGACCCCAGGAGCCAGCCCATCAACCTGAACCATTACGC | TCGA |
| GTGGATTTCCGGTGAAGGATGGGCGGAGAAAGATTCTGCCAGAAGAAGAG*GATGGGAGCAAAGTGTTTACGGCATCGTGTGATAAAACTGCCAAAATGTG | TCGA |
| ACATATTGGAACACAAATTTCAAGTGATTCATATGGAAGCATAGATAAAA*GGGTTGGCTGTGTTCCGGCCGCAGAGCACCGTCTGCGTGAGGAGATCCTG | TCGA |
| AGTGGGGGTCCTTCTACAGCCCAGCCCAGACGGCCAACAATCCCTGCAGT*GACTTGATTAGAGACCAAGGATTTCGTGGTGATGGAGGTAAGTAGTGATT | TCGA |
| AGCAGTCAGGTCAAAGAATATGGCCAGAAGAGGGCATTCTGCACAGATTG*ATCTGGGCAGTGAATTAGTTCGCTACGATGCAAGAGTACACACTCCTCAT | TCGA |
| GCTTTTCTTACCAGGTGGCAAAGGGCATGGCTTTCCTCGCCTCCAAGAAT*CACTATTGGAAATAATTATGAAAAATGGAAGCACGGCTGGAAATAGCCCT | TCGA |
| ATTGTTGACAGTTCTGAAGAATTCTAATAAAATGTACATATATAAATCAA*aaaaaaaaaaaaaaaaaaaccccaaaaaaaaaacttttttatttatattct | TCGA |
| CGCCTGGGATTTTCTCTGCGTTCTGCTCCTACTGCTTCGCGTCCAGACAG*CGCGCAGGGACCATGTCGGCGGAGACCGCGAGCGGCCCCACAGAGGACCA | TCGA |
| GTCTAGCCAGAGACATCAAGAATGATTCTAATTATGTGGTTAAAGGAAAC*GATGGAGGATGAAGCTGTCCTGGACAGAGGGGCTTCCTTCCTCAAGCATG | TCGA |
| CGCCTGGGATTTTCTCTGCGTTCTGCTCCTACTGCTTCGCGTCCAGACAG*GAAATGAAAGCATTTGTCCAGATCTCTGGCAAAGATGGCACTGTGGTGTT | TCGA |
| TACAGCTAATTCAGAATCATTTTGTGGACGAATATGATCCAACAATAGAG*GGTCTAAGAAAAAGAAGACATCTGAGTCACAAAAGCAATAAGCAGTCAAC | TCGA |
| TACAGCTAATTCAGAATCATTTTGTGGACGAATATGATCCAACAATAGAG*CCCTTACAGTGAGGTTTCTTACTAAGCGATTCATTGGAGAATATGCTTCT | TCGA |
| TTTGGAGAAAGGCATTTATTTCTCTGAAGAAGGAGGTCATGTCGTCTTGG*ATGGAGTGGATGTTGAAGATGATCCGACTTGCTCTTGGCCAGCTTCCTCA | TCGA |
| ACATGAGGAAACAGCTTGATCGTCTGGGCCTGTGTTTCAGCTGGGATAGG*GATGGTACCTTGTTTGATGGTCGACCAATAGAGTCTCTGTCCCTGATAGA | TCGA |
| CCATGTTGAAAAATCTATCCAGGATTCATCCCGTGTCAACAATGGTAAAG*ACTGGAAGGAAAGAAAAAGGAGATCCTCTCAACATTGCGATTGATAAGAT | TCGA |
| ACTTCCAGCGCAGAGTGCCCTACAACTACCTGCAGCGGGCCTACATCAAG*CCACAGTACACGAACCTGGGGCTCCTGAACAGCATGGACCAGCAGATTCA | TCGA |
| TCATACTGACCCCAAAGAGTCAAACTTCTCAGATGTTAATACCAACCGTT*CTGGTTCCCGAGCCTCATACAGCAGCCAGCATGGGCACCTGGGCCCAGAG | TCGA |
| TCCGGCTGTAAGGAGCCGCGGCGGGGGGAAAATGGAGCCCTTCACCAATG*CCATCCCCGACGGCTTCACCATCGAATCCAAGCCATCCGAGGAGCTACTA | TCGA |
| GAAACTTTCCTCCTCCACCACCTCTTGATGAAGAGGCTTTCAAAGTACAG*GTCAGTTATGCCTCCTCCAGCAGACTCCTCAGCAACAAGAATCAATTCAA | TCGA |
| AGCCATTCTATGCTGTGGAAAAGAAAGCATACTGCGAGCCCTGCTACATT*GATCCAGCTGGGGAAAACCGTGCATTACCTGCCCATCCTGTTCATCGACC | TCGA |
| TCATTGTTCCACTGGACGCTCATCAGAGGGAAGATCTTTTTCCTCAATTG*ATTTGTGACAAAGAGTGGCACTACTATGTCATCAATGTGGAGTTTCCTGT | TCGA |
| GTTGGCTGAACCTTGTGTCAACCATCCATTCCAGGAGCCAGCTATCTGAG*TGTTCCTGGCTCCTCAGCAAGCACGGTCGCTGCTCCAGCGGGTCCGGCGA | TCGA |
| TCATTGTTCCACTGGACGCTCATCAGAGGGAAGATCTTTTTCCTCAATTG*CATTGACAAGATATCTAAAGTCACCTCTCCTGTGTTGGTCATTCATGGTA | TCGA |
| TCATTGTTCCACTGGACGCTCATCAGAGGGAAGATCTTTTTCCTCAATTG*TTGTATATGAACACAGATCAAGATTAGAGAAATCCTTGCAAAAAGAAAGA | TCGA |
| GAAACTTTCCTCCTCCACCACCTCTTGATGAAGAGGCTTTCAAAGTACAG*ATGGCGTTTAATGATTGCTTCAGTTTGAACTACCCTGGCAACCCCTGCCC | TCGA |
| TCATTGTTCCACTGGACGCTCATCAGAGGGAAGATCTTTTTCCTCAATTG*ACAGCGTGGAAGATGAACTGGAGATGGCCACCGTCAGGCATCGGCCTGAA | TCGA |
| ATATCACAGATCCTGTTTCAGCCCCCTGTGCGCCACCATTGCAGCCAAAG*CTTAACCAGCTCGAAAAAGCAGTGGAAGCAGCTCACACATTTTTCGTGGC | TCGA |
| CAAAAAGATGCTGTATGACATGGAAAATCCACCTGCTGACGAATACTTTG*TATAATATGCTTTTTCTTAGCTTTCCTATGACTCTACTTAGCAACTTTTG | TCGA |
| TTCCCGGGATAAGTAGCTTTAGCGATCGGGAGACAGCCGGGCGCTGCAAG*TTGGTTACCACAGAGAAGCGCTTCCTCAAAGACAGTTTGTACAATGAAGG | TCGA |
| TCATTGTTCCACTGGACGCTCATCAGAGGGAAGATCTTTTTCCTCAATTG*GTTTCAAGTGCCCCATTTGCTCCAAGTCTGTGGCTTCTGACGAGATGGAA | TCGA |
| GATGGAATGCCTACCGGGTTATAAGCGGGTTAACAACACCTTTTGCCAAG*GTCATATTCTTCAATCTCCATCAGCCAATGTGCTTCCAACCCTTCCTTTC | TCGA |
| CAAGTTGGTGAAAAGGCTTGGCGCTGGGCAGTTTGGGGAAGTCTGGATGG*GTCCCGTGGCTGTCATCAGTGGTGAGGAGGACTCAGCCAGCCCACTGCAC | TCGA |
| TTGGACAAAGGGTGGATGAAATTGATGCTGCTATTCAGAGATCACAACAG*GACTTGATTAGAGACCAAGGATTTCGTGGTGATGGAGGTAAGTAGTGATT | TCGA |
| CTTTGTCTGATAGTGAAATTGAGACCAACTCTGCCACAAGCACCATCTTT*ATTGGAGCTGGAGGGAGCATCACTGGGCTGAAGTTTAACCCTCTCAATAC | TCGA |
| CAGCCAGTGGTCACAGCTGGATGTTTGCGACATCCCAGAGAGCTTCCAGA*GACCCACTGATCTCAGCATGAAGAGACAATTGGCGACTAGCTCAGGATCC | TCGA |
| AGCCGCCCGCTTCAACCCCAGGGGACCAGAGGAACTCAGCCCTGATTGCG*AATGGAATGACTCCTCTAATGCATGCAGCATATAAAGGAAAACTCGATAT | TCGA |
| ATCAGCCCATTGCAGGAATGGGATACCAAGTCTCCCAACAACAGAGACAG*AAGTCACCTGCACCGGTGCCCGTGACGTACGAGCTGCCCACACTGTATAG | TCGA |
| ATCAGCCCATTGCAGGAATGGGATACCAAGTCTCCCAACAACAGAGACAG*AAAGTGTGGACTATGGGCCAGTTTTTGTGCAAGAACCAGATGATATTATT | TCGA |
| AGCCGCCCGCTTCAACCCCAGGGGACCAGAGGAACTCAGCCCTGATTGCG*GAGTTAGCCCTGCGTAGCCAGTTACCAACACTGGAGCAGGATGGTGGGAC | TCGA |
| TCATCTGTGTGCTGGCCATCATCGTACCCATTCCCAAGTCCCACGAGGCG*GATGGAACAAAATTCATCTGTATTGGGGCTCTGTATTCTGAACTTCTGGC | TCGA |
| TCCTGGACGCGCTACTGGCCAGCAAGCAGCATTGGGACCGTGACCTGCAG*CTCCAGCTCCAGAAATCCCAGTACCTGCAACTGGGCCCCAGCCGAGGCCA | TCGA |

| | |
|---|---|
| ACGGGGCCTTCTACAGTGACGGGGAGATCAGCATTTGCATGGAACACATG*AAGCCCCTGAATGTGACAGTGATCCGCAGGGGGGAAAAACACCAGCTTAG | TCGA |
| ACGGGGCCTTCTACAGTGACGGGGAGATCAGCATTTGCATGGAACACATG*AAGCCCCTGAATGTGACAGTGATCCGCAGGGGGGAAAAACACCAGCTTAG | TCGA |
| CGCTACACCAACCTCTCGTACATCGGCGAGGGCGCCTACGGCATGGTGTG*GTCTACCCCGAGCCCCGGAGCGAGAGCGAGTGCCTGAGCAACATCCGCGA | TCGA |
| GTGACCTCAAGCCTTCCAACCTGCTGCTCAACACCACCTGTGATCTCAAG*GCCCTGCGTCTTCCCAGGTGACCACGCCGGCTTCAGGACATGCACGGACA | TCGA |
| CGCTACACCAACCTCTCGTACATCGGCGAGGGCGCCTACGGCATGGTGTG*CTGCTGTATGTTTCCAGGAAACTTCTAGGACAGAGGCTATTCAACAAGCT | TCGA |
| CGCTACACCAACCTCTCGTACATCGGCGAGGGCGCCTACGGCATGGTGTG*GCATGGATGAGTTGATGGAGGTGTCTTTCTCGCCCCTGGCGGCCACAGGC | TCGA |
| AGTTATTAAAGTCTGTTGGTGCACAAAAAGACACTTATACTATGAAAGAG*ATATCCACACAGAAGCAGTTCAGGCTGCACTGGCAAAGCATAAAGAACAG | TCGA |
| GAAAATATATACCATGATCTACAGGAACTTGGTAGTAGTCAATCAGCAGG*CCCGATGGCACCAGCAAGGAGTGTGTGTTCATCGAGAAGGTTCTGGAGAA | TCGA |
| ATTATAGCCTTAGTGAAGAAGGACAAGAACTCTCAGATGAAGATGATGAG*GCACAACGACTCTGTAAACGCTGTGGTCAAGCATGGAGAGCTGCAACACT | TCGA |
| ATTATAGCCTTAGTGAAGAAGGACAAGAACTCTCAGATGAAGATGATGAG*GATGCCAAAGAAGACCATGAAAGAACACATCAAATGGTCTTACTGAGAAA | TCGA |
| TTGAGGGACCCCCGACTCCAAGCGCGAAAACCCCGGATGGTGAGGAGCAG*GAGTGGCTTTGGAGAGATATCATCCCCTGTAATCCGGGAGGCAGAGGTGA | TCGA |
| AGTTATTAAAGTCTGTTGGTGCACAAAAAGACACTTATACTATGAAAGAG*ATCCCCTCAGCAGAGTGTGTTTAAGACCAAAAAATGTAACCGATTCATTG | TCGA |
| GGGAGAGTGATACAGATTCATTTGAAGAAGATCCTGAAATTTCCTTAGCT*GCAGCCAAAGGAGACTCTGGAGATGGTGTGGATCCGAGGAAGTGGTTGCG | TCGA |
| CATGAAGAGCGAAGACTATCCCCATGAAACTATGGCGCCGGATATCCACG*CTCAAAGCTTGGATTCTGCCGTGCCACTCACGAATGGCGACACAGAAGAC | TCGA |
| ATGTGGGAGAGCAGAGGTCAAACCTGAAAGACCCCAGTTATGGATGGGAG*GGCCATTCACATGTTAAACTGCAGCTTGCAGCCATGTTTTGTATATCAAA | TCGA |
| GACTGGGCACAGCATGAGATCCAAAGGCAGGGCAAGGAAACTGGCCACAA*CACTATGGGACTGAGTAACATTCTCTTTGTGATGGCCTTCCTGCTCTCTG | TCGA |
| ATTGGGAACAGCAACCATGGCAGCCAATCTCCCAGGAATGTGGAGGAGAG*GAAAAGAAAATTATTCTTACATATTTTGCTCCAACTCCTGAAGCGTGTAA | TCGA |
| GACTGGGCACAGCATGAGATCCAAAGGCAGGGCAAGGAAACTGGCCACAA*CCATGTTTCCTGAGAGACTGGGAGTTGCAGGTGCACTTCAAAATCCATGG | TCGA |
| GACTGGGCACAGCATGAGATCCAAAGGCAGGGCAAGGAAACTGGCCACAA*AATCTCCAGAATCTGGTCTCCAGAAACATTATTCTAATCTGTGTATGGAA | TCGA |
| ATGTGGGAGAGCAGAGGTCAAACCTGAAAGACCCCAGTTATGGATGGGAG*AAGAATCGGACTCTGAAATACCTAAGAATGACTGGAAACAAAATTGAAAA | TCGA |
| GACTGGGCACAGCATGAGATCCAAAGGCAGGGCAAGGAAACTGGCCACAA*AAATCAGGTATTATGATGAGCTGTGTACAGCAATGGCTGAGGAATTGAAT | TCGA |
| GACTGGGCACAGCATGAGATCCAAAGGCAGGGCAAGGAAACTGGCCACAA*GCGTCTGCTATGCAGAGTTTGGAGTTCGAGTCCCCAAGACCACAGGATCT | TCGA |
| CCGCCATCCCGTCAAGCTCCATCCCTTTGGGCAGACAGCCCATGGCACAG*CTGCCTGTGTCAGGGAATCTGCTTGATGTGTACAGTAGTCAAGGCGTGGC | TCGA |
| AGAAATGGTTTCAAATGAATCTGTAGACTACCGAGCTACTTTTCCAGAAG*AATCCAAAGGAACAAGAAGGGATTTCAGAAGCTATGATAAACCTTGGCAA | TCGA |
| GCCTGAGCGGGACATGGACTCAACAGATCTGTCTGCCTGCAATCTACAAG*ATTGACTTTGAAGATGTGATTGCAGAACCAGAAGGGACACACAGTTTTGA | TCGA |
| GTGTGGTCCTTGCGCCGCTGACTTCTCCACTGGTTCCTGGGCACCGAAAG*ATTGACTTTGAAGATGTGATTGCAGAACCAGAAGGGACACACAGTTTTGA | TCGA |
| GATGTCTCCAGCATTTTTACGGACCCAATCATGAGCACTGCTTTAATAGG*ATTGACTTTGAAGATGTGATTGCAGAACCAGAAGGGACACACAGTTTTGA | TCGA |
| GAGACCTCACCATAGCTAATCTTGGGACATCAGAGGGTCGCTTCATGCAG*GAAACCAACATGTCTGACAAAAGTGACTTAAAAGCTGAGCTAGAGCGCAA | TCGA |
| AGAAATGGTTTCAAATGAATCTGTAGACTACCGAGCTACTTTTCCAGAAG*GGCCACCCAGTGCTCCTGCAGAAGATCGTTCAGGAACACCCGACAGCATT | TCGA |
| GTTGGACAATGATGGCAAGAAAATTCACTGTGCTGTGAAATCCTTGAACA*GCTGTAAAGCGGTTCTTGAAACAAGAGTGCAAGTGCCACGGGGTGAGCGG | TCGA |
| GAAGAGCCCAAAACCTATTACGAACTCAAAAGTCAACCGCTGAAGAGCAG*ACTAAGAAAAATGGCTTTGTCAGCCCAACAGATACCCAGATGGTTTAACT | TCGA |
| GAAGAGCCCAAAACCTATTACGAACTCAAAAGTCAACCGCTGAAGAGCAG*GCCTCTCTGCTGATGGTGGTGCCAAGCGCCAGGAGCACCTATCCCGATTT | TCGA |
| CTCCTTGCAGTCCAGTCAAAACTTACCCCCTCTGAACGAAAGGAAAAATG*GCACCCCTGAGGAGCCGCCAGGTGGTGAGTCTGCAGAAGAGGAAGAGAAC | TCGA |
| TGCAGGTCCTGCATCCAATGGATGCTGCCCAGAGATCGCAGCATATCAAA*GACTTGATTAGAGACCAAGGATTTCGTGGTGATGGAGGTAAGTAGTGATT | TCGA |
| AGCAAAAAATTCCAGCAGATGGAGTCCACAGGATCAGAGTGGACTTTAAG*ACTTATACAAGCACTAGCAACAACTCTATATCTGGATCATTGAAGCGCTT | TCGA |
| AAGTGGCTCCCCGCCCAAGTATCCCTGTAAAACAAAAACCAAAAGAAAAG*AGATGTGAACTTTGTCCCCATAAGGATGGAGCTTTAAAAAGAACAGATAA | TCGA |
| TAACACCCAGGGTGGTTTGCTTTCTCTGTGCCAGTAGTGGGCATGTAGAG*AAATATAAAGAGAAGGACAAACACAAACAGAAACACAAGAAGCAGCCAGA | TCGA |
| GAATCTACAATGGATGCCTTCCAAAGCCTACCTGCAGAAGCAAGCTAAAG*GTATTTATAACAGCAATGATGTAGCAGTATCGTTTCCAAATGTAGTATCT | TCGA |
| AGCAAAAAATTCCAGCAGATGGAGTCCACAGGATCAGAGTGGACTTTAAG*GATTTGGAGTTCCATGGAGTGATGAGATTTTATTTTCAAGATAAAGCTGC | TCGA |
| AAGTGGCTCCCCGCCCAAGTATCCCTGTAAAACAAAAACCAAAAGAAAAG*GATTTGGAGTTCCATGGAGTGATGAGATTTTATTTTCAAGATAAAGCTGC | TCGA |
| GTGATTCTTCCTCTCACTCTCAGGATAAACATCATGAGAAAGAGAAAAAA*GCTACAAGAATTAGTTGATCGAGTGCTGGAACGTTTTCAGGCATCTGGAC | TCGA |
| GTGATTCTTCCTCTCACTCTCAGGATAAACATCATGAGAAAGAGAAAAAA*GTTCCCAGTAGCTGCAGCAGTGAAAGACAGTGATTGGCTCCAGTGCTCCC | TCGA |
| GACCGTGGTTTTGCAGGAAATGTGAATCTCAGGAGAGAGCAGCCAGAGTG*CTAACGTCCTGGAGGGCGACTCCATGGATCAGGACGTCGAAAGCCCAGTG | TCGA |

| | |
|---|---|
| CCCGCTGGTTTATTGCGACGGGCACGGCTGCAGCGTCGCGGTGCATCAAG*GTCTCAGCCAATTTCTTCTCCAGTCATCCTCCAGTTTGGTCATGCAGAGA | TCGA |
| CCCGCTGGTTTATTGCGACGGGCACGGCTGCAGCGTCGCGGTGCATCAAG*ATATTTGGATTTGACCTGCATTTTGGAATTTATCTACACTTAAAATGCCA | TCGA |
| TTCATCAACAGCAGTTTCAGCAGTTGTTAAATTCTCAACAGCTCACACCA*GTTCAGGTTCTTCGGAATGCTGGAGAAGAAGTGACTCTAACAGTGTCATT | TCGA |
| TTGATTTGGGAGGAGATATTCATAGTGGGACAGCATTACCGACAAGCAAG*GCCCAGCAGCAAAGAAACCTGGATAACATCGTCTTGCAACAACCCAGAAT | TCGA |
| ACTGGAACGCCAACCGGCTGGACCTGTTCGAGATCAGCCAGCCGACCGAG*GCTATGTGGTTGATGTTGCAGAATGATGAGCCGGAGGACTTCGTTATAGC | TCGA |
| GGGAAGGCAGATTTGTTCTTAAGAATGAGAATGACGCCATTCCTCCTAAG*GAAACATCCCAGTTATCTGGACTCCCTGAGTTTGTTAAAATAGTAGAAGT | TCGA |
| ACTGGAACGCCAACCGGCTGGACCTGTTCGAGATCAGCCAGCCGACCGAG*AAATTGTCCGAGCCATGACACATGTGATAAACCAAGGCATGGCGATGTAC | TCGA |
| ACTGGAACGCCAACCGGCTGGACCTGTTCGAGATCAGCCAGCCGACCGAG*CTGTGAAAAAGAAAGAGAAAAAGTCTAAGACCAGTGAAAAGAAAGACAGC | TCGA |
| AGAAGTATGGTCTGGAAAAAGAAAACCCTAAGGATTACTGCATCGCCCGG*GACGGTTCACTGGGAAACATCGATGACCTGGCGCAGCAGTATGCAGATTA | TCGA |
| ACTGGAACGCCAACCGGCTGGACCTGTTCGAGATCAGCCAGCCGACCGAG*ATCTTTGAGAACCACCACCCCAGCCTTTACCCTTAACATTCCATCCGAGG | TCGA |
| ACTGGAACGCCAACCGGCTGGACCTGTTCGAGATCAGCCAGCCGACCGAG*GGAGTGGTGTCCTGGCTGTCCTGATGATAGCTGCGTTCCTCCAACCCATA | TCGA |
| TATCGGAACCCCTCCTTGAAGCTGATGGTGAATGTGTGCGGACACACTCT*GGAGAACACCGAAAACAATAACTCCTCCTCCAACAAGCAGAACCAACTCT | TCGA |
| TCATCGATAGGTTTCTTCGACTTCATTGTGGCCTTTGGGTGAGGAGAAAG*GTTGCCAATTTGGCCTGTTCCATCTCCAACAATGAAGAAGGGGTGAAATT | TCGA |
| GGTGGAAGATGCAATGCTGATGTTTGATAAAACTACCAACAGGCACAGAG*GTGTAATTACTTCAAGCCTCCAGGATGGCAATCCAGTTCCGTTCGCTCTT | TCGA |
| GGCTTATGGACCAGTGGCAGCAGCGGCGGTGGCGGCAGCAAGAGGATCAG*GTGCTCAAAAATGTGATAACTTTGCTGAAAAAAGACCCCTCCTTGGTGTT | TCGA |
| GGTGGAAGATGCAATGCTGATGTTTGATAAAACTACCAACAGGCACAGAG*TCATTGCCTGAAGATGAAGGCCATACTAGGACACTTGACACAGCAAAAGA | TCGA |
| TTTCCCAGATTGACCCCAAAGTTGCATTTCCTCGTCGAGCGCAACCCAAG*GTTTAACATGTCAGGAAGTAAAGGCTCTCAGAGAGAAGGCATGGTCAAGG | TCGA |
| TTTCCCAGATTGACCCCAAAGTTGCATTTCCTCGTCGAGCGCAACCCAAG*TTCCTTCTTTCTGGGGATTGGTGAACTCAGCTTGGAATCTTTGCTCTGTG | TCGA |
| GGTGGAAGATGCAATGCTGATGTTTGATAAAACTACCAACAGGCACAGAG*CTCAACAGAATTCACCAAAAATCCATGAAGGCTGGTGGGCATACAAGGAG | TCGA |
| CGAACACAGTAGTGGAAGATGTAAAGCAATATTTCGAGCAGTTTGGCAAG*GACCTGAACCCGCGGCGGGACATCTCCTCCTGGCTGGTGAGTCCCCCCAT | TCGA |
| GGTGGAAGATGCAATGCTGATGTTTGATAAAACTACCAACAGGCACAGAG*GTAGGGTTTTTGGATCGGCAGAGGACATTGGCCCTGCTGGAATTGTTCTA | TCGA |
| CGAACACAGTAGTGGAAGATGTAAAGCAATATTTCGAGCAGTTTGGCAAG*GAGTGCGCTACAGCCAGCAGGGAAACAATGAGATCACTTCCTCCTCAACA | TCGA |
| TGGCCGTGGCTACCCCGGATTTGCTCCAAGCTATGGCTATCAGTTCCCAG*AGCCTCCAAGGCAGGTCATCCTGACACTGCAACCCACTTTGGTGGCTGTG | TCGA |
| TTTCCCAGATTGACCCCAAAGTTGCATTTCCTCGTCGAGCGCAACCCAAG*AAGTTGGGATATCTAAAGCAGAAGCCTTAGAAACTCTGCAAATTATCAGA | TCGA |
| GGTGGAAGATGCAATGCTGATGTTTGATAAAACTACCAACAGGCACAGAG*GAAGGAAAAGCTGATGCGATGCTCTCAGTGCCGCGTCGCCAAATACTGTA | TCGA |
| CGAACACAGTAGTGGAAGATGTAAAGCAATATTTCGAGCAGTTTGGCAAG*ATTCGAAATCAGAGCAAAGCCTCTGGGTCTGGGCTCTGTGAGGGAGATGA | TCGA |
| GGTGGAAGATGCAATGCTGATGTTTGATAAAACTACCAACAGGCACAGAG*GGATGGATGAATGAGATTTACAACTATGATCCAGAAACCTACCATGCGAC | TCGA |
| TGGCCGTGGCTACCCCGGATTTGCTCCAAGCTATGGCTATCAGTTCCCAG*ACTTTCCCCTTTTCCCAGTCCGAGTTTGACGTCGGTCTCCTGGATACCAG | TCGA |
| CGAACACAGTAGTGGAAGATGTAAAGCAATATTTCGAGCAGTTTGGCAAG*GTTGCACCATTACTCAGACTTCTAAAGAGACTATTTGAAGTAAGAAAACC | TCGA |
| TTTCCCAGATTGACCCCAAAGTTGCATTTCCTCGTCGAGCGCAACCCAAG*GCTGCTGAGAACTAGCCCTAGACCTCTGCGTGAGGGTTCTTCTGCCGAAG | TCGA |
| GGTGGAAGATGCAATGCTGATGTTTGATAAAACTACCAACAGGCACAGAG*GCTGCTGAGAACTAGCCCTAGACCTCTGCGTGAGGGTTCTTCTGCCGAAG | TCGA |
| CATCAGATTCCATTTCAACTTCTGCCGACAACTTTTCTCCTGATTTGAGG*CTCCAGGGTTCCTTGAAAAGAAAACAGGTAGTTAACCTATCTCCTGCCAA | TCGA |
| TGCCCACGTTCCTTAACGTCATTCGAGTCTGTGATGGGGCCATCCGGGAA*ATACCGACTCATTGGTTCCTCGTCTGCCACATGCATCATCTCAGGCAACA | TCGA |
| CCAGAGCTGATGCTGGGCCGCATGCGCTGCCTCGAGGCCTTGGGGGAATG*GTGAAATTCAATCAAAAAATCTGTACAAGATTCCACTTAGAAACCTCGTG | TCGA |
| AGGCAGCGGCCGGAGTGTTAGAATATGCCATGAAACACTTTGGAGAGCTG*GTAGCGGGAAGGATGACCACGCTCACACGACAAGACCTCAACTTTGGCCA | TCGA |
| GGTGTCCCTGAGCTGCCTGTGTCTCTGCCTCCTTCCGCATGTGGTCCCAG*ggaatgaactccattgcctttgtgtgcaggaagccggaagcacgaccact | TCGA |
| TGGGACGTGGCCCAGCGCTCAGGCAACCCGGTGCTCATGGGCTTCTCTAG*GTGAGTGTGGCTGTGAAGTGCCTGAAGCCCGATGTCCTGAGCCAGCCAGA | TCGA |
| CTGAGTTTCAAGAAAATGGACCACCCTTACTGAAGAAAATCAAACAAGAG*GATGAATTTCACCCATTCATCGAGGCACTTCTTCCACATGTCCGTGCAAT | TCGA |
| TGGACGGAAATGGCTATATCAGCAAGGCAGAGATGCTAGAGATCGTGCAG*GTTTACTTTTGGACATCAAGCCCATCACTGCCAGCCAGTGAAGAAGGATT | TCGA |
| GGTCAGCGCCGACGGCAGCCGGCACCTGACGGCGTGACCGACCCGAGCCG*AGGAAATAAAAGCGGAAACTGAAAAACAGAGGTTTGTGGGATTTCTTTTC | TCGA |
| GGTCAGCGCCGACGGCAGCCGGCACCTGACGGCGTGACCGACCCGAGCCG*GTACCAGGATCATCTATGACCGGAAATTCCTGATGGAGTGTCGGAACTCA | TCGA |
| TTGGATTTGGCTACACACTTATAGATCTTCTGCACTGTTTACAGGCACAG*AAATATTCCCAAGAGACTCTAACTTAAAAGACAAATTCATAAAGCATTTC | TCGA |
| TGCAATCTTAAAAGAAACTGTGAAGCAAATTCGTCAGATCAAAGAACAAG*CTGAATTCTTGCCGCCAGGATGGGGAAACAGAACAGCAAGCTGCGCCCGG | TCGA |

| | |
|---|---|
| GGTCAGCGCCGACGGCAGCCGGCACCTGACGGCGTGACCGACCCGAGCCG*ggcagatgattctggaccagatgaagcctgaggagccttccagctctaag | TCGA |
| TTGGATTTGGCTACACACTTATAGATCTTCTGCACTGTTTACAGGCACAG*AGCCCCCAATTTTGTGAGTGTCCCTGAAGCATTGTGACAATTCCATTTTA | TCGA |
| ACCAGCTGCCTGGAATGGATATGATTAAGCAGGAGGGAGACACAACACGG*TGAAGAAGAAGTAAAATATTCACAAGATGAAGATTTTTCCAGAAGGGACT | TCGA |
| CGGGAAGGGCCCTGAGCCAGAGTCCGTCATCGGTTACTCCGGAGAAGATT*ACACTAACAGCACATCTGGAGACCCGGTGGAGAAGAAGGACGAAACACCT | TCGA |
| GCAGCCGAGGGGCTGTCAAAGCTCCAACTGGAGGCCACCCTGTGGTTCAG*ATTTATCAGCGATGCTGGTTAGTATTCAAGAAAGCTTCAAGCAAAGGTCC | TCGA |
| GGTGACCATTAAACAGGAGCAGAACTTGGACCAGACCTACTTGGATGATG*GCAGCAATTCGTAAAGAATTAAATGAGTACAAAAGTAATGAAATGGAGGT | TCGA |
| GTTTACAGCCAGAGGTTCCACCATCCAAACAACTTTTGTCAACCATAATG*ATCGATCAGCTAAAGCACCGGTTGAATAAGATGGAGGAGGAATGTAAGCT | TCGA |
| GGCCACGTGCCTCGACCGCATAGGCCAGTTCACCTGTATCTGTATGGCAG*GTCCTGCCTAATCATTGGACACGGACTCTTAATAAAACGGTCTTCAGTTC | TCGA |
| TCCAGCTAATCCAGAACCACTTTGTAGATGAATATGATCCCACCATAGAG*GGCTCCGGCTGGCACCTGAGGAGCGGCGTGACCCCGAGGGCCCAGGGAGC | TCGA |
| GGAGCCAGGAATTCCCAGTTTGACACCACAGGCTGAGCTCCCTGAACCAG*CTTTGACAGATCCAAATGCTTCTGCTGCCCAGCAGGCTGTTCTCCAGCAG | TCGA |
| AACATGGATAAGCATGCCCTGAAAAAGTATCGTCGAGAAGCCTATCATCG*GATATGGAGCAAATTCTACAGATCAGACCCACGGATTGCCCTTGGGAAAT | TCGA |
| AATGGAAAAGATAAAGTGTTTTGGTTTTGATATGGATTATACCCTTGCTG*TGATTCTACCAGAAAGGAATGAAGAACAGAACCTTCAGGAATTGAGTCAC | TCGA |
| AATGGAAAAGATAAAGTGTTTTGGTTTTGATATGGATTATACCCTTGCTG*GGCTACAGGACAACCCCTGGGCATGTGACTGCCGACTCTATGACCTGGTT | TCGA |
| AACTGGTAGACCAGAGCCTACGGTTACTTGGAGACACATCTCTCCCAAAG*GTGAGACGATGCGGATCGCCTCCTCCGAGTTTGCAGATGACCCTTGCTCG | TCGA |
| CTTCGTCATGATCAACAAATATGGTCGACGGTCCAAATTTGGAATGAAGG*ATGAGCTCACAGTGCCTGCACTTTACCCCAGTAGCCCTGAAGTGTGGGCC | TCGA |
| ATTTGCTTTATAAAAGACCACTGGATCCCAAGAGTGAAGCTCAGCTTCAG*GAGGACACCATGGAGGTGGAAGAGTTCTTGAAAGAAGCTGCAGTCATGAA | TCGA |
| ACCAGCACCTGGGACTCTTGGAACTGGGCTTGGTGCAGGATTTGGAACAG*AGTCTCAGCATGTTTCAGCAGCAGCTAGAGCAAAGCAAGCTCCTATACAA | TCGA |
| TTTCCTTTGGTAATACCAGCACCATAGGACAGCCAAGCACCAACACCATG*TGGGAAGTTGGTGTCTCCAAAGTGGAAGAATTTCAAGGGCCTGAAGCTAC | TCGA |
| ACCAGCACCTGGGACTCTTGGAACTGGGCTTGGTGCAGGATTTGGAACAG*CTGTGCGGTCAGAGAAGAAACGCCTTAGGAAGCCAAGCAAGTGGCTTTTG | TCGA |
| GGCCAAGTAGTTTTACAGCTGCTCCTACTGGGACTACTATTAAATTTAAC*GTCACTGCAGCCAGAGGGGTCCAGAAGAGAGAGGAGGCACTGCCTCCACT | TCGA |
| ACCAGCACCTGGGACTCTTGGAACTGGGCTTGGTGCAGGATTTGGAACAG*GACTGGGAATTCCCACATTTCATGGGAGATGTTGATGTAAATCTCCCTGG | TCGA |
| GACTGACCCCTCAGTCAGGCCTATGCATGAAAGAGTGCCAAAAAAAGAAG*CAAAACCAGAAATCCTGACTTACGACAGGCTCGTGAATGGCATGCTCCAA | TCGA |
| CGGTCCCGCCGCCGCTGCTGCTGCTGCTGTGCTCAGGCCTGGCCGGACAG*CTCCAGCTCCAGAAATCCCAGTACCTGCAACTGGGCCCCAGCCGAGGCCA | TCGA |
| CTGAAACAATTGAGGATTCTAGTAATCAAGCAGCAGCCAGAGAATGGGAG*TGGAAGCAAGCAATTTCTTCAACCGTCCTTGGAAAAGTAATAGTTCAACC | TCGA |
| AACCTCAGCCCACCACTTCCAGAACACTGATCATGACAACCAACAATCAG*AGGCCTCGTGGACAGAGAGATTCAAGCTATTATTGGGAAATAGAAGCCAG | TCGA |
| TTACCAAGCAACAAGTGAAATTGAGGTCACACCTTCAGTCCTTCGAGCAA*ATGAAGGACTGAACCATGAATGCAAACTCTGCAGCCAGACCTTTGACTCT | TCGA |
| AGCACTACCCAGAGGCCTATGCCTCCCCCAGCCACACCAAAGGCGAGCAG*GTTTCTTCGGCTACGTTTCAGTCACTTGTTCCTGATATTCCCGGTCACAT | TCGA |
| AATGAAGCCTGCCTTGTTTAATGTGTTGTGTGAAATCAAAGAAAAAACAG*TTGCAGGCTCCTTTAGCTGCACCATGAAGTTTACAGTCCGGGACTGTGAC | TCGA |
| GTCAGCCCATGGAAGCCAAGCTAACTCGCCCTCAACTCCCAACTCGGCTG*GCTGGAAAAAAAGAAAGGAGTAATACCCTGAATATTGCTTTAGACAACAT | TCGA |
| AATGAAGCCTGCCTTGTTTAATGTGTTGTGTGAAATCAAAGAAAAAACAG*CAAAGTGATTTCCCAAAAGAGACCTTTGAAGGCCCTAAACATGAAGTACT | TCGA |
| AATGAAGCCTGCCTTGTTTAATGTGTTGTGTGAAATCAAAGAAAAAACAG*AAACAAGTGTGAAGGAAAATACTATGAAGTACATGGAAACACTTAAGAAT | TCGA |
| AATGAAGCCTGCCTTGTTTAATGTGTTGTGTGAAATCAAAGAAAAAACAG*CCATGGTCGAAGGCCTTCAGGTCACAGTGCCCGACAAGAAGAAGGTGGCC | TCGA |
| AATGAAGCCTGCCTTGTTTAATGTGTTGTGTGAAATCAAAGAAAAAACAG*AGCCATGGGGGTGTTGATGTCCAAGCGGCAGACAGTGGAGCAGGTGCAGA | TCGA |
| AATGAAGCCTGCCTTGTTTAATGTGTTGTGTGAAATCAAAGAAAAAACAG*ATGATCTAAATGCCACATGTGTGGAGCCCACAGAGCTGACAGGGTGGCCC | TCGA |
| AATGAAGCCTGCCTTGTTTAATGTGTTGTGTGAAATCAAAGAAAAAACAG*ATCACTGTGTCCACTACTATGATCTTCGTAACACTAAACAGCCAATCATG | TCGA |
| GTCAGCCCATGGAAGCCAAGCTAACTCGCCCTCAACTCCCAACTCGGCTG*GGCTCCTGCTGAAGCTACCCTTTCTGGACTCCCCCCCAGACTCCCAGCA | TCGA |
| CGGTGTTCTATAGGCAGGTGTTGGGTGCAGAGAGCGCCCCTCCCGGACAG*GAGGAGCAAAATAGAGGCAAGCCCAATTGGGAGCATCTAAATGAAGATTT | TCGA |
| CAGGAGTGGAGTGATATCAGACACCGAGCTTCAGCAAGCTCTCTCCAACG*GGGTTGGCTGTGTTCCGGCCGCAGAGCACCGTCTGCGTGAGGAGATCCTG | TCGA |
| GGCTTTGGAGAAATTGAGGAAACTGAGGAAAGAGGCTGCAGCGAGGAAAG*ATTCTGGAAGTTACGGTCAGTCTGGGGGTGAGCAGCAAAGGTAAAGTATA | TCGA |
| AGCTGCACGGCCTGTGACCCGAAGCCCGGGGGGTTCCCAGGAGCAGCGAG*AGCTGCGGTGCAACCCTGGGCAGTTTGCGTGTCGCAGCGGCACCATCCAG | TCGA |
| ACTTTTTTTTAAGGGAAAGATTTTAATATTAAACCTGGTGCTTCTCACTC*aaaaaaaaaaaaaaaaaaaccccaaaaaaaaaactttttttatttatattct | TCGA |
| GCTTCCTCCACCTGCCGCGGCGTCTCGTGGCCCAGCTGCATCTGCTGCAG*GATATGATGTTTCAGCTTCTCCGAGGTCTGGACTTTCTTCATTCACACCG | TCGA |
| TCGTGCCAGCCTCCTCCCCCATGTACCACACCTGTGTGGCCTTCGCCTGG*GAAGCATACGTGATGGCTGGTGTGGGCTCCCCATATGTCTCCCGCCTTCT | TCGA |

| | |
|---|---|
| CGACACGCCGTACCTCGCAGAGGCCAGGATCTCCTTTAACCTGGGGGCAG*GTTCTTCACACACCCCCATTCGGCGTAGTACCCAGAGAGCTCAAGATGTG | TCGA |
| AAATTGTCTTGGTTAAGGGAAATAAACTTAACACAGAGGAAACTGTAAAA*CAACAGGCGGAGGTGGCCCACATGTCCCAGACCCAGGAGGAGAAGCTGGC | TCGA |
| ATATCCAGTTGGAGGTACCTCGGGAAGCTACCATTTCTTATATTAAGCAG*CCTCTGCGACTTGATATCAAAAGAAAGCTAACTGCTAGATCTGATCGAGT | TCGA |
| ATGTTGCGCTTGATGGATTTACTCTGGAAAGAAGCTGGTTTGGATCTTCG*TGTCCCAGGGTCAACGGTGTCAAGGCACACACTAATACGTGCAACCGAAG | TCGA |
| TGACCCAGGGGAAAAATTAGACTCAAAAATTGGAGTCCTTATAGGAAAAG*ATAGAAGACATGCAGTGGGGTAAAGGAGTCCGAGAGATACCCGCCTCAGT | TCGA |
| ATATCCAGTTGGAGGTACCTCGGGAAGCTACCATTTCTTATATTAAGCAG*CATGACTAATATGTTCTGCACAAGACATGAAGGCACAGACAGCACTGTCT | TCGA |
| CCAGACCAGTACGTTCGAGAATATGCTGTAGGCTGCCTGCGACAGATGAG*GTTGCTGAACGGCTGATGACCATCGCCTATGAAAGTGGTGTTAACCTCTT | TCGA |
| TAATGCCATCTGTTTTCCTAAAAGACTGAAATGGGGGCGACCATTCTTAG*AGGAATGTTACGGCATCCGGAGTCATCCCTCAGATTTCTCTGATCATGGG | TCGA |
| GCCTTCAACAAAGATGCCCTTCTGAACTGGCTTAAAGAATACAACTCTGG*CCCATTTTTCAGTGAAGAATTTTACTTTGAGATTCCAAGAACTTTCCAGT | TCGA |
| AAATTGTCTTGGTTAAGGGAAATAAACTTAACACAGAGGAAACTGTAAAA*GTCTTCTCCCCTGTCTGCCTCCCGGAGCTAGGACTGCAGAGGGGCCTATC | TCGA |
| AGGACTGGGTGAAAAAGATCCGTGAAGTGGCCCAGACAGCAGACGCCAGG*GATTGTGACCAGATCCACGTTGATGACGTCTCATCAGATGACAATGGCCA | TCGA |
| GACGCGCCAGATCACGTGGAGCCGGGGCGCCGACAAGATCGAGGGGGCCA*GAAACTATGGAAAACGTGAAGAAATGTAAAAATTTCCTATCTACGTTAAT | TCGA |
| CTCCTGCACGTGCAACTGCAAACGCTCTTTGTTCCAGAGCATGGAGATCA*CTCAGGTACCAGACAGTGATGAGCAGTTTGTTCCTGATTTCCATTCAGAA | TCGA |
| TAGCAAAATACCCACACAAAATAAAGAGTGGAGCTGAAGCTAAGAAATTG*GCCCCAGAGCTACTGGAGCAGCAGAAGTACACAGTGACCGTCGACTACTG | TCGA |
| CAGGAGAAGTGCCTGAGACCCGAGGACATCCTGCGCTTCATGGAAACAAG*AGGCAAAGGCCCATTGAAAAATACATCTGATGTCATTAATGCTGCCAAGA | TCGA |
| ACTGGAGGATGCCTATGTATTCCCTGGGGATGGCGCATCACACACCAAAG*GAGCTATGCCTGTTCCAGACCAGCCTTCATCAGCCTCAGAGAAGACGAGT | TCGA |
| CCTCCTCAGTGTTGACCTATGCCTCTCCGCCACTCATCACCAATGTCACG*GTCCATAATTCAGCAGCCAAATCTTTCCCTCGAAGGTTTGTTTTGCCTTT | TCGA |
| TTCAAAGAGAAAAGGCAACATCCTGTCACAGGCCATGCTCTGGCAAAAAC*ACCTCCTTATCTATTTTTCTTGAATCATGAGAGATGAAATTGCAACAACA | TCGA |
| CATCCGCAAAGGCTTTCTCGCTTGTCACCTTGCCATGTGGAAGAAACTGG*GTGACATTCTGCGCCTCGGATGCGCCGGAGAGGTCCTTGCTGGCCGACCG | TCGA |
| AAGATGCCATCTCAATGTGCCAGGACCAAGAGGAGAAAAAATACCTGATG*GTGACCATCTTGGTGCTGCAGGGCCGGCTGGATGAGGCCCGACAGATGCT | TCGA |
| GCGAGCCGACACCATGCGATCCAAGGCGAGGGCGAGGAAGCTAGCCAAAA*GGATGGATTCGTTCTGTCCGATCCCAGAAGGAAGTCTTGTTCCTGCATGT | TCGA |
| GAACAGGACTGACTCCAGTGATGACGCAGGCCTTAAGGAGAAAGTTTCAG*AGGCTGCCTCCATTCGAAACATTTTCTCAGGGACCTACGTTGCAGTTCAC | TCGA |
| CAGTTGACATAACTACTCCAAAAGCTGCCAGAAGGGGGAGAAAGAGAAAG*GAGCCCTGGCCTCCTTGCAGAAACATCGAGGTGACATGTAGCATGACCGA | TCGA |
| TCAAAGGCAAGaaaacaaaaacaaaaatagatataaaaacaTCCTGCCCT*CCTACCTGAAACGTCGCTATGGCCTCATCAGCACCGGCTCAGACAGTGAG | TCGA |
| TAGCTGAGACCACAGATAAAGTCAAACAAGGCTTTTGGGAAGAATTTGAG*ATTGAAATAGCCACAGTGATGCCTTCAGGGAACAGAGAGCTCCTCACTCC | TCGA |
| AGCGGGTCCGTCGCGGAGCCGGAGGGCGGGAGGAACATGACATCGCGGAG*GATTGCTGTCACGTCTAATGTGGCTGCTGCCTCGTGTCACATCTGAAACT | TCGA |
| CAGATGTTCCAGCTAATTGACATTGCCCGGCAGACGGCTCAGGGAATGGA*GAATTAGAGAATAAAATCCCAGCACGATGTTCCTCACTGGAGACACCAGT | TCGA |
| GAGAAGCTGCCGCCGAACGACAGGACGTTGGGGCGGCCTGGCTCCCTCAG*GAGGCCTGGAGAACCCAGGAGTTTCCAAGCTGTGCTGCCACCCGAGCTCT | TCGA |
| GAGAAGCTGCCGCCGAACGACAGGACGTTGGGGCGGCCTGGCTCCCTCAG*CGTCGAGGGCGAGGCCCCCAGCAGTGAGACTGGCACATCCCTGGACAGCC | TCGA |
| GAGAAGCTGCCGCCGAACGACAGGACGTTGGGGCGGCCTGGCTCCCTCAG*GTCACATGTGCCAATTTAACGAACGGTGGAAAGTCAGAACTTCTGAAATC | TCGA |
| TCGGATTGGGTCAGGCTCTTTTGGAACTGTTTATAAGGGTAAATGGCACG*GCTGCCAGGATATCGCAGATGAATTCAGAGCACAGGAGATTGATGGACAG | TCGA |
| GAGAAGCTGCCGCCGAACGACAGGACGTTGGGGCGGCCTGGCTCCCTCAG*GTCCTGGGCATGGGGAGCCTGACGTTTTGAAGGATGAGCTTCAACTCTAT | TCGA |
| ATGTCCACATGGTCAGCACCACCCTGCCTGTGGACAGCAGGATGATTGAG*CTGCGTGAGCTGGAGGACAAGTACGCAGAGTGCATGGAGATGCTGCATGA | TCGA |
| GGGAGTCAAACAGACTGCTGTCTGCCCTTATACGACACAGTAAATCAAAA*AGACCACAGAGCTGGGCAGCAAGAAGGAGCTCAAGTCCATGCCCTTCATC | TCGA |
| AGTGCTGCTTCAAACGGGCAGGGCTCTAGGAAACATATGTTACGATAGCC*GAAAACAGTGCATGTGCTTGGTGGATCTGGATCGAGCAAGAGCTGCAGAA | TCGA |
| TGATAAGAGAGAAATGATTTTTGAAGTTCTTGCTCCATTGGCAGAAAATG*GTACTATAAGATCTCCGTGGTGCTCATGTGCTTTGTGGTCCCCACGCTGG | TCGA |
| TCTCCAGCACCAGCTTCCAGTTAGTGGATATAGCACACCATCCCCAGCCA*GGACAGAGGGTCTCACGGATGGCCCCGCTGCCAGGAGCAGAACTGGTTCG | TCGA |
| AGACGCTGTGCTCGCTGGCACCCCGAACGGGTTGCTTCCCCCGCTGCGAG*CTCAACAGAATTCACCAAAAATCCATGAAGGCTGGTGGGCATACAAGGAG | TCGA |
| CGGTTCTCCTGGGAACCCCATCGGCCCCCTGCCAGCACACACCTGAGCAG*GTAAATCACCTTCTGAGCCTCTGATTCCCAAGAGGGTTGGTCTTATTCAC | TCGA |
| CCATCGCCGACCAGATCACCCTCCTCAAGGCTGCCTGCCTGGACATCCTG*GAGGGGAGATTTGTCGCCTGCCGCTCGCTCTGGGGCTCGATGTGAATATA | TCGA |
| AGACGCTGTGCTCGCTGGCACCCCGAACGGGTTGCTTCCCCCGCTGCGAG*CTTATCATGGATACAACCAAATGATGGAATGCCTACCGGGTTATAAGCGG | TCGA |
| TCTCCAGCACCAGCTTCCAGTTAGTGGATATAGCACACCATCCCCAGCCA*CTCCTTGCTCGGATTGAACGTATGGAAAGGCGGATGCAGCTGGTAAAGAA | TCGA |
| AGACGCTGTGCTCGCTGGCACCCCGAACGGGTTGCTTCCCCCGCTGCGAG*AACTCTATGAACAAGAATGCAACGGAGAGACTGTGGTCTACTGGGAAGTG | TCGA |

| | |
|---|---|
| AGACGCTGTGCTCGCTGGCACCCCGAACGGGTTGCTTCCCCCGCTGCGAG*GTGTCCCTCAATGCATGGTTCTGGTCCACAGTTTTCCACACCAGGGACAC | TCGA |
| AGACGCTGTGCTCGCTGGCACCCCGAACGGGTTGCTTCCCCCGCTGCGAG*AAATCATGCCCAAGTTCAAACAACGAAGACGAAAGCTAAAAGCCAAAGCC | TCGA |
| TCTCCAGCACCAGCTTCCAGTTAGTGGATATAGCACACCATCCCCAGCCA*GTGCACAAGCTTCTCATCGTGGTGGAGCTGTTGCTGGGGGAGATCCCTGA | TCGA |
| TCTCCAGCACCAGCTTCCAGTTAGTGGATATAGCACACCATCCCCAGCCA*GAGTATAACATGTATGGCTGGTGGGTGGGAGAACTGAACAGCCTCGTTGG | TCGA |
| CCATCGCCGACCAGATCACCCTCCTCAAGGCTGCCTGCCTGGACATCCTG*CGCGAGCTTCGGCCTCGGCTCTGTACCATGAAGAAGGGCCCCAGTGGCTA | TCGA |
| AGACGCTGTGCTCGCTGGCACCCCGAACGGGTTGCTTCCCCCGCTGCGAG*CTCCAGCGATTCAAGCGCTCCCTCTCCCTCAAGACCATCCTCCGAAGTAA | TCGA |
| CAGTACTCCTCTGCCCAACACTGTACCTCAGTTCATTGCCAGAGAGCCAT*GTCCCGTGGCTGTCATCAGTGGTGAGGAGGACTCAGCCAGCCCACTGCAC | TCGA |
| CCATCGCCTGTCACCTGGACCCGCGCGTGTTCGTGGACGGCCTGTGCCGG*GTGGTGGCCCTAGGGGATGTTCCAGATGGCACTCTGGTCACTGTGATGGC | TCGA |
| TCTTGGCATGTTGGTCTGGTCACCCAATCAAAATCTGTCAGAAGCAAAGT*GCTCTTCTCAACCATCTGTGAGTCCAGGGGAACCGTCTCCACCATCCATC | TCGA |
| AAGACCTGGTCTCCTCGGTTCAATTGGAGAAGGAAGATACTTCAAAAAAG*ATCATACATCTGTCGGTGGCCTGGGAGACAGTTTTTATGAATACTTACTG | TCGA |
| TTTGACGACTGCTCTTCCTCCTGTTGTCTCGAACCCAATTTATGACAACC*GGGAAGAATGGCTGGTCACCACAGTAGGGGCGTACCTCCCAGCGGTGTTT | TCGA |
| CCAGTACCTACTCCCTCTCCGTGAGCAGGAGGGCTCGCCGATTTGCCCAG*GATCATAAGTTTTTACTGCCAGTATTGAGAAATTCGTGGAAGAAATGTCA | TCGA |
| AAAGAATTAAAGGGGGAAAAAGTAATTCGACCAGAATGGATTGTGGAAAG*GCCTCGCAGTGCCGATTATTTTATGCAAGAAGCTAAACGAATGAAGCATA | TCGA |
| CCCTCACCGAGGGGGAGATGAAGAAGGGTCTCGGGTCCCTAAGCCACGGG*GGTGCTGGTCAAGATAAACTAGGAATCTATGTGAAGTCGGTTGTGAAAGG | TCGA |
| ACCATCAACAGTTTGACTACATAATAAGGATGATGAATTGTACTCTACAG*GTTCTTTTTTATCTTGGCCAGTATATTATGACTAAACGATTATATGATGA | TCGA |
| TGAAAAAGCTGGGCTCAAAGAAGCCACAGAAACCTATTCCCCGCCCCTTG*GTATCATCTTTATCAGAAAGTGAGGAGTCCCAGGACTCATCCGACAGCAT | TCGA |
| TCACAGCCAGTGACCTTGCAGCCAGTGACCTCAAAGGATTTCAGCCACAG*ATTTTAAATGGAGGGACTCCAGACATTCCTTCCAGTGGCCTACTATCAGG | TCGA |
| TCACAGCCAGTGACCTTGCAGCCAGTGACCTCAAAGGATTTCAGCCACAG*TTGTCGAGATGTCCACAGAAGGAGGATTTGGTGGTACTAGCAGCAGTGAT | TCGA |
| GAGAACTAAGACTTGGCACAAAGGCACCCTTATTGCCATCCAGACAGTTG*ACGTGCCAAATACTGGTGTAAGCAAACCTAGAGTTTCTGACGCTGTCCAG | TCGA |
| AATAATTGAGATTCCTGATGAAGATGATGATGTCCTCAGTATTGATTCAG*GTAAAAGATGAGAAAAATATTCAAGAGGTATTTGACCTGAGTGACTATGA | TCGA |
| TTCCAGTGCTCGTAAAAACAGATGGGATGAAACCCCCAAAACAGAGAGAG*GCTAAAGGTTGGTCTTCAGGTTGTGGCAGTCAAGGCTCCAGGGTTTGGTG | TCGA |
| GCAGCATGTCGGTGGCGGGGCTGAAGAAGCAGTTCTACAAGGCGAGCCAG*GCCCAACCGCTTGCTTATCCTCCACGGCTTCCTGGACGAAAACGTGCACT | TCGA |
| CCTTCCCCACTCCATGGCGCCAGCCTCCTCCCCCGCTCCCCAACTTCTCG*CCCGCGCCGGCTGTGCTGCACAGGGGGAGGAGAGGGAACCCCAGGCGCGA | TCGA |
| GCAGCATGTCGGTGGCGGGGCTGAAGAAGCAGTTCTACAAGGCGAGCCAG*GACACATTGGCGTGAGACCTGGGAGTACGTTGTGCCAAATCATTGCCACT | TCGA |
| GCAGCATGTCGGTGGCGGGGCTGAAGAAGCAGTTCTACAAGGCGAGCCAG*GAAGATGAGAATGTGCCTACTGCTCCTGATCCTCCAAGTCAACATTTACG | TCGA |
| GGAGAGTGCTGGCTACATGGAGCCCTATGAGGCACAGAGGATCATGACAG*GACATGGAGGAGTGAATCAGCTTGGGGGGGTTTTTGTGAATGGACGGCCA | TCGA |
| TGAAGGAGAAATTCGACTATGGCAACAAGTACAAGCGGCGGGTGCCCCGG*GACAAAGAACAACCCTGACTGGCCTTGACGTTCAGGACATGCTGCCCAGG | TCGA |
| TGAAGGAGAAATTCGACTATGGCAACAAGTACAAGCGGCGGGTGCCCCGG*AGTGATCAGCAGCTGGACTGTGCCTTGGACCTAATGAGGCGCCTGCCTCC | TCGA |
| TGAAGGAGAAATTCGACTATGGCAACAAGTACAAGCGGCGGGTGCCCCGG*GATATTGACTGGGTACAGACAGAGAAGCACGTGTTTGAGCAGGCATCCAG | TCGA |
| TGAAGGAGAAATTCGACTATGGCAACAAGTACAAGCGGCGGGTGCCCCGG*CAAACGGGGATTCAGCGTCCGATCCTTTGGAACAGGGACTCACGTGAAGC | TCGA |
| TACGCCCTGCTGCGCGTGGAGCACGGTCCCCCCCACACCAAGAACTGGAG*GCTGCTCCTGCGTTTGGTGGATGATTTCTTGTTGGTGACACCTCACCTCA | TCGA |
| TGGTGATCGCCTCCAACATCGGGCTTCCAGTCAGCACCACGCACTGTAAG*GTGAAAGCTGATAAGGACAGCCCCAACCTAGCCCAGCTGCAGCAGGCCTC | TCGA |
| tttattagataatataGTTGGATCAAACAAAAACAACACAATTTGTCCCG*GTCATAGCTCCTTGGAATCACCAACAAACATGCCTTCTCCTTCTCCTGAT | TCGA |
| AGACTCGGACTCCCAGGAGGAAGCGCACGGCAAGCTCGTGCTCTGGTCGG*GTGATTTCCATGGTGATGGTGGCTGTGGGTGTCTACGCTCGGCTAATGAA | TCGA |
| ACATGTTTGAGGCCCGGGAATTTCTTCGAAAAAAGCTTATTGGGAAGAAG*AATGAAATTTTTGGTTCGACGGTGGCATCGAGTCACAGGTTCTGTTTCCA | TCGA |
| TTCACCTGTCCAGCATCCGACCACCGAGGCTGGAGGGGGAGAACACCCAG*GTGGTGTTTGCATTGCTCAGTCACAGAAAATCCCACGTGAACCAAGACCT | TCGA |
| CAAACTCTATTTGCCAAAGGAAACTTGCCTTATCACCTTCTTGCTTGCAG*GTTTCCCTAGGACTTCTGCATGCCCATGCCACACATATCCTGTGGCCCCC | TCGA |
| TCTGAGCAGCCTGTTACCTTCTGCCTCAGTGCCAGAGTCCATGACAATTA*GTGTGACCAATTCAGTGAAAGGAACGAATGCGATTCTCTGGACCTGTTTG | TCGA |
| CTCCGACATCCTCTTCCCTGTCAGTGGCGTGGGCTCCTACAGCCCAGCAG*GTGTGACCAATTCAGTGAAAGGAACGAATGCGATTCTCTGGACCTGTTTG | TCGA |
| TTTCCTGAAGAACGTTGGGGAGAGTGTGGCAGCTGCCCTTAGCCCTCTGG*ACACTAACAGCACATCTGGAGACCCGGTGGAGAAGAAGGACGAAACACCT | TCGA |
| TCCAGGAGGTGGAGGGCCACCAGGAACACCCATCATGCCTAGTCCAGCAG*GCCCGGCTGTGCTGGCTCCAGAGGATGGGCTGGCCATGTCCCTGCATTTC | TCGA |
| CAGTAGAGGCCATTCATAGAGCTGTGGAATTCAATCCTCATGTGCCAAAA*ACACTTCTGAGAAATTCATCAGGCTGTGAAGCGCGCCGTGATGAATATCG | TCGA |
| TCCCCGGCACACGCGCAGCCCCGGCCTCTCGGCCTCTGCCGGAGAAACAG*TCCAGTGACACAGGATGACCTTCAGTATCACAACCTCAGCAAGCAGCAGA | TCGA |

| | |
|---|---|
| TCTGGGACAAGTTGCATGAAGAGCATATCAATGCAGGACGTACAGTTCAG*TACTGTCCCACTGACCAGGCTGCTGCAGGCACAGATGCTGAGCATGTGCA | TCGA |
| GAGGAGCGGGTGGCCCCGCGCTGCGCCCGCCCTCGCCTCACCTGGCGCAG*CTAATCTTCCAGTACCAACAATTGCAGCAATAGATGGACTCGCTTTAGGT | TCGA |
| GGGCGTGGGGGATATGACAAGGATGGAAGAGGTCCTATGACAGGATCAAG*GTATGAAAGTATCATCGCCACTCTGTGTGAGAACTTAGACTCGCTGGATG | TCGA |
| CTGAAAAGGAAAATCTGCAAAGAACTTTCCTGCTTACAACCTCAAGTAAA*AAAACACTTGGTAGACGGGACTCGAGTGATGATTGGGAGATTCCTGATGG | TCGA |
| GTAGACAAGCCCTCAAGGATGCCCGGTCCCCATCACCGGCACACATTGTC*CATTCCCCGTCGGAGCCCTTTCTAGAGAAACCAGTGCCGGATATGACTCA | TCGA |
| GTAGACAAGCCCTCAAGGATGCCCGGTCCCCATCACCGGCACACATTGTC*GAGTTCCGATTCTGAAGAGGCATTTGAGACCCCGGAGTCAACGACCCCTG | TCGA |
| GCACACATTACTCTGCGTACAAAACGATTGAACACCAGATTGCAGTTCAG*AATCATCCAGAACCGCATCCTGCTCGTCATCCTAGGGATCATCGTGGTCA | TCGA |
| TTCAGACACCCCTACCCCACAGCTCTGACCGTCAATGCTTCCATGTCCAG*AATCATCCAGAACCGCATCCTGCTCGTCATCCTAGGGATCATCGTGGTCA | TCGA |
| GTAGACAAGCCCTCAAGGATGCCCGGTCCCCATCACCGGCACACATTGTC*AATCATCCAGAACCGCATCCTGCTCGTCATCCTAGGGATCATCGTGGTCA | TCGA |
| GTAGACAAGCCCTCAAGGATGCCCGGTCCCCATCACCGGCACACATTGTC*AGCAAATTGGTCAGGAGATGTTGGAAAACCTTAGTCATGACAGAGAAAAG | TCGA |
| CTGCACCTCGCAGTCTCCCCAGGTCCGCCCAGCAGCCGCGCTTCAGCCAG*AGCCTGGACGGCCGTATCGAAGTCAAAGGGCAGCATGGAAGCTCATCACT | TCGA |
| CTGCACCTCGCAGTCTCCCCAGGTCCGCCCAGCAGCCGCGCTTCAGCCAG*GCTGAGATTGTGGGCAAAAAATTATTAAGCCTTCTGCCTGATGAAGAGAA | TCGA |
| CCTGACAGCTTCTACGAGCTCAAAAGCCAACCCTTACCCCTTCGCTCAAG*ACGTCATCCTTCAGCAGCGTCACAGATTCCACAATGTCTCTCAATATCAT | TCGA |
| TGCTCAGCTATCTGCCCGGAGGCACCACAGGACTGCAGCTCCCCAGCACG*GTCAGCCAGAGCAGCCTCCCCATGCTGTCCTCGCCGTCACCGGGCCAGCA | TCGA |
| ACACATTGAGCCGTCCACCCCCTGCTCAGGTGCCCAGGGAGGTGCTCAAG*TCAGATTCTGAGGAAGATGAACCCACAAAGAAGAAAACTATCCTTCAGGT | TCGA |
| ACACATTGAGCCGTCCACCCCCTGCTCAGGTGCCCAGGGAGGTGCTCAAG*GGATCCAGTGAGGGGACTGGTTTGTCTGCCTTGCTTCCCCAACCTAAAAA | TCGA |
| TGCCGGAgccgcggggcgcgggcggcggACAGATTGACCTTCAGAGCGAG*GAATCCTAAAACCAAAATATTAGAACGAAAACAGAAACATGGCTCACTAT | TCGA |
| AAGGAGCTCAGACAATTTCTGTCCAGCAACATGACAGCAAGCTCAGGCTG*CCCCCACCTTCTGGCCCTTTCACCTCCTCCTTGGGAGGGGCAGGATTCTC | TCGA |
| TGCCGGAgccgcggggcgcgggcggcggACAGATTGACCTTCAGAGCGAG*GCCAGCCTCAGGGGAACCCCTTGGGCTGCACCCCACTTCTGCCGAATGAC | TCGA |
| TGCCGGAgccgcggggcgcgggcggcggACAGATTGACCTTCAGAGCGAG*CCAGTCACTCCCGCTTCAACCCCAGCGAGTCGTCCACCTACTCCACCAAT | TCGA |
| TGCCGGAgccgcggggcgcgggcggcggACAGATTGACCTTCAGAGCGAG*AGTGCCCCTGAAGAAATTTAAGTCTATCCGTGAGACCATGAAGGAGAAGG | TCGA |
| TGGACGATGTCCTTGGCTACATCAATCCTGAAATGCAGATGCCCAACACG*AATGCCTACCGCAGGAATGGGATGGCTTTTCTAGCCTGCTGATGTGACAA | TCGA |
| CAATAAAAATGTTATGTCAGCGTTTGGCTTAACAGATGATCAGGTTTCAG*ATCAGTTTCCTAATTCATCTCAGAACGGTTCATGCCGACAAGTGCAGTAT | TCGA |
| ACAGCAGCCACCATATACAGGAGCTCAGACTCAAGCAGGTCAGATTGAAG*ACACTAACAGCACATCTGGAGACCCGGTGGAGAAGAAGGACGAAACACCT | TCGA |
| TGTATTTGGGCTTCGGAGCATTTACTCTCTGATTCTGGGCCAAGATAATG*AAAACATCCCAGAAAAGTGGACCCCAGAAGTCAAGCATTTCTGTCCCAAC | TCGA |
| AAGATTATACCTCACTATTCTCTTGCTCGATGGCTACTCTGTAATGGCAG*GAATGTTAGTTCGGCCGGAGAGGAGGCCCGCAGAAGGATGAGAGAGTGTG | TCGA |
| AAAGAAAATTCTGGGCATATTTATAATAAGAACATATCTCAGAAAGATTG*TGCTTTTTTAGGACCAAAGGATATATTTCCTTACTCAGAAAATAAGGAAA | TCGA |
| CGTCGTCTGCACGCAGCCCAAATCCCCATCCGGGACAGTGTGCACCTCAA*ATGTCACCGGGTGCGCATCAATGTACCTCCACACAGAGGGCTTCTCTGGG | TCGA |
| GATCTTTCTTTCCAAAACATTTCTGGACAGTACCTGATTGTATTTTTTTT*TGCGTTTTAAaaattgtctttatttacattttacagaaagttgagaaagt | TCGA |
| AGCTGTCTGGCATGGGCCTCTCCACCGTGCCTGACCTGCTGCTGCCACTG*CTCTTAGAACAAGCTCTGGTGATTGAGGAACAGCTGCGCCGGGCTGCTTA | TCGA |
| CCTGGCCCTGAAGGTGCAGGAGGTTCCCGCGTCTGCTCAGAGTCGACCAG*GGAAACCAATGAAAAGCGTGCTGGTGGTGGCTCTCCTTGTCATTTTCCAG | TCGA |
| TGCGGAGAGGTCAGTAACTAAATTGGAGAAAAGCATTGATGACTTAGAAG*TGTACCGCCGGAAGCACCAGGAGCTGCAAGCCATGCAGATGGAGCTGCAG | TCGA |
| CGAAGATCCGCATCACCTCCAGCCCCATTTTTATCACCTTCTACATCCTG*AATACTCTGGAGCAGTGCAATGTGTGTTCCAAGCCCATCATGGAGCGGAT | TCGA |
| TCCAGCATCTGGGGGCTGCTAAGGATGCCCAGCGGCAGCTCACAGCCGAG*GATGCAATTCGAAGTCACAGCGAATCAGGTACTTTTCCATAGTCATTTAG | TCGA |
| GGTTATTTTTTTTCCTTTCTCTTGCAGGACATTGGGGACACATTGAGCAG*TGCAGGAATAGGTAGAACAGGGTGTTTTATTGCTACATCCATTGGCTGTC | TCGA |
| ACAGCCGAATTTTATGCACTGAAGGGAATGTTCTTGGCTCAGATCAACAA*AATGTGAAAGATGTTGCAGAAGTGTTCCAGAAGTGGCTGAAGATAGAAGG | TCGA |
| CCGACTTCCTCTACGACCACATCCAGCCGGTGCGCGCAGAGCTCATGCAG*TATGTGCAGAAGCTTATAATCCTGATGAAGAAGAAGATGATGCAGAGTCC | TCGA |
| AAGGCCTGATTGGGCTGTTGCAGGAAGGAAACACAACTGTATTAGTAGAG*CCACAACAAGTTGTTCAGAAGAAGCCTGCTCAGGTAAGACATAGTCATTA | TCGA |
| TAGCTATGCTATACAAAGGTTTAACTTTCTTTCACAGAGGACTTCTGAAG*GAAATTGAACTTAGCTCATTAAGGGAAGCTTTGTCTTTCGTGTCATTAAT | TCGA |
| TGTGGGAAGATGTTGTCATTTGTTATGAAAGAGCCGGGCAGCACGGAAAG*GTTTGTTTAAGCATCTTAAACACGTGGCATGGAAGACCAGAAGAGAAGTG | TCGA |
| ggggccgccgccacccgcctccCACCATGGCTCTGAAGAGAATCCACAAG*GTGGAGGTGGAGGTGGAGAGCATGGACAAGGCCGGCAACTTTATCGGCTG | TCGA |
| CATTTCGTGGTTCACCCGCTGCCGGGCACCGAGGACCAGCTCAATGACAG*ACTGAGAAGAACAACAAATAGAATTCTGGCTTCCTCCTGCTGTAGCAGTA | TCGA |
| TTTTCAGTGGAACCACTACCACCACGACCATCATCTGATCAGTCTAGCAG*AAGGGGACAGCCGGAGGCTGAAGGGGGCCATCCAGAGGAGCACGGAGACG | TCGA |

| | |
|---|---|
| GGTTCTGGTCAATAGTAGAGAAGATGAGCATGACAGAACGACAAGATCTT*AAACACGAAGACACCGACTGTCCCTGCGTGGTGGTGTCCTGCCCTCACAA | TCGA |
| GACAAAGAAGTATGCTGATGTGATCATCCCTAGAGGTGCAGATAATCTGG*TTTTGAGTATCCGAGGAGCCCAGGAGGAGGAACCCACAGACCCCCAGCTG | TCGA |
| GAAGGGAAAACAGTCCAGATCCCCGTGTATGACTTTGTCTCCCATTCCCG*AAAACATGCTTTAAACTGCCACAGAATGAAGCCTGCCTTGTTTAATGTGT | TCGA |
| CCTTAACAAGAAGAGAATCGCCTACTTCACTTTGCCCAAGACTGACTCTG*TCTTACTGTTAACAGTCCCAGGAATGGAAGAGACCACATCAGAGGCAGAC | TCGA |
| CAGATCACTATGAAATAGCACTACCAAATATTGAGGAGTTACCAGCCCTG*GTACAGATGAAGTTTTTAGTTGAGCAAATGAGGCGACCAGATTTCATGGA | TCGA |
| AAAAAAGCTGTCGAAGAAGCTGAAAGACTCTCTGAAAGCCTTAAATTAAG*TGTCCTGGACGTTGCATGGGCCGTGCTGTGAGGATGCAGCAGCGTCACAC | TCGA |
| ATGTTTCACCCATAATTCAGCCAGTGCCTAGTATAAAGAATGTTCCACAG*ATAAATGTCAGGCTATAAGGCCTGAGCTAAAGGAAGGTGAATGCAGTAAG | TCGA |
| ATTTACAGATTGGAACAACTCTCCGGAGTCTGAAAGATGCACTTTTCAAG*GGGGCAGCTGGCATTTTATGCTATGTGGGAGCCTATGTCTTCATCACTTA | TCGA |
| CTATTGGAGGAGATGTTCCAGCCTTACAGCTCCAGTATGACCATTGTAAG*AAAGCAGAAGCCACTGGAGAAAAACGGCCAAGAGGCAGACCTAGGAAATG | TCGA |
| CCGGCTCCATCGACCTCAAGGACATCAACTTCCGGCCGCAGATGTCCCAG*AGGAAGATGGATGGCCGAGAGTACCCAGACGCACAGGGCTTTGCTGCTGA | TCGA |
| TCACCGCGCCCGTGGCCGCCGTGCAGGCGGCCGTCAGCAACCTCGTCCGG*GAATACTTGGACCTCAGCCAACCTCTCGAACAGTATTCACCTAGTTACCC | TCGA |
| TCTACAAGGTCCTAGAGCTGTATGAGGAGGCCGTGGACCTGGCCCTGCAG*GTGGGAGGACACACCATGCTCCCCATTCGCTGGATGCCTCCTGAAAGCAT | TCGA |
| GGGCAGCCGCTACCAGGGGGTCAGAGGGATCGGAAGAGTCTTCAAAAACG*gttgcctcacggactgtttcaccaagtatgttgccatgacatagcacggc | TCGA |
| GGGCAGCCGCTACCAGGGGGTCAGAGGGATCGGAAGAGTCTTCAAAAACG*GTCAAAAGGCTGAGTTTCAAATTCTGGATAATGTAGACTCAACGGGAGAG | TCGA |
| TTGTCGAAGCAGCTCTTGTGTCTAAGATCTCAAGTCCTTCAGATAAAAAG*GTGGTGGTCAATGCATCATCAAGATTAATGCTCAGTTATGACCTCAAGAC | TCGA |
| TTGTCGAAGCAGCTCTTGTGTCTAAGATCTCAAGTCCTTCAGATAAAAAG*ATTGGAGCATATGATCAACAAATATGGGAAAAATCTGTTGAACAGAGAGA | TCGA |
| ccagcctgccgctccGGAGAGCCGCCCGCCGAGGATGCGACGCACCGCAG*CTTGACAGAGCCAATTCGCTATTAAACCAGACTCAACAGCCTTACAGGTA | TCGA |
| GAATGGAGATGTTCGAGTGGGACTATTTGCTCTCTGTGATATTCCTGCAG*TAATTCATCCGTTGTATCAGAAAGCGATGACGTGGGACATTAATATTGCA | TCGA |
| AAAGCCAACGCAGAGTGTATCATCTCCTGAAGCAACATCTGGTTCTACAG*CATCTGCATTGCCGGGACCGGATATGAGCATGAAACCTAGTGCCGCCCCG | TCGA |
| CAGAGAAATCCACTGAGGTTGTGCCAAAGAAGAAGATCAAAAAGGAGCAG*CTTATGGTGCATTACCAGCCAGTTGCAGGGTCTGATGATTGGGGTTCAAC | TCGA |
| ATTGGCATCACTTCCTGATAGCAAGTTCATCTGCATGGAATGTAAAACTG*AGAGCGAAATGTCATCAGTGCAGTCACAACAGGAGCAGTTGTCCCAGTCA | TCGA |
| CAGTAATTCTTCTGCTGTAAATGTAGGCTTTTGTTTCGTTTGTGCCAGAG*CAGCAGCTACCTGGCAACTGAACCCATCATCACCACAGCCACTCCTGCAG | TCGA |
| CCTGTGATCGCACTGACACGGCCGGGGGGTTAGAATGGAACAAACTGAAG*GCCTGGAAGAAACGCTGGTTTATCCTGCGGAGTGGCCGGATGAGCGGTGA | TCGA |
| CCTGTGATCGCACTGACACGGCCGGGGGGTTAGAATGGAACAAACTGAAG*GTCCCTATGTTAACTACTTGAGACAAAAATAAATATGGCCTTCTACAGTT | TCGA |
| CCTGTGATCGCACTGACACGGCCGGGGGGTTAGAATGGAACAAACTGAAG*AGAATCCGACAACAGCTGCTCCAGCTGACACGTATCCAGCTAGTGAGTCT | TCGA |
| CCTGTGATCGCACTGACACGGCCGGGGGGTTAGAATGGAACAAACTGAAG*TTCCTCCTAAGATATATGACATCTCAAATGATATGACCGTCAATGAAGGA | TCGA |
| AAACGGAGCGGAGAGGCTGGGGCCTCAGGACCAAAAGGAGCATTAAGAAG*GAAACGTCCTTTTCTAGATCCATTTCTTTGACCCGACCTGGAAGCTCCAG | TCGA |
| AAACGGAGCGGAGAGGCTGGGGCCTCAGGACCAAAAGGAGCATTAAGAAG*GCTCTGTACAACTCAATCAAGAATGAGAAGCTTGAATGGGCAGTGTAAGT | TCGA |
| CCTGTGATCGCACTGACACGGCCGGGGGGTTAGAATGGAACAAACTGAAG*CTCACAAATTACCCCAAATCCAGAGAGAATATCAACCTTTTTCACTGTTG | TCGA |
| CCTGTGATCGCACTGACACGGCCGGGGGGTTAGAATGGAACAAACTGAAG*CCTTGGAGCATGGCAAGTCCAGAGCACCCTGGGAGCCCTGGCTGCATGGG | TCGA |
| ACAGGTTGAAACAGTTCCTCAGGCTACAGTGAAGACTGGATTACAGAAAG*AACCAAGAAATTGAAGAACTGACAAAAATCTGTGATGAGCTGATTGCAAA | TCGA |
| ATTGGCATCACTTCCTGATAGCAAGTTCATCTGCATGGAATGTAAAACTG*CCTGAAGAATAAGTAGTGAAAGCATCAATCAAGACATAAGAACAAAACAT | TCGA |
| CCTGTGATCGCACTGACACGGCCGGGGGGTTAGAATGGAACAAACTGAAG*CTATGTGTTCTCATTTCACCCAAGACTTTTTGCCAGTGCAGGGGATAGAA | TCGA |
| TGTTTGGATGCAATGTTCTTAGGATCCTCCTCCCGTCCTTCTCATCCTTC*GAAGCCTTATCAGTTGTGAGTGAGGACCAGTCGTTGTTTGAGTGTGCCTA | TCGA |
| TTCCACACCAGTGCCTCAGAGGTCCATGGCAGTATCCCAGCCAAATCTCG*GGGTGCGTCTTGACAGAGTACGTGGAGGTCGGCAGAAGTACAAGCGCAGG | TCGA |
| AAATGTGATAAAAACAAGGTGGAAGATTCTTCCAAGGAACCAGTGCGAAG*TTTCCAAGGCAGGAAAGGTAAAGAGACAAAGAAAGTGCCTTGTACAGATG | TCGA |
| CCCTGGGATATATTTTTGAGACCAAATTAATCAGCACATTGATTTATAAG*GTTGAATTAAAGAAAATACTTTATCAGAAGAAGATGGCCACTGCCCAGTT | TCGA |
| AGATGGAGAAGGAGAGGCTGCGGCTGAAACAGCAAGAACTGCTTCGGCAG*CTCCAGGGTTCCTTGAAAAGAAAACAGGTAGTTAACCTATCTCCTGCCAA | TCGA |
| TTTGGTATTGGAAGAATGAATCTGGCACATGGATTCAGTATGGAGAAGAG*AAAACACTTGGTAGACGGGACTCGAGTGATGATTGGGAGATTCCTGATGG | TCGA |
| ATGGACTTGAGTGGCCAGTAAGAGAAGGCTGTCACCAATACAGCTTAACA*TCTAACAAAGTGCCAGTGGTGCAGCACCCTCACCATGTCCACCCCCTCAC | TCGA |
| TGGCCCCAGGTGTAAGCGCAGAAAGCAGGCGAACCCGCGGCGCAATAACG*GCTGCCCCTCCAAGAGCGGAGGCAGCGAGAGTACGCGTGTGCCTCGCGCC | TCGA |
| TGGCCCCAGGTGTAAGCGCAGAAAGCAGGCGAACCCGCGGCGCAATAACG*GAAAGATACTATTCAAGTCCAGTACATCATAATGCAGAATTCTGTATCAG | TCGA |
| GGATTCACCCACAGCGGggcggcggcggcggcggctgcggcCCAATATAG*AAACATTGCTGAGGCTCTTGTCAGCAAAGGTCTAGCCACAGTGATCAGAT | TCGA |

| | |
|---|---|
| GAAGGAGCCCTGGATGGTTGTGAGGGAAGAGACAGGAAGATGGTGCCCAG*GGGAGTACATCAAGACCTGGCGGCCACGCTACTTCCTCCTCAAGAATGAT | TCGA |

Supplementary Table S2: Gene short-read count

| Gene | 1N | 3N | 1T | 2T | 3T | 1M | 2M | 3M |
|---|---|---|---|---|---|---|---|---|
| A*01:01:01:01 | 0 | 0 | 20 | 11215 | 0 | 0 | 5295 | 0 |
| A*03:01:0:01 | 10833 | 0 | 26501 | 0 | 232 | 586 | 0 | 0 |
| A1BG | 0 | 0 | 0 | 0 | 0 | 256 | 0 | 0 |
| A1CF | 0 | 376 | 1290 | 0 | 0 | 0 | 0 | 0 |
| A2M | 62 | 506 | 2050 | 0 | 556 | 2216 | 0 | 0 |
| AAAS | 0 | 696 | 0 | 1272 | 984 | 0 | 422 | 0 |
| AACS | 0 | 0 | 0 | 248 | 692 | 0 | 0 | 0 |
| AADACP1 | 0 | 0 | 0 | 0 | 0 | 0 | 404 | 0 |
| AADAT | 0 | 0 | 0 | 430 | 0 | 0 | 198 | 0 |
| AAGAB | 63484 | 102354 | 207844 | 201647 | 285872 | 87123 | 145793 | 7374 |
| AAMDC | 0 | 590 | 1352 | 0 | 525 | 0 | 0 | 0 |
| AAMP | 118 | 882 | 940 | 1415 | 5292 | 0 | 1414 | 802 |
| AAR2 | 432 | 0 | 0 | 0 | 0 | 0 | 212 | 0 |
| AARS | 0 | 0 | 0 | 0 | 278 | 0 | 0 | 0 |
| AARS2 | 0 | 0 | 0 | 0 | 650 | 0 | 312 | 0 |
| AARSD1 | 43513 | 118501 | 343099 | 275753 | 413986 | 250572 | 541266 | 139284 |
| AASDHPPT | 0 | 787 | 0 | 0 | 746 | 0 | 276 | 1036 |
| AATF | 0 | 0 | 0 | 1242 | 1520 | 0 | 0 | 0 |
| ABCA10 | 0 | 0 | 0 | 0 | 0 | 0 | 212 | 0 |
| ABCA11P | 210 | 846 | 0 | 0 | 3550 | 0 | 2458 | 0 |
| ABCA17P | 0 | 0 | 0 | 0 | 488 | 0 | 0 | 772 |
| ABCA2 | 0 | 222 | 0 | 0 | 152 | 0 | 170 | 0 |
| ABCA3 | 0 | 0 | 0 | 0 | 0 | 0 | 0 | 518 |
| ABCA5 | 700 | 678 | 0 | 0 | 0 | 0 | 0 | 0 |
| ABCA7 | 0 | 100 | 0 | 0 | 0 | 0 | 0 | 0 |
| ABCA8 | 402 | 0 | 0 | 0 | 0 | 0 | 0 | 0 |
| ABCB1 | 0 | 0 | 1746 | 0 | 0 | 0 | 0 | 0 |
| ABCB10 | 0 | 0 | 0 | 0 | 246 | 0 | 0 | 0 |
| ABCB6 | 0 | 0 | 0 | 1274 | 0 | 0 | 174 | 0 |
| ABCB7 | 12210 | 21782 | 20916 | 27893 | 88246 | 32715 | 63619 | 23420 |
| ABCB8 | 676 | 736 | 6806 | 962 | 5525 | 0 | 3060 | 816 |
| ABCB9 | 0 | 0 | 0 | 0 | 18 | 0 | 0 | 0 |
| ABCC1 | 0 | 0 | 298 | 788 | 0 | 0 | 487 | 0 |
| ABCC10 | 0 | 0 | 0 | 0 | 0 | 0 | 0 | 2254 |
| ABCC13 | 0 | 0 | 0 | 0 | 288 | 0 | 0 | 0 |
| ABCC3 | 0 | 0 | 0 | 0 | 1460 | 0 | 0 | 1040 |
| ABCC4 | 438 | 1212 | 0 | 842 | 706 | 0 | 0 | 0 |
| ABCC6 | 0 | 0 | 0 | 0 | 0 | 0 | 176 | 1200 |
| ABCC6P2 | 652 | 0 | 0 | 0 | 0 | 0 | 0 | 0 |
| ABCD1 | 0 | 0 | 0 | 0 | 10 | 0 | 0 | 0 |
| ABCD3 | 482 | 0 | 1560 | 0 | 324 | 0 | 0 | 0 |

| | | | | | | | | |
|---|---|---|---|---|---|---|---|---|
| ABCD4 | 0 | 0 | 1132 | 0 | 880 | 0 | 286 | 0 |
| ABCE1 | 0 | 0 | 0 | 0 | 554 | 0 | 150 | 0 |
| ABCF1 | 0 | 0 | 0 | 0 | 1230 | 1950 | 0 | 0 |
| ABCF1-DT | 0 | 0 | 0 | 0 | 0 | 0 | 0 | 388 |
| ABCF2 | 0 | 0 | 0 | 1056 | 0 | 0 | 0 | 1768 |
| ABCF3 | 0 | 400 | 1348 | 0 | 0 | 0 | 0 | 986 |
| ABCG1 | 0 | 0 | 0 | 0 | 0 | 2278 | 0 | 0 |
| ABCG2 | 390 | 0 | 790 | 0 | 0 | 0 | 0 | 134 |
| ABHD10 | 0 | 192 | 0 | 0 | 502 | 0 | 0 | 0 |
| ABHD11 | 590 | 1940 | 634 | 737 | 942 | 0 | 0 | 0 |
| ABHD11-AS1 | 0 | 212 | 986 | 0 | 0 | 0 | 0 | 0 |
| ABHD12 | 570 | 292 | 1804 | 2068 | 3016 | 0 | 0 | 0 |
| ABHD13 | 618 | 0 | 1809 | 0 | 964 | 0 | 0 | 671 |
| ABHD14A | 614 | 0 | 0 | 0 | 0 | 1932 | 0 | 0 |
| ABHD14A-ACY1 | 610 | 0 | 6301 | 611 | 538 | 0 | 616 | 458 |
| ABHD14B | 0 | 2172 | 3716 | 0 | 3292 | 0 | 658 | 888 |
| ABHD16A | 1778 | 398 | 3330 | 2040 | 788 | 0 | 880 | 0 |
| ABHD16B | 0 | 0 | 0 | 0 | 154 | 0 | 0 | 0 |
| ABHD17A | 1984 | 0 | 0 | 0 | 174 | 2054 | 0 | 0 |
| ABHD17B | 0 | 182 | 0 | 342 | 0 | 0 | 0 | 0 |
| ABHD17C | 68435 | 92048 | 146962 | 100917 | 159452 | 59314 | 191831 | 51085 |
| ABHD18 | 0 | 0 | 0 | 0 | 0 | 0 | 450 | 0 |
| ABHD2 | 5366 | 10974 | 10906 | 45812 | 62154 | 22148 | 138377 | 21207 |
| ABHD3 | 0 | 306 | 482 | 0 | 1214 | 0 | 816 | 680 |
| ABHD4 | 0 | 298 | 0 | 0 | 1688 | 0 | 306 | 0 |
| ABHD5 | 0 | 0 | 1006 | 0 | 614 | 0 | 192 | 0 |
| ABHD6 | 11188 | 7060 | 13390 | 4372 | 10438 | 360 | 40559 | 0 |
| ABI1 | 98897 | 184144 | 252945 | 145951 | 276130 | 239687 | 402638 | 138179 |
| ABI2 | 0 | 0 | 0 | 515 | 1508 | 0 | 0 | 0 |
| ABI3 | 0 | 252 | 0 | 0 | 0 | 0 | 0 | 0 |
| ABI3BP | 0 | 554 | 0 | 0 | 522 | 0 | 296 | 0 |
| ABL1 | 4171 | 3458 | 3344 | 2636 | 8891 | 2309 | 9297 | 936 |
| ABLIM1 | 710 | 0 | 0 | 0 | 228 | 0 | 574 | 0 |
| ABLIM3 | 0 | 140 | 0 | 0 | 0 | 0 | 0 | 0 |
| ABR | 3393 | 2240 | 4881 | 5874 | 12413 | 2294 | 5187 | 2208 |
| ABRACL | 1782 | 2784 | 9034 | 7730 | 8148 | 0 | 5634 | 3718 |
| ABRAXAS1 | 364 | 0 | 0 | 206 | 1870 | 0 | 0 | 0 |
| ABRAXAS2 | 0 | 0 | 0 | 0 | 112 | 0 | 0 | 0 |
| ABT1 | 0 | 1042 | 756 | 2122 | 1316 | 0 | 1522 | 1022 |
| ABTB1 | 864 | 0 | 0 | 0 | 648 | 0 | 0 | 0 |
| ABTB2 | 0 | 0 | 126 | 0 | 492 | 0 | 0 | 0 |
| ACAA1 | 0 | 0 | 4386 | 2254 | 3809 | 394 | 1206 | 0 |
| ACAA2 | 1917 | 3478 | 16027 | 1678 | 3056 | 0 | 740 | 1104 |

| | | | | | | | | |
|---|---|---|---|---|---|---|---|---|
| ACACA | 0 | 76 | 0 | 0 | 3170 | 0 | 260 | 1912 |
| ACAD10 | 0 | 0 | 1036 | 0 | 0 | 0 | 0 | 0 |
| ACAD11 | 0 | 0 | 0 | 0 | 0 | 0 | 1323 | 0 |
| ACAD8 | 0 | 770 | 0 | 0 | 1170 | 0 | 0 | 0 |
| ACAD9 | 0 | 0 | 0 | 942 | 1392 | 0 | 0 | 1516 |
| ACADM | 57581 | 95509 | 109529 | 94915 | 68377 | 46430 | 331895 | 25299 |
| ACADS | 720 | 0 | 0 | 0 | 0 | 0 | 1382 | 0 |
| ACADVL | 376 | 322 | 1696 | 3249 | 3458 | 5428 | 1016 | 0 |
| ACAP1 | 0 | 62 | 0 | 0 | 0 | 0 | 0 | 0 |
| ACAP2 | 0 | 0 | 0 | 0 | 1834 | 0 | 0 | 0 |
| ACAP3 | 322 | 0 | 0 | 0 | 600 | 0 | 0 | 0 |
| ACAT1 | 862 | 897 | 6332 | 0 | 3955 | 0 | 1190 | 1077 |
| ACAT2 | 527 | 0 | 5582 | 776 | 428 | 0 | 0 | 2716 |
| ACBD3 | 0 | 0 | 0 | 0 | 130 | 0 | 330 | 692 |
| ACBD5 | 204 | 0 | 0 | 0 | 1004 | 0 | 178 | 0 |
| ACBD6 | 0 | 0 | 0 | 0 | 2627 | 0 | 0 | 0 |
| ACBD7 | 0 | 0 | 0 | 0 | 0 | 0 | 46 | 0 |
| ACCS | 0 | 0 | 382 | 0 | 0 | 0 | 0 | 0 |
| ACD | 0 | 0 | 0 | 1809 | 1260 | 0 | 370 | 0 |
| ACE | 0 | 0 | 0 | 0 | 0 | 0 | 732 | 0 |
| ACER2 | 0 | 0 | 1554 | 0 | 0 | 0 | 0 | 0 |
| ACER3 | 0 | 0 | 2060 | 0 | 0 | 0 | 0 | 0 |
| ACHE | 28 | 0 | 1052 | 0 | 0 | 0 | 0 | 0 |
| ACIN1 | 0 | 943 | 0 | 0 | 134 | 0 | 254 | 1496 |
| ACKR1 | 0 | 796 | 0 | 34 | 0 | 0 | 0 | 0 |
| ACKR3 | 0 | 0 | 0 | 0 | 52 | 0 | 0 | 0 |
| ACLY | 2980 | 3294 | 18152 | 17285 | 23745 | 11552 | 10750 | 11788 |
| ACO2 | 1146 | 864 | 1296 | 0 | 344 | 0 | 0 | 696 |
| ACOT11 | 832 | 0 | 1028 | 728 | 0 | 0 | 0 | 891 |
| ACOT13 | 1632 | 92 | 5642 | 2409 | 0 | 2217 | 1642 | 0 |
| ACOT2 | 0 | 0 | 236 | 0 | 2773 | 0 | 490 | 0 |
| ACOT4 | 0 | 0 | 4595 | 0 | 0 | 0 | 1032 | 0 |
| ACOT7 | 104535 | 42035 | 405240 | 231220 | 163205 | 192549 | 57019 | 82751 |
| ACOT8 | 0 | 678 | 2860 | 2410 | 5652 | 2222 | 1391 | 2145 |
| ACOT9 | 38274 | 69358 | 71929 | 107599 | 458062 | 189422 | 142992 | 131898 |
| ACOX1 | 578 | 0 | 1832 | 0 | 298 | 0 | 176 | 0 |
| ACOX3 | 0 | 0 | 0 | 0 | 872 | 0 | 0 | 0 |
| ACP1 | 1824 | 2696 | 0 | 9183 | 12920 | 7764 | 6271 | 3792 |
| ACP2 | 0 | 518 | 1559 | 0 | 122 | 0 | 0 | 420 |
| ACP5 | 0 | 0 | 0 | 1341 | 298 | 0 | 0 | 0 |
| ACP6 | 0 | 184 | 0 | 10 | 0 | 0 | 1530 | 0 |
| ACPP | 1654 | 22864 | 9286 | 7410 | 24943 | 2722 | 33759 | 12530 |
| ACSF2 | 766 | 0 | 0 | 0 | 0 | 0 | 1008 | 0 |

| | | | | | | | | |
|---|---|---|---|---|---|---|---|---|
| ACSF3 | 5645 | 454 | 16634 | 5856 | 6172 | 2514 | 7707 | 1406 |
| ACSL1 | 2356 | 730 | 11227 | 3706 | 22654 | 24101 | 22451 | 6223 |
| ACSL3 | 0 | 2094 | 990 | 0 | 4246 | 0 | 1768 | 0 |
| ACSL5 | 1361 | 762 | 752 | 1742 | 2576 | 0 | 1762 | 0 |
| ACSM3 | 446 | 1044 | 0 | 0 | 0 | 0 | 0 | 0 |
| ACSS1 | 0 | 0 | 0 | 738 | 1436 | 0 | 0 | 0 |
| ACSS2 | 652 | 922 | 964 | 868 | 1249 | 0 | 0 | 1094 |
| ACTA2 | 2215 | 1379 | 4416 | 0 | 15455 | 10504 | 400 | 3026 |
| ACTA2-AS1 | 0 | 0 | 0 | 0 | 0 | 0 | 146 | 0 |
| ACTB | 114210 | 113628 | 156985 | 241111 | 753089 | 136463 | 230901 | 269731 |
| ACTG1 | 7616383 | 9179296 | 14204177 | 9794132 | 20692705 | 13857421 | 13032478 | 7591924 |
| ACTG2 | 0 | 2486 | 4204 | 1246 | 4902 | 0 | 0 | 1138 |
| ACTL6A | 0 | 780 | 3224 | 166 | 2272 | 0 | 536 | 2784 |
| ACTN1 | 6446 | 18667 | 26175 | 21209 | 110073 | 18666 | 41308 | 17174 |
| ACTN4 | 5560 | 4024 | 29197 | 8039 | 26751 | 9018 | 10791 | 5556 |
| ACTR10 | 216 | 138 | 3504 | 1010 | 1512 | 0 | 0 | 529 |
| ACTR1A | 400 | 436 | 3992 | 0 | 950 | 0 | 0 | 997 |
| ACTR1B | 0 | 0 | 5201 | 532 | 658 | 1435 | 0 | 0 |
| ACTR2 | 280 | 222 | 2850 | 458 | 1258 | 0 | 264 | 0 |
| ACTR3 | 694 | 1586 | 2606 | 2354 | 1348 | 0 | 0 | 732 |
| ACTR3B | 2912 | 6824 | 4897 | 12329 | 32069 | 19025 | 31053 | 14854 |
| ACTR3C | 0 | 0 | 1162 | 0 | 630 | 0 | 334 | 0 |
| ACTR5 | 0 | 0 | 896 | 0 | 0 | 0 | 0 | 1264 |
| ACTR6 | 0 | 406 | 778 | 0 | 564 | 0 | 474 | 0 |
| ACTR8 | 0 | 0 | 0 | 0 | 504 | 0 | 0 | 0 |
| ACVR1 | 8664 | 12150 | 17150 | 9943 | 41381 | 33930 | 46291 | 10976 |
| ACVR1B | 0 | 522 | 0 | 0 | 970 | 0 | 0 | 0 |
| ACVR2A | 28502 | 25908 | 72169 | 27458 | 59533 | 37451 | 62930 | 24090 |
| ACVRL1 | 0 | 0 | 470 | 0 | 0 | 0 | 124 | 0 |
| ACY1 | 1056 | 0 | 0 | 0 | 3002 | 502 | 3312 | 0 |
| ACY3 | 0 | 0 | 0 | 0 | 964 | 1306 | 0 | 0 |
| ACYP1 | 1676 | 78 | 672 | 1577 | 3497 | 0 | 542 | 840 |
| ACYP2 | 0 | 162 | 0 | 2322 | 462 | 0 | 0 | 0 |
| ADA | 0 | 374 | 0 | 0 | 408 | 0 | 0 | 2042 |
| ADA2 | 528 | 0 | 0 | 0 | 0 | 0 | 0 | 0 |
| ADAM10 | 0 | 0 | 0 | 0 | 2152 | 0 | 248 | 0 |
| ADAM12 | 0 | 0 | 0 | 0 | 742 | 0 | 0 | 0 |
| ADAM15 | 1710 | 2148 | 2242 | 0 | 9075 | 0 | 1930 | 2288 |
| ADAM17 | 0 | 0 | 0 | 0 | 0 | 0 | 0 | 412 |
| ADAM28 | 0 | 262 | 0 | 0 | 0 | 0 | 0 | 0 |
| ADAM33 | 0 | 0 | 0 | 0 | 292 | 0 | 0 | 0 |
| ADAM8 | 0 | 184 | 0 | 0 | 0 | 0 | 330 | 0 |
| ADAM9 | 0 | 124 | 536 | 0 | 619 | 2167 | 338 | 0 |

| | | | | | | | | |
|---|---|---|---|---|---|---|---|---|
| ADAMDEC1 | 134 | 1028 | 1364 | 0 | 0 | 0 | 0 | 0 |
| ADAMTS10 | 0 | 0 | 0 | 0 | 190 | 0 | 0 | 0 |
| ADAMTS12 | 0 | 0 | 0 | 0 | 1012 | 0 | 0 | 0 |
| ADAMTS16 | 0 | 0 | 0 | 0 | 0 | 134 | 0 | 0 |
| ADAMTS2 | 0 | 0 | 0 | 0 | 715 | 0 | 0 | 0 |
| ADAMTS9 | 0 | 140 | 0 | 0 | 0 | 0 | 0 | 0 |
| ADAMTSL1 | 520 | 0 | 0 | 0 | 0 | 0 | 0 | 0 |
| ADAMTSL2 | 0 | 0 | 0 | 0 | 0 | 302 | 194 | 0 |
| ADAP1 | 14355 | 8432 | 82236 | 10848 | 14113 | 31552 | 38315 | 648 |
| ADAP2 | 1874 | 2926 | 14750 | 1808 | 14957 | 16494 | 5898 | 6591 |
| ADAT1 | 0 | 314 | 0 | 0 | 0 | 934 | 0 | 0 |
| ADAT2 | 0 | 238 | 0 | 0 | 0 | 0 | 0 | 0 |
| ADAT3 | 0 | 0 | 1495 | 0 | 0 | 0 | 118 | 0 |
| ADCK2 | 396 | 0 | 0 | 0 | 418 | 0 | 0 | 0 |
| ADCK5 | 460 | 200 | 0 | 0 | 2886 | 0 | 416 | 0 |
| ADCY3 | 0 | 0 | 0 | 0 | 960 | 0 | 0 | 0 |
| ADCY6 | 0 | 0 | 0 | 0 | 582 | 902 | 0 | 0 |
| ADCY7 | 0 | 0 | 0 | 0 | 270 | 0 | 0 | 0 |
| ADCY9 | 0 | 196 | 0 | 0 | 0 | 0 | 808 | 0 |
| ADD1 | 5650 | 11776 | 29405 | 7169 | 13816 | 9935 | 24279 | 6495 |
| ADD2 | 0 | 370 | 0 | 0 | 0 | 0 | 0 | 0 |
| ADD3 | 0 | 328 | 916 | 0 | 114 | 0 | 3930 | 0 |
| ADGB | 0 | 164 | 0 | 0 | 0 | 0 | 0 | 0 |
| ADGRA3 | 0 | 184 | 548 | 0 | 0 | 0 | 0 | 0 |
| ADGRE2 | 0 | 0 | 0 | 0 | 410 | 0 | 0 | 0 |
| ADGRE5 | 0 | 0 | 0 | 0 | 2660 | 38 | 0 | 486 |
| ADGRF1 | 0 | 0 | 0 | 0 | 804 | 0 | 0 | 0 |
| ADGRF4 | 0 | 118 | 0 | 0 | 0 | 0 | 0 | 0 |
| ADGRG1 | 0 | 350 | 0 | 0 | 358 | 0 | 1374 | 0 |
| ADGRL1 | 0 | 0 | 0 | 0 | 322 | 0 | 0 | 0 |
| ADGRL3 | 0 | 336 | 0 | 0 | 0 | 0 | 0 | 0 |
| ADH1B | 443 | 0 | 1375 | 0 | 0 | 2668 | 0 | 0 |
| ADH1C | 1934 | 6430 | 16657 | 993 | 0 | 0 | 0 | 0 |
| ADH5 | 578 | 1992 | 2056 | 116 | 1661 | 2696 | 3026 | 712 |
| ADH6 | 17744 | 12718 | 30153 | 1006 | 1042 | 0 | 9835 | 1694 |
| ADI1 | 2312 | 2076 | 3960 | 3941 | 11969 | 2356 | 2852 | 0 |
| ADIPOR1 | 506 | 358 | 0 | 2654 | 3256 | 0 | 1998 | 796 |
| ADIPOR2 | 0 | 0 | 1180 | 0 | 526 | 0 | 0 | 0 |
| ADIRF | 1368 | 0 | 2190 | 4880 | 1994 | 5065 | 7242 | 3294 |
| ADK | 42132 | 95311 | 138751 | 350391 | 644317 | 293210 | 486020 | 256012 |
| ADM | 0 | 316 | 7800 | 0 | 2098 | 0 | 410 | 544 |
| ADNP2 | 0 | 0 | 0 | 0 | 0 | 0 | 192 | 528 |
| ADO | 0 | 0 | 0 | 0 | 0 | 0 | 738 | 0 |

| | | | | | | | | |
|---|---|---|---|---|---|---|---|---|
| ADORA2B | 0 | 240 | 0 | 0 | 0 | 0 | 0 | 0 |
| ADORA3 | 0 | 48 | 0 | 0 | 0 | 0 | 0 | 0 |
| ADPGK | 0 | 184 | 0 | 939 | 176 | 1860 | 848 | 0 |
| ADPRHL2 | 630 | 0 | 3394 | 2531 | 688 | 0 | 630 | 0 |
| ADPRM | 0 | 0 | 0 | 1118 | 1344 | 0 | 0 | 0 |
| ADRM1 | 0 | 1056 | 1048 | 2726 | 6570 | 2836 | 4559 | 0 |
| ADSL | 0 | 438 | 0 | 142 | 1436 | 0 | 458 | 0 |
| ADSS | 7988 | 10210 | 10288 | 9329 | 38227 | 12723 | 15402 | 3316 |
| ADSSL1 | 0 | 326 | 0 | 0 | 0 | 0 | 0 | 0 |
| ADTRP | 1223 | 956 | 5598 | 0 | 0 | 0 | 0 | 0 |
| AEBP1 | 478 | 0 | 0 | 0 | 3560 | 0 | 0 | 728 |
| AEBP2 | 0 | 258 | 728 | 0 | 134 | 0 | 0 | 0 |
| AES | 28656 | 28640 | 88588 | 31658 | 53282 | 32309 | 43945 | 13520 |
| AFAP1 | 0 | 0 | 0 | 0 | 1408 | 0 | 0 | 0 |
| AFAP1-AS1 | 0 | 0 | 0 | 0 | 0 | 0 | 0 | 11073 |
| AFDN | 192083 | 92184 | 131299 | 136806 | 550442 | 135821 | 338748 | 90623 |
| AFDN-DT | 0 | 0 | 0 | 0 | 700 | 0 | 228 | 0 |
| AFF1 | 1930 | 1322 | 2306 | 0 | 5430 | 2074 | 5506 | 0 |
| AFF3 | 320 | 3548 | 1764 | 0 | 0 | 2018 | 0 | 502 |
| AFF4 | 6459 | 5748 | 15777 | 3456 | 5842 | 7210 | 6955 | 1274 |
| AFG1L | 7006 | 9434 | 20555 | 12417 | 38631 | 14620 | 10050 | 5213 |
| AFG3L1P | 0 | 0 | 0 | 0 | 136 | 2386 | 0 | 0 |
| AFG3L2 | 0 | 1021 | 4850 | 0 | 450 | 0 | 234 | 0 |
| AFMID | 208 | 338 | 0 | 0 | 0 | 0 | 1048 | 626 |
| AFTPH | 0 | 232 | 1116 | 0 | 0 | 0 | 0 | 0 |
| AGA | 0 | 0 | 1882 | 0 | 0 | 0 | 0 | 0 |
| AGAP1 | 2212 | 1496 | 4952 | 5458 | 6714 | 0 | 3406 | 1344 |
| AGAP12P | 0 | 0 | 1662 | 0 | 0 | 0 | 0 | 0 |
| AGAP2-AS1 | 0 | 0 | 0 | 0 | 982 | 0 | 0 | 0 |
| AGAP3 | 762 | 0 | 0 | 0 | 0 | 0 | 0 | 0 |
| AGAP6 | 0 | 0 | 0 | 0 | 554 | 0 | 212 | 0 |
| AGBL4 | 0 | 1959 | 2377 | 1071 | 6101 | 0 | 0 | 4924 |
| AGFG1 | 0 | 0 | 0 | 0 | 0 | 0 | 612 | 0 |
| AGFG2 | 310 | 218 | 0 | 0 | 0 | 0 | 0 | 0 |
| AGK | 5588 | 21529 | 35073 | 30319 | 61388 | 19645 | 58062 | 17136 |
| AGMAT | 0 | 0 | 0 | 0 | 0 | 0 | 508 | 0 |
| AGO2 | 0 | 0 | 0 | 0 | 0 | 0 | 110 | 0 |
| AGO3 | 0 | 0 | 0 | 0 | 196 | 0 | 0 | 0 |
| AGPAT1 | 0 | 0 | 36 | 0 | 134 | 0 | 200 | 0 |
| AGPAT2 | 812 | 368 | 3740 | 0 | 3248 | 0 | 0 | 880 |
| AGPAT3 | 4100 | 1856 | 25837 | 7509 | 4104 | 3805 | 13452 | 4803 |
| AGPAT4 | 0 | 0 | 0 | 0 | 238 | 0 | 0 | 0 |
| AGPAT5 | 0 | 134 | 0 | 1490 | 0 | 0 | 0 | 0 |

| | | | | | | | | |
|---|---|---|---|---|---|---|---|---|
| AGPS | 6439 | 5707 | 7373 | 13023 | 45287 | 4252 | 19854 | 19139 |
| AGR2 | 6509 | 33396 | 45341 | 44342 | 148724 | 10423 | 6108 | 25743 |
| AGR3 | 2724 | 15835 | 10523 | 1941 | 8789 | 1738 | 602 | 2945 |
| AGRN | 238 | 162 | 0 | 0 | 1202 | 0 | 0 | 964 |
| AGTPBP1 | 0 | 0 | 0 | 0 | 392 | 0 | 0 | 0 |
| AGTRAP | 0 | 0 | 0 | 2287 | 2300 | 1590 | 0 | 1060 |
| AGXT | 0 | 0 | 0 | 0 | 0 | 0 | 678 | 0 |
| AHCTF1 | 0 | 0 | 0 | 0 | 310 | 22 | 0 | 0 |
| AHCY | 0 | 114 | 0 | 3256 | 3540 | 1872 | 1472 | 1068 |
| AHCYL1 | 0 | 0 | 0 | 0 | 0 | 0 | 388 | 0 |
| AHCYL2 | 30772 | 24706 | 95244 | 5282 | 7031 | 0 | 5772 | 1366 |
| AHDC1 | 0 | 0 | 0 | 0 | 304 | 938 | 1532 | 742 |
| AHI1 | 260 | 650 | 0 | 4404 | 4874 | 0 | 1287 | 3634 |
| AHNAK | 1308 | 282 | 499 | 0 | 660 | 1557 | 0 | 1878 |
| AHNAK2 | 0 | 0 | 0 | 0 | 1540 | 0 | 0 | 0 |
| AHR | 0 | 0 | 0 | 0 | 0 | 116 | 0 | 0 |
| AHRR | 70 | 0 | 0 | 0 | 0 | 0 | 0 | 0 |
| AHSA1 | 0 | 1545 | 3382 | 0 | 9064 | 4088 | 2464 | 1742 |
| AHSA2P | 0 | 124 | 0 | 0 | 516 | 0 | 306 | 0 |
| AHSG | 0 | 0 | 0 | 0 | 906 | 0 | 0 | 0 |
| AIDA | 0 | 330 | 0 | 0 | 278 | 2110 | 0 | 0 |
| AIF1 | 0 | 660 | 1088 | 410 | 1906 | 1645 | 280 | 814 |
| AIF1L | 0 | 0 | 0 | 0 | 0 | 0 | 470 | 0 |
| AIFM1 | 706 | 0 | 0 | 1026 | 1152 | 0 | 866 | 1212 |
| AIFM2 | 0 | 0 | 0 | 0 | 388 | 0 | 0 | 1284 |
| AIG1 | 71723 | 108186 | 113718 | 219461 | 586461 | 165834 | 650355 | 249327 |
| AIM2 | 0 | 460 | 0 | 0 | 0 | 0 | 0 | 0 |
| AIMP1 | 942 | 1088 | 7125 | 2652 | 2044 | 1500 | 820 | 1670 |
| AIMP2 | 77675 | 207645 | 253878 | 343017 | 858786 | 246986 | 387185 | 266276 |
| AIP | 0 | 552 | 826 | 0 | 2045 | 0 | 1980 | 1382 |
| AJUBA | 0 | 0 | 0 | 0 | 510 | 0 | 0 | 0 |
| AK1 | 884 | 416 | 2158 | 1004 | 828 | 2901 | 462 | 1402 |
| AK2 | 1892 | 3272 | 3153 | 3475 | 8094 | 2602 | 2076 | 3878 |
| AK3 | 1144 | 702 | 1240 | 0 | 1124 | 0 | 256 | 0 |
| AK4 | 0 | 0 | 1595 | 930 | 696 | 0 | 34 | 977 |
| AK6 | 678 | 1635 | 1500 | 1199 | 1836 | 0 | 0 | 0 |
| AK9 | 0 | 0 | 0 | 0 | 0 | 0 | 0 | 388 |
| AKAP1 | 2108 | 2445 | 8573 | 938 | 5679 | 7928 | 7984 | 432 |
| AKAP11 | 0 | 0 | 0 | 476 | 249 | 0 | 0 | 0 |
| AKAP12 | 0 | 0 | 0 | 0 | 134 | 0 | 0 | 0 |
| AKAP13 | 50720 | 55718 | 52469 | 8275 | 63063 | 20115 | 36371 | 13191 |
| AKAP17A | 0 | 560 | 0 | 1275 | 0 | 0 | 568 | 0 |
| AKAP7 | 0 | 318 | 0 | 0 | 457 | 0 | 0 | 0 |

| | | | | | | | | |
|---|---|---|---|---|---|---|---|---|
| AKAP8 | 478 | 0 | 0 | 0 | 0 | 0 | 188 | 0 |
| AKAP8L | 7701 | 5730 | 23044 | 15228 | 30827 | 7102 | 19567 | 4907 |
| AKAP9 | 132357 | 99778 | 195836 | 29273 | 27160 | 21215 | 44582 | 8173 |
| AKIP1 | 0 | 0 | 0 | 0 | 1098 | 0 | 0 | 2016 |
| AKIRIN1 | 402 | 0 | 0 | 1705 | 204 | 0 | 196 | 0 |
| AKIRIN2 | 772 | 0 | 0 | 1106 | 390 | 536 | 220 | 1438 |
| AKR1A1 | 750 | 272 | 5114 | 1252 | 7050 | 4400 | 7276 | 3594 |
| AKR1B1 | 0 | 0 | 0 | 0 | 0 | 3064 | 0 | 694 |
| AKR1B10 | 676 | 228 | 1846 | 0 | 188 | 0 | 0 | 0 |
| AKR1C1 | 838 | 0 | 0 | 0 | 1484 | 0 | 0 | 0 |
| AKR1C2 | 0 | 282 | 1598 | 0 | 0 | 2701 | 518 | 0 |
| AKR1C3 | 484 | 3194 | 8785 | 2673 | 3266 | 9965 | 1058 | 2268 |
| AKR1E2 | 0 | 0 | 0 | 0 | 0 | 0 | 2112 | 0 |
| AKR7A2 | 0 | 1522 | 1126 | 0 | 4830 | 0 | 612 | 376 |
| AKR7A3 | 1552 | 840 | 0 | 2800 | 602 | 0 | 0 | 1020 |
| AKR7L | 0 | 0 | 0 | 0 | 10 | 0 | 0 | 0 |
| AKT1 | 8937 | 17830 | 28261 | 49603 | 102435 | 34432 | 38034 | 31291 |
| AKT1S1 | 0 | 0 | 0 | 304 | 28 | 0 | 0 | 0 |
| AKT2 | 2428 | 2201 | 8534 | 1176 | 7860 | 2815 | 4641 | 2487 |
| AKT3 | 3064 | 1941 | 2751 | 2522 | 4680 | 2009 | 1316 | 3995 |
| AKTIP | 0 | 222 | 1608 | 20 | 498 | 0 | 972 | 866 |
| ALAD | 0 | 38 | 0 | 0 | 0 | 0 | 1010 | 0 |
| ALAS1 | 0 | 1016 | 0 | 0 | 2881 | 0 | 14 | 956 |
| ALB | 0 | 666 | 0 | 0 | 0 | 0 | 306 | 0 |
| ALCAM | 0 | 0 | 0 | 0 | 604 | 0 | 0 | 0 |
| ALDH16A1 | 0 | 0 | 0 | 1285 | 0 | 0 | 0 | 0 |
| ALDH18A1 | 62 | 40 | 2018 | 0 | 0 | 0 | 1434 | 484 |
| ALDH1A1 | 0 | 0 | 0 | 0 | 0 | 0 | 0 | 3384 |
| ALDH1B1 | 0 | 0 | 0 | 0 | 0 | 0 | 560 | 0 |
| ALDH1L1 | 1144 | 7812 | 6026 | 10844 | 2412 | 0 | 35111 | 10554 |
| ALDH1L2 | 0 | 0 | 0 | 0 | 0 | 2745 | 0 | 0 |
| ALDH2 | 1364 | 976 | 2435 | 10422 | 2238 | 2227 | 4412 | 1736 |
| ALDH3A2 | 0 | 410 | 1970 | 0 | 1862 | 0 | 582 | 1851 |
| ALDH3B1 | 0 | 0 | 0 | 0 | 0 | 0 | 0 | 742 |
| ALDH3B2 | 0 | 0 | 0 | 0 | 408 | 0 | 0 | 0 |
| ALDH5A1 | 0 | 0 | 1506 | 0 | 0 | 0 | 0 | 0 |
| ALDH6A1 | 0 | 0 | 1118 | 0 | 0 | 0 | 360 | 0 |
| ALDH7A1 | 676 | 0 | 1808 | 2393 | 716 | 2333 | 274 | 0 |
| ALDH9A1 | 0 | 0 | 1838 | 0 | 978 | 0 | 266 | 0 |
| ALDOA | 1537672 | 1505717 | 5382577 | 3202839 | 10180417 | 4030883 | 3700820 | 3530438 |
| ALDOC | 0 | 0 | 1608 | 0 | 0 | 1952 | 0 | 0 |
| ALG1 | 0 | 276 | 0 | 1010 | 0 | 0 | 12 | 0 |
| ALG10 | 0 | 0 | 0 | 0 | 490 | 0 | 0 | 0 |

| | | | | | | | | |
|---|---|---|---|---|---|---|---|---|
| ALG10B | 0 | 0 | 0 | 0 | 0 | 0 | 246 | 0 |
| ALG11 | 0 | 0 | 0 | 0 | 1478 | 0 | 0 | 0 |
| ALG13 | 2214 | 190 | 184 | 0 | 0 | 0 | 440 | 0 |
| ALG14 | 148243 | 340046 | 341252 | 283100 | 514084 | 363100 | 395480 | 228799 |
| ALG1L | 0 | 0 | 0 | 0 | 520 | 0 | 0 | 0 |
| ALG2 | 0 | 372 | 0 | 0 | 482 | 0 | 0 | 0 |
| ALG3 | 570 | 798 | 0 | 1212 | 2266 | 2128 | 0 | 0 |
| ALG5 | 960 | 1422 | 1696 | 1623 | 5862 | 1812 | 502 | 1280 |
| ALG6 | 0 | 442 | 0 | 0 | 186 | 0 | 0 | 0 |
| ALG8 | 0 | 592 | 0 | 1036 | 0 | 0 | 474 | 0 |
| ALG9 | 0 | 540 | 0 | 0 | 0 | 0 | 546 | 0 |
| ALK | 198 | 0 | 0 | 0 | 0 | 0 | 272 | 0 |
| ALKAL1 | 0 | 583 | 0 | 2197 | 936 | 0 | 740 | 0 |
| ALKBH1 | 0 | 0 | 0 | 0 | 392 | 0 | 0 | 0 |
| ALKBH2 | 0 | 0 | 138 | 0 | 1308 | 0 | 0 | 1195 |
| ALKBH3 | 34060 | 21725 | 54400 | 47702 | 75985 | 50422 | 52778 | 18126 |
| ALKBH5 | 0 | 0 | 399 | 1132 | 1078 | 0 | 488 | 0 |
| ALKBH6 | 84 | 0 | 0 | 0 | 1438 | 0 | 0 | 0 |
| ALKBH7 | 790 | 502 | 1812 | 1003 | 4540 | 0 | 604 | 44 |
| ALMS1 | 0 | 0 | 0 | 0 | 748 | 0 | 0 | 0 |
| ALOX15 | 0 | 0 | 0 | 0 | 218 | 0 | 0 | 0 |
| ALOX5 | 0 | 0 | 0 | 0 | 934 | 0 | 0 | 0 |
| ALOX5AP | 0 | 0 | 0 | 1020 | 444 | 0 | 0 | 1052 |
| ALS2 | 0 | 0 | 0 | 510 | 392 | 0 | 0 | 0 |
| ALS2CL | 304 | 370 | 72 | 0 | 584 | 0 | 146 | 0 |
| ALYREF | 0 | 488 | 1564 | 554 | 1082 | 0 | 548 | 0 |
| AMACR | 22537 | 15648 | 132390 | 39991 | 39605 | 39531 | 203820 | 5040 |
| AMBRA1 | 1508 | 1178 | 4090 | 3106 | 5766 | 1666 | 6961 | 1992 |
| AMD1 | 36 | 730 | 0 | 700 | 1992 | 0 | 98 | 0 |
| AMFR | 0 | 0 | 0 | 0 | 348 | 0 | 0 | 0 |
| AMIGO2 | 0 | 0 | 2444 | 650 | 738 | 984 | 0 | 0 |
| AMMECR1 | 0 | 524 | 574 | 0 | 2718 | 234 | 1583 | 0 |
| AMMECR1L | 0 | 0 | 0 | 919 | 0 | 0 | 0 | 0 |
| AMN | 830 | 681 | 742 | 0 | 750 | 0 | 0 | 0 |
| AMN1 | 0 | 1140 | 0 | 0 | 0 | 0 | 0 | 0 |
| AMOTL1 | 848 | 934 | 3336 | 0 | 3490 | 0 | 3598 | 6068 |
| AMPD2 | 0 | 0 | 0 | 0 | 0 | 2088 | 0 | 0 |
| AMT | 104 | 520 | 0 | 0 | 800 | 2158 | 1774 | 0 |
| AMY2B | 0 | 0 | 0 | 0 | 0 | 550 | 0 | 0 |
| AMZ2 | 798 | 0 | 0 | 0 | 2329 | 1882 | 1593 | 0 |
| AMZ2P1 | 0 | 0 | 0 | 1122 | 0 | 1876 | 0 | 0 |
| ANAPC1 | 0 | 0 | 0 | 774 | 2459 | 0 | 0 | 0 |
| ANAPC11 | 3500 | 3134 | 7451 | 6260 | 5820 | 1780 | 3550 | 3288 |

| | | | | | | | | |
|---|---|---|---|---|---|---|---|---|
| ANAPC13 | 306 | 860 | 3195 | 4597 | 5278 | 2328 | 2622 | 2606 |
| ANAPC15 | 1166 | 472 | 2164 | 1344 | 4538 | 0 | 0 | 1239 |
| ANAPC16 | 0 | 2272 | 4275 | 415 | 4452 | 5928 | 3426 | 1554 |
| ANAPC2 | 856 | 0 | 0 | 970 | 812 | 0 | 0 | 0 |
| ANAPC4 | 0 | 0 | 256 | 0 | 706 | 0 | 0 | 0 |
| ANAPC5 | 472 | 204 | 3710 | 2408 | 2296 | 0 | 422 | 0 |
| ANAPC7 | 1674 | 494 | 0 | 0 | 1416 | 0 | 296 | 0 |
| ANG | 0 | 1182 | 1476 | 0 | 2027 | 0 | 1518 | 342 |
| ANGEL1 | 484 | 0 | 0 | 0 | 0 | 0 | 0 | 0 |
| ANGEL2 | 0 | 0 | 0 | 0 | 554 | 0 | 0 | 0 |
| ANGPTL2 | 0 | 0 | 0 | 852 | 0 | 0 | 0 | 0 |
| ANGPTL4 | 728 | 942 | 3587 | 1098 | 17403 | 0 | 0 | 4096 |
| ANK2 | 0 | 196 | 0 | 0 | 0 | 0 | 0 | 0 |
| ANK3 | 30393 | 21465 | 111658 | 11153 | 31606 | 3104 | 23544 | 6541 |
| ANKFN1 | 0 | 246 | 0 | 0 | 1668 | 0 | 0 | 0 |
| ANKH | 348 | 0 | 334 | 0 | 174 | 0 | 0 | 0 |
| ANKHD1 | 0 | 640 | 1044 | 0 | 0 | 140 | 0 | 0 |
| ANKHD1-EIF4EBP3 | 0 | 240 | 0 | 0 | 0 | 0 | 386 | 0 |
| ANKIB1 | 0 | 278 | 3444 | 540 | 3665 | 1885 | 1760 | 1290 |
| ANKLE2 | 0 | 0 | 1370 | 0 | 0 | 0 | 0 | 3325 |
| ANKMY1 | 0 | 38 | 2777 | 0 | 0 | 0 | 160 | 900 |
| ANKMY2 | 12200 | 10302 | 19622 | 982 | 47810 | 42514 | 15736 | 10463 |
| ANKRA2 | 138 | 0 | 0 | 658 | 0 | 0 | 0 | 0 |
| ANKRD10 | 17651 | 48209 | 57802 | 58851 | 145414 | 68162 | 53329 | 22760 |
| ANKRD11 | 751 | 1582 | 4634 | 1552 | 1156 | 2654 | 3298 | 1294 |
| ANKRD12 | 0 | 0 | 0 | 0 | 372 | 0 | 0 | 0 |
| ANKRD13A | 0 | 0 | 0 | 0 | 554 | 0 | 0 | 0 |
| ANKRD13B | 0 | 0 | 0 | 0 | 0 | 0 | 448 | 0 |
| ANKRD13D | 0 | 0 | 0 | 0 | 0 | 0 | 280 | 0 |
| ANKRD16 | 0 | 0 | 0 | 590 | 486 | 0 | 0 | 0 |
| ANKRD17 | 274 | 0 | 268 | 0 | 302 | 0 | 0 | 0 |
| ANKRD20A4-ANKRD20A20P | 0 | 0 | 0 | 0 | 1216 | 0 | 0 | 0 |
| ANKRD22 | 0 | 0 | 0 | 2023 | 1896 | 448 | 1153 | 74 |
| ANKRD27 | 0 | 440 | 0 | 0 | 0 | 0 | 158 | 0 |
| ANKRD28 | 0 | 0 | 0 | 0 | 968 | 0 | 0 | 0 |
| ANKRD29 | 0 | 0 | 0 | 0 | 0 | 0 | 1036 | 0 |
| ANKRD36 | 278 | 0 | 0 | 0 | 170 | 0 | 0 | 0 |
| ANKRD36B | 0 | 0 | 0 | 294 | 0 | 0 | 422 | 0 |
| ANKRD36BP2 | 159 | 0 | 0 | 0 | 0 | 0 | 0 | 0 |
| ANKRD36C | 230 | 0 | 0 | 0 | 1550 | 0 | 206 | 0 |
| ANKRD37 | 0 | 0 | 0 | 0 | 2612 | 0 | 1064 | 1698 |
| ANKRD39 | 878 | 62 | 0 | 214 | 396 | 0 | 0 | 0 |
| ANKRD40 | 240 | 0 | 0 | 0 | 0 | 0 | 12 | 0 |

| | | | | | | | | |
|---|---|---|---|---|---|---|---|---|
| ANKRD40CL | 534 | 0 | 1997 | 0 | 0 | 0 | 0 | 0 |
| ANKRD44 | 0 | 0 | 0 | 0 | 286 | 0 | 0 | 772 |
| ANKRD46 | 0 | 360 | 0 | 0 | 714 | 0 | 0 | 498 |
| ANKRD49 | 0 | 0 | 0 | 0 | 356 | 0 | 0 | 0 |
| ANKRD50 | 0 | 0 | 0 | 0 | 0 | 0 | 354 | 0 |
| ANKRD52 | 0 | 0 | 0 | 0 | 188 | 0 | 0 | 0 |
| ANKRD53 | 286 | 0 | 0 | 0 | 422 | 0 | 0 | 0 |
| ANKRD54 | 0 | 0 | 0 | 1299 | 0 | 3344 | 0 | 0 |
| ANKRD55 | 0 | 634 | 0 | 0 | 0 | 852 | 0 | 0 |
| ANKRD61 | 0 | 0 | 0 | 0 | 974 | 0 | 0 | 0 |
| ANKRD65 | 0 | 0 | 0 | 0 | 568 | 0 | 0 | 0 |
| ANKRD9 | 1483 | 228 | 1472 | 0 | 356 | 0 | 1506 | 0 |
| ANKS1B | 1226 | 0 | 0 | 0 | 0 | 0 | 862 | 0 |
| ANKS3 | 0 | 0 | 0 | 1068 | 0 | 0 | 0 | 0 |
| ANKS4B | 0 | 0 | 0 | 0 | 1500 | 0 | 0 | 0 |
| ANKS6 | 0 | 0 | 0 | 0 | 0 | 1440 | 0 | 0 |
| ANKZF1 | 0 | 678 | 0 | 52 | 1148 | 0 | 0 | 0 |
| ANLN | 0 | 0 | 0 | 0 | 884 | 0 | 0 | 0 |
| ANO10 | 0 | 0 | 4628 | 1306 | 0 | 0 | 0 | 0 |
| ANO6 | 0 | 0 | 0 | 112 | 1154 | 1432 | 0 | 0 |
| ANO7L1 | 0 | 0 | 0 | 0 | 0 | 1546 | 0 | 0 |
| ANO9 | 830 | 0 | 800 | 380 | 485 | 0 | 234 | 0 |
| ANOS1 | 0 | 540 | 0 | 0 | 2666 | 0 | 0 | 0 |
| ANP32A | 812 | 1750 | 1703 | 1651 | 4367 | 1592 | 1470 | 146 |
| ANP32B | 861 | 260 | 3248 | 8071 | 3791 | 4525 | 524 | 818 |
| ANP32E | 0 | 144 | 0 | 0 | 542 | 0 | 160 | 0 |
| ANPEP | 141253 | 18820 | 443925 | 4591 | 15950 | 4042 | 6760 | 5782 |
| ANTXR1 | 0 | 0 | 0 | 0 | 1889 | 0 | 0 | 0 |
| ANTXR2 | 470 | 0 | 0 | 0 | 0 | 0 | 362 | 0 |
| ANXA1 | 488 | 776 | 7469 | 0 | 13700 | 1268 | 572 | 8852 |
| ANXA11 | 300 | 1288 | 3870 | 0 | 1706 | 0 | 1618 | 2677 |
| ANXA13 | 0 | 148 | 0 | 0 | 1680 | 0 | 0 | 0 |
| ANXA2 | 10253 | 23137 | 63621 | 69537 | 72406 | 32556 | 12356 | 33698 |
| ANXA2R | 0 | 0 | 0 | 742 | 1624 | 0 | 0 | 0 |
| ANXA3 | 0 | 922 | 7000 | 4120 | 5356 | 0 | 716 | 1232 |
| ANXA4 | 810078 | 838735 | 2186504 | 1435070 | 3871985 | 986485 | 1759286 | 1662056 |
| ANXA5 | 556 | 706 | 0 | 1803 | 9690 | 5482 | 4862 | 7768 |
| ANXA6 | 0 | 0 | 0 | 0 | 983 | 0 | 1042 | 742 |
| ANXA7 | 0 | 696 | 3362 | 980 | 986 | 246 | 1758 | 0 |
| AOAH | 18556 | 16769 | 54428 | 84259 | 30257 | 178746 | 180415 | 34868 |
| AOC1 | 1088 | 966 | 2080 | 0 | 0 | 0 | 592 | 805 |
| AP1AR | 0 | 56 | 672 | 0 | 0 | 1174 | 198 | 0 |
| AP1B1 | 0 | 216 | 1458 | 0 | 178 | 0 | 0 | 504 |

| | | | | | | | | |
|---|---|---|---|---|---|---|---|---|
| AP1G2 | 0 | 282 | 816 | 584 | 1430 | 0 | 0 | 624 |
| AP1M1 | 0 | 0 | 1334 | 0 | 608 | 0 | 0 | 0 |
| AP1M2 | 0 | 1560 | 3851 | 0 | 2992 | 0 | 244 | 1184 |
| AP1S1 | 638 | 398 | 602 | 926 | 1185 | 0 | 3847 | 0 |
| AP1S2 | 0 | 0 | 0 | 1274 | 0 | 0 | 0 | 0 |
| AP2A1 | 484 | 82 | 1430 | 2357 | 2180 | 1662 | 1592 | 684 |
| AP2A2 | 534 | 0 | 0 | 0 | 0 | 0 | 0 | 374 |
| AP2B1 | 4466 | 14162 | 10025 | 15809 | 58432 | 11493 | 14757 | 12745 |
| AP2M1 | 171747 | 246973 | 496477 | 459945 | 1169089 | 613091 | 837446 | 630866 |
| AP2S1 | 430 | 1234 | 1844 | 3428 | 4441 | 2790 | 2014 | 1833 |
| AP3B1 | 16895 | 37367 | 19648 | 29747 | 64213 | 16537 | 26258 | 8870 |
| AP3D1 | 0 | 0 | 0 | 822 | 4948 | 0 | 0 | 0 |
| AP3M1 | 392 | 0 | 0 | 0 | 1351 | 0 | 386 | 0 |
| AP3M2 | 0 | 0 | 0 | 0 | 710 | 0 | 606 | 0 |
| AP3S1 | 84397 | 119856 | 133729 | 131665 | 343825 | 207525 | 181579 | 112282 |
| AP3S2 | 900 | 0 | 864 | 2366 | 0 | 0 | 0 | 0 |
| AP4B1 | 440 | 426 | 0 | 0 | 0 | 0 | 0 | 0 |
| AP4B1-AS1 | 0 | 0 | 0 | 0 | 0 | 0 | 0 | 1315 |
| AP4M1 | 0 | 778 | 410 | 1506 | 1916 | 0 | 16 | 0 |
| AP5M1 | 308 | 94 | 358 | 0 | 664 | 0 | 0 | 798 |
| AP5Z1 | 0 | 0 | 0 | 552 | 366 | 0 | 224 | 0 |
| APBA3 | 758 | 0 | 0 | 0 | 804 | 0 | 0 | 0 |
| APBB2 | 0 | 0 | 0 | 1282 | 0 | 0 | 58 | 486 |
| APC | 21328 | 17962 | 15299 | 1382 | 11090 | 6272 | 4118 | 2252 |
| APCDD1 | 0 | 350 | 0 | 0 | 0 | 1130 | 3992 | 0 |
| APEH | 0 | 354 | 454 | 0 | 1349 | 0 | 594 | 0 |
| APEX1 | 0 | 430 | 0 | 0 | 6625 | 0 | 2442 | 1828 |
| APEX2 | 0 | 0 | 0 | 0 | 0 | 0 | 0 | 584 |
| APH1A | 1050 | 3472 | 1392 | 2985 | 4459 | 2156 | 4120 | 2304 |
| APH1B | 0 | 0 | 0 | 0 | 528 | 0 | 0 | 0 |
| API5 | 0 | 0 | 0 | 617 | 0 | 0 | 290 | 0 |
| APIP | 90620 | 197543 | 362900 | 222612 | 679356 | 249067 | 442008 | 223243 |
| APLF | 0 | 0 | 0 | 0 | 178 | 0 | 0 | 0 |
| APLP2 | 144069 | 114014 | 216559 | 95041 | 291341 | 182140 | 338847 | 97321 |
| APMAP | 1266 | 380 | 2704 | 0 | 0 | 2877 | 0 | 0 |
| APOA1 | 0 | 0 | 0 | 0 | 372 | 0 | 0 | 0 |
| APOA2 | 0 | 0 | 0 | 1426 | 2834 | 0 | 1584 | 0 |
| APOBEC1 | 0 | 0 | 258 | 0 | 1905 | 2364 | 0 | 2134 |
| APOBEC3A_B | 0 | 0 | 0 | 0 | 0 | 0 | 0 | 1020 |
| APOBEC3D | 0 | 0 | 0 | 0 | 0 | 0 | 0 | 964 |
| APOBEC3F | 0 | 0 | 0 | 0 | 222 | 0 | 0 | 0 |
| APOBEC3G | 0 | 0 | 0 | 0 | 0 | 0 | 308 | 0 |
| APOBR | 568 | 328 | 2181 | 0 | 0 | 0 | 0 | 0 |

| | | | | | | | | |
|---|---|---|---|---|---|---|---|---|
| APOC1 | 1498 | 333 | 867 | 550 | 1560 | 2158 | 2796 | 2540 |
| APOC4-APOC2 | 864 | 0 | 1066 | 0 | 1322 | 0 | 498 | 0 |
| APOD | 0 | 0 | 0 | 0 | 1764 | 1730 | 0 | 988 |
| APOE | 7548 | 3054 | 18077 | 8908 | 34727 | 35213 | 74919 | 35998 |
| APOL1 | 17027 | 10644 | 25908 | 6781 | 43411 | 20499 | 16196 | 16256 |
| APOL2 | 618 | 170 | 0 | 0 | 0 | 0 | 0 | 220 |
| APOL3 | 0 | 526 | 0 | 0 | 642 | 0 | 0 | 0 |
| APOL4 | 530 | 0 | 0 | 940 | 332 | 1204 | 3113 | 0 |
| APOL6 | 0 | 0 | 0 | 0 | 570 | 0 | 152 | 0 |
| APOLD1 | 0 | 0 | 0 | 0 | 1811 | 0 | 0 | 0 |
| APOO | 84746 | 172423 | 373605 | 175702 | 464801 | 169740 | 241869 | 124628 |
| APOOL | 0 | 0 | 0 | 0 | 1630 | 0 | 0 | 0 |
| APOPT1 | 542 | 560 | 1668 | 0 | 4264 | 0 | 308 | 1100 |
| APP | 446 | 1510 | 0 | 3150 | 2286 | 0 | 1258 | 0 |
| APPBP2 | 5947 | 9860 | 14490 | 4008 | 18001 | 9834 | 14548 | 3335 |
| APPL1 | 0 | 0 | 0 | 0 | 612 | 0 | 0 | 0 |
| APPL2 | 0 | 0 | 0 | 0 | 776 | 0 | 0 | 0 |
| APRT | 3163 | 1972 | 10832 | 5338 | 12742 | 2224 | 5972 | 1642 |
| APTR | 768 | 656 | 1922 | 1071 | 1694 | 0 | 516 | 498 |
| APTX | 0 | 0 | 0 | 978 | 2459 | 0 | 0 | 0 |
| AQP1 | 0 | 0 | 0 | 0 | 0 | 0 | 0 | 762 |
| AQP12A | 0 | 0 | 262 | 0 | 0 | 0 | 0 | 0 |
| AQP12B | 636 | 0 | 4768 | 0 | 0 | 0 | 0 | 0 |
| AQP3 | 0 | 498 | 0 | 0 | 0 | 0 | 0 | 0 |
| AQP5 | 0 | 0 | 0 | 0 | 1104 | 0 | 0 | 0 |
| AQP7 | 0 | 0 | 0 | 0 | 0 | 4008 | 737 | 0 |
| AQP7P3 | 0 | 0 | 0 | 0 | 0 | 0 | 1262 | 0 |
| AQP8 | 8816 | 850 | 26331 | 0 | 0 | 0 | 0 | 0 |
| AQR | 0 | 0 | 0 | 0 | 422 | 0 | 0 | 0 |
| AR | 696 | 0 | 0 | 0 | 0 | 0 | 422 | 0 |
| ARAF | 0 | 0 | 0 | 0 | 1616 | 0 | 0 | 0 |
| ARAP1 | 0 | 0 | 3524 | 0 | 558 | 0 | 611 | 0 |
| ARAP3 | 0 | 264 | 0 | 0 | 0 | 0 | 0 | 0 |
| ARCN1 | 0 | 0 | 0 | 1011 | 1042 | 0 | 0 | 654 |
| AREG | 896 | 0 | 8393 | 9726 | 701 | 2026 | 0 | 534 |
| AREL1 | 0 | 466 | 0 | 0 | 0 | 0 | 0 | 0 |
| ARF1 | 2276 | 3304 | 8624 | 6601 | 10499 | 5937 | 5150 | 2622 |
| ARF3 | 200 | 212 | 3521 | 3884 | 1524 | 4101 | 1942 | 0 |
| ARF4 | 1456 | 1570 | 4961 | 1490 | 7299 | 1727 | 2766 | 762 |
| ARF4-AS1 | 0 | 0 | 0 | 502 | 0 | 0 | 0 | 0 |
| ARF5 | 1681 | 2464 | 5258 | 4204 | 6765 | 2020 | 4050 | 3053 |
| ARF6 | 770 | 1204 | 2784 | 1150 | 5200 | 0 | 590 | 840 |
| ARFGAP1 | 628 | 224 | 0 | 1556 | 0 | 0 | 0 | 0 |

| | | | | | | | | |
|---|---|---|---|---|---|---|---|---|
| ARFGAP2 | 12313 | 14045 | 15006 | 19747 | 39430 | 15995 | 58386 | 18407 |
| ARFGAP3 | 1108 | 1136 | 1541 | 0 | 379 | 2570 | 232 | 734 |
| ARFGEF1 | 0 | 0 | 0 | 0 | 234 | 0 | 0 | 0 |
| ARFGEF2 | 0 | 0 | 0 | 836 | 0 | 0 | 0 | 0 |
| ARFIP1 | 910 | 178 | 734 | 0 | 0 | 0 | 0 | 0 |
| ARFIP2 | 0 | 764 | 0 | 0 | 0 | 0 | 320 | 0 |
| ARFRP1 | 514 | 0 | 0 | 1250 | 724 | 0 | 328 | 0 |
| ARGLU1 | 312 | 299 | 3920 | 0 | 1498 | 1540 | 1214 | 0 |
| ARHGAP1 | 1754 | 3580 | 5582 | 4036 | 5797 | 5339 | 5236 | 1390 |
| ARHGAP10 | 432 | 870 | 4711 | 4140 | 3844 | 3666 | 6930 | 2398 |
| ARHGAP11A | 0 | 0 | 0 | 594 | 0 | 0 | 0 | 0 |
| ARHGAP12 | 0 | 640 | 0 | 0 | 0 | 0 | 0 | 662 |
| ARHGAP15 | 0 | 384 | 0 | 0 | 16 | 0 | 0 | 0 |
| ARHGAP17 | 0 | 450 | 0 | 0 | 0 | 0 | 0 | 0 |
| ARHGAP18 | 22803 | 20012 | 45392 | 51908 | 132428 | 104108 | 320169 | 61412 |
| ARHGAP21 | 0 | 36 | 934 | 0 | 388 | 0 | 0 | 0 |
| ARHGAP23 | 0 | 0 | 0 | 0 | 674 | 0 | 0 | 0 |
| ARHGAP24 | 1440 | 3114 | 0 | 1756 | 2073 | 3989 | 3154 | 5617 |
| ARHGAP25 | 2828 | 2444 | 7642 | 0 | 4989 | 0 | 120 | 472 |
| ARHGAP26 | 4836 | 2894 | 7796 | 1543 | 7780 | 2710 | 3704 | 2064 |
| ARHGAP27 | 196 | 0 | 0 | 0 | 0 | 0 | 550 | 310 |
| ARHGAP27P1 | 0 | 504 | 0 | 0 | 380 | 0 | 0 | 0 |
| ARHGAP31 | 1558 | 0 | 0 | 0 | 0 | 0 | 0 | 0 |
| ARHGAP32 | 0 | 0 | 0 | 0 | 1185 | 0 | 0 | 0 |
| ARHGAP39 | 0 | 0 | 0 | 0 | 112 | 0 | 0 | 0 |
| ARHGAP4 | 380 | 0 | 634 | 0 | 1118 | 0 | 0 | 0 |
| ARHGAP42 | 0 | 0 | 0 | 0 | 0 | 0 | 0 | 514 |
| ARHGAP44 | 0 | 0 | 0 | 0 | 0 | 0 | 632 | 0 |
| ARHGAP45 | 536 | 0 | 1620 | 0 | 0 | 1120 | 38 | 0 |
| ARHGAP5 | 12842 | 10948 | 14782 | 4614 | 37122 | 2579 | 6781 | 6574 |
| ARHGAP6 | 1008 | 244 | 1476 | 0 | 4152 | 0 | 1204 | 10 |
| ARHGAP8 | 0 | 896 | 0 | 0 | 0 | 0 | 530 | 0 |
| ARHGAP9 | 0 | 0 | 0 | 0 | 390 | 0 | 0 | 0 |
| ARHGDIA | 77995 | 70878 | 176051 | 87821 | 148838 | 119669 | 190756 | 48952 |
| ARHGDIB | 1758 | 964 | 0 | 3584 | 3290 | 0 | 427 | 2754 |
| ARHGDIG | 0 | 0 | 0 | 0 | 578 | 0 | 0 | 0 |
| ARHGEF1 | 1022 | 184 | 595 | 580 | 650 | 544 | 1340 | 1259 |
| ARHGEF10 | 0 | 216 | 0 | 0 | 1714 | 0 | 252 | 1804 |
| ARHGEF10L | 8608 | 9179 | 19562 | 6654 | 15975 | 19549 | 3170 | 6365 |
| ARHGEF12 | 24234 | 11376 | 29819 | 17044 | 21757 | 12940 | 39669 | 9440 |
| ARHGEF16 | 0 | 296 | 1222 | 432 | 0 | 0 | 0 | 0 |
| ARHGEF18 | 584 | 0 | 0 | 0 | 516 | 0 | 0 | 0 |
| ARHGEF19 | 0 | 0 | 0 | 0 | 1380 | 0 | 0 | 0 |

| | | | | | | | | |
|---|---|---|---|---|---|---|---|---|
| ARHGEF2 | 304 | 0 | 0 | 0 | 0 | 0 | 0 | 0 |
| ARHGEF3 | 0 | 0 | 0 | 0 | 0 | 0 | 238 | 0 |
| ARHGEF35 | 0 | 0 | 1219 | 0 | 0 | 0 | 720 | 0 |
| ARHGEF37 | 678 | 0 | 0 | 0 | 0 | 0 | 0 | 0 |
| ARHGEF38 | 0 | 0 | 0 | 0 | 584 | 0 | 0 | 0 |
| ARHGEF39 | 0 | 0 | 0 | 0 | 482 | 0 | 224 | 0 |
| ARHGEF4 | 0 | 0 | 0 | 0 | 0 | 0 | 206 | 0 |
| ARHGEF40 | 0 | 0 | 0 | 0 | 218 | 0 | 408 | 0 |
| ARHGEF5 | 0 | 0 | 1120 | 0 | 0 | 0 | 0 | 0 |
| ARHGEF7 | 10230 | 9840 | 33748 | 17038 | 33847 | 17641 | 23035 | 9026 |
| ARHGEF9 | 10343 | 13198 | 31568 | 5710 | 29181 | 3582 | 8358 | 7096 |
| ARID1A | 3482 | 1394 | 1232 | 5022 | 4050 | 748 | 2800 | 1478 |
| ARID1B | 5654 | 5204 | 13136 | 5187 | 26314 | 3954 | 12647 | 5125 |
| ARID2 | 5672 | 5918 | 6626 | 1807 | 7857 | 4054 | 13822 | 148 |
| ARID3A | 0 | 0 | 0 | 0 | 384 | 0 | 0 | 0 |
| ARID4A | 0 | 0 | 0 | 0 | 308 | 0 | 0 | 0 |
| ARID4B | 0 | 0 | 0 | 0 | 208 | 0 | 20 | 0 |
| ARID5A | 0 | 0 | 1282 | 0 | 752 | 0 | 0 | 0 |
| ARID5B | 0 | 0 | 0 | 0 | 390 | 0 | 0 | 0 |
| ARIH2 | 0 | 502 | 856 | 0 | 0 | 0 | 0 | 0 |
| ARIH2OS | 0 | 0 | 0 | 0 | 1618 | 0 | 0 | 0 |
| ARL1 | 228 | 1990 | 0 | 5572 | 6887 | 5460 | 2580 | 0 |
| ARL11 | 0 | 0 | 0 | 0 | 0 | 0 | 330 | 0 |
| ARL14 | 748 | 1420 | 7395 | 1078 | 2624 | 0 | 1058 | 256 |
| ARL14EP | 0 | 900 | 0 | 0 | 570 | 0 | 298 | 0 |
| ARL15 | 340 | 0 | 0 | 0 | 698 | 0 | 0 | 0 |
| ARL16 | 0 | 40 | 1132 | 1248 | 2672 | 1762 | 0 | 924 |
| ARL17B | 0 | 0 | 2166 | 970 | 0 | 0 | 0 | 0 |
| ARL2 | 846 | 522 | 916 | 56 | 558 | 0 | 4093 | 1128 |
| ARL2BP | 204 | 0 | 0 | 1884 | 190 | 0 | 0 | 0 |
| ARL3 | 176 | 0 | 0 | 0 | 1996 | 0 | 1098 | 1274 |
| ARL4A | 1470 | 408 | 6200 | 1286 | 188 | 2528 | 0 | 0 |
| ARL4D | 3252 | 1770 | 7784 | 0 | 6202 | 1777 | 767 | 2402 |
| ARL5A | 0 | 340 | 2504 | 748 | 1244 | 0 | 0 | 484 |
| ARL5B | 0 | 214 | 0 | 0 | 0 | 0 | 0 | 0 |
| ARL6 | 0 | 0 | 0 | 0 | 222 | 0 | 0 | 0 |
| ARL6IP1 | 0 | 590 | 3290 | 1350 | 4492 | 6789 | 3984 | 0 |
| ARL6IP4 | 2278 | 2094 | 5521 | 3560 | 6092 | 1926 | 2746 | 938 |
| ARL6IP5 | 0 | 430 | 1312 | 2713 | 5447 | 799 | 796 | 0 |
| ARL6IP6 | 380 | 0 | 1076 | 1340 | 1698 | 0 | 564 | 0 |
| ARL8A | 1684 | 706 | 806 | 0 | 1640 | 1936 | 2850 | 450 |
| ARL8B | 25401 | 36813 | 134593 | 50201 | 122190 | 88719 | 115348 | 53503 |
| ARL9 | 0 | 0 | 0 | 0 | 394 | 0 | 0 | 0 |

| | | | | | | | | |
|---|---|---|---|---|---|---|---|---|
| ARMC1 | 0 | 370 | 0 | 0 | 0 | 0 | 0 | 0 |
| ARMC10 | 0 | 0 | 0 | 680 | 2374 | 0 | 752 | 0 |
| ARMC2 | 0 | 0 | 0 | 798 | 0 | 0 | 0 | 0 |
| ARMC7 | 0 | 0 | 0 | 1076 | 1070 | 0 | 0 | 0 |
| ARMC8 | 15081 | 33173 | 36974 | 31925 | 84537 | 26071 | 69704 | 43890 |
| ARMCX2 | 0 | 0 | 0 | 340 | 0 | 0 | 0 | 0 |
| ARMCX3 | 0 | 0 | 650 | 660 | 0 | 1964 | 212 | 0 |
| ARMCX5 | 0 | 258 | 0 | 0 | 0 | 0 | 0 | 0 |
| ARMCX6 | 0 | 1036 | 0 | 0 | 1646 | 2022 | 0 | 0 |
| ARMH3 | 0 | 0 | 0 | 0 | 0 | 0 | 0 | 1008 |
| ARMH4 | 0 | 0 | 0 | 0 | 3680 | 0 | 0 | 0 |
| ARMT1 | 0 | 0 | 0 | 0 | 352 | 0 | 0 | 0 |
| ARNT | 3786 | 8679 | 25776 | 7310 | 27266 | 9207 | 21295 | 9773 |
| ARNTL | 8654 | 5262 | 12432 | 9148 | 10308 | 15339 | 9756 | 11102 |
| ARNTL2 | 0 | 0 | 0 | 0 | 272 | 0 | 0 | 0 |
| ARPC1A | 926 | 1056 | 3356 | 7320 | 5546 | 1788 | 682 | 0 |
| ARPC1B | 895 | 534 | 2894 | 5042 | 4792 | 3234 | 124 | 2582 |
| ARPC2 | 2994 | 3204 | 18199 | 8170 | 10946 | 1768 | 7780 | 4397 |
| ARPC3 | 3850 | 3108 | 7021 | 9451 | 19278 | 7520 | 5194 | 3226 |
| ARPC4 | 788 | 1360 | 4129 | 4384 | 11131 | 0 | 5430 | 1892 |
| ARPC5 | 1594 | 931 | 7044 | 6600 | 5450 | 4268 | 1852 | 4773 |
| ARPC5L | 1284 | 0 | 0 | 4583 | 7305 | 3591 | 955 | 2980 |
| ARPP19 | 0 | 686 | 0 | 1770 | 1888 | 0 | 0 | 0 |
| ARRB1 | 24337 | 12206 | 120413 | 30331 | 33917 | 11826 | 39513 | 15424 |
| ARRB2 | 0 | 0 | 568 | 0 | 458 | 0 | 212 | 0 |
| ARRDC1 | 30 | 360 | 1534 | 0 | 2302 | 0 | 1406 | 908 |
| ARRDC1-AS1 | 0 | 0 | 0 | 0 | 72 | 0 | 0 | 0 |
| ARRDC2 | 344 | 646 | 1472 | 806 | 202 | 0 | 0 | 0 |
| ARRDC3 | 0 | 0 | 754 | 0 | 0 | 0 | 0 | 0 |
| ARRDC3-AS1 | 0 | 636 | 0 | 0 | 0 | 0 | 0 | 0 |
| ARRDC4 | 504 | 376 | 0 | 0 | 608 | 0 | 0 | 0 |
| ARSA | 648 | 96 | 0 | 0 | 0 | 0 | 0 | 706 |
| ARSB | 0 | 0 | 0 | 0 | 0 | 0 | 0 | 568 |
| ARSD | 0 | 0 | 0 | 0 | 1330 | 1644 | 0 | 0 |
| ARSE | 0 | 874 | 0 | 0 | 1158 | 0 | 692 | 0 |
| ARSG | 0 | 0 | 0 | 0 | 0 | 1437 | 0 | 0 |
| ARSJ | 0 | 0 | 0 | 0 | 58 | 0 | 0 | 0 |
| ARSK | 0 | 0 | 0 | 0 | 880 | 0 | 0 | 0 |
| ARV1 | 0 | 394 | 0 | 0 | 1422 | 0 | 0 | 0 |
| ARVCF | 0 | 0 | 0 | 0 | 876 | 0 | 0 | 0 |
| AS3MT | 0 | 0 | 0 | 3293 | 6682 | 0 | 4048 | 882 |
| ASAH1 | 1738 | 1575 | 0 | 0 | 4746 | 1795 | 0 | 2336 |
| ASAH2B | 0 | 0 | 2024 | 0 | 0 | 0 | 0 | 0 |

| | | | | | | | | |
|---|---|---|---|---|---|---|---|---|
| ASB13 | 0 | 654 | 3189 | 0 | 330 | 0 | 590 | 0 |
| ASB3 | 0 | 0 | 556 | 1072 | 0 | 0 | 0 | 0 |
| ASB9 | 0 | 574 | 0 | 0 | 1016 | 0 | 0 | 0 |
| ASCC1 | 9210 | 27726 | 30918 | 27987 | 76295 | 46143 | 123484 | 56356 |
| ASCC2 | 0 | 0 | 0 | 0 | 996 | 0 | 0 | 1032 |
| ASCC3 | 0 | 0 | 0 | 0 | 248 | 0 | 174 | 0 |
| ASCL2 | 0 | 0 | 0 | 5935 | 3504 | 1082 | 13981 | 520 |
| ASDURF | 0 | 0 | 0 | 886 | 0 | 0 | 458 | 0 |
| ASF1A | 670 | 0 | 0 | 0 | 1372 | 0 | 0 | 702 |
| ASF1B | 616 | 298 | 0 | 1038 | 780 | 0 | 856 | 0 |
| ASGR1 | 0 | 0 | 0 | 0 | 0 | 0 | 24 | 0 |
| ASH1L | 0 | 300 | 0 | 0 | 0 | 0 | 0 | 290 |
| ASH1L-AS1 | 0 | 0 | 1206 | 0 | 390 | 1346 | 0 | 0 |
| ASH2L | 12248 | 7299 | 18996 | 14846 | 24823 | 23385 | 23105 | 6922 |
| ASIC2 | 0 | 324 | 929 | 1104 | 0 | 0 | 0 | 0 |
| ASIP | 0 | 0 | 0 | 0 | 0 | 0 | 382 | 0 |
| ASL | 1428 | 416 | 1816 | 2973 | 728 | 94 | 178 | 0 |
| ASNA1 | 0 | 0 | 0 | 0 | 560 | 0 | 1042 | 0 |
| ASNS | 0 | 0 | 1384 | 1649 | 0 | 0 | 0 | 1236 |
| ASNSD1 | 540 | 638 | 0 | 0 | 0 | 0 | 792 | 0 |
| ASPG | 168 | 0 | 0 | 0 | 0 | 0 | 0 | 0 |
| ASPH | 1100 | 6448 | 4367 | 8240 | 48976 | 22682 | 45623 | 17956 |
| ASPHD1 | 0 | 0 | 0 | 0 | 0 | 0 | 300 | 0 |
| ASPM | 0 | 0 | 0 | 0 | 1574 | 0 | 0 | 0 |
| ASPN | 0 | 0 | 800 | 0 | 0 | 0 | 0 | 0 |
| ASPSCR1 | 0 | 0 | 0 | 0 | 325 | 3765 | 7320 | 427 |
| ASS1 | 2882 | 2030 | 4159 | 5673 | 24424 | 0 | 1874 | 9307 |
| ASTE1 | 0 | 0 | 0 | 1307 | 0 | 0 | 0 | 0 |
| ASTN2 | 2990 | 4364 | 9745 | 3764 | 11396 | 3480 | 8693 | 1352 |
| ASXL1 | 9982 | 18495 | 16560 | 23175 | 84521 | 15100 | 142504 | 23590 |
| ASXL2 | 3216 | 4342 | 20504 | 6378 | 26511 | 5093 | 28534 | 5874 |
| ATAD1 | 0 | 1768 | 784 | 0 | 1632 | 0 | 0 | 0 |
| ATAD2 | 76 | 1690 | 974 | 2337 | 28995 | 10590 | 11504 | 5555 |
| ATAD3A | 0 | 0 | 853 | 0 | 0 | 0 | 0 | 0 |
| ATAD3B | 0 | 0 | 0 | 0 | 1242 | 0 | 0 | 0 |
| ATAD5 | 2046 | 1568 | 0 | 3910 | 8503 | 0 | 508 | 1112 |
| ATAT1 | 0 | 0 | 0 | 0 | 426 | 0 | 0 | 0 |
| ATE1 | 15148 | 12837 | 18086 | 5065 | 24268 | 11941 | 28741 | 7464 |
| ATF1 | 20232 | 41194 | 38078 | 25037 | 50944 | 24644 | 116739 | 14413 |
| ATF2 | 0 | 296 | 0 | 0 | 0 | 0 | 0 | 0 |
| ATF3 | 0 | 0 | 6797 | 0 | 714 | 2772 | 0 | 1230 |
| ATF4 | 1241 | 1042 | 4426 | 5362 | 2340 | 3542 | 1912 | 1044 |
| ATF5 | 518 | 0 | 0 | 744 | 510 | 1446 | 0 | 0 |

| | | | | | | | | |
|---|---|---|---|---|---|---|---|---|
| ATF6 | 44125 | 82789 | 85760 | 68016 | 218255 | 39896 | 130799 | 47851 |
| ATF6B | 412 | 288 | 824 | 609 | 2424 | 0 | 566 | 672 |
| ATF7 | 24222 | 43326 | 28570 | 23838 | 49020 | 17645 | 81242 | 24369 |
| ATF7IP | 218 | 0 | 0 | 0 | 0 | 0 | 0 | 0 |
| ATG10 | 4968 | 4948 | 5300 | 7580 | 22698 | 5697 | 8811 | 6984 |
| ATG101 | 896 | 0 | 0 | 0 | 770 | 0 | 0 | 0 |
| ATG12 | 0 | 784 | 1622 | 3033 | 3075 | 0 | 0 | 2561 |
| ATG13 | 0 | 0 | 0 | 0 | 628 | 356 | 0 | 0 |
| ATG16L1 | 0 | 0 | 0 | 0 | 300 | 0 | 660 | 330 |
| ATG16L2 | 0 | 0 | 2026 | 0 | 0 | 0 | 0 | 508 |
| ATG2A | 386 | 0 | 0 | 0 | 22 | 0 | 182 | 0 |
| ATG3 | 0 | 2094 | 942 | 3222 | 3000 | 0 | 776 | 1148 |
| ATG4A | 328 | 542 | 0 | 0 | 0 | 0 | 0 | 0 |
| ATG4B | 506 | 382 | 1376 | 0 | 1854 | 0 | 0 | 0 |
| ATG4D | 15040 | 9108 | 60308 | 9483 | 18228 | 17014 | 8995 | 6404 |
| ATG5 | 13944 | 19243 | 13203 | 35877 | 37890 | 27940 | 17408 | 11890 |
| ATG7 | 22202 | 34923 | 100861 | 54446 | 189165 | 72575 | 93376 | 50820 |
| ATG9A | 0 | 0 | 0 | 0 | 432 | 0 | 0 | 0 |
| ATG9B | 0 | 0 | 0 | 0 | 0 | 0 | 320 | 0 |
| ATIC | 0 | 0 | 2689 | 0 | 1154 | 2383 | 2134 | 0 |
| ATL2 | 0 | 840 | 1618 | 1266 | 5428 | 1522 | 928 | 1510 |
| ATL3 | 0 | 370 | 0 | 0 | 1332 | 0 | 0 | 334 |
| ATM | 0 | 478 | 0 | 0 | 0 | 0 | 639 | 0 |
| ATOH1 | 376 | 0 | 0 | 0 | 0 | 0 | 0 | 0 |
| ATOH8 | 0 | 0 | 0 | 0 | 142 | 0 | 0 | 0 |
| ATOX1 | 726 | 2456 | 1810 | 3409 | 586 | 924 | 1656 | 664 |
| ATP10B | 0 | 0 | 0 | 0 | 0 | 0 | 196 | 86 |
| ATP10D | 0 | 676 | 0 | 412 | 0 | 4842 | 88 | 0 |
| ATP11A | 0 | 0 | 0 | 0 | 1100 | 0 | 104 | 290 |
| ATP11B | 0 | 82 | 890 | 0 | 248 | 0 | 0 | 0 |
| ATP13A1 | 0 | 668 | 0 | 0 | 0 | 0 | 0 | 0 |
| ATP13A3 | 0 | 0 | 0 | 0 | 1240 | 0 | 464 | 0 |
| ATP1A1 | 550975 | 973971 | 949355 | 650858 | 822752 | 693191 | 806188 | 148549 |
| ATP1A1-AS1 | 0 | 0 | 2245 | 0 | 450 | 0 | 0 | 0 |
| ATP1B1 | 1028 | 1792 | 2241 | 3151 | 1746 | 200 | 60 | 0 |
| ATP1B3 | 3783 | 0 | 17814 | 4316 | 2436 | 0 | 3335 | 1120 |
| ATP23 | 8875 | 9226 | 20423 | 11597 | 7760 | 3738 | 16709 | 4315 |
| ATP2A1-AS1 | 0 | 0 | 0 | 0 | 420 | 0 | 0 | 0 |
| ATP2A2 | 1263 | 0 | 0 | 0 | 372 | 0 | 0 | 0 |
| ATP2A3 | 1048 | 444 | 1126 | 470 | 0 | 0 | 524 | 0 |
| ATP2B1 | 212 | 0 | 561 | 0 | 1208 | 0 | 0 | 0 |
| ATP2B1-AS1 | 0 | 0 | 0 | 950 | 0 | 0 | 0 | 0 |
| ATP2B4 | 0 | 0 | 1000 | 0 | 700 | 0 | 0 | 0 |

| | | | | | | | | |
|---|---|---|---|---|---|---|---|---|
| ATP2C1 | 426 | 168 | 926 | 0 | 1244 | 0 | 0 | 0 |
| ATP2C2 | 0 | 0 | 1863 | 0 | 60 | 1282 | 0 | 0 |
| ATP5F1A | 4625 | 5992 | 11960 | 4517 | 6944 | 5794 | 5059 | 1580 |
| ATP5F1B | 6531 | 7638 | 23789 | 17242 | 15312 | 9387 | 4467 | 10688 |
| ATP5F1C | 4656 | 10570 | 16987 | 7248 | 22140 | 6395 | 4674 | 4091 |
| ATP5F1D | 242 | 1771 | 3514 | 2406 | 7645 | 3884 | 0 | 2962 |
| ATP5F1E | 4450 | 15560 | 14902 | 24035 | 33068 | 8214 | 24073 | 10843 |
| ATP5IF1 | 2646 | 9180 | 10237 | 7004 | 9651 | 5196 | 3945 | 1378 |
| ATP5MC1 | 1518 | 2944 | 4608 | 5058 | 4884 | 3164 | 3593 | 1516 |
| ATP5MC2 | 12758 | 12331 | 9308 | 13981 | 22571 | 8346 | 10874 | 4056 |
| ATP5MC3 | 5532 | 14900 | 19556 | 24601 | 17833 | 10671 | 8193 | 5602 |
| ATP5MD | 7352 | 11486 | 5667 | 12939 | 15214 | 2941 | 8888 | 2000 |
| ATP5ME | 4016 | 6422 | 2163 | 3406 | 10480 | 1766 | 6315 | 1590 |
| ATP5MF | 2011 | 6300 | 4356 | 6108 | 15544 | 9295 | 5788 | 1566 |
| ATP5MG | 5448 | 9969 | 21926 | 12878 | 24286 | 7484 | 12874 | 10378 |
| ATP5MPL | 642215 | 682840 | 294399 | 197789 | 1511680 | 209128 | 855419 | 156433 |
| ATP5PB | 0 | 6824 | 16832 | 9421 | 8375 | 11893 | 4572 | 1232 |
| ATP5PD | 2362 | 5176 | 6532 | 2758 | 8380 | 9146 | 4756 | 2391 |
| ATP5PF | 3998 | 9522 | 12360 | 8780 | 6414 | 4470 | 4322 | 6462 |
| ATP5PO | 2829 | 6639 | 14698 | 18890 | 15373 | 4614 | 12292 | 9500 |
| ATP5S | 0 | 702 | 0 | 0 | 1142 | 3014 | 0 | 568 |
| ATP6AP1 | 950 | 644 | 54 | 926 | 5400 | 0 | 2456 | 1491 |
| ATP6AP2 | 508 | 456 | 0 | 0 | 2236 | 762 | 318 | 1954 |
| ATP6V0A1 | 0 | 558 | 0 | 494 | 0 | 0 | 218 | 0 |
| ATP6V0A2 | 714 | 0 | 0 | 0 | 426 | 0 | 0 | 0 |
| ATP6V0B | 0 | 4624 | 5813 | 8577 | 7177 | 9464 | 3384 | 2611 |
| ATP6V0C | 700 | 1830 | 1302 | 3500 | 2748 | 1992 | 1532 | 998 |
| ATP6V0D1 | 2487 | 1748 | 8737 | 1036 | 3051 | 3060 | 2693 | 1037 |
| ATP6V0E1 | 3480 | 1176 | 5509 | 9049 | 13105 | 0 | 2170 | 2064 |
| ATP6V0E2 | 0 | 0 | 0 | 0 | 634 | 0 | 0 | 0 |
| ATP6V1A | 0 | 888 | 1206 | 0 | 0 | 0 | 428 | 0 |
| ATP6V1B2 | 0 | 304 | 214 | 0 | 0 | 422 | 0 | 0 |
| ATP6V1C1 | 0 | 0 | 0 | 931 | 492 | 0 | 2240 | 844 |
| ATP6V1D | 1384 | 400 | 1830 | 698 | 1975 | 0 | 168 | 1294 |
| ATP6V1E1 | 608 | 82 | 4328 | 2627 | 920 | 2092 | 1750 | 0 |
| ATP6V1F | 58 | 302 | 2035 | 10538 | 6220 | 2835 | 3575 | 3094 |
| ATP6V1G1 | 0 | 7698 | 3736 | 4030 | 8680 | 6106 | 3729 | 616 |
| ATP6V1G2-DDX39B | 1489 | 108 | 1282 | 944 | 3238 | 2226 | 1973 | 1034 |
| ATP6V1H | 38334 | 33887 | 58075 | 78508 | 68134 | 264135 | 647062 | 31356 |
| ATP7B | 0 | 0 | 0 | 0 | 1442 | 0 | 0 | 0 |
| ATP8B1 | 358 | 354 | 0 | 0 | 828 | 0 | 655 | 0 |
| ATP9A | 0 | 0 | 0 | 0 | 436 | 0 | 116 | 0 |
| ATP9B | 682 | 742 | 3508 | 672 | 13149 | 540 | 3710 | 1182 |

| | | | | | | | | |
|---|---|---|---|---|---|---|---|---|
| ATPAF1 | 0 | 494 | 700 | 2333 | 962 | 0 | 864 | 0 |
| ATPAF2 | 0 | 512 | 0 | 0 | 2366 | 1774 | 0 | 0 |
| ATPSCKMT | 0 | 0 | 0 | 0 | 218 | 0 | 0 | 0 |
| ATR | 4836 | 5260 | 4384 | 12284 | 28032 | 7586 | 16400 | 5236 |
| ATRAID | 2264 | 5784 | 7657 | 0 | 14100 | 2101 | 4813 | 8985 |
| ATRIP-TREX1 | 514 | 368 | 762 | 1324 | 124 | 0 | 334 | 0 |
| ATRN | 790 | 348 | 0 | 0 | 372 | 0 | 0 | 722 |
| ATRX | 246836 | 306086 | 418534 | 297530 | 1326508 | 438888 | 694545 | 282350 |
| ATXN1 | 0 | 0 | 0 | 0 | 0 | 922 | 0 | 1022 |
| ATXN10 | 62 | 0 | 554 | 998 | 2288 | 0 | 456 | 0 |
| ATXN1L | 0 | 822 | 0 | 0 | 0 | 0 | 0 | 0 |
| ATXN2 | 2600 | 0 | 0 | 0 | 2026 | 0 | 0 | 0 |
| ATXN2L | 0 | 258 | 0 | 0 | 1581 | 0 | 0 | 0 |
| ATXN3 | 0 | 0 | 0 | 0 | 876 | 3112 | 0 | 0 |
| ATXN7L1 | 0 | 0 | 0 | 0 | 368 | 0 | 0 | 0 |
| ATXN7L2 | 1164 | 0 | 0 | 0 | 0 | 0 | 0 | 0 |
| ATXN7L3 | 512 | 0 | 0 | 0 | 0 | 0 | 0 | 0 |
| ATXN7L3B | 0 | 0 | 596 | 580 | 730 | 1648 | 246 | 0 |
| AUH | 41796 | 39684 | 95713 | 40202 | 124682 | 61738 | 68489 | 22094 |
| AUNIP | 0 | 550 | 0 | 1410 | 0 | 0 | 0 | 0 |
| AUP1 | 2107 | 1424 | 3510 | 3069 | 6082 | 1837 | 2788 | 1640 |
| AURKA | 0 | 0 | 0 | 0 | 0 | 0 | 798 | 0 |
| AURKAIP1 | 0 | 886 | 4949 | 2113 | 2798 | 0 | 0 | 912 |
| AURKB | 0 | 332 | 0 | 852 | 5518 | 0 | 0 | 0 |
| AUTS2 | 0 | 454 | 450 | 0 | 596 | 0 | 52 | 0 |
| AVPI1 | 420 | 0 | 1993 | 0 | 0 | 0 | 0 | 0 |
| AXIN1 | 6540 | 2536 | 9683 | 11063 | 10753 | 4819 | 14124 | 1560 |
| AXIN2 | 2356 | 912 | 3938 | 4113 | 5004 | 16172 | 54899 | 1864 |
| AXL | 0 | 0 | 0 | 0 | 152 | 0 | 0 | 0 |
| AZGP1 | 0 | 0 | 0 | 4738 | 1388 | 1628 | 7102 | 1219 |
| AZI2 | 0 | 0 | 0 | 0 | 0 | 0 | 1274 | 0 |
| AZIN1 | 554 | 0 | 276 | 1488 | 1944 | 658 | 0 | 1084 |
| AZIN1-AS1 | 0 | 562 | 0 | 0 | 0 | 0 | 0 | 1008 |
| B2M | 37488930 | 30556355 | 144472759 | 21137703 | 37768162 | 26399669 | 28426319 | 19521321 |
| B3GALT5 | 0 | 1272 | 0 | 0 | 0 | 0 | 0 | 0 |
| B3GALT6 | 0 | 0 | 0 | 0 | 486 | 0 | 158 | 0 |
| B3GAT3 | 0 | 972 | 0 | 0 | 0 | 0 | 0 | 673 |
| B3GLCT | 0 | 0 | 0 | 0 | 410 | 0 | 0 | 0 |
| B3GNT2 | 0 | 340 | 606 | 0 | 462 | 0 | 0 | 0 |
| B3GNT3 | 0 | 204 | 1978 | 1038 | 278 | 0 | 0 | 0 |
| B3GNT6 | 228 | 256 | 0 | 0 | 0 | 0 | 0 | 0 |
| B3GNT7 | 0 | 1228 | 1038 | 0 | 0 | 0 | 0 | 0 |
| B3GNT8 | 0 | 460 | 2386 | 0 | 0 | 0 | 0 | 0 |

| | | | | | | | | |
|---|---|---|---|---|---|---|---|---|
| B3GNTL1 | 0 | 0 | 0 | 0 | 534 | 0 | 0 | 0 |
| B4GALNT2 | 15275 | 1695 | 34785 | 0 | 0 | 0 | 280 | 0 |
| B4GALT1 | 10018 | 3454 | 7854 | 1146 | 104 | 1396 | 1904 | 1277 |
| B4GALT1-AS1 | 0 | 0 | 1094 | 0 | 727 | 0 | 0 | 0 |
| B4GALT2 | 0 | 0 | 0 | 1068 | 0 | 0 | 824 | 0 |
| B4GALT3 | 0 | 356 | 0 | 0 | 616 | 0 | 916 | 710 |
| B4GALT4 | 0 | 1428 | 1776 | 0 | 2797 | 0 | 0 | 0 |
| B4GALT5 | 410 | 1454 | 0 | 3597 | 822 | 0 | 2344 | 558 |
| B4GALT7 | 0 | 0 | 736 | 0 | 334 | 0 | 0 | 0 |
| B4GAT1 | 0 | 0 | 0 | 873 | 0 | 0 | 366 | 0 |
| B9D1 | 0 | 198 | 0 | 1187 | 3322 | 0 | 0 | 0 |
| B9D2 | 0 | 1083 | 0 | 0 | 0 | 0 | 0 | 0 |
| BAALC | 0 | 0 | 0 | 0 | 734 | 1012 | 0 | 0 |
| BAAT | 0 | 362 | 0 | 0 | 0 | 0 | 876 | 0 |
| BABAM1 | 314 | 20 | 1520 | 0 | 4008 | 0 | 290 | 0 |
| BABAM2 | 676 | 0 | 0 | 0 | 1345 | 0 | 674 | 0 |
| BACE2 | 0 | 0 | 0 | 1128 | 688 | 0 | 544 | 0 |
| BAD | 306 | 130 | 0 | 2060 | 6516 | 0 | 0 | 1386 |
| BAG1 | 14645 | 19711 | 45746 | 24697 | 35262 | 36076 | 29990 | 6873 |
| BAG2 | 0 | 780 | 0 | 0 | 1166 | 276 | 0 | 0 |
| BAG3 | 596 | 338 | 0 | 0 | 558 | 0 | 0 | 0 |
| BAG4 | 900 | 2226 | 8252 | 0 | 20581 | 0 | 5130 | 7104 |
| BAG5 | 312 | 202 | 0 | 0 | 0 | 0 | 0 | 0 |
| BAG6 | 0 | 40 | 2674 | 0 | 1333 | 2990 | 0 | 0 |
| BAHCC1 | 0 | 0 | 758 | 0 | 870 | 0 | 0 | 0 |
| BAHD1 | 0 | 0 | 0 | 0 | 162 | 0 | 0 | 0 |
| BAIAP2 | 0 | 0 | 0 | 0 | 0 | 3490 | 134 | 0 |
| BAIAP2L1 | 822 | 1240 | 0 | 0 | 1220 | 1828 | 0 | 744 |
| BAIAP2L2 | 1176 | 210 | 548 | 0 | 1348 | 0 | 0 | 478 |
| BAK1 | 0 | 290 | 924 | 2473 | 1306 | 0 | 812 | 0 |
| BAMBI | 0 | 0 | 0 | 1304 | 1816 | 2076 | 2254 | 0 |
| BANF1 | 566 | 2272 | 3461 | 13262 | 7262 | 9949 | 8289 | 1891 |
| BANK1 | 0 | 768 | 0 | 0 | 0 | 1553 | 0 | 0 |
| BANP | 0 | 0 | 0 | 0 | 1247 | 0 | 626 | 0 |
| BAP1 | 18782 | 19622 | 37463 | 30030 | 99102 | 23039 | 110621 | 13093 |
| BARD1 | 1942 | 4828 | 0 | 3719 | 12449 | 0 | 1424 | 3365 |
| BARX2 | 0 | 0 | 508 | 0 | 688 | 0 | 0 | 0 |
| BASP1 | 470 | 0 | 0 | 0 | 0 | 0 | 0 | 0 |
| BATF | 0 | 0 | 0 | 0 | 612 | 0 | 386 | 974 |
| BATF2 | 482 | 0 | 0 | 0 | 0 | 0 | 0 | 0 |
| BAX | 134038 | 99895 | 444088 | 134300 | 329888 | 138910 | 187995 | 67015 |
| BAZ1A | 175603 | 116640 | 160895 | 86538 | 360478 | 64427 | 152197 | 64402 |
| BAZ1B | 0 | 0 | 0 | 0 | 0 | 0 | 1004 | 0 |

| | | | | | | | | |
|---|---|---|---|---|---|---|---|---|
| BAZ2A | 0 | 0 | 0 | 0 | 448 | 0 | 0 | 0 |
| BAZ2B | 0 | 734 | 0 | 0 | 0 | 0 | 0 | 296 |
| BBC3 | 0 | 0 | 0 | 457 | 0 | 0 | 310 | 0 |
| BBIP1 | 1986 | 0 | 0 | 0 | 0 | 0 | 631 | 0 |
| BBOF1 | 292 | 0 | 0 | 0 | 0 | 0 | 0 | 0 |
| BBOX1 | 0 | 0 | 1602 | 0 | 0 | 0 | 0 | 0 |
| BBOX1-AS1 | 0 | 0 | 0 | 0 | 340 | 0 | 670 | 0 |
| BBS1 | 0 | 50 | 0 | 0 | 646 | 0 | 1308 | 0 |
| BBS10 | 0 | 0 | 752 | 0 | 0 | 0 | 0 | 0 |
| BBS4 | 0 | 0 | 0 | 0 | 1469 | 0 | 138 | 0 |
| BBS5 | 226 | 0 | 0 | 770 | 0 | 0 | 0 | 0 |
| BBS7 | 0 | 0 | 0 | 0 | 860 | 0 | 0 | 0 |
| BBS9 | 0 | 0 | 0 | 0 | 0 | 230 | 0 | 0 |
| BBX | 72989 | 74458 | 47173 | 62974 | 310075 | 93959 | 247610 | 57449 |
| BCAM | 0 | 0 | 0 | 0 | 0 | 0 | 86 | 0 |
| BCAP29 | 360 | 1215 | 2730 | 1568 | 624 | 0 | 874 | 0 |
| BCAP31 | 3714 | 2186 | 5891 | 8781 | 14855 | 3058 | 4110 | 6357 |
| BCAR1 | 0 | 1016 | 0 | 0 | 450 | 0 | 0 | 0 |
| BCAR3 | 612 | 150 | 0 | 0 | 520 | 806 | 0 | 0 |
| BCAS1 | 0 | 424 | 0 | 0 | 230 | 0 | 0 | 0 |
| BCAS2 | 0 | 434 | 2144 | 1526 | 1579 | 0 | 0 | 0 |
| BCAS3 | 1084 | 1712 | 7512 | 348 | 4999 | 2373 | 5678 | 0 |
| BCAS4 | 2828 | 7303 | 2962 | 580 | 22384 | 4719 | 19911 | 8118 |
| BCAT2 | 970 | 0 | 1532 | 0 | 2302 | 0 | 0 | 1392 |
| BCCIP | 692 | 522 | 0 | 1000 | 3097 | 0 | 0 | 2918 |
| BCKDHA | 0 | 136 | 1425 | 0 | 1786 | 1832 | 1062 | 0 |
| BCKDHB | 4188 | 2974 | 16085 | 6396 | 14922 | 2050 | 3196 | 1516 |
| BCKDK | 0 | 0 | 0 | 0 | 1064 | 0 | 312 | 1764 |
| BCL10 | 23348 | 16294 | 56695 | 6966 | 12728 | 11048 | 12254 | 4514 |
| BCL11A | 3904 | 15894 | 13223 | 61266 | 93470 | 19261 | 96563 | 15849 |
| BCL2 | 0 | 174 | 0 | 624 | 464 | 946 | 582 | 0 |
| BCL2A1 | 23422 | 47083 | 12078 | 4594 | 116184 | 16918 | 17013 | 31596 |
| BCL2L1 | 0 | 0 | 0 | 1893 | 1002 | 1280 | 1362 | 0 |
| BCL2L11 | 1500 | 928 | 0 | 190 | 10665 | 4890 | 3557 | 5166 |
| BCL2L12 | 3260 | 3766 | 20091 | 9751 | 28338 | 10817 | 5496 | 9026 |
| BCL2L13 | 0 | 440 | 0 | 0 | 306 | 0 | 0 | 0 |
| BCL2L14 | 50 | 460 | 0 | 0 | 0 | 0 | 0 | 0 |
| BCL2L15 | 0 | 2782 | 2432 | 0 | 1070 | 0 | 0 | 0 |
| BCL2L2 | 602 | 0 | 0 | 0 | 0 | 0 | 0 | 0 |
| BCL3 | 210 | 0 | 1468 | 0 | 930 | 0 | 508 | 526 |
| BCL7A | 0 | 0 | 0 | 0 | 110 | 0 | 222 | 0 |
| BCL7B | 0 | 1230 | 0 | 0 | 848 | 3973 | 484 | 0 |
| BCL7C | 0 | 0 | 3558 | 1305 | 856 | 3025 | 1339 | 0 |

| | | | | | | | | |
|---|---|---|---|---|---|---|---|---|
| BCL9 | 1006 | 178 | 0 | 0 | 0 | 0 | 3929 | 0 |
| BCL9L | 0 | 0 | 0 | 0 | 72 | 0 | 0 | 0 |
| BCLAF1 | 0 | 0 | 0 | 1036 | 0 | 0 | 0 | 0 |
| BCO1 | 0 | 446 | 0 | 0 | 0 | 0 | 0 | 0 |
| BCO2 | 646 | 1168 | 1240 | 0 | 602 | 1620 | 708 | 966 |
| BCOR | 1472 | 4144 | 3497 | 1198 | 5723 | 2273 | 8962 | 0 |
| BCORL1 | 296 | 22 | 1193 | 46 | 662 | 326 | 4337 | 308 |
| BCR | 3556 | 4966 | 5284 | 5898 | 10591 | 3241 | 21738 | 4606 |
| BCRP2 | 134 | 0 | 0 | 0 | 0 | 0 | 0 | 0 |
| BCS1L | 0 | 0 | 0 | 0 | 1777 | 2358 | 0 | 0 |
| BDH1 | 1130 | 946 | 3473 | 4235 | 442 | 0 | 1013 | 604 |
| BDH2 | 622 | 1092 | 2057 | 584 | 870 | 0 | 866 | 0 |
| BDKRB1 | 0 | 0 | 1966 | 0 | 0 | 0 | 0 | 0 |
| BDP1 | 6872 | 7512 | 6150 | 2510 | 22039 | 11528 | 7198 | 6458 |
| BECN1 | 0 | 198 | 2962 | 880 | 1782 | 1458 | 1864 | 0 |
| BEND5 | 8794 | 9735 | 16086 | 3902 | 24203 | 5972 | 0 | 10497 |
| BEND7 | 0 | 0 | 962 | 0 | 0 | 0 | 0 | 0 |
| BEST2 | 0 | 226 | 0 | 0 | 0 | 0 | 0 | 0 |
| BEST3 | 0 | 0 | 0 | 0 | 0 | 0 | 2537 | 0 |
| BEST4 | 0 | 0 | 934 | 0 | 0 | 0 | 396 | 0 |
| BET1 | 0 | 0 | 0 | 0 | 1848 | 0 | 1986 | 708 |
| BET1L | 0 | 552 | 0 | 126 | 786 | 0 | 340 | 0 |
| BEX3 | 0 | 0 | 1474 | 0 | 2684 | 0 | 0 | 0 |
| BEX4 | 940 | 0 | 854 | 0 | 1407 | 0 | 0 | 0 |
| BFAR | 318 | 1562 | 764 | 2917 | 542 | 0 | 310 | 0 |
| BFSP2 | 4704 | 1268 | 4146 | 1193 | 0 | 1948 | 0 | 0 |
| BGN | 11602 | 17728 | 14738 | 38374 | 546826 | 227913 | 112573 | 232670 |
| BHLHE40 | 0 | 0 | 840 | 230 | 0 | 1144 | 573 | 0 |
| BHLHE41 | 747 | 0 | 0 | 0 | 650 | 0 | 0 | 0 |
| BICC1 | 1130 | 882 | 3601 | 0 | 4921 | 2596 | 0 | 1768 |
| BICDL2 | 0 | 490 | 260 | 1632 | 0 | 3670 | 1336 | 0 |
| BICRA | 0 | 0 | 28 | 0 | 0 | 0 | 0 | 0 |
| BID | 0 | 902 | 1656 | 3786 | 3102 | 0 | 728 | 1778 |
| BIK | 0 | 988 | 0 | 4022 | 710 | 2590 | 1554 | 0 |
| BIN1 | 0 | 596 | 0 | 0 | 892 | 0 | 466 | 0 |
| BIN2 | 0 | 0 | 0 | 0 | 0 | 0 | 372 | 0 |
| BIN3 | 210 | 558 | 1586 | 0 | 198 | 2185 | 350 | 0 |
| BIRC2 | 0 | 102 | 0 | 0 | 586 | 0 | 578 | 0 |
| BIRC3 | 0 | 226 | 998 | 0 | 0 | 709 | 0 | 0 |
| BIRC5 | 0 | 0 | 0 | 1457 | 4668 | 0 | 0 | 0 |
| BIRC6 | 23666 | 34569 | 45769 | 28772 | 85813 | 33179 | 66659 | 11313 |
| BISPR | 3524 | 1582 | 11548 | 2568 | 71341 | 6849 | 4159 | 15415 |
| BIVM | 0 | 0 | 0 | 524 | 332 | 0 | 0 | 0 |

| | | | | | | | | |
|---|---|---|---|---|---|---|---|---|
| BIVM-ERCC5 | 580 | 0 | 0 | 0 | 0 | 1515 | 0 | 0 |
| BLCAP | 1040 | 894 | 3140 | 3632 | 4665 | 1666 | 3122 | 1700 |
| BLM | 4180 | 3284 | 11563 | 18701 | 33991 | 9858 | 8226 | 8046 |
| BLMH | 0 | 0 | 0 | 1096 | 0 | 0 | 0 | 0 |
| BLNK | 0 | 296 | 1770 | 0 | 0 | 0 | 0 | 616 |
| BLOC1S1 | 6567 | 3724 | 5232 | 2776 | 7634 | 1147 | 3832 | 746 |
| BLOC1S1-RDH5 | 0 | 0 | 1884 | 0 | 22 | 0 | 1246 | 0 |
| BLOC1S2 | 0 | 1050 | 0 | 2076 | 1642 | 0 | 0 | 2120 |
| BLOC1S3 | 0 | 0 | 0 | 0 | 0 | 0 | 456 | 0 |
| BLOC1S4 | 0 | 952 | 2948 | 0 | 1158 | 0 | 1718 | 748 |
| BLOC1S5 | 0 | 0 | 0 | 0 | 0 | 1854 | 0 | 0 |
| BLOC1S5-TXNDC5 | 288 | 384 | 0 | 0 | 419 | 75 | 1568 | 458 |
| BLOC1S6 | 0 | 0 | 0 | 0 | 616 | 0 | 280 | 0 |
| BLVRB | 1368 | 1028 | 4669 | 0 | 10246 | 3990 | 994 | 3572 |
| BLZF1 | 0 | 312 | 210 | 0 | 1400 | 0 | 130 | 0 |
| BMI1 | 0 | 0 | 1077 | 0 | 274 | 1089 | 54 | 750 |
| BMP1 | 0 | 0 | 0 | 0 | 268 | 0 | 278 | 0 |
| BMP2 | 0 | 114 | 1238 | 0 | 0 | 0 | 0 | 0 |
| BMP2K | 0 | 0 | 0 | 0 | 774 | 0 | 910 | 641 |
| BMP4 | 0 | 90 | 0 | 0 | 3628 | 0 | 0 | 944 |
| BMP5 | 10186 | 3326 | 16327 | 1091 | 0 | 0 | 0 | 0 |
| BMP7 | 0 | 0 | 0 | 1380 | 0 | 0 | 1269 | 0 |
| BMP8B | 0 | 0 | 0 | 0 | 226 | 0 | 0 | 0 |
| BMPR1A | 4968 | 10194 | 13620 | 10090 | 3064 | 10036 | 32136 | 1626 |
| BMPR2 | 0 | 92 | 0 | 0 | 590 | 0 | 464 | 0 |
| BMS1 | 0 | 0 | 1262 | 0 | 0 | 0 | 0 | 0 |
| BMS1P4 | 0 | 0 | 0 | 0 | 0 | 0 | 0 | 682 |
| BMT2 | 546 | 0 | 0 | 0 | 580 | 0 | 0 | 0 |
| BNC2 | 0 | 554 | 0 | 0 | 1755 | 0 | 0 | 0 |
| BNIP1 | 0 | 0 | 0 | 1284 | 0 | 0 | 0 | 160 |
| BNIP2 | 0 | 482 | 0 | 0 | 0 | 0 | 296 | 832 |
| BNIP3 | 0 | 0 | 1138 | 0 | 624 | 0 | 0 | 0 |
| BNIP3L | 0 | 0 | 1631 | 0 | 844 | 1328 | 0 | 0 |
| BOC | 544 | 0 | 0 | 0 | 0 | 0 | 0 | 0 |
| BOD1 | 0 | 0 | 0 | 1089 | 1084 | 0 | 0 | 967 |
| BOK | 0 | 0 | 0 | 0 | 1280 | 0 | 0 | 894 |
| BOLA1 | 1260 | 0 | 799 | 0 | 1430 | 0 | 470 | 1753 |
| BOLA2 | 0 | 0 | 0 | 0 | 318 | 0 | 0 | 0 |
| BOLA2-SMG1P6 | 0 | 610 | 0 | 1635 | 1076 | 0 | 0 | 470 |
| BOLA2B | 1846 | 1316 | 2346 | 1136 | 2542 | 838 | 828 | 1112 |
| BOLA3 | 800 | 776 | 598 | 3440 | 1664 | 0 | 332 | 1500 |
| BOP1 | 0 | 244 | 0 | 1232 | 832 | 0 | 1230 | 0 |
| BORA | 0 | 0 | 0 | 0 | 0 | 0 | 0 | 1190 |

| | | | | | | | | |
|---|---|---|---|---|---|---|---|---|
| BORCS5 | 0 | 0 | 0 | 902 | 0 | 0 | 0 | 0 |
| BORCS6 | 0 | 0 | 698 | 0 | 1086 | 0 | 0 | 0 |
| BORCS7 | 66592 | 109640 | 82986 | 73253 | 149625 | 96218 | 194955 | 39560 |
| BORCS7-ASMT | 2036 | 0 | 0 | 10 | 3910 | 0 | 3654 | 0 |
| BORCS8 | 0 | 0 | 0 | 1195 | 208 | 0 | 0 | 0 |
| BPHL | 26132 | 42565 | 372008 | 39611 | 202888 | 85787 | 135979 | 72210 |
| BPI | 0 | 0 | 0 | 0 | 860 | 0 | 0 | 1036 |
| BPNT1 | 0 | 0 | 2190 | 0 | 370 | 0 | 0 | 0 |
| BPTF | 3232 | 2122 | 5438 | 0 | 8686 | 5168 | 1266 | 1545 |
| BRAF | 8543 | 19236 | 20895 | 23233 | 39009 | 20428 | 68412 | 17113 |
| BRAP | 0 | 0 | 0 | 0 | 194 | 0 | 0 | 0 |
| BRAT1 | 576 | 0 | 0 | 2339 | 0 | 0 | 0 | 0 |
| BRCA1 | 0 | 980 | 6743 | 3426 | 9710 | 2334 | 2274 | 1930 |
| BRCA2 | 7392 | 2054 | 2306 | 29366 | 41347 | 20344 | 28134 | 7454 |
| BRCC3 | 0 | 0 | 1650 | 0 | 354 | 832 | 0 | 0 |
| BRD1 | 0 | 0 | 1170 | 0 | 250 | 0 | 0 | 535 |
| BRD2 | 70190 | 82973 | 44388 | 53963 | 285200 | 61138 | 289955 | 60584 |
| BRD3 | 4284 | 6119 | 4379 | 3115 | 18984 | 5019 | 10756 | 4039 |
| BRD4 | 1720 | 2147 | 6850 | 1442 | 14802 | 1320 | 8768 | 1538 |
| BRD7 | 29109 | 33708 | 48696 | 56803 | 127394 | 64027 | 201889 | 25413 |
| BRD8 | 0 | 0 | 0 | 0 | 22 | 32 | 652 | 0 |
| BRD9 | 8482 | 12376 | 35075 | 13661 | 79697 | 24583 | 29329 | 9344 |
| BRF1 | 0 | 0 | 0 | 0 | 1214 | 0 | 0 | 0 |
| BRF2 | 0 | 0 | 0 | 0 | 38 | 0 | 316 | 0 |
| BRI3 | 2105 | 1615 | 7306 | 3807 | 4894 | 4790 | 4122 | 2742 |
| BRINP3 | 0 | 0 | 0 | 0 | 320 | 0 | 0 | 0 |
| BRIP1 | 0 | 0 | 0 | 396 | 6226 | 1570 | 1054 | 1064 |
| BRIX1 | 0 | 524 | 2420 | 0 | 2948 | 2758 | 1548 | 3041 |
| BRK1 | 0 | 3834 | 6694 | 7953 | 14114 | 4901 | 3673 | 2422 |
| BRMS1 | 0 | 0 | 0 | 182 | 3320 | 0 | 0 | 0 |
| BRMS1L | 0 | 0 | 0 | 0 | 0 | 0 | 0 | 1348 |
| BROX | 562 | 0 | 0 | 0 | 1426 | 0 | 0 | 28 |
| BRPF1 | 529 | 0 | 0 | 0 | 0 | 0 | 474 | 0 |
| BRPF3 | 0 | 0 | 0 | 776 | 1927 | 0 | 0 | 0 |
| BRWD1 | 0 | 0 | 2009 | 0 | 34 | 0 | 0 | 0 |
| BRWD1-AS2 | 598 | 590 | 0 | 0 | 0 | 0 | 0 | 0 |
| BRWD3 | 0 | 395 | 0 | 0 | 0 | 0 | 0 | 0 |
| BSCL2 | 0 | 976 | 1988 | 4196 | 0 | 460 | 1261 | 0 |
| BSDC1 | 320 | 280 | 964 | 0 | 1448 | 0 | 0 | 0 |
| BSG | 1204 | 1991 | 18124 | 2425 | 10326 | 4326 | 1204 | 6251 |
| BSPRY | 15411 | 41572 | 64674 | 44197 | 101675 | 57021 | 52449 | 24497 |
| BST1 | 0 | 0 | 0 | 956 | 488 | 0 | 0 | 0 |
| BST2 | 0 | 134 | 1180 | 1891 | 6522 | 0 | 0 | 5466 |

| | | | | | | | | |
|---|---|---|---|---|---|---|---|---|
| BTAF1 | 272 | 404 | 1760 | 0 | 0 | 0 | 0 | 0 |
| BTBD1 | 0 | 0 | 60 | 32 | 38 | 0 | 408 | 0 |
| BTBD10 | 0 | 0 | 0 | 952 | 0 | 0 | 0 | 2154 |
| BTBD2 | 0 | 0 | 1580 | 296 | 0 | 0 | 0 | 0 |
| BTBD3 | 0 | 0 | 0 | 0 | 146 | 0 | 0 | 0 |
| BTBD6 | 218 | 362 | 1848 | 1632 | 0 | 0 | 174 | 246 |
| BTBD7 | 0 | 0 | 0 | 0 | 296 | 0 | 0 | 0 |
| BTBD9 | 333 | 0 | 2032 | 0 | 606 | 2633 | 1842 | 2267 |
| BTD | 236 | 494 | 1294 | 134 | 1220 | 0 | 0 | 0 |
| BTF3 | 4800 | 13055 | 15839 | 23200 | 32420 | 16332 | 4135 | 8500 |
| BTF3L4 | 0 | 1186 | 1306 | 1930 | 1376 | 1588 | 146 | 844 |
| BTG1 | 3530 | 1118 | 8703 | 2257 | 1212 | 3654 | 730 | 1120 |
| BTG2 | 585 | 806 | 702 | 0 | 0 | 0 | 0 | 0 |
| BTG3 | 388 | 2156 | 0 | 2422 | 980 | 0 | 766 | 3214 |
| BTN2A1 | 0 | 0 | 0 | 0 | 286 | 0 | 0 | 0 |
| BTN2A2 | 0 | 0 | 0 | 0 | 0 | 0 | 398 | 0 |
| BTN3A1 | 0 | 606 | 0 | 0 | 358 | 0 | 288 | 0 |
| BTN3A2 | 0 | 0 | 0 | 0 | 0 | 0 | 278 | 0 |
| BTNL3 | 896 | 412 | 7057 | 0 | 540 | 0 | 0 | 0 |
| BTNL8 | 692 | 1108 | 0 | 0 | 564 | 0 | 0 | 0 |
| BTRC | 3600 | 1936 | 5138 | 4493 | 14408 | 4992 | 20428 | 4359 |
| BUB1 | 0 | 0 | 0 | 0 | 108 | 0 | 0 | 0 |
| BUB1B | 5901 | 14670 | 12743 | 48140 | 142057 | 19844 | 48428 | 39938 |
| BUB3 | 0 | 674 | 2101 | 4564 | 1010 | 0 | 1434 | 0 |
| BUD13 | 0 | 0 | 0 | 1290 | 0 | 0 | 0 | 0 |
| BUD23 | 632 | 1156 | 0 | 1435 | 2168 | 1702 | 390 | 784 |
| BUD31 | 824 | 506 | 8212 | 4746 | 508 | 0 | 3687 | 2818 |
| BYSL | 0 | 0 | 0 | 0 | 868 | 0 | 0 | 0 |
| BZW1 | 0 | 314 | 3356 | 944 | 2010 | 0 | 280 | 0 |
| BZW2 | 0 | 472 | 0 | 1430 | 5744 | 0 | 968 | 2463 |
| C10orf143 | 742 | 0 | 0 | 0 | 0 | 0 | 636 | 0 |
| C10orf25 | 46 | 2336 | 0 | 1130 | 19204 | 0 | 3852 | 4840 |
| C10orf88 | 0 | 126 | 0 | 0 | 0 | 0 | 0 | 0 |
| C10orf95 | 0 | 0 | 0 | 0 | 0 | 0 | 192 | 0 |
| C10orf99 | 7587 | 4352 | 20108 | 1382 | 0 | 0 | 5061 | 0 |
| C11orf1 | 0 | 542 | 296 | 1315 | 1230 | 1888 | 2678 | 0 |
| C11orf24 | 0 | 0 | 0 | 757 | 365 | 0 | 0 | 1160 |
| C11orf45 | 0 | 0 | 0 | 0 | 838 | 0 | 0 | 0 |
| C11orf49 | 41290 | 53284 | 139217 | 120464 | 236502 | 135169 | 133242 | 79204 |
| C11orf53 | 0 | 0 | 0 | 0 | 1091 | 0 | 0 | 0 |
| C11orf54 | 0 | 356 | 0 | 0 | 0 | 0 | 370 | 350 |
| C11orf58 | 1128 | 2444 | 1502 | 3071 | 3233 | 1440 | 3582 | 3463 |
| C11orf68 | 0 | 0 | 0 | 810 | 924 | 0 | 0 | 1250 |

| | | | | | | | | |
|---|---|---|---|---|---|---|---|---|
| C11orf71 | 0 | 0 | 3022 | 286 | 3524 | 0 | 246 | 0 |
| C11orf72 | 0 | 0 | 0 | 0 | 0 | 2946 | 0 | 0 |
| C11orf74 | 0 | 0 | 0 | 446 | 488 | 0 | 0 | 0 |
| C11orf80 | 488 | 0 | 0 | 0 | 0 | 0 | 0 | 0 |
| C11orf86 | 960 | 0 | 2986 | 0 | 0 | 0 | 0 | 0 |
| C11orf96 | 2134 | 334 | 2838 | 0 | 808 | 3180 | 0 | 2473 |
| C11orf98 | 2621 | 502 | 12127 | 1184 | 7850 | 2684 | 2830 | 2615 |
| C12orf10 | 0 | 0 | 0 | 0 | 0 | 0 | 0 | 2723 |
| C12orf29 | 0 | 0 | 48 | 1596 | 2697 | 0 | 0 | 0 |
| C12orf4 | 562 | 0 | 1800 | 0 | 0 | 0 | 0 | 0 |
| C12orf43 | 0 | 258 | 0 | 0 | 0 | 0 | 0 | 0 |
| C12orf45 | 0 | 514 | 0 | 0 | 300 | 310 | 0 | 294 |
| C12orf49 | 3554 | 5730 | 15474 | 2836 | 9886 | 2843 | 7472 | 1322 |
| C12orf50 | 0 | 402 | 2103 | 0 | 824 | 0 | 2518 | 0 |
| C12orf56 | 738 | 0 | 0 | 0 | 0 | 0 | 222 | 0 |
| C12orf57 | 0 | 2398 | 2831 | 457 | 1601 | 4363 | 960 | 534 |
| C12orf65 | 0 | 0 | 2070 | 0 | 0 | 0 | 154 | 0 |
| C12orf73 | 0 | 0 | 0 | 0 | 1460 | 0 | 0 | 944 |
| C12orf75 | 602 | 1822 | 2050 | 6185 | 3866 | 1578 | 2230 | 1560 |
| C12orf76 | 0 | 0 | 0 | 0 | 0 | 0 | 130 | 0 |
| C14orf119 | 782 | 3156 | 1094 | 2207 | 8248 | 0 | 2812 | 1894 |
| C14orf93 | 0 | 0 | 0 | 0 | 0 | 0 | 202 | 0 |
| C15orf39 | 0 | 0 | 0 | 0 | 484 | 0 | 0 | 0 |
| C15orf40 | 0 | 566 | 0 | 642 | 938 | 0 | 0 | 0 |
| C15orf41 | 380 | 0 | 0 | 0 | 0 | 0 | 878 | 0 |
| C15orf48 | 29091 | 36280 | 85908 | 6030 | 22036 | 0 | 0 | 3074 |
| C15orf61 | 0 | 472 | 0 | 266 | 1506 | 0 | 166 | 86 |
| C16orf45 | 2084 | 1992 | 6405 | 0 | 8576 | 5472 | 326 | 2138 |
| C16orf54 | 0 | 0 | 1001 | 0 | 0 | 0 | 0 | 0 |
| C16orf58 | 0 | 350 | 0 | 1452 | 52 | 336 | 296 | 0 |
| C16orf72 | 505 | 0 | 0 | 0 | 432 | 0 | 170 | 0 |
| C16orf74 | 694 | 0 | 0 | 0 | 244 | 0 | 0 | 0 |
| C16orf87 | 0 | 0 | 0 | 0 | 0 | 0 | 755 | 786 |
| C16orf91 | 0 | 644 | 1969 | 2010 | 150 | 0 | 507 | 0 |
| C16orf95 | 0 | 0 | 0 | 0 | 0 | 0 | 106 | 0 |
| C17orf49 | 0 | 140 | 2010 | 0 | 2550 | 0 | 0 | 2052 |
| C17orf58 | 0 | 0 | 0 | 626 | 0 | 0 | 162 | 0 |
| C17orf64 | 0 | 0 | 0 | 0 | 66 | 0 | 0 | 0 |
| C17orf75 | 0 | 482 | 0 | 0 | 876 | 0 | 0 | 0 |
| C17orf80 | 0 | 0 | 0 | 1204 | 0 | 0 | 0 | 0 |
| C18orf21 | 0 | 0 | 0 | 1290 | 0 | 0 | 2384 | 0 |
| C18orf32 | 0 | 2606 | 1408 | 1726 | 2020 | 982 | 0 | 882 |
| C19orf12 | 0 | 602 | 0 | 0 | 358 | 0 | 0 | 0 |

| | | | | | | | | |
|---|---|---|---|---|---|---|---|---|
| C19orf24 | 768 | 568 | 0 | 1824 | 2644 | 2228 | 0 | 0 |
| C19orf25 | 0 | 298 | 0 | 1006 | 1608 | 0 | 204 | 364 |
| C19orf33 | 3759 | 6552 | 5725 | 1162 | 13033 | 1064 | 794 | 1428 |
| C19orf38 | 5623 | 2910 | 24067 | 0 | 25131 | 8887 | 2192 | 22397 |
| C19orf44 | 0 | 0 | 0 | 0 | 22 | 0 | 0 | 0 |
| C19orf48 | 100 | 398 | 1695 | 2148 | 8116 | 2109 | 662 | 1227 |
| C19orf53 | 2092 | 3148 | 4158 | 744 | 7796 | 2040 | 2354 | 1664 |
| C19orf54 | 0 | 0 | 1144 | 0 | 0 | 0 | 0 | 0 |
| C19orf66 | 12 | 1140 | 2044 | 0 | 6072 | 0 | 1226 | 4754 |
| C19orf70 | 0 | 0 | 0 | 0 | 948 | 1680 | 0 | 0 |
| C19orf73 | 0 | 0 | 0 | 0 | 988 | 0 | 0 | 858 |
| C19orf81 | 0 | 0 | 0 | 0 | 446 | 0 | 0 | 0 |
| C1D | 0 | 398 | 0 | 512 | 298 | 1302 | 352 | 586 |
| C1GALT1 | 0 | 0 | 697 | 0 | 392 | 0 | 0 | 0 |
| C1GALT1C1 | 0 | 706 | 3951 | 764 | 616 | 0 | 1036 | 2376 |
| C1orf105 | 0 | 0 | 0 | 0 | 0 | 0 | 765 | 0 |
| C1orf109 | 0 | 192 | 0 | 0 | 164 | 0 | 0 | 0 |
| C1orf115 | 0 | 0 | 640 | 406 | 2262 | 0 | 0 | 0 |
| C1orf122 | 776 | 48 | 1334 | 0 | 914 | 0 | 1600 | 654 |
| C1orf131 | 0 | 0 | 1675 | 0 | 370 | 0 | 0 | 0 |
| C1orf159 | 328 | 1550 | 5230 | 0 | 2676 | 3058 | 2138 | 0 |
| C1orf162 | 0 | 0 | 0 | 0 | 666 | 1646 | 0 | 1142 |
| C1orf174 | 0 | 630 | 0 | 944 | 0 | 0 | 0 | 0 |
| C1orf198 | 0 | 0 | 0 | 1380 | 1076 | 1112 | 96 | 0 |
| C1orf21 | 157869 | 215617 | 276468 | 112747 | 550848 | 183975 | 294999 | 123860 |
| C1orf210 | 998 | 342 | 1562 | 0 | 790 | 1014 | 2282 | 0 |
| C1orf226 | 1154 | 882 | 2684 | 0 | 1924 | 146 | 1232 | 2862 |
| C1orf35 | 0 | 490 | 1666 | 1402 | 0 | 0 | 0 | 0 |
| C1orf43 | 790 | 1311 | 476 | 0 | 5878 | 2283 | 3160 | 3254 |
| C1orf50 | 0 | 822 | 0 | 0 | 440 | 0 | 893 | 1048 |
| C1orf52 | 0 | 0 | 0 | 656 | 548 | 0 | 664 | 0 |
| C1orf53 | 0 | 0 | 0 | 0 | 598 | 0 | 0 | 0 |
| C1orf54 | 0 | 0 | 1476 | 0 | 0 | 882 | 398 | 504 |
| C1orf56 | 0 | 0 | 0 | 1380 | 1640 | 0 | 0 | 0 |
| C1QA | 1562 | 182 | 0 | 98 | 204 | 3180 | 0 | 0 |
| C1QB | 0 | 448 | 3949 | 968 | 258 | 9504 | 1512 | 0 |
| C1QBP | 0 | 7018 | 2825 | 9182 | 17207 | 4562 | 3268 | 3017 |
| C1QC | 0 | 876 | 4512 | 0 | 1968 | 2504 | 0 | 2270 |
| C1QL1 | 0 | 0 | 0 | 0 | 240 | 0 | 0 | 0 |
| C1QTNF1 | 0 | 0 | 0 | 0 | 1144 | 2010 | 0 | 0 |
| C1QTNF12 | 1390 | 0 | 4197 | 0 | 0 | 0 | 0 | 0 |
| C1QTNF3 | 0 | 0 | 0 | 0 | 556 | 0 | 0 | 0 |
| C1QTNF3-AMACR | 3362 | 340 | 1254 | 40 | 2978 | 2815 | 15503 | 0 |

| | | | | | | | | |
|---|---|---|---|---|---|---|---|---|
| C1QTNF6 | 0 | 0 | 0 | 0 | 754 | 0 | 0 | 0 |
| C1R | 740 | 672 | 2651 | 0 | 386 | 0 | 0 | 0 |
| C1RL | 0 | 0 | 0 | 0 | 14 | 0 | 0 | 0 |
| C1S | 786 | 0 | 0 | 292 | 1296 | 0 | 304 | 0 |
| C2 | 0 | 0 | 0 | 1694 | 165 | 0 | 0 | 0 |
| C20orf202 | 0 | 0 | 0 | 0 | 1062 | 0 | 0 | 0 |
| C20orf204 | 0 | 384 | 0 | 0 | 0 | 0 | 0 | 0 |
| C20orf27 | 0 | 0 | 0 | 292 | 2778 | 0 | 0 | 0 |
| C20orf96 | 0 | 0 | 0 | 0 | 68 | 0 | 0 | 0 |
| C21orf2 | 0 | 0 | 624 | 272 | 0 | 0 | 0 | 1044 |
| C21orf58 | 0 | 508 | 0 | 0 | 0 | 0 | 0 | 0 |
| C21orf59-TCP10L | 0 | 0 | 0 | 0 | 698 | 0 | 0 | 0 |
| C21orf91 | 0 | 0 | 0 | 0 | 0 | 0 | 1214 | 0 |
| C22orf31 | 0 | 2446 | 462 | 0 | 0 | 0 | 92 | 0 |
| C22orf39 | 0 | 1052 | 158 | 0 | 2696 | 0 | 1652 | 0 |
| C2CD2 | 0 | 0 | 0 | 0 | 625 | 0 | 0 | 0 |
| C2CD2L | 3628 | 1528 | 10074 | 388 | 1432 | 2150 | 4012 | 88 |
| C2CD3 | 0 | 0 | 0 | 0 | 0 | 0 | 556 | 0 |
| C2CD4B | 0 | 0 | 0 | 0 | 472 | 0 | 1580 | 0 |
| C2CD4D | 0 | 0 | 0 | 0 | 554 | 0 | 0 | 0 |
| C2CD5 | 0 | 0 | 0 | 388 | 0 | 0 | 0 | 0 |
| C2orf27A | 0 | 0 | 0 | 0 | 146 | 0 | 0 | 0 |
| C2orf49 | 324 | 1089 | 0 | 14 | 586 | 0 | 0 | 0 |
| C2orf54 | 148 | 0 | 0 | 0 | 0 | 0 | 0 | 0 |
| C2orf68 | 0 | 220 | 0 | 0 | 494 | 1022 | 0 | 0 |
| C2orf69 | 0 | 0 | 0 | 0 | 0 | 0 | 82 | 0 |
| C2orf72 | 0 | 568 | 0 | 0 | 0 | 0 | 0 | 0 |
| C2orf74 | 23487 | 36839 | 32312 | 30101 | 52337 | 95115 | 32740 | 42523 |
| C2orf76 | 0 | 316 | 0 | 548 | 0 | 0 | 508 | 0 |
| C2orf88 | 570 | 594 | 9053 | 0 | 0 | 0 | 0 | 0 |
| C3orf14 | 0 | 0 | 0 | 920 | 0 | 0 | 0 | 0 |
| C3orf38 | 0 | 374 | 0 | 0 | 1352 | 1247 | 74 | 0 |
| C3orf52 | 0 | 0 | 2992 | 0 | 0 | 0 | 0 | 0 |
| C3orf62 | 194 | 1060 | 0 | 0 | 0 | 0 | 0 | 0 |
| C3orf67 | 1222 | 1309 | 0 | 1386 | 18284 | 0 | 4872 | 0 |
| C3orf70 | 0 | 262 | 0 | 0 | 0 | 0 | 0 | 0 |
| C3orf85 | 394 | 0 | 0 | 0 | 1218 | 0 | 0 | 0 |
| C4A | 0 | 0 | 0 | 0 | 0 | 1274 | 0 | 0 |
| C4BPB | 0 | 0 | 0 | 1056 | 0 | 0 | 0 | 890 |
| C4orf19 | 456 | 830 | 602 | 0 | 0 | 0 | 0 | 0 |
| C4orf3 | 499 | 1554 | 1116 | 2222 | 8341 | 0 | 3810 | 2332 |
| C4orf33 | 0 | 0 | 0 | 194 | 0 | 0 | 0 | 1078 |
| C4orf36 | 0 | 0 | 0 | 0 | 150 | 0 | 0 | 0 |

| | | | | | | | | |
|---|---|---|---|---|---|---|---|---|
| C4orf47 | 0 | 0 | 0 | 0 | 278 | 0 | 0 | 0 |
| C4orf48 | 1988 | 0 | 2004 | 3256 | 4690 | 1266 | 1373 | 1503 |
| C5AR1 | 0 | 0 | 0 | 0 | 0 | 0 | 160 | 0 |
| C5orf15 | 126 | 698 | 1582 | 752 | 1140 | 1052 | 310 | 0 |
| C5orf22 | 0 | 0 | 0 | 0 | 0 | 0 | 216 | 0 |
| C5orf24 | 0 | 340 | 624 | 0 | 0 | 0 | 0 | 604 |
| C5orf30 | 6146 | 7940 | 7400 | 1056 | 2752 | 1544 | 158 | 2062 |
| C5orf34 | 0 | 0 | 0 | 0 | 0 | 0 | 0 | 104 |
| C5orf46 | 0 | 0 | 0 | 0 | 1215 | 1300 | 0 | 0 |
| C5orf51 | 6340 | 10200 | 7048 | 5295 | 38569 | 11162 | 12382 | 5335 |
| C5orf52 | 0 | 0 | 0 | 0 | 0 | 0 | 540 | 0 |
| C5orf56 | 608 | 0 | 0 | 0 | 836 | 0 | 0 | 0 |
| C5orf66-AS1 | 0 | 0 | 0 | 0 | 552 | 0 | 0 | 0 |
| C6orf106 | 0 | 320 | 0 | 564 | 0 | 1566 | 0 | 0 |
| C6orf120 | 380 | 134 | 0 | 0 | 0 | 0 | 278 | 0 |
| C6orf132 | 806 | 430 | 0 | 0 | 0 | 0 | 488 | 0 |
| C6orf136 | 0 | 0 | 3883 | 1066 | 294 | 428 | 276 | 0 |
| C6orf141 | 0 | 0 | 0 | 0 | 358 | 0 | 0 | 0 |
| C6orf15 | 0 | 0 | 0 | 0 | 442 | 0 | 0 | 0 |
| C6orf203 | 0 | 0 | 0 | 622 | 2152 | 0 | 1843 | 0 |
| C6orf222 | 1168 | 0 | 4326 | 822 | 4104 | 0 | 7806 | 1048 |
| C6orf223 | 0 | 0 | 0 | 0 | 134 | 0 | 0 | 0 |
| C6orf226 | 0 | 0 | 0 | 0 | 2005 | 0 | 592 | 0 |
| C6orf48 | 0 | 1894 | 1808 | 4485 | 5262 | 5890 | 8322 | 2516 |
| C6orf52 | 0 | 0 | 0 | 0 | 0 | 0 | 0 | 610 |
| C6orf62 | 0 | 0 | 0 | 0 | 644 | 1209 | 0 | 172 |
| C6orf89 | 0 | 534 | 794 | 1522 | 54 | 0 | 0 | 0 |
| C7 | 0 | 0 | 0 | 0 | 0 | 0 | 0 | 856 |
| C7orf13 | 446 | 0 | 0 | 0 | 0 | 0 | 186 | 0 |
| C7orf25 | 0 | 0 | 0 | 0 | 812 | 0 | 0 | 0 |
| C7orf26 | 0 | 0 | 0 | 0 | 426 | 0 | 0 | 928 |
| C7orf31 | 0 | 0 | 1927 | 0 | 0 | 0 | 0 | 0 |
| C7orf43 | 0 | 270 | 0 | 0 | 198 | 0 | 0 | 0 |
| C7orf50 | 1108 | 620 | 13959 | 10010 | 26760 | 6466 | 9694 | 4406 |
| C7orf61 | 0 | 0 | 0 | 0 | 877 | 0 | 0 | 0 |
| C8orf33 | 0 | 0 | 2420 | 796 | 2128 | 0 | 2501 | 636 |
| C8orf34 | 0 | 0 | 0 | 0 | 1976 | 1878 | 0 | 0 |
| C8orf37 | 0 | 0 | 1256 | 0 | 194 | 0 | 0 | 0 |
| C8orf44-SGK3 | 0 | 0 | 0 | 0 | 340 | 0 | 0 | 0 |
| C8orf59 | 0 | 1420 | 968 | 2304 | 11355 | 1705 | 5760 | 946 |
| C8orf76 | 0 | 0 | 0 | 5237 | 1643 | 2921 | 358 | 0 |
| C8orf82 | 492 | 0 | 1030 | 1054 | 1230 | 0 | 532 | 0 |
| C9orf152 | 0 | 732 | 729 | 2284 | 0 | 0 | 266 | 0 |

| | | | | | | | | |
|---|---|---|---|---|---|---|---|---|
| C9orf16 | 2166 | 478 | 2084 | 758 | 5152 | 1617 | 1793 | 764 |
| C9orf163 | 0 | 144 | 0 | 0 | 0 | 0 | 0 | 0 |
| C9orf3 | 21680 | 22005 | 58109 | 31429 | 127063 | 57798 | 260611 | 11677 |
| C9orf40 | 702 | 300 | 0 | 0 | 710 | 0 | 456 | 0 |
| C9orf64 | 4454 | 11628 | 23122 | 20280 | 54176 | 17126 | 35051 | 18908 |
| C9orf72 | 0 | 118 | 0 | 0 | 0 | 0 | 0 | 0 |
| C9orf78 | 562 | 462 | 1290 | 1761 | 2902 | 0 | 0 | 0 |
| C9orf85 | 0 | 0 | 0 | 0 | 562 | 0 | 0 | 0 |
| CA1 | 4882204 | 1497006 | 14424081 | 9237 | 65479 | 0 | 2656 | 1424 |
| CA10 | 0 | 0 | 2802 | 0 | 0 | 0 | 0 | 0 |
| CA11 | 516 | 0 | 0 | 114 | 0 | 0 | 0 | 0 |
| CA12 | 644 | 412 | 0 | 0 | 0 | 1699 | 1176 | 0 |
| CA13 | 0 | 0 | 0 | 0 | 0 | 1704 | 0 | 0 |
| CA2 | 13288 | 8544 | 48747 | 5211 | 0 | 0 | 0 | 0 |
| CA4 | 8142 | 488 | 37085 | 0 | 0 | 0 | 0 | 0 |
| CA6 | 0 | 0 | 0 | 0 | 0 | 13150 | 0 | 0 |
| CA7 | 25625 | 1225 | 77010 | 0 | 0 | 0 | 0 | 0 |
| CA9 | 0 | 0 | 0 | 3163 | 2206 | 0 | 0 | 0 |
| CAAP1 | 400 | 0 | 908 | 0 | 0 | 0 | 172 | 0 |
| CAB39 | 0 | 422 | 3747 | 0 | 288 | 1830 | 252 | 0 |
| CAB39L | 0 | 80 | 0 | 0 | 0 | 3504 | 334 | 0 |
| CABIN1 | 464 | 326 | 894 | 0 | 598 | 3121 | 314 | 0 |
| CABLES1 | 0 | 0 | 0 | 0 | 2240 | 0 | 1024 | 1924 |
| CABLES2 | 0 | 0 | 0 | 0 | 0 | 0 | 360 | 0 |
| CABP4 | 0 | 0 | 704 | 0 | 0 | 0 | 0 | 0 |
| CACFD1 | 326 | 0 | 1244 | 0 | 352 | 0 | 1124 | 0 |
| CACNA1D | 710 | 0 | 0 | 1956 | 1182 | 2195 | 3460 | 0 |
| CACNA2D1 | 1336 | 0 | 0 | 0 | 0 | 3160 | 0 | 0 |
| CACNA2D3 | 0 | 0 | 2509 | 0 | 0 | 0 | 0 | 0 |
| CACNA2D4 | 0 | 0 | 0 | 0 | 0 | 0 | 0 | 366 |
| CACNB3 | 356 | 0 | 0 | 0 | 456 | 0 | 0 | 0 |
| CACTIN | 0 | 0 | 0 | 0 | 30 | 0 | 0 | 0 |
| CACUL1 | 0 | 0 | 0 | 0 | 410 | 0 | 0 | 0 |
| CACYBP | 816 | 1674 | 1012 | 3113 | 7274 | 4946 | 6901 | 2780 |
| CAD | 62 | 0 | 0 | 0 | 0 | 0 | 228 | 0 |
| CADM1 | 0 | 192 | 0 | 0 | 0 | 0 | 0 | 0 |
| CADPS | 0 | 0 | 0 | 0 | 6275 | 0 | 802 | 0 |
| CAHM | 0 | 0 | 0 | 0 | 126 | 0 | 0 | 0 |
| CALB2 | 0 | 0 | 2130 | 0 | 0 | 202 | 0 | 0 |
| CALCOCO1 | 0 | 626 | 0 | 552 | 916 | 0 | 0 | 0 |
| CALCOCO2 | 52 | 812 | 6241 | 152 | 1422 | 0 | 1082 | 0 |
| CALD1 | 62522 | 27777 | 118225 | 22538 | 47380 | 39035 | 3495 | 14857 |
| CALHM6 | 0 | 0 | 0 | 906 | 0 | 0 | 0 | 0 |

| | | | | | | | | |
|---|---|---|---|---|---|---|---|---|
| CALM1 | 2418 | 4592 | 11152 | 8247 | 12899 | 5007 | 3113 | 2769 |
| CALM2 | 8134 | 13656 | 52445 | 36120 | 57168 | 25715 | 7696 | 24073 |
| CALM3 | 718 | 1370 | 2040 | 1165 | 6991 | 0 | 0 | 2260 |
| CALML4 | 5780 | 5582 | 14756 | 6480 | 10143 | 0 | 3044 | 2030 |
| CALR | 10443 | 12484 | 10991 | 23199 | 66389 | 7832 | 44498 | 35345 |
| CALU | 332 | 154 | 0 | 0 | 1293 | 1196 | 368 | 0 |
| CAMK1 | 0 | 0 | 0 | 0 | 1082 | 0 | 0 | 0 |
| CAMK1D | 0 | 56 | 0 | 0 | 0 | 0 | 0 | 0 |
| CAMK2D | 0 | 0 | 0 | 0 | 668 | 0 | 0 | 0 |
| CAMK2G | 362 | 0 | 0 | 0 | 0 | 0 | 0 | 0 |
| CAMK2N1 | 1766 | 1340 | 8738 | 1074 | 2655 | 0 | 2480 | 670 |
| CAMK4 | 342 | 1156 | 146 | 0 | 2132 | 0 | 0 | 1742 |
| CAMKK2 | 0 | 350 | 0 | 0 | 0 | 0 | 0 | 0 |
| CAMKMT | 0 | 0 | 0 | 0 | 1950 | 0 | 0 | 0 |
| CAMLG | 372 | 996 | 0 | 0 | 1526 | 2677 | 0 | 0 |
| CAMSAP1 | 0 | 0 | 0 | 0 | 448 | 0 | 0 | 0 |
| CAMSAP2 | 776 | 334 | 1688 | 748 | 4534 | 1682 | 3409 | 126 |
| CAMSAP3 | 0 | 772 | 0 | 0 | 1678 | 0 | 478 | 0 |
| CAMTA1 | 387655 | 427056 | 1107393 | 399293 | 743557 | 303619 | 326708 | 175304 |
| CAMTA2 | 896 | 908 | 532 | 166 | 2856 | 3489 | 2232 | 22 |
| CAND1 | 0 | 0 | 0 | 394 | 270 | 0 | 0 | 0 |
| CANT1 | 18855 | 27503 | 54937 | 18660 | 39510 | 37466 | 57809 | 10942 |
| CANX | 0 | 862 | 0 | 1086 | 1380 | 0 | 330 | 1313 |
| CAP1 | 0 | 342 | 1072 | 2841 | 1012 | 0 | 880 | 0 |
| CAPG | 668 | 1134 | 2481 | 722 | 2552 | 3644 | 1494 | 8131 |
| CAPN1 | 0 | 276 | 0 | 2308 | 1970 | 0 | 1564 | 2316 |
| CAPN10 | 0 | 0 | 0 | 0 | 714 | 0 | 0 | 0 |
| CAPN10-DT | 0 | 0 | 0 | 0 | 0 | 0 | 0 | 288 |
| CAPN12 | 204 | 250 | 0 | 951 | 0 | 0 | 318 | 0 |
| CAPN15 | 0 | 0 | 0 | 0 | 0 | 1642 | 0 | 0 |
| CAPN2 | 1383 | 300 | 4014 | 0 | 1638 | 0 | 206 | 0 |
| CAPN3 | 474 | 0 | 0 | 0 | 0 | 0 | 0 | 0 |
| CAPN5 | 882 | 204 | 3368 | 0 | 0 | 0 | 0 | 0 |
| CAPN8 | 0 | 736 | 0 | 3008 | 1185 | 0 | 0 | 868 |
| CAPNS1 | 0 | 722 | 4580 | 4391 | 3430 | 0 | 842 | 881 |
| CAPRIN1 | 0 | 330 | 1112 | 0 | 0 | 238 | 526 | 0 |
| CAPS | 0 | 0 | 0 | 0 | 804 | 0 | 668 | 0 |
| CAPZA1 | 0 | 749 | 1728 | 1478 | 4311 | 0 | 1320 | 0 |
| CAPZA2 | 535391 | 789187 | 1435767 | 775472 | 1857962 | 1433929 | 1756233 | 778823 |
| CAPZB | 378220 | 366255 | 1231599 | 653964 | 1288662 | 625901 | 439115 | 462699 |
| CARD10 | 1236 | 773 | 0 | 0 | 0 | 0 | 0 | 0 |
| CARD11 | 0 | 946 | 0 | 752 | 5948 | 0 | 444 | 680 |
| CARD14 | 0 | 0 | 0 | 0 | 492 | 0 | 0 | 0 |

| | | | | | | | | |
|---|---|---|---|---|---|---|---|---|
| CARD16 | 2898 | 1014 | 1664 | 966 | 244 | 0 | 0 | 0 |
| CARD19 | 22637 | 17483 | 86564 | 38648 | 74711 | 43246 | 53781 | 19244 |
| CARD6 | 0 | 0 | 0 | 0 | 216 | 0 | 0 | 0 |
| CARD8 | 0 | 0 | 0 | 828 | 0 | 0 | 0 | 0 |
| CARHSP1 | 198 | 0 | 0 | 0 | 3514 | 0 | 1072 | 0 |
| CARM1 | 1056 | 1160 | 4688 | 3485 | 9571 | 938 | 8272 | 1256 |
| CARMIL1 | 0 | 0 | 0 | 0 | 722 | 828 | 584 | 0 |
| CARMIL2 | 0 | 0 | 0 | 0 | 0 | 0 | 186 | 0 |
| CARS | 100629 | 170010 | 509435 | 382103 | 678988 | 517904 | 529757 | 572049 |
| CARS2 | 0 | 0 | 2032 | 2863 | 670 | 896 | 0 | 0 |
| CASC1 | 0 | 0 | 0 | 1504 | 0 | 0 | 0 | 0 |
| CASC3 | 1748 | 596 | 0 | 0 | 1155 | 0 | 2160 | 0 |
| CASC4 | 304 | 0 | 0 | 0 | 0 | 1139 | 128 | 0 |
| CASC8 | 0 | 0 | 0 | 502 | 1072 | 0 | 0 | 0 |
| CASC9 | 0 | 454 | 0 | 0 | 0 | 0 | 306 | 0 |
| CASK | 0 | 0 | 0 | 0 | 854 | 0 | 0 | 0 |
| CASKIN1 | 1096 | 808 | 328 | 784 | 1926 | 1773 | 3618 | 0 |
| CASKIN2 | 0 | 0 | 0 | 0 | 0 | 0 | 872 | 40 |
| CASP1 | 206 | 814 | 2086 | 4604 | 456 | 0 | 0 | 1536 |
| CASP10 | 0 | 0 | 1029 | 0 | 1424 | 0 | 126 | 0 |
| CASP3 | 0 | 0 | 4143 | 1162 | 456 | 0 | 202 | 0 |
| CASP4 | 0 | 0 | 908 | 830 | 374 | 0 | 3444 | 0 |
| CASP5 | 0 | 0 | 0 | 0 | 0 | 0 | 468 | 0 |
| CASP6 | 0 | 646 | 0 | 0 | 704 | 1933 | 2101 | 0 |
| CASP7 | 450 | 630 | 6850 | 0 | 1210 | 1238 | 0 | 0 |
| CASP8 | 26391 | 22988 | 40469 | 37881 | 64054 | 14640 | 43178 | 13398 |
| CASP8AP2 | 0 | 316 | 0 | 0 | 0 | 0 | 148 | 0 |
| CASR | 454 | 0 | 0 | 0 | 0 | 0 | 0 | 0 |
| CAST | 958 | 0 | 1964 | 904 | 1942 | 0 | 640 | 886 |
| CASTOR1 | 0 | 0 | 2068 | 0 | 364 | 0 | 0 | 0 |
| CASZ1 | 0 | 100 | 0 | 0 | 0 | 0 | 0 | 0 |
| CAT | 0 | 522 | 734 | 0 | 1212 | 0 | 684 | 828 |
| CATSPER1 | 0 | 0 | 1666 | 0 | 0 | 0 | 0 | 0 |
| CATSPERB | 0 | 316 | 0 | 0 | 216 | 0 | 0 | 0 |
| CAV1 | 248590 | 69698 | 161679 | 79596 | 447023 | 224224 | 32149 | 110024 |
| CAV2 | 0 | 0 | 0 | 785 | 1615 | 2428 | 0 | 656 |
| CAVIN1 | 0 | 0 | 0 | 0 | 946 | 0 | 0 | 0 |
| CAVIN3 | 0 | 0 | 408 | 1122 | 4511 | 0 | 0 | 4604 |
| CBFA2T2 | 0 | 0 | 0 | 0 | 614 | 0 | 0 | 0 |
| CBFB | 41087 | 38112 | 40527 | 131866 | 191269 | 109110 | 160542 | 70938 |
| CBL | 2853 | 2110 | 2814 | 1478 | 8362 | 2428 | 1450 | 3359 |
| CBLB | 12856 | 30315 | 29092 | 14047 | 80408 | 13241 | 28408 | 38863 |
| CBLC | 26129 | 14995 | 53138 | 23621 | 76469 | 16279 | 29374 | 38986 |

| | | | | | | | | |
|---|---|---|---|---|---|---|---|---|
| CBLL1 | 0 | 0 | 0 | 0 | 776 | 0 | 0 | 634 |
| CBLN2 | 0 | 0 | 0 | 1330 | 0 | 0 | 0 | 0 |
| CBLN3 | 0 | 0 | 0 | 0 | 294 | 0 | 0 | 0 |
| CBR1 | 1419 | 320 | 8184 | 3234 | 3460 | 2096 | 346 | 1473 |
| CBR3 | 0 | 0 | 0 | 1441 | 0 | 0 | 0 | 0 |
| CBR3-AS1 | 344 | 0 | 0 | 0 | 0 | 0 | 122 | 0 |
| CBR4 | 0 | 266 | 2329 | 0 | 0 | 0 | 0 | 0 |
| CBWD1 | 0 | 2124 | 2326 | 1264 | 1060 | 1240 | 870 | 0 |
| CBWD2 | 0 | 592 | 2870 | 2058 | 1004 | 1342 | 1212 | 0 |
| CBWD3 | 0 | 0 | 3671 | 0 | 0 | 0 | 446 | 0 |
| CBWD5 | 880 | 4448 | 0 | 7327 | 2444 | 4531 | 3882 | 4710 |
| CBWD6 | 0 | 0 | 0 | 0 | 0 | 0 | 158 | 0 |
| CBX1 | 0 | 0 | 0 | 0 | 0 | 0 | 494 | 0 |
| CBX2 | 0 | 0 | 0 | 0 | 318 | 0 | 0 | 0 |
| CBX3 | 20 | 1386 | 1133 | 7658 | 7648 | 2016 | 2072 | 1526 |
| CBX4 | 0 | 0 | 0 | 0 | 312 | 0 | 0 | 0 |
| CBX5 | 0 | 0 | 0 | 792 | 296 | 0 | 1136 | 492 |
| CBX7 | 332 | 0 | 0 | 0 | 0 | 1596 | 0 | 0 |
| CBX8 | 0 | 0 | 0 | 0 | 0 | 0 | 26 | 0 |
| CBY1 | 934 | 430 | 1678 | 1420 | 590 | 0 | 0 | 244 |
| CC2D1A | 0 | 198 | 664 | 0 | 1070 | 0 | 0 | 0 |
| CC2D1B | 0 | 0 | 0 | 0 | 0 | 0 | 524 | 0 |
| CC2D2B | 0 | 0 | 0 | 0 | 280 | 0 | 0 | 0 |
| CCAR1 | 66338 | 54931 | 64518 | 59519 | 239943 | 108156 | 215006 | 69489 |
| CCAR2 | 4213 | 2684 | 11160 | 5849 | 9092 | 11951 | 7465 | 574 |
| CCDC102B | 0 | 0 | 0 | 806 | 0 | 0 | 0 | 0 |
| CCDC106 | 0 | 0 | 0 | 1678 | 0 | 0 | 0 | 0 |
| CCDC107 | 0 | 10 | 4436 | 1300 | 3058 | 0 | 0 | 0 |
| CCDC112 | 0 | 0 | 0 | 0 | 0 | 0 | 0 | 1258 |
| CCDC115 | 1098 | 0 | 0 | 0 | 433 | 0 | 830 | 684 |
| CCDC12 | 944 | 2470 | 0 | 3478 | 7505 | 0 | 1164 | 0 |
| CCDC120 | 0 | 344 | 0 | 0 | 0 | 0 | 0 | 0 |
| CCDC121 | 0 | 0 | 0 | 0 | 322 | 0 | 0 | 0 |
| CCDC122 | 0 | 0 | 1034 | 0 | 0 | 0 | 0 | 794 |
| CCDC124 | 0 | 594 | 1796 | 1384 | 3568 | 0 | 944 | 1460 |
| CCDC126 | 178 | 0 | 0 | 0 | 546 | 0 | 0 | 0 |
| CCDC127 | 13712 | 38755 | 77503 | 29213 | 235407 | 89052 | 120288 | 45947 |
| CCDC13 | 0 | 0 | 0 | 0 | 1116 | 0 | 0 | 0 |
| CCDC130 | 0 | 0 | 0 | 0 | 814 | 0 | 0 | 446 |
| CCDC136 | 0 | 0 | 0 | 0 | 42 | 0 | 0 | 0 |
| CCDC137 | 0 | 0 | 2172 | 1040 | 0 | 0 | 0 | 0 |
| CCDC14 | 284 | 394 | 0 | 303 | 998 | 0 | 0 | 0 |
| CCDC142 | 0 | 0 | 2297 | 0 | 1668 | 0 | 874 | 0 |

| | | | | | | | | |
|---|---|---|---|---|---|---|---|---|
| CCDC144A | 368 | 0 | 0 | 0 | 0 | 0 | 0 | 0 |
| CCDC146 | 0 | 0 | 0 | 0 | 186 | 0 | 0 | 0 |
| CCDC148 | 0 | 0 | 0 | 0 | 658 | 0 | 0 | 0 |
| CCDC152 | 446 | 158 | 228 | 0 | 0 | 0 | 0 | 0 |
| CCDC159 | 0 | 682 | 0 | 268 | 748 | 0 | 0 | 784 |
| CCDC167 | 0 | 486 | 0 | 777 | 402 | 0 | 1166 | 666 |
| CCDC170 | 10420 | 984 | 9976 | 40777 | 7377 | 32159 | 215699 | 1034 |
| CCDC171 | 0 | 0 | 772 | 0 | 0 | 0 | 0 | 0 |
| CCDC174 | 0 | 0 | 0 | 0 | 0 | 0 | 692 | 0 |
| CCDC178 | 0 | 290 | 0 | 0 | 0 | 0 | 0 | 0 |
| CCDC18-AS1 | 0 | 254 | 0 | 0 | 0 | 0 | 0 | 0 |
| CCDC183 | 0 | 0 | 0 | 0 | 508 | 0 | 0 | 0 |
| CCDC186 | 918 | 332 | 0 | 0 | 0 | 0 | 1792 | 0 |
| CCDC24 | 0 | 0 | 0 | 0 | 422 | 0 | 0 | 0 |
| CCDC25 | 0 | 0 | 1392 | 0 | 566 | 0 | 0 | 0 |
| CCDC28A | 0 | 256 | 2969 | 0 | 146 | 0 | 1422 | 0 |
| CCDC30 | 0 | 0 | 1424 | 0 | 0 | 0 | 0 | 0 |
| CCDC32 | 0 | 431 | 1442 | 0 | 30 | 0 | 448 | 0 |
| CCDC34 | 0 | 0 | 0 | 618 | 482 | 0 | 864 | 0 |
| CCDC43 | 0 | 564 | 0 | 0 | 0 | 0 | 1064 | 0 |
| CCDC47 | 360 | 0 | 0 | 0 | 0 | 0 | 376 | 0 |
| CCDC50 | 174 | 623 | 0 | 0 | 1342 | 0 | 1184 | 0 |
| CCDC51 | 644 | 0 | 856 | 0 | 568 | 2801 | 0 | 0 |
| CCDC57 | 271 | 450 | 1566 | 0 | 0 | 880 | 918 | 0 |
| CCDC58 | 0 | 914 | 0 | 908 | 1652 | 0 | 554 | 1365 |
| CCDC59 | 0 | 496 | 0 | 1106 | 1146 | 0 | 988 | 758 |
| CCDC6 | 44313 | 66985 | 91371 | 64241 | 193537 | 69750 | 170577 | 56540 |
| CCDC61 | 0 | 170 | 0 | 866 | 0 | 0 | 288 | 0 |
| CCDC68 | 0 | 356 | 1132 | 0 | 0 | 0 | 0 | 0 |
| CCDC69 | 0 | 0 | 0 | 0 | 0 | 0 | 0 | 24 |
| CCDC7 | 13721 | 3140 | 3332 | 0 | 4892 | 5034 | 3350 | 1794 |
| CCDC71L | 496 | 0 | 0 | 0 | 232 | 0 | 260 | 0 |
| CCDC74A | 0 | 0 | 0 | 0 | 2014 | 0 | 0 | 1874 |
| CCDC77 | 2036 | 1701 | 1074 | 10401 | 17434 | 10626 | 22020 | 8284 |
| CCDC82 | 0 | 0 | 0 | 0 | 1450 | 0 | 0 | 0 |
| CCDC84 | 0 | 0 | 0 | 0 | 1150 | 0 | 212 | 1262 |
| CCDC85B | 0 | 0 | 1882 | 0 | 892 | 5280 | 1119 | 0 |
| CCDC85C | 0 | 0 | 0 | 0 | 326 | 0 | 0 | 0 |
| CCDC88A | 402 | 0 | 1836 | 0 | 1464 | 0 | 444 | 852 |
| CCDC88B | 436 | 0 | 0 | 0 | 552 | 0 | 0 | 844 |
| CCDC9 | 0 | 0 | 1498 | 0 | 550 | 0 | 0 | 0 |
| CCDC90B | 0 | 504 | 0 | 0 | 1577 | 0 | 0 | 0 |
| CCDC91 | 358 | 34 | 0 | 0 | 0 | 0 | 310 | 0 |

| | | | | | | | | |
|---|---|---|---|---|---|---|---|---|
| CCDC92 | 0 | 204 | 0 | 0 | 0 | 0 | 0 | 0 |
| CCDC93 | 0 | 0 | 0 | 0 | 0 | 1847 | 0 | 164 |
| CCDC97 | 0 | 0 | 748 | 68 | 0 | 1192 | 1242 | 0 |
| CCDC9B | 0 | 270 | 0 | 0 | 30 | 0 | 136 | 0 |
| CCHCR1 | 400 | 0 | 0 | 0 | 102 | 0 | 0 | 776 |
| CCK | 0 | 0 | 0 | 0 | 860 | 0 | 0 | 0 |
| CCL11 | 0 | 0 | 2080 | 0 | 0 | 0 | 0 | 0 |
| CCL13 | 0 | 0 | 2082 | 0 | 540 | 0 | 0 | 0 |
| CCL14 | 0 | 0 | 0 | 0 | 0 | 1533 | 0 | 0 |
| CCL15 | 3984 | 2182 | 5004 | 1246 | 642 | 0 | 632 | 0 |
| CCL15-CCL14 | 0 | 36 | 3418 | 0 | 992 | 0 | 736 | 0 |
| CCL18 | 0 | 418 | 0 | 0 | 1990 | 0 | 0 | 0 |
| CCL19 | 1750 | 624 | 0 | 0 | 0 | 2184 | 0 | 1268 |
| CCL2 | 698 | 0 | 880 | 0 | 1194 | 4099 | 0 | 2594 |
| CCL20 | 2008 | 378 | 1496 | 0 | 1484 | 0 | 0 | 792 |
| CCL21 | 854 | 1622 | 3990 | 1242 | 0 | 0 | 0 | 0 |
| CCL22 | 0 | 0 | 0 | 1202 | 0 | 0 | 0 | 0 |
| CCL23 | 0 | 290 | 0 | 0 | 0 | 0 | 0 | 0 |
| CCL24 | 0 | 328 | 0 | 0 | 0 | 1248 | 520 | 0 |
| CCL28 | 492 | 0 | 8372 | 664 | 286 | 0 | 236 | 0 |
| CCL3 | 0 | 0 | 0 | 0 | 804 | 1260 | 0 | 1192 |
| CCL4 | 0 | 0 | 1352 | 0 | 0 | 0 | 0 | 416 |
| CCL4L1 | 0 | 0 | 0 | 0 | 0 | 1200 | 0 | 0 |
| CCL4L2 | 0 | 0 | 0 | 0 | 0 | 14 | 0 | 0 |
| CCL5 | 774 | 0 | 0 | 852 | 0 | 0 | 502 | 1160 |
| CCL8 | 588 | 0 | 869 | 0 | 0 | 0 | 0 | 0 |
| CCM2 | 0 | 0 | 0 | 1126 | 1708 | 0 | 592 | 0 |
| CCM2L | 0 | 0 | 0 | 718 | 0 | 0 | 0 | 880 |
| CCN1 | 0 | 630 | 2572 | 0 | 0 | 472 | 0 | 1050 |
| CCN2 | 0 | 596 | 980 | 0 | 708 | 3463 | 558 | 760 |
| CCN3 | 0 | 394 | 0 | 0 | 0 | 0 | 0 | 0 |
| CCNA2 | 2332 | 11182 | 13482 | 39385 | 118694 | 20264 | 42614 | 29387 |
| CCNB1 | 0 | 0 | 0 | 6065 | 3724 | 0 | 1262 | 2500 |
| CCNB1IP1 | 0 | 444 | 0 | 0 | 688 | 2298 | 294 | 1144 |
| CCNB2 | 0 | 460 | 0 | 1389 | 2858 | 0 | 0 | 2678 |
| CCNC | 53166 | 135843 | 290405 | 257191 | 600207 | 194636 | 428770 | 287135 |
| CCND1 | 0 | 0 | 174 | 0 | 1636 | 0 | 520 | 0 |
| CCND2 | 548 | 0 | 0 | 0 | 418 | 0 | 268 | 0 |
| CCND2-AS1 | 362 | 0 | 0 | 0 | 0 | 0 | 0 | 0 |
| CCND3 | 9231 | 13999 | 53287 | 24342 | 18728 | 7519 | 37640 | 12793 |
| CCNDBP1 | 0 | 2756 | 0 | 0 | 1544 | 1944 | 406 | 1250 |
| CCNE1 | 0 | 0 | 0 | 0 | 504 | 0 | 0 | 0 |
| CCNE2 | 0 | 0 | 0 | 0 | 36 | 0 | 0 | 516 |

| | | | | | | | | |
|---|---|---|---|---|---|---|---|---|
| CCNF | 0 | 0 | 0 | 0 | 70 | 0 | 0 | 1470 |
| CCNG1 | 0 | 2042 | 0 | 586 | 2428 | 1321 | 694 | 0 |
| CCNG2 | 397 | 416 | 1776 | 0 | 560 | 0 | 266 | 0 |
| CCNH | 32070 | 56335 | 50264 | 117926 | 199582 | 71227 | 110921 | 57248 |
| CCNI | 220 | 0 | 0 | 386 | 0 | 1430 | 784 | 0 |
| CCNI2 | 0 | 0 | 0 | 0 | 904 | 0 | 0 | 0 |
| CCNK | 1280 | 604 | 2384 | 6133 | 8985 | 3948 | 3158 | 1080 |
| CCNL1 | 876 | 0 | 0 | 0 | 1928 | 1482 | 946 | 442 |
| CCNL2 | 15637 | 9456 | 19813 | 23578 | 30729 | 42399 | 20003 | 9660 |
| CCNO | 0 | 0 | 0 | 1046 | 256 | 0 | 0 | 0 |
| CCNQ | 0 | 228 | 0 | 1483 | 2683 | 0 | 488 | 1108 |
| CCNT2 | 196 | 0 | 0 | 0 | 0 | 0 | 0 | 0 |
| CCNY | 136 | 830 | 2142 | 0 | 6457 | 1848 | 3010 | 1077 |
| CCPG1 | 171907 | 278550 | 210661 | 60593 | 240604 | 96482 | 278077 | 69179 |
| CCR7 | 25479 | 25178 | 10362 | 12866 | 5771 | 8020 | 3772 | 66590 |
| CCRL2 | 34519 | 54086 | 122164 | 51359 | 69653 | 30662 | 148512 | 21370 |
| CCS | 634 | 0 | 3390 | 0 | 2885 | 0 | 0 | 2070 |
| CCSER1 | 2008 | 1210 | 0 | 0 | 2808 | 2694 | 2520 | 0 |
| CCSER2 | 0 | 0 | 1946 | 0 | 198 | 0 | 0 | 0 |
| CCT2 | 336 | 42 | 0 | 3987 | 4496 | 2372 | 3654 | 0 |
| CCT3 | 414 | 832 | 1592 | 4163 | 9169 | 5378 | 4037 | 2883 |
| CCT4 | 38430 | 107026 | 117573 | 344891 | 673549 | 326077 | 662408 | 324873 |
| CCT5 | 0 | 830 | 2102 | 5254 | 14382 | 0 | 2839 | 672 |
| CCT6A | 1152 | 722 | 1655 | 2616 | 5488 | 0 | 2576 | 2104 |
| CCT6P1 | 0 | 354 | 0 | 0 | 0 | 0 | 0 | 0 |
| CCT6P3 | 0 | 0 | 0 | 0 | 0 | 0 | 600 | 0 |
| CCT7 | 0 | 1392 | 0 | 4627 | 3831 | 0 | 772 | 1326 |
| CCT8 | 0 | 728 | 0 | 4291 | 2462 | 5065 | 2745 | 6152 |
| CCZ1 | 36 | 0 | 3434 | 662 | 4053 | 2434 | 0 | 1236 |
| CCZ1B | 0 | 0 | 0 | 2324 | 568 | 0 | 292 | 0 |
| CCZ1P-OR7E38P | 0 | 0 | 0 | 790 | 306 | 0 | 0 | 0 |
| CD101 | 1226 | 0 | 0 | 1140 | 0 | 0 | 0 | 706 |
| CD14 | 946 | 994 | 1844 | 1578 | 726 | 2557 | 0 | 0 |
| CD151 | 1283 | 2550 | 6408 | 5070 | 7447 | 2998 | 2954 | 2602 |
| CD163 | 0 | 408 | 864 | 0 | 0 | 0 | 0 | 0 |
| CD163L1 | 0 | 0 | 0 | 0 | 378 | 0 | 0 | 0 |
| CD164 | 1816 | 2854 | 9095 | 4865 | 11718 | 432 | 2878 | 1970 |
| CD177 | 6869 | 2124 | 16770 | 0 | 1446 | 0 | 0 | 0 |
| CD1A | 1012 | 1318 | 1324 | 0 | 2204 | 0 | 392 | 0 |
| CD1D | 0 | 0 | 0 | 1072 | 0 | 0 | 0 | 0 |
| CD2 | 0 | 110 | 0 | 0 | 0 | 0 | 0 | 0 |
| CD200 | 0 | 0 | 0 | 0 | 778 | 0 | 0 | 0 |
| CD209 | 0 | 0 | 0 | 0 | 0 | 0 | 0 | 662 |

| | | | | | | | | |
|---|---|---|---|---|---|---|---|---|
| CD226 | 512 | 674 | 6969 | 0 | 1002 | 0 | 284 | 0 |
| CD24 | 11651 | 9026 | 19920 | 10735 | 28116 | 4875 | 3496 | 1052 |
| CD27 | 0 | 238 | 0 | 0 | 0 | 0 | 0 | 0 |
| CD27-AS1 | 0 | 0 | 0 | 0 | 0 | 22 | 0 | 0 |
| CD274 | 508 | 1218 | 587 | 0 | 840 | 0 | 134 | 532 |
| CD276 | 0 | 0 | 0 | 0 | 0 | 0 | 0 | 2088 |
| CD28 | 0 | 0 | 1218 | 0 | 0 | 0 | 0 | 0 |
| CD2AP | 348 | 0 | 4238 | 0 | 0 | 0 | 0 | 0 |
| CD2BP2 | 738 | 208 | 0 | 550 | 0 | 1840 | 0 | 1076 |
| CD300LF | 630 | 0 | 768 | 0 | 0 | 0 | 0 | 0 |
| CD302 | 690 | 0 | 0 | 0 | 0 | 0 | 0 | 812 |
| CD320 | 700 | 480 | 0 | 1532 | 4354 | 1968 | 1518 | 0 |
| CD33 | 242 | 0 | 0 | 0 | 0 | 0 | 0 | 0 |
| CD36 | 534 | 0 | 292 | 0 | 300 | 0 | 0 | 0 |
| CD37 | 59048 | 103352 | 60840 | 14374 | 34745 | 69595 | 19127 | 73416 |
| CD38 | 3665 | 7830 | 10355 | 2220 | 2622 | 22780 | 59971 | 30054 |
| CD3D | 824 | 0 | 0 | 0 | 0 | 0 | 0 | 0 |
| CD3E | 708 | 0 | 1371 | 0 | 134 | 0 | 0 | 0 |
| CD3G | 0 | 560 | 0 | 0 | 0 | 0 | 0 | 0 |
| CD40 | 0 | 0 | 0 | 0 | 0 | 0 | 0 | 818 |
| CD44 | 60888 | 171610 | 355650 | 356015 | 1404062 | 495126 | 1024646 | 517446 |
| CD46 | 38657 | 87160 | 117755 | 162181 | 344516 | 464492 | 589328 | 100357 |
| CD47 | 870 | 1068 | 0 | 2241 | 6696 | 0 | 830 | 6624 |
| CD48 | 0 | 444 | 1426 | 0 | 0 | 0 | 0 | 0 |
| CD52 | 842 | 1270 | 2370 | 0 | 0 | 3378 | 358 | 566 |
| CD53 | 620 | 0 | 0 | 0 | 0 | 0 | 0 | 0 |
| CD55 | 0 | 0 | 0 | 354 | 3294 | 1664 | 640 | 2742 |
| CD58 | 784 | 426 | 0 | 1254 | 2496 | 0 | 658 | 0 |
| CD59 | 702 | 1712 | 11861 | 2811 | 9712 | 2214 | 2567 | 4250 |
| CD63 | 8695 | 23131 | 28288 | 24768 | 50058 | 36638 | 34746 | 32906 |
| CD68 | 612 | 612 | 1470 | 1248 | 4237 | 8936 | 2332 | 2220 |
| CD7 | 0 | 262 | 1144 | 0 | 0 | 0 | 0 | 0 |
| CD74 | 2966348 | 5587665 | 8728227 | 1238515 | 3425026 | 11945996 | 4350483 | 4877855 |
| CD79A | 0 | 232 | 0 | 0 | 0 | 0 | 0 | 817 |
| CD79B | 35049 | 90947 | 18410 | 17526 | 3970 | 42981 | 4118 | 91678 |
| CD81 | 2369 | 898 | 3030 | 4233 | 6676 | 1500 | 3156 | 2266 |
| CD82 | 0 | 0 | 0 | 0 | 518 | 0 | 0 | 734 |
| CD86 | 5558 | 5382 | 16317 | 0 | 30080 | 39568 | 5144 | 26408 |
| CD8B | 730 | 0 | 0 | 0 | 0 | 0 | 0 | 0 |
| CD9 | 6283 | 9041 | 35155 | 20033 | 25284 | 10741 | 18766 | 8832 |
| CD96 | 376 | 0 | 0 | 0 | 0 | 0 | 0 | 0 |
| CD99 | 3092 | 1345 | 10909 | 7145 | 17115 | 11845 | 1689 | 4597 |
| CD99L2 | 0 | 0 | 0 | 958 | 0 | 0 | 0 | 0 |

| | | | | | | | | |
|---|---|---|---|---|---|---|---|---|
| CD99P1 | 0 | 0 | 0 | 886 | 554 | 0 | 0 | 0 |
| CDA | 790 | 1454 | 3673 | 1282 | 3812 | 0 | 2190 | 0 |
| CDAN1 | 0 | 0 | 0 | 0 | 0 | 4851 | 0 | 0 |
| CDC123 | 0 | 462 | 0 | 2566 | 976 | 7746 | 442 | 0 |
| CDC14A | 0 | 146 | 0 | 0 | 0 | 0 | 0 | 0 |
| CDC14B | 0 | 0 | 0 | 0 | 100 | 0 | 0 | 0 |
| CDC16 | 41768 | 56587 | 57108 | 120222 | 215353 | 276891 | 180460 | 29505 |
| CDC20 | 0 | 610 | 0 | 1886 | 3732 | 0 | 818 | 1152 |
| CDC23 | 0 | 0 | 1428 | 0 | 0 | 661 | 0 | 0 |
| CDC25B | 0 | 0 | 0 | 0 | 740 | 0 | 0 | 0 |
| CDC26 | 0 | 1490 | 2648 | 0 | 3788 | 0 | 352 | 0 |
| CDC27 | 202 | 730 | 1708 | 1104 | 6706 | 2586 | 2444 | 1915 |
| CDC34 | 0 | 0 | 0 | 0 | 154 | 0 | 1190 | 0 |
| CDC37 | 118 | 1614 | 620 | 3884 | 5443 | 0 | 928 | 1824 |
| CDC37L1 | 0 | 0 | 1496 | 0 | 0 | 0 | 0 | 0 |
| CDC40 | 0 | 0 | 0 | 0 | 0 | 0 | 652 | 0 |
| CDC42 | 405 | 2056 | 7011 | 1132 | 3069 | 0 | 712 | 684 |
| CDC42BPB | 384 | 698 | 0 | 0 | 1574 | 0 | 238 | 0 |
| CDC42BPG | 210 | 430 | 1908 | 0 | 484 | 600 | 1610 | 0 |
| CDC42EP1 | 0 | 350 | 0 | 0 | 1632 | 1230 | 446 | 0 |
| CDC42EP2 | 0 | 0 | 0 | 0 | 1096 | 0 | 588 | 913 |
| CDC42EP4 | 2422 | 831 | 5447 | 2342 | 4837 | 0 | 5471 | 1771 |
| CDC42EP5 | 1096 | 2658 | 4270 | 908 | 4504 | 874 | 334 | 1142 |
| CDC42SE1 | 0 | 240 | 0 | 1058 | 2282 | 0 | 1258 | 1950 |
| CDC42SE2 | 314 | 668 | 1880 | 0 | 2013 | 0 | 38 | 556 |
| CDC45 | 0 | 0 | 0 | 1096 | 370 | 0 | 0 | 0 |
| CDC5L | 0 | 0 | 0 | 0 | 0 | 0 | 0 | 348 |
| CDC6 | 568 | 3320 | 5665 | 18827 | 53458 | 15760 | 17020 | 18417 |
| CDC73 | 34139 | 43742 | 47384 | 47079 | 133613 | 43317 | 121391 | 42633 |
| CDCA3 | 0 | 0 | 0 | 0 | 2158 | 0 | 522 | 0 |
| CDCA4 | 0 | 0 | 0 | 902 | 0 | 0 | 0 | 0 |
| CDCA5 | 0 | 0 | 0 | 3124 | 590 | 0 | 332 | 434 |
| CDCA7 | 212 | 652 | 0 | 2026 | 4821 | 0 | 596 | 0 |
| CDCA7L | 0 | 422 | 804 | 826 | 242 | 0 | 0 | 0 |
| CDCA8 | 0 | 0 | 0 | 0 | 0 | 0 | 644 | 895 |
| CDCP1 | 0 | 0 | 144 | 0 | 176 | 0 | 672 | 0 |
| CDH1 | 102232 | 111939 | 273136 | 116458 | 197525 | 114958 | 249220 | 40119 |
| CDH11 | 3130 | 1896 | 1232 | 610 | 11004 | 9979 | 2044 | 2650 |
| CDH13 | 0 | 0 | 0 | 0 | 50 | 0 | 0 | 0 |
| CDH17 | 272502 | 397292 | 808557 | 331579 | 873123 | 815477 | 1378433 | 381247 |
| CDH24 | 70 | 0 | 0 | 0 | 0 | 0 | 0 | 0 |
| CDH3 | 0 | 0 | 0 | 0 | 0 | 0 | 550 | 0 |
| CDH5 | 0 | 298 | 0 | 0 | 0 | 0 | 0 | 0 |

| | | | | | | | | |
|---|---|---|---|---|---|---|---|---|
| CDHR1 | 874 | 254 | 0 | 0 | 0 | 0 | 342 | 0 |
| CDHR2 | 384 | 402 | 828 | 0 | 0 | 0 | 0 | 0 |
| CDHR5 | 716 | 702 | 4226 | 0 | 0 | 0 | 752 | 0 |
| CDIP1 | 0 | 0 | 1598 | 0 | 0 | 0 | 0 | 0 |
| CDIPT | 0 | 340 | 440 | 0 | 200 | 0 | 0 | 0 |
| CDK1 | 0 | 0 | 0 | 1899 | 2827 | 0 | 660 | 708 |
| CDK10 | 374 | 854 | 0 | 682 | 402 | 0 | 612 | 0 |
| CDK11A | 0 | 0 | 1038 | 744 | 778 | 0 | 736 | 0 |
| CDK11B | 218 | 242 | 1346 | 0 | 216 | 0 | 0 | 0 |
| CDK12 | 10630 | 6505 | 8199 | 7396 | 17666 | 7124 | 22290 | 4186 |
| CDK14 | 878 | 1194 | 1024 | 0 | 8404 | 1812 | 598 | 1114 |
| CDK16 | 0 | 0 | 0 | 0 | 418 | 0 | 0 | 0 |
| CDK17 | 164 | 0 | 1094 | 0 | 0 | 0 | 0 | 0 |
| CDK18 | 0 | 0 | 0 | 0 | 390 | 0 | 0 | 0 |
| CDK19 | 250 | 0 | 0 | 0 | 590 | 0 | 0 | 0 |
| CDK2 | 0 | 0 | 0 | 0 | 0 | 0 | 260 | 0 |
| CDK2AP1 | 674 | 750 | 1158 | 2619 | 1185 | 0 | 1002 | 972 |
| CDK2AP2 | 0 | 454 | 1560 | 1882 | 2468 | 2217 | 2136 | 3165 |
| CDK4 | 680 | 614 | 1482 | 5732 | 4323 | 3266 | 930 | 2379 |
| CDK5 | 0 | 0 | 0 | 0 | 1825 | 0 | 362 | 0 |
| CDK5R1 | 0 | 70 | 0 | 0 | 0 | 0 | 0 | 0 |
| CDK5RAP1 | 0 | 0 | 0 | 0 | 98 | 0 | 1474 | 0 |
| CDK5RAP3 | 23836 | 43172 | 50023 | 56685 | 82524 | 56459 | 126117 | 42779 |
| CDK6 | 2444 | 919 | 1532 | 11098 | 5199 | 9432 | 11616 | 4132 |
| CDK7 | 0 | 0 | 3487 | 2800 | 452 | 0 | 0 | 0 |
| CDK8 | 540 | 0 | 0 | 572 | 188 | 0 | 306 | 0 |
| CDK9 | 0 | 606 | 0 | 412 | 572 | 0 | 834 | 0 |
| CDKL3 | 948 | 1394 | 0 | 938 | 3746 | 2466 | 1110 | 1098 |
| CDKN1A | 386 | 410 | 5848 | 0 | 588 | 0 | 2326 | 0 |
| CDKN1C | 0 | 0 | 0 | 706 | 1596 | 0 | 0 | 1052 |
| CDKN2A | 6684 | 2696 | 2820 | 15038 | 158012 | 5348 | 26687 | 41708 |
| CDKN2AIP | 0 | 0 | 0 | 0 | 76 | 0 | 0 | 902 |
| CDKN2AIPNL | 636 | 0 | 0 | 1280 | 1678 | 0 | 0 | 1542 |
| CDKN2B | 288 | 216 | 945 | 0 | 0 | 0 | 0 | 0 |
| CDKN2B-AS1 | 4440 | 2388 | 10890 | 0 | 1168 | 0 | 0 | 0 |
| CDKN2C | 0 | 0 | 0 | 0 | 380 | 586 | 0 | 656 |
| CDKN3 | 0 | 0 | 928 | 1126 | 9893 | 0 | 0 | 1905 |
| CDPF1 | 0 | 0 | 0 | 0 | 1164 | 0 | 0 | 0 |
| CDS1 | 0 | 0 | 1288 | 0 | 0 | 0 | 0 | 0 |
| CDS2 | 0 | 0 | 0 | 0 | 484 | 0 | 0 | 454 |
| CDSN | 80 | 0 | 0 | 0 | 0 | 0 | 0 | 0 |
| CDT1 | 0 | 0 | 0 | 0 | 406 | 0 | 0 | 0 |
| CDV3 | 0 | 0 | 0 | 528 | 0 | 0 | 0 | 0 |

| | | | | | | | | |
|---|---|---|---|---|---|---|---|---|
| CDX1 | 3195 | 3798 | 3277 | 1489 | 1944 | 0 | 276 | 0 |
| CDX2 | 2686 | 700 | 644 | 0 | 1180 | 0 | 2356 | 668 |
| CDYL | 0 | 0 | 0 | 188 | 0 | 0 | 0 | 0 |
| CEACAM1 | 1824 | 798 | 6081 | 0 | 5164 | 0 | 0 | 2322 |
| CEACAM3 | 0 | 0 | 1064 | 0 | 0 | 0 | 0 | 0 |
| CEACAM5 | 3662 | 7818 | 21689 | 13256 | 23837 | 1646 | 564 | 963 |
| CEACAM6 | 0 | 692 | 2822 | 5643 | 2718 | 0 | 0 | 4190 |
| CEACAM7 | 13324 | 2570 | 48732 | 0 | 646 | 0 | 0 | 0 |
| CEBPA | 0 | 0 | 0 | 0 | 594 | 0 | 900 | 0 |
| CEBPA-DT | 98 | 0 | 0 | 0 | 0 | 0 | 0 | 0 |
| CEBPB | 314 | 164 | 1606 | 1978 | 2471 | 704 | 1980 | 2513 |
| CEBPD | 1652 | 3908 | 2589 | 989 | 8442 | 0 | 1006 | 2285 |
| CEBPG | 0 | 948 | 5590 | 0 | 6460 | 3526 | 354 | 2580 |
| CEBPZ | 0 | 0 | 0 | 0 | 0 | 0 | 562 | 714 |
| CEBPZOS | 0 | 1054 | 1560 | 2270 | 898 | 1814 | 866 | 0 |
| CECR7 | 0 | 663 | 0 | 0 | 0 | 0 | 0 | 0 |
| CEL | 0 | 0 | 0 | 14108 | 490 | 28163 | 48472 | 0 |
| CELA3A | 0 | 0 | 0 | 0 | 0 | 0 | 678 | 0 |
| CELF1 | 234 | 390 | 0 | 0 | 0 | 0 | 0 | 0 |
| CELF2 | 96 | 0 | 0 | 0 | 864 | 0 | 0 | 0 |
| CELF3 | 0 | 0 | 0 | 0 | 300 | 0 | 0 | 0 |
| CELP | 740 | 0 | 2094 | 20842 | 0 | 40615 | 42121 | 0 |
| CELSR1 | 0 | 0 | 0 | 70 | 516 | 0 | 0 | 0 |
| CELSR3 | 0 | 0 | 0 | 0 | 248 | 0 | 0 | 0 |
| CEMIP | 0 | 0 | 0 | 0 | 0 | 0 | 1084 | 0 |
| CEMIP2 | 0 | 480 | 0 | 0 | 0 | 1982 | 842 | 0 |
| CENPA | 0 | 0 | 0 | 728 | 1732 | 0 | 0 | 1363 |
| CENPB | 0 | 0 | 0 | 0 | 482 | 0 | 0 | 308 |
| CENPBD1 | 0 | 294 | 0 | 0 | 266 | 0 | 0 | 0 |
| CENPE | 0 | 0 | 0 | 1151 | 0 | 0 | 0 | 0 |
| CENPF | 0 | 0 | 1202 | 0 | 0 | 0 | 0 | 796 |
| CENPH | 0 | 144 | 500 | 2128 | 472 | 1602 | 0 | 0 |
| CENPI | 0 | 0 | 0 | 0 | 0 | 0 | 0 | 975 |
| CENPJ | 574 | 0 | 0 | 592 | 0 | 0 | 0 | 0 |
| CENPK | 0 | 608 | 0 | 0 | 579 | 0 | 122 | 0 |
| CENPM | 0 | 0 | 0 | 1338 | 598 | 0 | 0 | 970 |
| CENPN | 0 | 0 | 0 | 2342 | 584 | 0 | 0 | 868 |
| CENPO | 0 | 578 | 0 | 0 | 0 | 0 | 0 | 0 |
| CENPP | 11882 | 9502 | 29889 | 77438 | 218361 | 51047 | 59150 | 36339 |
| CENPQ | 0 | 0 | 0 | 0 | 730 | 0 | 0 | 720 |
| CENPS | 0 | 0 | 2735 | 0 | 0 | 884 | 857 | 0 |
| CENPS-CORT | 0 | 0 | 0 | 728 | 0 | 0 | 0 | 0 |
| CENPT | 606 | 0 | 0 | 0 | 798 | 0 | 0 | 556 |

| | | | | | | | | |
|---|---|---|---|---|---|---|---|---|
| CENPU | 0 | 0 | 0 | 0 | 144 | 0 | 0 | 0 |
| CENPV | 1166 | 498 | 1930 | 1416 | 4772 | 0 | 0 | 1854 |
| CENPW | 0 | 660 | 0 | 1597 | 14108 | 1501 | 894 | 1760 |
| CENPX | 2088 | 830 | 5263 | 1462 | 2694 | 7023 | 6404 | 0 |
| CEP104 | 754 | 0 | 0 | 0 | 1110 | 12 | 0 | 0 |
| CEP112 | 0 | 0 | 0 | 0 | 650 | 0 | 0 | 0 |
| CEP120 | 0 | 0 | 0 | 576 | 0 | 0 | 0 | 0 |
| CEP131 | 0 | 0 | 0 | 0 | 494 | 0 | 194 | 0 |
| CEP164 | 17654 | 7929 | 19831 | 9271 | 39200 | 19738 | 35336 | 13149 |
| CEP170 | 1834 | 582 | 0 | 0 | 7154 | 5984 | 3432 | 6164 |
| CEP170B | 640 | 0 | 0 | 0 | 280 | 0 | 194 | 0 |
| CEP19 | 476 | 0 | 0 | 0 | 0 | 0 | 0 | 0 |
| CEP250 | 0 | 0 | 0 | 0 | 790 | 0 | 0 | 0 |
| CEP41 | 0 | 0 | 0 | 0 | 0 | 0 | 374 | 0 |
| CEP55 | 0 | 0 | 0 | 0 | 417 | 0 | 0 | 0 |
| CEP57 | 0 | 0 | 158 | 0 | 284 | 0 | 372 | 0 |
| CEP70 | 5148 | 2940 | 12675 | 4926 | 43194 | 1910 | 16445 | 7388 |
| CEP72 | 0 | 0 | 0 | 0 | 20 | 0 | 0 | 0 |
| CEP76 | 2952 | 2492 | 0 | 2614 | 12755 | 1542 | 4017 | 7361 |
| CEP78 | 0 | 0 | 0 | 0 | 536 | 0 | 0 | 0 |
| CEP83 | 0 | 0 | 0 | 530 | 0 | 0 | 0 | 0 |
| CEP85 | 0 | 0 | 0 | 0 | 344 | 0 | 0 | 0 |
| CEP85L | 1250 | 62 | 0 | 0 | 1830 | 0 | 0 | 0 |
| CEP89 | 0 | 0 | 0 | 662 | 0 | 0 | 0 | 0 |
| CEPT1 | 0 | 378 | 1492 | 0 | 386 | 1365 | 230 | 1244 |
| CERCAM | 0 | 0 | 0 | 0 | 706 | 0 | 0 | 0 |
| CERK | 0 | 0 | 0 | 1130 | 0 | 0 | 0 | 0 |
| CERKL | 452 | 0 | 0 | 0 | 0 | 0 | 0 | 0 |
| CERNA2 | 0 | 0 | 0 | 126 | 0 | 0 | 0 | 0 |
| CERS2 | 0 | 894 | 0 | 0 | 634 | 0 | 2684 | 2316 |
| CERS5 | 260 | 352 | 0 | 972 | 1216 | 2497 | 1166 | 1052 |
| CERS6 | 652 | 0 | 0 | 0 | 426 | 0 | 624 | 0 |
| CES1 | 0 | 0 | 0 | 0 | 0 | 0 | 1908 | 0 |
| CES2 | 81440 | 192951 | 632188 | 45531 | 50562 | 86032 | 24448 | 7257 |
| CES3 | 684 | 228 | 2887 | 0 | 0 | 0 | 0 | 0 |
| CETN2 | 0 | 0 | 96 | 242 | 1826 | 2192 | 466 | 1922 |
| CETN3 | 848 | 764 | 1828 | 913 | 638 | 2539 | 0 | 0 |
| CFAP20 | 0 | 0 | 0 | 1092 | 3052 | 4246 | 0 | 0 |
| CFAP298 | 0 | 1558 | 3174 | 1692 | 1682 | 5141 | 1934 | 0 |
| CFAP300 | 0 | 0 | 0 | 0 | 1074 | 0 | 0 | 0 |
| CFAP36 | 0 | 446 | 0 | 0 | 350 | 0 | 582 | 0 |
| CFAP410 | 0 | 0 | 0 | 0 | 1170 | 0 | 0 | 0 |
| CFAP65 | 0 | 0 | 0 | 0 | 230 | 0 | 0 | 0 |

| | | | | | | | | |
|---|---|---|---|---|---|---|---|---|
| CFAP69 | 0 | 0 | 0 | 0 | 14 | 0 | 0 | 0 |
| CFAP97 | 832 | 5098 | 0 | 3912 | 12142 | 4135 | 17303 | 1860 |
| CFB | 71513 | 38019 | 183544 | 110162 | 76887 | 70331 | 277228 | 32768 |
| CFD | 0 | 0 | 2532 | 0 | 8695 | 0 | 118 | 1072 |
| CFDP1 | 3056 | 320 | 5842 | 0 | 1854 | 856 | 1689 | 0 |
| CFH | 670 | 592 | 1283 | 0 | 0 | 0 | 0 | 0 |
| CFI | 0 | 0 | 0 | 0 | 0 | 0 | 1080 | 0 |
| CFL1 | 17024 | 21314 | 78508 | 47946 | 98431 | 28609 | 18537 | 27220 |
| CFLAR | 104231 | 135424 | 196666 | 166939 | 429278 | 190299 | 363444 | 398135 |
| CFTR | 470 | 0 | 0 | 0 | 0 | 516 | 0 | 0 |
| CGGBP1 | 0 | 654 | 224 | 0 | 2290 | 0 | 267 | 548 |
| CGN | 2906 | 4182 | 10267 | 0 | 6348 | 1155 | 7143 | 2068 |
| CH17-125A10.2 | 0 | 0 | 296 | 0 | 0 | 0 | 0 | 0 |
| CH17-340M24.3 | 0 | 0 | 0 | 0 | 1296 | 0 | 216 | 0 |
| CHAC1 | 0 | 0 | 620 | 0 | 0 | 0 | 0 | 0 |
| CHAC2 | 0 | 0 | 0 | 0 | 1230 | 0 | 0 | 0 |
| CHAF1A | 1211 | 0 | 2961 | 0 | 1192 | 0 | 0 | 0 |
| CHCHD1 | 543 | 1990 | 1238 | 696 | 2827 | 0 | 1373 | 1096 |
| CHCHD10 | 3400 | 1738 | 2470 | 864 | 1188 | 0 | 1052 | 312 |
| CHCHD2 | 6687 | 5656 | 15669 | 29001 | 41616 | 22011 | 6887 | 15971 |
| CHCHD3 | 0 | 620 | 535 | 1202 | 2650 | 0 | 918 | 1068 |
| CHCHD4 | 818 | 568 | 0 | 0 | 400 | 0 | 0 | 0 |
| CHCHD5 | 868 | 1442 | 0 | 1248 | 3070 | 0 | 0 | 1215 |
| CHCHD6 | 0 | 0 | 0 | 0 | 2118 | 2576 | 0 | 0 |
| CHCHD7 | 1646 | 1040 | 0 | 1041 | 3305 | 4068 | 2313 | 908 |
| CHD1 | 0 | 340 | 0 | 0 | 784 | 0 | 0 | 0 |
| CHD1L | 516 | 8844 | 7854 | 9550 | 20776 | 9323 | 20553 | 12518 |
| CHD2 | 85150 | 60194 | 93382 | 78586 | 181027 | 110847 | 264345 | 52457 |
| CHD3 | 1348 | 958 | 1118 | 852 | 3238 | 4486 | 3178 | 0 |
| CHD4 | 175127 | 98070 | 104145 | 97312 | 360206 | 116231 | 289824 | 54858 |
| CHD5 | 0 | 0 | 0 | 342 | 0 | 0 | 0 | 0 |
| CHD6 | 0 | 18 | 0 | 0 | 0 | 0 | 714 | 0 |
| CHD7 | 151826 | 98404 | 121994 | 147955 | 759895 | 396491 | 685119 | 101651 |
| CHD8 | 0 | 0 | 0 | 0 | 696 | 0 | 0 | 0 |
| CHD9 | 306 | 0 | 0 | 0 | 0 | 0 | 0 | 0 |
| CHDH | 36 | 214 | 0 | 0 | 640 | 0 | 108 | 0 |
| CHEK1 | 0 | 0 | 0 | 1778 | 0 | 0 | 1982 | 0 |
| CHEK2 | 7311 | 15180 | 21849 | 27991 | 118789 | 35889 | 42356 | 30519 |
| CHEK2P2 | 0 | 0 | 0 | 0 | 192 | 0 | 0 | 0 |
| CHFR | 706 | 0 | 0 | 0 | 0 | 0 | 0 | 0 |
| CHGA | 0 | 2576 | 0 | 0 | 126 | 0 | 0 | 0 |
| CHGB | 788 | 36303 | 3552 | 0 | 1456 | 0 | 0 | 0 |
| CHI3L1 | 0 | 192 | 0 | 836 | 0 | 0 | 964 | 0 |

| | | | | | | | | |
|---|---|---|---|---|---|---|---|---|
| CHIC1 | 0 | 0 | 0 | 0 | 82 | 0 | 0 | 0 |
| CHIC2 | 0 | 2424 | 2371 | 2054 | 2867 | 0 | 578 | 904 |
| CHID1 | 0 | 550 | 0 | 2200 | 1014 | 4714 | 0 | 508 |
| CHKA | 0 | 403 | 0 | 0 | 370 | 0 | 0 | 842 |
| CHKB | 1075 | 190 | 0 | 0 | 0 | 0 | 0 | 1622 |
| CHM | 0 | 0 | 0 | 0 | 0 | 0 | 160 | 0 |
| CHMP1A | 110574 | 69301 | 331246 | 79837 | 186395 | 116732 | 194268 | 54246 |
| CHMP1B | 366 | 726 | 2689 | 0 | 3488 | 2346 | 0 | 0 |
| CHMP2A | 4968 | 3486 | 11108 | 2725 | 4799 | 2144 | 952 | 1082 |
| CHMP2B | 0 | 418 | 1909 | 912 | 5912 | 1692 | 0 | 1000 |
| CHMP3 | 664 | 514 | 552 | 1055 | 2311 | 0 | 454 | 134 |
| CHMP4A | 765 | 1743 | 2682 | 1120 | 5874 | 3726 | 438 | 968 |
| CHMP4B | 3352 | 1386 | 11702 | 8417 | 2679 | 3884 | 2062 | 2800 |
| CHMP4C | 0 | 0 | 0 | 1990 | 482 | 0 | 3384 | 0 |
| CHMP5 | 1804 | 2568 | 14194 | 2336 | 2468 | 1478 | 3976 | 1702 |
| CHMP6 | 0 | 0 | 0 | 0 | 0 | 1999 | 0 | 0 |
| CHMP7 | 600 | 0 | 0 | 0 | 0 | 1037 | 0 | 0 |
| CHN1 | 0 | 0 | 0 | 0 | 842 | 0 | 0 | 0 |
| CHN2 | 0 | 0 | 0 | 0 | 794 | 0 | 2982 | 0 |
| CHODL | 0 | 400 | 0 | 0 | 0 | 0 | 0 | 0 |
| CHORDC1 | 0 | 478 | 0 | 0 | 455 | 720 | 0 | 0 |
| CHP1 | 832 | 488 | 1832 | 0 | 38 | 0 | 208 | 390 |
| CHP2 | 0 | 282 | 3446 | 0 | 946 | 0 | 0 | 950 |
| CHPF | 0 | 0 | 0 | 1588 | 868 | 0 | 0 | 0 |
| CHPF2 | 664 | 0 | 0 | 0 | 424 | 946 | 0 | 0 |
| CHPT1 | 596 | 694 | 0 | 0 | 110 | 1866 | 1982 | 0 |
| CHRAC1 | 0 | 0 | 0 | 710 | 576 | 0 | 0 | 0 |
| CHRNA5 | 0 | 0 | 0 | 0 | 440 | 0 | 0 | 0 |
| CHRNB1 | 0 | 652 | 0 | 0 | 0 | 0 | 0 | 0 |
| CHRNB2 | 0 | 350 | 0 | 0 | 880 | 0 | 0 | 952 |
| CHST10 | 0 | 0 | 0 | 0 | 516 | 0 | 0 | 0 |
| CHST11 | 4250 | 2944 | 9025 | 1980 | 21719 | 6898 | 15049 | 16645 |
| CHST12 | 0 | 0 | 0 | 0 | 822 | 0 | 0 | 0 |
| CHST3 | 0 | 0 | 0 | 0 | 486 | 0 | 0 | 0 |
| CHST5 | 612 | 32 | 2384 | 0 | 0 | 0 | 0 | 0 |
| CHST6 | 0 | 514 | 0 | 0 | 0 | 0 | 0 | 0 |
| CHST7 | 456 | 0 | 0 | 0 | 0 | 0 | 0 | 0 |
| CHTF18 | 0 | 0 | 0 | 506 | 0 | 0 | 0 | 0 |
| CHTF8 | 0 | 0 | 1216 | 1877 | 672 | 0 | 254 | 38 |
| CHTOP | 0 | 380 | 2740 | 0 | 2268 | 0 | 712 | 704 |
| CHUK | 0 | 0 | 0 | 0 | 0 | 16 | 0 | 0 |
| CHURC1 | 926 | 4776 | 2139 | 1358 | 9402 | 1879 | 1604 | 255 |
| CIAO1 | 304 | 698 | 4738 | 2684 | 2790 | 5602 | 900 | 986 |

| | | | | | | | | |
|---|---|---|---|---|---|---|---|---|
| CIAO2A | 1946 | 2035 | 6745 | 7544 | 2842 | 0 | 740 | 1778 |
| CIAO2B | 1652 | 1798 | 3731 | 4658 | 6537 | 1437 | 4784 | 1420 |
| CIAO3 | 566 | 450 | 0 | 0 | 0 | 0 | 0 | 0 |
| CIAPIN1 | 0 | 0 | 0 | 1366 | 2873 | 0 | 446 | 0 |
| CIB1 | 2672 | 6240 | 14346 | 7999 | 10846 | 532 | 1494 | 2692 |
| CIC | 1213 | 706 | 1950 | 1699 | 5320 | 0 | 990 | 775 |
| CIDEA | 2274 | 0 | 15351 | 0 | 0 | 40164 | 0 | 0 |
| CIDEB | 0 | 0 | 0 | 0 | 1022 | 0 | 0 | 0 |
| CIDEC | 724 | 0 | 2414 | 0 | 0 | 0 | 0 | 0 |
| CIDECP1 | 0 | 0 | 1677 | 0 | 0 | 0 | 0 | 0 |
| CIITA | 774 | 262 | 0 | 0 | 0 | 692 | 1748 | 0 |
| CINP | 0 | 0 | 2634 | 2584 | 3837 | 0 | 1098 | 0 |
| CIP2A | 330 | 3814 | 4358 | 15315 | 24734 | 2917 | 15270 | 9264 |
| CIPC | 0 | 328 | 0 | 0 | 0 | 0 | 0 | 0 |
| CIR1 | 0 | 252 | 0 | 0 | 1254 | 3097 | 0 | 0 |
| CIRBP | 2516 | 876 | 5535 | 2605 | 8904 | 0 | 1366 | 1775 |
| CISD1 | 0 | 0 | 1142 | 2006 | 4082 | 72 | 0 | 0 |
| CISD2 | 588 | 846 | 0 | 3972 | 2462 | 0 | 728 | 634 |
| CISD3 | 68112 | 138404 | 278782 | 122533 | 170703 | 71968 | 181812 | 53404 |
| CISH | 0 | 0 | 728 | 0 | 612 | 1396 | 894 | 0 |
| CIT | 0 | 0 | 0 | 668 | 384 | 0 | 0 | 0 |
| CITED2 | 0 | 716 | 1085 | 0 | 619 | 720 | 258 | 0 |
| CITED4 | 0 | 0 | 0 | 768 | 0 | 0 | 0 | 0 |
| CIZ1 | 0 | 0 | 0 | 1640 | 504 | 1338 | 0 | 0 |
| CKAP2 | 0 | 382 | 0 | 2594 | 680 | 0 | 0 | 70 |
| CKAP4 | 0 | 336 | 0 | 0 | 726 | 0 | 0 | 0 |
| CKAP5 | 2136 | 1930 | 2848 | 4766 | 7761 | 2992 | 10279 | 4640 |
| CKB | 30876 | 13928 | 99868 | 2014 | 12860 | 0 | 7412 | 3870 |
| CKLF | 910 | 460 | 3660 | 1824 | 3022 | 1668 | 462 | 1120 |
| CKMT1A | 0 | 1850 | 3792 | 0 | 790 | 0 | 324 | 2918 |
| CKMT1B | 2376 | 2968 | 4030 | 5343 | 2584 | 0 | 414 | 0 |
| CKMT2 | 0 | 0 | 0 | 1428 | 0 | 0 | 420 | 0 |
| CKS1B | 737 | 978 | 2882 | 8520 | 10671 | 0 | 3866 | 5886 |
| CKS2 | 0 | 350 | 2907 | 15155 | 23336 | 0 | 2366 | 3326 |
| CLASP1 | 2088 | 2852 | 18 | 1695 | 3744 | 682 | 7619 | 718 |
| CLASP2 | 1644 | 1082 | 918 | 1640 | 10181 | 605 | 7422 | 2163 |
| CLASRP | 1010 | 0 | 1427 | 1176 | 0 | 0 | 0 | 0 |
| CLC | 0 | 0 | 0 | 0 | 0 | 1683 | 0 | 0 |
| CLCA1 | 892 | 1696 | 17437 | 934 | 0 | 0 | 0 | 0 |
| CLCA4 | 3109 | 274 | 9036 | 0 | 0 | 0 | 0 | 0 |
| CLCA4-AS1 | 0 | 0 | 1726 | 0 | 0 | 0 | 0 | 0 |
| CLCF1 | 1106 | 804 | 3410 | 0 | 0 | 0 | 0 | 0 |
| CLCN2 | 0 | 0 | 986 | 0 | 0 | 1028 | 532 | 0 |

| | | | | | | | | |
|---|---|---|---|---|---|---|---|---|
| CLCN3 | 0 | 1122 | 0 | 0 | 0 | 0 | 1042 | 0 |
| CLCN6 | 2408 | 588 | 122 | 0 | 2444 | 1302 | 404 | 500 |
| CLCN7 | 2404 | 2998 | 7395 | 362 | 7345 | 1282 | 19620 | 5066 |
| CLDN1 | 0 | 0 | 0 | 0 | 0 | 2351 | 0 | 0 |
| CLDN10 | 0 | 1584 | 0 | 0 | 0 | 10700 | 0 | 0 |
| CLDN12 | 220 | 848 | 0 | 0 | 0 | 0 | 0 | 0 |
| CLDN15 | 0 | 140 | 0 | 960 | 418 | 0 | 264 | 0 |
| CLDN2 | 0 | 0 | 0 | 0 | 40 | 0 | 1866 | 1358 |
| CLDN23 | 0 | 0 | 3352 | 0 | 0 | 0 | 0 | 0 |
| CLDN3 | 3446 | 3957 | 8451 | 890 | 4667 | 0 | 5038 | 0 |
| CLDN4 | 11547 | 6428 | 46900 | 9736 | 18159 | 11548 | 7067 | 850 |
| CLDN5 | 1100 | 416 | 0 | 0 | 0 | 0 | 0 | 0 |
| CLDN7 | 8928 | 11629 | 38333 | 7763 | 10375 | 1826 | 3077 | 6973 |
| CLDN8 | 424 | 1194 | 1276 | 0 | 0 | 0 | 0 | 0 |
| CLDND1 | 0 | 0 | 0 | 784 | 2065 | 0 | 754 | 0 |
| CLDND2 | 150 | 0 | 0 | 0 | 0 | 0 | 804 | 0 |
| CLEC16A | 244 | 354 | 1160 | 1260 | 1508 | 1862 | 2870 | 0 |
| CLEC18B | 0 | 0 | 0 | 0 | 706 | 0 | 0 | 0 |
| CLEC18C | 0 | 0 | 0 | 0 | 208 | 0 | 0 | 0 |
| CLEC2B | 0 | 1128 | 0 | 1354 | 2016 | 1942 | 122 | 2350 |
| CLEC2D | 4368 | 2556 | 0 | 4181 | 2347 | 2328 | 3865 | 2632 |
| CLEC3B | 652 | 388 | 3972 | 0 | 0 | 0 | 0 | 0 |
| CLEC4A | 0 | 0 | 1952 | 0 | 0 | 0 | 0 | 0 |
| CLEC4M | 0 | 0 | 1704 | 0 | 0 | 0 | 0 | 0 |
| CLECL1 | 0 | 0 | 0 | 0 | 398 | 1366 | 0 | 0 |
| CLIC1 | 4127 | 6194 | 29201 | 25436 | 30139 | 10863 | 15136 | 16583 |
| CLIC3 | 0 | 0 | 470 | 0 | 1080 | 0 | 0 | 0 |
| CLIC4 | 0 | 918 | 0 | 0 | 1234 | 0 | 0 | 0 |
| CLIC5 | 638 | 466 | 2056 | 0 | 0 | 0 | 414 | 0 |
| CLINT1 | 328 | 206 | 0 | 0 | 457 | 0 | 0 | 454 |
| CLIP1 | 11927 | 16971 | 23903 | 8036 | 37912 | 13611 | 21662 | 4796 |
| CLIP2 | 0 | 0 | 0 | 1661 | 170 | 0 | 386 | 0 |
| CLK1 | 296 | 620 | 2593 | 0 | 312 | 1918 | 740 | 0 |
| CLK2 | 0 | 0 | 0 | 728 | 1030 | 0 | 0 | 0 |
| CLK3 | 0 | 0 | 1236 | 0 | 634 | 0 | 574 | 0 |
| CLK4 | 0 | 494 | 0 | 0 | 204 | 0 | 0 | 956 |
| CLMN | 158398 | 88227 | 84008 | 16982 | 271069 | 3440 | 29302 | 14399 |
| CLN3 | 846 | 407 | 3076 | 2257 | 264 | 1214 | 1430 | 1018 |
| CLN5 | 0 | 86 | 879 | 1984 | 586 | 2097 | 460 | 716 |
| CLN6 | 0 | 410 | 0 | 0 | 0 | 0 | 202 | 0 |
| CLN8 | 0 | 0 | 0 | 0 | 492 | 0 | 0 | 542 |
| CLNK | 0 | 0 | 2573 | 0 | 0 | 1704 | 0 | 0 |
| CLNS1A | 650 | 1552 | 1881 | 1286 | 5884 | 2054 | 0 | 3916 |

| | | | | | | | | |
|---|---|---|---|---|---|---|---|---|
| CLOCK | 0 | 0 | 0 | 0 | 952 | 0 | 0 | 0 |
| CLP1 | 0 | 0 | 0 | 0 | 410 | 0 | 1170 | 0 |
| CLPP | 848 | 0 | 0 | 0 | 3018 | 0 | 560 | 1457 |
| CLPTM1 | 584 | 2504 | 2554 | 0 | 2458 | 1720 | 3121 | 4612 |
| CLPTM1L | 10795 | 20968 | 39831 | 59537 | 129949 | 40738 | 66619 | 29080 |
| CLPX | 0 | 0 | 0 | 0 | 0 | 0 | 338 | 0 |
| CLRN3 | 1074 | 1602 | 8479 | 2964 | 0 | 2081 | 3869 | 2536 |
| CLSTN1 | 340 | 1820 | 1270 | 0 | 64 | 0 | 0 | 0 |
| CLSTN2 | 0 | 406 | 0 | 0 | 0 | 0 | 0 | 0 |
| CLTA | 1840 | 3924 | 15547 | 5173 | 12613 | 7262 | 3114 | 3950 |
| CLTB | 5263 | 3047 | 21444 | 2576 | 6640 | 2318 | 3953 | 3128 |
| CLTC | 83320 | 139376 | 201419 | 121172 | 260557 | 164909 | 262068 | 37714 |
| CLTCL1 | 0 | 620 | 0 | 1222 | 1570 | 0 | 234 | 0 |
| CLU | 0 | 1142 | 1629 | 0 | 0 | 8173 | 512 | 2071 |
| CLUAP1 | 0 | 0 | 0 | 0 | 690 | 0 | 0 | 506 |
| CLUH | 0 | 0 | 850 | 0 | 684 | 0 | 0 | 0 |
| CLVS1 | 0 | 0 | 0 | 0 | 0 | 0 | 200 | 0 |
| CLYBL | 0 | 0 | 2419 | 0 | 378 | 0 | 624 | 0 |
| CMAHP | 0 | 318 | 0 | 0 | 0 | 0 | 0 | 0 |
| CMAS | 62339 | 88079 | 123397 | 81696 | 86865 | 69858 | 129805 | 38154 |
| CMBL | 1172 | 732 | 1534 | 1377 | 3496 | 0 | 0 | 2548 |
| CMC1 | 622 | 2712 | 5289 | 6845 | 6330 | 3569 | 2220 | 1606 |
| CMC2 | 0 | 434 | 1279 | 2300 | 3012 | 3474 | 950 | 832 |
| CMC4 | 0 | 0 | 2346 | 1542 | 1038 | 0 | 0 | 0 |
| CMIP | 0 | 0 | 0 | 0 | 418 | 0 | 0 | 0 |
| CMKLR1 | 456 | 0 | 0 | 0 | 0 | 0 | 0 | 0 |
| CMPK1 | 2408 | 2412 | 5026 | 660 | 4673 | 4200 | 386 | 1082 |
| CMPK2 | 0 | 0 | 0 | 534 | 0 | 0 | 0 | 0 |
| CMSS1 | 0 | 370 | 0 | 0 | 2900 | 0 | 558 | 0 |
| CMTM3 | 0 | 0 | 0 | 0 | 230 | 0 | 1024 | 0 |
| CMTM4 | 250 | 0 | 688 | 0 | 476 | 0 | 0 | 0 |
| CMTM6 | 0 | 0 | 1062 | 694 | 1004 | 0 | 754 | 0 |
| CMTM7 | 308 | 0 | 0 | 0 | 3174 | 0 | 0 | 0 |
| CMTM8 | 608 | 0 | 0 | 1760 | 1596 | 0 | 1180 | 578 |
| CNBP | 516339 | 1067257 | 7430044 | 2030897 | 3912581 | 2714535 | 4703035 | 2194352 |
| CNDP2 | 156 | 962 | 3662 | 0 | 1228 | 0 | 0 | 0 |
| CNEP1R1 | 0 | 10 | 0 | 830 | 180 | 0 | 0 | 0 |
| CNFN | 0 | 142 | 0 | 0 | 430 | 0 | 0 | 0 |
| CNIH1 | 282 | 448 | 1144 | 1764 | 4603 | 1496 | 0 | 548 |
| CNIH3 | 0 | 0 | 0 | 1362 | 2306 | 0 | 0 | 0 |
| CNIH4 | 1498 | 0 | 3286 | 1636 | 1694 | 3741 | 1590 | 694 |
| CNN1 | 0 | 0 | 490 | 0 | 0 | 200 | 0 | 0 |
| CNN2 | 0 | 1428 | 1559 | 0 | 1616 | 0 | 1091 | 2591 |

| | | | | | | | | |
|---|---|---|---|---|---|---|---|---|
| CNN3 | 12095 | 17095 | 62073 | 28635 | 52843 | 41978 | 31027 | 24287 |
| CNNM2 | 0 | 0 | 0 | 0 | 0 | 0 | 1152 | 0 |
| CNNM3 | 0 | 108 | 0 | 0 | 1862 | 0 | 462 | 0 |
| CNNM4 | 0 | 326 | 0 | 0 | 548 | 0 | 0 | 0 |
| CNOT1 | 0 | 0 | 0 | 0 | 0 | 0 | 196 | 0 |
| CNOT11 | 0 | 116 | 0 | 0 | 1986 | 1682 | 0 | 1020 |
| CNOT2 | 322 | 0 | 360 | 0 | 724 | 0 | 0 | 0 |
| CNOT3 | 408 | 0 | 0 | 0 | 490 | 0 | 264 | 0 |
| CNOT6 | 2252 | 1256 | 3028 | 1000 | 5407 | 962 | 2854 | 640 |
| CNOT7 | 0 | 678 | 0 | 0 | 866 | 0 | 0 | 1102 |
| CNOT8 | 0 | 0 | 1885 | 130 | 156 | 1815 | 312 | 0 |
| CNOT9 | 0 | 322 | 0 | 0 | 2070 | 0 | 0 | 72 |
| CNP | 482 | 62 | 0 | 0 | 234 | 0 | 138 | 0 |
| CNPPD1 | 0 | 0 | 1351 | 0 | 1289 | 1764 | 128 | 0 |
| CNPY2 | 682 | 1774 | 3314 | 7659 | 6239 | 8056 | 2926 | 2212 |
| CNPY3 | 1078 | 400 | 2128 | 1449 | 4753 | 2628 | 1256 | 794 |
| CNPY3-GNMT | 950 | 0 | 0 | 0 | 0 | 0 | 0 | 0 |
| CNPY4 | 0 | 0 | 1500 | 0 | 422 | 0 | 0 | 0 |
| CNRIP1 | 658 | 0 | 0 | 1981 | 0 | 0 | 0 | 0 |
| CNST | 0 | 0 | 0 | 0 | 0 | 0 | 0 | 454 |
| CNTLN | 0 | 0 | 0 | 0 | 450 | 0 | 0 | 0 |
| CNTN5 | 0 | 0 | 0 | 0 | 0 | 0 | 278 | 0 |
| CNTROB | 0 | 0 | 0 | 0 | 50 | 0 | 0 | 300 |
| COA1 | 0 | 0 | 2628 | 1478 | 4960 | 0 | 56 | 1084 |
| COA3 | 3658 | 3901 | 7567 | 2192 | 6677 | 4297 | 8845 | 584 |
| COA4 | 0 | 2728 | 0 | 957 | 4846 | 3387 | 4254 | 2504 |
| COA5 | 0 | 492 | 3016 | 1938 | 4107 | 1699 | 1145 | 44 |
| COA6 | 0 | 0 | 1266 | 1435 | 2012 | 1548 | 852 | 1550 |
| COA6-AS1 | 0 | 186 | 0 | 382 | 0 | 0 | 404 | 0 |
| COASY | 0 | 0 | 0 | 0 | 494 | 0 | 0 | 600 |
| COBL | 0 | 331 | 0 | 0 | 816 | 0 | 360 | 0 |
| COCH | 0 | 0 | 0 | 1006 | 0 | 0 | 0 | 0 |
| COG1 | 0 | 0 | 0 | 0 | 492 | 0 | 510 | 0 |
| COG2 | 0 | 0 | 0 | 0 | 906 | 0 | 741 | 0 |
| COG3 | 136 | 0 | 0 | 346 | 0 | 0 | 0 | 0 |
| COG4 | 11021 | 14303 | 30104 | 39003 | 37165 | 30298 | 52963 | 19895 |
| COG5 | 0 | 0 | 0 | 0 | 686 | 0 | 0 | 0 |
| COG6 | 18227 | 28345 | 17077 | 28606 | 76596 | 83673 | 115673 | 27505 |
| COG7 | 0 | 0 | 0 | 0 | 0 | 0 | 692 | 0 |
| COG8 | 0 | 106 | 0 | 0 | 972 | 0 | 1500 | 0 |
| COL10A1 | 0 | 0 | 0 | 0 | 690 | 0 | 0 | 0 |
| COL11A1 | 0 | 0 | 0 | 0 | 862 | 0 | 0 | 0 |
| COL12A1 | 0 | 0 | 0 | 0 | 1756 | 2160 | 0 | 0 |

| | | | | | | | | |
|---|---|---|---|---|---|---|---|---|
| COL15A1 | 0 | 0 | 0 | 0 | 562 | 0 | 0 | 0 |
| COL16A1 | 14878 | 10925 | 3259 | 9295 | 32453 | 23197 | 7970 | 17920 |
| COL17A1 | 1784 | 228 | 0 | 0 | 0 | 0 | 0 | 0 |
| COL18A1 | 0 | 0 | 1194 | 1236 | 1568 | 0 | 0 | 888 |
| COL1A1 | 117 | 30 | 38 | 1122 | 7185 | 0 | 0 | 618 |
| COL1A2 | 228 | 532 | 0 | 1486 | 17387 | 1092 | 0 | 1150 |
| COL21A1 | 1056 | 614 | 0 | 0 | 2790 | 0 | 1314 | 0 |
| COL24A1 | 818 | 0 | 0 | 0 | 0 | 0 | 0 | 0 |
| COL3A1 | 868 | 518 | 0 | 0 | 11699 | 2918 | 662 | 2538 |
| COL4A1 | 0 | 0 | 40 | 182 | 0 | 1072 | 0 | 553 |
| COL4A2 | 0 | 0 | 0 | 752 | 822 | 0 | 0 | 0 |
| COL4A3BP | 0 | 0 | 0 | 632 | 0 | 1182 | 0 | 0 |
| COL4A5 | 362 | 0 | 0 | 0 | 0 | 0 | 0 | 0 |
| COL5A1 | 0 | 0 | 0 | 0 | 2212 | 0 | 0 | 0 |
| COL5A2 | 0 | 0 | 0 | 0 | 1782 | 0 | 0 | 0 |
| COL6A1 | 606 | 0 | 0 | 982 | 2013 | 1494 | 0 | 0 |
| COL6A2 | 296 | 814 | 0 | 0 | 3284 | 480 | 0 | 0 |
| COL6A3 | 0 | 0 | 0 | 0 | 5079 | 1238 | 0 | 0 |
| COL7A1 | 0 | 0 | 0 | 0 | 796 | 0 | 0 | 0 |
| COL8A1 | 0 | 0 | 0 | 0 | 522 | 0 | 0 | 0 |
| COL9A2 | 782 | 988 | 4737 | 0 | 0 | 0 | 744 | 0 |
| COL9A3 | 0 | 0 | 0 | 0 | 316 | 0 | 2878 | 0 |
| COLCA1 | 0 | 0 | 0 | 0 | 684 | 0 | 0 | 0 |
| COLCA2 | 0 | 0 | 0 | 0 | 0 | 0 | 0 | 1034 |
| COLGALT1 | 0 | 274 | 1810 | 0 | 1076 | 2376 | 520 | 0 |
| COMMD1 | 0 | 510 | 0 | 1350 | 1077 | 2114 | 508 | 3020 |
| COMMD10 | 43942 | 147081 | 124850 | 205093 | 493789 | 242317 | 107908 | 157253 |
| COMMD2 | 0 | 162 | 0 | 0 | 0 | 0 | 0 | 0 |
| COMMD3 | 0 | 1704 | 5083 | 2532 | 1605 | 2263 | 1490 | 2594 |
| COMMD4 | 0 | 592 | 3300 | 2195 | 3175 | 2475 | 1728 | 1116 |
| COMMD5 | 0 | 230 | 0 | 0 | 434 | 4267 | 664 | 0 |
| COMMD6 | 3139 | 3398 | 10902 | 12240 | 17068 | 5131 | 8804 | 9665 |
| COMMD7 | 0 | 400 | 50 | 3580 | 5321 | 0 | 1598 | 5016 |
| COMMD8 | 0 | 0 | 0 | 0 | 1953 | 324 | 324 | 1774 |
| COMMD9 | 0 | 0 | 0 | 0 | 1078 | 2566 | 322 | 2288 |
| COMP | 0 | 0 | 0 | 0 | 982 | 1524 | 0 | 0 |
| COMT | 18872 | 51559 | 201922 | 105360 | 364358 | 266504 | 267790 | 85517 |
| COMTD1 | 774 | 712 | 1162 | 192 | 564 | 0 | 1658 | 1078 |
| COP1 | 34534 | 68347 | 87509 | 96692 | 238581 | 71392 | 134538 | 77260 |
| COPA | 0 | 0 | 0 | 1202 | 870 | 0 | 58 | 0 |
| COPB1 | 928 | 1280 | 0 | 0 | 1196 | 0 | 0 | 0 |
| COPB2 | 46696 | 152972 | 198145 | 145493 | 367666 | 230341 | 358567 | 199808 |
| COPE | 1712 | 1614 | 11962 | 4324 | 9404 | 6998 | 452 | 938 |

| | | | | | | | | |
|---|---|---|---|---|---|---|---|---|
| COPG1 | 0 | 548 | 0 | 0 | 0 | 0 | 2274 | 1118 |
| COPG2 | 2518 | 4774 | 7253 | 6470 | 10850 | 12986 | 11076 | 3597 |
| COPG2IT1 | 0 | 0 | 0 | 0 | 0 | 0 | 1949 | 0 |
| COPRS | 912 | 598 | 0 | 2453 | 5703 | 0 | 1652 | 0 |
| COPS2 | 234 | 68 | 0 | 770 | 2762 | 916 | 0 | 848 |
| COPS3 | 44454 | 97622 | 66975 | 101203 | 236856 | 123698 | 124120 | 91643 |
| COPS4 | 726 | 673 | 2306 | 0 | 1992 | 1122 | 0 | 362 |
| COPS5 | 820 | 0 | 1942 | 1322 | 3583 | 2858 | 2034 | 3750 |
| COPS6 | 0 | 410 | 0 | 6491 | 5515 | 0 | 2990 | 1174 |
| COPS7A | 0 | 440 | 0 | 0 | 1766 | 0 | 558 | 552 |
| COPS7B | 492 | 338 | 0 | 715 | 3053 | 0 | 0 | 0 |
| COPS8 | 0 | 648 | 12 | 0 | 2802 | 0 | 0 | 662 |
| COPS9 | 0 | 1318 | 0 | 2310 | 5514 | 0 | 454 | 1536 |
| COPZ1 | 2194 | 2864 | 2881 | 5090 | 5268 | 2076 | 2055 | 1277 |
| COPZ2 | 0 | 524 | 0 | 954 | 0 | 0 | 0 | 0 |
| COQ10A | 6532 | 9651 | 9508 | 10074 | 23545 | 17770 | 9320 | 5993 |
| COQ10B | 25552 | 30109 | 79262 | 31196 | 90848 | 30877 | 74570 | 60539 |
| COQ2 | 0 | 732 | 0 | 0 | 60 | 1726 | 0 | 0 |
| COQ3 | 0 | 0 | 0 | 2306 | 702 | 0 | 0 | 0 |
| COQ4 | 154 | 320 | 958 | 1288 | 830 | 0 | 523 | 1990 |
| COQ5 | 0 | 970 | 1574 | 0 | 690 | 0 | 0 | 1264 |
| COQ6 | 0 | 1052 | 0 | 1380 | 576 | 0 | 678 | 0 |
| COQ7 | 892 | 0 | 0 | 824 | 754 | 0 | 0 | 0 |
| COQ8A | 0 | 0 | 1946 | 0 | 580 | 1240 | 0 | 404 |
| COQ9 | 3049 | 0 | 0 | 1456 | 1356 | 0 | 1332 | 0 |
| CORO1A | 1004 | 0 | 0 | 926 | 416 | 0 | 0 | 0 |
| CORO1B | 488 | 132 | 3305 | 960 | 1936 | 0 | 0 | 810 |
| CORO1C | 0 | 184 | 0 | 114 | 1470 | 0 | 0 | 0 |
| CORO2A | 2888 | 0 | 10275 | 4167 | 5076 | 8470 | 4440 | 1082 |
| CORO7 | 0 | 0 | 0 | 0 | 0 | 0 | 0 | 506 |
| COTL1 | 0 | 708 | 0 | 5843 | 1850 | 0 | 0 | 1538 |
| COX10 | 0 | 0 | 1104 | 0 | 0 | 0 | 0 | 0 |
| COX10-AS1 | 0 | 0 | 1722 | 0 | 0 | 0 | 0 | 0 |
| COX11 | 0 | 434 | 2198 | 0 | 0 | 0 | 974 | 814 |
| COX14 | 2516 | 2048 | 7486 | 832 | 320 | 0 | 514 | 462 |
| COX15 | 0 | 434 | 0 | 0 | 665 | 0 | 0 | 0 |
| COX16 | 0 | 2067 | 1495 | 3602 | 5692 | 726 | 1296 | 978 |
| COX17 | 688 | 2484 | 2807 | 4085 | 4002 | 1076 | 1758 | 1388 |
| COX19 | 290 | 164 | 0 | 1393 | 2094 | 0 | 2805 | 1253 |
| COX20 | 2016 | 1772 | 766 | 3733 | 5110 | 358 | 1185 | 0 |
| COX4I1 | 16351 | 21450 | 54518 | 37444 | 25670 | 20584 | 15623 | 10695 |
| COX4I2 | 10883 | 0 | 4496 | 656 | 1684 | 2900 | 648 | 0 |
| COX5A | 8901 | 12033 | 16170 | 13513 | 18120 | 7393 | 7928 | 2904 |

| | | | | | | | | |
|---|---|---|---|---|---|---|---|---|
| COX5B | 31391 | 37696 | 53473 | 28133 | 57459 | 14335 | 37738 | 8838 |
| COX6A1 | 13766 | 23437 | 23512 | 18778 | 22841 | 6387 | 13600 | 11196 |
| COX6B1 | 6946 | 13922 | 13297 | 12094 | 24226 | 8647 | 6586 | 5129 |
| COX6B2 | 0 | 0 | 0 | 0 | 316 | 0 | 0 | 0 |
| COX6C | 12015 | 20620 | 18448 | 13185 | 34246 | 19134 | 22462 | 13921 |
| COX7A1 | 0 | 424 | 0 | 0 | 0 | 0 | 0 | 276 |
| COX7A2 | 6356 | 20298 | 15030 | 16975 | 26393 | 4857 | 9766 | 4018 |
| COX7A2L | 0 | 2220 | 4769 | 9466 | 7357 | 1034 | 4664 | 5173 |
| COX7B | 7137 | 15697 | 6623 | 6239 | 17199 | 4426 | 5664 | 3487 |
| COX7C | 6510 | 17997 | 14461 | 10430 | 24310 | 9192 | 7956 | 6776 |
| COX8A | 7910 | 7243 | 5724 | 3038 | 14207 | 0 | 4966 | 5891 |
| CPA2 | 5068 | 580 | 35333 | 0 | 0 | 0 | 1208 | 0 |
| CPAMD8 | 0 | 82 | 0 | 0 | 0 | 0 | 0 | 0 |
| CPB1 | 0 | 662 | 0 | 0 | 0 | 0 | 0 | 0 |
| CPD | 480 | 634 | 0 | 176 | 3032 | 0 | 664 | 0 |
| CPE | 0 | 1170 | 0 | 0 | 276 | 1208 | 0 | 0 |
| CPEB2 | 0 | 0 | 0 | 0 | 210 | 0 | 0 | 0 |
| CPEB3 | 0 | 0 | 0 | 0 | 206 | 0 | 0 | 0 |
| CPED1 | 0 | 0 | 0 | 0 | 476 | 0 | 0 | 0 |
| CPLANE1 | 76 | 360 | 0 | 0 | 548 | 0 | 96 | 0 |
| CPLX1 | 0 | 102 | 0 | 844 | 3132 | 0 | 0 | 0 |
| CPM | 40464 | 131314 | 181577 | 6078 | 28012 | 5115 | 9096 | 14636 |
| CPNE1 | 0 | 744 | 0 | 0 | 7570 | 0 | 1426 | 0 |
| CPNE2 | 544 | 0 | 0 | 0 | 968 | 0 | 0 | 0 |
| CPNE3 | 284 | 0 | 0 | 0 | 878 | 0 | 454 | 0 |
| CPNE7 | 0 | 0 | 0 | 850 | 0 | 776 | 728 | 0 |
| CPOX | 242 | 0 | 0 | 0 | 1509 | 0 | 0 | 668 |
| CPPED1 | 0 | 0 | 0 | 0 | 704 | 932 | 344 | 0 |
| CPQ | 0 | 0 | 2134 | 0 | 438 | 1906 | 0 | 0 |
| CPSF1 | 130 | 106 | 1668 | 0 | 722 | 0 | 0 | 0 |
| CPSF2 | 0 | 0 | 0 | 0 | 590 | 0 | 0 | 0 |
| CPSF3 | 1184 | 0 | 2180 | 1308 | 522 | 0 | 0 | 0 |
| CPSF4 | 0 | 52 | 0 | 830 | 1232 | 0 | 776 | 0 |
| CPSF6 | 108415 | 263216 | 389639 | 321008 | 912661 | 327133 | 573734 | 287207 |
| CPSF7 | 148 | 0 | 0 | 954 | 0 | 0 | 0 | 0 |
| CPT1A | 7262 | 11531 | 56273 | 3514 | 18551 | 2682 | 14297 | 7401 |
| CPTP | 468 | 0 | 0 | 0 | 636 | 0 | 0 | 0 |
| CPVL | 0 | 0 | 0 | 0 | 538 | 0 | 0 | 1114 |
| CPXM1 | 0 | 0 | 0 | 0 | 1027 | 0 | 0 | 0 |
| CPXM2 | 0 | 0 | 0 | 0 | 0 | 1332 | 0 | 0 |
| CR1 | 2037 | 3491 | 0 | 1313 | 2167 | 7190 | 1266 | 0 |
| CR1L | 548 | 350 | 0 | 0 | 0 | 0 | 0 | 0 |
| CR2 | 0 | 240 | 0 | 0 | 0 | 0 | 0 | 0 |

| | | | | | | | | |
|---|---|---|---|---|---|---|---|---|
| CRABP2 | 0 | 0 | 0 | 0 | 0 | 0 | 978 | 0 |
| CRACR2A | 1007 | 2906 | 3615 | 0 | 2324 | 1820 | 1286 | 0 |
| CRACR2B | 422 | 0 | 392 | 0 | 108 | 0 | 150 | 0 |
| CRADD | 87135 | 66911 | 382042 | 58822 | 79735 | 69431 | 156776 | 41078 |
| CRAT | 274 | 0 | 0 | 0 | 0 | 1906 | 528 | 0 |
| CRB3 | 7292 | 8302 | 20581 | 15934 | 30674 | 8794 | 21061 | 10042 |
| CRBN | 0 | 274 | 994 | 0 | 146 | 1546 | 290 | 0 |
| CRCP | 0 | 286 | 0 | 324 | 0 | 0 | 1860 | 0 |
| CREB1 | 510 | 2572 | 2238 | 1542 | 4232 | 2290 | 2808 | 4935 |
| CREB3 | 0 | 0 | 0 | 0 | 0 | 0 | 1214 | 0 |
| CREB3L1 | 0 | 0 | 420 | 962 | 0 | 0 | 0 | 1087 |
| CREB3L2 | 808 | 1694 | 9099 | 1281 | 4956 | 1690 | 1194 | 4135 |
| CREBBP | 11364 | 7332 | 11973 | 2910 | 12732 | 4645 | 23869 | 4355 |
| CREBL2 | 0 | 102 | 970 | 0 | 54 | 0 | 0 | 0 |
| CREBRF | 0 | 0 | 0 | 0 | 528 | 0 | 0 | 0 |
| CREBZF | 142 | 0 | 1889 | 0 | 1088 | 1854 | 40 | 0 |
| CREG1 | 2670 | 988 | 3328 | 3958 | 2354 | 2636 | 2338 | 2002 |
| CREG2 | 0 | 0 | 1402 | 0 | 0 | 0 | 0 | 0 |
| CRELD1 | 306 | 0 | 0 | 0 | 984 | 0 | 506 | 0 |
| CRELD2 | 0 | 1690 | 3341 | 388 | 730 | 2504 | 604 | 0 |
| CREM | 0 | 498 | 2620 | 0 | 0 | 0 | 0 | 1064 |
| CRIM1 | 0 | 0 | 0 | 0 | 772 | 0 | 628 | 0 |
| CRIM1-DT | 0 | 0 | 0 | 0 | 506 | 0 | 0 | 0 |
| CRIP1 | 3366 | 5631 | 4144 | 3359 | 9203 | 768 | 378 | 1318 |
| CRIP2 | 0 | 452 | 0 | 470 | 118 | 0 | 0 | 1575 |
| CRIPAK | 666 | 0 | 0 | 0 | 0 | 0 | 0 | 0 |
| CRIPT | 0 | 938 | 0 | 944 | 1506 | 0 | 302 | 0 |
| CRISPLD2 | 382 | 0 | 0 | 0 | 0 | 0 | 0 | 0 |
| CRK | 908 | 552 | 0 | 0 | 1046 | 0 | 0 | 1316 |
| CRKL | 398 | 2554 | 1350 | 3176 | 9186 | 1358 | 4400 | 2066 |
| CRLF3 | 0 | 354 | 0 | 0 | 0 | 0 | 0 | 0 |
| CRLS1 | 158 | 1088 | 0 | 1002 | 5374 | 96 | 452 | 1032 |
| CRNDE | 0 | 0 | 0 | 2784 | 746 | 1440 | 534 | 0 |
| CRNKL1 | 0 | 0 | 0 | 52 | 0 | 0 | 66 | 0 |
| CROCCP2 | 584 | 0 | 0 | 0 | 0 | 0 | 0 | 0 |
| CROT | 0 | 168 | 0 | 0 | 0 | 0 | 0 | 0 |
| CRTAP | 0 | 0 | 0 | 980 | 1310 | 0 | 1386 | 516 |
| CRTC1 | 1004 | 1492 | 4669 | 597 | 4716 | 578 | 5630 | 0 |
| CRTC2 | 0 | 0 | 0 | 696 | 0 | 0 | 0 | 0 |
| CRTC3 | 0 | 0 | 0 | 1321 | 646 | 0 | 0 | 0 |
| CRY1 | 0 | 132 | 0 | 0 | 1174 | 0 | 0 | 0 |
| CRY2 | 0 | 548 | 0 | 0 | 0 | 0 | 0 | 0 |
| CRYAB | 0 | 0 | 3352 | 0 | 0 | 0 | 0 | 0 |

| | | | | | | | | |
|---|---|---|---|---|---|---|---|---|
| CRYBB1 | 0 | 1920 | 11203 | 1469 | 3616 | 16570 | 4690 | 0 |
| CRYBB2P1 | 0 | 0 | 0 | 1132 | 599 | 0 | 0 | 0 |
| CRYBB3 | 652 | 0 | 0 | 0 | 0 | 0 | 0 | 0 |
| CRYBG2 | 0 | 0 | 0 | 814 | 0 | 0 | 0 | 0 |
| CRYL1 | 0 | 1833 | 5130 | 7774 | 2993 | 3820 | 3262 | 1536 |
| CRYM | 0 | 0 | 0 | 0 | 250 | 0 | 0 | 0 |
| CRYZ | 0 | 854 | 0 | 0 | 788 | 0 | 1960 | 0 |
| CRYZL1 | 0 | 0 | 0 | 0 | 3744 | 0 | 384 | 0 |
| CRYZL2P | 0 | 0 | 0 | 0 | 94 | 0 | 138 | 0 |
| CS | 0 | 276 | 0 | 920 | 298 | 0 | 0 | 0 |
| CSAD | 0 | 0 | 0 | 0 | 386 | 0 | 0 | 0 |
| CSDE1 | 178164 | 364110 | 344756 | 386298 | 990047 | 485847 | 1569807 | 208450 |
| CSE1L | 0 | 166 | 0 | 634 | 1132 | 2299 | 352 | 0 |
| CSF1 | 0 | 399 | 0 | 0 | 402 | 0 | 0 | 0 |
| CSF2 | 0 | 0 | 0 | 540 | 0 | 0 | 0 | 0 |
| CSF2RA | 0 | 0 | 0 | 0 | 0 | 1690 | 0 | 0 |
| CSF2RB | 0 | 112 | 0 | 0 | 0 | 0 | 246 | 0 |
| CSF3R | 0 | 0 | 0 | 0 | 30 | 0 | 0 | 0 |
| CSGALNACT2 | 0 | 0 | 904 | 0 | 0 | 0 | 0 | 0 |
| CSK | 402 | 0 | 0 | 330 | 494 | 0 | 0 | 0 |
| CSKMT | 122 | 170 | 40 | 396 | 436 | 0 | 0 | 246 |
| CSMD1 | 0 | 576 | 0 | 0 | 0 | 0 | 0 | 0 |
| CSMD3 | 0 | 190 | 0 | 0 | 0 | 0 | 0 | 0 |
| CSNK1A1 | 886 | 140 | 2409 | 576 | 0 | 0 | 2036 | 0 |
| CSNK1D | 0 | 0 | 0 | 1758 | 860 | 0 | 0 | 720 |
| CSNK1E | 1590 | 788 | 318 | 943 | 5338 | 0 | 962 | 0 |
| CSNK2A1 | 0 | 0 | 1052 | 962 | 339 | 0 | 0 | 1386 |
| CSNK2B | 1283 | 3820 | 1300 | 6035 | 8363 | 3726 | 5876 | 822 |
| CSRNP1 | 0 | 0 | 0 | 0 | 0 | 0 | 452 | 0 |
| CSRNP3 | 0 | 232 | 0 | 0 | 0 | 0 | 0 | 1004 |
| CSRP1 | 528 | 378 | 4136 | 3599 | 1394 | 0 | 1224 | 3058 |
| CSRP2 | 976 | 0 | 0 | 0 | 0 | 0 | 0 | 0 |
| CST1 | 0 | 0 | 0 | 736 | 0 | 1204 | 0 | 0 |
| CST3 | 10774 | 12820 | 46689 | 10179 | 25064 | 6573 | 5192 | 7501 |
| CST6 | 0 | 0 | 0 | 0 | 612 | 0 | 0 | 0 |
| CST7 | 0 | 418 | 0 | 0 | 0 | 0 | 0 | 0 |
| CSTA | 0 | 1434 | 0 | 0 | 0 | 0 | 402 | 0 |
| CSTB | 1366 | 1280 | 5737 | 5517 | 26808 | 7620 | 6291 | 23808 |
| CSTF1 | 0 | 332 | 1359 | 0 | 340 | 0 | 0 | 0 |
| CSTF2 | 662 | 470 | 0 | 0 | 0 | 584 | 0 | 0 |
| CSTF2T | 0 | 0 | 0 | 0 | 1164 | 0 | 260 | 0 |
| CSTF3 | 2302 | 10810 | 7090 | 27113 | 58434 | 14905 | 36125 | 12828 |
| CTB-49A3.2 | 0 | 226 | 0 | 0 | 0 | 0 | 0 | 0 |

| | | | | | | | | |
|---|---|---|---|---|---|---|---|---|
| CTBP1 | 380 | 436 | 843 | 820 | 642 | 542 | 154 | 0 |
| CTBP2 | 1050 | 1110 | 0 | 780 | 0 | 0 | 182 | 176 |
| CTBS | 0 | 422 | 526 | 0 | 0 | 2240 | 328 | 0 |
| CTC-338M12.4 | 0 | 0 | 0 | 0 | 0 | 0 | 1000 | 0 |
| CTC1 | 0 | 0 | 0 | 0 | 0 | 0 | 0 | 1164 |
| CTCF | 27292 | 33923 | 53552 | 48225 | 85563 | 38994 | 89176 | 17330 |
| CTDNEP1 | 204 | 0 | 0 | 787 | 644 | 1214 | 110 | 626 |
| CTDSP1 | 0 | 638 | 3154 | 0 | 1664 | 0 | 310 | 650 |
| CTDSP2 | 658 | 0 | 1656 | 1502 | 914 | 0 | 96 | 0 |
| CTDSPL | 8522 | 1240 | 9421 | 9843 | 13758 | 4734 | 11647 | 4174 |
| CTDSPL2 | 38 | 0 | 0 | 0 | 0 | 0 | 298 | 0 |
| CTHRC1 | 0 | 542 | 0 | 0 | 5090 | 2691 | 3102 | 1510 |
| CTIF | 0 | 0 | 0 | 0 | 276 | 0 | 0 | 0 |
| CTNNA1 | 976 | 2146 | 1944 | 2694 | 1662 | 4678 | 1782 | 0 |
| CTNNA2 | 530 | 0 | 308 | 910 | 1503 | 3780 | 1906 | 2032 |
| CTNNA3 | 0 | 0 | 0 | 0 | 1442 | 0 | 288 | 2558 |
| CTNNAL1 | 282 | 0 | 444 | 0 | 0 | 0 | 0 | 498 |
| CTNNB1 | 99526 | 128371 | 213956 | 365382 | 848716 | 429031 | 434072 | 143678 |
| CTNNBIP1 | 418 | 0 | 0 | 720 | 2820 | 0 | 948 | 0 |
| CTNNBL1 | 0 | 484 | 1884 | 172 | 0 | 0 | 2348 | 0 |
| CTNND1 | 15356 | 14109 | 39946 | 20461 | 47978 | 9062 | 25850 | 11974 |
| CTNND2 | 434 | 0 | 3568 | 0 | 0 | 772 | 0 | 0 |
| CTNS | 1092 | 370 | 0 | 0 | 0 | 0 | 504 | 398 |
| CTPS1 | 0 | 292 | 0 | 929 | 254 | 0 | 0 | 821 |
| CTPS2 | 70 | 0 | 0 | 0 | 828 | 0 | 142 | 0 |
| CTSA | 3528 | 1790 | 13654 | 7006 | 5199 | 5300 | 20927 | 2890 |
| CTSB | 27456 | 32833 | 70591 | 51521 | 108584 | 94589 | 150762 | 34777 |
| CTSC | 650 | 720 | 1812 | 1051 | 1814 | 1487 | 308 | 2936 |
| CTSD | 3094 | 3798 | 9444 | 2246 | 7324 | 814 | 2728 | 9359 |
| CTSE | 0 | 0 | 0 | 0 | 816 | 0 | 0 | 1700 |
| CTSH | 0 | 352 | 0 | 0 | 7218 | 3950 | 0 | 3591 |
| CTSK | 0 | 630 | 1686 | 915 | 1468 | 2832 | 284 | 0 |
| CTSL | 0 | 258 | 4034 | 0 | 2490 | 0 | 362 | 2910 |
| CTSO | 0 | 0 | 0 | 0 | 294 | 0 | 0 | 0 |
| CTSS | 4538 | 5251 | 10346 | 2156 | 5162 | 1639 | 1922 | 3046 |
| CTSV | 0 | 0 | 0 | 0 | 0 | 0 | 1120 | 942 |
| CTSZ | 0 | 0 | 592 | 912 | 1364 | 0 | 1556 | 1132 |
| CTTN | 25236 | 23032 | 187787 | 45243 | 133881 | 43260 | 65458 | 65921 |
| CTTNBP2 | 0 | 442 | 0 | 0 | 0 | 0 | 264 | 0 |
| CTTNBP2NL | 0 | 0 | 0 | 0 | 128 | 0 | 0 | 0 |
| CTU1 | 1320 | 238 | 0 | 0 | 0 | 0 | 0 | 0 |
| CTU2 | 0 | 0 | 0 | 0 | 972 | 0 | 0 | 0 |
| CTXN1 | 0 | 0 | 0 | 488 | 0 | 0 | 0 | 0 |

| | | | | | | | | |
|---|---|---|---|---|---|---|---|---|
| CUBN | 0 | 0 | 0 | 0 | 0 | 855 | 0 | 0 |
| CUEDC1 | 0 | 0 | 0 | 0 | 492 | 0 | 934 | 0 |
| CUEDC2 | 0 | 0 | 4744 | 952 | 386 | 0 | 1496 | 1270 |
| CUL1 | 9082 | 9858 | 6721 | 15162 | 25277 | 11782 | 22679 | 1638 |
| CUL2 | 0 | 0 | 1275 | 0 | 981 | 1508 | 0 | 0 |
| CUL3 | 23501 | 26672 | 39932 | 29972 | 82244 | 8149 | 73381 | 17909 |
| CUL4A | 0 | 0 | 0 | 1246 | 775 | 3708 | 144 | 1569 |
| CUL4B | 758 | 0 | 0 | 0 | 532 | 0 | 0 | 0 |
| CUL5 | 0 | 296 | 0 | 0 | 0 | 0 | 0 | 0 |
| CUL7 | 280 | 0 | 0 | 0 | 0 | 0 | 0 | 0 |
| CUL9 | 0 | 250 | 0 | 0 | 540 | 0 | 0 | 0 |
| CUTA | 1396 | 2650 | 6785 | 8552 | 9570 | 2951 | 6606 | 7230 |
| CUTALP | 0 | 186 | 0 | 0 | 0 | 0 | 152 | 0 |
| CUTC | 0 | 556 | 1496 | 1094 | 0 | 0 | 0 | 0 |
| CUX1 | 140442 | 107854 | 154469 | 146757 | 298886 | 307277 | 571336 | 68394 |
| CWC15 | 1384 | 1176 | 0 | 2490 | 8208 | 0 | 2514 | 3120 |
| CWC22 | 0 | 0 | 0 | 0 | 1078 | 0 | 0 | 0 |
| CWC25 | 352109 | 219525 | 894102 | 191669 | 508178 | 198988 | 511971 | 92935 |
| CWC27 | 0 | 0 | 0 | 0 | 270 | 596 | 1524 | 0 |
| CWF19L1 | 0 | 290 | 0 | 0 | 0 | 1688 | 1495 | 0 |
| CWH43 | 30813 | 29725 | 69369 | 2818 | 7888 | 0 | 1718 | 1032 |
| CXADR | 0 | 696 | 0 | 0 | 1931 | 0 | 0 | 1282 |
| CXCL1 | 0 | 0 | 0 | 608 | 2940 | 0 | 16 | 0 |
| CXCL10 | 0 | 0 | 1046 | 0 | 1120 | 0 | 866 | 0 |
| CXCL11 | 0 | 396 | 0 | 0 | 0 | 0 | 0 | 0 |
| CXCL12 | 0 | 818 | 0 | 0 | 0 | 0 | 0 | 0 |
| CXCL14 | 1637 | 1052 | 1970 | 1116 | 108 | 3613 | 0 | 0 |
| CXCL16 | 0 | 0 | 0 | 0 | 0 | 0 | 588 | 420 |
| CXCL17 | 0 | 0 | 0 | 0 | 2156 | 0 | 0 | 1130 |
| CXCL2 | 112 | 0 | 0 | 0 | 462 | 0 | 0 | 0 |
| CXCL3 | 712 | 254 | 0 | 0 | 942 | 0 | 0 | 1040 |
| CXCL8 | 0 | 0 | 0 | 368 | 1182 | 0 | 0 | 850 |
| CXCR3 | 0 | 0 | 0 | 0 | 0 | 0 | 294 | 0 |
| CXCR4 | 155604 | 274257 | 120003 | 41362 | 132179 | 389441 | 75559 | 295374 |
| CXCR5 | 2176 | 1724 | 0 | 0 | 0 | 0 | 0 | 3559 |
| CXorf38 | 0 | 0 | 0 | 0 | 0 | 0 | 0 | 902 |
| CXorf40A | 0 | 0 | 0 | 1360 | 0 | 0 | 0 | 0 |
| CXorf40B | 0 | 0 | 0 | 572 | 0 | 0 | 0 | 0 |
| CXorf56 | 0 | 582 | 0 | 328 | 618 | 0 | 0 | 0 |
| CXorf65 | 522 | 0 | 0 | 0 | 0 | 0 | 0 | 0 |
| CXXC1 | 666 | 304 | 0 | 0 | 644 | 0 | 0 | 0 |
| CXXC4 | 0 | 0 | 0 | 0 | 0 | 0 | 370 | 0 |
| CXXC5 | 0 | 0 | 0 | 0 | 698 | 1502 | 274 | 0 |

| | | | | | | | | |
|---|---|---|---|---|---|---|---|---|
| CYB561 | 0 | 350 | 0 | 0 | 1538 | 0 | 344 | 0 |
| CYB561A3 | 0 | 0 | 1624 | 0 | 614 | 0 | 712 | 0 |
| CYB561D2 | 622 | 0 | 2412 | 980 | 1526 | 0 | 0 | 0 |
| CYB5A | 0 | 1630 | 8594 | 9270 | 7785 | 5485 | 788 | 4186 |
| CYB5B | 0 | 826 | 3175 | 1566 | 2112 | 0 | 1384 | 0 |
| CYB5D2 | 0 | 0 | 0 | 0 | 1240 | 0 | 0 | 0 |
| CYB5R1 | 0 | 396 | 0 | 0 | 0 | 0 | 0 | 0 |
| CYB5R2 | 0 | 0 | 1910 | 0 | 310 | 0 | 0 | 0 |
| CYB5R3 | 0 | 342 | 0 | 0 | 204 | 0 | 1136 | 1172 |
| CYB5R4 | 0 | 0 | 88 | 806 | 488 | 1788 | 0 | 0 |
| CYB5RL | 0 | 0 | 0 | 0 | 966 | 0 | 0 | 0 |
| CYBA | 3326 | 4130 | 9347 | 7492 | 13846 | 4024 | 2822 | 2644 |
| CYBB | 0 | 0 | 0 | 0 | 264 | 0 | 0 | 260 |
| CYBC1 | 0 | 0 | 2230 | 0 | 1900 | 0 | 0 | 0 |
| CYBRD1 | 0 | 0 | 0 | 0 | 808 | 204 | 0 | 0 |
| CYC1 | 1382 | 3992 | 8432 | 8694 | 12026 | 11211 | 9541 | 9874 |
| CYCS | 3974 | 9356 | 6498 | 9262 | 11004 | 3786 | 3452 | 3229 |
| CYFIP1 | 840 | 44 | 1681 | 0 | 372 | 0 | 2372 | 0 |
| CYFIP2 | 0 | 0 | 0 | 0 | 0 | 0 | 330 | 0 |
| CYGB | 2146 | 1338 | 2362 | 912 | 1300 | 3030 | 384 | 0 |
| CYHR1 | 1550 | 954 | 827 | 1488 | 4156 | 1236 | 1890 | 0 |
| CYLD | 4228 | 5826 | 8961 | 3436 | 8107 | 8900 | 8605 | 2383 |
| CYP17A1 | 0 | 0 | 0 | 0 | 0 | 1300 | 0 | 0 |
| CYP21A1P | 192 | 0 | 0 | 0 | 0 | 0 | 0 | 0 |
| CYP24A1 | 0 | 0 | 0 | 0 | 0 | 0 | 806 | 0 |
| CYP26A1 | 0 | 0 | 0 | 0 | 0 | 20 | 232 | 0 |
| CYP27A1 | 768 | 798 | 0 | 0 | 0 | 0 | 408 | 0 |
| CYP2B6 | 186 | 0 | 0 | 0 | 678 | 0 | 276 | 0 |
| CYP2C18 | 31837 | 23155 | 160206 | 6342 | 39478 | 0 | 2976 | 4363 |
| CYP2C19 | 40 | 0 | 0 | 0 | 0 | 0 | 0 | 0 |
| CYP2C9 | 0 | 224 | 2954 | 0 | 62 | 0 | 0 | 0 |
| CYP2D6 | 602 | 0 | 0 | 0 | 0 | 0 | 0 | 0 |
| CYP2J2 | 0 | 0 | 1910 | 0 | 0 | 0 | 210 | 1094 |
| CYP2R1 | 0 | 0 | 0 | 0 | 866 | 0 | 0 | 0 |
| CYP2S1 | 0 | 0 | 0 | 796 | 2895 | 0 | 144 | 0 |
| CYP2T1P | 0 | 0 | 0 | 0 | 0 | 1308 | 0 | 0 |
| CYP2W1 | 0 | 0 | 0 | 0 | 0 | 0 | 102 | 0 |
| CYP39A1 | 0 | 0 | 0 | 1070 | 0 | 1782 | 0 | 0 |
| CYP3A5 | 148 | 760 | 8913 | 0 | 638 | 0 | 0 | 0 |
| CYP4F11 | 0 | 0 | 0 | 0 | 0 | 0 | 324 | 0 |
| CYP4F12 | 0 | 368 | 2893 | 0 | 794 | 0 | 162 | 0 |
| CYP4F2 | 486 | 0 | 472 | 0 | 0 | 2354 | 0 | 0 |
| CYP4V2 | 0 | 0 | 1128 | 0 | 0 | 0 | 0 | 0 |

| | | | | | | | | |
|---|---|---|---|---|---|---|---|---|
| CYP4X1 | 0 | 0 | 0 | 910 | 0 | 0 | 0 | 0 |
| CYP51A1 | 354 | 0 | 1998 | 0 | 424 | 0 | 0 | 0 |
| CYP7B1 | 768 | 296 | 0 | 0 | 0 | 0 | 0 | 0 |
| CYREN | 0 | 0 | 1020 | 298 | 0 | 0 | 0 | 0 |
| CYSLTR1 | 0 | 0 | 0 | 0 | 0 | 0 | 0 | 218 |
| CYSRT1 | 0 | 0 | 0 | 0 | 410 | 0 | 0 | 792 |
| CYSTM1 | 3388 | 6326 | 10544 | 2743 | 10059 | 4838 | 2078 | 4108 |
| CYTH2 | 0 | 0 | 3308 | 1044 | 282 | 0 | 652 | 0 |
| CYTH3 | 0 | 0 | 0 | 0 | 0 | 0 | 482 | 0 |
| CYTH4 | 0 | 0 | 0 | 0 | 38 | 0 | 0 | 0 |
| CYTIP | 0 | 0 | 0 | 0 | 0 | 0 | 0 | 886 |
| CYTL1 | 0 | 0 | 0 | 0 | 356 | 0 | 0 | 0 |
| CYTOR | 0 | 0 | 0 | 226 | 2766 | 0 | 0 | 498 |
| CZ1P-ASNS | 0 | 0 | 0 | 0 | 0 | 0 | 312 | 0 |
| CZIB | 0 | 466 | 3865 | 4340 | 894 | 0 | 0 | 0 |
| D2HGDH | 0 | 122 | 0 | 0 | 0 | 316 | 0 | 0 |
| DAB2 | 0 | 0 | 0 | 0 | 336 | 0 | 468 | 0 |
| DAB2IP | 0 | 0 | 900 | 0 | 0 | 614 | 0 | 0 |
| DACH1 | 0 | 0 | 0 | 856 | 0 | 450 | 314 | 0 |
| DAD1 | 2330 | 4646 | 7829 | 6091 | 16632 | 4109 | 2820 | 5786 |
| DAG1 | 616 | 580 | 4372 | 0 | 990 | 2308 | 2172 | 0 |
| DAGLA | 214 | 0 | 0 | 0 | 0 | 0 | 283 | 0 |
| DAGLB | 0 | 176 | 1260 | 0 | 0 | 0 | 0 | 0 |
| DALRD3 | 332 | 0 | 0 | 0 | 0 | 0 | 0 | 0 |
| DANCR | 226 | 1622 | 2634 | 3174 | 5917 | 1146 | 762 | 4433 |
| DAND5 | 0 | 0 | 2642 | 0 | 918 | 0 | 0 | 0 |
| DAP | 15276 | 31035 | 39074 | 75772 | 149452 | 56461 | 132912 | 49465 |
| DAP3 | 0 | 0 | 0 | 0 | 796 | 3640 | 894 | 818 |
| DAPK1 | 0 | 0 | 0 | 0 | 206 | 0 | 0 | 0 |
| DAPK2 | 418 | 126 | 0 | 0 | 0 | 0 | 178 | 0 |
| DAPK3 | 0 | 0 | 0 | 0 | 84 | 0 | 0 | 0 |
| DARS | 326 | 0 | 0 | 0 | 640 | 0 | 0 | 0 |
| DARS2 | 0 | 0 | 0 | 0 | 0 | 0 | 236 | 0 |
| DAXX | 836 | 282 | 0 | 410 | 204 | 0 | 1098 | 0 |
| DAZAP1 | 534 | 692 | 0 | 482 | 1446 | 0 | 492 | 736 |
| DAZAP2 | 1030 | 2911 | 8059 | 3299 | 4294 | 2205 | 4677 | 680 |
| DBF4 | 0 | 312 | 0 | 0 | 410 | 0 | 0 | 0 |
| DBI | 4343 | 3724 | 13728 | 14598 | 27982 | 1707 | 8171 | 8365 |
| DBNDD1 | 0 | 0 | 0 | 0 | 398 | 0 | 0 | 0 |
| DBNDD2 | 0 | 2724 | 2452 | 1208 | 3915 | 356 | 1274 | 0 |
| DBNL | 0 | 1010 | 244 | 403 | 903 | 2082 | 490 | 954 |
| DBP | 0 | 0 | 0 | 0 | 768 | 0 | 0 | 0 |
| DBR1 | 1620 | 3178 | 1979 | 2391 | 5786 | 0 | 4794 | 2412 |

| | | | | | | | | |
|---|---|---|---|---|---|---|---|---|
| DBT | 0 | 0 | 0 | 0 | 0 | 0 | 134 | 0 |
| DCAF1 | 0 | 0 | 0 | 0 | 0 | 1050 | 0 | 0 |
| DCAF10 | 1054 | 2659 | 11302 | 3577 | 3398 | 260 | 9876 | 2728 |
| DCAF11 | 678 | 42 | 1576 | 0 | 578 | 2009 | 538 | 0 |
| DCAF12 | 0 | 58 | 0 | 0 | 0 | 2522 | 0 | 0 |
| DCAF13 | 0 | 1076 | 1319 | 3075 | 3155 | 10131 | 2232 | 398 |
| DCAF15 | 884 | 0 | 0 | 0 | 0 | 0 | 0 | 0 |
| DCAF16 | 0 | 0 | 0 | 1028 | 0 | 0 | 1462 | 0 |
| DCAF4 | 654 | 0 | 0 | 0 | 604 | 0 | 0 | 0 |
| DCAF5 | 0 | 18 | 0 | 0 | 0 | 2306 | 0 | 0 |
| DCAF6 | 0 | 94 | 0 | 0 | 554 | 0 | 78 | 1088 |
| DCAF7 | 11065 | 15675 | 14659 | 28469 | 74588 | 46753 | 106105 | 24839 |
| DCAF8 | 258 | 0 | 526 | 592 | 42 | 0 | 1041 | 0 |
| DCAKD | 0 | 0 | 0 | 0 | 0 | 0 | 714 | 0 |
| DCBLD1 | 0 | 84 | 0 | 370 | 506 | 0 | 0 | 444 |
| DCK | 0 | 0 | 0 | 0 | 378 | 0 | 0 | 246 |
| DCLRE1B | 0 | 0 | 0 | 0 | 902 | 0 | 0 | 0 |
| DCN | 0 | 1576 | 5068 | 0 | 3025 | 11033 | 0 | 1208 |
| DCP1A | 0 | 0 | 0 | 0 | 120 | 0 | 574 | 0 |
| DCP1B | 0 | 338 | 0 | 0 | 1184 | 0 | 0 | 0 |
| DCP2 | 0 | 596 | 0 | 0 | 0 | 0 | 0 | 0 |
| DCPS | 0 | 0 | 0 | 0 | 2532 | 0 | 0 | 1210 |
| DCST1-AS1 | 0 | 136 | 0 | 0 | 0 | 0 | 0 | 0 |
| DCST2 | 0 | 0 | 0 | 0 | 0 | 0 | 148 | 0 |
| DCTD | 0 | 292 | 1206 | 3186 | 360 | 3487 | 590 | 0 |
| DCTN1 | 0 | 0 | 3108 | 0 | 954 | 0 | 928 | 1023 |
| DCTN2 | 0 | 356 | 8216 | 0 | 1066 | 2191 | 386 | 0 |
| DCTN3 | 0 | 1422 | 0 | 98 | 1806 | 0 | 0 | 2038 |
| DCTN4 | 0 | 0 | 0 | 0 | 0 | 0 | 480 | 0 |
| DCTN5 | 278 | 428 | 1430 | 0 | 1274 | 0 | 136 | 660 |
| DCTN6 | 0 | 746 | 26 | 0 | 2286 | 0 | 1799 | 0 |
| DCTPP1 | 1670 | 0 | 1022 | 7294 | 3125 | 0 | 366 | 0 |
| DCUN1D1 | 14732 | 15748 | 11486 | 23508 | 48243 | 12007 | 39852 | 39819 |
| DCUN1D2 | 0 | 0 | 0 | 0 | 468 | 0 | 120 | 0 |
| DCUN1D4 | 10608 | 10742 | 9127 | 15979 | 53926 | 20105 | 58568 | 18662 |
| DCUN1D5 | 0 | 402 | 0 | 1094 | 488 | 0 | 0 | 1298 |
| DCXR | 1030 | 890 | 4354 | 3685 | 2348 | 2222 | 3822 | 0 |
| DDA1 | 0 | 0 | 0 | 742 | 466 | 0 | 0 | 0 |
| DDAH1 | 0 | 0 | 872 | 0 | 1818 | 1318 | 0 | 0 |
| DDAH2 | 0 | 0 | 0 | 406 | 50 | 1122 | 1234 | 1011 |
| DDB1 | 0 | 0 | 2490 | 1542 | 456 | 1572 | 314 | 0 |
| DDB2 | 6592 | 19460 | 30123 | 19594 | 57678 | 16136 | 23762 | 20377 |
| DDC | 718 | 560 | 50 | 732 | 0 | 0 | 1572 | 0 |

| | | | | | | | | |
|---|---|---|---|---|---|---|---|---|
| DDHD2 | 6723 | 6131 | 8981 | 5135 | 29572 | 6980 | 6650 | 15484 |
| DDIT3 | 192952 | 175665 | 1087806 | 341499 | 900817 | 442098 | 1394648 | 760094 |
| DDIT4 | 1190 | 0 | 2767 | 2275 | 5677 | 7989 | 164 | 1934 |
| DDN-AS1 | 0 | 0 | 0 | 0 | 284 | 0 | 0 | 0 |
| DDOST | 50 | 354 | 8558 | 8095 | 10702 | 846 | 1536 | 4074 |
| DDR1 | 1718 | 1195 | 0 | 1856 | 748 | 998 | 0 | 0 |
| DDR2 | 2720 | 836 | 2964 | 0 | 5461 | 2334 | 538 | 1360 |
| DDRGK1 | 0 | 948 | 4350 | 1410 | 3204 | 0 | 806 | 0 |
| DDT | 1520 | 2352 | 4709 | 3255 | 950 | 2874 | 832 | 1620 |
| DDTL | 0 | 0 | 1376 | 0 | 0 | 0 | 0 | 0 |
| DDX1 | 894 | 322 | 2606 | 888 | 1158 | 748 | 0 | 0 |
| DDX10 | 23459 | 39336 | 18116 | 60756 | 140603 | 52673 | 109825 | 56062 |
| DDX11 | 0 | 0 | 0 | 804 | 788 | 0 | 0 | 0 |
| DDX17 | 214 | 182 | 2732 | 216 | 1357 | 0 | 1148 | 0 |
| DDX18 | 726 | 604 | 0 | 0 | 326 | 0 | 268 | 0 |
| DDX19A | 1020 | 1356 | 2786 | 1237 | 1784 | 4136 | 1354 | 0 |
| DDX19B | 16434 | 38691 | 51706 | 56181 | 79272 | 76488 | 90858 | 22811 |
| DDX20 | 0 | 0 | 0 | 0 | 0 | 0 | 316 | 0 |
| DDX21 | 0 | 0 | 0 | 0 | 0 | 0 | 0 | 552 |
| DDX23 | 15266 | 23742 | 77869 | 18436 | 42067 | 24224 | 66616 | 12874 |
| DDX24 | 0 | 0 | 0 | 384 | 1314 | 0 | 700 | 450 |
| DDX27 | 0 | 338 | 0 | 0 | 1645 | 0 | 1298 | 0 |
| DDX28 | 0 | 134 | 0 | 1161 | 996 | 0 | 0 | 0 |
| DDX31 | 1840 | 8056 | 9225 | 8633 | 31552 | 13133 | 28480 | 15333 |
| DDX39A | 0 | 732 | 1528 | 5333 | 890 | 2499 | 0 | 1172 |
| DDX39B | 0 | 963 | 0 | 0 | 3304 | 0 | 0 | 822 |
| DDX3X | 838 | 1744 | 0 | 0 | 1255 | 0 | 0 | 0 |
| DDX3Y | 0 | 180 | 0 | 530 | 1598 | 0 | 0 | 0 |
| DDX41 | 274 | 0 | 0 | 1329 | 346 | 0 | 0 | 1288 |
| DDX42 | 657 | 630 | 1566 | 2139 | 0 | 0 | 382 | 656 |
| DDX46 | 748 | 0 | 0 | 1246 | 0 | 0 | 0 | 0 |
| DDX47 | 0 | 382 | 1278 | 0 | 332 | 0 | 1311 | 0 |
| DDX49 | 0 | 0 | 0 | 814 | 384 | 0 | 0 | 0 |
| DDX5 | 289216 | 463857 | 507423 | 398226 | 1068515 | 917641 | 830323 | 369934 |
| DDX50 | 0 | 0 | 0 | 0 | 900 | 0 | 176 | 0 |
| DDX51 | 0 | 0 | 0 | 0 | 450 | 0 | 408 | 0 |
| DDX52 | 21017 | 45405 | 48855 | 34945 | 113186 | 47303 | 48895 | 25802 |
| DDX54 | 0 | 26 | 0 | 0 | 1030 | 0 | 0 | 0 |
| DDX55 | 0 | 0 | 0 | 0 | 906 | 0 | 0 | 0 |
| DDX56 | 0 | 244 | 1118 | 0 | 642 | 0 | 0 | 0 |
| DDX58 | 0 | 560 | 0 | 0 | 0 | 0 | 0 | 0 |
| DDX59 | 0 | 0 | 0 | 0 | 286 | 0 | 0 | 0 |
| DDX6 | 25487 | 15920 | 20409 | 23501 | 55384 | 11199 | 61985 | 22618 |

| | | | | | | | | |
|---|---|---|---|---|---|---|---|---|
| DDX60 | 390 | 0 | 4508 | 0 | 0 | 0 | 0 | 0 |
| DDX60L | 0 | 0 | 0 | 0 | 136 | 0 | 106 | 0 |
| DEAF1 | 440 | 0 | 2703 | 0 | 0 | 0 | 0 | 0 |
| DECR1 | 74 | 2615 | 10607 | 2823 | 6782 | 6181 | 10800 | 2052 |
| DECR2 | 0 | 580 | 0 | 0 | 1134 | 0 | 1364 | 0 |
| DEDD | 0 | 148 | 0 | 0 | 894 | 0 | 310 | 0 |
| DEDD2 | 0 | 370 | 0 | 0 | 1030 | 0 | 0 | 0 |
| DEF8 | 480 | 486 | 3687 | 0 | 646 | 0 | 0 | 0 |
| DEFA5 | 0 | 0 | 0 | 350 | 0 | 0 | 0 | 0 |
| DEFB1 | 0 | 480 | 1034 | 396 | 472 | 0 | 322 | 0 |
| DEGS1 | 0 | 314 | 0 | 0 | 2510 | 0 | 0 | 1008 |
| DEGS2 | 0 | 346 | 0 | 0 | 1030 | 0 | 0 | 0 |
| DEK | 97674 | 126106 | 116655 | 277241 | 721956 | 187421 | 423395 | 266811 |
| DELE1 | 0 | 0 | 1404 | 0 | 0 | 1656 | 912 | 874 |
| DENND1A | 4632 | 2061 | 8241 | 7008 | 8385 | 5842 | 31579 | 10635 |
| DENND2A | 416 | 900 | 2277 | 0 | 0 | 0 | 362 | 0 |
| DENND2D | 0 | 0 | 772 | 0 | 0 | 0 | 0 | 0 |
| DENND3 | 0 | 542 | 0 | 0 | 362 | 3637 | 412 | 932 |
| DENND4A | 0 | 0 | 0 | 0 | 256 | 0 | 0 | 500 |
| DENND4B | 420 | 0 | 92 | 0 | 0 | 0 | 0 | 0 |
| DENND4C | 582 | 0 | 0 | 1204 | 584 | 0 | 200 | 0 |
| DENND5A | 0 | 0 | 0 | 0 | 0 | 639 | 0 | 0 |
| DENND5B | 0 | 0 | 0 | 0 | 0 | 0 | 204 | 0 |
| DENND6A | 0 | 0 | 0 | 966 | 0 | 0 | 0 | 0 |
| DENND6B | 0 | 0 | 0 | 0 | 988 | 0 | 0 | 717 |
| DENR | 0 | 632 | 0 | 990 | 0 | 0 | 660 | 0 |
| DEPDC1 | 0 | 0 | 0 | 0 | 936 | 0 | 0 | 0 |
| DEPDC1B | 2828 | 17142 | 24904 | 64280 | 187975 | 53587 | 36493 | 64996 |
| DEPDC7 | 44 | 0 | 0 | 0 | 470 | 0 | 0 | 0 |
| DEPP1 | 0 | 0 | 2334 | 0 | 1306 | 1954 | 1065 | 0 |
| DEPTOR | 0 | 2508 | 5090 | 3953 | 3600 | 0 | 2900 | 4648 |
| DERA | 0 | 1260 | 24 | 3282 | 1272 | 0 | 1958 | 0 |
| DERL1 | 791 | 0 | 0 | 1048 | 1738 | 2279 | 1369 | 702 |
| DERL2 | 0 | 422 | 3102 | 0 | 0 | 6947 | 854 | 606 |
| DERL3 | 0 | 0 | 0 | 0 | 390 | 0 | 0 | 0 |
| DES | 0 | 0 | 3667 | 0 | 0 | 0 | 0 | 0 |
| DESI1 | 0 | 633 | 0 | 2338 | 762 | 0 | 933 | 1690 |
| DESI2 | 0 | 214 | 0 | 0 | 0 | 0 | 0 | 0 |
| DEUP1 | 0 | 0 | 0 | 1870 | 284 | 0 | 0 | 0 |
| DEXI | 8925 | 18184 | 24589 | 20358 | 47003 | 27820 | 54458 | 11942 |
| DFFA | 0 | 0 | 0 | 1472 | 1520 | 0 | 548 | 1340 |
| DFFB | 0 | 0 | 0 | 1170 | 0 | 0 | 732 | 0 |
| DGAT1 | 0 | 1006 | 3959 | 2228 | 502 | 0 | 1644 | 830 |

| | | | | | | | | |
|---|---|---|---|---|---|---|---|---|
| DGAT2 | 0 | 0 | 0 | 788 | 0 | 0 | 630 | 0 |
| DGCR11 | 0 | 0 | 0 | 0 | 216 | 0 | 0 | 0 |
| DGCR2 | 2704 | 2068 | 1484 | 0 | 3364 | 4998 | 3898 | 776 |
| DGCR5 | 0 | 0 | 0 | 0 | 268 | 0 | 0 | 0 |
| DGCR6L | 1400 | 0 | 0 | 0 | 1289 | 0 | 0 | 1108 |
| DGCR8 | 968 | 2304 | 4854 | 3720 | 4212 | 2396 | 3886 | 768 |
| DGKA | 0 | 0 | 1375 | 408 | 0 | 0 | 0 | 0 |
| DGKD | 0 | 0 | 0 | 0 | 1513 | 0 | 0 | 0 |
| DGKE | 656 | 0 | 0 | 0 | 0 | 0 | 0 | 0 |
| DGKG | 0 | 810 | 2360 | 0 | 34 | 5509 | 1762 | 754 |
| DGKQ | 134 | 0 | 0 | 0 | 0 | 0 | 1786 | 0 |
| DGKZ | 0 | 538 | 0 | 910 | 870 | 0 | 1092 | 1112 |
| DGLUCY | 0 | 0 | 0 | 0 | 1746 | 0 | 0 | 0 |
| DGUOK | 1860 | 3436 | 2068 | 3697 | 9285 | 1591 | 2756 | 4202 |
| DGUOK-AS1 | 0 | 0 | 0 | 732 | 0 | 0 | 0 | 0 |
| DHCR24 | 117346 | 26747 | 161303 | 71312 | 192467 | 52060 | 92365 | 52079 |
| DHCR7 | 0 | 0 | 0 | 0 | 770 | 0 | 0 | 0 |
| DHDDS | 26298 | 29024 | 91593 | 28506 | 49394 | 37844 | 33761 | 46248 |
| DHFR | 290 | 0 | 0 | 0 | 0 | 2016 | 62 | 0 |
| DHODH | 0 | 0 | 672 | 0 | 0 | 0 | 366 | 318 |
| DHPS | 566 | 1910 | 2156 | 0 | 1744 | 1438 | 0 | 1048 |
| DHRS1 | 768 | 364 | 0 | 0 | 1506 | 0 | 0 | 0 |
| DHRS11 | 1366 | 760 | 2538 | 0 | 0 | 0 | 0 | 0 |
| DHRS12 | 0 | 0 | 2162 | 0 | 0 | 0 | 1428 | 0 |
| DHRS13 | 0 | 0 | 0 | 0 | 370 | 0 | 0 | 890 |
| DHRS3 | 21616 | 25284 | 101111 | 35237 | 32189 | 73617 | 52988 | 23152 |
| DHRS4 | 0 | 1290 | 7121 | 0 | 1894 | 2836 | 1340 | 0 |
| DHRS4-AS1 | 0 | 832 | 0 | 0 | 1513 | 0 | 577 | 0 |
| DHRS4L2 | 0 | 1288 | 2078 | 0 | 0 | 0 | 0 | 0 |
| DHRS7 | 1042 | 0 | 5218 | 0 | 6859 | 0 | 482 | 134 |
| DHRS7B | 468 | 796 | 1082 | 0 | 237 | 0 | 0 | 1047 |
| DHRS9 | 0 | 0 | 6269 | 0 | 0 | 0 | 0 | 1256 |
| DHRSX | 742 | 304 | 0 | 2348 | 430 | 4956 | 210 | 0 |
| DHTKD1 | 0 | 0 | 0 | 0 | 0 | 0 | 190 | 0 |
| DHX15 | 55887 | 104964 | 156648 | 116585 | 267439 | 130784 | 250304 | 86966 |
| DHX16 | 0 | 220 | 0 | 1591 | 1120 | 0 | 0 | 0 |
| DHX29 | 0 | 2472 | 0 | 0 | 4090 | 0 | 614 | 0 |
| DHX30 | 0 | 234 | 0 | 452 | 1444 | 0 | 1678 | 0 |
| DHX32 | 1248 | 0 | 1674 | 0 | 644 | 0 | 0 | 0 |
| DHX33 | 0 | 0 | 0 | 0 | 0 | 0 | 0 | 762 |
| DHX35 | 0 | 0 | 0 | 0 | 808 | 0 | 1228 | 0 |
| DHX36 | 0 | 0 | 0 | 0 | 0 | 0 | 802 | 0 |
| DHX37 | 1549 | 2334 | 2074 | 470 | 2934 | 2056 | 1038 | 0 |

| | | | | | | | | |
|---|---|---|---|---|---|---|---|---|
| DHX38 | 0 | 0 | 0 | 0 | 1274 | 0 | 0 | 0 |
| DHX40 | 5884 | 19528 | 10978 | 5796 | 60928 | 15764 | 38563 | 9972 |
| DHX58 | 0 | 0 | 0 | 0 | 0 | 0 | 658 | 0 |
| DHX8 | 18141 | 18843 | 26603 | 11046 | 46231 | 11285 | 41749 | 15503 |
| DHX9 | 0 | 182 | 0 | 777 | 2240 | 0 | 1256 | 0 |
| DIABLO | 0 | 980 | 3647 | 1462 | 1612 | 0 | 1426 | 1060 |
| DIAPH1 | 252 | 206 | 1120 | 0 | 0 | 0 | 0 | 812 |
| DICER1 | 17819 | 18690 | 7197 | 7945 | 30449 | 7908 | 9322 | 4153 |
| DICER1-AS1 | 2762 | 0 | 0 | 0 | 0 | 0 | 0 | 0 |
| DIDO1 | 846 | 106 | 270 | 1970 | 1312 | 1253 | 766 | 0 |
| DIEXF | 0 | 0 | 0 | 0 | 264 | 0 | 0 | 0 |
| DIMT1 | 410 | 1144 | 0 | 0 | 1016 | 0 | 240 | 722 |
| DIO3OS | 0 | 140 | 0 | 0 | 814 | 2658 | 0 | 0 |
| DIP2A | 106 | 0 | 0 | 0 | 1710 | 0 | 532 | 0 |
| DIP2B | 3334 | 0 | 0 | 710 | 2681 | 0 | 1218 | 0 |
| DIPK1A | 0 | 0 | 2352 | 0 | 270 | 0 | 602 | 0 |
| DIRAS3 | 1042 | 6274 | 5182 | 1158 | 4006 | 1862 | 6848 | 0 |
| DIRC2 | 0 | 0 | 1235 | 0 | 0 | 0 | 0 | 0 |
| DIS3 | 0 | 0 | 0 | 720 | 1070 | 1096 | 0 | 0 |
| DISP1 | 0 | 0 | 0 | 0 | 1044 | 0 | 0 | 0 |
| DKC1 | 0 | 190 | 288 | 894 | 2612 | 1779 | 0 | 0 |
| DKK3 | 0 | 0 | 1953 | 1000 | 976 | 0 | 0 | 0 |
| DKK4 | 0 | 0 | 0 | 0 | 0 | 2008 | 510 | 1004 |
| DLAT | 0 | 0 | 0 | 0 | 220 | 0 | 0 | 0 |
| DLD | 390 | 432 | 959 | 0 | 1074 | 1436 | 1084 | 774 |
| DLEU1 | 20912 | 55612 | 36579 | 110730 | 171154 | 359516 | 259610 | 43266 |
| DLEU2 | 0 | 0 | 0 | 0 | 778 | 0 | 0 | 0 |
| DLG1 | 0 | 62 | 0 | 391 | 218 | 0 | 0 | 0 |
| DLG3 | 17733 | 24752 | 15957 | 32235 | 159953 | 18424 | 42059 | 46212 |
| DLG5 | 102 | 0 | 0 | 0 | 1528 | 0 | 90 | 376 |
| DLGAP1-AS1 | 0 | 46 | 0 | 788 | 0 | 0 | 718 | 0 |
| DLGAP1-AS2 | 0 | 354 | 0 | 842 | 538 | 0 | 570 | 1313 |
| DLGAP4 | 860 | 9047 | 8556 | 32451 | 31383 | 22017 | 60783 | 18972 |
| DLGAP5 | 0 | 0 | 0 | 0 | 114 | 0 | 0 | 0 |
| DLL1 | 440 | 1638 | 2666 | 0 | 2198 | 0 | 0 | 936 |
| DLL4 | 0 | 0 | 0 | 0 | 0 | 0 | 84 | 0 |
| DLST | 0 | 0 | 0 | 0 | 498 | 0 | 696 | 0 |
| DMAC1 | 494 | 1942 | 1407 | 2370 | 2958 | 706 | 2172 | 342 |
| DMAC2 | 0 | 0 | 0 | 3334 | 328 | 0 | 0 | 854 |
| DMAP1 | 438 | 308 | 2076 | 1038 | 268 | 0 | 0 | 0 |
| DMBT1 | 0 | 0 | 934 | 3702 | 624 | 0 | 0 | 0 |
| DMD | 0 | 0 | 0 | 0 | 0 | 0 | 2530 | 0 |
| DMKN | 0 | 0 | 1254 | 0 | 7222 | 0 | 0 | 0 |

| | | | | | | | | |
|---|---|---|---|---|---|---|---|---|
| DMPK | 0 | 0 | 0 | 900 | 0 | 0 | 0 | 0 |
| DMRT2 | 0 | 0 | 0 | 0 | 1578 | 0 | 0 | 0 |
| DMTF1 | 24519 | 20282 | 20129 | 14430 | 53091 | 18694 | 47539 | 16801 |
| DMTN | 0 | 0 | 582 | 0 | 324 | 0 | 536 | 0 |
| DMWD | 0 | 0 | 0 | 772 | 0 | 0 | 0 | 0 |
| DMXL2 | 0 | 0 | 0 | 0 | 708 | 0 | 0 | 0 |
| DNAAF2 | 0 | 94 | 0 | 0 | 674 | 0 | 0 | 682 |
| DNAAF4-CCPG1 | 8375 | 27220 | 4733 | 4295 | 25468 | 3848 | 10802 | 1971 |
| DNAAF5 | 0 | 0 | 0 | 0 | 289 | 0 | 0 | 408 |
| DNAH1 | 0 | 0 | 0 | 0 | 18 | 0 | 184 | 0 |
| DNAH10 | 0 | 0 | 0 | 1030 | 0 | 0 | 0 | 0 |
| DNAH14 | 0 | 0 | 1235 | 0 | 0 | 0 | 0 | 0 |
| DNAH6 | 0 | 0 | 0 | 0 | 0 | 0 | 0 | 918 |
| DNAH9 | 1224 | 0 | 0 | 0 | 0 | 0 | 0 | 0 |
| DNAJA1 | 1440 | 0 | 3635 | 1148 | 260 | 6477 | 0 | 3258 |
| DNAJA2 | 0 | 322 | 2595 | 0 | 6524 | 1572 | 312 | 0 |
| DNAJA3 | 0 | 0 | 1724 | 964 | 0 | 0 | 2962 | 0 |
| DNAJA4 | 0 | 0 | 1110 | 0 | 0 | 0 | 0 | 0 |
| DNAJB1 | 649 | 0 | 2972 | 0 | 2178 | 6381 | 552 | 2726 |
| DNAJB11 | 575 | 1066 | 0 | 1247 | 2444 | 0 | 0 | 4456 |
| DNAJB12 | 0 | 902 | 2094 | 0 | 38 | 0 | 2188 | 0 |
| DNAJB14 | 18710 | 16412 | 25710 | 19828 | 26662 | 20016 | 46999 | 15843 |
| DNAJB2 | 60 | 322 | 0 | 0 | 1918 | 0 | 0 | 1094 |
| DNAJB4 | 23223 | 17875 | 52496 | 10401 | 26044 | 40163 | 61384 | 35408 |
| DNAJB6 | 575 | 799 | 1358 | 1164 | 1452 | 2002 | 244 | 648 |
| DNAJB9 | 0 | 0 | 1322 | 508 | 0 | 0 | 1462 | 1508 |
| DNAJC1 | 0 | 228 | 0 | 0 | 48 | 0 | 0 | 1196 |
| DNAJC10 | 0 | 340 | 0 | 0 | 500 | 0 | 0 | 1019 |
| DNAJC11 | 19276 | 34088 | 79688 | 47399 | 79033 | 30580 | 40832 | 17589 |
| DNAJC12 | 0 | 990 | 0 | 0 | 1274 | 0 | 0 | 0 |
| DNAJC13 | 0 | 166 | 0 | 0 | 0 | 0 | 0 | 0 |
| DNAJC15 | 358 | 832 | 2537 | 9538 | 7525 | 5450 | 3196 | 2892 |
| DNAJC17 | 778 | 150 | 783 | 0 | 0 | 0 | 378 | 998 |
| DNAJC19 | 612 | 4482 | 514 | 4460 | 2810 | 0 | 3058 | 5943 |
| DNAJC2 | 0 | 686 | 0 | 0 | 310 | 0 | 0 | 0 |
| DNAJC21 | 0 | 226 | 2398 | 357 | 1692 | 0 | 0 | 0 |
| DNAJC22 | 160 | 0 | 0 | 0 | 1038 | 0 | 0 | 0 |
| DNAJC24 | 0 | 0 | 0 | 809 | 0 | 0 | 0 | 0 |
| DNAJC25 | 0 | 450 | 0 | 0 | 0 | 0 | 0 | 714 |
| DNAJC25-GNG10 | 298 | 0 | 0 | 0 | 0 | 0 | 96 | 0 |
| DNAJC28 | 0 | 526 | 0 | 0 | 0 | 0 | 0 | 0 |
| DNAJC3 | 52564 | 46910 | 75482 | 59905 | 68135 | 98049 | 153438 | 26730 |
| DNAJC4 | 372 | 836 | 0 | 0 | 962 | 0 | 1550 | 0 |

| | | | | | | | | |
|---|---|---|---|---|---|---|---|---|
| DNAJC5 | 0 | 0 | 854 | 0 | 62 | 0 | 842 | 320 |
| DNAJC7 | 134245 | 203378 | 256842 | 223403 | 436637 | 285043 | 394922 | 166014 |
| DNAJC8 | 1678 | 1718 | 3685 | 1266 | 6623 | 3228 | 1168 | 6466 |
| DNAJC9 | 0 | 366 | 3891 | 3069 | 2117 | 0 | 0 | 0 |
| DNAJC9-AS1 | 0 | 0 | 0 | 0 | 554 | 76 | 0 | 0 |
| DNAL1 | 0 | 0 | 0 | 0 | 1428 | 0 | 0 | 938 |
| DNAL4 | 0 | 0 | 0 | 0 | 0 | 0 | 378 | 12 |
| DNALI1 | 0 | 426 | 0 | 0 | 0 | 0 | 0 | 0 |
| DNASE1 | 2470 | 3540 | 2758 | 2365 | 7539 | 14410 | 18616 | 2320 |
| DNASE1L1 | 508 | 0 | 0 | 0 | 502 | 0 | 0 | 972 |
| DNASE1L3 | 0 | 0 | 1910 | 0 | 0 | 0 | 0 | 0 |
| DNASE2 | 0 | 0 | 0 | 1118 | 900 | 3037 | 0 | 1408 |
| DNHD1 | 206 | 0 | 0 | 0 | 0 | 0 | 0 | 0 |
| DNLZ | 0 | 0 | 0 | 0 | 0 | 0 | 348 | 0 |
| DNM1 | 0 | 0 | 0 | 0 | 24 | 0 | 0 | 0 |
| DNM1P41 | 0 | 0 | 0 | 0 | 448 | 0 | 0 | 0 |
| DNM2 | 243669 | 175504 | 791104 | 136905 | 854254 | 128539 | 200599 | 187396 |
| DNM3 | 0 | 0 | 0 | 0 | 428 | 0 | 1696 | 0 |
| DNM3OS | 0 | 0 | 0 | 0 | 0 | 0 | 872 | 0 |
| DNMBP | 0 | 0 | 0 | 1332 | 296 | 0 | 0 | 1086 |
| DNMT1 | 0 | 0 | 0 | 0 | 1016 | 0 | 276 | 802 |
| DNMT3A | 1920 | 3119 | 2020 | 5204 | 10069 | 1387 | 9775 | 214 |
| DNPEP | 358 | 478 | 3650 | 0 | 1994 | 0 | 970 | 0 |
| DNPH1 | 2252 | 1916 | 2314 | 0 | 8856 | 0 | 2889 | 1675 |
| DNTTIP1 | 0 | 668 | 0 | 2706 | 3228 | 2596 | 6448 | 6140 |
| DNTTIP2 | 1132 | 0 | 0 | 0 | 0 | 2378 | 348 | 0 |
| DOCK1 | 11456 | 3956 | 5813 | 1663 | 5822 | 4226 | 12914 | 0 |
| DOCK5 | 0 | 246 | 0 | 0 | 406 | 0 | 0 | 562 |
| DOCK6 | 2964 | 4846 | 6494 | 3822 | 26205 | 8473 | 6411 | 5002 |
| DOCK7 | 0 | 510 | 224 | 1684 | 2856 | 0 | 868 | 0 |
| DOCK8 | 10605 | 11117 | 3286 | 1536 | 2762 | 6212 | 6932 | 1481 |
| DOCK9 | 272 | 0 | 0 | 0 | 0 | 0 | 0 | 0 |
| DOHH | 0 | 0 | 0 | 0 | 1484 | 0 | 0 | 0 |
| DOK1 | 0 | 408 | 0 | 0 | 0 | 0 | 0 | 0 |
| DOK2 | 828 | 0 | 0 | 0 | 0 | 0 | 0 | 0 |
| DOK3 | 110 | 0 | 0 | 0 | 0 | 0 | 0 | 0 |
| DOK4 | 489 | 60 | 0 | 0 | 424 | 0 | 916 | 254 |
| DOK5 | 0 | 376 | 910 | 0 | 3372 | 1388 | 0 | 5031 |
| DOK6 | 0 | 0 | 3132 | 0 | 0 | 0 | 0 | 0 |
| DOK7 | 0 | 0 | 0 | 0 | 0 | 0 | 324 | 0 |
| DOLK | 0 | 0 | 0 | 0 | 0 | 0 | 390 | 0 |
| DOLPP1 | 15130 | 10031 | 51807 | 11886 | 15590 | 5313 | 19541 | 4938 |
| DONSON | 0 | 356 | 0 | 0 | 582 | 0 | 0 | 0 |

| | | | | | | | | |
|---|---|---|---|---|---|---|---|---|
| DOPEY2 | 0 | 0 | 298 | 0 | 448 | 0 | 0 | 0 |
| DPAGT1 | 560 | 0 | 922 | 974 | 790 | 0 | 0 | 0 |
| DPCD | 0 | 72 | 0 | 1360 | 2040 | 0 | 1974 | 0 |
| DPEP1 | 0 | 0 | 0 | 4375 | 0 | 1142 | 10639 | 0 |
| DPF2 | 0 | 0 | 0 | 176 | 1444 | 0 | 738 | 558 |
| DPH1 | 0 | 0 | 0 | 1861 | 0 | 0 | 894 | 0 |
| DPH2 | 0 | 0 | 1698 | 804 | 0 | 1594 | 0 | 0 |
| DPH3 | 0 | 200 | 784 | 2405 | 3690 | 0 | 592 | 0 |
| DPH5 | 722 | 0 | 0 | 0 | 0 | 0 | 596 | 951 |
| DPH6 | 0 | 0 | 1710 | 0 | 0 | 0 | 398 | 0 |
| DPM1 | 50931 | 74955 | 103634 | 213884 | 446811 | 134266 | 650744 | 154724 |
| DPM2 | 0 | 1160 | 0 | 2516 | 1350 | 18 | 0 | 646 |
| DPM3 | 580 | 1086 | 0 | 862 | 3748 | 0 | 280 | 0 |
| DPP10-AS1 | 0 | 0 | 0 | 1147 | 0 | 0 | 0 | 0 |
| DPP3 | 0 | 364 | 0 | 912 | 1180 | 0 | 922 | 1265 |
| DPP4 | 0 | 0 | 962 | 0 | 2044 | 0 | 0 | 0 |
| DPP7 | 686 | 526 | 0 | 198 | 3860 | 1442 | 816 | 562 |
| DPP8 | 0 | 0 | 0 | 0 | 220 | 82 | 0 | 1136 |
| DPP9 | 2437 | 2264 | 6417 | 270 | 7160 | 2110 | 4046 | 1778 |
| DPT | 0 | 510 | 0 | 0 | 0 | 0 | 0 | 0 |
| DPY19L1 | 0 | 0 | 548 | 0 | 42 | 0 | 0 | 0 |
| DPY19L2P3 | 0 | 0 | 0 | 571 | 0 | 0 | 157 | 0 |
| DPY19L3 | 0 | 0 | 0 | 0 | 0 | 0 | 814 | 0 |
| DPY19L4 | 0 | 0 | 0 | 0 | 916 | 302 | 0 | 0 |
| DPY30 | 0 | 1532 | 4572 | 2555 | 7693 | 0 | 5129 | 2145 |
| DPYD | 2714 | 1648 | 4838 | 318 | 2002 | 5034 | 392 | 1094 |
| DPYSL2 | 0 | 0 | 1149 | 0 | 0 | 0 | 396 | 0 |
| DPYSL3 | 0 | 516 | 0 | 0 | 0 | 0 | 0 | 0 |
| DQX1 | 638 | 0 | 2184 | 0 | 0 | 0 | 428 | 0 |
| DR1 | 768 | 750 | 0 | 2702 | 2434 | 1849 | 492 | 0 |
| DRAM1 | 0 | 0 | 0 | 0 | 0 | 0 | 0 | 880 |
| DRAM2 | 0 | 1378 | 0 | 2832 | 568 | 0 | 0 | 0 |
| DRAP1 | 0 | 1452 | 1108 | 692 | 3654 | 4364 | 2254 | 3956 |
| DRG1 | 0 | 666 | 0 | 2664 | 4172 | 0 | 1028 | 1270 |
| DRG2 | 0 | 198 | 0 | 0 | 0 | 0 | 300 | 2182 |
| DROSHA | 198 | 0 | 0 | 0 | 0 | 0 | 0 | 1331 |
| DRP2 | 0 | 0 | 0 | 0 | 0 | 0 | 0 | 834 |
| DSC2 | 200 | 584 | 902 | 0 | 1258 | 0 | 0 | 0 |
| DSCC1 | 0 | 0 | 0 | 1240 | 1488 | 0 | 0 | 0 |
| DSG2 | 0 | 712 | 2099 | 1211 | 3438 | 1568 | 1324 | 2780 |
| DSN1 | 0 | 0 | 0 | 0 | 1092 | 0 | 304 | 838 |
| DSP | 27176 | 17321 | 24762 | 11894 | 58740 | 9713 | 76205 | 11817 |
| DST | 0 | 204 | 1342 | 0 | 498 | 0 | 0 | 0 |

| | | | | | | | | |
|---|---|---|---|---|---|---|---|---|
| DSTN | 2950 | 5146 | 12333 | 7369 | 8102 | 3129 | 1522 | 3230 |
| DSTNP2 | 0 | 416 | 0 | 2156 | 0 | 1854 | 3148 | 0 |
| DSTYK | 0 | 0 | 2728 | 0 | 922 | 0 | 514 | 0 |
| DTD1 | 24153 | 50161 | 82888 | 109034 | 506319 | 35423 | 71303 | 80165 |
| DTD2 | 0 | 0 | 0 | 0 | 0 | 0 | 0 | 462 |
| DTL | 776 | 1666 | 1888 | 15230 | 25051 | 7559 | 9154 | 4086 |
| DTNA | 1846 | 482 | 0 | 6652 | 1390 | 1440 | 3889 | 918 |
| DTNB | 1960 | 6316 | 13029 | 9237 | 58013 | 30061 | 60914 | 12624 |
| DTNBP1 | 774 | 992 | 1870 | 0 | 388 | 0 | 1184 | 0 |
| DTWD1 | 0 | 0 | 0 | 982 | 0 | 0 | 0 | 1102 |
| DTX2 | 0 | 838 | 660 | 0 | 998 | 0 | 0 | 1071 |
| DTX3 | 144 | 0 | 0 | 0 | 0 | 0 | 0 | 0 |
| DTX3L | 616 | 0 | 1242 | 0 | 602 | 0 | 0 | 0 |
| DTX4 | 1779 | 2230 | 1506 | 0 | 4614 | 994 | 3544 | 2236 |
| DTYMK | 0 | 0 | 3765 | 0 | 3442 | 2306 | 2294 | 388 |
| DUOXA2 | 0 | 0 | 0 | 494 | 1560 | 0 | 0 | 0 |
| DUS1L | 0 | 554 | 0 | 3972 | 2164 | 1803 | 0 | 0 |
| DUS3L | 0 | 0 | 0 | 0 | 810 | 0 | 0 | 0 |
| DUS4L | 0 | 0 | 0 | 729 | 0 | 0 | 0 | 0 |
| DUSP1 | 5684 | 1732 | 1634 | 860 | 3110 | 1674 | 0 | 2554 |
| DUSP11 | 0 | 60 | 674 | 1300 | 0 | 0 | 0 | 0 |
| DUSP12 | 0 | 266 | 0 | 0 | 974 | 0 | 328 | 1010 |
| DUSP14 | 0 | 0 | 0 | 0 | 1606 | 0 | 0 | 0 |
| DUSP16 | 2158 | 822 | 8606 | 2108 | 3878 | 3189 | 8929 | 2020 |
| DUSP22 | 520 | 0 | 0 | 0 | 1628 | 0 | 1062 | 0 |
| DUSP23 | 0 | 1292 | 2910 | 2233 | 2832 | 4467 | 2730 | 2020 |
| DUSP26 | 5160 | 0 | 4998 | 0 | 576 | 0 | 0 | 0 |
| DUSP27 | 0 | 0 | 0 | 0 | 348 | 0 | 0 | 0 |
| DUSP28 | 0 | 0 | 0 | 0 | 510 | 0 | 0 | 0 |
| DUSP3 | 0 | 0 | 116 | 490 | 0 | 0 | 1022 | 0 |
| DUSP4 | 0 | 0 | 0 | 0 | 136 | 0 | 0 | 0 |
| DUSP5 | 0 | 0 | 2899 | 0 | 0 | 0 | 0 | 0 |
| DUSP6 | 150 | 210 | 0 | 0 | 0 | 1010 | 406 | 0 |
| DUSP8 | 0 | 380 | 0 | 0 | 128 | 0 | 348 | 0 |
| DUT | 640 | 1664 | 2086 | 1095 | 2992 | 1914 | 2690 | 3260 |
| DVL1 | 0 | 0 | 0 | 692 | 472 | 0 | 0 | 0 |
| DVL2 | 0 | 104 | 0 | 0 | 2048 | 2606 | 362 | 0 |
| DVL3 | 0 | 0 | 0 | 740 | 406 | 0 | 0 | 0 |
| DWORF | 0 | 0 | 0 | 0 | 0 | 0 | 2354 | 0 |
| DXO | 0 | 0 | 0 | 1277 | 560 | 0 | 92 | 0 |
| DYM | 7613 | 11467 | 7435 | 7459 | 19924 | 16407 | 20338 | 6843 |
| DYNC1H1 | 386 | 722 | 1323 | 1208 | 340 | 0 | 546 | 0 |
| DYNC1I1 | 0 | 0 | 0 | 0 | 0 | 0 | 993 | 0 |

| | | | | | | | | |
|---|---|---|---|---|---|---|---|---|
| DYNC1I2 | 0 | 0 | 0 | 0 | 1408 | 0 | 1626 | 0 |
| DYNC1LI1 | 0 | 386 | 3516 | 0 | 176 | 0 | 990 | 0 |
| DYNC1LI2 | 670 | 286 | 884 | 0 | 486 | 0 | 28 | 0 |
| DYNC2H1 | 0 | 0 | 2641 | 0 | 3442 | 0 | 1292 | 2124 |
| DYNC2LI1 | 0 | 318 | 0 | 0 | 0 | 0 | 458 | 0 |
| DYNLL1 | 2882 | 4320 | 16815 | 11374 | 14887 | 8839 | 4966 | 3062 |
| DYNLL2 | 0 | 0 | 960 | 0 | 0 | 0 | 0 | 0 |
| DYNLRB1 | 300 | 1250 | 2360 | 8632 | 7346 | 1746 | 5136 | 2862 |
| DYNLT1 | 1658 | 640 | 2848 | 6228 | 6181 | 1888 | 1130 | 4015 |
| DYNLT3 | 0 | 0 | 0 | 0 | 1013 | 0 | 0 | 784 |
| DYRK1A | 318 | 1235 | 0 | 0 | 222 | 0 | 1522 | 294 |
| DYRK2 | 0 | 0 | 0 | 0 | 1032 | 0 | 0 | 0 |
| DYRK4 | 0 | 473 | 0 | 0 | 1846 | 0 | 0 | 0 |
| DZIP3 | 942 | 0 | 0 | 0 | 0 | 0 | 130 | 0 |
| E2F3 | 0 | 0 | 0 | 0 | 1078 | 0 | 0 | 0 |
| E2F4 | 6340 | 27824 | 20408 | 53273 | 70367 | 41969 | 75926 | 17021 |
| E2F5 | 3990 | 2675 | 0 | 8818 | 13136 | 13796 | 24590 | 9865 |
| E2F6 | 0 | 0 | 0 | 0 | 1236 | 0 | 0 | 0 |
| E2F8 | 0 | 0 | 0 | 0 | 0 | 0 | 228 | 0 |
| E4F1 | 0 | 0 | 0 | 0 | 1460 | 0 | 0 | 0 |
| EAF2 | 0 | 0 | 0 | 1456 | 0 | 0 | 0 | 0 |
| EAPP | 562 | 750 | 3777 | 1879 | 788 | 2238 | 486 | 1408 |
| EARS2 | 0 | 0 | 0 | 0 | 734 | 0 | 782 | 0 |
| EBAG9 | 468 | 0 | 0 | 0 | 296 | 0 | 0 | 0 |
| EBF4 | 236 | 0 | 0 | 0 | 0 | 0 | 0 | 0 |
| EBI3 | 722 | 0 | 0 | 0 | 0 | 0 | 0 | 0 |
| EBLN3P | 0 | 0 | 0 | 0 | 742 | 0 | 0 | 0 |
| EBNA1BP2 | 0 | 432 | 2454 | 3099 | 4106 | 0 | 3088 | 1430 |
| EBP | 0 | 2040 | 7134 | 4913 | 3728 | 3549 | 0 | 3075 |
| EBPL | 1496 | 1138 | 1162 | 4239 | 2304 | 1814 | 3346 | 4996 |
| ECE1 | 1510 | 1901 | 12134 | 1431 | 1961 | 1850 | 1242 | 3934 |
| ECH1 | 1984 | 2872 | 9498 | 5526 | 7120 | 5070 | 2892 | 4740 |
| ECHDC1 | 115617 | 227077 | 256401 | 293978 | 874285 | 254524 | 641034 | 446451 |
| ECHDC2 | 1548 | 252 | 72 | 0 | 1190 | 0 | 594 | 0 |
| ECHDC3 | 0 | 0 | 0 | 0 | 0 | 0 | 1128 | 0 |
| ECHS1 | 116 | 2618 | 13805 | 7256 | 7017 | 4243 | 4321 | 3296 |
| ECI1 | 0 | 742 | 4667 | 3416 | 1024 | 2132 | 2275 | 1217 |
| ECI2 | 0 | 0 | 0 | 910 | 306 | 0 | 0 | 0 |
| ECM2 | 208 | 0 | 0 | 0 | 0 | 0 | 0 | 0 |
| ECPAS | 656 | 0 | 0 | 362 | 0 | 0 | 0 | 0 |
| ECSCR | 0 | 0 | 0 | 0 | 0 | 0 | 0 | 1026 |
| ECSIT | 904 | 6262 | 13468 | 3650 | 7246 | 2280 | 1796 | 3354 |
| ECT2 | 0 | 0 | 0 | 0 | 0 | 0 | 0 | 732 |

| | | | | | | | | |
|---|---|---|---|---|---|---|---|---|
| EDARADD | 0 | 0 | 0 | 0 | 0 | 0 | 196 | 0 |
| EDC4 | 0 | 304 | 2089 | 0 | 964 | 2122 | 0 | 0 |
| EDDM13 | 0 | 0 | 272 | 0 | 0 | 0 | 0 | 0 |
| EDEM2 | 0 | 1610 | 1938 | 1178 | 1074 | 0 | 198 | 522 |
| EDEM3 | 0 | 0 | 880 | 0 | 0 | 0 | 0 | 0 |
| EDF1 | 11687 | 6606 | 13147 | 22248 | 23555 | 3846 | 9442 | 5822 |
| EDIL3 | 0 | 615 | 0 | 0 | 650 | 0 | 0 | 0 |
| EDN2 | 730 | 0 | 0 | 0 | 0 | 0 | 0 | 0 |
| EDN3 | 0 | 0 | 1318 | 0 | 0 | 0 | 0 | 0 |
| EDNRB | 0 | 0 | 1910 | 0 | 0 | 0 | 348 | 0 |
| EDRF1 | 570 | 226 | 0 | 0 | 0 | 0 | 0 | 0 |
| EEA1 | 0 | 0 | 0 | 0 | 0 | 0 | 0 | 200 |
| EED | 0 | 0 | 2290 | 0 | 0 | 0 | 0 | 774 |
| EEF1A1 | 14473 | 33664 | 63304 | 104929 | 138769 | 58672 | 51100 | 74467 |
| EEF1AKMT1 | 0 | 0 | 1944 | 2502 | 3006 | 0 | 866 | 980 |
| EEF1AKMT2 | 0 | 0 | 2467 | 0 | 1182 | 0 | 306 | 0 |
| EEF1AKMT4 | 0 | 0 | 0 | 2626 | 448 | 1679 | 0 | 0 |
| EEF1B2 | 4984 | 5508 | 23087 | 29365 | 45763 | 10374 | 19765 | 19846 |
| EEF1D | 2580 | 4901 | 14232 | 16959 | 39113 | 18135 | 19056 | 11243 |
| EEF1E1 | 0 | 936 | 0 | 2508 | 2406 | 2242 | 484 | 2862 |
| EEF1G | 5456 | 15058 | 36462 | 52588 | 104213 | 33557 | 74754 | 34619 |
| EEF2 | 969594 | 986788 | 1156905 | 1705779 | 4265912 | 2862108 | 2314049 | 880572 |
| EEF2KMT | 986 | 594 | 0 | 1370 | 920 | 0 | 276 | 1308 |
| EEFSEC | 0 | 714 | 0 | 0 | 694 | 0 | 0 | 0 |
| EEPD1 | 0 | 0 | 0 | 0 | 0 | 4258 | 0 | 0 |
| EFCAB11 | 0 | 0 | 0 | 1246 | 519 | 0 | 0 | 992 |
| EFCAB14 | 0 | 442 | 0 | 0 | 174 | 0 | 596 | 0 |
| EFCAB2 | 0 | 0 | 0 | 0 | 0 | 0 | 330 | 0 |
| EFCAB5 | 1306 | 0 | 3840 | 0 | 1190 | 0 | 0 | 0 |
| EFCC1 | 0 | 0 | 0 | 0 | 296 | 0 | 0 | 0 |
| EFEMP1 | 0 | 0 | 1882 | 1048 | 2286 | 4578 | 0 | 1600 |
| EFEMP2 | 0 | 0 | 26 | 0 | 596 | 3678 | 0 | 1122 |
| EFHC1 | 0 | 152 | 0 | 0 | 0 | 0 | 0 | 0 |
| EFHC2 | 176 | 0 | 1440 | 0 | 166 | 0 | 0 | 1320 |
| EFHD2 | 39683 | 34770 | 123258 | 44291 | 184236 | 37307 | 38643 | 38001 |
| EFL1 | 0 | 0 | 0 | 621 | 302 | 1767 | 402 | 0 |
| EFNA1 | 318 | 896 | 0 | 0 | 2524 | 2051 | 4516 | 2302 |
| EFNA2 | 0 | 0 | 0 | 616 | 0 | 0 | 0 | 0 |
| EFNA3 | 0 | 0 | 0 | 0 | 0 | 1050 | 956 | 0 |
| EFNA4 | 688 | 404 | 0 | 470 | 1132 | 0 | 557 | 0 |
| EFNA5 | 0 | 0 | 0 | 0 | 246 | 0 | 0 | 0 |
| EFNB1 | 782 | 168 | 0 | 0 | 2344 | 0 | 0 | 0 |
| EFNB2 | 0 | 0 | 0 | 0 | 0 | 374 | 0 | 0 |

| | | | | | | | | |
|---|---|---|---|---|---|---|---|---|
| EFR3A | 0 | 340 | 0 | 0 | 0 | 0 | 0 | 0 |
| EFTUD2 | 0 | 0 | 0 | 0 | 2048 | 718 | 290 | 508 |
| EGFL7 | 0 | 0 | 870 | 0 | 0 | 0 | 0 | 0 |
| EGFR | 4332 | 2300 | 5482 | 2170 | 12216 | 1092 | 3948 | 3865 |
| EGLN1 | 732 | 0 | 0 | 0 | 1241 | 0 | 0 | 0 |
| EGLN2 | 426 | 196 | 0 | 0 | 0 | 1914 | 0 | 0 |
| EGLN3 | 0 | 1357 | 1584 | 1026 | 1943 | 0 | 0 | 0 |
| EGR1 | 744 | 0 | 0 | 0 | 0 | 0 | 0 | 0 |
| EHBP1L1 | 1096 | 0 | 2958 | 0 | 2098 | 1250 | 1162 | 458 |
| EHD1 | 0 | 0 | 0 | 0 | 0 | 1597 | 454 | 290 |
| EHD4 | 0 | 0 | 0 | 926 | 0 | 1362 | 888 | 0 |
| EHF | 0 | 1024 | 935 | 0 | 3548 | 1760 | 0 | 0 |
| EHHADH | 0 | 0 | 0 | 0 | 0 | 0 | 0 | 1448 |
| EHMT1 | 0 | 0 | 0 | 0 | 430 | 0 | 0 | 0 |
| EHMT2 | 7900 | 3410 | 3201 | 9532 | 27083 | 10643 | 35146 | 10240 |
| EI24 | 1011 | 0 | 648 | 0 | 2220 | 0 | 1074 | 1228 |
| EID1 | 0 | 346 | 0 | 1779 | 2318 | 642 | 918 | 0 |
| EID2 | 0 | 0 | 1360 | 0 | 398 | 0 | 0 | 0 |
| EIF1 | 9718 | 12137 | 21346 | 24589 | 34075 | 8384 | 23395 | 7823 |
| EIF1AD | 0 | 0 | 1450 | 1327 | 610 | 0 | 0 | 1175 |
| EIF1AX | 0 | 682 | 0 | 1030 | 3112 | 0 | 206 | 0 |
| EIF1AY | 0 | 470 | 1942 | 1262 | 5465 | 2636 | 2454 | 1144 |
| EIF1B | 1512 | 480 | 0 | 2600 | 2856 | 948 | 2282 | 2624 |
| EIF2A | 370 | 682 | 0 | 1278 | 572 | 1735 | 884 | 223 |
| EIF2AK1 | 14502 | 32156 | 34294 | 35373 | 112010 | 41550 | 85895 | 20253 |
| EIF2AK2 | 0 | 0 | 0 | 0 | 270 | 0 | 0 | 0 |
| EIF2AK3 | 0 | 496 | 0 | 0 | 0 | 0 | 0 | 0 |
| EIF2AK4 | 812 | 964 | 0 | 530 | 1076 | 578 | 1257 | 404 |
| EIF2B1 | 0 | 0 | 568 | 0 | 1128 | 0 | 0 | 0 |
| EIF2B2 | 0 | 0 | 0 | 1314 | 164 | 0 | 122 | 0 |
| EIF2B3 | 0 | 0 | 0 | 0 | 992 | 2365 | 0 | 0 |
| EIF2B4 | 0 | 532 | 264 | 1362 | 2884 | 442 | 468 | 0 |
| EIF2B5 | 0 | 404 | 1792 | 0 | 754 | 1436 | 0 | 0 |
| EIF2D | 588 | 0 | 0 | 0 | 0 | 0 | 34 | 0 |
| EIF2S1 | 0 | 362 | 0 | 998 | 2298 | 0 | 418 | 0 |
| EIF2S2 | 0 | 0 | 702 | 7839 | 8836 | 2312 | 3895 | 7274 |
| EIF2S3 | 378 | 86 | 0 | 1614 | 758 | 0 | 2046 | 0 |
| EIF3A | 0 | 118 | 0 | 0 | 334 | 0 | 0 | 0 |
| EIF3B | 0 | 484 | 1574 | 1258 | 1470 | 1224 | 318 | 654 |
| EIF3C | 374 | 1150 | 3048 | 1190 | 664 | 2913 | 3228 | 746 |
| EIF3CL | 474 | 1050 | 958 | 0 | 806 | 0 | 1390 | 542 |
| EIF3D | 193960 | 443100 | 1140386 | 628141 | 1220675 | 1280193 | 1294176 | 527501 |
| EIF3E | 0 | 2800 | 3619 | 5547 | 4188 | 25222 | 9449 | 4211 |

| | | | | | | | | |
|---|---|---|---|---|---|---|---|---|
| EIF3F | 3236 | 1386 | 6305 | 10852 | 8436 | 3932 | 5738 | 4441 |
| EIF3G | 528 | 1200 | 0 | 4026 | 3378 | 4572 | 1626 | 0 |
| EIF3H | 485612 | 1042596 | 984864 | 2005213 | 6132166 | 5506142 | 10410180 | 2258833 |
| EIF3I | 718 | 2568 | 5952 | 11961 | 10428 | 7640 | 1484 | 2327 |
| EIF3J | 0 | 1048 | 0 | 0 | 582 | 0 | 558 | 0 |
| EIF3J-DT | 0 | 0 | 0 | 0 | 194 | 0 | 0 | 0 |
| EIF3K | 1332516 | 2014265 | 1814575 | 1893272 | 4719461 | 2381245 | 2024378 | 1622095 |
| EIF3L | 741 | 1548 | 1921 | 1932 | 3706 | 6010 | 2902 | 4002 |
| EIF3M | 404 | 750 | 3872 | 5598 | 4813 | 0 | 482 | 2080 |
| EIF4A1 | 0 | 0 | 940 | 2378 | 528 | 0 | 0 | 0 |
| EIF4A2 | 1544 | 3448 | 16561 | 6159 | 13933 | 4155 | 9115 | 10049 |
| EIF4A3 | 996 | 918 | 3517 | 0 | 3364 | 2048 | 0 | 0 |
| EIF4B | 0 | 2678 | 0 | 2856 | 6324 | 0 | 1660 | 324 |
| EIF4E | 582 | 0 | 1286 | 518 | 2268 | 2244 | 1544 | 876 |
| EIF4E2 | 422 | 978 | 0 | 1526 | 1868 | 0 | 512 | 3817 |
| EIF4E3 | 2284 | 4416 | 12225 | 1050 | 4238 | 1412 | 598 | 0 |
| EIF4EBP1 | 3378 | 6472 | 9676 | 33198 | 17402 | 13415 | 22984 | 12500 |
| EIF4EBP2 | 144 | 1210 | 220 | 452 | 0 | 2657 | 368 | 570 |
| EIF4EBP3 | 1038 | 1008 | 1881 | 0 | 0 | 1452 | 412 | 584 |
| EIF4G1 | 0 | 634 | 0 | 1275 | 1258 | 0 | 502 | 0 |
| EIF4G2 | 360 | 924 | 0 | 4519 | 3336 | 2112 | 988 | 0 |
| EIF4G3 | 1820 | 4574 | 10492 | 236 | 5142 | 5360 | 5642 | 3007 |
| EIF4H | 0 | 2394 | 3440 | 2078 | 2431 | 0 | 952 | 1619 |
| EIF5 | 430 | 304 | 0 | 0 | 4512 | 660 | 0 | 792 |
| EIF5A | 1724 | 2622 | 3565 | 7354 | 4848 | 1104 | 3128 | 3717 |
| EIF5B | 914 | 366 | 0 | 156 | 1800 | 0 | 1092 | 0 |
| EIF6 | 1932 | 4012 | 2620 | 18734 | 7119 | 11361 | 6053 | 6824 |
| EIPR1 | 516 | 370 | 626 | 0 | 0 | 0 | 0 | 1258 |
| ELAC1 | 0 | 472 | 0 | 1652 | 0 | 5444 | 0 | 0 |
| ELAC2 | 0 | 210 | 0 | 0 | 980 | 0 | 982 | 0 |
| ELAVL1 | 0 | 0 | 0 | 0 | 512 | 0 | 1070 | 0 |
| ELF1 | 83167 | 88765 | 51492 | 85358 | 144099 | 147759 | 235820 | 35989 |
| ELF2 | 0 | 0 | 0 | 0 | 0 | 0 | 174 | 0 |
| ELF3 | 715916 | 453303 | 1479175 | 754576 | 2761979 | 644419 | 1395785 | 505501 |
| ELF3-AS1 | 0 | 430 | 434 | 424 | 0 | 0 | 0 | 0 |
| ELF4 | 0 | 674 | 0 | 0 | 0 | 0 | 0 | 0 |
| ELFN1-AS1 | 0 | 398 | 18 | 930 | 3736 | 0 | 496 | 0 |
| ELK2AP | 30 | 0 | 0 | 0 | 0 | 0 | 0 | 34 |
| ELK3 | 2410 | 5528 | 66 | 8031 | 15524 | 3840 | 5702 | 4857 |
| ELK4 | 1250 | 1080 | 5359 | 3146 | 3352 | 3311 | 5602 | 2584 |
| ELL | 6992 | 11579 | 28992 | 5424 | 22894 | 20961 | 10782 | 12883 |
| ELL2 | 0 | 0 | 1622 | 0 | 286 | 0 | 0 | 0 |
| ELMO1 | 0 | 0 | 0 | 599 | 0 | 0 | 0 | 592 |

| | | | | | | | | |
|---|---|---|---|---|---|---|---|---|
| ELMO2 | 0 | 350 | 0 | 0 | 0 | 0 | 0 | 0 |
| ELMO3 | 0 | 0 | 2251 | 0 | 1156 | 0 | 0 | 0 |
| ELMOD1 | 0 | 1608 | 478 | 0 | 1718 | 0 | 1232 | 0 |
| ELMOD2 | 574 | 766 | 827 | 0 | 0 | 0 | 50 | 626 |
| ELMOD3 | 0 | 0 | 0 | 0 | 0 | 0 | 0 | 869 |
| ELMSAN1 | 0 | 30 | 765 | 0 | 1202 | 0 | 0 | 0 |
| ELN | 0 | 0 | 0 | 0 | 406 | 0 | 0 | 0 |
| ELOA-AS1 | 0 | 378 | 0 | 0 | 1212 | 0 | 1010 | 0 |
| ELOB | 9311 | 7924 | 9524 | 9686 | 21729 | 7146 | 7693 | 4652 |
| ELOC | 1016 | 2072 | 2113 | 5582 | 9428 | 3764 | 6019 | 4780 |
| ELOF1 | 926 | 0 | 0 | 1158 | 3308 | 400 | 0 | 1963 |
| ELOVL1 | 2258 | 0 | 2998 | 3524 | 1942 | 5541 | 1260 | 1640 |
| ELOVL5 | 354 | 0 | 0 | 0 | 0 | 0 | 0 | 0 |
| ELOVL6 | 0 | 540 | 0 | 0 | 986 | 0 | 0 | 0 |
| ELP1 | 0 | 140 | 0 | 0 | 0 | 0 | 0 | 0 |
| ELP2 | 0 | 0 | 2213 | 0 | 456 | 0 | 416 | 0 |
| ELP3 | 688 | 0 | 0 | 0 | 0 | 0 | 0 | 0 |
| ELP5 | 1694 | 249 | 3655 | 798 | 3756 | 0 | 0 | 2202 |
| ELP6 | 0 | 0 | 1586 | 0 | 3103 | 0 | 0 | 0 |
| EMC1 | 0 | 0 | 0 | 660 | 0 | 1108 | 0 | 0 |
| EMC1-AS1 | 0 | 0 | 0 | 0 | 598 | 0 | 848 | 0 |
| EMC10 | 0 | 804 | 2836 | 2910 | 2904 | 4218 | 546 | 2210 |
| EMC2 | 0 | 1489 | 2448 | 1554 | 1194 | 0 | 6191 | 516 |
| EMC3 | 425887 | 626657 | 1915957 | 734797 | 2093721 | 1018496 | 1585528 | 540242 |
| EMC4 | 0 | 546 | 1787 | 4069 | 1345 | 0 | 928 | 2696 |
| EMC6 | 880 | 1396 | 3618 | 936 | 772 | 0 | 1190 | 0 |
| EMC7 | 0 | 1494 | 0 | 2448 | 0 | 1734 | 244 | 0 |
| EMC8 | 298 | 428 | 1064 | 3544 | 0 | 0 | 1362 | 1052 |
| EMC9 | 0 | 0 | 558 | 1108 | 1610 | 0 | 0 | 3167 |
| EMCN | 362 | 0 | 0 | 0 | 0 | 0 | 0 | 0 |
| EMD | 0 | 258 | 778 | 0 | 0 | 0 | 880 | 0 |
| EMG1 | 1068 | 578 | 0 | 1579 | 3088 | 1692 | 526 | 2060 |
| EMID1 | 0 | 0 | 0 | 0 | 0 | 0 | 0 | 146 |
| EMILIN1 | 1394 | 0 | 0 | 0 | 640 | 0 | 0 | 0 |
| EMILIN2 | 0 | 0 | 0 | 0 | 3514 | 0 | 0 | 0 |
| EML2 | 0 | 166 | 1036 | 0 | 0 | 0 | 0 | 0 |
| EML3 | 450 | 0 | 0 | 0 | 366 | 1180 | 0 | 0 |
| EML4 | 28226 | 30109 | 21364 | 22558 | 109138 | 50057 | 58677 | 21301 |
| EML5 | 0 | 274 | 0 | 0 | 0 | 0 | 0 | 0 |
| EML6 | 686 | 1468 | 0 | 0 | 60 | 4843 | 1170 | 0 |
| EMP1 | 486 | 610 | 3991 | 0 | 1521 | 0 | 0 | 0 |
| EMP2 | 252 | 0 | 1370 | 0 | 2146 | 1714 | 646 | 0 |
| EMP3 | 644 | 0 | 0 | 0 | 1922 | 1332 | 0 | 0 |

| | | | | | | | | |
|---|---|---|---|---|---|---|---|---|
| EMX1 | 0 | 0 | 724 | 0 | 286 | 0 | 0 | 0 |
| EN1 | 0 | 0 | 0 | 0 | 0 | 1220 | 0 | 0 |
| ENAH | 556 | 300 | 0 | 0 | 0 | 0 | 328 | 0 |
| ENC1 | 330 | 368 | 0 | 0 | 868 | 0 | 0 | 0 |
| ENDOD1 | 0 | 0 | 0 | 0 | 1206 | 0 | 0 | 786 |
| ENDOG | 0 | 426 | 0 | 1334 | 492 | 0 | 580 | 0 |
| ENG | 0 | 0 | 0 | 0 | 330 | 0 | 0 | 0 |
| ENGASE | 142 | 362 | 0 | 441 | 308 | 1636 | 950 | 0 |
| ENKD1 | 0 | 0 | 358 | 0 | 0 | 0 | 738 | 0 |
| ENO1 | 1212499 | 1576248 | 2644223 | 5120014 | 14464603 | 5083894 | 3111528 | 4453096 |
| ENO2 | 0 | 0 | 0 | 0 | 108 | 0 | 0 | 0 |
| ENOPH1 | 0 | 1252 | 0 | 1774 | 6379 | 0 | 8725 | 2132 |
| ENOSF1 | 0 | 0 | 1024 | 0 | 452 | 0 | 858 | 0 |
| ENPP2 | 2972 | 15690 | 12076 | 7170 | 16137 | 59603 | 5118 | 12616 |
| ENPP4 | 4282 | 7122 | 8167 | 5600 | 9493 | 1883 | 11614 | 3834 |
| ENSA | 770 | 372 | 4042 | 2576 | 2571 | 1440 | 1524 | 0 |
| ENTPD1 | 9300 | 9090 | 13335 | 5898 | 24982 | 45620 | 17221 | 22264 |
| ENTPD2 | 0 | 0 | 0 | 0 | 0 | 0 | 0 | 426 |
| ENTPD4 | 0 | 1038 | 0 | 0 | 0 | 0 | 0 | 0 |
| ENTPD5 | 1430 | 350 | 280 | 0 | 0 | 0 | 0 | 0 |
| ENTPD6 | 1586 | 894 | 0 | 1192 | 1320 | 0 | 1878 | 886 |
| ENTPD8 | 460 | 128 | 0 | 0 | 0 | 0 | 0 | 0 |
| ENTR1 | 0 | 0 | 0 | 0 | 404 | 0 | 528 | 0 |
| ENY2 | 223751 | 452254 | 245478 | 357139 | 2155634 | 744607 | 1760425 | 468554 |
| EOGT | 0 | 384 | 0 | 0 | 238 | 0 | 0 | 350 |
| EOMES | 0 | 0 | 0 | 1147 | 0 | 0 | 0 | 0 |
| EP300 | 14122 | 10191 | 36333 | 8829 | 32314 | 18041 | 21685 | 7692 |
| EPAS1 | 942 | 3190 | 2064 | 1170 | 6232 | 2132 | 2818 | 2345 |
| EPB41 | 268 | 0 | 0 | 700 | 0 | 0 | 0 | 0 |
| EPB41L2 | 20188 | 28028 | 41398 | 59960 | 139236 | 41038 | 34054 | 34174 |
| EPB41L3 | 614 | 0 | 0 | 0 | 0 | 0 | 0 | 0 |
| EPB41L4A | 0 | 124 | 0 | 0 | 164 | 1838 | 0 | 0 |
| EPB41L4A-AS1 | 817 | 556 | 0 | 522 | 2292 | 2700 | 804 | 564 |
| EPB41L4B | 724 | 0 | 0 | 0 | 280 | 0 | 346 | 0 |
| EPB41L5 | 0 | 0 | 0 | 0 | 414 | 0 | 0 | 0 |
| EPCAM | 4289056 | 4060834 | 7539871 | 4613523 | 10633008 | 3820558 | 9530872 | 2739381 |
| EPDR1 | 0 | 0 | 0 | 0 | 540 | 0 | 0 | 0 |
| EPG5 | 0 | 0 | 0 | 0 | 0 | 0 | 242 | 0 |
| EPHA1 | 0 | 0 | 0 | 0 | 346 | 0 | 452 | 0 |
| EPHA2 | 398 | 0 | 0 | 0 | 0 | 0 | 0 | 244 |
| EPHB2 | 3704 | 9754 | 18322 | 20553 | 8054 | 38157 | 66217 | 9780 |
| EPHB3 | 0 | 0 | 0 | 616 | 747 | 0 | 0 | 0 |
| EPHB4 | 0 | 344 | 0 | 0 | 302 | 0 | 0 | 0 |

| | | | | | | | | |
|---|---|---|---|---|---|---|---|---|
| EPHX1 | 2598 | 18696 | 34364 | 8419 | 36385 | 44534 | 76413 | 4938 |
| EPHX2 | 35916 | 44048 | 153712 | 11635 | 13172 | 50254 | 46493 | 5293 |
| EPM2AIP1 | 0 | 0 | 0 | 0 | 0 | 1134 | 0 | 0 |
| EPN1 | 367 | 381 | 890 | 12 | 2732 | 0 | 0 | 0 |
| EPN3 | 0 | 0 | 0 | 0 | 656 | 0 | 0 | 0 |
| EPOR | 0 | 0 | 0 | 0 | 0 | 0 | 302 | 0 |
| EPPK1 | 0 | 402 | 0 | 500 | 0 | 1588 | 452 | 0 |
| EPRS | 702 | 488 | 0 | 0 | 0 | 0 | 0 | 816 |
| EPS15 | 46108 | 72497 | 81033 | 61990 | 105035 | 72538 | 172357 | 34471 |
| EPS15L1 | 1738 | 1034 | 4774 | 716 | 10275 | 3010 | 642 | 748 |
| EPS8 | 0 | 372 | 3376 | 2072 | 670 | 0 | 366 | 0 |
| EPS8L1 | 0 | 0 | 0 | 704 | 454 | 0 | 0 | 0 |
| EPS8L2 | 0 | 308 | 1374 | 1002 | 1094 | 1585 | 874 | 0 |
| EPS8L3 | 1550 | 930 | 2184 | 1388 | 1436 | 0 | 1739 | 710 |
| EPSTI1 | 1910 | 0 | 1469 | 246 | 300 | 0 | 3216 | 872 |
| EPYC | 0 | 0 | 0 | 0 | 186440 | 88437 | 1608 | 876 |
| ERAL1 | 168 | 160 | 1387 | 1229 | 450 | 0 | 859 | 0 |
| ERAP1 | 496 | 0 | 2070 | 0 | 30 | 0 | 0 | 0 |
| ERBB2 | 68949 | 41134 | 141577 | 43448 | 138417 | 35350 | 160842 | 29290 |
| ERBB3 | 63953 | 42985 | 203364 | 57452 | 125502 | 37797 | 177588 | 25859 |
| ERBIN | 1322 | 0 | 1098 | 0 | 154 | 1001 | 0 | 0 |
| ERC1 | 18410 | 13766 | 11227 | 8444 | 27602 | 13183 | 17484 | 11486 |
| ERCC1 | 0 | 52 | 532 | 0 | 1674 | 1543 | 0 | 0 |
| ERCC2 | 5783 | 11172 | 18579 | 19383 | 76869 | 9944 | 26361 | 28057 |
| ERCC3 | 7583 | 6576 | 16585 | 6905 | 35292 | 7922 | 49974 | 5973 |
| ERCC5 | 0 | 0 | 0 | 0 | 450 | 0 | 0 | 0 |
| ERCC6 | 0 | 0 | 0 | 0 | 0 | 0 | 266 | 0 |
| ERCC8 | 0 | 0 | 0 | 1608 | 624 | 0 | 0 | 0 |
| EREG | 0 | 0 | 0 | 754 | 0 | 0 | 0 | 0 |
| ERG | 6787 | 11481 | 11759 | 6477 | 25700 | 15822 | 7586 | 8416 |
| ERG28 | 0 | 514 | 8336 | 3015 | 2824 | 0 | 1000 | 1820 |
| ERGIC1 | 13857 | 37900 | 78321 | 45418 | 70299 | 64426 | 112279 | 24702 |
| ERGIC2 | 0 | 764 | 0 | 1682 | 1587 | 0 | 920 | 0 |
| ERGIC3 | 920 | 3172 | 1536 | 15945 | 11537 | 5962 | 10866 | 3842 |
| ERH | 760 | 2890 | 3078 | 3028 | 10205 | 2698 | 2262 | 954 |
| ERI1 | 0 | 0 | 0 | 0 | 748 | 2209 | 0 | 0 |
| ERI3 | 368 | 248 | 102 | 0 | 585 | 0 | 0 | 0 |
| ERICH1 | 711 | 0 | 0 | 0 | 0 | 0 | 0 | 0 |
| ERICH2 | 492 | 0 | 0 | 0 | 0 | 0 | 156 | 0 |
| ERICH5 | 0 | 0 | 0 | 0 | 0 | 0 | 588 | 0 |
| ERLEC1 | 544 | 776 | 1248 | 1046 | 388 | 1526 | 0 | 718 |
| ERLIN1 | 0 | 0 | 860 | 0 | 836 | 0 | 0 | 172 |
| ERLIN2 | 9846 | 17954 | 15222 | 8425 | 15299 | 8282 | 12640 | 7105 |

| | | | | | | | | |
|---|---|---|---|---|---|---|---|---|
| ERMAP | 0 | 0 | 0 | 652 | 620 | 0 | 0 | 422 |
| ERN2 | 56 | 1636 | 0 | 0 | 418 | 0 | 0 | 382 |
| ERO1A | 1012 | 254 | 1017 | 548 | 8713 | 0 | 1309 | 3272 |
| ERO1B | 0 | 104 | 0 | 0 | 0 | 0 | 0 | 0 |
| ERP29 | 0 | 2150 | 336 | 1278 | 4446 | 0 | 3150 | 2729 |
| ERP44 | 103606 | 231497 | 257237 | 283882 | 581466 | 392504 | 568635 | 204874 |
| ERRFI1 | 0 | 0 | 706 | 0 | 0 | 0 | 0 | 462 |
| ERV3-1 | 0 | 0 | 0 | 0 | 1390 | 0 | 0 | 426 |
| ERVK3-1 | 0 | 886 | 0 | 1431 | 644 | 0 | 0 | 0 |
| ESAM | 0 | 154 | 1038 | 776 | 494 | 0 | 0 | 48 |
| ESD | 0 | 2872 | 3962 | 4067 | 2046 | 2606 | 1916 | 1152 |
| ESF1 | 0 | 0 | 0 | 0 | 793 | 0 | 0 | 0 |
| ESM1 | 0 | 0 | 0 | 0 | 222 | 0 | 0 | 0 |
| ESPL1 | 0 | 0 | 0 | 926 | 0 | 0 | 0 | 0 |
| ESPN | 386 | 284 | 1233 | 0 | 0 | 0 | 0 | 0 |
| ESRG | 0 | 0 | 0 | 0 | 1264 | 0 | 0 | 0 |
| ESRP1 | 386 | 484 | 0 | 0 | 0 | 0 | 0 | 742 |
| ESRP2 | 0 | 254 | 0 | 0 | 692 | 0 | 532 | 0 |
| ESRRA | 222 | 422 | 5215 | 504 | 1002 | 0 | 1222 | 0 |
| ESRRG | 1000 | 0 | 0 | 1850 | 196 | 0 | 5479 | 0 |
| ESS2 | 1640 | 2551 | 10566 | 6290 | 15940 | 7100 | 5164 | 8745 |
| ESYT1 | 0 | 338 | 1996 | 0 | 0 | 0 | 0 | 0 |
| ESYT2 | 808 | 518 | 0 | 1004 | 0 | 0 | 0 | 0 |
| ESYT3 | 0 | 0 | 0 | 0 | 0 | 2144 | 0 | 0 |
| ETAA1 | 82 | 194 | 0 | 0 | 0 | 0 | 0 | 0 |
| ETF1 | 414 | 0 | 0 | 0 | 0 | 0 | 224 | 0 |
| ETFA | 1240 | 2406 | 7724 | 15946 | 6634 | 0 | 2888 | 1262 |
| ETFB | 0 | 1670 | 1890 | 3605 | 7459 | 1976 | 252 | 0 |
| ETFDH | 0 | 0 | 0 | 0 | 194 | 0 | 0 | 0 |
| ETFRF1 | 668 | 0 | 484 | 0 | 270 | 0 | 0 | 146 |
| ETHE1 | 3176 | 4298 | 28841 | 2875 | 11862 | 0 | 372 | 2109 |
| ETNK1 | 22554 | 23219 | 100187 | 20131 | 55295 | 25070 | 45541 | 25560 |
| ETS1 | 98 | 0 | 0 | 302 | 0 | 0 | 0 | 0 |
| ETS2 | 1494 | 734 | 644 | 1562 | 4136 | 16 | 4809 | 733 |
| ETV1 | 3617 | 4220 | 5324 | 5833 | 43206 | 4237 | 340 | 4979 |
| ETV3 | 0 | 0 | 0 | 0 | 0 | 0 | 608 | 0 |
| ETV4 | 214 | 4262 | 582 | 102098 | 52629 | 59492 | 332486 | 22191 |
| ETV5 | 2198 | 262 | 0 | 1500 | 4210 | 3643 | 4419 | 2224 |
| ETV6 | 25340 | 6492 | 43099 | 26524 | 65198 | 28065 | 30986 | 15341 |
| EVA1C | 350 | 318 | 1790 | 0 | 0 | 0 | 0 | 1662 |
| EVADR | 1108 | 0 | 0 | 1644 | 0 | 2556 | 2838 | 0 |
| EVI2B | 0 | 500 | 0 | 0 | 0 | 608 | 0 | 0 |
| EVI5L | 0 | 302 | 0 | 0 | 0 | 0 | 0 | 0 |

| | | | | | | | | |
|---|---|---|---|---|---|---|---|---|
| EVL | 0 | 366 | 0 | 0 | 78 | 1236 | 554 | 620 |
| EVPL | 0 | 0 | 0 | 620 | 212 | 0 | 296 | 206 |
| EWSR1 | 46216 | 107421 | 190353 | 166563 | 427859 | 281924 | 317145 | 159165 |
| EXD2 | 2484 | 11753 | 23082 | 10325 | 34345 | 7195 | 19340 | 6616 |
| EXD3 | 522 | 334 | 0 | 0 | 0 | 0 | 0 | 535 |
| EXO5 | 0 | 404 | 0 | 0 | 0 | 0 | 0 | 0 |
| EXOC1 | 0 | 0 | 1464 | 0 | 0 | 1922 | 0 | 0 |
| EXOC2 | 0 | 0 | 0 | 0 | 574 | 0 | 0 | 0 |
| EXOC3 | 0 | 0 | 0 | 0 | 1065 | 0 | 962 | 902 |
| EXOC3L4 | 102 | 0 | 0 | 0 | 0 | 0 | 0 | 0 |
| EXOC4 | 6304 | 13996 | 14779 | 20719 | 27740 | 21768 | 40494 | 13723 |
| EXOC6B | 1286 | 1534 | 2510 | 0 | 6206 | 1483 | 1346 | 1918 |
| EXOC7 | 0 | 0 | 0 | 0 | 178 | 0 | 0 | 0 |
| EXOC8 | 0 | 0 | 0 | 0 | 0 | 0 | 310 | 0 |
| EXOSC1 | 0 | 1602 | 0 | 0 | 3246 | 2914 | 1746 | 0 |
| EXOSC10 | 16511 | 54217 | 83542 | 106961 | 143810 | 69117 | 97697 | 60402 |
| EXOSC2 | 5673 | 16840 | 13980 | 75756 | 139204 | 75536 | 185930 | 42074 |
| EXOSC3 | 880 | 8096 | 922 | 4755 | 3127 | 5502 | 2882 | 1088 |
| EXOSC4 | 0 | 0 | 2150 | 3685 | 4998 | 0 | 466 | 0 |
| EXOSC5 | 0 | 810 | 1758 | 1618 | 3369 | 10768 | 2342 | 790 |
| EXOSC6 | 0 | 0 | 0 | 0 | 0 | 0 | 316 | 0 |
| EXOSC7 | 0 | 510 | 0 | 5212 | 3042 | 0 | 1128 | 0 |
| EXOSC8 | 0 | 338 | 1759 | 13376 | 5120 | 6601 | 526 | 2580 |
| EXOSC9 | 18459 | 41443 | 32768 | 48220 | 128095 | 51039 | 82859 | 40800 |
| EXPH5 | 330 | 236 | 0 | 0 | 0 | 0 | 0 | 0 |
| EXT1 | 94835 | 48939 | 154104 | 43885 | 275959 | 81971 | 119538 | 79067 |
| EXT2 | 72629 | 113112 | 155956 | 104568 | 419224 | 130888 | 193537 | 131604 |
| EXTL2 | 400 | 0 | 0 | 0 | 714 | 0 | 0 | 0 |
| EYA1 | 0 | 0 | 0 | 0 | 0 | 0 | 522 | 0 |
| EYA2 | 3011 | 4372 | 7018 | 0 | 1750 | 6854 | 0 | 0 |
| EYA3 | 2548 | 8102 | 9919 | 11133 | 20013 | 9558 | 7842 | 11014 |
| EYS | 0 | 0 | 0 | 0 | 766 | 0 | 0 | 0 |
| EZH1 | 0 | 0 | 0 | 828 | 0 | 0 | 564 | 0 |
| EZH2 | 8279 | 13178 | 20296 | 32884 | 42060 | 19577 | 15885 | 17715 |
| EZR | 724885 | 682794 | 2029258 | 491119 | 1860502 | 483071 | 782502 | 1029549 |
| F11R | 584 | 1208 | 2493 | 1498 | 1078 | 2136 | 0 | 883 |
| F12 | 0 | 0 | 0 | 1376 | 1314 | 0 | 0 | 0 |
| F13A1 | 0 | 0 | 0 | 0 | 0 | 0 | 513 | 0 |
| F2 | 0 | 0 | 0 | 1816 | 0 | 0 | 0 | 0 |
| F2RL1 | 448 | 712 | 0 | 656 | 1008 | 0 | 0 | 0 |
| F2RL2 | 0 | 0 | 0 | 0 | 42 | 0 | 0 | 0 |
| F3 | 266 | 0 | 0 | 524 | 634 | 0 | 0 | 600 |
| F5 | 1074 | 548 | 0 | 0 | 390 | 2604 | 0 | 5202 |

| | | | | | | | | |
|---|---|---|---|---|---|---|---|---|
| F8 | 0 | 0 | 0 | 1056 | 0 | 0 | 0 | 0 |
| F8A1 | 0 | 0 | 3318 | 0 | 2144 | 0 | 0 | 516 |
| FA2H | 610 | 244 | 0 | 0 | 0 | 0 | 0 | 0 |
| FAAH | 0 | 450 | 1484 | 0 | 1038 | 0 | 2252 | 0 |
| FAAH2 | 0 | 0 | 0 | 0 | 812 | 0 | 0 | 0 |
| FAAHP1 | 0 | 0 | 804 | 0 | 0 | 0 | 0 | 0 |
| FAAP100 | 0 | 322 | 0 | 0 | 1392 | 0 | 1067 | 0 |
| FAAP20 | 960 | 0 | 1984 | 2480 | 1226 | 0 | 648 | 1036 |
| FAAP24 | 0 | 0 | 0 | 0 | 0 | 0 | 98 | 962 |
| FABP1 | 195363 | 42815 | 344562 | 21785 | 15349 | 2746 | 3904 | 648 |
| FABP2 | 0 | 466 | 998 | 280 | 1126 | 0 | 578 | 0 |
| FABP3 | 0 | 0 | 1144 | 0 | 1370 | 0 | 0 | 0 |
| FABP4 | 0 | 662 | 1624 | 0 | 262 | 9506 | 0 | 0 |
| FABP5 | 2500 | 3566 | 7278 | 5394 | 7217 | 10202 | 2296 | 2748 |
| FABP6 | 0 | 0 | 0 | 0 | 1372 | 0 | 0 | 0 |
| FADD | 512 | 3134 | 2235 | 2380 | 7901 | 1912 | 6133 | 2268 |
| FADS1 | 0 | 0 | 0 | 596 | 0 | 0 | 0 | 0 |
| FADS2 | 0 | 0 | 0 | 0 | 1742 | 1594 | 286 | 484 |
| FADS3 | 0 | 0 | 0 | 884 | 74 | 0 | 0 | 0 |
| FAF1 | 16613 | 38061 | 35720 | 100002 | 152312 | 60731 | 192943 | 53132 |
| FAH | 0 | 496 | 0 | 0 | 2377 | 754 | 0 | 0 |
| FAHD1 | 210 | 976 | 0 | 904 | 646 | 0 | 1242 | 0 |
| FAHD2A | 482 | 418 | 0 | 300 | 904 | 3456 | 0 | 894 |
| FAHD2B | 368 | 0 | 196 | 0 | 1224 | 0 | 0 | 0 |
| FAIM | 0 | 430 | 0 | 0 | 1414 | 0 | 38 | 0 |
| FAM102A | 0 | 228 | 1064 | 0 | 1402 | 0 | 0 | 0 |
| FAM102B | 0 | 0 | 0 | 144 | 38 | 0 | 0 | 0 |
| FAM104A | 3712 | 8930 | 38690 | 7632 | 18730 | 19461 | 20773 | 9687 |
| FAM104B | 0 | 1026 | 40 | 0 | 1085 | 1477 | 0 | 0 |
| FAM107B | 616 | 886 | 1322 | 0 | 338 | 0 | 0 | 0 |
| FAM110A | 0 | 0 | 0 | 0 | 0 | 0 | 0 | 660 |
| FAM110C | 0 | 260 | 0 | 0 | 0 | 0 | 0 | 0 |
| FAM114A1 | 0 | 0 | 0 | 0 | 0 | 0 | 926 | 0 |
| FAM114A2 | 15542 | 24126 | 46036 | 45153 | 44091 | 47338 | 112034 | 28771 |
| FAM117A | 3421 | 2434 | 6480 | 5443 | 524 | 1822 | 6030 | 2044 |
| FAM117B | 0 | 0 | 0 | 246 | 0 | 0 | 0 | 0 |
| FAM118A | 0 | 0 | 0 | 0 | 0 | 0 | 0 | 1288 |
| FAM118B | 0 | 794 | 2714 | 1246 | 2452 | 1770 | 0 | 1046 |
| FAM120A | 0 | 292 | 550 | 0 | 514 | 0 | 0 | 0 |
| FAM120AOS | 496 | 162 | 1620 | 0 | 366 | 0 | 0 | 0 |
| FAM120B | 0 | 428 | 727 | 0 | 212 | 0 | 0 | 0 |
| FAM122A | 758 | 0 | 0 | 0 | 860 | 0 | 766 | 0 |
| FAM122B | 0 | 0 | 0 | 0 | 1411 | 580 | 0 | 0 |

| | | | | | | | | |
|---|---|---|---|---|---|---|---|---|
| FAM122C | 0 | 0 | 1086 | 0 | 0 | 0 | 0 | 0 |
| FAM124B | 0 | 0 | 0 | 0 | 280 | 0 | 0 | 0 |
| FAM126A | 988 | 1576 | 0 | 0 | 3374 | 1680 | 0 | 428 |
| FAM129A | 0 | 0 | 0 | 0 | 0 | 0 | 0 | 274 |
| FAM129B | 308 | 156 | 0 | 867 | 1174 | 0 | 252 | 0 |
| FAM129C | 414 | 0 | 0 | 0 | 0 | 0 | 0 | 0 |
| FAM133B | 0 | 448 | 0 | 0 | 774 | 0 | 0 | 0 |
| FAM135A | 0 | 0 | 298 | 0 | 374 | 0 | 0 | 0 |
| FAM136A | 0 | 410 | 3949 | 1298 | 2652 | 1234 | 2718 | 0 |
| FAM153C | 1018 | 0 | 0 | 0 | 0 | 0 | 0 | 0 |
| FAM155A | 0 | 0 | 0 | 0 | 768 | 0 | 0 | 0 |
| FAM156A | 0 | 0 | 0 | 0 | 0 | 0 | 0 | 651 |
| FAM156B | 0 | 0 | 0 | 526 | 192 | 4272 | 0 | 100 |
| FAM160A2 | 554 | 32 | 678 | 0 | 332 | 0 | 0 | 0 |
| FAM160B1 | 0 | 0 | 0 | 0 | 0 | 0 | 338 | 0 |
| FAM162A | 1338 | 6329 | 12375 | 6284 | 10526 | 2649 | 3196 | 4805 |
| FAM168A | 658 | 194 | 0 | 0 | 1712 | 396 | 268 | 1646 |
| FAM171A1 | 656 | 0 | 0 | 0 | 428 | 0 | 0 | 0 |
| FAM172A | 0 | 308 | 0 | 0 | 0 | 0 | 0 | 816 |
| FAM173A | 814 | 748 | 32 | 1222 | 2198 | 0 | 488 | 1180 |
| FAM173B | 0 | 0 | 0 | 0 | 518 | 1910 | 0 | 0 |
| FAM174A | 0 | 0 | 0 | 644 | 0 | 0 | 0 | 0 |
| FAM174B | 0 | 348 | 0 | 0 | 0 | 0 | 0 | 0 |
| FAM177A1 | 368451 | 441795 | 398328 | 164147 | 520650 | 204964 | 476057 | 152210 |
| FAM177B | 0 | 0 | 0 | 638 | 0 | 0 | 0 | 0 |
| FAM185A | 0 | 232 | 0 | 0 | 0 | 0 | 0 | 0 |
| FAM192A | 0 | 260 | 2561 | 2064 | 0 | 0 | 1576 | 0 |
| FAM193A | 992 | 976 | 3619 | 624 | 280 | 2272 | 3244 | 0 |
| FAM193B | 0 | 136 | 0 | 0 | 0 | 0 | 0 | 0 |
| FAM198B | 80 | 0 | 0 | 0 | 0 | 0 | 0 | 0 |
| FAM19A5 | 0 | 0 | 0 | 0 | 592 | 0 | 0 | 0 |
| FAM200A | 0 | 360 | 0 | 0 | 0 | 0 | 0 | 0 |
| FAM200B | 636 | 184 | 1904 | 0 | 1982 | 0 | 1140 | 870 |
| FAM204A | 0 | 1048 | 0 | 0 | 2780 | 2067 | 0 | 934 |
| FAM206A | 0 | 0 | 0 | 956 | 1050 | 0 | 470 | 0 |
| FAM207A | 0 | 0 | 0 | 0 | 3630 | 1508 | 1746 | 864 |
| FAM208A | 0 | 0 | 0 | 0 | 1148 | 268 | 0 | 0 |
| FAM20A | 6580 | 17494 | 1720 | 6129 | 15912 | 6405 | 43053 | 5964 |
| FAM20B | 0 | 0 | 0 | 518 | 0 | 0 | 0 | 0 |
| FAM20C | 0 | 446 | 0 | 0 | 0 | 0 | 0 | 0 |
| FAM210A | 0 | 380 | 0 | 0 | 1040 | 0 | 0 | 0 |
| FAM210B | 0 | 328 | 0 | 0 | 1788 | 0 | 418 | 0 |
| FAM213A | 0 | 0 | 1177 | 0 | 484 | 0 | 0 | 0 |

| | | | | | | | | |
|---|---|---|---|---|---|---|---|---|
| FAM213B | 313 | 0 | 1460 | 0 | 936 | 0 | 0 | 0 |
| FAM214A | 0 | 0 | 0 | 0 | 200 | 0 | 496 | 0 |
| FAM214B | 0 | 0 | 0 | 1818 | 0 | 0 | 0 | 0 |
| FAM216A | 0 | 0 | 0 | 0 | 916 | 0 | 0 | 150 |
| FAM219A | 0 | 0 | 0 | 0 | 934 | 0 | 0 | 0 |
| FAM219B | 600 | 702 | 1992 | 0 | 0 | 0 | 758 | 0 |
| FAM21EP | 0 | 0 | 0 | 0 | 0 | 0 | 408 | 1229 |
| FAM221A | 0 | 0 | 577 | 0 | 588 | 0 | 0 | 732 |
| FAM222B | 0 | 0 | 0 | 0 | 396 | 0 | 0 | 0 |
| FAM226A | 0 | 424 | 0 | 0 | 0 | 0 | 0 | 0 |
| FAM228B | 0 | 0 | 0 | 0 | 1414 | 0 | 0 | 0 |
| FAM229A | 0 | 0 | 0 | 0 | 618 | 0 | 0 | 0 |
| FAM229B | 0 | 0 | 0 | 0 | 0 | 0 | 0 | 845 |
| FAM234A | 0 | 0 | 0 | 2012 | 0 | 0 | 966 | 0 |
| FAM241B | 0 | 0 | 0 | 0 | 1056 | 0 | 0 | 0 |
| FAM24B | 0 | 318 | 0 | 0 | 0 | 570 | 0 | 0 |
| FAM27B | 0 | 0 | 0 | 0 | 168 | 0 | 0 | 0 |
| FAM30A | 0 | 508 | 0 | 0 | 0 | 0 | 0 | 0 |
| FAM32A | 884 | 659 | 5268 | 0 | 4406 | 0 | 1204 | 1271 |
| FAM35A | 0 | 0 | 0 | 0 | 370 | 0 | 0 | 0 |
| FAM3A | 0 | 348 | 1520 | 0 | 1714 | 0 | 284 | 0 |
| FAM3B | 0 | 0 | 0 | 0 | 968 | 0 | 0 | 0 |
| FAM3C | 202 | 754 | 1780 | 0 | 0 | 0 | 0 | 2686 |
| FAM3D | 5812 | 4348 | 12995 | 9910 | 26054 | 348 | 1685 | 618 |
| FAM43A | 80 | 0 | 0 | 0 | 0 | 0 | 0 | 0 |
| FAM45A | 0 | 280 | 0 | 0 | 800 | 0 | 0 | 0 |
| FAM47E | 242 | 0 | 0 | 0 | 308 | 0 | 0 | 0 |
| FAM49A | 310 | 0 | 0 | 1282 | 156 | 0 | 0 | 0 |
| FAM49B | 10234 | 6296 | 13285 | 34631 | 106279 | 37087 | 102164 | 41057 |
| FAM50A | 1022 | 0 | 948 | 1677 | 5164 | 0 | 722 | 2302 |
| FAM50B | 0 | 244 | 0 | 0 | 1373 | 0 | 22 | 2438 |
| FAM57A | 0 | 16 | 0 | 0 | 0 | 0 | 0 | 824 |
| FAM71E1 | 0 | 0 | 0 | 0 | 602 | 0 | 0 | 0 |
| FAM76B | 0 | 0 | 0 | 0 | 0 | 0 | 0 | 384 |
| FAM78B | 0 | 0 | 0 | 0 | 1348 | 0 | 0 | 0 |
| FAM83B | 0 | 58 | 0 | 0 | 0 | 0 | 0 | 0 |
| FAM83D | 0 | 0 | 0 | 0 | 244 | 0 | 0 | 0 |
| FAM83E | 0 | 1039 | 0 | 0 | 1976 | 0 | 0 | 0 |
| FAM83F | 966 | 1074 | 991 | 694 | 1871 | 0 | 1974 | 0 |
| FAM83G | 206 | 0 | 0 | 0 | 96 | 0 | 0 | 0 |
| FAM83H | 0 | 0 | 0 | 1366 | 1596 | 0 | 0 | 0 |
| FAM83H-AS1 | 0 | 0 | 0 | 0 | 0 | 0 | 350 | 0 |
| FAM84A | 0 | 0 | 1251 | 0 | 248 | 0 | 56 | 0 |

| | | | | | | | | |
|---|---|---|---|---|---|---|---|---|
| FAM84B | 0 | 202 | 0 | 0 | 936 | 0 | 144 | 0 |
| FAM86B1 | 0 | 0 | 1472 | 0 | 0 | 0 | 0 | 0 |
| FAM86C1 | 616 | 0 | 0 | 0 | 908 | 0 | 550 | 0 |
| FAM86C2P | 0 | 0 | 0 | 0 | 38 | 0 | 0 | 0 |
| FAM86DP | 0 | 0 | 0 | 1298 | 1058 | 0 | 206 | 0 |
| FAM86JP | 0 | 0 | 0 | 1066 | 910 | 0 | 0 | 0 |
| FAM89A | 0 | 0 | 0 | 528 | 52 | 0 | 1272 | 0 |
| FAM89B | 0 | 344 | 0 | 0 | 1482 | 2270 | 0 | 0 |
| FAM8A1 | 0 | 0 | 718 | 0 | 374 | 0 | 0 | 0 |
| FAM91A1 | 0 | 0 | 0 | 0 | 974 | 0 | 0 | 712 |
| FAM92A | 0 | 0 | 0 | 1178 | 484 | 0 | 0 | 492 |
| FAM95C | 62 | 0 | 0 | 0 | 0 | 0 | 0 | 0 |
| FAM98A | 0 | 354 | 0 | 0 | 994 | 0 | 0 | 390 |
| FAM98C | 2040 | 0 | 0 | 0 | 842 | 0 | 0 | 0 |
| FAN1 | 0 | 546 | 0 | 0 | 0 | 0 | 92 | 0 |
| FANCA | 1028 | 876 | 694 | 5462 | 8816 | 386 | 4734 | 3068 |
| FANCB | 0 | 0 | 0 | 0 | 2518 | 0 | 744 | 0 |
| FANCC | 6056 | 6056 | 20909 | 23485 | 42529 | 10562 | 22699 | 13104 |
| FANCD2 | 2726 | 2688 | 3171 | 21748 | 45729 | 2830 | 16204 | 4532 |
| FANCD2OS | 0 | 0 | 0 | 0 | 244 | 0 | 0 | 0 |
| FANCE | 0 | 0 | 3625 | 810 | 0 | 0 | 0 | 0 |
| FANCF | 0 | 0 | 0 | 0 | 706 | 0 | 306 | 0 |
| FANCG | 126 | 0 | 0 | 0 | 0 | 0 | 0 | 0 |
| FANCI | 0 | 0 | 0 | 0 | 42 | 0 | 0 | 0 |
| FANCL | 0 | 0 | 0 | 1916 | 1010 | 0 | 0 | 0 |
| FAP | 0 | 0 | 0 | 0 | 0 | 1394 | 0 | 0 |
| FAR1 | 706 | 0 | 0 | 0 | 432 | 0 | 0 | 0 |
| FAR2 | 0 | 0 | 713 | 0 | 330 | 0 | 144 | 0 |
| FARP1 | 0 | 126 | 0 | 812 | 0 | 0 | 1566 | 1098 |
| FARP2 | 990 | 2470 | 2736 | 1044 | 8877 | 192 | 2000 | 3826 |
| FARSA | 22794 | 48188 | 130217 | 84093 | 165671 | 85325 | 106594 | 57271 |
| FARSB | 0 | 0 | 0 | 1128 | 3426 | 0 | 0 | 2028 |
| FAS | 74398 | 211637 | 235878 | 24218 | 45012 | 31688 | 26556 | 32693 |
| FASN | 0 | 0 | 0 | 1140 | 0 | 0 | 536 | 0 |
| FASTK | 0 | 0 | 2961 | 1228 | 662 | 0 | 0 | 30 |
| FASTKD1 | 0 | 0 | 0 | 0 | 212 | 2396 | 270 | 0 |
| FASTKD2 | 0 | 0 | 0 | 4085 | 428 | 0 | 0 | 0 |
| FASTKD3 | 19851 | 30631 | 24829 | 18905 | 143379 | 15305 | 32630 | 13538 |
| FAT1 | 23012 | 13395 | 25990 | 11607 | 36161 | 4991 | 197590 | 8829 |
| FAT4 | 0 | 688 | 0 | 0 | 0 | 0 | 0 | 0 |
| FAU | 9722 | 18189 | 19072 | 32873 | 45616 | 20429 | 17728 | 20680 |
| FBH1 | 0 | 0 | 10 | 0 | 0 | 0 | 0 | 0 |
| FBL | 782 | 696 | 1923 | 3594 | 5026 | 2321 | 1386 | 755 |

| | | | | | | | | |
|---|---|---|---|---|---|---|---|---|
| FBLIM1 | 7988 | 15884 | 35762 | 12956 | 13689 | 6000 | 11100 | 6976 |
| FBLN1 | 3636 | 7226 | 139006 | 2519 | 28313 | 45804 | 2932 | 4980 |
| FBLN2 | 770 | 2442 | 17966 | 1106 | 27509 | 22201 | 8122 | 9857 |
| FBLN5 | 1742 | 4804 | 24990 | 4508 | 7400 | 35561 | 6762 | 18153 |
| FBN1 | 0 | 0 | 0 | 0 | 817 | 0 | 0 | 0 |
| FBP1 | 392 | 0 | 2138 | 3586 | 5847 | 0 | 2286 | 3319 |
| FBRS | 0 | 0 | 0 | 0 | 0 | 0 | 114 | 0 |
| FBRSL1 | 0 | 0 | 0 | 0 | 844 | 0 | 0 | 0 |
| FBXL12 | 464 | 238 | 0 | 0 | 528 | 0 | 0 | 0 |
| FBXL15 | 0 | 0 | 0 | 1674 | 0 | 0 | 274 | 0 |
| FBXL20 | 6161 | 2144 | 3658 | 864 | 4469 | 5670 | 7164 | 4283 |
| FBXL22 | 1084 | 132 | 0 | 0 | 168 | 0 | 0 | 0 |
| FBXL3 | 0 | 0 | 794 | 641 | 0 | 0 | 0 | 0 |
| FBXL5 | 0 | 234 | 1050 | 686 | 1343 | 0 | 586 | 1591 |
| FBXL6 | 0 | 0 | 480 | 72 | 1198 | 1618 | 2080 | 0 |
| FBXO11 | 10040 | 9406 | 4854 | 7266 | 47131 | 10862 | 30763 | 6245 |
| FBXO21 | 0 | 0 | 0 | 340 | 290 | 0 | 0 | 0 |
| FBXO22 | 0 | 326 | 0 | 0 | 812 | 0 | 0 | 546 |
| FBXO25 | 558 | 0 | 1982 | 0 | 708 | 0 | 0 | 0 |
| FBXO28 | 2444 | 5434 | 3158 | 5029 | 11402 | 9591 | 15695 | 1914 |
| FBXO3 | 14908 | 39800 | 65627 | 40123 | 80951 | 57086 | 87174 | 40702 |
| FBXO32 | 0 | 0 | 3256 | 0 | 2631 | 0 | 464 | 0 |
| FBXO34 | 560 | 0 | 0 | 0 | 0 | 0 | 150 | 0 |
| FBXO36 | 0 | 0 | 0 | 0 | 688 | 0 | 0 | 144 |
| FBXO38 | 532 | 0 | 0 | 0 | 810 | 0 | 0 | 408 |
| FBXO4 | 0 | 814 | 1194 | 0 | 2584 | 0 | 0 | 0 |
| FBXO41 | 0 | 0 | 0 | 0 | 0 | 0 | 220 | 0 |
| FBXO44 | 146 | 0 | 0 | 0 | 0 | 0 | 0 | 0 |
| FBXO45 | 814 | 0 | 0 | 0 | 0 | 0 | 0 | 0 |
| FBXO48 | 0 | 0 | 0 | 0 | 98 | 0 | 0 | 0 |
| FBXO5 | 0 | 1182 | 0 | 1672 | 3708 | 0 | 3306 | 1432 |
| FBXO6 | 0 | 0 | 0 | 1104 | 728 | 0 | 0 | 0 |
| FBXO7 | 0 | 0 | 0 | 904 | 762 | 0 | 0 | 0 |
| FBXO8 | 0 | 0 | 0 | 0 | 0 | 0 | 786 | 0 |
| FBXO9 | 4821 | 7874 | 4565 | 2614 | 10996 | 0 | 13773 | 6792 |
| FBXW11 | 0 | 0 | 302 | 0 | 196 | 0 | 0 | 260 |
| FBXW2 | 0 | 114 | 1731 | 1130 | 296 | 0 | 360 | 228 |
| FBXW4 | 382 | 0 | 0 | 0 | 0 | 0 | 316 | 0 |
| FBXW5 | 0 | 118 | 3405 | 0 | 1850 | 2997 | 922 | 0 |
| FBXW7 | 19546 | 15020 | 14210 | 3891 | 38688 | 7172 | 19775 | 8123 |
| FBXW8 | 0 | 0 | 0 | 0 | 670 | 0 | 0 | 0 |
| FCER1G | 0 | 454 | 4485 | 0 | 5466 | 356 | 784 | 1804 |
| FCER2 | 0 | 370 | 0 | 0 | 0 | 0 | 0 | 0 |

| | | | | | | | | |
|---|---|---|---|---|---|---|---|---|
| FCF1 | 0 | 0 | 0 | 0 | 1325 | 0 | 0 | 0 |
| FCGBP | 7010 | 12441 | 10456 | 2344 | 204 | 0 | 0 | 0 |
| FCGR1A | 0 | 692 | 0 | 0 | 0 | 0 | 0 | 1474 |
| FCGR1B | 0 | 0 | 0 | 0 | 0 | 2104 | 0 | 0 |
| FCGR2A | 0 | 114 | 0 | 0 | 0 | 0 | 0 | 0 |
| FCGR2B | 166 | 338 | 0 | 0 | 0 | 0 | 0 | 0 |
| FCGR3A | 0 | 0 | 0 | 0 | 408 | 1482 | 0 | 0 |
| FCGRT | 1216 | 3762 | 16729 | 5875 | 2266 | 1830 | 2270 | 1170 |
| FCHO1 | 0 | 0 | 2296 | 0 | 676 | 0 | 0 | 0 |
| FCHO2 | 0 | 168 | 0 | 0 | 0 | 0 | 0 | 0 |
| FCHSD1 | 434 | 0 | 0 | 0 | 0 | 0 | 0 | 0 |
| FCHSD2 | 8284 | 2008 | 0 | 284 | 8579 | 5188 | 3424 | 3846 |
| FCMR | 0 | 0 | 0 | 0 | 814 | 0 | 0 | 0 |
| FCN3 | 0 | 0 | 0 | 0 | 0 | 2114 | 0 | 844 |
| FDCSP | 952 | 3234 | 0 | 874 | 0 | 0 | 0 | 1246 |
| FDFT1 | 507390 | 380207 | 3840419 | 869028 | 2643807 | 1067595 | 711850 | 1045063 |
| FDPS | 2867 | 352 | 4248 | 8389 | 8372 | 0 | 0 | 794 |
| FDX1 | 514 | 1244 | 2724 | 970 | 3438 | 1823 | 176 | 1172 |
| FDX2 | 0 | 152 | 0 | 0 | 0 | 0 | 0 | 0 |
| FEM1B | 0 | 0 | 150 | 118 | 0 | 0 | 0 | 0 |
| FEM1C | 0 | 0 | 0 | 0 | 436 | 0 | 0 | 0 |
| FEN1 | 13882 | 33037 | 111202 | 165426 | 325113 | 72607 | 117137 | 112534 |
| FER | 0 | 2582 | 0 | 1398 | 3926 | 2441 | 0 | 2312 |
| FER1L4 | 0 | 0 | 0 | 0 | 491 | 0 | 0 | 0 |
| FERMT1 | 0 | 0 | 0 | 810 | 326 | 0 | 0 | 976 |
| FERMT2 | 148 | 312 | 0 | 0 | 492 | 0 | 0 | 0 |
| FERMT3 | 666 | 0 | 0 | 0 | 760 | 0 | 0 | 0 |
| FES | 0 | 442 | 0 | 0 | 0 | 0 | 0 | 0 |
| FETUB | 0 | 1186 | 0 | 0 | 480 | 0 | 0 | 0 |
| FEZ1 | 0 | 0 | 0 | 0 | 896 | 0 | 0 | 0 |
| FEZ2 | 0 | 96 | 0 | 0 | 1630 | 0 | 0 | 1312 |
| FEZF1-AS1 | 0 | 0 | 0 | 0 | 900 | 0 | 0 | 0 |
| FFAR4 | 458 | 0 | 0 | 0 | 0 | 0 | 0 | 0 |
| FGD2 | 0 | 0 | 0 | 0 | 0 | 0 | 310 | 0 |
| FGD4 | 0 | 506 | 0 | 0 | 0 | 0 | 0 | 0 |
| FGD5 | 0 | 0 | 0 | 0 | 0 | 2028 | 0 | 0 |
| FGD5-AS1 | 0 | 0 | 0 | 0 | 930 | 0 | 126 | 0 |
| FGF12 | 0 | 574 | 0 | 0 | 0 | 0 | 0 | 0 |
| FGF18 | 0 | 0 | 870 | 1120 | 868 | 8645 | 13928 | 1050 |
| FGF7 | 0 | 0 | 0 | 0 | 0 | 0 | 0 | 738 |
| FGF7P3 | 0 | 0 | 0 | 0 | 662 | 0 | 0 | 0 |
| FGFBP1 | 778 | 12 | 0 | 0 | 986 | 0 | 0 | 0 |
| FGFR1 | 4688 | 5416 | 4340 | 1635 | 11968 | 9148 | 322 | 3080 |

| | | | | | | | | |
|---|---|---|---|---|---|---|---|---|
| FGFR1OP | 850 | 130 | 956 | 0 | 948 | 2440 | 766 | 4724 |
| FGFR1OP2 | 0 | 0 | 0 | 0 | 188 | 1570 | 338 | 0 |
| FGFR2 | 27177 | 9844 | 27593 | 482 | 26243 | 2698 | 0 | 4420 |
| FGFR3 | 776 | 484 | 432 | 348 | 0 | 0 | 1376 | 0 |
| FGFR4 | 430 | 0 | 0 | 0 | 0 | 0 | 294 | 0 |
| FGFRL1 | 0 | 0 | 0 | 0 | 0 | 0 | 380 | 0 |
| FGGY | 6902 | 22783 | 48837 | 57067 | 34586 | 31131 | 204903 | 16634 |
| FGL2 | 670 | 40 | 0 | 0 | 0 | 0 | 0 | 0 |
| FGR | 948 | 1244 | 1042 | 906 | 2546 | 3335 | 2083 | 9516 |
| FH | 1216 | 1616 | 821 | 1353 | 874 | 0 | 0 | 0 |
| FHAD1 | 216 | 0 | 0 | 0 | 1002 | 0 | 1788 | 0 |
| FHDC1 | 0 | 588 | 0 | 0 | 128 | 0 | 0 | 0 |
| FHIT | 88653 | 102707 | 171002 | 79254 | 298723 | 147773 | 193838 | 29289 |
| FHL1 | 0 | 0 | 802 | 0 | 0 | 0 | 0 | 0 |
| FHL2 | 0 | 1354 | 1194 | 3979 | 7210 | 0 | 800 | 4321 |
| FHL3 | 0 | 0 | 0 | 0 | 18 | 0 | 0 | 0 |
| FHOD1 | 0 | 818 | 0 | 0 | 132 | 0 | 0 | 756 |
| FIBP | 836 | 506 | 3064 | 2781 | 4257 | 0 | 0 | 1567 |
| FIG4 | 0 | 0 | 2444 | 1160 | 0 | 0 | 0 | 0 |
| FIGNL1 | 0 | 0 | 0 | 0 | 1178 | 0 | 280 | 0 |
| FIP1L1 | 88381 | 112398 | 161146 | 100200 | 292647 | 174427 | 329064 | 94169 |
| FIS1 | 1548 | 2406 | 3859 | 6218 | 6638 | 5079 | 6588 | 1881 |
| FITM1 | 0 | 270 | 0 | 0 | 0 | 0 | 0 | 0 |
| FITM2 | 0 | 42 | 0 | 0 | 0 | 0 | 550 | 0 |
| FKBP10 | 0 | 0 | 0 | 0 | 616 | 0 | 0 | 0 |
| FKBP11 | 0 | 262 | 2958 | 869 | 1800 | 2574 | 2802 | 878 |
| FKBP14 | 0 | 0 | 0 | 0 | 0 | 0 | 0 | 982 |
| FKBP15 | 2032 | 1498 | 3684 | 3028 | 6434 | 3690 | 5275 | 2368 |
| FKBP1A | 1790 | 1966 | 5116 | 1410 | 9354 | 2845 | 654 | 1354 |
| FKBP1A-SDCBP2 | 0 | 186 | 2690 | 0 | 1362 | 0 | 0 | 0 |
| FKBP1B | 0 | 0 | 0 | 1070 | 0 | 0 | 0 | 0 |
| FKBP2 | 526 | 0 | 2136 | 304 | 2110 | 1439 | 0 | 0 |
| FKBP3 | 806 | 1076 | 1486 | 1008 | 4989 | 0 | 890 | 0 |
| FKBP4 | 774 | 436 | 692 | 1669 | 562 | 1481 | 1282 | 0 |
| FKBP5 | 0 | 668 | 0 | 0 | 72 | 0 | 0 | 0 |
| FKBP7 | 0 | 0 | 0 | 932 | 0 | 0 | 0 | 0 |
| FKBP8 | 25212 | 33196 | 150352 | 44064 | 142477 | 59192 | 85193 | 48888 |
| FKBP9 | 1132 | 1198 | 0 | 628 | 1800 | 0 | 1976 | 0 |
| FKBP9P1 | 0 | 0 | 0 | 0 | 162 | 0 | 0 | 0 |
| FKBPL | 0 | 0 | 0 | 1684 | 0 | 0 | 0 | 0 |
| FLAD1 | 82 | 0 | 0 | 1168 | 1208 | 0 | 0 | 0 |
| FLCN | 0 | 80 | 1330 | 0 | 0 | 1450 | 0 | 1438 |
| FLI1 | 0 | 0 | 0 | 350 | 0 | 0 | 0 | 0 |

| | | | | | | | | |
|---|---|---|---|---|---|---|---|---|
| FLII | 1536 | 12 | 630 | 0 | 502 | 0 | 262 | 0 |
| FLJ20021 | 798 | 0 | 0 | 830 | 1224 | 0 | 254 | 0 |
| FLJ23867 | 0 | 458 | 0 | 0 | 940 | 0 | 0 | 0 |
| FLJ37453 | 0 | 0 | 1206 | 0 | 386 | 0 | 428 | 704 |
| FLJ42627 | 0 | 0 | 0 | 0 | 0 | 0 | 58 | 0 |
| FLJ44635 | 0 | 0 | 0 | 1296 | 2534 | 3878 | 0 | 0 |
| FLJ46875 | 0 | 0 | 0 | 0 | 0 | 0 | 198 | 0 |
| FLNA | 0 | 212 | 0 | 0 | 2612 | 874 | 0 | 882 |
| FLNB | 1108 | 2049 | 7788 | 2548 | 4540 | 3094 | 422 | 1128 |
| FLOT1 | 600 | 380 | 286 | 680 | 4498 | 0 | 2378 | 4464 |
| FLOT2 | 0 | 722 | 0 | 0 | 0 | 0 | 0 | 0 |
| FLT3LG | 5195 | 1922 | 6058 | 2468 | 8857 | 9118 | 5655 | 1959 |
| FLT4 | 0 | 0 | 0 | 0 | 0 | 0 | 0 | 1148 |
| FLVCR1 | 0 | 0 | 0 | 0 | 601 | 0 | 0 | 0 |
| FLVCR2 | 0 | 262 | 0 | 0 | 0 | 0 | 0 | 0 |
| FLYWCH1 | 1040 | 0 | 0 | 0 | 108 | 0 | 0 | 0 |
| FLYWCH2 | 0 | 0 | 0 | 2642 | 1048 | 0 | 0 | 0 |
| FMC1 | 752 | 1092 | 0 | 1380 | 2530 | 1305 | 582 | 0 |
| FMNL1 | 784 | 0 | 0 | 0 | 704 | 0 | 0 | 0 |
| FMNL2 | 0 | 0 | 0 | 0 | 350 | 0 | 0 | 0 |
| FMNL3 | 0 | 0 | 0 | 256 | 1290 | 0 | 0 | 0 |
| FMO4 | 13451 | 9893 | 37569 | 3003 | 8286 | 4835 | 19267 | 2392 |
| FMO5 | 0 | 0 | 988 | 134 | 492 | 0 | 0 | 0 |
| FMOD | 0 | 0 | 1257 | 0 | 0 | 0 | 0 | 0 |
| FMR1 | 0 | 0 | 0 | 0 | 1908 | 0 | 0 | 0 |
| FN1 | 0 | 4648 | 840 | 984 | 38217 | 45134 | 3280 | 33917 |
| FN3KRP | 0 | 0 | 742 | 0 | 800 | 0 | 0 | 0 |
| FNBP1 | 0 | 362 | 1052 | 0 | 432 | 0 | 396 | 0 |
| FNBP1L | 0 | 0 | 0 | 0 | 0 | 10 | 18 | 823 |
| FNBP4 | 0 | 202 | 0 | 0 | 510 | 0 | 0 | 0 |
| FNDC1 | 0 | 0 | 0 | 0 | 662 | 0 | 0 | 0 |
| FNDC3A | 0 | 176 | 0 | 0 | 0 | 0 | 0 | 0 |
| FNDC3B | 0 | 0 | 0 | 0 | 364 | 0 | 0 | 0 |
| FNIP1 | 0 | 0 | 0 | 0 | 0 | 0 | 0 | 1186 |
| FNTA | 0 | 462 | 4487 | 3821 | 4002 | 1346 | 3130 | 4127 |
| FNTB | 798 | 236 | 0 | 0 | 1861 | 2214 | 0 | 0 |
| FOCAD | 120 | 258 | 0 | 380 | 1984 | 0 | 200 | 1076 |
| FOLR1 | 0 | 3236 | 2216 | 19939 | 23826 | 154238 | 1567484 | 12753 |
| FOLR2 | 750 | 0 | 0 | 0 | 374 | 1186 | 0 | 0 |
| FOPNL | 234 | 412 | 0 | 746 | 1224 | 0 | 1600 | 0 |
| FOS | 6676 | 2144 | 16204 | 2733 | 2394 | 1586 | 0 | 2162 |
| FOSB | 224 | 0 | 0 | 0 | 668 | 0 | 0 | 0 |
| FOSL1 | 0 | 0 | 0 | 2108 | 667 | 0 | 0 | 0 |

| | | | | | | | | |
|---|---|---|---|---|---|---|---|---|
| FOSL2 | 1640 | 0 | 1345 | 0 | 0 | 0 | 2653 | 842 |
| FOXA1 | 0 | 304 | 0 | 0 | 0 | 0 | 0 | 0 |
| FOXA2 | 0 | 214 | 0 | 0 | 0 | 0 | 182 | 0 |
| FOXA3 | 0 | 328 | 1220 | 0 | 1238 | 0 | 0 | 0 |
| FOXC1 | 0 | 0 | 0 | 0 | 186 | 0 | 0 | 0 |
| FOXD1 | 0 | 0 | 0 | 0 | 534 | 0 | 0 | 0 |
| FOXD2 | 0 | 0 | 0 | 0 | 152 | 0 | 0 | 0 |
| FOXD2-AS1 | 0 | 0 | 0 | 0 | 927 | 0 | 0 | 0 |
| FOXF1 | 0 | 226 | 0 | 0 | 0 | 0 | 0 | 0 |
| FOXJ1 | 0 | 0 | 1102 | 604 | 0 | 0 | 0 | 0 |
| FOXJ3 | 336 | 0 | 1842 | 180 | 316 | 0 | 0 | 0 |
| FOXK1 | 1370 | 166 | 0 | 2920 | 4102 | 1840 | 1884 | 476 |
| FOXK2 | 566 | 0 | 0 | 0 | 0 | 0 | 440 | 0 |
| FOXN2 | 0 | 136 | 0 | 0 | 0 | 0 | 0 | 0 |
| FOXN3 | 0 | 0 | 0 | 0 | 278 | 0 | 0 | 0 |
| FOXO1 | 26 | 4364 | 8517 | 6234 | 8214 | 3264 | 10186 | 594 |
| FOXO3 | 542 | 1137 | 8370 | 4280 | 11678 | 3445 | 9862 | 3444 |
| FOXO3B | 0 | 362 | 0 | 0 | 0 | 0 | 166 | 0 |
| FOXP1 | 105000 | 117752 | 98451 | 48938 | 281945 | 44268 | 64498 | 28929 |
| FOXP4 | 0 | 0 | 0 | 0 | 0 | 0 | 921 | 0 |
| FOXP4-AS1 | 0 | 0 | 0 | 502 | 0 | 0 | 590 | 638 |
| FOXQ1 | 0 | 0 | 0 | 624 | 0 | 0 | 0 | 0 |
| FOXRED1 | 314 | 0 | 0 | 2334 | 710 | 0 | 0 | 0 |
| FOXRED2 | 0 | 413 | 0 | 0 | 304 | 0 | 0 | 0 |
| FOXS1 | 0 | 0 | 0 | 0 | 0 | 0 | 0 | 560 |
| FPGS | 0 | 0 | 0 | 494 | 626 | 0 | 0 | 0 |
| FPGT | 0 | 222 | 0 | 0 | 266 | 0 | 332 | 0 |
| FPR1 | 0 | 568 | 0 | 0 | 0 | 2112 | 0 | 0 |
| FRA10AC1 | 0 | 0 | 0 | 1192 | 0 | 0 | 606 | 1020 |
| FRAS1 | 0 | 426 | 0 | 0 | 1140 | 0 | 0 | 0 |
| FRAT2 | 0 | 0 | 0 | 0 | 296 | 0 | 328 | 158 |
| FREM1 | 0 | 0 | 0 | 164 | 0 | 0 | 250 | 0 |
| FRG1 | 0 | 1056 | 3809 | 3110 | 1486 | 0 | 174 | 1130 |
| FRG1BP | 0 | 0 | 0 | 0 | 1468 | 0 | 0 | 916 |
| FRG1FP | 0 | 668 | 0 | 0 | 0 | 0 | 0 | 0 |
| FRG1HP | 0 | 0 | 0 | 0 | 1041 | 0 | 0 | 0 |
| FRG1JP | 0 | 0 | 0 | 704 | 0 | 0 | 0 | 0 |
| FRK | 0 | 360 | 0 | 0 | 0 | 0 | 0 | 0 |
| FRMD1 | 0 | 0 | 0 | 0 | 0 | 0 | 894 | 0 |
| FRMD3 | 0 | 0 | 0 | 0 | 268 | 0 | 0 | 0 |
| FRMD4B | 5358 | 7574 | 12926 | 1274 | 30917 | 5912 | 6304 | 5726 |
| FRMD5 | 0 | 0 | 0 | 1078 | 2139 | 1016 | 1426 | 842 |
| FRMD6-AS1 | 0 | 0 | 0 | 0 | 0 | 0 | 364 | 0 |

| | | | | | | | | |
|---|---|---|---|---|---|---|---|---|
| FRMD7 | 0 | 267 | 0 | 0 | 248 | 0 | 0 | 0 |
| FRMD8 | 0 | 0 | 0 | 0 | 0 | 0 | 0 | 460 |
| FRS2 | 0 | 0 | 0 | 0 | 428 | 0 | 0 | 0 |
| FRY | 270 | 0 | 0 | 0 | 0 | 0 | 0 | 0 |
| FRYL | 30859 | 21692 | 21254 | 3980 | 8455 | 0 | 11425 | 0 |
| FRZB | 506 | 0 | 0 | 0 | 1214 | 0 | 0 | 0 |
| FSBP | 0 | 0 | 0 | 0 | 0 | 0 | 414 | 0 |
| FSD1L | 0 | 0 | 0 | 710 | 0 | 0 | 0 | 0 |
| FSD2 | 0 | 0 | 0 | 0 | 236 | 0 | 0 | 0 |
| FSTL1 | 0 | 0 | 0 | 161 | 122 | 0 | 298 | 0 |
| FSTL3 | 1042 | 0 | 0 | 0 | 996 | 0 | 883 | 766 |
| FSTL4 | 0 | 390 | 0 | 4432 | 0 | 0 | 488 | 0 |
| FTCDNL1 | 0 | 0 | 0 | 0 | 772 | 0 | 0 | 0 |
| FTH1 | 64457 | 50112 | 258448 | 128867 | 188074 | 51972 | 35652 | 92519 |
| FTL | 24157 | 23889 | 112510 | 146003 | 208232 | 93162 | 85006 | 83832 |
| FTO | 0 | 0 | 3035 | 0 | 232 | 0 | 0 | 0 |
| FTSJ1 | 0 | 0 | 194 | 990 | 1198 | 0 | 0 | 0 |
| FTSJ3 | 0 | 0 | 0 | 0 | 774 | 0 | 0 | 962 |
| FUBP1 | 91293 | 180051 | 261292 | 247904 | 412458 | 399636 | 512729 | 219999 |
| FUBP3 | 6812 | 9997 | 25303 | 14302 | 38467 | 25362 | 46702 | 7295 |
| FUCA1 | 548 | 444 | 2674 | 0 | 2172 | 0 | 466 | 0 |
| FUCA2 | 372 | 0 | 2376 | 2565 | 0 | 0 | 1456 | 4846 |
| FUNDC1 | 24783 | 45747 | 46697 | 45055 | 220829 | 40220 | 137742 | 36106 |
| FUNDC2 | 140 | 222 | 2250 | 4005 | 3210 | 2340 | 1540 | 2885 |
| FUOM | 0 | 0 | 1146 | 0 | 2124 | 0 | 1514 | 0 |
| FURIN | 0 | 0 | 593 | 0 | 0 | 0 | 330 | 0 |
| FUS | 62946 | 53788 | 128032 | 119863 | 297920 | 128057 | 275422 | 81978 |
| FUT2 | 586 | 0 | 0 | 0 | 0 | 0 | 0 | 0 |
| FUT3 | 1904 | 276 | 0 | 0 | 820 | 3170 | 1984 | 840 |
| FUT4 | 0 | 0 | 0 | 0 | 430 | 0 | 644 | 0 |
| FUT6 | 1340 | 594 | 1256 | 524 | 0 | 2220 | 266 | 0 |
| FUT8 | 0 | 0 | 1743 | 0 | 1238 | 0 | 0 | 0 |
| FUT8-AS1 | 0 | 0 | 0 | 0 | 358 | 0 | 0 | 0 |
| FXN | 2127 | 3990 | 3879 | 13264 | 26805 | 6959 | 14536 | 1894 |
| FXR1 | 0 | 698 | 0 | 0 | 0 | 0 | 378 | 563 |
| FXR2 | 2858 | 92 | 3154 | 1540 | 4951 | 56 | 2618 | 1748 |
| FXYD1 | 264 | 0 | 0 | 0 | 0 | 0 | 0 | 0 |
| FXYD3 | 52034 | 58112 | 160591 | 12283 | 33971 | 1022 | 11383 | 8540 |
| FXYD5 | 792 | 172 | 0 | 4362 | 5570 | 6020 | 1332 | 3064 |
| FYN | 2656 | 4344 | 7383 | 1728 | 13760 | 3910 | 522 | 16971 |
| FYTTD1 | 0 | 398 | 0 | 0 | 618 | 0 | 0 | 0 |
| FZD1 | 114 | 0 | 0 | 0 | 0 | 0 | 0 | 0 |
| FZD5 | 612 | 452 | 0 | 0 | 0 | 1650 | 0 | 0 |

| | | | | | | | | |
|---|---|---|---|---|---|---|---|---|
| FZD6 | 508 | 0 | 0 | 0 | 396 | 637 | 512 | 0 |
| FZD8 | 0 | 0 | 0 | 0 | 188 | 0 | 0 | 0 |
| FZR1 | 1064 | 1636 | 1824 | 0 | 9013 | 3918 | 1965 | 0 |
| G0S2 | 54 | 810 | 0 | 1068 | 754 | 3222 | 0 | 805 |
| G2E3 | 0 | 94 | 0 | 0 | 328 | 0 | 166 | 0 |
| G3BP1 | 0 | 828 | 1204 | 424 | 2238 | 784 | 410 | 0 |
| G3BP2 | 0 | 0 | 0 | 0 | 897 | 0 | 0 | 512 |
| G6PC3 | 0 | 0 | 990 | 0 | 1384 | 0 | 22 | 0 |
| G6PD | 416 | 0 | 0 | 0 | 0 | 0 | 702 | 1950 |
| GAA | 500 | 288 | 0 | 0 | 1644 | 0 | 0 | 0 |
| GAB2 | 0 | 0 | 1665 | 0 | 0 | 0 | 0 | 0 |
| GABARAP | 4492 | 6400 | 11848 | 4736 | 18716 | 11231 | 7418 | 5865 |
| GABARAPL1 | 0 | 0 | 1938 | 0 | 572 | 0 | 150 | 0 |
| GABARAPL2 | 34711 | 40010 | 47569 | 46359 | 88244 | 63641 | 142192 | 37727 |
| GABPB1 | 0 | 0 | 0 | 54 | 336 | 1162 | 0 | 0 |
| GABPB1-AS1 | 0 | 294 | 0 | 0 | 0 | 0 | 0 | 0 |
| GABPB1-IT1 | 0 | 0 | 1616 | 0 | 768 | 0 | 330 | 0 |
| GABRE | 0 | 0 | 0 | 891 | 0 | 0 | 0 | 540 |
| GADD45A | 930 | 838 | 3376 | 1032 | 1176 | 0 | 0 | 698 |
| GADD45B | 0 | 0 | 0 | 0 | 2778 | 2910 | 0 | 0 |
| GADD45G | 850 | 902 | 3770 | 0 | 1006 | 2590 | 580 | 1156 |
| GADD45GIP1 | 4574 | 9254 | 16660 | 6620 | 42614 | 19728 | 6891 | 12980 |
| GADL1 | 0 | 0 | 1031 | 0 | 0 | 0 | 0 | 0 |
| GAK | 0 | 0 | 0 | 0 | 1270 | 0 | 870 | 0 |
| GAL3ST1 | 0 | 0 | 2455 | 0 | 0 | 0 | 0 | 0 |
| GAL3ST2 | 0 | 262 | 0 | 0 | 0 | 0 | 0 | 0 |
| GALC | 0 | 242 | 0 | 0 | 843 | 0 | 0 | 540 |
| GALE | 585 | 456 | 5069 | 0 | 212 | 304 | 2427 | 1173 |
| GALK1 | 0 | 0 | 0 | 0 | 534 | 0 | 0 | 0 |
| GALM | 0 | 0 | 0 | 0 | 1330 | 0 | 0 | 42 |
| GALNS | 482 | 0 | 0 | 0 | 0 | 0 | 0 | 0 |
| GALNT1 | 0 | 1494 | 776 | 148 | 4393 | 0 | 0 | 0 |
| GALNT10 | 0 | 0 | 0 | 1714 | 98 | 0 | 0 | 0 |
| GALNT11 | 620 | 0 | 0 | 0 | 304 | 0 | 0 | 0 |
| GALNT12 | 0 | 726 | 0 | 0 | 198 | 0 | 0 | 0 |
| GALNT14 | 1076 | 316 | 0 | 0 | 0 | 0 | 0 | 0 |
| GALNT15 | 0 | 54 | 0 | 0 | 0 | 0 | 0 | 0 |
| GALNT18 | 794 | 1770 | 733 | 1148 | 1176 | 2306 | 0 | 3199 |
| GALNT2 | 0 | 0 | 0 | 0 | 0 | 0 | 502 | 0 |
| GALNT3 | 172 | 382 | 698 | 2094 | 0 | 0 | 90 | 765 |
| GALNT5 | 0 | 0 | 0 | 266 | 0 | 0 | 0 | 0 |
| GALNT6 | 0 | 0 | 0 | 0 | 1166 | 0 | 0 | 0 |
| GALT | 0 | 240 | 0 | 0 | 0 | 0 | 0 | 1066 |

| | | | | | | | | |
|---|---|---|---|---|---|---|---|---|
| GANAB | 1102 | 0 | 0 | 2158 | 698 | 0 | 578 | 0 |
| GAPDH | 2559375 | 6016409 | 5811438 | 12475894 | 31391588 | 5747788 | 17171340 | 9618554 |
| GAPLINC | 0 | 614 | 0 | 0 | 0 | 0 | 0 | 0 |
| GAPVD1 | 6706 | 7082 | 951 | 5782 | 20485 | 5236 | 22538 | 4564 |
| GAR1 | 0 | 390 | 1956 | 1261 | 890 | 0 | 432 | 1012 |
| GARNL3 | 0 | 0 | 0 | 0 | 2276 | 0 | 706 | 594 |
| GARS | 0 | 0 | 0 | 6082 | 3697 | 0 | 1542 | 5132 |
| GART | 322 | 256 | 982 | 606 | 530 | 0 | 292 | 1326 |
| GAS2 | 0 | 0 | 0 | 0 | 0 | 0 | 0 | 996 |
| GAS2L1 | 0 | 0 | 0 | 0 | 152 | 0 | 0 | 0 |
| GAS2L3 | 3217 | 4473 | 8544 | 8449 | 15588 | 7280 | 12056 | 5769 |
| GAS5 | 3318 | 7016 | 8048 | 15216 | 18966 | 18698 | 24217 | 17432 |
| GAS6 | 1610 | 3978 | 19270 | 11613 | 6274 | 18496 | 9046 | 1180 |
| GAS8 | 0 | 0 | 0 | 881 | 588 | 4258 | 178 | 0 |
| GATA1 | 0 | 0 | 0 | 0 | 776 | 0 | 0 | 0 |
| GATA2 | 692 | 0 | 0 | 0 | 616 | 0 | 226 | 0 |
| GATA3 | 886 | 204 | 0 | 0 | 200 | 0 | 2336 | 602 |
| GATA3-AS1 | 0 | 0 | 0 | 0 | 264 | 0 | 0 | 0 |
| GATA6 | 568 | 0 | 0 | 0 | 280 | 0 | 74 | 0 |
| GATA6-AS1 | 401 | 0 | 0 | 0 | 480 | 0 | 114 | 0 |
| GATAD1 | 0 | 340 | 0 | 0 | 0 | 0 | 0 | 1108 |
| GATAD2A | 128 | 0 | 442 | 0 | 656 | 0 | 0 | 418 |
| GATAD2B | 0 | 36 | 1768 | 0 | 966 | 2754 | 1577 | 2584 |
| GATB | 0 | 0 | 1948 | 0 | 0 | 0 | 0 | 0 |
| GATC | 0 | 100 | 0 | 0 | 752 | 0 | 0 | 0 |
| GATD1 | 0 | 612 | 0 | 0 | 420 | 0 | 242 | 857 |
| GATD3A | 0 | 0 | 782 | 0 | 635 | 0 | 122 | 0 |
| GATD3B | 1295 | 566 | 0 | 0 | 0 | 0 | 0 | 0 |
| GATM | 0 | 348 | 0 | 0 | 0 | 0 | 0 | 0 |
| GBA | 602 | 404 | 0 | 0 | 1404 | 0 | 1066 | 808 |
| GBA2 | 0 | 762 | 1095 | 0 | 799 | 0 | 0 | 0 |
| GBAT2 | 0 | 0 | 0 | 0 | 0 | 0 | 0 | 62 |
| GBE1 | 0 | 0 | 0 | 0 | 1226 | 272 | 0 | 0 |
| GBF1 | 6172 | 4394 | 15894 | 3388 | 13236 | 7573 | 16336 | 4736 |
| GBGT1 | 0 | 230 | 0 | 0 | 0 | 0 | 0 | 0 |
| GBP1 | 360 | 512 | 0 | 0 | 0 | 0 | 0 | 0 |
| GBP1P1 | 0 | 0 | 0 | 0 | 0 | 2380 | 0 | 0 |
| GBP2 | 0 | 1262 | 3072 | 0 | 0 | 0 | 1026 | 0 |
| GBP3 | 0 | 0 | 858 | 0 | 453 | 0 | 0 | 0 |
| GCA | 0 | 354 | 0 | 0 | 942 | 1026 | 757 | 0 |
| GCAT | 0 | 0 | 0 | 0 | 0 | 0 | 730 | 0 |
| GCC2 | 306 | 0 | 0 | 0 | 0 | 0 | 304 | 0 |
| GCDH | 0 | 0 | 0 | 0 | 1402 | 0 | 24 | 0 |

| | | | | | | | | |
|---|---|---|---|---|---|---|---|---|
| GCFC2 | 0 | 124 | 0 | 0 | 406 | 0 | 1018 | 0 |
| GCG | 0 | 3632 | 1646 | 0 | 0 | 0 | 0 | 0 |
| GCH1 | 0 | 0 | 1130 | 0 | 0 | 0 | 0 | 0 |
| GCHFR | 0 | 0 | 0 | 2270 | 30 | 0 | 2346 | 0 |
| GCLC | 592 | 0 | 0 | 0 | 142 | 0 | 0 | 0 |
| GCLM | 0 | 240 | 0 | 0 | 0 | 0 | 0 | 1351 |
| GCN1 | 741 | 0 | 0 | 0 | 318 | 0 | 0 | 0 |
| GCNT1 | 0 | 948 | 0 | 0 | 746 | 0 | 400 | 0 |
| GCNT3 | 0 | 2294 | 9073 | 3912 | 1000 | 0 | 0 | 340 |
| GCSH | 0 | 1048 | 0 | 3296 | 2458 | 0 | 1353 | 732 |
| GCSHP3 | 0 | 0 | 0 | 0 | 350 | 0 | 0 | 0 |
| GDA | 0 | 254 | 0 | 0 | 142 | 0 | 0 | 0 |
| GDAP2 | 0 | 0 | 0 | 0 | 0 | 0 | 702 | 0 |
| GDE1 | 0 | 388 | 0 | 0 | 670 | 2018 | 0 | 0 |
| GDF11 | 0 | 1756 | 0 | 792 | 1900 | 734 | 5032 | 1220 |
| GDF15 | 0 | 419 | 3371 | 5278 | 4455 | 2656 | 1113 | 992 |
| GDI1 | 542 | 0 | 0 | 2793 | 374 | 1275 | 85 | 768 |
| GDI2 | 0 | 1272 | 3696 | 992 | 4621 | 0 | 2364 | 646 |
| GDPD1 | 0 | 276 | 0 | 0 | 0 | 0 | 0 | 0 |
| GDPD3 | 0 | 392 | 674 | 1192 | 54 | 0 | 0 | 0 |
| GDPD5 | 0 | 0 | 0 | 0 | 0 | 0 | 1340 | 608 |
| GEM | 0 | 0 | 1088 | 0 | 448 | 0 | 730 | 0 |
| GEMIN2 | 0 | 0 | 1927 | 0 | 3090 | 0 | 0 | 0 |
| GEMIN4 | 0 | 308 | 0 | 0 | 0 | 0 | 0 | 0 |
| GEMIN5 | 0 | 0 | 0 | 0 | 0 | 0 | 52 | 0 |
| GEMIN6 | 0 | 1576 | 0 | 0 | 1212 | 1838 | 0 | 440 |
| GEMIN7 | 1694 | 0 | 4684 | 0 | 1162 | 0 | 0 | 4814 |
| GEMIN8 | 198 | 0 | 0 | 0 | 0 | 0 | 0 | 0 |
| GEMIN8P4 | 0 | 342 | 0 | 1026 | 0 | 0 | 0 | 0 |
| GET4 | 238 | 418 | 0 | 418 | 600 | 0 | 542 | 816 |
| GFER | 2490 | 448 | 0 | 0 | 0 | 0 | 238 | 0 |
| GFM1 | 0 | 0 | 4387 | 0 | 602 | 0 | 0 | 0 |
| GFM2 | 0 | 0 | 0 | 1310 | 0 | 0 | 0 | 0 |
| GFOD2 | 1582 | 1640 | 7371 | 2558 | 4720 | 5106 | 4481 | 2570 |
| GFPT1 | 61998 | 159413 | 105067 | 262782 | 576300 | 107541 | 448441 | 156388 |
| GFRA1 | 0 | 0 | 752 | 0 | 0 | 0 | 0 | 0 |
| GGA1 | 0 | 416 | 0 | 1134 | 86 | 0 | 0 | 0 |
| GGA2 | 688 | 0 | 288 | 0 | 470 | 0 | 712 | 0 |
| GGA3 | 0 | 0 | 627 | 0 | 222 | 0 | 0 | 0 |
| GGACT | 0 | 376 | 0 | 0 | 316 | 0 | 0 | 0 |
| GGCT | 0 | 0 | 5202 | 3954 | 8018 | 1357 | 1216 | 2700 |
| GGCX | 0 | 0 | 0 | 0 | 548 | 1795 | 0 | 0 |
| GGH | 1338 | 1894 | 3002 | 11281 | 1928 | 8184 | 7852 | 2606 |

| | | | | | | | | |
|---|---|---|---|---|---|---|---|---|
| GGNBP2 | 1024 | 678 | 3227 | 0 | 702 | 0 | 2614 | 987 |
| GGPS1 | 0 | 0 | 0 | 0 | 0 | 0 | 336 | 1865 |
| GGT1 | 0 | 270 | 0 | 0 | 0 | 0 | 1066 | 0 |
| GGT6 | 1240 | 0 | 0 | 1766 | 548 | 0 | 0 | 0 |
| GGTA1P | 0 | 0 | 164 | 0 | 0 | 0 | 0 | 0 |
| GHDC | 0 | 0 | 0 | 1386 | 0 | 0 | 214 | 0 |
| GHITM | 1830 | 2108 | 10957 | 6213 | 2332 | 7631 | 992 | 1881 |
| GID4 | 50 | 0 | 3097 | 0 | 3562 | 0 | 1163 | 842 |
| GID8 | 464 | 1274 | 2576 | 2890 | 3390 | 514 | 2089 | 1921 |
| GIGYF1 | 306 | 0 | 0 | 0 | 498 | 0 | 570 | 0 |
| GIGYF2 | 302 | 0 | 0 | 380 | 320 | 0 | 0 | 0 |
| GIMAP1 | 0 | 274 | 0 | 0 | 0 | 2118 | 0 | 0 |
| GIMAP2 | 0 | 0 | 0 | 0 | 358 | 0 | 0 | 0 |
| GIMAP4 | 0 | 0 | 0 | 0 | 496 | 0 | 0 | 1510 |
| GIMAP5 | 0 | 0 | 2156 | 0 | 0 | 0 | 0 | 0 |
| GIMAP7 | 0 | 596 | 0 | 0 | 0 | 0 | 696 | 0 |
| GIN1 | 0 | 0 | 0 | 0 | 428 | 0 | 0 | 0 |
| GINM1 | 0 | 270 | 2860 | 0 | 2442 | 174 | 0 | 0 |
| GINS2 | 0 | 0 | 0 | 1470 | 1454 | 0 | 0 | 0 |
| GINS3 | 0 | 0 | 0 | 0 | 1022 | 0 | 0 | 0 |
| GINS4 | 0 | 0 | 0 | 0 | 0 | 0 | 0 | 1106 |
| GIPC1 | 1444 | 876 | 3110 | 3509 | 6566 | 0 | 0 | 870 |
| GIPC2 | 0 | 0 | 660 | 810 | 762 | 0 | 0 | 922 |
| GIT1 | 566 | 0 | 0 | 0 | 0 | 0 | 0 | 0 |
| GIT2 | 0 | 0 | 0 | 1571 | 0 | 0 | 0 | 0 |
| GJA1 | 106 | 0 | 0 | 0 | 252 | 0 | 0 | 0 |
| GJA4 | 0 | 0 | 0 | 1214 | 0 | 0 | 0 | 0 |
| GJA9-MYCBP | 198 | 62 | 1618 | 0 | 0 | 0 | 12 | 0 |
| GJB1 | 0 | 610 | 1564 | 779 | 1000 | 0 | 434 | 0 |
| GJB2 | 556 | 0 | 0 | 0 | 534 | 1678 | 0 | 0 |
| GJB3 | 0 | 0 | 0 | 472 | 0 | 0 | 0 | 0 |
| GJB4 | 0 | 0 | 0 | 0 | 912 | 0 | 0 | 0 |
| GJC1 | 0 | 0 | 0 | 1264 | 0 | 0 | 0 | 0 |
| GJC2 | 0 | 0 | 0 | 0 | 0 | 0 | 656 | 596 |
| GJC3 | 1228 | 0 | 0 | 0 | 0 | 0 | 23389 | 0 |
| GJD3 | 0 | 0 | 0 | 0 | 298 | 0 | 0 | 0 |
| GK | 0 | 0 | 472 | 346 | 222 | 856 | 0 | 0 |
| GKAP1 | 0 | 0 | 0 | 0 | 1518 | 0 | 0 | 0 |
| GLA | 918 | 0 | 0 | 0 | 722 | 0 | 896 | 0 |
| GLB1 | 39498 | 66984 | 133826 | 120981 | 263834 | 112058 | 228842 | 59727 |
| GLB1L | 0 | 0 | 0 | 0 | 0 | 2140 | 0 | 0 |
| GLCCI1 | 0 | 252 | 1350 | 0 | 220 | 0 | 0 | 0 |
| GLCE | 0 | 264 | 0 | 0 | 604 | 0 | 108 | 0 |

| | | | | | | | | |
|---|---|---|---|---|---|---|---|---|
| GLDC | 422 | 0 | 590 | 0 | 760 | 2459 | 0 | 0 |
| GLE1 | 184 | 0 | 878 | 0 | 844 | 0 | 0 | 0 |
| GLG1 | 0 | 0 | 0 | 1368 | 601 | 1794 | 0 | 0 |
| GLI1 | 570 | 0 | 0 | 0 | 0 | 0 | 0 | 0 |
| GLI3 | 0 | 0 | 0 | 0 | 138 | 0 | 0 | 0 |
| GLI4 | 0 | 0 | 0 | 0 | 256 | 1636 | 970 | 0 |
| GLIDR | 0 | 214 | 1258 | 892 | 700 | 0 | 0 | 0 |
| GLIPR1 | 0 | 0 | 0 | 0 | 0 | 0 | 242 | 1352 |
| GLIPR2 | 0 | 1458 | 1448 | 0 | 0 | 0 | 0 | 0 |
| GLIS1 | 0 | 198 | 0 | 0 | 0 | 0 | 0 | 0 |
| GLIS2 | 0 | 0 | 0 | 0 | 0 | 0 | 1740 | 676 |
| GLIS3 | 1116 | 354 | 0 | 0 | 0 | 0 | 0 | 0 |
| GLMN | 0 | 0 | 0 | 0 | 0 | 0 | 502 | 0 |
| GLMP | 627 | 318 | 0 | 1233 | 0 | 0 | 0 | 2046 |
| GLO1 | 0 | 1411 | 0 | 1480 | 4671 | 0 | 2086 | 652 |
| GLOD4 | 88104 | 172853 | 328610 | 118784 | 347423 | 161723 | 233752 | 107166 |
| GLOD5 | 0 | 414 | 0 | 0 | 670 | 0 | 0 | 0 |
| GLRX | 1648 | 498 | 6451 | 4029 | 1101 | 2627 | 542 | 1358 |
| GLRX2 | 646 | 504 | 1292 | 1492 | 1616 | 0 | 0 | 896 |
| GLRX3 | 1692 | 1096 | 4320 | 3909 | 6588 | 7196 | 1234 | 2123 |
| GLRX5 | 1416 | 1064 | 3862 | 0 | 4090 | 0 | 523 | 782 |
| GLS | 0 | 0 | 0 | 0 | 0 | 198 | 136 | 0 |
| GLT1D1 | 0 | 0 | 0 | 0 | 0 | 30 | 0 | 0 |
| GLT8D1 | 114 | 3780 | 552 | 0 | 2146 | 0 | 3976 | 1108 |
| GLT8D2 | 534 | 0 | 0 | 0 | 0 | 0 | 0 | 0 |
| GLTP | 342 | 0 | 824 | 0 | 36 | 1959 | 1065 | 0 |
| GLUD1 | 738 | 316 | 1167 | 0 | 310 | 208 | 464 | 0 |
| GLUL | 0 | 1634 | 2196 | 986 | 2925 | 1078 | 4066 | 740 |
| GLYCTK | 0 | 0 | 0 | 0 | 1613 | 0 | 0 | 0 |
| GLYR1 | 18454 | 17319 | 19356 | 25524 | 48158 | 16056 | 63105 | 11181 |
| GM2A | 0 | 800 | 0 | 0 | 0 | 0 | 0 | 666 |
| GMCL1 | 1010 | 2998 | 17609 | 2508 | 14892 | 2841 | 14880 | 357 |
| GMDS | 54174 | 75338 | 156976 | 198464 | 425548 | 101254 | 100514 | 141494 |
| GMDS-DT | 0 | 0 | 0 | 0 | 158 | 0 | 0 | 0 |
| GMFB | 0 | 682 | 0 | 0 | 0 | 0 | 0 | 0 |
| GMFG | 0 | 0 | 2094 | 0 | 326 | 4072 | 750 | 755 |
| GMIP | 890 | 0 | 0 | 0 | 0 | 0 | 0 | 0 |
| GMNN | 989 | 1806 | 3726 | 3817 | 3469 | 0 | 504 | 0 |
| GMPPA | 432 | 484 | 0 | 1933 | 162 | 0 | 2739 | 624 |
| GMPPB | 884 | 1342 | 54 | 1324 | 1434 | 0 | 1717 | 0 |
| GMPR2 | 0 | 874 | 1775 | 2351 | 860 | 0 | 0 | 762 |
| GMPS | 0 | 122 | 0 | 1348 | 562 | 0 | 100 | 0 |
| GNA11 | 8670 | 392 | 50353 | 0 | 4761 | 2435 | 4272 | 1500 |

| | | | | | | | | |
|---|---|---|---|---|---|---|---|---|
| GNA13 | 0 | 0 | 0 | 0 | 308 | 0 | 0 | 0 |
| GNA14 | 0 | 310 | 0 | 0 | 0 | 0 | 0 | 0 |
| GNAI2 | 0 | 0 | 0 | 0 | 424 | 0 | 1627 | 638 |
| GNAI3 | 3130 | 5228 | 6959 | 9967 | 16860 | 8672 | 17289 | 5818 |
| GNAO1 | 0 | 0 | 0 | 0 | 354 | 0 | 0 | 0 |
| GNAQ | 16756 | 7785 | 31284 | 3813 | 38477 | 9633 | 8396 | 6356 |
| GNAS | 399862 | 376022 | 832369 | 721857 | 1978578 | 912112 | 2679659 | 577971 |
| GNB1 | 440 | 426 | 2904 | 0 | 4557 | 926 | 250 | 2125 |
| GNB1L | 0 | 0 | 0 | 1110 | 0 | 0 | 0 | 688 |
| GNB2 | 0 | 350 | 1950 | 1450 | 1846 | 2427 | 678 | 0 |
| GNE | 0 | 0 | 0 | 476 | 956 | 0 | 1832 | 0 |
| GNG10 | 0 | 2236 | 3764 | 2110 | 4276 | 0 | 442 | 0 |
| GNG11 | 1091 | 1614 | 1230 | 2993 | 656 | 5584 | 0 | 875 |
| GNG12 | 104832 | 98852 | 746564 | 66764 | 178990 | 66365 | 201620 | 39485 |
| GNG2 | 0 | 0 | 0 | 0 | 0 | 0 | 410 | 0 |
| GNG4 | 0 | 0 | 0 | 1736 | 0 | 0 | 0 | 0 |
| GNG5 | 4484 | 8465 | 12392 | 8510 | 10613 | 3402 | 4318 | 1818 |
| GNG7 | 0 | 0 | 0 | 0 | 0 | 0 | 0 | 1061 |
| GNL2 | 0 | 0 | 0 | 0 | 0 | 0 | 402 | 0 |
| GNL3 | 58789 | 322672 | 123642 | 520797 | 1838305 | 516014 | 1326603 | 380986 |
| GNL3L | 0 | 0 | 0 | 0 | 724 | 0 | 290 | 0 |
| GNLY | 698 | 0 | 0 | 0 | 0 | 0 | 0 | 0 |
| GNPAT | 0 | 510 | 0 | 0 | 1576 | 0 | 0 | 0 |
| GNPDA1 | 0 | 0 | 0 | 3306 | 76 | 0 | 795 | 0 |
| GNPNAT1 | 0 | 0 | 0 | 0 | 544 | 0 | 0 | 0 |
| GNPTAB | 0 | 0 | 0 | 0 | 256 | 0 | 0 | 0 |
| GNPTG | 0 | 566 | 1967 | 2074 | 1984 | 0 | 2744 | 0 |
| GNS | 6162 | 11929 | 12637 | 17370 | 30082 | 34536 | 59663 | 28089 |
| GOLGA1 | 472 | 0 | 0 | 0 | 0 | 0 | 0 | 0 |
| GOLGA2 | 346 | 0 | 0 | 0 | 0 | 0 | 0 | 0 |
| GOLGA2P10 | 1727 | 0 | 0 | 0 | 386 | 0 | 0 | 196 |
| GOLGA2P5 | 780 | 0 | 0 | 0 | 0 | 0 | 0 | 0 |
| GOLGA2P7 | 462 | 0 | 0 | 0 | 202 | 0 | 0 | 672 |
| GOLGA3 | 0 | 0 | 0 | 0 | 414 | 0 | 0 | 0 |
| GOLGA4 | 63919 | 64027 | 61371 | 74943 | 260513 | 73859 | 156863 | 40142 |
| GOLGA5 | 0 | 373 | 0 | 0 | 324 | 0 | 136 | 519 |
| GOLGA6L9 | 0 | 0 | 0 | 0 | 122 | 0 | 0 | 0 |
| GOLGA7 | 0 | 60 | 0 | 0 | 3190 | 0 | 0 | 2820 |
| GOLGA8B | 482 | 0 | 0 | 0 | 0 | 210 | 332 | 0 |
| GOLGA8J | 408 | 0 | 0 | 0 | 0 | 0 | 0 | 0 |
| GOLGA8O | 524 | 0 | 0 | 0 | 0 | 0 | 0 | 0 |
| GOLGB1 | 622 | 704 | 0 | 0 | 282 | 1056 | 0 | 1606 |
| GOLIM4 | 79404 | 56173 | 49305 | 23154 | 41435 | 41764 | 92189 | 14148 |

| | | | | | | | | |
|---|---|---|---|---|---|---|---|---|
| GOLM1 | 0 | 556 | 4530 | 3294 | 6238 | 0 | 452 | 344 |
| GOLPH3 | 0 | 626 | 1498 | 924 | 2175 | 0 | 682 | 408 |
| GOLPH3L | 60232 | 69670 | 163085 | 51748 | 250922 | 38381 | 111924 | 62812 |
| GOLT1A | 0 | 0 | 0 | 554 | 528 | 0 | 958 | 1718 |
| GOLT1B | 20453 | 52021 | 50386 | 75952 | 147096 | 68743 | 95854 | 112158 |
| GON4L | 0 | 0 | 0 | 0 | 1478 | 0 | 0 | 0 |
| GON7 | 0 | 0 | 1606 | 0 | 0 | 0 | 0 | 0 |
| GOPC | 788 | 496 | 1814 | 0 | 68 | 0 | 0 | 0 |
| GORAB | 0 | 0 | 0 | 0 | 162 | 1543 | 0 | 0 |
| GORASP1 | 0 | 0 | 0 | 0 | 0 | 0 | 314 | 858 |
| GORASP2 | 676 | 286 | 1944 | 172 | 416 | 1778 | 586 | 0 |
| GOSR1 | 632 | 300 | 1329 | 0 | 608 | 0 | 512 | 1568 |
| GOSR2 | 0 | 0 | 0 | 776 | 980 | 1750 | 0 | 0 |
| GOT1 | 0 | 0 | 0 | 2896 | 1301 | 0 | 304 | 0 |
| GOT2 | 338 | 746 | 0 | 0 | 220 | 0 | 0 | 0 |
| GP1BA | 816 | 0 | 0 | 0 | 0 | 0 | 0 | 0 |
| GP6 | 0 | 0 | 0 | 0 | 108 | 0 | 0 | 0 |
| GPA33 | 2104 | 864 | 13105 | 438 | 790 | 0 | 598 | 0 |
| GPAA1 | 2386 | 372 | 0 | 876 | 2288 | 0 | 0 | 868 |
| GPALPP1 | 0 | 0 | 0 | 0 | 214 | 0 | 310 | 0 |
| GPANK1 | 0 | 0 | 0 | 0 | 0 | 0 | 0 | 248 |
| GPAT2 | 0 | 314 | 1408 | 0 | 0 | 0 | 0 | 0 |
| GPAT3 | 0 | 0 | 1442 | 0 | 0 | 0 | 0 | 0 |
| GPAT4 | 0 | 0 | 0 | 294 | 0 | 0 | 124 | 0 |
| GPATCH1 | 12057 | 8338 | 28903 | 13215 | 51140 | 21026 | 23126 | 11985 |
| GPATCH11 | 0 | 0 | 0 | 287 | 0 | 0 | 0 | 0 |
| GPATCH2 | 0 | 288 | 0 | 0 | 0 | 0 | 0 | 0 |
| GPATCH2L | 0 | 0 | 0 | 0 | 0 | 0 | 418 | 0 |
| GPATCH4 | 0 | 0 | 0 | 1270 | 498 | 0 | 854 | 0 |
| GPATCH8 | 0 | 0 | 0 | 0 | 0 | 0 | 1030 | 0 |
| GPBP1 | 1246 | 0 | 0 | 1182 | 472 | 1921 | 0 | 0 |
| GPBP1L1 | 0 | 0 | 22 | 0 | 244 | 1851 | 596 | 0 |
| GPC1 | 0 | 0 | 0 | 0 | 0 | 0 | 0 | 468 |
| GPC3 | 1312 | 2418 | 21159 | 5238 | 120242 | 8226 | 5740 | 1230 |
| GPC4 | 770 | 1382 | 598 | 846 | 12350 | 2706 | 2600 | 136 |
| GPC5 | 314 | 0 | 2788 | 0 | 0 | 0 | 0 | 0 |
| GPC6 | 7138 | 1716 | 7219 | 0 | 580 | 2836 | 0 | 1282 |
| GPCPD1 | 0 | 54 | 0 | 0 | 1606 | 0 | 0 | 0 |
| GPD1L | 6700 | 9742 | 18247 | 3082 | 14703 | 1426 | 16593 | 2108 |
| GPER1 | 0 | 0 | 1382 | 0 | 0 | 0 | 0 | 0 |
| GPHN | 7721 | 7643 | 10698 | 11894 | 46439 | 8752 | 38879 | 6644 |
| GPI | 94 | 0 | 2996 | 1326 | 6534 | 0 | 1876 | 1034 |
| GPLD1 | 0 | 498 | 0 | 0 | 0 | 0 | 0 | 0 |

| | | | | | | | | |
|---|---|---|---|---|---|---|---|---|
| GPN1 | 0 | 402 | 0 | 1244 | 1676 | 0 | 456 | 0 |
| GPN2 | 0 | 0 | 0 | 0 | 1594 | 0 | 0 | 720 |
| GPN3 | 0 | 0 | 0 | 1199 | 0 | 0 | 0 | 0 |
| GPNMB | 0 | 0 | 1517 | 0 | 0 | 0 | 0 | 1966 |
| GPR107 | 0 | 0 | 0 | 0 | 16 | 0 | 0 | 0 |
| GPR108 | 0 | 0 | 1814 | 220 | 778 | 0 | 602 | 0 |
| GPR132 | 0 | 0 | 0 | 0 | 0 | 0 | 374 | 0 |
| GPR137B | 0 | 0 | 0 | 0 | 578 | 0 | 0 | 0 |
| GPR146 | 0 | 398 | 712 | 0 | 884 | 0 | 0 | 2608 |
| GPR150 | 0 | 0 | 0 | 0 | 0 | 0 | 752 | 0 |
| GPR153 | 0 | 0 | 0 | 22 | 0 | 0 | 0 | 0 |
| GPR160 | 408 | 0 | 2859 | 1125 | 256 | 1036 | 3466 | 534 |
| GPR161 | 0 | 0 | 0 | 0 | 1016 | 0 | 0 | 0 |
| GPR180 | 0 | 0 | 0 | 0 | 150 | 1215 | 0 | 0 |
| GPR183 | 0 | 160 | 0 | 0 | 0 | 0 | 0 | 68 |
| GPR19 | 0 | 0 | 0 | 0 | 0 | 0 | 0 | 1028 |
| GPR27 | 0 | 0 | 0 | 588 | 0 | 0 | 0 | 0 |
| GPR34 | 0 | 0 | 0 | 0 | 0 | 0 | 366 | 0 |
| GPR35 | 14238 | 7289 | 19795 | 22307 | 36739 | 24530 | 54914 | 14650 |
| GPR39 | 0 | 0 | 0 | 0 | 54 | 0 | 0 | 0 |
| GPR89B | 0 | 0 | 0 | 0 | 0 | 0 | 138 | 514 |
| GPRC5A | 0 | 868 | 1712 | 970 | 1362 | 3364 | 0 | 1349 |
| GPRC5C | 0 | 0 | 476 | 0 | 1312 | 0 | 714 | 1026 |
| GPRIN2 | 108 | 0 | 0 | 0 | 390 | 0 | 1094 | 0 |
| GPRIN3 | 0 | 0 | 0 | 0 | 0 | 460 | 0 | 0 |
| GPS1 | 222 | 420 | 0 | 2436 | 1338 | 1919 | 0 | 192 |
| GPS2 | 348 | 0 | 2096 | 0 | 2672 | 0 | 802 | 0 |
| GPSM2 | 0 | 0 | 1304 | 0 | 0 | 1362 | 1162 | 1084 |
| GPSM3 | 0 | 124 | 0 | 0 | 0 | 0 | 0 | 0 |
| GPT | 426 | 230 | 0 | 0 | 0 | 0 | 0 | 0 |
| GPT2 | 570 | 0 | 0 | 0 | 428 | 0 | 0 | 0 |
| GPX1 | 1680 | 2248 | 4406 | 16165 | 25358 | 6361 | 9874 | 8798 |
| GPX2 | 11310 | 31454 | 23351 | 43104 | 51790 | 88037 | 17590 | 6127 |
| GPX3 | 26061 | 23690 | 125647 | 21663 | 17640 | 160410 | 55861 | 10452 |
| GPX4 | 2051 | 1938 | 5817 | 3389 | 14376 | 5105 | 4689 | 1868 |
| GPX7 | 0 | 0 | 0 | 0 | 0 | 0 | 176 | 1088 |
| GRAMD1A | 0 | 0 | 1598 | 0 | 670 | 0 | 1878 | 0 |
| GRAMD2B | 378 | 0 | 1042 | 150 | 0 | 1874 | 128 | 0 |
| GRAMD4 | 0 | 0 | 1018 | 0 | 320 | 878 | 0 | 0 |
| GRAP2 | 1180 | 1504 | 11680 | 1688 | 0 | 3120 | 0 | 0 |
| GRB10 | 1296 | 522 | 76 | 4160 | 8905 | 1474 | 2390 | 1594 |
| GRB2 | 814 | 450 | 0 | 218 | 766 | 0 | 1048 | 954 |
| GRB7 | 1738 | 4044 | 10312 | 29710 | 31878 | 14638 | 69938 | 7835 |

| | | | | | | | | |
|---|---|---|---|---|---|---|---|---|
| GRHL2 | 0 | 0 | 0 | 0 | 0 | 0 | 326 | 0 |
| GRHL3 | 0 | 0 | 0 | 254 | 242 | 0 | 0 | 0 |
| GRHPR | 0 | 800 | 1867 | 2585 | 1334 | 3563 | 2344 | 1322 |
| GRIA1 | 0 | 0 | 0 | 0 | 534 | 0 | 0 | 0 |
| GRIA3 | 1060 | 500 | 0 | 0 | 934 | 0 | 0 | 262 |
| GRID1 | 515 | 0 | 0 | 0 | 0 | 0 | 0 | 0 |
| GRIK2 | 0 | 0 | 2074 | 0 | 0 | 0 | 0 | 0 |
| GRINA | 0 | 0 | 567 | 832 | 902 | 0 | 3188 | 1008 |
| GRIP1 | 0 | 0 | 0 | 0 | 64 | 0 | 308 | 0 |
| GRIP2 | 0 | 0 | 0 | 0 | 0 | 0 | 636 | 0 |
| GRIPAP1 | 544 | 0 | 694 | 528 | 660 | 0 | 0 | 0 |
| GRK2 | 0 | 552 | 2082 | 1064 | 268 | 112 | 315 | 0 |
| GRK3 | 0 | 0 | 0 | 0 | 1142 | 706 | 2536 | 1715 |
| GRK6 | 573 | 356 | 0 | 0 | 0 | 66 | 0 | 0 |
| GRM8 | 444 | 274 | 0 | 4540 | 0 | 0 | 7579 | 0 |
| GRN | 3804 | 2386 | 20426 | 5878 | 8615 | 2900 | 6920 | 5152 |
| GRPEL1 | 0 | 0 | 0 | 3325 | 2410 | 0 | 716 | 2126 |
| GRSF1 | 0 | 2436 | 192 | 2056 | 3446 | 0 | 994 | 1834 |
| GRTP1 | 0 | 328 | 798 | 2683 | 1224 | 2670 | 2644 | 0 |
| GRWD1 | 0 | 0 | 0 | 1224 | 0 | 0 | 0 | 0 |
| GS1-124K5.11 | 0 | 0 | 0 | 1194 | 0 | 0 | 0 | 0 |
| GS1-124K5.4 | 0 | 0 | 0 | 0 | 1130 | 670 | 0 | 0 |
| GS1-594A7.3 | 0 | 0 | 1725 | 0 | 0 | 0 | 0 | 0 |
| GSAP | 0 | 198 | 0 | 0 | 0 | 0 | 0 | 0 |
| GSDMB | 63982 | 39529 | 161636 | 44545 | 22750 | 15845 | 37266 | 6519 |
| GSDMD | 0 | 144 | 0 | 1198 | 937 | 350 | 739 | 0 |
| GSE1 | 0 | 0 | 0 | 2044 | 418 | 0 | 0 | 0 |
| GSEC | 0 | 0 | 0 | 0 | 0 | 0 | 0 | 600 |
| GSK3A | 0 | 0 | 904 | 0 | 88 | 1034 | 0 | 482 |
| GSK3B | 0 | 0 | 1060 | 0 | 312 | 0 | 350 | 78 |
| GSKIP | 558 | 1357 | 6103 | 2348 | 1486 | 0 | 242 | 0 |
| GSN | 3380 | 1776 | 10729 | 1674 | 7524 | 1663 | 0 | 1712 |
| GSPT1 | 0 | 1094 | 0 | 1032 | 968 | 0 | 396 | 714 |
| GSR | 50 | 168 | 310 | 338 | 276 | 0 | 0 | 0 |
| GSS | 82349 | 180773 | 357142 | 374194 | 574883 | 253177 | 466556 | 180215 |
| GSTA1 | 0 | 0 | 2120 | 0 | 0 | 0 | 0 | 0 |
| GSTCD | 1106 | 726 | 1186 | 1812 | 5317 | 459 | 2274 | 0 |
| GSTK1 | 3881 | 8195 | 25496 | 23022 | 6300 | 10120 | 23156 | 2431 |
| GSTM2 | 0 | 324 | 0 | 0 | 0 | 0 | 0 | 0 |
| GSTM3 | 0 | 0 | 1039 | 0 | 0 | 0 | 876 | 1368 |
| GSTM4 | 800 | 410 | 1286 | 0 | 1088 | 0 | 644 | 0 |
| GSTO1 | 0 | 1564 | 5799 | 5076 | 2868 | 3591 | 1804 | 2593 |
| GSTO2 | 0 | 0 | 0 | 0 | 668 | 0 | 616 | 480 |

| | | | | | | | | |
|---|---|---|---|---|---|---|---|---|
| GSTP1 | 4628 | 9195 | 18018 | 37568 | 64977 | 8358 | 27321 | 29544 |
| GSTT1 | 1601 | 616 | 3914 | 0 | 3832 | 4330 | 1590 | 972 |
| GSTZ1 | 660 | 682 | 1332 | 0 | 124 | 0 | 0 | 0 |
| GTF2A1 | 0 | 350 | 0 | 0 | 0 | 0 | 0 | 0 |
| GTF2A2 | 989 | 2236 | 1288 | 7039 | 4108 | 0 | 624 | 2356 |
| GTF2B | 0 | 436 | 0 | 4845 | 2067 | 0 | 1240 | 0 |
| GTF2E2 | 1918 | 360 | 0 | 768 | 1213 | 0 | 0 | 0 |
| GTF2F1 | 0 | 374 | 306 | 1495 | 1777 | 0 | 406 | 0 |
| GTF2F2 | 0 | 340 | 0 | 2888 | 4949 | 2557 | 1297 | 0 |
| GTF2H1 | 0 | 0 | 0 | 947 | 0 | 0 | 542 | 0 |
| GTF2H2 | 0 | 0 | 0 | 0 | 0 | 0 | 304 | 734 |
| GTF2H2B | 0 | 0 | 0 | 0 | 574 | 0 | 0 | 0 |
| GTF2H3 | 2360 | 7414 | 4292 | 13972 | 28539 | 5764 | 22351 | 9535 |
| GTF2H4 | 0 | 0 | 0 | 0 | 492 | 0 | 0 | 0 |
| GTF2H5 | 0 | 0 | 2774 | 568 | 4137 | 1512 | 1322 | 1346 |
| GTF2I | 0 | 0 | 0 | 0 | 842 | 2410 | 166 | 0 |
| GTF2IP12 | 0 | 0 | 0 | 0 | 0 | 62 | 0 | 0 |
| GTF2IP20 | 0 | 0 | 0 | 0 | 20 | 0 | 702 | 0 |
| GTF2IP4 | 959 | 776 | 0 | 0 | 368 | 0 | 0 | 0 |
| GTF2IRD1 | 410 | 1370 | 0 | 3449 | 1722 | 6488 | 6573 | 286 |
| GTF2IRD2 | 0 | 0 | 1738 | 0 | 1418 | 784 | 1070 | 707 |
| GTF2IRD2B | 0 | 0 | 0 | 0 | 978 | 0 | 0 | 0 |
| GTF3A | 0 | 0 | 2078 | 3909 | 6003 | 4920 | 4059 | 1476 |
| GTF3C1 | 1404 | 0 | 0 | 0 | 149 | 0 | 0 | 0 |
| GTF3C2 | 16941 | 25866 | 108441 | 37554 | 105217 | 51522 | 109324 | 32727 |
| GTF3C3 | 0 | 0 | 0 | 544 | 1774 | 0 | 0 | 0 |
| GTF3C5 | 0 | 682 | 0 | 0 | 1058 | 2044 | 58 | 0 |
| GTF3C6 | 1678 | 808 | 2208 | 2014 | 9859 | 204 | 1278 | 1378 |
| GTPBP10 | 0 | 0 | 1916 | 1226 | 1668 | 0 | 110 | 0 |
| GTPBP2 | 0 | 0 | 0 | 0 | 0 | 102 | 492 | 0 |
| GTPBP3 | 0 | 0 | 0 | 0 | 318 | 0 | 0 | 0 |
| GTPBP4 | 0 | 284 | 0 | 1184 | 2420 | 2314 | 276 | 998 |
| GTPBP6 | 0 | 582 | 0 | 108 | 1582 | 0 | 0 | 0 |
| GTPBP8 | 0 | 352 | 0 | 1190 | 0 | 0 | 0 | 0 |
| GTSE1 | 0 | 0 | 0 | 0 | 410 | 1834 | 0 | 0 |
| GTSF1 | 0 | 0 | 836 | 0 | 0 | 0 | 0 | 0 |
| GUCA2A | 24810 | 10020 | 71949 | 1442 | 1816 | 0 | 0 | 0 |
| GUCA2B | 3150 | 0 | 10004 | 0 | 0 | 0 | 0 | 0 |
| GUCD1 | 1420 | 0 | 0 | 0 | 56 | 0 | 464 | 0 |
| GUCY1B1 | 0 | 0 | 0 | 0 | 346 | 0 | 0 | 0 |
| GUCY2C | 302 | 0 | 0 | 0 | 0 | 1464 | 504 | 0 |
| GUK1 | 0 | 3699 | 6204 | 10173 | 7274 | 9254 | 12825 | 7601 |
| GULP1 | 0 | 0 | 1052 | 0 | 0 | 0 | 0 | 0 |

| | | | | | | | | |
|---|---|---|---|---|---|---|---|---|
| GUSB | 0 | 312 | 1264 | 0 | 0 | 0 | 0 | 0 |
| GUSBP11 | 756 | 0 | 0 | 0 | 0 | 0 | 0 | 0 |
| GUSBP5 | 0 | 0 | 0 | 118 | 0 | 0 | 0 | 0 |
| GVQW3 | 0 | 0 | 0 | 0 | 0 | 0 | 0 | 452 |
| GXYLT1 | 0 | 0 | 0 | 0 | 506 | 0 | 0 | 0 |
| GXYLT2 | 0 | 0 | 0 | 0 | 0 | 0 | 2245 | 1046 |
| GYG1 | 0 | 0 | 0 | 414 | 0 | 0 | 0 | 0 |
| GYG2 | 0 | 0 | 0 | 0 | 354 | 410 | 0 | 0 |
| GYPC | 0 | 0 | 0 | 0 | 0 | 1833 | 0 | 0 |
| GYS1 | 0 | 0 | 0 | 362 | 0 | 0 | 0 | 0 |
| GZF1 | 426 | 0 | 0 | 0 | 180 | 0 | 0 | 0 |
| GZMB | 0 | 0 | 814 | 4484 | 0 | 0 | 4384 | 1192 |
| H1F0 | 982 | 426 | 1648 | 1125 | 2251 | 1977 | 2268 | 1012 |
| H1FX | 656 | 0 | 0 | 1092 | 4436 | 1800 | 442 | 2402 |
| H2AFJ | 1505 | 2094 | 0 | 1498 | 3286 | 1426 | 2834 | 1034 |
| H2AFV | 536 | 722 | 1975 | 4870 | 5208 | 0 | 1436 | 776 |
| H2AFX | 688 | 522 | 0 | 0 | 1378 | 0 | 0 | 0 |
| H2AFY | 98879 | 173924 | 202692 | 465561 | 685590 | 617821 | 400689 | 231067 |
| H2AFZ | 1122 | 3226 | 11845 | 32651 | 24304 | 6279 | 5677 | 13056 |
| H3F3A | 7596 | 4732 | 21633 | 17617 | 29974 | 8481 | 13612 | 5994 |
| H3F3AP4 | 916 | 3924 | 438 | 4111 | 6491 | 12 | 1644 | 2504 |
| H3F3B | 21613 | 18056 | 33973 | 26770 | 39985 | 57490 | 42463 | 10067 |
| H6PD | 290 | 736 | 0 | 0 | 2578 | 748 | 170 | 0 |
| HAAO | 0 | 0 | 0 | 2284 | 242 | 0 | 474 | 0 |
| HABP4 | 0 | 0 | 0 | 0 | 628 | 0 | 370 | 1260 |
| HACD1 | 3358 | 0 | 0 | 0 | 8243 | 2786 | 4660 | 1200 |
| HACD2 | 0 | 30 | 0 | 0 | 0 | 0 | 0 | 0 |
| HACD3 | 0 | 0 | 0 | 0 | 1973 | 0 | 0 | 0 |
| HACL1 | 0 | 302 | 1824 | 0 | 892 | 0 | 528 | 0 |
| HADH | 460 | 916 | 4090 | 1251 | 2982 | 0 | 3113 | 0 |
| HADHA | 153327 | 246301 | 349636 | 231780 | 573722 | 281706 | 417108 | 135321 |
| HADHB | 0 | 2898 | 4316 | 226 | 3354 | 0 | 1169 | 0 |
| HAGH | 550 | 0 | 1230 | 1267 | 0 | 0 | 444 | 1018 |
| HAGHL | 0 | 0 | 0 | 0 | 142 | 0 | 406 | 0 |
| HAGLR | 0 | 0 | 537 | 0 | 0 | 0 | 0 | 0 |
| HAGLROS | 0 | 0 | 0 | 0 | 556 | 0 | 0 | 0 |
| HAND1 | 0 | 0 | 0 | 0 | 0 | 0 | 350 | 0 |
| HAPLN1 | 0 | 422 | 0 | 0 | 0 | 0 | 0 | 0 |
| HAPLN3 | 0 | 0 | 0 | 0 | 0 | 1600 | 0 | 0 |
| HARBI1 | 8156 | 14011 | 22454 | 21397 | 26763 | 36536 | 35014 | 28863 |
| HARS | 532 | 0 | 652 | 348 | 0 | 0 | 0 | 0 |
| HARS2 | 806 | 210 | 3278 | 0 | 0 | 676 | 0 | 0 |
| HAS2 | 610 | 0 | 0 | 0 | 0 | 0 | 0 | 0 |

| | | | | | | | | |
|---|---|---|---|---|---|---|---|---|
| HAS2-AS1 | 0 | 0 | 0 | 0 | 0 | 0 | 266 | 0 |
| HAT1 | 0 | 506 | 1066 | 0 | 852 | 0 | 522 | 1196 |
| HAUS1 | 0 | 0 | 1865 | 1525 | 2068 | 0 | 0 | 0 |
| HAUS2 | 0 | 0 | 1744 | 1722 | 2878 | 0 | 1014 | 0 |
| HAUS3 | 0 | 0 | 0 | 726 | 650 | 0 | 0 | 0 |
| HAUS4 | 0 | 0 | 2188 | 0 | 1762 | 0 | 664 | 0 |
| HAUS7 | 0 | 186 | 0 | 0 | 1598 | 0 | 0 | 0 |
| HAUS8 | 17964 | 20414 | 140182 | 34167 | 107833 | 26845 | 22882 | 21637 |
| HAX1 | 1616 | 0 | 0 | 0 | 3658 | 2077 | 842 | 3086 |
| HBA1 | 0 | 0 | 1612 | 0 | 0 | 0 | 0 | 0 |
| HBA2 | 0 | 524 | 0 | 0 | 1058 | 0 | 0 | 0 |
| HBB | 630 | 0 | 0 | 0 | 996 | 0 | 0 | 0 |
| HBEGF | 402 | 0 | 1334 | 0 | 366 | 0 | 0 | 0 |
| HBP1 | 276 | 374 | 0 | 1996 | 720 | 0 | 0 | 64 |
| HBS1L | 518 | 0 | 0 | 1116 | 568 | 0 | 306 | 0 |
| HCAR1 | 0 | 0 | 0 | 0 | 182 | 0 | 0 | 0 |
| HCCS | 0 | 0 | 0 | 2332 | 1983 | 0 | 0 | 0 |
| HCFC1 | 0 | 138 | 0 | 0 | 0 | 0 | 0 | 0 |
| HCFC1R1 | 0 | 0 | 446 | 0 | 0 | 0 | 576 | 1056 |
| HCFC2 | 4405 | 4611 | 9478 | 2423 | 5284 | 10492 | 2592 | 3599 |
| HCG18 | 0 | 0 | 0 | 1418 | 736 | 0 | 0 | 756 |
| HCK | 0 | 0 | 0 | 0 | 0 | 0 | 0 | 1016 |
| HCLS1 | 0 | 150 | 0 | 0 | 0 | 1370 | 0 | 0 |
| HCN3 | 0 | 0 | 0 | 0 | 624 | 0 | 0 | 0 |
| HCP5 | 0 | 1012 | 0 | 0 | 0 | 3651 | 0 | 0 |
| HCST | 0 | 0 | 0 | 0 | 0 | 0 | 0 | 562 |
| HDAC1 | 1042 | 1526 | 2904 | 3646 | 3978 | 2564 | 580 | 0 |
| HDAC10 | 570 | 0 | 1140 | 590 | 616 | 0 | 0 | 278 |
| HDAC11 | 0 | 0 | 0 | 0 | 0 | 0 | 0 | 1176 |
| HDAC2 | 444 | 342 | 1064 | 3518 | 3108 | 0 | 1376 | 2136 |
| HDAC3 | 0 | 394 | 942 | 0 | 795 | 0 | 340 | 0 |
| HDAC5 | 0 | 0 | 1384 | 0 | 80 | 1652 | 1932 | 0 |
| HDAC6 | 0 | 0 | 3421 | 0 | 0 | 0 | 686 | 0 |
| HDAC7 | 0 | 188 | 0 | 0 | 918 | 0 | 0 | 0 |
| HDAC8 | 0 | 0 | 0 | 0 | 80 | 0 | 0 | 1104 |
| HDDC2 | 626 | 480 | 0 | 0 | 5708 | 0 | 2076 | 0 |
| HDDC3 | 0 | 0 | 0 | 2466 | 0 | 0 | 1054 | 0 |
| HDGF | 0 | 266 | 0 | 866 | 4158 | 1448 | 504 | 1297 |
| HDGFL2 | 456 | 332 | 0 | 0 | 750 | 926 | 584 | 1520 |
| HDGFL3 | 0 | 0 | 0 | 480 | 0 | 0 | 0 | 858 |
| HDHD2 | 0 | 360 | 1963 | 0 | 0 | 0 | 748 | 0 |
| HDHD3 | 1764 | 786 | 7772 | 0 | 1730 | 0 | 1548 | 1174 |
| HDHD5 | 772 | 374 | 0 | 0 | 3755 | 2356 | 0 | 2282 |

| | | | | | | | | |
|---|---|---|---|---|---|---|---|---|
| HDHD5-AS1 | 0 | 0 | 0 | 0 | 1340 | 0 | 0 | 0 |
| HDLBP | 0 | 192 | 0 | 1330 | 2498 | 0 | 1544 | 685 |
| HEATR1 | 3466 | 6679 | 3681 | 11619 | 28096 | 2506 | 23630 | 6024 |
| HEATR3 | 460 | 0 | 0 | 0 | 0 | 0 | 0 | 0 |
| HEATR5A | 0 | 0 | 0 | 0 | 814 | 0 | 0 | 0 |
| HEATR5B | 0 | 260 | 0 | 0 | 0 | 0 | 0 | 0 |
| HEBP1 | 1496 | 0 | 0 | 0 | 88 | 0 | 1262 | 0 |
| HEBP2 | 3212 | 3538 | 14506 | 3711 | 10912 | 6620 | 1455 | 4233 |
| HECA | 0 | 0 | 0 | 0 | 0 | 0 | 843 | 0 |
| HECTD1 | 0 | 450 | 792 | 0 | 792 | 0 | 996 | 0 |
| HECTD2-AS1 | 0 | 708 | 0 | 0 | 486 | 0 | 0 | 208 |
| HECTD3 | 622 | 0 | 0 | 0 | 0 | 0 | 230 | 0 |
| HECTD4 | 0 | 0 | 0 | 0 | 1246 | 0 | 0 | 0 |
| HECW2 | 0 | 0 | 0 | 0 | 164 | 0 | 0 | 0 |
| HEIH | 0 | 254 | 0 | 0 | 1250 | 0 | 190 | 0 |
| HELLS | 0 | 0 | 0 | 924 | 828 | 0 | 0 | 0 |
| HEMK1 | 18389 | 29966 | 78297 | 39736 | 130167 | 75654 | 106920 | 20562 |
| HENMT1 | 0 | 738 | 0 | 1205 | 2228 | 0 | 0 | 0 |
| HEPACAM2 | 63242 | 228893 | 436771 | 8140 | 24929 | 0 | 1090 | 754 |
| HEPH | 0 | 474 | 1110 | 644 | 682 | 0 | 876 | 0 |
| HERC1 | 0 | 348 | 0 | 0 | 196 | 0 | 348 | 0 |
| HERC2 | 0 | 0 | 0 | 0 | 174 | 0 | 590 | 0 |
| HERC2P2 | 222 | 0 | 0 | 578 | 0 | 654 | 56 | 0 |
| HERC2P9 | 0 | 0 | 0 | 0 | 314 | 0 | 0 | 774 |
| HERC4 | 8410 | 4771 | 9064 | 10904 | 16048 | 3796 | 20847 | 4372 |
| HERC6 | 0 | 0 | 0 | 0 | 0 | 0 | 32 | 518 |
| HERPUD1 | 938 | 566 | 2704 | 2324 | 462 | 0 | 2660 | 1414 |
| HERPUD2 | 3044 | 6835 | 14458 | 2677 | 23888 | 12172 | 27283 | 8738 |
| HES1 | 1228 | 560 | 3798 | 860 | 2148 | 468 | 808 | 0 |
| HES4 | 68 | 216 | 2328 | 1224 | 3107 | 1690 | 834 | 2134 |
| HES6 | 386 | 0 | 0 | 2666 | 1542 | 0 | 524 | 1482 |
| HEXA | 224 | 1029 | 0 | 1362 | 1462 | 0 | 600 | 0 |
| HEXB | 702 | 1368 | 7960 | 2239 | 4370 | 4733 | 238 | 2702 |
| HEXIM1 | 0 | 326 | 0 | 0 | 830 | 0 | 116 | 0 |
| HEXIM2 | 0 | 0 | 0 | 0 | 0 | 1760 | 0 | 0 |
| HEYL | 0 | 328 | 0 | 104 | 184 | 0 | 346 | 420 |
| HGD | 21412 | 58780 | 73605 | 24703 | 66694 | 11548 | 32423 | 79285 |
| HGH1 | 0 | 0 | 0 | 0 | 632 | 0 | 254 | 0 |
| HGS | 0 | 0 | 0 | 0 | 632 | 1936 | 844 | 0 |
| HGSNAT | 0 | 0 | 0 | 482 | 0 | 0 | 0 | 0 |
| HHAT | 0 | 0 | 694 | 0 | 0 | 0 | 0 | 0 |
| HHEX | 0 | 338 | 0 | 0 | 0 | 0 | 0 | 0 |
| HHLA2 | 157877 | 73840 | 350013 | 19122 | 39675 | 2978 | 29403 | 20323 |

| | | | | | | | | |
|---|---|---|---|---|---|---|---|---|
| HHLA3 | 164 | 0 | 0 | 0 | 0 | 0 | 0 | 0 |
| HIBADH | 51216 | 107460 | 190460 | 105873 | 403694 | 223726 | 308907 | 88997 |
| HIBCH | 6974 | 1906 | 2671 | 7900 | 8247 | 2684 | 8943 | 4947 |
| HIC1 | 0 | 0 | 1004 | 0 | 0 | 0 | 0 | 0 |
| HID1 | 0 | 0 | 0 | 0 | 792 | 402 | 0 | 0 |
| HIF1A | 10814 | 37990 | 27789 | 19884 | 88515 | 25098 | 15047 | 19503 |
| HIF1A-AS2 | 2178 | 10910 | 52 | 34 | 22807 | 190 | 4790 | 220 |
| HIF1AN | 0 | 744 | 0 | 0 | 0 | 194 | 0 | 0 |
| HIGD1A | 4150 | 7692 | 21461 | 3344 | 12544 | 3703 | 0 | 3112 |
| HIGD2A | 1644 | 1752 | 5472 | 3492 | 6364 | 6286 | 3348 | 146 |
| HIKESHI | 0 | 0 | 0 | 1238 | 4717 | 0 | 859 | 0 |
| HILPDA | 0 | 0 | 0 | 0 | 1210 | 0 | 178 | 0 |
| HINFP | 0 | 0 | 1912 | 0 | 866 | 0 | 530 | 864 |
| HINT1 | 7012 | 14138 | 29137 | 21054 | 22000 | 13144 | 10526 | 9926 |
| HINT2 | 0 | 1840 | 4490 | 1314 | 1760 | 654 | 1212 | 0 |
| HINT3 | 3454 | 6760 | 646 | 3302 | 17293 | 1838 | 27134 | 2794 |
| HIP1 | 506 | 0 | 0 | 0 | 1441 | 0 | 526 | 358 |
| HIP1R | 0 | 0 | 2980 | 0 | 398 | 0 | 0 | 0 |
| HIPK1 | 854 | 0 | 0 | 0 | 0 | 0 | 0 | 0 |
| HIPK2 | 0 | 370 | 0 | 0 | 0 | 0 | 0 | 0 |
| HIPK3 | 0 | 0 | 0 | 0 | 0 | 0 | 264 | 20 |
| HIRA | 2524 | 274 | 4014 | 438 | 1438 | 1506 | 1942 | 2066 |
| HIRIP3 | 0 | 0 | 0 | 1327 | 0 | 0 | 0 | 0 |
| HIST1H1C | 0 | 0 | 7885 | 0 | 1818 | 0 | 0 | 260 |
| HIST1H2AC | 0 | 0 | 0 | 0 | 1470 | 0 | 42 | 0 |
| HIST1H2AE | 0 | 0 | 0 | 0 | 752 | 830 | 0 | 0 |
| HIST1H2AH | 0 | 0 | 0 | 0 | 0 | 808 | 0 | 0 |
| HIST1H2AJ | 0 | 0 | 0 | 0 | 586 | 0 | 0 | 0 |
| HIST1H2AM | 0 | 0 | 0 | 0 | 666 | 0 | 0 | 0 |
| HIST1H2BD | 0 | 116 | 1954 | 2537 | 0 | 0 | 1226 | 1420 |
| HIST1H2BF | 0 | 0 | 0 | 0 | 970 | 0 | 0 | 0 |
| HIST1H2BG | 0 | 0 | 0 | 0 | 330 | 0 | 586 | 0 |
| HIST1H2BH | 930 | 0 | 0 | 0 | 0 | 0 | 0 | 0 |
| HIST1H2BJ | 0 | 0 | 0 | 0 | 478 | 0 | 0 | 0 |
| HIST1H2BK | 0 | 812 | 1658 | 3412 | 4446 | 1820 | 3812 | 1068 |
| HIST1H3D | 0 | 0 | 0 | 0 | 1060 | 0 | 0 | 0 |
| HIST1H4J | 0 | 478 | 0 | 0 | 1000 | 0 | 0 | 556 |
| HIST2H2AA3 | 0 | 1366 | 2542 | 1780 | 10023 | 0 | 1144 | 3189 |
| HIST2H2BC | 0 | 0 | 0 | 158 | 0 | 0 | 0 | 0 |
| HIST2H2BE | 0 | 242 | 0 | 0 | 1184 | 0 | 0 | 0 |
| HIST3H2A | 0 | 0 | 1436 | 820 | 2008 | 1768 | 0 | 0 |
| HJURP | 68 | 0 | 0 | 32 | 0 | 0 | 0 | 0 |
| HK1 | 0 | 394 | 0 | 0 | 0 | 0 | 50 | 112 |

| | | | | | | | | |
|---|---|---|---|---|---|---|---|---|
| HK2 | 564 | 296 | 1430 | 0 | 1310 | 0 | 0 | 0 |
| HKDC1 | 0 | 0 | 0 | 0 | 0 | 0 | 0 | 586 |
| HKR1 | 0 | 0 | 1058 | 0 | 0 | 0 | 0 | 0 |
| HLA-A | 21522 | 15446 | 54064 | 1926 | 22226 | 5477 | 2294 | 11793 |
| HLA-B | 3667082 | 2131484 | 6012380 | 1214478 | 3463670 | 2279314 | 1628592 | 2015858 |
| HLA-C | 173656 | 276864 | 333170 | 101903 | 474536 | 136231 | 168701 | 240072 |
| HLA-DMA | 1032 | 298 | 2778 | 0 | 0 | 0 | 66 | 0 |
| HLA-DMB | 0 | 0 | 0 | 0 | 32 | 0 | 588 | 0 |
| HLA-DOA | 0 | 0 | 400 | 0 | 0 | 0 | 454 | 0 |
| HLA-DPA1 | 1458 | 1082 | 6682 | 0 | 206 | 10553 | 482 | 7046 |
| HLA-DPB1 | 1812 | 1826 | 0 | 0 | 0 | 7888 | 910 | 2775 |
| HLA-DQA1 | 0 | 928 | 2518 | 0 | 0 | 6445 | 278 | 0 |
| HLA-DQB1 | 842 | 1506 | 3994 | 1688 | 0 | 3643 | 1054 | 0 |
| HLA-DRA | 3802 | 7548 | 9352 | 1834 | 7209 | 19340 | 1516 | 9894 |
| HLA-DRB1 | 1805 | 0 | 3325 | 522 | 0 | 4030 | 674 | 0 |
| HLA-DRB3 | 0 | 0 | 1677 | 0 | 0 | 0 | 0 | 0 |
| HLA-DRB4 | 0 | 286 | 0 | 0 | 1452 | 0 | 0 | 1892 |
| HLA-E | 2518 | 2246 | 5940 | 410 | 3774 | 10743 | 3880 | 2766 |
| HLA-F | 484 | 558 | 5079 | 830 | 1502 | 0 | 480 | 0 |
| HLA-F-AS1 | 1214 | 0 | 0 | 1239 | 0 | 0 | 0 | 0 |
| HLA-G | 0 | 440 | 0 | 0 | 0 | 0 | 0 | 0 |
| HLA-H | 0 | 0 | 92 | 0 | 0 | 0 | 0 | 0 |
| HLF | 738 | 0 | 0 | 0 | 0 | 0 | 0 | 0 |
| HLTF | 0 | 474 | 0 | 1424 | 7808 | 0 | 390 | 1808 |
| HM13 | 0 | 1142 | 2002 | 5623 | 4449 | 0 | 1204 | 0 |
| HMBS | 0 | 0 | 0 | 0 | 1320 | 0 | 356 | 1724 |
| HMCES | 990 | 0 | 0 | 0 | 0 | 1780 | 1040 | 956 |
| HMCN1 | 0 | 240 | 0 | 0 | 0 | 0 | 0 | 0 |
| HMCN2 | 0 | 0 | 0 | 0 | 444 | 0 | 0 | 0 |
| HMG20B | 0 | 532 | 2758 | 1591 | 450 | 0 | 0 | 0 |
| HMGA1 | 6593 | 21494 | 16533 | 65106 | 256302 | 74150 | 136505 | 111248 |
| HMGA2 | 560 | 2135 | 0 | 1511 | 4888 | 0 | 13116 | 6636 |
| HMGB1 | 1776 | 3976 | 5480 | 19325 | 29117 | 21812 | 15513 | 6214 |
| HMGB2 | 240 | 1660 | 8503 | 4614 | 13757 | 4166 | 4224 | 4686 |
| HMGB3 | 0 | 1436 | 0 | 1488 | 6225 | 1856 | 0 | 0 |
| HMGCL | 208 | 1300 | 5268 | 1346 | 1476 | 0 | 0 | 1060 |
| HMGCLL1 | 986 | 0 | 0 | 0 | 0 | 0 | 0 | 0 |
| HMGCR | 40 | 0 | 0 | 30 | 522 | 0 | 198 | 663 |
| HMGCS1 | 1504 | 476 | 2199 | 1138 | 1954 | 1138 | 0 | 0 |
| HMGCS2 | 1067 | 1720 | 14636 | 1631 | 492 | 0 | 6148 | 0 |
| HMGN1 | 2234 | 4318 | 1542 | 5386 | 14791 | 5484 | 5443 | 9308 |
| HMGN2 | 2605 | 6264 | 12340 | 25069 | 38260 | 8388 | 5984 | 4239 |
| HMGN3 | 222 | 1254 | 7322 | 900 | 5112 | 0 | 316 | 1689 |

| | | | | | | | | |
|---|---|---|---|---|---|---|---|---|
| HMGN4 | 518 | 30 | 4846 | 1468 | 1910 | 0 | 0 | 1028 |
| HMGXB3 | 0 | 0 | 0 | 1426 | 148 | 0 | 298 | 0 |
| HMOX1 | 1214 | 0 | 554 | 0 | 778 | 0 | 0 | 1180 |
| HMOX2 | 0 | 2778 | 2774 | 2222 | 1878 | 0 | 2436 | 826 |
| HMSD | 0 | 0 | 0 | 0 | 0 | 772 | 0 | 0 |
| HNF1A | 0 | 0 | 0 | 0 | 296 | 0 | 176 | 0 |
| HNF1B | 14211 | 16080 | 23182 | 6938 | 28040 | 2396 | 11440 | 4313 |
| HNF4A | 0 | 412 | 682 | 0 | 1090 | 0 | 882 | 0 |
| HNF4G | 0 | 0 | 0 | 0 | 794 | 0 | 0 | 0 |
| HNMT | 1114 | 1064 | 1544 | 0 | 1846 | 1478 | 1342 | 1800 |
| HNRNPA0 | 0 | 322 | 1098 | 898 | 3954 | 0 | 1242 | 0 |
| HNRNPA1 | 2634 | 7016 | 13753 | 27168 | 29118 | 19110 | 5698 | 10213 |
| HNRNPA2B1 | 304286 | 564429 | 827055 | 1100110 | 3099608 | 1381876 | 1451177 | 915126 |
| HNRNPA3 | 3874 | 3316 | 4628 | 4356 | 10973 | 9713 | 13166 | 4019 |
| HNRNPAB | 950 | 1870 | 3321 | 2082 | 2028 | 0 | 1448 | 1206 |
| HNRNPC | 2436 | 2064 | 1158 | 8779 | 16440 | 762 | 2810 | 5404 |
| HNRNPD | 412 | 158 | 2678 | 842 | 9832 | 2692 | 1289 | 2959 |
| HNRNPDL | 2240 | 2252 | 6336 | 4874 | 8120 | 2036 | 6080 | 0 |
| HNRNPF | 1289 | 1076 | 4941 | 562 | 1454 | 0 | 1258 | 699 |
| HNRNPH1 | 21814 | 39143 | 58187 | 107352 | 200275 | 121760 | 111243 | 71867 |
| HNRNPH2 | 394 | 118 | 2814 | 1054 | 2842 | 1370 | 1001 | 0 |
| HNRNPH3 | 0 | 664 | 1404 | 3420 | 2174 | 2088 | 0 | 0 |
| HNRNPK | 2288 | 2318 | 4546 | 4137 | 8656 | 1344 | 3396 | 4329 |
| HNRNPL | 200 | 252 | 1752 | 406 | 2964 | 1300 | 314 | 1138 |
| HNRNPLL | 28242 | 39081 | 46350 | 56412 | 158260 | 42304 | 110310 | 41298 |
| HNRNPM | 0 | 1730 | 708 | 3325 | 3472 | 0 | 2240 | 1587 |
| HNRNPR | 1084 | 0 | 1436 | 1371 | 3532 | 0 | 387 | 0 |
| HNRNPU | 332 | 854 | 0 | 2381 | 5143 | 0 | 1836 | 0 |
| HNRNPUL1 | 0 | 0 | 0 | 610 | 862 | 0 | 364 | 0 |
| HNRNPUL2 | 0 | 732 | 0 | 0 | 840 | 0 | 0 | 0 |
| HOGA1 | 0 | 0 | 0 | 0 | 0 | 0 | 668 | 0 |
| HOMER2 | 728 | 0 | 0 | 0 | 1124 | 306 | 1982 | 1104 |
| HOMEZ | 632 | 460 | 0 | 0 | 308 | 0 | 0 | 0 |
| HOOK2 | 0 | 450 | 0 | 0 | 1765 | 0 | 1273 | 0 |
| HOOK3 | 11881 | 6832 | 2330 | 17228 | 71000 | 45743 | 33772 | 20187 |
| HOPX | 12496 | 43174 | 122004 | 43658 | 887541 | 281526 | 21705 | 97562 |
| HORMAD1 | 0 | 0 | 0 | 0 | 720 | 0 | 0 | 0 |
| HOTAIRM1 | 752 | 0 | 0 | 0 | 1390 | 0 | 0 | 0 |
| HOTTIP | 0 | 0 | 0 | 304 | 794 | 1593 | 336 | 0 |
| HOXA-AS3 | 0 | 0 | 0 | 62 | 68 | 0 | 864 | 0 |
| HOXA10 | 612 | 122 | 0 | 893 | 1056 | 0 | 466 | 0 |
| HOXA10-AS | 0 | 0 | 0 | 0 | 688 | 1704 | 710 | 0 |
| HOXA10-HOXA9 | 0 | 156 | 0 | 0 | 1026 | 0 | 222 | 0 |

| | | | | | | | | |
|---|---|---|---|---|---|---|---|---|
| HOXA11 | 0 | 0 | 0 | 0 | 366 | 0 | 0 | 0 |
| HOXA11-AS | 0 | 0 | 0 | 888 | 78 | 0 | 298 | 0 |
| HOXA13 | 0 | 0 | 0 | 406 | 0 | 0 | 400 | 0 |
| HOXA3 | 0 | 0 | 0 | 1634 | 0 | 0 | 558 | 0 |
| HOXA4 | 942 | 0 | 0 | 0 | 0 | 0 | 408 | 0 |
| HOXA5 | 0 | 0 | 0 | 0 | 0 | 0 | 0 | 1021 |
| HOXA7 | 0 | 0 | 0 | 0 | 498 | 0 | 0 | 0 |
| HOXA9 | 0 | 450 | 0 | 1504 | 2018 | 0 | 1184 | 0 |
| HOXB-AS3 | 0 | 404 | 0 | 0 | 288 | 0 | 304 | 0 |
| HOXB13 | 0 | 1316 | 0 | 0 | 3166 | 0 | 0 | 1056 |
| HOXB2 | 0 | 0 | 0 | 0 | 0 | 0 | 0 | 598 |
| HOXB3 | 358 | 600 | 860 | 1428 | 3258 | 8155 | 2807 | 0 |
| HOXB4 | 0 | 0 | 626 | 0 | 714 | 0 | 0 | 0 |
| HOXB5 | 215 | 0 | 0 | 916 | 0 | 0 | 0 | 0 |
| HOXB6 | 1116 | 154 | 0 | 1784 | 2324 | 0 | 388 | 0 |
| HOXB7 | 106 | 0 | 0 | 1111 | 680 | 0 | 1336 | 0 |
| HOXB8 | 0 | 0 | 0 | 0 | 474 | 616 | 578 | 0 |
| HOXB9 | 0 | 0 | 322 | 0 | 1064 | 0 | 432 | 0 |
| HOXD1 | 458 | 0 | 0 | 0 | 0 | 0 | 0 | 0 |
| HOXD11 | 0 | 408 | 0 | 0 | 0 | 0 | 0 | 0 |
| HOXD13 | 0 | 358 | 0 | 0 | 0 | 0 | 0 | 0 |
| HP08777 | 0 | 0 | 0 | 0 | 0 | 458 | 0 | 0 |
| HPCAL1 | 314 | 160 | 0 | 946 | 502 | 0 | 0 | 962 |
| HPDL | 0 | 152 | 0 | 0 | 116 | 0 | 412 | 0 |
| HPF1 | 0 | 852 | 0 | 1260 | 0 | 0 | 0 | 298 |
| HPGD | 770 | 0 | 1756 | 914 | 0 | 0 | 764 | 220 |
| HPGDS | 0 | 490 | 0 | 0 | 1562 | 0 | 0 | 0 |
| HPRT1 | 0 | 862 | 587 | 0 | 1428 | 3324 | 0 | 2610 |
| HPS1 | 0 | 0 | 0 | 1096 | 996 | 774 | 540 | 0 |
| HPS3 | 0 | 0 | 0 | 914 | 0 | 0 | 0 | 448 |
| HPS4 | 854 | 162 | 1484 | 0 | 0 | 0 | 1342 | 0 |
| HPS5 | 0 | 0 | 0 | 0 | 0 | 0 | 398 | 0 |
| HPS6 | 726 | 482 | 1346 | 0 | 992 | 0 | 0 | 0 |
| HPSE | 0 | 0 | 0 | 0 | 0 | 1574 | 0 | 0 |
| HR | 292 | 0 | 0 | 0 | 0 | 0 | 0 | 0 |
| HRAS | 0 | 0 | 0 | 0 | 1066 | 0 | 766 | 0 |
| HRASLS | 1136 | 0 | 0 | 0 | 0 | 0 | 0 | 0 |
| HRASLS2 | 114922 | 7673 | 1093150 | 0 | 17170 | 9265 | 5148 | 28343 |
| HRCT1 | 904 | 262 | 60 | 0 | 374 | 0 | 0 | 0 |
| HRH1 | 5658 | 7491 | 21531 | 6310 | 18490 | 14352 | 23864 | 4208 |
| HS1BP3 | 1120 | 0 | 0 | 0 | 0 | 0 | 0 | 0 |
| HS6ST1 | 0 | 0 | 0 | 0 | 290 | 0 | 0 | 0 |
| HSBP1 | 2343 | 1642 | 5718 | 10590 | 9058 | 3285 | 3714 | 2744 |

| | | | | | | | | |
|---|---|---|---|---|---|---|---|---|
| HSBP1L1 | 858 | 1970 | 1560 | 0 | 3226 | 4498 | 2622 | 744 |
| HSCB | 0 | 628 | 0 | 2642 | 1368 | 0 | 1284 | 0 |
| HSD11B1 | 462 | 3254 | 9975 | 4783 | 6185 | 47160 | 4186 | 20492 |
| HSD11B1-AS1 | 0 | 0 | 0 | 0 | 0 | 0 | 0 | 540 |
| HSD11B2 | 16501 | 24697 | 35009 | 7269 | 13858 | 2342 | 21939 | 4539 |
| HSD17B10 | 0 | 2002 | 1548 | 3456 | 5047 | 2020 | 3743 | 0 |
| HSD17B11 | 0 | 1021 | 12839 | 888 | 2394 | 1651 | 2617 | 1828 |
| HSD17B12 | 12052 | 150117 | 86576 | 243817 | 311670 | 73943 | 231349 | 123744 |
| HSD17B14 | 0 | 420 | 10427 | 0 | 736 | 3111 | 5308 | 2013 |
| HSD17B2 | 2358 | 872 | 14780 | 0 | 736 | 0 | 0 | 1522 |
| HSD17B4 | 356 | 1492 | 0 | 2758 | 1520 | 1823 | 1524 | 0 |
| HSD17B6 | 0 | 0 | 0 | 0 | 142 | 0 | 0 | 0 |
| HSD17B7 | 0 | 0 | 1300 | 1100 | 1190 | 2478 | 0 | 0 |
| HSD17B7P2 | 0 | 278 | 0 | 0 | 0 | 0 | 0 | 0 |
| HSD17B8 | 448 | 442 | 1082 | 806 | 938 | 0 | 0 | 0 |
| HSD3B1 | 0 | 0 | 0 | 0 | 0 | 2326 | 0 | 0 |
| HSD3B2 | 0 | 0 | 1325 | 0 | 0 | 0 | 0 | 0 |
| HSDL2 | 0 | 0 | 0 | 0 | 338 | 0 | 422 | 579 |
| HSF1 | 0 | 0 | 0 | 674 | 1862 | 2091 | 3888 | 0 |
| HSF2 | 0 | 472 | 0 | 0 | 458 | 0 | 0 | 0 |
| HSF4 | 0 | 0 | 0 | 150 | 584 | 0 | 286 | 0 |
| HSF5 | 0 | 0 | 302 | 0 | 320 | 0 | 0 | 0 |
| HSFX2 | 0 | 0 | 1522 | 0 | 0 | 0 | 0 | 0 |
| HSH2D | 0 | 366 | 1116 | 0 | 612 | 0 | 0 | 0 |
| HSP90AA1 | 125278 | 228230 | 231562 | 237558 | 1604454 | 341457 | 516614 | 370681 |
| HSP90AB1 | 3348 | 3678 | 2486 | 9452 | 21455 | 14213 | 16612 | 13171 |
| HSP90B1 | 702 | 2363 | 1472 | 7494 | 5643 | 7967 | 4586 | 4283 |
| HSPA13 | 0 | 532 | 0 | 0 | 0 | 0 | 226 | 0 |
| HSPA14 | 0 | 0 | 0 | 0 | 740 | 0 | 0 | 0 |
| HSPA1A | 0 | 0 | 0 | 719 | 754 | 1408 | 0 | 176 |
| HSPA1B | 580 | 0 | 0 | 1648 | 5387 | 3363 | 1662 | 1762 |
| HSPA5 | 800 | 1302 | 3850 | 3927 | 12127 | 2206 | 1362 | 3246 |
| HSPA6 | 0 | 0 | 0 | 0 | 0 | 0 | 0 | 1190 |
| HSPA8 | 0 | 5852 | 4791 | 17247 | 7398 | 15553 | 6415 | 2040 |
| HSPA9 | 396 | 1601 | 3246 | 3098 | 1154 | 0 | 0 | 922 |
| HSPB1 | 2202 | 2083 | 8533 | 5224 | 16476 | 10274 | 6991 | 8711 |
| HSPB11 | 194 | 578 | 1466 | 852 | 1032 | 0 | 1695 | 1765 |
| HSPB2-C11orf52 | 0 | 0 | 0 | 0 | 560 | 0 | 0 | 0 |
| HSPB3 | 0 | 0 | 1936 | 0 | 0 | 0 | 1220 | 0 |
| HSPB7 | 0 | 0 | 0 | 0 | 386 | 1384 | 0 | 0 |
| HSPB8 | 0 | 0 | 1388 | 0 | 0 | 0 | 0 | 0 |
| HSPBAP1 | 0 | 0 | 0 | 0 | 678 | 0 | 432 | 998 |
| HSPBP1 | 0 | 334 | 1138 | 0 | 0 | 3724 | 0 | 958 |

| | | | | | | | | |
|---|---|---|---|---|---|---|---|---|
| HSPD1 | 347400 | 1053391 | 1598406 | 3230796 | 6972668 | 4550579 | 9363937 | 2191949 |
| HSPE1 | 182 | 9241 | 4502 | 19605 | 25938 | 4663 | 16648 | 6789 |
| HSPG2 | 0 | 0 | 0 | 0 | 1436 | 0 | 192 | 458 |
| HSPH1 | 2230 | 7233 | 3106 | 25085 | 31777 | 47184 | 44612 | 12897 |
| HTATIP2 | 1562 | 1224 | 9438 | 4039 | 3530 | 4666 | 410 | 1638 |
| HTATSF1 | 17715 | 29077 | 46811 | 32896 | 175432 | 71144 | 102880 | 19001 |
| HTD2 | 862 | 0 | 0 | 0 | 0 | 0 | 0 | 0 |
| HTR7P1 | 0 | 0 | 0 | 0 | 254 | 0 | 0 | 0 |
| HTRA1 | 648 | 0 | 0 | 0 | 1191 | 0 | 0 | 0 |
| HTRA2 | 0 | 0 | 0 | 0 | 0 | 0 | 0 | 242 |
| HUNK | 0 | 0 | 0 | 0 | 0 | 0 | 110 | 0 |
| HUS1 | 0 | 480 | 0 | 1262 | 0 | 0 | 0 | 0 |
| HUWE1 | 7510 | 8897 | 5830 | 3622 | 19979 | 3024 | 18481 | 9834 |
| HYAL2 | 0 | 0 | 3618 | 2107 | 1130 | 0 | 0 | 0 |
| HYI | 756 | 606 | 0 | 3858 | 430 | 0 | 1362 | 0 |
| HYOU1 | 848 | 236 | 696 | 0 | 434 | 578 | 952 | 446 |
| IAH1 | 826 | 904 | 1586 | 890 | 949 | 4014 | 170 | 1137 |
| IARS | 0 | 214 | 0 | 0 | 2200 | 0 | 0 | 0 |
| IARS2 | 58 | 1396 | 0 | 0 | 1554 | 0 | 0 | 2263 |
| ICA1 | 362 | 0 | 2373 | 574 | 0 | 0 | 320 | 1168 |
| ICAM1 | 1384 | 0 | 0 | 0 | 0 | 1464 | 0 | 0 |
| ICAM2 | 33973 | 33629 | 52784 | 35816 | 53545 | 109580 | 60309 | 35012 |
| ICAM3 | 0 | 0 | 0 | 528 | 0 | 0 | 0 | 1096 |
| ICAM4 | 0 | 488 | 0 | 0 | 0 | 0 | 0 | 0 |
| ICE1 | 166 | 0 | 0 | 1175 | 472 | 0 | 1043 | 0 |
| ICE2 | 0 | 0 | 0 | 0 | 310 | 0 | 0 | 0 |
| ICMT | 0 | 0 | 1206 | 0 | 1042 | 0 | 0 | 0 |
| ID1 | 8450 | 2928 | 21822 | 7285 | 986 | 1080 | 9804 | 0 |
| ID2 | 1052 | 1126 | 1358 | 0 | 3190 | 1342 | 1420 | 896 |
| ID2-AS1 | 0 | 0 | 0 | 0 | 0 | 0 | 414 | 839 |
| ID3 | 7562 | 184 | 4536 | 2946 | 1478 | 5318 | 2314 | 2556 |
| ID4 | 0 | 0 | 0 | 0 | 644 | 772 | 2284 | 0 |
| IDH1 | 190 | 1276 | 3317 | 4594 | 4486 | 1888 | 864 | 0 |
| IDH1-AS1 | 0 | 0 | 0 | 0 | 358 | 0 | 0 | 0 |
| IDH2 | 0 | 610 | 0 | 2906 | 13741 | 0 | 5978 | 3128 |
| IDH3A | 830 | 208 | 2616 | 602 | 1406 | 0 | 0 | 0 |
| IDH3B | 598 | 1314 | 5370 | 2949 | 3228 | 6554 | 1562 | 1874 |
| IDH3G | 698 | 830 | 3800 | 0 | 3150 | 0 | 278 | 1214 |
| IDI1 | 122 | 0 | 2517 | 1200 | 3176 | 0 | 416 | 260 |
| IDNK | 0 | 0 | 262 | 0 | 602 | 0 | 0 | 0 |
| IDO1 | 42272 | 6786 | 65555 | 5162 | 18648 | 52537 | 11343 | 27314 |
| IDS | 0 | 54 | 0 | 0 | 596 | 0 | 0 | 0 |
| IDUA | 310 | 0 | 0 | 0 | 186 | 0 | 0 | 0 |

| | | | | | | | | |
|---|---|---|---|---|---|---|---|---|
| IER2 | 1723 | 826 | 3913 | 2691 | 5074 | 0 | 0 | 0 |
| IER3 | 1455 | 1060 | 4536 | 2179 | 1554 | 2160 | 7022 | 2943 |
| IER3IP1 | 360 | 648 | 1486 | 1125 | 1154 | 0 | 0 | 545 |
| IER5L | 0 | 0 | 0 | 704 | 2138 | 0 | 854 | 218 |
| IFFO1 | 0 | 0 | 0 | 0 | 232 | 0 | 0 | 0 |
| IFI16 | 558 | 266 | 0 | 0 | 0 | 0 | 0 | 418 |
| IFI27 | 19813 | 9502 | 71029 | 20924 | 49392 | 15155 | 10165 | 11370 |
| IFI27L1 | 838 | 0 | 1764 | 0 | 0 | 1424 | 589 | 0 |
| IFI27L2 | 0 | 0 | 0 | 954 | 830 | 1232 | 0 | 512 |
| IFI30 | 3189 | 554 | 2275 | 1146 | 4876 | 1956 | 2288 | 6362 |
| IFI35 | 0 | 0 | 436 | 0 | 552 | 2088 | 312 | 1234 |
| IFI44 | 0 | 0 | 0 | 0 | 24 | 0 | 0 | 0 |
| IFI44L | 0 | 0 | 0 | 0 | 0 | 0 | 0 | 224 |
| IFI6 | 1956 | 1802 | 9331 | 906 | 10710 | 0 | 878 | 6506 |
| IFIH1 | 0 | 0 | 0 | 0 | 1024 | 0 | 0 | 0 |
| IFIT1 | 0 | 0 | 2818 | 0 | 278 | 0 | 0 | 0 |
| IFIT2 | 0 | 0 | 0 | 0 | 286 | 0 | 0 | 0 |
| IFIT3 | 298 | 0 | 1526 | 770 | 0 | 0 | 0 | 0 |
| IFIT5 | 0 | 0 | 0 | 0 | 0 | 0 | 0 | 232 |
| IFITM1 | 5809 | 4094 | 13916 | 28214 | 8204 | 21424 | 38013 | 5561 |
| IFITM2 | 476128 | 614349 | 1248564 | 917818 | 951080 | 2229386 | 1957643 | 437771 |
| IFITM3 | 6536 | 7141 | 12587 | 13072 | 39273 | 31337 | 13321 | 24827 |
| IFNAR1 | 5898 | 8729 | 5414 | 4318 | 26522 | 14782 | 24542 | 8228 |
| IFNAR2 | 0 | 0 | 814 | 1610 | 917 | 4335 | 0 | 1208 |
| IFNGR1 | 510 | 1104 | 0 | 880 | 1608 | 404 | 0 | 1988 |
| IFNGR2 | 638 | 688 | 4666 | 1684 | 3730 | 1602 | 2568 | 2462 |
| IFNLR1 | 0 | 0 | 0 | 0 | 0 | 0 | 368 | 0 |
| IFRD1 | 0 | 0 | 994 | 0 | 3520 | 1120 | 1134 | 742 |
| IFRD2 | 0 | 244 | 848 | 1236 | 1454 | 0 | 982 | 0 |
| IFT140 | 1156 | 2050 | 4835 | 5504 | 1950 | 0 | 8492 | 2138 |
| IFT172 | 0 | 0 | 3232 | 1140 | 68 | 0 | 450 | 0 |
| IFT20 | 1772 | 94 | 1464 | 0 | 3174 | 1890 | 0 | 0 |
| IFT22 | 0 | 0 | 0 | 0 | 166 | 2269 | 374 | 0 |
| IFT27 | 0 | 280 | 0 | 0 | 662 | 0 | 656 | 0 |
| IFT43 | 0 | 2041 | 1834 | 0 | 2657 | 2094 | 0 | 3172 |
| IFT46 | 0 | 0 | 0 | 0 | 0 | 0 | 0 | 554 |
| IFT52 | 0 | 0 | 2280 | 1262 | 580 | 0 | 496 | 0 |
| IFT57 | 1778 | 406 | 1535 | 0 | 2447 | 0 | 1416 | 0 |
| IFT74 | 0 | 0 | 1270 | 0 | 0 | 0 | 380 | 0 |
| IFT80 | 700 | 756 | 3186 | 3385 | 9170 | 0 | 1078 | 4072 |
| IFT81 | 0 | 0 | 0 | 0 | 0 | 0 | 0 | 1822 |
| IFT88 | 0 | 0 | 2046 | 0 | 0 | 0 | 540 | 0 |
| IGBP1 | 810 | 506 | 1078 | 0 | 4506 | 612 | 2482 | 1882 |

| | | | | | | | | |
|---|---|---|---|---|---|---|---|---|
| IGF1 | 0 | 0 | 0 | 0 | 400 | 0 | 0 | 0 |
| IGF1R | 326 | 0 | 0 | 0 | 0 | 0 | 0 | 0 |
| IGF2 | 0 | 0 | 4062 | 576 | 15473 | 6756 | 224 | 0 |
| IGF2BP2 | 10649 | 20875 | 17715 | 11037 | 39723 | 21853 | 59118 | 23774 |
| IGF2R | 3000 | 2256 | 2466 | 7117 | 12005 | 6124 | 22200 | 4648 |
| IGFBP2 | 0 | 0 | 740 | 5819 | 1467 | 1896 | 2901 | 0 |
| IGFBP3 | 0 | 282 | 0 | 0 | 248 | 0 | 0 | 254 |
| IGFBP4 | 0 | 330 | 2586 | 0 | 850 | 0 | 1322 | 1204 |
| IGFBP5 | 736 | 430 | 0 | 672 | 522 | 1170 | 0 | 0 |
| IGFBP6 | 0 | 0 | 0 | 0 | 0 | 1340 | 0 | 0 |
| IGFBP7 | 2334 | 3353 | 3610 | 9211 | 26042 | 13685 | 931 | 19082 |
| IGFL2 | 0 | 0 | 0 | 0 | 1430 | 0 | 0 | 0 |
| IGFL2-AS1 | 0 | 0 | 0 | 0 | 364 | 0 | 0 | 0 |
| IGLL1 | 0 | 0 | 0 | 0 | 0 | 0 | 787 | 0 |
| IGLL5 | 26275 | 33152 | 97382 | 14918 | 4444 | 58577 | 402 | 127386 |
| IGSF22 | 0 | 0 | 0 | 1134 | 0 | 0 | 0 | 0 |
| IGSF3 | 1146 | 0 | 0 | 0 | 902 | 0 | 0 | 0 |
| IGSF8 | 0 | 222 | 0 | 0 | 564 | 3533 | 1300 | 0 |
| IGSF9 | 0 | 136 | 0 | 0 | 0 | 0 | 0 | 0 |
| IHH | 336 | 204 | 1076 | 0 | 942 | 1454 | 874 | 0 |
| IK | 0 | 1046 | 7214 | 4168 | 4986 | 0 | 712 | 1423 |
| IKBIP | 0 | 0 | 0 | 0 | 582 | 0 | 0 | 0 |
| IKBKB | 9028 | 17308 | 17764 | 21181 | 84978 | 65031 | 76156 | 27279 |
| IKBKE | 0 | 0 | 0 | 0 | 646 | 0 | 0 | 0 |
| IKBKG | 0 | 0 | 0 | 0 | 1662 | 0 | 0 | 0 |
| IKZF1 | 3526 | 3878 | 2560 | 0 | 1280 | 1330 | 216 | 2164 |
| IKZF3 | 194 | 234 | 0 | 0 | 5517 | 0 | 0 | 0 |
| IKZF5 | 0 | 0 | 0 | 0 | 580 | 0 | 0 | 0 |
| IL10RB | 498 | 254 | 3000 | 0 | 3982 | 0 | 302 | 646 |
| IL13RA1 | 0 | 606 | 1294 | 0 | 674 | 1406 | 0 | 0 |
| IL15RA | 1654 | 0 | 216 | 419 | 0 | 0 | 0 | 436 |
| IL16 | 0 | 258 | 0 | 0 | 0 | 0 | 820 | 0 |
| IL17RA | 0 | 226 | 0 | 0 | 278 | 0 | 0 | 0 |
| IL17RB | 0 | 0 | 0 | 1472 | 0 | 0 | 934 | 0 |
| IL17RC | 0 | 0 | 1007 | 0 | 0 | 0 | 0 | 0 |
| IL17RD | 0 | 0 | 0 | 0 | 0 | 0 | 0 | 504 |
| IL17RE | 36989 | 41852 | 167718 | 29631 | 175417 | 40334 | 156780 | 25490 |
| IL18 | 922 | 1318 | 3448 | 1442 | 0 | 1958 | 0 | 1468 |
| IL1B | 0 | 0 | 0 | 1191 | 0 | 0 | 0 | 0 |
| IL1R1 | 0 | 0 | 0 | 0 | 322 | 1327 | 0 | 0 |
| IL1R2 | 0 | 0 | 4074 | 0 | 782 | 0 | 0 | 0 |
| IL1RAP | 0 | 0 | 0 | 0 | 274 | 0 | 0 | 0 |
| IL20RA | 0 | 3072 | 4019 | 10477 | 21178 | 6415 | 25246 | 6037 |

| | | | | | | | | |
|---|---|---|---|---|---|---|---|---|
| IL20RB | 0 | 370 | 3089 | 1408 | 0 | 0 | 0 | 0 |
| IL20RB-AS1 | 0 | 0 | 0 | 0 | 20 | 0 | 0 | 0 |
| IL22RA1 | 0 | 0 | 1401 | 0 | 0 | 0 | 1182 | 0 |
| IL27RA | 2854 | 1566 | 1268 | 2252 | 12715 | 1614 | 1756 | 0 |
| IL2RB | 0 | 0 | 0 | 434 | 0 | 0 | 0 | 0 |
| IL2RG | 30 | 1140 | 4965 | 0 | 782 | 1756 | 1260 | 832 |
| IL32 | 5494 | 1878 | 24835 | 8881 | 13627 | 13834 | 0 | 4590 |
| IL33 | 0 | 216 | 0 | 533 | 0 | 0 | 0 | 0 |
| IL34 | 732 | 0 | 0 | 0 | 0 | 0 | 0 | 0 |
| IL3RA | 0 | 0 | 0 | 0 | 320 | 0 | 0 | 0 |
| IL4I1 | 0 | 512 | 0 | 0 | 0 | 0 | 0 | 0 |
| IL4R | 330 | 0 | 0 | 0 | 0 | 0 | 0 | 0 |
| IL6R | 468 | 0 | 0 | 0 | 0 | 0 | 0 | 0 |
| IL6ST | 6880 | 12736 | 10072 | 3388 | 8704 | 4200 | 2632 | 2725 |
| IL7 | 290 | 0 | 0 | 0 | 0 | 0 | 0 | 0 |
| IL7R | 654 | 0 | 0 | 0 | 0 | 0 | 0 | 0 |
| ILDR1 | 0 | 0 | 0 | 244 | 0 | 0 | 0 | 0 |
| ILF2 | 574 | 2358 | 3646 | 7853 | 6580 | 3449 | 6282 | 1217 |
| ILF3 | 24128 | 18177 | 15833 | 29271 | 71542 | 27831 | 32359 | 13201 |
| ILF3-DT | 0 | 52 | 0 | 0 | 0 | 0 | 0 | 0 |
| ILK | 0 | 0 | 0 | 1316 | 3500 | 1430 | 980 | 3758 |
| ILKAP | 910 | 188 | 872 | 208 | 384 | 0 | 0 | 0 |
| ILVBL | 306 | 500 | 530 | 520 | 2110 | 3586 | 0 | 0 |
| IMMP1L | 0 | 0 | 0 | 1642 | 2288 | 0 | 446 | 0 |
| IMMP2L | 23514 | 44665 | 95274 | 87890 | 160803 | 204624 | 369964 | 42174 |
| IMMT | 352 | 0 | 2482 | 0 | 1112 | 0 | 686 | 1006 |
| IMP3 | 2464 | 1544 | 6833 | 757 | 4298 | 0 | 2343 | 0 |
| IMP4 | 0 | 1001 | 0 | 2264 | 4557 | 1402 | 0 | 1180 |
| IMPA1 | 0 | 300 | 0 | 0 | 720 | 2403 | 0 | 0 |
| IMPA2 | 0 | 1176 | 1206 | 0 | 5548 | 0 | 0 | 0 |
| IMPACT | 0 | 0 | 0 | 0 | 546 | 999 | 0 | 0 |
| IMPAD1 | 0 | 0 | 0 | 0 | 926 | 0 | 448 | 0 |
| IMPDH1 | 0 | 208 | 0 | 0 | 192 | 0 | 186 | 1241 |
| IMPDH2 | 204 | 1418 | 604 | 880 | 9949 | 448 | 1808 | 1452 |
| INAVA | 0 | 326 | 1469 | 248 | 1854 | 0 | 0 | 0 |
| INCA1 | 0 | 432 | 0 | 0 | 0 | 0 | 0 | 0 |
| INCENP | 0 | 0 | 0 | 0 | 0 | 0 | 1092 | 0 |
| INF2 | 0 | 354 | 2228 | 808 | 1604 | 0 | 0 | 0 |
| ING1 | 0 | 0 | 0 | 320 | 0 | 0 | 0 | 882 |
| ING2 | 0 | 0 | 0 | 0 | 1926 | 0 | 0 | 0 |
| ING3 | 694 | 0 | 0 | 0 | 1424 | 0 | 0 | 0 |
| ING4 | 0 | 296 | 2908 | 0 | 1506 | 0 | 252 | 0 |
| ING5 | 0 | 348 | 0 | 0 | 1689 | 576 | 312 | 0 |

| | | | | | | | | |
|---|---|---|---|---|---|---|---|---|
| INHBA | 0 | 0 | 0 | 0 | 622 | 0 | 0 | 0 |
| INIP | 66 | 0 | 0 | 0 | 0 | 0 | 0 | 0 |
| INMT | 0 | 216 | 0 | 0 | 0 | 0 | 0 | 0 |
| INO80 | 0 | 0 | 0 | 0 | 0 | 0 | 152 | 0 |
| INO80B | 0 | 0 | 0 | 0 | 310 | 0 | 0 | 0 |
| INO80B-WBP1 | 0 | 0 | 2176 | 0 | 694 | 0 | 0 | 0 |
| INO80C | 0 | 472 | 0 | 1796 | 2014 | 0 | 0 | 1044 |
| INO80E | 0 | 194 | 1700 | 540 | 1548 | 0 | 416 | 0 |
| INPP1 | 0 | 0 | 0 | 0 | 0 | 0 | 0 | 792 |
| INPP4B | 3469 | 4888 | 11397 | 3046 | 17369 | 2167 | 33580 | 7660 |
| INPP5A | 0 | 76 | 268 | 378 | 310 | 0 | 0 | 0 |
| INPP5D | 544 | 1548 | 0 | 1670 | 0 | 5847 | 3194 | 0 |
| INPP5E | 0 | 0 | 0 | 0 | 228 | 0 | 0 | 0 |
| INPP5F | 0 | 194 | 3088 | 0 | 0 | 0 | 0 | 0 |
| INPP5K | 4362 | 2576 | 20360 | 5368 | 4942 | 3772 | 5434 | 6584 |
| INPPL1 | 0 | 0 | 0 | 0 | 110 | 0 | 0 | 0 |
| INS-IGF2 | 2640 | 0 | 2761 | 440 | 16524 | 3367 | 962 | 0 |
| INSC | 1720 | 0 | 328 | 0 | 0 | 0 | 0 | 216 |
| INSIG1 | 756 | 0 | 1344 | 282 | 642 | 0 | 0 | 1042 |
| INSIG2 | 640 | 314 | 1592 | 0 | 750 | 0 | 0 | 0 |
| INSL5 | 0 | 4425 | 0 | 0 | 370 | 0 | 0 | 0 |
| INSR | 256 | 370 | 2330 | 0 | 0 | 0 | 374 | 0 |
| INTS1 | 0 | 0 | 0 | 0 | 110 | 0 | 0 | 0 |
| INTS10 | 706 | 0 | 0 | 0 | 1206 | 0 | 1505 | 0 |
| INTS11 | 476 | 0 | 0 | 338 | 1816 | 0 | 0 | 0 |
| INTS12 | 0 | 326 | 561 | 0 | 0 | 0 | 0 | 0 |
| INTS13 | 0 | 0 | 0 | 0 | 508 | 0 | 0 | 0 |
| INTS14 | 16253 | 45374 | 68699 | 65956 | 95621 | 28249 | 157501 | 30617 |
| INTS3 | 140 | 0 | 0 | 0 | 0 | 0 | 454 | 572 |
| INTS4 | 0 | 0 | 0 | 0 | 738 | 0 | 0 | 0 |
| INTS5 | 0 | 470 | 0 | 0 | 0 | 0 | 0 | 0 |
| INTS6 | 0 | 268 | 0 | 555 | 0 | 0 | 0 | 0 |
| INTS6L | 0 | 0 | 1670 | 0 | 0 | 828 | 0 | 0 |
| INTS7 | 5600 | 9675 | 24857 | 20876 | 34578 | 15517 | 20000 | 9919 |
| INTS8 | 0 | 0 | 0 | 881 | 358 | 0 | 286 | 0 |
| INTS9 | 1790 | 13004 | 18607 | 9676 | 33843 | 22684 | 22050 | 5480 |
| INTU | 0 | 0 | 0 | 0 | 290 | 0 | 0 | 0 |
| IP6K1 | 0 | 666 | 0 | 0 | 418 | 0 | 0 | 0 |
| IP6K2 | 882 | 0 | 1458 | 0 | 8587 | 0 | 1148 | 0 |
| IPMK | 0 | 0 | 0 | 0 | 180 | 0 | 0 | 0 |
| IPO11-LRRC70 | 0 | 0 | 0 | 0 | 114 | 1914 | 0 | 0 |
| IPO4 | 0 | 0 | 0 | 0 | 478 | 2394 | 314 | 236 |
| IPO5 | 0 | 0 | 510 | 1014 | 710 | 0 | 0 | 1531 |

| | | | | | | | | |
|---|---|---|---|---|---|---|---|---|
| IPO7 | 982 | 9240 | 6606 | 3498 | 17798 | 12641 | 19706 | 3226 |
| IPO8 | 80 | 150 | 0 | 0 | 0 | 0 | 280 | 0 |
| IPO9 | 0 | 0 | 0 | 380 | 250 | 0 | 0 | 0 |
| IPPK | 3422 | 3969 | 11558 | 4569 | 12122 | 10788 | 8349 | 1452 |
| IQCB1 | 0 | 0 | 0 | 590 | 758 | 0 | 0 | 2462 |
| IQCC | 0 | 0 | 0 | 0 | 60 | 0 | 0 | 0 |
| IQCE | 0 | 0 | 0 | 0 | 176 | 0 | 0 | 0 |
| IQCG | 0 | 146 | 0 | 0 | 0 | 0 | 0 | 0 |
| IQCH | 938 | 1996 | 1054 | 3156 | 11781 | 0 | 2104 | 0 |
| IQCH-AS1 | 0 | 0 | 0 | 0 | 300 | 0 | 406 | 0 |
| IQCK | 0 | 0 | 0 | 0 | 404 | 0 | 0 | 0 |
| IQGAP1 | 51729 | 43062 | 72604 | 45577 | 100132 | 37165 | 93393 | 25073 |
| IQGAP2 | 216 | 0 | 0 | 0 | 0 | 0 | 0 | 0 |
| IQGAP3 | 0 | 0 | 0 | 0 | 322 | 0 | 0 | 0 |
| IQSEC1 | 0 | 0 | 0 | 0 | 0 | 0 | 770 | 0 |
| IQSEC2 | 0 | 0 | 0 | 0 | 642 | 0 | 0 | 0 |
| IRAIN | 0 | 0 | 0 | 0 | 0 | 0 | 436 | 0 |
| IRAK1 | 306 | 0 | 0 | 299 | 1992 | 3190 | 0 | 380 |
| IREB2 | 0 | 350 | 0 | 0 | 0 | 0 | 0 | 0 |
| IRF1 | 1296 | 380 | 1650 | 0 | 986 | 1552 | 0 | 0 |
| IRF2 | 77424 | 82153 | 326240 | 47228 | 154150 | 101337 | 198630 | 48257 |
| IRF2BP1 | 0 | 354 | 0 | 0 | 482 | 0 | 190 | 0 |
| IRF2BP2 | 2242 | 3782 | 654 | 8142 | 10773 | 1925 | 13061 | 1700 |
| IRF3 | 302 | 0 | 0 | 1048 | 710 | 0 | 0 | 0 |
| IRF6 | 406 | 324 | 0 | 340 | 476 | 0 | 0 | 0 |
| IRF7 | 374 | 0 | 188 | 0 | 0 | 0 | 504 | 0 |
| IRF8 | 0 | 224 | 0 | 1500 | 889 | 428 | 0 | 0 |
| IRF9 | 1108 | 546 | 5933 | 0 | 4846 | 0 | 732 | 672 |
| IRGQ | 0 | 0 | 0 | 0 | 162 | 0 | 282 | 0 |
| IRS1 | 1293 | 1962 | 3341 | 4751 | 37316 | 0 | 2804 | 4838 |
| IRS2 | 0 | 0 | 0 | 0 | 488 | 0 | 0 | 0 |
| IRX2 | 0 | 150 | 0 | 0 | 220 | 0 | 0 | 0 |
| ISCA1 | 872 | 0 | 0 | 0 | 1490 | 2583 | 1426 | 0 |
| ISCA2 | 2456 | 396 | 3390 | 2400 | 4284 | 0 | 401 | 2761 |
| ISCU | 2506 | 3442 | 7422 | 9009 | 3382 | 7242 | 5476 | 0 |
| ISG15 | 3481 | 1916 | 10730 | 0 | 4638 | 2813 | 0 | 200 |
| ISG20 | 0 | 1726 | 7586 | 3881 | 0 | 0 | 1038 | 0 |
| ISG20L2 | 490 | 0 | 0 | 0 | 502 | 0 | 386 | 0 |
| ISLR | 0 | 478 | 958 | 0 | 0 | 0 | 0 | 0 |
| ISOC1 | 0 | 0 | 364 | 958 | 1859 | 0 | 0 | 0 |
| ISOC2 | 802 | 1474 | 2000 | 0 | 1880 | 0 | 755 | 0 |
| IST1 | 0 | 448 | 582 | 1094 | 1610 | 0 | 571 | 350 |
| ISX | 2097 | 0 | 1598 | 0 | 0 | 0 | 0 | 0 |

| | | | | | | | | |
|---|---|---|---|---|---|---|---|---|
| ISY1 | 994 | 340 | 1751 | 1588 | 2671 | 0 | 1360 | 0 |
| ISY1-RAB43 | 0 | 0 | 0 | 482 | 0 | 0 | 0 | 0 |
| ISYNA1 | 0 | 0 | 0 | 0 | 1200 | 0 | 0 | 0 |
| ITCH | 0 | 270 | 0 | 0 | 0 | 0 | 0 | 0 |
| ITFG1 | 0 | 508 | 1010 | 0 | 586 | 0 | 212 | 568 |
| ITFG2 | 1390 | 624 | 0 | 0 | 0 | 0 | 170 | 0 |
| ITFG2-AS1 | 0 | 0 | 0 | 0 | 552 | 0 | 0 | 0 |
| ITGA1 | 0 | 0 | 556 | 0 | 478 | 0 | 0 | 0 |
| ITGA11 | 0 | 0 | 0 | 0 | 748 | 0 | 0 | 0 |
| ITGA2 | 602 | 284 | 1584 | 2000 | 5060 | 1374 | 2302 | 872 |
| ITGA3 | 80 | 0 | 468 | 684 | 0 | 0 | 0 | 0 |
| ITGA5 | 0 | 0 | 0 | 0 | 1782 | 0 | 0 | 0 |
| ITGA6 | 558 | 18 | 0 | 0 | 631 | 0 | 20 | 78 |
| ITGA7 | 0 | 0 | 0 | 0 | 0 | 0 | 594 | 0 |
| ITGA9 | 0 | 0 | 0 | 0 | 0 | 1532 | 0 | 0 |
| ITGA9-AS1 | 0 | 0 | 0 | 0 | 514 | 0 | 238 | 0 |
| ITGAE | 1026 | 1960 | 1462 | 0 | 3586 | 3199 | 0 | 1908 |
| ITGAV | 214 | 342 | 578 | 1582 | 3344 | 206 | 1122 | 944 |
| ITGB1 | 0 | 622 | 1319 | 2218 | 2072 | 0 | 0 | 172 |
| ITGB1-DT | 0 | 0 | 0 | 0 | 886 | 0 | 0 | 0 |
| ITGB1BP1 | 302 | 544 | 751 | 1185 | 4058 | 1903 | 372 | 0 |
| ITGB2 | 114 | 0 | 0 | 0 | 2972 | 1538 | 0 | 550 |
| ITGB3 | 0 | 0 | 0 | 0 | 0 | 0 | 386 | 0 |
| ITGB3BP | 0 | 0 | 0 | 1762 | 2262 | 0 | 0 | 0 |
| ITGB4 | 0 | 216 | 0 | 0 | 0 | 1072 | 0 | 390 |
| ITGB5 | 3532 | 2218 | 5250 | 4774 | 18635 | 9432 | 5145 | 7144 |
| ITGB6 | 6256 | 11312 | 15384 | 9330 | 54797 | 8595 | 25134 | 14695 |
| ITGB8 | 0 | 0 | 0 | 0 | 0 | 0 | 68 | 0 |
| ITIH3 | 158 | 0 | 0 | 0 | 0 | 0 | 0 | 0 |
| ITIH5 | 0 | 0 | 0 | 522 | 0 | 0 | 0 | 0 |
| ITK | 0 | 0 | 0 | 0 | 476 | 0 | 0 | 0 |
| ITLN1 | 223059 | 2623282 | 1991972 | 146621 | 122868 | 0 | 2572 | 0 |
| ITM2A | 0 | 0 | 322 | 0 | 0 | 0 | 0 | 0 |
| ITM2B | 7953 | 6075 | 21438 | 6522 | 13843 | 15953 | 7785 | 3258 |
| ITM2C | 6690 | 7533 | 15223 | 4384 | 2944 | 0 | 620 | 0 |
| ITPA | 832 | 622 | 0 | 6004 | 1788 | 0 | 630 | 2356 |
| ITPKA | 62 | 0 | 2794 | 1047 | 0 | 0 | 0 | 0 |
| ITPKC | 876 | 856 | 4524 | 0 | 3260 | 4983 | 880 | 1258 |
| ITPR1 | 1096 | 0 | 0 | 3581 | 430 | 1176 | 932 | 0 |
| ITPR2 | 12254 | 12148 | 2153 | 8484 | 8504 | 17991 | 12168 | 4875 |
| ITPR3 | 0 | 0 | 0 | 971 | 1062 | 0 | 0 | 0 |
| ITPRID2 | 0 | 0 | 0 | 862 | 0 | 0 | 0 | 0 |
| ITPRIP | 276 | 0 | 0 | 0 | 0 | 0 | 0 | 0 |

| | | | | | | | | |
|---|---|---|---|---|---|---|---|---|
| ITSN2 | 0 | 0 | 870 | 0 | 0 | 0 | 0 | 494 |
| IVD | 0 | 582 | 0 | 0 | 795 | 1780 | 0 | 1524 |
| IVNS1ABP | 1326 | 762 | 1332 | 0 | 1566 | 1246 | 0 | 2145 |
| IWS1 | 0 | 0 | 0 | 1050 | 0 | 1902 | 633 | 0 |
| IYD | 0 | 0 | 1180 | 0 | 590 | 1405 | 0 | 0 |
| JADE1 | 0 | 0 | 0 | 0 | 0 | 0 | 670 | 0 |
| JADE3 | 0 | 0 | 0 | 898 | 2966 | 0 | 1372 | 1212 |
| JAG1 | 0 | 374 | 0 | 0 | 0 | 1257 | 0 | 504 |
| JAG2 | 0 | 0 | 0 | 0 | 438 | 0 | 0 | 0 |
| JAGN1 | 0 | 898 | 6088 | 2248 | 2173 | 2454 | 2649 | 944 |
| JAK1 | 13839 | 24921 | 37720 | 26497 | 39996 | 26261 | 44537 | 11577 |
| JAK2 | 6403 | 7881 | 15684 | 3822 | 9602 | 2918 | 5225 | 6344 |
| JAML | 0 | 0 | 0 | 0 | 336 | 0 | 0 | 0 |
| JAZF1 | 0 | 0 | 0 | 0 | 0 | 0 | 586 | 0 |
| JCHAIN | 1774 | 8170 | 20680 | 0 | 0 | 2169 | 0 | 900 |
| JDP2 | 0 | 0 | 160 | 0 | 370 | 0 | 0 | 522 |
| JKAMP | 0 | 250 | 1031 | 0 | 2644 | 0 | 0 | 1900 |
| JMJD1C | 0 | 148 | 0 | 0 | 0 | 0 | 0 | 0 |
| JMJD1C-AS1 | 0 | 0 | 500 | 0 | 0 | 0 | 0 | 0 |
| JMJD4 | 0 | 0 | 0 | 132 | 1396 | 0 | 0 | 0 |
| JMJD6 | 0 | 0 | 1767 | 0 | 1042 | 0 | 666 | 0 |
| JMJD7 | 0 | 0 | 532 | 1254 | 0 | 0 | 1238 | 28 |
| JMJD7-PLA2G4B | 490 | 0 | 0 | 0 | 288 | 0 | 0 | 0 |
| JMJD8 | 0 | 0 | 1746 | 818 | 0 | 0 | 312 | 0 |
| JMY | 0 | 0 | 0 | 0 | 231 | 0 | 0 | 0 |
| JOSD1 | 0 | 445 | 750 | 710 | 0 | 0 | 334 | 0 |
| JOSD2 | 472 | 0 | 0 | 0 | 578 | 0 | 0 | 0 |
| JPH1 | 0 | 0 | 0 | 0 | 504 | 904 | 0 | 0 |
| JPH4 | 0 | 0 | 0 | 0 | 0 | 398 | 0 | 0 |
| JPT1 | 2048 | 834 | 8831 | 7545 | 4844 | 0 | 952 | 2706 |
| JPT2 | 1126 | 410 | 0 | 0 | 758 | 1679 | 838 | 0 |
| JPX | 0 | 0 | 0 | 0 | 516 | 0 | 0 | 182 |
| JRK | 414 | 0 | 0 | 0 | 0 | 0 | 0 | 0 |
| JRKL | 0 | 124 | 0 | 532 | 482 | 600 | 0 | 0 |
| JSRP1 | 0 | 0 | 2170 | 0 | 0 | 0 | 0 | 0 |
| JTB | 1815 | 3764 | 12206 | 10083 | 11161 | 2376 | 2586 | 1994 |
| JUN | 1622 | 1986 | 9126 | 322 | 1326 | 0 | 1584 | 736 |
| JUNB | 5284 | 2582 | 16483 | 1888 | 1434 | 2864 | 572 | 2700 |
| JUND | 1304 | 390 | 6150 | 0 | 744 | 0 | 0 | 0 |
| JUP | 0 | 352 | 2519 | 892 | 1717 | 0 | 702 | 0 |
| KALRN | 0 | 0 | 0 | 0 | 0 | 0 | 524 | 36 |
| KANK1 | 1566 | 3272 | 4689 | 7277 | 8470 | 17797 | 26102 | 830 |
| KANSL1 | 0 | 0 | 658 | 0 | 0 | 0 | 0 | 0 |

| | | | | | | | | |
|---|---|---|---|---|---|---|---|---|
| KANSL1-AS1 | 0 | 0 | 1152 | 0 | 0 | 0 | 0 | 0 |
| KANSL1L | 342 | 0 | 0 | 0 | 192 | 0 | 0 | 0 |
| KANSL2 | 530 | 666 | 1539 | 0 | 100 | 0 | 0 | 0 |
| KANSL3 | 838 | 90 | 0 | 0 | 0 | 0 | 0 | 138 |
| KANTR | 0 | 704 | 0 | 0 | 66 | 0 | 422 | 0 |
| KARS | 0 | 544 | 2266 | 4468 | 3782 | 0 | 404 | 0 |
| KAT14 | 4810 | 4344 | 18010 | 3354 | 32561 | 12490 | 20878 | 3433 |
| KAT2A | 4688 | 3912 | 6479 | 13010 | 32310 | 6001 | 31604 | 6714 |
| KAT5 | 0 | 442 | 0 | 0 | 456 | 0 | 1278 | 1460 |
| KAT6A | 0 | 0 | 0 | 0 | 22 | 0 | 1346 | 62 |
| KAT7 | 0 | 596 | 0 | 0 | 0 | 0 | 152 | 0 |
| KAT8 | 544 | 0 | 487 | 988 | 0 | 0 | 1606 | 0 |
| KATNA1 | 10267 | 13264 | 23164 | 40973 | 117087 | 44948 | 71609 | 28820 |
| KATNB1 | 14 | 0 | 0 | 0 | 0 | 0 | 0 | 0 |
| KATNBL1 | 0 | 320 | 1968 | 0 | 0 | 0 | 0 | 0 |
| KAZALD1 | 0 | 0 | 0 | 0 | 0 | 3542 | 244 | 0 |
| KAZN | 0 | 0 | 0 | 700 | 582 | 0 | 0 | 0 |
| KBTBD2 | 728 | 0 | 0 | 0 | 618 | 0 | 0 | 0 |
| KBTBD3 | 0 | 144 | 1624 | 1222 | 860 | 0 | 0 | 0 |
| KBTBD6 | 0 | 132 | 0 | 0 | 0 | 0 | 0 | 0 |
| KBTBD7 | 0 | 0 | 0 | 0 | 0 | 1280 | 0 | 0 |
| KCMF1 | 700 | 938 | 2076 | 0 | 1974 | 0 | 1160 | 812 |
| KCNA6 | 618 | 0 | 0 | 0 | 0 | 0 | 0 | 0 |
| KCNAB1 | 452 | 114 | 2514 | 0 | 3982 | 488 | 1042 | 1110 |
| KCNE3 | 0 | 0 | 0 | 214 | 410 | 0 | 947 | 672 |
| KCNE4 | 0 | 0 | 0 | 0 | 446 | 0 | 0 | 0 |
| KCNH2 | 60 | 0 | 0 | 0 | 0 | 0 | 585 | 0 |
| KCNIP2-AS1 | 0 | 0 | 0 | 0 | 0 | 0 | 883 | 0 |
| KCNIP4 | 5058 | 394 | 7760 | 986 | 0 | 0 | 0 | 0 |
| KCNJ6 | 0 | 1158 | 702 | 0 | 0 | 0 | 0 | 0 |
| KCNK1 | 490 | 234 | 668 | 0 | 0 | 0 | 688 | 0 |
| KCNK12 | 0 | 0 | 819 | 0 | 0 | 0 | 0 | 0 |
| KCNK15 | 0 | 328 | 0 | 0 | 0 | 0 | 0 | 0 |
| KCNK17 | 0 | 0 | 0 | 0 | 0 | 0 | 358 | 0 |
| KCNK5 | 1024 | 0 | 1294 | 316 | 881 | 0 | 0 | 0 |
| KCNK6 | 0 | 54 | 0 | 0 | 0 | 0 | 0 | 1184 |
| KCNMA1 | 294 | 0 | 0 | 0 | 0 | 0 | 0 | 0 |
| KCNN3 | 3040 | 4826 | 8480 | 970 | 2087 | 1396 | 1274 | 5102 |
| KCNN4 | 1186 | 0 | 394 | 428 | 2868 | 0 | 308 | 986 |
| KCNQ1 | 0 | 0 | 0 | 357 | 512 | 0 | 1186 | 0 |
| KCNQ1OT1 | 0 | 74 | 0 | 1188 | 2050 | 0 | 2174 | 3492 |
| KCNQ4 | 0 | 166 | 0 | 0 | 0 | 0 | 0 | 0 |
| KCNRG | 1622 | 2618 | 0 | 2271 | 17220 | 3839 | 3764 | 1185 |

| | | | | | | | | |
|---|---|---|---|---|---|---|---|---|
| KCNS3 | 0 | 0 | 3142 | 788 | 0 | 0 | 0 | 0 |
| KCTD10 | 0 | 0 | 0 | 324 | 1624 | 0 | 0 | 0 |
| KCTD12 | 0 | 124 | 0 | 0 | 0 | 0 | 0 | 0 |
| KCTD13 | 0 | 196 | 0 | 0 | 280 | 0 | 0 | 0 |
| KCTD14 | 0 | 0 | 0 | 556 | 1444 | 0 | 0 | 1296 |
| KCTD15 | 0 | 0 | 0 | 0 | 330 | 0 | 0 | 0 |
| KCTD17 | 0 | 0 | 0 | 0 | 642 | 1193 | 0 | 0 |
| KCTD18 | 0 | 0 | 0 | 0 | 326 | 0 | 0 | 0 |
| KCTD20 | 0 | 0 | 0 | 1429 | 1094 | 0 | 0 | 972 |
| KCTD3 | 0 | 72 | 0 | 0 | 158 | 0 | 0 | 0 |
| KCTD5 | 0 | 0 | 2165 | 576 | 330 | 0 | 398 | 0 |
| KCTD6 | 0 | 448 | 0 | 0 | 418 | 0 | 0 | 828 |
| KCTD9 | 780 | 0 | 1378 | 0 | 0 | 0 | 550 | 0 |
| KDELC2 | 0 | 0 | 0 | 0 | 194 | 0 | 0 | 930 |
| KDELR1 | 1036 | 1402 | 0 | 3633 | 2650 | 1252 | 1555 | 2324 |
| KDELR2 | 137 | 2252 | 2550 | 7333 | 11362 | 3298 | 5482 | 4160 |
| KDELR3 | 0 | 0 | 0 | 0 | 2612 | 0 | 1134 | 1144 |
| KDF1 | 0 | 430 | 1448 | 0 | 1348 | 0 | 0 | 0 |
| KDM1A | 198 | 271 | 0 | 2674 | 2129 | 0 | 0 | 0 |
| KDM2A | 3002 | 3667 | 4403 | 1070 | 6229 | 5493 | 8091 | 5869 |
| KDM3A | 0 | 62 | 0 | 0 | 674 | 0 | 192 | 0 |
| KDM4A | 8328 | 1814 | 1802 | 2722 | 8437 | 6608 | 13722 | 1532 |
| KDM4B | 1190 | 1798 | 12939 | 294 | 9150 | 2832 | 2202 | 0 |
| KDM4C | 0 | 0 | 1010 | 0 | 346 | 0 | 0 | 0 |
| KDM5A | 127584 | 60272 | 72736 | 35500 | 128998 | 44912 | 140933 | 23166 |
| KDM5C | 1929 | 2779 | 3619 | 4275 | 16114 | 5611 | 8797 | 5058 |
| KDM5D | 606 | 1016 | 1368 | 999 | 1948 | 4240 | 1746 | 412 |
| KDM6A | 11432 | 11797 | 7880 | 4750 | 27485 | 5292 | 20111 | 4056 |
| KDM7A | 7866 | 4204 | 8160 | 4560 | 12228 | 8504 | 38569 | 0 |
| KDM7A-DT | 950 | 0 | 0 | 0 | 1686 | 0 | 390 | 0 |
| KDSR | 0 | 86 | 0 | 0 | 302 | 0 | 58 | 0 |
| KEAP1 | 0 | 0 | 866 | 0 | 1426 | 0 | 0 | 0 |
| KHDC4 | 0 | 322 | 1773 | 0 | 1096 | 0 | 1297 | 1156 |
| KHDRBS1 | 0 | 262 | 0 | 976 | 1264 | 1733 | 0 | 0 |
| KHDRBS3 | 5590 | 7724 | 5227 | 17446 | 40984 | 27030 | 68302 | 3858 |
| KHK | 0 | 0 | 0 | 0 | 0 | 0 | 630 | 0 |
| KHNYN | 0 | 0 | 0 | 0 | 182 | 0 | 226 | 0 |
| KHSRP | 0 | 0 | 0 | 0 | 470 | 0 | 0 | 0 |
| KIAA0232 | 694 | 132 | 0 | 0 | 324 | 0 | 0 | 0 |
| KIAA0319L | 0 | 0 | 0 | 0 | 88 | 0 | 432 | 0 |
| KIAA0391 | 18568 | 42074 | 42367 | 38788 | 99688 | 27722 | 33183 | 16191 |
| KIAA0753 | 954 | 506 | 0 | 1289 | 1398 | 0 | 1074 | 1288 |
| KIAA0895L | 606 | 132 | 0 | 0 | 0 | 0 | 0 | 0 |

| | | | | | | | | |
|---|---|---|---|---|---|---|---|---|
| KIAA0930 | 0 | 0 | 0 | 0 | 702 | 0 | 0 | 0 |
| KIAA1109 | 3414 | 2383 | 7086 | 3530 | 4428 | 0 | 5810 | 0 |
| KIAA1143 | 0 | 0 | 1132 | 998 | 3146 | 0 | 278 | 0 |
| KIAA1147 | 0 | 0 | 0 | 0 | 838 | 0 | 0 | 0 |
| KIAA1191 | 0 | 122 | 0 | 936 | 1124 | 0 | 0 | 1040 |
| KIAA1211 | 0 | 94 | 0 | 0 | 0 | 0 | 0 | 0 |
| KIAA1211L | 8162 | 3270 | 14469 | 7360 | 10946 | 760 | 16050 | 6073 |
| KIAA1257 | 0 | 0 | 0 | 0 | 0 | 0 | 0 | 932 |
| KIAA1324 | 0 | 244 | 0 | 860 | 726 | 194 | 0 | 0 |
| KIAA1324L | 0 | 172 | 0 | 0 | 0 | 0 | 0 | 0 |
| KIAA1328 | 0 | 0 | 0 | 0 | 850 | 0 | 0 | 0 |
| KIAA1468 | 114 | 0 | 0 | 1006 | 0 | 0 | 0 | 0 |
| KIAA1522 | 0 | 0 | 752 | 0 | 0 | 0 | 0 | 0 |
| KIAA1549 | 0 | 0 | 0 | 0 | 974 | 1761 | 1004 | 0 |
| KIAA1551 | 0 | 446 | 0 | 0 | 510 | 0 | 0 | 0 |
| KIAA1586 | 0 | 0 | 0 | 0 | 0 | 0 | 452 | 0 |
| KIAA1841 | 0 | 0 | 0 | 0 | 1362 | 0 | 0 | 0 |
| KIAA1958 | 896 | 0 | 0 | 0 | 0 | 608 | 3062 | 728 |
| KIAA2013 | 0 | 492 | 1048 | 0 | 0 | 0 | 0 | 772 |
| KIAA2026 | 0 | 0 | 0 | 0 | 1606 | 0 | 0 | 0 |
| KIDINS220 | 0 | 0 | 0 | 0 | 682 | 0 | 0 | 0 |
| KIF11 | 0 | 0 | 0 | 0 | 0 | 0 | 140 | 0 |
| KIF12 | 732 | 0 | 0 | 0 | 0 | 0 | 0 | 0 |
| KIF13A | 0 | 0 | 758 | 702 | 0 | 0 | 260 | 0 |
| KIF13B | 958 | 0 | 0 | 0 | 498 | 0 | 0 | 0 |
| KIF16B | 0 | 0 | 0 | 0 | 0 | 0 | 450 | 0 |
| KIF18A | 0 | 0 | 0 | 0 | 0 | 0 | 0 | 610 |
| KIF18B | 0 | 0 | 470 | 0 | 0 | 0 | 0 | 0 |
| KIF1BP | 0 | 0 | 0 | 0 | 1808 | 1350 | 0 | 0 |
| KIF1C | 454 | 0 | 1654 | 0 | 0 | 0 | 498 | 0 |
| KIF20A | 0 | 0 | 0 | 0 | 902 | 0 | 0 | 0 |
| KIF20B | 2514 | 5938 | 7319 | 15932 | 35915 | 5585 | 10600 | 2426 |
| KIF22 | 14910 | 24209 | 61450 | 58283 | 113862 | 17111 | 40830 | 26283 |
| KIF23 | 0 | 0 | 0 | 735 | 230 | 0 | 0 | 0 |
| KIF25 | 0 | 468 | 0 | 0 | 0 | 0 | 3122 | 0 |
| KIF26B | 0 | 0 | 0 | 0 | 0 | 0 | 180 | 0 |
| KIF27 | 0 | 0 | 0 | 0 | 770 | 0 | 0 | 0 |
| KIF2A | 0 | 0 | 0 | 0 | 0 | 0 | 400 | 0 |
| KIF2C | 0 | 0 | 0 | 910 | 1622 | 0 | 1606 | 188 |
| KIF3C | 0 | 780 | 0 | 1110 | 1018 | 0 | 674 | 0 |
| KIF4A | 0 | 0 | 0 | 0 | 628 | 0 | 0 | 0 |
| KIF5B | 10210 | 17096 | 14264 | 16723 | 60204 | 16721 | 85225 | 12017 |
| KIF9 | 204 | 1756 | 2155 | 2043 | 3874 | 909 | 1824 | 3761 |

| | | | | | | | | |
|---|---|---|---|---|---|---|---|---|
| KIFAP3 | 0 | 0 | 0 | 854 | 0 | 0 | 0 | 0 |
| KIFC1 | 0 | 0 | 0 | 0 | 424 | 0 | 0 | 0 |
| KIFC2 | 0 | 0 | 0 | 0 | 0 | 0 | 510 | 900 |
| KIFC3 | 0 | 304 | 0 | 0 | 5818 | 875 | 776 | 1481 |
| KIN | 0 | 0 | 0 | 0 | 733 | 0 | 330 | 1002 |
| KIT | 2105 | 10799 | 10228 | 1114 | 5936 | 0 | 4709 | 1158 |
| KLB | 0 | 96 | 0 | 0 | 0 | 0 | 0 | 0 |
| KLC1 | 202 | 682 | 0 | 0 | 0 | 0 | 0 | 0 |
| KLC2 | 0 | 0 | 0 | 0 | 394 | 0 | 98 | 0 |
| KLC4 | 0 | 0 | 582 | 0 | 416 | 824 | 0 | 0 |
| KLF10 | 0 | 0 | 1930 | 0 | 894 | 470 | 136 | 0 |
| KLF11 | 0 | 0 | 0 | 0 | 1462 | 0 | 0 | 0 |
| KLF13 | 0 | 0 | 1780 | 0 | 0 | 0 | 688 | 0 |
| KLF16 | 0 | 0 | 0 | 0 | 60 | 0 | 0 | 0 |
| KLF2 | 0 | 0 | 0 | 0 | 1706 | 1084 | 0 | 0 |
| KLF3 | 0 | 0 | 1426 | 0 | 0 | 0 | 0 | 0 |
| KLF3-AS1 | 0 | 0 | 284 | 0 | 0 | 976 | 0 | 0 |
| KLF4 | 1953 | 266 | 2525 | 0 | 146 | 0 | 0 | 0 |
| KLF5 | 4318 | 5598 | 4905 | 3176 | 13294 | 2524 | 5625 | 4256 |
| KLF6 | 1586 | 0 | 1226 | 0 | 596 | 0 | 104 | 0 |
| KLF8 | 0 | 0 | 0 | 0 | 0 | 538 | 0 | 0 |
| KLF9 | 0 | 0 | 0 | 0 | 0 | 0 | 0 | 38 |
| KLHDC1 | 0 | 254 | 0 | 0 | 0 | 0 | 0 | 0 |
| KLHDC2 | 882 | 938 | 7002 | 2384 | 2976 | 3378 | 1284 | 0 |
| KLHDC3 | 0 | 219 | 1262 | 1786 | 634 | 0 | 448 | 0 |
| KLHDC4 | 0 | 0 | 0 | 0 | 288 | 0 | 0 | 166 |
| KLHDC8B | 0 | 0 | 0 | 0 | 0 | 0 | 1641 | 0 |
| KLHDC9 | 0 | 372 | 0 | 0 | 0 | 0 | 0 | 0 |
| KLHL12 | 0 | 202 | 0 | 0 | 1418 | 0 | 0 | 0 |
| KLHL17 | 388 | 0 | 0 | 0 | 870 | 0 | 0 | 0 |
| KLHL2 | 0 | 374 | 0 | 0 | 578 | 0 | 452 | 0 |
| KLHL20 | 0 | 0 | 0 | 0 | 580 | 1188 | 154 | 0 |
| KLHL21 | 498 | 0 | 0 | 0 | 0 | 0 | 0 | 0 |
| KLHL23 | 0 | 0 | 0 | 0 | 314 | 0 | 0 | 0 |
| KLHL24 | 0 | 0 | 0 | 0 | 1020 | 0 | 0 | 0 |
| KLHL25 | 0 | 0 | 0 | 0 | 614 | 0 | 0 | 0 |
| KLHL26 | 0 | 124 | 0 | 0 | 0 | 0 | 0 | 0 |
| KLHL3 | 0 | 0 | 0 | 678 | 0 | 0 | 0 | 0 |
| KLHL31 | 0 | 0 | 0 | 0 | 14 | 0 | 0 | 262 |
| KLHL35 | 0 | 0 | 0 | 0 | 4308 | 2066 | 0 | 1082 |
| KLHL36 | 0 | 0 | 0 | 0 | 216 | 0 | 0 | 0 |
| KLHL42 | 0 | 916 | 0 | 0 | 0 | 0 | 488 | 0 |
| KLHL6 | 0 | 414 | 1715 | 0 | 0 | 0 | 0 | 0 |

| | | | | | | | | |
|---|---|---|---|---|---|---|---|---|
| KLHL7 | 880 | 0 | 0 | 0 | 0 | 0 | 0 | 38 |
| KLK1 | 1693 | 5952 | 5052 | 3187 | 4459 | 0 | 0 | 658 |
| KLK10 | 0 | 0 | 0 | 3507 | 730 | 0 | 0 | 2975 |
| KLK11 | 0 | 5392 | 1558 | 5970 | 1284 | 0 | 2587 | 7698 |
| KLK12 | 0 | 0 | 0 | 0 | 0 | 1696 | 0 | 780 |
| KLK3 | 1340 | 1300 | 2992 | 0 | 254 | 0 | 0 | 0 |
| KLK6 | 0 | 0 | 0 | 1362 | 1092 | 2114 | 292 | 1142 |
| KLKB1 | 0 | 0 | 0 | 0 | 714 | 0 | 0 | 0 |
| KLRC3 | 340 | 1040 | 0 | 0 | 2108 | 0 | 3804 | 940 |
| KMT2A | 20328 | 5456 | 18265 | 8100 | 32354 | 18250 | 25818 | 12330 |
| KMT2B | 0 | 0 | 0 | 0 | 330 | 0 | 0 | 0 |
| KMT2C | 6290 | 8566 | 32 | 6263 | 8862 | 1572 | 10566 | 3994 |
| KMT2D | 478 | 0 | 0 | 0 | 268 | 0 | 0 | 0 |
| KMT2E | 549934 | 390486 | 467866 | 137977 | 685259 | 288274 | 728153 | 103423 |
| KMT2E-AS1 | 0 | 0 | 0 | 500 | 0 | 0 | 0 | 0 |
| KMT5A | 0 | 426 | 0 | 566 | 248 | 0 | 0 | 0 |
| KMT5B | 0 | 106 | 0 | 0 | 274 | 0 | 0 | 0 |
| KMT5C | 516 | 0 | 0 | 0 | 0 | 0 | 0 | 0 |
| KNL1 | 6759 | 4520 | 9949 | 17318 | 37793 | 8390 | 15792 | 1920 |
| KNOP1 | 0 | 0 | 0 | 1129 | 0 | 0 | 951 | 0 |
| KNSTRN | 0 | 0 | 0 | 0 | 1432 | 0 | 0 | 1213 |
| KPNA1 | 0 | 0 | 0 | 762 | 1072 | 0 | 0 | 624 |
| KPNA2 | 148 | 0 | 0 | 932 | 7236 | 0 | 596 | 1179 |
| KPNA3 | 0 | 0 | 0 | 0 | 0 | 0 | 0 | 540 |
| KPNA4 | 0 | 0 | 0 | 0 | 607 | 0 | 236 | 0 |
| KPNA6 | 0 | 0 | 0 | 0 | 0 | 0 | 0 | 786 |
| KPNA7 | 580 | 0 | 0 | 0 | 0 | 0 | 0 | 0 |
| KPNB1 | 0 | 538 | 0 | 0 | 0 | 0 | 178 | 0 |
| KRAS | 38984 | 30277 | 29510 | 12560 | 42540 | 5179 | 37260 | 7108 |
| KRBOX4 | 3898 | 632 | 0 | 0 | 13616 | 2950 | 6700 | 4174 |
| KRCC1 | 526 | 860 | 1100 | 0 | 1576 | 0 | 946 | 0 |
| KREMEN1 | 0 | 0 | 546 | 0 | 0 | 0 | 0 | 0 |
| KRIT1 | 0 | 0 | 0 | 0 | 668 | 0 | 0 | 0 |
| KRR1 | 0 | 522 | 0 | 0 | 0 | 0 | 0 | 588 |
| KRT10 | 686 | 658 | 2602 | 2784 | 7888 | 0 | 0 | 1282 |
| KRT12 | 0 | 0 | 400 | 0 | 0 | 0 | 0 | 0 |
| KRT14 | 10667 | 5782 | 8256 | 868 | 880 | 2035 | 416 | 8206 |
| KRT15 | 766 | 430 | 0 | 0 | 0 | 0 | 0 | 0 |
| KRT17 | 0 | 0 | 0 | 0 | 2044 | 0 | 0 | 0 |
| KRT17P5 | 0 | 0 | 0 | 0 | 362 | 0 | 0 | 0 |
| KRT18 | 4488 | 12627 | 57886 | 52413 | 19914 | 45730 | 19881 | 34266 |
| KRT19 | 4501492 | 6020253 | 8530825 | 3072591 | 12934865 | 3069469 | 2591884 | 1195596 |
| KRT20 | 7390 | 8161 | 60384 | 6724 | 2516 | 2202 | 0 | 0 |

| | | | | | | | | |
|---|---|---|---|---|---|---|---|---|
| KRT222 | 5100 | 2682 | 0 | 1692 | 0 | 9291 | 0 | 1142 |
| KRT23 | 0 | 0 | 0 | 0 | 0 | 0 | 4896 | 0 |
| KRT6A | 0 | 0 | 0 | 2716 | 4598 | 1782 | 0 | 1264 |
| KRT6B | 0 | 0 | 0 | 0 | 1886 | 0 | 0 | 0 |
| KRT8 | 28018 | 27644 | 141312 | 41201 | 44289 | 21575 | 17077 | 31458 |
| KRT80 | 0 | 0 | 0 | 0 | 1486 | 0 | 0 | 0 |
| KRT86 | 0 | 0 | 0 | 0 | 478 | 0 | 0 | 0 |
| KRTAP13-2 | 0 | 472 | 2042 | 0 | 0 | 0 | 0 | 0 |
| KRTCAP2 | 2396 | 1264 | 1787 | 4399 | 8230 | 0 | 4767 | 2769 |
| KRTCAP3 | 0 | 876 | 18 | 0 | 4228 | 0 | 3786 | 0 |
| KTN1 | 1362 | 682 | 609 | 0 | 2064 | 0 | 1180 | 0 |
| KXD1 | 326 | 1196 | 270 | 1265 | 1162 | 0 | 246 | 1444 |
| KYAT1 | 120 | 936 | 6126 | 0 | 2458 | 9077 | 1985 | 1849 |
| KYAT3 | 0 | 0 | 0 | 1252 | 1172 | 0 | 0 | 0 |
| KYNU | 2394 | 7306 | 10808 | 690 | 71894 | 28543 | 7666 | 71375 |
| L2HGDH | 598 | 694 | 8946 | 3592 | 27148 | 5077 | 3646 | 4433 |
| L3HYPDH | 0 | 0 | 732 | 0 | 0 | 0 | 0 | 0 |
| L3MBTL2 | 2926 | 6553 | 16063 | 3476 | 27935 | 5510 | 7930 | 2768 |
| LACTB | 0 | 344 | 0 | 0 | 466 | 2126 | 0 | 0 |
| LACTB2 | 0 | 590 | 956 | 1142 | 1964 | 6610 | 3900 | 1046 |
| LAD1 | 1002 | 0 | 2195 | 4246 | 3377 | 0 | 38 | 1326 |
| LAGE3 | 76 | 294 | 768 | 1534 | 4077 | 1880 | 1232 | 1644 |
| LAIR1 | 0 | 10 | 1022 | 0 | 0 | 0 | 0 | 183 |
| LAMA1 | 0 | 1080 | 3728 | 0 | 0 | 0 | 0 | 0 |
| LAMA3 | 10478 | 18271 | 15845 | 12501 | 52043 | 1228 | 5562 | 15876 |
| LAMA4 | 0 | 0 | 0 | 0 | 1576 | 0 | 0 | 0 |
| LAMA5 | 456 | 0 | 0 | 1340 | 1886 | 0 | 0 | 0 |
| LAMB1 | 0 | 0 | 0 | 1190 | 0 | 0 | 0 | 1732 |
| LAMB2 | 0 | 0 | 1894 | 0 | 336 | 0 | 0 | 0 |
| LAMB3 | 0 | 300 | 1396 | 2092 | 2136 | 0 | 234 | 0 |
| LAMC1 | 0 | 216 | 0 | 1132 | 343 | 2604 | 0 | 0 |
| LAMC2 | 2174 | 2168 | 5767 | 3212 | 10162 | 6235 | 2092 | 2448 |
| LAMP1 | 284 | 0 | 104 | 2655 | 824 | 1858 | 830 | 0 |
| LAMP2 | 510 | 714 | 1910 | 0 | 4516 | 0 | 180 | 402 |
| LAMP3 | 1119 | 28 | 0 | 969 | 1876 | 5289 | 11403 | 4970 |
| LAMTOR1 | 1410 | 3282 | 1812 | 2666 | 2290 | 1874 | 874 | 3246 |
| LAMTOR2 | 4250 | 594 | 4388 | 762 | 8716 | 792 | 1914 | 2384 |
| LAMTOR3 | 1217 | 622 | 724 | 1996 | 1332 | 1430 | 440 | 950 |
| LAMTOR4 | 5158 | 5903 | 9801 | 4849 | 14782 | 14026 | 6296 | 4192 |
| LAMTOR5 | 3046 | 6467 | 10285 | 5753 | 11502 | 6654 | 7610 | 3974 |
| LANCL1 | 1024 | 0 | 0 | 0 | 0 | 0 | 490 | 0 |
| LANCL2 | 466 | 0 | 0 | 0 | 0 | 0 | 1058 | 0 |
| LAP3 | 0 | 1218 | 0 | 1236 | 630 | 0 | 448 | 852 |

| | | | | | | | | |
|---|---|---|---|---|---|---|---|---|
| LAPTM4A | 3950 | 3532 | 8920 | 9597 | 7972 | 5670 | 7048 | 3255 |
| LAPTM4B | 75292 | 181334 | 216533 | 601089 | 110036 | 1689817 | 2820839 | 59799 |
| LAPTM5 | 2542 | 1648 | 1914 | 0 | 806 | 0 | 0 | 1042 |
| LARGE2 | 0 | 0 | 0 | 0 | 474 | 574 | 230 | 0 |
| LARP1 | 3162 | 2645 | 13592 | 5252 | 12138 | 7488 | 22117 | 1454 |
| LARP1B | 0 | 0 | 0 | 0 | 0 | 1588 | 336 | 0 |
| LARP4 | 0 | 364 | 0 | 0 | 0 | 0 | 0 | 0 |
| LARP4B | 11118 | 16961 | 21238 | 8270 | 31882 | 12074 | 32132 | 14452 |
| LARP6 | 0 | 0 | 0 | 0 | 0 | 0 | 1162 | 0 |
| LARP7 | 141824 | 217081 | 201272 | 101328 | 396511 | 124231 | 371575 | 75690 |
| LARS | 0 | 246 | 54 | 588 | 2366 | 1034 | 0 | 0 |
| LARS2 | 1676 | 6320 | 5150 | 4888 | 20907 | 6685 | 9064 | 1498 |
| LAS1L | 0 | 0 | 0 | 0 | 3824 | 0 | 0 | 778 |
| LASP1 | 0 | 116 | 0 | 612 | 0 | 0 | 0 | 0 |
| LAT2 | 0 | 278 | 0 | 0 | 0 | 0 | 0 | 0 |
| LATS1 | 1470 | 1688 | 1742 | 1078 | 5569 | 3385 | 4452 | 4426 |
| LATS2 | 0 | 0 | 0 | 0 | 554 | 0 | 108 | 0 |
| LAYN | 0 | 0 | 0 | 0 | 0 | 0 | 0 | 912 |
| LBH | 0 | 198 | 0 | 0 | 1142 | 48 | 0 | 0 |
| LBHD1 | 0 | 0 | 0 | 0 | 0 | 0 | 324 | 0 |
| LBP | 0 | 0 | 0 | 1242 | 0 | 5081 | 0 | 3216 |
| LBR | 0 | 0 | 0 | 865 | 2335 | 0 | 246 | 0 |
| LCA5L | 0 | 0 | 0 | 1366 | 2104 | 0 | 4526 | 0 |
| LCMT1 | 0 | 1752 | 5534 | 4415 | 0 | 0 | 0 | 0 |
| LCN10 | 1474 | 1182 | 1850 | 0 | 0 | 0 | 0 | 976 |
| LCN12 | 0 | 0 | 1130 | 0 | 0 | 0 | 0 | 0 |
| LCN2 | 803481 | 1352788 | 1328670 | 1336260 | 1666338 | 788588 | 3178528 | 1002543 |
| LCN6 | 0 | 0 | 0 | 820 | 0 | 0 | 0 | 0 |
| LCP1 | 0 | 0 | 0 | 0 | 538 | 0 | 0 | 0 |
| LCP2 | 0 | 0 | 0 | 0 | 0 | 0 | 328 | 0 |
| LDAH | 0 | 0 | 0 | 0 | 796 | 0 | 0 | 0 |
| LDB1 | 438 | 0 | 0 | 0 | 0 | 0 | 0 | 0 |
| LDHA | 2876 | 2981 | 21752 | 10320 | 34355 | 9405 | 6890 | 19613 |
| LDHB | 242036 | 803183 | 783905 | 2524057 | 4984994 | 540027 | 5419359 | 2283560 |
| LDHC | 0 | 0 | 0 | 790 | 0 | 0 | 0 | 0 |
| LDHD | 326 | 0 | 0 | 0 | 0 | 0 | 0 | 0 |
| LDLR | 13550 | 8478 | 78833 | 8486 | 54013 | 16551 | 14245 | 27439 |
| LDLRAD1 | 0 | 1072 | 5060 | 66 | 5034 | 2838 | 1393 | 1666 |
| LDLRAP1 | 0 | 358 | 0 | 626 | 666 | 0 | 0 | 0 |
| LEAP2 | 480 | 0 | 774 | 0 | 0 | 0 | 0 | 0 |
| LEFTY1 | 408 | 2216 | 694 | 282 | 2147 | 0 | 0 | 0 |
| LEMD1 | 0 | 0 | 0 | 0 | 0 | 4135 | 0 | 1550 |
| LEMD2 | 0 | 610 | 1636 | 0 | 0 | 0 | 0 | 0 |

| | | | | | | | | |
|---|---|---|---|---|---|---|---|---|
| LEMD3 | 0 | 348 | 0 | 0 | 0 | 0 | 0 | 0 |
| LENG1 | 0 | 0 | 1608 | 0 | 0 | 0 | 0 | 0 |
| LENG8 | 492 | 0 | 0 | 0 | 1028 | 572 | 0 | 0 |
| LEO1 | 518 | 0 | 0 | 0 | 0 | 0 | 0 | 0 |
| LEPR | 0 | 0 | 0 | 0 | 0 | 0 | 202 | 0 |
| LEPROT | 0 | 786 | 1124 | 0 | 654 | 0 | 0 | 0 |
| LEPROTL1 | 120203 | 129287 | 127877 | 61956 | 201930 | 80790 | 198649 | 63710 |
| LETM1 | 0 | 14 | 1342 | 0 | 0 | 0 | 0 | 0 |
| LETM2 | 0 | 820 | 2188 | 0 | 3152 | 0 | 0 | 2627 |
| LETMD1 | 370 | 1820 | 1744 | 1250 | 1302 | 0 | 1405 | 710 |
| LFNG | 486 | 0 | 0 | 644 | 1430 | 100 | 0 | 0 |
| LGALS1 | 808 | 1042 | 4288 | 3037 | 21335 | 5082 | 0 | 4018 |
| LGALS2 | 1186 | 1736 | 3410 | 0 | 0 | 0 | 88 | 0 |
| LGALS3 | 46834 | 60285 | 163285 | 42370 | 133439 | 10871 | 9293 | 23132 |
| LGALS3BP | 920 | 1344 | 11483 | 2462 | 3788 | 6244 | 1146 | 400 |
| LGALS4 | 29347 | 21785 | 94969 | 17582 | 62768 | 7616 | 4210 | 15760 |
| LGALS8 | 5392 | 5136 | 5491 | 0 | 12182 | 6241 | 15542 | 2398 |
| LGALS8-AS1 | 0 | 0 | 0 | 1098 | 0 | 0 | 0 | 0 |
| LGALS9 | 0 | 0 | 1171 | 572 | 1190 | 0 | 0 | 0 |
| LGALSL | 0 | 0 | 0 | 0 | 0 | 0 | 372 | 0 |
| LGMN | 148 | 128 | 3223 | 332 | 2331 | 0 | 0 | 0 |
| LGR4 | 0 | 0 | 614 | 0 | 136 | 0 | 0 | 0 |
| LGR5 | 0 | 466 | 0 | 0 | 422 | 0 | 144 | 0 |
| LGR6 | 0 | 0 | 0 | 0 | 204 | 0 | 0 | 0 |
| LHFPL2 | 1088 | 0 | 0 | 0 | 0 | 0 | 0 | 0 |
| LHFPL3 | 1368 | 636 | 1824 | 0 | 0 | 0 | 0 | 0 |
| LHFPL6 | 0 | 0 | 0 | 338 | 0 | 0 | 0 | 0 |
| LHPP | 280 | 0 | 0 | 0 | 2066 | 104 | 0 | 0 |
| LIAS | 0 | 460 | 0 | 0 | 94 | 0 | 0 | 0 |
| LIG3 | 0 | 0 | 0 | 0 | 552 | 0 | 0 | 0 |
| LILRA2 | 0 | 0 | 1694 | 0 | 0 | 0 | 0 | 0 |
| LILRA5 | 0 | 354 | 0 | 0 | 0 | 0 | 0 | 0 |
| LILRB2 | 0 | 0 | 0 | 0 | 0 | 934 | 0 | 0 |
| LILRB3 | 0 | 0 | 1422 | 0 | 0 | 0 | 0 | 0 |
| LILRB4 | 0 | 0 | 0 | 2453 | 0 | 0 | 0 | 0 |
| LIMA1 | 0 | 172 | 0 | 432 | 846 | 0 | 0 | 0 |
| LIMCH1 | 15162 | 6124 | 9048 | 8493 | 71564 | 19711 | 29143 | 14449 |
| LIMD1 | 138 | 0 | 0 | 0 | 370 | 0 | 0 | 0 |
| LIME1 | 0 | 0 | 0 | 0 | 0 | 0 | 2423 | 884 |
| LIMK2 | 674 | 0 | 0 | 0 | 302 | 0 | 0 | 0 |
| LIMS1 | 0 | 0 | 0 | 0 | 528 | 0 | 104 | 0 |
| LIMS2 | 354 | 0 | 1330 | 0 | 0 | 0 | 0 | 0 |
| LIN37 | 0 | 0 | 888 | 0 | 0 | 0 | 208 | 0 |

| | | | | | | | | |
|---|---|---|---|---|---|---|---|---|
| LIN54 | 0 | 0 | 0 | 0 | 1130 | 0 | 0 | 434 |
| LIN7A | 0 | 0 | 0 | 0 | 0 | 0 | 756 | 646 |
| LIN7B | 0 | 0 | 1310 | 0 | 132 | 1296 | 456 | 0 |
| LIN7C | 0 | 0 | 1613 | 0 | 700 | 0 | 0 | 1066 |
| LIN9 | 0 | 0 | 0 | 0 | 1232 | 0 | 0 | 0 |
| LINC-PINT | 0 | 442 | 0 | 0 | 0 | 1132 | 766 | 0 |
| LINC00114 | 282 | 0 | 0 | 0 | 0 | 0 | 0 | 0 |
| LINC00167 | 0 | 0 | 170 | 0 | 0 | 0 | 0 | 0 |
| LINC00239 | 0 | 0 | 0 | 0 | 502 | 0 | 993 | 0 |
| LINC00261 | 0 | 0 | 0 | 0 | 308 | 0 | 1338 | 0 |
| LINC00265 | 0 | 276 | 0 | 0 | 0 | 0 | 568 | 0 |
| LINC00339 | 0 | 0 | 1431 | 0 | 0 | 0 | 690 | 0 |
| LINC00379 | 0 | 0 | 0 | 0 | 414 | 0 | 0 | 0 |
| LINC00467 | 0 | 250 | 874 | 0 | 0 | 0 | 282 | 0 |
| LINC00476 | 0 | 0 | 2214 | 2258 | 0 | 0 | 0 | 748 |
| LINC00482 | 0 | 320 | 0 | 0 | 0 | 0 | 0 | 0 |
| LINC00504 | 572 | 16 | 0 | 0 | 198 | 0 | 2950 | 0 |
| LINC00506 | 0 | 792 | 2206 | 1382 | 5374 | 0 | 3005 | 0 |
| LINC00511 | 0 | 0 | 0 | 0 | 250 | 0 | 0 | 0 |
| LINC00513 | 0 | 0 | 0 | 336 | 0 | 0 | 872 | 0 |
| LINC00526 | 0 | 0 | 0 | 0 | 0 | 0 | 0 | 1184 |
| LINC00539 | 0 | 0 | 0 | 0 | 0 | 0 | 598 | 0 |
| LINC00543 | 0 | 0 | 1296 | 0 | 350 | 0 | 0 | 642 |
| LINC00623 | 0 | 206 | 0 | 0 | 588 | 1633 | 644 | 0 |
| LINC00630 | 0 | 0 | 0 | 0 | 254 | 0 | 0 | 0 |
| LINC00639 | 0 | 334 | 0 | 0 | 0 | 0 | 0 | 0 |
| LINC00649 | 0 | 0 | 0 | 0 | 600 | 2000 | 1787 | 0 |
| LINC00667 | 0 | 0 | 0 | 632 | 1458 | 0 | 0 | 1960 |
| LINC00668 | 0 | 0 | 0 | 324 | 0 | 0 | 0 | 0 |
| LINC00673 | 260 | 0 | 0 | 0 | 0 | 0 | 0 | 0 |
| LINC00674 | 3730 | 0 | 0 | 2197 | 1862 | 0 | 1340 | 959 |
| LINC00680 | 0 | 22 | 0 | 0 | 0 | 0 | 0 | 0 |
| LINC00842 | 0 | 666 | 0 | 0 | 0 | 0 | 406 | 0 |
| LINC00847 | 0 | 0 | 1821 | 0 | 496 | 0 | 0 | 0 |
| LINC00852 | 0 | 0 | 0 | 0 | 368 | 0 | 0 | 0 |
| LINC00857 | 0 | 0 | 0 | 158 | 1012 | 0 | 0 | 0 |
| LINC00863 | 0 | 154 | 0 | 0 | 474 | 0 | 0 | 0 |
| LINC00869 | 0 | 260 | 0 | 1596 | 1294 | 0 | 1245 | 1648 |
| LINC00884 | 0 | 0 | 0 | 0 | 0 | 0 | 0 | 1388 |
| LINC00894 | 0 | 0 | 0 | 0 | 0 | 2772 | 0 | 0 |
| LINC00896 | 0 | 192 | 0 | 0 | 418 | 0 | 114 | 0 |
| LINC00909 | 0 | 398 | 1720 | 0 | 0 | 0 | 0 | 0 |
| LINC00920 | 0 | 0 | 0 | 0 | 418 | 0 | 0 | 0 |

| | | | | | | | | |
|---|---|---|---|---|---|---|---|---|
| LINC00921 | 0 | 0 | 0 | 0 | 0 | 0 | 280 | 0 |
| LINC00923 | 0 | 36 | 0 | 0 | 0 | 0 | 0 | 0 |
| LINC00926 | 0 | 256 | 0 | 0 | 508 | 692 | 0 | 0 |
| LINC00938 | 0 | 0 | 0 | 0 | 0 | 1250 | 0 | 0 |
| LINC00960 | 0 | 0 | 0 | 0 | 602 | 0 | 0 | 1260 |
| LINC00963 | 0 | 0 | 0 | 0 | 0 | 0 | 0 | 682 |
| LINC01000 | 1468 | 0 | 0 | 0 | 0 | 0 | 62 | 0 |
| LINC01003 | 0 | 0 | 0 | 0 | 1032 | 1288 | 510 | 0 |
| LINC01004 | 0 | 56 | 0 | 0 | 238 | 0 | 0 | 0 |
| LINC01006 | 0 | 0 | 0 | 0 | 1068 | 0 | 0 | 0 |
| LINC01011 | 0 | 0 | 0 | 0 | 0 | 1864 | 0 | 0 |
| LINC01023 | 0 | 0 | 0 | 0 | 384 | 0 | 372 | 0 |
| LINC01082 | 0 | 0 | 0 | 0 | 466 | 0 | 0 | 0 |
| LINC01089 | 0 | 272 | 0 | 48 | 0 | 0 | 0 | 0 |
| LINC01123 | 0 | 0 | 0 | 702 | 942 | 0 | 294 | 0 |
| LINC01124 | 0 | 0 | 0 | 0 | 266 | 0 | 0 | 0 |
| LINC01133 | 0 | 0 | 6316 | 0 | 384 | 0 | 0 | 0 |
| LINC01137 | 932 | 0 | 0 | 0 | 1290 | 0 | 0 | 914 |
| LINC01138 | 0 | 0 | 0 | 0 | 992 | 0 | 0 | 0 |
| LINC01176 | 0 | 0 | 0 | 0 | 0 | 0 | 0 | 792 |
| LINC01184 | 0 | 40 | 0 | 0 | 0 | 0 | 0 | 0 |
| LINC01224 | 0 | 0 | 0 | 0 | 376 | 0 | 0 | 0 |
| LINC01232 | 0 | 0 | 0 | 0 | 0 | 1864 | 0 | 0 |
| LINC01278 | 0 | 0 | 0 | 0 | 404 | 0 | 0 | 0 |
| LINC01279 | 0 | 222 | 0 | 0 | 0 | 0 | 0 | 0 |
| LINC01311 | 0 | 0 | 572 | 0 | 0 | 0 | 0 | 0 |
| LINC01315 | 0 | 0 | 0 | 0 | 0 | 1797 | 2363 | 0 |
| LINC01389 | 640 | 0 | 0 | 0 | 0 | 0 | 0 | 0 |
| LINC01424 | 0 | 0 | 0 | 0 | 278 | 0 | 0 | 0 |
| LINC01431 | 0 | 0 | 574 | 0 | 142 | 0 | 0 | 0 |
| LINC01481 | 0 | 0 | 522 | 878 | 494 | 0 | 312 | 1300 |
| LINC01504 | 1080 | 0 | 0 | 0 | 0 | 0 | 0 | 0 |
| LINC01534 | 0 | 0 | 0 | 0 | 1238 | 0 | 0 | 0 |
| LINC01547 | 554 | 0 | 0 | 0 | 0 | 0 | 0 | 0 |
| LINC01550 | 422 | 0 | 0 | 0 | 0 | 0 | 0 | 0 |
| LINC01551 | 0 | 0 | 0 | 0 | 0 | 0 | 48 | 0 |
| LINC01558 | 0 | 0 | 0 | 2340 | 0 | 0 | 0 | 0 |
| LINC01559 | 0 | 0 | 1517 | 0 | 0 | 0 | 0 | 0 |
| LINC01560 | 0 | 0 | 0 | 0 | 735 | 0 | 0 | 0 |
| LINC01578 | 1472 | 1916 | 8792 | 6507 | 6163 | 0 | 4913 | 940 |
| LINC01590 | 346 | 0 | 0 | 0 | 0 | 0 | 0 | 0 |
| LINC01612 | 0 | 956 | 2756 | 0 | 496 | 0 | 334 | 0 |
| LINC01637 | 0 | 170 | 0 | 0 | 0 | 0 | 0 | 0 |

| | | | | | | | | |
|---|---|---|---|---|---|---|---|---|
| LINC01645 | 0 | 0 | 0 | 0 | 538 | 0 | 0 | 0 |
| LINC01715 | 0 | 0 | 0 | 0 | 385 | 0 | 0 | 0 |
| LINC01748 | 0 | 0 | 0 | 54 | 0 | 0 | 0 | 0 |
| LINC01752 | 0 | 1144 | 0 | 1045 | 0 | 0 | 0 | 0 |
| LINC01781 | 0 | 394 | 0 | 0 | 0 | 0 | 0 | 0 |
| LINC01814 | 0 | 0 | 0 | 0 | 210 | 0 | 0 | 0 |
| LINC01816 | 0 | 0 | 0 | 0 | 494 | 0 | 0 | 0 |
| LINC01843 | 0 | 0 | 0 | 1494 | 0 | 0 | 0 | 0 |
| LINC01857 | 0 | 0 | 0 | 0 | 0 | 0 | 0 | 1170 |
| LINC01871 | 522 | 0 | 0 | 0 | 0 | 0 | 0 | 0 |
| LINC01873 | 1028 | 1362 | 0 | 0 | 2657 | 0 | 4116 | 0 |
| LINC01881 | 0 | 0 | 0 | 3321 | 3660 | 0 | 994 | 0 |
| LINC01889 | 0 | 0 | 0 | 0 | 44 | 0 | 0 | 0 |
| LINC01943 | 0 | 0 | 0 | 0 | 60 | 0 | 0 | 0 |
| LINC02001 | 0 | 418 | 0 | 990 | 368 | 0 | 0 | 0 |
| LINC02014 | 0 | 0 | 0 | 830 | 0 | 0 | 0 | 0 |
| LINC02018 | 0 | 0 | 0 | 0 | 208 | 0 | 0 | 0 |
| LINC02019 | 582 | 376 | 0 | 700 | 1192 | 2365 | 1052 | 0 |
| LINC02023 | 0 | 288 | 0 | 0 | 0 | 0 | 0 | 0 |
| LINC02037 | 0 | 0 | 0 | 0 | 1186 | 0 | 0 | 0 |
| LINC02043 | 1522 | 0 | 0 | 0 | 448 | 0 | 1988 | 0 |
| LINC02080 | 0 | 0 | 652 | 0 | 0 | 0 | 0 | 0 |
| LINC02086 | 0 | 0 | 0 | 0 | 1084 | 0 | 142 | 0 |
| LINC02108 | 1030 | 142 | 0 | 0 | 0 | 0 | 0 | 0 |
| LINC02193 | 630 | 0 | 0 | 0 | 364 | 0 | 1018 | 0 |
| LINC02253 | 0 | 0 | 0 | 1242 | 0 | 0 | 496 | 0 |
| LINC02298 | 0 | 0 | 0 | 0 | 92 | 0 | 0 | 0 |
| LINC02313 | 0 | 0 | 0 | 0 | 600 | 0 | 0 | 0 |
| LINC02321 | 0 | 0 | 0 | 16 | 0 | 0 | 0 | 0 |
| LINC02328 | 0 | 0 | 2101 | 0 | 0 | 0 | 0 | 0 |
| LINC02332 | 0 | 0 | 0 | 0 | 204 | 0 | 0 | 0 |
| LINC02385 | 0 | 130 | 0 | 0 | 0 | 0 | 0 | 0 |
| LINC02397 | 0 | 0 | 0 | 0 | 0 | 0 | 0 | 140 |
| LINC02441 | 0 | 474 | 3216 | 0 | 0 | 0 | 0 | 0 |
| LINC02449 | 0 | 0 | 0 | 0 | 580 | 0 | 0 | 0 |
| LINC02453 | 0 | 0 | 0 | 0 | 0 | 0 | 576 | 0 |
| LINC02482 | 0 | 0 | 0 | 734 | 0 | 0 | 848 | 0 |
| LINC02535 | 0 | 320 | 0 | 0 | 0 | 0 | 0 | 0 |
| LINC02542 | 0 | 0 | 0 | 0 | 0 | 0 | 168 | 0 |
| LINC02562 | 0 | 0 | 0 | 1050 | 186 | 0 | 0 | 808 |
| LINC02585 | 0 | 0 | 0 | 0 | 406 | 0 | 0 | 0 |
| LINC02588 | 0 | 354 | 0 | 0 | 0 | 0 | 0 | 0 |
| LINGO1 | 0 | 80 | 0 | 0 | 0 | 0 | 0 | 0 |

| | | | | | | | | |
|---|---|---|---|---|---|---|---|---|
| LINS1 | 0 | 0 | 0 | 0 | 1890 | 0 | 0 | 0 |
| LIPA | 0 | 40 | 0 | 0 | 0 | 0 | 218 | 0 |
| LIPC | 1144 | 0 | 2480 | 0 | 1360 | 0 | 0 | 0 |
| LIPH | 0 | 1084 | 1850 | 0 | 600 | 0 | 0 | 0 |
| LIPT1 | 0 | 0 | 0 | 0 | 936 | 0 | 664 | 0 |
| LIPT2 | 0 | 1398 | 0 | 0 | 0 | 0 | 0 | 0 |
| LITAF | 1286 | 0 | 1107 | 1048 | 2266 | 0 | 434 | 1320 |
| LL21NC02-21A1.1 | 0 | 0 | 0 | 0 | 0 | 0 | 0 | 360 |
| LLGL1 | 0 | 0 | 0 | 0 | 300 | 0 | 0 | 0 |
| LLGL2 | 0 | 562 | 0 | 368 | 4192 | 0 | 3406 | 0 |
| LLPH | 711 | 482 | 3026 | 2357 | 6149 | 0 | 0 | 3194 |
| LMAN1 | 0 | 0 | 0 | 0 | 1516 | 0 | 588 | 1068 |
| LMAN2 | 946 | 140 | 984 | 1459 | 1711 | 957 | 2678 | 1432 |
| LMAN2L | 7802 | 7545 | 14796 | 15494 | 42606 | 27399 | 35871 | 9622 |
| LMBR1L | 0 | 0 | 932 | 0 | 0 | 1522 | 0 | 0 |
| LMBRD1 | 740 | 864 | 4802 | 0 | 1422 | 1160 | 0 | 0 |
| LMBRD2 | 3772 | 2908 | 5106 | 1028 | 7574 | 1585 | 1752 | 230 |
| LMCD1 | 0 | 364 | 0 | 0 | 400 | 0 | 0 | 212 |
| LMF1 | 178 | 0 | 1056 | 0 | 0 | 0 | 0 | 0 |
| LMF2 | 0 | 0 | 0 | 0 | 440 | 0 | 689 | 0 |
| LMNA | 3434 | 6320 | 6647 | 8728 | 12434 | 3824 | 5196 | 4614 |
| LMNB1 | 0 | 0 | 0 | 0 | 828 | 0 | 0 | 0 |
| LMNB2 | 770 | 0 | 0 | 1040 | 1640 | 0 | 0 | 0 |
| LMO2 | 0 | 0 | 0 | 0 | 0 | 1592 | 0 | 0 |
| LMO4 | 0 | 322 | 0 | 0 | 790 | 0 | 0 | 650 |
| LMO7 | 0 | 40 | 838 | 620 | 1050 | 646 | 0 | 1115 |
| LMO7-AS1 | 0 | 0 | 0 | 0 | 0 | 0 | 360 | 0 |
| LMOD1 | 0 | 446 | 0 | 0 | 0 | 0 | 0 | 0 |
| LMTK2 | 244 | 0 | 414 | 624 | 0 | 0 | 0 | 0 |
| LMTK3 | 0 | 0 | 0 | 0 | 248 | 0 | 0 | 204 |
| LNPEP | 1592 | 4928 | 8825 | 6072 | 8664 | 6800 | 12977 | 4128 |
| LNPK | 0 | 0 | 0 | 0 | 1410 | 400 | 0 | 0 |
| LNX1 | 0 | 680 | 60 | 30 | 532 | 0 | 818 | 78 |
| LNX2 | 0 | 0 | 0 | 1002 | 0 | 0 | 0 | 0 |
| LOC100049716 | 0 | 0 | 0 | 0 | 474 | 0 | 828 | 0 |
| LOC100128108 | 166 | 0 | 0 | 0 | 0 | 0 | 0 | 0 |
| LOC100128361 | 0 | 0 | 0 | 0 | 694 | 0 | 44 | 0 |
| LOC100129034 | 0 | 542 | 0 | 0 | 0 | 0 | 0 | 0 |
| LOC100129434 | 0 | 1452 | 0 | 260 | 10296 | 0 | 15321 | 1534 |
| LOC100130298 | 0 | 448 | 0 | 0 | 0 | 0 | 0 | 0 |
| LOC100130370 | 0 | 0 | 0 | 0 | 706 | 0 | 0 | 0 |
| LOC100130449 | 0 | 0 | 0 | 0 | 380 | 0 | 0 | 0 |
| LOC100130872 | 225 | 0 | 826 | 0 | 0 | 0 | 0 | 0 |

| | | | | | | | | |
|---|---|---|---|---|---|---|---|---|
| LOC100130899 | 0 | 0 | 0 | 0 | 0 | 0 | 38 | 0 |
| LOC100131257 | 0 | 0 | 0 | 768 | 0 | 0 | 0 | 0 |
| LOC100131289 | 0 | 0 | 0 | 0 | 0 | 0 | 172 | 0 |
| LOC100132062 | 0 | 172 | 0 | 0 | 0 | 0 | 0 | 0 |
| LOC100132154 | 0 | 0 | 0 | 0 | 0 | 0 | 658 | 1602 |
| LOC100132215 | 0 | 0 | 0 | 0 | 590 | 0 | 0 | 0 |
| LOC100132249 | 0 | 0 | 0 | 0 | 0 | 0 | 206 | 0 |
| LOC100133315 | 0 | 0 | 0 | 0 | 506 | 0 | 0 | 0 |
| LOC100133331 | 86 | 0 | 450 | 0 | 1391 | 0 | 0 | 0 |
| LOC100134368 | 0 | 0 | 0 | 0 | 72 | 0 | 0 | 0 |
| LOC100190986 | 0 | 0 | 714 | 0 | 330 | 0 | 0 | 0 |
| LOC100233156 | 0 | 0 | 0 | 0 | 250 | 0 | 0 | 0 |
| LOC100270746 | 0 | 0 | 0 | 0 | 0 | 0 | 494 | 0 |
| LOC100272217 | 0 | 0 | 0 | 0 | 1140 | 0 | 0 | 0 |
| LOC100287042 | 0 | 0 | 0 | 0 | 0 | 0 | 388 | 0 |
| LOC100287896 | 0 | 0 | 0 | 720 | 0 | 0 | 0 | 0 |
| LOC100288203 | 0 | 0 | 0 | 0 | 762 | 0 | 0 | 0 |
| LOC100288748 | 0 | 0 | 0 | 1166 | 0 | 0 | 0 | 0 |
| LOC100288798 | 0 | 506 | 0 | 0 | 1758 | 0 | 0 | 0 |
| LOC100288846 | 0 | 368 | 0 | 0 | 0 | 0 | 0 | 0 |
| LOC100289230 | 0 | 222 | 0 | 0 | 0 | 0 | 0 | 0 |
| LOC100289361 | 0 | 0 | 0 | 0 | 0 | 0 | 384 | 432 |
| LOC100289561 | 0 | 0 | 1197 | 984 | 656 | 0 | 0 | 0 |
| LOC100310756 | 0 | 0 | 0 | 0 | 172 | 0 | 0 | 0 |
| LOC100379224 | 0 | 0 | 0 | 0 | 0 | 0 | 94 | 0 |
| LOC100419583 | 238 | 436 | 0 | 1221 | 950 | 0 | 0 | 0 |
| LOC100505501 | 486 | 0 | 0 | 0 | 1104 | 0 | 0 | 0 |
| LOC100505549 | 570 | 390 | 0 | 0 | 0 | 0 | 0 | 0 |
| LOC100505938 | 0 | 0 | 988 | 0 | 0 | 0 | 388 | 0 |
| LOC100506023 | 0 | 0 | 1108 | 0 | 0 | 0 | 0 | 0 |
| LOC100506098 | 0 | 0 | 0 | 0 | 0 | 200 | 0 | 0 |
| LOC100506123 | 0 | 542 | 0 | 0 | 0 | 0 | 0 | 0 |
| LOC100506235 | 0 | 0 | 0 | 630 | 0 | 0 | 0 | 0 |
| LOC100506253 | 714 | 0 | 714 | 0 | 0 | 0 | 0 | 0 |
| LOC100506302 | 632 | 0 | 46 | 0 | 0 | 0 | 402 | 580 |
| LOC100506472 | 662 | 0 | 0 | 0 | 0 | 0 | 0 | 0 |
| LOC100506476 | 220 | 300 | 0 | 0 | 0 | 0 | 0 | 0 |
| LOC100506544 | 874 | 0 | 0 | 0 | 0 | 0 | 1570 | 0 |
| LOC100506548 | 0 | 398 | 0 | 0 | 364 | 0 | 0 | 0 |
| LOC100506585 | 0 | 0 | 0 | 0 | 0 | 0 | 368 | 0 |
| LOC100506639 | 0 | 0 | 0 | 0 | 0 | 0 | 442 | 0 |
| LOC100506691 | 0 | 0 | 0 | 0 | 1708 | 0 | 0 | 0 |
| LOC100506804 | 0 | 182 | 0 | 0 | 0 | 0 | 192 | 0 |

| | | | | | | | | |
|---|---|---|---|---|---|---|---|---|
| LOC100506922 | 571 | 0 | 942 | 0 | 86 | 0 | 584 | 0 |
| LOC100507002 | 0 | 0 | 0 | 0 | 736 | 0 | 0 | 0 |
| LOC100507291 | 0 | 0 | 0 | 0 | 386 | 0 | 0 | 0 |
| LOC100507336 | 0 | 0 | 0 | 0 | 593 | 0 | 0 | 0 |
| LOC100507373 | 0 | 0 | 0 | 0 | 0 | 0 | 0 | 918 |
| LOC100507412 | 442 | 337 | 0 | 552 | 268 | 0 | 142 | 0 |
| LOC100507507 | 0 | 0 | 0 | 0 | 192 | 0 | 0 | 0 |
| LOC100507547 | 0 | 0 | 0 | 0 | 736 | 0 | 0 | 0 |
| LOC100507557 | 0 | 0 | 0 | 0 | 0 | 0 | 0 | 841 |
| LOC100507599 | 0 | 0 | 0 | 0 | 0 | 0 | 1145 | 0 |
| LOC100507634 | 0 | 0 | 0 | 0 | 0 | 0 | 0 | 366 |
| LOC100630923 | 316 | 202 | 1764 | 0 | 0 | 0 | 0 | 0 |
| LOC100652758 | 0 | 252 | 0 | 0 | 556 | 0 | 84 | 0 |
| LOC100652768 | 1240 | 0 | 0 | 0 | 852 | 0 | 0 | 0 |
| LOC100652901 | 0 | 0 | 0 | 0 | 302 | 0 | 0 | 0 |
| LOC100996318 | 0 | 0 | 0 | 0 | 0 | 0 | 0 | 478 |
| LOC100996442 | 0 | 0 | 0 | 0 | 0 | 0 | 306 | 0 |
| LOC100996447 | 0 | 128 | 16 | 0 | 754 | 0 | 560 | 0 |
| LOC100996709 | 0 | 0 | 0 | 1014 | 0 | 0 | 236 | 0 |
| LOC100996717 | 976 | 1674 | 2995 | 0 | 0 | 3211 | 812 | 432 |
| LOC100996732 | 0 | 0 | 54 | 484 | 0 | 0 | 0 | 0 |
| LOC100996740 | 446 | 0 | 1466 | 0 | 0 | 2058 | 868 | 0 |
| LOC100996747 | 0 | 976 | 344 | 250 | 804 | 0 | 1170 | 398 |
| LOC100996756 | 0 | 0 | 1457 | 0 | 576 | 0 | 0 | 0 |
| LOC100996763 | 2380 | 5158 | 454 | 0 | 3596 | 0 | 962 | 11414 |
| LOC100996842 | 0 | 0 | 0 | 205 | 1348 | 0 | 0 | 0 |
| LOC101059954 | 0 | 0 | 0 | 0 | 0 | 0 | 0 | 1159 |
| LOC101060400 | 1264 | 0 | 932 | 2126 | 864 | 0 | 2289 | 0 |
| LOC101409256 | 0 | 0 | 0 | 0 | 0 | 0 | 16 | 0 |
| LOC101926913 | 0 | 0 | 0 | 0 | 0 | 938 | 0 | 0 |
| LOC101926935 | 0 | 0 | 0 | 0 | 548 | 692 | 0 | 0 |
| LOC101926944 | 0 | 116 | 0 | 0 | 0 | 0 | 0 | 0 |
| LOC101926967 | 362 | 0 | 0 | 0 | 0 | 0 | 0 | 0 |
| LOC101927021 | 0 | 0 | 0 | 0 | 1680 | 0 | 0 | 0 |
| LOC101927069 | 0 | 368 | 0 | 0 | 0 | 0 | 0 | 0 |
| LOC101927100 | 970 | 0 | 0 | 0 | 0 | 0 | 0 | 0 |
| LOC101927241 | 42 | 0 | 0 | 0 | 0 | 0 | 0 | 0 |
| LOC101927245 | 0 | 0 | 0 | 0 | 724 | 0 | 0 | 0 |
| LOC101927278 | 0 | 0 | 0 | 0 | 919 | 0 | 0 | 0 |
| LOC101927300 | 0 | 0 | 0 | 0 | 320 | 0 | 0 | 0 |
| LOC101927318 | 0 | 0 | 0 | 0 | 596 | 0 | 0 | 0 |
| LOC101927345 | 0 | 0 | 0 | 0 | 0 | 1150 | 0 | 0 |
| LOC101927417 | 0 | 0 | 0 | 0 | 0 | 0 | 718 | 0 |

| | | | | | | | | |
|---|---|---|---|---|---|---|---|---|
| LOC101927420 | 0 | 0 | 0 | 0 | 1004 | 0 | 0 | 0 |
| LOC101927436 | 0 | 0 | 0 | 0 | 0 | 0 | 0 | 602 |
| LOC101927476 | 0 | 0 | 0 | 0 | 0 | 0 | 78 | 0 |
| LOC101927479 | 0 | 130 | 0 | 0 | 998 | 0 | 0 | 872 |
| LOC101927503 | 0 | 0 | 0 | 0 | 0 | 1812 | 0 | 0 |
| LOC101927506 | 152 | 0 | 0 | 0 | 0 | 0 | 0 | 0 |
| LOC101927511 | 249 | 0 | 0 | 0 | 0 | 0 | 0 | 0 |
| LOC101927556 | 0 | 0 | 0 | 0 | 0 | 0 | 314 | 0 |
| LOC101927610 | 0 | 0 | 0 | 0 | 0 | 0 | 342 | 0 |
| LOC101927627 | 0 | 0 | 0 | 0 | 0 | 3295 | 0 | 0 |
| LOC101927635 | 0 | 0 | 0 | 0 | 478 | 0 | 0 | 0 |
| LOC101927752 | 0 | 0 | 0 | 0 | 960 | 0 | 346 | 0 |
| LOC101927809 | 0 | 0 | 0 | 0 | 2462 | 0 | 0 | 758 |
| LOC101927811 | 0 | 0 | 0 | 0 | 38 | 0 | 674 | 0 |
| LOC101927884 | 0 | 0 | 0 | 0 | 614 | 0 | 0 | 0 |
| LOC101927898 | 0 | 0 | 1962 | 0 | 0 | 0 | 0 | 0 |
| LOC101927963 | 0 | 0 | 0 | 0 | 124 | 0 | 0 | 0 |
| LOC101928069 | 0 | 0 | 0 | 696 | 457 | 0 | 120 | 0 |
| LOC101928076 | 0 | 382 | 0 | 0 | 0 | 0 | 0 | 0 |
| LOC101928092 | 0 | 0 | 0 | 0 | 1434 | 0 | 0 | 0 |
| LOC101928095 | 0 | 418 | 0 | 746 | 0 | 0 | 0 | 0 |
| LOC101928096 | 0 | 0 | 0 | 0 | 0 | 0 | 190 | 0 |
| LOC101928100 | 0 | 0 | 0 | 0 | 1132 | 0 | 0 | 0 |
| LOC101928156 | 0 | 0 | 1472 | 0 | 616 | 0 | 0 | 0 |
| LOC101928177 | 0 | 0 | 0 | 0 | 0 | 0 | 0 | 414 |
| LOC101928215 | 0 | 1115 | 0 | 0 | 0 | 0 | 0 | 0 |
| LOC101928217 | 164 | 0 | 0 | 0 | 0 | 0 | 0 | 0 |
| LOC101928323 | 0 | 0 | 0 | 0 | 194 | 0 | 0 | 0 |
| LOC101928343 | 0 | 0 | 0 | 0 | 0 | 2081 | 0 | 0 |
| LOC101928347 | 0 | 0 | 232 | 0 | 0 | 0 | 0 | 0 |
| LOC101928433 | 0 | 0 | 0 | 0 | 116 | 0 | 536 | 0 |
| LOC101928487 | 0 | 0 | 0 | 0 | 491 | 0 | 0 | 0 |
| LOC101928504 | 0 | 0 | 0 | 516 | 0 | 0 | 0 | 0 |
| LOC101928659 | 0 | 0 | 0 | 0 | 390 | 0 | 0 | 0 |
| LOC101928669 | 0 | 0 | 0 | 0 | 0 | 294 | 0 | 0 |
| LOC101928764 | 0 | 0 | 0 | 0 | 0 | 0 | 310 | 0 |
| LOC101928837 | 0 | 0 | 0 | 1991 | 0 | 0 | 0 | 0 |
| LOC101928881 | 0 | 0 | 0 | 569 | 920 | 0 | 782 | 0 |
| LOC101928963 | 0 | 0 | 0 | 0 | 0 | 0 | 32 | 0 |
| LOC101929128 | 0 | 0 | 0 | 0 | 0 | 0 | 0 | 975 |
| LOC101929185 | 438 | 0 | 0 | 0 | 232 | 0 | 0 | 0 |
| LOC101929243 | 0 | 0 | 474 | 0 | 301 | 0 | 558 | 0 |
| LOC101929272 | 0 | 0 | 746 | 0 | 0 | 0 | 0 | 0 |

| | | | | | | | | |
|---|---|---|---|---|---|---|---|---|
| LOC101929340 | 810 | 362 | 1012 | 0 | 0 | 0 | 0 | 0 |
| LOC101929408 | 0 | 0 | 0 | 0 | 0 | 1122 | 0 | 0 |
| LOC101929444 | 0 | 500 | 616 | 0 | 0 | 0 | 0 | 0 |
| LOC101929703 | 0 | 0 | 0 | 0 | 0 | 0 | 590 | 0 |
| LOC101929705 | 0 | 0 | 0 | 0 | 634 | 0 | 0 | 0 |
| LOC101929709 | 322 | 0 | 0 | 0 | 0 | 0 | 0 | 0 |
| LOC101929756 | 0 | 0 | 1462 | 0 | 0 | 0 | 0 | 0 |
| LOC101929796 | 0 | 802 | 0 | 0 | 0 | 0 | 664 | 0 |
| LOC101929898 | 0 | 0 | 0 | 0 | 506 | 0 | 0 | 0 |
| LOC101929937 | 0 | 82 | 0 | 0 | 0 | 0 | 0 | 0 |
| LOC101929951 | 0 | 0 | 602 | 0 | 0 | 0 | 0 | 0 |
| LOC101929959 | 250 | 0 | 0 | 0 | 1223 | 0 | 0 | 0 |
| LOC101930010 | 0 | 0 | 0 | 586 | 0 | 0 | 0 | 0 |
| LOC101930071 | 0 | 0 | 0 | 0 | 188 | 292 | 0 | 0 |
| LOC101930085 | 0 | 540 | 0 | 0 | 478 | 0 | 0 | 0 |
| LOC101930665 | 0 | 0 | 0 | 0 | 0 | 0 | 336 | 0 |
| LOC102546294 | 0 | 0 | 1584 | 0 | 0 | 0 | 0 | 0 |
| LOC102606465 | 0 | 0 | 86 | 0 | 132 | 1618 | 0 | 0 |
| LOC102723342 | 3634 | 5250 | 2342 | 2545 | 23724 | 1109 | 6973 | 1698 |
| LOC102723564 | 0 | 0 | 0 | 548 | 0 | 0 | 762 | 0 |
| LOC102723575 | 0 | 0 | 0 | 0 | 0 | 788 | 0 | 0 |
| LOC102723708 | 0 | 0 | 0 | 0 | 0 | 0 | 0 | 954 |
| LOC102723724 | 0 | 0 | 0 | 0 | 0 | 0 | 878 | 0 |
| LOC102723728 | 366 | 0 | 0 | 0 | 0 | 0 | 0 | 0 |
| LOC102723878 | 0 | 0 | 0 | 389 | 0 | 0 | 40 | 0 |
| LOC102723897 | 324 | 0 | 418 | 0 | 0 | 0 | 410 | 0 |
| LOC102723906 | 0 | 0 | 0 | 0 | 180 | 0 | 0 | 0 |
| LOC102723983 | 0 | 0 | 0 | 0 | 656 | 0 | 466 | 0 |
| LOC102724015 | 0 | 0 | 0 | 0 | 989 | 0 | 0 | 0 |
| LOC102724093 | 0 | 0 | 0 | 0 | 732 | 0 | 0 | 0 |
| LOC102724156 | 0 | 0 | 236 | 0 | 0 | 0 | 0 | 0 |
| LOC102724200 | 0 | 0 | 0 | 0 | 0 | 0 | 338 | 0 |
| LOC102724297 | 0 | 0 | 0 | 0 | 288 | 0 | 0 | 0 |
| LOC102724330 | 0 | 0 | 0 | 0 | 0 | 0 | 1014 | 0 |
| LOC102724364 | 0 | 680 | 0 | 0 | 1912 | 0 | 458 | 0 |
| LOC102724397 | 0 | 0 | 0 | 1118 | 646 | 0 | 0 | 0 |
| LOC102724479 | 0 | 0 | 0 | 0 | 0 | 0 | 0 | 1022 |
| LOC102724517 | 1120 | 366 | 1134 | 1032 | 1707 | 1499 | 242 | 0 |
| LOC102724532 | 0 | 594 | 0 | 0 | 694 | 0 | 0 | 0 |
| LOC102724615 | 0 | 0 | 0 | 770 | 0 | 0 | 0 | 0 |
| LOC102724701 | 434 | 366 | 2193 | 0 | 848 | 0 | 158 | 0 |
| LOC102724715 | 6416 | 2498 | 25193 | 0 | 14560 | 0 | 3549 | 1208 |
| LOC102724720 | 0 | 0 | 0 | 0 | 210 | 0 | 0 | 0 |

| | | | | | | | | |
|---|---|---|---|---|---|---|---|---|
| LOC102724740 | 0 | 0 | 0 | 0 | 0 | 738 | 0 | 0 |
| LOC102724760 | 0 | 912 | 0 | 0 | 0 | 0 | 0 | 0 |
| LOC102724805 | 0 | 0 | 0 | 0 | 590 | 0 | 0 | 0 |
| LOC102724985 | 0 | 368 | 400 | 2167 | 398 | 0 | 854 | 974 |
| LOC102724993 | 0 | 0 | 1482 | 0 | 0 | 0 | 0 | 0 |
| LOC102725021 | 0 | 0 | 0 | 770 | 0 | 0 | 0 | 0 |
| LOC102725044 | 0 | 0 | 0 | 0 | 126 | 0 | 0 | 0 |
| LOC102725231 | 0 | 0 | 0 | 0 | 300 | 0 | 0 | 0 |
| LOC103091866 | 0 | 580 | 0 | 0 | 0 | 0 | 0 | 0 |
| LOC105369140 | 0 | 0 | 0 | 0 | 0 | 0 | 334 | 0 |
| LOC105369171 | 0 | 0 | 625 | 0 | 0 | 0 | 0 | 0 |
| LOC105369194 | 0 | 0 | 0 | 0 | 0 | 0 | 0 | 220 |
| LOC105369203 | 0 | 0 | 0 | 492 | 0 | 0 | 0 | 0 |
| LOC105369205 | 562 | 2372 | 110 | 0 | 3496 | 0 | 3690 | 116 |
| LOC105369230 | 0 | 2383 | 0 | 0 | 374 | 0 | 0 | 2113 |
| LOC105369315 | 194 | 0 | 0 | 0 | 300 | 0 | 0 | 0 |
| LOC105369339 | 0 | 0 | 0 | 622 | 0 | 0 | 0 | 0 |
| LOC105369347 | 0 | 608 | 0 | 0 | 0 | 0 | 0 | 0 |
| LOC105369360 | 596 | 0 | 0 | 0 | 0 | 0 | 0 | 0 |
| LOC105369370 | 0 | 356 | 0 | 0 | 0 | 0 | 0 | 0 |
| LOC105369388 | 0 | 0 | 0 | 0 | 0 | 0 | 0 | 256 |
| LOC105369486 | 0 | 0 | 0 | 0 | 174 | 0 | 0 | 0 |
| LOC105369517 | 0 | 0 | 0 | 0 | 628 | 0 | 0 | 0 |
| LOC105369526 | 0 | 0 | 0 | 0 | 526 | 0 | 0 | 0 |
| LOC105369606 | 0 | 0 | 0 | 1204 | 0 | 0 | 0 | 0 |
| LOC105369656 | 0 | 264 | 0 | 0 | 0 | 0 | 0 | 0 |
| LOC105369690 | 0 | 268 | 0 | 0 | 0 | 0 | 0 | 0 |
| LOC105369699 | 0 | 0 | 0 | 0 | 0 | 1990 | 0 | 0 |
| LOC105369724 | 0 | 0 | 592 | 0 | 0 | 0 | 0 | 0 |
| LOC105369748 | 0 | 0 | 0 | 0 | 0 | 1796 | 0 | 651 |
| LOC105369755 | 0 | 496 | 272 | 0 | 0 | 0 | 0 | 0 |
| LOC105369779 | 600 | 0 | 0 | 402 | 826 | 0 | 990 | 0 |
| LOC105369782 | 2658 | 2050 | 2284 | 0 | 3480 | 658 | 13138 | 48 |
| LOC105369819 | 618 | 0 | 0 | 0 | 0 | 0 | 0 | 0 |
| LOC105369828 | 0 | 0 | 0 | 0 | 0 | 0 | 340 | 0 |
| LOC105369885 | 0 | 0 | 0 | 496 | 0 | 0 | 0 | 0 |
| LOC105370047 | 0 | 0 | 0 | 0 | 66 | 0 | 0 | 0 |
| LOC105370190 | 0 | 0 | 1076 | 0 | 0 | 0 | 0 | 0 |
| LOC105370203 | 0 | 590 | 0 | 0 | 208 | 0 | 0 | 0 |
| LOC105370333 | 0 | 0 | 0 | 1296 | 0 | 0 | 180 | 0 |
| LOC105370532 | 0 | 0 | 0 | 0 | 590 | 0 | 0 | 0 |
| LOC105370546 | 0 | 0 | 628 | 1112 | 0 | 0 | 0 | 0 |
| LOC105370624 | 0 | 0 | 0 | 0 | 0 | 0 | 496 | 856 |

| | | | | | | | | |
|---|---|---|---|---|---|---|---|---|
| LOC105370647 | 0 | 0 | 0 | 0 | 46 | 0 | 0 | 0 |
| LOC105370792 | 0 | 0 | 0 | 0 | 0 | 1894 | 0 | 0 |
| LOC105370858 | 0 | 378 | 0 | 0 | 0 | 0 | 0 | 0 |
| LOC105370969 | 0 | 384 | 918 | 302 | 558 | 0 | 644 | 0 |
| LOC105371049 | 0 | 0 | 0 | 640 | 0 | 0 | 0 | 0 |
| LOC105371056 | 0 | 0 | 0 | 654 | 0 | 0 | 0 | 0 |
| LOC105371058 | 0 | 0 | 0 | 0 | 0 | 0 | 422 | 0 |
| LOC105371074 | 0 | 0 | 697 | 0 | 0 | 0 | 0 | 0 |
| LOC105371081 | 0 | 0 | 0 | 0 | 0 | 0 | 270 | 0 |
| LOC105371114 | 0 | 0 | 0 | 0 | 0 | 2200 | 0 | 0 |
| LOC105371115 | 0 | 0 | 0 | 1180 | 0 | 0 | 0 | 0 |
| LOC105371257 | 0 | 552 | 0 | 0 | 0 | 0 | 0 | 0 |
| LOC105371264 | 0 | 0 | 0 | 0 | 366 | 0 | 0 | 474 |
| LOC105371413 | 0 | 0 | 1522 | 0 | 0 | 0 | 0 | 0 |
| LOC105371448 | 0 | 0 | 474 | 0 | 252 | 0 | 0 | 420 |
| LOC105371452 | 0 | 0 | 0 | 0 | 0 | 0 | 230 | 0 |
| LOC105371472 | 0 | 0 | 0 | 0 | 0 | 0 | 202 | 0 |
| LOC105371554 | 0 | 0 | 0 | 0 | 194 | 0 | 0 | 0 |
| LOC105371622 | 0 | 0 | 0 | 1125 | 0 | 0 | 0 | 0 |
| LOC105371755 | 0 | 50 | 12 | 0 | 44 | 0 | 0 | 0 |
| LOC105371762 | 1738 | 532 | 2825 | 58 | 1034 | 0 | 0 | 0 |
| LOC105371763 | 626 | 0 | 0 | 0 | 0 | 1750 | 678 | 0 |
| LOC105371811 | 0 | 0 | 0 | 0 | 332 | 0 | 406 | 0 |
| LOC105371839 | 0 | 0 | 0 | 0 | 1058 | 0 | 268 | 0 |
| LOC105371870 | 206 | 0 | 0 | 0 | 0 | 0 | 0 | 0 |
| LOC105371887 | 24 | 0 | 1294 | 0 | 584 | 0 | 419 | 0 |
| LOC105371893 | 0 | 100 | 0 | 0 | 0 | 0 | 0 | 0 |
| LOC105371984 | 0 | 0 | 0 | 0 | 0 | 1458 | 0 | 0 |
| LOC105372217 | 0 | 166 | 0 | 0 | 0 | 0 | 0 | 0 |
| LOC105372234 | 0 | 0 | 0 | 0 | 326 | 0 | 0 | 0 |
| LOC105372246 | 0 | 302 | 0 | 0 | 0 | 0 | 0 | 0 |
| LOC105372295 | 0 | 0 | 1150 | 0 | 0 | 0 | 0 | 0 |
| LOC105372401 | 0 | 290 | 0 | 0 | 0 | 0 | 0 | 0 |
| LOC105372426 | 0 | 0 | 0 | 136 | 0 | 0 | 0 | 0 |
| LOC105372432 | 0 | 0 | 1134 | 0 | 0 | 0 | 0 | 0 |
| LOC105372441 | 2343 | 4254 | 3953 | 835 | 2266 | 0 | 0 | 0 |
| LOC105372480 | 0 | 0 | 1162 | 0 | 771 | 0 | 0 | 0 |
| LOC105372578 | 0 | 0 | 1924 | 0 | 558 | 0 | 0 | 0 |
| LOC105372582 | 0 | 0 | 0 | 0 | 158 | 888 | 0 | 0 |
| LOC105372629 | 0 | 0 | 0 | 496 | 1146 | 0 | 1415 | 0 |
| LOC105372675 | 0 | 0 | 0 | 0 | 0 | 0 | 246 | 0 |
| LOC105372803 | 0 | 0 | 0 | 0 | 0 | 2023 | 2635 | 0 |
| LOC105372809 | 0 | 0 | 0 | 0 | 0 | 0 | 0 | 1112 |

| | | | | | | | | |
|---|---|---|---|---|---|---|---|---|
| LOC105372825 | 0 | 0 | 0 | 0 | 130 | 0 | 0 | 0 |
| LOC105372962 | 348 | 0 | 0 | 0 | 0 | 0 | 0 | 0 |
| LOC105373159 | 0 | 0 | 0 | 0 | 0 | 1068 | 0 | 0 |
| LOC105373195 | 0 | 0 | 0 | 0 | 709 | 0 | 0 | 0 |
| LOC105373335 | 0 | 0 | 0 | 0 | 0 | 0 | 462 | 0 |
| LOC105373383 | 0 | 0 | 714 | 0 | 0 | 822 | 0 | 0 |
| LOC105373428 | 0 | 0 | 0 | 0 | 272 | 0 | 0 | 0 |
| LOC105373733 | 0 | 0 | 0 | 0 | 260 | 0 | 0 | 0 |
| LOC105373759 | 0 | 624 | 0 | 0 | 0 | 0 | 0 | 0 |
| LOC105373785 | 0 | 530 | 0 | 0 | 0 | 0 | 366 | 0 |
| LOC105373830 | 0 | 0 | 0 | 0 | 0 | 0 | 886 | 0 |
| LOC105373902 | 0 | 460 | 22 | 0 | 0 | 0 | 0 | 0 |
| LOC105373933 | 0 | 0 | 1582 | 0 | 0 | 0 | 0 | 0 |
| LOC105374041 | 0 | 0 | 222 | 0 | 0 | 0 | 0 | 0 |
| LOC105374080 | 328 | 0 | 0 | 0 | 0 | 0 | 0 | 0 |
| LOC105374101 | 0 | 62 | 0 | 0 | 0 | 0 | 0 | 0 |
| LOC105374148 | 884 | 0 | 0 | 0 | 0 | 0 | 0 | 0 |
| LOC105374187 | 0 | 0 | 0 | 0 | 0 | 0 | 182 | 0 |
| LOC105374205 | 0 | 22 | 0 | 0 | 0 | 0 | 0 | 0 |
| LOC105374264 | 0 | 0 | 0 | 0 | 0 | 0 | 0 | 1050 |
| LOC105374298 | 108 | 0 | 0 | 1108 | 2691 | 0 | 2929 | 1192 |
| LOC105374363 | 0 | 0 | 0 | 1054 | 0 | 0 | 0 | 0 |
| LOC105374367 | 0 | 0 | 0 | 0 | 0 | 0 | 86 | 0 |
| LOC105374588 | 0 | 356 | 1168 | 0 | 0 | 0 | 0 | 0 |
| LOC105374836 | 0 | 0 | 0 | 738 | 0 | 0 | 0 | 0 |
| LOC105374852 | 0 | 0 | 0 | 0 | 0 | 1488 | 0 | 0 |
| LOC105374879 | 0 | 0 | 0 | 1168 | 0 | 0 | 0 | 0 |
| LOC105374888 | 0 | 108 | 0 | 0 | 0 | 0 | 0 | 0 |
| LOC105374898 | 0 | 0 | 0 | 0 | 0 | 0 | 214 | 0 |
| LOC105374947 | 0 | 0 | 0 | 0 | 0 | 2412 | 0 | 0 |
| LOC105374952 | 0 | 0 | 0 | 679 | 0 | 0 | 0 | 0 |
| LOC105375170 | 0 | 0 | 0 | 0 | 0 | 0 | 0 | 426 |
| LOC105375304 | 0 | 0 | 0 | 0 | 358 | 0 | 0 | 0 |
| LOC105375431 | 418 | 0 | 1274 | 0 | 532 | 0 | 0 | 391 |
| LOC105375546 | 0 | 0 | 0 | 0 | 0 | 0 | 14 | 0 |
| LOC105375566 | 0 | 454 | 0 | 0 | 54 | 0 | 0 | 0 |
| LOC105375666 | 0 | 0 | 0 | 0 | 361 | 0 | 318 | 142 |
| LOC105375683 | 0 | 0 | 668 | 0 | 0 | 0 | 0 | 210 |
| LOC105375740 | 518 | 1502 | 3074 | 3172 | 2524 | 1958 | 3024 | 0 |
| LOC105375767 | 0 | 0 | 0 | 0 | 0 | 0 | 1060 | 0 |
| LOC105375771 | 0 | 0 | 0 | 0 | 56 | 0 | 0 | 0 |
| LOC105375918 | 0 | 0 | 1684 | 0 | 0 | 0 | 0 | 0 |
| LOC105376037 | 0 | 4017 | 0 | 3570 | 0 | 1566 | 7244 | 0 |

| | | | | | | | | |
|---|---|---|---|---|---|---|---|---|
| LOC105376095 | 304 | 0 | 0 | 0 | 0 | 0 | 0 | 0 |
| LOC105376106 | 0 | 586 | 0 | 0 | 398 | 0 | 0 | 0 |
| LOC105376156 | 0 | 0 | 0 | 0 | 1244 | 0 | 0 | 0 |
| LOC105376159 | 1152 | 0 | 0 | 0 | 0 | 0 | 1535 | 0 |
| LOC105376289 | 0 | 0 | 0 | 457 | 0 | 0 | 0 | 0 |
| LOC105376294 | 0 | 0 | 0 | 0 | 494 | 0 | 0 | 0 |
| LOC105376352 | 0 | 0 | 0 | 0 | 122 | 0 | 0 | 0 |
| LOC105376512 | 0 | 214 | 0 | 0 | 0 | 0 | 524 | 0 |
| LOC105376526 | 622 | 2712 | 0 | 0 | 1304 | 0 | 0 | 0 |
| LOC105376567 | 1658 | 0 | 1660 | 0 | 0 | 0 | 0 | 0 |
| LOC105376748 | 0 | 0 | 0 | 0 | 0 | 998 | 0 | 0 |
| LOC105376805 | 606 | 468 | 0 | 412 | 0 | 0 | 380 | 0 |
| LOC105376810 | 110 | 0 | 0 | 0 | 0 | 0 | 0 | 0 |
| LOC105376938 | 0 | 0 | 0 | 0 | 978 | 0 | 0 | 0 |
| LOC105376952 | 0 | 0 | 0 | 0 | 0 | 0 | 284 | 0 |
| LOC105377094 | 0 | 414 | 0 | 1078 | 424 | 0 | 0 | 0 |
| LOC105377115 | 0 | 0 | 0 | 0 | 0 | 2418 | 0 | 0 |
| LOC105377205 | 46 | 0 | 0 | 0 | 0 | 0 | 0 | 0 |
| LOC105377290 | 0 | 0 | 1934 | 0 | 0 | 0 | 0 | 542 |
| LOC105377297 | 0 | 0 | 0 | 0 | 0 | 0 | 606 | 0 |
| LOC105377442 | 496 | 330 | 0 | 0 | 3622 | 0 | 566 | 0 |
| LOC105377621 | 0 | 0 | 0 | 0 | 644 | 0 | 0 | 0 |
| LOC105377668 | 0 | 0 | 750 | 0 | 0 | 0 | 0 | 0 |
| LOC105377763 | 512 | 0 | 0 | 0 | 0 | 0 | 0 | 0 |
| LOC105377809 | 0 | 0 | 0 | 0 | 0 | 0 | 334 | 0 |
| LOC105377921 | 0 | 34 | 0 | 0 | 0 | 0 | 24 | 0 |
| LOC105378077 | 0 | 0 | 44 | 0 | 0 | 0 | 0 | 0 |
| LOC105378083 | 0 | 0 | 0 | 0 | 0 | 0 | 124 | 0 |
| LOC105378100 | 1116 | 126 | 374 | 0 | 0 | 0 | 0 | 0 |
| LOC105378120 | 0 | 0 | 0 | 1275 | 0 | 0 | 0 | 0 |
| LOC105378177 | 0 | 0 | 0 | 0 | 0 | 0 | 384 | 0 |
| LOC105378179 | 236 | 0 | 0 | 0 | 0 | 0 | 0 | 0 |
| LOC105378309 | 0 | 0 | 0 | 0 | 374 | 0 | 0 | 0 |
| LOC105378539 | 296 | 0 | 0 | 0 | 0 | 0 | 0 | 0 |
| LOC105378589 | 0 | 0 | 0 | 0 | 0 | 0 | 0 | 460 |
| LOC105378604 | 0 | 0 | 0 | 1572 | 0 | 0 | 0 | 0 |
| LOC105378669 | 0 | 0 | 0 | 794 | 0 | 0 | 0 | 0 |
| LOC105378720 | 0 | 452 | 0 | 0 | 0 | 0 | 0 | 0 |
| LOC105378939 | 0 | 0 | 0 | 0 | 0 | 0 | 0 | 1018 |
| LOC105378948 | 758 | 0 | 0 | 0 | 0 | 0 | 0 | 0 |
| LOC105378954 | 0 | 0 | 88 | 0 | 836 | 0 | 0 | 0 |
| LOC105378980 | 0 | 0 | 0 | 0 | 170 | 0 | 0 | 0 |
| LOC105379151 | 0 | 0 | 0 | 0 | 0 | 0 | 474 | 0 |

| | | | | | | | | |
|---|---|---|---|---|---|---|---|---|
| LOC105379172 | 798 | 0 | 0 | 0 | 0 | 0 | 278 | 0 |
| LOC105379173 | 0 | 152 | 0 | 0 | 198 | 0 | 0 | 0 |
| LOC105379257 | 0 | 0 | 0 | 0 | 0 | 0 | 514 | 0 |
| LOC105379280 | 0 | 0 | 0 | 0 | 0 | 1453 | 378 | 0 |
| LOC105379395 | 0 | 0 | 0 | 0 | 0 | 0 | 0 | 508 |
| LOC105379407 | 470 | 0 | 0 | 0 | 508 | 0 | 0 | 0 |
| LOC105379499 | 0 | 0 | 0 | 134 | 0 | 12 | 0 | 0 |
| LOC105379508 | 0 | 0 | 795 | 0 | 500 | 0 | 0 | 0 |
| LOC105379524 | 354 | 0 | 0 | 0 | 0 | 0 | 0 | 0 |
| LOC105379561 | 0 | 0 | 0 | 0 | 2324 | 0 | 0 | 0 |
| LOC106780825 | 0 | 0 | 376 | 0 | 0 | 0 | 732 | 714 |
| LOC107983998 | 0 | 0 | 0 | 0 | 0 | 0 | 266 | 0 |
| LOC107984158 | 0 | 0 | 0 | 0 | 592 | 0 | 0 | 0 |
| LOC107984251 | 0 | 0 | 1988 | 0 | 0 | 0 | 0 | 0 |
| LOC107984262 | 0 | 0 | 0 | 0 | 468 | 2052 | 0 | 0 |
| LOC107984329 | 0 | 342 | 0 | 0 | 0 | 0 | 0 | 0 |
| LOC107984347 | 0 | 744 | 0 | 0 | 0 | 0 | 1714 | 826 |
| LOC107984421 | 0 | 0 | 0 | 868 | 0 | 0 | 0 | 0 |
| LOC107984522 | 0 | 0 | 0 | 0 | 86 | 0 | 121 | 0 |
| LOC107984638 | 0 | 0 | 0 | 0 | 0 | 0 | 0 | 761 |
| LOC107984658 | 0 | 0 | 0 | 0 | 972 | 0 | 36 | 0 |
| LOC107984659 | 0 | 0 | 0 | 0 | 848 | 0 | 0 | 0 |
| LOC107984709 | 0 | 0 | 1567 | 0 | 0 | 0 | 0 | 0 |
| LOC107984763 | 0 | 0 | 0 | 0 | 388 | 0 | 0 | 0 |
| LOC107984813 | 0 | 0 | 0 | 0 | 218 | 0 | 0 | 0 |
| LOC107984834 | 0 | 0 | 0 | 1062 | 0 | 0 | 0 | 0 |
| LOC107984875 | 474 | 0 | 264 | 288 | 526 | 0 | 748 | 0 |
| LOC107984895 | 0 | 0 | 0 | 0 | 824 | 0 | 464 | 0 |
| LOC107984916 | 0 | 0 | 0 | 0 | 416 | 0 | 604 | 0 |
| LOC107984948 | 0 | 498 | 0 | 0 | 0 | 0 | 0 | 0 |
| LOC107984950 | 0 | 0 | 1297 | 0 | 0 | 0 | 0 | 0 |
| LOC107984974 | 0 | 54 | 0 | 0 | 0 | 0 | 52 | 0 |
| LOC107985000 | 0 | 0 | 0 | 0 | 162 | 0 | 0 | 0 |
| LOC107985009 | 0 | 0 | 0 | 0 | 0 | 0 | 0 | 970 |
| LOC107985120 | 0 | 0 | 0 | 581 | 0 | 0 | 0 | 0 |
| LOC107985200 | 0 | 88 | 0 | 2395 | 1926 | 0 | 838 | 1551 |
| LOC107985208 | 986 | 0 | 0 | 0 | 0 | 0 | 0 | 0 |
| LOC107985216 | 242 | 0 | 0 | 0 | 0 | 0 | 0 | 0 |
| LOC107985236 | 5305 | 4586 | 290 | 0 | 6914 | 28 | 4388 | 525 |
| LOC107985249 | 0 | 604 | 0 | 0 | 0 | 0 | 0 | 0 |
| LOC107985250 | 0 | 0 | 0 | 0 | 330 | 0 | 0 | 0 |
| LOC107985303 | 0 | 210 | 0 | 0 | 0 | 0 | 0 | 0 |
| LOC107985334 | 0 | 0 | 0 | 0 | 0 | 520 | 0 | 0 |

| | | | | | | | | |
|---|---|---|---|---|---|---|---|---|
| LOC107985367 | 0 | 344 | 0 | 0 | 0 | 0 | 0 | 0 |
| LOC107985403 | 0 | 0 | 0 | 0 | 0 | 0 | 514 | 0 |
| LOC107985404 | 0 | 0 | 0 | 0 | 0 | 0 | 934 | 326 |
| LOC107985428 | 0 | 0 | 0 | 42 | 0 | 0 | 0 | 0 |
| LOC107985433 | 828 | 0 | 0 | 0 | 0 | 0 | 0 | 0 |
| LOC107985452 | 0 | 0 | 0 | 0 | 0 | 0 | 446 | 0 |
| LOC107985480 | 230 | 0 | 0 | 0 | 0 | 1252 | 0 | 0 |
| LOC107985554 | 778 | 0 | 0 | 0 | 0 | 0 | 0 | 0 |
| LOC107985567 | 0 | 0 | 0 | 873 | 1134 | 1832 | 0 | 0 |
| LOC107985733 | 0 | 627 | 1298 | 0 | 1149 | 0 | 0 | 0 |
| LOC107985770 | 0 | 0 | 0 | 0 | 936 | 0 | 0 | 0 |
| LOC107985788 | 0 | 0 | 0 | 0 | 34 | 0 | 0 | 0 |
| LOC107985803 | 0 | 98 | 1224 | 0 | 0 | 1848 | 0 | 976 |
| LOC107985875 | 0 | 0 | 0 | 0 | 286 | 0 | 0 | 0 |
| LOC107985910 | 388 | 0 | 0 | 0 | 0 | 0 | 0 | 0 |
| LOC107985911 | 0 | 0 | 0 | 0 | 92 | 0 | 0 | 0 |
| LOC107985917 | 274 | 0 | 0 | 0 | 0 | 0 | 0 | 0 |
| LOC107985923 | 0 | 0 | 168 | 0 | 0 | 0 | 0 | 0 |
| LOC107985961 | 0 | 0 | 0 | 0 | 0 | 0 | 922 | 0 |
| LOC107986003 | 0 | 0 | 0 | 156 | 0 | 0 | 0 | 0 |
| LOC107986084 | 0 | 1716 | 4746 | 2750 | 11218 | 4762 | 3488 | 7530 |
| LOC107986114 | 0 | 0 | 0 | 0 | 0 | 0 | 40 | 0 |
| LOC107986152 | 688 | 2118 | 0 | 0 | 2024 | 0 | 1290 | 0 |
| LOC107986167 | 0 | 0 | 0 | 0 | 0 | 0 | 708 | 0 |
| LOC107986282 | 0 | 202 | 0 | 0 | 540 | 0 | 496 | 0 |
| LOC107986298 | 336 | 0 | 0 | 0 | 0 | 0 | 0 | 0 |
| LOC107986348 | 0 | 558 | 0 | 0 | 0 | 0 | 0 | 0 |
| LOC107986396 | 0 | 0 | 0 | 0 | 420 | 0 | 0 | 0 |
| LOC107986411 | 0 | 0 | 0 | 0 | 744 | 0 | 0 | 0 |
| LOC107986432 | 0 | 0 | 0 | 0 | 1134 | 0 | 0 | 0 |
| LOC107986483 | 0 | 0 | 0 | 0 | 0 | 716 | 1681 | 0 |
| LOC107986554 | 0 | 0 | 0 | 0 | 0 | 0 | 164 | 0 |
| LOC107986596 | 0 | 0 | 0 | 0 | 370 | 0 | 0 | 0 |
| LOC107986632 | 0 | 0 | 0 | 0 | 1164 | 0 | 0 | 0 |
| LOC107986742 | 0 | 0 | 0 | 0 | 1367 | 0 | 0 | 0 |
| LOC107986762 | 0 | 0 | 0 | 0 | 0 | 0 | 0 | 222 |
| LOC107986824 | 0 | 0 | 0 | 0 | 0 | 0 | 314 | 0 |
| LOC107986847 | 0 | 12 | 0 | 0 | 0 | 0 | 0 | 0 |
| LOC107986852 | 0 | 0 | 0 | 810 | 0 | 0 | 0 | 0 |
| LOC107986854 | 0 | 498 | 0 | 0 | 0 | 0 | 0 | 0 |
| LOC107986898 | 0 | 0 | 0 | 34 | 544 | 0 | 0 | 0 |
| LOC107986962 | 0 | 0 | 0 | 0 | 0 | 64 | 0 | 0 |
| LOC107986973 | 0 | 0 | 0 | 0 | 0 | 0 | 564 | 0 |

| | | | | | | | | |
|---|---|---|---|---|---|---|---|---|
| LOC107986983 | 0 | 0 | 0 | 0 | 1146 | 0 | 0 | 4078 |
| LOC107986991 | 478 | 0 | 0 | 0 | 0 | 0 | 0 | 0 |
| LOC107986997 | 1076 | 0 | 0 | 0 | 0 | 0 | 0 | 0 |
| LOC107987009 | 0 | 0 | 16 | 0 | 0 | 0 | 0 | 0 |
| LOC107987043 | 634 | 0 | 0 | 0 | 0 | 0 | 0 | 0 |
| LOC107987063 | 0 | 0 | 0 | 0 | 0 | 396 | 0 | 0 |
| LOC107987072 | 538 | 0 | 0 | 0 | 0 | 0 | 148 | 0 |
| LOC107987102 | 0 | 544 | 0 | 0 | 0 | 0 | 608 | 0 |
| LOC107987108 | 0 | 0 | 0 | 0 | 1256 | 0 | 0 | 0 |
| LOC107987135 | 660 | 0 | 0 | 0 | 0 | 0 | 0 | 0 |
| LOC107987136 | 0 | 0 | 0 | 318 | 424 | 0 | 0 | 0 |
| LOC107987142 | 0 | 0 | 0 | 0 | 68 | 0 | 0 | 0 |
| LOC107987206 | 0 | 0 | 880 | 0 | 1556 | 0 | 0 | 214 |
| LOC107987223 | 0 | 0 | 0 | 792 | 0 | 0 | 0 | 0 |
| LOC107987251 | 386 | 0 | 0 | 0 | 0 | 0 | 0 | 0 |
| LOC107987272 | 0 | 0 | 0 | 0 | 0 | 1546 | 0 | 0 |
| LOC107987276 | 0 | 0 | 0 | 0 | 262 | 0 | 0 | 0 |
| LOC107987294 | 570 | 0 | 0 | 0 | 0 | 0 | 0 | 0 |
| LOC107987299 | 0 | 712 | 0 | 0 | 0 | 0 | 0 | 0 |
| LOC107987344 | 0 | 0 | 0 | 0 | 754 | 0 | 0 | 0 |
| LOC107987373 | 212 | 0 | 1216 | 1863 | 0 | 2448 | 1452 | 0 |
| LOC107987423 | 0 | 0 | 0 | 0 | 0 | 0 | 60 | 0 |
| LOC107987457 | 0 | 0 | 0 | 0 | 0 | 0 | 466 | 0 |
| LOC107987545 | 0 | 116 | 0 | 0 | 0 | 0 | 0 | 0 |
| LOC108783654 | 640 | 0 | 0 | 0 | 0 | 0 | 0 | 0 |
| LOC110384692 | 0 | 706 | 0 | 0 | 0 | 0 | 0 | 0 |
| LOC112267857 | 142 | 0 | 0 | 0 | 0 | 0 | 0 | 0 |
| LOC112267874 | 0 | 0 | 0 | 862 | 0 | 0 | 0 | 0 |
| LOC112267877 | 0 | 0 | 0 | 0 | 0 | 0 | 1008 | 0 |
| LOC112267883 | 672 | 0 | 0 | 0 | 0 | 0 | 0 | 242 |
| LOC112267915 | 0 | 0 | 0 | 0 | 0 | 0 | 262 | 0 |
| LOC112267941 | 0 | 0 | 0 | 0 | 362 | 0 | 0 | 0 |
| LOC112267943 | 0 | 0 | 0 | 0 | 0 | 0 | 360 | 0 |
| LOC112267983 | 0 | 0 | 0 | 0 | 972 | 0 | 0 | 0 |
| LOC112268035 | 0 | 0 | 0 | 0 | 238 | 0 | 0 | 0 |
| LOC112268037 | 0 | 0 | 0 | 0 | 240 | 0 | 0 | 0 |
| LOC112268045 | 0 | 0 | 2232 | 0 | 0 | 0 | 0 | 0 |
| LOC112268063 | 326 | 0 | 0 | 0 | 0 | 0 | 0 | 0 |
| LOC112268064 | 0 | 0 | 0 | 0 | 0 | 0 | 481 | 0 |
| LOC112268088 | 228 | 0 | 0 | 0 | 0 | 0 | 0 | 0 |
| LOC112268098 | 0 | 0 | 676 | 414 | 0 | 0 | 0 | 0 |
| LOC112268105 | 0 | 0 | 0 | 0 | 684 | 0 | 0 | 0 |
| LOC112268109 | 0 | 0 | 218 | 640 | 198 | 0 | 0 | 0 |

| | | | | | | | | |
|---|---|---|---|---|---|---|---|---|
| LOC112268124 | 0 | 0 | 0 | 0 | 238 | 0 | 0 | 0 |
| LOC112268146 | 0 | 460 | 0 | 0 | 0 | 0 | 0 | 0 |
| LOC112268155 | 250 | 0 | 0 | 1032 | 1700 | 0 | 3891 | 472 |
| LOC112268184 | 734 | 262 | 0 | 0 | 0 | 0 | 0 | 0 |
| LOC112268186 | 0 | 0 | 0 | 0 | 404 | 0 | 258 | 0 |
| LOC112268196 | 0 | 0 | 0 | 0 | 94 | 0 | 0 | 0 |
| LOC112268218 | 0 | 0 | 0 | 0 | 0 | 792 | 0 | 0 |
| LOC112268233 | 0 | 0 | 0 | 0 | 0 | 0 | 832 | 0 |
| LOC112268238 | 0 | 0 | 0 | 0 | 1534 | 0 | 0 | 0 |
| LOC112268260 | 0 | 0 | 0 | 0 | 0 | 0 | 0 | 604 |
| LOC112268277 | 0 | 0 | 0 | 0 | 1158 | 2568 | 1472 | 835 |
| LOC112268284 | 2718 | 2202 | 2913 | 930 | 11900 | 4904 | 8347 | 1818 |
| LOC112268295 | 0 | 0 | 0 | 0 | 0 | 0 | 1022 | 0 |
| LOC112268313 | 0 | 0 | 0 | 752 | 1329 | 0 | 0 | 244 |
| LOC112268317 | 0 | 0 | 0 | 0 | 0 | 1588 | 0 | 0 |
| LOC112268389 | 0 | 390 | 360 | 0 | 0 | 0 | 0 | 0 |
| LOC112268398 | 0 | 238 | 0 | 358 | 0 | 0 | 0 | 0 |
| LOC112268426 | 18 | 66 | 0 | 0 | 40 | 18 | 0 | 0 |
| LOC112268432 | 712 | 152 | 0 | 0 | 158 | 0 | 0 | 0 |
| LOC112268439 | 730 | 366 | 786 | 1280 | 0 | 0 | 1286 | 0 |
| LOC112268458 | 0 | 0 | 0 | 12 | 942 | 0 | 708 | 0 |
| LOC143666 | 0 | 0 | 0 | 0 | 0 | 0 | 0 | 450 |
| LOC148709 | 0 | 0 | 0 | 0 | 0 | 0 | 740 | 0 |
| LOC150776 | 0 | 0 | 366 | 0 | 116 | 0 | 366 | 0 |
| LOC154761 | 0 | 0 | 0 | 0 | 294 | 0 | 0 | 0 |
| LOC171391 | 0 | 278 | 0 | 0 | 0 | 0 | 0 | 0 |
| LOC220729 | 720 | 0 | 0 | 0 | 656 | 0 | 272 | 0 |
| LOC257396 | 0 | 650 | 0 | 0 | 0 | 0 | 0 | 0 |
| LOC283177 | 0 | 0 | 0 | 0 | 0 | 1536 | 0 | 0 |
| LOC283710 | 0 | 220 | 0 | 0 | 0 | 0 | 0 | 0 |
| LOC283788 | 0 | 0 | 1606 | 0 | 1052 | 0 | 0 | 1362 |
| LOC284454 | 0 | 0 | 220 | 949 | 58 | 0 | 0 | 440 |
| LOC284581 | 0 | 0 | 0 | 518 | 434 | 0 | 0 | 0 |
| LOC285074 | 0 | 580 | 0 | 0 | 974 | 0 | 0 | 0 |
| LOC339803 | 0 | 0 | 0 | 0 | 0 | 0 | 0 | 880 |
| LOC374443 | 0 | 0 | 0 | 1016 | 0 | 0 | 0 | 0 |
| LOC389602 | 0 | 0 | 0 | 0 | 0 | 0 | 236 | 1188 |
| LOC389641 | 0 | 0 | 0 | 0 | 0 | 1566 | 0 | 0 |
| LOC389831 | 0 | 0 | 1582 | 0 | 702 | 0 | 0 | 0 |
| LOC389834 | 0 | 0 | 410 | 0 | 79 | 0 | 0 | 0 |
| LOC389906 | 0 | 430 | 0 | 0 | 0 | 0 | 0 | 0 |
| LOC400706 | 0 | 0 | 0 | 550 | 0 | 0 | 0 | 0 |
| LOC440434 | 0 | 0 | 0 | 0 | 134 | 0 | 0 | 0 |

| | | | | | | | | |
|---|---|---|---|---|---|---|---|---|
| LOC441179 | 0 | 0 | 0 | 0 | 0 | 0 | 0 | 748 |
| LOC441239 | 0 | 0 | 0 | 0 | 0 | 0 | 42 | 0 |
| LOC554206 | 0 | 0 | 0 | 0 | 284 | 0 | 0 | 0 |
| LOC643072 | 0 | 0 | 1280 | 0 | 0 | 0 | 0 | 0 |
| LOC645513 | 0 | 0 | 1050 | 0 | 0 | 0 | 0 | 0 |
| LOC646214 | 938 | 0 | 704 | 0 | 712 | 0 | 1753 | 0 |
| LOC646626 | 1278 | 0 | 1570 | 0 | 5702 | 0 | 0 | 0 |
| LOC652276 | 0 | 0 | 0 | 1042 | 0 | 0 | 304 | 1040 |
| LOC728392 | 0 | 406 | 0 | 0 | 204 | 0 | 0 | 0 |
| LOC728613 | 2862 | 246 | 1358 | 710 | 2097 | 0 | 3510 | 1268 |
| LOC728743 | 854 | 0 | 0 | 0 | 0 | 0 | 0 | 1004 |
| LOC729737 | 0 | 0 | 0 | 0 | 202 | 0 | 0 | 0 |
| LOC729966 | 596 | 550 | 0 | 1627 | 1632 | 0 | 0 | 662 |
| LOC730202 | 0 | 0 | 0 | 0 | 408 | 0 | 122 | 0 |
| LOC91548 | 0 | 0 | 24 | 0 | 434 | 0 | 0 | 0 |
| LOC93622 | 0 | 0 | 1804 | 0 | 218 | 0 | 142 | 0 |
| LONP1 | 306 | 0 | 0 | 390 | 264 | 0 | 0 | 0 |
| LONP2 | 0 | 0 | 0 | 1172 | 1414 | 0 | 0 | 0 |
| LONRF1 | 444 | 0 | 0 | 0 | 0 | 0 | 0 | 0 |
| LOXL1 | 0 | 0 | 0 | 0 | 1548 | 0 | 0 | 1224 |
| LOXL2 | 0 | 0 | 0 | 0 | 524 | 0 | 0 | 0 |
| LOXL3 | 0 | 0 | 0 | 0 | 716 | 0 | 0 | 0 |
| LPAR4 | 0 | 0 | 0 | 922 | 0 | 0 | 0 | 0 |
| LPAR6 | 270 | 154 | 0 | 905 | 636 | 2958 | 870 | 732 |
| LPCAT1 | 0 | 0 | 0 | 570 | 268 | 0 | 0 | 0 |
| LPCAT2 | 0 | 0 | 0 | 0 | 0 | 0 | 274 | 0 |
| LPCAT3 | 670 | 338 | 0 | 346 | 1768 | 0 | 96 | 0 |
| LPCAT4 | 0 | 0 | 0 | 0 | 420 | 1514 | 0 | 0 |
| LPGAT1 | 0 | 0 | 0 | 0 | 456 | 0 | 0 | 0 |
| LPIN1 | 0 | 0 | 0 | 0 | 368 | 0 | 0 | 0 |
| LPIN2 | 0 | 0 | 0 | 0 | 40 | 0 | 0 | 372 |
| LPIN3 | 488 | 0 | 0 | 0 | 0 | 0 | 90 | 0 |
| LPP | 40465 | 35005 | 62945 | 19166 | 78949 | 27214 | 47232 | 27389 |
| LPP-AS2 | 1174 | 1654 | 4114 | 0 | 0 | 0 | 0 | 296 |
| LPXN | 0 | 0 | 0 | 869 | 0 | 0 | 428 | 0 |
| LRBA | 5004 | 7668 | 15397 | 4130 | 11432 | 4118 | 28166 | 2584 |
| LRCH1 | 0 | 0 | 0 | 0 | 0 | 0 | 0 | 380 |
| LRCH3 | 0 | 0 | 0 | 0 | 0 | 0 | 494 | 0 |
| LRCH4 | 841 | 0 | 0 | 0 | 0 | 0 | 0 | 768 |
| LRFN4 | 398 | 0 | 0 | 0 | 0 | 0 | 396 | 0 |
| LRG1 | 0 | 0 | 1338 | 0 | 0 | 0 | 0 | 688 |
| LRIF1 | 0 | 0 | 0 | 1154 | 0 | 0 | 0 | 0 |
| LRIG1 | 0 | 0 | 0 | 0 | 0 | 0 | 188 | 0 |

| | | | | | | | | |
|---|---|---|---|---|---|---|---|---|
| LRIG2 | 218 | 0 | 0 | 0 | 0 | 0 | 0 | 0 |
| LRIG3 | 23803 | 28213 | 22143 | 12949 | 51919 | 10548 | 30045 | 5515 |
| LRMDA | 27161 | 90140 | 108403 | 60692 | 53302 | 89005 | 58769 | 32428 |
| LRP1 | 554 | 0 | 1586 | 520 | 104 | 0 | 146 | 0 |
| LRP10 | 484 | 0 | 0 | 0 | 995 | 0 | 316 | 0 |
| LRP11 | 0 | 0 | 0 | 0 | 646 | 0 | 0 | 0 |
| LRP12 | 1366 | 2238 | 5940 | 582 | 5354 | 0 | 3766 | 4563 |
| LRP5 | 0 | 0 | 0 | 146 | 1106 | 0 | 0 | 0 |
| LRP5L | 0 | 0 | 0 | 0 | 655 | 0 | 0 | 0 |
| LRP6 | 359 | 0 | 0 | 0 | 662 | 0 | 0 | 0 |
| LRPAP1 | 0 | 784 | 1206 | 3240 | 126 | 0 | 2451 | 1054 |
| LRPPRC | 0 | 1108 | 0 | 0 | 2571 | 990 | 520 | 0 |
| LRR1 | 0 | 0 | 0 | 1042 | 418 | 2096 | 254 | 1126 |
| LRRC1 | 24131 | 12634 | 57606 | 9401 | 33951 | 7588 | 12427 | 15462 |
| LRRC14 | 0 | 0 | 0 | 369 | 1258 | 0 | 0 | 0 |
| LRRC19 | 0 | 216 | 0 | 0 | 0 | 0 | 0 | 0 |
| LRRC2 | 0 | 0 | 1728 | 0 | 0 | 0 | 0 | 0 |
| LRRC23 | 1650 | 6724 | 12976 | 10021 | 15592 | 22969 | 38690 | 9446 |
| LRRC26 | 0 | 490 | 0 | 1376 | 922 | 852 | 0 | 0 |
| LRRC28 | 0 | 0 | 0 | 0 | 522 | 0 | 0 | 0 |
| LRRC32 | 0 | 370 | 0 | 0 | 0 | 0 | 398 | 0 |
| LRRC34 | 0 | 0 | 0 | 924 | 12827 | 1881 | 604 | 4243 |
| LRRC37A2 | 0 | 0 | 0 | 0 | 452 | 0 | 0 | 0 |
| LRRC40 | 0 | 0 | 0 | 0 | 882 | 0 | 0 | 0 |
| LRRC41 | 0 | 0 | 0 | 0 | 0 | 0 | 0 | 893 |
| LRRC42 | 0 | 550 | 0 | 0 | 0 | 274 | 0 | 0 |
| LRRC45 | 0 | 0 | 0 | 0 | 318 | 1176 | 0 | 0 |
| LRRC47 | 0 | 564 | 0 | 0 | 690 | 0 | 0 | 0 |
| LRRC49 | 0 | 0 | 0 | 728 | 0 | 0 | 0 | 0 |
| LRRC57 | 0 | 0 | 1122 | 0 | 0 | 0 | 1002 | 0 |
| LRRC58 | 0 | 0 | 0 | 0 | 292 | 0 | 398 | 0 |
| LRRC59 | 17205 | 41922 | 106241 | 85111 | 143018 | 67932 | 89394 | 66565 |
| LRRC6 | 13094 | 3534 | 36361 | 20426 | 133243 | 15284 | 50208 | 14840 |
| LRRC66 | 0 | 0 | 1806 | 0 | 0 | 0 | 0 | 0 |
| LRRC75A | 0 | 364 | 0 | 0 | 0 | 2508 | 0 | 0 |
| LRRC75B | 0 | 0 | 0 | 0 | 348 | 0 | 0 | 0 |
| LRRC8A | 0 | 668 | 1418 | 0 | 0 | 431 | 0 | 0 |
| LRRC8B | 0 | 0 | 0 | 922 | 0 | 0 | 0 | 0 |
| LRRC8D | 0 | 0 | 0 | 0 | 1008 | 0 | 0 | 1012 |
| LRRCC1 | 0 | 330 | 0 | 0 | 206 | 0 | 230 | 0 |
| LRRFIP1 | 272 | 0 | 0 | 0 | 538 | 0 | 0 | 0 |
| LRRFIP2 | 786 | 212 | 1684 | 0 | 558 | 0 | 0 | 0 |
| LRTOMT | 0 | 0 | 1601 | 0 | 0 | 0 | 1069 | 0 |

| | | | | | | | | |
|---|---|---|---|---|---|---|---|---|
| LSAMP | 1236 | 966 | 2460 | 0 | 13367 | 2513 | 0 | 7863 |
| LSG1 | 0 | 0 | 0 | 0 | 36 | 2109 | 483 | 1557 |
| LSM1 | 844 | 1032 | 3557 | 0 | 564 | 0 | 0 | 506 |
| LSM10 | 0 | 454 | 14 | 0 | 0 | 0 | 484 | 0 |
| LSM11 | 0 | 0 | 0 | 0 | 0 | 0 | 926 | 0 |
| LSM12 | 540 | 288 | 1194 | 772 | 1412 | 850 | 1592 | 482 |
| LSM14A | 0 | 0 | 1447 | 312 | 2394 | 0 | 44 | 0 |
| LSM14B | 0 | 0 | 0 | 0 | 672 | 0 | 456 | 0 |
| LSM2 | 0 | 530 | 2831 | 1808 | 4046 | 0 | 2520 | 1086 |
| LSM3 | 1296 | 4664 | 3794 | 10168 | 9149 | 1028 | 2984 | 3695 |
| LSM4 | 1102 | 1262 | 6586 | 4679 | 16752 | 6684 | 2560 | 3578 |
| LSM5 | 2588 | 2364 | 2170 | 2274 | 10978 | 2013 | 1280 | 2215 |
| LSM6 | 0 | 1478 | 2117 | 778 | 4427 | 0 | 1394 | 1390 |
| LSM7 | 1378 | 748 | 1634 | 3123 | 7300 | 2778 | 454 | 1296 |
| LSM8 | 916 | 150 | 2096 | 2568 | 3898 | 1698 | 3234 | 1958 |
| LSP1 | 0 | 0 | 0 | 0 | 1150 | 0 | 694 | 0 |
| LSP1P5 | 0 | 0 | 0 | 0 | 418 | 946 | 0 | 0 |
| LSR | 2432 | 1754 | 5286 | 2579 | 4954 | 2880 | 430 | 2072 |
| LSS | 356 | 0 | 0 | 0 | 602 | 0 | 0 | 0 |
| LST1 | 954 | 0 | 422 | 0 | 0 | 0 | 0 | 0 |
| LTA4H | 0 | 164 | 0 | 1888 | 1740 | 0 | 704 | 482 |
| LTB | 1032 | 0 | 0 | 0 | 0 | 0 | 0 | 0 |
| LTBP1 | 5909 | 2838 | 2238 | 363 | 13435 | 0 | 1625 | 1547 |
| LTBP2 | 0 | 0 | 0 | 0 | 469 | 0 | 0 | 0 |
| LTBP3 | 0 | 0 | 0 | 0 | 298 | 0 | 352 | 920 |
| LTBP4 | 530 | 402 | 614 | 0 | 640 | 0 | 0 | 1928 |
| LTBR | 1084 | 338 | 0 | 974 | 942 | 0 | 532 | 0 |
| LTC4S | 0 | 384 | 0 | 582 | 0 | 0 | 0 | 0 |
| LTF | 11252 | 8490 | 2267 | 1439 | 0 | 4892 | 1509 | 634 |
| LTN1 | 0 | 0 | 858 | 0 | 0 | 2163 | 0 | 0 |
| LTO1 | 3611 | 2448 | 46 | 3001 | 14539 | 8665 | 9673 | 4231 |
| LTV1 | 0 | 0 | 3603 | 802 | 1136 | 0 | 466 | 0 |
| LUC7L | 52865 | 37471 | 115752 | 58304 | 140072 | 49172 | 76222 | 30210 |
| LUC7L2 | 20565 | 49470 | 103941 | 49293 | 71478 | 66065 | 105998 | 21466 |
| LUC7L3 | 1628 | 1242 | 661 | 1415 | 962 | 0 | 1272 | 0 |
| LUM | 0 | 1070 | 4695 | 668 | 24716 | 6747 | 0 | 4350 |
| LURAP1L | 494 | 248 | 946 | 0 | 1955 | 0 | 492 | 0 |
| LUZP1 | 906 | 226 | 264 | 0 | 1074 | 0 | 851 | 0 |
| LXN | 852 | 792 | 0 | 2666 | 602 | 0 | 0 | 1190 |
| LY6E | 0 | 770 | 0 | 736 | 7528 | 3782 | 4718 | 3166 |
| LY6E-DT | 0 | 0 | 0 | 0 | 0 | 0 | 0 | 882 |
| LY6G6C | 0 | 0 | 0 | 0 | 0 | 2348 | 0 | 0 |
| LY6G6D | 0 | 0 | 0 | 755 | 0 | 0 | 13032 | 0 |

| | | | | | | | | |
|---|---|---|---|---|---|---|---|---|
| LY6G6E | 0 | 0 | 0 | 976 | 0 | 0 | 46 | 0 |
| LY6G6F-LY6G6D | 0 | 0 | 0 | 2024 | 0 | 1082 | 3324 | 0 |
| LY75 | 426 | 592 | 0 | 0 | 1068 | 0 | 0 | 0 |
| LY86 | 0 | 0 | 1584 | 0 | 0 | 0 | 0 | 0 |
| LY96 | 0 | 0 | 1810 | 0 | 64 | 0 | 0 | 0 |
| LYAR | 35309 | 111485 | 107251 | 240335 | 570484 | 217703 | 459440 | 166010 |
| LYL1 | 0 | 618 | 0 | 0 | 486 | 1045 | 0 | 960 |
| LYN | 10208 | 14800 | 15131 | 27152 | 32043 | 41703 | 47061 | 13071 |
| LYPD3 | 0 | 0 | 0 | 0 | 314 | 0 | 0 | 0 |
| LYPD6B | 0 | 0 | 0 | 0 | 936 | 0 | 0 | 0 |
| LYPD8 | 6222 | 3885 | 38018 | 0 | 0 | 0 | 0 | 0 |
| LYPLA1 | 0 | 0 | 2184 | 0 | 0 | 628 | 214 | 0 |
| LYPLA2 | 524 | 814 | 1556 | 2197 | 1034 | 885 | 442 | 0 |
| LYPLAL1 | 0 | 542 | 2659 | 0 | 1462 | 0 | 1094 | 0 |
| LYRM1 | 0 | 606 | 0 | 0 | 0 | 0 | 0 | 0 |
| LYRM2 | 644 | 0 | 162 | 0 | 0 | 0 | 0 | 1121 |
| LYRM4 | 0 | 610 | 0 | 5421 | 1256 | 0 | 375 | 0 |
| LYRM7 | 0 | 104 | 0 | 0 | 0 | 0 | 0 | 507 |
| LYRM9 | 12157 | 16748 | 47640 | 5921 | 25336 | 58453 | 42038 | 8290 |
| LYSMD2 | 0 | 459 | 0 | 0 | 1075 | 0 | 0 | 0 |
| LYSMD4 | 366 | 0 | 0 | 0 | 1189 | 1290 | 0 | 0 |
| LYST | 318 | 1814 | 2962 | 0 | 2408 | 0 | 1422 | 1500 |
| LYVE1 | 318 | 0 | 0 | 0 | 0 | 0 | 0 | 0 |
| LYZ | 309019 | 173559 | 670004 | 234261 | 5652000 | 1421411 | 406492 | 5486069 |
| LZIC | 0 | 0 | 748 | 0 | 2149 | 0 | 1112 | 0 |
| LZTFL1 | 0 | 0 | 0 | 991 | 84 | 0 | 870 | 1080 |
| LZTR1 | 1158 | 0 | 0 | 0 | 74 | 0 | 0 | 0 |
| LZTS2 | 0 | 0 | 0 | 0 | 0 | 0 | 0 | 672 |
| M6PR | 0 | 302 | 1321 | 1160 | 782 | 1894 | 460 | 0 |
| MAB21L3 | 0 | 0 | 0 | 0 | 452 | 0 | 0 | 0 |
| MACC1 | 0 | 0 | 0 | 0 | 0 | 0 | 0 | 208 |
| MACF1 | 4540 | 1414 | 3035 | 810 | 3116 | 1210 | 1218 | 286 |
| MACROD1 | 0 | 490 | 0 | 497 | 478 | 0 | 2893 | 1798 |
| MACROD2 | 0 | 0 | 0 | 0 | 620 | 0 | 168 | 0 |
| MAD1L1 | 1062 | 5316 | 7834 | 3278 | 14001 | 9004 | 10474 | 21486 |
| MAD2L1 | 0 | 28 | 0 | 0 | 2438 | 1828 | 1104 | 360 |
| MAD2L1BP | 1660 | 1168 | 0 | 0 | 0 | 0 | 640 | 0 |
| MAD2L2 | 0 | 1624 | 5637 | 7183 | 17335 | 24510 | 3260 | 9665 |
| MADD | 1224 | 590 | 2590 | 4324 | 2088 | 2863 | 4430 | 1156 |
| MAEA | 658 | 606 | 2322 | 0 | 324 | 636 | 1718 | 0 |
| MAF | 0 | 0 | 330 | 0 | 0 | 0 | 0 | 0 |
| MAF1 | 1728 | 0 | 0 | 0 | 2411 | 0 | 920 | 0 |
| MAFB | 0 | 474 | 0 | 0 | 0 | 0 | 0 | 0 |

| | | | | | | | | |
|---|---|---|---|---|---|---|---|---|
| MAFF | 0 | 0 | 1812 | 804 | 0 | 0 | 0 | 0 |
| MAFG | 0 | 0 | 0 | 0 | 396 | 0 | 0 | 0 |
| MAFIP | 0 | 804 | 3108 | 0 | 5794 | 0 | 0 | 114 |
| MAFK | 622 | 368 | 1334 | 0 | 0 | 0 | 0 | 0 |
| MAGED1 | 0 | 0 | 1716 | 249 | 1680 | 0 | 430 | 0 |
| MAGED2 | 0 | 0 | 1576 | 4698 | 1758 | 2209 | 4087 | 0 |
| MAGEF1 | 0 | 0 | 1573 | 0 | 636 | 1960 | 0 | 1910 |
| MAGEH1 | 0 | 348 | 0 | 0 | 0 | 0 | 0 | 1189 |
| MAGI2-AS3 | 0 | 0 | 0 | 0 | 0 | 0 | 0 | 1134 |
| MAGI3 | 0 | 0 | 0 | 0 | 0 | 0 | 92 | 0 |
| MAGIX | 0 | 438 | 0 | 0 | 0 | 0 | 0 | 0 |
| MAGOH | 30 | 1474 | 2594 | 1103 | 264 | 3668 | 1066 | 2482 |
| MAGOHB | 0 | 328 | 0 | 1212 | 2668 | 0 | 618 | 1156 |
| MAGT1 | 49275 | 84347 | 90721 | 74435 | 298426 | 83326 | 198941 | 74216 |
| MAIP1 | 0 | 0 | 0 | 952 | 1040 | 0 | 0 | 0 |
| MAL2 | 1408 | 1578 | 3528 | 3294 | 15240 | 4846 | 5587 | 7446 |
| MALAT1 | 2730 | 3722 | 10699 | 9951 | 19404 | 2873 | 16716 | 6121 |
| MALL | 1314 | 216 | 2963 | 1322 | 520 | 0 | 776 | 3138 |
| MALSU1 | 960 | 2036 | 0 | 848 | 0 | 0 | 724 | 984 |
| MALT1 | 5278 | 11972 | 11357 | 7204 | 23040 | 7187 | 8636 | 6970 |
| MAMDC4 | 324 | 0 | 0 | 0 | 0 | 0 | 0 | 0 |
| MAML2 | 2198 | 3280 | 606 | 2809 | 5939 | 3382 | 8278 | 3106 |
| MAML3 | 0 | 0 | 0 | 1268 | 38 | 0 | 704 | 0 |
| MAN1A1 | 0 | 0 | 0 | 0 | 430 | 0 | 0 | 0 |
| MAN1A2 | 1960 | 2548 | 3028 | 1267 | 7974 | 3800 | 7043 | 1032 |
| MAN1B1 | 0 | 0 | 0 | 0 | 200 | 0 | 0 | 0 |
| MAN2A1 | 6301 | 7872 | 5664 | 5028 | 5753 | 2316 | 5735 | 928 |
| MAN2B1 | 0 | 0 | 1922 | 0 | 0 | 0 | 518 | 0 |
| MAN2B2 | 0 | 0 | 226 | 0 | 588 | 0 | 0 | 0 |
| MAN2C1 | 280 | 60 | 3024 | 1333 | 476 | 0 | 0 | 0 |
| MANBA | 0 | 0 | 0 | 0 | 421 | 0 | 0 | 0 |
| MANBAL | 142023 | 178596 | 441474 | 277093 | 803105 | 279439 | 1144209 | 224012 |
| MANEA-DT | 0 | 0 | 0 | 0 | 0 | 0 | 0 | 434 |
| MANEAL | 0 | 100 | 0 | 1392 | 940 | 0 | 0 | 0 |
| MANF | 0 | 1102 | 5454 | 1225 | 3528 | 0 | 1710 | 1826 |
| MANSC1 | 0 | 0 | 0 | 0 | 524 | 0 | 1392 | 0 |
| MAOA | 344 | 2146 | 0 | 0 | 0 | 0 | 0 | 0 |
| MAOB | 0 | 0 | 0 | 0 | 148 | 0 | 0 | 0 |
| MAP1LC3A | 432 | 492 | 0 | 1036 | 470 | 0 | 1052 | 0 |
| MAP1LC3B | 2092 | 920 | 4058 | 4568 | 2174 | 0 | 4369 | 2885 |
| MAP1S | 1230 | 4294 | 8469 | 2348 | 12692 | 2500 | 3981 | 234 |
| MAP2K1 | 1663 | 1722 | 924 | 1884 | 7336 | 620 | 3224 | 4550 |
| MAP2K2 | 45565 | 38569 | 115210 | 43710 | 161958 | 78456 | 67528 | 43256 |

| | | | | | | | | |
|---|---|---|---|---|---|---|---|---|
| MAP2K3 | 0 | 284 | 0 | 944 | 784 | 0 | 0 | 931 |
| MAP2K4 | 25087 | 39783 | 44062 | 24057 | 87174 | 35888 | 31506 | 35502 |
| MAP2K5 | 0 | 490 | 773 | 730 | 0 | 0 | 1700 | 0 |
| MAP2K6 | 660 | 0 | 0 | 1252 | 1371 | 0 | 0 | 762 |
| MAP2K7 | 0 | 0 | 866 | 0 | 248 | 0 | 0 | 0 |
| MAP3K1 | 1074 | 0 | 446 | 0 | 2010 | 1570 | 762 | 420 |
| MAP3K11 | 0 | 168 | 0 | 0 | 872 | 0 | 0 | 0 |
| MAP3K12 | 7338 | 7003 | 3711 | 562 | 9530 | 426 | 16094 | 0 |
| MAP3K13 | 107598 | 214194 | 186433 | 123231 | 638795 | 160458 | 228639 | 177308 |
| MAP3K14 | 0 | 0 | 0 | 0 | 290 | 0 | 0 | 0 |
| MAP3K2 | 4594 | 1672 | 6425 | 2362 | 15476 | 4908 | 13546 | 4724 |
| MAP3K20 | 0 | 0 | 0 | 0 | 530 | 0 | 0 | 0 |
| MAP3K21 | 0 | 0 | 0 | 0 | 256 | 0 | 740 | 0 |
| MAP3K3 | 0 | 0 | 0 | 0 | 0 | 0 | 184 | 0 |
| MAP3K4 | 878 | 0 | 0 | 0 | 0 | 0 | 452 | 0 |
| MAP3K5 | 0 | 350 | 0 | 0 | 0 | 0 | 0 | 0 |
| MAP3K6 | 0 | 272 | 0 | 0 | 0 | 0 | 0 | 0 |
| MAP3K7 | 0 | 0 | 26 | 0 | 158 | 0 | 0 | 0 |
| MAP3K8 | 0 | 0 | 0 | 0 | 0 | 0 | 0 | 750 |
| MAP4 | 120353 | 86736 | 249432 | 91896 | 510605 | 157603 | 401210 | 99633 |
| MAP4K3-DT | 0 | 0 | 0 | 0 | 118 | 0 | 118 | 0 |
| MAP4K4 | 0 | 264 | 0 | 0 | 0 | 0 | 0 | 0 |
| MAP4K5 | 0 | 320 | 0 | 0 | 0 | 0 | 0 | 348 |
| MAP7 | 0 | 1320 | 0 | 0 | 426 | 1588 | 326 | 0 |
| MAP7D1 | 0 | 0 | 0 | 342 | 272 | 1485 | 546 | 0 |
| MAPK1 | 61472 | 94010 | 133213 | 73520 | 224589 | 110722 | 114645 | 111841 |
| MAPK10 | 2875 | 3568 | 0 | 836 | 4606 | 5505 | 0 | 0 |
| MAPK12 | 0 | 144 | 0 | 0 | 0 | 0 | 0 | 2142 |
| MAPK13 | 226 | 340 | 1392 | 0 | 1296 | 0 | 0 | 2010 |
| MAPK14 | 0 | 0 | 0 | 0 | 570 | 0 | 208 | 868 |
| MAPK15 | 0 | 0 | 0 | 0 | 0 | 0 | 526 | 0 |
| MAPK1IP1L | 0 | 1225 | 1610 | 538 | 2118 | 0 | 0 | 0 |
| MAPK3 | 1282 | 474 | 182 | 569 | 0 | 1520 | 430 | 0 |
| MAPK6 | 0 | 0 | 0 | 0 | 172 | 0 | 0 | 0 |
| MAPK7 | 0 | 60 | 0 | 0 | 156 | 0 | 412 | 0 |
| MAPK8IP1 | 0 | 0 | 0 | 0 | 408 | 0 | 0 | 0 |
| MAPK8IP3 | 626 | 424 | 1026 | 0 | 1678 | 1491 | 268 | 2472 |
| MAPK9 | 0 | 0 | 0 | 1324 | 0 | 0 | 0 | 0 |
| MAPKAP1 | 0 | 342 | 0 | 0 | 2293 | 0 | 302 | 0 |
| MAPKAPK2 | 0 | 446 | 0 | 618 | 1220 | 0 | 0 | 0 |
| MAPKAPK3 | 5064 | 6140 | 5458 | 2632 | 26799 | 5291 | 18095 | 0 |
| MAPKAPK5 | 0 | 218 | 0 | 0 | 638 | 0 | 1086 | 0 |
| MAPKAPK5-AS1 | 985 | 1926 | 3715 | 1115 | 3406 | 4596 | 734 | 2840 |

| | | | | | | | | |
|---|---|---|---|---|---|---|---|---|
| MAPRE1 | 0 | 0 | 0 | 527 | 2332 | 0 | 620 | 558 |
| MAPRE2 | 0 | 318 | 0 | 714 | 294 | 0 | 158 | 0 |
| MAPRE3 | 0 | 0 | 0 | 0 | 0 | 0 | 618 | 0 |
| 2-Mar | 0 | 402 | 0 | 0 | 0 | 0 | 0 | 0 |
| 2-Mar | 0 | 0 | 0 | 0 | 404 | 274 | 0 | 0 |
| 3-Mar | 0 | 0 | 0 | 0 | 0 | 492 | 0 | 0 |
| 5-Mar | 0 | 218 | 980 | 0 | 1052 | 0 | 576 | 0 |
| 6-Mar | 0 | 0 | 256 | 0 | 956 | 414 | 366 | 528 |
| 7-Mar | 0 | 124 | 0 | 64 | 0 | 0 | 0 | 0 |
| 9-Mar | 0 | 0 | 0 | 0 | 0 | 0 | 0 | 818 |
| MARCKS | 514 | 280 | 0 | 0 | 504 | 0 | 0 | 239 |
| MARCKSL1 | 290 | 1928 | 1365 | 7981 | 8828 | 0 | 2606 | 2191 |
| MARF1 | 0 | 0 | 2020 | 0 | 1012 | 0 | 0 | 0 |
| MARK1 | 582 | 0 | 0 | 0 | 0 | 0 | 0 | 0 |
| MARK2 | 3254 | 1758 | 2014 | 2310 | 2390 | 776 | 1828 | 876 |
| MARK3 | 704 | 810 | 0 | 0 | 572 | 0 | 0 | 0 |
| MARK4 | 0 | 0 | 0 | 0 | 0 | 0 | 22 | 0 |
| MARS | 0 | 0 | 0 | 624 | 629 | 0 | 324 | 2285 |
| MARVELD1 | 370 | 0 | 0 | 450 | 0 | 0 | 0 | 0 |
| MARVELD2 | 0 | 0 | 1002 | 0 | 698 | 0 | 0 | 0 |
| MARVELD3 | 0 | 0 | 798 | 0 | 882 | 0 | 0 | 0 |
| MAST2 | 196 | 648 | 1688 | 0 | 330 | 0 | 0 | 0 |
| MAST3 | 594 | 520 | 935 | 0 | 884 | 0 | 64 | 0 |
| MAT2A | 1548 | 1210 | 3030 | 2282 | 722 | 96 | 0 | 1310 |
| MAT2B | 470 | 718 | 566 | 4371 | 1438 | 0 | 1626 | 0 |
| MATN1-AS1 | 0 | 182 | 0 | 0 | 0 | 0 | 0 | 0 |
| MATN2 | 242 | 0 | 1683 | 0 | 0 | 0 | 96 | 0 |
| MATN3 | 110 | 0 | 0 | 0 | 0 | 2451 | 0 | 0 |
| MATR3 | 0 | 344 | 1524 | 60 | 790 | 0 | 180 | 0 |
| MAU2 | 588 | 0 | 0 | 190 | 0 | 0 | 216 | 0 |
| MAVS | 0 | 0 | 0 | 938 | 202 | 0 | 0 | 0 |
| MAX | 281437 | 246818 | 745094 | 202549 | 717510 | 291007 | 329485 | 164508 |
| MAZ | 0 | 240 | 0 | 0 | 3000 | 0 | 0 | 608 |
| MB | 0 | 619 | 0 | 0 | 0 | 0 | 0 | 0 |
| MBD1 | 18 | 186 | 0 | 0 | 568 | 0 | 1018 | 526 |
| MBD2 | 622 | 540 | 0 | 0 | 246 | 0 | 0 | 0 |
| MBD6 | 322 | 186 | 0 | 0 | 0 | 0 | 1366 | 0 |
| MBIP | 526 | 812 | 0 | 0 | 402 | 0 | 274 | 0 |
| MBLAC2 | 0 | 0 | 0 | 0 | 604 | 0 | 0 | 0 |
| MBNL1 | 416 | 420 | 0 | 464 | 490 | 0 | 1192 | 0 |
| MBNL1-AS1 | 0 | 0 | 821 | 0 | 0 | 0 | 0 | 0 |
| MBNL2 | 532 | 202 | 0 | 861 | 222 | 0 | 985 | 0 |
| MBOAT1 | 1118 | 760 | 0 | 0 | 746 | 0 | 0 | 713 |

| | | | | | | | | |
|---|---|---|---|---|---|---|---|---|
| MBOAT2 | 0 | 0 | 0 | 0 | 150 | 0 | 0 | 720 |
| MBOAT4 | 652 | 0 | 0 | 0 | 0 | 0 | 0 | 0 |
| MBOAT7 | 0 | 476 | 0 | 556 | 1672 | 0 | 198 | 0 |
| MBP | 0 | 0 | 0 | 0 | 836 | 0 | 0 | 0 |
| MBTD1 | 0 | 0 | 0 | 0 | 0 | 0 | 394 | 0 |
| MBTPS1 | 260 | 0 | 0 | 0 | 1302 | 0 | 0 | 0 |
| MBTPS2 | 212 | 320 | 0 | 0 | 0 | 0 | 0 | 0 |
| MC1R | 0 | 0 | 0 | 0 | 0 | 0 | 134 | 0 |
| MCAM | 0 | 0 | 0 | 0 | 56 | 0 | 52 | 0 |
| MCAT | 0 | 0 | 0 | 1576 | 0 | 0 | 0 | 0 |
| MCCC1 | 0 | 0 | 0 | 0 | 242 | 0 | 278 | 902 |
| MCCC2 | 874 | 624 | 0 | 794 | 870 | 0 | 1174 | 604 |
| MCEE | 0 | 0 | 2249 | 634 | 356 | 0 | 536 | 0 |
| MCF2L | 0 | 948 | 0 | 906 | 0 | 0 | 0 | 0 |
| MCFD2 | 21449 | 71185 | 101428 | 97564 | 322149 | 152894 | 254657 | 145058 |
| MCL1 | 276 | 844 | 3012 | 0 | 1231 | 2204 | 0 | 0 |
| MCM2 | 0 | 0 | 0 | 0 | 0 | 0 | 0 | 760 |
| MCM3 | 0 | 532 | 0 | 0 | 732 | 1732 | 1329 | 0 |
| MCM4 | 0 | 132 | 0 | 426 | 894 | 786 | 0 | 0 |
| MCM5 | 0 | 160 | 0 | 0 | 244 | 0 | 0 | 848 |
| MCM6 | 0 | 0 | 0 | 0 | 0 | 2037 | 0 | 0 |
| MCM7 | 790 | 256 | 806 | 840 | 570 | 0 | 0 | 1269 |
| MCM8 | 0 | 0 | 0 | 0 | 416 | 0 | 0 | 0 |
| MCM9 | 276 | 0 | 0 | 0 | 352 | 0 | 0 | 0 |
| MCMBP | 0 | 0 | 0 | 1178 | 1706 | 0 | 0 | 870 |
| MCMDC2 | 0 | 0 | 0 | 0 | 0 | 676 | 0 | 0 |
| MCOLN1 | 0 | 0 | 822 | 0 | 0 | 0 | 0 | 0 |
| MCOLN2 | 416 | 0 | 1706 | 0 | 0 | 0 | 0 | 0 |
| MCPH1 | 0 | 0 | 0 | 0 | 334 | 0 | 0 | 0 |
| MCRIP1 | 0 | 0 | 1162 | 910 | 2428 | 0 | 1388 | 1006 |
| MCRIP2 | 682 | 0 | 0 | 0 | 2868 | 3216 | 714 | 0 |
| MCRS1 | 660 | 306 | 868 | 128 | 2204 | 0 | 302 | 0 |
| MCTP1 | 1054 | 4468 | 1903 | 1100 | 4870 | 1712 | 4706 | 2817 |
| MCTS1 | 0 | 906 | 1828 | 3443 | 3710 | 1894 | 518 | 2802 |
| MCU | 428 | 0 | 0 | 0 | 644 | 0 | 943 | 0 |
| MCUB | 0 | 0 | 0 | 210 | 1586 | 0 | 728 | 0 |
| MCUR1 | 0 | 0 | 0 | 0 | 1230 | 0 | 0 | 0 |
| MDFI | 0 | 0 | 0 | 0 | 446 | 0 | 680 | 0 |
| MDFIC | 0 | 0 | 0 | 0 | 208 | 0 | 0 | 0 |
| MDH1 | 0 | 3847 | 7404 | 3137 | 8772 | 0 | 1220 | 2637 |
| MDH2 | 0 | 3760 | 6647 | 11677 | 7264 | 2712 | 8178 | 2371 |
| MDK | 4220 | 970 | 10082 | 2024 | 1100 | 3806 | 7931 | 6000 |
| MDM1 | 0 | 248 | 0 | 0 | 0 | 0 | 0 | 0 |

| | | | | | | | | |
|---|---|---|---|---|---|---|---|---|
| MDM2 | 17246 | 25329 | 37947 | 15827 | 33481 | 35380 | 45365 | 18246 |
| MDM4 | 0 | 0 | 697 | 0 | 0 | 0 | 390 | 0 |
| MDP1 | 26 | 0 | 1614 | 942 | 2686 | 0 | 0 | 0 |
| ME2 | 597 | 0 | 1032 | 0 | 0 | 0 | 508 | 0 |
| ME3 | 0 | 340 | 726 | 0 | 964 | 0 | 0 | 0 |
| MEA1 | 3029 | 1726 | 5362 | 1304 | 3192 | 2106 | 3758 | 0 |
| MEAF6 | 450 | 1396 | 0 | 1850 | 1599 | 0 | 991 | 0 |
| MECOM | 88223 | 210284 | 137873 | 111125 | 178507 | 42317 | 242418 | 44437 |
| MECP2 | 0 | 366 | 1976 | 1870 | 1248 | 0 | 1792 | 0 |
| MECR | 0 | 0 | 0 | 132 | 348 | 0 | 0 | 1568 |
| MED1 | 6464 | 6626 | 620 | 3456 | 22160 | 7889 | 25209 | 4159 |
| MED10 | 1340 | 0 | 1099 | 959 | 7919 | 2056 | 0 | 2032 |
| MED11 | 0 | 0 | 0 | 0 | 474 | 0 | 0 | 0 |
| MED12 | 0 | 0 | 0 | 832 | 0 | 0 | 0 | 0 |
| MED13 | 0 | 0 | 0 | 0 | 74 | 0 | 326 | 0 |
| MED13L | 1608 | 1134 | 0 | 1276 | 170 | 0 | 2574 | 0 |
| MED14 | 0 | 0 | 150 | 0 | 0 | 0 | 0 | 0 |
| MED15 | 1326 | 1382 | 1372 | 0 | 9449 | 4930 | 1915 | 0 |
| MED16 | 0 | 0 | 0 | 668 | 0 | 0 | 0 | 0 |
| MED17 | 0 | 0 | 0 | 0 | 374 | 0 | 556 | 0 |
| MED18 | 0 | 214 | 0 | 1252 | 0 | 0 | 0 | 0 |
| MED19 | 0 | 0 | 0 | 0 | 1336 | 0 | 0 | 0 |
| MED20 | 0 | 0 | 0 | 0 | 426 | 0 | 0 | 0 |
| MED21 | 0 | 0 | 0 | 0 | 352 | 0 | 1430 | 0 |
| MED22 | 0 | 0 | 0 | 0 | 750 | 0 | 0 | 0 |
| MED23 | 0 | 0 | 0 | 0 | 0 | 0 | 231 | 0 |
| MED24 | 0 | 358 | 1350 | 0 | 1400 | 0 | 352 | 0 |
| MED25 | 0 | 436 | 0 | 0 | 0 | 0 | 0 | 0 |
| MED26 | 0 | 0 | 477 | 0 | 0 | 0 | 0 | 0 |
| MED27 | 0 | 0 | 539 | 1410 | 1224 | 0 | 0 | 1252 |
| MED28 | 0 | 0 | 0 | 2285 | 2372 | 0 | 1698 | 692 |
| MED29 | 0 | 0 | 0 | 120 | 0 | 0 | 0 | 0 |
| MED30 | 838 | 322 | 864 | 0 | 2452 | 4098 | 1338 | 904 |
| MED31 | 0 | 152 | 0 | 852 | 1108 | 0 | 0 | 561 |
| MED4 | 247684 | 329172 | 433001 | 824504 | 1217511 | 1610555 | 1626752 | 320070 |
| MED6 | 0 | 0 | 0 | 1380 | 605 | 0 | 0 | 0 |
| MED7 | 804 | 770 | 0 | 698 | 976 | 684 | 346 | 0 |
| MED8 | 0 | 818 | 0 | 3021 | 1808 | 2312 | 796 | 2620 |
| MED9 | 772 | 0 | 0 | 0 | 1801 | 924 | 268 | 0 |
| MEF2D | 424 | 0 | 0 | 0 | 0 | 0 | 436 | 0 |
| MEG3 | 458 | 0 | 0 | 0 | 72 | 0 | 0 | 0 |
| MEGF6 | 0 | 0 | 0 | 0 | 259 | 0 | 0 | 0 |
| MEIS1 | 0 | 388 | 1764 | 0 | 0 | 0 | 0 | 0 |

| | | | | | | | | |
|---|---|---|---|---|---|---|---|---|
| MELK | 0 | 0 | 1920 | 0 | 1032 | 0 | 0 | 0 |
| MELTF | 688 | 0 | 0 | 0 | 0 | 0 | 0 | 0 |
| MELTF-AS1 | 0 | 0 | 0 | 0 | 0 | 0 | 18 | 0 |
| MEMO1 | 0 | 0 | 2224 | 0 | 1430 | 0 | 1086 | 0 |
| MEN1 | 446 | 0 | 0 | 0 | 834 | 0 | 1776 | 0 |
| MEP1A | 1490 | 974 | 4651 | 0 | 0 | 0 | 428 | 0 |
| MEPCE | 184 | 0 | 0 | 0 | 0 | 0 | 248 | 0 |
| MERTK | 322 | 512 | 4440 | 0 | 1625 | 0 | 6258 | 1102 |
| MESD | 0 | 224 | 788 | 2298 | 792 | 0 | 0 | 2056 |
| MESP1 | 874 | 0 | 0 | 0 | 0 | 0 | 0 | 768 |
| MEST | 0 | 396 | 0 | 0 | 192 | 0 | 820 | 0 |
| MET | 18541 | 32558 | 32666 | 46979 | 99306 | 49304 | 202098 | 74154 |
| METAP1 | 0 | 0 | 0 | 0 | 501 | 0 | 550 | 432 |
| METAP2 | 0 | 0 | 1134 | 0 | 344 | 0 | 1436 | 985 |
| METRN | 0 | 0 | 0 | 0 | 1178 | 0 | 522 | 0 |
| METRNL | 0 | 0 | 0 | 0 | 426 | 0 | 650 | 950 |
| METTL1 | 0 | 0 | 938 | 2403 | 232 | 2574 | 0 | 0 |
| METTL13 | 0 | 396 | 0 | 624 | 0 | 0 | 0 | 0 |
| METTL14 | 0 | 0 | 0 | 0 | 0 | 0 | 266 | 0 |
| METTL17 | 404 | 0 | 0 | 0 | 1096 | 0 | 802 | 832 |
| METTL18 | 0 | 0 | 1946 | 0 | 1653 | 0 | 318 | 0 |
| METTL21A | 0 | 550 | 0 | 0 | 0 | 1288 | 636 | 0 |
| METTL21EP | 0 | 0 | 0 | 0 | 0 | 0 | 1784 | 0 |
| METTL22 | 212 | 244 | 0 | 0 | 464 | 0 | 0 | 0 |
| METTL23 | 432 | 0 | 0 | 1822 | 1246 | 1180 | 2394 | 0 |
| METTL25 | 0 | 368 | 0 | 0 | 0 | 0 | 0 | 0 |
| METTL26 | 1880 | 0 | 847 | 3968 | 3824 | 2355 | 2678 | 204 |
| METTL27 | 0 | 398 | 0 | 0 | 1330 | 0 | 0 | 0 |
| METTL2A | 0 | 0 | 0 | 0 | 724 | 0 | 1274 | 0 |
| METTL2B | 0 | 428 | 0 | 0 | 0 | 830 | 298 | 0 |
| METTL3 | 0 | 96 | 0 | 1602 | 480 | 675 | 0 | 0 |
| METTL4 | 0 | 331 | 0 | 0 | 0 | 0 | 0 | 0 |
| METTL5 | 0 | 902 | 1638 | 5187 | 8058 | 0 | 5730 | 1706 |
| METTL6 | 0 | 0 | 0 | 0 | 1730 | 0 | 0 | 0 |
| METTL7A | 0 | 344 | 0 | 0 | 1132 | 0 | 1074 | 0 |
| METTL7B | 0 | 462 | 0 | 0 | 0 | 0 | 0 | 0 |
| METTL9 | 0 | 1678 | 0 | 1647 | 440 | 0 | 736 | 0 |
| MEX3B | 0 | 0 | 0 | 0 | 164 | 0 | 0 | 0 |
| MEX3C | 690 | 1066 | 0 | 0 | 0 | 0 | 1390 | 0 |
| MFAP1 | 0 | 0 | 1258 | 0 | 372 | 0 | 0 | 0 |
| MFAP2 | 0 | 514 | 0 | 0 | 2930 | 0 | 0 | 0 |
| MFAP4 | 0 | 0 | 816 | 0 | 2822 | 1832 | 0 | 622 |
| MFAP5 | 0 | 0 | 3684 | 0 | 352 | 0 | 0 | 0 |

| | | | | | | | | |
|---|---|---|---|---|---|---|---|---|
| MFF | 0 | 748 | 0 | 2042 | 5297 | 448 | 703 | 0 |
| MFGE8 | 0 | 0 | 0 | 634 | 1037 | 0 | 0 | 0 |
| MFHAS1 | 0 | 0 | 0 | 0 | 0 | 0 | 0 | 616 |
| MFN1 | 0 | 0 | 0 | 0 | 340 | 0 | 0 | 0 |
| MFN2 | 0 | 264 | 0 | 1340 | 0 | 0 | 0 | 0 |
| MFNG | 0 | 0 | 1450 | 0 | 0 | 0 | 0 | 0 |
| MFRP | 0 | 0 | 2028 | 254 | 3858 | 0 | 272 | 0 |
| MFSD1 | 194 | 0 | 3257 | 0 | 368 | 118 | 0 | 0 |
| MFSD10 | 0 | 0 | 1820 | 844 | 1136 | 0 | 0 | 420 |
| MFSD11 | 0 | 0 | 3386 | 502 | 1308 | 0 | 528 | 972 |
| MFSD13A | 0 | 180 | 0 | 0 | 302 | 0 | 0 | 0 |
| MFSD14A | 0 | 0 | 0 | 1361 | 1762 | 0 | 690 | 0 |
| MFSD14B | 264 | 84 | 468 | 0 | 1636 | 0 | 0 | 0 |
| MFSD14C | 0 | 0 | 0 | 0 | 724 | 0 | 201 | 0 |
| MFSD2A | 658 | 0 | 0 | 0 | 0 | 0 | 1136 | 0 |
| MFSD3 | 142 | 0 | 724 | 826 | 812 | 1816 | 1655 | 994 |
| MFSD5 | 356 | 0 | 0 | 0 | 346 | 774 | 526 | 0 |
| MFSD6 | 0 | 430 | 0 | 0 | 0 | 0 | 0 | 0 |
| MFSD6L | 554 | 0 | 0 | 0 | 0 | 0 | 0 | 84 |
| MFSD9 | 430 | 0 | 0 | 0 | 482 | 0 | 0 | 0 |
| MGA | 0 | 0 | 0 | 157 | 338 | 0 | 1486 | 0 |
| MGAM2 | 0 | 0 | 0 | 0 | 116 | 0 | 0 | 0 |
| MGARP | 0 | 0 | 0 | 1598 | 0 | 0 | 0 | 0 |
| MGAT1 | 0 | 0 | 0 | 0 | 192 | 0 | 1262 | 275 |
| MGAT2 | 264 | 386 | 532 | 803 | 364 | 0 | 0 | 0 |
| MGAT3 | 0 | 0 | 0 | 0 | 948 | 0 | 0 | 0 |
| MGAT4A | 11852 | 6400 | 34051 | 6276 | 25085 | 14794 | 8422 | 16658 |
| MGAT4B | 560 | 2330 | 6837 | 2994 | 1328 | 462 | 1501 | 1344 |
| MGAT5 | 1014 | 0 | 0 | 96 | 0 | 1169 | 0 | 0 |
| MGC32805 | 0 | 0 | 0 | 0 | 0 | 0 | 514 | 0 |
| MGEA5 | 0 | 0 | 0 | 0 | 44 | 0 | 486 | 0 |
| MGLL | 1144 | 32 | 2792 | 306 | 510 | 410 | 966 | 0 |
| MGME1 | 27291 | 46371 | 72597 | 59641 | 211809 | 70765 | 81759 | 34936 |
| MGMT | 170379 | 193595 | 687295 | 22218 | 763429 | 332709 | 15875 | 245028 |
| MGP | 958 | 0 | 2792 | 4314 | 3065 | 14512 | 2281 | 6951 |
| MGRN1 | 0 | 142 | 0 | 0 | 0 | 0 | 0 | 0 |
| MGST1 | 4066 | 2354 | 9333 | 8618 | 11122 | 767 | 0 | 2304 |
| MGST2 | 3444 | 3478 | 1728 | 2680 | 1688 | 3324 | 2826 | 0 |
| MGST3 | 10644 | 9893 | 14519 | 7957 | 19868 | 11011 | 6121 | 6939 |
| MHENCR | 934 | 0 | 0 | 1062 | 644 | 24 | 0 | 0 |
| MIA | 0 | 0 | 0 | 0 | 0 | 0 | 0 | 1720 |
| MIA-RAB4B | 0 | 0 | 0 | 0 | 1226 | 0 | 0 | 0 |
| MIA2 | 564 | 194 | 486 | 0 | 356 | 0 | 0 | 0 |

| | | | | | | | | |
|---|---|---|---|---|---|---|---|---|
| MIA3 | 0 | 0 | 0 | 1308 | 932 | 0 | 720 | 0 |
| MIAT | 652 | 0 | 0 | 0 | 0 | 0 | 0 | 0 |
| MIB1 | 0 | 0 | 0 | 0 | 0 | 942 | 0 | 0 |
| MICA | 0 | 0 | 0 | 0 | 994 | 0 | 0 | 718 |
| MICAL1 | 1517 | 2948 | 1512 | 0 | 3321 | 1258 | 3642 | 3242 |
| MICAL2 | 0 | 272 | 1144 | 0 | 0 | 1978 | 0 | 0 |
| MICAL3 | 734 | 0 | 0 | 0 | 554 | 0 | 0 | 0 |
| MICALL2 | 1043 | 0 | 566 | 1275 | 940 | 0 | 0 | 0 |
| MICB-DT | 0 | 0 | 0 | 0 | 0 | 3281 | 0 | 0 |
| MICOS10 | 0 | 0 | 0 | 2092 | 652 | 0 | 358 | 0 |
| MICOS10-NBL1 | 0 | 0 | 1202 | 0 | 0 | 0 | 0 | 0 |
| MICOS13 | 0 | 50 | 2222 | 3361 | 468 | 1189 | 1014 | 838 |
| MICU1 | 0 | 0 | 0 | 0 | 1782 | 0 | 1796 | 0 |
| MICU2 | 115732 | 157549 | 222711 | 243705 | 354389 | 341860 | 337658 | 108176 |
| MID1IP1 | 0 | 0 | 0 | 0 | 976 | 0 | 397 | 568 |
| MID2 | 268 | 938 | 0 | 0 | 5072 | 1932 | 984 | 1708 |
| MIDN | 0 | 0 | 96 | 0 | 0 | 0 | 0 | 0 |
| MIEN1 | 4581 | 3166 | 2917 | 3993 | 5860 | 2863 | 2148 | 386 |
| MIER1 | 0 | 0 | 3134 | 0 | 0 | 0 | 0 | 587 |
| MIER2 | 0 | 0 | 0 | 0 | 542 | 0 | 0 | 0 |
| MIER3 | 476 | 0 | 926 | 0 | 0 | 0 | 0 | 0 |
| MIF | 3335 | 3520 | 11646 | 4678 | 33731 | 14143 | 10321 | 11269 |
| MIF4GD | 0 | 930 | 0 | 1146 | 0 | 0 | 0 | 0 |
| MIGA1 | 0 | 0 | 1457 | 0 | 290 | 0 | 312 | 0 |
| MIGA2 | 0 | 0 | 0 | 0 | 554 | 0 | 0 | 0 |
| MIIP | 0 | 228 | 0 | 0 | 252 | 0 | 0 | 0 |
| MILR1 | 0 | 0 | 0 | 0 | 0 | 0 | 642 | 0 |
| MINCR | 0 | 398 | 0 | 0 | 1156 | 3562 | 1188 | 0 |
| MINDY1 | 0 | 1044 | 1176 | 0 | 2292 | 0 | 724 | 0 |
| MINDY2 | 0 | 0 | 0 | 0 | 0 | 2026 | 422 | 0 |
| MINDY3 | 762 | 0 | 0 | 0 | 638 | 0 | 0 | 0 |
| MINDY4 | 0 | 0 | 0 | 0 | 888 | 0 | 0 | 0 |
| MINK1 | 0 | 608 | 0 | 0 | 610 | 0 | 584 | 0 |
| MINOS1 | 1756 | 5470 | 2682 | 8541 | 10936 | 1198 | 2484 | 2334 |
| MINPP1 | 33562 | 87797 | 182711 | 70453 | 146681 | 89424 | 234635 | 35064 |
| MIOS | 0 | 0 | 0 | 0 | 1594 | 0 | 0 | 0 |
| MIPEP | 34963 | 54363 | 150009 | 167515 | 286978 | 173329 | 309036 | 94893 |
| MIPEPP3 | 0 | 0 | 0 | 812 | 624 | 0 | 0 | 0 |
| MIPOL1 | 0 | 98 | 1290 | 1687 | 4841 | 0 | 0 | 0 |
| MIR155HG | 0 | 0 | 0 | 0 | 412 | 0 | 0 | 0 |
| MIR17HG | 0 | 0 | 0 | 0 | 328 | 0 | 0 | 0 |
| MIR193BHG | 0 | 0 | 0 | 0 | 0 | 0 | 1045 | 0 |
| MIR194-2HG | 0 | 1144 | 5204 | 1114 | 440 | 712 | 1432 | 476 |

| | | | | | | | | |
|---|---|---|---|---|---|---|---|---|
| MIR200CHG | 270 | 282 | 735 | 0 | 514 | 0 | 0 | 0 |
| MIR210HG | 0 | 302 | 0 | 0 | 1106 | 32 | 0 | 654 |
| MIR22HG | 0 | 0 | 3316 | 0 | 0 | 0 | 0 | 0 |
| MIR29B2CHG | 0 | 0 | 1003 | 0 | 0 | 0 | 196 | 0 |
| MIR34AHG | 0 | 0 | 0 | 0 | 164 | 0 | 0 | 0 |
| MIR4435-2HG | 0 | 238 | 653 | 1660 | 2768 | 0 | 338 | 1444 |
| MIR4458HG | 0 | 310 | 0 | 747 | 1016 | 0 | 0 | 811 |
| MIR762HG | 0 | 138 | 0 | 0 | 720 | 0 | 0 | 0 |
| MIRLET7BHG | 384 | 0 | 0 | 0 | 24 | 0 | 0 | 0 |
| MIS12 | 534 | 0 | 0 | 0 | 0 | 0 | 0 | 0 |
| MIS18A | 0 | 0 | 0 | 996 | 144 | 565 | 788 | 0 |
| MIS18BP1 | 0 | 0 | 0 | 0 | 0 | 0 | 424 | 0 |
| MISP | 1216 | 782 | 5200 | 264 | 1554 | 0 | 0 | 748 |
| MISP3 | 0 | 768 | 734 | 0 | 518 | 0 | 0 | 0 |
| MITD1 | 0 | 0 | 4393 | 0 | 4808 | 0 | 2158 | 0 |
| MITF | 1074 | 4294 | 9006 | 1843 | 32507 | 11086 | 3299 | 4026 |
| MKI67 | 0 | 0 | 0 | 0 | 80 | 0 | 0 | 0 |
| MKKS | 1742 | 1220 | 3108 | 4823 | 9373 | 0 | 840 | 1080 |
| MKL1 | 3974 | 2184 | 7591 | 3428 | 7006 | 1746 | 1831 | 3239 |
| MKL2 | 1988 | 1178 | 1499 | 1854 | 5584 | 3432 | 6912 | 0 |
| MKLN1 | 0 | 0 | 1000 | 0 | 0 | 0 | 0 | 0 |
| MKNK1 | 0 | 0 | 0 | 484 | 0 | 0 | 0 | 690 |
| MKNK2 | 7008 | 5821 | 66745 | 12661 | 69382 | 11172 | 7464 | 9869 |
| MKRN1 | 47008 | 60739 | 154785 | 68433 | 155815 | 68235 | 201694 | 28325 |
| MKRN2 | 10639 | 35180 | 62793 | 53569 | 174042 | 57493 | 123754 | 24717 |
| MKRN2OS | 0 | 0 | 0 | 0 | 0 | 0 | 0 | 1132 |
| MKS1 | 0 | 0 | 0 | 1094 | 0 | 0 | 0 | 0 |
| MLEC | 328 | 552 | 1546 | 3234 | 4148 | 2830 | 1502 | 574 |
| MLF1 | 9806 | 46604 | 14304 | 26981 | 135879 | 85789 | 118558 | 168707 |
| MLF2 | 1374 | 544 | 2970 | 868 | 8037 | 1254 | 4098 | 2222 |
| MLH1 | 28191 | 47928 | 72410 | 91011 | 205414 | 101416 | 151707 | 54037 |
| MLH3 | 0 | 0 | 0 | 54 | 0 | 0 | 0 | 0 |
| MLLT1 | 0 | 0 | 0 | 0 | 2011 | 0 | 368 | 0 |
| MLLT10 | 43094 | 32435 | 129118 | 102178 | 135163 | 86561 | 194986 | 70145 |
| MLLT3 | 802 | 0 | 0 | 0 | 0 | 0 | 218 | 0 |
| MLLT6 | 454 | 0 | 904 | 0 | 366 | 0 | 404 | 0 |
| MLPH | 0 | 354 | 0 | 768 | 1034 | 0 | 0 | 0 |
| MLST8 | 0 | 0 | 0 | 804 | 1342 | 0 | 300 | 0 |
| MLX | 558 | 0 | 1278 | 1770 | 1030 | 918 | 1830 | 0 |
| MLXIP | 224 | 996 | 0 | 1376 | 5066 | 2032 | 6943 | 920 |
| MLXIPL | 0 | 0 | 0 | 0 | 0 | 0 | 0 | 706 |
| MLYCD | 0 | 0 | 3052 | 0 | 0 | 0 | 0 | 0 |
| MMAA | 0 | 0 | 1100 | 0 | 0 | 0 | 0 | 0 |

| | | | | | | | | |
|---|---|---|---|---|---|---|---|---|
| MMAB | 0 | 0 | 0 | 0 | 2108 | 5253 | 440 | 3082 |
| MMACHC | 120 | 0 | 0 | 914 | 0 | 0 | 0 | 0 |
| MMADHC | 1168 | 2315 | 6856 | 4590 | 2456 | 0 | 3329 | 4090 |
| MMD | 202 | 0 | 0 | 0 | 238 | 1010 | 206 | 0 |
| MME | 0 | 0 | 0 | 0 | 0 | 0 | 550 | 0 |
| MMEL1 | 0 | 0 | 0 | 0 | 0 | 0 | 240 | 910 |
| MMP11 | 0 | 0 | 0 | 0 | 7362 | 214 | 362 | 0 |
| MMP12 | 102 | 0 | 0 | 0 | 1670 | 0 | 0 | 0 |
| MMP14 | 0 | 0 | 540 | 0 | 630 | 0 | 0 | 0 |
| MMP15 | 0 | 0 | 0 | 0 | 0 | 0 | 536 | 0 |
| MMP17 | 576 | 0 | 0 | 0 | 0 | 0 | 0 | 0 |
| MMP19 | 58 | 2952 | 498 | 0 | 5188 | 5025 | 0 | 1142 |
| MMP2 | 0 | 136 | 0 | 515 | 7622 | 0 | 0 | 0 |
| MMP24 | 0 | 0 | 54 | 0 | 0 | 0 | 0 | 0 |
| MMP24-AS1-EDEM2 | 574 | 0 | 1831 | 0 | 0 | 0 | 1080 | 620 |
| MMP24OS | 242 | 1076 | 370 | 930 | 0 | 462 | 1114 | 0 |
| MMP25 | 0 | 0 | 0 | 0 | 232 | 0 | 0 | 0 |
| MMP28 | 1230 | 2884 | 5978 | 0 | 0 | 0 | 0 | 0 |
| MMP7 | 0 | 0 | 0 | 0 | 44547 | 1522 | 0 | 1598 |
| MMP9 | 0 | 0 | 0 | 504 | 0 | 0 | 0 | 0 |
| MMRN2 | 0 | 0 | 2400 | 0 | 0 | 0 | 0 | 0 |
| MMS19 | 0 | 0 | 0 | 0 | 0 | 0 | 166 | 0 |
| MMS22L | 0 | 0 | 0 | 0 | 298 | 990 | 0 | 0 |
| MMUT | 0 | 310 | 0 | 0 | 0 | 874 | 0 | 0 |
| MNAT1 | 66818 | 57494 | 72136 | 48994 | 600986 | 170535 | 163324 | 114366 |
| MND1 | 0 | 0 | 352 | 1242 | 0 | 0 | 0 | 0 |
| MNDA | 4084 | 3980 | 34703 | 932 | 22331 | 155877 | 13076 | 20857 |
| MNT | 0 | 0 | 0 | 627 | 448 | 0 | 0 | 0 |
| MNX1 | 0 | 0 | 0 | 216 | 134 | 530 | 256 | 0 |
| MNX1-AS1 | 0 | 0 | 0 | 0 | 536 | 0 | 0 | 160 |
| MOAP1 | 0 | 92 | 0 | 0 | 0 | 0 | 366 | 0 |
| MOB1A | 5826 | 11852 | 33706 | 5760 | 26382 | 8046 | 22360 | 7776 |
| MOB2 | 878 | 0 | 0 | 0 | 0 | 0 | 0 | 0 |
| MOB3A | 0 | 464 | 2642 | 0 | 0 | 0 | 1104 | 0 |
| MOB3B | 0 | 238 | 0 | 0 | 0 | 0 | 0 | 0 |
| MOB4 | 732 | 475 | 2666 | 0 | 952 | 4726 | 1580 | 500 |
| MOCS1 | 0 | 0 | 906 | 0 | 0 | 0 | 0 | 0 |
| MOCS2 | 332 | 0 | 1105 | 2344 | 0 | 0 | 0 | 0 |
| MOCS3 | 0 | 0 | 0 | 0 | 0 | 0 | 174 | 0 |
| MOGAT2 | 0 | 0 | 1437 | 0 | 0 | 0 | 0 | 0 |
| MOGAT3 | 0 | 0 | 0 | 0 | 0 | 0 | 1328 | 0 |
| MOGS | 0 | 350 | 266 | 0 | 0 | 0 | 212 | 788 |
| MON1A | 0 | 0 | 0 | 0 | 478 | 0 | 0 | 0 |

| | | | | | | | | |
|---|---|---|---|---|---|---|---|---|
| MON2 | 0 | 166 | 0 | 0 | 646 | 0 | 0 | 0 |
| MORC2 | 0 | 0 | 0 | 0 | 1098 | 0 | 360 | 366 |
| MORC3 | 2420 | 6550 | 6276 | 3410 | 8251 | 5855 | 10396 | 3625 |
| MORC4 | 0 | 0 | 0 | 0 | 238 | 0 | 0 | 0 |
| MORF4L1 | 482 | 1274 | 3436 | 2686 | 4152 | 1810 | 2534 | 7874 |
| MORF4L2 | 0 | 432 | 2425 | 3267 | 6044 | 1804 | 2510 | 1570 |
| MORN2 | 745 | 0 | 0 | 726 | 5477 | 0 | 0 | 0 |
| MORN3 | 0 | 0 | 0 | 0 | 739 | 0 | 32941 | 1160 |
| MORN4 | 600 | 0 | 0 | 0 | 0 | 0 | 0 | 134 |
| MOSPD1 | 0 | 0 | 1218 | 846 | 538 | 0 | 0 | 0 |
| MOSPD3 | 0 | 290 | 0 | 0 | 462 | 0 | 0 | 0 |
| MOV10 | 0 | 202 | 0 | 0 | 454 | 1907 | 0 | 0 |
| MPC1 | 2934 | 2107 | 11858 | 4872 | 2684 | 3541 | 972 | 698 |
| MPC2 | 1618 | 3460 | 5531 | 3890 | 8679 | 0 | 1318 | 2096 |
| MPDU1 | 842 | 454 | 3224 | 0 | 0 | 0 | 0 | 302 |
| MPEG1 | 0 | 0 | 332 | 0 | 0 | 0 | 0 | 0 |
| MPG | 730 | 14 | 1038 | 0 | 3626 | 0 | 0 | 0 |
| MPHOSPH10 | 526 | 352 | 1674 | 0 | 0 | 0 | 0 | 1012 |
| MPHOSPH6 | 0 | 0 | 1256 | 1305 | 2216 | 0 | 1078 | 0 |
| MPHOSPH8 | 0 | 0 | 0 | 0 | 0 | 5784 | 44 | 0 |
| MPI | 108 | 0 | 0 | 850 | 296 | 0 | 870 | 0 |
| MPLKIP | 0 | 952 | 1712 | 4479 | 2100 | 599 | 1771 | 870 |
| MPND | 1212 | 596 | 508 | 0 | 766 | 0 | 0 | 0 |
| MPP1 | 0 | 0 | 838 | 0 | 0 | 0 | 0 | 892 |
| MPP2 | 0 | 0 | 0 | 0 | 0 | 0 | 812 | 0 |
| MPP3 | 0 | 204 | 0 | 0 | 0 | 0 | 0 | 0 |
| MPP5 | 510 | 1548 | 0 | 1932 | 4232 | 1488 | 2980 | 0 |
| MPP6 | 281 | 0 | 0 | 0 | 0 | 0 | 0 | 0 |
| MPP7 | 0 | 0 | 0 | 0 | 320 | 0 | 0 | 0 |
| MPPE1 | 0 | 662 | 0 | 1114 | 0 | 0 | 732 | 0 |
| MPPED2 | 1696 | 280 | 0 | 0 | 0 | 0 | 1076 | 0 |
| MPRIP | 686 | 0 | 0 | 840 | 0 | 0 | 450 | 0 |
| MPST | 684 | 1150 | 3922 | 3544 | 5138 | 1772 | 1490 | 0 |
| MPV17 | 0 | 836 | 60 | 0 | 2083 | 0 | 2156 | 2438 |
| MPV17L | 0 | 0 | 0 | 0 | 0 | 0 | 230 | 0 |
| MPV17L2 | 0 | 0 | 0 | 0 | 0 | 0 | 0 | 1254 |
| MPZL1 | 0 | 568 | 0 | 2196 | 616 | 0 | 378 | 0 |
| MPZL2 | 0 | 870 | 896 | 1312 | 3763 | 0 | 0 | 0 |
| MPZL3 | 216 | 0 | 804 | 0 | 308 | 0 | 0 | 0 |
| MRE11 | 0 | 0 | 0 | 0 | 0 | 0 | 266 | 0 |
| MREG | 0 | 0 | 0 | 0 | 250 | 0 | 0 | 0 |
| MRFAP1 | 0 | 1940 | 580 | 1110 | 4244 | 0 | 4058 | 870 |
| MRFAP1L1 | 0 | 584 | 2010 | 1084 | 3154 | 0 | 0 | 0 |

| | | | | | | | | |
|---|---|---|---|---|---|---|---|---|
| MRGBP | 0 | 286 | 0 | 1384 | 1834 | 1568 | 470 | 0 |
| MRM1 | 0 | 0 | 0 | 2177 | 0 | 0 | 0 | 0 |
| MRM2 | 0 | 370 | 0 | 620 | 1212 | 0 | 0 | 1158 |
| MRM3 | 0 | 0 | 0 | 0 | 734 | 0 | 0 | 0 |
| MRNIP | 0 | 0 | 22 | 0 | 0 | 0 | 0 | 0 |
| MROH1 | 410 | 0 | 0 | 0 | 602 | 0 | 0 | 0 |
| MROH3P | 0 | 0 | 149 | 0 | 0 | 0 | 0 | 0 |
| MROH6 | 0 | 0 | 0 | 0 | 480 | 0 | 274 | 0 |
| MRPL1 | 0 | 843 | 2844 | 0 | 1470 | 0 | 332 | 1324 |
| MRPL10 | 0 | 0 | 691 | 2470 | 2111 | 0 | 0 | 1057 |
| MRPL11 | 338 | 2646 | 0 | 3845 | 9241 | 0 | 5996 | 188 |
| MRPL12 | 1498 | 2558 | 3116 | 2396 | 2081 | 0 | 1526 | 0 |
| MRPL13 | 1326 | 2334 | 2967 | 6000 | 8060 | 2414 | 6157 | 4944 |
| MRPL14 | 0 | 1096 | 3902 | 3622 | 6511 | 0 | 1400 | 356 |
| MRPL15 | 1426 | 1066 | 0 | 2948 | 1197 | 7854 | 7488 | 2568 |
| MRPL16 | 0 | 1692 | 4784 | 5937 | 1560 | 0 | 532 | 498 |
| MRPL17 | 826 | 0 | 0 | 6074 | 2572 | 1208 | 2766 | 2786 |
| MRPL18 | 2968 | 800 | 1112 | 4296 | 5771 | 2030 | 1580 | 1360 |
| MRPL19 | 0 | 0 | 0 | 0 | 1240 | 0 | 360 | 0 |
| MRPL2 | 0 | 218 | 0 | 1592 | 2960 | 0 | 712 | 2756 |
| MRPL20 | 954 | 1864 | 2528 | 5561 | 5751 | 2126 | 876 | 1433 |
| MRPL20-AS1 | 0 | 0 | 0 | 1008 | 311 | 0 | 0 | 0 |
| MRPL21 | 0 | 1442 | 3169 | 5308 | 8007 | 0 | 2324 | 1690 |
| MRPL22 | 830 | 2322 | 1295 | 1392 | 3540 | 4530 | 2509 | 1198 |
| MRPL23 | 1012 | 4230 | 0 | 0 | 1605 | 2418 | 0 | 995 |
| MRPL24 | 0 | 1544 | 0 | 624 | 2038 | 0 | 866 | 0 |
| MRPL27 | 68 | 2164 | 5802 | 4941 | 3303 | 0 | 1094 | 3076 |
| MRPL28 | 806 | 440 | 1082 | 2696 | 530 | 0 | 0 | 0 |
| MRPL3 | 0 | 476 | 0 | 1922 | 1678 | 0 | 126 | 734 |
| MRPL30 | 0 | 0 | 776 | 0 | 382 | 1680 | 398 | 660 |
| MRPL32 | 0 | 1056 | 0 | 1083 | 1878 | 0 | 909 | 1704 |
| MRPL33 | 0 | 2090 | 3082 | 2644 | 6099 | 2412 | 1502 | 1063 |
| MRPL34 | 2662 | 0 | 3090 | 958 | 7348 | 1989 | 740 | 1736 |
| MRPL35 | 98377 | 195322 | 157806 | 95804 | 392621 | 96013 | 169417 | 140583 |
| MRPL36 | 0 | 1104 | 1374 | 4788 | 9113 | 5603 | 3072 | 2796 |
| MRPL37 | 224 | 956 | 3191 | 1254 | 3228 | 0 | 0 | 0 |
| MRPL38 | 1034 | 382 | 0 | 1119 | 2492 | 1390 | 538 | 1224 |
| MRPL39 | 0 | 982 | 0 | 0 | 1006 | 0 | 1032 | 2044 |
| MRPL4 | 4360 | 4332 | 25746 | 17681 | 26018 | 8678 | 6173 | 10981 |
| MRPL40 | 0 | 0 | 0 | 1142 | 4456 | 0 | 1122 | 447 |
| MRPL41 | 1506 | 1046 | 2336 | 2788 | 2398 | 0 | 690 | 0 |
| MRPL42 | 124009 | 267070 | 160817 | 166916 | 556148 | 165629 | 480419 | 153535 |
| MRPL43 | 570 | 296 | 0 | 2284 | 2210 | 1754 | 1320 | 2062 |

| | | | | | | | | |
|---|---|---|---|---|---|---|---|---|
| MRPL44 | 438 | 0 | 3540 | 2162 | 556 | 1338 | 0 | 664 |
| MRPL45 | 0 | 460 | 1916 | 978 | 2968 | 0 | 872 | 1130 |
| MRPL45P2 | 0 | 0 | 0 | 0 | 0 | 0 | 498 | 0 |
| MRPL46 | 0 | 0 | 0 | 2575 | 598 | 0 | 2068 | 0 |
| MRPL47 | 1100 | 1152 | 0 | 2351 | 4842 | 0 | 1260 | 1234 |
| MRPL48 | 128336 | 251275 | 270750 | 561827 | 868881 | 534398 | 1062462 | 430869 |
| MRPL49 | 370 | 44 | 0 | 1944 | 430 | 32 | 386 | 1102 |
| MRPL50 | 154 | 1330 | 1712 | 8495 | 2946 | 0 | 512 | 0 |
| MRPL51 | 160 | 2418 | 3438 | 8426 | 7182 | 2254 | 7341 | 1620 |
| MRPL52 | 956 | 0 | 2968 | 2202 | 5020 | 10 | 2096 | 2558 |
| MRPL53 | 0 | 572 | 2286 | 1292 | 4580 | 0 | 978 | 1582 |
| MRPL54 | 0 | 754 | 6064 | 890 | 0 | 3802 | 0 | 1990 |
| MRPL55 | 0 | 492 | 0 | 1412 | 3776 | 0 | 454 | 0 |
| MRPL57 | 1966 | 2164 | 2079 | 6728 | 2936 | 4101 | 7824 | 1492 |
| MRPL58 | 504 | 0 | 3578 | 968 | 594 | 0 | 0 | 1453 |
| MRPL9 | 560 | 0 | 0 | 5462 | 2135 | 1682 | 3062 | 4000 |
| MRPS10 | 0 | 0 | 0 | 1968 | 2538 | 0 | 668 | 3611 |
| MRPS11 | 744 | 1254 | 0 | 2796 | 0 | 2750 | 412 | 4423 |
| MRPS12 | 0 | 580 | 2980 | 590 | 2820 | 0 | 652 | 2628 |
| MRPS14 | 0 | 470 | 3162 | 3291 | 3930 | 254 | 1146 | 1722 |
| MRPS15 | 0 | 1786 | 0 | 4062 | 4716 | 0 | 0 | 2843 |
| MRPS16 | 1110 | 1738 | 4860 | 1017 | 1420 | 0 | 1132 | 1285 |
| MRPS17 | 0 | 0 | 1630 | 2370 | 2032 | 0 | 924 | 465 |
| MRPS18A | 0 | 3042 | 4868 | 1469 | 2334 | 0 | 638 | 0 |
| MRPS18B | 0 | 0 | 948 | 2257 | 2404 | 1110 | 1622 | 1142 |
| MRPS18C | 624 | 3514 | 3278 | 2571 | 3935 | 1260 | 1810 | 442 |
| MRPS2 | 0 | 296 | 1887 | 2616 | 0 | 2743 | 428 | 0 |
| MRPS21 | 762 | 1130 | 0 | 4594 | 4140 | 5634 | 2033 | 604 |
| MRPS22 | 426 | 1192 | 0 | 0 | 920 | 1904 | 0 | 2330 |
| MRPS23 | 87935 | 213736 | 219940 | 354987 | 680502 | 402815 | 782238 | 303352 |
| MRPS24 | 1046 | 422 | 3765 | 6080 | 6312 | 5280 | 4884 | 894 |
| MRPS25 | 0 | 1014 | 3587 | 1647 | 4233 | 0 | 326 | 0 |
| MRPS26 | 720 | 1956 | 3716 | 956 | 6981 | 2275 | 486 | 2433 |
| MRPS27 | 70 | 376 | 0 | 0 | 156 | 0 | 0 | 0 |
| MRPS28 | 506 | 466 | 0 | 578 | 1170 | 1404 | 742 | 894 |
| MRPS30 | 0 | 0 | 2010 | 0 | 736 | 0 | 0 | 0 |
| MRPS31 | 16421 | 19582 | 28988 | 76363 | 92744 | 74863 | 139181 | 35934 |
| MRPS31P5 | 0 | 0 | 0 | 0 | 1010 | 0 | 194 | 0 |
| MRPS33 | 266745 | 463934 | 265908 | 299494 | 544200 | 475431 | 1041216 | 107942 |
| MRPS34 | 186 | 358 | 3068 | 2071 | 4420 | 10302 | 1314 | 0 |
| MRPS35 | 1454 | 2818 | 432 | 1232 | 2826 | 0 | 982 | 2154 |
| MRPS36 | 662 | 2366 | 1524 | 1542 | 4248 | 2046 | 1044 | 645 |
| MRPS5 | 1510 | 1187 | 0 | 1569 | 880 | 0 | 0 | 1233 |

| | | | | | | | | |
|---|---|---|---|---|---|---|---|---|
| MRPS6 | 1966 | 0 | 1717 | 1213 | 2358 | 1884 | 304 | 997 |
| MRPS7 | 728 | 3404 | 4590 | 4130 | 5116 | 1749 | 1728 | 1178 |
| MRPS9 | 952 | 1446 | 0 | 0 | 2923 | 0 | 402 | 0 |
| MRRF | 0 | 218 | 0 | 0 | 0 | 0 | 398 | 0 |
| MRS2 | 0 | 1326 | 0 | 0 | 0 | 0 | 672 | 1883 |
| MRTFA | 0 | 6790 | 36490 | 316 | 9149 | 10598 | 4282 | 7019 |
| MRTFA-AS1 | 0 | 0 | 0 | 0 | 0 | 0 | 36 | 0 |
| MRTFB | 1116 | 1535 | 3030 | 2458 | 4542 | 3514 | 5370 | 1592 |
| MRTO4 | 420 | 0 | 1756 | 1546 | 739 | 0 | 0 | 0 |
| MS4A1 | 0 | 0 | 1336 | 232 | 0 | 0 | 0 | 1730 |
| MS4A12 | 11608 | 5460 | 73039 | 0 | 1540 | 0 | 0 | 0 |
| MS4A4A | 0 | 0 | 0 | 0 | 912 | 108 | 0 | 1350 |
| MS4A4E | 0 | 152 | 0 | 0 | 444 | 0 | 0 | 0 |
| MS4A6A | 142 | 0 | 386 | 0 | 0 | 0 | 1392 | 2458 |
| MS4A7 | 0 | 0 | 748 | 0 | 588 | 0 | 0 | 0 |
| MS4A8 | 0 | 466 | 980 | 0 | 0 | 0 | 0 | 0 |
| MSANTD2 | 224 | 0 | 0 | 0 | 0 | 0 | 0 | 0 |
| MSANTD3 | 0 | 0 | 0 | 732 | 0 | 0 | 0 | 0 |
| MSANTD4 | 0 | 0 | 0 | 0 | 98 | 0 | 0 | 0 |
| MSC | 0 | 0 | 0 | 0 | 426 | 0 | 0 | 0 |
| MSC-AS1 | 0 | 0 | 0 | 0 | 1080 | 0 | 0 | 0 |
| MSH2 | 26809 | 57497 | 94641 | 160220 | 394269 | 101289 | 175155 | 114225 |
| MSH5-SAPCD1 | 0 | 340 | 0 | 0 | 0 | 0 | 0 | 0 |
| MSI2 | 9045 | 23459 | 17391 | 42106 | 133057 | 63172 | 113554 | 45447 |
| MSL1 | 2114 | 3240 | 1232 | 1445 | 7348 | 1110 | 11826 | 910 |
| MSL2 | 1496 | 3414 | 3210 | 3143 | 6309 | 3270 | 7859 | 2630 |
| MSL3 | 0 | 0 | 0 | 0 | 472 | 0 | 0 | 0 |
| MSL3P1 | 0 | 0 | 0 | 934 | 0 | 0 | 0 | 0 |
| MSLN | 0 | 0 | 0 | 0 | 654 | 0 | 2311 | 3907 |
| MSMB | 0 | 0 | 0 | 182 | 1713 | 0 | 616 | 0 |
| MSMO1 | 0 | 0 | 0 | 0 | 1200 | 0 | 100 | 743 |
| MSN | 0 | 482 | 1152 | 0 | 522 | 0 | 0 | 0 |
| MSRA | 0 | 0 | 0 | 0 | 642 | 0 | 0 | 0 |
| MSRB1 | 1634 | 700 | 3900 | 1016 | 856 | 2232 | 0 | 2194 |
| MSRB2 | 2122 | 568 | 0 | 2586 | 2033 | 1644 | 1246 | 832 |
| MSRB3 | 0 | 0 | 0 | 190 | 0 | 0 | 0 | 0 |
| MST1L | 364 | 0 | 1712 | 0 | 0 | 0 | 0 | 0 |
| MST1R | 0 | 932 | 0 | 494 | 54 | 0 | 428 | 0 |
| MSX2 | 0 | 0 | 0 | 0 | 0 | 0 | 552 | 0 |
| MT1A | 0 | 376 | 0 | 0 | 0 | 0 | 0 | 0 |
| MT1DP | 0 | 0 | 786 | 0 | 176 | 0 | 0 | 0 |
| MT1E | 14792 | 14753 | 11895 | 588 | 3716 | 0 | 780 | 292 |
| MT1F | 4868 | 252 | 9175 | 0 | 218 | 0 | 514 | 0 |

| | | | | | | | | |
|---|---|---|---|---|---|---|---|---|
| MT1G | 18630 | 7301 | 23445 | 606 | 2282 | 698 | 3422 | 0 |
| MT1H | 2480 | 1846 | 5943 | 0 | 0 | 0 | 0 | 0 |
| MT1L | 585 | 0 | 0 | 0 | 420 | 0 | 424 | 0 |
| MT1M | 5180 | 966 | 3185 | 0 | 0 | 0 | 0 | 0 |
| MT1X | 5084 | 4666 | 4416 | 2864 | 2951 | 0 | 1886 | 976 |
| MT2A | 14148 | 19816 | 24848 | 3656 | 6954 | 0 | 1016 | 4718 |
| MT3 | 0 | 0 | 0 | 0 | 0 | 0 | 1812 | 0 |
| MTA1 | 0 | 0 | 478 | 0 | 364 | 0 | 782 | 116 |
| MTA2 | 0 | 0 | 0 | 0 | 34 | 310 | 0 | 0 |
| MTA3 | 0 | 230 | 1710 | 1072 | 0 | 0 | 0 | 0 |
| MTAP | 28376 | 50713 | 50404 | 127942 | 311494 | 139025 | 235030 | 166315 |
| MTCH1 | 0 | 330 | 5516 | 214 | 5482 | 890 | 1832 | 944 |
| MTCH2 | 470 | 2402 | 4102 | 5600 | 5867 | 0 | 3521 | 1120 |
| MTDH | 770 | 0 | 745 | 0 | 1732 | 0 | 1634 | 0 |
| MTERF2 | 0 | 0 | 0 | 1523 | 0 | 0 | 306 | 0 |
| MTERF3 | 3776 | 16640 | 9827 | 24390 | 83452 | 50576 | 71935 | 22039 |
| MTERF4 | 0 | 534 | 2384 | 668 | 540 | 0 | 554 | 0 |
| MTF2 | 34158 | 35500 | 63410 | 70900 | 112014 | 43764 | 74322 | 19718 |
| MTFMT | 0 | 346 | 1584 | 1192 | 0 | 1819 | 736 | 0 |
| MTFP1 | 0 | 12 | 0 | 1574 | 482 | 2263 | 0 | 0 |
| MTFR1 | 0 | 0 | 0 | 0 | 420 | 1115 | 0 | 0 |
| MTFR1L | 0 | 0 | 0 | 0 | 1042 | 0 | 0 | 0 |
| MTFR2 | 0 | 0 | 0 | 844 | 296 | 0 | 568 | 0 |
| MTG1 | 0 | 0 | 0 | 0 | 458 | 0 | 0 | 0 |
| MTG2 | 0 | 0 | 0 | 0 | 1964 | 0 | 0 | 669 |
| MTHFD1 | 0 | 0 | 0 | 0 | 554 | 0 | 0 | 1080 |
| MTHFD1L | 1030 | 5134 | 7011 | 30250 | 134838 | 68942 | 46971 | 45188 |
| MTHFD2 | 0 | 0 | 0 | 2630 | 2377 | 948 | 448 | 2502 |
| MTHFD2L | 852 | 0 | 0 | 0 | 2118 | 874 | 52 | 0 |
| MTHFR | 246 | 412 | 0 | 0 | 0 | 0 | 0 | 0 |
| MTHFS | 104074 | 177377 | 351095 | 92876 | 713360 | 169185 | 309932 | 230303 |
| MTHFSD | 0 | 0 | 0 | 0 | 348 | 0 | 0 | 0 |
| MTIF2 | 0 | 0 | 490 | 0 | 1326 | 0 | 0 | 0 |
| MTIF3 | 0 | 1148 | 5232 | 5070 | 4686 | 1984 | 4220 | 2305 |
| MTM1 | 0 | 0 | 768 | 0 | 0 | 0 | 0 | 0 |
| MTMR10 | 0 | 0 | 0 | 790 | 508 | 0 | 0 | 0 |
| MTMR11 | 450 | 1018 | 678 | 0 | 953 | 2540 | 1178 | 981 |
| MTMR12 | 0 | 1338 | 4011 | 0 | 12439 | 2111 | 4510 | 1360 |
| MTMR14 | 20533 | 27386 | 50848 | 43234 | 139052 | 57296 | 107030 | 32302 |
| MTMR2 | 5130 | 11286 | 6336 | 13864 | 39101 | 19428 | 23304 | 13427 |
| MTMR4 | 308 | 272 | 0 | 0 | 0 | 0 | 0 | 0 |
| MTMR9 | 0 | 0 | 0 | 0 | 0 | 0 | 672 | 0 |
| MTMR9LP | 0 | 0 | 588 | 0 | 0 | 0 | 0 | 0 |

| | | | | | | | | |
|---|---|---|---|---|---|---|---|---|
| MTO1 | 0 | 52 | 0 | 0 | 0 | 0 | 0 | 0 |
| MTOR | 308 | 3806 | 4562 | 1442 | 4853 | 0 | 4770 | 1360 |
| MTPAP | 378 | 0 | 0 | 1282 | 0 | 0 | 0 | 0 |
| MTPN | 60 | 0 | 0 | 0 | 0 | 442 | 209 | 0 |
| MTR | 0 | 0 | 0 | 0 | 240 | 0 | 0 | 0 |
| MTREX | 0 | 0 | 0 | 0 | 1976 | 0 | 0 | 0 |
| MTRF1 | 0 | 398 | 0 | 0 | 2120 | 1192 | 0 | 0 |
| MTRF1L | 0 | 408 | 0 | 0 | 0 | 0 | 0 | 944 |
| MTRNR2L1 | 92 | 0 | 0 | 0 | 0 | 0 | 0 | 0 |
| MTRNR2L10 | 0 | 0 | 0 | 0 | 286 | 0 | 0 | 0 |
| MTRNR2L2 | 19961 | 4592 | 13531 | 1539 | 7625 | 5040 | 11216 | 2822 |
| MTRNR2L6 | 434 | 0 | 0 | 0 | 0 | 0 | 0 | 0 |
| MTRNR2L8 | 12908 | 3310 | 15041 | 2520 | 4464 | 2552 | 3792 | 1200 |
| MTRR | 0 | 40 | 0 | 618 | 106 | 0 | 1127 | 0 |
| MTSS1 | 3056 | 0 | 2346 | 0 | 1156 | 0 | 5217 | 1049 |
| MTURN | 0 | 0 | 1554 | 0 | 600 | 0 | 314 | 0 |
| MTUS1 | 12 | 0 | 0 | 0 | 272 | 0 | 0 | 0 |
| MTX1 | 0 | 0 | 0 | 0 | 1972 | 1895 | 0 | 2706 |
| MTX2 | 0 | 514 | 1245 | 1294 | 1466 | 0 | 0 | 553 |
| MUC1 | 889 | 334 | 650 | 0 | 1334 | 0 | 0 | 392 |
| MUC12 | 1944 | 6291 | 4590 | 1694 | 0 | 0 | 0 | 0 |
| MUC13 | 5660 | 1888 | 9665 | 2441 | 3615 | 0 | 1072 | 0 |
| MUC17 | 0 | 0 | 0 | 0 | 672 | 0 | 0 | 0 |
| MUC2 | 766 | 2094 | 5070 | 1468 | 0 | 0 | 0 | 0 |
| MUC20 | 2114 | 272 | 6846 | 750 | 4508 | 993 | 538 | 988 |
| MUC3A | 520 | 948 | 856 | 1255 | 2494 | 0 | 1248 | 346 |
| MUC4 | 3672 | 3290 | 5226 | 1052 | 1246 | 0 | 0 | 652 |
| MUC5B | 0 | 1749 | 1408 | 0 | 316 | 0 | 0 | 0 |
| MUCL1 | 0 | 0 | 0 | 0 | 0 | 2222 | 0 | 0 |
| MUL1 | 0 | 386 | 1828 | 562 | 0 | 0 | 0 | 0 |
| MUM1 | 2928 | 3022 | 1899 | 2000 | 16904 | 13418 | 4542 | 1878 |
| MUS81 | 0 | 0 | 0 | 0 | 638 | 0 | 146 | 0 |
| MUSTN1 | 0 | 0 | 1400 | 0 | 244 | 0 | 0 | 0 |
| MUT | 0 | 924 | 0 | 0 | 110 | 0 | 158 | 0 |
| MUTYH | 0 | 0 | 0 | 0 | 0 | 0 | 186 | 0 |
| MVB12A | 68 | 676 | 0 | 786 | 5488 | 2222 | 1432 | 0 |
| MVB12B | 0 | 0 | 0 | 1012 | 0 | 0 | 0 | 0 |
| MVD | 0 | 0 | 1388 | 0 | 0 | 0 | 0 | 0 |
| MVK | 0 | 486 | 0 | 580 | 652 | 0 | 200 | 62 |
| MVP | 16821 | 22238 | 76374 | 17347 | 25319 | 19951 | 28095 | 12682 |
| MX1 | 104 | 340 | 2150 | 0 | 1470 | 0 | 204 | 748 |
| MXD1 | 1319 | 0 | 1270 | 0 | 624 | 0 | 0 | 0 |
| MXD3 | 0 | 0 | 0 | 432 | 1078 | 0 | 0 | 0 |

| | | | | | | | | |
|---|---|---|---|---|---|---|---|---|
| MXD4 | 0 | 298 | 0 | 0 | 0 | 1561 | 0 | 0 |
| MXI1 | 378 | 388 | 640 | 376 | 0 | 0 | 726 | 0 |
| MXRA8 | 0 | 0 | 0 | 0 | 2776 | 0 | 0 | 504 |
| MYADM | 0 | 234 | 0 | 0 | 0 | 0 | 200 | 0 |
| MYB | 21879 | 95790 | 58053 | 77266 | 248942 | 66984 | 146755 | 26471 |
| MYBL2 | 0 | 0 | 0 | 0 | 1804 | 0 | 0 | 934 |
| MYC | 3704 | 6492 | 15025 | 19327 | 30694 | 24060 | 57656 | 12568 |
| MYCBP | 458 | 728 | 1338 | 3907 | 4136 | 0 | 0 | 0 |
| MYD88 | 0 | 0 | 1198 | 738 | 1442 | 0 | 0 | 0 |
| MYDGF | 1876 | 1114 | 0 | 4718 | 3792 | 0 | 2431 | 652 |
| MYEF2 | 0 | 0 | 508 | 0 | 0 | 0 | 0 | 0 |
| MYEOV | 0 | 0 | 0 | 904 | 150 | 0 | 0 | 782 |
| MYH10 | 0 | 0 | 0 | 0 | 0 | 0 | 794 | 0 |
| MYH11 | 10429 | 3913 | 76321 | 1040 | 6306 | 9306 | 2352 | 0 |
| MYH14 | 2150 | 1452 | 7882 | 3466 | 12332 | 3606 | 1654 | 2655 |
| MYH15 | 0 | 0 | 0 | 0 | 0 | 0 | 556 | 0 |
| MYH9 | 336860 | 270048 | 468266 | 225311 | 1128298 | 313671 | 480680 | 216814 |
| MYL12A | 5622 | 10510 | 38930 | 17671 | 42507 | 15994 | 6104 | 28323 |
| MYL12B | 9811 | 11213 | 50657 | 13190 | 56339 | 17994 | 9521 | 23747 |
| MYL6 | 30726 | 40786 | 139153 | 39766 | 106985 | 43796 | 22773 | 29984 |
| MYL6B | 0 | 0 | 1786 | 0 | 1308 | 0 | 364 | 0 |
| MYL9 | 460 | 452 | 1222 | 810 | 3008 | 0 | 1276 | 0 |
| MYLIP | 0 | 468 | 0 | 0 | 876 | 0 | 0 | 0 |
| MYLK | 6412 | 3914 | 24924 | 6264 | 5380 | 2544 | 2828 | 2942 |
| MYNN | 0 | 240 | 0 | 0 | 0 | 0 | 0 | 0 |
| MYO10 | 0 | 0 | 0 | 140 | 0 | 0 | 0 | 0 |
| MYO15B | 268 | 270 | 1810 | 724 | 0 | 0 | 334 | 0 |
| MYO19 | 0 | 0 | 0 | 0 | 0 | 0 | 0 | 340 |
| MYO1A | 1440 | 512 | 0 | 0 | 654 | 0 | 72 | 0 |
| MYO1B | 392 | 0 | 0 | 0 | 966 | 0 | 0 | 0 |
| MYO1C | 150 | 0 | 1350 | 124 | 0 | 0 | 212 | 0 |
| MYO1D | 6554 | 6009 | 7224 | 3855 | 5445 | 3628 | 2894 | 0 |
| MYO1E | 230 | 0 | 0 | 0 | 0 | 0 | 0 | 0 |
| MYO1G | 0 | 0 | 0 | 0 | 218 | 0 | 0 | 0 |
| MYO5B | 1334 | 140 | 0 | 2524 | 1257 | 0 | 0 | 0 |
| MYO6 | 91850 | 106610 | 595751 | 193371 | 713125 | 182466 | 261940 | 179160 |
| MYO7B | 0 | 0 | 1152 | 0 | 1270 | 0 | 940 | 0 |
| MYO9A | 0 | 0 | 0 | 0 | 0 | 0 | 0 | 638 |
| MYO9B | 0 | 490 | 462 | 1248 | 1400 | 2466 | 0 | 104 |
| MYOF | 0 | 410 | 0 | 0 | 1634 | 0 | 0 | 0 |
| MYOM1 | 530 | 0 | 0 | 0 | 0 | 0 | 0 | 0 |
| MYOM3 | 0 | 0 | 0 | 1462 | 0 | 0 | 0 | 0 |
| MYOZ2 | 1112 | 0 | 0 | 0 | 0 | 0 | 0 | 0 |

| | | | | | | | | |
|---|---|---|---|---|---|---|---|---|
| MYPOP | 0 | 0 | 0 | 0 | 432 | 0 | 0 | 366 |
| MYRF | 96 | 0 | 0 | 0 | 416 | 0 | 0 | 0 |
| MYRIP | 0 | 1194 | 0 | 340 | 0 | 2016 | 8144 | 0 |
| MYSM1 | 0 | 0 | 0 | 0 | 554 | 0 | 0 | 0 |
| MYZAP | 0 | 0 | 114 | 815 | 0 | 0 | 0 | 0 |
| MZB1 | 1008 | 590 | 1553 | 0 | 0 | 0 | 0 | 2081 |
| MZT1 | 0 | 46 | 2203 | 3826 | 3342 | 1150 | 106 | 0 |
| MZT2A | 1104 | 562 | 3051 | 3785 | 3296 | 0 | 2453 | 846 |
| MZT2B | 3342 | 4476 | 4390 | 3117 | 10709 | 1704 | 4271 | 1827 |
| N4BP1 | 0 | 190 | 0 | 1303 | 1827 | 0 | 318 | 0 |
| N4BP2 | 0 | 0 | 0 | 0 | 0 | 0 | 350 | 0 |
| N4BP2L2 | 0 | 444 | 0 | 4906 | 976 | 2756 | 2248 | 1380 |
| N4BP3 | 0 | 368 | 790 | 0 | 0 | 0 | 0 | 0 |
| N6AMT1 | 0 | 294 | 0 | 1094 | 128 | 0 | 358 | 0 |
| NAA10 | 0 | 232 | 3826 | 7494 | 3348 | 0 | 0 | 3668 |
| NAA15 | 0 | 0 | 0 | 538 | 0 | 0 | 0 | 0 |
| NAA16 | 352 | 0 | 0 | 0 | 128 | 0 | 18 | 0 |
| NAA20 | 1400 | 2094 | 3535 | 86 | 3750 | 1292 | 446 | 2058 |
| NAA25 | 0 | 0 | 117 | 0 | 0 | 0 | 0 | 0 |
| NAA35 | 0 | 0 | 0 | 0 | 0 | 0 | 976 | 0 |
| NAA38 | 504 | 408 | 2076 | 1640 | 960 | 1632 | 416 | 1666 |
| NAA40 | 0 | 0 | 0 | 0 | 704 | 0 | 0 | 0 |
| NAA50 | 0 | 404 | 0 | 1097 | 326 | 0 | 78 | 367 |
| NAA60 | 1452 | 314 | 582 | 0 | 982 | 0 | 0 | 762 |
| NAAA | 0 | 278 | 3730 | 0 | 0 | 0 | 2126 | 0 |
| NAB1 | 422 | 0 | 0 | 0 | 0 | 0 | 924 | 0 |
| NAB2 | 0 | 0 | 0 | 0 | 0 | 1526 | 0 | 0 |
| NABP1 | 0 | 0 | 866 | 0 | 286 | 0 | 0 | 0 |
| NABP2 | 0 | 0 | 0 | 0 | 698 | 1051 | 1418 | 929 |
| NACA | 1695443 | 3448509 | 3137457 | 6056045 | 8822564 | 7816126 | 15816763 | 3695048 |
| NACC1 | 0 | 0 | 0 | 926 | 0 | 0 | 0 | 0 |
| NACC2 | 198 | 0 | 0 | 0 | 532 | 0 | 0 | 0 |
| NADK | 0 | 582 | 0 | 0 | 442 | 0 | 0 | 0 |
| NADK2 | 0 | 235 | 0 | 0 | 1210 | 0 | 0 | 0 |
| NADSYN1 | 0 | 458 | 2492 | 0 | 0 | 0 | 334 | 0 |
| NAE1 | 83694 | 196219 | 233136 | 336064 | 613011 | 309721 | 591629 | 192312 |
| NAF1 | 0 | 0 | 0 | 0 | 266 | 0 | 0 | 0 |
| NAGA | 0 | 0 | 0 | 0 | 930 | 0 | 676 | 0 |
| NAGK | 0 | 0 | 1248 | 0 | 0 | 0 | 0 | 1038 |
| NAGLU | 0 | 314 | 0 | 1218 | 0 | 0 | 0 | 0 |
| NAGPA | 164 | 0 | 0 | 0 | 0 | 0 | 762 | 0 |
| NAGS | 0 | 0 | 0 | 0 | 0 | 0 | 0 | 952 |
| NAMPT | 178 | 1441 | 3118 | 1565 | 2268 | 0 | 1436 | 936 |

| | | | | | | | | |
|---|---|---|---|---|---|---|---|---|
| NANS | 1890 | 8643 | 3036 | 6650 | 8548 | 1932 | 0 | 1215 |
| NAP1L1 | 0 | 549 | 0 | 2760 | 4372 | 0 | 3354 | 1276 |
| NAP1L4 | 63922 | 58207 | 58147 | 122678 | 158281 | 131523 | 225334 | 85128 |
| NAPA | 646 | 1350 | 0 | 1795 | 502 | 2632 | 1470 | 598 |
| NAPA-AS1 | 0 | 350 | 0 | 0 | 682 | 0 | 0 | 0 |
| NAPEPLD | 0 | 0 | 568 | 0 | 0 | 0 | 0 | 0 |
| NAPG | 0 | 0 | 0 | 0 | 574 | 0 | 278 | 0 |
| NAPRT | 390 | 1150 | 4128 | 636 | 1984 | 0 | 2826 | 1796 |
| NARF | 0 | 0 | 1922 | 0 | 1732 | 4249 | 734 | 0 |
| NARS | 1010 | 876 | 3061 | 1458 | 1308 | 0 | 278 | 0 |
| NARS2 | 2190 | 11739 | 6656 | 19048 | 46643 | 9404 | 36456 | 15462 |
| NASP | 66755 | 117343 | 106393 | 236871 | 493648 | 120323 | 269283 | 114065 |
| NAT1 | 0 | 0 | 0 | 614 | 0 | 0 | 272 | 0 |
| NAT10 | 0 | 230 | 0 | 1184 | 1062 | 0 | 0 | 0 |
| NAT2 | 0 | 860 | 0 | 0 | 0 | 0 | 0 | 0 |
| NAT9 | 0 | 748 | 0 | 0 | 0 | 0 | 0 | 0 |
| NAV1 | 0 | 0 | 0 | 0 | 382 | 0 | 0 | 0 |
| NAXD | 454 | 0 | 0 | 502 | 782 | 0 | 672 | 0 |
| NAXE | 0 | 1134 | 3318 | 0 | 5121 | 0 | 2292 | 1924 |
| NBAS | 1182 | 218 | 0 | 0 | 1664 | 0 | 1198 | 0 |
| NBDY | 970 | 0 | 1600 | 1244 | 4692 | 0 | 748 | 3130 |
| NBEAL1 | 108 | 0 | 657 | 0 | 0 | 0 | 0 | 1376 |
| NBEAL2 | 0 | 0 | 0 | 0 | 502 | 0 | 96 | 0 |
| NBL1 | 648 | 392 | 3210 | 990 | 4087 | 0 | 806 | 698 |
| NBN | 100090 | 185929 | 57058 | 121512 | 494520 | 160696 | 502604 | 147630 |
| NBPF10 | 0 | 0 | 0 | 0 | 378 | 0 | 0 | 0 |
| NBPF14 | 602 | 274 | 0 | 0 | 0 | 0 | 758 | 700 |
| NBPF15 | 0 | 0 | 0 | 0 | 0 | 0 | 146 | 0 |
| NBPF19 | 1330 | 0 | 0 | 0 | 0 | 0 | 0 | 0 |
| NBPF6 | 0 | 214 | 0 | 0 | 0 | 0 | 0 | 0 |
| NBPF8 | 0 | 0 | 0 | 0 | 1150 | 0 | 0 | 0 |
| NBR1 | 0 | 0 | 1140 | 1170 | 1881 | 0 | 1470 | 0 |
| NCALD | 5988 | 3948 | 19442 | 25870 | 8260 | 11562 | 43275 | 1564 |
| NCAM2 | 334 | 438 | 1192 | 786 | 1429 | 0 | 482 | 0 |
| NCAPD2 | 0 | 0 | 0 | 0 | 176 | 0 | 0 | 0 |
| NCAPD3 | 0 | 0 | 0 | 0 | 618 | 0 | 0 | 0 |
| NCAPG2 | 0 | 0 | 0 | 0 | 190 | 0 | 0 | 0 |
| NCAPH | 0 | 526 | 0 | 0 | 1118 | 0 | 0 | 410 |
| NCAPH2 | 0 | 0 | 0 | 0 | 224 | 0 | 0 | 0 |
| NCBP1 | 0 | 0 | 2030 | 0 | 874 | 0 | 0 | 0 |
| NCBP2 | 1286 | 1420 | 1690 | 740 | 3040 | 0 | 0 | 2102 |
| NCBP2-AS2 | 0 | 1412 | 1815 | 2714 | 2600 | 0 | 2530 | 2208 |
| NCBP3 | 0 | 232 | 0 | 0 | 382 | 0 | 0 | 0 |

| | | | | | | | | |
|---|---|---|---|---|---|---|---|---|
| NCEH1 | 3571 | 5404 | 15241 | 6224 | 18190 | 3133 | 6708 | 10828 |
| NCF1 | 0 | 0 | 0 | 0 | 0 | 0 | 0 | 252 |
| NCF1B | 0 | 0 | 0 | 592 | 0 | 0 | 0 | 0 |
| NCF1C | 0 | 0 | 0 | 0 | 290 | 0 | 0 | 0 |
| NCF4 | 0 | 280 | 1512 | 0 | 0 | 0 | 0 | 0 |
| NCK1 | 17200 | 45632 | 62480 | 38975 | 106052 | 50112 | 105743 | 88227 |
| NCK1-DT | 0 | 0 | 0 | 0 | 1310 | 0 | 0 | 0 |
| NCK2 | 0 | 0 | 0 | 144 | 652 | 0 | 230 | 0 |
| NCKAP1 | 560 | 0 | 642 | 0 | 1419 | 0 | 0 | 0 |
| NCKAP5 | 36196 | 7042 | 27644 | 680 | 632 | 4369 | 13745 | 1824 |
| NCKIPSD | 0 | 0 | 0 | 0 | 312 | 1534 | 621 | 0 |
| NCL | 46996 | 77799 | 67440 | 133545 | 383710 | 163974 | 368774 | 70872 |
| NCLN | 0 | 252 | 0 | 881 | 250 | 0 | 0 | 968 |
| NCOA1 | 268 | 72 | 1352 | 278 | 0 | 0 | 0 | 0 |
| NCOA2 | 3588 | 5340 | 19755 | 4239 | 29902 | 8576 | 42361 | 19420 |
| NCOA3 | 1312 | 2946 | 360 | 0 | 10726 | 0 | 12958 | 4078 |
| NCOA4 | 387100 | 779068 | 1868687 | 524420 | 878455 | 733464 | 1140729 | 386914 |
| NCOA5 | 0 | 338 | 0 | 0 | 0 | 0 | 0 | 0 |
| NCOA6 | 0 | 266 | 0 | 1244 | 262 | 0 | 50 | 0 |
| NCOA7 | 0 | 347 | 1376 | 490 | 1262 | 0 | 414 | 0 |
| NCOR1 | 577117 | 589889 | 1001444 | 442340 | 1390075 | 452096 | 980787 | 308451 |
| NCOR2 | 13272 | 6662 | 14332 | 11025 | 41015 | 12967 | 26627 | 6447 |
| NCSTN | 0 | 328 | 626 | 0 | 230 | 0 | 718 | 0 |
| NDC1 | 0 | 0 | 0 | 1934 | 164 | 0 | 0 | 0 |
| NDE1 | 0 | 0 | 0 | 0 | 646 | 0 | 0 | 0 |
| NDEL1 | 482 | 0 | 612 | 1510 | 306 | 0 | 204 | 0 |
| NDFIP1 | 0 | 0 | 1033 | 2513 | 2002 | 2126 | 402 | 0 |
| NDFIP2 | 18 | 978 | 990 | 7758 | 3788 | 3289 | 1004 | 1478 |
| NDN | 0 | 804 | 0 | 0 | 0 | 0 | 0 | 0 |
| NDOR1 | 0 | 426 | 0 | 0 | 502 | 0 | 1120 | 0 |
| NDRG1 | 116132 | 40489 | 148740 | 12824 | 546649 | 121385 | 136640 | 134193 |
| NDRG2 | 0 | 446 | 0 | 0 | 1176 | 0 | 1377 | 0 |
| NDRG3 | 0 | 0 | 0 | 0 | 2286 | 0 | 550 | 0 |
| NDST1 | 0 | 896 | 0 | 0 | 0 | 0 | 0 | 0 |
| NDST2 | 0 | 40 | 0 | 0 | 460 | 0 | 0 | 0 |
| NDUFA1 | 3732 | 6846 | 8404 | 5509 | 11975 | 4035 | 3572 | 974 |
| NDUFA10 | 170237 | 348931 | 431874 | 392399 | 914970 | 312007 | 625483 | 308644 |
| NDUFA11 | 4054 | 1676 | 7346 | 3699 | 11386 | 2469 | 440 | 1546 |
| NDUFA12 | 2194 | 2858 | 3737 | 3898 | 5782 | 3326 | 2392 | 1820 |
| NDUFA13 | 7174 | 16163 | 14257 | 11788 | 21440 | 8182 | 7963 | 5777 |
| NDUFA2 | 4233 | 4488 | 3118 | 4011 | 5284 | 4094 | 1172 | 1978 |
| NDUFA3 | 496 | 1572 | 1038 | 1241 | 4404 | 1380 | 578 | 154 |
| NDUFA4 | 7268 | 13329 | 14443 | 19042 | 26128 | 27474 | 17112 | 9256 |

| | | | | | | | | |
|---|---|---|---|---|---|---|---|---|
| NDUFA4L2 | 0 | 0 | 0 | 0 | 178 | 0 | 0 | 0 |
| NDUFA5 | 0 | 4660 | 4250 | 379 | 3868 | 1352 | 0 | 0 |
| NDUFA6 | 4097 | 2495 | 5830 | 998 | 7713 | 4051 | 2073 | 1619 |
| NDUFA6-DT | 0 | 0 | 0 | 1001 | 0 | 0 | 0 | 0 |
| NDUFA7 | 1254 | 814 | 912 | 3756 | 5768 | 1284 | 2870 | 1338 |
| NDUFA8 | 1116 | 2896 | 9202 | 2246 | 3004 | 4890 | 3080 | 1072 |
| NDUFA9 | 1704 | 3456 | 4114 | 10556 | 3111 | 0 | 1396 | 982 |
| NDUFAB1 | 3944 | 6594 | 14325 | 2618 | 7650 | 6200 | 4860 | 3969 |
| NDUFAF1 | 0 | 0 | 0 | 0 | 0 | 0 | 452 | 0 |
| NDUFAF2 | 0 | 274 | 0 | 3802 | 1460 | 3142 | 726 | 0 |
| NDUFAF3 | 0 | 948 | 2487 | 2476 | 5286 | 0 | 2692 | 648 |
| NDUFAF4 | 0 | 0 | 0 | 4320 | 2950 | 0 | 2194 | 0 |
| NDUFAF5 | 642 | 0 | 0 | 0 | 4582 | 0 | 0 | 0 |
| NDUFAF6 | 0 | 0 | 0 | 468 | 9046 | 2182 | 304 | 1271 |
| NDUFAF7 | 0 | 396 | 0 | 0 | 791 | 0 | 0 | 0 |
| NDUFAF8 | 132 | 1262 | 0 | 5043 | 2686 | 0 | 573 | 382 |
| NDUFB1 | 1404 | 4994 | 4210 | 1130 | 11498 | 560 | 1138 | 1712 |
| NDUFB10 | 2094 | 4132 | 2206 | 3412 | 4834 | 3111 | 1738 | 1932 |
| NDUFB11 | 888 | 2320 | 4140 | 4210 | 8974 | 3534 | 3957 | 1776 |
| NDUFB2 | 2924 | 8044 | 10748 | 4991 | 8966 | 5610 | 5178 | 3016 |
| NDUFB3 | 2120 | 3570 | 2957 | 7541 | 8032 | 4761 | 3631 | 2584 |
| NDUFB4 | 1812 | 3906 | 3967 | 2346 | 8474 | 402 | 6064 | 4758 |
| NDUFB5 | 1212 | 1162 | 3004 | 1600 | 1334 | 0 | 1028 | 748 |
| NDUFB6 | 1232 | 4182 | 3860 | 5857 | 8739 | 3324 | 1822 | 2635 |
| NDUFB7 | 41603 | 34634 | 212495 | 46149 | 95563 | 60305 | 29072 | 28416 |
| NDUFB8 | 4660 | 2954 | 9812 | 4719 | 10144 | 4008 | 3211 | 5627 |
| NDUFB9 | 4441031 | 5311629 | 11220456 | 4489010 | 18798449 | 10347016 | 23468049 | 4594743 |
| NDUFC1 | 1026 | 4586 | 3969 | 4067 | 8619 | 3099 | 9618 | 818 |
| NDUFC2 | 706 | 2380 | 6235 | 5886 | 12863 | 2422 | 4449 | 7585 |
| NDUFS1 | 782 | 550 | 982 | 2119 | 386 | 0 | 180 | 0 |
| NDUFS2 | 0 | 3388 | 3354 | 3270 | 4039 | 1799 | 492 | 3013 |
| NDUFS3 | 274 | 4472 | 9484 | 1289 | 1612 | 3394 | 0 | 1194 |
| NDUFS4 | 2402 | 2242 | 5300 | 1124 | 4900 | 5302 | 746 | 1256 |
| NDUFS5 | 667304 | 1012119 | 523285 | 792211 | 1500780 | 604955 | 1567176 | 276180 |
| NDUFS6 | 2442 | 2094 | 4322 | 2233 | 10278 | 0 | 2500 | 1384 |
| NDUFS7 | 1422 | 2238 | 6017 | 1270 | 2546 | 5530 | 0 | 3612 |
| NDUFS8 | 826 | 952 | 1462 | 8301 | 8615 | 1508 | 1280 | 964 |
| NDUFV1 | 2224 | 1006 | 7247 | 6272 | 3976 | 1204 | 660 | 1542 |
| NDUFV2 | 2764 | 1750 | 13879 | 8311 | 17406 | 3300 | 1960 | 4245 |
| NDUFV3 | 522 | 810 | 2697 | 0 | 442 | 2322 | 0 | 1264 |
| NEAT1 | 648 | 614 | 2052 | 422 | 1580 | 1534 | 1868 | 62 |
| NEBL | 2228 | 1950 | 2698 | 2912 | 9569 | 12059 | 31240 | 996 |
| NECAP1 | 506 | 0 | 1671 | 640 | 1526 | 0 | 410 | 946 |

| | | | | | | | | |
|---|---|---|---|---|---|---|---|---|
| NECAP2 | 0 | 0 | 1848 | 0 | 1714 | 0 | 348 | 0 |
| NECTIN1 | 0 | 0 | 0 | 0 | 330 | 0 | 0 | 0 |
| NECTIN2 | 274 | 0 | 0 | 0 | 2650 | 224 | 456 | 1664 |
| NECTIN3 | 0 | 0 | 0 | 0 | 436 | 0 | 574 | 0 |
| NECTIN4 | 0 | 0 | 0 | 0 | 0 | 0 | 528 | 0 |
| NEDD4 | 0 | 0 | 0 | 0 | 0 | 0 | 460 | 0 |
| NEDD4L | 80 | 296 | 2609 | 0 | 0 | 0 | 210 | 10 |
| NEDD8 | 2592 | 2808 | 2959 | 5404 | 18060 | 5058 | 2704 | 5297 |
| NEDD8-MDP1 | 0 | 0 | 0 | 0 | 0 | 0 | 116 | 0 |
| NEDD9 | 0 | 0 | 284 | 0 | 1886 | 0 | 0 | 0 |
| NEFL | 346 | 0 | 0 | 0 | 0 | 0 | 0 | 0 |
| NEGR1 | 0 | 464 | 0 | 0 | 1064 | 0 | 0 | 0 |
| NEIL1 | 0 | 452 | 0 | 0 | 0 | 0 | 0 | 0 |
| NEIL2 | 0 | 0 | 0 | 0 | 122 | 0 | 0 | 0 |
| NEK2 | 0 | 0 | 0 | 0 | 1366 | 0 | 0 | 0 |
| NEK3 | 0 | 0 | 423 | 0 | 0 | 3968 | 0 | 0 |
| NEK4 | 2703 | 2916 | 5975 | 9088 | 34349 | 13768 | 20156 | 3191 |
| NEK5 | 0 | 0 | 0 | 0 | 0 | 0 | 1126 | 0 |
| NEK6 | 0 | 524 | 0 | 1784 | 4277 | 0 | 1674 | 6639 |
| NEK7 | 0 | 342 | 0 | 0 | 0 | 0 | 0 | 738 |
| NEK9 | 0 | 0 | 0 | 0 | 670 | 0 | 0 | 0 |
| NELFA | 0 | 0 | 0 | 0 | 0 | 0 | 1750 | 0 |
| NELFB | 0 | 214 | 779 | 0 | 1646 | 1928 | 0 | 738 |
| NELFCD | 0 | 552 | 844 | 0 | 294 | 774 | 178 | 0 |
| NELFE | 574 | 564 | 2639 | 2425 | 6998 | 3066 | 1506 | 478 |
| NELL2 | 856 | 1700 | 1055 | 0 | 3004 | 0 | 2662 | 1054 |
| NEMF | 0 | 0 | 0 | 0 | 0 | 606 | 0 | 0 |
| NEMP1 | 0 | 0 | 0 | 0 | 772 | 0 | 0 | 0 |
| NEMP2 | 0 | 776 | 0 | 0 | 3158 | 1084 | 546 | 794 |
| NENF | 0 | 2058 | 1197 | 1640 | 4661 | 4146 | 3310 | 1555 |
| NEO1 | 0 | 278 | 0 | 0 | 0 | 0 | 0 | 0 |
| NEPRO | 398 | 0 | 0 | 0 | 0 | 0 | 0 | 0 |
| NET1 | 944 | 216 | 380 | 0 | 590 | 0 | 272 | 0 |
| NEU1 | 362 | 0 | 0 | 0 | 1430 | 0 | 2564 | 588 |
| NEURL2 | 0 | 0 | 0 | 1140 | 0 | 0 | 200 | 0 |
| NEURL4 | 0 | 0 | 1206 | 0 | 380 | 0 | 0 | 0 |
| NEXN | 0 | 0 | 0 | 0 | 0 | 410 | 398 | 0 |
| NF1 | 37655 | 24609 | 28814 | 25414 | 94593 | 16097 | 62734 | 24023 |
| NF2 | 3370 | 3854 | 5708 | 6379 | 12644 | 2089 | 5802 | 5757 |
| NFASC | 0 | 1453 | 1428 | 0 | 518 | 0 | 0 | 0 |
| NFAT5 | 0 | 0 | 840 | 0 | 0 | 0 | 398 | 0 |
| NFATC1 | 0 | 0 | 0 | 0 | 786 | 0 | 0 | 632 |
| NFATC2 | 0 | 696 | 0 | 0 | 662 | 584 | 1454 | 0 |

| | | | | | | | | |
|---|---|---|---|---|---|---|---|---|
| NFATC4 | 0 | 0 | 0 | 1114 | 0 | 0 | 564 | 0 |
| NFE2L1 | 510 | 0 | 0 | 1336 | 212 | 0 | 750 | 0 |
| NFE2L2 | 245516 | 365402 | 658423 | 364203 | 771941 | 510765 | 1605811 | 278558 |
| NFE2L3 | 0 | 0 | 0 | 1536 | 0 | 0 | 226 | 596 |
| NFIA | 0 | 0 | 0 | 0 | 0 | 0 | 1039 | 0 |
| NFIB | 8733 | 12720 | 13075 | 33092 | 36648 | 22337 | 30265 | 8410 |
| NFIC | 3931 | 1956 | 2594 | 563 | 10742 | 4301 | 2420 | 1573 |
| NFIL3 | 0 | 0 | 0 | 0 | 1240 | 2012 | 0 | 0 |
| NFIX | 0 | 0 | 0 | 3936 | 5041 | 954 | 5145 | 800 |
| NFKB1 | 0 | 510 | 0 | 0 | 0 | 0 | 80 | 0 |
| NFKB2 | 3594 | 6023 | 4032 | 2934 | 10101 | 8431 | 12447 | 5692 |
| NFKBIA | 998 | 1252 | 5478 | 0 | 1348 | 0 | 738 | 1762 |
| NFKBIB | 0 | 0 | 0 | 308 | 586 | 0 | 0 | 0 |
| NFKBIE | 7178 | 10274 | 18329 | 27351 | 43777 | 15357 | 33267 | 27395 |
| NFKBIZ | 0 | 122 | 0 | 0 | 0 | 0 | 0 | 460 |
| NFS1 | 0 | 250 | 0 | 964 | 432 | 0 | 0 | 420 |
| NFU1 | 798 | 1072 | 3212 | 34 | 2928 | 0 | 1820 | 0 |
| NFX1 | 0 | 412 | 0 | 0 | 0 | 0 | 142 | 0 |
| NFYA | 0 | 0 | 1978 | 0 | 0 | 0 | 1480 | 0 |
| NFYB | 0 | 0 | 0 | 0 | 1733 | 0 | 0 | 0 |
| NFYC | 20675 | 43530 | 84564 | 53442 | 73244 | 35292 | 145847 | 33816 |
| NGDN | 544 | 482 | 2114 | 0 | 3300 | 0 | 1366 | 2708 |
| NGEF | 478 | 0 | 1964 | 1284 | 990 | 0 | 294 | 0 |
| NGLY1 | 0 | 298 | 0 | 862 | 2271 | 0 | 522 | 0 |
| NGRN | 0 | 818 | 0 | 761 | 4019 | 0 | 3050 | 1149 |
| NHEJ1 | 0 | 1034 | 1578 | 1110 | 0 | 0 | 632 | 934 |
| NHLRC1 | 0 | 0 | 0 | 834 | 1584 | 0 | 0 | 0 |
| NHLRC2 | 0 | 194 | 0 | 0 | 0 | 0 | 0 | 0 |
| NHLRC3 | 0 | 0 | 0 | 0 | 0 | 0 | 1836 | 0 |
| NHP2 | 974 | 1142 | 2282 | 7764 | 11979 | 6740 | 4521 | 2018 |
| NHSL1 | 23171 | 17270 | 36658 | 14117 | 37982 | 15036 | 6593 | 16217 |
| NICN1 | 6062 | 4448 | 5097 | 7971 | 18089 | 5411 | 26111 | 0 |
| NIF3L1 | 0 | 0 | 0 | 0 | 820 | 0 | 1497 | 0 |
| NIFK | 0 | 18 | 0 | 1093 | 3404 | 0 | 64 | 0 |
| NIFK-AS1 | 0 | 0 | 2141 | 0 | 0 | 0 | 356 | 0 |
| NINJ1 | 0 | 348 | 0 | 0 | 956 | 606 | 0 | 1328 |
| NINJ2 | 17958 | 36043 | 111605 | 50845 | 107843 | 88079 | 131767 | 24272 |
| NIP7 | 604 | 20 | 0 | 1432 | 742 | 0 | 382 | 0 |
| NIPA1 | 484 | 0 | 0 | 0 | 300 | 0 | 0 | 0 |
| NIPA2 | 468 | 578 | 1626 | 1296 | 1409 | 0 | 1216 | 0 |
| NIPAL1 | 126 | 0 | 0 | 662 | 566 | 0 | 156 | 0 |
| NIPBL | 37998 | 17092 | 5887 | 2798 | 91073 | 9967 | 41879 | 2460 |
| NIPSNAP1 | 0 | 0 | 0 | 0 | 0 | 0 | 0 | 502 |

| | | | | | | | | |
|---|---|---|---|---|---|---|---|---|
| NIPSNAP2 | 0 | 536 | 1314 | 1008 | 2462 | 0 | 1316 | 402 |
| NIPSNAP3A | 974 | 1058 | 1289 | 2580 | 26 | 0 | 0 | 0 |
| NISCH | 2024 | 4143 | 11154 | 3011 | 9392 | 3714 | 10790 | 8469 |
| NIT1 | 662 | 0 | 2844 | 0 | 991 | 0 | 226 | 0 |
| NIT2 | 2282 | 1024 | 2090 | 910 | 11096 | 8894 | 140 | 416 |
| NKAP | 0 | 454 | 0 | 0 | 0 | 0 | 0 | 0 |
| NKAPD1 | 0 | 0 | 0 | 0 | 382 | 0 | 0 | 0 |
| NKD1 | 0 | 0 | 0 | 1564 | 128 | 0 | 3426 | 0 |
| NKD2 | 0 | 0 | 0 | 0 | 1102 | 0 | 998 | 0 |
| NKG7 | 0 | 0 | 1064 | 0 | 0 | 0 | 0 | 0 |
| NKIRAS1 | 0 | 0 | 0 | 0 | 396 | 0 | 0 | 0 |
| NKIRAS2 | 0 | 0 | 0 | 0 | 1690 | 0 | 310 | 0 |
| NKRF | 0 | 0 | 0 | 0 | 96 | 0 | 0 | 0 |
| NKTR | 0 | 0 | 0 | 0 | 0 | 364 | 0 | 0 |
| NKX2-3 | 0 | 0 | 708 | 0 | 0 | 0 | 0 | 0 |
| NLK | 1473 | 2092 | 6275 | 4222 | 6176 | 4529 | 5736 | 3746 |
| NLN | 970 | 0 | 1392 | 0 | 548 | 0 | 0 | 0 |
| NLRC4 | 0 | 0 | 0 | 1060 | 0 | 0 | 0 | 0 |
| NLRC5 | 0 | 0 | 1292 | 0 | 0 | 0 | 0 | 0 |
| NLRP1 | 0 | 916 | 881 | 0 | 0 | 0 | 0 | 82 |
| NLRP11 | 0 | 980 | 0 | 0 | 6727 | 0 | 0 | 0 |
| NLRP2 | 1022 | 0 | 0 | 0 | 0 | 0 | 0 | 0 |
| NLRX1 | 0 | 258 | 0 | 0 | 0 | 0 | 0 | 0 |
| NMB | 0 | 0 | 0 | 0 | 0 | 0 | 0 | 700 |
| NMD3 | 2372 | 5354 | 4873 | 9626 | 24510 | 18466 | 30450 | 16280 |
| NME1 | 0 | 3273 | 3664 | 3277 | 7342 | 23376 | 6420 | 3685 |
| NME1-NME2 | 1860 | 0 | 0 | 1582 | 2341 | 2113 | 1308 | 0 |
| NME2 | 3806 | 14800 | 9013 | 15198 | 24060 | 14927 | 14148 | 17231 |
| NME3 | 1605 | 828 | 0 | 3159 | 0 | 1082 | 1484 | 0 |
| NME4 | 0 | 0 | 0 | 988 | 1630 | 0 | 754 | 0 |
| NME6 | 0 | 0 | 0 | 865 | 3082 | 0 | 478 | 0 |
| NME7 | 0 | 0 | 0 | 0 | 0 | 0 | 450 | 0 |
| NME8 | 0 | 154 | 0 | 0 | 0 | 0 | 0 | 0 |
| NME9 | 1092 | 4268 | 0 | 0 | 2256 | 0 | 3432 | 0 |
| NMI | 0 | 0 | 468 | 0 | 1414 | 4256 | 0 | 0 |
| NMNAT3 | 0 | 0 | 0 | 0 | 0 | 0 | 196 | 0 |
| NMRAL1 | 0 | 82 | 0 | 0 | 4313 | 0 | 1354 | 0 |
| NMRK1 | 0 | 0 | 1648 | 0 | 1102 | 0 | 230 | 0 |
| NMT1 | 0 | 0 | 0 | 0 | 472 | 212 | 1624 | 0 |
| NMU | 0 | 0 | 0 | 0 | 792 | 0 | 0 | 2568 |
| NNMT | 0 | 494 | 3166 | 3582 | 1030 | 6382 | 180 | 5650 |
| NNT | 396 | 0 | 0 | 0 | 420 | 0 | 254 | 0 |
| NNT-AS1 | 0 | 0 | 0 | 0 | 560 | 0 | 156 | 0 |

| | | | | | | | | |
|---|---|---|---|---|---|---|---|---|
| NOA1 | 190 | 0 | 2996 | 0 | 470 | 0 | 58 | 0 |
| NOB1 | 484 | 758 | 0 | 0 | 208 | 2000 | 276 | 0 |
| NOC2L | 0 | 0 | 0 | 2626 | 0 | 0 | 898 | 0 |
| NOC4L | 2110 | 4242 | 21217 | 21428 | 9754 | 6267 | 23295 | 9263 |
| NOL10 | 410 | 344 | 0 | 1423 | 0 | 0 | 0 | 0 |
| NOL11 | 16539 | 49095 | 39570 | 38621 | 127722 | 65243 | 127769 | 35299 |
| NOL12 | 686 | 1016 | 882 | 0 | 1104 | 1632 | 0 | 0 |
| NOL3 | 0 | 1388 | 928 | 956 | 0 | 0 | 0 | 0 |
| NOL4 | 0 | 1132 | 0 | 0 | 1105 | 0 | 810 | 1242 |
| NOL4L | 348 | 0 | 0 | 788 | 0 | 0 | 1388 | 648 |
| NOL7 | 1222 | 92 | 7528 | 2552 | 9028 | 0 | 2966 | 16 |
| NOL8 | 0 | 536 | 0 | 0 | 1142 | 0 | 0 | 0 |
| NOL9 | 0 | 0 | 0 | 478 | 0 | 0 | 0 | 0 |
| NOLC1 | 12326 | 74292 | 30565 | 131535 | 268701 | 118448 | 324905 | 81188 |
| NOMO1 | 0 | 0 | 0 | 766 | 230 | 0 | 0 | 0 |
| NOMO2 | 0 | 0 | 0 | 1000 | 0 | 0 | 514 | 818 |
| NOMO3 | 998 | 0 | 0 | 0 | 0 | 0 | 0 | 0 |
| NONO | 748 | 2134 | 0 | 2264 | 4891 | 0 | 3468 | 1011 |
| NOP10 | 3718 | 2832 | 5544 | 3290 | 4588 | 1248 | 3658 | 2651 |
| NOP14-AS1 | 0 | 0 | 0 | 1213 | 0 | 0 | 0 | 1180 |
| NOP16 | 794 | 440 | 0 | 1708 | 2338 | 2446 | 418 | 796 |
| NOP2 | 0 | 142 | 0 | 0 | 0 | 0 | 0 | 0 |
| NOP53 | 856 | 2056 | 202 | 5309 | 6260 | 2284 | 796 | 812 |
| NOP56 | 0 | 0 | 2055 | 1161 | 8373 | 0 | 532 | 0 |
| NOP58 | 0 | 423 | 1634 | 2594 | 3896 | 0 | 376 | 0 |
| NOP9 | 0 | 0 | 0 | 0 | 998 | 0 | 428 | 0 |
| NORAD | 0 | 510 | 0 | 0 | 1538 | 2180 | 2532 | 298 |
| NOS1AP | 3460 | 574 | 1022 | 0 | 2118 | 2074 | 1822 | 964 |
| NOS2 | 5567 | 433 | 11305 | 791 | 4628 | 2042 | 0 | 0 |
| NOS3 | 0 | 0 | 1386 | 0 | 0 | 0 | 1626 | 0 |
| NOSIP | 1084 | 1576 | 4275 | 2902 | 4076 | 0 | 380 | 1074 |
| NOTCH1 | 0 | 100 | 66 | 0 | 0 | 924 | 508 | 0 |
| NOTCH2 | 1528 | 354 | 0 | 0 | 1498 | 0 | 528 | 82 |
| NOTCH2NLA | 2768 | 5136 | 8090 | 1867 | 7287 | 3446 | 2604 | 16925 |
| NOTCH3 | 514 | 0 | 0 | 0 | 194 | 0 | 0 | 0 |
| NOTCH4 | 0 | 0 | 0 | 0 | 0 | 0 | 0 | 1120 |
| NOTUM | 0 | 0 | 0 | 468 | 0 | 0 | 11503 | 0 |
| NOX1 | 40 | 864 | 1628 | 447 | 1776 | 0 | 636 | 0 |
| NOXA1 | 0 | 226 | 686 | 0 | 0 | 0 | 0 | 540 |
| NOXO1 | 0 | 0 | 448 | 0 | 524 | 0 | 0 | 830 |
| NOXRED1 | 0 | 0 | 0 | 0 | 722 | 0 | 0 | 0 |
| NPAS2 | 2172 | 2134 | 6382 | 4051 | 19716 | 6449 | 14539 | 2527 |
| NPAS3 | 0 | 0 | 706 | 0 | 766 | 0 | 0 | 0 |

| | | | | | | | | |
|---|---|---|---|---|---|---|---|---|
| NPC2 | 3423 | 5036 | 9407 | 6638 | 16908 | 9342 | 2646 | 5672 |
| NPDC1 | 1230 | 1180 | 0 | 0 | 2561 | 0 | 724 | 0 |
| NPEPL1 | 0 | 0 | 0 | 1078 | 0 | 0 | 0 | 0 |
| NPEPPS | 0 | 0 | 0 | 0 | 278 | 908 | 0 | 0 |
| NPFFR2 | 0 | 590 | 1135 | 0 | 544 | 0 | 0 | 0 |
| NPIPA1 | 0 | 0 | 0 | 0 | 430 | 0 | 0 | 0 |
| NPIPA2 | 0 | 0 | 2018 | 0 | 0 | 0 | 0 | 0 |
| NPIPA3 | 0 | 342 | 0 | 0 | 0 | 0 | 0 | 0 |
| NPIPA5 | 1138 | 162 | 0 | 0 | 0 | 0 | 0 | 0 |
| NPIPA7 | 0 | 0 | 0 | 0 | 400 | 0 | 0 | 0 |
| NPIPA8 | 0 | 0 | 0 | 0 | 484 | 0 | 0 | 0 |
| NPIPB12 | 0 | 0 | 0 | 0 | 390 | 0 | 0 | 0 |
| NPIPB3 | 0 | 0 | 0 | 0 | 628 | 0 | 0 | 0 |
| NPIPB4 | 0 | 0 | 438 | 0 | 2067 | 2280 | 604 | 0 |
| NPL | 0 | 0 | 1940 | 962 | 1898 | 0 | 0 | 0 |
| NPLOC4 | 286 | 72 | 0 | 200 | 412 | 2122 | 492 | 934 |
| NPM1 | 474095 | 1174476 | 1037756 | 3189343 | 5500597 | 4441100 | 6941800 | 2012334 |
| NPM2 | 0 | 0 | 0 | 0 | 1174 | 0 | 0 | 0 |
| NPM3 | 0 | 410 | 0 | 2260 | 1144 | 0 | 250 | 478 |
| NPNT | 0 | 330 | 0 | 0 | 0 | 0 | 461 | 0 |
| NPPA-AS1 | 0 | 264 | 0 | 0 | 0 | 0 | 0 | 0 |
| NPPC | 0 | 374 | 0 | 0 | 0 | 0 | 0 | 0 |
| NPR3 | 0 | 0 | 0 | 0 | 316 | 0 | 0 | 0 |
| NPRL2 | 0 | 0 | 1720 | 0 | 2270 | 604 | 0 | 0 |
| NPRL3 | 0 | 0 | 0 | 851 | 950 | 0 | 0 | 0 |
| NPTN | 592 | 348 | 1280 | 152 | 362 | 972 | 1380 | 0 |
| NPW | 0 | 0 | 0 | 0 | 1528 | 0 | 0 | 0 |
| NPY1R | 1758 | 0 | 0 | 0 | 0 | 0 | 0 | 586 |
| NQO1 | 484 | 0 | 5934 | 13743 | 8844 | 4668 | 11586 | 2054 |
| NQO2 | 0 | 456 | 0 | 0 | 764 | 2756 | 12 | 0 |
| NR1D2 | 0 | 0 | 275 | 0 | 0 | 0 | 440 | 1040 |
| NR1H2 | 0 | 240 | 1446 | 0 | 1052 | 0 | 0 | 0 |
| NR1H3 | 0 | 0 | 0 | 404 | 0 | 0 | 490 | 0 |
| NR1H4 | 298 | 0 | 0 | 0 | 0 | 0 | 0 | 0 |
| NR1I2 | 0 | 0 | 0 | 0 | 0 | 0 | 1282 | 0 |
| NR2C1 | 1722 | 10815 | 3403 | 5398 | 23071 | 8046 | 15300 | 7058 |
| NR2C2 | 194 | 0 | 0 | 0 | 1020 | 0 | 3080 | 986 |
| NR2C2AP | 2170 | 0 | 0 | 684 | 1672 | 0 | 152 | 0 |
| NR2F1 | 378 | 0 | 0 | 0 | 0 | 0 | 0 | 0 |
| NR2F2 | 0 | 0 | 642 | 0 | 867 | 0 | 346 | 0 |
| NR2F6 | 0 | 0 | 0 | 2324 | 3275 | 0 | 0 | 0 |
| NR3C1 | 0 | 0 | 0 | 0 | 108 | 0 | 0 | 0 |
| NR3C2 | 378 | 326 | 0 | 0 | 0 | 0 | 0 | 258 |

| | | | | | | | | |
|---|---|---|---|---|---|---|---|---|
| NR4A1 | 30 | 0 | 0 | 0 | 502 | 1428 | 360 | 580 |
| NR4A2 | 0 | 0 | 0 | 0 | 654 | 0 | 0 | 0 |
| NR5A2 | 0 | 0 | 0 | 0 | 0 | 1418 | 0 | 0 |
| NRARP | 0 | 666 | 0 | 0 | 558 | 0 | 0 | 0 |
| NRAS | 23104 | 41612 | 38700 | 31823 | 82749 | 23372 | 105767 | 9515 |
| NRAV | 788 | 0 | 0 | 0 | 0 | 0 | 0 | 0 |
| NRBF2 | 0 | 470 | 2136 | 0 | 1977 | 0 | 1416 | 0 |
| NRBP1 | 37994 | 44044 | 116666 | 43542 | 158474 | 90428 | 120514 | 64070 |
| NRBP2 | 0 | 118 | 0 | 0 | 621 | 0 | 1264 | 0 |
| NRDC | 0 | 0 | 0 | 0 | 1082 | 1276 | 0 | 0 |
| NREP | 0 | 8982 | 13693 | 17099 | 114540 | 11949 | 4847 | 9908 |
| NRF1 | 0 | 76 | 0 | 0 | 0 | 0 | 0 | 0 |
| NRG1 | 9600 | 866 | 6818 | 9585 | 2814 | 2162 | 163 | 2573 |
| NRG4 | 0 | 0 | 0 | 0 | 0 | 0 | 126 | 446 |
| NRM | 0 | 0 | 0 | 1005 | 0 | 0 | 0 | 0 |
| NRN1 | 0 | 0 | 0 | 0 | 1292 | 0 | 0 | 0 |
| NRP1 | 864 | 3930 | 8610 | 11736 | 18971 | 10278 | 3605 | 18458 |
| NRP2 | 6111 | 1160 | 0 | 4156 | 21968 | 4415 | 4397 | 6722 |
| NRSN2 | 0 | 280 | 0 | 0 | 0 | 0 | 508 | 0 |
| NSA2 | 614 | 154 | 2728 | 3350 | 2056 | 1912 | 452 | 406 |
| NSD1 | 2770 | 1586 | 1180 | 1968 | 2771 | 606 | 4713 | 3174 |
| NSD2 | 33661 | 10752 | 5678 | 14632 | 40483 | 10563 | 24246 | 12738 |
| NSD3 | 88394 | 79730 | 69680 | 38467 | 397174 | 29977 | 110423 | 50247 |
| NSDHL | 0 | 923 | 0 | 1338 | 812 | 0 | 0 | 1077 |
| NSF | 0 | 512 | 0 | 2050 | 326 | 0 | 0 | 0 |
| NSFL1C | 0 | 0 | 0 | 1390 | 2054 | 0 | 642 | 1576 |
| NSG1 | 0 | 30 | 0 | 668 | 0 | 0 | 0 | 0 |
| NSL1 | 0 | 532 | 0 | 0 | 1199 | 0 | 0 | 0 |
| NSMAF | 0 | 420 | 0 | 0 | 342 | 0 | 0 | 0 |
| NSMCE1 | 564 | 624 | 2144 | 2460 | 3960 | 0 | 412 | 2951 |
| NSMCE2 | 25254 | 25345 | 36269 | 50565 | 217438 | 109266 | 187187 | 85299 |
| NSMCE3 | 0 | 0 | 0 | 0 | 928 | 0 | 0 | 696 |
| NSMCE4A | 436 | 796 | 3786 | 3706 | 2158 | 0 | 466 | 0 |
| NSMF | 200 | 0 | 0 | 0 | 628 | 0 | 406 | 0 |
| NSUN2 | 3322 | 10405 | 5830 | 17953 | 54344 | 16488 | 34153 | 15343 |
| NSUN4 | 0 | 0 | 0 | 0 | 0 | 0 | 68 | 0 |
| NSUN5 | 0 | 338 | 0 | 940 | 72 | 0 | 224 | 856 |
| NSUN5P1 | 0 | 0 | 665 | 0 | 734 | 0 | 0 | 0 |
| NSUN5P2 | 438 | 0 | 0 | 0 | 0 | 0 | 0 | 0 |
| NSUN6 | 0 | 0 | 0 | 1466 | 0 | 0 | 0 | 0 |
| NSUN7 | 276 | 328 | 0 | 0 | 913 | 196 | 778 | 936 |
| NT5C | 1268 | 594 | 8423 | 180 | 3888 | 0 | 2495 | 0 |
| NT5C2 | 43931 | 60710 | 78393 | 54308 | 188480 | 61608 | 143320 | 36434 |

| | | | | | | | | |
|---|---|---|---|---|---|---|---|---|
| NT5C3A | 902 | 736 | 3236 | 2098 | 1024 | 0 | 424 | 804 |
| NT5C3B | 0 | 452 | 258 | 0 | 0 | 0 | 1148 | 0 |
| NT5DC1 | 0 | 402 | 0 | 2738 | 1038 | 0 | 58 | 0 |
| NT5DC2 | 0 | 0 | 0 | 0 | 1122 | 0 | 0 | 1366 |
| NT5DC3 | 0 | 0 | 0 | 0 | 704 | 0 | 0 | 0 |
| NT5E | 0 | 0 | 0 | 2110 | 1238 | 0 | 0 | 764 |
| NTAN1 | 0 | 0 | 64 | 1548 | 352 | 0 | 0 | 2050 |
| NTHL1 | 2166 | 6874 | 41002 | 33247 | 19747 | 18537 | 36347 | 14099 |
| NTM | 2608 | 2810 | 0 | 4202 | 38208 | 11985 | 0 | 0 |
| NTMT1 | 0 | 360 | 0 | 1170 | 3566 | 0 | 1908 | 1052 |
| NTPCR | 0 | 676 | 0 | 0 | 828 | 1970 | 496 | 1988 |
| NTRK1 | 0 | 976 | 0 | 0 | 624 | 0 | 0 | 0 |
| NTRK3 | 3894 | 2832 | 3446 | 0 | 1161 | 0 | 0 | 4468 |
| NTS | 0 | 0 | 0 | 0 | 0 | 0 | 0 | 924 |
| NUB1 | 600 | 596 | 0 | 0 | 0 | 1688 | 0 | 344 |
| NUBP1 | 14557 | 34656 | 28283 | 27454 | 50113 | 40390 | 45432 | 33478 |
| NUBP2 | 46 | 0 | 0 | 720 | 1816 | 0 | 264 | 0 |
| NUBPL | 132 | 0 | 0 | 124 | 2222 | 0 | 280 | 0 |
| NUCB1 | 0 | 860 | 0 | 720 | 810 | 2729 | 618 | 0 |
| NUCB2 | 0 | 1320 | 0 | 0 | 1427 | 184 | 2696 | 982 |
| NUCKS1 | 0 | 686 | 0 | 0 | 2262 | 0 | 0 | 0 |
| NUDC | 0 | 518 | 935 | 3648 | 4099 | 6121 | 2298 | 48 |
| NUDCD1 | 0 | 0 | 0 | 2330 | 0 | 0 | 666 | 744 |
| NUDCD2 | 518 | 556 | 0 | 816 | 3018 | 0 | 0 | 982 |
| NUDCD3 | 0 | 0 | 0 | 0 | 2719 | 0 | 2242 | 556 |
| NUDT1 | 0 | 662 | 0 | 530 | 990 | 0 | 0 | 0 |
| NUDT13 | 0 | 302 | 0 | 0 | 0 | 0 | 0 | 0 |
| NUDT14 | 1120 | 0 | 2272 | 736 | 2530 | 0 | 0 | 934 |
| NUDT15 | 0 | 0 | 0 | 932 | 92 | 0 | 374 | 0 |
| NUDT16 | 550 | 0 | 770 | 1346 | 2604 | 0 | 0 | 1128 |
| NUDT16L1 | 0 | 988 | 6115 | 1262 | 392 | 0 | 278 | 1516 |
| NUDT17 | 0 | 0 | 0 | 0 | 702 | 0 | 0 | 0 |
| NUDT19 | 0 | 0 | 0 | 602 | 0 | 0 | 0 | 0 |
| NUDT2 | 0 | 0 | 0 | 0 | 0 | 0 | 918 | 0 |
| NUDT21 | 0 | 0 | 1926 | 0 | 298 | 552 | 1046 | 292 |
| NUDT22 | 1252 | 454 | 1724 | 658 | 2754 | 0 | 1252 | 0 |
| NUDT3 | 0 | 0 | 0 | 870 | 1451 | 0 | 1016 | 654 |
| NUDT4 | 0 | 250 | 56 | 527 | 865 | 1712 | 226 | 0 |
| NUDT4B | 282 | 0 | 0 | 0 | 0 | 0 | 0 | 0 |
| NUDT5 | 0 | 466 | 0 | 0 | 2246 | 2231 | 915 | 1356 |
| NUDT6 | 620 | 840 | 0 | 0 | 1919 | 0 | 0 | 0 |
| NUDT7 | 0 | 344 | 3696 | 0 | 0 | 0 | 0 | 0 |
| NUDT8 | 0 | 0 | 4012 | 0 | 408 | 0 | 1211 | 0 |

| | | | | | | | | |
|---|---|---|---|---|---|---|---|---|
| NUDT9 | 1680 | 442 | 1340 | 1254 | 516 | 0 | 0 | 0 |
| NUF2 | 0 | 0 | 0 | 0 | 268 | 0 | 200 | 0 |
| NUFIP1 | 0 | 0 | 0 | 250 | 420 | 0 | 0 | 0 |
| NUFIP2 | 622 | 0 | 0 | 0 | 306 | 0 | 0 | 0 |
| NUMA1 | 1064 | 300 | 902 | 174 | 1719 | 0 | 868 | 0 |
| NUMB | 23693 | 41574 | 65331 | 23210 | 100671 | 34586 | 46885 | 23309 |
| NUMBL | 0 | 0 | 0 | 0 | 0 | 0 | 272 | 0 |
| NUP107 | 53323 | 118839 | 116734 | 189732 | 405952 | 164258 | 309695 | 139718 |
| NUP133 | 0 | 0 | 0 | 0 | 80 | 0 | 0 | 0 |
| NUP153 | 0 | 0 | 0 | 0 | 108 | 0 | 0 | 0 |
| NUP155 | 0 | 0 | 0 | 0 | 404 | 0 | 0 | 0 |
| NUP160 | 0 | 0 | 0 | 0 | 282 | 0 | 0 | 350 |
| NUP205 | 0 | 578 | 0 | 1188 | 0 | 2128 | 0 | 0 |
| NUP210 | 4042 | 0 | 3480 | 0 | 1822 | 0 | 140 | 782 |
| NUP214 | 0 | 342 | 0 | 2903 | 986 | 0 | 2673 | 0 |
| NUP35 | 0 | 414 | 0 | 1402 | 524 | 0 | 328 | 0 |
| NUP37 | 22904 | 69892 | 47807 | 180588 | 295206 | 98694 | 268008 | 132975 |
| NUP43 | 0 | 0 | 0 | 0 | 268 | 0 | 0 | 0 |
| NUP50 | 0 | 0 | 0 | 0 | 1280 | 0 | 0 | 562 |
| NUP50-DT | 116 | 0 | 0 | 756 | 0 | 0 | 0 | 0 |
| NUP54 | 0 | 318 | 0 | 534 | 1251 | 0 | 0 | 0 |
| NUP58 | 0 | 0 | 0 | 0 | 1024 | 0 | 0 | 0 |
| NUP62 | 0 | 0 | 0 | 0 | 1300 | 0 | 186 | 0 |
| NUP62CL | 0 | 16 | 0 | 0 | 0 | 0 | 186 | 526 |
| NUP85 | 12589 | 22646 | 82621 | 42914 | 72439 | 64589 | 61617 | 31402 |
| NUP88 | 0 | 352 | 0 | 0 | 592 | 0 | 358 | 0 |
| NUP93 | 912 | 196 | 0 | 0 | 1710 | 0 | 0 | 772 |
| NUP98 | 19002 | 67905 | 65692 | 47101 | 97263 | 56594 | 125116 | 50918 |
| NUPL2 | 58 | 0 | 1116 | 0 | 0 | 0 | 0 | 0 |
| NUPR1 | 3260 | 1092 | 1385 | 6790 | 12546 | 4478 | 6263 | 7035 |
| NUS1 | 0 | 0 | 690 | 0 | 1806 | 0 | 424 | 0 |
| NUSAP1 | 0 | 0 | 0 | 0 | 1626 | 0 | 298 | 0 |
| NUTF2 | 0 | 1822 | 1712 | 7161 | 6246 | 2274 | 4010 | 5504 |
| NUTM2A-AS1 | 0 | 0 | 710 | 0 | 1130 | 0 | 0 | 0 |
| NUTM2B-AS1 | 0 | 0 | 0 | 746 | 0 | 0 | 1084 | 526 |
| NVL | 0 | 0 | 1359 | 132 | 734 | 0 | 0 | 0 |
| NXF1 | 1192 | 0 | 0 | 1005 | 652 | 0 | 831 | 698 |
| NXN | 0 | 0 | 0 | 0 | 442 | 0 | 0 | 0 |
| NXPE1 | 948 | 802 | 0 | 0 | 0 | 0 | 0 | 0 |
| NXPE2 | 7665 | 5212 | 7319 | 1908 | 3549 | 1840 | 708 | 1272 |
| NXPE3 | 424 | 0 | 0 | 0 | 0 | 0 | 34 | 159 |
| NXPE4 | 666 | 1095 | 1450 | 0 | 822 | 0 | 0 | 0 |
| NXT1 | 0 | 1141 | 0 | 1380 | 2251 | 0 | 0 | 0 |

| | | | | | | | | |
|---|---|---|---|---|---|---|---|---|
| NXT2 | 0 | 0 | 664 | 0 | 0 | 0 | 0 | 0 |
| NYNRIN | 176 | 0 | 0 | 334 | 238 | 0 | 0 | 0 |
| OAF | 4614 | 6232 | 12807 | 3658 | 3740 | 2026 | 6732 | 1511 |
| OARD1 | 736 | 434 | 0 | 0 | 904 | 2526 | 634 | 1000 |
| OAS1 | 2160 | 436 | 1310 | 1373 | 3389 | 0 | 1720 | 0 |
| OAS2 | 0 | 400 | 0 | 0 | 240 | 0 | 0 | 0 |
| OAS3 | 0 | 0 | 0 | 0 | 156 | 0 | 0 | 1180 |
| OASL | 0 | 0 | 710 | 0 | 0 | 0 | 0 | 0 |
| OAT | 0 | 866 | 3982 | 3009 | 1338 | 0 | 2532 | 836 |
| OAZ1 | 1682 | 5996 | 18069 | 18787 | 28267 | 16101 | 8641 | 4101 |
| OAZ2 | 1912 | 438 | 0 | 0 | 204 | 0 | 312 | 0 |
| OAZ3 | 0 | 0 | 0 | 0 | 506 | 0 | 0 | 0 |
| OBP2B | 0 | 0 | 0 | 0 | 0 | 5287 | 0 | 0 |
| OCEL1 | 938 | 312 | 0 | 0 | 1288 | 0 | 0 | 0 |
| OCIAD1 | 303917 | 464171 | 498627 | 304732 | 721886 | 370355 | 1334769 | 216671 |
| OCIAD2 | 1782 | 1978 | 8156 | 10747 | 16330 | 4178 | 4006 | 3588 |
| OCLN | 0 | 0 | 882 | 0 | 2161 | 0 | 0 | 0 |
| OCRL | 0 | 360 | 0 | 0 | 496 | 0 | 424 | 0 |
| ODAM | 0 | 0 | 0 | 0 | 0 | 1527 | 0 | 0 |
| ODC1 | 0 | 424 | 812 | 2476 | 5003 | 1752 | 2016 | 2146 |
| ODF1 | 0 | 0 | 0 | 0 | 386 | 0 | 214 | 0 |
| ODF2 | 0 | 0 | 0 | 0 | 0 | 0 | 122 | 988 |
| ODF2L | 0 | 0 | 4650 | 0 | 936 | 2676 | 0 | 1688 |
| ODF3L2 | 0 | 0 | 0 | 0 | 0 | 0 | 0 | 298 |
| ODR4 | 0 | 0 | 0 | 0 | 816 | 0 | 0 | 0 |
| OFD1 | 23698 | 17405 | 29901 | 17265 | 35357 | 29876 | 42448 | 4470 |
| OGA | 0 | 0 | 0 | 0 | 0 | 1635 | 0 | 990 |
| OGDH | 0 | 172 | 482 | 404 | 760 | 324 | 0 | 0 |
| OGFOD1 | 0 | 326 | 0 | 0 | 448 | 0 | 0 | 0 |
| OGFOD2 | 0 | 0 | 0 | 0 | 0 | 0 | 824 | 0 |
| OGFOD3 | 1410 | 362 | 0 | 0 | 1226 | 0 | 0 | 0 |
| OGFR | 0 | 0 | 0 | 0 | 0 | 0 | 536 | 0 |
| OGN | 0 | 538 | 0 | 0 | 0 | 0 | 0 | 0 |
| OGT | 0 | 374 | 0 | 1855 | 2523 | 1227 | 2008 | 0 |
| OIP5 | 0 | 0 | 0 | 0 | 1120 | 0 | 384 | 1023 |
| OIP5-AS1 | 520 | 642 | 1568 | 1391 | 292 | 0 | 0 | 0 |
| OIT3 | 0 | 490 | 0 | 0 | 0 | 0 | 0 | 0 |
| OLA1 | 976 | 352 | 831 | 1841 | 1798 | 0 | 118 | 1294 |
| OLFM1 | 0 | 420 | 0 | 0 | 1223 | 0 | 0 | 0 |
| OLFM2 | 346 | 0 | 0 | 0 | 0 | 0 | 0 | 0 |
| OLFM4 | 0 | 7023 | 0 | 2224 | 2090 | 0 | 0 | 0 |
| OLFML1 | 0 | 0 | 0 | 0 | 366 | 0 | 0 | 0 |
| OLFML2B | 0 | 0 | 492 | 0 | 0 | 979 | 0 | 0 |

| | | | | | | | | |
|---|---|---|---|---|---|---|---|---|
| OLFML3 | 496 | 0 | 0 | 0 | 1088 | 0 | 0 | 0 |
| OLMALINC | 0 | 0 | 1244 | 0 | 0 | 0 | 0 | 0 |
| OLR1 | 0 | 384 | 0 | 905 | 10895 | 20659 | 5310 | 10631 |
| OMA1 | 0 | 0 | 3040 | 0 | 494 | 0 | 0 | 0 |
| OPA1 | 0 | 0 | 0 | 0 | 0 | 1341 | 0 | 0 |
| OPLAH | 0 | 158 | 0 | 0 | 270 | 0 | 625 | 0 |
| OPN3 | 972 | 0 | 0 | 0 | 0 | 0 | 0 | 0 |
| OPTN | 446 | 0 | 0 | 1112 | 1316 | 0 | 1998 | 0 |
| OR7E14P | 0 | 0 | 0 | 0 | 0 | 1456 | 0 | 0 |
| OR7E37P | 0 | 0 | 0 | 1037 | 0 | 0 | 0 | 0 |
| OR7E91P | 0 | 0 | 0 | 0 | 524 | 0 | 0 | 0 |
| ORAI1 | 0 | 0 | 0 | 0 | 1216 | 0 | 0 | 0 |
| ORAI3 | 0 | 0 | 0 | 0 | 0 | 1411 | 0 | 1185 |
| ORAOV1 | 6068 | 6658 | 4550 | 5914 | 15954 | 7878 | 4572 | 1716 |
| ORC1 | 0 | 0 | 0 | 514 | 164 | 0 | 0 | 0 |
| ORC2 | 0 | 0 | 0 | 0 | 2092 | 0 | 0 | 598 |
| ORC3 | 0 | 0 | 0 | 0 | 0 | 1721 | 0 | 948 |
| ORC4 | 812 | 426 | 0 | 2483 | 788 | 2124 | 276 | 0 |
| ORC6 | 0 | 0 | 0 | 1125 | 0 | 0 | 0 | 0 |
| ORM1 | 0 | 0 | 0 | 0 | 3797 | 0 | 0 | 0 |
| ORM2 | 0 | 0 | 0 | 0 | 0 | 0 | 500 | 0 |
| ORMDL1 | 754 | 1928 | 2028 | 2604 | 1254 | 0 | 770 | 0 |
| ORMDL2 | 0 | 2996 | 1623 | 10 | 3061 | 832 | 1632 | 0 |
| ORMDL3 | 0 | 0 | 0 | 0 | 0 | 0 | 830 | 0 |
| OS9 | 1784 | 894 | 1988 | 2151 | 266 | 1772 | 1246 | 686 |
| OSBP | 0 | 0 | 0 | 0 | 462 | 0 | 0 | 0 |
| OSBPL10 | 0 | 268 | 0 | 0 | 0 | 0 | 0 | 0 |
| OSBPL1A | 0 | 0 | 0 | 0 | 442 | 476 | 0 | 78 |
| OSBPL2 | 324 | 356 | 1696 | 1264 | 126 | 0 | 0 | 869 |
| OSBPL3 | 682 | 0 | 0 | 806 | 0 | 0 | 0 | 0 |
| OSBPL5 | 1270 | 2170 | 628 | 584 | 258 | 1332 | 2771 | 1222 |
| OSBPL7 | 446 | 0 | 0 | 746 | 578 | 0 | 293 | 0 |
| OSBPL8 | 0 | 0 | 0 | 0 | 0 | 0 | 0 | 1122 |
| OSBPL9 | 19115 | 44285 | 71274 | 68786 | 140899 | 77852 | 157476 | 57239 |
| OSCAR | 0 | 0 | 0 | 168 | 0 | 0 | 0 | 0 |
| OSCP1 | 0 | 0 | 0 | 0 | 618 | 0 | 0 | 0 |
| OSER1 | 0 | 1054 | 2205 | 0 | 1572 | 0 | 3240 | 2170 |
| OSER1-DT | 0 | 0 | 0 | 0 | 0 | 0 | 262 | 0 |
| OSGEP | 0 | 184 | 0 | 1272 | 3060 | 0 | 0 | 316 |
| OSGIN1 | 0 | 0 | 1769 | 0 | 0 | 0 | 0 | 0 |
| OSGIN2 | 0 | 772 | 0 | 0 | 3818 | 0 | 778 | 198 |
| OSMR | 0 | 0 | 0 | 0 | 210 | 0 | 0 | 0 |
| OST4 | 3073 | 4262 | 5336 | 4985 | 15049 | 4158 | 9253 | 5666 |

| | | | | | | | | |
|---|---|---|---|---|---|---|---|---|
| OSTC | 2640 | 608 | 6687 | 8428 | 5898 | 1454 | 3301 | 2162 |
| OSTF1 | 1723 | 904 | 6788 | 2404 | 3085 | 2133 | 0 | 0 |
| OSTM1 | 0 | 0 | 0 | 0 | 418 | 0 | 0 | 0 |
| OTOP2 | 402 | 0 | 0 | 0 | 0 | 0 | 0 | 0 |
| OTUB1 | 486 | 722 | 2094 | 1822 | 3568 | 884 | 266 | 4014 |
| OTUD5 | 0 | 0 | 1166 | 0 | 604 | 0 | 612 | 0 |
| OTUD6B-AS1 | 0 | 558 | 438 | 0 | 0 | 4852 | 2298 | 0 |
| OTUD7B | 160 | 0 | 0 | 0 | 0 | 0 | 0 | 0 |
| OTULIN | 0 | 0 | 0 | 0 | 280 | 0 | 0 | 0 |
| OTULINL | 928 | 496 | 1404 | 3098 | 960 | 4550 | 3989 | 0 |
| OVCA2 | 0 | 1426 | 1624 | 787 | 1592 | 0 | 0 | 872 |
| OVGP1 | 0 | 0 | 0 | 1828 | 0 | 2316 | 0 | 0 |
| OVOL1 | 0 | 0 | 0 | 0 | 348 | 0 | 0 | 0 |
| OVOL2 | 0 | 0 | 0 | 1044 | 70 | 0 | 0 | 956 |
| OXA1L | 0 | 2455 | 0 | 0 | 5279 | 0 | 0 | 1784 |
| OXLD1 | 0 | 282 | 511 | 2155 | 814 | 1934 | 3578 | 298 |
| OXNAD1 | 0 | 468 | 1252 | 0 | 0 | 0 | 0 | 0 |
| OXR1 | 22167 | 29948 | 34184 | 24956 | 115954 | 104347 | 189187 | 41093 |
| OXSM | 644 | 384 | 471 | 260 | 306 | 2228 | 0 | 0 |
| P2RX1 | 548 | 0 | 0 | 0 | 0 | 0 | 0 | 0 |
| P2RX4 | 476 | 0 | 5116 | 42 | 1345 | 0 | 210 | 0 |
| P2RX5 | 638 | 2860 | 0 | 0 | 3156 | 0 | 0 | 622 |
| P2RX5-TAX1BP3 | 2358 | 470 | 0 | 104 | 3144 | 1997 | 830 | 0 |
| P2RX7 | 62 | 0 | 0 | 0 | 0 | 0 | 0 | 0 |
| P2RY1 | 570 | 0 | 0 | 0 | 0 | 0 | 0 | 0 |
| P2RY13 | 4684 | 7138 | 5232 | 0 | 420 | 8368 | 1772 | 4356 |
| P2RY14 | 0 | 0 | 0 | 0 | 0 | 0 | 0 | 554 |
| P3H1 | 426 | 356 | 0 | 0 | 112 | 0 | 306 | 0 |
| P3H2 | 0 | 18658 | 440 | 0 | 17022 | 1769 | 0 | 1036 |
| P3H3 | 0 | 0 | 0 | 0 | 2030 | 0 | 0 | 0 |
| P3H4 | 0 | 0 | 0 | 506 | 0 | 0 | 1156 | 461 |
| P4HA1 | 0 | 106 | 0 | 0 | 923 | 0 | 0 | 0 |
| P4HA2 | 50356 | 44753 | 195404 | 60759 | 235832 | 229136 | 108655 | 68957 |
| P4HA3 | 0 | 0 | 0 | 0 | 472 | 0 | 0 | 0 |
| P4HB | 441708 | 404995 | 516031 | 572043 | 1138012 | 942566 | 1752172 | 456327 |
| P4HTM | 0 | 192 | 0 | 0 | 1140 | 1002 | 0 | 1046 |
| PA2G4 | 644 | 450 | 1744 | 2648 | 2532 | 0 | 2691 | 0 |
| PAAF1 | 0 | 664 | 0 | 0 | 54 | 2588 | 424 | 0 |
| PABPC1 | 711671 | 1240345 | 1041522 | 1664173 | 9133627 | 5858904 | 9398304 | 1423296 |
| PABPC1L | 946 | 0 | 0 | 736 | 346 | 0 | 138 | 0 |
| PABPC3 | 146 | 186 | 0 | 534 | 5230 | 1858 | 0 | 1466 |
| PABPC4 | 0 | 0 | 0 | 0 | 728 | 0 | 1036 | 0 |
| PABPN1 | 0 | 212 | 0 | 1209 | 504 | 0 | 378 | 0 |

| | | | | | | | | |
|---|---|---|---|---|---|---|---|---|
| PACRG | 1934 | 3300 | 5440 | 814 | 5408 | 2478 | 1168 | 0 |
| PACRGL | 0 | 0 | 944 | 0 | 138 | 0 | 0 | 0 |
| PACS1 | 0 | 0 | 590 | 130 | 0 | 0 | 0 | 0 |
| PACS2 | 470 | 0 | 586 | 0 | 524 | 0 | 0 | 0 |
| PACSIN2 | 39952 | 88328 | 298668 | 17192 | 78113 | 36715 | 66273 | 25975 |
| PACSIN3 | 0 | 0 | 0 | 0 | 318 | 0 | 0 | 0 |
| PADI2 | 1138 | 762 | 4929 | 0 | 0 | 0 | 0 | 0 |
| PADI4 | 0 | 854 | 2651 | 0 | 1208 | 2038 | 0 | 0 |
| PAF1 | 0 | 0 | 0 | 0 | 1760 | 0 | 0 | 0 |
| PAFAH1B1 | 440 | 0 | 1644 | 0 | 1584 | 0 | 0 | 0 |
| PAFAH1B2 | 0 | 1658 | 0 | 0 | 1468 | 0 | 344 | 0 |
| PAFAH1B3 | 0 | 0 | 0 | 2026 | 3378 | 210 | 434 | 858 |
| PAFAH2 | 0 | 0 | 2716 | 0 | 0 | 0 | 0 | 0 |
| PAG1 | 376 | 166 | 0 | 0 | 456 | 0 | 0 | 0 |
| PAGR1 | 0 | 0 | 0 | 1090 | 2940 | 0 | 675 | 694 |
| PAICS | 24206 | 126349 | 93664 | 299604 | 709747 | 270683 | 711623 | 216559 |
| PAIP1 | 0 | 840 | 1929 | 3930 | 5245 | 1240 | 2471 | 3872 |
| PAIP2 | 0 | 701 | 3886 | 1924 | 2078 | 1890 | 786 | 2299 |
| PAIP2B | 0 | 0 | 1190 | 0 | 352 | 0 | 0 | 0 |
| PAK1 | 0 | 0 | 2545 | 798 | 1684 | 2326 | 262 | 900 |
| PAK1IP1 | 0 | 0 | 0 | 0 | 494 | 0 | 1004 | 0 |
| PAK2 | 0 | 426 | 0 | 0 | 302 | 0 | 0 | 0 |
| PAK4 | 1286 | 2024 | 2932 | 770 | 4624 | 448 | 4060 | 0 |
| PALD1 | 0 | 240 | 0 | 0 | 0 | 0 | 0 | 0 |
| PALLD | 0 | 0 | 0 | 0 | 138 | 0 | 240 | 0 |
| PALM | 48 | 0 | 0 | 0 | 0 | 0 | 0 | 0 |
| PALM3 | 0 | 0 | 0 | 0 | 0 | 0 | 0 | 150 |
| PALMD | 790 | 1166 | 3882 | 368 | 884 | 22399 | 2192 | 3099 |
| PAM | 0 | 0 | 0 | 0 | 1106 | 0 | 0 | 0 |
| PAM16 | 0 | 404 | 1726 | 1939 | 1876 | 0 | 1784 | 0 |
| PAMR1 | 1722 | 388 | 4818 | 850 | 8186 | 0 | 0 | 782 |
| PAN2 | 0 | 0 | 0 | 0 | 1156 | 0 | 0 | 0 |
| PAN3 | 0 | 338 | 562 | 416 | 1036 | 1406 | 0 | 0 |
| PANK1 | 0 | 0 | 0 | 0 | 651 | 0 | 0 | 0 |
| PANK2 | 0 | 0 | 0 | 676 | 526 | 0 | 0 | 554 |
| PANK3 | 0 | 0 | 973 | 1202 | 0 | 0 | 0 | 0 |
| PAOX | 0 | 360 | 0 | 0 | 0 | 0 | 0 | 0 |
| PAPD4 | 128 | 0 | 0 | 0 | 230 | 0 | 0 | 144 |
| PAPD7 | 0 | 200 | 0 | 0 | 3749 | 86 | 1330 | 1008 |
| PAPLN | 0 | 0 | 0 | 0 | 112 | 0 | 0 | 0 |
| PAPOLA | 25093 | 34771 | 38402 | 24785 | 139273 | 28009 | 70784 | 26132 |
| PAPSS1 | 0 | 0 | 2168 | 0 | 0 | 324 | 316 | 0 |
| PAPSS2 | 0 | 936 | 0 | 2492 | 610 | 0 | 52 | 0 |

| | | | | | | | | |
|---|---|---|---|---|---|---|---|---|
| PAQR3 | 0 | 0 | 0 | 0 | 746 | 0 | 0 | 0 |
| PAQR5 | 610 | 0 | 0 | 0 | 0 | 0 | 0 | 0 |
| PAQR8 | 0 | 526 | 1140 | 0 | 460 | 0 | 534 | 0 |
| PARD3 | 9311 | 7012 | 17113 | 9512 | 31102 | 14929 | 18832 | 13219 |
| PARD3-AS1 | 0 | 0 | 0 | 0 | 128 | 198 | 0 | 0 |
| PARD3B | 0 | 0 | 512 | 0 | 1600 | 0 | 0 | 0 |
| PARD6A | 0 | 0 | 0 | 2044 | 924 | 0 | 0 | 0 |
| PARD6B | 0 | 0 | 0 | 0 | 40 | 0 | 96 | 326 |
| PARD6G | 0 | 0 | 0 | 1186 | 0 | 0 | 0 | 0 |
| PARK7 | 3154 | 7543 | 7498 | 16830 | 26787 | 8470 | 1848 | 2816 |
| PARL | 928 | 1164 | 0 | 0 | 748 | 0 | 1134 | 2257 |
| PARM1 | 1228 | 1210 | 2531 | 0 | 654 | 1416 | 363 | 0 |
| PARN | 4902 | 15034 | 9578 | 9149 | 37474 | 23553 | 31179 | 12034 |
| PARP1 | 0 | 0 | 1978 | 716 | 592 | 0 | 0 | 0 |
| PARP10 | 0 | 0 | 1440 | 375 | 688 | 0 | 164 | 0 |
| PARP11 | 0 | 0 | 0 | 0 | 664 | 0 | 0 | 0 |
| PARP12 | 0 | 0 | 1417 | 0 | 0 | 1386 | 0 | 0 |
| PARP14 | 0 | 0 | 0 | 0 | 0 | 0 | 308 | 0 |
| PARP15 | 0 | 286 | 42 | 0 | 0 | 0 | 0 | 0 |
| PARP2 | 0 | 0 | 0 | 950 | 1130 | 2184 | 0 | 0 |
| PARP3 | 0 | 152 | 0 | 0 | 474 | 0 | 296 | 0 |
| PARP4 | 0 | 382 | 0 | 0 | 866 | 0 | 974 | 0 |
| PARP6 | 0 | 412 | 0 | 0 | 516 | 0 | 182 | 0 |
| PARP8 | 0 | 0 | 0 | 0 | 198 | 0 | 414 | 0 |
| PARPBP | 0 | 0 | 0 | 0 | 1328 | 0 | 0 | 0 |
| PART1 | 0 | 0 | 0 | 0 | 0 | 0 | 1308 | 0 |
| PARTICL | 150 | 0 | 0 | 0 | 172 | 0 | 0 | 0 |
| PARVA | 47017 | 84098 | 265911 | 91440 | 137707 | 122890 | 185197 | 103119 |
| PARVB | 0 | 0 | 1455 | 3724 | 0 | 0 | 1086 | 914 |
| PARVG | 0 | 0 | 0 | 0 | 744 | 1502 | 0 | 0 |
| PASK | 0 | 0 | 0 | 0 | 296 | 0 | 0 | 0 |
| PATJ | 774 | 1292 | 5947 | 1873 | 6168 | 6151 | 12320 | 2447 |
| PATL1 | 0 | 0 | 0 | 0 | 0 | 0 | 268 | 0 |
| PATL2 | 0 | 268 | 0 | 0 | 0 | 0 | 0 | 0 |
| PATZ1 | 0 | 0 | 0 | 0 | 0 | 0 | 0 | 617 |
| PAWR | 314 | 128 | 0 | 1034 | 1150 | 882 | 342 | 0 |
| PAX5 | 0 | 982 | 0 | 0 | 0 | 0 | 0 | 572 |
| PAX8 | 5824 | 4805 | 31171 | 1402 | 2443 | 13151 | 186 | 1178 |
| PAX8-AS1 | 0 | 186 | 0 | 0 | 0 | 0 | 0 | 0 |
| PAXIP1 | 0 | 0 | 0 | 0 | 428 | 0 | 142 | 0 |
| PAXIP1-AS1 | 0 | 432 | 0 | 1260 | 34 | 0 | 232 | 130 |
| PAXX | 0 | 234 | 780 | 2661 | 4304 | 0 | 1052 | 2940 |
| PBDC1 | 0 | 494 | 1928 | 1526 | 1736 | 0 | 544 | 1295 |

| | | | | | | | | |
|---|---|---|---|---|---|---|---|---|
| PBK | 0 | 728 | 0 | 0 | 0 | 0 | 0 | 0 |
| PBLD | 114 | 0 | 0 | 0 | 0 | 0 | 1126 | 0 |
| PBRM1 | 27797 | 30886 | 43983 | 63512 | 175721 | 57769 | 89941 | 27099 |
| PBX1 | 12612 | 3180 | 23838 | 3206 | 15248 | 4669 | 12654 | 4408 |
| PBX3 | 0 | 0 | 0 | 0 | 0 | 467 | 0 | 0 |
| PBX4 | 0 | 0 | 0 | 0 | 634 | 0 | 0 | 0 |
| PBXIP1 | 158 | 272 | 0 | 0 | 204 | 0 | 0 | 722 |
| PC | 410 | 0 | 0 | 0 | 0 | 0 | 0 | 0 |
| PCAT6 | 0 | 0 | 0 | 0 | 672 | 0 | 0 | 0 |
| PCAT7 | 0 | 0 | 0 | 0 | 0 | 215 | 0 | 0 |
| PCBD1 | 844 | 1724 | 678 | 3097 | 2755 | 8797 | 3656 | 1202 |
| PCBD2 | 588 | 0 | 0 | 1052 | 0 | 0 | 0 | 0 |
| PCBP1 | 966 | 1000 | 4092 | 2512 | 4715 | 1938 | 2502 | 1972 |
| PCBP2 | 220283 | 300199 | 2128329 | 838374 | 910833 | 769964 | 1874128 | 419607 |
| PCBP4 | 0 | 0 | 0 | 0 | 642 | 0 | 0 | 0 |
| PCCA | 0 | 400 | 1076 | 0 | 712 | 0 | 1696 | 863 |
| PCCB | 23713 | 42800 | 97954 | 77970 | 255110 | 47182 | 71042 | 92931 |
| PCDH1 | 738 | 0 | 0 | 926 | 0 | 0 | 0 | 0 |
| PCDHB13 | 0 | 228 | 0 | 0 | 0 | 0 | 0 | 0 |
| PCDHB14 | 0 | 0 | 0 | 0 | 0 | 0 | 168 | 0 |
| PCDHGC3 | 0 | 1200 | 0 | 0 | 374 | 0 | 0 | 0 |
| PCED1A | 300 | 0 | 0 | 0 | 226 | 0 | 0 | 0 |
| PCED1B | 0 | 0 | 0 | 0 | 374 | 0 | 0 | 0 |
| PCGF1 | 0 | 0 | 1452 | 1317 | 2498 | 1454 | 812 | 0 |
| PCGF2 | 318 | 566 | 2593 | 0 | 544 | 0 | 1914 | 0 |
| PCGF3 | 0 | 0 | 0 | 0 | 498 | 2138 | 0 | 0 |
| PCGF5 | 810 | 206 | 2656 | 898 | 0 | 1670 | 0 | 2345 |
| PCGF6 | 0 | 0 | 0 | 0 | 0 | 0 | 316 | 0 |
| PCID2 | 38 | 0 | 0 | 474 | 2414 | 0 | 0 | 1896 |
| PCIF1 | 0 | 776 | 0 | 0 | 856 | 1796 | 0 | 0 |
| PCK1 | 452 | 2276 | 2228 | 0 | 0 | 0 | 0 | 0 |
| PCK2 | 566 | 384 | 0 | 0 | 2176 | 0 | 346 | 816 |
| PCLAF | 0 | 0 | 0 | 3626 | 2848 | 0 | 1778 | 1142 |
| PCM1 | 3122 | 5750 | 8938 | 3250 | 8792 | 8260 | 7036 | 3364 |
| PCMT1 | 0 | 1194 | 668 | 2684 | 2444 | 2556 | 2451 | 1318 |
| PCMTD1 | 50924 | 79155 | 236184 | 81555 | 116264 | 277561 | 667047 | 45324 |
| PCMTD2 | 0 | 0 | 0 | 0 | 732 | 0 | 38 | 614 |
| PCNA | 626 | 750 | 1062 | 4603 | 3643 | 2130 | 0 | 86 |
| PCNP | 354 | 0 | 5537 | 2137 | 1010 | 1170 | 674 | 0 |
| PCNT | 748 | 0 | 190 | 0 | 700 | 0 | 80 | 0 |
| PCNX1 | 0 | 260 | 0 | 0 | 0 | 0 | 0 | 0 |
| PCNX3 | 418 | 0 | 0 | 1136 | 1629 | 0 | 296 | 620 |
| PCNX4 | 0 | 30 | 0 | 0 | 184 | 0 | 0 | 0 |

| | | | | | | | | |
|---|---|---|---|---|---|---|---|---|
| PCOLCE | 0 | 0 | 0 | 0 | 1684 | 0 | 0 | 0 |
| PCOTH | 0 | 0 | 0 | 1146 | 0 | 1766 | 0 | 0 |
| PCP2 | 0 | 0 | 0 | 0 | 929 | 0 | 0 | 0 |
| PCSK1N | 168 | 1610 | 4376 | 0 | 0 | 0 | 0 | 0 |
| PCSK4 | 0 | 0 | 0 | 0 | 164 | 0 | 0 | 0 |
| PCSK5 | 0 | 0 | 0 | 0 | 392 | 0 | 0 | 0 |
| PCSK7 | 460 | 26 | 2390 | 42 | 243 | 0 | 0 | 0 |
| PCSK9 | 0 | 0 | 0 | 0 | 170 | 0 | 0 | 0 |
| PCTP | 9580 | 18242 | 30740 | 24448 | 40706 | 43478 | 36827 | 8171 |
| PCYOX1 | 0 | 0 | 0 | 0 | 0 | 0 | 0 | 1034 |
| PCYOX1L | 0 | 666 | 0 | 0 | 0 | 0 | 0 | 0 |
| PCYT1A | 27391 | 32333 | 196599 | 43636 | 113600 | 86135 | 38429 | 64551 |
| PCYT2 | 0 | 0 | 1102 | 0 | 721 | 0 | 1308 | 0 |
| PDAP1 | 558 | 0 | 765 | 1820 | 1805 | 0 | 596 | 74 |
| PDCD10 | 712 | 982 | 1458 | 1388 | 6290 | 0 | 4100 | 3569 |
| PDCD2 | 0 | 318 | 2334 | 954 | 4212 | 0 | 702 | 1406 |
| PDCD2L | 0 | 0 | 570 | 0 | 1088 | 0 | 0 | 950 |
| PDCD4 | 3090 | 3856 | 16528 | 914 | 5941 | 0 | 3092 | 0 |
| PDCD4-AS1 | 0 | 216 | 0 | 0 | 0 | 0 | 0 | 0 |
| PDCD5 | 0 | 1143 | 6440 | 4989 | 9402 | 1794 | 866 | 5966 |
| PDCD6 | 71763 | 95331 | 310432 | 122733 | 476510 | 126406 | 219572 | 91096 |
| PDCD6IP | 532 | 732 | 0 | 0 | 1234 | 0 | 352 | 1864 |
| PDCD7 | 28181 | 19964 | 7000 | 33116 | 51714 | 16340 | 75416 | 9405 |
| PDCL | 0 | 248 | 12 | 0 | 481 | 2800 | 204 | 0 |
| PDCL3 | 0 | 720 | 1761 | 934 | 4444 | 2005 | 1040 | 0 |
| PDE12 | 1496 | 1355 | 0 | 682 | 6938 | 0 | 2954 | 114 |
| PDE4A | 0 | 0 | 0 | 0 | 412 | 0 | 202 | 0 |
| PDE4D | 956 | 946 | 0 | 0 | 726 | 0 | 1984 | 1032 |
| PDE4DIP | 0 | 0 | 0 | 0 | 0 | 0 | 381 | 0 |
| PDE6A | 802 | 0 | 0 | 0 | 0 | 0 | 0 | 0 |
| PDE6B | 628 | 0 | 0 | 0 | 0 | 0 | 0 | 0 |
| PDE6D | 0 | 0 | 1808 | 0 | 2390 | 0 | 0 | 930 |
| PDE7B | 1894 | 2648 | 4593 | 0 | 572 | 0 | 0 | 892 |
| PDE8A | 0 | 0 | 658 | 0 | 180 | 0 | 0 | 0 |
| PDE9A | 572 | 763 | 378 | 1204 | 0 | 0 | 166 | 0 |
| PDF | 0 | 1150 | 0 | 2076 | 494 | 0 | 1148 | 0 |
| PDGFB | 0 | 0 | 816 | 0 | 714 | 1498 | 0 | 448 |
| PDGFRA | 3610 | 1382 | 2388 | 1410 | 4249 | 1554 | 134 | 0 |
| PDGFRB | 638 | 236 | 0 | 0 | 2934 | 5210 | 1386 | 0 |
| PDHA1 | 1638 | 1111 | 5884 | 2367 | 3970 | 2743 | 0 | 0 |
| PDHB | 2465 | 1144 | 5961 | 2242 | 3321 | 2118 | 2186 | 2345 |
| PDHX | 29021 | 77230 | 205113 | 105933 | 363386 | 100472 | 258298 | 118240 |
| PDIA3 | 822 | 1216 | 856 | 4240 | 5557 | 3162 | 3067 | 1900 |

| | | | | | | | | |
|---|---|---|---|---|---|---|---|---|
| PDIA4 | 0 | 422 | 0 | 254 | 3662 | 0 | 1514 | 4265 |
| PDIA5 | 0 | 40 | 0 | 0 | 336 | 0 | 962 | 0 |
| PDIA6 | 688 | 2870 | 0 | 7908 | 9563 | 0 | 2946 | 3052 |
| PDK1 | 0 | 0 | 0 | 862 | 498 | 1544 | 0 | 698 |
| PDK2 | 0 | 0 | 0 | 0 | 0 | 0 | 1062 | 0 |
| PDK3 | 711 | 292 | 0 | 2985 | 2789 | 3006 | 4257 | 1584 |
| PDK4 | 0 | 390 | 0 | 0 | 0 | 0 | 0 | 0 |
| PDLIM1 | 4646 | 5430 | 10660 | 4277 | 9024 | 1992 | 2174 | 0 |
| PDLIM2 | 712 | 222 | 1848 | 0 | 674 | 2192 | 106 | 0 |
| PDLIM4 | 0 | 0 | 0 | 0 | 1086 | 1010 | 0 | 0 |
| PDLIM5 | 710 | 0 | 0 | 0 | 0 | 0 | 0 | 0 |
| PDLIM7 | 0 | 0 | 632 | 0 | 0 | 0 | 0 | 0 |
| PDP1 | 0 | 0 | 0 | 0 | 0 | 0 | 364 | 0 |
| PDP2 | 0 | 402 | 0 | 0 | 0 | 0 | 0 | 0 |
| PDPK1 | 160 | 406 | 0 | 0 | 0 | 0 | 0 | 0 |
| PDPN | 0 | 176 | 0 | 0 | 1060 | 1748 | 0 | 0 |
| PDRG1 | 28306 | 36069 | 63532 | 186137 | 291939 | 170865 | 503128 | 82384 |
| PDS5A | 924 | 0 | 0 | 0 | 0 | 0 | 0 | 0 |
| PDS5B | 0 | 1196 | 0 | 2954 | 6227 | 662 | 2356 | 1846 |
| PDSS1 | 0 | 66 | 0 | 0 | 582 | 0 | 1352 | 0 |
| PDSS2 | 0 | 0 | 0 | 1156 | 0 | 0 | 438 | 0 |
| PDXDC1 | 576 | 0 | 1860 | 0 | 604 | 0 | 0 | 0 |
| PDXK | 7096 | 8144 | 7789 | 34295 | 88605 | 12631 | 42218 | 58778 |
| PDXP | 0 | 424 | 0 | 0 | 1236 | 0 | 0 | 0 |
| PDZD11 | 0 | 591 | 0 | 0 | 6708 | 0 | 470 | 0 |
| PDZD3 | 4992 | 1700 | 11841 | 472 | 1917 | 0 | 6774 | 0 |
| PDZD7 | 0 | 218 | 0 | 0 | 0 | 0 | 0 | 0 |
| PDZD8 | 0 | 528 | 0 | 0 | 587 | 0 | 0 | 0 |
| PDZK1 | 0 | 0 | 846 | 0 | 0 | 0 | 0 | 0 |
| PDZK1IP1 | 1016 | 412 | 4340 | 3228 | 4352 | 288 | 5196 | 2948 |
| PDZRN3 | 0 | 0 | 0 | 0 | 0 | 0 | 0 | 830 |
| PEA15 | 0 | 154 | 0 | 0 | 200 | 1219 | 778 | 0 |
| PEAK3 | 0 | 0 | 0 | 0 | 28 | 0 | 0 | 0 |
| PEBP1 | 650 | 130 | 3734 | 6789 | 7025 | 1973 | 434 | 1330 |
| PECAM1 | 726 | 964 | 0 | 0 | 0 | 0 | 0 | 0 |
| PECR | 0 | 0 | 1338 | 0 | 0 | 0 | 0 | 714 |
| PEF1 | 0 | 408 | 0 | 1434 | 2046 | 0 | 16 | 0 |
| PEG10 | 0 | 0 | 0 | 0 | 288 | 0 | 0 | 0 |
| PELI1 | 0 | 0 | 0 | 0 | 635 | 0 | 0 | 0 |
| PELO | 0 | 0 | 0 | 0 | 1678 | 0 | 0 | 0 |
| PELP1 | 0 | 0 | 0 | 0 | 30 | 0 | 0 | 0 |
| PEMT | 0 | 0 | 698 | 2075 | 1782 | 0 | 0 | 0 |
| PEPD | 0 | 0 | 0 | 1410 | 4202 | 1378 | 0 | 780 |

| | | | | | | | | |
|---|---|---|---|---|---|---|---|---|
| PER1 | 3418 | 1286 | 0 | 0 | 0 | 0 | 1165 | 0 |
| PER2 | 0 | 0 | 0 | 0 | 0 | 0 | 152 | 0 |
| PERP | 1576 | 2134 | 9875 | 9930 | 14238 | 4323 | 3528 | 866 |
| PES1 | 0 | 160 | 0 | 1606 | 836 | 0 | 1904 | 0 |
| PET100 | 816 | 1208 | 2102 | 1581 | 6782 | 1460 | 992 | 274 |
| PET117 | 708 | 880 | 0 | 0 | 3012 | 2087 | 0 | 2076 |
| PEX1 | 832 | 6006 | 15483 | 3648 | 7070 | 14417 | 13520 | 432 |
| PEX10 | 700 | 284 | 0 | 0 | 0 | 0 | 0 | 0 |
| PEX11B | 782 | 92 | 368 | 0 | 0 | 0 | 853 | 1058 |
| PEX13 | 0 | 0 | 0 | 1116 | 0 | 0 | 0 | 0 |
| PEX14 | 0 | 0 | 0 | 0 | 850 | 0 | 0 | 0 |
| PEX16 | 0 | 1002 | 1939 | 1180 | 1128 | 0 | 614 | 0 |
| PEX19 | 0 | 0 | 0 | 982 | 464 | 0 | 0 | 0 |
| PEX2 | 566 | 210 | 0 | 1580 | 4280 | 2430 | 594 | 1650 |
| PEX26 | 780 | 0 | 1298 | 0 | 568 | 0 | 0 | 0 |
| PEX3 | 0 | 0 | 0 | 457 | 1370 | 0 | 132 | 0 |
| PEX5 | 0 | 242 | 0 | 0 | 0 | 0 | 154 | 0 |
| PEX6 | 0 | 568 | 0 | 0 | 0 | 0 | 551 | 0 |
| PEX7 | 0 | 0 | 162 | 1097 | 692 | 0 | 0 | 0 |
| PF4 | 0 | 264 | 0 | 0 | 0 | 0 | 0 | 0 |
| PFAS | 0 | 124 | 0 | 534 | 338 | 0 | 288 | 0 |
| PFDN1 | 1054 | 38 | 3347 | 3728 | 2112 | 2600 | 486 | 1255 |
| PFDN2 | 0 | 404 | 826 | 1046 | 5380 | 1807 | 2552 | 560 |
| PFDN4 | 94158 | 159243 | 42575 | 167764 | 661507 | 83293 | 1300477 | 105509 |
| PFDN5 | 5513 | 8738 | 14215 | 18270 | 25198 | 7316 | 13596 | 6432 |
| PFDN6 | 780 | 1772 | 0 | 1566 | 2800 | 0 | 592 | 878 |
| PFKFB2 | 0 | 520 | 0 | 0 | 0 | 0 | 0 | 0 |
| PFKFB3 | 0 | 24 | 726 | 498 | 1868 | 0 | 730 | 698 |
| PFKL | 1316 | 956 | 0 | 0 | 4145 | 0 | 1432 | 1853 |
| PFKM | 0 | 0 | 0 | 1204 | 0 | 0 | 680 | 0 |
| PFKP | 61488 | 29106 | 67498 | 34924 | 175321 | 34040 | 34048 | 65807 |
| PFN1 | 16697 | 17274 | 46307 | 48563 | 59600 | 26189 | 14261 | 28365 |
| PFN2 | 0 | 0 | 1488 | 918 | 1273 | 0 | 0 | 0 |
| PGAM1 | 608 | 1162 | 1122 | 1044 | 5832 | 6002 | 1938 | 722 |
| PGAM5 | 0 | 0 | 0 | 816 | 0 | 0 | 0 | 1184 |
| PGAP1 | 0 | 0 | 0 | 0 | 0 | 0 | 262 | 0 |
| PGAP2 | 272 | 306 | 0 | 0 | 124 | 0 | 828 | 0 |
| PGAP3 | 18131 | 10867 | 54472 | 5320 | 15938 | 13592 | 21544 | 8487 |
| PGBD2 | 0 | 254 | 0 | 0 | 0 | 0 | 0 | 0 |
| PGBD4 | 0 | 206 | 0 | 0 | 0 | 0 | 0 | 0 |
| PGC | 0 | 0 | 0 | 0 | 2359 | 0 | 982 | 609 |
| PGD | 22012 | 55288 | 62204 | 141754 | 207955 | 87624 | 62978 | 82803 |
| PGF | 706 | 0 | 0 | 0 | 0 | 0 | 0 | 0 |

| | | | | | | | | |
|---|---|---|---|---|---|---|---|---|
| PGGHG | 120 | 0 | 0 | 1198 | 222 | 1112 | 0 | 1678 |
| PGGT1B | 442 | 358 | 932 | 0 | 0 | 0 | 0 | 0 |
| PGK1 | 1384 | 1232 | 2639 | 5885 | 29404 | 1614 | 3879 | 12840 |
| PGLS | 168 | 1910 | 1796 | 0 | 2064 | 0 | 302 | 36 |
| PGM1 | 0 | 0 | 3272 | 0 | 676 | 1823 | 0 | 0 |
| PGM2 | 0 | 320 | 0 | 0 | 0 | 0 | 0 | 0 |
| PGM2L1 | 0 | 0 | 0 | 0 | 0 | 1484 | 0 | 0 |
| PGM3 | 0 | 0 | 0 | 0 | 1176 | 0 | 820 | 2326 |
| PGM5 | 112 | 0 | 0 | 0 | 0 | 0 | 0 | 0 |
| PGP | 638 | 0 | 0 | 466 | 0 | 0 | 0 | 0 |
| PGPEP1 | 0 | 186 | 2616 | 986 | 158 | 0 | 1420 | 1255 |
| PGRMC1 | 638 | 978 | 3330 | 24 | 3638 | 1978 | 1846 | 1066 |
| PGRMC2 | 0 | 700 | 536 | 0 | 582 | 1048 | 2123 | 0 |
| PGS1 | 0 | 402 | 0 | 0 | 0 | 0 | 0 | 602 |
| PHACTR1 | 0 | 0 | 0 | 0 | 636 | 0 | 610 | 888 |
| PHACTR2 | 240 | 1204 | 0 | 3160 | 438 | 0 | 570 | 2636 |
| PHACTR3 | 0 | 19969 | 1474 | 3878 | 16942 | 4408 | 7918 | 6244 |
| PHACTR4 | 11713 | 17569 | 43276 | 13353 | 26019 | 16873 | 18826 | 7736 |
| PHAX | 0 | 0 | 0 | 0 | 676 | 0 | 357 | 1460 |
| PHB | 1666 | 3360 | 5274 | 10275 | 6370 | 4822 | 8056 | 2514 |
| PHB2 | 62 | 1028 | 5648 | 4787 | 3402 | 6694 | 1518 | 1918 |
| PHC1 | 596 | 0 | 0 | 0 | 0 | 0 | 0 | 0 |
| PHC3 | 1170 | 1614 | 2578 | 1158 | 9665 | 0 | 8487 | 1964 |
| PHETA2 | 0 | 0 | 0 | 0 | 0 | 0 | 518 | 720 |
| PHEX | 68 | 94 | 0 | 0 | 0 | 0 | 0 | 0 |
| PHF1 | 0 | 0 | 931 | 544 | 0 | 1912 | 286 | 0 |
| PHF10 | 0 | 0 | 0 | 974 | 0 | 0 | 246 | 1114 |
| PHF11 | 528 | 0 | 3596 | 0 | 338 | 0 | 522 | 0 |
| PHF12 | 0 | 526 | 0 | 0 | 0 | 0 | 0 | 0 |
| PHF13 | 0 | 0 | 0 | 0 | 136 | 0 | 0 | 0 |
| PHF14 | 0 | 230 | 0 | 0 | 634 | 0 | 1074 | 0 |
| PHF19 | 0 | 0 | 0 | 766 | 2200 | 0 | 0 | 1014 |
| PHF2 | 0 | 0 | 870 | 0 | 0 | 0 | 0 | 0 |
| PHF20 | 0 | 0 | 0 | 0 | 82 | 0 | 0 | 0 |
| PHF20L1 | 0 | 392 | 0 | 0 | 0 | 0 | 230 | 0 |
| PHF21A | 11852 | 5077 | 3486 | 1670 | 8886 | 8040 | 6050 | 5949 |
| PHF23 | 0 | 0 | 0 | 0 | 380 | 0 | 0 | 1046 |
| PHF5A | 0 | 302 | 986 | 514 | 4863 | 1460 | 0 | 0 |
| PHF6 | 0 | 0 | 28 | 0 | 518 | 0 | 0 | 0 |
| PHF7 | 0 | 0 | 2214 | 0 | 0 | 0 | 0 | 0 |
| PHGDH | 0 | 0 | 0 | 3678 | 0 | 1676 | 60 | 0 |
| PHGR1 | 48972 | 35771 | 58871 | 11354 | 12937 | 2620 | 8791 | 0 |
| PHIP | 0 | 0 | 0 | 0 | 356 | 1620 | 0 | 0 |

| | | | | | | | | |
|---|---|---|---|---|---|---|---|---|
| PHKA1 | 0 | 0 | 0 | 0 | 0 | 0 | 0 | 350 |
| PHKB | 496 | 0 | 2280 | 0 | 302 | 0 | 0 | 0 |
| PHKG2 | 0 | 0 | 0 | 2084 | 0 | 0 | 172 | 2094 |
| PHLDA1 | 0 | 312 | 0 | 1454 | 3673 | 0 | 0 | 594 |
| PHLDA2 | 654 | 0 | 1492 | 2950 | 2754 | 3748 | 456 | 836 |
| PHLDA3 | 0 | 0 | 0 | 0 | 1384 | 0 | 0 | 0 |
| PHLDB1 | 1122 | 1727 | 1556 | 720 | 3542 | 5048 | 250 | 726 |
| PHLDB3 | 418 | 0 | 0 | 0 | 986 | 0 | 0 | 0 |
| PHLPP1 | 0 | 0 | 1414 | 0 | 0 | 0 | 42 | 0 |
| PHOSPHO2 | 0 | 0 | 0 | 0 | 1448 | 0 | 0 | 0 |
| PHOSPHO2-KLHL23 | 0 | 0 | 0 | 0 | 338 | 0 | 194 | 638 |
| PHPT1 | 2658 | 3336 | 4878 | 4860 | 7887 | 1533 | 3564 | 4630 |
| PHRF1 | 0 | 0 | 0 | 2032 | 269 | 0 | 212 | 0 |
| PHTF2 | 344 | 6941 | 7794 | 2119 | 23339 | 11375 | 9921 | 4282 |
| PHYH | 2287 | 1974 | 496 | 2714 | 5527 | 0 | 968 | 1140 |
| PHYHD1 | 0 | 0 | 420 | 0 | 0 | 0 | 0 | 0 |
| PHYKPL | 302 | 222 | 702 | 0 | 0 | 1328 | 0 | 0 |
| PI16 | 0 | 318 | 0 | 0 | 0 | 0 | 0 | 0 |
| PI3 | 3857 | 2346 | 6722 | 3360 | 514 | 0 | 0 | 624 |
| PI4K2A | 0 | 0 | 1198 | 0 | 0 | 0 | 0 | 0 |
| PI4K2B | 0 | 0 | 0 | 0 | 0 | 0 | 84 | 0 |
| PI4KA | 2384 | 1368 | 596 | 686 | 7107 | 1354 | 1677 | 2678 |
| PI4KAP1 | 0 | 0 | 0 | 372 | 0 | 0 | 0 | 0 |
| PI4KB | 0 | 0 | 0 | 1284 | 0 | 1520 | 0 | 0 |
| PIAS1 | 39477 | 48979 | 108769 | 45038 | 159338 | 68141 | 128560 | 40257 |
| PIAS2 | 238 | 0 | 0 | 0 | 340 | 0 | 0 | 0 |
| PIAS3 | 0 | 0 | 0 | 100 | 0 | 0 | 0 | 0 |
| PIBF1 | 5969 | 13558 | 9980 | 28311 | 26576 | 33152 | 41674 | 14573 |
| PICALM | 33145 | 25315 | 66734 | 37980 | 132645 | 53676 | 92441 | 45773 |
| PID1 | 444 | 392 | 1094 | 0 | 908 | 0 | 0 | 1491 |
| PIDD1 | 513 | 0 | 0 | 0 | 0 | 0 | 0 | 0 |
| PIEZO1 | 0 | 620 | 38 | 0 | 232 | 1176 | 300 | 666 |
| PIF1 | 0 | 0 | 0 | 0 | 378 | 0 | 0 | 0 |
| PIFO | 945 | 0 | 0 | 0 | 0 | 0 | 0 | 0 |
| PIGA | 0 | 0 | 0 | 0 | 454 | 0 | 0 | 0 |
| PIGBOS1 | 0 | 0 | 3140 | 166 | 682 | 0 | 1162 | 0 |
| PIGC | 0 | 450 | 0 | 1330 | 3829 | 0 | 402 | 0 |
| PIGF | 1670 | 634 | 2706 | 0 | 874 | 1449 | 2068 | 0 |
| PIGH | 0 | 0 | 1728 | 0 | 1362 | 0 | 0 | 0 |
| PIGK | 0 | 0 | 0 | 0 | 648 | 0 | 0 | 0 |
| PIGL | 15027 | 18445 | 36994 | 27393 | 60621 | 38002 | 37430 | 30295 |
| PIGM | 0 | 0 | 0 | 0 | 576 | 0 | 646 | 0 |
| PIGN | 0 | 250 | 0 | 0 | 42 | 0 | 0 | 0 |

| | | | | | | | | |
|---|---|---|---|---|---|---|---|---|
| PIGO | 0 | 0 | 0 | 0 | 0 | 0 | 636 | 0 |
| PIGP | 1140 | 0 | 0 | 0 | 2260 | 0 | 730 | 0 |
| PIGQ | 310 | 0 | 961 | 0 | 62 | 0 | 0 | 0 |
| PIGR | 19220 | 24838 | 35692 | 5713 | 1254 | 0 | 0 | 0 |
| PIGS | 0 | 848 | 938 | 0 | 484 | 288 | 0 | 740 |
| PIGT | 582 | 1104 | 1756 | 974 | 1620 | 290 | 570 | 144 |
| PIGU | 0 | 296 | 0 | 4075 | 0 | 3126 | 0 | 0 |
| PIGV | 1411 | 0 | 0 | 0 | 0 | 0 | 836 | 0 |
| PIGW | 0 | 0 | 0 | 0 | 0 | 914 | 0 | 0 |
| PIGX | 184 | 0 | 0 | 1102 | 1532 | 0 | 0 | 0 |
| PIGY-DT | 0 | 0 | 334 | 0 | 0 | 0 | 0 | 0 |
| PIGZ | 470 | 440 | 2589 | 0 | 1122 | 0 | 982 | 0 |
| PIH1D1 | 0 | 654 | 1932 | 0 | 2557 | 2020 | 1309 | 3831 |
| PIH1D2 | 576 | 0 | 0 | 0 | 0 | 0 | 0 | 0 |
| PIK3C2A | 0 | 0 | 0 | 0 | 262 | 0 | 0 | 0 |
| PIK3C2B | 383 | 0 | 2446 | 0 | 660 | 0 | 480 | 0 |
| PIK3C3 | 0 | 306 | 0 | 0 | 502 | 0 | 0 | 0 |
| PIK3CA | 1190 | 526 | 0 | 2012 | 6474 | 3086 | 6665 | 1612 |
| PIK3CB | 28040 | 19376 | 34186 | 26384 | 64069 | 23617 | 66417 | 27332 |
| PIK3IP1 | 0 | 0 | 0 | 1359 | 0 | 1767 | 0 | 0 |
| PIK3R1 | 7018 | 17203 | 8008 | 15307 | 29805 | 25235 | 11779 | 4926 |
| PIK3R2 | 0 | 1288 | 0 | 0 | 0 | 752 | 462 | 0 |
| PIK3R3 | 274 | 0 | 0 | 0 | 0 | 0 | 0 | 0 |
| PIK3R4 | 0 | 0 | 0 | 0 | 456 | 0 | 0 | 0 |
| PIK3R5 | 0 | 0 | 0 | 0 | 0 | 0 | 294 | 0 |
| PIM1 | 0 | 0 | 0 | 0 | 350 | 2940 | 70 | 0 |
| PIM2 | 406 | 0 | 0 | 0 | 0 | 0 | 0 | 1210 |
| PIM3 | 790 | 562 | 986 | 860 | 668 | 0 | 0 | 216 |
| PIN1 | 0 | 366 | 2286 | 1998 | 2982 | 1356 | 336 | 0 |
| PIN4 | 800 | 0 | 0 | 100 | 3124 | 0 | 1542 | 1687 |
| PINK1 | 0 | 0 | 0 | 0 | 1468 | 0 | 0 | 0 |
| PINK1-AS | 0 | 0 | 0 | 0 | 0 | 0 | 96 | 0 |
| PINLYP | 948 | 0 | 0 | 0 | 0 | 0 | 0 | 0 |
| PINX1 | 454 | 0 | 1940 | 1482 | 0 | 0 | 0 | 0 |
| PIP4K2B | 654 | 1932 | 3460 | 1700 | 3993 | 0 | 3624 | 1622 |
| PIP4K2C | 6266 | 5444 | 17597 | 5110 | 23133 | 5781 | 16969 | 5438 |
| PIP4P1 | 0 | 0 | 0 | 726 | 258 | 0 | 0 | 0 |
| PIP4P2 | 0 | 0 | 0 | 0 | 1057 | 0 | 0 | 0 |
| PIP5K1B | 314 | 328 | 0 | 202 | 534 | 0 | 0 | 0 |
| PIP5K1C | 0 | 752 | 0 | 0 | 0 | 0 | 0 | 0 |
| PIR | 0 | 0 | 0 | 0 | 202 | 0 | 0 | 0 |
| PISD | 0 | 262 | 0 | 0 | 690 | 0 | 1452 | 0 |
| PITHD1 | 0 | 0 | 0 | 2152 | 1760 | 0 | 598 | 3372 |

| | | | | | | | | |
|---|---|---|---|---|---|---|---|---|
| PITPNA | 9924 | 10638 | 19565 | 35664 | 47954 | 24556 | 30357 | 25345 |
| PITPNA-AS1 | 318 | 0 | 0 | 0 | 936 | 0 | 0 | 494 |
| PITPNB | 0 | 0 | 0 | 0 | 490 | 0 | 554 | 0 |
| PITPNC1 | 4154 | 4484 | 1448 | 7256 | 19739 | 4450 | 7885 | 11822 |
| PITPNM2 | 568 | 0 | 890 | 0 | 498 | 0 | 0 | 0 |
| PITRM1 | 0 | 174 | 1504 | 0 | 378 | 0 | 0 | 0 |
| PITX1 | 0 | 0 | 0 | 764 | 148 | 0 | 0 | 0 |
| PITX2 | 0 | 0 | 0 | 766 | 0 | 0 | 1182 | 0 |
| PJA1 | 0 | 0 | 0 | 0 | 0 | 0 | 0 | 124 |
| PJA2 | 0 | 138 | 1000 | 1800 | 68 | 0 | 0 | 62 |
| PKD1P3-NPIPA1 | 0 | 0 | 0 | 0 | 592 | 0 | 0 | 0 |
| PKD1P6-NPIPP1 | 0 | 0 | 0 | 0 | 130 | 0 | 1332 | 0 |
| PKD2 | 0 | 0 | 0 | 0 | 0 | 0 | 0 | 932 |
| PKDCC | 0 | 0 | 0 | 0 | 754 | 794 | 0 | 746 |
| PKHD1L1 | 502 | 488 | 0 | 0 | 0 | 0 | 0 | 0 |
| PKIB | 1782 | 504 | 14895 | 0 | 248 | 0 | 0 | 0 |
| PKIG | 564 | 340 | 3030 | 0 | 2258 | 2137 | 995 | 0 |
| PKM | 335991 | 316282 | 373096 | 1053791 | 2961365 | 756374 | 1337245 | 1014461 |
| PKMYT1 | 0 | 0 | 368 | 532 | 0 | 0 | 0 | 0 |
| PKN1 | 0 | 560 | 0 | 0 | 1070 | 1940 | 0 | 0 |
| PKNOX1 | 0 | 0 | 0 | 0 | 0 | 0 | 352 | 0 |
| PKP1 | 0 | 0 | 0 | 0 | 124 | 0 | 0 | 0 |
| PKP2 | 0 | 331 | 360 | 0 | 0 | 0 | 0 | 0 |
| PKP3 | 816 | 0 | 924 | 708 | 1277 | 0 | 0 | 412 |
| PKP4 | 104 | 1520 | 1860 | 2300 | 5732 | 0 | 5798 | 5366 |
| PLA1A | 0 | 0 | 0 | 0 | 0 | 0 | 762 | 0 |
| PLA2G10 | 4233 | 706 | 1672 | 0 | 0 | 0 | 0 | 744 |
| PLA2G12A | 524 | 0 | 1607 | 1214 | 1175 | 3098 | 1918 | 0 |
| PLA2G15 | 0 | 394 | 0 | 1220 | 0 | 0 | 0 | 98 |
| PLA2G16 | 1560 | 0 | 1992 | 1475 | 13635 | 2431 | 0 | 9276 |
| PLA2G1B | 1376 | 1540 | 3722 | 0 | 686 | 1996 | 4964 | 1052 |
| PLA2G2A | 0 | 11712 | 7410 | 5107 | 10620 | 1056 | 0 | 0 |
| PLA2G2F | 0 | 0 | 0 | 0 | 36 | 0 | 0 | 0 |
| PLA2G6 | 2416 | 1346 | 10658 | 1390 | 5322 | 1112 | 6286 | 1056 |
| PLA2R1 | 0 | 0 | 0 | 0 | 0 | 0 | 12 | 0 |
| PLAA | 0 | 510 | 0 | 0 | 0 | 0 | 0 | 0 |
| PLAC8 | 10098 | 2152 | 36163 | 1909 | 8376 | 0 | 0 | 902 |
| PLAC9 | 0 | 356 | 565 | 0 | 0 | 0 | 0 | 0 |
| PLAGL1 | 620 | 685 | 1102 | 0 | 860 | 876 | 1834 | 0 |
| PLAGL2 | 0 | 0 | 0 | 826 | 0 | 0 | 0 | 0 |
| PLAT | 272 | 716 | 0 | 1070 | 916 | 0 | 0 | 0 |
| PLAU | 0 | 0 | 0 | 0 | 44 | 302 | 0 | 0 |
| PLAUR | 3137 | 0 | 1560 | 0 | 4491 | 0 | 0 | 2052 |

| | | | | | | | | |
|---|---|---|---|---|---|---|---|---|
| PLBD1 | 0 | 704 | 0 | 1222 | 3340 | 0 | 1438 | 1040 |
| PLCB2 | 0 | 0 | 0 | 0 | 384 | 0 | 0 | 0 |
| PLCB3 | 244 | 0 | 0 | 146 | 1314 | 0 | 64 | 668 |
| PLCB4 | 434 | 212 | 0 | 0 | 0 | 0 | 0 | 0 |
| PLCD1 | 0 | 442 | 0 | 990 | 0 | 0 | 0 | 0 |
| PLCD4 | 0 | 266 | 0 | 0 | 0 | 0 | 0 | 0 |
| PLCE1 | 0 | 604 | 2234 | 0 | 362 | 0 | 0 | 0 |
| PLCG1 | 2682 | 1818 | 5924 | 862 | 9334 | 3904 | 9754 | 1890 |
| PLCXD1 | 0 | 0 | 362 | 0 | 486 | 0 | 0 | 0 |
| PLD1 | 0 | 0 | 507 | 983 | 0 | 0 | 0 | 0 |
| PLD3 | 1270 | 0 | 0 | 0 | 1960 | 0 | 836 | 0 |
| PLD6 | 0 | 0 | 0 | 0 | 758 | 0 | 0 | 0 |
| PLEC | 832 | 0 | 642 | 504 | 0 | 1626 | 968 | 0 |
| PLEK2 | 420 | 0 | 754 | 2922 | 3078 | 0 | 0 | 0 |
| PLEKHA1 | 0 | 0 | 0 | 0 | 694 | 0 | 0 | 558 |
| PLEKHA2 | 0 | 426 | 0 | 0 | 188 | 2130 | 758 | 0 |
| PLEKHA3 | 0 | 0 | 0 | 0 | 342 | 0 | 58 | 0 |
| PLEKHA4 | 0 | 448 | 0 | 0 | 0 | 2222 | 0 | 0 |
| PLEKHA5 | 0 | 0 | 0 | 0 | 136 | 0 | 0 | 0 |
| PLEKHA6 | 494 | 70 | 0 | 0 | 408 | 0 | 0 | 0 |
| PLEKHA7 | 0 | 0 | 0 | 0 | 730 | 0 | 554 | 0 |
| PLEKHA8 | 0 | 0 | 0 | 0 | 314 | 0 | 0 | 0 |
| PLEKHB1 | 0 | 190 | 0 | 0 | 22 | 0 | 0 | 0 |
| PLEKHB2 | 162 | 552 | 0 | 0 | 1486 | 1882 | 1028 | 1618 |
| PLEKHF1 | 240 | 974 | 458 | 1346 | 1460 | 2368 | 152 | 74 |
| PLEKHF2 | 0 | 318 | 0 | 0 | 260 | 0 | 300 | 1192 |
| PLEKHG1 | 0 | 0 | 0 | 0 | 550 | 0 | 0 | 0 |
| PLEKHG3 | 0 | 0 | 0 | 0 | 752 | 0 | 0 | 0 |
| PLEKHG4 | 0 | 0 | 0 | 0 | 0 | 0 | 420 | 0 |
| PLEKHG5 | 0 | 0 | 510 | 0 | 458 | 0 | 0 | 0 |
| PLEKHG6 | 3944 | 3177 | 15939 | 1378 | 4344 | 0 | 4926 | 1960 |
| PLEKHH1 | 292 | 0 | 962 | 0 | 1080 | 0 | 444 | 0 |
| PLEKHH3 | 0 | 0 | 0 | 0 | 0 | 0 | 542 | 0 |
| PLEKHJ1 | 2712 | 1262 | 4904 | 3495 | 4248 | 3010 | 1512 | 1344 |
| PLEKHM1 | 0 | 0 | 0 | 0 | 0 | 0 | 680 | 0 |
| PLEKHM2 | 886 | 228 | 0 | 2347 | 4316 | 0 | 1676 | 384 |
| PLEKHM3 | 0 | 26 | 993 | 0 | 0 | 0 | 0 | 0 |
| PLEKHO1 | 2486 | 418 | 1546 | 0 | 4732 | 5106 | 1320 | 1315 |
| PLEKHO2 | 0 | 614 | 0 | 0 | 0 | 0 | 0 | 0 |
| PLGRKT | 606 | 724 | 5663 | 1766 | 422 | 3976 | 1342 | 0 |
| PLIN2 | 0 | 0 | 0 | 0 | 4896 | 0 | 2346 | 0 |
| PLIN3 | 0 | 594 | 3221 | 0 | 840 | 0 | 0 | 0 |
| PLK1 | 0 | 406 | 0 | 508 | 1338 | 0 | 0 | 0 |

| | | | | | | | | |
|---|---|---|---|---|---|---|---|---|
| PLK2 | 0 | 198 | 0 | 0 | 372 | 0 | 0 | 0 |
| PLK3 | 0 | 688 | 0 | 1240 | 0 | 0 | 0 | 0 |
| PLK4 | 0 | 0 | 0 | 0 | 0 | 0 | 252 | 0 |
| PLLP | 0 | 382 | 0 | 0 | 0 | 0 | 0 | 0 |
| PLN | 20465 | 12239 | 33725 | 3148 | 28074 | 23729 | 8265 | 6167 |
| PLOD1 | 0 | 0 | 0 | 0 | 1728 | 0 | 0 | 0 |
| PLOD2 | 0 | 124 | 1454 | 0 | 1113 | 0 | 0 | 0 |
| PLOD3 | 0 | 0 | 0 | 1206 | 0 | 0 | 0 | 832 |
| PLP1 | 0 | 0 | 1239 | 0 | 0 | 0 | 0 | 0 |
| PLP2 | 0 | 1884 | 358 | 6946 | 10759 | 0 | 2600 | 2048 |
| PLPBP | 27984 | 28899 | 79906 | 27296 | 48870 | 30970 | 54229 | 14284 |
| PLPP1 | 249855 | 127399 | 1870222 | 24621 | 77070 | 120772 | 80675 | 43623 |
| PLPP2 | 710 | 0 | 2712 | 3050 | 5964 | 0 | 680 | 0 |
| PLPP3 | 0 | 76 | 0 | 0 | 0 | 0 | 216 | 0 |
| PLPP4 | 0 | 0 | 0 | 0 | 50198 | 9250 | 488 | 5438 |
| PLPP5 | 820 | 2024 | 556 | 650 | 4102 | 0 | 1967 | 1232 |
| PLPP6 | 0 | 0 | 0 | 0 | 0 | 0 | 654 | 0 |
| PLPPR2 | 0 | 350 | 0 | 0 | 0 | 0 | 0 | 0 |
| PLRG1 | 0 | 0 | 1512 | 0 | 266 | 0 | 0 | 0 |
| PLS1 | 232 | 336 | 550 | 1478 | 1385 | 0 | 633 | 478 |
| PLS3 | 0 | 0 | 1222 | 0 | 1630 | 0 | 0 | 548 |
| PLSCR1 | 1672 | 1404 | 3098 | 2800 | 4514 | 0 | 220 | 750 |
| PLSCR2 | 0 | 0 | 0 | 0 | 532 | 0 | 0 | 0 |
| PLSCR3 | 0 | 0 | 0 | 0 | 0 | 3419 | 0 | 0 |
| PLSCR4 | 0 | 0 | 558 | 0 | 544 | 0 | 0 | 0 |
| PLTP | 0 | 550 | 494 | 1068 | 1374 | 494 | 2126 | 896 |
| PLUT | 0 | 0 | 0 | 1024 | 0 | 0 | 0 | 0 |
| PLVAP | 572 | 0 | 0 | 0 | 0 | 0 | 0 | 744 |
| PLXDC2 | 192 | 0 | 0 | 0 | 84 | 0 | 0 | 0 |
| PLXNA2 | 456 | 390 | 2585 | 0 | 64 | 0 | 582 | 0 |
| PLXNA3 | 0 | 298 | 0 | 0 | 1230 | 0 | 542 | 0 |
| PLXNB1 | 0 | 244 | 550 | 0 | 1586 | 2518 | 698 | 0 |
| PLXNB2 | 206 | 0 | 0 | 1348 | 414 | 2952 | 538 | 1222 |
| PLXND1 | 0 | 0 | 0 | 0 | 0 | 0 | 0 | 564 |
| PM20D2 | 0 | 0 | 568 | 0 | 0 | 0 | 0 | 0 |
| PMAIP1 | 0 | 338 | 0 | 0 | 747 | 0 | 0 | 408 |
| PMEL | 0 | 0 | 0 | 0 | 0 | 0 | 158 | 0 |
| PMEPA1 | 11850 | 13264 | 33928 | 27557 | 70550 | 93696 | 135302 | 12915 |
| PMF1 | 138 | 560 | 2666 | 0 | 3944 | 4081 | 0 | 0 |
| PMF1-BGLAP | 0 | 0 | 0 | 1290 | 0 | 0 | 0 | 0 |
| PML | 1918 | 2294 | 2608 | 6588 | 7495 | 4841 | 8930 | 3086 |
| PMM1 | 1454 | 958 | 3592 | 1372 | 806 | 0 | 384 | 548 |
| PMM2 | 400 | 846 | 415 | 0 | 56 | 0 | 34 | 0 |

| | | | | | | | | |
|---|---|---|---|---|---|---|---|---|
| PMP22 | 0 | 188 | 400 | 0 | 244 | 1444 | 0 | 0 |
| PMPCA | 0 | 0 | 46 | 0 | 3268 | 0 | 260 | 0 |
| PMPCB | 0 | 2196 | 4589 | 0 | 3456 | 0 | 1050 | 0 |
| PMS1 | 0 | 0 | 962 | 0 | 0 | 0 | 0 | 0 |
| PMS2 | 5682 | 11245 | 27516 | 22180 | 47215 | 15116 | 49796 | 12900 |
| PMS2CL | 0 | 0 | 0 | 0 | 0 | 0 | 46 | 0 |
| PMS2P1 | 31598 | 77394 | 81370 | 131158 | 337564 | 218606 | 307237 | 115330 |
| PMS2P2 | 100 | 0 | 0 | 1032 | 1930 | 4560 | 3283 | 0 |
| PMS2P3 | 0 | 784 | 0 | 0 | 1420 | 0 | 0 | 0 |
| PMS2P4 | 4647 | 7764 | 9500 | 3332 | 10398 | 6553 | 12109 | 4754 |
| PMS2P5 | 1998 | 7374 | 0 | 8218 | 16878 | 7012 | 26445 | 8028 |
| PMS2P7 | 5522 | 3812 | 2114 | 11101 | 15862 | 14577 | 16723 | 3430 |
| PMS2P9 | 0 | 0 | 0 | 3648 | 763 | 2996 | 732 | 2194 |
| PMVK | 0 | 0 | 988 | 674 | 616 | 2110 | 1190 | 1224 |
| PNISR | 0 | 676 | 0 | 722 | 369 | 435 | 379 | 736 |
| PNKD | 1044 | 736 | 3601 | 3423 | 2991 | 1734 | 2087 | 2575 |
| PNKP | 0 | 228 | 0 | 0 | 248 | 0 | 0 | 0 |
| PNMA1 | 0 | 0 | 0 | 0 | 1100 | 0 | 0 | 1246 |
| PNMA2 | 0 | 0 | 0 | 0 | 0 | 0 | 54 | 0 |
| PNN | 594 | 0 | 833 | 172 | 614 | 0 | 320 | 570 |
| PNO1 | 0 | 988 | 0 | 1158 | 882 | 0 | 0 | 0 |
| PNP | 0 | 1166 | 0 | 0 | 622 | 2253 | 0 | 816 |
| PNPLA2 | 692 | 74 | 0 | 0 | 286 | 1011 | 1886 | 214 |
| PNPLA4 | 0 | 0 | 0 | 1140 | 602 | 0 | 1644 | 0 |
| PNPLA6 | 696 | 0 | 0 | 772 | 244 | 0 | 0 | 0 |
| PNPLA8 | 0 | 340 | 0 | 0 | 0 | 0 | 0 | 726 |
| PNPO | 0 | 0 | 0 | 0 | 0 | 0 | 621 | 388 |
| PNPT1 | 0 | 0 | 0 | 0 | 318 | 0 | 570 | 0 |
| PNRC1 | 706 | 920 | 0 | 0 | 1690 | 0 | 0 | 1056 |
| PNRC2 | 216 | 746 | 2720 | 2557 | 1618 | 0 | 0 | 1288 |
| POC1B | 0 | 276 | 138 | 0 | 0 | 1046 | 0 | 0 |
| POC1B-AS1 | 0 | 0 | 0 | 0 | 688 | 996 | 0 | 0 |
| PODXL | 0 | 0 | 0 | 0 | 2084 | 0 | 0 | 0 |
| PODXL2 | 0 | 0 | 1642 | 994 | 544 | 0 | 0 | 0 |
| POF1B | 0 | 388 | 0 | 0 | 514 | 0 | 0 | 0 |
| POFUT1 | 0 | 0 | 0 | 624 | 346 | 1328 | 338 | 0 |
| POFUT2 | 0 | 0 | 0 | 0 | 0 | 0 | 124 | 0 |
| POGLUT1 | 0 | 368 | 0 | 0 | 0 | 0 | 0 | 0 |
| POLA2 | 0 | 0 | 0 | 0 | 298 | 0 | 0 | 0 |
| POLB | 11536 | 27728 | 27443 | 84924 | 284994 | 202789 | 263121 | 145534 |
| POLD1 | 0 | 0 | 0 | 0 | 1036 | 0 | 0 | 0 |
| POLD2 | 0 | 448 | 1354 | 2268 | 3896 | 2016 | 440 | 1106 |
| POLD4 | 3207 | 2958 | 9317 | 1818 | 8020 | 7395 | 1550 | 6982 |

| | | | | | | | | |
|---|---|---|---|---|---|---|---|---|
| POLDIP2 | 738 | 0 | 0 | 1526 | 0 | 0 | 0 | 946 |
| POLDIP3 | 944 | 256 | 0 | 480 | 0 | 1264 | 652 | 0 |
| POLE | 0 | 178 | 1052 | 954 | 392 | 0 | 1108 | 0 |
| POLE3 | 0 | 0 | 0 | 1470 | 670 | 2067 | 1159 | 0 |
| POLE4 | 3136 | 1434 | 0 | 3320 | 2086 | 0 | 3894 | 0 |
| POLG | 2756 | 1328 | 5468 | 1188 | 9680 | 7832 | 7740 | 2206 |
| POLG2 | 0 | 340 | 0 | 0 | 1310 | 0 | 0 | 0 |
| POLH | 0 | 0 | 0 | 0 | 322 | 0 | 0 | 0 |
| POLI | 0 | 308 | 0 | 30 | 0 | 0 | 0 | 0 |
| POLM | 0 | 0 | 0 | 914 | 0 | 0 | 0 | 0 |
| POLQ | 0 | 0 | 0 | 454 | 0 | 0 | 872 | 0 |
| POLR1A | 490 | 72 | 1398 | 777 | 4550 | 1598 | 1624 | 824 |
| POLR1B | 0 | 1020 | 0 | 0 | 0 | 0 | 0 | 0 |
| POLR1C | 0 | 1328 | 1500 | 0 | 1928 | 2007 | 818 | 2360 |
| POLR1D | 0 | 1082 | 5822 | 3960 | 5060 | 6690 | 1582 | 5403 |
| POLR2A | 158 | 0 | 0 | 0 | 0 | 0 | 268 | 0 |
| POLR2B | 81122 | 97541 | 47241 | 69982 | 199674 | 45492 | 195594 | 31661 |
| POLR2C | 0 | 490 | 0 | 1168 | 1184 | 0 | 1822 | 0 |
| POLR2D | 0 | 0 | 0 | 4158 | 944 | 0 | 0 | 0 |
| POLR2E | 604 | 356 | 2893 | 3171 | 5210 | 5093 | 1244 | 2126 |
| POLR2F | 1178 | 1520 | 0 | 3082 | 3154 | 1750 | 1430 | 1078 |
| POLR2G | 1654 | 462 | 0 | 5467 | 6550 | 0 | 3278 | 7222 |
| POLR2H | 812 | 576 | 1329 | 1989 | 4148 | 0 | 2605 | 2374 |
| POLR2I | 724 | 830 | 0 | 3046 | 5091 | 0 | 1314 | 480 |
| POLR2J | 802 | 2745 | 3270 | 2368 | 3938 | 3093 | 1850 | 816 |
| POLR2J2 | 0 | 380 | 0 | 68 | 366 | 0 | 354 | 964 |
| POLR2J3 | 10 | 336 | 0 | 0 | 0 | 0 | 0 | 0 |
| POLR2J4 | 0 | 0 | 0 | 0 | 0 | 0 | 360 | 0 |
| POLR2K | 434 | 3919 | 1302 | 1656 | 6465 | 1124 | 5514 | 3108 |
| POLR2L | 724 | 5963 | 866 | 4248 | 9127 | 3235 | 1478 | 2438 |
| POLR2M | 0 | 0 | 0 | 1536 | 0 | 0 | 0 | 0 |
| POLR3A | 624 | 0 | 0 | 0 | 0 | 0 | 316 | 0 |
| POLR3C | 0 | 0 | 0 | 4002 | 674 | 648 | 0 | 0 |
| POLR3D | 0 | 0 | 0 | 0 | 0 | 0 | 476 | 0 |
| POLR3E | 0 | 0 | 0 | 442 | 1270 | 0 | 0 | 0 |
| POLR3F | 0 | 510 | 0 | 0 | 248 | 0 | 0 | 0 |
| POLR3G | 0 | 166 | 0 | 0 | 760 | 0 | 0 | 0 |
| POLR3GL | 0 | 0 | 0 | 0 | 0 | 0 | 0 | 366 |
| POLR3H | 7666 | 27823 | 19782 | 33940 | 110639 | 78160 | 56497 | 30701 |
| POLR3K | 944 | 0 | 0 | 2241 | 1162 | 0 | 1532 | 0 |
| POLRMT | 374 | 0 | 0 | 0 | 226 | 0 | 390 | 0 |
| POM121 | 0 | 0 | 0 | 635 | 572 | 0 | 920 | 0 |
| POM121C | 0 | 0 | 0 | 0 | 0 | 1742 | 0 | 0 |

| | | | | | | | | |
|---|---|---|---|---|---|---|---|---|
| POMGNT1 | 0 | 36 | 0 | 866 | 388 | 0 | 1900 | 0 |
| POMGNT2 | 0 | 0 | 0 | 0 | 798 | 0 | 0 | 0 |
| POMK | 0 | 0 | 0 | 754 | 0 | 0 | 0 | 0 |
| POMP | 2048 | 5591 | 7365 | 25469 | 13717 | 12508 | 4922 | 8120 |
| POMT1 | 0 | 372 | 0 | 0 | 690 | 0 | 0 | 0 |
| POMZP3 | 0 | 0 | 1374 | 814 | 0 | 0 | 0 | 0 |
| PON2 | 0 | 240 | 0 | 1076 | 998 | 2690 | 3278 | 1236 |
| PON3 | 0 | 0 | 0 | 0 | 0 | 0 | 1319 | 0 |
| POP4 | 0 | 0 | 0 | 0 | 2003 | 0 | 0 | 1312 |
| POP5 | 1090 | 460 | 922 | 1266 | 3492 | 2594 | 544 | 1406 |
| POP7 | 15537 | 14076 | 48225 | 40122 | 47200 | 23845 | 49621 | 16654 |
| POPDC3 | 0 | 0 | 0 | 0 | 1042 | 0 | 0 | 0 |
| POR | 0 | 0 | 1056 | 0 | 580 | 0 | 336 | 646 |
| PORCN | 0 | 0 | 0 | 0 | 376 | 0 | 0 | 0 |
| POSTN | 0 | 0 | 0 | 642 | 1884 | 0 | 226 | 1076 |
| POT1 | 2394 | 4066 | 5415 | 7648 | 30938 | 250 | 12475 | 9098 |
| POU2AF1 | 1048 | 936 | 0 | 0 | 0 | 0 | 0 | 0 |
| POU2F1 | 718 | 0 | 0 | 0 | 0 | 0 | 0 | 0 |
| POU2F3 | 0 | 494 | 1590 | 1421 | 2150 | 4108 | 3938 | 2188 |
| PP7080 | 110 | 0 | 0 | 0 | 506 | 0 | 338 | 0 |
| PPA1 | 2216 | 4660 | 4188 | 13707 | 15000 | 4118 | 7547 | 2814 |
| PPA2 | 1688 | 1104 | 4386 | 3992 | 7109 | 2882 | 942 | 2564 |
| PPAN | 0 | 0 | 0 | 0 | 3070 | 0 | 0 | 1104 |
| PPARA | 1233 | 0 | 0 | 0 | 0 | 0 | 0 | 0 |
| PPARD | 0 | 0 | 0 | 0 | 0 | 0 | 1155 | 110 |
| PPARG | 773616 | 251030 | 1955732 | 422282 | 2713305 | 182152 | 1801788 | 246955 |
| PPARGC1B | 1090 | 74 | 2000 | 0 | 1450 | 0 | 1206 | 0 |
| PPCDC | 0 | 376 | 0 | 0 | 0 | 0 | 466 | 0 |
| PPCS | 374 | 2594 | 6898 | 2536 | 8409 | 0 | 2287 | 2153 |
| PPDPF | 7833 | 4403 | 11506 | 14537 | 20240 | 3225 | 16109 | 2528 |
| PPFIA1 | 11960 | 8132 | 18054 | 13873 | 25500 | 7168 | 21876 | 6935 |
| PPFIA2 | 0 | 714 | 0 | 0 | 2472 | 0 | 0 | 0 |
| PPFIA3 | 686 | 0 | 0 | 0 | 0 | 0 | 0 | 621 |
| PPFIBP1 | 0 | 0 | 0 | 1104 | 0 | 0 | 0 | 0 |
| PPFIBP2 | 470 | 0 | 0 | 0 | 592 | 0 | 378 | 0 |
| PPHLN1 | 0 | 0 | 0 | 0 | 316 | 1431 | 312 | 768 |
| PPIA | 14814 | 21441 | 53473 | 70670 | 163977 | 65887 | 36435 | 55491 |
| PPIB | 250 | 7792 | 15776 | 16186 | 11270 | 2792 | 18939 | 7304 |
| PPIC | 1152 | 1182 | 3149 | 1122 | 1936 | 0 | 302 | 0 |
| PPID | 0 | 0 | 0 | 884 | 0 | 0 | 622 | 0 |
| PPIE | 720 | 454 | 0 | 4941 | 4950 | 230 | 332 | 0 |
| PPIF | 0 | 846 | 1049 | 398 | 1502 | 0 | 190 | 1256 |
| PPIG | 0 | 736 | 0 | 0 | 0 | 0 | 726 | 0 |

| | | | | | | | | |
|---|---|---|---|---|---|---|---|---|
| PPIH | 915 | 390 | 1062 | 1326 | 2101 | 0 | 1528 | 0 |
| PPIL1 | 0 | 776 | 0 | 0 | 0 | 0 | 0 | 587 |
| PPIL2 | 468 | 0 | 0 | 0 | 324 | 0 | 0 | 0 |
| PPIL3 | 0 | 460 | 0 | 0 | 1592 | 0 | 290 | 0 |
| PPIL4 | 0 | 0 | 0 | 700 | 0 | 1862 | 386 | 0 |
| PPIL6 | 0 | 0 | 0 | 0 | 692 | 0 | 304 | 0 |
| PPIP5K1 | 0 | 0 | 276 | 0 | 0 | 0 | 0 | 0 |
| PPIP5K2 | 0 | 480 | 0 | 0 | 0 | 0 | 0 | 0 |
| PPL | 0 | 0 | 0 | 452 | 1402 | 0 | 178 | 1050 |
| PPM1A | 0 | 0 | 1412 | 0 | 180 | 0 | 0 | 0 |
| PPM1B | 306 | 1054 | 1190 | 0 | 851 | 0 | 0 | 0 |
| PPM1D | 6781 | 7634 | 16786 | 3296 | 11174 | 7954 | 18695 | 3074 |
| PPM1F | 0 | 0 | 0 | 0 | 1274 | 0 | 0 | 0 |
| PPM1G | 712 | 1748 | 0 | 1925 | 2048 | 0 | 0 | 0 |
| PPM1H | 0 | 326 | 0 | 0 | 1408 | 0 | 2974 | 0 |
| PPM1K | 0 | 0 | 0 | 0 | 0 | 1608 | 0 | 0 |
| PPM1L | 0 | 0 | 0 | 0 | 0 | 0 | 0 | 481 |
| PPM1M | 0 | 0 | 0 | 0 | 390 | 0 | 0 | 0 |
| PPME1 | 0 | 0 | 2096 | 758 | 358 | 0 | 0 | 0 |
| PPOX | 0 | 0 | 0 | 0 | 0 | 0 | 550 | 0 |
| PPP1CA | 125812 | 183145 | 1175559 | 406357 | 870709 | 365698 | 486849 | 345448 |
| PPP1CB | 896 | 1432 | 0 | 860 | 2288 | 166 | 1118 | 1760 |
| PPP1CC | 528 | 686 | 5960 | 1495 | 2534 | 1382 | 1346 | 2722 |
| PPP1R10 | 698 | 0 | 3383 | 0 | 1200 | 986 | 206 | 0 |
| PPP1R11 | 0 | 0 | 2428 | 574 | 434 | 0 | 0 | 902 |
| PPP1R12B | 0 | 372 | 0 | 0 | 0 | 0 | 0 | 0 |
| PPP1R12C | 0 | 764 | 0 | 0 | 882 | 0 | 0 | 0 |
| PPP1R13B | 0 | 1632 | 1444 | 0 | 1070 | 0 | 0 | 120 |
| PPP1R14A | 0 | 0 | 1407 | 0 | 0 | 0 | 0 | 0 |
| PPP1R14B | 996 | 1052 | 1779 | 6116 | 8674 | 1060 | 2788 | 5919 |
| PPP1R14C | 0 | 400 | 0 | 1241 | 0 | 0 | 336 | 0 |
| PPP1R14D | 4888 | 1074 | 7544 | 1925 | 1700 | 0 | 0 | 1158 |
| PPP1R15A | 898 | 0 | 1010 | 0 | 480 | 0 | 1130 | 1149 |
| PPP1R15B | 0 | 192 | 0 | 0 | 0 | 1318 | 0 | 0 |
| PPP1R16A | 148 | 344 | 1678 | 645 | 1916 | 1420 | 1862 | 0 |
| PPP1R18 | 0 | 0 | 0 | 0 | 184 | 0 | 0 | 0 |
| PPP1R1B | 160200 | 332061 | 672316 | 211077 | 531590 | 204778 | 903725 | 126457 |
| PPP1R2 | 584 | 590 | 0 | 3248 | 2057 | 2019 | 2836 | 1866 |
| PPP1R21 | 0 | 0 | 0 | 0 | 464 | 0 | 286 | 0 |
| PPP1R26 | 0 | 0 | 0 | 0 | 0 | 0 | 310 | 0 |
| PPP1R27 | 0 | 0 | 122 | 0 | 0 | 0 | 0 | 0 |
| PPP1R32 | 0 | 858 | 0 | 2626 | 0 | 0 | 732 | 0 |
| PPP1R35 | 0 | 330 | 2999 | 2661 | 2066 | 0 | 360 | 882 |

| | | | | | | | | |
|---|---|---|---|---|---|---|---|---|
| PPP1R37 | 0 | 0 | 128 | 0 | 536 | 0 | 462 | 0 |
| PPP1R3B | 0 | 0 | 1460 | 0 | 0 | 0 | 406 | 0 |
| PPP1R3E | 0 | 260 | 0 | 0 | 0 | 0 | 0 | 0 |
| PPP1R3G | 0 | 302 | 0 | 0 | 668 | 0 | 690 | 0 |
| PPP1R7 | 0 | 2095 | 2354 | 1638 | 2016 | 0 | 1485 | 1187 |
| PPP1R9B | 0 | 356 | 0 | 0 | 0 | 0 | 504 | 0 |
| PPP2CA | 1486 | 84 | 2230 | 4312 | 2604 | 1860 | 352 | 3118 |
| PPP2CB | 0 | 564 | 2888 | 0 | 0 | 0 | 470 | 762 |
| PPP2R1A | 43919 | 137246 | 374388 | 247487 | 531602 | 407128 | 384919 | 163524 |
| PPP2R1B | 0 | 0 | 2041 | 0 | 0 | 1036 | 0 | 0 |
| PPP2R2A | 20931 | 26379 | 21602 | 27423 | 99572 | 41245 | 38590 | 24278 |
| PPP2R2C | 0 | 0 | 0 | 0 | 0 | 0 | 538 | 0 |
| PPP2R2D | 0 | 366 | 0 | 0 | 738 | 32 | 0 | 458 |
| PPP2R3A | 0 | 0 | 0 | 0 | 138 | 0 | 266 | 0 |
| PPP2R3B | 0 | 0 | 0 | 0 | 256 | 0 | 0 | 0 |
| PPP2R3C | 0 | 418 | 0 | 0 | 674 | 0 | 0 | 0 |
| PPP2R5A | 0 | 1046 | 0 | 0 | 202 | 1398 | 0 | 0 |
| PPP2R5B | 0 | 0 | 0 | 0 | 246 | 0 | 0 | 0 |
| PPP2R5C | 0 | 494 | 0 | 0 | 1412 | 0 | 0 | 0 |
| PPP2R5D | 0 | 206 | 0 | 0 | 740 | 1168 | 0 | 0 |
| PPP3CA | 0 | 0 | 0 | 0 | 1092 | 144 | 0 | 0 |
| PPP3CC | 0 | 0 | 0 | 0 | 0 | 836 | 0 | 0 |
| PPP3R1 | 0 | 242 | 0 | 0 | 218 | 0 | 0 | 0 |
| PPP4C | 0 | 194 | 8224 | 3824 | 1440 | 0 | 1356 | 1100 |
| PPP4R1 | 8122 | 14756 | 19704 | 8040 | 27347 | 2463 | 9318 | 9857 |
| PPP4R1L | 0 | 0 | 0 | 0 | 0 | 0 | 256 | 0 |
| PPP4R2 | 162 | 0 | 0 | 0 | 723 | 220 | 0 | 0 |
| PPP4R3A | 1608 | 1396 | 3339 | 1924 | 4358 | 0 | 704 | 1558 |
| PPP4R3B | 0 | 0 | 1440 | 1100 | 1311 | 1916 | 1380 | 0 |
| PPP5C | 0 | 568 | 0 | 816 | 922 | 0 | 234 | 2302 |
| PPP6C | 60589 | 96942 | 100527 | 94009 | 246612 | 112410 | 210315 | 50840 |
| PPP6R1 | 0 | 0 | 0 | 0 | 0 | 1326 | 0 | 0 |
| PPP6R2 | 0 | 380 | 0 | 0 | 424 | 0 | 0 | 1188 |
| PPP6R3 | 0 | 0 | 0 | 1034 | 258 | 0 | 444 | 704 |
| PPRC1 | 0 | 0 | 0 | 0 | 0 | 0 | 518 | 0 |
| PPT1 | 414 | 0 | 1832 | 0 | 736 | 0 | 322 | 2368 |
| PPT2-EGFL8 | 0 | 0 | 0 | 0 | 779 | 0 | 0 | 0 |
| PPWD1 | 22320 | 60376 | 55001 | 71905 | 119342 | 77722 | 64591 | 41707 |
| PPY | 385 | 12915 | 16 | 0 | 572 | 0 | 0 | 0 |
| PQBP1 | 0 | 0 | 1822 | 4729 | 2622 | 0 | 0 | 0 |
| PQLC1 | 666 | 360 | 2681 | 716 | 952 | 0 | 0 | 0 |
| PQLC3 | 698 | 368 | 0 | 0 | 986 | 0 | 0 | 0 |
| PRAC1 | 1118 | 1634 | 50 | 0 | 6992 | 0 | 0 | 282 |

| | | | | | | | | |
|---|---|---|---|---|---|---|---|---|
| PRAC2 | 0 | 364 | 956 | 0 | 0 | 0 | 0 | 0 |
| PRADC1 | 24 | 306 | 4051 | 0 | 1604 | 0 | 0 | 0 |
| PRAF2 | 0 | 0 | 0 | 0 | 968 | 0 | 0 | 0 |
| PRAP1 | 7340 | 74 | 22720 | 4404 | 3673 | 4715 | 4174 | 3965 |
| PRC1 | 2324 | 11066 | 8223 | 48203 | 123471 | 13385 | 48364 | 41059 |
| PRCC | 4778 | 16385 | 35753 | 27238 | 53313 | 23266 | 35850 | 18375 |
| PRCP | 0 | 0 | 586 | 1252 | 0 | 0 | 424 | 16 |
| PRDM1 | 2532 | 842 | 1176 | 66 | 1540 | 90 | 158 | 1298 |
| PRDM10 | 0 | 0 | 0 | 607 | 0 | 0 | 0 | 0 |
| PRDM11 | 706 | 0 | 0 | 0 | 0 | 0 | 0 | 0 |
| PRDM16 | 0 | 0 | 0 | 0 | 468 | 0 | 0 | 0 |
| PRDM2 | 1354 | 4282 | 5419 | 10534 | 24084 | 12899 | 10764 | 5737 |
| PRDX1 | 3415 | 3242 | 14646 | 23426 | 24470 | 8709 | 13595 | 6380 |
| PRDX2 | 795 | 3488 | 7153 | 18433 | 17002 | 2600 | 4954 | 1454 |
| PRDX3 | 366 | 2106 | 3418 | 5444 | 5885 | 0 | 4654 | 5096 |
| PRDX4 | 769 | 2042 | 1850 | 9629 | 15616 | 6259 | 1884 | 5771 |
| PRDX5 | 6642 | 12884 | 31926 | 64329 | 29852 | 10507 | 46310 | 13808 |
| PRDX6 | 4304 | 7137 | 28588 | 3393 | 14724 | 4613 | 3434 | 5364 |
| PREB | 0 | 450 | 0 | 0 | 768 | 2727 | 0 | 0 |
| PRELID1 | 9705 | 5292 | 17427 | 9547 | 2802 | 15262 | 12643 | 2728 |
| PRELID2 | 710 | 680 | 690 | 1743 | 4453 | 0 | 1202 | 954 |
| PRELID3A | 0 | 0 | 0 | 0 | 0 | 1526 | 0 | 0 |
| PRELID3B | 0 | 380 | 4048 | 4727 | 12316 | 1798 | 977 | 2815 |
| PRELP | 422 | 1512 | 3968 | 1186 | 2236 | 20746 | 1198 | 4502 |
| PREP | 19723 | 54777 | 49904 | 55053 | 171364 | 33409 | 70102 | 45233 |
| PREPL | 26 | 454 | 1448 | 0 | 512 | 0 | 662 | 0 |
| PREX1 | 548 | 0 | 0 | 0 | 924 | 0 | 0 | 0 |
| PRF1 | 0 | 0 | 0 | 0 | 0 | 1532 | 0 | 0 |
| PRH1-PRR4 | 0 | 0 | 0 | 0 | 0 | 332 | 0 | 0 |
| PRICKLE2 | 398 | 0 | 0 | 0 | 26 | 0 | 72 | 0 |
| PRICKLE3 | 0 | 0 | 0 | 0 | 220 | 0 | 0 | 0 |
| PRIM1 | 0 | 0 | 0 | 1925 | 1218 | 0 | 0 | 1490 |
| PRIMPOL | 0 | 0 | 664 | 0 | 0 | 0 | 0 | 0 |
| PRKAA1 | 0 | 770 | 0 | 0 | 704 | 0 | 0 | 0 |
| PRKAB1 | 0 | 552 | 3408 | 0 | 424 | 0 | 1028 | 0 |
| PRKAB2 | 456 | 0 | 0 | 0 | 0 | 0 | 0 | 0 |
| PRKACA | 0 | 406 | 0 | 0 | 44 | 0 | 0 | 0 |
| PRKACB | 574 | 718 | 225 | 0 | 0 | 0 | 0 | 0 |
| PRKAG1 | 1070 | 340 | 5405 | 0 | 1330 | 0 | 0 | 0 |
| PRKAG2 | 165 | 832 | 0 | 0 | 620 | 0 | 0 | 0 |
| PRKAG2-AS1 | 0 | 890 | 0 | 0 | 1340 | 0 | 0 | 0 |
| PRKAR1A | 12973 | 12167 | 26771 | 21706 | 24649 | 59425 | 20891 | 16147 |
| PRKAR1B | 0 | 0 | 658 | 0 | 0 | 0 | 0 | 0 |

| | | | | | | | | |
|---|---|---|---|---|---|---|---|---|
| PRKAR2A | 0 | 0 | 0 | 0 | 1098 | 0 | 0 | 0 |
| PRKAR2A-AS1 | 0 | 0 | 0 | 0 | 496 | 0 | 0 | 0 |
| PRKAR2B | 2066 | 1274 | 3526 | 416 | 0 | 890 | 0 | 0 |
| PRKCA | 0 | 480 | 0 | 0 | 0 | 0 | 0 | 0 |
| PRKCD | 40759 | 40880 | 100007 | 23732 | 117800 | 45836 | 53215 | 13551 |
| PRKCH | 4234 | 7744 | 10138 | 1524 | 20115 | 11884 | 14260 | 9246 |
| PRKCI | 0 | 0 | 0 | 724 | 694 | 0 | 250 | 0 |
| PRKCQ | 0 | 0 | 0 | 1184 | 0 | 0 | 0 | 0 |
| PRKCQ-AS1 | 0 | 302 | 0 | 0 | 0 | 0 | 0 | 0 |
| PRKCSH | 672 | 314 | 1836 | 2350 | 1236 | 1534 | 176 | 2746 |
| PRKCZ | 18826 | 11098 | 42904 | 38371 | 41727 | 19485 | 42956 | 10151 |
| PRKD2 | 262 | 0 | 1244 | 0 | 598 | 0 | 150 | 0 |
| PRKDC | 236 | 184 | 0 | 0 | 658 | 0 | 0 | 744 |
| PRKN | 0 | 1134 | 1728 | 0 | 2778 | 1416 | 964 | 0 |
| PRKRA | 0 | 1706 | 0 | 946 | 2526 | 0 | 730 | 0 |
| PRKRA-AS1 | 534 | 0 | 0 | 0 | 0 | 0 | 0 | 0 |
| PRKRIP1 | 0 | 518 | 0 | 0 | 2028 | 0 | 278 | 0 |
| PRLR | 0 | 0 | 138 | 0 | 0 | 0 | 0 | 0 |
| PRMT1 | 0 | 878 | 2576 | 2952 | 7035 | 0 | 0 | 940 |
| PRMT2 | 788 | 758 | 3825 | 1294 | 2312 | 0 | 282 | 1740 |
| PRMT3 | 0 | 258 | 0 | 0 | 0 | 0 | 0 | 614 |
| PRMT5 | 0 | 0 | 0 | 0 | 706 | 0 | 0 | 614 |
| PRMT6 | 0 | 0 | 0 | 0 | 0 | 0 | 322 | 0 |
| PRMT7 | 640 | 218 | 0 | 0 | 728 | 0 | 0 | 0 |
| PRMT9 | 1040 | 0 | 0 | 1394 | 866 | 0 | 3486 | 0 |
| PRNP | 0 | 248 | 0 | 0 | 1164 | 0 | 0 | 0 |
| PROCA1 | 0 | 0 | 0 | 0 | 0 | 0 | 0 | 726 |
| PROCR | 0 | 118 | 0 | 3552 | 1937 | 2016 | 510 | 891 |
| PRODH | 0 | 0 | 0 | 0 | 360 | 0 | 0 | 0 |
| PROM1 | 0 | 0 | 0 | 0 | 310 | 0 | 0 | 0 |
| PROS1 | 322 | 6952 | 3311 | 5054 | 7823 | 1399 | 7743 | 3082 |
| PROSER1 | 0 | 0 | 0 | 0 | 292 | 1722 | 0 | 0 |
| PROSER2 | 0 | 0 | 0 | 0 | 202 | 0 | 0 | 0 |
| PRPF18 | 500 | 1308 | 0 | 788 | 0 | 0 | 770 | 0 |
| PRPF19 | 0 | 322 | 2171 | 1188 | 1037 | 0 | 0 | 0 |
| PRPF3 | 0 | 364 | 80 | 642 | 638 | 0 | 0 | 1220 |
| PRPF31 | 0 | 0 | 0 | 1583 | 0 | 0 | 0 | 0 |
| PRPF38A | 0 | 484 | 0 | 400 | 818 | 5377 | 1221 | 0 |
| PRPF39 | 0 | 0 | 402 | 0 | 236 | 0 | 0 | 0 |
| PRPF4 | 0 | 0 | 0 | 0 | 372 | 0 | 0 | 0 |
| PRPF40A | 0 | 0 | 0 | 624 | 1116 | 0 | 0 | 0 |
| PRPF40B | 0 | 0 | 0 | 0 | 192 | 0 | 0 | 0 |
| PRPF4B | 0 | 0 | 96 | 0 | 468 | 0 | 0 | 0 |

| | | | | | | | | |
|---|---|---|---|---|---|---|---|---|
| PRPF6 | 0 | 0 | 3362 | 1702 | 1246 | 0 | 2012 | 0 |
| PRPF8 | 500 | 368 | 0 | 0 | 1543 | 0 | 626 | 0 |
| PRPS1 | 0 | 0 | 0 | 0 | 844 | 617 | 0 | 444 |
| PRPS2 | 0 | 320 | 1231 | 496 | 1998 | 740 | 0 | 534 |
| PRPSAP1 | 0 | 0 | 0 | 1244 | 418 | 0 | 0 | 0 |
| PRPSAP2 | 434 | 320 | 2502 | 1334 | 498 | 0 | 252 | 0 |
| PRR11 | 2304 | 6654 | 0 | 19250 | 35664 | 16856 | 21072 | 16969 |
| PRR12 | 0 | 440 | 0 | 0 | 1482 | 0 | 100 | 2122 |
| PRR13 | 4005 | 1916 | 20595 | 4761 | 768 | 4743 | 2953 | 2220 |
| PRR14L | 0 | 0 | 2306 | 0 | 0 | 0 | 0 | 0 |
| PRR15 | 1084 | 626 | 0 | 3007 | 1861 | 0 | 1946 | 435 |
| PRR15L | 1140 | 430 | 1496 | 3232 | 5982 | 2169 | 1976 | 1226 |
| PRR22 | 0 | 164 | 0 | 0 | 0 | 0 | 0 | 0 |
| PRR26 | 0 | 0 | 0 | 0 | 814 | 0 | 0 | 0 |
| PRR3 | 464 | 0 | 0 | 0 | 0 | 0 | 0 | 0 |
| PRR34-AS1 | 0 | 0 | 0 | 0 | 192 | 0 | 0 | 0 |
| PRR5 | 572 | 0 | 544 | 0 | 0 | 0 | 302 | 0 |
| PRR5L | 0 | 124 | 832 | 0 | 0 | 0 | 0 | 275 |
| PRR7 | 0 | 1048 | 0 | 0 | 0 | 0 | 0 | 0 |
| PRRC1 | 343 | 362 | 0 | 0 | 0 | 0 | 0 | 0 |
| PRRC2A | 0 | 0 | 0 | 0 | 734 | 0 | 72 | 0 |
| PRRC2B | 0 | 0 | 0 | 0 | 16 | 0 | 0 | 0 |
| PRRC2C | 0 | 0 | 0 | 0 | 1118 | 0 | 0 | 0 |
| PRRG2 | 3368 | 4542 | 32042 | 5840 | 15474 | 6176 | 9793 | 0 |
| PRRG4 | 0 | 0 | 0 | 0 | 0 | 0 | 726 | 0 |
| PRRT3-AS1 | 0 | 0 | 0 | 833 | 0 | 0 | 0 | 0 |
| PRRX1 | 0 | 0 | 0 | 0 | 0 | 502 | 0 | 0 |
| PRRX2 | 0 | 0 | 0 | 0 | 993 | 0 | 0 | 0 |
| PRSS1 | 0 | 0 | 0 | 0 | 718 | 0 | 0 | 1086 |
| PRSS12 | 0 | 534 | 0 | 0 | 0 | 0 | 0 | 0 |
| PRSS16 | 648 | 0 | 0 | 0 | 570 | 0 | 0 | 0 |
| PRSS2 | 0 | 0 | 0 | 0 | 2024 | 0 | 0 | 7009 |
| PRSS22 | 0 | 276 | 0 | 0 | 442 | 0 | 366 | 0 |
| PRSS23 | 872 | 424 | 1550 | 0 | 3378 | 3094 | 3336 | 52 |
| PRSS3 | 562 | 3292 | 11464 | 0 | 5487 | 3497 | 2300 | 1903 |
| PRSS33 | 0 | 0 | 0 | 0 | 0 | 1816 | 0 | 0 |
| PRSS8 | 2200 | 1728 | 11954 | 4634 | 706 | 0 | 1084 | 1222 |
| PRTFDC1 | 0 | 0 | 0 | 1300 | 1388 | 0 | 0 | 0 |
| PRUNE1 | 0 | 376 | 0 | 0 | 675 | 0 | 250 | 0 |
| PRXL2A | 0 | 0 | 0 | 0 | 2190 | 0 | 1194 | 0 |
| PRXL2C | 0 | 434 | 0 | 0 | 0 | 0 | 0 | 0 |
| PSAP | 138665 | 183459 | 745010 | 249172 | 526630 | 573543 | 772166 | 351570 |
| PSAT1 | 0 | 0 | 0 | 1328 | 3121 | 0 | 306 | 0 |

| | | | | | | | | |
|---|---|---|---|---|---|---|---|---|
| PSD3 | 2080 | 428 | 446 | 0 | 2522 | 0 | 1068 | 446 |
| PSD4 | 0 | 178 | 626 | 0 | 572 | 0 | 0 | 0 |
| PSEN1 | 0 | 0 | 0 | 0 | 588 | 0 | 0 | 0 |
| PSEN2 | 0 | 36 | 0 | 0 | 276 | 0 | 0 | 650 |
| PSENEN | 3702 | 3752 | 7375 | 739 | 5452 | 3000 | 1388 | 2143 |
| PSIP1 | 96064 | 111438 | 58320 | 115974 | 316842 | 104969 | 313418 | 91375 |
| PSKH1 | 0 | 0 | 991 | 0 | 524 | 0 | 0 | 0 |
| PSMA1 | 3316 | 2880 | 7361 | 4168 | 2786 | 9169 | 2524 | 11779 |
| PSMA2 | 600 | 4431 | 6974 | 15540 | 29815 | 13673 | 8658 | 5189 |
| PSMA3 | 1334 | 1562 | 5698 | 3875 | 12189 | 2672 | 3344 | 2012 |
| PSMA3-AS1 | 828 | 416 | 0 | 0 | 808 | 2370 | 410 | 1616 |
| PSMA4 | 3230 | 946 | 9353 | 4683 | 4917 | 0 | 6521 | 1190 |
| PSMA5 | 283480 | 502673 | 477052 | 818316 | 1358069 | 684934 | 1555554 | 496355 |
| PSMA6 | 4746 | 2594 | 6992 | 1214 | 16212 | 156 | 2612 | 2784 |
| PSMA7 | 388 | 706 | 15211 | 32398 | 18263 | 3867 | 7342 | 7666 |
| PSMB1 | 3578 | 4580 | 4440 | 8319 | 17704 | 4208 | 3253 | 8234 |
| PSMB10 | 786 | 2782 | 1832 | 4977 | 5799 | 0 | 1556 | 0 |
| PSMB2 | 1169 | 1490 | 0 | 5890 | 4678 | 7192 | 812 | 1908 |
| PSMB3 | 624096 | 828595 | 2455992 | 1356045 | 2548249 | 1346219 | 2611317 | 764806 |
| PSMB4 | 2174 | 918 | 4644 | 3614 | 1962 | 2804 | 544 | 3034 |
| PSMB5 | 0 | 445 | 3812 | 4260 | 6821 | 1980 | 4673 | 864 |
| PSMB6 | 0 | 1928 | 7497 | 1142 | 4618 | 4004 | 470 | 2626 |
| PSMB7 | 1542 | 994 | 6775 | 6431 | 9676 | 0 | 5859 | 2024 |
| PSMB8 | 1712 | 3656 | 17062 | 4427 | 4880 | 0 | 6900 | 5855 |
| PSMB8-AS1 | 0 | 0 | 762 | 0 | 0 | 0 | 0 | 0 |
| PSMB9 | 0 | 212 | 4444 | 2865 | 170 | 5525 | 1844 | 4879 |
| PSMC1 | 0 | 1288 | 4913 | 1268 | 8350 | 0 | 1032 | 4372 |
| PSMC2 | 0 | 1092 | 0 | 1654 | 2222 | 0 | 1822 | 0 |
| PSMC3 | 176 | 1836 | 1998 | 4017 | 2773 | 3652 | 3372 | 480 |
| PSMC3IP | 0 | 0 | 0 | 914 | 0 | 0 | 0 | 0 |
| PSMC4 | 402 | 908 | 1719 | 3646 | 3074 | 1943 | 488 | 1165 |
| PSMC5 | 0 | 635 | 5966 | 3200 | 5468 | 304 | 616 | 1258 |
| PSMC6 | 0 | 932 | 3505 | 4404 | 9948 | 0 | 300 | 1376 |
| PSMD1 | 376 | 304 | 0 | 0 | 1462 | 0 | 334 | 0 |
| PSMD10 | 0 | 766 | 0 | 0 | 2986 | 0 | 340 | 0 |
| PSMD11 | 156105 | 220525 | 636261 | 293967 | 968818 | 311838 | 339618 | 358866 |
| PSMD12 | 7928 | 17192 | 27608 | 25937 | 44747 | 49048 | 55349 | 21321 |
| PSMD13 | 2972 | 958 | 0 | 2000 | 4037 | 2277 | 1205 | 2924 |
| PSMD14 | 1834 | 794 | 0 | 3342 | 7639 | 0 | 0 | 0 |
| PSMD2 | 0 | 0 | 2987 | 1612 | 1098 | 0 | 0 | 1222 |
| PSMD3 | 17395 | 18085 | 55904 | 43631 | 83025 | 24650 | 68843 | 26786 |
| PSMD4 | 0 | 546 | 5907 | 4127 | 9924 | 6995 | 3843 | 5983 |
| PSMD5 | 0 | 0 | 0 | 0 | 0 | 0 | 534 | 0 |

| | | | | | | | | |
|---|---|---|---|---|---|---|---|---|
| PSMD6 | 1288 | 0 | 1258 | 6567 | 7718 | 0 | 1004 | 680 |
| PSMD7 | 0 | 2074 | 2356 | 1505 | 4678 | 4180 | 1134 | 1302 |
| PSMD8 | 2051 | 2906 | 1836 | 2516 | 13671 | 4698 | 210 | 1966 |
| PSMD9 | 104015 | 208954 | 360307 | 215310 | 447568 | 202663 | 276109 | 129066 |
| PSME1 | 10777 | 4122 | 17526 | 8398 | 19044 | 11622 | 6913 | 6830 |
| PSME2 | 5318 | 4598 | 13773 | 7234 | 14542 | 14458 | 2364 | 2849 |
| PSME3 | 304 | 0 | 0 | 0 | 2300 | 0 | 544 | 0 |
| PSME4 | 0 | 0 | 0 | 0 | 208 | 0 | 0 | 0 |
| PSMF1 | 0 | 26 | 0 | 1326 | 3174 | 0 | 528 | 918 |
| PSMG1 | 0 | 0 | 0 | 0 | 2289 | 0 | 30 | 1156 |
| PSMG2 | 1058 | 840 | 4593 | 1392 | 3040 | 5048 | 666 | 0 |
| PSMG3 | 0 | 0 | 0 | 1072 | 3580 | 0 | 2857 | 0 |
| PSMG4 | 694 | 1186 | 0 | 0 | 2412 | 1366 | 1957 | 750 |
| PSORS1C1 | 0 | 340 | 0 | 1500 | 248 | 0 | 0 | 0 |
| PSORS1C3 | 0 | 0 | 0 | 0 | 0 | 0 | 0 | 442 |
| PSPC1 | 0 | 402 | 1820 | 1101 | 1028 | 2676 | 1381 | 0 |
| PSPH | 0 | 0 | 0 | 996 | 544 | 1717 | 436 | 0 |
| PSRC1 | 0 | 0 | 0 | 0 | 0 | 0 | 0 | 690 |
| PSTK | 0 | 0 | 0 | 0 | 763 | 1300 | 0 | 582 |
| PSTPIP1 | 0 | 456 | 0 | 0 | 2067 | 0 | 0 | 0 |
| PSTPIP2 | 0 | 0 | 0 | 0 | 0 | 1560 | 0 | 0 |
| PTAFR | 0 | 0 | 0 | 0 | 0 | 0 | 0 | 780 |
| PTAR1 | 0 | 344 | 0 | 0 | 0 | 0 | 0 | 0 |
| PTBP1 | 660 | 408 | 0 | 352 | 2492 | 0 | 1348 | 972 |
| PTBP3 | 0 | 0 | 0 | 0 | 0 | 0 | 0 | 1284 |
| PTCD2 | 0 | 336 | 0 | 0 | 1198 | 0 | 0 | 0 |
| PTCD3 | 698 | 106 | 0 | 2111 | 841 | 576 | 288 | 0 |
| PTCH1 | 582 | 0 | 56 | 1446 | 1886 | 1932 | 1856 | 172 |
| PTDSS1 | 0 | 0 | 0 | 844 | 1242 | 1317 | 22 | 1120 |
| PTEN | 240458 | 289552 | 388276 | 286965 | 235915 | 584036 | 942011 | 109374 |
| PTENP1 | 0 | 708 | 0 | 0 | 580 | 0 | 0 | 0 |
| PTER | 116 | 0 | 0 | 0 | 0 | 0 | 826 | 0 |
| PTGDS | 36 | 244 | 0 | 0 | 0 | 0 | 0 | 292 |
| PTGER3 | 598 | 0 | 0 | 0 | 0 | 0 | 0 | 0 |
| PTGER4 | 0 | 318 | 0 | 0 | 328 | 0 | 0 | 0 |
| PTGES2 | 0 | 898 | 954 | 2222 | 858 | 0 | 1096 | 0 |
| PTGES2-AS1 | 0 | 0 | 0 | 0 | 0 | 0 | 0 | 382 |
| PTGES3 | 144623 | 238535 | 1126278 | 652892 | 663023 | 530874 | 810930 | 276356 |
| PTGES3L | 274 | 0 | 0 | 1038 | 0 | 0 | 454 | 0 |
| PTGES3L-AARSD1 | 3248 | 5128 | 22333 | 20173 | 8101 | 30263 | 47254 | 10107 |
| PTGIR | 0 | 0 | 0 | 0 | 0 | 0 | 380 | 0 |
| PTGR1 | 0 | 398 | 5018 | 0 | 2182 | 0 | 894 | 0 |
| PTGR2 | 0 | 0 | 1620 | 0 | 554 | 0 | 0 | 0 |

| | | | | | | | | |
|---|---|---|---|---|---|---|---|---|
| PTGS2 | 0 | 0 | 792 | 586 | 0 | 1601 | 0 | 0 |
| PTHLH | 0 | 0 | 0 | 0 | 0 | 1602 | 0 | 628 |
| PTK2 | 0 | 1098 | 0 | 0 | 160 | 0 | 0 | 0 |
| PTK2B | 0 | 0 | 298 | 0 | 244 | 0 | 0 | 0 |
| PTK6 | 0 | 0 | 0 | 0 | 948 | 0 | 408 | 412 |
| PTK7 | 0 | 0 | 0 | 0 | 502 | 0 | 302 | 0 |
| PTMA | 6434 | 10167 | 26144 | 36139 | 84241 | 14767 | 13910 | 26455 |
| PTMS | 208 | 1994 | 0 | 558 | 2734 | 1367 | 390 | 0 |
| PTN | 0 | 304 | 0 | 0 | 0 | 0 | 0 | 0 |
| PTOV1 | 0 | 0 | 0 | 1774 | 2193 | 0 | 0 | 0 |
| PTP4A1 | 662 | 292 | 2609 | 0 | 384 | 0 | 390 | 0 |
| PTP4A2 | 790 | 974 | 8280 | 2304 | 3428 | 0 | 0 | 1556 |
| PTP4A3 | 486 | 0 | 0 | 1899 | 0 | 208 | 8355 | 0 |
| PTPA | 552 | 0 | 1062 | 338 | 1310 | 0 | 1044 | 0 |
| PTPMT1 | 0 | 1168 | 0 | 0 | 5008 | 2024 | 1348 | 468 |
| PTPN11 | 30993 | 47777 | 87810 | 97965 | 184346 | 61759 | 185099 | 55394 |
| PTPN12 | 0 | 0 | 0 | 0 | 520 | 996 | 0 | 0 |
| PTPN13 | 784 | 562 | 0 | 5176 | 4536 | 5515 | 0 | 0 |
| PTPN14 | 0 | 0 | 0 | 0 | 0 | 0 | 0 | 394 |
| PTPN18 | 0 | 450 | 0 | 911 | 1734 | 844 | 642 | 0 |
| PTPN2 | 0 | 2118 | 1791 | 0 | 1450 | 0 | 0 | 1094 |
| PTPN23 | 0 | 554 | 0 | 0 | 0 | 0 | 0 | 0 |
| PTPN3 | 0 | 182 | 0 | 0 | 0 | 0 | 0 | 0 |
| PTPN6 | 48606 | 51229 | 139283 | 40045 | 74942 | 69159 | 139587 | 46905 |
| PTPN7 | 0 | 0 | 0 | 0 | 1106 | 0 | 0 | 0 |
| PTPRA | 2152 | 478 | 0 | 565 | 306 | 0 | 0 | 0 |
| PTPRB | 294 | 1118 | 1857 | 2807 | 6117 | 12161 | 10643 | 732 |
| PTPRC | 0 | 0 | 0 | 1464 | 0 | 0 | 0 | 0 |
| PTPRCAP | 2390 | 0 | 0 | 0 | 0 | 1715 | 366 | 812 |
| PTPRD | 10996 | 1508 | 3722 | 3314 | 0 | 0 | 966 | 0 |
| PTPRE | 0 | 0 | 0 | 0 | 362 | 0 | 362 | 0 |
| PTPRF | 0 | 608 | 0 | 0 | 0 | 0 | 998 | 0 |
| PTPRG | 686 | 550 | 1542 | 841 | 3014 | 2974 | 1322 | 1104 |
| PTPRG-AS1 | 0 | 0 | 0 | 0 | 0 | 0 | 214 | 0 |
| PTPRH | 656 | 0 | 0 | 0 | 0 | 0 | 0 | 0 |
| PTPRJ | 0 | 801 | 2109 | 0 | 456 | 0 | 0 | 444 |
| PTPRK | 137881 | 171103 | 256150 | 120461 | 478726 | 56928 | 324722 | 191684 |
| PTPRM | 370 | 0 | 0 | 0 | 0 | 0 | 0 | 0 |
| PTPRN2 | 0 | 0 | 0 | 0 | 502 | 1501 | 722 | 776 |
| PTPRO | 0 | 424 | 0 | 0 | 0 | 0 | 0 | 0 |
| PTPRR | 127079 | 7996 | 579702 | 1858 | 97662 | 20641 | 5250 | 10793 |
| PTPRZ1 | 0 | 0 | 0 | 0 | 0 | 0 | 750 | 0 |
| PTRH1 | 0 | 0 | 0 | 0 | 1316 | 0 | 854 | 0 |

| | | | | | | | | |
|---|---|---|---|---|---|---|---|---|
| PTRH2 | 59005 | 190361 | 119328 | 149225 | 268525 | 174839 | 297810 | 138841 |
| PTRHD1 | 0 | 1498 | 1308 | 1040 | 2214 | 0 | 588 | 866 |
| PTS | 518 | 1362 | 1140 | 5827 | 1235 | 0 | 198 | 1131 |
| PTTG1 | 0 | 0 | 1362 | 7184 | 7018 | 2106 | 1822 | 3418 |
| PTTG1IP | 0 | 384 | 6237 | 1631 | 3292 | 433 | 320 | 636 |
| PUDP | 0 | 0 | 1432 | 0 | 728 | 1934 | 342 | 0 |
| PUF60 | 1190 | 442 | 0 | 3354 | 4084 | 2832 | 3678 | 4655 |
| PUM1 | 0 | 0 | 0 | 1080 | 416 | 0 | 0 | 0 |
| PUM2 | 0 | 0 | 0 | 0 | 236 | 0 | 430 | 0 |
| PUM3 | 0 | 0 | 0 | 2512 | 354 | 0 | 1049 | 0 |
| PURA | 0 | 0 | 0 | 0 | 0 | 0 | 316 | 0 |
| PUS1 | 0 | 274 | 0 | 416 | 0 | 0 | 0 | 0 |
| PUS10 | 0 | 0 | 0 | 1087 | 1076 | 0 | 0 | 0 |
| PUS3 | 0 | 0 | 0 | 914 | 0 | 0 | 318 | 0 |
| PUS7 | 0 | 594 | 974 | 662 | 8412 | 5599 | 6579 | 1390 |
| PUS7L | 0 | 936 | 0 | 0 | 0 | 0 | 40 | 0 |
| PUSL1 | 0 | 508 | 0 | 0 | 296 | 0 | 0 | 0 |
| PVALB | 1106 | 794 | 0 | 0 | 0 | 1050 | 0 | 0 |
| PVR | 0 | 270 | 1586 | 1326 | 2003 | 0 | 272 | 1098 |
| PVT1 | 686 | 0 | 0 | 0 | 848 | 0 | 466 | 0 |
| PWP1 | 0 | 414 | 0 | 894 | 0 | 0 | 0 | 0 |
| PWWP2A | 0 | 0 | 927 | 0 | 0 | 0 | 0 | 0 |
| PWWP2B | 1408 | 0 | 0 | 0 | 0 | 0 | 82 | 0 |
| PWWP3A | 520 | 616 | 0 | 398 | 560 | 0 | 540 | 0 |
| PXDC1 | 0 | 474 | 0 | 0 | 614 | 0 | 948 | 0 |
| PXDN | 0 | 0 | 0 | 0 | 0 | 0 | 1053 | 0 |
| PXDNL | 0 | 0 | 0 | 0 | 0 | 0 | 172 | 0 |
| PXK | 0 | 0 | 1326 | 0 | 266 | 0 | 0 | 0 |
| PXMP2 | 1190 | 3034 | 286 | 2734 | 0 | 0 | 0 | 0 |
| PXMP4 | 0 | 0 | 0 | 0 | 0 | 2043 | 0 | 1624 |
| PXN | 0 | 772 | 0 | 992 | 4018 | 0 | 2160 | 848 |
| PXYLP1 | 2650 | 8660 | 2671 | 6818 | 27593 | 9613 | 9114 | 5068 |
| PYCARD | 288 | 1256 | 2668 | 3030 | 1212 | 0 | 0 | 1084 |
| PYCR1 | 0 | 0 | 0 | 1592 | 782 | 0 | 400 | 528 |
| PYCR2 | 1273 | 0 | 0 | 2120 | 1162 | 0 | 554 | 1155 |
| PYCR3 | 696 | 0 | 0 | 0 | 56 | 0 | 0 | 0 |
| PYGB | 8888 | 14643 | 29063 | 6628 | 39043 | 9939 | 72847 | 6726 |
| PYGL | 0 | 0 | 0 | 0 | 144 | 0 | 0 | 0 |
| PYGO2 | 152 | 0 | 1468 | 0 | 0 | 0 | 512 | 617 |
| PYM1 | 384 | 474 | 2876 | 0 | 378 | 1290 | 910 | 1926 |
| PYROXD1 | 0 | 0 | 0 | 0 | 662 | 0 | 0 | 0 |
| PYURF | 1486 | 1998 | 9487 | 4724 | 11494 | 0 | 1906 | 2194 |
| PYY | 0 | 6560 | 909 | 0 | 486 | 0 | 0 | 0 |

| | | | | | | | | |
|---|---|---|---|---|---|---|---|---|
| QARS | 1658 | 1262 | 0 | 0 | 1406 | 1311 | 2115 | 836 |
| QDPR | 0 | 328 | 0 | 2470 | 4821 | 0 | 2416 | 0 |
| QKI | 4388 | 4103 | 13071 | 16932 | 18164 | 24221 | 30773 | 14333 |
| QPCT | 0 | 0 | 1528 | 0 | 434 | 1138 | 1712 | 1124 |
| QPCTL | 0 | 0 | 0 | 0 | 930 | 0 | 0 | 0 |
| QPRT | 0 | 170 | 662 | 3325 | 602 | 0 | 3751 | 716 |
| QRICH1 | 9898 | 9249 | 19928 | 20170 | 57066 | 22648 | 30960 | 11835 |
| QRSL1 | 29174 | 59612 | 38284 | 54969 | 214304 | 82982 | 163238 | 57075 |
| QSOX1 | 724 | 0 | 430 | 0 | 1699 | 0 | 208 | 1558 |
| QSOX2 | 0 | 0 | 0 | 0 | 0 | 0 | 582 | 0 |
| QTRT1 | 12535 | 20693 | 44260 | 28593 | 86878 | 29931 | 19208 | 17863 |
| QTRT2 | 0 | 0 | 0 | 0 | 0 | 0 | 14 | 0 |
| R3HCC1 | 0 | 0 | 3420 | 0 | 0 | 0 | 0 | 0 |
| R3HCC1L | 0 | 0 | 0 | 0 | 0 | 0 | 66 | 976 |
| R3HDM1 | 0 | 0 | 0 | 0 | 871 | 0 | 0 | 0 |
| R3HDM2 | 1390 | 2804 | 1448 | 4661 | 4971 | 3491 | 15688 | 536 |
| R3HDM4 | 0 | 498 | 0 | 1957 | 1278 | 1822 | 816 | 1158 |
| RAB10 | 910 | 1452 | 0 | 986 | 979 | 0 | 652 | 0 |
| RAB11A | 500 | 158 | 2062 | 1448 | 642 | 0 | 1086 | 160 |
| RAB11B | 1490 | 424 | 2885 | 0 | 3387 | 0 | 0 | 1342 |
| RAB11B-AS1 | 0 | 0 | 0 | 0 | 0 | 1739 | 0 | 0 |
| RAB11FIP1 | 0 | 518 | 270 | 0 | 330 | 0 | 145 | 198 |
| RAB13 | 0 | 1076 | 402 | 2748 | 2180 | 2885 | 0 | 1290 |
| RAB14 | 830 | 692 | 0 | 266 | 1672 | 0 | 242 | 0 |
| RAB17 | 744 | 0 | 0 | 0 | 0 | 0 | 90 | 0 |
| RAB18 | 0 | 1242 | 144 | 684 | 240 | 2738 | 2080 | 0 |
| RAB1A | 0 | 912 | 3389 | 1931 | 4557 | 762 | 738 | 5787 |
| RAB1B | 1494 | 2240 | 1108 | 2536 | 4064 | 2654 | 1478 | 1331 |
| RAB20 | 734 | 360 | 1545 | 1138 | 1074 | 0 | 0 | 164 |
| RAB21 | 94 | 234 | 0 | 374 | 358 | 0 | 0 | 320 |
| RAB22A | 652 | 420 | 0 | 2044 | 1176 | 0 | 1982 | 1817 |
| RAB23 | 0 | 0 | 0 | 630 | 210 | 0 | 0 | 0 |
| RAB24 | 492 | 850 | 0 | 0 | 1212 | 0 | 1722 | 430 |
| RAB25 | 1438 | 3362 | 4566 | 6380 | 11462 | 1153 | 3312 | 3482 |
| RAB26 | 0 | 0 | 400 | 0 | 0 | 0 | 0 | 602 |
| RAB27A | 248 | 0 | 0 | 0 | 0 | 0 | 0 | 0 |
| RAB2A | 0 | 1824 | 1859 | 1362 | 6140 | 4380 | 5777 | 2461 |
| RAB30 | 0 | 0 | 1480 | 412 | 588 | 0 | 278 | 0 |
| RAB30-AS1 | 854 | 0 | 0 | 0 | 762 | 0 | 0 | 0 |
| RAB31 | 0 | 0 | 0 | 0 | 1168 | 0 | 0 | 484 |
| RAB32 | 0 | 838 | 4096 | 1220 | 2041 | 0 | 0 | 0 |
| RAB33A | 0 | 664 | 0 | 0 | 0 | 0 | 0 | 0 |
| RAB34 | 782 | 0 | 0 | 0 | 534 | 0 | 0 | 0 |

| | | | | | | | | |
|---|---|---|---|---|---|---|---|---|
| RAB35 | 724 | 0 | 0 | 0 | 0 | 0 | 600 | 0 |
| RAB36 | 0 | 0 | 0 | 0 | 374 | 0 | 0 | 0 |
| RAB37 | 328 | 0 | 0 | 0 | 0 | 0 | 0 | 0 |
| RAB38 | 0 | 0 | 0 | 314 | 0 | 0 | 0 | 0 |
| RAB3B | 0 | 0 | 0 | 0 | 702 | 0 | 340 | 0 |
| RAB3D | 0 | 0 | 1722 | 0 | 720 | 0 | 328 | 0 |
| RAB3GAP1 | 14986 | 28424 | 38489 | 10842 | 57068 | 16942 | 47059 | 10524 |
| RAB3GAP2 | 0 | 0 | 0 | 0 | 142 | 0 | 0 | 0 |
| RAB3IL1 | 0 | 0 | 0 | 0 | 300 | 0 | 0 | 644 |
| RAB3IP | 0 | 0 | 0 | 0 | 0 | 0 | 354 | 0 |
| RAB40B | 0 | 1128 | 0 | 0 | 426 | 845 | 1790 | 0 |
| RAB40C | 0 | 0 | 0 | 658 | 502 | 944 | 432 | 0 |
| RAB43 | 88 | 0 | 0 | 0 | 0 | 0 | 0 | 818 |
| RAB4A | 0 | 2684 | 1914 | 2414 | 6664 | 0 | 974 | 988 |
| RAB4B | 524 | 48 | 0 | 0 | 0 | 0 | 0 | 0 |
| RAB4B-EGLN2 | 524 | 376 | 0 | 0 | 0 | 0 | 536 | 1150 |
| RAB5A | 0 | 34 | 651 | 0 | 1014 | 1972 | 812 | 0 |
| RAB5B | 0 | 0 | 660 | 0 | 0 | 0 | 0 | 0 |
| RAB5C | 2840 | 1172 | 4180 | 2715 | 1945 | 2445 | 894 | 2050 |
| RAB5IF | 1380 | 1346 | 5657 | 16301 | 8731 | 6086 | 7414 | 4070 |
| RAB6A | 82509 | 134038 | 296549 | 246223 | 503692 | 223107 | 580174 | 194064 |
| RAB6B | 0 | 0 | 0 | 0 | 0 | 1606 | 0 | 0 |
| RAB6D | 2182 | 2938 | 9946 | 2450 | 2484 | 0 | 13750 | 706 |
| RAB7A | 178 | 1100 | 0 | 3300 | 3204 | 0 | 2272 | 3759 |
| RAB8A | 232 | 606 | 1560 | 1028 | 2814 | 0 | 0 | 0 |
| RAB8B | 0 | 486 | 0 | 0 | 290 | 0 | 0 | 0 |
| RAB9A | 680 | 1082 | 0 | 0 | 2250 | 0 | 200 | 0 |
| RABAC1 | 2239 | 992 | 1554 | 0 | 1887 | 3342 | 1616 | 4434 |
| RABEP1 | 23298 | 21389 | 9832 | 14008 | 40470 | 34367 | 58800 | 11351 |
| RABEP2 | 0 | 0 | 0 | 376 | 0 | 0 | 810 | 0 |
| RABEPK | 0 | 0 | 0 | 760 | 0 | 0 | 0 | 0 |
| RABGAP1 | 0 | 0 | 605 | 0 | 0 | 0 | 0 | 0 |
| RABGAP1L | 0 | 0 | 0 | 1896 | 0 | 0 | 1332 | 0 |
| RABGEF1 | 0 | 828 | 874 | 292 | 1090 | 0 | 624 | 402 |
| RABGGTA | 8362 | 12816 | 38138 | 13257 | 38896 | 10979 | 20843 | 11038 |
| RABGGTB | 0 | 1208 | 0 | 1012 | 3683 | 1842 | 4128 | 902 |
| RABIF | 0 | 0 | 3268 | 1238 | 0 | 0 | 292 | 0 |
| RABL2A | 314 | 0 | 0 | 0 | 220 | 0 | 0 | 1092 |
| RABL2B | 0 | 328 | 0 | 0 | 0 | 0 | 0 | 0 |
| RABL3 | 0 | 0 | 0 | 0 | 324 | 0 | 536 | 932 |
| RABL6 | 254 | 828 | 2670 | 176 | 2920 | 0 | 536 | 0 |
| RAC1 | 736 | 892 | 15894 | 5163 | 13114 | 0 | 3436 | 5497 |
| RAC2 | 72091 | 98285 | 174784 | 62146 | 113499 | 120988 | 118248 | 100658 |

| | | | | | | | | |
|---|---|---|---|---|---|---|---|---|
| RACGAP1 | 0 | 246 | 0 | 896 | 0 | 0 | 376 | 0 |
| RACK1 | 4911115 | 8857359 | 15087205 | 16570291 | 34773135 | 36431859 | 54126485 | 10101667 |
| RAD1 | 0 | 0 | 0 | 1482 | 3154 | 0 | 0 | 0 |
| RAD17 | 28462 | 62520 | 42125 | 49860 | 132976 | 54720 | 106634 | 32020 |
| RAD21 | 73525 | 97910 | 103325 | 128412 | 666429 | 265452 | 625666 | 161206 |
| RAD23A | 0 | 456 | 380 | 1078 | 3328 | 0 | 1390 | 630 |
| RAD23B | 0 | 84 | 0 | 134 | 1785 | 0 | 506 | 0 |
| RAD50 | 622 | 0 | 0 | 0 | 0 | 0 | 0 | 0 |
| RAD51 | 0 | 0 | 0 | 888 | 394 | 0 | 0 | 850 |
| RAD51AP1 | 0 | 0 | 0 | 849 | 288 | 0 | 0 | 0 |
| RAD51B | 0 | 0 | 0 | 0 | 1094 | 0 | 0 | 0 |
| RAD51C | 28505 | 25774 | 21388 | 23826 | 105460 | 48568 | 57157 | 22222 |
| RAD51L3-RFFL | 0 | 0 | 0 | 1004 | 0 | 0 | 0 | 0 |
| RAD54B | 1128 | 1958 | 3319 | 11501 | 42568 | 6337 | 31377 | 13854 |
| RAD54L2 | 0 | 0 | 0 | 0 | 24 | 0 | 0 | 0 |
| RAD9A | 0 | 0 | 28 | 0 | 334 | 0 | 0 | 0 |
| RAE1 | 27409 | 59420 | 59484 | 256943 | 373212 | 194439 | 549664 | 143062 |
| RAET1E-AS1 | 0 | 0 | 0 | 0 | 0 | 0 | 116 | 0 |
| RAF1 | 52461 | 76131 | 171459 | 106127 | 280110 | 100850 | 250859 | 75643 |
| RAI14 | 0 | 0 | 0 | 0 | 114 | 0 | 0 | 0 |
| RALA | 0 | 0 | 34 | 775 | 1939 | 0 | 332 | 0 |
| RALB | 0 | 620 | 0 | 314 | 4091 | 2182 | 0 | 0 |
| RALBP1 | 0 | 304 | 0 | 0 | 402 | 0 | 548 | 788 |
| RALGAPA2 | 0 | 0 | 0 | 0 | 0 | 0 | 487 | 0 |
| RALGAPB | 0 | 0 | 0 | 1293 | 0 | 0 | 0 | 0 |
| RALGDS | 0 | 0 | 0 | 990 | 376 | 0 | 0 | 0 |
| RALGPS1 | 0 | 322 | 0 | 997 | 0 | 1284 | 480 | 0 |
| RALGPS2 | 0 | 138 | 1335 | 0 | 1272 | 0 | 530 | 0 |
| RALY | 1268 | 1890 | 6326 | 3398 | 7292 | 2196 | 1373 | 1242 |
| RALYL | 708 | 0 | 0 | 0 | 476 | 0 | 0 | 0 |
| RAMAC | 664 | 470 | 1773 | 0 | 0 | 0 | 628 | 828 |
| RAMP1 | 0 | 0 | 0 | 208 | 2570 | 0 | 0 | 1464 |
| RAMP2 | 0 | 0 | 0 | 0 | 1688 | 1293 | 0 | 0 |
| RAMP3 | 0 | 318 | 0 | 0 | 842 | 0 | 0 | 0 |
| RAN | 3326 | 4806 | 1658 | 32387 | 30189 | 10178 | 8619 | 7476 |
| RANBP1 | 0 | 2409 | 4391 | 8867 | 8569 | 0 | 2863 | 7544 |
| RANBP17 | 2094 | 1198 | 1204 | 1439 | 4444 | 0 | 628 | 513 |
| RANBP2 | 0 | 2194 | 938 | 1270 | 1312 | 408 | 2858 | 0 |
| RANBP3 | 8063 | 19619 | 19875 | 36072 | 76529 | 36996 | 34852 | 23234 |
| RANGRF | 0 | 0 | 0 | 0 | 2821 | 0 | 72 | 0 |
| RAP1A | 304 | 384 | 0 | 386 | 1538 | 0 | 1129 | 0 |
| RAP1B | 0 | 1368 | 6223 | 2046 | 1726 | 0 | 1676 | 0 |
| RAP1GAP | 0 | 0 | 0 | 0 | 356 | 0 | 0 | 0 |

| | | | | | | | | |
|---|---|---|---|---|---|---|---|---|
| RAP1GAP2 | 0 | 178 | 0 | 0 | 238 | 0 | 0 | 0 |
| RAP1GDS1 | 23672 | 59084 | 58871 | 85822 | 196000 | 82103 | 212270 | 53255 |
| RAP2A | 256 | 0 | 0 | 0 | 852 | 0 | 48 | 0 |
| RAP2B | 450 | 0 | 0 | 736 | 0 | 975 | 0 | 0 |
| RAP2C | 0 | 214 | 0 | 0 | 0 | 0 | 0 | 0 |
| RAPGEF1 | 0 | 0 | 0 | 0 | 0 | 396 | 424 | 0 |
| RAPGEF5 | 0 | 0 | 356 | 0 | 0 | 0 | 0 | 0 |
| RAPGEFL1 | 0 | 252 | 470 | 772 | 600 | 0 | 302 | 0 |
| RARA | 22650 | 15920 | 42297 | 12137 | 36305 | 28649 | 63388 | 11721 |
| RARA-AS1 | 0 | 0 | 0 | 1072 | 0 | 1601 | 0 | 0 |
| RARG | 0 | 0 | 0 | 0 | 588 | 0 | 0 | 0 |
| RARRES1 | 84722 | 103794 | 208558 | 58139 | 790093 | 65006 | 71082 | 537769 |
| RARRES2 | 0 | 562 | 8479 | 899 | 3546 | 8595 | 0 | 2193 |
| RARRES3 | 1934 | 807 | 8266 | 0 | 1576 | 0 | 0 | 0 |
| RARS | 214 | 96 | 0 | 0 | 850 | 2180 | 450 | 0 |
| RARS2 | 0 | 440 | 2138 | 0 | 1350 | 0 | 692 | 0 |
| RASA1 | 0 | 0 | 0 | 0 | 1416 | 0 | 514 | 0 |
| RASA2 | 9926 | 10706 | 5911 | 3270 | 7190 | 5937 | 8767 | 4906 |
| RASA3 | 0 | 0 | 0 | 0 | 0 | 0 | 0 | 696 |
| RASD1 | 0 | 298 | 1172 | 0 | 0 | 884 | 0 | 0 |
| RASEF | 276 | 627 | 0 | 0 | 0 | 0 | 0 | 0 |
| RASL11A | 564 | 0 | 964 | 1008 | 0 | 0 | 702 | 0 |
| RASSF1 | 0 | 0 | 0 | 4501 | 3393 | 1804 | 0 | 0 |
| RASSF1-AS1 | 0 | 0 | 0 | 0 | 546 | 0 | 0 | 0 |
| RASSF10 | 0 | 0 | 0 | 0 | 0 | 0 | 812 | 0 |
| RASSF2 | 0 | 0 | 0 | 0 | 0 | 0 | 794 | 0 |
| RASSF3 | 626 | 0 | 0 | 1926 | 376 | 0 | 0 | 0 |
| RASSF4 | 398 | 0 | 0 | 0 | 456 | 1592 | 0 | 0 |
| RASSF6 | 140 | 0 | 0 | 0 | 0 | 0 | 0 | 0 |
| RASSF7 | 1734 | 2097 | 1124 | 2622 | 5264 | 142 | 148 | 1644 |
| RASSF8 | 0 | 0 | 0 | 0 | 226 | 0 | 0 | 0 |
| RAVER1 | 0 | 0 | 0 | 0 | 274 | 0 | 0 | 0 |
| RAVER2 | 264 | 0 | 0 | 0 | 0 | 0 | 0 | 0 |
| RB1 | 26221 | 26164 | 30300 | 63463 | 111133 | 117967 | 102216 | 39316 |
| RB1CC1 | 0 | 0 | 0 | 0 | 0 | 0 | 90 | 0 |
| RBAK | 0 | 220 | 0 | 0 | 0 | 0 | 0 | 0 |
| RBBP4 | 0 | 706 | 2048 | 5494 | 1382 | 1286 | 714 | 726 |
| RBBP5 | 0 | 0 | 0 | 0 | 242 | 0 | 0 | 0 |
| RBBP6 | 0 | 0 | 0 | 1467 | 948 | 0 | 532 | 0 |
| RBBP7 | 0 | 1532 | 0 | 1358 | 5200 | 0 | 1038 | 1380 |
| RBBP8 | 728 | 0 | 0 | 0 | 0 | 0 | 0 | 0 |
| RBBP8NL | 632 | 0 | 0 | 0 | 230 | 0 | 0 | 0 |
| RBCK1 | 2893 | 1020 | 3490 | 4524 | 4712 | 2412 | 1507 | 666 |

| | | | | | | | | |
|---|---|---|---|---|---|---|---|---|
| RBFA | 768 | 0 | 0 | 0 | 2276 | 0 | 0 | 0 |
| RBFOX2 | 640 | 422 | 0 | 4628 | 1448 | 0 | 284 | 0 |
| RBKS | 610 | 0 | 0 | 0 | 504 | 0 | 1052 | 0 |
| RBM10 | 9354 | 7814 | 7987 | 1881 | 19379 | 11934 | 3631 | 7519 |
| RBM12 | 0 | 0 | 1392 | 0 | 1445 | 0 | 210 | 0 |
| RBM14 | 0 | 0 | 0 | 0 | 816 | 0 | 36 | 0 |
| RBM14-RBM4 | 0 | 0 | 0 | 1249 | 0 | 0 | 0 | 0 |
| RBM15 | 0 | 0 | 0 | 0 | 344 | 0 | 0 | 0 |
| RBM15B | 0 | 0 | 0 | 0 | 0 | 0 | 608 | 0 |
| RBM17 | 0 | 0 | 0 | 510 | 4064 | 1418 | 2583 | 0 |
| RBM18 | 0 | 246 | 1380 | 0 | 1764 | 0 | 0 | 858 |
| RBM19 | 0 | 0 | 1110 | 0 | 1229 | 2150 | 0 | 0 |
| RBM22 | 0 | 0 | 0 | 0 | 122 | 0 | 939 | 0 |
| RBM23 | 520 | 320 | 1806 | 0 | 922 | 0 | 0 | 0 |
| RBM24 | 0 | 496 | 0 | 0 | 0 | 0 | 0 | 0 |
| RBM25 | 271648 | 121797 | 165702 | 100886 | 399098 | 110818 | 174902 | 106008 |
| RBM26 | 0 | 0 | 0 | 0 | 838 | 0 | 759 | 0 |
| RBM27 | 0 | 0 | 0 | 0 | 214 | 0 | 0 | 0 |
| RBM28 | 0 | 0 | 0 | 0 | 0 | 1204 | 0 | 0 |
| RBM3 | 0 | 1216 | 4996 | 338 | 10502 | 1803 | 482 | 3479 |
| RBM33 | 1048 | 0 | 20 | 0 | 904 | 0 | 0 | 94 |
| RBM34 | 0 | 464 | 0 | 1378 | 960 | 0 | 384 | 0 |
| RBM38 | 0 | 0 | 0 | 10 | 0 | 0 | 0 | 0 |
| RBM39 | 914 | 340 | 2056 | 1854 | 8621 | 3583 | 5668 | 2874 |
| RBM4 | 0 | 446 | 902 | 0 | 8086 | 1486 | 4305 | 978 |
| RBM42 | 1186 | 609 | 3433 | 1235 | 6964 | 0 | 336 | 1600 |
| RBM45 | 0 | 576 | 0 | 1355 | 1852 | 0 | 0 | 0 |
| RBM47 | 4766 | 4362 | 7600 | 4955 | 18825 | 5686 | 8905 | 1603 |
| RBM48 | 0 | 0 | 0 | 0 | 0 | 0 | 28 | 0 |
| RBM4B | 0 | 358 | 0 | 0 | 0 | 0 | 0 | 734 |
| RBM5 | 1586 | 0 | 0 | 0 | 846 | 0 | 288 | 0 |
| RBM6 | 15184 | 4196 | 12963 | 16477 | 33170 | 10277 | 15344 | 4886 |
| RBM7 | 0 | 356 | 0 | 364 | 2286 | 0 | 1625 | 1500 |
| RBM8A | 0 | 928 | 1770 | 5978 | 6884 | 0 | 2887 | 8975 |
| RBMS1 | 5894 | 11484 | 12125 | 3300 | 36320 | 15986 | 19068 | 18236 |
| RBMS2 | 0 | 0 | 0 | 0 | 36 | 0 | 114 | 0 |
| RBMS3 | 0 | 0 | 1012 | 0 | 0 | 0 | 0 | 0 |
| RBMX | 0 | 86 | 0 | 1620 | 2132 | 0 | 772 | 1121 |
| RBMX2 | 0 | 0 | 0 | 692 | 0 | 0 | 0 | 0 |
| RBMXL1 | 664 | 0 | 0 | 0 | 0 | 1322 | 0 | 0 |
| RBP1 | 0 | 0 | 0 | 2756 | 0 | 7734 | 1190 | 0 |
| RBP2 | 0 | 156 | 0 | 0 | 924 | 0 | 0 | 0 |
| RBP4 | 258 | 0 | 1488 | 0 | 3091 | 0 | 0 | 0 |

| | | | | | | | | |
|---|---|---|---|---|---|---|---|---|
| RBP5 | 200 | 0 | 0 | 0 | 0 | 0 | 0 | 0 |
| RBPJ | 0 | 0 | 0 | 0 | 440 | 0 | 0 | 0 |
| RBPMS | 32462 | 47909 | 41335 | 71392 | 289335 | 186836 | 153522 | 68057 |
| RBPMS-AS1 | 0 | 0 | 0 | 0 | 468 | 0 | 0 | 0 |
| RBSN | 0 | 0 | 0 | 1334 | 0 | 0 | 0 | 0 |
| RBX1 | 428986 | 630138 | 468770 | 335995 | 1299724 | 462148 | 676221 | 330930 |
| RC3H1 | 594 | 0 | 296 | 0 | 262 | 0 | 0 | 0 |
| RC3H2 | 0 | 480 | 0 | 0 | 0 | 0 | 0 | 0 |
| RCAN1 | 4738 | 3368 | 15803 | 3678 | 6752 | 0 | 1328 | 919 |
| RCAN2 | 3172 | 2562 | 1646 | 864 | 6122 | 2114 | 2084 | 0 |
| RCAN3 | 304 | 0 | 0 | 0 | 0 | 0 | 0 | 0 |
| RCBTB1 | 0 | 236 | 0 | 0 | 0 | 0 | 0 | 0 |
| RCBTB2 | 0 | 0 | 0 | 0 | 328 | 2397 | 0 | 0 |
| RCC1 | 0 | 612 | 640 | 0 | 378 | 0 | 0 | 0 |
| RCC1L | 0 | 272 | 0 | 0 | 584 | 0 | 1266 | 0 |
| RCC2 | 0 | 28 | 0 | 578 | 82 | 0 | 0 | 0 |
| RCCD1 | 0 | 0 | 0 | 0 | 106 | 0 | 0 | 0 |
| RCE1 | 464 | 826 | 2200 | 0 | 1782 | 0 | 396 | 0 |
| RCHY1 | 514 | 0 | 0 | 1502 | 1076 | 2496 | 0 | 0 |
| RCL1 | 6186 | 26456 | 29127 | 41855 | 73470 | 55445 | 153877 | 32889 |
| RCN1 | 0 | 1212 | 4653 | 3332 | 1310 | 3167 | 6758 | 1931 |
| RCN2 | 0 | 2400 | 0 | 3547 | 3788 | 1742 | 170 | 620 |
| RCN3 | 0 | 0 | 1987 | 921 | 0 | 0 | 0 | 0 |
| RCOR1 | 3080 | 1366 | 2998 | 1460 | 9784 | 1720 | 302 | 0 |
| RCOR2 | 0 | 0 | 0 | 0 | 0 | 0 | 126 | 0 |
| RCOR3 | 1008 | 0 | 128 | 0 | 222 | 1660 | 0 | 0 |
| RCSD1 | 0 | 576 | 0 | 0 | 0 | 0 | 0 | 0 |
| RDH10 | 0 | 424 | 0 | 0 | 310 | 0 | 0 | 1208 |
| RDH11 | 0 | 696 | 0 | 0 | 303 | 0 | 0 | 0 |
| RDH13 | 0 | 1125 | 0 | 1022 | 0 | 0 | 0 | 0 |
| RDH14 | 0 | 1106 | 0 | 1094 | 2678 | 1792 | 0 | 705 |
| REC8 | 0 | 0 | 0 | 0 | 404 | 0 | 0 | 0 |
| RECQL4 | 0 | 0 | 0 | 0 | 502 | 0 | 0 | 0 |
| RECQL5 | 0 | 0 | 0 | 0 | 1042 | 0 | 0 | 0 |
| REEP1 | 0 | 0 | 0 | 1176 | 0 | 0 | 7662 | 0 |
| REEP3 | 0 | 771 | 4025 | 0 | 1324 | 0 | 907 | 0 |
| REEP4 | 0 | 0 | 722 | 0 | 1605 | 1588 | 0 | 0 |
| REEP5 | 1770 | 1452 | 2120 | 3835 | 7010 | 1592 | 1998 | 1706 |
| REEP6 | 0 | 0 | 0 | 0 | 0 | 1708 | 0 | 0 |
| REG1A | 0 | 0 | 0 | 28167 | 538 | 0 | 0 | 0 |
| REG1B | 0 | 0 | 0 | 2118 | 0 | 0 | 0 | 0 |
| REG3A | 0 | 0 | 0 | 1316 | 0 | 0 | 0 | 0 |
| REG4 | 0 | 4294 | 0 | 7530 | 16379 | 0 | 0 | 7134 |

| | | | | | | | | |
|---|---|---|---|---|---|---|---|---|
| REL | 3292 | 4416 | 4848 | 4074 | 23173 | 14153 | 13526 | 5628 |
| RELA | 0 | 384 | 0 | 0 | 1056 | 0 | 437 | 752 |
| RELB | 1314 | 2336 | 13988 | 490 | 4732 | 0 | 956 | 1594 |
| RELCH | 0 | 0 | 0 | 0 | 0 | 0 | 0 | 694 |
| RELL1 | 0 | 424 | 1761 | 0 | 0 | 960 | 0 | 0 |
| RENBP | 0 | 0 | 1666 | 0 | 564 | 0 | 0 | 0 |
| REP15 | 0 | 2073 | 4207 | 0 | 0 | 0 | 0 | 0 |
| REPIN1 | 816 | 472 | 2264 | 1105 | 1264 | 1220 | 512 | 0 |
| REPS1 | 0 | 0 | 0 | 0 | 894 | 0 | 0 | 0 |
| REPS2 | 0 | 0 | 0 | 0 | 462 | 0 | 0 | 0 |
| RER1 | 1082 | 2734 | 4227 | 4732 | 8464 | 2016 | 1364 | 2310 |
| RERE | 0 | 0 | 0 | 0 | 879 | 0 | 0 | 0 |
| RERGL | 27160 | 0 | 11284 | 0 | 1062 | 5136 | 2220 | 1402 |
| REST | 1148 | 0 | 0 | 0 | 0 | 0 | 436 | 0 |
| RET | 1096 | 0 | 0 | 0 | 0 | 0 | 0 | 0 |
| RETNLB | 0 | 2617 | 1678 | 0 | 822 | 0 | 0 | 0 |
| RETREG1 | 100696 | 165129 | 233649 | 12330 | 93700 | 8636 | 69955 | 15173 |
| RETREG2 | 896 | 700 | 2922 | 0 | 528 | 0 | 116 | 0 |
| RETREG3 | 0 | 358 | 0 | 0 | 162 | 0 | 0 | 0 |
| RETSAT | 876 | 0 | 2748 | 0 | 0 | 2172 | 0 | 0 |
| REV1 | 5008 | 4746 | 3242 | 10279 | 7874 | 3764 | 11952 | 6280 |
| REV3L | 2276 | 970 | 6028 | 4606 | 6845 | 0 | 5212 | 568 |
| REX1BD | 2516 | 1012 | 0 | 2796 | 7072 | 4451 | 540 | 0 |
| REXO2 | 694 | 1904 | 0 | 0 | 3505 | 1663 | 522 | 2596 |
| REXO4 | 0 | 0 | 0 | 0 | 0 | 0 | 786 | 0 |
| RFC1 | 0 | 0 | 0 | 0 | 0 | 486 | 0 | 120 |
| RFC2 | 550 | 0 | 0 | 0 | 1118 | 0 | 0 | 0 |
| RFC3 | 0 | 296 | 0 | 0 | 612 | 0 | 0 | 0 |
| RFC4 | 0 | 0 | 0 | 1206 | 4432 | 0 | 0 | 907 |
| RFC5 | 0 | 622 | 0 | 0 | 690 | 0 | 0 | 0 |
| RFFL | 0 | 0 | 0 | 0 | 936 | 0 | 0 | 0 |
| RFK | 0 | 222 | 4266 | 0 | 987 | 0 | 891 | 0 |
| RFLNA | 0 | 352 | 1279 | 0 | 0 | 0 | 0 | 0 |
| RFNG | 0 | 0 | 0 | 510 | 0 | 0 | 213 | 0 |
| RFPL3S | 0 | 0 | 0 | 0 | 0 | 0 | 0 | 958 |
| RFT1 | 0 | 420 | 0 | 0 | 1376 | 0 | 440 | 1046 |
| RFTN1 | 0 | 0 | 0 | 0 | 406 | 0 | 0 | 0 |
| RFTN2 | 586 | 422 | 864 | 0 | 3432 | 2380 | 0 | 1912 |
| RFWD3 | 846 | 1738 | 6051 | 7133 | 5220 | 6547 | 3688 | 4540 |
| RFX1 | 0 | 364 | 362 | 0 | 856 | 0 | 244 | 0 |
| RFX2 | 588 | 194 | 0 | 0 | 200 | 0 | 0 | 0 |
| RFX7 | 172 | 0 | 0 | 0 | 0 | 0 | 0 | 0 |
| RFXANK | 0 | 410 | 935 | 0 | 2180 | 0 | 0 | 0 |

| | | | | | | | | |
|---|---|---|---|---|---|---|---|---|
| RGCC | 0 | 0 | 0 | 756 | 1778 | 6863 | 1935 | 0 |
| RGL1 | 0 | 0 | 0 | 0 | 0 | 0 | 346 | 0 |
| RGL2 | 0 | 0 | 0 | 0 | 1226 | 0 | 692 | 1590 |
| RGMB | 0 | 0 | 1692 | 0 | 0 | 0 | 330 | 0 |
| RGN | 0 | 0 | 1771 | 0 | 0 | 0 | 0 | 0 |
| RGP1 | 0 | 0 | 0 | 0 | 0 | 0 | 0 | 1046 |
| RGPD2 | 0 | 284 | 0 | 0 | 0 | 0 | 0 | 0 |
| RGPD5 | 0 | 0 | 0 | 0 | 132 | 0 | 0 | 0 |
| RGPD8 | 0 | 0 | 0 | 0 | 170 | 0 | 0 | 0 |
| RGS1 | 0 | 0 | 6469 | 0 | 1048 | 6016 | 254 | 1748 |
| RGS10 | 0 | 194 | 992 | 0 | 942 | 0 | 582 | 1178 |
| RGS13 | 0 | 460 | 0 | 0 | 232 | 0 | 0 | 0 |
| RGS14 | 0 | 658 | 0 | 0 | 682 | 0 | 0 | 0 |
| RGS16 | 0 | 0 | 0 | 0 | 0 | 1775 | 0 | 0 |
| RGS19 | 232 | 0 | 0 | 0 | 0 | 0 | 0 | 0 |
| RGS2 | 830 | 1087 | 1686 | 1125 | 718 | 2846 | 0 | 2040 |
| RGS3 | 0 | 116 | 1630 | 0 | 0 | 1070 | 0 | 0 |
| RGS7 | 0 | 0 | 0 | 0 | 2876 | 0 | 0 | 0 |
| RHBDD1 | 634 | 0 | 0 | 0 | 0 | 0 | 398 | 0 |
| RHBDD2 | 0 | 332 | 4852 | 0 | 1038 | 325 | 0 | 0 |
| RHBDD3 | 0 | 0 | 0 | 1156 | 0 | 0 | 330 | 0 |
| RHBDF1 | 374 | 0 | 0 | 0 | 0 | 0 | 0 | 0 |
| RHBDF2 | 0 | 0 | 0 | 0 | 0 | 1958 | 0 | 0 |
| RHBDL1 | 0 | 0 | 0 | 0 | 0 | 0 | 1117 | 640 |
| RHBDL2 | 0 | 0 | 0 | 0 | 1930 | 0 | 0 | 0 |
| RHBG | 0 | 678 | 0 | 0 | 0 | 0 | 548 | 0 |
| RHCE | 0 | 136 | 0 | 0 | 414 | 0 | 0 | 0 |
| RHEB | 0 | 1039 | 3141 | 4672 | 4332 | 3211 | 3304 | 2658 |
| RHEBL1 | 0 | 0 | 0 | 0 | 0 | 0 | 0 | 1040 |
| RHEX | 0 | 0 | 1142 | 0 | 0 | 0 | 0 | 0 |
| RHNO1 | 0 | 0 | 0 | 0 | 0 | 0 | 852 | 0 |
| RHOA | 2299571 | 3392354 | 12175714 | 4947774 | 14718874 | 6048049 | 8728758 | 3764658 |
| RHOB | 684 | 352 | 0 | 0 | 1320 | 0 | 458 | 0 |
| RHOBTB3 | 0 | 172 | 0 | 758 | 284 | 0 | 0 | 443 |
| RHOC | 11766 | 5954 | 27679 | 7879 | 10263 | 0 | 4284 | 10710 |
| RHOD | 3480 | 5628 | 8215 | 3233 | 18714 | 5167 | 11673 | 12131 |
| RHOF | 0 | 0 | 988 | 0 | 0 | 0 | 0 | 0 |
| RHOG | 14320 | 10351 | 35242 | 12790 | 10635 | 190 | 11988 | 14550 |
| RHOH | 0 | 0 | 506 | 0 | 0 | 0 | 0 | 0 |
| RHOJ | 0 | 0 | 0 | 0 | 0 | 0 | 0 | 500 |
| RHOT1 | 0 | 578 | 0 | 0 | 224 | 0 | 0 | 0 |
| RHOT2 | 594 | 0 | 2165 | 471 | 164 | 1036 | 0 | 638 |
| RHOU | 4066 | 256 | 15095 | 1450 | 12050 | 986 | 3539 | 4859 |

| | | | | | | | | |
|---|---|---|---|---|---|---|---|---|
| RHOV | 0 | 0 | 0 | 0 | 878 | 0 | 0 | 0 |
| RHPN1 | 0 | 306 | 0 | 0 | 0 | 0 | 274 | 0 |
| RHPN1-AS1 | 0 | 0 | 0 | 0 | 0 | 0 | 0 | 884 |
| RHPN2 | 0 | 480 | 0 | 0 | 0 | 0 | 0 | 0 |
| RIBC2 | 0 | 0 | 0 | 0 | 862 | 0 | 0 | 0 |
| RIC3 | 0 | 0 | 0 | 0 | 0 | 0 | 0 | 260 |
| RIC8A | 0 | 0 | 0 | 0 | 0 | 0 | 0 | 566 |
| RIDA | 0 | 118 | 0 | 1015 | 3022 | 0 | 98 | 1104 |
| RIIAD1 | 0 | 0 | 0 | 0 | 0 | 0 | 518 | 0 |
| RILP | 1760 | 1332 | 3829 | 1248 | 214 | 2982 | 0 | 3018 |
| RILPL1 | 0 | 0 | 0 | 0 | 500 | 0 | 0 | 0 |
| RILPL2 | 0 | 0 | 0 | 257 | 0 | 0 | 0 | 0 |
| RIMKLB | 0 | 0 | 0 | 0 | 0 | 0 | 362 | 0 |
| RIMS3 | 0 | 334 | 0 | 0 | 0 | 0 | 0 | 0 |
| RIN1 | 0 | 0 | 5472 | 0 | 0 | 0 | 454 | 0 |
| RIN2 | 0 | 336 | 0 | 1688 | 1626 | 0 | 0 | 0 |
| RING1 | 0 | 0 | 0 | 0 | 2098 | 1742 | 0 | 0 |
| RINT1 | 0 | 0 | 0 | 0 | 1102 | 0 | 0 | 0 |
| RIOK1 | 64964 | 91409 | 88384 | 122321 | 350236 | 155626 | 264222 | 117853 |
| RIOK2 | 386 | 0 | 1916 | 0 | 0 | 0 | 0 | 0 |
| RIOK3 | 378 | 0 | 220 | 0 | 867 | 0 | 372 | 450 |
| RIOX1 | 0 | 298 | 0 | 0 | 0 | 0 | 0 | 0 |
| RIOX2 | 0 | 0 | 378 | 0 | 1236 | 0 | 220 | 595 |
| RIPK1 | 0 | 0 | 0 | 0 | 478 | 0 | 0 | 0 |
| RIPK2 | 8269 | 30813 | 14380 | 64716 | 287054 | 201102 | 146466 | 102005 |
| RIPK3 | 276 | 0 | 2068 | 0 | 316 | 0 | 0 | 570 |
| RIT1 | 0 | 0 | 0 | 0 | 720 | 0 | 0 | 1222 |
| RITA1 | 0 | 0 | 0 | 0 | 1320 | 0 | 0 | 0 |
| RLIM | 3857 | 1056 | 2635 | 813 | 2597 | 0 | 1396 | 290 |
| RLN3 | 0 | 0 | 0 | 0 | 0 | 0 | 0 | 1174 |
| RMC1 | 310 | 0 | 0 | 0 | 0 | 0 | 0 | 0 |
| RMDN1 | 0 | 828 | 2188 | 1378 | 1550 | 0 | 0 | 908 |
| RMDN2 | 0 | 0 | 2360 | 0 | 1240 | 0 | 416 | 0 |
| RMDN3 | 0 | 0 | 1054 | 1249 | 0 | 0 | 0 | 0 |
| RMI1 | 329 | 310 | 0 | 0 | 0 | 0 | 0 | 0 |
| RMI2 | 0 | 0 | 0 | 1834 | 1566 | 0 | 0 | 0 |
| RMND1 | 0 | 202 | 0 | 0 | 0 | 0 | 0 | 0 |
| RMND5A | 0 | 0 | 692 | 0 | 1216 | 0 | 0 | 944 |
| RMND5B | 0 | 200 | 0 | 0 | 0 | 0 | 0 | 0 |
| RNA18SN3 | 69024 | 23710 | 44728 | 19257 | 75422 | 66533 | 66879 | 16687 |
| RNA28SN1 | 7218 | 808 | 11983 | 1488 | 13123 | 12423 | 5146 | 4457 |
| RNA28SN2 | 3034 | 1156 | 3411 | 390 | 1148 | 1088 | 1490 | 894 |
| RNA28SN3 | 33214 | 21107 | 29375 | 18611 | 68676 | 22421 | 57774 | 14678 |

| | | | | | | | | |
|---|---|---|---|---|---|---|---|---|
| RNA28SN4 | 17126 | 11543 | 10064 | 11141 | 35401 | 12292 | 22617 | 9979 |
| RNA45SN3 | 948 | 0 | 0 | 426 | 2022 | 0 | 600 | 344 |
| RNA45SN4 | 0 | 954 | 0 | 118 | 942 | 0 | 0 | 0 |
| RNASE1 | 1086 | 1762 | 11049 | 3101 | 5659 | 5524 | 3254 | 1464 |
| RNASE4 | 210 | 2830 | 6921 | 1186 | 6906 | 0 | 440 | 2574 |
| RNASE6 | 0 | 1552 | 0 | 834 | 0 | 1712 | 548 | 0 |
| RNASEH1 | 756 | 670 | 0 | 758 | 38 | 0 | 1048 | 232 |
| RNASEH1-AS1 | 0 | 0 | 0 | 0 | 1944 | 0 | 504 | 0 |
| RNASEH2A | 0 | 0 | 0 | 2818 | 2362 | 0 | 1116 | 0 |
| RNASEH2B | 0 | 354 | 0 | 2816 | 4828 | 0 | 948 | 0 |
| RNASEH2C | 660 | 892 | 0 | 758 | 932 | 58 | 1520 | 1476 |
| RNASEK | 1982 | 5861 | 8142 | 2674 | 4942 | 6848 | 3612 | 1232 |
| RNASEK-C17orf49 | 644 | 630 | 3265 | 928 | 4326 | 2404 | 1958 | 590 |
| RNASET2 | 1316 | 682 | 6001 | 5694 | 11471 | 3143 | 4606 | 3132 |
| RND1 | 4266 | 0 | 18902 | 6235 | 5427 | 5363 | 3658 | 0 |
| RND3 | 0 | 430 | 0 | 0 | 412 | 0 | 0 | 0 |
| RNF103 | 0 | 306 | 1412 | 570 | 384 | 0 | 0 | 0 |
| RNF11 | 0 | 0 | 0 | 0 | 1112 | 1048 | 0 | 460 |
| RNF111 | 0 | 500 | 0 | 0 | 0 | 0 | 0 | 0 |
| RNF113A | 0 | 0 | 0 | 1534 | 1004 | 6310 | 0 | 960 |
| RNF114 | 0 | 884 | 0 | 2332 | 5684 | 2346 | 3788 | 1027 |
| RNF115 | 2072 | 1187 | 820 | 3328 | 6406 | 0 | 6699 | 2914 |
| RNF121 | 0 | 488 | 0 | 0 | 962 | 0 | 0 | 0 |
| RNF125 | 674 | 0 | 0 | 0 | 0 | 0 | 0 | 0 |
| RNF126 | 0 | 816 | 0 | 0 | 270 | 1783 | 0 | 0 |
| RNF128 | 0 | 96 | 0 | 0 | 1118 | 0 | 148 | 0 |
| RNF13 | 0 | 0 | 3436 | 182 | 0 | 0 | 1340 | 0 |
| RNF130 | 450 | 538 | 294 | 1811 | 3288 | 4241 | 2276 | 0 |
| RNF135 | 23875 | 16206 | 64423 | 16368 | 50118 | 13253 | 49892 | 18266 |
| RNF138 | 212 | 0 | 0 | 0 | 0 | 0 | 0 | 0 |
| RNF138P1 | 0 | 0 | 792 | 0 | 0 | 0 | 0 | 0 |
| RNF139 | 0 | 0 | 1120 | 0 | 332 | 0 | 632 | 0 |
| RNF14 | 136 | 0 | 0 | 0 | 0 | 0 | 996 | 0 |
| RNF141 | 0 | 0 | 970 | 0 | 2020 | 0 | 0 | 0 |
| RNF144A | 0 | 0 | 0 | 632 | 0 | 0 | 0 | 0 |
| RNF145 | 688 | 812 | 0 | 78 | 1694 | 1420 | 0 | 975 |
| RNF146 | 1924 | 8060 | 322 | 1295 | 28964 | 118 | 32889 | 526 |
| RNF149 | 482 | 428 | 0 | 1346 | 3194 | 1612 | 214 | 0 |
| RNF152 | 0 | 388 | 0 | 0 | 0 | 0 | 0 | 0 |
| RNF167 | 17934 | 31548 | 54752 | 24189 | 68560 | 26226 | 53939 | 29061 |
| RNF168 | 0 | 266 | 336 | 10 | 0 | 0 | 442 | 0 |
| RNF170 | 0 | 0 | 1460 | 72 | 536 | 0 | 358 | 0 |
| RNF181 | 1594 | 3440 | 1386 | 4499 | 6249 | 0 | 2784 | 1162 |

| | | | | | | | | |
|---|---|---|---|---|---|---|---|---|
| RNF185 | 0 | 0 | 0 | 0 | 12 | 0 | 0 | 0 |
| RNF186 | 658 | 0 | 0 | 0 | 2654 | 0 | 0 | 334 |
| RNF187 | 580 | 116 | 0 | 3355 | 1234 | 0 | 536 | 0 |
| RNF19B | 1130 | 0 | 1034 | 0 | 0 | 0 | 0 | 0 |
| RNF2 | 0 | 220 | 0 | 0 | 0 | 950 | 0 | 174 |
| RNF20 | 0 | 274 | 0 | 0 | 910 | 0 | 0 | 0 |
| RNF207 | 0 | 0 | 0 | 0 | 274 | 0 | 0 | 0 |
| RNF208 | 0 | 0 | 0 | 0 | 246 | 0 | 0 | 251 |
| RNF213 | 1244 | 0 | 0 | 1832 | 1386 | 0 | 668 | 0 |
| RNF216 | 17084 | 5886 | 6735 | 7392 | 31363 | 9054 | 18438 | 8060 |
| RNF216P1 | 7063 | 5830 | 1279 | 3816 | 16217 | 4414 | 11732 | 1886 |
| RNF217 | 0 | 298 | 5317 | 0 | 4434 | 1702 | 2612 | 546 |
| RNF219 | 0 | 0 | 122 | 254 | 328 | 0 | 0 | 0 |
| RNF220 | 0 | 544 | 0 | 0 | 1484 | 0 | 946 | 0 |
| RNF227 | 0 | 238 | 0 | 0 | 0 | 0 | 0 | 0 |
| RNF25 | 0 | 1328 | 1742 | 484 | 0 | 0 | 0 | 0 |
| RNF26 | 0 | 450 | 0 | 0 | 744 | 0 | 0 | 0 |
| RNF31 | 1392 | 1161 | 8632 | 2700 | 8600 | 0 | 2082 | 1154 |
| RNF32 | 0 | 0 | 0 | 0 | 0 | 0 | 512 | 0 |
| RNF34 | 0 | 0 | 0 | 938 | 0 | 0 | 0 | 0 |
| RNF38 | 660 | 0 | 503 | 904 | 0 | 0 | 588 | 0 |
| RNF39 | 0 | 0 | 0 | 0 | 317 | 0 | 426 | 0 |
| RNF4 | 0 | 0 | 1288 | 0 | 0 | 0 | 1290 | 0 |
| RNF40 | 318 | 556 | 0 | 0 | 0 | 0 | 396 | 0 |
| RNF41 | 0 | 0 | 0 | 1142 | 0 | 0 | 0 | 0 |
| RNF43 | 2308 | 5780 | 6800 | 16655 | 18469 | 45550 | 229024 | 4057 |
| RNF44 | 0 | 0 | 0 | 0 | 194 | 0 | 196 | 0 |
| RNF5 | 136 | 292 | 1557 | 1078 | 1610 | 0 | 2164 | 1064 |
| RNF6 | 0 | 402 | 930 | 898 | 278 | 0 | 0 | 384 |
| RNF7 | 3684 | 3590 | 11438 | 6676 | 8504 | 7076 | 8235 | 3115 |
| RNF8 | 868 | 556 | 0 | 0 | 356 | 0 | 0 | 0 |
| RNFT1 | 0 | 110 | 284 | 0 | 308 | 0 | 352 | 0 |
| RNGTT | 0 | 386 | 0 | 0 | 0 | 0 | 0 | 0 |
| RNH1 | 698 | 1366 | 894 | 1264 | 1860 | 0 | 190 | 1845 |
| RNLS | 12078 | 27499 | 35708 | 6324 | 53742 | 20665 | 46792 | 22409 |
| RNMT | 0 | 0 | 0 | 0 | 34 | 0 | 0 | 0 |
| RNPC3 | 0 | 0 | 0 | 0 | 0 | 0 | 0 | 404 |
| RNPEP | 0 | 786 | 316 | 1989 | 1858 | 0 | 0 | 0 |
| RNPEPL1 | 706 | 0 | 0 | 0 | 0 | 0 | 0 | 0 |
| RNPS1 | 590 | 3238 | 2652 | 2118 | 5335 | 2160 | 1779 | 2928 |
| ROBO1 | 330 | 0 | 0 | 0 | 0 | 0 | 0 | 0 |
| ROBO2 | 496 | 3538 | 0 | 2002 | 18796 | 0 | 0 | 0 |
| ROCK1 | 30422 | 25343 | 24373 | 20975 | 111866 | 47393 | 34770 | 34675 |

| | | | | | | | | |
|---|---|---|---|---|---|---|---|---|
| ROCK2 | 454 | 376 | 0 | 0 | 560 | 0 | 250 | 0 |
| ROGDI | 0 | 404 | 0 | 0 | 2647 | 0 | 0 | 0 |
| ROM1 | 0 | 0 | 386 | 0 | 0 | 1584 | 948 | 0 |
| ROMO1 | 1970 | 2004 | 3197 | 6146 | 6403 | 800 | 7148 | 1624 |
| ROR1 | 0 | 0 | 0 | 0 | 410 | 0 | 0 | 0 |
| ROR2 | 0 | 326 | 0 | 0 | 174 | 2876 | 0 | 0 |
| RP9 | 0 | 0 | 0 | 1380 | 4048 | 0 | 394 | 0 |
| RPA1 | 224 | 116 | 0 | 524 | 0 | 0 | 1342 | 588 |
| RPA2 | 0 | 232 | 0 | 0 | 1418 | 2454 | 1272 | 0 |
| RPA3 | 492 | 1886 | 3792 | 6992 | 3777 | 97 | 728 | 776 |
| RPAIN | 0 | 696 | 828 | 3666 | 1772 | 0 | 1282 | 516 |
| RPAP1 | 0 | 0 | 0 | 0 | 672 | 0 | 0 | 0 |
| RPAP2 | 0 | 0 | 0 | 0 | 654 | 0 | 700 | 0 |
| RPAP3 | 0 | 202 | 0 | 566 | 1380 | 0 | 478 | 0 |
| RPARP-AS1 | 94 | 0 | 0 | 0 | 306 | 0 | 562 | 0 |
| RPE | 0 | 334 | 358 | 616 | 386 | 0 | 0 | 1385 |
| RPF1 | 702 | 462 | 0 | 0 | 1861 | 0 | 444 | 0 |
| RPF2 | 686 | 1704 | 0 | 3374 | 1737 | 0 | 568 | 0 |
| RPGRIP1L | 0 | 0 | 0 | 605 | 346 | 0 | 0 | 0 |
| RPIA | 0 | 414 | 632 | 1084 | 1916 | 1292 | 0 | 666 |
| RPL10 | 17024 | 54826 | 83605 | 104200 | 242903 | 145460 | 72409 | 109266 |
| RPL10A | 4565 | 28714 | 28566 | 59897 | 73401 | 38972 | 71724 | 52883 |
| RPL10L | 0 | 276 | 0 | 0 | 0 | 0 | 0 | 0 |
| RPL11 | 13344 | 37834 | 35627 | 99789 | 87647 | 61077 | 24946 | 40820 |
| RPL12 | 13308 | 14062 | 35928 | 72090 | 84056 | 75660 | 73588 | 25620 |
| RPL13 | 26729 | 33393 | 40408 | 119772 | 131220 | 80051 | 112567 | 44528 |
| RPL13A | 35743 | 66039 | 122329 | 129170 | 213177 | 146319 | 74460 | 69392 |
| RPL14 | 10533 | 24362 | 27006 | 59869 | 160286 | 48942 | 94482 | 25258 |
| RPL15 | 12266 | 28797 | 40813 | 60612 | 143304 | 58760 | 60487 | 40711 |
| RPL17 | 17241 | 42662 | 56051 | 49964 | 62432 | 63841 | 42868 | 39856 |
| RPL17-C18orf32 | 466 | 0 | 14 | 0 | 0 | 0 | 176 | 0 |
| RPL18 | 82206 | 190800 | 115893 | 193692 | 611410 | 313577 | 260301 | 134614 |
| RPL18A | 18477 | 36658 | 43937 | 65492 | 138595 | 35576 | 26076 | 25047 |
| RPL19 | 11900 | 30752 | 43458 | 47064 | 75390 | 35814 | 49769 | 29710 |
| RPL21 | 30322 | 38967 | 62753 | 208048 | 141474 | 220676 | 114056 | 74014 |
| RPL22 | 4341392 | 7330982 | 4929557 | 7608069 | 20104480 | 8921036 | 11989567 | 4467709 |
| RPL22L1 | 0 | 3486 | 1418 | 1998 | 3272 | 1857 | 6542 | 3378 |
| RPL23 | 3436 | 5540 | 4370 | 9406 | 15427 | 4622 | 13022 | 2423 |
| RPL23A | 12203 | 28830 | 31508 | 23920 | 82154 | 26323 | 34830 | 23062 |
| RPL23AP53 | 0 | 0 | 0 | 0 | 1104 | 0 | 0 | 0 |
| RPL23AP7 | 0 | 0 | 0 | 1211 | 934 | 0 | 720 | 0 |
| RPL23AP82 | 0 | 0 | 0 | 0 | 0 | 0 | 0 | 16 |
| RPL24 | 2678 | 11939 | 16694 | 33919 | 54872 | 17757 | 24899 | 25623 |

| | | | | | | | | |
|---|---|---|---|---|---|---|---|---|
| RPL26 | 14664 | 39098 | 33813 | 53811 | 100195 | 54640 | 53484 | 41748 |
| RPL26L1 | 638 | 874 | 1306 | 2346 | 2558 | 0 | 1018 | 1634 |
| RPL27 | 6172 | 12688 | 14959 | 23619 | 43456 | 15106 | 21355 | 11099 |
| RPL27A | 10359 | 22120 | 15284 | 35134 | 49578 | 23081 | 49870 | 23868 |
| RPL28 | 9976 | 21163 | 4890 | 15243 | 66687 | 14667 | 20443 | 11191 |
| RPL29 | 16283 | 20673 | 25685 | 69253 | 121651 | 59634 | 55971 | 24827 |
| RPL3 | 14261 | 23858 | 58436 | 89480 | 112824 | 113973 | 51107 | 42917 |
| RPL30 | 10244 | 16397 | 21473 | 47027 | 134840 | 84150 | 85891 | 38621 |
| RPL31 | 10738 | 26570 | 21234 | 43632 | 86544 | 38139 | 46359 | 27645 |
| RPL32 | 14132 | 49064 | 38434 | 80800 | 166152 | 68597 | 64801 | 35900 |
| RPL32P3 | 0 | 0 | 0 | 0 | 330 | 0 | 0 | 0 |
| RPL34 | 22450 | 34661 | 21612 | 44152 | 81843 | 32258 | 54798 | 13851 |
| RPL35 | 11999 | 20504 | 14304 | 29064 | 95245 | 20924 | 35399 | 15486 |
| RPL35A | 10708 | 22821 | 20279 | 38480 | 75210 | 28501 | 55568 | 29081 |
| RPL36 | 10768 | 25922 | 8143 | 29873 | 75141 | 18888 | 28867 | 17627 |
| RPL36A | 3030 | 24167 | 14572 | 38331 | 105723 | 25894 | 41422 | 14605 |
| RPL36AL | 3788 | 10211 | 9904 | 3119 | 15746 | 862 | 5284 | 6106 |
| RPL37 | 8662 | 19108 | 6732 | 19094 | 95301 | 11949 | 12786 | 11077 |
| RPL37A | 10782 | 19910 | 5159 | 23155 | 74022 | 18095 | 37388 | 14791 |
| RPL38 | 4653 | 5771 | 1724 | 8810 | 29927 | 5976 | 15865 | 4189 |
| RPL39 | 7868 | 26684 | 5136 | 14002 | 88414 | 11732 | 45888 | 5063 |
| RPL39L | 0 | 496 | 0 | 0 | 942 | 0 | 0 | 0 |
| RPL4 | 6964 | 13770 | 24376 | 67973 | 52037 | 25905 | 31893 | 23257 |
| RPL41 | 47449 | 120589 | 79289 | 119955 | 196098 | 81011 | 131232 | 62015 |
| RPL5 | 2629520 | 4489868 | 3552908 | 7457350 | 12776159 | 11244216 | 21141486 | 4516647 |
| RPL6 | 13991 | 18603 | 37795 | 108976 | 75611 | 62359 | 49725 | 29118 |
| RPL7 | 14909 | 44087 | 84705 | 148244 | 332150 | 424900 | 283670 | 151879 |
| RPL7A | 6922 | 10042 | 17997 | 48688 | 62535 | 67121 | 43267 | 17664 |
| RPL7L1 | 0 | 512 | 0 | 1052 | 64 | 0 | 972 | 88 |
| RPL8 | 6877 | 10839 | 32753 | 47716 | 125996 | 119918 | 59817 | 51391 |
| RPL9 | 16660 | 23819 | 38785 | 46895 | 79687 | 49794 | 52294 | 37904 |
| RPLP0 | 12238 | 27324 | 56308 | 160734 | 134873 | 86925 | 93071 | 46313 |
| RPLP1 | 29896 | 39266 | 53123 | 109429 | 138969 | 38945 | 106457 | 37925 |
| RPLP2 | 12993 | 22467 | 8448 | 34018 | 50040 | 25575 | 33430 | 21682 |
| RPN1 | 0 | 554 | 2871 | 3694 | 3790 | 1570 | 308 | 3291 |
| RPN2 | 1044 | 1318 | 1238 | 5516 | 3040 | 4140 | 2554 | 2753 |
| RPP14 | 0 | 0 | 0 | 88 | 414 | 0 | 0 | 0 |
| RPP21 | 680 | 1990 | 3524 | 2105 | 4462 | 2470 | 2262 | 918 |
| RPP25 | 0 | 0 | 0 | 1288 | 1103 | 0 | 2205 | 0 |
| RPP25L | 822 | 240 | 1646 | 2778 | 2834 | 2046 | 0 | 0 |
| RPP30 | 556 | 422 | 5479 | 1212 | 822 | 0 | 0 | 0 |
| RPP38 | 0 | 0 | 0 | 2797 | 4080 | 0 | 0 | 0 |
| RPP40 | 0 | 0 | 0 | 1112 | 608 | 0 | 0 | 0 |

| | | | | | | | | |
|---|---|---|---|---|---|---|---|---|
| RPRD1A | 0 | 0 | 0 | 1044 | 274 | 0 | 196 | 0 |
| RPRD1B | 0 | 488 | 0 | 0 | 1322 | 0 | 0 | 0 |
| RPRD2 | 0 | 0 | 0 | 0 | 690 | 0 | 1032 | 0 |
| RPRM | 0 | 0 | 0 | 497 | 0 | 0 | 0 | 0 |
| RPS10 | 5673 | 10178 | 10890 | 13028 | 30946 | 13356 | 19468 | 12676 |
| RPS10-NUDT3 | 0 | 0 | 0 | 0 | 0 | 0 | 24 | 12 |
| RPS11 | 6840 | 7312 | 9457 | 16602 | 33215 | 10083 | 9695 | 10333 |
| RPS12 | 17648 | 50383 | 37448 | 59887 | 158361 | 48005 | 85513 | 55269 |
| RPS13 | 14032 | 28144 | 24189 | 53930 | 66880 | 30240 | 43598 | 41609 |
| RPS14 | 14514 | 36755 | 54193 | 79349 | 80787 | 66790 | 81498 | 37531 |
| RPS14P3 | 160 | 0 | 0 | 0 | 0 | 0 | 0 | 0 |
| RPS15 | 11557 | 23845 | 24072 | 31532 | 80097 | 36920 | 25839 | 15916 |
| RPS15A | 9194 | 40220 | 30252 | 72430 | 94108 | 46908 | 74461 | 42948 |
| RPS16 | 11669 | 10784 | 14271 | 36159 | 67439 | 20436 | 19079 | 25527 |
| RPS17 | 12124 | 33460 | 20617 | 47712 | 78482 | 22277 | 40686 | 24040 |
| RPS18 | 17773 | 46425 | 41823 | 114631 | 191535 | 98378 | 109021 | 75438 |
| RPS19 | 18498 | 42217 | 43677 | 73870 | 219672 | 63162 | 36432 | 51363 |
| RPS19BP1 | 982 | 702 | 650 | 3126 | 40 | 5506 | 1502 | 2358 |
| RPS2 | 4500 | 22335 | 35984 | 103179 | 91946 | 59355 | 63652 | 40607 |
| RPS20 | 6953 | 13094 | 12924 | 25958 | 22268 | 30981 | 45867 | 13548 |
| RPS21 | 8350 | 8720 | 7127 | 15205 | 30127 | 11872 | 28631 | 5128 |
| RPS23 | 27011 | 54490 | 34476 | 70895 | 138217 | 76417 | 61077 | 36989 |
| RPS24 | 14738 | 21706 | 16724 | 18814 | 49644 | 19342 | 32973 | 11126 |
| RPS25 | 10688 | 28936 | 29015 | 37216 | 83232 | 36153 | 43471 | 27309 |
| RPS26 | 8993 | 14628 | 9335 | 14148 | 26299 | 5322 | 17125 | 2746 |
| RPS27 | 19156 | 49060 | 16986 | 35951 | 112231 | 26036 | 73992 | 27040 |
| RPS27A | 22383 | 45626 | 39461 | 88656 | 168953 | 53559 | 70865 | 54715 |
| RPS27L | 4588 | 9165 | 15585 | 10122 | 14380 | 5991 | 7870 | 3820 |
| RPS28 | 16760 | 27968 | 16254 | 27420 | 108470 | 21228 | 34918 | 11237 |
| RPS29 | 9202 | 31793 | 11292 | 12083 | 82426 | 6112 | 17381 | 9256 |
| RPS3 | 17042 | 20616 | 46969 | 102614 | 162316 | 72084 | 81142 | 82997 |
| RPS3A | 18059 | 45445 | 88241 | 137258 | 168722 | 125617 | 90555 | 100425 |
| RPS4X | 9339 | 28629 | 49485 | 96405 | 153468 | 105231 | 75221 | 57861 |
| RPS4Y1 | 92 | 3226 | 7666 | 0 | 33375 | 1900 | 0 | 5542 |
| RPS5 | 3768 | 8702 | 28033 | 36778 | 71033 | 55595 | 21150 | 19786 |
| RPS6 | 2696172 | 4369503 | 5418912 | 8991734 | 14410053 | 16150074 | 23387771 | 4942195 |
| RPS6KA1 | 22406 | 13731 | 72396 | 24049 | 31009 | 12231 | 16224 | 3252 |
| RPS6KA2 | 0 | 0 | 0 | 0 | 0 | 0 | 0 | 674 |
| RPS6KA3 | 4856 | 5728 | 6418 | 6361 | 19574 | 3812 | 5606 | 1540 |
| RPS6KA4 | 0 | 0 | 0 | 556 | 284 | 706 | 652 | 0 |
| RPS6KB1 | 0 | 0 | 0 | 0 | 941 | 0 | 0 | 0 |
| RPS6KB2 | 500 | 0 | 0 | 998 | 0 | 0 | 400 | 0 |
| RPS6KL1 | 0 | 0 | 0 | 0 | 226 | 0 | 0 | 0 |

| | | | | | | | | |
|---|---|---|---|---|---|---|---|---|
| RPS7 | 10418 | 20271 | 40116 | 65485 | 105856 | 50826 | 59328 | 52967 |
| RPS8 | 5080 | 31704 | 37625 | 73019 | 124450 | 86026 | 74839 | 38815 |
| RPS9 | 6304 | 11799 | 17347 | 30998 | 44054 | 29592 | 17944 | 15733 |
| RPSA | 4894 | 12545 | 10263 | 47351 | 69036 | 26877 | 19297 | 19282 |
| RPSAP52 | 0 | 0 | 0 | 0 | 0 | 2042 | 750 | 0 |
| RPSAP58 | 0 | 48 | 2470 | 2531 | 1854 | 4452 | 3178 | 0 |
| RPSAP9 | 714 | 0 | 0 | 0 | 0 | 0 | 0 | 0 |
| RPTOR | 0 | 1254 | 1713 | 270 | 5970 | 0 | 3794 | 690 |
| RPUSD1 | 0 | 390 | 0 | 898 | 242 | 0 | 0 | 0 |
| RPUSD3 | 0 | 462 | 0 | 0 | 1778 | 0 | 1722 | 0 |
| RPUSD4 | 638 | 0 | 0 | 0 | 700 | 0 | 150 | 0 |
| RRAGA | 0 | 1514 | 0 | 0 | 2738 | 38 | 3867 | 0 |
| RRAGB | 0 | 0 | 0 | 0 | 842 | 184 | 0 | 0 |
| RRAGC | 0 | 0 | 0 | 0 | 264 | 0 | 0 | 0 |
| RRAS | 1250 | 324 | 2454 | 1096 | 274 | 0 | 1050 | 3070 |
| RRAS2 | 252 | 0 | 1392 | 646 | 292 | 0 | 388 | 0 |
| RRBP1 | 408 | 1316 | 438 | 1426 | 3292 | 0 | 928 | 868 |
| RREB1 | 0 | 468 | 0 | 0 | 0 | 0 | 0 | 0 |
| RRM1 | 0 | 0 | 0 | 0 | 2056 | 0 | 0 | 0 |
| RRM2 | 0 | 538 | 0 | 2134 | 228 | 0 | 0 | 0 |
| RRM2B | 0 | 0 | 0 | 0 | 724 | 0 | 0 | 1384 |
| RRN3 | 0 | 0 | 0 | 80 | 464 | 0 | 234 | 0 |
| RRNAD1 | 0 | 220 | 0 | 0 | 418 | 0 | 438 | 462 |
| RRP1 | 0 | 0 | 0 | 0 | 552 | 0 | 0 | 2145 |
| RRP12 | 0 | 278 | 0 | 0 | 0 | 0 | 0 | 0 |
| RRP15 | 0 | 240 | 0 | 0 | 1520 | 0 | 0 | 0 |
| RRP1B | 0 | 0 | 0 | 0 | 1048 | 0 | 162 | 0 |
| RRP36 | 0 | 0 | 0 | 0 | 22 | 2511 | 0 | 0 |
| RRP7A | 0 | 1038 | 5095 | 0 | 2630 | 0 | 2178 | 554 |
| RRP8 | 0 | 0 | 1293 | 0 | 0 | 754 | 0 | 0 |
| RRP9 | 0 | 148 | 0 | 1268 | 1244 | 0 | 0 | 284 |
| RRS1 | 0 | 0 | 0 | 392 | 1922 | 741 | 1282 | 0 |
| RSAD1 | 0 | 0 | 0 | 438 | 160 | 0 | 0 | 0 |
| RSAD2 | 0 | 0 | 828 | 0 | 72 | 0 | 0 | 0 |
| RSBN1 | 0 | 596 | 0 | 0 | 0 | 0 | 0 | 0 |
| RSF1 | 100043 | 78972 | 59328 | 52192 | 176753 | 45729 | 150580 | 43135 |
| RSL1D1 | 0 | 128 | 0 | 0 | 4340 | 2953 | 1444 | 4208 |
| RSL24D1 | 0 | 564 | 1200 | 3046 | 2718 | 2016 | 3533 | 1712 |
| RSPH1 | 0 | 422 | 0 | 1102 | 0 | 0 | 0 | 0 |
| RSPH14 | 0 | 0 | 0 | 0 | 0 | 0 | 354 | 0 |
| RSPH3 | 0 | 0 | 1024 | 0 | 314 | 0 | 0 | 0 |
| RSPO2 | 0 | 0 | 0 | 0 | 170 | 0 | 0 | 0 |
| RSPO3 | 8848 | 19657 | 16402 | 3702 | 11923 | 2568 | 584 | 3585 |

| | | | | | | | | |
|---|---|---|---|---|---|---|---|---|
| RSPRY1 | 0 | 232 | 0 | 0 | 0 | 0 | 198 | 420 |
| RSRC1 | 4926 | 16856 | 24355 | 46013 | 129025 | 21569 | 36959 | 51596 |
| RSRC2 | 68927 | 126628 | 152841 | 153107 | 415425 | 192182 | 332978 | 137124 |
| RSRP1 | 0 | 856 | 843 | 414 | 1454 | 1578 | 1542 | 0 |
| RSU1 | 290222 | 283305 | 1154185 | 618404 | 818267 | 637541 | 1329614 | 316994 |
| RTCA | 354 | 644 | 2141 | 0 | 476 | 0 | 2566 | 1192 |
| RTCB | 662 | 0 | 2410 | 1970 | 2695 | 0 | 1370 | 670 |
| RTEL1 | 0 | 0 | 617 | 0 | 1192 | 0 | 0 | 692 |
| RTEL1-TNFRSF6B | 0 | 248 | 0 | 0 | 90 | 0 | 0 | 0 |
| RTF1 | 0 | 0 | 1812 | 0 | 892 | 0 | 1051 | 0 |
| RTF2 | 364 | 794 | 4044 | 6598 | 6290 | 670 | 1693 | 1108 |
| RTKN | 0 | 420 | 0 | 0 | 1310 | 0 | 324 | 0 |
| RTKN2 | 58 | 392 | 0 | 4513 | 5381 | 40 | 2048 | 4550 |
| RTL10 | 0 | 0 | 0 | 0 | 674 | 0 | 0 | 0 |
| RTL5 | 1788 | 0 | 0 | 0 | 930 | 4416 | 324 | 0 |
| RTL8A | 1406 | 328 | 1340 | 1894 | 0 | 0 | 194 | 456 |
| RTL8C | 0 | 0 | 580 | 0 | 1156 | 0 | 0 | 0 |
| RTN3 | 782 | 474 | 32 | 2324 | 770 | 0 | 1414 | 1074 |
| RTN4 | 2746 | 1386 | 3054 | 0 | 2006 | 0 | 1409 | 0 |
| RTN4IP1 | 680 | 386 | 0 | 0 | 1002 | 0 | 0 | 0 |
| RTN4RL2 | 0 | 126 | 0 | 0 | 0 | 0 | 0 | 0 |
| RTRAF | 744 | 3106 | 4935 | 5740 | 7248 | 0 | 1942 | 3364 |
| RUBCN | 0 | 0 | 0 | 0 | 404 | 0 | 0 | 0 |
| RUBCNL | 58 | 474 | 0 | 1016 | 1404 | 1506 | 492 | 0 |
| RUFY1 | 32873 | 25130 | 112568 | 54534 | 87576 | 83480 | 105242 | 18933 |
| RUFY2 | 428 | 0 | 0 | 0 | 0 | 0 | 0 | 0 |
| RUFY3 | 0 | 0 | 0 | 388 | 0 | 0 | 0 | 0 |
| RUNDC1 | 0 | 0 | 0 | 0 | 778 | 0 | 0 | 0 |
| RUNDC3A | 0 | 0 | 0 | 0 | 418 | 0 | 0 | 0 |
| RUNX1 | 1572 | 2400 | 1044 | 8966 | 28255 | 26269 | 26380 | 7767 |
| RUNX1T1 | 446 | 188 | 0 | 0 | 2712 | 2937 | 772 | 2200 |
| RUNX2 | 0 | 608 | 2242 | 540 | 932 | 997 | 214 | 1996 |
| RUSC1 | 0 | 0 | 3442 | 764 | 1448 | 0 | 0 | 0 |
| RUVBL1 | 646 | 0 | 0 | 0 | 816 | 2242 | 0 | 2358 |
| RUVBL2 | 0 | 0 | 0 | 2280 | 2471 | 0 | 454 | 0 |
| RWDD1 | 0 | 580 | 1776 | 1312 | 1734 | 0 | 1233 | 464 |
| RWDD2A | 1100 | 498 | 0 | 0 | 1320 | 0 | 0 | 0 |
| RWDD2B | 572 | 0 | 0 | 0 | 0 | 0 | 358 | 1088 |
| RWDD3 | 488 | 196 | 0 | 0 | 1190 | 0 | 380 | 0 |
| RWDD4 | 0 | 214 | 446 | 0 | 1721 | 1639 | 470 | 0 |
| RXRA | 0 | 0 | 1190 | 0 | 0 | 0 | 0 | 730 |
| RXRB | 410 | 440 | 0 | 634 | 0 | 0 | 164 | 0 |
| RXYLT1 | 0 | 0 | 0 | 0 | 0 | 0 | 290 | 0 |

| | | | | | | | | |
|---|---|---|---|---|---|---|---|---|
| RYBP | 5045 | 2758 | 4273 | 2696 | 10270 | 2247 | 6783 | 4516 |
| RYK | 0 | 386 | 0 | 0 | 0 | 0 | 0 | 0 |
| S100A10 | 29322 | 58695 | 70347 | 39696 | 143730 | 32057 | 12653 | 77196 |
| S100A11 | 3554 | 9917 | 3449 | 12064 | 52577 | 12515 | 11149 | 27286 |
| S100A12 | 0 | 0 | 0 | 0 | 0 | 0 | 0 | 120 |
| S100A13 | 294096 | 428575 | 499102 | 153616 | 1558937 | 404791 | 283271 | 399705 |
| S100A14 | 3282 | 1324 | 8095 | 1066 | 6092 | 0 | 2476 | 3856 |
| S100A16 | 2316 | 1262 | 4460 | 7143 | 17093 | 2773 | 1526 | 13791 |
| S100A2 | 1184 | 0 | 0 | 832 | 3138 | 0 | 194 | 565 |
| S100A4 | 462073 | 255859 | 297157 | 176436 | 2495102 | 567292 | 268761 | 292873 |
| S100A5 | 0 | 0 | 0 | 0 | 332 | 0 | 0 | 0 |
| S100A6 | 1773667 | 1126913 | 837540 | 866894 | 8603948 | 950523 | 1846456 | 1401375 |
| S100A7 | 0 | 0 | 0 | 0 | 0 | 0 | 0 | 796 |
| S100A8 | 0 | 0 | 1358 | 684 | 846 | 0 | 0 | 828 |
| S100A9 | 0 | 0 | 1426 | 0 | 618 | 0 | 0 | 80 |
| S100B | 0 | 0 | 0 | 0 | 0 | 0 | 0 | 528 |
| S100P | 0 | 5611 | 1148 | 10025 | 12880 | 4737 | 13010 | 6412 |
| S100PBP | 0 | 0 | 0 | 0 | 0 | 0 | 228 | 0 |
| SAA1 | 0 | 0 | 0 | 0 | 0 | 3643 | 0 | 0 |
| SAA2 | 0 | 0 | 0 | 0 | 0 | 2596 | 0 | 0 |
| SAA2-SAA4 | 0 | 0 | 0 | 0 | 0 | 1724 | 0 | 0 |
| SAAL1 | 1553 | 0 | 0 | 893 | 0 | 0 | 0 | 0 |
| SAC3D1 | 0 | 0 | 2288 | 0 | 0 | 0 | 168 | 0 |
| SACM1L | 0 | 0 | 1230 | 0 | 0 | 0 | 0 | 0 |
| SAE1 | 0 | 0 | 0 | 2057 | 6108 | 1884 | 2104 | 0 |
| SAFB | 732 | 0 | 1296 | 0 | 2346 | 0 | 364 | 0 |
| SAFB2 | 274 | 0 | 0 | 742 | 1092 | 0 | 74 | 562 |
| SAMD10 | 0 | 76 | 0 | 0 | 308 | 1096 | 0 | 0 |
| SAMD12 | 5712 | 8216 | 14425 | 16909 | 92760 | 23161 | 83306 | 29190 |
| SAMD13 | 0 | 0 | 1298 | 0 | 1202 | 0 | 0 | 0 |
| SAMD3 | 1180 | 508 | 1446 | 755 | 15623 | 3020 | 696 | 4792 |
| SAMD4A | 0 | 0 | 0 | 0 | 116 | 0 | 0 | 0 |
| SAMD4B | 0 | 0 | 0 | 0 | 434 | 0 | 0 | 0 |
| SAMD5 | 0 | 104 | 0 | 0 | 2927 | 2354 | 2415 | 1150 |
| SAMD8 | 0 | 0 | 0 | 0 | 0 | 1474 | 0 | 0 |
| SAMD9 | 0 | 928 | 658 | 0 | 0 | 0 | 0 | 0 |
| SAMM50 | 0 | 420 | 1384 | 354 | 312 | 0 | 1571 | 1961 |
| SAMSN1 | 0 | 0 | 0 | 541 | 0 | 1090 | 0 | 0 |
| SAP130 | 0 | 136 | 0 | 0 | 624 | 1377 | 0 | 0 |
| SAP18 | 816 | 2236 | 3961 | 6580 | 9280 | 12621 | 8795 | 6158 |
| SAP30 | 0 | 0 | 0 | 0 | 50 | 0 | 0 | 1502 |
| SAP30BP | 926 | 340 | 0 | 4264 | 408 | 0 | 0 | 0 |
| SAP30L | 0 | 448 | 1836 | 0 | 0 | 0 | 804 | 0 |

| | | | | | | | | |
|---|---|---|---|---|---|---|---|---|
| SAPCD2 | 0 | 0 | 0 | 0 | 852 | 0 | 0 | 0 |
| SAR1A | 0 | 1302 | 1452 | 0 | 124 | 3814 | 1518 | 3584 |
| SAR1B | 0 | 0 | 3414 | 0 | 3557 | 0 | 0 | 242 |
| SARAF | 706 | 2923 | 850 | 2180 | 2304 | 2444 | 1806 | 824 |
| SARNP | 271153 | 443499 | 1053435 | 727840 | 1513517 | 897651 | 1072025 | 580910 |
| SARS | 1438 | 1132 | 4798 | 10202 | 24 | 2676 | 2982 | 1744 |
| SARS2 | 0 | 0 | 1218 | 0 | 1679 | 0 | 0 | 0 |
| SART1 | 0 | 778 | 0 | 0 | 74 | 0 | 362 | 0 |
| SART3 | 0 | 0 | 0 | 0 | 962 | 0 | 0 | 0 |
| SASH1 | 128 | 1322 | 0 | 338 | 3776 | 2434 | 1692 | 2100 |
| SASH3 | 0 | 302 | 0 | 0 | 0 | 0 | 0 | 0 |
| SAT1 | 9744 | 21863 | 68881 | 12245 | 85864 | 19586 | 12512 | 34837 |
| SAT2 | 994 | 404 | 1770 | 472 | 908 | 0 | 484 | 0 |
| SATB1 | 0 | 0 | 0 | 0 | 690 | 0 | 0 | 0 |
| SATB1-AS1 | 0 | 0 | 0 | 0 | 308 | 0 | 0 | 0 |
| SATB2 | 0 | 0 | 0 | 636 | 0 | 0 | 0 | 0 |
| SATB2-AS1 | 0 | 0 | 0 | 0 | 620 | 0 | 0 | 0 |
| SAV1 | 0 | 0 | 0 | 0 | 0 | 0 | 248 | 564 |
| SAYSD1 | 0 | 580 | 0 | 0 | 1960 | 0 | 766 | 0 |
| SBDS | 760 | 694 | 1534 | 0 | 1622 | 767 | 2853 | 1005 |
| SBDSP1 | 0 | 340 | 926 | 0 | 0 | 1828 | 0 | 0 |
| SBF1 | 0 | 618 | 0 | 0 | 778 | 0 | 0 | 0 |
| SBF2 | 0 | 462 | 3677 | 282 | 352 | 0 | 3027 | 132 |
| SBNO1 | 0 | 258 | 1094 | 1389 | 980 | 0 | 124 | 0 |
| SBNO2 | 530 | 0 | 0 | 1082 | 746 | 0 | 688 | 0 |
| SC5D | 10876 | 8430 | 22030 | 14478 | 65792 | 20037 | 42850 | 42475 |
| SCAF1 | 0 | 0 | 0 | 0 | 272 | 0 | 0 | 0 |
| SCAF11 | 0 | 106 | 0 | 0 | 402 | 0 | 1196 | 0 |
| SCAF8 | 0 | 0 | 0 | 334 | 776 | 0 | 0 | 0 |
| SCAI | 0 | 150 | 0 | 0 | 470 | 0 | 0 | 0 |
| SCAMP1 | 0 | 160 | 0 | 0 | 202 | 1216 | 0 | 288 |
| SCAMP1-AS1 | 0 | 0 | 74 | 1150 | 1244 | 0 | 0 | 0 |
| SCAMP2 | 1486 | 1240 | 1444 | 0 | 946 | 0 | 344 | 0 |
| SCAMP3 | 0 | 1044 | 78 | 5084 | 3819 | 2487 | 480 | 988 |
| SCAMP4 | 0 | 206 | 0 | 96 | 1108 | 0 | 0 | 0 |
| SCAND1 | 674 | 806 | 0 | 5679 | 6659 | 0 | 2228 | 0 |
| SCAND2P | 826 | 0 | 0 | 0 | 0 | 0 | 0 | 0 |
| SCAP | 12355 | 10222 | 18475 | 14073 | 45696 | 15243 | 45639 | 8766 |
| SCARB1 | 70093 | 49876 | 211632 | 75916 | 322191 | 105271 | 165256 | 69218 |
| SCARB2 | 0 | 700 | 0 | 0 | 0 | 0 | 0 | 1226 |
| SCARF1 | 0 | 0 | 0 | 0 | 0 | 0 | 304 | 0 |
| SCARF2 | 0 | 0 | 0 | 0 | 236 | 0 | 0 | 0 |
| SCART1 | 0 | 0 | 18 | 0 | 0 | 0 | 0 | 0 |

| | | | | | | | | |
|---|---|---|---|---|---|---|---|---|
| SCCPDH | 0 | 0 | 0 | 130 | 1122 | 0 | 1656 | 3083 |
| SCD | 0 | 240 | 0 | 470 | 1730 | 0 | 2432 | 0 |
| SCD5 | 0 | 1320 | 0 | 1106 | 1422 | 1202 | 1088 | 188 |
| SCEL | 290 | 276 | 0 | 0 | 8504 | 0 | 0 | 6850 |
| SCFD1 | 0 | 0 | 0 | 1352 | 0 | 0 | 1156 | 0 |
| SCFD2 | 0 | 0 | 0 | 0 | 0 | 1550 | 0 | 832 |
| SCG2 | 0 | 0 | 0 | 0 | 310 | 0 | 0 | 0 |
| SCG5 | 0 | 1142 | 0 | 0 | 0 | 0 | 0 | 0 |
| SCGB1C2 | 0 | 426 | 2116 | 0 | 0 | 0 | 0 | 0 |
| SCGB2A1 | 0 | 462 | 1022 | 0 | 0 | 0 | 0 | 0 |
| SCHIP1 | 0 | 0 | 0 | 0 | 838 | 0 | 0 | 0 |
| SCIMP | 0 | 0 | 0 | 886 | 0 | 0 | 0 | 0 |
| SCIN | 1544 | 0 | 8018 | 0 | 1654 | 0 | 0 | 1499 |
| SCLY | 0 | 0 | 0 | 0 | 0 | 0 | 0 | 1080 |
| SCMH1 | 0 | 0 | 0 | 0 | 0 | 0 | 190 | 0 |
| SCML1 | 0 | 0 | 0 | 0 | 0 | 0 | 398 | 532 |
| SCN1B | 0 | 0 | 0 | 0 | 684 | 0 | 0 | 0 |
| SCN8A | 0 | 0 | 0 | 0 | 0 | 0 | 116 | 0 |
| SCNM1 | 208 | 1506 | 7422 | 2588 | 4600 | 5434 | 1782 | 0 |
| SCNN1A | 420 | 1226 | 1712 | 952 | 0 | 2724 | 0 | 0 |
| SCNN1B | 822 | 678 | 0 | 0 | 0 | 0 | 0 | 0 |
| SCNN1G | 0 | 301 | 0 | 0 | 0 | 0 | 0 | 0 |
| SCO1 | 0 | 0 | 724 | 0 | 908 | 0 | 0 | 1032 |
| SCO2 | 968 | 0 | 2020 | 1200 | 1070 | 0 | 546 | 1040 |
| SCOC | 0 | 358 | 0 | 1293 | 832 | 4406 | 976 | 645 |
| SCP2 | 1932 | 1796 | 3597 | 3408 | 3742 | 2604 | 3012 | 156 |
| SCPEP1 | 0 | 510 | 1730 | 0 | 36 | 0 | 0 | 0 |
| SCRIB | 0 | 0 | 18 | 0 | 1002 | 3289 | 302 | 314 |
| SCRN1 | 0 | 0 | 0 | 0 | 902 | 0 | 0 | 0 |
| SCRN2 | 448 | 0 | 1650 | 0 | 0 | 0 | 0 | 0 |
| SCRN3 | 22 | 0 | 0 | 0 | 0 | 0 | 0 | 0 |
| SCT | 0 | 0 | 0 | 0 | 122 | 0 | 204 | 0 |
| SCUBE2 | 0 | 0 | 1756 | 0 | 0 | 0 | 0 | 0 |
| SCX | 0 | 0 | 0 | 138 | 0 | 0 | 0 | 0 |
| SCYL1 | 979 | 0 | 0 | 734 | 706 | 0 | 490 | 502 |
| SCYL2 | 0 | 426 | 0 | 0 | 172 | 0 | 0 | 0 |
| SCYL3 | 0 | 0 | 0 | 0 | 0 | 0 | 330 | 0 |
| SDAD1 | 0 | 162 | 0 | 0 | 0 | 0 | 0 | 128 |
| SDC1 | 0 | 428 | 4652 | 1998 | 2086 | 0 | 1212 | 764 |
| SDC2 | 0 | 0 | 632 | 1501 | 224 | 0 | 0 | 992 |
| SDC4 | 604 | 420 | 1156 | 330 | 2062 | 1058 | 890 | 478 |
| SDCBP | 558 | 1264 | 2651 | 1909 | 3884 | 2134 | 1442 | 3926 |
| SDCBP2 | 3592 | 1990 | 15741 | 0 | 2192 | 0 | 0 | 1870 |

| | | | | | | | | |
|---|---|---|---|---|---|---|---|---|
| SDCCAG3 | 0 | 0 | 0 | 0 | 0 | 0 | 240 | 0 |
| SDCCAG8 | 384 | 314 | 0 | 0 | 1116 | 0 | 728 | 150 |
| SDE2 | 0 | 0 | 0 | 0 | 116 | 0 | 398 | 0 |
| SDF2 | 606 | 154 | 4588 | 0 | 2586 | 1586 | 0 | 0 |
| SDF2L1 | 0 | 600 | 1588 | 1836 | 585 | 0 | 464 | 444 |
| SDF4 | 2822 | 990 | 713 | 1014 | 1104 | 1536 | 260 | 558 |
| SDHA | 402 | 134 | 2435 | 1450 | 9182 | 1604 | 1254 | 282 |
| SDHAF1 | 332 | 1260 | 1738 | 1396 | 1485 | 0 | 0 | 1625 |
| SDHAF2 | 343528 | 497644 | 633998 | 484942 | 1424515 | 644547 | 1391325 | 694130 |
| SDHAF3 | 684 | 134 | 1515 | 1545 | 1526 | 0 | 0 | 0 |
| SDHAF4 | 0 | 276 | 652 | 0 | 562 | 0 | 0 | 0 |
| SDHAP2 | 0 | 0 | 882 | 0 | 1398 | 0 | 0 | 0 |
| SDHB | 403506 | 576653 | 4341273 | 446563 | 921647 | 577885 | 418080 | 362944 |
| SDHC | 168 | 862 | 2854 | 1254 | 5493 | 2230 | 1678 | 792 |
| SDHD | 1918 | 2020 | 10982 | 7702 | 3906 | 6069 | 2056 | 942 |
| SDK2 | 692 | 0 | 0 | 0 | 0 | 0 | 0 | 0 |
| SDR16C5 | 0 | 0 | 2155 | 2706 | 0 | 0 | 368 | 0 |
| SDR39U1 | 296 | 562 | 542 | 1830 | 1056 | 0 | 0 | 0 |
| SDS | 0 | 0 | 0 | 0 | 456 | 0 | 0 | 0 |
| SDSL | 0 | 416 | 0 | 1158 | 0 | 0 | 0 | 0 |
| SEC11A | 238257 | 443427 | 470200 | 540964 | 1051650 | 504676 | 900226 | 427536 |
| SEC11C | 0 | 3048 | 4588 | 2983 | 3606 | 0 | 2460 | 2830 |
| SEC13 | 203518 | 428057 | 1349017 | 762589 | 1703127 | 1052352 | 1467266 | 476541 |
| SEC14L1 | 0 | 0 | 0 | 0 | 230 | 0 | 0 | 0 |
| SEC16A | 1330 | 2175 | 4032 | 3343 | 7802 | 0 | 6354 | 2006 |
| SEC16B | 0 | 0 | 0 | 0 | 226 | 0 | 258 | 0 |
| SEC22B | 0 | 1764 | 976 | 2318 | 3019 | 0 | 1015 | 402 |
| SEC22C | 0 | 0 | 0 | 0 | 2014 | 0 | 496 | 0 |
| SEC23A | 654 | 0 | 0 | 0 | 626 | 1456 | 0 | 0 |
| SEC23A-AS1 | 0 | 0 | 36 | 0 | 0 | 0 | 0 | 0 |
| SEC23B | 11936 | 42464 | 43937 | 72512 | 154396 | 46224 | 84435 | 30247 |
| SEC23IP | 0 | 0 | 0 | 0 | 0 | 0 | 280 | 0 |
| SEC24A | 5362 | 6604 | 6273 | 6118 | 15364 | 13422 | 7534 | 5789 |
| SEC24B | 0 | 34 | 0 | 0 | 474 | 0 | 0 | 0 |
| SEC24C | 614 | 268 | 764 | 864 | 0 | 0 | 0 | 934 |
| SEC24D | 0 | 151 | 0 | 0 | 442 | 672 | 0 | 0 |
| SEC31A | 751 | 0 | 0 | 0 | 0 | 0 | 1684 | 0 |
| SEC31B | 0 | 0 | 0 | 0 | 116 | 0 | 0 | 0 |
| SEC61A1 | 760 | 0 | 0 | 0 | 734 | 0 | 1804 | 0 |
| SEC61B | 2672 | 1834 | 5923 | 5242 | 13291 | 7022 | 6237 | 4587 |
| SEC61G | 1510 | 4380 | 538 | 2770 | 12852 | 1164 | 5230 | 730 |
| SEC62 | 0 | 1058 | 0 | 0 | 374 | 1544 | 1228 | 426 |
| SEC63 | 3035 | 5006 | 2292 | 3437 | 16786 | 2412 | 10560 | 9769 |

| | | | | | | | | |
|---|---|---|---|---|---|---|---|---|
| SECISBP2 | 8962 | 17308 | 7931 | 11318 | 35342 | 33279 | 33752 | 6418 |
| SECTM1 | 730 | 0 | 0 | 0 | 0 | 0 | 0 | 0 |
| SEH1L | 0 | 0 | 72 | 0 | 228 | 0 | 142 | 955 |
| SEL1L | 0 | 0 | 0 | 0 | 222 | 0 | 0 | 0 |
| SEL1L3 | 0 | 120 | 0 | 1060 | 0 | 0 | 326 | 0 |
| SELENBP1 | 8825 | 2920 | 18731 | 3783 | 3356 | 0 | 11628 | 910 |
| SELENOF | 0 | 2228 | 42 | 3241 | 3398 | 0 | 3632 | 3290 |
| SELENOH | 3382 | 1832 | 6611 | 11195 | 15064 | 3488 | 7674 | 9153 |
| SELENOI | 0 | 0 | 0 | 0 | 1698 | 0 | 0 | 0 |
| SELENOK | 1480 | 0 | 1362 | 4633 | 6370 | 6417 | 1461 | 1776 |
| SELENOM | 1586 | 941 | 1430 | 530 | 3354 | 1692 | 416 | 1732 |
| SELENON | 0 | 610 | 0 | 496 | 538 | 0 | 612 | 0 |
| SELENOO | 0 | 0 | 2044 | 0 | 0 | 0 | 0 | 0 |
| SELENOP | 9958 | 8214 | 83260 | 1676 | 12798 | 3866 | 236 | 0 |
| SELENOS | 1366 | 1386 | 3245 | 7703 | 4230 | 0 | 5290 | 1720 |
| SELENOT | 1208 | 710 | 1440 | 2520 | 3173 | 0 | 1462 | 3265 |
| SELENOW | 5166 | 7490 | 25486 | 18269 | 16277 | 8716 | 8750 | 8116 |
| SELP | 0 | 0 | 0 | 842 | 0 | 0 | 0 | 0 |
| SELPLG | 0 | 0 | 210 | 0 | 0 | 1547 | 0 | 1628 |
| SEM1 | 252 | 6403 | 5208 | 9387 | 15883 | 5573 | 4307 | 2071 |
| SEMA3A | 0 | 0 | 0 | 0 | 0 | 0 | 0 | 1648 |
| SEMA3B | 506 | 162 | 26 | 0 | 188 | 1048 | 446 | 0 |
| SEMA3C | 76 | 152 | 102 | 0 | 0 | 0 | 0 | 0 |
| SEMA3D | 904 | 728 | 0 | 0 | 582 | 708 | 188 | 0 |
| SEMA3F | 0 | 0 | 0 | 0 | 858 | 0 | 0 | 0 |
| SEMA4A | 1108 | 224 | 1112 | 1034 | 4067 | 1717 | 1686 | 2985 |
| SEMA4B | 388 | 746 | 240 | 1114 | 6225 | 0 | 883 | 1856 |
| SEMA4D | 0 | 0 | 0 | 0 | 176 | 0 | 0 | 0 |
| SEMA4G | 1112 | 194 | 0 | 922 | 168 | 0 | 0 | 0 |
| SEMA5A | 572 | 558 | 0 | 0 | 1148 | 0 | 0 | 0 |
| SEMA5B | 0 | 0 | 0 | 0 | 528 | 0 | 0 | 0 |
| SEMA6A | 504 | 496 | 0 | 0 | 0 | 0 | 0 | 0 |
| SEMA6D | 514 | 0 | 0 | 0 | 0 | 0 | 0 | 0 |
| SEMA7A | 62 | 0 | 0 | 0 | 0 | 0 | 0 | 0 |
| SENP2 | 0 | 0 | 0 | 0 | 504 | 0 | 0 | 0 |
| SENP3-EIF4A1 | 808 | 2764 | 10458 | 8938 | 16662 | 7527 | 5458 | 9995 |
| SENP6 | 8228 | 5683 | 14143 | 11045 | 27727 | 10807 | 21905 | 11939 |
| SENP7 | 0 | 292 | 0 | 0 | 0 | 0 | 0 | 0 |
| SENP8 | 2218 | 7841 | 22256 | 11364 | 15924 | 5781 | 31332 | 9259 |
| SEPHS1 | 0 | 316 | 1938 | 629 | 0 | 3309 | 566 | 1270 |
| SEPHS2 | 1120 | 486 | 1496 | 786 | 2318 | 0 | 1500 | 0 |
| SEPSECS | 0 | 0 | 0 | 0 | 0 | 0 | 146 | 0 |
| 1-Sep | 629 | 0 | 998 | 1032 | 0 | 0 | 414 | 0 |

| | | | | | | | | |
|---|---|---|---|---|---|---|---|---|
| 10-Sep | 0 | 26 | 0 | 1537 | 319 | 1102 | 284 | 0 |
| 11-Sep | 566 | 0 | 0 | 0 | 1352 | 0 | 0 | 0 |
| 14-Sep | 0 | 0 | 0 | 0 | 750 | 0 | 884 | 0 |
| 2-Sep | 270 | 1466 | 164 | 3648 | 3628 | 0 | 646 | 1384 |
| 4-Sep | 0 | 0 | 1640 | 0 | 1580 | 0 | 0 | 0 |
| SEPT5-GP1BB | 568 | 342 | 0 | 0 | 0 | 0 | 0 | 0 |
| 7-Sep | 874 | 884 | 2720 | 0 | 1782 | 0 | 1300 | 671 |
| 8-Sep | 0 | 0 | 704 | 0 | 152 | 0 | 0 | 0 |
| 9-Sep | 418 | 208 | 0 | 0 | 1732 | 0 | 0 | 0 |
| SERAC1 | 0 | 362 | 0 | 0 | 0 | 0 | 0 | 0 |
| SERBP1 | 886 | 354 | 2263 | 1329 | 1322 | 2159 | 2920 | 3212 |
| SERF1A | 1172 | 1334 | 2730 | 3051 | 1690 | 0 | 426 | 758 |
| SERF2 | 13954 | 21258 | 19322 | 24188 | 42856 | 13523 | 18992 | 10258 |
| SERF2-C15ORF63 | 2763 | 3951 | 712 | 1974 | 4581 | 1852 | 2659 | 1237 |
| SERGEF | 0 | 584 | 0 | 0 | 716 | 0 | 0 | 0 |
| SERINC1 | 342 | 346 | 844 | 0 | 1244 | 0 | 606 | 1320 |
| SERINC2 | 85579 | 94571 | 299340 | 72973 | 120395 | 53637 | 73223 | 18739 |
| SERINC3 | 0 | 1394 | 1629 | 3512 | 280 | 1925 | 5022 | 878 |
| SERINC5 | 170 | 0 | 0 | 0 | 448 | 0 | 410 | 0 |
| SERP1 | 1186 | 1486 | 3959 | 5022 | 2162 | 2065 | 4069 | 4394 |
| SERP2 | 0 | 0 | 0 | 0 | 0 | 1198 | 0 | 0 |
| SERPINA1 | 85515 | 113217 | 1035241 | 167611 | 71240 | 191475 | 93997 | 67301 |
| SERPINB1 | 644 | 5764 | 1456 | 3279 | 17868 | 4560 | 1047 | 7608 |
| SERPINB5 | 0 | 420 | 0 | 948 | 1330 | 0 | 0 | 0 |
| SERPINB6 | 1440 | 1494 | 7558 | 5930 | 6598 | 5784 | 727 | 6780 |
| SERPINB8 | 0 | 554 | 0 | 0 | 0 | 0 | 0 | 1362 |
| SERPINB9 | 0 | 0 | 0 | 0 | 214 | 0 | 282 | 608 |
| SERPINB9P1 | 0 | 0 | 0 | 0 | 0 | 0 | 46 | 0 |
| SERPIND1 | 0 | 0 | 0 | 0 | 666 | 0 | 0 | 0 |
| SERPINE1 | 0 | 0 | 0 | 0 | 302 | 0 | 0 | 0 |
| SERPINE2 | 0 | 0 | 0 | 0 | 1176 | 0 | 290 | 0 |
| SERPINF1 | 0 | 0 | 0 | 1171 | 542 | 0 | 0 | 3411 |
| SERPINF2 | 0 | 0 | 0 | 0 | 0 | 0 | 428 | 0 |
| SERPING1 | 118 | 230 | 1112 | 0 | 1978 | 2204 | 0 | 404 |
| SERPINH1 | 188 | 0 | 1922 | 0 | 3134 | 0 | 0 | 790 |
| SERPINI1 | 0 | 296 | 0 | 0 | 0 | 0 | 678 | 0 |
| SERTAD1 | 172 | 0 | 1852 | 0 | 912 | 2168 | 0 | 0 |
| SERTAD3 | 602 | 470 | 456 | 0 | 0 | 0 | 0 | 852 |
| SERTAD4 | 0 | 0 | 1442 | 0 | 0 | 0 | 52 | 0 |
| SERTAD4-AS1 | 0 | 0 | 1404 | 0 | 0 | 0 | 0 | 0 |
| SESN1 | 300 | 0 | 0 | 0 | 2324 | 0 | 316 | 260 |
| SESN2 | 0 | 0 | 0 | 0 | 448 | 0 | 0 | 0 |
| SESTD1 | 0 | 0 | 0 | 326 | 1106 | 0 | 0 | 0 |

| | | | | | | | | |
|---|---|---|---|---|---|---|---|---|
| SET | 1514 | 2064 | 0 | 1342 | 8598 | 0 | 1906 | 812 |
| SETBP1 | 2011 | 458 | 0 | 0 | 0 | 0 | 1680 | 1171 |
| SETD1A | 0 | 0 | 0 | 0 | 592 | 0 | 0 | 0 |
| SETD1B | 0 | 0 | 0 | 0 | 0 | 0 | 226 | 0 |
| SETD2 | 48124 | 41607 | 66193 | 31294 | 118983 | 42580 | 104084 | 10912 |
| SETD3 | 0 | 316 | 0 | 0 | 1166 | 0 | 0 | 0 |
| SETD5 | 0 | 0 | 0 | 0 | 642 | 0 | 0 | 0 |
| SETD6 | 0 | 0 | 0 | 270 | 212 | 0 | 550 | 0 |
| SETD9 | 770 | 0 | 372 | 0 | 0 | 0 | 0 | 0 |
| SETDB1 | 20078 | 14997 | 12323 | 9609 | 50188 | 21805 | 41957 | 19923 |
| SETDB2 | 0 | 0 | 0 | 0 | 1414 | 0 | 218 | 0 |
| SETMAR | 0 | 0 | 666 | 0 | 0 | 0 | 0 | 1085 |
| SETX | 0 | 0 | 0 | 0 | 1320 | 0 | 0 | 0 |
| SEZ6L2 | 0 | 0 | 0 | 0 | 306 | 0 | 0 | 0 |
| SF1 | 0 | 0 | 0 | 766 | 632 | 0 | 218 | 0 |
| SF3A1 | 350 | 704 | 0 | 0 | 648 | 0 | 140 | 0 |
| SF3A2 | 0 | 88 | 0 | 518 | 780 | 1086 | 256 | 0 |
| SF3A3 | 0 | 0 | 10 | 3392 | 0 | 352 | 1094 | 0 |
| SF3B1 | 87382 | 139558 | 108571 | 111233 | 427817 | 138344 | 250941 | 86673 |
| SF3B2 | 1556 | 1652 | 2154 | 1508 | 3822 | 0 | 3908 | 730 |
| SF3B3 | 3914 | 13934 | 14113 | 17575 | 55739 | 17342 | 67951 | 19254 |
| SF3B4 | 484 | 0 | 0 | 1756 | 1090 | 0 | 2148 | 1474 |
| SF3B5 | 1784 | 1380 | 0 | 3192 | 8195 | 1352 | 4164 | 1443 |
| SF3B6 | 700 | 2300 | 7247 | 11788 | 14758 | 11381 | 8362 | 6502 |
| SFI1 | 0 | 0 | 0 | 0 | 260 | 0 | 0 | 0 |
| SFMBT1 | 0 | 0 | 0 | 0 | 0 | 0 | 96 | 0 |
| SFMBT2 | 0 | 322 | 0 | 0 | 0 | 0 | 0 | 0 |
| SFN | 236051 | 169389 | 506505 | 170530 | 338591 | 172940 | 162654 | 79122 |
| SFPQ | 185734 | 357058 | 278771 | 380094 | 819647 | 445003 | 566296 | 211355 |
| SFR1 | 0 | 0 | 0 | 0 | 320 | 0 | 0 | 0 |
| SFRP2 | 0 | 0 | 0 | 0 | 2116 | 1026 | 0 | 0 |
| SFRP4 | 0 | 0 | 0 | 0 | 0 | 2952 | 0 | 0 |
| SFSWAP | 0 | 0 | 732 | 0 | 418 | 0 | 0 | 0 |
| SFT2D1 | 570 | 2306 | 4653 | 4943 | 5076 | 3453 | 1028 | 2140 |
| SFT2D2 | 0 | 0 | 1483 | 0 | 0 | 0 | 414 | 0 |
| SFT2D3 | 484 | 208 | 781 | 1742 | 276 | 0 | 0 | 0 |
| SFTA2 | 0 | 0 | 0 | 0 | 748 | 0 | 222 | 0 |
| SFTPA2 | 0 | 0 | 1348 | 0 | 0 | 0 | 0 | 0 |
| SFXN1 | 1284 | 0 | 560 | 0 | 1442 | 934 | 0 | 1550 |
| SFXN2 | 0 | 0 | 0 | 1643 | 366 | 0 | 0 | 0 |
| SFXN4 | 980 | 808 | 0 | 2335 | 660 | 0 | 0 | 1096 |
| SFXN5 | 0 | 0 | 0 | 0 | 0 | 0 | 546 | 0 |
| SGCB | 0 | 506 | 0 | 1214 | 672 | 2339 | 0 | 0 |

| | | | | | | | | |
|---|---|---|---|---|---|---|---|---|
| SGF29 | 0 | 0 | 0 | 0 | 1766 | 0 | 0 | 0 |
| SGK1 | 1634 | 1016 | 4454 | 0 | 0 | 0 | 0 | 0 |
| SGK2 | 0 | 0 | 1196 | 0 | 544 | 0 | 790 | 0 |
| SGK3 | 0 | 272 | 0 | 0 | 0 | 0 | 0 | 0 |
| SGMS1 | 0 | 0 | 0 | 0 | 278 | 0 | 0 | 510 |
| SGMS2 | 0 | 0 | 0 | 0 | 0 | 0 | 0 | 810 |
| SGO1 | 0 | 0 | 0 | 0 | 0 | 0 | 0 | 1145 |
| SGPL1 | 0 | 654 | 0 | 0 | 332 | 0 | 0 | 0 |
| SGPP2 | 16298 | 14871 | 37489 | 17619 | 26619 | 6505 | 26478 | 12239 |
| SGSH | 0 | 0 | 481 | 0 | 0 | 0 | 0 | 0 |
| SGSM1 | 232 | 0 | 0 | 0 | 854 | 0 | 324 | 0 |
| SGSM2 | 0 | 0 | 0 | 0 | 0 | 0 | 696 | 0 |
| SGSM3 | 24032 | 37403 | 78721 | 23557 | 53277 | 34728 | 35188 | 29053 |
| SGTA | 4296 | 6231 | 16512 | 10470 | 22352 | 8769 | 12425 | 5776 |
| SGTB | 0 | 0 | 0 | 0 | 1974 | 0 | 0 | 0 |
| SH2B1 | 864 | 0 | 0 | 0 | 668 | 0 | 658 | 0 |
| SH2D2A | 0 | 0 | 0 | 0 | 454 | 0 | 30 | 0 |
| SH2D4A | 194 | 0 | 0 | 0 | 0 | 0 | 834 | 0 |
| SH3BGR | 0 | 0 | 0 | 0 | 0 | 0 | 272 | 0 |
| SH3BGRL | 0 | 2248 | 9710 | 0 | 5098 | 0 | 462 | 522 |
| SH3BGRL2 | 314 | 190 | 434 | 0 | 910 | 0 | 182 | 0 |
| SH3BGRL3 | 12542 | 7864 | 32075 | 18599 | 23146 | 4256 | 7867 | 13709 |
| SH3BP1 | 38 | 358 | 0 | 0 | 0 | 0 | 0 | 0 |
| SH3BP2 | 0 | 134 | 0 | 0 | 0 | 0 | 0 | 0 |
| SH3BP5 | 0 | 508 | 0 | 0 | 564 | 0 | 0 | 0 |
| SH3BP5L | 0 | 0 | 0 | 714 | 0 | 0 | 0 | 0 |
| SH3D19 | 352 | 866 | 0 | 1522 | 1366 | 0 | 2235 | 0 |
| SH3GL1 | 3312 | 5084 | 16236 | 5176 | 17936 | 5679 | 9968 | 5549 |
| SH3GLB1 | 0 | 846 | 0 | 0 | 3156 | 0 | 866 | 3223 |
| SH3GLB2 | 864 | 194 | 2457 | 788 | 218 | 0 | 1106 | 0 |
| SH3KBP1 | 0 | 978 | 0 | 0 | 0 | 0 | 0 | 532 |
| SH3PXD2A | 0 | 312 | 0 | 1237 | 278 | 2180 | 0 | 0 |
| SH3PXD2B | 0 | 0 | 0 | 0 | 1440 | 0 | 0 | 0 |
| SH3RF1 | 0 | 124 | 0 | 0 | 192 | 0 | 0 | 0 |
| SH3RF2 | 7046 | 3684 | 17126 | 14248 | 19160 | 4895 | 24433 | 10094 |
| SH3TC1 | 0 | 358 | 0 | 0 | 224 | 0 | 408 | 0 |
| SH3YL1 | 600 | 166 | 0 | 0 | 2206 | 0 | 1264 | 0 |
| SHANK2 | 0 | 0 | 0 | 0 | 0 | 0 | 296 | 1182 |
| SHARPIN | 0 | 274 | 84 | 0 | 6405 | 1219 | 1190 | 1540 |
| SHB | 6340 | 17860 | 34540 | 27585 | 27067 | 27155 | 56810 | 18529 |
| SHC1 | 1801 | 3115 | 11169 | 3464 | 5084 | 2769 | 6736 | 2804 |
| SHC2 | 0 | 0 | 0 | 0 | 0 | 0 | 326 | 678 |
| SHCBP1 | 0 | 0 | 0 | 0 | 0 | 0 | 290 | 0 |

| | | | | | | | | |
|---|---|---|---|---|---|---|---|---|
| SHD | 554 | 0 | 0 | 0 | 572 | 0 | 0 | 0 |
| SHISA5 | 808 | 74 | 2673 | 2080 | 1394 | 0 | 3029 | 2917 |
| SHISA8 | 0 | 144 | 0 | 0 | 0 | 0 | 0 | 0 |
| SHISA9 | 246 | 0 | 1062 | 1090 | 0 | 0 | 2242 | 0 |
| SHKBP1 | 882 | 0 | 0 | 0 | 260 | 0 | 0 | 0 |
| SHLD1 | 0 | 0 | 0 | 0 | 524 | 0 | 0 | 1094 |
| SHLD2 | 0 | 0 | 728 | 0 | 0 | 0 | 0 | 0 |
| SHMT1 | 0 | 0 | 0 | 0 | 132 | 0 | 558 | 0 |
| SHMT2 | 0 | 572 | 0 | 1150 | 1470 | 0 | 1148 | 0 |
| SHOC2 | 0 | 0 | 928 | 0 | 1100 | 0 | 225 | 0 |
| SHOX | 0 | 0 | 0 | 316 | 0 | 0 | 0 | 0 |
| SHPRH | 452 | 0 | 0 | 0 | 0 | 0 | 0 | 0 |
| SHQ1 | 0 | 332 | 1008 | 0 | 0 | 0 | 0 | 0 |
| SHROOM1 | 0 | 164 | 188 | 0 | 0 | 0 | 0 | 0 |
| SHTN1 | 0 | 478 | 0 | 208 | 312 | 0 | 0 | 0 |
| SI | 0 | 300 | 0 | 0 | 0 | 0 | 0 | 0 |
| SIAE | 400 | 0 | 0 | 0 | 0 | 0 | 0 | 0 |
| SIAH1 | 0 | 458 | 2392 | 0 | 754 | 0 | 238 | 0 |
| SIAH2 | 0 | 0 | 2198 | 892 | 0 | 0 | 352 | 0 |
| SIDT1 | 0 | 222 | 0 | 0 | 0 | 0 | 0 | 0 |
| SIDT2 | 0 | 0 | 0 | 0 | 212 | 0 | 0 | 0 |
| SIGIRR | 1012 | 0 | 1940 | 1332 | 828 | 0 | 2364 | 2335 |
| SIGLEC10 | 0 | 0 | 328 | 0 | 0 | 0 | 0 | 0 |
| SIGLEC9 | 0 | 0 | 0 | 0 | 0 | 2054 | 0 | 0 |
| SIGMAR1 | 0 | 476 | 0 | 900 | 1000 | 1541 | 476 | 996 |
| SIK1B | 582 | 0 | 0 | 0 | 924 | 0 | 0 | 0 |
| SIK2 | 0 | 0 | 0 | 0 | 614 | 0 | 0 | 0 |
| SIK3 | 1920 | 0 | 0 | 0 | 0 | 0 | 0 | 0 |
| SIKE1 | 0 | 256 | 1874 | 2442 | 0 | 0 | 0 | 0 |
| SIL1 | 236 | 0 | 0 | 0 | 0 | 0 | 1429 | 0 |
| SIMC1 | 0 | 0 | 0 | 0 | 336 | 0 | 422 | 0 |
| SIN3A | 0 | 0 | 0 | 0 | 304 | 0 | 244 | 0 |
| SINHCAF | 0 | 0 | 0 | 2852 | 556 | 0 | 959 | 0 |
| SIPA1 | 240 | 0 | 0 | 1304 | 1222 | 0 | 472 | 0 |
| SIPA1L1 | 0 | 0 | 0 | 0 | 1004 | 0 | 0 | 0 |
| SIPA1L3 | 0 | 0 | 1668 | 0 | 90 | 0 | 0 | 0 |
| SIRT1 | 0 | 0 | 0 | 0 | 290 | 0 | 0 | 0 |
| SIRT2 | 0 | 0 | 0 | 0 | 0 | 2024 | 0 | 0 |
| SIRT3 | 0 | 0 | 0 | 1172 | 1590 | 0 | 0 | 1495 |
| SIRT5 | 0 | 0 | 0 | 0 | 0 | 0 | 418 | 1242 |
| SIRT6 | 484 | 0 | 0 | 0 | 0 | 0 | 0 | 0 |
| SIRT7 | 0 | 0 | 823 | 0 | 0 | 0 | 1050 | 0 |
| SIVA1 | 1526 | 642 | 1794 | 1446 | 4522 | 0 | 310 | 1019 |

| | | | | | | | | |
|---|---|---|---|---|---|---|---|---|
| SIX1 | 0 | 0 | 0 | 0 | 548 | 0 | 1018 | 0 |
| SIX5 | 0 | 0 | 0 | 0 | 832 | 0 | 0 | 0 |
| SKA2 | 0 | 0 | 0 | 602 | 2018 | 0 | 130 | 722 |
| SKA3 | 0 | 0 | 0 | 1246 | 850 | 0 | 90 | 0 |
| SKAP1 | 10668 | 10000 | 19396 | 49202 | 83181 | 169189 | 174612 | 43367 |
| SKAP2 | 340 | 0 | 344 | 0 | 154 | 0 | 718 | 0 |
| SKI | 0 | 212 | 0 | 411 | 0 | 0 | 280 | 0 |
| SKIDA1 | 0 | 12 | 0 | 0 | 138 | 2007 | 284 | 257 |
| SKIL | 196 | 0 | 1066 | 0 | 390 | 0 | 0 | 772 |
| SKIV2L | 0 | 0 | 0 | 0 | 0 | 0 | 580 | 0 |
| SKP1 | 4002 | 7040 | 15644 | 22817 | 17058 | 25578 | 11577 | 3502 |
| SLAIN2 | 0 | 0 | 0 | 0 | 608 | 0 | 0 | 0 |
| SLAMF1 | 7736 | 7429 | 8849 | 1356 | 3361 | 26968 | 1144 | 13129 |
| SLAMF9 | 0 | 582 | 0 | 0 | 39927 | 4133 | 4064 | 36829 |
| SLBP | 0 | 1447 | 1324 | 0 | 1324 | 0 | 480 | 1126 |
| SLC10A3 | 0 | 0 | 0 | 1054 | 448 | 0 | 0 | 0 |
| SLC10A5 | 0 | 458 | 0 | 0 | 1004 | 0 | 100 | 0 |
| SLC11A1 | 0 | 0 | 0 | 0 | 620 | 0 | 0 | 0 |
| SLC11A2 | 550 | 240 | 0 | 0 | 156 | 0 | 0 | 0 |
| SLC12A2 | 160 | 0 | 0 | 2812 | 1088 | 3346 | 0 | 0 |
| SLC12A7 | 260 | 452 | 1344 | 1410 | 5941 | 0 | 1608 | 670 |
| SLC12A9 | 0 | 588 | 0 | 620 | 680 | 0 | 672 | 0 |
| SLC13A2 | 0 | 0 | 0 | 1824 | 0 | 0 | 0 | 0 |
| SLC13A3 | 0 | 0 | 0 | 0 | 0 | 0 | 1006 | 0 |
| SLC15A1 | 0 | 0 | 1167 | 0 | 0 | 0 | 0 | 0 |
| SLC15A2 | 0 | 0 | 0 | 992 | 0 | 0 | 0 | 0 |
| SLC15A3 | 0 | 492 | 0 | 0 | 0 | 0 | 0 | 0 |
| SLC16A1 | 20282 | 4188 | 26141 | 19419 | 42986 | 14217 | 144169 | 10702 |
| SLC16A1-AS1 | 0 | 0 | 0 | 0 | 0 | 0 | 262 | 0 |
| SLC16A14 | 192 | 0 | 0 | 0 | 3576 | 0 | 0 | 0 |
| SLC16A3 | 0 | 322 | 0 | 0 | 854 | 1896 | 479 | 294 |
| SLC16A5 | 0 | 496 | 0 | 0 | 756 | 0 | 0 | 0 |
| SLC16A7 | 1728 | 2372 | 1425 | 2056 | 0 | 0 | 0 | 0 |
| SLC16A9 | 766 | 374 | 0 | 0 | 426 | 0 | 0 | 0 |
| SLC17A4 | 0 | 0 | 1187 | 0 | 0 | 0 | 0 | 0 |
| SLC17A5 | 0 | 0 | 0 | 0 | 26 | 0 | 0 | 0 |
| SLC17A9 | 0 | 420 | 0 | 0 | 751 | 0 | 0 | 0 |
| SLC18A1 | 0 | 300 | 0 | 0 | 0 | 0 | 0 | 0 |
| SLC18B1 | 0 | 0 | 1566 | 0 | 344 | 0 | 0 | 0 |
| SLC19A2 | 0 | 0 | 0 | 0 | 0 | 414 | 0 | 0 |
| SLC19A3 | 0 | 0 | 0 | 0 | 0 | 0 | 36 | 0 |
| SLC1A1 | 0 | 176 | 0 | 0 | 0 | 0 | 0 | 0 |
| SLC1A3 | 0 | 0 | 0 | 0 | 600 | 0 | 0 | 0 |

| | | | | | | | | |
|---|---|---|---|---|---|---|---|---|
| SLC1A5 | 13556 | 32911 | 28765 | 50626 | 84707 | 103824 | 121418 | 32953 |
| SLC1A7 | 0 | 0 | 0 | 0 | 0 | 0 | 1020 | 0 |
| SLC20A1 | 412 | 0 | 1576 | 2079 | 260 | 0 | 222 | 0 |
| SLC20A2 | 18097 | 14526 | 53581 | 33419 | 54867 | 20701 | 47682 | 19429 |
| SLC22A16 | 842 | 0 | 0 | 0 | 2407 | 0 | 0 | 1252 |
| SLC22A17 | 0 | 0 | 0 | 0 | 366 | 0 | 286 | 0 |
| SLC22A18 | 462 | 200 | 0 | 1474 | 2529 | 1933 | 1322 | 792 |
| SLC22A18AS | 510 | 470 | 735 | 0 | 0 | 0 | 0 | 0 |
| SLC22A23 | 4452 | 958 | 4328 | 1206 | 4266 | 3022 | 6280 | 1153 |
| SLC23A3 | 0 | 0 | 0 | 0 | 0 | 0 | 0 | 470 |
| SLC24A3 | 0 | 0 | 0 | 0 | 690 | 0 | 0 | 0 |
| SLC25A1 | 602 | 600 | 950 | 0 | 2572 | 0 | 864 | 802 |
| SLC25A10 | 62 | 0 | 892 | 0 | 484 | 0 | 0 | 0 |
| SLC25A11 | 1268 | 340 | 3113 | 0 | 1769 | 0 | 1438 | 0 |
| SLC25A12 | 0 | 0 | 0 | 0 | 160 | 0 | 0 | 0 |
| SLC25A13 | 0 | 252 | 0 | 0 | 0 | 0 | 0 | 0 |
| SLC25A14 | 4065 | 10193 | 8908 | 1370 | 59920 | 16888 | 2948 | 11096 |
| SLC25A17 | 24399 | 63414 | 78766 | 67468 | 236339 | 118935 | 119291 | 93010 |
| SLC25A18 | 0 | 0 | 0 | 900 | 0 | 0 | 0 | 0 |
| SLC25A19 | 0 | 0 | 0 | 0 | 0 | 0 | 320 | 0 |
| SLC25A20 | 0 | 276 | 2220 | 0 | 1148 | 0 | 364 | 0 |
| SLC25A21 | 0 | 0 | 22 | 0 | 342 | 0 | 0 | 0 |
| SLC25A24 | 100 | 774 | 0 | 936 | 817 | 852 | 0 | 512 |
| SLC25A25 | 0 | 0 | 0 | 0 | 0 | 0 | 0 | 1172 |
| SLC25A26 | 0 | 450 | 2572 | 2807 | 4415 | 0 | 0 | 1356 |
| SLC25A27 | 0 | 0 | 0 | 0 | 70 | 0 | 0 | 0 |
| SLC25A28 | 78 | 0 | 0 | 0 | 0 | 0 | 0 | 740 |
| SLC25A3 | 4491 | 10503 | 21771 | 19781 | 20720 | 4634 | 2730 | 12988 |
| SLC25A32 | 2718 | 7120 | 3760 | 12726 | 38832 | 39981 | 44606 | 15262 |
| SLC25A33 | 0 | 0 | 0 | 1318 | 774 | 0 | 248 | 0 |
| SLC25A34 | 0 | 550 | 0 | 0 | 0 | 0 | 0 | 0 |
| SLC25A35 | 0 | 0 | 0 | 1116 | 1362 | 0 | 0 | 0 |
| SLC25A36 | 0 | 0 | 0 | 1006 | 168 | 0 | 0 | 0 |
| SLC25A37 | 0 | 466 | 0 | 0 | 150 | 0 | 0 | 0 |
| SLC25A38 | 0 | 1219 | 2575 | 154 | 1053 | 1480 | 0 | 0 |
| SLC25A39 | 1116 | 340 | 0 | 1134 | 5058 | 0 | 2041 | 0 |
| SLC25A4 | 0 | 1202 | 2216 | 486 | 666 | 1398 | 490 | 1624 |
| SLC25A40 | 0 | 224 | 0 | 0 | 0 | 1638 | 0 | 0 |
| SLC25A43 | 0 | 0 | 0 | 0 | 0 | 0 | 578 | 0 |
| SLC25A44 | 2278 | 7113 | 31512 | 8460 | 13441 | 7725 | 7391 | 4028 |
| SLC25A45 | 360 | 0 | 0 | 0 | 0 | 0 | 0 | 0 |
| SLC25A46 | 0 | 0 | 0 | 346 | 0 | 0 | 0 | 38 |
| SLC25A5 | 8258 | 10610 | 31834 | 17853 | 23527 | 4215 | 4565 | 3031 |

| | | | | | | | | |
|---|---|---|---|---|---|---|---|---|
| SLC25A6 | 86 | 3286 | 13653 | 15177 | 20669 | 15806 | 9315 | 7371 |
| SLC26A1 | 0 | 0 | 0 | 0 | 660 | 0 | 0 | 0 |
| SLC26A11 | 864 | 0 | 2618 | 0 | 2106 | 1810 | 1386 | 0 |
| SLC26A2 | 1356 | 466 | 1403 | 0 | 0 | 0 | 0 | 0 |
| SLC26A3 | 3290 | 6332 | 40775 | 1734 | 1702 | 0 | 1418 | 0 |
| SLC26A4 | 0 | 112 | 0 | 0 | 0 | 0 | 0 | 0 |
| SLC26A6 | 0 | 1212 | 0 | 0 | 380 | 0 | 0 | 0 |
| SLC27A2 | 0 | 1026 | 0 | 1864 | 424 | 0 | 0 | 0 |
| SLC27A3 | 0 | 420 | 0 | 0 | 862 | 0 | 0 | 0 |
| SLC27A6 | 716 | 0 | 0 | 0 | 0 | 0 | 0 | 0 |
| SLC28A2 | 494 | 0 | 0 | 0 | 0 | 0 | 414 | 0 |
| SLC29A1 | 2820 | 8704 | 22864 | 90025 | 174384 | 75875 | 209879 | 41459 |
| SLC29A2 | 0 | 146 | 0 | 0 | 0 | 0 | 814 | 0 |
| SLC29A3 | 2609 | 5692 | 7188 | 5429 | 10427 | 12581 | 24884 | 2920 |
| SLC2A1 | 0 | 0 | 0 | 292 | 2619 | 1268 | 0 | 1720 |
| SLC2A10 | 0 | 0 | 0 | 0 | 0 | 0 | 686 | 0 |
| SLC2A13 | 6138 | 3443 | 12256 | 2444 | 5951 | 0 | 3858 | 52 |
| SLC2A4RG | 694 | 0 | 636 | 0 | 1389 | 0 | 1154 | 1418 |
| SLC2A5 | 0 | 0 | 0 | 0 | 0 | 1280 | 0 | 0 |
| SLC2A8 | 714 | 0 | 0 | 984 | 256 | 0 | 286 | 0 |
| SLC2A9 | 0 | 0 | 0 | 1099 | 0 | 0 | 152 | 0 |
| SLC30A1 | 0 | 0 | 0 | 806 | 0 | 0 | 0 | 0 |
| SLC30A10 | 0 | 0 | 0 | 0 | 0 | 0 | 0 | 12 |
| SLC30A4 | 0 | 0 | 0 | 22 | 0 | 0 | 0 | 0 |
| SLC30A5 | 444 | 0 | 0 | 1466 | 250 | 0 | 658 | 406 |
| SLC30A6 | 1356 | 10464 | 6973 | 17040 | 32009 | 6230 | 41810 | 7740 |
| SLC30A9 | 802 | 632 | 1310 | 0 | 372 | 1501 | 540 | 0 |
| SLC31A1 | 0 | 0 | 0 | 0 | 1724 | 0 | 494 | 0 |
| SLC33A1 | 0 | 318 | 0 | 0 | 0 | 0 | 0 | 0 |
| SLC35A1 | 1028 | 34 | 1662 | 1124 | 1980 | 0 | 0 | 892 |
| SLC35A2 | 0 | 442 | 0 | 742 | 602 | 0 | 0 | 0 |
| SLC35A3 | 894 | 574 | 0 | 894 | 146 | 0 | 410 | 0 |
| SLC35A4 | 268 | 158 | 0 | 0 | 0 | 0 | 0 | 842 |
| SLC35A5 | 0 | 0 | 0 | 787 | 0 | 0 | 0 | 0 |
| SLC35B1 | 152639 | 247643 | 397591 | 326581 | 652082 | 386439 | 832759 | 282346 |
| SLC35B2 | 0 | 0 | 1170 | 0 | 886 | 1528 | 422 | 540 |
| SLC35B3 | 622 | 420 | 0 | 0 | 1606 | 0 | 822 | 0 |
| SLC35C1 | 0 | 0 | 1787 | 900 | 0 | 0 | 0 | 0 |
| SLC35C2 | 0 | 856 | 3210 | 24 | 1366 | 0 | 530 | 0 |
| SLC35D1 | 306 | 284 | 0 | 0 | 236 | 0 | 302 | 0 |
| SLC35D2 | 0 | 0 | 1398 | 0 | 1194 | 0 | 624 | 0 |
| SLC35E2A | 656 | 0 | 0 | 0 | 306 | 0 | 0 | 0 |
| SLC35E2B | 0 | 0 | 0 | 0 | 1518 | 0 | 0 | 0 |

| | | | | | | | | |
|---|---|---|---|---|---|---|---|---|
| SLC35E3 | 0 | 314 | 0 | 0 | 0 | 0 | 0 | 0 |
| SLC35E4 | 0 | 0 | 0 | 736 | 920 | 0 | 0 | 0 |
| SLC35F2 | 0 | 584 | 0 | 0 | 1328 | 0 | 0 | 0 |
| SLC35F5 | 954 | 422 | 0 | 0 | 1898 | 1200 | 502 | 0 |
| SLC35F6 | 0 | 0 | 0 | 0 | 496 | 0 | 0 | 0 |
| SLC35G1 | 608 | 0 | 0 | 0 | 0 | 0 | 0 | 0 |
| SLC37A1 | 348 | 544 | 0 | 0 | 902 | 0 | 0 | 0 |
| SLC37A3 | 0 | 636 | 0 | 2164 | 608 | 74 | 0 | 0 |
| SLC37A4 | 0 | 0 | 1482 | 0 | 1228 | 0 | 0 | 0 |
| SLC38A1 | 600 | 248 | 0 | 0 | 1068 | 0 | 288 | 0 |
| SLC38A10 | 0 | 2200 | 3050 | 1722 | 5834 | 0 | 2148 | 662 |
| SLC38A11 | 0 | 0 | 646 | 0 | 0 | 0 | 0 | 0 |
| SLC38A2 | 0 | 262 | 0 | 0 | 296 | 0 | 468 | 742 |
| SLC38A3 | 0 | 0 | 0 | 0 | 0 | 1014 | 300 | 0 |
| SLC38A5 | 1104 | 6046 | 4558 | 4059 | 58041 | 0 | 19115 | 2903 |
| SLC38A6 | 0 | 0 | 0 | 0 | 516 | 0 | 0 | 0 |
| SLC38A9 | 2939 | 6698 | 14489 | 12737 | 15348 | 3617 | 9572 | 7220 |
| SLC39A1 | 0 | 0 | 0 | 3542 | 374 | 0 | 830 | 0 |
| SLC39A10 | 0 | 0 | 0 | 0 | 316 | 0 | 0 | 0 |
| SLC39A11 | 19258 | 21229 | 85504 | 26940 | 45161 | 48829 | 52478 | 25462 |
| SLC39A13 | 0 | 0 | 0 | 676 | 400 | 0 | 0 | 0 |
| SLC39A14 | 390 | 396 | 1276 | 0 | 196 | 0 | 220 | 0 |
| SLC39A3 | 12684 | 14470 | 41146 | 20815 | 77838 | 18423 | 20483 | 14961 |
| SLC39A4 | 1638 | 390 | 1725 | 3766 | 5954 | 4889 | 15176 | 4940 |
| SLC39A5 | 590 | 1078 | 1400 | 0 | 0 | 0 | 444 | 0 |
| SLC39A7 | 890 | 1150 | 0 | 0 | 1595 | 1608 | 1149 | 0 |
| SLC39A8 | 0 | 628 | 0 | 0 | 0 | 0 | 0 | 0 |
| SLC3A1 | 81281 | 12362 | 320752 | 10203 | 24884 | 0 | 60668 | 3082 |
| SLC3A2 | 0 | 854 | 1882 | 1332 | 1910 | 3293 | 2955 | 1621 |
| SLC40A1 | 0 | 168 | 806 | 2332 | 0 | 0 | 1166 | 176 |
| SLC41A2 | 0 | 0 | 0 | 0 | 208 | 0 | 0 | 0 |
| SLC41A3 | 0 | 0 | 0 | 894 | 510 | 0 | 0 | 0 |
| SLC43A1 | 0 | 480 | 0 | 1178 | 0 | 0 | 0 | 0 |
| SLC43A2 | 934 | 1928 | 1891 | 596 | 1390 | 0 | 2092 | 2175 |
| SLC43A3 | 0 | 0 | 0 | 0 | 1704 | 0 | 0 | 768 |
| SLC44A1 | 0 | 0 | 0 | 0 | 1828 | 0 | 0 | 0 |
| SLC44A2 | 41056 | 59198 | 172327 | 64682 | 156290 | 65771 | 52286 | 50795 |
| SLC44A3 | 0 | 0 | 0 | 634 | 0 | 0 | 920 | 0 |
| SLC44A3-AS1 | 890 | 320 | 0 | 0 | 2574 | 0 | 396 | 0 |
| SLC44A4 | 2509 | 2464 | 730 | 0 | 2948 | 580 | 664 | 902 |
| SLC44A5 | 1786 | 2870 | 1266 | 1292 | 5916 | 2751 | 6072 | 224 |
| SLC45A2 | 0 | 0 | 2095 | 0 | 0 | 0 | 0 | 0 |
| SLC45A3 | 1040 | 442 | 0 | 915 | 1844 | 4000 | 788 | 594 |

| | | | | | | | | |
|---|---|---|---|---|---|---|---|---|
| SLC45A4 | 0 | 0 | 110 | 132 | 0 | 0 | 0 | 0 |
| SLC46A3 | 0 | 0 | 0 | 0 | 835 | 0 | 0 | 0 |
| SLC47A1 | 864 | 726 | 0 | 0 | 640 | 0 | 0 | 492 |
| SLC48A1 | 0 | 0 | 0 | 974 | 956 | 0 | 342 | 0 |
| SLC49A4 | 0 | 0 | 0 | 0 | 1192 | 0 | 0 | 0 |
| SLC4A11 | 0 | 0 | 0 | 0 | 332 | 0 | 0 | 0 |
| SLC4A1AP | 0 | 0 | 0 | 0 | 0 | 0 | 0 | 1230 |
| SLC4A2 | 0 | 306 | 0 | 2952 | 156 | 0 | 0 | 0 |
| SLC4A4 | 1852 | 1918 | 68 | 0 | 38 | 0 | 372 | 0 |
| SLC4A7 | 0 | 106 | 0 | 0 | 0 | 0 | 0 | 0 |
| SLC50A1 | 0 | 1220 | 5601 | 1802 | 4121 | 0 | 2066 | 5372 |
| SLC51B | 2948 | 540 | 2392 | 0 | 0 | 0 | 0 | 0 |
| SLC52A2 | 920 | 194 | 0 | 0 | 154 | 0 | 1160 | 0 |
| SLC52A3 | 706 | 0 | 38 | 0 | 578 | 0 | 0 | 670 |
| SLC5A1 | 0 | 366 | 3050 | 0 | 816 | 0 | 0 | 0 |
| SLC5A3 | 0 | 0 | 0 | 0 | 694 | 0 | 0 | 0 |
| SLC5A6 | 0 | 0 | 1158 | 2604 | 0 | 0 | 800 | 0 |
| SLC6A12 | 0 | 0 | 0 | 936 | 0 | 0 | 0 | 0 |
| SLC6A14 | 0 | 0 | 28 | 0 | 0 | 0 | 0 | 0 |
| SLC6A19 | 1104 | 0 | 0 | 0 | 0 | 0 | 0 | 0 |
| SLC6A20 | 0 | 0 | 0 | 0 | 796 | 0 | 0 | 0 |
| SLC6A4 | 0 | 0 | 0 | 0 | 0 | 28 | 0 | 0 |
| SLC6A8 | 556 | 136 | 0 | 1022 | 1118 | 0 | 0 | 0 |
| SLC7A1 | 0 | 0 | 0 | 0 | 270 | 0 | 808 | 0 |
| SLC7A11 | 0 | 0 | 0 | 0 | 0 | 0 | 0 | 264 |
| SLC7A14 | 0 | 0 | 247 | 0 | 0 | 0 | 0 | 0 |
| SLC7A5 | 0 | 0 | 0 | 112 | 90 | 0 | 462 | 0 |
| SLC7A5P2 | 0 | 0 | 0 | 0 | 0 | 0 | 393 | 0 |
| SLC7A6 | 0 | 0 | 0 | 0 | 0 | 0 | 1066 | 0 |
| SLC7A6OS | 0 | 452 | 0 | 0 | 0 | 0 | 0 | 0 |
| SLC7A7 | 0 | 0 | 1028 | 0 | 0 | 0 | 946 | 0 |
| SLC7A8 | 0 | 394 | 0 | 0 | 0 | 0 | 901 | 0 |
| SLC9A2 | 0 | 0 | 968 | 0 | 0 | 0 | 0 | 534 |
| SLC9A3-AS1 | 0 | 0 | 0 | 0 | 292 | 0 | 0 | 0 |
| SLC9A3R1 | 3114 | 0 | 5686 | 3772 | 1716 | 1072 | 1344 | 704 |
| SLC9A3R2 | 552 | 234 | 0 | 0 | 442 | 0 | 138 | 0 |
| SLC9A4 | 0 | 0 | 0 | 0 | 272 | 0 | 0 | 0 |
| SLC9A7 | 0 | 88 | 0 | 0 | 0 | 0 | 0 | 0 |
| SLC9A8 | 0 | 0 | 0 | 0 | 0 | 0 | 334 | 0 |
| SLCO1B3 | 0 | 0 | 0 | 0 | 1205 | 0 | 718 | 122 |
| SLCO1C1 | 550 | 0 | 0 | 0 | 0 | 0 | 0 | 0 |
| SLCO2A1 | 566 | 536 | 112 | 0 | 628 | 0 | 0 | 0 |
| SLCO2B1 | 0 | 0 | 0 | 0 | 0 | 0 | 466 | 0 |

| | | | | | | | | |
|---|---|---|---|---|---|---|---|---|
| SLCO3A1 | 0 | 0 | 0 | 0 | 622 | 0 | 0 | 0 |
| SLCO4A1 | 0 | 0 | 818 | 1186 | 446 | 0 | 252 | 1264 |
| SLF2 | 0 | 880 | 0 | 931 | 212 | 1104 | 0 | 0 |
| SLIRP | 2112 | 6214 | 1467 | 4030 | 11390 | 0 | 342 | 1480 |
| SLIT3 | 0 | 0 | 0 | 0 | 0 | 830 | 0 | 0 |
| SLK | 0 | 0 | 0 | 0 | 888 | 0 | 0 | 0 |
| SLMAP | 0 | 0 | 0 | 0 | 70 | 0 | 0 | 0 |
| SLMO2-ATP5E | 64 | 0 | 0 | 0 | 470 | 0 | 0 | 1158 |
| SLPI | 0 | 1698 | 0 | 2157 | 4768 | 0 | 276 | 716 |
| SLTM | 0 | 0 | 0 | 0 | 330 | 0 | 0 | 0 |
| SLX1A-SULT1A3 | 204 | 582 | 0 | 0 | 0 | 0 | 126 | 0 |
| SLX1B | 879 | 70 | 3572 | 2995 | 844 | 0 | 944 | 850 |
| SLX1B-SULT1A4 | 0 | 0 | 0 | 0 | 0 | 623 | 0 | 0 |
| SLX4IP | 0 | 128 | 0 | 0 | 1120 | 0 | 246 | 0 |
| SMAD2 | 33166 | 84941 | 123112 | 77361 | 98987 | 73994 | 113446 | 65600 |
| SMAD3 | 5742 | 6308 | 14114 | 4388 | 16432 | 0 | 6757 | 2434 |
| SMAD4 | 3078 | 5128 | 14767 | 4777 | 2766 | 4928 | 16347 | 3778 |
| SMAD6 | 0 | 0 | 0 | 1634 | 294 | 2802 | 2358 | 0 |
| SMAD7 | 0 | 0 | 1150 | 0 | 0 | 0 | 456 | 0 |
| SMAGP | 0 | 826 | 2138 | 3728 | 2640 | 2224 | 2498 | 878 |
| SMAP1 | 0 | 424 | 0 | 960 | 0 | 3899 | 0 | 0 |
| SMAP2 | 9476 | 13410 | 13367 | 11873 | 14850 | 20140 | 16620 | 13104 |
| SMARCA2 | 0 | 230 | 0 | 0 | 1519 | 0 | 0 | 0 |
| SMARCA4 | 86393 | 56522 | 72603 | 38861 | 326769 | 58697 | 83429 | 29859 |
| SMARCA5 | 0 | 94 | 0 | 0 | 0 | 0 | 148 | 0 |
| SMARCB1 | 76891 | 124248 | 260716 | 117527 | 351446 | 198881 | 234726 | 103922 |
| SMARCC1 | 430 | 1138 | 6325 | 6160 | 32151 | 5876 | 16507 | 4520 |
| SMARCC2 | 0 | 0 | 0 | 192 | 0 | 0 | 0 | 0 |
| SMARCD1 | 0 | 0 | 0 | 0 | 252 | 0 | 0 | 0 |
| SMARCD2 | 0 | 0 | 2350 | 334 | 1300 | 1448 | 212 | 0 |
| SMARCE1 | 61851 | 168055 | 150421 | 221884 | 632324 | 186005 | 542255 | 157117 |
| SMC1A | 0 | 0 | 0 | 0 | 0 | 0 | 32 | 0 |
| SMC3 | 2210 | 50 | 4279 | 0 | 5054 | 0 | 4637 | 0 |
| SMC4 | 0 | 0 | 1786 | 1536 | 0 | 0 | 0 | 1346 |
| SMCO3 | 0 | 0 | 0 | 0 | 0 | 0 | 796 | 0 |
| SMCO4 | 210 | 0 | 7066 | 2649 | 3835 | 532 | 0 | 0 |
| SMCR8 | 0 | 262 | 296 | 0 | 0 | 0 | 0 | 0 |
| SMDT1 | 4626 | 2586 | 3606 | 588 | 4258 | 3055 | 442 | 98 |
| SMG1 | 0 | 560 | 0 | 0 | 0 | 0 | 380 | 0 |
| SMG1P2 | 0 | 0 | 0 | 0 | 592 | 0 | 0 | 0 |
| SMG1P5 | 0 | 0 | 2331 | 88 | 0 | 0 | 0 | 0 |
| SMG5 | 126 | 382 | 0 | 510 | 858 | 0 | 0 | 0 |
| SMG6 | 276 | 646 | 0 | 1360 | 2414 | 0 | 2366 | 0 |

| | | | | | | | | |
|---|---|---|---|---|---|---|---|---|
| SMG7 | 386 | 0 | 0 | 0 | 0 | 0 | 994 | 0 |
| SMG8 | 0 | 0 | 0 | 0 | 753 | 0 | 0 | 0 |
| SMG9 | 0 | 0 | 0 | 0 | 438 | 0 | 0 | 0 |
| SMIM1 | 0 | 0 | 0 | 0 | 0 | 0 | 0 | 298 |
| SMIM10L1 | 0 | 508 | 0 | 732 | 1432 | 0 | 626 | 0 |
| SMIM11A | 0 | 0 | 0 | 1082 | 972 | 0 | 258 | 0 |
| SMIM12 | 0 | 702 | 5307 | 0 | 1266 | 1601 | 202 | 1064 |
| SMIM13 | 0 | 0 | 0 | 476 | 0 | 0 | 0 | 0 |
| SMIM14 | 0 | 1550 | 5128 | 0 | 2382 | 0 | 1533 | 696 |
| SMIM15 | 0 | 46 | 2742 | 0 | 4192 | 1513 | 66 | 0 |
| SMIM19 | 1552 | 502 | 1893 | 3204 | 3826 | 2458 | 5008 | 2310 |
| SMIM2-AS1 | 0 | 0 | 1532 | 0 | 0 | 0 | 88 | 0 |
| SMIM20 | 0 | 1798 | 1602 | 1528 | 0 | 2945 | 2020 | 0 |
| SMIM22 | 6841 | 3619 | 6671 | 4446 | 8254 | 0 | 6510 | 2477 |
| SMIM24 | 0 | 196 | 1748 | 0 | 0 | 0 | 0 | 0 |
| SMIM25 | 0 | 0 | 0 | 0 | 0 | 0 | 400 | 0 |
| SMIM26 | 1856 | 1232 | 2650 | 2648 | 8560 | 3978 | 1592 | 1122 |
| SMIM27 | 0 | 22 | 0 | 0 | 0 | 572 | 184 | 0 |
| SMIM29 | 1034 | 0 | 0 | 632 | 0 | 0 | 758 | 0 |
| SMIM3 | 0 | 0 | 0 | 1110 | 1226 | 1598 | 732 | 464 |
| SMIM30 | 0 | 14 | 3198 | 0 | 872 | 0 | 296 | 44 |
| SMIM31 | 0 | 740 | 0 | 0 | 1252 | 0 | 0 | 418 |
| SMIM35 | 450 | 0 | 0 | 0 | 0 | 0 | 0 | 0 |
| SMIM37 | 2600 | 2094 | 2226 | 374 | 4050 | 0 | 692 | 304 |
| SMIM4 | 0 | 1268 | 0 | 533 | 2278 | 456 | 426 | 0 |
| SMIM6 | 840 | 0 | 3424 | 0 | 0 | 0 | 462 | 0 |
| SMIM7 | 236941 | 377989 | 885576 | 309509 | 1253851 | 498306 | 497765 | 282740 |
| SMIM8 | 0 | 0 | 1584 | 0 | 1660 | 0 | 198 | 0 |
| SMN1 | 0 | 0 | 0 | 3287 | 1598 | 840 | 514 | 828 |
| SMN2 | 0 | 1248 | 0 | 767 | 1899 | 0 | 888 | 380 |
| SMNDC1 | 16360 | 29461 | 20679 | 28551 | 83943 | 55475 | 85421 | 29049 |
| SMOC2 | 3207 | 5430 | 11895 | 14299 | 14383 | 34007 | 6770 | 3268 |
| SMPD1 | 374 | 740 | 1354 | 1204 | 804 | 0 | 312 | 0 |
| SMPD2 | 688 | 0 | 151 | 0 | 1742 | 0 | 0 | 0 |
| SMPD4 | 0 | 0 | 0 | 0 | 0 | 0 | 0 | 932 |
| SMPDL3A | 140 | 1036 | 8520 | 512 | 1190 | 0 | 848 | 0 |
| SMPDL3B | 0 | 350 | 2300 | 0 | 5746 | 0 | 1588 | 0 |
| SMPX | 0 | 0 | 0 | 0 | 0 | 1442 | 0 | 0 |
| SMS | 0 | 616 | 2176 | 20 | 3557 | 0 | 518 | 808 |
| SMTN | 1042 | 0 | 0 | 0 | 316 | 0 | 724 | 0 |
| SMTNL1 | 0 | 0 | 0 | 0 | 0 | 0 | 0 | 844 |
| SMU1 | 0 | 0 | 0 | 0 | 118 | 0 | 991 | 0 |
| SMUG1 | 700 | 196 | 0 | 0 | 2757 | 0 | 1329 | 0 |

| | | | | | | | | |
|---|---|---|---|---|---|---|---|---|
| SMURF1 | 0 | 0 | 0 | 1228 | 320 | 0 | 2010 | 1114 |
| SMURF2 | 420 | 0 | 0 | 0 | 0 | 0 | 0 | 0 |
| SMURF2P1-LRRC37BP1 | 0 | 0 | 0 | 0 | 350 | 0 | 0 | 0 |
| SMYD2 | 0 | 390 | 0 | 1794 | 544 | 0 | 0 | 0 |
| SMYD3 | 43434 | 112398 | 217341 | 240846 | 719981 | 524807 | 372439 | 267714 |
| SMYD5 | 0 | 196 | 0 | 0 | 0 | 0 | 0 | 0 |
| SNAI1 | 0 | 0 | 0 | 1177 | 7404 | 0 | 4754 | 2996 |
| SNAI2 | 0 | 0 | 0 | 0 | 852 | 0 | 0 | 0 |
| SNAP23 | 270 | 512 | 1832 | 238 | 2002 | 0 | 1344 | 0 |
| SNAP29 | 854 | 490 | 1148 | 3778 | 382 | 0 | 558 | 0 |
| SNAP47 | 0 | 0 | 0 | 793 | 0 | 0 | 992 | 0 |
| SNAPC1 | 0 | 358 | 0 | 0 | 0 | 0 | 0 | 0 |
| SNAPC2 | 0 | 0 | 0 | 2314 | 0 | 0 | 0 | 0 |
| SNAPC3 | 0 | 0 | 0 | 556 | 0 | 0 | 186 | 0 |
| SNAPC4 | 0 | 0 | 0 | 0 | 330 | 0 | 0 | 0 |
| SNAPC5 | 1026 | 374 | 0 | 1911 | 1681 | 0 | 622 | 0 |
| SNAPIN | 0 | 1274 | 0 | 0 | 3346 | 0 | 468 | 1054 |
| SNCA | 0 | 216 | 0 | 0 | 0 | 0 | 0 | 0 |
| SNCG | 0 | 0 | 0 | 0 | 0 | 0 | 0 | 946 |
| SND1 | 56642 | 120673 | 261737 | 316525 | 420526 | 621082 | 759939 | 184507 |
| SND1-IT1 | 0 | 0 | 0 | 0 | 22 | 0 | 0 | 0 |
| SNF8 | 710 | 1394 | 4687 | 1446 | 2429 | 2294 | 1555 | 2042 |
| SNHG1 | 0 | 306 | 0 | 1454 | 3556 | 806 | 1746 | 0 |
| SNHG10 | 0 | 0 | 406 | 0 | 2664 | 0 | 0 | 844 |
| SNHG11 | 0 | 0 | 0 | 50 | 22 | 0 | 460 | 1130 |
| SNHG12 | 0 | 0 | 1848 | 798 | 490 | 0 | 686 | 524 |
| SNHG15 | 0 | 0 | 0 | 644 | 4085 | 0 | 0 | 0 |
| SNHG16 | 302 | 0 | 0 | 0 | 3164 | 2472 | 1722 | 1504 |
| SNHG17 | 0 | 0 | 0 | 0 | 3494 | 4401 | 1368 | 0 |
| SNHG18 | 0 | 1428 | 0 | 2113 | 2038 | 1794 | 1450 | 0 |
| SNHG19 | 338 | 388 | 0 | 0 | 1938 | 844 | 586 | 0 |
| SNHG21 | 0 | 0 | 647 | 0 | 0 | 0 | 0 | 0 |
| SNHG29 | 3641 | 9254 | 11276 | 32388 | 40661 | 18860 | 14616 | 16709 |
| SNHG3 | 862 | 2700 | 0 | 2806 | 3585 | 2449 | 0 | 0 |
| SNHG5 | 5824 | 4392 | 6577 | 17483 | 19390 | 16128 | 11622 | 12439 |
| SNHG6 | 458 | 1960 | 1268 | 5943 | 9401 | 3454 | 16453 | 1783 |
| SNHG7 | 0 | 968 | 0 | 1830 | 8761 | 0 | 1708 | 2825 |
| SNHG8 | 1850 | 5374 | 210 | 2068 | 6818 | 1862 | 4072 | 346 |
| SNHG9 | 0 | 134 | 0 | 84 | 250 | 0 | 0 | 0 |
| SNIP1 | 0 | 0 | 604 | 0 | 0 | 0 | 0 | 0 |
| SNORC | 320 | 1374 | 1054 | 0 | 1802 | 0 | 699 | 0 |
| SNORD15B | 0 | 0 | 0 | 0 | 0 | 0 | 90 | 0 |
| SNORD3A | 0 | 0 | 0 | 0 | 220 | 0 | 0 | 0 |

| | | | | | | | | |
|---|---|---|---|---|---|---|---|---|
| SNRK | 0 | 0 | 0 | 0 | 260 | 0 | 0 | 0 |
| SNRNP200 | 0 | 0 | 0 | 1776 | 3052 | 2332 | 1386 | 0 |
| SNRNP25 | 2014 | 0 | 2100 | 3414 | 2302 | 0 | 0 | 0 |
| SNRNP27 | 0 | 430 | 0 | 74 | 1632 | 1875 | 0 | 1256 |
| SNRNP35 | 562 | 654 | 1308 | 0 | 1428 | 0 | 758 | 0 |
| SNRNP40 | 0 | 1088 | 0 | 2601 | 504 | 0 | 0 | 0 |
| SNRNP70 | 11931 | 10756 | 20199 | 11053 | 69861 | 16748 | 31929 | 15138 |
| SNRPA | 0 | 142 | 1570 | 1352 | 4378 | 0 | 336 | 1662 |
| SNRPA1 | 0 | 472 | 1458 | 3939 | 5843 | 0 | 5228 | 4200 |
| SNRPB | 618 | 2437 | 1280 | 4031 | 17562 | 0 | 3656 | 1482 |
| SNRPB2 | 0 | 588 | 0 | 0 | 3398 | 0 | 567 | 1098 |
| SNRPC | 1044 | 4878 | 2928 | 3691 | 13580 | 4534 | 6069 | 2968 |
| SNRPD1 | 1046 | 3506 | 842 | 9098 | 12889 | 6777 | 3146 | 6197 |
| SNRPD2 | 1772 | 3706 | 6590 | 13883 | 16875 | 8780 | 4074 | 7811 |
| SNRPD3 | 574 | 1680 | 4164 | 3462 | 9032 | 3135 | 2220 | 1520 |
| SNRPE | 858 | 5449 | 901 | 5565 | 14360 | 630 | 2254 | 3464 |
| SNRPF | 584 | 1778 | 1166 | 3793 | 5474 | 0 | 4126 | 372 |
| SNRPG | 1898 | 6470 | 2355 | 6478 | 13341 | 2845 | 5646 | 4332 |
| SNTB1 | 0 | 0 | 0 | 0 | 0 | 0 | 226 | 0 |
| SNTB2 | 4323 | 5194 | 30306 | 5488 | 18690 | 33943 | 23493 | 11438 |
| SNTG2 | 2214 | 1158 | 0 | 0 | 0 | 0 | 0 | 0 |
| SNU13 | 511700 | 1029328 | 901544 | 737025 | 2492164 | 1324489 | 2490862 | 575109 |
| SNUPN | 0 | 0 | 0 | 3450 | 1389 | 0 | 1288 | 0 |
| SNURF | 0 | 276 | 0 | 0 | 0 | 0 | 0 | 0 |
| SNW1 | 40275 | 54305 | 74469 | 63185 | 214135 | 77860 | 108070 | 59912 |
| SNX1 | 544 | 326 | 0 | 420 | 2280 | 1752 | 2752 | 0 |
| SNX10 | 0 | 0 | 0 | 0 | 350 | 0 | 0 | 0 |
| SNX12 | 0 | 36 | 0 | 0 | 0 | 0 | 0 | 0 |
| SNX13 | 525 | 258 | 0 | 0 | 404 | 0 | 536 | 0 |
| SNX14 | 0 | 0 | 0 | 0 | 0 | 1725 | 0 | 0 |
| SNX15 | 0 | 0 | 0 | 916 | 930 | 0 | 872 | 0 |
| SNX17 | 0 | 360 | 914 | 2096 | 1630 | 3859 | 1800 | 1075 |
| SNX18 | 0 | 0 | 422 | 0 | 0 | 0 | 0 | 0 |
| SNX2 | 976 | 596 | 2403 | 0 | 1660 | 0 | 0 | 1154 |
| SNX20 | 1366 | 478 | 0 | 1722 | 0 | 0 | 0 | 0 |
| SNX21 | 0 | 0 | 0 | 0 | 534 | 0 | 0 | 0 |
| SNX22 | 0 | 0 | 0 | 0 | 0 | 0 | 162 | 0 |
| SNX24 | 0 | 0 | 0 | 0 | 0 | 2163 | 0 | 0 |
| SNX25 | 0 | 0 | 0 | 0 | 576 | 0 | 0 | 0 |
| SNX29 | 2073 | 1628 | 1096 | 808 | 6140 | 2202 | 5544 | 1533 |
| SNX3 | 3330 | 1642 | 8141 | 3931 | 7258 | 8105 | 3611 | 886 |
| SNX30 | 5988 | 500 | 6443 | 2452 | 13300 | 488 | 16164 | 0 |
| SNX32 | 478 | 0 | 0 | 124 | 1272 | 0 | 0 | 0 |

| | | | | | | | | |
|---|---|---|---|---|---|---|---|---|
| SNX33 | 0 | 0 | 0 | 0 | 372 | 0 | 0 | 0 |
| SNX4 | 0 | 376 | 0 | 0 | 1264 | 0 | 394 | 0 |
| SNX5 | 438 | 0 | 0 | 1288 | 3218 | 420 | 0 | 0 |
| SNX6 | 0 | 1238 | 0 | 0 | 1996 | 0 | 0 | 1642 |
| SNX7 | 0 | 386 | 1196 | 0 | 0 | 0 | 1062 | 654 |
| SNX8 | 0 | 0 | 0 | 1302 | 542 | 0 | 0 | 2710 |
| SNX9 | 1342 | 1448 | 1568 | 320 | 7976 | 0 | 4547 | 1360 |
| SOAT1 | 204 | 0 | 574 | 775 | 0 | 384 | 0 | 0 |
| SOBP | 0 | 0 | 0 | 0 | 0 | 0 | 480 | 0 |
| SOCS1 | 0 | 0 | 370 | 0 | 0 | 0 | 0 | 0 |
| SOCS2 | 0 | 518 | 0 | 0 | 128 | 0 | 0 | 0 |
| SOCS6 | 0 | 0 | 0 | 0 | 0 | 1336 | 0 | 0 |
| SOCS7 | 0 | 0 | 0 | 0 | 0 | 0 | 58 | 0 |
| SOD1 | 9806 | 4932 | 28390 | 14558 | 20695 | 7961 | 7136 | 15505 |
| SOD2 | 0 | 1438 | 0 | 820 | 5057 | 4738 | 2693 | 1056 |
| SOD2-OT1 | 0 | 0 | 0 | 0 | 1176 | 0 | 0 | 0 |
| SOD3 | 301 | 252 | 0 | 768 | 1254 | 0 | 1094 | 0 |
| SOGA1 | 0 | 0 | 0 | 0 | 146 | 0 | 0 | 0 |
| SON | 681167 | 485291 | 301244 | 380485 | 1067163 | 386401 | 993670 | 252039 |
| SORBS2 | 4760 | 5588 | 8174 | 3628 | 24745 | 5165 | 9896 | 2734 |
| SORBS3 | 560 | 0 | 0 | 432 | 0 | 0 | 546 | 0 |
| SORD | 0 | 206 | 546 | 0 | 2019 | 0 | 0 | 0 |
| SORL1 | 1224 | 0 | 0 | 0 | 746 | 0 | 544 | 0 |
| SORT1 | 0 | 0 | 392 | 0 | 1470 | 0 | 0 | 988 |
| SOS1 | 2417 | 308 | 6684 | 1289 | 3406 | 0 | 2679 | 0 |
| SOS2 | 0 | 0 | 842 | 0 | 470 | 0 | 0 | 0 |
| SOWAHA | 0 | 0 | 1776 | 0 | 514 | 0 | 0 | 0 |
| SOWAHC | 0 | 0 | 0 | 0 | 452 | 0 | 304 | 0 |
| SOX12 | 0 | 0 | 0 | 0 | 192 | 0 | 0 | 0 |
| SOX13 | 0 | 164 | 0 | 0 | 100 | 0 | 0 | 0 |
| SOX15 | 0 | 0 | 0 | 290 | 0 | 0 | 0 | 0 |
| SOX4 | 0 | 0 | 508 | 770 | 1460 | 0 | 168 | 0 |
| SOX6 | 3224 | 196 | 0 | 0 | 744 | 0 | 2162 | 0 |
| SOX9 | 0 | 0 | 0 | 248 | 0 | 0 | 256 | 0 |
| SOX9-AS1 | 0 | 0 | 0 | 248 | 0 | 0 | 0 | 0 |
| SP1 | 0 | 0 | 0 | 0 | 0 | 0 | 1260 | 0 |
| SP100 | 0 | 182 | 676 | 1678 | 1730 | 3688 | 542 | 0 |
| SP110 | 430 | 0 | 0 | 0 | 0 | 0 | 0 | 0 |
| SP2 | 396 | 180 | 704 | 0 | 248 | 0 | 0 | 0 |
| SP3 | 0 | 0 | 0 | 844 | 659 | 0 | 512 | 0 |
| SP4 | 0 | 0 | 1110 | 0 | 0 | 0 | 0 | 0 |
| SP5 | 0 | 0 | 0 | 692 | 270 | 0 | 1100 | 0 |
| SP6 | 0 | 0 | 0 | 0 | 0 | 0 | 124 | 0 |

| | | | | | | | | |
|---|---|---|---|---|---|---|---|---|
| SPA17 | 2134 | 546 | 7004 | 2663 | 15460 | 2555 | 0 | 2710 |
| SPACA3 | 3712 | 0 | 2162 | 0 | 1800 | 0 | 6049 | 2765 |
| SPACA6 | 0 | 0 | 0 | 0 | 790 | 0 | 0 | 0 |
| SPACA9 | 0 | 0 | 0 | 0 | 0 | 0 | 0 | 940 |
| SPAG1 | 0 | 0 | 1948 | 0 | 554 | 0 | 1094 | 0 |
| SPAG16 | 43435 | 72882 | 71158 | 62343 | 167602 | 17671 | 168888 | 68921 |
| SPAG4 | 0 | 0 | 0 | 0 | 638 | 0 | 0 | 0 |
| SPAG7 | 153354 | 254250 | 2039993 | 202662 | 551195 | 358091 | 581360 | 247568 |
| SPAG9 | 12417 | 8268 | 1978 | 7857 | 8764 | 13630 | 22783 | 5189 |
| SPARC | 154 | 430 | 0 | 3086 | 10564 | 3872 | 1180 | 4805 |
| SPARCL1 | 115214 | 131788 | 117399 | 78681 | 101554 | 128252 | 48735 | 31521 |
| SPART | 0 | 150 | 0 | 0 | 0 | 0 | 0 | 0 |
| SPAST | 5598 | 9977 | 5860 | 8688 | 30737 | 7234 | 27668 | 7578 |
| SPATA12 | 0 | 482 | 335 | 5860 | 8451 | 11996 | 6442 | 2286 |
| SPATA13 | 0 | 0 | 0 | 0 | 186 | 0 | 0 | 0 |
| SPATA2 | 0 | 0 | 0 | 956 | 0 | 0 | 0 | 0 |
| SPATA20 | 1224 | 0 | 0 | 0 | 454 | 0 | 0 | 0 |
| SPATA21 | 206 | 0 | 0 | 0 | 0 | 0 | 224 | 0 |
| SPATA24 | 0 | 0 | 0 | 0 | 494 | 0 | 0 | 0 |
| SPATA33 | 760 | 0 | 0 | 1562 | 0 | 0 | 270 | 0 |
| SPATA5L1 | 0 | 0 | 0 | 0 | 950 | 0 | 234 | 0 |
| SPATA6L | 0 | 354 | 0 | 0 | 200 | 0 | 0 | 0 |
| SPATA7 | 0 | 232 | 0 | 0 | 0 | 0 | 0 | 694 |
| SPATC1L | 0 | 0 | 0 | 0 | 0 | 468 | 0 | 998 |
| SPATS2 | 0 | 0 | 0 | 0 | 0 | 0 | 360 | 0 |
| SPATS2L | 56 | 0 | 1721 | 638 | 2314 | 0 | 242 | 0 |
| SPC24 | 672 | 0 | 0 | 0 | 1292 | 0 | 0 | 0 |
| SPC25 | 0 | 0 | 0 | 0 | 2052 | 1395 | 0 | 1940 |
| SPCS1 | 1974 | 6188 | 8773 | 19542 | 21247 | 11507 | 9526 | 4051 |
| SPCS2 | 1972 | 1108 | 8824 | 4607 | 4248 | 1337 | 4016 | 2544 |
| SPCS3 | 684 | 0 | 4531 | 2640 | 4011 | 0 | 820 | 1818 |
| SPDEF | 498 | 1142 | 1889 | 0 | 810 | 0 | 448 | 2016 |
| SPDL1 | 0 | 0 | 0 | 2191 | 320 | 0 | 0 | 0 |
| SPDYA | 0 | 0 | 0 | 0 | 0 | 0 | 670 | 0 |
| SPDYE11 | 412 | 0 | 0 | 0 | 429 | 0 | 0 | 0 |
| SPDYE3 | 910 | 344 | 376 | 364 | 6593 | 0 | 4843 | 1106 |
| SPECC1 | 80 | 0 | 0 | 0 | 0 | 0 | 0 | 0 |
| SPECC1L | 404 | 0 | 0 | 0 | 0 | 0 | 0 | 0 |
| SPECC1L-ADORA2A | 106 | 0 | 0 | 0 | 0 | 0 | 0 | 0 |
| SPEN | 20279 | 21491 | 5700 | 27590 | 33030 | 27844 | 36498 | 7409 |
| SPG11 | 0 | 0 | 0 | 0 | 0 | 0 | 1310 | 0 |
| SPG21 | 0 | 780 | 0 | 3239 | 0 | 0 | 664 | 0 |
| SPG7 | 1000 | 96 | 0 | 0 | 0 | 0 | 401 | 0 |

| | | | | | | | | |
|---|---|---|---|---|---|---|---|---|
| SPHK2 | 0 | 406 | 0 | 0 | 0 | 0 | 0 | 0 |
| SPI1 | 0 | 0 | 0 | 0 | 0 | 1416 | 0 | 0 |
| SPIB | 638 | 296 | 1584 | 0 | 0 | 0 | 0 | 0 |
| SPICE1 | 0 | 0 | 0 | 0 | 140 | 0 | 0 | 0 |
| SPIDR | 13204 | 19088 | 61332 | 36213 | 198805 | 84185 | 167059 | 41348 |
| SPIN1 | 0 | 0 | 0 | 0 | 322 | 0 | 174 | 0 |
| SPIN2B | 0 | 0 | 0 | 0 | 1070 | 0 | 156 | 0 |
| SPIN3 | 0 | 198 | 0 | 0 | 0 | 0 | 0 | 0 |
| SPIN4 | 322 | 0 | 0 | 0 | 369 | 0 | 0 | 0 |
| SPINDOC | 0 | 0 | 0 | 444 | 570 | 582 | 2450 | 0 |
| SPINK1 | 972 | 11155 | 3987 | 1932 | 3218 | 14 | 11492 | 0 |
| SPINK2 | 0 | 540 | 0 | 0 | 0 | 0 | 0 | 0 |
| SPINK4 | 0 | 16123 | 851 | 6663 | 1816 | 0 | 0 | 0 |
| SPINK5 | 0 | 2578 | 0 | 0 | 1108 | 0 | 0 | 0 |
| SPINT1 | 1596 | 2430 | 2755 | 830 | 4077 | 0 | 1567 | 1616 |
| SPINT1-AS1 | 842 | 546 | 2522 | 0 | 930 | 1898 | 287 | 10 |
| SPINT2 | 3524 | 6838 | 35037 | 26937 | 42093 | 25134 | 13478 | 18347 |
| SPIRE2 | 286 | 968 | 0 | 2356 | 939 | 1208 | 724 | 0 |
| SPN | 306 | 0 | 0 | 0 | 0 | 0 | 0 | 0 |
| SPNS1 | 0 | 306 | 0 | 0 | 576 | 0 | 0 | 492 |
| SPNS2 | 0 | 0 | 0 | 0 | 76 | 0 | 0 | 0 |
| SPOCK1 | 0 | 0 | 0 | 0 | 3957 | 0 | 0 | 2439 |
| SPOCK2 | 0 | 0 | 0 | 0 | 502 | 0 | 710 | 0 |
| SPON1 | 0 | 0 | 0 | 0 | 340 | 2902 | 0 | 656 |
| SPON2 | 50 | 0 | 0 | 0 | 446 | 0 | 0 | 0 |
| SPOP | 247087 | 305981 | 1247700 | 277915 | 716473 | 426511 | 1009337 | 242907 |
| SPOUT1 | 0 | 340 | 0 | 0 | 0 | 3301 | 292 | 0 |
| SPP1 | 0 | 470 | 0 | 988 | 23147 | 3699 | 10 | 21577 |
| SPPL2A | 958 | 418 | 1498 | 0 | 1638 | 0 | 974 | 0 |
| SPPL2B | 2152 | 0 | 968 | 0 | 760 | 0 | 132 | 0 |
| SPPL3 | 856 | 0 | 0 | 174 | 2259 | 0 | 0 | 0 |
| SPR | 80 | 196 | 1938 | 0 | 0 | 0 | 2140 | 1256 |
| SPRED1 | 0 | 0 | 0 | 0 | 0 | 2218 | 188 | 0 |
| SPRR1A | 0 | 0 | 0 | 0 | 592 | 0 | 0 | 0 |
| SPRY1 | 0 | 0 | 0 | 0 | 0 | 0 | 44 | 0 |
| SPRY2 | 324 | 322 | 2990 | 0 | 146 | 1280 | 432 | 676 |
| SPRY4 | 456 | 0 | 0 | 0 | 0 | 0 | 68 | 0 |
| SPRYD3 | 0 | 0 | 0 | 0 | 112 | 0 | 0 | 0 |
| SPRYD4 | 0 | 0 | 1612 | 0 | 0 | 0 | 0 | 144 |
| SPRYD7 | 646 | 736 | 0 | 1012 | 946 | 0 | 0 | 0 |
| SPSB1 | 1034 | 1510 | 10389 | 0 | 2972 | 840 | 2486 | 0 |
| SPSB2 | 0 | 444 | 0 | 0 | 666 | 0 | 0 | 0 |
| SPSB3 | 1214 | 156 | 3717 | 1368 | 2495 | 0 | 0 | 0 |

| | | | | | | | | |
|---|---|---|---|---|---|---|---|---|
| SPTAN1 | 1128 | 438 | 2944 | 572 | 1306 | 660 | 0 | 0 |
| SPTBN1 | 340 | 406 | 1352 | 2123 | 4298 | 0 | 0 | 0 |
| SPTBN2 | 0 | 0 | 0 | 0 | 292 | 0 | 0 | 0 |
| SPTLC1 | 90537 | 185094 | 257938 | 203462 | 470428 | 206525 | 378565 | 67270 |
| SPTLC2 | 0 | 0 | 0 | 0 | 538 | 1204 | 876 | 1179 |
| SPTLC3 | 0 | 0 | 1394 | 0 | 0 | 0 | 0 | 0 |
| SPTSSA | 1520 | 1130 | 3717 | 1005 | 1562 | 202 | 192 | 378 |
| SPTY2D1 | 60 | 0 | 0 | 0 | 0 | 0 | 0 | 0 |
| SPX | 0 | 210 | 0 | 0 | 0 | 0 | 0 | 0 |
| SQLE | 0 | 0 | 0 | 0 | 214 | 0 | 0 | 0 |
| SQOR | 398 | 2226 | 10064 | 5488 | 0 | 0 | 648 | 2192 |
| SQSTM1 | 268959 | 256760 | 2064913 | 609410 | 580931 | 711211 | 1627647 | 260810 |
| SRA1 | 900 | 440 | 0 | 5432 | 1281 | 0 | 0 | 0 |
| SRBD1 | 4028 | 9606 | 11276 | 7587 | 5594 | 4602 | 7679 | 4530 |
| SRC | 0 | 0 | 274 | 0 | 898 | 0 | 1887 | 0 |
| SRCAP | 0 | 0 | 0 | 0 | 0 | 0 | 0 | 1340 |
| SRD5A1 | 0 | 0 | 0 | 0 | 0 | 454 | 0 | 0 |
| SRD5A3 | 0 | 644 | 1710 | 0 | 2659 | 0 | 0 | 1840 |
| SREBF1 | 0 | 0 | 1408 | 0 | 312 | 0 | 238 | 0 |
| SREBF2 | 560 | 0 | 0 | 0 | 502 | 58 | 286 | 712 |
| SREK1 | 0 | 506 | 2174 | 0 | 0 | 0 | 0 | 0 |
| SREK1IP1 | 0 | 0 | 1878 | 892 | 742 | 847 | 124 | 0 |
| SRF | 0 | 0 | 0 | 0 | 0 | 1249 | 114 | 0 |
| SRFBP1 | 23837 | 25616 | 8632 | 33881 | 62879 | 25033 | 32287 | 12001 |
| SRGAP1 | 0 | 0 | 892 | 0 | 0 | 0 | 0 | 0 |
| SRGAP2 | 0 | 0 | 0 | 0 | 174 | 0 | 234 | 1166 |
| SRGAP2B | 0 | 0 | 534 | 0 | 0 | 0 | 0 | 0 |
| SRGAP2C | 0 | 0 | 0 | 0 | 0 | 1026 | 0 | 0 |
| SRGN | 726 | 1406 | 3708 | 788 | 688 | 1884 | 346 | 1552 |
| SRI | 7193 | 18738 | 62802 | 8942 | 18696 | 5246 | 6948 | 2697 |
| SRM | 0 | 894 | 0 | 418 | 3988 | 0 | 718 | 968 |
| SRMS | 0 | 0 | 0 | 0 | 44 | 0 | 0 | 0 |
| SRP14 | 6198 | 13557 | 31044 | 26191 | 21211 | 9796 | 20905 | 9654 |
| SRP19 | 948 | 576 | 0 | 3727 | 112 | 0 | 1134 | 1236 |
| SRP54 | 942 | 1026 | 0 | 1294 | 2032 | 0 | 430 | 0 |
| SRP54-AS1 | 0 | 0 | 0 | 0 | 452 | 0 | 0 | 0 |
| SRP68 | 0 | 0 | 0 | 0 | 244 | 0 | 1068 | 0 |
| SRP72 | 0 | 0 | 486 | 0 | 0 | 2334 | 0 | 0 |
| SRP9 | 1348 | 3390 | 1067 | 7421 | 15445 | 5987 | 1666 | 4830 |
| SRPK1 | 0 | 84 | 1452 | 446 | 4498 | 0 | 848 | 0 |
| SRPK2 | 7425 | 13330 | 30429 | 14957 | 27484 | 21883 | 29908 | 6222 |
| SRPK3 | 0 | 0 | 0 | 530 | 0 | 0 | 0 | 0 |
| SRPRA | 0 | 0 | 0 | 0 | 1232 | 0 | 752 | 0 |

| | | | | | | | | |
|---|---|---|---|---|---|---|---|---|
| SRPRB | 0 | 620 | 0 | 1128 | 1366 | 0 | 4050 | 478 |
| SRPX | 0 | 0 | 0 | 0 | 0 | 0 | 406 | 0 |
| SRPX2 | 0 | 0 | 0 | 0 | 294 | 0 | 0 | 0 |
| SRR | 0 | 0 | 0 | 0 | 0 | 0 | 22 | 0 |
| SRRD | 0 | 0 | 0 | 0 | 990 | 0 | 0 | 0 |
| SRRM1 | 0 | 458 | 0 | 0 | 1350 | 0 | 780 | 0 |
| SRRM2 | 6285 | 3290 | 20313 | 7401 | 12205 | 2781 | 15248 | 2346 |
| SRRM5 | 0 | 0 | 0 | 0 | 0 | 1452 | 0 | 0 |
| SRRT | 0 | 0 | 1833 | 1080 | 688 | 0 | 0 | 0 |
| SRSF1 | 648 | 464 | 0 | 2901 | 1594 | 1706 | 1774 | 636 |
| SRSF10 | 0 | 160 | 0 | 1322 | 0 | 0 | 873 | 640 |
| SRSF11 | 790 | 1272 | 0 | 870 | 1090 | 0 | 0 | 0 |
| SRSF2 | 562 | 3444 | 2513 | 4927 | 4934 | 1215 | 236 | 1828 |
| SRSF3 | 34 | 2130 | 8992 | 8325 | 8753 | 5180 | 5556 | 5247 |
| SRSF4 | 0 | 410 | 1460 | 0 | 2251 | 0 | 0 | 916 |
| SRSF5 | 3140 | 1130 | 9558 | 2647 | 10788 | 593 | 688 | 1649 |
| SRSF6 | 1286 | 246 | 0 | 1693 | 2172 | 0 | 1498 | 130 |
| SRSF7 | 866 | 838 | 44 | 1988 | 4295 | 3910 | 1244 | 3542 |
| SRSF8 | 0 | 62 | 814 | 1209 | 204 | 0 | 416 | 854 |
| SRSF9 | 2932 | 2186 | 1290 | 2587 | 7227 | 3470 | 1716 | 1376 |
| SRXN1 | 0 | 0 | 0 | 0 | 1336 | 958 | 0 | 0 |
| SS18 | 26021 | 42266 | 51726 | 18290 | 171261 | 40297 | 40013 | 32150 |
| SS18L1 | 0 | 0 | 0 | 0 | 0 | 0 | 164 | 800 |
| SS18L2 | 0 | 0 | 1732 | 3790 | 4876 | 1382 | 1732 | 0 |
| SSB | 0 | 1976 | 1658 | 2579 | 1480 | 3907 | 1996 | 0 |
| SSBP1 | 1026 | 4640 | 2740 | 7339 | 15125 | 12106 | 8105 | 2463 |
| SSBP2 | 20974 | 18912 | 89304 | 11790 | 37264 | 39363 | 17063 | 15958 |
| SSBP3 | 320 | 0 | 0 | 0 | 768 | 0 | 0 | 0 |
| SSBP4 | 202 | 528 | 0 | 0 | 1224 | 0 | 298 | 1329 |
| SSC4D | 0 | 0 | 0 | 0 | 1004 | 0 | 0 | 0 |
| SSFA2 | 0 | 506 | 0 | 0 | 532 | 0 | 0 | 0 |
| SSH3 | 0 | 0 | 1300 | 247 | 403 | 978 | 382 | 0 |
| SSNA1 | 558 | 936 | 2303 | 7110 | 4504 | 972 | 0 | 0 |
| SSR1 | 0 | 420 | 0 | 0 | 1604 | 0 | 1068 | 2298 |
| SSR2 | 2108 | 3808 | 6569 | 5871 | 9714 | 4891 | 3730 | 6602 |
| SSR3 | 0 | 386 | 4958 | 2754 | 2461 | 1676 | 1126 | 5512 |
| SSR4 | 2068 | 4490 | 2071 | 7983 | 20934 | 7652 | 5230 | 11892 |
| SSRP1 | 0 | 0 | 1073 | 236 | 5964 | 940 | 1008 | 2006 |
| SSSCA1 | 902 | 0 | 1288 | 0 | 2926 | 0 | 744 | 0 |
| SSSCA1-AS1 | 0 | 0 | 0 | 0 | 0 | 0 | 0 | 872 |
| SST | 0 | 0 | 0 | 0 | 676 | 0 | 0 | 0 |
| SSU72 | 60685 | 109878 | 276072 | 73733 | 281645 | 122291 | 54212 | 64749 |
| SSX2IP | 0 | 0 | 0 | 0 | 0 | 0 | 350 | 0 |

| | | | | | | | | |
|---|---|---|---|---|---|---|---|---|
| SSX6P | 0 | 0 | 0 | 0 | 506 | 0 | 0 | 0 |
| ST13 | 1512 | 1168 | 709 | 0 | 7490 | 13508 | 4558 | 1704 |
| ST14 | 0 | 2084 | 4988 | 2997 | 3981 | 2174 | 1001 | 884 |
| ST18 | 0 | 1310 | 0 | 0 | 4068 | 0 | 0 | 292 |
| ST20 | 2016 | 422 | 1192 | 0 | 2125 | 0 | 502 | 1044 |
| ST20-MTHFS | 652 | 460 | 872 | 3697 | 0 | 0 | 3492 | 360 |
| ST3GAL1 | 926 | 2080 | 1174 | 2340 | 6097 | 2114 | 65709 | 11718 |
| ST3GAL2 | 0 | 0 | 0 | 0 | 0 | 0 | 212 | 0 |
| ST3GAL3 | 0 | 0 | 0 | 0 | 2160 | 0 | 0 | 0 |
| ST3GAL4 | 0 | 1224 | 1122 | 888 | 519 | 0 | 0 | 0 |
| ST3GAL5 | 72 | 0 | 0 | 0 | 0 | 0 | 0 | 0 |
| ST5 | 0 | 0 | 0 | 0 | 34 | 0 | 0 | 272 |
| ST6GAL1 | 2008 | 6780 | 1687 | 3312 | 4988 | 0 | 10784 | 3546 |
| ST6GALNAC1 | 500 | 1950 | 1254 | 1592 | 944 | 0 | 0 | 0 |
| ST6GALNAC2 | 580 | 764 | 0 | 0 | 3675 | 46 | 0 | 0 |
| ST6GALNAC4 | 0 | 664 | 0 | 846 | 0 | 0 | 0 | 0 |
| ST6GALNAC6 | 1050 | 72 | 3076 | 0 | 0 | 0 | 950 | 0 |
| ST7 | 16093 | 46462 | 82376 | 83933 | 116289 | 131660 | 116903 | 32326 |
| ST7-AS1 | 606 | 0 | 0 | 0 | 344 | 0 | 430 | 0 |
| ST7-OT4 | 0 | 0 | 0 | 0 | 770 | 0 | 488 | 0 |
| ST7L | 0 | 1132 | 0 | 758 | 496 | 2068 | 108 | 0 |
| ST8SIA6 | 0 | 0 | 0 | 0 | 0 | 0 | 660 | 0 |
| STAB1 | 0 | 166 | 522 | 0 | 0 | 0 | 0 | 0 |
| STAG1 | 32785 | 34617 | 62430 | 32266 | 71127 | 22509 | 84545 | 27792 |
| STAG2 | 72419 | 51275 | 78246 | 44013 | 164277 | 47117 | 183352 | 16487 |
| STAG3 | 0 | 934 | 0 | 0 | 0 | 0 | 0 | 0 |
| STAG3L4 | 0 | 0 | 0 | 0 | 738 | 2602 | 536 | 0 |
| STAG3L5P-PVRIG2P-PILRB | 0 | 0 | 974 | 0 | 1026 | 0 | 0 | 0 |
| STAM | 348 | 498 | 0 | 0 | 0 | 0 | 0 | 0 |
| STAM2 | 0 | 0 | 2444 | 0 | 0 | 0 | 0 | 0 |
| STAMBP | 0 | 0 | 1189 | 230 | 766 | 0 | 0 | 0 |
| STAMBPL1 | 0 | 430 | 1730 | 1076 | 0 | 0 | 372 | 0 |
| STAP2 | 5668 | 2190 | 8507 | 1240 | 6859 | 304 | 0 | 3350 |
| STARD10 | 0 | 0 | 2125 | 3738 | 3871 | 2440 | 3616 | 2936 |
| STARD13 | 0 | 0 | 0 | 0 | 0 | 1285 | 0 | 0 |
| STARD3 | 0 | 0 | 0 | 1148 | 0 | 0 | 2434 | 0 |
| STARD3NL | 272 | 0 | 0 | 2448 | 726 | 1774 | 836 | 0 |
| STARD5 | 864 | 286 | 0 | 1078 | 0 | 0 | 0 | 1926 |
| STARD7 | 1150 | 990 | 178 | 0 | 1757 | 0 | 650 | 1298 |
| STARD7-AS1 | 0 | 0 | 0 | 594 | 170 | 0 | 0 | 0 |
| STAT1 | 0 | 0 | 1751 | 0 | 1335 | 0 | 198 | 2376 |
| STAT2 | 0 | 50 | 0 | 0 | 0 | 484 | 0 | 0 |
| STAT3 | 22359 | 37865 | 53953 | 29942 | 65241 | 30489 | 90874 | 22630 |

| | | | | | | | | |
|---|---|---|---|---|---|---|---|---|
| STAT5A | 2696 | 3020 | 2646 | 5768 | 19767 | 2178 | 14216 | 6838 |
| STAT5B | 5750 | 8786 | 11690 | 11707 | 37728 | 12917 | 22841 | 11547 |
| STAT6 | 7414 | 2750 | 9430 | 9603 | 9216 | 9584 | 18089 | 1338 |
| STAU1 | 0 | 0 | 1028 | 0 | 896 | 0 | 484 | 712 |
| STAU2 | 490 | 432 | 0 | 0 | 992 | 0 | 0 | 0 |
| STBD1 | 946 | 376 | 0 | 0 | 0 | 864 | 0 | 0 |
| STEAP1 | 654 | 0 | 0 | 1394 | 774 | 0 | 0 | 0 |
| STEAP3 | 0 | 0 | 0 | 0 | 652 | 0 | 0 | 0 |
| STIL | 132 | 0 | 0 | 0 | 0 | 0 | 768 | 0 |
| STIM1 | 0 | 0 | 0 | 0 | 810 | 0 | 0 | 0 |
| STIM2 | 0 | 0 | 0 | 0 | 200 | 1134 | 0 | 0 |
| STIMATE | 0 | 0 | 0 | 0 | 68 | 0 | 0 | 0 |
| STIP1 | 0 | 0 | 0 | 3628 | 4721 | 0 | 2384 | 0 |
| STK10 | 0 | 360 | 0 | 0 | 0 | 0 | 0 | 0 |
| STK11 | 10266 | 5099 | 13670 | 20695 | 25604 | 10825 | 13916 | 7513 |
| STK11IP | 0 | 0 | 0 | 0 | 143 | 0 | 0 | 1022 |
| STK16 | 0 | 0 | 0 | 0 | 0 | 0 | 470 | 1002 |
| STK17A | 1436 | 0 | 2860 | 102 | 170 | 0 | 0 | 0 |
| STK17B | 0 | 404 | 0 | 0 | 0 | 0 | 0 | 734 |
| STK19 | 0 | 0 | 746 | 0 | 454 | 0 | 0 | 676 |
| STK24 | 3164 | 1369 | 15700 | 12091 | 11630 | 4736 | 10945 | 0 |
| STK25 | 848 | 390 | 1826 | 0 | 4439 | 0 | 322 | 2299 |
| STK3 | 0 | 0 | 0 | 0 | 0 | 0 | 933 | 0 |
| STK31 | 0 | 420 | 0 | 0 | 80971 | 0 | 0 | 13904 |
| STK32C | 906 | 480 | 0 | 0 | 172 | 0 | 0 | 0 |
| STK35 | 1752 | 0 | 1093 | 1734 | 3951 | 0 | 0 | 82 |
| STK38 | 18826 | 23171 | 24568 | 25785 | 47347 | 22903 | 111267 | 10359 |
| STK38L | 0 | 252 | 252 | 0 | 378 | 0 | 0 | 0 |
| STK39 | 3430 | 3031 | 25078 | 9021 | 33966 | 8154 | 12286 | 7092 |
| STK4 | 0 | 0 | 0 | 0 | 310 | 0 | 870 | 1956 |
| STK40 | 0 | 0 | 0 | 2236 | 0 | 0 | 828 | 0 |
| STMN1 | 0 | 0 | 1002 | 18441 | 12770 | 5082 | 1158 | 7789 |
| STMN2 | 64830 | 40274 | 92922 | 1720 | 199956 | 19545 | 886 | 13237 |
| STMN3 | 0 | 0 | 0 | 0 | 232 | 290 | 0 | 0 |
| STMP1 | 0 | 1504 | 2690 | 1922 | 4256 | 0 | 950 | 0 |
| STN1 | 8868 | 7166 | 8189 | 10355 | 18285 | 11465 | 24855 | 10984 |
| STOM | 0 | 573 | 0 | 894 | 1232 | 0 | 0 | 0 |
| STOML2 | 0 | 822 | 2914 | 7778 | 1438 | 0 | 0 | 424 |
| STON2 | 0 | 0 | 0 | 0 | 128 | 0 | 0 | 0 |
| STOX1 | 458 | 0 | 0 | 0 | 0 | 0 | 0 | 0 |
| STPG1 | 0 | 0 | 0 | 0 | 0 | 0 | 488 | 0 |
| STRADB | 0 | 0 | 0 | 0 | 0 | 0 | 0 | 906 |
| STRAP | 0 | 1932 | 1304 | 3646 | 2522 | 1250 | 1168 | 0 |

| | | | | | | | | |
|---|---|---|---|---|---|---|---|---|
| STRBP | 160 | 1216 | 4419 | 0 | 3006 | 244 | 10031 | 440 |
| STRIP1 | 0 | 0 | 1070 | 0 | 0 | 0 | 0 | 0 |
| STRN | 8040 | 3358 | 4122 | 4640 | 7422 | 12163 | 8493 | 0 |
| STRN4 | 0 | 0 | 0 | 0 | 1604 | 1044 | 506 | 0 |
| STT3A | 0 | 1056 | 0 | 1540 | 1299 | 0 | 3180 | 736 |
| STT3B | 1610 | 600 | 9772 | 10980 | 16623 | 12562 | 27314 | 7551 |
| STUB1 | 0 | 422 | 7722 | 5913 | 4878 | 0 | 1392 | 2395 |
| STX10 | 212 | 0 | 0 | 0 | 1796 | 0 | 0 | 1388 |
| STX16 | 0 | 454 | 0 | 0 | 484 | 0 | 626 | 0 |
| STX16-NPEPL1 | 0 | 624 | 2237 | 0 | 84 | 0 | 736 | 0 |
| STX17 | 53534 | 53220 | 39318 | 28396 | 149590 | 44370 | 143020 | 28829 |
| STX17-AS1 | 0 | 596 | 0 | 0 | 0 | 0 | 62 | 0 |
| STX18 | 0 | 662 | 585 | 0 | 2398 | 180 | 0 | 742 |
| STX19 | 0 | 212 | 4153 | 0 | 0 | 0 | 0 | 0 |
| STX3 | 0 | 408 | 130 | 0 | 1130 | 0 | 440 | 0 |
| STX4 | 0 | 0 | 0 | 0 | 1368 | 0 | 0 | 0 |
| STX5 | 48798 | 69519 | 393950 | 90957 | 245554 | 128634 | 227125 | 166183 |
| STX6 | 0 | 318 | 0 | 0 | 0 | 0 | 594 | 0 |
| STX7 | 514 | 774 | 2604 | 0 | 774 | 2886 | 0 | 0 |
| STX8 | 0 | 66 | 0 | 1206 | 2601 | 0 | 1116 | 2620 |
| STXBP1 | 196 | 0 | 0 | 814 | 152 | 958 | 0 | 0 |
| STXBP2 | 0 | 644 | 3498 | 1054 | 1296 | 0 | 2296 | 2432 |
| STXBP3 | 0 | 0 | 1644 | 0 | 0 | 0 | 266 | 0 |
| STXBP5 | 314 | 0 | 0 | 0 | 454 | 0 | 0 | 0 |
| STXBP6 | 30649 | 115757 | 74507 | 67413 | 304532 | 21959 | 37675 | 18643 |
| STYX | 0 | 0 | 0 | 0 | 176 | 0 | 0 | 0 |
| STYXL1 | 0 | 0 | 0 | 0 | 708 | 1502 | 830 | 0 |
| SUB1 | 2840 | 5163 | 10791 | 13183 | 39160 | 4441 | 8805 | 12888 |
| SUCLA2 | 136 | 646 | 0 | 5208 | 2294 | 0 | 444 | 2188 |
| SUCLG1 | 1358 | 6124 | 13743 | 5978 | 9301 | 0 | 1824 | 0 |
| SUCLG2 | 2008 | 828 | 0 | 1404 | 1514 | 0 | 0 | 0 |
| SUCNR1 | 0 | 0 | 0 | 0 | 186 | 3360 | 488 | 506 |
| SUCO | 666 | 0 | 0 | 0 | 0 | 1757 | 0 | 0 |
| SUDS3 | 0 | 0 | 0 | 0 | 538 | 0 | 0 | 1410 |
| SUFU | 4295 | 3068 | 15009 | 8863 | 8505 | 4422 | 11055 | 4290 |
| SUGCT | 22796 | 9678 | 54249 | 1383 | 43122 | 62022 | 4284 | 7038 |
| SUGP1 | 1142 | 854 | 0 | 926 | 982 | 0 | 676 | 0 |
| SUGP2 | 0 | 0 | 126 | 0 | 436 | 0 | 0 | 0 |
| SUGT1 | 0 | 824 | 0 | 2769 | 2449 | 2670 | 1208 | 3069 |
| SUGT1P3 | 0 | 0 | 0 | 0 | 372 | 0 | 0 | 0 |
| SULF1 | 0 | 0 | 0 | 0 | 252 | 1022 | 0 | 746 |
| SULF2 | 0 | 528 | 0 | 0 | 2540 | 1906 | 284 | 0 |
| SULT1A1 | 737 | 882 | 8004 | 0 | 0 | 0 | 0 | 0 |

| | | | | | | | | |
|---|---|---|---|---|---|---|---|---|
| SULT1A2 | 788 | 290 | 3372 | 0 | 0 | 0 | 0 | 0 |
| SULT1A3 | 0 | 1008 | 0 | 2028 | 0 | 0 | 0 | 0 |
| SULT1B1 | 0 | 428 | 40 | 0 | 0 | 0 | 0 | 0 |
| SULT1C2 | 0 | 0 | 0 | 0 | 168 | 0 | 0 | 0 |
| SULT2B1 | 0 | 0 | 116 | 0 | 446 | 0 | 0 | 0 |
| SUMF1 | 0 | 0 | 660 | 0 | 920 | 0 | 0 | 0 |
| SUMF2 | 0 | 1408 | 456 | 1539 | 1410 | 802 | 762 | 2008 |
| SUMO1 | 282868 | 514491 | 632268 | 600272 | 1813015 | 598971 | 1399029 | 602415 |
| SUMO1P3 | 0 | 0 | 0 | 0 | 0 | 2530 | 0 | 0 |
| SUMO2 | 2146 | 3252 | 8079 | 8411 | 10058 | 6769 | 3103 | 11093 |
| SUMO3 | 0 | 922 | 3862 | 3139 | 2890 | 288 | 771 | 732 |
| SUN1 | 118 | 0 | 2046 | 0 | 0 | 0 | 0 | 0 |
| SUN2 | 328 | 0 | 676 | 428 | 0 | 0 | 0 | 0 |
| SUOX | 6298 | 7606 | 30297 | 15901 | 25642 | 8945 | 47415 | 5838 |
| SUPT16H | 0 | 0 | 454 | 0 | 200 | 0 | 234 | 0 |
| SUPT20H | 0 | 26 | 0 | 0 | 305 | 1708 | 692 | 0 |
| SUPT3H | 2166 | 20462 | 21569 | 20140 | 100973 | 24516 | 59694 | 40931 |
| SUPT4H1 | 0 | 947 | 690 | 886 | 1218 | 78 | 2920 | 1392 |
| SUPT5H | 0 | 214 | 0 | 442 | 606 | 0 | 822 | 0 |
| SUPT6H | 460 | 0 | 312 | 164 | 560 | 0 | 0 | 0 |
| SUPT7L | 0 | 454 | 1267 | 0 | 694 | 0 | 0 | 0 |
| SUPV3L1 | 0 | 0 | 0 | 0 | 1102 | 0 | 684 | 0 |
| SURF1 | 816 | 416 | 6409 | 6857 | 6066 | 5526 | 1404 | 0 |
| SURF2 | 0 | 988 | 0 | 0 | 1974 | 5716 | 0 | 1420 |
| SURF4 | 0 | 0 | 1423 | 1213 | 0 | 0 | 804 | 0 |
| SURF6 | 0 | 662 | 0 | 1128 | 0 | 0 | 1920 | 0 |
| SUSD3 | 0 | 1452 | 3680 | 5746 | 0 | 23908 | 4110 | 1056 |
| SUSD6 | 226 | 400 | 0 | 0 | 1473 | 0 | 0 | 0 |
| SUV39H1 | 0 | 0 | 0 | 0 | 586 | 0 | 0 | 0 |
| SUZ12 | 72766 | 75004 | 98880 | 57258 | 255334 | 74305 | 102309 | 52868 |
| SUZ12P1 | 2622 | 3546 | 1972 | 3713 | 6925 | 5232 | 2156 | 4641 |
| SVBP | 0 | 0 | 0 | 1062 | 3094 | 1834 | 0 | 0 |
| SVIL | 0 | 302 | 0 | 0 | 0 | 0 | 0 | 0 |
| SVIL-AS1 | 0 | 0 | 0 | 0 | 812 | 842 | 0 | 0 |
| SVIP | 0 | 0 | 0 | 0 | 2214 | 0 | 974 | 843 |
| SVOP | 0 | 12 | 0 | 0 | 0 | 0 | 0 | 0 |
| SVOPL | 1374 | 2994 | 3876 | 932 | 0 | 2331 | 128 | 0 |
| SWI5 | 0 | 0 | 1868 | 784 | 1302 | 0 | 1450 | 0 |
| SWSAP1 | 585 | 0 | 0 | 0 | 0 | 0 | 0 | 0 |
| SWT1 | 0 | 0 | 0 | 0 | 0 | 0 | 338 | 0 |
| SYAP1 | 0 | 662 | 0 | 0 | 2265 | 0 | 0 | 852 |
| SYBU | 750 | 5051 | 2231 | 1594 | 12532 | 2426 | 7044 | 4800 |
| SYCE1L | 0 | 0 | 0 | 0 | 0 | 0 | 246 | 0 |

| | | | | | | | | |
|---|---|---|---|---|---|---|---|---|
| SYDE2 | 0 | 182 | 0 | 0 | 0 | 0 | 0 | 0 |
| SYF2 | 1266 | 364 | 2904 | 2842 | 3437 | 1593 | 554 | 1900 |
| SYK | 5334 | 7728 | 7359 | 12420 | 21881 | 21143 | 29486 | 4255 |
| SYMPK | 0 | 508 | 0 | 0 | 0 | 0 | 672 | 0 |
| SYN2 | 0 | 0 | 220 | 0 | 0 | 0 | 0 | 0 |
| SYN3 | 0 | 0 | 0 | 0 | 1128 | 0 | 0 | 0 |
| SYNC | 0 | 0 | 0 | 0 | 372 | 0 | 107 | 0 |
| SYNCRIP | 0 | 0 | 0 | 0 | 768 | 3061 | 2164 | 1158 |
| SYNDIG1 | 0 | 0 | 0 | 0 | 1376 | 0 | 0 | 412 |
| SYNE1 | 1658 | 3412 | 5926 | 0 | 8891 | 1108 | 0 | 5890 |
| SYNE2 | 0 | 482 | 0 | 0 | 592 | 0 | 478 | 838 |
| SYNE3 | 0 | 0 | 0 | 0 | 410 | 0 | 0 | 0 |
| SYNE4 | 0 | 0 | 0 | 0 | 0 | 0 | 1234 | 0 |
| SYNGR1 | 0 | 0 | 0 | 0 | 0 | 1298 | 0 | 0 |
| SYNGR2 | 106 | 1478 | 11565 | 8331 | 9018 | 9455 | 5631 | 88 |
| SYNJ2 | 0 | 0 | 0 | 0 | 0 | 1466 | 0 | 528 |
| SYNJ2BP | 0 | 358 | 0 | 0 | 1913 | 0 | 0 | 0 |
| SYNJ2BP-COX16 | 0 | 0 | 0 | 0 | 372 | 0 | 0 | 0 |
| SYNPO | 0 | 0 | 1486 | 1393 | 904 | 0 | 0 | 0 |
| SYNPO2 | 1546 | 248 | 1558 | 0 | 276 | 0 | 164 | 0 |
| SYNPR | 0 | 0 | 0 | 0 | 0 | 831 | 0 | 0 |
| SYNRG | 0 | 168 | 0 | 0 | 418 | 0 | 0 | 0 |
| SYPL1 | 140 | 2032 | 3235 | 3058 | 2621 | 1549 | 2910 | 2145 |
| SYS1 | 0 | 0 | 0 | 1088 | 1218 | 0 | 0 | 0 |
| SYS1-DBNDD2 | 0 | 0 | 0 | 0 | 954 | 0 | 78 | 0 |
| SYT1 | 932 | 1078 | 6770 | 0 | 0 | 0 | 506 | 0 |
| SYT11 | 0 | 414 | 0 | 0 | 0 | 0 | 0 | 0 |
| SYT13 | 0 | 348 | 0 | 0 | 0 | 0 | 0 | 0 |
| SYT7 | 0 | 0 | 0 | 908 | 978 | 0 | 3372 | 542 |
| SYTL2 | 0 | 100 | 18 | 0 | 0 | 0 | 0 | 0 |
| SYTL3 | 725 | 1470 | 214 | 110 | 2610 | 24 | 1482 | 116 |
| SYVN1 | 0 | 0 | 0 | 0 | 0 | 0 | 998 | 0 |
| SZRD1 | 710 | 0 | 0 | 0 | 276 | 0 | 0 | 0 |
| SZT2 | 0 | 126 | 0 | 925 | 1405 | 0 | 0 | 450 |
| TAB2 | 0 | 0 | 0 | 0 | 0 | 0 | 60 | 0 |
| TAB3 | 0 | 0 | 0 | 0 | 824 | 0 | 0 | 0 |
| TACC1 | 12575 | 5904 | 15493 | 14331 | 21108 | 7546 | 25509 | 7616 |
| TACC2 | 7092 | 14063 | 14834 | 14581 | 23185 | 10060 | 34570 | 8889 |
| TACC3 | 12531 | 11054 | 31188 | 37325 | 86757 | 10701 | 24080 | 33278 |
| TACO1 | 0 | 0 | 3606 | 0 | 2278 | 0 | 0 | 0 |
| TACSTD2 | 0 | 0 | 0 | 812 | 746 | 2949 | 0 | 386 |
| TADA1 | 0 | 0 | 1035 | 0 | 410 | 0 | 1118 | 0 |
| TADA3 | 0 | 0 | 0 | 1966 | 492 | 0 | 0 | 0 |

| | | | | | | | | |
|---|---|---|---|---|---|---|---|---|
| TAF1 | 0 | 0 | 0 | 0 | 174 | 0 | 0 | 0 |
| TAF10 | 2992 | 560 | 3274 | 2286 | 5208 | 1454 | 1686 | 4351 |
| TAF11 | 0 | 266 | 1531 | 0 | 1952 | 0 | 648 | 406 |
| TAF12 | 0 | 492 | 0 | 0 | 600 | 0 | 0 | 970 |
| TAF15 | 70913 | 74713 | 83764 | 59059 | 221310 | 127958 | 132925 | 42326 |
| TAF1A-AS1 | 0 | 452 | 0 | 0 | 806 | 0 | 470 | 0 |
| TAF1C | 0 | 204 | 0 | 0 | 262 | 0 | 0 | 0 |
| TAF1D | 0 | 338 | 0 | 2833 | 2901 | 0 | 312 | 0 |
| TAF2 | 0 | 0 | 0 | 0 | 869 | 0 | 276 | 568 |
| TAF3 | 0 | 1637 | 0 | 0 | 1168 | 0 | 950 | 44 |
| TAF4 | 0 | 1654 | 258 | 2644 | 1308 | 1285 | 2399 | 0 |
| TAF4B | 0 | 0 | 616 | 0 | 0 | 0 | 0 | 0 |
| TAF5L | 0 | 0 | 0 | 0 | 0 | 0 | 164 | 0 |
| TAF6 | 0 | 0 | 1798 | 0 | 686 | 0 | 0 | 0 |
| TAF6L | 0 | 0 | 742 | 0 | 418 | 0 | 0 | 520 |
| TAF7 | 638 | 0 | 584 | 1244 | 598 | 3358 | 1380 | 0 |
| TAF8 | 11000 | 14677 | 10503 | 22542 | 46526 | 6660 | 30890 | 27160 |
| TAF9 | 108 | 984 | 4236 | 3506 | 3508 | 2184 | 408 | 2978 |
| TAF9B | 0 | 0 | 0 | 0 | 104 | 0 | 0 | 0 |
| TAGLN | 688 | 696 | 1408 | 0 | 11381 | 7055 | 0 | 2746 |
| TAGLN2 | 0 | 2630 | 8215 | 10868 | 12859 | 16818 | 2684 | 4952 |
| TALAM1 | 16 | 108 | 3458 | 0 | 508 | 0 | 0 | 0 |
| TALDO1 | 834 | 2798 | 2185 | 10967 | 13045 | 9063 | 4439 | 7673 |
| TAMM41 | 0 | 396 | 0 | 0 | 0 | 914 | 404 | 0 |
| TANC1 | 144 | 0 | 0 | 0 | 0 | 0 | 0 | 0 |
| TANC2 | 0 | 0 | 0 | 0 | 428 | 0 | 0 | 0 |
| TANGO2 | 0 | 34 | 0 | 0 | 1030 | 2230 | 0 | 0 |
| TANGO6 | 668 | 718 | 1170 | 1565 | 1054 | 0 | 358 | 1324 |
| TANK | 826 | 0 | 0 | 784 | 0 | 0 | 462 | 0 |
| TAOK1 | 13361 | 12190 | 8436 | 5972 | 17687 | 4719 | 9788 | 5071 |
| TAOK2 | 0 | 0 | 216 | 0 | 0 | 0 | 0 | 0 |
| TAOK3 | 0 | 0 | 462 | 0 | 0 | 0 | 0 | 0 |
| TAP1 | 716 | 388 | 0 | 0 | 1554 | 2112 | 1418 | 0 |
| TAP2 | 0 | 0 | 0 | 420 | 0 | 0 | 0 | 0 |
| TAPBP | 1564 | 342 | 1130 | 0 | 1564 | 0 | 18 | 818 |
| TAPBPL | 0 | 0 | 1150 | 834 | 0 | 0 | 0 | 0 |
| TAPT1 | 0 | 0 | 0 | 0 | 738 | 0 | 0 | 0 |
| TARBP1 | 458 | 0 | 0 | 0 | 458 | 0 | 0 | 0 |
| TARBP2 | 0 | 0 | 0 | 0 | 170 | 0 | 382 | 0 |
| TARDBP | 0 | 488 | 0 | 784 | 1824 | 0 | 1184 | 0 |
| TARS | 0 | 0 | 0 | 0 | 2264 | 0 | 0 | 658 |
| TARS2 | 0 | 0 | 398 | 0 | 772 | 0 | 194 | 0 |
| TARSL2 | 0 | 516 | 0 | 0 | 0 | 0 | 0 | 0 |

| | | | | | | | | |
|---|---|---|---|---|---|---|---|---|
| TASP1 | 1846 | 19026 | 24270 | 11185 | 89475 | 10265 | 19965 | 16810 |
| TATDN1 | 800 | 1724 | 1332 | 3643 | 5424 | 0 | 1098 | 1186 |
| TATDN2 | 0 | 0 | 0 | 0 | 0 | 2472 | 0 | 0 |
| TATDN3 | 342 | 0 | 0 | 0 | 360 | 1548 | 388 | 0 |
| TAX1BP1 | 149471 | 267140 | 308620 | 166264 | 1011927 | 349914 | 395957 | 348041 |
| TAX1BP3 | 1392 | 0 | 3755 | 971 | 2180 | 0 | 1010 | 1096 |
| TAZ | 0 | 408 | 1278 | 0 | 468 | 0 | 0 | 0 |
| TBC1D10B | 0 | 0 | 0 | 522 | 188 | 855 | 0 | 0 |
| TBC1D12 | 0 | 0 | 0 | 800 | 0 | 0 | 0 | 0 |
| TBC1D13 | 0 | 0 | 0 | 0 | 0 | 0 | 570 | 0 |
| TBC1D14 | 896 | 0 | 0 | 1176 | 0 | 0 | 516 | 704 |
| TBC1D16 | 0 | 0 | 0 | 800 | 0 | 0 | 0 | 0 |
| TBC1D17 | 3260 | 3823 | 4060 | 4070 | 13964 | 968 | 3992 | 3458 |
| TBC1D19 | 0 | 0 | 0 | 0 | 294 | 0 | 0 | 0 |
| TBC1D2 | 0 | 192 | 0 | 0 | 645 | 0 | 0 | 0 |
| TBC1D20 | 156 | 70 | 0 | 302 | 624 | 0 | 382 | 0 |
| TBC1D22A | 0 | 728 | 0 | 0 | 0 | 0 | 0 | 0 |
| TBC1D22B | 0 | 0 | 0 | 0 | 66 | 0 | 0 | 0 |
| TBC1D23 | 0 | 0 | 0 | 512 | 878 | 0 | 0 | 0 |
| TBC1D24 | 574 | 0 | 0 | 0 | 0 | 0 | 0 | 0 |
| TBC1D2B | 0 | 220 | 0 | 825 | 0 | 0 | 0 | 0 |
| TBC1D31 | 0 | 0 | 0 | 0 | 272 | 0 | 0 | 0 |
| TBC1D3L | 0 | 0 | 0 | 0 | 608 | 60 | 0 | 0 |
| TBC1D4 | 0 | 0 | 0 | 0 | 156 | 0 | 576 | 0 |
| TBC1D7 | 684 | 0 | 1996 | 2454 | 1022 | 622 | 580 | 0 |
| TBC1D7-LOC100130357 | 0 | 0 | 0 | 0 | 0 | 0 | 0 | 324 |
| TBC1D8 | 586 | 0 | 0 | 0 | 324 | 0 | 0 | 0 |
| TBC1D9B | 708 | 316 | 0 | 0 | 1500 | 0 | 612 | 0 |
| TBCA | 3603 | 2537 | 3065 | 6171 | 9441 | 2218 | 3512 | 6017 |
| TBCB | 0 | 0 | 1260 | 1288 | 4664 | 0 | 1260 | 664 |
| TBCC | 1694 | 0 | 1106 | 1406 | 488 | 0 | 0 | 632 |
| TBCD | 0 | 0 | 0 | 0 | 0 | 1486 | 0 | 0 |
| TBCE | 0 | 20 | 0 | 0 | 0 | 0 | 0 | 1412 |
| TBCK | 0 | 0 | 0 | 0 | 120 | 0 | 0 | 0 |
| TBK1 | 0 | 0 | 0 | 668 | 742 | 0 | 0 | 0 |
| TBL1X | 1536 | 470 | 0 | 2658 | 1912 | 0 | 2444 | 1546 |
| TBL1XR1 | 58016 | 58843 | 128564 | 95551 | 259518 | 86887 | 204603 | 97984 |
| TBL2 | 0 | 356 | 1578 | 0 | 1052 | 3028 | 0 | 0 |
| TBL3 | 0 | 0 | 0 | 0 | 0 | 0 | 266 | 794 |
| TBP | 0 | 0 | 0 | 0 | 1998 | 0 | 0 | 0 |
| TBPL1 | 0 | 0 | 82 | 0 | 994 | 0 | 26 | 1178 |
| TBRG1 | 588 | 0 | 0 | 1106 | 1066 | 0 | 0 | 0 |
| TBRG4 | 970 | 636 | 0 | 0 | 3122 | 0 | 318 | 846 |

| | | | | | | | | |
|---|---|---|---|---|---|---|---|---|
| TBX10 | 0 | 1024 | 0 | 1112 | 0 | 0 | 0 | 0 |
| TBX18 | 0 | 0 | 0 | 0 | 1229 | 0 | 0 | 0 |
| TBX3 | 0 | 0 | 0 | 0 | 396 | 0 | 1616 | 0 |
| TBXAS1 | 0 | 132 | 0 | 892 | 634 | 0 | 0 | 775 |
| TC2N | 5132 | 22173 | 7161 | 772 | 40608 | 1932 | 3556 | 3986 |
| TCAF1 | 0 | 0 | 0 | 0 | 0 | 0 | 0 | 572 |
| TCAIM | 0 | 0 | 0 | 1557 | 794 | 0 | 402 | 0 |
| TCAM1P | 0 | 0 | 0 | 0 | 276 | 0 | 0 | 0 |
| TCEA1 | 0 | 0 | 1495 | 3278 | 1518 | 598 | 3422 | 1028 |
| TCEA3 | 149638 | 212145 | 777060 | 137216 | 56332 | 49437 | 319976 | 18666 |
| TCEAL1 | 430 | 0 | 0 | 0 | 514 | 0 | 0 | 914 |
| TCEAL2 | 0 | 344 | 0 | 0 | 0 | 0 | 0 | 0 |
| TCEAL3 | 0 | 0 | 0 | 0 | 690 | 0 | 0 | 0 |
| TCEAL4 | 0 | 494 | 2510 | 1623 | 6315 | 0 | 642 | 0 |
| TCEAL7 | 354 | 0 | 0 | 0 | 0 | 0 | 0 | 0 |
| TCEAL8 | 0 | 574 | 0 | 1392 | 1398 | 0 | 414 | 0 |
| TCEAL9 | 594 | 460 | 2800 | 1228 | 5451 | 1864 | 0 | 1646 |
| TCEANC | 0 | 0 | 0 | 0 | 430 | 0 | 0 | 0 |
| TCERG1 | 0 | 464 | 0 | 792 | 732 | 0 | 0 | 0 |
| TCF12 | 0 | 426 | 0 | 0 | 220 | 0 | 168 | 0 |
| TCF19 | 0 | 0 | 0 | 1564 | 0 | 0 | 0 | 0 |
| TCF20 | 790 | 274 | 460 | 618 | 1624 | 0 | 730 | 750 |
| TCF25 | 0 | 388 | 0 | 568 | 1526 | 0 | 0 | 40 |
| TCF3 | 816 | 618 | 10 | 2426 | 4717 | 1410 | 1084 | 1120 |
| TCF4 | 0 | 0 | 0 | 0 | 500 | 0 | 0 | 0 |
| TCF7L2 | 258078 | 248195 | 194353 | 54730 | 360273 | 40516 | 320303 | 25757 |
| TCFL5 | 0 | 0 | 0 | 0 | 0 | 1316 | 278 | 0 |
| TCHH | 0 | 0 | 2070 | 0 | 0 | 0 | 0 | 0 |
| TCHP | 0 | 0 | 1557 | 0 | 218 | 0 | 0 | 0 |
| TCIM | 1978 | 1130 | 4108 | 2639 | 6038 | 8991 | 314 | 2074 |
| TCIRG1 | 370 | 484 | 0 | 0 | 0 | 0 | 742 | 0 |
| TCN1 | 0 | 522 | 0 | 0 | 8076 | 0 | 0 | 1365 |
| TCN2 | 0 | 0 | 6509 | 0 | 648 | 0 | 0 | 0 |
| TCOF1 | 0 | 0 | 0 | 200 | 2350 | 1524 | 1106 | 0 |
| TCP1 | 0 | 1304 | 0 | 1308 | 3893 | 2820 | 1304 | 3017 |
| TCP10L2 | 0 | 0 | 0 | 0 | 756 | 0 | 0 | 0 |
| TCP11L1 | 0 | 0 | 0 | 1286 | 0 | 0 | 426 | 0 |
| TCTA | 0 | 416 | 0 | 0 | 1968 | 0 | 0 | 0 |
| TCTE3 | 0 | 208 | 0 | 0 | 476 | 0 | 0 | 0 |
| TCTEX1D1 | 0 | 414 | 0 | 0 | 0 | 0 | 0 | 0 |
| TCTEX1D2 | 22214 | 20890 | 29680 | 17622 | 126231 | 40644 | 13589 | 24010 |
| TCTEX1D4 | 0 | 0 | 0 | 0 | 0 | 0 | 204 | 0 |
| TCTN3 | 458 | 376 | 0 | 0 | 0 | 0 | 0 | 0 |

| | | | | | | | | |
|---|---|---|---|---|---|---|---|---|
| TDG | 0 | 0 | 732 | 902 | 238 | 0 | 0 | 836 |
| TDGF1 | 0 | 0 | 0 | 1941 | 0 | 0 | 1593 | 0 |
| TDO2 | 0 | 0 | 0 | 0 | 0 | 0 | 0 | 946 |
| TDP1 | 0 | 0 | 0 | 0 | 556 | 0 | 0 | 0 |
| TDP2 | 1308 | 480 | 16430 | 554 | 1848 | 0 | 1352 | 0 |
| TDRD3 | 0 | 0 | 0 | 1036 | 708 | 0 | 818 | 1620 |
| TDRD7 | 0 | 0 | 0 | 744 | 0 | 0 | 0 | 0 |
| TDRG1 | 0 | 0 | 0 | 0 | 506 | 0 | 0 | 0 |
| TDRKH | 62 | 344 | 0 | 3295 | 9020 | 6580 | 13464 | 8238 |
| TDRP | 541 | 0 | 0 | 0 | 0 | 0 | 0 | 0 |
| TEAD1 | 0 | 0 | 0 | 658 | 814 | 0 | 496 | 1052 |
| TEAD2 | 1074 | 0 | 80 | 1882 | 1978 | 0 | 2142 | 398 |
| TEAD3 | 1070 | 0 | 592 | 0 | 1032 | 0 | 0 | 0 |
| TEAD4 | 0 | 0 | 0 | 0 | 0 | 0 | 626 | 0 |
| TEC | 13516 | 12077 | 18958 | 9383 | 17006 | 13613 | 27372 | 1762 |
| TECR | 960 | 992 | 2158 | 6827 | 7110 | 7095 | 1540 | 2446 |
| TEDC1 | 0 | 1078 | 0 | 1354 | 316 | 0 | 0 | 0 |
| TEFM | 0 | 0 | 1994 | 0 | 634 | 0 | 0 | 0 |
| TEKT4P2 | 0 | 380 | 0 | 212 | 860 | 0 | 0 | 0 |
| TEN1 | 0 | 0 | 0 | 0 | 0 | 2893 | 0 | 0 |
| TENM3 | 0 | 0 | 0 | 0 | 1392 | 1413 | 0 | 0 |
| TENT2 | 0 | 354 | 0 | 0 | 0 | 0 | 0 | 0 |
| TENT5C | 0 | 382 | 0 | 0 | 0 | 0 | 0 | 0 |
| TERF1 | 0 | 288 | 0 | 0 | 840 | 0 | 0 | 0 |
| TERF2 | 0 | 0 | 0 | 1114 | 0 | 0 | 0 | 0 |
| TERF2IP | 416 | 294 | 0 | 0 | 0 | 0 | 230 | 0 |
| TERT | 0 | 0 | 0 | 0 | 594 | 0 | 0 | 0 |
| TES | 782 | 0 | 0 | 1676 | 288 | 0 | 0 | 0 |
| TESC | 0 | 0 | 0 | 2828 | 1312 | 5798 | 0 | 1080 |
| TESK1 | 0 | 0 | 1118 | 0 | 0 | 0 | 30 | 0 |
| TESK2 | 0 | 0 | 0 | 0 | 0 | 1144 | 0 | 0 |
| TESMIN | 0 | 0 | 0 | 1002 | 883 | 0 | 334 | 0 |
| TET2 | 25618 | 13064 | 21654 | 2391 | 17093 | 2620 | 12796 | 6229 |
| TEX10 | 5997 | 9932 | 1505 | 14370 | 33667 | 9113 | 36874 | 3071 |
| TEX11 | 0 | 114 | 0 | 0 | 0 | 0 | 0 | 0 |
| TEX2 | 2546 | 2630 | 0 | 4512 | 2671 | 3156 | 6830 | 0 |
| TEX261 | 0 | 602 | 0 | 0 | 0 | 0 | 0 | 0 |
| TEX264 | 0 | 872 | 0 | 4836 | 2075 | 2080 | 316 | 418 |
| TEX30 | 0 | 458 | 0 | 3546 | 2360 | 0 | 0 | 0 |
| TFAM | 0 | 0 | 0 | 0 | 0 | 0 | 280 | 0 |
| TFAP2A-AS2 | 0 | 0 | 0 | 0 | 0 | 0 | 0 | 690 |
| TFAP4 | 0 | 516 | 0 | 471 | 598 | 0 | 0 | 0 |
| TFB1M | 38289 | 104643 | 160976 | 160712 | 319714 | 215312 | 270078 | 146286 |

| | | | | | | | | |
|---|---|---|---|---|---|---|---|---|
| TFB2M | 0 | 0 | 1710 | 1048 | 0 | 0 | 772 | 0 |
| TFCP2 | 0 | 0 | 0 | 342 | 0 | 0 | 492 | 0 |
| TFCP2L1 | 998 | 910 | 1592 | 0 | 0 | 0 | 0 | 0 |
| TFDP1 | 0 | 0 | 0 | 114 | 2524 | 686 | 0 | 1840 |
| TFDP2 | 11758 | 22640 | 20250 | 44945 | 110090 | 46111 | 146881 | 63895 |
| TFE3 | 4194 | 3526 | 9969 | 6777 | 16608 | 3727 | 9450 | 4070 |
| TFEB | 5684 | 4616 | 7939 | 6785 | 5181 | 4056 | 10644 | 4585 |
| TFEC | 0 | 0 | 256 | 0 | 0 | 0 | 0 | 0 |
| TFF1 | 630 | 3569 | 8945 | 2134 | 10795 | 0 | 0 | 12192 |
| TFF2 | 0 | 0 | 116 | 0 | 2362 | 1620 | 0 | 13172 |
| TFF3 | 26874 | 53906 | 68438 | 16762 | 58246 | 36860 | 10042 | 14751 |
| TFG | 250608 | 509219 | 983402 | 789119 | 1731835 | 715159 | 1625950 | 852721 |
| TFIP11 | 0 | 260 | 0 | 0 | 0 | 0 | 0 | 0 |
| TFPI | 0 | 844 | 0 | 0 | 0 | 0 | 0 | 724 |
| TFPI2 | 0 | 444 | 0 | 0 | 0 | 0 | 0 | 0 |
| TFPT | 292 | 0 | 0 | 0 | 967 | 0 | 0 | 0 |
| TFR2 | 0 | 414 | 0 | 0 | 1264 | 0 | 0 | 0 |
| TFRC | 722 | 390 | 1298 | 0 | 1254 | 0 | 384 | 0 |
| TGDS | 456 | 0 | 0 | 474 | 764 | 0 | 0 | 0 |
| TGFA | 0 | 0 | 1773 | 0 | 0 | 0 | 0 | 0 |
| TGFB1 | 500 | 0 | 0 | 0 | 0 | 0 | 0 | 960 |
| TGFB1I1 | 348 | 0 | 0 | 0 | 0 | 0 | 0 | 740 |
| TGFB3 | 0 | 0 | 0 | 56 | 0 | 0 | 0 | 0 |
| TGFBI | 0 | 0 | 0 | 6672 | 5978 | 5300 | 3998 | 804 |
| TGFBR2 | 2212 | 1810 | 4102 | 1888 | 8992 | 1450 | 19784 | 3523 |
| TGFBR3 | 0 | 380 | 1334 | 0 | 0 | 0 | 0 | 0 |
| TGFBR3L | 0 | 0 | 0 | 0 | 708 | 0 | 0 | 0 |
| TGFBRAP1 | 0 | 296 | 0 | 0 | 196 | 0 | 673 | 0 |
| TGIF1 | 0 | 612 | 7314 | 7684 | 5108 | 7267 | 10913 | 0 |
| TGM1 | 0 | 0 | 0 | 0 | 720 | 0 | 0 | 0 |
| TGM2 | 0 | 0 | 0 | 0 | 502 | 0 | 556 | 0 |
| TGOLN2 | 1134 | 0 | 1004 | 740 | 1902 | 0 | 3191 | 0 |
| TH | 0 | 0 | 0 | 0 | 340 | 0 | 0 | 0 |
| THADA | 0 | 0 | 0 | 0 | 0 | 354 | 0 | 0 |
| THAP1 | 0 | 0 | 0 | 0 | 1260 | 0 | 300 | 786 |
| THAP10 | 0 | 0 | 0 | 0 | 364 | 0 | 0 | 0 |
| THAP11 | 490 | 0 | 1476 | 0 | 468 | 1434 | 1806 | 0 |
| THAP12 | 0 | 0 | 2398 | 0 | 1214 | 0 | 0 | 1566 |
| THAP2 | 0 | 0 | 0 | 1228 | 0 | 0 | 0 | 0 |
| THAP3 | 17455 | 24010 | 114644 | 29527 | 67436 | 74360 | 58172 | 26701 |
| THAP4 | 1170 | 1034 | 5648 | 0 | 490 | 0 | 0 | 0 |
| THAP6 | 0 | 0 | 1546 | 0 | 1054 | 0 | 0 | 0 |
| THAP7 | 11459 | 13873 | 43494 | 11801 | 65412 | 31699 | 34689 | 25267 |

| | | | | | | | | |
|---|---|---|---|---|---|---|---|---|
| THAP7-AS1 | 0 | 360 | 0 | 0 | 1447 | 0 | 814 | 0 |
| THAP9 | 0 | 0 | 1720 | 0 | 0 | 0 | 0 | 0 |
| THAP9-AS1 | 0 | 284 | 0 | 2185 | 486 | 0 | 0 | 750 |
| THBD | 0 | 0 | 0 | 1287 | 0 | 0 | 0 | 0 |
| THBS1 | 1088 | 1408 | 2486 | 0 | 2158 | 1512 | 0 | 0 |
| THBS2 | 0 | 0 | 0 | 0 | 2878 | 0 | 0 | 0 |
| THBS3 | 0 | 316 | 0 | 0 | 0 | 0 | 0 | 0 |
| THBS4 | 0 | 0 | 0 | 0 | 264 | 0 | 0 | 0 |
| THEM4 | 0 | 358 | 0 | 0 | 3002 | 0 | 0 | 0 |
| THEM6 | 422 | 0 | 0 | 0 | 1866 | 1716 | 2341 | 0 |
| THEMIS2 | 0 | 202 | 0 | 0 | 796 | 0 | 334 | 798 |
| THG1L | 0 | 574 | 0 | 1410 | 0 | 0 | 0 | 0 |
| THNSL2 | 0 | 0 | 0 | 0 | 0 | 0 | 816 | 0 |
| THOC1 | 7886 | 26636 | 14973 | 14128 | 111768 | 28262 | 25694 | 28081 |
| THOC2 | 940 | 0 | 0 | 0 | 108 | 0 | 0 | 0 |
| THOC3 | 0 | 454 | 0 | 2717 | 2837 | 2204 | 4639 | 614 |
| THOC5 | 0 | 180 | 0 | 0 | 0 | 0 | 0 | 0 |
| THOC6 | 0 | 618 | 1400 | 1249 | 506 | 0 | 1360 | 0 |
| THOC7 | 2679 | 3106 | 1838 | 2824 | 13606 | 3082 | 5596 | 3334 |
| THOP1 | 956 | 378 | 0 | 765 | 0 | 0 | 0 | 0 |
| THRA | 3916 | 4840 | 7502 | 3648 | 2466 | 3026 | 15302 | 2804 |
| THRAP3 | 0 | 372 | 0 | 364 | 1889 | 862 | 0 | 0 |
| THRB | 3674 | 3188 | 9393 | 1070 | 13229 | 0 | 4832 | 2218 |
| THSD4 | 334 | 890 | 330 | 0 | 901 | 0 | 0 | 0 |
| THSD7A | 3564 | 0 | 0 | 0 | 1214 | 0 | 414 | 0 |
| THTPA | 0 | 0 | 0 | 0 | 1556 | 0 | 0 | 0 |
| THUMPD1 | 0 | 0 | 0 | 0 | 0 | 0 | 194 | 0 |
| THUMPD2 | 0 | 364 | 1754 | 824 | 0 | 0 | 527 | 0 |
| THUMPD3 | 0 | 0 | 0 | 822 | 0 | 2436 | 0 | 0 |
| THUMPD3-AS1 | 0 | 0 | 0 | 0 | 0 | 0 | 154 | 350 |
| THY1 | 0 | 0 | 0 | 0 | 1628 | 0 | 0 | 0 |
| THYN1 | 0 | 658 | 2422 | 4530 | 3997 | 1884 | 1644 | 2358 |
| TIA1 | 32 | 0 | 0 | 0 | 346 | 0 | 214 | 0 |
| TIAF1 | 1610 | 0 | 0 | 0 | 0 | 0 | 0 | 0 |
| TIAL1 | 442 | 128 | 0 | 0 | 452 | 0 | 1712 | 0 |
| TICAM1 | 0 | 0 | 0 | 658 | 0 | 0 | 0 | 0 |
| TIFA | 0 | 0 | 0 | 0 | 414 | 0 | 0 | 0 |
| TIGAR | 0 | 0 | 0 | 0 | 312 | 0 | 0 | 0 |
| TIGD1 | 0 | 0 | 0 | 0 | 1514 | 0 | 1048 | 0 |
| TIGD2 | 0 | 0 | 0 | 0 | 0 | 0 | 292 | 0 |
| TIGD6 | 0 | 0 | 0 | 0 | 1390 | 0 | 0 | 0 |
| TIMELESS | 0 | 0 | 0 | 0 | 670 | 0 | 0 | 586 |
| TIMM10 | 1522 | 2026 | 0 | 1456 | 5322 | 0 | 2526 | 2367 |

| | | | | | | | | |
|---|---|---|---|---|---|---|---|---|
| TIMM10B | 0 | 0 | 164 | 3089 | 2348 | 0 | 792 | 1678 |
| TIMM13 | 2298 | 2246 | 6984 | 1702 | 11149 | 2403 | 3529 | 1769 |
| TIMM17A | 225 | 0 | 0 | 9187 | 1170 | 2033 | 0 | 2268 |
| TIMM17B | 0 | 1932 | 1080 | 1066 | 4541 | 5626 | 1226 | 990 |
| TIMM21 | 148 | 0 | 1808 | 0 | 1884 | 2576 | 0 | 0 |
| TIMM22 | 0 | 1402 | 2053 | 0 | 0 | 0 | 0 | 1380 |
| TIMM23 | 164 | 604 | 5457 | 1358 | 1488 | 0 | 612 | 1283 |
| TIMM23B | 0 | 66 | 0 | 0 | 0 | 0 | 0 | 0 |
| TIMM29 | 0 | 0 | 0 | 0 | 376 | 0 | 0 | 0 |
| TIMM44 | 1410 | 378 | 0 | 1162 | 358 | 0 | 0 | 0 |
| TIMM50 | 982 | 416 | 1818 | 2238 | 1397 | 1640 | 682 | 2124 |
| TIMM8A | 0 | 786 | 0 | 0 | 0 | 0 | 0 | 0 |
| TIMM8B | 972 | 3104 | 3572 | 598 | 3835 | 0 | 3530 | 1609 |
| TIMM9 | 0 | 0 | 0 | 1180 | 3912 | 0 | 1404 | 0 |
| TIMMDC1 | 0 | 1312 | 3945 | 0 | 492 | 0 | 3676 | 4196 |
| TIMP1 | 912 | 2728 | 1851 | 7032 | 19337 | 21805 | 5901 | 38206 |
| TIMP2 | 1000 | 122 | 0 | 0 | 1301 | 98 | 0 | 0 |
| TIMP3 | 0 | 0 | 0 | 0 | 1154 | 0 | 0 | 628 |
| TINAG | 1104 | 1180 | 4479 | 1272 | 0 | 0 | 542 | 0 |
| TINAGL1 | 0 | 880 | 658 | 2980 | 4362 | 4148 | 1902 | 1036 |
| TINCR | 690 | 346 | 0 | 0 | 0 | 0 | 0 | 0 |
| TINF2 | 0 | 368 | 1316 | 0 | 0 | 0 | 816 | 0 |
| TIPIN | 0 | 374 | 0 | 0 | 0 | 0 | 0 | 1186 |
| TIPRL | 0 | 878 | 1774 | 0 | 3589 | 0 | 0 | 1194 |
| TIRAP | 0 | 0 | 0 | 976 | 0 | 0 | 0 | 0 |
| TJAP1 | 612 | 0 | 1420 | 0 | 214 | 0 | 0 | 0 |
| TJP1 | 540 | 394 | 0 | 1100 | 0 | 40 | 792 | 387 |
| TJP2 | 2494 | 3018 | 4400 | 3614 | 5442 | 7098 | 15980 | 1057 |
| TJP3 | 932 | 626 | 0 | 0 | 2512 | 0 | 0 | 0 |
| TK1 | 0 | 0 | 0 | 0 | 3620 | 0 | 434 | 154 |
| TK2 | 0 | 0 | 0 | 0 | 0 | 0 | 218 | 882 |
| TKFC | 795 | 0 | 0 | 0 | 1008 | 0 | 0 | 0 |
| TKT | 1686 | 0 | 2981 | 2248 | 5224 | 6108 | 2807 | 3156 |
| TLCD1 | 0 | 180 | 0 | 0 | 252 | 0 | 0 | 0 |
| TLCD2 | 0 | 0 | 907 | 406 | 0 | 0 | 0 | 0 |
| TLDC2 | 0 | 0 | 1648 | 0 | 1188 | 0 | 0 | 0 |
| TLE1 | 0 | 454 | 0 | 0 | 2062 | 0 | 0 | 0 |
| TLE2 | 0 | 308 | 0 | 0 | 0 | 0 | 382 | 0 |
| TLE4 | 0 | 0 | 0 | 0 | 0 | 0 | 920 | 0 |
| TLE6 | 0 | 0 | 0 | 0 | 150 | 0 | 0 | 0 |
| TLK1 | 654 | 0 | 0 | 0 | 0 | 0 | 0 | 1008 |
| TLK2 | 236 | 0 | 0 | 0 | 0 | 40 | 0 | 0 |
| TLL2 | 0 | 0 | 0 | 1066 | 0 | 0 | 0 | 0 |

| | | | | | | | | |
|---|---|---|---|---|---|---|---|---|
| TLN1 | 0 | 0 | 110 | 0 | 194 | 0 | 196 | 0 |
| TLN2 | 0 | 0 | 0 | 0 | 14 | 0 | 0 | 0 |
| TLNRD1 | 0 | 0 | 836 | 42 | 720 | 0 | 358 | 0 |
| TLR3 | 76 | 0 | 0 | 0 | 0 | 0 | 0 | 0 |
| TM2D1 | 586 | 378 | 2468 | 774 | 1588 | 3055 | 0 | 808 |
| TM2D2 | 0 | 348 | 1704 | 0 | 1000 | 5285 | 26 | 0 |
| TM2D3 | 572 | 696 | 1736 | 2098 | 366 | 3986 | 2388 | 0 |
| TM4SF1 | 2408 | 1178 | 3205 | 630 | 14185 | 2552 | 0 | 19629 |
| TM4SF19 | 0 | 0 | 0 | 1231 | 0 | 0 | 0 | 0 |
| TM4SF20 | 884 | 0 | 1544 | 0 | 0 | 484 | 330 | 0 |
| TM4SF4 | 398 | 1850 | 1310 | 0 | 15758 | 7801 | 0 | 102770 |
| TM4SF5 | 0 | 0 | 194 | 1022 | 0 | 0 | 482 | 0 |
| TM7SF3 | 0 | 0 | 0 | 1414 | 0 | 1351 | 244 | 503 |
| TM9SF1 | 0 | 0 | 1250 | 0 | 0 | 1425 | 570 | 959 |
| TM9SF2 | 296 | 2485 | 2598 | 3395 | 2340 | 3473 | 1292 | 0 |
| TM9SF3 | 146 | 1138 | 1813 | 746 | 1366 | 0 | 0 | 403 |
| TM9SF4 | 12886 | 25310 | 32231 | 66334 | 93934 | 39361 | 184491 | 23504 |
| TMA16 | 0 | 0 | 0 | 0 | 758 | 0 | 0 | 0 |
| TMA7 | 2252 | 10198 | 10113 | 14316 | 37303 | 3152 | 10114 | 2184 |
| TMBIM1 | 0 | 1724 | 1478 | 0 | 3192 | 0 | 1092 | 1222 |
| TMBIM4 | 914 | 2104 | 4207 | 2946 | 2370 | 1510 | 1714 | 930 |
| TMBIM6 | 7224 | 9530 | 14189 | 10542 | 25484 | 15710 | 38099 | 6196 |
| TMC3-AS1 | 0 | 0 | 0 | 0 | 0 | 0 | 980 | 0 |
| TMC4 | 2480 | 772 | 2852 | 0 | 594 | 1132 | 596 | 1028 |
| TMC5 | 212 | 0 | 0 | 0 | 758 | 1839 | 0 | 860 |
| TMC6 | 514 | 422 | 0 | 0 | 0 | 0 | 0 | 0 |
| TMCC1 | 0 | 0 | 0 | 0 | 0 | 0 | 824 | 0 |
| TMCC1-AS1 | 682 | 244 | 0 | 0 | 0 | 0 | 0 | 0 |
| TMCC3 | 0 | 294 | 0 | 0 | 0 | 0 | 0 | 0 |
| TMCO1 | 1134 | 3703 | 17307 | 5579 | 4254 | 1656 | 3804 | 954 |
| TMCO3 | 20004 | 26883 | 85918 | 57770 | 60928 | 87317 | 66286 | 15875 |
| TMCO4 | 896 | 0 | 1238 | 0 | 0 | 0 | 0 | 0 |
| TMCO6 | 0 | 0 | 0 | 0 | 444 | 0 | 0 | 0 |
| TMED1 | 7232 | 6802 | 1338 | 2028 | 17975 | 0 | 7397 | 1254 |
| TMED10 | 0 | 1668 | 3958 | 184 | 4772 | 0 | 278 | 2439 |
| TMED2 | 0 | 3204 | 4138 | 5820 | 7001 | 0 | 3918 | 3955 |
| TMED3 | 0 | 1836 | 1649 | 5771 | 4186 | 0 | 504 | 1282 |
| TMED4 | 1480 | 1504 | 3068 | 7241 | 1776 | 1320 | 672 | 708 |
| TMED5 | 0 | 426 | 1186 | 1627 | 0 | 0 | 212 | 0 |
| TMED7 | 0 | 0 | 0 | 0 | 528 | 0 | 0 | 1496 |
| TMED8 | 0 | 0 | 0 | 0 | 1210 | 0 | 0 | 0 |
| TMED9 | 792 | 1484 | 4184 | 7278 | 4669 | 3287 | 2364 | 2580 |
| TMEM101 | 0 | 0 | 0 | 0 | 1130 | 0 | 456 | 325 |

| | | | | | | | | |
|---|---|---|---|---|---|---|---|---|
| TMEM102 | 0 | 0 | 0 | 0 | 30 | 0 | 0 | 0 |
| TMEM105 | 0 | 180 | 0 | 0 | 0 | 0 | 0 | 0 |
| TMEM106A | 0 | 0 | 0 | 0 | 0 | 0 | 454 | 0 |
| TMEM106B | 0 | 826 | 1494 | 2078 | 0 | 0 | 360 | 0 |
| TMEM106C | 1374 | 1900 | 3626 | 8992 | 3032 | 9718 | 4200 | 1114 |
| TMEM107 | 0 | 0 | 1618 | 2300 | 426 | 0 | 0 | 1483 |
| TMEM109 | 0 | 284 | 0 | 0 | 1376 | 0 | 0 | 0 |
| TMEM11 | 0 | 1106 | 1144 | 1186 | 1614 | 0 | 0 | 0 |
| TMEM115 | 264 | 0 | 0 | 0 | 0 | 0 | 0 | 0 |
| TMEM116 | 0 | 0 | 0 | 0 | 0 | 0 | 558 | 688 |
| TMEM117 | 4590 | 3768 | 8420 | 6007 | 11784 | 15162 | 7555 | 2594 |
| TMEM120A | 1232 | 272 | 3825 | 2298 | 2119 | 3739 | 1164 | 2654 |
| TMEM120B | 0 | 338 | 0 | 0 | 182 | 0 | 0 | 0 |
| TMEM123 | 0 | 1322 | 1244 | 0 | 5665 | 0 | 984 | 3689 |
| TMEM125 | 576 | 492 | 0 | 758 | 1684 | 0 | 1128 | 0 |
| TMEM126A | 0 | 1398 | 1774 | 776 | 7750 | 0 | 352 | 1562 |
| TMEM126B | 84394 | 163321 | 119769 | 220779 | 511520 | 155506 | 395577 | 245003 |
| TMEM127 | 1616 | 0 | 0 | 0 | 956 | 0 | 346 | 0 |
| TMEM128 | 270 | 200 | 0 | 0 | 426 | 0 | 0 | 0 |
| TMEM129 | 0 | 242 | 0 | 500 | 354 | 0 | 0 | 0 |
| TMEM131 | 584 | 0 | 0 | 542 | 0 | 0 | 0 | 0 |
| TMEM131L | 664 | 0 | 0 | 0 | 0 | 0 | 0 | 0 |
| TMEM134 | 906 | 598 | 1264 | 416 | 3294 | 0 | 842 | 0 |
| TMEM135 | 36829 | 35818 | 78781 | 36669 | 147696 | 41458 | 63348 | 39119 |
| TMEM136 | 0 | 0 | 0 | 0 | 1816 | 0 | 0 | 0 |
| TMEM138 | 0 | 664 | 972 | 1108 | 2539 | 0 | 0 | 3455 |
| TMEM139 | 0 | 420 | 1444 | 2344 | 1178 | 0 | 572 | 0 |
| TMEM140 | 350 | 0 | 998 | 0 | 0 | 0 | 0 | 0 |
| TMEM141 | 8530 | 3922 | 23204 | 6086 | 14276 | 2040 | 10937 | 2601 |
| TMEM147 | 1104 | 1294 | 0 | 7921 | 6625 | 4666 | 1983 | 3291 |
| TMEM14A | 0 | 0 | 3440 | 2932 | 1352 | 584 | 852 | 1154 |
| TMEM14B | 1262 | 2028 | 0 | 7749 | 10767 | 2250 | 4727 | 3218 |
| TMEM14C | 742 | 2287 | 4622 | 8908 | 9023 | 0 | 3598 | 1018 |
| TMEM150A | 0 | 0 | 0 | 1300 | 168 | 0 | 182 | 0 |
| TMEM150B | 0 | 0 | 0 | 0 | 770 | 0 | 0 | 0 |
| TMEM154 | 608 | 3930 | 2409 | 2620 | 4466 | 8826 | 2430 | 534 |
| TMEM158 | 0 | 0 | 0 | 0 | 380 | 0 | 0 | 0 |
| TMEM159 | 0 | 0 | 2052 | 548 | 450 | 0 | 0 | 354 |
| TMEM160 | 1004 | 484 | 1754 | 784 | 508 | 3428 | 0 | 0 |
| TMEM161A | 0 | 0 | 0 | 2110 | 1926 | 0 | 0 | 0 |
| TMEM161B | 11605 | 10672 | 21473 | 13007 | 16427 | 12322 | 9238 | 11675 |
| TMEM161B-AS1 | 0 | 0 | 958 | 248 | 474 | 0 | 0 | 0 |
| TMEM163 | 0 | 0 | 0 | 242 | 0 | 0 | 0 | 0 |

| | | | | | | | | |
|---|---|---|---|---|---|---|---|---|
| TMEM164 | 0 | 0 | 0 | 0 | 0 | 0 | 632 | 0 |
| TMEM165 | 112 | 368 | 4065 | 2515 | 1414 | 2038 | 210 | 2298 |
| TMEM167A | 0 | 2774 | 1764 | 2006 | 3971 | 0 | 0 | 0 |
| TMEM167B | 788 | 0 | 0 | 0 | 1226 | 0 | 360 | 0 |
| TMEM168 | 0 | 14 | 0 | 0 | 566 | 566 | 0 | 0 |
| TMEM170A | 0 | 0 | 952 | 0 | 0 | 0 | 322 | 0 |
| TMEM171 | 1015 | 578 | 1854 | 1302 | 548 | 0 | 0 | 0 |
| TMEM173 | 0 | 0 | 0 | 1120 | 436 | 0 | 594 | 0 |
| TMEM175 | 750 | 526 | 0 | 0 | 0 | 0 | 0 | 0 |
| TMEM176A | 762 | 2900 | 8842 | 3334 | 2753 | 9281 | 6666 | 3050 |
| TMEM176B | 1531039 | 1635375 | 4748111 | 1446447 | 2991804 | 2805711 | 5374627 | 907129 |
| TMEM177 | 0 | 824 | 0 | 2522 | 1972 | 354 | 0 | 0 |
| TMEM179B | 1730 | 2718 | 5658 | 5652 | 8038 | 4858 | 6813 | 2008 |
| TMEM18 | 0 | 324 | 212 | 2046 | 0 | 2114 | 0 | 308 |
| TMEM181 | 428 | 172 | 0 | 0 | 1232 | 898 | 126 | 710 |
| TMEM182 | 2255 | 5285 | 1392 | 10424 | 34694 | 484 | 28546 | 22393 |
| TMEM183A | 1876 | 0 | 4928 | 0 | 664 | 0 | 4078 | 2066 |
| TMEM184A | 0 | 0 | 2438 | 0 | 402 | 0 | 0 | 0 |
| TMEM184B | 0 | 110 | 0 | 710 | 0 | 0 | 0 | 0 |
| TMEM185A | 476 | 0 | 2622 | 0 | 470 | 0 | 0 | 0 |
| TMEM185B | 0 | 0 | 0 | 0 | 900 | 0 | 0 | 0 |
| TMEM186 | 0 | 0 | 0 | 0 | 0 | 0 | 1198 | 0 |
| TMEM187 | 0 | 0 | 0 | 0 | 0 | 0 | 0 | 584 |
| TMEM189 | 535 | 0 | 0 | 918 | 1693 | 0 | 0 | 0 |
| TMEM19 | 0 | 314 | 0 | 500 | 0 | 0 | 344 | 0 |
| TMEM191B | 868 | 0 | 1340 | 0 | 0 | 0 | 510 | 802 |
| TMEM191C | 0 | 0 | 1190 | 1072 | 0 | 0 | 0 | 0 |
| TMEM192 | 0 | 0 | 0 | 0 | 0 | 0 | 719 | 0 |
| TMEM199 | 0 | 0 | 0 | 296 | 0 | 0 | 0 | 0 |
| TMEM200A | 614 | 0 | 534 | 0 | 999 | 0 | 0 | 0 |
| TMEM200B | 0 | 0 | 0 | 466 | 0 | 0 | 0 | 0 |
| TMEM203 | 0 | 510 | 3823 | 1126 | 2554 | 0 | 1524 | 881 |
| TMEM204 | 476 | 0 | 0 | 0 | 0 | 0 | 0 | 0 |
| TMEM205 | 140 | 476 | 4175 | 930 | 6340 | 0 | 252 | 862 |
| TMEM206 | 0 | 0 | 0 | 0 | 1083 | 0 | 0 | 0 |
| TMEM208 | 0 | 666 | 2735 | 3931 | 2486 | 3207 | 4300 | 2045 |
| TMEM209 | 0 | 0 | 0 | 512 | 0 | 0 | 0 | 0 |
| TMEM211 | 0 | 0 | 0 | 0 | 0 | 0 | 1634 | 0 |
| TMEM214 | 0 | 0 | 0 | 0 | 1026 | 0 | 456 | 0 |
| TMEM216 | 1006 | 478 | 3322 | 0 | 2079 | 0 | 2880 | 0 |
| TMEM218 | 3514 | 4138 | 9140 | 3235 | 9797 | 2555 | 7360 | 4213 |
| TMEM219 | 2654 | 3412 | 11330 | 9181 | 7318 | 5597 | 2145 | 5319 |
| TMEM220 | 670 | 894 | 0 | 0 | 862 | 0 | 0 | 0 |

| | | | | | | | | |
|---|---|---|---|---|---|---|---|---|
| TMEM222 | 18718 | 43411 | 112412 | 89830 | 108909 | 75299 | 118073 | 39911 |
| TMEM223 | 0 | 0 | 0 | 0 | 1271 | 0 | 2610 | 0 |
| TMEM230 | 0 | 1456 | 1365 | 939 | 2903 | 0 | 814 | 1520 |
| TMEM234 | 0 | 0 | 1297 | 0 | 0 | 1580 | 0 | 1048 |
| TMEM236 | 0 | 0 | 216 | 0 | 0 | 0 | 0 | 0 |
| TMEM237 | 0 | 0 | 0 | 1318 | 562 | 0 | 0 | 0 |
| TMEM238 | 138 | 0 | 1981 | 2120 | 1812 | 672 | 760 | 0 |
| TMEM238L | 698 | 386 | 1687 | 0 | 2426 | 0 | 354 | 0 |
| TMEM241 | 0 | 0 | 0 | 0 | 0 | 1590 | 0 | 0 |
| TMEM242 | 36688 | 60169 | 71144 | 75965 | 168196 | 61585 | 107661 | 38565 |
| TMEM243 | 14130 | 41249 | 20126 | 31561 | 50909 | 39250 | 113251 | 14242 |
| TMEM246 | 572 | 0 | 0 | 0 | 0 | 0 | 770 | 604 |
| TMEM248 | 0 | 876 | 0 | 0 | 816 | 0 | 1378 | 812 |
| TMEM250 | 0 | 306 | 0 | 0 | 0 | 0 | 0 | 0 |
| TMEM251 | 1012 | 672 | 2892 | 1556 | 1032 | 0 | 0 | 0 |
| TMEM253 | 0 | 494 | 0 | 0 | 0 | 954 | 0 | 0 |
| TMEM254 | 0 | 0 | 2144 | 0 | 0 | 882 | 0 | 0 |
| TMEM256 | 2274 | 2402 | 2928 | 0 | 3122 | 0 | 2006 | 174 |
| TMEM256-PLSCR3 | 0 | 266 | 0 | 0 | 494 | 0 | 0 | 0 |
| TMEM258 | 4260 | 10205 | 6126 | 5895 | 23374 | 6212 | 14887 | 6155 |
| TMEM259 | 734 | 210 | 0 | 0 | 3053 | 540 | 1467 | 0 |
| TMEM26-AS1 | 0 | 358 | 0 | 0 | 0 | 0 | 0 | 0 |
| TMEM265 | 0 | 170 | 0 | 898 | 2756 | 0 | 0 | 0 |
| TMEM267 | 304 | 0 | 0 | 0 | 0 | 0 | 0 | 0 |
| TMEM268 | 0 | 528 | 0 | 0 | 0 | 0 | 0 | 0 |
| TMEM30A | 80 | 410 | 484 | 1307 | 1008 | 0 | 787 | 0 |
| TMEM30B | 0 | 0 | 1768 | 0 | 576 | 0 | 0 | 0 |
| TMEM33 | 0 | 1118 | 0 | 20 | 1028 | 0 | 0 | 0 |
| TMEM35B | 1038 | 0 | 1328 | 0 | 2346 | 1392 | 1622 | 18 |
| TMEM37 | 0 | 450 | 3408 | 0 | 0 | 0 | 0 | 0 |
| TMEM38A | 0 | 0 | 0 | 0 | 0 | 0 | 0 | 162 |
| TMEM38B | 38 | 858 | 0 | 0 | 334 | 0 | 1087 | 282 |
| TMEM39A | 0 | 0 | 0 | 0 | 0 | 0 | 330 | 574 |
| TMEM40 | 0 | 0 | 0 | 4719 | 9372 | 0 | 8953 | 72 |
| TMEM41A | 142 | 0 | 0 | 780 | 411 | 0 | 1310 | 0 |
| TMEM41B | 1996 | 992 | 2414 | 0 | 3080 | 0 | 734 | 3455 |
| TMEM42 | 0 | 0 | 0 | 0 | 304 | 0 | 0 | 0 |
| TMEM44 | 0 | 266 | 0 | 0 | 436 | 0 | 0 | 0 |
| TMEM45B | 3852 | 2476 | 12285 | 0 | 4877 | 1594 | 2488 | 552 |
| TMEM47 | 0 | 0 | 0 | 534 | 0 | 0 | 0 | 0 |
| TMEM50A | 694 | 322 | 1540 | 3612 | 2080 | 1661 | 1290 | 2188 |
| TMEM50B | 628 | 460 | 3552 | 1126 | 616 | 0 | 1162 | 2343 |
| TMEM51 | 2334 | 1646 | 6445 | 2207 | 4973 | 5009 | 7994 | 4550 |

| | | | | | | | | |
|---|---|---|---|---|---|---|---|---|
| TMEM53 | 280 | 0 | 0 | 556 | 0 | 0 | 1122 | 1665 |
| TMEM54 | 7149 | 3282 | 33941 | 12308 | 12925 | 0 | 1154 | 2328 |
| TMEM59 | 6258 | 11313 | 29082 | 9869 | 12787 | 12152 | 9155 | 4383 |
| TMEM60 | 822 | 1155 | 0 | 1058 | 1516 | 1919 | 2019 | 921 |
| TMEM61 | 0 | 0 | 0 | 0 | 0 | 0 | 862 | 0 |
| TMEM62 | 0 | 0 | 0 | 2608 | 0 | 0 | 1358 | 0 |
| TMEM63A | 526 | 374 | 0 | 0 | 1392 | 3115 | 64 | 0 |
| TMEM63B | 0 | 0 | 0 | 0 | 92 | 0 | 250 | 0 |
| TMEM64 | 0 | 0 | 0 | 0 | 188 | 1182 | 0 | 0 |
| TMEM65 | 1791 | 594 | 3105 | 0 | 7382 | 12384 | 6171 | 3454 |
| TMEM68 | 0 | 0 | 0 | 678 | 0 | 1128 | 1738 | 0 |
| TMEM69 | 0 | 896 | 646 | 0 | 1418 | 0 | 0 | 0 |
| TMEM70 | 802 | 544 | 1120 | 1286 | 4945 | 2079 | 952 | 3067 |
| TMEM72 | 0 | 0 | 392 | 0 | 0 | 0 | 0 | 0 |
| TMEM74B | 0 | 0 | 0 | 0 | 1760 | 0 | 0 | 0 |
| TMEM80 | 0 | 1320 | 0 | 0 | 1518 | 0 | 726 | 0 |
| TMEM81 | 0 | 0 | 0 | 0 | 194 | 0 | 0 | 0 |
| TMEM86B | 0 | 0 | 0 | 0 | 0 | 0 | 0 | 336 |
| TMEM87A | 956 | 1534 | 2570 | 1074 | 1363 | 0 | 3082 | 0 |
| TMEM87B | 2732 | 6716 | 14985 | 3274 | 9242 | 4438 | 4796 | 3379 |
| TMEM8A | 0 | 118 | 4232 | 790 | 134 | 0 | 3102 | 0 |
| TMEM8B | 432 | 0 | 0 | 0 | 0 | 0 | 212 | 0 |
| TMEM9 | 0 | 594 | 0 | 0 | 950 | 0 | 3352 | 0 |
| TMEM91 | 188 | 0 | 0 | 0 | 0 | 0 | 0 | 0 |
| TMEM92 | 0 | 0 | 0 | 1062 | 0 | 0 | 0 | 0 |
| TMEM94 | 0 | 1758 | 3554 | 1011 | 1078 | 1204 | 1759 | 0 |
| TMEM97 | 0 | 0 | 1688 | 0 | 0 | 2969 | 0 | 0 |
| TMEM98 | 567 | 606 | 514 | 0 | 1966 | 0 | 986 | 0 |
| TMEM99 | 0 | 0 | 0 | 646 | 1134 | 0 | 0 | 0 |
| TMEM9B | 206503 | 363138 | 1008921 | 189593 | 333730 | 256120 | 313607 | 171053 |
| TMF1 | 0 | 262 | 0 | 0 | 0 | 742 | 0 | 0 |
| TMIGD1 | 93315 | 14732 | 120086 | 0 | 790 | 0 | 0 | 0 |
| TMIGD2 | 0 | 256 | 0 | 0 | 0 | 0 | 0 | 0 |
| TMLHE | 0 | 628 | 816 | 0 | 0 | 1864 | 0 | 342 |
| TMOD3 | 0 | 960 | 3126 | 172 | 0 | 0 | 0 | 0 |
| TMPO | 0 | 424 | 0 | 0 | 374 | 0 | 0 | 1032 |
| TMPO-AS1 | 0 | 0 | 0 | 0 | 686 | 0 | 0 | 0 |
| TMPRSS2 | 158885 | 208753 | 892190 | 96440 | 243768 | 111022 | 200589 | 126123 |
| TMPRSS3 | 0 | 0 | 0 | 1242 | 0 | 2486 | 0 | 0 |
| TMPRSS4 | 0 | 474 | 2388 | 0 | 1700 | 2462 | 1244 | 0 |
| TMPRSS7 | 0 | 0 | 0 | 0 | 0 | 0 | 382 | 0 |
| TMSB10 | 42260 | 52900 | 60481 | 117830 | 285715 | 53261 | 70283 | 76296 |
| TMSB4X | 90705 | 81442 | 334127 | 166974 | 323140 | 139267 | 40146 | 101988 |

| | | | | | | | | |
|---|---|---|---|---|---|---|---|---|
| TMSB4Y | 88 | 0 | 0 | 0 | 1059 | 0 | 0 | 0 |
| TMTC3 | 0 | 0 | 0 | 0 | 0 | 0 | 364 | 0 |
| TMTC4 | 0 | 136 | 0 | 386 | 0 | 0 | 0 | 0 |
| TMUB1 | 384 | 0 | 4405 | 480 | 260 | 2379 | 1298 | 0 |
| TMUB2 | 466 | 766 | 350 | 0 | 0 | 0 | 204 | 977 |
| TMX1 | 0 | 742 | 3757 | 780 | 860 | 0 | 0 | 0 |
| TMX2 | 117522 | 296745 | 270628 | 513653 | 707878 | 493395 | 843395 | 316533 |
| TMX2-CTNND1 | 8993 | 16707 | 494 | 10424 | 44780 | 4347 | 24589 | 6901 |
| TMX4 | 0 | 0 | 0 | 0 | 428 | 0 | 0 | 906 |
| TNC | 166 | 810 | 1328 | 2954 | 16472 | 1834 | 250 | 4048 |
| TNFAIP2 | 0 | 0 | 0 | 0 | 2050 | 0 | 196 | 0 |
| TNFAIP3 | 1734 | 4252 | 7847 | 984 | 6224 | 1616 | 6286 | 6246 |
| TNFAIP8 | 0 | 0 | 0 | 0 | 0 | 0 | 244 | 0 |
| TNFAIP8L1 | 0 | 0 | 0 | 0 | 838 | 0 | 310 | 0 |
| TNFAIP8L3 | 336 | 0 | 0 | 0 | 0 | 0 | 0 | 0 |
| TNFRSF10A | 0 | 0 | 0 | 0 | 1002 | 0 | 0 | 0 |
| TNFRSF10B | 0 | 0 | 0 | 1105 | 0 | 0 | 1246 | 962 |
| TNFRSF11A | 0 | 390 | 0 | 0 | 290 | 0 | 186 | 0 |
| TNFRSF11B | 0 | 398 | 0 | 0 | 50 | 0 | 0 | 0 |
| TNFRSF12A | 0 | 530 | 2210 | 1723 | 5258 | 0 | 1064 | 2023 |
| TNFRSF14 | 11204 | 11130 | 5969 | 12140 | 17800 | 13758 | 11395 | 3115 |
| TNFRSF14-AS1 | 1360 | 216 | 3306 | 0 | 1040 | 0 | 0 | 0 |
| TNFRSF17 | 21412 | 59348 | 83723 | 2966 | 1840 | 19080 | 0 | 33049 |
| TNFRSF18 | 0 | 0 | 1380 | 0 | 0 | 0 | 0 | 0 |
| TNFRSF1A | 96475 | 67062 | 553378 | 56982 | 114626 | 77717 | 155802 | 53855 |
| TNFRSF1B | 648 | 0 | 2274 | 0 | 644 | 0 | 800 | 0 |
| TNFRSF21 | 0 | 0 | 0 | 0 | 1034 | 0 | 1038 | 0 |
| TNFRSF6B | 0 | 0 | 1191 | 1094 | 0 | 0 | 0 | 0 |
| TNFSF10 | 622011 | 259062 | 986659 | 151268 | 326257 | 157470 | 218971 | 95815 |
| TNFSF13 | 860 | 874 | 1560 | 0 | 0 | 0 | 842 | 776 |
| TNFSF15 | 0 | 0 | 0 | 0 | 604 | 0 | 0 | 0 |
| TNIK | 608 | 0 | 0 | 0 | 1598 | 0 | 0 | 0 |
| TNIP1 | 1802 | 1686 | 2266 | 0 | 1050 | 0 | 366 | 1205 |
| TNIP2 | 0 | 168 | 0 | 0 | 1362 | 0 | 364 | 1928 |
| TNK2 | 434 | 0 | 0 | 0 | 340 | 0 | 88 | 70 |
| TNKS1BP1 | 602 | 378 | 876 | 0 | 376 | 0 | 398 | 0 |
| TNKS2 | 0 | 300 | 0 | 0 | 0 | 0 | 0 | 0 |
| TNNC1 | 0 | 2053 | 0 | 0 | 1382 | 2066 | 1570 | 834 |
| TNNC2 | 610 | 0 | 1147 | 500 | 1450 | 0 | 1644 | 790 |
| TNNT2 | 0 | 0 | 2176 | 0 | 0 | 0 | 0 | 0 |
| TNNT3 | 0 | 570 | 0 | 0 | 0 | 0 | 0 | 0 |
| TNPO1 | 0 | 0 | 0 | 0 | 0 | 0 | 0 | 200 |
| TNPO2 | 0 | 0 | 374 | 0 | 176 | 0 | 1036 | 0 |

| | | | | | | | | |
|---|---|---|---|---|---|---|---|---|
| TNRC18 | 334 | 402 | 0 | 0 | 1266 | 993 | 74 | 0 |
| TNRC18P1 | 0 | 0 | 0 | 0 | 418 | 0 | 0 | 0 |
| TNRC6A | 0 | 0 | 0 | 0 | 434 | 0 | 222 | 0 |
| TNRC6B | 146922 | 61093 | 110944 | 13083 | 125532 | 44272 | 106686 | 14237 |
| TNS1 | 0 | 449 | 0 | 0 | 70 | 0 | 0 | 0 |
| TNS3 | 554 | 1526 | 771 | 456 | 874 | 0 | 358 | 478 |
| TNS4 | 158 | 0 | 5296 | 5837 | 2754 | 670 | 14043 | 5578 |
| TNXB | 0 | 0 | 479 | 0 | 0 | 0 | 0 | 0 |
| TOB1 | 584 | 1806 | 1844 | 940 | 4220 | 2366 | 1480 | 1707 |
| TOB1-AS1 | 0 | 2261 | 0 | 498 | 2678 | 5509 | 5496 | 144 |
| TOB2 | 496 | 376 | 0 | 0 | 0 | 0 | 718 | 0 |
| TOE1 | 0 | 304 | 0 | 554 | 0 | 0 | 0 | 0 |
| TOLLIP | 0 | 104 | 1273 | 612 | 0 | 0 | 0 | 0 |
| TOLLIP-AS1 | 0 | 0 | 0 | 0 | 1048 | 0 | 468 | 0 |
| TOM1 | 252 | 658 | 0 | 0 | 572 | 0 | 0 | 0 |
| TOM1L1 | 0 | 0 | 0 | 0 | 1360 | 0 | 1333 | 0 |
| TOM1L2 | 0 | 0 | 0 | 906 | 0 | 0 | 0 | 0 |
| TOMM20 | 0 | 886 | 708 | 3223 | 4312 | 3527 | 2326 | 2647 |
| TOMM22 | 1693 | 1096 | 2898 | 3471 | 6567 | 2336 | 0 | 1820 |
| TOMM34 | 0 | 0 | 0 | 3164 | 1870 | 0 | 3724 | 0 |
| TOMM40 | 0 | 0 | 0 | 0 | 706 | 0 | 0 | 562 |
| TOMM5 | 1234 | 2226 | 2519 | 8999 | 5906 | 854 | 4402 | 1520 |
| TOMM6 | 1106 | 2372 | 9137 | 9162 | 14454 | 5334 | 5891 | 2926 |
| TOMM7 | 2754 | 5785 | 1956 | 3416 | 13191 | 1264 | 2606 | 2861 |
| TOMM70 | 0 | 0 | 0 | 810 | 0 | 0 | 516 | 0 |
| TONSL | 0 | 0 | 0 | 0 | 860 | 0 | 0 | 0 |
| TOP1 | 0 | 0 | 924 | 228 | 286 | 0 | 34 | 0 |
| TOP1MT | 0 | 0 | 0 | 0 | 96 | 1152 | 1474 | 0 |
| TOP2A | 0 | 0 | 0 | 1312 | 570 | 0 | 0 | 0 |
| TOP2B | 102 | 582 | 0 | 0 | 0 | 0 | 0 | 0 |
| TOP3A | 368 | 2018 | 6641 | 2272 | 3934 | 2742 | 2229 | 2038 |
| TOP3B | 1220 | 946 | 3614 | 1010 | 7710 | 0 | 3300 | 1232 |
| TOPBP1 | 622 | 1572 | 4870 | 6618 | 13836 | 3122 | 6693 | 2442 |
| TOPORS | 0 | 668 | 448 | 0 | 0 | 0 | 0 | 0 |
| TOR1A | 0 | 444 | 4850 | 0 | 616 | 0 | 342 | 1658 |
| TOR1AIP1 | 0 | 0 | 0 | 499 | 0 | 0 | 0 | 0 |
| TOR1AIP2 | 0 | 0 | 0 | 0 | 0 | 0 | 188 | 0 |
| TOR1B | 0 | 586 | 0 | 0 | 226 | 0 | 1170 | 408 |
| TOR2A | 0 | 422 | 1012 | 1228 | 0 | 0 | 0 | 0 |
| TOR3A | 600 | 646 | 0 | 1182 | 1594 | 0 | 702 | 0 |
| TOX | 9109 | 11956 | 15219 | 5178 | 11142 | 48452 | 528 | 1610 |
| TOX3 | 0 | 0 | 1319 | 0 | 118 | 0 | 0 | 0 |
| TOX4 | 0 | 0 | 36 | 0 | 524 | 0 | 187 | 0 |

| | | | | | | | | |
|---|---|---|---|---|---|---|---|---|
| TP53 | 11206 | 26247 | 45314 | 10181 | 10306 | 12545 | 8347 | 1117 |
| TP53BP1 | 3665 | 5828 | 2210 | 2339 | 10668 | 6389 | 19930 | 698 |
| TP53I11 | 0 | 0 | 0 | 1418 | 742 | 0 | 0 | 0 |
| TP53I13 | 0 | 264 | 0 | 0 | 0 | 0 | 0 | 498 |
| TP53I3 | 650 | 748 | 1800 | 238 | 536 | 0 | 0 | 0 |
| TP53INP1 | 254 | 232 | 0 | 0 | 0 | 0 | 0 | 0 |
| TP53INP2 | 242 | 0 | 0 | 0 | 0 | 0 | 0 | 0 |
| TP53RK | 0 | 226 | 0 | 2130 | 0 | 2198 | 3582 | 1086 |
| TP53TG1 | 0 | 0 | 2588 | 0 | 0 | 0 | 510 | 0 |
| TP73 | 0 | 0 | 0 | 0 | 162 | 0 | 764 | 0 |
| TPBG | 0 | 0 | 0 | 632 | 830 | 0 | 0 | 0 |
| TPCN1 | 322 | 0 | 0 | 0 | 0 | 0 | 646 | 0 |
| TPCN2 | 0 | 0 | 0 | 644 | 0 | 0 | 0 | 0 |
| TPD52 | 1444 | 1096 | 2794 | 4023 | 4872 | 2262 | 372 | 1984 |
| TPD52L1 | 0 | 0 | 0 | 1077 | 4055 | 2057 | 1270 | 0 |
| TPD52L2 | 0 | 0 | 0 | 989 | 402 | 0 | 2252 | 806 |
| TPGS1 | 0 | 2050 | 2761 | 0 | 584 | 0 | 644 | 2050 |
| TPGS2 | 9074 | 33099 | 48176 | 43241 | 201690 | 30008 | 68929 | 75931 |
| TPH1 | 858 | 0 | 0 | 0 | 0 | 0 | 0 | 0 |
| TPI1 | 7514 | 9289 | 44953 | 21187 | 39047 | 14606 | 5754 | 15744 |
| TPK1 | 0 | 0 | 1970 | 0 | 0 | 0 | 0 | 0 |
| TPM1 | 283385 | 332436 | 892181 | 286680 | 881914 | 389706 | 372097 | 242729 |
| TPM2 | 994 | 80 | 2366 | 3292 | 15647 | 1457 | 844 | 7880 |
| TPM3 | 569482 | 810464 | 1355500 | 1140712 | 2517375 | 991028 | 1482633 | 1222259 |
| TPM3P9 | 558 | 0 | 0 | 1582 | 4028 | 0 | 1776 | 1282 |
| TPM4 | 742 | 1838 | 3058 | 2619 | 3569 | 4654 | 1628 | 2806 |
| TPMT | 80404 | 115567 | 186246 | 131804 | 356440 | 81178 | 174358 | 144932 |
| TPP1 | 0 | 0 | 620 | 0 | 0 | 0 | 270 | 0 |
| TPP2 | 0 | 0 | 414 | 750 | 0 | 1032 | 0 | 0 |
| TPPP | 0 | 0 | 0 | 0 | 154 | 0 | 0 | 0 |
| TPPP3 | 0 | 0 | 0 | 0 | 1234 | 0 | 0 | 1022 |
| TPR | 0 | 636 | 0 | 0 | 1116 | 702 | 0 | 878 |
| TPRA1 | 6512 | 5527 | 18468 | 13634 | 28506 | 35745 | 28156 | 5118 |
| TPRG1 | 1202 | 586 | 1336 | 1552 | 2374 | 1798 | 7050 | 818 |
| TPRG1L | 0 | 474 | 314 | 1538 | 336 | 0 | 434 | 0 |
| TPRKB | 738 | 826 | 1703 | 1556 | 5246 | 0 | 664 | 611 |
| TPRN | 712 | 0 | 0 | 0 | 500 | 0 | 538 | 0 |
| TPSAB1 | 0 | 692 | 0 | 0 | 830 | 0 | 0 | 0 |
| TPSB2 | 0 | 1006 | 0 | 0 | 0 | 0 | 0 | 0 |
| TPSG1 | 0 | 0 | 1150 | 0 | 0 | 0 | 0 | 0 |
| TPST1 | 0 | 0 | 0 | 0 | 366 | 0 | 0 | 940 |
| TPST2 | 15831 | 17374 | 38022 | 34872 | 42713 | 129892 | 161261 | 27406 |
| TPT1 | 4061502 | 5507734 | 4378929 | 8496685 | 15316522 | 17584559 | 31057160 | 6926669 |

| | | | | | | | | |
|---|---|---|---|---|---|---|---|---|
| TPTEP1 | 0 | 0 | 0 | 250 | 814 | 0 | 0 | 0 |
| TPTEP2-CSNK1E | 0 | 284 | 3758 | 0 | 2492 | 0 | 0 | 0 |
| TPX2 | 3316 | 14644 | 12373 | 104059 | 283840 | 32467 | 120041 | 74936 |
| TRA2A | 53680 | 59024 | 103917 | 61710 | 215544 | 97159 | 125238 | 63268 |
| TRA2B | 1072 | 0 | 936 | 1036 | 3160 | 0 | 514 | 0 |
| TRABD | 1766 | 1314 | 0 | 834 | 106 | 0 | 396 | 654 |
| TRABD2A | 678 | 712 | 3206 | 1934 | 3252 | 1536 | 760 | 0 |
| TRADD | 1396 | 438 | 0 | 956 | 914 | 2473 | 1554 | 0 |
| TRAF2 | 160 | 0 | 0 | 0 | 0 | 2478 | 548 | 0 |
| TRAF3 | 5704 | 6642 | 17701 | 4404 | 25081 | 8123 | 8099 | 8139 |
| TRAF3IP2 | 0 | 0 | 0 | 593 | 0 | 0 | 0 | 0 |
| TRAF3IP3 | 0 | 0 | 1436 | 0 | 0 | 0 | 0 | 0 |
| TRAF4 | 52 | 618 | 2494 | 0 | 1436 | 0 | 648 | 640 |
| TRAF5 | 0 | 452 | 0 | 0 | 622 | 0 | 422 | 1008 |
| TRAF6 | 0 | 0 | 0 | 0 | 0 | 0 | 218 | 0 |
| TRAF7 | 12655 | 9594 | 34948 | 20588 | 26448 | 20886 | 39826 | 14321 |
| TRAFD1 | 0 | 0 | 0 | 844 | 0 | 0 | 0 | 0 |
| TRAIP | 0 | 0 | 0 | 0 | 150 | 0 | 0 | 0 |
| TRAK1 | 5188 | 4431 | 25510 | 1848 | 37933 | 6543 | 10508 | 8631 |
| TRAM1 | 0 | 572 | 1786 | 184 | 1500 | 0 | 474 | 2428 |
| TRAM1L1 | 0 | 484 | 0 | 0 | 0 | 0 | 0 | 0 |
| TRAM2-AS1 | 0 | 0 | 0 | 0 | 0 | 0 | 364 | 0 |
| TRANK1 | 198 | 0 | 783 | 0 | 236 | 0 | 0 | 0 |
| TRAP1 | 2560 | 10358 | 10362 | 20082 | 27065 | 4790 | 35078 | 15959 |
| TRAPPC1 | 0 | 1038 | 1796 | 2632 | 4186 | 248 | 1546 | 554 |
| TRAPPC10 | 0 | 0 | 0 | 0 | 164 | 0 | 0 | 0 |
| TRAPPC12 | 318 | 236 | 0 | 1556 | 208 | 0 | 0 | 0 |
| TRAPPC2 | 4646 | 8451 | 7425 | 15259 | 46215 | 15244 | 21036 | 6929 |
| TRAPPC2B | 56968 | 62655 | 40698 | 50221 | 272516 | 93741 | 101590 | 87954 |
| TRAPPC2L | 0 | 0 | 1078 | 1626 | 3753 | 1525 | 4155 | 1854 |
| TRAPPC3 | 859 | 1804 | 946 | 0 | 3271 | 2302 | 572 | 1078 |
| TRAPPC4 | 235221 | 410036 | 668262 | 270052 | 635162 | 388105 | 977917 | 520797 |
| TRAPPC5 | 0 | 0 | 3186 | 1362 | 4056 | 0 | 1174 | 1158 |
| TRAPPC6A | 0 | 612 | 1751 | 3393 | 0 | 863 | 0 | 844 |
| TRAPPC6B | 0 | 0 | 0 | 0 | 908 | 0 | 0 | 712 |
| TRAPPC8 | 0 | 0 | 0 | 0 | 242 | 0 | 0 | 0 |
| TRAPPC9 | 0 | 0 | 0 | 0 | 0 | 0 | 468 | 0 |
| TRDMT1 | 0 | 404 | 0 | 0 | 0 | 0 | 0 | 0 |
| TREH | 0 | 0 | 0 | 0 | 1406 | 0 | 0 | 0 |
| TREM2 | 0 | 0 | 0 | 0 | 0 | 0 | 68 | 0 |
| TRERF1 | 0 | 516 | 0 | 222 | 2136 | 0 | 0 | 1280 |
| TRHDE | 2144 | 996 | 6270 | 852 | 0 | 0 | 19341 | 0 |
| TRIAP1 | 0 | 0 | 0 | 1088 | 0 | 1464 | 0 | 982 |

| | | | | | | | | |
|---|---|---|---|---|---|---|---|---|
| TRIB1 | 0 | 0 | 0 | 0 | 0 | 0 | 368 | 252 |
| TRIB2 | 0 | 0 | 0 | 0 | 98 | 0 | 0 | 0 |
| TRIB3 | 0 | 0 | 0 | 0 | 514 | 0 | 0 | 665 |
| TRIM13 | 0 | 1368 | 0 | 0 | 1500 | 3429 | 772 | 0 |
| TRIM14 | 0 | 468 | 1544 | 0 | 0 | 0 | 0 | 0 |
| TRIM15 | 734 | 0 | 656 | 844 | 369 | 0 | 0 | 0 |
| TRIM16 | 0 | 0 | 0 | 900 | 936 | 0 | 0 | 1383 |
| TRIM16L | 0 | 50 | 0 | 0 | 0 | 0 | 0 | 0 |
| TRIM2 | 0 | 716 | 1234 | 1315 | 118 | 1414 | 1104 | 488 |
| TRIM21 | 508 | 0 | 1688 | 2318 | 0 | 1058 | 1142 | 0 |
| TRIM22 | 0 | 0 | 800 | 0 | 0 | 0 | 410 | 0 |
| TRIM23 | 448 | 0 | 0 | 0 | 0 | 0 | 0 | 0 |
| TRIM24 | 13765 | 12798 | 23933 | 38620 | 90667 | 20257 | 76467 | 20114 |
| TRIM26 | 1029 | 0 | 700 | 704 | 386 | 0 | 512 | 788 |
| TRIM27 | 13779 | 18797 | 55993 | 37871 | 67032 | 41796 | 123870 | 27577 |
| TRIM28 | 0 | 716 | 0 | 3049 | 2772 | 0 | 0 | 0 |
| TRIM29 | 0 | 424 | 0 | 768 | 802 | 1677 | 0 | 746 |
| TRIM3 | 0 | 252 | 0 | 0 | 0 | 0 | 0 | 0 |
| TRIM31 | 1336 | 672 | 5963 | 907 | 4985 | 0 | 2714 | 0 |
| TRIM33 | 24839 | 16893 | 32709 | 11408 | 52106 | 16539 | 71855 | 9565 |
| TRIM34 | 0 | 0 | 1484 | 0 | 830 | 0 | 0 | 0 |
| TRIM35 | 0 | 0 | 1806 | 0 | 0 | 0 | 0 | 0 |
| TRIM4 | 3229 | 3102 | 4065 | 3116 | 3354 | 12461 | 11246 | 1784 |
| TRIM40 | 0 | 0 | 1764 | 0 | 0 | 0 | 0 | 0 |
| TRIM41 | 0 | 0 | 0 | 0 | 28 | 0 | 0 | 0 |
| TRIM44 | 372 | 236 | 0 | 0 | 1904 | 3598 | 374 | 0 |
| TRIM45 | 0 | 0 | 0 | 0 | 202 | 0 | 0 | 0 |
| TRIM46 | 0 | 0 | 0 | 0 | 0 | 0 | 0 | 1090 |
| TRIM52 | 0 | 368 | 0 | 0 | 816 | 0 | 0 | 0 |
| TRIM52-AS1 | 0 | 0 | 0 | 0 | 764 | 0 | 0 | 0 |
| TRIM54 | 0 | 0 | 0 | 0 | 0 | 0 | 1126 | 0 |
| TRIM56 | 0 | 0 | 0 | 0 | 344 | 0 | 0 | 0 |
| TRIM59 | 0 | 0 | 0 | 0 | 402 | 0 | 0 | 0 |
| TRIM59-IFT80 | 1286 | 2376 | 1495 | 6235 | 14140 | 1672 | 7373 | 4344 |
| TRIM65 | 610 | 0 | 0 | 0 | 0 | 0 | 0 | 0 |
| TRIM66 | 108 | 0 | 0 | 0 | 0 | 0 | 0 | 0 |
| TRIM7 | 0 | 0 | 0 | 0 | 684 | 0 | 0 | 0 |
| TRIM74 | 258 | 340 | 0 | 0 | 0 | 0 | 0 | 0 |
| TRIM8 | 0 | 38 | 0 | 0 | 160 | 264 | 208 | 0 |
| TRIO | 0 | 0 | 0 | 0 | 1970 | 1723 | 2192 | 0 |
| TRIOBP | 0 | 432 | 0 | 0 | 146 | 0 | 0 | 946 |
| TRIP10 | 1122 | 0 | 1270 | 0 | 0 | 0 | 0 | 630 |
| TRIP11 | 0 | 0 | 1064 | 0 | 0 | 0 | 0 | 0 |

| | | | | | | | | |
|---|---|---|---|---|---|---|---|---|
| TRIP12 | 5208 | 3948 | 4831 | 3956 | 12856 | 4274 | 8466 | 6065 |
| TRIP13 | 0 | 282 | 0 | 1556 | 400 | 0 | 262 | 0 |
| TRIP4 | 908 | 0 | 0 | 0 | 580 | 0 | 1506 | 0 |
| TRIP6 | 0 | 262 | 0 | 0 | 456 | 0 | 0 | 0 |
| TRIQK | 0 | 998 | 0 | 0 | 104 | 0 | 430 | 0 |
| TRIR | 726 | 42 | 892 | 4225 | 7690 | 0 | 542 | 2852 |
| TRIT1 | 0 | 0 | 0 | 0 | 0 | 0 | 340 | 0 |
| TRMO | 174 | 0 | 714 | 0 | 0 | 0 | 0 | 0 |
| TRMT1 | 0 | 214 | 0 | 0 | 626 | 0 | 238 | 828 |
| TRMT10B | 0 | 0 | 0 | 0 | 660 | 0 | 0 | 0 |
| TRMT10C | 0 | 394 | 0 | 1276 | 6342 | 992 | 2622 | 962 |
| TRMT11 | 35218 | 65973 | 150129 | 113832 | 332709 | 142487 | 195600 | 102276 |
| TRMT112 | 1196 | 1204 | 4356 | 6670 | 10132 | 2046 | 11616 | 5656 |
| TRMT12 | 0 | 0 | 0 | 0 | 258 | 0 | 0 | 0 |
| TRMT13 | 0 | 0 | 0 | 0 | 718 | 0 | 210 | 0 |
| TRMT1L | 1322 | 0 | 0 | 1078 | 0 | 0 | 0 | 0 |
| TRMT2B | 0 | 300 | 0 | 0 | 360 | 0 | 0 | 0 |
| TRMT44 | 0 | 0 | 718 | 0 | 0 | 0 | 538 | 0 |
| TRMT6 | 0 | 894 | 0 | 0 | 675 | 0 | 0 | 1152 |
| TRMT61A | 820 | 0 | 0 | 0 | 0 | 0 | 0 | 0 |
| TRMT61B | 13777 | 26881 | 36941 | 40418 | 86341 | 50128 | 97670 | 24393 |
| TRMT9B | 388 | 0 | 0 | 0 | 0 | 0 | 0 | 0 |
| TRMU | 0 | 346 | 0 | 0 | 500 | 0 | 226 | 928 |
| TRNAU1AP | 0 | 0 | 0 | 1408 | 0 | 0 | 0 | 1428 |
| TRNP1 | 0 | 0 | 0 | 0 | 0 | 24 | 0 | 0 |
| TRNT1 | 0 | 917 | 0 | 0 | 312 | 0 | 396 | 0 |
| TROAP | 0 | 370 | 0 | 0 | 1048 | 0 | 606 | 0 |
| TROVE2 | 3195 | 6022 | 8901 | 5733 | 11366 | 2297 | 13187 | 2236 |
| TRPC1 | 0 | 0 | 0 | 0 | 0 | 664 | 0 | 0 |
| TRPC4AP | 0 | 154 | 0 | 0 | 346 | 0 | 230 | 0 |
| TRPM2 | 0 | 0 | 0 | 1234 | 0 | 0 | 0 | 0 |
| TRPM2-AS | 0 | 0 | 0 | 0 | 1216 | 0 | 0 | 0 |
| TRPM4 | 2648 | 1038 | 4415 | 1480 | 1852 | 0 | 150 | 0 |
| TRPM6 | 480 | 0 | 1888 | 0 | 0 | 0 | 0 | 0 |
| TRPM7 | 0 | 0 | 0 | 296 | 0 | 0 | 0 | 0 |
| TRPT1 | 368 | 0 | 3039 | 1240 | 1740 | 1988 | 1180 | 1058 |
| TRPV2 | 0 | 0 | 0 | 0 | 0 | 0 | 274 | 0 |
| TRRAP | 610 | 266 | 0 | 647 | 1885 | 1184 | 2210 | 0 |
| TRUB1 | 0 | 0 | 0 | 0 | 0 | 0 | 0 | 1136 |
| TRUB2 | 0 | 0 | 0 | 2342 | 408 | 2256 | 350 | 0 |
| TSC1 | 584 | 11053 | 6003 | 8965 | 28035 | 11184 | 15839 | 7661 |
| TSC2 | 7498 | 8260 | 19025 | 6816 | 22507 | 17621 | 34374 | 2584 |
| TSC22D1 | 2402 | 2216 | 18195 | 4250 | 7864 | 11413 | 8859 | 568 |

| | | | | | | | | |
|---|---|---|---|---|---|---|---|---|
| TSC22D1-AS1 | 0 | 0 | 0 | 0 | 0 | 1524 | 0 | 0 |
| TSC22D2 | 0 | 0 | 1790 | 0 | 0 | 0 | 0 | 0 |
| TSC22D3 | 550 | 834 | 32 | 0 | 1326 | 0 | 0 | 1050 |
| TSC22D4 | 0 | 146 | 0 | 1930 | 194 | 0 | 648 | 214 |
| TSEN15 | 0 | 0 | 0 | 0 | 174 | 0 | 0 | 741 |
| TSEN2 | 0 | 372 | 0 | 0 | 2430 | 0 | 0 | 0 |
| TSEN34 | 594 | 0 | 90 | 1272 | 1450 | 1510 | 0 | 2220 |
| TSEN54 | 548 | 286 | 0 | 0 | 348 | 0 | 286 | 803 |
| TSFM | 58255 | 139105 | 100003 | 143543 | 333758 | 134609 | 361677 | 95810 |
| TSG101 | 436 | 234 | 1092 | 3502 | 1682 | 2014 | 782 | 3204 |
| TSGA10 | 0 | 0 | 0 | 0 | 642 | 0 | 0 | 0 |
| TSHR | 0 | 0 | 0 | 0 | 0 | 0 | 0 | 224 |
| TSHZ2 | 1072 | 0 | 818 | 0 | 2410 | 5563 | 0 | 0 |
| TSHZ3 | 376 | 0 | 0 | 0 | 0 | 0 | 0 | 0 |
| TSKU | 0 | 0 | 0 | 408 | 464 | 0 | 615 | 0 |
| TSN | 0 | 310 | 0 | 3295 | 246 | 0 | 590 | 380 |
| TSNARE1 | 0 | 132 | 0 | 426 | 8732 | 0 | 454 | 50593 |
| TSNAX | 0 | 300 | 0 | 124 | 0 | 0 | 0 | 0 |
| TSNAX-DISC1 | 0 | 0 | 0 | 0 | 628 | 0 | 0 | 0 |
| TSPAN1 | 10495 | 11690 | 80870 | 4428 | 5273 | 0 | 0 | 0 |
| TSPAN12 | 0 | 462 | 580 | 1757 | 720 | 1232 | 882 | 0 |
| TSPAN13 | 638 | 902 | 4083 | 1884 | 3836 | 1638 | 1968 | 1654 |
| TSPAN14 | 0 | 0 | 0 | 0 | 50 | 0 | 778 | 0 |
| TSPAN15 | 500 | 928 | 592 | 1024 | 1738 | 348 | 0 | 422 |
| TSPAN17 | 0 | 0 | 0 | 224 | 1887 | 0 | 0 | 1068 |
| TSPAN18 | 0 | 0 | 0 | 0 | 274 | 0 | 0 | 0 |
| TSPAN2 | 0 | 0 | 0 | 0 | 428 | 0 | 0 | 908 |
| TSPAN3 | 355118 | 402567 | 1092590 | 255798 | 669022 | 349061 | 331082 | 207714 |
| TSPAN31 | 188 | 62 | 1720 | 0 | 496 | 0 | 1038 | 0 |
| TSPAN33 | 110 | 542 | 886 | 2172 | 990 | 2198 | 13577 | 2046 |
| TSPAN4 | 0 | 0 | 682 | 0 | 0 | 1983 | 0 | 0 |
| TSPAN5 | 0 | 0 | 0 | 689 | 0 | 0 | 0 | 0 |
| TSPAN6 | 738 | 266 | 3709 | 1422 | 1330 | 3222 | 2944 | 0 |
| TSPAN7 | 380 | 846 | 4161 | 0 | 236 | 0 | 0 | 0 |
| TSPAN8 | 13982 | 21184 | 68534 | 30837 | 24372 | 8706 | 11911 | 8855 |
| TSPAN9 | 0 | 0 | 0 | 0 | 522 | 0 | 0 | 0 |
| TSPO | 662 | 5002 | 16699 | 10120 | 10258 | 6626 | 7137 | 3144 |
| TSPYL1 | 1844 | 330 | 0 | 762 | 392 | 530 | 0 | 676 |
| TSPYL2 | 0 | 0 | 0 | 0 | 110 | 0 | 0 | 0 |
| TSPYL4 | 0 | 0 | 0 | 353 | 0 | 0 | 0 | 0 |
| TSR1 | 750 | 478 | 0 | 0 | 0 | 0 | 0 | 0 |
| TSR2 | 770 | 0 | 1532 | 1314 | 5437 | 1395 | 1188 | 0 |
| TSR3 | 730 | 1374 | 1716 | 1199 | 378 | 5219 | 1556 | 1013 |

| | | | | | | | | |
|---|---|---|---|---|---|---|---|---|
| TSSC4 | 0 | 0 | 1772 | 0 | 572 | 0 | 0 | 0 |
| TST | 3444 | 412 | 2814 | 2584 | 568 | 0 | 2046 | 0 |
| TSTA3 | 564 | 1322 | 2346 | 1614 | 10398 | 9709 | 1689 | 1532 |
| TSTD1 | 0 | 4791 | 7630 | 2479 | 4938 | 5539 | 4218 | 1426 |
| TSTD2 | 210 | 0 | 0 | 0 | 0 | 526 | 0 | 0 |
| TSTD3 | 0 | 80 | 0 | 0 | 0 | 0 | 0 | 0 |
| TTC1 | 0 | 0 | 0 | 0 | 2684 | 0 | 1767 | 0 |
| TTC13 | 478 | 1705 | 3484 | 6564 | 15941 | 3713 | 6984 | 4501 |
| TTC14 | 0 | 0 | 1280 | 0 | 0 | 0 | 0 | 0 |
| TTC17 | 0 | 0 | 0 | 0 | 658 | 0 | 0 | 0 |
| TTC19 | 0 | 0 | 0 | 0 | 0 | 0 | 56 | 0 |
| TTC22 | 0 | 414 | 1380 | 0 | 0 | 0 | 0 | 0 |
| TTC27 | 8056 | 20553 | 20340 | 15908 | 82043 | 35567 | 86771 | 14016 |
| TTC28 | 1490 | 0 | 3668 | 0 | 0 | 2436 | 1618 | 0 |
| TTC28-AS1 | 0 | 684 | 0 | 1006 | 278 | 0 | 0 | 0 |
| TTC3 | 0 | 388 | 0 | 0 | 4689 | 0 | 214 | 0 |
| TTC32 | 0 | 0 | 0 | 0 | 0 | 0 | 342 | 0 |
| TTC33 | 0 | 320 | 0 | 0 | 526 | 0 | 0 | 0 |
| TTC36 | 0 | 0 | 0 | 0 | 0 | 455 | 0 | 0 |
| TTC38 | 0 | 564 | 0 | 144 | 0 | 0 | 80 | 762 |
| TTC39A | 332 | 709 | 0 | 0 | 1178 | 0 | 0 | 1888 |
| TTC39B | 0 | 336 | 0 | 0 | 0 | 0 | 0 | 0 |
| TTC39C | 0 | 0 | 0 | 0 | 546 | 1946 | 424 | 0 |
| TTC4 | 0 | 440 | 0 | 0 | 1122 | 0 | 0 | 0 |
| TTC5 | 0 | 0 | 0 | 0 | 756 | 0 | 748 | 0 |
| TTC7A | 1702 | 378 | 2855 | 0 | 1966 | 0 | 366 | 1050 |
| TTC7B | 0 | 24 | 0 | 0 | 1886 | 0 | 1260 | 746 |
| TTC8 | 0 | 0 | 0 | 0 | 266 | 0 | 0 | 0 |
| TTC9C | 0 | 0 | 0 | 0 | 1444 | 0 | 1066 | 0 |
| TTF1 | 0 | 0 | 0 | 0 | 0 | 0 | 498 | 0 |
| TTK | 0 | 0 | 0 | 0 | 1668 | 0 | 0 | 0 |
| TTLL1 | 4310 | 4342 | 16036 | 8898 | 27710 | 9756 | 15162 | 7691 |
| TTLL12 | 0 | 0 | 0 | 0 | 0 | 2088 | 0 | 0 |
| TTLL2 | 0 | 0 | 0 | 0 | 70 | 0 | 0 | 0 |
| TTLL3 | 362 | 0 | 0 | 0 | 0 | 0 | 0 | 0 |
| TTLL4 | 0 | 0 | 0 | 808 | 0 | 0 | 0 | 0 |
| TTLL5 | 0 | 384 | 0 | 0 | 0 | 0 | 0 | 0 |
| TTLL6 | 436 | 0 | 0 | 0 | 0 | 0 | 0 | 0 |
| TTLL9 | 0 | 0 | 0 | 0 | 0 | 0 | 644 | 0 |
| TTN-AS1 | 0 | 296 | 0 | 0 | 0 | 0 | 0 | 0 |
| TTTY14 | 0 | 222 | 0 | 0 | 374 | 0 | 0 | 0 |
| TTTY15 | 0 | 272 | 0 | 0 | 0 | 0 | 0 | 0 |
| TTYH2 | 0 | 0 | 0 | 0 | 0 | 0 | 875 | 0 |

| | | | | | | | | |
|---|---|---|---|---|---|---|---|---|
| TTYH3 | 0 | 297 | 140 | 454 | 2345 | 192 | 0 | 0 |
| TUBA1A | 0 | 0 | 1567 | 0 | 3268 | 1660 | 0 | 0 |
| TUBA1B | 1362 | 1730 | 9256 | 21720 | 25808 | 8763 | 3528 | 3740 |
| TUBA1C | 1606 | 0 | 6630 | 2110 | 3312 | 1696 | 1552 | 2931 |
| TUBA4A | 0 | 468 | 2066 | 0 | 1912 | 0 | 810 | 0 |
| TUBB | 712 | 710 | 2010 | 9835 | 9195 | 0 | 2014 | 8235 |
| TUBB1 | 0 | 0 | 0 | 0 | 920 | 0 | 0 | 0 |
| TUBB2A | 0 | 0 | 872 | 0 | 1062 | 0 | 0 | 0 |
| TUBB2B | 28 | 0 | 0 | 0 | 1128 | 0 | 0 | 0 |
| TUBB3 | 0 | 0 | 0 | 1190 | 0 | 0 | 0 | 0 |
| TUBB4B | 1234 | 424 | 3135 | 7245 | 5896 | 0 | 628 | 1032 |
| TUBB6 | 216 | 0 | 0 | 0 | 0 | 0 | 0 | 4770 |
| TUBD1 | 4036 | 4318 | 5786 | 4686 | 25207 | 7064 | 20446 | 4560 |
| TUBE1 | 0 | 336 | 0 | 0 | 0 | 0 | 0 | 0 |
| TUBG1 | 0 | 498 | 0 | 1066 | 1506 | 0 | 0 | 2377 |
| TUBG2 | 0 | 0 | 0 | 0 | 764 | 0 | 650 | 0 |
| TUBGCP2 | 0 | 0 | 0 | 0 | 646 | 0 | 650 | 0 |
| TUBGCP4 | 0 | 0 | 0 | 0 | 896 | 0 | 470 | 0 |
| TUBGCP5 | 718 | 0 | 0 | 0 | 0 | 0 | 0 | 0 |
| TUBGCP6 | 0 | 0 | 0 | 0 | 450 | 0 | 0 | 0 |
| TUFM | 2150 | 1580 | 3596 | 4423 | 7795 | 2164 | 3657 | 2326 |
| TULP3 | 0 | 0 | 0 | 0 | 609 | 0 | 192 | 0 |
| TULP4 | 1146 | 0 | 12 | 1000 | 1910 | 0 | 426 | 908 |
| TUSC1 | 0 | 0 | 1714 | 0 | 1428 | 192 | 0 | 0 |
| TUSC2 | 874 | 220 | 1481 | 584 | 858 | 0 | 742 | 0 |
| TUSC3 | 0 | 514 | 1684 | 0 | 0 | 0 | 0 | 0 |
| TUT1 | 0 | 0 | 0 | 2348 | 0 | 0 | 0 | 0 |
| TUT7 | 488 | 0 | 0 | 0 | 0 | 0 | 0 | 0 |
| TVP23B | 602 | 1164 | 1054 | 0 | 0 | 0 | 0 | 774 |
| TWF1 | 0 | 574 | 1614 | 0 | 1046 | 0 | 0 | 586 |
| TWF2 | 0 | 0 | 0 | 1218 | 976 | 0 | 392 | 0 |
| TWIST1 | 0 | 0 | 0 | 0 | 0 | 0 | 0 | 766 |
| TWIST2 | 1290 | 0 | 0 | 1120 | 0 | 0 | 0 | 0 |
| TWISTNB | 0 | 0 | 0 | 1590 | 424 | 0 | 0 | 0 |
| TWSG1 | 0 | 0 | 0 | 0 | 868 | 0 | 0 | 0 |
| TXK | 0 | 1010 | 0 | 0 | 0 | 618 | 276 | 0 |
| TXLNA | 112 | 930 | 502 | 548 | 4646 | 4284 | 2594 | 836 |
| TXLNGY | 0 | 190 | 0 | 0 | 138 | 0 | 484 | 0 |
| TXN | 8126 | 17854 | 21613 | 34071 | 63154 | 17731 | 19222 | 10670 |
| TXN2 | 771513 | 980102 | 3789307 | 1051043 | 1675439 | 1498327 | 2339999 | 636256 |
| TXNDC11 | 582 | 224 | 1827 | 0 | 352 | 0 | 1592 | 0 |
| TXNDC12 | 0 | 0 | 3640 | 2563 | 714 | 2890 | 1614 | 0 |
| TXNDC15 | 0 | 564 | 0 | 1082 | 0 | 1382 | 686 | 0 |

| | | | | | | | | |
|---|---|---|---|---|---|---|---|---|
| TXNDC16 | 0 | 206 | 0 | 0 | 0 | 0 | 0 | 0 |
| TXNDC17 | 3344 | 8046 | 7714 | 3136 | 11434 | 4532 | 1652 | 4382 |
| TXNDC5 | 0 | 0 | 1354 | 580 | 1710 | 0 | 582 | 724 |
| TXNDC9 | 0 | 832 | 4273 | 736 | 3139 | 0 | 1088 | 1146 |
| TXNIP | 672 | 1796 | 6987 | 2243 | 2578 | 0 | 264 | 806 |
| TXNL1 | 223271 | 397230 | 1920430 | 387647 | 618155 | 624757 | 786510 | 277418 |
| TXNL4A | 2151 | 1932 | 1774 | 1456 | 12338 | 1394 | 786 | 586 |
| TXNL4B | 0 | 0 | 0 | 0 | 106 | 0 | 0 | 0 |
| TXNRD1 | 19526 | 32451 | 46375 | 56805 | 76253 | 31752 | 75173 | 38352 |
| TXNRD2 | 0 | 0 | 0 | 0 | 1308 | 1114 | 0 | 0 |
| TYK2 | 0 | 0 | 0 | 0 | 0 | 0 | 482 | 0 |
| TYMP | 964 | 0 | 184 | 574 | 2512 | 1096 | 178 | 522 |
| TYMS | 0 | 0 | 0 | 1254 | 2673 | 0 | 0 | 2566 |
| TYMSOS | 0 | 0 | 1402 | 408 | 0 | 0 | 0 | 0 |
| TYROBP | 0 | 710 | 506 | 0 | 1342 | 0 | 640 | 2507 |
| TYRP1 | 466 | 0 | 0 | 0 | 0 | 0 | 0 | 0 |
| TYSND1 | 1026 | 0 | 1969 | 0 | 0 | 610 | 594 | 444 |
| TYW1 | 0 | 0 | 0 | 1488 | 646 | 0 | 0 | 0 |
| TYW5 | 0 | 0 | 0 | 0 | 534 | 0 | 0 | 0 |
| U2AF1 | 546 | 1366 | 1920 | 5079 | 5093 | 1876 | 2161 | 3234 |
| U2AF1L4 | 0 | 0 | 0 | 1112 | 2092 | 0 | 478 | 0 |
| U2AF1L5 | 0 | 0 | 3891 | 1124 | 758 | 5424 | 0 | 0 |
| U2AF2 | 0 | 934 | 0 | 0 | 3194 | 515 | 0 | 0 |
| U2SURP | 0 | 0 | 0 | 0 | 0 | 0 | 744 | 0 |
| UACA | 30554 | 29474 | 67081 | 34316 | 77212 | 107014 | 59267 | 47416 |
| UAP1 | 660 | 1448 | 4178 | 2067 | 6244 | 574 | 276 | 0 |
| UBA1 | 478 | 0 | 0 | 585 | 1630 | 0 | 1112 | 0 |
| UBA2 | 0 | 0 | 580 | 2291 | 1426 | 1888 | 192 | 0 |
| UBA3 | 21928 | 24770 | 55548 | 60547 | 178353 | 51055 | 100246 | 41556 |
| UBA5 | 0 | 646 | 0 | 0 | 1658 | 0 | 0 | 3115 |
| UBA52 | 5494 | 10472 | 14235 | 18469 | 41040 | 12303 | 8135 | 11943 |
| UBA6-AS1 | 0 | 0 | 0 | 0 | 0 | 0 | 608 | 0 |
| UBAC1 | 4401 | 3160 | 42441 | 5340 | 4398 | 2774 | 5189 | 1882 |
| UBAC2 | 0 | 424 | 0 | 3680 | 998 | 1480 | 0 | 0 |
| UBAC2-AS1 | 0 | 0 | 0 | 0 | 564 | 0 | 0 | 0 |
| UBALD1 | 0 | 0 | 0 | 0 | 838 | 1486 | 0 | 836 |
| UBALD2 | 367 | 422 | 0 | 994 | 2394 | 700 | 0 | 0 |
| UBAP1 | 0 | 0 | 0 | 0 | 1426 | 0 | 0 | 698 |
| UBAP1L | 0 | 0 | 0 | 0 | 0 | 0 | 36 | 0 |
| UBAP2 | 766 | 0 | 0 | 0 | 540 | 0 | 928 | 0 |
| UBAP2L | 0 | 0 | 0 | 0 | 1016 | 0 | 940 | 0 |
| UBB | 9330 | 5647 | 21931 | 10927 | 35353 | 9980 | 6256 | 5626 |
| UBC | 4649 | 5337 | 29756 | 12356 | 24383 | 20209 | 8390 | 10566 |

| | | | | | | | | |
|---|---|---|---|---|---|---|---|---|
| UBD | 4103 | 0 | 0 | 0 | 2504 | 3584 | 1392 | 2068 |
| UBE2A | 1512 | 0 | 0 | 0 | 4817 | 1214 | 840 | 0 |
| UBE2B | 732 | 106 | 494 | 758 | 1592 | 1812 | 722 | 588 |
| UBE2C | 0 | 412 | 0 | 9304 | 17476 | 0 | 2858 | 2528 |
| UBE2D1 | 6766 | 21240 | 27268 | 9318 | 29810 | 10047 | 20103 | 18214 |
| UBE2D2 | 25427 | 36817 | 89654 | 39190 | 68289 | 17874 | 51940 | 13034 |
| UBE2D3 | 2648 | 1944 | 9768 | 6045 | 3152 | 4392 | 4071 | 2786 |
| UBE2D3-AS1 | 0 | 0 | 0 | 0 | 416 | 1365 | 0 | 0 |
| UBE2D4 | 0 | 0 | 0 | 1424 | 460 | 0 | 0 | 0 |
| UBE2E1 | 0 | 550 | 1890 | 0 | 616 | 0 | 268 | 0 |
| UBE2E2 | 0 | 0 | 0 | 0 | 500 | 0 | 0 | 0 |
| UBE2E3 | 0 | 0 | 1502 | 5077 | 3593 | 0 | 460 | 1330 |
| UBE2F | 788 | 390 | 0 | 1046 | 2763 | 0 | 0 | 0 |
| UBE2F-SCLY | 0 | 0 | 0 | 0 | 0 | 0 | 426 | 0 |
| UBE2G1 | 0 | 0 | 0 | 0 | 0 | 0 | 290 | 226 |
| UBE2G2 | 0 | 330 | 0 | 0 | 568 | 0 | 0 | 0 |
| UBE2H | 426 | 0 | 0 | 0 | 0 | 0 | 0 | 592 |
| UBE2I | 560 | 2000 | 2792 | 3990 | 3628 | 1350 | 2567 | 1246 |
| UBE2J1 | 0 | 202 | 1212 | 0 | 0 | 2411 | 14 | 1316 |
| UBE2J2 | 614 | 0 | 2295 | 1789 | 1454 | 0 | 650 | 0 |
| UBE2K | 1220 | 0 | 1716 | 972 | 1192 | 1843 | 618 | 0 |
| UBE2L3 | 3784 | 1918 | 2010 | 2632 | 4720 | 0 | 406 | 1148 |
| UBE2L6 | 858 | 0 | 2346 | 0 | 4698 | 0 | 1103 | 0 |
| UBE2M | 978 | 0 | 0 | 0 | 0 | 0 | 0 | 0 |
| UBE2N | 0 | 1090 | 1642 | 3054 | 5928 | 0 | 428 | 796 |
| UBE2O | 0 | 476 | 0 | 1002 | 274 | 0 | 0 | 0 |
| UBE2Q1 | 410 | 0 | 0 | 0 | 364 | 0 | 0 | 0 |
| UBE2Q2 | 0 | 0 | 1526 | 736 | 226 | 0 | 0 | 0 |
| UBE2R2 | 0 | 318 | 1398 | 3322 | 1661 | 0 | 0 | 1886 |
| UBE2S | 0 | 322 | 806 | 1656 | 6004 | 2102 | 0 | 3719 |
| UBE2T | 584 | 788 | 1508 | 3247 | 4791 | 0 | 0 | 880 |
| UBE2V1 | 294 | 1256 | 3724 | 3043 | 4274 | 3550 | 2942 | 1968 |
| UBE2V2 | 0 | 524 | 1680 | 1104 | 518 | 4891 | 1884 | 928 |
| UBE2W | 0 | 0 | 1218 | 0 | 812 | 3405 | 626 | 0 |
| UBE2Z | 0 | 448 | 0 | 974 | 0 | 0 | 598 | 0 |
| UBE3A | 0 | 0 | 0 | 0 | 90 | 0 | 266 | 0 |
| UBE3B | 654 | 0 | 0 | 0 | 1652 | 0 | 0 | 0 |
| UBE3C | 0 | 454 | 0 | 0 | 2186 | 0 | 0 | 0 |
| UBE3D | 434 | 0 | 0 | 0 | 0 | 0 | 0 | 0 |
| UBE4A | 0 | 0 | 0 | 0 | 1097 | 0 | 0 | 438 |
| UBE4B | 0 | 0 | 0 | 200 | 984 | 0 | 0 | 0 |
| UBFD1 | 0 | 24 | 0 | 0 | 304 | 0 | 0 | 0 |
| UBIAD1 | 0 | 318 | 0 | 268 | 0 | 0 | 426 | 0 |

| | | | | | | | | |
|---|---|---|---|---|---|---|---|---|
| UBL3 | 0 | 340 | 950 | 0 | 0 | 489 | 0 | 292 |
| UBL4A | 0 | 0 | 446 | 0 | 1538 | 0 | 294 | 0 |
| UBL5 | 4461 | 4431 | 3681 | 2949 | 12722 | 1120 | 3216 | 1045 |
| UBL7 | 1014 | 286 | 0 | 0 | 220 | 2382 | 0 | 0 |
| UBL7-AS1 | 0 | 0 | 0 | 0 | 0 | 0 | 546 | 0 |
| UBLCP1 | 0 | 236 | 0 | 0 | 294 | 0 | 202 | 0 |
| UBN2 | 342 | 340 | 0 | 0 | 0 | 0 | 0 | 0 |
| UBOX5 | 0 | 0 | 0 | 654 | 0 | 0 | 0 | 0 |
| UBP1 | 70 | 0 | 0 | 0 | 680 | 0 | 128 | 0 |
| UBQLN1 | 0 | 336 | 0 | 293 | 770 | 3037 | 841 | 0 |
| UBQLN2 | 0 | 200 | 1056 | 0 | 0 | 0 | 326 | 0 |
| UBQLN4 | 0 | 0 | 0 | 0 | 688 | 0 | 0 | 0 |
| UBR1 | 0 | 0 | 450 | 0 | 0 | 2098 | 1344 | 0 |
| UBR2 | 718 | 0 | 588 | 0 | 0 | 0 | 0 | 0 |
| UBR3 | 2446 | 1934 | 3064 | 2151 | 3820 | 0 | 4808 | 450 |
| UBR4 | 0 | 0 | 1452 | 752 | 1730 | 0 | 402 | 2010 |
| UBR5 | 15168 | 16817 | 16269 | 19016 | 129998 | 71553 | 115186 | 26090 |
| UBR7 | 0 | 414 | 1003 | 0 | 540 | 1690 | 0 | 0 |
| UBTD2 | 0 | 0 | 0 | 0 | 0 | 0 | 192 | 0 |
| UBTF | 14170 | 12112 | 52484 | 23315 | 55269 | 20913 | 51335 | 13853 |
| UBXN1 | 1048 | 2776 | 2286 | 7150 | 9137 | 3024 | 9910 | 3274 |
| UBXN10 | 0 | 0 | 0 | 0 | 536 | 0 | 0 | 0 |
| UBXN11 | 0 | 0 | 2772 | 116 | 1421 | 0 | 330 | 0 |
| UBXN2A | 0 | 0 | 0 | 0 | 694 | 0 | 0 | 758 |
| UBXN2B | 0 | 270 | 0 | 0 | 0 | 0 | 0 | 0 |
| UBXN4 | 1732 | 332 | 0 | 0 | 214 | 0 | 584 | 0 |
| UBXN6 | 0 | 0 | 0 | 276 | 502 | 0 | 1440 | 0 |
| UBXN7 | 0 | 0 | 0 | 0 | 236 | 0 | 0 | 0 |
| UCA1 | 0 | 0 | 0 | 1557 | 704 | 0 | 3932 | 1170 |
| UCHL1 | 312 | 0 | 0 | 0 | 0 | 0 | 0 | 1208 |
| UCHL3 | 0 | 1126 | 3759 | 7975 | 3066 | 8396 | 2320 | 0 |
| UCHL5 | 0 | 28 | 1250 | 1900 | 2930 | 0 | 864 | 1086 |
| UCK1 | 0 | 0 | 0 | 0 | 570 | 0 | 0 | 0 |
| UCK2 | 81059 | 76712 | 117058 | 137012 | 381827 | 99512 | 200243 | 151478 |
| UCKL1 | 7684 | 6212 | 25407 | 11262 | 25620 | 9616 | 20123 | 3400 |
| UCP2 | 0 | 257 | 0 | 2140 | 270 | 0 | 246 | 560 |
| UEVLD | 0 | 0 | 0 | 0 | 294 | 0 | 0 | 0 |
| UFC1 | 988 | 2572 | 5524 | 3665 | 4021 | 2974 | 3099 | 5406 |
| UFD1 | 854 | 676 | 222 | 0 | 1262 | 2250 | 1750 | 2861 |
| UFL1 | 0 | 0 | 0 | 0 | 230 | 808 | 0 | 322 |
| UFL1-AS1 | 0 | 0 | 0 | 0 | 254 | 562 | 184 | 0 |
| UFM1 | 20 | 1370 | 1050 | 1044 | 2132 | 1838 | 4382 | 647 |
| UFSP1 | 0 | 0 | 0 | 0 | 0 | 0 | 0 | 464 |

| | | | | | | | | |
|---|---|---|---|---|---|---|---|---|
| UFSP2 | 0 | 0 | 3402 | 0 | 0 | 0 | 518 | 0 |
| UGCG | 0 | 0 | 0 | 0 | 166 | 0 | 0 | 0 |
| UGDH | 340 | 1068 | 952 | 1426 | 406 | 0 | 196 | 0 |
| UGDH-AS1 | 112 | 174 | 0 | 0 | 402 | 0 | 637 | 0 |
| UGGT1 | 238 | 3645 | 3026 | 7585 | 12951 | 4563 | 18705 | 4122 |
| UGGT2 | 688 | 0 | 984 | 0 | 2548 | 3977 | 4333 | 0 |
| UGP2 | 3082 | 1194 | 216 | 392 | 3012 | 0 | 987 | 720 |
| UGT1A10 | 0 | 0 | 0 | 0 | 0 | 0 | 260 | 0 |
| UGT1A6 | 622 | 0 | 1206 | 90 | 0 | 0 | 0 | 0 |
| UGT1A8 | 0 | 358 | 1662 | 0 | 0 | 0 | 0 | 0 |
| UGT2A3 | 216 | 200 | 490 | 0 | 0 | 0 | 164 | 0 |
| UGT2B17 | 2298 | 898 | 11429 | 988 | 0 | 0 | 0 | 0 |
| UGT2B7 | 0 | 420 | 0 | 0 | 0 | 0 | 0 | 0 |
| UGT8 | 0 | 510 | 1626 | 172 | 0 | 0 | 48 | 764 |
| UHRF1 | 0 | 0 | 0 | 1584 | 186 | 0 | 0 | 0 |
| UHRF1BP1 | 0 | 0 | 0 | 0 | 376 | 0 | 0 | 0 |
| UHRF1BP1L | 710 | 1648 | 426 | 0 | 3137 | 0 | 4248 | 346 |
| UIMC1 | 0 | 0 | 0 | 0 | 266 | 0 | 0 | 0 |
| ULBP2 | 0 | 0 | 0 | 0 | 920 | 0 | 0 | 0 |
| ULK1 | 0 | 0 | 0 | 0 | 416 | 0 | 0 | 0 |
| ULK3 | 274 | 285 | 0 | 574 | 288 | 1528 | 0 | 0 |
| ULK4 | 1582 | 386 | 7272 | 0 | 3894 | 0 | 0 | 1014 |
| UMAD1 | 0 | 0 | 2371 | 0 | 1788 | 4969 | 0 | 422 |
| UMPS | 0 | 418 | 0 | 1702 | 1676 | 0 | 860 | 2506 |
| UNC119 | 0 | 0 | 1875 | 0 | 344 | 0 | 824 | 1839 |
| UNC119B | 268 | 338 | 0 | 0 | 580 | 0 | 458 | 0 |
| UNC13B | 362 | 1276 | 8650 | 1770 | 5056 | 4674 | 2208 | 976 |
| UNC45A | 6961 | 5803 | 13574 | 8421 | 22305 | 15234 | 26796 | 5467 |
| UNC50 | 530 | 1164 | 348 | 0 | 6298 | 844 | 2264 | 2063 |
| UNC5A | 0 | 0 | 0 | 500 | 0 | 0 | 0 | 0 |
| UNC5B | 362 | 0 | 0 | 0 | 0 | 0 | 0 | 0 |
| UNC5CL | 0 | 0 | 0 | 0 | 274 | 1358 | 0 | 0 |
| UNC93A | 0 | 2112 | 726 | 10526 | 79365 | 24517 | 4883 | 29816 |
| UNC93B1 | 0 | 218 | 0 | 0 | 738 | 3238 | 0 | 0 |
| UNG | 28545 | 39947 | 128613 | 115287 | 171108 | 115008 | 170332 | 49061 |
| UNK | 9116 | 18114 | 84935 | 12700 | 20399 | 25741 | 19953 | 4943 |
| UPF1 | 808 | 722 | 0 | 3067 | 2044 | 0 | 3588 | 0 |
| UPF2 | 0 | 0 | 1486 | 0 | 434 | 0 | 20 | 0 |
| UPF3A | 190733 | 208002 | 224704 | 333421 | 666595 | 723720 | 562645 | 76570 |
| UPK1A | 0 | 0 | 0 | 0 | 0 | 0 | 940 | 0 |
| UPK3B | 0 | 0 | 0 | 0 | 22 | 0 | 0 | 0 |
| UPK3BL2 | 0 | 0 | 0 | 0 | 0 | 0 | 0 | 712 |
| UPP1 | 650 | 930 | 1708 | 0 | 1772 | 2740 | 0 | 374 |

| | | | | | | | | |
|---|---|---|---|---|---|---|---|---|
| UQCC1 | 12090 | 22729 | 47131 | 57230 | 44847 | 26868 | 134432 | 23225 |
| UQCC2 | 111038 | 184478 | 84656 | 176123 | 908440 | 124481 | 408135 | 188321 |
| UQCC3 | 1172 | 454 | 436 | 4448 | 2728 | 1362 | 2032 | 0 |
| UQCR10 | 7620 | 10049 | 4492 | 7878 | 9026 | 2034 | 3694 | 2518 |
| UQCR11 | 5726 | 9610 | 6409 | 8188 | 18089 | 9536 | 2670 | 3282 |
| UQCRB | 2475540 | 4419209 | 683395 | 867705 | 9621149 | 1527482 | 12345447 | 660795 |
| UQCRC1 | 1556 | 3209 | 7291 | 2536 | 17254 | 4231 | 4766 | 4695 |
| UQCRC2 | 259 | 1148 | 3904 | 3160 | 4148 | 0 | 922 | 1556 |
| UQCRFS1 | 1660 | 8750 | 13015 | 11493 | 9831 | 0 | 1674 | 2486 |
| UQCRH | 5412 | 9334 | 12644 | 16389 | 16046 | 8613 | 12357 | 7660 |
| UQCRQ | 7677 | 14459 | 14164 | 18740 | 19642 | 6130 | 6373 | 3698 |
| URB1-AS1 | 0 | 312 | 0 | 0 | 0 | 0 | 0 | 0 |
| URGCP | 296 | 452 | 0 | 0 | 1882 | 0 | 388 | 0 |
| URGCP-MRPS24 | 462 | 172 | 0 | 606 | 0 | 0 | 0 | 0 |
| URI1 | 28691 | 49906 | 51362 | 50622 | 201840 | 55481 | 88731 | 36883 |
| URM1 | 0 | 1358 | 0 | 3256 | 3948 | 1430 | 3023 | 0 |
| UROD | 0 | 1260 | 2214 | 0 | 568 | 0 | 848 | 1176 |
| UROS | 752 | 302 | 0 | 0 | 2696 | 2509 | 2394 | 0 |
| USE1 | 0 | 1336 | 566 | 2653 | 1042 | 0 | 890 | 906 |
| USF1 | 420 | 0 | 0 | 0 | 0 | 0 | 0 | 0 |
| USF2 | 0 | 344 | 868 | 0 | 1888 | 0 | 0 | 1608 |
| USH1C | 74 | 748 | 0 | 2898 | 592 | 0 | 0 | 278 |
| USHBP1 | 460 | 0 | 0 | 0 | 0 | 0 | 0 | 0 |
| USO1 | 0 | 0 | 1629 | 0 | 388 | 0 | 0 | 0 |
| USP10 | 7782 | 27183 | 24263 | 58326 | 57830 | 23808 | 72957 | 30073 |
| USP11 | 0 | 0 | 1382 | 0 | 1134 | 0 | 0 | 0 |
| USP12 | 0 | 0 | 1880 | 0 | 0 | 0 | 0 | 0 |
| USP13 | 1640 | 668 | 4284 | 3613 | 33634 | 4008 | 16186 | 12556 |
| USP14 | 0 | 0 | 0 | 0 | 462 | 0 | 0 | 0 |
| USP15 | 0 | 0 | 0 | 0 | 0 | 0 | 0 | 630 |
| USP16 | 0 | 0 | 0 | 0 | 0 | 0 | 786 | 0 |
| USP18 | 0 | 0 | 1650 | 0 | 0 | 0 | 0 | 0 |
| USP19 | 0 | 148 | 0 | 0 | 550 | 0 | 0 | 0 |
| USP22 | 2227 | 2906 | 9272 | 2684 | 11085 | 1860 | 5182 | 3936 |
| USP24 | 0 | 0 | 0 | 0 | 218 | 0 | 0 | 474 |
| USP3 | 0 | 0 | 1451 | 0 | 1070 | 0 | 0 | 0 |
| USP3-AS1 | 156 | 0 | 0 | 0 | 0 | 0 | 0 | 0 |
| USP30 | 0 | 0 | 0 | 0 | 0 | 0 | 289 | 780 |
| USP32 | 760 | 406 | 1462 | 0 | 2084 | 0 | 0 | 0 |
| USP33 | 232 | 324 | 0 | 0 | 300 | 0 | 98 | 0 |
| USP34 | 8113 | 3790 | 6010 | 1244 | 14493 | 4698 | 12168 | 2208 |
| USP36 | 0 | 0 | 0 | 0 | 0 | 668 | 0 | 0 |
| USP38 | 0 | 186 | 0 | 0 | 0 | 0 | 0 | 0 |

| | | | | | | | | |
|---|---|---|---|---|---|---|---|---|
| USP39 | 0 | 0 | 0 | 0 | 534 | 0 | 0 | 0 |
| USP4 | 19846 | 27629 | 77963 | 35493 | 122505 | 26750 | 48507 | 24562 |
| USP40 | 2453 | 4840 | 4941 | 6142 | 11380 | 10842 | 30456 | 1986 |
| USP42 | 0 | 272 | 0 | 2905 | 1106 | 0 | 2390 | 0 |
| USP43 | 0 | 0 | 0 | 0 | 0 | 0 | 0 | 568 |
| USP45 | 0 | 0 | 0 | 852 | 0 | 0 | 0 | 0 |
| USP47 | 0 | 616 | 0 | 0 | 0 | 0 | 0 | 0 |
| USP49 | 0 | 0 | 0 | 0 | 780 | 0 | 0 | 0 |
| USP5 | 0 | 0 | 0 | 0 | 196 | 0 | 1001 | 554 |
| USP53 | 696 | 804 | 0 | 3829 | 642 | 0 | 16234 | 0 |
| USP54 | 830 | 178 | 0 | 0 | 0 | 0 | 0 | 0 |
| USP7 | 0 | 0 | 0 | 0 | 1623 | 0 | 268 | 0 |
| USP8 | 398663 | 390674 | 347615 | 294672 | 682866 | 269219 | 1013532 | 141052 |
| USP9X | 2964 | 1292 | 0 | 0 | 4398 | 3273 | 3332 | 332 |
| USP9Y | 0 | 0 | 0 | 0 | 186 | 0 | 0 | 0 |
| USPL1 | 0 | 0 | 0 | 0 | 194 | 0 | 0 | 0 |
| UTP11 | 0 | 442 | 0 | 1414 | 2364 | 2822 | 0 | 0 |
| UTP14A | 17922 | 41282 | 40262 | 85709 | 173337 | 144886 | 217423 | 31658 |
| UTP14C | 0 | 0 | 0 | 0 | 226 | 0 | 0 | 0 |
| UTP15 | 0 | 0 | 0 | 0 | 790 | 0 | 0 | 0 |
| UTP18 | 1326 | 292 | 0 | 3116 | 2139 | 1212 | 3822 | 3028 |
| UTP23 | 0 | 0 | 0 | 0 | 692 | 0 | 0 | 0 |
| UTP3 | 0 | 0 | 0 | 0 | 0 | 0 | 474 | 966 |
| UTP4 | 764 | 0 | 1232 | 0 | 1086 | 0 | 0 | 0 |
| UTP6 | 0 | 368 | 0 | 1022 | 428 | 777 | 458 | 756 |
| UTRN | 6206 | 3956 | 0 | 1242 | 1986 | 3156 | 2194 | 544 |
| UTS2 | 676 | 0 | 0 | 0 | 0 | 0 | 0 | 0 |
| UTS2B | 0 | 0 | 0 | 0 | 518 | 0 | 0 | 0 |
| UTY | 1790 | 486 | 1010 | 910 | 1440 | 0 | 1018 | 0 |
| UVRAG | 0 | 0 | 0 | 0 | 0 | 0 | 762 | 1050 |
| UVSSA | 0 | 0 | 0 | 0 | 0 | 842 | 0 | 0 |
| UXS1 | 390 | 636 | 0 | 0 | 1080 | 1539 | 1000 | 896 |
| UXT | 0 | 2716 | 2155 | 4554 | 10238 | 4235 | 212 | 1674 |
| VAC14 | 399 | 0 | 0 | 0 | 2564 | 0 | 0 | 0 |
| VAMP2 | 400 | 650 | 0 | 0 | 482 | 0 | 0 | 0 |
| VAMP3 | 556 | 366 | 0 | 1267 | 0 | 2166 | 0 | 0 |
| VAMP4 | 0 | 0 | 0 | 0 | 464 | 0 | 0 | 776 |
| VAMP5 | 0 | 474 | 0 | 0 | 0 | 1585 | 0 | 778 |
| VAMP7 | 0 | 0 | 0 | 564 | 262 | 2294 | 0 | 0 |
| VAMP8 | 6908 | 3910 | 19561 | 2426 | 8854 | 2732 | 4909 | 3166 |
| VANGL1 | 5302 | 9214 | 54560 | 40600 | 45067 | 19949 | 56405 | 29486 |
| VAPA | 1750 | 2472 | 12327 | 4174 | 5082 | 0 | 1250 | 4243 |
| VAPB | 370 | 124 | 0 | 494 | 0 | 0 | 0 | 0 |

| | | | | | | | | |
|---|---|---|---|---|---|---|---|---|
| VARS | 644 | 0 | 0 | 0 | 188 | 0 | 393 | 0 |
| VARS2 | 0 | 0 | 0 | 0 | 369 | 0 | 0 | 0 |
| VASP | 1276 | 248 | 0 | 2045 | 1214 | 0 | 411 | 324 |
| VAT1 | 76 | 22 | 0 | 252 | 0 | 0 | 156 | 0 |
| VAV1 | 0 | 328 | 0 | 0 | 0 | 0 | 0 | 0 |
| VAV2 | 1964 | 562 | 0 | 2836 | 5079 | 1773 | 7914 | 1126 |
| VAV3 | 454 | 372 | 0 | 0 | 0 | 0 | 2422 | 0 |
| VBP1 | 0 | 0 | 0 | 958 | 4339 | 0 | 1812 | 834 |
| VCAM1 | 0 | 0 | 0 | 544 | 0 | 0 | 0 | 0 |
| VCAN | 0 | 0 | 0 | 0 | 1328 | 0 | 0 | 0 |
| VCL | 18891 | 19348 | 65571 | 13605 | 73572 | 18050 | 10092 | 20065 |
| VCP | 0 | 274 | 3326 | 1714 | 2125 | 0 | 508 | 0 |
| VCPKMT | 0 | 0 | 0 | 0 | 312 | 0 | 0 | 0 |
| VDAC1 | 532 | 246 | 1520 | 3746 | 6556 | 4953 | 2846 | 914 |
| VDAC2 | 366 | 1126 | 11279 | 7145 | 6126 | 4877 | 4463 | 684 |
| VDAC3 | 214 | 3217 | 0 | 9831 | 4396 | 8293 | 2062 | 1532 |
| VDR | 0 | 0 | 1426 | 0 | 0 | 0 | 222 | 0 |
| VEGFA | 16390 | 3928 | 33744 | 7589 | 88591 | 27877 | 27673 | 22031 |
| VEGFB | 0 | 0 | 1512 | 292 | 0 | 0 | 0 | 0 |
| VENTX | 0 | 0 | 0 | 0 | 0 | 0 | 1016 | 0 |
| VEZF1 | 0 | 330 | 1282 | 0 | 596 | 0 | 0 | 212 |
| VEZT | 0 | 0 | 0 | 0 | 292 | 0 | 0 | 0 |
| VGF | 0 | 270 | 0 | 568 | 1667 | 436 | 2946 | 0 |
| VGLL4 | 0 | 212 | 622 | 417 | 774 | 530 | 566 | 0 |
| VHL | 4074 | 6560 | 13044 | 8136 | 37845 | 5581 | 22803 | 5805 |
| VIL1 | 0 | 0 | 0 | 762 | 1642 | 0 | 1820 | 0 |
| VILL | 0 | 0 | 1361 | 0 | 244 | 0 | 0 | 0 |
| VIM | 387085 | 423242 | 474537 | 344305 | 2046530 | 1058245 | 367766 | 876741 |
| VIP | 22 | 0 | 2287 | 0 | 0 | 0 | 0 | 0 |
| VIPAS39 | 0 | 0 | 0 | 1562 | 306 | 0 | 0 | 0 |
| VIPR1 | 1220 | 0 | 864 | 234 | 0 | 0 | 0 | 0 |
| VIRMA | 59882 | 35869 | 36053 | 34360 | 149901 | 56421 | 280119 | 29771 |
| VKORC1 | 870 | 424 | 0 | 0 | 1226 | 2460 | 1410 | 777 |
| VKORC1L1 | 0 | 14 | 1372 | 226 | 1409 | 650 | 0 | 544 |
| VMA21 | 0 | 96 | 0 | 0 | 1976 | 0 | 0 | 494 |
| VMAC | 468 | 0 | 0 | 0 | 0 | 0 | 0 | 850 |
| VMO1 | 0 | 0 | 1213 | 0 | 802 | 0 | 0 | 0 |
| VMP1 | 2802 | 4802 | 6576 | 11111 | 9834 | 3832 | 7240 | 2888 |
| VNN1 | 0 | 0 | 1572 | 0 | 0 | 0 | 0 | 58 |
| VNN3 | 0 | 0 | 0 | 0 | 0 | 0 | 0 | 1008 |
| VOPP1 | 0 | 168 | 0 | 1639 | 1533 | 0 | 996 | 0 |
| VPREB3 | 0 | 1122 | 0 | 0 | 0 | 0 | 0 | 0 |
| VPS11 | 0 | 0 | 0 | 0 | 424 | 0 | 226 | 0 |

| | | | | | | | | |
|---|---|---|---|---|---|---|---|---|
| VPS13A | 0 | 284 | 0 | 0 | 0 | 0 | 0 | 0 |
| VPS13B | 2781 | 714 | 1156 | 790 | 7202 | 1439 | 7196 | 796 |
| VPS13C | 1000 | 388 | 0 | 0 | 0 | 0 | 0 | 0 |
| VPS13D | 0 | 0 | 1394 | 0 | 258 | 0 | 0 | 0 |
| VPS16 | 582 | 0 | 1488 | 748 | 460 | 0 | 0 | 0 |
| VPS18 | 1634 | 1282 | 5353 | 3833 | 3160 | 0 | 6536 | 844 |
| VPS25 | 394 | 0 | 1152 | 2539 | 3004 | 0 | 2745 | 1066 |
| VPS26A | 816 | 466 | 968 | 2558 | 1418 | 0 | 236 | 1240 |
| VPS26B | 0 | 0 | 0 | 0 | 1360 | 0 | 1220 | 0 |
| VPS26C | 0 | 120 | 0 | 0 | 0 | 0 | 0 | 0 |
| VPS28 | 770 | 2382 | 8660 | 5597 | 5742 | 8341 | 15195 | 5962 |
| VPS29 | 220 | 3466 | 4682 | 3711 | 10271 | 0 | 2924 | 3442 |
| VPS33A | 0 | 0 | 656 | 1078 | 0 | 0 | 0 | 0 |
| VPS33B | 0 | 220 | 0 | 0 | 0 | 0 | 284 | 636 |
| VPS33B-DT | 0 | 0 | 0 | 0 | 0 | 0 | 178 | 0 |
| VPS35 | 474 | 268 | 1686 | 1136 | 0 | 44 | 460 | 0 |
| VPS36 | 0 | 542 | 0 | 0 | 1418 | 586 | 440 | 0 |
| VPS37B | 536 | 0 | 0 | 1120 | 0 | 0 | 0 | 434 |
| VPS37C | 0 | 0 | 0 | 666 | 492 | 0 | 0 | 0 |
| VPS39 | 0 | 48 | 50 | 0 | 0 | 0 | 410 | 0 |
| VPS45 | 0 | 0 | 12 | 0 | 578 | 2338 | 1058 | 234 |
| VPS4A | 2138 | 0 | 0 | 0 | 1020 | 0 | 592 | 786 |
| VPS4B | 0 | 370 | 114 | 942 | 0 | 956 | 0 | 520 |
| VPS50 | 4263 | 5258 | 3439 | 3080 | 10173 | 3770 | 14070 | 966 |
| VPS51 | 742 | 492 | 582 | 0 | 766 | 1736 | 354 | 0 |
| VPS52 | 15018 | 26458 | 43621 | 30129 | 63056 | 22806 | 111109 | 33532 |
| VPS53 | 0 | 522 | 412 | 0 | 1316 | 0 | 1160 | 1227 |
| VPS54 | 214 | 0 | 1382 | 0 | 0 | 0 | 0 | 0 |
| VPS72 | 0 | 0 | 1046 | 1117 | 720 | 4502 | 822 | 0 |
| VPS8 | 2618 | 6476 | 21408 | 2854 | 20447 | 4110 | 3167 | 2162 |
| VPS9D1 | 904 | 0 | 0 | 0 | 0 | 0 | 0 | 0 |
| VRK1 | 18718 | 43104 | 75460 | 116164 | 415906 | 55856 | 103666 | 111804 |
| VRK2 | 0 | 0 | 0 | 370 | 308 | 0 | 194 | 0 |
| VRK3 | 0 | 828 | 0 | 1398 | 2404 | 0 | 412 | 3082 |
| VSIG1 | 0 | 474 | 0 | 2760 | 0 | 0 | 0 | 0 |
| VSIG10 | 0 | 0 | 0 | 0 | 318 | 0 | 0 | 0 |
| VSIG2 | 2696 | 4668 | 9915 | 396 | 1194 | 0 | 887 | 0 |
| VSIG4 | 0 | 0 | 1280 | 0 | 0 | 6441 | 0 | 0 |
| VSIR | 700 | 394 | 194 | 0 | 1462 | 0 | 0 | 0 |
| VSNL1 | 0 | 0 | 0 | 0 | 1410 | 1302 | 0 | 0 |
| VSTM4 | 0 | 0 | 0 | 0 | 172 | 0 | 0 | 0 |
| VTA1 | 496 | 514 | 1244 | 0 | 624 | 0 | 1350 | 1560 |
| VTI1A | 44960 | 55104 | 127933 | 95162 | 190824 | 63579 | 164702 | 82736 |

| | | | | | | | | |
|---|---|---|---|---|---|---|---|---|
| VTI1B | 576 | 430 | 1907 | 0 | 4612 | 0 | 512 | 1239 |
| VWA1 | 0 | 0 | 0 | 0 | 2376 | 0 | 406 | 0 |
| VWA2 | 0 | 0 | 0 | 0 | 0 | 0 | 206 | 0 |
| VWA5A | 55468 | 40019 | 153534 | 29479 | 24757 | 22079 | 58503 | 5901 |
| VWA7 | 0 | 0 | 811 | 0 | 0 | 0 | 0 | 0 |
| VWF | 0 | 0 | 0 | 0 | 1282 | 2184 | 0 | 0 |
| WAC | 8776 | 12962 | 20391 | 15781 | 34705 | 25454 | 40054 | 6194 |
| WAC-AS1 | 0 | 0 | 0 | 0 | 0 | 0 | 550 | 0 |
| WAPL | 0 | 0 | 0 | 0 | 0 | 0 | 0 | 1340 |
| WARS | 0 | 0 | 0 | 2663 | 1808 | 0 | 0 | 0 |
| WARS2 | 0 | 178 | 0 | 1342 | 1110 | 0 | 0 | 0 |
| WAS | 44 | 0 | 0 | 0 | 0 | 262 | 0 | 0 |
| WASF2 | 30810 | 23976 | 444713 | 72979 | 97686 | 102138 | 50834 | 42633 |
| WASH3P | 0 | 0 | 0 | 0 | 162 | 0 | 358 | 0 |
| WASH7P | 392 | 0 | 736 | 890 | 560 | 0 | 0 | 0 |
| WASH8P | 0 | 259 | 0 | 0 | 0 | 0 | 696 | 0 |
| WASHC1 | 604 | 1246 | 1754 | 0 | 0 | 0 | 1080 | 0 |
| WASHC2A | 0 | 0 | 434 | 0 | 1492 | 0 | 0 | 0 |
| WASHC3 | 522 | 120 | 1514 | 2244 | 1142 | 2486 | 710 | 545 |
| WASHC4 | 0 | 0 | 0 | 0 | 118 | 0 | 0 | 0 |
| WASHC5 | 0 | 0 | 0 | 1290 | 574 | 0 | 546 | 1108 |
| WASL | 306 | 326 | 1076 | 0 | 0 | 0 | 0 | 0 |
| WBP1 | 284 | 0 | 3518 | 186 | 1275 | 0 | 1520 | 0 |
| WBP11 | 98 | 56 | 0 | 2076 | 204 | 368 | 0 | 0 |
| WBP1L | 1092 | 1370 | 5667 | 1872 | 4130 | 3514 | 3706 | 2685 |
| WBP2 | 410 | 0 | 122 | 0 | 938 | 2529 | 500 | 648 |
| WBP4 | 0 | 943 | 0 | 0 | 0 | 0 | 0 | 1128 |
| WDCP | 612 | 950 | 0 | 1351 | 5464 | 2219 | 2170 | 2902 |
| WDFY1 | 0 | 102 | 0 | 0 | 1116 | 0 | 0 | 0 |
| WDFY2 | 3974 | 4204 | 3848 | 10583 | 25719 | 17342 | 9743 | 10956 |
| WDFY3 | 0 | 0 | 0 | 0 | 0 | 0 | 94 | 0 |
| WDR1 | 198 | 626 | 5039 | 4165 | 1452 | 1848 | 948 | 1596 |
| WDR11 | 0 | 0 | 0 | 0 | 520 | 0 | 0 | 688 |
| WDR11-AS1 | 0 | 450 | 0 | 0 | 0 | 0 | 0 | 0 |
| WDR12 | 0 | 0 | 0 | 0 | 2802 | 0 | 1052 | 0 |
| WDR13 | 538 | 354 | 0 | 0 | 643 | 344 | 376 | 0 |
| WDR18 | 0 | 0 | 1645 | 2124 | 3000 | 0 | 470 | 0 |
| WDR20 | 450 | 0 | 0 | 0 | 156 | 0 | 0 | 0 |
| WDR24 | 810 | 1270 | 1818 | 1126 | 2008 | 2617 | 1612 | 440 |
| WDR25 | 0 | 0 | 0 | 0 | 368 | 0 | 0 | 0 |
| WDR26 | 0 | 0 | 0 | 0 | 850 | 0 | 0 | 799 |
| WDR27 | 0 | 0 | 0 | 0 | 502 | 0 | 0 | 0 |
| WDR3 | 0 | 0 | 0 | 0 | 504 | 0 | 0 | 0 |

| | | | | | | | | |
|---|---|---|---|---|---|---|---|---|
| WDR31 | 0 | 0 | 0 | 0 | 40 | 0 | 0 | 0 |
| WDR33 | 220 | 306 | 3064 | 1864 | 4366 | 2462 | 48 | 0 |
| WDR34 | 0 | 826 | 0 | 323 | 74 | 0 | 0 | 0 |
| WDR35 | 0 | 0 | 0 | 0 | 526 | 0 | 244 | 0 |
| WDR36 | 0 | 0 | 852 | 968 | 0 | 0 | 0 | 0 |
| WDR4 | 0 | 326 | 0 | 0 | 280 | 0 | 378 | 1206 |
| WDR41 | 0 | 440 | 0 | 0 | 192 | 0 | 0 | 0 |
| WDR43 | 0 | 0 | 26 | 0 | 870 | 0 | 0 | 302 |
| WDR45 | 320 | 766 | 3402 | 0 | 724 | 0 | 374 | 0 |
| WDR45B | 646 | 1040 | 0 | 770 | 1518 | 2760 | 332 | 0 |
| WDR46 | 0 | 0 | 0 | 0 | 0 | 1923 | 0 | 392 |
| WDR47 | 392 | 0 | 0 | 0 | 0 | 0 | 0 | 0 |
| WDR5 | 0 | 236 | 0 | 0 | 752 | 0 | 0 | 584 |
| WDR53 | 0 | 0 | 0 | 0 | 0 | 0 | 0 | 1432 |
| WDR54 | 0 | 0 | 0 | 0 | 2148 | 0 | 0 | 1010 |
| WDR55 | 0 | 446 | 0 | 0 | 898 | 0 | 0 | 0 |
| WDR59 | 0 | 0 | 2262 | 1554 | 1876 | 2110 | 2882 | 1076 |
| WDR5B | 0 | 240 | 0 | 0 | 0 | 0 | 668 | 0 |
| WDR6 | 374 | 12 | 0 | 0 | 2748 | 0 | 264 | 0 |
| WDR61 | 0 | 0 | 1560 | 4669 | 702 | 4496 | 480 | 1268 |
| WDR62 | 0 | 0 | 0 | 0 | 847 | 0 | 0 | 0 |
| WDR70 | 9802 | 24500 | 22948 | 18838 | 167589 | 26830 | 40753 | 41493 |
| WDR73 | 22 | 0 | 3393 | 0 | 412 | 0 | 0 | 0 |
| WDR74 | 1006 | 400 | 2892 | 1362 | 3464 | 828 | 3050 | 506 |
| WDR75 | 0 | 156 | 0 | 0 | 476 | 0 | 0 | 0 |
| WDR77 | 0 | 0 | 0 | 2218 | 2082 | 1634 | 1466 | 0 |
| WDR78 | 0 | 0 | 0 | 0 | 0 | 2089 | 0 | 0 |
| WDR82 | 1104 | 350 | 182 | 0 | 324 | 0 | 80 | 0 |
| WDR83OS | 2346 | 2241 | 10485 | 36 | 4433 | 8434 | 4311 | 4898 |
| WDR90 | 0 | 0 | 0 | 0 | 258 | 0 | 0 | 0 |
| WDR91 | 0 | 0 | 0 | 0 | 446 | 0 | 0 | 0 |
| WDR92 | 411 | 0 | 0 | 0 | 302 | 0 | 0 | 0 |
| WDYHV1 | 6487 | 0 | 8653 | 34933 | 5165 | 92612 | 88211 | 2693 |
| WEE1 | 0 | 0 | 0 | 0 | 0 | 0 | 0 | 264 |
| WFDC2 | 862 | 8175 | 1498 | 830 | 1048 | 0 | 0 | 0 |
| WFDC3 | 0 | 0 | 0 | 458 | 1068 | 422 | 0 | 0 |
| WFS1 | 0 | 0 | 116 | 640 | 0 | 762 | 2444 | 124 |
| WHAMM | 0 | 0 | 558 | 0 | 430 | 0 | 0 | 0 |
| WHRN | 0 | 0 | 0 | 0 | 242 | 0 | 614 | 390 |
| WI2-87327B8.2 | 0 | 0 | 0 | 0 | 0 | 2042 | 0 | 0 |
| WIF1 | 0 | 0 | 0 | 1010 | 0 | 106780 | 81926 | 0 |
| WIPF2 | 0 | 554 | 0 | 0 | 0 | 0 | 0 | 0 |
| WIPI1 | 0 | 838 | 1958 | 0 | 0 | 2294 | 0 | 0 |

| | | | | | | | | |
|---|---|---|---|---|---|---|---|---|
| WIPI2 | 0 | 0 | 680 | 0 | 352 | 1858 | 404 | 0 |
| WLS | 35196 | 30915 | 50280 | 38164 | 94136 | 100336 | 117394 | 21576 |
| WNK1 | 0 | 488 | 966 | 0 | 898 | 0 | 0 | 0 |
| WNK2 | 1026 | 518 | 209 | 640 | 1460 | 0 | 460 | 0 |
| WNK4 | 0 | 0 | 1048 | 0 | 0 | 0 | 530 | 0 |
| WNT11 | 0 | 0 | 0 | 0 | 1280 | 0 | 1176 | 635 |
| WNT2 | 0 | 0 | 892 | 4034 | 37462 | 53781 | 0 | 0 |
| WRAP53 | 4088 | 3388 | 5988 | 8169 | 15604 | 2688 | 6255 | 9954 |
| WRAP73 | 488 | 136 | 598 | 824 | 0 | 0 | 0 | 0 |
| WRB | 28456 | 43325 | 86780 | 84963 | 152858 | 183882 | 154408 | 88983 |
| WRB-SH3BGR | 0 | 0 | 0 | 0 | 7083 | 3368 | 0 | 3495 |
| WRN | 14260 | 12846 | 21761 | 6950 | 31343 | 6660 | 10766 | 6795 |
| WRNIP1 | 0 | 26 | 778 | 0 | 496 | 64 | 746 | 1176 |
| WSB1 | 0 | 346 | 0 | 0 | 108 | 0 | 0 | 0 |
| WSB2 | 514 | 0 | 0 | 0 | 420 | 1010 | 178 | 0 |
| WSCD1 | 2376 | 426 | 2618 | 0 | 0 | 0 | 544 | 0 |
| WT1 | 0 | 0 | 0 | 0 | 0 | 3934 | 0 | 0 |
| WTAP | 1162 | 754 | 860 | 2452 | 2796 | 0 | 0 | 718 |
| WWC1 | 0 | 0 | 0 | 0 | 0 | 0 | 0 | 798 |
| WWC3 | 0 | 702 | 0 | 782 | 504 | 0 | 0 | 0 |
| WWP1 | 0 | 0 | 0 | 621 | 0 | 0 | 506 | 0 |
| WWP2 | 5320 | 11388 | 21049 | 12674 | 26729 | 3888 | 26933 | 8528 |
| WWTR1 | 1012 | 1208 | 1642 | 1116 | 6647 | 3389 | 4854 | 3809 |
| XAB2 | 0 | 0 | 0 | 0 | 1470 | 0 | 0 | 2382 |
| XBP1 | 278 | 3210 | 5864 | 1138 | 2636 | 1684 | 3803 | 1959 |
| XCL1 | 0 | 0 | 0 | 0 | 0 | 2089 | 0 | 0 |
| XDH | 0 | 0 | 1516 | 0 | 0 | 0 | 0 | 0 |
| XIAP | 0 | 268 | 0 | 0 | 2563 | 0 | 502 | 246 |
| XK | 0 | 464 | 0 | 0 | 0 | 0 | 160 | 0 |
| XKR8 | 0 | 0 | 0 | 0 | 0 | 0 | 1423 | 0 |
| XKR9 | 2564 | 1916 | 80 | 3478 | 47260 | 0 | 34737 | 26297 |
| XPA | 25300 | 21548 | 16828 | 21265 | 102829 | 50711 | 54470 | 17620 |
| XPC | 17010 | 28800 | 27391 | 24789 | 63844 | 37864 | 61749 | 9369 |
| XPNPEP1 | 0 | 1550 | 1502 | 1840 | 754 | 0 | 0 | 942 |
| XPNPEP3 | 0 | 0 | 0 | 1180 | 0 | 0 | 0 | 0 |
| XPO1 | 6794 | 8785 | 13493 | 30864 | 53488 | 18893 | 60291 | 16856 |
| XPO6 | 0 | 0 | 0 | 0 | 0 | 0 | 116 | 0 |
| XPO7 | 0 | 178 | 0 | 0 | 862 | 0 | 0 | 0 |
| XPOT | 0 | 334 | 0 | 1615 | 212 | 0 | 286 | 0 |
| XRCC1 | 0 | 142 | 0 | 1046 | 264 | 0 | 0 | 0 |
| XRCC3 | 0 | 164 | 0 | 0 | 0 | 0 | 0 | 0 |
| XRCC4 | 0 | 0 | 0 | 0 | 0 | 0 | 274 | 0 |
| XRCC5 | 0 | 1192 | 2284 | 2087 | 3584 | 0 | 528 | 1715 |

| | | | | | | | | |
|---|---|---|---|---|---|---|---|---|
| XRCC6 | 1560 | 578 | 4124 | 1007 | 6372 | 3274 | 1078 | 0 |
| XRN1 | 0 | 0 | 0 | 1174 | 284 | 0 | 0 | 0 |
| XRN2 | 82 | 0 | 0 | 0 | 792 | 338 | 504 | 0 |
| XRRA1 | 232 | 74 | 1720 | 1268 | 1694 | 1024 | 612 | 0 |
| XXYLT1 | 196 | 576 | 0 | 0 | 932 | 0 | 0 | 0 |
| XYLB | 0 | 194 | 0 | 0 | 284 | 0 | 706 | 0 |
| XYLT2 | 602 | 0 | 1428 | 0 | 657 | 0 | 1054 | 646 |
| YAE1 | 0 | 602 | 0 | 0 | 1342 | 0 | 2198 | 2639 |
| YAF2 | 0 | 0 | 0 | 0 | 0 | 1176 | 0 | 1195 |
| YAP1 | 5278 | 6870 | 7375 | 25995 | 36682 | 19650 | 57159 | 29514 |
| YARS | 0 | 0 | 0 | 2010 | 566 | 0 | 0 | 0 |
| YARS2 | 668 | 0 | 0 | 1218 | 2203 | 0 | 0 | 0 |
| YBEY | 1198 | 0 | 1999 | 0 | 218 | 0 | 0 | 0 |
| YBX1 | 576 | 2420 | 6350 | 11074 | 12933 | 9714 | 1810 | 3670 |
| YBX2 | 0 | 0 | 0 | 0 | 350 | 0 | 0 | 0 |
| YBX3 | 354 | 186 | 92 | 2443 | 1730 | 0 | 2772 | 186 |
| YDJC | 0 | 0 | 0 | 0 | 5695 | 0 | 840 | 2080 |
| YEATS2 | 0 | 826 | 0 | 0 | 3343 | 0 | 1662 | 286 |
| YEATS4 | 23379 | 88794 | 38689 | 108185 | 333226 | 109403 | 239672 | 87082 |
| YES1 | 552 | 528 | 0 | 0 | 732 | 0 | 0 | 0 |
| YIF1A | 702 | 0 | 2608 | 1412 | 5314 | 0 | 4627 | 4026 |
| YIF1B | 8918 | 8963 | 60796 | 34551 | 39841 | 32104 | 15828 | 29812 |
| YIPF1 | 0 | 0 | 0 | 1696 | 1341 | 0 | 468 | 0 |
| YIPF2 | 1712 | 1086 | 0 | 18 | 1916 | 0 | 1830 | 816 |
| YIPF3 | 1091 | 2114 | 5610 | 3825 | 2197 | 2061 | 869 | 2756 |
| YIPF4 | 15166 | 23474 | 30696 | 37225 | 84067 | 29940 | 80261 | 37495 |
| YIPF5 | 0 | 0 | 993 | 0 | 102 | 1133 | 912 | 2039 |
| YIPF6 | 0 | 0 | 4561 | 0 | 2070 | 540 | 408 | 0 |
| YJEFN3 | 0 | 0 | 1050 | 0 | 0 | 0 | 0 | 0 |
| YJU2 | 0 | 0 | 960 | 0 | 2334 | 0 | 0 | 0 |
| YKT6 | 0 | 612 | 0 | 0 | 1486 | 1108 | 1062 | 648 |
| YME1L1 | 290 | 304 | 0 | 1288 | 0 | 0 | 52 | 642 |
| YPEL2 | 0 | 0 | 3217 | 0 | 130 | 1522 | 1442 | 0 |
| YPEL3 | 0 | 0 | 4955 | 0 | 1604 | 3866 | 1562 | 1082 |
| YPEL5 | 111236 | 131093 | 243415 | 80202 | 294879 | 178735 | 327582 | 134227 |
| YRDC | 0 | 0 | 0 | 904 | 268 | 0 | 0 | 0 |
| YTHDC1 | 0 | 392 | 0 | 0 | 440 | 0 | 0 | 0 |
| YTHDC2 | 0 | 0 | 0 | 0 | 588 | 0 | 0 | 0 |
| YTHDF1 | 0 | 0 | 0 | 0 | 1040 | 0 | 208 | 0 |
| YTHDF2 | 0 | 0 | 0 | 0 | 184 | 0 | 0 | 0 |
| YTHDF3 | 0 | 156 | 0 | 94 | 414 | 0 | 0 | 0 |
| YTHDF3-AS1 | 0 | 398 | 0 | 0 | 1052 | 0 | 1260 | 626 |
| YWHAB | 582 | 3033 | 2063 | 8201 | 2262 | 672 | 5380 | 3320 |

| | | | | | | | | |
|---|---|---|---|---|---|---|---|---|
| YWHAE | 1129385 | 2422227 | 8261723 | 2492335 | 9184105 | 3132271 | 4665264 | 2728109 |
| YWHAEP1 | 0 | 0 | 0 | 0 | 3686 | 0 | 0 | 1994 |
| YWHAG | 310 | 1126 | 0 | 396 | 0 | 0 | 0 | 0 |
| YWHAH | 898 | 1038 | 6300 | 3142 | 3082 | 0 | 0 | 0 |
| YWHAQ | 536 | 1475 | 10375 | 1852 | 6180 | 1998 | 2672 | 902 |
| YWHAZ | 956 | 4236 | 9781 | 13037 | 23636 | 7615 | 9650 | 7133 |
| YY1 | 890 | 822 | 0 | 268 | 2298 | 0 | 805 | 0 |
| YY1AP1 | 0 | 0 | 0 | 0 | 726 | 836 | 0 | 0 |
| ZADH2 | 110 | 0 | 0 | 0 | 386 | 726 | 296 | 0 |
| ZAR1 | 2994 | 0 | 1236 | 0 | 0 | 0 | 90 | 0 |
| ZBED1 | 0 | 0 | 0 | 950 | 1956 | 0 | 0 | 0 |
| ZBED5 | 228 | 0 | 1208 | 0 | 0 | 394 | 582 | 0 |
| ZBED5-AS1 | 0 | 394 | 3201 | 0 | 166 | 0 | 468 | 0 |
| ZBED8 | 0 | 532 | 0 | 0 | 0 | 0 | 0 | 0 |
| ZBP1 | 0 | 442 | 0 | 0 | 516 | 0 | 0 | 0 |
| ZBTB11 | 0 | 0 | 0 | 1140 | 0 | 0 | 0 | 0 |
| ZBTB16 | 1478 | 9610 | 2494 | 0 | 1574 | 0 | 0 | 3280 |
| ZBTB2 | 0 | 144 | 0 | 0 | 0 | 0 | 0 | 0 |
| ZBTB25 | 0 | 0 | 0 | 0 | 0 | 0 | 486 | 0 |
| ZBTB38 | 28432 | 46328 | 16798 | 47010 | 61101 | 11510 | 133196 | 13946 |
| ZBTB4 | 0 | 0 | 0 | 0 | 454 | 0 | 0 | 128 |
| ZBTB43 | 0 | 270 | 0 | 0 | 142 | 0 | 0 | 864 |
| ZBTB44 | 0 | 0 | 0 | 0 | 0 | 0 | 208 | 0 |
| ZBTB47 | 0 | 0 | 0 | 0 | 0 | 0 | 780 | 0 |
| ZBTB48 | 0 | 0 | 0 | 704 | 872 | 0 | 0 | 0 |
| ZBTB6 | 0 | 384 | 0 | 0 | 0 | 0 | 0 | 0 |
| ZBTB7A | 0 | 0 | 0 | 0 | 837 | 0 | 510 | 0 |
| ZBTB7B | 0 | 0 | 0 | 0 | 880 | 0 | 0 | 0 |
| ZBTB7C | 0 | 0 | 270 | 0 | 0 | 0 | 0 | 0 |
| ZBTB8OS | 1336 | 2094 | 756 | 1302 | 3132 | 1330 | 0 | 0 |
| ZBTB9 | 0 | 0 | 0 | 0 | 1256 | 0 | 0 | 0 |
| ZC2HC1A | 10608 | 10850 | 17746 | 12026 | 28748 | 9926 | 47122 | 7996 |
| ZC3H11A | 610 | 200 | 0 | 302 | 1384 | 995 | 140 | 1472 |
| ZC3H12A | 0 | 0 | 2018 | 222 | 0 | 418 | 42 | 0 |
| ZC3H12D | 0 | 0 | 406 | 144 | 0 | 0 | 0 | 0 |
| ZC3H13 | 25121 | 12522 | 10649 | 20158 | 72511 | 49176 | 52181 | 9224 |
| ZC3H14 | 0 | 258 | 0 | 0 | 1804 | 0 | 0 | 0 |
| ZC3H15 | 99679 | 136591 | 91274 | 132013 | 451748 | 101095 | 331675 | 132626 |
| ZC3H18 | 2862 | 4932 | 24259 | 9719 | 13523 | 7374 | 20581 | 3345 |
| ZC3H4 | 0 | 340 | 0 | 0 | 136 | 0 | 0 | 0 |
| ZC3H7A | 0 | 0 | 0 | 280 | 0 | 0 | 0 | 0 |
| ZC3H7B | 0 | 0 | 0 | 0 | 570 | 0 | 0 | 0 |
| ZC3H8 | 0 | 720 | 0 | 1440 | 2862 | 0 | 0 | 908 |

| | | | | | | | | |
|---|---|---|---|---|---|---|---|---|
| ZC3HAV1 | 3458 | 3118 | 6027 | 2170 | 7215 | 4034 | 9133 | 2635 |
| ZC3HAV1L | 0 | 0 | 0 | 1484 | 0 | 0 | 0 | 0 |
| ZC3HC1 | 722 | 590 | 0 | 0 | 0 | 0 | 1122 | 0 |
| ZCCHC10 | 742 | 486 | 1144 | 3298 | 1156 | 0 | 150 | 0 |
| ZCCHC11 | 0 | 0 | 0 | 0 | 568 | 0 | 260 | 0 |
| ZCCHC14 | 312 | 0 | 0 | 150 | 0 | 0 | 0 | 0 |
| ZCCHC17 | 0 | 456 | 0 | 0 | 0 | 1028 | 328 | 0 |
| ZCCHC18 | 0 | 0 | 0 | 398 | 0 | 0 | 0 | 0 |
| ZCCHC24 | 0 | 0 | 0 | 0 | 372 | 2952 | 350 | 0 |
| ZCCHC3 | 0 | 0 | 328 | 0 | 0 | 0 | 0 | 0 |
| ZCCHC6 | 0 | 0 | 0 | 0 | 934 | 0 | 0 | 0 |
| ZCCHC7 | 1220 | 440 | 0 | 0 | 0 | 0 | 0 | 392 |
| ZCCHC8 | 3004 | 3096 | 1330 | 3653 | 4207 | 0 | 8839 | 1898 |
| ZCCHC9 | 72 | 0 | 0 | 1196 | 12 | 0 | 0 | 556 |
| ZCRB1 | 0 | 0 | 0 | 1907 | 1946 | 0 | 752 | 446 |
| ZDHHC1 | 0 | 400 | 0 | 0 | 516 | 0 | 0 | 0 |
| ZDHHC11B | 0 | 0 | 0 | 0 | 0 | 818 | 0 | 1852 |
| ZDHHC12 | 570 | 396 | 0 | 688 | 1166 | 2276 | 0 | 0 |
| ZDHHC14 | 144 | 850 | 2400 | 0 | 3046 | 0 | 602 | 0 |
| ZDHHC16 | 0 | 0 | 0 | 0 | 500 | 0 | 0 | 0 |
| ZDHHC19 | 0 | 656 | 0 | 0 | 0 | 0 | 0 | 0 |
| ZDHHC2 | 222 | 540 | 0 | 0 | 600 | 0 | 461 | 0 |
| ZDHHC20 | 6158 | 5960 | 12091 | 9113 | 32442 | 29806 | 19314 | 16434 |
| ZDHHC21 | 353 | 0 | 0 | 0 | 0 | 0 | 0 | 0 |
| ZDHHC23 | 0 | 38 | 0 | 0 | 0 | 0 | 0 | 0 |
| ZDHHC24 | 0 | 0 | 0 | 0 | 630 | 0 | 0 | 0 |
| ZDHHC3 | 0 | 606 | 0 | 1026 | 4125 | 0 | 0 | 0 |
| ZDHHC4 | 0 | 1712 | 1088 | 0 | 128 | 0 | 2060 | 0 |
| ZDHHC5 | 0 | 988 | 0 | 48 | 0 | 0 | 0 | 0 |
| ZDHHC6 | 29188 | 66158 | 75352 | 96742 | 184181 | 78910 | 215091 | 50999 |
| ZDHHC7 | 752 | 466 | 617 | 0 | 398 | 0 | 404 | 0 |
| ZDHHC8 | 0 | 334 | 0 | 0 | 0 | 0 | 0 | 0 |
| ZDHHC9 | 678 | 0 | 0 | 0 | 0 | 0 | 645 | 0 |
| ZEB1 | 17027 | 13112 | 48532 | 4938 | 70910 | 25911 | 3984 | 4571 |
| ZEB2 | 226 | 0 | 0 | 0 | 0 | 0 | 0 | 0 |
| ZER1 | 346 | 0 | 0 | 0 | 1135 | 0 | 0 | 0 |
| ZFAND1 | 0 | 328 | 0 | 4881 | 2862 | 2030 | 1971 | 0 |
| ZFAND2A | 2085 | 484 | 4762 | 880 | 2308 | 0 | 0 | 4183 |
| ZFAND2B | 334 | 310 | 0 | 3310 | 1306 | 0 | 434 | 1266 |
| ZFAND3 | 0 | 0 | 0 | 0 | 0 | 0 | 65 | 0 |
| ZFAND4 | 0 | 0 | 0 | 0 | 0 | 0 | 282 | 0 |
| ZFAND5 | 284 | 270 | 0 | 2745 | 4765 | 1628 | 2296 | 0 |
| ZFAND6 | 358 | 1100 | 1286 | 0 | 730 | 0 | 684 | 732 |

| | | | | | | | | |
|---|---|---|---|---|---|---|---|---|
| ZFAS1 | 800 | 2932 | 2060 | 10885 | 13018 | 8312 | 13964 | 6401 |
| ZFC3H1 | 0 | 0 | 0 | 0 | 468 | 0 | 0 | 0 |
| ZFHX3 | 294 | 624 | 791 | 0 | 2512 | 152 | 2046 | 0 |
| ZFP36 | 3082 | 910 | 7826 | 1658 | 3423 | 711 | 358 | 1082 |
| ZFP36L1 | 0 | 0 | 0 | 0 | 2076 | 0 | 0 | 0 |
| ZFP36L2 | 1788 | 1258 | 4401 | 0 | 1370 | 0 | 450 | 250 |
| ZFP62 | 0 | 570 | 0 | 0 | 0 | 0 | 290 | 0 |
| ZFP64 | 0 | 0 | 0 | 0 | 2374 | 0 | 370 | 0 |
| ZFP69B | 0 | 0 | 0 | 120 | 0 | 0 | 0 | 0 |
| ZFP82 | 386 | 0 | 0 | 0 | 0 | 0 | 0 | 0 |
| ZFP90 | 0 | 0 | 0 | 0 | 0 | 0 | 966 | 0 |
| ZFP91 | 0 | 0 | 1416 | 0 | 0 | 0 | 600 | 0 |
| ZFP91-CNTF | 0 | 0 | 0 | 0 | 622 | 0 | 0 | 0 |
| ZFPL1 | 28 | 0 | 1420 | 0 | 2260 | 0 | 0 | 0 |
| ZFPM1 | 0 | 0 | 0 | 286 | 162 | 0 | 0 | 0 |
| ZFR | 12598 | 12856 | 339421 | 16135 | 79635 | 25479 | 48813 | 15655 |
| ZFX | 0 | 0 | 0 | 0 | 0 | 0 | 36 | 0 |
| ZFYVE1 | 526 | 0 | 0 | 0 | 0 | 0 | 0 | 0 |
| ZFYVE16 | 397 | 0 | 0 | 1014 | 614 | 0 | 490 | 0 |
| ZFYVE19 | 0 | 0 | 0 | 0 | 0 | 0 | 1002 | 0 |
| ZFYVE21 | 690 | 972 | 3412 | 0 | 1058 | 2258 | 0 | 896 |
| ZFYVE27 | 76 | 154 | 714 | 1160 | 2066 | 0 | 0 | 864 |
| ZFYVE28 | 936 | 0 | 0 | 0 | 0 | 0 | 70 | 0 |
| ZG16 | 30664 | 32181 | 168520 | 4530 | 3678 | 0 | 0 | 0 |
| ZG16B | 0 | 890 | 899 | 2582 | 4268 | 0 | 0 | 1062 |
| ZGLP1 | 0 | 0 | 0 | 1206 | 414 | 0 | 0 | 0 |
| ZGPAT | 0 | 0 | 4990 | 1787 | 158 | 0 | 380 | 0 |
| ZHX1 | 0 | 82 | 0 | 0 | 0 | 0 | 0 | 0 |
| ZHX2 | 0 | 0 | 0 | 0 | 400 | 0 | 0 | 0 |
| ZKSCAN1 | 0 | 0 | 0 | 1046 | 0 | 0 | 184 | 0 |
| ZKSCAN3 | 0 | 0 | 0 | 0 | 392 | 0 | 0 | 0 |
| ZKSCAN5 | 0 | 0 | 0 | 982 | 0 | 0 | 0 | 0 |
| ZKSCAN8 | 264 | 0 | 0 | 0 | 0 | 0 | 0 | 0 |
| ZMAT2 | 1648 | 1180 | 3602 | 0 | 5277 | 1893 | 928 | 802 |
| ZMAT3 | 2976 | 708 | 6689 | 2054 | 2538 | 2842 | 1748 | 2008 |
| ZMAT4 | 0 | 0 | 1751 | 0 | 0 | 0 | 0 | 1374 |
| ZMAT5 | 0 | 0 | 0 | 0 | 0 | 0 | 66 | 1213 |
| ZMIZ1 | 0 | 0 | 0 | 1006 | 733 | 0 | 0 | 0 |
| ZMIZ2 | 1276 | 219 | 0 | 0 | 1532 | 0 | 0 | 0 |
| ZMPSTE24 | 25208 | 82627 | 71414 | 57124 | 174580 | 57815 | 110961 | 51453 |
| ZMYM2 | 0 | 110 | 0 | 0 | 0 | 0 | 186 | 1358 |
| ZMYM3 | 0 | 178 | 0 | 0 | 238 | 0 | 0 | 0 |
| ZMYM4 | 20 | 270 | 1824 | 0 | 5606 | 2462 | 886 | 0 |

| | | | | | | | | |
|---|---|---|---|---|---|---|---|---|
| ZMYM5 | 94 | 0 | 0 | 0 | 62 | 0 | 290 | 0 |
| ZMYND11 | 10464 | 24260 | 15306 | 16641 | 38915 | 16385 | 53838 | 15953 |
| ZMYND19 | 0 | 242 | 0 | 1124 | 1052 | 0 | 2272 | 0 |
| ZMYND8 | 13110 | 12192 | 8398 | 18232 | 46645 | 26381 | 51805 | 2572 |
| ZNF100 | 0 | 0 | 0 | 0 | 142 | 0 | 0 | 0 |
| ZNF101 | 0 | 468 | 0 | 0 | 0 | 0 | 0 | 0 |
| ZNF107 | 0 | 0 | 0 | 353 | 0 | 0 | 560 | 0 |
| ZNF117 | 0 | 0 | 0 | 0 | 0 | 0 | 370 | 0 |
| ZNF12 | 0 | 516 | 0 | 0 | 520 | 0 | 0 | 0 |
| ZNF121 | 0 | 522 | 0 | 0 | 0 | 0 | 0 | 0 |
| ZNF124 | 0 | 0 | 0 | 460 | 74 | 0 | 0 | 0 |
| ZNF131 | 0 | 164 | 0 | 0 | 0 | 0 | 0 | 0 |
| ZNF133 | 0 | 0 | 0 | 0 | 1554 | 0 | 0 | 0 |
| ZNF134 | 778 | 0 | 0 | 0 | 0 | 0 | 0 | 0 |
| ZNF136 | 0 | 302 | 1298 | 0 | 0 | 0 | 0 | 0 |
| ZNF138 | 0 | 370 | 774 | 0 | 344 | 0 | 0 | 0 |
| ZNF140 | 0 | 0 | 652 | 0 | 1415 | 0 | 1362 | 766 |
| ZNF141 | 0 | 0 | 1958 | 0 | 0 | 0 | 0 | 0 |
| ZNF143 | 0 | 0 | 0 | 0 | 0 | 0 | 726 | 0 |
| ZNF146 | 0 | 0 | 0 | 462 | 1246 | 0 | 180 | 0 |
| ZNF148 | 0 | 0 | 0 | 0 | 0 | 0 | 0 | 708 |
| ZNF154 | 0 | 0 | 0 | 0 | 126 | 0 | 0 | 0 |
| ZNF16 | 0 | 0 | 0 | 0 | 0 | 0 | 232 | 0 |
| ZNF165 | 0 | 0 | 0 | 0 | 766 | 0 | 0 | 0 |
| ZNF169 | 0 | 483 | 0 | 0 | 0 | 0 | 0 | 0 |
| ZNF182 | 0 | 0 | 0 | 0 | 0 | 0 | 394 | 0 |
| ZNF184 | 0 | 0 | 0 | 0 | 256 | 0 | 0 | 0 |
| ZNF185 | 0 | 0 | 0 | 0 | 0 | 0 | 410 | 0 |
| ZNF189 | 3700 | 2378 | 6076 | 2650 | 10186 | 4521 | 9213 | 1136 |
| ZNF195 | 0 | 242 | 10 | 0 | 94 | 0 | 0 | 0 |
| ZNF197 | 0 | 0 | 0 | 0 | 0 | 0 | 0 | 1009 |
| ZNF2 | 0 | 0 | 0 | 0 | 562 | 0 | 0 | 0 |
| ZNF20 | 0 | 0 | 762 | 0 | 698 | 0 | 0 | 0 |
| ZNF200 | 0 | 0 | 0 | 0 | 342 | 0 | 0 | 0 |
| ZNF202 | 0 | 0 | 0 | 1037 | 0 | 0 | 0 | 0 |
| ZNF205 | 0 | 0 | 554 | 0 | 0 | 0 | 0 | 0 |
| ZNF207 | 0 | 270 | 2472 | 402 | 3125 | 0 | 235 | 0 |
| ZNF213 | 0 | 0 | 0 | 0 | 0 | 0 | 184 | 0 |
| ZNF214 | 742 | 426 | 0 | 1082 | 6146 | 0 | 2729 | 1472 |
| ZNF217 | 0 | 0 | 0 | 885 | 346 | 0 | 0 | 0 |
| ZNF219 | 0 | 0 | 924 | 1364 | 0 | 0 | 0 | 0 |
| ZNF22 | 0 | 0 | 0 | 1080 | 434 | 0 | 312 | 0 |
| ZNF221 | 0 | 0 | 0 | 0 | 98 | 0 | 0 | 0 |

| | | | | | | | | |
|---|---|---|---|---|---|---|---|---|
| ZNF222 | 0 | 0 | 0 | 0 | 1744 | 0 | 0 | 0 |
| ZNF224 | 0 | 0 | 1345 | 0 | 0 | 414 | 0 | 0 |
| ZNF226 | 0 | 0 | 1866 | 0 | 0 | 0 | 0 | 0 |
| ZNF227 | 0 | 0 | 0 | 0 | 906 | 0 | 0 | 0 |
| ZNF23 | 0 | 0 | 0 | 0 | 162 | 0 | 0 | 0 |
| ZNF230 | 0 | 0 | 0 | 0 | 1694 | 1846 | 0 | 0 |
| ZNF234 | 0 | 0 | 0 | 0 | 96 | 0 | 0 | 0 |
| ZNF236 | 0 | 2438 | 2392 | 0 | 4264 | 6668 | 766 | 1338 |
| ZNF239 | 0 | 0 | 0 | 0 | 0 | 0 | 0 | 1078 |
| ZNF24 | 0 | 0 | 0 | 0 | 0 | 0 | 276 | 0 |
| ZNF248 | 0 | 0 | 0 | 0 | 588 | 0 | 0 | 1244 |
| ZNF25 | 0 | 0 | 0 | 0 | 422 | 0 | 0 | 0 |
| ZNF250 | 0 | 0 | 0 | 736 | 486 | 1168 | 0 | 0 |
| ZNF251 | 0 | 562 | 0 | 0 | 0 | 0 | 0 | 0 |
| ZNF252P | 0 | 376 | 0 | 0 | 634 | 0 | 0 | 0 |
| ZNF253 | 22292 | 29936 | 28840 | 16284 | 124796 | 11045 | 37714 | 11406 |
| ZNF254 | 798 | 2156 | 7292 | 2580 | 12297 | 2104 | 8743 | 316 |
| ZNF256 | 0 | 0 | 0 | 0 | 120 | 0 | 0 | 0 |
| ZNF26 | 0 | 0 | 0 | 0 | 226 | 0 | 0 | 0 |
| ZNF263 | 0 | 288 | 0 | 0 | 0 | 0 | 422 | 0 |
| ZNF264 | 0 | 372 | 0 | 0 | 0 | 0 | 0 | 0 |
| ZNF266 | 1604 | 4332 | 4957 | 1750 | 13046 | 156 | 1176 | 3762 |
| ZNF267 | 0 | 0 | 0 | 0 | 905 | 0 | 0 | 320 |
| ZNF271P | 0 | 0 | 0 | 175 | 254 | 0 | 458 | 750 |
| ZNF274 | 0 | 0 | 1534 | 0 | 0 | 0 | 0 | 0 |
| ZNF277 | 0 | 0 | 0 | 0 | 428 | 0 | 1288 | 0 |
| ZNF28 | 0 | 0 | 0 | 0 | 508 | 0 | 0 | 0 |
| ZNF280D | 0 | 422 | 48 | 0 | 2217 | 0 | 688 | 0 |
| ZNF282 | 0 | 0 | 0 | 0 | 0 | 0 | 432 | 1128 |
| ZNF284 | 0 | 0 | 0 | 0 | 416 | 0 | 0 | 0 |
| ZNF292 | 818 | 2305 | 6026 | 1408 | 8175 | 5360 | 3712 | 916 |
| ZNF296 | 0 | 316 | 0 | 0 | 0 | 0 | 0 | 0 |
| ZNF3 | 0 | 0 | 1240 | 0 | 0 | 1052 | 0 | 0 |
| ZNF304 | 0 | 200 | 0 | 0 | 0 | 0 | 0 | 530 |
| ZNF311 | 0 | 0 | 0 | 907 | 0 | 0 | 0 | 591 |
| ZNF317 | 0 | 0 | 0 | 0 | 170 | 0 | 0 | 0 |
| ZNF32 | 0 | 314 | 0 | 0 | 3896 | 0 | 0 | 0 |
| ZNF320 | 0 | 0 | 0 | 0 | 0 | 0 | 396 | 0 |
| ZNF321P | 0 | 162 | 0 | 0 | 1780 | 0 | 0 | 0 |
| ZNF322 | 0 | 0 | 0 | 646 | 0 | 0 | 0 | 662 |
| ZNF326 | 0 | 364 | 1690 | 1334 | 0 | 0 | 0 | 0 |
| ZNF329 | 0 | 0 | 1304 | 0 | 0 | 0 | 0 | 0 |
| ZNF330 | 436 | 0 | 0 | 956 | 1192 | 0 | 0 | 0 |

| | | | | | | | | |
|---|---|---|---|---|---|---|---|---|
| ZNF331 | 1334 | 4102 | 6581 | 0 | 15311 | 26045 | 4054 | 7394 |
| ZNF335 | 0 | 478 | 0 | 367 | 514 | 0 | 0 | 0 |
| ZNF337 | 0 | 0 | 0 | 0 | 389 | 0 | 0 | 0 |
| ZNF33A | 0 | 82 | 1182 | 1234 | 538 | 0 | 680 | 1106 |
| ZNF33B | 0 | 0 | 0 | 0 | 0 | 0 | 1016 | 0 |
| ZNF34 | 0 | 0 | 0 | 0 | 977 | 14 | 370 | 0 |
| ZNF346 | 0 | 0 | 0 | 807 | 0 | 0 | 158 | 0 |
| ZNF350 | 5851 | 25346 | 25436 | 12336 | 9394 | 13926 | 43846 | 7843 |
| ZNF354A | 0 | 0 | 0 | 0 | 1778 | 0 | 0 | 0 |
| ZNF358 | 402 | 0 | 908 | 0 | 1816 | 0 | 122 | 0 |
| ZNF362 | 0 | 0 | 468 | 0 | 426 | 0 | 0 | 0 |
| ZNF366 | 0 | 324 | 0 | 0 | 0 | 0 | 0 | 0 |
| ZNF37A | 0 | 0 | 246 | 0 | 0 | 0 | 0 | 1056 |
| ZNF384 | 0 | 470 | 0 | 0 | 180 | 0 | 236 | 406 |
| ZNF391 | 0 | 0 | 0 | 0 | 0 | 0 | 114 | 0 |
| ZNF398 | 0 | 0 | 0 | 0 | 540 | 0 | 0 | 0 |
| ZNF407 | 0 | 384 | 402 | 0 | 0 | 0 | 0 | 0 |
| ZNF408 | 0 | 320 | 0 | 0 | 0 | 0 | 0 | 0 |
| ZNF41 | 0 | 336 | 0 | 0 | 580 | 0 | 0 | 0 |
| ZNF410 | 0 | 0 | 2750 | 0 | 412 | 0 | 0 | 0 |
| ZNF414 | 0 | 0 | 0 | 0 | 192 | 0 | 0 | 0 |
| ZNF426 | 388 | 0 | 0 | 0 | 864 | 0 | 0 | 0 |
| ZNF426-DT | 0 | 218 | 0 | 0 | 196 | 0 | 0 | 0 |
| ZNF428 | 0 | 0 | 0 | 0 | 488 | 0 | 867 | 817 |
| ZNF431 | 1168 | 0 | 0 | 0 | 1655 | 0 | 0 | 0 |
| ZNF432 | 0 | 0 | 0 | 0 | 687 | 0 | 0 | 0 |
| ZNF433 | 0 | 0 | 0 | 0 | 0 | 0 | 544 | 0 |
| ZNF438 | 604 | 1344 | 566 | 831 | 3164 | 0 | 734 | 230 |
| ZNF439 | 0 | 0 | 160 | 0 | 0 | 0 | 0 | 0 |
| ZNF44 | 0 | 998 | 1490 | 0 | 332 | 0 | 0 | 0 |
| ZNF440 | 0 | 0 | 0 | 0 | 182 | 0 | 0 | 0 |
| ZNF443 | 0 | 0 | 0 | 0 | 878 | 6064 | 0 | 0 |
| ZNF444 | 0 | 554 | 0 | 0 | 0 | 0 | 0 | 0 |
| ZNF445 | 0 | 0 | 0 | 1384 | 1583 | 0 | 284 | 0 |
| ZNF446 | 0 | 0 | 802 | 0 | 0 | 0 | 0 | 0 |
| ZNF45 | 0 | 0 | 0 | 0 | 0 | 0 | 328 | 0 |
| ZNF451 | 0 | 0 | 0 | 1086 | 721 | 0 | 508 | 0 |
| ZNF454 | 550 | 0 | 0 | 0 | 0 | 0 | 0 | 0 |
| ZNF461 | 1494 | 4162 | 9800 | 3795 | 5884 | 2210 | 6250 | 3040 |
| ZNF462 | 0 | 0 | 0 | 0 | 1196 | 0 | 860 | 0 |
| ZNF468 | 0 | 362 | 0 | 0 | 0 | 0 | 0 | 0 |
| ZNF471 | 0 | 0 | 0 | 0 | 0 | 0 | 677 | 0 |
| ZNF480 | 596 | 44 | 0 | 204 | 178 | 0 | 1458 | 751 |

| | | | | | | | | |
|---|---|---|---|---|---|---|---|---|
| ZNF485 | 516 | 94 | 0 | 3008 | 0 | 0 | 0 | 0 |
| ZNF486 | 0 | 18 | 0 | 0 | 90 | 0 | 758 | 0 |
| ZNF487 | 0 | 0 | 0 | 0 | 0 | 0 | 0 | 320 |
| ZNF490 | 0 | 0 | 0 | 996 | 0 | 0 | 0 | 0 |
| ZNF492 | 74 | 0 | 0 | 0 | 0 | 0 | 0 | 0 |
| ZNF493 | 0 | 0 | 0 | 0 | 0 | 1460 | 0 | 0 |
| ZNF496 | 0 | 0 | 0 | 0 | 212 | 0 | 2797 | 0 |
| ZNF503-AS1 | 0 | 658 | 0 | 0 | 0 | 0 | 0 | 0 |
| ZNF503-AS2 | 0 | 0 | 0 | 0 | 718 | 0 | 206 | 0 |
| ZNF511 | 0 | 0 | 3346 | 3418 | 3775 | 1624 | 1048 | 0 |
| ZNF512 | 0 | 358 | 0 | 0 | 0 | 0 | 0 | 0 |
| ZNF512B | 0 | 0 | 0 | 0 | 468 | 0 | 0 | 0 |
| ZNF513 | 0 | 402 | 0 | 0 | 0 | 0 | 272 | 450 |
| ZNF514 | 0 | 0 | 0 | 436 | 0 | 0 | 200 | 0 |
| ZNF521 | 2095 | 126 | 8040 | 0 | 13837 | 7385 | 372 | 438 |
| ZNF524 | 46 | 0 | 172 | 0 | 660 | 0 | 0 | 0 |
| ZNF527 | 0 | 384 | 0 | 0 | 0 | 0 | 0 | 0 |
| ZNF528 | 0 | 316 | 0 | 0 | 0 | 0 | 0 | 0 |
| ZNF532 | 0 | 0 | 0 | 0 | 278 | 0 | 0 | 0 |
| ZNF542P | 0 | 0 | 0 | 0 | 308 | 0 | 0 | 0 |
| ZNF547 | 364 | 0 | 0 | 0 | 0 | 0 | 0 | 0 |
| ZNF552 | 0 | 0 | 966 | 0 | 0 | 0 | 0 | 0 |
| ZNF557 | 658 | 0 | 0 | 0 | 0 | 0 | 0 | 0 |
| ZNF561 | 0 | 0 | 0 | 0 | 0 | 0 | 780 | 0 |
| ZNF562 | 0 | 62 | 0 | 0 | 0 | 0 | 0 | 0 |
| ZNF563 | 0 | 564 | 0 | 0 | 98 | 0 | 0 | 0 |
| ZNF564 | 0 | 0 | 1032 | 0 | 0 | 0 | 0 | 0 |
| ZNF565 | 0 | 0 | 0 | 0 | 62 | 0 | 0 | 0 |
| ZNF567 | 0 | 0 | 1142 | 0 | 168 | 0 | 0 | 0 |
| ZNF57 | 0 | 0 | 0 | 997 | 554 | 0 | 0 | 0 |
| ZNF573 | 0 | 0 | 0 | 0 | 0 | 0 | 0 | 368 |
| ZNF574 | 334 | 0 | 0 | 0 | 0 | 0 | 0 | 0 |
| ZNF575 | 0 | 0 | 1040 | 0 | 0 | 0 | 0 | 0 |
| ZNF576 | 20462 | 29251 | 59561 | 48088 | 112067 | 48626 | 48107 | 28328 |
| ZNF577 | 0 | 0 | 1548 | 0 | 0 | 0 | 0 | 0 |
| ZNF579 | 0 | 0 | 0 | 0 | 596 | 0 | 0 | 0 |
| ZNF580 | 0 | 0 | 0 | 1632 | 546 | 0 | 0 | 0 |
| ZNF581 | 0 | 0 | 2150 | 0 | 683 | 0 | 0 | 0 |
| ZNF582-AS1 | 0 | 330 | 0 | 0 | 0 | 0 | 0 | 0 |
| ZNF585A | 0 | 0 | 828 | 838 | 20 | 0 | 0 | 0 |
| ZNF585B | 282 | 0 | 0 | 0 | 0 | 0 | 0 | 0 |
| ZNF587 | 0 | 26 | 0 | 0 | 560 | 0 | 147 | 0 |
| ZNF589 | 0 | 0 | 0 | 0 | 0 | 0 | 196 | 0 |

| | | | | | | | | |
|---|---|---|---|---|---|---|---|---|
| ZNF592 | 0 | 0 | 0 | 44 | 858 | 0 | 0 | 0 |
| ZNF593 | 0 | 0 | 2028 | 2172 | 3036 | 1966 | 2092 | 902 |
| ZNF595 | 530 | 124 | 0 | 0 | 0 | 0 | 0 | 0 |
| ZNF598 | 0 | 0 | 0 | 24 | 0 | 0 | 152 | 0 |
| ZNF599 | 416 | 0 | 0 | 0 | 0 | 0 | 0 | 0 |
| ZNF613 | 0 | 0 | 0 | 0 | 346 | 0 | 0 | 0 |
| ZNF614 | 0 | 0 | 0 | 0 | 38 | 0 | 0 | 0 |
| ZNF615 | 0 | 0 | 680 | 0 | 0 | 0 | 0 | 0 |
| ZNF616 | 0 | 0 | 0 | 0 | 252 | 0 | 0 | 0 |
| ZNF621 | 0 | 0 | 0 | 872 | 0 | 0 | 0 | 0 |
| ZNF622 | 0 | 356 | 30 | 0 | 1044 | 150 | 0 | 0 |
| ZNF623 | 0 | 0 | 0 | 0 | 442 | 0 | 82 | 0 |
| ZNF625 | 502 | 0 | 0 | 0 | 0 | 0 | 0 | 0 |
| ZNF625-ZNF20 | 0 | 0 | 0 | 0 | 86 | 0 | 0 | 0 |
| ZNF626 | 0 | 398 | 0 | 0 | 0 | 0 | 0 | 0 |
| ZNF627 | 0 | 0 | 1400 | 0 | 784 | 0 | 0 | 0 |
| ZNF628 | 0 | 0 | 0 | 0 | 148 | 0 | 0 | 478 |
| ZNF630 | 0 | 0 | 0 | 0 | 0 | 0 | 0 | 822 |
| ZNF638 | 0 | 362 | 0 | 0 | 0 | 0 | 0 | 0 |
| ZNF639 | 0 | 792 | 0 | 1279 | 0 | 0 | 276 | 0 |
| ZNF649 | 0 | 0 | 0 | 0 | 2158 | 0 | 0 | 394 |
| ZNF652 | 0 | 0 | 738 | 986 | 638 | 0 | 0 | 0 |
| ZNF653 | 0 | 580 | 0 | 0 | 176 | 0 | 0 | 0 |
| ZNF655 | 216 | 0 | 2032 | 0 | 0 | 0 | 0 | 0 |
| ZNF662 | 0 | 172 | 0 | 0 | 0 | 0 | 0 | 0 |
| ZNF664 | 0 | 296 | 0 | 1006 | 0 | 272 | 0 | 0 |
| ZNF665 | 0 | 0 | 1348 | 0 | 0 | 0 | 0 | 0 |
| ZNF667-AS1 | 0 | 0 | 0 | 0 | 0 | 0 | 0 | 756 |
| ZNF668 | 0 | 390 | 0 | 0 | 0 | 0 | 0 | 0 |
| ZNF670 | 0 | 0 | 1416 | 0 | 0 | 0 | 0 | 0 |
| ZNF670-ZNF695 | 0 | 1240 | 0 | 1198 | 8252 | 0 | 2280 | 0 |
| ZNF672 | 0 | 0 | 0 | 0 | 2544 | 2161 | 0 | 0 |
| ZNF674 | 0 | 1516 | 638 | 0 | 7575 | 0 | 1824 | 0 |
| ZNF675 | 0 | 0 | 0 | 0 | 458 | 0 | 0 | 0 |
| ZNF681 | 0 | 146 | 0 | 0 | 772 | 0 | 0 | 0 |
| ZNF688 | 0 | 0 | 1392 | 0 | 0 | 0 | 0 | 981 |
| ZNF69 | 0 | 450 | 0 | 0 | 1438 | 0 | 0 | 0 |
| ZNF692 | 780 | 0 | 0 | 0 | 314 | 0 | 0 | 0 |
| ZNF695 | 942 | 0 | 124 | 3076 | 40705 | 0 | 1610 | 9341 |
| ZNF7 | 0 | 836 | 0 | 1445 | 1340 | 0 | 718 | 1540 |
| ZNF700 | 1158 | 0 | 0 | 0 | 0 | 0 | 158 | 590 |
| ZNF701 | 0 | 0 | 0 | 0 | 0 | 446 | 0 | 0 |
| ZNF703 | 0 | 0 | 0 | 140 | 664 | 250 | 2565 | 0 |

| | | | | | | | | |
|---|---|---|---|---|---|---|---|---|
| ZNF704 | 0 | 0 | 0 | 0 | 766 | 1529 | 0 | 0 |
| ZNF706 | 2411 | 2336 | 4542 | 2655 | 24361 | 3464 | 11354 | 3526 |
| ZNF709 | 508 | 1090 | 0 | 0 | 7648 | 3013 | 1370 | 595 |
| ZNF71 | 0 | 70 | 0 | 0 | 1008 | 0 | 0 | 0 |
| ZNF710 | 0 | 0 | 680 | 0 | 488 | 0 | 894 | 0 |
| ZNF721 | 7190 | 12156 | 8100 | 10480 | 35706 | 21363 | 108941 | 12742 |
| ZNF724 | 0 | 0 | 0 | 0 | 584 | 0 | 0 | 0 |
| ZNF726 | 0 | 0 | 14 | 0 | 0 | 0 | 0 | 0 |
| ZNF736 | 0 | 0 | 0 | 0 | 0 | 0 | 404 | 0 |
| ZNF738 | 0 | 0 | 0 | 0 | 866 | 0 | 0 | 0 |
| ZNF74 | 0 | 306 | 0 | 0 | 0 | 0 | 0 | 0 |
| ZNF740 | 0 | 0 | 0 | 1478 | 406 | 0 | 0 | 0 |
| ZNF746 | 524 | 0 | 0 | 0 | 0 | 0 | 0 | 0 |
| ZNF75A | 588 | 0 | 0 | 0 | 614 | 42 | 0 | 0 |
| ZNF75D | 0 | 338 | 0 | 0 | 0 | 0 | 0 | 0 |
| ZNF76 | 0 | 0 | 0 | 998 | 0 | 0 | 0 | 0 |
| ZNF766 | 0 | 0 | 0 | 0 | 264 | 0 | 0 | 0 |
| ZNF767P | 0 | 0 | 0 | 0 | 0 | 0 | 516 | 0 |
| ZNF768 | 0 | 0 | 0 | 498 | 0 | 0 | 0 | 672 |
| ZNF77 | 0 | 0 | 1438 | 0 | 0 | 0 | 0 | 0 |
| ZNF770 | 0 | 394 | 0 | 0 | 0 | 0 | 0 | 0 |
| ZNF774 | 1324 | 6460 | 3920 | 3450 | 8637 | 0 | 5631 | 546 |
| ZNF775 | 0 | 0 | 0 | 0 | 486 | 0 | 0 | 0 |
| ZNF780A | 0 | 0 | 0 | 0 | 720 | 0 | 212 | 0 |
| ZNF780B | 0 | 0 | 0 | 0 | 307 | 0 | 0 | 0 |
| ZNF782 | 0 | 58 | 0 | 0 | 0 | 0 | 552 | 0 |
| ZNF784 | 0 | 0 | 0 | 0 | 604 | 0 | 0 | 82 |
| ZNF787 | 0 | 0 | 0 | 0 | 0 | 0 | 0 | 376 |
| ZNF79 | 0 | 0 | 0 | 0 | 250 | 0 | 0 | 0 |
| ZNF791 | 0 | 676 | 0 | 0 | 1333 | 0 | 272 | 0 |
| ZNF793 | 176 | 0 | 0 | 0 | 0 | 0 | 0 | 0 |
| ZNF799 | 0 | 0 | 0 | 0 | 458 | 0 | 406 | 0 |
| ZNF8-ERVK3-1 | 1854 | 0 | 0 | 44 | 2198 | 0 | 336 | 184 |
| ZNF800 | 0 | 0 | 114 | 0 | 676 | 0 | 284 | 0 |
| ZNF805 | 0 | 0 | 0 | 0 | 714 | 0 | 0 | 0 |
| ZNF808 | 0 | 0 | 0 | 0 | 0 | 0 | 0 | 722 |
| ZNF81 | 0 | 0 | 0 | 0 | 0 | 0 | 194 | 0 |
| ZNF814 | 0 | 144 | 0 | 0 | 0 | 0 | 292 | 0 |
| ZNF815P | 0 | 0 | 0 | 0 | 598 | 0 | 0 | 0 |
| ZNF816 | 0 | 0 | 0 | 0 | 438 | 0 | 0 | 0 |
| ZNF821 | 6172 | 3172 | 11732 | 0 | 3516 | 5365 | 6905 | 1504 |
| ZNF823 | 0 | 0 | 6299 | 0 | 0 | 0 | 0 | 0 |
| ZNF83 | 0 | 0 | 0 | 0 | 1340 | 0 | 0 | 1194 |

| | | | | | | | | |
|---|---|---|---|---|---|---|---|---|
| ZNF830 | 0 | 0 | 0 | 0 | 194 | 0 | 84 | 0 |
| ZNF836 | 0 | 0 | 0 | 0 | 194 | 0 | 0 | 0 |
| ZNF84 | 0 | 0 | 0 | 462 | 0 | 0 | 0 | 0 |
| ZNF846 | 780 | 0 | 0 | 0 | 944 | 0 | 0 | 0 |
| ZNF85 | 76 | 1272 | 0 | 904 | 0 | 2460 | 1956 | 916 |
| ZNF852 | 0 | 0 | 0 | 0 | 392 | 0 | 0 | 0 |
| ZNF865 | 0 | 360 | 0 | 0 | 426 | 0 | 0 | 0 |
| ZNF880 | 5212 | 10963 | 2760 | 0 | 3502 | 6929 | 0 | 2154 |
| ZNF888 | 0 | 0 | 0 | 0 | 0 | 0 | 0 | 212 |
| ZNF891 | 0 | 0 | 0 | 0 | 447 | 0 | 0 | 0 |
| ZNF91 | 60462 | 43142 | 49455 | 23730 | 90516 | 28748 | 55577 | 15218 |
| ZNF93 | 0 | 380 | 0 | 1270 | 9452 | 1504 | 528 | 0 |
| ZNFX1 | 690 | 40 | 0 | 1160 | 1606 | 0 | 2894 | 0 |
| ZNHIT1 | 594 | 3858 | 2898 | 8317 | 2562 | 12477 | 2240 | 808 |
| ZNHIT2 | 1084 | 364 | 0 | 1400 | 0 | 1800 | 0 | 0 |
| ZNHIT3 | 0 | 851 | 0 | 0 | 1051 | 0 | 1810 | 0 |
| ZNRD1 | 155897 | 231303 | 392777 | 255512 | 950243 | 278724 | 742008 | 336433 |
| ZNRD1ASP | 0 | 0 | 0 | 0 | 0 | 1612 | 0 | 0 |
| ZNRF1 | 7614 | 3242 | 11081 | 5220 | 16485 | 2948 | 13761 | 4758 |
| ZNRF2 | 590 | 0 | 0 | 1458 | 0 | 324 | 0 | 0 |
| ZNRF3 | 166 | 214 | 0 | 0 | 1405 | 1638 | 6850 | 420 |
| ZP3 | 0 | 0 | 0 | 0 | 636 | 0 | 0 | 0 |
| ZPR1 | 0 | 0 | 0 | 1022 | 1980 | 0 | 0 | 2363 |
| ZRANB2 | 768 | 184 | 4476 | 330 | 1612 | 0 | 1050 | 1504 |
| ZRSR2 | 0 | 424 | 0 | 0 | 1354 | 0 | 0 | 0 |
| ZSCAN16 | 0 | 24 | 0 | 0 | 0 | 0 | 718 | 1108 |
| ZSCAN16-AS1 | 0 | 0 | 0 | 0 | 326 | 0 | 0 | 0 |
| ZSCAN18 | 1110 | 274 | 278 | 0 | 0 | 0 | 0 | 0 |
| ZSCAN2 | 0 | 0 | 0 | 0 | 0 | 0 | 444 | 0 |
| ZSCAN22 | 372 | 0 | 0 | 0 | 0 | 0 | 0 | 0 |
| ZSCAN25 | 0 | 0 | 0 | 1046 | 0 | 1218 | 0 | 0 |
| ZSCAN32 | 0 | 0 | 0 | 0 | 0 | 0 | 334 | 0 |
| ZSCAN9 | 0 | 244 | 0 | 0 | 0 | 0 | 0 | 0 |
| ZSWIM1 | 0 | 0 | 0 | 0 | 438 | 0 | 0 | 0 |
| ZSWIM3 | 0 | 0 | 2048 | 0 | 154 | 0 | 0 | 0 |
| ZSWIM4 | 0 | 0 | 0 | 0 | 0 | 0 | 0 | 482 |
| ZSWIM6 | 0 | 0 | 0 | 0 | 284 | 0 | 0 | 0 |
| ZSWIM7 | 0 | 644 | 0 | 1122 | 1039 | 0 | 1831 | 0 |
| ZUP1 | 0 | 464 | 0 | 0 | 0 | 0 | 36 | 0 |
| ZW10 | 0 | 0 | 0 | 0 | 422 | 0 | 206 | 0 |
| ZWILCH | 506 | 0 | 0 | 1146 | 0 | 0 | 0 | 0 |
| ZWINT | 0 | 0 | 0 | 0 | 730 | 0 | 0 | 0 |
| ZXDC | 494 | 0 | 88 | 0 | 0 | 0 | 0 | 0 |

| | | | | | | | | |
|---|---|---|---|---|---|---|---|---|
| ZYG11B | 592 | 338 | 0 | 0 | 0 | 0 | 0 | 0 |
| ZYX | 224 | 0 | 0 | 2725 | 992 | 0 | 226 | 0 |
| ZZZ3 | 0 | 322 | 0 | 0 | 0 | 0 | 0 | 0 |

Supplementary Table S3: Differentially expressed genes

| Gene | 1N | 3N | 1T | 2T | 3T | 1M | 2M | 3M | padj_NT | padj_TL |
|---|---|---|---|---|---|---|---|---|---|---|
| AACS | 0 | 0 | 0 | 7.963225934 | 8.21279698 | 0 | 0 | 0 | 0.102136532 | 0.037815738 |
| AAMDC | 0 | 9.670244415 | 10.06397233 | 0 | 7.815888334 | 0 | 0 | 0 | 0.999660881 | 0.012558249 |
| AAR2 | 9.68252076 | 0 | 0 | 0 | 0 | 0 | 7.178500772 | 0 | 7.42E-08 | 1 |
| AATF | 0 | 0 | 0 | 10.28284497 | 9.345373041 | 0 | 0 | 0 | 0.027402944 | 0.010003595 |
| ABCA5 | 10.37817204 | 9.870584892 | 0 | 0 | 0 | 0 | 0 | 0 | 0.000888042 | 1 |
| ABCA8 | 9.578815964 | 0 | 0 | 0 | 0 | 0 | 0 | 0 | 8.04E-08 | 1 |
| ABCB10 | 0 | 0 | 0 | 0 | 6.729472521 | 0 | 0 | 0 | 0.000125541 | 1 |
| ABCC4 | 9.702396267 | 10.70793825 | 0 | 9.722621973 | 8.241596695 | 0 | 0 | 0 | 0.852692235 | 0.01420877 |
| ABCC6 | 0 | 0 | 0 | 0 | 0 | 0 | 6.912047636 | 10.66136277 | 1 | 0.009245646 |
| ABCC6P2 | 10.27576888 | 0 | 0 | 0 | 0 | 0 | 0 | 0 | 7.07E-08 | 1 |
| ABCD3 | 9.840340458 | 0 | 10.27024349 | 0 | 7.123531262 | 0 | 0 | 0 | 0.999660881 | 0.012307739 |
| ABHD11 | 10.13173416 | 11.3862815 | 8.972957461 | 9.530709613 | 8.656460903 | 0 | 0 | 0 | 0.601259282 | 0.002374802 |
| ABHD12 | 10.08202624 | 8.657303766 | 10.47973879 | 11.01795335 | 10.33283761 | 0 | 0 | 0 | 0.800431515 | 0.000291056 |
| ABHD14A | 10.1892076 | 0 | 0 | 0 | 0 | 10.86881043 | 0 | 0 | 7.07E-08 | 1 |
| ABI2 | 0 | 0 | 0 | 9.014457977 | 9.333955795 | 0 | 0 | 0 | 0.058968573 | 0.013778143 |
| ABI3 | 0 | 8.445326205 | 0 | 0 | 0 | 0 | 0 | 0 | 1.44E-07 | 1 |
| ABRAXAS1 | 9.435755845 | 0 | 0 | 7.696708325 | 9.643924818 | 0 | 0 | 0 | 0.999483812 | 0.01616544 |
| ACAA1 | 0 | 0 | 11.76085037 | 11.14214727 | 10.66938057 | 8.577991331 | 9.678362459 | 0 | 7.07E-08 | 0.999854694 |
| ACAD9 | 0 | 0 | 0 | 9.88434752 | 9.218664857 | 0 | 0 | 10.99841242 | 0.039663385 | 0.999854694 |
| ACADS | 10.41878391 | 0 | 0 | 0 | 0 | 0 | 9.874665916 | 0 | 7.07E-08 | 1 |
| ACD | 0 | 0 | 0 | 10.82500918 | 9.075183036 | 0 | 7.97770259 | 0 | 0.016219305 | 0.999854694 |
| ACLY | 12.46722793 | 12.14984385 | 13.80968586 | 14.08055325 | 13.30877617 | 13.44814347 | 12.83283324 | 13.95677507 | 0.045923577 | 0.999854694 |
| ACOT2 | 0 | 0 | 7.552095279 | 0 | 10.21174682 | 0 | 8.381556762 | 0 | 0.046967639 | 0.999854694 |
| ACP5 | 0 | 0 | 0 | 10.39340353 | 7.00375218 | 0 | 0 | 0 | 0.046818355 | 0.012259484 |
| ACSF2 | 10.5080681 | 0 | 0 | 0 | 0 | 0 | 9.41997401 | 0 | 7.07E-08 | 1 |
| ACSM3 | 9.728478045 | 10.49280905 | 0 | 0 | 0 | 0 | 0 | 0 | 0.000804484 | 1 |
| ACSS1 | 0 | 0 | 0 | 9.532663167 | 9.263487197 | 0 | 0 | 0 | 0.048029488 | 0.012111456 |
| ACTR1B | 0 | 0 | 12.00666845 | 9.061222644 | 8.140363768 | 10.44003325 | 0 | 0 | 3.02E-05 | 0.999854694 |
| ACYP2 | 0 | 7.810193807 | 0 | 11.18500903 | 7.632336583 | 0 | 0 | 0 | 0.816268987 | 0.008991934 |
| ADA2 | 9.971708127 | 0 | 0 | 0 | 0 | 0 | 0 | 0 | 7.07E-08 | 1 |
| ADAM28 | 0 | 8.501311304 | 0 | 0 | 0 | 0 | 0 | 0 | 1.38E-07 | 1 |
| ADAMTS10 | 0 | 0 | 0 | 0 | 6.360815222 | 0 | 0 | 0 | 6.12E-05 | 1 |
| ADAMTSL1 | 9.949703925 | 0 | 0 | 0 | 0 | 0 | 0 | 0 | 7.07E-08 | 1 |
| ADAMTSL2 | 0 | 0 | 0 | 0 | 0 | 8.195493859 | 7.051416926 | 0 | 1 | 0.039241395 |
| ADAT1 | 0 | 8.761849577 | 0 | 0 | 0 | 9.821033926 | 0 | 0 | 1.16E-07 | 1 |
| ADAT2 | 0 | 8.363107485 | 0 | 0 | 0 | 0 | 0 | 0 | 1.55E-07 | 1 |
| ADCY7 | 0 | 0 | 0 | 0 | 6.8625646 | 0 | 0 | 0 | 4.09E-05 | 1 |
| ADD2 | 0 | 8.998107186 | 0 | 0 | 0 | 0 | 0 | 0 | 2.64E-07 | 1 |
| ADGRL3 | 0 | 8.859328625 | 0 | 0 | 0 | 0 | 0 | 0 | 1.21E-07 | 1 |
| ADH1C | 11.84364118 | 13.11466719 | 13.68569495 | 9.960335776 | 0 | 0 | 0 | 0 | 0.988867264 | 3.43E-12 |
| ADIPOR2 | 0 | 0 | 9.867860481 | 0 | 7.818621538 | 0 | 0 | 0 | 0.064693636 | 0.013910154 |

| | | | | | | | | | | |
|---|---|---|---|---|---|---|---|---|---|---|
| ADNP2 | 0 | 0 | 0 | 0 | 0 | 0 | 7.036579876 | 9.478071506 | 1 | 0.015061471 |
| ADORA2B | 0 | 8.375143804 | 0 | 0 | 0 | 0 | 0 | 0 | 1.55E-07 | 1 |
| ADPRM | 0 | 0 | 0 | 10.13122842 | 9.168125263 | 0 | 0 | 0 | 0.033829888 | 0.010542114 |
| ADSSL1 | 0 | 8.81583464 | 0 | 0 | 0 | 0 | 0 | 0 | 1.15E-07 | 1 |
| AEBP2 | 0 | 8.479177282 | 9.172042307 | 0 | 5.864366253 | 0 | 0 | 0 | 0.998077553 | 0.031655119 |
| AFMID | 8.62996247 | 8.867872257 | 0 | 0 | 0 | 0 | 9.4760367 | 9.723379502 | 0.019707134 | 0.009209122 |
| AGAP3 | 10.50051991 | 0 | 0 | 0 | 0 | 0 | 0 | 0 | 7.07E-08 | 1 |
| AGFG2 | 9.204448442 | 8.236875968 | 0 | 0 | 0 | 0 | 0 | 0 | 0.019179867 | 1 |
| AGO3 | 0 | 0 | 0 | 0 | 6.405131987 | 0 | 0 | 0 | 5.97E-05 | 1 |
| AGPAT4 | 0 | 0 | 0 | 0 | 6.682232705 | 0 | 0 | 0 | 4.74E-05 | 1 |
| AGTRAP | 0 | 0 | 0 | 11.16310691 | 9.942183234 | 10.58790804 | 0 | 10.48251027 | 0.030796109 | 0.999854694 |
| AIM2 | 0 | 9.311663645 | 0 | 0 | 0 | 0 | 0 | 0 | 8.05E-08 | 1 |
| AK4 | 0 | 0 | 10.30222825 | 9.865870872 | 8.221084302 | 0 | 4.589263136 | 10.36496214 | 0.002147873 | 0.999854694 |
| AK6 | 10.33213756 | 11.13961603 | 10.21370669 | 10.23205298 | 9.617486 | 0 | 0 | 0 | 0.910605604 | 0.000631775 |
| AKAP11 | 0 | 0 | 0 | 8.901075665 | 6.746796135 | 0 | 0 | 0 | 0.099427067 | 0.036143846 |
| AKAP13 | 16.55615861 | 16.22977612 | 15.34095693 | 13.01795453 | 14.71785757 | 14.2482176 | 14.59114587 | 14.11900039 | 0.020623482 | 0.999854694 |
| AKAP8 | 9.828331099 | 0 | 0 | 0 | 0 | 0 | 7.006440014 | 0 | 7.17E-08 | 1 |
| AKR1B1 | 0 | 0 | 0 | 0 | 0 | 11.53384658 | 0 | 9.871985276 | 1 | 0.004031828 |
| AKR1B10 | 10.32787885 | 8.301371897 | 10.51291902 | 0 | 6.345735199 | 0 | 0 | 0 | 0.980815574 | 0.011685672 |
| ALB | 0 | 9.844849563 | 0 | 0 | 0 | 0 | 7.704905507 | 0 | 7.07E-08 | 1 |
| ALDH9A1 | 0 | 0 | 10.50665753 | 0 | 8.710436694 | 0 | 7.503839514 | 0 | 0.030796109 | 0.999854694 |
| ALG9 | 0 | 9.54265281 | 0 | 0 | 0 | 0 | 8.537232216 | 0 | 7.53E-08 | 1 |
| ALK | 8.559063282 | 0 | 0 | 0 | 0 | 0 | 7.535844571 | 0 | 1.38E-07 | 1 |
| ALKBH5 | 0 | 0 | 8.306553364 | 10.14916628 | 8.850567979 | 0 | 8.37567389 | 0 | 0.006236283 | 0.999854694 |
| ALOX15 | 0 | 0 | 0 | 0 | 6.556887521 | 0 | 0 | 0 | 9.34E-05 | 1 |
| ALS2 | 0 | 0 | 0 | 9.000410141 | 7.396595249 | 0 | 0 | 0 | 0.089646608 | 0.027725114 |
| AMIGO2 | 0 | 0 | 10.91751914 | 9.349745754 | 8.305342632 | 9.896188644 | 0 | 0 | 0.001067109 | 0.999854694 |
| AMN | 10.62375866 | 9.876947628 | 9.199475854 | 0 | 8.328539441 | 0 | 0 | 0 | 0.789943424 | 0.017341809 |
| AMN1 | 0 | 10.61963685 | 0 | 0 | 0 | 0 | 0 | 0 | 7.07E-08 | 1 |
| ANAPC1 | 0 | 0 | 0 | 9.601285316 | 10.03852628 | 0 | 0 | 0 | 0.028305576 | 0.010097347 |
| ANAPC2 | 10.66823035 | 0 | 0 | 9.926561143 | 8.442785867 | 0 | 0 | 0 | 0.983546955 | 0.012440088 |
| ANAPC4 | 0 | 0 | 7.668851543 | 0 | 8.241596695 | 0 | 0 | 0 | 0.109086237 | 0.041430247 |
| ANGEL1 | 9.846307856 | 0 | 0 | 0 | 0 | 0 | 0 | 0 | 7.17E-08 | 1 |
| ANKHD1-EIF4EBP3 | 0 | 8.375143804 | 0 | 0 | 0 | 0 | 8.038540912 | 0 | 1.55E-07 | 1 |
| ANKRD16 | 0 | 0 | 0 | 9.210245837 | 7.705040924 | 0 | 0 | 0 | 0.080740233 | 0.021776607 |
| ANKRD22 | 0 | 0 | 0 | 10.98622896 | 9.663820783 | 8.762839314 | 9.613605999 | 6.655488901 | 0.034161488 | 0.999854694 |
| ANKRD27 | 0 | 9.2476365 | 0 | 0 | 0 | 0 | 6.757760881 | 0 | 8.25E-08 | 1 |
| ANKRD36B | 0 | 0 | 0 | 8.207797171 | 0 | 0 | 8.166714905 | 0 | 1.16E-05 | 1 |
| ANKRD40 | 8.835927714 | 0 | 0 | 0 | 0 | 0 | 3.192657415 | 0 | 1.15E-07 | 0.999854694 |
| ANKRD52 | 0 | 0 | 0 | 0 | 6.345735199 | 0 | 0 | 0 | 6.22E-05 | 1 |
| ANKRD55 | 0 | 9.773889409 | 0 | 0 | 0 | 9.688618303 | 0 | 0 | 7.17E-08 | 1 |
| ANKS1B | 11.18623903 | 0 | 0 | 0 | 0 | 0 | 9.19459482 | 0 | 4.18E-08 | 1 |
| ANKZF1 | 0 | 9.870584892 | 0 | 5.731096434 | 8.941142865 | 0 | 0 | 0 | 0.961310672 | 0.040504575 |

| | | | | | | | | | | |
|---|---|---|---|---|---|---|---|---|---|---|
| ANO10 | 0 | 0 | 11.83831169 | 10.35527794 | 0 | 0 | 0 | 0 | 0.023102837 | 0.004757269 |
| ANTXR2 | 9.804008249 | 0 | 0 | 0 | 0 | 0 | 7.946293493 | 0 | 7.17E-08 | 1 |
| ANXA2R | 0 | 0 | 0 | 9.540451037 | 9.440711312 | 0 | 0 | 0 | 0.043619256 | 0.011493473 |
| AP1M1 | 0 | 0 | 10.04465403 | 0 | 8.026767928 | 0 | 0 | 0 | 0.055080694 | 0.012307739 |
| AP2A2 | 9.987993796 | 0 | 0 | 0 | 0 | 0 | 0 | 8.981404596 | 7.07E-08 | 1 |
| AP3D1 | 0 | 0 | 0 | 9.687981679 | 11.04660984 | 0 | 0 | 0 | 0.032597368 | 0.007353911 |
| AP3S2 | 10.74050121 | 0 | 9.418741439 | 11.21207961 | 0 | 0 | 0 | 0 | 0.996280769 | 0.007654893 |
| AP4B1 | 9.708961049 | 9.201064401 | 0 | 0 | 0 | 0 | 0 | 0 | 0.006606788 | 1 |
| AP4M1 | 0 | 10.06887167 | 8.345666268 | 10.56069854 | 9.678940814 | 0 | 3.567697193 | 0 | 0.972184991 | 0.023347883 |
| APC | 15.30637318 | 14.59662011 | 13.56301344 | 10.43682009 | 12.21057402 | 12.56710695 | 11.44883075 | 11.56911899 | 0.041252274 | 0.999854694 |
| APCDD1 | 0 | 8.918098054 | 0 | 0 | 0 | 10.09558557 | 11.40401493 | 0 | 1.08E-07 | 0.003597041 |
| APOA2 | 0 | 0 | 0 | 10.48200434 | 10.24311278 | 0 | 10.07128465 | 0 | 0.03566582 | 0.999854694 |
| APOBEC3F | 0 | 0 | 0 | 0 | 6.582842939 | 0 | 0 | 0 | 0.000136511 | 1 |
| APOL2 | 10.19856778 | 7.879432217 | 0 | 0 | 0 | 0 | 0 | 8.21786649 | 0.007583189 | 1 |
| APTX | 0 | 0 | 0 | 9.938398735 | 10.03852628 | 0 | 0 | 0 | 0.022283634 | 0.009547202 |
| AQP12A | 0 | 0 | 7.702112179 | 0 | 0 | 0 | 0 | 0 | 2.25E-05 | 1 |
| AQP3 | 0 | 9.426002279 | 0 | 0 | 0 | 0 | 0 | 0 | 7.87E-08 | 1 |
| AQP7 | 0 | 0 | 0 | 0 | 0 | 11.92119818 | 8.968989117 | 0 | 1 | 3.43E-12 |
| AR | 10.36991065 | 0 | 0 | 0 | 0 | 0 | 8.166714905 | 0 | 7.07E-08 | 1 |
| ARAP1 | 0 | 0 | 11.44526017 | 0 | 7.903457334 | 0 | 8.699090458 | 0 | 0.034167337 | 0.999854694 |
| ARAP3 | 0 | 8.51225226 | 0 | 0 | 0 | 0 | 0 | 0 | 1.38E-07 | 1 |
| ARCN1 | 0 | 0 | 0 | 9.986227367 | 8.801674049 | 0 | 0 | 9.786434411 | 0.043827245 | 0.999854694 |
| AREL1 | 0 | 9.330330507 | 0 | 0 | 0 | 0 | 0 | 0 | 8.05E-08 | 1 |
| ARFGEF1 | 0 | 0 | 0 | 0 | 6.658019782 | 0 | 0 | 0 | 4.84E-05 | 1 |
| ARFIP2 | 0 | 10.04269877 | 0 | 0 | 0 | 0 | 7.769143214 | 0 | 7.07E-08 | 1 |
| ARHGAP12 | 0 | 9.787463024 | 0 | 0 | 0 | 0 | 0 | 9.80395525 | 7.17E-08 | 1 |
| ARHGAP17 | 0 | 9.280005236 | 0 | 0 | 0 | 0 | 0 | 0 | 8.10E-08 | 1 |
| ARHGAP21 | 0 | 5.662592985 | 9.530974688 | 0 | 7.381886513 | 0 | 0 | 0 | 0.704660194 | 0.018029099 |
| ARHGAP27 | 8.544455536 | 0 | 0 | 0 | 0 | 0 | 8.54773464 | 8.711223699 | 1.39E-07 | 0.015141771 |
| ARHGAP31 | 11.53184335 | 0 | 0 | 0 | 0 | 0 | 0 | 0 | 7.07E-08 | 1 |
| ARHGAP4 | 9.497729097 | 0 | 8.972957461 | 0 | 8.903019168 | 0 | 0 | 0 | 0.999483812 | 0.015785316 |
| ARHGAP8 | 0 | 10.27242335 | 0 | 0 | 0 | 0 | 8.494440854 | 0 | 7.07E-08 | 1 |
| ARHGEF16 | 0 | 8.676884284 | 9.918264838 | 8.761452735 | 0 | 0 | 0 | 0 | 0.968776858 | 0.012259484 |
| ARHGEF2 | 9.176299815 | 0 | 0 | 0 | 0 | 0 | 0 | 0 | 1.15E-07 | 1 |
| ARHGEF37 | 10.33213756 | 0 | 0 | 0 | 0 | 0 | 0 | 0 | 7.07E-08 | 1 |
| ARHGEF40 | 0 | 0 | 0 | 0 | 6.556887521 | 0 | 8.11821332 | 0 | 9.34E-05 | 1 |
| ARID4B | 0 | 0 | 0 | 0 | 6.489879427 | 0 | 3.865083322 | 0 | 5.56E-05 | 1 |
| ARID5A | 0 | 0 | 9.987347031 | 0 | 8.332369572 | 0 | 0 | 0 | 0.053103275 | 0.012259484 |
| ARL17B | 0 | 0 | 10.74340383 | 9.926561143 | 0 | 0 | 0 | 0 | 0.012916396 | 0.00825262 |
| ARL2BP | 8.602019497 | 0 | 0 | 10.88358408 | 6.360815222 | 0 | 0 | 0 | 0.912935917 | 2.29E-10 |
| ARL6 | 0 | 0 | 0 | 0 | 6.582842939 | 0 | 0 | 0 | 0.000136511 | 1 |
| ARMC1 | 0 | 8.998107186 | 0 | 0 | 0 | 0 | 0 | 0 | 2.64E-07 | 1 |
| ARMC10 | 0 | 0 | 0 | 9.414743228 | 9.987823595 | 0 | 8.997999732 | 0 | 0.032597368 | 0.999854694 |

| | | | | | | | | | | |
|---|---|---|---|---|---|---|---|---|---|---|
| ARMC7 | 0 | 0 | 0 | 10.07603652 | 8.839844964 | 0 | 0 | 0 | 0.040469999 | 0.011067641 |
| ARMCX2 | 0 | 0 | 0 | 8.416855435 | 0 | 0 | 0 | 0 | 9.33E-06 | 1 |
| ARMCX5 | 0 | 8.479177282 | 0 | 0 | 0 | 0 | 0 | 0 | 1.39E-07 | 1 |
| ARNTL2 | 0 | 0 | 0 | 0 | 6.873120683 | 0 | 0 | 0 | 4.05E-05 | 1 |
| ARPP19 | 0 | 9.887490223 | 0 | 10.79358366 | 9.65772812 | 0 | 0 | 0 | 0.981170768 | 0.00825262 |
| ARRDC2 | 9.35434707 | 9.800910121 | 10.18654497 | 9.659657844 | 6.448127902 | 0 | 0 | 0 | 0.99353678 | 0.00208311 |
| ARRDC3-AS1 | 0 | 9.778428152 | 0 | 0 | 0 | 0 | 0 | 0 | 7.17E-08 | 1 |
| ARSA | 10.26689792 | 7.05971876 | 0 | 0 | 0 | 0 | 0 | 9.896691627 | 0.008952142 | 1 |
| AS3MT | 0 | 0 | 0 | 11.68886085 | 11.47986732 | 0 | 11.42410513 | 10.21750006 | 0.017818159 | 0.999854694 |
| ASB3 | 0 | 0 | 8.783962225 | 10.07066834 | 0 | 0 | 0 | 0 | 0.045270943 | 0.011625223 |
| ASCC3 | 0 | 0 | 0 | 0 | 6.741044679 | 0 | 6.895697203 | 0 | 0.000123975 | 1 |
| ASCL2 | 0 | 0 | 0 | 12.53851182 | 10.54904816 | 10.03302179 | 13.21191844 | 9.456076322 | 1.35E-07 | 0.999854694 |
| ASH1L | 0 | 8.696202607 | 0 | 0 | 0 | 0 | 0 | 8.615245757 | 1.18E-07 | 1 |
| ASIC2 | 0 | 8.80697625 | 9.52324123 | 10.11306472 | 0 | 0 | 0 | 0 | 0.946338504 | 0.01037888 |
| ASNS | 0 | 0 | 10.09768996 | 10.69148534 | 0 | 0 | 0 | 10.70398116 | 0.035967833 | 0.999854694 |
| ASNSD1 | 10.00409768 | 9.78295266 | 0 | 0 | 0 | 0 | 9.072624984 | 0 | 0.002143819 | 1 |
| ASPG | 8.322706971 | 0 | 0 | 0 | 0 | 0 | 0 | 0 | 2.56E-07 | 1 |
| ATAD1 | 0 | 11.25239556 | 9.278778857 | 0 | 9.44779056 | 0 | 0 | 0 | 0.958210869 | 0.012061641 |
| ATF2 | 0 | 8.676884284 | 0 | 0 | 0 | 0 | 0 | 0 | 1.20E-07 | 1 |
| ATF3 | 0 | 0 | 12.39269521 | 0 | 8.257799178 | 11.38940879 | 0 | 10.69696495 | 1.14E-07 | 0.999854694 |
| ATF7 | 15.48993823 | 15.86687107 | 14.46402175 | 14.54427295 | 14.35444959 | 14.05921567 | 15.75055075 | 15.00445631 | 0.03884807 | 0.999854694 |
| ATF7IP | 8.697540019 | 0 | 0 | 0 | 0 | 0 | 0 | 0 | 1.24E-07 | 1 |
| ATG4A | 9.285741834 | 9.547979116 | 0 | 0 | 0 | 0 | 0 | 0 | 0.007258162 | 1 |
| ATG4B | 9.91037005 | 9.044065918 | 10.08933414 | 0 | 9.631543354 | 0 | 0 | 0 | 0.99499976 | 0.009818636 |
| ATIC | 0 | 0 | 11.05527665 | 0 | 8.948648186 | 11.17134827 | 10.50092684 | 0 | 0.013148052 | 0.999854694 |
| ATM | 0 | 9.366954907 | 0 | 0 | 0 | 0 | 8.763581896 | 0 | 7.88E-08 | 1 |
| ATOH1 | 9.48248357 | 0 | 0 | 0 | 0 | 0 | 0 | 0 | 7.87E-08 | 1 |
| ATP11B | 0 | 6.83415338 | 9.461453882 | 0 | 6.741044679 | 0 | 0 | 0 | 0.889993558 | 0.02144636 |
| ATP13A1 | 0 | 9.849170791 | 0 | 0 | 0 | 0 | 0 | 0 | 7.07E-08 | 1 |
| ATP1A1 | 19.99750522 | 20.35742186 | 19.51833455 | 19.31522249 | 18.42340006 | 19.3550549 | 19.06134586 | 17.61223956 | 0.041441572 | 0.999854694 |
| ATP1A1-AS1 | 0 | 0 | 10.79505642 | 0 | 7.594562621 | 0 | 0 | 0 | 0.030016686 | 0.010003595 |
| ATP2B1 | 8.657374495 | 0 | 8.796848942 | 0 | 9.014494887 | 0 | 0 | 0 | 0.983995642 | 0.016168618 |
| ATP2B4 | 0 | 0 | 9.629351518 | 0 | 8.22932429 | 0 | 0 | 0 | 0.06727895 | 0.014388177 |
| ATP2C1 | 9.662367605 | 7.862431639 | 9.518581179 | 0 | 9.056780188 | 0 | 0 | 0 | 0.99353678 | 0.012348695 |
| ATP6V1F | 6.796891173 | 8.705765621 | 10.65345355 | 13.36668704 | 11.37653944 | 11.42181832 | 11.24490855 | 12.02725614 | 0.001047501 | 0.999854694 |
| ATPSCKMT | 0 | 0 | 0 | 0 | 6.556887521 | 0 | 0 | 0 | 9.34E-05 | 1 |
| ATXN1 | 0 | 0 | 0 | 0 | 0 | 9.802398885 | 0 | 10.4298787 | 1 | 0.007255458 |
| ATXN1L | 0 | 10.14816801 | 0 | 0 | 0 | 0 | 0 | 0 | 7.07E-08 | 1 |
| ATXN7L2 | 11.11140409 | 0 | 0 | 0 | 0 | 0 | 0 | 0 | 5.99E-08 | 1 |
| ATXN7L3 | 9.927358907 | 0 | 0 | 0 | 0 | 0 | 0 | 0 | 7.07E-08 | 1 |
| ATXN7L3B | 0 | 0 | 8.883969995 | 9.185625776 | 8.289668258 | 10.63956453 | 7.391717672 | 0 | 0.016964947 | 0.999854694 |
| AURKB | 0 | 8.842088059 | 0 | 9.739635128 | 11.20383934 | 0 | 0 | 0 | 0.886363058 | 0.006817884 |
| AZGP1 | 0 | 0 | 0 | 12.21360325 | 9.214520192 | 10.6219601 | 12.23489542 | 10.6840126 | 1.14E-07 | 0.999854694 |

| | | | | | | | | | | |
|---|---|---|---|---|---|---|---|---|---|---|
| AZIN1-AS1 | 0 | 9.600187807 | 0 | 0 | 0 | 0 | 0 | 10.40999367 | 7.44E-08 | 1 |
| B3GALT5 | 0 | 10.77760654 | 0 | 0 | 0 | 0 | 0 | 0 | 7.07E-08 | 1 |
| B3GAT3 | 0 | 10.38978972 | 0 | 0 | 0 | 0 | 0 | 9.827704158 | 7.07E-08 | 1 |
| B3GNT2 | 0 | 8.876365592 | 8.907925066 | 0 | 7.632336583 | 0 | 0 | 0 | 0.999483812 | 0.026344013 |
| B3GNT3 | 0 | 8.141445139 | 10.612493 | 10.02421381 | 6.904333267 | 0 | 0 | 0 | 0.679782208 | 0.001190665 |
| B3GNT6 | 8.762093289 | 8.467981611 | 0 | 0 | 0 | 0 | 0 | 0 | 0.022683929 | 1 |
| B4GALT1-AS1 | 0 | 0 | 9.758807731 | 0 | 8.283746183 | 0 | 0 | 0 | 0.062335861 | 0.013610425 |
| B4GALT4 | 0 | 10.94441406 | 10.45718696 | 0 | 10.22416902 | 0 | 0 | 0 | 0.997434996 | 0.007909465 |
| B4GALT7 | 0 | 0 | 9.187782438 | 0 | 7.167075754 | 0 | 0 | 0 | 0.088635954 | 0.024398886 |
| B9D1 | 0 | 8.098531233 | 0 | 10.21755339 | 10.47215022 | 0 | 0 | 0 | 0.838607701 | 0.008009116 |
| B9D2 | 0 | 10.54568453 | 0 | 0 | 0 | 0 | 0 | 0 | 7.07E-08 | 1 |
| BAAT | 0 | 8.966633964 | 0 | 0 | 0 | 0 | 9.217798469 | 0 | 1.07E-07 | 1 |
| BACE2 | 0 | 0 | 0 | 10.14406389 | 8.204461778 | 0 | 8.531952193 | 0 | 0.046749858 | 0.999854694 |
| BAG5 | 9.21371058 | 8.127281865 | 0 | 0 | 0 | 0 | 0 | 0 | 0.019856263 | 1 |
| BAHCC1 | 0 | 0 | 9.230202718 | 0 | 8.542045106 | 0 | 0 | 0 | 0.072776828 | 0.016054013 |
| BAMBI | 0 | 0 | 0 | 10.3530686 | 9.601704371 | 10.97246827 | 10.57980117 | 0 | 0.021534056 | 0.999854694 |
| BANK1 | 0 | 10.05022532 | 0 | 0 | 0 | 10.55396144 | 0 | 0 | 7.07E-08 | 1 |
| BARX2 | 0 | 0 | 8.65401506 | 0 | 8.204461778 | 0 | 0 | 0 | 0.089278262 | 0.024814074 |
| BASP1 | 9.804008249 | 0 | 0 | 0 | 0 | 0 | 0 | 0 | 7.17E-08 | 1 |
| BATF2 | 9.840340458 | 0 | 0 | 0 | 0 | 0 | 0 | 0 | 7.17E-08 | 1 |
| BAZ2B | 0 | 9.984962099 | 0 | 0 | 0 | 0 | 0 | 8.644715457 | 7.07E-08 | 1 |
| BBIP1 | 11.88190873 | 0 | 0 | 0 | 0 | 0 | 8.745448055 | 0 | 5.99E-08 | 1 |
| BBOF1 | 9.118299335 | 0 | 0 | 0 | 0 | 0 | 0 | 0 | 9.99E-08 | 1 |
| BCAS2 | 0 | 9.227860829 | 10.72868413 | 10.57971922 | 9.400230015 | 0 | 0 | 0 | 0.726059685 | 0.000468734 |
| BCL2L1 | 0 | 0 | 0 | 10.8904559 | 8.745330337 | 10.27525208 | 9.853657569 | 0 | 0.017066863 | 0.999854694 |
| BCL2L14 | 6.584840871 | 9.311663645 | 0 | 0 | 0 | 0 | 0 | 0 | 0.026424156 | 1 |
| BCL2L15 | 0 | 11.90618414 | 10.91042177 | 0 | 8.839844964 | 0 | 0 | 0 | 0.963126669 | 0.008668791 |
| BCL2L2 | 10.16075706 | 0 | 0 | 0 | 0 | 0 | 0 | 0 | 7.17E-08 | 1 |
| BCL7C | 0 | 0 | 11.45910781 | 10.35417369 | 8.518703802 | 11.51537169 | 9.82911361 | 0 | 4.06E-05 | 0.999854694 |
| BCL9 | 10.90104575 | 7.945499326 | 0 | 0 | 0 | 0 | 11.38107391 | 0 | 0.001520628 | 1 |
| BCO1 | 0 | 9.267144757 | 0 | 0 | 0 | 0 | 0 | 0 | 8.18E-08 | 1 |
| BEST2 | 0 | 8.288701315 | 0 | 0 | 0 | 0 | 0 | 0 | 1.78E-07 | 1 |
| BEX3 | 0 | 0 | 10.18850215 | 0 | 10.16472422 | 0 | 0 | 0 | 0.016438003 | 0.008621456 |
| BEX4 | 10.80320108 | 0 | 9.401970879 | 0 | 9.234102159 | 0 | 0 | 0 | 0.978907185 | 0.012283188 |
| BIRC5 | 0 | 0 | 0 | 10.51300977 | 10.96260982 | 0 | 0 | 0 | 0.029164006 | 0.006244708 |
| BIVM | 0 | 0 | 0 | 9.039404442 | 7.158471365 | 0 | 0 | 0 | 0.090827893 | 0.028731957 |
| BIVM-ERCC5 | 10.10709428 | 0 | 0 | 0 | 0 | 10.51824547 | 0 | 0 | 7.07E-08 | 1 |
| BMP1 | 0 | 0 | 0 | 0 | 6.85193071 | 0 | 7.567155003 | 0 | 4.12E-05 | 1 |
| BMP5 | 14.24025106 | 12.16378841 | 13.6568283 | 10.09599117 | 0 | 0 | 0 | 0 | 0.932679569 | 3.43E-12 |
| BMP8B | 0 | 0 | 0 | 0 | 6.608339636 | 0 | 0 | 0 | 5.07E-05 | 1 |
| BNIP2 | 0 | 9.378959303 | 0 | 0 | 0 | 0 | 7.657204612 | 10.13337775 | 7.88E-08 | 0.010542114 |
| BNIP3 | 0 | 0 | 9.815631163 | 0 | 8.064100782 | 0 | 0 | 0 | 0.06347494 | 0.013733031 |
| BNIP3L | 0 | 0 | 10.33440339 | 0 | 8.498391931 | 10.32832134 | 0 | 0 | 0.038486768 | 0.999854694 |

| | | | | | | | | | | |
|---|---|---|---|---|---|---|---|---|---|---|
| BOC | 10.01473459 | 0 | 0 | 0 | 0 | 0 | 0 | 0 | 7.07E-08 | 1 |
| BOD1 | 0 | 0 | 0 | 10.09334644 | 8.858558259 | 0 | 0 | 10.35013078 | 0.039663385 | 0.999854694 |
| BORCS6 | 0 | 0 | 9.111438383 | 0 | 8.861211882 | 0 | 0 | 0 | 0.069590538 | 0.015141771 |
| BORCS8 | 0 | 0 | 0 | 10.22723596 | 6.489879427 | 0 | 0 | 0 | 0.05760153 | 0.01390329 |
| BPNT1 | 0 | 0 | 10.75929223 | 0 | 7.31377587 | 0 | 0 | 0 | 0.032736381 | 0.01037888 |
| BRAP | 0 | 0 | 0 | 0 | 6.39051047 | 0 | 0 | 0 | 5.98E-05 | 1 |
| BRCC3 | 0 | 0 | 10.35109977 | 0 | 7.250409752 | 9.654390426 | 0 | 0 | 0.049574671 | 0.999854694 |
| BRIP1 | 0 | 0 | 0 | 8.63622404 | 11.37792991 | 10.56965778 | 9.484261318 | 10.48794037 | 0.031432954 | 0.999854694 |
| BRMS1 | 0 | 0 | 0 | 7.518919157 | 10.471282 | 0 | 0 | 0 | 0.036529844 | 0.011034216 |
| BRPF1 | 9.974435204 | 0 | 0 | 0 | 0 | 0 | 8.333808086 | 0 | 7.07E-08 | 1 |
| BRPF3 | 0 | 0 | 0 | 9.605003615 | 9.687189778 | 0 | 0 | 0 | 0.036171464 | 0.010829604 |
| BRWD1-AS2 | 10.15114749 | 9.670244415 | 0 | 0 | 0 | 0 | 0 | 0 | 0.001949994 | 1 |
| BRWD3 | 0 | 9.092255983 | 0 | 0 | 0 | 0 | 0 | 0 | 1.16E-07 | 1 |
| BSDC1 | 9.25017571 | 8.596915356 | 9.576524594 | 0 | 9.275473597 | 0 | 0 | 0 | 0.999483812 | 0.011578584 |
| BST1 | 0 | 0 | 0 | 9.905608724 | 7.710937422 | 0 | 0 | 0 | 0.0622522 | 0.014264276 |
| BTBD2 | 0 | 0 | 10.28860724 | 8.217545222 | 0 | 0 | 0 | 0 | 0.046457532 | 0.011578584 |
| BTD | 8.811733678 | 9.414384561 | 10.00077521 | 7.080028634 | 9.028728141 | 0 | 0 | 0 | 0.999485949 | 0.002802269 |
| BTNL3 | 10.7340787 | 9.152938599 | 12.44684221 | 0 | 7.856352457 | 0 | 0 | 0 | 0.979247546 | 9.11E-12 |
| C10orf143 | 10.46217494 | 0 | 0 | 0 | 0 | 0 | 8.756808388 | 0 | 7.07E-08 | 1 |
| C11orf54 | 0 | 8.942570108 | 0 | 0 | 0 | 0 | 7.97770259 | 8.885916959 | 1.07E-07 | 0.017215256 |
| C11orf71 | 0 | 0 | 11.22363585 | 8.168132648 | 10.55725385 | 0 | 7.391717672 | 0 | 6.26E-05 | 0.613591685 |
| C11orf74 | 0 | 0 | 0 | 8.807360775 | 7.710937422 | 0 | 0 | 0 | 0.091379336 | 0.028721926 |
| C11orf80 | 9.85816911 | 0 | 0 | 0 | 0 | 0 | 0 | 0 | 7.17E-08 | 1 |
| C12orf29 | 0 | 0 | 5.284213006 | 10.64438356 | 10.17168901 | 0 | 0 | 0 | 0.001540288 | 0.001491158 |
| C12orf43 | 0 | 8.479177282 | 0 | 0 | 0 | 0 | 0 | 0 | 1.39E-07 | 1 |
| C12orf56 | 10.45438212 | 0 | 0 | 0 | 0 | 0 | 7.244547495 | 0 | 7.07E-08 | 1 |
| C15orf40 | 0 | 9.610406593 | 0 | 9.331906887 | 8.650337012 | 0 | 0 | 0 | 0.998873521 | 0.015215749 |
| C15orf41 | 9.497729097 | 0 | 0 | 0 | 0 | 0 | 9.22108302 | 0 | 7.87E-08 | 1 |
| C16orf87 | 0 | 0 | 0 | 0 | 0 | 0 | 9.003732502 | 10.05139871 | 1 | 0.009209122 |
| C1GALT1 | 0 | 0 | 9.109373744 | 0 | 7.396595249 | 0 | 0 | 0 | 0.088798079 | 0.024404781 |
| C1orf115 | 0 | 0 | 8.986519609 | 8.672114028 | 9.918172938 | 0 | 0 | 0 | 0.006061566 | 0.001805281 |
| C1orf131 | 0 | 0 | 10.37277835 | 0 | 7.31377587 | 0 | 0 | 0 | 0.048159116 | 0.011625223 |
| C1orf198 | 0 | 0 | 0 | 10.43473225 | 8.847894688 | 10.07244093 | 6.047527104 | 0 | 0.029719049 | 0.999854694 |
| C1orf35 | 0 | 9.402672528 | 10.36501154 | 10.45753398 | 0 | 0 | 0 | 0 | 0.961746291 | 0.008180039 |
| C1orf56 | 0 | 0 | 0 | 10.43473225 | 9.45483524 | 0 | 0 | 0 | 0.021732998 | 0.009470925 |
| C1QL1 | 0 | 0 | 0 | 0 | 6.694188464 | 0 | 0 | 0 | 0.000126205 | 1 |
| C1R | 10.45828379 | 9.857774611 | 11.03475328 | 0 | 7.374475525 | 0 | 0 | 0 | 0.975550004 | 0.009361372 |
| C2 | 0 | 0 | 0 | 10.73030464 | 6.15993909 | 0 | 0 | 0 | 0.035967833 | 2.04E-10 |
| C20orf204 | 0 | 9.051585356 | 0 | 0 | 0 | 0 | 0 | 0 | 1.18E-07 | 1 |
| C20orf27 | 0 | 0 | 0 | 8.197982807 | 10.21434362 | 0 | 0 | 0 | 0.04139993 | 0.011464572 |
| C21orf58 | 0 | 9.454643748 | 0 | 0 | 0 | 0 | 0 | 0 | 7.73E-08 | 1 |
| C2CD5 | 0 | 0 | 0 | 8.606855021 | 0 | 0 | 0 | 0 | 7.56E-06 | 1 |
| C2orf72 | 0 | 9.615488971 | 0 | 0 | 0 | 0 | 0 | 0 | 7.42E-08 | 1 |

| | | | | | | | | | | |
|---|---|---|---|---|---|---|---|---|---|---|
| C3orf62 | 8.529698369 | 10.51473649 | 0 | 0 | 0 | 0 | 0 | 0 | 0.002635642 | 1 |
| C3orf70 | 0 | 8.501311304 | 0 | 0 | 0 | 0 | 0 | 0 | 1.38E-07 | 1 |
| C4orf33 | 0 | 0 | 0 | 7.61055072 | 0 | 0 | 0 | 10.50678635 | 2.15E-05 | 1 |
| C6orf120 | 9.497729097 | 7.537775521 | 0 | 0 | 0 | 0 | 7.567155003 | 0 | 0.01936014 | 1 |
| C6orf132 | 10.58145439 | 9.214524828 | 0 | 0 | 0 | 0 | 8.37567389 | 0 | 0.001363749 | 1 |
| C6orf136 | 0 | 0 | 11.58517012 | 10.06257842 | 6.984408946 | 8.697106567 | 7.556793512 | 0 | 0.000105947 | 0.999854694 |
| C6orf203 | 0 | 0 | 0 | 9.286320151 | 9.846328303 | 0 | 10.28957992 | 0 | 0.038478525 | 0.999854694 |
| C6orf89 | 0 | 9.52655488 | 9.297034957 | 10.57593509 | 4.589393553 | 0 | 0 | 0 | 0.963126669 | 0.002663527 |
| C7orf13 | 9.728478045 | 0 | 0 | 0 | 0 | 0 | 6.991130623 | 0 | 7.27E-08 | 1 |
| C7orf50 | 11.04030387 | 9.741711995 | 13.43078312 | 13.29253514 | 13.48121451 | 12.61104776 | 12.68368211 | 12.53714952 | 0.000134196 | 0.999854694 |
| C8orf33 | 0 | 0 | 10.9032893 | 9.641668845 | 9.830166051 | 0 | 10.72972562 | 9.746216776 | 0.000161601 | 0.999854694 |
| C8orf37 | 0 | 0 | 9.957816655 | 0 | 6.39051047 | 0 | 0 | 0 | 0.07064749 | 0.015771922 |
| C8orf76 | 0 | 0 | 0 | 12.35803671 | 9.457468152 | 11.46491646 | 7.930328737 | 0 | 1.07E-07 | 0.999854694 |
| C9orf78 | 10.0616534 | 9.317912807 | 9.99631302 | 10.78623336 | 10.27729265 | 0 | 0 | 0 | 0.937966741 | 0.000418169 |
| CA12 | 10.25797206 | 9.152938599 | 0 | 0 | 0 | 10.68350699 | 9.642065528 | 0 | 0.002961748 | 0.006244708 |
| CA2 | 14.62377001 | 13.52471985 | 15.23480772 | 12.35085774 | 0 | 0 | 0 | 0 | 0.990582117 | 5.02E-13 |
| CA9 | 0 | 0 | 0 | 11.63076985 | 9.882044651 | 0 | 0 | 0 | 0.026043162 | 0.00558734 |
| CAB39L | 0 | 6.79884554 | 0 | 0 | 0 | 11.72737195 | 7.830642173 | 0 | 0.123472343 | 6.18E-12 |
| CACNA2D1 | 11.31014909 | 0 | 0 | 0 | 0 | 11.57834006 | 0 | 0 | 2.76E-08 | 1 |
| CALCOCO1 | 0 | 9.755590285 | 0 | 9.114366811 | 8.616182922 | 0 | 0 | 0 | 0.991384446 | 0.017127937 |
| CAMK2G | 9.427818601 | 0 | 0 | 0 | 0 | 0 | 0 | 0 | 7.88E-08 | 1 |
| CAMKK2 | 0 | 8.918098054 | 0 | 0 | 0 | 0 | 0 | 0 | 1.08E-07 | 1 |
| CAND1 | 0 | 0 | 0 | 8.628937645 | 6.8625646 | 0 | 0 | 0 | 0.108732536 | 0.044192116 |
| CAPN3 | 9.816220932 | 0 | 0 | 0 | 0 | 0 | 0 | 0 | 7.17E-08 | 1 |
| CARD10 | 11.19795379 | 10.05957862 | 0 | 0 | 0 | 0 | 0 | 0 | 4.51E-05 | 1 |
| CARD16 | 12.4269804 | 10.45077461 | 10.36327988 | 9.920605724 | 6.717806789 | 0 | 0 | 0 | 0.333712949 | 0.001521549 |
| CARD6 | 0 | 0 | 0 | 0 | 6.543732574 | 0 | 0 | 0 | 5.33E-05 | 1 |
| CARS2 | 0 | 0 | 10.65132648 | 11.48705164 | 8.166345699 | 9.76117775 | 0 | 0 | 2.11E-05 | 0.999854694 |
| CASC4 | 9.176299815 | 0 | 0 | 0 | 0 | 10.10702012 | 6.457101373 | 0 | 1.15E-07 | 0.011412104 |
| CASC8 | 0 | 0 | 0 | 8.977645149 | 8.842533199 | 0 | 0 | 0 | 0.071806411 | 0.016596849 |
| CASC9 | 0 | 9.292752085 | 0 | 0 | 0 | 0 | 7.704905507 | 0 | 8.08E-08 | 1 |
| CASKIN2 | 0 | 0 | 0 | 0 | 0 | 0 | 9.211206849 | 5.78007706 | 1 | 0.021057037 |
| CASP10 | 0 | 0 | 9.670543162 | 0 | 9.251400376 | 0 | 6.43464192 | 0 | 0.04606091 | 0.999854694 |
| CASP3 | 0 | 0 | 11.67864476 | 10.18686959 | 7.613573228 | 0 | 7.109284752 | 0 | 4.34E-05 | 0.475335351 |
| CASP4 | 0 | 0 | 9.490300275 | 9.701937764 | 7.329191883 | 0 | 11.19107306 | 0 | 0.010249298 | 1 |
| CASP8AP2 | 0 | 8.770988541 | 0 | 0 | 0 | 0 | 6.664334008 | 0 | 1.15E-07 | 1 |
| CASR | 9.754096665 | 0 | 0 | 0 | 0 | 0 | 0 | 0 | 7.17E-08 | 1 |
| CASTOR1 | 0 | 0 | 10.67664665 | 0 | 7.290338514 | 0 | 0 | 0 | 0.035912867 | 0.010542114 |
| CAV2 | 0 | 0 | 0 | 9.621618376 | 9.432705414 | 11.19832615 | 0 | 9.79083461 | 0.041903927 | 0.999854694 |
| CAVIN3 | 0 | 0 | 8.33863325 | 10.13637632 | 10.9132777 | 0 | 0 | 12.60055742 | 0.000254149 | 0.999854694 |
| CBR3-AS1 | 9.35434707 | 0 | 0 | 0 | 0 | 0 | 6.388646063 | 0 | 8.08E-08 | 1 |
| CBX7 | 9.303201411 | 0 | 0 | 0 | 0 | 10.5933384 | 0 | 0 | 8.36E-08 | 1 |
| CC2D1A | 0 | 8.098531233 | 9.039528139 | 0 | 8.839844964 | 0 | 0 | 0 | 0.963126669 | 0.015774309 |

| | | | | | | | | | | |
|---|---|---|---|---|---|---|---|---|---|---|
| CCDC107 | 0 | 3.886810325 | 11.77719924 | 10.34863975 | 10.35277423 | 0 | 0 | 0 | 0.000127551 | 0.000193605 |
| CCDC120 | 0 | 8.893203714 | 0 | 0 | 0 | 0 | 0 | 0 | 1.09E-07 | 1 |
| CCDC137 | 0 | 0 | 10.74739236 | 10.02698824 | 0 | 0 | 0 | 0 | 0.03605376 | 0.008085065 |
| CCDC14 | 9.078295019 | 9.088605668 | 0 | 8.251153865 | 8.739573023 | 0 | 0 | 0 | 0.960445231 | 0.023914707 |
| CCDC142 | 0 | 0 | 10.82807345 | 0 | 9.479224203 | 0 | 9.214506424 | 0 | 0.013810945 | 0.999854694 |
| CCDC144A | 9.451500517 | 0 | 0 | 0 | 0 | 0 | 0 | 0 | 7.88E-08 | 1 |
| CCDC146 | 0 | 0 | 0 | 0 | 6.330495883 | 0 | 0 | 0 | 6.26E-05 | 1 |
| CCDC178 | 0 | 8.64741294 | 0 | 0 | 0 | 0 | 0 | 0 | 1.23E-07 | 1 |
| CCDC18-AS1 | 0 | 8.456698379 | 0 | 0 | 0 | 0 | 0 | 0 | 1.42E-07 | 1 |
| CCDC186 | 10.76905382 | 8.842088059 | 0 | 0 | 0 | 0 | 10.24912729 | 0 | 0.001199611 | 1 |
| CCDC25 | 0 | 0 | 10.10599766 | 0 | 7.923909096 | 0 | 0 | 0 | 0.053844504 | 0.012259484 |
| CCDC43 | 0 | 9.605306248 | 0 | 0 | 0 | 0 | 9.49786567 | 0 | 7.44E-08 | 1 |
| CCDC47 | 9.419837447 | 0 | 0 | 0 | 0 | 0 | 8.000818649 | 0 | 7.91E-08 | 1 |
| CCDC85B | 0 | 0 | 10.5407642 | 0 | 8.577977952 | 12.31876398 | 9.570479476 | 0 | 0.031058569 | 0.999854694 |
| CCDC9 | 0 | 0 | 10.21178344 | 0 | 7.882711472 | 0 | 0 | 0 | 0.049865925 | 0.011703791 |
| CCDC91 | 9.411811895 | 5.581805241 | 0 | 0 | 0 | 0 | 7.723552846 | 0 | 0.028619301 | 1 |
| CCDC93 | 0 | 0 | 0 | 0 | 0 | 10.80393471 | 0 | 7.795712438 | 1 | 0.008549078 |
| CCL13 | 0 | 0 | 10.68637461 | 0 | 7.856352457 | 0 | 0 | 0 | 0.032276595 | 0.01024397 |
| CCL19 | 11.69944958 | 9.750979006 | 0 | 0 | 0 | 11.04559917 | 0 | 10.74083533 | 5.83E-06 | 0.003988377 |
| CCL21 | 10.6648577 | 11.12810433 | 11.62437457 | 10.28284497 | 0 | 0 | 0 | 0 | 0.988867264 | 0.005430957 |
| CCL23 | 0 | 8.64741294 | 0 | 0 | 0 | 0 | 0 | 0 | 1.23E-07 | 1 |
| CCL24 | 0 | 8.824638969 | 0 | 0 | 0 | 10.23875605 | 8.467037044 | 0 | 1.36E-07 | 0.009237175 |
| CCM2 | 0 | 0 | 0 | 10.14150592 | 9.51336555 | 0 | 8.653626659 | 0 | 0.028071845 | 0.999854694 |
| CCN3 | 0 | 9.088605668 | 0 | 0 | 0 | 0 | 0 | 0 | 1.16E-07 | 1 |
| CCNB1 | 0 | 0 | 0 | 12.56976624 | 10.63684157 | 0 | 9.743766328 | 11.71979316 | 8.08E-08 | 0.999854694 |
| CCND2-AS1 | 9.427818601 | 0 | 0 | 0 | 0 | 0 | 0 | 0 | 7.88E-08 | 1 |
| CCNO | 0 | 0 | 0 | 10.03527963 | 6.786426836 | 0 | 0 | 0 | 0.064713996 | 0.014818059 |
| CCNT2 | 8.544455536 | 0 | 0 | 0 | 0 | 0 | 0 | 0 | 1.39E-07 | 1 |
| CCPG1 | 18.31714969 | 18.55148204 | 17.34631687 | 15.89012426 | 16.64961299 | 16.510147 | 17.52571899 | 16.50971499 | 0.001703133 | 0.999854694 |
| CCR7 | 15.56292735 | 15.08381911 | 13.00093537 | 13.65462484 | 11.26848841 | 12.92173181 | 11.32226394 | 16.45468697 | 0.001334367 | 0.999854694 |
| CCSER2 | 0 | 0 | 10.58897743 | 0 | 6.419606803 | 0 | 0 | 0 | 0.043827245 | 0.011326554 |
| CCT6P1 | 0 | 8.934458793 | 0 | 0 | 0 | 0 | 0 | 0 | 1.07E-07 | 1 |
| CCZ1 | 6.116742619 | 0 | 11.40794985 | 9.376097171 | 10.75890512 | 11.20188538 | 0 | 10.70398116 | 0.039354497 | 0.999854694 |
| CCZ1B | 0 | 0 | 0 | 11.1862506 | 7.928977054 | 0 | 7.637673707 | 0 | 0.01361219 | 0.999854694 |
| CCZ1P-OR7E38P | 0 | 0 | 0 | 9.630766785 | 7.041677569 | 0 | 0 | 0 | 0.074522731 | 0.018718621 |
| CD177 | 13.67187142 | 11.51696193 | 13.69544831 | 0 | 9.273482767 | 0 | 0 | 0 | 1 | 3.43E-12 |
| CD27 | 0 | 8.363107485 | 0 | 0 | 0 | 0 | 0 | 0 | 1.55E-07 | 1 |
| CD302 | 10.35742919 | 0 | 0 | 0 | 0 | 0 | 0 | 10.09830559 | 7.07E-08 | 1 |
| CD33 | 8.847874263 | 0 | 0 | 0 | 0 | 0 | 0 | 0 | 1.15E-07 | 1 |
| CD37 | 16.77548994 | 17.1211184 | 15.554507 | 13.8145111 | 13.85791278 | 16.03887682 | 13.66402058 | 16.59547445 | 0.01175574 | 0.999854694 |
| CD3D | 10.61329832 | 0 | 0 | 0 | 0 | 0 | 0 | 0 | 7.07E-08 | 1 |
| CD3E | 10.39455423 | 0 | 10.08408707 | 0 | 5.864366253 | 0 | 0 | 0 | 0.980815574 | 0.015141771 |
| CD3G | 0 | 9.595051142 | 0 | 0 | 0 | 0 | 0 | 0 | 7.73E-08 | 1 |

| | | | | | | | | | | |
|---|---|---|---|---|---|---|---|---|---|---|
| CD53 | 10.2032252 | 0 | 0 | 0 | 0 | 0 | 0 | 0 | 7.07E-08 | 1 |
| CD55 | 0 | 0 | 0 | 8.474903094 | 10.45994733 | 10.65349502 | 8.765832683 | 11.85305589 | 0.030396424 | 0.999854694 |
| CD79A | 0 | 8.326384029 | 0 | 0 | 0 | 0 | 0 | 10.10715388 | 1.61E-07 | 1 |
| CD79B | 16.02298384 | 16.93665143 | 13.83004556 | 14.10052826 | 10.72907136 | 15.34360547 | 11.44883075 | 16.91595261 | 0.02149044 | 0.999854694 |
| CD8B | 10.43866903 | 0 | 0 | 0 | 0 | 0 | 0 | 0 | 7.07E-08 | 1 |
| CD96 | 9.48248357 | 0 | 0 | 0 | 0 | 0 | 0 | 0 | 7.87E-08 | 1 |
| CD99P1 | 0 | 0 | 0 | 9.796023637 | 7.893121693 | 0 | 0 | 0 | 0.064375435 | 0.01457526 |
| CDC45 | 0 | 0 | 0 | 10.10258185 | 7.31377587 | 0 | 0 | 0 | 0.05760153 | 0.013697851 |
| CDC6 | 10.07695994 | 12.16118405 | 12.12992665 | 14.20382895 | 14.47947925 | 13.89623389 | 13.49565477 | 14.60046152 | 0.044790293 | 0.999854694 |
| CDCA5 | 0 | 0 | 0 | 11.61287643 | 7.98358032 | 0 | 7.822015489 | 9.195666708 | 0.030438756 | 0.999854694 |
| CDCA7L | 0 | 9.187477203 | 9.315062926 | 9.694976597 | 6.706045958 | 0 | 0 | 0 | 0.975665177 | 0.003381284 |
| CDCA8 | 0 | 0 | 0 | 0 | 0 | 0 | 8.77480088 | 10.23859149 | 1 | 0.009118992 |
| CDH5 | 0 | 8.68657578 | 0 | 0 | 0 | 0 | 0 | 0 | 1.19E-07 | 1 |
| CDHR1 | 10.69823457 | 8.456698379 | 0 | 0 | 0 | 0 | 7.864642143 | 0 | 0.001969598 | 1 |
| CDK1 | 0 | 0 | 0 | 10.89501899 | 10.23954785 | 0 | 8.810126365 | 9.900768529 | 0.031881853 | 0.999854694 |
| CDK11A | 0 | 0 | 9.683091274 | 9.544329263 | 8.381257477 | 0 | 8.967034175 | 0 | 0.006502772 | 0.999854694 |
| CDK11B | 8.697540019 | 8.387080535 | 10.0575616 | 0 | 6.543732574 | 0 | 0 | 0 | 0.999660881 | 0.01453734 |
| CDK7 | 0 | 0 | 11.4300381 | 11.4549621 | 7.600927365 | 0 | 0 | 0 | 2.94E-06 | 0.000317935 |
| CDKN1C | 0 | 0 | 0 | 9.468798819 | 9.415656752 | 0 | 0 | 10.47158838 | 0.04602085 | 1 |
| CDKN2B-AS1 | 13.04239149 | 11.68592657 | 13.0726283 | 0 | 8.966010238 | 0 | 0 | 0 | 0.961825223 | 4.43E-12 |
| CDKN3 | 0 | 0 | 9.521689552 | 10.14150592 | 12.04583146 | 0 | 0 | 11.32778965 | 3.06E-07 | 0.999854694 |
| CDYL | 0 | 0 | 0 | 7.565462304 | 0 | 0 | 0 | 0 | 2.38E-05 | 1 |
| CEBPZ | 0 | 0 | 0 | 0 | 0 | 0 | 8.578790831 | 9.91293056 | 1 | 0.010278758 |
| CECR7 | 0 | 9.838343355 | 0 | 0 | 0 | 0 | 0 | 0 | 7.07E-08 | 1 |
| CELF1 | 8.799482788 | 9.073911341 | 0 | 0 | 0 | 0 | 0 | 0 | 0.01586022 | 1 |
| CELSR3 | 0 | 0 | 0 | 0 | 6.741044679 | 0 | 0 | 0 | 0.000123975 | 1 |
| CEMIP2 | 0 | 9.37296959 | 0 | 0 | 0 | 10.90565284 | 9.160785668 | 0 | 7.88E-08 | 0.006244708 |
| CENPA | 0 | 0 | 0 | 9.513007557 | 9.53346915 | 0 | 0 | 10.84500739 | 0.042013317 | 1 |
| CENPM | 0 | 0 | 0 | 10.39017482 | 8.002934593 | 0 | 0 | 10.35459622 | 0.039544447 | 0.999854694 |
| CENPN | 0 | 0 | 0 | 11.19737685 | 7.968892295 | 0 | 0 | 10.19443599 | 0.013148052 | 0.999854694 |
| CENPO | 0 | 9.640635714 | 0 | 0 | 0 | 0 | 0 | 0 | 7.38E-08 | 1 |
| CENPP | 14.46243049 | 13.67802707 | 14.52913252 | 16.24400965 | 16.509669 | 15.59172712 | 15.29271293 | 15.58091352 | 0.009644909 | 0.999854694 |
| CEP19 | 9.822288722 | 0 | 0 | 0 | 0 | 0 | 0 | 0 | 7.17E-08 | 1 |
| CERKL | 9.747734533 | 0 | 0 | 0 | 0 | 0 | 0 | 0 | 7.17E-08 | 1 |
| CERNA2 | 0 | 0 | 0 | 6.991896215 | 0 | 0 | 0 | 0 | 4.01E-05 | 1 |
| CETN2 | 0 | 0 | 6.265581877 | 7.928036185 | 9.609616765 | 11.05087162 | 8.309327743 | 11.34060202 | 0.027508034 | 0.999854694 |
| CFAP20 | 0 | 0 | 0 | 10.09731172 | 10.34994295 | 12.00439919 | 0 | 0 | 0.013847966 | 0.999854694 |
| CFAP65 | 0 | 0 | 0 | 0 | 6.633393544 | 0 | 0 | 0 | 4.95E-05 | 1 |
| CFD | 0 | 0 | 10.96852634 | 0 | 11.85965642 | 0 | 6.341135342 | 10.49873963 | 0.019179867 | 0.999854694 |
| CFTR | 9.804008249 | 0 | 0 | 0 | 0 | 8.966273901 | 0 | 0 | 7.17E-08 | 1 |
| CH17-125A10.2 | 0 | 0 | 7.87734654 | 0 | 0 | 0 | 0 | 0 | 1.77E-05 | 1 |
| CHAF1A | 11.16848659 | 0 | 11.19422911 | 0 | 8.995296112 | 0 | 0 | 0 | 0.996280769 | 0.007609416 |
| CHD5 | 0 | 0 | 0 | 8.425292327 | 0 | 0 | 0 | 0 | 9.22E-06 | 1 |

| | | | | | | | | | | |
|---|---|---|---|---|---|---|---|---|---|---|
| CHD9 | 9.185743846 | 0 | 0 | 0 | 0 | 0 | 0 | 0 | 1.15E-07 | 1 |
| CHEK2P2 | 0 | 0 | 0 | 0 | 6.375739247 | 0 | 0 | 0 | 6.06E-05 | 1 |
| CHFR | 10.39047609 | 0 | 0 | 0 | 0 | 0 | 0 | 0 | 7.07E-08 | 1 |
| CHGB | 10.54889168 | 15.61173147 | 11.45667375 | 0 | 9.283409559 | 0 | 0 | 0 | 0.340088361 | 0.006310119 |
| CHKB | 10.99670368 | 8.039251825 | 0 | 0 | 0 | 0 | 0 | 11.0958704 | 0.001071743 | 1 |
| CHMP4C | 0 | 0 | 0 | 10.96251287 | 7.693175133 | 0 | 11.16572843 | 0 | 0.021320809 | 0.999854694 |
| CHMP7 | 10.15596027 | 0 | 0 | 0 | 0 | 9.971796937 | 0 | 0 | 7.07E-08 | 1 |
| CHODL | 0 | 9.11037029 | 0 | 0 | 0 | 0 | 0 | 0 | 1.15E-07 | 1 |
| CHPF | 0 | 0 | 0 | 10.63713836 | 8.538733665 | 0 | 0 | 0 | 0.026684226 | 0.010003595 |
| CHRAC1 | 0 | 0 | 0 | 9.47693819 | 7.949072713 | 0 | 0 | 0 | 0.072776828 | 0.017225749 |
| CHRNB1 | 0 | 9.814233037 | 0 | 0 | 0 | 0 | 0 | 0 | 7.15E-08 | 1 |
| CHST6 | 0 | 9.471559609 | 0 | 0 | 0 | 0 | 0 | 0 | 7.73E-08 | 1 |
| CHST7 | 9.760430865 | 0 | 0 | 0 | 0 | 0 | 0 | 0 | 7.17E-08 | 1 |
| CHTF8 | 0 | 0 | 9.911171138 | 10.87821661 | 8.170630891 | 0 | 7.437617244 | 5.707457615 | 0.000590594 | 0.723933139 |
| CIAO3 | 10.07187578 | 9.280005236 | 0 | 0 | 0 | 0 | 0 | 0 | 0.003631747 | 1 |
| CIAPIN1 | 0 | 0 | 0 | 10.42003215 | 10.26281485 | 0 | 8.246245428 | 0 | 0.035967833 | 1 |
| CIITA | 10.52304704 | 8.501311304 | 0 | 0 | 0 | 9.388941079 | 10.21329168 | 0 | 0.002925422 | 0.007912372 |
| CINP | 0 | 0 | 11.0254764 | 11.3391843 | 10.67994058 | 0 | 9.543183784 | 0 | 1.27E-07 | 0.999854694 |
| CIPC | 0 | 8.824638969 | 0 | 0 | 0 | 0 | 0 | 0 | 1.36E-07 | 1 |
| CISD1 | 0 | 0 | 9.820687648 | 10.97406028 | 10.76918522 | 6.142650091 | 0 | 0 | 1.10E-05 | 0.17185335 |
| CIT | 0 | 0 | 0 | 9.389094555 | 7.367026271 | 0 | 0 | 0 | 0.078825993 | 0.021040057 |
| CIZ1 | 0 | 0 | 0 | 10.68359454 | 7.757261396 | 10.33913592 | 0 | 0 | 0.030796109 | 0.999854694 |
| CLASRP | 10.90676775 | 0 | 10.14179168 | 10.20413285 | 0 | 0 | 0 | 0 | 0.995827349 | 0.009118992 |
| CLCA1 | 10.72762748 | 11.19243856 | 13.75171331 | 9.872056084 | 0 | 0 | 0 | 0 | 0.947522648 | 8.28E-12 |
| CLCN3 | 0 | 10.59669041 | 0 | 0 | 0 | 0 | 9.467764926 | 0 | 7.07E-08 | 1 |
| CLDN10 | 0 | 11.09391831 | 0 | 0 | 0 | 13.33762189 | 0 | 0 | 4.69E-08 | 1 |
| CLDN12 | 8.710683817 | 10.1930549 | 0 | 0 | 0 | 0 | 0 | 0 | 0.004558524 | 1 |
| CLDN5 | 11.0298545 | 9.166853421 | 0 | 0 | 0 | 0 | 0 | 0 | 0.000364041 | 1 |
| CLDND1 | 0 | 0 | 0 | 9.619781713 | 9.786858034 | 0 | 9.001824109 | 0 | 0.033826141 | 0.999854694 |
| CLEC18C | 0 | 0 | 0 | 0 | 6.489879427 | 0 | 0 | 0 | 5.56E-05 | 1 |
| CLIC3 | 0 | 0 | 8.542136853 | 0 | 8.853236326 | 0 | 0 | 0 | 0.078817485 | 0.019130243 |
| CLIP2 | 0 | 0 | 0 | 10.70193971 | 6.20241426 | 0 | 8.038540912 | 0 | 0.036873615 | 0.999854694 |
| CLK2 | 0 | 0 | 0 | 9.513007557 | 8.785000775 | 0 | 0 | 0 | 0.059387286 | 0.013787886 |
| CLN6 | 0 | 9.145930534 | 0 | 0 | 0 | 0 | 7.109284752 | 0 | 1.07E-07 | 1 |
| CLSTN2 | 0 | 9.131811442 | 0 | 0 | 0 | 0 | 0 | 0 | 1.10E-07 | 1 |
| CLUH | 0 | 0 | 9.395207686 | 0 | 8.19607814 | 0 | 0 | 0 | 0.073814976 | 0.01630416 |
| CLYBL | 0 | 0 | 10.90269333 | 0 | 7.344444907 | 0 | 8.729391788 | 0 | 0.027486506 | 0.999854694 |
| CMAHP | 0 | 8.780069978 | 0 | 0 | 0 | 0 | 0 | 0 | 1.15E-07 | 1 |
| CMC4 | 0 | 0 | 10.85850903 | 10.59475724 | 8.796137674 | 0 | 0 | 0 | 0.000119569 | 0.000525098 |
| CMKLR1 | 9.760430865 | 0 | 0 | 0 | 0 | 0 | 0 | 0 | 7.17E-08 | 1 |
| CMTM3 | 0 | 0 | 0 | 0 | 6.633393544 | 0 | 9.442661178 | 0 | 4.95E-05 | 1 |
| CMTM4 | 8.894695111 | 0 | 9.090657822 | 0 | 7.675191427 | 0 | 0 | 0 | 0.999483812 | 0.022769437 |
| CMTM6 | 0 | 0 | 9.716028935 | 9.444101503 | 8.748200401 | 0 | 9.001824109 | 0 | 0.005531672 | 0.999854694 |

| | | | | | | | | | | |
|---|---|---|---|---|---|---|---|---|---|---|
| CNDP2 | 8.21613834 | 10.37488148 | 11.50065854 | 0 | 9.038139559 | 0 | 0 | 0 | 0.985757273 | 0.006428177 |
| CNEP1R1 | 0 | 3.886810325 | 0 | 9.701937764 | 6.283787608 | 0 | 0 | 0 | 0.410447612 | 0.019900018 |
| CNIH3 | 0 | 0 | 0 | 10.41580446 | 9.94593807 | 0 | 0 | 0 | 0.015107937 | 0.008668791 |
| CNOT2 | 9.259149778 | 0 | 8.158655085 | 0 | 8.277799698 | 0 | 0 | 0 | 0.98939231 | 0.031616492 |
| CNOT8 | 0 | 0 | 10.54306056 | 7.036635307 | 6.08018266 | 10.77873462 | 7.732786904 | 0 | 0.015019105 | 0.999854694 |
| CNPPD1 | 0 | 0 | 10.06290585 | 0 | 9.107951389 | 10.73763937 | 6.457101373 | 0 | 0.038169102 | 0.999854694 |
| CNPY3-GNMT | 10.81845934 | 0 | 0 | 0 | 0 | 0 | 0 | 0 | 7.07E-08 | 1 |
| CNPY4 | 0 | 0 | 10.21370669 | 0 | 7.502375789 | 0 | 0 | 0 | 0.053686747 | 0.012259484 |
| COA1 | 0 | 0 | 11.02218791 | 10.53364114 | 11.05010281 | 0 | 5.285415716 | 10.51478844 | 5.99E-07 | 0.999854694 |
| COA6 | 0 | 0 | 9.969246136 | 10.49107477 | 9.749389578 | 10.54931218 | 9.17778928 | 11.03039541 | 0.00028877 | 0.999854694 |
| COL17A1 | 11.72720201 | 8.301371897 | 0 | 0 | 0 | 0 | 0 | 0 | 3.66E-05 | 1 |
| COL18A1 | 0 | 0 | 9.884858354 | 10.27586416 | 9.390159382 | 0 | 0 | 10.22727289 | 0.00081351 | 0.999854694 |
| COL24A1 | 10.60276158 | 0 | 0 | 0 | 0 | 0 | 0 | 0 | 7.07E-08 | 1 |
| COL4A2 | 0 | 0 | 0 | 9.559738748 | 8.460394093 | 0 | 0 | 0 | 0.064284256 | 0.01453734 |
| COL4A5 | 9.427818601 | 0 | 0 | 0 | 0 | 0 | 0 | 0 | 7.88E-08 | 1 |
| COMT | 15.12987645 | 16.11785845 | 17.2851921 | 16.68821838 | 17.24830426 | 17.9759626 | 17.47133695 | 16.81558912 | 0.016730737 | 0.999854694 |
| COPA | 0 | 0 | 0 | 10.23565523 | 8.542045106 | 0 | 5.334765644 | 0 | 0.039395275 | 0.617752072 |
| COPG1 | 0 | 9.563841056 | 0 | 0 | 0 | 0 | 10.59253762 | 10.55931388 | 7.58E-08 | 0.005049454 |
| COPS7B | 9.86993364 | 8.867872257 | 0 | 9.487048249 | 10.35041522 | 0 | 0 | 0 | 0.999483812 | 0.009547202 |
| COQ3 | 0 | 0 | 0 | 11.17503787 | 8.233426702 | 0 | 0 | 0 | 0.036442685 | 0.008637628 |
| COQ7 | 10.72762748 | 0 | 0 | 9.691483378 | 8.336189561 | 0 | 0 | 0 | 0.975665177 | 0.01420877 |
| COQ8A | 0 | 0 | 10.58897743 | 0 | 7.959016539 | 10.22948594 | 0 | 9.092509649 | 0.034883265 | 0.999854694 |
| CORO1A | 10.89817622 | 0 | 0 | 9.859659028 | 7.481831074 | 0 | 0 | 0 | 0.963126669 | 0.015083909 |
| CORO1C | 0 | 7.993136997 | 0 | 6.848698922 | 9.297193309 | 0 | 0 | 0 | 0.975665177 | 0.024260309 |
| COX7A1 | 0 | 9.194286798 | 0 | 0 | 0 | 0 | 0 | 8.544047727 | 8.58E-08 | 1 |
| CPB1 | 0 | 9.836168082 | 0 | 0 | 0 | 0 | 0 | 0 | 7.07E-08 | 1 |
| CPEB2 | 0 | 0 | 0 | 0 | 6.503532343 | 0 | 0 | 0 | 5.50E-05 | 1 |
| CPEB3 | 0 | 0 | 0 | 0 | 6.476096071 | 0 | 0 | 0 | 5.63E-05 | 1 |
| CPLX1 | 0 | 7.146545342 | 0 | 9.726040692 | 10.38724397 | 0 | 0 | 0 | 0.648972745 | 0.009118992 |
| CPQ | 0 | 0 | 10.72194338 | 0 | 7.555772974 | 10.84927398 | 0 | 0 | 0.032791209 | 0.999854694 |
| CPSF1 | 7.954073896 | 7.201656162 | 10.36674111 | 0 | 8.273821715 | 0 | 0 | 0 | 0.930569319 | 0.010829604 |
| CPSF3 | 11.1359711 | 0 | 10.75269332 | 10.3574839 | 7.807657514 | 0 | 0 | 0 | 0.999483812 | 0.000779052 |
| CR1L | 10.02529365 | 8.918098054 | 0 | 0 | 0 | 0 | 0 | 0 | 0.005477696 | 1 |
| CR2 | 0 | 8.375143804 | 0 | 0 | 0 | 0 | 0 | 0 | 1.55E-07 | 1 |
| CRAT | 9.026677385 | 0 | 0 | 0 | 0 | 10.84927398 | 8.489001577 | 0 | 1.07E-07 | 0.007255458 |
| CRIPAK | 10.30639463 | 0 | 0 | 0 | 0 | 0 | 0 | 0 | 7.07E-08 | 1 |
| CRISPLD2 | 9.505291868 | 0 | 0 | 0 | 0 | 0 | 0 | 0 | 7.80E-08 | 1 |
| CRLF3 | 0 | 8.934458793 | 0 | 0 | 0 | 0 | 0 | 0 | 1.07E-07 | 1 |
| CRNDE | 0 | 0 | 0 | 11.44669744 | 8.320848539 | 10.44504772 | 8.505258271 | 0 | 0.032339976 | 0.999854694 |
| CROCCP2 | 10.11700079 | 0 | 0 | 0 | 0 | 0 | 0 | 0 | 7.07E-08 | 1 |
| CRTAP | 0 | 0 | 0 | 9.941343018 | 9.131224023 | 0 | 9.878831122 | 9.444951687 | 0.039670009 | 0.999854694 |
| CRTC3 | 0 | 0 | 0 | 10.371741 | 8.113905325 | 0 | 0 | 0 | 0.039224551 | 0.011034216 |
| CRY2 | 0 | 9.563841056 | 0 | 0 | 0 | 0 | 0 | 0 | 7.58E-08 | 1 |

| | | | | | | | | | | |
|---|---|---|---|---|---|---|---|---|---|---|
| CRYBB2P1 | 0 | 0 | 0 | 10.14916628 | 8.005335723 | 0 | 0 | 0 | 0.048589202 | 0.012259484 |
| CRYBB3 | 10.27576888 | 0 | 0 | 0 | 0 | 0 | 0 | 0 | 7.07E-08 | 1 |
| CRYM | 0 | 0 | 0 | 0 | 6.752524753 | 0 | 0 | 0 | 0.000121249 | 1 |
| CS | 0 | 8.576210795 | 0 | 9.850290823 | 7.00375218 | 0 | 0 | 0 | 0.980815574 | 0.01630416 |
| CSK | 9.578815964 | 0 | 0 | 8.373914626 | 7.728483681 | 0 | 0 | 0 | 0.976136384 | 0.038552483 |
| CSMD1 | 0 | 9.635641309 | 0 | 0 | 0 | 0 | 0 | 0 | 7.38E-08 | 1 |
| CSNK1D | 0 | 0 | 0 | 10.78377492 | 8.525411369 | 0 | 0 | 9.924990921 | 0.021928345 | 0.999854694 |
| CSNK2A1 | 0 | 0 | 9.702396179 | 9.914625618 | 7.188364856 | 0 | 0 | 10.86913607 | 0.006885203 | 1 |
| CSRNP3 | 0 | 8.326384029 | 0 | 0 | 0 | 0 | 0 | 10.40426152 | 1.61E-07 | 1 |
| CSRP2 | 10.85739169 | 0 | 0 | 0 | 0 | 0 | 0 | 0 | 7.07E-08 | 1 |
| CST7 | 0 | 9.173760825 | 0 | 0 | 0 | 0 | 0 | 0 | 8.78E-08 | 1 |
| CSTA | 0 | 10.95046004 | 0 | 0 | 0 | 0 | 8.096917163 | 0 | 6.72E-08 | 1 |
| CSTF1 | 0 | 8.842088059 | 10.07141569 | 0 | 7.192585236 | 0 | 0 | 0 | 0.983546955 | 0.013697851 |
| CSTF2 | 10.29771056 | 9.342642235 | 0 | 0 | 0 | 9.144535309 | 0 | 0 | 0.002234858 | 1 |
| CTB-49A3.2 | 0 | 8.288701315 | 0 | 0 | 0 | 0 | 0 | 0 | 1.78E-07 | 1 |
| CTNS | 11.01932888 | 8.998107186 | 0 | 0 | 0 | 0 | 8.422078578 | 9.070962627 | 0.000458407 | 0.013809342 |
| CTSV | 0 | 0 | 0 | 0 | 0 | 0 | 9.571766478 | 10.31237128 | 1 | 0.007595775 |
| CTSZ | 0 | 0 | 8.874275474 | 9.837704496 | 9.189398997 | 0 | 10.0455785 | 10.57725583 | 0.00523953 | 0.999854694 |
| CTTNBP2 | 0 | 9.254168606 | 0 | 0 | 0 | 0 | 7.493011414 | 0 | 8.22E-08 | 1 |
| CTU1 | 11.29277389 | 8.363107485 | 0 | 0 | 0 | 0 | 0 | 0 | 0.000279094 | 1 |
| CUEDC2 | 0 | 0 | 11.8740172 | 9.899565999 | 7.374475525 | 0 | 9.988901365 | 10.74310776 | 3.38E-05 | 0.999854694 |
| CUL2 | 0 | 0 | 9.979455819 | 0 | 8.714844832 | 10.51156867 | 0 | 0 | 0.047389518 | 0.999854694 |
| CUL4A | 0 | 0 | 0 | 10.28748014 | 8.375700381 | 11.80898701 | 6.62520069 | 11.04796419 | 0.039544447 | 0.999854694 |
| CUL5 | 0 | 8.676884284 | 0 | 0 | 0 | 0 | 0 | 0 | 1.20E-07 | 1 |
| CUL7 | 9.057869043 | 0 | 0 | 0 | 0 | 0 | 0 | 0 | 1.07E-07 | 1 |
| CUTC | 0 | 9.584722618 | 10.20985762 | 10.09994919 | 0 | 0 | 0 | 0 | 0.980815574 | 0.009160167 |
| CWC27 | 0 | 0 | 0 | 0 | 6.8625646 | 9.173827945 | 10.01562801 | 0 | 4.09E-05 | 0.999854694 |
| CWF19L1 | 0 | 8.64741294 | 0 | 0 | 0 | 10.67414176 | 9.987937624 | 0 | 1.23E-07 | 0.005709433 |
| CXCL1 | 0 | 0 | 0 | 9.253530096 | 10.29604626 | 0 | 3.567697193 | 0 | 0.026997065 | 0.132463079 |
| CXCL11 | 0 | 9.095897086 | 0 | 0 | 0 | 0 | 0 | 0 | 1.15E-07 | 1 |
| CXCL12 | 0 | 10.14113668 | 0 | 0 | 0 | 0 | 0 | 0 | 7.07E-08 | 1 |
| CXCL16 | 0 | 0 | 0 | 0 | 0 | 0 | 8.643870008 | 9.148443004 | 1 | 0.012440088 |
| CXCR4 | 18.17340101 | 18.52907423 | 16.53447064 | 15.33929012 | 15.78545686 | 18.52320843 | 15.64593057 | 18.60383993 | 1.11E-06 | 0.999854694 |
| CXCR5 | 12.01368829 | 11.21605215 | 0 | 0 | 0 | 0 | 0 | 12.22920972 | 3.38E-08 | 1 |
| CXorf56 | 0 | 9.650572952 | 0 | 8.365170934 | 8.050213926 | 0 | 0 | 0 | 0.976055111 | 0.033156588 |
| CXorf65 | 9.955236519 | 0 | 0 | 0 | 0 | 0 | 0 | 0 | 7.07E-08 | 1 |
| CYB561A3 | 0 | 0 | 10.32820306 | 0 | 8.040881201 | 0 | 8.919302823 | 0 | 0.043675612 | 0.999854694 |
| CYB561D2 | 10.20786763 | 0 | 10.89851466 | 9.941343018 | 9.35104796 | 0 | 0 | 0 | 0.939819551 | 0.000566119 |
| CYB5R1 | 0 | 9.095897086 | 0 | 0 | 0 | 0 | 0 | 0 | 1.15E-07 | 1 |
| CYB5R2 | 0 | 0 | 10.56205602 | 0 | 7.060272833 | 0 | 0 | 0 | 0.041866744 | 0.011034216 |
| CYB5R4 | 0 | 0 | 6.141754711 | 9.659657844 | 7.710937422 | 10.75712421 | 0 | 0 | 0.029923551 | 0.999854694 |
| CYBB | 0 | 0 | 0 | 0 | 6.830424858 | 0 | 0 | 8.458128915 | 4.17E-05 | 1 |
| CYBC1 | 0 | 0 | 10.78539014 | 0 | 9.666857493 | 0 | 0 | 0 | 0.012769246 | 0.008002596 |

| | | | | | | | | | |
|---|---|---|---|---|---|---|---|---|---|
| CYP21A1P | 8.514788691 | 0 | 0 | 0 | 0 | 0 | 0 | 0 | 1.43E-07 | 1 |
| CYP27A1 | 10.51182744 | 10.1054566 | 0 | 0 | 0 | 0 | 8.11821332 | 0 | 0.000403392 | 1 |
| CYP2D6 | 10.16075706 | 0 | 0 | 0 | 0 | 0 | 0 | 0 | 7.17E-08 | 1 |
| CYP2S1 | 0 | 0 | 0 | 9.641668845 | 10.27381129 | 0 | 6.62520069 | 0 | 0.021913665 | 0.929301027 |
| CYP3A5 | 8.140451703 | 10.03513275 | 12.7836445 | 0 | 8.095992883 | 0 | 0 | 0 | 0.869935364 | 1.40E-11 |
| CYP51A1 | 9.395625601 | 0 | 10.62699793 | 0 | 7.509159523 | 0 | 0 | 0 | 0.981160878 | 0.010542114 |
| CYP7B1 | 10.51182744 | 8.676884284 | 0 | 0 | 0 | 0 | 0 | 0 | 0.002639452 | 1 |
| CYREN | 0 | 0 | 9.657884952 | 8.227227848 | 0 | 0 | 0 | 0 | 0.069590538 | 0.015147164 |
| CYTH2 | 0 | 0 | 11.35403926 | 10.03252112 | 6.924772702 | 0 | 8.792571741 | 0 | 0.00025824 | 0.999854694 |
| CYTOR | 0 | 0 | 0 | 7.829771231 | 10.20810344 | 0 | 0 | 9.393801179 | 0.044978193 | 0.999854694 |
| CZIB | 0 | 9.330330507 | 11.57846901 | 12.08704795 | 8.581200627 | 0 | 0 | 0 | 0.536632258 | 0.000158964 |
| DAGLA | 8.670887633 | 0 | 0 | 0 | 0 | 0 | 7.592737761 | 0 | 1.28E-07 | 1 |
| DALRD3 | 9.303201411 | 0 | 0 | 0 | 0 | 0 | 0 | 0 | 8.36E-08 | 1 |
| DAND5 | 0 | 0 | 11.02984942 | 0 | 8.619321467 | 0 | 0 | 0 | 0.015956891 | 0.008551774 |
| DAPK1 | 0 | 0 | 0 | 0 | 6.476096071 | 0 | 0 | 0 | 5.63E-05 | 1 |
| DAPK2 | 9.635051191 | 7.449459114 | 0 | 0 | 0 | 0 | 6.928214841 | 0 | 0.017544661 | 1 |
| DCAF15 | 10.71463784 | 0 | 0 | 0 | 0 | 0 | 0 | 0 | 7.07E-08 | 1 |
| DCP2 | 0 | 9.684823965 | 0 | 0 | 0 | 0 | 0 | 0 | 7.17E-08 | 1 |
| DCTN1 | 0 | 0 | 11.264102 | 0 | 8.674678135 | 0 | 9.300856631 | 10.43128862 | 0.035359158 | 0.999854694 |
| DDA1 | 0 | 0 | 0 | 9.540451037 | 7.644711271 | 0 | 0 | 0 | 0.073459279 | 0.017687269 |
| DDAH1 | 0 | 0 | 9.432018923 | 0 | 9.603290326 | 10.31742507 | 0 | 0 | 0.042985444 | 0.999854694 |
| DDB1 | 0 | 0 | 10.94440682 | 10.59475724 | 7.613573228 | 10.57149323 | 7.741962234 | 0 | 0.000218668 | 0.999854694 |
| DDOST | 6.584840871 | 8.934458793 | 12.72501556 | 12.98623016 | 12.15920612 | 9.678434935 | 10.02693243 | 12.42414578 | 1.55E-05 | 0.999854694 |
| DDX11 | 0 | 0 | 0 | 9.656077944 | 8.399628042 | 0 | 0 | 0 | 0.062244365 | 0.01420877 |
| DDX28 | 0 | 7.537775521 | 0 | 10.18562856 | 8.736685727 | 0 | 0 | 0 | 0.825192631 | 0.011034216 |
| DDX3Y | 0 | 7.961553963 | 0 | 9.055798949 | 9.417460865 | 0 | 0 | 0 | 0.930207109 | 0.013215896 |
| DDX49 | 0 | 0 | 0 | 9.67388927 | 7.367026271 | 0 | 0 | 0 | 0.072286065 | 0.017226216 |
| DDX56 | 0 | 8.398919312 | 9.790079432 | 0 | 8.104976904 | 0 | 0 | 0 | 0.968436438 | 0.013787886 |
| DDX58 | 0 | 9.595051142 | 0 | 0 | 0 | 0 | 0 | 0 | 7.73E-08 | 1 |
| DEF8 | 9.834348275 | 9.390864636 | 11.51047079 | 0 | 8.113905325 | 0 | 0 | 0 | 0.98939231 | 7.83E-11 |
| DEFA5 | 0 | 0 | 0 | 8.458554996 | 0 | 0 | 0 | 0 | 1.20E-05 | 1 |
| DENND4A | 0 | 0 | 0 | 0 | 6.786426836 | 0 | 0 | 9.399574953 | 4.30E-05 | 1 |
| DEPP1 | 0 | 0 | 10.85111457 | 0 | 9.126819988 | 10.88513712 | 9.499219076 | 0 | 0.016576946 | 0.999854694 |
| DEUP1 | 0 | 0 | 0 | 10.8728291 | 6.934884845 | 0 | 0 | 0 | 0.027910066 | 0.01037888 |
| DFFA | 0 | 0 | 0 | 10.52777651 | 9.345373041 | 0 | 8.542492985 | 10.82046829 | 0.020815667 | 0.999854694 |
| DGCR11 | 0 | 0 | 0 | 0 | 6.543732574 | 0 | 0 | 0 | 5.33E-05 | 1 |
| DGCR5 | 0 | 0 | 0 | 0 | 6.85193071 | 0 | 0 | 0 | 4.12E-05 | 1 |
| DGKA | 0 | 0 | 10.08828625 | 8.679186115 | 0 | 0 | 0 | 0 | 0.047848441 | 0.011671668 |
| DGKE | 10.28458564 | 0 | 0 | 0 | 0 | 0 | 0 | 0 | 7.07E-08 | 1 |
| DHFR | 9.108401769 | 0 | 0 | 0 | 0 | 10.93017883 | 5.428672801 | 0 | 1.03E-07 | 1 |
| DHX16 | 0 | 8.250007882 | 0 | 10.63985958 | 8.905592331 | 0 | 0 | 0 | 0.854032916 | 0.009634076 |
| DHX32 | 11.21188707 | 0 | 10.37191743 | 0 | 8.109448021 | 0 | 0 | 0 | 0.971036351 | 0.010943033 |
| DICER1 | 15.04704788 | 14.65393641 | 12.47517784 | 12.95924966 | 13.66751596 | 12.90144508 | 12.62723829 | 12.45184868 | 0.018508422 | 0.999854694 |

| | | | | | | | | | |
|---|---|---|---|---|---|---|---|---|---|
| DICER1-AS1 | 12.35764902 | 0 | 0 | 0 | 0 | 0 | 0 | 0 | 4.41E-08 | 1 |
| DIEXF | 0 | 0 | 0 | 0 | 6.830424858 | 0 | 0 | 0 | 4.17E-05 | 1 |
| DIPK1A | 0 | 0 | 10.86219209 | 0 | 6.8625646 | 0 | 8.67773346 | 0 | 0.030946452 | 0.999854694 |
| DKK3 | 0 | 0 | 10.59415432 | 9.970460016 | 8.707490434 | 0 | 0 | 0 | 0.000774342 | 0.000795451 |
| DKK4 | 0 | 0 | 0 | 0 | 0 | 10.92444541 | 8.439102608 | 10.40426152 | 1 | 0.000336322 |
| DLAT | 0 | 0 | 0 | 0 | 6.569923599 | 0 | 0 | 0 | 5.22E-05 | 1 |
| DLG1 | 0 | 6.434870687 | 0 | 8.617938583 | 6.556887521 | 0 | 0 | 0 | 0.925765556 | 0.048671455 |
| DMAC2 | 0 | 0 | 0 | 11.70670706 | 7.141107086 | 0 | 0 | 10.1709972 | 1.94E-07 | 1 |
| DMAP1 | 9.702396267 | 8.734080111 | 10.68221351 | 10.02421381 | 6.85193071 | 0 | 0 | 0 | 0.975665177 | 0.00116937 |
| DMBT1 | 0 | 0 | 9.530974688 | 11.85771495 | 8.064100782 | 0 | 0 | 0 | 2.95E-05 | 0.000503943 |
| DMKN | 0 | 0 | 9.955519852 | 0 | 11.59194799 | 0 | 0 | 0 | 0.025644743 | 0.005387024 |
| DNAH9 | 11.18388462 | 0 | 0 | 0 | 0 | 0 | 0 | 0 | 4.18E-08 | 1 |
| DNAJA3 | 0 | 0 | 10.41434611 | 9.917618769 | 0 | 0 | 10.97365997 | 0 | 0.01941848 | 0.999854694 |
| DNAJB9 | 0 | 0 | 10.03162994 | 8.994752481 | 0 | 0 | 9.955767518 | 10.99078283 | 0.044733564 | 0.999854694 |
| DNAJC21 | 0 | 8.288701315 | 10.89012082 | 8.487043733 | 9.499805811 | 0 | 0 | 0 | 0.657500903 | 0.000835188 |
| DNAJC25 | 0 | 9.280005236 | 0 | 0 | 0 | 0 | 0 | 9.91293056 | 8.10E-08 | 1 |
| DNAJC25-GNG10 | 9.147591029 | 0 | 0 | 0 | 0 | 0 | 6.047527104 | 0 | 1.17E-07 | 1 |
| DNAJC28 | 0 | 9.504807688 | 0 | 0 | 0 | 0 | 0 | 0 | 7.60E-08 | 1 |
| DNAJC9 | 0 | 8.982456399 | 11.58813843 | 11.58725887 | 9.822697405 | 0 | 0 | 0 | 0.425877135 | 0.000148447 |
| DNALI1 | 0 | 9.201064401 | 0 | 0 | 0 | 0 | 0 | 0 | 8.49E-08 | 1 |
| DNASE2 | 0 | 0 | 0 | 10.13122842 | 8.590825685 | 11.52108151 | 0 | 10.89184409 | 0.042080155 | 0.999854694 |
| DNHD1 | 8.616058634 | 0 | 0 | 0 | 0 | 0 | 0 | 0 | 1.33E-07 | 1 |
| DNMBP | 0 | 0 | 0 | 10.38369563 | 6.994112981 | 0 | 0 | 10.51744597 | 0.047315079 | 0.999854694 |
| DNTTIP2 | 11.07120542 | 0 | 0 | 0 | 0 | 11.16831935 | 7.889626398 | 0 | 6.01E-08 | 0.006569304 |
| DOCK1 | 14.40975857 | 12.41399259 | 12.16712541 | 10.70367476 | 11.28117679 | 11.99758925 | 13.09739938 | 0 | 0.024417503 | 0.999854694 |
| DOCK8 | 14.29840511 | 13.90447548 | 11.3444162 | 10.58913633 | 10.20601737 | 12.55324152 | 12.19994905 | 10.96473097 | 2.20E-06 | 0.999854694 |
| DOCK9 | 9.016128482 | 0 | 0 | 0 | 0 | 0 | 0 | 0 | 1.07E-07 | 1 |
| DOK1 | 0 | 9.13888826 | 0 | 0 | 0 | 0 | 0 | 0 | 1.09E-07 | 1 |
| DOK2 | 10.6202803 | 0 | 0 | 0 | 0 | 0 | 0 | 0 | 7.07E-08 | 1 |
| DPAGT1 | 10.05651492 | 0 | 9.51234427 | 9.93249208 | 8.40327425 | 0 | 0 | 0 | 0.989756518 | 0.001559115 |
| DPH2 | 0 | 0 | 10.39243898 | 9.656077944 | 0 | 10.59153055 | 0 | 0 | 0.024220922 | 0.999854694 |
| DPH5 | 10.42278292 | 0 | 0 | 0 | 0 | 0 | 8.66331777 | 10.32607883 | 7.07E-08 | 0.008991934 |
| DPP4 | 0 | 0 | 9.57353227 | 0 | 9.772128228 | 0 | 0 | 0 | 0.035892743 | 0.010542114 |
| DPP8 | 0 | 0 | 0 | 0 | 6.569923599 | 6.327784716 | 0 | 10.58234138 | 5.22E-05 | 0.999854694 |
| DPT | 0 | 9.460304435 | 0 | 0 | 0 | 0 | 0 | 0 | 7.73E-08 | 1 |
| DPYSL3 | 0 | 9.477154439 | 0 | 0 | 0 | 0 | 0 | 0 | 7.73E-08 | 1 |
| DRAM2 | 0 | 10.89302053 | 0 | 11.47135073 | 7.928977054 | 0 | 0 | 0 | 0.999483812 | 5.22E-11 |
| DRG2 | 0 | 8.098531233 | 0 | 0 | 0 | 0 | 7.67647464 | 11.5235785 | 0.069224624 | 5.81E-12 |
| DROSHA | 8.559063282 | 0 | 0 | 0 | 0 | 0 | 0 | 10.81075125 | 1.38E-07 | 1 |
| DSC2 | 8.5735246 | 9.655516021 | 9.480748754 | 0 | 9.072895476 | 0 | 0 | 0 | 0.987185781 | 0.012440088 |
| DSCC1 | 0 | 0 | 0 | 10.28052178 | 9.31472394 | 0 | 0 | 0 | 0.027792408 | 0.010003595 |
| DST | 0 | 8.141445139 | 10.05327189 | 0 | 7.740063742 | 0 | 0 | 0 | 0.945194724 | 0.012777646 |
| DSTYK | 0 | 0 | 11.07604085 | 0 | 8.625578148 | 0 | 8.450341371 | 0 | 0.014705735 | 0.999854694 |

| | | | | | | | | | |
|---|---|---|---|---|---|---|---|---|---|
| DTX3L | 10.19389528 | 0 | 9.941661716 | 0 | 8.012515226 | 0 | 0 | 0 | 0.989756518 | 0.013007083 |
| DTYMK | 0 | 0 | 11.54066299 | 0 | 10.52330983 | 11.12398272 | 10.60516261 | 9.034319984 | 0.024898456 | 0.999854694 |
| DUOXA2 | 0 | 0 | 0 | 8.954515171 | 9.382790877 | 0 | 0 | 0 | 0.058823139 | 0.013775418 |
| DUSP1 | 13.39870094 | 11.22272851 | 10.33705254 | 9.75310266 | 10.37708196 | 10.66213377 | 0 | 11.75061455 | 0.047910271 | 0.999854694 |
| DUSP11 | 0 | 6.388120724 | 9.061052812 | 10.34863975 | 0 | 0 | 0 | 0 | 0.60839395 | 0.010541 |
| DUSP26 | 13.25917902 | 0 | 11.94924456 | 0 | 7.949072713 | 0 | 0 | 0 | 0.932679569 | 1.40E-11 |
| DUSP6 | 8.159748856 | 8.183119329 | 0 | 0 | 0 | 9.933774742 | 8.111149473 | 0 | 0.03230555 | 0.010547287 |
| DVL1 | 0 | 0 | 0 | 9.439943864 | 7.66307653 | 0 | 0 | 0 | 0.075280036 | 0.018766577 |
| DVL3 | 0 | 0 | 0 | 9.536562357 | 7.446926049 | 0 | 0 | 0 | 0.074720419 | 0.018560008 |
| DXO | 0 | 0 | 0 | 10.32290658 | 7.908597519 | 0 | 5.987074578 | 0 | 0.042806302 | 0.868809265 |
| DYNC2H1 | 0 | 0 | 11.02930352 | 0 | 10.52330983 | 0 | 9.777621415 | 11.48472455 | 0.029493366 | 0.999854694 |
| DYNC2LI1 | 0 | 8.780069978 | 0 | 0 | 0 | 0 | 8.284424824 | 0 | 1.15E-07 | 1 |
| DZIP3 | 10.80626567 | 0 | 0 | 0 | 0 | 0 | 6.479216537 | 0 | 7.07E-08 | 1 |
| EBF4 | 8.811733678 | 0 | 0 | 0 | 0 | 0 | 0 | 0 | 1.17E-07 | 1 |
| EBI3 | 10.42278292 | 0 | 0 | 0 | 0 | 0 | 0 | 0 | 7.07E-08 | 1 |
| ECI2 | 0 | 0 | 0 | 9.834540682 | 7.041677569 | 0 | 0 | 0 | 0.070017371 | 0.016321748 |
| ECM2 | 8.62996247 | 0 | 0 | 0 | 0 | 0 | 0 | 0 | 1.33E-07 | 1 |
| EDDM13 | 0 | 0 | 7.75589729 | 0 | 0 | 0 | 0 | 0 | 2.01E-05 | 1 |
| EDN2 | 10.43866903 | 0 | 0 | 0 | 0 | 0 | 0 | 0 | 7.07E-08 | 1 |
| EDRF1 | 10.08202624 | 8.288701315 | 0 | 0 | 0 | 0 | 0 | 0 | 0.00751471 | 1 |
| EEF1AKMT1 | 0 | 0 | 10.58749491 | 11.29267827 | 10.32804992 | 0 | 9.201262603 | 10.36938198 | 1.92E-06 | 0.999854694 |
| EEF1AKMT2 | 0 | 0 | 10.93102562 | 0 | 8.983165828 | 0 | 7.704905507 | 0 | 0.015871109 | 0.999854694 |
| EEF1AKMT4 | 0 | 0 | 0 | 11.36243624 | 7.588169674 | 10.66643382 | 0 | 0 | 0.034710928 | 0.999854694 |
| EEF1B2 | 13.20911722 | 12.89140423 | 14.15661659 | 14.84509602 | 14.25526564 | 13.29298775 | 13.71135434 | 14.70826752 | 0.033963012 | 0.999854694 |
| EEF1G | 13.33964361 | 14.34221351 | 14.81590059 | 15.68571037 | 15.44250598 | 14.98652991 | 15.63047804 | 15.51096032 | 0.034883265 | 0.999854694 |
| EFCAB11 | 0 | 0 | 0 | 10.28748014 | 7.799379467 | 0 | 0 | 10.38692716 | 0.045270943 | 0.999854694 |
| EFCAB5 | 11.27739703 | 0 | 11.56910998 | 0 | 8.992878201 | 0 | 0 | 0 | 0.999483812 | 0.00624926 |
| EFEMP1 | 0 | 0 | 10.5407642 | 10.03803287 | 9.933383758 | 12.11298682 | 0 | 11.07617755 | 0.000180407 | 0.999854694 |
| EFNA3 | 0 | 0 | 0 | 0 | 0 | 9.989752583 | 9.343675448 | 0 | 1 | 0.008668791 |
| EFNA5 | 0 | 0 | 0 | 0 | 6.729472521 | 0 | 0 | 0 | 0.000125541 | 1 |
| EFR3A | 0 | 8.876365592 | 0 | 0 | 0 | 0 | 0 | 0 | 1.10E-07 | 1 |
| EGLN2 | 9.662367605 | 8.083938166 | 0 | 0 | 0 | 10.85531342 | 0 | 0 | 0.013963785 | 1 |
| EGLN3 | 0 | 10.87087711 | 10.2922521 | 10.00745431 | 9.699104696 | 0 | 0 | 0 | 0.997434996 | 0.000646553 |
| EGR1 | 10.46605563 | 0 | 0 | 0 | 0 | 0 | 0 | 0 | 7.07E-08 | 1 |
| EHD1 | 0 | 0 | 0 | 0 | 0 | 10.59424148 | 8.271810291 | 8.615245757 | 1 | 0.000877642 |
| EID2 | 0 | 0 | 10.0724759 | 0 | 7.418380944 | 0 | 0 | 0 | 0.060476799 | 0.013309664 |
| EIF1AD | 0 | 0 | 10.16483899 | 10.37827398 | 8.03148771 | 0 | 0 | 10.63100807 | 0.001334033 | 0.999854694 |
| EIF2AK2 | 0 | 0 | 0 | 0 | 6.8625646 | 0 | 0 | 0 | 4.09E-05 | 1 |
| EIF2AK3 | 0 | 9.420205115 | 0 | 0 | 0 | 0 | 0 | 0 | 7.87E-08 | 1 |
| EIF2B1 | 0 | 0 | 8.814699123 | 0 | 8.91583933 | 0 | 0 | 0 | 0.074178145 | 0.016683828 |
| EIF2D | 10.12683973 | 0 | 0 | 0 | 0 | 0 | 4.589263136 | 0 | 7.07E-08 | 0.68580637 |
| EIF2S2 | 0 | 0 | 9.119667514 | 12.93987451 | 11.88285762 | 11.12772993 | 11.3685398 | 13.26033324 | 6.71E-08 | 0.999854694 |
| EIF3J-DT | 0 | 0 | 0 | 0 | 6.39051047 | 0 | 0 | 0 | 5.98E-05 | 1 |

| | | | | | | | | | | |
|---|---|---|---|---|---|---|---|---|---|---|
| EIF4A1 | 0 | 0 | 9.540200447 | 11.21937519 | 7.824072462 | 0 | 0 | 0 | 0.000426311 | 0.000795451 |
| ELF3-AS1 | 0 | 9.214524828 | 8.427492135 | 8.734548411 | 0 | 0 | 0 | 0 | 0.996280769 | 0.023752146 |
| ELF4 | 0 | 9.862057356 | 0 | 0 | 0 | 0 | 0 | 0 | 7.07E-08 | 1 |
| ELL2 | 0 | 0 | 10.32642663 | 0 | 6.944926602 | 0 | 0 | 0 | 0.053003076 | 0.012259484 |
| ELMO2 | 0 | 8.918098054 | 0 | 0 | 0 | 0 | 0 | 0 | 1.08E-07 | 1 |
| ELMO3 | 0 | 0 | 10.79890486 | 0 | 8.951141308 | 0 | 0 | 0 | 0.019573795 | 0.008954152 |
| ELMSAN1 | 0 | 5.405243681 | 9.243442636 | 0 | 9.007325253 | 0 | 0 | 0 | 0.591073813 | 0.013910154 |
| ELOVL5 | 9.395625601 | 0 | 0 | 0 | 0 | 0 | 0 | 0 | 8.04E-08 | 1 |
| ELP2 | 0 | 0 | 10.77435616 | 0 | 7.613573228 | 0 | 8.146127844 | 0 | 0.030665635 | 0.999854694 |
| ELP3 | 10.35324459 | 0 | 0 | 0 | 0 | 0 | 0 | 0 | 7.07E-08 | 1 |
| ELP6 | 0 | 0 | 10.29407108 | 0 | 10.37383352 | 0 | 0 | 0 | 0.011496749 | 0.007912009 |
| EMC9 | 0 | 0 | 8.789130733 | 10.11827772 | 9.42823842 | 0 | 0 | 12.06089185 | 0.002974244 | 0.999854694 |
| EMCN | 9.427818601 | 0 | 0 | 0 | 0 | 0 | 0 | 0 | 7.88E-08 | 1 |
| EML5 | 0 | 8.565746011 | 0 | 0 | 0 | 0 | 0 | 0 | 1.33E-07 | 1 |
| EML6 | 10.34904781 | 10.9842501 | 0 | 0 | 4.73539137 | 12.19415254 | 9.634695257 | 0 | 0.044784409 | 0.110469387 |
| EMP1 | 9.852250672 | 9.718280646 | 11.62473598 | 0 | 9.346320413 | 0 | 0 | 0 | 0.981170768 | 0.005907036 |
| EMX1 | 0 | 0 | 9.164107372 | 0 | 6.944926602 | 0 | 0 | 0 | 0.091190689 | 0.026198065 |
| ENAH | 10.04618272 | 8.696202607 | 0 | 0 | 0 | 0 | 7.80460581 | 0 | 0.006248654 | 1 |
| ENTPD4 | 0 | 10.48449957 | 0 | 0 | 0 | 0 | 0 | 0 | 7.07E-08 | 1 |
| ENTPD8 | 9.773016437 | 7.472050216 | 0 | 0 | 0 | 0 | 0 | 0 | 0.015258864 | 1 |
| EPB41L3 | 10.1892076 | 0 | 0 | 0 | 0 | 0 | 0 | 0 | 7.07E-08 | 1 |
| EPHA2 | 9.564407853 | 0 | 0 | 0 | 0 | 0 | 0 | 8.366767416 | 7.60E-08 | 1 |
| EPHB3 | 0 | 0 | 0 | 9.272358426 | 8.322775112 | 0 | 0 | 0 | 0.073373752 | 0.017341809 |
| EPN1 | 9.447580428 | 9.040291451 | 9.461453882 | 3.703427965 | 10.19027495 | 0 | 0 | 0 | 0.998873521 | 0.003658514 |
| EPRS | 10.38228506 | 9.396780662 | 0 | 0 | 0 | 0 | 0 | 10.10538856 | 0.001754985 | 1 |
| EPS8L1 | 0 | 0 | 0 | 9.464711849 | 7.607264152 | 0 | 0 | 0 | 0.075193869 | 0.018753555 |
| ERC1 | 15.09411976 | 14.21279928 | 13.11659179 | 13.04711831 | 13.52590536 | 13.63866356 | 13.5344557 | 13.91933496 | 0.040416394 | 0.999854694 |
| ERCC8 | 0 | 0 | 0 | 10.65518359 | 8.064100782 | 0 | 0 | 0 | 0.029866903 | 0.01037888 |
| ERICH1 | 10.4006499 | 0 | 0 | 0 | 0 | 0 | 0 | 0 | 9.27E-08 | 1 |
| ERICH2 | 9.86993364 | 0 | 0 | 0 | 0 | 0 | 6.739553261 | 0 | 7.15E-08 | 1 |
| ERVK3-1 | 0 | 10.25624448 | 0 | 10.48705051 | 8.109448021 | 0 | 0 | 0 | 0.999483812 | 0.010829604 |
| ESM1 | 0 | 0 | 0 | 0 | 6.582842939 | 0 | 0 | 0 | 0.000136511 | 1 |
| ESRP1 | 9.520299506 | 9.38492425 | 0 | 0 | 0 | 0 | 0 | 9.968369199 | 0.006858453 | 1 |
| ETAA1 | 7.292659623 | 8.069195977 | 0 | 0 | 0 | 0 | 0 | 0 | 0.042080155 | 1 |
| ETF1 | 9.621196547 | 0 | 0 | 0 | 0 | 0 | 7.257401605 | 0 | 8.05E-08 | 1 |
| ETFDH | 0 | 0 | 0 | 0 | 6.39051047 | 0 | 0 | 0 | 5.98E-05 | 1 |
| EVI2B | 0 | 9.431776243 | 0 | 0 | 0 | 9.202537636 | 0 | 0 | 7.88E-08 | 1 |
| EVI5L | 0 | 8.705765621 | 0 | 0 | 0 | 0 | 0 | 0 | 1.18E-07 | 1 |
| EXD3 | 9.955236519 | 8.850734095 | 0 | 0 | 0 | 0 | 0 | 9.497045998 | 0.006398527 | 1 |
| EXO5 | 0 | 9.124699738 | 0 | 0 | 0 | 0 | 0 | 0 | 1.13E-07 | 1 |
| EXOSC4 | 0 | 0 | 10.73271351 | 11.85107647 | 11.06110842 | 0 | 8.309327743 | 0 | 7.07E-08 | 0.645223676 |
| EXPH5 | 9.294498035 | 8.350969903 | 0 | 0 | 0 | 0 | 0 | 0 | 0.017061299 | 1 |
| F12 | 0 | 0 | 0 | 10.43054748 | 9.135614655 | 0 | 0 | 0 | 0.026410201 | 0.009927907 |

| | | | | | | | | | | |
|---|---|---|---|---|---|---|---|---|---|---|
| F2RL1 | 9.734925475 | 9.941103266 | 0 | 9.362981627 | 8.753923456 | 0 | 0 | 0 | 0.931652833 | 0.014577658 |
| F8A1 | 0 | 0 | 11.35839225 | 0 | 9.840960978 | 0 | 0 | 9.444951687 | 0.030003792 | 0.999854694 |
| FA2H | 10.17978629 | 8.398919312 | 0 | 0 | 0 | 0 | 0 | 0 | 0.006097458 | 1 |
| FAAP24 | 0 | 0 | 0 | 0 | 0 | 0 | 6.076829244 | 10.34265753 | 1 | 0.010829604 |
| FABP3 | 0 | 0 | 9.823209261 | 0 | 9.195720423 | 0 | 0 | 0 | 0.043083237 | 0.011034216 |
| FADS3 | 0 | 0 | 0 | 9.792766979 | 5.027671069 | 0 | 0 | 0 | 0.075053315 | 0.02109466 |
| FAHD2B | 9.451500517 | 0 | 7.285730034 | 0 | 9.033441525 | 0 | 0 | 0 | 0.990582117 | 0.026344013 |
| FAM102A | 0 | 8.301371897 | 9.718740095 | 0 | 9.228974724 | 0 | 0 | 0 | 0.937677193 | 0.011246701 |
| FAM107B | 10.19389528 | 10.25624448 | 10.03162994 | 0 | 7.184132094 | 0 | 0 | 0 | 0.864814725 | 0.013910154 |
| FAM110C | 0 | 8.49028674 | 0 | 0 | 0 | 0 | 0 | 0 | 1.38E-07 | 1 |
| FAM117B | 0 | 0 | 0 | 7.951591129 | 0 | 0 | 0 | 0 | 1.54E-05 | 1 |
| FAM120A | 0 | 8.657303766 | 8.768344665 | 0 | 7.785476285 | 0 | 0 | 0 | 0.999483812 | 0.027487602 |
| FAM120AOS | 9.881603011 | 7.810193807 | 10.32464801 | 0 | 7.298193348 | 0 | 0 | 0 | 0.992101433 | 0.011872284 |
| FAM120B | 0 | 9.207810313 | 9.170062661 | 0 | 6.517057266 | 0 | 0 | 0 | 0.990582117 | 0.028360154 |
| FAM129C | 9.621196547 | 0 | 0 | 0 | 0 | 0 | 0 | 0 | 8.05E-08 | 1 |
| FAM153C | 10.91814411 | 0 | 0 | 0 | 0 | 0 | 0 | 0 | 7.07E-08 | 1 |
| FAM160A2 | 10.04098874 | 5.496223797 | 9.069573559 | 0 | 7.158471365 | 0 | 0 | 0 | 0.949627741 | 0.026826243 |
| FAM172A | 0 | 8.734080111 | 0 | 0 | 0 | 0 | 0 | 10.10538856 | 1.17E-07 | 1 |
| FAM174B | 0 | 8.909847578 | 0 | 0 | 0 | 0 | 0 | 0 | 1.08E-07 | 1 |
| FAM177A1 | 19.41699181 | 19.21692143 | 18.26534644 | 17.32787461 | 17.76325868 | 17.59717763 | 18.30136571 | 17.64736364 | 0.001156691 | 0.999854694 |
| FAM185A | 0 | 8.326384029 | 0 | 0 | 0 | 0 | 0 | 0 | 1.61E-07 | 1 |
| FAM200A | 0 | 8.958657193 | 0 | 0 | 0 | 0 | 0 | 0 | 1.07E-07 | 1 |
| FAM206A | 0 | 0 | 0 | 9.905608724 | 8.812683467 | 0 | 8.321619838 | 0 | 0.046353196 | 0.999854694 |
| FAM20C | 0 | 9.267144757 | 0 | 0 | 0 | 0 | 0 | 0 | 8.18E-08 | 1 |
| FAM213A | 0 | 0 | 9.864191877 | 0 | 7.699120227 | 0 | 0 | 0 | 0.066119036 | 0.01420877 |
| FAM213B | 9.218319445 | 0 | 10.17474585 | 0 | 8.647265291 | 0 | 0 | 0 | 0.980815574 | 0.010965748 |
| FAM214A | 0 | 0 | 0 | 0 | 6.433937832 | 0 | 8.399062803 | 0 | 5.81E-05 | 1 |
| FAM21EP | 0 | 0 | 0 | 0 | 0 | 0 | 8.11821332 | 10.69579225 | 1 | 0.008359571 |
| FAM226A | 0 | 9.194286798 | 0 | 0 | 0 | 0 | 0 | 0 | 8.58E-08 | 1 |
| FAM24B | 0 | 8.780069978 | 0 | 0 | 0 | 9.109591469 | 0 | 0 | 1.15E-07 | 1 |
| FAM30A | 0 | 9.454643748 | 0 | 0 | 0 | 0 | 0 | 0 | 7.73E-08 | 1 |
| FAM49A | 9.204448442 | 0 | 0 | 10.32853992 | 6.08018266 | 0 | 0 | 0 | 0.985757273 | 0.013610425 |
| FAM83D | 0 | 0 | 0 | 0 | 6.717806789 | 0 | 0 | 0 | 0.000126205 | 1 |
| FAM83H | 0 | 0 | 0 | 10.42003215 | 9.415656752 | 0 | 0 | 0 | 0.022668776 | 0.009547202 |
| FAM86DP | 0 | 0 | 0 | 10.34642021 | 8.823609506 | 0 | 7.137370893 | 0 | 0.032447379 | 0.999854694 |
| FAM86JP | 0 | 0 | 0 | 10.06257842 | 8.60672611 | 0 | 0 | 0 | 0.04436568 | 0.011578584 |
| FAM8A1 | 0 | 0 | 9.152122534 | 0 | 7.329191883 | 0 | 0 | 0 | 0.088199879 | 0.024124185 |
| FAM92A | 0 | 0 | 0 | 10.20658225 | 7.699120227 | 0 | 0 | 9.376339907 | 0.049413137 | 0.999854694 |
| FAN1 | 0 | 9.558573097 | 0 | 0 | 0 | 0 | 5.987074578 | 0 | 7.53E-08 | 1 |
| FANCD2OS | 0 | 0 | 0 | 0 | 6.717806789 | 0 | 0 | 0 | 0.000126205 | 1 |
| FANCE | 0 | 0 | 11.48601283 | 9.666791105 | 0 | 0 | 0 | 0 | 0.029719049 | 0.006327432 |
| FANCL | 0 | 0 | 0 | 10.90786992 | 8.756776493 | 0 | 0 | 0 | 0.016576946 | 0.009001108 |
| FARSB | 0 | 0 | 0 | 10.14406389 | 10.51659246 | 0 | 0 | 11.41802226 | 0.035907399 | 0.999854694 |

| | | | | | | | | | | |
|---|---|---|---|---|---|---|---|---|---|---|
| FASTK | 0 | 0 | 11.19422911 | 10.26650356 | 8.149076506 | 0 | 0 | 5.373764509 | 0.00015087 | 0.112010874 |
| FASTKD1 | 0 | 0 | 0 | 0 | 6.517057266 | 11.17919383 | 7.525254913 | 0 | 5.50E-05 | 0.80968273 |
| FASTKD2 | 0 | 0 | 0 | 11.99970969 | 7.52263204 | 0 | 0 | 0 | 1.55E-07 | 1.68E-11 |
| FAT4 | 0 | 9.891685783 | 0 | 0 | 0 | 0 | 0 | 0 | 7.07E-08 | 1 |
| FBXL3 | 0 | 0 | 9.297034957 | 9.329661438 | 0 | 0 | 0 | 0 | 0.058621566 | 0.013309664 |
| FBXL6 | 0 | 0 | 8.572429871 | 6.193017253 | 9.002525626 | 10.61307666 | 10.46397604 | 0 | 0.033484176 | 0.999854694 |
| FBXO25 | 10.05135807 | 0 | 10.61540567 | 0 | 8.245664407 | 0 | 0 | 0 | 0.997434996 | 0.010099462 |
| FBXO32 | 0 | 0 | 11.33118953 | 0 | 10.13597611 | 0 | 8.303142196 | 0 | 0.028871635 | 0.999854694 |
| FBXO34 | 10.05651492 | 0 | 0 | 0 | 0 | 0 | 6.683509672 | 0 | 7.07E-08 | 1 |
| FBXO4 | 0 | 10.13407091 | 9.884858354 | 0 | 10.10999426 | 0 | 0 | 0 | 0.996280769 | 0.009236883 |
| FBXO45 | 10.59569409 | 0 | 0 | 0 | 0 | 0 | 0 | 0 | 7.07E-08 | 1 |
| FBXO6 | 0 | 0 | 0 | 10.11306472 | 8.28572291 | 0 | 0 | 0 | 0.046892553 | 0.011957177 |
| FBXO7 | 0 | 0 | 0 | 9.825007397 | 8.351369167 | 0 | 0 | 0 | 0.056863878 | 0.013425237 |
| FBXW4 | 9.505291868 | 0 | 0 | 0 | 0 | 0 | 7.751079579 | 0 | 7.80E-08 | 1 |
| FCER2 | 0 | 8.998107186 | 0 | 0 | 0 | 0 | 0 | 0 | 2.64E-07 | 1 |
| FCGBP | 13.70118356 | 14.06680041 | 13.01396234 | 11.19860782 | 6.46217976 | 0 | 0 | 0 | 0.435737206 | 0.000158964 |
| FCGR1A | 0 | 9.900040478 | 0 | 0 | 0 | 0 | 0 | 10.95789928 | 7.07E-08 | 1 |
| FCGR2B | 8.305483268 | 8.867872257 | 0 | 0 | 0 | 0 | 0 | 0 | 0.022739103 | 1 |
| FCHO1 | 0 | 0 | 10.82744558 | 0 | 8.179163277 | 0 | 0 | 0 | 0.025282695 | 0.009547202 |
| FCHSD1 | 9.689176399 | 0 | 0 | 0 | 0 | 0 | 0 | 0 | 7.88E-08 | 1 |
| FCN3 | 0 | 0 | 0 | 0 | 0 | 10.99862429 | 0 | 10.15401896 | 1 | 0.005217323 |
| FEN1 | 14.68685902 | 15.47572823 | 16.4245841 | 17.33907216 | 17.08388996 | 16.10000091 | 16.27844192 | 17.21166392 | 0.00075391 | 0.999854694 |
| FES | 0 | 9.254168606 | 0 | 0 | 0 | 0 | 0 | 0 | 8.22E-08 | 1 |
| FFAR4 | 9.766737375 | 0 | 0 | 0 | 0 | 0 | 0 | 0 | 7.17E-08 | 1 |
| FGD4 | 0 | 9.448960762 | 0 | 0 | 0 | 0 | 0 | 0 | 7.80E-08 | 1 |
| FGF12 | 0 | 9.630629553 | 0 | 0 | 0 | 0 | 0 | 0 | 1.36E-07 | 1 |
| FGF18 | 0 | 0 | 9.42871099 | 10.13380467 | 8.538733665 | 13.02998209 | 13.20643957 | 10.46884494 | 0.003651797 | 0.666516277 |
| FGFR1OP2 | 0 | 0 | 0 | 0 | 6.345735199 | 10.56965778 | 7.847742315 | 0 | 6.22E-05 | 0.910945216 |
| FGFR4 | 9.675834273 | 0 | 0 | 0 | 0 | 0 | 7.64747221 | 0 | 7.44E-08 | 1 |
| FGL2 | 10.31502675 | 5.811745105 | 0 | 0 | 0 | 0 | 0 | 0 | 0.010675182 | 1 |
| FH | 11.17442838 | 11.12276011 | 9.345202746 | 10.40624662 | 8.54864527 | 0 | 0 | 0 | 0.621522984 | 0.001216795 |
| FIG4 | 0 | 0 | 10.91751914 | 10.18438646 | 0 | 0 | 0 | 0 | 0.033443771 | 0.007355649 |
| FITM1 | 0 | 8.54458593 | 0 | 0 | 0 | 0 | 0 | 0 | 1.34E-07 | 1 |
| FKBP1A-SDCBP2 | 0 | 8.008672964 | 11.05581282 | 0 | 9.187285685 | 0 | 0 | 0 | 0.831624692 | 0.007900733 |
| FLAD1 | 7.292659623 | 0 | 0 | 10.19429344 | 9.014494887 | 0 | 0 | 0 | 0.765361891 | 0.010590691 |
| FLI1 | 0 | 0 | 0 | 8.458554996 | 0 | 0 | 0 | 0 | 1.20E-05 | 1 |
| FLJ44635 | 0 | 0 | 0 | 10.34419726 | 10.08183047 | 11.87364095 | 0 | 0 | 0.014410414 | 1 |
| FLOT2 | 0 | 9.961204535 | 0 | 0 | 0 | 0 | 0 | 0 | 7.07E-08 | 1 |
| FLVCR2 | 0 | 8.501311304 | 0 | 0 | 0 | 0 | 0 | 0 | 1.38E-07 | 1 |
| FLYWCH2 | 0 | 0 | 0 | 11.37119647 | 8.809938979 | 0 | 0 | 0 | 0.032339976 | 0.007427763 |
| FMNL3 | 0 | 0 | 0 | 8.008848956 | 9.109068164 | 0 | 0 | 0 | 0.078777618 | 0.020852829 |
| FMO5 | 0 | 0 | 9.611956589 | 7.080028634 | 7.722658607 | 0 | 0 | 0 | 0.028079188 | 0.005940842 |
| FN3KRP | 0 | 0 | 9.199475854 | 0 | 8.42136833 | 0 | 0 | 0 | 0.074923244 | 0.016902113 |

| | | | | | | | | | | |
|---|---|---|---|---|---|---|---|---|---|---|
| FOSL1 | 0 | 0 | 0 | 11.04557883 | 8.159893952 | 0 | 0 | 0 | 0.016451241 | 0.009118992 |
| FOXA1 | 0 | 8.715265663 | 0 | 0 | 0 | 0 | 0 | 0 | 1.18E-07 | 1 |
| FOXA3 | 0 | 8.824638969 | 9.915904145 | 0 | 9.049818146 | 0 | 0 | 0 | 0.967841956 | 0.011034216 |
| FOXC1 | 0 | 0 | 0 | 0 | 6.330495883 | 0 | 0 | 0 | 6.26E-05 | 1 |
| FOXF1 | 0 | 8.288701315 | 0 | 0 | 0 | 0 | 0 | 0 | 1.78E-07 | 1 |
| FOXJ1 | 0 | 0 | 9.769307127 | 9.244022975 | 0 | 0 | 0 | 0 | 0.047797648 | 0.011678653 |
| FOXK2 | 10.07187578 | 0 | 0 | 0 | 0 | 0 | 8.22677003 | 0 | 7.07E-08 | 1 |
| FOXO3B | 0 | 8.966633964 | 0 | 0 | 0 | 0 | 6.828376929 | 0 | 1.07E-07 | 1 |
| FOXRED1 | 9.222913633 | 0 | 0 | 11.19244244 | 8.249720682 | 0 | 0 | 0 | 0.946046188 | 0.008551774 |
| FPGS | 0 | 0 | 0 | 8.954515171 | 8.068700188 | 0 | 0 | 0 | 0.082850386 | 0.022769437 |
| FPR1 | 0 | 9.615488971 | 0 | 0 | 0 | 10.99725942 | 0 | 0 | 7.42E-08 | 1 |
| FREM1 | 0 | 0 | 0 | 7.369539606 | 0 | 0 | 7.414849988 | 0 | 2.74E-05 | 1 |
| FRG1FP | 0 | 9.849170791 | 0 | 0 | 0 | 0 | 0 | 0 | 7.07E-08 | 1 |
| FRK | 0 | 8.958657193 | 0 | 0 | 0 | 0 | 0 | 0 | 1.07E-07 | 1 |
| FRMD3 | 0 | 0 | 0 | 0 | 6.85193071 | 0 | 0 | 0 | 4.12E-05 | 1 |
| FRMD5 | 0 | 0 | 0 | 10.07871314 | 9.837596232 | 9.94231114 | 9.919834864 | 10.1505992 | 0.02359436 | 0.999854694 |
| FRY | 9.005501878 | 0 | 0 | 0 | 0 | 0 | 0 | 0 | 1.07E-07 | 1 |
| FSD2 | 0 | 0 | 0 | 0 | 6.670177039 | 0 | 0 | 0 | 4.78E-05 | 1 |
| FTL | 15.48606162 | 15.00800413 | 16.44145441 | 17.15888533 | 16.44114633 | 16.45962935 | 15.81588851 | 16.78687918 | 0.007395872 | 0.999854694 |
| FTO | 0 | 0 | 11.22982612 | 0 | 6.645759207 | 0 | 0 | 0 | 7.07E-08 | 9.61E-11 |
| FTSJ1 | 0 | 0 | 7.271028348 | 9.955974972 | 9.002525626 | 0 | 0 | 0 | 0.010538501 | 0.002827437 |
| FUOM | 0 | 0 | 9.825726473 | 0 | 9.827454651 | 0 | 10.00613952 | 0 | 0.029981024 | 1 |
| FUT2 | 10.12192865 | 0 | 0 | 0 | 0 | 0 | 0 | 0 | 7.07E-08 | 1 |
| FUT8 | 0 | 0 | 10.43014738 | 0 | 9.049818146 | 0 | 0 | 0 | 0.028970911 | 0.009818636 |
| FXR1 | 0 | 9.912482497 | 0 | 0 | 0 | 0 | 8.008442421 | 9.570542731 | 7.07E-08 | 0.012259484 |
| FXYD1 | 8.973144195 | 0 | 0 | 0 | 0 | 0 | 0 | 0 | 1.08E-07 | 1 |
| FZD5 | 10.18450463 | 9.286392738 | 0 | 0 | 0 | 10.64131321 | 0 | 0 | 0.00296033 | 1 |
| FZD8 | 0 | 0 | 0 | 0 | 6.345735199 | 0 | 0 | 0 | 6.22E-05 | 1 |
| G6PC3 | 0 | 0 | 9.614870348 | 0 | 9.210363586 | 0 | 3.993557961 | 0 | 0.04828143 | 0.269144954 |
| G6PD | 9.6281405 | 0 | 0 | 0 | 0 | 0 | 8.89893906 | 11.36145982 | 7.53E-08 | 0.005427968 |
| GABARAPL1 | 0 | 0 | 10.58303816 | 0 | 7.939059873 | 0 | 6.683509672 | 0 | 0.035248332 | 0.999854694 |
| GABPB1-AS1 | 0 | 8.667127243 | 0 | 0 | 0 | 0 | 0 | 0 | 1.21E-07 | 1 |
| GABPB1-IT1 | 0 | 0 | 10.32108418 | 0 | 8.36264997 | 0 | 7.813336911 | 0 | 0.040562762 | 0.999854694 |
| GAL3ST2 | 0 | 8.501311304 | 0 | 0 | 0 | 0 | 0 | 0 | 1.38E-07 | 1 |
| GALNS | 9.840340458 | 0 | 0 | 0 | 0 | 0 | 0 | 0 | 7.17E-08 | 1 |
| GALNT1 | 0 | 11.00956589 | 9.2640058 | 7.222383808 | 10.87505701 | 0 | 0 | 0 | 0.995827349 | 0.001245844 |
| GALNT14 | 10.99804444 | 8.770988541 | 0 | 0 | 0 | 0 | 0 | 0 | 0.000622074 | 1 |
| GALNT5 | 0 | 0 | 0 | 8.063920809 | 0 | 0 | 0 | 0 | 1.35E-05 | 1 |
| GALT | 0 | 8.375143804 | 0 | 0 | 0 | 0 | 0 | 10.49064777 | 1.55E-07 | 1 |
| GAPLINC | 0 | 9.727698902 | 0 | 0 | 0 | 0 | 0 | 0 | 7.17E-08 | 1 |
| GARS | 0 | 0 | 0 | 12.57380375 | 10.62635016 | 0 | 10.0325516 | 12.75716685 | 7.91E-08 | 0.999854694 |
| GATA3-AS1 | 0 | 0 | 0 | 0 | 6.830424858 | 0 | 0 | 0 | 4.17E-05 | 1 |
| GATAD1 | 0 | 8.876365592 | 0 | 0 | 0 | 0 | 0 | 10.5463602 | 1.10E-07 | 1 |

| | | | | | | | | | | |
|---|---|---|---|---|---|---|---|---|---|---|
| GATD3B | 11.26519916 | 9.610406593 | 0 | 0 | 0 | 0 | 0 | 0 | 7.51E-05 | 1 |
| GATM | 0 | 8.909847578 | 0 | 0 | 0 | 0 | 0 | 0 | 1.08E-07 | 1 |
| GBA2 | 0 | 10.03892072 | 9.760124341 | 0 | 8.4195691 | 0 | 0 | 0 | 0.990582117 | 0.01316597 |
| GBAT2 | 0 | 0 | 0 | 0 | 0 | 0 | 0 | 6.402999079 | 1 | 2.48E-06 |
| GBGT1 | 0 | 8.313932167 | 0 | 0 | 0 | 0 | 0 | 0 | 1.64E-07 | 1 |
| GBP1 | 9.419837447 | 9.465942998 | 0 | 0 | 0 | 0 | 0 | 0 | 0.006979146 | 1 |
| GBP3 | 0 | 0 | 9.408702515 | 0 | 7.604099238 | 0 | 0 | 0 | 0.07870342 | 0.01869339 |
| GCC2 | 9.185743846 | 0 | 0 | 0 | 0 | 0 | 7.695490668 | 0 | 1.15E-07 | 1 |
| GCLM | 0 | 8.375143804 | 0 | 0 | 0 | 0 | 0 | 10.83225647 | 1.55E-07 | 1 |
| GDPD1 | 0 | 8.576210795 | 0 | 0 | 0 | 0 | 0 | 0 | 1.31E-07 | 1 |
| GDPD3 | 0 | 9.081277213 | 9.061052812 | 10.22361261 | 4.589393553 | 0 | 0 | 0 | 0.960239563 | 0.003840265 |
| GDPD5 | 0 | 0 | 0 | 0 | 0 | 0 | 9.830189467 | 9.681338694 | 1 | 0.008582302 |
| GEMIN2 | 0 | 0 | 10.57483153 | 0 | 10.36778123 | 0 | 0 | 0 | 0.034478854 | 0.007255458 |
| GEMIN4 | 0 | 8.734080111 | 0 | 0 | 0 | 0 | 0 | 0 | 1.17E-07 | 1 |
| GEMIN8 | 8.559063282 | 0 | 0 | 0 | 0 | 0 | 0 | 0 | 1.38E-07 | 1 |
| GFER | 12.20811147 | 9.273589327 | 0 | 0 | 0 | 0 | 7.344309675 | 0 | 1.55E-07 | 1 |
| GFM1 | 0 | 0 | 11.76117917 | 0 | 8.012515226 | 0 | 0 | 0 | 5.65E-07 | 1.76E-11 |
| GGA1 | 0 | 9.166853421 | 0 | 10.15171072 | 5.238297909 | 0 | 0 | 0 | 1 | 0.015810782 |
| GGA3 | 0 | 0 | 8.956972117 | 0 | 6.582842939 | 0 | 0 | 0 | 0.101543344 | 0.033895535 |
| GGCT | 0 | 0 | 12.00694575 | 11.95269815 | 11.74274559 | 10.35946293 | 9.6902613 | 11.83079279 | 4.99E-10 | 0.999854694 |
| GGPS1 | 0 | 0 | 0 | 0 | 0 | 0 | 7.83921758 | 11.29718632 | 1 | 0.00625024 |
| GGT1 | 0 | 8.54458593 | 0 | 0 | 0 | 0 | 9.500571214 | 0 | 1.34E-07 | 1 |
| GGT6 | 11.20261319 | 0 | 0 | 10.79032148 | 7.877478056 | 0 | 0 | 0 | 0.985686044 | 0.010065115 |
| GGTA1P | 0 | 0 | 7.030375154 | 0 | 0 | 0 | 0 | 0 | 5.74E-05 | 1 |
| GIGYF2 | 9.166793555 | 0 | 0 | 8.576875689 | 7.105738675 | 0 | 0 | 0 | 0.987285143 | 0.042743578 |
| GIMAP1 | 0 | 8.565746011 | 0 | 0 | 0 | 11.00135018 | 0 | 0 | 1.33E-07 | 1 |
| GIMAP7 | 0 | 9.684823965 | 0 | 0 | 0 | 0 | 8.886581389 | 0 | 7.17E-08 | 1 |
| GINS2 | 0 | 0 | 0 | 10.52581632 | 9.281429657 | 0 | 0 | 0 | 0.02172948 | 0.009471665 |
| GIPC2 | 0 | 0 | 9.030827538 | 9.666791105 | 8.351369167 | 0 | 0 | 10.28143559 | 0.01011564 | 0.999854694 |
| GIT1 | 10.07187578 | 0 | 0 | 0 | 0 | 0 | 0 | 0 | 7.07E-08 | 1 |
| GJC2 | 0 | 0 | 0 | 0 | 0 | 0 | 8.801375753 | 9.652615079 | 1 | 0.010542114 |
| GJC3 | 11.18858961 | 0 | 0 | 0 | 0 | 0 | 13.9542199 | 0 | 4.18E-08 | 1 |
| GK | 0 | 0 | 8.548246557 | 8.442019522 | 6.582842939 | 9.6953675 | 0 | 0 | 0.041448065 | 1 |
| GLCCI1 | 0 | 8.445326205 | 10.06183858 | 0 | 6.569923599 | 0 | 0 | 0 | 0.971198277 | 0.014472868 |
| GLE1 | 8.453559626 | 0 | 9.441897455 | 0 | 8.498391931 | 0 | 0 | 0 | 0.974069502 | 0.014691949 |
| GLG1 | 0 | 0 | 0 | 10.42214136 | 8.010126026 | 10.76195457 | 0 | 0 | 0.038217032 | 0.999854694 |
| GLI1 | 10.08202624 | 0 | 0 | 0 | 0 | 0 | 0 | 0 | 7.07E-08 | 1 |
| GLI4 | 0 | 0 | 0 | 0 | 6.786426836 | 10.62902767 | 9.364617528 | 0 | 4.30E-05 | 0.929188366 |
| GLIDR | 0 | 8.210248014 | 9.960109806 | 9.805749732 | 8.22932429 | 0 | 0 | 0 | 0.778511001 | 0.00143388 |
| GLIPR1 | 0 | 0 | 0 | 0 | 0 | 0 | 7.368208397 | 10.83332336 | 1 | 0.008519862 |
| GLIS2 | 0 | 0 | 0 | 0 | 0 | 0 | 10.20667939 | 9.834113854 | 1 | 0.007255458 |
| GLIS3 | 11.05067811 | 8.934458793 | 0 | 0 | 0 | 0 | 0 | 0 | 0.000438176 | 1 |
| GLT8D2 | 9.987993796 | 0 | 0 | 0 | 0 | 0 | 0 | 0 | 7.07E-08 | 1 |

| | | | | | | | | | | |
|---|---|---|---|---|---|---|---|---|---|---|
| GM2A | 0 | 10.10906458 | 0 | 0 | 0 | 0 | 0 | 9.812636517 | 7.07E-08 | 1 |
| GMFB | 0 | 9.879062319 | 0 | 0 | 0 | 0 | 0 | 0 | 7.07E-08 | 1 |
| GMFG | 0 | 0 | 10.69466096 | 0 | 7.132345939 | 11.94404739 | 8.99416519 | 9.99340186 | 0.035967833 | 0.999854694 |
| GMIP | 10.72439102 | 0 | 0 | 0 | 0 | 0 | 0 | 0 | 7.07E-08 | 1 |
| GNA14 | 0 | 8.743396117 | 0 | 0 | 0 | 0 | 0 | 0 | 1.17E-07 | 1 |
| GNLY | 10.37404726 | 0 | 0 | 0 | 0 | 0 | 0 | 0 | 7.07E-08 | 1 |
| GNPTAB | 0 | 0 | 0 | 0 | 6.786426836 | 0 | 0 | 0 | 4.30E-05 | 1 |
| GOLGA1 | 9.810127513 | 0 | 0 | 0 | 0 | 0 | 0 | 0 | 7.25E-08 | 1 |
| GOLGA2 | 9.362697802 | 0 | 0 | 0 | 0 | 0 | 0 | 0 | 8.05E-08 | 1 |
| GOLGA2P5 | 10.53418006 | 0 | 0 | 0 | 0 | 0 | 0 | 0 | 7.07E-08 | 1 |
| GOLGA8B | 9.840340458 | 0 | 0 | 0 | 0 | 7.673489052 | 7.822015489 | 0 | 7.17E-08 | 0.037167969 |
| GOLGA8J | 9.600161871 | 0 | 0 | 0 | 0 | 0 | 0 | 0 | 7.54E-08 | 1 |
| GOLGA8O | 9.960747977 | 0 | 0 | 0 | 0 | 0 | 0 | 0 | 7.07E-08 | 1 |
| GOLIM4 | 17.20281014 | 16.24150935 | 15.25122782 | 14.50227298 | 14.11194164 | 15.30216733 | 15.9329165 | 14.2200391 | 0.00027591 | 0.999854694 |
| GORASP1 | 0 | 0 | 0 | 0 | 0 | 0 | 7.741962234 | 10.17773295 | 1 | 0.01037888 |
| GOT1 | 0 | 0 | 0 | 11.50357984 | 9.121295971 | 0 | 7.695490668 | 0 | 0.029901269 | 0.999854694 |
| GP1BA | 10.59923216 | 0 | 0 | 0 | 0 | 0 | 0 | 0 | 7.07E-08 | 1 |
| GPALPP1 | 0 | 0 | 0 | 0 | 6.530456572 | 0 | 7.723552846 | 0 | 5.38E-05 | 1 |
| GPAT4 | 0 | 0 | 0 | 8.207797171 | 0 | 0 | 6.411827288 | 0 | 1.16E-05 | 1 |
| GPATCH11 | 0 | 0 | 0 | 8.173150762 | 0 | 0 | 0 | 0 | 1.21E-05 | 1 |
| GPATCH2 | 0 | 8.637453836 | 0 | 0 | 0 | 0 | 0 | 0 | 1.24E-07 | 1 |
| GPATCH4 | 0 | 0 | 0 | 10.31498276 | 7.740063742 | 0 | 9.181166085 | 0 | 0.044779417 | 0.999854694 |
| GPC3 | 11.28400719 | 11.70393255 | 14.03081451 | 12.35831212 | 15.64890732 | 12.95831599 | 11.92779181 | 10.69696495 | 0.025248586 | 0.999854694 |
| GPLD1 | 0 | 9.426002279 | 0 | 0 | 0 | 0 | 0 | 0 | 7.87E-08 | 1 |
| GPR108 | 0 | 0 | 10.48770834 | 7.791124916 | 8.381257477 | 0 | 8.67773346 | 0 | 0.006682609 | 0.999854694 |
| GPR183 | 0 | 7.792352246 | 0 | 0 | 0 | 0 | 0 | 6.534760568 | 0.077643549 | 2.11E-06 |
| GPR89B | 0 | 0 | 0 | 0 | 0 | 0 | 6.564435428 | 9.439357035 | 1 | 0.016586748 |
| GPT | 9.662367605 | 8.313932167 | 0 | 0 | 0 | 0 | 0 | 0 | 0.012658427 | 1 |
| GPX7 | 0 | 0 | 0 | 0 | 0 | 0 | 6.912047636 | 10.52009861 | 1 | 0.009818636 |
| GRAMD1A | 0 | 0 | 10.30493709 | 0 | 8.166345699 | 0 | 10.31669943 | 0 | 0.043227351 | 0.999854694 |
| GRB7 | 11.68952574 | 12.44572741 | 12.9939579 | 14.86194642 | 13.73367726 | 13.78969206 | 15.53440576 | 13.36750706 | 0.018813146 | 0.999854694 |
| GRID1 | 9.935778898 | 0 | 0 | 0 | 0 | 0 | 0 | 0 | 7.07E-08 | 1 |
| GRINA | 0 | 0 | 8.81216258 | 9.705405792 | 8.59401982 | 0 | 11.0796891 | 10.40999367 | 0.009767771 | 0.999854694 |
| GRIPAP1 | 10.01473459 | 0 | 9.103162042 | 9.050354788 | 8.144726714 | 0 | 0 | 0 | 0.997434996 | 0.003648618 |
| GRK6 | 10.0895925 | 8.942570108 | 0 | 0 | 0 | 6.018974366 | 0 | 0 | 0.004849511 | 0.360056801 |
| GRPEL1 | 0 | 0 | 0 | 11.70280847 | 10.00951558 | 0 | 8.927368522 | 11.4860819 | 0.024711474 | 0.999854694 |
| GSE1 | 0 | 0 | 0 | 11.00112053 | 7.488711871 | 0 | 0 | 0 | 0.02091984 | 0.009697633 |
| GSR | 6.584840871 | 7.862431639 | 7.943740461 | 8.408368913 | 6.894003922 | 0 | 0 | 0 | 0.985757273 | 0.009904995 |
| GSTM2 | 0 | 8.80697625 | 0 | 0 | 0 | 0 | 0 | 0 | 1.15E-07 | 1 |
| GSTZ1 | 10.29334884 | 9.879062319 | 10.0424915 | 0 | 5.754469106 | 0 | 0 | 0 | 0.878789877 | 0.015774309 |
| GTF2A1 | 0 | 8.918098054 | 0 | 0 | 0 | 0 | 0 | 0 | 1.08E-07 | 1 |
| GTF2E2 | 11.83165938 | 8.958657193 | 0 | 9.590072571 | 9.020442484 | 0 | 0 | 0 | 0.676663915 | 0.012467263 |
| GTF2H2 | 0 | 0 | 0 | 0 | 0 | 0 | 7.695490668 | 9.952745771 | 1 | 0.011034216 |

| | | | | | | | | | | |
|---|---|---|---|---|---|---|---|---|---|---|
| GTF2H5 | 0 | 0 | 11.10015391 | 9.155516148 | 10.788483 | 10.51538777 | 9.810700222 | 10.82691014 | 2.11E-05 | 0.999854694 |
| GTF2IRD2 | 0 | 0 | 10.4260059 | 0 | 9.245318778 | 9.568770155 | 9.505967127 | 9.898731518 | 0.026612843 | 0.999854694 |
| GTF3A | 0 | 0 | 10.68360188 | 11.93618905 | 11.32532814 | 12.21690503 | 11.42801876 | 10.9598545 | 5.73E-08 | 0.999854694 |
| GTF3C3 | 0 | 0 | 0 | 9.093343475 | 9.567990139 | 0 | 0 | 0 | 0.050164663 | 0.012307739 |
| GTPBP10 | 0 | 0 | 10.56657795 | 10.26415389 | 9.479224203 | 0 | 6.241145621 | 0 | 0.000222712 | 0.297997125 |
| GTPBP2 | 0 | 0 | 0 | 0 | 0 | 6.639132057 | 8.387415743 | 0 | 1 | 0.03673018 |
| GTPBP6 | 0 | 9.650572952 | 0 | 6.771391648 | 9.402964394 | 0 | 0 | 0 | 0.98939231 | 0.02260752 |
| GUCY2C | 9.166793555 | 0 | 0 | 0 | 0 | 10.4688775 | 8.422078578 | 0 | 1.16E-07 | 0.008668791 |
| GUSBP11 | 10.48912305 | 0 | 0 | 0 | 0 | 0 | 0 | 0 | 7.07E-08 | 1 |
| GUSBP5 | 0 | 0 | 0 | 6.89802758 | 0 | 0 | 0 | 0 | 0.000113293 | 1 |
| GXYLT2 | 0 | 0 | 0 | 0 | 0 | 0 | 10.57403288 | 10.46334236 | 1 | 0.005291109 |
| GYG1 | 0 | 0 | 0 | 8.700196723 | 0 | 0 | 0 | 0 | 6.87E-06 | 1 |
| GYS1 | 0 | 0 | 0 | 8.507053818 | 0 | 0 | 0 | 0 | 8.45E-06 | 1 |
| GZMB | 0 | 0 | 9.332868393 | 12.13412854 | 0 | 0 | 11.5391034 | 10.65171857 | 3.11E-07 | 0.999854694 |
| H2AFZ | 11.05841012 | 12.11975642 | 13.19388904 | 14.99812063 | 13.34234292 | 12.56871594 | 11.91187393 | 14.10416026 | 0.030947978 | 0.999854694 |
| HAAO | 0 | 0 | 0 | 11.16121402 | 6.706045958 | 0 | 8.333808086 | 0 | 7.17E-08 | 0.999854694 |
| HAPLN1 | 0 | 9.187477203 | 0 | 0 | 0 | 0 | 0 | 0 | 8.58E-08 | 1 |
| HARS | 9.98258564 | 0 | 9.01326732 | 8.450310949 | 0 | 0 | 0 | 0 | 0.981170768 | 0.020144701 |
| HAS2 | 10.17978629 | 0 | 0 | 0 | 0 | 0 | 0 | 0 | 7.07E-08 | 1 |
| HAUS1 | 0 | 0 | 10.52768203 | 10.57877412 | 9.788950068 | 0 | 0 | 0 | 7.99E-05 | 0.000450503 |
| HAUS2 | 0 | 0 | 10.43097425 | 10.7539421 | 10.26532141 | 0 | 9.428523561 | 0 | 2.67E-05 | 0.999854694 |
| HAUS3 | 0 | 0 | 0 | 9.509044094 | 8.122778831 | 0 | 0 | 0 | 0.070464448 | 0.016256925 |
| HAUS4 | 0 | 0 | 10.75797486 | 0 | 9.558210998 | 0 | 8.81882422 | 0 | 0.014419516 | 0.999854694 |
| HBEGF | 9.578815964 | 0 | 10.04465403 | 0 | 7.298193348 | 0 | 0 | 0 | 0.999483812 | 0.013697851 |
| HCAR1 | 0 | 0 | 0 | 0 | 6.299525656 | 0 | 0 | 0 | 9.81E-05 | 1 |
| HCCS | 0 | 0 | 0 | 11.19120619 | 9.728468463 | 0 | 0 | 0 | 0.031489831 | 0.007044839 |
| HCG18 | 0 | 0 | 0 | 10.47389358 | 8.301439975 | 0 | 0 | 9.995309578 | 0.033856317 | 0.999854694 |
| HCP5 | 0 | 10.44792829 | 0 | 0 | 0 | 11.78664372 | 0 | 0 | 7.07E-08 | 1 |
| HEATR3 | 9.773016437 | 0 | 0 | 0 | 0 | 0 | 0 | 0 | 7.17E-08 | 1 |
| HEATR5B | 0 | 8.49028674 | 0 | 0 | 0 | 0 | 0 | 0 | 1.38E-07 | 1 |
| HECTD3 | 10.20786763 | 0 | 0 | 0 | 0 | 0 | 7.295290731 | 0 | 7.07E-08 | 1 |
| HELLS | 0 | 0 | 0 | 9.856543048 | 8.470856787 | 0 | 0 | 0 | 0.053731692 | 0.012777646 |
| HENMT1 | 0 | 9.992795136 | 0 | 10.2392485 | 9.896345996 | 0 | 0 | 0 | 0.990582117 | 0.009159898 |
| HHEX | 0 | 8.867872257 | 0 | 0 | 0 | 0 | 0 | 0 | 1.10E-07 | 1 |
| HIF1AN | 0 | 10.00446552 | 0 | 0 | 0 | 7.559739089 | 0 | 0 | 7.07E-08 | 1 |
| HIKESHI | 0 | 0 | 0 | 10.27819485 | 10.97766735 | 0 | 9.189573681 | 0 | 0.030414641 | 0.999854694 |
| HINFP | 0 | 0 | 10.5635649 | 0 | 8.535414607 | 0 | 8.494440854 | 10.18777796 | 0.030796109 | 0.999854694 |
| HIP1R | 0 | 0 | 11.20345302 | 0 | 7.418380944 | 0 | 0 | 0 | 0.016730737 | 4.84E-11 |
| HIPK1 | 10.6648577 | 0 | 0 | 0 | 0 | 0 | 0 | 0 | 7.07E-08 | 1 |
| HIPK2 | 0 | 8.998107186 | 0 | 0 | 0 | 0 | 0 | 0 | 2.64E-07 | 1 |
| HIST1H1C | 0 | 0 | 12.60687082 | 0 | 9.603290326 | 0 | 0 | 8.458128915 | 1.88E-07 | 0.968895524 |
| HIST1H2BH | 10.78777972 | 0 | 0 | 0 | 0 | 0 | 0 | 0 | 7.07E-08 | 1 |
| HIST2H2BC | 0 | 0 | 0 | 7.316099607 | 0 | 0 | 0 | 0 | 2.89E-05 | 1 |

| | | | | | | | | | |
|---|---|---|---|---|---|---|---|---|---|
| HIST3H2A | 0 | 0 | 10.15085409 | 9.684471461 | 9.746521881 | 10.74090518 | 0 | 0 | 0.000861115 | 1 |
| HK1 | 0 | 9.088605668 | 0 | 0 | 0 | 0 | 5.126349189 | 7.248526719 | 1.16E-07 | 0.13921824 |
| HK2 | 10.06677365 | 8.676884284 | 10.14481881 | 0 | 9.131224023 | 0 | 0 | 0 | 0.990582117 | 0.01037888 |
| HLA-DPB1 | 11.74966277 | 11.29894527 | 0 | 0 | 0 | 12.89779224 | 9.272643615 | 11.87031045 | 7.07E-08 | 2.37E-05 |
| HLA-G | 0 | 9.2476365 | 0 | 0 | 0 | 0 | 0 | 0 | 8.25E-08 | 1 |
| HLF | 10.45438212 | 0 | 0 | 0 | 0 | 0 | 0 | 0 | 7.07E-08 | 1 |
| HMCES | 10.87792807 | 0 | 0 | 0 | 0 | 10.75065846 | 9.464997095 | 10.33363823 | 7.07E-08 | 0.000283858 |
| HMCN1 | 0 | 8.375143804 | 0 | 0 | 0 | 0 | 0 | 0 | 1.55E-07 | 1 |
| HMG20B | 0 | 9.521148737 | 11.09181239 | 10.63985958 | 7.594562621 | 0 | 0 | 0 | 0.802737132 | 0.000590012 |
| HMGCLL1 | 10.8720903 | 0 | 0 | 0 | 0 | 0 | 0 | 0 | 7.07E-08 | 1 |
| HMGXB3 | 0 | 0 | 0 | 10.48200434 | 6.005385965 | 0 | 7.666871799 | 0 | 0.047632903 | 0.999854694 |
| HOXA13 | 0 | 0 | 0 | 8.672114028 | 0 | 0 | 8.089748008 | 0 | 7.06E-06 | 1 |
| HOXA4 | 10.80626567 | 0 | 0 | 0 | 0 | 0 | 8.11821332 | 0 | 7.07E-08 | 1 |
| HOXB4 | 0 | 0 | 8.954673969 | 0 | 8.257799178 | 0 | 0 | 0 | 0.081940324 | 0.020497277 |
| HOXD1 | 9.766737375 | 0 | 0 | 0 | 0 | 0 | 0 | 0 | 7.17E-08 | 1 |
| HOXD11 | 0 | 9.13888826 | 0 | 0 | 0 | 0 | 0 | 0 | 1.09E-07 | 1 |
| HOXD13 | 0 | 8.950636073 | 0 | 0 | 0 | 0 | 0 | 0 | 1.07E-07 | 1 |
| HPS1 | 0 | 0 | 0 | 10.10258185 | 8.736685727 | 9.550274611 | 8.52133382 | 0 | 0.041219944 | 0.999854694 |
| HPS6 | 10.43074784 | 9.378959303 | 10.0575616 | 0 | 8.730893742 | 0 | 0 | 0 | 0.950589633 | 0.011067641 |
| HR | 9.118299335 | 0 | 0 | 0 | 0 | 0 | 0 | 0 | 9.99E-08 | 1 |
| HRASLS | 11.07629194 | 0 | 0 | 0 | 0 | 0 | 0 | 0 | 6.01E-08 | 1 |
| HS1BP3 | 11.05583738 | 0 | 0 | 0 | 0 | 0 | 0 | 0 | 6.24E-08 | 1 |
| HSD17B7 | 0 | 0 | 10.00744271 | 10.1078328 | 8.992878201 | 11.22772153 | 0 | 0 | 0.001368275 | 1 |
| HSD17B7P2 | 0 | 8.586600217 | 0 | 0 | 0 | 0 | 0 | 0 | 1.30E-07 | 1 |
| HSD17B8 | 9.734925475 | 9.254168606 | 9.74291396 | 9.659657844 | 8.650337012 | 0 | 0 | 0 | 0.999483812 | 0.00150475 |
| HSF1 | 0 | 0 | 0 | 9.40197589 | 9.637747368 | 10.98284974 | 11.36594567 | 0 | 0.041864252 | 0.999854694 |
| HSH2D | 0 | 8.982456399 | 9.787499192 | 0 | 8.036192101 | 0 | 0 | 0 | 0.98939231 | 0.013932059 |
| HSPA13 | 0 | 9.521148737 | 0 | 0 | 0 | 0 | 7.270142197 | 0 | 7.58E-08 | 1 |
| HTD2 | 10.67830125 | 0 | 0 | 0 | 0 | 0 | 0 | 0 | 7.07E-08 | 1 |
| HTR7P1 | 0 | 0 | 0 | 0 | 6.775214428 | 0 | 0 | 0 | 4.34E-05 | 1 |
| HYAL2 | 0 | 0 | 11.48322521 | 11.04489461 | 8.918389748 | 0 | 0 | 0 | 3.11E-06 | 0.000283858 |
| ICAM1 | 11.36105332 | 0 | 0 | 0 | 0 | 10.4688775 | 0 | 0 | 2.76E-08 | 1 |
| ICAM4 | 0 | 9.396780662 | 0 | 0 | 0 | 0 | 0 | 0 | 7.88E-08 | 1 |
| ICMT | 0 | 0 | 9.899270241 | 0 | 8.801674049 | 0 | 0 | 0 | 0.04828143 | 0.011578584 |
| ID2-AS1 | 0 | 0 | 0 | 0 | 0 | 0 | 8.139199685 | 10.14545432 | 1 | 0.010075248 |
| IDH3A | 10.62375866 | 8.169361257 | 11.01558836 | 9.239245818 | 9.233078129 | 0 | 0 | 0 | 0.990582117 | 0.000676953 |
| IDNK | 0 | 0 | 7.702112179 | 0 | 8.012515226 | 0 | 0 | 0 | 0.115807388 | 0.047557087 |
| IER2 | 11.67702416 | 10.15516524 | 11.59626992 | 11.39769848 | 11.08287093 | 0 | 0 | 0 | 0.971036351 | 8.85E-05 |
| IER5L | 0 | 0 | 0 | 9.464711849 | 9.83692234 | 0 | 9.181166085 | 8.204735556 | 0.035488892 | 0.999854694 |
| IFFO1 | 0 | 0 | 0 | 0 | 6.645759207 | 0 | 0 | 0 | 7.73E-05 | 1 |
| IFI16 | 10.05135807 | 8.523110867 | 0 | 0 | 0 | 0 | 0 | 9.141568782 | 0.006926823 | 1 |
| IFIT1 | 0 | 0 | 11.12284747 | 0 | 6.904333267 | 0 | 0 | 0 | 0.021132864 | 6.70E-11 |
| IFIT3 | 9.147591029 | 0 | 10.23847846 | 9.593819844 | 0 | 0 | 0 | 0 | 0.959397646 | 0.010003595 |

| | | | | | | | | | | |
|---|---|---|---|---|---|---|---|---|---|---|
| IFNAR2 | 0 | 0 | 9.332868393 | 10.65697576 | 8.617753048 | 12.03431961 | 0 | 10.67094291 | 0.001446947 | 0.999854694 |
| IFRD1 | 0 | 0 | 9.62068027 | 0 | 10.55561644 | 10.0827733 | 9.589664981 | 9.968369199 | 0.016158732 | 0.999854694 |
| IFT172 | 0 | 0 | 11.32052019 | 10.15931723 | 4.909577724 | 0 | 8.259084486 | 0 | 0.000450922 | 0.999854694 |
| IFT52 | 0 | 0 | 10.81736233 | 10.30587335 | 7.959016539 | 0 | 8.399062803 | 0 | 0.000444451 | 0.999854694 |
| IGF1R | 9.276932164 | 0 | 0 | 0 | 0 | 0 | 0 | 0 | 8.39E-08 | 1 |
| IGF2 | 0 | 0 | 11.65016796 | 9.175658893 | 12.69098137 | 12.67433369 | 7.257401605 | 0 | 2.35E-08 | 0.999854694 |
| IGFBP2 | 0 | 0 | 9.19558857 | 12.51003987 | 9.294250714 | 10.84168895 | 10.94365371 | 0 | 1.54E-07 | 1 |
| IL16 | 0 | 8.479177282 | 0 | 0 | 0 | 0 | 9.122656968 | 0 | 1.39E-07 | 1 |
| IL1R2 | 0 | 0 | 11.65442238 | 0 | 8.388633797 | 0 | 0 | 0 | 0.030396424 | 2.26E-11 |
| IL20RB | 0 | 8.998107186 | 11.25525897 | 10.46369059 | 0 | 0 | 0 | 0 | 0.888160297 | 0.006058239 |
| IL34 | 10.44261338 | 0 | 0 | 0 | 0 | 0 | 0 | 0 | 7.07E-08 | 1 |
| IL4I1 | 0 | 9.465942998 | 0 | 0 | 0 | 0 | 0 | 0 | 7.73E-08 | 1 |
| IL4R | 9.294498035 | 0 | 0 | 0 | 0 | 0 | 0 | 0 | 8.36E-08 | 1 |
| IL6R | 9.797862919 | 0 | 0 | 0 | 0 | 0 | 0 | 0 | 7.17E-08 | 1 |
| IL6ST | 13.67417973 | 14.10060825 | 12.95998813 | 11.72987996 | 11.86114855 | 11.988688 | 10.80333776 | 11.84408598 | 0.049081679 | 0.999854694 |
| IL7 | 9.108401769 | 0 | 0 | 0 | 0 | 0 | 0 | 0 | 1.03E-07 | 1 |
| IL7R | 10.280184 | 0 | 0 | 0 | 0 | 0 | 0 | 0 | 7.07E-08 | 1 |
| ILDR1 | 0 | 0 | 0 | 7.939861729 | 0 | 0 | 0 | 0 | 1.53E-05 | 1 |
| ILK | 0 | 0 | 0 | 10.36627416 | 10.54740141 | 10.4350013 | 9.379392199 | 12.30768694 | 0.033921414 | 0.999854694 |
| ILKAP | 10.75643348 | 8.024043411 | 9.432018923 | 7.710580649 | 7.367026271 | 0 | 0 | 0 | 0.847289161 | 0.006244708 |
| IMMP1L | 0 | 0 | 0 | 10.68535178 | 9.934644116 | 0 | 8.246245428 | 0 | 0.035962426 | 0.999854694 |
| IMPA2 | 0 | 10.66446301 | 9.899270241 | 0 | 11.21165837 | 0 | 0 | 0 | 0.991196379 | 0.006270106 |
| INAVA | 0 | 8.81583464 | 10.18360423 | 7.963225934 | 9.631543354 | 0 | 0 | 0 | 0.873009502 | 0.001461729 |
| INCA1 | 0 | 9.221208238 | 0 | 0 | 0 | 0 | 0 | 0 | 8.39E-08 | 1 |
| INF2 | 0 | 8.934458793 | 10.7840964 | 9.663228883 | 9.422859707 | 0 | 0 | 0 | 0.7550754 | 0.000631775 |
| ING1 | 0 | 0 | 0 | 8.329656402 | 0 | 0 | 0 | 10.21750006 | 1.02E-05 | 1 |
| INMT | 0 | 8.223623425 | 0 | 0 | 0 | 0 | 0 | 0 | 2.64E-07 | 1 |
| INO80B-WBP1 | 0 | 0 | 10.75004528 | 0 | 8.216946592 | 0 | 0 | 0 | 0.027523413 | 0.009724498 |
| INPP5A | 0 | 6.725526766 | 7.734623255 | 8.569282495 | 7.060272833 | 0 | 0 | 0 | 0.847043566 | 0.009658338 |
| INPP5E | 0 | 0 | 0 | 0 | 6.620920975 | 0 | 0 | 0 | 5.02E-05 | 1 |
| INSIG2 | 10.24899064 | 8.761849577 | 10.29951432 | 0 | 8.328539441 | 0 | 0 | 0 | 0.980815574 | 0.01094825 |
| INTS11 | 9.822288722 | 0 | 0 | 8.408368913 | 9.601704371 | 0 | 0 | 0 | 0.997434996 | 0.014264276 |
| INTS3 | 8.060573482 | 0 | 0 | 0 | 0 | 0 | 8.271810291 | 9.593393101 | 0.071806411 | 0.01159012 |
| INTS5 | 0 | 9.342642235 | 0 | 0 | 0 | 0 | 0 | 0 | 8.01E-08 | 1 |
| IPMK | 0 | 0 | 0 | 0 | 6.283787608 | 0 | 0 | 0 | 0.000155241 | 1 |
| IPO5 | 0 | 0 | 8.659669766 | 9.990497814 | 8.249720682 | 0 | 0 | 11.01261004 | 0.009179992 | 1 |
| IPO8 | 7.257265736 | 7.699676638 | 0 | 0 | 0 | 0 | 7.577442608 | 0 | 0.048207122 | 1 |
| IPO9 | 0 | 0 | 0 | 8.576875689 | 6.752524753 | 0 | 0 | 0 | 0.112134614 | 0.047762148 |
| IQGAP2 | 8.684275372 | 0 | 0 | 0 | 0 | 0 | 0 | 0 | 1.25E-07 | 1 |
| IREB2 | 0 | 8.918098054 | 0 | 0 | 0 | 0 | 0 | 0 | 1.08E-07 | 1 |
| IRF3 | 9.166793555 | 0 | 0 | 10.03803287 | 8.249720682 | 0 | 0 | 0 | 0.983546955 | 0.012307739 |
| IRF6 | 9.593081603 | 8.80697625 | 0 | 8.416855435 | 7.675191427 | 0 | 0 | 0 | 0.899315913 | 0.038392165 |
| ISOC1 | 0 | 0 | 8.174541148 | 9.908620621 | 9.635423988 | 0 | 0 | 0 | 0.004502943 | 0.001588084 |

| | | | | | | | | | | |
|---|---|---|---|---|---|---|---|---|---|---|
| ITCH | 0 | 8.54458593 | 0 | 0 | 0 | 0 | 0 | 0 | 1.34E-07 | 1 |
| ITFG2 | 11.36729189 | 9.750979006 | 0 | 0 | 0 | 0 | 6.8624297 | 0 | 4.01E-05 | 1 |
| ITGA1 | 0 | 0 | 8.783962225 | 0 | 7.681210938 | 0 | 0 | 0 | 0.094767502 | 0.028219261 |
| ITGA3 | 7.257265736 | 0 | 8.536001166 | 9.423192445 | 0 | 0 | 0 | 0 | 0.880392942 | 0.015978244 |
| ITGB3BP | 0 | 0 | 0 | 10.78705191 | 9.918172938 | 0 | 0 | 0 | 0.034989165 | 0.007894598 |
| ITGB4 | 0 | 8.223623425 | 0 | 0 | 0 | 10.01963904 | 0 | 9.041723345 | 2.64E-07 | 0.009554591 |
| ITLN1 | 18.69294575 | 21.78684337 | 20.58751134 | 17.16497903 | 15.68007503 | 0 | 10.77008775 | 0 | 0.588193933 | 0.012592755 |
| ITM2A | 0 | 0 | 7.998314546 | 0 | 0 | 0 | 0 | 0 | 1.57E-05 | 1 |
| ITPKA | 6.89226915 | 0 | 11.11051344 | 10.0366569 | 0 | 0 | 0 | 0 | 0.626699816 | 0.007178231 |
| ITPR2 | 14.50690361 | 14.03241879 | 10.73472399 | 13.05393555 | 11.82762071 | 14.08723 | 13.0115656 | 12.68305889 | 0.019967737 | 0.999854694 |
| ITPR3 | 0 | 0 | 0 | 9.928046165 | 8.829041651 | 0 | 0 | 0 | 0.045270943 | 0.011671668 |
| ITPRIP | 9.037149714 | 0 | 0 | 0 | 0 | 0 | 0 | 0 | 1.07E-07 | 1 |
| JADE3 | 0 | 0 | 0 | 9.815410697 | 10.30873864 | 0 | 9.864199982 | 10.67570924 | 0.018571474 | 0.999854694 |
| JAG1 | 0 | 9.013590009 | 0 | 0 | 0 | 10.24911422 | 0 | 9.411053639 | 1.04E-07 | 0.008419794 |
| JMJD4 | 0 | 0 | 0 | 7.058495112 | 9.222797649 | 0 | 0 | 0 | 0.083400879 | 0.024631349 |
| JMJD6 | 0 | 0 | 10.44986264 | 0 | 8.801674049 | 0 | 8.823153561 | 0 | 0.031432954 | 0.999854694 |
| JMJD7 | 0 | 0 | 8.720451232 | 10.29670606 | 0 | 0 | 9.716098353 | 5.276712016 | 0.038370584 | 0.999854694 |
| JMJD8 | 0 | 0 | 10.43262657 | 9.680952681 | 0 | 0 | 7.732786904 | 0 | 0.022750895 | 0.999854694 |
| JMY | 0 | 0 | 0 | 0 | 6.639589624 | 0 | 0 | 0 | 5.07E-05 | 1 |
| JRK | 9.621196547 | 0 | 0 | 0 | 0 | 0 | 0 | 0 | 8.05E-08 | 1 |
| JUND | 11.27518689 | 9.073911341 | 12.24841172 | 0 | 8.316987658 | 0 | 0 | 0 | 0.997434996 | 9.11E-12 |
| KALRN | 0 | 0 | 0 | 0 | 0 | 0 | 8.47806111 | 5.63098815 | 1 | 0.043736931 |
| KANSL2 | 9.977157135 | 9.844849563 | 10.25070664 | 0 | 5.450528608 | 0 | 0 | 0 | 1 | 0.01420877 |
| KANSL3 | 10.63758884 | 6.96733011 | 0 | 0 | 0 | 0 | 0 | 7.547907651 | 0.004493938 | 1 |
| KAT7 | 0 | 9.684823965 | 0 | 0 | 0 | 0 | 6.702433802 | 0 | 7.17E-08 | 1 |
| KAZALD1 | 0 | 0 | 0 | 0 | 0 | 11.74292883 | 7.38001092 | 0 | 1 | 4.71E-12 |
| KAZN | 0 | 0 | 0 | 9.456503009 | 7.963962867 | 0 | 0 | 0 | 0.073048044 | 0.017339178 |
| KBTBD3 | 0 | 7.641072074 | 10.32820306 | 10.25944303 | 8.525411369 | 0 | 0 | 0 | 0.524320932 | 0.000835188 |
| KCNA6 | 10.19856778 | 0 | 0 | 0 | 0 | 0 | 0 | 0 | 7.07E-08 | 1 |
| KCNIP4 | 13.23037796 | 9.088605668 | 12.58382045 | 9.950139986 | 0 | 0 | 0 | 0 | 0.958210869 | 9.52E-12 |
| KCNK15 | 0 | 8.824638969 | 0 | 0 | 0 | 0 | 0 | 0 | 1.36E-07 | 1 |
| KCNK5 | 10.9266179 | 0 | 10.00077521 | 8.311565818 | 8.560123397 | 0 | 0 | 0 | 0.973004436 | 0.002355872 |
| KCNMA1 | 9.128129462 | 0 | 0 | 0 | 0 | 0 | 0 | 0 | 9.93E-08 | 1 |
| KCNS3 | 0 | 0 | 11.27979233 | 9.62711438 | 0 | 0 | 0 | 0 | 0.032289461 | 0.007236136 |
| KCTD10 | 0 | 0 | 0 | 8.347522947 | 9.440711312 | 0 | 0 | 0 | 0.067902631 | 0.01566498 |
| KCTD20 | 0 | 0 | 0 | 10.48503416 | 8.87177783 | 0 | 0 | 10.35756552 | 0.027870278 | 0.999854694 |
| KCTD5 | 0 | 0 | 10.742738 | 9.175658893 | 7.14981535 | 0 | 8.08254305 | 0 | 0.003144577 | 0.999854694 |
| KDELC2 | 0 | 0 | 0 | 0 | 6.39051047 | 0 | 0 | 10.29388958 | 5.98E-05 | 1 |
| KDF1 | 0 | 9.214524828 | 10.16284942 | 0 | 9.172405179 | 0 | 0 | 0 | 0.975665177 | 0.01037888 |
| KDM1A | 8.559063282 | 8.549905117 | 0 | 11.38855896 | 9.830843106 | 0 | 0 | 0 | 0.832701413 | 0.006244708 |
| KDM4C | 0 | 0 | 9.643688775 | 0 | 7.21765149 | 0 | 0 | 0 | 0.075193869 | 0.017328364 |
| KDM5A | 17.88697035 | 16.34311917 | 15.81215154 | 15.11880894 | 15.7503132 | 15.4070059 | 16.54525147 | 14.9314205 | 0.007330965 | 0.999854694 |
| KEAP1 | 0 | 0 | 9.422072283 | 0 | 9.253421891 | 0 | 0 | 0 | 0.052536874 | 0.012259484 |

| | | | | | | | | | | |
|---|---|---|---|---|---|---|---|---|---|---|
| KHNYN | 0 | 0 | 0 | 0 | 6.299525656 | 0 | 7.270142197 | 0 | 9.81E-05 | 1 |
| KIAA0895L | 10.17030305 | 7.51619808 | 0 | 0 | 0 | 0 | 0 | 0 | 0.009107867 | 1 |
| KIAA1143 | 0 | 0 | 9.808013049 | 9.967574619 | 10.39367363 | 0 | 7.567155003 | 0 | 0.000301204 | 0.80968273 |
| KIAA1958 | 10.7340787 | 0 | 0 | 0 | 0 | 9.202537636 | 11.02153918 | 9.940916157 | 7.07E-08 | 0.000299696 |
| KIF12 | 10.44261338 | 0 | 0 | 0 | 0 | 0 | 0 | 0 | 7.07E-08 | 1 |
| KIF23 | 0 | 0 | 0 | 9.526794551 | 6.633393544 | 0 | 0 | 0 | 0.079076279 | 0.021760252 |
| KIF25 | 0 | 9.336499504 | 0 | 0 | 0 | 0 | 11.0495221 | 0 | 8.04E-08 | 1 |
| KIF2C | 0 | 0 | 0 | 9.834540682 | 9.438936059 | 0 | 10.09116584 | 7.99192018 | 0.03643578 | 0.999854694 |
| KIFC2 | 0 | 0 | 0 | 0 | 0 | 0 | 8.439102608 | 10.24662217 | 1 | 0.009470925 |
| KLC1 | 8.587842398 | 9.879062319 | 0 | 0 | 0 | 0 | 0 | 0 | 0.008061964 | 1 |
| KLF10 | 0 | 0 | 10.57707434 | 0 | 8.581200627 | 8.831845869 | 6.543597963 | 0 | 0.029981024 | 0.999854694 |
| KLF3-AS1 | 0 | 0 | 7.817899521 | 0 | 0 | 9.884423886 | 0 | 0 | 1.89E-05 | 1 |
| KLHDC1 | 0 | 8.456698379 | 0 | 0 | 0 | 0 | 0 | 0 | 1.42E-07 | 1 |
| KLHDC9 | 0 | 9.005869367 | 0 | 0 | 0 | 0 | 0 | 0 | 1.04E-07 | 1 |
| KLHL21 | 9.88740249 | 0 | 0 | 0 | 0 | 0 | 0 | 0 | 7.14E-08 | 1 |
| KLHL36 | 0 | 0 | 0 | 0 | 6.543732574 | 0 | 0 | 0 | 5.33E-05 | 1 |
| KLHL42 | 0 | 10.30424674 | 0 | 0 | 0 | 0 | 8.37567389 | 0 | 7.07E-08 | 1 |
| KLHL7 | 10.70809889 | 0 | 0 | 0 | 0 | 0 | 0 | 5.707457615 | 7.07E-08 | 0.42410514 |
| KLK10 | 0 | 0 | 0 | 11.7796691 | 8.289668258 | 0 | 0 | 11.97068644 | 1.59E-07 | 0.999854694 |
| KLK12 | 0 | 0 | 0 | 0 | 0 | 10.68095886 | 0 | 10.04035398 | 1 | 0.006112506 |
| KLK3 | 11.31446038 | 10.80900179 | 11.20924841 | 0 | 6.775214428 | 0 | 0 | 0 | 1 | 6.66E-11 |
| KLK6 | 0 | 0 | 0 | 10.41580446 | 8.869143589 | 10.99862429 | 7.637673707 | 10.58993625 | 0.029981024 | 0.999854694 |
| KMT2E | 19.99477685 | 19.03881548 | 18.49748426 | 17.07731569 | 18.15959262 | 18.08924572 | 18.91447146 | 17.08986087 | 0.026774159 | 0.999854694 |
| KMT5A | 0 | 9.201064401 | 0 | 9.15043621 | 6.741044679 | 0 | 0 | 0 | 0.99236975 | 0.028988986 |
| KMT5C | 9.938574676 | 0 | 0 | 0 | 0 | 0 | 0 | 0 | 7.07E-08 | 1 |
| KPNA7 | 10.10709428 | 0 | 0 | 0 | 0 | 0 | 0 | 0 | 7.07E-08 | 1 |
| KPNB1 | 0 | 9.537306767 | 0 | 0 | 0 | 0 | 6.928214841 | 0 | 7.53E-08 | 1 |
| KRAS | 16.17649049 | 15.34987069 | 14.5107226 | 13.61990043 | 14.14990963 | 12.29090512 | 14.62598356 | 13.22703143 | 0.007276607 | 0.999854694 |
| KRR1 | 0 | 9.493809912 | 0 | 0 | 0 | 0 | 0 | 9.63314329 | 8.05E-08 | 1 |
| KRT15 | 10.5080681 | 9.214524828 | 0 | 0 | 0 | 0 | 0 | 0 | 0.001626223 | 1 |
| KRT6A | 0 | 0 | 0 | 11.41103465 | 10.94082273 | 10.75227762 | 0 | 10.73627972 | 0.022850244 | 0.999854694 |
| KYAT3 | 0 | 0 | 0 | 10.29440511 | 8.970932687 | 0 | 0 | 0 | 0.032197685 | 0.010433427 |
| L2HGDH | 10.15114749 | 9.904199751 | 12.7889754 | 11.81420931 | 13.50198051 | 12.26221361 | 11.27326829 | 12.54596191 | 0.00246704 | 0.999854694 |
| LAMA5 | 9.760430865 | 0 | 0 | 10.3923281 | 9.656200926 | 0 | 0 | 0 | 0.983546955 | 0.009209122 |
| LAMB2 | 0 | 0 | 10.54992777 | 0 | 7.17562913 | 0 | 0 | 0 | 0.041720285 | 0.011034216 |
| LANCL1 | 10.9266179 | 0 | 0 | 0 | 0 | 0 | 8.381556762 | 0 | 7.07E-08 | 1 |
| LANCL2 | 9.7916913 | 0 | 0 | 0 | 0 | 0 | 9.489718463 | 0 | 7.44E-08 | 1 |
| LARP1B | 0 | 0 | 0 | 0 | 0 | 10.58609337 | 7.83921758 | 0 | 1 | 0.008817225 |
| LARP4 | 0 | 8.974566872 | 0 | 0 | 0 | 0 | 0 | 0 | 1.07E-07 | 1 |
| LAT2 | 0 | 8.586600217 | 0 | 0 | 0 | 0 | 0 | 0 | 1.30E-07 | 1 |
| LBR | 0 | 0 | 0 | 9.761456469 | 9.963949941 | 0 | 7.391717672 | 0 | 0.027385701 | 0.999854694 |
| LCA5L | 0 | 0 | 0 | 10.42003215 | 9.813820683 | 0 | 11.58507698 | 0 | 0.01687064 | 0.999854694 |
| LCMT1 | 0 | 11.23928546 | 12.09618096 | 12.11176071 | 0 | 0 | 0 | 0 | 0.9690195 | 1.40E-11 |

| | | | | | | | | | |
|---|---|---|---|---|---|---|---|---|---|
| LDB1 | 9.702396267 | 0 | 0 | 0 | 0 | 0 | 0 | 0 | 7.38E-08 | 1 |
| LDHA | 12.41598848 | 12.005833 | 14.07068877 | 13.33653181 | 13.84162859 | 13.15153014 | 12.19118325 | 14.69123013 | 0.031348886 | 0.999854694 |
| LDHD | 9.276932164 | 0 | 0 | 0 | 0 | 0 | 0 | 0 | 8.39E-08 | 1 |
| LDLRAP1 | 0 | 8.950636073 | 0 | 9.295553464 | 8.157736942 | 0 | 0 | 0 | 0.99499976 | 0.017977768 |
| LEFTY1 | 9.600161871 | 11.5781156 | 9.103162042 | 8.147883805 | 9.842976066 | 0 | 0 | 0 | 0.64695618 | 0.00219522 |
| LEMD1 | 0 | 0 | 0 | 0 | 0 | 11.96619158 | 0 | 11.03039541 | 1 | 0.002088375 |
| LEMD3 | 0 | 8.909847578 | 0 | 0 | 0 | 0 | 0 | 0 | 1.08E-07 | 1 |
| LEO1 | 9.944150032 | 0 | 0 | 0 | 0 | 0 | 0 | 0 | 7.07E-08 | 1 |
| LEPROT | 0 | 10.08361716 | 9.797792581 | 0 | 8.131598092 | 0 | 0 | 0 | 0.98939231 | 0.013697851 |
| LEPROTL1 | 17.8009962 | 17.44412728 | 16.62615603 | 15.9222166 | 16.39681031 | 16.25406692 | 17.04045915 | 16.390902 | 0.014512125 | 0.999854694 |
| LGALS9 | 0 | 0 | 9.856826564 | 9.165622674 | 8.992878201 | 0 | 0 | 0 | 0.005044036 | 0.001538174 |
| LGMN | 8.140451703 | 7.472050216 | 11.31649875 | 8.382605645 | 9.961478874 | 0 | 0 | 0 | 0.482408615 | 0.000566119 |
| LGR4 | 0 | 0 | 8.926806794 | 0 | 5.885375679 | 0 | 0 | 0 | 0.10818032 | 0.039883202 |
| LGR6 | 0 | 0 | 0 | 0 | 6.46217976 | 0 | 0 | 0 | 5.69E-05 | 1 |
| LHFPL2 | 11.01403714 | 0 | 0 | 0 | 0 | 0 | 0 | 0 | 7.07E-08 | 1 |
| LHFPL6 | 0 | 0 | 0 | 8.408368913 | 0 | 0 | 0 | 0 | 9.42E-06 | 1 |
| LILRA5 | 0 | 8.934458793 | 0 | 0 | 0 | 0 | 0 | 0 | 1.07E-07 | 1 |
| LIMA1 | 0 | 7.896234792 | 0 | 8.761452735 | 8.501797164 | 0 | 0 | 0 | 0.968436438 | 0.021306839 |
| LIME1 | 0 | 0 | 0 | 0 | 0 | 0 | 10.68404231 | 10.22076503 | 1 | 0.005441159 |
| LIN7A | 0 | 0 | 0 | 0 | 0 | 0 | 9.005638375 | 9.768698172 | 1 | 0.010003595 |
| LIN7C | 0 | 0 | 10.31840552 | 0 | 8.22932429 | 0 | 0 | 10.49064777 | 0.042053591 | 0.999854694 |
| LINC-PINT | 0 | 9.254168606 | 0 | 0 | 0 | 10.09813443 | 9.02455989 | 0 | 8.22E-08 | 0.008991934 |
| LINC00114 | 9.06811818 | 0 | 0 | 0 | 0 | 0 | 0 | 0 | 1.09E-07 | 1 |
| LINC00167 | 0 | 0 | 7.081824519 | 0 | 0 | 0 | 0 | 0 | 0.000144173 | 1 |
| LINC00265 | 0 | 8.576210795 | 0 | 0 | 0 | 0 | 8.594071773 | 0 | 1.31E-07 | 1 |
| LINC00476 | 0 | 0 | 10.77500756 | 11.14470411 | 0 | 0 | 0 | 9.97997673 | 0.027413221 | 0.999854694 |
| LINC00482 | 0 | 8.789094606 | 0 | 0 | 0 | 0 | 0 | 0 | 1.15E-07 | 1 |
| LINC00511 | 0 | 0 | 0 | 0 | 6.752524753 | 0 | 0 | 0 | 0.000121249 | 1 |
| LINC00513 | 0 | 0 | 0 | 8.399832174 | 0 | 0 | 9.211206849 | 0 | 2.12E-05 | 1 |
| LINC00630 | 0 | 0 | 0 | 0 | 6.775214428 | 0 | 0 | 0 | 4.34E-05 | 1 |
| LINC00639 | 0 | 8.850734095 | 0 | 0 | 0 | 0 | 0 | 0 | 1.17E-07 | 1 |
| LINC00668 | 0 | 0 | 0 | 8.347522947 | 0 | 0 | 0 | 0 | 1.03E-05 | 1 |
| LINC00673 | 8.951162053 | 0 | 0 | 0 | 0 | 0 | 0 | 0 | 1.10E-07 | 1 |
| LINC00842 | 0 | 9.844849563 | 0 | 0 | 0 | 0 | 8.111149473 | 0 | 7.07E-08 | 1 |
| LINC00847 | 0 | 0 | 10.49326094 | 0 | 7.73428533 | 0 | 0 | 0 | 0.040172136 | 0.010845902 |
| LINC00857 | 0 | 0 | 0 | 7.316099607 | 8.759623899 | 0 | 0 | 0 | 0.095930888 | 0.033412241 |
| LINC01000 | 11.44602996 | 0 | 0 | 0 | 0 | 0 | 5.428672801 | 0 | 7.07E-08 | 0.467810605 |
| LINC01124 | 0 | 0 | 0 | 0 | 6.841217856 | 0 | 0 | 0 | 4.14E-05 | 1 |
| LINC01133 | 0 | 0 | 12.28682869 | 0 | 7.367026271 | 0 | 0 | 0 | 2.27E-07 | 1.17E-11 |
| LINC01279 | 0 | 8.263021342 | 0 | 0 | 0 | 0 | 0 | 0 | 2.55E-07 | 1 |
| LINC01315 | 0 | 0 | 0 | 0 | 0 | 10.76436371 | 10.64788981 | 0 | 1 | 0.00418531 |
| LINC01389 | 10.24899064 | 0 | 0 | 0 | 0 | 0 | 0 | 0 | 7.07E-08 | 1 |
| LINC01431 | 0 | 0 | 8.829825439 | 0 | 5.946628341 | 0 | 0 | 0 | 0.112166591 | 0.043375133 |

| | | | | | | | | | | |
|---|---|---|---|---|---|---|---|---|---|---|
| LINC01481 | 0 | 0 | 8.693140313 | 9.782952664 | 7.728483681 | 0 | 7.732786904 | 10.77677146 | 0.01418249 | 0.999854694 |
| LINC01504 | 11.00339506 | 0 | 0 | 0 | 0 | 0 | 0 | 0 | 7.07E-08 | 1 |
| LINC01547 | 10.04098874 | 0 | 0 | 0 | 0 | 0 | 0 | 0 | 7.07E-08 | 1 |
| LINC01550 | 9.648774049 | 0 | 0 | 0 | 0 | 0 | 0 | 0 | 7.53E-08 | 1 |
| LINC01590 | 9.362697802 | 0 | 0 | 0 | 0 | 0 | 0 | 0 | 8.05E-08 | 1 |
| LINC01781 | 0 | 9.088605668 | 0 | 0 | 0 | 0 | 0 | 0 | 1.16E-07 | 1 |
| LINC01814 | 0 | 0 | 0 | 0 | 6.503532343 | 0 | 0 | 0 | 5.50E-05 | 1 |
| LINC01871 | 9.955236519 | 0 | 0 | 0 | 0 | 0 | 0 | 0 | 7.07E-08 | 1 |
| LINC01881 | 0 | 0 | 0 | 11.70107237 | 10.61184799 | 0 | 9.399825791 | 0 | 0.022156595 | 0.999854694 |
| LINC02001 | 0 | 9.173760825 | 0 | 9.955974972 | 7.306005647 | 0 | 0 | 0 | 0.991384446 | 0.01463487 |
| LINC02018 | 0 | 0 | 0 | 0 | 6.489879427 | 0 | 0 | 0 | 5.56E-05 | 1 |
| LINC02023 | 0 | 8.637453836 | 0 | 0 | 0 | 0 | 0 | 0 | 1.24E-07 | 1 |
| LINC02108 | 10.9350422 | 7.620995983 | 0 | 0 | 0 | 0 | 0 | 0 | 0.001607806 | 1 |
| LINC02332 | 0 | 0 | 0 | 0 | 6.46217976 | 0 | 0 | 0 | 5.69E-05 | 1 |
| LINC02535 | 0 | 8.789094606 | 0 | 0 | 0 | 0 | 0 | 0 | 1.15E-07 | 1 |
| LINC02588 | 0 | 8.934458793 | 0 | 0 | 0 | 0 | 0 | 0 | 1.07E-07 | 1 |
| LIPC | 11.08641148 | 0 | 10.93860415 | 0 | 9.185169272 | 0 | 0 | 0 | 0.994561213 | 0.00825262 |
| LIPH | 0 | 10.54701515 | 10.5160396 | 0 | 8.007732862 | 0 | 0 | 0 | 0.991384446 | 0.010542114 |
| LIPT2 | 0 | 10.91379815 | 0 | 0 | 0 | 0 | 0 | 0 | 7.07E-08 | 1 |
| LMNB2 | 10.51557701 | 0 | 0 | 10.02698824 | 9.45483524 | 0 | 0 | 0 | 0.997434996 | 0.01037888 |
| LMOD1 | 0 | 9.267144757 | 0 | 0 | 0 | 0 | 0 | 0 | 8.18E-08 | 1 |
| LMTK2 | 8.859722698 | 0 | 8.359630277 | 9.290944194 | 0 | 0 | 0 | 0 | 0.990582117 | 0.017871642 |
| LMTK3 | 0 | 0 | 0 | 0 | 6.741044679 | 0 | 0 | 8.109312118 | 0.000123975 | 1 |
| LOC100128108 | 8.305483268 | 0 | 0 | 0 | 0 | 0 | 0 | 0 | 2.57E-07 | 1 |
| LOC100129034 | 0 | 9.547979116 | 0 | 0 | 0 | 0 | 0 | 0 | 7.53E-08 | 1 |
| LOC100130298 | 0 | 9.273589327 | 0 | 0 | 0 | 0 | 0 | 0 | 8.16E-08 | 1 |
| LOC100132154 | 0 | 0 | 0 | 0 | 0 | 0 | 8.805757694 | 11.07797895 | 1 | 0.006244708 |
| LOC100133331 | 7.360944326 | 0 | 8.479573097 | 0 | 9.217629807 | 0 | 0 | 0 | 0.904635539 | 0.016321748 |
| LOC100190986 | 0 | 0 | 9.14407697 | 0 | 7.14981535 | 0 | 0 | 0 | 0.090045423 | 0.025297739 |
| LOC100233156 | 0 | 0 | 0 | 0 | 6.752524753 | 0 | 0 | 0 | 0.000121249 | 1 |
| LOC100288846 | 0 | 8.990303015 | 0 | 0 | 0 | 0 | 0 | 0 | 1.05E-07 | 1 |
| LOC100289230 | 0 | 8.263021342 | 0 | 0 | 0 | 0 | 0 | 0 | 2.55E-07 | 1 |
| LOC100289361 | 0 | 0 | 0 | 0 | 0 | 0 | 8.031074953 | 9.189014369 | 1 | 0.014458334 |
| LOC100289561 | 0 | 0 | 9.888474845 | 9.947213619 | 8.135987587 | 0 | 0 | 0 | 0.003651797 | 0.001429087 |
| LOC100419583 | 8.823881412 | 9.234482884 | 0 | 10.25826291 | 8.668631248 | 0 | 0 | 0 | 0.999483812 | 0.010883809 |
| LOC100505549 | 10.08202624 | 9.073911341 | 0 | 0 | 0 | 0 | 0 | 0 | 0.004378017 | 1 |
| LOC100506123 | 0 | 9.547979116 | 0 | 0 | 0 | 0 | 0 | 0 | 7.53E-08 | 1 |
| LOC100506472 | 10.29771056 | 0 | 0 | 0 | 0 | 0 | 0 | 0 | 7.07E-08 | 1 |
| LOC100506476 | 8.710683817 | 8.696202607 | 0 | 0 | 0 | 0 | 0 | 0 | 0.020815667 | 1 |
| LOC100506544 | 10.69823457 | 0 | 0 | 0 | 0 | 0 | 10.05848883 | 0 | 7.07E-08 | 1 |
| LOC100507507 | 0 | 0 | 0 | 0 | 6.375739247 | 0 | 0 | 0 | 6.06E-05 | 1 |
| LOC100996732 | 0 | 0 | 5.450018499 | 8.925071258 | 0 | 0 | 0 | 0 | 0.107878425 | 0.047456091 |
| LOC100996756 | 0 | 0 | 10.17178093 | 0 | 7.949072713 | 0 | 0 | 0 | 0.050752178 | 0.011852991 |

| | | | | | | | | | | |
|---|---|---|---|---|---|---|---|---|---|---|
| LOC100996842 | 0 | 0 | 0 | 7.689721819 | 9.172405179 | 0 | 0 | 0 | 0.080078756 | 0.02191077 |
| LOC101926967 | 9.427818601 | 0 | 0 | 0 | 0 | 0 | 0 | 0 | 7.88E-08 | 1 |
| LOC101927069 | 0 | 8.990303015 | 0 | 0 | 0 | 0 | 0 | 0 | 1.05E-07 | 1 |
| LOC101927100 | 10.8485001 | 0 | 0 | 0 | 0 | 0 | 0 | 0 | 7.07E-08 | 1 |
| LOC101927511 | 8.888924932 | 0 | 0 | 0 | 0 | 0 | 0 | 0 | 1.14E-07 | 1 |
| LOC101928076 | 0 | 9.044065918 | 0 | 0 | 0 | 0 | 0 | 0 | 1.01E-07 | 1 |
| LOC101928156 | 0 | 0 | 10.18654497 | 0 | 8.04555511 | 0 | 0 | 0 | 0.049042449 | 0.011625223 |
| LOC101928215 | 0 | 10.58766729 | 0 | 0 | 0 | 0 | 0 | 0 | 7.07E-08 | 1 |
| LOC101928323 | 0 | 0 | 0 | 0 | 6.39051047 | 0 | 0 | 0 | 5.98E-05 | 1 |
| LOC101928347 | 0 | 0 | 7.527565756 | 0 | 0 | 0 | 0 | 0 | 2.53E-05 | 1 |
| LOC101929709 | 9.259149778 | 0 | 0 | 0 | 0 | 0 | 0 | 0 | 8.58E-08 | 1 |
| LOC101929796 | 0 | 10.11266356 | 0 | 0 | 0 | 0 | 8.81882422 | 0 | 7.07E-08 | 1 |
| LOC101930071 | 0 | 0 | 0 | 0 | 6.345735199 | 8.147082208 | 0 | 0 | 6.22E-05 | 1 |
| LOC102723728 | 9.443649659 | 0 | 0 | 0 | 0 | 0 | 0 | 0 | 7.88E-08 | 1 |
| LOC102723878 | 0 | 0 | 0 | 8.610559011 | 0 | 0 | 4.814710258 | 0 | 7.54E-06 | 1 |
| LOC102723906 | 0 | 0 | 0 | 0 | 6.283787608 | 0 | 0 | 0 | 0.000155241 | 1 |
| LOC102724156 | 0 | 0 | 7.552095279 | 0 | 0 | 0 | 0 | 0 | 2.82E-05 | 1 |
| LOC102724397 | 0 | 0 | 0 | 10.13122842 | 8.113905325 | 0 | 0 | 0 | 0.048152031 | 0.012259484 |
| LOC102724720 | 0 | 0 | 0 | 0 | 6.503532343 | 0 | 0 | 0 | 5.50E-05 | 1 |
| LOC102724760 | 0 | 10.29793797 | 0 | 0 | 0 | 0 | 0 | 0 | 7.07E-08 | 1 |
| LOC103091866 | 0 | 9.645612889 | 0 | 0 | 0 | 0 | 0 | 0 | 7.38E-08 | 1 |
| LOC105369347 | 0 | 9.713548361 | 0 | 0 | 0 | 0 | 0 | 0 | 7.17E-08 | 1 |
| LOC105369360 | 10.14631859 | 0 | 0 | 0 | 0 | 0 | 0 | 0 | 7.07E-08 | 1 |
| LOC105369370 | 0 | 8.942570108 | 0 | 0 | 0 | 0 | 0 | 0 | 1.07E-07 | 1 |
| LOC105369656 | 0 | 8.51225226 | 0 | 0 | 0 | 0 | 0 | 0 | 1.38E-07 | 1 |
| LOC105369690 | 0 | 8.533888356 | 0 | 0 | 0 | 0 | 0 | 0 | 1.35E-07 | 1 |
| LOC105369748 | 0 | 0 | 0 | 0 | 0 | 10.76356111 | 0 | 9.779808847 | 1 | 0.006244708 |
| LOC105369819 | 10.19856778 | 0 | 0 | 0 | 0 | 0 | 0 | 0 | 7.07E-08 | 1 |
| LOC105370546 | 0 | 0 | 8.95926661 | 10.12347196 | 0 | 0 | 0 | 0 | 0.041111287 | 0.011070952 |
| LOC105370624 | 0 | 0 | 0 | 0 | 0 | 0 | 8.399062803 | 10.17436901 | 1 | 0.009724498 |
| LOC105370858 | 0 | 9.028908435 | 0 | 0 | 0 | 0 | 0 | 0 | 1.03E-07 | 1 |
| LOC105371257 | 0 | 9.574319617 | 0 | 0 | 0 | 0 | 0 | 0 | 7.58E-08 | 1 |
| LOC105371554 | 0 | 0 | 0 | 0 | 6.39051047 | 0 | 0 | 0 | 5.98E-05 | 1 |
| LOC105371762 | 11.68952574 | 9.521148737 | 11.12642513 | 5.885825233 | 8.790579971 | 0 | 0 | 0 | 0.837104821 | 0.001538174 |
| LOC105371763 | 10.21710791 | 0 | 0 | 0 | 0 | 10.7261505 | 8.848860281 | 0 | 7.07E-08 | 0.007255458 |
| LOC105371870 | 8.616058634 | 0 | 0 | 0 | 0 | 0 | 0 | 0 | 1.33E-07 | 1 |
| LOC105372246 | 0 | 8.705765621 | 0 | 0 | 0 | 0 | 0 | 0 | 1.18E-07 | 1 |
| LOC105372401 | 0 | 8.64741294 | 0 | 0 | 0 | 0 | 0 | 0 | 1.23E-07 | 1 |
| LOC105372426 | 0 | 0 | 0 | 7.101245469 | 0 | 0 | 0 | 0 | 3.59E-05 | 1 |
| LOC105372441 | 12.12034182 | 12.51875168 | 11.61093805 | 9.710592252 | 9.920719238 | 0 | 0 | 0 | 0.257087869 | 0.00031612 |
| LOC105372480 | 0 | 0 | 9.845707608 | 0 | 8.368257464 | 0 | 0 | 0 | 0.057737276 | 0.012722919 |
| LOC105372578 | 0 | 0 | 10.57258523 | 0 | 7.903457334 | 0 | 0 | 0 | 0.035847856 | 0.010542114 |
| LOC105372803 | 0 | 0 | 0 | 0 | 0 | 10.93517695 | 10.80498032 | 0 | 1 | 0.003610405 |

| | | | | | | | | | | |
|---|---|---|---|---|---|---|---|---|---|---|
| LOC105372962 | 9.371000476 | 0 | 0 | 0 | 0 | 0 | 0 | 0 | 8.05E-08 | 1 |
| LOC105373428 | 0 | 0 | 0 | 0 | 6.873120683 | 0 | 0 | 0 | 4.05E-05 | 1 |
| LOC105373733 | 0 | 0 | 0 | 0 | 6.808593567 | 0 | 0 | 0 | 6.32E-05 | 1 |
| LOC105373759 | 0 | 9.750979006 | 0 | 0 | 0 | 0 | 0 | 0 | 7.17E-08 | 1 |
| LOC105373785 | 0 | 9.515722261 | 0 | 0 | 0 | 0 | 7.962083516 | 0 | 7.58E-08 | 1 |
| LOC105374041 | 0 | 0 | 7.464352792 | 0 | 0 | 0 | 0 | 0 | 2.70E-05 | 1 |
| LOC105374080 | 9.285741834 | 0 | 0 | 0 | 0 | 0 | 0 | 0 | 8.38E-08 | 1 |
| LOC105374148 | 10.71463784 | 0 | 0 | 0 | 0 | 0 | 0 | 0 | 7.07E-08 | 1 |
| LOC105376095 | 9.176299815 | 0 | 0 | 0 | 0 | 0 | 0 | 0 | 1.15E-07 | 1 |
| LOC105376159 | 11.09646054 | 0 | 0 | 0 | 0 | 0 | 10.02599377 | 0 | 5.99E-08 | 1 |
| LOC105377094 | 0 | 9.159912786 | 0 | 10.07871314 | 7.509159523 | 0 | 0 | 0 | 0.98939231 | 0.013610425 |
| LOC105377763 | 9.927358907 | 0 | 0 | 0 | 0 | 0 | 0 | 0 | 7.07E-08 | 1 |
| LOC105378179 | 8.811733678 | 0 | 0 | 0 | 0 | 0 | 0 | 0 | 1.17E-07 | 1 |
| LOC105378539 | 9.137893062 | 0 | 0 | 0 | 0 | 0 | 0 | 0 | 1.17E-07 | 1 |
| LOC105378720 | 0 | 9.286392738 | 0 | 0 | 0 | 0 | 0 | 0 | 8.08E-08 | 1 |
| LOC105378948 | 10.49293202 | 0 | 0 | 0 | 0 | 0 | 0 | 0 | 7.07E-08 | 1 |
| LOC105379172 | 10.56707273 | 0 | 0 | 0 | 0 | 0 | 7.567155003 | 0 | 7.07E-08 | 1 |
| LOC105379280 | 0 | 0 | 0 | 0 | 0 | 10.45800436 | 8.008442421 | 0 | 1 | 0.009118992 |
| LOC105379499 | 0 | 0 | 0 | 7.080028634 | 0 | 3.656335968 | 0 | 0 | 3.68E-05 | 1 |
| LOC105379508 | 0 | 0 | 9.298847925 | 0 | 7.745819102 | 0 | 0 | 0 | 0.080058324 | 0.019369457 |
| LOC105379524 | 9.395625601 | 0 | 0 | 0 | 0 | 0 | 0 | 0 | 8.04E-08 | 1 |
| LOC107984329 | 0 | 8.884809218 | 0 | 0 | 0 | 0 | 0 | 0 | 1.09E-07 | 1 |
| LOC107984347 | 0 | 10.00446552 | 0 | 0 | 0 | 0 | 10.18497771 | 10.12294533 | 7.07E-08 | 0.00670453 |
| LOC107984813 | 0 | 0 | 0 | 0 | 6.556887521 | 0 | 0 | 0 | 9.34E-05 | 1 |
| LOC107984948 | 0 | 9.426002279 | 0 | 0 | 0 | 0 | 0 | 0 | 7.87E-08 | 1 |
| LOC107985208 | 10.8720903 | 0 | 0 | 0 | 0 | 0 | 0 | 0 | 7.07E-08 | 1 |
| LOC107985216 | 8.847874263 | 0 | 0 | 0 | 0 | 0 | 0 | 0 | 1.15E-07 | 1 |
| LOC107985249 | 0 | 9.704036967 | 0 | 0 | 0 | 0 | 0 | 0 | 7.37E-08 | 1 |
| LOC107985367 | 0 | 8.893203714 | 0 | 0 | 0 | 0 | 0 | 0 | 1.09E-07 | 1 |
| LOC107985404 | 0 | 0 | 0 | 0 | 0 | 0 | 9.310139681 | 8.783658497 | 1 | 0.01143424 |
| LOC107985433 | 10.6202803 | 0 | 0 | 0 | 0 | 0 | 0 | 0 | 7.07E-08 | 1 |
| LOC107985480 | 8.77466443 | 0 | 0 | 0 | 0 | 10.24336887 | 0 | 0 | 1.18E-07 | 1 |
| LOC107985554 | 10.53047859 | 0 | 0 | 0 | 0 | 0 | 0 | 0 | 7.07E-08 | 1 |
| LOC107985567 | 0 | 0 | 0 | 9.774722758 | 8.923477098 | 10.79217695 | 0 | 0 | 0.04828143 | 0.999854694 |
| LOC107985733 | 0 | 9.757890409 | 10.00522363 | 0 | 8.942396466 | 0 | 0 | 0 | 0.997434996 | 0.011034216 |
| LOC107985910 | 9.527745184 | 0 | 0 | 0 | 0 | 0 | 0 | 0 | 7.73E-08 | 1 |
| LOC107985917 | 9.026677385 | 0 | 0 | 0 | 0 | 0 | 0 | 0 | 1.07E-07 | 1 |
| LOC107985923 | 0 | 0 | 7.064877781 | 0 | 0 | 0 | 0 | 0 | 4.02E-05 | 1 |
| LOC107986003 | 0 | 0 | 0 | 7.297837142 | 0 | 0 | 0 | 0 | 2.94E-05 | 1 |
| LOC107986298 | 9.320452216 | 0 | 0 | 0 | 0 | 0 | 0 | 0 | 8.22E-08 | 1 |
| LOC107986348 | 0 | 9.589896123 | 0 | 0 | 0 | 0 | 0 | 0 | 7.53E-08 | 1 |
| LOC107986483 | 0 | 0 | 0 | 0 | 0 | 9.438056497 | 10.15695466 | 0 | 1 | 0.008002596 |
| LOC107986854 | 0 | 9.426002279 | 0 | 0 | 0 | 0 | 0 | 0 | 7.87E-08 | 1 |

| | | | | | | | | | | |
|---|---|---|---|---|---|---|---|---|---|---|
| LOC107986991 | 9.828331099 | 0 | 0 | 0 | 0 | 0 | 0 | 0 | 7.17E-08 | 1 |
| LOC107986997 | 10.99804444 | 0 | 0 | 0 | 0 | 0 | 0 | 0 | 7.07E-08 | 1 |
| LOC107987043 | 10.2354128 | 0 | 0 | 0 | 0 | 0 | 0 | 0 | 7.07E-08 | 1 |
| LOC107987072 | 9.998749664 | 0 | 0 | 0 | 0 | 0 | 6.664334008 | 0 | 7.07E-08 | 1 |
| LOC107987102 | 0 | 9.553285831 | 0 | 0 | 0 | 0 | 8.69200653 | 0 | 7.53E-08 | 1 |
| LOC107987135 | 10.29334884 | 0 | 0 | 0 | 0 | 0 | 0 | 0 | 7.07E-08 | 1 |
| LOC107987136 | 0 | 0 | 0 | 8.320639466 | 7.509159523 | 0 | 0 | 0 | 0.109442361 | 0.044296831 |
| LOC107987206 | 0 | 0 | 9.445175324 | 0 | 9.379092464 | 0 | 0 | 8.178109615 | 0.04846603 | 1 |
| LOC107987251 | 9.520299506 | 0 | 0 | 0 | 0 | 0 | 0 | 0 | 7.73E-08 | 1 |
| LOC107987276 | 0 | 0 | 0 | 0 | 6.819550507 | 0 | 0 | 0 | 4.21E-05 | 1 |
| LOC107987294 | 10.08202624 | 0 | 0 | 0 | 0 | 0 | 0 | 0 | 7.07E-08 | 1 |
| LOC107987299 | 0 | 9.941103266 | 0 | 0 | 0 | 0 | 0 | 0 | 7.07E-08 | 1 |
| LOC108783654 | 10.24899064 | 0 | 0 | 0 | 0 | 0 | 0 | 0 | 7.07E-08 | 1 |
| LOC110384692 | 0 | 9.92890668 | 0 | 0 | 0 | 0 | 0 | 0 | 7.07E-08 | 1 |
| LOC112267883 | 10.31932352 | 0 | 0 | 0 | 0 | 0 | 0 | 8.354929434 | 7.07E-08 | 1 |
| LOC112268035 | 0 | 0 | 0 | 0 | 6.682232705 | 0 | 0 | 0 | 4.74E-05 | 1 |
| LOC112268037 | 0 | 0 | 0 | 0 | 6.694188464 | 0 | 0 | 0 | 0.000126205 | 1 |
| LOC112268063 | 9.276932164 | 0 | 0 | 0 | 0 | 0 | 0 | 0 | 8.39E-08 | 1 |
| LOC112268088 | 8.762093289 | 0 | 0 | 0 | 0 | 0 | 0 | 0 | 1.18E-07 | 1 |
| LOC112268098 | 0 | 0 | 9.065319476 | 8.700196723 | 0 | 0 | 0 | 0 | 0.074990497 | 0.017717402 |
| LOC112268109 | 0 | 0 | 7.438271136 | 9.32741249 | 6.419606803 | 0 | 0 | 0 | 0.037972528 | 0.008378564 |
| LOC112268124 | 0 | 0 | 0 | 0 | 6.682232705 | 0 | 0 | 0 | 4.74E-05 | 1 |
| LOC112268146 | 0 | 9.311663645 | 0 | 0 | 0 | 0 | 0 | 0 | 8.05E-08 | 1 |
| LOC112268184 | 10.44654697 | 8.501311304 | 0 | 0 | 0 | 0 | 0 | 0 | 0.003455535 | 1 |
| LOC112268313 | 0 | 0 | 0 | 9.559738748 | 9.151961534 | 0 | 0 | 8.366767416 | 0.049865925 | 0.999854694 |
| LOC171391 | 0 | 8.586600217 | 0 | 0 | 0 | 0 | 0 | 0 | 1.30E-07 | 1 |
| LOC257396 | 0 | 9.809805721 | 0 | 0 | 0 | 0 | 0 | 0 | 7.15E-08 | 1 |
| LOC283710 | 0 | 8.250007882 | 0 | 0 | 0 | 0 | 0 | 0 | 2.64E-07 | 1 |
| LOC283788 | 0 | 0 | 10.3121359 | 0 | 8.815422744 | 0 | 0 | 10.8439491 | 0.035320365 | 0.999854694 |
| LOC284454 | 0 | 0 | 7.451370902 | 9.895017287 | 4.688348153 | 0 | 0 | 9.215441638 | 0.030866583 | 0.999854694 |
| LOC284581 | 0 | 0 | 0 | 9.022821486 | 7.542607814 | 0 | 0 | 0 | 0.087401939 | 0.026167363 |
| LOC389602 | 0 | 0 | 0 | 0 | 0 | 0 | 7.332210197 | 10.64687219 | 1 | 0.009118992 |
| LOC389831 | 0 | 0 | 10.29043082 | 0 | 8.233426702 | 0 | 0 | 0 | 0.043012141 | 0.011034216 |
| LOC389906 | 0 | 9.214524828 | 0 | 0 | 0 | 0 | 0 | 0 | 8.39E-08 | 1 |
| LOC646626 | 11.2461427 | 0 | 10.27945458 | 0 | 11.25114219 | 0 | 0 | 0 | 0.999483812 | 0.005764538 |
| LOC728743 | 10.6648577 | 0 | 0 | 0 | 0 | 0 | 0 | 10.40426152 | 7.07E-08 | 1 |
| LOC729737 | 0 | 0 | 0 | 0 | 6.448127902 | 0 | 0 | 0 | 5.74E-05 | 1 |
| LOC93622 | 0 | 0 | 10.47973879 | 0 | 6.556887521 | 0 | 6.605228635 | 0 | 0.048100936 | 0.999854694 |
| LONP1 | 9.185743846 | 0 | 0 | 8.614253516 | 6.830424858 | 0 | 0 | 0 | 0.985757273 | 0.045174485 |
| LONP2 | 0 | 0 | 0 | 10.19922153 | 9.241250091 | 0 | 0 | 0 | 0.030946452 | 0.01037888 |
| LONRF1 | 9.722001672 | 0 | 0 | 0 | 0 | 0 | 0 | 0 | 7.30E-08 | 1 |
| LPCAT1 | 0 | 0 | 0 | 9.16057826 | 6.85193071 | 0 | 0 | 0 | 0.089445503 | 0.028091929 |
| LPIN3 | 9.85816911 | 0 | 0 | 0 | 0 | 0 | 5.955871074 | 0 | 7.17E-08 | 1 |

| | | | | | | | | | | |
|---|---|---|---|---|---|---|---|---|---|---|
| LRCH4 | 10.64274116 | 0 | 0 | 0 | 0 | 0 | 0 | 10.01800757 | 7.07E-08 | 1 |
| LRFN4 | 9.564407853 | 0 | 0 | 0 | 0 | 0 | 8.075301928 | 0 | 7.60E-08 | 1 |
| LRIG2 | 8.697540019 | 0 | 0 | 0 | 0 | 0 | 0 | 0 | 1.24E-07 | 1 |
| LRP5 | 0 | 0 | 0 | 7.202887337 | 8.887483063 | 0 | 0 | 0 | 0.092900759 | 0.031153436 |
| LRRC14 | 0 | 0 | 0 | 8.534609709 | 9.072895476 | 0 | 0 | 0 | 0.073972838 | 0.017687269 |
| LRRC19 | 0 | 8.223623425 | 0 | 0 | 0 | 0 | 0 | 0 | 2.64E-07 | 1 |
| LRRC32 | 0 | 8.998107186 | 0 | 0 | 0 | 0 | 8.08254305 | 0 | 2.64E-07 | 1 |
| LRRC34 | 0 | 0 | 0 | 9.856543048 | 12.42045722 | 10.83023611 | 8.682506857 | 12.48277403 | 1.15E-07 | 0.999854694 |
| LRRC42 | 0 | 9.56908985 | 0 | 0 | 0 | 8.055624028 | 0 | 0 | 7.53E-08 | 1 |
| LRRC75A | 0 | 8.974566872 | 0 | 0 | 0 | 11.24507549 | 0 | 0 | 1.07E-07 | 1 |
| LRRFIP2 | 10.54522782 | 8.196747437 | 10.38050358 | 0 | 7.903457334 | 0 | 0 | 0 | 0.971198277 | 0.011034216 |
| LSG1 | 0 | 0 | 0 | 0 | 4.034088102 | 10.99520968 | 8.360860894 | 11.03689304 | 0.611721328 | 0.019369457 |
| LSM14A | 0 | 0 | 10.16185361 | 8.293245504 | 9.999914943 | 0 | 4.947545984 | 0 | 0.00194559 | 0.130219786 |
| LTB | 10.93783941 | 0 | 0 | 0 | 0 | 0 | 0 | 0 | 7.07E-08 | 1 |
| LTV1 | 0 | 0 | 11.47723355 | 9.652489139 | 8.926014062 | 0 | 8.309327743 | 0 | 7.97E-05 | 0.99475784 |
| LYRM1 | 0 | 9.708800502 | 0 | 0 | 0 | 0 | 0 | 0 | 7.17E-08 | 1 |
| LYVE1 | 9.24114547 | 0 | 0 | 0 | 0 | 0 | 0 | 0 | 1.07E-07 | 1 |
| LZIC | 0 | 0 | 9.211075254 | 0 | 9.844317895 | 0 | 9.561438172 | 0 | 0.04059308 | 0.999854694 |
| MAD2L1BP | 11.62330142 | 10.65462132 | 0 | 0 | 0 | 0 | 8.765832683 | 0 | 5.29E-07 | 1 |
| MAF | 0 | 0 | 8.033583097 | 0 | 0 | 0 | 0 | 0 | 1.51E-05 | 1 |
| MAFB | 0 | 9.354849786 | 0 | 0 | 0 | 0 | 0 | 0 | 7.91E-08 | 1 |
| MAFF | 0 | 0 | 10.48611794 | 9.656077944 | 0 | 0 | 0 | 0 | 0.021854315 | 0.009237175 |
| MAGED1 | 0 | 0 | 10.40764081 | 7.969008339 | 9.489551709 | 0 | 8.193715158 | 0 | 0.00299894 | 0.999854694 |
| MAGED2 | 0 | 0 | 10.28495314 | 12.20137435 | 9.554936499 | 11.06201201 | 11.43793308 | 0 | 1.38E-07 | 0.999854694 |
| MAGEF1 | 0 | 0 | 10.28220648 | 0 | 8.091479805 | 10.88955797 | 0 | 11.33156982 | 0.044978193 | 0.999854694 |
| MAGEH1 | 0 | 8.909847578 | 0 | 0 | 0 | 0 | 0 | 10.64808532 | 1.08E-07 | 1 |
| MAGIX | 0 | 9.241074683 | 0 | 0 | 0 | 0 | 0 | 0 | 8.30E-08 | 1 |
| MAIP1 | 0 | 0 | 0 | 9.899565999 | 8.798908517 | 0 | 0 | 0 | 0.046802627 | 0.011852991 |
| MAMDC4 | 9.268068368 | 0 | 0 | 0 | 0 | 0 | 0 | 0 | 8.58E-08 | 1 |
| MAN1B1 | 0 | 0 | 0 | 0 | 6.433937832 | 0 | 0 | 0 | 5.81E-05 | 1 |
| MAN2C1 | 9.057869043 | 6.388120724 | 11.22458993 | 10.38477752 | 7.675191427 | 0 | 0 | 0 | 0.624632107 | 0.000592153 |
| MANEAL | 0 | 7.118179823 | 0 | 10.44721421 | 8.653402207 | 0 | 0 | 0 | 0.727712695 | 0.01037888 |
| MAOA | 9.35434707 | 11.53182319 | 0 | 0 | 0 | 0 | 0 | 0 | 2.60E-05 | 1 |
| MAP2K7 | 0 | 0 | 9.422072283 | 0 | 6.741044679 | 0 | 0 | 0 | 0.084044243 | 0.022155303 |
| MAP3K12 | 13.76715118 | 13.23780823 | 11.51982818 | 9.140222357 | 11.9919126 | 8.69036552 | 13.41495386 | 0 | 0.045408205 | 0.999854694 |
| MAP3K21 | 0 | 0 | 0 | 0 | 6.786426836 | 0 | 8.974838097 | 0 | 4.30E-05 | 1 |
| MAP3K4 | 10.70481827 | 0 | 0 | 0 | 0 | 0 | 8.26546142 | 0 | 7.07E-08 | 1 |
| MAP3K5 | 0 | 8.918098054 | 0 | 0 | 0 | 0 | 0 | 0 | 1.08E-07 | 1 |
| MAP3K6 | 0 | 8.555204765 | 0 | 0 | 0 | 0 | 0 | 0 | 1.34E-07 | 1 |
| MAP4K4 | 0 | 8.51225226 | 0 | 0 | 0 | 0 | 0 | 0 | 1.38E-07 | 1 |
| MAP4K5 | 0 | 8.789094606 | 0 | 0 | 0 | 0 | 0 | 8.87766687 | 1.15E-07 | 1 |
| MAPK1IP1L | 0 | 10.72332115 | 10.31572187 | 9.077372452 | 9.823377972 | 0 | 0 | 0 | 0.999483812 | 0.000859037 |
| MAPKAPK2 | 0 | 9.267144757 | 0 | 9.277027363 | 9.028728141 | 0 | 0 | 0 | 0.99353678 | 0.014003311 |

| | | | | | | | | | | |
|---|---|---|---|---|---|---|---|---|---|---|
| MAPRE1 | 0 | 0 | 0 | 9.047624983 | 9.962097038 | 0 | 8.720135904 | 9.557689932 | 0.039224551 | 0.999854694 |
| 2-Mar | 0 | 9.117552805 | 0 | 0 | 0 | 0 | 0 | 0 | 1.14E-07 | 1 |
| MARF1 | 0 | 0 | 10.6427867 | 0 | 8.759623899 | 0 | 0 | 0 | 0.026296017 | 0.009547202 |
| MARK1 | 10.11205603 | 0 | 0 | 0 | 0 | 0 | 0 | 0 | 7.07E-08 | 1 |
| MARVELD2 | 0 | 0 | 9.632230391 | 0 | 8.225210179 | 0 | 0 | 0 | 0.067266939 | 0.014388177 |
| MARVELD3 | 0 | 0 | 9.3042732 | 0 | 8.5617557 | 0 | 0 | 0 | 0.070604497 | 0.015338243 |
| MAST2 | 8.544455536 | 9.805364777 | 10.38392378 | 0 | 7.14981535 | 0 | 0 | 0 | 0.98939231 | 0.011685672 |
| MAVS | 0 | 0 | 0 | 9.878214892 | 6.448127902 | 0 | 0 | 0 | 0.070604497 | 0.017185058 |
| MB | 0 | 9.739385911 | 0 | 0 | 0 | 0 | 0 | 0 | 7.25E-08 | 1 |
| MBD6 | 9.259149778 | 8.008672964 | 0 | 0 | 0 | 0 | 9.857883783 | 0 | 0.020086812 | 1 |
| MBOAT4 | 10.27576888 | 0 | 0 | 0 | 0 | 0 | 0 | 0 | 7.07E-08 | 1 |
| MBTPS2 | 8.657374495 | 8.789094606 | 0 | 0 | 0 | 0 | 0 | 0 | 0.020329865 | 1 |
| MCCC1 | 0 | 0 | 0 | 0 | 6.706045958 | 0 | 7.567155003 | 10.24982197 | 0.000126325 | 0.999854694 |
| MCEE | 0 | 0 | 10.79762319 | 9.313844677 | 7.258484401 | 0 | 8.510636715 | 0 | 0.002389299 | 0.999854694 |
| MCMBP | 0 | 0 | 0 | 10.20658225 | 9.511677536 | 0 | 0 | 10.19775352 | 0.026684226 | 0.999854694 |
| MCRIP1 | 0 | 0 | 9.845707608 | 9.834540682 | 10.02024048 | 0 | 9.880909225 | 10.40713044 | 0.000724658 | 0.999854694 |
| MDFIC | 0 | 0 | 0 | 0 | 6.489879427 | 0 | 0 | 0 | 5.56E-05 | 1 |
| MDM1 | 0 | 8.422309346 | 0 | 0 | 0 | 0 | 0 | 0 | 1.48E-07 | 1 |
| MDP1 | 5.655231357 | 0 | 10.31929896 | 9.88434752 | 10.16579792 | 0 | 0 | 0 | 0.092132211 | 0.000566119 |
| ME3 | 0 | 8.876365592 | 9.168080295 | 0 | 8.689685392 | 0 | 0 | 0 | 0.990582117 | 0.015771922 |
| MECOM | 17.35475232 | 18.14588603 | 16.73473815 | 16.76507383 | 16.21893789 | 15.32114432 | 17.32773251 | 15.87115086 | 0.026913455 | 0.999854694 |
| MED14 | 0 | 0 | 6.902671521 | 0 | 0 | 0 | 0 | 0 | 4.61E-05 | 1 |
| MED25 | 0 | 9.234482884 | 0 | 0 | 0 | 0 | 0 | 0 | 8.36E-08 | 1 |
| MED27 | 0 | 0 | 8.73926584 | 10.46573697 | 9.033441525 | 0 | 0 | 10.72252592 | 0.002477655 | 1 |
| MED28 | 0 | 0 | 0 | 11.16184526 | 9.986608868 | 0 | 10.17145874 | 9.867826101 | 0.030621622 | 0.999854694 |
| MED29 | 0 | 0 | 0 | 6.922073503 | 0 | 0 | 0 | 0 | 0.000113876 | 1 |
| MED6 | 0 | 0 | 0 | 10.43473225 | 8.019659178 | 0 | 0 | 0 | 0.037622992 | 0.011034216 |
| MEF2D | 9.655586838 | 0 | 0 | 0 | 0 | 0 | 8.213638823 | 0 | 7.44E-08 | 1 |
| MEGF6 | 0 | 0 | 0 | 0 | 6.803083732 | 0 | 0 | 0 | 6.79E-05 | 1 |
| MELK | 0 | 0 | 10.56958472 | 0 | 8.78779307 | 0 | 0 | 0 | 0.028041796 | 0.009724498 |
| MELTF | 10.35324459 | 0 | 0 | 0 | 0 | 0 | 0 | 0 | 7.07E-08 | 1 |
| MEMO1 | 0 | 0 | 10.78150543 | 0 | 9.257456445 | 0 | 9.527351199 | 0 | 0.016961701 | 0.999854694 |
| MEPCE | 8.453559626 | 0 | 0 | 0 | 0 | 0 | 7.403330192 | 0 | 1.53E-07 | 1 |
| MESP1 | 10.69823457 | 0 | 0 | 0 | 0 | 0 | 0 | 10.01800757 | 7.07E-08 | 1 |
| METTL1 | 0 | 0 | 9.537131744 | 11.23445682 | 6.645759207 | 11.28253495 | 0 | 0 | 0.000676929 | 0.999854694 |
| METTL18 | 0 | 0 | 10.58897743 | 0 | 9.466209982 | 0 | 7.760139668 | 0 | 0.019215938 | 0.999854694 |
| METTL21A | 0 | 9.56908985 | 0 | 0 | 0 | 10.28423363 | 8.756808388 | 0 | 7.53E-08 | 0.008817225 |
| METTL25 | 0 | 8.990303015 | 0 | 0 | 0 | 0 | 0 | 0 | 1.05E-07 | 1 |
| METTL2B | 0 | 9.207810313 | 0 | 0 | 0 | 9.650922548 | 7.666871799 | 0 | 8.43E-08 | 0.012288713 |
| METTL4 | 0 | 8.837745532 | 0 | 0 | 0 | 0 | 0 | 0 | 1.13E-07 | 1 |
| METTL7B | 0 | 9.317912807 | 0 | 0 | 0 | 0 | 0 | 0 | 8.05E-08 | 1 |
| MEX3C | 10.35742919 | 10.52287411 | 0 | 0 | 0 | 0 | 9.882984338 | 0 | 0.000212422 | 1 |
| MFAP1 | 0 | 0 | 9.960109806 | 0 | 7.321504468 | 0 | 0 | 0 | 0.066119036 | 0.01420877 |

| | | | | | | | | | | |
|---|---|---|---|---|---|---|---|---|---|---|
| MFAP4 | 0 | 0 | 9.336403268 | 0 | 10.23699607 | 10.79217695 | 0 | 9.714142401 | 0.027908204 | 0.999854694 |
| MFAP5 | 0 | 0 | 11.50929684 | 0 | 7.242289655 | 0 | 0 | 0 | 9.19E-07 | 3.14E-11 |
| MFGE8 | 0 | 0 | 0 | 9.313844677 | 8.794750254 | 0 | 0 | 0 | 0.06536001 | 0.014692722 |
| MFRP | 0 | 0 | 10.6484855 | 7.997577745 | 10.68781017 | 0 | 7.535844571 | 0 | 0.000238169 | 1 |
| MFSD10 | 0 | 0 | 10.49246902 | 9.726040692 | 8.926014062 | 0 | 0 | 9.148443004 | 0.001114205 | 0.999854694 |
| MFSD11 | 0 | 0 | 11.38764931 | 8.977645149 | 9.129023686 | 0 | 8.489001577 | 10.35756552 | 0.000191393 | 0.999854694 |
| MFSD14A | 0 | 0 | 0 | 10.4147456 | 9.558210998 | 0 | 8.874116951 | 0 | 0.020704959 | 0.999854694 |
| MFSD14B | 8.973144195 | 6.868617716 | 8.536001166 | 0 | 9.4513172 | 0 | 0 | 0 | 0.99396342 | 0.014458334 |
| MFSD2A | 10.28897389 | 0 | 0 | 0 | 0 | 0 | 9.592203879 | 0 | 7.07E-08 | 1 |
| MFSD6 | 0 | 9.214524828 | 0 | 0 | 0 | 0 | 0 | 0 | 8.39E-08 | 1 |
| MFSD6L | 10.04098874 | 0 | 0 | 0 | 0 | 0 | 0 | 6.83664825 | 7.07E-08 | 1 |
| MGAT1 | 0 | 0 | 0 | 0 | 6.375739247 | 0 | 9.743766328 | 8.538825133 | 6.06E-05 | 0.999854694 |
| MGAT2 | 8.973144195 | 9.059065806 | 8.720451232 | 9.654284657 | 7.290338514 | 0 | 0 | 0 | 0.996280769 | 0.003988377 |
| MIA2 | 10.06677365 | 8.069195977 | 8.59030499 | 0 | 7.258484401 | 0 | 0 | 0 | 0.865943987 | 0.038158619 |
| MIA3 | 0 | 0 | 0 | 10.3574839 | 8.641102159 | 0 | 8.935389379 | 0 | 0.034170548 | 0.999854694 |
| MIAT | 10.27576888 | 0 | 0 | 0 | 0 | 0 | 0 | 0 | 7.07E-08 | 1 |
| MICALL2 | 10.95312784 | 0 | 8.809621568 | 10.32064707 | 8.653402207 | 0 | 0 | 0 | 0.987285143 | 0.001521549 |
| MICOS10 | 0 | 0 | 0 | 11.03459204 | 8.1271952 | 0 | 7.930328737 | 0 | 0.016865451 | 0.999854694 |
| MICOS13 | 0 | 6.128527206 | 10.78020819 | 11.71834004 | 7.650859037 | 10.16894608 | 9.428523561 | 10.14373527 | 0.043619256 | 0.999854694 |
| MINDY2 | 0 | 0 | 0 | 0 | 0 | 10.93731372 | 8.166714905 | 0 | 1 | 0.007255458 |
| MIPEPP3 | 0 | 0 | 0 | 9.670344553 | 8.064100782 | 0 | 0 | 0 | 0.06658548 | 0.015061471 |
| MIPOL1 | 0 | 7.089245392 | 9.99631302 | 10.72433426 | 11.01508451 | 0 | 0 | 0 | 0.118670292 | 0.000291056 |
| MIR200CHG | 9.005501878 | 8.607157265 | 9.185824288 | 0 | 7.785476285 | 0 | 0 | 0 | 0.980815574 | 0.020809157 |
| MIS12 | 9.987993796 | 0 | 0 | 0 | 0 | 0 | 0 | 0 | 7.07E-08 | 1 |
| MISP3 | 0 | 10.05022532 | 9.183863476 | 0 | 7.796609528 | 0 | 0 | 0 | 0.971198277 | 0.020809157 |
| MITD1 | 0 | 0 | 11.7631504 | 0 | 11.0052211 | 0 | 10.51705045 | 0 | 0.020363098 | 0.999854694 |
| MKNK2 | 13.70077192 | 12.97113248 | 15.68814396 | 13.63145441 | 14.85562032 | 13.39989269 | 12.30660453 | 13.70044956 | 0.045024163 | 0.999854694 |
| MLLT3 | 10.57428148 | 0 | 0 | 0 | 0 | 0 | 7.218490506 | 0 | 7.07E-08 | 1 |
| MLPH | 0 | 8.934458793 | 0 | 9.590072571 | 8.790579971 | 0 | 0 | 0 | 0.983546955 | 0.013425237 |
| MLST8 | 0 | 0 | 0 | 9.656077944 | 9.165980533 | 0 | 7.67647464 | 0 | 0.046970352 | 0.999854694 |
| MMEL1 | 0 | 0 | 0 | 0 | 0 | 0 | 7.356308522 | 10.26255067 | 1 | 0.01037888 |
| MMP14 | 0 | 0 | 8.741933722 | 0 | 8.077855242 | 0 | 0 | 0 | 0.089462396 | 0.024827605 |
| MMP17 | 10.09711927 | 0 | 0 | 0 | 0 | 0 | 0 | 0 | 7.07E-08 | 1 |
| MMP2 | 0 | 7.559034993 | 0 | 9.014457977 | 11.66969465 | 0 | 0 | 0 | 0.682271578 | 0.006115638 |
| MMP25 | 0 | 0 | 0 | 0 | 6.645759207 | 0 | 0 | 0 | 7.73E-05 | 1 |
| MMUT | 0 | 8.743396117 | 0 | 0 | 0 | 9.725354145 | 0 | 0 | 1.17E-07 | 1 |
| MND1 | 0 | 0 | 8.126348344 | 10.28284497 | 0 | 0 | 0 | 0 | 0.045270943 | 0.011814381 |
| MNT | 0 | 0 | 0 | 9.297852588 | 7.588169674 | 0 | 0 | 0 | 0.079549852 | 0.021202278 |
| MOB2 | 10.70481827 | 0 | 0 | 0 | 0 | 0 | 0 | 0 | 7.07E-08 | 1 |
| MOB3B | 0 | 8.363107485 | 0 | 0 | 0 | 0 | 0 | 0 | 1.55E-07 | 1 |
| MOCS2 | 9.303201411 | 0 | 9.773224785 | 11.19860782 | 0 | 0 | 0 | 0 | 0.935616984 | 0.007255458 |
| MORC4 | 0 | 0 | 0 | 0 | 6.682232705 | 0 | 0 | 0 | 4.74E-05 | 1 |
| MORN2 | 10.46799206 | 0 | 0 | 9.509044094 | 11.19308435 | 0 | 0 | 0 | 0.98867228 | 0.007236136 |

| | | | | | | | | | | |
|---|---|---|---|---|---|---|---|---|---|---|
| MORN4 | 10.15596027 | 0 | 0 | 0 | 0 | 0 | 0 | 7.505702501 | 7.07E-08 | 1 |
| MOSPD1 | 0 | 0 | 9.913539582 | 9.72945133 | 7.851022366 | 0 | 0 | 0 | 0.00531921 | 0.001661226 |
| MPEG1 | 0 | 0 | 8.042267146 | 0 | 0 | 0 | 0 | 0 | 1.50E-05 | 1 |
| MPG | 10.43866903 | 4.344099474 | 9.683091274 | 0 | 10.59839191 | 0 | 0 | 0 | 0.99353678 | 0.008532333 |
| MPHOSPH6 | 0 | 0 | 9.957816655 | 10.35417369 | 9.888562842 | 0 | 9.516698783 | 0 | 0.000314807 | 0.999854694 |
| MPND | 11.16967691 | 9.684823965 | 8.65401506 | 0 | 8.358899494 | 0 | 0 | 0 | 0.626823469 | 0.023100282 |
| MPP6 | 9.063002713 | 0 | 0 | 0 | 0 | 0 | 0 | 0 | 1.05E-07 | 1 |
| MPPED2 | 11.65424464 | 8.596915356 | 0 | 0 | 0 | 0 | 9.514023343 | 0 | 4.19E-05 | 1 |
| MPZL2 | 0 | 10.22997487 | 9.471133574 | 10.36188573 | 10.6518624 | 0 | 0 | 0 | 0.936409549 | 0.000525098 |
| MPZL3 | 8.684275372 | 0 | 9.315062926 | 0 | 7.051005161 | 0 | 0 | 0 | 0.995442919 | 0.02266389 |
| MREG | 0 | 0 | 0 | 0 | 6.752524753 | 0 | 0 | 0 | 0.000121249 | 1 |
| MRFAP1L1 | 0 | 9.655516021 | 10.6356314 | 10.08671334 | 10.3973349 | 0 | 0 | 0 | 0.800923714 | 0.000413289 |
| MROH3P | 0 | 0 | 6.893102272 | 0 | 0 | 0 | 0 | 0 | 5.95E-05 | 1 |
| MRPL10 | 0 | 0 | 9.096923479 | 11.27411497 | 9.818607242 | 0 | 0 | 10.47842425 | 8.60E-05 | 0.999854694 |
| MRPL20-AS1 | 0 | 0 | 0 | 9.981944242 | 7.064884439 | 0 | 0 | 0 | 0.065076325 | 0.014818059 |
| MRPL28 | 10.58145439 | 9.2476365 | 9.74291396 | 11.4003756 | 7.829502868 | 0 | 0 | 0 | 0.98939231 | 0.000646553 |
| MRPL37 | 8.736617533 | 10.36586202 | 11.30210885 | 10.29670606 | 10.4307683 | 0 | 0 | 0 | 0.838607701 | 0.00026433 |
| MRPL40 | 0 | 0 | 0 | 10.16184385 | 10.89558888 | 0 | 9.574337042 | 9.23817494 | 0.031968155 | 0.999854694 |
| MRPL46 | 0 | 0 | 0 | 11.33415261 | 8.002934593 | 0 | 10.45563462 | 0 | 0.034540642 | 0.999854694 |
| MRPS10 | 0 | 0 | 0 | 10.94648274 | 10.08410392 | 0 | 8.82746995 | 12.25013188 | 0.03230827 | 0.999854694 |
| MRPS17 | 0 | 0 | 10.33351926 | 11.21451557 | 9.763643177 | 0 | 9.294634579 | 9.295038347 | 1.96E-05 | 0.999854694 |
| MRPS18B | 0 | 0 | 9.552410397 | 11.14406533 | 10.00592283 | 10.06984623 | 10.10545472 | 10.58993625 | 5.88E-05 | 0.999854694 |
| MRPS30 | 0 | 0 | 10.6356314 | 0 | 8.301439975 | 0 | 0 | 0 | 0.030634721 | 0.010003595 |
| MRPS33 | 18.95098106 | 19.28746381 | 17.68231901 | 18.19540937 | 17.82708149 | 18.81103893 | 19.43042644 | 17.1515597 | 0.010019796 | 0.999854694 |
| MRRF | 0 | 8.236875968 | 0 | 0 | 0 | 0 | 8.08254305 | 0 | 2.64E-07 | 1 |
| MRS2 | 0 | 10.83755517 | 0 | 0 | 0 | 0 | 8.836064177 | 11.31103821 | 7.07E-08 | 0.00558734 |
| MRTO4 | 9.641928937 | 0 | 10.44085991 | 10.59849238 | 8.307290009 | 0 | 0 | 0 | 0.880392942 | 0.000712114 |
| MS4A7 | 0 | 0 | 9.211075254 | 0 | 7.978700908 | 0 | 0 | 0 | 0.079733579 | 0.01919178 |
| MSANTD2 | 8.736617533 | 0 | 0 | 0 | 0 | 0 | 0 | 0 | 1.20E-07 | 1 |
| MSH5-SAPCD1 | 0 | 8.876365592 | 0 | 0 | 0 | 0 | 0 | 0 | 1.10E-07 | 1 |
| MSN | 0 | 9.378959303 | 9.833251866 | 0 | 7.807657514 | 0 | 0 | 0 | 0.999483812 | 0.01420877 |
| MSRB3 | 0 | 0 | 0 | 7.580648887 | 0 | 0 | 0 | 0 | 2.21E-05 | 1 |
| MT1A | 0 | 9.021269553 | 0 | 0 | 0 | 0 | 0 | 0 | 1.03E-07 | 1 |
| MT1DP | 0 | 0 | 9.282448605 | 0 | 6.251786905 | 0 | 0 | 0 | 0.091489372 | 0.027012222 |
| MTA3 | 0 | 8.313932167 | 10.40259131 | 10.07066834 | 0 | 0 | 0 | 0 | 0.885848337 | 0.008817225 |
| MTHFD1L | 10.9350422 | 12.7899728 | 12.43740913 | 14.88793209 | 15.8141905 | 16.02527651 | 14.96011439 | 15.89532875 | 0.045923577 | 0.999854694 |
| MTHFD2 | 0 | 0 | 0 | 11.36463129 | 9.98964377 | 9.84247488 | 8.252679239 | 11.72094651 | 0.028355055 | 0.999854694 |
| MTHFR | 8.871474618 | 9.152938599 | 0 | 0 | 0 | 0 | 0 | 0 | 0.014355457 | 1 |
| MTIF2 | 0 | 0 | 8.602099869 | 0 | 9.148706942 | 0 | 0 | 0 | 0.072311509 | 0.01630416 |
| MTMR10 | 0 | 0 | 0 | 9.630766785 | 7.768613653 | 0 | 0 | 0 | 0.070807764 | 0.016324977 |
| MTMR4 | 9.195126457 | 8.555204765 | 0 | 0 | 0 | 0 | 0 | 0 | 0.016711674 | 1 |
| MTPN | 6.84536821 | 0 | 0 | 0 | 0 | 8.743432035 | 7.158082399 | 0 | 0.124501839 | 0.024297017 |
| MTR | 0 | 0 | 0 | 0 | 6.694188464 | 0 | 0 | 0 | 0.000126205 | 1 |

| | | | | | | | | | | |
|---|---|---|---|---|---|---|---|---|---|---|
| MTRF1L | 0 | 9.13888826 | 0 | 0 | 0 | 0 | 0 | 10.31542868 | 1.09E-07 | 1 |
| MTRNR2L6 | 9.689176399 | 0 | 0 | 0 | 0 | 0 | 0 | 0 | 7.88E-08 | 1 |
| MTURN | 0 | 0 | 10.26468848 | 0 | 8.007732862 | 0 | 7.741962234 | 0 | 0.04658487 | 0.999854694 |
| MUC12 | 11.85107959 | 13.0831414 | 11.82642024 | 10.73030464 | 0 | 0 | 0 | 0 | 0.843321238 | 0.00418531 |
| MUC2 | 10.5080681 | 11.49644666 | 11.96987422 | 10.52385347 | 0 | 0 | 0 | 0 | 0.991384446 | 0.004098578 |
| MUC5B | 0 | 11.236814 | 10.12247091 | 0 | 7.087723911 | 0 | 0 | 0 | 0.946589028 | 0.0135152 |
| MUL1 | 0 | 9.059065806 | 10.49879226 | 9.140222357 | 0 | 0 | 0 | 0 | 0.958261149 | 0.010003595 |
| MUSTN1 | 0 | 0 | 10.1142578 | 0 | 6.717806789 | 0 | 0 | 0 | 0.064369108 | 0.013938369 |
| MXD1 | 11.29168097 | 0 | 9.973792696 | 0 | 8.064100782 | 0 | 0 | 0 | 0.952260332 | 0.012659623 |
| MXD3 | 0 | 0 | 0 | 8.761452735 | 8.850567979 | 0 | 0 | 0 | 0.074720419 | 0.018035133 |
| MXD4 | 0 | 8.68657578 | 0 | 0 | 0 | 10.56136923 | 0 | 0 | 1.19E-07 | 1 |
| MYADM | 0 | 8.338729338 | 0 | 0 | 0 | 0 | 7.095033944 | 0 | 1.59E-07 | 1 |
| MYCBP | 9.766737375 | 9.97313222 | 10.0489694 | 11.93545091 | 10.78813443 | 0 | 0 | 0 | 0.63523793 | 0.000158964 |
| MYD88 | 0 | 0 | 9.88967833 | 9.532663167 | 9.269492846 | 0 | 0 | 0 | 0.002648196 | 0.00116937 |
| MYL6B | 0 | 0 | 10.46528171 | 0 | 9.129023686 | 0 | 7.954210107 | 0 | 0.027073951 | 0.999854694 |
| MYNN | 0 | 8.375143804 | 0 | 0 | 0 | 0 | 0 | 0 | 1.55E-07 | 1 |
| MYO10 | 0 | 0 | 0 | 7.142765408 | 0 | 0 | 0 | 0 | 3.45E-05 | 1 |
| MYO1E | 8.77466443 | 0 | 0 | 0 | 0 | 0 | 0 | 0 | 1.18E-07 | 1 |
| MYO1G | 0 | 0 | 0 | 0 | 6.556887521 | 0 | 0 | 0 | 9.34E-05 | 1 |
| MYO5B | 11.3079886 | 7.600636572 | 0 | 11.30530338 | 9.071750335 | 0 | 0 | 0 | 0.990582117 | 0.007383648 |
| MYO7B | 0 | 0 | 9.833251866 | 0 | 9.086566738 | 0 | 9.319363382 | 0 | 0.045069962 | 0.999854694 |
| MYOM1 | 9.977157135 | 0 | 0 | 0 | 0 | 0 | 0 | 0 | 7.07E-08 | 1 |
| MYOZ2 | 11.04550031 | 0 | 0 | 0 | 0 | 0 | 0 | 0 | 6.54E-08 | 1 |
| MYZAP | 0 | 0 | 6.510545513 | 9.675658367 | 0 | 0 | 0 | 0 | 0.075723274 | 0.020809157 |
| MZT1 | 0 | 6.010024966 | 10.76782595 | 11.90523433 | 10.4808038 | 10.12087373 | 6.188425891 | 0 | 0.003944049 | 0.999854694 |
| NAA25 | 0 | 0 | 6.547614429 | 0 | 0 | 0 | 0 | 0 | 6.12E-05 | 1 |
| NAB1 | 9.648774049 | 0 | 0 | 0 | 0 | 0 | 9.294634579 | 0 | 7.53E-08 | 1 |
| NABP1 | 0 | 0 | 9.422072283 | 0 | 6.944926602 | 0 | 0 | 0 | 0.083005621 | 0.021337307 |
| NAF1 | 0 | 0 | 0 | 0 | 6.841217856 | 0 | 0 | 0 | 4.14E-05 | 1 |
| NARF | 0 | 0 | 10.57108576 | 0 | 9.53346915 | 12.00541792 | 8.963116328 | 0 | 0.018786523 | 0.999854694 |
| NAT10 | 0 | 8.313932167 | 0 | 10.2139056 | 8.829041651 | 0 | 0 | 0 | 0.915944843 | 0.010829604 |
| NAT2 | 0 | 10.2133101 | 0 | 0 | 0 | 0 | 0 | 0 | 7.07E-08 | 1 |
| NAT9 | 0 | 10.01219366 | 0 | 0 | 0 | 0 | 0 | 0 | 7.07E-08 | 1 |
| NBPF14 | 10.16075706 | 8.565746011 | 0 | 0 | 0 | 0 | 9.009442583 | 9.884391338 | 0.005672999 | 0.009724498 |
| NBPF19 | 11.30365789 | 0 | 0 | 0 | 0 | 0 | 0 | 0 | 2.76E-08 | 1 |
| NBR1 | 0 | 0 | 9.818161621 | 10.19675959 | 9.652375851 | 0 | 9.963632456 | 0 | 0.00073984 | 1 |
| NCAPG2 | 0 | 0 | 0 | 0 | 6.360815222 | 0 | 0 | 0 | 6.12E-05 | 1 |
| NCAPH2 | 0 | 0 | 0 | 0 | 6.595647612 | 0 | 0 | 0 | 5.10E-05 | 1 |
| NCBP1 | 0 | 0 | 10.64990669 | 0 | 8.54864527 | 0 | 0 | 0 | 0.027969853 | 0.009724498 |
| NCKAP1 | 10.05651492 | 0 | 8.991012141 | 0 | 9.24633416 | 0 | 0 | 0 | 0.990582117 | 0.013910154 |
| NCOA1 | 8.99479642 | 6.64828144 | 10.06397233 | 8.12734671 | 0 | 0 | 0 | 0 | 0.987819958 | 0.012777646 |
| NCOA5 | 0 | 8.867872257 | 0 | 0 | 0 | 0 | 0 | 0 | 1.10E-07 | 1 |
| NDC1 | 0 | 0 | 0 | 10.92135317 | 6.151291903 | 0 | 0 | 0 | 1.98E-07 | 1.58E-10 |

| | | | | | | | | | | |
|---|---|---|---|---|---|---|---|---|---|---|
| NDFIP1 | 0 | 0 | 9.676133579 | 11.29900464 | 9.742209619 | 11.00678653 | 8.096917163 | 0 | 4.67E-05 | 0.999854694 |
| NDN | 0 | 10.11625358 | 0 | 0 | 0 | 0 | 0 | 0 | 7.07E-08 | 1 |
| NDST1 | 0 | 10.27242335 | 0 | 0 | 0 | 0 | 0 | 0 | 7.07E-08 | 1 |
| NDUFAF4 | 0 | 0 | 0 | 12.08038576 | 10.30094117 | 0 | 10.54090296 | 0 | 0.019775179 | 0.999854694 |
| NDUFAF6 | 0 | 0 | 0 | 8.876674206 | 11.91673533 | 11.04427804 | 7.695490668 | 10.74424263 | 1.28E-07 | 0.999854694 |
| NDUFS5 | 20.27386227 | 20.41285003 | 18.65898536 | 19.59876401 | 19.29058118 | 19.15863012 | 20.02032358 | 18.50690619 | 0.032419163 | 0.999854694 |
| NECAP2 | 0 | 0 | 10.51448015 | 0 | 9.518417774 | 0 | 7.889626398 | 0 | 0.020256571 | 0.999854694 |
| NEDD9 | 0 | 0 | 7.817899521 | 0 | 9.656200926 | 0 | 0 | 0 | 0.067529586 | 0.015215749 |
| NEFL | 9.362697802 | 0 | 0 | 0 | 0 | 0 | 0 | 0 | 8.05E-08 | 1 |
| NEIL1 | 0 | 9.286392738 | 0 | 0 | 0 | 0 | 0 | 0 | 8.08E-08 | 1 |
| NEK7 | 0 | 8.884809218 | 0 | 0 | 0 | 0 | 0 | 9.960578634 | 1.09E-07 | 1 |
| NEO1 | 0 | 8.586600217 | 0 | 0 | 0 | 0 | 0 | 0 | 1.30E-07 | 1 |
| NEPRO | 9.564407853 | 0 | 0 | 0 | 0 | 0 | 0 | 0 | 7.60E-08 | 1 |
| NEURL4 | 0 | 0 | 9.899270241 | 0 | 7.352011368 | 0 | 0 | 0 | 0.067902631 | 0.01453734 |
| NEXN | 0 | 0 | 0 | 0 | 0 | 8.635272278 | 8.08254305 | 0 | 1 | 0.018552193 |
| NFASC | 0 | 10.96944019 | 10.14280143 | 0 | 7.796609528 | 0 | 0 | 0 | 0.969395783 | 0.012259484 |
| NFIX | 0 | 0 | 0 | 11.94611716 | 11.07346172 | 9.851567204 | 11.76995497 | 10.0768456 | 0.018119434 | 0.999854694 |
| NFKB1 | 0 | 9.460304435 | 0 | 0 | 0 | 0 | 5.788848436 | 0 | 7.73E-08 | 1 |
| NFKBIB | 0 | 0 | 0 | 8.274689552 | 7.973804938 | 0 | 0 | 0 | 0.100902568 | 0.036419143 |
| NFX1 | 0 | 9.152938599 | 0 | 0 | 0 | 0 | 6.605228635 | 0 | 8.91E-08 | 1 |
| NHLRC1 | 0 | 0 | 0 | 9.708865503 | 9.404784439 | 0 | 0 | 0 | 0.040416394 | 0.011067641 |
| NIPSNAP3A | 10.85443391 | 10.51201372 | 9.995195314 | 11.33695016 | 3.598081693 | 0 | 0 | 0 | 0.976114635 | 0.001349516 |
| NKAP | 0 | 9.292752085 | 0 | 0 | 0 | 0 | 0 | 0 | 8.08E-08 | 1 |
| NKD1 | 0 | 0 | 0 | 10.61518186 | 5.799437418 | 0 | 11.18351631 | 0 | 0.042080155 | 1 |
| NLN | 10.8485001 | 0 | 10.10599766 | 0 | 7.877478056 | 0 | 0 | 0 | 0.976114635 | 0.012283188 |
| NLRP2 | 10.92379883 | 0 | 0 | 0 | 0 | 0 | 0 | 0 | 7.07E-08 | 1 |
| NLRX1 | 0 | 8.479177282 | 0 | 0 | 0 | 0 | 0 | 0 | 1.39E-07 | 1 |
| NME4 | 0 | 0 | 0 | 9.953060429 | 9.446024002 | 0 | 9.001824109 | 0 | 0.033711114 | 0.999854694 |
| NME6 | 0 | 0 | 0 | 9.761456469 | 10.36404409 | 0 | 8.345894226 | 0 | 0.018117483 | 0.999854694 |
| NMRK1 | 0 | 0 | 10.34935133 | 0 | 8.882266959 | 0 | 7.295290731 | 0 | 0.033420487 | 0.999854694 |
| NOL3 | 0 | 10.90344675 | 9.521689552 | 9.905608724 | 0 | 0 | 0 | 0 | 0.983546955 | 0.010829604 |
| NOMO1 | 0 | 0 | 0 | 9.58631554 | 6.633393544 | 0 | 0 | 0 | 0.077165773 | 0.020809157 |
| NOMO3 | 10.88953321 | 0 | 0 | 0 | 0 | 0 | 0 | 0 | 7.07E-08 | 1 |
| NOP56 | 0 | 0 | 10.66755444 | 10.18562856 | 11.80522995 | 0 | 8.499859701 | 0 | 1.18E-07 | 0.81310999 |
| NPAS3 | 0 | 0 | 9.127849973 | 0 | 8.358899494 | 0 | 0 | 0 | 0.07678267 | 0.017804903 |
| NPFFR2 | 0 | 9.670244415 | 9.811827134 | 0 | 7.866953924 | 0 | 0 | 0 | 0.997434996 | 0.01420877 |
| NPIPA3 | 0 | 8.884809218 | 0 | 0 | 0 | 0 | 0 | 0 | 1.09E-07 | 1 |
| NPIPA5 | 11.07882849 | 7.810193807 | 0 | 0 | 0 | 0 | 0 | 0 | 0.000921291 | 1 |
| NPL | 0 | 0 | 10.58452527 | 9.914625618 | 9.665339937 | 0 | 0 | 0 | 0.000303666 | 0.000592153 |
| NPNT | 0 | 8.833389894 | 0 | 0 | 0 | 0 | 8.293813865 | 0 | 1.14E-07 | 1 |
| NPPA-AS1 | 0 | 8.51225226 | 0 | 0 | 0 | 0 | 0 | 0 | 1.38E-07 | 1 |
| NPPC | 0 | 9.013590009 | 0 | 0 | 0 | 0 | 0 | 0 | 1.04E-07 | 1 |
| NPRL2 | 0 | 0 | 10.41099735 | 0 | 9.923261051 | 9.193031079 | 0 | 0 | 0.016556811 | 0.999854694 |

| | | | | | | | | | | |
|---|---|---|---|---|---|---|---|---|---|---|
| NPRL3 | 0 | 0 | 0 | 9.737942813 | 8.668631248 | 0 | 0 | 0 | 0.054485639 | 0.012947783 |
| NPY1R | 11.70602775 | 0 | 0 | 0 | 0 | 0 | 0 | 9.628234 | 6.24E-08 | 1 |
| NR1D2 | 0 | 0 | 7.771649443 | 0 | 0 | 0 | 8.22677003 | 10.45504893 | 1.98E-05 | 0.999854694 |
| NR1H2 | 0 | 8.375143804 | 10.16085711 | 0 | 8.815422744 | 0 | 0 | 0 | 0.932679569 | 0.010829604 |
| NR1H3 | 0 | 0 | 0 | 8.665007102 | 0 | 0 | 8.381556762 | 0 | 7.09E-06 | 1 |
| NR1H4 | 9.147591029 | 0 | 0 | 0 | 0 | 0 | 0 | 0 | 1.17E-07 | 1 |
| NR2F1 | 9.490126471 | 0 | 0 | 0 | 0 | 0 | 0 | 0 | 7.87E-08 | 1 |
| NR2F6 | 0 | 0 | 0 | 11.1862506 | 10.45160762 | 0 | 0 | 0 | 0.027792408 | 0.005927524 |
| NR3C2 | 9.490126471 | 8.81583464 | 0 | 0 | 0 | 0 | 0 | 8.447020158 | 0.011329815 | 1 |
| NRAV | 10.54889168 | 0 | 0 | 0 | 0 | 0 | 0 | 0 | 7.07E-08 | 1 |
| NRG4 | 0 | 0 | 0 | 0 | 0 | 0 | 6.43464192 | 9.234949175 | 1 | 0.019761711 |
| NRSN2 | 0 | 8.596915356 | 0 | 0 | 0 | 0 | 8.433450222 | 0 | 1.27E-07 | 1 |
| NSF | 0 | 9.465942998 | 0 | 11.00534719 | 7.132345939 | 0 | 0 | 0 | 0.977268168 | 0.00982437 |
| NSFL1C | 0 | 0 | 0 | 10.44514137 | 9.77916118 | 0 | 8.77032375 | 11.05438335 | 0.016868965 | 0.999854694 |
| NSUN5P1 | 0 | 0 | 9.041695116 | 0 | 8.297526732 | 0 | 0 | 0 | 0.079539815 | 0.019111579 |
| NSUN5P2 | 9.702396267 | 0 | 0 | 0 | 0 | 0 | 0 | 0 | 7.38E-08 | 1 |
| NT5E | 0 | 0 | 0 | 11.04694632 | 9.049818146 | 0 | 0 | 10.01048118 | 0.035972209 | 0.999854694 |
| NTAN1 | 0 | 0 | 5.689965016 | 10.60035633 | 7.242289655 | 0 | 0 | 11.43358286 | 0.012350769 | 1 |
| NTHL1 | 12.00704458 | 13.21098779 | 14.98519564 | 15.02421683 | 13.0428147 | 14.13035996 | 14.59019361 | 14.21503407 | 0.028305576 | 0.999854694 |
| NUB1 | 10.15596027 | 9.684823965 | 0 | 0 | 0 | 10.67414176 | 0 | 8.861023792 | 0.001894414 | 0.007912009 |
| NUDT1 | 0 | 9.836168082 | 0 | 9.055798949 | 8.727989007 | 0 | 0 | 0 | 0.990582117 | 0.016825783 |
| NUDT13 | 0 | 8.705765621 | 0 | 0 | 0 | 0 | 0 | 0 | 1.18E-07 | 1 |
| NUDT21 | 0 | 0 | 10.57408315 | 0 | 7.00375218 | 9.063382976 | 9.473284708 | 8.625136027 | 0.04170989 | 0.999854694 |
| NUDT3 | 0 | 0 | 0 | 9.769762184 | 9.278454701 | 0 | 9.43136219 | 9.786434411 | 0.041548195 | 0.999854694 |
| NUDT4B | 9.06811818 | 0 | 0 | 0 | 0 | 0 | 0 | 0 | 1.09E-07 | 1 |
| NUDT8 | 0 | 0 | 11.63230492 | 0 | 7.453974941 | 0 | 9.684324147 | 0 | 6.10E-07 | 0.999854694 |
| NUDT9 | 11.64057397 | 9.254168606 | 10.05112225 | 10.29670606 | 7.791053646 | 0 | 0 | 0 | 0.818016271 | 0.00116937 |
| NUF2 | 0 | 0 | 0 | 0 | 6.85193071 | 0 | 7.095033944 | 0 | 4.12E-05 | 1 |
| NUP43 | 0 | 0 | 0 | 0 | 6.85193071 | 0 | 0 | 0 | 4.12E-05 | 1 |
| NUP54 | 0 | 8.780069978 | 0 | 9.066626025 | 9.064860334 | 0 | 0 | 0 | 0.983546955 | 0.014738878 |
| NUP62CL | 0 | 4.527837782 | 0 | 0 | 0 | 0 | 6.991130623 | 9.472604067 | 0.283148217 | 0.015176609 |
| NUTM2A-AS1 | 0 | 0 | 9.135986286 | 0 | 8.918389748 | 0 | 0 | 0 | 0.067902631 | 0.014692722 |
| NVL | 0 | 0 | 10.07141569 | 7.058495112 | 8.297526732 | 0 | 0 | 0 | 0.015471258 | 0.003597041 |
| NXPE1 | 10.81542057 | 10.11266356 | 0 | 0 | 0 | 0 | 0 | 0 | 0.000159813 | 1 |
| NXPE3 | 9.655586838 | 0 | 0 | 0 | 0 | 0 | 4.589263136 | 7.75124755 | 7.44E-08 | 0.111025767 |
| NXPE4 | 10.30639463 | 10.56157158 | 10.16483899 | 0 | 8.460394093 | 0 | 0 | 0 | 0.870196658 | 0.01107436 |
| NXT1 | 0 | 10.62090101 | 0 | 10.43473225 | 9.911147352 | 0 | 0 | 0 | 0.999660881 | 0.008668791 |
| OCLN | 0 | 0 | 9.448445763 | 0 | 9.85234277 | 0 | 0 | 0 | 0.035992573 | 0.010542114 |
| ODF2 | 0 | 0 | 0 | 0 | 0 | 0 | 6.388646063 | 10.38110244 | 1 | 0.010542114 |
| ODF2L | 0 | 0 | 11.84515167 | 0 | 8.647265291 | 11.33857894 | 0 | 11.15338571 | 0.027450171 | 0.999854694 |
| OGA | 0 | 0 | 0 | 0 | 0 | 10.62814611 | 0 | 10.38401774 | 1 | 0.005637576 |
| OGN | 0 | 9.537306767 | 0 | 0 | 0 | 0 | 0 | 0 | 7.53E-08 | 1 |
| OIP5-AS1 | 9.949703925 | 9.79195933 | 10.27761706 | 10.44617816 | 6.974639195 | 0 | 0 | 0 | 0.999483812 | 0.00116937 |

| | | | | | | | | | | |
|---|---|---|---|---|---|---|---|---|---|---|
| OIT3 | 0 | 9.402672528 | 0 | 0 | 0 | 0 | 0 | 0 | 7.88E-08 | 1 |
| OLFM2 | 9.362697802 | 0 | 0 | 0 | 0 | 0 | 0 | 0 | 8.05E-08 | 1 |
| OLFM4 | 0 | 13.24192215 | 0 | 11.12282516 | 9.804199658 | 0 | 0 | 0 | 0.894736815 | 0.007159623 |
| OMA1 | 0 | 0 | 11.23219994 | 0 | 7.728483681 | 0 | 0 | 0 | 0.014805854 | 0.008551774 |
| OPN3 | 10.85147005 | 0 | 0 | 0 | 0 | 0 | 0 | 0 | 7.07E-08 | 1 |
| ORAI3 | 0 | 0 | 0 | 0 | 0 | 10.41571817 | 0 | 10.64322669 | 1 | 0.00558734 |
| ORC1 | 0 | 0 | 0 | 9.011659331 | 6.151291903 | 0 | 0 | 0 | 0.099453785 | 0.037016979 |
| ORC3 | 0 | 0 | 0 | 0 | 0 | 10.70205702 | 0 | 10.3215241 | 1 | 0.00558734 |
| OSBPL10 | 0 | 8.533888356 | 0 | 0 | 0 | 0 | 0 | 0 | 1.35E-07 | 1 |
| OSCAR | 0 | 0 | 0 | 7.404097292 | 0 | 0 | 0 | 0 | 2.66E-05 | 1 |
| OSMR | 0 | 0 | 0 | 0 | 6.503532343 | 0 | 0 | 0 | 5.50E-05 | 1 |
| OTOP2 | 9.578815964 | 0 | 0 | 0 | 0 | 0 | 0 | 0 | 8.04E-08 | 1 |
| P2RX1 | 10.02529365 | 0 | 0 | 0 | 0 | 0 | 0 | 0 | 7.07E-08 | 1 |
| P2RY1 | 10.08202624 | 0 | 0 | 0 | 0 | 0 | 0 | 0 | 7.07E-08 | 1 |
| PACRGL | 0 | 0 | 9.546318339 | 0 | 5.906083538 | 0 | 0 | 0 | 0.083625397 | 0.02260752 |
| PACS1 | 0 | 0 | 8.869403674 | 7.036635307 | 0 | 0 | 0 | 0 | 0.10149484 | 0.035568676 |
| PACS2 | 9.804008249 | 0 | 8.85961044 | 0 | 7.813149942 | 0 | 0 | 0 | 0.979109272 | 0.025558844 |
| PAFAH1B1 | 9.708961049 | 0 | 10.34584807 | 0 | 9.404784439 | 0 | 0 | 0 | 0.987185781 | 0.009547202 |
| PAFAH1B3 | 0 | 0 | 0 | 10.98836577 | 10.49625063 | 7.673489052 | 8.207028126 | 10.17773295 | 0.02932436 | 0.999854694 |
| PAGR1 | 0 | 0 | 0 | 10.09466941 | 10.29604626 | 0 | 8.842476416 | 9.871985276 | 0.014778777 | 0.999854694 |
| PAIP2B | 0 | 0 | 9.88002222 | 0 | 7.242289655 | 0 | 0 | 0 | 0.069475653 | 0.014818059 |
| PAK1 | 0 | 0 | 10.97591091 | 9.645284634 | 9.492977848 | 11.13643575 | 7.482101429 | 10.24662217 | 0.000214871 | 0.999854694 |
| PALD1 | 0 | 8.375143804 | 0 | 0 | 0 | 0 | 0 | 0 | 1.55E-07 | 1 |
| PANK3 | 0 | 0 | 9.589913775 | 10.23565523 | 0 | 0 | 0 | 0 | 0.028391554 | 0.010003595 |
| PAOX | 0 | 8.958657193 | 0 | 0 | 0 | 0 | 0 | 0 | 1.07E-07 | 1 |
| PAQR5 | 10.17978629 | 0 | 0 | 0 | 0 | 0 | 0 | 0 | 7.07E-08 | 1 |
| PARD3B | 0 | 0 | 8.665302393 | 0 | 9.419262725 | 0 | 0 | 0 | 0.064245808 | 0.01420877 |
| PARD6A | 0 | 0 | 0 | 11.00112053 | 8.628696343 | 0 | 0 | 0 | 0.014998869 | 0.008817225 |
| PARP1 | 0 | 0 | 10.612493 | 9.489061787 | 7.988443284 | 0 | 0 | 0 | 0.002168344 | 0.00116937 |
| PARP10 | 0 | 0 | 10.15486362 | 8.55781724 | 8.204461778 | 0 | 6.811044309 | 0 | 0.008479619 | 0.847545326 |
| PARP2 | 0 | 0 | 0 | 9.896535119 | 8.918389748 | 11.04559917 | 0 | 0 | 0.044784409 | 0.999854694 |
| PARP8 | 0 | 0 | 0 | 0 | 6.419606803 | 0 | 8.139199685 | 0 | 5.88E-05 | 1 |
| PARVB | 0 | 0 | 10.16980092 | 11.86626083 | 0 | 0 | 9.527351199 | 10.26887315 | 0.022889485 | 0.999854694 |
| PATL2 | 0 | 8.533888356 | 0 | 0 | 0 | 0 | 0 | 0 | 1.35E-07 | 1 |
| PAX5 | 0 | 10.40454548 | 0 | 0 | 0 | 0 | 0 | 9.593393101 | 7.07E-08 | 1 |
| PBK | 0 | 9.97313222 | 0 | 0 | 0 | 0 | 0 | 0 | 7.07E-08 | 1 |
| PC | 9.607207561 | 0 | 0 | 0 | 0 | 0 | 0 | 0 | 7.53E-08 | 1 |
| PCDHB13 | 0 | 8.301371897 | 0 | 0 | 0 | 0 | 0 | 0 | 1.66E-07 | 1 |
| PCGF1 | 0 | 0 | 10.16682581 | 10.36736919 | 10.06120661 | 10.45899622 | 9.108538284 | 0 | 0.000161186 | 0.999854694 |
| PCLAF | 0 | 0 | 0 | 11.82779713 | 10.25021632 | 0 | 10.23782131 | 10.58993625 | 0.022422238 | 0.999854694 |
| PCNX1 | 0 | 8.49028674 | 0 | 0 | 0 | 0 | 0 | 0 | 1.38E-07 | 1 |
| PCSK7 | 9.773016437 | 5.204020779 | 10.88530233 | 5.429426546 | 6.711938358 | 0 | 0 | 0 | 0.981170768 | 0.003527496 |
| PCYOX1L | 0 | 9.844849563 | 0 | 0 | 0 | 0 | 0 | 0 | 7.07E-08 | 1 |

| | | | | | | | | | |
|---|---|---|---|---|---|---|---|---|---|
| PDCD4-AS1 | 0 | 8.223623425 | 0 | 0 | 0 | 0 | 0 | 0 | 2.64E-07 | 1 |
| PDE6A | 10.57428148 | 0 | 0 | 0 | 0 | 0 | 0 | 0 | 7.07E-08 | 1 |
| PDE6B | 10.22170595 | 0 | 0 | 0 | 0 | 0 | 0 | 0 | 7.07E-08 | 1 |
| PDE6D | 0 | 0 | 10.48293189 | 0 | 9.997504767 | 0 | 0 | 10.29388958 | 0.014194385 | 0.999854694 |
| PDE8A | 0 | 0 | 9.026457481 | 0 | 6.283787608 | 0 | 0 | 0 | 0.101138461 | 0.033871406 |
| PDK4 | 0 | 9.073911341 | 0 | 0 | 0 | 0 | 0 | 0 | 1.17E-07 | 1 |
| PDLIM5 | 10.39862087 | 0 | 0 | 0 | 0 | 0 | 0 | 0 | 7.07E-08 | 1 |
| PDP2 | 0 | 9.117552805 | 0 | 0 | 0 | 0 | 0 | 0 | 1.14E-07 | 1 |
| PDPK1 | 8.252542925 | 9.131811442 | 0 | 0 | 0 | 0 | 0 | 0 | 0.019674583 | 1 |
| PDS5A | 10.77844715 | 0 | 0 | 0 | 0 | 0 | 0 | 0 | 7.07E-08 | 1 |
| PDXDC1 | 10.09711927 | 0 | 10.52381165 | 0 | 8.017281789 | 0 | 0 | 0 | 0.999483812 | 0.010542114 |
| PDZD7 | 0 | 8.236875968 | 0 | 0 | 0 | 0 | 0 | 0 | 2.64E-07 | 1 |
| PECAM1 | 10.43074784 | 10.37787548 | 0 | 0 | 0 | 0 | 0 | 0 | 0.000261972 | 1 |
| PEMT | 0 | 0 | 9.111438383 | 11.02282616 | 9.574472933 | 0 | 0 | 0 | 0.000269597 | 0.000631775 |
| PEPD | 0 | 0 | 0 | 10.46573697 | 10.81096159 | 10.38160136 | 0 | 10.04035398 | 0.030808339 | 0.999854694 |
| PER1 | 12.66503534 | 10.79338956 | 0 | 0 | 0 | 0 | 9.628524471 | 0 | 4.99E-10 | 1 |
| PEX10 | 10.37817204 | 8.617326978 | 0 | 0 | 0 | 0 | 0 | 0 | 0.003674003 | 1 |
| PEX19 | 0 | 0 | 0 | 9.944281305 | 7.638537195 | 0 | 0 | 0 | 0.061401022 | 0.01420877 |
| PEX26 | 10.53418006 | 0 | 10.00522363 | 0 | 7.928977054 | 0 | 0 | 0 | 0.983546955 | 0.012725218 |
| PEX5 | 0 | 8.387080535 | 0 | 0 | 0 | 0 | 6.72111291 | 0 | 1.53E-07 | 1 |
| PEX6 | 0 | 9.615488971 | 0 | 0 | 0 | 0 | 8.550348343 | 0 | 7.42E-08 | 1 |
| PEX7 | 0 | 0 | 7.012809396 | 10.10389638 | 8.21279698 | 0 | 0 | 0 | 0.014483031 | 0.003676766 |
| PF4 | 0 | 8.51225226 | 0 | 0 | 0 | 0 | 0 | 0 | 1.38E-07 | 1 |
| PFKFB2 | 0 | 9.488279425 | 0 | 0 | 0 | 0 | 0 | 0 | 7.73E-08 | 1 |
| PFN2 | 0 | 0 | 10.20212852 | 9.847154521 | 9.089964407 | 0 | 0 | 0 | 0.001368275 | 0.000893674 |
| PGBD2 | 0 | 8.456698379 | 0 | 0 | 0 | 0 | 0 | 0 | 1.42E-07 | 1 |
| PGF | 10.39047609 | 0 | 0 | 0 | 0 | 0 | 0 | 0 | 7.07E-08 | 1 |
| PGM1 | 0 | 0 | 11.33825884 | 0 | 8.179163277 | 10.78507603 | 0 | 0 | 0.035379258 | 0.999854694 |
| PGM2 | 0 | 8.789094606 | 0 | 0 | 0 | 0 | 0 | 0 | 1.15E-07 | 1 |
| PGS1 | 0 | 9.117552805 | 0 | 0 | 0 | 0 | 0 | 9.66704837 | 1.14E-07 | 1 |
| PHC1 | 10.14631859 | 0 | 0 | 0 | 0 | 0 | 0 | 0 | 7.07E-08 | 1 |
| PHETA2 | 0 | 0 | 0 | 0 | 0 | 0 | 8.46149326 | 9.924990921 | 1 | 0.01037888 |
| PHF12 | 0 | 9.504807688 | 0 | 0 | 0 | 0 | 0 | 0 | 7.60E-08 | 1 |
| PHF19 | 0 | 0 | 0 | 9.58631554 | 9.878119554 | 0 | 0 | 10.41854941 | 0.032339976 | 0.999854694 |
| PHF20L1 | 0 | 9.081277213 | 0 | 0 | 0 | 0 | 7.295290731 | 0 | 1.17E-07 | 1 |
| PHF21A | 14.45878349 | 12.77386805 | 11.42962446 | 10.70973107 | 11.89099619 | 12.92532461 | 12.00365724 | 12.97026424 | 0.007847049 | 0.999854694 |
| PHKB | 9.881603011 | 0 | 10.81736233 | 0 | 7.022839493 | 0 | 0 | 0 | 0.991757787 | 0.01037888 |
| PHRF1 | 0 | 0 | 0 | 10.9926299 | 6.857257452 | 0 | 7.178500772 | 0 | 0.023731195 | 0.999854694 |
| PI16 | 0 | 8.780069978 | 0 | 0 | 0 | 0 | 0 | 0 | 1.15E-07 | 1 |
| PI4KAP1 | 0 | 0 | 0 | 8.546260139 | 0 | 0 | 0 | 0 | 2.46E-05 | 1 |
| PIAS3 | 0 | 0 | 0 | 6.661416457 | 0 | 0 | 0 | 0 | 5.29E-05 | 1 |
| PIDD1 | 9.930171034 | 0 | 0 | 0 | 0 | 0 | 0 | 0 | 7.07E-08 | 1 |
| PIFO | 10.81085038 | 0 | 0 | 0 | 0 | 0 | 0 | 0 | 7.07E-08 | 1 |

| | | | | | | | | | | |
|---|---|---|---|---|---|---|---|---|---|---|
| PIGBOS1 | 0 | 0 | 11.27887408 | 7.386921919 | 8.19186798 | 0 | 9.624809291 | 0 | 0.001444857 | 0.999854694 |
| PIGH | 0 | 0 | 10.4176871 | 0 | 9.187285685 | 0 | 0 | 0 | 0.027508034 | 0.009724498 |
| PIGQ | 9.204448442 | 0 | 9.572033777 | 0 | 4.780948925 | 0 | 0 | 0 | 1 | 0.024766336 |
| PIGR | 15.15623673 | 15.064205 | 14.78510872 | 12.48352174 | 9.068309447 | 0 | 0 | 0 | 0.576918806 | 6.69E-06 |
| PIGV | 11.38891686 | 0 | 0 | 0 | 0 | 0 | 9.150486467 | 0 | 2.97E-08 | 1 |
| PIGX | 8.453559626 | 0 | 0 | 10.11045113 | 9.356700643 | 0 | 0 | 0 | 0.91063389 | 0.01037888 |
| PIGY-DT | 0 | 0 | 8.050899235 | 0 | 0 | 0 | 0 | 0 | 1.49E-05 | 1 |
| PIH1D2 | 10.09711927 | 0 | 0 | 0 | 0 | 0 | 0 | 0 | 7.07E-08 | 1 |
| PIK3C2A | 0 | 0 | 0 | 0 | 6.819550507 | 0 | 0 | 0 | 4.21E-05 | 1 |
| PIK3R2 | 0 | 10.79563025 | 0 | 0 | 0 | 9.508729999 | 8.296930015 | 0 | 7.07E-08 | 0.011685672 |
| PIK3R3 | 9.026677385 | 0 | 0 | 0 | 0 | 0 | 0 | 0 | 1.07E-07 | 1 |
| PIM2 | 9.593081603 | 0 | 0 | 0 | 0 | 0 | 0 | 10.67332804 | 7.58E-08 | 1 |
| PINLYP | 10.81542057 | 0 | 0 | 0 | 0 | 0 | 0 | 0 | 7.07E-08 | 1 |
| PINX1 | 9.754096665 | 0 | 10.58452527 | 10.53753769 | 0 | 0 | 0 | 0 | 0.968729704 | 0.007496493 |
| PIP4P1 | 0 | 0 | 0 | 9.509044094 | 6.797552774 | 0 | 0 | 0 | 0.078825993 | 0.02144636 |
| PIP5K1C | 0 | 10.01988062 | 0 | 0 | 0 | 0 | 0 | 0 | 7.07E-08 | 1 |
| PIR | 0 | 0 | 0 | 0 | 6.448127902 | 0 | 0 | 0 | 5.74E-05 | 1 |
| PITHD1 | 0 | 0 | 0 | 11.07536809 | 9.556574678 | 0 | 8.668139023 | 12.15135899 | 0.033678295 | 0.999854694 |
| PITPNM2 | 10.07695994 | 0 | 9.461453882 | 0 | 7.740063742 | 0 | 0 | 0 | 0.981170768 | 0.017505868 |
| PITRM1 | 0 | 7.912843925 | 10.21754553 | 0 | 7.344444907 | 0 | 0 | 0 | 0.922337827 | 0.012348695 |
| PITX1 | 0 | 0 | 0 | 9.582548699 | 6.005385965 | 0 | 0 | 0 | 0.079549852 | 0.022671312 |
| PKHD1L1 | 9.898931975 | 9.396780662 | 0 | 0 | 0 | 0 | 0 | 0 | 0.00420142 | 1 |
| PKMYT1 | 0 | 0 | 8.190254187 | 9.061222644 | 0 | 0 | 0 | 0 | 0.081046379 | 0.021739634 |
| PLAA | 0 | 9.460304435 | 0 | 0 | 0 | 0 | 0 | 0 | 7.73E-08 | 1 |
| PLAT | 9.016128482 | 9.949177413 | 0 | 10.06797673 | 8.616182922 | 0 | 0 | 0 | 0.981769365 | 0.011578584 |
| PLCB4 | 9.689176399 | 8.196747437 | 0 | 0 | 0 | 0 | 0 | 0 | 0.012978692 | 1 |
| PLCD4 | 0 | 8.523110867 | 0 | 0 | 0 | 0 | 0 | 0 | 1.36E-07 | 1 |
| PLCE1 | 0 | 9.704036967 | 10.78797416 | 0 | 7.282440679 | 0 | 0 | 0 | 0.990582117 | 0.010278758 |
| PLCXD1 | 0 | 0 | 8.166619982 | 0 | 7.705040924 | 0 | 0 | 0 | 0.112232418 | 0.043055332 |
| PLD1 | 0 | 0 | 8.651179375 | 9.945748207 | 0 | 0 | 0 | 0 | 0.051802237 | 0.012417553 |
| PLEK2 | 9.641928937 | 0 | 9.222582136 | 11.51647 | 10.36217188 | 0 | 0 | 0 | 0.679525733 | 0.000334306 |
| PLEKHA4 | 0 | 9.273589327 | 0 | 0 | 0 | 11.07047346 | 0 | 0 | 8.16E-08 | 1 |
| PLEKHG5 | 0 | 0 | 8.659669766 | 0 | 7.619854833 | 0 | 0 | 0 | 0.099345499 | 0.03164999 |
| PLEKHO2 | 0 | 9.727698902 | 0 | 0 | 0 | 0 | 0 | 0 | 7.17E-08 | 1 |
| PLIN3 | 0 | 9.679980467 | 11.31560358 | 0 | 8.491557257 | 0 | 0 | 0 | 0.971036351 | 0.007647289 |
| PLK1 | 0 | 9.131811442 | 0 | 8.994752481 | 9.161681486 | 0 | 0 | 0 | 0.991384446 | 0.01457526 |
| PLLP | 0 | 9.044065918 | 0 | 0 | 0 | 0 | 0 | 0 | 1.01E-07 | 1 |
| PLOD2 | 0 | 7.426508624 | 10.1688099 | 0 | 8.896566111 | 0 | 0 | 0 | 0.813725912 | 0.010595256 |
| PLPPR2 | 0 | 8.918098054 | 0 | 0 | 0 | 0 | 0 | 0 | 1.08E-07 | 1 |
| PLRG1 | 0 | 0 | 10.22519269 | 0 | 6.841217856 | 0 | 0 | 0 | 0.058312937 | 0.013007083 |
| PLS3 | 0 | 0 | 9.918264838 | 0 | 9.446024002 | 0 | 0 | 9.531635636 | 0.03569984 | 0.999854694 |
| PLSCR4 | 0 | 0 | 8.789130733 | 0 | 7.866953924 | 0 | 0 | 0 | 0.091489372 | 0.026317017 |
| PLVAP | 10.08707482 | 0 | 0 | 0 | 0 | 0 | 0 | 9.972248762 | 7.07E-08 | 1 |

| | | | | | | | | | | |
|---|---|---|---|---|---|---|---|---|---|---|
| PMS2P9 | 0 | 0 | 0 | 11.83652154 | 8.353255437 | 11.50147895 | 8.959187811 | 11.53148824 | 1.51E-07 | 0.999854694 |
| PMVK | 0 | 0 | 9.611956589 | 9.40197589 | 8.04555511 | 10.99589325 | 9.659117799 | 10.68991445 | 0.009413337 | 0.999854694 |
| PNO1 | 0 | 10.41332703 | 0 | 10.18189905 | 8.5617557 | 0 | 0 | 0 | 0.992483691 | 0.011154849 |
| PNPLA4 | 0 | 0 | 0 | 10.15931723 | 8.012515226 | 0 | 10.12487367 | 0 | 0.048159116 | 0.999854694 |
| PNPLA6 | 10.36991065 | 0 | 0 | 9.597557409 | 6.717806789 | 0 | 0 | 0 | 0.966245748 | 0.020371116 |
| PNPLA8 | 0 | 8.876365592 | 0 | 0 | 0 | 0 | 0 | 9.936951298 | 1.10E-07 | 1 |
| PNPO | 0 | 0 | 0 | 0 | 0 | 0 | 8.722455448 | 9.034319984 | 1 | 0.012722919 |
| PODXL2 | 0 | 0 | 10.34409325 | 9.961786454 | 7.866953924 | 0 | 0 | 0 | 0.002122301 | 0.00116937 |
| POGLUT1 | 0 | 8.990303015 | 0 | 0 | 0 | 0 | 0 | 0 | 1.05E-07 | 1 |
| POLE3 | 0 | 0 | 0 | 10.52581632 | 8.166345699 | 10.96620334 | 9.621084518 | 0 | 0.033365155 | 0.999854694 |
| POLR1B | 0 | 10.45928005 | 0 | 0 | 0 | 0 | 0 | 0 | 7.07E-08 | 1 |
| POLR2D | 0 | 0 | 0 | 12.02525718 | 8.659513127 | 0 | 0 | 0 | 1.30E-07 | 3.12E-11 |
| POLR2J3 | 4.321811331 | 8.859328625 | 0 | 0 | 0 | 0 | 0 | 0 | 0.042057457 | 1 |
| POLR3A | 10.21249517 | 0 | 0 | 0 | 0 | 0 | 7.751079579 | 0 | 7.07E-08 | 1 |
| POLR3C | 0 | 0 | 0 | 11.97010208 | 8.174903392 | 9.294308964 | 0 | 0 | 4.57E-07 | 0.999854694 |
| POLR3E | 0 | 0 | 0 | 8.794392577 | 9.086566738 | 0 | 0 | 0 | 0.070094441 | 0.016168618 |
| POMZP3 | 0 | 0 | 10.0872376 | 9.67388927 | 0 | 0 | 0 | 0 | 0.031058569 | 0.01024397 |
| POSTN | 0 | 0 | 0 | 9.331906887 | 9.654672113 | 0 | 7.270142197 | 10.50410909 | 0.042753786 | 0.999854694 |
| POU2AF1 | 10.96002393 | 10.33538329 | 0 | 0 | 0 | 0 | 0 | 0 | 5.81E-05 | 1 |
| POU2F1 | 10.41477378 | 0 | 0 | 0 | 0 | 0 | 0 | 0 | 7.07E-08 | 1 |
| PPARA | 11.19444934 | 0 | 0 | 0 | 0 | 0 | 0 | 0 | 4.18E-08 | 1 |
| PPARD | 0 | 0 | 0 | 0 | 0 | 0 | 9.616103148 | 7.222704 | 1 | 0.013289501 |
| PPCDC | 0 | 9.021269553 | 0 | 0 | 0 | 0 | 8.309327743 | 0 | 1.03E-07 | 1 |
| PPFIA3 | 10.34904781 | 0 | 0 | 0 | 0 | 0 | 0 | 9.711823855 | 7.07E-08 | 1 |
| PPIG | 0 | 9.988883934 | 0 | 0 | 0 | 0 | 8.947337666 | 0 | 7.07E-08 | 1 |
| PPIL1 | 0 | 10.06516163 | 0 | 0 | 0 | 0 | 0 | 9.630690734 | 7.07E-08 | 1 |
| PPIP5K1 | 0 | 0 | 7.776862171 | 0 | 0 | 0 | 0 | 0 | 1.97E-05 | 1 |
| PPIP5K2 | 0 | 9.37296959 | 0 | 0 | 0 | 0 | 0 | 0 | 7.88E-08 | 1 |
| PPM1A | 0 | 0 | 10.12656 | 0 | 6.283787608 | 0 | 0 | 0 | 0.066125078 | 0.014309205 |
| PPM1B | 9.185743846 | 10.50655271 | 9.88002222 | 0 | 8.510275248 | 0 | 0 | 0 | 0.930569319 | 0.012259484 |
| PPM1G | 10.40267609 | 11.23598924 | 0 | 10.9146273 | 9.774945524 | 0 | 0 | 0 | 0.958196509 | 0.007674926 |
| PPME1 | 0 | 0 | 10.69603741 | 9.571188803 | 7.266514108 | 0 | 0 | 0 | 0.002283735 | 0.00126979 |
| PPP1CA | 17.86679255 | 17.946534 | 19.82666156 | 18.63562875 | 18.50513284 | 18.43245664 | 18.33370566 | 18.82976512 | 0.028529356 | 0.999854694 |
| PPP1R11 | 0 | 0 | 10.90804819 | 9.17064951 | 7.542607814 | 0 | 0 | 10.24982197 | 0.001900482 | 0.999854694 |
| PPP1R12B | 0 | 9.005869367 | 0 | 0 | 0 | 0 | 0 | 0 | 1.04E-07 | 1 |
| PPP1R18 | 0 | 0 | 0 | 0 | 6.315093872 | 0 | 0 | 0 | 6.33E-05 | 1 |
| PPP1R27 | 0 | 0 | 6.607354851 | 0 | 0 | 0 | 0 | 0 | 5.86E-05 | 1 |
| PPP1R3E | 0 | 8.49028674 | 0 | 0 | 0 | 0 | 0 | 0 | 1.38E-07 | 1 |
| PPP1R9B | 0 | 8.942570108 | 0 | 0 | 0 | 0 | 8.422078578 | 0 | 1.07E-07 | 1 |
| PPP2R3B | 0 | 0 | 0 | 0 | 6.786426836 | 0 | 0 | 0 | 4.30E-05 | 1 |
| PPP2R5B | 0 | 0 | 0 | 0 | 6.729472521 | 0 | 0 | 0 | 0.000125541 | 1 |
| PPP4R3B | 0 | 0 | 10.15486362 | 10.1078328 | 9.132322934 | 10.85681934 | 9.872578794 | 0 | 0.000960251 | 0.999854694 |
| PQBP1 | 0 | 0 | 10.49405243 | 12.21086077 | 10.13103695 | 0 | 0 | 0 | 7.07E-08 | 0.000148727 |

| | | | | | | | | | | |
|---|---|---|---|---|---|---|---|---|---|---|
| PQLC1 | 10.30639463 | 8.958657193 | 11.05098014 | 9.489061787 | 8.671657859 | 0 | 0 | 0 | 0.988867264 | 0.000726381 |
| PRADC1 | 5.542137777 | 8.724703557 | 11.64625702 | 0 | 9.422859707 | 0 | 0 | 0 | 0.694043898 | 0.005727576 |
| PRCP | 0 | 0 | 8.85961044 | 10.29440511 | 0 | 0 | 8.173512489 | 4.497002282 | 0.036573995 | 0.999854694 |
| PRDM11 | 10.39047609 | 0 | 0 | 0 | 0 | 0 | 0 | 0 | 7.07E-08 | 1 |
| PRICKLE3 | 0 | 0 | 0 | 0 | 6.569923599 | 0 | 0 | 0 | 5.22E-05 | 1 |
| PRIM1 | 0 | 0 | 0 | 10.9146273 | 9.026365663 | 0 | 0 | 10.9734673 | 0.01443822 | 0.999854694 |
| PRKAB2 | 9.760430865 | 0 | 0 | 0 | 0 | 0 | 0 | 0 | 7.17E-08 | 1 |
| PRKAG1 | 10.98998111 | 8.876365592 | 12.0621608 | 0 | 9.153044768 | 0 | 0 | 0 | 0.990582117 | 9.09E-12 |
| PRKCA | 0 | 9.37296959 | 0 | 0 | 0 | 0 | 0 | 0 | 7.88E-08 | 1 |
| PRKCQ-AS1 | 0 | 8.705765621 | 0 | 0 | 0 | 0 | 0 | 0 | 1.18E-07 | 1 |
| PRKRA-AS1 | 9.987993796 | 0 | 0 | 0 | 0 | 0 | 0 | 0 | 7.07E-08 | 1 |
| PRLR | 0 | 0 | 6.783425399 | 0 | 0 | 0 | 0 | 0 | 5.07E-05 | 1 |
| PRMT3 | 0 | 8.479177282 | 0 | 0 | 0 | 0 | 0 | 9.695488855 | 1.39E-07 | 1 |
| PROSER2 | 0 | 0 | 0 | 0 | 6.448127902 | 0 | 0 | 0 | 5.74E-05 | 1 |
| PRPF19 | 0 | 8.798063133 | 10.74672837 | 10.21876727 | 8.794750254 | 0 | 0 | 0 | 0.713367231 | 0.000631775 |
| PRPF40A | 0 | 0 | 0 | 9.290944194 | 8.900441408 | 0 | 0 | 0 | 0.063546946 | 0.01439849 |
| PRPF40B | 0 | 0 | 0 | 0 | 6.375739247 | 0 | 0 | 0 | 6.06E-05 | 1 |
| PRPF6 | 0 | 0 | 11.3773909 | 10.73709781 | 9.059093422 | 0 | 10.41605733 | 0 | 1.17E-05 | 0.999854694 |
| PRPSAP1 | 0 | 0 | 0 | 10.28516442 | 7.488711871 | 0 | 0 | 0 | 0.048007264 | 0.012259484 |
| PRR3 | 9.785493166 | 0 | 0 | 0 | 0 | 0 | 0 | 0 | 7.17E-08 | 1 |
| PRR34-AS1 | 0 | 0 | 0 | 0 | 6.375739247 | 0 | 0 | 0 | 6.06E-05 | 1 |
| PRR7 | 0 | 10.49832224 | 0 | 0 | 0 | 0 | 0 | 0 | 7.07E-08 | 1 |
| PRRC1 | 9.350153507 | 8.966633964 | 0 | 0 | 0 | 0 | 0 | 0 | 0.011613136 | 1 |
| PRSS12 | 0 | 9.52655488 | 0 | 0 | 0 | 0 | 0 | 0 | 7.60E-08 | 1 |
| PRTFDC1 | 0 | 0 | 0 | 10.34863975 | 9.214520192 | 0 | 0 | 0 | 0.027484689 | 0.010003595 |
| PRXL2C | 0 | 9.227860829 | 0 | 0 | 0 | 0 | 0 | 0 | 8.36E-08 | 1 |
| PSAT1 | 0 | 0 | 0 | 10.37935994 | 10.38217191 | 0 | 7.704905507 | 0 | 0.035365565 | 1 |
| PSD4 | 0 | 7.945499326 | 8.954673969 | 0 | 7.939059873 | 0 | 0 | 0 | 0.979109272 | 0.022881398 |
| PSKH1 | 0 | 0 | 9.616325024 | 0 | 7.813149942 | 0 | 0 | 0 | 0.072543146 | 0.015785316 |
| PSMA7 | 9.527745184 | 9.92890668 | 13.55469177 | 14.98689855 | 12.93011887 | 11.869544 | 12.28283339 | 13.3360508 | 7.07E-08 | 0.999854694 |
| PSMD14 | 11.76706843 | 10.09821346 | 0 | 11.71016366 | 11.67290785 | 0 | 0 | 0 | 0.999485949 | 0.002827437 |
| PSMD2 | 0 | 0 | 11.20683649 | 10.65876571 | 8.877031927 | 0 | 0 | 10.68755661 | 3.55E-05 | 0.999854694 |
| PSME4 | 0 | 0 | 0 | 0 | 6.489879427 | 0 | 0 | 0 | 5.56E-05 | 1 |
| PSMG3 | 0 | 0 | 0 | 10.07066834 | 10.57998453 | 0 | 10.92161573 | 0 | 0.035781052 | 0.999854694 |
| PSORS1C1 | 0 | 8.876365592 | 0 | 10.5549431 | 6.741044679 | 0 | 0 | 0 | 0.963126669 | 0.011578584 |
| PTAR1 | 0 | 8.893203714 | 0 | 0 | 0 | 0 | 0 | 0 | 1.09E-07 | 1 |
| PTDSS1 | 0 | 0 | 0 | 9.726040692 | 9.054463238 | 10.31633091 | 3.993557961 | 10.56189072 | 0.047359454 | 0.999854694 |
| PTGDS | 6.116742619 | 8.398919312 | 0 | 0 | 0 | 0 | 0 | 8.625136027 | 0.045024163 | 1 |
| PTGER3 | 10.15114749 | 0 | 0 | 0 | 0 | 0 | 0 | 0 | 7.07E-08 | 1 |
| PTGR2 | 0 | 0 | 10.32464801 | 0 | 7.893121693 | 0 | 0 | 0 | 0.045270943 | 0.011235459 |
| PTHLH | 0 | 0 | 0 | 0 | 0 | 10.5987484 | 0 | 9.727975969 | 1 | 0.006854959 |
| PTN | 0 | 8.715265663 | 0 | 0 | 0 | 0 | 0 | 0 | 1.18E-07 | 1 |
| PTOV1 | 0 | 0 | 0 | 10.79683848 | 9.873526736 | 0 | 0 | 0 | 0.035207169 | 0.007910112 |

| | | | | | | | | | | |
|---|---|---|---|---|---|---|---|---|---|---|
| PTPN23 | 0 | 9.579530494 | 0 | 0 | 0 | 0 | 0 | 0 | 7.44E-08 | 1 |
| PTPRA | 11.9976917 | 9.366954907 | 0 | 9.147889518 | 7.041677569 | 0 | 0 | 0 | 0.387746571 | 0.027192267 |
| PTPRCAP | 12.14898911 | 0 | 0 | 0 | 0 | 10.69702153 | 7.962083516 | 10.09830559 | 7.07E-08 | 0.000532763 |
| PTPRF | 0 | 9.713548361 | 0 | 0 | 0 | 0 | 9.405611195 | 0 | 7.17E-08 | 1 |
| PTPRH | 10.28458564 | 0 | 0 | 0 | 0 | 0 | 0 | 0 | 7.07E-08 | 1 |
| PTPRM | 9.459308883 | 0 | 0 | 0 | 0 | 0 | 0 | 0 | 7.88E-08 | 1 |
| PTPRO | 0 | 9.194286798 | 0 | 0 | 0 | 0 | 0 | 0 | 8.58E-08 | 1 |
| PTTG1 | 0 | 0 | 10.07459399 | 12.81400898 | 11.5506231 | 10.99315703 | 10.2730601 | 12.17090258 | 4.52E-09 | 0.999854694 |
| PUDP | 0 | 0 | 10.14683338 | 0 | 8.28572291 | 10.87030234 | 7.864642143 | 0 | 0.047780591 | 0.999854694 |
| PUM1 | 0 | 0 | 0 | 10.0813848 | 7.481831074 | 0 | 0 | 0 | 0.056985669 | 0.013610425 |
| PUM2 | 0 | 0 | 0 | 0 | 6.670177039 | 0 | 8.193715158 | 0 | 4.78E-05 | 1 |
| PUM3 | 0 | 0 | 0 | 11.29843066 | 7.250409752 | 0 | 9.47741073 | 0 | 0.035972209 | 0.999854694 |
| PUS10 | 0 | 0 | 0 | 10.09069686 | 8.847894688 | 0 | 0 | 0 | 0.039913639 | 0.011034216 |
| PUS7L | 0 | 10.33538329 | 0 | 0 | 0 | 0 | 4.814710258 | 0 | 7.17E-08 | 0.625785924 |
| PVALB | 11.03769861 | 10.09821346 | 0 | 0 | 0 | 9.989752583 | 0 | 0 | 7.51E-05 | 1 |
| PWWP2B | 11.38584736 | 0 | 0 | 0 | 0 | 0 | 5.823835742 | 0 | 2.97E-08 | 1 |
| PXK | 0 | 0 | 10.03598438 | 0 | 6.841217856 | 0 | 0 | 0 | 0.066636219 | 0.014309205 |
| PXMP2 | 11.14326036 | 12.03125158 | 7.827979026 | 11.42056092 | 0 | 0 | 0 | 0 | 0.873275474 | 1.56E-10 |
| PXMP4 | 0 | 0 | 0 | 0 | 0 | 10.94936263 | 0 | 11.0976474 | 1 | 0.003589484 |
| PYCR1 | 0 | 0 | 0 | 10.64076551 | 8.388633797 | 0 | 8.089748008 | 9.478071506 | 0.027747177 | 0.999854694 |
| PYY | 0 | 13.14354104 | 9.491886065 | 0 | 7.705040924 | 0 | 0 | 0 | 0.665116211 | 0.017339178 |
| R3HCC1L | 0 | 0 | 0 | 0 | 0 | 0 | 5.516839174 | 10.36348585 | 1 | 0.011034216 |
| RAB17 | 10.46605563 | 0 | 0 | 0 | 0 | 0 | 5.955871074 | 0 | 7.07E-08 | 1 |
| RAB23 | 0 | 0 | 0 | 9.304728059 | 6.503532343 | 0 | 0 | 0 | 0.086638287 | 0.026700293 |
| RAB27A | 8.883131582 | 0 | 0 | 0 | 0 | 0 | 0 | 0 | 1.14E-07 | 1 |
| RAB30 | 0 | 0 | 10.19435779 | 8.69322713 | 7.978700908 | 0 | 7.567155003 | 0 | 0.008236263 | 0.999854694 |
| RAB32 | 0 | 10.17595559 | 11.66218971 | 10.25708182 | 9.770011639 | 0 | 0 | 0 | 0.854138757 | 0.000273087 |
| RAB33A | 0 | 9.840515353 | 0 | 0 | 0 | 0 | 0 | 0 | 7.07E-08 | 1 |
| RAB35 | 10.42677088 | 0 | 0 | 0 | 0 | 0 | 8.672944217 | 0 | 7.07E-08 | 1 |
| RAB37 | 9.285741834 | 0 | 0 | 0 | 0 | 0 | 0 | 0 | 8.38E-08 | 1 |
| RAB38 | 0 | 0 | 0 | 8.302434741 | 0 | 0 | 0 | 0 | 1.05E-05 | 1 |
| RAB3D | 0 | 0 | 10.4126727 | 0 | 8.269832732 | 0 | 7.80460581 | 0 | 0.038303276 | 0.999854694 |
| RAB4B | 9.960747977 | 6.070492458 | 0 | 0 | 0 | 0 | 0 | 0 | 0.016548457 | 1 |
| RAB4B-EGLN2 | 9.960747977 | 9.021269553 | 0 | 0 | 0 | 0 | 8.510636715 | 10.60000095 | 0.005537628 | 0.00825262 |
| RAB8A | 8.787126977 | 9.708800502 | 10.27024349 | 10.01026111 | 10.23290381 | 0 | 0 | 0 | 0.922337827 | 0.000538325 |
| RABEP1 | 15.43382759 | 14.84853181 | 12.92519919 | 13.77730311 | 14.07794651 | 15.02093904 | 15.28415114 | 13.902279 | 0.041776164 | 0.999854694 |
| RABEP2 | 0 | 0 | 0 | 8.561649124 | 0 | 0 | 9.104986918 | 0 | 1.67E-05 | 1 |
| RABIF | 0 | 0 | 11.33649476 | 10.27819485 | 0 | 0 | 7.637673707 | 0 | 0.02858177 | 0.999005823 |
| RABL2B | 0 | 8.824638969 | 0 | 0 | 0 | 0 | 0 | 0 | 1.36E-07 | 1 |
| RAC1 | 10.45046986 | 10.26597355 | 13.61805387 | 12.33750965 | 12.45237547 | 0 | 11.18771939 | 12.85627633 | 0.004788156 | 0.999854694 |
| RAD1 | 0 | 0 | 0 | 10.53753769 | 10.3973349 | 0 | 0 | 0 | 0.033856317 | 0.007496493 |
| RAD50 | 10.20786763 | 0 | 0 | 0 | 0 | 0 | 0 | 0 | 7.07E-08 | 1 |
| RAD51AP1 | 0 | 0 | 0 | 9.734552215 | 6.954898948 | 0 | 0 | 0 | 0.072543146 | 0.017622038 |

| | | | | | | | | | | |
|---|---|---|---|---|---|---|---|---|---|---|
| RAD54B | 11.06610091 | 11.39960066 | 11.35882683 | 13.49283383 | 14.15085886 | 12.58197897 | 14.378074 | 14.18974515 | 0.030796109 | 0.999854694 |
| RALA | 0 | 0 | 4.801878001 | 9.603145663 | 9.696135179 | 0 | 7.822015489 | 0 | 0.01298805 | 0.999854694 |
| RALGDS | 0 | 0 | 0 | 9.955974972 | 7.336838553 | 0 | 0 | 0 | 0.063916801 | 0.01457526 |
| RAPGEF1 | 0 | 0 | 0 | 0 | 0 | 8.585277065 | 8.173512489 | 0 | 1 | 0.018299451 |
| RARRES3 | 11.84364118 | 10.12162192 | 12.67493877 | 0 | 9.397490443 | 0 | 0 | 0 | 0.999485949 | 4.76E-12 |
| RASA2 | 14.20294975 | 13.85013113 | 12.19123952 | 11.67875204 | 11.58554342 | 12.48792892 | 12.53869646 | 12.69220252 | 0.001597874 | 0.999854694 |
| RASEF | 9.037149714 | 9.757890409 | 0 | 0 | 0 | 0 | 0 | 0 | 0.006977122 | 1 |
| RASSF1 | 0 | 0 | 0 | 12.13958661 | 10.50263832 | 10.76996942 | 0 | 0 | 0.018676813 | 0.999854694 |
| RASSF3 | 10.21710791 | 0 | 0 | 10.91537617 | 7.336838553 | 0 | 0 | 0 | 0.996280769 | 0.010003595 |
| RASSF8 | 0 | 0 | 0 | 0 | 6.608339636 | 0 | 0 | 0 | 5.07E-05 | 1 |
| RAVER2 | 8.973144195 | 0 | 0 | 0 | 0 | 0 | 0 | 0 | 1.08E-07 | 1 |
| RBAK | 0 | 8.250007882 | 0 | 0 | 0 | 0 | 0 | 0 | 2.64E-07 | 1 |
| RBBP5 | 0 | 0 | 0 | 0 | 6.706045958 | 0 | 0 | 0 | 0.000126325 | 1 |
| RBBP6 | 0 | 0 | 0 | 10.52287104 | 8.665598273 | 0 | 8.499859701 | 0 | 0.028897473 | 0.999854694 |
| RBBP8 | 10.43471387 | 0 | 0 | 0 | 0 | 0 | 0 | 0 | 7.07E-08 | 1 |
| RBM12 | 0 | 0 | 10.10599766 | 0 | 9.272486321 | 0 | 7.164920683 | 0 | 0.034222004 | 0.999854694 |
| RBM17 | 0 | 0 | 0 | 9.000410141 | 10.7628131 | 10.4228525 | 10.77624123 | 0 | 0.016596934 | 0.999854694 |
| RBM19 | 0 | 0 | 9.779730665 | 0 | 9.039311682 | 11.02297377 | 0 | 0 | 0.047632903 | 0.999854694 |
| RBM23 | 9.949703925 | 8.789094606 | 10.48133622 | 0 | 8.625578148 | 0 | 0 | 0 | 0.993570664 | 0.010189075 |
| RBM24 | 0 | 9.420205115 | 0 | 0 | 0 | 0 | 0 | 0 | 7.87E-08 | 1 |
| RBM27 | 0 | 0 | 0 | 0 | 6.530456572 | 0 | 0 | 0 | 5.38E-05 | 1 |
| RBM45 | 0 | 9.635641309 | 0 | 10.40837607 | 9.629988173 | 0 | 0 | 0 | 0.981170768 | 0.009209122 |
| RBM4B | 0 | 8.950636073 | 0 | 0 | 0 | 0 | 0 | 9.952745771 | 1.07E-07 | 1 |
| RBMXL1 | 10.30205913 | 0 | 0 | 0 | 0 | 10.32179346 | 0 | 0 | 7.07E-08 | 1 |
| RBP4 | 8.940044091 | 0 | 10.20212852 | 0 | 10.3682477 | 0 | 0 | 0 | 0.932679569 | 0.008092529 |
| RBP5 | 8.5735246 | 0 | 0 | 0 | 0 | 0 | 0 | 0 | 1.55E-07 | 1 |
| RC3H2 | 0 | 9.37296959 | 0 | 0 | 0 | 0 | 0 | 0 | 7.88E-08 | 1 |
| RCAN3 | 9.176299815 | 0 | 0 | 0 | 0 | 0 | 0 | 0 | 1.15E-07 | 1 |
| RCBTB1 | 0 | 8.350969903 | 0 | 0 | 0 | 0 | 0 | 0 | 1.57E-07 | 1 |
| RCC1 | 0 | 9.72299746 | 8.986519609 | 0 | 7.344444907 | 0 | 0 | 0 | 0.976136384 | 0.027192267 |
| RCC2 | 0 | 5.308137638 | 0 | 9.180650941 | 5.171448341 | 0 | 0 | 0 | 0.741955828 | 0.036356873 |
| RCN3 | 0 | 0 | 10.61903827 | 9.851856421 | 0 | 0 | 0 | 0 | 0.016192794 | 0.008668791 |
| RCSD1 | 0 | 9.635641309 | 0 | 0 | 0 | 0 | 0 | 0 | 7.38E-08 | 1 |
| RENBP | 0 | 0 | 10.36501154 | 0 | 7.918823272 | 0 | 0 | 0 | 0.0434622 | 0.011034216 |
| REST | 11.09144476 | 0 | 0 | 0 | 0 | 0 | 8.213638823 | 0 | 5.99E-08 | 1 |
| RET | 11.02460129 | 0 | 0 | 0 | 0 | 0 | 0 | 0 | 6.72E-08 | 1 |
| RETNLB | 0 | 11.81799934 | 10.37535803 | 0 | 8.460394093 | 0 | 0 | 0 | 0.937966741 | 0.010542114 |
| RFC1 | 0 | 0 | 0 | 0 | 0 | 8.880037188 | 0 | 7.347429756 | 1 | 0.022348702 |
| RFC4 | 0 | 0 | 0 | 10.24044427 | 10.88780163 | 0 | 0 | 10.25779056 | 0.031509902 | 0.999854694 |
| RFX7 | 8.356549509 | 0 | 0 | 0 | 0 | 0 | 0 | 0 | 2.53E-07 | 1 |
| RFXANK | 0 | 9.145930534 | 9.532516417 | 0 | 9.86495823 | 0 | 0 | 0 | 0.971198277 | 0.01037888 |
| RGCC | 0 | 0 | 0 | 9.567382206 | 9.571235177 | 12.69700027 | 10.35980261 | 0 | 0.039956712 | 0.999854694 |
| RGPD2 | 0 | 8.617326978 | 0 | 0 | 0 | 0 | 0 | 0 | 1.27E-07 | 1 |

| | | | | | | | | | | |
|---|---|---|---|---|---|---|---|---|---|---|
| RGS1 | 0 | 0 | 12.3213534 | 0 | 8.809938979 | 12.50699607 | 7.437617244 | 11.20375425 | 1.15E-07 | 0.999854694 |
| RGS19 | 8.787126977 | 0 | 0 | 0 | 0 | 0 | 0 | 0 | 1.22E-07 | 1 |
| RHBDD1 | 10.2354128 | 0 | 0 | 0 | 0 | 0 | 8.08254305 | 0 | 7.07E-08 | 1 |
| RHBDF1 | 9.474799963 | 0 | 0 | 0 | 0 | 0 | 0 | 0 | 7.88E-08 | 1 |
| RHBDL1 | 0 | 0 | 0 | 0 | 0 | 0 | 9.567902023 | 9.755251417 | 1 | 0.008991934 |
| RHBG | 0 | 9.870584892 | 0 | 0 | 0 | 0 | 8.542492985 | 0 | 7.07E-08 | 1 |
| RHPN1 | 0 | 8.724703557 | 0 | 0 | 0 | 0 | 7.546357065 | 0 | 1.17E-07 | 1 |
| RHPN2 | 0 | 9.37296959 | 0 | 0 | 0 | 0 | 0 | 0 | 7.88E-08 | 1 |
| RILPL2 | 0 | 0 | 0 | 8.014451711 | 0 | 0 | 0 | 0 | 3.02E-05 | 1 |
| RIMS3 | 0 | 8.850734095 | 0 | 0 | 0 | 0 | 0 | 0 | 1.17E-07 | 1 |
| RIN2 | 0 | 8.859328625 | 0 | 10.72518869 | 9.442484383 | 0 | 0 | 0 | 0.935616984 | 0.008668791 |
| RIOX1 | 0 | 8.68657578 | 0 | 0 | 0 | 0 | 0 | 0 | 1.19E-07 | 1 |
| RMC1 | 9.204448442 | 0 | 0 | 0 | 0 | 0 | 0 | 0 | 1.14E-07 | 1 |
| RMDN2 | 0 | 0 | 10.86708826 | 0 | 9.052142562 | 0 | 8.146127844 | 0 | 0.016826944 | 0.999854694 |
| RMDN3 | 0 | 0 | 9.705133056 | 10.29094678 | 0 | 0 | 0 | 0 | 0.025282695 | 0.009724498 |
| RMI1 | 9.290126577 | 8.743396117 | 0 | 0 | 0 | 0 | 0 | 0 | 0.014096311 | 1 |
| RMI2 | 0 | 0 | 0 | 10.84479957 | 9.388320781 | 0 | 0 | 0 | 0.013056079 | 0.008532333 |
| RNA45SN4 | 0 | 10.36284296 | 0 | 6.89802758 | 8.656460903 | 0 | 0 | 0 | 0.913328892 | 0.041287884 |
| RNASEH2A | 0 | 0 | 0 | 11.4642036 | 9.980519843 | 0 | 9.566611567 | 0 | 0.027450171 | 0.999854694 |
| RNF103 | 0 | 8.724703557 | 10.12656 | 9.16057826 | 7.367026271 | 0 | 0 | 0 | 0.915944843 | 0.002232985 |
| RNF111 | 0 | 9.431776243 | 0 | 0 | 0 | 0 | 0 | 0 | 7.88E-08 | 1 |
| RNF113A | 0 | 0 | 0 | 10.58725782 | 8.748200401 | 12.57581996 | 0 | 10.33965736 | 0.026271629 | 0.999854694 |
| RNF125 | 10.32360752 | 0 | 0 | 0 | 0 | 0 | 0 | 0 | 7.07E-08 | 1 |
| RNF13 | 0 | 0 | 11.40878953 | 7.518919157 | 0 | 0 | 9.830189467 | 0 | 0.035423319 | 0.999854694 |
| RNF138 | 8.657374495 | 0 | 0 | 0 | 0 | 0 | 0 | 0 | 1.30E-07 | 1 |
| RNF141 | 0 | 0 | 9.585464507 | 0 | 9.755107927 | 0 | 0 | 0 | 0.035967833 | 0.010542114 |
| RNF152 | 0 | 9.066507669 | 0 | 0 | 0 | 0 | 0 | 0 | 1.17E-07 | 1 |
| RNF170 | 0 | 0 | 10.17474585 | 6.193017253 | 7.845672509 | 0 | 7.930328737 | 0 | 0.019052053 | 0.999854694 |
| RNF2 | 0 | 8.250007882 | 0 | 0 | 0 | 9.845512026 | 0 | 7.880730732 | 2.64E-07 | 0.011578584 |
| RNF208 | 0 | 0 | 0 | 0 | 6.729472521 | 0 | 0 | 8.407451742 | 0.000125541 | 1 |
| RNF219 | 0 | 0 | 6.607354851 | 7.997577745 | 7.141107086 | 0 | 0 | 0 | 0.066299881 | 0.014258614 |
| RNF227 | 0 | 8.363107485 | 0 | 0 | 0 | 0 | 0 | 0 | 1.55E-07 | 1 |
| RNF25 | 0 | 10.83972836 | 10.42932003 | 8.925071258 | 0 | 0 | 0 | 0 | 0.988867264 | 0.01037888 |
| RNF40 | 9.24114547 | 9.584722618 | 0 | 0 | 0 | 0 | 8.075301928 | 0 | 0.007252428 | 1 |
| RNF44 | 0 | 0 | 0 | 0 | 6.39051047 | 0 | 7.06610294 | 0 | 5.98E-05 | 1 |
| RNGTT | 0 | 9.059065806 | 0 | 0 | 0 | 0 | 0 | 0 | 1.18E-07 | 1 |
| RNPEP | 0 | 10.08361716 | 7.971285541 | 10.96178808 | 9.634648696 | 0 | 0 | 0 | 0.958728107 | 0.000870107 |
| RNPEPL1 | 10.39047609 | 0 | 0 | 0 | 0 | 0 | 0 | 0 | 7.07E-08 | 1 |
| ROBO1 | 9.294498035 | 0 | 0 | 0 | 0 | 0 | 0 | 0 | 8.36E-08 | 1 |
| ROBO2 | 9.881603011 | 12.25291545 | 0 | 10.97118208 | 12.97161553 | 0 | 0 | 0 | 0.99353678 | 3.43E-12 |
| RP9 | 0 | 0 | 0 | 10.43473225 | 10.75712527 | 0 | 8.068024279 | 0 | 0.031510127 | 0.999854694 |
| RPGRIP1L | 0 | 0 | 0 | 9.246405636 | 7.21765149 | 0 | 0 | 0 | 0.083902631 | 0.02416948 |
| RPL10L | 0 | 8.576210795 | 0 | 0 | 0 | 0 | 0 | 0 | 1.31E-07 | 1 |

| | | | | | | | | | | |
|---|---|---|---|---|---|---|---|---|---|---|
| RPL23AP7 | 0 | 0 | 0 | 10.2464083 | 8.644187016 | 0 | 8.935389379 | 0 | 0.037622992 | 0.999854694 |
| RPL7 | 14.78982332 | 15.89199091 | 16.03192553 | 17.18086089 | 17.11478341 | 18.64892668 | 17.554448 | 17.64422291 | 0.032597368 | 0.999854694 |
| RPL8 | 13.67355056 | 13.86794204 | 14.66113961 | 15.54545264 | 15.71634306 | 16.82386049 | 15.30888992 | 16.08090191 | 0.010984532 | 0.999854694 |
| RPP25 | 0 | 0 | 0 | 10.33527103 | 8.883572754 | 0 | 10.54811326 | 0 | 0.032119053 | 0.999854694 |
| RPP30 | 10.04618272 | 9.187477203 | 12.08177423 | 10.24759816 | 8.460394093 | 0 | 0 | 0 | 0.758223081 | 0.000291056 |
| RPP38 | 0 | 0 | 0 | 11.45341608 | 10.76847859 | 0 | 0 | 0 | 0.023570644 | 0.00480484 |
| RPP40 | 0 | 0 | 0 | 10.12347196 | 8.026767928 | 0 | 0 | 0 | 0.049413137 | 0.012283188 |
| RPS5 | 12.80566128 | 13.551153 | 14.43664808 | 15.16983153 | 14.88954718 | 15.71485369 | 13.80905701 | 14.70389941 | 0.011658031 | 0.999854694 |
| RPS6KL1 | 0 | 0 | 0 | 0 | 6.608339636 | 0 | 0 | 0 | 5.07E-05 | 1 |
| RPSAP52 | 0 | 0 | 0 | 0 | 0 | 10.94865665 | 8.99416519 | 0 | 1 | 0.00624926 |
| RPSAP58 | 0 | 6.070492458 | 10.93277805 | 11.3092974 | 9.631543354 | 12.07273217 | 11.07515869 | 0 | 0.032842841 | 0.999854694 |
| RPSAP9 | 10.40671994 | 0 | 0 | 0 | 0 | 0 | 0 | 0 | 7.07E-08 | 1 |
| RPUSD1 | 0 | 9.073911341 | 0 | 9.815410697 | 6.706045958 | 0 | 0 | 0 | 0.995442919 | 0.017334618 |
| RRAGC | 0 | 0 | 0 | 0 | 6.830424858 | 0 | 0 | 0 | 4.17E-05 | 1 |
| RREB1 | 0 | 9.336499504 | 0 | 0 | 0 | 0 | 0 | 0 | 8.04E-08 | 1 |
| RRM2 | 0 | 9.537306767 | 0 | 11.06325583 | 6.620920975 | 0 | 0 | 0 | 0.980430955 | 2.04E-10 |
| RRP12 | 0 | 8.586600217 | 0 | 0 | 0 | 0 | 0 | 0 | 1.30E-07 | 1 |
| RSAD1 | 0 | 0 | 0 | 8.781306752 | 6.116175352 | 0 | 0 | 0 | 0.109513557 | 0.046371395 |
| RSAD2 | 0 | 0 | 9.357432539 | 0 | 4.98937073 | 0 | 0 | 0 | 0.092970915 | 0.029546388 |
| RSBN1 | 0 | 9.684823965 | 0 | 0 | 0 | 0 | 0 | 0 | 7.17E-08 | 1 |
| RSF1 | 17.5361448 | 16.73296835 | 15.51820076 | 15.67480563 | 16.20469213 | 15.43301371 | 16.6407712 | 15.82824912 | 0.027500278 | 0.999854694 |
| RSPH3 | 0 | 0 | 9.663524539 | 0 | 7.078631465 | 0 | 0 | 0 | 0.075280036 | 0.017436678 |
| RSPRY1 | 0 | 8.326384029 | 0 | 0 | 0 | 0 | 7.080640961 | 9.148443004 | 1.61E-07 | 0.018594788 |
| RTF1 | 0 | 0 | 10.48611794 | 0 | 8.577977952 | 0 | 9.48015487 | 0 | 0.032736381 | 0.999854694 |
| RTL8C | 0 | 0 | 8.844794804 | 0 | 8.951141308 | 0 | 0 | 0 | 0.072776828 | 0.01630416 |
| RUFY2 | 9.669116652 | 0 | 0 | 0 | 0 | 0 | 0 | 0 | 7.44E-08 | 1 |
| RUFY3 | 0 | 0 | 0 | 8.606855021 | 0 | 0 | 0 | 0 | 7.56E-06 | 1 |
| RUSC1 | 0 | 0 | 11.41130567 | 9.582548699 | 9.275473597 | 0 | 0 | 0 | 7.65E-05 | 0.000485974 |
| RUVBL2 | 0 | 0 | 0 | 11.1586863 | 10.04554289 | 0 | 8.271810291 | 0 | 0.030353545 | 0.999854694 |
| RWDD2B | 10.08707482 | 0 | 0 | 0 | 0 | 0 | 7.930328737 | 10.52009861 | 7.07E-08 | 0.009118992 |
| RYK | 0 | 9.059065806 | 0 | 0 | 0 | 0 | 0 | 0 | 1.18E-07 | 1 |
| S100A8 | 0 | 0 | 10.0703547 | 9.423192445 | 8.501797164 | 0 | 0 | 10.1264312 | 0.004291968 | 0.999854694 |
| SAE1 | 0 | 0 | 0 | 11.01026268 | 11.35033487 | 10.83253396 | 10.48051556 | 0 | 0.022735435 | 0.999854694 |
| SAMD13 | 0 | 0 | 10.00522363 | 0 | 9.007325253 | 0 | 0 | 0 | 0.041618605 | 0.010943033 |
| SAP30BP | 10.78156472 | 8.876365592 | 0 | 12.06156626 | 7.453974941 | 0 | 0 | 0 | 1 | 1.61E-11 |
| SAR1B | 0 | 0 | 11.39952597 | 0 | 10.57069201 | 0 | 0 | 8.354929434 | 0.026011992 | 0.999854694 |
| SARS2 | 0 | 0 | 9.913539582 | 0 | 9.488693902 | 0 | 0 | 0 | 0.034989165 | 0.01037888 |
| SASH3 | 0 | 8.705765621 | 0 | 0 | 0 | 0 | 0 | 0 | 1.18E-07 | 1 |
| SAV1 | 0 | 0 | 0 | 0 | 0 | 0 | 7.403330192 | 9.573099607 | 1 | 0.013697851 |
| SCAF1 | 0 | 0 | 0 | 0 | 6.873120683 | 0 | 0 | 0 | 4.05E-05 | 1 |
| SCAF8 | 0 | 0 | 0 | 8.391244621 | 8.377555125 | 0 | 0 | 0 | 0.089646608 | 0.027439132 |
| SCAMP1-AS1 | 0 | 0 | 5.895636902 | 10.1719063 | 9.056780188 | 0 | 0 | 0 | 0.010249298 | 0.003243461 |
| SCAMP4 | 0 | 8.155470719 | 0 | 6.603116496 | 8.890084061 | 0 | 0 | 0 | 0.990582117 | 0.036233885 |

| | | | | | | | | | |
|---|---|---|---|---|---|---|---|---|---|
| SCAND2P | 10.61679353 | 0 | 0 | 0 | 0 | 0 | 0 | 0 | 7.07E-08 | 1 |
| SCARB2 | 0 | 9.916606105 | 0 | 0 | 0 | 0 | 0 | 10.69226845 | 7.07E-08 | 1 |
| SCARF2 | 0 | 0 | 0 | 0 | 6.670177039 | 0 | 0 | 0 | 4.78E-05 | 1 |
| SCFD2 | 0 | 0 | 0 | 0 | 0 | 10.55117368 | 0 | 10.13337775 | 1 | 0.006244708 |
| SCG5 | 0 | 10.62216407 | 0 | 0 | 0 | 0 | 0 | 0 | 7.07E-08 | 1 |
| SCML1 | 0 | 0 | 0 | 0 | 0 | 0 | 8.08254305 | 9.488944611 | 1 | 0.012440088 |
| SCNN1B | 10.60979461 | 9.870584892 | 0 | 0 | 0 | 0 | 0 | 0 | 0.00049104 | 1 |
| SCNN1G | 0 | 8.700992037 | 0 | 0 | 0 | 0 | 0 | 0 | 1.18E-07 | 1 |
| SCX | 0 | 0 | 0 | 7.122154799 | 0 | 0 | 0 | 0 | 3.56E-05 | 1 |
| SDC2 | 0 | 0 | 8.968408265 | 10.55590393 | 6.595647612 | 0 | 0 | 10.38692716 | 0.005639397 | 0.999854694 |
| SDHAF3 | 10.3448388 | 7.537775521 | 10.22804996 | 10.5975595 | 9.35104796 | 0 | 0 | 0 | 0.960318461 | 0.000590012 |
| SDHAF4 | 0 | 8.576210795 | 9.01326732 | 0 | 7.913719455 | 0 | 0 | 0 | 0.99353678 | 0.022345897 |
| SDHAP2 | 0 | 0 | 9.448445763 | 0 | 9.224859614 | 0 | 0 | 0 | 0.052572637 | 0.012259484 |
| SDK2 | 10.36160169 | 0 | 0 | 0 | 0 | 0 | 0 | 0 | 7.07E-08 | 1 |
| SDR16C5 | 0 | 0 | 10.73606275 | 11.40571497 | 0 | 0 | 7.96991419 | 0 | 0.025267978 | 0.999854694 |
| SDR39U1 | 9.137893062 | 9.600187807 | 8.747254732 | 10.8416513 | 8.820885744 | 0 | 0 | 0 | 0.981095742 | 0.001042905 |
| SEC14L1 | 0 | 0 | 0 | 0 | 6.633393544 | 0 | 0 | 0 | 4.95E-05 | 1 |
| SEC16B | 0 | 0 | 0 | 0 | 6.608339636 | 0 | 7.460030783 | 0 | 5.07E-05 | 1 |
| SEC31A | 10.47955641 | 0 | 0 | 0 | 0 | 0 | 10.15952482 | 0 | 7.07E-08 | 1 |
| SECTM1 | 10.43866903 | 0 | 0 | 0 | 0 | 0 | 0 | 0 | 7.07E-08 | 1 |
| SEL1L | 0 | 0 | 0 | 0 | 6.582842939 | 0 | 0 | 0 | 0.000136511 | 1 |
| SELPLG | 0 | 0 | 7.38464918 | 0 | 0 | 10.54838053 | 0 | 11.10119485 | 2.91E-05 | 0.912844529 |
| SEMA4G | 11.04550031 | 8.069195977 | 0 | 9.853420322 | 6.185573957 | 0 | 0 | 0 | 0.850876376 | 0.01786378 |
| SEMA6A | 9.904662348 | 9.420205115 | 0 | 0 | 0 | 0 | 0 | 0 | 0.004054775 | 1 |
| SEMA6D | 9.932977691 | 0 | 0 | 0 | 0 | 0 | 0 | 0 | 7.07E-08 | 1 |
| SENP7 | 0 | 8.657303766 | 0 | 0 | 0 | 0 | 0 | 0 | 1.22E-07 | 1 |
| 4-Sep | 0 | 0 | 10.34233629 | 0 | 9.401142051 | 0 | 0 | 0 | 0.026504187 | 0.009551658 |
| SEPT5-GP1BB | 10.07695994 | 8.884809218 | 0 | 0 | 0 | 0 | 0 | 0 | 0.005186306 | 1 |
| 8-Sep | 0 | 0 | 9.123764545 | 0 | 6.043268989 | 0 | 0 | 0 | 0.098772605 | 0.032241775 |
| SERAC1 | 0 | 8.966633964 | 0 | 0 | 0 | 0 | 0 | 0 | 1.07E-07 | 1 |
| SERPINB5 | 0 | 9.180635315 | 0 | 9.893497858 | 9.153044768 | 0 | 0 | 0 | 0.980815574 | 0.011067641 |
| SERPINB8 | 0 | 9.579530494 | 0 | 0 | 0 | 0 | 0 | 10.8439491 | 7.44E-08 | 1 |
| SERPINB9 | 0 | 0 | 0 | 0 | 6.530456572 | 0 | 7.587657373 | 9.681338694 | 5.38E-05 | 0.999854694 |
| SERPINF1 | 0 | 0 | 0 | 10.19799108 | 7.861662929 | 0 | 0 | 12.16794558 | 0.048178463 | 0.999854694 |
| SERPINI1 | 0 | 8.676884284 | 0 | 0 | 0 | 0 | 8.848860281 | 0 | 1.20E-07 | 1 |
| SESTD1 | 0 | 0 | 0 | 8.356373925 | 8.887483063 | 0 | 0 | 0 | 0.079549852 | 0.021057037 |
| SETBP1 | 11.89995144 | 9.305387296 | 0 | 0 | 0 | 0 | 10.15609692 | 10.6260915 | 3.02E-06 | 0.005705522 |
| SFI1 | 0 | 0 | 0 | 0 | 6.808593567 | 0 | 0 | 0 | 6.32E-05 | 1 |
| SFMBT2 | 0 | 8.798063133 | 0 | 0 | 0 | 0 | 0 | 0 | 1.15E-07 | 1 |
| SFSWAP | 0 | 0 | 9.179933839 | 0 | 7.488711871 | 0 | 0 | 0 | 0.085828317 | 0.022637767 |
| SFT2D3 | 9.846307856 | 8.169361257 | 9.273256673 | 10.77059199 | 6.894003922 | 0 | 0 | 0 | 0.981170768 | 0.001496582 |
| SFXN2 | 0 | 0 | 0 | 10.6862296 | 7.298193348 | 0 | 0 | 0 | 0.033255093 | 0.010789277 |
| SGK3 | 0 | 8.555204765 | 0 | 0 | 0 | 0 | 0 | 0 | 1.34E-07 | 1 |

| | | | | | | | | | | |
|---|---|---|---|---|---|---|---|---|---|---|
| SH2D4A | 8.529698369 | 0 | 0 | 0 | 0 | 0 | 9.147036996 | 0 | 1.41E-07 | 1 |
| SH3BP1 | 6.193650512 | 8.950636073 | 0 | 0 | 0 | 0 | 0 | 0 | 0.034157652 | 1 |
| SH3KBP1 | 0 | 10.39866127 | 0 | 0 | 0 | 0 | 0 | 9.488944611 | 7.07E-08 | 1 |
| SHANK2 | 0 | 0 | 0 | 0 | 0 | 0 | 7.657204612 | 10.63957196 | 1 | 0.008968837 |
| SHC2 | 0 | 0 | 0 | 0 | 0 | 0 | 7.795821548 | 9.838371219 | 1 | 0.011395116 |
| SHOX | 0 | 0 | 0 | 8.311565818 | 0 | 0 | 0 | 0 | 1.04E-05 | 1 |
| SHPRH | 9.747734533 | 0 | 0 | 0 | 0 | 0 | 0 | 0 | 7.17E-08 | 1 |
| SI | 0 | 8.696202607 | 0 | 0 | 0 | 0 | 0 | 0 | 1.18E-07 | 1 |
| SIAE | 9.571629895 | 0 | 0 | 0 | 0 | 0 | 0 | 0 | 7.60E-08 | 1 |
| SIAH2 | 0 | 0 | 10.76454972 | 9.805749732 | 0 | 0 | 7.906045388 | 0 | 0.013855366 | 0.999854694 |
| SIDT1 | 0 | 8.263021342 | 0 | 0 | 0 | 0 | 0 | 0 | 2.55E-07 | 1 |
| SIDT2 | 0 | 0 | 0 | 0 | 6.517057266 | 0 | 0 | 0 | 5.50E-05 | 1 |
| SIGLEC10 | 0 | 0 | 8.024846459 | 0 | 0 | 0 | 0 | 0 | 1.52E-05 | 1 |
| SIK3 | 11.83316256 | 0 | 0 | 0 | 0 | 0 | 0 | 0 | 6.01E-08 | 1 |
| SIKE1 | 0 | 8.467981611 | 10.53462266 | 11.25767381 | 0 | 0 | 0 | 0 | 0.821026538 | 0.005940842 |
| SIL1 | 8.811733678 | 0 | 0 | 0 | 0 | 0 | 9.922863671 | 0 | 1.17E-07 | 1 |
| SINHCAF | 0 | 0 | 0 | 11.48149989 | 7.898298769 | 0 | 9.348188699 | 0 | 0.032447379 | 0.999854694 |
| SIRT3 | 0 | 0 | 0 | 10.19922153 | 9.410230838 | 0 | 0 | 10.97829805 | 0.028344895 | 0.999854694 |
| SIRT5 | 0 | 0 | 0 | 0 | 0 | 0 | 8.153022891 | 10.71096341 | 1 | 0.00825262 |
| SIRT6 | 9.846307856 | 0 | 0 | 0 | 0 | 0 | 0 | 0 | 7.17E-08 | 1 |
| SKA2 | 0 | 0 | 0 | 9.239245818 | 9.753680464 | 0 | 6.479216537 | 9.928988741 | 0.041925642 | 0.999854694 |
| SKA3 | 0 | 0 | 0 | 10.28748014 | 8.508583613 | 0 | 5.955871074 | 0 | 0.037958306 | 0.826575509 |
| SLC10A3 | 0 | 0 | 0 | 10.04626121 | 7.588169674 | 0 | 0 | 0 | 0.057561092 | 0.013690456 |
| SLC15A3 | 0 | 9.40854043 | 0 | 0 | 0 | 0 | 0 | 0 | 7.88E-08 | 1 |
| SLC16A7 | 11.6812034 | 11.6762307 | 10.13977005 | 11.00956149 | 0 | 0 | 0 | 0 | 0.867466701 | 0.007255458 |
| SLC18A1 | 0 | 8.696202607 | 0 | 0 | 0 | 0 | 0 | 0 | 1.18E-07 | 1 |
| SLC18B1 | 0 | 0 | 10.2757772 | 0 | 7.209344369 | 0 | 0 | 0 | 0.053379301 | 0.012259484 |
| SLC25A10 | 6.89226915 | 0 | 9.464687668 | 0 | 7.699120227 | 0 | 0 | 0 | 0.864901994 | 0.017622038 |
| SLC25A13 | 0 | 8.445326205 | 0 | 0 | 0 | 0 | 0 | 0 | 1.44E-07 | 1 |
| SLC25A33 | 0 | 0 | 0 | 10.36846339 | 8.373843248 | 0 | 7.403330192 | 0 | 0.036503789 | 0.999854694 |
| SLC25A34 | 0 | 9.56908985 | 0 | 0 | 0 | 0 | 0 | 0 | 7.53E-08 | 1 |
| SLC25A35 | 0 | 0 | 0 | 10.12864756 | 9.187285685 | 0 | 0 | 0 | 0.033678295 | 0.010542114 |
| SLC25A36 | 0 | 0 | 0 | 9.979081745 | 6.185573957 | 0 | 0 | 0 | 0.068915188 | 0.016357977 |
| SLC25A40 | 0 | 8.275918465 | 0 | 0 | 0 | 10.63078916 | 0 | 0 | 1.73E-07 | 1 |
| SLC25A45 | 9.419837447 | 0 | 0 | 0 | 0 | 0 | 0 | 0 | 7.91E-08 | 1 |
| SLC25A46 | 0 | 0 | 0 | 8.442019522 | 0 | 0 | 0 | 5.707457615 | 1.20E-05 | 1 |
| SLC27A2 | 0 | 10.46773564 | 0 | 10.86819517 | 7.509159523 | 0 | 0 | 0 | 0.999483812 | 0.010065115 |
| SLC27A6 | 10.41075248 | 0 | 0 | 0 | 0 | 0 | 0 | 0 | 7.07E-08 | 1 |
| SLC28A2 | 9.875780124 | 0 | 0 | 0 | 0 | 0 | 8.139199685 | 0 | 7.15E-08 | 1 |
| SLC29A1 | 12.38762521 | 13.55148451 | 14.14261446 | 16.46129105 | 16.18522536 | 16.16351599 | 17.11979487 | 15.77107649 | 0.002284458 | 0.999854694 |
| SLC2A1 | 0 | 0 | 0 | 8.197982807 | 10.1293868 | 10.26167403 | 0 | 11.18046758 | 0.044587169 | 0.999854694 |
| SLC33A1 | 0 | 8.780069978 | 0 | 0 | 0 | 0 | 0 | 0 | 1.15E-07 | 1 |
| SLC35A2 | 0 | 9.254168606 | 0 | 9.540451037 | 8.012515226 | 0 | 0 | 0 | 0.999483812 | 0.016321748 |

| | | | | | | | | | | |
|---|---|---|---|---|---|---|---|---|---|---|
| SLC35A4 | 8.99479642 | 7.774287275 | 0 | 0 | 0 | 0 | 0 | 10.1505992 | 0.025853276 | 1 |
| SLC35C1 | 0 | 0 | 10.4660887 | 9.818616695 | 0 | 0 | 0 | 0 | 0.019839019 | 0.009118992 |
| SLC35D2 | 0 | 0 | 10.11219719 | 0 | 8.997709978 | 0 | 8.729391788 | 0 | 0.038582958 | 0.999854694 |
| SLC35E3 | 0 | 8.761849577 | 0 | 0 | 0 | 0 | 0 | 0 | 1.16E-07 | 1 |
| SLC35E4 | 0 | 0 | 0 | 9.52875341 | 8.622453199 | 0 | 0 | 0 | 0.06237984 | 0.01420877 |
| SLC35G1 | 10.17505246 | 0 | 0 | 0 | 0 | 0 | 0 | 0 | 7.07E-08 | 1 |
| SLC37A4 | 0 | 0 | 10.1963044 | 0 | 9.038139559 | 0 | 0 | 0 | 0.035525927 | 0.01037888 |
| SLC38A3 | 0 | 0 | 0 | 0 | 0 | 9.939471283 | 7.67647464 | 0 | 1 | 0.011034216 |
| SLC39A1 | 0 | 0 | 0 | 11.79399183 | 7.329191883 | 0 | 9.140113213 | 0 | 1.73E-07 | 0.999854694 |
| SLC39A13 | 0 | 0 | 0 | 9.406244235 | 7.425570344 | 0 | 0 | 0 | 0.077991075 | 0.020497277 |
| SLC39A8 | 0 | 9.760186872 | 0 | 0 | 0 | 0 | 0 | 0 | 7.17E-08 | 1 |
| SLC41A2 | 0 | 0 | 0 | 0 | 6.489879427 | 0 | 0 | 0 | 5.56E-05 | 1 |
| SLC41A3 | 0 | 0 | 0 | 9.808977247 | 7.774256458 | 0 | 0 | 0 | 0.065339989 | 0.014756957 |
| SLC48A1 | 0 | 0 | 0 | 9.93249208 | 8.677692101 | 0 | 7.864642143 | 0 | 0.047685799 | 0.999854694 |
| SLC4A4 | 11.78115486 | 11.36983375 | 5.775783019 | 0 | 4.107448263 | 0 | 7.98544917 | 0 | 0.001979659 | 0.999854694 |
| SLC5A1 | 0 | 8.982456399 | 11.23693589 | 0 | 8.449854967 | 0 | 0 | 0 | 0.935518721 | 0.007978891 |
| SLC5A6 | 0 | 0 | 9.840738208 | 11.3503034 | 0 | 0 | 9.087097759 | 0 | 0.029981024 | 0.999854694 |
| SLC6A19 | 11.03508864 | 0 | 0 | 0 | 0 | 0 | 0 | 0 | 6.71E-08 | 1 |
| SLC6A8 | 10.04618272 | 7.559034993 | 0 | 10.00182426 | 8.903019168 | 0 | 0 | 0 | 0.994897187 | 0.011255646 |
| SLC7A1 | 0 | 0 | 0 | 0 | 6.8625646 | 0 | 9.101426788 | 0 | 4.09E-05 | 1 |
| SLC7A14 | 0 | 0 | 7.617477076 | 0 | 0 | 0 | 0 | 0 | 2.31E-05 | 1 |
| SLC7A6OS | 0 | 9.286392738 | 0 | 0 | 0 | 0 | 0 | 0 | 8.08E-08 | 1 |
| SLC7A8 | 0 | 9.088605668 | 0 | 0 | 0 | 0 | 9.258327475 | 0 | 1.16E-07 | 1 |
| SLC9A4 | 0 | 0 | 0 | 0 | 6.873120683 | 0 | 0 | 0 | 4.05E-05 | 1 |
| SLCO1C1 | 10.03054434 | 0 | 0 | 0 | 0 | 0 | 0 | 0 | 7.07E-08 | 1 |
| SLCO4A1 | 0 | 0 | 9.339929502 | 10.21633848 | 7.581748272 | 0 | 7.426278527 | 10.73627972 | 0.005596508 | 0.999854694 |
| SLX1A-SULT1A3 | 8.602019497 | 9.650572952 | 0 | 0 | 0 | 0 | 6.43464192 | 0 | 0.010635804 | 1 |
| SMAD6 | 0 | 0 | 0 | 10.67830993 | 6.984408946 | 11.40493273 | 10.6448358 | 0 | 0.034989165 | 0.999854694 |
| SMARCC1 | 9.675834273 | 10.61710519 | 12.28888259 | 12.59218528 | 13.74597885 | 12.47303181 | 13.45150556 | 12.57399667 | 0.002434762 | 0.999854694 |
| SMARCC2 | 0 | 0 | 0 | 7.595677272 | 0 | 0 | 0 | 0 | 2.19E-05 | 1 |
| SMARCD1 | 0 | 0 | 0 | 0 | 6.763914196 | 0 | 0 | 0 | 0.000116718 | 1 |
| SMARCD2 | 0 | 0 | 10.86096545 | 8.391244621 | 9.120188625 | 10.4530348 | 7.178500772 | 0 | 0.001394328 | 0.999854694 |
| SMC4 | 0 | 0 | 10.46528171 | 10.58913633 | 0 | 0 | 0 | 10.82691014 | 0.034324231 | 1 |
| SMG1 | 0 | 9.595051142 | 0 | 0 | 0 | 0 | 8.016026117 | 0 | 7.73E-08 | 1 |
| SMG1P5 | 0 | 0 | 10.84926002 | 6.478934088 | 0 | 0 | 0 | 0 | 0.034671354 | 1.90E-10 |
| SMG5 | 7.909170687 | 9.044065918 | 0 | 9.000410141 | 8.522061484 | 0 | 0 | 0 | 0.991384446 | 0.01865314 |
| SMG7 | 9.520299506 | 0 | 0 | 0 | 0 | 0 | 9.399825791 | 0 | 7.73E-08 | 1 |
| SMIM11A | 0 | 0 | 0 | 10.08405153 | 8.701579802 | 0 | 7.460030783 | 0 | 0.042080155 | 0.999854694 |
| SMIM27 | 0 | 4.970111344 | 0 | 0 | 0 | 9.114635565 | 6.975657029 | 0 | 0.257504232 | 0.019111579 |
| SMIM3 | 0 | 0 | 0 | 10.12087718 | 9.035792454 | 10.59514399 | 8.959187811 | 9.291937386 | 0.035972209 | 0.999854694 |
| SMIM35 | 9.74134422 | 0 | 0 | 0 | 0 | 0 | 0 | 0 | 7.22E-08 | 1 |
| SMIM8 | 0 | 0 | 10.2922521 | 0 | 9.472297898 | 0 | 7.080640961 | 0 | 0.026612843 | 0.999854694 |
| SMN1 | 0 | 0 | 0 | 11.68623059 | 9.417460865 | 9.668179176 | 8.450341371 | 10.1264312 | 0.026801204 | 0.999854694 |

| | | | | | | | | | |
|---|---|---|---|---|---|---|---|---|---|
| SMPD2 | 10.35324459 | 0 | 6.912177715 | 0 | 9.541763669 | 0 | 0 | 0 | 0.970266847 | 0.019243012 |
| SMURF2 | 9.641928937 | 0 | 0 | 0 | 0 | 0 | 0 | 0 | 7.44E-08 | 1 |
| SMYD2 | 0 | 9.073911341 | 0 | 10.81300332 | 7.866953924 | 0 | 0 | 0 | 0.952260332 | 0.010003595 |
| SNAI1 | 0 | 0 | 0 | 10.20535807 | 11.62784298 | 0 | 11.65595978 | 11.98083187 | 0.024019715 | 0.999854694 |
| SNAPC1 | 0 | 8.950636073 | 0 | 0 | 0 | 0 | 0 | 0 | 1.07E-07 | 1 |
| SNCA | 0 | 8.223623425 | 0 | 0 | 0 | 0 | 0 | 0 | 2.64E-07 | 1 |
| SNHG10 | 0 | 0 | 8.331565778 | 0 | 10.15394307 | 0 | 0 | 10.15401896 | 0.042132385 | 0.999854694 |
| SNHG12 | 0 | 0 | 10.51448015 | 9.645284634 | 7.716809918 | 0 | 8.865747094 | 9.46711583 | 0.002499376 | 0.999854694 |
| SNHG15 | 0 | 0 | 0 | 9.336387325 | 10.7702445 | 0 | 0 | 0 | 0.013779569 | 0.008668791 |
| SNORD3A | 0 | 0 | 0 | 0 | 6.569923599 | 0 | 0 | 0 | 5.22E-05 | 1 |
| SNRK | 0 | 0 | 0 | 0 | 6.808593567 | 0 | 0 | 0 | 6.32E-05 | 1 |
| SNRNP200 | 0 | 0 | 0 | 10.79846313 | 10.34994295 | 11.14015079 | 9.878831122 | 0 | 0.032011714 | 0.999854694 |
| SNRNP25 | 11.90210148 | 0 | 10.69878636 | 11.7409059 | 9.943435932 | 0 | 0 | 0 | 0.997540019 | 0.000201395 |
| SNRNP40 | 0 | 10.55232541 | 0 | 11.34864099 | 7.757261396 | 0 | 0 | 0 | 0.99574929 | 0.008359571 |
| SNTG2 | 12.03865896 | 10.64222403 | 0 | 0 | 0 | 0 | 0 | 0 | 7.07E-08 | 1 |
| SNUPN | 0 | 0 | 0 | 11.75603481 | 9.215557475 | 0 | 9.773153043 | 0 | 0.026504187 | 0.999854694 |
| SNURF | 0 | 8.576210795 | 0 | 0 | 0 | 0 | 0 | 0 | 1.31E-07 | 1 |
| SNX32 | 9.828331099 | 0 | 0 | 6.968995405 | 9.088832739 | 0 | 0 | 0 | 0.980815574 | 0.028010277 |
| SNX8 | 0 | 0 | 0 | 10.35085587 | 7.861662929 | 0 | 0 | 11.83612477 | 0.042080155 | 0.999854694 |
| SON | 20.30352664 | 19.35239423 | 17.86232353 | 18.54072082 | 18.79865014 | 18.51190251 | 19.36299587 | 18.3749447 | 0.008342202 | 0.999854694 |
| SORD | 0 | 8.155470719 | 8.757838236 | 0 | 9.754394372 | 0 | 0 | 0 | 0.937031857 | 0.012259484 |
| SOS2 | 0 | 0 | 9.381585435 | 0 | 7.656980716 | 0 | 0 | 0 | 0.078825993 | 0.018761292 |
| SOWAHA | 0 | 0 | 10.45718696 | 0 | 7.785476285 | 0 | 0 | 0 | 0.041219944 | 0.010943033 |
| SOX12 | 0 | 0 | 0 | 0 | 6.375739247 | 0 | 0 | 0 | 6.06E-05 | 1 |
| SOX15 | 0 | 0 | 0 | 8.188101219 | 0 | 0 | 0 | 0 | 3.96E-05 | 1 |
| SOX4 | 0 | 0 | 8.65401506 | 9.593819844 | 9.28736123 | 0 | 6.845503783 | 0 | 0.007232675 | 0.855584422 |
| SOX9 | 0 | 0 | 0 | 7.963225934 | 0 | 0 | 7.44886754 | 0 | 1.50E-05 | 1 |
| SOX9-AS1 | 0 | 0 | 0 | 7.963225934 | 0 | 0 | 0 | 0 | 1.50E-05 | 1 |
| SP110 | 9.675834273 | 0 | 0 | 0 | 0 | 0 | 0 | 0 | 7.44E-08 | 1 |
| SP2 | 9.557149476 | 7.961553963 | 9.123764545 | 0 | 6.741044679 | 0 | 0 | 0 | 0.956864147 | 0.028219261 |
| SPAG1 | 0 | 0 | 10.59045844 | 0 | 7.893121693 | 0 | 9.537925538 | 0 | 0.035320365 | 0.999854694 |
| SPAG9 | 14.5259667 | 13.47735068 | 10.612493 | 12.94318303 | 11.87105682 | 13.6867667 | 13.91634987 | 12.77309993 | 0.040321937 | 0.999854694 |
| SPARCL1 | 17.73983961 | 17.47176892 | 16.50282062 | 16.26698296 | 15.40521864 | 16.92079277 | 15.01330069 | 15.37571261 | 0.005130171 | 0.999854694 |
| SPATA13 | 0 | 0 | 0 | 0 | 6.330495883 | 0 | 0 | 0 | 6.26E-05 | 1 |
| SPATA21 | 8.616058634 | 0 | 0 | 0 | 0 | 0 | 7.257401605 | 0 | 1.33E-07 | 1 |
| SPATA7 | 0 | 8.326384029 | 0 | 0 | 0 | 0 | 0 | 9.871985276 | 1.61E-07 | 1 |
| SPATC1L | 0 | 0 | 0 | 0 | 0 | 8.825707172 | 0 | 10.39562037 | 1 | 0.008817225 |
| SPDL1 | 0 | 0 | 0 | 11.1012676 | 7.105738675 | 0 | 0 | 0 | 0.019035111 | 6.34E-10 |
| SPECC1L | 9.585966416 | 0 | 0 | 0 | 0 | 0 | 0 | 0 | 7.58E-08 | 1 |
| SPG7 | 10.89241997 | 7.05971876 | 0 | 0 | 0 | 0 | 8.093337038 | 0 | 0.002267519 | 1 |
| SPHK2 | 0 | 9.131811442 | 0 | 0 | 0 | 0 | 0 | 0 | 1.10E-07 | 1 |
| SPINK2 | 0 | 9.54265281 | 0 | 0 | 0 | 0 | 0 | 0 | 7.53E-08 | 1 |
| SPINT2 | 12.70909019 | 13.20341317 | 14.75838818 | 14.72059199 | 14.13467078 | 14.56957135 | 13.15906299 | 14.59496785 | 0.01298805 | 0.999854694 |

| | | | | | | | | | | |
|---|---|---|---|---|---|---|---|---|---|---|
| SPN | 9.185743846 | 0 | 0 | 0 | 0 | 0 | 0 | 0 | 1.15E-07 | 1 |
| SPOUT1 | 0 | 8.876365592 | 0 | 0 | 0 | 11.64129848 | 7.637673707 | 0 | 1.10E-07 | 4.71E-12 |
| SPPL3 | 10.66823035 | 0 | 0 | 7.454429623 | 9.91626026 | 0 | 0 | 0 | 0.980815574 | 0.014358084 |
| SPRED1 | 0 | 0 | 0 | 0 | 0 | 11.06787522 | 7.006440014 | 0 | 1 | 9.52E-12 |
| SPRY4 | 9.760430865 | 0 | 0 | 0 | 0 | 0 | 5.558980851 | 0 | 7.17E-08 | 0.437359002 |
| SPRYD7 | 10.26244189 | 9.988883934 | 0 | 9.987652255 | 8.662558908 | 0 | 0 | 0 | 0.932604529 | 0.011753849 |
| SPSB3 | 11.1720546 | 7.75599323 | 11.52215807 | 10.42214136 | 10.05947457 | 0 | 0 | 0 | 0.958210869 | 0.000260424 |
| SPTBN1 | 9.337499182 | 9.131811442 | 10.06397233 | 11.05580352 | 10.84353297 | 0 | 0 | 0 | 0.641936512 | 0.000273087 |
| SQLE | 0 | 0 | 0 | 0 | 6.530456572 | 0 | 0 | 0 | 5.38E-05 | 1 |
| SRA1 | 10.74050121 | 9.2476365 | 0 | 12.41076972 | 9.09898593 | 0 | 0 | 0 | 0.971198277 | 7.68E-12 |
| SREK1IP1 | 0 | 0 | 10.5376967 | 9.805749732 | 8.313116418 | 9.680137163 | 6.411827288 | 0 | 0.001443324 | 0.999854694 |
| SRF | 0 | 0 | 0 | 0 | 0 | 10.23991064 | 6.292006564 | 0 | 1 | 0.011034216 |
| SRP68 | 0 | 0 | 0 | 0 | 6.717806789 | 0 | 9.503271693 | 0 | 0.000126205 | 1 |
| SRRT | 0 | 0 | 10.50273025 | 10.0813848 | 8.204461778 | 0 | 0 | 0 | 0.00111349 | 0.000893674 |
| SRSF11 | 10.55254627 | 10.77760654 | 0 | 9.769762184 | 8.866504529 | 0 | 0 | 0 | 0.798581521 | 0.012259484 |
| SS18L1 | 0 | 0 | 0 | 0 | 0 | 0 | 6.811044309 | 10.0768456 | 1 | 0.011412104 |
| SS18L2 | 0 | 0 | 10.42102038 | 11.89159888 | 11.02547254 | 10.38577996 | 10.20003665 | 0 | 7.07E-08 | 0.999854694 |
| SSH3 | 0 | 0 | 10.00744271 | 7.95742026 | 7.436287711 | 9.887374082 | 8.023570157 | 0 | 0.017362915 | 0.999854694 |
| SSRP1 | 0 | 0 | 9.730877659 | 7.891966594 | 11.31592841 | 9.830261952 | 9.41997401 | 11.402292 | 0.000174798 | 0.999854694 |
| ST3GAL4 | 0 | 10.72214365 | 9.795226111 | 9.799272959 | 7.799379467 | 0 | 0 | 0 | 0.98939231 | 0.001749423 |
| ST6GALNAC1 | 9.89317875 | 11.39369621 | 9.955519852 | 10.64076551 | 8.659513127 | 0 | 0 | 0 | 0.878789877 | 0.000795451 |
| STAG3 | 0 | 10.33229971 | 0 | 0 | 0 | 0 | 0 | 0 | 7.07E-08 | 1 |
| STAG3L5P-PVRIG2P-PILRB | 0 | 0 | 9.59139382 | 0 | 8.779399919 | 0 | 0 | 0 | 0.058683489 | 0.013007083 |
| STAM | 9.371000476 | 9.426002279 | 0 | 0 | 0 | 0 | 0 | 0 | 0.007614398 | 1 |
| STAMBP | 0 | 0 | 9.878810649 | 7.854972032 | 8.358899494 | 0 | 0 | 0 | 0.014846685 | 0.003330216 |
| STARD10 | 0 | 0 | 10.71584966 | 11.87167288 | 10.6926604 | 11.20543585 | 11.26135324 | 11.95165351 | 7.07E-08 | 0.999854694 |
| STARD7-AS1 | 0 | 0 | 0 | 9.219977412 | 6.20241426 | 0 | 0 | 0 | 0.091190689 | 0.030225307 |
| STAT1 | 0 | 0 | 10.43674911 | 0 | 9.158448773 | 0 | 7.080640961 | 11.64642222 | 0.027450171 | 0.999854694 |
| STBD1 | 10.81237538 | 9.021269553 | 0 | 0 | 0 | 9.708771903 | 0 | 0 | 0.000888393 | 1 |
| STEAP1 | 10.280184 | 0 | 0 | 10.44928408 | 8.373843248 | 0 | 0 | 0 | 0.999485949 | 0.010595256 |
| STIM2 | 0 | 0 | 0 | 0 | 6.433937832 | 10.10067879 | 0 | 0 | 5.81E-05 | 1 |
| STIP1 | 0 | 0 | 0 | 11.82859244 | 10.97888962 | 0 | 10.66064649 | 0 | 0.01941848 | 0.999854694 |
| STK10 | 0 | 8.958657193 | 0 | 0 | 0 | 0 | 0 | 0 | 1.07E-07 | 1 |
| STK16 | 0 | 0 | 0 | 0 | 0 | 0 | 8.321619838 | 10.40138689 | 1 | 0.009118992 |
| STK17B | 0 | 9.124699738 | 0 | 0 | 0 | 0 | 0 | 9.952745771 | 1.13E-07 | 1 |
| STMN1 | 0 | 0 | 9.632230391 | 14.17394431 | 12.41403313 | 12.26363343 | 9.619840787 | 13.35901271 | 7.07E-08 | 0.999854694 |
| STMN3 | 0 | 0 | 0 | 0 | 6.645759207 | 8.137201838 | 0 | 0 | 7.73E-05 | 1 |
| STOM | 0 | 9.628117131 | 0 | 9.808977247 | 9.042822345 | 0 | 0 | 0 | 0.9955195 | 0.011625223 |
| STOX1 | 9.766737375 | 0 | 0 | 0 | 0 | 0 | 0 | 0 | 7.17E-08 | 1 |
| STT3B | 11.57919307 | 9.694462449 | 12.91636927 | 13.42595851 | 12.7943941 | 13.56905661 | 14.17801726 | 13.31424664 | 0.024300757 | 0.999854694 |
| STX17-AS1 | 0 | 9.684823965 | 0 | 0 | 0 | 0 | 5.428672801 | 0 | 7.17E-08 | 0.467810605 |
| STX6 | 0 | 8.780069978 | 0 | 0 | 0 | 0 | 8.658480352 | 0 | 1.15E-07 | 1 |
| SUCLG2 | 11.89779819 | 10.15865117 | 0 | 10.4595891 | 9.339675713 | 0 | 0 | 0 | 0.832701413 | 0.009547202 |

| | | | | | | | | | | |
|---|---|---|---|---|---|---|---|---|---|---|
| SUCNR1 | 0 | 0 | 0 | 0 | 6.330495883 | 11.66684865 | 8.37567389 | 9.416758914 | 6.26E-05 | 0.273913501 |
| SUCO | 10.30639463 | 0 | 0 | 0 | 0 | 10.73190637 | 0 | 0 | 7.07E-08 | 1 |
| SULF1 | 0 | 0 | 0 | 0 | 6.763914196 | 9.950797325 | 0 | 9.976117921 | 0.000116718 | 1 |
| SUMF1 | 0 | 0 | 9.030827538 | 0 | 8.622453199 | 0 | 0 | 0 | 0.075193869 | 0.01717411 |
| SUN2 | 9.285741834 | 0 | 9.065319476 | 8.748063288 | 0 | 0 | 0 | 0 | 0.999483812 | 0.017443763 |
| SUPT6H | 9.773016437 | 0 | 7.952980712 | 7.369539606 | 7.908597519 | 0 | 0 | 0 | 0.970998255 | 0.01037888 |
| SUPT7L | 0 | 9.292752085 | 9.97038412 | 0 | 8.216946592 | 0 | 0 | 0 | 0.991384446 | 0.012348695 |
| SURF4 | 0 | 0 | 10.13774559 | 10.24878703 | 0 | 0 | 9.094280063 | 0 | 0.018857296 | 1 |
| SVBP | 0 | 0 | 0 | 10.05715983 | 10.36964618 | 10.7937502 | 0 | 0 | 0.014031743 | 0.999854694 |
| SVIL | 0 | 8.705765621 | 0 | 0 | 0 | 0 | 0 | 0 | 1.18E-07 | 1 |
| SWI5 | 0 | 0 | 10.52999929 | 9.619781713 | 9.122402468 | 0 | 9.94388913 | 0 | 0.00098632 | 0.999854694 |
| SWSAP1 | 10.11946682 | 0 | 0 | 0 | 0 | 0 | 0 | 0 | 7.07E-08 | 1 |
| SYMPK | 0 | 9.454643748 | 0 | 0 | 0 | 0 | 8.836064177 | 0 | 7.73E-08 | 1 |
| SYN2 | 0 | 0 | 7.451370902 | 0 | 0 | 0 | 0 | 0 | 4.73E-05 | 1 |
| SYNGR2 | 7.660943177 | 10.99403958 | 13.15937984 | 13.02768372 | 11.912264 | 13.15917881 | 11.90013939 | 6.903188591 | 0.031988575 | 0.999854694 |
| SYNJ2 | 0 | 0 | 0 | 0 | 0 | 10.47084566 | 0 | 9.478071506 | 1 | 0.007640511 |
| SYNPO | 0 | 0 | 10.20018975 | 10.44824951 | 8.597206899 | 0 | 0 | 0 | 0.00076353 | 0.000795451 |
| SYS1 | 0 | 0 | 0 | 10.09202226 | 9.026365663 | 0 | 0 | 0 | 0.03689453 | 0.01086644 |
| SYT11 | 0 | 9.159912786 | 0 | 0 | 0 | 0 | 0 | 0 | 8.91E-08 | 1 |
| SYT13 | 0 | 8.909847578 | 0 | 0 | 0 | 0 | 0 | 0 | 1.08E-07 | 1 |
| TACO1 | 0 | 0 | 11.47843388 | 0 | 9.928331283 | 0 | 0 | 0 | 0.028273623 | 0.005705522 |
| TADA3 | 0 | 0 | 0 | 10.94501659 | 7.722658607 | 0 | 0 | 0 | 0.021773259 | 0.009724498 |
| TAF6 | 0 | 0 | 10.47493584 | 0 | 8.200276049 | 0 | 0 | 0 | 0.036589339 | 0.010542114 |
| TALAM1 | 4.972573658 | 7.228441702 | 11.41799399 | 0 | 7.768613653 | 0 | 0 | 0 | 0.548037712 | 2.99E-11 |
| TAMM41 | 0 | 9.095897086 | 0 | 0 | 0 | 9.789840441 | 8.104050868 | 0 | 1.15E-07 | 0.011034216 |
| TAOK1 | 14.63167371 | 14.03739781 | 12.70430406 | 12.54747645 | 12.88389048 | 12.1567408 | 12.69760202 | 12.73991847 | 0.019035111 | 0.999854694 |
| TAOK2 | 0 | 0 | 7.425051332 | 0 | 0 | 0 | 0 | 0 | 2.80E-05 | 1 |
| TAPBPL | 0 | 0 | 9.830747761 | 9.708865503 | 0 | 0 | 0 | 0 | 0.035967833 | 0.010590691 |
| TARSL2 | 0 | 9.477154439 | 0 | 0 | 0 | 0 | 0 | 0 | 7.73E-08 | 1 |
| TAZ | 0 | 9.13888826 | 9.982843054 | 0 | 7.650859037 | 0 | 0 | 0 | 0.990582117 | 0.013603802 |
| TBC1D22A | 0 | 9.97313222 | 0 | 0 | 0 | 0 | 0 | 0 | 7.07E-08 | 1 |
| TBC1D23 | 0 | 0 | 0 | 9.0060457 | 8.555215378 | 0 | 0 | 0 | 0.075193869 | 0.018340322 |
| TBC1D24 | 10.09210579 | 0 | 0 | 0 | 0 | 0 | 0 | 0 | 7.07E-08 | 1 |
| TBC1D31 | 0 | 0 | 0 | 0 | 6.873120683 | 0 | 0 | 0 | 4.05E-05 | 1 |
| TBCB | 0 | 0 | 9.962399319 | 10.33527103 | 10.96137366 | 0 | 9.741480823 | 9.808302414 | 2.76E-05 | 0.999854694 |
| TBK1 | 0 | 0 | 0 | 9.389094555 | 8.313116418 | 0 | 0 | 0 | 0.071163453 | 0.01634826 |
| TBL3 | 0 | 0 | 0 | 0 | 0 | 0 | 7.503839514 | 10.0659947 | 1 | 0.010883809 |
| TBRG1 | 10.12683973 | 0 | 0 | 10.11567357 | 8.83445342 | 0 | 0 | 0 | 0.999660881 | 0.011034216 |
| TCAIM | 0 | 0 | 0 | 10.60871443 | 8.410539137 | 0 | 8.096917163 | 0 | 0.028532957 | 0.999854694 |
| TCEA1 | 0 | 0 | 10.20889374 | 11.68227619 | 9.343476431 | 9.178652744 | 11.18183165 | 10.43831766 | 5.91E-06 | 0.999854694 |
| TCEAL2 | 0 | 8.893203714 | 0 | 0 | 0 | 0 | 0 | 0 | 1.09E-07 | 1 |
| TCEAL7 | 9.395625601 | 0 | 0 | 0 | 0 | 0 | 0 | 0 | 8.04E-08 | 1 |
| TCERG1 | 0 | 9.324135017 | 0 | 9.634409966 | 8.293602846 | 0 | 0 | 0 | 0.997434996 | 0.01457526 |

| | | | | | | | | | | |
|---|---|---|---|---|---|---|---|---|---|---|
| TCF7L2 | 18.90332709 | 18.38502024 | 17.23007373 | 15.74330752 | 17.23203823 | 15.25840028 | 17.72967087 | 15.08437147 | 0.002182759 | 0.999854694 |
| TCFL5 | 0 | 0 | 0 | 0 | 0 | 10.31523591 | 7.567155003 | 0 | 1 | 0.010003595 |
| TCHP | 0 | 0 | 10.26746866 | 0 | 6.556887521 | 0 | 0 | 0 | 0.0581952 | 0.013021032 |
| TCIRG1 | 9.459308883 | 9.38492425 | 0 | 0 | 0 | 0 | 8.978724284 | 0 | 0.007294269 | 1 |
| TCN2 | 0 | 0 | 12.33024487 | 0 | 8.1183489 | 0 | 0 | 0 | 1.69E-07 | 9.11E-12 |
| TCTEX1D1 | 0 | 9.159912786 | 0 | 0 | 0 | 0 | 0 | 0 | 8.91E-08 | 1 |
| TCTN3 | 9.766737375 | 9.021269553 | 0 | 0 | 0 | 0 | 0 | 0 | 0.007196113 | 1 |
| TDG | 0 | 0 | 9.179933839 | 9.821815585 | 6.682232705 | 0 | 0 | 10.14029102 | 0.013592164 | 0.999854694 |
| TDRP | 10.00676427 | 0 | 0 | 0 | 0 | 0 | 0 | 0 | 7.07E-08 | 1 |
| TEAD3 | 10.98998111 | 0 | 8.874275474 | 0 | 8.78779307 | 0 | 0 | 0 | 0.943204702 | 0.017222122 |
| TEDC1 | 0 | 10.53901294 | 0 | 10.40731174 | 7.087723911 | 0 | 0 | 0 | 0.98939231 | 0.012072966 |
| TEFM | 0 | 0 | 10.62410859 | 0 | 8.086952566 | 0 | 0 | 0 | 0.032674287 | 0.010274509 |
| TEKT4P2 | 0 | 9.036507082 | 0 | 7.737931428 | 8.525411369 | 0 | 0 | 0 | 0.992483691 | 0.034075145 |
| TENT2 | 0 | 8.934458793 | 0 | 0 | 0 | 0 | 0 | 0 | 1.07E-07 | 1 |
| TENT5C | 0 | 9.044065918 | 0 | 0 | 0 | 0 | 0 | 0 | 1.01E-07 | 1 |
| TERF2IP | 9.6281405 | 8.667127243 | 0 | 0 | 0 | 0 | 7.295290731 | 0 | 0.010819872 | 1 |
| TES | 10.53787205 | 0 | 0 | 10.71490203 | 6.954898948 | 0 | 0 | 0 | 0.997434996 | 0.010829604 |
| TESC | 0 | 0 | 0 | 11.46931231 | 9.133421009 | 12.45375616 | 0 | 10.50945865 | 0.030240507 | 0.999854694 |
| TESMIN | 0 | 0 | 0 | 9.973339655 | 8.563386158 | 0 | 7.830642173 | 0 | 0.048007264 | 0.999854694 |
| TEX261 | 0 | 9.699257651 | 0 | 0 | 0 | 0 | 0 | 0 | 7.17E-08 | 1 |
| TEX30 | 0 | 9.305387296 | 0 | 11.79561969 | 9.979298948 | 0 | 0 | 0 | 0.887232895 | 0.005049454 |
| TFAP4 | 0 | 9.477154439 | 0 | 8.885873187 | 8.002934593 | 0 | 0 | 0 | 0.989756518 | 0.024406584 |
| TFB2M | 0 | 0 | 10.40259131 | 10.03803287 | 0 | 0 | 9.035794815 | 0 | 0.017713866 | 0.999854694 |
| TFCP2 | 0 | 0 | 0 | 8.425292327 | 0 | 0 | 8.387415743 | 0 | 9.22E-06 | 1 |
| TFEC | 0 | 0 | 7.668851543 | 0 | 0 | 0 | 0 | 0 | 2.19E-05 | 1 |
| TFIP11 | 0 | 8.49028674 | 0 | 0 | 0 | 0 | 0 | 0 | 1.38E-07 | 1 |
| TFPI | 0 | 10.18623948 | 0 | 0 | 0 | 0 | 0 | 9.932975513 | 7.07E-08 | 1 |
| TFPI2 | 0 | 9.260671271 | 0 | 0 | 0 | 0 | 0 | 0 | 8.22E-08 | 1 |
| TGDS | 9.760430865 | 0 | 0 | 8.895013884 | 8.355139244 | 0 | 0 | 0 | 0.98867228 | 0.021151099 |
| TGFB1 | 9.89317875 | 0 | 0 | 0 | 0 | 0 | 0 | 10.33965736 | 7.10E-08 | 1 |
| TGFB1I1 | 9.371000476 | 0 | 0 | 0 | 0 | 0 | 0 | 9.964479175 | 8.05E-08 | 1 |
| TGFBI | 0 | 0 | 0 | 12.70735629 | 11.31930972 | 12.32421735 | 11.40618088 | 10.08403446 | 7.38E-08 | 0.999854694 |
| THAP12 | 0 | 0 | 10.89012082 | 0 | 9.021629067 | 0 | 0 | 11.04520436 | 0.016559096 | 0.999854694 |
| THAP4 | 11.11881822 | 10.47893321 | 12.12559176 | 0 | 7.716809918 | 0 | 0 | 0 | 1 | 1.31E-11 |
| THAP6 | 0 | 0 | 10.25724836 | 0 | 8.81815683 | 0 | 0 | 0 | 0.036709899 | 0.010542114 |
| THBS3 | 0 | 8.770988541 | 0 | 0 | 0 | 0 | 0 | 0 | 1.15E-07 | 1 |
| THBS4 | 0 | 0 | 0 | 0 | 6.830424858 | 0 | 0 | 0 | 4.17E-05 | 1 |
| THSD4 | 9.311852598 | 10.26273781 | 8.033583097 | 0 | 8.592423637 | 0 | 0 | 0 | 0.815498455 | 0.027721629 |
| THUMPD3-AS1 | 0 | 0 | 0 | 0 | 0 | 0 | 6.72111291 | 8.885916959 | 1 | 0.024827605 |
| TIAF1 | 11.57919307 | 0 | 0 | 0 | 0 | 0 | 0 | 0 | 6.86E-08 | 1 |
| TIMM10B | 0 | 0 | 7.030375154 | 11.59662706 | 9.971951802 | 0 | 9.072624984 | 11.14481729 | 7.60E-05 | 0.999854694 |
| TIMM44 | 11.38789442 | 9.028908435 | 0 | 10.18686959 | 7.266514108 | 0 | 0 | 0 | 0.812976922 | 0.01316597 |
| TIMM8A | 0 | 10.08361716 | 0 | 0 | 0 | 0 | 0 | 0 | 7.07E-08 | 1 |

| | | | | | | | | | | |
|---|---|---|---|---|---|---|---|---|---|---|
| TIMM9 | 0 | 0 | 0 | 10.2090275 | 10.70785133 | 0 | 9.897427155 | 0 | 0.033562922 | 0.999854694 |
| TINCR | 10.35742919 | 8.901549648 | 0 | 0 | 0 | 0 | 0 | 0 | 0.003067322 | 1 |
| TIPIN | 0 | 9.013590009 | 0 | 0 | 0 | 0 | 0 | 10.64444288 | 1.04E-07 | 1 |
| TJAP1 | 10.18450463 | 0 | 10.13470357 | 0 | 6.530456572 | 0 | 0 | 0 | 0.98939231 | 0.01404322 |
| TK2 | 0 | 0 | 0 | 0 | 0 | 0 | 7.218490506 | 10.21750006 | 1 | 0.010552942 |
| TLCD2 | 0 | 0 | 9.48871274 | 8.672114028 | 0 | 0 | 0 | 0 | 0.067902631 | 0.014756957 |
| TLDC2 | 0 | 0 | 10.34935133 | 0 | 8.990456231 | 0 | 0 | 0 | 0.032011714 | 0.010097347 |
| TLE2 | 0 | 8.734080111 | 0 | 0 | 0 | 0 | 8.023570157 | 0 | 1.17E-07 | 1 |
| TLK1 | 10.280184 | 0 | 0 | 0 | 0 | 0 | 0 | 10.40999367 | 7.07E-08 | 1 |
| TLK2 | 8.811733678 | 0 | 0 | 0 | 0 | 5.310897272 | 0 | 0 | 1.17E-07 | 0.518196643 |
| TLNRD1 | 0 | 0 | 9.371283667 | 5.429426546 | 8.269832732 | 0 | 7.930328737 | 0 | 0.034110874 | 0.999854694 |
| TMC6 | 9.932977691 | 9.187477203 | 0 | 0 | 0 | 0 | 0 | 0 | 0.004961468 | 1 |
| TMCC1-AS1 | 10.34061747 | 8.398919312 | 0 | 0 | 0 | 0 | 0 | 0 | 0.004573034 | 1 |
| TMCC3 | 0 | 8.667127243 | 0 | 0 | 0 | 0 | 0 | 0 | 1.21E-07 | 1 |
| TMEM107 | 0 | 0 | 10.3228672 | 11.17128085 | 7.515911508 | 0 | 0 | 10.96667695 | 0.000192696 | 0.999854694 |
| TMEM11 | 0 | 10.5759826 | 9.823209261 | 10.21633848 | 9.431813121 | 0 | 0 | 0 | 0.991384446 | 0.000810113 |
| TMEM115 | 8.973144195 | 0 | 0 | 0 | 0 | 0 | 0 | 0 | 1.08E-07 | 1 |
| TMEM116 | 0 | 0 | 0 | 0 | 0 | 0 | 8.56851287 | 9.859471605 | 1 | 0.01037888 |
| TMEM129 | 0 | 8.387080535 | 0 | 8.971897327 | 7.250409752 | 0 | 0 | 0 | 0.992766244 | 0.029546388 |
| TMEM131L | 10.30205913 | 0 | 0 | 0 | 0 | 0 | 0 | 0 | 7.07E-08 | 1 |
| TMEM14A | 0 | 0 | 11.41046745 | 11.52139724 | 9.176672437 | 9.144535309 | 9.17778928 | 10.60500709 | 3.60E-07 | 0.999854694 |
| TMEM159 | 0 | 0 | 10.66544807 | 9.103893437 | 7.594562621 | 0 | 0 | 8.902276938 | 0.003317549 | 0.999854694 |
| TMEM161A | 0 | 0 | 0 | 11.04694632 | 9.686441818 | 0 | 0 | 0 | 0.033419045 | 0.007428164 |
| TMEM161B-AS1 | 0 | 0 | 9.567528938 | 7.963225934 | 7.669146695 | 0 | 0 | 0 | 0.024019715 | 0.005132109 |
| TMEM163 | 0 | 0 | 0 | 7.928036185 | 0 | 0 | 0 | 0 | 1.55E-05 | 1 |
| TMEM167A | 0 | 11.90203059 | 10.44741292 | 10.97406028 | 10.7294345 | 0 | 0 | 0 | 0.996458807 | 0.000263179 |
| TMEM171 | 10.91388848 | 9.640635714 | 10.51915345 | 10.35085587 | 7.877478056 | 0 | 0 | 0 | 0.975665177 | 0.000855798 |
| TMEM175 | 10.47763544 | 9.504807688 | 0 | 0 | 0 | 0 | 0 | 0 | 0.001232699 | 1 |
| TMEM184A | 0 | 0 | 10.91397482 | 0 | 7.432724094 | 0 | 0 | 0 | 0.026743343 | 0.009724498 |
| TMEM185A | 9.822288722 | 0 | 11.01889191 | 0 | 7.656980716 | 0 | 0 | 0 | 0.985757273 | 0.009227496 |
| TMEM189 | 9.99069029 | 0 | 0 | 9.847154521 | 9.500657039 | 0 | 0 | 0 | 0.997434996 | 0.010650095 |
| TMEM191C | 0 | 0 | 9.88002222 | 10.07066834 | 0 | 0 | 0 | 0 | 0.027450171 | 0.009904995 |
| TMEM199 | 0 | 0 | 0 | 8.217545222 | 0 | 0 | 0 | 0 | 5.08E-05 | 1 |
| TMEM200A | 10.1892076 | 0 | 8.725851918 | 0 | 8.741014507 | 0 | 0 | 0 | 0.974263258 | 0.01869339 |
| TMEM204 | 9.822288722 | 0 | 0 | 0 | 0 | 0 | 0 | 0 | 7.17E-08 | 1 |
| TMEM236 | 0 | 0 | 7.425051332 | 0 | 0 | 0 | 0 | 0 | 2.80E-05 | 1 |
| TMEM237 | 0 | 0 | 0 | 10.36846339 | 7.913719455 | 0 | 0 | 0 | 0.041219216 | 0.011326554 |
| TMEM246 | 10.08707482 | 0 | 0 | 0 | 0 | 0 | 9.032059553 | 9.671827556 | 7.07E-08 | 0.010189075 |
| TMEM250 | 0 | 8.724703557 | 0 | 0 | 0 | 0 | 0 | 0 | 1.17E-07 | 1 |
| TMEM251 | 10.90962026 | 9.857774611 | 11.16022686 | 10.60778814 | 8.78779307 | 0 | 0 | 0 | 0.999660881 | 0.000440129 |
| TMEM253 | 0 | 9.414384561 | 0 | 0 | 0 | 9.851567204 | 0 | 0 | 7.88E-08 | 1 |
| TMEM26-AS1 | 0 | 8.950636073 | 0 | 0 | 0 | 0 | 0 | 0 | 1.07E-07 | 1 |
| TMEM265 | 0 | 7.879432217 | 0 | 9.815410697 | 10.2028826 | 0 | 0 | 0 | 0.796336567 | 0.009360533 |

| | | | | | | | | | | |
|---|---|---|---|---|---|---|---|---|---|---|
| TMEM267 | 9.176299815 | 0 | 0 | 0 | 0 | 0 | 0 | 0 | 1.15E-07 | 1 |
| TMEM268 | 0 | 9.510275296 | 0 | 0 | 0 | 0 | 0 | 0 | 7.88E-08 | 1 |
| TMEM30B | 0 | 0 | 10.4506783 | 0 | 7.949072713 | 0 | 0 | 0 | 0.039989728 | 0.010829604 |
| TMEM39A | 0 | 0 | 0 | 0 | 0 | 0 | 7.813336911 | 9.598422184 | 1 | 0.012440088 |
| TMEM40 | 0 | 0 | 0 | 12.20780745 | 11.96779931 | 0 | 12.56897936 | 6.616358012 | 0.012849367 | 0.999854694 |
| TMEM64 | 0 | 0 | 0 | 0 | 6.345735199 | 10.16043482 | 0 | 0 | 6.22E-05 | 1 |
| TMEM69 | 0 | 10.27242335 | 8.999955452 | 0 | 9.245318778 | 0 | 0 | 0 | 0.981769365 | 0.013910154 |
| TMEM81 | 0 | 0 | 0 | 0 | 6.39051047 | 0 | 0 | 0 | 5.98E-05 | 1 |
| TMEM8B | 9.68252076 | 0 | 0 | 0 | 0 | 0 | 7.178500772 | 0 | 7.42E-08 | 1 |
| TMEM91 | 8.48449896 | 0 | 0 | 0 | 0 | 0 | 0 | 0 | 1.47E-07 | 1 |
| TMEM99 | 0 | 0 | 0 | 9.340853892 | 8.923477098 | 0 | 0 | 0 | 0.061711916 | 0.014200185 |
| TMF1 | 0 | 8.501311304 | 0 | 0 | 0 | 9.489443214 | 0 | 0 | 1.38E-07 | 1 |
| TMIGD2 | 0 | 8.467981611 | 0 | 0 | 0 | 0 | 0 | 0 | 1.41E-07 | 1 |
| TMOD3 | 0 | 10.37188125 | 11.2724299 | 7.437846525 | 0 | 0 | 0 | 0 | 0.9955195 | 6.63E-11 |
| TMX1 | 0 | 10.00058587 | 11.53759528 | 9.612411586 | 8.525411369 | 0 | 0 | 0 | 0.879507658 | 0.000538325 |
| TNFAIP8L3 | 9.320452216 | 0 | 0 | 0 | 0 | 0 | 0 | 0 | 8.22E-08 | 1 |
| TNFRSF14-AS1 | 11.3358257 | 8.223623425 | 11.35316709 | 0 | 8.798908517 | 0 | 0 | 0 | 0.985757273 | 0.007255458 |
| TNFRSF6B | 0 | 0 | 9.881232774 | 10.09994919 | 0 | 0 | 0 | 0 | 0.026801204 | 0.009822772 |
| TNKS2 | 0 | 8.696202607 | 0 | 0 | 0 | 0 | 0 | 0 | 1.18E-07 | 1 |
| TNNT3 | 0 | 9.620553507 | 0 | 0 | 0 | 0 | 0 | 0 | 7.42E-08 | 1 |
| TOB2 | 9.881603011 | 9.021269553 | 0 | 0 | 0 | 0 | 8.931384524 | 0 | 0.006202938 | 1 |
| TOLLIP | 0 | 7.174363889 | 9.977193237 | 9.262974977 | 0 | 0 | 0 | 0 | 0.786425151 | 0.011034216 |
| TOMM34 | 0 | 0 | 0 | 11.63122575 | 9.643924818 | 0 | 11.3037946 | 0 | 0.026801204 | 0.999854694 |
| TOP1 | 0 | 0 | 9.515466095 | 7.842426656 | 6.944926602 | 0 | 4.589263136 | 0 | 0.029981024 | 0.306674545 |
| TOP2A | 0 | 0 | 0 | 10.36188573 | 7.934027272 | 0 | 0 | 0 | 0.041246238 | 0.011326554 |
| TOP2B | 7.605727588 | 9.650572952 | 0 | 0 | 0 | 0 | 0 | 0 | 0.015925185 | 1 |
| TOR2A | 0 | 9.187477203 | 9.646539208 | 10.26650356 | 0 | 0 | 0 | 0 | 0.961310672 | 0.009904995 |
| TOX3 | 0 | 0 | 10.02835546 | 0 | 5.684274069 | 0 | 0 | 0 | 0.070256089 | 0.015994804 |
| TP53I11 | 0 | 0 | 0 | 10.47389358 | 8.313116418 | 0 | 0 | 0 | 0.033815753 | 0.010590691 |
| TP53I13 | 0 | 8.51225226 | 0 | 0 | 0 | 0 | 0 | 9.393801179 | 1.38E-07 | 1 |
| TP53I3 | 10.27134022 | 10.01219366 | 10.4765386 | 7.904090267 | 7.845672509 | 0 | 0 | 0 | 0.937966741 | 0.00233029 |
| TP53INP1 | 8.917547778 | 8.326384029 | 0 | 0 | 0 | 0 | 0 | 0 | 0.022156595 | 1 |
| TP53INP2 | 8.847874263 | 0 | 0 | 0 | 0 | 0 | 0 | 0 | 1.15E-07 | 1 |
| TPBG | 0 | 0 | 0 | 9.309293569 | 8.474327557 | 0 | 0 | 0 | 0.070665441 | 0.01630416 |
| TPCN1 | 9.259149778 | 0 | 0 | 0 | 0 | 0 | 8.779264159 | 0 | 8.58E-08 | 1 |
| TPD52L1 | 0 | 0 | 0 | 10.07737545 | 10.75961645 | 10.95921025 | 9.752872315 | 0 | 0.033826141 | 0.999854694 |
| TPH1 | 10.67159513 | 0 | 0 | 0 | 0 | 0 | 0 | 0 | 7.07E-08 | 1 |
| TPSB2 | 0 | 10.43935546 | 0 | 0 | 0 | 0 | 0 | 0 | 7.07E-08 | 1 |
| TPTEP1 | 0 | 0 | 0 | 7.97476766 | 8.446324747 | 0 | 0 | 0 | 0.095884584 | 0.032381737 |
| TPTEP2-CSNK1E | 0 | 8.617326978 | 11.5379791 | 0 | 10.05774045 | 0 | 0 | 0 | 0.835596588 | 0.00543027 |
| TPX2 | 12.62133379 | 14.30199498 | 13.25679962 | 16.67029304 | 16.88802814 | 14.93889174 | 16.31377187 | 16.62503861 | 0.042132385 | 0.999854694 |
| TRAF2 | 8.252542925 | 0 | 0 | 0 | 0 | 11.22772153 | 8.542492985 | 0 | 0.066636219 | 0.005938081 |
| TRAM1L1 | 0 | 9.38492425 | 0 | 0 | 0 | 0 | 0 | 0 | 7.88E-08 | 1 |

| | | | | | | | | | | |
|---|---|---|---|---|---|---|---|---|---|---|
| TRANK1 | 8.559063282 | 0 | 9.276940477 | 0 | 6.670177039 | 0 | 0 | 0 | 0.994141362 | 0.025185565 |
| TRAPPC12 | 9.24114547 | 8.350969903 | 0 | 10.60778814 | 6.489879427 | 0 | 0 | 0 | 0.99353678 | 0.011488338 |
| TRAPPC2L | 0 | 0 | 9.737576887 | 10.67123354 | 10.64802579 | 10.52773047 | 11.4617308 | 11.28865534 | 4.56E-05 | 0.999854694 |
| TRAPPC5 | 0 | 0 | 11.2998474 | 10.41580446 | 10.75997198 | 0 | 9.639612952 | 10.60999592 | 8.56E-07 | 0.999854694 |
| TRAPPC8 | 0 | 0 | 0 | 0 | 6.706045958 | 0 | 0 | 0 | 0.000126325 | 1 |
| TRDMT1 | 0 | 9.124699738 | 0 | 0 | 0 | 0 | 0 | 0 | 1.13E-07 | 1 |
| TRIB1 | 0 | 0 | 0 | 0 | 0 | 0 | 7.96991419 | 8.41317125 | 1 | 0.022645335 |
| TRIM15 | 10.44654697 | 0 | 9.022074146 | 9.726040692 | 7.30989599 | 0 | 0 | 0 | 0.98939231 | 0.003330216 |
| TRIM23 | 9.734925475 | 0 | 0 | 0 | 0 | 0 | 0 | 0 | 7.25E-08 | 1 |
| TRIM28 | 0 | 9.949177413 | 0 | 11.57782945 | 10.2112269 | 0 | 0 | 0 | 0.955300467 | 0.005384503 |
| TRIM3 | 0 | 8.445326205 | 0 | 0 | 0 | 0 | 0 | 0 | 1.44E-07 | 1 |
| TRIM34 | 0 | 0 | 10.19824838 | 0 | 8.474327557 | 0 | 0 | 0 | 0.043421293 | 0.011034216 |
| TRIM45 | 0 | 0 | 0 | 0 | 6.448127902 | 0 | 0 | 0 | 5.74E-05 | 1 |
| TRIM65 | 10.17978629 | 0 | 0 | 0 | 0 | 0 | 0 | 0 | 7.07E-08 | 1 |
| TRIM74 | 8.940044091 | 8.876365592 | 0 | 0 | 0 | 0 | 0 | 0 | 0.016575383 | 1 |
| TRMT12 | 0 | 0 | 0 | 0 | 6.797552774 | 0 | 0 | 0 | 5.66E-05 | 1 |
| TRMT61A | 10.60628238 | 0 | 0 | 0 | 0 | 0 | 0 | 0 | 7.07E-08 | 1 |
| TRMT9B | 9.527745184 | 0 | 0 | 0 | 0 | 0 | 0 | 0 | 7.73E-08 | 1 |
| TRPM7 | 0 | 0 | 0 | 8.217545222 | 0 | 0 | 0 | 0 | 5.08E-05 | 1 |
| TRUB2 | 0 | 0 | 0 | 11.19737685 | 7.453974941 | 11.0923716 | 7.89785925 | 0 | 0.014918833 | 0.999854694 |
| TSHZ3 | 9.48248357 | 0 | 0 | 0 | 0 | 0 | 0 | 0 | 7.87E-08 | 1 |
| TSPYL4 | 0 | 0 | 0 | 8.470833404 | 0 | 0 | 0 | 0 | 8.80E-06 | 1 |
| TSR1 | 10.47763544 | 9.366954907 | 0 | 0 | 0 | 0 | 0 | 0 | 0.001466198 | 1 |
| TSSC4 | 0 | 0 | 10.4539363 | 0 | 7.939059873 | 0 | 0 | 0 | 0.039966309 | 0.010829604 |
| TSTD2 | 8.643733587 | 0 | 0 | 0 | 0 | 8.993910797 | 0 | 0 | 1.31E-07 | 1 |
| TTC28-AS1 | 0 | 9.883282425 | 0 | 9.979081745 | 6.904333267 | 0 | 0 | 0 | 0.99353678 | 0.015165481 |
| TTC39B | 0 | 8.859328625 | 0 | 0 | 0 | 0 | 0 | 0 | 1.21E-07 | 1 |
| TTC8 | 0 | 0 | 0 | 0 | 6.841217856 | 0 | 0 | 0 | 4.14E-05 | 1 |
| TTLL3 | 9.427818601 | 0 | 0 | 0 | 0 | 0 | 0 | 0 | 7.88E-08 | 1 |
| TTLL5 | 0 | 9.051585356 | 0 | 0 | 0 | 0 | 0 | 0 | 1.18E-07 | 1 |
| TTLL6 | 9.695801475 | 0 | 0 | 0 | 0 | 0 | 0 | 0 | 7.38E-08 | 1 |
| TTN-AS1 | 0 | 8.676884284 | 0 | 0 | 0 | 0 | 0 | 0 | 1.20E-07 | 1 |
| TTTY15 | 0 | 8.555204765 | 0 | 0 | 0 | 0 | 0 | 0 | 1.34E-07 | 1 |
| TUBA1A | 0 | 0 | 10.27669743 | 0 | 10.44852291 | 10.65002499 | 0 | 0 | 0.036007876 | 0.999854694 |
| TUBA1B | 11.33794493 | 11.22106231 | 12.83811468 | 14.41003974 | 13.4289594 | 13.04953867 | 11.22582379 | 12.30076151 | 0.001568217 | 0.999854694 |
| TUBB2A | 0 | 0 | 9.432018923 | 0 | 8.829041651 | 0 | 0 | 0 | 0.062606601 | 0.013697851 |
| TUBB6 | 8.684275372 | 0 | 0 | 0 | 0 | 0 | 0 | 12.65165077 | 1.25E-07 | 1 |
| TUBE1 | 0 | 8.859328625 | 0 | 0 | 0 | 0 | 0 | 0 | 1.21E-07 | 1 |
| TUBGCP5 | 10.41477378 | 0 | 0 | 0 | 0 | 0 | 0 | 0 | 7.07E-08 | 1 |
| TUSC1 | 0 | 0 | 10.40595961 | 0 | 9.255440578 | 7.544868397 | 0 | 0 | 0.026952792 | 0.999854694 |
| TUT7 | 9.85816911 | 0 | 0 | 0 | 0 | 0 | 0 | 0 | 7.17E-08 | 1 |
| TWF2 | 0 | 0 | 0 | 10.25471675 | 8.707490434 | 0 | 8.060709732 | 0 | 0.036442685 | 0.999854694 |
| TWISTNB | 0 | 0 | 0 | 10.63895308 | 7.509159523 | 0 | 0 | 0 | 0.033826141 | 0.010821525 |

| | | | | | | | | | | |
|---|---|---|---|---|---|---|---|---|---|---|
| TXK | 0 | 10.44507634 | 0 | 0 | 0 | 9.226033527 | 7.556793512 | 0 | 7.07E-08 | 0.015652521 |
| TXNDC12 | 0 | 0 | 11.49196821 | 11.32741627 | 8.257799178 | 11.44952907 | 10.09832797 | 0 | 2.11E-06 | 0.999854694 |
| TXNDC5 | 0 | 0 | 10.06610293 | 9.185625776 | 9.515051591 | 0 | 8.629110188 | 9.932975513 | 0.002261343 | 0.999854694 |
| TYMS | 0 | 0 | 0 | 10.29670606 | 10.15880456 | 0 | 0 | 11.75737523 | 0.01402602 | 1 |
| TYMSOS | 0 | 0 | 10.11631546 | 8.679186115 | 0 | 0 | 0 | 0 | 0.046892553 | 0.011578584 |
| TYRP1 | 9.7916913 | 0 | 0 | 0 | 0 | 0 | 0 | 0 | 7.44E-08 | 1 |
| TYW1 | 0 | 0 | 0 | 10.54336285 | 8.113905325 | 0 | 0 | 0 | 0.033169536 | 0.010552942 |
| U2AF1L4 | 0 | 0 | 0 | 10.12347196 | 9.805578025 | 0 | 8.345894226 | 0 | 0.023220489 | 0.999854694 |
| U2AF1L5 | 0 | 0 | 11.58813843 | 10.1389434 | 8.343799328 | 12.35757574 | 0 | 0 | 4.21E-05 | 0.999854694 |
| UBA2 | 0 | 0 | 8.844794804 | 11.16562691 | 9.253421891 | 10.83559208 | 7.036579876 | 0 | 0.000323925 | 0.999854694 |
| UBE2D4 | 0 | 0 | 0 | 10.47998092 | 7.626109207 | 0 | 0 | 0 | 0.039141627 | 0.011070952 |
| UBE2E3 | 0 | 0 | 10.21562739 | 12.31328103 | 10.58521047 | 0 | 8.290690969 | 10.80966753 | 7.07E-08 | 0.999854694 |
| UBE2F | 10.54889168 | 9.073911341 | 0 | 10.03527963 | 10.20653917 | 0 | 0 | 0 | 0.98939231 | 0.008991934 |
| UBE2G1 | 0 | 0 | 0 | 0 | 0 | 0 | 7.627808199 | 8.256557088 | 1 | 0.029399892 |
| UBE2H | 9.662367605 | 0 | 0 | 0 | 0 | 0 | 0 | 9.642912035 | 7.44E-08 | 1 |
| UBE2M | 10.86034341 | 0 | 0 | 0 | 0 | 0 | 0 | 0 | 7.07E-08 | 1 |
| UBE2O | 0 | 9.360915042 | 0 | 9.973339655 | 6.883600088 | 0 | 0 | 0 | 0.997434996 | 0.015230279 |
| UBE2Q2 | 0 | 0 | 10.23847846 | 9.52875341 | 6.608339636 | 0 | 0 | 0 | 0.006398527 | 0.00208311 |
| UBE3D | 9.689176399 | 0 | 0 | 0 | 0 | 0 | 0 | 0 | 7.88E-08 | 1 |
| UBE4B | 0 | 0 | 0 | 7.654272592 | 8.719239543 | 0 | 0 | 0 | 0.092975531 | 0.030614355 |
| UBN2 | 9.34594772 | 8.876365592 | 0 | 0 | 0 | 0 | 0 | 0 | 0.012412073 | 1 |
| UBR4 | 0 | 0 | 10.16682581 | 9.559738748 | 9.531804509 | 0 | 8.096917163 | 11.40516483 | 0.001299662 | 0.999854694 |
| UBXN11 | 0 | 0 | 11.09911385 | 6.873574072 | 9.248362782 | 0 | 7.813336911 | 0 | 0.001328126 | 0.999854694 |
| UBXN2B | 0 | 8.54458593 | 0 | 0 | 0 | 0 | 0 | 0 | 1.34E-07 | 1 |
| UBXN7 | 0 | 0 | 0 | 0 | 6.670177039 | 0 | 0 | 0 | 4.78E-05 | 1 |
| UCA1 | 0 | 0 | 0 | 10.60871443 | 8.237517482 | 0 | 11.38217465 | 10.62485973 | 0.030003792 | 0.999854694 |
| UCHL1 | 9.21371058 | 0 | 0 | 0 | 0 | 0 | 0 | 10.67094291 | 1.13E-07 | 1 |
| UCHL5 | 0 | 5.308137638 | 9.950915249 | 10.8957781 | 10.29113468 | 0 | 9.197932563 | 10.51744597 | 0.035972209 | 0.999854694 |
| UFL1 | 0 | 0 | 0 | 0 | 6.633393544 | 9.612215366 | 0 | 8.765887886 | 4.95E-05 | 0.999854694 |
| UFL1-AS1 | 0 | 0 | 0 | 0 | 6.775214428 | 9.089236856 | 6.975657029 | 0 | 4.34E-05 | 0.999854694 |
| UGT1A6 | 10.20786763 | 0 | 9.899270241 | 6.510996096 | 0 | 0 | 0 | 0 | 0.981170768 | 0.017222122 |
| UGT2B17 | 12.09237 | 10.27563746 | 13.14231562 | 9.953060429 | 0 | 0 | 0 | 0 | 0.996280769 | 4.43E-12 |
| UGT2B7 | 0 | 9.180635315 | 0 | 0 | 0 | 0 | 0 | 0 | 8.54E-08 | 1 |
| UHRF1 | 0 | 0 | 0 | 10.63350207 | 6.330495883 | 0 | 0 | 0 | 0.039556325 | 0.011464572 |
| UIMC1 | 0 | 0 | 0 | 0 | 6.841217856 | 0 | 0 | 0 | 4.14E-05 | 1 |
| UMAD1 | 0 | 0 | 10.87379348 | 0 | 9.579315983 | 12.23119927 | 0 | 9.155284626 | 0.011919704 | 0.999854694 |
| UNC119 | 0 | 0 | 10.53539179 | 0 | 7.209344369 | 0 | 9.129664832 | 11.27694028 | 0.042034455 | 0.999854694 |
| UNC5B | 9.427818601 | 0 | 0 | 0 | 0 | 0 | 0 | 0 | 7.88E-08 | 1 |
| URB1-AS1 | 0 | 8.752652351 | 0 | 0 | 0 | 0 | 0 | 0 | 1.16E-07 | 1 |
| USF1 | 9.641928937 | 0 | 0 | 0 | 0 | 0 | 0 | 0 | 7.44E-08 | 1 |
| USHBP1 | 9.773016437 | 0 | 0 | 0 | 0 | 0 | 0 | 0 | 7.17E-08 | 1 |
| USO1 | 0 | 0 | 10.33263459 | 0 | 7.381886513 | 0 | 0 | 0 | 0.049397317 | 0.011703791 |
| USP11 | 0 | 0 | 10.09560554 | 0 | 8.923477098 | 0 | 0 | 0 | 0.040321937 | 0.010829604 |

| | | | | | | | | | | |
|---|---|---|---|---|---|---|---|---|---|---|
| USP24 | 0 | 0 | 0 | 0 | 6.556887521 | 0 | 0 | 9.322651082 | 9.34E-05 | 1 |
| USP3 | 0 | 0 | 10.16583274 | 0 | 8.839844964 | 0 | 0 | 0 | 0.03951089 | 0.010789277 |
| USP30 | 0 | 0 | 0 | 0 | 0 | 0 | 7.62285003 | 10.04035398 | 1 | 0.01086644 |
| USP32 | 10.49673095 | 9.131811442 | 10.17671908 | 0 | 9.80005664 | 0 | 0 | 0 | 0.983311717 | 0.009237175 |
| USP47 | 0 | 9.732385072 | 0 | 0 | 0 | 0 | 0 | 0 | 7.17E-08 | 1 |
| USP5 | 0 | 0 | 0 | 0 | 6.405131987 | 0 | 9.409935073 | 9.547324608 | 5.97E-05 | 1 |
| USP54 | 10.62375866 | 7.945499326 | 0 | 0 | 0 | 0 | 0 | 0 | 0.003190254 | 1 |
| USP8 | 19.5306886 | 19.0395099 | 18.06888055 | 18.17199233 | 18.15454577 | 17.99058559 | 19.39154875 | 17.53752817 | 0.009164425 | 0.999854694 |
| USP9Y | 0 | 0 | 0 | 0 | 6.330495883 | 0 | 0 | 0 | 6.26E-05 | 1 |
| USPL1 | 0 | 0 | 0 | 0 | 6.39051047 | 0 | 0 | 0 | 5.98E-05 | 1 |
| UTP14C | 0 | 0 | 0 | 0 | 6.608339636 | 0 | 0 | 0 | 5.07E-05 | 1 |
| UTP3 | 0 | 0 | 0 | 0 | 0 | 0 | 8.333808086 | 10.34863922 | 1 | 0.009209122 |
| UTP4 | 10.50429894 | 0 | 9.930010706 | 0 | 8.861211882 | 0 | 0 | 0 | 0.989756518 | 0.011354734 |
| UTS2 | 10.32787885 | 0 | 0 | 0 | 0 | 0 | 0 | 0 | 7.07E-08 | 1 |
| UVRAG | 0 | 0 | 0 | 0 | 0 | 0 | 9.017021037 | 10.46884494 | 1 | 0.00801604 |
| VAMP5 | 0 | 9.354849786 | 0 | 0 | 0 | 10.58336707 | 0 | 10.03665353 | 7.91E-08 | 0.00624926 |
| VANGL1 | 13.29834068 | 13.6336269 | 15.39733394 | 15.31246455 | 14.23315657 | 14.23626291 | 15.22415957 | 15.27943177 | 0.014932011 | 0.999854694 |
| VAV1 | 0 | 8.824638969 | 0 | 0 | 0 | 0 | 0 | 0 | 1.36E-07 | 1 |
| VAV3 | 9.754096665 | 9.005869367 | 0 | 0 | 0 | 0 | 10.68344714 | 0 | 0.007370522 | 1 |
| VBP1 | 0 | 0 | 0 | 9.908620621 | 10.85722266 | 0 | 10.26512653 | 10.13683853 | 0.033778303 | 0.999854694 |
| VDAC2 | 9.443649659 | 10.60182125 | 13.12325773 | 12.80615673 | 11.35457855 | 12.20424336 | 11.56486083 | 9.851068447 | 0.027523413 | 0.999854694 |
| VEGFB | 0 | 0 | 10.22519269 | 8.197982807 | 0 | 0 | 0 | 0 | 0.049016692 | 0.011852991 |
| VIL1 | 0 | 0 | 0 | 9.578771997 | 9.456591048 | 0 | 10.27147687 | 0 | 0.042180622 | 1 |
| VILL | 0 | 0 | 10.07353533 | 0 | 6.717806789 | 0 | 0 | 0 | 0.066119036 | 0.01420877 |
| VIPAS39 | 0 | 0 | 0 | 10.61333698 | 7.041677569 | 0 | 0 | 0 | 0.037089642 | 0.011034216 |
| VIPR1 | 11.17916425 | 0 | 9.418741439 | 7.879740175 | 0 | 0 | 0 | 0 | 0.926424839 | 0.018766577 |
| VMAC | 9.797862919 | 0 | 0 | 0 | 0 | 0 | 0 | 10.16422986 | 7.17E-08 | 1 |
| VMO1 | 0 | 0 | 9.907611166 | 0 | 8.424960073 | 0 | 0 | 0 | 0.054622676 | 0.012283188 |
| VPREB3 | 0 | 10.59669041 | 0 | 0 | 0 | 0 | 0 | 0 | 7.07E-08 | 1 |
| VPS13A | 0 | 8.617326978 | 0 | 0 | 0 | 0 | 0 | 0 | 1.27E-07 | 1 |
| VPS13C | 10.89241997 | 9.066507669 | 0 | 0 | 0 | 0 | 0 | 0 | 0.000659735 | 1 |
| VPS13D | 0 | 0 | 10.10806713 | 0 | 6.797552774 | 0 | 0 | 0 | 0.064178982 | 0.013910154 |
| VPS16 | 10.11205603 | 0 | 10.20212852 | 9.552054579 | 7.626109207 | 0 | 0 | 0 | 0.990582117 | 0.001612925 |
| VPS33A | 0 | 0 | 9.022074146 | 10.07871314 | 0 | 0 | 0 | 0 | 0.041548195 | 0.011106965 |
| VPS33B | 0 | 8.250007882 | 0 | 0 | 0 | 0 | 7.597800322 | 9.746216776 | 2.64E-07 | 0.012259484 |
| VPS37C | 0 | 0 | 0 | 9.384775091 | 7.722658607 | 0 | 0 | 0 | 0.076102473 | 0.019205096 |
| VPS72 | 0 | 0 | 9.694154259 | 10.12993857 | 8.269832732 | 12.08884091 | 9.126165155 | 0 | 0.00328272 | 0.999854694 |
| VPS9D1 | 10.74689525 | 0 | 0 | 0 | 0 | 0 | 0 | 0 | 7.07E-08 | 1 |
| VSIR | 10.37817204 | 9.088605668 | 7.271028348 | 0 | 9.289333013 | 0 | 0 | 0 | 0.865762967 | 0.021776607 |
| WARS | 0 | 0 | 0 | 11.38261414 | 9.595343063 | 0 | 0 | 0 | 0.029901269 | 0.006444802 |
| WARS2 | 0 | 7.945499326 | 0 | 10.39447817 | 8.892680377 | 0 | 0 | 0 | 0.846923553 | 0.01037888 |
| WASH7P | 9.542522242 | 0 | 9.187782438 | 9.80251498 | 7.908597519 | 0 | 0 | 0 | 0.981170768 | 0.002374387 |
| WASH8P | 0 | 8.484742705 | 0 | 0 | 0 | 0 | 8.886581389 | 0 | 1.58E-07 | 1 |

| | | | | | | | | | | |
|---|---|---|---|---|---|---|---|---|---|---|
| WASHC2A | 0 | 0 | 8.427492135 | 0 | 9.318590875 | 0 | 0 | 0 | 0.070483102 | 0.015810782 |
| WASHC5 | 0 | 0 | 0 | 10.33750777 | 7.944074979 | 0 | 8.537232216 | 10.5463602 | 0.041988354 | 0.999854694 |
| WBP4 | 0 | 10.34612425 | 0 | 0 | 0 | 0 | 0 | 10.57215229 | 7.07E-08 | 1 |
| WDR11-AS1 | 0 | 9.280005236 | 0 | 0 | 0 | 0 | 0 | 0 | 8.10E-08 | 1 |
| WDR18 | 0 | 0 | 10.34672468 | 11.05648259 | 10.32516965 | 0 | 8.321619838 | 0 | 1.02E-05 | 0.907268934 |
| WDR36 | 0 | 0 | 9.398593245 | 9.923586506 | 0 | 0 | 0 | 0 | 0.039709783 | 0.011034216 |
| WDR46 | 0 | 0 | 0 | 0 | 0 | 10.86207771 | 0 | 9.049088908 | 1 | 0.007159623 |
| WDR47 | 9.542522242 | 0 | 0 | 0 | 0 | 0 | 0 | 0 | 7.73E-08 | 1 |
| WDR59 | 0 | 0 | 10.8059338 | 10.60593377 | 9.648540608 | 10.99589325 | 10.93417858 | 10.50410909 | 4.96E-05 | 0.999854694 |
| WDR5B | 0 | 8.375143804 | 0 | 0 | 0 | 0 | 8.82746995 | 0 | 1.55E-07 | 1 |
| WDR61 | 0 | 0 | 10.27024349 | 12.19244313 | 8.233426702 | 12.08691733 | 8.351899538 | 10.74083533 | 8.86E-07 | 0.999854694 |
| WDR73 | 5.419418853 | 0 | 11.39062765 | 0 | 7.467970239 | 0 | 0 | 0 | 0.422878711 | 1.27E-10 |
| WDR77 | 0 | 0 | 0 | 11.11892948 | 9.798672985 | 10.62726402 | 9.959705347 | 0 | 0.032011714 | 0.999854694 |
| WDR90 | 0 | 0 | 0 | 0 | 6.797552774 | 0 | 0 | 0 | 5.66E-05 | 1 |
| WFDC2 | 10.67830125 | 13.46103246 | 10.21178344 | 9.701937764 | 8.809938979 | 0 | 0 | 0 | 0.024974767 | 0.001091209 |
| WHAMM | 0 | 0 | 8.789130733 | 0 | 7.52932141 | 0 | 0 | 0 | 0.096729883 | 0.029661957 |
| WHRN | 0 | 0 | 0 | 0 | 6.706045958 | 0 | 8.706139773 | 9.041723345 | 0.000126325 | 0.999854694 |
| WIPF2 | 0 | 9.579530494 | 0 | 0 | 0 | 0 | 0 | 0 | 7.44E-08 | 1 |
| WNK1 | 0 | 9.396780662 | 9.579510725 | 0 | 8.587624462 | 0 | 0 | 0 | 0.998393647 | 0.013715581 |
| WNT2 | 0 | 0 | 9.464687668 | 11.98158914 | 13.96652828 | 15.66699604 | 0 | 0 | 7.07E-08 | 0.999854694 |
| WRAP73 | 9.85816911 | 7.559034993 | 8.888792936 | 9.691483378 | 0 | 0 | 0 | 0 | 0.991384446 | 0.013122865 |
| WWC3 | 0 | 9.920717961 | 0 | 9.616101356 | 7.757261396 | 0 | 0 | 0 | 0.98939231 | 0.016536489 |
| XK | 0 | 9.324135017 | 0 | 0 | 0 | 0 | 6.775741571 | 0 | 8.05E-08 | 1 |
| XRCC1 | 0 | 7.620995983 | 0 | 10.03527963 | 6.830424858 | 0 | 0 | 0 | 0.911081797 | 0.014775816 |
| XRN1 | 0 | 0 | 0 | 10.20167928 | 6.934884845 | 0 | 0 | 0 | 0.056135231 | 0.013610425 |
| YAF2 | 0 | 0 | 0 | 0 | 0 | 10.15309928 | 0 | 10.6553427 | 1 | 0.005940842 |
| YARS | 0 | 0 | 0 | 10.97693275 | 7.923909096 | 0 | 0 | 0 | 0.0196795 | 0.009470925 |
| YARS2 | 10.31071715 | 0 | 0 | 10.25471675 | 9.880083437 | 0 | 0 | 0 | 0.996280769 | 0.009159898 |
| YBEY | 11.15292243 | 0 | 10.62771936 | 0 | 6.556887521 | 0 | 0 | 0 | 0.976114635 | 0.011034216 |
| YIPF1 | 0 | 0 | 0 | 10.73200593 | 9.164906972 | 0 | 8.315486882 | 0 | 0.017657163 | 0.999854694 |
| YIPF6 | 0 | 0 | 11.81727878 | 0 | 9.790343075 | 9.031734041 | 8.11821332 | 0 | 0.02506161 | 0.999854694 |
| YJU2 | 0 | 0 | 9.570533726 | 0 | 9.963332571 | 0 | 0 | 0 | 0.031408765 | 0.010189075 |
| YPEL3 | 0 | 0 | 11.93678188 | 0 | 9.422859707 | 11.86917098 | 10.05112565 | 10.512126 | 0.024675797 | 0.999854694 |
| YRDC | 0 | 0 | 0 | 9.825007397 | 6.85193071 | 0 | 0 | 0 | 0.070665441 | 0.016902113 |
| YTHDF2 | 0 | 0 | 0 | 0 | 6.315093872 | 0 | 0 | 0 | 6.33E-05 | 1 |
| YWHAH | 10.73729353 | 10.48449957 | 12.28317008 | 11.62116252 | 10.36404409 | 0 | 0 | 0 | 0.670671829 | 6.80E-05 |
| ZBED1 | 0 | 0 | 0 | 9.896535119 | 9.708713631 | 0 | 0 | 0 | 0.03000204 | 0.010278758 |
| ZBED8 | 0 | 9.521148737 | 0 | 0 | 0 | 0 | 0 | 0 | 7.58E-08 | 1 |
| ZBTB48 | 0 | 0 | 0 | 9.464711849 | 8.545348962 | 0 | 0 | 0 | 0.065905321 | 0.014775816 |
| ZBTB6 | 0 | 9.051585356 | 0 | 0 | 0 | 0 | 0 | 0 | 1.18E-07 | 1 |
| ZBTB7C | 0 | 0 | 7.745299485 | 0 | 0 | 0 | 0 | 0 | 2.03E-05 | 1 |
| ZC3H12A | 0 | 0 | 10.64135848 | 7.804122388 | 0 | 8.663081869 | 4.882656432 | 0 | 0.036199827 | 0.999854694 |
| ZC3H7A | 0 | 0 | 0 | 8.137651801 | 0 | 0 | 0 | 0 | 1.27E-05 | 1 |

| | | | | | | | | | | |
|---|---|---|---|---|---|---|---|---|---|---|
| ZC3HC1 | 10.42278292 | 9.670244415 | 0 | 0 | 0 | 0 | 9.574337042 | 0 | 0.001108626 | 1 |
| ZCCHC17 | 0 | 9.299083523 | 0 | 0 | 0 | 9.959233885 | 7.80460581 | 0 | 8.05E-08 | 0.010829815 |
| ZCCHC18 | 0 | 0 | 0 | 8.64347382 | 0 | 0 | 0 | 0 | 7.29E-06 | 1 |
| ZCCHC3 | 0 | 0 | 8.024846459 | 0 | 0 | 0 | 0 | 0 | 1.52E-05 | 1 |
| ZCCHC7 | 11.17916425 | 9.2476365 | 0 | 0 | 0 | 0 | 0 | 9.049088908 | 0.000183438 | 1 |
| ZCRB1 | 0 | 0 | 0 | 10.90108075 | 9.70132783 | 0 | 8.997999732 | 9.234949175 | 0.034989165 | 0.999854694 |
| ZDHHC11B | 0 | 0 | 0 | 0 | 0 | 9.629938381 | 0 | 11.28709881 | 1 | 0.005132109 |
| ZDHHC19 | 0 | 9.823047116 | 0 | 0 | 0 | 0 | 0 | 0 | 7.12E-08 | 1 |
| ZDHHC21 | 9.391550492 | 0 | 0 | 0 | 0 | 0 | 0 | 0 | 8.04E-08 | 1 |
| ZDHHC3 | 0 | 9.708800502 | 0 | 10.00745431 | 10.78429454 | 0 | 0 | 0 | 0.969794791 | 0.007615607 |
| ZDHHC8 | 0 | 8.850734095 | 0 | 0 | 0 | 0 | 0 | 0 | 1.17E-07 | 1 |
| ZDHHC9 | 10.33213756 | 0 | 0 | 0 | 0 | 0 | 8.777034246 | 0 | 7.07E-08 | 1 |
| ZEB2 | 8.749411643 | 0 | 0 | 0 | 0 | 0 | 0 | 0 | 1.19E-07 | 1 |
| ZFP62 | 0 | 9.620553507 | 0 | 0 | 0 | 0 | 7.627808199 | 0 | 7.42E-08 | 1 |
| ZFP69B | 0 | 0 | 0 | 6.922073503 | 0 | 0 | 0 | 0 | 0.000113876 | 1 |
| ZFP82 | 9.520299506 | 0 | 0 | 0 | 0 | 0 | 0 | 0 | 7.73E-08 | 1 |
| ZFPL1 | 5.760100304 | 0 | 10.13470357 | 0 | 9.916898101 | 0 | 0 | 0 | 0.421993138 | 0.009159898 |
| ZFYVE1 | 9.96623846 | 0 | 0 | 0 | 0 | 0 | 0 | 0 | 7.07E-08 | 1 |
| ZFYVE28 | 10.7970523 | 0 | 0 | 0 | 0 | 0 | 5.599926409 | 0 | 7.07E-08 | 1 |
| ZGLP1 | 0 | 0 | 0 | 10.24044427 | 7.474917303 | 0 | 0 | 0 | 0.04999624 | 0.012440088 |
| ZGPAT | 0 | 0 | 11.94693405 | 10.8073662 | 6.098291247 | 0 | 8.016026117 | 0 | 6.87E-06 | 0.72489419 |
| ZKSCAN8 | 8.973144195 | 0 | 0 | 0 | 0 | 0 | 0 | 0 | 1.08E-07 | 1 |
| ZMIZ1 | 0 | 0 | 0 | 9.979081745 | 8.295566123 | 0 | 0 | 0 | 0.051802237 | 0.012473101 |
| ZMYM2 | 0 | 7.254738985 | 0 | 0 | 0 | 0 | 6.991130623 | 10.83970819 | 0.099427067 | 0.008668791 |
| ZNF101 | 0 | 9.336499504 | 0 | 0 | 0 | 0 | 0 | 0 | 8.04E-08 | 1 |
| ZNF107 | 0 | 0 | 0 | 8.470833404 | 0 | 0 | 8.573661004 | 0 | 8.80E-06 | 1 |
| ZNF121 | 0 | 9.493809912 | 0 | 0 | 0 | 0 | 0 | 0 | 8.05E-08 | 1 |
| ZNF134 | 10.53047859 | 0 | 0 | 0 | 0 | 0 | 0 | 0 | 7.07E-08 | 1 |
| ZNF138 | 0 | 8.998107186 | 9.260288777 | 0 | 7.209344369 | 0 | 0 | 0 | 0.999483812 | 0.022769437 |
| ZNF169 | 0 | 9.381944859 | 0 | 0 | 0 | 0 | 0 | 0 | 7.88E-08 | 1 |
| ZNF184 | 0 | 0 | 0 | 0 | 6.786426836 | 0 | 0 | 0 | 4.30E-05 | 1 |
| ZNF20 | 0 | 0 | 9.237783257 | 0 | 8.225210179 | 0 | 0 | 0 | 0.07622068 | 0.017524835 |
| ZNF217 | 0 | 0 | 0 | 9.794396227 | 7.21765149 | 0 | 0 | 0 | 0.07021841 | 0.016321748 |
| ZNF219 | 0 | 0 | 9.515466095 | 10.41791985 | 0 | 0 | 0 | 0 | 0.024898456 | 0.009724498 |
| ZNF251 | 0 | 9.600187807 | 0 | 0 | 0 | 0 | 0 | 0 | 7.44E-08 | 1 |
| ZNF26 | 0 | 0 | 0 | 0 | 6.608339636 | 0 | 0 | 0 | 5.07E-05 | 1 |
| ZNF263 | 0 | 8.637453836 | 0 | 0 | 0 | 0 | 8.166714905 | 0 | 1.24E-07 | 1 |
| ZNF264 | 0 | 9.005869367 | 0 | 0 | 0 | 0 | 0 | 0 | 1.04E-07 | 1 |
| ZNF282 | 0 | 0 | 0 | 0 | 0 | 0 | 8.200386998 | 10.57215229 | 1 | 0.008668791 |
| ZNF296 | 0 | 8.770988541 | 0 | 0 | 0 | 0 | 0 | 0 | 1.15E-07 | 1 |
| ZNF326 | 0 | 8.974566872 | 10.38563085 | 10.38585859 | 0 | 0 | 0 | 0 | 0.937031857 | 0.00825262 |
| ZNF330 | 9.695801475 | 0 | 0 | 9.905608724 | 8.995296112 | 0 | 0 | 0 | 0.99353678 | 0.01143424 |
| ZNF335 | 0 | 9.366954907 | 0 | 8.526790157 | 7.785476285 | 0 | 0 | 0 | 0.985285296 | 0.033895535 |

| | | | | | | | | | | |
|---|---|---|---|---|---|---|---|---|---|---|
| ZNF362 | 0 | 0 | 8.536001166 | 0 | 7.515911508 | 0 | 0 | 0 | 0.104856829 | 0.036201267 |
| ZNF366 | 0 | 8.80697625 | 0 | 0 | 0 | 0 | 0 | 0 | 1.15E-07 | 1 |
| ZNF37A | 0 | 0 | 7.611654213 | 0 | 0 | 0 | 0 | 10.47705966 | 2.33E-05 | 1 |
| ZNF408 | 0 | 8.789094606 | 0 | 0 | 0 | 0 | 0 | 0 | 1.15E-07 | 1 |
| ZNF410 | 0 | 0 | 11.08762348 | 0 | 7.467970239 | 0 | 0 | 0 | 0.020248687 | 0.009159898 |
| ZNF414 | 0 | 0 | 0 | 0 | 6.375739247 | 0 | 0 | 0 | 6.06E-05 | 1 |
| ZNF439 | 0 | 0 | 6.995027124 | 0 | 0 | 0 | 0 | 0 | 4.21E-05 | 1 |
| ZNF44 | 0 | 10.42784521 | 10.20406468 | 0 | 7.158471365 | 0 | 0 | 0 | 0.985757273 | 0.012705982 |
| ZNF440 | 0 | 0 | 0 | 0 | 6.299525656 | 0 | 0 | 0 | 9.81E-05 | 1 |
| ZNF444 | 0 | 9.579530494 | 0 | 0 | 0 | 0 | 0 | 0 | 7.44E-08 | 1 |
| ZNF445 | 0 | 0 | 0 | 10.43890491 | 9.403874704 | 0 | 7.597800322 | 0 | 0.022330739 | 0.999854694 |
| ZNF451 | 0 | 0 | 0 | 10.08937024 | 8.271828602 | 0 | 8.433450222 | 0 | 0.047894608 | 0.999854694 |
| ZNF454 | 10.03054434 | 0 | 0 | 0 | 0 | 0 | 0 | 0 | 7.07E-08 | 1 |
| ZNF468 | 0 | 8.966633964 | 0 | 0 | 0 | 0 | 0 | 0 | 1.07E-07 | 1 |
| ZNF496 | 0 | 0 | 0 | 0 | 6.517057266 | 0 | 10.8910109 | 0 | 5.50E-05 | 1 |
| ZNF503-AS1 | 0 | 9.827434044 | 0 | 0 | 0 | 0 | 0 | 0 | 7.10E-08 | 1 |
| ZNF511 | 0 | 0 | 11.37051122 | 11.74259475 | 10.6564529 | 10.61841328 | 9.4760367 | 0 | 7.07E-08 | 0.999854694 |
| ZNF512 | 0 | 8.950636073 | 0 | 0 | 0 | 0 | 0 | 0 | 1.07E-07 | 1 |
| ZNF513 | 0 | 9.117552805 | 0 | 0 | 0 | 0 | 7.535844571 | 9.247809184 | 1.14E-07 | 0.015785316 |
| ZNF527 | 0 | 9.051585356 | 0 | 0 | 0 | 0 | 0 | 0 | 1.18E-07 | 1 |
| ZNF528 | 0 | 8.770988541 | 0 | 0 | 0 | 0 | 0 | 0 | 1.15E-07 | 1 |
| ZNF547 | 9.435755845 | 0 | 0 | 0 | 0 | 0 | 0 | 0 | 7.88E-08 | 1 |
| ZNF557 | 10.28897389 | 0 | 0 | 0 | 0 | 0 | 0 | 0 | 7.07E-08 | 1 |
| ZNF567 | 0 | 0 | 9.820687648 | 0 | 6.185573957 | 0 | 0 | 0 | 0.074522731 | 0.017524835 |
| ZNF57 | 0 | 0 | 0 | 9.966129753 | 7.893121693 | 0 | 0 | 0 | 0.057561092 | 0.013610425 |
| ZNF574 | 9.311852598 | 0 | 0 | 0 | 0 | 0 | 0 | 0 | 8.29E-08 | 1 |
| ZNF580 | 0 | 0 | 0 | 10.67654408 | 7.872225587 | 0 | 0 | 0 | 0.030353545 | 0.01037888 |
| ZNF581 | 0 | 0 | 10.73271351 | 0 | 8.193974596 | 0 | 0 | 0 | 0.028273623 | 0.009818636 |
| ZNF582-AS1 | 0 | 8.833389894 | 0 | 0 | 0 | 0 | 0 | 0 | 1.14E-07 | 1 |
| ZNF585A | 0 | 0 | 9.357432539 | 9.715760134 | 3.254875415 | 0 | 0 | 0 | 0.020482047 | 0.005661908 |
| ZNF585B | 9.06811818 | 0 | 0 | 0 | 0 | 0 | 0 | 0 | 1.09E-07 | 1 |
| ZNF593 | 0 | 0 | 10.6484855 | 11.08870796 | 10.34236561 | 10.89396532 | 10.4722695 | 10.24982197 | 3.74E-06 | 0.999854694 |
| ZNF595 | 9.977157135 | 7.426508624 | 0 | 0 | 0 | 0 | 0 | 0 | 0.012301623 | 1 |
| ZNF599 | 9.6281405 | 0 | 0 | 0 | 0 | 0 | 0 | 0 | 7.53E-08 | 1 |
| ZNF616 | 0 | 0 | 0 | 0 | 6.763914196 | 0 | 0 | 0 | 0.000116718 | 1 |
| ZNF625 | 9.898931975 | 0 | 0 | 0 | 0 | 0 | 0 | 0 | 7.07E-08 | 1 |
| ZNF626 | 0 | 9.103151837 | 0 | 0 | 0 | 0 | 0 | 0 | 1.15E-07 | 1 |
| ZNF627 | 0 | 0 | 10.1142578 | 0 | 8.392307862 | 0 | 0 | 0 | 0.047605081 | 0.011464572 |
| ZNF638 | 0 | 8.966633964 | 0 | 0 | 0 | 0 | 0 | 0 | 1.07E-07 | 1 |
| ZNF652 | 0 | 0 | 9.191690784 | 9.950139986 | 8.095992883 | 0 | 0 | 0 | 0.007337719 | 0.002049112 |
| ZNF668 | 0 | 9.073911341 | 0 | 0 | 0 | 0 | 0 | 0 | 1.17E-07 | 1 |
| ZNF700 | 11.10395166 | 0 | 0 | 0 | 0 | 0 | 6.757760881 | 9.638035931 | 5.99E-08 | 0.014264276 |
| ZNF74 | 0 | 8.724703557 | 0 | 0 | 0 | 0 | 0 | 0 | 1.17E-07 | 1 |

| | | | | | | | | | | |
|---|---|---|---|---|---|---|---|---|---|---|
| ZNF740 | 0 | 0 | 0 | 10.53364114 | 7.446926049 | 0 | 0 | 0 | 0.038156624 | 0.011034216 |
| ZNF746 | 9.960747977 | 0 | 0 | 0 | 0 | 0 | 0 | 0 | 7.07E-08 | 1 |
| ZNF75D | 0 | 8.867872257 | 0 | 0 | 0 | 0 | 0 | 0 | 1.10E-07 | 1 |
| ZNF766 | 0 | 0 | 0 | 0 | 6.830424858 | 0 | 0 | 0 | 4.17E-05 | 1 |
| ZNF770 | 0 | 9.088605668 | 0 | 0 | 0 | 0 | 0 | 0 | 1.16E-07 | 1 |
| ZNF79 | 0 | 0 | 0 | 0 | 6.752524753 | 0 | 0 | 0 | 0.000121249 | 1 |
| ZNF793 | 8.389616335 | 0 | 0 | 0 | 0 | 0 | 0 | 0 | 1.62E-07 | 1 |
| ZNF830 | 0 | 0 | 0 | 0 | 6.39051047 | 0 | 5.857994609 | 0 | 5.98E-05 | 1 |
| ZNF836 | 0 | 0 | 0 | 0 | 6.39051047 | 0 | 0 | 0 | 5.98E-05 | 1 |
| ZNF91 | 16.80963015 | 15.86073118 | 15.25561014 | 14.53772213 | 15.23922036 | 14.76338519 | 15.20282504 | 14.32521442 | 0.035525927 | 0.999854694 |
| ZPR1 | 0 | 0 | 0 | 10.00182426 | 9.726286792 | 0 | 0 | 11.63850949 | 0.027467126 | 1 |
| ZSCAN16 | 0 | 5.091801452 | 0 | 0 | 0 | 0 | 8.931384524 | 10.5463602 | 0.252336971 | 0.007912372 |
| ZSCAN22 | 9.467075215 | 0 | 0 | 0 | 0 | 0 | 0 | 0 | 7.88E-08 | 1 |
| ZSCAN9 | 0 | 8.398919312 | 0 | 0 | 0 | 0 | 0 | 0 | 1.51E-07 | 1 |
| ZSWIM3 | 0 | 0 | 10.66263479 | 0 | 6.061843883 | 0 | 0 | 0 | 0.041988354 | 2.04E-10 |
| ZUP1 | 0 | 9.324135017 | 0 | 0 | 0 | 0 | 4.668391817 | 0 | 8.05E-08 | 0.663581498 |
| ZYG11B | 10.13661204 | 8.867872257 | 0 | 0 | 0 | 0 | 0 | 0 | 0.004770209 | 1 |
| ZZZ3 | 0 | 8.798063133 | 0 | 0 | 0 | 0 | 0 | 0 | 1.15E-07 | 1 |

### Supplementary Table S4: Differentially expressed isoforms

| Gene | Isoform | 1N | 3N | 1T | 2T | 3T | 1M | 2M | 3M | padj_NT | padj_TL |
|---|---|---|---|---|---|---|---|---|---|---|---|
| A1CF | XM_005269720 | 0 | 9.006487637 | 0 | 0 | 0 | 0 | 0 | 0 | 6.80E-08 | 1 |
| AAAS | XM_011538780 | 0 | 8.475517571 | 0 | 0 | 0 | 0 | 0 | 0 | 1.09E-07 | 1 |
| AACS | NM_001319839 | 0 | 0 | 0 | 8.029665411 | 0 | 0 | 0 | 0 | 1.02E-05 | 1 |
| AAGAB | XM_024450052 | 0 | 0 | 0 | 0 | 0 | 11.27285901 | 9.592700938 | 0 | 1 | 0.02072721 |
| AAMP | NM_001302545 | 7.782511257 | 0 | 0 | 10.53749502 | 9.603429314 | 0 | 0 | 0 | 0.827857426 | 0.02072721 |
| AAR2 | XM_011528763 | 9.649985154 | 0 | 0 | 0 | 0 | 0 | 6.953230848 | 0 | 2.95E-08 | 1 |
| ABCA2 | NM_212533 | 0 | 8.248259235 | 0 | 0 | 0 | 0 | 0 | 0 | 2.02E-07 | 1 |
| ABCA5 | NM_172232 | 10.34562109 | 9.855790228 | 0 | 0 | 0 | 0 | 0 | 0 | 0.000742596 | 1 |
| ABCA8 | XR_002957945 | 9.546283346 | 0 | 0 | 0 | 0 | 0 | 0 | 0 | 3.25E-08 | 1 |
| ABCB10 | XR_001737084 | 0 | 0 | 0 | 0 | 6.690841957 | 0 | 0 | 0 | 3.72E-05 | 1 |
| ABCB8 | NM_001282293 | 0 | 0 | 0 | 0 | 0 | 0 | 10.29425491 | 10.12983217 | 1 | 0.02072721 |
| ABCC1 | XM_017023242 | 0 | 0 | 7.936379488 | 9.693732641 | 0 | 0 | 8.042470426 | 0 | 0.029881811 | 0.999932046 |
| ABCC4 | XM_017020322 | 9.669860113 | 8.20886266 | 0 | 0 | 0 | 0 | 0 | 0 | 0.002935751 | 1 |
| ABCC6P2 | NR_023387 | 10.24321976 | 0 | 0 | 0 | 0 | 0 | 0 | 0 | 1.59E-08 | 1 |
| ABCD3 | XM_011541877 | 9.807800698 | 0 | 9.022425311 | 0 | 7.084811668 | 0 | 0 | 0 | 0.999953634 | 0.03159928 |
| ABCF3 | XM_024453625 | 0 | 9.095586658 | 0 | 0 | 0 | 0 | 0 | 10.40262864 | 5.54E-08 | 1 |
| ABCG2 | XM_011532420 | 9.502621339 | 0 | 0 | 0 | 0 | 0 | 0 | 0 | 3.44E-08 | 1 |
| ABHD11 | NM_148912 | 0 | 11.3714765 | 9.022425311 | 0 | 7.44304914 | 0 | 0 | 0 | 0.902319944 | 0.029357153 |
| ABHD11 | XM_024446949 | 10.09918783 | 0 | 0 | 0 | 0 | 0 | 0 | 0 | 1.74E-08 | 1 |
| ABHD12 | NM_001042472 | 8.678186754 | 0 | 0 | 9.875605271 | 8.876918096 | 0 | 0 | 0 | 0.999953634 | 0.02072721 |
| ABHD12 | NM_015600 | 0 | 8.642530071 | 0 | 10.26832462 | 8.473070111 | 0 | 0 | 0 | 0.999953634 | 0.02072721 |
| ABHD12 | XM_017027796 | 0 | 0 | 10.52926948 | 0 | 8.754447516 | 0 | 0 | 0 | 0.016251078 | 0.02072721 |
| ABHD12 | XR_002958467 | 9.346729415 | 0 | 0 | 0 | 0 | 0 | 0 | 0 | 4.43E-08 | 1 |
| ABHD13 | NM_032859 | 10.16602012 | 0 | 10.53325985 | 0 | 8.650778048 | 0 | 0 | 0 | 0.999953634 | 0.020768938 |
| ABHD14A | NM_015407 | 10.15666012 | 0 | 0 | 0 | 0 | 10.84242745 | 0 | 0 | 1.62E-08 | 1 |
| ABHD14B | NM_001254753 | 0 | 8.943876548 | 10.40411791 | 0 | 9.170386259 | 0 | 0 | 0 | 0.999953634 | 0.02072721 |
| ABHD16A | NR_033489 | 10.05452941 | 9.08836834 | 10.42316685 | 9.957086203 | 0 | 0 | 0 | 0 | 0.999953634 | 0.02072721 |
| ABHD17A | NM_001130111 | 11.84788856 | 0 | 0 | 0 | 0 | 10.93072186 | 0 | 0 | 1.34E-08 | 1 |
| ABHD17B | XM_024447567 | 0 | 0 | 0 | 8.491803396 | 0 | 0 | 0 | 0 | 6.26E-06 | 1 |
| ABHD2 | XM_024449829 | 10.35792493 | 0 | 0 | 0 | 0 | 0 | 0 | 0 | 1.48E-08 | 1 |
| ABHD3 | XR_935209 | 0 | 0 | 8.627850236 | 0 | 6.351978551 | 0 | 0 | 0 | 0.071180577 | 0.046384936 |
| ABHD5 | NM_016006 | 0 | 0 | 9.68747462 | 0 | 8.002028225 | 0 | 0 | 0 | 0.028264333 | 0.02072721 |
| ABI1 | NM_001348031 | 0 | 0 | 10.39362058 | 0 | 6.97463031 | 0 | 6.514885902 | 0 | 0.022307656 | 0.999932046 |
| ABI1 | XM_005252332 | 9.682959584 | 0 | 0 | 0 | 0 | 0 | 0 | 10.72375851 | 2.84E-08 | 1 |
| ABI3 | NM_016428 | 0 | 8.430558346 | 0 | 0 | 0 | 0 | 0 | 0 | 1.13E-07 | 1 |
| ABLIM1 | XM_006717846 | 10.36606958 | 0 | 0 | 0 | 0 | 0 | 0 | 0 | 1.48E-08 | 1 |
| ABLIM1 | XM_024448022 | 0 | 0 | 0 | 0 | 6.582319518 | 0 | 8.382856136 | 0 | 4.04E-05 | 1 |
| ABR | NM_001092 | 0 | 0 | 9.357366688 | 0 | 7.44304914 | 0 | 0 | 0 | 0.038107772 | 0.0247531 |
| ABRAXAS1 | XM_005263280 | 0 | 0 | 0 | 7.763094795 | 0 | 0 | 0 | 0 | 2.57E-05 | 1 |
| ABTB1 | XR_924182 | 10.64908701 | 0 | 0 | 0 | 0 | 0 | 0 | 0 | 1.34E-08 | 1 |
| ACAA1 | NM_001607 | 0 | 0 | 10.70086101 | 10.43073285 | 9.845694196 | 0 | 8.493752915 | 0 | 0.000764956 | 0.999932046 |

| | | | | | | | | | | | |
|---|---|---|---|---|---|---|---|---|---|---|---|
| ACAA1 | XM_011533650 | 0 | 0 | 0 | 9.947916041 | 7.983181653 | 0 | 0 | 0 | 0.024330175 | 0.02072721 |
| ACAD8 | XM_024448438 | 0 | 8.935855595 | 0 | 0 | 0 | 0 | 0 | 0 | 7.69E-08 | 1 |
| ACAD9 | NM_014049 | 0 | 0 | 0 | 9.950979243 | 9.179728125 | 0 | 0 | 11.02286619 | 0.018782969 | 0.999932046 |
| ACADS | NM_001302554 | 10.38623228 | 0 | 0 | 0 | 0 | 0 | 0 | 0 | 1.70E-08 | 1 |
| ACADVL | XR_934021 | 0 | 0 | 0 | 0 | 6.758905041 | 0 | 8.058247234 | 0 | 9.53E-05 | 1 |
| ACADVL | XR_934023 | 0 | 0 | 0 | 0 | 6.667421491 | 4.321702656 | 0 | 0 | 3.78E-05 | 1 |
| ACAP3 | XM_011540609 | 9.226627846 | 0 | 0 | 0 | 0 | 0 | 0 | 0 | 5.40E-08 | 1 |
| ACD | XR_429728 | 0 | 0 | 0 | 9.116932532 | 7.012300182 | 0 | 0 | 0 | 0.046582167 | 0.032030331 |
| ACIN1 | NM_014977 | 0 | 9.819195269 | 0 | 0 | 0 | 0 | 0 | 0 | 2.01E-08 | 1 |
| ACLY | NM_001096 | 0 | 0 | 12.30964602 | 11.74851699 | 10.00716038 | 11.70180555 | 9.740828669 | 10.61185893 | 0.00016744 | 0.999932046 |
| ACOT2 | NM_006821 | 0 | 0 | 7.601400457 | 0 | 0 | 0 | 8.15532787 | 0 | 1.81E-05 | 1 |
| ACOT8 | XR_244130 | 0 | 0 | 9.396446213 | 0 | 9.058930383 | 0 | 0 | 10.45995827 | 0.024305675 | 0.999932046 |
| ACP1 | NR_024080 | 9.734199496 | 0 | 0 | 0 | 0 | 0 | 0 | 10.35506601 | 5.08E-08 | 1 |
| ACP2 | NM_001610 | 0 | 0 | 10.31884448 | 0 | 5.693189014 | 0 | 0 | 9.172865883 | 0.026022619 | 1 |
| ACP2 | XR_001747908 | 0 | 9.467937896 | 0 | 0 | 0 | 0 | 0 | 0 | 3.10E-08 | 1 |
| ACP5 | XM_011528069 | 0 | 0 | 0 | 10.46005576 | 6.965057125 | 0 | 0 | 0 | 0.0209853 | 0.02072721 |
| ACSF2 | NM_025149 | 10.47551503 | 0 | 0 | 0 | 0 | 0 | 0 | 0 | 1.44E-08 | 1 |
| ACSF3 | NR_147929 | 0 | 0 | 7.526510252 | 0 | 0 | 0 | 0 | 0 | 1.86E-05 | 1 |
| ACSF3 | XM_017023019 | 0 | 0 | 0 | 10.48668527 | 7.415197319 | 0 | 9.8554239 | 0 | 0.019753354 | 0.999932046 |
| ACSF3 | XR_933238 | 0 | 0 | 0 | 10.12921816 | 8.002028225 | 0 | 0 | 0 | 0.022058827 | 0.02072721 |
| ACSL1 | XM_005262828 | 0 | 0 | 10.39010443 | 10.22342977 | 8.712153435 | 0 | 0 | 11.24134951 | 0.00139908 | 0.999932046 |
| ACSM3 | NM_005622 | 0 | 9.405416277 | 0 | 0 | 0 | 0 | 0 | 0 | 3.31E-08 | 1 |
| ACSM3 | XM_011545911 | 0 | 8.818611963 | 0 | 0 | 0 | 0 | 0 | 0 | 8.12E-08 | 1 |
| ACSM3 | XM_024450370 | 9.695941183 | 0 | 0 | 0 | 0 | 0 | 0 | 0 | 2.66E-08 | 1 |
| ACSS1 | XR_937170 | 0 | 0 | 0 | 9.59927585 | 9.224548436 | 0 | 0 | 0 | 0.021181076 | 0.02072721 |
| ACSS2 | XM_011528912 | 10.24321976 | 9.872695374 | 9.626025595 | 9.833071484 | 7.983181653 | 0 | 0 | 0 | 0.999953634 | 0.004381107 |
| ACTA2 | NM_001320855 | 0 | 9.799439006 | 0 | 0 | 0 | 11.26619798 | 0 | 0 | 2.04E-08 | 1 |
| ACTL6A | NM_177989 | 0 | 0 | 10.42660343 | 0 | 8.800928156 | 0 | 0 | 10.17161555 | 0.016912939 | 0.999932046 |
| ACTN4 | NM_001322033 | 10.22989302 | 9.411213355 | 8.456887945 | 0 | 7.801472153 | 0 | 0 | 0 | 0.877337094 | 0.035282533 |
| ACTR1B | NM_005735 | 0 | 0 | 10.97470853 | 9.127801312 | 0 | 10.4136552 | 0 | 0 | 0.013156658 | 0.999932046 |
| ACTR1B | XM_017003116 | 0 | 0 | 10.36345654 | 0 | 8.10150079 | 0 | 0 | 0 | 0.019686112 | 0.02072721 |
| ACTR3 | NR_102318 | 0 | 8.737876298 | 10.72199233 | 9.334269269 | 8.234946357 | 0 | 0 | 0 | 0.932308674 | 0.003016079 |
| ACTR3C | NM_001164458 | 0 | 0 | 9.89521947 | 0 | 8.038998467 | 0 | 7.604755783 | 0 | 0.025216192 | 0.999932046 |
| ACTR6 | NR_048569 | 0 | 9.117027412 | 9.317199039 | 0 | 7.305685067 | 0 | 0 | 0 | 0.999953634 | 0.025754319 |
| ACVR1 | NM_001347664 | 0 | 0 | 10.17608123 | 9.389502959 | 0 | 11.42613238 | 9.777507907 | 0 | 0.017830139 | 0.999932046 |
| ACVR1 | XM_006712825 | 0 | 0 | 9.197584296 | 0 | 9.690238446 | 0 | 0 | 9.871287726 | 0.020568316 | 0.999932046 |
| ACY1 | NM_001198897 | 10.93843027 | 0 | 0 | 0 | 0 | 0 | 0 | 0 | 1.34E-08 | 1 |
| ACYP2 | NM_138448 | 0 | 0 | 0 | 10.83890988 | 7.593532742 | 0 | 0 | 0 | 0.016012631 | 0.02072721 |
| ADA | NM_001322050 | 0 | 8.998808245 | 0 | 0 | 0 | 0 | 0 | 0 | 6.89E-08 | 1 |
| ADA2 | XM_011546133 | 9.939165239 | 0 | 0 | 0 | 0 | 0 | 0 | 0 | 2.01E-08 | 1 |
| ADAM15 | NM_003815 | 0 | 0 | 7.826204809 | 0 | 10.49933402 | 0 | 0 | 10.18187502 | 0.018094055 | 0.999932046 |
| ADAM15 | NM_207191 | 0 | 9.585396421 | 0 | 0 | 0 | 0 | 0 | 0 | 2.61E-08 | 1 |
| ADAM28 | XR_949375 | 0 | 8.486541819 | 0 | 0 | 0 | 0 | 0 | 0 | 1.09E-07 | 1 |

| | | | | | | | | | | | |
|---|---|---|---|---|---|---|---|---|---|---|---|
| ADAMTS10 | XM_017027340 | 0 | 0 | 0 | 0 | 6.3222931 | 0 | 0 | 0 | 0.000244538 | 1 |
| ADAMTSL1 | XM_017015312 | 9.917161541 | 0 | 0 | 0 | 0 | 0 | 0 | 0 | 2.04E-08 | 1 |
| ADAMTSL2 | XM_011519242 | 0 | 0 | 0 | 0 | 0 | 8.169187498 | 6.826303056 | 0 | 1 | 0.043708568 |
| ADAP2 | NM_001346716 | 0 | 0 | 9.514183525 | 0 | 8.66562982 | 0 | 0 | 8.868599317 | 0.026087146 | 0.999932046 |
| ADAT1 | NM_012091 | 0 | 8.747073305 | 0 | 0 | 0 | 9.794666143 | 0 | 0 | 8.61E-08 | 1 |
| ADAT2 | XM_011535439 | 0 | 8.348342131 | 0 | 0 | 0 | 0 | 0 | 0 | 1.24E-07 | 1 |
| ADCK5 | XM_011516910 | 9.740478394 | 0 | 0 | 0 | 0 | 0 | 0 | 0 | 4.01E-08 | 1 |
| ADD1 | NM_001354756 | 10.31649738 | 0 | 0 | 0 | 0 | 0 | 0 | 0 | 1.52E-08 | 1 |
| ADD1 | NM_001354757 | 0 | 8.903318262 | 0 | 0 | 0 | 0 | 0 | 0 | 7.75E-08 | 1 |
| ADD1 | NM_014189 | 0 | 0 | 0 | 0 | 0 | 0 | 9.511367848 | 9.724611366 | 1 | 0.02072721 |
| ADD1 | XR_002959706 | 0 | 9.081113725 | 0 | 10.8487973 | 9.08125658 | 0 | 0 | 0 | 0.999953634 | 0.02072721 |
| ADD2 | XM_011532502 | 0 | 8.983325733 | 0 | 0 | 0 | 0 | 0 | 0 | 7.17E-08 | 1 |
| ADGRL3 | NM_015236 | 0 | 8.844550112 | 0 | 0 | 0 | 0 | 0 | 0 | 8.06E-08 | 1 |
| ADH1C | NM_000669 | 11.47312432 | 12.95061281 | 13.73421643 | 10.02697104 | 0 | 0 | 0 | 0 | 0.999953634 | 1.37E-09 |
| ADH1C | NR_133005 | 9.553433586 | 9.754542834 | 3.436126667 | 0 | 0 | 0 | 0 | 0 | 0.044077791 | 0.999932046 |
| ADIPOR1 | XM_024447427 | 0 | 0 | 0 | 7.520758627 | 8.232953421 | 0 | 0 | 0 | 0.064584444 | 0.044792007 |
| ADIPOR2 | NM_024551 | 0 | 0 | 0 | 0 | 6.725274845 | 0 | 0 | 0 | 9.53E-05 | 1 |
| ADK | NM_001202449 | 0 | 0 | 0 | 9.907491 | 9.392867275 | 0 | 9.829916618 | 10.57600511 | 0.018050422 | 0.999932046 |
| ADK | XM_017015701 | 0 | 0 | 0 | 11.91344764 | 12.55001872 | 10.62016598 | 9.615273323 | 13.94605799 | 1.34E-08 | 0.999932046 |
| ADK | XM_017015704 | 0 | 0 | 0 | 10.58065506 | 8.977950098 | 10.6672736 | 10.24960523 | 11.76714537 | 0.014954989 | 0.999932046 |
| ADORA2B | NM_000676 | 0 | 8.360378074 | 0 | 0 | 0 | 0 | 0 | 0 | 1.21E-07 | 1 |
| ADPRHL2 | XM_011541636 | 0 | 0 | 10.23997979 | 0 | 8.165592712 | 0 | 7.704370312 | 0 | 0.020731579 | 0.999932046 |
| ADPRM | XR_243561 | 0 | 0 | 0 | 10.19787099 | 9.129190896 | 0 | 0 | 0 | 0.017209362 | 0.02072721 |
| ADRM1 | NM_175573 | 0 | 0 | 9.74641291 | 9.310616166 | 8.2973088 | 0 | 9.837250547 | 0 | 0.003264094 | 0.999932046 |
| ADSSL1 | NM_152328 | 0 | 8.801057108 | 0 | 0 | 0 | 0 | 0 | 0 | 8.24E-08 | 1 |
| ADTRP | XR_926322 | 10.82925902 | 9.277965255 | 0 | 0 | 0 | 0 | 0 | 0 | 0.000675612 | 1 |
| AFDN | XM_017010883 | 10.2867736 | 0 | 0 | 0 | 0 | 10.45621996 | 9.224463138 | 0 | 1.54E-08 | 0.02072721 |
| AFDN | XM_017010887 | 0 | 0 | 0 | 0 | 6.702411139 | 0 | 0 | 0 | 3.69E-05 | 1 |
| AFF1 | NM_001313960 | 0 | 0 | 0 | 0 | 6.30721813 | 9.60357477 | 0 | 0 | 4.99E-05 | 1 |
| AFF1 | XM_005263013 | 0 | 0 | 10.79825593 | 0 | 7.752229049 | 0 | 7.543303808 | 0 | 0.016458389 | 0.999932046 |
| AFF1 | XM_011531973 | 0 | 9.405416277 | 0 | 0 | 0 | 0 | 4.596451906 | 0 | 3.31E-08 | 0.706383321 |
| AFF3 | XM_011511175 | 0 | 11.43403091 | 0 | 0 | 0 | 0 | 0 | 0 | 1.34E-08 | 1 |
| AFF3 | XM_011511177 | 0 | 9.006487637 | 0 | 0 | 0 | 0 | 0 | 0 | 6.80E-08 | 1 |
| AFF3 | XM_024452883 | 9.217654114 | 9.868487622 | 0 | 0 | 0 | 0 | 0 | 9.429755563 | 0.001490299 | 1 |
| AFG1L | XM_011535657 | 11.33057265 | 9.527862205 | 10.86373457 | 10.78843004 | 8.765519627 | 0 | 0 | 0 | 0.999953634 | 0.001669278 |
| AFG1L | XM_017010647 | 10.62553334 | 9.411213355 | 11.56431529 | 0 | 10.29338713 | 0 | 0 | 0 | 0.999953634 | 0.02072721 |
| AFG3L2 | NM_006796 | 0 | 0 | 11.67969586 | 0 | 7.012300182 | 0 | 7.09458011 | 0 | 3.44E-08 | 0.999932046 |
| AFG3L2 | XM_011525601 | 0 | 10.44589276 | 9.434495016 | 0 | 5.908266127 | 0 | 0 | 0 | 0.999953634 | 0.02912589 |
| AFMID | NM_001010982 | 0 | 8.853093555 | 0 | 0 | 0 | 0 | 0 | 0 | 7.91E-08 | 1 |
| AFMID | XM_011524332 | 8.597469932 | 0 | 0 | 0 | 0 | 0 | 0 | 0 | 1.05E-07 | 1 |
| AFTPH | XR_244941 | 0 | 8.31161984 | 0 | 0 | 0 | 0 | 0 | 0 | 1.27E-07 | 1 |
| AGAP1 | XM_006712235 | 8.951500292 | 10.07978126 | 0 | 10.99099956 | 9.175583651 | 0 | 0 | 0 | 0.999953634 | 0.02072721 |
| AGAP1 | XM_024452676 | 0 | 0 | 9.420774911 | 0 | 6.130081192 | 0 | 0 | 0 | 0.045065368 | 0.028715875 |

| | | | | | | | | | | | |
|---|---|---|---|---|---|---|---|---|---|---|---|
| AGAP3 | XM_017011735 | 10.46796695 | 0 | 0 | 0 | 0 | 0 | 0 | 0 | 1.45E-08 | 1 |
| AGBL4 | NM_001323574 | 0 | 9.849399377 | 0 | 0 | 0 | 0 | 0 | 10.01210526 | 1.99E-08 | 1 |
| AGFG2 | XM_017012114 | 9.171928589 | 0 | 0 | 0 | 0 | 0 | 0 | 0 | 7.86E-08 | 1 |
| AGO3 | XM_017000523 | 0 | 0 | 0 | 0 | 6.366595318 | 0 | 0 | 0 | 4.76E-05 | 1 |
| AGPAT3 | XM_005261160 | 0 | 0 | 10.31699395 | 0 | 8.320016823 | 0 | 0 | 0 | 0.019455171 | 0.02072721 |
| AGPAT4 | XR_001743516 | 0 | 0 | 0 | 0 | 6.643614539 | 0 | 0 | 0 | 3.86E-05 | 1 |
| AGRN | NM_198576 | 0 | 0 | 0 | 0 | 6.582319518 | 0 | 0 | 0 | 4.04E-05 | 1 |
| AGRN | XM_005244749 | 0 | 0 | 0 | 0 | 6.758905041 | 0 | 0 | 0 | 9.53E-05 | 1 |
| AGTRAP | NM_020350 | 0 | 0 | 0 | 10.06705036 | 9.551601796 | 10.56152812 | 0 | 10.50695895 | 0.016206733 | 0.999932046 |
| AHCY | NM_000687 | 0 | 0 | 0 | 9.897999058 | 7.97366523 | 10.79693797 | 0 | 10.5177989 | 0.02494212 | 0.999932046 |
| AHCY | XM_017027708 | 0 | 0 | 0 | 10.22089417 | 8.691983703 | 0 | 0 | 0 | 0.018725083 | 0.02072721 |
| AHCYL2 | NM_015328 | 0 | 9.277965255 | 0 | 0 | 0 | 0 | 0 | 0 | 4.01E-08 | 1 |
| AHCYL2 | XM_005250237 | 14.05514731 | 13.42363683 | 14.43567508 | 11.64781618 | 7.910229599 | 0 | 0 | 0 | 0.999953634 | 0.000126798 |
| AHDC1 | XM_024446461 | 0 | 0 | 0 | 0 | 0 | 9.800824589 | 8.259993762 | 0 | 1 | 0.02072721 |
| AHI1 | NM_017651 | 0 | 0 | 0 | 9.146626846 | 9.277717614 | 0 | 0 | 0 | 0.024375168 | 0.02072721 |
| AHI1 | XR_001743486 | 0 | 0 | 0 | 9.733411602 | 7.992635714 | 0 | 7.84197419 | 0 | 0.027125273 | 0.999932046 |
| AHNAK | NM_024060 | 8.638392769 | 0 | 0 | 0 | 0 | 0 | 0 | 0 | 1.01E-07 | 1 |
| AIDA | NM_022831 | 0 | 8.818611963 | 0 | 0 | 0 | 10.96950906 | 0 | 0 | 8.12E-08 | 1 |
| AIF1 | NM_001318970 | 0 | 0 | 9.800390998 | 0 | 8.919671304 | 0 | 0 | 10.12629497 | 0.021611375 | 0.999932046 |
| AIFM1 | NM_004208 | 0 | 0 | 0 | 9.916920899 | 8.864098664 | 0 | 0 | 10.70016004 | 0.020600679 | 0.999932046 |
| AIG1 | XM_017010928 | 0 | 0 | 0 | 0 | 6.505153187 | 0 | 6.781384747 | 0 | 4.28E-05 | 1 |
| AIM2 | NM_004833 | 0 | 9.296876505 | 0 | 0 | 0 | 0 | 0 | 0 | 4.01E-08 | 1 |
| AIMP2 | XM_006715772 | 0 | 0 | 0 | 0 | 0 | 11.12730581 | 0 | 10.80343984 | 1 | 0.02072721 |
| AIMP2 | XM_011515519 | 0 | 0 | 10.71568536 | 11.82438792 | 9.608049874 | 0 | 0 | 0 | 0.000391783 | 0.000940769 |
| AIP | NM_003977 | 0 | 0 | 9.403439234 | 0 | 8.287740609 | 0 | 8.172825027 | 0 | 0.030452016 | 0.999932046 |
| AK1 | NM_001318122 | 0 | 0 | 0 | 10.04284953 | 7.24369152 | 0 | 0 | 0 | 0.025746385 | 0.02072721 |
| AK2 | NR_134976 | 10.57020707 | 0 | 0 | 0 | 0 | 0 | 0 | 0 | 1.34E-08 | 1 |
| AK2 | XR_001737036 | 0 | 0 | 0 | 10.01091583 | 9.181795906 | 0 | 0 | 0 | 0.018373202 | 0.02072721 |
| AK4 | NM_203464 | 0 | 0 | 10.35175448 | 9.932501706 | 6.42362501 | 0 | 0 | 0 | 0.002689497 | 0.004096053 |
| AK6 | NM_016283 | 10.29958742 | 11.12481206 | 0 | 10.29869948 | 9.578533242 | 0 | 0 | 0 | 0.999953634 | 0.02072721 |
| AKAP1 | NM_003488 | 0 | 0 | 10.65764653 | 0 | 9.383914218 | 12.26562681 | 9.765064406 | 0 | 0.013317244 | 0.999932046 |
| AKAP1 | XM_024450960 | 0 | 0 | 9.757371304 | 9.485579489 | 9.337368818 | 0 | 0 | 0 | 0.002104881 | 0.003193095 |
| AKAP11 | XM_017020383 | 0 | 0 | 0 | 8.967640006 | 6.708161125 | 0 | 0 | 0 | 0.054045385 | 0.037047176 |
| AKAP8 | XR_001753582 | 9.79579164 | 0 | 0 | 0 | 0 | 0 | 6.781384747 | 0 | 2.47E-08 | 1 |
| AKAP8L | NM_001291478 | 0 | 9.138154184 | 0 | 0 | 0 | 0 | 0 | 0 | 5.16E-08 | 1 |
| AKAP8L | XR_002958295 | 0 | 0 | 0 | 0 | 6.59478863 | 0 | 0 | 0 | 4.02E-05 | 1 |
| AKAP9 | XM_017011658 | 11.59420723 | 0 | 0 | 0 | 0 | 0 | 0 | 0 | 1.34E-08 | 1 |
| AKR1A1 | NM_153326 | 0 | 0 | 11.2921433 | 0 | 10.22032751 | 11.13463829 | 10.50001952 | 10.54719553 | 0.00770263 | 0.999932046 |
| AKR1A1 | XM_011540492 | 0 | 0 | 0 | 0 | 0 | 10.91661263 | 9.149853398 | 0 | 1 | 0.02072721 |
| AKR1B10 | XR_927491 | 0 | 0 | 0 | 0 | 6.30721813 | 0 | 0 | 0 | 4.99E-05 | 1 |
| AKR7A3 | NM_012067 | 11.49371403 | 10.16459397 | 0 | 11.52163957 | 7.97366523 | 0 | 0 | 0 | 0.999953634 | 7.83E-09 |
| AKT1 | NM_001014432 | 0 | 0 | 9.753271654 | 9.357540824 | 9.220530915 | 0 | 0 | 11.36054924 | 0.00232869 | 1 |
| AKT3 | NM_001206729 | 0 | 0 | 0 | 10.3702361 | 8.674468046 | 0 | 0 | 11.02286619 | 0.017534783 | 0.999932046 |

| | | | | | | | | | | | |
|---|---|---|---|---|---|---|---|---|---|---|---|
| AKT3 | XM_024446892 | 11.17237551 | 0 | 0 | 0 | 0 | 0 | 0 | 0 | 1.34E-08 | 1 |
| AKTIP | NM_022476 | 0 | 0 | 0 | 0 | 6.725274845 | 0 | 0 | 0 | 9.53E-05 | 1 |
| AKTIP | XM_017023564 | 0 | 0 | 0 | 0 | 6.690841957 | 0 | 0 | 0 | 3.72E-05 | 1 |
| ALB | NM_000477 | 0 | 9.830055185 | 0 | 0 | 0 | 0 | 7.479117364 | 0 | 2.01E-08 | 1 |
| ALDH18A1 | NM_002860 | 0 | 0 | 0 | 0 | 0 | 0 | 7.743928512 | 9.377153042 | 1 | 0.02072721 |
| ALDH2 | NM_000690 | 0 | 0 | 10.96173892 | 13.02430717 | 7.555764066 | 11.04732982 | 10.66365905 | 11.21827551 | 0.000252816 | 0.999932046 |
| ALDH9A1 | NM_000696 | 0 | 0 | 10.55618885 | 0 | 8.153000089 | 0 | 7.27822739 | 0 | 0.017771359 | 0.999932046 |
| ALDOA | NM_001355562 | 0 | 0 | 10.4274613 | 0 | 11.6876554 | 10.93632715 | 8.032920857 | 11.62211152 | 0.005678281 | 0.999932046 |
| ALG13 | NR_136735 | 9.025355161 | 0 | 0 | 0 | 0 | 0 | 0 | 0 | 7.69E-08 | 1 |
| ALG13 | XM_011531031 | 10.19374119 | 0 | 0 | 0 | 0 | 0 | 0 | 0 | 1.60E-08 | 1 |
| ALG13 | XM_017029855 | 9.190393069 | 7.795449567 | 0 | 0 | 0 | 0 | 0 | 0 | 0.005416432 | 1 |
| ALG3 | XM_024453296 | 0 | 0 | 0 | 6.698604544 | 0 | 0 | 0 | 0 | 3.86E-05 | 1 |
| ALG8 | XR_950044 | 0 | 9.660328297 | 0 | 0 | 0 | 0 | 0 | 0 | 2.46E-08 | 1 |
| ALG9 | NM_001077690 | 0 | 9.527862205 | 0 | 0 | 0 | 0 | 0 | 0 | 2.90E-08 | 1 |
| ALK | XR_001738688 | 8.526574932 | 0 | 0 | 0 | 0 | 0 | 0 | 0 | 1.12E-07 | 1 |
| ALKBH2 | XM_005253835 | 0 | 0 | 6.832547844 | 0 | 0 | 0 | 0 | 0 | 3.72E-05 | 1 |
| ALKBH5 | NM_017758 | 0 | 0 | 8.35596428 | 10.21580957 | 8.811650533 | 0 | 8.149448007 | 0 | 0.002892298 | 0.999932046 |
| ALOX15 | NM_001140 | 0 | 0 | 0 | 0 | 6.518304288 | 0 | 0 | 0 | 4.23E-05 | 1 |
| ALOX5AP | NM_001629 | 0 | 0 | 0 | 10.06563796 | 7.536502354 | 0 | 0 | 10.49603692 | 0.02446904 | 0.999932046 |
| ALS2CL | NR_135622 | 0 | 0 | 0 | 0 | 6.505153187 | 0 | 0 | 0 | 4.28E-05 | 1 |
| ALS2CL | XR_001740097 | 0 | 8.983325733 | 0 | 0 | 0 | 0 | 0 | 0 | 7.17E-08 | 1 |
| AMD1 | NM_001634 | 0 | 8.519117955 | 0 | 9.523110922 | 9.04309389 | 0 | 0 | 0 | 0.999953634 | 0.02072721 |
| AMIGO2 | XM_005268837 | 0 | 0 | 10.96705877 | 9.416346541 | 7.162254726 | 0 | 0 | 0 | 0.001978261 | 0.003067596 |
| AMN | XM_024449714 | 10.26950953 | 9.862152894 | 9.248957794 | 0 | 8.28965933 | 0 | 0 | 0 | 0.977787243 | 0.022064246 |
| AMN1 | NR_103523 | 0 | 8.407542174 | 0 | 0 | 0 | 0 | 0 | 0 | 1.16E-07 | 1 |
| AMN1 | XM_006719054 | 0 | 8.959785899 | 0 | 0 | 0 | 0 | 0 | 0 | 7.69E-08 | 1 |
| AMN1 | XM_017018965 | 0 | 9.495485143 | 0 | 0 | 0 | 0 | 0 | 0 | 2.95E-08 | 1 |
| AMOTL1 | XM_011542626 | 0 | 0 | 10.99510467 | 0 | 9.609586778 | 0 | 0 | 0 | 0.010630445 | 0.02072721 |
| AMT | NM_001164712 | 0 | 9.124104101 | 0 | 0 | 0 | 0 | 0 | 0 | 5.29E-08 | 1 |
| AMZ2 | NR_144442 | 10.53451875 | 0 | 0 | 0 | 0 | 0 | 0 | 0 | 1.36E-08 | 1 |
| ANAPC1 | NM_022662 | 0 | 0 | 0 | 9.667902088 | 8.893414856 | 0 | 0 | 0 | 0.022674865 | 0.02072721 |
| ANAPC11 | NM_001289415 | 0 | 9.517150383 | 9.88276327 | 0 | 8.278108536 | 0 | 0 | 0 | 0.999953634 | 0.02072721 |
| ANAPC11 | NM_001289418 | 9.616239393 | 0 | 0 | 0 | 0 | 0 | 0 | 0 | 2.91E-08 | 1 |
| ANAPC11 | NM_016476 | 10.76449512 | 9.759095847 | 0 | 0 | 0 | 0 | 0 | 0 | 0.000568271 | 1 |
| ANAPC13 | NM_001242374 | 0 | 0 | 0 | 10.03565369 | 10.00014679 | 0 | 0 | 10.43444146 | 0.014191273 | 1 |
| ANAPC15 | NM_001278485 | 0 | 9.333971188 | 0 | 0 | 0 | 0 | 0 | 0 | 3.79E-08 | 1 |
| ANAPC15 | NM_014042 | 11.08131887 | 0 | 10.79160823 | 10.46327738 | 8.442351687 | 0 | 0 | 0 | 0.999953634 | 0.002089223 |
| ANAPC2 | XR_929779 | 10.63567489 | 0 | 0 | 0 | 0 | 0 | 0 | 0 | 2.23E-08 | 1 |
| ANAPC5 | NM_001330489 | 0 | 0 | 10.59793683 | 10.10741968 | 8.153000089 | 0 | 7.940604293 | 0 | 0.001533648 | 0.999932046 |
| ANAPC5 | NM_016237 | 9.777588513 | 0 | 0 | 10.47821718 | 8.614497302 | 0 | 0 | 0 | 0.999953634 | 0.021157255 |
| ANAPC7 | XM_017019428 | 0 | 9.399595812 | 0 | 0 | 0 | 0 | 0 | 0 | 3.36E-08 | 1 |
| ANGEL1 | NM_015305 | 9.813767948 | 0 | 0 | 0 | 0 | 0 | 0 | 0 | 2.49E-08 | 1 |
| ANGEL2 | XR_001737532 | 0 | 0 | 0 | 0 | 6.30721813 | 0 | 0 | 0 | 4.99E-05 | 1 |

| | | | | | | | | | | | |
|---|---|---|---|---|---|---|---|---|---|---|---|
| ANGPTL4 | NM_001039667 | 0 | 0 | 0 | 10.17185123 | 8.8089774 | 0 | 0 | 0 | 0.018697628 | 0.02072721 |
| ANK3 | NM_001320874 | 0 | 0 | 10.29554044 | 0 | 8.772396791 | 0 | 0 | 0 | 0.01808998 | 0.02072721 |
| ANK3 | NM_020987 | 9.296476405 | 6.05590339 | 0 | 0 | 0 | 0 | 0 | 0 | 0.00832905 | 1 |
| ANK3 | XM_005269715 | 0 | 0 | 11.1371659 | 9.637803806 | 8.320016823 | 0 | 0 | 0 | 0.001291781 | 0.002366752 |
| ANKH | NM_054027 | 0 | 0 | 8.100280313 | 0 | 0 | 0 | 0 | 0 | 1.03E-05 | 1 |
| ANKHD1 | NM_024668 | 0 | 9.772669303 | 0 | 0 | 0 | 0 | 0 | 0 | 4.12E-08 | 1 |
| ANKHD1-EIF4EBP3 | NM_020690 | 0 | 8.360378074 | 0 | 0 | 0 | 0 | 7.812509721 | 0 | 1.21E-07 | 1 |
| ANKRD10 | NR_104587 | 0 | 0 | 0 | 10.60613892 | 9.734579646 | 0 | 7.234460372 | 0 | 0.012278029 | 0.999932046 |
| ANKRD10 | XM_024449382 | 0 | 0 | 0 | 9.862651006 | 7.864619359 | 0 | 0 | 0 | 0.025919438 | 0.02072721 |
| ANKRD10 | XR_002957462 | 0 | 0 | 0 | 9.79947801 | 9.181795906 | 0 | 0 | 0 | 0.019914518 | 0.02072721 |
| ANKRD10 | XR_002957463 | 0 | 0 | 9.665827761 | 0 | 10.52765603 | 11.39695608 | 0 | 0 | 0.012767294 | 0.999932046 |
| ANKRD10 | XR_002957465 | 0 | 0 | 9.256701524 | 7.326879247 | 9.528226765 | 11.22328188 | 0 | 10.48226708 | 0.004707012 | 0.999932046 |
| ANKRD11 | NM_001256183 | 0 | 9.905922754 | 0 | 0 | 0 | 0 | 0 | 0 | 1.87E-08 | 1 |
| ANKRD11 | NR_045839 | 0 | 8.809861237 | 0 | 0 | 0 | 0 | 0 | 0 | 8.24E-08 | 1 |
| ANKRD13A | NM_033121 | 0 | 0 | 0 | 0 | 6.725274845 | 0 | 0 | 0 | 9.53E-05 | 1 |
| ANKRD17 | XM_017008017 | 8.994164854 | 0 | 0 | 0 | 0 | 0 | 0 | 0 | 7.73E-08 | 1 |
| ANKRD22 | NM_144590 | 0 | 0 | 0 | 11.05289752 | 9.624866434 | 8.736503407 | 9.386954008 | 6.679713864 | 0.010158825 | 0.999932046 |
| ANKRD27 | XM_017027349 | 0 | 9.232850423 | 0 | 0 | 0 | 0 | 6.533064721 | 0 | 4.38E-08 | 1 |
| ANKRD36 | XR_001738741 | 9.015033136 | 0 | 0 | 0 | 0 | 0 | 0 | 0 | 7.69E-08 | 1 |
| ANKRD36B | XR_002959321 | 0 | 0 | 0 | 8.162444728 | 0 | 0 | 0 | 0 | 8.86E-06 | 1 |
| ANKRD40 | XM_017025296 | 8.803424112 | 0 | 0 | 0 | 0 | 0 | 2.992327696 | 0 | 8.89E-08 | 0.999932046 |
| ANKRD46 | NM_198401 | 0 | 8.943876548 | 0 | 0 | 0 | 0 | 0 | 0 | 7.69E-08 | 1 |
| ANKRD52 | XM_017019184 | 0 | 0 | 0 | 0 | 6.30721813 | 0 | 0 | 0 | 4.99E-05 | 1 |
| ANKRD55 | XM_017009853 | 0 | 9.759095847 | 0 | 0 | 0 | 9.662253352 | 0 | 0 | 2.12E-08 | 1 |
| ANKS1B | NM_001204080 | 10.88841554 | 0 | 0 | 0 | 0 | 0 | 0 | 0 | 1.34E-08 | 1 |
| ANKS1B | NM_181670 | 8.583566901 | 0 | 0 | 0 | 0 | 0 | 0 | 0 | 1.06E-07 | 1 |
| ANKZF1 | XR_922953 | 0 | 9.855790228 | 0 | 0 | 0 | 0 | 0 | 0 | 1.98E-08 | 1 |
| ANO10 | NM_018075 | 0 | 0 | 11.8878632 | 10.42192887 | 0 | 0 | 0 | 0 | 0.005954954 | 0.02072721 |
| ANTXR2 | NM_058172 | 9.771469405 | 0 | 0 | 0 | 0 | 0 | 7.720323987 | 0 | 2.47E-08 | 1 |
| ANXA11 | NM_145868 | 0 | 9.830055185 | 0 | 0 | 0 | 0 | 9.126098068 | 11.45733604 | 2.01E-08 | 0.02072721 |
| ANXA2R | NM_001014279 | 0 | 0 | 0 | 9.607064194 | 9.401765114 | 0 | 0 | 0 | 0.020013101 | 0.02072721 |
| ANXA7 | NM_004034 | 0 | 8.728620286 | 0 | 0 | 0 | 0 | 7.58746354 | 0 | 8.80E-08 | 1 |
| AOAH | NM_001177507 | 0 | 9.595615072 | 0 | 0 | 0 | 0 | 0 | 0 | 2.59E-08 | 1 |
| AOAH | XM_011515335 | 0 | 0 | 7.576866482 | 0 | 0 | 0 | 9.146905285 | 0 | 3.23E-05 | 1 |
| AOAH | XM_011515341 | 0 | 0 | 7.487554998 | 10.70923559 | 8.274237629 | 9.459194651 | 9.843637293 | 10.54719553 | 0.002970093 | 0.999932046 |
| AP1AR | XM_017008380 | 0 | 6.275194576 | 0 | 0 | 0 | 9.016014309 | 6.85548998 | 0 | 0.149373223 | 0.026465236 |
| AP1G2 | XR_001750584 | 0 | 0 | 0 | 9.26211379 | 7.187172039 | 0 | 0 | 0 | 0.041269221 | 0.028702352 |
| AP1M1 | NM_032493 | 0 | 0 | 10.09417272 | 0 | 7.987916427 | 0 | 0 | 0 | 0.022985197 | 0.02072721 |
| AP2A2 | XM_011519929 | 9.95545054 | 0 | 0 | 0 | 0 | 0 | 0 | 9.005822227 | 2.01E-08 | 1 |
| AP2B1 | NM_001030006 | 0 | 0 | 0 | 11.43349902 | 8.323766986 | 10.98175813 | 9.365556577 | 10.18187502 | 0.010311917 | 0.999932046 |
| AP2B1 | NM_001282 | 0 | 0 | 9.539797346 | 10.50555865 | 11.17526887 | 11.82664884 | 0 | 9.729282335 | 0.000625761 | 0.999932046 |
| AP2B1 | XM_011524450 | 0 | 9.290600259 | 0 | 0 | 0 | 0 | 0 | 0 | 4.07E-08 | 1 |
| AP2S1 | XM_011526424 | 0 | 0 | 0 | 9.0384682 | 8.258649294 | 0 | 0 | 0 | 0.035901725 | 0.025004616 |

| | | | | | | | | | | |
|---|---|---|---|---|---|---|---|---|---|---|
| AP3D1 | NM_003938 | 0 | 0 | 0 | 9.754603346 | 7.401067144 | 0 | 0 | 0 | 0.029708212 | 0.021805825 |
| AP3S2 | NR_037582 | 10.70794476 | 0 | 9.468235064 | 10.15335408 | 0 | 0 | 0 | 0 | 0.999953634 | 0.02096331 |
| AP4B1 | NM_006594 | 0 | 9.081113725 | 0 | 0 | 0 | 0 | 0 | 0 | 5.70E-08 | 1 |
| AP4B1 | XR_002958807 | 9.676424716 | 0 | 0 | 0 | 0 | 0 | 0 | 0 | 2.73E-08 | 1 |
| AP4M1 | NM_004722 | 0 | 9.102769039 | 0 | 10.62735608 | 7.075944552 | 0 | 0 | 0 | 0.999953634 | 0.02072721 |
| AP5M1 | XM_011536940 | 9.162606966 | 7.014878551 | 8.200039322 | 0 | 8.114548983 | 0 | 0 | 0 | 0.999953634 | 0.035282533 |
| APBA3 | XM_006722950 | 10.46037918 | 0 | 0 | 0 | 0 | 0 | 0 | 0 | 1.45E-08 | 1 |
| APC | NM_001354905 | 0 | 9.416987232 | 0 | 0 | 0 | 0 | 0 | 0 | 3.28E-08 | 1 |
| APCDD1 | NM_153000 | 0 | 8.903318262 | 0 | 0 | 0 | 10.06921268 | 11.17713992 | 0 | 7.75E-08 | 0.02072721 |
| APH1A | NM_001243771 | 0 | 0 | 0 | 9.551641627 | 7.752229049 | 0 | 8.441777415 | 0 | 0.031303306 | 0.999932046 |
| APH1A | NR_045033 | 0 | 0 | 0 | 7.124706719 | 0 | 0 | 0 | 0 | 2.57E-05 | 1 |
| APH1A | NR_045035 | 0 | 0 | 9.474889408 | 0 | 5.597297131 | 0 | 0 | 0 | 0.045943966 | 0.029253745 |
| APLP2 | NM_001142276 | 0 | 0 | 14.3876643 | 13.33437334 | 10.35790399 | 14.83860715 | 11.83423572 | 12.56128364 | 2.15E-05 | 0.999932046 |
| APLP2 | NM_001328682 | 0 | 0 | 12.478712 | 11.09296612 | 7.587306227 | 11.04408862 | 10.75603587 | 10.38940937 | 0.00036816 | 0.999932046 |
| APLP2 | NM_001328684 | 0 | 0 | 0 | 10.74143689 | 7.964085617 | 0 | 0 | 10.8276001 | 0.016182345 | 0.999932046 |
| APOA2 | NM_001643 | 0 | 0 | 0 | 10.54865946 | 10.20414231 | 0 | 9.844547382 | 0 | 0.01082976 | 0.999932046 |
| APOBEC1 | NM_005889 | 0 | 0 | 7.729351051 | 0 | 9.631689835 | 11.13341879 | 0 | 11.5159558 | 0.029252715 | 0.999932046 |
| APOBEC3F | XR_002958675 | 0 | 0 | 0 | 0 | 6.544252221 | 0 | 0 | 0 | 4.20E-05 | 1 |
| APOC1 | NM_001321066 | 10.89124009 | 0 | 0 | 0 | 0 | 0 | 0 | 0 | 1.34E-08 | 1 |
| APOL1 | NM_001136541 | 9.643298854 | 8.168359982 | 0 | 10.37480538 | 7.40814953 | 0 | 0 | 0 | 0.999953634 | 0.02072721 |
| APOL2 | NM_145637 | 10.16602012 | 0 | 0 | 0 | 0 | 0 | 0 | 0 | 1.60E-08 | 1 |
| APOL3 | XM_024452276 | 0 | 9.490017613 | 0 | 0 | 0 | 0 | 0 | 0 | 3.40E-08 | 1 |
| APOL4 | NM_145660 | 9.944614124 | 0 | 0 | 0 | 0 | 10.16064276 | 10.81850969 | 0 | 2.01E-08 | 0.02072721 |
| APP | NM_001136130 | 0 | 8.728620286 | 0 | 0 | 0 | 0 | 8.195829284 | 0 | 8.80E-08 | 1 |
| APP | NM_201413 | 0 | 0 | 0 | 10.23855048 | 6.544252221 | 0 | 0 | 0 | 0.024736775 | 0.02072721 |
| AQP12A | NM_198998 | 0 | 0 | 7.751443001 | 0 | 0 | 0 | 0 | 0 | 1.49E-05 | 1 |
| AQP12B | XR_922993 | 10.20740451 | 0 | 0 | 0 | 0 | 0 | 0 | 0 | 1.60E-08 | 1 |
| AQP3 | XR_001746289 | 0 | 9.411213355 | 0 | 0 | 0 | 0 | 0 | 0 | 3.28E-08 | 1 |
| AR | NM_000044 | 10.33735985 | 0 | 0 | 0 | 0 | 0 | 7.940604293 | 0 | 1.58E-08 | 1 |
| ARAP3 | XM_011537677 | 0 | 8.497482465 | 0 | 0 | 0 | 0 | 0 | 0 | 1.06E-07 | 1 |
| ARCN1 | XM_005271542 | 0 | 0 | 0 | 10.05286379 | 8.762759555 | 0 | 0 | 0 | 0.019870902 | 0.02072721 |
| AREL1 | NM_001039479 | 0 | 9.315543066 | 0 | 0 | 0 | 0 | 0 | 0 | 3.90E-08 | 1 |
| ARFGAP1 | NR_104023 | 10.18915784 | 0 | 0 | 0 | 0 | 0 | 0 | 0 | 1.60E-08 | 1 |
| ARFGAP2 | NM_001242832 | 0 | 10.47247452 | 0 | 0 | 0 | 0 | 0 | 0 | 1.34E-08 | 1 |
| ARFGAP2 | XM_006718346 | 11.37765889 | 0 | 0 | 0 | 0 | 0 | 9.104986362 | 0 | 1.34E-08 | 1 |
| ARFGEF1 | XM_005251135 | 0 | 0 | 0 | 0 | 6.61940813 | 0 | 0 | 0 | 5.85E-05 | 1 |
| ARFIP2 | XM_005252840 | 0 | 9.689244249 | 0 | 0 | 0 | 0 | 7.543303808 | 0 | 2.34E-08 | 1 |
| ARFRP1 | NM_001267544 | 9.900435696 | 0 | 0 | 0 | 0 | 0 | 0 | 0 | 2.07E-08 | 1 |
| ARHGAP10 | XM_005263215 | 0 | 0 | 11.08429488 | 9.802872777 | 9.777598559 | 11.46736833 | 9.790789648 | 9.933328324 | 0.000823432 | 0.999932046 |
| ARHGAP10 | XM_017008602 | 9.649985154 | 0 | 0 | 0 | 0 | 0 | 9.146905285 | 0 | 2.95E-08 | 1 |
| ARHGAP12 | XM_017016954 | 0 | 9.772669303 | 0 | 0 | 0 | 0 | 0 | 9.828393739 | 4.12E-08 | 1 |
| ARHGAP15 | NM_018460 | 0 | 9.036802844 | 0 | 0 | 0 | 0 | 0 | 0 | 6.38E-08 | 1 |
| ARHGAP17 | XR_950811 | 0 | 8.33620493 | 0 | 0 | 0 | 0 | 0 | 0 | 1.25E-07 | 1 |

| | | | | | | | | | | | |
|---|---|---|---|---|---|---|---|---|---|---|---|
| ARHGAP21 | XR_930513 | 0 | 0 | 9.580473641 | 0 | 7.343120441 | 0 | 0 | 0 | 0.034088034 | 0.022619671 |
| ARHGAP24 | NM_001287805 | 0 | 8.497482465 | 0 | 0 | 0 | 0 | 0 | 0 | 1.06E-07 | 1 |
| ARHGAP24 | NM_001346093 | 9.509990722 | 0 | 0 | 0 | 0 | 0 | 8.680665279 | 0 | 3.47E-08 | 1 |
| ARHGAP25 | NM_001166276 | 0 | 0 | 12.04704062 | 0 | 7.290434628 | 0 | 0 | 0 | 1.47E-08 | 5.45E-09 |
| ARHGAP25 | NM_001166277 | 0 | 8.774317692 | 0 | 0 | 0 | 0 | 0 | 0 | 8.46E-08 | 1 |
| ARHGAP25 | NM_014882 | 10.47927431 | 0 | 9.81094843 | 0 | 8.194554972 | 0 | 0 | 0 | 0.999953634 | 0.02072721 |
| ARHGAP25 | XM_005264676 | 0 | 9.500932031 | 0 | 0 | 0 | 0 | 0 | 0 | 2.95E-08 | 1 |
| ARHGAP25 | XM_011533206 | 10.50900103 | 7.881486713 | 0 | 0 | 0 | 0 | 0 | 0 | 0.001490299 | 1 |
| ARHGAP26 | NM_001135608 | 0 | 0 | 10.72199233 | 0 | 7.660307374 | 0 | 0 | 10.07216922 | 0.017370381 | 0.999932046 |
| ARHGAP26 | NR_146198 | 10.821876 | 0 | 0 | 0 | 0 | 0 | 0 | 0 | 1.34E-08 | 1 |
| ARHGAP26 | XM_017009248 | 0 | 10.3051001 | 0 | 0 | 0 | 0 | 0 | 0 | 1.47E-08 | 1 |
| ARHGAP27 | XR_934417 | 8.511968076 | 0 | 0 | 0 | 0 | 0 | 8.321425601 | 8.735631445 | 1.14E-07 | 0.021870855 |
| ARHGAP31 | XM_017006955 | 11.49927878 | 0 | 0 | 0 | 0 | 0 | 0 | 0 | 1.34E-08 | 1 |
| ARHGAP4 | NM_001666 | 9.46519897 | 0 | 9.022425311 | 0 | 8.864098664 | 0 | 0 | 0 | 0.999953634 | 0.020942118 |
| ARHGAP45 | XM_024451435 | 9.960838376 | 0 | 0 | 0 | 0 | 10.05640063 | 4.525586624 | 0 | 2.01E-08 | 1 |
| ARHGAP6 | NR_109776 | 0 | 0 | 9.795083168 | 0 | 7.724127604 | 0 | 9.123100959 | 3.8795549 | 0.028179353 | 0.999932046 |
| ARHGAP8 | NM_181335 | 0 | 10.25762482 | 0 | 0 | 0 | 0 | 8.268156521 | 0 | 1.47E-08 | 1 |
| ARHGEF10L | NM_001328124 | 0 | 0 | 10.70227943 | 0 | 7.806841281 | 0 | 0 | 8.893783085 | 0.017233205 | 0.999932046 |
| ARHGEF10L | XM_006710728 | 0 | 8.582143237 | 0 | 0 | 0 | 0 | 0 | 0 | 9.91E-08 | 1 |
| ARHGEF10L | XM_017001620 | 0 | 0 | 0 | 8.378061333 | 0 | 0 | 0 | 0 | 2.07E-05 | 1 |
| ARHGEF10L | XM_017001622 | 9.663265502 | 0 | 0 | 0 | 0 | 9.607093313 | 0 | 0 | 2.79E-08 | 1 |
| ARHGEF10L | XR_946690 | 11.49557133 | 0 | 0 | 0 | 0 | 11.05443495 | 0 | 0 | 1.34E-08 | 1 |
| ARHGEF16 | NM_014448 | 0 | 8.662110092 | 9.967779298 | 8.82800322 | 0 | 0 | 0 | 0 | 0.999953634 | 0.02072721 |
| ARHGEF2 | XM_024450823 | 9.143781062 | 0 | 0 | 0 | 0 | 0 | 0 | 0 | 7.26E-08 | 1 |
| ARHGEF37 | XM_017009461 | 10.29958742 | 0 | 0 | 0 | 0 | 0 | 0 | 0 | 1.54E-08 | 1 |
| ARHGEF40 | XM_017021437 | 0 | 0 | 0 | 0 | 6.518304288 | 0 | 7.892131936 | 0 | 4.23E-05 | 1 |
| ARHGEF7 | NM_001320851 | 0 | 0 | 0 | 11.09296612 | 7.636381755 | 0 | 8.032920857 | 0 | 0.013785183 | 0.999932046 |
| ARHGEF7 | NM_001354053 | 0 | 0 | 10.37950427 | 10.15069217 | 9.008561607 | 0 | 9.346409285 | 0 | 0.001335097 | 0.999932046 |
| ARHGEF7 | NM_145735 | 0 | 9.321711965 | 0 | 0 | 0 | 0 | 0 | 0 | 3.84E-08 | 1 |
| ARHGEF7 | XR_001749712 | 0 | 0 | 10.35356099 | 0 | 7.536502354 | 0 | 0 | 0 | 0.021438448 | 0.02072721 |
| ARHGEF9 | NM_001330495 | 0 | 8.571828374 | 0 | 0 | 0 | 0 | 0 | 0 | 9.97E-08 | 1 |
| ARHGEF9 | NM_001353923 | 0 | 0 | 10.66493943 | 10.49720094 | 0 | 0 | 8.095422193 | 0 | 0.010382649 | 0.999932046 |
| ARHGEF9 | NM_001353927 | 0 | 0 | 0 | 9.083826095 | 8.707855025 | 0 | 0 | 0 | 0.030452016 | 0.021934047 |
| ARHGEF9 | XM_005262250 | 0 | 0 | 10.41626903 | 0 | 8.907227812 | 0 | 0 | 0 | 0.016626721 | 0.020808337 |
| ARID1B | XM_017011103 | 0 | 0 | 0 | 11.52266935 | 9.038537064 | 0 | 9.058679075 | 0 | 0.008746237 | 0.999932046 |
| ARID2 | NM_001347839 | 0 | 0 | 0 | 0 | 0 | 10.03304826 | 6.043023068 | 0 | 1 | 0.02072721 |
| ARID4B | XR_002956802 | 0 | 0 | 0 | 0 | 6.451316153 | 0 | 3.654887771 | 0 | 4.51E-05 | 1 |
| ARID5A | NM_001319096 | 0 | 0 | 10.03686385 | 0 | 8.293489135 | 0 | 0 | 0 | 0.022381568 | 0.02072721 |
| ARIH2 | XM_024453311 | 0 | 9.422738093 | 0 | 0 | 0 | 0 | 0 | 0 | 3.26E-08 | 1 |
| ARL1 | NM_001301068 | 8.729593473 | 8.261155927 | 0 | 0 | 0 | 0 | 0 | 0 | 0.00625909 | 1 |
| ARL15 | XM_017009598 | 9.304974408 | 0 | 0 | 0 | 0 | 0 | 0 | 0 | 4.80E-08 | 1 |
| ARL17B | NM_001039083 | 0 | 0 | 10.79294022 | 9.993194855 | 0 | 0 | 0 | 0 | 0.011637903 | 0.02072721 |
| ARL2BP | NM_012106 | 8.569528585 | 0 | 0 | 10.95025027 | 6.3222931 | 0 | 0 | 0 | 0.998900485 | 1.68E-08 |

| | | | | | | | | | | | |
|---|---|---|---|---|---|---|---|---|---|---|---|
| ARL5A | NM_012097 | 0 | 0 | 10.57489281 | 9.618668438 | 9.017851335 | 0 | 0 | 0 | 0.001497522 | 0.002589366 |
| ARL6 | NR_136601 | 0 | 0 | 0 | 0 | 6.544252221 | 0 | 0 | 0 | 4.20E-05 | 1 |
| ARL6IP4 | NM_001002251 | 0 | 10.3630761 | 10.17403807 | 11.43130726 | 8.999211672 | 0 | 0 | 0 | 0.999953634 | 0.001535028 |
| ARL6IP4 | NM_001278379 | 0 | 8.453213086 | 0 | 0 | 0 | 0 | 0 | 0 | 1.11E-07 | 1 |
| ARL6IP4 | NR_103512 | 9.567628621 | 0 | 0 | 0 | 0 | 0 | 0 | 0 | 3.16E-08 | 1 |
| ARL6IP6 | NM_152522 | 9.46519897 | 0 | 9.784408563 | 10.45898029 | 8.60528274 | 0 | 0 | 0 | 0.999953634 | 0.002835114 |
| ARMC1 | XM_017013605 | 0 | 8.983325733 | 0 | 0 | 0 | 0 | 0 | 0 | 7.17E-08 | 1 |
| ARMC10 | XM_017012687 | 0 | 0 | 0 | 9.481348416 | 8.904726182 | 0 | 0 | 0 | 0.024405981 | 0.02072721 |
| ARMC7 | XR_002958068 | 0 | 0 | 0 | 10.14267682 | 8.136037099 | 0 | 0 | 0 | 0.021472144 | 0.02072721 |
| ARMC8 | XM_006713565 | 0 | 0 | 12.22927919 | 9.792664417 | 7.075944552 | 6.526460115 | 10.3733185 | 11.27448425 | 0.000743273 | 0.999932046 |
| ARMC8 | XM_017006098 | 10.27384495 | 0 | 0 | 0 | 0 | 0 | 0 | 9.667347319 | 1.81E-08 | 1 |
| ARMC8 | XM_017006099 | 9.124706245 | 0 | 0 | 0 | 0 | 0 | 0 | 0 | 6.80E-08 | 1 |
| ARMCX2 | XM_024452484 | 0 | 0 | 0 | 8.483365391 | 0 | 0 | 0 | 0 | 2.83E-05 | 1 |
| ARMCX3 | XM_005262141 | 0 | 0 | 9.058313897 | 9.438340856 | 0 | 0 | 6.953230848 | 0 | 0.025733678 | 0.999932046 |
| ARMCX5 | NM_001168485 | 0 | 8.464408432 | 0 | 0 | 0 | 0 | 0 | 0 | 1.11E-07 | 1 |
| ARNT | NM_001668 | 0 | 0 | 9.917373151 | 9.695560103 | 11.23224748 | 0 | 9.296734012 | 0 | 0.000723177 | 0.999932046 |
| ARNT | XM_005245157 | 0 | 0 | 0 | 9.127801312 | 7.729791854 | 0 | 0 | 0 | 0.039670524 | 0.02750692 |
| ARNT | XM_011509546 | 8.873659666 | 0 | 0 | 0 | 0 | 0 | 0 | 0 | 8.39E-08 | 1 |
| ARNTL | NM_001030273 | 0 | 0 | 0 | 0 | 0 | 11.4750909 | 0 | 10.77209341 | 1 | 0.019987819 |
| ARNTL | NM_001178 | 0 | 0 | 11.33437758 | 9.32956957 | 0 | 10.71859535 | 9.199070987 | 0 | 0.010408209 | 0.999932046 |
| ARNTL | NM_001297719 | 0 | 0 | 9.946391943 | 9.872377593 | 9.277717614 | 9.779154025 | 8.849746381 | 9.815272851 | 0.001689179 | 0.999932046 |
| ARNTL | NM_001351804 | 9.502621339 | 0 | 8.522592574 | 0 | 7.267252374 | 0 | 0 | 0 | 0.999953634 | 0.040276199 |
| ARNTL | NM_001351807 | 0 | 0 | 0 | 9.999126067 | 8.659707446 | 0 | 0 | 8.885437246 | 0.020792098 | 0.999932046 |
| ARNTL | NM_001351808 | 9.449953927 | 0 | 0 | 0 | 0 | 0 | 0 | 0 | 3.80E-08 | 1 |
| ARNTL | NM_001351814 | 0 | 0 | 9.886511435 | 8.937070225 | 0 | 0 | 0 | 0 | 0.022393848 | 0.02072721 |
| ARNTL | NM_001351823 | 0 | 8.861586703 | 0 | 0 | 0 | 0 | 0 | 0 | 7.86E-08 | 1 |
| ARNTL | NM_001351824 | 11.35123402 | 0 | 0 | 0 | 0 | 0 | 9.4516967 | 0 | 1.34E-08 | 1 |
| ARNTL | NR_147788 | 9.911607777 | 0 | 0 | 0 | 0 | 0 | 0 | 10.47672214 | 2.05E-08 | 1 |
| ARNTL | XM_006718234 | 0 | 8.612652582 | 0 | 0 | 0 | 0 | 0 | 0 | 9.67E-08 | 1 |
| ARNTL | XM_017017744 | 0 | 9.864267562 | 0 | 0 | 0 | 0 | 0 | 0 | 1.96E-08 | 1 |
| ARNTL | XR_001747876 | 0 | 8.78328601 | 0 | 0 | 0 | 0 | 8.251784555 | 0 | 8.40E-08 | 1 |
| ARPC2 | XM_017003114 | 0 | 0 | 0 | 0 | 0 | 0 | 7.720323987 | 9.441189048 | 1 | 0.02072721 |
| ARPC2 | XM_024452509 | 0 | 0 | 10.39712818 | 9.985746385 | 0 | 0 | 9.961582686 | 0 | 0.013762639 | 0.999932046 |
| ARPC5L | XM_011519067 | 0 | 0 | 0 | 0 | 6.758905041 | 9.455295751 | 7.038418754 | 10.23539302 | 9.53E-05 | 0.868924005 |
| ARPP19 | NM_006628 | 0 | 9.872695374 | 0 | 10.81892939 | 8.917191177 | 0 | 0 | 0 | 0.999953634 | 0.02072721 |
| ARRDC2 | NM_015683 | 9.321821705 | 9.786116244 | 7.33498256 | 9.726277944 | 6.409577539 | 0 | 0 | 0 | 0.999953634 | 0.015827107 |
| ARRDC3-AS1 | NR_027435 | 0 | 9.763634537 | 0 | 0 | 0 | 0 | 0 | 0 | 2.11E-08 | 1 |
| ARSA | XM_011530690 | 10.23434896 | 7.045019777 | 0 | 0 | 0 | 0 | 0 | 9.921131806 | 0.00244652 | 1 |
| ARSE | NM_001282631 | 0 | 8.765293275 | 0 | 0 | 0 | 0 | 0 | 0 | 8.55E-08 | 1 |
| ARV1 | XR_002957381 | 0 | 0 | 0 | 0 | 6.381065478 | 0 | 0 | 0 | 4.71E-05 | 1 |
| ARVCF | XM_006724248 | 0 | 0 | 0 | 0 | 6.667421491 | 0 | 0 | 0 | 3.78E-05 | 1 |
| AS3MT | NM_020682 | 0 | 0 | 0 | 11.75554179 | 11.44087808 | 0 | 11.19722887 | 10.24194531 | 0.004030609 | 0.999932046 |
| ASB3 | NM_016115 | 0 | 0 | 8.833416502 | 10.13730841 | 0 | 0 | 0 | 0 | 0.020212395 | 0.02072721 |

| | | | | | | | | | | | |
|---|---|---|---|---|---|---|---|---|---|---|---|
| ASB9 | XM_017029285 | 0 | 9.615837767 | 0 | 0 | 0 | 0 | 0 | 0 | 2.54E-08 | 1 |
| ASCC1 | XM_017016291 | 0 | 0 | 0 | 9.756355353 | 9.712579392 | 0 | 10.47320203 | 0 | 0.017233205 | 1 |
| ASCC3 | NM_022091 | 0 | 0 | 0 | 0 | 6.702411139 | 0 | 6.67079425 | 0 | 3.69E-05 | 1 |
| ASCL2 | NM_005170 | 0 | 0 | 0 | 12.60520156 | 10.51007146 | 10.00664998 | 12.98497863 | 9.48050741 | 1.34E-08 | 0.999932046 |
| ASH1L | XM_017001788 | 0 | 8.681427931 | 0 | 0 | 0 | 0 | 0 | 8.639649521 | 9.11E-08 | 1 |
| ASH1L-AS1 | NR_147963 | 0 | 0 | 9.948784034 | 0 | 7.350492346 | 0 | 0 | 0 | 0.027469095 | 0.02072721 |
| ASIC2 | NM_183377 | 0 | 8.792198922 | 9.57273983 | 10.17970655 | 0 | 0 | 0 | 0 | 0.999953634 | 0.020808337 |
| ASNS | NM_001673 | 0 | 0 | 10.14721031 | 10.75814661 | 0 | 0 | 0 | 10.72843224 | 0.011138656 | 0.999932046 |
| ASNSD1 | NM_019048 | 9.97155406 | 9.768158992 | 0 | 0 | 0 | 0 | 8.846115701 | 0 | 0.001020126 | 1 |
| ASPH | NM_004318 | 0 | 0 | 0 | 0 | 0 | 10.21468908 | 10.40759247 | 0 | 1 | 0.02072721 |
| ASPH | NM_032466 | 0 | 0 | 0 | 9.433968736 | 8.973182448 | 9.877123777 | 8.470359551 | 9.63286261 | 0.02436216 | 0.999932046 |
| ASPH | NM_032468 | 0 | 8.261155927 | 0 | 0 | 0 | 0 | 0 | 0 | 1.93E-07 | 1 |
| ASPH | XM_005251242 | 0 | 9.495485143 | 0 | 0 | 0 | 0 | 0 | 0 | 2.95E-08 | 1 |
| ASPH | XM_005251247 | 0 | 0 | 0 | 9.961649519 | 7.735433951 | 0 | 0 | 0 | 0.025103979 | 0.02072721 |
| ASPSCR1 | NR_045351 | 0 | 0 | 0 | 0 | 0 | 0 | 8.920504771 | 9.196671647 | 1 | 0.02072721 |
| ASPSCR1 | XR_001752618 | 0 | 0 | 0 | 0 | 0 | 11.42563431 | 10.88857421 | 0 | 1 | 0.019357428 |
| ASTN2 | NM_198187 | 0 | 9.790570849 | 0 | 0 | 0 | 0 | 8.195829284 | 0 | 2.05E-08 | 1 |
| ASTN2 | NM_198188 | 0 | 0 | 9.620034462 | 11.41476212 | 7.939854543 | 0 | 6.953230848 | 10.22219849 | 0.001145381 | 0.999932046 |
| ASXL1 | XM_006723727 | 0 | 0 | 9.844736088 | 0 | 9.726103171 | 0 | 9.863527812 | 0 | 0.01671739 | 1 |
| ASXL2 | XM_011532950 | 0 | 0 | 4.765355971 | 7.946164089 | 10.25167108 | 0 | 8.618176364 | 0 | 0.006780809 | 0.999932046 |
| ATAD1 | NM_001321967 | 0 | 0 | 9.328265229 | 0 | 8.499836882 | 0 | 0 | 0 | 0.029713165 | 0.020903961 |
| ATAD1 | XM_011540303 | 0 | 10.07978126 | 0 | 0 | 0 | 0 | 0 | 0 | 1.60E-08 | 1 |
| ATAD3B | XM_005244806 | 0 | 0 | 0 | 0 | 6.337212177 | 0 | 0 | 0 | 4.86E-05 | 1 |
| ATAD5 | XM_011525274 | 0 | 0 | 0 | 10.29267545 | 8.034428667 | 0 | 7.299622854 | 5.654964161 | 0.020208087 | 0.999932046 |
| ATE1 | XM_005269459 | 0 | 0 | 8.742587407 | 8.086482355 | 0 | 0 | 0 | 0 | 0.048122487 | 0.031450357 |
| ATF2 | NR_045774 | 0 | 8.662110092 | 0 | 0 | 0 | 0 | 0 | 0 | 9.15E-08 | 1 |
| ATF3 | XR_001737203 | 0 | 0 | 10.80090645 | 0 | 8.218925247 | 9.960633697 | 0 | 0 | 0.015453755 | 0.999932046 |
| ATF4 | NM_001675 | 10.65075479 | 0 | 0 | 0 | 0 | 0 | 0 | 0 | 1.34E-08 | 1 |
| ATF4 | NM_182810 | 0 | 0 | 10.72059316 | 0 | 6.945718048 | 0 | 7.728235151 | 0 | 0.018684502 | 0.999932046 |
| ATF5 | NM_001193646 | 9.911607777 | 0 | 0 | 0 | 0 | 0 | 0 | 0 | 2.05E-08 | 1 |
| ATF7 | NR_073163 | 9.834461071 | 0 | 0 | 0 | 0 | 0 | 8.357577191 | 0 | 2.30E-08 | 1 |
| ATF7 | XM_005268587 | 10.87420921 | 0 | 0 | 0 | 0 | 0 | 10.40512709 | 0 | 1.34E-08 | 1 |
| ATF7 | XM_017018722 | 9.480284601 | 9.095586658 | 0 | 0 | 0 | 0 | 8.039294254 | 0 | 0.002156021 | 1 |
| ATF7IP | XR_001748808 | 8.665043676 | 0 | 0 | 0 | 0 | 0 | 0 | 0 | 9.91E-08 | 1 |
| ATG10 | XM_011543660 | 10.34973404 | 0 | 0 | 0 | 0 | 0 | 8.04563962 | 0 | 1.51E-08 | 1 |
| ATG101 | NM_001098673 | 10.70152235 | 0 | 0 | 0 | 0 | 0 | 0 | 0 | 1.34E-08 | 1 |
| ATG4A | NM_052936 | 9.25321892 | 0 | 0 | 0 | 0 | 0 | 0 | 0 | 5.16E-08 | 1 |
| ATG4A | XM_011530842 | 0 | 9.533188437 | 0 | 0 | 0 | 0 | 0 | 0 | 2.83E-08 | 1 |
| ATG4B | XM_005246996 | 0 | 0 | 10.13885423 | 0 | 9.045366917 | 0 | 0 | 0 | 0.01822676 | 0.02072721 |
| ATG4B | XM_017003639 | 9.877828587 | 0 | 0 | 0 | 0 | 0 | 0 | 0 | 2.13E-08 | 1 |
| ATG5 | NR_104403 | 0 | 9.625843798 | 0 | 0 | 0 | 0 | 0 | 0 | 2.50E-08 | 1 |
| ATG5 | XM_024446590 | 0 | 0 | 4.580269178 | 12.17288205 | 8.375270711 | 10.18203309 | 8.867763981 | 5.80410115 | 0.001468274 | 0.999932046 |
| ATL2 | NM_001135673 | 0 | 0 | 0 | 10.09085191 | 8.629725311 | 10.49851236 | 0 | 10.27106783 | 0.02006963 | 0.999932046 |

| | | | | | | | | | | | |
|---|---|---|---|---|---|---|---|---|---|---|---|
| ATL2 | NM_001308076 | 0 | 9.352166886 | 0 | 0 | 0 | 0 | 0 | 0 | 3.68E-08 | 1 |
| ATL2 | NM_001330464 | 0 | 0 | 10.11562229 | 0 | 10.008326 | 0 | 9.074280573 | 0 | 0.013786995 | 1 |
| ATL2 | NM_022374 | 0 | 0 | 7.762363398 | 7.908843317 | 9.101058952 | 0 | 0 | 9.71050703 | 0.008885977 | 1 |
| ATL3 | NM_015459 | 0 | 8.983325733 | 0 | 0 | 0 | 0 | 0 | 0 | 7.17E-08 | 1 |
| ATM | NM_001351834 | 0 | 9.352166886 | 0 | 0 | 0 | 0 | 8.537181625 | 0 | 3.68E-08 | 1 |
| ATOH1 | NM_005172 | 9.449953927 | 0 | 0 | 0 | 0 | 0 | 0 | 0 | 3.80E-08 | 1 |
| ATP10D | XM_017008472 | 0 | 0 | 0 | 8.422875715 | 0 | 11.15039863 | 4.37247236 | 0 | 6.71E-06 | 1 |
| ATP13A1 | NM_020410 | 0 | 9.834376365 | 0 | 0 | 0 | 0 | 0 | 0 | 2.00E-08 | 1 |
| ATP1A1 | NM_000701 | 0 | 0 | 17.19927666 | 17.25219577 | 14.36491468 | 17.51062961 | 14.5164283 | 15.55224607 | 8.96E-08 | 0.999932046 |
| ATP1A1-AS1 | NR_024124 | 0 | 0 | 10.84459381 | 0 | 7.555764066 | 0 | 0 | 0 | 0.01640737 | 0.02072721 |
| ATP2A2 | NM_170665 | 10.14013255 | 0 | 0 | 0 | 0 | 0 | 0 | 0 | 2.06E-08 | 1 |
| ATP2A2 | XR_243009 | 9.854861577 | 0 | 0 | 0 | 0 | 0 | 0 | 0 | 2.23E-08 | 1 |
| ATP2B1 | XR_002957330 | 8.624880391 | 0 | 8.846304202 | 0 | 8.975568243 | 0 | 0 | 0 | 0.999953634 | 0.021119768 |
| ATP2B4 | NM_001684 | 0 | 0 | 9.678854813 | 0 | 8.190452933 | 0 | 0 | 0 | 0.027295981 | 0.02072721 |
| ATP2C1 | NM_014382 | 9.629832564 | 7.847685041 | 9.568079565 | 0 | 8.539078316 | 0 | 0 | 0 | 0.999953634 | 0.02072721 |
| ATP5ME | NR_033743 | 0 | 0 | 0 | 7.831174272 | 0 | 0 | 6.781384747 | 0 | 1.26E-05 | 1 |
| ATP5MF | NM_001039178 | 0 | 0 | 0 | 7.80432675 | 0 | 8.978473625 | 0 | 0 | 3.71E-05 | 1 |
| ATP5MG | NR_033759 | 0 | 0 | 9.981863397 | 6.608362223 | 7.618173501 | 0 | 0 | 9.043817182 | 0.007409743 | 0.999932046 |
| ATP5PF | NM_001003701 | 0 | 8.959785899 | 0 | 0 | 0 | 0 | 0 | 0 | 7.69E-08 | 1 |
| ATP5PF | NM_001320266 | 0 | 0 | 0 | 8.14114623 | 0 | 0 | 8.207195305 | 0 | 9.06E-06 | 1 |
| ATP5S | XM_005267537 | 0 | 9.585396421 | 0 | 0 | 0 | 0 | 0 | 0 | 2.61E-08 | 1 |
| ATP6V0B | NM_001039457 | 0 | 10.77858738 | 0 | 9.804567169 | 9.286391711 | 0 | 0 | 0 | 0.999953634 | 0.02072721 |
| ATP6V1C1 | NM_001695 | 0 | 0 | 0 | 9.934050572 | 7.683842675 | 0 | 10.34401416 | 10.17846329 | 0.025729408 | 0.999932046 |
| ATP6V1H | XM_006716455 | 0 | 0 | 0 | 10.37708461 | 7.997339613 | 0 | 11.62028408 | 0 | 0.01946381 | 0.999932046 |
| ATP9B | NM_198531 | 0 | 0 | 8.879283175 | 0 | 7.724127604 | 0 | 8.820441837 | 0 | 0.046548677 | 0.999932046 |
| ATP9B | XM_024451169 | 0 | 0 | 0 | 9.46429893 | 8.131765004 | 0 | 0 | 0 | 0.030166088 | 0.021907747 |
| ATP9B | XM_024451172 | 0 | 0 | 9.735370642 | 0 | 8.509747759 | 0 | 8.251784555 | 0 | 0.024623293 | 0.999932046 |
| ATP9B | XR_002958169 | 0 | 0 | 0 | 0 | 0 | 0 | 7.696326825 | 10.37308625 | 1 | 0.02072721 |
| ATPAF2 | XR_001752677 | 0 | 8.681427931 | 0 | 0 | 0 | 0 | 0 | 0 | 9.11E-08 | 1 |
| ATPSCKMT | NM_001258388 | 0 | 0 | 0 | 0 | 6.518304288 | 0 | 0 | 0 | 4.23E-05 | 1 |
| ATR | XR_001740179 | 0 | 0 | 0 | 8.466339867 | 9.472728608 | 0 | 9.963259532 | 0 | 0.027360166 | 0.999932046 |
| ATRAID | NM_016085 | 11.32222453 | 10.54808691 | 0 | 0 | 0 | 0 | 9.423956922 | 11.67391214 | 0.000235392 | 0.02072721 |
| ATRX | XM_005262155 | 0 | 0 | 11.60051722 | 13.39944712 | 11.59118797 | 14.10909143 | 11.84768864 | 11.95533117 | 5.12E-05 | 0.999932046 |
| ATXN1 | NM_001357857 | 0 | 0 | 0 | 0 | 0 | 9.776031486 | 0 | 10.45432674 | 1 | 0.02072721 |
| ATXN1L | NM_001137675 | 0 | 10.13337057 | 0 | 0 | 0 | 0 | 0 | 0 | 1.58E-08 | 1 |
| ATXN2 | NM_001310121 | 11.53403233 | 0 | 0 | 0 | 0 | 0 | 0 | 0 | 1.34E-08 | 1 |
| ATXN2 | NM_002973 | 0 | 0 | 0 | 0 | 6.3222931 | 0 | 0 | 0 | 0.000244538 | 1 |
| ATXN7L2 | XM_017000268 | 11.07884328 | 0 | 0 | 0 | 0 | 0 | 0 | 0 | 1.34E-08 | 1 |
| ATXN7L3 | XM_011525046 | 9.894817043 | 0 | 0 | 0 | 0 | 0 | 0 | 0 | 2.09E-08 | 1 |
| ATXN7L3B | NM_001136262 | 0 | 0 | 8.933431676 | 9.252214525 | 8.250791506 | 10.61318399 | 7.166214885 | 0 | 0.004950428 | 0.999932046 |
| AUH | XM_006717150 | 0 | 8.529815232 | 8.994910573 | 0 | 8.589793133 | 0 | 0 | 0 | 0.999953634 | 0.022438445 |
| AUP1 | NR_126510 | 0 | 0 | 10.87446932 | 0 | 7.82812018 | 0 | 9.383200978 | 0 | 0.015643013 | 0.999932046 |
| AURKAIP1 | XM_024447924 | 0 | 0 | 0 | 10.01531228 | 7.581052723 | 0 | 0 | 0 | 0.024946167 | 0.02072721 |

| | | | | | | | | | | | |
|---|---|---|---|---|---|---|---|---|---|---|---|
| AURKB | NM_004217 | 0 | 0 | 0 | 9.806259575 | 10.44871393 | 0 | 0 | 0 | 0.012773773 | 0.02072721 |
| AVPI1 | XM_017016494 | 9.609394475 | 0 | 0 | 0 | 0 | 0 | 0 | 0 | 2.97E-08 | 1 |
| AXIN1 | NM_003502 | 8.815370067 | 8.195487713 | 0 | 0 | 0 | 0 | 9.068060205 | 0 | 0.006089482 | 1 |
| AXIN2 | XM_011525319 | 0 | 0 | 9.583555595 | 0 | 8.840732662 | 0 | 9.750571874 | 0 | 0.024156081 | 0.999932046 |
| AXIN2 | XM_017025193 | 10.21644202 | 0 | 0 | 0 | 0 | 0 | 0 | 0 | 1.59E-08 | 1 |
| AZGP1 | NM_001185 | 0 | 0 | 0 | 12.28029022 | 9.175583651 | 10.59557977 | 12.0079807 | 10.70846348 | 1.34E-08 | 0.999932046 |
| AZIN1-AS1 | NR_126339 | 0 | 9.585396421 | 0 | 0 | 0 | 0 | 0 | 10.43444146 | 2.61E-08 | 1 |
| B3GALT5 | NM_001356336 | 0 | 10.76280445 | 0 | 0 | 0 | 0 | 0 | 0 | 1.34E-08 | 1 |
| B3GAT3 | NM_012200 | 0 | 9.885245492 | 0 | 0 | 0 | 0 | 0 | 9.85214309 | 2.00E-08 | 1 |
| B3GAT3 | NR_109991 | 0 | 8.582143237 | 0 | 0 | 0 | 0 | 0 | 0 | 9.91E-08 | 1 |
| B3GNT2 | NM_006577 | 0 | 8.861586703 | 8.957388444 | 0 | 7.593532742 | 0 | 0 | 0 | 0.999953634 | 0.029357153 |
| B3GNT3 | XM_011527626 | 0 | 8.126687292 | 0 | 10.09085191 | 6.865660185 | 0 | 0 | 0 | 0.999953634 | 0.02072721 |
| B3GNT6 | NM_138706 | 8.729593473 | 8.453213086 | 0 | 0 | 0 | 0 | 0 | 0 | 0.005548459 | 1 |
| B3GNT7 | NM_145236 | 0 | 9.19973932 | 0 | 0 | 0 | 0 | 0 | 0 | 4.67E-08 | 1 |
| B3GNT8 | XM_011526934 | 0 | 9.296876505 | 0 | 0 | 0 | 0 | 0 | 0 | 4.01E-08 | 1 |
| B4GALNT2 | XM_011524314 | 8.850625659 | 0 | 0 | 0 | 0 | 0 | 0 | 0 | 8.53E-08 | 1 |
| B4GALT1-AS1 | NR_108109 | 0 | 0 | 9.808316306 | 0 | 8.24486995 | 0 | 0 | 0 | 0.025256332 | 0.02072721 |
| B4GALT3 | NM_001199874 | 0 | 8.927789798 | 0 | 0 | 0 | 0 | 0 | 0 | 7.69E-08 | 1 |
| B4GALT3 | XM_017002714 | 0 | 0 | 0 | 0 | 6.544252221 | 0 | 0 | 0 | 4.20E-05 | 1 |
| B4GALT4 | NM_212543 | 0 | 9.490017613 | 10.50671712 | 0 | 9.170386259 | 0 | 0 | 0 | 0.999953634 | 0.02072721 |
| B4GALT4 | XM_006713800 | 0 | 8.818611963 | 0 | 0 | 0 | 0 | 0 | 0 | 8.12E-08 | 1 |
| B4GALT4 | XM_024453807 | 0 | 9.036802844 | 0 | 0 | 0 | 0 | 0 | 0 | 6.38E-08 | 1 |
| B9D1 | NM_015681 | 0 | 8.083774979 | 0 | 10.28419933 | 10.43317496 | 0 | 0 | 0 | 0.85973045 | 0.02072721 |
| B9D2 | NM_030578 | 0 | 10.53088392 | 0 | 0 | 0 | 0 | 0 | 0 | 1.34E-08 | 1 |
| BAAT | NM_001701 | 0 | 8.951853153 | 0 | 0 | 0 | 0 | 8.991245499 | 0 | 7.60E-08 | 1 |
| BABAM1 | NM_014173 | 9.190393069 | 4.822913417 | 10.28232386 | 0 | 9.688785145 | 0 | 0 | 0 | 0.999953634 | 0.02072721 |
| BABAM2 | NR_137438 | 10.29532878 | 0 | 0 | 0 | 0 | 0 | 0 | 0 | 1.54E-08 | 1 |
| BACE2 | NM_138992 | 0 | 0 | 0 | 10.21070698 | 8.165592712 | 0 | 0 | 0 | 0.020641811 | 0.02072721 |
| BAG2 | XM_024446596 | 0 | 8.756212053 | 0 | 0 | 0 | 8.039779276 | 0 | 0 | 8.55E-08 | 1 |
| BAG3 | XM_005270287 | 10.11377196 | 0 | 0 | 0 | 0 | 0 | 0 | 0 | 1.70E-08 | 1 |
| BAG5 | NM_004873 | 9.181190369 | 0 | 0 | 0 | 0 | 0 | 0 | 0 | 7.80E-08 | 1 |
| BAG6 | XM_017011297 | 0 | 0 | 9.022425311 | 0 | 9.117355799 | 11.47219978 | 0 | 0 | 0.027138717 | 1 |
| BAIAP2L2 | XM_011530379 | 10.17531978 | 0 | 0 | 0 | 0 | 0 | 0 | 0 | 1.60E-08 | 1 |
| BAMBI | NM_012342 | 0 | 0 | 0 | 10.41971946 | 9.562752167 | 10.94608429 | 10.35299605 | 0 | 0.013993253 | 0.999932046 |
| BANF1 | XM_017018514 | 0 | 0 | 0 | 11.34312291 | 8.74608731 | 0 | 8.207195305 | 0 | 0.01025918 | 0.999932046 |
| BANF1 | XM_017018515 | 0 | 0 | 0 | 8.841270918 | 8.513036313 | 0 | 8.402765142 | 0 | 0.036022978 | 0.999932046 |
| BANK1 | XM_017008337 | 0 | 10.03542879 | 0 | 0 | 0 | 0 | 0 | 0 | 1.60E-08 | 1 |
| BAP1 | XM_011534149 | 0 | 0 | 9.366341809 | 0 | 7.854284905 | 10.15584598 | 0 | 0 | 0.034649792 | 0.999932046 |
| BARD1 | NM_000465 | 10.55959296 | 11.09900498 | 0 | 11.0131504 | 10.86563873 | 0 | 0 | 0 | 0.999953634 | 0.02072721 |
| BASP1 | NM_006317 | 9.771469405 | 0 | 0 | 0 | 0 | 0 | 0 | 0 | 2.47E-08 | 1 |
| BATF2 | NM_138456 | 9.807800698 | 0 | 0 | 0 | 0 | 0 | 0 | 0 | 2.49E-08 | 1 |
| BAZ2B | XM_017003936 | 0 | 9.684464992 | 0 | 0 | 0 | 0 | 0 | 0 | 2.36E-08 | 1 |
| BBIP1 | NM_001195304 | 10.24321976 | 0 | 0 | 0 | 0 | 0 | 0 | 0 | 1.59E-08 | 1 |

| | | | | | | | | | | | |
|---|---|---|---|---|---|---|---|---|---|---|---|
| BBIP1 | NM_001195305 | 9.866390785 | 0 | 0 | 0 | 0 | 0 | 8.519054934 | 0 | 2.17E-08 | 1 |
| BBIP1 | NM_001243783 | 10.59467378 | 0 | 0 | 0 | 0 | 0 | 0 | 0 | 1.34E-08 | 1 |
| BBOF1 | XM_017021663 | 9.085782918 | 0 | 0 | 0 | 0 | 0 | 0 | 0 | 7.29E-08 | 1 |
| BBX | NM_001142568 | 0 | 0 | 11.45539534 | 10.60224751 | 9.707564817 | 11.44345748 | 0 | 11.08977591 | 0.000504685 | 0.999932046 |
| BBX | XM_024453644 | 0 | 9.993538654 | 0 | 0 | 0 | 0 | 8.801819545 | 9.941402427 | 1.68E-08 | 0.02072721 |
| BCAP29 | XM_011516408 | 9.387309849 | 9.387883956 | 10.19836703 | 0 | 8.025245409 | 0 | 0 | 0 | 0.999953634 | 0.02072721 |
| BCAP31 | XR_002958759 | 11.71390931 | 0 | 10.31699395 | 0 | 6.147113942 | 0 | 0 | 0 | 0.999953634 | 0.02072721 |
| BCAP31 | XR_002958761 | 10.39819601 | 4.51367241 | 0 | 10.00797741 | 8.218925247 | 0 | 0 | 0 | 0.999953634 | 0.02072721 |
| BCAR1 | XM_005256260 | 0 | 8.774317692 | 0 | 0 | 0 | 0 | 0 | 0 | 8.46E-08 | 1 |
| BCAS1 | XM_011529090 | 0 | 9.179501644 | 0 | 0 | 0 | 0 | 0 | 0 | 4.86E-08 | 1 |
| BCAS1 | XM_017028112 | 0 | 0 | 0 | 0 | 6.59478863 | 0 | 0 | 0 | 4.02E-05 | 1 |
| BCAS2 | NM_005872 | 0 | 9.213075091 | 10.77822023 | 10.64637732 | 9.361285436 | 0 | 0 | 0 | 0.894771227 | 0.001586242 |
| BCAS3 | NM_001353145 | 0 | 9.451153482 | 0 | 0 | 0 | 0 | 7.863684239 | 0 | 3.15E-08 | 1 |
| BCAT2 | NM_001190 | 0 | 0 | 10.29365975 | 0 | 9.46934677 | 0 | 0 | 10.06479657 | 0.015309434 | 0.999932046 |
| BCL11A | XM_011532912 | 0 | 0 | 8.300853839 | 0 | 9.761097899 | 0 | 9.319168841 | 10.37457782 | 0.024576224 | 0.999932046 |
| BCL11A | XM_017004333 | 0 | 9.124104101 | 0 | 0 | 0 | 0 | 7.974256665 | 0 | 5.29E-08 | 1 |
| BCL2L11 | NM_207002 | 0 | 0 | 0 | 0 | 0 | 12.18169144 | 10.38712595 | 9.929274263 | 1 | 0.000493446 |
| BCL2L11 | NM_207003 | 0 | 8.975521721 | 0 | 0 | 0 | 0 | 0 | 8.998101666 | 7.27E-08 | 1 |
| BCL2L14 | NM_138722 | 0 | 9.296876505 | 0 | 0 | 0 | 0 | 0 | 0 | 4.01E-08 | 1 |
| BCL2L15 | NM_001010922 | 0 | 11.89137745 | 10.95996127 | 0 | 8.800928156 | 0 | 0 | 0 | 0.999953634 | 0.02072721 |
| BCL2L2 | NM_004050 | 10.12821014 | 0 | 0 | 0 | 0 | 0 | 0 | 0 | 1.68E-08 | 1 |
| BCL3 | XM_011527197 | 0 | 0 | 9.614018345 | 0 | 7.587306227 | 0 | 0 | 9.497035549 | 0.032005189 | 0.999932046 |
| BCL3 | XM_017027110 | 8.611240259 | 0 | 0 | 0 | 0 | 0 | 0 | 0 | 1.04E-07 | 1 |
| BCL7B | NR_036682 | 0 | 8.602554319 | 0 | 0 | 0 | 11.02317065 | 8.137615898 | 0 | 9.73E-08 | 0.02072721 |
| BCL7C | NM_004765 | 0 | 0 | 10.52926948 | 10.42082459 | 8.479808505 | 11.48898356 | 9.602418094 | 0 | 0.001265952 | 0.999932046 |
| BCL9 | XM_005277418 | 10.06457365 | 0 | 0 | 0 | 0 | 0 | 10.81061397 | 0 | 1.80E-08 | 1 |
| BCL9 | XM_017002026 | 9.643298854 | 0 | 0 | 0 | 0 | 0 | 8.686767846 | 0 | 2.88E-08 | 1 |
| BCO1 | XM_017023289 | 0 | 9.252358352 | 0 | 0 | 0 | 0 | 0 | 0 | 4.28E-08 | 1 |
| BDH2 | XM_017008462 | 10.17531978 | 0 | 0 | 0 | 0 | 0 | 0 | 0 | 1.60E-08 | 1 |
| BDP1 | NM_018429 | 10.10406561 | 9.346127117 | 0 | 0 | 0 | 0 | 0 | 11.3162193 | 0.001185107 | 1 |
| BECN1 | NM_001313998 | 0 | 0 | 10.29741868 | 0 | 8.608360805 | 10.39240229 | 9.360475428 | 0 | 0.018675558 | 0.999932046 |
| BEND5 | XM_017002331 | 10.74276593 | 5.390784359 | 0 | 0 | 0 | 11.12975411 | 0 | 0 | 0.001948997 | 1 |
| BEND5 | XR_001737408 | 0 | 11.0459632 | 11.14865646 | 0 | 9.624866434 | 0 | 0 | 0 | 0.999953634 | 0.02072721 |
| BEST2 | XM_005259963 | 0 | 8.273938353 | 0 | 0 | 0 | 0 | 0 | 0 | 2.02E-07 | 1 |
| BET1L | NM_016526 | 0 | 0 | 0 | 7.058084711 | 0 | 0 | 0 | 0 | 2.75E-05 | 1 |
| BEX3 | NM_206917 | 0 | 0 | 10.23802521 | 0 | 10.12575557 | 0 | 0 | 0 | 0.012636617 | 0.02072721 |
| BFSP2 | XM_024453788 | 10.94115867 | 0 | 0 | 0 | 0 | 0 | 0 | 0 | 1.34E-08 | 1 |
| BHLHE40 | XM_024453798 | 0 | 0 | 9.427651273 | 7.921391152 | 0 | 10.08696053 | 8.380348081 | 0 | 0.035569763 | 0.999932046 |
| BICDL2 | XM_005255135 | 0 | 0 | 7.740439313 | 10.74320495 | 0 | 10.57236835 | 0 | 0 | 0.017612901 | 0.999932046 |
| BICDL2 | XM_011522391 | 0 | 9.387883956 | 0 | 0 | 0 | 10.94051686 | 7.892131936 | 0 | 3.40E-08 | 0.02072721 |
| BID | NM_001244570 | 0 | 0 | 0 | 11.35241958 | 9.315876167 | 0 | 0 | 0 | 0.00918515 | 0.02072721 |
| BIN1 | XM_017003828 | 0 | 9.145128245 | 0 | 0 | 0 | 0 | 0 | 0 | 5.08E-08 | 1 |
| BIN3 | XM_011544586 | 0 | 8.273938353 | 0 | 0 | 0 | 0 | 0 | 0 | 2.02E-07 | 1 |

| | | | | | | | | | | | |
|---|---|---|---|---|---|---|---|---|---|---|---|
| BIN3 | XR_949444 | 8.611240259 | 0 | 0 | 0 | 0 | 0 | 0 | 0 | 1.04E-07 | 1 |
| BIRC3 | XM_024448467 | 0 | 8.273938353 | 0 | 0 | 0 | 9.397544622 | 0 | 0 | 2.02E-07 | 1 |
| BIRC5 | NM_001168 | 0 | 0 | 0 | 10.57966586 | 10.7252512 | 0 | 0 | 0 | 0.008755979 | 0.02072721 |
| BIRC6 | NM_016252 | 0 | 0 | 8.615858226 | 9.656688688 | 9.845694196 | 0 | 8.465635012 | 9.677050566 | 0.002355992 | 0.999932046 |
| BIVM | NM_017693 | 0 | 0 | 0 | 9.10598125 | 7.119744988 | 0 | 0 | 0 | 0.045985196 | 0.031617224 |
| BIVM-ERCC5 | NM_001204425 | 10.07454845 | 0 | 0 | 0 | 0 | 10.4918664 | 0 | 0 | 1.79E-08 | 1 |
| BLCAP | XM_017027599 | 0 | 0 | 0 | 10.54967014 | 8.993337041 | 0 | 8.692844709 | 0 | 0.015130311 | 0.999932046 |
| BLM | NM_001287247 | 0 | 0 | 12.76561202 | 12.04110495 | 10.05419515 | 10.96950906 | 9.912086445 | 10.71671941 | 0.000109488 | 0.999932046 |
| BLNK | NM_001114094 | 0 | 8.475517571 | 0 | 0 | 0 | 0 | 0 | 0 | 1.09E-07 | 1 |
| BLZF1 | NM_001320973 | 0 | 0 | 7.433922402 | 0 | 9.187981525 | 0 | 6.255003263 | 0 | 0.039981685 | 0.999932046 |
| BLZF1 | NM_003666 | 0 | 8.737876298 | 0 | 0 | 0 | 0 | 0 | 0 | 9.11E-08 | 1 |
| BMI1 | NM_005180 | 0 | 0 | 9.785747216 | 0 | 6.844931782 | 10.01594442 | 5.013866823 | 10.00826692 | 0.032458111 | 0.999932046 |
| BMP2K | XM_017008382 | 0 | 0 | 0 | 0 | 0 | 0 | 7.954159584 | 9.781938865 | 1 | 0.02072721 |
| BMP5 | NM_021073 | 14.12220041 | 11.9027384 | 13.21888365 | 10.1626323 | 0 | 0 | 0 | 0 | 0.999953634 | 1.92E-09 |
| BMP8B | XM_024449299 | 0 | 0 | 0 | 0 | 6.569741695 | 0 | 0 | 0 | 4.12E-05 | 1 |
| BNIP2 | NM_001320674 | 0 | 9.364171095 | 0 | 0 | 0 | 0 | 0 | 0 | 3.61E-08 | 1 |
| BNIP2 | NM_004330 | 0 | 0 | 0 | 0 | 0 | 0 | 7.431456038 | 10.15782178 | 1 | 0.02072721 |
| BNIP3 | NM_004052 | 0 | 0 | 9.86514191 | 0 | 8.025245409 | 0 | 0 | 0 | 0.025713421 | 0.02072721 |
| BNIP3L | NM_004331 | 0 | 0 | 10.38393047 | 0 | 8.459498162 | 10.30194482 | 0 | 0 | 0.018443387 | 0.999932046 |
| BOC | XM_017007443 | 9.982190738 | 0 | 0 | 0 | 0 | 0 | 0 | 0 | 1.98E-08 | 1 |
| BOD1 | NM_138369 | 0 | 0 | 0 | 10.15998747 | 8.81964034 | 0 | 0 | 0 | 0.018740683 | 0.02072721 |
| BOLA2-SMG1P6 | NM_001320623 | 0 | 9.703487753 | 0 | 0 | 0 | 0 | 0 | 9.334871374 | 2.30E-08 | 1 |
| BOLA2B | NM_001320607 | 0 | 8.895067962 | 0 | 8.359738397 | 8.066117405 | 0 | 0 | 0 | 0.999953634 | 0.035225234 |
| BOLA3 | NM_212552 | 10.53812757 | 9.422738093 | 8.93825496 | 11.03352211 | 8.327507425 | 0 | 0 | 0 | 0.999953634 | 0.002874229 |
| BORCS6 | NM_017622 | 0 | 0 | 9.160915107 | 0 | 8.822293806 | 0 | 0 | 0 | 0.028726223 | 0.020781046 |
| BORCS8 | NM_001145783 | 0 | 0 | 0 | 10.29388227 | 6.451316153 | 0 | 0 | 0 | 0.024213691 | 0.02072721 |
| BPNT1 | NM_006085 | 0 | 0 | 10.80882894 | 0 | 5.716198176 | 0 | 0 | 0 | 3.42E-08 | 1 |
| BPNT1 | XM_005273000 | 0 | 0 | 0 | 0 | 6.690841957 | 0 | 0 | 0 | 3.72E-05 | 1 |
| BPTF | XM_005257151 | 0 | 9.411213355 | 0 | 0 | 0 | 11.42014412 | 0 | 0 | 3.28E-08 | 1 |
| BPTF | XM_024450646 | 9.595606435 | 0 | 0 | 0 | 0 | 0 | 0 | 0 | 3.08E-08 | 1 |
| BRAP | XM_017019994 | 0 | 0 | 0 | 0 | 6.351978551 | 0 | 0 | 0 | 4.81E-05 | 1 |
| BRAT1 | XM_011515183 | 10.06457365 | 0 | 0 | 0 | 0 | 0 | 0 | 0 | 1.80E-08 | 1 |
| BRCA1 | NM_007299 | 0 | 0 | 9.313491365 | 0 | 7.587306227 | 0 | 0 | 0 | 0.037907308 | 0.024639803 |
| BRCC3 | XM_005274751 | 0 | 0 | 10.40062728 | 0 | 7.211666293 | 0 | 0 | 0 | 0.021703066 | 0.02072721 |
| BRD1 | XR_244360 | 0 | 0 | 0 | 0 | 6.713888284 | 0 | 0 | 9.521478054 | 3.66E-05 | 1 |
| BRD8 | XR_948230 | 0 | 0 | 0 | 0 | 0 | 4.972462907 | 8.566160223 | 0 | 1 | 0.043192544 |
| BRD9 | XM_024446193 | 0 | 0 | 0 | 11.09850178 | 11.38946867 | 11.42663028 | 8.479762465 | 10.31833588 | 0.005396476 | 0.999932046 |
| BRD9 | XM_024446195 | 0 | 9.265218616 | 0 | 0 | 0 | 0 | 7.655422644 | 0 | 4.18E-08 | 1 |
| BRD9 | XM_024446198 | 0 | 0 | 11.64249798 | 0 | 11.1015469 | 11.15641475 | 10.71332949 | 9.607716039 | 0.004824193 | 0.999932046 |
| BRD9 | XR_002956174 | 0 | 0 | 9.77095327 | 10.34367813 | 9.545191248 | 0 | 10.27345508 | 0 | 0.001321369 | 1 |
| BRIP1 | NM_032043 | 0 | 0 | 0 | 0 | 6.381065478 | 0 | 0 | 0 | 4.71E-05 | 1 |
| BRIP1 | XM_017025201 | 0 | 0 | 0 | 8.702760903 | 7.905232421 | 0 | 0 | 0 | 0.047238212 | 0.032030331 |
| BRMS1 | XR_002957138 | 0 | 0 | 0 | 7.585264412 | 9.31399202 | 0 | 0 | 0 | 0.036190965 | 0.02496249 |

| | | | | | | | | | | | |
|---|---|---|---|---|---|---|---|---|---|---|---|
| BRPF1 | XR_001740258 | 9.941892254 | 0 | 0 | 0 | 0 | 0 | 8.107603982 | 0 | 2.01E-08 | 1 |
| BRWD1-AS2 | NR_111942 | 10.11860076 | 9.655452119 | 0 | 0 | 0 | 0 | 0 | 0 | 0.000986254 | 1 |
| BRWD3 | XM_005262113 | 0 | 9.077472692 | 0 | 0 | 0 | 0 | 0 | 0 | 7.73E-08 | 1 |
| BSCL2 | NR_037949 | 0 | 8.78328601 | 0 | 0 | 0 | 0 | 0 | 0 | 8.40E-08 | 1 |
| BSDC1 | NR_125352 | 5.510268966 | 8.582143237 | 0 | 0 | 0 | 0 | 0 | 0 | 0.017521547 | 1 |
| BSG | XM_017027173 | 0 | 0 | 11.69829667 | 0 | 9.918180981 | 0 | 7.154520047 | 9.793135989 | 0.006710754 | 0.999932046 |
| BTBD2 | XM_011528127 | 0 | 0 | 10.3381331 | 8.284026919 | 0 | 0 | 0 | 0 | 0.020792098 | 0.02072721 |
| BTBD6 | NM_033271 | 0 | 0 | 8.791384731 | 10.29990127 | 0 | 0 | 6.67079425 | 8.402901782 | 0.018919595 | 0.999932046 |
| BTBD6 | XM_005268188 | 8.665043676 | 8.951853153 | 10.06580283 | 8.82800322 | 0 | 0 | 0 | 0 | 0.999953634 | 0.02072721 |
| BTBD9 | NM_001099272 | 0 | 0 | 0 | 0 | 0 | 11.28883067 | 7.960889765 | 10.02735731 | 1 | 0.001535028 |
| BTD | NM_001281725 | 0 | 0 | 0 | 0 | 6.395391943 | 0 | 0 | 0 | 4.65E-05 | 1 |
| BTD | NM_001281726 | 0 | 9.399595812 | 10.02330906 | 7.146247488 | 8.732045066 | 0 | 0 | 0 | 0.999953634 | 0.006675853 |
| BTD | XM_024453724 | 8.779231295 | 0 | 0 | 0 | 0 | 0 | 0 | 0 | 9.02E-08 | 1 |
| BTF3L4 | NM_001136497 | 0 | 0 | 0 | 10.14535356 | 7.944733425 | 0 | 0 | 0 | 0.022059726 | 0.02072721 |
| BTN3A1 | NM_007048 | 0 | 9.694007725 | 0 | 0 | 0 | 0 | 0 | 0 | 2.55E-08 | 1 |
| BTNL3 | NM_197975 | 10.70152235 | 9.138154184 | 12.4963983 | 0 | 7.817519962 | 0 | 0 | 0 | 0.999953634 | 3.33E-09 |
| BTNL8 | NM_024850 | 10.11377196 | 0 | 0 | 0 | 0 | 0 | 0 | 0 | 1.70E-08 | 1 |
| BTNL8 | XM_011534649 | 0 | 10.14036773 | 0 | 0 | 0 | 0 | 0 | 0 | 1.56E-08 | 1 |
| BTNL8 | XM_011534653 | 7.486331536 | 8.592384875 | 0 | 0 | 0 | 0 | 0 | 0 | 0.010150947 | 1 |
| BUB3 | NM_001007793 | 0 | 0 | 10.7490083 | 10.91460632 | 0 | 0 | 0 | 0 | 0.008515228 | 0.02072721 |
| BUD23 | XM_011515778 | 0 | 0 | 0 | 7.528981351 | 0 | 0 | 0 | 0 | 1.76E-05 | 1 |
| BZW1 | NM_001207067 | 0 | 8.747073305 | 10.04359378 | 9.954035954 | 0 | 0 | 0 | 0 | 0.999953634 | 0.02072721 |
| BZW1 | NM_014670 | 0 | 0 | 10.23410809 | 0 | 8.66562982 | 0 | 4.790286182 | 0 | 0.018956978 | 0.509456447 |
| BZW1 | XM_017005393 | 0 | 0 | 8.93825496 | 0 | 8.754447516 | 0 | 7.09458011 | 0 | 0.032133817 | 0.999932046 |
| BZW2 | NR_027624 | 0 | 0 | 0 | 10.55269793 | 7.843875888 | 0 | 9.135052218 | 0 | 0.01818298 | 0.999932046 |
| C10orf143 | NM_001355042 | 10.4296226 | 0 | 0 | 0 | 0 | 0 | 8.530410777 | 0 | 1.47E-08 | 1 |
| C11orf24 | NM_022338 | 0 | 0 | 0 | 9.63590165 | 7.255520044 | 0 | 0 | 0 | 0.032567382 | 0.023358895 |
| C11orf49 | NR_103471 | 0 | 0 | 7.275241756 | 0 | 0 | 0 | 0 | 0 | 5.92E-05 | 1 |
| C11orf54 | XM_011542782 | 0 | 8.927789798 | 0 | 0 | 0 | 0 | 0 | 0 | 7.69E-08 | 1 |
| C11orf68 | NM_001135635 | 0 | 0 | 0 | 0 | 6.607150896 | 0 | 0 | 9.871287726 | 0.00011086 | 1 |
| C11orf68 | NM_031450 | 0 | 0 | 0 | 9.733411602 | 8.173927142 | 0 | 0 | 9.607716039 | 0.026115516 | 0.999932046 |
| C11orf71 | NM_001271562 | 0 | 0 | 11.27318029 | 8.234606714 | 10.518277 | 0 | 7.166214885 | 0 | 0.000855364 | 0.630780089 |
| C11orf74 | NM_138787 | 0 | 0 | 0 | 8.873915965 | 7.672123016 | 0 | 0 | 0 | 0.046268417 | 0.03159928 |
| C11orf80 | NM_024650 | 9.825628908 | 0 | 0 | 0 | 0 | 0 | 0 | 0 | 2.30E-08 | 1 |
| C11orf86 | NM_001136485 | 10.80100026 | 0 | 0 | 0 | 0 | 0 | 0 | 0 | 1.34E-08 | 1 |
| C12orf29 | NM_001009894 | 0 | 0 | 5.33252694 | 10.71104353 | 10.1327202 | 0 | 0 | 0 | 0.002268724 | 0.003656388 |
| C12orf4 | XM_006718992 | 10.02910853 | 0 | 0 | 0 | 0 | 0 | 0 | 0 | 1.87E-08 | 1 |
| C12orf43 | NM_022895 | 0 | 8.464408432 | 0 | 0 | 0 | 0 | 0 | 0 | 1.11E-07 | 1 |
| C12orf49 | NM_001353625 | 0 | 0 | 9.974838534 | 0 | 9.052164589 | 0 | 0 | 0 | 0.019425648 | 0.02072721 |
| C12orf56 | XR_944486 | 10.4218299 | 0 | 0 | 0 | 0 | 0 | 0 | 0 | 1.47E-08 | 1 |
| C14orf119 | XM_017021390 | 0 | 9.346127117 | 0 | 0 | 0 | 0 | 0 | 0 | 3.72E-08 | 1 |
| C15orf40 | NM_001160114 | 0 | 9.186279128 | 0 | 9.39850643 | 7.523516842 | 0 | 0 | 0 | 0.999953634 | 0.025070991 |
| C15orf41 | XR_931924 | 9.46519897 | 0 | 0 | 0 | 0 | 0 | 0 | 0 | 3.72E-08 | 1 |

| | | | | | | | | | | | |
|---|---|---|---|---|---|---|---|---|---|---|---|
| C16orf72 | XM_011522462 | 9.874977627 | 0 | 0 | 0 | 0 | 0 | 0 | 0 | 2.15E-08 | 1 |
| C19orf12 | XM_024451736 | 0 | 8.835955774 | 0 | 0 | 0 | 0 | 0 | 0 | 2.22E-07 | 1 |
| C19orf12 | XM_024451738 | 0 | 8.519117955 | 0 | 0 | 0 | 0 | 0 | 0 | 1.04E-07 | 1 |
| C19orf48 | NM_001290152 | 0 | 9.08836834 | 0 | 0 | 0 | 0 | 0 | 0 | 5.61E-08 | 1 |
| C1GALT1 | XM_024446838 | 0 | 0 | 9.158850343 | 0 | 7.357826773 | 0 | 0 | 0 | 0.04371157 | 0.02788959 |
| C1orf115 | NM_024709 | 0 | 0 | 9.035988366 | 8.738654916 | 9.879210711 | 0 | 0 | 0 | 0.002882974 | 0.004325677 |
| C1orf131 | NM_152379 | 0 | 0 | 10.42230642 | 0 | 7.275021256 | 0 | 0 | 0 | 0.021277588 | 0.02072721 |
| C1orf159 | NM_017891 | 0 | 0 | 10.15965437 | 0 | 9.564338065 | 0 | 9.828077295 | 0 | 0.015691656 | 0.999932046 |
| C1orf159 | XM_005244765 | 0 | 0 | 9.093331342 | 0 | 4.270485932 | 0 | 0 | 0 | 0.064834412 | 0.04268834 |
| C1orf159 | XM_024447912 | 0 | 0 | 0 | 0 | 6.464964927 | 0 | 0 | 0 | 4.45E-05 | 1 |
| C1orf198 | XM_017002599 | 0 | 0 | 0 | 10.50138585 | 6.736572239 | 0 | 5.824272679 | 0 | 0.020924225 | 0.90102082 |
| C1orf35 | NM_024319 | 0 | 0 | 10.41453941 | 10.52418831 | 0 | 0 | 0 | 0 | 0.011262512 | 0.02072721 |
| C1orf35 | NR_130975 | 0 | 9.387883956 | 0 | 0 | 0 | 0 | 0 | 0 | 3.40E-08 | 1 |
| C1orf43 | NM_001297718 | 0 | 9.186279128 | 0 | 0 | 0 | 0 | 0 | 0 | 4.80E-08 | 1 |
| C1orf43 | NM_015449 | 0 | 0 | 8.609824632 | 0 | 10.06655763 | 0 | 9.672770285 | 12.1244403 | 0.019980007 | 0.999932046 |
| C1orf52 | NR_024113 | 0 | 0 | 0 | 9.429583326 | 7.838643081 | 0 | 0 | 0 | 0.032823165 | 0.023358895 |
| C1orf56 | NM_017860 | 0 | 0 | 0 | 10.50138585 | 9.415888488 | 0 | 0 | 0 | 0.014019265 | 0.02072721 |
| C1QL1 | NM_006688 | 0 | 0 | 0 | 0 | 6.655567121 | 0 | 0 | 0 | 3.82E-05 | 1 |
| C1QTNF3 | NR_146599 | 0 | 0 | 0 | 0 | 6.505153187 | 0 | 0 | 0 | 4.28E-05 | 1 |
| C1R | NM_001733 | 10.42573151 | 9.842980089 | 11.08429488 | 0 | 7.335710674 | 0 | 0 | 0 | 0.999953634 | 0.02072721 |
| C20orf204 | XM_024451876 | 0 | 9.036802844 | 0 | 0 | 0 | 0 | 0 | 0 | 6.38E-08 | 1 |
| C20orf27 | NM_001258429 | 0 | 0 | 0 | 8.264461514 | 9.212462155 | 0 | 0 | 0 | 0.032607408 | 0.02296717 |
| C21orf2 | XR_937571 | 0 | 0 | 8.999532959 | 8.162444728 | 0 | 0 | 0 | 0 | 0.04155345 | 0.027403095 |
| C21orf58 | NM_058180 | 0 | 9.439854398 | 0 | 0 | 0 | 0 | 0 | 0 | 3.19E-08 | 1 |
| C2CD5 | XM_017020278 | 0 | 0 | 0 | 8.673388514 | 0 | 0 | 0 | 0 | 5.13E-06 | 1 |
| C2orf49 | XM_024453136 | 9.235546106 | 0 | 0 | 0 | 0 | 0 | 0 | 0 | 5.31E-08 | 1 |
| C2orf72 | NM_001144994 | 0 | 9.600697383 | 0 | 0 | 0 | 0 | 0 | 0 | 2.56E-08 | 1 |
| C3orf38 | XM_005264745 | 0 | 8.998808245 | 0 | 0 | 0 | 0 | 5.456313559 | 0 | 6.89E-08 | 1 |
| C3orf62 | NM_198562 | 8.497211815 | 10.4999361 | 0 | 0 | 0 | 0 | 0 | 0 | 0.001174853 | 1 |
| C3orf70 | NM_001025266 | 0 | 8.486541819 | 0 | 0 | 0 | 0 | 0 | 0 | 1.09E-07 | 1 |
| C4orf19 | NM_018302 | 0 | 10.14733113 | 0 | 0 | 0 | 0 | 0 | 0 | 1.55E-08 | 1 |
| C4orf33 | XM_005262734 | 0 | 0 | 0 | 7.676917851 | 0 | 0 | 0 | 0 | 1.47E-05 | 1 |
| C4orf48 | XM_011513469 | 0 | 0 | 0 | 10.55068011 | 5.646039634 | 0 | 0 | 0 | 0.022059726 | 5.45E-09 |
| C5orf51 | XM_005248289 | 0 | 0 | 0 | 10.38616567 | 10.26489835 | 0 | 0 | 10.45432674 | 0.011298501 | 1 |
| C6orf120 | NM_001317342 | 9.46519897 | 7.523045066 | 0 | 0 | 0 | 0 | 7.341484785 | 0 | 0.00465859 | 1 |
| C6orf132 | NM_001164446 | 10.54890019 | 9.19973932 | 0 | 0 | 0 | 0 | 8.149448007 | 0 | 0.000888245 | 1 |
| C6orf136 | NM_001109938 | 0 | 0 | 11.63471908 | 9.481348416 | 0 | 0 | 0 | 0 | 0.008495729 | 0.02072721 |
| C6orf203 | NM_016487 | 0 | 0 | 0 | 9.352916447 | 9.807368162 | 0 | 9.756386369 | 0 | 0.018817304 | 0.999932046 |
| C6orf48 | NM_001287488 | 0 | 9.006487637 | 0 | 0 | 0 | 0 | 0 | 0 | 6.80E-08 | 1 |
| C7orf13 | NR_026865 | 9.695941183 | 0 | 0 | 0 | 0 | 0 | 6.766095723 | 0 | 2.66E-08 | 1 |
| C7orf50 | NR_134537 | 0 | 0 | 0 | 9.726277944 | 10.40866557 | 0 | 8.058247234 | 0 | 0.013319883 | 0.999932046 |
| C7orf50 | NR_146967 | 0 | 0 | 0 | 9.531320288 | 8.034428667 | 0 | 0 | 0 | 0.029825232 | 0.02173026 |
| C7orf50 | XM_017012720 | 0 | 0 | 0 | 7.662040946 | 0 | 0 | 6.750642936 | 0 | 1.49E-05 | 1 |

| | | | | | | | | | | | |
|---|---|---|---|---|---|---|---|---|---|---|---|
| C8orf33 | NM_023080 | 0 | 0 | 10.95282868 | 9.708287933 | 9.202312328 | 0 | 9.690088235 | 9.770654157 | 0.001110043 | 0.999932046 |
| C8orf34 | XR_928756 | 0 | 0 | 0 | 0 | 6.42362501 | 0 | 0 | 0 | 4.54E-05 | 1 |
| C8orf37 | XM_005250799 | 0 | 0 | 10.00733248 | 0 | 6.351978551 | 0 | 0 | 0 | 0.029881811 | 0.020910753 |
| C8orf76 | NM_032847 | 0 | 0 | 0 | 12.42472499 | 9.418521297 | 11.43852865 | 7.704370312 | 0 | 1.58E-08 | 0.999932046 |
| C9orf16 | XM_011519004 | 0 | 0 | 7.945990394 | 0 | 0 | 0 | 7.604755783 | 0 | 1.21E-05 | 1 |
| C9orf3 | XM_005252284 | 0 | 8.853093555 | 0 | 0 | 0 | 0 | 0 | 0 | 7.91E-08 | 1 |
| C9orf3 | XM_017015234 | 0 | 9.918188339 | 0 | 0 | 0 | 0 | 0 | 0 | 1.84E-08 | 1 |
| C9orf3 | XR_929852 | 0 | 0 | 9.017875808 | 9.07261961 | 8.529367569 | 9.311738383 | 13.02827577 | 0 | 0.00472914 | 0.999932046 |
| C9orf78 | NM_016520 | 10.02910853 | 9.303125566 | 10.04583013 | 10.85289714 | 10.23832142 | 0 | 0 | 0 | 0.999953634 | 0.001524206 |
| CA1 | NM_001164830 | 23.01000981 | 20.88750487 | 23.43763904 | 13.24329512 | 14.69222216 | 0 | 10.29691705 | 0 | 1 | 0.016557995 |
| CA12 | NR_135511 | 10.22542327 | 9.138154184 | 0 | 0 | 0 | 10.65712596 | 9.41540741 | 0 | 0.001197431 | 0.02072721 |
| CA2 | NM_000067 | 14.43977792 | 13.41067312 | 15.20704636 | 12.41754595 | 0 | 0 | 0 | 0 | 0.999953634 | 8.87E-10 |
| CA4 | XR_001752609 | 9.922694008 | 0 | 0 | 0 | 0 | 0 | 0 | 0 | 2.04E-08 | 1 |
| CA9 | NM_001216 | 0 | 0 | 0 | 11.69744999 | 9.843083461 | 0 | 0 | 0 | 0.006732503 | 0.02072721 |
| CAAP1 | XR_001746381 | 9.539097493 | 0 | 0 | 0 | 0 | 0 | 0 | 0 | 3.28E-08 | 1 |
| CACNA2D1 | NM_000722 | 9.279328737 | 0 | 0 | 0 | 0 | 11.55195153 | 0 | 0 | 4.99E-08 | 1 |
| CACNA2D1 | NM_001302890 | 10.86274256 | 0 | 0 | 0 | 0 | 0 | 0 | 0 | 1.34E-08 | 1 |
| CADPS | XM_024453800 | 0 | 0 | 0 | 0 | 6.736572239 | 0 | 0 | 0 | 9.53E-05 | 1 |
| CALCOCO1 | XM_017019709 | 0 | 9.740796941 | 0 | 0 | 0 | 0 | 0 | 0 | 2.17E-08 | 1 |
| CALD1 | NM_033157 | 0 | 0 | 8.833416502 | 0 | 6.823901211 | 0 | 0 | 0 | 0.058416043 | 0.037082673 |
| CALM2 | NM_001305624 | 0 | 10.15080026 | 9.476548209 | 10.33783939 | 0 | 0 | 0 | 0 | 0.999953634 | 0.02072721 |
| CALML4 | NM_001286694 | 0 | 10.06146648 | 10.64221037 | 10.47821718 | 10.69009157 | 0 | 0 | 0 | 0.999953634 | 0.001228497 |
| CAMK2G | XR_945824 | 9.395290737 | 0 | 0 | 0 | 0 | 0 | 0 | 0 | 4.12E-08 | 1 |
| CAMKK2 | XR_001748558 | 0 | 8.903318262 | 0 | 0 | 0 | 0 | 0 | 0 | 7.75E-08 | 1 |
| CAMSAP2 | NM_203459 | 0 | 0 | 0 | 0 | 0 | 10.08191196 | 8.246285684 | 0 | 1 | 0.02072721 |
| CAMTA1 | XR_001737063 | 0 | 0 | 10.34177806 | 0 | 7.695467896 | 0 | 0 | 0 | 0.02111548 | 0.02072721 |
| CAND1 | NM_018448 | 0 | 0 | 0 | 8.695473678 | 6.823901211 | 0 | 0 | 0 | 0.061119289 | 0.042994552 |
| CANT1 | XM_005257022 | 0 | 9.029283553 | 10.87195067 | 10.02261004 | 0 | 0 | 0 | 0 | 0.999953634 | 0.02072721 |
| CANT1 | XM_006721683 | 0 | 11.38332176 | 10.65325302 | 10.51490355 | 9.045366917 | 0 | 0 | 0 | 0.999953634 | 0.001924802 |
| CAPN1 | NM_005186 | 0 | 0 | 0 | 9.981258774 | 7.343120441 | 0 | 8.40769976 | 10.71200753 | 0.026303837 | 0.999932046 |
| CAPN1 | XM_011545292 | 0 | 0 | 0 | 10.46542113 | 9.118435738 | 0 | 8.609636205 | 10.55247661 | 0.015315707 | 0.999932046 |
| CAPN12 | XR_001753611 | 8.569528585 | 8.419096158 | 0 | 0 | 0 | 0 | 7.534307311 | 0 | 0.006351647 | 1 |
| CAPN2 | NM_001748 | 11.03481839 | 0 | 11.26648391 | 0 | 9.204348014 | 0 | 0 | 0 | 0.999953634 | 0.02072721 |
| CAPN3 | NM_024344 | 9.783681778 | 0 | 0 | 0 | 0 | 0 | 0 | 0 | 2.45E-08 | 1 |
| CAPN5 | NM_004055 | 10.67881602 | 0 | 0 | 0 | 0 | 0 | 0 | 0 | 1.34E-08 | 1 |
| CAPNS1 | XM_005259296 | 0 | 0 | 0 | 7.749084719 | 0 | 0 | 0 | 0 | 1.37E-05 | 1 |
| CAPRIN1 | XR_930869 | 0 | 8.818611963 | 0 | 0 | 0 | 7.826947636 | 7.913105533 | 0 | 8.12E-08 | 0.034581849 |
| CAPZB | NM_001206541 | 0 | 7.606261038 | 10.96528766 | 9.893229565 | 7.757784245 | 0 | 0 | 0 | 0.564327024 | 0.002588426 |
| CAPZB | XM_011542228 | 9.588662682 | 0 | 0 | 0 | 0 | 0 | 0 | 0 | 3.15E-08 | 1 |
| CAPZB | XM_017002429 | 0 | 9.08836834 | 0 | 0 | 0 | 0 | 0 | 0 | 5.61E-08 | 1 |
| CARD10 | NM_014550 | 11.16539211 | 10.04478201 | 0 | 0 | 0 | 0 | 0 | 0 | 0.000351441 | 1 |
| CARD16 | NM_052889 | 0 | 0 | 10.40062728 | 9.987239158 | 0 | 0 | 0 | 0 | 0.013742945 | 0.02072721 |
| CARD19 | NM_001318010 | 0 | 0 | 0 | 10.21580957 | 9.787137504 | 0 | 9.173223903 | 0 | 0.014251827 | 0.999932046 |

| | | | | | | | | | | | |
|---|---|---|---|---|---|---|---|---|---|---|---|
| CARD19 | NR_134467 | 0 | 0 | 11.41097785 | 8.713623186 | 10.26099178 | 0 | 11.43331103 | 0 | 0.000782981 | 0.999932046 |
| CARD6 | XM_017009990 | 0 | 0 | 0 | 0 | 6.505153187 | 0 | 0 | 0 | 4.28E-05 | 1 |
| CARM1 | NM_199141 | 0 | 0 | 0 | 8.893154207 | 6.97463031 | 0 | 0 | 0 | 0.053874509 | 0.036824496 |
| CARS | XR_001747997 | 0 | 0 | 10.15551827 | 0 | 9.426391018 | 10.3084432 | 10.4440764 | 7.921513153 | 0.016372233 | 0.999932046 |
| CARS2 | XR_243051 | 0 | 0 | 10.70086101 | 10.29508808 | 7.393949817 | 9.734811215 | 0 | 0 | 0.001572701 | 0.999932046 |
| CASC4 | XR_001751066 | 9.143781062 | 0 | 0 | 0 | 0 | 0 | 6.232930575 | 0 | 7.26E-08 | 1 |
| CASC8 | NR_117100 | 0 | 0 | 0 | 9.044216539 | 8.803616231 | 0 | 0 | 0 | 0.029939523 | 0.021614814 |
| CASC9 | NR_103850 | 0 | 9.277965255 | 0 | 0 | 0 | 0 | 7.479117364 | 0 | 4.01E-08 | 1 |
| CASKIN2 | NM_020753 | 0 | 0 | 0 | 0 | 0 | 0 | 8.984655768 | 5.80410115 | 1 | 0.030130471 |
| CASP1 | NM_001257118 | 8.583566901 | 9.914111386 | 0 | 0 | 0 | 0 | 0 | 0 | 0.00193281 | 1 |
| CASP1 | NM_033292 | 0 | 0 | 10.63108253 | 11.18429994 | 7.574771995 | 0 | 0 | 11.04176562 | 0.000990121 | 0.999932046 |
| CASP3 | NM_032991 | 0 | 0 | 11.72819472 | 10.25351436 | 7.574771995 | 0 | 6.884098118 | 0 | 0.000833682 | 0.499074315 |
| CASP4 | NM_001225 | 0 | 0 | 9.539797346 | 9.768560191 | 6.631562102 | 0 | 10.78524292 | 0 | 0.004262501 | 1 |
| CASP8 | XM_005246885 | 0 | 0 | 0 | 10.3656523 | 8.95395271 | 0 | 8.149448007 | 0 | 0.016615172 | 0.999932046 |
| CASP8 | XM_005246886 | 11.16071767 | 10.7964161 | 0 | 0 | 0 | 0 | 9.709200726 | 0 | 0.000229177 | 1 |
| CASP8 | XM_005246894 | 0 | 4.822913417 | 0 | 7.035175705 | 9.139866469 | 0 | 0 | 0 | 0.549414983 | 0.030130471 |
| CASP8AP2 | NM_001137667 | 0 | 8.756212053 | 0 | 0 | 0 | 0 | 0 | 0 | 8.55E-08 | 1 |
| CASR | NM_000388 | 9.721559119 | 0 | 0 | 0 | 0 | 0 | 0 | 0 | 2.60E-08 | 1 |
| CAST | NM_001042442 | 0 | 0 | 0 | 9.891636224 | 7.683842675 | 0 | 0 | 0 | 0.026299336 | 0.02072721 |
| CAST | NR_104285 | 0 | 0 | 0 | 0 | 6.702411139 | 0 | 0 | 10.20553379 | 3.69E-05 | 1 |
| CASTOR1 | NM_001037666 | 0 | 0 | 10.7261817 | 0 | 7.251587968 | 0 | 0 | 0 | 0.018094055 | 0.020835214 |
| CAV2 | NM_001233 | 0 | 0 | 0 | 9.688236323 | 9.393759533 | 11.17194025 | 0 | 9.815272851 | 0.019490764 | 0.999932046 |
| CAVIN3 | NM_145040 | 0 | 0 | 8.388047551 | 10.2030191 | 10.87429511 | 0 | 0 | 12.62501914 | 0.001145381 | 0.999932046 |
| CBLB | NM_001321788 | 0 | 0 | 10.63257121 | 10.77206939 | 9.090091207 | 0 | 0 | 9.988920522 | 0.000857784 | 0.999932046 |
| CBLB | NM_001321791 | 0 | 0 | 0 | 9.993194855 | 8.410964939 | 10.55243195 | 0 | 0 | 0.021944559 | 0.999932046 |
| CBLB | NM_001321807 | 0 | 0 | 0 | 9.71551088 | 7.581052723 | 0 | 0 | 0 | 0.029493346 | 0.021614814 |
| CBLB | NM_001321820 | 0 | 9.014126367 | 0 | 0 | 0 | 0 | 0 | 0 | 6.71E-08 | 1 |
| CBLB | NM_170662 | 0 | 0 | 9.693192679 | 0 | 9.003894214 | 11.35676488 | 0 | 11.41374956 | 0.022015058 | 0.999932046 |
| CBLB | NR_135807 | 0 | 9.595615072 | 0 | 0 | 0 | 0 | 0 | 0 | 2.59E-08 | 1 |
| CBLB | XM_017007395 | 0 | 0 | 10.37417483 | 12.6025256 | 10.18158696 | 9.721887413 | 0 | 12.65785323 | 0.000220285 | 0.999932046 |
| CBLB | XM_017007396 | 0 | 0 | 0 | 10.55269793 | 8.499836882 | 0 | 8.668382154 | 10.78336691 | 0.016529494 | 0.999932046 |
| CBLB | XR_001740338 | 0 | 0 | 6.893420866 | 0 | 0 | 0 | 0 | 0 | 3.53E-05 | 1 |
| CBLC | XM_011526690 | 9.609394475 | 0 | 0 | 0 | 0 | 0 | 0 | 0 | 2.97E-08 | 1 |
| CBR3-AS1 | NR_038892 | 9.321821705 | 0 | 0 | 0 | 0 | 0 | 0 | 0 | 4.64E-08 | 1 |
| CBR4 | XR_001741341 | 0 | 8.508340767 | 0 | 0 | 0 | 0 | 0 | 0 | 1.06E-07 | 1 |
| CBWD1 | XR_001746352 | 0 | 9.81482831 | 10.89570214 | 0 | 7.542951541 | 0 | 0 | 0 | 0.999953634 | 0.02072721 |
| CBWD2 | XM_024452717 | 0 | 0 | 0 | 11.07763264 | 8.709289252 | 0 | 8.013629967 | 0 | 0.012051755 | 0.999932046 |
| CBWD5 | NR_104600 | 0 | 0 | 0 | 0 | 0 | 0 | 7.892131936 | 9.316364312 | 1 | 0.02072721 |
| CBX5 | NM_012117 | 0 | 0 | 0 | 9.701028643 | 6.955419991 | 0 | 9.365556577 | 9.400769032 | 0.03270441 | 0.999932046 |
| CBX7 | NM_001346744 | 9.270677863 | 0 | 0 | 0 | 0 | 0 | 0 | 0 | 5.05E-08 | 1 |
| CCAR1 | NM_001282959 | 0 | 9.193024922 | 0 | 10.59443305 | 9.024779586 | 0 | 0 | 0 | 0.999953634 | 0.02072721 |
| CCDC107 | NM_174923 | 0 | 3.873005774 | 9.272065393 | 10.41529045 | 0 | 0 | 0 | 0 | 0.15697939 | 0.02072721 |
| CCDC12 | XM_024453364 | 0 | 8.747073305 | 0 | 0 | 0 | 0 | 8.736643032 | 0 | 8.61E-08 | 1 |

| | | | | | | | | | | | |
|---|---|---|---|---|---|---|---|---|---|---|---|
| CCDC120 | NM_033626 | 0 | 8.878424457 | 0 | 0 | 0 | 0 | 0 | 0 | 9.15E-08 | 1 |
| CCDC124 | NM_001136203 | 0 | 0 | 10.52286184 | 10.50555865 | 10.07992424 | 0 | 9.098897264 | 10.9685914 | 0.000735886 | 0.999932046 |
| CCDC126 | XM_024446993 | 8.373391248 | 0 | 0 | 0 | 0 | 0 | 0 | 0 | 1.86E-07 | 1 |
| CCDC14 | XM_011513081 | 0 | 9.073822446 | 0 | 0 | 0 | 0 | 0 | 0 | 7.73E-08 | 1 |
| CCDC14 | XM_017007061 | 0 | 0 | 0 | 0 | 6.631562102 | 0 | 0 | 0 | 3.88E-05 | 1 |
| CCDC14 | XR_001740237 | 9.045780268 | 0 | 0 | 8.317640605 | 7.219739493 | 0 | 0 | 0 | 0.999953634 | 0.04832555 |
| CCDC142 | NM_032779 | 0 | 0 | 10.87761146 | 0 | 9.440276507 | 0 | 8.987954396 | 0 | 0.01177658 | 0.999932046 |
| CCDC144A | XR_001752703 | 9.418971874 | 0 | 0 | 0 | 0 | 0 | 0 | 0 | 4.01E-08 | 1 |
| CCDC159 | XM_017026259 | 0 | 0 | 0 | 8.14114623 | 0 | 0 | 0 | 0 | 9.06E-06 | 1 |
| CCDC159 | XM_017026260 | 0 | 9.451153482 | 0 | 0 | 0 | 0 | 0 | 0 | 3.15E-08 | 1 |
| CCDC178 | XM_017025722 | 0 | 8.632639499 | 0 | 0 | 0 | 0 | 0 | 0 | 9.44E-08 | 1 |
| CCDC18-AS1 | NR_034089 | 0 | 8.441930184 | 0 | 0 | 0 | 0 | 0 | 0 | 1.12E-07 | 1 |
| CCDC186 | NM_153249 | 0 | 8.827309931 | 0 | 0 | 0 | 0 | 0 | 0 | 8.09E-08 | 1 |
| CCDC186 | XM_011539916 | 10.736497 | 0 | 0 | 0 | 0 | 0 | 0 | 0 | 1.34E-08 | 1 |
| CCDC32 | NM_001080792 | 0 | 9.203084837 | 0 | 0 | 0 | 0 | 7.044767963 | 0 | 4.65E-08 | 1 |
| CCDC34 | NM_030771 | 0 | 0 | 0 | 9.343622985 | 7.654363064 | 0 | 8.971385315 | 0 | 0.035817219 | 0.999932046 |
| CCDC43 | NM_144609 | 0 | 9.590514794 | 0 | 0 | 0 | 0 | 9.271239882 | 0 | 2.90E-08 | 1 |
| CCDC47 | NM_020198 | 9.387309849 | 0 | 0 | 0 | 0 | 0 | 7.774812207 | 0 | 4.17E-08 | 1 |
| CCDC51 | NM_024661 | 10.22542327 | 0 | 0 | 0 | 0 | 0 | 0 | 0 | 1.59E-08 | 1 |
| CCDC57 | XR_001752488 | 8.978313131 | 0 | 0 | 0 | 0 | 0 | 0 | 0 | 7.76E-08 | 1 |
| CCDC57 | XR_001752491 | 0 | 9.265218616 | 0 | 0 | 0 | 8.305369235 | 8.479762465 | 0 | 4.18E-08 | 0.023358895 |
| CCDC7 | NM_145023 | 10.36606958 | 9.970166223 | 0 | 0 | 0 | 11.57459279 | 0 | 0 | 0.000683901 | 1 |
| CCDC7 | XM_006717506 | 11.43491862 | 0 | 0 | 0 | 0 | 0 | 8.347340272 | 9.486037855 | 1.34E-08 | 0.02072721 |
| CCDC7 | XM_011519674 | 0 | 9.029283553 | 0 | 0 | 0 | 0 | 0 | 0 | 6.49E-08 | 1 |
| CCDC7 | XM_011519678 | 10.75677221 | 0 | 0 | 0 | 0 | 0 | 0 | 0 | 1.34E-08 | 1 |
| CCDC7 | XM_024448167 | 10.66897426 | 0 | 0 | 0 | 0 | 0 | 0 | 0 | 1.34E-08 | 1 |
| CCDC85B | NM_006848 | 0 | 0 | 10.59029631 | 0 | 8.539078316 | 12.29237197 | 9.343837004 | 0 | 0.016406281 | 0.999932046 |
| CCDC88A | XM_005264421 | 9.546283346 | 0 | 0 | 0 | 0 | 0 | 0 | 0 | 3.25E-08 | 1 |
| CCDC9 | XM_017026578 | 0 | 0 | 10.26130717 | 0 | 7.843875888 | 0 | 0 | 0 | 0.021598639 | 0.02072721 |
| CCDC91 | XM_024449039 | 9.379284565 | 5.567305463 | 0 | 0 | 0 | 0 | 0 | 0 | 0.008364887 | 1 |
| CCL13 | NM_005408 | 0 | 0 | 10.73590986 | 0 | 7.817519962 | 0 | 0 | 0 | 0.016902631 | 0.02072721 |
| CCL19 | NM_006274 | 11.66688379 | 9.736185717 | 0 | 0 | 0 | 11.01921454 | 0 | 10.76528678 | 0.000258274 | 0.02072721 |
| CCL20 | NM_004591 | 11.16539211 | 9.014126367 | 10.25938129 | 0 | 7.636381755 | 0 | 0 | 0 | 0.999953634 | 0.02072721 |
| CCL21 | NM_002989 | 10.63230229 | 11.11330042 | 11.67392395 | 10.34949334 | 0 | 0 | 0 | 0 | 0.999953634 | 0.02072721 |
| CCL23 | NM_005064 | 0 | 8.632639499 | 0 | 0 | 0 | 0 | 0 | 0 | 9.44E-08 | 1 |
| CCL24 | XM_011516459 | 0 | 8.809861237 | 0 | 0 | 0 | 0 | 6.796513442 | 0 | 8.24E-08 | 1 |
| CCL24 | XM_011516460 | 0 | 0 | 0 | 0 | 0 | 10.21238087 | 7.58746354 | 0 | 1 | 0.02072721 |
| CCM2 | NM_031443 | 0 | 0 | 0 | 10.2081489 | 9.474416558 | 0 | 8.229662188 | 0 | 0.015672839 | 0.999932046 |
| CCN3 | NM_002514 | 0 | 9.073822446 | 0 | 0 | 0 | 0 | 0 | 0 | 7.73E-08 | 1 |
| CCNB1 | NM_031966 | 0 | 0 | 0 | 12.63645621 | 10.59786332 | 0 | 9.517087276 | 11.74425159 | 1.34E-08 | 0.999932046 |
| CCND1 | NM_053056 | 0 | 0 | 7.164345561 | 0 | 9.412370586 | 0 | 8.240765774 | 0 | 0.036548105 | 0.999932046 |
| CCND2-AS1 | NR_149146 | 9.395290737 | 0 | 0 | 0 | 0 | 0 | 0 | 0 | 4.12E-08 | 1 |
| CCND3 | NM_001136125 | 0 | 0 | 10.7613086 | 0 | 8.73486447 | 0 | 7.234460372 | 0 | 0.014525643 | 0.999932046 |

| | | | | | | | | | | | |
|---|---|---|---|---|---|---|---|---|---|---|---|
| CCND3 | NM_001287427 | 10.27816739 | 0 | 0 | 0 | 0 | 0 | 9.475392161 | 0 | 1.60E-08 | 1 |
| CCNDBP1 | XR_001751188 | 0 | 9.387883956 | 0 | 0 | 0 | 0 | 0 | 0 | 3.40E-08 | 1 |
| CCNG2 | NM_004354 | 0 | 0 | 10.50671712 | 0 | 7.477124411 | 0 | 7.27822739 | 0 | 0.019870902 | 0.999932046 |
| CCNH | XM_011543706 | 10.71593279 | 0 | 0 | 0 | 0 | 0 | 0 | 0 | 1.34E-08 | 1 |
| CCNK | XM_005268154 | 0 | 0 | 0 | 11.56826469 | 7.030772579 | 0 | 0 | 9.592414981 | 1.71E-08 | 0.999932046 |
| CCNL2 | NM_001320155 | 9.663265502 | 8.286608519 | 0 | 10.59834557 | 8.798235064 | 0 | 0 | 0 | 0.999953634 | 0.02072721 |
| CCNL2 | NM_001350498 | 10.19374119 | 9.522516235 | 11.05239527 | 0 | 6.229385857 | 0 | 0 | 0 | 1 | 1.34E-08 |
| CCNO | NR_125348 | 0 | 0 | 0 | 10.1019182 | 6.747781854 | 0 | 0 | 0 | 0.026303998 | 0.020730817 |
| CCNT2 | XR_001739027 | 8.511968076 | 0 | 0 | 0 | 0 | 0 | 0 | 0 | 1.14E-07 | 1 |
| CCPG1 | NM_001204450 | 0 | 0 | 13.96672311 | 9.765083618 | 10.68134883 | 12.51261428 | 10.29425491 | 12.04891617 | 8.34E-05 | 0.999932046 |
| CCSER1 | XM_017008197 | 9.676424716 | 0 | 0 | 0 | 0 | 0 | 10.03358004 | 0 | 2.73E-08 | 1 |
| CCSER2 | XR_001747113 | 0 | 0 | 0 | 0 | 6.381065478 | 0 | 0 | 0 | 4.71E-05 | 1 |
| CCT4 | NM_001256721 | 0 | 0 | 11.05739168 | 10.1560111 | 8.118872287 | 8.865496475 | 0 | 10.61059366 | 0.001172608 | 0.999932046 |
| CCT6P1 | NR_003110 | 0 | 8.919678654 | 0 | 0 | 0 | 0 | 0 | 0 | 7.73E-08 | 1 |
| CCT7 | NM_001166285 | 0 | 0 | 0 | 8.254578401 | 0 | 0 | 0 | 0 | 8.02E-06 | 1 |
| CCT7 | XM_011532479 | 0 | 0 | 0 | 8.979688824 | 7.971276284 | 0 | 7.082287382 | 7.347646499 | 0.040242734 | 0.999932046 |
| CCZ1B | NM_198097 | 0 | 0 | 0 | 11.25292332 | 7.890136184 | 0 | 7.411941715 | 0 | 0.012165562 | 0.999932046 |
| CCZ1P-OR7E38P | NR_148008 | 0 | 0 | 0 | 9.697385254 | 7.002974523 | 0 | 0 | 0 | 0.032571513 | 0.023358895 |
| CD151 | NM_001039490 | 10.12581379 | 0 | 0 | 0 | 0 | 0 | 0 | 0 | 1.68E-08 | 1 |
| CD151 | NM_139030 | 0 | 0 | 10.13675763 | 11.64828805 | 9.399989931 | 8.956574905 | 8.930825448 | 10.97647107 | 0.000578159 | 0.999932046 |
| CD164 | NM_001142402 | 0 | 0 | 10.03235972 | 0 | 9.374905252 | 0 | 0 | 0 | 0.017458727 | 0.02072721 |
| CD177 | NM_020406 | 13.23079404 | 11.50215644 | 13.54273745 | 0 | 8.806299306 | 0 | 0 | 0 | 1 | 1.80E-09 |
| CD177 | XM_017027022 | 11.61997741 | 0 | 10.81146013 | 0 | 7.282748526 | 0 | 0 | 0 | 0.999953634 | 0.02072721 |
| CD1A | NM_001320652 | 10.87706169 | 7.434733716 | 0 | 0 | 0 | 0 | 0 | 0 | 0.001160953 | 1 |
| CD226 | NM_006566 | 0 | 0 | 10.84908275 | 0 | 7.87488031 | 0 | 0 | 0 | 0.01576065 | 0.02072721 |
| CD226 | XM_017025527 | 0 | 9.847262786 | 0 | 0 | 0 | 0 | 7.372102888 | 0 | 2.00E-08 | 1 |
| CD27 | XM_017020234 | 0 | 8.348342131 | 0 | 0 | 0 | 0 | 0 | 0 | 1.24E-07 | 1 |
| CD274 | NM_014143 | 0 | 10.70025673 | 0 | 0 | 0 | 0 | 0 | 9.513376477 | 1.34E-08 | 1 |
| CD302 | NM_014880 | 10.3248786 | 0 | 0 | 0 | 0 | 0 | 0 | 10.12274908 | 1.52E-08 | 1 |
| CD33 | XM_017027510 | 8.815370067 | 0 | 0 | 0 | 0 | 0 | 0 | 0 | 8.84E-08 | 1 |
| CD36 | XM_024447002 | 0 | 0 | 7.907156541 | 0 | 0 | 0 | 0 | 0 | 1.26E-05 | 1 |
| CD36 | XM_024447003 | 9.95545054 | 0 | 0 | 0 | 0 | 0 | 0 | 0 | 2.01E-08 | 1 |
| CD37 | XM_005259436 | 10.97882423 | 9.416987232 | 0 | 0 | 0 | 0 | 0 | 0 | 0.00055775 | 1 |
| CD3D | NM_000732 | 10.58074365 | 0 | 0 | 0 | 0 | 0 | 0 | 0 | 1.34E-08 | 1 |
| CD3E | NM_000733 | 10.362003 | 0 | 10.133607 | 0 | 5.826041635 | 0 | 0 | 0 | 0.999953634 | 0.020781046 |
| CD3G | NM_000073 | 0 | 9.580259825 | 0 | 0 | 0 | 0 | 0 | 0 | 2.61E-08 | 1 |
| CD44 | NM_001001392 | 0 | 0 | 10.7128734 | 8.064023368 | 0 | 0 | 0 | 9.929274263 | 0.017830139 | 0.999932046 |
| CD44 | XM_005253232 | 0 | 0 | 0 | 0 | 0 | 0 | 12.65497307 | 10.50150827 | 1 | 1.37E-09 |
| CD44 | XM_006718388 | 0 | 0 | 0 | 10.21070698 | 7.393949817 | 11.00261298 | 10.87972254 | 0 | 0.022995441 | 0.999932046 |
| CD46 | NM_172352 | 0 | 0 | 10.02330906 | 14.78664745 | 11.46272646 | 15.26369939 | 15.53647707 | 12.57594454 | 3.32E-05 | 0.999932046 |
| CD46 | XM_017001310 | 0 | 9.059128398 | 0 | 0 | 0 | 8.296679657 | 8.289700746 | 9.945422596 | 7.80E-08 | 0.004246182 |
| CD47 | XM_005247908 | 0 | 0 | 0 | 10.45359087 | 8.94667496 | 0 | 8.913583073 | 0 | 0.015963133 | 0.999932046 |
| CD47 | XM_017007536 | 8.691211177 | 0 | 0 | 0 | 0 | 0 | 0 | 0 | 9.73E-08 | 1 |

| | | | | | | | | | | | |
|---|---|---|---|---|---|---|---|---|---|---|---|
| CD48 | XM_017002867 | 0 | 9.245884974 | 0 | 0 | 0 | 0 | 0 | 0 | 4.33E-08 | 1 |
| CD53 | XM_024451057 | 10.17067744 | 0 | 0 | 0 | 0 | 0 | 0 | 0 | 1.60E-08 | 1 |
| CD55 | NM_000574 | 0 | 0 | 0 | 8.541420573 | 9.781694404 | 0 | 7.361968883 | 0 | 0.023250768 | 0.999932046 |
| CD55 | XM_017000467 | 0 | 0 | 0 | 0 | 0 | 10.62711433 | 7.704370312 | 0 | 1 | 0.02072721 |
| CD63 | NM_001267698 | 0 | 10.64966162 | 0 | 10.80376031 | 8.304917924 | 0 | 0 | 0 | 0.999953634 | 0.02072721 |
| CD68 | NM_001040059 | 0 | 0 | 0 | 9.02108447 | 7.885068797 | 0 | 7.245527232 | 0 | 0.040355468 | 0.999932046 |
| CD79A | NM_001783 | 0 | 8.31161984 | 0 | 0 | 0 | 0 | 0 | 0 | 1.27E-07 | 1 |
| CD86 | NM_006889 | 12.0623103 | 11.99633855 | 0 | 0 | 0 | 0 | 8.372797648 | 11.14500615 | 5.39E-05 | 0.02072721 |
| CD8B | NM_172101 | 10.40611707 | 0 | 0 | 0 | 0 | 0 | 0 | 0 | 1.47E-08 | 1 |
| CD96 | XR_924090 | 9.449953927 | 0 | 0 | 0 | 0 | 0 | 0 | 0 | 3.80E-08 | 1 |
| CD99 | NM_001321368 | 0 | 0 | 0 | 10.49720094 | 9.872825412 | 11.04018953 | 0 | 0 | 0.012358485 | 0.999932046 |
| CD99P1 | NR_033381 | 0 | 0 | 0 | 9.862651006 | 7.854284905 | 0 | 0 | 0 | 0.025964912 | 0.02072721 |
| CDC16 | NM_001318518 | 0 | 0 | 0 | 11.48354058 | 9.210437894 | 10.52293272 | 8.484441039 | 0 | 0.008684252 | 0.999932046 |
| CDC16 | XR_001749713 | 0 | 0 | 0 | 0 | 6.531336591 | 11.14919239 | 0 | 0 | 4.23E-05 | 1 |
| CDC26 | XM_017014575 | 0 | 9.456770012 | 0 | 0 | 0 | 0 | 0 | 0 | 3.14E-08 | 1 |
| CDC42BPG | XM_011545157 | 0 | 0 | 7.40633941 | 0 | 0 | 0 | 8.816736559 | 0 | 2.10E-05 | 1 |
| CDC42SE1 | NM_020239 | 0 | 0 | 0 | 9.704662854 | 9.029380004 | 0 | 9.512513549 | 10.4315782 | 0.021564719 | 0.999932046 |
| CDC42SE1 | XM_017001847 | 0 | 8.360378074 | 0 | 8.119528586 | 8.743289774 | 0 | 0 | 0 | 0.999953634 | 0.029099217 |
| CDCA5 | NM_080668 | 0 | 0 | 0 | 10.93021201 | 7.944733425 | 0 | 7.596135569 | 9.220090964 | 0.014621629 | 0.999932046 |
| CDCA7L | NM_018719 | 0 | 9.17269217 | 0 | 9.761598646 | 6.667421491 | 0 | 0 | 0 | 0.999953634 | 0.023492144 |
| CDH1 | NM_001317186 | 0 | 0 | 9.68747462 | 10.45682694 | 0 | 0 | 8.521333265 | 0 | 0.014508243 | 0.999932046 |
| CDH11 | NM_001308392 | 0 | 0 | 0 | 0 | 0 | 11.97766871 | 0 | 9.043817182 | 1 | 1.92E-09 |
| CDH5 | XM_024450133 | 0 | 8.671801345 | 0 | 0 | 0 | 0 | 0 | 0 | 9.15E-08 | 1 |
| CDHR1 | XM_011540339 | 10.6656787 | 8.441930184 | 0 | 0 | 0 | 0 | 7.638730623 | 0 | 0.001079203 | 1 |
| CDHR5 | NM_001171968 | 0 | 8.508340767 | 0 | 0 | 0 | 0 | 0 | 0 | 1.06E-07 | 1 |
| CDHR5 | XM_024448584 | 0 | 9.219697032 | 0 | 0 | 0 | 0 | 0 | 0 | 4.50E-08 | 1 |
| CDK1 | NM_001786 | 0 | 0 | 0 | 10.96168545 | 9.955522041 | 0 | 7.899157068 | 9.925208779 | 0.009735518 | 0.999932046 |
| CDK10 | NM_052988 | 0 | 9.073822446 | 0 | 0 | 0 | 0 | 0 | 0 | 7.73E-08 | 1 |
| CDK10 | XM_017023808 | 9.442270566 | 0 | 0 | 0 | 0 | 0 | 7.543303808 | 0 | 3.84E-08 | 1 |
| CDK10 | XM_017023810 | 0 | 9.296876505 | 0 | 0 | 0 | 0 | 0 | 0 | 4.01E-08 | 1 |
| CDK11A | NR_132740 | 0 | 0 | 8.86415573 | 9.610942656 | 8.342372954 | 0 | 8.740559545 | 0 | 0.004093436 | 0.999932046 |
| CDK11B | XR_002958171 | 8.665043676 | 0 | 0 | 0 | 0 | 0 | 0 | 0 | 9.91E-08 | 1 |
| CDK14 | NM_012395 | 0 | 0 | 9.713029274 | 0 | 7.999685824 | 0 | 0 | 0 | 0.027908952 | 0.02072721 |
| CDK19 | XM_024446377 | 8.862188632 | 0 | 0 | 0 | 0 | 0 | 0 | 0 | 8.55E-08 | 1 |
| CDK7 | NM_001324069 | 0 | 0 | 9.781727525 | 0 | 7.562127909 | 0 | 0 | 0 | 0.029218393 | 0.020785849 |
| CDK7 | NM_001799 | 0 | 0 | 10.94865166 | 11.52163957 | 0 | 0 | 0 | 0 | 0.006049086 | 0.02072721 |
| CDK8 | XM_017020317 | 0 | 0 | 0 | 0 | 6.30721813 | 0 | 0 | 0 | 4.99E-05 | 1 |
| CDKL3 | XM_017009536 | 0 | 0 | 0 | 0 | 6.42362501 | 0 | 0 | 0 | 4.54E-05 | 1 |
| CDKL3 | XR_002956161 | 0 | 0 | 0 | 9.944846321 | 8.061632835 | 0 | 0 | 0 | 0.02404068 | 0.02072721 |
| CDKN1C | NM_001122630 | 0 | 0 | 0 | 9.535407519 | 8.748879432 | 0 | 0 | 10.49603692 | 0.024907781 | 0.999932046 |
| CDKN2A | NM_000077 | 0 | 10.36008212 | 0 | 0 | 0 | 11.3764851 | 10.86005557 | 0 | 1.43E-08 | 0.019790742 |
| CDKN2B-AS1 | NR_047536 | 10.12341345 | 0 | 0 | 0 | 0 | 0 | 0 | 0 | 1.68E-08 | 1 |
| CDKN2B-AS1 | NR_047543 | 12.80039366 | 11.67112052 | 13.12218746 | 0 | 8.927086206 | 0 | 0 | 0 | 1 | 1.92E-09 |

| | | | | | | | | | | | |
|---|---|---|---|---|---|---|---|---|---|---|---|
| CDKN3 | NM_005192 | 0 | 0 | 9.57118808 | 10.2081489 | 11.99218411 | 0 | 0 | 11.35224584 | 0.000416315 | 0.999932046 |
| CDX2 | XM_011534879 | 11.30961107 | 9.138154184 | 0 | 0 | 0 | 0 | 9.373144902 | 9.841396372 | 0.000458906 | 0.02072721 |
| CDYL | XM_011514997 | 0 | 0 | 0 | 7.631818844 | 0 | 0 | 0 | 0 | 1.70E-05 | 1 |
| CEACAM1 | NM_001024912 | 0 | 9.479019994 | 11.09405303 | 0 | 10.37743143 | 0 | 0 | 0 | 0.999953634 | 0.02072721 |
| CEACAM5 | XM_011526322 | 0 | 0 | 10.98581648 | 10.88448224 | 8.342372954 | 0 | 0 | 0 | 0.000832479 | 0.001669278 |
| CEACAM5 | XM_017026145 | 0 | 9.327854598 | 11.81696301 | 0 | 9.574604127 | 0 | 0 | 0 | 0.975136659 | 0.02072721 |
| CEBPZ | NM_005760 | 0 | 0 | 0 | 0 | 0 | 0 | 8.352467811 | 9.937371023 | 1 | 0.02072721 |
| CEBPZOS | NM_001322374 | 0 | 0 | 10.31976886 | 0 | 8.131765004 | 0 | 8.026519178 | 0 | 0.020034139 | 0.999932046 |
| CEBPZOS | XM_017003107 | 0 | 0 | 0 | 0 | 6.569741695 | 0 | 0 | 0 | 4.12E-05 | 1 |
| CECR7 | NR_015352 | 0 | 9.648106811 | 0 | 0 | 0 | 0 | 0 | 0 | 2.60E-08 | 1 |
| CELF1 | XM_024448335 | 8.766981031 | 9.059128398 | 0 | 0 | 0 | 0 | 0 | 0 | 0.003550553 | 1 |
| CELSR3 | NM_001407 | 0 | 0 | 0 | 0 | 6.702411139 | 0 | 0 | 0 | 3.69E-05 | 1 |
| CEMIP2 | NM_013390 | 0 | 9.358181476 | 0 | 0 | 0 | 0 | 8.934249332 | 0 | 3.64E-08 | 1 |
| CENPA | NM_001809 | 0 | 0 | 0 | 9.579619033 | 9.49451941 | 0 | 0 | 10.86945983 | 0.019593982 | 1 |
| CENPJ | NM_018451 | 10.05956027 | 0 | 0 | 0 | 0 | 0 | 0 | 0 | 5.01E-08 | 1 |
| CENPM | NM_024053 | 0 | 0 | 0 | 10.45682694 | 7.964085617 | 0 | 0 | 10.37904332 | 0.018782969 | 0.999932046 |
| CENPN | NM_018455 | 0 | 0 | 0 | 10.27567312 | 7.930046999 | 0 | 0 | 10.21888092 | 0.020696947 | 0.999932046 |
| CENPO | NR_136184 | 0 | 9.625843798 | 0 | 0 | 0 | 0 | 0 | 0 | 2.50E-08 | 1 |
| CENPX | NM_144998 | 0 | 0 | 11.39352333 | 10.0399755 | 8.190452933 | 12.05634967 | 10.8894564 | 0 | 0.000983427 | 0.999932046 |
| CENPX | XM_017024329 | 10.24321976 | 8.453213086 | 0 | 0 | 0 | 0 | 0 | 0 | 0.001585468 | 1 |
| CEP104 | XM_024451106 | 10.45275129 | 0 | 0 | 0 | 0 | 0 | 0 | 0 | 1.45E-08 | 1 |
| CEP164 | XM_017017365 | 0 | 0 | 9.554007759 | 10.7502556 | 10.39000635 | 10.78300665 | 11.43300853 | 10.30900536 | 0.000825021 | 0.999932046 |
| CEP164 | XM_017017377 | 11.75480525 | 0 | 0 | 0 | 0 | 0 | 9.010835996 | 0 | 1.34E-08 | 1 |
| CEP164 | XM_017017380 | 0 | 10.10145643 | 0 | 0 | 0 | 0 | 10.20423866 | 0 | 2.36E-08 | 1 |
| CEP164 | XM_017017381 | 0 | 9.914111386 | 10.91418094 | 0 | 8.371652232 | 0 | 0 | 0 | 0.999953634 | 0.02072721 |
| CEP170 | XM_011544337 | 10.5981354 | 0 | 0 | 0 | 0 | 0 | 0 | 0 | 1.34E-08 | 1 |
| CEP19 | XM_011513246 | 9.789749415 | 0 | 0 | 0 | 0 | 0 | 0 | 0 | 2.40E-08 | 1 |
| CEP57 | XM_024448778 | 0 | 0 | 7.026200938 | 0 | 0 | 0 | 0 | 0 | 3.08E-05 | 1 |
| CEP70 | NM_001320600 | 0 | 0 | 0 | 0 | 0 | 0 | 6.796513442 | 8.934803088 | 1 | 0.028217974 |
| CEP70 | XM_017007283 | 9.708807013 | 0 | 10.39537545 | 0 | 7.87488031 | 0 | 0 | 0 | 0.999953634 | 0.02072721 |
| CEP76 | XM_005258149 | 10.34149639 | 0 | 0 | 0 | 0 | 0 | 0 | 0 | 1.60E-08 | 1 |
| CEP76 | XM_017025982 | 9.502621339 | 9.439854398 | 0 | 0 | 0 | 0 | 0 | 0 | 0.001716408 | 1 |
| CEPT1 | NM_001330743 | 0 | 8.774317692 | 0 | 0 | 0 | 0 | 0 | 0 | 8.46E-08 | 1 |
| CEPT1 | XM_024450080 | 0 | 0 | 0 | 0 | 6.582319518 | 0 | 0 | 0 | 4.04E-05 | 1 |
| CERKL | NR_027690 | 9.715197156 | 0 | 0 | 0 | 0 | 0 | 0 | 0 | 2.61E-08 | 1 |
| CERNA2 | NR_134505 | 0 | 0 | 0 | 7.058084711 | 0 | 0 | 0 | 0 | 2.75E-05 | 1 |
| CERS2 | NM_181746 | 0 | 9.296876505 | 0 | 0 | 0 | 0 | 8.525879161 | 0 | 4.01E-08 | 1 |
| CERS2 | XM_011509452 | 0 | 0 | 0 | 0 | 6.366595318 | 0 | 8.655993551 | 0 | 4.76E-05 | 1 |
| CERS5 | NM_147190 | 0 | 0 | 0 | 9.996163509 | 7.449928885 | 9.250061503 | 9.403105259 | 0 | 0.025729408 | 0.999932046 |
| CETN2 | NM_004344 | 0 | 0 | 6.314513258 | 7.994469212 | 9.570664282 | 11.02448695 | 8.083136668 | 11.3650583 | 0.008785189 | 0.999932046 |
| CFAP20 | NM_013242 | 0 | 0 | 0 | 10.1639529 | 9.433350468 | 11.97800844 | 0 | 0 | 0.016139745 | 0.999932046 |
| CFAP65 | XM_017003755 | 0 | 0 | 0 | 0 | 6.59478863 | 0 | 0 | 0 | 4.02E-05 | 1 |
| CFD | NM_001928 | 0 | 0 | 11.01806684 | 0 | 11.82066397 | 0 | 6.117198234 | 10.52318849 | 0.004456296 | 0.999932046 |

| | | | | | | | | | | | |
|---|---|---|---|---|---|---|---|---|---|---|---|
| CFH | NM_001014975 | 10.28247691 | 9.660328297 | 0 | 0 | 0 | 0 | 0 | 0 | 0.000870102 | 1 |
| CFLAR | NM_001351590 | 0 | 0 | 0 | 12.883509 | 8.301118378 | 11.80727869 | 12.14264325 | 11.27687913 | 1.34E-08 | 0.999932046 |
| CFLAR | NM_001351594 | 0 | 0 | 0 | 11.3208002 | 8.258649294 | 0 | 9.779412801 | 0 | 0.011138299 | 0.999932046 |
| CFTR | NM_000492 | 9.771469405 | 0 | 0 | 0 | 0 | 8.939929929 | 0 | 0 | 2.47E-08 | 1 |
| CGGBP1 | NM_003663 | 0 | 9.803852726 | 7.526510252 | 0 | 8.360742019 | 0 | 0 | 0 | 0.999953634 | 0.040312936 |
| CH17-125A10.2 | XR_426817 | 0 | 0 | 7.926704127 | 0 | 0 | 0 | 0 | 0 | 1.25E-05 | 1 |
| CHAC2 | NM_001008708 | 0 | 0 | 0 | 0 | 6.42362501 | 0 | 0 | 0 | 4.54E-05 | 1 |
| CHAF1A | XR_936135 | 11.1359252 | 0 | 11.24377313 | 0 | 8.410964939 | 0 | 0 | 0 | 0.999953634 | 0.02072721 |
| CHCHD4 | NM_001098502 | 0 | 9.600697383 | 0 | 0 | 0 | 0 | 0 | 0 | 2.56E-08 | 1 |
| CHD1 | XR_427702 | 0 | 8.861586703 | 0 | 0 | 0 | 0 | 0 | 0 | 7.86E-08 | 1 |
| CHD1L | NR_046070 | 0 | 9.102769039 | 0 | 0 | 0 | 0 | 0 | 0 | 5.49E-08 | 1 |
| CHD3 | XM_017024064 | 0 | 0 | 9.839589214 | 0 | 8.620607807 | 0 | 0 | 0 | 0.022861747 | 0.02072721 |
| CHD4 | XM_017018733 | 0 | 9.08836834 | 0 | 0 | 0 | 0 | 9.775600494 | 0 | 5.61E-08 | 1 |
| CHD5 | NM_015557 | 0 | 0 | 0 | 8.491803396 | 0 | 0 | 0 | 0 | 6.26E-06 | 1 |
| CHD9 | XR_001751995 | 9.153224721 | 0 | 0 | 0 | 0 | 0 | 0 | 0 | 6.38E-08 | 1 |
| CHEK2 | NM_001005735 | 0 | 9.239382419 | 8.909070364 | 11.47877301 | 10.53049845 | 0 | 0 | 0 | 0.835097266 | 0.001394691 |
| CHEK2P2 | NR_038836 | 0 | 0 | 0 | 0 | 6.337212177 | 0 | 0 | 0 | 4.86E-05 | 1 |
| CHFR | NM_001161346 | 10.35792493 | 0 | 0 | 0 | 0 | 0 | 0 | 0 | 1.48E-08 | 1 |
| CHGA | NM_001275 | 0 | 10.48627065 | 0 | 0 | 0 | 0 | 0 | 0 | 1.34E-08 | 1 |
| CHGA | NM_001301690 | 0 | 8.835955774 | 0 | 0 | 0 | 0 | 0 | 0 | 2.22E-07 | 1 |
| CHGB | NM_001819 | 10.51633798 | 15.5969212 | 11.50622123 | 0 | 9.244469915 | 0 | 0 | 0 | 0.379609501 | 0.02072721 |
| CHIC2 | XM_006714037 | 0 | 9.138154184 | 0 | 0 | 0 | 0 | 0 | 0 | 5.16E-08 | 1 |
| CHID1 | NM_001142674 | 0 | 9.284296589 | 0 | 0 | 0 | 0 | 0 | 0 | 4.07E-08 | 1 |
| CHKA | XM_017017148 | 0 | 9.106346865 | 0 | 0 | 0 | 0 | 0 | 0 | 5.47E-08 | 1 |
| CHKB | NM_005198 | 10.9641441 | 8.024497849 | 0 | 0 | 0 | 0 | 0 | 11.12032494 | 0.000914795 | 1 |
| CHMP2B | NM_014043 | 0 | 0 | 10.61083355 | 9.904333953 | 11.05915161 | 10.65117337 | 0 | 10.42295417 | 0.000572294 | 0.999932046 |
| CHMP3 | NR_036454 | 9.524617511 | 9.456770012 | 0 | 0 | 0 | 0 | 8.04563962 | 0 | 0.001672333 | 1 |
| CHMP4C | NM_152284 | 0 | 0 | 0 | 11.0291809 | 7.654363064 | 0 | 10.93886971 | 0 | 0.014279378 | 0.999932046 |
| CHMP7 | NM_152272 | 10.12341345 | 0 | 0 | 0 | 0 | 9.945426233 | 0 | 0 | 1.68E-08 | 1 |
| CHODL | NM_024944 | 0 | 9.095586658 | 0 | 0 | 0 | 0 | 0 | 0 | 5.54E-08 | 1 |
| CHPF | NM_024536 | 0 | 0 | 0 | 10.70379812 | 8.499836882 | 0 | 0 | 0 | 0.015309434 | 0.02072721 |
| CHRAC1 | NM_017444 | 0 | 0 | 0 | 9.543547407 | 7.456775978 | 0 | 0 | 0 | 0.033216108 | 0.023669364 |
| CHRNB1 | NM_000747 | 0 | 9.799439006 | 0 | 0 | 0 | 0 | 0 | 0 | 2.04E-08 | 1 |
| CHST6 | XM_005255955 | 0 | 9.456770012 | 0 | 0 | 0 | 0 | 0 | 0 | 3.14E-08 | 1 |
| CHST7 | NM_019886 | 9.727893151 | 0 | 0 | 0 | 0 | 0 | 0 | 0 | 2.55E-08 | 1 |
| CHTF8 | XM_011523167 | 0 | 0 | 0 | 9.699208099 | 8.002028225 | 0 | 0 | 0 | 0.027517844 | 0.020869972 |
| CHTF8 | XM_011523168 | 0 | 0 | 9.96068535 | 10.1560111 | 0 | 0 | 0 | 5.731458906 | 0.015141066 | 0.793649388 |
| CHTOP | NM_001206612 | 0 | 0 | 9.040481197 | 0 | 7.593532742 | 0 | 7.797548711 | 9.917043278 | 0.044224465 | 0.999932046 |
| CHURC1 | NM_001204064 | 0 | 8.681427931 | 0 | 0 | 0 | 0 | 0 | 0 | 9.11E-08 | 1 |
| CIAO2B | NR_046109 | 0 | 0 | 9.053876344 | 8.14114623 | 6.30721813 | 0 | 0 | 0 | 0.012293095 | 0.019308333 |
| CIAO3 | NM_001304799 | 10.0393307 | 0 | 0 | 0 | 0 | 0 | 0 | 0 | 1.86E-08 | 1 |
| CIAO3 | NM_022493 | 0 | 9.265218616 | 0 | 0 | 0 | 0 | 0 | 0 | 4.18E-08 | 1 |
| CIAPIN1 | NM_020313 | 0 | 0 | 0 | 10.48668527 | 10.22384395 | 0 | 8.020088967 | 0 | 0.011034603 | 0.999932046 |

| | | | | | | | | | | | |
|---|---|---|---|---|---|---|---|---|---|---|---|
| CIB1 | NR_102428 | 0 | 0 | 9.718647135 | 9.32956957 | 7.779793563 | 0 | 8.864178418 | 0 | 0.003778934 | 0.999932046 |
| CIC | XM_024451433 | 9.724729612 | 0 | 8.496668691 | 0 | 8.558305894 | 0 | 0 | 0 | 0.999953634 | 0.026681724 |
| CIC | XR_002958287 | 0 | 0 | 7.945990394 | 0 | 0 | 0 | 0 | 0 | 1.21E-05 | 1 |
| CIITA | XR_932846 | 10.49049373 | 8.486541819 | 0 | 0 | 0 | 0 | 7.299622854 | 0 | 0.001248737 | 1 |
| CINP | NM_032630 | 0 | 0 | 11.07501784 | 11.40585984 | 10.64096159 | 0 | 9.316547485 | 0 | 0.000298565 | 0.999932046 |
| CIPC | NM_033426 | 0 | 8.809861237 | 0 | 0 | 0 | 0 | 0 | 0 | 8.24E-08 | 1 |
| CIRBP | NR_023313 | 10.10406561 | 8.248259235 | 0 | 0 | 0 | 0 | 0 | 0 | 0.0019392 | 1 |
| CIRBP | XM_006722637 | 4.755095516 | 8.911521649 | 0 | 9.083826095 | 8.479808505 | 0 | 0 | 0 | 0.999953634 | 0.023251362 |
| CIRBP | XR_001753598 | 0 | 0 | 9.877750519 | 0 | 5.295077683 | 0 | 0 | 0 | 0.035646895 | 0.02352709 |
| CIRBP | XR_001753600 | 9.434546067 | 6.509408673 | 9.784408563 | 9.257172648 | 7.093624618 | 0 | 0 | 0 | 0.999953634 | 0.006477692 |
| CISD1 | NM_018464 | 0 | 0 | 9.870198585 | 11.04072857 | 10.73020478 | 6.116629873 | 0 | 0 | 0.000552318 | 0.227466204 |
| CIT | XM_017018739 | 0 | 0 | 0 | 9.45569803 | 7.328262653 | 0 | 0 | 0 | 0.035784786 | 0.025156042 |
| CIZ1 | XM_024447497 | 0 | 0 | 0 | 10.7502556 | 6.582319518 | 10.31275925 | 0 | 0 | 0.018494796 | 1 |
| CKLF | NM_181641 | 0 | 0 | 9.824037495 | 0 | 8.956370491 | 9.708846791 | 0 | 8.105223474 | 0.021166613 | 0.999932046 |
| CLASP2 | XM_024453409 | 0 | 0 | 9.555578088 | 9.324854511 | 0 | 0 | 8.362668539 | 10.7698281 | 0.022759645 | 0.999932046 |
| CLASRP | NM_007056 | 10.87420921 | 0 | 0 | 0 | 0 | 0 | 0 | 0 | 1.34E-08 | 1 |
| CLASRP | NR_103529 | 0 | 0 | 10.19131336 | 10.27077828 | 0 | 0 | 0 | 0 | 0.013436527 | 0.02072721 |
| CLCA1 | NM_001285 | 10.69507121 | 11.17763435 | 13.8012746 | 9.938687217 | 0 | 0 | 0 | 0 | 0.999953634 | 1.37E-09 |
| CLCN3 | NM_173872 | 0 | 10.58188946 | 0 | 0 | 0 | 0 | 8.342194444 | 0 | 1.34E-08 | 1 |
| CLDN10 | NM_006984 | 0 | 11.07911456 | 0 | 0 | 0 | 13.31122723 | 0 | 0 | 1.34E-08 | 1 |
| CLDN12 | NM_012129 | 8.678186754 | 10.17825705 | 0 | 0 | 0 | 0 | 0 | 0 | 0.001475017 | 1 |
| CLDN5 | XM_017028930 | 10.99729455 | 9.152068755 | 0 | 0 | 0 | 0 | 0 | 0 | 0.000611815 | 1 |
| CLDN7 | NM_001185023 | 0 | 0 | 0 | 0 | 6.569741695 | 0 | 0 | 0 | 4.12E-05 | 1 |
| CLDND1 | NM_001040181 | 0 | 0 | 0 | 9.686399554 | 9.145174777 | 0 | 0 | 0 | 0.02099522 | 0.02072721 |
| CLEC16A | NM_001243403 | 0 | 0 | 9.892736819 | 0 | 8.368024654 | 0 | 10.02075404 | 0 | 0.023695558 | 0.999932046 |
| CLEC16A | XM_005255210 | 0 | 0 | 0 | 9.85612999 | 6.736572239 | 10.78921496 | 9.058679075 | 0 | 0.030538991 | 0.999932046 |
| CLEC16A | XR_932810 | 8.827217917 | 8.919678654 | 0 | 0 | 0 | 0 | 0 | 0 | 0.003780405 | 1 |
| CLEC18C | XM_024450239 | 0 | 0 | 0 | 0 | 6.451316153 | 0 | 0 | 0 | 4.51E-05 | 1 |
| CLEC2D | NM_001197318 | 0 | 8.861586703 | 0 | 0 | 0 | 0 | 0 | 0 | 7.86E-08 | 1 |
| CLIC1 | NM_001287594 | 0 | 0 | 6.811672367 | 11.48512628 | 0 | 0 | 0 | 0 | 2.06E-08 | 4.22E-08 |
| CLIC3 | NM_004669 | 0 | 0 | 8.554355783 | 0 | 8.63275172 | 0 | 0 | 0 | 0.03898352 | 0.025525363 |
| CLINT1 | NM_001195556 | 9.25321892 | 0 | 0 | 0 | 0 | 0 | 0 | 0 | 5.16E-08 | 1 |
| CLIP1 | XM_005253593 | 0 | 9.559528578 | 0 | 0 | 0 | 0 | 0 | 0 | 2.70E-08 | 1 |
| CLIP2 | XR_001744871 | 0 | 0 | 0 | 10.76860127 | 6.163947944 | 0 | 7.812509721 | 0 | 0.018822751 | 0.999932046 |
| CLK3 | NM_001130028 | 0 | 0 | 9.984197437 | 0 | 8.048094872 | 0 | 8.382856136 | 0 | 0.024116461 | 0.999932046 |
| CLMN | XM_017021646 | 0 | 11.10795622 | 10.62660726 | 0 | 11.12242091 | 0 | 0 | 0 | 0.999953634 | 0.02072721 |
| CLMN | XR_429330 | 0 | 0 | 0 | 0 | 6.61940813 | 0 | 0 | 0 | 5.85E-05 | 1 |
| CLN6 | NM_017882 | 0 | 9.131146246 | 0 | 0 | 0 | 0 | 6.884098118 | 0 | 5.23E-08 | 1 |
| CLPTM1L | XR_002956182 | 0 | 0 | 11.00892578 | 0 | 9.998974538 | 0 | 9.566463924 | 0 | 0.009469329 | 0.999932046 |
| CLSTN2 | XM_017007022 | 0 | 9.117027412 | 0 | 0 | 0 | 0 | 0 | 0 | 5.36E-08 | 1 |
| CLTB | XR_001741995 | 0 | 9.002653051 | 0 | 0 | 0 | 0 | 0 | 0 | 6.87E-08 | 1 |
| CLTC | NM_001288653 | 0 | 0 | 11.05295129 | 0 | 8.165592712 | 0 | 7.994077632 | 0 | 0.013535352 | 0.999932046 |
| CLTCL1 | XR_937923 | 0 | 9.19973932 | 0 | 0 | 0 | 0 | 0 | 0 | 4.67E-08 | 1 |

| | | | | | | | | | | | |
|---|---|---|---|---|---|---|---|---|---|---|---|
| CLUH | XM_024450680 | 0 | 0 | 9.44470014 | 0 | 7.195383085 | 0 | 0 | 0 | 0.037935462 | 0.024639803 |
| CMAHP | NR_002174 | 0 | 8.765293275 | 0 | 0 | 0 | 0 | 0 | 0 | 8.55E-08 | 1 |
| CMC1 | NM_001331186 | 0 | 9.376076245 | 0 | 0 | 0 | 0 | 0 | 0 | 3.47E-08 | 1 |
| CMC4 | NM_001018024 | 0 | 0 | 10.9080476 | 10.66141578 | 8.75722352 | 0 | 0 | 0 | 0.000868155 | 0.001737491 |
| CMKLR1 | XM_017018820 | 9.727893151 | 0 | 0 | 0 | 0 | 0 | 0 | 0 | 2.55E-08 | 1 |
| CMTM3 | XM_011522861 | 0 | 0 | 0 | 0 | 6.59478863 | 0 | 9.216048648 | 0 | 4.02E-05 | 1 |
| CMTM4 | XM_011522882 | 8.862188632 | 0 | 9.140133269 | 0 | 7.636381755 | 0 | 0 | 0 | 0.999953634 | 0.026465236 |
| CMTM6 | NM_017801 | 0 | 0 | 9.765535817 | 9.510708615 | 8.709289252 | 0 | 8.775337779 | 0 | 0.002606619 | 0.999932046 |
| CNDP2 | XM_011526073 | 8.183673428 | 8.632639499 | 10.06690435 | 0 | 8.452664007 | 0 | 0 | 0 | 0.999953634 | 0.02072721 |
| CNIH3 | NR_136297 | 0 | 0 | 0 | 10.48245744 | 7.843875888 | 0 | 0 | 0 | 0.018822751 | 0.02072721 |
| CNN2 | NM_201277 | 0 | 10.3389485 | 10.31884448 | 0 | 8.743289774 | 0 | 0 | 0 | 0.999953634 | 0.02072721 |
| CNOT6 | XM_011534606 | 0 | 9.970166223 | 0 | 8.78058117 | 8.88962462 | 0 | 0 | 0 | 0.999953634 | 0.022671467 |
| CNOT6 | XR_001742163 | 10.90248333 | 0 | 0 | 0 | 0 | 0 | 8.62242756 | 0 | 1.34E-08 | 1 |
| CNOT8 | NM_004779 | 0 | 0 | 0 | 6.783532865 | 0 | 0 | 0 | 0 | 3.65E-05 | 1 |
| CNOT9 | XM_017005249 | 0 | 0 | 0 | 0 | 0 | 0 | 0 | 6.640576357 | 1 | 2.07E-06 |
| CNPPD1 | NM_015680 | 0 | 0 | 10.11242512 | 0 | 9.023627185 | 10.71125774 | 0 | 0 | 0.018526857 | 0.999932046 |
| CNPY3 | NM_001318845 | 0 | 0 | 10.75106565 | 0 | 6.42362501 | 0 | 0 | 0 | 0.019130036 | 0.020942118 |
| CNPY3-GNMT | NR_134892 | 10.78590188 | 0 | 0 | 0 | 0 | 0 | 0 | 0 | 1.34E-08 | 1 |
| CNPY4 | NM_152755 | 0 | 0 | 10.26323048 | 0 | 7.463590728 | 0 | 0 | 0 | 0.02268042 | 0.02072721 |
| CNRIP1 | XM_006711984 | 10.25642453 | 0 | 0 | 0 | 0 | 0 | 0 | 0 | 1.60E-08 | 1 |
| COA1 | NM_001321198 | 0 | 0 | 10.10922084 | 0 | 8.048094872 | 0 | 0 | 0 | 0.022571784 | 0.02072721 |
| COA1 | XM_024446830 | 0 | 0 | 10.03461354 | 10.60029786 | 0 | 0 | 5.064738612 | 9.193294779 | 0.012363807 | 0.999932046 |
| COA6 | NM_001206641 | 0 | 0 | 10.01876234 | 10.55773017 | 9.710432422 | 10.52293272 | 8.951247914 | 10.25819701 | 0.001018211 | 0.999932046 |
| COA6-AS1 | NR_125962 | 0 | 0 | 0 | 8.650960001 | 0 | 0 | 7.877978204 | 0 | 5.26E-06 | 1 |
| COG6 | NM_001145079 | 11.11312079 | 0 | 0 | 0 | 0 | 0 | 10.10519542 | 0 | 1.34E-08 | 1 |
| COL16A1 | XM_005270481 | 0 | 8.612652582 | 0 | 0 | 0 | 0 | 0 | 0 | 9.67E-08 | 1 |
| COL16A1 | XM_011540722 | 0 | 0 | 0 | 0 | 0 | 10.74198805 | 0 | 9.84570467 | 1 | 0.02072721 |
| COL16A1 | XR_001736982 | 0 | 0 | 0 | 10.11836009 | 5.354946376 | 0 | 0 | 10.64437234 | 0.029273635 | 1 |
| COL17A1 | NM_000494 | 11.69463602 | 8.286608519 | 0 | 0 | 0 | 0 | 0 | 0 | 0.000405924 | 1 |
| COL18A1 | NM_130445 | 0 | 0 | 9.934371634 | 10.34251227 | 9.351215213 | 0 | 0 | 10.25171828 | 0.001328654 | 0.999932046 |
| COL21A1 | XM_006715223 | 0 | 9.712905893 | 0 | 0 | 0 | 0 | 0 | 0 | 2.55E-08 | 1 |
| COL24A1 | NR_146342 | 10.57020707 | 0 | 0 | 0 | 0 | 0 | 0 | 0 | 1.34E-08 | 1 |
| COL4A2 | NM_001846 | 0 | 0 | 0 | 9.626353068 | 8.421503242 | 0 | 0 | 0 | 0.026025838 | 0.02072721 |
| COL4A5 | XM_017029263 | 9.395290737 | 0 | 0 | 0 | 0 | 0 | 0 | 0 | 4.12E-08 | 1 |
| COL6A2 | NM_001849 | 0 | 0 | 0 | 0 | 6.544252221 | 0 | 0 | 0 | 4.20E-05 | 1 |
| COL9A2 | XM_017000332 | 0 | 9.817013442 | 0 | 0 | 0 | 0 | 8.756120132 | 0 | 2.01E-08 | 1 |
| COLGALT1 | NM_024656 | 0 | 0 | 10.53405661 | 0 | 7.672123016 | 11.14072035 | 8.240765774 | 0 | 0.019121873 | 0.999932046 |
| COMMD1 | XM_017003412 | 0 | 0 | 0 | 0 | 0 | 10.97224008 | 0 | 10.07216922 | 1 | 0.02072721 |
| COMMD10 | XR_001742100 | 11.68490196 | 10.63981999 | 9.473228697 | 11.56576907 | 6.667421491 | 3.39210272 | 0 | 0 | 0.999953634 | 0.022619671 |
| COMMD10 | XR_001742101 | 0 | 0 | 0 | 10.07830002 | 6.478485785 | 0 | 8.051957199 | 10.39676836 | 0.027433758 | 0.999932046 |
| COMMD5 | NM_001081004 | 0 | 8.29916838 | 0 | 0 | 0 | 0 | 7.759452999 | 0 | 1.31E-07 | 1 |
| COMMD9 | NM_014186 | 0 | 0 | 0 | 0 | 6.582319518 | 11.25165937 | 7.552244553 | 11.61644952 | 4.04E-05 | 0.338740357 |
| COMT | NM_001135162 | 0 | 0 | 10.00043084 | 10.40862134 | 0 | 10.64946811 | 9.086641422 | 0 | 0.013535352 | 0.999932046 |

| | | | | | | | | | | | |
|---|---|---|---|---|---|---|---|---|---|---|---|
| COMTD1 | XM_024447791 | 0 | 0 | 0 | 0 | 0 | 0 | 9.246664761 | 10.5312353 | 1 | 0.02072721 |
| COP1 | XM_017002066 | 10.62213692 | 0 | 0 | 0 | 0 | 0 | 7.944005075 | 0 | 1.34E-08 | 1 |
| COP1 | XM_017002079 | 0 | 0 | 10.90312209 | 10.31543415 | 10.57208211 | 0 | 9.532982091 | 12.63376518 | 0.00054663 | 0.999932046 |
| COPA | NM_004371 | 0 | 0 | 0 | 10.30230186 | 8.503148079 | 0 | 5.11387751 | 0 | 0.018681381 | 0.624273269 |
| COPB2 | XM_017007513 | 11.19541234 | 0 | 0 | 0 | 0 | 0 | 0 | 0 | 1.34E-08 | 1 |
| COPG1 | NM_016128 | 0 | 9.54905016 | 0 | 0 | 0 | 0 | 10.36573109 | 10.58376343 | 2.79E-08 | 0.02072721 |
| COPS2 | NM_004236 | 8.766981031 | 6.552011553 | 0 | 9.66043618 | 9.956126209 | 0 | 0 | 0 | 0.999953634 | 0.02096331 |
| COPS3 | NM_001199125 | 0 | 0 | 0 | 0 | 0 | 0 | 8.190112517 | 9.446871989 | 1 | 0.02072721 |
| COPS3 | NM_001316357 | 0 | 0 | 11.11284033 | 0 | 8.811650533 | 0 | 10.37709729 | 11.88851903 | 0.011828889 | 0.999932046 |
| COPS3 | NM_001316358 | 0 | 0 | 0 | 0 | 0 | 11.63357538 | 10.02557716 | 10.92344798 | 1 | 0.000537185 |
| COPS3 | XM_005256842 | 0 | 0 | 0 | 11.49565329 | 7.555764066 | 10.98175813 | 4.961135439 | 11.22490582 | 1.64E-08 | 0.999932046 |
| COPZ2 | XR_934474 | 0 | 9.484529283 | 0 | 0 | 0 | 0 | 0 | 0 | 2.99E-08 | 1 |
| COQ6 | XM_024449619 | 0 | 0 | 0 | 0 | 6.725274845 | 0 | 0 | 0 | 9.53E-05 | 1 |
| COQ6 | XR_943466 | 0 | 9.428466121 | 0 | 0 | 0 | 0 | 0 | 0 | 3.25E-08 | 1 |
| COQ8A | XM_024448518 | 0 | 0 | 8.843735856 | 0 | 7.920172326 | 0 | 0 | 9.116930837 | 0.044749943 | 0.999932046 |
| CORO1A | NM_007074 | 10.02910853 | 0 | 0 | 9.926289561 | 7.44304914 | 0 | 0 | 0 | 0.999953634 | 0.020781046 |
| CORO1A | XM_017022885 | 9.682959584 | 0 | 0 | 0 | 0 | 0 | 0 | 0 | 2.84E-08 | 1 |
| CORO1B | NM_001018070 | 0 | 0 | 0 | 0 | 6.725274845 | 0 | 0 | 0 | 9.53E-05 | 1 |
| COX15 | XM_011539298 | 0 | 9.213075091 | 0 | 0 | 0 | 0 | 0 | 0 | 4.56E-08 | 1 |
| COX20 | NM_001312872 | 0 | 0 | 0 | 8.491803396 | 0 | 0 | 0 | 0 | 6.26E-06 | 1 |
| COX20 | NR_132421 | 10.52727391 | 0 | 0 | 0 | 0 | 0 | 0 | 0 | 1.38E-08 | 1 |
| COX7A1 | NM_001864 | 0 | 9.179501644 | 0 | 0 | 0 | 0 | 0 | 8.56844836 | 4.86E-08 | 1 |
| COX7A2 | NR_029466 | 10.63903963 | 0 | 0 | 0 | 0 | 0 | 0 | 0 | 1.34E-08 | 1 |
| CPB1 | NM_001871 | 0 | 9.821373802 | 0 | 0 | 0 | 0 | 0 | 0 | 2.01E-08 | 1 |
| CPEB2 | XM_005248135 | 0 | 0 | 0 | 0 | 6.464964927 | 0 | 0 | 0 | 4.45E-05 | 1 |
| CPEB3 | XR_001747064 | 0 | 0 | 0 | 0 | 6.437537018 | 0 | 0 | 0 | 4.51E-05 | 1 |
| CPLX1 | XM_011513391 | 0 | 0 | 0 | 9.792664417 | 8.316256887 | 0 | 0 | 0 | 0.024623293 | 0.02072721 |
| CPM | XR_001748579 | 10.54172739 | 11.05339628 | 0 | 0 | 0 | 0 | 0 | 0 | 0.000286167 | 1 |
| CPM | XR_001748580 | 10.63903963 | 11.70218941 | 11.64913239 | 0 | 8.020631767 | 0 | 0 | 0 | 0.999953634 | 6.49E-09 |
| CPM | XR_001748582 | 0 | 11.78823528 | 9.696043232 | 10.70016173 | 0 | 0 | 0 | 0 | 0.999953634 | 0.02072721 |
| CPM | XR_001748583 | 10.73963487 | 10.46691892 | 0 | 0 | 0 | 0 | 0 | 0 | 0.000383723 | 1 |
| CPOX | XR_001740025 | 8.815370067 | 0 | 0 | 0 | 0 | 0 | 0 | 0 | 8.84E-08 | 1 |
| CPQ | NM_016134 | 0 | 0 | 10.77147935 | 0 | 7.516979992 | 10.82289119 | 0 | 0 | 0.017186294 | 0.999932046 |
| CPSF1 | NM_013291 | 7.92163105 | 0 | 10.41626903 | 0 | 5.572294089 | 0 | 0 | 0 | 0.996110883 | 5.45E-09 |
| CPSF3 | NM_016207 | 11.10341004 | 0 | 10.80222989 | 10.42413491 | 7.768830875 | 0 | 0 | 0 | 0.999953634 | 0.002417476 |
| CPT1A | NM_001031847 | 0 | 7.962680405 | 12.24054988 | 0 | 9.665330992 | 0 | 0 | 0 | 0.607485361 | 3.65E-09 |
| CR1L | NM_175710 | 9.992749568 | 8.903318262 | 0 | 0 | 0 | 0 | 0 | 0 | 0.001621727 | 1 |
| CR2 | NM_001877 | 0 | 8.360378074 | 0 | 0 | 0 | 0 | 0 | 0 | 1.21E-07 | 1 |
| CRACR2B | XM_017017596 | 9.616239393 | 0 | 0 | 0 | 0 | 0 | 0 | 0 | 2.91E-08 | 1 |
| CRADD | NM_001330126 | 11.27326218 | 0 | 0 | 0 | 0 | 0 | 0 | 0 | 1.34E-08 | 1 |
| CRADD | XR_001748908 | 10.94115867 | 0 | 0 | 0 | 0 | 0 | 8.15532787 | 0 | 1.34E-08 | 1 |
| CRAT | NM_004003 | 8.994164854 | 0 | 0 | 0 | 0 | 10.82289119 | 8.262719816 | 0 | 7.73E-08 | 0.02072721 |
| CRB3 | NM_139161 | 0 | 0 | 11.95989559 | 0 | 11.38812609 | 10.64605153 | 9.953169096 | 10.46416753 | 0.00374296 | 0.999932046 |

| | | | | | | | | | | | |
|---|---|---|---|---|---|---|---|---|---|---|---|
| CRBN | XM_011533793 | 0 | 8.550974732 | 0 | 0 | 0 | 0 | 0 | 0 | 1.01E-07 | 1 |
| CRCP | NM_014478 | 0 | 0 | 0 | 8.414023513 | 0 | 0 | 7.899157068 | 0 | 3.17E-05 | 1 |
| CRCP | NR_024548 | 0 | 8.612652582 | 0 | 0 | 0 | 0 | 9.717172953 | 0 | 9.67E-08 | 1 |
| CREBBP | XM_011522381 | 0 | 9.219697032 | 0 | 0 | 0 | 10.62364434 | 9.514802225 | 0 | 4.50E-08 | 0.02072721 |
| CREBZF | NR_028027 | 0 | 0 | 9.906338833 | 0 | 8.567824433 | 0 | 0 | 0 | 0.022465028 | 0.02072721 |
| CRELD1 | NM_015513 | 9.153224721 | 0 | 0 | 0 | 0 | 0 | 8.201523487 | 0 | 6.38E-08 | 1 |
| CRELD2 | NR_104295 | 0 | 8.895067962 | 0 | 0 | 0 | 0 | 0 | 0 | 7.75E-08 | 1 |
| CREM | NM_182721 | 0 | 9.411213355 | 0 | 0 | 0 | 0 | 0 | 10.5123891 | 3.28E-08 | 1 |
| CRIPAK | NM_175918 | 10.27384495 | 0 | 0 | 0 | 0 | 0 | 0 | 0 | 1.81E-08 | 1 |
| CRISPLD2 | XM_005256190 | 9.472761503 | 0 | 0 | 0 | 0 | 0 | 0 | 0 | 3.68E-08 | 1 |
| CRK | NM_005206 | 10.39421912 | 8.430558346 | 0 | 0 | 0 | 0 | 0 | 10.81886129 | 0.001393331 | 1 |
| CRLF3 | XR_001752525 | 0 | 8.919678654 | 0 | 0 | 0 | 0 | 0 | 0 | 7.73E-08 | 1 |
| CROCCP2 | NR_026752 | 10.08445476 | 0 | 0 | 0 | 0 | 0 | 0 | 0 | 1.77E-08 | 1 |
| CRTAP | NM_006371 | 0 | 0 | 0 | 10.00797741 | 9.092291437 | 0 | 9.652126454 | 9.469382508 | 0.018777731 | 0.999932046 |
| CRTC1 | XM_017026538 | 0 | 0 | 11.41271174 | 0 | 8.89215255 | 0 | 10.55108918 | 0 | 0.009783949 | 0.999932046 |
| CRY2 | NM_021117 | 0 | 9.54905016 | 0 | 0 | 0 | 0 | 0 | 0 | 2.79E-08 | 1 |
| CRYBB3 | XM_017028599 | 10.24321976 | 0 | 0 | 0 | 0 | 0 | 0 | 0 | 1.59E-08 | 1 |
| CRYL1 | XM_005266415 | 0 | 9.37013595 | 0 | 0 | 0 | 0 | 8.026519178 | 0 | 3.50E-08 | 1 |
| CRYL1 | XM_005266416 | 0 | 8.911521649 | 0 | 0 | 0 | 4.88241379 | 0 | 0 | 7.73E-08 | 0.662531298 |
| CRYM | XM_024450157 | 0 | 0 | 0 | 0 | 6.713888284 | 0 | 0 | 0 | 3.66E-05 | 1 |
| CRYZ | XM_005270491 | 0 | 9.37013595 | 0 | 0 | 0 | 0 | 0 | 0 | 3.50E-08 | 1 |
| CS | NM_004077 | 0 | 8.561439232 | 0 | 9.916920899 | 6.965057125 | 0 | 0 | 0 | 0.999953634 | 0.021292807 |
| CSF1 | XM_017000369 | 0 | 9.091982013 | 0 | 0 | 0 | 0 | 0 | 0 | 5.59E-08 | 1 |
| CSK | NM_004383 | 9.546283346 | 0 | 0 | 8.44041882 | 7.689666995 | 0 | 0 | 0 | 0.999953634 | 0.038974575 |
| CSMD1 | XM_011534754 | 0 | 9.620849457 | 0 | 0 | 0 | 0 | 0 | 0 | 2.53E-08 | 1 |
| CSNK1D | NM_139062 | 0 | 0 | 0 | 9.352916447 | 8.486515572 | 0 | 0 | 0 | 0.029044836 | 0.021217141 |
| CSNK1D | XM_005256337 | 0 | 0 | 0 | 7.600950069 | 0 | 0 | 0 | 0 | 1.59E-05 | 1 |
| CSNK1E | NM_001894 | 10.62553334 | 0 | 0 | 0 | 0 | 0 | 0 | 0 | 1.34E-08 | 1 |
| CSNK2A1 | NM_001895 | 0 | 0 | 9.751902512 | 9.981258774 | 0 | 0 | 0 | 10.89358872 | 0.017193829 | 1 |
| CSRNP3 | NM_001172173 | 0 | 8.31161984 | 0 | 0 | 0 | 0 | 0 | 0 | 1.27E-07 | 1 |
| CSRP2 | NM_001300965 | 10.82483375 | 0 | 0 | 0 | 0 | 0 | 0 | 0 | 1.34E-08 | 1 |
| CST7 | NM_003650 | 0 | 9.158976035 | 0 | 0 | 0 | 0 | 0 | 0 | 5.01E-08 | 1 |
| CSTA | NM_005213 | 0 | 10.935657 | 0 | 0 | 0 | 0 | 7.870848924 | 0 | 1.34E-08 | 1 |
| CSTF1 | NM_001324 | 0 | 0 | 10.12093523 | 0 | 7.153852394 | 0 | 0 | 0 | 0.025559486 | 0.02072721 |
| CSTF1 | XM_011528600 | 0 | 8.827309931 | 0 | 0 | 0 | 0 | 0 | 0 | 8.09E-08 | 1 |
| CSTF2 | NM_001325 | 10.26516103 | 9.327854598 | 0 | 0 | 0 | 0 | 0 | 0 | 0.00105779 | 1 |
| CTB-49A3.2 | XR_001742887 | 0 | 8.273938353 | 0 | 0 | 0 | 0 | 0 | 0 | 2.02E-07 | 1 |
| CTBP1 | NM_001012614 | 9.46519897 | 0 | 0 | 0 | 0 | 0 | 0 | 0 | 3.72E-08 | 1 |
| CTBP1 | XM_017007762 | 0 | 0 | 0 | 7.20900477 | 0 | 0 | 0 | 0 | 2.36E-05 | 1 |
| CTBP2 | NM_022802 | 10.32905103 | 0 | 0 | 0 | 0 | 0 | 0 | 0 | 1.51E-08 | 1 |
| CTBP2 | XM_024447831 | 9.379284565 | 10.56638663 | 0 | 0 | 0 | 0 | 6.735022839 | 7.921513153 | 0.000780688 | 0.052083002 |
| CTCF | NM_001191022 | 10.6885911 | 0 | 0 | 0 | 0 | 0 | 0 | 0 | 1.34E-08 | 1 |
| CTDSP2 | NM_005730 | 0 | 0 | 10.40586006 | 0 | 7.796082968 | 0 | 0 | 0 | 0.020129384 | 0.02072721 |

| | | | | | | | | | | |
|---|---|---|---|---|---|---|---|---|---|---|
| CTNNA2 | NM_001282600 | 0 | 0 | 0 | 9.901169983 | 8.764140251 | 11.8103344 | 9.230045657 | 11.06781567 | 0.021221788 | 0.999932046 |
| CTNNA2 | XM_017003405 | 0 | 0 | 7.983806275 | 0 | 0 | 0 | 0 | 0 | 1.18E-05 | 1 |
| CTNNAL1 | NM_003798 | 9.035603861 | 0 | 0 | 0 | 0 | 0 | 0 | 9.418230744 | 7.68E-08 | 1 |
| CTNNB1 | NM_001098209 | 0 | 0 | 14.30634654 | 14.42796744 | 13.15999615 | 14.50313505 | 12.36602042 | 14.0907855 | 4.69E-06 | 0.999932046 |
| CTNNB1 | NM_001098210 | 0 | 0 | 14.53404053 | 16.17862451 | 14.24303055 | 16.96489593 | 13.8302112 | 14.8385217 | 7.84E-07 | 0.999932046 |
| CTNNB1 | NM_001904 | 0 | 0 | 14.09124412 | 13.32436884 | 12.16362362 | 15.64352026 | 10.56661575 | 11.93626706 | 1.44E-05 | 0.999932046 |
| CTNNB1 | XM_017005738 | 0 | 0 | 0 | 11.43130726 | 9.751384634 | 10.39443992 | 9.80395023 | 11.26244988 | 0.007996442 | 0.999932046 |
| CTNNB1 | XM_024453356 | 0 | 0 | 12.38400458 | 11.50454124 | 10.77814382 | 11.83829702 | 9.798324675 | 9.116930837 | 0.000138295 | 0.999932046 |
| CTNNBIP1 | XM_017001855 | 9.602516927 | 0 | 0 | 9.563698439 | 7.87488031 | 0 | 0 | 0 | 0.999953634 | 0.022027568 |
| CTNND1 | NM_001206889 | 9.418971874 | 0 | 0 | 0 | 0 | 0 | 0 | 0 | 4.01E-08 | 1 |
| CTNS | XM_017024258 | 10.98676906 | 8.983325733 | 0 | 0 | 0 | 0 | 0 | 9.095383146 | 0.000659892 | 1 |
| CTPS1 | NR_125440 | 0 | 0 | 0 | 0 | 6.736572239 | 0 | 0 | 10.13863739 | 9.53E-05 | 1 |
| CTSA | NM_001167594 | 8.815370067 | 0 | 0 | 0 | 0 | 0 | 9.586186281 | 0 | 8.84E-08 | 1 |
| CTSV | NM_001333 | 0 | 0 | 0 | 0 | 0 | 0 | 9.345123718 | 10.33681783 | 1 | 0.02072721 |
| CTSZ | NM_001336 | 0 | 0 | 8.923736459 | 9.904333953 | 9.150463625 | 0 | 9.818845344 | 10.60170558 | 0.002578041 | 0.999932046 |
| CTTN | NM_001184740 | 0 | 7.962680405 | 0 | 10.51593815 | 7.490532315 | 0 | 0 | 0 | 0.972866145 | 0.02072721 |
| CTTNBP2 | NM_033427 | 0 | 9.239382419 | 0 | 0 | 0 | 0 | 0 | 0 | 4.33E-08 | 1 |
| CTU1 | NM_145232 | 11.26021133 | 8.348342131 | 0 | 0 | 0 | 0 | 0 | 0 | 0.000622718 | 1 |
| CUEDC2 | NM_024040 | 0 | 0 | 11.92356903 | 9.966198446 | 7.335710674 | 0 | 9.762177523 | 10.76755923 | 0.000840442 | 0.999932046 |
| CUL3 | XM_011511994 | 0 | 9.131146246 | 10.79825593 | 0 | 7.251587968 | 0 | 0 | 0 | 0.999953634 | 0.02072721 |
| CUL4A | NM_001354943 | 0 | 0 | 0 | 8.834652321 | 8.336816315 | 9.546078736 | 0 | 10.52453274 | 0.038439475 | 0.999932046 |
| CUL4A | XM_011537523 | 0 | 0 | 0 | 0 | 0 | 0 | 6.400722758 | 9.412433625 | 1 | 0.02296717 |
| CUL4B | XM_011531400 | 10.46037918 | 0 | 0 | 0 | 0 | 0 | 0 | 0 | 1.45E-08 | 1 |
| CUL5 | NM_003478 | 0 | 8.662110092 | 0 | 0 | 0 | 0 | 0 | 0 | 9.15E-08 | 1 |
| CUL7 | XM_017011539 | 9.025355161 | 0 | 0 | 0 | 0 | 0 | 0 | 0 | 7.69E-08 | 1 |
| CUTC | NM_015960 | 0 | 9.569931439 | 10.25938129 | 10.16659049 | 0 | 0 | 0 | 0 | 0.999953634 | 0.02072721 |
| CWF19L1 | NM_018294 | 0 | 8.632639499 | 0 | 0 | 0 | 10.64776083 | 9.761213943 | 0 | 9.44E-08 | 0.02072721 |
| CWH43 | XM_011513755 | 10.35792493 | 9.352166886 | 11.6612168 | 0 | 9.200273765 | 0 | 0 | 0 | 0.999953634 | 0.02072721 |
| CWH43 | XM_011513757 | 0 | 9.517150383 | 0 | 0 | 0 | 0 | 7.534307311 | 0 | 2.90E-08 | 1 |
| CXCL1 | NM_001511 | 0 | 0 | 0 | 9.320123993 | 10.07325642 | 0 | 0 | 0 | 0.017142442 | 0.02072721 |
| CXCL10 | NM_001565 | 0 | 0 | 9.743660257 | 0 | 8.86667168 | 0 | 8.97471439 | 0 | 0.022397485 | 0.999932046 |
| CXCL11 | NM_005409 | 0 | 9.081113725 | 0 | 0 | 0 | 0 | 0 | 0 | 5.70E-08 | 1 |
| CXCL12 | NM_199168 | 0 | 10.1263393 | 0 | 0 | 0 | 0 | 0 | 0 | 1.58E-08 | 1 |
| CXCL16 | NM_022059 | 0 | 0 | 0 | 0 | 0 | 0 | 8.417518646 | 9.172865883 | 1 | 0.02072721 |
| CXCL8 | NM_000584 | 0 | 0 | 0 | 8.597229662 | 8.944240862 | 0 | 0 | 10.18867435 | 0.033357567 | 0.999932046 |
| CXCR4 | NM_001008540 | 0 | 10.38680682 | 0 | 0 | 0 | 0 | 0 | 9.418230744 | 1.38E-08 | 1 |
| CXCR5 | NM_001716 | 10.41399488 | 8.286608519 | 0 | 0 | 0 | 0 | 0 | 10.45432674 | 0.00143772 | 1 |
| CXCR5 | NM_032966 | 11.3876948 | 10.99669162 | 0 | 0 | 0 | 0 | 0 | 11.76544089 | 0.000170553 | 1 |
| CXorf56 | NM_022101 | 0 | 9.635780908 | 0 | 8.431673933 | 8.011359982 | 0 | 0 | 0 | 0.999953634 | 0.034880283 |
| CXorf65 | XM_005262244 | 9.922694008 | 0 | 0 | 0 | 0 | 0 | 0 | 0 | 2.04E-08 | 1 |
| CXXC1 | NM_014593 | 10.27384495 | 0 | 0 | 0 | 0 | 0 | 0 | 0 | 1.81E-08 | 1 |
| CYB561A3 | NM_001300763 | 0 | 0 | 10.37772997 | 0 | 7.925118111 | 0 | 0 | 0 | 0.020062723 | 0.02072721 |
| CYB561D2 | NM_001291284 | 10.17531978 | 0 | 10.14512582 | 0 | 9.31210541 | 0 | 0 | 0 | 0.999953634 | 0.02096331 |

| | | | | | | | | | | | |
|---|---|---|---|---|---|---|---|---|---|---|---|
| CYB561D2 | NM_007022 | 0 | 0 | 9.721447883 | 10.00797741 | 0 | 0 | 0 | 0 | 0.017187999 | 0.020910753 |
| CYB5R1 | NM_016243 | 0 | 9.081113725 | 0 | 0 | 0 | 0 | 0 | 0 | 5.70E-08 | 1 |
| CYB5R3 | NM_000398 | 0 | 8.870030144 | 0 | 0 | 0 | 0 | 7.671923744 | 10.6517725 | 7.86E-08 | 0.02072721 |
| CYB5R3 | NM_007326 | 0 | 0 | 0 | 0 | 6.42362501 | 0 | 8.835168514 | 0 | 4.54E-05 | 1 |
| CYB5R4 | NM_016230 | 0 | 0 | 6.190629309 | 9.726277944 | 7.672123016 | 10.73074237 | 0 | 0 | 0.008999766 | 0.999932046 |
| CYBA | XM_011522905 | 9.531875668 | 0 | 0 | 0 | 0 | 0 | 0 | 0 | 3.28E-08 | 1 |
| CYBC1 | XM_017025078 | 0 | 0 | 8.899209478 | 0 | 9.627903042 | 0 | 0 | 0 | 0.022792691 | 0.02072721 |
| CYC1 | XM_017013102 | 0 | 0 | 10.54516517 | 0 | 7.219739493 | 0 | 7.351763189 | 0 | 0.020013257 | 0.999932046 |
| CYLD | XM_011522907 | 0 | 9.559528578 | 0 | 0 | 0 | 0 | 8.9780358 | 0 | 2.70E-08 | 1 |
| CYP21A1P | NR_040090 | 8.482303063 | 0 | 0 | 0 | 0 | 0 | 0 | 0 | 1.17E-07 | 1 |
| CYP27A1 | NM_000784 | 0 | 8.690990708 | 0 | 0 | 0 | 0 | 0 | 0 | 9.02E-08 | 1 |
| CYP27A1 | XM_017003488 | 10.47927431 | 9.405416277 | 0 | 0 | 0 | 0 | 7.892131936 | 0 | 0.000854273 | 1 |
| CYP2C18 | NM_001128925 | 11.30114042 | 10.71907374 | 6.893420866 | 10.61582178 | 8.966001299 | 0 | 0 | 0 | 0.984801075 | 0.00458654 |
| CYP2D6 | NM_000106 | 10.12821014 | 0 | 0 | 0 | 0 | 0 | 0 | 0 | 1.68E-08 | 1 |
| CYP2S1 | NM_030622 | 0 | 0 | 0 | 9.708287933 | 8.246846504 | 0 | 0 | 0 | 0.026031127 | 0.02072721 |
| CYP3A5 | NM_000777 | 0 | 8.870030144 | 11.80842676 | 0 | 8.057134281 | 0 | 0 | 0 | 0.933304018 | 5.45E-09 |
| CYP4F12 | NM_023944 | 0 | 8.975521721 | 0 | 0 | 0 | 0 | 0 | 0 | 7.27E-08 | 1 |
| CYP4F12 | NR_117085 | 0 | 0 | 10.05917594 | 0 | 8.371652232 | 0 | 0 | 0 | 0.021809154 | 0.02072721 |
| CYP4F12 | XM_017027172 | 0 | 0 | 6.811672367 | 0 | 0 | 0 | 0 | 0 | 3.79E-05 | 1 |
| CYP51A1 | NM_000786 | 0 | 0 | 10.08550279 | 0 | 7.470373439 | 0 | 0 | 0 | 0.024974675 | 0.02072721 |
| CYP7B1 | XM_017014002 | 10.47927431 | 8.662110092 | 0 | 0 | 0 | 0 | 0 | 0 | 0.001182901 | 1 |
| CYREN | XM_024446930 | 0 | 0 | 0 | 8.29371101 | 0 | 0 | 0 | 0 | 7.78E-06 | 1 |
| CYTH2 | NM_004228 | 0 | 0 | 9.787084628 | 0 | 6.886094979 | 0 | 8.178610571 | 0 | 0.032286982 | 0.999932046 |
| CYTH2 | NM_017457 | 0 | 0 | 9.15263826 | 10.09915958 | 0 | 0 | 0 | 0 | 0.019177062 | 0.02072721 |
| CYTOR | NR_024206 | 0 | 0 | 0 | 7.896185389 | 10.11389214 | 0 | 0 | 9.418230744 | 0.02211931 | 0.999932046 |
| CZIB | NM_017887 | 0 | 9.315543066 | 11.6280179 | 12.10986863 | 8.542300761 | 0 | 0 | 0 | 0.675684994 | 0.000688496 |
| CZIB | NR_130897 | 0 | 0 | 0 | 7.102839446 | 0 | 0 | 0 | 0 | 3.69E-05 | 1 |
| DACH1 | NM_080760 | 0 | 0 | 0 | 0 | 0 | 8.742914643 | 7.516144242 | 0 | 1 | 0.026465236 |
| DAG1 | NM_001177636 | 9.906032551 | 0 | 0 | 0 | 0 | 0 | 0 | 0 | 2.06E-08 | 1 |
| DAG1 | NM_001177638 | 0 | 9.630820909 | 0 | 0 | 0 | 0 | 0 | 0 | 4.30E-08 | 1 |
| DAG1 | NM_001177639 | 0 | 0 | 0 | 0 | 0 | 10.91093007 | 7.863684239 | 0 | 1 | 0.02072721 |
| DAGLA | XM_017018239 | 8.638392769 | 0 | 0 | 0 | 0 | 0 | 0 | 0 | 1.01E-07 | 1 |
| DALRD3 | XM_011533891 | 9.270677863 | 0 | 0 | 0 | 0 | 0 | 0 | 0 | 5.05E-08 | 1 |
| DAND5 | NM_152654 | 0 | 0 | 11.07939094 | 0 | 8.580418909 | 0 | 0 | 0 | 0.012541147 | 0.02072721 |
| DAP3 | XM_024449697 | 0 | 0 | 0 | 0 | 0 | 11.75590206 | 7.856483795 | 0 | 1 | 2.51E-09 |
| DAPK1 | XM_017014360 | 0 | 0 | 0 | 0 | 6.437537018 | 0 | 0 | 0 | 4.51E-05 | 1 |
| DAPK2 | NM_014326 | 9.602516927 | 0 | 0 | 0 | 0 | 0 | 6.703265946 | 0 | 3.14E-08 | 1 |
| DAXX | XM_005248860 | 0 | 8.592384875 | 0 | 0 | 0 | 0 | 9.039731805 | 0 | 9.87E-08 | 1 |
| DAZAP1 | NM_170711 | 0 | 0 | 0 | 0 | 0 | 0 | 7.212068669 | 9.981108733 | 1 | 0.02072721 |
| DAZAP2 | NM_001136266 | 0 | 9.352166886 | 0 | 0 | 0 | 0 | 0 | 0 | 3.68E-08 | 1 |
| DBI | NM_001282634 | 0 | 8.453213086 | 0 | 0 | 0 | 0 | 0 | 0 | 1.11E-07 | 1 |
| DBNDD2 | NM_001048221 | 0 | 0 | 9.754639499 | 0 | 7.95444197 | 0 | 0 | 0 | 0.027566145 | 0.02072721 |
| DBNL | NM_001284315 | 0 | 0 | 0 | 0 | 6.3222931 | 0 | 0 | 0 | 0.000244538 | 1 |

| | | | | | | | | | | | |
|---|---|---|---|---|---|---|---|---|---|---|---|
| DCAF10 | XM_005251577 | 0 | 0 | 10.65764653 | 9.71551088 | 8.342372954 | 0 | 7.759452999 | 10.55247661 | 0.001664377 | 0.999932046 |
| DCAF10 | XM_017015129 | 0 | 8.642530071 | 0 | 0 | 0 | 0 | 7.680104054 | 0 | 9.87E-08 | 1 |
| DCAF10 | XR_002956810 | 10.50900103 | 0 | 0 | 0 | 0 | 0 | 0 | 0 | 1.40E-08 | 1 |
| DCAF15 | NM_138353 | 10.68208174 | 0 | 0 | 0 | 0 | 0 | 0 | 0 | 1.34E-08 | 1 |
| DCAF4 | XM_017021207 | 0 | 0 | 0 | 0 | 6.655567121 | 0 | 0 | 0 | 3.82E-05 | 1 |
| DCAF4 | XM_017021209 | 10.24763479 | 0 | 0 | 0 | 0 | 0 | 0 | 0 | 1.58E-08 | 1 |
| DCP2 | NM_152624 | 0 | 9.670031486 | 0 | 0 | 0 | 0 | 0 | 0 | 2.40E-08 | 1 |
| DCTD | NM_001012732 | 0 | 0 | 9.948784034 | 10.27322778 | 7.235751614 | 11.69396828 | 7.569961514 | 0 | 0.002375746 | 0.999932046 |
| DCTD | XM_017007822 | 0 | 8.642530071 | 0 | 0 | 0 | 0 | 0 | 0 | 9.87E-08 | 1 |
| DCTN1 | NR_033935 | 0 | 0 | 11.313647 | 0 | 8.635771793 | 0 | 8.15532787 | 10.45573668 | 0.010823484 | 0.999932046 |
| DCTN5 | XR_001752007 | 9.015033136 | 9.193024922 | 0 | 0 | 0 | 0 | 0 | 0 | 0.002768448 | 1 |
| DCUN1D4 | XM_017007912 | 0 | 9.17269217 | 0 | 0 | 0 | 0 | 0 | 0 | 4.92E-08 | 1 |
| DCUN1D4 | XR_245254 | 0 | 0 | 9.995811331 | 0 | 6.935950419 | 9.852136402 | 9.754450806 | 0 | 0.028217352 | 0.999932046 |
| DCUN1D5 | NM_001318739 | 0 | 9.102769039 | 0 | 0 | 0 | 0 | 0 | 0 | 5.49E-08 | 1 |
| DCUN1D5 | NM_032299 | 0 | 0 | 0 | 10.16659049 | 7.672123016 | 0 | 0 | 10.79900326 | 0.022698714 | 0.999932046 |
| DDA1 | NM_024050 | 0 | 0 | 0 | 9.607064194 | 6.478485785 | 0 | 0 | 0 | 0.036850336 | 0.026003313 |
| DDA1 | XM_024451701 | 0 | 0 | 0 | 0 | 6.736572239 | 0 | 0 | 0 | 9.53E-05 | 1 |
| DDAH1 | XM_024446130 | 0 | 0 | 9.4815132 | 0 | 7.305685067 | 0 | 0 | 0 | 0.036362422 | 0.023763327 |
| DDAH2 | NM_013974 | 0 | 0 | 0 | 8.738654916 | 0 | 10.05897216 | 0 | 8.712227954 | 4.77E-06 | 0.999932046 |
| DDB1 | NM_001923 | 0 | 0 | 10.99394691 | 10.66141578 | 7.574771995 | 10.5451135 | 7.516144242 | 0 | 0.001095549 | 0.999932046 |
| DDC | NM_001242886 | 0 | 9.580259825 | 0 | 0 | 0 | 0 | 7.933778586 | 0 | 2.61E-08 | 1 |
| DDC | XM_011515161 | 10.38222222 | 0 | 5.389987782 | 9.587513928 | 0 | 0 | 0 | 0 | 0.999953634 | 0.030953422 |
| DDHD2 | XM_005273454 | 0 | 0 | 10.68516559 | 0 | 9.73879928 | 11.06726439 | 8.058247234 | 0 | 0.011938998 | 0.999932046 |
| DDHD2 | XR_001745506 | 0 | 0 | 0 | 10.5766942 | 6.544252221 | 0 | 0 | 10.43729905 | 0.020393518 | 1 |
| DDR1 | NM_013994 | 0 | 9.358181476 | 0 | 10.92865902 | 6.491881102 | 0 | 0 | 0 | 0.999953634 | 1.70E-08 |
| DDX19A | NM_018332 | 0 | 0 | 0 | 10.34367813 | 8.079488076 | 0 | 7.421731871 | 0 | 0.019565878 | 0.999932046 |
| DDX24 | NM_020414 | 0 | 0 | 0 | 8.658474981 | 9.096681855 | 0 | 8.668382154 | 9.272234908 | 0.030616778 | 0.999932046 |
| DDX28 | NM_018380 | 0 | 7.523045066 | 0 | 10.25227328 | 8.697775315 | 0 | 0 | 0 | 0.887723585 | 0.02072721 |
| DDX31 | NM_022779 | 0 | 0 | 11.66849173 | 12.67633195 | 10.2472349 | 10.42067049 | 10.80688334 | 12.51435695 | 0.000127205 | 0.999932046 |
| DDX31 | XM_011518921 | 0 | 0 | 11.73756353 | 0 | 8.489857449 | 0 | 10.60364173 | 0 | 0.008489868 | 0.999932046 |
| DDX39A | XM_011527620 | 0 | 8.384152849 | 0 | 0 | 0 | 0 | 0 | 0 | 1.18E-07 | 1 |
| DDX3X | XM_011543892 | 0 | 10.03167046 | 0 | 0 | 0 | 0 | 0 | 0 | 1.96E-08 | 1 |
| DDX3X | XM_017029313 | 10.34562109 | 0 | 0 | 0 | 0 | 0 | 0 | 0 | 2.06E-08 | 1 |
| DDX3Y | NM_004660 | 0 | 0 | 0 | 9.122377157 | 8.476443242 | 0 | 0 | 0 | 0.032342569 | 0.02296717 |
| DDX46 | XR_948314 | 10.44123329 | 0 | 0 | 0 | 0 | 0 | 0 | 0 | 1.46E-08 | 1 |
| DDX52 | NM_001291476 | 9.860637698 | 0 | 0 | 0 | 0 | 0 | 0 | 0 | 3.34E-08 | 1 |
| DDX56 | NM_019082 | 0 | 8.384152849 | 9.839589214 | 0 | 8.066117405 | 0 | 0 | 0 | 0.999953634 | 0.02072721 |
| DDX58 | NM_014314 | 0 | 9.580259825 | 0 | 0 | 0 | 0 | 0 | 0 | 2.61E-08 | 1 |
| DEF8 | NM_001242822 | 0 | 9.17269217 | 0 | 0 | 0 | 0 | 0 | 0 | 4.92E-08 | 1 |
| DEF8 | XM_017023369 | 9.801808664 | 0 | 0 | 0 | 0 | 0 | 0 | 0 | 2.38E-08 | 1 |
| DEFA5 | NM_021010 | 0 | 0 | 0 | 8.525070387 | 0 | 0 | 0 | 0 | 6.60E-06 | 1 |
| DEGS2 | XM_006720043 | 0 | 8.886770211 | 0 | 0 | 0 | 0 | 0 | 0 | 7.80E-08 | 1 |
| DELE1 | NM_014773 | 0 | 0 | 0 | 0 | 0 | 10.62016598 | 9.049236545 | 10.22881084 | 1 | 0.001535028 |

| | | | | | | | | | | | |
|---|---|---|---|---|---|---|---|---|---|---|---|
| DENND1A | XM_017014950 | 0 | 0 | 0 | 0 | 0 | 0 | 10.28019752 | 11.49964423 | 1 | 0.02072721 |
| DENND3 | XR_928310 | 0 | 0 | 0 | 0 | 0 | 11.75471288 | 0 | 10.3214327 | 1 | 0.019618292 |
| DENND4A | XR_002957625 | 0 | 0 | 0 | 0 | 6.747781854 | 0 | 0 | 0 | 9.53E-05 | 1 |
| DEPDC1B | XM_011543510 | 0 | 0 | 11.86556463 | 10.53851354 | 7.203547662 | 0 | 7.877978204 | 0 | 0.000738401 | 0.803660423 |
| DEPP1 | NM_007021 | 0 | 0 | 10.90065301 | 0 | 9.087887617 | 10.85875397 | 9.272592969 | 0 | 0.012611131 | 0.999932046 |
| DERL2 | NR_130906 | 0 | 9.17269217 | 0 | 0 | 0 | 0 | 0 | 0 | 4.92E-08 | 1 |
| DEUP1 | XR_947800 | 0 | 0 | 0 | 10.93949503 | 6.896204849 | 0 | 0 | 0 | 0.016176358 | 0.02072721 |
| DFFA | NM_004401 | 0 | 0 | 0 | 0 | 0 | 0 | 8.316186335 | 10.8449205 | 1 | 0.02072721 |
| DFFA | NM_213566 | 0 | 0 | 0 | 10.59443305 | 9.30643073 | 0 | 0 | 0 | 0.01379215 | 0.02072721 |
| DGAT2 | NM_032564 | 0 | 0 | 0 | 7.844412973 | 0 | 0 | 0 | 0 | 3.66E-05 | 1 |
| DGCR11 | NR_024157 | 0 | 0 | 0 | 0 | 6.505153187 | 0 | 0 | 0 | 4.28E-05 | 1 |
| DGCR2 | NM_005137 | 10.34562109 | 0 | 0 | 0 | 0 | 0 | 0 | 0 | 2.06E-08 | 1 |
| DGKE | XR_001752670 | 10.25203635 | 0 | 0 | 0 | 0 | 0 | 0 | 0 | 1.58E-08 | 1 |
| DGUOK | NM_001318861 | 10.59120384 | 0 | 0 | 0 | 0 | 0 | 0 | 0 | 1.34E-08 | 1 |
| DGUOK | NM_080918 | 0 | 0 | 0 | 10.2572312 | 8.693433787 | 0 | 5.11387751 | 0 | 0.01844115 | 0.618053445 |
| DGUOK | XR_244926 | 0 | 0 | 0 | 7.058084711 | 0 | 0 | 0 | 0 | 2.75E-05 | 1 |
| DHDDS | NM_001243565 | 10.33321142 | 9.265218616 | 9.601910322 | 10.56474595 | 9.945212287 | 0 | 0 | 0 | 0.999953634 | 0.002089223 |
| DHFR | NR_110936 | 9.07588576 | 0 | 0 | 0 | 0 | 0 | 5.207402423 | 0 | 7.35E-08 | 0.538759989 |
| DHODH | NM_001361 | 0 | 0 | 0 | 0 | 0 | 0 | 7.73610317 | 8.772304849 | 1 | 0.02496249 |
| DHPS | XM_024451406 | 0 | 9.399595812 | 0 | 0 | 0 | 0 | 0 | 0 | 3.36E-08 | 1 |
| DHRS11 | XM_011525233 | 0 | 0 | 7.625524183 | 0 | 0 | 0 | 0 | 0 | 1.68E-05 | 1 |
| DHRS4 | XM_006720005 | 0 | 7.501468832 | 10.71779075 | 0 | 8.60528274 | 0 | 0 | 0 | 0.826891385 | 0.02072721 |
| DHRS7B | XM_024450685 | 9.765324233 | 0 | 0 | 0 | 0 | 0 | 0 | 0 | 2.50E-08 | 1 |
| DHX16 | XR_001743678 | 0 | 0 | 0 | 10.70651942 | 8.86667168 | 0 | 0 | 0 | 0.014323767 | 0.02072721 |
| DHX32 | NM_018180 | 11.17932527 | 0 | 10.42144548 | 0 | 8.070588078 | 0 | 0 | 0 | 0.999953634 | 0.02072721 |
| DHX37 | XM_005253590 | 0 | 0 | 0 | 0 | 6.59478863 | 0 | 7.223307962 | 0 | 4.02E-05 | 1 |
| DHX40 | XM_006722083 | 0 | 0 | 10.97646811 | 0 | 8.066117405 | 0 | 0 | 0 | 0.014325624 | 0.02072721 |
| DHX8 | NM_001322218 | 9.676424716 | 0 | 0 | 0 | 0 | 0 | 0 | 0 | 2.73E-08 | 1 |
| DHX8 | NM_001322220 | 0 | 0 | 0 | 8.886769922 | 5.847041254 | 0 | 0 | 0 | 0.062748172 | 0.044872901 |
| DIABLO | NM_138930 | 0 | 0 | 0 | 0 | 6.702411139 | 0 | 0 | 0 | 3.69E-05 | 1 |
| DICER1 | XM_011536600 | 10.54890019 | 0 | 0 | 0 | 0 | 0 | 0 | 0 | 1.36E-08 | 1 |
| DICER1 | XM_011536605 | 0 | 0 | 8.02972346 | 0 | 0 | 0 | 0 | 0 | 1.13E-05 | 1 |
| DICER1-AS1 | NR_015415 | 12.3250796 | 0 | 0 | 0 | 0 | 0 | 0 | 0 | 1.34E-08 | 1 |
| DIDO1 | NM_080796 | 0 | 0 | 7.794637209 | 9.897999058 | 8.8249424 | 0 | 0 | 0 | 0.004042324 | 0.006119986 |
| DIDO1 | NM_080797 | 10.61873248 | 0 | 0 | 0 | 0 | 0 | 0 | 0 | 1.34E-08 | 1 |
| DIP2A | NM_015151 | 0 | 0 | 0 | 0 | 6.607150896 | 0 | 7.361968883 | 0 | 0.00011086 | 1 |
| DIPK1A | NM_001252271 | 0 | 0 | 10.39537545 | 0 | 6.823901211 | 0 | 8.451367849 | 0 | 0.022624736 | 0.999932046 |
| DIS3 | NM_014953 | 0 | 0 | 0 | 9.563698439 | 8.800928156 | 10.02517937 | 0 | 0 | 0.024330175 | 0.999932046 |
| DKK4 | XM_017013316 | 0 | 0 | 0 | 0 | 0 | 10.89806188 | 8.212844911 | 10.42870925 | 1 | 0.001535028 |
| DLAT | XM_011542647 | 0 | 0 | 0 | 0 | 6.531336591 | 0 | 0 | 0 | 4.23E-05 | 1 |
| DLG1 | XM_017005822 | 0 | 0 | 0 | 0 | 6.518304288 | 0 | 0 | 0 | 4.23E-05 | 1 |
| DLG1 | XR_001740044 | 0 | 0 | 0 | 8.684473357 | 0 | 0 | 0 | 0 | 5.06E-06 | 1 |
| DLG3 | NM_021120 | 0 | 7.585902702 | 9.831834309 | 10.24355571 | 9.514345989 | 0 | 0 | 0 | 0.574454921 | 0.002477101 |

| | | | | | | | | | | | |
|---|---|---|---|---|---|---|---|---|---|---|---|
| DLG3 | XM_011530883 | 0 | 8.475517571 | 0 | 0 | 0 | 0 | 0 | 0 | 1.09E-07 | 1 |
| DLG5 | NM_004747 | 0 | 0 | 0 | 0 | 6.758905041 | 0 | 2.992327696 | 0 | 9.53E-05 | 1 |
| DLGAP4 | XM_011528692 | 0 | 0 | 9.953556348 | 10.74849617 | 10.87141542 | 11.13707419 | 12.47170453 | 0 | 0.000578268 | 0.999932046 |
| DLGAP4 | XR_001754205 | 0 | 0 | 0 | 10.77639284 | 10.43750816 | 0 | 11.61202368 | 12.0903529 | 0.009048863 | 0.999932046 |
| DLL1 | XM_005266934 | 0 | 10.42455562 | 9.217560411 | 0 | 8.904726182 | 0 | 0 | 0 | 0.999953634 | 0.02072721 |
| DMAC1 | NM_001318058 | 0 | 0 | 0 | 8.75977034 | 0 | 0 | 0 | 0 | 4.66E-06 | 1 |
| DMAC1 | NM_001318059 | 0 | 0 | 9.322742744 | 0 | 7.689666995 | 0 | 0 | 0 | 0.03684379 | 0.024114988 |
| DMAC2 | NR_135478 | 0 | 0 | 0 | 7.691642912 | 0 | 0 | 0 | 0 | 5.63E-05 | 1 |
| DMBT1 | XM_024447854 | 0 | 0 | 9.580473641 | 10.56674421 | 8.025245409 | 0 | 0 | 0 | 0.001959786 | 0.003125591 |
| DMKN | NM_001352329 | 0 | 0 | 10.0050356 | 0 | 8.020631767 | 0 | 0 | 0 | 0.023961656 | 0.02072721 |
| DMTF1 | NM_001142327 | 0 | 0 | 11.87659996 | 10.80545366 | 10.9894427 | 12.10910372 | 10.29158785 | 12.31632449 | 0.000227971 | 0.999932046 |
| DMTF1 | XM_017012870 | 11.09117895 | 9.970166223 | 0 | 0 | 0 | 0 | 8.639308172 | 0 | 0.00038889 | 1 |
| DMTF1 | XM_017012875 | 0 | 0 | 10.09633208 | 0 | 7.849089784 | 11.20945984 | 0 | 10.8276001 | 0.023535365 | 0.999932046 |
| DMTF1 | XM_024447021 | 10.7333523 | 0 | 0 | 0 | 0 | 0 | 0 | 0 | 1.34E-08 | 1 |
| DMTF1 | XM_024447022 | 0 | 0 | 10.24582768 | 10.55269793 | 8.105863313 | 0 | 0 | 10.52453274 | 0.001483327 | 0.999932046 |
| DNAH9 | NM_004662 | 11.15132309 | 0 | 0 | 0 | 0 | 0 | 0 | 0 | 1.34E-08 | 1 |
| DNAJB1 | NM_001300914 | 0 | 0 | 7.684125614 | 0 | 0 | 0 | 0 | 0 | 1.68E-05 | 1 |
| DNAJB4 | NM_001317100 | 9.860637698 | 8.582143237 | 0 | 0 | 0 | 0 | 8.77915078 | 0 | 0.002095514 | 1 |
| DNAJB6 | NM_058246 | 0 | 0 | 9.96068535 | 0 | 8.338670907 | 0 | 7.154520047 | 0 | 0.023004047 | 0.999932046 |
| DNAJB9 | NM_012328 | 0 | 0 | 10.08114821 | 9.061325395 | 0 | 0 | 9.729049295 | 11.01523654 | 0.02004138 | 0.999932046 |
| DNAJC19 | NR_033723 | 0 | 9.684464992 | 0 | 0 | 0 | 0 | 8.432122801 | 0 | 2.36E-08 | 1 |
| DNAJC2 | NM_014377 | 0 | 9.872695374 | 0 | 0 | 0 | 0 | 0 | 0 | 1.94E-08 | 1 |
| DNAJC21 | XM_011513965 | 0 | 0 | 0 | 8.553562748 | 8.274237629 | 0 | 0 | 0 | 0.044583687 | 0.030260794 |
| DNAJC25 | NM_001015882 | 0 | 9.265218616 | 0 | 0 | 0 | 0 | 0 | 9.937371023 | 4.18E-08 | 1 |
| DNAJC25-GNG10 | NM_004125 | 9.11507342 | 0 | 0 | 0 | 0 | 0 | 5.824272679 | 0 | 6.89E-08 | 1 |
| DNAJC28 | NM_001040192 | 0 | 9.490017613 | 0 | 0 | 0 | 0 | 0 | 0 | 3.40E-08 | 1 |
| DNAJC4 | NM_001307980 | 0 | 8.592384875 | 0 | 0 | 0 | 0 | 0 | 0 | 9.87E-08 | 1 |
| DNAJC4 | NM_001307981 | 0 | 9.165850403 | 0 | 0 | 0 | 0 | 0 | 0 | 4.97E-08 | 1 |
| DNAJC7 | NM_001144766 | 0 | 10.2988596 | 0 | 0 | 0 | 11.20482291 | 8.240765774 | 0 | 2.61E-08 | 0.02072721 |
| DNAJC7 | NR_029431 | 0 | 10.78752928 | 10.85801896 | 10.81725179 | 8.206791567 | 0 | 0 | 0 | 0.999953634 | 0.001876635 |
| DNAJC9 | NM_015190 | 0 | 8.967675264 | 11.63768742 | 11.65393838 | 9.783737974 | 0 | 0 | 0 | 0.600730987 | 0.00064493 |
| DNALI1 | NM_003462 | 0 | 9.186279128 | 0 | 0 | 0 | 0 | 0 | 0 | 4.80E-08 | 1 |
| DNASE1 | NM_001351825 | 0 | 9.585396421 | 0 | 0 | 0 | 0 | 8.736643032 | 0 | 2.61E-08 | 1 |
| DNASE1 | NM_005223 | 0 | 11.70926512 | 0 | 0 | 0 | 12.20915519 | 12.50028314 | 0 | 1.34E-08 | 1.37E-09 |
| DNASE1 | XM_017022992 | 11.268925 | 0 | 0 | 0 | 0 | 0 | 0 | 0 | 1.34E-08 | 1 |
| DNASE1L1 | NM_001303620 | 9.883513662 | 0 | 0 | 0 | 0 | 0 | 0 | 0 | 2.12E-08 | 1 |
| DNASE2 | NM_001375 | 0 | 0 | 0 | 10.19787099 | 8.551925132 | 11.49469334 | 0 | 10.91629695 | 0.019411172 | 0.999932046 |
| DNHD1 | NM_144666 | 8.583566901 | 0 | 0 | 0 | 0 | 0 | 0 | 0 | 1.06E-07 | 1 |
| DNPEP | NM_001319121 | 0 | 0 | 10.57179227 | 0 | 8.697775315 | 0 | 0 | 0 | 0.016097447 | 0.02072721 |
| DNPEP | NM_001319122 | 0 | 0 | 10.50833974 | 0 | 8.61755579 | 0 | 0 | 0 | 0.016856584 | 0.020808337 |
| DNTTIP1 | NM_052951 | 0 | 0 | 0 | 11.47239163 | 9.623345731 | 11.26842175 | 11.55894967 | 13.04031283 | 0.008045082 | 0.999932046 |
| DNTTIP2 | NM_014597 | 11.03864504 | 0 | 0 | 0 | 0 | 11.14193369 | 7.663696786 | 0 | 1.34E-08 | 0.02072721 |
| DOCK8 | XM_011518046 | 11.12756546 | 10.95165675 | 0 | 0 | 0 | 0 | 9.53974091 | 0 | 0.000212059 | 1 |

| | | | | | | | | | | | |
|---|---|---|---|---|---|---|---|---|---|---|---|
| DOCK8 | XM_011518047 | 10.32905103 | 0 | 0 | 0 | 0 | 0 | 0 | 0 | 1.51E-08 | 1 |
| DOCK8 | XM_011518048 | 0 | 10.15426107 | 0 | 0 | 0 | 0 | 0 | 10.98918446 | 1.54E-08 | 1 |
| DOCK8 | XM_017015174 | 11.1299589 | 9.490017613 | 0 | 10.65579471 | 8.596008947 | 0 | 0 | 0 | 0.999953634 | 0.02072721 |
| DOCK9 | XM_017020520 | 8.983616415 | 0 | 0 | 0 | 0 | 0 | 0 | 0 | 7.75E-08 | 1 |
| DOK1 | NM_001318867 | 0 | 9.124104101 | 0 | 0 | 0 | 0 | 0 | 0 | 5.29E-08 | 1 |
| DOK2 | NM_201349 | 10.58772553 | 0 | 0 | 0 | 0 | 0 | 0 | 0 | 1.34E-08 | 1 |
| DOPEY2 | XR_001754923 | 0 | 0 | 7.936379488 | 0 | 0 | 0 | 0 | 0 | 1.22E-05 | 1 |
| DPAGT1 | NM_001382 | 0 | 0 | 9.561842367 | 9.999126067 | 7.82812018 | 0 | 0 | 0 | 0.002879534 | 0.004331325 |
| DPAGT1 | XM_017017295 | 0 | 0 | 0 | 0 | 6.690841957 | 0 | 0 | 0 | 3.72E-05 | 1 |
| DPAGT1 | XR_001747785 | 10.02397016 | 0 | 0 | 0 | 0 | 0 | 0 | 0 | 1.88E-08 | 1 |
| DPH3 | NM_001047434 | 0 | 0 | 9.328265229 | 11.30233023 | 9.730347633 | 0 | 8.427271157 | 0 | 0.000834239 | 0.999932046 |
| DPH5 | NM_015958 | 10.39023122 | 0 | 0 | 0 | 0 | 0 | 0 | 10.35052556 | 1.47E-08 | 1 |
| DPP4 | XM_005246371 | 0 | 0 | 9.623033138 | 0 | 9.094488316 | 0 | 0 | 0 | 0.022023199 | 0.02072721 |
| DPP7 | XM_017014652 | 0 | 7.847685041 | 0 | 0 | 0 | 0 | 5.161397696 | 9.592414981 | 0.065565564 | 0.023251362 |
| DPP7 | XM_024447525 | 10.31649738 | 8.453213086 | 0 | 7.706219197 | 8.771023979 | 0 | 0 | 0 | 0.979637466 | 0.032030331 |
| DPP8 | XR_001751338 | 0 | 0 | 0 | 0 | 6.531336591 | 6.301719098 | 0 | 0 | 4.23E-05 | 1 |
| DPP9 | XM_005259673 | 9.623051988 | 7.830482506 | 0 | 0 | 0 | 0 | 0 | 0 | 0.003569457 | 1 |
| DPT | NM_001937 | 0 | 9.445515002 | 0 | 0 | 0 | 0 | 0 | 0 | 3.16E-08 | 1 |
| DPYD | XM_006710397 | 10.34973404 | 9.131146246 | 9.796411957 | 0 | 6.780897397 | 0 | 0 | 0 | 0.999953634 | 0.021907747 |
| DPYD | XM_017000507 | 9.321821705 | 0 | 0 | 0 | 0 | 0 | 0 | 0 | 4.64E-08 | 1 |
| DPYSL3 | NM_001387 | 0 | 9.46236476 | 0 | 0 | 0 | 0 | 0 | 0 | 3.12E-08 | 1 |
| DQX1 | XM_017003520 | 10.21193034 | 0 | 0 | 0 | 0 | 0 | 0 | 0 | 1.60E-08 | 1 |
| DRAM2 | XM_011540708 | 0 | 10.03542879 | 0 | 0 | 0 | 0 | 0 | 0 | 1.60E-08 | 1 |
| DRG2 | NM_001388 | 0 | 8.083774979 | 0 | 0 | 0 | 0 | 7.450709923 | 9.981108733 | 0.058343624 | 0.02072721 |
| DROSHA | XM_024446035 | 8.526574932 | 0 | 0 | 0 | 0 | 0 | 0 | 0 | 1.12E-07 | 1 |
| DSC2 | NM_004949 | 0 | 9.640723913 | 0 | 0 | 0 | 0 | 0 | 0 | 2.47E-08 | 1 |
| DSC2 | XM_005258206 | 8.541035379 | 0 | 9.530245374 | 0 | 9.033965799 | 0 | 0 | 0 | 0.999953634 | 0.02072721 |
| DSCC1 | NM_024094 | 0 | 0 | 0 | 10.34717007 | 8.853760467 | 0 | 0 | 0 | 0.01712436 | 0.02072721 |
| DTL | NM_001286230 | 0 | 9.165850403 | 0 | 0 | 0 | 0 | 0 | 0 | 4.97E-08 | 1 |
| DTNA | NM_001198938 | 10.62553334 | 0 | 0 | 0 | 0 | 0 | 0 | 0 | 1.34E-08 | 1 |
| DTNA | NM_001198940 | 0 | 8.827309931 | 0 | 0 | 0 | 0 | 0 | 0 | 8.09E-08 | 1 |
| DTNA | XM_017025591 | 9.442270566 | 0 | 0 | 0 | 0 | 0 | 0 | 0 | 3.84E-08 | 1 |
| DTNA | XM_017025601 | 0 | 0 | 0 | 12.28636607 | 7.925118111 | 0 | 10.58688 | 0 | 1.34E-08 | 0.999932046 |
| DTNA | XM_017025602 | 9.279328737 | 0 | 0 | 0 | 0 | 0 | 9.489424851 | 10.29961409 | 4.99E-08 | 0.02072721 |
| DTNA | XR_001753159 | 0 | 0 | 0 | 8.491803396 | 0 | 0 | 0 | 0 | 6.26E-06 | 1 |
| DTNA | XR_001753161 | 9.065920236 | 0 | 0 | 0 | 0 | 0 | 0 | 0 | 7.80E-08 | 1 |
| DTNB | NM_001256303 | 0 | 0 | 12.06919708 | 11.16925437 | 12.04723506 | 13.47193948 | 10.88415517 | 9.38309338 | 0.00010839 | 0.999932046 |
| DTNB | NM_001320936 | 0 | 0 | 0 | 9.664173964 | 7.012300182 | 0 | 0 | 9.761562403 | 0.0331797 | 0.999932046 |
| DTNB | NM_001351394 | 0 | 0 | 0 | 0 | 0 | 11.14979564 | 7.751711637 | 0 | 1 | 0.02072721 |
| DTNB | NM_021907 | 0 | 0 | 0 | 10.33080157 | 5.804731829 | 0 | 0 | 0 | 0.024847467 | 0.02072721 |
| DTNB | NM_183360 | 0 | 0 | 10.63405836 | 9.792664417 | 7.817519962 | 0 | 0 | 0 | 0.001853534 | 0.003016079 |
| DTNBP1 | NM_032122 | 0 | 9.165850403 | 0 | 0 | 0 | 0 | 8.507608997 | 0 | 4.97E-08 | 1 |
| DTNBP1 | XM_005249447 | 10.49049373 | 0 | 0 | 0 | 0 | 0 | 0 | 0 | 1.43E-08 | 1 |

| | | | | | | | | | | | |
|---|---|---|---|---|---|---|---|---|---|---|---|
| DTNBP1 | XM_011514937 | 0 | 9.610808606 | 0 | 0 | 0 | 0 | 0 | 0 | 2.54E-08 | 1 |
| DTX2 | NM_020892 | 0 | 10.16115789 | 0 | 0 | 0 | 0 | 0 | 0 | 1.54E-08 | 1 |
| DTX3L | NM_138287 | 10.16134771 | 0 | 9.991176986 | 0 | 7.97366523 | 0 | 0 | 0 | 0.999953634 | 0.02072721 |
| DTYMK | NM_012145 | 0 | 0 | 11.24912055 | 0 | 9.900711739 | 11.09759742 | 9.49175042 | 9.058739343 | 0.008625002 | 0.999932046 |
| DUOXA2 | NM_207581 | 0 | 0 | 0 | 9.02108447 | 9.34384701 | 0 | 0 | 0 | 0.024716486 | 0.02072721 |
| DUS1L | NM_022156 | 0 | 0 | 0 | 11.66470673 | 8.323766986 | 0 | 0 | 0 | 0.008885257 | 1.73E-08 |
| DUS1L | XM_005256393 | 0 | 0 | 0 | 0 | 6.544252221 | 0 | 0 | 0 | 4.20E-05 | 1 |
| DUSP11 | NM_003584 | 0 | 6.373487887 | 9.110526406 | 10.41529045 | 0 | 0 | 0 | 0 | 0.584713146 | 0.020835214 |
| DUSP16 | NM_030640 | 0 | 0 | 11.73132442 | 0 | 8.262562196 | 10.065381 | 9.083561124 | 0 | 0.008779317 | 0.999932046 |
| DUSP22 | NM_020185 | 0 | 0 | 0 | 0 | 6.758905041 | 0 | 9.268529894 | 0 | 9.53E-05 | 1 |
| DUSP22 | NR_104473 | 9.917161541 | 0 | 0 | 0 | 0 | 0 | 0 | 0 | 2.04E-08 | 1 |
| DUSP26 | NM_001305115 | 11.66523456 | 0 | 10.02330906 | 0 | 7.910229599 | 0 | 0 | 0 | 0.999953634 | 0.02072721 |
| DUSP26 | NM_024025 | 11.57149456 | 0 | 0 | 0 | 0 | 0 | 0 | 0 | 1.34E-08 | 1 |
| DUSP6 | NM_022652 | 8.12728836 | 8.168359982 | 0 | 0 | 0 | 0 | 7.885072427 | 0 | 0.010039832 | 1 |
| DUT | XR_931760 | 0 | 0 | 0 | 0 | 6.42362501 | 0 | 7.534307311 | 0 | 4.54E-05 | 1 |
| DVL3 | XM_011512513 | 0 | 0 | 0 | 9.603175278 | 7.40814953 | 0 | 0 | 0 | 0.032402276 | 0.023251362 |
| DYM | NM_001353211 | 0 | 0 | 0 | 9.523110922 | 8.858938826 | 11.19316495 | 0 | 10.59148026 | 0.024330175 | 0.999932046 |
| DYM | XM_006722488 | 0 | 9.186279128 | 0 | 0 | 0 | 0 | 7.56113023 | 0 | 4.80E-08 | 1 |
| DYM | XM_006722492 | 0 | 0 | 0 | 8.254578401 | 8.641793038 | 0 | 0 | 0 | 0.043003113 | 0.029148463 |
| DYM | XM_011526038 | 10.22765987 | 0 | 0 | 0 | 0 | 0 | 0 | 0 | 1.59E-08 | 1 |
| DYM | XM_017025796 | 10.56313968 | 9.376076245 | 0 | 0 | 0 | 0 | 9.857228707 | 0 | 0.000809073 | 1 |
| DYNC1LI2 | NR_136658 | 10.28247691 | 8.612652582 | 0 | 0 | 0 | 0 | 4.107213043 | 0 | 0.001437855 | 0.840351602 |
| DYNC2H1 | XM_017018292 | 0 | 0 | 11.07884502 | 0 | 10.48433361 | 0 | 9.550935715 | 11.50918171 | 0.007824428 | 0.999932046 |
| DYNC2LI1 | NM_001193464 | 0 | 8.765293275 | 0 | 0 | 0 | 0 | 0 | 0 | 8.55E-08 | 1 |
| DYNLL1 | NM_001037495 | 0 | 0 | 10.65031657 | 10.89724281 | 9.653459362 | 0 | 9.190506642 | 0 | 0.000673026 | 0.999932046 |
| DYNLRB1 | NM_001281727 | 9.124706245 | 0 | 0 | 0 | 0 | 0 | 0 | 0 | 6.80E-08 | 1 |
| DYNLRB1 | NM_177953 | 0 | 0 | 10.91662699 | 10.23100994 | 0 | 0 | 0 | 0 | 0.010257629 | 0.02072721 |
| DYRK1A | NM_130438 | 8.716912497 | 0 | 0 | 0 | 0 | 0 | 6.826303056 | 0 | 9.50E-08 | 1 |
| DYRK4 | NM_001282286 | 0 | 9.337019786 | 0 | 0 | 0 | 0 | 0 | 0 | 3.77E-08 | 1 |
| DZIP3 | XM_005247918 | 10.77370837 | 0 | 0 | 0 | 0 | 0 | 0 | 0 | 1.34E-08 | 1 |
| EBAG9 | NM_198120 | 9.765324233 | 0 | 0 | 0 | 0 | 0 | 0 | 0 | 2.50E-08 | 1 |
| EBF4 | XM_017027985 | 8.779231295 | 0 | 0 | 0 | 0 | 0 | 0 | 0 | 9.02E-08 | 1 |
| EBI3 | XM_011527619 | 10.39023122 | 0 | 0 | 0 | 0 | 0 | 0 | 0 | 1.47E-08 | 1 |
| ECE1 | NM_001113348 | 11.0437315 | 9.095586658 | 0 | 0 | 0 | 0 | 0 | 10.53657502 | 0.000599243 | 1 |
| ECHDC1 | XM_005267050 | 0 | 0 | 0 | 12.59103571 | 11.24218959 | 11.40708366 | 11.66759216 | 12.834069 | 0.002828364 | 0.999932046 |
| ECHDC2 | XR_002957014 | 11.16305678 | 0 | 0 | 0 | 0 | 0 | 0 | 0 | 1.34E-08 | 1 |
| ECM2 | XM_024447437 | 8.597469932 | 0 | 0 | 0 | 0 | 0 | 0 | 0 | 1.05E-07 | 1 |
| ECSIT | NM_001142464 | 0 | 9.258802813 | 0 | 0 | 0 | 0 | 0 | 0 | 4.23E-08 | 1 |
| EDDM13 | NM_001354658 | 0 | 0 | 7.805236676 | 0 | 0 | 0 | 0 | 0 | 1.40E-05 | 1 |
| EDN2 | NM_001956 | 10.29105753 | 0 | 0 | 0 | 0 | 0 | 0 | 0 | 1.54E-08 | 1 |
| EDRF1 | XM_017016066 | 0 | 8.273938353 | 0 | 0 | 0 | 0 | 0 | 0 | 2.02E-07 | 1 |
| EDRF1 | XR_945662 | 10.04948094 | 0 | 0 | 0 | 0 | 0 | 0 | 0 | 5.08E-08 | 1 |
| EEF1AKMT1 | NM_001318939 | 0 | 0 | 10.63702806 | 10.32136409 | 10.28907758 | 0 | 8.97471439 | 10.39382927 | 0.000695881 | 0.999932046 |

| | | | | | | | | | | | |
|---|---|---|---|---|---|---|---|---|---|---|---|
| EEF1AKMT2 | NM_212554 | 0 | 0 | 10.98056548 | 0 | 7.864619359 | 0 | 0 | 0 | 0.014680579 | 0.02072721 |
| EEF1AKMT4 | NM_032331 | 0 | 0 | 0 | 11.42911218 | 7.549372027 | 10.64005298 | 0 | 0 | 0.011298501 | 0.999932046 |
| EEF1B2 | NM_001037663 | 0 | 0 | 0 | 10.29749668 | 9.429874939 | 0 | 8.326645909 | 0 | 0.015306964 | 0.999932046 |
| EEF1D | XM_024447088 | 11.15132309 | 5.481743011 | 0 | 0 | 0 | 6.193537681 | 0 | 0 | 0.00125465 | 1 |
| EEF1D | XM_024447089 | 0 | 0 | 0 | 11.3419566 | 10.55104023 | 0 | 10.21342682 | 10.28064615 | 0.006731982 | 0.999932046 |
| EEF2KMT | NM_201400 | 10.83953218 | 9.665188049 | 0 | 0 | 0 | 0 | 0 | 0 | 0.000559903 | 1 |
| EEF2KMT | NM_201598 | 0 | 0 | 0 | 10.49090074 | 8.583550423 | 0 | 7.331132628 | 10.81006923 | 0.016811367 | 0.999932046 |
| EEFSEC | NM_021937 | 0 | 9.930350523 | 0 | 0 | 0 | 0 | 0 | 0 | 1.81E-08 | 1 |
| EFCAB11 | NM_145231 | 0 | 0 | 0 | 10.35412868 | 7.760553842 | 0 | 0 | 10.41137467 | 0.020345289 | 0.999932046 |
| EFEMP1 | NM_001039349 | 0 | 0 | 0 | 10.10467156 | 9.349376687 | 12.0865956 | 0 | 10.51509654 | 0.016909521 | 0.999932046 |
| EFEMP1 | XM_024452757 | 0 | 0 | 10.06580283 | 0 | 8.230957729 | 0 | 0 | 8.649540214 | 0.022331735 | 0.999932046 |
| EFL1 | XM_011522000 | 0 | 0 | 0 | 9.350598688 | 6.98414039 | 0 | 7.870848924 | 0 | 0.040460065 | 0.999932046 |
| EFNA3 | NM_004952 | 0 | 0 | 0 | 0 | 0 | 9.96338155 | 9.117087997 | 0 | 1 | 0.02072721 |
| EFNA5 | XM_017009205 | 0 | 0 | 0 | 0 | 6.690841957 | 0 | 0 | 0 | 3.72E-05 | 1 |
| EFR3A | XR_002956625 | 0 | 8.861586703 | 0 | 0 | 0 | 0 | 0 | 0 | 7.86E-08 | 1 |
| EGLN2 | NM_080732 | 9.629832564 | 8.069182463 | 0 | 0 | 0 | 10.82893057 | 0 | 0 | 0.00322741 | 1 |
| EGLN3 | NM_022073 | 0 | 10.85607449 | 10.34177806 | 0 | 9.66014917 | 0 | 0 | 0 | 0.999953634 | 0.02072721 |
| EGR1 | NM_001964 | 10.43350322 | 0 | 0 | 0 | 0 | 0 | 0 | 0 | 1.47E-08 | 1 |
| EHBP1L1 | NM_001351087 | 0 | 0 | 0 | 0 | 6.631562102 | 0 | 0 | 0 | 3.88E-05 | 1 |
| EHBP1L1 | XM_005273868 | 0 | 0 | 9.4815132 | 0 | 7.752229049 | 0 | 8.618176364 | 0 | 0.03328205 | 0.999932046 |
| EHD1 | XM_011544739 | 0 | 0 | 0 | 0 | 0 | 0 | 8.04563962 | 8.639649521 | 1 | 0.024311363 |
| EHF | XM_024448434 | 0 | 10.45012258 | 9.582015441 | 0 | 9.017851335 | 0 | 0 | 0 | 0.999953634 | 0.02072721 |
| EID2 | NM_153232 | 0 | 0 | 10.12199547 | 0 | 7.379608952 | 0 | 0 | 0 | 0.024784435 | 0.02072721 |
| EIF2A | XM_011513224 | 0 | 8.642530071 | 0 | 0 | 0 | 0 | 0 | 8.261726724 | 9.87E-08 | 1 |
| EIF2AK3 | NM_004836 | 0 | 8.497482465 | 0 | 0 | 0 | 0 | 0 | 0 | 1.06E-07 | 1 |
| EIF2AK3 | XM_017005376 | 0 | 8.31161984 | 0 | 0 | 0 | 0 | 0 | 0 | 1.27E-07 | 1 |
| EIF2AK4 | XM_011521600 | 7.486331536 | 8.671801345 | 0 | 0 | 0 | 0 | 7.351763189 | 9.116930837 | 0.009454429 | 0.023251362 |
| EIF2AK4 | XM_017022219 | 10.07951011 | 0 | 0 | 0 | 0 | 0 | 4.906403331 | 0 | 1.78E-08 | 0.614577643 |
| EIF2B1 | NM_001414 | 0 | 0 | 8.86415573 | 0 | 8.876918096 | 0 | 0 | 0 | 0.031496943 | 0.02158975 |
| EIF2B2 | NM_014239 | 0 | 0 | 0 | 10.43073285 | 6.112844945 | 0 | 6.164610941 | 0 | 0.022896462 | 0.999932046 |
| EIF2B5 | NM_003907 | 0 | 9.10991584 | 0 | 0 | 0 | 10.41465947 | 0 | 0 | 5.42E-08 | 1 |
| EIF2B5 | XR_924208 | 0 | 0 | 10.51964732 | 0 | 8.2973088 | 0 | 0 | 0 | 0.017712464 | 0.02096331 |
| EIF2D | XM_006711196 | 10.0942935 | 0 | 0 | 0 | 0 | 0 | 0 | 0 | 1.80E-08 | 1 |
| EIF2S2 | NM_003908 | 0 | 0 | 0 | 12.83687083 | 11.61433825 | 11.1013446 | 9.942157556 | 13.04219111 | 1.34E-08 | 0.999932046 |
| EIF2S2 | XM_017028118 | 0 | 0 | 9.16914474 | 9.836388206 | 6.197037886 | 0 | 9.866219032 | 0 | 0.005208197 | 0.999932046 |
| EIF3B | XM_017012752 | 0 | 4.676562808 | 10.33264833 | 0 | 7.10238406 | 0 | 0 | 0 | 0.342061359 | 0.02072721 |
| EIF3I | XM_024450518 | 0 | 0 | 9.484813727 | 12.01206825 | 6.30721813 | 11.31757107 | 0 | 7.837479392 | 0.001039056 | 0.999932046 |
| EIF3J-DT | NR_034171 | 0 | 0 | 0 | 0 | 6.351978551 | 0 | 0 | 0 | 4.81E-05 | 1 |
| EIF3L | XM_006724260 | 0 | 0 | 0 | 6.810766492 | 0 | 0 | 5.824272679 | 0 | 3.53E-05 | 1 |
| EIF4A1 | NM_001416 | 0 | 0 | 9.58969982 | 11.28604854 | 7.785243826 | 0 | 0 | 0 | 0.001303194 | 0.002469438 |
| EIF4A3 | XM_011525522 | 10.85408237 | 0 | 0 | 0 | 0 | 0 | 0 | 0 | 1.34E-08 | 1 |
| EIF4E2 | NM_001276336 | 0 | 9.500932031 | 0 | 0 | 0 | 0 | 0 | 0 | 2.95E-08 | 1 |
| EIF4G3 | NM_001198802 | 0 | 9.117027412 | 0 | 0 | 0 | 0 | 0 | 0 | 5.36E-08 | 1 |

| | | | | | | | | | | | |
|---|---|---|---|---|---|---|---|---|---|---|---|
| EIF4G3 | XM_017002684 | 0 | 8.261155927 | 0 | 0 | 0 | 0 | 0 | 10.01593341 | 1.93E-07 | 1 |
| EIF4G3 | XM_017002700 | 0 | 0 | 11.28648052 | 0 | 8.8511643 | 0 | 0 | 0 | 0.01062117 | 0.02072721 |
| EIF4G3 | XM_017002704 | 0 | 9.934381906 | 9.382357652 | 7.958392843 | 8.912218095 | 0 | 0 | 0 | 0.999953634 | 0.007428226 |
| EIF5A | NM_001143761 | 0 | 0 | 9.004140582 | 0 | 8.148778003 | 0 | 8.212844911 | 9.61278054 | 0.038432671 | 0.999932046 |
| EIF5A | XM_005256509 | 0 | 8.453213086 | 0 | 0 | 0 | 0 | 0 | 0 | 1.11E-07 | 1 |
| EIF5A | XR_002957981 | 0 | 0 | 0 | 0 | 6.769943125 | 0 | 0 | 10.26464677 | 9.43E-05 | 1 |
| EIF6 | NM_181468 | 0 | 0 | 0 | 10.25351436 | 7.328262653 | 8.853696497 | 7.223307962 | 8.966799504 | 0.022638468 | 0.999932046 |
| EIF6 | NR_052023 | 0 | 0 | 10.21833239 | 0 | 5.520949241 | 0 | 8.032920857 | 0 | 0.028033481 | 1 |
| ELF3-AS1 | NR_146472 | 0 | 9.19973932 | 8.476915428 | 8.801096068 | 0 | 0 | 0 | 0 | 0.999953634 | 0.027373137 |
| ELF4 | XM_011531308 | 0 | 9.847262786 | 0 | 0 | 0 | 0 | 0 | 0 | 2.00E-08 | 1 |
| ELFN1-AS1 | NR_120508 | 0 | 0 | 0 | 0 | 6.351978551 | 0 | 0 | 0 | 4.81E-05 | 1 |
| ELFN1-AS1 | NR_120510 | 0 | 9.08836834 | 3.975960667 | 9.932501706 | 9.845694196 | 0 | 0 | 0 | 0.999953634 | 0.007638895 |
| ELMO2 | XM_024451961 | 0 | 8.903318262 | 0 | 0 | 0 | 0 | 0 | 0 | 7.75E-08 | 1 |
| ELMO3 | NM_024712 | 0 | 0 | 10.23704693 | 0 | 8.768274428 | 0 | 0 | 0 | 0.018583294 | 0.02072721 |
| ELMO3 | XM_024450447 | 0 | 0 | 9.317199039 | 0 | 5.54685007 | 0 | 0 | 0 | 0.051572072 | 0.032485916 |
| ELMOD2 | NM_153702 | 0 | 10.03167046 | 0 | 0 | 0 | 0 | 0 | 9.747816432 | 1.96E-08 | 1 |
| ELMOD2 | XR_001741193 | 10.05956027 | 0 | 0 | 0 | 0 | 0 | 0 | 0 | 5.01E-08 | 1 |
| ELMSAN1 | XM_005268206 | 0 | 5.390784359 | 9.292927063 | 0 | 7.510413387 | 0 | 0 | 0 | 0.646598684 | 0.025156042 |
| ELOC | NM_001204859 | 0 | 0 | 0 | 0 | 0 | 11.26619798 | 8.878467512 | 9.550800565 | 1 | 0.001516752 |
| ELOC | NM_001204863 | 0 | 0 | 0 | 0 | 0 | 8.690808823 | 8.925674339 | 0 | 1 | 0.02072721 |
| ELOVL5 | NM_021814 | 9.363098816 | 0 | 0 | 0 | 0 | 0 | 0 | 0 | 4.33E-08 | 1 |
| ELP3 | XM_024447185 | 10.32069408 | 0 | 0 | 0 | 0 | 0 | 0 | 0 | 1.52E-08 | 1 |
| ELP5 | NM_015362 | 8.791378415 | 0 | 0 | 0 | 0 | 0 | 0 | 0 | 9.38E-08 | 1 |
| ELP6 | NM_001031703 | 0 | 0 | 10.34359709 | 0 | 7.895185835 | 0 | 0 | 0 | 0.020534682 | 0.02072721 |
| EMC2 | NM_001329494 | 0 | 8.690990708 | 0 | 0 | 0 | 0 | 0 | 0 | 9.02E-08 | 1 |
| EMC2 | NR_138033 | 0 | 0 | 0 | 0 | 6.409577539 | 0 | 4.961135439 | 0 | 4.59E-05 | 1 |
| EMC9 | XM_024449620 | 0 | 0 | 0 | 10.18491977 | 8.564658559 | 0 | 0 | 0 | 0.019475972 | 0.02072721 |
| EMCN | NM_016242 | 9.395290737 | 0 | 0 | 0 | 0 | 0 | 0 | 0 | 4.12E-08 | 1 |
| EMD | XM_024452349 | 0 | 8.464408432 | 0 | 0 | 0 | 0 | 0 | 0 | 1.11E-07 | 1 |
| EML4 | XM_006711991 | 10.85408237 | 0 | 0 | 0 | 0 | 0 | 0 | 0 | 1.34E-08 | 1 |
| EML5 | XM_017021065 | 0 | 8.550974732 | 0 | 0 | 0 | 0 | 0 | 0 | 1.01E-07 | 1 |
| EML6 | XR_002959298 | 0 | 10.24144608 | 0 | 0 | 0 | 12.16776099 | 9.408038715 | 0 | 1.48E-08 | 1.92E-09 |
| EMP1 | NM_001423 | 0 | 0 | 10.86816444 | 0 | 8.678866937 | 0 | 0 | 0 | 0.013868375 | 0.02072721 |
| EMP1 | XM_024448877 | 9.819710617 | 9.703487753 | 10.45127681 | 0 | 7.812190501 | 0 | 0 | 0 | 0.999953634 | 0.02072721 |
| ENAH | XM_024448319 | 10.01363818 | 8.681427931 | 0 | 0 | 0 | 0 | 7.578739068 | 0 | 0.001769865 | 1 |
| ENC1 | NR_046318 | 0 | 8.975521721 | 0 | 0 | 0 | 0 | 0 | 0 | 7.27E-08 | 1 |
| ENGASE | XR_001752594 | 0 | 8.951853153 | 0 | 0 | 0 | 0 | 0 | 0 | 7.60E-08 | 1 |
| ENPP2 | XM_017013572 | 0 | 0 | 0 | 0 | 0 | 11.3867569 | 0 | 11.12387678 | 1 | 0.018973637 |
| ENSA | NM_207046 | 0 | 0 | 10.6656667 | 6.914833961 | 7.24369152 | 0 | 7.630311627 | 0 | 0.004513359 | 1 |
| ENTPD1 | NM_001164179 | 0 | 0 | 10.88074677 | 9.02108447 | 9.296923046 | 12.84779877 | 0 | 11.95383523 | 0.001400773 | 0.999932046 |
| ENTPD1 | NM_001320916 | 0 | 9.610808606 | 0 | 0 | 0 | 0 | 9.808621494 | 0 | 2.54E-08 | 1 |
| ENTPD1 | XM_011540370 | 0 | 0 | 0 | 9.960130017 | 7.735433951 | 0 | 0 | 10.96265327 | 0.025123737 | 0.999932046 |
| ENTPD1 | XM_017016960 | 0 | 0 | 0 | 0 | 6.667421491 | 0 | 0 | 0 | 3.78E-05 | 1 |

| | | | | | | | | | | | |
|---|---|---|---|---|---|---|---|---|---|---|---|
| ENTPD1 | XM_017016964 | 0 | 0 | 11.02034308 | 0 | 10.11983605 | 11.20366134 | 0 | 0 | 0.009069669 | 0.999932046 |
| ENTPD4 | NM_004901 | 0 | 10.4696994 | 0 | 0 | 0 | 0 | 0 | 0 | 1.34E-08 | 1 |
| ENTPD5 | NM_001330189 | 10.3538353 | 0 | 0 | 0 | 0 | 0 | 0 | 0 | 2.01E-08 | 1 |
| ENTPD5 | XM_006720325 | 0 | 8.903318262 | 0 | 0 | 0 | 0 | 0 | 0 | 7.75E-08 | 1 |
| ENTPD8 | NM_001033113 | 9.676424716 | 0 | 0 | 0 | 0 | 0 | 0 | 0 | 2.73E-08 | 1 |
| EPB41L2 | NM_001135555 | 0 | 8.602554319 | 0 | 0 | 0 | 0 | 0 | 0 | 9.73E-08 | 1 |
| EPB41L2 | NM_001199388 | 0 | 0 | 0 | 9.0384682 | 7.790673576 | 0 | 7.767153042 | 0 | 0.041029487 | 0.999932046 |
| EPB41L2 | NM_001350306 | 0 | 0 | 11.7274983 | 12.53242786 | 11.45312823 | 12.26507012 | 9.826235625 | 12.74577126 | 8.91E-05 | 0.999932046 |
| EPB41L2 | NM_001350310 | 0 | 0 | 10.87823907 | 8.378061333 | 10.63983387 | 11.39695608 | 9.483594476 | 0 | 0.000971241 | 0.999932046 |
| EPB41L3 | XM_017025656 | 10.15666012 | 0 | 0 | 0 | 0 | 0 | 0 | 0 | 1.62E-08 | 1 |
| EPHA2 | NM_004431 | 9.531875668 | 0 | 0 | 0 | 0 | 0 | 0 | 8.391159551 | 3.28E-08 | 1 |
| EPHB2 | NM_001309193 | 0 | 8.662110092 | 0 | 0 | 0 | 0 | 0 | 0 | 9.15E-08 | 1 |
| EPHB3 | NM_004443 | 0 | 0 | 0 | 9.338953707 | 8.283895493 | 0 | 0 | 0 | 0.03087447 | 0.022212956 |
| EPN1 | XM_011526880 | 0 | 0 | 9.510949584 | 0 | 7.905232421 | 0 | 0 | 0 | 0.031697942 | 0.021463205 |
| EPRS | NM_004446 | 0 | 9.38199218 | 0 | 0 | 0 | 0 | 0 | 10.12983217 | 3.43E-08 | 1 |
| EPRS | XM_017000614 | 10.34973404 | 0 | 0 | 0 | 0 | 0 | 0 | 0 | 1.51E-08 | 1 |
| EPS15L1 | XM_017027090 | 0 | 0 | 0 | 9.55567177 | 5.467709404 | 0 | 0 | 0 | 0.042342975 | 0.029924055 |
| EPS15L1 | XM_017027092 | 0 | 0 | 8.09164712 | 0 | 8.768274428 | 0 | 0 | 0 | 0.042956148 | 0.028000789 |
| EPS8L1 | XM_011527050 | 0 | 0 | 0 | 9.531320288 | 7.568463804 | 0 | 0 | 0 | 0.032774176 | 0.023358895 |
| EPS8L3 | NM_133181 | 0 | 0 | 9.993496019 | 8.466339867 | 0 | 0 | 8.822290914 | 9.929274263 | 0.023834661 | 0.999932046 |
| EPS8L3 | XR_001737407 | 0 | 8.407542174 | 0 | 0 | 0 | 0 | 0 | 0 | 1.16E-07 | 1 |
| ERC1 | NR_027948 | 10.54172739 | 11.93226809 | 0 | 0 | 0 | 0 | 8.268156521 | 10.77661339 | 0.000136417 | 0.02072721 |
| ERC1 | XM_017019058 | 0 | 0 | 0 | 10.42633757 | 9.392867275 | 0 | 9.427605546 | 0 | 0.014599385 | 0.999932046 |
| ERC1 | XM_017019068 | 9.403227718 | 0 | 0 | 0 | 0 | 10.95853301 | 8.412617556 | 0 | 4.07E-08 | 0.02072721 |
| ERC1 | XM_017019075 | 0 | 0 | 10.93363422 | 0 | 8.407434995 | 0 | 9.731019215 | 0 | 0.013988632 | 0.999932046 |
| ERC1 | XR_001748636 | 0 | 9.930350523 | 0 | 0 | 0 | 0 | 0 | 0 | 1.81E-08 | 1 |
| ERG | NM_001136154 | 0 | 0 | 10.10708069 | 0 | 8.659707446 | 0 | 0 | 10.88314745 | 0.020111354 | 0.999932046 |
| ERG | NR_111949 | 0 | 0 | 0 | 7.167471359 | 0 | 0 | 0 | 0 | 2.47E-05 | 1 |
| ERI3 | XM_017002303 | 9.418971874 | 8.407542174 | 0 | 0 | 0 | 0 | 0 | 0 | 0.003324612 | 1 |
| ERICH1 | XM_017013116 | 10.36809858 | 0 | 0 | 0 | 0 | 0 | 0 | 0 | 1.48E-08 | 1 |
| ERICH2 | NM_001290031 | 9.83739315 | 0 | 0 | 0 | 0 | 0 | 6.514885902 | 0 | 2.27E-08 | 1 |
| ERLIN1 | NM_001100626 | 0 | 0 | 9.461549885 | 0 | 7.746652379 | 0 | 0 | 0 | 0.033703373 | 0.022480743 |
| ERN2 | NM_033266 | 0 | 9.834376365 | 0 | 0 | 0 | 0 | 0 | 0 | 2.00E-08 | 1 |
| ERO1A | NM_014584 | 0 | 0 | 9.683171154 | 9.170475661 | 10.73619707 | 0 | 9.250789795 | 10.26464677 | 0.00123663 | 0.999932046 |
| ERP29 | XM_017018720 | 0 | 0 | 7.589185621 | 0 | 0 | 0 | 9.135052218 | 0 | 1.75E-05 | 1 |
| ERVK3-1 | NR_144445 | 0 | 10.24144608 | 0 | 10.55370578 | 8.070588078 | 0 | 0 | 0 | 0.999953634 | 0.02072721 |
| ESM1 | NM_007036 | 0 | 0 | 0 | 0 | 6.544252221 | 0 | 0 | 0 | 4.20E-05 | 1 |
| ESRP1 | XM_005250992 | 8.624880391 | 9.37013595 | 0 | 0 | 0 | 0 | 0 | 0 | 0.003015025 | 1 |
| ESS2 | XM_006724330 | 0 | 0 | 7.974444826 | 0 | 8.218925247 | 0 | 0 | 0 | 0.057403437 | 0.036962457 |
| ESYT2 | XM_024446848 | 0 | 9.467937896 | 0 | 0 | 0 | 0 | 0 | 0 | 3.10E-08 | 1 |
| ETAA1 | XM_017004377 | 7.260294043 | 8.054440838 | 0 | 0 | 0 | 0 | 0 | 0 | 0.016923608 | 1 |
| ETF1 | XM_005271921 | 9.588662682 | 0 | 0 | 0 | 0 | 0 | 0 | 0 | 3.15E-08 | 1 |
| ETFA | XR_931766 | 0 | 8.039547025 | 0 | 10.48139854 | 6.945718048 | 0 | 0 | 0 | 0.992240069 | 0.02072721 |

| | | | | | | | | | | | |
|---|---|---|---|---|---|---|---|---|---|---|---|
| ETFDH | XM_024453935 | 0 | 0 | 0 | 0 | 6.351978551 | 0 | 0 | 0 | 4.81E-05 | 1 |
| ETNK1 | NM_001039481 | 0 | 9.416987232 | 12.06037529 | 0 | 7.251587968 | 0 | 0 | 0 | 0.990228069 | 5.16E-09 |
| ETNK1 | XM_017019581 | 0 | 8.632639499 | 0 | 0 | 0 | 10.67651331 | 0 | 0 | 9.44E-08 | 1 |
| ETV1 | NM_001163150 | 10.20286443 | 9.284296589 | 10.00043084 | 0 | 6.351978551 | 0 | 0 | 0 | 0.999953634 | 0.02092873 |
| ETV1 | NM_004956 | 0 | 0 | 0 | 10.11016257 | 11.77961153 | 0 | 0 | 11.10423254 | 0.005920625 | 0.999932046 |
| ETV6 | XM_011520612 | 0 | 0 | 7.89728252 | 0 | 0 | 0 | 0 | 0 | 3.19E-05 | 1 |
| EVA1C | XM_024452126 | 9.346729415 | 0 | 0 | 0 | 0 | 0 | 0 | 0 | 4.43E-08 | 1 |
| EVA1C | XR_001754885 | 0 | 8.765293275 | 0 | 0 | 0 | 0 | 0 | 0 | 8.55E-08 | 1 |
| EVI2B | NM_006495 | 0 | 9.416987232 | 0 | 0 | 0 | 0 | 0 | 0 | 3.28E-08 | 1 |
| EVI5L | NM_145245 | 0 | 8.690990708 | 0 | 0 | 0 | 0 | 0 | 0 | 9.02E-08 | 1 |
| EVL | NM_016337 | 0 | 0 | 0 | 0 | 0 | 10.19845348 | 8.331847395 | 0 | 1 | 0.02072721 |
| EVL | XM_005267749 | 0 | 8.967675264 | 0 | 0 | 0 | 0 | 0 | 0 | 7.35E-08 | 1 |
| EVPL | NM_001988 | 0 | 0 | 0 | 9.348277199 | 6.478485785 | 0 | 7.431456038 | 8.147715207 | 0.043770831 | 0.999932046 |
| EWSR1 | NM_001163287 | 0 | 0 | 10.67364253 | 11.2171125 | 11.16824643 | 12.61577161 | 11.0440269 | 11.41302394 | 0.00030731 | 0.999932046 |
| EWSR1 | NM_013986 | 0 | 0 | 10.50346638 | 9.498198764 | 10.31097016 | 11.11252808 | 10.07138775 | 0 | 0.000995733 | 0.999932046 |
| EWSR1 | XM_005261390 | 0 | 0 | 0 | 0 | 0 | 11.20366134 | 8.246285684 | 0 | 1 | 0.02072721 |
| EWSR1 | XM_011529995 | 0 | 0 | 0 | 8.331810132 | 0 | 0 | 7.200741128 | 9.534880393 | 7.48E-06 | 0.999932046 |
| EWSR1 | XM_017028646 | 0 | 0 | 10.52446642 | 0 | 7.838643081 | 0 | 6.884098118 | 0 | 0.018822751 | 0.999932046 |
| EXD2 | NM_001193361 | 0 | 0 | 12.22410446 | 11.21838419 | 9.818042958 | 0 | 10.900876 | 9.657578368 | 0.000237536 | 0.999932046 |
| EXD3 | NR_104599 | 9.922694008 | 8.835955774 | 0 | 0 | 0 | 0 | 0 | 9.521478054 | 0.001776413 | 1 |
| EXO5 | XM_017002102 | 0 | 9.10991584 | 0 | 0 | 0 | 0 | 0 | 0 | 5.42E-08 | 1 |
| EXOSC10 | XR_002956821 | 9.055885396 | 0 | 0 | 0 | 0 | 0 | 0 | 0 | 7.54E-08 | 1 |
| EXOSC2 | NM_001282708 | 0 | 0 | 0 | 11.95752109 | 10.18158696 | 11.41312626 | 12.11405828 | 11.78070926 | 0.005340378 | 0.999932046 |
| EXOSC4 | NM_019037 | 0 | 0 | 10.78224969 | 11.91775951 | 11.02212384 | 0 | 8.083136668 | 0 | 0.000218422 | 0.668933693 |
| EXOSC8 | XR_429212 | 0 | 0 | 0 | 11.37312061 | 9.435085097 | 0 | 0 | 0 | 0.008865284 | 0.02072721 |
| EXOSC8 | XR_941480 | 0 | 0 | 0 | 9.744046385 | 8.006701649 | 0 | 0 | 0 | 0.026880182 | 0.020781046 |
| EXPH5 | XM_017017404 | 9.261974802 | 8.33620493 | 0 | 0 | 0 | 0 | 0 | 0 | 0.003926103 | 1 |
| EXT2 | XM_011519950 | 0 | 0 | 10.1709679 | 9.645387449 | 0 | 0 | 0 | 0 | 0.016630459 | 0.02072721 |
| EXTL2 | XM_017000651 | 9.539097493 | 0 | 0 | 0 | 0 | 0 | 0 | 0 | 3.28E-08 | 1 |
| EYA2 | XM_017027721 | 0 | 10.60483574 | 0 | 0 | 0 | 0 | 0 | 0 | 1.34E-08 | 1 |
| EYA3 | NM_001282562 | 0 | 0 | 11.59670885 | 0 | 10.02684903 | 12.21924894 | 0 | 0 | 0.006913248 | 0.999932046 |
| EYA3 | XM_011541003 | 0 | 9.812639863 | 0 | 0 | 0 | 0 | 0 | 0 | 2.01E-08 | 1 |
| EYA3 | XM_024454014 | 0 | 8.31161984 | 0 | 0 | 0 | 0 | 0 | 0 | 1.27E-07 | 1 |
| EZH2 | XM_011515887 | 0 | 0 | 0 | 10.47821718 | 8.368024654 | 0 | 0 | 0 | 0.017538747 | 0.02072721 |
| EZH2 | XM_017011818 | 9.07588576 | 10.19851207 | 0 | 0 | 0 | 0 | 0 | 0 | 0.001223046 | 1 |
| EZH2 | XM_017011820 | 0 | 0 | 0 | 8.814612372 | 8.985072162 | 0 | 0 | 0 | 0.030452016 | 0.021907747 |
| F2RL1 | NM_005242 | 0 | 0 | 0 | 9.429583326 | 8.513036313 | 0 | 0 | 0 | 0.027842286 | 0.020903961 |
| F8A1 | NM_012151 | 0 | 0 | 11.40793851 | 0 | 9.802000998 | 0 | 0 | 9.469382508 | 0.00811514 | 0.999932046 |
| FA2H | XM_011523319 | 10.14723899 | 8.384152849 | 0 | 0 | 0 | 0 | 0 | 0 | 0.001778148 | 1 |
| FABP3 | NM_004102 | 0 | 0 | 9.872720291 | 0 | 9.156784755 | 0 | 0 | 0 | 0.019666085 | 0.02072721 |
| FADS3 | NM_021727 | 0 | 0 | 0 | 9.859394182 | 4.989879473 | 0 | 0 | 0 | 0.035609697 | 0.02562326 |
| FAF1 | XM_017000136 | 10.47927431 | 0 | 0 | 0 | 0 | 0 | 0 | 0 | 1.44E-08 | 1 |
| FAHD2A | XM_024452939 | 0 | 0 | 0 | 8.30333053 | 0 | 0 | 0 | 0 | 7.64E-06 | 1 |

| | | | | | | | | | | | |
|---|---|---|---|---|---|---|---|---|---|---|---|
| FAIM | NM_018147 | 0 | 9.19973932 | 0 | 0 | 0 | 0 | 0 | 0 | 4.67E-08 | 1 |
| FAM102A | NM_203305 | 0 | 8.286608519 | 9.768247087 | 0 | 9.190037519 | 0 | 0 | 0 | 0.999953634 | 0.02072721 |
| FAM104A | NM_001289410 | 0 | 9.490017613 | 0 | 0 | 0 | 9.825198805 | 0 | 10.44867315 | 3.40E-08 | 0.02072721 |
| FAM104B | NM_001166702 | 0 | 9.754542834 | 0 | 0 | 0 | 0 | 0 | 0 | 2.13E-08 | 1 |
| FAM110C | XR_001738890 | 0 | 8.475517571 | 0 | 0 | 0 | 0 | 0 | 0 | 1.09E-07 | 1 |
| FAM114A2 | NM_001317993 | 0 | 0 | 8.957388444 | 0 | 8.958784227 | 12.13797054 | 8.951247914 | 0 | 0.02949256 | 0.999932046 |
| FAM114A2 | NM_001317994 | 0 | 0 | 0 | 9.645387449 | 9.703252693 | 0 | 10.04787429 | 0 | 0.017946574 | 0.999932046 |
| FAM117B | NM_173511 | 0 | 0 | 0 | 8.01802849 | 0 | 0 | 0 | 0 | 1.03E-05 | 1 |
| FAM118B | NM_001330446 | 0 | 0 | 10.36882564 | 10.35412868 | 7.084811668 | 0 | 0 | 0 | 0.001922313 | 0.003057899 |
| FAM118B | XM_011542977 | 0 | 9.165850403 | 0 | 0 | 0 | 10.71615363 | 0 | 0 | 4.97E-08 | 1 |
| FAM120AOS | NR_136235 | 9.849062238 | 0 | 0 | 0 | 0 | 0 | 0 | 0 | 2.23E-08 | 1 |
| FAM120B | NM_032448 | 0 | 9.193024922 | 0 | 0 | 0 | 0 | 0 | 0 | 4.74E-08 | 1 |
| FAM120B | XR_001743682 | 0 | 0 | 0 | 0 | 6.478485785 | 0 | 0 | 0 | 4.36E-05 | 1 |
| FAM129B | NM_022833 | 9.162606966 | 0 | 0 | 0 | 0 | 0 | 0 | 0 | 7.27E-08 | 1 |
| FAM129C | NM_001321827 | 9.588662682 | 0 | 0 | 0 | 0 | 0 | 0 | 0 | 3.15E-08 | 1 |
| FAM133B | NR_109929 | 0 | 9.258802813 | 0 | 0 | 0 | 0 | 0 | 0 | 4.23E-08 | 1 |
| FAM135A | XM_017011136 | 0 | 0 | 7.936379488 | 0 | 0 | 0 | 0 | 0 | 1.22E-05 | 1 |
| FAM136A | NM_032822 | 0 | 0 | 11.28032066 | 10.41307084 | 10.10846726 | 0 | 9.647962002 | 0 | 0.000523393 | 1 |
| FAM153C | NR_149722 | 10.88558544 | 0 | 0 | 0 | 0 | 0 | 0 | 0 | 1.34E-08 | 1 |
| FAM156B | XM_024452434 | 0 | 0 | 0 | 0 | 6.337212177 | 0 | 0 | 7.110453114 | 4.86E-05 | 1 |
| FAM160A2 | XR_001747999 | 10.00844432 | 0 | 0 | 0 | 0 | 0 | 0 | 0 | 1.93E-08 | 1 |
| FAM171A1 | XM_017015905 | 10.25203635 | 0 | 0 | 0 | 0 | 0 | 0 | 0 | 1.58E-08 | 1 |
| FAM172A | XR_001742304 | 0 | 8.719304505 | 0 | 0 | 0 | 0 | 0 | 10.12983217 | 8.96E-08 | 1 |
| FAM174B | XR_931815 | 0 | 8.895067962 | 0 | 0 | 0 | 0 | 0 | 0 | 7.75E-08 | 1 |
| FAM185A | NM_001350987 | 0 | 8.31161984 | 0 | 0 | 0 | 0 | 0 | 0 | 1.27E-07 | 1 |
| FAM192A | NM_024946 | 0 | 0 | 11.03448881 | 10.06846137 | 0 | 0 | 8.058247234 | 0 | 0.010201494 | 0.999932046 |
| FAM193A | NR_046335 | 0 | 0 | 0 | 0 | 0 | 11.0761776 | 9.199070987 | 0 | 1 | 0.02072721 |
| FAM200A | NM_145111 | 0 | 8.943876548 | 0 | 0 | 0 | 0 | 0 | 0 | 7.69E-08 | 1 |
| FAM206A | NM_017832 | 0 | 0 | 0 | 9.972241456 | 8.773768299 | 0 | 8.095422193 | 0 | 0.020545819 | 0.999932046 |
| FAM208A | XM_005264999 | 0 | 0 | 0 | 0 | 6.758905041 | 7.99750765 | 0 | 0 | 9.53E-05 | 1 |
| FAM20C | XR_242097 | 0 | 9.252358352 | 0 | 0 | 0 | 0 | 0 | 0 | 4.28E-08 | 1 |
| FAM213A | XM_017016775 | 0 | 0 | 9.913704414 | 0 | 7.660307374 | 0 | 0 | 0 | 0.026709285 | 0.02072721 |
| FAM213B | XM_011540665 | 9.185799057 | 0 | 10.22426851 | 0 | 8.608360805 | 0 | 0 | 0 | 0.999953634 | 0.02072721 |
| FAM214A | XM_011521790 | 0 | 0 | 0 | 0 | 6.395391943 | 0 | 0 | 0 | 4.65E-05 | 1 |
| FAM219B | NR_135844 | 0 | 9.213075091 | 0 | 0 | 0 | 0 | 0 | 0 | 4.56E-08 | 1 |
| FAM219B | XR_002957668 | 0 | 8.519117955 | 0 | 0 | 0 | 0 | 0 | 0 | 1.04E-07 | 1 |
| FAM219B | XR_429468 | 10.12341345 | 0 | 0 | 0 | 0 | 0 | 8.782953729 | 0 | 1.68E-08 | 1 |
| FAM21EP | NR_038275 | 0 | 0 | 0 | 0 | 0 | 0 | 7.892131936 | 10.72024326 | 1 | 0.02072721 |
| FAM226A | NR_026595 | 0 | 9.179501644 | 0 | 0 | 0 | 0 | 0 | 0 | 4.86E-08 | 1 |
| FAM24B | NM_152644 | 0 | 8.765293275 | 0 | 0 | 0 | 9.08324246 | 0 | 0 | 8.55E-08 | 1 |
| FAM30A | NR_026800 | 0 | 9.439854398 | 0 | 0 | 0 | 0 | 0 | 0 | 3.19E-08 | 1 |
| FAM3C | XM_011515737 | 8.555352323 | 10.00891253 | 0 | 0 | 0 | 0 | 0 | 0 | 0.00180424 | 1 |
| FAM3D | XM_006712967 | 0 | 0 | 10.31421369 | 11.87525943 | 7.393949817 | 0 | 0 | 0 | 0.00074474 | 0.001555546 |

| | | | | | | | | | | | |
|---|---|---|---|---|---|---|---|---|---|---|---|
| FAM45A | XM_017016261 | 0 | 8.582143237 | 0 | 0 | 0 | 0 | 0 | 0 | 9.91E-08 | 1 |
| FAM49B | NM_001353247 | 0 | 8.983325733 | 0 | 0 | 0 | 9.052602861 | 9.959903889 | 11.07517295 | 7.17E-08 | 0.001394691 |
| FAM49B | NM_001353249 | 0 | 0 | 9.047194317 | 10.27811433 | 12.57678305 | 11.16061116 | 12.72682088 | 13.89045089 | 0.000305676 | 0.999932046 |
| FAM49B | NM_001353271 | 0 | 0 | 0 | 10.1560111 | 10.23085121 | 11.08440513 | 10.9178377 | 0 | 0.01248512 | 0.999932046 |
| FAM49B | NM_001353298 | 0 | 8.886770211 | 0 | 0 | 0 | 0 | 7.933778586 | 0 | 7.80E-08 | 1 |
| FAM49B | NM_001353300 | 0 | 8.911521649 | 0 | 0 | 0 | 0 | 0 | 0 | 7.73E-08 | 1 |
| FAM49B | NM_016623 | 0 | 0 | 10.21833239 | 11.19079287 | 9.314934401 | 0 | 8.6351065 | 8.63467863 | 0.000759752 | 0.999932046 |
| FAM49B | XM_017013554 | 11.27109522 | 0 | 0 | 0 | 0 | 0 | 0 | 0 | 1.34E-08 | 1 |
| FAM83D | XM_017028088 | 0 | 0 | 0 | 0 | 6.679179249 | 0 | 0 | 0 | 3.78E-05 | 1 |
| FAM83G | XM_017024953 | 8.583566901 | 0 | 0 | 0 | 0 | 0 | 0 | 0 | 1.06E-07 | 1 |
| FAM84A | NM_145175 | 0 | 0 | 10.0015834 | 0 | 6.702411139 | 0 | 0 | 0 | 0.028915101 | 0.020781046 |
| FAM86DP | NR_024241 | 0 | 0 | 0 | 10.41307084 | 8.784693675 | 0 | 6.91214998 | 0 | 0.01681908 | 0.999932046 |
| FAM86JP | NR_024251 | 0 | 0 | 0 | 10.12921816 | 8.567824433 | 0 | 0 | 0 | 0.01998239 | 0.02072721 |
| FAM89B | NM_152832 | 0 | 0 | 0 | 0 | 6.505153187 | 11.07490765 | 0 | 0 | 4.28E-05 | 1 |
| FAM8A1 | NM_016255 | 0 | 0 | 9.201601708 | 0 | 7.290434628 | 0 | 0 | 0 | 0.043222808 | 0.02754963 |
| FAM92A | NM_145269 | 0 | 0 | 0 | 10.27322778 | 7.660307374 | 0 | 0 | 9.400769032 | 0.021535116 | 0.999932046 |
| FAN1 | XR_001751151 | 0 | 9.543782272 | 0 | 0 | 0 | 0 | 0 | 0 | 2.90E-08 | 1 |
| FANCA | XR_933244 | 0 | 0 | 0 | 9.99022008 | 6.531336591 | 0 | 0 | 0 | 0.028784129 | 0.02137193 |
| FANCA | XR_933247 | 0 | 0 | 0 | 10.08528677 | 10.34642811 | 0 | 0 | 8.11239291 | 0.01222617 | 0.999932046 |
| FANCC | NM_001243744 | 0 | 0 | 9.754639499 | 11.84758452 | 9.934215172 | 11.51496981 | 12.0962112 | 10.71436539 | 0.000486183 | 0.999932046 |
| FANCC | XM_017014453 | 0 | 0 | 10.88762057 | 0 | 7.71273195 | 0 | 8.113656519 | 0 | 0.01572837 | 0.999932046 |
| FANCD2OS | NM_173472 | 0 | 0 | 0 | 0 | 6.679179249 | 0 | 0 | 0 | 3.78E-05 | 1 |
| FANCE | NM_021922 | 0 | 0 | 11.53556066 | 9.733411602 | 0 | 0 | 0 | 0 | 0.008531783 | 0.02072721 |
| FANCL | NM_018062 | 0 | 0 | 0 | 9.79947801 | 7.401067144 | 0 | 0 | 0 | 0.028938309 | 0.021342962 |
| FANCL | XR_002959309 | 0 | 0 | 0 | 10.13191995 | 7.71273195 | 0 | 0 | 0 | 0.023001802 | 0.02072721 |
| FAR2 | NM_001271783 | 0 | 0 | 9.191537123 | 0 | 7.111090638 | 0 | 0 | 0 | 0.045053826 | 0.028655532 |
| FARSB | NM_005687 | 0 | 0 | 0 | 10.21070698 | 10.01645911 | 0 | 0 | 11.3991674 | 0.013240365 | 1 |
| FAS | NR_028036 | 9.035603861 | 0 | 0 | 0 | 0 | 0 | 0 | 10.67858625 | 7.68E-08 | 1 |
| FAS | NR_135315 | 0 | 11.04409893 | 9.229414648 | 0 | 7.075944552 | 0 | 0 | 0 | 0.974126475 | 0.028240248 |
| FASTK | NM_033015 | 0 | 0 | 0 | 0 | 6.3222931 | 0 | 0 | 0 | 0.000244538 | 1 |
| FASTKD1 | NR_104020 | 0 | 0 | 0 | 0 | 6.478485785 | 11.15280809 | 0 | 0 | 4.36E-05 | 1 |
| FASTKD2 | NM_014929 | 0 | 0 | 0 | 12.06639446 | 7.483843939 | 0 | 0 | 0 | 1.47E-08 | 5.17E-09 |
| FASTKD3 | XR_001742253 | 0 | 8.709928179 | 9.580473641 | 10.04284953 | 8.551925132 | 0 | 0 | 0 | 0.972972452 | 0.003533857 |
| FAT4 | NM_001291285 | 0 | 9.876890889 | 0 | 0 | 0 | 0 | 0 | 0 | 1.94E-08 | 1 |
| FBL | XM_005258651 | 0 | 0 | 8.330507817 | 10.82895422 | 0 | 0 | 8.720869785 | 0 | 0.015672839 | 0.999932046 |
| FBLIM1 | NM_001350151 | 0 | 0 | 10.23214554 | 10.39406497 | 7.530024208 | 0 | 0 | 0 | 0.001842713 | 0.003016079 |
| FBLIM1 | XM_011541617 | 0 | 0 | 0 | 9.79947801 | 7.618173501 | 0 | 0 | 0 | 0.027943555 | 0.020947511 |
| FBLIM1 | XM_017001520 | 9.279328737 | 0 | 0 | 0 | 0 | 0 | 0 | 0 | 4.99E-08 | 1 |
| FBLN5 | XM_011536357 | 0 | 8.486541819 | 9.461549885 | 0 | 8.052621656 | 0 | 0 | 0 | 0.999953634 | 0.021342962 |
| FBP1 | NM_000507 | 0 | 0 | 9.729817651 | 10.47183335 | 9.843083461 | 0 | 10.36699843 | 12.15296835 | 0.001124846 | 0.999932046 |
| FBP1 | XM_006717005 | 9.509990722 | 0 | 9.81881611 | 11.19596625 | 9.315876167 | 0 | 0 | 0 | 0.954763034 | 0.001765436 |
| FBXL12 | NM_001316937 | 9.752954799 | 8.348342131 | 0 | 0 | 0 | 0 | 0 | 0 | 0.002562092 | 1 |
| FBXL22 | XM_005254320 | 10.9761662 | 0 | 0 | 0 | 0 | 0 | 0 | 0 | 1.34E-08 | 1 |

| | | | | | | | | | | | |
|---|---|---|---|---|---|---|---|---|---|---|---|
| FBXL3 | NM_012158 | 0 | 0 | 9.346522316 | 9.396260824 | 0 | 0 | 0 | 0 | 0.023916106 | 0.02072721 |
| FBXL6 | NM_012162 | 0 | 0 | 8.621866691 | 0 | 8.795536935 | 0 | 8.032920857 | 0 | 0.035510717 | 0.999932046 |
| FBXO11 | NM_001190274 | 0 | 0 | 9.067148263 | 11.03352211 | 11.38924499 | 12.32798886 | 10.05260774 | 11.45240068 | 0.000502741 | 0.999932046 |
| FBXO11 | XM_017005015 | 0 | 0 | 0 | 8.312886334 | 8.057134281 | 0 | 0 | 0 | 0.053733299 | 0.036097409 |
| FBXO21 | XM_017019039 | 0 | 0 | 0 | 8.483365391 | 6.926116208 | 0 | 0 | 0 | 0.06703217 | 0.047689182 |
| FBXO32 | NM_058229 | 0 | 0 | 10.43515924 | 0 | 9.655692746 | 0 | 8.07695445 | 0 | 0.013668266 | 0.999932046 |
| FBXO32 | NM_148177 | 0 | 0 | 10.32530271 | 0 | 8.17807637 | 0 | 0 | 0 | 0.019839422 | 0.02072721 |
| FBXO34 | XM_017021393 | 10.02397016 | 0 | 0 | 0 | 0 | 0 | 6.458933278 | 0 | 1.88E-08 | 1 |
| FBXO38 | XR_001742284 | 9.950042506 | 0 | 0 | 0 | 0 | 0 | 0 | 9.131119222 | 2.01E-08 | 1 |
| FBXO4 | NM_012176 | 0 | 8.919678654 | 9.934371634 | 0 | 9.015534505 | 0 | 0 | 0 | 0.999953634 | 0.02072721 |
| FBXO4 | XM_011514026 | 0 | 8.550974732 | 0 | 0 | 0 | 0 | 0 | 0 | 1.01E-07 | 1 |
| FBXO45 | NM_001105573 | 10.56313968 | 0 | 0 | 0 | 0 | 0 | 0 | 0 | 1.34E-08 | 1 |
| FBXO6 | NM_018438 | 0 | 0 | 0 | 10.17970655 | 8.246846504 | 0 | 0 | 0 | 0.020677201 | 0.02072721 |
| FBXO9 | NM_012347 | 11.22705814 | 0 | 0 | 0 | 0 | 0 | 0 | 0 | 1.34E-08 | 1 |
| FBXW2 | NM_012164 | 0 | 0 | 0 | 7.504171232 | 0 | 0 | 0 | 0 | 1.75E-05 | 1 |
| FBXW4 | NM_022039 | 9.472761503 | 0 | 0 | 0 | 0 | 0 | 7.525254359 | 0 | 3.68E-08 | 1 |
| FBXW5 | XM_005266089 | 0 | 0 | 9.331935177 | 0 | 7.581052723 | 0 | 0 | 0 | 0.0375737 | 0.024490333 |
| FBXW7 | NM_001349798 | 0 | 0 | 10.03686385 | 0 | 7.774322632 | 0 | 11.27422371 | 0 | 0.024546536 | 0.999932046 |
| FBXW7 | XM_011532086 | 0 | 0 | 0 | 0 | 0 | 0 | 0 | 6.427178007 | 1 | 2.72E-06 |
| FBXW7 | XM_024454121 | 0 | 9.893551993 | 10.947456 | 0 | 8.853760467 | 0 | 0 | 0 | 0.999953634 | 0.02072721 |
| FBXW7 | XM_024454125 | 12.08098591 | 0 | 0 | 0 | 0 | 0 | 9.342549142 | 0 | 1.34E-08 | 1 |
| FCER2 | NM_001220500 | 0 | 8.983325733 | 0 | 0 | 0 | 0 | 0 | 0 | 7.17E-08 | 1 |
| FCGBP | NM_003890 | 13.66861034 | 14.05199071 | 13.06352127 | 11.26528077 | 6.42362501 | 0 | 0 | 0 | 0.85973045 | 0.000650464 |
| FCGR1A | NM_000566 | 0 | 9.885245492 | 0 | 0 | 0 | 0 | 0 | 10.98235271 | 2.00E-08 | 1 |
| FCGR2B | NM_004001 | 0 | 8.853093555 | 0 | 0 | 0 | 0 | 0 | 0 | 7.91E-08 | 1 |
| FCHO1 | XR_002958285 | 0 | 0 | 10.87698358 | 0 | 8.140296582 | 0 | 0 | 0 | 0.015034736 | 0.02072721 |
| FCHSD1 | XR_002956197 | 9.656640609 | 0 | 0 | 0 | 0 | 0 | 0 | 0 | 3.16E-08 | 1 |
| FCHSD2 | XM_011545409 | 0 | 9.179501644 | 0 | 0 | 0 | 0 | 0 | 0 | 4.86E-08 | 1 |
| FDPS | NM_001135822 | 10.72070446 | 0 | 8.065433177 | 12.52627995 | 11.37756574 | 0 | 0 | 0 | 0.966161772 | 0.000657675 |
| FDPS | NM_001242824 | 0 | 0 | 10.31421369 | 9.869142678 | 8.110212684 | 0 | 0 | 10.0904377 | 0.001991244 | 0.999932046 |
| FDPS | XM_005244963 | 11.82959893 | 8.911521649 | 10.92150669 | 9.003488718 | 9.097777375 | 0 | 0 | 0 | 0.999953634 | 0.002548758 |
| FER | NM_001308031 | 0 | 11.10973982 | 0 | 0 | 0 | 0 | 0 | 0 | 1.34E-08 | 1 |
| FERMT2 | XM_005267285 | 0 | 8.737876298 | 0 | 0 | 0 | 0 | 0 | 0 | 9.11E-08 | 1 |
| FES | XR_243206 | 0 | 9.239382419 | 0 | 0 | 0 | 0 | 0 | 0 | 4.33E-08 | 1 |
| FETUB | XM_011512681 | 0 | 10.66187 | 0 | 0 | 0 | 0 | 0 | 0 | 1.34E-08 | 1 |
| FFAR4 | NM_181745 | 9.734199496 | 0 | 0 | 0 | 0 | 0 | 0 | 0 | 5.08E-08 | 1 |
| FGD4 | XM_024448840 | 0 | 9.434171496 | 0 | 0 | 0 | 0 | 0 | 0 | 3.24E-08 | 1 |
| FGF12 | NM_004113 | 0 | 9.615837767 | 0 | 0 | 0 | 0 | 0 | 0 | 2.54E-08 | 1 |
| FGF18 | NM_003862 | 0 | 0 | 9.478205106 | 10.20044734 | 8.499836882 | 13.00358805 | 12.97949987 | 10.49329345 | 0.002225842 | 0.882519284 |
| FGFR1 | NM_001354368 | 0 | 0 | 0 | 9.15727502 | 7.869758957 | 0 | 0 | 9.667347319 | 0.037710982 | 0.999932046 |
| FGFR1 | XM_006716314 | 8.237937428 | 7.93074915 | 0 | 0 | 0 | 0 | 0 | 10.4315782 | 0.010830367 | 1 |
| FGFR1 | XM_017013226 | 0 | 8.719304505 | 0 | 0 | 0 | 0 | 0 | 0 | 8.96E-08 | 1 |
| FGFR1 | XM_024447097 | 8.555352323 | 9.740796941 | 10.02330906 | 0 | 8.712153435 | 0 | 0 | 0 | 0.999953634 | 0.02072721 |

| | | | | | | | | | | | |
|---|---|---|---|---|---|---|---|---|---|---|---|
| FGFR1 | XR_001745496 | 11.42325679 | 8.709928179 | 0 | 0 | 0 | 0 | 0 | 0 | 0.000474527 | 1 |
| FGFR1OP2 | NM_015633 | 0 | 0 | 0 | 0 | 6.30721813 | 0 | 7.621843212 | 0 | 4.99E-05 | 1 |
| FGFR2 | NM_001144915 | 0 | 8.835955774 | 0 | 0 | 0 | 0 | 0 | 0 | 2.22E-07 | 1 |
| FGFR2 | NM_022970 | 10.41399488 | 0 | 10.00962571 | 0 | 8.118872287 | 0 | 0 | 0 | 0.999953634 | 0.02072721 |
| FGFR2 | XM_017015920 | 0 | 0 | 8.709114096 | 8.985675707 | 0 | 0 | 0 | 0 | 0.034647391 | 0.023763327 |
| FGFR2 | XM_024447890 | 10.95202098 | 0 | 11.11177334 | 0 | 9.365839676 | 0 | 0 | 0 | 0.999953634 | 0.02072721 |
| FGFR3 | NM_022965 | 0 | 0 | 8.470270444 | 8.516825278 | 0 | 0 | 8.15532787 | 0 | 0.046046611 | 1 |
| FGFR3 | NR_148971 | 8.611240259 | 8.168359982 | 0 | 0 | 0 | 0 | 0 | 0 | 0.0072133 | 1 |
| FGFR3 | XM_006713871 | 0 | 8.550974732 | 0 | 0 | 0 | 0 | 6.232930575 | 0 | 1.01E-07 | 1 |
| FGFR3 | XM_011513422 | 10.0393307 | 0 | 0 | 0 | 0 | 0 | 8.427271157 | 0 | 1.86E-08 | 1 |
| FGFR4 | NM_213647 | 9.643298854 | 0 | 0 | 0 | 0 | 0 | 7.421731871 | 0 | 2.88E-08 | 1 |
| FGGY | NM_001113411 | 0 | 9.772669303 | 0 | 0 | 0 | 0 | 10.53989557 | 0 | 4.12E-08 | 1 |
| FGGY | NM_001350797 | 0 | 0 | 8.450149855 | 9.744046385 | 0 | 0 | 10.04787429 | 11.76259559 | 0.026283267 | 0.999932046 |
| FGGY | XM_017001673 | 0 | 0 | 0 | 0 | 0 | 10.38011571 | 8.273572814 | 0 | 1 | 0.02072721 |
| FGGY | XM_017001679 | 0 | 0 | 10.43515924 | 0 | 10.24624723 | 0 | 5.493922725 | 0 | 0.011198731 | 0.659352223 |
| FGGY | XM_024448227 | 0 | 0 | 0 | 10.70470579 | 9.295013967 | 0 | 11.89808989 | 0 | 0.013120978 | 0.999932046 |
| FGL2 | NM_006682 | 10.28247691 | 5.797199543 | 0 | 0 | 0 | 0 | 0 | 0 | 0.002996015 | 1 |
| FGR | NM_001042729 | 0 | 8.248259235 | 0 | 0 | 0 | 0 | 7.728235151 | 10.57859585 | 2.02E-07 | 0.02072721 |
| FGR | XM_011541010 | 0 | 0 | 0 | 0 | 0 | 0 | 7.720323987 | 10.37009846 | 1 | 0.02072721 |
| FGR | XR_946583 | 0 | 9.277965255 | 0 | 0 | 0 | 0 | 0 | 9.888094489 | 4.01E-08 | 1 |
| FH | NM_000143 | 11.14186693 | 11.10795622 | 9.394692648 | 10.47289928 | 8.400349091 | 0 | 0 | 0 | 0.983589543 | 0.003174145 |
| FHDC1 | XM_011532389 | 0 | 9.650559405 | 0 | 0 | 0 | 0 | 0 | 0 | 2.46E-08 | 1 |
| FHIT | NM_001354590 | 0 | 0 | 12.17973519 | 9.929398977 | 11.06812222 | 0 | 9.049236545 | 10.13863739 | 0.000249307 | 0.999932046 |
| FHIT | XM_017005882 | 0 | 10.46134184 | 10.03686385 | 11.23607105 | 9.977709443 | 0 | 0 | 0 | 0.999953634 | 0.001394691 |
| FHL1 | XM_024452355 | 0 | 0 | 7.259911857 | 0 | 0 | 0 | 0 | 0 | 6.79E-05 | 1 |
| FHL2 | NM_001450 | 0 | 0 | 0 | 9.089396862 | 9.251377948 | 0 | 0 | 10.29647003 | 0.025031088 | 0.999932046 |
| FHOD1 | NM_013241 | 0 | 9.439854398 | 0 | 0 | 0 | 0 | 0 | 0 | 3.19E-08 | 1 |
| FHOD1 | XR_933285 | 0 | 8.728620286 | 0 | 0 | 0 | 0 | 0 | 10.01975144 | 8.80E-08 | 1 |
| FIBP | NM_198897 | 0 | 0 | 10.6100781 | 0 | 8.626692541 | 0 | 0 | 0 | 0.015991489 | 0.02072721 |
| FIP1L1 | XM_005265779 | 0 | 0 | 10.05696019 | 10.80291289 | 0 | 9.998635381 | 0 | 0 | 0.011205574 | 0.999932046 |
| FITM1 | NM_203402 | 0 | 8.529815232 | 0 | 0 | 0 | 0 | 0 | 0 | 1.04E-07 | 1 |
| FKBP1A-SDCBP2 | NR_037661 | 0 | 7.9939202 | 11.10535474 | 0 | 9.148350412 | 0 | 0 | 0 | 0.842663899 | 0.02072721 |
| FKBP2 | NM_004470 | 0 | 0 | 10.77283005 | 8.322379261 | 8.110212684 | 10.41766811 | 0 | 0 | 0.002549147 | 1 |
| FKBP3 | XM_011536565 | 0 | 8.911521649 | 0 | 0 | 0 | 0 | 0 | 0 | 7.73E-08 | 1 |
| FKBP8 | XM_011527863 | 9.727893151 | 0 | 0 | 8.893154207 | 7.470373439 | 0 | 0 | 0 | 0.999953634 | 0.032833213 |
| FLAD1 | NM_201398 | 7.260294043 | 0 | 0 | 10.26093849 | 8.583550423 | 0 | 0 | 0 | 0.821917016 | 0.02072721 |
| FLI1 | NM_002017 | 0 | 0 | 0 | 8.525070387 | 0 | 0 | 0 | 0 | 6.60E-06 | 1 |
| FLJ37453 | NR_024279 | 0 | 0 | 9.948784034 | 0 | 7.335710674 | 0 | 7.960889765 | 9.917043278 | 0.0275216 | 0.999932046 |
| FLJ44635 | NM_207422 | 0 | 0 | 0 | 10.41084781 | 10.04286386 | 11.84725081 | 0 | 0 | 0.012132852 | 0.999932046 |
| FLNB | NM_001457 | 9.849062238 | 0 | 10.51803737 | 0 | 9.31399202 | 0 | 0 | 0 | 0.999953634 | 0.02072721 |
| FLNB | XM_005264978 | 0 | 0 | 0 | 0 | 6.3222931 | 0 | 0 | 10.47949727 | 0.000244538 | 1 |
| FLOT1 | XM_006714947 | 10.12341345 | 0 | 0 | 0 | 0 | 0 | 7.402084669 | 0 | 1.68E-08 | 1 |
| FLOT2 | NM_004475 | 0 | 9.265218616 | 0 | 0 | 0 | 0 | 0 | 0 | 4.18E-08 | 1 |

| | | | | | | | | | | | |
|---|---|---|---|---|---|---|---|---|---|---|---|
| FLOT2 | XM_017024395 | 0 | 8.540433774 | 0 | 0 | 0 | 0 | 0 | 0 | 1.02E-07 | 1 |
| FLT3LG | NM_001204502 | 10.61018607 | 9.694007725 | 0 | 0 | 0 | 0 | 7.696326825 | 0 | 0.000665677 | 1 |
| FLT3LG | XM_011526675 | 10.41399488 | 0 | 9.838299621 | 10.46542113 | 8.043553839 | 0 | 0 | 0 | 0.999953634 | 0.00302572 |
| FLT3LG | XM_011526680 | 0 | 0 | 9.733984397 | 10.2030191 | 8.548724137 | 11.07744643 | 8.596730422 | 0 | 0.001978549 | 0.999932046 |
| FLT3LG | XM_017026534 | 10.73020072 | 0 | 0 | 0 | 0 | 0 | 9.713192346 | 0 | 1.34E-08 | 1 |
| FLT3LG | XR_935782 | 0 | 8.550974732 | 0 | 0 | 0 | 0 | 0 | 0 | 1.01E-07 | 1 |
| FLVCR2 | NM_017791 | 0 | 8.486541819 | 0 | 0 | 0 | 0 | 0 | 0 | 1.09E-07 | 1 |
| FLYWCH1 | XR_001752004 | 10.91641528 | 0 | 0 | 0 | 0 | 0 | 0 | 0 | 1.34E-08 | 1 |
| FLYWCH2 | NM_138439 | 0 | 0 | 0 | 11.43787256 | 8.771023979 | 0 | 0 | 0 | 0.009630342 | 0.02072721 |
| FMNL3 | XM_011538969 | 0 | 0 | 0 | 8.075296565 | 0 | 0 | 0 | 0 | 9.81E-06 | 1 |
| FMO4 | XM_005245045 | 0 | 0 | 9.364551246 | 0 | 7.574771995 | 0 | 7.84197419 | 0 | 0.036909647 | 0.999932046 |
| FMO4 | XM_005245048 | 11.61315024 | 10.57674034 | 0 | 0 | 0 | 0 | 9.798324675 | 10.71436539 | 0.000182674 | 0.02072721 |
| FMO5 | XM_006711245 | 0 | 0 | 0 | 7.146247488 | 0 | 0 | 0 | 0 | 3.03E-05 | 1 |
| FMO5 | XM_011509350 | 0 | 0 | 9.661459138 | 0 | 7.683842675 | 0 | 0 | 0 | 0.030540956 | 0.02096331 |
| FN1 | NM_001306132 | 0 | 0 | 0 | 0 | 0 | 8.994682249 | 0 | 9.507950041 | 1 | 0.02072721 |
| FN1 | XM_017003692 | 0 | 0 | 0 | 0 | 0 | 10.83943903 | 0 | 8.67881184 | 1 | 0.02072721 |
| FN1 | XM_017003695 | 0 | 9.059128398 | 0 | 0 | 0 | 0 | 7.954159584 | 8.563225531 | 7.80E-08 | 0.02562326 |
| FN3KRP | NM_024619 | 0 | 0 | 9.248957794 | 0 | 8.382480556 | 0 | 0 | 0 | 0.031919759 | 0.02164958 |
| FNBP1 | XM_024447457 | 0 | 8.951853153 | 0 | 0 | 0 | 0 | 0 | 0 | 7.60E-08 | 1 |
| FOCAD | NM_017794 | 0 | 0 | 0 | 8.643405671 | 6.557053251 | 0 | 0 | 0 | 0.065809018 | 0.046965326 |
| FOLR1 | NM_016729 | 0 | 0 | 0 | 13.11517734 | 12.65997942 | 16.31256856 | 19.40624433 | 13.58068084 | 1.34E-08 | 0.300510615 |
| FOPNL | NM_144600 | 0 | 0 | 0 | 9.614810718 | 8.281969086 | 0 | 0 | 0 | 0.027081236 | 0.020781046 |
| FOPNL | NR_130757 | 8.766981031 | 8.774317692 | 0 | 0 | 0 | 0 | 7.913105533 | 0 | 0.004361196 | 1 |
| FOSL1 | NM_005438 | 0 | 0 | 0 | 11.11224869 | 8.12102909 | 0 | 0 | 0 | 0.012856698 | 0.02072721 |
| FOXA1 | XM_017021246 | 0 | 8.700490516 | 0 | 0 | 0 | 0 | 0 | 0 | 9.01E-08 | 1 |
| FOXA3 | NM_004497 | 0 | 8.809861237 | 9.965418522 | 0 | 9.010889653 | 0 | 0 | 0 | 0.999953634 | 0.02072721 |
| FOXF1 | NM_001451 | 0 | 8.273938353 | 0 | 0 | 0 | 0 | 0 | 0 | 2.02E-07 | 1 |
| FOXJ1 | NM_001454 | 0 | 0 | 9.81881611 | 9.310616166 | 0 | 0 | 0 | 0 | 0.020792172 | 0.02072721 |
| FOXJ3 | NM_014947 | 0 | 0 | 0 | 7.569406338 | 0 | 0 | 0 | 0 | 1.69E-05 | 1 |
| FOXJ3 | XM_024454163 | 9.287928048 | 0 | 10.55932307 | 0 | 7.049011441 | 0 | 0 | 0 | 0.999953634 | 0.02072721 |
| FOXK2 | XR_933935 | 10.0393307 | 0 | 0 | 0 | 0 | 0 | 8.000624564 | 0 | 1.86E-08 | 1 |
| FOXO1 | XM_011535008 | 0 | 0 | 0 | 10.4845729 | 7.187172039 | 0 | 0 | 0 | 0.020252726 | 0.02072721 |
| FOXO3B | NR_026718 | 0 | 8.951853153 | 0 | 0 | 0 | 0 | 6.603572437 | 0 | 7.60E-08 | 1 |
| FOXP1 | NM_001349342 | 0 | 0 | 10.70227943 | 10.82395052 | 0 | 0 | 0 | 0 | 0.008996652 | 0.02072721 |
| FOXP1 | XM_017006167 | 11.40657281 | 10.79419952 | 11.19523667 | 0 | 10.56813537 | 0 | 0 | 0 | 0.999953634 | 0.02072721 |
| FOXRED1 | XM_017018000 | 9.190393069 | 0 | 0 | 0 | 0 | 0 | 0 | 0 | 7.73E-08 | 1 |
| FOXRED2 | XM_017028942 | 0 | 9.141645428 | 0 | 0 | 0 | 0 | 0 | 0 | 5.13E-08 | 1 |
| FPR1 | NM_002029 | 0 | 9.600697383 | 0 | 0 | 0 | 10.97087522 | 0 | 0 | 2.56E-08 | 1 |
| FREM1 | XR_001746195 | 0 | 0 | 0 | 7.43584608 | 0 | 0 | 7.189323943 | 0 | 1.87E-05 | 1 |
| FRG1 | NM_004477 | 0 | 0 | 11.06623168 | 11.67307841 | 8.811650533 | 0 | 0 | 10.59915604 | 0.000464259 | 0.999932046 |
| FRG1 | XR_002959721 | 0 | 10.49448544 | 9.930745925 | 0 | 7.415197319 | 0 | 0 | 0 | 0.999953634 | 0.02072721 |
| FRG1FP | NR_132320 | 0 | 9.834376365 | 0 | 0 | 0 | 0 | 0 | 0 | 2.00E-08 | 1 |
| FRK | XM_011535656 | 0 | 8.943876548 | 0 | 0 | 0 | 0 | 0 | 0 | 7.69E-08 | 1 |

| | | | | | | | | | | | |
|---|---|---|---|---|---|---|---|---|---|---|---|
| FRMD4B | XM_005264723 | 0 | 0 | 9.740902341 | 0 | 8.17807637 | 0 | 9.033360344 | 10.82323731 | 0.026499275 | 0.999932046 |
| FRMD5 | NM_001322950 | 0 | 0 | 0 | 0 | 0 | 9.915940983 | 7.899157068 | 10.12274908 | 1 | 0.002651192 |
| FRY | XM_017020307 | 8.972990281 | 0 | 0 | 0 | 0 | 0 | 0 | 0 | 7.80E-08 | 1 |
| FSD2 | NM_001281806 | 0 | 0 | 0 | 0 | 6.631562102 | 0 | 0 | 0 | 3.88E-05 | 1 |
| FSTL4 | XM_011543283 | 0 | 9.059128398 | 0 | 0 | 0 | 0 | 0 | 0 | 7.80E-08 | 1 |
| FTO | XM_024450437 | 0 | 0 | 0 | 0 | 6.607150896 | 0 | 0 | 0 | 0.00011086 | 1 |
| FTSJ1 | NM_012280 | 0 | 0 | 7.320277675 | 0 | 0 | 0 | 0 | 0 | 2.29E-05 | 1 |
| FTSJ1 | XM_005272595 | 0 | 0 | 0 | 10.02261004 | 8.522857234 | 0 | 0 | 0 | 0.021166613 | 0.02072721 |
| FUBP1 | XM_017002739 | 8.918652917 | 0 | 0 | 0 | 0 | 0 | 0 | 0 | 8.06E-08 | 1 |
| FUBP1 | XM_024450628 | 0 | 0 | 0 | 7.994469212 | 8.748879432 | 10.38011571 | 10.82728348 | 10.99015779 | 0.044463488 | 0.999932046 |
| FUBP1 | XR_001737521 | 0 | 0 | 9.4173244 | 8.831331601 | 7.187172039 | 9.95512224 | 7.899157068 | 0 | 0.006406752 | 0.999932046 |
| FUBP1 | XR_946790 | 0 | 8.286608519 | 0 | 0 | 0 | 0 | 8.827823966 | 0 | 2.03E-07 | 1 |
| FUBP3 | XM_011519173 | 0 | 0 | 0 | 0 | 0 | 8.717096959 | 8.676582513 | 0 | 1 | 0.02072721 |
| FUBP3 | XM_017015269 | 0 | 0 | 8.79670616 | 9.0384682 | 9.641489231 | 10.00931165 | 9.845456897 | 0 | 0.003198028 | 0.999932046 |
| FUBP3 | XM_017015273 | 0 | 0 | 9.879005341 | 11.77770777 | 0 | 0 | 9.930204221 | 9.165991457 | 0.006985032 | 0.999932046 |
| FUCA1 | XM_017000905 | 0 | 9.245884974 | 9.675970061 | 0 | 8.386072012 | 0 | 0 | 0 | 0.999953634 | 0.02072721 |
| FUS | NM_001170937 | 0 | 0 | 0 | 0 | 6.758905041 | 0 | 0 | 0 | 9.53E-05 | 1 |
| FUS | XM_011545781 | 0 | 0 | 11.57325751 | 0 | 9.627903042 | 0 | 10.15886318 | 4.683316329 | 0.007701018 | 0.999932046 |
| FUT2 | NM_000511 | 10.08938252 | 0 | 0 | 0 | 0 | 0 | 0 | 0 | 1.76E-08 | 1 |
| FUT3 | NM_001097639 | 9.825628908 | 0 | 0 | 0 | 0 | 10.41063817 | 0 | 10.17161555 | 2.30E-08 | 0.02072721 |
| FUT3 | NM_001097641 | 0 | 8.561439232 | 0 | 0 | 0 | 0 | 8.660134917 | 0 | 1.00E-07 | 1 |
| FUT8 | XM_017021140 | 0 | 0 | 10.47967688 | 0 | 8.516317387 | 0 | 0 | 0 | 0.017422361 | 0.020942118 |
| FXR1 | XM_017007299 | 0 | 9.897687377 | 0 | 0 | 0 | 0 | 0 | 0 | 1.88E-08 | 1 |
| FXYD1 | NM_021902 | 8.940634053 | 0 | 0 | 0 | 0 | 0 | 0 | 0 | 7.88E-08 | 1 |
| FXYD3 | NM_001136012 | 12.58003697 | 10.88030906 | 10.38481408 | 0 | 10.29434307 | 0 | 0 | 0 | 0.910753494 | 0.02072721 |
| FYN | XM_005266892 | 0 | 9.245884974 | 0 | 0 | 0 | 0 | 0 | 0 | 4.33E-08 | 1 |
| FZD5 | XM_024453130 | 10.15195724 | 9.271606013 | 0 | 0 | 0 | 10.61493266 | 0 | 0 | 0.001187467 | 1 |
| FZD6 | NR_133921 | 6.49435489 | 0 | 0 | 0 | 0 | 9.243294129 | 8.21847248 | 0 | 0.139461426 | 0.02072721 |
| FZD8 | NM_031866 | 0 | 0 | 0 | 0 | 6.30721813 | 0 | 0 | 0 | 4.99E-05 | 1 |
| G3BP1 | NM_005754 | 0 | 0 | 9.946391943 | 0 | 9.856090149 | 9.542408 | 0 | 0 | 0.015437659 | 1 |
| G6PC3 | NM_138387 | 0 | 0 | 8.159308007 | 0 | 8.583550423 | 0 | 0 | 0 | 0.04596385 | 0.029686259 |
| G6PC3 | XM_017025335 | 0 | 0 | 9.040481197 | 0 | 7.599732499 | 0 | 3.781941135 | 0 | 0.044175735 | 0.52090538 |
| G6PD | NM_001360016 | 9.595606435 | 0 | 0 | 0 | 0 | 0 | 8.67248816 | 11.38591623 | 3.08E-08 | 0.02072721 |
| GAA | XM_005257194 | 9.860637698 | 8.622680652 | 0 | 0 | 0 | 0 | 0 | 0 | 0.002059396 | 1 |
| GABARAPL1 | NM_031412 | 0 | 0 | 10.63257121 | 0 | 7.900217873 | 0 | 6.458933278 | 0 | 0.017693164 | 0.999932046 |
| GABPB1-AS1 | NR_024490 | 0 | 8.652353298 | 0 | 0 | 0 | 0 | 0 | 0 | 9.28E-08 | 1 |
| GABPB1-IT1 | NR_026891 | 0 | 0 | 10.37061091 | 0 | 8.323766986 | 0 | 7.58746354 | 0 | 0.01892723 | 0.999932046 |
| GAK | XM_005272270 | 0 | 0 | 0 | 0 | 6.736572239 | 0 | 0 | 0 | 9.53E-05 | 1 |
| GAL3ST2 | NM_022134 | 0 | 8.486541819 | 0 | 0 | 0 | 0 | 0 | 0 | 1.09E-07 | 1 |
| GALE | NM_001008216 | 0 | 0 | 9.81094843 | 0 | 6.478485785 | 8.178677299 | 0 | 0 | 0.033383857 | 0.999932046 |
| GALNS | XM_017023113 | 9.807800698 | 0 | 0 | 0 | 0 | 0 | 0 | 0 | 2.49E-08 | 1 |
| GALNT1 | NM_020474 | 0 | 10.99476254 | 9.313491365 | 7.288647943 | 10.83607497 | 0 | 0 | 0 | 0.999953634 | 0.003193095 |
| GALNT14 | NM_024572 | 10.96548485 | 0 | 0 | 0 | 0 | 0 | 0 | 0 | 1.34E-08 | 1 |

| | | | | | | | | | | | |
|---|---|---|---|---|---|---|---|---|---|---|---|
| GALNT14 | XM_017004906 | 0 | 8.756212053 | 0 | 0 | 0 | 0 | 0 | 0 | 8.55E-08 | 1 |
| GALNT3 | NM_004482 | 0 | 7.86468487 | 9.160915107 | 10.92865902 | 0 | 0 | 0 | 0 | 0.842259755 | 0.02072721 |
| GALNT5 | XM_017003237 | 0 | 0 | 0 | 8.130377898 | 0 | 0 | 0 | 0 | 9.41E-06 | 1 |
| GALT | NM_001258332 | 0 | 8.360378074 | 0 | 0 | 0 | 0 | 0 | 4.126159536 | 1.21E-07 | 0.868419696 |
| GAPDH | NM_001256799 | 0 | 0 | 0 | 11.5491888 | 8.452664007 | 0 | 10.27952469 | 6.516359403 | 0.009414775 | 0.999932046 |
| GAPLINC | NR_110428 | 0 | 9.712905893 | 0 | 0 | 0 | 0 | 0 | 0 | 2.55E-08 | 1 |
| GAPVD1 | NM_001354294 | 0 | 0 | 0 | 10.95864492 | 8.066117405 | 0 | 9.033360344 | 10.27746044 | 0.014170397 | 0.999932046 |
| GAPVD1 | NM_001354295 | 0 | 0 | 0 | 0 | 0 | 10.40761482 | 0 | 10.56820483 | 1 | 0.02072721 |
| GAPVD1 | XM_017014600 | 11.09608384 | 0 | 0 | 0 | 0 | 0 | 7.106768981 | 9.657578368 | 1.34E-08 | 0.02072721 |
| GAR1 | NM_018983 | 0 | 0 | 10.64590065 | 10.37137978 | 8.535848657 | 0 | 7.974256665 | 10.44015099 | 0.00120534 | 0.999932046 |
| GAR1 | NM_032993 | 0 | 9.059128398 | 0 | 0 | 0 | 0 | 0 | 0 | 7.80E-08 | 1 |
| GARS | NM_001316772 | 0 | 0 | 0 | 11.39521653 | 10.41175211 | 0 | 0 | 12.23777706 | 0.006840384 | 0.999932046 |
| GARS | XM_006715686 | 0 | 0 | 0 | 11.17056893 | 7.470373439 | 0 | 8.032920857 | 9.35314403 | 0.013394216 | 0.999932046 |
| GART | NM_001136006 | 0 | 0 | 0 | 0 | 6.491881102 | 0 | 0 | 0 | 4.33E-05 | 1 |
| GAS5 | NR_152525 | 0 | 0 | 0 | 0 | 0 | 10.9979301 | 0 | 10.35506601 | 1 | 0.02072721 |
| GAS5 | NR_152526 | 0 | 0 | 0 | 8.474877744 | 0 | 0 | 0 | 0 | 6.35E-06 | 1 |
| GAS5 | NR_152530 | 0 | 0 | 0 | 0 | 0 | 10.00931165 | 9.829916618 | 0 | 1 | 0.02072721 |
| GAS5 | NR_152531 | 9.721559119 | 0 | 0 | 0 | 0 | 0 | 0 | 10.15608825 | 2.60E-08 | 1 |
| GAS5 | NR_152533 | 0 | 8.475517571 | 0 | 0 | 0 | 3.837722533 | 0 | 0 | 1.09E-07 | 0.976243727 |
| GAS5 | NR_152534 | 0 | 0 | 0 | 7.720649682 | 0 | 0 | 0 | 0 | 1.41E-05 | 1 |
| GATAD1 | NR_052016 | 0 | 8.861586703 | 0 | 0 | 0 | 0 | 0 | 0 | 7.86E-08 | 1 |
| GATAD2A | XM_024451560 | 7.899356205 | 0 | 8.503193464 | 0 | 8.097125035 | 0 | 0 | 0 | 0.999953634 | 0.031459623 |
| GATD1 | NM_182612 | 0 | 9.708204508 | 0 | 0 | 0 | 0 | 0 | 0 | 2.47E-08 | 1 |
| GATD3B | XM_006723901 | 10.43737343 | 0 | 0 | 0 | 0 | 0 | 0 | 0 | 1.47E-08 | 1 |
| GATD3B | XM_024452142 | 9.995377245 | 9.595615072 | 0 | 0 | 0 | 0 | 0 | 0 | 0.001115345 | 1 |
| GATM | NM_001482 | 0 | 8.895067962 | 0 | 0 | 0 | 0 | 0 | 0 | 7.75E-08 | 1 |
| GBA2 | NM_020944 | 0 | 0 | 9.809632968 | 0 | 8.38068147 | 0 | 0 | 0 | 0.024535363 | 0.02072721 |
| GBA2 | XM_017014946 | 0 | 10.02412431 | 0 | 0 | 0 | 0 | 0 | 0 | 1.63E-08 | 1 |
| GBAT2 | NR_131972 | 0 | 0 | 0 | 0 | 0 | 0 | 0 | 6.427178007 | 1 | 2.72E-06 |
| GBGT1 | NM_021996 | 0 | 8.29916838 | 0 | 0 | 0 | 0 | 0 | 0 | 1.31E-07 | 1 |
| GBP1 | XM_024446351 | 9.387309849 | 9.451153482 | 0 | 0 | 0 | 0 | 0 | 0 | 0.001832515 | 1 |
| GCC2 | XR_923067 | 9.153224721 | 0 | 0 | 0 | 0 | 0 | 7.469710231 | 0 | 6.38E-08 | 1 |
| GCLM | XM_017001057 | 0 | 8.360378074 | 0 | 0 | 0 | 0 | 0 | 10.85670879 | 1.21E-07 | 1 |
| GDE1 | NM_001324066 | 0 | 9.051724868 | 0 | 0 | 0 | 0 | 0 | 0 | 7.86E-08 | 1 |
| GDF15 | NM_004864 | 0 | 0 | 0 | 10.53239157 | 9.168302049 | 0 | 0 | 0 | 0.014680579 | 0.02072721 |
| GDPD1 | NM_182569 | 0 | 8.561439232 | 0 | 0 | 0 | 0 | 0 | 0 | 1.00E-07 | 1 |
| GDPD3 | NM_024307 | 0 | 9.06649413 | 9.110526406 | 10.29025878 | 4.552031775 | 0 | 0 | 0 | 0.999953634 | 0.007991001 |
| GEM | NM_005261 | 0 | 0 | 9.800390998 | 0 | 7.549372027 | 0 | 8.728777965 | 0 | 0.028975418 | 0.999932046 |
| GEMIN2 | NM_003616 | 0 | 0 | 10.6243644 | 0 | 9.630933271 | 0 | 0 | 0 | 0.012658311 | 0.02072721 |
| GEMIN4 | NM_015721 | 0 | 8.719304505 | 0 | 0 | 0 | 0 | 0 | 0 | 8.96E-08 | 1 |
| GEMIN7 | NM_001007269 | 10.54531825 | 0 | 0 | 0 | 0 | 0 | 0 | 0 | 1.36E-08 | 1 |
| GEMIN7 | NM_001007270 | 0 | 0 | 10.62361601 | 0 | 8.919671304 | 0 | 0 | 10.46556789 | 0.015059301 | 0.999932046 |
| GEMIN8 | NM_017856 | 8.526574932 | 0 | 0 | 0 | 0 | 0 | 0 | 0 | 1.12E-07 | 1 |

| | | | | | | | | | | | |
|---|---|---|---|---|---|---|---|---|---|---|---|
| GFER | NM_005262 | 12.17554274 | 9.258802813 | 0 | 0 | 0 | 0 | 7.118855734 | 0 | 0.000186164 | 1 |
| GFOD2 | NM_001243650 | 0 | 0 | 10.06359724 | 10.59639063 | 0 | 9.875665851 | 9.669692479 | 0 | 0.012282915 | 0.999932046 |
| GGA1 | NM_013365 | 0 | 9.152068755 | 0 | 0 | 0 | 0 | 0 | 0 | 5.06E-08 | 1 |
| GGA1 | XR_002958684 | 0 | 0 | 0 | 10.21835411 | 5.200341806 | 0 | 0 | 0 | 0.027779243 | 1 |
| GGA3 | XM_017024385 | 0 | 0 | 0 | 0 | 6.544252221 | 0 | 0 | 0 | 4.20E-05 | 1 |
| GGCT | NM_024051 | 0 | 0 | 11.28364078 | 12.0193824 | 10.85790033 | 0 | 9.319168841 | 11.85525176 | 0.000180274 | 0.999932046 |
| GGCT | NR_037669 | 0 | 0 | 10.78760486 | 0 | 9.984832692 | 0 | 6.092894468 | 0 | 0.01057057 | 0.802086624 |
| GGNBP2 | XM_011525266 | 0 | 0 | 0 | 0 | 0 | 0 | 8.964704031 | 10.40409 | 1 | 0.02072721 |
| GGPS1 | NM_001037277 | 0 | 0 | 0 | 0 | 0 | 0 | 7.613324795 | 10.54056688 | 1 | 0.02072721 |
| GGT1 | NM_013430 | 0 | 8.529815232 | 0 | 0 | 0 | 0 | 8.107603982 | 0 | 1.04E-07 | 1 |
| GGT6 | NM_153338 | 11.17005147 | 0 | 0 | 10.85698537 | 7.838643081 | 0 | 0 | 0 | 0.999953634 | 0.02072721 |
| GGTA1P | NR_045211 | 0 | 0 | 7.079567203 | 0 | 0 | 0 | 0 | 0 | 2.94E-05 | 1 |
| GID4 | NM_024052 | 0 | 0 | 11.3085339 | 0 | 10.14336978 | 0 | 8.930825448 | 0 | 0.00781162 | 0.999932046 |
| GIMAP1 | NM_130759 | 0 | 8.550974732 | 0 | 0 | 0 | 10.97496594 | 0 | 0 | 1.01E-07 | 1 |
| GIMAP7 | NM_153236 | 0 | 9.670031486 | 0 | 0 | 0 | 0 | 8.660134917 | 0 | 2.40E-08 | 1 |
| GINS2 | NM_016095 | 0 | 0 | 0 | 10.5924728 | 9.2424901 | 0 | 0 | 0 | 0.014014528 | 0.02072721 |
| GIPC1 | NM_005716 | 0 | 0 | 10.89322032 | 9.57366938 | 10.84360453 | 0 | 0 | 10.22219849 | 0.000627847 | 1 |
| GIPC1 | NM_202468 | 10.25642453 | 8.348342131 | 9.144313654 | 11.41697913 | 8.131765004 | 0 | 0 | 0 | 0.999953634 | 0.002417476 |
| GIPC1 | XM_017026148 | 0 | 9.768158992 | 6.343890623 | 0 | 8.907227812 | 0 | 0 | 0 | 0.999953634 | 0.037810409 |
| GIPC2 | NM_017655 | 0 | 0 | 9.0802992 | 9.733411602 | 8.312487126 | 0 | 0 | 10.30588172 | 0.003558135 | 0.999932046 |
| GIT1 | XM_024450734 | 10.0393307 | 0 | 0 | 0 | 0 | 0 | 0 | 0 | 1.86E-08 | 1 |
| GJA9-MYCBP | NR_037637 | 8.526574932 | 6.420232184 | 0 | 0 | 0 | 0 | 2.992327696 | 0 | 0.016118645 | 0.999932046 |
| GJB1 | XM_011530907 | 0 | 0 | 10.32346045 | 9.677180466 | 0 | 0 | 7.980893958 | 0 | 0.015536466 | 0.999932046 |
| GJC2 | NM_020435 | 0 | 0 | 0 | 0 | 0 | 0 | 8.574960864 | 9.677050566 | 1 | 0.02072721 |
| GJC3 | NM_181538 | 11.15602802 | 0 | 0 | 0 | 0 | 0 | 13.72726968 | 0 | 1.34E-08 | 1 |
| GK | NM_001205019 | 0 | 0 | 0 | 0 | 6.544252221 | 0 | 0 | 0 | 4.20E-05 | 1 |
| GK | XM_017029409 | 0 | 0 | 0 | 8.508532777 | 0 | 0 | 0 | 0 | 6.12E-06 | 1 |
| GLA | XR_938397 | 10.736497 | 0 | 0 | 0 | 0 | 0 | 7.402084669 | 0 | 1.34E-08 | 1 |
| GLCCI1 | NM_138426 | 0 | 8.430558346 | 10.11135781 | 0 | 6.531336591 | 0 | 0 | 0 | 0.999953634 | 0.02072721 |
| GLCE | XM_017022074 | 0 | 8.497482465 | 0 | 0 | 0 | 0 | 5.991365783 | 0 | 1.06E-07 | 1 |
| GLE1 | XM_024447519 | 8.421077904 | 0 | 9.491392213 | 0 | 7.618173501 | 0 | 0 | 0 | 0.999953634 | 0.022619671 |
| GLG1 | NM_012201 | 0 | 0 | 0 | 10.48879454 | 7.278890064 | 0 | 0 | 0 | 0.02002499 | 0.02072721 |
| GLG1 | NR_027265 | 0 | 0 | 0 | 0 | 6.59478863 | 10.73557268 | 0 | 0 | 4.02E-05 | 1 |
| GLI1 | NM_005269 | 9.581685348 | 0 | 0 | 0 | 0 | 0 | 0 | 0 | 3.13E-08 | 1 |
| GLI4 | NM_138465 | 0 | 0 | 0 | 0 | 6.747781854 | 10.60264726 | 9.138024627 | 0 | 9.53E-05 | 0.999932046 |
| GLIPR2 | NR_104641 | 0 | 10.95959057 | 0 | 0 | 0 | 0 | 0 | 0 | 1.34E-08 | 1 |
| GLIS2 | NM_001318918 | 0 | 0 | 0 | 0 | 0 | 0 | 9.077380728 | 9.858552905 | 1 | 0.02072721 |
| GLIS3 | XM_017014361 | 11.01811794 | 0 | 0 | 0 | 0 | 0 | 0 | 0 | 1.34E-08 | 1 |
| GLIS3 | XR_929206 | 0 | 8.919678654 | 0 | 0 | 0 | 0 | 0 | 0 | 7.73E-08 | 1 |
| GLRX2 | XM_017001397 | 10.22989302 | 0 | 0 | 0 | 0 | 0 | 0 | 0 | 1.59E-08 | 1 |
| GLT8D2 | NM_031302 | 9.95545054 | 0 | 0 | 0 | 0 | 0 | 0 | 0 | 2.01E-08 | 1 |
| GLYR1 | NM_001308096 | 0 | 0 | 10.83621996 | 0 | 8.238923989 | 10.53036427 | 0 | 0 | 0.015146083 | 0.999932046 |
| GLYR1 | NM_001324098 | 0 | 0 | 0 | 0 | 0 | 0 | 8.436958184 | 9.400769032 | 1 | 0.02072721 |

| | | | | | | | | | | | |
|---|---|---|---|---|---|---|---|---|---|---|---|
| GLYR1 | NR_136698 | 9.560548562 | 0 | 0 | 0 | 0 | 0 | 0 | 0 | 3.19E-08 | 1 |
| GLYR1 | XM_011522716 | 0 | 0 | 0 | 9.257172648 | 6.464964927 | 0 | 9.089715157 | 9.63286261 | 0.046489006 | 0.999932046 |
| GLYR1 | XM_017023786 | 11.56796835 | 0 | 0 | 0 | 0 | 0 | 6.796513442 | 10.4711558 | 1.34E-08 | 0.02072721 |
| GM2A | NM_001167607 | 0 | 10.09426749 | 0 | 0 | 0 | 0 | 0 | 9.837075169 | 1.59E-08 | 1 |
| GMCL1 | XM_011533033 | 0 | 0 | 0 | 0 | 6.491881102 | 0 | 6.586265919 | 8.938841664 | 4.33E-05 | 0.999932046 |
| GMFB | NM_004124 | 0 | 9.864267562 | 0 | 0 | 0 | 0 | 0 | 0 | 1.96E-08 | 1 |
| GMFG | NM_004877 | 0 | 0 | 10.74419638 | 0 | 7.093624618 | 11.91765692 | 8.767681412 | 10.01784369 | 0.018196212 | 0.999932046 |
| GMIP | XM_017026861 | 10.69183479 | 0 | 0 | 0 | 0 | 0 | 0 | 0 | 2.15E-08 | 1 |
| GMNN | NM_001251989 | 0 | 9.620849457 | 0 | 11.34777873 | 8.973182448 | 0 | 0 | 0 | 0.999953634 | 0.02072721 |
| GMPR2 | NM_001002001 | 0 | 0 | 10.50590512 | 10.25599332 | 7.320775981 | 0 | 0 | 10.03114526 | 0.001804162 | 0.999932046 |
| GMPR2 | NM_016576 | 0 | 10.22178903 | 0 | 0 | 0 | 0 | 0 | 0 | 1.51E-08 | 1 |
| GMPS | NM_003875 | 0 | 0 | 0 | 10.4675617 | 7.87488031 | 0 | 0 | 0 | 0.018884777 | 0.02072721 |
| GNA14 | NM_004297 | 0 | 8.728620286 | 0 | 0 | 0 | 0 | 0 | 0 | 8.80E-08 | 1 |
| GNAS | NM_001309883 | 0 | 0 | 10.21436136 | 0 | 8.555119041 | 0 | 8.207195305 | 0 | 0.019565878 | 0.999932046 |
| GNAS | XM_017027815 | 0 | 0 | 11.5693764 | 0 | 10.74146379 | 0 | 11.88510093 | 9.75241299 | 0.005681098 | 0.999932046 |
| GNG10 | NM_001198664 | 0 | 9.946408903 | 0 | 10.65767084 | 9.812715433 | 0 | 0 | 0 | 0.999953634 | 0.02072721 |
| GNLY | NM_006433 | 10.34149639 | 0 | 0 | 0 | 0 | 0 | 0 | 0 | 1.60E-08 | 1 |
| GNPNAT1 | XM_006720238 | 0 | 0 | 0 | 0 | 6.437537018 | 0 | 0 | 0 | 4.51E-05 | 1 |
| GNPTAB | XM_017019962 | 0 | 0 | 0 | 0 | 6.747781854 | 0 | 0 | 0 | 9.53E-05 | 1 |
| GOLGA1 | NM_002077 | 9.777588513 | 0 | 0 | 0 | 0 | 0 | 0 | 0 | 2.46E-08 | 1 |
| GOLGA2 | NM_004486 | 9.330172147 | 0 | 0 | 0 | 0 | 0 | 0 | 0 | 4.56E-08 | 1 |
| GOLGA2P5 | NR_024261 | 10.50162657 | 0 | 0 | 0 | 0 | 0 | 0 | 0 | 1.41E-08 | 1 |
| GOLGA4 | XM_024453456 | 0 | 0 | 11.59518268 | 0 | 9.91072005 | 0 | 9.36301824 | 0 | 0.007122137 | 0.999932046 |
| GOLGA8B | XM_024449931 | 9.807800698 | 0 | 0 | 0 | 0 | 7.647222305 | 7.596135569 | 0 | 2.49E-08 | 0.043035185 |
| GOLGA8J | XM_006720651 | 9.567628621 | 0 | 0 | 0 | 0 | 0 | 0 | 0 | 3.16E-08 | 1 |
| GOLGA8O | NM_001277308 | 9.928205339 | 0 | 0 | 0 | 0 | 0 | 0 | 0 | 2.03E-08 | 1 |
| GOLGB1 | XR_001740105 | 10.17531978 | 0 | 0 | 0 | 0 | 0 | 0 | 0 | 1.60E-08 | 1 |
| GOLM1 | NM_016548 | 0 | 5.077427354 | 11.4209194 | 10.99693978 | 10.50595113 | 0 | 7.720323987 | 8.885437246 | 0.042718283 | 0.999932046 |
| GOLPH3 | NM_022130 | 0 | 0 | 10.24582768 | 9.371325678 | 8.967200643 | 0 | 8.382856136 | 9.131119222 | 0.00198569 | 0.999932046 |
| GOSR1 | XM_017025376 | 0 | 8.681427931 | 0 | 0 | 0 | 0 | 0 | 0 | 9.11E-08 | 1 |
| GOT1 | NM_002079 | 0 | 0 | 0 | 11.57025808 | 9.082363872 | 0 | 7.469710231 | 0 | 0.008435145 | 0.999932046 |
| GP1BA | NM_000173 | 10.5666777 | 0 | 0 | 0 | 0 | 0 | 0 | 0 | 1.34E-08 | 1 |
| GPALPP1 | XR_941622 | 0 | 0 | 0 | 0 | 6.491881102 | 0 | 7.497749587 | 0 | 4.33E-05 | 1 |
| GPAT2 | NM_207328 | 0 | 8.747073305 | 0 | 0 | 0 | 0 | 0 | 0 | 8.61E-08 | 1 |
| GPAT4 | XR_001745474 | 0 | 0 | 0 | 8.274277384 | 0 | 0 | 6.187745501 | 0 | 7.87E-06 | 1 |
| GPATCH11 | XM_017003748 | 0 | 0 | 0 | 8.239625615 | 0 | 0 | 0 | 0 | 1.80E-05 | 1 |
| GPATCH2 | NM_001297754 | 0 | 8.622680652 | 0 | 0 | 0 | 0 | 0 | 0 | 9.58E-08 | 1 |
| GPBP1 | XM_024446176 | 9.894817043 | 0 | 0 | 0 | 0 | 0 | 0 | 0 | 2.09E-08 | 1 |
| GPBP1 | XR_002956169 | 0 | 0 | 0 | 0 | 6.464964927 | 0 | 0 | 0 | 4.45E-05 | 1 |
| GPBP1L1 | XM_024448810 | 0 | 0 | 0 | 0 | 6.679179249 | 0 | 8.262719816 | 0 | 3.78E-05 | 1 |
| GPC6 | XM_017020300 | 0 | 10.01653847 | 0 | 0 | 0 | 0 | 0 | 0 | 2.27E-08 | 1 |
| GPHN | XM_011536345 | 0 | 0 | 0 | 9.74051016 | 8.280040102 | 0 | 0 | 0 | 0.025455996 | 0.02072721 |
| GPI | NM_000175 | 0 | 0 | 11.26071931 | 9.981258774 | 10.58620167 | 0 | 0 | 10.4711558 | 0.000513649 | 0.999932046 |

| | | | | | | | | | | | |
|---|---|---|---|---|---|---|---|---|---|---|---|
| GPLD1 | XM_024446403 | 0 | 9.411213355 | 0 | 0 | 0 | 0 | 0 | 0 | 3.28E-08 | 1 |
| GPR108 | XM_024451618 | 0 | 0 | 10.53723922 | 7.857531294 | 6.906244365 | 0 | 8.451367849 | 0 | 0.004392043 | 0.999932046 |
| GPR160 | XM_024453450 | 0 | 0 | 0 | 0 | 6.747781854 | 0 | 0 | 0 | 9.53E-05 | 1 |
| GPR160 | XM_024453451 | 0 | 0 | 0 | 0 | 0 | 0 | 8.964704031 | 9.51878258 | 1 | 0.02072721 |
| GPR183 | XM_017020405 | 0 | 7.777608832 | 0 | 0 | 0 | 0 | 0 | 6.558964522 | 0.067978813 | 2.30E-06 |
| GPS1 | NM_001330539 | 8.691211177 | 0 | 0 | 0 | 0 | 0 | 0 | 0 | 9.73E-08 | 1 |
| GPS1 | XM_017024534 | 0 | 9.165850403 | 0 | 0 | 0 | 0 | 0 | 0 | 4.97E-08 | 1 |
| GPSM2 | XM_017001098 | 0 | 0 | 0 | 0 | 0 | 0 | 8.981349581 | 10.53923749 | 1 | 0.02072721 |
| GPT | NM_005309 | 9.629832564 | 8.29916838 | 0 | 0 | 0 | 0 | 0 | 0 | 0.002921259 | 1 |
| GPT2 | NM_133443 | 10.04948094 | 0 | 0 | 0 | 0 | 0 | 0 | 0 | 5.08E-08 | 1 |
| GPX1 | NM_001329455 | 0 | 9.044283149 | 0 | 0 | 0 | 0 | 0 | 0 | 6.33E-08 | 1 |
| GPX4 | NM_001039847 | 0 | 9.484529283 | 0 | 0 | 0 | 0 | 0 | 0 | 2.99E-08 | 1 |
| GPX7 | NM_015696 | 0 | 0 | 0 | 0 | 0 | 0 | 6.687121454 | 10.54454772 | 1 | 0.02072721 |
| GRAMD2B | XM_017009785 | 9.457596586 | 0 | 0 | 0 | 0 | 0 | 0 | 0 | 3.78E-08 | 1 |
| GRAMD2B | XM_024446201 | 0 | 0 | 0 | 7.307890232 | 0 | 0 | 0 | 0 | 2.14E-05 | 1 |
| GRAP2 | NM_001291824 | 9.045780268 | 0 | 0 | 0 | 0 | 0 | 0 | 0 | 7.60E-08 | 1 |
| GRB10 | XM_024446730 | 0 | 0 | 5.982240563 | 0 | 9.289271525 | 5.75998802 | 6.811485137 | 0 | 0.046268417 | 0.999932046 |
| GRB2 | NM_002086 | 10.56313968 | 0 | 0 | 0 | 0 | 0 | 8.207195305 | 0 | 1.34E-08 | 1 |
| GRB7 | NM_001242443 | 0 | 0 | 0 | 0 | 0 | 0 | 10.42046706 | 10.24846798 | 1 | 0.02072721 |
| GRB7 | XM_024450729 | 0 | 0 | 9.26440391 | 11.22599084 | 10.8011927 | 0 | 11.82734684 | 11.70151026 | 0.00058198 | 0.999932046 |
| GRID1 | NM_017551 | 9.903236837 | 0 | 0 | 0 | 0 | 0 | 0 | 0 | 2.07E-08 | 1 |
| GRINA | NM_001009184 | 0 | 0 | 8.861618996 | 8.224516227 | 8.555119041 | 0 | 10.6595278 | 10.43444146 | 0.007053794 | 0.999932046 |
| GRIPAP1 | XM_024452410 | 9.982190738 | 0 | 9.15263826 | 9.116932532 | 2.367080301 | 0 | 0 | 0 | 0.999953634 | 0.02072721 |
| GRK2 | XR_001747779 | 0 | 0 | 10.73590986 | 0 | 6.813269833 | 0 | 0 | 0 | 0.018733627 | 0.020920478 |
| GRK3 | NM_005160 | 0 | 0 | 0 | 0 | 0 | 9.391436267 | 9.501015388 | 11.20072455 | 1 | 0.001394691 |
| GRK6 | XM_006714859 | 0 | 8.927789798 | 0 | 0 | 0 | 0 | 0 | 0 | 7.69E-08 | 1 |
| GRK6 | XM_024446032 | 10.05704703 | 0 | 0 | 0 | 0 | 0 | 0 | 0 | 5.05E-08 | 1 |
| GRM8 | XM_017012079 | 9.502621339 | 8.550974732 | 0 | 0 | 0 | 0 | 9.792677098 | 0 | 0.002884596 | 1 |
| GRPEL1 | NM_025196 | 0 | 0 | 0 | 11.76948961 | 9.970550849 | 0 | 8.700907578 | 11.51053907 | 0.006275517 | 0.999932046 |
| GRSF1 | XM_011531897 | 0 | 10.25440355 | 7.305421363 | 10.5262435 | 9.626385537 | 0 | 0 | 0 | 0.999953634 | 0.003589884 |
| GRTP1 | XM_017020756 | 0 | 0 | 0 | 0 | 6.42362501 | 0 | 0 | 0 | 4.54E-05 | 1 |
| GSDMD | NM_024736 | 0 | 0 | 0 | 10.28541326 | 8.609897378 | 8.381306358 | 0 | 0 | 0.018491109 | 0.999932046 |
| GSE1 | XM_017023083 | 0 | 0 | 0 | 11.06778942 | 7.449928885 | 0 | 0 | 0 | 0.014265821 | 0.02072721 |
| GSK3B | NM_002093 | 0 | 0 | 0 | 0 | 0 | 0 | 0 | 6.754940855 | 1 | 1.78E-06 |
| GSKIP | XM_024449626 | 0 | 0 | 9.468235064 | 0 | 5.354946376 | 0 | 0 | 0 | 0.047165139 | 0.030070338 |
| GSN | NM_000177 | 0 | 0 | 11.32381917 | 0 | 7.350492346 | 0 | 0 | 0 | 0.012650499 | 9.42E-09 |
| GSN | XR_001746283 | 11.92433918 | 9.439854398 | 10.44874387 | 0 | 8.89719516 | 0 | 0 | 0 | 0.975550828 | 0.020781046 |
| GSPT1 | NM_001130006 | 0 | 8.878424457 | 0 | 0 | 0 | 0 | 0 | 0 | 9.15E-08 | 1 |
| GSR | NM_000637 | 0 | 7.103476409 | 7.99310737 | 8.474877744 | 0 | 0 | 0 | 0 | 0.999953634 | 0.036829179 |
| GSS | NM_001322494 | 0 | 0 | 0 | 10.35412868 | 7.910229599 | 0 | 7.856483795 | 0 | 0.019954389 | 0.999932046 |
| GSTM2 | NM_000848 | 0 | 8.792198922 | 0 | 0 | 0 | 0 | 0 | 0 | 8.40E-08 | 1 |
| GSTO1 | NM_001191002 | 0 | 0 | 0 | 10.27077828 | 8.057134281 | 0 | 0 | 0 | 0.020372859 | 0.02072721 |
| GSTT1 | NM_001293809 | 0 | 9.352166886 | 0 | 0 | 0 | 0 | 0 | 0 | 3.68E-08 | 1 |

| | | | | | | | | | | | |
|---|---|---|---|---|---|---|---|---|---|---|---|
| GSTT1 | NM_001293814 | 10.40414088 | 0 | 0 | 0 | 0 | 0 | 0 | 0 | 1.47E-08 | 1 |
| GSTZ1 | XM_024449552 | 10.26079939 | 0 | 0 | 0 | 0 | 0 | 0 | 0 | 1.58E-08 | 1 |
| GTF2A1 | NM_201595 | 0 | 8.903318262 | 0 | 0 | 0 | 0 | 0 | 0 | 7.75E-08 | 1 |
| GTF2E2 | NM_002095 | 0 | 8.943876548 | 0 | 9.656688688 | 8.981515525 | 0 | 0 | 0 | 0.999953634 | 0.02072721 |
| GTF2E2 | XM_011544510 | 11.43878504 | 0 | 0 | 0 | 0 | 0 | 0 | 0 | 1.34E-08 | 1 |
| GTF2E2 | XM_024447138 | 9.623051988 | 0 | 0 | 0 | 0 | 0 | 0 | 0 | 2.93E-08 | 1 |
| GTF2F1 | XM_024451475 | 0 | 0 | 7.974444826 | 10.6167865 | 9.056678642 | 0 | 7.885072427 | 0 | 0.002270731 | 0.999932046 |
| GTF2H3 | NM_001271867 | 0 | 8.622680652 | 0 | 0 | 0 | 0 | 7.863684239 | 0 | 9.58E-08 | 1 |
| GTF2H5 | NM_207118 | 0 | 0 | 0 | 9.222102536 | 10.74950227 | 0 | 9.584008175 | 10.85136241 | 0.012811187 | 0.999932046 |
| GTF2IRD1 | XM_006716183 | 0 | 9.006487637 | 0 | 0 | 0 | 0 | 0 | 0 | 6.80E-08 | 1 |
| GTF2IRD1 | XM_011516713 | 0 | 0 | 0 | 0 | 0 | 10.34367032 | 6.603572437 | 0 | 1 | 0.02072721 |
| GTF2IRD2 | NM_001281447 | 0 | 0 | 10.4755353 | 0 | 9.206380832 | 9.542408 | 9.279339435 | 9.923171732 | 0.015146083 | 0.999932046 |
| GTF3A | NM_002097 | 0 | 0 | 10.73313707 | 12.00287311 | 11.28634046 | 12.1905134 | 11.20114226 | 10.98430795 | 0.000188918 | 0.999932046 |
| GTF3C2 | NM_001318909 | 0 | 0 | 12.11171422 | 0 | 10.16495871 | 0 | 9.889335013 | 11.15371953 | 0.004953244 | 0.999932046 |
| GTF3C3 | XM_005246965 | 0 | 0 | 0 | 9.159924829 | 9.529039137 | 0 | 0 | 0 | 0.022059726 | 0.02072721 |
| GTF3C5 | XR_002956827 | 0 | 9.303125566 | 0 | 0 | 0 | 0 | 5.11387751 | 0 | 4.01E-08 | 0.561568989 |
| GTPBP10 | NM_033107 | 0 | 0 | 10.61611064 | 10.33080157 | 9.440276507 | 0 | 6.017425619 | 0 | 0.000960006 | 0.32133666 |
| GTPBP2 | XM_024446478 | 0 | 0 | 0 | 0 | 0 | 6.613002074 | 8.161183866 | 0 | 1 | 0.044141775 |
| GUCY2C | XM_011520631 | 9.134275178 | 0 | 0 | 0 | 0 | 10.44249906 | 8.195829284 | 0 | 6.66E-08 | 0.02072721 |
| GUK1 | NM_000858 | 0 | 0 | 12.31057631 | 12.17288205 | 9.375808684 | 11.0963462 | 10.30685654 | 11.89372971 | 0.000162304 | 0.999932046 |
| GUSB | XR_927461 | 0 | 8.737876298 | 0 | 0 | 0 | 0 | 0 | 0 | 9.11E-08 | 1 |
| GUSBP11 | NR_024448 | 10.45657028 | 0 | 0 | 0 | 0 | 0 | 0 | 0 | 1.45E-08 | 1 |
| GUSBP5 | NR_003675 | 0 | 0 | 0 | 6.964181636 | 0 | 0 | 0 | 0 | 3.03E-05 | 1 |
| GYG1 | NM_001184720 | 0 | 0 | 0 | 8.766740692 | 0 | 0 | 0 | 0 | 4.64E-06 | 1 |
| GYS1 | NR_027763 | 0 | 0 | 0 | 8.573575335 | 0 | 0 | 0 | 0 | 2.47E-05 | 1 |
| GZMB | NM_004131 | 0 | 0 | 9.382357652 | 12.20081473 | 0 | 0 | 11.20504506 | 10.67616912 | 1.34E-08 | 0.999932046 |
| H2AFJ | NR_027716 | 0 | 0 | 0 | 0 | 6.518304288 | 0 | 8.626666267 | 0 | 4.23E-05 | 1 |
| H2AFV | NM_138635 | 0 | 0 | 9.371700203 | 0 | 6.736572239 | 0 | 0 | 0 | 0.042956148 | 0.027403095 |
| H2AFV | NM_201516 | 0 | 9.411213355 | 0 | 0 | 0 | 0 | 0 | 0 | 3.28E-08 | 1 |
| H2AFY | NM_138609 | 0 | 0 | 10.63257121 | 10.76251193 | 0 | 0 | 0 | 0 | 0.009464101 | 0.02072721 |
| H2AFY | XM_011543731 | 0 | 8.78328601 | 10.22229251 | 9.451378274 | 9.382116919 | 0 | 0 | 0 | 0.941836129 | 0.002835114 |
| H2AFY | XR_948309 | 0 | 8.286608519 | 0 | 0 | 0 | 0 | 6.551017323 | 0 | 2.03E-07 | 1 |
| H6PD | XM_006711052 | 0 | 0 | 0 | 0 | 6.366595318 | 0 | 0 | 0 | 4.76E-05 | 1 |
| HAAO | XM_024452774 | 0 | 0 | 0 | 10.25846802 | 6.667421491 | 0 | 0 | 0 | 0.024242028 | 0.02072721 |
| HABP4 | NM_014282 | 0 | 0 | 0 | 0 | 6.769943125 | 0 | 7.751711637 | 0 | 9.43E-05 | 1 |
| HADHB | NM_001281513 | 0 | 9.131146246 | 0 | 0 | 0 | 0 | 0 | 0 | 5.23E-08 | 1 |
| HAPLN1 | XM_017009054 | 0 | 9.17269217 | 0 | 0 | 0 | 0 | 0 | 0 | 4.92E-08 | 1 |
| HARBI1 | XM_017017605 | 0 | 8.39589491 | 0 | 0 | 0 | 11.0349743 | 8.759984195 | 0 | 1.17E-07 | 0.02072721 |
| HARS | NM_002109 | 9.950042506 | 0 | 9.062737842 | 8.516825278 | 0 | 0 | 0 | 0 | 0.999953634 | 0.024319498 |
| HAS2 | NM_005328 | 10.14723899 | 0 | 0 | 0 | 0 | 0 | 0 | 0 | 1.64E-08 | 1 |
| HAT1 | XM_024453187 | 0 | 0 | 0 | 0 | 6.557053251 | 0 | 0 | 0 | 4.16E-05 | 1 |
| HAUS1 | NM_138443 | 0 | 0 | 0 | 10.64543219 | 9.33458352 | 0 | 0 | 0 | 0.013378444 | 0.02072721 |
| HAUS1 | NR_026978 | 0 | 0 | 10.57721384 | 0 | 7.757784245 | 0 | 0 | 0 | 0.018540178 | 0.02072721 |

| | | | | | | | | | | | |
|---|---|---|---|---|---|---|---|---|---|---|---|
| HAUS2 | NM_018097 | 0 | 0 | 0 | 9.711903927 | 9.857384392 | 0 | 8.240765774 | 0 | 0.016626721 | 0.999932046 |
| HAUS3 | NM_001303143 | 0 | 0 | 0 | 9.575655325 | 8.083917571 | 0 | 0 | 0 | 0.028827511 | 0.021190083 |
| HAUS4 | NM_017815 | 0 | 0 | 10.80751154 | 0 | 9.283506136 | 0 | 7.671923744 | 0 | 0.012666004 | 0.999932046 |
| HBEGF | NM_001945 | 9.546283346 | 0 | 10.09417272 | 0 | 7.259441431 | 0 | 0 | 0 | 0.999953634 | 0.02072721 |
| HBP1 | XR_001744628 | 0 | 8.998808245 | 0 | 0 | 0 | 0 | 0 | 0 | 6.89E-08 | 1 |
| HBP1 | XR_002956420 | 0 | 0 | 0 | 9.857763009 | 7.40814953 | 0 | 0 | 6.472457698 | 0.027943555 | 0.999932046 |
| HBS1L | XR_245499 | 0 | 0 | 0 | 0 | 6.655567121 | 0 | 0 | 0 | 3.82E-05 | 1 |
| HCCS | NM_005333 | 0 | 0 | 0 | 10.52213019 | 8.970792701 | 0 | 0 | 0 | 0.015381692 | 0.02072721 |
| HCP5 | NR_040662 | 0 | 10.43312838 | 0 | 0 | 0 | 11.76025402 | 0 | 0 | 1.34E-08 | 1 |
| HDAC5 | XM_017023996 | 0 | 0 | 10.03235972 | 0 | 5.098945058 | 10.61667921 | 9.626957495 | 0 | 0.032603039 | 0.999932046 |
| HDGF | XR_001737133 | 0 | 0 | 0 | 7.418246942 | 9.418521297 | 6.496419406 | 6.953230848 | 9.513376477 | 0.035188886 | 0.999932046 |
| HDGFL2 | NM_001001520 | 9.727893151 | 8.827309931 | 0 | 0 | 0 | 0 | 0 | 0 | 0.002078809 | 1 |
| HDGFL2 | NM_032631 | 0 | 0 | 0 | 0 | 6.702411139 | 9.78226982 | 8.40769976 | 11.02666593 | 3.69E-05 | 0.594061793 |
| HDHD2 | NM_001318765 | 0 | 8.943876548 | 0 | 0 | 0 | 0 | 0 | 0 | 7.69E-08 | 1 |
| HDHD5-AS1 | NR_024482 | 0 | 0 | 0 | 0 | 6.464964927 | 0 | 0 | 0 | 4.45E-05 | 1 |
| HEATR3 | XM_017023386 | 9.740478394 | 0 | 0 | 0 | 0 | 0 | 0 | 0 | 4.01E-08 | 1 |
| HEATR5B | XM_006712035 | 0 | 8.475517571 | 0 | 0 | 0 | 0 | 0 | 0 | 1.09E-07 | 1 |
| HEBP2 | NM_001326381 | 0 | 0 | 0 | 9.954035954 | 9.977709443 | 0 | 0 | 0 | 0.014723845 | 0.02072721 |
| HECTD3 | XM_024449803 | 10.17531978 | 0 | 0 | 0 | 0 | 0 | 0 | 0 | 1.60E-08 | 1 |
| HENMT1 | NM_001102592 | 0 | 8.886770211 | 0 | 10.30589526 | 8.592904388 | 0 | 0 | 0 | 0.999953634 | 0.02072721 |
| HENMT1 | XM_017000179 | 0 | 9.06649413 | 0 | 0 | 0 | 0 | 0 | 0 | 7.75E-08 | 1 |
| HEPACAM2 | NM_001288804 | 0 | 12.00115118 | 9.795083168 | 0 | 8.910972142 | 0 | 0 | 0 | 0.97128531 | 0.02072721 |
| HERPUD2 | XM_006715765 | 0 | 0 | 0 | 9.583571881 | 8.516317387 | 0 | 9.158661762 | 9.452532632 | 0.025939694 | 0.999932046 |
| HERPUD2 | XR_001744846 | 0 | 9.284296589 | 0 | 0 | 0 | 0 | 8.913583073 | 10.98039483 | 4.07E-08 | 0.02072721 |
| HES4 | NM_021170 | 0 | 0 | 10.3123572 | 0 | 8.301118378 | 0 | 8.920504771 | 10.66645992 | 0.019562833 | 0.999932046 |
| HEYL | NM_014571 | 0 | 8.809861237 | 0 | 0 | 0 | 0 | 0 | 0 | 8.24E-08 | 1 |
| HHEX | NM_002729 | 0 | 8.853093555 | 0 | 0 | 0 | 0 | 0 | 0 | 7.91E-08 | 1 |
| HHLA2 | XM_011512363 | 0 | 9.554298882 | 0 | 0 | 0 | 0 | 0 | 0 | 2.73E-08 | 1 |
| HHLA2 | XM_011512364 | 11.90350606 | 11.53172734 | 10.53644422 | 0 | 8.671527983 | 0 | 0 | 0 | 0.845329783 | 0.020730817 |
| HIBCH | NM_198047 | 11.97047595 | 9.014126367 | 9.39994696 | 10.73611967 | 9.300733642 | 0 | 0 | 0 | 0.999953634 | 0.002469438 |
| HIF1AN | NM_017902 | 0 | 9.989669449 | 0 | 0 | 0 | 0 | 0 | 0 | 2.68E-08 | 1 |
| HIGD1A | NM_001099668 | 0 | 0 | 11.15799022 | 0 | 7.562127909 | 0 | 0 | 0 | 0.013689899 | 0.02072721 |
| HIGD1A | NM_001099669 | 0 | 0 | 10.23997979 | 0 | 7.523516842 | 0 | 0 | 0 | 0.022781092 | 0.02072721 |
| HIKESHI | NM_016401 | 0 | 0 | 0 | 10.34484305 | 10.1439002 | 0 | 7.206416016 | 0 | 0.012020784 | 1 |
| HINFP | NM_198971 | 0 | 0 | 10.18219339 | 0 | 8.496518068 | 0 | 0 | 10.21222279 | 0.020077459 | 0.999932046 |
| HIP1R | XM_024449270 | 0 | 0 | 11.25299718 | 0 | 7.379608952 | 0 | 0 | 0 | 0.013184477 | 1.04E-08 |
| HIPK1 | XM_024453908 | 10.63230229 | 0 | 0 | 0 | 0 | 0 | 0 | 0 | 1.34E-08 | 1 |
| HIPK2 | XM_017012069 | 0 | 8.983325733 | 0 | 0 | 0 | 0 | 0 | 0 | 7.17E-08 | 1 |
| HIST1H1C | NM_005319 | 0 | 0 | 12.65642783 | 0 | 9.564338065 | 0 | 0 | 8.482525561 | 1.34E-08 | 0.999932046 |
| HIST1H2BD | NM_138720 | 0 | 0 | 10.64442567 | 11.37938713 | 0 | 0 | 9.475392161 | 10.92853424 | 0.007139217 | 0.999932046 |
| HIST1H2BH | NM_003524 | 10.75522266 | 0 | 0 | 0 | 0 | 0 | 0 | 0 | 1.34E-08 | 1 |
| HIST2H2BC | NR_036461 | 0 | 0 | 0 | 7.382391203 | 0 | 0 | 0 | 0 | 1.98E-05 | 1 |
| HIST3H2A | NM_033445 | 0 | 0 | 10.20037605 | 9.751092935 | 9.707564817 | 10.71452351 | 0 | 0 | 0.001348695 | 1 |

| | | | | | | | | | | | |
|---|---|---|---|---|---|---|---|---|---|---|---|
| HK1 | XM_024447969 | 0 | 9.073822446 | 0 | 0 | 0 | 0 | 4.906403331 | 7.272832804 | 7.73E-08 | 0.126943288 |
| HLA-DPB1 | NM_002121 | 11.71709664 | 11.28414061 | 0 | 0 | 0 | 12.87139849 | 9.046075251 | 11.8947696 | 0.000110735 | 0.000199228 |
| HLA-F | NM_018950 | 9.813767948 | 9.575104875 | 11.57325751 | 9.768560191 | 0 | 0 | 0 | 0 | 0.999953634 | 0.02072721 |
| HLA-F | XM_017010815 | 0 | 0 | 10.1198742 | 0 | 7.436136431 | 0 | 8.125685946 | 0 | 0.024636832 | 0.999932046 |
| HLA-F-AS1 | NR_026972 | 10.76757274 | 0 | 0 | 0 | 0 | 0 | 0 | 0 | 1.34E-08 | 1 |
| HLA-G | NM_002127 | 0 | 9.232850423 | 0 | 0 | 0 | 0 | 0 | 0 | 4.38E-08 | 1 |
| HLF | NM_002126 | 10.4218299 | 0 | 0 | 0 | 0 | 0 | 0 | 0 | 1.47E-08 | 1 |
| HMCES | NM_020187 | 0 | 0 | 0 | 0 | 0 | 0 | 9.238379139 | 10.35808505 | 1 | 0.02072721 |
| HMCES | XM_005247636 | 10.84536988 | 0 | 0 | 0 | 0 | 10.72427669 | 0 | 0 | 1.34E-08 | 1 |
| HMCN1 | NM_031935 | 0 | 8.360378074 | 0 | 0 | 0 | 0 | 0 | 0 | 1.21E-07 | 1 |
| HMG20B | NM_006339 | 0 | 8.919678654 | 11.14135488 | 10.70651942 | 7.555764066 | 0 | 0 | 0 | 0.827857426 | 0.001934792 |
| HMGA1 | NM_001319079 | 0 | 0 | 0 | 0 | 0 | 11.60570521 | 8.039294254 | 9.761562403 | 1 | 0.001394691 |
| HMGA1 | NM_001319081 | 0 | 0 | 0 | 0 | 0 | 0 | 9.36301824 | 11.60505841 | 1 | 0.02072721 |
| HMGA1 | NM_145903 | 0 | 0 | 0 | 10.11016257 | 11.20123904 | 10.39443992 | 9.815135958 | 10.92853424 | 0.008138588 | 0.999932046 |
| HMGB1 | XM_024449342 | 0 | 0 | 0 | 7.662040946 | 0 | 0 | 8.427271157 | 0 | 1.49E-05 | 1 |
| HMGB1 | XM_024449343 | 9.560548562 | 8.441930184 | 0 | 8.533268642 | 8.396793045 | 0 | 0 | 0 | 0.999953634 | 0.029173008 |
| HMGCLL1 | NM_001042406 | 10.83953218 | 0 | 0 | 0 | 0 | 0 | 0 | 0 | 1.34E-08 | 1 |
| HMGCS1 | XM_024446038 | 0 | 0 | 0 | 0 | 6.582319518 | 0 | 0 | 0 | 4.04E-05 | 1 |
| HMGCS2 | XM_011541313 | 10.95337303 | 10.716735 | 0 | 0 | 0 | 0 | 10.15299774 | 0 | 0.000279608 | 1 |
| HMGN1 | XM_024452071 | 0 | 9.942411034 | 0 | 0 | 0 | 0 | 0 | 10.24846798 | 5.01E-08 | 1 |
| HMGN3 | NM_004242 | 8.691211177 | 9.014126367 | 0 | 9.885245203 | 6.643614539 | 0 | 0 | 0 | 0.999953634 | 0.022212956 |
| HNF1B | NM_001304286 | 0 | 0 | 9.604946868 | 0 | 9.66385237 | 0 | 5.064738612 | 0 | 0.018491109 | 0.589596487 |
| HNMT | XR_002959286 | 9.623051988 | 9.258802813 | 0 | 0 | 0 | 0 | 8.125685946 | 9.587278352 | 0.001775117 | 0.02072721 |
| HNRNPA1 | XM_005268826 | 9.801808664 | 0 | 0 | 0 | 0 | 0 | 0 | 0 | 2.38E-08 | 1 |
| HNRNPA3 | NM_001330248 | 0 | 0 | 0 | 9.782383308 | 8.884555432 | 0 | 0 | 0 | 0.021703066 | 0.02072721 |
| HNRNPA3 | NM_001330249 | 0 | 0 | 0 | 9.74051016 | 6.758905041 | 0 | 8.181494666 | 0 | 0.032730902 | 0.999932046 |
| HNRNPC | NM_001077442 | 0 | 0 | 9.797739524 | 0 | 10.63229312 | 0 | 0 | 0 | 0.011724071 | 0.02072721 |
| HNRNPC | NM_031314 | 9.411121273 | 0 | 0 | 11.91109026 | 7.111090638 | 0 | 0 | 0 | 0.988370305 | 6.49E-09 |
| HNRNPC | XM_024449558 | 0 | 8.844550112 | 0 | 8.814612372 | 8.410964939 | 0 | 0 | 0 | 0.999953634 | 0.026003313 |
| HNRNPD | NM_001003810 | 0 | 0 | 11.09890747 | 0 | 7.093624618 | 0 | 7.684176881 | 11.26244988 | 0.014876792 | 0.999932046 |
| HNRNPF | NM_001098204 | 9.363098816 | 0 | 10.9468578 | 9.206807527 | 7.328262653 | 0 | 0 | 0 | 0.999953634 | 0.003174145 |
| HNRNPH1 | XM_005265901 | 0 | 0 | 0 | 0 | 0 | 0 | 8.525879161 | 10.11919446 | 1 | 0.02072721 |
| HNRNPH1 | XM_005265902 | 0 | 9.445515002 | 9.814887633 | 10.11563275 | 0 | 0 | 0 | 0 | 0.999953634 | 0.02072721 |
| HNRNPH1 | XM_006714862 | 0 | 0 | 0 | 11.83975597 | 9.105422809 | 0 | 9.889335013 | 10.46136272 | 0.007094609 | 0.999932046 |
| HNRNPH1 | XM_011534545 | 0 | 8.719304505 | 10.11349163 | 0 | 7.084811668 | 0 | 0 | 0 | 0.999953634 | 0.02072721 |
| HNRNPH1 | XM_017009414 | 0 | 0 | 0 | 10.12379934 | 10.08545732 | 0 | 10.22185657 | 0 | 0.013333863 | 1 |
| HNRNPH1 | XM_017009415 | 0 | 0 | 0 | 0 | 0 | 0 | 6.340064421 | 10.38793306 | 1 | 0.02072721 |
| HNRNPH1 | XM_017009417 | 0 | 0 | 0 | 9.910641153 | 9.164124572 | 0 | 0 | 0 | 0.019139493 | 0.02072721 |
| HNRNPH3 | NM_021644 | 0 | 8.140712363 | 0 | 11.81012011 | 9.365839676 | 0 | 0 | 0 | 0.728137993 | 0.02072721 |
| HNRNPK | NM_001318188 | 9.055885396 | 8.998808245 | 0 | 0 | 0 | 0 | 0 | 9.664911275 | 0.003084663 | 1 |
| HNRNPK | NM_031263 | 0 | 0 | 11.07392249 | 10.49510393 | 7.71273195 | 0 | 9.510221236 | 9.941402427 | 0.001100269 | 0.999932046 |
| HNRNPL | NM_001533 | 0 | 0 | 9.140133269 | 8.738654916 | 7.707000189 | 10.27122592 | 6.766095723 | 10.54983848 | 0.006762078 | 0.999932046 |
| HNRNPL | XR_243927 | 0 | 0 | 9.696043232 | 0 | 7.833391225 | 0 | 0 | 0 | 0.029138196 | 0.020781046 |

| | | | | | | | | | | | |
|---|---|---|---|---|---|---|---|---|---|---|---|
| HNRNPM | NM_005968 | 0 | 0 | 9.181401832 | 11.76948961 | 9.154680788 | 0 | 7.06988901 | 0 | 0.000756598 | 0.573976879 |
| HNRNPM | XM_005272481 | 0 | 8.453213086 | 0 | 0 | 0 | 0 | 0 | 0 | 1.11E-07 | 1 |
| HNRNPM | XM_017026832 | 0 | 9.014126367 | 0 | 0 | 0 | 0 | 8.937665109 | 7.530034637 | 6.71E-08 | 0.023727932 |
| HNRNPR | NM_001102397 | 0 | 0 | 0 | 8.214354668 | 7.997339613 | 0 | 0 | 0 | 0.056732441 | 0.038608076 |
| HNRNPUL1 | NM_144732 | 0 | 0 | 0 | 9.324854511 | 7.415197319 | 0 | 0 | 0 | 0.038020836 | 0.026484475 |
| HOMER2 | XM_005272448 | 10.40216198 | 0 | 0 | 0 | 0 | 0 | 0 | 0 | 1.47E-08 | 1 |
| HOMER2 | XM_005272449 | 0 | 0 | 0 | 0 | 0 | 0 | 9.378181702 | 10.56559534 | 1 | 0.02072721 |
| HOOK2 | NM_013312 | 0 | 8.323964754 | 0 | 0 | 0 | 0 | 0 | 0 | 1.27E-07 | 1 |
| HOPX | XM_017008729 | 0 | 0 | 12.42409884 | 0 | 9.103242531 | 0 | 8.574960864 | 0 | 1.34E-08 | 0.999932046 |
| HOTTIP | NR_037843 | 0 | 0 | 0 | 8.322379261 | 8.371652232 | 10.56424582 | 7.613324795 | 0 | 0.047165139 | 0.999932046 |
| HOXA11-AS | NR_002795 | 0 | 0 | 0 | 9.865900493 | 5.063498159 | 0 | 7.4411151 | 0 | 0.035275508 | 1 |
| HOXA13 | NM_000522 | 0 | 0 | 0 | 8.738654916 | 0 | 0 | 7.863684239 | 0 | 4.77E-06 | 1 |
| HOXA4 | NM_002141 | 10.77370837 | 0 | 0 | 0 | 0 | 0 | 7.892131936 | 0 | 1.34E-08 | 1 |
| HOXB-AS3 | NR_110331 | 0 | 9.10991584 | 0 | 0 | 0 | 0 | 0 | 0 | 5.42E-08 | 1 |
| HOXB3 | NM_002146 | 0 | 0 | 9.461549885 | 9.711903927 | 0 | 8.780795026 | 7.578739068 | 0 | 0.020589676 | 0.999932046 |
| HOXB4 | NM_024015 | 0 | 0 | 9.004140582 | 0 | 8.218925247 | 0 | 0 | 0 | 0.037593808 | 0.024607173 |
| HOXB6 | NM_018952 | 0 | 0 | 0 | 9.227165049 | 6.713888284 | 0 | 0 | 0 | 0.045742205 | 0.031617224 |
| HOXB6 | XM_011524727 | 0 | 7.722723415 | 0 | 10.31662209 | 9.504466758 | 0 | 0 | 0 | 0.866161272 | 0.02072721 |
| HOXB8 | XM_005257286 | 0 | 0 | 0 | 0 | 6.351978551 | 0 | 7.402084669 | 0 | 4.81E-05 | 1 |
| HOXB9 | NM_024017 | 0 | 0 | 8.047688802 | 0 | 8.792833751 | 0 | 7.974256665 | 0 | 0.042982435 | 0.999932046 |
| HOXD1 | NM_024501 | 9.734199496 | 0 | 0 | 0 | 0 | 0 | 0 | 0 | 5.08E-08 | 1 |
| HOXD11 | NM_021192 | 0 | 9.124104101 | 0 | 0 | 0 | 0 | 0 | 0 | 5.29E-08 | 1 |
| HOXD13 | NM_000523 | 0 | 8.935855595 | 0 | 0 | 0 | 0 | 0 | 0 | 7.69E-08 | 1 |
| HPS6 | NM_024747 | 10.39819601 | 9.364171095 | 10.10708069 | 0 | 8.691983703 | 0 | 0 | 0 | 0.999953634 | 0.02072721 |
| HR | XM_006716367 | 9.085782918 | 0 | 0 | 0 | 0 | 0 | 0 | 0 | 7.29E-08 | 1 |
| HRASLS | NM_020386 | 11.0437315 | 0 | 0 | 0 | 0 | 0 | 0 | 0 | 1.34E-08 | 1 |
| HRH1 | XM_011533652 | 0 | 0 | 11.21772675 | 0 | 7.568463804 | 0 | 9.544229262 | 0 | 0.013207232 | 0.999932046 |
| HS1BP3 | XM_017004700 | 11.02327716 | 0 | 0 | 0 | 0 | 0 | 0 | 0 | 1.34E-08 | 1 |
| HSD17B11 | XM_011532021 | 0 | 8.261155927 | 0 | 0 | 0 | 0 | 0 | 0 | 1.93E-07 | 1 |
| HSD17B14 | XM_005258970 | 0 | 0 | 10.15551827 | 0 | 8.262562196 | 11.52941294 | 10.22325675 | 0 | 0.021259426 | 0.999932046 |
| HSD17B7 | NM_016371 | 0 | 0 | 10.05696019 | 0 | 8.95395271 | 11.2013354 | 0 | 0 | 0.019226475 | 0.999932046 |
| HSD17B7P2 | NR_003086 | 0 | 8.571828374 | 0 | 0 | 0 | 0 | 0 | 0 | 9.97E-08 | 1 |
| HSD17B8 | NM_014234 | 9.70238844 | 9.239382419 | 9.792421912 | 9.726277944 | 8.611432316 | 0 | 0 | 0 | 0.999953634 | 0.003648567 |
| HSF1 | NM_005526 | 0 | 0 | 0 | 9.468580228 | 7.959271851 | 0 | 11.06916074 | 0 | 0.031334336 | 0.999932046 |
| HSF1 | XM_017013377 | 0 | 0 | 0 | 0 | 6.351978551 | 0 | 6.750642936 | 0 | 4.81E-05 | 1 |
| HSH2D | NM_001352265 | 0 | 8.818611963 | 0 | 0 | 0 | 0 | 0 | 0 | 8.12E-08 | 1 |
| HSPA13 | NM_006948 | 0 | 9.506358431 | 0 | 0 | 0 | 0 | 7.044767963 | 0 | 2.91E-08 | 1 |
| HSPA1A | NM_005345 | 0 | 0 | 0 | 9.561695954 | 8.2973088 | 10.38627208 | 0 | 7.921513153 | 0.027633976 | 0.999932046 |
| HSPBP1 | NM_012267 | 0 | 0 | 0 | 0 | 0 | 10.80461986 | 0 | 10.3610978 | 1 | 0.02072721 |
| HSPBP1 | XM_017026545 | 0 | 8.835955774 | 0 | 0 | 0 | 10.7736438 | 0 | 0 | 2.22E-07 | 1 |
| HSPH1 | NM_001286503 | 0 | 0 | 0 | 0 | 0 | 10.61929507 | 0 | 10.62570413 | 1 | 0.02072721 |
| HSPH1 | NM_001286504 | 0 | 0 | 9.201601708 | 13.28970407 | 0 | 12.06373075 | 10.29824628 | 12.36553508 | 1.34E-08 | 0.999932046 |
| HSPH1 | XM_017020363 | 0 | 0 | 0 | 10.40862134 | 8.944240862 | 0 | 9.922467304 | 10.8166683 | 0.016311267 | 0.999932046 |

| | | | | | | | | | | | |
|---|---|---|---|---|---|---|---|---|---|---|---|
| HTATIP2 | NM_001098522 | 10.22542327 | 0 | 10.21535514 | 10.31424523 | 8.486515572 | 0 | 0 | 0 | 0.999953634 | 0.002651192 |
| HTD2 | NM_001348712 | 10.48302382 | 0 | 0 | 0 | 0 | 0 | 0 | 0 | 1.47E-08 | 1 |
| HTR7P1 | NR_002774 | 0 | 0 | 0 | 0 | 6.736572239 | 0 | 0 | 0 | 9.53E-05 | 1 |
| HUS1 | XR_001744682 | 0 | 9.358181476 | 0 | 0 | 0 | 0 | 0 | 0 | 3.64E-08 | 1 |
| HUWE1 | XM_005261965 | 0 | 9.315543066 | 10.93725269 | 0 | 8.048094872 | 0 | 0 | 0 | 0.999953634 | 0.02072721 |
| HYAL2 | NM_033158 | 0 | 0 | 10.21237174 | 10.34134547 | 8.879468369 | 0 | 0 | 0 | 0.001369273 | 0.002477101 |
| HYAL2 | XM_005265525 | 0 | 0 | 9.093331342 | 9.83969732 | 0 | 0 | 0 | 0 | 0.021701799 | 0.02072721 |
| HYOU1 | NM_006389 | 0 | 8.33620493 | 0 | 0 | 0 | 0 | 8.525879161 | 0 | 1.25E-07 | 1 |
| IAH1 | XR_002959279 | 10.58423881 | 0 | 0 | 0 | 0 | 0 | 0 | 0 | 1.34E-08 | 1 |
| ICA1 | XM_011515351 | 9.395290737 | 0 | 0 | 0 | 0 | 0 | 0 | 0 | 4.12E-08 | 1 |
| ICAM1 | NM_000201 | 11.32849014 | 0 | 0 | 0 | 0 | 10.44249906 | 0 | 0 | 1.34E-08 | 1 |
| ICAM4 | NM_001544 | 0 | 9.38199218 | 0 | 0 | 0 | 0 | 0 | 0 | 3.43E-08 | 1 |
| ICMT | NM_012405 | 0 | 0 | 9.948784034 | 0 | 8.623653382 | 0 | 0 | 0 | 0.021776748 | 0.02072721 |
| ID2-AS1 | NR_110153 | 0 | 0 | 0 | 0 | 0 | 0 | 7.913105533 | 10.16989853 | 1 | 0.02072721 |
| IDH2 | NM_001290114 | 0 | 6.16971154 | 0 | 10.24105527 | 9.706847024 | 0 | 0 | 0 | 0.501061225 | 0.02072721 |
| IDH3A | NM_005530 | 8.873659666 | 0 | 11.06512964 | 9.305838651 | 7.10238406 | 0 | 0 | 0 | 0.920720885 | 0.00302572 |
| IDH3A | XM_024449912 | 10.06956967 | 8.1546024 | 0 | 0 | 0 | 0 | 0 | 0 | 0.00207295 | 1 |
| IDH3B | NM_006899 | 0 | 0 | 11.51754536 | 11.48459791 | 8.689079155 | 12.6041538 | 7.782430925 | 10.10488773 | 0.000428414 | 0.999932046 |
| IDH3B | XR_001754266 | 0 | 0 | 0 | 0 | 6.505153187 | 0 | 0 | 0 | 4.28E-05 | 1 |
| IDH3G | NM_174869 | 0 | 0 | 10.40760011 | 0 | 8.19864538 | 0 | 0 | 0 | 0.018959561 | 0.02072721 |
| IDI1 | NR_134301 | 0 | 0 | 11.0094988 | 0 | 7.849089784 | 0 | 0 | 8.482525561 | 0.0144596 | 0.999932046 |
| IDNK | NM_001001551 | 0 | 0 | 7.751443001 | 0 | 0 | 0 | 0 | 0 | 1.49E-05 | 1 |
| IDS | NM_006123 | 0 | 0 | 0 | 0 | 6.725274845 | 0 | 0 | 0 | 9.53E-05 | 1 |
| IER2 | NM_004907 | 11.64445852 | 10.14036773 | 11.645819 | 11.46437501 | 11.04388607 | 0 | 0 | 0 | 0.999953634 | 0.000448935 |
| IER5L | NM_203434 | 0 | 0 | 0 | 9.531320288 | 9.797962481 | 0 | 8.954623728 | 8.229118956 | 0.017971914 | 0.999932046 |
| IFFO1 | XM_017019173 | 0 | 0 | 0 | 0 | 6.607150896 | 0 | 0 | 0 | 0.00011086 | 1 |
| IFI16 | XM_017001149 | 10.01881342 | 8.508340767 | 0 | 0 | 0 | 0 | 0 | 0 | 0.001893423 | 1 |
| IFI27 | NM_001288954 | 0 | 0 | 0 | 10.10604627 | 7.746652379 | 0 | 0 | 0 | 0.023223652 | 0.02072721 |
| IFI27L1 | XM_017020961 | 10.5666777 | 0 | 0 | 0 | 0 | 0 | 0 | 0 | 1.34E-08 | 1 |
| IFI27L2 | NM_032036 | 0 | 0 | 0 | 9.969223115 | 8.435435627 | 10.19378098 | 0 | 9.458171152 | 0.022067013 | 0.999932046 |
| IFIT1 | NM_001548 | 0 | 0 | 11.17239044 | 0 | 6.865660185 | 0 | 0 | 0 | 0.014511831 | 1.11E-08 |
| IFIT3 | NM_001031683 | 9.11507342 | 0 | 10.28800294 | 9.66043618 | 0 | 0 | 0 | 0 | 0.999953634 | 0.02072721 |
| IFNAR2 | NM_207584 | 0 | 0 | 0 | 10.72363608 | 8.578850599 | 8.24341352 | 0 | 10.69539366 | 0.014968221 | 0.999932046 |
| IFRD1 | NM_001550 | 0 | 0 | 9.670183195 | 0 | 10.51663962 | 10.05640063 | 9.36301824 | 9.992810611 | 0.012811187 | 0.999932046 |
| IFT140 | XM_005255726 | 0 | 0 | 0 | 10.45034753 | 6.607150896 | 0 | 8.862382289 | 0 | 0.021780147 | 0.999932046 |
| IFT140 | XM_017023911 | 0 | 0 | 6.124455375 | 10.07409167 | 8.848563454 | 0 | 10.6281605 | 10.56559534 | 0.004974277 | 0.999932046 |
| IFT172 | XM_006711987 | 0 | 0 | 10.49366998 | 10.22596092 | 0 | 0 | 8.032920857 | 0 | 0.012235628 | 0.999932046 |
| IFT80 | NM_001190241 | 0 | 0 | 10.14303832 | 10.15069217 | 0 | 0 | 9.290068587 | 0 | 0.014267268 | 1 |
| IGBP1 | XR_938398 | 9.004636726 | 0 | 0 | 0 | 0 | 9.185629648 | 9.477740442 | 0 | 7.73E-08 | 0.02072721 |
| IGF1R | XM_017022138 | 9.244409574 | 0 | 0 | 0 | 0 | 0 | 0 | 0 | 5.26E-08 | 1 |
| IGF2 | NM_000612 | 0 | 0 | 10.58261512 | 9.242246866 | 10.1364565 | 11.49943429 | 0 | 0 | 0.001124753 | 0.999932046 |
| IGF2 | NM_001291861 | 0 | 0 | 9.403439234 | 0 | 11.43135016 | 11.04603421 | 0 | 0 | 0.008312731 | 0.999932046 |
| IGF2 | NM_001291862 | 0 | 0 | 10.12622866 | 0 | 11.31676714 | 10.46206063 | 7.032041479 | 0 | 0.007542278 | 0.999932046 |

| | | | | | | | | | | | |
|---|---|---|---|---|---|---|---|---|---|---|---|
| IGF2BP2 | NM_001291874 | 0 | 0 | 0 | 0 | 0 | 0 | 6.939666787 | 9.38309338 | 1 | 0.021929828 |
| IGF2BP2 | NM_006548 | 0 | 0 | 9.326426749 | 10.31899504 | 10.78155655 | 11.43011079 | 11.62452908 | 12.52950004 | 0.000892813 | 0.999932046 |
| IGFBP2 | NM_000597 | 0 | 0 | 9.245070286 | 12.57672939 | 9.255310595 | 10.81530624 | 10.70783868 | 0 | 0.00039722 | 1 |
| IGSF8 | NM_052868 | 0 | 0 | 0 | 0 | 0 | 11.71286999 | 8.137615898 | 0 | 1 | 5.45E-09 |
| IKBKB | NM_001190720 | 0 | 0 | 0 | 0 | 6.769943125 | 0 | 0 | 0 | 9.43E-05 | 1 |
| IKBKB | NM_001556 | 0 | 0 | 11.86904198 | 11.58856757 | 10.35653169 | 13.64910872 | 11.86864261 | 11.47482577 | 0.000206545 | 0.999932046 |
| IKBKB | NR_033819 | 0 | 0 | 0 | 9.442699766 | 7.523516842 | 11.00127656 | 0 | 0 | 0.034804797 | 0.999932046 |
| IKZF1 | NM_001291846 | 0 | 10.31131371 | 0 | 0 | 0 | 0 | 0 | 0 | 1.45E-08 | 1 |
| IKZF1 | NM_001291847 | 0 | 8.818611963 | 0 | 0 | 0 | 0 | 0 | 10.38497589 | 8.12E-08 | 1 |
| IKZF1 | XM_011515068 | 0 | 9.422738093 | 0 | 0 | 0 | 0 | 0 | 0 | 3.26E-08 | 1 |
| IKZF1 | XM_017011670 | 11.83327545 | 10.88656474 | 0 | 0 | 0 | 0 | 0 | 0 | 0.000128128 | 1 |
| IL15RA | NM_002189 | 0 | 0 | 7.474332608 | 0 | 0 | 0 | 0 | 0 | 1.96E-05 | 1 |
| IL15RA | NM_172200 | 0 | 0 | 0 | 8.784020642 | 0 | 0 | 0 | 0 | 4.55E-06 | 1 |
| IL15RA | XM_017016198 | 10.33321142 | 0 | 0 | 0 | 0 | 0 | 0 | 0 | 1.51E-08 | 1 |
| IL15RA | XM_024447983 | 10.80100026 | 0 | 0 | 0 | 0 | 0 | 0 | 9.226712959 | 1.34E-08 | 1 |
| IL16 | NM_172217 | 0 | 8.464408432 | 0 | 0 | 0 | 0 | 0 | 0 | 1.11E-07 | 1 |
| IL17RE | XM_006712975 | 0 | 0 | 11.58060299 | 8.905838641 | 10.81191312 | 11.58444347 | 9.87515368 | 11.62775139 | 0.000537047 | 0.999932046 |
| IL1R2 | XM_006712734 | 0 | 0 | 10.57179227 | 0 | 7.624268488 | 0 | 0 | 0 | 0.018852492 | 0.02072721 |
| IL1R2 | XM_011511804 | 0 | 0 | 10.82584651 | 0 | 7.021565945 | 0 | 0 | 0 | 0.017517226 | 0.02072721 |
| IL20RA | NM_001278722 | 0 | 0 | 0 | 10.53136871 | 7.763318133 | 0 | 9.460034453 | 0 | 0.018567968 | 0.999932046 |
| IL20RA | XM_017010954 | 0 | 9.265218616 | 9.806998441 | 0 | 6.993588191 | 0 | 0 | 0 | 0.999953634 | 0.021326758 |
| IL20RB | NM_144717 | 0 | 0 | 11.30480386 | 10.53034512 | 0 | 0 | 0 | 0 | 0.007778678 | 0.02072721 |
| IL20RB | XM_017006632 | 0 | 8.983325733 | 0 | 0 | 0 | 0 | 0 | 0 | 7.17E-08 | 1 |
| IL32 | NM_001012633 | 10.39819601 | 0 | 0 | 0 | 0 | 0 | 0 | 0 | 1.60E-08 | 1 |
| IL32 | NM_001012634 | 10.07951011 | 8.959785899 | 0 | 9.920050551 | 9.692415656 | 0 | 0 | 0 | 0.999953634 | 0.021060749 |
| IL34 | NM_152456 | 10.41006135 | 0 | 0 | 0 | 0 | 0 | 0 | 0 | 1.59E-08 | 1 |
| IL4I1 | NR_047577 | 0 | 9.451153482 | 0 | 0 | 0 | 0 | 0 | 0 | 3.15E-08 | 1 |
| IL4R | NM_001257997 | 9.261974802 | 0 | 0 | 0 | 0 | 0 | 0 | 0 | 5.08E-08 | 1 |
| IL6R | XM_017001201 | 9.765324233 | 0 | 0 | 0 | 0 | 0 | 0 | 0 | 2.50E-08 | 1 |
| IL7 | NM_001199886 | 9.07588576 | 0 | 0 | 0 | 0 | 0 | 0 | 0 | 7.35E-08 | 1 |
| IL7R | XM_005248299 | 10.24763479 | 0 | 0 | 0 | 0 | 0 | 0 | 0 | 1.58E-08 | 1 |
| ILDR1 | XM_011512739 | 0 | 0 | 0 | 8.006296941 | 0 | 0 | 0 | 0 | 1.04E-05 | 1 |
| ILF3 | NM_001137673 | 0 | 0 | 11.09134905 | 9.645387449 | 7.959271851 | 0 | 8.184373006 | 11.4481569 | 0.001438434 | 0.999932046 |
| ILF3 | XM_006722742 | 0 | 0 | 0 | 0 | 6.702411139 | 10.06793659 | 0 | 9.850000141 | 3.69E-05 | 0.999932046 |
| ILF3 | XM_017026763 | 0 | 0 | 0 | 0 | 0 | 0 | 10.0367688 | 9.627868238 | 1 | 0.02072721 |
| ILK | NM_001014794 | 0 | 0 | 0 | 9.726277944 | 8.250791506 | 10.40862331 | 9.152795499 | 10.92446666 | 0.025798264 | 0.999932046 |
| ILK | NM_004517 | 0 | 0 | 0 | 0 | 6.544252221 | 0 | 0 | 11.57553248 | 4.20E-05 | 1 |
| ILK | XM_011520065 | 0 | 0 | 0 | 9.066983555 | 8.614497302 | 0 | 0 | 7.347646499 | 0.031669825 | 0.999932046 |
| ILKAP | NM_030768 | 10.72387683 | 0 | 0 | 0 | 0 | 0 | 0 | 0 | 1.34E-08 | 1 |
| ILKAP | XR_923033 | 0 | 0 | 0 | 7.776970127 | 0 | 0 | 0 | 0 | 1.33E-05 | 1 |
| IMMP1L | XM_011519945 | 0 | 0 | 0 | 10.27322778 | 8.85635197 | 0 | 0 | 0 | 0.01770313 | 0.02072721 |
| IMMP2L | NM_001350960 | 0 | 0 | 11.34349641 | 0 | 7.178913994 | 0 | 11.14794731 | 0 | 0.012705018 | 1 |
| IMMT | XM_017003195 | 9.354937332 | 0 | 0 | 0 | 0 | 0 | 0 | 10.4315782 | 4.37E-08 | 1 |

| | | | | | | | | | | | |
|---|---|---|---|---|---|---|---|---|---|---|---|
| IMP4 | XM_024453222 | 0 | 8.486541819 | 0 | 0 | 0 | 0 | 0 | 0 | 1.09E-07 | 1 |
| IMPA2 | NM_014214 | 0 | 10.1816527 | 9.948784034 | 0 | 11.17267195 | 0 | 0 | 0 | 0.999953634 | 0.02072721 |
| IMPA2 | XM_011525659 | 0 | 8.801057108 | 0 | 0 | 0 | 0 | 0 | 0 | 8.24E-08 | 1 |
| IMPDH1 | XM_024446758 | 0 | 0 | 0 | 0 | 6.337212177 | 0 | 0 | 7.323137152 | 4.86E-05 | 1 |
| INAVA | NM_001142569 | 0 | 8.801057108 | 0 | 0 | 0 | 0 | 0 | 0 | 8.24E-08 | 1 |
| INAVA | NM_018265 | 0 | 0 | 9.924682753 | 0 | 8.555119041 | 0 | 0 | 0 | 0.022348432 | 0.02072721 |
| INAVA | XM_011509755 | 0 | 0 | 7.862181347 | 8.029665411 | 8.63275172 | 0 | 0 | 0 | 0.010645402 | 0.016659858 |
| INCA1 | XM_024450746 | 0 | 9.206422614 | 0 | 0 | 0 | 0 | 0 | 0 | 4.63E-08 | 1 |
| INF2 | NM_032714 | 0 | 0 | 0 | 9.729849182 | 6.42362501 | 0 | 0 | 0 | 0.034274416 | 0.024568095 |
| ING1 | NM_198219 | 0 | 0 | 0 | 8.396154474 | 0 | 0 | 0 | 0 | 1.37E-05 | 1 |
| ING3 | NM_198267 | 10.33321142 | 0 | 0 | 0 | 0 | 0 | 0 | 0 | 1.51E-08 | 1 |
| ING4 | XM_017019393 | 0 | 0 | 10.08550279 | 0 | 8.342372954 | 0 | 0 | 0 | 0.021701799 | 0.02072721 |
| INO80B-WBP1 | NR_037849 | 0 | 0 | 10.7995818 | 0 | 8.17807637 | 0 | 0 | 0 | 0.015557448 | 0.02072721 |
| INO80C | NM_001098817 | 0 | 0 | 0 | 0 | 6.643614539 | 0 | 0 | 0 | 3.86E-05 | 1 |
| INO80E | NM_173618 | 0 | 0 | 10.44366461 | 8.79429013 | 7.806841281 | 0 | 0 | 0 | 0.003001449 | 0.004380867 |
| INPP4B | XM_017008794 | 9.849062238 | 0 | 0 | 0 | 0 | 0 | 9.420299048 | 0 | 2.23E-08 | 1 |
| INPP4B | XM_017008796 | 0 | 0 | 11.68436859 | 0 | 9.561164523 | 0 | 0 | 0 | 0.00730753 | 0.02072721 |
| INPP5A | XR_945735 | 0 | 0 | 0 | 7.921391152 | 0 | 0 | 0 | 0 | 1.14E-05 | 1 |
| INPP5D | NM_001017915 | 0 | 0 | 0 | 0 | 0 | 10.50229635 | 7.856483795 | 0 | 1 | 0.02072721 |
| INPP5E | NM_019892 | 0 | 0 | 0 | 0 | 6.582319518 | 0 | 0 | 0 | 4.04E-05 | 1 |
| INPP5K | XM_024450802 | 0 | 0 | 9.821429165 | 10.02842178 | 8.19864538 | 0 | 7.331132628 | 0 | 0.002298335 | 0.999932046 |
| INSC | XR_930868 | 10.38222222 | 0 | 0 | 0 | 0 | 0 | 0 | 0 | 1.47E-08 | 1 |
| INSIG2 | NM_001321331 | 10.21644202 | 0 | 0 | 0 | 0 | 0 | 0 | 0 | 1.59E-08 | 1 |
| INSIG2 | NM_016133 | 0 | 8.747073305 | 10.34904047 | 0 | 8.28965933 | 0 | 0 | 0 | 0.999953634 | 0.02072721 |
| INTS10 | XR_002956637 | 10.35792493 | 0 | 0 | 0 | 0 | 0 | 9.474216586 | 0 | 1.48E-08 | 1 |
| INTS11 | XM_017001558 | 9.789749415 | 0 | 0 | 8.474877744 | 8.89719516 | 0 | 0 | 0 | 0.999953634 | 0.024631639 |
| INTS14 | XM_006720711 | 0 | 0 | 10.7163875 | 0 | 8.532611753 | 0 | 8.899639246 | 0 | 0.015398614 | 0.999932046 |
| INTS3 | NM_023015 | 8.028121183 | 0 | 0 | 0 | 0 | 0 | 8.04563962 | 9.617827325 | 0.062457626 | 0.02072721 |
| INTS5 | NM_030628 | 0 | 9.327854598 | 0 | 0 | 0 | 0 | 0 | 0 | 3.82E-08 | 1 |
| INTS7 | NM_001199812 | 0 | 0 | 9.501204039 | 0 | 9.748597361 | 0 | 0 | 10.35959221 | 0.018494796 | 0.999932046 |
| INTS9 | NM_001145159 | 0 | 0 | 0 | 9.211923901 | 7.618173501 | 11.39695608 | 9.164504268 | 10.0368086 | 0.038869166 | 0.999932046 |
| INTS9 | XM_006716356 | 0 | 0 | 0 | 10.05428874 | 7.483843939 | 10.02517937 | 0 | 0 | 0.024756429 | 0.999932046 |
| IPO4 | NR_051979 | 0 | 0 | 0 | 0 | 0 | 11.15160386 | 7.516144242 | 8.343210923 | 1 | 0.002622729 |
| IPO5 | XM_024449346 | 0 | 0 | 8.709114096 | 0 | 8.210847477 | 0 | 0 | 11.03706392 | 0.043295862 | 0.999932046 |
| IPO8 | XM_017018692 | 7.224905374 | 7.684937679 | 0 | 0 | 0 | 0 | 7.351763189 | 0 | 0.02193917 | 1 |
| IPO9 | NM_018085 | 0 | 0 | 0 | 8.643405671 | 6.713888284 | 0 | 0 | 0 | 0.064035647 | 0.045375566 |
| IPPK | XM_017015041 | 0 | 0 | 6.190629309 | 8.234606714 | 6.834454817 | 0 | 0 | 0 | 0.027379751 | 0.02072721 |
| IQCH | NM_001284348 | 0 | 0 | 0 | 0 | 6.758905041 | 0 | 6.796513442 | 0 | 9.53E-05 | 1 |
| IQCH | NM_001322475 | 0 | 0 | 9.754639499 | 0 | 8.89215255 | 0 | 8.295036906 | 0 | 0.022125423 | 0.999932046 |
| IQCH | XM_017022494 | 0 | 8.39589491 | 0 | 0 | 0 | 0 | 0 | 0 | 1.17E-07 | 1 |
| IQCH | XM_017022504 | 0 | 0 | 0 | 8.967640006 | 9.064544407 | 0 | 0 | 0 | 0.027986508 | 0.020903961 |
| IQGAP2 | XM_024454337 | 8.651779762 | 0 | 0 | 0 | 0 | 0 | 0 | 0 | 1.00E-07 | 1 |
| IREB2 | NM_004136 | 0 | 8.903318262 | 0 | 0 | 0 | 0 | 0 | 0 | 7.75E-08 | 1 |

| | | | | | | | | | | | |
|---|---|---|---|---|---|---|---|---|---|---|---|
| IRF3 | NR_045568 | 9.134275178 | 0 | 0 | 0 | 0 | 0 | 0 | 0 | 6.66E-08 | 1 |
| IRF3 | XR_002958312 | 0 | 0 | 0 | 0 | 6.451316153 | 0 | 0 | 0 | 4.51E-05 | 1 |
| IRF6 | NM_006147 | 9.560548562 | 8.792198922 | 0 | 8.483365391 | 7.636381755 | 0 | 0 | 0 | 0.999953634 | 0.038833809 |
| IRF7 | NM_001572 | 9.442270566 | 0 | 0 | 0 | 0 | 0 | 0 | 0 | 3.84E-08 | 1 |
| IRF7 | XM_017017674 | 0 | 0 | 7.275241756 | 0 | 0 | 0 | 0 | 0 | 5.92E-05 | 1 |
| IRF8 | NM_002163 | 0 | 0 | 0 | 9.498198764 | 8.447517061 | 8.670773519 | 0 | 0 | 0.027454051 | 0.999932046 |
| ISG20 | NM_001303237 | 0 | 0 | 0 | 8.167720583 | 0 | 0 | 0 | 0 | 8.82E-06 | 1 |
| ISG20 | XM_005254899 | 0 | 9.428466121 | 0 | 0 | 0 | 0 | 0 | 0 | 3.25E-08 | 1 |
| ISG20 | XM_017022147 | 0 | 8.998808245 | 0 | 0 | 0 | 0 | 0 | 0 | 6.89E-08 | 1 |
| ISOC1 | NM_016048 | 0 | 0 | 8.223937317 | 9.975253495 | 9.596470608 | 0 | 0 | 0 | 0.002532437 | 0.003932411 |
| ITCH | XM_024452005 | 0 | 8.529815232 | 0 | 0 | 0 | 0 | 0 | 0 | 1.04E-07 | 1 |
| ITFG2 | NR_147202 | 10.23434896 | 0 | 0 | 0 | 0 | 0 | 0 | 0 | 1.59E-08 | 1 |
| ITFG2 | XM_005253709 | 10.4296226 | 0 | 0 | 0 | 0 | 0 | 0 | 0 | 1.47E-08 | 1 |
| ITFG2 | XM_024449057 | 0 | 9.517150383 | 0 | 0 | 0 | 0 | 0 | 0 | 2.90E-08 | 1 |
| ITGA1 | NM_181501 | 0 | 0 | 8.833416502 | 0 | 7.642400461 | 0 | 0 | 0 | 0.048833995 | 0.03091407 |
| ITGA3 | XM_005257308 | 7.224905374 | 0 | 8.585434713 | 9.489798191 | 0 | 0 | 0 | 0 | 0.93568295 | 0.02096331 |
| ITGA6 | XM_017004008 | 10.01881342 | 4.676562808 | 0 | 0 | 0 | 0 | 3.654887771 | 6.754940855 | 0.004743045 | 0.218648763 |
| ITGAE | XM_024450740 | 0 | 9.271606013 | 0 | 0 | 0 | 10.28556965 | 0 | 0 | 4.15E-08 | 1 |
| ITGB1 | NM_002211 | 0 | 0 | 9.544549719 | 10.7502556 | 5.167328428 | 0 | 0 | 0 | 0.002960326 | 0.004603096 |
| ITGB1BP1 | NR_134957 | 9.134275178 | 0 | 0 | 0 | 0 | 0 | 0 | 0 | 6.66E-08 | 1 |
| ITGB1BP1 | XM_005246183 | 0 | 0 | 9.266323099 | 0 | 8.816981984 | 0 | 7.759452999 | 0 | 0.027592147 | 0.999932046 |
| ITGB3BP | NM_014288 | 0 | 0 | 0 | 10.85371572 | 9.481148681 | 0 | 0 | 0 | 0.011643831 | 0.02072721 |
| ITLN1 | NM_017625 | 18.66037013 | 21.77203281 | 20.63707613 | 17.23167928 | 15.6410727 | 0 | 10.54326279 | 0 | 0.937178729 | 0.02072721 |
| ITM2A | NM_004867 | 0 | 0 | 8.047688802 | 0 | 0 | 0 | 0 | 0 | 1.25E-05 | 1 |
| ITPKA | NM_002220 | 0 | 0 | 11.16005622 | 10.10329554 | 0 | 0 | 0 | 0 | 0.009480747 | 0.02072721 |
| ITPR3 | NM_002224 | 0 | 0 | 0 | 8.364340994 | 7.536502354 | 0 | 0 | 0 | 0.061298604 | 0.042673774 |
| ITPR3 | XM_011514577 | 0 | 0 | 0 | 9.433968736 | 8.011359982 | 0 | 0 | 0 | 0.03153386 | 0.022619671 |
| ITPRIP | XM_005270257 | 9.004636726 | 0 | 0 | 0 | 0 | 0 | 0 | 0 | 7.73E-08 | 1 |
| IVD | NR_148923 | 0 | 9.635780908 | 0 | 0 | 0 | 0 | 0 | 0 | 2.48E-08 | 1 |
| JADE3 | NM_014735 | 0 | 0 | 0 | 9.882039044 | 10.26976672 | 0 | 9.637497974 | 10.70016004 | 0.013477571 | 0.999932046 |
| JAG1 | NM_000214 | 0 | 8.998808245 | 0 | 0 | 0 | 10.22273888 | 0 | 9.435483632 | 6.89E-08 | 0.02072721 |
| JAK2 | NM_001322198 | 0 | 0 | 8.05658827 | 0 | 9.916940169 | 0 | 0 | 0 | 0.02379327 | 0.02072721 |
| JKAMP | NM_001284201 | 0 | 8.419096158 | 0 | 0 | 0 | 0 | 0 | 0 | 1.14E-07 | 1 |
| JKAMP | NM_016475 | 0 | 0 | 9.722846221 | 0 | 9.635466715 | 0 | 0 | 10.25819701 | 0.017971914 | 1 |
| JMJD4 | XM_011544262 | 0 | 0 | 0 | 7.124706719 | 8.2973088 | 0 | 0 | 0 | 0.069628491 | 0.048679581 |
| JMJD6 | XR_934429 | 0 | 0 | 10.49939263 | 0 | 8.656737116 | 0 | 0 | 0 | 0.016811352 | 0.020808337 |
| JMJD7 | NM_001114632 | 0 | 0 | 8.769900534 | 10.36335493 | 0 | 0 | 9.489424851 | 5.300551716 | 0.018494796 | 0.999932046 |
| JMY | XM_011543155 | 0 | 0 | 0 | 0 | 6.600983004 | 0 | 0 | 0 | 4.00E-05 | 1 |
| JRK | XM_006716678 | 9.588662682 | 0 | 0 | 0 | 0 | 0 | 0 | 0 | 3.15E-08 | 1 |
| JRKL | NM_003772 | 0 | 7.411784592 | 0 | 9.127801312 | 7.654363064 | 0 | 0 | 0 | 0.999953634 | 0.028011146 |
| JUND | NM_005354 | 11.24262448 | 9.059128398 | 12.29796653 | 0 | 8.278108536 | 0 | 0 | 0 | 0.999953634 | 3.87E-09 |
| KANSL2 | NM_017822 | 9.944614124 | 9.830055185 | 10.30023146 | 0 | 5.412429324 | 0 | 0 | 0 | 1 | 0.02072721 |
| KANSL3 | XR_002959316 | 10.60503382 | 0 | 0 | 0 | 0 | 0 | 0 | 0 | 1.34E-08 | 1 |

| | | | | | | | | | | | |
|---|---|---|---|---|---|---|---|---|---|---|---|
| KARS | NM_001130089 | 0 | 0 | 0 | 10.81976746 | 8.902220207 | 0 | 7.877978204 | 0 | 0.013425772 | 0.999932046 |
| KARS | NM_005548 | 0 | 9.538495079 | 0 | 11.24608133 | 9.641489231 | 0 | 0 | 0 | 0.999953634 | 0.02072721 |
| KARS | XM_017023217 | 0 | 0 | 10.85801896 | 8.834652321 | 8.222947235 | 0 | 0 | 0 | 0.00203168 | 0.003110083 |
| KAT5 | XM_006718421 | 0 | 9.239382419 | 0 | 0 | 0 | 0 | 0 | 0 | 4.33E-08 | 1 |
| KAT7 | NM_007067 | 0 | 9.615837767 | 0 | 0 | 0 | 0 | 0 | 0 | 2.54E-08 | 1 |
| KAT8 | NM_182958 | 9.982190738 | 0 | 0 | 0 | 0 | 0 | 0 | 0 | 1.98E-08 | 1 |
| KATNA1 | XM_017010207 | 0 | 0 | 0 | 10.59050989 | 8.773768299 | 0 | 0 | 0 | 0.015451077 | 0.02072721 |
| KBTBD2 | XM_011515257 | 10.40216198 | 0 | 0 | 0 | 0 | 0 | 0 | 0 | 1.47E-08 | 1 |
| KBTBD3 | NM_198439 | 0 | 0 | 10.37772997 | 10.32609055 | 8.486515572 | 0 | 0 | 0 | 0.001423802 | 0.002548758 |
| KCNA6 | XM_005253686 | 10.16602012 | 0 | 0 | 0 | 0 | 0 | 0 | 0 | 1.60E-08 | 1 |
| KCNIP4 | NM_025221 | 0 | 0 | 10.17812149 | 10.01677478 | 0 | 0 | 0 | 0 | 0.01478954 | 0.02072721 |
| KCNK1 | XM_011544184 | 0 | 8.323964754 | 0 | 0 | 0 | 0 | 0 | 0 | 1.27E-07 | 1 |
| KCNK15 | NM_022358 | 0 | 8.809861237 | 0 | 0 | 0 | 0 | 0 | 0 | 8.24E-08 | 1 |
| KCNK5 | NM_003740 | 10.89405913 | 0 | 0 | 8.378061333 | 8.228959271 | 0 | 0 | 0 | 0.986247317 | 0.032843069 |
| KCNK5 | XM_006715235 | 0 | 0 | 10.05029247 | 0 | 6.095400279 | 0 | 0 | 0 | 0.02981136 | 0.020903961 |
| KCNMA1 | NM_001271522 | 9.095612642 | 0 | 0 | 0 | 0 | 0 | 0 | 0 | 7.18E-08 | 1 |
| KCNN3 | NM_001204087 | 0 | 8.983325733 | 0 | 0 | 0 | 0 | 0 | 11.65009295 | 7.17E-08 | 1 |
| KCNN3 | NM_002249 | 0 | 0 | 11.2364491 | 9.993194855 | 7.328262653 | 0 | 0 | 0 | 0.001321339 | 0.002417476 |
| KCNN4 | XR_935823 | 11.10584386 | 0 | 8.337827042 | 0 | 8.882014143 | 0 | 0 | 0 | 0.996706997 | 0.024841872 |
| KCNS3 | NM_002252 | 0 | 0 | 11.32933755 | 9.693732641 | 0 | 0 | 0 | 0 | 0.009655334 | 0.02072721 |
| KCTD10 | NR_133898 | 0 | 0 | 0 | 8.414023513 | 7.523516842 | 0 | 0 | 0 | 0.060224177 | 0.041757194 |
| KCTD13 | NR_110933 | 0 | 0 | 0 | 0 | 6.643614539 | 0 | 0 | 0 | 3.86E-05 | 1 |
| KCTD14 | NM_023930 | 0 | 0 | 0 | 9.191348626 | 9.232550068 | 0 | 0 | 10.79677985 | 0.024409102 | 1 |
| KCTD20 | XR_001743256 | 0 | 0 | 0 | 10.55168937 | 8.061632835 | 0 | 0 | 0 | 0.017693164 | 0.02072721 |
| KCTD5 | NM_018992 | 0 | 0 | 10.79227438 | 9.242246866 | 7.111090638 | 0 | 7.856483795 | 0 | 0.002378921 | 0.999932046 |
| KDELC2 | XM_011542623 | 0 | 0 | 0 | 0 | 6.351978551 | 0 | 0 | 0 | 4.81E-05 | 1 |
| KDM1A | XM_005245786 | 8.526574932 | 8.535134272 | 0 | 10.72184386 | 8.628209723 | 0 | 0 | 0 | 0.999953634 | 0.02072721 |
| KDM1A | XM_006710474 | 0 | 0 | 0 | 10.12921816 | 6.276587234 | 0 | 0 | 0 | 0.027111383 | 0.020869972 |
| KDM1A | XM_017000717 | 0 | 0 | 0 | 0 | 6.544252221 | 0 | 0 | 0 | 4.20E-05 | 1 |
| KDM2A | NR_027473 | 0 | 0 | 10.30956799 | 0 | 8.869240114 | 12.18198637 | 10.28422791 | 11.04364202 | 0.017612901 | 0.999932046 |
| KDM4B | NM_015015 | 0 | 0 | 11.42264139 | 0 | 9.561164523 | 0 | 0 | 0 | 0.00851906 | 0.02072721 |
| KDM4B | XM_011527814 | 0 | 9.059128398 | 0 | 0 | 0 | 0 | 0 | 0 | 7.80E-08 | 1 |
| KDM4B | XM_011527819 | 0 | 0 | 9.607977036 | 0 | 7.235751614 | 0 | 0 | 0 | 0.034150253 | 0.022650952 |
| KDM4B | XM_017026505 | 0 | 0 | 10.1017164 | 0 | 7.483843939 | 0 | 7.392159812 | 0 | 0.024727596 | 0.999932046 |
| KDM5C | NR_148672 | 0 | 0 | 0 | 9.242246866 | 6.582319518 | 11.1401133 | 0 | 0 | 0.046183495 | 0.999932046 |
| KDM5C | XM_011530824 | 0 | 0 | 0 | 8.01802849 | 0 | 0 | 0 | 0 | 1.03E-05 | 1 |
| KDM5C | XR_001755737 | 0 | 0 | 7.8773294 | 0 | 8.790125492 | 0 | 6.477826297 | 0 | 0.045177439 | 0.999932046 |
| KDM5D | NM_004653 | 0 | 7.914514525 | 0 | 0 | 0 | 10.62364434 | 8.84247586 | 9.145169427 | 0.063339158 | 0.002297095 |
| KDM5D | XR_001756012 | 0 | 0 | 0 | 0 | 6.582319518 | 9.531339422 | 0 | 0 | 4.04E-05 | 1 |
| KDM6A | NM_001291418 | 0 | 9.073822446 | 0 | 0 | 0 | 0 | 0 | 0 | 7.73E-08 | 1 |
| KEAP1 | NM_203500 | 0 | 0 | 9.471566073 | 0 | 7.067022599 | 0 | 0 | 0 | 0.038143142 | 0.024755256 |
| KHDRBS1 | NR_073498 | 0 | 8.486541819 | 0 | 10.00208255 | 8.479808505 | 0 | 0 | 0 | 0.999953634 | 0.02072721 |
| KHDRBS3 | XM_005250757 | 0 | 0 | 0 | 8.108597066 | 0 | 0 | 0 | 0 | 9.41E-06 | 1 |

| | | | | | | | | | | | |
|---|---|---|---|---|---|---|---|---|---|---|---|
| KIAA0232 | XM_024454291 | 10.33321142 | 0 | 0 | 0 | 0 | 0 | 0 | 0 | 1.51E-08 | 1 |
| KIAA0391 | NM_001256678 | 0 | 0 | 10.46052638 | 11.59347645 | 11.55355668 | 0 | 0 | 11.4481569 | 0.000228483 | 1 |
| KIAA0895L | NM_001040715 | 10.13775594 | 7.501468832 | 0 | 0 | 0 | 0 | 0 | 0 | 0.002395984 | 1 |
| KIAA1109 | XM_011532327 | 0 | 0 | 0 | 10.94257615 | 8.509747759 | 0 | 8.724829293 | 0 | 0.013428611 | 0.999932046 |
| KIAA1143 | NM_020696 | 0 | 0 | 9.85752351 | 10.03421021 | 10.35469993 | 0 | 7.341484785 | 0 | 0.001048615 | 0.81105098 |
| KIAA1191 | XM_024446134 | 0 | 7.388464503 | 0 | 9.941770056 | 8.871803984 | 0 | 0 | 0 | 0.893878189 | 0.02072721 |
| KIAA1324 | NM_020775 | 0 | 0 | 0 | 9.664173964 | 8.242890685 | 0 | 0 | 0 | 0.02662564 | 0.02072721 |
| KIAA1958 | NM_001287036 | 10.70152235 | 0 | 0 | 0 | 0 | 0 | 0 | 0 | 1.34E-08 | 1 |
| KIAA1958 | XM_011518311 | 0 | 0 | 0 | 0 | 0 | 9.176185619 | 10.79469171 | 9.965357103 | 1 | 0.001516752 |
| KIF12 | XM_024447408 | 10.41006135 | 0 | 0 | 0 | 0 | 0 | 0 | 0 | 1.59E-08 | 1 |
| KIF13A | NM_001243423 | 0 | 0 | 9.279686404 | 9.527221444 | 0 | 0 | 0 | 0 | 0.023250768 | 0.02072721 |
| KIF23 | XM_005254799 | 0 | 0 | 0 | 0 | 6.59478863 | 0 | 0 | 0 | 4.02E-05 | 1 |
| KIF25 | XM_011535802 | 0 | 9.321711965 | 0 | 0 | 0 | 0 | 9.378181702 | 0 | 3.84E-08 | 1 |
| KIF2C | XM_011540541 | 0 | 0 | 0 | 9.901169983 | 9.399989931 | 0 | 9.864425444 | 8.01629051 | 0.018053568 | 0.999932046 |
| KIF9 | NM_001134878 | 0 | 9.059128398 | 0 | 0 | 0 | 0 | 0 | 0 | 7.80E-08 | 1 |
| KIF9 | XM_011534003 | 0 | 9.08836834 | 0 | 0 | 0 | 0 | 0 | 0 | 5.61E-08 | 1 |
| KIT | XM_005265742 | 0 | 0 | 10.55775681 | 0 | 8.066117405 | 0 | 8.566160223 | 0 | 0.017979898 | 0.999932046 |
| KLC1 | NM_182923 | 8.555352323 | 9.864267562 | 0 | 0 | 0 | 0 | 0 | 0 | 0.002042243 | 1 |
| KLC4 | NM_201523 | 0 | 0 | 8.899209478 | 0 | 7.44304914 | 0 | 0 | 0 | 0.049719673 | 0.031380494 |
| KLF10 | NM_005655 | 0 | 0 | 9.335595812 | 0 | 8.131765004 | 0 | 0 | 0 | 0.032811359 | 0.022064246 |
| KLF10 | NR_103760 | 0 | 0 | 9.870198585 | 0 | 6.544252221 | 8.805507098 | 6.31926467 | 0 | 0.031959438 | 0.999932046 |
| KLF3-AS1 | NR_026804 | 0 | 0 | 7.867248391 | 0 | 0 | 9.858054838 | 0 | 0 | 1.32E-05 | 1 |
| KLF6 | NM_001300 | 11.52496752 | 0 | 0 | 0 | 0 | 0 | 0 | 0 | 1.34E-08 | 1 |
| KLF6 | NR_027653 | 0 | 0 | 9.972489291 | 0 | 7.959271851 | 0 | 5.937789944 | 0 | 0.024605612 | 0.999932046 |
| KLHDC1 | XR_002957530 | 0 | 8.441930184 | 0 | 0 | 0 | 0 | 0 | 0 | 1.12E-07 | 1 |
| KLHDC2 | XM_006720094 | 0 | 9.158976035 | 0 | 0 | 0 | 0 | 8.184373006 | 0 | 5.01E-08 | 1 |
| KLHDC2 | XM_011536610 | 0 | 0 | 9.233344514 | 0 | 8.61755579 | 0 | 8.831500914 | 0 | 0.029868098 | 0.999932046 |
| KLHDC3 | XM_024446320 | 0 | 0 | 10.01420126 | 0 | 8.048094872 | 0 | 0 | 0 | 0.02376504 | 0.02072721 |
| KLHDC9 | NR_033385 | 0 | 8.991087758 | 0 | 0 | 0 | 0 | 0 | 0 | 7.02E-08 | 1 |
| KLHL2 | XM_024453881 | 0 | 8.998808245 | 0 | 0 | 0 | 0 | 0 | 0 | 6.89E-08 | 1 |
| KLHL21 | NM_014851 | 9.854861577 | 0 | 0 | 0 | 0 | 0 | 0 | 0 | 2.23E-08 | 1 |
| KLHL24 | XM_017006661 | 0 | 0 | 0 | 0 | 6.582319518 | 0 | 0 | 0 | 4.04E-05 | 1 |
| KLHL36 | XM_005256149 | 0 | 0 | 0 | 0 | 6.505153187 | 0 | 0 | 0 | 4.28E-05 | 1 |
| KLHL42 | XM_017019698 | 0 | 10.28944795 | 0 | 0 | 0 | 0 | 0 | 0 | 1.47E-08 | 1 |
| KLHL6 | XR_924209 | 0 | 9.145128245 | 0 | 0 | 0 | 0 | 0 | 0 | 5.08E-08 | 1 |
| KLHL7 | NM_001172428 | 10.67554289 | 0 | 0 | 0 | 0 | 0 | 0 | 5.731458906 | 1.34E-08 | 0.436498398 |
| KLK1 | NM_002257 | 0 | 12.28543031 | 12.01429702 | 10.18751933 | 9.384812028 | 0 | 0 | 0 | 0.999953634 | 0.001042882 |
| KLK11 | NM_006853 | 0 | 9.495485143 | 0 | 0 | 0 | 0 | 0 | 10.26303703 | 2.95E-08 | 1 |
| KLK11 | XM_005258439 | 0 | 9.645650041 | 0 | 0 | 0 | 0 | 0 | 0 | 2.48E-08 | 1 |
| KLK12 | NM_145894 | 0 | 0 | 0 | 0 | 0 | 10.65457785 | 0 | 10.06479657 | 1 | 0.02072721 |
| KLK3 | NM_001030047 | 8.526574932 | 7.626336084 | 0 | 0 | 0 | 0 | 0 | 0 | 0.010348295 | 1 |
| KLK3 | NM_001648 | 11.05132772 | 10.6249305 | 11.25879265 | 0 | 6.736572239 | 0 | 0 | 0 | 1 | 1.04E-08 |
| KLK6 | NM_001012964 | 0 | 0 | 0 | 10.48245744 | 8.830225047 | 0 | 0 | 0 | 0.016122834 | 0.020903961 |

| | | | | | | | | | | | |
|---|---|---|---|---|---|---|---|---|---|---|---|
| KLK6 | NM_002774 | 0 | 0 | 0 | 0 | 0 | 10.97224008 | 7.411941715 | 0 | 1 | 0.02072721 |
| KMT2A | XM_006718839 | 11.74077977 | 0 | 11.89159815 | 0 | 8.088333507 | 0 | 0 | 0 | 0.999953634 | 5.45E-09 |
| KMT2A | XM_011542829 | 0 | 9.352166886 | 0 | 0 | 0 | 0 | 0 | 0 | 3.68E-08 | 1 |
| KMT2C | XM_005250025 | 0 | 9.352166886 | 0 | 0 | 0 | 0 | 0 | 0 | 3.68E-08 | 1 |
| KMT2C | XM_017012480 | 0 | 10.05407497 | 0 | 0 | 0 | 0 | 8.605347092 | 0 | 1.60E-08 | 1 |
| KMT2C | XM_017012484 | 0 | 8.756212053 | 0 | 0 | 0 | 0 | 0 | 0 | 8.55E-08 | 1 |
| KMT5A | XM_006719394 | 0 | 9.186279128 | 0 | 9.217022195 | 6.702411139 | 0 | 0 | 0 | 0.999953634 | 0.031862586 |
| KMT5C | XM_024451742 | 9.906032551 | 0 | 0 | 0 | 0 | 0 | 0 | 0 | 2.06E-08 | 1 |
| KPNA7 | XM_011516215 | 10.07454845 | 0 | 0 | 0 | 0 | 0 | 0 | 0 | 1.79E-08 | 1 |
| KPNB1 | NM_002265 | 0 | 9.522516235 | 0 | 0 | 0 | 0 | 6.703265946 | 0 | 2.88E-08 | 1 |
| KRR1 | NM_007043 | 0 | 9.479019994 | 0 | 0 | 0 | 0 | 0 | 9.657578368 | 3.04E-08 | 1 |
| KRT10 | XM_005257343 | 0 | 9.014126367 | 0 | 9.671620603 | 8.274237629 | 0 | 0 | 0 | 0.999953634 | 0.02072721 |
| KRT15 | XM_011524784 | 0 | 9.19973932 | 0 | 0 | 0 | 0 | 0 | 0 | 4.67E-08 | 1 |
| KRT15 | XM_017024614 | 10.47551503 | 0 | 0 | 0 | 0 | 0 | 0 | 0 | 1.44E-08 | 1 |
| KRT6A | NM_005554 | 0 | 0 | 0 | 11.4777114 | 10.90183975 | 10.72589583 | 0 | 10.76073112 | 0.005595512 | 0.999932046 |
| KRTCAP3 | NM_001168364 | 0 | 9.416987232 | 0 | 0 | 0 | 0 | 7.805048609 | 0 | 3.28E-08 | 1 |
| KTN1 | NM_004986 | 0 | 0 | 0 | 0 | 6.351978551 | 0 | 0 | 0 | 4.81E-05 | 1 |
| KYAT1 | NR_109829 | 0 | 8.690990708 | 0 | 0 | 0 | 0 | 0 | 0 | 9.02E-08 | 1 |
| KYNU | NM_001032998 | 10.21644202 | 0 | 0 | 0 | 0 | 0 | 0 | 0 | 1.59E-08 | 1 |
| L2HGDH | XM_017021657 | 0 | 0 | 10.57334337 | 0 | 10.28715809 | 0 | 0 | 10.76073112 | 0.01046073 | 0.999932046 |
| LAMA5 | XR_936532 | 9.727893151 | 0 | 0 | 10.45898029 | 7.859461386 | 0 | 0 | 0 | 0.999953634 | 0.02072721 |
| LAMP2 | NM_002294 | 0 | 6.984094144 | 10.6115886 | 0 | 8.161407372 | 0 | 0 | 0 | 0.744967879 | 0.020810615 |
| LAMP3 | XM_011512688 | 0 | 0 | 0 | 0 | 0 | 9.052602861 | 6.31926467 | 0 | 1 | 0.028374914 |
| LAMTOR3 | NM_021970 | 11.14305235 | 9.731559707 | 0 | 0 | 0 | 10.40862331 | 0 | 0 | 0.000417442 | 1 |
| LAMTOR3 | NR_024170 | 0 | 0 | 9.213587255 | 11.03352211 | 9.116275051 | 0 | 8.000624564 | 10.34900889 | 0.001242785 | 0.999932046 |
| LAMTOR4 | NR_134526 | 0 | 11.06355464 | 8.856532107 | 0 | 7.386797204 | 0 | 0 | 0 | 0.93494198 | 0.03243257 |
| LAMTOR4 | NR_134528 | 10.25203635 | 10.25117508 | 6.072753641 | 0 | 8.914706779 | 0 | 0 | 0 | 0.78971698 | 0.039163676 |
| LANCL1 | XM_005246243 | 10.89405913 | 0 | 0 | 0 | 0 | 0 | 8.15532787 | 0 | 1.34E-08 | 1 |
| LANCL2 | NM_018697 | 9.759152772 | 0 | 0 | 0 | 0 | 0 | 9.2630946 | 0 | 2.49E-08 | 1 |
| LARP1 | XM_005268404 | 0 | 0 | 10.92576293 | 9.180949892 | 0 | 9.968861605 | 0 | 0 | 0.013339002 | 0.999932046 |
| LARP4 | XM_024448827 | 0 | 8.959785899 | 0 | 0 | 0 | 0 | 0 | 0 | 7.69E-08 | 1 |
| LARP4B | XM_005252435 | 0 | 10.27043843 | 0 | 0 | 0 | 0 | 0 | 0 | 1.47E-08 | 1 |
| LARP4B | XM_017015988 | 0 | 0 | 10.72129291 | 11.163324 | 7.654363064 | 0 | 10.16616161 | 10.94268111 | 0.000947645 | 0.999932046 |
| LARP7 | XM_024454084 | 0 | 9.726918817 | 0 | 0 | 0 | 0 | 0 | 0 | 2.23E-08 | 1 |
| LAT2 | XM_011516558 | 0 | 8.571828374 | 0 | 0 | 0 | 0 | 0 | 0 | 9.97E-08 | 1 |
| LATS1 | NM_001270519 | 0 | 0 | 0 | 10.14535356 | 9.739501354 | 10.81378445 | 0 | 12.56814394 | 0.014882197 | 0.999932046 |
| LBR | XR_001737168 | 0 | 0 | 0 | 9.828082059 | 9.237528645 | 0 | 7.166214885 | 0 | 0.019385121 | 0.999932046 |
| LCA5L | XM_024452056 | 0 | 0 | 0 | 10.48668527 | 9.774861522 | 0 | 11.35819127 | 0 | 0.012779132 | 0.999932046 |
| LCMT1 | NM_016309 | 0 | 10.35106274 | 11.38736401 | 11.75904141 | 0 | 0 | 0 | 0 | 0.999953634 | 0.019059177 |
| LCMT1 | XM_005255354 | 0 | 9.620849457 | 0 | 0 | 0 | 0 | 0 | 0 | 2.53E-08 | 1 |
| LDB1 | NM_003893 | 9.669860113 | 0 | 0 | 0 | 0 | 0 | 0 | 0 | 2.76E-08 | 1 |
| LDHB | NM_001174097 | 0 | 0 | 0 | 10.71465263 | 9.101058952 | 0 | 11.11249595 | 12.0139312 | 0.013616444 | 0.999932046 |
| LDHD | XM_024450179 | 9.244409574 | 0 | 0 | 0 | 0 | 0 | 0 | 0 | 5.26E-08 | 1 |

| | | | | | | | | | | | |
|---|---|---|---|---|---|---|---|---|---|---|---|
| LDLR | NM_000527 | 0 | 0 | 10.73035894 | 0 | 9.641489231 | 11.68067166 | 0 | 0 | 0.012020784 | 0.999932046 |
| LDLR | NM_001195799 | 0 | 0 | 0 | 10.38390075 | 8.410964939 | 9.108287582 | 0 | 0 | 0.018247043 | 0.999932046 |
| LDLR | XM_011528010 | 10.15195724 | 10.23818845 | 0 | 0 | 0 | 0 | 0 | 0 | 0.000664527 | 1 |
| LEFTY1 | NM_020997 | 9.567628621 | 11.56330991 | 9.15263826 | 8.214354668 | 9.804016025 | 0 | 0 | 0 | 0.933304018 | 0.004942065 |
| LEMD1 | NM_001199052 | 0 | 0 | 0 | 0 | 0 | 10.65882219 | 0 | 11.05484944 | 1 | 0.02072721 |
| LEMD3 | NM_014319 | 0 | 8.895067962 | 0 | 0 | 0 | 0 | 0 | 0 | 7.75E-08 | 1 |
| LEO1 | NM_138792 | 9.911607777 | 0 | 0 | 0 | 0 | 0 | 0 | 0 | 2.05E-08 | 1 |
| LEPROT | NM_017526 | 0 | 9.543782272 | 9.847302656 | 0 | 6.451316153 | 0 | 0 | 0 | 0.999953634 | 0.022027568 |
| LETMD1 | XR_944516 | 0 | 0 | 0 | 7.616467016 | 0 | 0 | 0 | 0 | 1.56E-05 | 1 |
| LFNG | NM_002304 | 0 | 0 | 0 | 9.402987182 | 8.88962462 | 0 | 0 | 0 | 0.025255996 | 0.02072721 |
| LGALS3 | NM_001357678 | 10.54531825 | 8.967675264 | 10.75790236 | 10.38616567 | 7.111090638 | 0 | 0 | 0 | 0.999953634 | 0.002651192 |
| LGALS4 | XM_011526973 | 11.85803037 | 10.17144169 | 10.88574914 | 9.59927585 | 8.016003323 | 0 | 0 | 0 | 0.999953634 | 0.002751024 |
| LGALS9 | NM_009587 | 0 | 0 | 9.88401374 | 9.23220986 | 0 | 0 | 0 | 0 | 0.020743471 | 0.02072721 |
| LGMN | NM_005606 | 0 | 0 | 10.93242604 | 0 | 9.827979349 | 0 | 0 | 0 | 0.010348295 | 0.02072721 |
| LGR4 | NM_018490 | 0 | 0 | 8.976271492 | 0 | 5.847041254 | 0 | 0 | 0 | 0.061554247 | 0.03962103 |
| LGR5 | NR_110596 | 0 | 9.315543066 | 0 | 0 | 0 | 0 | 6.400722758 | 0 | 3.90E-08 | 1 |
| LGR6 | XM_017001998 | 0 | 0 | 0 | 0 | 6.42362501 | 0 | 0 | 0 | 4.54E-05 | 1 |
| LHFPL2 | XM_024454325 | 10.98147737 | 0 | 0 | 0 | 0 | 0 | 0 | 0 | 1.34E-08 | 1 |
| LHFPL6 | NM_005780 | 0 | 0 | 0 | 8.474877744 | 0 | 0 | 0 | 0 | 6.35E-06 | 1 |
| LIAS | XM_017007666 | 0 | 9.296876505 | 0 | 0 | 0 | 0 | 0 | 0 | 4.01E-08 | 1 |
| LILRA5 | NM_021250 | 0 | 8.919678654 | 0 | 0 | 0 | 0 | 0 | 0 | 7.73E-08 | 1 |
| LIMA1 | NM_016357 | 0 | 0 | 0 | 8.82800322 | 7.630337835 | 0 | 0 | 0 | 0.047984681 | 0.032586253 |
| LIMCH1 | NM_001330790 | 10.17994723 | 0 | 0 | 0 | 0 | 0 | 0 | 0 | 1.60E-08 | 1 |
| LIME1 | NM_017806 | 0 | 0 | 0 | 0 | 0 | 0 | 9.955698329 | 10.24521033 | 1 | 0.02072721 |
| LIMK2 | NM_016733 | 10.29105753 | 0 | 0 | 0 | 0 | 0 | 0 | 0 | 1.54E-08 | 1 |
| LIMS2 | NM_001161403 | 9.363098816 | 0 | 0 | 0 | 0 | 0 | 0 | 0 | 4.33E-08 | 1 |
| LIN7A | XM_011538928 | 0 | 0 | 0 | 0 | 0 | 0 | 7.671923744 | 9.793135989 | 1 | 0.02072721 |
| LIN7C | NM_018362 | 0 | 0 | 10.36793218 | 0 | 8.190452933 | 0 | 0 | 10.51509654 | 0.019383327 | 0.999932046 |
| LINC-PINT | NR_109851 | 0 | 9.239382419 | 0 | 0 | 0 | 10.0717615 | 0 | 0 | 4.33E-08 | 1 |
| LINC00114 | NR_027066 | 9.035603861 | 0 | 0 | 0 | 0 | 0 | 0 | 0 | 7.68E-08 | 1 |
| LINC00167 | NR_024233 | 0 | 0 | 7.131029629 | 0 | 0 | 0 | 0 | 0 | 2.78E-05 | 1 |
| LINC00265 | NR_026999 | 0 | 8.561439232 | 0 | 0 | 0 | 0 | 8.367741983 | 0 | 1.00E-07 | 1 |
| LINC00476 | NR_023389 | 0 | 0 | 10.82454457 | 11.21137601 | 0 | 0 | 0 | 10.00441834 | 0.007283323 | 0.999932046 |
| LINC00482 | NR_038080 | 0 | 8.774317692 | 0 | 0 | 0 | 0 | 0 | 0 | 8.46E-08 | 1 |
| LINC00511 | NR_033876 | 0 | 0 | 0 | 0 | 6.713888284 | 0 | 0 | 0 | 3.66E-05 | 1 |
| LINC00513 | NR_109780 | 0 | 0 | 0 | 8.466339867 | 0 | 0 | 8.984655768 | 0 | 6.39E-06 | 1 |
| LINC00543 | NR_135254 | 0 | 0 | 10.05251847 | 0 | 7.195383085 | 0 | 0 | 9.784185253 | 0.026422607 | 0.999932046 |
| LINC00630 | NR_146589 | 0 | 0 | 0 | 0 | 6.736572239 | 0 | 0 | 0 | 9.53E-05 | 1 |
| LINC00639 | NR_039982 | 0 | 8.835955774 | 0 | 0 | 0 | 0 | 0 | 0 | 2.22E-07 | 1 |
| LINC00667 | NR_015389 | 0 | 0 | 0 | 9.375891515 | 9.246447016 | 0 | 0 | 11.39329301 | 0.022762091 | 1 |
| LINC00668 | NR_034100 | 0 | 0 | 0 | 8.414023513 | 0 | 0 | 0 | 0 | 3.17E-05 | 1 |
| LINC00673 | NR_152516 | 8.918652917 | 0 | 0 | 0 | 0 | 0 | 0 | 0 | 8.06E-08 | 1 |
| LINC00842 | NR_033957 | 0 | 9.830055185 | 0 | 0 | 0 | 0 | 7.885072427 | 0 | 2.01E-08 | 1 |

| | | | | | | | | | | | |
|---|---|---|---|---|---|---|---|---|---|---|---|
| LINC00847 | NR_045679 | 0 | 0 | 10.54279195 | 0 | 7.695467896 | 0 | 0 | 0 | 0.01896696 | 0.02072721 |
| LINC00857 | NR_038464 | 0 | 0 | 0 | 7.382391203 | 8.720712025 | 0 | 0 | 0 | 0.053417031 | 0.035576279 |
| LINC00869 | NR_111952 | 0 | 0 | 0 | 10.71104353 | 6.79176912 | 0 | 9.497547994 | 0 | 0.018573259 | 0.999932046 |
| LINC01000 | NR_024368 | 11.41346607 | 0 | 0 | 0 | 0 | 0 | 5.207402423 | 0 | 1.34E-08 | 0.538759989 |
| LINC01123 | NR_046111 | 0 | 0 | 0 | 7.921391152 | 8.61755579 | 0 | 0 | 0 | 0.048527622 | 0.032537612 |
| LINC01133 | NR_038849 | 0 | 0 | 12.33638376 | 0 | 7.328262653 | 0 | 0 | 0 | 3.15E-08 | 1 |
| LINC01279 | NR_109779 | 0 | 8.248259235 | 0 | 0 | 0 | 0 | 0 | 0 | 2.02E-07 | 1 |
| LINC01315 | NR_120596 | 0 | 0 | 0 | 0 | 0 | 10.73798179 | 10.42107728 | 0 | 1 | 0.02072721 |
| LINC01389 | NR_126355 | 10.21644202 | 0 | 0 | 0 | 0 | 0 | 0 | 0 | 1.59E-08 | 1 |
| LINC01431 | NR_109884 | 0 | 0 | 8.879283175 | 0 | 5.908266127 | 0 | 0 | 0 | 0.06476835 | 0.04202927 |
| LINC01481 | NR_120459 | 0 | 0 | 8.742587407 | 9.849579366 | 7.689666995 | 0 | 7.506976231 | 10.80122326 | 0.004474148 | 0.999932046 |
| LINC01504 | NR_110953 | 10.97083541 | 0 | 0 | 0 | 0 | 0 | 0 | 0 | 1.34E-08 | 1 |
| LINC01547 | NR_027128 | 10.00844432 | 0 | 0 | 0 | 0 | 0 | 0 | 0 | 1.93E-08 | 1 |
| LINC01550 | NR_152747 | 9.616239393 | 0 | 0 | 0 | 0 | 0 | 0 | 0 | 2.91E-08 | 1 |
| LINC01578 | NR_037602 | 0 | 0 | 0 | 9.19652002 | 7.219739493 | 0 | 6.703265946 | 0 | 0.042740325 | 0.999932046 |
| LINC01590 | NR_026784 | 9.330172147 | 0 | 0 | 0 | 0 | 0 | 0 | 0 | 4.56E-08 | 1 |
| LINC01781 | NR_125942 | 0 | 9.073822446 | 0 | 0 | 0 | 0 | 0 | 0 | 7.73E-08 | 1 |
| LINC01814 | NR_110257 | 0 | 0 | 0 | 0 | 6.464964927 | 0 | 0 | 0 | 4.45E-05 | 1 |
| LINC01871 | XR_241262 | 9.922694008 | 0 | 0 | 0 | 0 | 0 | 0 | 0 | 2.04E-08 | 1 |
| LINC01881 | NR_130699 | 0 | 0 | 0 | 10.91695796 | 9.545191248 | 0 | 0 | 0 | 0.011096107 | 0.02072721 |
| LINC01881 | NR_130701 | 0 | 0 | 0 | 10.60224751 | 9.601885833 | 0 | 9.173223903 | 0 | 0.012752199 | 0.999932046 |
| LINC02001 | NR_135479 | 0 | 9.158976035 | 0 | 10.02261004 | 7.267252374 | 0 | 0 | 0 | 0.999953634 | 0.02072721 |
| LINC02018 | NR_151705 | 0 | 0 | 0 | 0 | 6.451316153 | 0 | 0 | 0 | 4.51E-05 | 1 |
| LINC02023 | XR_241610 | 0 | 8.622680652 | 0 | 0 | 0 | 0 | 0 | 0 | 9.58E-08 | 1 |
| LINC02108 | NR_109904 | 10.90248333 | 7.606261038 | 0 | 0 | 0 | 0 | 0 | 0 | 0.001086601 | 1 |
| LINC02332 | NR_146543 | 0 | 0 | 0 | 0 | 6.42362501 | 0 | 0 | 0 | 4.54E-05 | 1 |
| LINC02535 | NR_134633 | 0 | 8.774317692 | 0 | 0 | 0 | 0 | 0 | 0 | 8.46E-08 | 1 |
| LINC02562 | NR_149051 | 0 | 0 | 0 | 10.10741968 | 6.291983974 | 0 | 0 | 10.11563106 | 0.027421515 | 0.999932046 |
| LINC02588 | NR_147061 | 0 | 8.919678654 | 0 | 0 | 0 | 0 | 0 | 0 | 7.73E-08 | 1 |
| LIPH | XM_017005852 | 0 | 8.943876548 | 10.359866 | 0 | 7.968883375 | 0 | 0 | 0 | 0.999953634 | 0.02072721 |
| LIPT2 | NM_001144869 | 0 | 10.54017255 | 0 | 0 | 0 | 0 | 0 | 0 | 1.34E-08 | 1 |
| LIPT2 | NM_001329942 | 0 | 8.719304505 | 0 | 0 | 0 | 0 | 0 | 0 | 8.96E-08 | 1 |
| LITAF | NM_001136473 | 0 | 0 | 9.825339894 | 9.362150426 | 0 | 0 | 0 | 0 | 0.020466596 | 0.02072721 |
| LITAF | NR_024320 | 10.37012473 | 0 | 0 | 0 | 0 | 0 | 0 | 0 | 3.14E-08 | 1 |
| LLGL2 | XM_017024627 | 0 | 0 | 0 | 8.597229662 | 0 | 0 | 8.78674668 | 0 | 5.56E-06 | 1 |
| LLGL2 | XM_017024630 | 0 | 8.652353298 | 0 | 0 | 0 | 0 | 0 | 0 | 9.28E-08 | 1 |
| LLGL2 | XR_002958000 | 0 | 8.519117955 | 0 | 0 | 0 | 0 | 8.530410777 | 0 | 1.04E-07 | 1 |
| LMBRD2 | NM_001007527 | 0 | 0 | 11.23154567 | 0 | 8.548724137 | 10.15464429 | 0 | 0 | 0.0115288 | 0.999932046 |
| LMCD1 | NM_014583 | 0 | 8.959785899 | 0 | 0 | 0 | 0 | 0 | 0 | 7.69E-08 | 1 |
| LMF1 | XM_006720929 | 8.373391248 | 0 | 0 | 0 | 0 | 0 | 0 | 0 | 1.86E-07 | 1 |
| LMNA | NM_170707 | 0 | 0 | 0 | 7.831174272 | 8.278108536 | 0 | 0 | 0 | 0.058023297 | 0.039557029 |
| LMNA | NM_170708 | 10.14250526 | 9.817013442 | 0 | 10.73256394 | 7.939854543 | 0 | 0 | 0 | 0.999953634 | 0.020785849 |
| LMNB2 | NM_032737 | 10.48302382 | 0 | 0 | 10.09362645 | 9.415888488 | 0 | 0 | 0 | 0.999953634 | 0.02072721 |

| | | | | | | | | | | | |
|---|---|---|---|---|---|---|---|---|---|---|---|
| LMO4 | NM_006769 | 0 | 8.78328601 | 0 | 0 | 0 | 0 | 0 | 9.802031535 | 8.40E-08 | 1 |
| LMOD1 | NM_012134 | 0 | 9.252358352 | 0 | 0 | 0 | 0 | 0 | 0 | 4.28E-08 | 1 |
| LMTK2 | XM_011515981 | 8.827217917 | 0 | 8.409046753 | 9.357540824 | 0 | 0 | 0 | 0 | 0.999953634 | 0.022611547 |
| LMTK3 | XM_011526412 | 0 | 0 | 0 | 0 | 6.702411139 | 0 | 0 | 8.133689895 | 3.69E-05 | 1 |
| LNX1 | XM_024454262 | 0 | 9.428466121 | 0 | 0 | 0 | 0 | 0 | 0 | 3.25E-08 | 1 |
| LOC100129034 | NR_027406 | 0 | 9.533188437 | 0 | 0 | 0 | 0 | 0 | 0 | 2.83E-08 | 1 |
| LOC100130298 | NR_034003 | 0 | 9.258802813 | 0 | 0 | 0 | 0 | 0 | 0 | 4.23E-08 | 1 |
| LOC100132154 | XR_001746678 | 0 | 0 | 0 | 0 | 0 | 0 | 8.579341135 | 11.10243336 | 1 | 0.02072721 |
| LOC100133331 | NR_028327 | 7.328569034 | 0 | 8.529001408 | 0 | 9.178693122 | 0 | 0 | 0 | 0.979593971 | 0.021342962 |
| LOC100190986 | NR_024456 | 0 | 0 | 9.193555665 | 0 | 7.111090638 | 0 | 0 | 0 | 0.045002636 | 0.028645392 |
| LOC100233156 | NR_037872 | 0 | 0 | 0 | 0 | 6.713888284 | 0 | 0 | 0 | 3.66E-05 | 1 |
| LOC100288798 | NR_125380 | 0 | 0 | 0 | 0 | 6.451316153 | 0 | 0 | 0 | 4.51E-05 | 1 |
| LOC100288846 | NR_038935 | 0 | 8.975521721 | 0 | 0 | 0 | 0 | 0 | 0 | 7.27E-08 | 1 |
| LOC100289230 | NR_036530 | 0 | 8.248259235 | 0 | 0 | 0 | 0 | 0 | 0 | 2.02E-07 | 1 |
| LOC100289361 | NR_034032 | 0 | 0 | 0 | 0 | 0 | 0 | 7.805048609 | 9.213438433 | 1 | 0.02092873 |
| LOC100289561 | NM_001347880 | 0 | 0 | 9.937988255 | 0 | 8.097125035 | 0 | 0 | 0 | 0.02442701 | 0.02072721 |
| LOC100419583 | NR_125376 | 8.791378415 | 9.219697032 | 0 | 10.32491038 | 8.629725311 | 0 | 0 | 0 | 0.999953634 | 0.02072721 |
| LOC100505549 | NM_001242804 | 10.04948094 | 9.059128398 | 0 | 0 | 0 | 0 | 0 | 0 | 0.001434603 | 1 |
| LOC100506123 | NR_130704 | 0 | 9.533188437 | 0 | 0 | 0 | 0 | 0 | 0 | 2.83E-08 | 1 |
| LOC100506472 | NR_040535 | 10.26516103 | 0 | 0 | 0 | 0 | 0 | 0 | 0 | 4.25E-08 | 1 |
| LOC100506476 | NR_109995 | 8.678186754 | 8.681427931 | 0 | 0 | 0 | 0 | 0 | 0 | 0.004929085 | 1 |
| LOC100506544 | NR_148200 | 10.6656787 | 0 | 0 | 0 | 0 | 0 | 9.831753598 | 0 | 1.34E-08 | 1 |
| LOC100506922 | XR_001739698 | 0 | 0 | 9.592762148 | 0 | 5.200341806 | 0 | 8.40769976 | 0 | 0.04382872 | 1 |
| LOC100506922 | XR_001739700 | 10.05200738 | 0 | 0 | 0 | 0 | 0 | 0 | 0 | 5.08E-08 | 1 |
| LOC100507507 | NM_001322263 | 0 | 0 | 0 | 0 | 6.337212177 | 0 | 0 | 0 | 4.86E-05 | 1 |
| LOC100996717 | XM_024452061 | 10.82483375 | 0 | 0 | 0 | 0 | 0 | 0 | 9.213438433 | 1.34E-08 | 1 |
| LOC100996756 | XR_159169 | 0 | 0 | 10.2213035 | 0 | 7.910229599 | 0 | 0 | 0 | 0.02179728 | 0.02072721 |
| LOC100996763 | XM_011510256 | 0 | 10.46413307 | 0 | 0 | 0 | 0 | 0 | 0 | 1.34E-08 | 1 |
| LOC100996763 | XM_024452035 | 0 | 10.23492345 | 0 | 0 | 0 | 0 | 0 | 0 | 1.48E-08 | 1 |
| LOC101926967 | XR_243706 | 9.395290737 | 0 | 0 | 0 | 0 | 0 | 0 | 0 | 4.12E-08 | 1 |
| LOC101927069 | NR_110647 | 0 | 8.975521721 | 0 | 0 | 0 | 0 | 0 | 0 | 7.27E-08 | 1 |
| LOC101927100 | NR_109875 | 10.81594227 | 0 | 0 | 0 | 0 | 0 | 0 | 0 | 1.34E-08 | 1 |
| LOC101927511 | XR_001753362 | 8.85641873 | 0 | 0 | 0 | 0 | 0 | 0 | 0 | 8.52E-08 | 1 |
| LOC101928069 | NR_120586 | 0 | 0 | 0 | 9.514854575 | 7.577915777 | 0 | 6.14109935 | 0 | 0.033023468 | 0.999932046 |
| LOC101928076 | XR_241971 | 0 | 9.029283553 | 0 | 0 | 0 | 0 | 0 | 0 | 6.49E-08 | 1 |
| LOC101928156 | XR_923872 | 0 | 0 | 10.23606798 | 0 | 8.006701649 | 0 | 0 | 0 | 0.021328674 | 0.02072721 |
| LOC101928215 | XR_950968 | 0 | 10.5728664 | 0 | 0 | 0 | 0 | 0 | 0 | 1.34E-08 | 1 |
| LOC101928323 | NR_125400 | 0 | 0 | 0 | 0 | 6.351978551 | 0 | 0 | 0 | 4.81E-05 | 1 |
| LOC101928347 | XR_245041 | 0 | 0 | 7.576866482 | 0 | 0 | 0 | 0 | 0 | 3.23E-05 | 1 |
| LOC101928881 | NR_136325 | 0 | 0 | 0 | 9.224636013 | 8.583550423 | 0 | 8.827823966 | 0 | 0.029881811 | 0.999932046 |
| LOC101929243 | NR_134626 | 0 | 0 | 8.603765699 | 0 | 6.979393187 | 0 | 0 | 0 | 0.063902367 | 0.041354729 |
| LOC101929709 | NR_125822 | 9.226627846 | 0 | 0 | 0 | 0 | 0 | 0 | 0 | 5.40E-08 | 1 |
| LOC101929796 | XM_017003012 | 0 | 10.09786644 | 0 | 0 | 0 | 0 | 8.59240271 | 0 | 1.59E-08 | 1 |

| | | | | | | | | | | | |
|---|---|---|---|---|---|---|---|---|---|---|---|
| LOC101930071 | NR_126041 | 0 | 0 | 0 | 0 | 6.30721813 | 8.120778949 | 0 | 0 | 4.99E-05 | 1 |
| LOC102723728 | XM_006720996 | 9.411121273 | 0 | 0 | 0 | 0 | 0 | 0 | 0 | 4.01E-08 | 1 |
| LOC102723878 | XR_950194 | 0 | 0 | 0 | 8.677092933 | 0 | 0 | 4.596451906 | 0 | 7.46E-06 | 1 |
| LOC102724156 | NR_135137 | 0 | 0 | 7.601400457 | 0 | 0 | 0 | 0 | 0 | 1.81E-05 | 1 |
| LOC102724397 | XR_430055 | 0 | 0 | 0 | 10.19787099 | 8.07504494 | 0 | 0 | 0 | 0.02107584 | 0.02072721 |
| LOC102724715 | XR_002958137 | 12.21327678 | 0 | 12.63263359 | 0 | 8.218925247 | 0 | 0 | 0 | 0.999953634 | 2.81E-09 |
| LOC102724720 | XM_017010129 | 0 | 0 | 0 | 0 | 6.464964927 | 0 | 0 | 0 | 4.45E-05 | 1 |
| LOC102724760 | NR_135695 | 0 | 10.28313923 | 0 | 0 | 0 | 0 | 0 | 0 | 1.47E-08 | 1 |
| LOC102724985 | XM_024452518 | 0 | 8.612652582 | 0 | 0 | 0 | 0 | 7.341484785 | 0 | 9.67E-08 | 1 |
| LOC103091866 | NR_111957 | 0 | 9.630820909 | 0 | 0 | 0 | 0 | 0 | 0 | 4.30E-08 | 1 |
| LOC105369315 | XR_950132 | 8.497211815 | 0 | 0 | 0 | 0 | 0 | 0 | 0 | 1.15E-07 | 1 |
| LOC105369347 | XR_002957252 | 0 | 9.698755526 | 0 | 0 | 0 | 0 | 0 | 0 | 2.40E-08 | 1 |
| LOC105369360 | XR_950233 | 10.11377196 | 0 | 0 | 0 | 0 | 0 | 0 | 0 | 1.70E-08 | 1 |
| LOC105369370 | XR_950269 | 0 | 8.927789798 | 0 | 0 | 0 | 0 | 0 | 0 | 7.69E-08 | 1 |
| LOC105369656 | XR_001749002 | 0 | 8.497482465 | 0 | 0 | 0 | 0 | 0 | 0 | 1.06E-07 | 1 |
| LOC105369690 | XR_931424 | 0 | 8.519117955 | 0 | 0 | 0 | 0 | 0 | 0 | 1.04E-07 | 1 |
| LOC105369819 | XR_945057 | 10.16602012 | 0 | 0 | 0 | 0 | 0 | 0 | 0 | 1.60E-08 | 1 |
| LOC105370624 | XR_944148 | 0 | 0 | 0 | 0 | 0 | 0 | 8.172825027 | 10.19881364 | 1 | 0.02072721 |
| LOC105370858 | XR_932357 | 0 | 9.014126367 | 0 | 0 | 0 | 0 | 0 | 0 | 6.71E-08 | 1 |
| LOC105370969 | XR_001751655 | 0 | 9.036802844 | 9.555578088 | 8.312886334 | 3.920337561 | 0 | 0 | 0 | 0.999953634 | 0.019255725 |
| LOC105371257 | XR_001752182 | 0 | 9.559528578 | 0 | 0 | 0 | 0 | 0 | 0 | 2.70E-08 | 1 |
| LOC105371554 | XR_001753080 | 0 | 0 | 0 | 0 | 6.351978551 | 0 | 0 | 0 | 4.81E-05 | 1 |
| LOC105371762 | XR_951997 | 11.65696001 | 9.506358431 | 11.17596815 | 5.951423154 | 8.75166616 | 0 | 0 | 0 | 0.999953634 | 0.00377824 |
| LOC105371763 | XM_017030220 | 10.18455989 | 0 | 0 | 0 | 0 | 10.69976899 | 8.62242756 | 0 | 1.60E-08 | 0.02072721 |
| LOC105371870 | XR_001752982 | 8.583566901 | 0 | 0 | 0 | 0 | 0 | 0 | 0 | 1.06E-07 | 1 |
| LOC105372246 | XR_936266 | 0 | 8.690990708 | 0 | 0 | 0 | 0 | 0 | 0 | 9.02E-08 | 1 |
| LOC105372401 | XR_935968 | 0 | 8.632639499 | 0 | 0 | 0 | 0 | 0 | 0 | 9.44E-08 | 1 |
| LOC105372426 | XR_936001 | 0 | 0 | 0 | 7.167471359 | 0 | 0 | 0 | 0 | 2.47E-05 | 1 |
| LOC105372441 | NR_131203 | 9.759152772 | 11.09900498 | 9.662916816 | 9.777215148 | 0 | 0 | 0 | 0 | 0.999953634 | 0.02072721 |
| LOC105372441 | NR_131205 | 11.76792749 | 11.82017737 | 11.24523348 | 0 | 9.881756939 | 0 | 0 | 0 | 0.999055803 | 0.02072721 |
| LOC105372480 | NR_144444 | 0 | 0 | 9.89521947 | 0 | 8.329374014 | 0 | 0 | 0 | 0.023851196 | 0.02072721 |
| LOC105372578 | XR_937399 | 0 | 0 | 10.62211805 | 0 | 7.864619359 | 0 | 0 | 0 | 0.017870966 | 0.02092873 |
| LOC105372629 | XR_936745 | 0 | 0 | 0 | 9.026902352 | 8.899709872 | 0 | 9.681964501 | 0 | 0.029138196 | 1 |
| LOC105372803 | XR_937713 | 0 | 0 | 0 | 0 | 0 | 10.90879333 | 10.57815199 | 0 | 1 | 0.02072721 |
| LOC105372962 | XR_938084 | 9.338474533 | 0 | 0 | 0 | 0 | 0 | 0 | 0 | 4.50E-08 | 1 |
| LOC105373733 | XR_923558 | 0 | 0 | 0 | 0 | 6.769943125 | 0 | 0 | 0 | 9.43E-05 | 1 |
| LOC105373759 | XR_923620 | 0 | 9.736185717 | 0 | 0 | 0 | 0 | 0 | 0 | 2.22E-08 | 1 |
| LOC105373785 | XR_923673 | 0 | 9.500932031 | 0 | 0 | 0 | 0 | 7.73610317 | 0 | 2.95E-08 | 1 |
| LOC105374041 | XR_924334 | 0 | 0 | 7.513641688 | 0 | 0 | 0 | 0 | 0 | 1.89E-05 | 1 |
| LOC105374080 | XR_924427 | 9.25321892 | 0 | 0 | 0 | 0 | 0 | 0 | 0 | 5.16E-08 | 1 |
| LOC105374148 | XR_924567 | 10.68208174 | 0 | 0 | 0 | 0 | 0 | 0 | 0 | 1.34E-08 | 1 |
| LOC105375566 | XR_928164 | 0 | 9.277965255 | 0 | 0 | 0 | 0 | 0 | 0 | 4.01E-08 | 1 |
| LOC105376095 | XR_002956896 | 9.143781062 | 0 | 0 | 0 | 0 | 0 | 0 | 0 | 7.26E-08 | 1 |

| | | | | | | | | | | | |
|---|---|---|---|---|---|---|---|---|---|---|---|
| LOC105376159 | XR_930143 | 11.06389989 | 0 | 0 | 0 | 0 | 0 | 9.799263792 | 0 | 1.34E-08 | 1 |
| LOC105376567 | XR_931068 | 10.41791771 | 0 | 0 | 0 | 0 | 0 | 0 | 0 | 1.47E-08 | 1 |
| LOC105377763 | NR_134259 | 9.894817043 | 0 | 0 | 0 | 0 | 0 | 0 | 0 | 2.09E-08 | 1 |
| LOC105378179 | XR_943839 | 8.779231295 | 0 | 0 | 0 | 0 | 0 | 0 | 0 | 9.02E-08 | 1 |
| LOC105378539 | XR_946432 | 9.105375844 | 0 | 0 | 0 | 0 | 0 | 0 | 0 | 7.04E-08 | 1 |
| LOC105378720 | XR_001737639 | 0 | 9.271606013 | 0 | 0 | 0 | 0 | 0 | 0 | 4.15E-08 | 1 |
| LOC105378948 | XR_002958512 | 10.46037918 | 0 | 0 | 0 | 0 | 0 | 0 | 0 | 1.45E-08 | 1 |
| LOC105379172 | XR_948782 | 10.53451875 | 0 | 0 | 0 | 0 | 0 | 7.341484785 | 0 | 1.36E-08 | 1 |
| LOC105379280 | XR_001756103 | 0 | 0 | 0 | 0 | 0 | 10.43162606 | 7.782430925 | 0 | 1 | 0.02072721 |
| LOC105379499 | XR_001755120 | 0 | 0 | 0 | 7.146247488 | 0 | 3.63205005 | 0 | 0 | 3.03E-05 | 1 |
| LOC105379508 | XR_951151 | 0 | 0 | 9.348335382 | 0 | 7.707000189 | 0 | 0 | 0 | 0.036173147 | 0.023711603 |
| LOC105379524 | XR_951301 | 9.363098816 | 0 | 0 | 0 | 0 | 0 | 0 | 0 | 4.33E-08 | 1 |
| LOC107984329 | XR_001748206 | 0 | 8.870030144 | 0 | 0 | 0 | 0 | 0 | 0 | 7.86E-08 | 1 |
| LOC107984347 | XR_001748289 | 0 | 9.989669449 | 0 | 0 | 0 | 0 | 9.958223136 | 10.14738921 | 2.68E-08 | 0.02072721 |
| LOC107984813 | XM_024450502 | 0 | 0 | 0 | 0 | 6.518304288 | 0 | 0 | 0 | 4.23E-05 | 1 |
| LOC107984948 | XR_001738029 | 0 | 9.411213355 | 0 | 0 | 0 | 0 | 0 | 0 | 3.28E-08 | 1 |
| LOC107985200 | XR_001738207 | 0 | 0 | 0 | 10.19529003 | 8.1822137 | 0 | 8.92739342 | 10.46136272 | 0.020731579 | 0.999932046 |
| LOC107985208 | XR_001738245 | 10.83953218 | 0 | 0 | 0 | 0 | 0 | 0 | 0 | 1.34E-08 | 1 |
| LOC107985216 | XR_001738263 | 8.815370067 | 0 | 0 | 0 | 0 | 0 | 0 | 0 | 8.84E-08 | 1 |
| LOC107985249 | XR_001738390 | 0 | 9.689244249 | 0 | 0 | 0 | 0 | 0 | 0 | 2.34E-08 | 1 |
| LOC107985367 | XR_001738546 | 0 | 8.878424457 | 0 | 0 | 0 | 0 | 0 | 0 | 9.15E-08 | 1 |
| LOC107985404 | XR_001754634 | 0 | 0 | 0 | 0 | 0 | 0 | 9.083561124 | 8.808069076 | 1 | 0.02072721 |
| LOC107985433 | XM_017028192 | 10.58772553 | 0 | 0 | 0 | 0 | 0 | 0 | 0 | 1.34E-08 | 1 |
| LOC107985480 | XR_001755102 | 8.742163956 | 0 | 0 | 0 | 0 | 10.21699361 | 0 | 0 | 9.27E-08 | 1 |
| LOC107985554 | XR_001755448 | 10.49792516 | 0 | 0 | 0 | 0 | 0 | 0 | 0 | 1.51E-08 | 1 |
| LOC107985567 | XR_001755555 | 0 | 0 | 0 | 9.841349036 | 8.884555432 | 10.76579474 | 0 | 0 | 0.021150973 | 0.999932046 |
| LOC107985733 | XR_001737880 | 0 | 9.743097037 | 10.05474104 | 0 | 8.903473739 | 0 | 0 | 0 | 0.999953634 | 0.02072721 |
| LOC107985910 | XR_001739586 | 9.495214119 | 0 | 0 | 0 | 0 | 0 | 0 | 0 | 3.47E-08 | 1 |
| LOC107985917 | XR_002959409 | 8.994164854 | 0 | 0 | 0 | 0 | 0 | 0 | 0 | 7.73E-08 | 1 |
| LOC107985923 | XR_001739606 | 0 | 0 | 7.114078641 | 0 | 0 | 0 | 0 | 0 | 2.84E-05 | 1 |
| LOC107986003 | XR_001739969 | 0 | 0 | 0 | 7.364123525 | 0 | 0 | 0 | 0 | 2.02E-05 | 1 |
| LOC107986298 | XR_001741783 | 9.287928048 | 0 | 0 | 0 | 0 | 0 | 0 | 0 | 4.93E-08 | 1 |
| LOC107986348 | XR_001742411 | 0 | 9.575104875 | 0 | 0 | 0 | 0 | 0 | 0 | 2.75E-08 | 1 |
| LOC107986483 | XR_001743008 | 0 | 0 | 0 | 0 | 0 | 9.411697668 | 9.930204221 | 0 | 1 | 0.02072721 |
| LOC107986854 | XR_001745386 | 0 | 9.411213355 | 0 | 0 | 0 | 0 | 0 | 0 | 3.28E-08 | 1 |
| LOC107986991 | XR_001746442 | 9.79579164 | 0 | 0 | 0 | 0 | 0 | 0 | 0 | 2.47E-08 | 1 |
| LOC107986997 | XR_001746463 | 10.96548485 | 0 | 0 | 0 | 0 | 0 | 0 | 0 | 1.34E-08 | 1 |
| LOC107987043 | XR_001746601 | 10.20286443 | 0 | 0 | 0 | 0 | 0 | 0 | 0 | 1.60E-08 | 1 |
| LOC107987072 | XR_001746688 | 9.966206167 | 0 | 0 | 0 | 0 | 0 | 6.439789557 | 0 | 2.00E-08 | 1 |
| LOC107987102 | XR_001746847 | 0 | 9.538495079 | 0 | 0 | 0 | 0 | 8.465635012 | 0 | 2.95E-08 | 1 |
| LOC107987135 | XR_001746960 | 10.26079939 | 0 | 0 | 0 | 0 | 0 | 0 | 0 | 1.58E-08 | 1 |
| LOC107987136 | XR_001746961 | 0 | 0 | 0 | 8.387136267 | 7.470373439 | 0 | 0 | 0 | 0.061794915 | 0.04303999 |
| LOC107987206 | XR_001750762 | 0 | 0 | 9.494670242 | 0 | 9.340148749 | 0 | 0 | 8.202491483 | 0.021348124 | 0.999932046 |

| | | | | | | | | | | | |
|---|---|---|---|---|---|---|---|---|---|---|---|
| LOC107987251 | XR_001753050 | 9.487768672 | 0 | 0 | 0 | 0 | 0 | 0 | 0 | 3.53E-08 | 1 |
| LOC107987294 | XR_001754939 | 10.04948094 | 0 | 0 | 0 | 0 | 0 | 0 | 0 | 5.08E-08 | 1 |
| LOC107987299 | XR_002958640 | 0 | 9.926307843 | 0 | 0 | 0 | 0 | 0 | 0 | 1.81E-08 | 1 |
| LOC108783654 | NR_144402 | 10.21644202 | 0 | 0 | 0 | 0 | 0 | 0 | 0 | 1.59E-08 | 1 |
| LOC110384692 | NM_001352000 | 0 | 9.914111386 | 0 | 0 | 0 | 0 | 0 | 0 | 1.85E-08 | 1 |
| LOC112267883 | XR_002959130 | 10.2867736 | 0 | 0 | 0 | 0 | 0 | 0 | 8.379320965 | 1.54E-08 | 1 |
| LOC112268035 | XR_002956842 | 0 | 0 | 0 | 0 | 6.643614539 | 0 | 0 | 0 | 3.86E-05 | 1 |
| LOC112268037 | XR_002956860 | 0 | 0 | 0 | 0 | 6.655567121 | 0 | 0 | 0 | 3.82E-05 | 1 |
| LOC112268063 | XR_002957092 | 9.244409574 | 0 | 0 | 0 | 0 | 0 | 0 | 0 | 5.26E-08 | 1 |
| LOC112268088 | XR_002957396 | 8.729593473 | 0 | 0 | 0 | 0 | 0 | 0 | 0 | 9.38E-08 | 1 |
| LOC112268098 | XR_002957416 | 0 | 0 | 9.114793339 | 8.766740692 | 0 | 0 | 0 | 0 | 0.032403902 | 0.022421975 |
| LOC112268109 | XR_002957492 | 0 | 0 | 7.487554998 | 9.394011718 | 6.381065478 | 0 | 0 | 0 | 0.01163522 | 0.019308333 |
| LOC112268124 | XM_024449797 | 0 | 0 | 0 | 0 | 6.643614539 | 0 | 0 | 0 | 3.86E-05 | 1 |
| LOC112268146 | XR_002957708 | 0 | 9.296876505 | 0 | 0 | 0 | 0 | 0 | 0 | 4.01E-08 | 1 |
| LOC112268184 | XM_024451054 | 10.41399488 | 8.486541819 | 0 | 0 | 0 | 0 | 0 | 0 | 0.001341012 | 1 |
| LOC112268277 | XR_002958637 | 0 | 0 | 0 | 0 | 0 | 11.25278294 | 9.210411598 | 0 | 1 | 0.02072721 |
| LOC112268313 | XR_002958845 | 0 | 0 | 0 | 9.626353068 | 9.113027941 | 0 | 0 | 8.391159551 | 0.021703066 | 0.999932046 |
| LOC112268389 | XR_002959139 | 0 | 9.059128398 | 0 | 0 | 0 | 0 | 0 | 0 | 7.80E-08 | 1 |
| LOC171391 | NR_126343 | 0 | 8.571828374 | 0 | 0 | 0 | 0 | 0 | 0 | 9.97E-08 | 1 |
| LOC257396 | NR_034107 | 0 | 9.795011742 | 0 | 0 | 0 | 0 | 0 | 0 | 2.04E-08 | 1 |
| LOC283788 | NR_027436 | 0 | 0 | 10.36166239 | 0 | 8.77650741 | 0 | 0 | 10.86840153 | 0.017534783 | 0.999932046 |
| LOC284454 | NR_036515 | 0 | 0 | 7.500657305 | 9.961649519 | 4.650877629 | 0 | 0 | 9.239866457 | 0.009849908 | 0.999932046 |
| LOC389602 | NR_144629 | 0 | 0 | 0 | 0 | 0 | 0 | 7.106768981 | 10.67132269 | 1 | 0.02072721 |
| LOC389831 | NM_001347681 | 0 | 0 | 10.33995673 | 0 | 8.194554972 | 0 | 0 | 0 | 0.019641255 | 0.02072721 |
| LOC389906 | NR_034031 | 0 | 9.19973932 | 0 | 0 | 0 | 0 | 0 | 0 | 4.67E-08 | 1 |
| LOC646626 | NR_045484 | 11.21358056 | 0 | 10.3289802 | 0 | 11.21215532 | 0 | 0 | 0 | 0.999953634 | 0.02072721 |
| LOC728743 | NR_027237 | 10.63230229 | 0 | 0 | 0 | 0 | 0 | 0 | 10.42870925 | 1.34E-08 | 1 |
| LOC729737 | NR_039983 | 0 | 0 | 0 | 0 | 6.409577539 | 0 | 0 | 0 | 4.59E-05 | 1 |
| LOC93622 | NR_015433 | 0 | 0 | 10.52926948 | 0 | 6.518304288 | 0 | 6.380785351 | 0 | 0.021572016 | 0.999932046 |
| LONP1 | NR_076392 | 0 | 0 | 0 | 8.680787864 | 0 | 0 | 0 | 0 | 5.10E-06 | 1 |
| LONP2 | NM_031490 | 0 | 0 | 0 | 10.26586678 | 9.202312328 | 0 | 0 | 0 | 0.016407821 | 0.021032806 |
| LONRF1 | NR_138256 | 9.689464985 | 0 | 0 | 0 | 0 | 0 | 0 | 0 | 3.15E-08 | 1 |
| LPCAT1 | XM_017009879 | 0 | 0 | 0 | 9.227165049 | 6.197037886 | 0 | 0 | 0 | 0.049139065 | 0.034037301 |
| LPCAT3 | NM_005768 | 10.28247691 | 0 | 0 | 8.508532777 | 8.762759555 | 0 | 0 | 0 | 0.999953634 | 0.02561723 |
| LPIN3 | XM_011529002 | 9.825628908 | 0 | 0 | 0 | 0 | 0 | 5.732859748 | 0 | 2.30E-08 | 1 |
| LPP | XM_005247453 | 0 | 0 | 10.75926582 | 0 | 9.047636369 | 0 | 9.167416673 | 0 | 0.013678815 | 0.999932046 |
| LPP | XM_011512828 | 11.53403233 | 0 | 10.70793922 | 0 | 7.785243826 | 0 | 0 | 0 | 0.999953634 | 0.02074727 |
| LPP | XM_011512834 | 0 | 0 | 8.216015222 | 0 | 9.30643073 | 0 | 10.56108971 | 10.95369999 | 0.031510534 | 0.999932046 |
| LPP | XM_017006377 | 10.7333523 | 10.60230411 | 0 | 0 | 0 | 0 | 8.684736524 | 10.0940638 | 0.000352182 | 0.02072721 |
| LPP | XM_017006380 | 9.304974408 | 9.303125566 | 0 | 0 | 0 | 0 | 7.460241356 | 0 | 0.002125851 | 1 |
| LRCH4 | NM_002319 | 10.61018607 | 0 | 0 | 0 | 0 | 0 | 0 | 10.0424498 | 1.34E-08 | 1 |
| LRFN4 | XM_005274239 | 9.531875668 | 0 | 0 | 0 | 0 | 0 | 7.849247234 | 0 | 3.28E-08 | 1 |
| LRIG2 | XM_024451231 | 8.665043676 | 0 | 0 | 0 | 0 | 0 | 0 | 0 | 9.91E-08 | 1 |

| | | | | | | | | | | | |
|---|---|---|---|---|---|---|---|---|---|---|---|
| LRP5 | XR_949926 | 0 | 0 | 0 | 0 | 6.679179249 | 0 | 0 | 0 | 3.78E-05 | 1 |
| LRP6 | NM_002336 | 9.383302787 | 0 | 0 | 0 | 0 | 0 | 0 | 0 | 4.18E-08 | 1 |
| LRPAP1 | NR_110005 | 0 | 9.095586658 | 0 | 0 | 0 | 0 | 9.733969061 | 0 | 5.54E-08 | 1 |
| LRPAP1 | XR_002959730 | 0 | 0 | 8.720358382 | 10.01091583 | 0 | 0 | 0 | 0 | 0.021907134 | 0.02072721 |
| LRRC14 | NM_014665 | 0 | 0 | 0 | 8.601134616 | 8.662671672 | 0 | 0 | 0 | 0.037488995 | 0.025754319 |
| LRRC23 | NM_001135217 | 0 | 0 | 0 | 10.6349947 | 8.78741214 | 11.3764851 | 9.512513549 | 0 | 0.015112298 | 0.999932046 |
| LRRC32 | XM_017017527 | 0 | 8.983325733 | 0 | 0 | 0 | 0 | 7.856483795 | 0 | 7.17E-08 | 1 |
| LRRC34 | XM_024453362 | 0 | 0 | 0 | 9.923173429 | 9.42813403 | 0 | 0 | 0 | 0.017768441 | 0.02072721 |
| LRRC40 | NM_017768 | 0 | 0 | 0 | 0 | 6.679179249 | 0 | 0 | 0 | 3.78E-05 | 1 |
| LRRC42 | NM_052940 | 0 | 8.809861237 | 0 | 0 | 0 | 8.029326918 | 0 | 0 | 8.24E-08 | 1 |
| LRRC42 | XM_006710328 | 0 | 8.248259235 | 0 | 0 | 0 | 0 | 0 | 0 | 2.02E-07 | 1 |
| LRRC6 | XM_011516950 | 0 | 0 | 8.663239538 | 0 | 8.210847477 | 0 | 0 | 0 | 0.044198995 | 0.028374914 |
| LRRC6 | XM_017013296 | 0 | 0 | 0 | 10.88768298 | 8.323766986 | 9.435641615 | 12.56838317 | 0 | 0.014241889 | 0.999932046 |
| LRRC75A | NM_207387 | 0 | 8.959785899 | 0 | 0 | 0 | 0 | 0 | 0 | 7.69E-08 | 1 |
| LRRC8A | NM_019594 | 0 | 9.834376365 | 0 | 0 | 0 | 8.68082595 | 0 | 0 | 2.00E-08 | 1 |
| LRRFIP2 | NM_017724 | 0 | 0 | 9.637933735 | 0 | 7.864619359 | 0 | 0 | 0 | 0.029881302 | 0.020903961 |
| LSG1 | NM_018385 | 0 | 0 | 0 | 0 | 0 | 10.9688255 | 8.134642646 | 0 | 1 | 0.02072721 |
| LSM1 | NM_014462 | 10.61531999 | 10.46134184 | 11.50824993 | 0 | 7.879983547 | 0 | 0 | 0 | 0.999953634 | 7.47E-09 |
| LSM14A | NM_015578 | 0 | 0 | 10.2113759 | 8.359738397 | 9.707564817 | 0 | 0 | 0 | 0.002095514 | 0.003174145 |
| LSM6 | XM_017007673 | 0 | 9.736185717 | 0 | 0 | 0 | 0 | 0 | 0 | 2.22E-08 | 1 |
| LSR | NM_001260489 | 0 | 0 | 9.927111081 | 9.645387449 | 0 | 0 | 0 | 0 | 0.018173581 | 0.02072721 |
| LSR | NM_015925 | 10.4296226 | 10.42168667 | 11.71244355 | 0 | 7.328262653 | 0 | 0 | 0 | 1 | 6.49E-09 |
| LSR | XM_005258980 | 9.771469405 | 0 | 0 | 0 | 0 | 0 | 0 | 0 | 2.47E-08 | 1 |
| LSS | XM_011529564 | 9.371214389 | 0 | 0 | 0 | 0 | 0 | 0 | 0 | 4.28E-08 | 1 |
| LST1 | NM_205838 | 10.79196021 | 0 | 0 | 0 | 0 | 0 | 0 | 0 | 1.34E-08 | 1 |
| LTA4H | NM_000895 | 0 | 0 | 0 | 10.95330853 | 9.275782933 | 0 | 7.856483795 | 9.371188143 | 0.011593205 | 0.999932046 |
| LTA4H | XM_005268871 | 0 | 0 | 0 | 0 | 6.725274845 | 0 | 0 | 0 | 9.53E-05 | 1 |
| LTB | NM_002341 | 10.88558544 | 0 | 0 | 0 | 0 | 0 | 0 | 0 | 1.34E-08 | 1 |
| LTBP1 | NM_001166265 | 0 | 0 | 9.071545242 | 0 | 8.548724137 | 0 | 0 | 0 | 0.032717173 | 0.022107613 |
| LTBP1 | XM_017004109 | 0 | 9.059128398 | 0 | 0 | 0 | 0 | 0 | 0 | 7.80E-08 | 1 |
| LTBP1 | XM_017004110 | 12.27405319 | 9.277965255 | 0 | 0 | 0 | 0 | 0 | 0 | 0.000167345 | 1 |
| LTBP3 | XM_011545032 | 0 | 0 | 0 | 0 | 6.569741695 | 0 | 0 | 0 | 4.12E-05 | 1 |
| LTBP4 | XM_011527379 | 0 | 9.102769039 | 0 | 0 | 0 | 0 | 0 | 0 | 5.49E-08 | 1 |
| LTF | NM_001199149 | 9.777588513 | 0 | 0 | 0 | 0 | 0 | 0 | 9.766115442 | 2.46E-08 | 1 |
| LTV1 | NM_032860 | 0 | 0 | 11.52678128 | 9.719108837 | 8.887092252 | 0 | 8.083136668 | 0 | 0.000847929 | 0.987221099 |
| LUC7L3 | NM_001330330 | 10.87135108 | 0 | 0 | 0 | 0 | 0 | 0 | 0 | 1.34E-08 | 1 |
| LUC7L3 | NM_006107 | 0 | 0 | 9.082479415 | 10.53749502 | 8.020631767 | 0 | 8.9780358 | 0 | 0.002375768 | 0.999932046 |
| LUC7L3 | XM_005257449 | 0 | 10.72839101 | 0 | 0 | 0 | 0 | 0 | 0 | 1.34E-08 | 1 |
| LUC7L3 | XR_002958022 | 10.17067744 | 0 | 0 | 0 | 0 | 0 | 0 | 0 | 1.60E-08 | 1 |
| LURAP1L | NM_203403 | 9.843239492 | 8.407542174 | 9.598867372 | 0 | 8.882014143 | 0 | 0 | 0 | 0.999953634 | 0.02072721 |
| LYPLA2 | XM_005245728 | 0 | 0 | 0 | 8.78058117 | 0 | 0 | 0 | 0 | 4.59E-06 | 1 |
| LYPLAL1 | XR_002959345 | 0 | 0 | 10.14408245 | 0 | 8.830225047 | 0 | 9.311290442 | 0 | 0.019056111 | 0.999932046 |
| LYRM1 | NM_001128302 | 0 | 9.694007725 | 0 | 0 | 0 | 0 | 0 | 0 | 2.55E-08 | 1 |

| | | | | | | | | | | | |
|---|---|---|---|---|---|---|---|---|---|---|---|
| LYRM2 | NM_020466 | 0 | 0 | 7.061996878 | 0 | 0 | 0 | 0 | 0 | 2.99E-05 | 1 |
| LYRM2 | NR_028495 | 10.22542327 | 0 | 0 | 0 | 0 | 0 | 0 | 10.58762701 | 1.59E-08 | 1 |
| LYRM4 | NM_020408 | 0 | 8.1546024 | 0 | 11.79911343 | 9.031674724 | 0 | 0 | 0 | 0.746259121 | 0.02072721 |
| LYRM4 | NR_104418 | 0 | 9.102769039 | 0 | 0 | 0 | 0 | 0 | 0 | 5.49E-08 | 1 |
| LYSMD2 | XM_017022057 | 0 | 9.293741795 | 0 | 0 | 0 | 0 | 0 | 0 | 4.01E-08 | 1 |
| LYVE1 | NM_006691 | 9.208624214 | 0 | 0 | 0 | 0 | 0 | 0 | 0 | 5.58E-08 | 1 |
| LZTFL1 | XM_017006645 | 0 | 0 | 0 | 10.02406517 | 5.167328428 | 0 | 0 | 0 | 0.03153925 | 0.023194226 |
| M6PR | NM_002355 | 0 | 0 | 9.901407472 | 10.25103113 | 8.349748669 | 10.81378445 | 8.064509964 | 0 | 0.001900595 | 0.999932046 |
| MACF1 | XM_005270696 | 0 | 0 | 7.061996878 | 0 | 0 | 8.234341443 | 0 | 8.619662358 | 2.99E-05 | 0.999932046 |
| MAD1L1 | XM_017012690 | 0 | 0 | 10.39537545 | 0 | 7.877434185 | 10.37908708 | 0 | 0 | 0.02002499 | 0.999932046 |
| MAD1L1 | XM_017012691 | 0 | 0 | 9.721447883 | 0 | 9.491188327 | 11.34943104 | 0 | 10.74927864 | 0.01881306 | 0.999932046 |
| MAD2L1BP | NM_014628 | 11.59073616 | 10.63981999 | 0 | 0 | 0 | 0 | 8.53943153 | 0 | 0.000180012 | 1 |
| MADD | NM_001135944 | 0 | 0 | 0 | 11.08183069 | 7.605905728 | 11.40960449 | 0 | 10.30900536 | 0.013927877 | 0.999932046 |
| MADD | NM_130474 | 0 | 0 | 10.59335737 | 9.510708615 | 7.039920832 | 0 | 8.647674983 | 0 | 0.002493586 | 0.999932046 |
| MADD | NM_130475 | 0 | 0 | 0 | 0 | 6.451316153 | 0 | 6.939666787 | 0 | 4.51E-05 | 1 |
| MAF | NM_005360 | 0 | 0 | 8.082961956 | 0 | 0 | 0 | 0 | 0 | 1.05E-05 | 1 |
| MAFB | NM_005461 | 0 | 9.340061955 | 0 | 0 | 0 | 0 | 0 | 0 | 3.95E-08 | 1 |
| MAFF | NM_012323 | 0 | 0 | 10.53564879 | 9.722697844 | 0 | 0 | 0 | 0 | 0.014067333 | 0.02072721 |
| MAFIP | NR_046442 | 0 | 10.10145643 | 11.313647 | 0 | 11.08479324 | 0 | 0 | 0 | 0.999953634 | 0.02072721 |
| MAFK | XM_006715773 | 10.17531978 | 8.975521721 | 0 | 0 | 0 | 0 | 0 | 0 | 0.001347927 | 1 |
| MAGED1 | NM_006986 | 0 | 0 | 10.45716976 | 8.035448861 | 8.238923989 | 0 | 0 | 0 | 0.003304959 | 0.004762166 |
| MAGED2 | NM_014599 | 0 | 0 | 0 | 0 | 0 | 11.03562724 | 8.696881776 | 0 | 1 | 0.02072721 |
| MAGED2 | NM_177433 | 0 | 0 | 0 | 11.24545771 | 7.707000189 | 0 | 10.2869086 | 0 | 0.012484273 | 0.999932046 |
| MAGED2 | NM_201222 | 0 | 0 | 10.3344789 | 11.29089193 | 9.033965799 | 0 | 9.468324297 | 0 | 0.000742596 | 0.999932046 |
| MAGEF1 | NM_022149 | 0 | 0 | 10.33173217 | 0 | 8.052621656 | 10.86317478 | 0 | 11.35602604 | 0.020153035 | 0.999932046 |
| MAGEH1 | NM_014061 | 0 | 8.895067962 | 0 | 0 | 0 | 0 | 0 | 10.67253583 | 7.75E-08 | 1 |
| MAGIX | NM_001099682 | 0 | 9.226288718 | 0 | 0 | 0 | 0 | 0 | 0 | 4.45E-08 | 1 |
| MAGOHB | NR_135120 | 0 | 8.642530071 | 0 | 0 | 0 | 0 | 0 | 0 | 9.87E-08 | 1 |
| MAIP1 | NM_024520 | 0 | 0 | 0 | 9.966198446 | 8.759994193 | 0 | 0 | 0 | 0.020656552 | 0.02072721 |
| MALL | NM_005434 | 0 | 0 | 11.24474686 | 10.43948342 | 7.497189827 | 0 | 0 | 11.07425534 | 0.001068298 | 0.999932046 |
| MALT1 | XR_001753135 | 0 | 10.9396736 | 0 | 10.45898029 | 7.574771995 | 0 | 0 | 0 | 0.999953634 | 0.02072721 |
| MAMDC4 | NM_206920 | 9.235546106 | 0 | 0 | 0 | 0 | 0 | 0 | 0 | 5.31E-08 | 1 |
| MAN1B1 | XM_017014239 | 0 | 0 | 0 | 0 | 6.395391943 | 0 | 0 | 0 | 4.65E-05 | 1 |
| MAN2A1 | XM_017009472 | 0 | 9.522516235 | 0 | 0 | 0 | 0 | 0 | 0 | 2.88E-08 | 1 |
| MAN2B2 | XR_001741189 | 0 | 0 | 7.539265046 | 0 | 0 | 0 | 0 | 0 | 1.83E-05 | 1 |
| MAN2C1 | XM_017022188 | 0 | 0 | 10.02103749 | 0 | 7.497189827 | 0 | 0 | 0 | 0.025821961 | 0.02072721 |
| MAN2C1 | XR_002957639 | 9.025355161 | 6.373487887 | 0 | 0 | 0 | 0 | 0 | 0 | 0.010272288 | 1 |
| MAOA | NM_000240 | 0 | 10.83142586 | 0 | 0 | 0 | 0 | 0 | 0 | 1.34E-08 | 1 |
| MAOA | NM_001270458 | 9.321821705 | 10.11572772 | 0 | 0 | 0 | 0 | 0 | 0 | 0.001165059 | 1 |
| MAP1LC3A | XM_011529085 | 9.649985154 | 0 | 0 | 0 | 0 | 0 | 7.680104054 | 0 | 2.95E-08 | 1 |
| MAP1S | XM_017026930 | 0 | 0 | 0 | 10.41971946 | 10.78291938 | 0 | 9.138024627 | 0 | 0.009042552 | 0.999932046 |
| MAP2K6 | XM_011525027 | 0 | 0 | 0 | 10.36105389 | 8.594457502 | 0 | 0 | 10.03114526 | 0.017878668 | 0.999932046 |
| MAP2K7 | XM_006722800 | 0 | 0 | 0 | 0 | 6.702411139 | 0 | 0 | 0 | 3.69E-05 | 1 |

| | | | | | | | | | | |
|---|---|---|---|---|---|---|---|---|---|---|
| MAP3K20 | NM_133646 | 0 | 0 | 0 | 0 | 6.747781854 | 0 | 0 | 0 | 9.53E-05 | 1 |
| MAP3K21 | NM_032435 | 0 | 0 | 0 | 0 | 6.747781854 | 0 | 8.748360817 | 0 | 9.53E-05 | 1 |
| MAP3K4 | NR_120425 | 10.67226231 | 0 | 0 | 0 | 0 | 0 | 8.039294254 | 0 | 1.34E-08 | 1 |
| MAP3K5 | NM_005923 | 0 | 8.903318262 | 0 | 0 | 0 | 0 | 0 | 0 | 7.75E-08 | 1 |
| MAP3K6 | XR_946795 | 0 | 8.540433774 | 0 | 0 | 0 | 0 | 0 | 0 | 1.02E-07 | 1 |
| MAP4 | XM_017006402 | 0 | 0 | 0 | 0 | 0 | 10.02517937 | 8.763837937 | 0 | 1 | 0.02072721 |
| MAP4 | XM_017006412 | 0 | 0 | 9.715840939 | 10.59639063 | 0 | 9.949589646 | 10.80033142 | 10.98821048 | 0.013535352 | 0.999932046 |
| MAP4 | XM_017006414 | 10.29532878 | 0 | 0 | 0 | 0 | 0 | 0 | 0 | 1.54E-08 | 1 |
| MAP4 | XM_017006416 | 0 | 0 | 9.350146171 | 0 | 7.555764066 | 11.11993587 | 0 | 9.743205182 | 0.037386007 | 0.999932046 |
| MAP4 | XR_002959530 | 0 | 0 | 13.098818 | 0 | 11.8779019 | 0 | 10.28221413 | 0 | 1.34E-08 | 0.999932046 |
| MAP4K4 | XM_017005375 | 0 | 8.497482465 | 0 | 0 | 0 | 0 | 0 | 0 | 1.06E-07 | 1 |
| MAP4K5 | XM_024449461 | 0 | 8.774317692 | 0 | 0 | 0 | 0 | 0 | 8.902080922 | 8.46E-08 | 1 |
| MAP7 | XM_011536246 | 0 | 9.239382419 | 0 | 0 | 0 | 10.55971347 | 7.569961514 | 0 | 4.33E-08 | 0.02072721 |
| MAP7D1 | XM_006710761 | 0 | 0 | 0 | 8.491803396 | 0 | 10.46303178 | 8.310927973 | 0 | 6.26E-06 | 0.999932046 |
| MAPK10 | XM_005263131 | 0 | 9.245884974 | 0 | 0 | 0 | 0 | 0 | 0 | 4.33E-08 | 1 |
| MAPK10 | XM_005263135 | 10.01881342 | 0 | 0 | 0 | 0 | 0 | 0 | 0 | 1.89E-08 | 1 |
| MAPK10 | XM_017008433 | 0 | 9.117027412 | 0 | 0 | 0 | 0 | 0 | 0 | 5.36E-08 | 1 |
| MAPK10 | XM_017008452 | 10.22093963 | 0 | 0 | 0 | 0 | 0 | 0 | 0 | 1.59E-08 | 1 |
| MAPK10 | XM_024454142 | 10.49421423 | 0 | 0 | 0 | 0 | 0 | 0 | 0 | 1.42E-08 | 1 |
| MAPK10 | XM_024454143 | 8.791378415 | 0 | 0 | 0 | 0 | 0 | 0 | 0 | 9.38E-08 | 1 |
| MAPK10 | XM_024454147 | 0 | 10.75142427 | 0 | 0 | 0 | 0 | 0 | 0 | 1.34E-08 | 1 |
| MAPK1IP1L | NM_144578 | 0 | 0 | 10.36524846 | 9.143952478 | 9.116275051 | 0 | 0 | 0 | 0.001922607 | 0.00302572 |
| MAPK1IP1L | XM_017021789 | 0 | 8.486541819 | 0 | 0 | 0 | 0 | 0 | 0 | 1.09E-07 | 1 |
| MAPKAP1 | XM_017015126 | 0 | 8.870030144 | 0 | 0 | 0 | 0 | 0 | 0 | 7.86E-08 | 1 |
| MAPKAPK2 | XM_017002810 | 0 | 9.252358352 | 0 | 9.343622985 | 8.989800746 | 0 | 0 | 0 | 0.999953634 | 0.02072721 |
| MAPKAPK5-AS1 | NR_152605 | 0 | 8.612652582 | 0 | 0 | 0 | 11.02908455 | 0 | 6.860901615 | 9.67E-08 | 5.17E-09 |
| MAPKAPK5-AS1 | NR_152607 | 0 | 8.690990708 | 0 | 0 | 0 | 0 | 0 | 0 | 9.02E-08 | 1 |
| MAPRE1 | NM_012325 | 0 | 0 | 0 | 9.114202495 | 9.491188327 | 0 | 8.493752915 | 9.582123369 | 0.022690368 | 0.999932046 |
| MAPRE2 | NM_014268 | 0 | 8.765293275 | 0 | 0 | 0 | 0 | 0 | 0 | 8.55E-08 | 1 |
| 2-Mar | NM_017898 | 0 | 9.102769039 | 0 | 0 | 0 | 0 | 0 | 0 | 5.49E-08 | 1 |
| 5-Mar | NM_017824 | 0 | 0 | 9.649744388 | 0 | 8.77650741 | 0 | 8.387859204 | 0 | 0.02391827 | 0.999932046 |
| 6-Mar | XR_925576 | 0 | 0 | 7.718176908 | 0 | 0 | 0 | 0 | 0 | 1.53E-05 | 1 |
| MARF1 | XR_001752038 | 0 | 0 | 10.69232105 | 0 | 8.720712025 | 0 | 0 | 0 | 0.015117263 | 0.02072721 |
| MARK1 | XM_017001307 | 10.07951011 | 0 | 0 | 0 | 0 | 0 | 0 | 0 | 1.78E-08 | 1 |
| MARK2 | NM_001039469 | 0 | 0 | 10.1575878 | 8.731547201 | 7.335710674 | 0 | 0 | 0 | 0.004113735 | 0.005954559 |
| MARK2 | NM_017490 | 0 | 0 | 0 | 0 | 0 | 8.110899224 | 7.638730623 | 0 | 1 | 0.03435538 |
| MARK3 | NM_002376 | 0 | 10.11217311 | 0 | 0 | 0 | 0 | 0 | 0 | 1.59E-08 | 1 |
| MARK3 | XM_024449587 | 10.3538353 | 0 | 0 | 0 | 0 | 0 | 0 | 0 | 2.01E-08 | 1 |
| MARVELD3 | NM_001017967 | 0 | 0 | 9.353760947 | 0 | 7.763318133 | 0 | 0 | 0 | 0.035623064 | 0.023366125 |
| MATN2 | XM_017013418 | 8.815370067 | 0 | 0 | 0 | 0 | 0 | 0 | 0 | 8.84E-08 | 1 |
| MAVS | NR_037921 | 0 | 0 | 0 | 9.944846321 | 6.409577539 | 0 | 0 | 0 | 0.029945006 | 0.022064246 |
| MAX | XR_943450 | 0 | 0 | 0 | 10.9760411 | 9.629418952 | 0 | 9.696151157 | 0 | 0.010551563 | 0.999932046 |
| MAZ | XR_002957806 | 0 | 0 | 0 | 0 | 6.30721813 | 0 | 0 | 0 | 4.99E-05 | 1 |

| | | | | | | | | | | | |
|---|---|---|---|---|---|---|---|---|---|---|---|
| MB | XM_005261605 | 0 | 9.724592761 | 0 | 0 | 0 | 0 | 0 | 0 | 2.28E-08 | 1 |
| MBD6 | XR_429073 | 9.226627846 | 0 | 0 | 0 | 0 | 0 | 0 | 0 | 5.40E-08 | 1 |
| MBIP | XM_005267756 | 0 | 9.06649413 | 0 | 0 | 0 | 0 | 0 | 0 | 7.75E-08 | 1 |
| MBIP | XR_001750371 | 9.933695697 | 0 | 0 | 0 | 0 | 0 | 0 | 0 | 2.01E-08 | 1 |
| MBOAT4 | XM_017013726 | 10.24321976 | 0 | 0 | 0 | 0 | 0 | 0 | 0 | 1.59E-08 | 1 |
| MBP | NM_001025101 | 0 | 0 | 0 | 0 | 6.747781854 | 0 | 0 | 0 | 9.53E-05 | 1 |
| MBP | XM_017025780 | 0 | 0 | 0 | 0 | 6.544252221 | 0 | 0 | 0 | 4.20E-05 | 1 |
| MBTPS2 | NM_015884 | 8.624880391 | 8.774317692 | 0 | 0 | 0 | 0 | 0 | 0 | 0.00477578 | 1 |
| MCAT | NM_173467 | 0 | 0 | 0 | 7.631818844 | 0 | 0 | 0 | 0 | 1.70E-05 | 1 |
| MCCC1 | NM_020166 | 0 | 0 | 0 | 0 | 6.667421491 | 0 | 0 | 10.27426767 | 3.78E-05 | 1 |
| MCEE | NM_032601 | 0 | 0 | 9.623033138 | 9.380442947 | 7.219739493 | 0 | 8.284344777 | 0 | 0.004439593 | 0.999932046 |
| MCM7 | NM_005916 | 0 | 0 | 0 | 9.785818491 | 7.895185835 | 0 | 0 | 0 | 0.026861894 | 0.020781046 |
| MCM7 | XM_017012217 | 10.51999251 | 8.453213086 | 0 | 0 | 0 | 0 | 0 | 10.76642345 | 0.001229566 | 1 |
| MCM9 | NM_153255 | 9.004636726 | 0 | 0 | 0 | 0 | 0 | 0 | 0 | 7.73E-08 | 1 |
| MCOLN2 | XM_011541188 | 9.595606435 | 0 | 0 | 0 | 0 | 0 | 0 | 0 | 3.08E-08 | 1 |
| MCRIP1 | NM_207368 | 0 | 0 | 9.89521947 | 9.901169983 | 9.730347633 | 0 | 9.654204181 | 0 | 0.001434603 | 1 |
| MCRIP2 | NR_138610 | 10.30806717 | 0 | 0 | 0 | 0 | 0 | 0 | 0 | 1.54E-08 | 1 |
| MCRS1 | NM_006337 | 0 | 0 | 9.474889408 | 7.080635617 | 8.924618804 | 0 | 7.460241356 | 0 | 0.00603166 | 0.999932046 |
| MCRS1 | XM_017018690 | 10.26079939 | 8.709928179 | 0 | 0 | 0 | 0 | 0 | 0 | 0.001410695 | 1 |
| MCTP1 | XM_017009855 | 9.676424716 | 8.39589491 | 0 | 0 | 0 | 0 | 0 | 0 | 0.002684267 | 1 |
| MCTP1 | XM_017009859 | 0 | 9.868487622 | 0 | 0 | 0 | 0 | 6.979982279 | 9.779688974 | 1.95E-08 | 0.02072721 |
| MCTP1 | XM_017009860 | 0 | 0 | 9.1995944 | 0 | 9.867696746 | 0 | 9.138024627 | 0 | 0.01912717 | 0.999932046 |
| MCTP1 | XM_017009863 | 0 | 9.538495079 | 0 | 0 | 0 | 0 | 0 | 10.46976085 | 2.95E-08 | 1 |
| MCTP1 | XM_017009865 | 0 | 9.131146246 | 0 | 0 | 0 | 0 | 0 | 0 | 5.23E-08 | 1 |
| MCU | XM_017016882 | 9.636581421 | 0 | 0 | 0 | 0 | 0 | 7.189323943 | 0 | 2.90E-08 | 1 |
| MDFI | XM_011514625 | 0 | 0 | 0 | 0 | 6.725274845 | 0 | 7.288964784 | 0 | 9.53E-05 | 1 |
| MDFI | XM_017010867 | 0 | 0 | 0 | 0 | 6.351978551 | 0 | 7.906148158 | 0 | 4.81E-05 | 1 |
| MDFIC | NM_199072 | 0 | 0 | 0 | 0 | 6.451316153 | 0 | 0 | 0 | 4.51E-05 | 1 |
| MDM1 | NM_020128 | 0 | 8.407542174 | 0 | 0 | 0 | 0 | 0 | 0 | 1.16E-07 | 1 |
| MDM2 | XM_006719399 | 0 | 0 | 10.66202669 | 10.58460507 | 9.855442592 | 12.48562977 | 11.07188338 | 10.61438614 | 0.000725973 | 0.999932046 |
| MDP1 | NM_001199821 | 0 | 0 | 10.36882564 | 0 | 6.395391943 | 0 | 0 | 0 | 0.023928826 | 0.02072721 |
| MDP1 | NM_138476 | 5.623309261 | 0 | 0 | 9.950979243 | 10.01530004 | 0 | 0 | 0 | 0.380893159 | 0.02072721 |
| ME3 | XM_017017138 | 0 | 8.861586703 | 0 | 0 | 0 | 0 | 0 | 0 | 7.86E-08 | 1 |
| MEA1 | NM_014623 | 0 | 9.439854398 | 0 | 0 | 0 | 0 | 0 | 0 | 3.19E-08 | 1 |
| MEAF6 | NM_001270876 | 0 | 9.640723913 | 0 | 0 | 0 | 0 | 0 | 0 | 2.47E-08 | 1 |
| MECOM | NM_001105078 | 10.6557466 | 0 | 11.10213471 | 10.0313189 | 0 | 0 | 0 | 0 | 0.999953634 | 0.02072721 |
| MECR | NR_146214 | 0 | 0 | 0 | 7.124706719 | 0 | 0 | 0 | 0 | 2.57E-05 | 1 |
| MED1 | XM_005257465 | 0 | 0 | 0 | 10.69377591 | 6.42362501 | 0 | 9.833588242 | 0 | 0.019260992 | 1 |
| MED14 | XM_017029962 | 0 | 0 | 6.951829064 | 0 | 0 | 0 | 0 | 0 | 8.96E-05 | 1 |
| MED15 | NM_001293234 | 0 | 9.021724865 | 0 | 0 | 0 | 0 | 0 | 0 | 6.61E-08 | 1 |
| MED15 | NM_015889 | 10.18455989 | 9.117027412 | 9.781727525 | 0 | 9.83127633 | 0 | 0 | 0 | 0.999953634 | 0.02092873 |
| MED15 | XR_002958701 | 9.595606435 | 0 | 0 | 0 | 0 | 9.960633697 | 0 | 0 | 3.08E-08 | 1 |
| MED24 | NR_052017 | 0 | 8.935855595 | 10.11135781 | 0 | 9.187981525 | 0 | 0 | 0 | 0.999953634 | 0.02072721 |

| | | | | | | | | | | | |
|---|---|---|---|---|---|---|---|---|---|---|---|
| MED25 | NM_030973 | 0 | 9.219697032 | 0 | 0 | 0 | 0 | 0 | 0 | 4.50E-08 | 1 |
| MED27 | NM_004269 | 0 | 0 | 8.788716638 | 10.53239157 | 8.994513883 | 0 | 0 | 10.74697719 | 0.001980826 | 1 |
| MED28 | NM_025205 | 0 | 0 | 0 | 11.2285175 | 9.947644748 | 0 | 9.944706151 | 9.892265765 | 0.008512173 | 0.999932046 |
| MED29 | NM_017592 | 0 | 0 | 0 | 6.988236596 | 0 | 0 | 0 | 0 | 4.34E-05 | 1 |
| MED30 | XM_011517360 | 8.452015244 | 0 | 0 | 0 | 0 | 4.177693599 | 0 | 0 | 1.21E-07 | 0.862574616 |
| MED6 | NM_005466 | 0 | 0 | 0 | 10.50138585 | 7.980808426 | 0 | 0 | 0 | 0.018360815 | 0.02072721 |
| MED8 | NM_001001653 | 0 | 9.029283553 | 0 | 0 | 0 | 0 | 0 | 0 | 6.49E-08 | 1 |
| MEF2D | XM_017001315 | 9.623051988 | 0 | 0 | 0 | 0 | 0 | 0 | 0 | 2.93E-08 | 1 |
| MEG3 | NR_046469 | 9.734199496 | 0 | 0 | 0 | 0 | 0 | 0 | 0 | 5.08E-08 | 1 |
| MEGF6 | XM_017000533 | 0 | 0 | 0 | 0 | 6.76443464 | 0 | 0 | 0 | 9.51E-05 | 1 |
| MELTF | NM_033316 | 10.32069408 | 0 | 0 | 0 | 0 | 0 | 0 | 0 | 1.52E-08 | 1 |
| MEN1 | NM_130804 | 9.695941183 | 0 | 0 | 0 | 0 | 0 | 9.098897264 | 0 | 2.66E-08 | 1 |
| MEPCE | NM_001194991 | 8.421077904 | 0 | 0 | 0 | 0 | 0 | 0 | 0 | 1.25E-07 | 1 |
| MERTK | XM_005263565 | 0 | 9.245884974 | 0 | 0 | 0 | 0 | 9.184768702 | 0 | 4.33E-08 | 1 |
| MERTK | XM_005263568 | 0 | 0 | 8.966860862 | 0 | 7.401067144 | 0 | 0 | 0 | 0.048301188 | 0.030530376 |
| MESD | NR_126327 | 0 | 0 | 9.335595812 | 10.56674421 | 6.130081192 | 0 | 0 | 8.361378918 | 0.00310129 | 0.999932046 |
| MESP1 | NM_018670 | 10.6656787 | 0 | 0 | 0 | 0 | 0 | 0 | 10.0424498 | 1.34E-08 | 1 |
| MET | NM_001127500 | 0 | 0 | 0 | 0 | 0 | 0 | 6.840970326 | 10.52587575 | 1 | 0.02072721 |
| METTL1 | NM_005371 | 0 | 0 | 9.586630979 | 11.29270402 | 6.607150896 | 0 | 0 | 0 | 0.0016157 | 0.002835114 |
| METTL13 | XM_024447587 | 0 | 9.081113725 | 0 | 0 | 0 | 0 | 0 | 0 | 5.70E-08 | 1 |
| METTL17 | NM_001029991 | 0 | 0 | 0 | 0 | 6.381065478 | 0 | 0 | 0 | 4.71E-05 | 1 |
| METTL17 | XM_006720236 | 9.553433586 | 0 | 0 | 0 | 0 | 0 | 0 | 0 | 3.27E-08 | 1 |
| METTL21A | XM_024452722 | 0 | 9.554298882 | 0 | 0 | 0 | 0 | 0 | 0 | 2.73E-08 | 1 |
| METTL25 | XM_017020021 | 0 | 8.975521721 | 0 | 0 | 0 | 0 | 0 | 0 | 7.27E-08 | 1 |
| METTL26 | NM_001040161 | 0 | 0 | 9.439606601 | 10.87805938 | 0 | 0 | 7.960889765 | 0 | 0.012636617 | 0.999932046 |
| METTL26 | NM_001040162 | 0 | 0 | 0 | 9.324854511 | 7.689666995 | 0 | 0 | 0 | 0.035909283 | 0.025156042 |
| METTL26 | NR_109978 | 10.31228843 | 0 | 0 | 0 | 0 | 0 | 8.820441837 | 0 | 1.54E-08 | 1 |
| METTL2B | NM_018396 | 0 | 9.193024922 | 0 | 0 | 0 | 9.624558452 | 7.4411151 | 0 | 4.74E-08 | 0.02072721 |
| METTL3 | NM_019852 | 0 | 0 | 0 | 10.17970655 | 6.464964927 | 9.32675424 | 0 | 0 | 0.025821961 | 0.999932046 |
| METTL3 | XM_006720206 | 0 | 0 | 0 | 9.032696866 | 6.823901211 | 0 | 0 | 0 | 0.050889364 | 0.034829065 |
| METTL5 | NM_001293186 | 0 | 0 | 0 | 11.44874876 | 9.987199314 | 0 | 8.91010973 | 0 | 0.007477659 | 0.999932046 |
| METTL6 | NM_152396 | 0 | 0 | 0 | 0 | 6.366595318 | 0 | 0 | 0 | 4.76E-05 | 1 |
| METTL7B | NM_152637 | 0 | 9.303125566 | 0 | 0 | 0 | 0 | 0 | 0 | 4.01E-08 | 1 |
| METTL9 | XM_024450293 | 0 | 9.193024922 | 0 | 0 | 0 | 0 | 7.892131936 | 0 | 4.74E-08 | 1 |
| MEX3C | NM_016626 | 10.3248786 | 10.50807367 | 0 | 0 | 0 | 0 | 9.65627892 | 0 | 0.000492715 | 1 |
| MFAP1 | NM_005926 | 0 | 0 | 10.00962571 | 0 | 7.282748526 | 0 | 0 | 0 | 0.026792228 | 0.02072721 |
| MFAP4 | NM_002404 | 0 | 0 | 9.385892712 | 0 | 10.19802574 | 8.421828153 | 0 | 9.738579146 | 0.015929619 | 0.999932046 |
| MFF | NM_001277062 | 0 | 8.818611963 | 0 | 0 | 0 | 0 | 0 | 0 | 8.12E-08 | 1 |
| MFF | NM_001277065 | 0 | 0 | 0 | 9.79947801 | 10.14071475 | 0 | 0 | 0 | 0.01455307 | 0.02072721 |
| MFRP | NM_031433 | 0 | 0 | 10.69801997 | 8.064023368 | 10.64883105 | 0 | 7.310202764 | 0 | 0.001136564 | 1 |
| MFSD1 | NM_022736 | 8.497211815 | 0 | 11.38117829 | 0 | 7.267252374 | 0 | 0 | 0 | 0.93322348 | 9.62E-09 |
| MFSD10 | NM_001146069 | 0 | 0 | 10.54200001 | 0 | 8.002028225 | 0 | 0 | 0 | 0.018291955 | 0.02072721 |
| MFSD10 | XR_427465 | 0 | 0 | 0 | 9.792664417 | 7.768830875 | 0 | 0 | 0 | 0.027385771 | 0.020835214 |

| | | | | | | | | | | | |
|---|---|---|---|---|---|---|---|---|---|---|---|
| MFSD11 | NR_148242 | 0 | 0 | 0 | 9.044216539 | 7.282748526 | 0 | 0 | 0 | 0.046159167 | 0.031649306 |
| MFSD14A | NM_033055 | 0 | 0 | 0 | 10.48139854 | 9.022473864 | 0 | 8.647674983 | 0 | 0.015504134 | 0.999932046 |
| MFSD14B | XM_017015221 | 8.940634053 | 0 | 0 | 0 | 0 | 0 | 0 | 0 | 7.88E-08 | 1 |
| MFSD14B | XR_002956813 | 0 | 6.853934537 | 8.585434713 | 0 | 9.412370586 | 0 | 0 | 0 | 0.875696772 | 0.02072721 |
| MFSD2A | NM_032793 | 10.25642453 | 0 | 0 | 0 | 0 | 0 | 9.365556577 | 0 | 1.60E-08 | 1 |
| MFSD3 | XM_017013005 | 0 | 0 | 0 | 0 | 6.531336591 | 0 | 6.514885902 | 3.8795549 | 4.23E-05 | 0.999932046 |
| MFSD5 | NM_032889 | 0 | 0 | 0 | 0 | 0 | 9.523912909 | 8.257262546 | 0 | 1 | 0.02072721 |
| MFSD6 | XR_002959308 | 0 | 9.19973932 | 0 | 0 | 0 | 0 | 0 | 0 | 4.67E-08 | 1 |
| MFSD6L | NM_152599 | 10.00844432 | 0 | 0 | 0 | 0 | 0 | 0 | 6.860901615 | 1.93E-08 | 1 |
| MFSD9 | XM_011512007 | 9.643298854 | 0 | 0 | 0 | 0 | 0 | 0 | 0 | 2.88E-08 | 1 |
| MGA | XR_002957630 | 0 | 0 | 0 | 7.373286278 | 0 | 0 | 6.56874927 | 0 | 2.01E-05 | 1 |
| MGAT1 | NM_002406 | 0 | 0 | 0 | 0 | 0 | 0 | 8.732715858 | 8.563225531 | 1 | 0.020835214 |
| MGAT1 | XM_011534561 | 0 | 0 | 0 | 0 | 6.337212177 | 0 | 0 | 0 | 4.86E-05 | 1 |
| MGAT2 | NM_002408 | 8.940634053 | 9.044283149 | 8.769900534 | 9.720904456 | 7.251587968 | 0 | 0 | 0 | 0.999953634 | 0.008021236 |
| MGAT4B | XM_024454347 | 0 | 9.340061955 | 9.649744388 | 10.02551884 | 0 | 0 | 0 | 0 | 0.999953634 | 0.02096331 |
| MGST1 | NM_145792 | 11.34506661 | 8.747073305 | 9.346522316 | 0 | 7.24369152 | 0 | 0 | 0 | 0.836284875 | 0.025630112 |
| MGST2 | NM_001204367 | 9.944614124 | 0 | 0 | 0 | 0 | 0 | 8.798066067 | 0 | 2.01E-08 | 1 |
| MGST2 | XM_011531972 | 0 | 0 | 8.09164712 | 0 | 0 | 0 | 0 | 0 | 1.04E-05 | 1 |
| MIA2 | XM_024449592 | 8.497211815 | 0 | 0 | 0 | 0 | 0 | 0 | 0 | 1.15E-07 | 1 |
| MIAT | NR_033320 | 10.24321976 | 0 | 0 | 0 | 0 | 0 | 0 | 0 | 1.59E-08 | 1 |
| MICAL2 | NM_001282667 | 0 | 8.540433774 | 0 | 0 | 0 | 0 | 0 | 0 | 1.02E-07 | 1 |
| MICALL2 | NM_182924 | 0 | 0 | 8.859077794 | 0 | 8.614497302 | 0 | 0 | 0 | 0.034932228 | 0.023261296 |
| MID2 | NM_012216 | 0 | 0 | 0 | 0 | 0 | 10.72103295 | 0 | 9.985019915 | 1 | 0.02072721 |
| MIF4GD | XM_017024891 | 0 | 9.527862205 | 0 | 0 | 0 | 0 | 0 | 0 | 2.90E-08 | 1 |
| MIGA1 | XR_001737157 | 0 | 0 | 10.2213035 | 0 | 6.926116208 | 0 | 7.506976231 | 0 | 0.024736775 | 0.999932046 |
| MINDY1 | XM_024448340 | 0 | 9.559528578 | 0 | 0 | 0 | 0 | 0 | 0 | 2.70E-08 | 1 |
| MINDY3 | NM_024948 | 10.46796695 | 0 | 0 | 0 | 0 | 0 | 0 | 0 | 1.45E-08 | 1 |
| MINPP1 | NM_001178117 | 0 | 0 | 9.181401832 | 0 | 10.5764111 | 0 | 7.516144242 | 9.550800565 | 0.013993803 | 0.999932046 |
| MIPEPP3 | NR_046461 | 0 | 0 | 0 | 9.736965247 | 6.505153187 | 0 | 0 | 0 | 0.033812265 | 0.02430751 |
| MIR200CHG | NR_135032 | 8.972990281 | 8.592384875 | 9.235305439 | 0 | 7.746652379 | 0 | 0 | 0 | 0.999953634 | 0.024770715 |
| MIS12 | NM_001258218 | 9.95545054 | 0 | 0 | 0 | 0 | 0 | 0 | 0 | 2.01E-08 | 1 |
| MIS18A | NM_018944 | 0 | 0 | 0 | 10.0313189 | 5.928110332 | 9.070554946 | 8.838826814 | 0 | 0.029658808 | 1 |
| MISP | XM_011527686 | 11.14186693 | 9.296876505 | 0 | 8.119528586 | 9.338296057 | 0 | 0 | 0 | 0.917668892 | 0.022496982 |
| MISP3 | NM_001291291 | 0 | 9.014126367 | 9.233344514 | 0 | 7.555764066 | 0 | 0 | 0 | 0.999953634 | 0.025677608 |
| MITD1 | NM_001320418 | 0 | 0 | 11.26168168 | 0 | 8.864098664 | 0 | 9.181891152 | 0 | 0.010751756 | 0.999932046 |
| MITD1 | NM_138798 | 0 | 0 | 10.15862146 | 0 | 10.56576212 | 0 | 8.470359551 | 0 | 0.010839168 | 0.999932046 |
| MITF | NM_001354604 | 0 | 0 | 10.43430594 | 9.952508408 | 10.70670183 | 11.04279009 | 0 | 0 | 0.000725973 | 0.999932046 |
| MITF | NM_001354605 | 0 | 0 | 10.23214554 | 0 | 9.47103868 | 0 | 0 | 0 | 0.015674579 | 0.02072721 |
| MITF | NM_198158 | 0 | 10.6422867 | 0 | 0 | 0 | 11.13463829 | 0 | 0 | 1.34E-08 | 1 |
| MKL1 | XM_011530285 | 10.81296622 | 0 | 0 | 0 | 0 | 0 | 0 | 0 | 1.34E-08 | 1 |
| MKL2 | XM_017023502 | 10.54890019 | 0 | 0 | 0 | 0 | 0 | 0 | 0 | 1.36E-08 | 1 |
| MKRN1 | XM_011515997 | 0 | 0 | 11.11337353 | 0 | 8.136037099 | 0 | 8.91010973 | 0 | 0.013129332 | 0.999932046 |
| MLH1 | NM_001167617 | 0 | 0 | 0 | 10.83228042 | 8.486515572 | 11.13950599 | 11.14573387 | 11.23643606 | 0.014312157 | 0.999932046 |

| | | | | | | | | | | | |
|---|---|---|---|---|---|---|---|---|---|---|---|
| MLH1 | NM_001167619 | 10.10892695 | 0 | 10.81211718 | 11.32316629 | 9.940335027 | 0 | 0 | 0 | 0.979637466 | 0.001042882 |
| MLLT10 | NM_001195626 | 0 | 0 | 0 | 9.829747119 | 8.644794263 | 11.18494796 | 9.862629622 | 0 | 0.022498114 | 0.999932046 |
| MLLT10 | NM_001324297 | 0 | 0 | 11.74101807 | 12.13533789 | 0 | 0 | 10.95749782 | 11.62399393 | 0.003420003 | 0.999932046 |
| MLLT10 | XM_024448190 | 0 | 9.67966985 | 0 | 0 | 0 | 0 | 8.172825027 | 10.13688064 | 2.40E-08 | 0.02072721 |
| MLLT10 | XM_024448200 | 0 | 0 | 0 | 0 | 0 | 11.42314136 | 10.04866428 | 0 | 1 | 0.02072721 |
| MLLT10 | XM_024448202 | 0 | 0 | 0 | 11.65628617 | 9.423772545 | 11.34943104 | 11.31943915 | 10.74927864 | 0.00751387 | 0.999932046 |
| MLLT10 | XM_024448206 | 0 | 0 | 0 | 0 | 0 | 0 | 8.21847248 | 9.862810347 | 1 | 0.02072721 |
| MLLT3 | NM_004529 | 10.54172739 | 0 | 0 | 0 | 0 | 0 | 0 | 0 | 1.36E-08 | 1 |
| MLPH | NM_001042467 | 0 | 0 | 0 | 9.656688688 | 6.886094979 | 0 | 0 | 0 | 0.033928732 | 0.024302692 |
| MLPH | XM_006712740 | 0 | 8.919678654 | 0 | 0 | 0 | 0 | 0 | 0 | 7.73E-08 | 1 |
| MMEL1 | XM_017002315 | 0 | 0 | 0 | 0 | 0 | 0 | 7.130842066 | 10.28699654 | 1 | 0.02072721 |
| MMP14 | NM_004995 | 0 | 0 | 8.791384731 | 0 | 8.038998467 | 0 | 0 | 0 | 0.044222052 | 0.028340948 |
| MMP17 | NM_016155 | 10.06457365 | 0 | 0 | 0 | 0 | 0 | 0 | 0 | 1.80E-08 | 1 |
| MMP19 | XM_006719401 | 0 | 0 | 0 | 0 | 6.582319518 | 0 | 0 | 0 | 4.04E-05 | 1 |
| MMP19 | XM_017019309 | 0 | 8.911521649 | 0 | 0 | 0 | 0 | 0 | 0 | 7.73E-08 | 1 |
| MMP2 | NM_004530 | 0 | 7.544303366 | 0 | 9.081032624 | 11.6307037 | 0 | 0 | 0 | 0.623557501 | 0.02072721 |
| MMP24OS | NM_001355003 | 0 | 0 | 0 | 7.896185389 | 0 | 0 | 0 | 0 | 1.17E-05 | 1 |
| MMP24OS | NR_149148 | 8.815370067 | 0 | 0 | 0 | 0 | 0 | 8.871340655 | 0 | 8.84E-08 | 1 |
| MMP25 | XM_024450391 | 0 | 0 | 0 | 0 | 6.607150896 | 0 | 0 | 0 | 0.00011086 | 1 |
| MMP28 | XM_011525226 | 10.16602012 | 0 | 0 | 0 | 0 | 0 | 0 | 0 | 1.60E-08 | 1 |
| MMP28 | XM_017025064 | 0 | 10.18842011 | 0 | 0 | 0 | 0 | 0 | 0 | 1.52E-08 | 1 |
| MMUT | NM_000255 | 0 | 8.728620286 | 0 | 0 | 0 | 9.698988382 | 0 | 0 | 8.80E-08 | 1 |
| MNAT1 | XM_017021332 | 10.59467378 | 8.29916838 | 0 | 0 | 0 | 9.994611322 | 8.660134917 | 0 | 0.001207228 | 0.02072721 |
| MNAT1 | XM_017021334 | 0 | 8.441930184 | 0 | 9.257172648 | 9.111943945 | 0 | 0 | 0 | 0.999953634 | 0.02072721 |
| MOB1A | NM_001317110 | 0 | 9.396676749 | 0 | 0 | 0 | 0 | 7.460241356 | 0 | 3.36E-08 | 1 |
| MOB2 | NM_053005 | 10.67226231 | 0 | 0 | 0 | 0 | 0 | 0 | 0 | 1.34E-08 | 1 |
| MOB3B | NM_024761 | 0 | 8.348342131 | 0 | 0 | 0 | 0 | 0 | 0 | 1.24E-07 | 1 |
| MOCS2 | NM_176806 | 9.270677863 | 0 | 9.822733919 | 11.26528077 | 0 | 0 | 0 | 0 | 0.998900485 | 0.02072721 |
| MORC2 | XM_011530004 | 0 | 0 | 0 | 0 | 0 | 0 | 7.712369201 | 8.974688946 | 1 | 0.023084584 |
| MORC3 | NM_001320445 | 0 | 0 | 0 | 0 | 0 | 9.938460398 | 0 | 9.545513337 | 1 | 0.02072721 |
| MORC4 | XM_011531027 | 0 | 0 | 0 | 0 | 6.643614539 | 0 | 0 | 0 | 3.86E-05 | 1 |
| MORF4L2 | NM_001142424 | 0 | 0 | 0 | 10.19011422 | 8.334959336 | 0 | 0 | 0 | 0.020254269 | 0.02072721 |
| MORN2 | NM_001145450 | 10.43543963 | 0 | 0 | 9.575655325 | 11.15409815 | 0 | 0 | 0 | 0.999953634 | 0.02072721 |
| MORN4 | NM_178832 | 10.12341345 | 0 | 0 | 0 | 0 | 0 | 0 | 7.530034637 | 1.68E-08 | 1 |
| MOSPD1 | XM_017029654 | 0 | 0 | 9.963053877 | 9.796075236 | 7.812190501 | 0 | 0 | 0 | 0.002661621 | 0.004029843 |
| MOSPD3 | NM_001040097 | 0 | 8.632639499 | 0 | 0 | 0 | 0 | 0 | 0 | 9.44E-08 | 1 |
| MPC1 | XM_024446456 | 0 | 0 | 10.41020626 | 0 | 5.693189014 | 0 | 6.67079425 | 0 | 0.024620084 | 0.999932046 |
| MPDU1 | NM_004870 | 10.61189941 | 0 | 0 | 0 | 0 | 0 | 0 | 0 | 1.34E-08 | 1 |
| MPDU1 | XM_024451040 | 0 | 9.277965255 | 0 | 0 | 0 | 0 | 0 | 0 | 4.01E-08 | 1 |
| MPEG1 | NM_001039396 | 0 | 0 | 8.09164712 | 0 | 0 | 0 | 0 | 0 | 1.04E-05 | 1 |
| MPG | NM_001015052 | 0 | 0 | 9.732596818 | 0 | 10.49601404 | 0 | 0 | 0 | 0.012723719 | 0.02072721 |
| MPHOSPH6 | NM_005792 | 0 | 0 | 10.00733248 | 10.42082459 | 9.476102537 | 0 | 9.290068587 | 0 | 0.001187467 | 0.999932046 |
| MPND | NM_001300862 | 11.13711551 | 9.670031486 | 8.703458918 | 0 | 8.320016823 | 0 | 0 | 0 | 0.796420688 | 0.0268468 |

| | | | | | | | | | | | |
|---|---|---|---|---|---|---|---|---|---|---|---|
| MPP6 | XM_017012317 | 9.030488612 | 0 | 0 | 0 | 0 | 0 | 0 | 0 | 7.69E-08 | 1 |
| MPPE1 | XM_017025929 | 0 | 9.265218616 | 0 | 0 | 0 | 0 | 0 | 0 | 4.18E-08 | 1 |
| MPPED2 | XM_017018234 | 11.62167916 | 0 | 0 | 0 | 0 | 0 | 9.28739377 | 0 | 1.34E-08 | 1 |
| MPPED2 | XM_024448676 | 0 | 8.582143237 | 0 | 0 | 0 | 0 | 0 | 0 | 9.91E-08 | 1 |
| MPZL2 | NM_005797 | 0 | 8.878424457 | 9.520629739 | 0 | 8.803616231 | 0 | 0 | 0 | 0.999953634 | 0.02072721 |
| MPZL2 | NM_144765 | 0 | 9.490017613 | 0 | 10.42853688 | 10.12950993 | 0 | 0 | 0 | 0.999953634 | 0.02072721 |
| MPZL3 | XM_017017304 | 8.651779762 | 0 | 9.364551246 | 0 | 7.012300182 | 0 | 0 | 0 | 0.999953634 | 0.026342382 |
| MREG | XM_011511468 | 0 | 0 | 0 | 0 | 6.713888284 | 0 | 0 | 0 | 3.66E-05 | 1 |
| MRFAP1L1 | NM_203462 | 0 | 9.640723913 | 10.68516559 | 10.15335408 | 10.03602212 | 0 | 0 | 0 | 0.991101673 | 0.001616366 |
| MRGBP | NR_136405 | 0 | 0 | 0 | 8.650960001 | 0 | 10.5414403 | 0 | 0 | 5.26E-06 | 1 |
| MROH3P | NR_147176 | 0 | 0 | 6.942257105 | 0 | 0 | 0 | 0 | 0 | 8.99E-05 | 1 |
| MRPL10 | NM_145255 | 0 | 0 | 9.146399313 | 11.34078935 | 9.779647935 | 0 | 0 | 10.50287287 | 0.000844449 | 0.999932046 |
| MRPL14 | NM_001318768 | 0 | 0 | 0 | 10.01384828 | 8.410964939 | 0 | 8.007141921 | 0 | 0.021739874 | 0.999932046 |
| MRPL20-AS1 | NR_015434 | 0 | 0 | 0 | 10.04858048 | 7.026176606 | 0 | 0 | 0 | 0.026325448 | 0.020730817 |
| MRPL21 | NM_181515 | 0 | 0 | 7.695565578 | 10.07127927 | 9.295013967 | 0 | 0 | 0 | 0.003109711 | 0.00475718 |
| MRPL22 | NM_001014990 | 0 | 0 | 10.0514059 | 0 | 8.63275172 | 0 | 0 | 0 | 0.020742069 | 0.02072721 |
| MRPL28 | NM_006428 | 10.54890019 | 0 | 9.792421912 | 11.46705217 | 7.790673576 | 0 | 0 | 0 | 0.999953634 | 0.002129361 |
| MRPL28 | XM_011522351 | 0 | 9.232850423 | 0 | 0 | 0 | 0 | 0 | 0 | 4.38E-08 | 1 |
| MRPL34 | NM_023937 | 0 | 0 | 10.31791951 | 0 | 7.987916427 | 0 | 0 | 0 | 0.020527359 | 0.02072721 |
| MRPL37 | NM_016491 | 8.704119069 | 10.35106274 | 11.35165437 | 10.36335493 | 10.39179385 | 0 | 0 | 0 | 0.999953634 | 0.001042882 |
| MRPL40 | NM_003776 | 0 | 0 | 0 | 10.22848764 | 10.55941433 | 0 | 9.347693707 | 9.262600397 | 0.01064748 | 0.999932046 |
| MRPL42 | NR_038160 | 9.643298854 | 9.102769039 | 0 | 0 | 0 | 0 | 10.00129867 | 10.19544179 | 0.001912091 | 0.02072721 |
| MRPL43 | NM_176793 | 0 | 8.662110092 | 0 | 0 | 0 | 0 | 0 | 0 | 9.15E-08 | 1 |
| MRPL46 | NM_022163 | 0 | 0 | 0 | 11.40082806 | 7.964085617 | 0 | 10.22884393 | 0 | 0.010999766 | 0.999932046 |
| MRPL47 | NM_177988 | 0 | 0 | 0 | 9.996163509 | 9.800656081 | 0 | 8.456139264 | 0 | 0.015398614 | 1 |
| MRPL49 | NM_004927 | 9.426779986 | 0 | 0 | 10.18231551 | 7.490532315 | 0 | 0 | 0 | 0.999953634 | 0.02072721 |
| MRPL52 | NM_181306 | 0 | 0 | 0 | 9.429583326 | 7.925118111 | 0 | 8.07695445 | 0 | 0.032258822 | 0.999932046 |
| MRPL52 | NM_181307 | 0 | 0 | 10.06359724 | 0 | 7.473752874 | 0 | 0 | 0 | 0.025281678 | 0.02072721 |
| MRPL55 | NM_181463 | 0 | 0 | 0 | 8.75977034 | 0 | 0 | 0 | 0 | 4.66E-06 | 1 |
| MRPL55 | XM_005273059 | 0 | 0 | 0 | 10.03709574 | 7.636381755 | 0 | 0 | 0 | 0.024505355 | 0.02072721 |
| MRPL55 | XR_002959482 | 0 | 0 | 0 | 0 | 6.61940813 | 0 | 0 | 0 | 5.85E-05 | 1 |
| MRPL57 | XM_017020740 | 10.54531825 | 0 | 0 | 0 | 0 | 7.920414269 | 8.947864183 | 0 | 1.36E-08 | 0.021614814 |
| MRPL58 | NM_001545 | 0 | 0 | 11.51673943 | 9.99022008 | 7.95444197 | 0 | 0 | 10.9616612 | 0.000973622 | 0.999932046 |
| MRPL58 | XM_017024574 | 9.872121021 | 0 | 0 | 0 | 0 | 0 | 0 | 0 | 2.16E-08 | 1 |
| MRPL9 | NM_031420 | 0 | 0 | 0 | 12.48540289 | 9.604200436 | 10.64262684 | 10.71631569 | 11.84183488 | 1.34E-08 | 0.999932046 |
| MRPL9 | NR_125331 | 10.02397016 | 0 | 0 | 0 | 0 | 0 | 6.56874927 | 10.82868874 | 1.88E-08 | 0.02072721 |
| MRPS10 | NM_018141 | 0 | 0 | 0 | 11.0131504 | 9.796613792 | 0 | 0 | 12.27459253 | 0.009922086 | 0.999932046 |
| MRPS11 | NR_135918 | 10.43350322 | 0 | 0 | 0 | 0 | 0 | 0 | 0 | 1.47E-08 | 1 |
| MRPS12 | NM_021107 | 0 | 0 | 10.33630716 | 0 | 7.707000189 | 0 | 0 | 0 | 0.021142039 | 0.02072721 |
| MRPS17 | NM_015969 | 0 | 0 | 10.38304631 | 11.28118883 | 9.72468557 | 0 | 9.068060205 | 9.319465356 | 0.000596493 | 0.999932046 |
| MRPS18B | NM_014046 | 0 | 0 | 0 | 10.01384828 | 9.966958188 | 10.04347378 | 9.738872106 | 10.61438614 | 0.014463966 | 0.999932046 |
| MRPS18B | XM_024446408 | 0 | 0 | 9.601910322 | 10.38503365 | 0 | 0 | 6.458933278 | 0 | 0.015309434 | 0.999932046 |
| MRPS25 | NM_022497 | 0 | 8.571828374 | 0 | 0 | 0 | 0 | 0 | 0 | 9.97E-08 | 1 |

| | | | | | | | | | | | |
|---|---|---|---|---|---|---|---|---|---|---|---|
| MRPS30 | NM_016640 | 0 | 0 | 10.68516559 | 0 | 8.262562196 | 0 | 0 | 0 | 0.016337878 | 0.02072721 |
| MRPS35 | NM_001190864 | 10.53090088 | 0 | 0 | 0 | 0 | 0 | 0 | 0 | 1.37E-08 | 1 |
| MRPS5 | NM_001321996 | 0 | 8.709928179 | 0 | 0 | 0 | 0 | 0 | 0 | 8.96E-08 | 1 |
| MRPS9 | XR_001738903 | 10.78893422 | 0 | 0 | 0 | 0 | 0 | 0 | 0 | 1.34E-08 | 1 |
| MRS2 | NM_001286265 | 0 | 8.642530071 | 0 | 0 | 0 | 0 | 0 | 0 | 9.87E-08 | 1 |
| MRS2 | NM_020662 | 0 | 10.19515592 | 0 | 0 | 0 | 0 | 7.899157068 | 11.33549429 | 1.51E-08 | 0.02072721 |
| MRTFA | NM_001318139 | 0 | 11.72331328 | 0 | 0 | 0 | 0 | 0 | 10.64807716 | 1.34E-08 | 1 |
| MRTO4 | NM_016183 | 9.609394475 | 0 | 10.49038967 | 10.66515103 | 8.268411723 | 0 | 0 | 0 | 0.999953634 | 0.002294967 |
| MS4A1 | NM_152866 | 0 | 0 | 10.09633208 | 7.933830793 | 0 | 0 | 0 | 10.5312353 | 0.024847467 | 0.999932046 |
| MS4A7 | NM_206940 | 0 | 0 | 9.260557857 | 0 | 7.939854543 | 0 | 0 | 0 | 0.035854182 | 0.02352709 |
| MSANTD2 | XM_017018298 | 8.704119069 | 0 | 0 | 0 | 0 | 0 | 0 | 0 | 9.60E-08 | 1 |
| MSH2 | XM_005264332 | 0 | 0 | 0 | 10.03421021 | 8.641793038 | 0 | 9.535238551 | 0 | 0.020568316 | 0.999932046 |
| MSH5-SAPCD1 | NR_037846 | 0 | 8.861586703 | 0 | 0 | 0 | 0 | 0 | 0 | 7.86E-08 | 1 |
| MSI2 | NM_001322250 | 0 | 0 | 0 | 10.4845729 | 9.969354289 | 0 | 9.368090456 | 11.75745987 | 0.012054145 | 0.999932046 |
| MSI2 | NM_001322251 | 0 | 0 | 0 | 11.63263454 | 11.25255559 | 12.46929009 | 12.25643002 | 13.37176069 | 0.004578156 | 0.999932046 |
| MSI2 | XM_011524286 | 0 | 0 | 0 | 10.14133659 | 7.170608406 | 0 | 0 | 0 | 0.024563463 | 0.02072721 |
| MSLN | NM_013404 | 0 | 0 | 0 | 0 | 0 | 0 | 7.621843212 | 11.9726701 | 1 | 1.92E-09 |
| MSMB | NM_002443 | 0 | 0 | 0 | 7.585264412 | 9.478627815 | 0 | 8.484441039 | 0 | 0.032955018 | 0.999932046 |
| MSN | NM_002444 | 0 | 9.364171095 | 0 | 0 | 0 | 0 | 0 | 0 | 3.61E-08 | 1 |
| MSN | XM_017029546 | 0 | 0 | 9.88276327 | 0 | 7.768830875 | 0 | 0 | 0 | 0.026665054 | 0.02072721 |
| MSRB3 | XM_024448921 | 0 | 0 | 0 | 7.647009032 | 0 | 0 | 0 | 0 | 1.52E-05 | 1 |
| MST1R | NR_134919 | 0 | 9.321711965 | 0 | 0 | 0 | 0 | 0 | 0 | 3.84E-08 | 1 |
| MT1A | NM_005946 | 0 | 9.006487637 | 0 | 0 | 0 | 0 | 0 | 0 | 6.80E-08 | 1 |
| MT1DP | NR_027781 | 0 | 0 | 9.331935177 | 0 | 6.213302533 | 0 | 0 | 0 | 0.047356815 | 0.030063529 |
| MTA1 | XM_017021759 | 0 | 0 | 0 | 0 | 6.643614539 | 0 | 0 | 0 | 3.86E-05 | 1 |
| MTA3 | NM_020744 | 0 | 8.29916838 | 10.45212013 | 10.13730841 | 0 | 0 | 0 | 0 | 0.93155113 | 0.02072721 |
| MTCH1 | NM_001271641 | 0 | 0 | 11.60961651 | 7.817812962 | 9.545191248 | 0 | 0 | 10.31833588 | 0.001080108 | 0.999932046 |
| MTCH1 | NR_130739 | 0 | 0 | 0 | 0 | 6.582319518 | 0 | 0 | 0 | 4.04E-05 | 1 |
| MTCH1 | XM_005248979 | 0 | 8.818611963 | 0 | 0 | 0 | 9.725129234 | 8.21847248 | 0 | 8.12E-08 | 0.02072721 |
| MTCH2 | NM_001317232 | 0 | 9.479019994 | 10.61611064 | 0 | 7.320775981 | 0 | 0 | 0 | 0.999953634 | 0.02072721 |
| MTERF4 | XM_017003380 | 0 | 9.511764498 | 0 | 0 | 0 | 0 | 0 | 0 | 2.90E-08 | 1 |
| MTF2 | XM_011541019 | 0 | 9.511764498 | 0 | 0 | 0 | 0 | 0 | 0 | 2.90E-08 | 1 |
| MTF2 | XM_017000677 | 0 | 10.43597468 | 0 | 0 | 0 | 0 | 0 | 0 | 1.34E-08 | 1 |
| MTFR2 | NM_138419 | 0 | 0 | 0 | 9.792664417 | 6.955419991 | 0 | 0 | 0 | 0.030907891 | 0.022511788 |
| MTHFD1L | NM_001242767 | 0 | 0 | 0 | 11.78973404 | 11.57003027 | 12.50013161 | 11.02729161 | 13.49584803 | 0.00377125 | 0.999932046 |
| MTHFD1L | NM_001242768 | 0 | 0 | 0 | 0 | 0 | 12.61947835 | 9.427605546 | 11.32241536 | 1 | 0.000286686 |
| MTHFD1L | XM_017010703 | 0 | 0 | 0 | 9.768560191 | 9.952497401 | 0 | 7.310202764 | 0 | 0.01576347 | 0.999932046 |
| MTHFD1L | XR_002956274 | 0 | 0 | 0 | 10.2030191 | 10.36201305 | 10.53591292 | 8.436958184 | 6.062825694 | 0.011657592 | 0.999932046 |
| MTHFD2 | NM_006636 | 0 | 0 | 0 | 11.43130726 | 8.07504494 | 0 | 8.026519178 | 11.74540495 | 0.010645402 | 0.999932046 |
| MTHFR | XM_005263460 | 0 | 9.138154184 | 0 | 0 | 0 | 0 | 0 | 0 | 5.16E-08 | 1 |
| MTHFR | XR_002956640 | 8.838969261 | 0 | 0 | 0 | 0 | 0 | 0 | 0 | 8.56E-08 | 1 |
| MTIF3 | NM_152912 | 0 | 0 | 11.7382551 | 12.37797878 | 10.45342876 | 0 | 8.229662188 | 11.62712583 | 0.000138355 | 0.999932046 |
| MTMR11 | NM_001145862 | 9.708807013 | 0 | 0 | 0 | 0 | 0 | 0 | 0 | 2.61E-08 | 1 |

| | | | | | | | | | | | |
|---|---|---|---|---|---|---|---|---|---|---|---|
| MTMR11 | XR_426760 | 0 | 10.4416505 | 0 | 0 | 0 | 3.39210272 | 8.724829293 | 0 | 1.34E-08 | 1 |
| MTMR12 | NM_001040446 | 0 | 0 | 10.78760486 | 0 | 10.82056518 | 0 | 10.74486803 | 10.77886808 | 0.007769725 | 0.999932046 |
| MTMR12 | XM_005248313 | 0 | 9.17269217 | 10.56790721 | 0 | 10.10520251 | 0 | 0 | 0 | 0.999953634 | 0.02072721 |
| MTMR12 | XM_017009589 | 0 | 9.456770012 | 0 | 0 | 0 | 0 | 0 | 0 | 3.14E-08 | 1 |
| MTMR14 | XM_017007045 | 0 | 0 | 0 | 10.43730075 | 9.08125658 | 0 | 8.60104519 | 0 | 0.015637687 | 0.999932046 |
| MTMR2 | NM_001243571 | 0 | 0 | 0 | 0 | 0 | 10.03827044 | 8.862382289 | 0 | 1 | 0.02072721 |
| MTMR2 | NM_016156 | 0 | 0 | 10.88137302 | 9.752849208 | 9.998388054 | 12.98291667 | 10.76038246 | 9.784185253 | 0.000860948 | 0.999932046 |
| MTMR4 | XM_005257784 | 9.162606966 | 8.540433774 | 0 | 0 | 0 | 0 | 0 | 0 | 0.003799591 | 1 |
| MTOR | NM_004958 | 0 | 0 | 10.80355212 | 0 | 6.155555496 | 0 | 8.377835658 | 10.86628262 | 0.018948466 | 0.999932046 |
| MTPN | NM_145808 | 6.813078996 | 0 | 0 | 0 | 0 | 8.717096959 | 6.932836632 | 0 | 0.118377845 | 0.030915082 |
| MTR | XM_017001330 | 0 | 0 | 0 | 0 | 6.655567121 | 0 | 0 | 0 | 3.82E-05 | 1 |
| MTRF1 | XM_017020875 | 0 | 9.08836834 | 0 | 0 | 0 | 0 | 0 | 0 | 5.61E-08 | 1 |
| MTRF1L | NR_126056 | 0 | 9.124104101 | 0 | 0 | 0 | 0 | 0 | 0 | 5.29E-08 | 1 |
| MTRNR2L6 | NM_001190487 | 9.656640609 | 0 | 0 | 0 | 0 | 0 | 0 | 0 | 3.16E-08 | 1 |
| MTSS1 | NM_014751 | 9.928205339 | 0 | 9.776350455 | 0 | 7.74105407 | 0 | 0 | 0 | 0.999953634 | 0.02072721 |
| MTURN | NM_152793 | 0 | 0 | 10.31421369 | 0 | 7.968883375 | 0 | 7.516144242 | 0 | 0.020611584 | 0.999932046 |
| MTX2 | NM_001006635 | 0 | 0 | 9.99465414 | 0 | 7.854284905 | 0 | 0 | 0 | 0.024736775 | 0.02072721 |
| MUC12 | NM_001164462 | 11.81851281 | 13.06833255 | 11.87597164 | 10.79696696 | 0 | 0 | 0 | 0 | 0.993233175 | 0.020604072 |
| MUC2 | NM_002457 | 10.47551503 | 11.48164124 | 12.01942689 | 10.59050989 | 0 | 0 | 0 | 0 | 0.999953634 | 0.019987819 |
| MUC4 | NM_001322468 | 11.71550385 | 8.407542174 | 10.95936823 | 0 | 7.058045127 | 0 | 0 | 0 | 0.999953634 | 0.02072721 |
| MUC5B | NM_002458 | 0 | 11.2220096 | 10.17199201 | 0 | 7.049011441 | 0 | 0 | 0 | 0.999953634 | 0.02072721 |
| MUL1 | NM_024544 | 0 | 9.044283149 | 10.5483234 | 9.206807527 | 0 | 0 | 0 | 0 | 0.999953634 | 0.02072721 |
| MUSTN1 | NM_205853 | 0 | 0 | 10.16377865 | 0 | 6.679179249 | 0 | 0 | 0 | 0.026283267 | 0.02072721 |
| MXD1 | NM_002357 | 11.25911841 | 0 | 10.02330906 | 0 | 8.025245409 | 0 | 0 | 0 | 0.999953634 | 0.02072721 |
| MXD3 | NM_031300 | 0 | 0 | 0 | 8.82800322 | 8.811650533 | 0 | 0 | 0 | 0.032449977 | 0.02296717 |
| MXD4 | XR_427467 | 0 | 8.671801345 | 0 | 0 | 0 | 0 | 0 | 0 | 9.15E-08 | 1 |
| MXI1 | NM_005962 | 0 | 0 | 0 | 8.628177296 | 0 | 0 | 0 | 0 | 6.35E-06 | 1 |
| MYADM | NM_138373 | 0 | 8.323964754 | 0 | 0 | 0 | 0 | 6.869864959 | 0 | 1.27E-07 | 1 |
| MYB | NM_001161656 | 0 | 0 | 0 | 10.48879454 | 9.031674724 | 8.21602433 | 9.422738661 | 0 | 0.015422143 | 0.999932046 |
| MYB | NR_134958 | 0 | 0 | 0 | 11.19596625 | 7.223759215 | 9.829723478 | 7.812509721 | 0 | 0.013504702 | 0.999932046 |
| MYB | NR_134959 | 0 | 0 | 9.148481961 | 0 | 8.635771793 | 0 | 0 | 0 | 0.030738053 | 0.021157255 |
| MYCBP | NM_012333 | 9.734199496 | 9.958336465 | 10.09848822 | 11.36453125 | 10.7491537 | 0 | 0 | 0 | 0.999953634 | 0.001024038 |
| MYD88 | NM_002468 | 0 | 0 | 9.939191783 | 0 | 8.861521052 | 0 | 0 | 0 | 0.020681047 | 0.02072721 |
| MYL6B | NM_002475 | 0 | 0 | 10.49857649 | 0 | 9.090091207 | 0 | 7.728235151 | 0 | 0.015381692 | 0.999932046 |
| MYL9 | NM_181526 | 0 | 0 | 0 | 9.733411602 | 6.42362501 | 0 | 0 | 0 | 0.034200079 | 0.024533936 |
| MYLK | NM_053026 | 0 | 0 | 11.32197501 | 9.061325395 | 8.620607807 | 0 | 0 | 9.597533386 | 0.001267779 | 1 |
| MYLK | XM_017006472 | 0 | 0 | 0 | 0 | 6.725274845 | 0 | 0 | 0 | 9.53E-05 | 1 |
| MYNN | XM_017006868 | 0 | 8.360378074 | 0 | 0 | 0 | 0 | 0 | 0 | 1.21E-07 | 1 |
| MYO10 | XM_011514046 | 0 | 0 | 0 | 7.20900477 | 0 | 0 | 0 | 0 | 2.36E-05 | 1 |
| MYO1A | NM_005379 | 11.38569319 | 9.451153482 | 0 | 0 | 0 | 0 | 5.417697669 | 0 | 0.000378687 | 1 |
| MYO1D | NM_001303279 | 0 | 10.06330846 | 0 | 10.13865239 | 9.291188213 | 0 | 0 | 0 | 0.999953634 | 0.02072721 |
| MYO1E | XM_024449932 | 8.742163956 | 0 | 0 | 0 | 0 | 0 | 0 | 0 | 9.27E-08 | 1 |
| MYO1G | XM_017012504 | 0 | 0 | 0 | 0 | 6.518304288 | 0 | 0 | 0 | 4.23E-05 | 1 |

| | | | | | | | | | | | |
|---|---|---|---|---|---|---|---|---|---|---|---|
| MYO5B | NM_001080467 | 11.27542589 | 7.585902702 | 0 | 11.37197831 | 9.032820716 | 0 | 0 | 0 | 0.999953634 | 0.02072721 |
| MYO6 | XM_005248726 | 0 | 0 | 9.936783722 | 9.583571881 | 12.43076296 | 11.05700998 | 9.089715157 | 12.61968306 | 0.000330273 | 0.999932046 |
| MYO7B | XR_001738748 | 0 | 0 | 9.88276327 | 0 | 5.783102534 | 0 | 0 | 0 | 0.034150253 | 0.022766316 |
| MYOF | NM_133337 | 0 | 8.951853153 | 0 | 0 | 0 | 0 | 0 | 0 | 7.60E-08 | 1 |
| MYOM1 | XM_024451281 | 9.944614124 | 0 | 0 | 0 | 0 | 0 | 0 | 0 | 2.01E-08 | 1 |
| MYOZ2 | NM_016599 | 11.0129402 | 0 | 0 | 0 | 0 | 0 | 0 | 0 | 1.34E-08 | 1 |
| MYRIP | NR_104316 | 0 | 10.67156286 | 0 | 0 | 0 | 0 | 11.4242083 | 0 | 1.34E-08 | 1 |
| MYRIP | XM_011533575 | 0 | 0 | 0 | 8.483365391 | 0 | 10.90379525 | 10.94821383 | 0 | 2.83E-05 | 0.999932046 |
| MYZAP | NM_152451 | 0 | 0 | 6.559575832 | 0 | 0 | 0 | 0 | 0 | 4.56E-05 | 1 |
| MZT2A | NM_001085365 | 0 | 0 | 0 | 0 | 0 | 0 | 8.62242756 | 10.18187502 | 1 | 0.02072721 |
| N4BP1 | XR_001752040 | 0 | 0 | 0 | 10.41861348 | 9.571453112 | 0 | 7.534307311 | 0 | 0.013971626 | 0.999932046 |
| N4BP2L2 | XM_017020349 | 0 | 0 | 0 | 11.54209677 | 7.895185835 | 0 | 10.34915352 | 0 | 0.010153174 | 0.999932046 |
| N4BP3 | XM_006714834 | 0 | 8.975521721 | 0 | 0 | 0 | 0 | 0 | 0 | 7.27E-08 | 1 |
| NAA20 | NM_181527 | 10.15195724 | 8.323964754 | 0 | 0 | 0 | 0 | 0 | 0 | 0.001810461 | 1 |
| NAA25 | XR_243023 | 0 | 0 | 6.596658308 | 0 | 0 | 0 | 0 | 0 | 4.50E-05 | 1 |
| NAA38 | NM_032356 | 9.872121021 | 6.887591794 | 0 | 0 | 0 | 0 | 7.920029518 | 8.8168733 | 0.003716755 | 0.023358895 |
| NAB1 | XM_024452917 | 9.616239393 | 0 | 0 | 0 | 0 | 0 | 0 | 0 | 2.91E-08 | 1 |
| NABP2 | XM_005269147 | 0 | 0 | 0 | 0 | 0 | 9.964753517 | 0 | 10.31678498 | 1 | 0.02072721 |
| NACA | NM_001113203 | 0 | 0 | 9.81094843 | 0 | 8.729220141 | 0 | 9.752512644 | 0 | 0.022559014 | 0.999932046 |
| NADK | XM_017002146 | 0 | 9.635780908 | 0 | 0 | 0 | 0 | 0 | 0 | 2.48E-08 | 1 |
| NANS | XM_011518787 | 0 | 9.239382419 | 0 | 0 | 0 | 0 | 0 | 0 | 4.33E-08 | 1 |
| NANS | XM_024447574 | 11.44071437 | 0 | 10.15551827 | 0 | 7.718441029 | 0 | 0 | 0 | 0.999953634 | 0.02072721 |
| NAP1L1 | NM_139207 | 0 | 0 | 0 | 10.28298438 | 8.709289252 | 0 | 9.590532652 | 0 | 0.018141319 | 0.999932046 |
| NAP1L1 | XM_017019339 | 0 | 0 | 0 | 8.414023513 | 0 | 0 | 0 | 0 | 3.17E-05 | 1 |
| NAPA | NR_038456 | 0 | 0 | 0 | 0 | 0 | 11.28828285 | 9.020532381 | 0 | 1 | 0.02072721 |
| NAPA | XR_002958377 | 10.22989302 | 0 | 0 | 0 | 0 | 0 | 0 | 9.681877826 | 1.59E-08 | 1 |
| NAPG | XM_017026063 | 0 | 0 | 0 | 0 | 6.478485785 | 0 | 7.341484785 | 0 | 4.36E-05 | 1 |
| NAPRT | XM_017013976 | 0 | 0 | 0 | 0 | 6.725274845 | 0 | 8.268156521 | 10.06479657 | 9.53E-05 | 0.999932046 |
| NARF | NM_012336 | 0 | 0 | 10.62061854 | 0 | 9.398212561 | 11.97358566 | 8.736643032 | 0 | 0.013497063 | 0.999932046 |
| NARS2 | XM_011545253 | 0 | 0 | 0 | 7.933830793 | 7.879983547 | 0 | 0 | 0 | 0.065368181 | 0.045569868 |
| NARS2 | XR_001747964 | 0 | 8.975521721 | 10.59488547 | 0 | 9.405308941 | 0 | 0 | 0 | 0.999953634 | 0.02072721 |
| NAT10 | NM_024662 | 0 | 8.29916838 | 0 | 10.28055141 | 8.790125492 | 0 | 0 | 0 | 0.999953634 | 0.02072721 |
| NAT2 | NM_000015 | 0 | 8.903318262 | 0 | 0 | 0 | 0 | 0 | 0 | 7.75E-08 | 1 |
| NAT2 | XM_017012938 | 0 | 9.445515002 | 0 | 0 | 0 | 0 | 0 | 0 | 3.16E-08 | 1 |
| NAT9 | XR_934438 | 0 | 9.99739751 | 0 | 0 | 0 | 0 | 0 | 0 | 1.68E-08 | 1 |
| NAXD | NM_001242882 | 9.721559119 | 0 | 0 | 0 | 0 | 0 | 0 | 0 | 2.60E-08 | 1 |
| NAXD | NR_040104 | 0 | 0 | 0 | 9.044216539 | 8.349748669 | 0 | 8.609636205 | 0 | 0.034845362 | 0.999932046 |
| NBL1 | NM_005380 | 0 | 0 | 0 | 9.987239158 | 9.433350468 | 0 | 8.767681412 | 9.904707713 | 0.017313785 | 0.999932046 |
| NBPF14 | NM_015383 | 10.12821014 | 8.550974732 | 0 | 0 | 0 | 0 | 8.782953729 | 9.908831299 | 0.001691733 | 0.02072721 |
| NBPF19 | NM_001351365 | 11.27109522 | 0 | 0 | 0 | 0 | 0 | 0 | 0 | 1.34E-08 | 1 |
| NBR1 | XM_024450750 | 0 | 0 | 9.867672463 | 10.26340474 | 9.613421891 | 0 | 9.736912887 | 0 | 0.00129096 | 1 |
| NCALD | NM_001040627 | 0 | 0 | 12.02368789 | 0 | 8.982702045 | 11.60526544 | 0 | 0 | 0.006560853 | 0.999932046 |
| NCALD | NM_001040628 | 0 | 9.585396421 | 0 | 0 | 0 | 0 | 7.782430925 | 0 | 2.61E-08 | 1 |

| | | | | | | | | | | | |
|---|---|---|---|---|---|---|---|---|---|---|---|
| NCALD | NM_001040629 | 10.28247691 | 0 | 0 | 0 | 0 | 0 | 0 | 0 | 1.54E-08 | 1 |
| NCALD | XM_024447299 | 10.41006135 | 9.290600259 | 0 | 0 | 0 | 0 | 10.46494025 | 0 | 0.000958476 | 1 |
| NCALD | XM_024447301 | 0 | 0 | 10.80090645 | 0 | 8.304917924 | 0 | 0 | 0 | 0.015292503 | 0.02072721 |
| NCAM2 | NM_004540 | 0 | 9.226288718 | 9.931955507 | 0 | 9.123823339 | 0 | 0 | 0 | 0.999953634 | 0.02072721 |
| NCAPG2 | XM_011516362 | 0 | 0 | 0 | 0 | 6.3222931 | 0 | 0 | 0 | 0.000244538 | 1 |
| NCAPH | XM_006712388 | 0 | 9.490017613 | 0 | 0 | 0 | 0 | 0 | 0 | 3.40E-08 | 1 |
| NCAPH2 | XR_001755232 | 0 | 0 | 0 | 0 | 6.557053251 | 0 | 0 | 0 | 4.16E-05 | 1 |
| NCBP1 | XR_001746305 | 0 | 0 | 10.68803203 | 0 | 8.509747759 | 0 | 0 | 0 | 0.015680082 | 0.02072721 |
| NCBP3 | NM_001114118 | 0 | 8.31161984 | 0 | 0 | 0 | 0 | 0 | 0 | 1.27E-07 | 1 |
| NCKAP1 | XM_006712201 | 10.02397016 | 0 | 9.040481197 | 0 | 9.207396167 | 0 | 0 | 0 | 0.999953634 | 0.02072721 |
| NCKAP5 | NM_207481 | 10.39819601 | 9.554298882 | 0 | 0 | 0 | 0 | 0 | 0 | 0.000844449 | 1 |
| NCKAP5 | XM_005263659 | 11.32012993 | 0 | 0 | 0 | 0 | 0 | 0 | 0 | 1.34E-08 | 1 |
| NCKAP5 | XM_011511102 | 0 | 9.708204508 | 0 | 0 | 0 | 0 | 0 | 0 | 2.47E-08 | 1 |
| NCKAP5 | XM_017003975 | 11.37967166 | 0 | 0 | 0 | 0 | 0 | 0 | 0 | 1.34E-08 | 1 |
| NCOA1 | XM_017005168 | 8.9622853 | 6.633619279 | 10.11349163 | 8.193814277 | 0 | 0 | 0 | 0 | 0.999953634 | 0.02072721 |
| NCOA2 | NM_001321707 | 10.63903963 | 0 | 0 | 0 | 0 | 0 | 9.143951136 | 10.46836455 | 1.34E-08 | 0.02072721 |
| NCOA2 | NM_006540 | 0 | 0 | 12.22237541 | 9.926289561 | 5.027158347 | 9.399575006 | 10.52974653 | 10.04993725 | 0.001007891 | 0.999932046 |
| NCOA2 | XM_017012968 | 0 | 9.117027412 | 0 | 0 | 0 | 0 | 10.51724439 | 0 | 5.36E-08 | 1 |
| NCOA4 | NM_001145262 | 0 | 0 | 11.30246768 | 9.932501706 | 9.572241512 | 0 | 9.519368714 | 10.06479657 | 0.000742818 | 0.999932046 |
| NCOA5 | NM_020967 | 0 | 8.853093555 | 0 | 0 | 0 | 0 | 0 | 0 | 7.91E-08 | 1 |
| NCOA7 | NM_001199622 | 0 | 0 | 10.13885423 | 0 | 7.365124102 | 0 | 0 | 0 | 0.024623293 | 0.02072721 |
| NCOR1 | XM_005256875 | 0 | 9.585396421 | 0 | 0 | 0 | 0 | 0 | 0 | 2.61E-08 | 1 |
| NDEL1 | NM_030808 | 9.807800698 | 0 | 8.97157385 | 10.63118044 | 0 | 0 | 0 | 0 | 0.999953634 | 0.02072721 |
| NDFIP1 | NM_030571 | 0 | 0 | 9.725638837 | 11.36567946 | 9.703252693 | 10.98040225 | 7.870848924 | 0 | 0.000719561 | 0.999932046 |
| NDN | NM_002487 | 0 | 10.10145643 | 0 | 0 | 0 | 0 | 0 | 0 | 2.36E-08 | 1 |
| NDRG2 | NM_201538 | 0 | 9.252358352 | 0 | 0 | 0 | 0 | 0 | 0 | 4.28E-08 | 1 |
| NDST1 | XM_017009433 | 0 | 10.25762482 | 0 | 0 | 0 | 0 | 0 | 0 | 1.47E-08 | 1 |
| NDUFA10 | NM_001322019 | 0 | 0 | 0 | 9.761598646 | 9.754166532 | 11.63443771 | 9.620596044 | 10.39529956 | 0.016955028 | 0.999932046 |
| NDUFA11 | NR_034166 | 0 | 0 | 10.24193172 | 0 | 9.315876167 | 0 | 0 | 0 | 0.016289345 | 0.020808337 |
| NDUFA2 | NR_033697 | 9.998000145 | 0 | 0 | 9.802872777 | 9.008561607 | 0 | 0 | 0 | 0.999953634 | 0.02072721 |
| NDUFA3 | NM_004542 | 0 | 0 | 8.943062173 | 0 | 8.114548983 | 9.016014309 | 0 | 0 | 0.040017754 | 0.999932046 |
| NDUFAB1 | XM_011545856 | 10.03422867 | 9.206422614 | 0 | 0 | 0 | 0 | 8.020088967 | 0 | 0.001347692 | 1 |
| NDUFAF4 | NM_014165 | 0 | 0 | 0 | 12.1470714 | 10.26196942 | 0 | 10.31410223 | 0 | 0.004681095 | 0.999932046 |
| NDUFAF5 | XR_430269 | 10.22093963 | 0 | 0 | 0 | 0 | 0 | 0 | 0 | 1.59E-08 | 1 |
| NDUFAF6 | NR_148909 | 0 | 0 | 0 | 8.943236221 | 8.94180265 | 0 | 0 | 0 | 0.029613199 | 0.02137193 |
| NDUFAF7 | XM_024452977 | 0 | 0 | 0 | 0 | 6.409577539 | 0 | 0 | 0 | 4.59E-05 | 1 |
| NDUFB3 | XM_017004186 | 0 | 0 | 0 | 0 | 0 | 9.391436267 | 8.60104519 | 0 | 1 | 0.02072721 |
| NDUFB6 | NM_001199987 | 0 | 0 | 0 | 7.706219197 | 0 | 0 | 0 | 0 | 1.42E-05 | 1 |
| NDUFS1 | NM_005006 | 0 | 0 | 9.652682006 | 10.20044734 | 7.335710674 | 0 | 0 | 0 | 0.002767154 | 0.004246182 |
| NDUFS2 | XM_005245209 | 0 | 9.06649413 | 0 | 0 | 0 | 0 | 0 | 0 | 7.75E-08 | 1 |
| NDUFS4 | NR_134475 | 11.19083431 | 8.632639499 | 0 | 0 | 0 | 0 | 0 | 0 | 0.000611815 | 1 |
| NDUFV2 | XM_017025782 | 0 | 0 | 8.456887945 | 0 | 8.943022271 | 0 | 0 | 0 | 0.035160755 | 0.02348728 |
| NDUFV3 | NM_021075 | 9.922694008 | 8.991087758 | 0 | 0 | 0 | 0 | 0 | 0 | 0.00164307 | 1 |

| | | | | | | | | | | | |
|---|---|---|---|---|---|---|---|---|---|---|---|
| NEAT1 | NR_028272 | 0 | 0 | 7.983806275 | 0 | 0 | 0 | 0 | 0 | 1.18E-05 | 1 |
| NEBL | XR_001746995 | 0 | 9.145128245 | 0 | 0 | 0 | 0 | 6.703265946 | 0 | 5.08E-08 | 1 |
| NECAP1 | NM_015509 | 0 | 0 | 10.41885958 | 9.394011718 | 9.31210541 | 0 | 0 | 10.34292624 | 0.001604841 | 1 |
| NECAP1 | NR_024260 | 9.877828587 | 0 | 0 | 0 | 0 | 0 | 7.899157068 | 0 | 2.13E-08 | 1 |
| NECAP2 | NM_018090 | 0 | 0 | 10.56401166 | 0 | 6.896204849 | 0 | 7.663696786 | 0 | 0.020450162 | 0.999932046 |
| NEDD4L | NM_001144971 | 0 | 8.662110092 | 0 | 0 | 0 | 0 | 0 | 0 | 9.15E-08 | 1 |
| NEFL | NM_006158 | 9.330172147 | 0 | 0 | 0 | 0 | 0 | 0 | 0 | 4.56E-08 | 1 |
| NEIL1 | XM_011522003 | 0 | 9.271606013 | 0 | 0 | 0 | 0 | 0 | 0 | 4.15E-08 | 1 |
| NEK4 | XM_017007085 | 0 | 0 | 9.364551246 | 0 | 9.443727098 | 9.995953923 | 9.316547485 | 0 | 0.021505498 | 0.999932046 |
| NEK6 | NM_014397 | 0 | 0 | 0 | 10.12651129 | 10.74251483 | 0 | 9.924190208 | 12.59686179 | 0.010173246 | 0.999932046 |
| NEK7 | XM_017000346 | 0 | 8.870030144 | 0 | 0 | 0 | 0 | 0 | 9.985019915 | 7.86E-08 | 1 |
| NELFCD | XR_244142 | 0 | 8.348342131 | 0 | 0 | 0 | 9.523912909 | 0 | 0 | 1.24E-07 | 1 |
| NELL2 | NM_001145108 | 0 | 0 | 9.756006048 | 0 | 8.827586142 | 0 | 9.2630946 | 0 | 0.022512148 | 0.999932046 |
| NELL2 | XM_011538396 | 0 | 9.124104101 | 0 | 0 | 0 | 0 | 9.458846292 | 0 | 5.29E-08 | 1 |
| NELL2 | XM_017019344 | 10.63567489 | 10.785299 | 0 | 0 | 0 | 0 | 7.834664296 | 0 | 0.000330611 | 1 |
| NEMP2 | XM_017003098 | 0 | 10.05036496 | 0 | 0 | 0 | 0 | 8.310927973 | 0 | 1.60E-08 | 1 |
| NEO1 | XM_017022238 | 0 | 8.571828374 | 0 | 0 | 0 | 0 | 0 | 0 | 9.97E-08 | 1 |
| NEPRO | XM_005247276 | 9.531875668 | 0 | 0 | 0 | 0 | 0 | 0 | 0 | 3.28E-08 | 1 |
| NET1 | XM_024447755 | 9.524617511 | 0 | 0 | 0 | 0 | 0 | 0 | 0 | 3.32E-08 | 1 |
| NEURL4 | NM_032442 | 0 | 0 | 9.948784034 | 0 | 7.313250256 | 0 | 0 | 0 | 0.02760405 | 0.02072721 |
| NEXN | XM_005271327 | 0 | 0 | 0 | 0 | 0 | 8.608942041 | 7.856483795 | 0 | 1 | 0.02562326 |
| NFASC | NM_015090 | 0 | 9.970166223 | 0 | 0 | 0 | 0 | 0 | 0 | 1.75E-08 | 1 |
| NFASC | XM_017000740 | 0 | 9.940407937 | 0 | 0 | 0 | 0 | 0 | 0 | 5.01E-08 | 1 |
| NFIB | NM_001190737 | 0 | 0 | 0 | 10.41307084 | 8.226958042 | 10.66305408 | 8.305650376 | 0 | 0.018526066 | 0.999932046 |
| NFIB | XM_005251470 | 0 | 0 | 10.40586006 | 0 | 10.55941433 | 10.96128486 | 8.981349581 | 10.17846329 | 0.010014229 | 0.999932046 |
| NFIC | NM_005597 | 0 | 0 | 10.88325012 | 0 | 8.144543525 | 0 | 0 | 0 | 0.014967492 | 0.02072721 |
| NFIX | NM_001271044 | 0 | 0 | 0 | 10.47183335 | 8.131765004 | 8.529584284 | 10.38086621 | 0 | 0.018238253 | 0.999932046 |
| NFIX | NM_002501 | 0 | 0 | 0 | 0 | 0 | 0 | 9.049236545 | 10.10128877 | 1 | 0.02072721 |
| NFIX | XM_005259918 | 0 | 0 | 0 | 7.958392843 | 0 | 0 | 0 | 0 | 1.09E-05 | 1 |
| NFIX | XM_017026837 | 0 | 0 | 0 | 9.679029003 | 6.98414039 | 0 | 7.688238242 | 0 | 0.033013618 | 0.999932046 |
| NFKB1 | XM_024454069 | 0 | 9.445515002 | 0 | 0 | 0 | 0 | 5.566321721 | 0 | 3.16E-08 | 1 |
| NFKBIB | NM_002503 | 0 | 0 | 0 | 8.341179754 | 0 | 0 | 0 | 0 | 7.39E-06 | 1 |
| NFS1 | NM_021100 | 0 | 8.419096158 | 0 | 0 | 0 | 0 | 0 | 9.172865883 | 1.14E-07 | 1 |
| NFS1 | NR_037570 | 0 | 0 | 0 | 9.984252065 | 7.497189827 | 0 | 0 | 0 | 0.025729408 | 0.02072721 |
| NFU1 | XM_017003808 | 0 | 0 | 9.185464497 | 5.197268818 | 0 | 0 | 0 | 0 | 0.060502716 | 0.039621915 |
| NFX1 | XR_001746314 | 0 | 9.138154184 | 0 | 0 | 0 | 0 | 0 | 0 | 5.16E-08 | 1 |
| NFYC | NM_001308114 | 0 | 8.765293275 | 0 | 0 | 0 | 0 | 0 | 0 | 8.55E-08 | 1 |
| NFYC | XM_011541517 | 8.611240259 | 0 | 0 | 0 | 0 | 0 | 9.461221637 | 10.45432674 | 1.04E-07 | 0.02072721 |
| NFYC | XM_017001369 | 0 | 0 | 10.69232105 | 9.849579366 | 7.490532315 | 0 | 9.905124141 | 10.80786282 | 0.001878066 | 0.999932046 |
| NGEF | NM_019850 | 0 | 0 | 10.65178554 | 10.3974372 | 0 | 0 | 7.421731871 | 0 | 0.010814351 | 0.999932046 |
| NGEF | XM_011510925 | 9.79579164 | 0 | 0 | 0 | 0 | 0 | 0 | 0 | 2.47E-08 | 1 |
| NGLY1 | XR_940471 | 0 | 0 | 0 | 0 | 6.679179249 | 0 | 0 | 0 | 3.78E-05 | 1 |
| NHLRC1 | NM_198586 | 0 | 0 | 0 | 9.775488305 | 9.365839676 | 0 | 0 | 0 | 0.019039619 | 0.02072721 |

| | | | | | | | | | | | |
|---|---|---|---|---|---|---|---|---|---|---|---|
| NHSL1 | XM_017011089 | 10.67554289 | 0 | 10.83492735 | 10.65391614 | 10.29673013 | 0 | 0 | 0 | 0.999953634 | 0.001228497 |
| NINJ1 | XM_011518716 | 0 | 8.895067962 | 0 | 0 | 0 | 0 | 0 | 0 | 7.75E-08 | 1 |
| NIPA2 | XM_017022663 | 0 | 8.801057108 | 0 | 0 | 0 | 0 | 8.676582513 | 0 | 8.24E-08 | 1 |
| NIPA2 | XM_024450083 | 9.765324233 | 8.430558346 | 7.8773294 | 0 | 9.19721051 | 0 | 0 | 0 | 0.999953634 | 0.024319498 |
| NIPSNAP3A | NM_015469 | 10.821876 | 10.16115789 | 10.04471239 | 11.40362566 | 3.562339684 | 0 | 0 | 0 | 0.999953634 | 0.003286822 |
| NISCH | XM_005264839 | 0 | 0 | 0 | 0 | 0 | 11.02317065 | 7.647100774 | 0 | 1 | 0.02072721 |
| NIT1 | NM_005600 | 0 | 0 | 10.09201011 | 0 | 8.15089059 | 0 | 0 | 0 | 0.022367631 | 0.02072721 |
| NKAP | NM_024528 | 0 | 9.277965255 | 0 | 0 | 0 | 0 | 0 | 0 | 4.01E-08 | 1 |
| NLN | XM_017009673 | 10.81594227 | 0 | 10.15551827 | 0 | 7.838643081 | 0 | 0 | 0 | 0.999953634 | 0.02072721 |
| NLRP2 | NR_145325 | 10.89124009 | 0 | 0 | 0 | 0 | 0 | 0 | 0 | 1.34E-08 | 1 |
| NLRX1 | XM_011542980 | 0 | 8.464408432 | 0 | 0 | 0 | 0 | 0 | 0 | 1.11E-07 | 1 |
| NMD3 | XM_005247511 | 0 | 0 | 10.13885423 | 9.926289561 | 9.931759943 | 12.81755584 | 11.11664572 | 12.51841068 | 0.001181042 | 0.999932046 |
| NME3 | XM_011522504 | 11.18278763 | 0 | 0 | 0 | 0 | 9.803893982 | 0 | 0 | 1.34E-08 | 1 |
| NME4 | NM_005009 | 0 | 0 | 0 | 10.01969536 | 9.407077595 | 0 | 7.73610317 | 0 | 0.017218565 | 0.999932046 |
| NMI | NM_004688 | 0 | 0 | 8.585434713 | 0 | 9.202312328 | 11.98140138 | 0 | 0 | 0.029967258 | 0.999932046 |
| NMT1 | XR_002958013 | 0 | 0 | 0 | 0 | 0 | 7.660829348 | 8.525879161 | 0 | 1 | 0.02750692 |
| NOB1 | NR_074074 | 0 | 8.935855595 | 0 | 0 | 0 | 10.89230564 | 0 | 0 | 7.69E-08 | 1 |
| NOL3 | NM_003946 | 0 | 9.021724865 | 0 | 0 | 0 | 0 | 0 | 0 | 6.61E-08 | 1 |
| NOL3 | XM_017023843 | 0 | 10.42741887 | 9.57118808 | 9.972241456 | 0 | 0 | 0 | 0 | 0.999953634 | 0.02072721 |
| NOL4 | XM_017026059 | 0 | 9.219697032 | 0 | 0 | 0 | 0 | 0 | 0 | 4.50E-08 | 1 |
| NOL8 | XM_024447595 | 0 | 9.517150383 | 0 | 0 | 0 | 0 | 0 | 0 | 2.90E-08 | 1 |
| NOMO1 | NM_014287 | 0 | 0 | 0 | 9.652931436 | 6.59478863 | 0 | 0 | 0 | 0.035305959 | 0.025070991 |
| NOMO3 | XM_005255318 | 10.85697488 | 0 | 0 | 0 | 0 | 0 | 0 | 0 | 1.34E-08 | 1 |
| NOP56 | NM_006392 | 0 | 0 | 10.7170893 | 8.699121892 | 10.00949069 | 0 | 8.273572814 | 0 | 0.001293319 | 0.999932046 |
| NOSIP | NM_001270960 | 0 | 10.78752928 | 11.77343019 | 11.44005437 | 10.47424598 | 0 | 0 | 0 | 0.999953634 | 0.000527933 |
| NOSIP | XM_024451529 | 10.9761662 | 0 | 0 | 0 | 0 | 0 | 0 | 0 | 1.34E-08 | 1 |
| NOTCH1 | NM_017617 | 0 | 7.103476409 | 0 | 0 | 0 | 9.779154025 | 8.207195305 | 0 | 0.097206878 | 0.02072721 |
| NOX1 | NM_001271815 | 0 | 0 | 9.795083168 | 0 | 7.536502354 | 0 | 0 | 0 | 0.029138196 | 0.020781046 |
| NOX1 | NM_007052 | 0 | 0 | 8.802008033 | 0 | 9.116275051 | 0 | 8.530410777 | 0 | 0.029148751 | 0.999932046 |
| NOX1 | NM_013955 | 0 | 9.051724868 | 0 | 0 | 0 | 0 | 0 | 0 | 7.86E-08 | 1 |
| NOXA1 | XM_024447388 | 0 | 8.273938353 | 0 | 0 | 0 | 0 | 0 | 9.534880393 | 2.02E-07 | 1 |
| NPAS2 | NM_002518 | 0 | 0 | 10.37950427 | 0 | 9.394651239 | 10.45816948 | 7.892131936 | 0 | 0.015087452 | 0.999932046 |
| NPFFR2 | XM_011531554 | 0 | 9.655452119 | 0 | 0 | 0 | 0 | 0 | 0 | 2.46E-08 | 1 |
| NPIPA3 | XM_017023553 | 0 | 8.870030144 | 0 | 0 | 0 | 0 | 0 | 0 | 7.86E-08 | 1 |
| NPIPA5 | XM_024450132 | 11.04626802 | 0 | 0 | 0 | 0 | 0 | 0 | 0 | 1.34E-08 | 1 |
| NPIPB4 | XM_024450289 | 0 | 0 | 8.490114275 | 0 | 9.749294684 | 11.08124624 | 8.456139264 | 0 | 0.023834661 | 0.999932046 |
| NPL | NM_030769 | 0 | 0 | 10.63405836 | 9.981258774 | 9.626385537 | 0 | 0 | 0 | 0.001034241 | 0.001991505 |
| NPNT | XM_017007984 | 0 | 8.818611963 | 0 | 0 | 0 | 0 | 8.067631163 | 0 | 8.12E-08 | 1 |
| NPPA-AS1 | NR_037806 | 0 | 8.497482465 | 0 | 0 | 0 | 0 | 0 | 0 | 1.06E-07 | 1 |
| NPPC | NM_024409 | 0 | 8.998808245 | 0 | 0 | 0 | 0 | 0 | 0 | 6.89E-08 | 1 |
| NPTN | NM_017455 | 0 | 0 | 9.461549885 | 7.326879247 | 6.702411139 | 0 | 7.057383163 | 0 | 0.011278824 | 0.999932046 |
| NPY1R | NM_000909 | 11.67346192 | 0 | 0 | 0 | 0 | 0 | 0 | 9.652668974 | 1.34E-08 | 1 |
| NQO1 | NM_001025434 | 0 | 0 | 0 | 10.40639144 | 8.861521052 | 0 | 7.974256665 | 0 | 0.016615172 | 0.999932046 |

| | | | | | | | | | | | |
|---|---|---|---|---|---|---|---|---|---|---|---|
| NR1D2 | XM_006713451 | 0 | 0 | 7.820991277 | 0 | 0 | 0 | 8.000624564 | 0 | 1.38E-05 | 1 |
| NR1H2 | NM_007121 | 0 | 8.360378074 | 10.21037937 | 0 | 8.083917571 | 0 | 0 | 0 | 0.999953634 | 0.02072721 |
| NR1H3 | NM_005693 | 0 | 0 | 0 | 8.731547201 | 0 | 0 | 0 | 0 | 4.81E-06 | 1 |
| NR1H4 | NM_001206992 | 9.11507342 | 0 | 0 | 0 | 0 | 0 | 0 | 0 | 6.89E-08 | 1 |
| NR2C1 | XM_006719584 | 0 | 9.605761852 | 0 | 0 | 0 | 0 | 0 | 0 | 2.55E-08 | 1 |
| NR2C1 | XM_011538717 | 0 | 9.245884974 | 0 | 0 | 0 | 0 | 0 | 0 | 4.33E-08 | 1 |
| NR2C2 | NM_001291694 | 0 | 0 | 0 | 0 | 0 | 0 | 9.055538427 | 10.40262864 | 1 | 0.02072721 |
| NR2F1 | NM_005654 | 9.457596586 | 0 | 0 | 0 | 0 | 0 | 0 | 0 | 3.78E-08 | 1 |
| NR2F2 | NM_021005 | 0 | 0 | 9.040481197 | 0 | 8.49817843 | 0 | 7.655422644 | 0 | 0.033732162 | 0.999932046 |
| NR2F6 | NM_005234 | 0 | 0 | 0 | 11.25292332 | 10.41263277 | 0 | 0 | 0 | 0.007334558 | 0.02072721 |
| NR3C2 | NM_001166104 | 0 | 8.801057108 | 0 | 0 | 0 | 0 | 0 | 0 | 8.24E-08 | 1 |
| NR3C2 | XM_011531975 | 9.457596586 | 0 | 0 | 0 | 0 | 0 | 0 | 8.47141627 | 3.78E-08 | 1 |
| NRAV | NR_038854 | 10.51633798 | 0 | 0 | 0 | 0 | 0 | 0 | 0 | 1.39E-08 | 1 |
| NRBP1 | NM_001321358 | 0 | 8.407542174 | 0 | 0 | 0 | 10.83719366 | 0 | 9.797590618 | 1.16E-07 | 0.02072721 |
| NRBP1 | NM_001321363 | 0 | 0 | 11.04514756 | 0 | 6.095400279 | 0 | 0 | 0 | 2.70E-08 | 1 |
| NRP1 | NM_001024628 | 0 | 5.648076284 | 11.70891981 | 12.10506416 | 10.15183321 | 12.65603116 | 9.218858934 | 12.93153008 | 0.048818018 | 0.999932046 |
| NRP1 | XM_006717526 | 0 | 0 | 0 | 10.24105527 | 8.822293806 | 0 | 0 | 11.19651409 | 0.01808444 | 0.999932046 |
| NRSN2 | XM_017028074 | 0 | 8.582143237 | 0 | 0 | 0 | 0 | 0 | 0 | 9.91E-08 | 1 |
| NSD1 | XM_024446161 | 0 | 0 | 9.917373151 | 0 | 8.900965585 | 0 | 9.523920794 | 0 | 0.020674594 | 0.999932046 |
| NSD1 | XM_024446163 | 8.754625855 | 9.265218616 | 0 | 0 | 0 | 9.171440319 | 9.514802225 | 8.076184046 | 0.003056885 | 0.004150193 |
| NSD2 | XM_017008587 | 0 | 0 | 0 | 8.046946667 | 0 | 0 | 0 | 0 | 1.03E-05 | 1 |
| NSDHL | NM_001129765 | 0 | 9.506358431 | 0 | 0 | 0 | 0 | 0 | 0 | 2.91E-08 | 1 |
| NSFL1C | NM_016143 | 0 | 0 | 0 | 10.5117953 | 9.740203086 | 0 | 0 | 11.07883757 | 0.012773773 | 0.999932046 |
| NSMCE1 | XM_006721023 | 0 | 0 | 0 | 6.258826736 | 0 | 0 | 0 | 0 | 0.342041679 | 1.77E-05 |
| NSMCE2 | XM_017013331 | 0 | 0 | 0 | 10.36105389 | 9.190037519 | 0 | 10.30619604 | 0 | 0.015773073 | 0.999932046 |
| NSMCE4A | NM_017615 | 0 | 0 | 10.3759535 | 11.92595566 | 9.273845655 | 0 | 0 | 0 | 0.000459317 | 0.001056205 |
| NSMCE4A | XR_001747115 | 0 | 9.650559405 | 0 | 0 | 0 | 0 | 0 | 0 | 2.46E-08 | 1 |
| NSUN2 | NM_001193455 | 0 | 0 | 0 | 9.523110922 | 6.855333211 | 0 | 0 | 11.08431711 | 0.037019753 | 0.999932046 |
| NSUN5 | NM_001168347 | 0 | 8.853093555 | 0 | 0 | 0 | 0 | 0 | 0 | 7.91E-08 | 1 |
| NSUN5P1 | NR_104013 | 0 | 0 | 9.091167477 | 0 | 6.261024403 | 0 | 0 | 0 | 0.054995015 | 0.034593848 |
| NSUN5P2 | NR_033323 | 9.669860113 | 0 | 0 | 0 | 0 | 0 | 0 | 0 | 2.76E-08 | 1 |
| NT5C | NR_045513 | 0 | 0 | 0 | 7.569406338 | 0 | 0 | 0 | 0 | 1.69E-05 | 1 |
| NT5C | XR_429885 | 0 | 0 | 11.51593306 | 0 | 9.30643073 | 0 | 7.630311627 | 0 | 0.00851432 | 0.999932046 |
| NT5C2 | NM_001351187 | 0 | 0 | 10.36524846 | 0 | 7.599732499 | 0 | 0 | 0 | 0.02111548 | 0.02072721 |
| NT5C2 | NM_001351190 | 10.12341345 | 0 | 0 | 0 | 0 | 0 | 9.187640525 | 0 | 1.68E-08 | 1 |
| NT5C2 | NM_001351194 | 11.24704134 | 0 | 0 | 0 | 0 | 0 | 0 | 0 | 1.34E-08 | 1 |
| NT5C2 | XM_006717723 | 0 | 0 | 0 | 8.997575451 | 7.968883375 | 0 | 9.042907014 | 10.05366645 | 0.039913075 | 0.999932046 |
| NT5C3A | NM_001002009 | 0 | 9.186279128 | 9.181401832 | 9.993194855 | 7.350492346 | 0 | 0 | 0 | 0.999953634 | 0.005644895 |
| NT5C3B | NM_052935 | 0 | 9.213075091 | 0 | 0 | 0 | 0 | 8.954623728 | 0 | 4.56E-08 | 1 |
| NT5DC1 | NM_152729 | 0 | 9.102769039 | 0 | 11.48934631 | 8.75722352 | 0 | 0 | 0 | 0.989151139 | 0.02072721 |
| NT5E | NM_002526 | 0 | 0 | 0 | 11.1136162 | 9.010889653 | 0 | 0 | 10.03492329 | 0.011181154 | 0.999932046 |
| NTAN1 | NM_001270766 | 0 | 0 | 0 | 10.15335408 | 7.203547662 | 0 | 0 | 0 | 0.024330175 | 0.02072721 |
| NTM | NM_001048209 | 0 | 10.37795348 | 0 | 0 | 0 | 0 | 0 | 0 | 1.39E-08 | 1 |

| | | | | | | | | | | | |
|---|---|---|---|---|---|---|---|---|---|---|---|
| NTM | XM_017017854 | 0 | 0 | 0 | 10.74320495 | 8.114548983 | 0 | 0 | 0 | 0.015844586 | 0.02072721 |
| NTRK3 | NM_001320134 | 10.79196021 | 8.919678654 | 0 | 0 | 0 | 0 | 0 | 7.551293494 | 0.000806913 | 1 |
| NTRK3 | XM_006720544 | 10.07454845 | 0 | 0 | 0 | 0 | 0 | 0 | 9.429755563 | 1.79E-08 | 1 |
| NTRK3 | XM_017022247 | 10.25203635 | 9.245884974 | 0 | 0 | 0 | 0 | 0 | 0 | 0.001113543 | 1 |
| NTRK3 | XM_024449936 | 8.791378415 | 9.808252985 | 0 | 0 | 0 | 0 | 0 | 9.116930837 | 0.001928961 | 1 |
| NTRK3 | XR_001751293 | 0 | 8.719304505 | 0 | 0 | 0 | 0 | 0 | 0 | 8.96E-08 | 1 |
| NUB1 | XM_017012306 | 10.12341345 | 0 | 0 | 0 | 0 | 0 | 0 | 0 | 1.68E-08 | 1 |
| NUB1 | XM_024446799 | 0 | 9.670031486 | 0 | 0 | 0 | 10.64776083 | 0 | 8.885437246 | 2.40E-08 | 0.02072721 |
| NUCB1 | NM_006184 | 0 | 10.19851207 | 0 | 0 | 0 | 10.44741438 | 0 | 0 | 1.51E-08 | 1 |
| NUCB1 | XM_017026845 | 0 | 0 | 0 | 9.563698439 | 8.400349091 | 10.22617509 | 8.48910449 | 0 | 0.026969025 | 0.999932046 |
| NUCB2 | XM_024448529 | 0 | 9.239382419 | 0 | 0 | 0 | 0 | 0 | 0 | 4.33E-08 | 1 |
| NUDT13 | XM_024447926 | 0 | 8.690990708 | 0 | 0 | 0 | 0 | 0 | 0 | 9.02E-08 | 1 |
| NUDT16 | NM_152395 | 0 | 0 | 0 | 10.17970655 | 10.08214002 | 0 | 0 | 0 | 0.013082656 | 0.02072721 |
| NUDT16L1 | XM_005255633 | 0 | 8.959785899 | 0 | 0 | 0 | 0 | 0 | 9.715223813 | 7.69E-08 | 1 |
| NUDT21 | NM_007006 | 0 | 0 | 10.62361601 | 0 | 6.965057125 | 9.037035537 | 9.246664761 | 8.649540214 | 0.019641255 | 0.999932046 |
| NUDT3 | NM_006703 | 0 | 0 | 0 | 9.836388206 | 9.239515275 | 0 | 9.204752435 | 9.810872569 | 0.019304981 | 0.999932046 |
| NUDT4 | NM_199040 | 0 | 0 | 0 | 9.114202495 | 7.787961256 | 0 | 0 | 0 | 0.039404016 | 0.027341676 |
| NUDT4B | NM_001355407 | 9.035603861 | 0 | 0 | 0 | 0 | 0 | 0 | 0 | 7.68E-08 | 1 |
| NUDT8 | NM_001243750 | 0 | 0 | 11.68185439 | 0 | 7.415197319 | 0 | 0 | 0 | 1.59E-08 | 6.49E-09 |
| NUDT9 | NM_024047 | 0 | 0 | 10.10064114 | 0 | 7.752229049 | 0 | 0 | 0 | 0.023850113 | 0.02072721 |
| NUDT9 | XM_011532032 | 10.51999251 | 7.014878551 | 0 | 0 | 0 | 0 | 0 | 0 | 0.001836387 | 1 |
| NUDT9 | XR_002959734 | 10.69183479 | 0 | 0 | 0 | 0 | 0 | 0 | 0 | 2.15E-08 | 1 |
| NUFIP2 | NM_020772 | 10.17531978 | 0 | 0 | 0 | 0 | 0 | 0 | 0 | 1.60E-08 | 1 |
| NUMA1 | NR_104476 | 10.94931306 | 8.681427931 | 9.530245374 | 7.520758627 | 8.280040102 | 0 | 0 | 0 | 0.976761196 | 0.009871761 |
| NUP210 | XM_005264994 | 0 | 0 | 9.239219313 | 0 | 7.968883375 | 0 | 0 | 10.0684876 | 0.035966805 | 0.999932046 |
| NUP210 | XM_024453416 | 0 | 0 | 0 | 0 | 6.464964927 | 0 | 0 | 0 | 4.45E-05 | 1 |
| NUP35 | NM_138285 | 0 | 0 | 0 | 10.24355571 | 6.780897397 | 0 | 7.578739068 | 0 | 0.024191991 | 0.999932046 |
| NUP54 | XR_001741238 | 0 | 8.765293275 | 0 | 0 | 0 | 0 | 0 | 0 | 8.55E-08 | 1 |
| NUP85 | XR_002958072 | 0 | 0 | 0 | 10.11016257 | 7.944733425 | 0 | 5.207402423 | 10.40262864 | 0.022468178 | 0.999932046 |
| NUP88 | NM_002532 | 0 | 8.911521649 | 0 | 0 | 0 | 0 | 0 | 0 | 7.73E-08 | 1 |
| NUPR1 | NM_001042483 | 8.452015244 | 0 | 0 | 0 | 0 | 0 | 0 | 0 | 1.21E-07 | 1 |
| NUS1 | NM_138459 | 0 | 0 | 9.144313654 | 0 | 9.55479642 | 0 | 7.947397859 | 0 | 0.022002131 | 0.999932046 |
| NUTM2A-AS1 | NR_024397 | 0 | 0 | 9.185464497 | 0 | 8.879468369 | 0 | 0 | 0 | 0.027884198 | 0.02072721 |
| NVL | XM_024447367 | 0 | 0 | 10.12093523 | 7.124706719 | 0 | 0 | 0 | 0 | 0.027437274 | 0.02072721 |
| NXF1 | NM_001081491 | 10.19374119 | 0 | 0 | 0 | 0 | 0 | 8.21847248 | 0 | 1.60E-08 | 1 |
| NXF1 | NM_006362 | 0 | 0 | 0 | 10.04428441 | 8.088333507 | 0 | 7.538812572 | 9.904707713 | 0.02269095 | 0.999932046 |
| NXF1 | XR_001747724 | 10.02910853 | 0 | 0 | 0 | 0 | 0 | 0 | 0 | 1.87E-08 | 1 |
| NXPE1 | XR_947793 | 10.78286315 | 10.09786644 | 0 | 0 | 0 | 0 | 0 | 0 | 0.000465423 | 1 |
| NXPE2 | XM_011542604 | 12.83013688 | 11.89655261 | 11.60469485 | 10.14802533 | 8.511392973 | 0 | 0 | 0 | 0.886385869 | 0.001535028 |
| NXPE3 | NR_145993 | 9.623051988 | 0 | 0 | 0 | 0 | 0 | 4.37247236 | 7.775600578 | 2.93E-08 | 0.093283431 |
| NXPE4 | NM_001077639 | 0 | 9.470716411 | 0 | 0 | 0 | 0 | 0 | 0 | 3.08E-08 | 1 |
| NXPE4 | XM_011542883 | 10.05452941 | 0 | 0 | 0 | 0 | 0 | 0 | 0 | 5.06E-08 | 1 |
| NXT1 | NM_013248 | 0 | 10.6060999 | 0 | 10.50138585 | 9.872185325 | 0 | 0 | 0 | 0.999953634 | 0.02072721 |

| | | | | | | | | | | | |
|---|---|---|---|---|---|---|---|---|---|---|---|
| NYNRIN | XM_011537016 | 8.357138871 | 0 | 0 | 0 | 0 | 0 | 0 | 0 | 1.96E-07 | 1 |
| OARD1 | NM_001329686 | 10.41791771 | 0 | 0 | 0 | 0 | 0 | 0 | 10.42295417 | 1.47E-08 | 1 |
| OARD1 | NM_145063 | 0 | 9.213075091 | 0 | 0 | 0 | 11.22900221 | 8.525879161 | 0 | 4.56E-08 | 0.02072721 |
| OAS1 | NM_016816 | 0 | 0 | 8.72594783 | 10.49405428 | 0 | 0 | 9.080474236 | 0 | 0.017521547 | 0.999932046 |
| OAS1 | XM_006719434 | 0 | 9.219697032 | 0 | 0 | 0 | 0 | 0 | 0 | 4.50E-08 | 1 |
| OAS1 | XM_011538413 | 10.34149639 | 0 | 0 | 0 | 0 | 0 | 0 | 0 | 1.60E-08 | 1 |
| OAT | NM_001322971 | 0 | 0 | 0 | 0 | 6.758905041 | 0 | 0 | 0 | 9.53E-05 | 1 |
| OCEL1 | NM_024578 | 0 | 8.737876298 | 0 | 0 | 0 | 0 | 0 | 0 | 9.11E-08 | 1 |
| OCEL1 | XM_006722899 | 10.76757274 | 0 | 0 | 0 | 0 | 0 | 0 | 0 | 1.34E-08 | 1 |
| OCIAD2 | NM_001286774 | 0 | 0 | 10.43856741 | 10.20558628 | 8.790125492 | 0 | 8.566160223 | 0 | 0.001347927 | 0.999932046 |
| OCIAD2 | NM_152398 | 0 | 0 | 0 | 10.81389073 | 8.043553839 | 0 | 0 | 0 | 0.015398614 | 0.02072721 |
| OFD1 | XM_005274606 | 11.61486007 | 0 | 10.40237365 | 10.62735608 | 7.71273195 | 0 | 0 | 0 | 0.999953634 | 0.002588426 |
| OFD1 | XM_011545595 | 0 | 9.613325378 | 0 | 0 | 0 | 0 | 9.017307488 | 0 | 2.54E-08 | 1 |
| OFD1 | XM_011545597 | 0 | 0 | 9.431077203 | 0 | 8.493191603 | 0 | 10.01186955 | 0 | 0.028391218 | 0.999932046 |
| OGA | NM_012215 | 0 | 0 | 0 | 0 | 0 | 10.60176571 | 0 | 10.40846521 | 1 | 0.02072721 |
| OGDH | XM_024446783 | 0 | 0 | 8.535381898 | 8.731547201 | 8.308707489 | 8.270292406 | 0 | 0 | 0.007294856 | 0.999932046 |
| OGFOD3 | NR_033265 | 10.27816739 | 8.054440838 | 0 | 0 | 0 | 0 | 0 | 0 | 0.001771088 | 1 |
| OGN | NM_033014 | 0 | 9.522516235 | 0 | 0 | 0 | 0 | 0 | 0 | 2.88E-08 | 1 |
| OGT | XM_024452467 | 0 | 0 | 0 | 10.92788189 | 9.465956986 | 10.18791899 | 0 | 0 | 0.011239152 | 0.999932046 |
| OIP5-AS1 | NR_026757 | 0 | 0 | 0 | 10.51283213 | 6.935950419 | 0 | 0 | 0 | 0.020446648 | 0.02072721 |
| OIT3 | NR_130125 | 0 | 9.387883956 | 0 | 0 | 0 | 0 | 0 | 0 | 3.40E-08 | 1 |
| OLFM1 | NM_014279 | 0 | 9.165850403 | 0 | 0 | 0 | 0 | 0 | 0 | 4.97E-08 | 1 |
| OLFM2 | NM_058164 | 9.330172147 | 0 | 0 | 0 | 0 | 0 | 0 | 0 | 4.56E-08 | 1 |
| OLFM4 | NM_006418 | 0 | 13.22711312 | 0 | 11.18949662 | 9.765240791 | 0 | 0 | 0 | 0.982665931 | 0.02072721 |
| OLR1 | NM_001172633 | 0 | 0 | 0 | 9.603175278 | 8.105863313 | 0 | 6.719231771 | 0 | 0.028265655 | 0.999932046 |
| OMA1 | NM_145243 | 0 | 0 | 11.2817445 | 0 | 7.689666995 | 0 | 0 | 0 | 0.012531836 | 0.02072721 |
| OPN3 | NM_014322 | 10.81891219 | 0 | 0 | 0 | 0 | 0 | 0 | 0 | 1.34E-08 | 1 |
| ORAI3 | NM_152288 | 0 | 0 | 0 | 0 | 0 | 10.38934044 | 0 | 10.66767715 | 1 | 0.02072721 |
| ORC1 | XM_017001389 | 0 | 0 | 0 | 9.078233734 | 6.112844945 | 0 | 0 | 0 | 0.054667199 | 0.037973953 |
| ORMDL1 | NM_001128150 | 10.45275129 | 9.333971188 | 0 | 10.47609238 | 9.029380004 | 0 | 0 | 0 | 0.999953634 | 0.020771528 |
| OS9 | NM_001261423 | 10.54172739 | 0 | 0 | 0 | 0 | 0 | 0 | 0 | 1.36E-08 | 1 |
| OS9 | XM_006719200 | 10.41006135 | 0 | 0 | 0 | 0 | 0 | 0 | 0 | 1.59E-08 | 1 |
| OSBPL10 | NM_017784 | 0 | 8.519117955 | 0 | 0 | 0 | 0 | 0 | 0 | 1.04E-07 | 1 |
| OSBPL2 | XM_017028163 | 0 | 8.927789798 | 0 | 0 | 0 | 0 | 0 | 0 | 7.69E-08 | 1 |
| OSBPL3 | XR_428075 | 10.30806717 | 0 | 0 | 0 | 0 | 0 | 0 | 0 | 1.54E-08 | 1 |
| OSBPL7 | NM_145798 | 9.695941183 | 0 | 0 | 0 | 0 | 0 | 7.416845097 | 0 | 2.66E-08 | 1 |
| OSBPL9 | NM_001350210 | 0 | 0 | 0 | 11.42691374 | 7.357826773 | 0 | 0 | 0 | 1.80E-08 | 1.07E-08 |
| OSBPL9 | NM_148906 | 11.07884328 | 9.517150383 | 10.65325302 | 9.667902088 | 8.210847477 | 0 | 0 | 0 | 0.999953634 | 0.002881517 |
| OSBPL9 | XM_006710318 | 0 | 0 | 0 | 0 | 0 | 0 | 9.781315183 | 11.18125388 | 1 | 0.02072721 |
| OSBPL9 | XM_006710323 | 0 | 0 | 9.872720291 | 10.6762991 | 0 | 0 | 9.458846292 | 0 | 0.012529135 | 0.999932046 |
| OSBPL9 | XM_011540600 | 0 | 0 | 0 | 0 | 0 | 0 | 9.676863839 | 10.1971287 | 1 | 0.02072721 |
| OSCAR | NM_206818 | 0 | 0 | 0 | 7.470413098 | 0 | 0 | 0 | 0 | 1.82E-05 | 1 |
| OSER1 | XM_024451891 | 0 | 9.708204508 | 7.857096443 | 0 | 8.428486253 | 0 | 0 | 0 | 0.999953634 | 0.034941497 |

| | | | | | | | | | | | |
|---|---|---|---|---|---|---|---|---|---|---|---|
| OSMR | XM_005248384 | 0 | 0 | 0 | 0 | 6.464964927 | 0 | 0 | 0 | 4.45E-05 | 1 |
| OSTF1 | XM_017014622 | 9.708807013 | 0 | 0 | 0 | 0 | 0 | 0 | 0 | 2.61E-08 | 1 |
| OTOP2 | XM_011525479 | 9.546283346 | 0 | 0 | 0 | 0 | 0 | 0 | 0 | 3.25E-08 | 1 |
| OTUB1 | NR_003089 | 0 | 0 | 5.782160807 | 10.90199908 | 8.532611753 | 9.715381837 | 7.27822739 | 8.950889997 | 0.003212912 | 0.999932046 |
| OTUD5 | XM_017029628 | 0 | 0 | 0 | 0 | 6.747781854 | 0 | 0 | 0 | 9.53E-05 | 1 |
| OTUD6B-AS1 | NR_110439 | 0 | 0 | 0 | 0 | 0 | 10.38217075 | 7.506976231 | 0 | 1 | 0.02072721 |
| OXLD1 | NR_130938 | 0 | 0 | 0 | 9.496103202 | 6.725274845 | 0 | 9.216048648 | 0 | 0.03835619 | 0.999932046 |
| OXNAD1 | XM_017007492 | 0 | 9.321711965 | 0 | 0 | 0 | 0 | 0 | 0 | 3.84E-08 | 1 |
| OXR1 | NM_001198533 | 0 | 0 | 11.2276109 | 10.51593815 | 8.966001299 | 13.04454957 | 11.4125951 | 10.41427827 | 0.000739643 | 0.999932046 |
| OXR1 | NM_001198534 | 0 | 0 | 0 | 8.350488917 | 0 | 9.307419275 | 8.62454847 | 0 | 1.02E-05 | 0.999932046 |
| OXR1 | NM_018002 | 9.789749415 | 0 | 0 | 0 | 0 | 0 | 0 | 0 | 2.40E-08 | 1 |
| P2RX1 | XR_934030 | 9.992749568 | 0 | 0 | 0 | 0 | 0 | 0 | 0 | 2.01E-08 | 1 |
| P2RX4 | NR_046373 | 0 | 0 | 10.59793683 | 0 | 6.945718048 | 0 | 0 | 0 | 0.019977469 | 0.02072721 |
| P2RY1 | NM_002563 | 10.04948094 | 0 | 0 | 0 | 0 | 0 | 0 | 0 | 5.08E-08 | 1 |
| P2RY13 | XM_006713664 | 10.29958742 | 9.309347676 | 0 | 0 | 0 | 0 | 7.79000962 | 0 | 0.001038587 | 1 |
| P3H1 | XR_946739 | 0 | 8.927789798 | 0 | 0 | 0 | 0 | 0 | 0 | 7.69E-08 | 1 |
| P3H2 | NM_001134418 | 0 | 9.439854398 | 0 | 0 | 0 | 0 | 0 | 0 | 3.19E-08 | 1 |
| P4HA2 | XM_006714730 | 0 | 0 | 10.69801997 | 11.17908448 | 10.62243085 | 11.6885813 | 0 | 11.20156517 | 0.000396997 | 0.999932046 |
| PAAF1 | NM_025155 | 0 | 9.825721024 | 0 | 0 | 0 | 11.26397079 | 7.947397859 | 0 | 2.01E-08 | 0.02072721 |
| PABPC1L | NR_134987 | 10.77981801 | 0 | 0 | 0 | 0 | 0 | 0 | 0 | 1.34E-08 | 1 |
| PACRG | NM_152410 | 0 | 0 | 9.017875808 | 0 | 8.038998467 | 0 | 0 | 0 | 0.039491414 | 0.025566236 |
| PACRG | XM_011535468 | 0 | 0 | 0 | 9.74051016 | 7.305685067 | 0 | 0 | 0 | 0.03038219 | 0.02215675 |
| PACS1 | XM_011545164 | 0 | 0 | 8.918864307 | 7.102839446 | 0 | 0 | 0 | 0 | 0.056322481 | 0.036421251 |
| PACS2 | XR_001750200 | 9.771469405 | 0 | 8.909070364 | 0 | 7.774322632 | 0 | 0 | 0 | 0.999953634 | 0.028820499 |
| PACSIN2 | NM_001349969 | 0 | 9.799439006 | 0 | 0 | 0 | 0 | 8.688796313 | 9.793135989 | 2.04E-08 | 0.02072721 |
| PACSIN2 | XM_017028563 | 8.373391248 | 10.07248322 | 0 | 0 | 0 | 11.42314136 | 0 | 0 | 0.001826003 | 1 |
| PADI2 | NM_007365 | 0 | 8.508340767 | 0 | 0 | 0 | 0 | 0 | 0 | 1.06E-07 | 1 |
| PADI4 | NM_012387 | 0 | 0 | 10.26323048 | 0 | 8.975568243 | 10.91944554 | 0 | 0 | 0.017562673 | 0.999932046 |
| PADI4 | XM_011541157 | 0 | 10.18842011 | 0 | 0 | 0 | 0 | 0 | 0 | 1.52E-08 | 1 |
| PAFAH1B1 | XM_017024701 | 9.676424716 | 0 | 10.39537545 | 0 | 9.365839676 | 0 | 0 | 0 | 0.999953634 | 0.02072721 |
| PAFAH1B2 | NM_002572 | 0 | 9.905922754 | 0 | 0 | 0 | 0 | 0 | 0 | 1.87E-08 | 1 |
| PAFAH1B3 | NM_002573 | 0 | 0 | 0 | 10.90989151 | 9.472728608 | 7.647222305 | 7.980893958 | 10.20217763 | 0.011322603 | 0.999932046 |
| PAFAH1B3 | XM_017026848 | 0 | 0 | 0 | 7.676917851 | 0 | 0 | 0 | 0 | 1.47E-05 | 1 |
| PAG1 | XM_017013643 | 9.449953927 | 0 | 0 | 0 | 0 | 0 | 0 | 0 | 3.80E-08 | 1 |
| PAGR1 | NM_024516 | 0 | 0 | 0 | 10.16131049 | 10.25707461 | 0 | 8.616046059 | 9.896425015 | 0.012332386 | 0.999932046 |
| PAK1 | XM_024448564 | 0 | 0 | 0 | 9.711903927 | 6.80255953 | 0 | 7.256509845 | 10.27106783 | 0.03313868 | 0.999932046 |
| PAK4 | NM_001014834 | 0 | 9.095586658 | 0 | 0 | 0 | 0 | 0 | 0 | 5.54E-08 | 1 |
| PALD1 | NM_014431 | 0 | 8.360378074 | 0 | 0 | 0 | 0 | 0 | 0 | 1.21E-07 | 1 |
| PAMR1 | NM_001001991 | 11.2492447 | 0 | 9.85752351 | 9.802872777 | 8.638785557 | 0 | 0 | 0 | 0.999953634 | 0.003399121 |
| PAMR1 | NM_001282676 | 0 | 0 | 0 | 0 | 6.725274845 | 0 | 0 | 0 | 9.53E-05 | 1 |
| PAN3 | NM_175854 | 0 | 8.737876298 | 0 | 0 | 0 | 0 | 0 | 0 | 9.11E-08 | 1 |
| PANK3 | NM_024594 | 0 | 0 | 9.639415365 | 10.30230186 | 0 | 0 | 0 | 0 | 0.015672839 | 0.02072721 |
| PAOX | NR_109765 | 0 | 8.943876548 | 0 | 0 | 0 | 0 | 0 | 0 | 7.69E-08 | 1 |

| | | | | | | | | | | | |
|---|---|---|---|---|---|---|---|---|---|---|---|
| PAPOLA | NM_001252006 | 0 | 0 | 0 | 6.669146541 | 10.86756687 | 10.52665328 | 9.482425568 | 9.513376477 | 0.016708288 | 1 |
| PAQR5 | XM_017022368 | 10.14723899 | 0 | 0 | 0 | 0 | 0 | 0 | 0 | 1.64E-08 | 1 |
| PARD3B | NM_205863 | 0 | 0 | 8.714747194 | 0 | 7.666227291 | 0 | 0 | 0 | 0.051677854 | 0.032485916 |
| PARD6A | NM_001037281 | 0 | 0 | 0 | 11.06778942 | 7.790673576 | 0 | 0 | 0 | 0.013758272 | 0.02072721 |
| PARL | NM_018622 | 10.75211854 | 9.399595812 | 0 | 0 | 0 | 0 | 0 | 9.715223813 | 0.000683351 | 1 |
| PARN | NM_001134477 | 0 | 0 | 10.08114821 | 9.719108837 | 7.630337835 | 10.55425578 | 0 | 0 | 0.002715205 | 0.999932046 |
| PARP1 | NM_001618 | 0 | 0 | 10.66202669 | 9.55567177 | 7.949595864 | 0 | 0 | 0 | 0.001938334 | 0.00302572 |
| PARP10 | NM_032789 | 0 | 0 | 0 | 8.624344952 | 0 | 0 | 0 | 0 | 5.42E-06 | 1 |
| PARP10 | XM_011517336 | 0 | 0 | 10.2043857 | 0 | 7.187172039 | 0 | 0 | 0 | 0.024318018 | 0.02072721 |
| PARP6 | XR_002957661 | 0 | 8.384152849 | 0 | 0 | 0 | 0 | 6.735022839 | 0 | 1.18E-07 | 1 |
| PARP8 | XR_002956178 | 0 | 0 | 0 | 0 | 6.381065478 | 0 | 0 | 0 | 4.71E-05 | 1 |
| PATJ | XM_011540465 | 0 | 0 | 11.11070555 | 10.94180649 | 0 | 0 | 10.56440787 | 0 | 0.007315082 | 0.999932046 |
| PATL2 | XM_017022000 | 0 | 8.519117955 | 0 | 0 | 0 | 0 | 0 | 0 | 1.04E-07 | 1 |
| PAWR | NM_002583 | 9.190393069 | 0 | 0 | 0 | 0 | 0 | 0 | 0 | 7.73E-08 | 1 |
| PAX5 | NM_016734 | 0 | 9.321711965 | 0 | 0 | 0 | 0 | 0 | 9.617827325 | 3.84E-08 | 1 |
| PAX5 | NR_104000 | 0 | 9.456770012 | 0 | 0 | 0 | 0 | 0 | 0 | 3.14E-08 | 1 |
| PAX8 | NM_013952 | 10.38222222 | 11.18780178 | 11.66449506 | 0 | 9.49784282 | 0 | 0 | 0 | 0.999953634 | 0.02072721 |
| PAX8 | NM_013992 | 0 | 0 | 9.328265229 | 10.52418831 | 4.989879473 | 7.164671276 | 0 | 0 | 0.003795038 | 0.987221099 |
| PAXX | NR_138077 | 0 | 0 | 0 | 9.603175278 | 9.533903787 | 0 | 0 | 10.49603692 | 0.019183213 | 1 |
| PBK | XM_006716368 | 0 | 9.958336465 | 0 | 0 | 0 | 0 | 0 | 0 | 1.76E-08 | 1 |
| PBRM1 | XM_017006730 | 9.045780268 | 0 | 0 | 0 | 0 | 0 | 0 | 0 | 7.60E-08 | 1 |
| PBRM1 | XM_017006732 | 0 | 8.919678654 | 0 | 0 | 0 | 0 | 0 | 0 | 7.73E-08 | 1 |
| PBRM1 | XM_017006738 | 0 | 10.07248322 | 0 | 0 | 0 | 0 | 9.592700938 | 10.42870925 | 1.60E-08 | 0.02072721 |
| PBRM1 | XM_017006744 | 0 | 0 | 0 | 0 | 0 | 11.29101984 | 9.196221851 | 0 | 1 | 0.02072721 |
| PBRM1 | XM_017006745 | 0 | 0 | 10.20838425 | 9.996163509 | 0 | 0 | 9.004335343 | 0 | 0.014723845 | 0.999932046 |
| PC | XM_017017872 | 9.574674104 | 0 | 0 | 0 | 0 | 0 | 0 | 0 | 3.15E-08 | 1 |
| PCCA | XR_001749576 | 0 | 9.095586658 | 0 | 0 | 0 | 0 | 0 | 0 | 5.54E-08 | 1 |
| PCCA | XR_931615 | 0 | 0 | 0 | 0 | 0 | 0 | 8.732715858 | 10.21055345 | 1 | 0.02072721 |
| PCDHB13 | NM_018933 | 0 | 8.286608519 | 0 | 0 | 0 | 0 | 0 | 0 | 2.03E-07 | 1 |
| PCDHGC3 | NM_002588 | 0 | 9.165850403 | 0 | 0 | 0 | 0 | 0 | 0 | 4.97E-08 | 1 |
| PCED1A | NM_022760 | 9.124706245 | 0 | 0 | 0 | 0 | 0 | 0 | 0 | 6.80E-08 | 1 |
| PCED1A | XM_005260804 | 0 | 0 | 0 | 0 | 6.569741695 | 0 | 0 | 0 | 4.12E-05 | 1 |
| PCGF1 | NM_032673 | 0 | 0 | 10.21634824 | 10.43402054 | 9.336440983 | 0 | 0 | 0 | 0.001136102 | 0.002168299 |
| PCID2 | NM_001127202 | 0 | 0 | 0 | 8.961577651 | 8.671527983 | 0 | 0 | 0 | 0.032436255 | 0.02296717 |
| PCIF1 | NM_022104 | 0 | 9.081113725 | 0 | 0 | 0 | 0 | 0 | 0 | 5.70E-08 | 1 |
| PCIF1 | XM_017028013 | 0 | 9.021724865 | 0 | 0 | 0 | 0 | 0 | 0 | 6.61E-08 | 1 |
| PCK1 | XM_024451888 | 9.715197156 | 10.43597468 | 0 | 0 | 0 | 0 | 0 | 0 | 0.000738063 | 1 |
| PCLAF | NM_014736 | 0 | 0 | 0 | 10.95636031 | 10.21124569 | 0 | 10.01105915 | 10.61438614 | 0.009020639 | 0.999932046 |
| PCM1 | NM_006197 | 8.452015244 | 0 | 0 | 0 | 0 | 10.34893351 | 0 | 0 | 1.21E-07 | 1 |
| PCM1 | XM_006716340 | 0 | 0 | 10.43430594 | 11.62306424 | 0 | 0 | 0 | 0 | 0.006701127 | 0.02072721 |
| PCM1 | XM_017013481 | 0 | 10.66187 | 0 | 0 | 0 | 11.3619806 | 0 | 0 | 1.34E-08 | 1 |
| PCM1 | XM_024447167 | 9.663265502 | 9.296876505 | 0 | 0 | 0 | 0 | 8.412617556 | 0 | 0.001684343 | 1 |
| PCMT1 | NM_001252053 | 0 | 0 | 9.097649359 | 9.044216539 | 6.582319518 | 0 | 0 | 0 | 0.007781319 | 0.011824395 |

| | | | | | | | | | | | |
|---|---|---|---|---|---|---|---|---|---|---|---|
| PCNP | NR_135223 | 0 | 0 | 11.76225359 | 0 | 8.717864799 | 0 | 0 | 0 | 0.008079076 | 1.06E-08 |
| PCNP | NR_135232 | 9.363098816 | 0 | 0 | 0 | 0 | 0 | 0 | 0 | 4.33E-08 | 1 |
| PCNP | XM_017006922 | 0 | 0 | 10.0514059 | 8.766740692 | 0 | 0 | 7.331132628 | 0 | 0.021726192 | 0.999932046 |
| PCNT | NM_006031 | 0 | 0 | 7.290410474 | 0 | 0 | 0 | 0 | 0 | 2.36E-05 | 1 |
| PCNX1 | XM_024449513 | 0 | 8.475517571 | 0 | 0 | 0 | 0 | 0 | 0 | 1.09E-07 | 1 |
| PCSK7 | XR_947870 | 9.740478394 | 5.189614022 | 10.42488616 | 5.494613896 | 6.673312348 | 0 | 0 | 0 | 0.999953634 | 0.011373486 |
| PCTP | NM_001330377 | 0 | 0 | 0 | 10.34949334 | 8.435435627 | 11.09884756 | 8.493752915 | 10.57211024 | 0.018494796 | 0.999932046 |
| PCYOX1L | XM_024446205 | 0 | 9.830055185 | 0 | 0 | 0 | 0 | 0 | 0 | 2.01E-08 | 1 |
| PDCD10 | NM_145859 | 9.296476405 | 9.327854598 | 0 | 0 | 0 | 0 | 9.201914508 | 0 | 0.00210171 | 1 |
| PDCD10 | XM_005247086 | 0 | 9.059128398 | 0 | 0 | 0 | 0 | 0 | 0 | 7.80E-08 | 1 |
| PDCD2 | NM_002598 | 0 | 0 | 0 | 9.969223115 | 9.172467463 | 0 | 7.310202764 | 10.91424727 | 0.018707468 | 0.999932046 |
| PDCD2 | XM_017010926 | 0 | 8.765293275 | 10.6115886 | 0 | 7.683842675 | 0 | 0 | 0 | 0.999953634 | 0.021049696 |
| PDCD2L | NM_032346 | 0 | 0 | 8.869215857 | 0 | 8.8249424 | 0 | 0 | 10.34900889 | 0.032105363 | 1 |
| PDE4D | XM_011543471 | 0 | 8.756212053 | 0 | 0 | 0 | 0 | 0 | 0 | 8.55E-08 | 1 |
| PDE4D | XM_024446112 | 10.79497986 | 0 | 0 | 0 | 0 | 0 | 0 | 0 | 1.34E-08 | 1 |
| PDE6A | NM_000440 | 10.54172739 | 0 | 0 | 0 | 0 | 0 | 0 | 0 | 1.36E-08 | 1 |
| PDE6B | NM_001145291 | 10.18915784 | 0 | 0 | 0 | 0 | 0 | 0 | 0 | 1.60E-08 | 1 |
| PDE6D | NM_002601 | 0 | 0 | 10.53246266 | 0 | 9.958540354 | 0 | 0 | 10.31833588 | 0.011935756 | 0.999932046 |
| PDE8A | XR_001751313 | 0 | 0 | 9.075928861 | 0 | 6.245291858 | 0 | 0 | 0 | 0.055569846 | 0.03500036 |
| PDE9A | NM_001001567 | 0 | 8.951853153 | 0 | 0 | 0 | 0 | 0 | 0 | 7.60E-08 | 1 |
| PDGFB | NM_002608 | 0 | 0 | 0 | 0 | 0 | 9.123108582 | 0 | 9.265819055 | 1 | 0.02072721 |
| PDGFB | NM_033016 | 0 | 0 | 9.385892712 | 0 | 8.218925247 | 9.760316602 | 0 | 0 | 0.031303306 | 0.999932046 |
| PDGFRA | NM_006206 | 0 | 0 | 0 | 9.222102536 | 6.381065478 | 0 | 0 | 0 | 0.048132384 | 0.033268532 |
| PDHA1 | NM_001173456 | 10.37416851 | 9.672447121 | 10.40760011 | 0 | 9.522527317 | 0 | 0 | 0 | 0.999953634 | 0.02072721 |
| PDHB | NR_033384 | 9.387309849 | 0 | 0 | 9.969223115 | 8.274237629 | 0 | 0 | 0 | 0.999953634 | 0.02072721 |
| PDK1 | XR_427095 | 0 | 0 | 0 | 9.82307532 | 5.167328428 | 10.51920254 | 0 | 0 | 0.036008564 | 1 |
| PDK4 | NM_002612 | 0 | 9.059128398 | 0 | 0 | 0 | 0 | 0 | 0 | 7.80E-08 | 1 |
| PDLIM1 | XM_011540330 | 12.40334215 | 9.605761852 | 7.751443001 | 10.90831647 | 7.203547662 | 0 | 0 | 0 | 0.88125609 | 0.004799127 |
| PDLIM5 | XM_017007658 | 10.36606958 | 0 | 0 | 0 | 0 | 0 | 0 | 0 | 1.48E-08 | 1 |
| PDP2 | NM_020786 | 0 | 9.102769039 | 0 | 0 | 0 | 0 | 0 | 0 | 5.49E-08 | 1 |
| PDPK1 | XM_024450296 | 0 | 9.117027412 | 0 | 0 | 0 | 0 | 0 | 0 | 5.36E-08 | 1 |
| PDS5A | NM_001100400 | 9.83739315 | 0 | 0 | 0 | 0 | 0 | 0 | 0 | 2.27E-08 | 1 |
| PDS5A | XR_001741188 | 9.649985154 | 0 | 0 | 0 | 0 | 0 | 0 | 0 | 2.95E-08 | 1 |
| PDS5B | XM_017020449 | 0 | 8.33620493 | 0 | 0 | 0 | 0 | 7.106768981 | 0 | 1.25E-07 | 1 |
| PDS5B | XM_017020450 | 0 | 0 | 0 | 0 | 6.582319518 | 9.298742073 | 0 | 10.11563106 | 4.04E-05 | 0.999932046 |
| PDXK | XM_005261199 | 0 | 0 | 0 | 10.68368358 | 9.183860727 | 0 | 9.523920794 | 11.65009295 | 0.013585579 | 0.999932046 |
| PDXK | XM_011529762 | 9.849062238 | 0 | 0 | 0 | 0 | 0 | 7.267409483 | 10.83194973 | 2.23E-08 | 0.02072721 |
| PDXK | XM_017028483 | 0 | 0 | 0 | 0 | 0 | 11.12116681 | 8.521333265 | 0 | 1 | 0.02072721 |
| PDZD11 | XM_017029568 | 0 | 9.657892268 | 0 | 0 | 0 | 0 | 0 | 0 | 2.44E-08 | 1 |
| PDZD3 | XM_011543005 | 0 | 0 | 7.695565578 | 0 | 0 | 0 | 7.421731871 | 0 | 1.56E-05 | 1 |
| PDZD8 | NM_173791 | 0 | 9.495485143 | 0 | 0 | 0 | 0 | 0 | 0 | 2.95E-08 | 1 |
| PEA15 | XM_006711599 | 0 | 0 | 0 | 0 | 6.395391943 | 10.17848999 | 0 | 0 | 4.65E-05 | 1 |
| PECAM1 | NM_000442 | 10.39819601 | 0 | 0 | 0 | 0 | 0 | 0 | 0 | 1.60E-08 | 1 |

| | | | | | | | | | | | |
|---|---|---|---|---|---|---|---|---|---|---|---|
| PECAM1 | XM_017024739 | 0 | 9.232850423 | 0 | 0 | 0 | 0 | 0 | 0 | 4.38E-08 | 1 |
| PECAM1 | XM_017024741 | 0 | 9.484529283 | 0 | 0 | 0 | 0 | 0 | 0 | 2.99E-08 | 1 |
| PEMT | NM_148172 | 0 | 0 | 9.160915107 | 10.17053784 | 9.535521698 | 0 | 0 | 0 | 0.00179366 | 0.002996065 |
| PEPD | NM_001166057 | 0 | 0 | 0 | 10.53239157 | 9.561164523 | 0 | 0 | 0 | 0.013319883 | 0.02072721 |
| PER1 | NM_002616 | 12.48039309 | 10.07613685 | 0 | 0 | 0 | 0 | 9.401869254 | 0 | 0.000102758 | 1 |
| PER1 | XM_005256690 | 9.313422649 | 7.291221295 | 0 | 0 | 0 | 0 | 0 | 0 | 0.005875572 | 1 |
| PER1 | XM_024450803 | 0 | 9.029283553 | 0 | 0 | 0 | 0 | 0 | 0 | 6.49E-08 | 1 |
| PES1 | XM_017028678 | 0 | 7.777608832 | 0 | 10.72004941 | 6.544252221 | 0 | 0 | 0 | 0.922322894 | 0.02072721 |
| PET100 | NR_033242 | 0 | 8.054440838 | 9.058313897 | 0 | 7.593532742 | 0 | 0 | 0 | 0.999953634 | 0.027966656 |
| PEX10 | NM_002617 | 10.34562109 | 0 | 0 | 0 | 0 | 0 | 0 | 0 | 2.06E-08 | 1 |
| PEX10 | NM_153818 | 0 | 8.602554319 | 0 | 0 | 0 | 0 | 0 | 0 | 9.73E-08 | 1 |
| PEX16 | NM_057174 | 0 | 9.158976035 | 0 | 0 | 0 | 0 | 0 | 0 | 5.01E-08 | 1 |
| PEX19 | NM_002857 | 0 | 0 | 0 | 10.01091583 | 7.599732499 | 0 | 0 | 0 | 0.02494212 | 0.02072721 |
| PEX26 | NM_017929 | 10.50162657 | 0 | 10.05474104 | 0 | 7.890136184 | 0 | 0 | 0 | 0.999953634 | 0.02072721 |
| PEX3 | NM_003630 | 0 | 0 | 0 | 8.9089924 | 9.156784755 | 0 | 6.276743329 | 0 | 0.027557563 | 0.999932046 |
| PEX5 | XM_017019750 | 0 | 8.372314435 | 0 | 0 | 0 | 0 | 6.496475094 | 0 | 1.20E-07 | 1 |
| PEX6 | NR_133009 | 0 | 9.600697383 | 0 | 0 | 0 | 0 | 0 | 0 | 2.56E-08 | 1 |
| PEX7 | NM_000288 | 0 | 0 | 7.061996878 | 10.17053784 | 8.173927142 | 0 | 0 | 0 | 0.004902567 | 0.007638895 |
| PF4 | NM_002619 | 0 | 8.497482465 | 0 | 0 | 0 | 0 | 0 | 0 | 1.06E-07 | 1 |
| PFDN6 | NM_001185181 | 0 | 0 | 0 | 9.768560191 | 10.18674423 | 0 | 0 | 0 | 0.014428755 | 0.02072721 |
| PFDN6 | NM_001265596 | 10.50162657 | 0 | 0 | 0 | 0 | 0 | 8.427271157 | 10.23539302 | 1.41E-08 | 0.02072721 |
| PFKFB2 | XM_024447656 | 0 | 8.612652582 | 0 | 0 | 0 | 0 | 0 | 0 | 9.67E-08 | 1 |
| PFKFB2 | XM_024447657 | 0 | 8.323964754 | 0 | 0 | 0 | 0 | 0 | 0 | 1.27E-07 | 1 |
| PFKP | NM_001345944 | 9.854861577 | 0 | 0 | 0 | 0 | 0 | 0 | 9.622856517 | 2.23E-08 | 1 |
| PFN2 | NM_002628 | 0 | 0 | 10.25165197 | 9.913784443 | 9.051033866 | 0 | 0 | 0 | 0.001554654 | 0.002651192 |
| PGAP2 | XM_024448442 | 0 | 8.709928179 | 0 | 0 | 0 | 0 | 0 | 0 | 8.96E-08 | 1 |
| PGAP2 | XM_024448446 | 8.983616415 | 0 | 0 | 0 | 0 | 0 | 8.101525945 | 0 | 7.75E-08 | 1 |
| PGAP3 | NM_001291730 | 10.23434896 | 0 | 0 | 0 | 0 | 9.968861605 | 7.663696786 | 11.1274199 | 1.59E-08 | 0.001765436 |
| PGBD2 | XM_011544161 | 0 | 8.441930184 | 0 | 0 | 0 | 0 | 0 | 0 | 1.12E-07 | 1 |
| PGF | NM_001207012 | 10.35792493 | 0 | 0 | 0 | 0 | 0 | 0 | 0 | 1.48E-08 | 1 |
| PGGT1B | XR_001742121 | 0 | 8.935855595 | 0 | 0 | 0 | 0 | 0 | 0 | 7.69E-08 | 1 |
| PGLS | XM_011527897 | 0 | 8.519117955 | 0 | 0 | 0 | 0 | 0 | 5.654964161 | 1.04E-07 | 0.454496582 |
| PGM1 | NM_002633 | 0 | 0 | 11.38780484 | 0 | 8.140296582 | 10.75869389 | 0 | 0 | 0.011106879 | 0.999932046 |
| PGM2 | XM_011513711 | 0 | 8.774317692 | 0 | 0 | 0 | 0 | 0 | 0 | 8.46E-08 | 1 |
| PGM3 | XR_942478 | 0 | 0 | 0 | 0 | 0 | 0 | 8.456139264 | 11.64020606 | 1 | 2.51E-09 |
| PGS1 | XR_934609 | 0 | 9.102769039 | 0 | 0 | 0 | 0 | 0 | 0 | 5.49E-08 | 1 |
| PHACTR1 | XM_017010469 | 0 | 0 | 0 | 0 | 0 | 0 | 8.470359551 | 10.25171828 | 1 | 0.02072721 |
| PHACTR2 | NM_014721 | 0 | 10.68358807 | 0 | 0 | 0 | 0 | 0 | 0 | 1.34E-08 | 1 |
| PHACTR4 | XM_017002162 | 0 | 9.860035122 | 0 | 0 | 0 | 0 | 8.143544083 | 0 | 1.97E-08 | 1 |
| PHC1 | XM_017018958 | 10.11377196 | 0 | 0 | 0 | 0 | 0 | 0 | 0 | 1.70E-08 | 1 |
| PHC3 | XR_924181 | 0 | 0 | 0 | 10.24854363 | 5.646039634 | 0 | 4.596451906 | 0 | 0.026426152 | 0.594061793 |
| PHF1 | NR_027692 | 0 | 0 | 0 | 0 | 0 | 10.82742309 | 7.382166205 | 0 | 1 | 0.02072721 |
| PHF10 | NM_133325 | 0 | 0 | 0 | 8.01802849 | 0 | 0 | 0 | 0 | 1.03E-05 | 1 |

| | | | | | | | | | | | |
|---|---|---|---|---|---|---|---|---|---|---|---|
| PHF12 | NM_020889 | 0 | 9.08836834 | 0 | 0 | 0 | 0 | 0 | 0 | 5.61E-08 | 1 |
| PHF14 | XM_017012814 | 0 | 8.29916838 | 0 | 0 | 0 | 0 | 8.712918017 | 0 | 1.31E-07 | 1 |
| PHF19 | NM_001009936 | 0 | 0 | 0 | 9.652931436 | 9.648982193 | 0 | 0 | 10.44299731 | 0.018221818 | 1 |
| PHF20L1 | XR_001745536 | 0 | 9.06649413 | 0 | 0 | 0 | 0 | 3.188585034 | 0 | 7.75E-08 | 0.999932046 |
| PHF21A | NR_147890 | 10.58074365 | 9.686856599 | 9.552435718 | 0 | 9.139866469 | 0 | 0 | 0 | 0.999953634 | 0.02072721 |
| PHF21A | XM_011520179 | 0 | 9.855790228 | 0 | 0 | 0 | 0 | 0 | 0 | 1.98E-08 | 1 |
| PHF21A | XM_017017892 | 0 | 8.671801345 | 0 | 0 | 0 | 0 | 0 | 0 | 9.15E-08 | 1 |
| PHF21A | XM_017017893 | 9.623051988 | 0 | 0 | 0 | 0 | 0 | 0 | 0 | 2.93E-08 | 1 |
| PHF21A | XM_017017906 | 0 | 9.387883956 | 0 | 0 | 0 | 0 | 0 | 0 | 3.40E-08 | 1 |
| PHF21A | XM_017017907 | 10.48676361 | 0 | 0 | 0 | 0 | 0 | 0 | 0 | 1.44E-08 | 1 |
| PHF21A | XM_024448577 | 11.22929222 | 0 | 0 | 0 | 0 | 0 | 6.953230848 | 0 | 1.34E-08 | 1 |
| PHF21A | XR_001747902 | 11.07636344 | 8.991087758 | 0 | 0 | 0 | 0 | 8.704922178 | 0 | 0.000604515 | 1 |
| PHKB | XM_017023282 | 9.849062238 | 0 | 0 | 0 | 0 | 0 | 0 | 0 | 2.23E-08 | 1 |
| PHLDB1 | NM_001144759 | 8.850625659 | 9.051724868 | 0 | 0 | 0 | 0 | 0 | 0 | 0.003390542 | 1 |
| PHLDB1 | XR_947813 | 0 | 9.885245492 | 0 | 0 | 0 | 10.04087446 | 0 | 0 | 2.00E-08 | 1 |
| PHRF1 | XM_011520237 | 0 | 0 | 0 | 10.27811433 | 6.818595315 | 0 | 6.953230848 | 0 | 0.023629222 | 0.999932046 |
| PHTF2 | XM_024446841 | 0 | 0 | 11.68400969 | 0 | 9.146234099 | 9.645241301 | 0 | 0 | 0.007920949 | 0.999932046 |
| PHYH | NM_001323082 | 0 | 9.095586658 | 0 | 0 | 0 | 0 | 0 | 0 | 5.54E-08 | 1 |
| PHYKPL | XM_024446250 | 9.134275178 | 8.248259235 | 0 | 0 | 0 | 0 | 0 | 0 | 0.004572986 | 1 |
| PI16 | NM_153370 | 0 | 8.765293275 | 0 | 0 | 0 | 0 | 0 | 0 | 8.55E-08 | 1 |
| PI4KAP1 | NR_003563 | 0 | 0 | 0 | 8.61278646 | 0 | 0 | 0 | 0 | 5.47E-06 | 1 |
| PIAS1 | XM_011522127 | 10.10892695 | 10.58445714 | 10.24582768 | 0 | 7.920172326 | 0 | 0 | 0 | 0.999953634 | 0.02072721 |
| PIAS1 | XM_017022689 | 11.25583468 | 0 | 0 | 0 | 0 | 0 | 0 | 0 | 1.34E-08 | 1 |
| PIAS2 | NM_001324046 | 8.791378415 | 0 | 0 | 0 | 0 | 0 | 0 | 0 | 9.38E-08 | 1 |
| PIAS3 | NM_006099 | 0 | 0 | 0 | 6.727473068 | 0 | 0 | 0 | 0 | 3.78E-05 | 1 |
| PIBF1 | NM_001349655 | 0 | 0 | 0 | 9.301045263 | 7.768830875 | 0 | 7.994077632 | 0 | 0.03576428 | 0.999932046 |
| PICALM | NM_001206947 | 0 | 0 | 10.61761484 | 0 | 8.10150079 | 10.89086298 | 8.519054934 | 0 | 0.017348084 | 0.999932046 |
| PICALM | XM_005274337 | 0 | 8.33620493 | 0 | 0 | 0 | 0 | 0 | 0 | 1.25E-07 | 1 |
| PICALM | XM_017018382 | 0 | 9.271606013 | 0 | 0 | 0 | 0 | 0 | 0 | 4.15E-08 | 1 |
| PID1 | NM_001100818 | 0 | 0 | 9.808316306 | 0 | 8.564658559 | 0 | 0 | 0 | 0.023562616 | 0.02072721 |
| PID1 | XM_017004404 | 9.689464985 | 9.06649413 | 0 | 0 | 0 | 0 | 0 | 0 | 0.001886644 | 1 |
| PIDD1 | XR_001747922 | 9.897629105 | 0 | 0 | 0 | 0 | 0 | 0 | 0 | 2.08E-08 | 1 |
| PIFO | NM_181643 | 10.77829302 | 0 | 0 | 0 | 0 | 0 | 0 | 0 | 1.34E-08 | 1 |
| PIGBOS1 | NM_001308422 | 0 | 0 | 10.46554672 | 0 | 8.153000089 | 0 | 0 | 0 | 0.018596161 | 0.02072721 |
| PIGBOS1 | NM_001308423 | 0 | 0 | 0 | 7.453233115 | 0 | 0 | 9.398154874 | 0 | 1.84E-05 | 1 |
| PIGH | NM_004569 | 0 | 0 | 10.4672163 | 0 | 7.002974523 | 0 | 0 | 0 | 0.021362935 | 0.02072721 |
| PIGL | XM_017025350 | 10.70152235 | 0 | 10.89694145 | 9.575655325 | 7.313250256 | 0 | 0 | 0 | 0.999953634 | 0.00302572 |
| PIGQ | NM_004204 | 9.171928589 | 0 | 0 | 0 | 0 | 0 | 0 | 0 | 7.86E-08 | 1 |
| PIGQ | NM_148920 | 0 | 0 | 9.621534579 | 0 | 4.743383171 | 0 | 0 | 0 | 0.044437493 | 1 |
| PIGR | NM_002644 | 13.41061056 | 12.42877781 | 14.07703174 | 10.86512723 | 0 | 0 | 0 | 0 | 0.999953634 | 1.92E-09 |
| PIGR | XM_011509629 | 14.59877535 | 14.79343765 | 13.54333114 | 12.01316774 | 9.029380004 | 0 | 0 | 0 | 0.763613371 | 0.000184208 |
| PIGT | NR_047694 | 0 | 8.622680652 | 0 | 0 | 0 | 0 | 0 | 0 | 9.58E-08 | 1 |
| PIGV | NM_017837 | 11.35635345 | 0 | 0 | 0 | 0 | 0 | 8.923953207 | 0 | 1.34E-08 | 1 |

| | | | | | | | | | | | |
|---|---|---|---|---|---|---|---|---|---|---|---|
| PIGX | NM_017861 | 8.421077904 | 0 | 0 | 10.17709286 | 9.317757856 | 0 | 0 | 0 | 0.997913553 | 0.021157255 |
| PIGY-DT | NR_134663 | 0 | 0 | 8.100280313 | 0 | 0 | 0 | 0 | 0 | 1.03E-05 | 1 |
| PIH1D2 | XM_017017201 | 10.06457365 | 0 | 0 | 0 | 0 | 0 | 0 | 0 | 1.80E-08 | 1 |
| PIK3C2B | XR_241085 | 9.476527956 | 0 | 10.9682383 | 0 | 6.337212177 | 0 | 0 | 0 | 0.999953634 | 1.49E-08 |
| PIK3C3 | XM_011526031 | 0 | 8.709928179 | 0 | 0 | 0 | 0 | 0 | 0 | 8.96E-08 | 1 |
| PIK3CB | XM_024453596 | 0 | 0 | 9.732596818 | 11.42691374 | 0 | 11.58176358 | 9.218858934 | 11.25193827 | 0.008817413 | 0.999932046 |
| PIK3R2 | NR_073517 | 0 | 10.78082806 | 0 | 0 | 0 | 0 | 8.070745625 | 0 | 1.34E-08 | 1 |
| PIK3R3 | NM_001328654 | 8.994164854 | 0 | 0 | 0 | 0 | 0 | 0 | 0 | 7.73E-08 | 1 |
| PIM2 | NM_006875 | 9.560548562 | 0 | 0 | 0 | 0 | 0 | 0 | 10.69777882 | 3.19E-08 | 1 |
| PIN1 | XM_011528068 | 0 | 0 | 0 | 9.159924829 | 7.920172326 | 0 | 0 | 0 | 0.037163677 | 0.025805586 |
| PINLYP | NM_001321124 | 10.78286315 | 0 | 0 | 0 | 0 | 0 | 0 | 0 | 1.34E-08 | 1 |
| PINX1 | NM_017884 | 9.721559119 | 0 | 10.63405836 | 10.60419453 | 0 | 0 | 0 | 0 | 0.999953634 | 0.02072721 |
| PIP4K2C | XM_005269152 | 0 | 0 | 10.38127638 | 9.785818491 | 7.930046999 | 0 | 0 | 10.26625472 | 0.002079161 | 0.999932046 |
| PIP4P1 | NM_144568 | 0 | 0 | 0 | 9.575655325 | 6.758905041 | 0 | 0 | 0 | 0.03628733 | 0.025627121 |
| PIP5K1C | XM_017026540 | 0 | 10.00508439 | 0 | 0 | 0 | 0 | 0 | 0 | 1.67E-08 | 1 |
| PIR | NM_003662 | 0 | 0 | 0 | 0 | 6.409577539 | 0 | 0 | 0 | 4.59E-05 | 1 |
| PISD | NM_001326411 | 0 | 8.486541819 | 0 | 0 | 0 | 0 | 0 | 0 | 1.09E-07 | 1 |
| PITHD1 | NM_020362 | 0 | 0 | 0 | 11.14203857 | 9.51762409 | 0 | 8.441777415 | 12.17581928 | 0.00992961 | 0.999932046 |
| PITPNC1 | NM_012417 | 10.64239654 | 0 | 0 | 9.816372523 | 7.516979992 | 0 | 0 | 0 | 0.999953634 | 0.02096331 |
| PITPNM2 | XM_024449101 | 10.04441475 | 0 | 9.510949584 | 0 | 7.701245565 | 0 | 0 | 0 | 0.999953634 | 0.02218467 |
| PITX1 | NM_002653 | 0 | 0 | 0 | 9.649164374 | 5.96699818 | 0 | 0 | 0 | 0.037888838 | 0.0268468 |
| PKHD1L1 | NM_177531 | 9.866390785 | 9.38199218 | 0 | 0 | 0 | 0 | 0 | 0 | 0.00138583 | 1 |
| PKIG | XM_017027615 | 0 | 8.861586703 | 0 | 0 | 0 | 0 | 0 | 0 | 7.86E-08 | 1 |
| PKM | XM_006720570 | 0 | 0 | 0 | 9.180949892 | 7.503816758 | 0 | 7.56113023 | 10.14389485 | 0.0405894 | 0.999932046 |
| PKP4 | XM_017005134 | 0 | 0 | 0 | 0 | 6.505153187 | 0 | 7.856483795 | 0 | 4.28E-05 | 1 |
| PLA2G10 | NR_133651 | 8.827217917 | 0 | 0 | 0 | 0 | 0 | 0 | 0 | 8.68E-08 | 1 |
| PLA2G15 | NM_012320 | 0 | 9.073822446 | 0 | 0 | 0 | 0 | 0 | 7.081519821 | 7.73E-08 | 1 |
| PLA2G16 | NM_001128203 | 0 | 0 | 10.67219566 | 0 | 11.42154018 | 0 | 0 | 12.56226568 | 0.006111706 | 0.999932046 |
| PLA2G2A | NM_001161727 | 0 | 13.33378281 | 12.14521328 | 12.15073945 | 11.86041915 | 0 | 0 | 0 | 0.999953634 | 0.000162382 |
| PLA2G6 | XM_005261765 | 9.663265502 | 0 | 0 | 0 | 0 | 0 | 9.758319337 | 0 | 2.79E-08 | 1 |
| PLA2G6 | XM_006724332 | 0 | 0 | 0 | 6.964181636 | 0 | 0 | 0 | 0 | 3.03E-05 | 1 |
| PLA2G6 | XM_024452283 | 9.682959584 | 0 | 0 | 8.474877744 | 7.477124411 | 0 | 0 | 0 | 0.999953634 | 0.041050462 |
| PLA2G6 | XR_937940 | 8.873659666 | 7.213730404 | 10.21535514 | 0 | 8.202724223 | 0 | 0 | 0 | 0.999953634 | 0.02072721 |
| PLAA | XR_001746420 | 0 | 9.445515002 | 0 | 0 | 0 | 0 | 0 | 0 | 3.16E-08 | 1 |
| PLCB4 | XM_024451907 | 9.656640609 | 0 | 0 | 0 | 0 | 0 | 0 | 0 | 3.16E-08 | 1 |
| PLCD4 | XM_017005120 | 0 | 8.508340767 | 0 | 0 | 0 | 0 | 0 | 0 | 1.06E-07 | 1 |
| PLCG1 | NM_002660 | 0 | 0 | 8.709114096 | 0 | 8.110212684 | 10.19378098 | 10.21201706 | 10.49054475 | 0.044846006 | 0.999932046 |
| PLCG1 | XM_011528865 | 10.62213692 | 0 | 0 | 0 | 0 | 0 | 7.479117364 | 0 | 1.34E-08 | 1 |
| PLD1 | XR_001740175 | 0 | 0 | 8.700622994 | 10.0123828 | 0 | 0 | 0 | 0 | 0.021972203 | 0.02072721 |
| PLD3 | XM_017026546 | 10.00323168 | 0 | 0 | 0 | 0 | 0 | 7.382166205 | 0 | 1.94E-08 | 1 |
| PLEK2 | NM_016445 | 9.609394475 | 0 | 9.272065393 | 11.58314844 | 10.32319883 | 0 | 0 | 0 | 0.895127701 | 0.001334576 |
| PLEKHA2 | NM_021623 | 0 | 0 | 0 | 0 | 6.30721813 | 0 | 0 | 0 | 4.99E-05 | 1 |
| PLEKHA2 | XM_024447227 | 0 | 9.186279128 | 0 | 0 | 0 | 10.98311274 | 8.782953729 | 0 | 4.80E-08 | 0.02072721 |

| | | | | | | | | | | | |
|---|---|---|---|---|---|---|---|---|---|---|---|
| PLEKHA4 | NM_020904 | 0 | 9.258802813 | 0 | 0 | 0 | 11.04408862 | 0 | 0 | 4.23E-08 | 1 |
| PLEKHA7 | XM_017017242 | 0 | 0 | 0 | 0 | 6.478485785 | 0 | 8.331847395 | 0 | 4.36E-05 | 1 |
| PLEKHB2 | NM_001100623 | 0 | 0 | 0 | 0 | 0 | 10.80461986 | 0 | 10.38201266 | 1 | 0.02072721 |
| PLEKHG5 | NM_198681 | 0 | 0 | 8.709114096 | 0 | 7.581052723 | 0 | 0 | 0 | 0.053153696 | 0.033333108 |
| PLEKHG6 | XM_011520967 | 11.08625733 | 0 | 10.6488461 | 0 | 7.599732499 | 0 | 0 | 0 | 0.999953634 | 0.020942118 |
| PLEKHG6 | XM_017019555 | 0 | 0 | 11.12187807 | 0 | 7.806841281 | 0 | 9.614206418 | 0 | 0.0136072 | 0.999932046 |
| PLEKHH1 | XR_002957564 | 9.085782918 | 0 | 9.623033138 | 0 | 8.814318721 | 0 | 0 | 0 | 0.999953634 | 0.02072721 |
| PLEKHM2 | XM_005245790 | 10.68534009 | 0 | 0 | 10.2030191 | 9.148350412 | 0 | 0 | 0 | 0.999953634 | 0.02072721 |
| PLEKHM2 | XM_017000758 | 0 | 0 | 0 | 10.32962526 | 10.02339403 | 0 | 7.688238242 | 9.043817182 | 0.012612408 | 0.999932046 |
| PLEKHO2 | NM_025201 | 0 | 9.712905893 | 0 | 0 | 0 | 0 | 0 | 0 | 2.55E-08 | 1 |
| PLIN3 | NM_005817 | 0 | 9.665188049 | 11.36514928 | 0 | 8.452664007 | 0 | 0 | 0 | 0.999953634 | 0.02072721 |
| PLK1 | NM_005030 | 0 | 9.117027412 | 0 | 9.061325395 | 9.122747427 | 0 | 0 | 0 | 0.999953634 | 0.02072721 |
| PLLP | NM_015993 | 0 | 9.029283553 | 0 | 0 | 0 | 0 | 0 | 0 | 6.49E-08 | 1 |
| PLOD2 | XR_001740176 | 0 | 7.411784592 | 10.21833239 | 0 | 8.351586721 | 0 | 0 | 0 | 0.898301168 | 0.02072721 |
| PLPBP | NM_001349346 | 0 | 8.935855595 | 0 | 0 | 0 | 0 | 0 | 0 | 7.69E-08 | 1 |
| PLPBP | NM_001349348 | 10.57724 | 9.073822446 | 11.21176359 | 11.29451383 | 8.070588078 | 0 | 0 | 0 | 0.999953634 | 0.001394691 |
| PLPP2 | NM_003712 | 0 | 0 | 11.11710044 | 11.39465417 | 10.93654241 | 0 | 0 | 0 | 0.000260576 | 0.000628869 |
| PLPP5 | NM_001354456 | 0 | 10.19179195 | 0 | 0 | 0 | 0 | 0 | 0 | 2.01E-08 | 1 |
| PLPPR2 | XM_017027148 | 0 | 8.903318262 | 0 | 0 | 0 | 0 | 0 | 0 | 7.75E-08 | 1 |
| PLRG1 | NM_002669 | 0 | 0 | 10.2747168 | 0 | 6.80255953 | 0 | 0 | 0 | 0.024341057 | 0.02072721 |
| PLS3 | NM_005032 | 0 | 0 | 9.967779298 | 0 | 9.407077595 | 0 | 0 | 9.556068487 | 0.017720024 | 0.999932046 |
| PLSCR1 | XM_005247538 | 10.39819601 | 8.235246214 | 8.697781485 | 0 | 8.798235064 | 0 | 0 | 0 | 0.999953634 | 0.023174056 |
| PLSCR1 | XM_017006629 | 9.457596586 | 8.602554319 | 5.076635497 | 0 | 8.929549395 | 0 | 0 | 0 | 0.999953634 | 0.04359418 |
| PLTP | NM_006227 | 0 | 0 | 8.663239538 | 10.13191995 | 7.757784245 | 0 | 10.14415462 | 10.26464677 | 0.003775641 | 0.999932046 |
| PLVAP | NM_031310 | 10.05452941 | 0 | 0 | 0 | 0 | 0 | 0 | 9.996690239 | 5.06E-08 | 1 |
| PLXNA3 | XR_938510 | 0 | 8.671801345 | 0 | 0 | 0 | 0 | 0 | 0 | 9.15E-08 | 1 |
| PML | NM_033240 | 0 | 0 | 10.21634824 | 9.920050551 | 6.758905041 | 0 | 10.2148352 | 9.607716039 | 0.002744041 | 0.999932046 |
| PMM2 | NM_000303 | 9.539097493 | 0 | 0 | 0 | 0 | 0 | 0 | 0 | 3.28E-08 | 1 |
| PMP22 | NM_153321 | 0 | 0 | 0 | 0 | 6.679179249 | 0 | 0 | 0 | 3.78E-05 | 1 |
| PMS2 | NM_001322005 | 0 | 0 | 9.4173244 | 0 | 9.003894214 | 9.600047624 | 10.20494752 | 8.790297784 | 0.024510347 | 0.999932046 |
| PMS2 | NM_001322009 | 0 | 0 | 10.69801997 | 10.43948342 | 0 | 0 | 0 | 0 | 0.010465596 | 0.02072721 |
| PMS2 | NM_001322010 | 0 | 0 | 0 | 9.357540824 | 7.449928885 | 10.96950906 | 9.430032845 | 10.36410426 | 0.037036695 | 0.999932046 |
| PMS2P9 | NR_028058 | 0 | 0 | 0 | 11.90320441 | 8.314373238 | 11.4750909 | 8.732715858 | 11.55594568 | 1.49E-08 | 0.999932046 |
| PMVK | NM_006556 | 0 | 0 | 9.661459138 | 9.468580228 | 8.006701649 | 10.96950906 | 9.432456067 | 10.71436539 | 0.003441193 | 0.999932046 |
| PNKD | NM_022572 | 0 | 0 | 0 | 0 | 6.409577539 | 0 | 8.251784555 | 0 | 4.59E-05 | 1 |
| PNO1 | NM_001329917 | 0 | 8.652353298 | 0 | 0 | 0 | 0 | 0 | 0 | 9.28E-08 | 1 |
| PNO1 | NM_020143 | 0 | 9.277965255 | 0 | 10.24854363 | 8.522857234 | 0 | 0 | 0 | 0.999953634 | 0.02072721 |
| PNO1 | NR_138146 | 0 | 8.360378074 | 0 | 0 | 0 | 0 | 0 | 0 | 1.21E-07 | 1 |
| PNPLA2 | NM_020376 | 9.015033136 | 0 | 0 | 0 | 0 | 0 | 7.604755783 | 0 | 7.69E-08 | 1 |
| PNPLA2 | XM_024448618 | 0 | 6.672754896 | 0 | 0 | 0 | 9.908830964 | 7.044767963 | 0 | 0.121830059 | 0.02072721 |
| PNPLA4 | NM_004650 | 0 | 0 | 0 | 10.22596092 | 7.97366523 | 0 | 9.898128075 | 0 | 0.021106645 | 0.999932046 |
| PNPLA6 | NM_006702 | 10.33735985 | 0 | 0 | 9.664173964 | 6.679179249 | 0 | 0 | 0 | 0.999953634 | 0.0247531 |
| PNPLA8 | XM_011516274 | 0 | 8.861586703 | 0 | 0 | 0 | 0 | 0 | 0 | 7.86E-08 | 1 |

| | | | | | | | | | | | |
|---|---|---|---|---|---|---|---|---|---|---|---|
| PNPO | XM_017024813 | 0 | 0 | 0 | 0 | 0 | 0 | 8.496071524 | 9.058739343 | 1 | 0.02072721 |
| PNRC2 | XM_017001691 | 0 | 9.993538654 | 8.869215857 | 9.114202495 | 8.114548983 | 0 | 0 | 0 | 0.999953634 | 0.00838335 |
| PODXL2 | NM_015720 | 0 | 0 | 10.39362058 | 10.02842178 | 7.82812018 | 0 | 0 | 0 | 0.001907192 | 0.00302572 |
| POFUT1 | XR_001754218 | 0 | 0 | 0 | 9.357540824 | 7.178913994 | 10.30194482 | 7.621843212 | 0 | 0.038968883 | 0.999932046 |
| POGLUT1 | XR_001740212 | 0 | 8.975521721 | 0 | 0 | 0 | 0 | 0 | 0 | 7.27E-08 | 1 |
| POLB | XM_005273536 | 0 | 0 | 0 | 10.80714503 | 9.35488525 | 12.63291691 | 10.54662216 | 11.35224584 | 0.012287872 | 0.999932046 |
| POLD2 | XR_002956452 | 0 | 0 | 0 | 0 | 0 | 10.90379525 | 8.000624564 | 0 | 1 | 0.02072721 |
| POLDIP3 | XM_017029010 | 10.77676642 | 8.453213086 | 0 | 0 | 0 | 0 | 8.566160223 | 0 | 0.000967049 | 1 |
| POLE | XR_002957339 | 0 | 0 | 9.751902512 | 0 | 7.357826773 | 0 | 9.329606891 | 0 | 0.030725933 | 0.999932046 |
| POLE3 | NR_027261 | 0 | 0 | 0 | 10.5924728 | 8.127480221 | 10.93981942 | 9.394430906 | 0 | 0.017164245 | 0.999932046 |
| POLI | NM_007195 | 0 | 8.719304505 | 0 | 0 | 0 | 0 | 0 | 0 | 8.96E-08 | 1 |
| POLR1B | NM_019014 | 0 | 10.44448006 | 0 | 0 | 0 | 0 | 0 | 0 | 1.34E-08 | 1 |
| POLR2D | NM_004805 | 0 | 0 | 0 | 12.09194223 | 8.620607807 | 0 | 0 | 0 | 1.36E-08 | 4.99E-09 |
| POLR2E | NM_001316323 | 0 | 0 | 0 | 9.686399554 | 7.10238406 | 0 | 0 | 0 | 0.032357265 | 0.023261296 |
| POLR2E | XM_017026883 | 0 | 0 | 0 | 8.483365391 | 0 | 0 | 8.377835658 | 0 | 2.83E-05 | 1 |
| POLR2F | NR_125371 | 0 | 0 | 0 | 10.26093849 | 8.400349091 | 0 | 0 | 0 | 0.019361142 | 0.02072721 |
| POLR2J3 | XM_024446812 | 0 | 8.844550112 | 0 | 0 | 0 | 0 | 0 | 0 | 8.06E-08 | 1 |
| POLR3A | NM_007055 | 10.17994723 | 0 | 0 | 0 | 0 | 0 | 7.525254359 | 0 | 1.60E-08 | 1 |
| POLR3E | XM_017023443 | 0 | 0 | 0 | 8.860946453 | 7.251587968 | 0 | 0 | 0 | 0.051780798 | 0.035213518 |
| POLR3F | NM_006466 | 0 | 9.445515002 | 0 | 0 | 0 | 0 | 0 | 0 | 3.16E-08 | 1 |
| POLR3F | NR_104209 | 0 | 0 | 0 | 0 | 6.702411139 | 0 | 0 | 0 | 3.69E-05 | 1 |
| POLR3H | NM_001018052 | 0 | 9.674858717 | 0 | 0 | 0 | 10.33626944 | 0 | 0 | 2.40E-08 | 1 |
| POLRMT | NM_005035 | 0 | 0 | 0 | 0 | 6.569741695 | 0 | 7.827317174 | 0 | 4.12E-05 | 1 |
| POLRMT | XR_001753708 | 9.442270566 | 0 | 0 | 0 | 0 | 0 | 0 | 0 | 3.84E-08 | 1 |
| POMT1 | NM_001353200 | 0 | 8.991087758 | 0 | 0 | 0 | 0 | 0 | 0 | 7.02E-08 | 1 |
| POMZP3 | NM_012230 | 0 | 0 | 10.13675763 | 9.74051016 | 0 | 0 | 0 | 0 | 0.016372233 | 0.02072721 |
| PON2 | NM_000305 | 0 | 0 | 0 | 0 | 0 | 11.3197171 | 9.030164031 | 10.72843224 | 1 | 0.000964324 |
| POR | NM_000941 | 0 | 0 | 9.757371304 | 0 | 7.920172326 | 0 | 7.613324795 | 9.793135989 | 0.027713773 | 0.999932046 |
| POSTN | NM_001330517 | 0 | 0 | 0 | 9.39850643 | 7.415197319 | 0 | 0 | 0 | 0.036402051 | 0.025566236 |
| POT1 | NM_001042594 | 0 | 0 | 10.07239941 | 9.362150426 | 9.812715433 | 0 | 8.572765732 | 10.55247661 | 0.001582719 | 0.999932046 |
| POT1 | NM_015450 | 0 | 0 | 10.92028832 | 11.70018327 | 11.32450857 | 0 | 10.63974118 | 11.87751496 | 0.000204194 | 0.999932046 |
| POT1 | NR_003102 | 0 | 8.571828374 | 0 | 0 | 0 | 0 | 0 | 0 | 9.97E-08 | 1 |
| POT1 | XR_001744618 | 0 | 0 | 0 | 0 | 0 | 7.897609146 | 0 | 9.904707713 | 1 | 0.02072721 |
| POU2AF1 | XM_006718860 | 10.92746477 | 0 | 0 | 0 | 0 | 0 | 0 | 0 | 1.34E-08 | 1 |
| POU2AF1 | XM_017017932 | 0 | 10.32058425 | 0 | 0 | 0 | 0 | 0 | 0 | 1.45E-08 | 1 |
| POU2F1 | XM_024447727 | 10.38222222 | 0 | 0 | 0 | 0 | 0 | 0 | 0 | 1.47E-08 | 1 |
| PPA1 | XR_002956985 | 0 | 0 | 0 | 0 | 6.690841957 | 0 | 0 | 0 | 3.72E-05 | 1 |
| PPARA | XR_001755253 | 11.1618877 | 0 | 0 | 0 | 0 | 0 | 0 | 0 | 1.34E-08 | 1 |
| PPARD | XM_024446474 | 0 | 0 | 0 | 0 | 0 | 0 | 9.389450616 | 7.247007203 | 1 | 0.020835214 |
| PPARG | XM_011533843 | 0 | 0 | 11.20527557 | 10.71915135 | 12.14453468 | 13.19723285 | 13.0756502 | 11.08249291 | 0.000180274 | 0.999932046 |
| PPARG | XM_024453605 | 0 | 0 | 10.31049833 | 10.70923559 | 9.965158496 | 0 | 8.60104519 | 0 | 0.000763192 | 0.999932046 |
| PPCDC | NM_021823 | 0 | 9.006487637 | 0 | 0 | 0 | 0 | 8.083136668 | 0 | 6.80E-08 | 1 |
| PPDPF | NM_001353423 | 0 | 9.081113725 | 0 | 0 | 0 | 0 | 0 | 0 | 5.70E-08 | 1 |

| | | | | | | | | | | | |
|---|---|---|---|---|---|---|---|---|---|---|---|
| PPFIA1 | NR_045286 | 0 | 0 | 9.546130371 | 11.01095056 | 0 | 0 | 9.523920794 | 12.30502668 | 0.01152683 | 0.999932046 |
| PPFIA1 | XM_011545309 | 10.18915784 | 0 | 9.229414648 | 0 | 7.516979992 | 0 | 0 | 0 | 0.999953634 | 0.025915601 |
| PPFIA1 | XM_011545314 | 10.41791771 | 0 | 0 | 0 | 0 | 0 | 0 | 0 | 1.47E-08 | 1 |
| PPFIA2 | NM_001220477 | 0 | 9.830055185 | 0 | 0 | 0 | 0 | 0 | 0 | 2.01E-08 | 1 |
| PPFIA3 | XM_017027407 | 10.31649738 | 0 | 0 | 0 | 0 | 0 | 0 | 9.736260554 | 1.52E-08 | 1 |
| PPFIBP2 | NM_001256569 | 9.771469405 | 0 | 0 | 0 | 0 | 0 | 0 | 0 | 2.47E-08 | 1 |
| PPHLN1 | XR_001748751 | 0 | 0 | 0 | 0 | 0 | 0 | 6.21051493 | 10.0424498 | 1 | 0.02072721 |
| PPIE | XM_006710290 | 0 | 0 | 0 | 10.41529045 | 7.503816758 | 0 | 0 | 0 | 0.020327145 | 0.02072721 |
| PPIG | XM_005246967 | 0 | 9.974088018 | 0 | 0 | 0 | 0 | 8.720869785 | 0 | 1.72E-08 | 1 |
| PPIL1 | NM_016059 | 0 | 10.05036496 | 0 | 0 | 0 | 0 | 0 | 9.655125759 | 1.60E-08 | 1 |
| PPIL2 | XM_017028706 | 9.765324233 | 0 | 0 | 0 | 0 | 0 | 0 | 0 | 2.50E-08 | 1 |
| PPIL3 | XM_017004354 | 0 | 9.296876505 | 0 | 0 | 0 | 0 | 0 | 0 | 4.01E-08 | 1 |
| PPIP5K1 | XM_017022754 | 0 | 0 | 7.826204809 | 0 | 0 | 0 | 0 | 0 | 1.37E-05 | 1 |
| PPIP5K2 | XM_017009274 | 0 | 9.358181476 | 0 | 0 | 0 | 0 | 0 | 0 | 3.64E-08 | 1 |
| PPL | NM_002705 | 0 | 0 | 0 | 8.893154207 | 8.522857234 | 0 | 6.703265946 | 0 | 0.035170076 | 0.999932046 |
| PPM1B | NM_001033557 | 0 | 9.315543066 | 0 | 0 | 0 | 0 | 0 | 0 | 3.90E-08 | 1 |
| PPM1B | XM_017004395 | 9.153224721 | 8.747073305 | 0 | 0 | 0 | 0 | 0 | 0 | 0.003389097 | 1 |
| PPM1B | XM_017004397 | 0 | 8.550974732 | 9.929535327 | 0 | 7.263352189 | 0 | 0 | 0 | 0.999953634 | 0.02072721 |
| PPM1G | NM_177983 | 10.37012473 | 11.22118484 | 0 | 10.98129422 | 9.735987562 | 0 | 0 | 0 | 0.999953634 | 0.02072721 |
| PPM1H | XM_017019676 | 0 | 8.801057108 | 0 | 0 | 0 | 0 | 8.588061978 | 0 | 8.24E-08 | 1 |
| PPME1 | NM_016147 | 0 | 0 | 10.74557285 | 9.637803806 | 7.227767768 | 0 | 0 | 0 | 0.00207886 | 0.003174145 |
| PPP1CC | XM_011538505 | 0 | 0 | 9.458195643 | 9.227165049 | 0 | 10.35940264 | 0 | 0 | 0.02436216 | 0.999932046 |
| PPP1R11 | XM_006715174 | 0 | 0 | 10.95758765 | 0 | 7.503816758 | 0 | 0 | 0 | 0.015473203 | 0.02072721 |
| PPP1R12B | XR_001737200 | 0 | 8.991087758 | 0 | 0 | 0 | 0 | 0 | 0 | 7.02E-08 | 1 |
| PPP1R12C | XM_005259013 | 0 | 9.405416277 | 0 | 0 | 0 | 0 | 0 | 0 | 3.31E-08 | 1 |
| PPP1R13B | XM_017021116 | 0 | 10.97923632 | 10.20838425 | 0 | 8.800928156 | 0 | 0 | 0 | 0.999953634 | 0.02072721 |
| PPP1R27 | NM_001007533 | 0 | 0 | 6.656419865 | 0 | 0 | 0 | 0 | 0 | 4.26E-05 | 1 |
| PPP1R32 | NM_001170753 | 0 | 10.19515592 | 0 | 0 | 0 | 0 | 0 | 0 | 1.51E-08 | 1 |
| PPP1R35 | NM_001346938 | 0 | 8.818611963 | 0 | 0 | 0 | 0 | 7.712369201 | 0 | 8.12E-08 | 1 |
| PPP1R35 | NM_145030 | 0 | 0 | 11.26216262 | 11.37140683 | 7.483843939 | 0 | 0 | 10.24194531 | 0.000678214 | 1 |
| PPP1R35 | XM_011515914 | 0 | 0 | 0 | 7.188387521 | 9.414130609 | 0 | 0 | 0 | 0.036665092 | 0.02519635 |
| PPP1R3E | NM_001276318 | 0 | 8.475517571 | 0 | 0 | 0 | 0 | 0 | 0 | 1.09E-07 | 1 |
| PPP1R7 | NM_001282409 | 0 | 9.799439006 | 0 | 0 | 0 | 0 | 0 | 0 | 2.04E-08 | 1 |
| PPP1R7 | NM_001282413 | 0 | 0 | 0 | 10.74849617 | 8.230957729 | 0 | 0 | 0 | 0.01554975 | 0.02072721 |
| PPP1R9B | NM_032595 | 0 | 8.927789798 | 0 | 0 | 0 | 0 | 6.869864959 | 0 | 7.69E-08 | 1 |
| PPP2R2A | NM_001177591 | 10.29532878 | 0 | 10.71568536 | 0 | 11.98154058 | 0 | 0 | 0 | 0.999953634 | 0.019357428 |
| PPP2R2D | NM_018461 | 0 | 8.967675264 | 0 | 0 | 0 | 0 | 0 | 9.297616749 | 7.35E-08 | 1 |
| PPP2R3B | NM_013239 | 0 | 0 | 0 | 0 | 6.747781854 | 0 | 0 | 0 | 9.53E-05 | 1 |
| PPP2R3C | XM_005267782 | 0 | 9.158976035 | 0 | 0 | 0 | 0 | 0 | 0 | 5.01E-08 | 1 |
| PPP2R3C | XM_024449639 | 0 | 0 | 0 | 0 | 6.557053251 | 0 | 0 | 0 | 4.16E-05 | 1 |
| PPP2R5B | XM_011545134 | 0 | 0 | 0 | 0 | 6.690841957 | 0 | 0 | 0 | 3.72E-05 | 1 |
| PPP4C | NM_001303503 | 0 | 0 | 10.96587827 | 9.932501706 | 0 | 0 | 0 | 0 | 0.010933472 | 0.02072721 |
| PPP4C | NM_002720 | 0 | 0 | 12.016009 | 11.49039939 | 9.228554799 | 0 | 9.357928127 | 10.56036215 | 0.000281734 | 0.999932046 |

| | | | | | | | | | | | |
|---|---|---|---|---|---|---|---|---|---|---|---|
| PPP4R1 | NM_005134 | 0 | 0 | 11.51027577 | 0 | 8.811650533 | 0 | 0 | 0 | 0.009335806 | 0.02072721 |
| PPP4R1 | NR_052003 | 9.457596586 | 0 | 0 | 0 | 0 | 0 | 0 | 0 | 3.78E-08 | 1 |
| PPP4R1 | XM_011525776 | 0 | 0 | 11.18563458 | 10.07970008 | 0 | 0 | 0 | 0 | 0.009435887 | 0.02072721 |
| PPP4R3A | NM_001284280 | 0 | 9.428466121 | 0 | 9.267038066 | 10.10193035 | 0 | 0 | 0 | 0.999953634 | 0.02072721 |
| PPP4R3B | XR_001738855 | 0 | 0 | 10.2043857 | 0 | 8.759994193 | 0 | 8.192973732 | 0 | 0.018852492 | 0.999932046 |
| PPP6R2 | XM_024452307 | 0 | 9.021724865 | 0 | 0 | 0 | 0 | 0 | 0 | 6.61E-08 | 1 |
| PPP6R3 | XM_024448601 | 0 | 0 | 0 | 10.08528677 | 6.758905041 | 0 | 8.013629967 | 9.917043278 | 0.026537558 | 0.999932046 |
| PPT1 | NM_000310 | 0 | 0 | 10.4838066 | 0 | 7.415197319 | 0 | 7.552244553 | 11.66601602 | 0.020252726 | 0.999932046 |
| PQBP1 | XM_017029207 | 0 | 0 | 9.313491365 | 10.49195269 | 6.041763794 | 0 | 0 | 0 | 0.003333553 | 0.005121877 |
| PQLC1 | XR_002958186 | 0 | 7.881486713 | 0 | 9.55567177 | 8.63275172 | 0 | 0 | 0 | 0.999953634 | 0.02072721 |
| PRAC2 | NM_001282275 | 0 | 8.959785899 | 0 | 0 | 0 | 0 | 0 | 0 | 7.69E-08 | 1 |
| PRAC2 | XM_011524747 | 0 | 0 | 7.513641688 | 0 | 0 | 0 | 0 | 0 | 1.89E-05 | 1 |
| PRADC1 | NM_032319 | 0 | 8.709928179 | 11.69580664 | 0 | 9.383914218 | 0 | 0 | 0 | 0.886651935 | 0.02072721 |
| PRAP1 | NM_001145201 | 10.44508285 | 0 | 9.870198585 | 0 | 7.372384706 | 0 | 0 | 0 | 0.999953634 | 0.02072721 |
| PRC1 | NM_003981 | 0 | 0 | 0 | 12.94144883 | 9.400877796 | 0 | 10.1172953 | 11.14413191 | 1.34E-08 | 0.999932046 |
| PRC1 | XM_011522188 | 0 | 0 | 10.89694145 | 10.08807202 | 12.02033329 | 5.708506274 | 8.740559545 | 11.37104854 | 0.000272514 | 0.999932046 |
| PRC1 | XM_017022713 | 0 | 0 | 0 | 11.14404748 | 10.62850791 | 0 | 7.980893958 | 9.945422596 | 0.007195098 | 0.999932046 |
| PRDM11 | XM_011520228 | 10.35792493 | 0 | 0 | 0 | 0 | 0 | 0 | 0 | 1.48E-08 | 1 |
| PRDM2 | NM_001135610 | 0 | 0 | 10.39712818 | 11.43240356 | 9.212462155 | 0 | 0 | 0 | 0.000634066 | 0.001394691 |
| PRDM2 | NM_012231 | 0 | 9.962290505 | 0 | 0 | 0 | 0 | 8.11968377 | 0 | 1.76E-08 | 1 |
| PRDM2 | XM_017002263 | 0 | 0 | 0 | 10.97152315 | 11.00237015 | 12.85669397 | 10.32522901 | 10.84814505 | 0.006714027 | 0.999932046 |
| PRDX3 | NM_001302272 | 0 | 0 | 0 | 7.345621569 | 0 | 0 | 0 | 10.10128877 | 2.05E-05 | 1 |
| PRDX5 | NM_001358511 | 0 | 0 | 10.74419638 | 0 | 8.262562196 | 10.77833282 | 9.562044322 | 0 | 0.015833405 | 0.999932046 |
| PRELID2 | NM_138492 | 0 | 0 | 0 | 10.16790747 | 8.790125492 | 0 | 0 | 0 | 0.018782969 | 0.02072721 |
| PREX1 | XM_011528934 | 9.992749568 | 0 | 0 | 0 | 0 | 0 | 0 | 0 | 2.01E-08 | 1 |
| PRICKLE3 | NM_006150 | 0 | 0 | 0 | 0 | 6.531336591 | 0 | 0 | 0 | 4.23E-05 | 1 |
| PRIM1 | NM_000946 | 0 | 0 | 0 | 10.98129422 | 8.987438391 | 0 | 0 | 10.99792086 | 0.012112499 | 0.999932046 |
| PRKAA1 | XM_017009624 | 0 | 8.719304505 | 0 | 0 | 0 | 0 | 0 | 0 | 8.96E-08 | 1 |
| PRKAB1 | NM_006253 | 0 | 8.878424457 | 11.44653594 | 0 | 7.470373439 | 0 | 0 | 0 | 0.991435662 | 7.83E-09 |
| PRKAB2 | NM_005399 | 9.727893151 | 0 | 0 | 0 | 0 | 0 | 0 | 0 | 2.55E-08 | 1 |
| PRKACB | NM_001300916 | 0 | 8.801057108 | 0 | 0 | 0 | 0 | 0 | 0 | 8.24E-08 | 1 |
| PRKACB | NM_207578 | 0 | 0 | 7.532901745 | 0 | 0 | 0 | 0 | 0 | 1.85E-05 | 1 |
| PRKACB | XM_005271020 | 8.307253465 | 9.06649413 | 0 | 0 | 0 | 0 | 0 | 0 | 0.004571979 | 1 |
| PRKACB | XM_006710758 | 9.553433586 | 0 | 0 | 0 | 0 | 0 | 0 | 0 | 3.27E-08 | 1 |
| PRKAG1 | NM_002733 | 10.95742161 | 8.765293275 | 11.66048729 | 0 | 9.114111124 | 0 | 0 | 0 | 0.999953634 | 0.02072721 |
| PRKAG2 | NM_024429 | 0 | 9.511764498 | 0 | 0 | 0 | 0 | 0 | 0 | 2.90E-08 | 1 |
| PRKCA | XR_001752558 | 0 | 9.358181476 | 0 | 0 | 0 | 0 | 0 | 0 | 3.64E-08 | 1 |
| PRKCD | NM_001354678 | 10.14723899 | 0 | 8.42974468 | 0 | 8.848563454 | 0 | 0 | 0 | 0.999953634 | 0.024568095 |
| PRKCH | XM_017021458 | 0 | 0 | 10.34177806 | 0 | 9.503640427 | 0 | 0 | 0 | 0.014872081 | 0.02072721 |
| PRKCI | XR_001740206 | 0 | 0 | 0 | 9.571680697 | 7.24369152 | 0 | 0 | 0 | 0.033889169 | 0.02416307 |
| PRKCQ-AS1 | NR_036503 | 0 | 8.690990708 | 0 | 0 | 0 | 0 | 0 | 0 | 9.02E-08 | 1 |
| PRKCSH | XM_011528130 | 0 | 0 | 9.758735269 | 10.03709574 | 8.715011943 | 0 | 0 | 0 | 0.002032121 | 0.003174145 |
| PRKCZ | NM_001033582 | 10.2867736 | 0 | 0 | 0 | 0 | 0 | 9.559829433 | 0 | 1.54E-08 | 1 |

| | | | | | | | | | | | |
|---|---|---|---|---|---|---|---|---|---|---|---|
| PRKCZ | NM_001242874 | 0 | 0 | 0 | 12.12350816 | 8.144543525 | 0 | 11.34731678 | 0 | 1.49E-08 | 0.999932046 |
| PRKCZ | XM_017001790 | 0 | 0 | 7.044209926 | 0 | 0 | 0 | 8.570567255 | 0 | 3.03E-05 | 1 |
| PRKCZ | XR_002957130 | 10.81296622 | 0 | 0 | 0 | 0 | 0 | 0 | 0 | 1.34E-08 | 1 |
| PRKRA | XR_001739008 | 0 | 9.772669303 | 0 | 0 | 0 | 0 | 0 | 0 | 4.12E-08 | 1 |
| PRKRA-AS1 | NR_110206 | 9.95545054 | 0 | 0 | 0 | 0 | 0 | 0 | 0 | 2.01E-08 | 1 |
| PRLR | XM_024446132 | 0 | 0 | 6.832547844 | 0 | 0 | 0 | 0 | 0 | 3.72E-05 | 1 |
| PRMT1 | NM_198318 | 0 | 9.193024922 | 11.04291014 | 11.5978802 | 11.22424416 | 0 | 0 | 0 | 0.634898279 | 0.000515743 |
| PRMT2 | NM_001242866 | 0 | 0 | 8.38097934 | 0 | 9.527413936 | 0 | 0 | 0 | 0.027138955 | 0.02072721 |
| PRMT2 | XM_017028340 | 0 | 0 | 10.60856602 | 0 | 5.804731829 | 0 | 0 | 10.14389485 | 0.021722667 | 1 |
| PRMT2 | XR_001754841 | 0 | 9.698755526 | 0 | 0 | 0 | 0 | 7.361968883 | 0 | 2.40E-08 | 1 |
| PRMT3 | XM_011519836 | 0 | 8.464408432 | 0 | 0 | 0 | 0 | 0 | 0 | 1.11E-07 | 1 |
| PRMT9 | NM_138364 | 0 | 0 | 0 | 10.51593815 | 7.128347732 | 0 | 10.1734233 | 0 | 0.02002499 | 0.999932046 |
| PRMT9 | XM_017008823 | 10.91641528 | 0 | 0 | 0 | 0 | 0 | 0 | 0 | 1.34E-08 | 1 |
| PROCR | NM_006404 | 0 | 0 | 0 | 11.86474044 | 9.655692746 | 10.90379525 | 8.212844911 | 10.25658006 | 0.006337432 | 0.999932046 |
| PROSER2 | NM_153256 | 0 | 0 | 0 | 0 | 6.409577539 | 0 | 0 | 0 | 4.59E-05 | 1 |
| PRPF19 | NM_014502 | 0 | 8.78328601 | 10.79626483 | 10.28541326 | 8.755836186 | 0 | 0 | 0 | 0.845329783 | 0.002089223 |
| PRPF40A | XM_024452980 | 0 | 0 | 0 | 9.357540824 | 7.806841281 | 0 | 0 | 0 | 0.034380846 | 0.02431343 |
| PRPF40B | XM_017019137 | 0 | 0 | 0 | 0 | 6.337212177 | 0 | 0 | 0 | 4.86E-05 | 1 |
| PRPF6 | NM_012469 | 0 | 0 | 10.34904047 | 10.80376031 | 9.020164451 | 0 | 9.781315183 | 0 | 0.000978533 | 0.999932046 |
| PRPSAP1 | NM_002766 | 0 | 0 | 0 | 9.138588821 | 7.449928885 | 0 | 0 | 0 | 0.0421553 | 0.029103945 |
| PRPSAP2 | NM_001353107 | 8.951500292 | 0 | 0 | 0 | 0 | 0 | 0 | 0 | 7.86E-08 | 1 |
| PRPSAP2 | XM_017024866 | 0 | 8.774317692 | 0 | 0 | 0 | 0 | 0 | 0 | 8.46E-08 | 1 |
| PRPSAP2 | XM_017024875 | 0 | 0 | 0 | 10.45251057 | 7.701245565 | 0 | 0 | 0 | 0.01946381 | 0.02072721 |
| PRR15 | NM_001329996 | 0 | 8.818611963 | 0 | 0 | 0 | 0 | 0 | 0 | 8.12E-08 | 1 |
| PRR3 | NM_001077497 | 9.752954799 | 0 | 0 | 0 | 0 | 0 | 0 | 0 | 2.49E-08 | 1 |
| PRR34-AS1 | NR_027034 | 0 | 0 | 0 | 0 | 6.337212177 | 0 | 0 | 0 | 4.86E-05 | 1 |
| PRR7 | XM_017009896 | 0 | 10.48352196 | 0 | 0 | 0 | 0 | 0 | 0 | 1.34E-08 | 1 |
| PRRC1 | XM_006714529 | 9.317628289 | 8.951853153 | 0 | 0 | 0 | 0 | 0 | 0 | 0.002654219 | 1 |
| PRRC2A | XM_017011274 | 0 | 0 | 0 | 0 | 6.736572239 | 0 | 0 | 0 | 9.53E-05 | 1 |
| PRRG2 | XM_006723286 | 0 | 0 | 11.07775258 | 0 | 7.854284905 | 0 | 0 | 0 | 0.013889064 | 0.02072721 |
| PRSS12 | NM_003619 | 0 | 9.511764498 | 0 | 0 | 0 | 0 | 0 | 0 | 2.90E-08 | 1 |
| PRSS16 | XM_017010163 | 10.23434896 | 0 | 0 | 0 | 0 | 0 | 0 | 0 | 1.59E-08 | 1 |
| PRSS3 | XM_011517965 | 9.883513662 | 10.97141009 | 0 | 0 | 0 | 0 | 0 | 0 | 0.000436285 | 1 |
| PRTFDC1 | NM_020200 | 0 | 0 | 0 | 10.41529045 | 9.175583651 | 0 | 0 | 0 | 0.015451077 | 0.020768938 |
| PRUNE1 | XM_017001957 | 0 | 9.006487637 | 0 | 0 | 0 | 0 | 7.189323943 | 0 | 6.80E-08 | 1 |
| PRXL2C | NM_153698 | 0 | 9.213075091 | 0 | 0 | 0 | 0 | 0 | 0 | 4.56E-08 | 1 |
| PSAT1 | NM_058179 | 0 | 0 | 0 | 9.591445233 | 10.34319844 | 0 | 7.479117364 | 0 | 0.014236928 | 0.999932046 |
| PSD3 | XM_017013265 | 10.71433872 | 0 | 0 | 0 | 0 | 0 | 0 | 0 | 1.34E-08 | 1 |
| PSIP1 | NM_021144 | 0 | 0 | 0 | 10.16131049 | 8.884555432 | 0 | 0 | 10.63693401 | 0.018494796 | 0.999932046 |
| PSIP1 | XM_024447399 | 0 | 0 | 0 | 9.186158628 | 11.11918762 | 0 | 10.18352865 | 11.74770891 | 0.010567574 | 0.999932046 |
| PSKH1 | NM_006742 | 0 | 0 | 9.665827761 | 0 | 7.774322632 | 0 | 0 | 0 | 0.02994657 | 0.02091605 |
| PSMA1 | NM_001143937 | 0 | 0 | 0 | 0 | 6.437537018 | 0 | 0 | 0 | 4.51E-05 | 1 |
| PSMA3 | NR_038123 | 9.682959584 | 0 | 0 | 0 | 0 | 0 | 0 | 0 | 2.84E-08 | 1 |

| | | | | | | | | | | | |
|---|---|---|---|---|---|---|---|---|---|---|---|
| PSMA3-AS1 | NR_029434 | 0 | 8.652353298 | 0 | 0 | 0 | 0 | 0 | 10.24521033 | 9.28E-08 | 1 |
| PSMA6 | NM_001282233 | 11.10584386 | 0 | 10.11774981 | 0 | 9.160983508 | 0 | 0 | 0 | 0.999953634 | 0.02072721 |
| PSMB3 | NR_104194 | 0 | 9.258802813 | 0 | 0 | 0 | 0 | 0 | 0 | 4.23E-08 | 1 |
| PSMB5 | NM_001144932 | 0 | 9.155526529 | 0 | 0 | 0 | 0 | 0 | 0 | 5.04E-08 | 1 |
| PSMB5 | NM_002797 | 0 | 5.293702769 | 11.60810394 | 12.12689799 | 11.19306135 | 10.87781373 | 11.40428644 | 10.21222279 | 0.028431543 | 0.999932046 |
| PSMC4 | NM_153001 | 0 | 0 | 0 | 8.854417711 | 8.78741214 | 0 | 0 | 0 | 0.032435818 | 0.02296717 |
| PSMC6 | XR_943495 | 0 | 9.467937896 | 0 | 0 | 0 | 0 | 0 | 0 | 3.10E-08 | 1 |
| PSMD1 | NR_034059 | 9.449953927 | 8.700490516 | 0 | 0 | 0 | 0 | 7.604755783 | 0 | 0.002780768 | 1 |
| PSMD14 | NM_005805 | 11.73450218 | 10.08341647 | 0 | 11.7768449 | 11.63391687 | 0 | 0 | 0 | 0.999953634 | 0.016689853 |
| PSMD2 | NM_002808 | 0 | 0 | 11.2563807 | 10.72542608 | 8.838112924 | 0 | 0 | 10.71200753 | 0.000687058 | 0.999932046 |
| PSME4 | XR_001738679 | 0 | 0 | 0 | 0 | 6.451316153 | 0 | 0 | 0 | 4.51E-05 | 1 |
| PSMF1 | NM_006814 | 0 | 0 | 0 | 9.83969732 | 9.383914218 | 0 | 8.262719816 | 10.29961409 | 0.018549998 | 0.999932046 |
| PSMF1 | XM_024452019 | 0 | 5.189614022 | 0 | 8.899510364 | 9.353051398 | 0 | 0 | 0 | 0.468739024 | 0.02072721 |
| PSMG3 | NM_001134340 | 0 | 0 | 0 | 10.13730841 | 10.09096926 | 0 | 10.68870802 | 0 | 0.013246205 | 1 |
| PSMG4 | XM_011514593 | 10.33321142 | 0 | 0 | 0 | 0 | 0 | 0 | 0 | 1.51E-08 | 1 |
| PSORS1C1 | NM_014068 | 0 | 8.861586703 | 0 | 10.62160046 | 6.702411139 | 0 | 0 | 0 | 0.999953634 | 0.02072721 |
| PSPH | XR_002956470 | 0 | 0 | 0 | 10.0313189 | 7.82812018 | 0 | 7.987500854 | 0 | 0.023906615 | 0.999932046 |
| PSTPIP1 | NM_003978 | 0 | 9.284296589 | 0 | 0 | 0 | 0 | 0 | 0 | 4.07E-08 | 1 |
| PTAR1 | XM_011518640 | 0 | 8.878424457 | 0 | 0 | 0 | 0 | 0 | 0 | 9.15E-08 | 1 |
| PTBP1 | NM_002819 | 0 | 0 | 0 | 8.533268642 | 0 | 0 | 0 | 0 | 5.97E-06 | 1 |
| PTCD2 | NM_024754 | 0 | 8.844550112 | 0 | 0 | 0 | 0 | 0 | 0 | 8.06E-08 | 1 |
| PTDSS1 | NM_014754 | 0 | 0 | 0 | 9.792664417 | 9.015534505 | 10.28995457 | 3.781941135 | 10.5863403 | 0.020857704 | 0.999932046 |
| PTGDS | NM_000954 | 6.084641612 | 8.384152849 | 0 | 0 | 0 | 0 | 0 | 8.649540214 | 0.018713016 | 1 |
| PTGER3 | NM_198717 | 10.11860076 | 0 | 0 | 0 | 0 | 0 | 0 | 0 | 1.70E-08 | 1 |
| PTGES2 | NM_025072 | 0 | 9.808252985 | 9.611000853 | 10.88127438 | 7.503816758 | 0 | 0 | 0 | 0.999953634 | 0.003016079 |
| PTGES2 | XM_005252239 | 0 | 8.372314435 | 0 | 0 | 0 | 0 | 0 | 0 | 1.20E-07 | 1 |
| PTGES3L | NM_001261430 | 8.994164854 | 0 | 0 | 0 | 0 | 0 | 8.04563962 | 0 | 7.73E-08 | 1 |
| PTGR1 | NM_001146108 | 0 | 0 | 11.31318292 | 0 | 8.496518068 | 0 | 9.020532381 | 0 | 0.011061298 | 0.999932046 |
| PTGR2 | NM_152444 | 0 | 0 | 10.37417483 | 0 | 7.854284905 | 0 | 0 | 0 | 0.02031574 | 0.02072721 |
| PTGS2 | NM_000963 | 0 | 0 | 9.342889335 | 9.267038066 | 0 | 10.5714681 | 0 | 0 | 0.024974675 | 1 |
| PTK2 | NM_001352752 | 0 | 8.323964754 | 0 | 0 | 0 | 0 | 0 | 0 | 1.27E-07 | 1 |
| PTK2 | XM_024447210 | 0 | 9.376076245 | 0 | 0 | 0 | 0 | 0 | 0 | 3.47E-08 | 1 |
| PTK2 | XM_024447211 | 0 | 9.014126367 | 0 | 0 | 0 | 0 | 0 | 0 | 6.71E-08 | 1 |
| PTK2B | XM_017013215 | 0 | 0 | 7.936379488 | 0 | 0 | 0 | 0 | 0 | 1.22E-05 | 1 |
| PTK2B | XM_017013216 | 0 | 0 | 0 | 0 | 6.679179249 | 0 | 0 | 0 | 3.78E-05 | 1 |
| PTMA | XM_011511520 | 0 | 0 | 0 | 0 | 6.643614539 | 0 | 0 | 0 | 3.86E-05 | 1 |
| PTN | NM_001321387 | 0 | 8.700490516 | 0 | 0 | 0 | 0 | 0 | 0 | 9.01E-08 | 1 |
| PTOV1 | NM_001305105 | 0 | 0 | 0 | 10.07127927 | 8.989800746 | 0 | 0 | 0 | 0.018750373 | 0.02072721 |
| PTP4A1 | NM_003463 | 0 | 0 | 10.6273541 | 0 | 5.54685007 | 0 | 0 | 0 | 0.0218555 | 1 |
| PTP4A2 | XR_002957649 | 0 | 0 | 0 | 10.09915958 | 7.536502354 | 0 | 0 | 10.02735731 | 0.024048834 | 0.999932046 |
| PTPA | NM_178000 | 0 | 0 | 0 | 8.474877744 | 0 | 0 | 0 | 0 | 6.35E-06 | 1 |
| PTPN11 | XM_011538613 | 0 | 9.006487637 | 0 | 9.425184544 | 6.607150896 | 0 | 0 | 0 | 0.999953634 | 0.028645392 |
| PTPN13 | XM_017008511 | 0 | 0 | 0 | 0 | 6.478485785 | 0 | 0 | 0 | 4.36E-05 | 1 |

| | | | | | | | | | | | |
|---|---|---|---|---|---|---|---|---|---|---|---|
| PTPN13 | XM_017008514 | 10.50900103 | 8.943876548 | 0 | 10.91460632 | 9.486177237 | 0 | 0 | 0 | 0.999953634 | 0.02072721 |
| PTPN18 | XM_024452789 | 0 | 0 | 0 | 7.080635617 | 0 | 0 | 0 | 0 | 2.70E-05 | 1 |
| PTPN2 | XM_017025888 | 0 | 9.434171496 | 0 | 0 | 0 | 0 | 0 | 0 | 3.24E-08 | 1 |
| PTPN23 | XM_005265031 | 0 | 9.564739385 | 0 | 0 | 0 | 0 | 0 | 0 | 2.69E-08 | 1 |
| PTPRCAP | NM_005608 | 12.11642066 | 0 | 0 | 0 | 0 | 10.67064035 | 7.73610317 | 10.12274908 | 1.34E-08 | 0.002089223 |
| PTPRF | XM_017001944 | 0 | 9.698755526 | 0 | 0 | 0 | 0 | 0 | 0 | 2.40E-08 | 1 |
| PTPRH | XR_001753730 | 10.25203635 | 0 | 0 | 0 | 0 | 0 | 0 | 0 | 1.58E-08 | 1 |
| PTPRK | XM_006715537 | 11.36755267 | 0 | 0 | 0 | 0 | 11.23185389 | 9.7011842 | 0 | 1.34E-08 | 0.02072721 |
| PTPRM | XM_017025908 | 9.426779986 | 0 | 0 | 0 | 0 | 0 | 0 | 0 | 3.96E-08 | 1 |
| PTPRN2 | NM_130842 | 0 | 0 | 0 | 0 | 0 | 10.47848193 | 0 | 10.05738604 | 1 | 0.02072721 |
| PTPRO | NM_030669 | 0 | 9.179501644 | 0 | 0 | 0 | 0 | 0 | 0 | 4.86E-08 | 1 |
| PTPRR | NM_002849 | 10.97083541 | 0 | 0 | 0 | 0 | 0 | 0 | 0 | 1.34E-08 | 1 |
| PTTG1 | NM_004219 | 0 | 0 | 10.12411362 | 12.60398583 | 11.50834144 | 0 | 10.04629302 | 12.19536295 | 0.000145855 | 0.999932046 |
| PTTG1IP | NM_004339 | 0 | 0 | 11.87722801 | 0 | 9.597245458 | 0 | 0 | 9.770654157 | 0.006469986 | 0.999932046 |
| PUDP | NM_012080 | 0 | 0 | 10.19635521 | 0 | 8.246846504 | 10.84391933 | 7.638730623 | 0 | 0.020883497 | 0.999932046 |
| PUF60 | NM_001271099 | 0 | 9.051724868 | 0 | 0 | 0 | 0 | 8.032920857 | 0 | 7.86E-08 | 1 |
| PUM1 | NM_014676 | 0 | 0 | 0 | 10.14802533 | 7.44304914 | 0 | 0 | 0 | 0.023723013 | 0.02072721 |
| PUM2 | XM_024452773 | 0 | 0 | 0 | 0 | 6.631562102 | 0 | 7.967588696 | 0 | 3.88E-05 | 1 |
| PUM3 | NM_014878 | 0 | 0 | 0 | 11.36510547 | 7.211666293 | 0 | 9.250789795 | 0 | 1.89E-08 | 0.999932046 |
| PUS1 | NM_025215 | 0 | 8.550974732 | 0 | 0 | 0 | 0 | 0 | 0 | 1.01E-07 | 1 |
| PUS1 | XM_011538768 | 0 | 0 | 0 | 8.773677528 | 0 | 0 | 0 | 0 | 4.62E-06 | 1 |
| PUS10 | XM_011532574 | 0 | 0 | 0 | 10.15733777 | 8.8089774 | 0 | 0 | 0 | 0.018789492 | 0.02072721 |
| PUS7L | XM_024449217 | 0 | 10.32058425 | 0 | 0 | 0 | 0 | 0 | 0 | 1.45E-08 | 1 |
| PUSL1 | NM_153339 | 0 | 9.439854398 | 0 | 0 | 0 | 0 | 0 | 0 | 3.19E-08 | 1 |
| PVALB | NM_001315532 | 11.00513859 | 10.08341647 | 0 | 0 | 0 | 9.96338155 | 0 | 0 | 0.000393121 | 1 |
| PVR | NM_001135768 | 0 | 0 | 0 | 10.44383889 | 8.276174381 | 0 | 7.310202764 | 10.55773842 | 0.018094055 | 0.999932046 |
| PVR | NM_006505 | 0 | 8.529815232 | 10.34359709 | 0 | 9.036253242 | 0 | 0 | 0 | 0.999953634 | 0.020869972 |
| PWWP2B | XM_011539387 | 11.35328397 | 0 | 0 | 0 | 0 | 0 | 0 | 0 | 1.34E-08 | 1 |
| PXK | XM_005265250 | 0 | 0 | 10.08550279 | 0 | 6.80255953 | 0 | 0 | 0 | 0.027200218 | 0.02072721 |
| PXMP2 | NM_018663 | 11.11069923 | 12.01644457 | 7.8773294 | 11.48723784 | 0 | 0 | 0 | 0 | 0.999953634 | 7.83E-09 |
| PXMP4 | NM_007238 | 0 | 0 | 0 | 0 | 0 | 10.92297886 | 0 | 11.12210196 | 1 | 0.02072721 |
| PXN | NM_002859 | 0 | 8.540433774 | 0 | 0 | 0 | 0 | 4.37247236 | 0 | 1.02E-07 | 0.764734118 |
| PXYLP1 | NM_001282728 | 0 | 0 | 10.31606779 | 12.02229771 | 8.435435627 | 0 | 9.023750081 | 10.13688064 | 0.000541819 | 0.999932046 |
| PXYLP1 | XM_005247903 | 0 | 9.014126367 | 0 | 0 | 0 | 9.887288217 | 0 | 0 | 6.71E-08 | 1 |
| PYCR1 | NM_006907 | 0 | 0 | 0 | 8.937070225 | 7.735433951 | 0 | 0 | 0 | 0.044041324 | 0.030130471 |
| PYM1 | NM_001143853 | 0 | 0 | 9.808316306 | 0 | 7.305685067 | 10.26009441 | 0 | 0 | 0.029967258 | 0.999932046 |
| PYY | NM_004160 | 0 | 13.12873212 | 9.54138321 | 0 | 7.666227291 | 0 | 0 | 0 | 0.622417231 | 0.022027568 |
| QARS | NM_005051 | 0 | 9.640723913 | 0 | 0 | 0 | 10.28337218 | 0 | 10.16473515 | 2.47E-08 | 0.02072721 |
| QKI | NM_001301085 | 0 | 0 | 8.769900534 | 9.07261961 | 7.605905728 | 10.06921268 | 11.02689078 | 0 | 0.007052827 | 0.999932046 |
| QKI | NM_206853 | 0 | 8.951853153 | 0 | 0 | 0 | 11.80574841 | 0 | 9.961392177 | 7.60E-08 | 0.020188082 |
| QPCT | NM_012413 | 0 | 0 | 10.28989101 | 0 | 7.503816758 | 10.07938103 | 9.956540422 | 10.59148026 | 0.022245073 | 0.999932046 |
| QPRT | NM_014298 | 0 | 0 | 9.084656341 | 10.52521627 | 0 | 0 | 10.07294177 | 9.917043278 | 0.016144258 | 0.999932046 |
| QRICH1 | NM_017730 | 0 | 0 | 11.23693853 | 0 | 9.289271525 | 11.42812295 | 0 | 0 | 0.010065133 | 0.999932046 |

| | | | | | | | | | | | |
|---|---|---|---|---|---|---|---|---|---|---|---|
| RAB10 | XM_024452565 | 10.04948094 | 8.248259235 | 0 | 10.01677478 | 8.31813808 | 0 | 0 | 0 | 0.999953634 | 0.02072721 |
| RAB17 | XM_006712689 | 10.43350322 | 0 | 0 | 0 | 0 | 0 | 5.732859748 | 0 | 1.47E-08 | 1 |
| RAB18 | NM_021252 | 0 | 10.72839101 | 0 | 0 | 0 | 10.57416716 | 10.23718434 | 0 | 1.34E-08 | 0.02072721 |
| RAB18 | NR_046172 | 0 | 0 | 6.893420866 | 9.489798191 | 6.655567121 | 0 | 0 | 0 | 0.011783851 | 0.019618292 |
| RAB23 | NR_103822 | 0 | 0 | 0 | 0 | 6.464964927 | 0 | 0 | 0 | 4.45E-05 | 1 |
| RAB24 | NR_109790 | 9.83739315 | 0 | 0 | 0 | 0 | 0 | 6.56874927 | 0 | 2.27E-08 | 1 |
| RAB27A | XM_024450009 | 8.850625659 | 0 | 0 | 0 | 0 | 0 | 0 | 0 | 8.53E-08 | 1 |
| RAB30 | NM_014488 | 0 | 0 | 10.24388102 | 8.75977034 | 0 | 0 | 7.341484785 | 0 | 0.019978324 | 0.999932046 |
| RAB32 | NM_006834 | 0 | 10.16115789 | 11.71173949 | 10.32372925 | 9.731053831 | 0 | 0 | 0 | 0.996679353 | 0.001056205 |
| RAB33A | NM_004794 | 0 | 9.825721024 | 0 | 0 | 0 | 0 | 0 | 0 | 2.01E-08 | 1 |
| RAB34 | NM_001256277 | 10.50531851 | 0 | 0 | 0 | 0 | 0 | 0 | 0 | 1.41E-08 | 1 |
| RAB35 | XM_011537779 | 10.39421912 | 0 | 0 | 0 | 0 | 0 | 0 | 0 | 1.47E-08 | 1 |
| RAB37 | NM_175738 | 9.25321892 | 0 | 0 | 0 | 0 | 0 | 0 | 0 | 5.16E-08 | 1 |
| RAB38 | NM_022337 | 0 | 0 | 0 | 8.368928953 | 0 | 0 | 0 | 0 | 7.12E-06 | 1 |
| RAB3D | NM_004283 | 0 | 0 | 10.46220177 | 0 | 8.230957729 | 0 | 7.578739068 | 0 | 0.018439374 | 0.999932046 |
| RAB40B | XM_024450551 | 0 | 9.179501644 | 0 | 0 | 0 | 0 | 7.819932445 | 0 | 4.86E-08 | 1 |
| RAB40C | NM_021168 | 0 | 0 | 0 | 9.433968736 | 7.71273195 | 9.810013246 | 7.974256665 | 0 | 0.033649828 | 0.999932046 |
| RAB4A | NM_004578 | 0 | 11.39507056 | 10.61460487 | 11.30771688 | 11.32731336 | 0 | 0 | 0 | 0.999953634 | 0.000657675 |
| RAB4B | NM_016154 | 9.928205339 | 6.05590339 | 0 | 0 | 0 | 0 | 0 | 0 | 0.004195538 | 1 |
| RAB4B-EGLN2 | NR_037791 | 9.928205339 | 9.006487637 | 0 | 0 | 0 | 0 | 8.284344777 | 10.62445095 | 0.001622708 | 0.02072721 |
| RAB5A | NM_004162 | 0 | 0 | 9.060527565 | 0 | 7.178913994 | 10.87197597 | 8.882017781 | 0 | 0.048103307 | 0.999932046 |
| RAB5C | NM_201434 | 9.801808664 | 0 | 0 | 0 | 0 | 0 | 8.026519178 | 10.64931 | 2.38E-08 | 0.02072721 |
| RAB6A | NM_001243718 | 0 | 0 | 10.22525549 | 0 | 9.795263842 | 10.47174284 | 0 | 10.50695895 | 0.014251827 | 0.999932046 |
| RAB7A | XR_002959583 | 8.373391248 | 8.181987608 | 0 | 0 | 0 | 0 | 7.460241356 | 0 | 0.008467741 | 1 |
| RAB8A | NM_005370 | 8.754625855 | 9.694007725 | 10.31976886 | 10.0768986 | 10.19393357 | 0 | 0 | 0 | 0.999953634 | 0.001816902 |
| RAB8B | XM_017022312 | 0 | 9.376076245 | 0 | 0 | 0 | 0 | 0 | 0 | 3.47E-08 | 1 |
| RABEP2 | NM_024816 | 0 | 0 | 0 | 8.628177296 | 0 | 0 | 8.878467512 | 0 | 6.35E-06 | 1 |
| RABGEF1 | NM_014504 | 0 | 0 | 0 | 8.264461514 | 0 | 0 | 0 | 0 | 7.93E-06 | 1 |
| RABIF | NM_002871 | 0 | 0 | 11.38604073 | 10.34484305 | 0 | 0 | 7.411941715 | 0 | 0.007911222 | 0.999932046 |
| RABL2A | NM_001306158 | 9.190393069 | 0 | 0 | 0 | 0 | 0 | 0 | 0 | 7.73E-08 | 1 |
| RABL2A | XR_002959036 | 0 | 0 | 0 | 0 | 6.531336591 | 0 | 0 | 0 | 4.23E-05 | 1 |
| RABL2B | XM_024452147 | 0 | 8.809861237 | 0 | 0 | 0 | 0 | 0 | 0 | 8.24E-08 | 1 |
| RAD1 | NM_002853 | 0 | 0 | 0 | 10.60419453 | 9.772119282 | 0 | 0 | 0 | 0.012150779 | 0.02072721 |
| RAD17 | NM_133338 | 0 | 0 | 0 | 9.645387449 | 10.54100727 | 9.379141419 | 10.19427783 | 10.35204063 | 0.012815669 | 0.999932046 |
| RAD23A | NM_001270362 | 0 | 0 | 0 | 10.14535356 | 9.224548436 | 0 | 9.461221637 | 0 | 0.017193829 | 0.999932046 |
| RAD50 | NM_005732 | 10.17531978 | 0 | 0 | 0 | 0 | 0 | 0 | 0 | 1.60E-08 | 1 |
| RAD51 | NM_002875 | 0 | 0 | 0 | 9.865900493 | 7.365124102 | 0 | 0 | 10.18867435 | 0.027986508 | 0.999932046 |
| RAD51AP1 | NM_006479 | 0 | 0 | 0 | 9.801176392 | 6.916214501 | 0 | 0 | 0 | 0.030907891 | 0.022511788 |
| RAD51C | XM_006722001 | 0 | 0 | 10.85291933 | 10.59639063 | 10.27364963 | 0 | 9.697159171 | 0 | 0.000564265 | 0.999932046 |
| RAD51C | XM_011525094 | 10.92471032 | 0 | 0 | 0 | 0 | 0 | 0 | 0 | 1.34E-08 | 1 |
| RAD54B | NM_001205262 | 0 | 10.06146648 | 0 | 0 | 0 | 0 | 0 | 0 | 1.60E-08 | 1 |
| RAF1 | NM_001354690 | 0 | 0 | 0 | 10.95636031 | 7.859461386 | 0 | 8.475068668 | 0 | 0.014550507 | 0.999932046 |
| RAF1 | NM_001354692 | 0 | 0 | 12.03836268 | 11.2763127 | 10.83607497 | 10.10449269 | 8.820441837 | 11.50510193 | 0.000182422 | 0.999932046 |

| | | | | | | | | | | | |
|---|---|---|---|---|---|---|---|---|---|---|---|
| RAF1 | NR_148941 | 0 | 0 | 0 | 9.583571881 | 9.135605715 | 0 | 0 | 0 | 0.021890341 | 0.02072721 |
| RALA | NM_005402 | 0 | 0 | 0 | 9.669762543 | 9.400877796 | 0 | 0 | 0 | 0.019573075 | 0.02072721 |
| RALB | XM_011511574 | 0 | 9.726918817 | 0 | 8.368928953 | 8.242890685 | 0 | 0 | 0 | 0.999953634 | 0.032833213 |
| RALGDS | NM_006266 | 0 | 0 | 0 | 10.02261004 | 7.298079998 | 0 | 0 | 0 | 0.025856022 | 0.02072721 |
| RALGPS1 | XR_001746426 | 0 | 8.78328601 | 0 | 0 | 0 | 0 | 8.125685946 | 0 | 8.40E-08 | 1 |
| RALY | NM_007367 | 0 | 0 | 10.61911747 | 10.14802533 | 7.128347732 | 0 | 7.56113023 | 0 | 0.001850326 | 0.999932046 |
| RALYL | NM_001100391 | 10.362003 | 0 | 0 | 0 | 0 | 0 | 0 | 0 | 1.48E-08 | 1 |
| RAMP1 | XM_017003156 | 0 | 0 | 0 | 7.776970127 | 8.580418909 | 0 | 0 | 0 | 0.05174167 | 0.034581849 |
| RANBP1 | NM_001278640 | 0 | 0 | 0 | 9.907491 | 9.685874143 | 0 | 8.966377255 | 13.00174171 | 0.016498375 | 0.999932046 |
| RANBP1 | XM_011530291 | 0 | 0 | 0 | 8.414023513 | 0 | 0 | 0 | 0 | 3.17E-05 | 1 |
| RANBP1 | XM_017028890 | 0 | 9.271606013 | 0 | 0 | 0 | 0 | 7.200741128 | 0 | 4.15E-08 | 1 |
| RANBP17 | NM_022897 | 0 | 9.46236476 | 9.946391943 | 0 | 6.844931782 | 0 | 0 | 0 | 0.999953634 | 0.020810615 |
| RANBP17 | XM_017009747 | 11.92571742 | 0 | 0 | 10.56174337 | 9.256292084 | 0 | 0 | 0 | 0.999953634 | 0.02072721 |
| RANBP3 | XM_006722928 | 11.05132772 | 0 | 9.189515753 | 0 | 6.366595318 | 0 | 0 | 0 | 0.966161772 | 0.032047775 |
| RANBP3 | XM_011528393 | 11.02069986 | 0 | 0 | 0 | 0 | 0 | 0 | 0 | 1.34E-08 | 1 |
| RAP1A | XM_024448786 | 0 | 0 | 0 | 0 | 6.544252221 | 0 | 7.712369201 | 0 | 4.20E-05 | 1 |
| RAP1A | XR_946724 | 9.143781062 | 0 | 0 | 0 | 0 | 0 | 0 | 0 | 7.26E-08 | 1 |
| RAP1GDS1 | NM_001100428 | 0 | 0 | 11.71876467 | 10.12244145 | 11.65936781 | 12.76241468 | 10.54550324 | 10.32760645 | 0.000230526 | 0.999932046 |
| RAP1GDS1 | XM_024454167 | 11.23819407 | 0 | 0 | 0 | 0 | 10.42067049 | 10.23023735 | 0 | 1.34E-08 | 0.02072721 |
| RAP1GDS1 | XM_024454170 | 10.28247691 | 0 | 0 | 0 | 0 | 0 | 0 | 0 | 1.54E-08 | 1 |
| RAP1GDS1 | XM_024454172 | 0 | 0 | 0 | 9.792664417 | 7.768830875 | 0 | 8.605347092 | 11.870132 | 0.027385771 | 0.999932046 |
| RAPGEFL1 | NM_016339 | 0 | 0 | 0 | 9.664173964 | 7.968883375 | 0 | 7.460241356 | 0 | 0.02820214 | 0.999932046 |
| RARRES3 | NM_004585 | 11.81107445 | 10.10682472 | 12.72449614 | 0 | 9.358545975 | 0 | 0 | 0 | 0.999953634 | 2.45E-09 |
| RARS2 | NM_001350506 | 0 | 0 | 10.77417948 | 0 | 7.320775981 | 0 | 0 | 0 | 0.017517226 | 0.02072721 |
| RARS2 | NR_146743 | 0 | 9.232850423 | 0 | 0 | 0 | 0 | 0 | 0 | 4.38E-08 | 1 |
| RASA2 | XM_017006969 | 0 | 9.014126367 | 0 | 0 | 0 | 0 | 8.609636205 | 0 | 6.71E-08 | 1 |
| RASA2 | XM_017006970 | 9.162606966 | 9.54905016 | 0 | 0 | 0 | 11.02514465 | 0 | 0 | 0.001949936 | 1 |
| RASA2 | XM_017006971 | 0 | 9.590514794 | 0 | 0 | 0 | 0 | 0 | 0 | 2.90E-08 | 1 |
| RASEF | XM_005251731 | 9.004636726 | 9.743097037 | 0 | 0 | 0 | 0 | 0 | 0 | 0.001836387 | 1 |
| RASSF1 | XM_024453328 | 0 | 0 | 0 | 11.4622297 | 10.46366248 | 10.74358744 | 0 | 0 | 0.006517015 | 0.999932046 |
| RASSF4 | XR_001747228 | 9.531875668 | 0 | 0 | 0 | 0 | 0 | 0 | 0 | 3.28E-08 | 1 |
| RAVER2 | XR_946693 | 8.940634053 | 0 | 0 | 0 | 0 | 0 | 0 | 0 | 7.88E-08 | 1 |
| RBBP5 | NM_001193272 | 0 | 0 | 0 | 0 | 6.667421491 | 0 | 0 | 0 | 3.78E-05 | 1 |
| RBBP6 | NM_032626 | 0 | 0 | 0 | 10.58952743 | 7.379608952 | 0 | 8.273572814 | 0 | 0.018777731 | 0.999932046 |
| RBBP8 | XM_024451233 | 10.40216198 | 0 | 0 | 0 | 0 | 0 | 0 | 0 | 1.47E-08 | 1 |
| RBCK1 | NM_031229 | 9.97155406 | 0 | 0 | 0 | 0 | 0 | 0 | 0 | 2.00E-08 | 1 |
| RBM10 | NM_001204467 | 0 | 0 | 11.26408479 | 0 | 8.285819332 | 11.01921454 | 0 | 0 | 0.011751319 | 0.999932046 |
| RBM10 | NM_001204468 | 0 | 9.044283149 | 0 | 0 | 0 | 0 | 0 | 0 | 6.33E-08 | 1 |
| RBM10 | NM_005676 | 10.5981354 | 0 | 0 | 0 | 0 | 0 | 0 | 0 | 1.34E-08 | 1 |
| RBM12 | NM_152838 | 0 | 0 | 10.15551827 | 0 | 9.233547159 | 0 | 6.939666787 | 0 | 0.017259379 | 0.999932046 |
| RBM17 | XR_001747246 | 0 | 0 | 0 | 9.066983555 | 8.932008386 | 10.39647467 | 0 | 0 | 0.028348703 | 1 |
| RBM18 | NM_033117 | 0 | 0 | 10.14303832 | 0 | 9.079039444 | 0 | 0 | 10.20217763 | 0.018050422 | 0.999932046 |
| RBM18 | XM_006717319 | 0 | 8.39589491 | 0 | 0 | 0 | 0 | 0 | 0 | 1.17E-07 | 1 |

| | | | | | | | | | | | |
|---|---|---|---|---|---|---|---|---|---|---|---|
| RBM23 | NM_001077352 | 9.917161541 | 0 | 0 | 0 | 0 | 0 | 0 | 0 | 2.04E-08 | 1 |
| RBM23 | NM_018107 | 0 | 0 | 9.751902512 | 0 | 7.757784245 | 0 | 0 | 0 | 0.028687356 | 0.020730817 |
| RBM24 | XM_011514389 | 0 | 9.405416277 | 0 | 0 | 0 | 0 | 0 | 0 | 3.31E-08 | 1 |
| RBM26 | XM_017020699 | 0 | 0 | 0 | 0 | 6.59478863 | 0 | 0 | 0 | 4.02E-05 | 1 |
| RBM27 | XR_944325 | 0 | 0 | 0 | 0 | 6.491881102 | 0 | 0 | 0 | 4.33E-05 | 1 |
| RBM3 | XM_024452413 | 0 | 0 | 0 | 8.474877744 | 0 | 0 | 0 | 0 | 6.35E-06 | 1 |
| RBM3 | XM_024452414 | 0 | 9.158976035 | 10.9962615 | 0 | 8.2973088 | 0 | 0 | 0 | 0.999953634 | 0.02072721 |
| RBM33 | XM_005249535 | 10.92746477 | 0 | 0 | 0 | 0 | 0 | 0 | 0 | 1.34E-08 | 1 |
| RBM34 | NM_015014 | 0 | 9.309347676 | 0 | 10.49929491 | 8.644794263 | 0 | 0 | 0 | 0.999953634 | 0.02096331 |
| RBM39 | NM_004902 | 0 | 0 | 0 | 9.133205149 | 8.907227812 | 0 | 0 | 0 | 0.027904375 | 0.020903731 |
| RBM39 | NM_184234 | 0 | 0 | 9.987691433 | 0 | 9.594143561 | 10.63316681 | 0 | 0 | 0.016611626 | 0.999932046 |
| RBM39 | XR_001754439 | 0 | 0 | 0 | 8.491803396 | 8.882014143 | 0 | 9.869799534 | 0 | 0.035528459 | 0.999932046 |
| RBM42 | XM_005259257 | 0 | 8.195487713 | 10.34450575 | 0 | 8.214892016 | 0 | 0 | 0 | 0.999953634 | 0.02072721 |
| RBM45 | XM_017003322 | 0 | 9.620849457 | 0 | 10.4750288 | 9.087887617 | 0 | 0 | 0 | 0.999953634 | 0.02072721 |
| RBM47 | XM_017008308 | 0 | 9.06649413 | 0 | 0 | 0 | 0 | 0 | 0 | 7.75E-08 | 1 |
| RBM4B | XR_247214 | 0 | 8.935855595 | 0 | 0 | 0 | 0 | 0 | 0 | 7.69E-08 | 1 |
| RBM6 | NM_001167582 | 0 | 0 | 0 | 12.03390038 | 10.61479834 | 12.09539053 | 10.14784584 | 11.04551598 | 0.004581124 | 0.999932046 |
| RBM6 | NM_005777 | 0 | 0 | 9.958312928 | 11.99547457 | 0 | 11.52057786 | 9.762177523 | 9.513376477 | 0.006035185 | 0.999932046 |
| RBM6 | XM_006712916 | 0 | 9.847262786 | 10.19434059 | 9.26211379 | 9.283506136 | 0 | 0 | 0 | 0.999953634 | 0.00302572 |
| RBM6 | XR_001739975 | 11.04626802 | 9.717592007 | 0 | 0 | 0 | 0 | 0 | 0 | 0.000457725 | 1 |
| RBMS1 | XM_005246737 | 0 | 9.554298882 | 0 | 0 | 0 | 0 | 7.712369201 | 10.66402238 | 2.73E-08 | 0.02072721 |
| RBMS1 | XM_005246738 | 0 | 0 | 0 | 10.49720094 | 8.369839583 | 0 | 0 | 0 | 0.017377207 | 0.02072721 |
| RBMXL1 | NM_019610 | 10.26950953 | 0 | 0 | 0 | 0 | 10.29541704 | 0 | 0 | 1.57E-08 | 1 |
| RBP4 | NM_006744 | 8.907535469 | 0 | 10.25165197 | 0 | 10.32927452 | 0 | 0 | 0 | 0.999953634 | 0.02072721 |
| RBP5 | NM_031491 | 8.541035379 | 0 | 0 | 0 | 0 | 0 | 0 | 0 | 1.11E-07 | 1 |
| RBPMS | XM_017012988 | 0 | 0 | 11.21076733 | 11.31189264 | 12.03026518 | 11.22156135 | 12.65315619 | 9.389009359 | 0.000152024 | 0.999932046 |
| RBPMS | XM_017012989 | 0 | 0 | 0 | 0 | 6.409577539 | 0 | 0 | 0 | 4.59E-05 | 1 |
| RBPMS | XM_017012992 | 11.54483545 | 11.30608635 | 8.769900534 | 10.21835411 | 9.8339085 | 0 | 0 | 0 | 0.930063746 | 0.002937674 |
| RC3H2 | NM_018835 | 0 | 9.358181476 | 0 | 0 | 0 | 0 | 0 | 0 | 3.64E-08 | 1 |
| RCAN1 | XM_017028283 | 11.50297672 | 0 | 0 | 10.36105389 | 10.29386518 | 0 | 0 | 0 | 0.999953634 | 0.02072721 |
| RCAN3 | NM_001251977 | 9.143781062 | 0 | 0 | 0 | 0 | 0 | 0 | 0 | 7.26E-08 | 1 |
| RCBTB1 | XR_941613 | 0 | 8.33620493 | 0 | 0 | 0 | 0 | 0 | 0 | 1.25E-07 | 1 |
| RCC1 | NR_030726 | 0 | 9.708204508 | 9.035988366 | 0 | 7.305685067 | 0 | 0 | 0 | 0.999953634 | 0.03006982 |
| RCC2 | NM_018715 | 0 | 5.293702769 | 0 | 9.247239304 | 5.133541874 | 0 | 0 | 0 | 0.708119097 | 0.037810409 |
| RCE1 | NM_001032279 | 0 | 8.951853153 | 0 | 0 | 0 | 0 | 0 | 0 | 7.60E-08 | 1 |
| RCHY1 | NM_015436 | 0 | 0 | 0 | 10.62352155 | 8.8089774 | 0 | 0 | 0 | 0.015130311 | 0.02072721 |
| RCHY1 | XM_024453984 | 9.900435696 | 0 | 0 | 0 | 0 | 0 | 0 | 0 | 2.07E-08 | 1 |
| RCN3 | NM_020650 | 0 | 0 | 9.064944738 | 9.918486574 | 0 | 0 | 0 | 0 | 0.021123344 | 0.02072721 |
| RCOR3 | NR_146434 | 10.87135108 | 0 | 0 | 0 | 0 | 0 | 0 | 0 | 1.34E-08 | 1 |
| RCSD1 | NM_052862 | 0 | 9.620849457 | 0 | 0 | 0 | 0 | 0 | 0 | 2.53E-08 | 1 |
| RDH11 | NM_001252650 | 0 | 8.384152849 | 0 | 0 | 0 | 0 | 0 | 0 | 1.18E-07 | 1 |
| RDH13 | XM_024451339 | 0 | 9.293741795 | 0 | 0 | 0 | 0 | 0 | 0 | 4.01E-08 | 1 |
| REEP4 | NM_025232 | 0 | 0 | 9.209603126 | 0 | 9.384812028 | 10.55971347 | 0 | 0 | 0.023018067 | 0.999932046 |

| | | | | | | | | | | | |
|---|---|---|---|---|---|---|---|---|---|---|---|
| RELA | XM_011545207 | 0 | 0 | 0 | 0 | 0 | 0 | 7.990792991 | 10.01210526 | 1 | 0.02072721 |
| RELB | XM_005259128 | 8.201989148 | 10.30198322 | 9.88276327 | 0 | 9.572241512 | 0 | 0 | 0 | 0.999953634 | 0.02072721 |
| REPS1 | XM_011536202 | 0 | 0 | 0 | 0 | 6.569741695 | 0 | 0 | 0 | 4.12E-05 | 1 |
| RERGL | NR_104413 | 11.69139863 | 0 | 0 | 0 | 0 | 0 | 9.199070987 | 0 | 1.34E-08 | 1 |
| REST | XM_017008527 | 11.05888416 | 0 | 0 | 0 | 0 | 0 | 0 | 0 | 1.34E-08 | 1 |
| RET | NM_020975 | 10.9920414 | 0 | 0 | 0 | 0 | 0 | 0 | 0 | 1.34E-08 | 1 |
| RETNLB | NM_032579 | 0 | 11.8031929 | 10.42488616 | 0 | 8.421503242 | 0 | 0 | 0 | 0.999953634 | 0.02072721 |
| RETREG3 | NR_026697 | 0 | 8.935855595 | 0 | 0 | 0 | 0 | 0 | 0 | 7.69E-08 | 1 |
| REV1 | NM_016316 | 0 | 0 | 0 | 0 | 6.478485785 | 0 | 0 | 0 | 4.36E-05 | 1 |
| REV1 | NR_135652 | 0 | 9.158976035 | 0 | 0 | 0 | 0 | 8.465635012 | 9.028739063 | 5.01E-08 | 0.02072721 |
| REV1 | XM_017004313 | 0 | 0 | 9.371700203 | 0 | 8.331238192 | 0 | 0 | 0 | 0.030538205 | 0.021032806 |
| REV3L | XM_011536029 | 0 | 0 | 10.52686995 | 11.18690062 | 9.160983508 | 0 | 9.042907014 | 0 | 0.000705997 | 0.999932046 |
| REV3L | XM_017011153 | 0 | 9.620849457 | 0 | 0 | 0 | 0 | 0 | 0 | 2.53E-08 | 1 |
| REV3L | XM_017011154 | 0 | 9.073822446 | 0 | 0 | 0 | 0 | 8.412617556 | 9.607716039 | 7.73E-08 | 0.02072721 |
| RFC2 | NM_001278791 | 9.998000145 | 0 | 0 | 0 | 0 | 0 | 0 | 0 | 1.94E-08 | 1 |
| RFC4 | NM_181573 | 0 | 0 | 0 | 10.30709108 | 8.835488421 | 0 | 0 | 10.28223637 | 0.017517226 | 0.999932046 |
| RFC5 | XM_011538643 | 0 | 8.497482465 | 0 | 0 | 0 | 0 | 0 | 0 | 1.06E-07 | 1 |
| RFT1 | NM_052859 | 0 | 9.165850403 | 0 | 0 | 0 | 0 | 6.85548998 | 10.4877908 | 4.97E-08 | 0.02072721 |
| RFWD3 | XM_017023391 | 0 | 0 | 0 | 0 | 0 | 10.81073605 | 8.816736559 | 10.10847773 | 1 | 0.001555546 |
| RGCC | NM_014059 | 0 | 0 | 0 | 9.633996983 | 9.532284059 | 12.67060705 | 10.13302394 | 0 | 0.018994951 | 0.999932046 |
| RGPD2 | XM_017004845 | 0 | 8.602554319 | 0 | 0 | 0 | 0 | 0 | 0 | 9.73E-08 | 1 |
| RGS1 | NM_002922 | 0 | 0 | 12.3709087 | 0 | 8.771023979 | 12.48060342 | 7.212068669 | 11.22820959 | 1.34E-08 | 0.999932046 |
| RGS13 | NM_002927 | 0 | 9.296876505 | 0 | 0 | 0 | 0 | 0 | 0 | 4.01E-08 | 1 |
| RGS13 | NM_144766 | 0 | 0 | 0 | 0 | 6.607150896 | 0 | 0 | 0 | 0.00011086 | 1 |
| RGS14 | XM_005265794 | 0 | 9.812639863 | 0 | 0 | 0 | 0 | 0 | 0 | 2.01E-08 | 1 |
| RGS19 | XM_011528486 | 8.754625855 | 0 | 0 | 0 | 0 | 0 | 0 | 0 | 9.15E-08 | 1 |
| RHBDD1 | XR_001738982 | 10.20286443 | 0 | 0 | 0 | 0 | 0 | 0 | 0 | 1.60E-08 | 1 |
| RHBDF1 | XM_017023556 | 9.442270566 | 0 | 0 | 0 | 0 | 0 | 0 | 0 | 3.84E-08 | 1 |
| RHBG | XR_001737328 | 0 | 9.855790228 | 0 | 0 | 0 | 0 | 0 | 0 | 1.98E-08 | 1 |
| RHOG | XM_005252916 | 0 | 0 | 12.29444416 | 0 | 7.683842675 | 0 | 0 | 0 | 1.34E-08 | 3.87E-09 |
| RHOT1 | XM_017024825 | 0 | 0 | 0 | 0 | 6.557053251 | 0 | 0 | 0 | 4.16E-05 | 1 |
| RHOT1 | XR_934499 | 0 | 9.625843798 | 0 | 0 | 0 | 0 | 0 | 0 | 2.50E-08 | 1 |
| RHOU | NR_037962 | 0 | 0 | 10.59793683 | 9.531320288 | 10.67328784 | 0 | 8.856980446 | 10.04993725 | 0.000792562 | 0.999932046 |
| RHPN1 | XM_017013016 | 0 | 8.709928179 | 0 | 0 | 0 | 0 | 0 | 0 | 8.96E-08 | 1 |
| RHPN2 | NM_033103 | 0 | 9.358181476 | 0 | 0 | 0 | 0 | 0 | 0 | 3.64E-08 | 1 |
| RILP | XM_005256811 | 9.539097493 | 0 | 0 | 0 | 0 | 0 | 0 | 10.83844969 | 3.28E-08 | 1 |
| RILPL2 | NR_130703 | 0 | 0 | 0 | 8.080900301 | 0 | 0 | 0 | 0 | 9.71E-06 | 1 |
| RIMS3 | XM_011542479 | 0 | 8.835955774 | 0 | 0 | 0 | 0 | 0 | 0 | 2.22E-07 | 1 |
| RIN2 | XM_024451913 | 0 | 8.844550112 | 0 | 10.79185087 | 9.403538116 | 0 | 0 | 0 | 0.999953634 | 0.02072721 |
| RINT1 | NM_021930 | 0 | 0 | 0 | 0 | 6.725274845 | 0 | 0 | 0 | 9.53E-05 | 1 |
| RIOK2 | NM_018343 | 9.487768672 | 0 | 0 | 0 | 0 | 0 | 0 | 0 | 3.53E-08 | 1 |
| RIOX1 | NM_024644 | 0 | 8.671801345 | 0 | 0 | 0 | 0 | 0 | 0 | 9.15E-08 | 1 |
| RMC1 | NM_001318709 | 9.171928589 | 0 | 0 | 0 | 0 | 0 | 0 | 0 | 7.86E-08 | 1 |

| | | | | | | | | | | | |
|---|---|---|---|---|---|---|---|---|---|---|---|
| RMDN1 | XM_017013518 | 0 | 8.690990708 | 0 | 0 | 0 | 0 | 0 | 0 | 9.02E-08 | 1 |
| RMI1 | XM_017015140 | 9.257603503 | 8.728620286 | 0 | 0 | 0 | 0 | 0 | 0 | 0.003171989 | 1 |
| RMI2 | NM_152308 | 0 | 0 | 0 | 10.91146482 | 8.671527983 | 0 | 0 | 0 | 0.013317244 | 0.02072721 |
| RMND5A | NM_022780 | 0 | 0 | 9.148481961 | 0 | 8.985072162 | 0 | 0 | 10.33987527 | 0.027278922 | 1 |
| RNA45SN4 | NR_146117 | 0 | 10.3480437 | 0 | 6.964181636 | 8.61755579 | 0 | 0 | 0 | 0.999953634 | 0.041789917 |
| RNASE1 | NM_198234 | 0 | 0 | 7.244417312 | 0 | 0 | 0 | 0 | 0 | 3.21E-05 | 1 |
| RNASE6 | XM_017021566 | 0 | 9.10991584 | 0 | 0 | 0 | 0 | 0 | 0 | 5.42E-08 | 1 |
| RNASEH1 | NM_002936 | 0 | 0 | 0 | 0 | 0 | 0 | 9.023750081 | 8.31862546 | 1 | 0.02072721 |
| RNASEH1 | XR_922673 | 0 | 9.838684641 | 0 | 0 | 0 | 0 | 6.477826297 | 0 | 2.00E-08 | 1 |
| RNASEH2A | NM_006397 | 0 | 0 | 0 | 11.53088122 | 9.941555888 | 0 | 9.339969963 | 0 | 0.007233002 | 0.999932046 |
| RNASET2 | XM_017011397 | 0 | 0 | 9.135940735 | 0 | 9.960950465 | 0 | 0 | 0 | 0.018733627 | 0.02072721 |
| RND3 | NM_005168 | 0 | 9.19973932 | 0 | 0 | 0 | 0 | 0 | 0 | 4.67E-08 | 1 |
| RNF103 | NM_005667 | 0 | 8.709928179 | 10.17608123 | 9.227165049 | 7.328262653 | 0 | 0 | 0 | 0.999953634 | 0.004942065 |
| RNF111 | XM_017022343 | 0 | 9.416987232 | 0 | 0 | 0 | 0 | 0 | 0 | 3.28E-08 | 1 |
| RNF113A | NM_006978 | 0 | 0 | 0 | 10.65391614 | 8.709289252 | 12.54942709 | 0 | 10.36410426 | 0.015164505 | 0.999932046 |
| RNF121 | XM_024448604 | 0 | 9.38199218 | 0 | 0 | 0 | 0 | 0 | 0 | 3.43E-08 | 1 |
| RNF125 | NM_017831 | 10.29105753 | 0 | 0 | 0 | 0 | 0 | 0 | 0 | 1.54E-08 | 1 |
| RNF126 | NM_194460 | 0 | 8.31161984 | 0 | 0 | 0 | 10.72670472 | 0 | 0 | 1.27E-07 | 1 |
| RNF126 | XM_005259594 | 0 | 9.117027412 | 0 | 0 | 0 | 0 | 0 | 0 | 5.36E-08 | 1 |
| RNF13 | XM_024453332 | 0 | 0 | 0 | 7.585264412 | 0 | 0 | 6.67079425 | 0 | 1.62E-05 | 1 |
| RNF130 | NM_001280801 | 9.708807013 | 0 | 0 | 0 | 0 | 0 | 0 | 0 | 2.61E-08 | 1 |
| RNF135 | XM_024451000 | 0 | 0 | 0 | 10.25846802 | 9.793912627 | 10.30519767 | 0 | 0 | 0.013985854 | 0.999932046 |
| RNF138 | XR_935225 | 8.624880391 | 0 | 0 | 0 | 0 | 0 | 0 | 0 | 1.02E-07 | 1 |
| RNF139 | NM_007218 | 0 | 0 | 9.842164946 | 0 | 7.119744988 | 0 | 8.521333265 | 0 | 0.030210229 | 0.999932046 |
| RNF141 | NM_016422 | 0 | 0 | 9.634965902 | 0 | 9.71615059 | 0 | 0 | 0 | 0.017971914 | 0.02072721 |
| RNF145 | XM_017009138 | 0 | 9.081113725 | 0 | 0 | 0 | 0 | 0 | 0 | 5.70E-08 | 1 |
| RNF146 | NM_001242848 | 0 | 8.78328601 | 0 | 0 | 0 | 0 | 0 | 0 | 8.40E-08 | 1 |
| RNF149 | XM_005263921 | 0 | 0 | 0 | 0 | 6.409577539 | 0 | 0 | 0 | 4.59E-05 | 1 |
| RNF152 | XM_017025613 | 0 | 9.051724868 | 0 | 0 | 0 | 0 | 0 | 0 | 7.86E-08 | 1 |
| RNF170 | NM_030954 | 0 | 0 | 10.22426851 | 6.258826736 | 0 | 0 | 0 | 0 | 0.028092044 | 0.02072721 |
| RNF19B | NM_153341 | 9.502621339 | 0 | 0 | 0 | 0 | 0 | 0 | 0 | 3.44E-08 | 1 |
| RNF20 | NM_019592 | 0 | 8.550974732 | 0 | 0 | 0 | 0 | 0 | 0 | 1.01E-07 | 1 |
| RNF208 | NM_031297 | 0 | 0 | 0 | 0 | 6.690841957 | 0 | 0 | 8.431845921 | 3.72E-05 | 1 |
| RNF216 | XM_017012364 | 12.12124062 | 0 | 0 | 0 | 0 | 0 | 0 | 0 | 1.34E-08 | 1 |
| RNF216 | XM_024446806 | 0 | 0 | 0 | 0 | 0 | 11.89624654 | 9.713192346 | 0 | 1 | 0.019697931 |
| RNF216P1 | NR_023384 | 0 | 0 | 10.03348707 | 10.13730841 | 8.88962462 | 0 | 9.650045731 | 0 | 0.001633613 | 1 |
| RNF217 | XM_011535495 | 0 | 0 | 10.38304631 | 0 | 9.137737665 | 10.65966957 | 9.388202852 | 0 | 0.016022152 | 0.999932046 |
| RNF219 | XM_024449410 | 0 | 0 | 6.656419865 | 8.064023368 | 7.10238406 | 0 | 0 | 0 | 0.025216192 | 0.02072721 |
| RNF220 | NM_018150 | 0 | 8.983325733 | 0 | 0 | 0 | 0 | 0 | 0 | 7.17E-08 | 1 |
| RNF227 | NR_152444 | 0 | 8.348342131 | 0 | 0 | 0 | 0 | 0 | 0 | 1.24E-07 | 1 |
| RNF25 | NM_022453 | 0 | 10.82492591 | 10.47884951 | 8.991637848 | 0 | 0 | 0 | 0 | 0.999953634 | 0.02072721 |
| RNF38 | NM_194328 | 10.26079939 | 0 | 0 | 0 | 0 | 0 | 0 | 0 | 1.58E-08 | 1 |
| RNF40 | NM_014771 | 9.208624214 | 9.569931439 | 0 | 0 | 0 | 0 | 7.849247234 | 0 | 0.001874083 | 1 |

| | | | | | | | | | | | |
|---|---|---|---|---|---|---|---|---|---|---|---|
| RNF43 | NM_001305544 | 10.21644202 | 0 | 0 | 0 | 0 | 11.16479539 | 12.83747077 | 0 | 1.59E-08 | 1.92E-09 |
| RNF44 | XR_941091 | 0 | 0 | 0 | 0 | 6.351978551 | 0 | 6.840970326 | 0 | 4.81E-05 | 1 |
| RNF6 | XM_024449392 | 0 | 9.102769039 | 9.574289913 | 9.882039044 | 6.865660185 | 0 | 0 | 0 | 0.999953634 | 0.005617506 |
| RNF7 | NR_037702 | 0 | 8.870030144 | 0 | 0 | 0 | 0 | 0 | 0 | 7.86E-08 | 1 |
| RNF8 | XR_001743734 | 10.6557466 | 0 | 0 | 0 | 0 | 0 | 0 | 0 | 1.34E-08 | 1 |
| RNF8 | XR_427853 | 0 | 9.569931439 | 0 | 0 | 0 | 0 | 0 | 0 | 2.66E-08 | 1 |
| RNGTT | NM_003800 | 0 | 9.044283149 | 0 | 0 | 0 | 0 | 0 | 0 | 6.33E-08 | 1 |
| RNPEP | NM_020216 | 0 | 0 | 8.020656193 | 11.02845609 | 9.595695343 | 0 | 0 | 0 | 0.001440983 | 0.002618715 |
| RNPEP | XM_017002022 | 0 | 10.06882031 | 0 | 0 | 0 | 0 | 0 | 0 | 1.60E-08 | 1 |
| RNPEPL1 | NM_018226 | 10.35792493 | 0 | 0 | 0 | 0 | 0 | 0 | 0 | 1.48E-08 | 1 |
| RNPS1 | NM_006711 | 0 | 0 | 0 | 0 | 0 | 9.763473303 | 7.488463554 | 0 | 1 | 0.02072721 |
| RNPS1 | NM_080594 | 0 | 0 | 10.04583013 | 0 | 9.283506136 | 0 | 0 | 11.44957288 | 0.0178104 | 0.999932046 |
| ROBO1 | XM_017006985 | 9.261974802 | 0 | 0 | 0 | 0 | 0 | 0 | 0 | 5.08E-08 | 1 |
| ROBO2 | XM_017007006 | 9.849062238 | 12.00307174 | 0 | 11.0378503 | 12.72638377 | 0 | 0 | 0 | 0.999953634 | 1.92E-09 |
| ROGDI | NM_024589 | 0 | 9.10991584 | 0 | 0 | 0 | 0 | 0 | 0 | 5.42E-08 | 1 |
| RP9 | NM_203288 | 0 | 0 | 0 | 10.50138585 | 10.11713731 | 0 | 7.84197419 | 0 | 0.011378785 | 0.999932046 |
| RPA1 | XM_024450863 | 8.704119069 | 0 | 0 | 0 | 0 | 0 | 8.020088967 | 9.657578368 | 9.60E-08 | 0.02072721 |
| RPAIN | NM_001160266 | 0 | 0 | 0 | 8.086482355 | 0 | 0 | 0 | 0 | 9.60E-06 | 1 |
| RPAIN | XR_001752663 | 0 | 9.213075091 | 0 | 0 | 0 | 0 | 0 | 0 | 4.56E-08 | 1 |
| RPE | NM_199229 | 0 | 0 | 0 | 9.338953707 | 7.335710674 | 0 | 0 | 10.89254799 | 0.038286522 | 0.999932046 |
| RPF2 | NM_032194 | 10.31649738 | 11.1844206 | 0 | 10.69651615 | 9.498672477 | 0 | 0 | 0 | 0.999953634 | 0.02072721 |
| RPGRIP1L | NM_001328423 | 0 | 0 | 0 | 9.312999003 | 7.178913994 | 0 | 0 | 0 | 0.040029929 | 0.027966656 |
| RPL10 | NM_001303625 | 0 | 0 | 8.443380147 | 10.00797741 | 10.15236053 | 10.5414403 | 7.704370312 | 11.1718481 | 0.001810422 | 0.999932046 |
| RPL10L | NM_080746 | 0 | 8.561439232 | 0 | 0 | 0 | 0 | 0 | 0 | 1.00E-07 | 1 |
| RPL13A | NM_001270491 | 9.97155406 | 0 | 0 | 0 | 0 | 0 | 0 | 0 | 2.00E-08 | 1 |
| RPL14 | NM_001034996 | 0 | 0 | 10.85929105 | 10.42963528 | 10.5474364 | 0 | 10.65122964 | 0 | 0.000538775 | 0.999932046 |
| RPL22L1 | NM_001320451 | 0 | 8.975521721 | 0 | 0 | 0 | 0 | 8.300353401 | 0 | 7.27E-08 | 1 |
| RPL23AP7 | NR_000029 | 0 | 0 | 0 | 10.31305533 | 8.148778003 | 0 | 0 | 0 | 0.01966012 | 0.02072721 |
| RPL23AP7 | NR_024529 | 0 | 0 | 0 | 0 | 6.736572239 | 0 | 8.708925636 | 0 | 9.53E-05 | 1 |
| RPL31 | NM_001098577 | 0 | 0 | 0 | 0 | 0 | 12.12207309 | 8.058247234 | 0 | 1 | 1.92E-09 |
| RPL32 | NM_001007073 | 0 | 10.14036773 | 0 | 0 | 0 | 0 | 8.251784555 | 0 | 1.56E-08 | 1 |
| RPL34 | NM_000995 | 9.1949725 | 0 | 10.40237365 | 11.05574595 | 4.270485932 | 0 | 0 | 0 | 0.941836129 | 0.003193095 |
| RPL7L1 | NM_198486 | 0 | 9.451153482 | 0 | 0 | 0 | 0 | 8.484441039 | 6.927451524 | 3.15E-08 | 0.03429376 |
| RPL9 | NM_001024921 | 0 | 0 | 11.6612168 | 11.82772464 | 9.268018164 | 11.2628559 | 9.809553935 | 12.05729928 | 0.000283329 | 0.999932046 |
| RPN2 | NM_001324305 | 0 | 8.927789798 | 0 | 0 | 0 | 0 | 0 | 0 | 7.69E-08 | 1 |
| RPN2 | XM_006723852 | 0 | 8.844550112 | 0 | 0 | 0 | 0 | 8.039294254 | 0 | 8.06E-08 | 1 |
| RPP14 | NM_001098783 | 0 | 0 | 0 | 6.544903833 | 0 | 0 | 0 | 0 | 4.35E-05 | 1 |
| RPP25 | NM_017793 | 0 | 0 | 0 | 10.40192126 | 8.844653371 | 0 | 10.3213117 | 0 | 0.016708288 | 0.999932046 |
| RPP30 | NM_006413 | 10.01363818 | 9.17269217 | 12.13132781 | 10.31424523 | 7.503816758 | 0 | 0 | 0 | 0.999953634 | 0.001376968 |
| RPP38 | NM_006414 | 0 | 0 | 0 | 11.52009353 | 10.19700379 | 0 | 0 | 0 | 0.006816422 | 0.02072721 |
| RPP40 | NM_006638 | 0 | 0 | 0 | 10.19011422 | 7.987916427 | 0 | 0 | 0 | 0.021466641 | 0.02072721 |
| RPRD1A | XM_011526050 | 0 | 0 | 0 | 10.09915958 | 6.163947944 | 0 | 6.840970326 | 0 | 0.027842286 | 0.999932046 |
| RPS19BP1 | NR_130151 | 0 | 0 | 0 | 0 | 0 | 9.535038392 | 8.184373006 | 0 | 1 | 0.02072721 |

| | | | | | | | | | | | |
|---|---|---|---|---|---|---|---|---|---|---|---|
| RPS21 | XR_002958499 | 10.51267417 | 9.500932031 | 0 | 0 | 0 | 0 | 0 | 0 | 0.000792882 | 1 |
| RPS27 | NM_001349946 | 9.502621339 | 0 | 0 | 0 | 0 | 0 | 0 | 0 | 3.44E-08 | 1 |
| RPS3 | NM_001260506 | 0 | 0 | 0 | 9.27683648 | 7.42919044 | 3.63205005 | 5.417697669 | 0 | 0.03898352 | 0.999932046 |
| RPS3A | NM_001267699 | 0 | 0 | 0 | 9.066983555 | 8.567824433 | 0 | 0 | 0 | 0.032181828 | 0.022861828 |
| RPS6KA1 | NM_001330441 | 9.539097493 | 0 | 0 | 0 | 0 | 0 | 0 | 0 | 3.28E-08 | 1 |
| RPS6KA3 | NM_004586 | 0 | 0 | 11.67897563 | 0 | 10.20006748 | 0 | 0 | 0 | 0.006321535 | 0.02072721 |
| RPS6KA3 | XM_017029717 | 0 | 0 | 9.74641291 | 11.06355026 | 9.924369085 | 11.82249291 | 9.20333417 | 10.38497589 | 0.000790295 | 0.999932046 |
| RPS6KL1 | XM_024449718 | 0 | 0 | 0 | 0 | 6.569741695 | 0 | 0 | 0 | 4.12E-05 | 1 |
| RPS9 | NM_001321702 | 0 | 8.384152849 | 0 | 0 | 0 | 0 | 0 | 10.19206204 | 1.18E-07 | 1 |
| RPS9 | NM_001321705 | 0 | 0 | 0 | 0 | 6.667421491 | 0 | 6.586265919 | 0 | 3.78E-05 | 1 |
| RPS9 | NM_001321706 | 0 | 8.464408432 | 0 | 0 | 0 | 0 | 0 | 0 | 1.11E-07 | 1 |
| RPS9 | NR_135763 | 0 | 0 | 10.30210361 | 10.62544009 | 9.965758643 | 0 | 7.647100774 | 8.851562547 | 0.000797575 | 0.999932046 |
| RPSAP52 | NR_026825 | 0 | 0 | 0 | 0 | 0 | 10.92227289 | 8.767681412 | 0 | 1 | 0.02072721 |
| RPSAP58 | NM_001355287 | 0 | 0 | 10.98231794 | 11.37597241 | 8.161407372 | 12.04634112 | 10.04471001 | 0 | 0.000669983 | 0.999932046 |
| RPSAP9 | NR_026890 | 10.37416851 | 0 | 0 | 0 | 0 | 0 | 0 | 0 | 1.48E-08 | 1 |
| RPUSD1 | NM_058192 | 0 | 9.059128398 | 0 | 9.882039044 | 6.667421491 | 0 | 0 | 0 | 0.999953634 | 0.022212956 |
| RRBP1 | NM_001042576 | 0 | 9.722262949 | 0 | 0 | 0 | 0 | 0 | 0 | 2.23E-08 | 1 |
| RREB1 | NM_001168344 | 0 | 9.321711965 | 0 | 0 | 0 | 0 | 0 | 0 | 3.84E-08 | 1 |
| RRM2 | NM_001165931 | 0 | 9.522516235 | 0 | 11.12992606 | 6.582319518 | 0 | 0 | 0 | 0.999953634 | 1.31E-08 |
| RRP12 | NM_015179 | 0 | 8.571828374 | 0 | 0 | 0 | 0 | 0 | 0 | 9.97E-08 | 1 |
| RRS1 | NM_015169 | 0 | 0 | 0 | 8.688149458 | 9.644491088 | 9.461140157 | 9.53974091 | 0 | 0.02395319 | 0.999932046 |
| RSAD1 | NR_130911 | 0 | 0 | 0 | 8.84785929 | 6.077742093 | 0 | 0 | 0 | 0.062545636 | 0.044603371 |
| RSAD2 | XM_011510415 | 0 | 0 | 9.406923074 | 0 | 4.951611714 | 0 | 0 | 0 | 0.05097348 | 0.032284732 |
| RSBN1 | NM_018364 | 0 | 9.670031486 | 0 | 0 | 0 | 0 | 0 | 0 | 2.40E-08 | 1 |
| RSPH1 | XM_011529786 | 0 | 9.17269217 | 0 | 0 | 0 | 0 | 0 | 0 | 4.92E-08 | 1 |
| RSPO3 | XM_017011379 | 11.26675151 | 12.34796908 | 11.01920541 | 0 | 8.165592712 | 0 | 0 | 0 | 0.87797743 | 0.02072721 |
| RSPRY1 | XM_024450483 | 0 | 8.31161984 | 0 | 0 | 0 | 0 | 0 | 9.172865883 | 1.27E-07 | 1 |
| RSRC1 | XM_024453560 | 0 | 0 | 5.498468293 | 0 | 10.26733459 | 0 | 9.496390342 | 0 | 0.025231964 | 1 |
| RSRC2 | XM_005253602 | 0 | 10.7073419 | 0 | 10.5354558 | 10.1439002 | 0 | 0 | 0 | 0.999953634 | 0.02072721 |
| RSRC2 | XM_017019831 | 10.29958742 | 9.694007725 | 0 | 10.50555865 | 10.44871393 | 0 | 0 | 0 | 0.999953634 | 0.02072721 |
| RSRC2 | XR_001748847 | 0 | 0 | 0 | 0 | 6.478485785 | 0 | 10.65382795 | 9.652668974 | 4.36E-05 | 0.976243727 |
| RSRP1 | NR_135789 | 0 | 0 | 0 | 8.766740692 | 0 | 0 | 0 | 0 | 4.64E-06 | 1 |
| RTCA | XM_005271298 | 0 | 8.248259235 | 0 | 0 | 0 | 0 | 0 | 0 | 2.02E-07 | 1 |
| RTEL1 | NM_032957 | 0 | 0 | 8.983289397 | 0 | 8.956370491 | 0 | 0 | 9.892265765 | 0.029230292 | 1 |
| RTF1 | NM_015138 | 0 | 0 | 10.53564879 | 0 | 8.539078316 | 0 | 9.25353328 | 0 | 0.016865508 | 0.999932046 |
| RTKN | XM_017004635 | 0 | 0 | 0 | 0 | 6.557053251 | 0 | 7.56113023 | 0 | 4.16E-05 | 1 |
| RTKN2 | NM_145307 | 0 | 0 | 0 | 9.744046385 | 10.22835252 | 0 | 0 | 8.877042846 | 0.014286405 | 0.999932046 |
| RTL8A | NM_001078172 | 10.28247691 | 0 | 0 | 0 | 0 | 0 | 0 | 9.29131303 | 1.54E-08 | 1 |
| RTL8C | NM_001078171 | 0 | 0 | 8.894253645 | 0 | 8.912218095 | 0 | 0 | 0 | 0.030683752 | 0.021215703 |
| RTN3 | NM_201429 | 0 | 0 | 0 | 0 | 6.505153187 | 0 | 0 | 0 | 4.28E-05 | 1 |
| RTN3 | NR_049750 | 0 | 8.895067962 | 0 | 0 | 0 | 0 | 0 | 0 | 7.75E-08 | 1 |
| RTN4 | NR_135830 | 0 | 0 | 9.724243205 | 0 | 8.206791567 | 0 | 0 | 0 | 0.026551524 | 0.02072721 |
| RTN4IP1 | NM_032730 | 0 | 9.044283149 | 0 | 0 | 0 | 0 | 0 | 0 | 6.33E-08 | 1 |

| | | | | | | | | | | | |
|---|---|---|---|---|---|---|---|---|---|---|---|
| RUBCNL | NM_001349772 | 0 | 9.340061955 | 0 | 0 | 0 | 10.48327637 | 0 | 0 | 3.95E-08 | 1 |
| RUFY2 | XR_246097 | 9.636581421 | 0 | 0 | 0 | 0 | 0 | 0 | 0 | 2.90E-08 | 1 |
| RUFY3 | XR_938714 | 0 | 0 | 0 | 8.673388514 | 0 | 0 | 0 | 0 | 5.13E-06 | 1 |
| RUNX1 | NM_001122607 | 0 | 0 | 0 | 10.68368358 | 8.958784227 | 11.10383732 | 10.07914117 | 0 | 0.014247971 | 0.999932046 |
| RUNX1 | XM_011529770 | 0 | 0 | 0 | 11.8327153 | 9.960950465 | 12.10817286 | 0 | 11.48314612 | 0.006056596 | 0.999932046 |
| RUNX1T1 | NM_001198633 | 0 | 0 | 0 | 0 | 0 | 10.60264726 | 0 | 10.40262864 | 1 | 0.02072721 |
| RUSC1 | NM_014328 | 0 | 0 | 10.67797447 | 0 | 7.677994746 | 0 | 0 | 0 | 0.017748616 | 0.020835214 |
| RUSC1 | XR_921763 | 0 | 0 | 10.20638636 | 0 | 8.641793038 | 0 | 0 | 0 | 0.019299864 | 0.02072721 |
| RUVBL1 | XM_005247841 | 0 | 0 | 0 | 0 | 6.518304288 | 0 | 0 | 0 | 4.23E-05 | 1 |
| RUVBL2 | XM_011526330 | 0 | 0 | 0 | 11.22535848 | 8.85635197 | 0 | 0 | 0 | 0.010779876 | 0.02072721 |
| RWDD2A | XM_017010223 | 0 | 9.411213355 | 0 | 0 | 0 | 0 | 0 | 0 | 3.28E-08 | 1 |
| RWDD2B | NM_016940 | 10.05452941 | 0 | 0 | 0 | 0 | 0 | 7.704370312 | 0 | 5.06E-08 | 1 |
| RWDD3 | NM_001278248 | 9.825628908 | 0 | 0 | 0 | 0 | 0 | 0 | 0 | 2.30E-08 | 1 |
| RXRB | NM_021976 | 0 | 9.232850423 | 0 | 0 | 0 | 0 | 0 | 0 | 4.38E-08 | 1 |
| RXRB | XM_017011176 | 9.574674104 | 0 | 0 | 0 | 0 | 0 | 6.586265919 | 0 | 3.15E-08 | 1 |
| RYK | XR_924160 | 0 | 9.044283149 | 0 | 0 | 0 | 0 | 0 | 0 | 6.33E-08 | 1 |
| S100A13 | XM_005245434 | 10.97350327 | 0 | 0 | 0 | 0 | 0 | 0 | 0 | 1.34E-08 | 1 |
| S100A13 | XM_011509864 | 9.83739315 | 8.809861237 | 0 | 0 | 0 | 9.358414258 | 6.750642936 | 0 | 0.001922313 | 0.022611547 |
| S100A16 | NM_001317007 | 0 | 7.585902702 | 0 | 10.03709574 | 9.022473864 | 0 | 0 | 0 | 0.910217967 | 0.02072721 |
| S100A8 | NM_002964 | 0 | 0 | 10.1198742 | 9.489798191 | 8.462903138 | 0 | 0 | 10.15087512 | 0.002370246 | 0.999932046 |
| S100A9 | NM_002965 | 0 | 0 | 10.19030288 | 0 | 8.011359982 | 0 | 0 | 6.791132747 | 0.02179728 | 0.999932046 |
| SAE1 | NM_005500 | 0 | 0 | 0 | 11.07693177 | 10.38193507 | 10.80615134 | 0 | 0 | 0.00806635 | 0.999932046 |
| SAFB | XM_017027118 | 10.41006135 | 0 | 10.05251847 | 0 | 8.656737116 | 0 | 0 | 0 | 0.999953634 | 0.02072721 |
| SAMD12 | XM_017013412 | 0 | 0 | 0 | 11.02337221 | 7.701245565 | 0 | 0 | 0 | 0.014259619 | 0.02072721 |
| SAMD13 | NM_001134663 | 0 | 0 | 10.05474104 | 0 | 6.607150896 | 0 | 0 | 0 | 0.028261608 | 0.02072721 |
| SAMD13 | NM_001134664 | 0 | 0 | 0 | 0 | 6.3222931 | 0 | 0 | 0 | 0.000244538 | 1 |
| SAMD3 | NM_001258275 | 0 | 0 | 0 | 9.632089798 | 10.39714311 | 0 | 0 | 0 | 0.013745088 | 0.02072721 |
| SAP18 | XM_011534868 | 0 | 0 | 0 | 7.763094795 | 0 | 0 | 0 | 0 | 2.57E-05 | 1 |
| SAP30BP | NM_013260 | 0 | 8.861586703 | 0 | 11.90715272 | 7.415197319 | 0 | 0 | 0 | 0.911156323 | 5.45E-09 |
| SAP30BP | XM_011524693 | 10.74900774 | 0 | 0 | 0 | 0 | 0 | 0 | 0 | 1.34E-08 | 1 |
| SAP30L | NR_024084 | 0 | 9.258802813 | 0 | 0 | 0 | 0 | 0 | 0 | 4.23E-08 | 1 |
| SAR1B | NM_016103 | 0 | 0 | 11.44907275 | 0 | 10.53171492 | 0 | 0 | 8.379320965 | 0.006469986 | 0.999932046 |
| SARAF | NM_001284239 | 9.623051988 | 9.479019994 | 0 | 0 | 0 | 0 | 8.347340272 | 0 | 0.001544315 | 1 |
| SARS2 | NM_017827 | 0 | 0 | 9.963053877 | 0 | 9.449745843 | 0 | 0 | 0 | 0.017527994 | 0.02072721 |
| SART1 | XM_011545345 | 0 | 9.434171496 | 0 | 0 | 0 | 0 | 0 | 0 | 3.24E-08 | 1 |
| SASH1 | XM_017010600 | 0 | 10.27362413 | 0 | 0 | 0 | 0 | 0 | 0 | 1.47E-08 | 1 |
| SASH3 | XM_006724763 | 0 | 8.690990708 | 0 | 0 | 0 | 0 | 0 | 0 | 9.02E-08 | 1 |
| SAT2 | NM_001320845 | 10.85118405 | 9.10991584 | 10.50183826 | 0 | 6.631562102 | 0 | 0 | 0 | 0.999953634 | 0.02072721 |
| SAT2 | NM_133491 | 0 | 0 | 0 | 8.955489713 | 8.131765004 | 0 | 8.137615898 | 0 | 0.038848737 | 0.999932046 |
| SAV1 | XM_017021568 | 0 | 0 | 0 | 0 | 0 | 0 | 7.177815683 | 9.597533386 | 1 | 0.02072721 |
| SBDSP1 | NR_024111 | 0 | 8.861586703 | 0 | 0 | 0 | 10.76264312 | 0 | 0 | 7.86E-08 | 1 |
| SBF1 | XM_011530710 | 0 | 8.273938353 | 0 | 0 | 0 | 0 | 0 | 0 | 2.02E-07 | 1 |
| SBNO2 | NM_014963 | 0 | 0 | 0 | 8.573575335 | 0 | 0 | 7.431456038 | 0 | 2.47E-05 | 1 |

| | | | | | | | | | | | |
|---|---|---|---|---|---|---|---|---|---|---|---|
| SCAF8 | NM_014892 | 0 | 0 | 0 | 8.457751161 | 8.338670907 | 0 | 0 | 0 | 0.045265831 | 0.030697281 |
| SCAMP1-AS1 | NR_105014 | 0 | 0 | 5.944383075 | 10.23855048 | 9.017851335 | 0 | 0 | 0 | 0.004327114 | 0.006774797 |
| SCAMP2 | NM_005697 | 10.21644202 | 0 | 10.20838425 | 0 | 7.119744988 | 0 | 0 | 0 | 0.999953634 | 0.02072721 |
| SCAMP3 | NM_052837 | 0 | 0 | 6.019129988 | 11.56526942 | 10.24772849 | 0 | 0 | 0 | 0.001166592 | 0.002311272 |
| SCAMP4 | NM_079834 | 0 | 8.140712363 | 0 | 6.669146541 | 8.8511643 | 0 | 0 | 0 | 0.999953634 | 0.038184118 |
| SCAND1 | NM_016558 | 0 | 0 | 0 | 10.06280902 | 7.10238406 | 0 | 0 | 0 | 0.025886001 | 0.02072721 |
| SCAND2P | NR_004859 | 10.58423881 | 0 | 0 | 0 | 0 | 0 | 0 | 0 | 1.34E-08 | 1 |
| SCARB2 | NM_005506 | 0 | 9.901810942 | 0 | 0 | 0 | 0 | 0 | 10.71671941 | 1.87E-08 | 1 |
| SCARF2 | NM_182895 | 0 | 0 | 0 | 0 | 6.631562102 | 0 | 0 | 0 | 3.88E-05 | 1 |
| SCEL | XM_011535290 | 0 | 8.561439232 | 0 | 0 | 0 | 0 | 0 | 0 | 1.00E-07 | 1 |
| SCG5 | NM_001144757 | 0 | 10.60736295 | 0 | 0 | 0 | 0 | 0 | 0 | 1.34E-08 | 1 |
| SCIN | NM_001112706 | 10.10406561 | 0 | 12.19794865 | 0 | 7.718441029 | 0 | 0 | 0 | 0.999953634 | 6.14E-09 |
| SCIN | XR_926951 | 10.19374119 | 0 | 0 | 0 | 0 | 0 | 0 | 9.929274263 | 1.60E-08 | 1 |
| SCML1 | XM_017029723 | 0 | 0 | 0 | 0 | 0 | 0 | 7.856483795 | 9.513376477 | 1 | 0.02072721 |
| SCNM1 | NM_001204856 | 0 | 0 | 11.63286076 | 10.59050989 | 8.70354377 | 11.39543079 | 8.62242756 | 0 | 0.000607324 | 0.999932046 |
| SCNN1B | XM_011545913 | 0 | 9.186279128 | 0 | 0 | 0 | 0 | 0 | 0 | 4.80E-08 | 1 |
| SCNN1B | XM_017023525 | 0 | 8.430558346 | 0 | 0 | 0 | 0 | 0 | 0 | 1.13E-07 | 1 |
| SCNN1B | XM_017023526 | 10.57724 | 0 | 0 | 0 | 0 | 0 | 0 | 0 | 1.34E-08 | 1 |
| SCNN1G | NM_001039 | 0 | 8.686217243 | 0 | 0 | 0 | 0 | 0 | 0 | 9.07E-08 | 1 |
| SCO1 | XM_005256751 | 0 | 0 | 9.213587255 | 0 | 8.070588078 | 0 | 0 | 0 | 0.035495606 | 0.023358895 |
| SCX | XM_006716616 | 0 | 0 | 0 | 7.188387521 | 0 | 0 | 0 | 0 | 2.41E-05 | 1 |
| SCYL1 | XM_017018033 | 0 | 0 | 0 | 9.591445233 | 8.202724223 | 0 | 8.15532787 | 0 | 0.027842606 | 0.999932046 |
| SCYL1 | XM_024448620 | 10.82925902 | 0 | 0 | 0 | 0 | 0 | 0 | 0 | 1.34E-08 | 1 |
| SDC1 | NM_002997 | 0 | 0 | 11.89532346 | 10.64826572 | 7.357826773 | 0 | 0 | 0 | 0.00067514 | 0.001418508 |
| SDC2 | NM_002998 | 0 | 0 | 0 | 7.011897042 | 0 | 0 | 0 | 10.41137467 | 2.89E-05 | 1 |
| SDC2 | XM_024447228 | 0 | 0 | 9.017875808 | 10.50034076 | 6.557053251 | 0 | 0 | 0 | 0.003360109 | 0.005156764 |
| SDCBP | NM_001007067 | 0 | 9.051724868 | 0 | 0 | 0 | 0 | 0 | 0 | 7.86E-08 | 1 |
| SDF2 | NM_006923 | 0 | 0 | 10.5002083 | 0 | 8.250791506 | 10.55789653 | 0 | 0 | 0.018013892 | 0.999932046 |
| SDF2 | NR_045585 | 0 | 7.722723415 | 11.17341355 | 0 | 9.072367519 | 0 | 0 | 0 | 0.774030274 | 0.02072721 |
| SDF4 | NM_016547 | 0 | 8.024497849 | 0 | 9.591445233 | 7.227767768 | 0 | 0 | 0 | 0.999953634 | 0.023933914 |
| SDHAF3 | NM_020186 | 10.31228843 | 7.523045066 | 10.27757415 | 10.66421812 | 9.31210541 | 0 | 0 | 0 | 0.999953634 | 0.00194552 |
| SDHAF4 | NM_145267 | 0 | 8.561439232 | 9.062737842 | 0 | 7.87488031 | 0 | 0 | 0 | 0.999953634 | 0.026003313 |
| SDHAP2 | NR_003265 | 0 | 0 | 9.497940839 | 0 | 9.185922598 | 0 | 0 | 0 | 0.022406821 | 0.02072721 |
| SDK2 | XM_011524915 | 10.32905103 | 0 | 0 | 0 | 0 | 0 | 0 | 0 | 1.51E-08 | 1 |
| SDR16C5 | NM_138969 | 0 | 0 | 10.785599 | 11.47239163 | 0 | 0 | 7.743928512 | 0 | 0.006531798 | 0.999932046 |
| SDR39U1 | NM_020195 | 0 | 9.585396421 | 8.79670616 | 10.90831647 | 8.781970078 | 0 | 0 | 0 | 0.999953634 | 0.002874229 |
| SDR39U1 | XM_011536976 | 9.105375844 | 0 | 0 | 0 | 0 | 0 | 0 | 0 | 7.04E-08 | 1 |
| SEC11C | NM_001307941 | 0 | 0 | 0 | 8.688149458 | 0 | 0 | 0 | 0 | 5.05E-06 | 1 |
| SEC13 | XM_005265379 | 0 | 0 | 9.934371634 | 8.405116658 | 0 | 0 | 9.612070238 | 0 | 0.024736775 | 0.999932046 |
| SEC14L1 | NM_003003 | 0 | 0 | 0 | 0 | 6.59478863 | 0 | 0 | 0 | 4.02E-05 | 1 |
| SEC16A | XM_011519259 | 0 | 0 | 0 | 9.425184544 | 8.186339198 | 0 | 0 | 0 | 0.030371674 | 0.022014132 |
| SEC16A | XM_011519263 | 0 | 0 | 8.769900534 | 8.924658605 | 8.656737116 | 0 | 9.715184022 | 0 | 0.005333955 | 1 |
| SEC16B | NM_033127 | 0 | 0 | 0 | 0 | 6.569741695 | 0 | 7.234460372 | 0 | 4.12E-05 | 1 |

| | | | | | | | | | | | |
|---|---|---|---|---|---|---|---|---|---|---|---|
| SEC23A | XM_005267262 | 10.24763479 | 0 | 0 | 0 | 0 | 0 | 0 | 0 | 1.58E-08 | 1 |
| SEC24A | XM_017008961 | 0 | 0 | 11.34258712 | 10.95483523 | 7.282748526 | 0 | 10.35299605 | 12.59462368 | 0.000834494 | 0.999932046 |
| SEC31A | NM_016211 | 10.44700378 | 0 | 0 | 0 | 0 | 0 | 9.932774001 | 0 | 1.45E-08 | 1 |
| SEC62 | XM_011513114 | 0 | 10.49721335 | 0 | 0 | 0 | 0 | 8.470359551 | 0 | 1.34E-08 | 1 |
| SECISBP2 | NM_001354697 | 0 | 0 | 9.852422134 | 0 | 9.16934453 | 10.72670472 | 9.984883005 | 10.15435263 | 0.019753398 | 0.999932046 |
| SECISBP2 | XM_024447666 | 9.777588513 | 0 | 0 | 0 | 0 | 0 | 0 | 0 | 2.46E-08 | 1 |
| SECTM1 | XM_011523588 | 10.40611707 | 0 | 0 | 0 | 0 | 0 | 0 | 0 | 1.47E-08 | 1 |
| SEH1L | XR_935069 | 0 | 0 | 0 | 0 | 6.582319518 | 0 | 6.380785351 | 0 | 4.04E-05 | 1 |
| SEL1L | NM_005065 | 0 | 0 | 0 | 0 | 6.544252221 | 0 | 0 | 0 | 4.20E-05 | 1 |
| SELENOS | NM_203472 | 10.16134771 | 0 | 0 | 0 | 0 | 0 | 8.305650376 | 0 | 1.61E-08 | 1 |
| SELPLG | NM_003006 | 0 | 0 | 7.433922402 | 0 | 0 | 10.52200108 | 0 | 11.12564943 | 2.05E-05 | 0.999932046 |
| SEMA3B | NM_001005914 | 9.162606966 | 7.795449567 | 0 | 0 | 0 | 0 | 0 | 0 | 0.005556192 | 1 |
| SEMA3B | NR_110697 | 0 | 0 | 0 | 0 | 6.30721813 | 0 | 0 | 0 | 4.99E-05 | 1 |
| SEMA6A | XM_024446138 | 9.872121021 | 9.405416277 | 0 | 0 | 0 | 0 | 0 | 0 | 0.001360464 | 1 |
| SEMA6D | XM_024450076 | 9.900435696 | 0 | 0 | 0 | 0 | 0 | 0 | 0 | 2.07E-08 | 1 |
| SENP7 | XR_001740218 | 0 | 8.642530071 | 0 | 0 | 0 | 0 | 0 | 0 | 9.87E-08 | 1 |
| SENP8 | XM_005254157 | 0 | 9.543782272 | 0 | 0 | 0 | 0 | 9.956540422 | 0 | 2.90E-08 | 1 |
| SENP8 | XM_011521217 | 8.497211815 | 0 | 0 | 0 | 0 | 0 | 8.759984195 | 6.959609593 | 1.15E-07 | 0.029103945 |
| SEPHS1 | XM_017015944 | 0 | 8.756212053 | 0 | 0 | 0 | 10.59114494 | 7.044767963 | 0 | 8.55E-08 | 0.02072721 |
| SEPHS1 | XM_017015945 | 0 | 0 | 0 | 0 | 0 | 10.64605153 | 7.630311627 | 3.8795549 | 1 | 0.007079904 |
| 10-Sep | XM_011510699 | 0 | 0 | 0 | 8.680787864 | 0 | 0 | 0 | 0 | 5.10E-06 | 1 |
| 2-Sep | NM_004404 | 0 | 9.158976035 | 0 | 10.359902 | 7.490532315 | 0 | 0 | 0 | 0.999953634 | 0.02072721 |
| SEPT5-GP1BB | NR_037611 | 10.04441475 | 0 | 0 | 0 | 0 | 0 | 0 | 0 | 1.85E-08 | 1 |
| SEPT5-GP1BB | NR_037612 | 0 | 8.870030144 | 0 | 0 | 0 | 0 | 0 | 0 | 7.86E-08 | 1 |
| 7-Sep | XM_024447010 | 9.338474533 | 9.517150383 | 10.34541384 | 0 | 6.747781854 | 0 | 0 | 0 | 0.999953634 | 0.02072721 |
| 9-Sep | NM_001113492 | 9.134275178 | 0 | 0 | 0 | 0 | 0 | 0 | 0 | 6.66E-08 | 1 |
| SERAC1 | XR_942606 | 0 | 8.951853153 | 0 | 0 | 0 | 0 | 0 | 0 | 7.60E-08 | 1 |
| SERF2 | NM_001199875 | 10.97083541 | 0 | 0 | 0 | 0 | 0 | 0 | 0 | 1.34E-08 | 1 |
| SERF2 | NM_001199878 | 11.85224378 | 10.33590398 | 3.817450822 | 10.09915958 | 7.593532742 | 0 | 0 | 0 | 0.748372929 | 0.016592643 |
| SERGEF | NR_104041 | 0 | 9.640723913 | 0 | 0 | 0 | 0 | 0 | 0 | 2.47E-08 | 1 |
| SERINC2 | NM_001199037 | 0 | 0 | 9.053876344 | 0 | 8.8249424 | 0 | 0 | 0 | 0.029899378 | 0.020942118 |
| SERPINA1 | NM_001002236 | 0 | 0 | 11.87659996 | 0 | 8.647789258 | 10.80461986 | 10.0045595 | 0 | 0.007577711 | 0.999932046 |
| SERPINA1 | NM_001127703 | 10.46037918 | 10.16802189 | 13.25714517 | 11.28058022 | 9.267044622 | 0 | 0 | 0 | 0.912132403 | 0.0002946 |
| SERPINA1 | NM_001127705 | 10.62892178 | 10.34501833 | 10.38304631 | 0 | 8.016003323 | 0 | 0 | 0 | 0.999953634 | 0.02072721 |
| SERPINB5 | NM_002639 | 0 | 9.165850403 | 0 | 9.960130017 | 9.114111124 | 0 | 0 | 0 | 0.999953634 | 0.02072721 |
| SERPINB6 | XM_024446465 | 9.270677863 | 0 | 10.29177661 | 9.758105236 | 0 | 0 | 0 | 0 | 0.999953634 | 0.02072721 |
| SERPINB8 | NM_002640 | 0 | 9.564739385 | 0 | 0 | 0 | 0 | 0 | 10.86840153 | 2.69E-08 | 1 |
| SERPINB9 | XM_017010943 | 0 | 0 | 0 | 0 | 6.491881102 | 0 | 7.361968883 | 0 | 4.33E-05 | 1 |
| SERPINE2 | NR_073116 | 0 | 0 | 0 | 0 | 6.337212177 | 0 | 0 | 0 | 4.86E-05 | 1 |
| SERPINI1 | NM_001122752 | 0 | 8.662110092 | 0 | 0 | 0 | 0 | 0 | 0 | 9.15E-08 | 1 |
| SERTAD3 | NM_203344 | 10.12821014 | 9.327854598 | 0 | 0 | 0 | 0 | 0 | 10.19206204 | 0.001175124 | 1 |
| SESTD1 | NM_178123 | 0 | 0 | 0 | 8.422875715 | 0 | 0 | 0 | 0 | 6.71E-06 | 1 |
| SETBP1 | XM_024451153 | 10.82778544 | 0 | 0 | 0 | 0 | 0 | 0 | 0 | 1.34E-08 | 1 |

| | | | | | | | | | | | |
|---|---|---|---|---|---|---|---|---|---|---|---|
| SETBP1 | XM_024451155 | 10.90667706 | 0 | 0 | 0 | 0 | 0 | 0 | 0 | 1.34E-08 | 1 |
| SETBP1 | XM_024451157 | 0 | 8.248259235 | 0 | 0 | 0 | 0 | 0 | 0 | 2.02E-07 | 1 |
| SETBP1 | XM_024451158 | 0 | 8.33620493 | 0 | 0 | 0 | 0 | 0 | 0 | 1.25E-07 | 1 |
| SETD2 | XM_011533634 | 10.362003 | 0 | 0 | 0 | 0 | 0 | 8.997805267 | 0 | 1.48E-08 | 1 |
| SETD2 | XR_002959512 | 0 | 0 | 9.283481866 | 0 | 7.477124411 | 0 | 0 | 0 | 0.039501884 | 0.025497016 |
| SETD6 | XR_001751988 | 0 | 0 | 0 | 8.151834782 | 0 | 0 | 0 | 0 | 8.96E-06 | 1 |
| SETD6 | XR_933437 | 0 | 0 | 0 | 0 | 6.478485785 | 0 | 0 | 0 | 4.36E-05 | 1 |
| SETD9 | NR_136556 | 10.48302382 | 0 | 0 | 0 | 0 | 0 | 0 | 0 | 1.47E-08 | 1 |
| SF3B1 | XR_001738681 | 0 | 0 | 0 | 10.83311077 | 8.922147175 | 0 | 8.262719816 | 0 | 0.013285295 | 0.999932046 |
| SFI1 | XM_024452308 | 0 | 0 | 0 | 0 | 6.769943125 | 0 | 0 | 0 | 9.43E-05 | 1 |
| SFMBT2 | XM_017016466 | 0 | 8.78328601 | 0 | 0 | 0 | 0 | 0 | 0 | 8.40E-08 | 1 |
| SFPQ | XM_017002053 | 10.71433872 | 7.881486713 | 0 | 0 | 0 | 0 | 0 | 0 | 0.001218003 | 1 |
| SFSWAP | XM_024449125 | 0 | 0 | 9.229414648 | 0 | 7.449928885 | 0 | 0 | 0 | 0.041029487 | 0.026265059 |
| SFT2D3 | NM_032740 | 9.813767948 | 8.1546024 | 9.322742744 | 10.83725537 | 6.855333211 | 0 | 0 | 0 | 0.999953634 | 0.003637102 |
| SFXN1 | NM_001322981 | 9.574674104 | 0 | 0 | 0 | 0 | 0 | 0 | 0 | 3.15E-08 | 1 |
| SFXN2 | XM_024447792 | 0 | 0 | 0 | 10.75289074 | 7.259441431 | 0 | 0 | 0 | 0.017413184 | 0.020958228 |
| SFXN4 | NR_110305 | 10.8307311 | 0 | 0 | 0 | 0 | 0 | 0 | 0 | 1.34E-08 | 1 |
| SGK2 | NM_170693 | 0 | 0 | 9.936783722 | 0 | 7.82812018 | 0 | 8.84247586 | 0 | 0.025671337 | 0.999932046 |
| SGK3 | NM_170709 | 0 | 8.540433774 | 0 | 0 | 0 | 0 | 0 | 0 | 1.02E-07 | 1 |
| SGPL1 | NM_003901 | 0 | 9.803852726 | 0 | 0 | 0 | 0 | 0 | 0 | 2.02E-08 | 1 |
| SGSM3 | NM_001350039 | 0 | 8.441930184 | 0 | 0 | 0 | 0 | 0 | 10.21555569 | 1.12E-07 | 1 |
| SGSM3 | NM_001350040 | 0 | 9.754542834 | 0 | 0 | 0 | 0 | 0 | 9.965357103 | 2.13E-08 | 1 |
| SGSM3 | NM_001350041 | 0 | 0 | 9.98885422 | 0 | 8.922147175 | 0 | 0 | 0 | 0.019971794 | 0.02072721 |
| SGSM3 | NM_001350046 | 0 | 0 | 11.00892578 | 11.32670817 | 9.220530915 | 12.16567469 | 10.65745773 | 11.5533139 | 0.000520083 | 0.999932046 |
| SGSM3 | XR_001755217 | 6.992141016 | 0 | 11.4140108 | 0 | 7.939854543 | 0 | 0 | 0 | 0.579085065 | 7.83E-09 |
| SGSM3 | XR_001755221 | 0 | 0 | 11.25541479 | 9.796075236 | 7.343120441 | 0 | 0 | 0 | 0.001395429 | 0.002477101 |
| SGSM3 | XR_001755225 | 9.746730083 | 0 | 8.09164712 | 0 | 8.683252456 | 0 | 0 | 0 | 0.999953634 | 0.029082591 |
| SGSM3 | XR_002958686 | 0 | 0 | 0 | 10.76860127 | 9.008561607 | 0 | 0 | 0 | 0.013502454 | 0.02072721 |
| SH2B1 | NM_015503 | 10.16134771 | 0 | 0 | 0 | 0 | 0 | 0 | 0 | 1.61E-08 | 1 |
| SH2D4A | XR_001745571 | 8.497211815 | 0 | 0 | 0 | 0 | 0 | 0 | 0 | 1.15E-07 | 1 |
| SH3BP1 | NM_018957 | 6.161524867 | 8.935855595 | 0 | 0 | 0 | 0 | 0 | 0 | 0.011038233 | 1 |
| SH3D19 | XM_017007782 | 0 | 8.508340767 | 0 | 0 | 0 | 0 | 0 | 0 | 1.06E-07 | 1 |
| SH3GLB1 | XM_017001410 | 0 | 9.38199218 | 0 | 0 | 0 | 0 | 0 | 0 | 3.43E-08 | 1 |
| SH3KBP1 | NM_001184960 | 0 | 10.32058425 | 0 | 0 | 0 | 0 | 0 | 0 | 1.45E-08 | 1 |
| SH3RF2 | XM_006714757 | 0 | 0 | 0 | 0 | 0 | 0 | 8.997805267 | 10.11563106 | 1 | 0.02072721 |
| SH3RF2 | XM_011537567 | 0 | 0 | 10.81408653 | 10.11016257 | 8.623653382 | 0 | 0 | 9.6722071 | 0.001209117 | 0.999932046 |
| SH3TC1 | NM_018986 | 0 | 0 | 0 | 0 | 6.557053251 | 0 | 7.892131936 | 0 | 4.16E-05 | 1 |
| SH3TC1 | XR_002959735 | 0 | 8.935855595 | 0 | 0 | 0 | 0 | 0 | 0 | 7.69E-08 | 1 |
| SH3YL1 | NR_104226 | 0 | 0 | 0 | 0 | 6.769943125 | 0 | 0 | 0 | 9.43E-05 | 1 |
| SH3YL1 | NR_104227 | 10.12341345 | 7.830482506 | 0 | 0 | 0 | 0 | 0 | 0 | 0.002198605 | 1 |
| SHANK2 | XM_017017389 | 0 | 0 | 0 | 0 | 0 | 0 | 7.431456038 | 10.66402238 | 1 | 0.02072721 |
| SHC1 | XM_005245451 | 0 | 9.580259825 | 0 | 0 | 0 | 0 | 0 | 0 | 2.61E-08 | 1 |
| SHC2 | NM_012435 | 0 | 0 | 0 | 0 | 0 | 0 | 7.569961514 | 9.862810347 | 1 | 0.02072721 |

| | | | | | | | | | | | |
|---|---|---|---|---|---|---|---|---|---|---|---|
| SHISA5 | NM_001272082 | 0 | 0 | 10.3280617 | 11.09296612 | 8.653760659 | 0 | 10.03118383 | 11.96674877 | 0.000927978 | 0.999932046 |
| SHKBP1 | XM_017027473 | 0 | 0 | 0 | 0 | 6.769943125 | 0 | 0 | 0 | 9.43E-05 | 1 |
| SHKBP1 | XM_017027474 | 10.67881602 | 0 | 0 | 0 | 0 | 0 | 0 | 0 | 1.34E-08 | 1 |
| SHMT2 | NM_005412 | 0 | 0 | 0 | 10.23855048 | 8.061632835 | 0 | 0 | 0 | 0.020683846 | 0.02072721 |
| SHMT2 | NR_029417 | 0 | 9.239382419 | 0 | 0 | 0 | 0 | 9.249416094 | 0 | 4.33E-08 | 1 |
| SHOX | NM_000451 | 0 | 0 | 0 | 8.378061333 | 0 | 0 | 0 | 0 | 2.07E-05 | 1 |
| SHPRH | XR_942390 | 9.715197156 | 0 | 0 | 0 | 0 | 0 | 0 | 0 | 2.61E-08 | 1 |
| SHROOM1 | NM_001172700 | 0 | 0 | 7.275241756 | 0 | 0 | 0 | 0 | 0 | 5.92E-05 | 1 |
| SHTN1 | NM_001258298 | 0 | 0 | 0 | 7.776970127 | 0 | 0 | 0 | 0 | 1.33E-05 | 1 |
| SHTN1 | XM_006717931 | 0 | 9.352166886 | 0 | 0 | 0 | 0 | 0 | 0 | 3.68E-08 | 1 |
| SI | XM_011513078 | 0 | 8.681427931 | 0 | 0 | 0 | 0 | 0 | 0 | 9.11E-08 | 1 |
| SIAE | XM_017017930 | 9.539097493 | 0 | 0 | 0 | 0 | 0 | 0 | 0 | 3.28E-08 | 1 |
| SIAH1 | NM_003031 | 0 | 9.290600259 | 10.93604754 | 0 | 8.2973088 | 0 | 0 | 0 | 0.999953634 | 0.02072721 |
| SIAH2 | NM_005067 | 0 | 0 | 10.81408653 | 9.872377593 | 0 | 0 | 7.680104054 | 0 | 0.011938998 | 0.999932046 |
| SIDT1 | XR_001740185 | 0 | 8.248259235 | 0 | 0 | 0 | 0 | 0 | 0 | 2.02E-07 | 1 |
| SIDT2 | NM_001040455 | 0 | 0 | 0 | 0 | 6.478485785 | 0 | 0 | 0 | 4.36E-05 | 1 |
| SIGLEC10 | NM_033130 | 0 | 0 | 8.074224188 | 0 | 0 | 0 | 0 | 0 | 2.07E-05 | 1 |
| SIK3 | NM_001281748 | 10.47927431 | 0 | 0 | 0 | 0 | 0 | 0 | 0 | 1.44E-08 | 1 |
| SIK3 | XR_001747816 | 11.06389989 | 0 | 0 | 0 | 0 | 0 | 0 | 0 | 1.34E-08 | 1 |
| SIL1 | XM_024446164 | 8.779231295 | 0 | 0 | 0 | 0 | 0 | 9.118593588 | 0 | 9.02E-08 | 1 |
| SIRT3 | NM_012239 | 0 | 0 | 0 | 10.26586678 | 9.087887617 | 0 | 0 | 10.98528457 | 0.016866753 | 0.999932046 |
| SIRT6 | NM_016539 | 9.813767948 | 0 | 0 | 0 | 0 | 0 | 0 | 0 | 2.49E-08 | 1 |
| SKAP1 | XM_017025258 | 0 | 0 | 11.09243125 | 11.2215585 | 8.274237629 | 0 | 7.411941715 | 0 | 0.000674783 | 0.666516354 |
| SKAP1 | XM_017025259 | 0 | 0 | 10.89322032 | 0 | 9.341999064 | 0 | 10.69326195 | 0 | 0.011965813 | 0.999932046 |
| SLAMF1 | NR_104399 | 0 | 9.451153482 | 0 | 0 | 0 | 0 | 0 | 10.35204063 | 3.15E-08 | 1 |
| SLAMF1 | NR_104400 | 10.65906491 | 10.74799264 | 0 | 0 | 0 | 12.66365434 | 0 | 10.97647107 | 0.000334443 | 1.37E-09 |
| SLAMF1 | XM_011509905 | 7.573321022 | 0 | 0 | 0 | 0 | 11.03693222 | 0 | 9.474955682 | 0.079081949 | 0.02072721 |
| SLAMF1 | XM_017002131 | 8.4366295 | 10.05777546 | 0 | 0 | 0 | 0 | 0 | 9.152143548 | 0.001808267 | 1 |
| SLBP | NM_001306074 | 0 | 9.541141097 | 0 | 0 | 0 | 0 | 0 | 0 | 2.79E-08 | 1 |
| SLC15A3 | NM_016582 | 0 | 9.393751769 | 0 | 0 | 0 | 0 | 0 | 0 | 3.37E-08 | 1 |
| SLC16A3 | XM_024451023 | 0 | 8.78328601 | 0 | 0 | 0 | 0 | 0 | 0 | 8.40E-08 | 1 |
| SLC16A5 | XM_011525462 | 0 | 8.29916838 | 0 | 0 | 0 | 0 | 0 | 0 | 1.31E-07 | 1 |
| SLC16A5 | XM_017025293 | 0 | 0 | 0 | 0 | 6.631562102 | 0 | 0 | 0 | 3.88E-05 | 1 |
| SLC16A7 | NM_001270623 | 11.41739033 | 11.32463489 | 9.691765287 | 10.71284921 | 0 | 0 | 0 | 0 | 0.999953634 | 0.02072721 |
| SLC16A7 | XM_024449276 | 8.896331684 | 9.399595812 | 8.415979108 | 8.91213928 | 0 | 0 | 0 | 0 | 0.999953634 | 0.02662288 |
| SLC16A9 | XM_017015883 | 0 | 8.998808245 | 0 | 0 | 0 | 0 | 0 | 0 | 6.89E-08 | 1 |
| SLC17A4 | XM_011514214 | 0 | 0 | 7.420196825 | 0 | 0 | 0 | 0 | 0 | 2.07E-05 | 1 |
| SLC18A1 | NM_003053 | 0 | 8.681427931 | 0 | 0 | 0 | 0 | 0 | 0 | 9.11E-08 | 1 |
| SLC20A1 | NM_005415 | 9.581685348 | 0 | 10.3344789 | 9.926289561 | 0 | 0 | 0 | 0 | 0.999953634 | 0.02072721 |
| SLC20A1 | XM_024453052 | 0 | 0 | 0 | 0 | 6.769943125 | 0 | 7.01920173 | 0 | 9.43E-05 | 1 |
| SLC20A2 | NM_001257180 | 0 | 0 | 9.410398522 | 0 | 8.270356307 | 10.33414791 | 0 | 0 | 0.030481013 | 0.999932046 |
| SLC20A2 | NM_006749 | 0 | 0 | 0 | 10.07830002 | 7.42919044 | 0 | 0 | 10.27106783 | 0.024623293 | 0.999932046 |
| SLC20A2 | XM_017013750 | 0 | 0 | 9.816198315 | 8.918412523 | 9.839158477 | 10.95715511 | 9.787007325 | 8.962838545 | 0.002030012 | 0.999932046 |

| | | | | | | | | | | | |
|---|---|---|---|---|---|---|---|---|---|---|---|
| SLC20A2 | XM_024447235 | 0 | 0 | 11.8614442 | 0 | 8.088333507 | 0 | 8.775337779 | 0 | 1.45E-08 | 0.999932046 |
| SLC22A18 | NM_001315502 | 0 | 0 | 0 | 0 | 0 | 10.84317358 | 9.155731612 | 0 | 1 | 0.02072721 |
| SLC22A23 | NM_015482 | 0 | 0 | 11.19372479 | 0 | 8.876918096 | 0 | 0 | 0 | 0.011150339 | 0.02072721 |
| SLC22A23 | XM_017011181 | 0 | 0 | 0 | 0 | 0 | 11.48755258 | 7.782430925 | 0 | 1 | 3.33E-09 |
| SLC22A23 | XM_017011185 | 9.055885396 | 6.223416896 | 0 | 0 | 0 | 0 | 6.898192229 | 0 | 0.010315872 | 1 |
| SLC25A1 | NR_046298 | 0 | 8.372314435 | 0 | 0 | 0 | 0 | 6.620673805 | 0 | 1.20E-07 | 1 |
| SLC25A11 | NM_001165417 | 9.854861577 | 0 | 0 | 0 | 0 | 0 | 0 | 0 | 2.23E-08 | 1 |
| SLC25A11 | XM_024450994 | 10.48302382 | 0 | 0 | 0 | 0 | 0 | 0 | 0 | 1.47E-08 | 1 |
| SLC25A13 | XR_002956405 | 0 | 8.430558346 | 0 | 0 | 0 | 0 | 0 | 0 | 1.13E-07 | 1 |
| SLC25A14 | XM_005262489 | 0 | 0 | 10.95044329 | 0 | 8.428486253 | 10.24210339 | 0 | 0 | 0.013807038 | 0.999932046 |
| SLC25A26 | XR_001739995 | 0 | 0 | 6.534313836 | 0 | 0 | 0 | 0 | 0 | 0.000227503 | 1 |
| SLC25A34 | XM_017001083 | 0 | 9.554298882 | 0 | 0 | 0 | 0 | 0 | 0 | 2.73E-08 | 1 |
| SLC25A36 | XM_005247575 | 0 | 0 | 0 | 10.04571785 | 6.147113942 | 0 | 0 | 0 | 0.028827511 | 0.021463205 |
| SLC25A39 | NM_001143780 | 0 | 0 | 0 | 10.21835411 | 9.198232318 | 0 | 0 | 0 | 0.016774312 | 0.02072721 |
| SLC25A40 | NM_018843 | 0 | 8.261155927 | 0 | 0 | 0 | 10.60440873 | 0 | 0 | 1.93E-07 | 1 |
| SLC25A44 | XM_017002906 | 0 | 9.6676118 | 0 | 0 | 0 | 0 | 8.906628006 | 0 | 2.40E-08 | 1 |
| SLC25A45 | NR_147973 | 9.387309849 | 0 | 0 | 0 | 0 | 0 | 0 | 0 | 4.17E-08 | 1 |
| SLC25A46 | NR_138151 | 0 | 0 | 0 | 8.508532777 | 0 | 0 | 0 | 5.731458906 | 6.12E-06 | 1 |
| SLC26A2 | XM_017009191 | 11.29901496 | 9.315543066 | 0 | 0 | 0 | 0 | 0 | 0 | 0.000433343 | 1 |
| SLC26A6 | NM_134263 | 0 | 10.69313658 | 0 | 0 | 0 | 0 | 0 | 0 | 1.34E-08 | 1 |
| SLC27A2 | NM_003645 | 0 | 10.45293558 | 0 | 10.93486099 | 7.470373439 | 0 | 0 | 0 | 0.999953634 | 0.02072721 |
| SLC27A6 | NM_014031 | 10.37820098 | 0 | 0 | 0 | 0 | 0 | 0 | 0 | 1.48E-08 | 1 |
| SLC28A2 | XM_011522200 | 9.843239492 | 0 | 0 | 0 | 0 | 0 | 0 | 0 | 2.27E-08 | 1 |
| SLC29A1 | XM_005248876 | 0 | 0 | 0 | 10.19270445 | 12.31871762 | 11.21696321 | 10.10974477 | 10.11205883 | 1.34E-08 | 0.999932046 |
| SLC29A1 | XM_005248878 | 0 | 0 | 10.76877415 | 11.16925437 | 8.707855025 | 11.31165298 | 10.52748146 | 0 | 0.000727038 | 0.999932046 |
| SLC29A1 | XM_005248880 | 0 | 0 | 0 | 11.25726048 | 7.757784245 | 10.47944209 | 10.46848681 | 9.529534389 | 0.012318995 | 0.999932046 |
| SLC29A3 | NM_001174098 | 0 | 9.522516235 | 0 | 0 | 0 | 0 | 0 | 0 | 2.88E-08 | 1 |
| SLC2A1 | NM_006516 | 0 | 0 | 0 | 8.264461514 | 10.09041901 | 10.2352985 | 0 | 11.20492276 | 0.021150973 | 0.999932046 |
| SLC2A8 | NM_014580 | 0 | 0 | 0 | 0 | 6.747781854 | 0 | 0 | 0 | 9.53E-05 | 1 |
| SLC30A5 | XM_005248569 | 0 | 0 | 0 | 0 | 6.61940813 | 0 | 0 | 9.124042472 | 5.85E-05 | 1 |
| SLC30A5 | XM_006714672 | 9.689464985 | 0 | 0 | 0 | 0 | 0 | 0 | 0 | 3.15E-08 | 1 |
| SLC30A6 | NM_001330477 | 0 | 0 | 0 | 0 | 6.366595318 | 0 | 7.223307962 | 0 | 4.76E-05 | 1 |
| SLC30A6 | NM_001330479 | 0 | 0 | 8.869215857 | 10.66701503 | 0 | 11.09884756 | 8.944472497 | 0 | 0.015680082 | 0.999932046 |
| SLC33A1 | NM_004733 | 0 | 8.765293275 | 0 | 0 | 0 | 0 | 0 | 0 | 8.55E-08 | 1 |
| SLC35A1 | NM_001168398 | 0 | 0 | 0 | 0 | 6.531336591 | 0 | 0 | 10.25819701 | 4.23E-05 | 1 |
| SLC35A1 | NM_006416 | 10.89968072 | 5.567305463 | 10.41107393 | 10.20558628 | 9.51762409 | 0 | 0 | 0 | 0.999953634 | 0.002089223 |
| SLC35A3 | NM_012243 | 9.162606966 | 8.943876548 | 0 | 0 | 0 | 0 | 0 | 0 | 0.002976062 | 1 |
| SLC35A3 | XM_017000869 | 8.482303063 | 0 | 0 | 0 | 0 | 0 | 0 | 0 | 1.17E-07 | 1 |
| SLC35A4 | NM_080670 | 8.9622853 | 7.759544708 | 0 | 0 | 0 | 0 | 0 | 10.17504349 | 0.006751153 | 1 |
| SLC35B2 | NM_178148 | 0 | 0 | 9.905107572 | 0 | 8.529367569 | 10.50418463 | 7.940604293 | 9.534880393 | 0.022674865 | 0.999932046 |
| SLC35B3 | XM_006715103 | 0 | 0 | 0 | 0 | 6.437537018 | 0 | 0 | 0 | 4.51E-05 | 1 |
| SLC35C2 | NM_015945 | 0 | 0 | 10.73590986 | 0 | 7.372384706 | 0 | 0 | 0 | 0.017789013 | 0.020781046 |
| SLC35C2 | XM_011528833 | 0 | 8.844550112 | 0 | 0 | 0 | 0 | 0 | 0 | 8.06E-08 | 1 |

| | | | | | | | | | | | |
|---|---|---|---|---|---|---|---|---|---|---|---|
| SLC35D1 | NM_015139 | 9.153224721 | 8.602554319 | 0 | 0 | 0 | 0 | 7.460241356 | 0 | 0.0036961 | 1 |
| SLC35D1 | XR_001737058 | 0 | 0 | 0 | 0 | 6.631562102 | 0 | 0 | 0 | 3.88E-05 | 1 |
| SLC35D2 | NM_007001 | 0 | 0 | 10.08984426 | 0 | 7.779793563 | 0 | 0 | 0 | 0.023883416 | 0.02072721 |
| SLC35E3 | NM_018656 | 0 | 8.747073305 | 0 | 0 | 0 | 0 | 0 | 0 | 8.61E-08 | 1 |
| SLC35E4 | XM_017028795 | 0 | 0 | 0 | 9.595365854 | 8.583550423 | 0 | 0 | 0 | 0.025361203 | 0.02072721 |
| SLC35F5 | XM_011511922 | 0 | 0 | 0 | 0 | 6.679179249 | 0 | 0 | 0 | 3.78E-05 | 1 |
| SLC35F5 | XM_011511923 | 10.79196021 | 0 | 0 | 0 | 0 | 10.01196842 | 7.774812207 | 0 | 1.34E-08 | 0.02072721 |
| SLC35G1 | NM_153226 | 10.14250526 | 0 | 0 | 0 | 0 | 0 | 0 | 0 | 1.67E-08 | 1 |
| SLC37A4 | NM_001467 | 0 | 0 | 10.24582768 | 0 | 8.999211672 | 0 | 0 | 0 | 0.017608563 | 0.02072721 |
| SLC38A2 | XR_944589 | 0 | 8.486541819 | 0 | 0 | 0 | 0 | 8.089292508 | 9.992810611 | 1.09E-07 | 0.02072721 |
| SLC38A9 | NM_001258286 | 0 | 0 | 0 | 8.414023513 | 0 | 0 | 0 | 0 | 3.17E-05 | 1 |
| SLC38A9 | NM_001349385 | 0 | 0 | 10.63405836 | 10.31899504 | 8.93936031 | 0 | 0 | 11.21661316 | 0.001115345 | 0.999932046 |
| SLC38A9 | XM_017009082 | 10.05956027 | 10.48627065 | 0 | 0 | 0 | 0 | 0 | 10.7096458 | 0.00058717 | 1 |
| SLC38A9 | XR_001742002 | 0 | 9.158976035 | 0 | 0 | 0 | 0 | 0 | 0 | 5.01E-08 | 1 |
| SLC39A1 | NM_014437 | 0 | 0 | 0 | 11.47132532 | 7.290434628 | 0 | 7.234460372 | 0 | 1.76E-08 | 0.999932046 |
| SLC39A11 | NM_001352693 | 0 | 0 | 10.73452413 | 9.138588821 | 10.24920822 | 0 | 9.742782581 | 0 | 0.001023139 | 0.999932046 |
| SLC39A11 | XM_006721757 | 0 | 0 | 0 | 0 | 0 | 10.96950906 | 8.534928204 | 10.81447196 | 1 | 0.001228497 |
| SLC39A11 | XM_006721758 | 0 | 9.885245492 | 0 | 0 | 0 | 0 | 0 | 0 | 2.00E-08 | 1 |
| SLC39A13 | XR_001748028 | 0 | 0 | 0 | 9.472848859 | 7.386797204 | 0 | 0 | 0 | 0.035083581 | 0.024770715 |
| SLC39A14 | XM_017013293 | 9.502621339 | 9.081113725 | 10.03010237 | 0 | 6.366595318 | 0 | 0 | 0 | 0.999953634 | 0.020835214 |
| SLC39A4 | NM_130849 | 0 | 0 | 0 | 10.1639529 | 7.885068797 | 0 | 12.04645632 | 11.2232511 | 0.022058827 | 0.999932046 |
| SLC39A4 | XM_024447188 | 0 | 0 | 10.22722744 | 0 | 8.811650533 | 5.75998802 | 9.510221236 | 10.20217763 | 0.018502273 | 0.999932046 |
| SLC39A4 | XM_024447189 | 0 | 0 | 0 | 7.487390903 | 0 | 0 | 7.01920173 | 0 | 1.78E-05 | 1 |
| SLC39A5 | NM_001135195 | 10.09918783 | 9.868487622 | 0 | 0 | 0 | 0 | 8.013629967 | 0 | 0.000875308 | 1 |
| SLC39A5 | XM_011538201 | 0 | 9.073822446 | 0 | 0 | 0 | 0 | 0 | 0 | 7.73E-08 | 1 |
| SLC39A8 | XM_017008541 | 0 | 9.745393473 | 0 | 0 | 0 | 0 | 0 | 0 | 2.16E-08 | 1 |
| SLC3A2 | NM_001013251 | 0 | 10.18842011 | 10.14721031 | 0 | 9.248421412 | 0 | 0 | 0 | 0.999953634 | 0.021157255 |
| SLC3A2 | NM_002394 | 0 | 0 | 0 | 10.45034753 | 7.555764066 | 11.61141008 | 9.849089234 | 11.11943561 | 0.019833701 | 0.999932046 |
| SLC40A1 | NM_014585 | 0 | 0 | 9.368130153 | 11.25787901 | 0 | 0 | 8.981349581 | 7.921513153 | 0.01046073 | 0.999932046 |
| SLC41A2 | XM_024449219 | 0 | 0 | 0 | 0 | 6.451316153 | 0 | 0 | 0 | 4.51E-05 | 1 |
| SLC43A1 | XM_024448728 | 0 | 9.358181476 | 0 | 0 | 0 | 0 | 0 | 0 | 3.64E-08 | 1 |
| SLC43A2 | XM_017024178 | 0 | 0 | 9.769600813 | 0 | 6.464964927 | 0 | 0 | 9.743205182 | 0.034348203 | 0.999932046 |
| SLC44A5 | NM_001320287 | 0 | 9.749975406 | 10.01876234 | 0 | 9.144114677 | 0 | 0 | 0 | 0.999953634 | 0.02072721 |
| SLC48A1 | XM_024449045 | 0 | 0 | 0 | 9.999126067 | 7.718441029 | 0 | 7.638730623 | 0 | 0.024675447 | 0.999932046 |
| SLC4A2 | NM_003040 | 0 | 8.709928179 | 0 | 10.84056249 | 6.041763794 | 0 | 0 | 0 | 1 | 1.94E-08 |
| SLC4A4 | XM_024454271 | 11.11795177 | 0 | 0 | 0 | 0 | 0 | 0 | 0 | 1.34E-08 | 1 |
| SLC4A4 | XM_024454272 | 0 | 10.25117508 | 0 | 0 | 0 | 0 | 7.759452999 | 0 | 1.47E-08 | 1 |
| SLC50A1 | NM_001287591 | 0 | 9.768158992 | 0 | 0 | 0 | 0 | 0 | 0 | 2.09E-08 | 1 |
| SLC52A2 | XM_006716659 | 10.73963487 | 0 | 0 | 0 | 0 | 0 | 0 | 0 | 1.34E-08 | 1 |
| SLC6A19 | NM_001003841 | 11.00252865 | 0 | 0 | 0 | 0 | 0 | 0 | 0 | 1.34E-08 | 1 |
| SLC6A8 | NM_005629 | 10.01363818 | 7.544303366 | 0 | 10.06846137 | 8.864098664 | 0 | 0 | 0 | 0.999953634 | 0.02072721 |
| SLC7A14 | NM_020949 | 0 | 0 | 7.66679376 | 0 | 0 | 0 | 0 | 0 | 1.62E-05 | 1 |
| SLC7A6OS | NM_032178 | 0 | 9.271606013 | 0 | 0 | 0 | 0 | 0 | 0 | 4.15E-08 | 1 |

| | | | | | | | | | | | |
|---|---|---|---|---|---|---|---|---|---|---|---|
| SLC7A8 | NR_049767 | 0 | 9.073822446 | 0 | 0 | 0 | 0 | 8.781053508 | 0 | 7.73E-08 | 1 |
| SLC9A3R2 | NM_001130012 | 0 | 8.323964754 | 0 | 0 | 0 | 0 | 0 | 0 | 1.27E-07 | 1 |
| SLC9A3R2 | XM_024450492 | 10.00323168 | 0 | 0 | 0 | 0 | 0 | 0 | 0 | 1.94E-08 | 1 |
| SLCO1C1 | XR_001748771 | 9.998000145 | 0 | 0 | 0 | 0 | 0 | 0 | 0 | 1.94E-08 | 1 |
| SLCO4A1 | XR_244115 | 0 | 0 | 0 | 10.28298438 | 7.542951541 | 0 | 0 | 0 | 0.021722667 | 0.02072721 |
| SLF2 | XR_001747138 | 0 | 0 | 0 | 0 | 6.478485785 | 0 | 0 | 0 | 4.36E-05 | 1 |
| SLIRP | NR_052025 | 0 | 9.036802844 | 0 | 0 | 0 | 0 | 0 | 0 | 6.38E-08 | 1 |
| SLX1A-SULT1A3 | NR_037608 | 8.569528585 | 9.635780908 | 0 | 0 | 0 | 0 | 6.21051493 | 0 | 0.002478848 | 1 |
| SLX1B | NM_178044 | 0 | 0 | 0 | 8.75977034 | 0 | 0 | 0 | 0 | 4.66E-06 | 1 |
| SMAD2 | XM_017025745 | 0 | 11.05524858 | 10.83880171 | 9.789245515 | 7.900217873 | 0 | 0 | 0 | 0.999953634 | 0.00272352 |
| SMAD2 | XM_017025747 | 0 | 8.975521721 | 0 | 0 | 0 | 0 | 0 | 0 | 7.27E-08 | 1 |
| SMAP2 | XM_017002093 | 0 | 0 | 0 | 0 | 0 | 10.35313028 | 9.652126454 | 0 | 1 | 0.02072721 |
| SMARCA4 | NM_001128846 | 0 | 0 | 0 | 8.214354668 | 0 | 0 | 7.469710231 | 0 | 3.97E-05 | 1 |
| SMARCC1 | NM_003074 | 0 | 0 | 10.73590986 | 11.89447989 | 11.28778114 | 0 | 11.46915827 | 10.69539366 | 0.000200614 | 0.999932046 |
| SMARCC1 | XM_011534034 | 0 | 0 | 0 | 9.626353068 | 10.86209712 | 11.06150526 | 10.09452412 | 0 | 0.011023326 | 0.999932046 |
| SMARCC2 | XR_002957374 | 0 | 0 | 0 | 7.662040946 | 0 | 0 | 0 | 0 | 1.49E-05 | 1 |
| SMARCD1 | XR_944684 | 0 | 0 | 0 | 0 | 6.725274845 | 0 | 0 | 0 | 9.53E-05 | 1 |
| SMARCD2 | NM_001098426 | 0 | 0 | 10.91050407 | 8.457751161 | 8.400349091 | 0 | 6.953230848 | 0 | 0.002077062 | 0.801491164 |
| SMC4 | XM_017005491 | 0 | 0 | 10.51481206 | 9.667902088 | 0 | 0 | 0 | 0 | 0.014414922 | 0.02072721 |
| SMDT1 | NR_146715 | 9.418971874 | 0 | 9.789755738 | 0 | 9.289271525 | 0 | 0 | 0 | 0.999953634 | 0.02072721 |
| SMG1 | XM_024450199 | 0 | 9.580259825 | 0 | 0 | 0 | 0 | 7.79000962 | 0 | 2.61E-08 | 1 |
| SMG1P5 | NR_002453 | 0 | 0 | 10.89879842 | 6.544903833 | 0 | 0 | 0 | 0 | 0.018674369 | 1.64E-08 |
| SMG5 | NM_015327 | 7.876732041 | 9.029283553 | 0 | 9.066983555 | 8.483165936 | 0 | 0 | 0 | 0.999953634 | 0.023353992 |
| SMG6 | XM_005256571 | 0 | 9.165850403 | 0 | 0 | 0 | 0 | 0 | 0 | 4.97E-08 | 1 |
| SMG7 | XM_017002968 | 9.487768672 | 0 | 0 | 0 | 0 | 0 | 9.173223903 | 0 | 3.53E-08 | 1 |
| SMIM11A | NM_058182 | 0 | 0 | 0 | 10.15069217 | 8.662671672 | 0 | 7.234460372 | 0 | 0.019412996 | 0.999932046 |
| SMIM12 | NM_138428 | 0 | 0 | 12.08532217 | 0 | 9.04309389 | 10.5714681 | 6.884098118 | 10.5123891 | 0.006247795 | 0.999932046 |
| SMIM12 | XM_011540555 | 0 | 9.905922754 | 0 | 0 | 0 | 0 | 0 | 0 | 1.87E-08 | 1 |
| SMIM19 | NM_001135674 | 0 | 0 | 0 | 10.82060504 | 6.993588191 | 0 | 7.827317174 | 10.32452288 | 0.017186809 | 0.999932046 |
| SMIM22 | XM_011522501 | 10.0942935 | 0 | 0 | 0 | 0 | 0 | 7.331132628 | 0 | 1.80E-08 | 1 |
| SMIM26 | NM_001348957 | 10.50531851 | 8.774317692 | 9.690336482 | 11.42140296 | 10.63002319 | 0 | 0 | 0 | 0.999953634 | 0.001148745 |
| SMIM27 | XM_024447368 | 0 | 4.955775652 | 0 | 0 | 0 | 9.088286388 | 6.750642936 | 0 | 0.283848684 | 0.025858564 |
| SMIM29 | NM_178508 | 8.754625855 | 0 | 0 | 0 | 0 | 0 | 0 | 0 | 9.15E-08 | 1 |
| SMIM3 | NM_032947 | 0 | 0 | 0 | 10.18751933 | 8.99686469 | 10.56876398 | 8.732715858 | 9.316364312 | 0.017830406 | 0.999932046 |
| SMIM35 | XR_002957113 | 9.708807013 | 0 | 0 | 0 | 0 | 0 | 0 | 0 | 2.61E-08 | 1 |
| SMIM8 | NM_020425 | 0 | 0 | 10.34177806 | 0 | 9.433350468 | 0 | 6.85548998 | 0 | 0.015156737 | 0.999932046 |
| SMN1 | NM_000344 | 0 | 0 | 0 | 11.7529115 | 8.830225047 | 9.641814685 | 8.224078182 | 10.15087512 | 0.00782275 | 0.999932046 |
| SMN2 | NM_022875 | 0 | 10.1748534 | 0 | 0 | 0 | 0 | 7.994077632 | 9.028739063 | 1.54E-08 | 0.021326758 |
| SMOC2 | XM_011536065 | 0 | 0 | 8.09164712 | 0 | 0 | 0 | 0 | 0 | 1.04E-05 | 1 |
| SMPD1 | XR_002957158 | 0 | 9.38199218 | 0 | 0 | 0 | 0 | 0 | 0 | 3.43E-08 | 1 |
| SMPDL3A | XM_017010204 | 0 | 0 | 0 | 0 | 6.736572239 | 0 | 0 | 0 | 9.53E-05 | 1 |
| SMTN | XM_017028915 | 10.91918559 | 0 | 0 | 0 | 0 | 0 | 0 | 0 | 1.34E-08 | 1 |
| SMURF1 | XM_017012457 | 0 | 0 | 0 | 10.33315133 | 7.067022599 | 0 | 0 | 10.57859585 | 0.022284458 | 0.999932046 |

| | | | | | | | | | | | |
|---|---|---|---|---|---|---|---|---|---|---|---|
| SMURF2 | XM_005257585 | 9.609394475 | 0 | 0 | 0 | 0 | 0 | 0 | 0 | 2.97E-08 | 1 |
| SMYD2 | NM_020197 | 0 | 0 | 0 | 10.87966778 | 7.82812018 | 0 | 0 | 0 | 0.015273184 | 0.02072721 |
| SMYD2 | XR_426790 | 0 | 9.059128398 | 0 | 0 | 0 | 0 | 0 | 0 | 7.80E-08 | 1 |
| SMYD3 | XM_011544254 | 0 | 0 | 0 | 10.87644918 | 9.797962481 | 0 | 8.137615898 | 10.9616612 | 0.010609021 | 0.999932046 |
| SMYD3 | XM_011544257 | 0 | 0 | 11.84869164 | 10.67814877 | 11.75412947 | 0 | 6.21051493 | 10.00441834 | 0.000169828 | 0.999932046 |
| SNAI1 | NM_005985 | 0 | 0 | 0 | 10.27200355 | 11.58885238 | 0 | 11.42907024 | 12.0052915 | 0.006342967 | 0.999932046 |
| SNAPC1 | NM_003082 | 0 | 8.935855595 | 0 | 0 | 0 | 0 | 0 | 0 | 7.69E-08 | 1 |
| SNAPIN | NR_052019 | 0 | 8.261155927 | 0 | 0 | 0 | 0 | 0 | 10.49877519 | 1.93E-07 | 1 |
| SNHG10 | NR_003138 | 0 | 0 | 8.38097934 | 0 | 9.160983508 | 0 | 0 | 0 | 0.032435818 | 0.022209035 |
| SNHG15 | NR_003697 | 0 | 0 | 0 | 9.402987182 | 10.73126405 | 0 | 0 | 0 | 0.012438691 | 0.02072721 |
| SNHG17 | NR_152755 | 0 | 0 | 0 | 0 | 0 | 11.1790499 | 7.79000962 | 0 | 1 | 0.02072721 |
| SNHG21 | NR_110096 | 0 | 0 | 7.420196825 | 0 | 0 | 0 | 0 | 0 | 2.07E-05 | 1 |
| SNHG29 | NR_027159 | 0 | 0 | 0 | 10.24105527 | 8.242890685 | 0 | 0 | 0 | 0.020062723 | 0.02072721 |
| SNHG29 | NR_027167 | 0 | 0 | 0 | 10.80376031 | 7.774322632 | 0 | 0 | 10.14039201 | 0.016001739 | 0.999932046 |
| SNHG29 | NR_027179 | 9.153224721 | 0 | 0 | 0 | 0 | 0 | 0 | 0 | 6.38E-08 | 1 |
| SNHG6 | NR_152598 | 0 | 0 | 0 | 9.802872777 | 9.669757788 | 0 | 10.76999494 | 0 | 0.017209362 | 1 |
| SNHG7 | NR_024543 | 0 | 0 | 0 | 10.90831647 | 10.14124615 | 0 | 9.036549592 | 10.5312353 | 0.009435897 | 0.999932046 |
| SNORD3A | NR_006880 | 0 | 0 | 0 | 0 | 6.531336591 | 0 | 0 | 0 | 4.23E-05 | 1 |
| SNRK | XM_005265245 | 0 | 0 | 0 | 0 | 6.769943125 | 0 | 0 | 0 | 9.43E-05 | 1 |
| SNRNP200 | NM_014014 | 0 | 0 | 0 | 10.86512723 | 10.31097016 | 11.11376536 | 9.652126454 | 0 | 0.00909637 | 0.999932046 |
| SNRNP25 | NM_024571 | 11.86953439 | 0 | 10.74832186 | 11.80758755 | 9.904472996 | 0 | 0 | 0 | 0.999953634 | 0.000855759 |
| SNRNP35 | NR_104103 | 10.02910853 | 0 | 0 | 0 | 0 | 0 | 0 | 0 | 1.87E-08 | 1 |
| SNRNP40 | NM_004814 | 0 | 10.53752475 | 0 | 11.41531669 | 7.718441029 | 0 | 0 | 0 | 0.999953634 | 9.16E-09 |
| SNRPA1 | NM_003090 | 0 | 0 | 10.22229251 | 11.65909849 | 10.89964287 | 0 | 11.28833914 | 12.4925426 | 0.00032526 | 0.999932046 |
| SNRPC | NR_029472 | 0 | 0 | 3.199506948 | 10.50034076 | 8.202724223 | 0 | 7.506976231 | 0 | 0.006603094 | 0.999932046 |
| SNTG2 | NM_018968 | 11.2514447 | 0 | 0 | 0 | 0 | 0 | 0 | 0 | 1.34E-08 | 1 |
| SNTG2 | XM_017004363 | 10.71114528 | 0 | 0 | 0 | 0 | 0 | 0 | 0 | 1.34E-08 | 1 |
| SNTG2 | XM_017004368 | 0 | 10.62742277 | 0 | 0 | 0 | 0 | 0 | 0 | 1.34E-08 | 1 |
| SNUPN | NM_005701 | 0 | 0 | 0 | 11.82271666 | 9.176620886 | 0 | 9.546468212 | 0 | 0.007089982 | 0.999932046 |
| SNURF | NM_005678 | 0 | 8.561439232 | 0 | 0 | 0 | 0 | 0 | 0 | 1.00E-07 | 1 |
| SNW1 | NM_001318844 | 0 | 0 | 9.520629739 | 0 | 6.544252221 | 0 | 10.32848533 | 0 | 0.039981685 | 0.999932046 |
| SNX13 | XM_017011893 | 9.93095313 | 8.464408432 | 0 | 0 | 0 | 0 | 8.284344777 | 0 | 0.00207886 | 1 |
| SNX15 | NM_147777 | 0 | 0 | 0 | 9.910641153 | 8.59910684 | 0 | 0 | 0 | 0.021938407 | 0.02072721 |
| SNX20 | NM_153337 | 9.877828587 | 9.117027412 | 0 | 0 | 0 | 0 | 0 | 0 | 0.001592807 | 1 |
| SNX29 | XM_005255682 | 0 | 9.821373802 | 0 | 0 | 0 | 0 | 0 | 0 | 2.01E-08 | 1 |
| SNX29 | XM_011522744 | 0 | 0 | 0 | 0 | 6.667421491 | 0 | 0 | 8.838651627 | 3.78E-05 | 1 |
| SNX3 | NM_001300929 | 0 | 9.303125566 | 0 | 0 | 0 | 0 | 0 | 0 | 4.01E-08 | 1 |
| SNX3 | NM_152827 | 0 | 8.632639499 | 9.721447883 | 9.523110922 | 4.331362461 | 0 | 0 | 0 | 0.999953634 | 0.009691049 |
| SNX32 | XM_024448420 | 9.79579164 | 0 | 0 | 7.035175705 | 9.049902256 | 0 | 0 | 0 | 0.999953634 | 0.031708371 |
| SNX4 | XM_017007414 | 0 | 9.006487637 | 0 | 0 | 0 | 0 | 0 | 0 | 6.80E-08 | 1 |
| SNX5 | NM_014426 | 0 | 0 | 0 | 10.40192126 | 6.823901211 | 8.643619478 | 0 | 0 | 0.021956886 | 0.999932046 |
| SNX8 | NM_013321 | 0 | 0 | 0 | 10.41750665 | 7.822829806 | 0 | 0 | 10.88524176 | 0.019520563 | 0.999932046 |
| SOCS2 | XM_017020152 | 0 | 8.991087758 | 0 | 0 | 0 | 0 | 0 | 0 | 7.02E-08 | 1 |

| | | | | | | | | | | |
|---|---|---|---|---|---|---|---|---|---|---|
| SOD2 | NM_000636 | 0 | 0 | 0 | 0 | 0 | 11.11005035 | 8.192973732 | 0 | 1 | 0.02072721 |
| SOD3 | NM_003102 | 9.129498645 | 8.430558346 | 0 | 0 | 0 | 0 | 9.311290442 | 0 | 0.004155733 | 1 |
| SOD3 | XR_427488 | 0 | 0 | 0 | 9.656688688 | 9.029380004 | 0 | 0 | 0 | 0.021944552 | 0.02072721 |
| SON | NM_001291411 | 0 | 0 | 0 | 0 | 0 | 11.02711594 | 7.856483795 | 8.8168733 | 1 | 0.002469438 |
| SORBS2 | XM_017008770 | 0 | 0 | 10.61911747 | 10.26832462 | 7.211666293 | 0 | 0 | 0 | 0.001725988 | 0.002874229 |
| SORBS3 | NM_005775 | 10.02397016 | 0 | 0 | 0 | 0 | 0 | 0 | 0 | 1.88E-08 | 1 |
| SORD | NR_034039 | 0 | 8.140712363 | 8.807290493 | 0 | 7.920172326 | 0 | 0 | 0 | 0.999953634 | 0.029094394 |
| SOS2 | NM_006939 | 0 | 0 | 9.431077203 | 0 | 7.618173501 | 0 | 0 | 0 | 0.035247905 | 0.023251362 |
| SOWAHA | NM_175873 | 0 | 0 | 10.50671712 | 0 | 7.746652379 | 0 | 0 | 0 | 0.01921012 | 0.02072721 |
| SOX12 | NM_006943 | 0 | 0 | 0 | 0 | 6.337212177 | 0 | 0 | 0 | 4.86E-05 | 1 |
| SOX15 | NM_006942 | 0 | 0 | 0 | 8.254578401 | 0 | 0 | 0 | 0 | 8.02E-06 | 1 |
| SOX4 | NM_003107 | 0 | 0 | 8.703458918 | 9.66043618 | 9.248421412 | 0 | 6.620673805 | 0 | 0.003017132 | 0.841126572 |
| SOX6 | NM_001145811 | 10.64574565 | 0 | 0 | 0 | 0 | 0 | 0 | 0 | 1.34E-08 | 1 |
| SOX9 | NM_000346 | 0 | 0 | 0 | 8.029665411 | 0 | 0 | 7.223307962 | 0 | 1.02E-05 | 1 |
| SOX9-AS1 | NR_103738 | 0 | 0 | 0 | 8.029665411 | 0 | 0 | 0 | 0 | 1.02E-05 | 1 |
| SP100 | NM_001206702 | 0 | 0 | 9.114793339 | 10.78328353 | 9.08125658 | 11.07236439 | 0 | 0 | 0.001508139 | 1 |
| SP110 | NM_080424 | 9.643298854 | 0 | 0 | 0 | 0 | 0 | 0 | 0 | 2.88E-08 | 1 |
| SP2 | NM_003110 | 0 | 0 | 0 | 0 | 6.702411139 | 0 | 0 | 0 | 3.69E-05 | 1 |
| SP2 | XM_024450892 | 9.524617511 | 7.946803117 | 0 | 0 | 0 | 0 | 0 | 0 | 0.003738691 | 1 |
| SP3 | NM_003111 | 0 | 0 | 0 | 9.792664417 | 8.1036837 | 0 | 0 | 0 | 0.025729408 | 0.02072721 |
| SP5 | NM_001003845 | 0 | 0 | 0 | 9.506550706 | 6.823901211 | 0 | 9.319168841 | 0 | 0.037591072 | 0.999932046 |
| SPATA21 | XR_946646 | 8.583566901 | 0 | 0 | 0 | 0 | 0 | 7.032041479 | 0 | 1.06E-07 | 1 |
| SPATA6L | NM_001353484 | 0 | 0 | 0 | 0 | 6.395391943 | 0 | 0 | 0 | 4.65E-05 | 1 |
| SPATA6L | XR_001746339 | 0 | 8.919678654 | 0 | 0 | 0 | 0 | 0 | 0 | 7.73E-08 | 1 |
| SPATA7 | NM_018418 | 0 | 8.31161984 | 0 | 0 | 0 | 0 | 0 | 9.896425015 | 1.27E-07 | 1 |
| SPATS2L | XM_017003785 | 0 | 0 | 0 | 0 | 6.366595318 | 0 | 0 | 0 | 4.76E-05 | 1 |
| SPC24 | XM_011527702 | 0 | 0 | 0 | 0 | 6.702411139 | 0 | 0 | 0 | 3.69E-05 | 1 |
| SPDL1 | NM_017785 | 0 | 0 | 0 | 11.16793862 | 7.067022599 | 0 | 0 | 0 | 0.01394138 | 3.22E-08 |
| SPECC1L | NM_015330 | 9.553433586 | 0 | 0 | 0 | 0 | 0 | 0 | 0 | 3.27E-08 | 1 |
| SPG7 | XM_006721264 | 9.708807013 | 0 | 0 | 0 | 0 | 0 | 0 | 0 | 2.61E-08 | 1 |
| SPG7 | XR_001751972 | 9.998000145 | 0 | 0 | 0 | 0 | 0 | 7.867271029 | 0 | 1.94E-08 | 1 |
| SPHK2 | XM_017027010 | 0 | 9.117027412 | 0 | 0 | 0 | 0 | 0 | 0 | 5.36E-08 | 1 |
| SPIDR | NM_001352931 | 0 | 0 | 12.14181976 | 10.07830002 | 7.654363064 | 11.42961408 | 8.647674983 | 9.29131303 | 0.000655701 | 0.999932046 |
| SPIDR | NM_001352932 | 0 | 0 | 9.476548209 | 10.79611554 | 7.920172326 | 0 | 11.50298398 | 0 | 0.001812516 | 0.999932046 |
| SPIDR | NM_001352933 | 0 | 0 | 10.00733248 | 9.061325395 | 7.87488031 | 11.33730107 | 8.609636205 | 10.93056373 | 0.003518038 | 0.999932046 |
| SPIDR | NM_001352934 | 0 | 0 | 9.193555665 | 0 | 9.822025731 | 10.28337218 | 8.357577191 | 9.867055263 | 0.019520563 | 0.999932046 |
| SPIDR | NM_001352935 | 0 | 0 | 9.721447883 | 0 | 7.763318133 | 0 | 0 | 0 | 0.029138196 | 0.020781046 |
| SPIDR | NM_001352937 | 9.877828587 | 0 | 0 | 0 | 0 | 0 | 0 | 0 | 2.13E-08 | 1 |
| SPIDR | NM_001352940 | 0 | 0 | 9.181401832 | 0 | 9.101058952 | 11.07109108 | 8.720869785 | 10.76528678 | 0.025821961 | 0.999932046 |
| SPIDR | NM_001352941 | 0 | 0 | 0 | 9.904333953 | 7.82812018 | 0 | 0 | 0 | 0.025521303 | 0.02072721 |
| SPIDR | NM_001352943 | 0 | 0 | 10.39186358 | 0 | 8.555119041 | 0 | 7.479117364 | 0 | 0.018049709 | 0.999932046 |
| SPIDR | NM_001352945 | 0 | 0 | 0 | 0 | 0 | 0 | 9.216048648 | 10.15782178 | 1 | 0.02072721 |
| SPIDR | NM_001352948 | 0 | 0 | 0 | 0 | 0 | 0 | 7.272828576 | 9.981108733 | 1 | 0.02072721 |

| | | | | | | | | | | | |
|---|---|---|---|---|---|---|---|---|---|---|---|
| SPIDR | NM_001352949 | 0 | 0 | 9.097649359 | 0 | 7.145400839 | 0 | 0 | 0 | 0.047356815 | 0.030016964 |
| SPIDR | NM_001352961 | 0 | 0 | 9.787084628 | 12.60908502 | 12.88882845 | 10.7720774 | 12.79749385 | 11.52538641 | 8.05E-05 | 0.999932046 |
| SPIDR | XM_017013268 | 0 | 0 | 8.918864307 | 0 | 8.904726182 | 0 | 10.02557716 | 10.26464677 | 0.030505146 | 0.999932046 |
| SPIDR | XM_017013269 | 0 | 0 | 11.09675196 | 10.93253837 | 10.91867563 | 12.7114297 | 11.64400134 | 11.75115796 | 0.000329783 | 0.999932046 |
| SPIDR | XM_017013273 | 0 | 0 | 0 | 10.18361823 | 7.449928885 | 0 | 9.910349016 | 0 | 0.023184684 | 0.999932046 |
| SPINDOC | XR_001747777 | 0 | 0 | 0 | 8.867445783 | 7.895185835 | 9.113244852 | 9.437290336 | 0 | 0.043678575 | 0.999932046 |
| SPINK2 | NM_021114 | 0 | 9.527862205 | 0 | 0 | 0 | 0 | 0 | 0 | 2.90E-08 | 1 |
| SPINT1 | NM_003710 | 10.57020707 | 8.652353298 | 0 | 0 | 0 | 0 | 0 | 11.11498073 | 0.001095213 | 1 |
| SPINT1-AS1 | NR_146466 | 10.06956967 | 0 | 0 | 0 | 0 | 10.81682642 | 0 | 0 | 1.80E-08 | 1 |
| SPINT1-AS1 | NR_146468 | 8.940634053 | 0 | 10.30677337 | 0 | 7.843875888 | 0 | 0 | 0 | 0.999953634 | 0.02072721 |
| SPINT2 | NM_001166103 | 0 | 0 | 11.47378334 | 11.75247265 | 10.34965057 | 10.7705093 | 9.28739377 | 0 | 0.000231594 | 0.999932046 |
| SPIRE2 | NM_032451 | 0 | 8.765293275 | 0 | 0 | 0 | 0 | 0 | 0 | 8.55E-08 | 1 |
| SPIRE2 | XM_011523380 | 9.055885396 | 0 | 0 | 0 | 0 | 0 | 0 | 0 | 7.54E-08 | 1 |
| SPIRE2 | XM_017023780 | 0 | 8.671801345 | 0 | 0 | 0 | 0 | 0 | 0 | 9.15E-08 | 1 |
| SPIRE2 | XM_017023781 | 0 | 0 | 0 | 10.72184386 | 7.867191447 | 0 | 8.235224663 | 0 | 0.016556936 | 0.999932046 |
| SPN | NM_003123 | 9.153224721 | 0 | 0 | 0 | 0 | 0 | 0 | 0 | 6.38E-08 | 1 |
| SPNS1 | XM_024450467 | 0 | 8.709928179 | 0 | 0 | 0 | 0 | 0 | 0 | 8.96E-08 | 1 |
| SPOCK2 | XM_011540404 | 0 | 0 | 0 | 0 | 6.451316153 | 0 | 7.728235151 | 0 | 4.51E-05 | 1 |
| SPOUT1 | XM_017014804 | 0 | 8.861586703 | 0 | 0 | 0 | 11.61490959 | 7.411941715 | 0 | 7.86E-08 | 3.87E-09 |
| SPP1 | NM_000582 | 0 | 0 | 0 | 10.01969536 | 11.28104563 | 0 | 0 | 13.56753617 | 0.007996442 | 0.999932046 |
| SPPL3 | XM_011537925 | 10.63567489 | 0 | 0 | 7.520758627 | 8.975568243 | 0 | 0 | 0 | 0.999953634 | 0.030130471 |
| SPRED1 | XM_011521289 | 0 | 0 | 0 | 0 | 0 | 11.0414904 | 6.781384747 | 0 | 1 | 5.17E-09 |
| SPRY4 | XM_017009910 | 9.727893151 | 0 | 0 | 0 | 0 | 0 | 5.33721949 | 0 | 2.55E-08 | 0.507804484 |
| SPRYD7 | NM_020456 | 10.22989302 | 9.974088018 | 0 | 10.05428874 | 8.07281823 | 0 | 0 | 0 | 0.999953634 | 0.02072721 |
| SPSB3 | NM_080861 | 10.4296226 | 0 | 10.65325302 | 0 | 9.394651239 | 0 | 0 | 0 | 0.999953634 | 0.02072721 |
| SPSB3 | XM_005255673 | 9.777588513 | 7.74125153 | 10.48627878 | 10.48879454 | 8.51795513 | 0 | 0 | 0 | 0.999953634 | 0.002319834 |
| SPTAN1 | XM_017015061 | 10.24763479 | 9.226288718 | 10.00273505 | 9.23220986 | 7.436136431 | 0 | 0 | 0 | 0.999953634 | 0.005311885 |
| SPTBN1 | XR_002959325 | 9.304974408 | 9.117027412 | 10.11349163 | 11.12247359 | 9.68003446 | 0 | 0 | 0 | 0.999953634 | 0.001535028 |
| SPTLC1 | NM_178324 | 0 | 9.059128398 | 0 | 0 | 0 | 0 | 4.006722785 | 0 | 7.80E-08 | 0.870999832 |
| SPTLC1 | XM_024447379 | 0 | 0 | 0 | 11.06142599 | 10.02742406 | 8.906052537 | 8.557305568 | 9.917043278 | 0.009058342 | 0.999932046 |
| SQLE | XM_011517246 | 0 | 0 | 0 | 0 | 6.491881102 | 0 | 0 | 0 | 4.33E-05 | 1 |
| SRA1 | NM_001035235 | 0 | 8.690990708 | 0 | 12.47745845 | 9.060054937 | 0 | 0 | 0 | 0.737757269 | 5.45E-09 |
| SRA1 | NR_045587 | 10.70794476 | 7.565252966 | 0 | 0 | 0 | 0 | 0 | 0 | 0.001333781 | 1 |
| SRBD1 | XR_001738805 | 6.370278018 | 9.754542834 | 0 | 0 | 0 | 0 | 0 | 0 | 0.004419298 | 1 |
| SREBF2 | NR_103834 | 0 | 0 | 0 | 0 | 0 | 5.809695791 | 7.382166205 | 9.933328324 | 1 | 0.009674774 |
| SREK1 | NM_001077199 | 0 | 9.434171496 | 0 | 0 | 0 | 0 | 0 | 0 | 3.24E-08 | 1 |
| SREK1IP1 | NM_173829 | 0 | 0 | 10.58722874 | 9.872377593 | 8.274237629 | 9.653772402 | 6.187745501 | 0 | 0.001653878 | 0.999932046 |
| SRF | NM_003131 | 0 | 0 | 0 | 0 | 0 | 10.21353544 | 6.068174253 | 0 | 1 | 0.02072721 |
| SRGN | NM_001321054 | 0 | 9.06649413 | 0 | 0 | 0 | 0 | 0 | 0 | 7.75E-08 | 1 |
| SRI | XM_011516528 | 0 | 11.59294126 | 12.32675998 | 8.597229662 | 2.936277275 | 0 | 0 | 0 | 0.999953634 | 0.002769306 |
| SRM | XM_017002179 | 0 | 0 | 0 | 8.029665411 | 0 | 0 | 0 | 0 | 1.02E-05 | 1 |
| SRP68 | NM_014230 | 0 | 0 | 0 | 0 | 6.679179249 | 0 | 8.04563962 | 0 | 3.78E-05 | 1 |
| SRPK1 | NM_003137 | 0 | 0 | 10.21634824 | 0 | 9.35488525 | 0 | 8.89613213 | 0 | 0.016289345 | 0.999932046 |

| | | | | | | | | | | | |
|---|---|---|---|---|---|---|---|---|---|---|---|
| SRRM2 | XR_002957796 | 0 | 9.708204508 | 0 | 0 | 0 | 0 | 0 | 9.502503116 | 2.47E-08 | 1 |
| SRRM2 | XR_932815 | 0 | 0 | 12.50394132 | 0 | 10.71243477 | 0 | 11.08464991 | 10.9566906 | 0.00344269 | 0.999932046 |
| SRRT | NM_015908 | 0 | 0 | 0 | 8.491803396 | 7.477124411 | 0 | 0 | 0 | 0.058859562 | 0.040669105 |
| SRRT | XM_005250406 | 0 | 0 | 0 | 0 | 6.3222931 | 0 | 0 | 0 | 0.000244538 | 1 |
| SRSF10 | XM_017000102 | 0 | 0 | 0 | 0 | 0 | 0 | 6.788968925 | 9.779688974 | 1 | 0.02072721 |
| SRSF11 | NM_001350607 | 0 | 9.860035122 | 0 | 0 | 0 | 0 | 0 | 0 | 1.97E-08 | 1 |
| SRSF11 | NR_146810 | 9.807800698 | 0 | 0 | 0 | 0 | 0 | 0 | 0 | 2.49E-08 | 1 |
| SRSF2 | NR_036608 | 0 | 10.81621352 | 10.48956843 | 11.01168421 | 6.366595318 | 0 | 0 | 0 | 0.999953634 | 0.002589366 |
| SRSF2 | XR_002958053 | 0 | 9.640723913 | 0 | 0 | 0 | 0 | 0 | 0 | 2.47E-08 | 1 |
| SRSF4 | XM_011541951 | 0 | 7.830482506 | 10.22426851 | 0 | 7.145400839 | 0 | 0 | 0 | 0.998900485 | 0.02072721 |
| SRSF5 | NM_006925 | 9.124706245 | 8.919678654 | 11.75646198 | 10.07970008 | 6.823901211 | 0 | 0 | 0 | 0.966161772 | 0.001924802 |
| SRSF6 | NR_034009 | 0 | 0 | 0 | 9.334269269 | 8.486515572 | 0 | 0 | 7.486553413 | 0.029273635 | 0.999932046 |
| SRSF7 | XR_939708 | 9.449953927 | 0 | 0 | 0 | 0 | 0 | 9.033360344 | 0 | 3.80E-08 | 1 |
| SRSF8 | NR_103726 | 0 | 0 | 9.382357652 | 10.31067259 | 5.986055956 | 0 | 7.920029518 | 10.19544179 | 0.003689313 | 0.999932046 |
| SS18 | XM_011526149 | 0 | 0 | 0 | 9.026902352 | 9.407077595 | 0 | 7.827317174 | 0 | 0.024124651 | 0.999932046 |
| SS18L1 | XR_001754230 | 0 | 0 | 0 | 0 | 0 | 0 | 6.586265919 | 10.10128877 | 1 | 0.02072721 |
| SS18L2 | XM_017006548 | 0 | 0 | 10.47054966 | 11.95828241 | 10.98648842 | 10.35940264 | 9.973279884 | 0 | 0.000242177 | 0.999932046 |
| SSBP1 | NM_001256512 | 0 | 0 | 0 | 8.79429013 | 8.136037099 | 0 | 9.678906268 | 0 | 0.041977579 | 0.999932046 |
| SSBP2 | NM_001256733 | 0 | 0 | 11.21574177 | 0 | 7.695467896 | 0 | 0 | 0 | 0.01303131 | 0.02072721 |
| SSBP3 | XM_017000899 | 9.217654114 | 0 | 0 | 0 | 0 | 0 | 0 | 0 | 5.49E-08 | 1 |
| SSBP4 | XR_936164 | 8.555352323 | 0 | 0 | 0 | 0 | 0 | 0 | 0 | 1.09E-07 | 1 |
| SSRP1 | NM_003146 | 0 | 0 | 9.780385135 | 0 | 10.83344675 | 9.803893982 | 0 | 11.42674866 | 0.010659562 | 0.999932046 |
| SSRP1 | XM_024448666 | 0 | 0 | 0 | 7.958392843 | 9.361285436 | 0 | 9.193367076 | 0 | 0.032595365 | 0.999932046 |
| ST3GAL1 | NM_003033 | 0 | 0 | 9.910026324 | 10.29025878 | 8.949104958 | 0 | 9.567566713 | 11.75803141 | 0.001568096 | 0.999932046 |
| ST3GAL1 | XM_017013736 | 0 | 0 | 0 | 0 | 0 | 0 | 9.642739475 | 10.22219849 | 1 | 0.02072721 |
| ST3GAL4 | NM_006278 | 0 | 8.561439232 | 0 | 0 | 0 | 0 | 0 | 0 | 1.00E-07 | 1 |
| ST3GAL4 | XM_024448660 | 0 | 10.3389485 | 9.844736088 | 9.865900493 | 0 | 0 | 0 | 0 | 0.999953634 | 0.020942118 |
| ST6GALNAC1 | NM_001289107 | 0 | 9.303125566 | 0 | 0 | 0 | 0 | 0 | 0 | 4.01E-08 | 1 |
| ST6GALNAC1 | NM_018414 | 0 | 0 | 9.382357652 | 8.841270918 | 8.620607807 | 0 | 0 | 0 | 0.004274073 | 0.006180585 |
| ST6GALNAC1 | XR_001752559 | 9.860637698 | 10.02790232 | 8.496668691 | 10.24605182 | 0 | 0 | 0 | 0 | 0.999953634 | 0.02072721 |
| ST6GALNAC1 | XR_002958047 | 0 | 9.950395724 | 0 | 0 | 0 | 0 | 0 | 0 | 4.93E-08 | 1 |
| ST6GALNAC4 | XM_024447518 | 0 | 9.416987232 | 0 | 0 | 0 | 0 | 0 | 0 | 3.28E-08 | 1 |
| ST6GALNAC6 | NM_013443 | 10.18915784 | 0 | 0 | 0 | 0 | 0 | 9.108021301 | 0 | 1.60E-08 | 1 |
| ST7L | NM_017744 | 0 | 8.927789798 | 0 | 0 | 0 | 0 | 0 | 0 | 7.69E-08 | 1 |
| STAG1 | XM_011512329 | 0 | 0 | 12.8492965 | 9.396260824 | 6.79176912 | 0 | 12.8360987 | 10.33681783 | 0.000533462 | 0.999932046 |
| STAG2 | XM_006724728 | 9.976882202 | 0 | 0 | 0 | 0 | 0 | 9.603493749 | 0 | 1.99E-08 | 1 |
| STAG2 | XM_017029234 | 10.99729455 | 0 | 0 | 0 | 0 | 0 | 8.643497643 | 0 | 1.34E-08 | 1 |
| STAG3 | XM_011515742 | 0 | 10.31750069 | 0 | 0 | 0 | 0 | 0 | 0 | 1.45E-08 | 1 |
| STAG3L5P-PVRIG2P-PILRB | NR_036569 | 0 | 0 | 9.640895475 | 0 | 8.720712025 | 0 | 0 | 0 | 0.02436216 | 0.02072721 |
| STAM | NM_003473 | 9.338474533 | 9.411213355 | 0 | 0 | 0 | 0 | 0 | 0 | 0.001936436 | 1 |
| STAMBPL1 | NM_020799 | 0 | 0 | 10.46888394 | 10.14267682 | 0 | 0 | 0 | 0 | 0.01270693 | 0.02072721 |
| STAMBPL1 | XM_006717928 | 0 | 9.19973932 | 0 | 0 | 0 | 0 | 7.759452999 | 0 | 4.67E-08 | 1 |
| STARD10 | NM_006645 | 0 | 0 | 10.76538551 | 11.93835618 | 10.6536812 | 11.1790499 | 11.03448766 | 11.97611302 | 0.000250883 | 0.999932046 |

| | | | | | | | | | | | |
|---|---|---|---|---|---|---|---|---|---|---|---|
| STARD3 | XM_024450549 | 0 | 0 | 0 | 7.307890232 | 0 | 0 | 9.818845344 | 0 | 2.14E-05 | 1 |
| STARD7-AS1 | NR_046322 | 0 | 0 | 0 | 9.286568795 | 6.163947944 | 0 | 0 | 0 | 0.04738611 | 0.032833589 |
| STAT1 | XM_017004783 | 0 | 0 | 0 | 0 | 6.451316153 | 0 | 0 | 10.59915604 | 4.51E-05 | 1 |
| STAT1 | XR_001738915 | 0 | 0 | 10.48627878 | 0 | 8.875641266 | 0 | 6.85548998 | 10.74005074 | 0.016190273 | 0.999932046 |
| STAT5A | NM_001288719 | 0 | 0 | 10.20238227 | 0 | 9.687330378 | 0 | 0 | 0 | 0.014895076 | 0.02072721 |
| STAT5B | XM_024450897 | 0 | 0 | 0 | 0 | 6.337212177 | 0 | 0 | 0 | 4.86E-05 | 1 |
| STAT6 | NM_003153 | 0 | 0 | 9.634965902 | 10.3702361 | 0 | 11.69933522 | 8.137615898 | 0 | 0.015278471 | 0.999932046 |
| STAU2 | NM_001164383 | 0 | 9.206422614 | 0 | 0 | 0 | 0 | 0 | 0 | 4.63E-08 | 1 |
| STAU2 | NM_014393 | 9.83152302 | 0 | 0 | 0 | 0 | 0 | 0 | 0 | 2.28E-08 | 1 |
| STBD1 | NM_003943 | 10.77981801 | 9.006487637 | 0 | 0 | 0 | 9.682406504 | 0 | 0 | 0.00078888 | 1 |
| STEAP1 | NM_012449 | 10.24763479 | 0 | 0 | 10.51593815 | 8.334959336 | 0 | 0 | 0 | 0.999953634 | 0.02072721 |
| STIM2 | NM_020860 | 0 | 0 | 0 | 0 | 6.395391943 | 10.07430581 | 0 | 0 | 4.65E-05 | 1 |
| STIP1 | NM_006819 | 0 | 0 | 0 | 11.6942546 | 10.93990612 | 0 | 10.10215452 | 0 | 0.004967217 | 0.999932046 |
| STK10 | XM_017009788 | 0 | 8.943876548 | 0 | 0 | 0 | 0 | 0 | 0 | 7.69E-08 | 1 |
| STK11 | XM_005259617 | 0 | 0 | 0 | 9.257172648 | 6.747781854 | 0 | 0 | 0 | 0.044634758 | 0.030925278 |
| STK16 | NM_001008910 | 0 | 0 | 0 | 0 | 0 | 0 | 8.095422193 | 10.42583458 | 1 | 0.02072721 |
| STK17A | XM_017012792 | 0 | 0 | 0 | 6.755775244 | 0 | 0 | 0 | 0 | 3.72E-05 | 1 |
| STK17B | XM_011512171 | 0 | 9.10991584 | 0 | 0 | 0 | 0 | 0 | 0 | 5.42E-08 | 1 |
| STK19 | NR_026717 | 0 | 0 | 9.256701524 | 0 | 7.568463804 | 0 | 0 | 9.858552905 | 0.039325144 | 0.999932046 |
| STK24 | NM_001286649 | 0 | 0 | 0 | 0 | 6.464964927 | 0 | 5.566321721 | 0 | 4.45E-05 | 1 |
| STK25 | NM_001271977 | 0 | 0 | 9.287267368 | 0 | 8.784693675 | 0 | 7.552244553 | 0 | 0.027646859 | 0.999932046 |
| STK25 | NM_006374 | 9.487768672 | 8.582143237 | 0 | 0 | 0 | 0 | 0 | 0 | 0.002875166 | 1 |
| STK25 | XM_017003171 | 0 | 0 | 9.768247087 | 0 | 8.469689074 | 0 | 0 | 10.07216922 | 0.024510347 | 0.999932046 |
| STK32C | NM_001318881 | 0 | 9.358181476 | 0 | 0 | 0 | 0 | 0 | 0 | 3.64E-08 | 1 |
| STK32C | NM_001318883 | 10.7175251 | 0 | 0 | 0 | 0 | 0 | 0 | 0 | 1.34E-08 | 1 |
| STK35 | XM_011529174 | 0 | 0 | 0 | 9.352916447 | 9.257272906 | 0 | 0 | 0 | 0.022861747 | 0.02072721 |
| STK38 | NM_001305102 | 0 | 0 | 12.70517142 | 12.43843123 | 10.07436986 | 12.38878902 | 11.61229089 | 11.41229796 | 9.41E-05 | 0.999932046 |
| STK38L | NM_015000 | 0 | 0 | 0 | 0 | 6.464964927 | 0 | 0 | 0 | 4.45E-05 | 1 |
| STK38L | XM_005253342 | 0 | 8.430558346 | 0 | 0 | 0 | 0 | 0 | 0 | 1.13E-07 | 1 |
| STK39 | XM_017003813 | 0 | 0 | 10.99857238 | 0 | 6.945718048 | 0 | 7.704370312 | 0 | 0.015960877 | 0.999932046 |
| STK39 | XM_017003815 | 0 | 0 | 9.664373023 | 0 | 8.459498162 | 0 | 10.51381583 | 0 | 0.025774039 | 0.999932046 |
| STMN1 | NM_005563 | 0 | 0 | 9.681733808 | 14.09835771 | 12.37503729 | 12.23724162 | 8.732715858 | 13.13201589 | 4.32E-05 | 0.999932046 |
| STMN3 | NR_075070 | 0 | 0 | 0 | 0 | 6.607150896 | 8.110899224 | 0 | 0 | 0.00011086 | 1 |
| STOM | NM_004099 | 0 | 0 | 0 | 9.875605271 | 8.445797324 | 0 | 0 | 0 | 0.023018009 | 0.02072721 |
| STOX1 | NM_001130160 | 9.734199496 | 0 | 0 | 0 | 0 | 0 | 0 | 0 | 5.08E-08 | 1 |
| STRAP | XM_005253294 | 0 | 8.737876298 | 0 | 0 | 0 | 0 | 0 | 0 | 9.11E-08 | 1 |
| STRBP | XM_017014898 | 0 | 0 | 0 | 0 | 6.643614539 | 0 | 0 | 0 | 3.86E-05 | 1 |
| STRN4 | NM_013403 | 0 | 0 | 0 | 0 | 0 | 9.95512224 | 8.201523487 | 0 | 1 | 0.02072721 |
| STT3A | NM_152713 | 0 | 0 | 0 | 9.966198446 | 7.526774194 | 0 | 8.794302797 | 9.724611366 | 0.025856575 | 0.999932046 |
| STX17-AS1 | NR_038853 | 0 | 9.670031486 | 0 | 0 | 0 | 0 | 5.207402423 | 0 | 2.40E-08 | 0.538759989 |
| STX5 | NM_001330294 | 0 | 0 | 8.928592211 | 0 | 10.06319654 | 0 | 8.212844911 | 12.85024068 | 0.018777731 | 0.999932046 |
| STX6 | XM_017000007 | 0 | 8.765293275 | 0 | 0 | 0 | 0 | 8.432122801 | 0 | 8.55E-08 | 1 |
| STX8 | NM_004853 | 0 | 0 | 0 | 10.30709108 | 9.777598559 | 0 | 9.339969963 | 11.81187105 | 0.013783825 | 0.999932046 |

| | | | | | | | | | | | |
|---|---|---|---|---|---|---|---|---|---|---|---|
| STXBP6 | NM_001304476 | 10.39023122 | 12.24217905 | 10.86119709 | 11.52678113 | 11.92449052 | 0 | 0 | 0 | 0.999953634 | 0.000399462 |
| STXBP6 | NM_001351943 | 0 | 0 | 0 | 0 | 0 | 0 | 10.59285003 | 10.88314745 | 1 | 0.02072721 |
| SUCLG2 | NM_003848 | 11.86523112 | 10.14385363 | 0 | 10.50971941 | 8.656737116 | 0 | 0 | 0 | 0.969603763 | 0.020947361 |
| SUCO | NM_016227 | 10.27384495 | 0 | 0 | 0 | 0 | 10.7055248 | 0 | 0 | 1.81E-08 | 1 |
| SUFU | XM_011539860 | 0 | 0 | 10.41799658 | 0 | 7.510413387 | 0 | 9.3126065 | 0 | 0.020742069 | 0.999932046 |
| SUFU | XM_011539864 | 0 | 0 | 9.721447883 | 9.247239304 | 7.654363064 | 0 | 8.278968849 | 0 | 0.004042324 | 0.999932046 |
| SUGCT | XM_006715775 | 0 | 8.497482465 | 11.80513009 | 0 | 8.84595791 | 0 | 0 | 0 | 0.842663899 | 5.45E-09 |
| SUGCT | XM_011515525 | 10.20740451 | 0 | 0 | 0 | 0 | 0 | 0 | 0 | 1.60E-08 | 1 |
| SUGP2 | NM_014884 | 0 | 0 | 6.702508352 | 0 | 0 | 0 | 0 | 0 | 4.12E-05 | 1 |
| SUGT1 | NM_001130912 | 0 | 10.13687339 | 0 | 11.50558329 | 7.939854543 | 0 | 0 | 0 | 0.999953634 | 7.83E-09 |
| SULF1 | XR_928764 | 0 | 0 | 0 | 0 | 6.725274845 | 0 | 0 | 0 | 9.53E-05 | 1 |
| SULT1A1 | NM_001055 | 0 | 9.650559405 | 0 | 0 | 0 | 0 | 0 | 0 | 2.46E-08 | 1 |
| SULT1A2 | NM_001054 | 10.51633798 | 0 | 0 | 0 | 0 | 0 | 0 | 0 | 1.39E-08 | 1 |
| SUN2 | XM_024452206 | 9.25321892 | 0 | 0 | 0 | 0 | 0 | 0 | 0 | 5.16E-08 | 1 |
| SUOX | XM_017019905 | 0 | 0 | 10.89508209 | 0 | 7.42919044 | 0 | 9.731019215 | 0 | 0.016169398 | 0.999932046 |
| SUPT16H | NM_007192 | 0 | 0 | 0 | 0 | 6.395391943 | 0 | 0 | 0 | 4.65E-05 | 1 |
| SUPT3H | NM_001350325 | 0 | 10.08341647 | 0 | 0 | 0 | 0 | 0 | 0 | 1.59E-08 | 1 |
| SUPT3H | NM_001350326 | 0 | 8.419096158 | 0 | 0 | 0 | 0 | 0 | 0 | 1.14E-07 | 1 |
| SUPT3H | NM_001350327 | 0 | 9.46236476 | 0 | 0 | 0 | 0 | 0 | 0 | 3.12E-08 | 1 |
| SUPT3H | NM_181356 | 0 | 0 | 10.0514059 | 11.96587366 | 8.323766986 | 0 | 10.91740527 | 8.918534776 | 0.000629184 | 0.999932046 |
| SUPT6H | XM_017024980 | 9.740478394 | 0 | 8.002348885 | 7.43584608 | 7.869758957 | 0 | 0 | 0 | 0.999953634 | 0.02072721 |
| SUPT7L | NM_014860 | 0 | 0 | 10.01990037 | 0 | 7.145400839 | 0 | 0 | 0 | 0.027138955 | 0.02072721 |
| SUPT7L | XM_005264672 | 0 | 9.277965255 | 0 | 0 | 0 | 0 | 0 | 0 | 4.01E-08 | 1 |
| SURF4 | NM_033161 | 0 | 0 | 10.18726716 | 9.610942656 | 0 | 0 | 8.352467811 | 0 | 0.016705938 | 0.999932046 |
| SUSD3 | XM_011518358 | 0 | 0 | 10.77552766 | 10.26709622 | 0 | 11.23924199 | 9.939604452 | 0 | 0.010753988 | 0.999932046 |
| SUSD3 | XM_011518359 | 0 | 10.7964161 | 0 | 0 | 0 | 11.50651664 | 0 | 0 | 1.34E-08 | 1 |
| SUSD3 | XM_011518361 | 0 | 0 | 0 | 0 | 0 | 11.18494796 | 7.827317174 | 0 | 1 | 0.02072721 |
| SUSD3 | XM_017014449 | 0 | 0 | 10.2747168 | 8.943236221 | 0 | 11.31649686 | 9.055538427 | 10.50150827 | 0.0189536 | 0.999932046 |
| SVBP | NM_199342 | 0 | 0 | 0 | 10.12379934 | 10.33067298 | 10.76736798 | 0 | 0 | 0.012140445 | 0.999932046 |
| SVIL | NM_001323599 | 0 | 8.690990708 | 0 | 0 | 0 | 0 | 0 | 0 | 9.02E-08 | 1 |
| SWI5 | NM_001318089 | 0 | 0 | 10.57953115 | 9.686399554 | 9.083470314 | 0 | 9.717172953 | 0 | 0.001427309 | 0.999932046 |
| SWSAP1 | NM_175871 | 10.08692074 | 0 | 0 | 0 | 0 | 0 | 0 | 0 | 1.76E-08 | 1 |
| SYF2 | NM_207170 | 0 | 8.959785899 | 0 | 9.62251587 | 7.949595864 | 0 | 0 | 0 | 0.999953634 | 0.021282828 |
| SYK | NM_001174167 | 10.89124009 | 0 | 0 | 0 | 0 | 0 | 0 | 0 | 1.34E-08 | 1 |
| SYMPK | XR_243962 | 0 | 9.439854398 | 0 | 0 | 0 | 0 | 8.609636205 | 0 | 3.19E-08 | 1 |
| SYN2 | NM_133625 | 0 | 0 | 7.500657305 | 0 | 0 | 0 | 0 | 0 | 2.22E-05 | 1 |
| SYNE1 | XM_006715421 | 10.98676906 | 0 | 0 | 0 | 0 | 0 | 0 | 0 | 1.41E-08 | 1 |
| SYNE1 | XR_001743287 | 0 | 0 | 9.135940735 | 0 | 11.4990429 | 0 | 0 | 11.81187105 | 0.00836901 | 0.999932046 |
| SYNGR2 | NM_001320523 | 0 | 0 | 10.47221345 | 0 | 9.199253403 | 0 | 0 | 0 | 0.01519341 | 0.02072721 |
| SYNJ2 | XR_245556 | 0 | 0 | 0 | 0 | 0 | 10.4444672 | 0 | 9.502503116 | 1 | 0.02072721 |
| SYNPO | XM_024454351 | 0 | 0 | 10.24971315 | 10.51490355 | 8.558305894 | 0 | 0 | 0 | 0.001347927 | 0.002469438 |
| SYNPO2 | NM_001286755 | 11.02584987 | 0 | 0 | 0 | 0 | 0 | 0 | 0 | 1.34E-08 | 1 |
| SYNPO2 | NM_133477 | 0 | 8.407542174 | 0 | 0 | 0 | 0 | 6.586265919 | 0 | 1.16E-07 | 1 |

| | | | | | | | | | | | |
|---|---|---|---|---|---|---|---|---|---|---|---|
| SYS1 | NM_033542 | 0 | 0 | 0 | 10.15866323 | 8.987438391 | 0 | 0 | 0 | 0.01808998 | 0.02072721 |
| SYT11 | XM_017000759 | 0 | 9.145128245 | 0 | 0 | 0 | 0 | 0 | 0 | 5.08E-08 | 1 |
| SYT13 | NM_020826 | 0 | 8.895067962 | 0 | 0 | 0 | 0 | 0 | 0 | 7.75E-08 | 1 |
| SYT7 | NM_004200 | 0 | 0 | 0 | 9.897999058 | 8.368024654 | 0 | 10.15739905 | 0 | 0.023163283 | 0.999932046 |
| SZRD1 | NM_001114600 | 10.36606958 | 0 | 0 | 0 | 0 | 0 | 0 | 0 | 1.48E-08 | 1 |
| SZT2 | NM_015284 | 0 | 0 | 0 | 9.924732336 | 9.193116026 | 0 | 0 | 9.272234908 | 0.018891923 | 0.999932046 |
| TACC1 | NM_001352782 | 0 | 0 | 10.90312209 | 0 | 7.849089784 | 0 | 10.04629302 | 0 | 0.015369033 | 0.999932046 |
| TACC1 | NM_001352784 | 10.29532878 | 0 | 0 | 10.56674421 | 8.157209854 | 0 | 0 | 0 | 0.999953634 | 0.02072721 |
| TACC1 | NM_001352794 | 0 | 0 | 0 | 0 | 6.505153187 | 0 | 0 | 0 | 4.28E-05 | 1 |
| TACC1 | NM_001352799 | 0 | 0 | 0 | 10.29025878 | 9.742306238 | 0 | 0 | 0 | 0.014019265 | 0.02072721 |
| TACC1 | NM_001352804 | 0 | 0 | 0 | 0 | 0 | 11.29429739 | 9.624840127 | 0 | 1 | 0.02072721 |
| TACC1 | NR_148049 | 11.23375001 | 0 | 0 | 0 | 0 | 0 | 7.834664296 | 0 | 1.34E-08 | 1 |
| TACC2 | NM_001291877 | 0 | 0 | 8.823022803 | 9.667902088 | 0 | 10.47463488 | 8.708925636 | 0 | 0.024925775 | 0.999932046 |
| TACC2 | NM_206862 | 0 | 0 | 6.124455375 | 0 | 9.121670713 | 0 | 0 | 0 | 0.051161624 | 0.033843338 |
| TACC2 | XM_006717552 | 0 | 8.844550112 | 0 | 0 | 0 | 0 | 7.552244553 | 0 | 8.06E-08 | 1 |
| TACC2 | XM_011539134 | 0 | 0 | 0 | 0 | 0 | 10.3266979 | 9.484762437 | 0 | 1 | 0.02072721 |
| TACC2 | XM_017015499 | 11.04626802 | 9.296876505 | 0 | 0 | 0 | 0 | 9.622719646 | 0 | 0.000552318 | 1 |
| TACC2 | XM_024447766 | 10.9761662 | 9.745393473 | 0 | 0 | 0 | 0 | 0 | 11.01140653 | 0.000479568 | 1 |
| TACC3 | XM_005247930 | 10.61702724 | 0 | 0 | 10.71284921 | 11.11675788 | 0 | 0 | 0 | 0.999953634 | 0.02072721 |
| TACC3 | XM_011513386 | 0 | 0 | 11.44653594 | 0 | 10.16704775 | 0 | 9.711197916 | 0 | 0.007222629 | 0.999932046 |
| TACO1 | NM_016360 | 0 | 0 | 11.52798162 | 0 | 9.889368769 | 0 | 0 | 0 | 0.007428178 | 0.02072721 |
| TACSTD2 | NM_002353 | 0 | 0 | 0 | 9.736965247 | 8.281969086 | 11.45228718 | 0 | 9.051297555 | 0.02549138 | 0.999932046 |
| TADA1 | NM_053053 | 0 | 0 | 9.728426058 | 0 | 7.422210844 | 0 | 9.342549142 | 0 | 0.030822321 | 0.999932046 |
| TADA3 | NM_133480 | 0 | 0 | 0 | 0 | 6.531336591 | 0 | 0 | 0 | 4.23E-05 | 1 |
| TAF1D | NM_024116 | 0 | 0 | 0 | 10.32491038 | 9.364929978 | 0 | 0 | 0 | 0.015376159 | 0.02074727 |
| TAF6 | NM_139315 | 0 | 0 | 10.16171799 | 0 | 7.510413387 | 0 | 0 | 0 | 0.023894825 | 0.02072721 |
| TAF6L | NM_006473 | 0 | 0 | 0 | 0 | 6.437537018 | 0 | 0 | 9.48050741 | 4.51E-05 | 1 |
| TAF6L | XM_017017100 | 0 | 0 | 9.248957794 | 0 | 6.478485785 | 0 | 0 | 0 | 0.048299441 | 0.030529185 |
| TALAM1 | NR_145459 | 4.941046909 | 7.213730404 | 11.467541 | 0 | 7.729791854 | 0 | 0 | 0 | 0.600027199 | 2.22E-08 |
| TAMM41 | NR_104314 | 0 | 9.081113725 | 0 | 0 | 0 | 0 | 0 | 0 | 5.70E-08 | 1 |
| TANK | NM_001199135 | 10.58423881 | 0 | 0 | 0 | 0 | 0 | 0 | 0 | 1.34E-08 | 1 |
| TAOK2 | XM_011545983 | 0 | 0 | 7.474332608 | 0 | 0 | 0 | 0 | 0 | 1.96E-05 | 1 |
| TAP2 | NM_001290043 | 0 | 0 | 0 | 8.787451934 | 0 | 0 | 0 | 0 | 4.55E-06 | 1 |
| TARBP1 | XR_001737382 | 9.734199496 | 0 | 0 | 0 | 0 | 0 | 0 | 0 | 5.08E-08 | 1 |
| TARS2 | NR_073514 | 0 | 0 | 8.300853839 | 0 | 7.642400461 | 0 | 0 | 0 | 0.062287147 | 0.040312936 |
| TARSL2 | XR_001751084 | 0 | 9.46236476 | 0 | 0 | 0 | 0 | 0 | 0 | 3.12E-08 | 1 |
| TASP1 | NM_001323602 | 0 | 0 | 9.040481197 | 0 | 10.61479834 | 0 | 8.660134917 | 0 | 0.014130977 | 0.999932046 |
| TASP1 | NR_136631 | 0 | 0 | 10.50590512 | 0 | 8.629725311 | 0 | 0 | 0 | 0.016838301 | 0.020808337 |
| TASP1 | XR_001754319 | 0 | 0 | 0 | 10.26586678 | 9.120593193 | 0 | 7.431456038 | 11.42531009 | 0.016736682 | 0.999932046 |
| TASP1 | XR_001754320 | 0 | 0 | 0 | 10.31899504 | 8.577280582 | 0 | 0 | 0 | 0.018291955 | 0.02072721 |
| TATDN1 | XM_017013898 | 0 | 9.938402056 | 0 | 0 | 0 | 0 | 0 | 0 | 5.04E-08 | 1 |
| TATDN1 | XM_024447294 | 0 | 0 | 0 | 7.706219197 | 0 | 0 | 0 | 0 | 1.42E-05 | 1 |
| TATDN3 | NM_001146171 | 9.313422649 | 0 | 0 | 0 | 0 | 0 | 0 | 0 | 4.70E-08 | 1 |

| | | | | | | | | | | | |
|---|---|---|---|---|---|---|---|---|---|---|---|
| TAZ | NM_181313 | 0 | 9.124104101 | 0 | 0 | 0 | 0 | 0 | 0 | 5.29E-08 | 1 |
| TAZ | XM_024452431 | 0 | 0 | 7.447518626 | 0 | 0 | 0 | 0 | 0 | 3.62E-05 | 1 |
| TBC1D10B | XM_011545790 | 0 | 0 | 0 | 0 | 6.30721813 | 9.667318094 | 0 | 0 | 4.99E-05 | 1 |
| TBC1D14 | XR_241653 | 10.70152235 | 0 | 0 | 0 | 0 | 0 | 8.229662188 | 9.917043278 | 1.34E-08 | 0.02072721 |
| TBC1D2 | NM_018421 | 0 | 0 | 0 | 0 | 6.381065478 | 0 | 0 | 0 | 4.71E-05 | 1 |
| TBC1D22A | NR_104292 | 0 | 9.958336465 | 0 | 0 | 0 | 0 | 0 | 0 | 1.76E-08 | 1 |
| TBC1D23 | NM_018309 | 0 | 0 | 0 | 9.07261961 | 7.049011441 | 0 | 0 | 0 | 0.047551391 | 0.032586253 |
| TBC1D24 | XR_001751956 | 10.05956027 | 0 | 0 | 0 | 0 | 0 | 0 | 0 | 5.01E-08 | 1 |
| TBC1D7 | NM_001143965 | 0 | 0 | 10.67508796 | 0 | 8.686168747 | 0 | 8.397813588 | 0 | 0.015325386 | 0.999932046 |
| TBC1D8 | XM_017003199 | 10.08938252 | 0 | 0 | 0 | 0 | 0 | 0 | 0 | 1.76E-08 | 1 |
| TBCA | NM_001297738 | 9.854861577 | 0 | 0 | 0 | 0 | 0 | 0 | 0 | 2.23E-08 | 1 |
| TBCB | NM_001281 | 0 | 0 | 10.0119153 | 10.40192126 | 10.58229341 | 0 | 8.367741983 | 9.832740984 | 0.000759752 | 0.999932046 |
| TBK1 | XM_005268810 | 0 | 0 | 0 | 9.45569803 | 7.859461386 | 0 | 0 | 0 | 0.032273783 | 0.023084584 |
| TBL1X | XM_011545571 | 0 | 0 | 0 | 9.402987182 | 6.518304288 | 0 | 6.42038839 | 0 | 0.041977579 | 0.999932046 |
| TBL1XR1 | XM_011513143 | 0 | 0 | 10.73035894 | 8.854417711 | 10.19393357 | 0 | 0 | 10.03492329 | 0.001138524 | 0.999932046 |
| TBL2 | NM_012453 | 0 | 0 | 10.33630716 | 0 | 8.77650741 | 10.44052824 | 0 | 0 | 0.017748616 | 0.999932046 |
| TBL2 | XR_001744627 | 0 | 8.927789798 | 0 | 0 | 0 | 0 | 0 | 0 | 7.69E-08 | 1 |
| TBL3 | NM_006453 | 0 | 0 | 0 | 0 | 0 | 0 | 7.27822739 | 10.0904377 | 1 | 0.02072721 |
| TBRG1 | NR_016021 | 0 | 0 | 0 | 10.18231551 | 7.707000189 | 0 | 0 | 0 | 0.022406821 | 0.02072721 |
| TBRG1 | XR_001748012 | 10.0942935 | 0 | 0 | 0 | 0 | 0 | 0 | 0 | 1.80E-08 | 1 |
| TBX10 | XM_011544993 | 0 | 10.45012258 | 0 | 0 | 0 | 0 | 0 | 0 | 1.34E-08 | 1 |
| TBX18 | XM_017011472 | 0 | 0 | 0 | 0 | 6.537808859 | 0 | 0 | 0 | 4.23E-05 | 1 |
| TBXAS1 | XM_024446901 | 0 | 7.501468832 | 0 | 9.872377593 | 5.264185146 | 0 | 0 | 0 | 0.999953634 | 0.02496249 |
| TCEA1 | NM_006756 | 0 | 0 | 10.25841739 | 11.74895705 | 0 | 9.152301482 | 10.23162943 | 10.36410426 | 0.006531798 | 0.999932046 |
| TCEA3 | XM_011542053 | 0 | 10.03917736 | 0 | 0 | 0 | 0 | 9.682982473 | 0 | 1.60E-08 | 1 |
| TCEA3 | XM_017002201 | 9.531875668 | 0 | 0 | 0 | 0 | 0 | 0 | 0 | 3.28E-08 | 1 |
| TCEAL1 | NM_001006639 | 0 | 0 | 0 | 0 | 6.667421491 | 0 | 0 | 10.29331911 | 3.78E-05 | 1 |
| TCEAL2 | NM_080390 | 0 | 8.878424457 | 0 | 0 | 0 | 0 | 0 | 0 | 9.15E-08 | 1 |
| TCEAL7 | NM_152278 | 9.363098816 | 0 | 0 | 0 | 0 | 0 | 0 | 0 | 4.33E-08 | 1 |
| TCEAL9 | NM_001006613 | 0 | 9.296876505 | 11.16314967 | 0 | 9.553199992 | 0 | 0 | 0 | 0.999953634 | 0.02072721 |
| TCERG1 | XR_001741980 | 0 | 9.309347676 | 0 | 0 | 0 | 0 | 0 | 0 | 3.94E-08 | 1 |
| TCF12 | NM_207040 | 0 | 0 | 0 | 0 | 6.531336591 | 0 | 0 | 0 | 4.23E-05 | 1 |
| TCF12 | XM_017022520 | 0 | 9.186279128 | 0 | 0 | 0 | 0 | 6.620673805 | 0 | 4.80E-08 | 1 |
| TCF3 | XM_024451672 | 0 | 0 | 0 | 0 | 0 | 10.38831838 | 7.782430925 | 0 | 1 | 0.02072721 |
| TCF3 | XM_024451675 | 10.5666777 | 0 | 0 | 0 | 0 | 0 | 8.680665279 | 6.860901615 | 1.34E-08 | 0.031007738 |
| TCF3 | XM_024451679 | 0 | 0 | 0 | 10.23855048 | 7.145400839 | 0 | 0 | 0 | 0.023324588 | 0.02072721 |
| TCF7L2 | NM_001198530 | 0 | 10.35407547 | 9.320897208 | 0 | 9.844389419 | 0 | 0 | 0 | 0.999953634 | 0.02072721 |
| TCF7L2 | NM_001198531 | 9.383302787 | 0 | 9.368130153 | 9.310616166 | 8.118872287 | 0 | 0 | 0 | 0.999953634 | 0.006061516 |
| TCF7L2 | NM_030756 | 0 | 0 | 0 | 7.631818844 | 0 | 0 | 0 | 0 | 1.70E-05 | 1 |
| TCF7L2 | XM_005270078 | 0 | 0 | 10.83363358 | 0 | 9.080148438 | 0 | 8.704922178 | 0 | 0.01308243 | 0.999932046 |
| TCF7L2 | XM_005270102 | 0 | 9.451153482 | 0 | 0 | 0 | 11.12730581 | 8.278968849 | 0 | 3.15E-08 | 0.02072721 |
| TCF7L2 | XM_011540110 | 0 | 0 | 11.3507502 | 0 | 9.538752088 | 0 | 8.095422193 | 0 | 0.008919075 | 0.999932046 |
| TCF7L2 | XM_017016589 | 13.27147057 | 11.63442015 | 10.26515123 | 0 | 10.36110094 | 0 | 0 | 0 | 0.726385406 | 0.02072721 |

| | | | | | | | | | | | |
|---|---|---|---|---|---|---|---|---|---|---|---|
| TCFL5 | XM_024451811 | 0 | 0 | 0 | 0 | 0 | 10.28885959 | 6.551017323 | 0 | 1 | 0.02072721 |
| TCHP | XR_001748901 | 0 | 0 | 0 | 0 | 6.518304288 | 0 | 0 | 0 | 4.23E-05 | 1 |
| TCIRG1 | XR_002957115 | 9.426779986 | 8.592384875 | 0 | 0 | 0 | 0 | 7.704370312 | 0 | 0.003001449 | 1 |
| TCN2 | NM_000355 | 0 | 0 | 12.37713852 | 0 | 8.079488076 | 0 | 0 | 0 | 1.34E-08 | 3.65E-09 |
| TCOF1 | XM_017009794 | 0 | 0 | 0 | 7.720649682 | 9.365839676 | 10.5004056 | 7.647100774 | 0 | 0.034212516 | 0.999932046 |
| TCTEX1D1 | NM_152665 | 0 | 9.145128245 | 0 | 0 | 0 | 0 | 0 | 0 | 5.08E-08 | 1 |
| TCTEX1D2 | NM_001351628 | 0 | 9.625843798 | 0 | 0 | 0 | 0 | 0 | 0 | 2.50E-08 | 1 |
| TCTN3 | NM_015631 | 9.734199496 | 0 | 0 | 0 | 0 | 0 | 0 | 0 | 5.08E-08 | 1 |
| TCTN3 | XM_024447935 | 0 | 9.006487637 | 0 | 0 | 0 | 0 | 0 | 0 | 6.80E-08 | 1 |
| TDG | NM_003211 | 0 | 0 | 9.177327695 | 9.888444253 | 0 | 0 | 0 | 8.772304849 | 0.020791475 | 0.999932046 |
| TDRD3 | XM_017020777 | 0 | 0 | 0 | 10.08807202 | 8.206791567 | 0 | 0 | 0 | 0.021776748 | 0.02072721 |
| TDRKH | XM_017000122 | 0 | 0 | 0 | 9.806259575 | 7.71273195 | 11.02053445 | 0 | 9.491547182 | 0.027437274 | 0.999932046 |
| TDRKH | XM_017000125 | 0 | 8.878424457 | 0 | 0 | 0 | 0 | 10.44046925 | 0 | 9.15E-08 | 1 |
| TDRP | XM_006716232 | 9.97422059 | 0 | 0 | 0 | 0 | 0 | 0 | 0 | 2.00E-08 | 1 |
| TEAD1 | NM_021961 | 0 | 0 | 0 | 9.433968736 | 8.407434995 | 0 | 8.172825027 | 10.49603692 | 0.028577946 | 0.999932046 |
| TEAD3 | NM_003214 | 10.95742161 | 0 | 8.923736459 | 0 | 8.748879432 | 0 | 0 | 0 | 0.999953634 | 0.022014709 |
| TECR | NM_001321170 | 0 | 0 | 0 | 7.382391203 | 0 | 0 | 7.751711637 | 0 | 1.98E-05 | 1 |
| TECR | NR_038104 | 10.14723899 | 0 | 0 | 0 | 0 | 0 | 0 | 0 | 1.64E-08 | 1 |
| TECR | XM_024451796 | 0 | 0 | 8.463594711 | 9.910641153 | 0 | 0 | 0 | 0 | 0.02423108 | 0.02072721 |
| TECR | XM_024451797 | 0 | 0 | 0 | 6.988236596 | 0 | 0 | 0 | 0 | 4.34E-05 | 1 |
| TECR | XM_024451799 | 9.346729415 | 0 | 0 | 0 | 0 | 0 | 0 | 0 | 4.43E-08 | 1 |
| TEFM | NM_024683 | 0 | 0 | 10.67364253 | 0 | 8.048094872 | 0 | 0 | 0 | 0.016952901 | 0.020730817 |
| TEKT4P2 | NR_038327 | 0 | 0 | 0 | 7.80432675 | 0 | 0 | 0 | 0 | 3.71E-05 | 1 |
| TENT2 | NM_001114394 | 0 | 8.919678654 | 0 | 0 | 0 | 0 | 0 | 0 | 7.73E-08 | 1 |
| TENT5C | NM_017709 | 0 | 9.029283553 | 0 | 0 | 0 | 0 | 0 | 0 | 6.49E-08 | 1 |
| TERF1 | NM_003218 | 0 | 8.622680652 | 0 | 0 | 0 | 0 | 0 | 0 | 9.58E-08 | 1 |
| TERF2IP | NM_018975 | 9.595606435 | 8.652353298 | 0 | 0 | 0 | 0 | 7.06988901 | 0 | 0.00253626 | 1 |
| TESC | NM_017899 | 0 | 0 | 0 | 11.51957781 | 9.094488316 | 12.42736367 | 0 | 10.53390763 | 0.00867744 | 0.999932046 |
| TEX261 | NM_144582 | 0 | 9.684464992 | 0 | 0 | 0 | 0 | 0 | 0 | 2.36E-08 | 1 |
| TEX264 | NM_001278195 | 0 | 9.158976035 | 0 | 0 | 0 | 0 | 0 | 0 | 5.01E-08 | 1 |
| TEX264 | NM_015926 | 0 | 0 | 0 | 11.2215585 | 9.383914218 | 10.94885997 | 0 | 9.165991457 | 0.009782036 | 0.999932046 |
| TFAP4 | XM_011522635 | 0 | 8.747073305 | 0 | 8.952436083 | 7.449928885 | 0 | 0 | 0 | 0.999953634 | 0.032030331 |
| TFB2M | NM_022366 | 0 | 0 | 10.45212013 | 10.10467156 | 0 | 0 | 8.809297332 | 0 | 0.012949897 | 0.999932046 |
| TFCP2 | NM_005653 | 0 | 0 | 0 | 8.491803396 | 0 | 0 | 8.161183866 | 0 | 6.26E-06 | 1 |
| TFDP2 | XM_017007098 | 0 | 0 | 0 | 6.444154352 | 9.876021594 | 0 | 9.885802727 | 0 | 0.030265127 | 0.999932046 |
| TFEB | NM_001167827 | 0 | 8.453213086 | 0 | 0 | 0 | 9.930056757 | 9.949789873 | 0 | 1.11E-07 | 0.02072721 |
| TFEB | NM_001271943 | 0 | 0 | 0 | 0 | 0 | 0 | 7.767153042 | 11.03988678 | 1 | 0.02072721 |
| TFEB | XM_005249412 | 0 | 10.08341647 | 0 | 0 | 0 | 0 | 0 | 0 | 1.59E-08 | 1 |
| TFEB | XM_011514915 | 0 | 9.226288718 | 0 | 0 | 0 | 0 | 0 | 0 | 4.45E-08 | 1 |
| TFEC | XM_017011878 | 0 | 0 | 7.718176908 | 0 | 0 | 0 | 0 | 0 | 1.53E-05 | 1 |
| TFF2 | NM_005423 | 0 | 0 | 6.584403084 | 0 | 9.941555888 | 10.58847748 | 0 | 14.14138525 | 0.028265655 | 0.999932046 |
| TFIP11 | XM_024452195 | 0 | 8.475517571 | 0 | 0 | 0 | 0 | 0 | 0 | 1.09E-07 | 1 |
| TFPI | NM_006287 | 0 | 10.17144169 | 0 | 0 | 0 | 0 | 0 | 9.957416323 | 1.54E-08 | 1 |

| | | | | | | | | | | | |
|---|---|---|---|---|---|---|---|---|---|---|---|
| TFPI2 | NM_006528 | 0 | 9.245884974 | 0 | 0 | 0 | 0 | 0 | 0 | 4.33E-08 | 1 |
| TGFB1 | NM_000660 | 9.860637698 | 0 | 0 | 0 | 0 | 0 | 0 | 10.36410426 | 3.34E-08 | 1 |
| TGFB1I1 | NM_001042454 | 9.338474533 | 0 | 0 | 0 | 0 | 0 | 0 | 9.988920522 | 4.50E-08 | 1 |
| TGFBI | NM_000358 | 0 | 0 | 0 | 12.77404724 | 11.2803221 | 12.29782531 | 11.17930574 | 10.10847773 | 1.34E-08 | 0.999932046 |
| TGFBR2 | NM_001024847 | 0 | 0 | 9.517410232 | 9.954035954 | 7.463590728 | 0 | 10.40451009 | 0 | 0.003280213 | 0.999932046 |
| TGFBR3L | XM_011527612 | 0 | 0 | 0 | 0 | 6.655567121 | 0 | 0 | 0 | 3.82E-05 | 1 |
| TGFBRAP1 | NR_137328 | 0 | 8.662110092 | 0 | 0 | 0 | 0 | 0 | 0 | 9.15E-08 | 1 |
| TGFBRAP1 | XM_017005339 | 0 | 0 | 0 | 0 | 6.366595318 | 0 | 8.611775989 | 0 | 4.76E-05 | 1 |
| TGIF1 | NM_003244 | 0 | 0 | 10.77417948 | 9.591445233 | 9.496182072 | 0 | 7.920029518 | 0 | 0.001162192 | 0.999932046 |
| TGIF1 | XM_017025959 | 0 | 0 | 0 | 10.27322778 | 9.511060423 | 10.67651331 | 8.172825027 | 0 | 0.015130311 | 0.999932046 |
| THAP1 | NM_199003 | 0 | 0 | 0 | 0 | 6.582319518 | 0 | 0 | 0 | 4.04E-05 | 1 |
| THAP12 | NR_130898 | 0 | 0 | 9.824037495 | 0 | 8.982702045 | 0 | 0 | 11.06965851 | 0.021023081 | 0.999932046 |
| THAP3 | XM_011542400 | 0 | 0 | 10.80817039 | 0 | 6.865660185 | 0 | 0 | 0 | 0.017938842 | 0.02072721 |
| THAP4 | NM_015963 | 0 | 0 | 9.718647135 | 0 | 7.677994746 | 0 | 0 | 0 | 0.029662337 | 0.020868927 |
| THBS1 | NM_003246 | 9.752954799 | 0 | 0 | 0 | 0 | 0 | 0 | 0 | 2.49E-08 | 1 |
| THBS3 | NR_045553 | 0 | 8.756212053 | 0 | 0 | 0 | 0 | 0 | 0 | 8.55E-08 | 1 |
| THOC7 | NM_001285387 | 0 | 9.428466121 | 0 | 0 | 0 | 0 | 0 | 0 | 3.25E-08 | 1 |
| THOC7 | XM_006713339 | 0 | 0 | 0 | 0 | 0 | 5.904250755 | 9.235606657 | 0 | 1 | 0.024880486 |
| THOP1 | NM_003249 | 10.79497986 | 0 | 0 | 0 | 0 | 0 | 0 | 0 | 1.34E-08 | 1 |
| THRA | NM_001190918 | 0 | 0 | 0 | 0 | 0 | 0 | 9.887569951 | 10.59660198 | 1 | 0.02072721 |
| THRAP3 | NM_005119 | 0 | 0 | 0 | 8.581503282 | 7.859461386 | 0 | 0 | 0 | 0.050953094 | 0.034403718 |
| THRB | NM_000461 | 0 | 10.11572772 | 0 | 0 | 0 | 0 | 9.537491487 | 0 | 1.80E-08 | 1 |
| THRB | NM_001354708 | 0 | 9.645650041 | 0 | 0 | 0 | 0 | 0 | 0 | 2.48E-08 | 1 |
| THRB | NM_001354710 | 0 | 0 | 10.70864514 | 0 | 7.930046999 | 0 | 0 | 0 | 0.016907692 | 0.02072721 |
| THRB | NM_001354711 | 0 | 0 | 10.17608123 | 0 | 10.2876382 | 0 | 9.089715157 | 0 | 0.012139313 | 0.999932046 |
| THRB | XM_011534052 | 0 | 9.580259825 | 10.56245049 | 0 | 10.44871393 | 0 | 0 | 0 | 0.999953634 | 0.02072721 |
| THRB | XM_017007111 | 11.15014445 | 0 | 0 | 0 | 0 | 0 | 0 | 0 | 1.34E-08 | 1 |
| THRB | XM_017007112 | 10.49792516 | 0 | 0 | 0 | 0 | 0 | 0 | 0 | 1.51E-08 | 1 |
| THSD4 | XM_017022586 | 9.279328737 | 9.772669303 | 8.082961956 | 0 | 7.6513817 | 0 | 0 | 0 | 0.885891781 | 0.044603371 |
| THSD7A | XM_011515193 | 12.10308214 | 0 | 0 | 0 | 0 | 0 | 0 | 0 | 1.34E-08 | 1 |
| THYN1 | NM_199298 | 0 | 0 | 0 | 9.498198764 | 5.354946376 | 0 | 9.523920794 | 0 | 0.044469437 | 1 |
| TIAF1 | NM_004740 | 11.54662814 | 0 | 0 | 0 | 0 | 0 | 0 | 0 | 1.34E-08 | 1 |
| TIAL1 | XR_945808 | 9.682959584 | 0 | 0 | 0 | 0 | 0 | 0 | 0 | 2.84E-08 | 1 |
| TIMM10B | NM_012192 | 0 | 0 | 7.079567203 | 11.66330671 | 9.93298808 | 0 | 8.846115701 | 11.16927221 | 0.001017703 | 0.999932046 |
| TIMM17B | NM_001167947 | 0 | 0 | 0 | 0 | 0 | 11.02974016 | 8.62242756 | 0 | 1 | 0.02072721 |
| TIMM44 | NM_006351 | 11.35533102 | 9.014126367 | 0 | 10.25351436 | 7.227767768 | 0 | 0 | 0 | 0.997913553 | 0.02072721 |
| TIMM50 | XM_011527491 | 10.83367076 | 0 | 0 | 0 | 0 | 0 | 0 | 10.67616912 | 1.34E-08 | 1 |
| TIMM8A | NM_004085 | 0 | 10.06882031 | 0 | 0 | 0 | 0 | 0 | 0 | 1.60E-08 | 1 |
| TIMM8B | NR_028383 | 0 | 10.05777546 | 0 | 0 | 0 | 0 | 9.986533002 | 10.41572788 | 1.60E-08 | 0.02072721 |
| TIMM9 | NM_012460 | 0 | 0 | 0 | 10.27567312 | 10.2948208 | 0 | 9.670719144 | 0 | 0.011657592 | 1 |
| TINCR | NR_027064 | 10.3248786 | 8.886770211 | 0 | 0 | 0 | 0 | 0 | 0 | 0.001237872 | 1 |
| TINF2 | NM_012461 | 0 | 8.975521721 | 0 | 0 | 0 | 0 | 0 | 0 | 7.27E-08 | 1 |
| TIPIN | NM_017858 | 0 | 8.998808245 | 0 | 0 | 0 | 0 | 0 | 10.66889336 | 6.89E-08 | 1 |

| | | | | | | | | | | | |
|---|---|---|---|---|---|---|---|---|---|---|---|
| TJAP1 | NR_027394 | 0 | 0 | 0 | 0 | 6.491881102 | 0 | 0 | 0 | 4.33E-05 | 1 |
| TJP1 | XM_017022531 | 9.97155406 | 9.073822446 | 0 | 0 | 0 | 5.285170999 | 5.964826548 | 9.055023248 | 0.0015144 | 0.02072721 |
| TJP2 | NM_004817 | 0 | 9.736185717 | 0 | 0 | 0 | 0 | 9.284713985 | 0 | 2.22E-08 | 1 |
| TKT | NM_001064 | 0 | 0 | 10.59641196 | 11.20497527 | 10.06095145 | 12.27201344 | 8.740559545 | 10.70016004 | 0.000509311 | 0.999932046 |
| TLCD1 | NM_138463 | 0 | 0 | 0 | 0 | 6.725274845 | 0 | 0 | 0 | 9.53E-05 | 1 |
| TLCD2 | NM_001164407 | 0 | 0 | 9.538209736 | 8.738654916 | 0 | 0 | 0 | 0 | 0.027385771 | 0.02072721 |
| TLE1 | NM_001303103 | 0 | 9.277965255 | 0 | 0 | 0 | 0 | 0 | 0 | 4.01E-08 | 1 |
| TLE2 | XM_011528230 | 0 | 8.719304505 | 0 | 0 | 0 | 0 | 7.797548711 | 0 | 8.96E-08 | 1 |
| TLK1 | XR_001739085 | 10.24763479 | 0 | 0 | 0 | 0 | 0 | 0 | 0 | 1.58E-08 | 1 |
| TLK2 | XM_024450553 | 8.779231295 | 0 | 0 | 0 | 0 | 5.285170999 | 0 | 0 | 9.02E-08 | 0.555221121 |
| TLNRD1 | NM_022566 | 0 | 0 | 9.420774911 | 5.494613896 | 8.230957729 | 0 | 7.704370312 | 0 | 0.010779876 | 0.999932046 |
| TM2D1 | NR_135160 | 0 | 9.014126367 | 0 | 0 | 0 | 0 | 0 | 0 | 6.71E-08 | 1 |
| TM2D1 | NR_135162 | 9.217654114 | 0 | 0 | 0 | 0 | 0 | 0 | 0 | 5.49E-08 | 1 |
| TM2D3 | NM_078474 | 10.05452941 | 7.213730404 | 0 | 0 | 0 | 10.82137739 | 10.23579762 | 0 | 0.002828549 | 0.02072721 |
| TM4SF5 | NM_003963 | 0 | 0 | 7.320277675 | 10.06846137 | 0 | 0 | 8.131663253 | 0 | 0.026578239 | 0.999932046 |
| TMBIM1 | NR_135643 | 0 | 9.610808606 | 0 | 0 | 0 | 0 | 7.431456038 | 8.330970562 | 2.54E-08 | 0.033407166 |
| TMC6 | NM_001127198 | 0 | 9.17269217 | 0 | 0 | 0 | 0 | 0 | 0 | 4.92E-08 | 1 |
| TMC6 | XR_001752420 | 9.900435696 | 0 | 0 | 0 | 0 | 0 | 0 | 0 | 2.07E-08 | 1 |
| TMCC1-AS1 | NR_037893 | 10.30806717 | 8.384152849 | 0 | 0 | 0 | 0 | 0 | 0 | 0.001533131 | 1 |
| TMCC3 | NM_020698 | 0 | 8.652353298 | 0 | 0 | 0 | 0 | 0 | 0 | 9.28E-08 | 1 |
| TMCO3 | NM_001349742 | 0 | 0 | 0 | 0 | 0 | 9.810013246 | 8.794302797 | 0 | 1 | 0.02072721 |
| TMCO3 | NR_146221 | 0 | 0 | 0 | 9.747573962 | 9.690238446 | 0 | 7.032041479 | 0 | 0.017428499 | 1 |
| TMED1 | NM_006858 | 10.5981354 | 9.533188437 | 0 | 0 | 0 | 0 | 0 | 0 | 0.000729089 | 1 |
| TMED3 | NR_125394 | 0 | 8.870030144 | 0 | 0 | 0 | 0 | 7.655422644 | 0 | 7.86E-08 | 1 |
| TMED4 | NM_001303058 | 0 | 8.612652582 | 0 | 0 | 0 | 0 | 0 | 0 | 9.67E-08 | 1 |
| TMEM107 | NR_147092 | 0 | 0 | 10.37239397 | 10.28541326 | 7.477124411 | 0 | 0 | 0 | 0.001840907 | 0.003016079 |
| TMEM11 | NM_003876 | 0 | 9.422738093 | 9.872720291 | 10.28298438 | 9.392867275 | 0 | 0 | 0 | 0.999953634 | 0.002488287 |
| TMEM11 | NR_024547 | 0 | 9.689244249 | 0 | 0 | 0 | 0 | 0 | 0 | 2.34E-08 | 1 |
| TMEM115 | NM_007024 | 8.940634053 | 0 | 0 | 0 | 0 | 0 | 0 | 0 | 7.88E-08 | 1 |
| TMEM120A | XM_024446954 | 0 | 0 | 7.156088427 | 0 | 0 | 0 | 0 | 10.39970147 | 2.71E-05 | 1 |
| TMEM126B | NM_001256546 | 0 | 0 | 0 | 11.63692055 | 9.940335027 | 0 | 12.37848442 | 10.59148026 | 0.006815226 | 0.999932046 |
| TMEM131 | XR_001738682 | 10.08445476 | 0 | 0 | 0 | 0 | 0 | 0 | 0 | 1.77E-08 | 1 |
| TMEM131L | XM_024453958 | 10.26950953 | 0 | 0 | 0 | 0 | 0 | 0 | 0 | 1.57E-08 | 1 |
| TMEM134 | NR_073413 | 9.801808664 | 0 | 0 | 0 | 0 | 0 | 0 | 0 | 2.38E-08 | 1 |
| TMEM138 | NM_016464 | 0 | 0 | 0 | 10.18491977 | 9.642240281 | 0 | 0 | 12.21089297 | 0.015112298 | 0.999932046 |
| TMEM138 | NR_028473 | 0 | 9.825721024 | 0 | 0 | 0 | 0 | 0 | 0 | 2.01E-08 | 1 |
| TMEM139 | NM_001242773 | 0 | 9.044283149 | 0 | 0 | 0 | 0 | 0 | 0 | 6.33E-08 | 1 |
| TMEM147 | NM_001242597 | 0 | 0 | 0 | 9.792664417 | 8.421503242 | 0 | 0 | 0 | 0.024124651 | 0.02072721 |
| TMEM14A | NM_014051 | 0 | 0 | 11.44568934 | 11.58807576 | 8.532611753 | 9.118185147 | 8.951247914 | 10.62945714 | 0.000436448 | 0.999932046 |
| TMEM14B | NM_001127711 | 0 | 0 | 0 | 10.00503299 | 8.638785557 | 0 | 0 | 9.815272851 | 0.020821831 | 0.999932046 |
| TMEM150A | XM_017003307 | 0 | 0 | 0 | 10.41529045 | 6.147113942 | 0 | 0 | 0 | 0.023042822 | 0.02072721 |
| TMEM159 | NM_020422 | 0 | 0 | 10.71498289 | 9.170475661 | 0 | 0 | 0 | 8.926691863 | 0.014855254 | 0.999932046 |
| TMEM161A | NM_017814 | 0 | 0 | 0 | 10.05855517 | 8.476443242 | 0 | 0 | 0 | 0.021014004 | 0.02072721 |

| | | | | | | | | | | | |
|---|---|---|---|---|---|---|---|---|---|---|---|
| TMEM161A | XM_017026916 | 0 | 0 | 0 | 10.16790747 | 8.803616231 | 0 | 0 | 0 | 0.018740683 | 0.02072721 |
| TMEM161B | NM_001349403 | 0 | 0 | 0 | 7.676917851 | 0 | 0 | 0 | 0 | 1.47E-05 | 1 |
| TMEM161B | XM_017009093 | 0 | 9.91002288 | 11.05406267 | 0 | 7.153852394 | 0 | 0 | 0 | 0.999953634 | 0.02072721 |
| TMEM161B | XM_024454373 | 0 | 8.935855595 | 0 | 0 | 0 | 0 | 0 | 0 | 7.69E-08 | 1 |
| TMEM161B | XM_024454376 | 0 | 8.384152849 | 0 | 0 | 0 | 0 | 0 | 0 | 1.18E-07 | 1 |
| TMEM161B | XM_024454377 | 0 | 0 | 8.769900534 | 0 | 7.920172326 | 0 | 6.811485137 | 0 | 0.046397469 | 0.999932046 |
| TMEM161B | XR_001742013 | 9.715197156 | 0 | 0 | 0 | 0 | 0 | 7.899157068 | 9.138161428 | 2.61E-08 | 0.02096331 |
| TMEM161B-AS1 | NR_039994 | 0 | 0 | 9.617029539 | 0 | 7.630337835 | 0 | 0 | 0 | 0.031670037 | 0.021440042 |
| TMEM161B-AS1 | NR_105019 | 0 | 0 | 0 | 8.029665411 | 0 | 0 | 0 | 0 | 1.02E-05 | 1 |
| TMEM163 | XM_017005061 | 0 | 0 | 0 | 7.994469212 | 0 | 0 | 0 | 0 | 1.07E-05 | 1 |
| TMEM165 | XM_017008412 | 7.707574112 | 0 | 0 | 10.55873452 | 8.048094872 | 0 | 0 | 0 | 0.889593186 | 0.02072721 |
| TMEM167A | NM_174909 | 0 | 11.88722391 | 10.49694284 | 11.04072857 | 10.6904547 | 0 | 0 | 0 | 0.999953634 | 0.001042882 |
| TMEM171 | NM_173490 | 10.88132986 | 9.625843798 | 10.56868506 | 10.41750665 | 7.838643081 | 0 | 0 | 0 | 0.999953634 | 0.002581075 |
| TMEM173 | NM_198282 | 0 | 0 | 0 | 10.20044734 | 7.510413387 | 0 | 0 | 0 | 0.022772235 | 0.02072721 |
| TMEM175 | NM_001297424 | 0 | 9.490017613 | 0 | 0 | 0 | 0 | 0 | 0 | 3.40E-08 | 1 |
| TMEM175 | XM_024454255 | 10.44508285 | 0 | 0 | 0 | 0 | 0 | 0 | 0 | 1.45E-08 | 1 |
| TMEM176A | XM_011516377 | 0 | 0 | 7.974444826 | 0 | 0 | 10.40458512 | 0 | 0 | 1.18E-05 | 1 |
| TMEM182 | NM_001321344 | 0 | 0 | 0 | 10.50764053 | 10.10846726 | 0 | 9.707200767 | 10.14738921 | 0.011380616 | 0.999932046 |
| TMEM183A | NM_001322956 | 0 | 0 | 10.71498289 | 0 | 8.114548983 | 0 | 7.4411151 | 10.69777882 | 0.016425231 | 0.999932046 |
| TMEM183A | NR_136530 | 9.911607777 | 0 | 0 | 0 | 0 | 0 | 7.578739068 | 0 | 2.05E-08 | 1 |
| TMEM183A | NR_146286 | 10.29958742 | 0 | 0 | 0 | 0 | 0 | 9.337386166 | 0 | 1.54E-08 | 1 |
| TMEM183A | NR_146289 | 9.695941183 | 0 | 0 | 0 | 0 | 0 | 9.282029213 | 0 | 2.66E-08 | 1 |
| TMEM185A | NM_001282302 | 0 | 0 | 9.951172164 | 0 | 7.618173501 | 0 | 0 | 0 | 0.026325634 | 0.02072721 |
| TMEM185A | NR_104121 | 9.789749415 | 0 | 0 | 0 | 0 | 0 | 0 | 0 | 2.40E-08 | 1 |
| TMEM189 | NM_199129 | 9.958146973 | 0 | 0 | 9.913784443 | 8.761377537 | 0 | 0 | 0 | 0.999953634 | 0.02072721 |
| TMEM191B | NM_001242313 | 0 | 0 | 0 | 0 | 0 | 0 | 8.212844911 | 9.691484157 | 1 | 0.02072721 |
| TMEM191C | NM_001207052 | 0 | 0 | 9.929535327 | 10.13730841 | 0 | 0 | 0 | 0 | 0.015380705 | 0.02072721 |
| TMEM199 | NM_152464 | 0 | 0 | 0 | 8.284026919 | 0 | 0 | 0 | 0 | 7.78E-06 | 1 |
| TMEM200A | XM_017010230 | 10.15666012 | 0 | 8.775301651 | 0 | 8.702103817 | 0 | 0 | 0 | 0.999953634 | 0.023261296 |
| TMEM204 | NM_024600 | 9.789749415 | 0 | 0 | 0 | 0 | 0 | 0 | 0 | 2.40E-08 | 1 |
| TMEM208 | NM_001318217 | 0 | 8.719304505 | 0 | 0 | 0 | 0 | 0 | 0 | 8.96E-08 | 1 |
| TMEM216 | NM_001173991 | 0 | 0 | 10.53564879 | 0 | 8.136037099 | 0 | 10.18640294 | 0 | 0.018004498 | 0.999932046 |
| TMEM216 | XM_005274039 | 0 | 9.352166886 | 10.27280875 | 0 | 7.415197319 | 0 | 0 | 0 | 0.999953634 | 0.02072721 |
| TMEM218 | NM_001258239 | 0 | 0 | 0 | 0 | 6.3222931 | 11.24546404 | 8.064509964 | 0 | 0.000244538 | 0.903316045 |
| TMEM218 | NM_001258244 | 10.88841554 | 10.82275274 | 0 | 9.987239158 | 9.99603973 | 0 | 0 | 0 | 0.999953634 | 0.02072721 |
| TMEM218 | NM_001258245 | 0 | 8.998808245 | 0 | 0 | 0 | 0 | 0 | 0 | 6.89E-08 | 1 |
| TMEM218 | NR_047589 | 0 | 0 | 0 | 10.01823581 | 8.790125492 | 0 | 9.088179108 | 10.10847773 | 0.02004138 | 0.999932046 |
| TMEM218 | XM_006718783 | 0 | 10.09426749 | 0 | 0 | 0 | 0 | 9.350259127 | 0 | 1.59E-08 | 1 |
| TMEM218 | XM_024448388 | 9.025355161 | 0 | 0 | 0 | 0 | 0 | 0 | 0 | 7.69E-08 | 1 |
| TMEM218 | XM_024448389 | 0 | 0 | 10.54674515 | 7.364123525 | 8.180146518 | 0 | 0 | 0 | 0.003669917 | 0.005243741 |
| TMEM220 | XM_011523835 | 0 | 10.25440355 | 0 | 0 | 0 | 0 | 0 | 0 | 1.47E-08 | 1 |
| TMEM230 | NM_001330984 | 0 | 9.06649413 | 0 | 0 | 0 | 0 | 0 | 0 | 7.75E-08 | 1 |
| TMEM236 | NM_001098844 | 0 | 0 | 7.474332608 | 0 | 0 | 0 | 0 | 0 | 1.96E-05 | 1 |

| | | | | | | | | | | | |
|---|---|---|---|---|---|---|---|---|---|---|---|
| TMEM237 | NM_152388 | 0 | 0 | 0 | 10.43511477 | 7.87488031 | 0 | 0 | 0 | 0.019200388 | 0.02072721 |
| TMEM246 | NM_032342 | 10.05452941 | 0 | 0 | 0 | 0 | 0 | 8.805563283 | 9.696263442 | 5.06E-08 | 0.02072721 |
| TMEM250 | XM_024447712 | 0 | 8.709928179 | 0 | 0 | 0 | 0 | 0 | 0 | 8.96E-08 | 1 |
| TMEM251 | NM_015676 | 10.87706169 | 9.842980089 | 11.20977038 | 10.67444706 | 3.220007989 | 0 | 0 | 0 | 0.999953634 | 0.003106942 |
| TMEM253 | XM_011537083 | 0 | 9.399595812 | 0 | 0 | 0 | 0 | 0 | 0 | 3.36E-08 | 1 |
| TMEM26-AS1 | NR_120643 | 0 | 8.935855595 | 0 | 0 | 0 | 0 | 0 | 0 | 7.69E-08 | 1 |
| TMEM265 | NM_001256829 | 0 | 7.86468487 | 0 | 9.882039044 | 10.16391305 | 0 | 0 | 0 | 0.881784333 | 0.02072721 |
| TMEM267 | XM_011514075 | 9.143781062 | 0 | 0 | 0 | 0 | 0 | 0 | 0 | 7.26E-08 | 1 |
| TMEM268 | XR_001746226 | 0 | 9.495485143 | 0 | 0 | 0 | 0 | 0 | 0 | 2.95E-08 | 1 |
| TMEM30B | NM_001017970 | 0 | 0 | 10.5002083 | 0 | 7.910229599 | 0 | 0 | 0 | 0.018876503 | 0.02072721 |
| TMEM33 | XM_005248117 | 0 | 9.799439006 | 0 | 0 | 0 | 0 | 0 | 0 | 2.04E-08 | 1 |
| TMEM37 | XM_011510659 | 0 | 9.265218616 | 0 | 0 | 0 | 0 | 0 | 0 | 4.18E-08 | 1 |
| TMEM38B | XM_011518833 | 6.161524867 | 8.519117955 | 0 | 0 | 0 | 0 | 0 | 8.599394397 | 0.016372233 | 1 |
| TMEM40 | NM_001284407 | 0 | 0 | 0 | 10.30947974 | 10.53535819 | 0 | 10.15520008 | 6.640576357 | 0.010465839 | 0.999932046 |
| TMEM40 | NM_001284408 | 0 | 0 | 0 | 10.59639063 | 8.77650741 | 0 | 11.55894967 | 0 | 0.015398633 | 0.999932046 |
| TMEM40 | NM_018306 | 0 | 0 | 0 | 11.06284252 | 10.87141542 | 0 | 9.132073672 | 0 | 0.006813967 | 1 |
| TMEM40 | XM_011533937 | 0 | 0 | 0 | 0 | 6.725274845 | 0 | 5.763978756 | 0 | 9.53E-05 | 1 |
| TMEM41B | NR_028491 | 0 | 0 | 9.420774911 | 0 | 7.812190501 | 0 | 8.04563962 | 0 | 0.03396298 | 0.999932046 |
| TMEM41B | XR_001747881 | 10.80400107 | 9.665188049 | 0 | 0 | 0 | 0 | 7.351763189 | 8.542143059 | 0.000576817 | 0.030862825 |
| TMEM44 | XM_017007518 | 0 | 8.508340767 | 0 | 0 | 0 | 0 | 0 | 0 | 1.06E-07 | 1 |
| TMEM50A | NM_014313 | 0 | 0 | 10.30116784 | 11.88890032 | 8.999211672 | 10.62451262 | 9.548703693 | 11.5519962 | 0.000520537 | 0.999932046 |
| TMEM51 | XM_005245919 | 0 | 9.506358431 | 0 | 0 | 0 | 0 | 0 | 0 | 2.91E-08 | 1 |
| TMEM53 | NM_024587 | 0 | 0 | 0 | 0 | 0 | 0 | 9.347693707 | 11.15805656 | 1 | 0.02072721 |
| TMEM53 | XM_011542138 | 9.025355161 | 0 | 0 | 0 | 0 | 0 | 0 | 0 | 7.69E-08 | 1 |
| TMEM54 | NM_001329722 | 0 | 10.73764849 | 12.0785115 | 0 | 9.454884858 | 0 | 0 | 0 | 0.999953634 | 4.10E-09 |
| TMEM54 | NM_001329724 | 0 | 9.17269217 | 0 | 0 | 0 | 0 | 0 | 0 | 4.92E-08 | 1 |
| TMEM59 | NM_001305049 | 10.80998402 | 6.113929405 | 0 | 10.50451658 | 8.246846504 | 0 | 0 | 0 | 0.999953634 | 0.02072721 |
| TMEM59 | NM_001305051 | 0 | 9.416987232 | 0 | 0 | 0 | 0 | 8.207195305 | 0 | 3.28E-08 | 1 |
| TMEM62 | XM_024450071 | 0 | 0 | 0 | 8.405116658 | 0 | 0 | 5.882147525 | 0 | 6.82E-06 | 1 |
| TMEM63A | XM_011544332 | 0 | 0 | 0 | 0 | 6.655567121 | 0 | 0 | 0 | 3.82E-05 | 1 |
| TMEM64 | NM_001146273 | 0 | 0 | 0 | 0 | 6.30721813 | 10.13406087 | 0 | 0 | 4.99E-05 | 1 |
| TMEM65 | XM_011516847 | 0 | 0 | 9.368130153 | 0 | 7.801472153 | 11.91694833 | 7.479117364 | 0 | 0.035063012 | 0.999932046 |
| TMEM65 | XM_017013104 | 0 | 0 | 0 | 0 | 0 | 10.70306084 | 8.882017781 | 0 | 1 | 0.02072721 |
| TMEM65 | XM_017013105 | 0 | 0 | 0 | 0 | 0 | 0 | 8.521333265 | 9.908831299 | 1 | 0.02072721 |
| TMEM68 | XM_017013040 | 0 | 0 | 0 | 0 | 0 | 10.06665936 | 7.987500854 | 0 | 1 | 0.02072721 |
| TMEM80 | NM_001042463 | 0 | 10.61491811 | 0 | 0 | 0 | 0 | 0 | 0 | 1.34E-08 | 1 |
| TMEM81 | NM_203376 | 0 | 0 | 0 | 0 | 6.351978551 | 0 | 0 | 0 | 4.81E-05 | 1 |
| TMEM87A | XM_011521443 | 9.442270566 | 8.886770211 | 0 | 0 | 0 | 0 | 0 | 0 | 0.002516835 | 1 |
| TMEM8B | XR_002956793 | 9.649985154 | 0 | 0 | 0 | 0 | 0 | 0 | 0 | 2.95E-08 | 1 |
| TMEM9 | NM_016456 | 0 | 9.665188049 | 0 | 0 | 0 | 0 | 10.9251693 | 0 | 2.41E-08 | 1 |
| TMEM91 | NM_001098824 | 8.452015244 | 0 | 0 | 0 | 0 | 0 | 0 | 0 | 1.21E-07 | 1 |
| TMEM94 | XM_017025450 | 0 | 0 | 9.053876344 | 10.05286379 | 7.774322632 | 0 | 7.421731871 | 0 | 0.003453972 | 0.999932046 |
| TMEM98 | NM_001033504 | 0 | 8.801057108 | 0 | 0 | 0 | 0 | 8.89613213 | 0 | 8.24E-08 | 1 |

| | | | | | | | | | | | |
|---|---|---|---|---|---|---|---|---|---|---|---|
| TMEM99 | NM_145274 | 0 | 0 | 0 | 9.407454061 | 8.884555432 | 0 | 0 | 0 | 0.02525384 | 0.02072721 |
| TMF1 | XM_011534056 | 0 | 8.486541819 | 0 | 0 | 0 | 9.463083043 | 0 | 0 | 1.09E-07 | 1 |
| TMIGD2 | NM_001169126 | 0 | 8.453213086 | 0 | 0 | 0 | 0 | 0 | 0 | 1.11E-07 | 1 |
| TMLHE | NM_018196 | 0 | 9.745393473 | 0 | 0 | 0 | 10.79076288 | 0 | 8.877042846 | 2.16E-08 | 0.02072721 |
| TMOD3 | XM_017022088 | 0 | 10.35708192 | 9.530245374 | 7.504171232 | 0 | 0 | 0 | 0 | 0.999953634 | 0.024319498 |
| TMPO | XM_005269132 | 0 | 9.179501644 | 0 | 0 | 0 | 0 | 0 | 0 | 4.86E-08 | 1 |
| TMUB1 | NM_001136044 | 0 | 0 | 11.81663562 | 8.979688824 | 6.769943125 | 11.14253998 | 9.557611138 | 0 | 0.001314309 | 0.999932046 |
| TMUB1 | NM_031434 | 9.480284601 | 0 | 0 | 0 | 0 | 0 | 0 | 0 | 3.64E-08 | 1 |
| TMUB2 | NM_001353184 | 0 | 0 | 8.167546787 | 0 | 0 | 0 | 0 | 0 | 2.98E-05 | 1 |
| TMUB2 | XM_024450933 | 0 | 9.321711965 | 0 | 0 | 0 | 0 | 0 | 10.38940937 | 3.84E-08 | 1 |
| TMUB2 | XR_001752619 | 9.759152772 | 8.671801345 | 0 | 0 | 0 | 0 | 0 | 0 | 0.002190645 | 1 |
| TMX1 | NM_030755 | 0 | 9.985789839 | 11.5871437 | 9.679029003 | 8.486515572 | 0 | 0 | 0 | 0.999953634 | 0.001765436 |
| TNFAIP3 | XM_006715555 | 0 | 9.527862205 | 0 | 0 | 0 | 0 | 0 | 0 | 2.90E-08 | 1 |
| TNFAIP8L3 | NM_001311175 | 9.287928048 | 0 | 0 | 0 | 0 | 0 | 0 | 0 | 4.93E-08 | 1 |
| TNFRSF10B | NM_147187 | 0 | 0 | 0 | 0 | 0 | 0 | 7.720323987 | 10.36710447 | 1 | 0.02072721 |
| TNFRSF14-AS1 | NR_037844 | 11.30326275 | 8.20886266 | 11.40271328 | 0 | 8.759994193 | 0 | 0 | 0 | 0.999953634 | 0.02072721 |
| TNFRSF1B | XM_011542063 | 10.23434896 | 0 | 10.86310061 | 0 | 8.070588078 | 0 | 0 | 0 | 0.999953634 | 0.02072721 |
| TNFRSF6B | NM_003823 | 0 | 0 | 9.930745925 | 10.16659049 | 0 | 0 | 0 | 0 | 0.015220299 | 0.02072721 |
| TNFSF10 | NM_001190942 | 10.20058902 | 0 | 0 | 0 | 0 | 0 | 0 | 0 | 1.60E-08 | 1 |
| TNFSF10 | NM_001190943 | 0 | 9.239382419 | 0 | 0 | 0 | 0 | 0 | 0 | 4.33E-08 | 1 |
| TNIP1 | NM_006058 | 10.15666012 | 9.506358431 | 9.527047278 | 0 | 8.558305894 | 0 | 0 | 0 | 0.999953634 | 0.02072721 |
| TNIP1 | XM_017008950 | 0 | 9.473489586 | 0 | 0 | 0 | 0 | 7.73610317 | 0 | 3.08E-08 | 1 |
| TNIP1 | XM_017008951 | 9.771469405 | 9.456770012 | 0 | 0 | 0 | 0 | 0 | 10.69180852 | 0.001416814 | 1 |
| TNKS2 | XM_017016700 | 0 | 8.681427931 | 0 | 0 | 0 | 0 | 0 | 0 | 9.11E-08 | 1 |
| TNNT3 | XM_024448672 | 0 | 9.605761852 | 0 | 0 | 0 | 0 | 0 | 0 | 2.55E-08 | 1 |
| TNRC18 | XM_017012734 | 0 | 8.248259235 | 0 | 0 | 0 | 0 | 0 | 0 | 2.02E-07 | 1 |
| TNS4 | NM_032865 | 0 | 0 | 8.117393241 | 8.244627116 | 6.631562102 | 9.316044599 | 10.14119483 | 10.82868874 | 0.017088229 | 0.999932046 |
| TNS4 | XM_005257744 | 0 | 0 | 10.84009086 | 12.29482951 | 8.506451693 | 0 | 11.79992686 | 11.88381327 | 0.000366602 | 0.999932046 |
| TOB2 | NM_016272 | 0 | 9.006487637 | 0 | 0 | 0 | 0 | 0 | 0 | 6.80E-08 | 1 |
| TOB2 | XM_017028539 | 9.849062238 | 0 | 0 | 0 | 0 | 0 | 8.704922178 | 0 | 2.23E-08 | 1 |
| TOE1 | XR_426587 | 0 | 8.700490516 | 0 | 0 | 0 | 0 | 0 | 0 | 9.01E-08 | 1 |
| TOM1 | XM_017028530 | 8.873659666 | 9.812639863 | 0 | 0 | 0 | 0 | 0 | 0 | 0.001850451 | 1 |
| TOM1L1 | XM_017024002 | 0 | 0 | 0 | 0 | 6.381065478 | 0 | 8.40029149 | 0 | 4.71E-05 | 1 |
| TOMM34 | NM_006809 | 0 | 0 | 0 | 10.93021201 | 8.020631767 | 0 | 11.07692611 | 0 | 0.014479375 | 0.999932046 |
| TOMM34 | XM_011528501 | 0 | 0 | 0 | 7.229331528 | 0 | 0 | 0 | 0 | 6.73E-05 | 1 |
| TOMM34 | XR_244131 | 0 | 0 | 0 | 10.25599332 | 9.022473864 | 0 | 0 | 0 | 0.017203645 | 0.02072721 |
| TOP1 | NM_003286 | 0 | 0 | 9.564964337 | 7.908843317 | 6.906244365 | 0 | 4.37247236 | 0 | 0.008670965 | 0.30092499 |
| TOP2A | XM_011525165 | 0 | 0 | 0 | 10.42853688 | 7.895185835 | 0 | 0 | 0 | 0.01921012 | 0.02072721 |
| TOP2B | XR_001740241 | 7.573321022 | 9.635780908 | 0 | 0 | 0 | 0 | 0 | 0 | 0.003664734 | 1 |
| TOP3A | NM_004618 | 0 | 0 | 10.25358819 | 10.59834557 | 9.206380832 | 10.29541704 | 0 | 9.5769499 | 0.001074345 | 0.999932046 |
| TOR2A | NM_001085347 | 0 | 9.17269217 | 9.696043232 | 10.33315133 | 0 | 0 | 0 | 0 | 0.999953634 | 0.02072721 |
| TOX3 | XM_024450230 | 0 | 0 | 10.07787362 | 0 | 5.646039634 | 0 | 0 | 0 | 0.030398091 | 0.02096331 |
| TP53I13 | XM_024451017 | 0 | 8.497482465 | 0 | 0 | 0 | 0 | 0 | 0 | 1.06E-07 | 1 |

| | | | | | | | | | | | |
|---|---|---|---|---|---|---|---|---|---|---|---|
| TP53I3 | NM_004881 | 0 | 0 | 0 | 7.970518813 | 0 | 0 | 0 | 0 | 1.08E-05 | 1 |
| TP53I3 | XM_006712150 | 0 | 8.540433774 | 0 | 0 | 0 | 0 | 0 | 0 | 1.02E-07 | 1 |
| TP53INP1 | NM_033285 | 8.885040211 | 0 | 0 | 0 | 0 | 0 | 0 | 0 | 8.26E-08 | 1 |
| TP53INP1 | XM_011517386 | 0 | 8.31161984 | 0 | 0 | 0 | 0 | 0 | 0 | 1.27E-07 | 1 |
| TP53INP2 | XM_006723841 | 8.815370067 | 0 | 0 | 0 | 0 | 0 | 0 | 0 | 8.84E-08 | 1 |
| TPCN1 | XR_001748765 | 9.226627846 | 0 | 0 | 0 | 0 | 0 | 8.032920857 | 0 | 5.40E-08 | 1 |
| TPD52 | NM_001025252 | 9.509990722 | 0 | 9.427651273 | 10.06563796 | 7.920172326 | 0 | 0 | 0 | 0.999953634 | 0.004331325 |
| TPD52L2 | NR_045090 | 0 | 0 | 0 | 10.02115343 | 5.232616603 | 0 | 9.734951004 | 0 | 0.031470501 | 1 |
| TPGS2 | NM_001330572 | 0 | 0 | 0 | 0 | 6.518304288 | 10.93702628 | 0 | 0 | 4.23E-05 | 1 |
| TPH1 | NM_004179 | 10.63903963 | 0 | 0 | 0 | 0 | 0 | 0 | 0 | 1.34E-08 | 1 |
| TPI1 | NM_000365 | 10.54531825 | 0 | 0 | 0 | 0 | 0 | 0 | 0 | 1.36E-08 | 1 |
| TPM1 | NM_000366 | 0 | 0 | 0 | 10.18751933 | 9.220530915 | 0 | 0 | 0 | 0.016902631 | 0.02072721 |
| TPM1 | XM_005254650 | 0 | 0 | 0 | 0 | 0 | 12.03887029 | 8.382856136 | 0 | 1 | 1.92E-09 |
| TPM1 | XM_006720667 | 9.819710617 | 0 | 10.74694801 | 0 | 9.216502176 | 0 | 0 | 0 | 0.999953634 | 0.02072721 |
| TPM3 | NM_001278191 | 0 | 0 | 10.44705277 | 0 | 9.020164451 | 0 | 10.82451857 | 0 | 0.015991489 | 0.999932046 |
| TPM3 | NM_152263 | 10.86274256 | 0 | 0 | 0 | 0 | 0 | 0 | 0 | 1.34E-08 | 1 |
| TPP2 | XR_243046 | 0 | 0 | 8.409046753 | 9.62251587 | 0 | 9.938460398 | 0 | 0 | 0.028115448 | 0.999932046 |
| TPRA1 | NM_001353002 | 0 | 0 | 10.52286184 | 10.15998747 | 7.568463804 | 10.77442637 | 0 | 10.0424498 | 0.001778148 | 0.999932046 |
| TPRKB | NM_001330391 | 0 | 8.384152849 | 0 | 0 | 0 | 0 | 0 | 0 | 1.18E-07 | 1 |
| TPSB2 | NM_024164 | 0 | 10.42455562 | 0 | 0 | 0 | 0 | 0 | 0 | 1.34E-08 | 1 |
| TPST2 | NM_001008566 | 0 | 0 | 0 | 0 | 0 | 0 | 7.870848924 | 10.08315804 | 1 | 0.02072721 |
| TPTEP1 | NR_001591 | 0 | 0 | 0 | 8.041209218 | 8.407434995 | 0 | 0 | 0 | 0.051614541 | 0.034581849 |
| TPTEP2-CSNK1E | NM_001289912 | 0 | 8.602554319 | 11.58752753 | 0 | 10.01877445 | 0 | 0 | 0 | 0.851226931 | 0.02072721 |
| TPX2 | XM_011528697 | 0 | 0 | 0 | 11.93255671 | 11.68656361 | 0 | 12.30979638 | 12.01057746 | 0.00338361 | 0.999932046 |
| TRA2A | NM_001282757 | 0 | 0 | 10.9916286 | 9.711903927 | 10.5337401 | 11.86425846 | 10.57815199 | 11.23150581 | 0.000662244 | 0.999932046 |
| TRA2A | NM_001282759 | 0 | 0 | 10.86056202 | 0 | 10.05870286 | 0 | 9.593783859 | 0 | 0.010000443 | 0.999932046 |
| TRABD | NM_001320487 | 0 | 9.152068755 | 0 | 0 | 0 | 0 | 0 | 0 | 5.06E-08 | 1 |
| TRABD | XM_011530715 | 0 | 9.872695374 | 0 | 0 | 0 | 0 | 0 | 0 | 1.94E-08 | 1 |
| TRABD | XM_017028944 | 10.78590188 | 0 | 0 | 0 | 0 | 0 | 0 | 0 | 1.34E-08 | 1 |
| TRABD2A | NM_001080824 | 0 | 0 | 10.22031381 | 0 | 9.371285855 | 0 | 0 | 0 | 0.016190273 | 0.020781046 |
| TRABD2A | NM_001307978 | 0 | 0 | 0 | 0 | 0 | 10.51171313 | 8.051957199 | 0 | 1 | 0.02072721 |
| TRABD2A | XM_011532504 | 9.296476405 | 7.646135616 | 0 | 10.98802025 | 6.245291858 | 0 | 0 | 0 | 1 | 9.95E-08 |
| TRAF2 | NM_021138 | 0 | 0 | 0 | 0 | 0 | 11.2013354 | 8.316186335 | 0 | 1 | 0.02072721 |
| TRAF3 | XM_011537117 | 9.79579164 | 0 | 0 | 0 | 0 | 0 | 0 | 0 | 2.47E-08 | 1 |
| TRAF7 | XM_005255627 | 0 | 0 | 9.902641893 | 0 | 8.715011943 | 11.31703407 | 9.475392161 | 9.400769032 | 0.021767497 | 0.999932046 |
| TRAK1 | NM_001265608 | 0 | 0 | 10.44111826 | 0 | 8.127480221 | 0 | 0 | 8.119526894 | 0.018859362 | 0.999932046 |
| TRAK1 | NR_146089 | 0 | 0 | 0 | 0 | 6.409577539 | 0 | 0 | 0 | 4.59E-05 | 1 |
| TRAM1L1 | NM_152402 | 0 | 9.37013595 | 0 | 0 | 0 | 0 | 0 | 0 | 3.50E-08 | 1 |
| TRANK1 | XM_017007572 | 8.526574932 | 0 | 9.326426749 | 0 | 6.631562102 | 0 | 0 | 0 | 0.999953634 | 0.02861509 |
| TRAPPC1 | NM_021210 | 0 | 0 | 8.72594783 | 0 | 9.561164523 | 0 | 8.763837937 | 9.571757813 | 0.024563463 | 0.999932046 |
| TRAPPC12 | NM_016030 | 9.208624214 | 8.33620493 | 0 | 10.67444706 | 6.451316153 | 0 | 0 | 0 | 0.999953634 | 0.02072721 |
| TRAPPC2L | NM_001318525 | 0 | 0 | 6.019129988 | 9.802872777 | 9.410608412 | 10.50135128 | 10.1233074 | 11.31311127 | 0.004731654 | 0.999932046 |
| TRAPPC2L | NM_016209 | 0 | 0 | 9.678854813 | 9.671620603 | 9.065664595 | 0 | 10.33950216 | 0 | 0.002230559 | 1 |

| | | | | | | | | | | | |
|---|---|---|---|---|---|---|---|---|---|---|---|
| TRAPPC3 | NM_001270896 | 0 | 9.258802813 | 0 | 0 | 0 | 0 | 0 | 0 | 4.23E-08 | 1 |
| TRAPPC4 | NM_001318489 | 10.41791771 | 0 | 0 | 0 | 0 | 0 | 8.417518646 | 0 | 1.47E-08 | 1 |
| TRAPPC5 | NM_001042462 | 0 | 0 | 11.34939289 | 9.849579366 | 10.25707461 | 0 | 9.412955358 | 10.63444603 | 0.000584814 | 0.999932046 |
| TRAPPC8 | XR_001753169 | 0 | 0 | 0 | 0 | 6.667421491 | 0 | 0 | 0 | 3.78E-05 | 1 |
| TRDMT1 | NM_004412 | 0 | 9.10991584 | 0 | 0 | 0 | 0 | 0 | 0 | 5.42E-08 | 1 |
| TRERF1 | XM_017011049 | 0 | 8.323964754 | 0 | 0 | 0 | 0 | 0 | 0 | 1.27E-07 | 1 |
| TRERF1 | XM_017011055 | 0 | 0 | 0 | 7.870531406 | 8.773768299 | 0 | 0 | 10.77886808 | 0.045542194 | 0.999932046 |
| TRIB1 | XR_428373 | 0 | 0 | 0 | 0 | 0 | 0 | 7.743928512 | 8.437565712 | 1 | 0.028987618 |
| TRIM14 | XR_929880 | 0 | 8.430558346 | 0 | 0 | 0 | 0 | 0 | 0 | 1.13E-07 | 1 |
| TRIM15 | NM_033229 | 10.41399488 | 0 | 9.071545242 | 9.792664417 | 6.42362501 | 0 | 0 | 0 | 0.999953634 | 0.008099638 |
| TRIM16 | NM_006470 | 0 | 0 | 0 | 9.885245203 | 8.608360805 | 0 | 0 | 10.89046427 | 0.022115815 | 0.999932046 |
| TRIM23 | NM_001656 | 9.70238844 | 0 | 0 | 0 | 0 | 0 | 0 | 0 | 2.63E-08 | 1 |
| TRIM28 | NM_005762 | 0 | 9.934381906 | 0 | 11.64450882 | 9.901966581 | 0 | 0 | 0 | 0.999953634 | 0.02072721 |
| TRIM29 | XM_017017453 | 0 | 0 | 0 | 0 | 6.667421491 | 0 | 0 | 0 | 3.78E-05 | 1 |
| TRIM3 | XM_024448329 | 0 | 8.430558346 | 0 | 0 | 0 | 0 | 0 | 0 | 1.13E-07 | 1 |
| TRIM31 | NR_134871 | 0 | 8.959785899 | 11.19875824 | 0 | 9.504466758 | 0 | 0 | 0 | 0.989141962 | 0.02072721 |
| TRIM31 | XM_011514264 | 0 | 0 | 9.017875808 | 0 | 8.816981984 | 0 | 9.473040052 | 0 | 0.030410519 | 1 |
| TRIM31 | XM_011514265 | 10.34973404 | 6.784168672 | 0 | 0 | 0 | 0 | 0 | 0 | 0.002299792 | 1 |
| TRIM34 | NM_021616 | 0 | 0 | 10.24777173 | 0 | 8.435435627 | 0 | 0 | 0 | 0.019693301 | 0.02072721 |
| TRIM45 | NM_025188 | 0 | 0 | 0 | 0 | 6.409577539 | 0 | 0 | 0 | 4.59E-05 | 1 |
| TRIM65 | XM_017024344 | 10.14723899 | 0 | 0 | 0 | 0 | 0 | 0 | 0 | 1.64E-08 | 1 |
| TRIM74 | XR_001744736 | 0 | 8.861586703 | 0 | 0 | 0 | 0 | 0 | 0 | 7.86E-08 | 1 |
| TRIM74 | XR_002956443 | 8.907535469 | 0 | 0 | 0 | 0 | 0 | 0 | 0 | 2.11E-07 | 1 |
| TRIP12 | NM_001348327 | 0 | 8.927789798 | 0 | 0 | 0 | 0 | 8.472716031 | 0 | 7.69E-08 | 1 |
| TRIP12 | XM_017005283 | 0 | 0 | 0 | 8.773677528 | 0 | 0 | 0 | 0 | 4.62E-06 | 1 |
| TRIP12 | XM_024453224 | 10.72387683 | 0 | 0 | 0 | 0 | 0 | 0 | 0 | 1.34E-08 | 1 |
| TRIP12 | XM_024453242 | 0 | 0 | 0 | 10.91460632 | 9.844389419 | 0 | 7.834664296 | 10.0940638 | 0.010288468 | 0.999932046 |
| TRIP4 | NM_016213 | 10.72070446 | 0 | 0 | 0 | 0 | 0 | 9.771778086 | 0 | 1.34E-08 | 1 |
| TRIQK | NM_001171797 | 0 | 9.340061955 | 0 | 0 | 0 | 0 | 7.967588696 | 0 | 3.95E-08 | 1 |
| TRIR | NM_024038 | 0 | 0 | 9.514183525 | 11.90359972 | 11.53446756 | 0 | 6.898192229 | 11.67209379 | 0.000261841 | 0.999932046 |
| TRMT10B | XR_001746192 | 0 | 0 | 0 | 0 | 6.518304288 | 0 | 0 | 0 | 4.23E-05 | 1 |
| TRMT11 | NM_001350586 | 0 | 0 | 0 | 11.44114404 | 9.755555472 | 11.2849916 | 7.885072427 | 9.627868238 | 0.007944948 | 0.999932046 |
| TRMT11 | NM_001350592 | 0 | 0 | 10.17608123 | 10.01677478 | 0 | 0 | 0 | 9.73161216 | 0.014798905 | 0.999932046 |
| TRMT11 | NR_146803 | 0 | 9.117027412 | 0 | 0 | 0 | 0 | 0 | 0 | 5.36E-08 | 1 |
| TRMT12 | NM_017956 | 0 | 0 | 0 | 0 | 6.758905041 | 0 | 0 | 0 | 9.53E-05 | 1 |
| TRMT2B | XM_017029871 | 0 | 8.681427931 | 0 | 0 | 0 | 0 | 0 | 0 | 9.11E-08 | 1 |
| TRMT61A | NM_152307 | 10.57372782 | 0 | 0 | 0 | 0 | 0 | 0 | 0 | 1.34E-08 | 1 |
| TRMT9B | XM_024447219 | 9.495214119 | 0 | 0 | 0 | 0 | 0 | 0 | 0 | 3.47E-08 | 1 |
| TRMU | NM_018006 | 0 | 8.886770211 | 0 | 0 | 0 | 0 | 0 | 0 | 7.80E-08 | 1 |
| TRNT1 | XM_011533776 | 0 | 9.439854398 | 0 | 0 | 0 | 0 | 0 | 0 | 3.19E-08 | 1 |
| TROAP | NM_005480 | 0 | 8.983325733 | 0 | 0 | 0 | 0 | 0 | 0 | 7.17E-08 | 1 |
| TRPM4 | NM_017636 | 0 | 0 | 10.51238834 | 0 | 8.011359982 | 0 | 0 | 0 | 0.018540605 | 0.02072721 |
| TRPM7 | XR_001751328 | 0 | 0 | 0 | 8.284026919 | 0 | 0 | 0 | 0 | 7.78E-06 | 1 |

| | | | | | | | | | | | |
|---|---|---|---|---|---|---|---|---|---|---|---|
| TRUB2 | NM_015679 | 0 | 0 | 0 | 11.26404978 | 7.415197319 | 11.06598657 | 7.671923744 | 0 | 0.012714034 | 0.999932046 |
| TSC22D1 | NM_001243797 | 0 | 0 | 0 | 0 | 0 | 10.95093823 | 7.525254359 | 0 | 1 | 0.02072721 |
| TSC22D4 | NR_130119 | 0 | 0 | 0 | 10.11290025 | 6.351978551 | 0 | 8.557305568 | 0 | 0.02718775 | 0.999932046 |
| TSEN2 | XR_001740293 | 0 | 0 | 0 | 0 | 6.42362501 | 0 | 0 | 0 | 4.54E-05 | 1 |
| TSEN54 | NM_207346 | 9.992749568 | 8.612652582 | 0 | 0 | 0 | 0 | 0 | 0 | 0.001850451 | 1 |
| TSFM | NM_001172696 | 0 | 10.66187 | 0 | 0 | 0 | 11.60614485 | 11.67630825 | 0 | 1.34E-08 | 0.012818915 |
| TSHZ2 | XM_017027641 | 0 | 0 | 9.38941913 | 0 | 7.930046999 | 11.74596214 | 0 | 0 | 0.033597348 | 0.999932046 |
| TSHZ3 | NM_020856 | 9.449953927 | 0 | 0 | 0 | 0 | 0 | 0 | 0 | 3.80E-08 | 1 |
| TSPAN1 | XM_011540460 | 0 | 0 | 3.436126667 | 10.21580957 | 9.294058479 | 0 | 0 | 0 | 0.005577453 | 0.009119021 |
| TSPAN12 | XM_017011913 | 0 | 0 | 0 | 10.01823581 | 8.230957729 | 10.19378098 | 6.514885902 | 0 | 0.022417213 | 0.999932046 |
| TSPAN15 | NM_001351263 | 0 | 0 | 0 | 0 | 6.437537018 | 0 | 0 | 0 | 4.51E-05 | 1 |
| TSPAN33 | XM_006715960 | 0 | 8.222114748 | 0 | 0 | 0 | 0 | 10.40574383 | 9.095383146 | 0.054579863 | 0.02072721 |
| TSPYL4 | NM_021648 | 0 | 0 | 0 | 8.537350365 | 0 | 0 | 0 | 0 | 5.99E-06 | 1 |
| TSR1 | NM_018128 | 10.44508285 | 9.352166886 | 0 | 0 | 0 | 0 | 0 | 0 | 0.000901948 | 1 |
| TST | NM_001270483 | 0 | 0 | 10.38127638 | 10.42853688 | 7.859461386 | 0 | 0 | 0 | 0.001572701 | 0.002701234 |
| TSTD2 | XR_929723 | 8.611240259 | 0 | 0 | 0 | 0 | 8.967565814 | 0 | 0 | 1.04E-07 | 1 |
| TTC13 | NM_024525 | 0 | 0 | 10.41280771 | 11.82855761 | 10.70741969 | 0 | 10.03358004 | 10.80343984 | 0.000298554 | 0.999932046 |
| TTC13 | XM_005273260 | 0 | 0 | 10.51964732 | 9.527221444 | 8.551925132 | 0 | 8.801819545 | 10.63195374 | 0.001832267 | 0.999932046 |
| TTC13 | XM_005273262 | 0 | 0 | 0 | 10.0399755 | 8.357086869 | 10.80001563 | 9.393187444 | 0 | 0.021704832 | 0.999932046 |
| TTC13 | XM_017002323 | 0 | 0 | 0 | 8.924658605 | 10.04001708 | 0 | 8.475068668 | 0 | 0.019039077 | 0.999932046 |
| TTC22 | XM_017001582 | 0 | 9.145128245 | 0 | 0 | 0 | 0 | 0 | 0 | 5.08E-08 | 1 |
| TTC27 | NM_001193509 | 0 | 9.795011742 | 0 | 0 | 0 | 0 | 11.64138527 | 11.086139 | 2.04E-08 | 0.015827107 |
| TTC28-AS1 | NR_026962 | 0 | 9.868487622 | 0 | 10.04571785 | 6.865660185 | 0 | 0 | 0 | 0.999953634 | 0.020808337 |
| TTC3 | XM_017028462 | 0 | 9.051724868 | 0 | 0 | 0 | 0 | 0 | 0 | 7.86E-08 | 1 |
| TTC33 | XM_011514003 | 0 | 8.774317692 | 0 | 0 | 0 | 0 | 0 | 0 | 8.46E-08 | 1 |
| TTC38 | NM_017931 | 0 | 0 | 0 | 0 | 0 | 0 | 5.566321721 | 10.03114526 | 1 | 0.02072721 |
| TTC38 | XR_001755259 | 0 | 0 | 0 | 7.249375869 | 0 | 0 | 0 | 0 | 2.27E-05 | 1 |
| TTC38 | XR_244380 | 0 | 9.590514794 | 0 | 0 | 0 | 0 | 0 | 0 | 2.90E-08 | 1 |
| TTC39A | NM_001297663 | 9.270677863 | 0 | 0 | 0 | 0 | 0 | 0 | 0 | 5.05E-08 | 1 |
| TTC39A | XM_011541053 | 0 | 9.10991584 | 0 | 0 | 0 | 0 | 0 | 0 | 5.42E-08 | 1 |
| TTC39A | XM_017000715 | 0 | 8.705217065 | 0 | 0 | 0 | 0 | 0 | 0 | 8.97E-08 | 1 |
| TTC39B | XR_001746190 | 0 | 8.844550112 | 0 | 0 | 0 | 0 | 0 | 0 | 8.06E-08 | 1 |
| TTC7A | NM_020458 | 10.66897426 | 0 | 10.24971315 | 0 | 8.281969086 | 0 | 0 | 0 | 0.999953634 | 0.02072721 |
| TTC7A | XM_017004525 | 0 | 0 | 8.183884528 | 0 | 7.335710674 | 0 | 0 | 0 | 0.072254 | 0.047448988 |
| TTC7A | XM_024453013 | 9.304974408 | 9.014126367 | 0 | 0 | 0 | 0 | 7.73610317 | 0 | 0.002574573 | 1 |
| TTLL1 | XM_006724225 | 10.16602012 | 0 | 9.471566073 | 0 | 10.5916555 | 0 | 0 | 0 | 0.999953634 | 0.02072721 |
| TTLL1 | XM_011530110 | 0 | 0 | 9.542967333 | 0 | 10.01355969 | 0 | 0 | 0 | 0.016497002 | 0.020942118 |
| TTLL1 | XM_017028752 | 0 | 0 | 0 | 10.78156394 | 7.900217873 | 0 | 8.070745625 | 0 | 0.015963133 | 0.999932046 |
| TTLL3 | NR_037162 | 9.395290737 | 0 | 0 | 0 | 0 | 0 | 0 | 0 | 4.12E-08 | 1 |
| TTLL5 | NM_015072 | 0 | 9.036802844 | 0 | 0 | 0 | 0 | 0 | 0 | 6.38E-08 | 1 |
| TTLL6 | XM_017024496 | 9.663265502 | 0 | 0 | 0 | 0 | 0 | 0 | 0 | 2.79E-08 | 1 |
| TTN-AS1 | NR_038271 | 0 | 8.662110092 | 0 | 0 | 0 | 0 | 0 | 0 | 9.15E-08 | 1 |
| TTTY15 | NR_001545 | 0 | 8.540433774 | 0 | 0 | 0 | 0 | 0 | 0 | 1.02E-07 | 1 |

| | | | | | | | | | | | |
|---|---|---|---|---|---|---|---|---|---|---|---|
| TUBA1A | NM_006009 | 0 | 0 | 10.32622296 | 0 | 10.40954811 | 10.62364434 | 0 | 0 | 0.010945384 | 1 |
| TUBB | NR_120608 | 0 | 0 | 0 | 10.26709622 | 8.473070111 | 0 | 0 | 11.59485691 | 0.019039619 | 0.999932046 |
| TUBB2A | NM_001069 | 0 | 0 | 9.4815132 | 0 | 6.945718048 | 0 | 0 | 0 | 0.038677162 | 0.025004616 |
| TUBB6 | NM_001303530 | 8.651779762 | 0 | 0 | 0 | 0 | 0 | 0 | 0 | 1.00E-07 | 1 |
| TUBD1 | NM_001193612 | 0 | 0 | 10.52526804 | 0 | 8.876918096 | 0 | 9.668665083 | 0 | 0.015892489 | 0.999932046 |
| TUBE1 | NM_016262 | 0 | 8.844550112 | 0 | 0 | 0 | 0 | 0 | 0 | 8.06E-08 | 1 |
| TUBG1 | NM_001070 | 0 | 0 | 0 | 10.12921816 | 9.293102358 | 0 | 0 | 10.73309088 | 0.016995481 | 0.999932046 |
| TUBG1 | XM_024450904 | 0 | 9.411213355 | 0 | 0 | 0 | 0 | 0 | 10.60805979 | 3.28E-08 | 1 |
| TUBGCP5 | NM_052903 | 10.38222222 | 0 | 0 | 0 | 0 | 0 | 0 | 0 | 1.47E-08 | 1 |
| TUSC1 | NM_001004125 | 0 | 0 | 10.45548851 | 0 | 9.216502176 | 7.518613818 | 0 | 0 | 0.015250928 | 0.999932046 |
| TUSC3 | NM_001356429 | 0 | 9.456770012 | 0 | 0 | 0 | 0 | 0 | 0 | 3.14E-08 | 1 |
| TUT7 | NM_024617 | 9.825628908 | 0 | 0 | 0 | 0 | 0 | 0 | 0 | 2.30E-08 | 1 |
| TWF2 | NM_007284 | 0 | 0 | 0 | 10.32136409 | 8.668581916 | 0 | 7.834664296 | 0 | 0.017971914 | 0.999932046 |
| TWISTNB | NM_001002926 | 0 | 0 | 0 | 10.70561289 | 7.470373439 | 0 | 0 | 0 | 0.017504135 | 0.021060749 |
| TXK | XM_017008581 | 0 | 0 | 0 | 0 | 0 | 9.199680779 | 7.331132628 | 0 | 1 | 0.022389345 |
| TXK | XR_925166 | 0 | 10.43027645 | 0 | 0 | 0 | 0 | 0 | 0 | 1.34E-08 | 1 |
| TXNDC11 | XM_017023268 | 10.07951011 | 8.261155927 | 10.54753449 | 0 | 7.203547662 | 0 | 0 | 0 | 0.999953634 | 0.02072721 |
| TXNDC12 | NM_015913 | 0 | 0 | 11.54151611 | 11.3940916 | 0 | 11.42314136 | 9.871586458 | 0 | 0.005110104 | 0.999932046 |
| TXNDC5 | NM_030810 | 0 | 0 | 10.11562229 | 9.252214525 | 9.476102537 | 0 | 8.402765142 | 9.957416323 | 0.001850326 | 0.999932046 |
| TXNIP | XM_017000085 | 8.791378415 | 10.19515592 | 10.08984426 | 0 | 9.258253061 | 0 | 0 | 0 | 0.999953634 | 0.02072721 |
| TXNL1 | XM_024451289 | 0 | 9.543782272 | 9.320897208 | 10.74673458 | 0 | 0 | 0 | 0 | 0.999953634 | 0.02072721 |
| TXNRD1 | NM_001093771 | 0 | 0 | 12.37558358 | 12.18529113 | 9.252362115 | 11.19783944 | 0 | 11.80081895 | 0.000160797 | 0.999932046 |
| TYMS | NM_001071 | 0 | 0 | 0 | 10.36335493 | 9.557983985 | 0 | 0 | 11.78183385 | 0.014349865 | 0.999932046 |
| TYMSOS | NM_001012716 | 0 | 0 | 10.16583638 | 8.745727785 | 0 | 0 | 0 | 0 | 0.020731579 | 0.02072721 |
| TYRP1 | XR_001746372 | 9.759152772 | 0 | 0 | 0 | 0 | 0 | 0 | 0 | 2.49E-08 | 1 |
| TYSND1 | NR_073594 | 9.789749415 | 0 | 0 | 0 | 0 | 0 | 0 | 0 | 2.40E-08 | 1 |
| TYSND1 | NR_073595 | 9.998000145 | 0 | 0 | 0 | 0 | 9.180915361 | 8.432122801 | 9.252901113 | 1.94E-08 | 0.004251584 |
| TYW1 | XM_011516372 | 0 | 0 | 0 | 10.61001987 | 6.747781854 | 0 | 0 | 0 | 0.019666085 | 0.02072721 |
| U2AF1 | XM_024452131 | 0 | 0 | 0 | 9.647277147 | 7.944733425 | 0 | 0 | 0 | 0.028597706 | 0.021119768 |
| U2AF1L5 | NM_001320648 | 0 | 0 | 10.53962158 | 10.20558628 | 8.304917924 | 10.64690643 | 0 | 0 | 0.001454799 | 0.999932046 |
| UAP1 | NM_001324116 | 0 | 9.838684641 | 10.96705877 | 11.08392514 | 10.94691409 | 0 | 0 | 0 | 0.893878189 | 0.000757934 |
| UBA2 | NM_005499 | 0 | 0 | 8.894253645 | 9.819727814 | 8.580418909 | 10.80920943 | 0 | 0 | 0.003339676 | 1 |
| UBA2 | XM_006722962 | 0 | 0 | 0 | 10.55370578 | 7.729791854 | 0 | 6.811485137 | 0 | 0.018447224 | 0.999932046 |
| UBA3 | XM_011534210 | 0 | 0 | 0 | 10.45359087 | 9.876021594 | 0 | 0 | 10.14913322 | 0.012583806 | 0.999932046 |
| UBA5 | NM_198329 | 0 | 9.786116244 | 0 | 0 | 0 | 0 | 0 | 10.29016129 | 2.05E-08 | 1 |
| UBAC2 | NM_001144072 | 0 | 9.179501644 | 0 | 10.16131049 | 5.888145156 | 0 | 0 | 0 | 0.999953634 | 0.020942118 |
| UBAP2L | XM_017002990 | 0 | 0 | 0 | 0 | 6.747781854 | 0 | 8.240765774 | 0 | 9.53E-05 | 1 |
| UBB | NM_001281719 | 0 | 9.411213355 | 0 | 0 | 0 | 0 | 0 | 0 | 3.28E-08 | 1 |
| UBE2E1 | NM_003341 | 0 | 9.554298882 | 10.59641196 | 0 | 8.006701649 | 0 | 0 | 0 | 0.999953634 | 0.020942118 |
| UBE2E3 | NM_006357 | 0 | 0 | 10.26515123 | 11.57869941 | 10.54623312 | 0 | 0 | 10.83411964 | 0.000388966 | 0.999932046 |
| UBE2F | NM_080678 | 10.51633798 | 0 | 0 | 10.1019182 | 8.990980475 | 0 | 0 | 0 | 0.999953634 | 0.02072721 |
| UBE2G1 | NM_003342 | 0 | 0 | 0 | 0 | 0 | 0 | 7.402084669 | 8.280943389 | 1 | 0.034593848 |
| UBE2G2 | XM_024452132 | 0 | 8.818611963 | 0 | 0 | 0 | 0 | 0 | 0 | 8.12E-08 | 1 |

| | | | | | | | | | | | |
|---|---|---|---|---|---|---|---|---|---|---|---|
| UBE2H | NM_182697 | 9.629832564 | 0 | 0 | 0 | 0 | 0 | 0 | 9.667347319 | 2.91E-08 | 1 |
| UBE2J2 | NM_058167 | 0 | 0 | 10.87635543 | 10.87564341 | 9.2424901 | 0 | 8.561739689 | 0 | 0.000695881 | 0.999932046 |
| UBE2J2 | XM_005244719 | 10.15666012 | 0 | 0 | 0 | 0 | 0 | 0 | 0 | 1.62E-08 | 1 |
| UBE2L3 | NR_046082 | 10.35792493 | 9.554298882 | 0 | 0 | 0 | 0 | 0 | 0 | 0.00087082 | 1 |
| UBE2M | NM_003969 | 10.82778544 | 0 | 0 | 0 | 0 | 0 | 0 | 0 | 1.34E-08 | 1 |
| UBE2O | XM_024450867 | 0 | 9.346127117 | 0 | 10.0399755 | 6.844931782 | 0 | 0 | 0 | 0.999953634 | 0.020835214 |
| UBE2Q2 | XR_002957692 | 0 | 0 | 0 | 9.595365854 | 6.569741695 | 0 | 0 | 0 | 0.036721706 | 0.025915601 |
| UBE2R2 | XM_005251496 | 0 | 0 | 0 | 10.25846802 | 9.326195217 | 0 | 0 | 0 | 0.015963133 | 0.020903961 |
| UBE2V1 | NM_001282578 | 0 | 8.372314435 | 0 | 0 | 0 | 0 | 0 | 0 | 1.20E-07 | 1 |
| UBE2W | NM_018299 | 0 | 0 | 9.963053877 | 0 | 8.403896394 | 10.53314126 | 8.300353401 | 0 | 0.022674865 | 0.999932046 |
| UBE3B | XM_024449269 | 10.24763479 | 0 | 0 | 0 | 0 | 0 | 0 | 0 | 1.58E-08 | 1 |
| UBE3D | NR_146807 | 9.656640609 | 0 | 0 | 0 | 0 | 0 | 0 | 0 | 3.16E-08 | 1 |
| UBE4B | XM_017000036 | 0 | 0 | 0 | 7.720649682 | 8.680330258 | 0 | 0 | 0 | 0.049945306 | 0.033375792 |
| UBL4A | NM_014235 | 0 | 0 | 8.516155144 | 0 | 9.323388243 | 0 | 7.421731871 | 0 | 0.028915468 | 0.999932046 |
| UBL7 | NM_201265 | 10.87990854 | 0 | 0 | 0 | 0 | 0 | 0 | 0 | 1.34E-08 | 1 |
| UBN2 | NM_173569 | 9.313422649 | 8.861586703 | 0 | 0 | 0 | 0 | 0 | 0 | 0.002813852 | 1 |
| UBQLN1 | NM_013438 | 0 | 8.844550112 | 0 | 0 | 0 | 0 | 7.578739068 | 0 | 8.06E-08 | 1 |
| UBQLN1 | NM_053067 | 0 | 0 | 0 | 8.269377797 | 8.327507425 | 11.49469334 | 8.221278054 | 0 | 0.049053547 | 0.999932046 |
| UBR2 | XR_001743284 | 10.38222222 | 0 | 0 | 0 | 0 | 0 | 0 | 0 | 1.47E-08 | 1 |
| UBR3 | NM_172070 | 0 | 0 | 9.101954491 | 10.31780905 | 5.063498159 | 0 | 7.604755783 | 0 | 0.00466771 | 0.999932046 |
| UBR4 | XM_017000830 | 0 | 0 | 10.21634824 | 9.626353068 | 9.49285483 | 0 | 0 | 11.42962152 | 0.001526324 | 1 |
| UBR5 | NM_001282873 | 0 | 0 | 9.044960079 | 0 | 8.048094872 | 0 | 0 | 0 | 0.038856313 | 0.025156042 |
| UBTF | NM_001076683 | 0 | 0 | 11.2285956 | 10.77466501 | 10.60635568 | 10.21238087 | 11.25072999 | 11.34921454 | 0.000380352 | 0.999932046 |
| UBTF | NM_001076684 | 0 | 0 | 13.17000063 | 10.83394064 | 8.353422435 | 12.64452032 | 9.647962002 | 11.75000919 | 0.000207251 | 0.999932046 |
| UBTF | NM_014233 | 0 | 0 | 11.05683738 | 10.44383889 | 0 | 0 | 0 | 10.48503159 | 0.008967537 | 0.999932046 |
| UBTF | XM_017024996 | 0 | 9.284296589 | 0 | 0 | 0 | 0 | 0 | 0 | 4.07E-08 | 1 |
| UBTF | XM_017025000 | 9.574674104 | 6.952638507 | 0 | 0 | 0 | 0 | 8.543920843 | 0 | 0.004986392 | 1 |
| UBXN1 | NM_015853 | 0 | 0 | 0 | 9.999126067 | 8.712153435 | 0 | 11.34217086 | 0 | 0.020579536 | 0.999932046 |
| UBXN11 | NM_183008 | 0 | 0 | 11.14865646 | 6.939718781 | 0 | 0 | 0 | 0 | 0.015700644 | 1.31E-08 |
| UBXN2B | NM_001077619 | 0 | 8.529815232 | 0 | 0 | 0 | 0 | 0 | 0 | 1.04E-07 | 1 |
| UBXN6 | XM_017027325 | 0 | 0 | 0 | 8.183433365 | 7.71273195 | 0 | 0 | 0 | 0.062785321 | 0.043708568 |
| UBXN7 | XM_011512671 | 0 | 0 | 0 | 0 | 6.631562102 | 0 | 0 | 0 | 3.88E-05 | 1 |
| UCA1 | NR_015379 | 0 | 0 | 0 | 10.67537338 | 8.19864538 | 0 | 11.15530103 | 10.64931 | 0.016246119 | 0.999932046 |
| UCHL1 | NM_004181 | 9.181190369 | 0 | 0 | 0 | 0 | 0 | 0 | 10.69539366 | 7.80E-08 | 1 |
| UCHL5 | NM_001199261 | 0 | 0 | 10.00043084 | 9.996163509 | 7.890136184 | 0 | 0 | 10.54189505 | 0.002320736 | 1 |
| UCKL1 | NR_148434 | 0 | 8.709928179 | 0 | 0 | 0 | 0 | 0 | 0 | 8.96E-08 | 1 |
| UCKL1 | NR_148441 | 0 | 0 | 9.795083168 | 0 | 8.838112924 | 0 | 8.692844709 | 0 | 0.022074318 | 0.999932046 |
| UCKL1 | NR_148442 | 0 | 8.737876298 | 0 | 0 | 0 | 0 | 0 | 0 | 9.11E-08 | 1 |
| UCKL1 | XR_001754304 | 0 | 0 | 0 | 0 | 0 | 0 | 7.630311627 | 9.48050741 | 1 | 0.02072721 |
| UCKL1 | XR_430301 | 11.14186693 | 0 | 0 | 0 | 0 | 0 | 0 | 0 | 1.34E-08 | 1 |
| UCP2 | NM_003355 | 0 | 0 | 0 | 11.13397487 | 6.823901211 | 0 | 0 | 9.587278352 | 2.30E-08 | 0.999932046 |
| UCP2 | XM_024448674 | 0 | 8.458821619 | 0 | 0 | 0 | 0 | 7.166214885 | 0 | 1.10E-07 | 1 |
| UFL1 | NM_015323 | 0 | 0 | 0 | 0 | 6.59478863 | 9.58585217 | 0 | 8.790297784 | 4.02E-05 | 0.999932046 |

| | | | | | | | | | | | |
|---|---|---|---|---|---|---|---|---|---|---|---|
| UFL1-AS1 | XR_002956363 | 0 | 0 | 0 | 0 | 6.736572239 | 9.062888532 | 6.750642936 | 0 | 9.53E-05 | 0.999932046 |
| UFM1 | NM_001286706 | 0 | 0 | 0 | 8.491803396 | 7.203547662 | 0 | 0 | 0 | 0.062748172 | 0.044041714 |
| UGDH | NM_003359 | 0 | 8.747073305 | 9.607977036 | 10.54865946 | 0 | 0 | 0 | 0 | 0.999953634 | 0.02072721 |
| UGGT1 | XM_017004509 | 0 | 0 | 10.82193716 | 10.70107169 | 0 | 0 | 8.207195305 | 0 | 0.009023721 | 0.999932046 |
| UGP2 | NM_006759 | 10.33735985 | 10.27362413 | 7.474332608 | 0 | 8.927086206 | 0 | 0 | 0 | 0.836175819 | 0.030260794 |
| UGP2 | XM_005264537 | 10.11377196 | 0 | 0 | 0 | 0 | 0 | 0 | 0 | 1.70E-08 | 1 |
| UGP2 | XM_005264538 | 0 | 8.622680652 | 0 | 0 | 0 | 0 | 0 | 0 | 9.58E-08 | 1 |
| UGT1A6 | NM_205862 | 10.17531978 | 0 | 9.948784034 | 6.57698191 | 0 | 0 | 0 | 0 | 0.999953634 | 0.022055558 |
| UGT2A3 | XM_011532247 | 8.651779762 | 8.098221372 | 0 | 0 | 0 | 0 | 6.586265919 | 0 | 0.007289974 | 1 |
| UGT2B17 | XM_024454205 | 12.05980185 | 10.13687339 | 13.19187505 | 10.01969536 | 0 | 0 | 0 | 0 | 0.999953634 | 1.92E-09 |
| UGT2B7 | NM_001349568 | 0 | 9.165850403 | 0 | 0 | 0 | 0 | 0 | 0 | 4.97E-08 | 1 |
| UGT8 | NM_001128174 | 0 | 9.445515002 | 0 | 0 | 0 | 0 | 0 | 0 | 3.16E-08 | 1 |
| UGT8 | NM_003360 | 0 | 0 | 10.37950427 | 7.504171232 | 0 | 0 | 0 | 0 | 0.022674865 | 0.02072721 |
| UHRF1 | XM_011527942 | 0 | 0 | 0 | 9.711903927 | 6.291983974 | 0 | 0 | 0 | 0.035188886 | 0.025070991 |
| UHRF1BP1L | XM_005268738 | 0 | 9.803852726 | 0 | 0 | 0 | 0 | 0 | 0 | 2.02E-08 | 1 |
| ULK3 | XM_024449897 | 0 | 8.607612286 | 0 | 0 | 0 | 0 | 0 | 0 | 9.73E-08 | 1 |
| ULK4 | XM_024453613 | 0 | 0 | 11.64064966 | 0 | 7.313250256 | 0 | 0 | 0 | 1.60E-08 | 6.56E-09 |
| UMAD1 | NM_001302350 | 0 | 0 | 10.92333233 | 0 | 9.087887617 | 0 | 0 | 0 | 0.012459054 | 0.02072721 |
| UNC119 | NM_005148 | 0 | 0 | 10.58492377 | 0 | 7.170608406 | 0 | 8.903137858 | 11.30139613 | 0.019666085 | 0.999932046 |
| UNC45A | NM_001323619 | 0 | 0 | 10.40150073 | 0 | 7.683842675 | 0 | 0 | 9.91088868 | 0.020505868 | 0.999932046 |
| UNC5B | XR_945617 | 9.395290737 | 0 | 0 | 0 | 0 | 0 | 0 | 0 | 4.12E-08 | 1 |
| UNK | NM_001080419 | 0 | 0 | 8.456887945 | 8.064023368 | 9.685874143 | 0 | 6.340064421 | 0 | 0.004830709 | 0.858016099 |
| UNK | XM_017025248 | 5.623309261 | 9.081113725 | 0 | 0 | 0 | 0 | 0 | 0 | 0.010559366 | 1 |
| UPF2 | NM_080599 | 0 | 0 | 10.24971315 | 0 | 7.503816758 | 0 | 0 | 0 | 0.02272315 | 0.02072721 |
| UPF3A | XR_001749649 | 0 | 0 | 10.86690015 | 0 | 9.338296057 | 0 | 8.402765142 | 0 | 0.012139547 | 0.999932046 |
| UPP1 | XM_024446913 | 0 | 9.131146246 | 0 | 0 | 0 | 0 | 0 | 0 | 5.23E-08 | 1 |
| UQCC1 | NM_001184977 | 0 | 9.684464992 | 0 | 0 | 0 | 0 | 10.00618716 | 0 | 2.36E-08 | 1 |
| URB1-AS1 | NR_026845 | 0 | 8.737876298 | 0 | 0 | 0 | 0 | 0 | 0 | 9.11E-08 | 1 |
| URGCP | NM_001077663 | 0 | 0 | 0 | 0 | 6.690841957 | 0 | 0 | 0 | 3.72E-05 | 1 |
| URM1 | NM_001265582 | 0 | 0 | 0 | 8.745727785 | 0 | 0 | 0 | 0 | 9.20E-06 | 1 |
| UROD | XM_005271170 | 0 | 9.014126367 | 0 | 0 | 0 | 0 | 8.944472497 | 0 | 6.71E-08 | 1 |
| UROS | NR_136676 | 0 | 8.690990708 | 0 | 0 | 0 | 0 | 0 | 0 | 9.02E-08 | 1 |
| USF1 | NM_207005 | 9.609394475 | 0 | 0 | 0 | 0 | 0 | 0 | 0 | 2.97E-08 | 1 |
| USF2 | NM_003367 | 0 | 0 | 9.474889408 | 0 | 8.700662425 | 0 | 0 | 10.66402238 | 0.026279362 | 0.999932046 |
| USH1C | NM_005709 | 0 | 9.99739751 | 0 | 11.57125375 | 7.949595864 | 0 | 0 | 0 | 0.999953634 | 7.83E-09 |
| USHBP1 | NR_135632 | 9.740478394 | 0 | 0 | 0 | 0 | 0 | 0 | 0 | 4.01E-08 | 1 |
| USO1 | NM_003715 | 0 | 0 | 10.38216162 | 0 | 7.343120441 | 0 | 0 | 0 | 0.021608743 | 0.02072721 |
| USP10 | XM_006721332 | 0 | 0 | 9.760097945 | 9.170475661 | 0 | 0 | 0 | 0 | 0.022115815 | 0.02072721 |
| USP11 | NM_004651 | 0 | 0 | 10.14512582 | 0 | 8.17807637 | 0 | 0 | 0 | 0.021704832 | 0.02072721 |
| USP13 | NM_003940 | 0 | 0 | 0 | 11.57175132 | 11.7055517 | 9.553392305 | 10.32978581 | 12.00673505 | 0.003900528 | 0.999932046 |
| USP13 | XM_017007426 | 9.900435696 | 0 | 9.795083168 | 9.551641627 | 10.57365779 | 0 | 0 | 0 | 0.999953634 | 0.002186952 |
| USP24 | XM_017000834 | 0 | 0 | 0 | 0 | 6.518304288 | 0 | 0 | 9.347078824 | 4.23E-05 | 1 |
| USP3 | XM_017022768 | 0 | 0 | 9.79906587 | 0 | 8.800928156 | 0 | 0 | 0 | 0.022267881 | 0.02072721 |

| | | | | | | | | | | | |
|---|---|---|---|---|---|---|---|---|---|---|---|
| USP32 | XM_017025234 | 0 | 9.117027412 | 10.2262418 | 0 | 6.569741695 | 0 | 0 | 0 | 0.999953634 | 0.02072721 |
| USP33 | XM_017000726 | 0 | 8.792198922 | 0 | 0 | 0 | 0 | 0 | 0 | 8.40E-08 | 1 |
| USP42 | NM_032172 | 0 | 0 | 0 | 0 | 6.464964927 | 0 | 10.00618716 | 0 | 4.45E-05 | 1 |
| USP47 | XM_017017954 | 0 | 9.219697032 | 0 | 0 | 0 | 0 | 0 | 0 | 4.50E-08 | 1 |
| USP5 | NM_001098536 | 0 | 0 | 0 | 0 | 6.366595318 | 0 | 0 | 0 | 4.76E-05 | 1 |
| USP53 | XM_005263077 | 10.33735985 | 0 | 0 | 0 | 0 | 0 | 9.652126454 | 0 | 1.58E-08 | 1 |
| USP53 | XM_006714240 | 0 | 9.245884974 | 0 | 0 | 0 | 0 | 0 | 0 | 4.33E-08 | 1 |
| USP53 | XM_011532038 | 0 | 0 | 0 | 11.37312061 | 7.636381755 | 0 | 12.02814808 | 0 | 0.011617501 | 1 |
| USP54 | XR_002956962 | 10.59120384 | 7.93074915 | 0 | 0 | 0 | 0 | 0 | 0 | 0.00135572 | 1 |
| USPL1 | XM_017020315 | 0 | 0 | 0 | 0 | 6.351978551 | 0 | 0 | 0 | 4.81E-05 | 1 |
| UTP14C | NM_021645 | 0 | 0 | 0 | 0 | 6.569741695 | 0 | 0 | 0 | 4.12E-05 | 1 |
| UTP18 | XM_011524870 | 10.80998402 | 0 | 0 | 0 | 0 | 0 | 0 | 0 | 1.34E-08 | 1 |
| UTP3 | NM_020368 | 0 | 0 | 0 | 0 | 0 | 0 | 8.107603982 | 10.37308625 | 1 | 0.02072721 |
| UTP4 | NM_032830 | 0 | 0 | 9.979525575 | 0 | 8.822293806 | 0 | 0 | 0 | 0.020536039 | 0.02072721 |
| UTP4 | XR_001752011 | 10.47174593 | 0 | 0 | 0 | 0 | 0 | 0 | 0 | 1.45E-08 | 1 |
| UTS2 | XM_011540538 | 10.29532878 | 0 | 0 | 0 | 0 | 0 | 0 | 0 | 1.54E-08 | 1 |
| UVRAG | XR_001747951 | 0 | 0 | 0 | 0 | 0 | 0 | 8.790529686 | 10.49329345 | 1 | 0.02072721 |
| UXS1 | NM_025076 | 0 | 9.051724868 | 0 | 0 | 0 | 10.51452622 | 9.181891152 | 10.26464677 | 7.86E-08 | 0.001535028 |
| UXS1 | XM_024453157 | 9.502621339 | 8.407542174 | 0 | 0 | 0 | 0 | 0 | 0 | 0.003096844 | 1 |
| VAMP5 | NM_006634 | 0 | 9.340061955 | 0 | 0 | 0 | 10.5569872 | 0 | 10.06109606 | 3.95E-08 | 0.02072721 |
| VAMP7 | XM_017030063 | 0 | 0 | 0 | 9.211923901 | 6.780897397 | 0 | 0 | 0 | 0.045711879 | 0.03159928 |
| VAMP8 | XM_017005170 | 10.70152235 | 8.886770211 | 3.639350893 | 0 | 0 | 0 | 8.827823966 | 0 | 0.039634865 | 1 |
| VAPA | XM_017026079 | 0 | 0 | 4.248700506 | 10.29508808 | 6.464964927 | 0 | 0 | 0 | 0.009435887 | 0.016659858 |
| VAPB | NM_004738 | 9.426779986 | 0 | 0 | 0 | 0 | 0 | 0 | 0 | 3.96E-08 | 1 |
| VASP | XM_005259199 | 0 | 0 | 0 | 9.16520989 | 7.672123016 | 0 | 6.742854027 | 0 | 0.039409365 | 0.999932046 |
| VAV1 | NM_005428 | 0 | 8.809861237 | 0 | 0 | 0 | 0 | 0 | 0 | 8.24E-08 | 1 |
| VAV3 | XR_001736913 | 9.721559119 | 8.991087758 | 0 | 0 | 0 | 0 | 9.837250547 | 0 | 0.001918957 | 1 |
| VBP1 | NM_003372 | 0 | 0 | 0 | 9.975253495 | 10.81824088 | 0 | 10.03836054 | 10.16128261 | 0.010273035 | 0.999932046 |
| VDAC1 | NR_036625 | 0 | 0 | 10.28232386 | 11.65816166 | 11.14273184 | 11.6596472 | 9.959063757 | 0 | 0.000285675 | 0.999932046 |
| VDAC2 | NM_001324088 | 0 | 0 | 0 | 10.52213019 | 8.86667168 | 0 | 0 | 0 | 0.015691656 | 0.020781046 |
| VEGFB | NM_003377 | 0 | 0 | 10.2747168 | 8.264461514 | 0 | 0 | 0 | 0 | 0.021564719 | 0.02072721 |
| VEZF1 | NM_007146 | 0 | 8.818611963 | 0 | 0 | 0 | 0 | 0 | 0 | 8.12E-08 | 1 |
| VEZF1 | XM_017025018 | 0 | 0 | 10.03686385 | 0 | 7.959271851 | 0 | 0 | 8.188991154 | 0.023850113 | 0.999932046 |
| VGF | NM_003378 | 0 | 0 | 0 | 9.222102536 | 9.263143875 | 8.6974259 | 9.584008175 | 0 | 0.023918092 | 0.999932046 |
| VIL1 | NM_007127 | 0 | 0 | 0 | 9.645387449 | 9.417644228 | 0 | 10.04471001 | 0 | 0.019641255 | 1 |
| VILL | XM_011533770 | 0 | 0 | 0 | 0 | 6.679179249 | 0 | 0 | 0 | 3.78E-05 | 1 |
| VKORC1L1 | XM_011515831 | 0 | 0 | 10.13465797 | 0 | 8.885824399 | 9.272392864 | 0 | 0 | 0.018905382 | 0.999932046 |
| VMAC | NM_001017921 | 9.765324233 | 0 | 0 | 0 | 0 | 0 | 0 | 10.18867435 | 2.50E-08 | 1 |
| VMP1 | NM_001329398 | 0 | 0 | 9.696043232 | 11.15205523 | 9.681496598 | 0 | 8.512198274 | 10.22550845 | 0.000831581 | 0.999932046 |
| VPREB3 | NM_013378 | 0 | 10.58188946 | 0 | 0 | 0 | 0 | 0 | 0 | 1.34E-08 | 1 |
| VPS13A | XR_001746260 | 0 | 8.602554319 | 0 | 0 | 0 | 0 | 0 | 0 | 9.73E-08 | 1 |
| VPS13B | XM_024447074 | 0 | 0 | 0 | 0 | 6.667421491 | 0 | 0 | 0 | 3.78E-05 | 1 |
| VPS13C | XR_001751332 | 10.8598616 | 9.051724868 | 0 | 0 | 0 | 0 | 0 | 0 | 0.000722101 | 1 |

| | | | | | | | | | | | |
|---|---|---|---|---|---|---|---|---|---|---|---|
| VPS13D | NM_018156 | 0 | 0 | 10.1575878 | 0 | 6.758905041 | 0 | 0 | 0 | 0.026155876 | 0.02072721 |
| VPS16 | NM_022575 | 10.07951011 | 0 | 10.25165197 | 9.618668438 | 7.587306227 | 0 | 0 | 0 | 0.999953634 | 0.003932411 |
| VPS18 | XM_011521844 | 0 | 0 | 9.44470014 | 0 | 7.227767768 | 0 | 8.007141921 | 0 | 0.037715864 | 0.999932046 |
| VPS29 | NR_104099 | 0 | 0 | 0 | 10.02115343 | 7.587306227 | 0 | 0 | 9.91088868 | 0.024847467 | 0.999932046 |
| VPS29 | NR_104102 | 0 | 9.393751769 | 0 | 0 | 0 | 0 | 0 | 0 | 3.37E-08 | 1 |
| VPS29 | XM_017019457 | 8.678186754 | 7.86468487 | 0 | 0 | 0 | 0 | 0 | 0 | 0.00811514 | 1 |
| VPS33A | NM_022916 | 0 | 0 | 9.071545242 | 10.14535356 | 0 | 0 | 0 | 0 | 0.01914756 | 0.02072721 |
| VPS33B | XR_001751213 | 0 | 0 | 0 | 0 | 0 | 0 | 7.372102888 | 9.770654157 | 1 | 0.02072721 |
| VPS37C | XM_017017955 | 0 | 0 | 0 | 9.451378274 | 7.683842675 | 0 | 0 | 0 | 0.033522361 | 0.023763327 |
| VPS45 | NM_007259 | 0 | 0 | 0 | 0 | 0 | 11.11747083 | 9.010835996 | 0 | 1 | 0.02072721 |
| VPS52 | NM_001289176 | 0 | 9.684464992 | 0 | 0 | 0 | 0 | 9.243908171 | 0 | 2.36E-08 | 1 |
| VPS53 | NM_001128159 | 0 | 8.747073305 | 0 | 0 | 0 | 0 | 0 | 0 | 8.61E-08 | 1 |
| VPS54 | XR_244939 | 8.638392769 | 0 | 0 | 0 | 0 | 0 | 0 | 0 | 1.01E-07 | 1 |
| VPS72 | NM_005997 | 0 | 0 | 9.743660257 | 10.19658109 | 8.230957729 | 12.06244979 | 8.441777415 | 0 | 0.002153909 | 0.999932046 |
| VPS9D1 | XR_933495 | 10.71433872 | 0 | 0 | 0 | 0 | 0 | 0 | 0 | 1.34E-08 | 1 |
| VRK2 | XM_017004864 | 0 | 0 | 0 | 8.60502903 | 0 | 0 | 0 | 0 | 1.54E-05 | 1 |
| VSIG1 | NM_001170553 | 0 | 9.340061955 | 0 | 0 | 0 | 0 | 0 | 0 | 3.95E-08 | 1 |
| VSIG2 | NM_001329920 | 0 | 9.777165557 | 0 | 0 | 0 | 0 | 0 | 0 | 4.32E-08 | 1 |
| VSIR | NM_022153 | 10.34562109 | 9.073822446 | 7.320277675 | 0 | 9.250393109 | 0 | 0 | 0 | 0.986247317 | 0.026224312 |
| VTI1A | XM_005269544 | 0 | 0 | 9.993496019 | 10.24978792 | 9.986016488 | 0 | 11.11060574 | 10.71554288 | 0.001069465 | 0.999932046 |
| VTI1A | XR_001747028 | 0 | 9.772669303 | 0 | 0 | 0 | 0 | 0 | 0 | 4.12E-08 | 1 |
| VWA5A | NM_001130142 | 0 | 9.893551993 | 9.053876344 | 11.67770845 | 0 | 0 | 0 | 0 | 0.999953634 | 0.02072721 |
| WARS | XM_024449708 | 0 | 0 | 0 | 9.873992335 | 7.259441431 | 0 | 0 | 0 | 0.028265655 | 0.021060749 |
| WASH7P | NR_024540 | 9.509990722 | 0 | 9.237263703 | 9.869142678 | 7.869758957 | 0 | 0 | 0 | 0.999953634 | 0.005205109 |
| WASH8P | NR_130745 | 0 | 8.469973695 | 0 | 0 | 0 | 0 | 8.660134917 | 0 | 1.09E-07 | 1 |
| WBP4 | XM_017020380 | 0 | 10.33132512 | 0 | 0 | 0 | 0 | 0 | 0 | 1.45E-08 | 1 |
| WDFY2 | XR_941483 | 0 | 0 | 0 | 9.885245203 | 8.157209854 | 0 | 0 | 0 | 0.024330175 | 0.02072721 |
| WDR1 | NM_017491 | 0 | 0 | 8.980953887 | 11.00506786 | 0 | 10.77833282 | 0 | 9.696263442 | 0.012900615 | 0.999932046 |
| WDR11-AS1 | NR_033850 | 0 | 9.265218616 | 0 | 0 | 0 | 0 | 0 | 0 | 4.18E-08 | 1 |
| WDR13 | XM_024452424 | 9.966206167 | 8.919678654 | 0 | 0 | 0 | 0 | 0 | 0 | 0.001643707 | 1 |
| WDR18 | NM_024100 | 0 | 0 | 10.39625208 | 11.12315268 | 10.28619738 | 0 | 8.095422193 | 0 | 0.000529502 | 0.910706866 |
| WDR20 | NM_001320130 | 9.279328737 | 0 | 0 | 0 | 0 | 0 | 0 | 0 | 4.99E-08 | 1 |
| WDR34 | NM_052844 | 0 | 9.630820909 | 0 | 0 | 0 | 0 | 0 | 0 | 4.30E-08 | 1 |
| WDR36 | NM_139281 | 0 | 0 | 9.448085869 | 9.99022008 | 0 | 0 | 0 | 0 | 0.018674369 | 0.02072721 |
| WDR41 | XR_001742140 | 0 | 9.232850423 | 0 | 0 | 0 | 0 | 0 | 0 | 4.38E-08 | 1 |
| WDR41 | XR_001742141 | 0 | 0 | 0 | 0 | 6.337212177 | 0 | 0 | 0 | 4.86E-05 | 1 |
| WDR45 | NM_007075 | 8.08843558 | 9.309347676 | 0 | 0 | 0 | 0 | 0 | 0 | 0.004109168 | 1 |
| WDR45B | NM_019613 | 0 | 9.808252985 | 0 | 9.55567177 | 7.497189827 | 0 | 0 | 0 | 0.999953634 | 0.023358895 |
| WDR46 | XR_002956316 | 0 | 0 | 0 | 0 | 0 | 10.83569479 | 0 | 9.073508739 | 1 | 0.02072721 |
| WDR47 | NM_014969 | 9.509990722 | 0 | 0 | 0 | 0 | 0 | 0 | 0 | 3.47E-08 | 1 |
| WDR59 | XR_933430 | 0 | 0 | 0 | 9.291410442 | 9.609586778 | 10.96950906 | 9.734951004 | 9.429755563 | 0.020617455 | 0.999932046 |
| WDR5B | NM_019069 | 0 | 8.360378074 | 0 | 0 | 0 | 0 | 8.60104519 | 0 | 1.21E-07 | 1 |
| WDR61 | NM_025234 | 0 | 0 | 0 | 12.24297535 | 8.194554972 | 12.06052622 | 8.125685946 | 10.76528678 | 1.34E-08 | 0.999932046 |

| | | | | | | | | | | | |
|---|---|---|---|---|---|---|---|---|---|---|---|
| WDR61 | XM_011522094 | 0 | 0 | 10.31976886 | 5.796569454 | 0 | 0 | 0 | 0 | 0.027486381 | 0.02072721 |
| WDR74 | XM_024448585 | 0 | 0 | 0 | 0 | 0 | 0 | 7.142729632 | 7.969679987 | 1 | 0.044457133 |
| WDR77 | NM_024102 | 0 | 0 | 0 | 11.18560087 | 9.759714287 | 10.60088363 | 9.732986449 | 0 | 0.00912758 | 0.999932046 |
| WDR90 | XM_017023024 | 0 | 0 | 0 | 0 | 6.758905041 | 0 | 0 | 0 | 9.53E-05 | 1 |
| WDYHV1 | NM_001283027 | 0 | 0 | 11.93477455 | 13.05185323 | 8.864098664 | 13.77624715 | 13.39940404 | 0 | 0.000129545 | 0.999932046 |
| WFDC2 | NM_006103 | 10.64574565 | 13.44622321 | 10.26130717 | 9.768560191 | 8.771023979 | 0 | 0 | 0 | 0.540170928 | 0.002940381 |
| WFDC3 | XM_011528553 | 0 | 0 | 0 | 8.91213928 | 8.798235064 | 8.650456053 | 0 | 0 | 0.031596444 | 1 |
| WFS1 | XM_017008586 | 0 | 0 | 6.584403084 | 9.394011718 | 0 | 9.501401201 | 10.46966706 | 0 | 0.044274191 | 0.999932046 |
| WHRN | XM_011518486 | 0 | 0 | 0 | 0 | 6.667421491 | 0 | 0 | 0 | 3.78E-05 | 1 |
| WIPF2 | XM_011524413 | 0 | 9.564739385 | 0 | 0 | 0 | 0 | 0 | 0 | 2.69E-08 | 1 |
| WLS | NM_001193334 | 0 | 0 | 0 | 9.950979243 | 7.153852394 | 0 | 9.052390927 | 0 | 0.027454051 | 0.999932046 |
| WNK1 | XM_017019839 | 0 | 9.38199218 | 9.629011857 | 0 | 8.548724137 | 0 | 0 | 0 | 0.999953634 | 0.02072721 |
| WNK2 | XM_017015055 | 0 | 0 | 7.427075936 | 0 | 8.364387931 | 0 | 0 | 0 | 0.062629642 | 0.041497342 |
| WNK2 | XR_001746367 | 0 | 8.632639499 | 0 | 0 | 0 | 0 | 0 | 0 | 9.44E-08 | 1 |
| WNT2 | NM_003391 | 0 | 0 | 9.514183525 | 12.04827372 | 13.68860035 | 15.56646044 | 0 | 0 | 6.52E-05 | 0.999932046 |
| WRAP53 | XM_011523952 | 0 | 8.419096158 | 0 | 0 | 0 | 0 | 0 | 0 | 1.14E-07 | 1 |
| WRAP53 | XM_024450824 | 9.883513662 | 0 | 0 | 0 | 0 | 0 | 0 | 0 | 2.12E-08 | 1 |
| WTAP | NM_001270531 | 0 | 8.737876298 | 0 | 9.551641627 | 8.700662425 | 0 | 0 | 0 | 0.999953634 | 0.02072721 |
| WTAP | NM_004906 | 0 | 0 | 9.461549885 | 0 | 9.096681855 | 0 | 0 | 9.945422596 | 0.023411965 | 0.999932046 |
| WTAP | NM_152858 | 0 | 0 | 0 | 10.53647577 | 7.136899482 | 0 | 0 | 0 | 0.019785074 | 0.02072721 |
| WWC3 | NM_015691 | 0 | 9.905922754 | 0 | 9.682718985 | 7.718441029 | 0 | 0 | 0 | 0.999953634 | 0.021464771 |
| WWP2 | NM_001270454 | 0 | 0 | 9.929535327 | 0 | 7.672123016 | 9.95512224 | 10.08300232 | 11.4151997 | 0.026419775 | 0.999932046 |
| WWP2 | NM_007014 | 10.08938252 | 10.44730408 | 10.77485373 | 10.73700724 | 10.70849582 | 0 | 0 | 0 | 0.999953634 | 0.001042882 |
| WWTR1 | NM_015472 | 0 | 0 | 10.19030288 | 0 | 9.563545334 | 11.3676962 | 10.92860662 | 11.59293607 | 0.015512397 | 0.999932046 |
| XK | NM_021083 | 0 | 9.309347676 | 0 | 0 | 0 | 0 | 6.551017323 | 0 | 3.94E-08 | 1 |
| XKR9 | XM_011517525 | 0 | 0 | 0 | 10.11016257 | 10.12897419 | 0 | 11.98711537 | 12.68575729 | 0.013190416 | 0.999932046 |
| XKR9 | XM_017013404 | 0 | 0 | 6.055099627 | 0 | 10.51500038 | 0 | 9.984883005 | 0 | 0.020993955 | 1 |
| XKR9 | XM_017013405 | 0 | 0 | 0 | 9.206807527 | 9.548400083 | 0 | 9.311290442 | 11.20994454 | 0.021649437 | 0.999932046 |
| XPC | NR_148950 | 0 | 0 | 9.948784034 | 0 | 9.450603618 | 0 | 9.176118775 | 0 | 0.017612901 | 0.999932046 |
| XRCC1 | NM_006297 | 0 | 7.606261038 | 0 | 10.1019182 | 6.79176912 | 0 | 0 | 0 | 0.983589543 | 0.02072721 |
| XRN1 | XR_001740178 | 0 | 0 | 0 | 10.26832462 | 6.896204849 | 0 | 0 | 0 | 0.023573664 | 0.02072721 |
| XRRA1 | XM_017017232 | 8.754625855 | 0 | 0 | 0 | 0 | 0 | 0 | 0 | 9.15E-08 | 1 |
| XRRA1 | XM_017017234 | 0 | 0 | 10.46052638 | 10.37936023 | 0 | 0 | 0 | 0 | 0.011721841 | 0.02072721 |
| YAF2 | XR_002957280 | 0 | 0 | 0 | 0 | 0 | 10.12672544 | 0 | 9.537545985 | 1 | 0.02072721 |
| YAP1 | NM_001282097 | 0 | 0 | 8.05658827 | 10.95025027 | 10.5514401 | 12.08028052 | 10.35491349 | 12.08671745 | 0.001023139 | 0.999932046 |
| YAP1 | NM_001282101 | 0 | 0 | 0 | 10.78156394 | 10.051936 | 11.72385023 | 10.16543342 | 12.38231413 | 0.010308615 | 0.999932046 |
| YAP1 | NM_006106 | 0 | 0 | 8.856532107 | 8.688149458 | 8.73486447 | 0 | 8.039294254 | 0 | 0.005511385 | 1 |
| YARS2 | NM_001040436 | 0 | 0 | 0 | 10.32136409 | 9.454884858 | 0 | 0 | 0 | 0.015063156 | 0.02072721 |
| YBEY | NM_001314022 | 0 | 0 | 0 | 0 | 6.518304288 | 0 | 0 | 0 | 4.23E-05 | 1 |
| YBEY | NM_001314025 | 11.1203612 | 0 | 0 | 0 | 0 | 0 | 0 | 0 | 1.34E-08 | 1 |
| YBX2 | NM_015982 | 0 | 0 | 0 | 0 | 6.725274845 | 0 | 0 | 0 | 9.53E-05 | 1 |
| YBX3 | XR_001748906 | 9.363098816 | 0 | 6.253900611 | 9.527221444 | 6.875913762 | 0 | 0 | 0 | 0.999953634 | 0.020465354 |
| YEATS2 | NM_018023 | 0 | 0 | 0 | 0 | 0 | 0 | 8.54839623 | 8.619662358 | 1 | 0.021237439 |

| | | | | | | | | | | | |
|---|---|---|---|---|---|---|---|---|---|---|---|
| YIF1B | NM_001145463 | 0 | 0 | 9.294808705 | 0 | 7.801472153 | 0 | 0 | 0 | 0.036425639 | 0.023849651 |
| YIPF2 | XM_011528270 | 0 | 8.612652582 | 0 | 0 | 0 | 0 | 8.696881776 | 0 | 9.67E-08 | 1 |
| YIPF5 | NM_030799 | 0 | 0 | 9.668732844 | 0 | 5.44033412 | 10.07303422 | 9.049236545 | 11.45028035 | 0.040662567 | 0.999932046 |
| YIPF6 | NM_173834 | 0 | 0 | 9.624530142 | 0 | 9.587919688 | 0 | 0 | 0 | 0.01881306 | 0.02072721 |
| YIPF6 | XM_011530924 | 0 | 0 | 10.41107393 | 0 | 6.544252221 | 9.005387706 | 7.892131936 | 0 | 0.023001521 | 0.999932046 |
| YJU2 | NM_018074 | 0 | 0 | 9.620034462 | 0 | 9.924369085 | 0 | 0 | 0 | 0.016708288 | 0.02096331 |
| YME1L1 | XM_011519300 | 9.07588576 | 8.700490516 | 0 | 0 | 0 | 0 | 4.961135439 | 9.784185253 | 0.003691525 | 0.021326758 |
| YPEL3 | NM_001145524 | 0 | 0 | 11.79086793 | 0 | 8.8249424 | 11.13219827 | 9.313921358 | 0 | 0.007808446 | 0.999932046 |
| YPEL3 | NM_031477 | 0 | 0 | 9.008733536 | 0 | 7.752229049 | 10.4813605 | 8.083136668 | 10.53657502 | 0.043174099 | 0.999932046 |
| YPEL5 | XM_017004318 | 0 | 0 | 0 | 0 | 0 | 11.89156544 | 9.740828669 | 0 | 1 | 0.019697931 |
| YPEL5 | XM_017004321 | 0 | 9.981899769 | 0 | 0 | 0 | 0 | 8.867763981 | 10.8276001 | 1.70E-08 | 0.02072721 |
| YRDC | NM_024640 | 0 | 0 | 0 | 9.891636224 | 6.813269833 | 0 | 0 | 0 | 0.029658808 | 0.021893363 |
| YWHAB | NM_003404 | 0 | 9.580259825 | 0 | 0 | 0 | 0 | 0 | 0 | 2.61E-08 | 1 |
| YWHAH | NM_003405 | 10.70473713 | 10.4696994 | 12.33272512 | 11.68784252 | 10.325071 | 0 | 0 | 0 | 0.999953634 | 0.000354166 |
| YWHAZ | NM_003406 | 0 | 0 | 9.877750519 | 11.64639967 | 6.906244365 | 10.95715511 | 9.303368783 | 10.39088417 | 0.001100465 | 0.999932046 |
| YWHAZ | NM_145690 | 0 | 0 | 9.665827761 | 10.29388227 | 10.15341459 | 8.421828153 | 9.514802225 | 0 | 0.001101043 | 0.999932046 |
| YWHAZ | XM_005251063 | 0 | 0 | 10.30303877 | 10.15069217 | 0 | 0 | 0 | 0 | 0.013495105 | 0.02072721 |
| ZADH2 | NM_175907 | 0 | 0 | 0 | 0 | 0 | 9.431678452 | 7.431456038 | 0 | 1 | 0.02074727 |
| ZBED1 | NM_004729 | 0 | 0 | 0 | 9.963167422 | 9.669757788 | 0 | 0 | 0 | 0.016251236 | 0.02096331 |
| ZBED5 | NM_001143667 | 0 | 0 | 0 | 0 | 0 | 8.551663808 | 8.402765142 | 0 | 1 | 0.022291204 |
| ZBED8 | NM_022090 | 0 | 9.506358431 | 0 | 0 | 0 | 0 | 0 | 0 | 2.91E-08 | 1 |
| ZBP1 | NR_136660 | 0 | 9.239382419 | 0 | 0 | 0 | 0 | 0 | 0 | 4.33E-08 | 1 |
| ZBTB16 | XR_001747955 | 0 | 9.422738093 | 0 | 0 | 0 | 0 | 0 | 0 | 3.26E-08 | 1 |
| ZBTB38 | XM_005247257 | 11.09608384 | 0 | 0 | 0 | 0 | 0 | 0 | 0 | 1.34E-08 | 1 |
| ZBTB38 | XM_017006050 | 10.96280211 | 10.15080026 | 0 | 0 | 0 | 0 | 10.21342682 | 0 | 0.000392062 | 1 |
| ZBTB38 | XM_017006057 | 11.25583468 | 10.79419952 | 0 | 0 | 0 | 0 | 10.73213854 | 0 | 0.000213965 | 1 |
| ZBTB6 | NM_006626 | 0 | 9.036802844 | 0 | 0 | 0 | 0 | 0 | 0 | 6.38E-08 | 1 |
| ZBTB7C | NM_001318841 | 0 | 0 | 7.794637209 | 0 | 0 | 0 | 0 | 0 | 2.93E-05 | 1 |
| ZBTB8OS | XM_011541329 | 0 | 9.315543066 | 0 | 0 | 0 | 0 | 0 | 0 | 3.90E-08 | 1 |
| ZC2HC1A | XM_011517539 | 0 | 0 | 0 | 0 | 0 | 10.7736438 | 9.12009761 | 0 | 1 | 0.02072721 |
| ZC3H11A | NM_001350261 | 0 | 0 | 0 | 0 | 6.478485785 | 0 | 0 | 0 | 4.36E-05 | 1 |
| ZC3H11A | NM_014827 | 0 | 8.098221372 | 0 | 0 | 0 | 9.885840531 | 0 | 10.98039483 | 0.057926468 | 0.02072721 |
| ZC3H12A | NM_025079 | 0 | 0 | 10.69089279 | 7.870531406 | 0 | 8.636750353 | 0 | 0 | 0.018491109 | 0.999932046 |
| ZC3H12D | NM_207360 | 0 | 0 | 8.38097934 | 7.249375869 | 0 | 0 | 0 | 0 | 0.071612221 | 0.048305709 |
| ZC3H13 | XM_005266302 | 0 | 0 | 0 | 10.7343429 | 8.025245409 | 0 | 9.931061323 | 0 | 0.016122834 | 0.999932046 |
| ZC3H13 | XM_017020477 | 0 | 0 | 0 | 11.47239163 | 10.17329679 | 11.2628559 | 9.651086467 | 0 | 0.007041034 | 0.999932046 |
| ZC3H13 | XM_017020479 | 0 | 0 | 10.53723922 | 0 | 9.42114931 | 11.21177273 | 10.77525463 | 10.56820483 | 0.013944793 | 0.999932046 |
| ZC3H14 | NM_024824 | 0 | 8.464408432 | 0 | 0 | 0 | 0 | 0 | 0 | 1.11E-07 | 1 |
| ZC3H7A | XR_932825 | 0 | 0 | 0 | 8.204121028 | 0 | 0 | 0 | 0 | 8.47E-06 | 1 |
| ZC3H8 | XR_001738995 | 0 | 9.942411034 | 0 | 0 | 0 | 0 | 0 | 0 | 5.01E-08 | 1 |
| ZC3HC1 | NM_016478 | 10.39023122 | 0 | 0 | 0 | 0 | 0 | 0 | 0 | 1.47E-08 | 1 |
| ZC3HC1 | XM_017012287 | 0 | 9.655452119 | 0 | 0 | 0 | 0 | 9.347693707 | 0 | 2.46E-08 | 1 |
| ZCCHC10 | XM_011543503 | 9.517322653 | 9.376076245 | 0 | 0 | 0 | 0 | 6.458933278 | 0 | 0.001773798 | 1 |

| | | | | | | | | | | | |
|---|---|---|---|---|---|---|---|---|---|---|---|
| ZCCHC17 | NM_016505 | 0 | 9.284296589 | 0 | 0 | 0 | 9.932863413 | 0 | 0 | 4.07E-08 | 1 |
| ZCCHC18 | XM_011531012 | 0 | 0 | 0 | 8.710011504 | 0 | 0 | 0 | 0 | 4.92E-06 | 1 |
| ZCCHC3 | NM_033089 | 0 | 0 | 8.074224188 | 0 | 0 | 0 | 0 | 0 | 2.07E-05 | 1 |
| ZCCHC7 | NM_001289119 | 11.14660276 | 0 | 0 | 0 | 0 | 0 | 0 | 0 | 1.34E-08 | 1 |
| ZCCHC7 | XM_024447691 | 0 | 9.232850423 | 0 | 0 | 0 | 0 | 0 | 9.073508739 | 4.38E-08 | 1 |
| ZCRB1 | NM_033114 | 0 | 0 | 0 | 10.96774735 | 9.66237223 | 0 | 7.704370312 | 9.259374542 | 0.010506752 | 0.999932046 |
| ZDHHC1 | XM_024450245 | 0 | 9.095586658 | 0 | 0 | 0 | 0 | 0 | 0 | 5.54E-08 | 1 |
| ZDHHC11B | XM_024446189 | 0 | 0 | 0 | 0 | 0 | 9.60357477 | 0 | 10.83628629 | 1 | 0.02072721 |
| ZDHHC14 | XM_017011312 | 8.068609638 | 9.081113725 | 10.78359035 | 0 | 8.85635197 | 0 | 0 | 0 | 0.999953634 | 0.02072721 |
| ZDHHC19 | XM_011512417 | 0 | 9.808252985 | 0 | 0 | 0 | 0 | 0 | 0 | 2.01E-08 | 1 |
| ZDHHC2 | XM_011544544 | 0 | 9.527862205 | 0 | 0 | 0 | 0 | 8.067631163 | 0 | 2.90E-08 | 1 |
| ZDHHC2 | XM_024447177 | 8.691211177 | 0 | 0 | 0 | 0 | 0 | 0 | 0 | 9.73E-08 | 1 |
| ZDHHC20 | NM_001286638 | 0 | 0 | 0 | 11.37654209 | 7.992635714 | 0 | 0 | 0 | 0.011126981 | 0.02072721 |
| ZDHHC20 | NM_001330059 | 0 | 0 | 0 | 9.83969732 | 8.949104958 | 0 | 0 | 9.879715581 | 0.02081351 | 0.999932046 |
| ZDHHC20 | XR_429215 | 10.02910853 | 0 | 0 | 0 | 0 | 0 | 8.441777415 | 0 | 1.87E-08 | 1 |
| ZDHHC21 | XM_024447531 | 9.359023845 | 0 | 0 | 0 | 0 | 0 | 0 | 0 | 4.33E-08 | 1 |
| ZDHHC3 | XM_024453557 | 0 | 0 | 0 | 10.07409167 | 7.012300182 | 0 | 0 | 0 | 0.025988879 | 0.02072721 |
| ZDHHC5 | XM_017017498 | 0 | 9.930350523 | 0 | 0 | 0 | 0 | 0 | 0 | 1.81E-08 | 1 |
| ZDHHC6 | XM_017016567 | 0 | 11.18611218 | 0 | 0 | 0 | 0 | 0 | 0 | 1.34E-08 | 1 |
| ZDHHC8 | NM_013373 | 0 | 8.835955774 | 0 | 0 | 0 | 0 | 0 | 0 | 2.22E-07 | 1 |
| ZDHHC9 | XR_001755694 | 10.29958742 | 0 | 0 | 0 | 0 | 0 | 0 | 0 | 1.54E-08 | 1 |
| ZEB2 | NM_014795 | 8.716912497 | 0 | 0 | 0 | 0 | 0 | 0 | 0 | 9.50E-08 | 1 |
| ZER1 | XM_017014190 | 9.330172147 | 0 | 0 | 0 | 0 | 0 | 0 | 0 | 4.56E-08 | 1 |
| ZFAND1 | NR_033194 | 0 | 0 | 0 | 9.591445233 | 8.462903138 | 0 | 0 | 0 | 0.026195635 | 0.02072721 |
| ZFAND2B | XM_017003368 | 0 | 0 | 0 | 11.04503522 | 6.780897397 | 0 | 7.980893958 | 0 | 0.015375939 | 0.999932046 |
| ZFAND5 | NM_001102420 | 0 | 0 | 0 | 11.26835367 | 8.029844345 | 0 | 9.76024972 | 0 | 0.011851755 | 0.999932046 |
| ZFAS1 | NR_003605 | 0 | 0 | 10.15137027 | 7.326879247 | 0 | 0 | 0 | 8.942868965 | 0.026299336 | 0.999932046 |
| ZFAS1 | NR_036659 | 0 | 8.39589491 | 0 | 0 | 0 | 0 | 0 | 7.22071086 | 1.17E-07 | 1 |
| ZFP62 | XM_017009717 | 0 | 9.605761852 | 0 | 0 | 0 | 0 | 0 | 0 | 2.55E-08 | 1 |
| ZFP69B | XM_017002147 | 0 | 0 | 0 | 6.988236596 | 0 | 0 | 0 | 0 | 4.34E-05 | 1 |
| ZFP82 | NM_133466 | 9.487768672 | 0 | 0 | 0 | 0 | 0 | 0 | 0 | 3.53E-08 | 1 |
| ZFPL1 | NM_006782 | 5.728132393 | 0 | 10.18422504 | 0 | 9.87793591 | 0 | 0 | 0 | 0.394234789 | 0.02072721 |
| ZFYVE1 | XR_001750376 | 9.933695697 | 0 | 0 | 0 | 0 | 0 | 0 | 0 | 2.01E-08 | 1 |
| ZFYVE16 | NR_146174 | 0 | 0 | 0 | 10.05713443 | 6.59478863 | 0 | 8.15532787 | 0 | 0.027454051 | 0.999932046 |
| ZFYVE27 | XR_002956957 | 7.151412288 | 0 | 9.193555665 | 8.414023513 | 7.925118111 | 0 | 0 | 0 | 0.805720161 | 0.010308798 |
| ZFYVE28 | XR_001741314 | 10.76449512 | 0 | 0 | 0 | 0 | 0 | 0 | 0 | 1.34E-08 | 1 |
| ZGLP1 | NM_001103167 | 0 | 0 | 0 | 10.30709108 | 7.436136431 | 0 | 0 | 0 | 0.021722667 | 0.02072721 |
| ZGPAT | NM_181485 | 0 | 0 | 11.99648653 | 10.87403052 | 6.059865095 | 0 | 7.79000962 | 0 | 0.000732924 | 0.736446283 |
| ZKSCAN8 | XM_017011266 | 8.940634053 | 0 | 0 | 0 | 0 | 0 | 0 | 0 | 7.88E-08 | 1 |
| ZMIZ1 | XM_017016443 | 0 | 0 | 0 | 10.04571785 | 8.256688855 | 0 | 0 | 0 | 0.022023199 | 0.02072721 |
| ZMIZ2 | NM_174929 | 10.81891219 | 0 | 0 | 0 | 0 | 0 | 0 | 0 | 1.34E-08 | 1 |
| ZMYM3 | XM_011531062 | 0 | 0 | 0 | 0 | 6.643614539 | 0 | 0 | 0 | 3.86E-05 | 1 |
| ZMYM4 | NM_005095 | 0 | 0 | 6.3726817 | 0 | 9.514345989 | 0 | 7.696326825 | 0 | 0.038028333 | 0.999932046 |

| | | | | | | | | | | | |
|---|---|---|---|---|---|---|---|---|---|---|---|
| ZMYM4 | XM_024450838 | 0 | 0 | 10.46387522 | 0 | 10.64583822 | 11.19199396 | 8.268156521 | 0 | 0.009435897 | 0.999932046 |
| ZMYM4 | XR_001737542 | 5.25355136 | 8.529815232 | 0 | 0 | 0 | 0 | 0 | 0 | 0.019039619 | 1 |
| ZMYND11 | NM_001202464 | 9.623051988 | 11.42259638 | 0 | 0 | 0 | 0 | 11.05153432 | 10.81886129 | 0.000325769 | 0.02072721 |
| ZMYND11 | NM_212479 | 0 | 0 | 0 | 9.693732641 | 9.793912627 | 0 | 0 | 0 | 0.01712436 | 0.02072721 |
| ZMYND11 | XM_017015592 | 0 | 9.786116244 | 0 | 0 | 0 | 0 | 0 | 0 | 2.05E-08 | 1 |
| ZMYND8 | NM_001281776 | 0 | 0 | 0 | 0 | 6.679179249 | 0 | 0 | 0 | 3.78E-05 | 1 |
| ZMYND8 | NM_001281777 | 0 | 0 | 0 | 9.806259575 | 8.327507425 | 0 | 0 | 0 | 0.02442701 | 0.02072721 |
| ZMYND8 | XM_017027778 | 9.244409574 | 0 | 0 | 0 | 0 | 0 | 8.62242756 | 0 | 5.26E-08 | 1 |
| ZNF101 | NM_033204 | 0 | 9.321711965 | 0 | 0 | 0 | 0 | 0 | 0 | 3.84E-08 | 1 |
| ZNF107 | XM_017012286 | 0 | 0 | 0 | 8.537350365 | 0 | 0 | 8.347340272 | 0 | 5.99E-06 | 1 |
| ZNF121 | NM_001008727 | 0 | 9.479019994 | 0 | 0 | 0 | 0 | 0 | 0 | 3.04E-08 | 1 |
| ZNF134 | NM_003435 | 10.49792516 | 0 | 0 | 0 | 0 | 0 | 0 | 0 | 1.51E-08 | 1 |
| ZNF169 | NM_194320 | 0 | 9.367156605 | 0 | 0 | 0 | 0 | 0 | 0 | 3.68E-08 | 1 |
| ZNF184 | NM_007149 | 0 | 0 | 0 | 0 | 6.747781854 | 0 | 0 | 0 | 9.53E-05 | 1 |
| ZNF20 | NM_021143 | 0 | 0 | 9.287267368 | 0 | 8.186339198 | 0 | 0 | 0 | 0.033121503 | 0.022212956 |
| ZNF207 | NM_001098507 | 0 | 0 | 10.05474104 | 0 | 7.211666293 | 0 | 0 | 0 | 0.026325634 | 0.02072721 |
| ZNF207 | XR_002958060 | 0 | 0 | 0 | 8.724404295 | 8.719289114 | 0 | 0 | 0 | 0.034997793 | 0.024372606 |
| ZNF207 | XR_002958063 | 0 | 8.529815232 | 9.910026324 | 0 | 7.718441029 | 0 | 0 | 0 | 0.999953634 | 0.02072721 |
| ZNF214 | NM_013249 | 0 | 0 | 0 | 10.15069217 | 9.765240791 | 0 | 0 | 10.4711558 | 0.014723845 | 0.999932046 |
| ZNF217 | XM_024451998 | 0 | 0 | 0 | 9.861023513 | 7.178913994 | 0 | 0 | 0 | 0.028827511 | 0.021326758 |
| ZNF219 | XM_024449625 | 0 | 0 | 9.564964337 | 10.4845729 | 0 | 0 | 0 | 0 | 0.014801369 | 0.02072721 |
| ZNF22 | NM_006963 | 0 | 0 | 0 | 10.14802533 | 7.503816758 | 0 | 7.506976231 | 0 | 0.023521214 | 0.999932046 |
| ZNF250 | NM_001109689 | 0 | 0 | 0 | 9.595365854 | 7.666227291 | 0 | 0 | 0 | 0.031043881 | 0.022456606 |
| ZNF251 | XM_005272356 | 0 | 8.975521721 | 0 | 0 | 0 | 0 | 0 | 0 | 7.27E-08 | 1 |
| ZNF254 | NM_001278664 | 10.49421423 | 9.876890889 | 0 | 0 | 0 | 0 | 0 | 0 | 0.000659097 | 1 |
| ZNF254 | XM_017027515 | 0 | 0 | 0 | 8.5893879 | 0 | 0 | 0 | 0 | 5.96E-06 | 1 |
| ZNF26 | XR_001748866 | 0 | 0 | 0 | 0 | 6.569741695 | 0 | 0 | 0 | 4.12E-05 | 1 |
| ZNF263 | XM_006720834 | 0 | 8.622680652 | 0 | 0 | 0 | 0 | 0 | 0 | 9.58E-08 | 1 |
| ZNF264 | NM_003417 | 0 | 8.991087758 | 0 | 0 | 0 | 0 | 0 | 0 | 7.02E-08 | 1 |
| ZNF266 | XM_017026154 | 0 | 0 | 10.33081543 | 0 | 8.729220141 | 0 | 0 | 0 | 0.017971914 | 0.02072721 |
| ZNF266 | XM_017026160 | 0 | 0 | 0 | 9.498198764 | 7.145400839 | 0 | 0 | 11.37552496 | 0.035966805 | 0.999932046 |
| ZNF271P | NR_024565 | 0 | 0 | 0 | 0 | 6.736572239 | 0 | 0 | 10.00826692 | 9.53E-05 | 1 |
| ZNF271P | NR_024566 | 0 | 0 | 0 | 7.528981351 | 0 | 0 | 8.058247234 | 0 | 1.76E-05 | 1 |
| ZNF280D | NM_001288589 | 0 | 9.17269217 | 0 | 0 | 0 | 0 | 0 | 0 | 4.92E-08 | 1 |
| ZNF296 | NM_145288 | 0 | 8.756212053 | 0 | 0 | 0 | 0 | 0 | 0 | 8.55E-08 | 1 |
| ZNF326 | XM_005270779 | 0 | 8.959785899 | 0 | 0 | 0 | 0 | 0 | 0 | 7.69E-08 | 1 |
| ZNF330 | NM_014487 | 9.663265502 | 0 | 0 | 9.679029003 | 8.709289252 | 0 | 0 | 0 | 0.999953634 | 0.02072721 |
| ZNF350 | XM_011527196 | 0 | 8.886770211 | 0 | 0 | 0 | 0 | 9.458846292 | 10.93663514 | 7.80E-08 | 0.02072721 |
| ZNF350 | XM_017027094 | 0 | 8.602554319 | 0 | 0 | 0 | 0 | 9.790789648 | 0 | 9.73E-08 | 1 |
| ZNF350 | XM_017027099 | 0 | 9.029283553 | 0 | 0 | 0 | 11.96023537 | 0 | 0 | 6.49E-08 | 1 |
| ZNF350 | XM_017027100 | 9.819710617 | 10.02412431 | 0 | 0 | 0 | 0 | 9.098897264 | 0 | 0.000947155 | 1 |
| ZNF350 | XM_017027103 | 0 | 0 | 0 | 0 | 0 | 11.25838771 | 9.948097288 | 0 | 1 | 0.02072721 |
| ZNF358 | NM_018083 | 0 | 0 | 9.539797346 | 0 | 8.118872287 | 0 | 0 | 0 | 0.029724747 | 0.020903731 |

| | | | | | | | | | | | |
|---|---|---|---|---|---|---|---|---|---|---|---|
| ZNF358 | XM_005272460 | 9.546283346 | 0 | 0 | 0 | 0 | 0 | 0 | 0 | 3.25E-08 | 1 |
| ZNF362 | XM_017000415 | 0 | 0 | 8.585434713 | 0 | 7.477124411 | 0 | 0 | 0 | 0.057569302 | 0.036633555 |
| ZNF366 | NM_152625 | 0 | 8.792198922 | 0 | 0 | 0 | 0 | 0 | 0 | 8.40E-08 | 1 |
| ZNF37A | XM_017016622 | 0 | 0 | 7.660969894 | 0 | 0 | 0 | 0 | 0 | 1.62E-05 | 1 |
| ZNF384 | NM_133476 | 0 | 9.021724865 | 0 | 0 | 0 | 0 | 0 | 0 | 6.61E-08 | 1 |
| ZNF408 | NM_024741 | 0 | 8.774317692 | 0 | 0 | 0 | 0 | 0 | 0 | 8.46E-08 | 1 |
| ZNF41 | XM_017029817 | 0 | 8.844550112 | 0 | 0 | 0 | 0 | 0 | 0 | 8.06E-08 | 1 |
| ZNF410 | NM_021188 | 0 | 0 | 11.1371659 | 0 | 7.42919044 | 0 | 0 | 0 | 0.014047466 | 0.02072721 |
| ZNF414 | NM_001146175 | 0 | 0 | 0 | 0 | 6.337212177 | 0 | 0 | 0 | 4.86E-05 | 1 |
| ZNF431 | NR_138053 | 11.08379022 | 0 | 0 | 0 | 0 | 0 | 0 | 0 | 1.34E-08 | 1 |
| ZNF439 | NM_152262 | 0 | 0 | 7.044209926 | 0 | 0 | 0 | 0 | 0 | 3.03E-05 | 1 |
| ZNF44 | NM_001353549 | 0 | 9.17269217 | 0 | 0 | 0 | 0 | 0 | 0 | 4.92E-08 | 1 |
| ZNF44 | NM_016264 | 0 | 9.620849457 | 0 | 0 | 0 | 0 | 0 | 0 | 2.53E-08 | 1 |
| ZNF44 | XM_011528066 | 0 | 0 | 10.25358819 | 0 | 7.119744988 | 0 | 0 | 0 | 0.023843098 | 0.02072721 |
| ZNF444 | NM_018337 | 0 | 9.564739385 | 0 | 0 | 0 | 0 | 0 | 0 | 2.69E-08 | 1 |
| ZNF445 | XM_017006346 | 0 | 0 | 0 | 10.50555865 | 8.860230516 | 0 | 7.372102888 | 0 | 0.015843823 | 0.999932046 |
| ZNF454 | NM_182594 | 9.998000145 | 0 | 0 | 0 | 0 | 0 | 0 | 0 | 1.94E-08 | 1 |
| ZNF461 | NM_001322825 | 10.13775594 | 0 | 0 | 0 | 0 | 0 | 0 | 0 | 4.01E-08 | 1 |
| ZNF461 | XM_017027470 | 0 | 10.73764849 | 0 | 0 | 0 | 0 | 8.846115701 | 0 | 1.34E-08 | 1 |
| ZNF468 | XM_017027444 | 0 | 8.951853153 | 0 | 0 | 0 | 0 | 0 | 0 | 7.60E-08 | 1 |
| ZNF480 | XM_011526465 | 0 | 0 | 0 | 7.749084719 | 0 | 0 | 0 | 0 | 1.37E-05 | 1 |
| ZNF485 | NM_001318142 | 9.906032551 | 0 | 0 | 0 | 0 | 0 | 0 | 0 | 2.06E-08 | 1 |
| ZNF496 | XM_024450375 | 0 | 0 | 0 | 0 | 6.478485785 | 0 | 10.08762209 | 0 | 4.36E-05 | 1 |
| ZNF503-AS1 | NR_038226 | 0 | 9.812639863 | 0 | 0 | 0 | 0 | 0 | 0 | 2.01E-08 | 1 |
| ZNF511 | NM_145806 | 0 | 0 | 11.42005763 | 11.80927642 | 10.61747431 | 10.592033 | 9.249416094 | 0 | 0.000209174 | 0.999932046 |
| ZNF512 | NM_032434 | 0 | 8.935855595 | 0 | 0 | 0 | 0 | 0 | 0 | 7.69E-08 | 1 |
| ZNF513 | XM_005264143 | 0 | 9.102769039 | 0 | 0 | 0 | 0 | 7.310202764 | 9.272234908 | 5.49E-08 | 0.021907747 |
| ZNF521 | NM_015461 | 0 | 0 | 11.67247736 | 0 | 10.84751746 | 11.26731029 | 0 | 0 | 0.005206411 | 0.999932046 |
| ZNF521 | XM_017025697 | 0 | 0 | 10.59488547 | 0 | 9.623345731 | 0 | 0 | 0 | 0.012843867 | 0.02072721 |
| ZNF527 | XM_017027381 | 0 | 9.036802844 | 0 | 0 | 0 | 0 | 0 | 0 | 6.38E-08 | 1 |
| ZNF528 | NM_032423 | 0 | 8.756212053 | 0 | 0 | 0 | 0 | 0 | 0 | 8.55E-08 | 1 |
| ZNF547 | NM_173631 | 9.403227718 | 0 | 0 | 0 | 0 | 0 | 0 | 0 | 4.07E-08 | 1 |
| ZNF557 | NM_001044388 | 10.25642453 | 0 | 0 | 0 | 0 | 0 | 0 | 0 | 1.60E-08 | 1 |
| ZNF563 | NM_145276 | 0 | 9.590514794 | 0 | 0 | 0 | 0 | 0 | 0 | 2.90E-08 | 1 |
| ZNF567 | XM_017026421 | 0 | 0 | 9.870198585 | 0 | 6.147113942 | 0 | 0 | 0 | 0.033279154 | 0.022291204 |
| ZNF57 | NM_173480 | 0 | 0 | 0 | 10.03276528 | 7.854284905 | 0 | 0 | 0 | 0.023808508 | 0.02072721 |
| ZNF574 | XM_011527230 | 9.279328737 | 0 | 0 | 0 | 0 | 0 | 0 | 0 | 4.99E-08 | 1 |
| ZNF580 | NM_207115 | 0 | 0 | 0 | 10.74320495 | 7.833391225 | 0 | 0 | 0 | 0.016425231 | 0.020808337 |
| ZNF581 | NM_016535 | 0 | 0 | 10.78224969 | 0 | 8.155106507 | 0 | 0 | 0 | 0.015753713 | 0.02072721 |
| ZNF582-AS1 | NR_037161 | 0 | 8.818611963 | 0 | 0 | 0 | 0 | 0 | 0 | 8.12E-08 | 1 |
| ZNF585A | NR_110152 | 0 | 0 | 9.406923074 | 9.782383308 | 3.220007989 | 0 | 0 | 0 | 0.00730753 | 0.011791742 |
| ZNF585B | NM_152279 | 9.035603861 | 0 | 0 | 0 | 0 | 0 | 0 | 0 | 7.68E-08 | 1 |
| ZNF593 | NM_015871 | 0 | 0 | 10.69801997 | 11.15537872 | 10.30339297 | 10.86758209 | 10.24547681 | 10.27426767 | 0.000457725 | 0.999932046 |

| | | | | | | | | | | | |
|---|---|---|---|---|---|---|---|---|---|---|---|
| ZNF595 | NM_182524 | 9.944614124 | 0 | 0 | 0 | 0 | 0 | 0 | 0 | 2.01E-08 | 1 |
| ZNF599 | NM_001007248 | 9.595606435 | 0 | 0 | 0 | 0 | 0 | 0 | 0 | 3.08E-08 | 1 |
| ZNF616 | NR_135823 | 0 | 0 | 0 | 0 | 6.725274845 | 0 | 0 | 0 | 9.53E-05 | 1 |
| ZNF625 | NM_145233 | 9.866390785 | 0 | 0 | 0 | 0 | 0 | 0 | 0 | 2.17E-08 | 1 |
| ZNF626 | NM_145297 | 0 | 9.08836834 | 0 | 0 | 0 | 0 | 0 | 0 | 5.61E-08 | 1 |
| ZNF638 | XM_017003810 | 0 | 8.951853153 | 0 | 0 | 0 | 0 | 0 | 0 | 7.60E-08 | 1 |
| ZNF639 | NM_001303425 | 0 | 9.073822446 | 0 | 0 | 0 | 0 | 0 | 0 | 7.73E-08 | 1 |
| ZNF639 | XM_017006552 | 0 | 9.08836834 | 0 | 0 | 0 | 0 | 0 | 0 | 5.61E-08 | 1 |
| ZNF652 | XM_024450656 | 0 | 0 | 9.241172275 | 10.01677478 | 8.057134281 | 0 | 0 | 0 | 0.003062913 | 0.004603096 |
| ZNF655 | XM_017012607 | 8.651779762 | 0 | 0 | 0 | 0 | 0 | 0 | 0 | 1.00E-07 | 1 |
| ZNF668 | NM_024706 | 0 | 9.059128398 | 0 | 0 | 0 | 0 | 0 | 0 | 7.80E-08 | 1 |
| ZNF674 | NM_001190417 | 0 | 7.411784592 | 9.031481501 | 0 | 9.974134586 | 0 | 0 | 0 | 0.88332723 | 0.02072721 |
| ZNF674 | XM_011543941 | 0 | 8.998808245 | 0 | 0 | 0 | 0 | 0 | 0 | 6.89E-08 | 1 |
| ZNF674 | XM_017029728 | 0 | 8.878424457 | 0 | 0 | 0 | 0 | 8.664264428 | 0 | 9.15E-08 | 1 |
| ZNF69 | NM_001321163 | 0 | 9.265218616 | 0 | 0 | 0 | 0 | 0 | 0 | 4.18E-08 | 1 |
| ZNF692 | NM_017865 | 10.18455989 | 0 | 0 | 0 | 0 | 0 | 0 | 0 | 1.60E-08 | 1 |
| ZNF695 | NM_020394 | 9.181190369 | 0 | 0 | 0 | 0 | 0 | 0 | 0 | 7.80E-08 | 1 |
| ZNF7 | XM_017013815 | 0 | 10.1577136 | 0 | 0 | 0 | 0 | 7.774812207 | 0 | 1.54E-08 | 1 |
| ZNF700 | NM_144566 | 11.07139093 | 0 | 0 | 0 | 0 | 0 | 6.533064721 | 9.662471112 | 1.34E-08 | 0.020781046 |
| ZNF738 | NM_001355241 | 0 | 0 | 0 | 0 | 6.366595318 | 0 | 0 | 0 | 4.76E-05 | 1 |
| ZNF74 | NR_046282 | 0 | 8.709928179 | 0 | 0 | 0 | 0 | 0 | 0 | 8.96E-08 | 1 |
| ZNF746 | XM_011515838 | 9.928205339 | 0 | 0 | 0 | 0 | 0 | 0 | 0 | 2.03E-08 | 1 |
| ZNF75A | XM_017023645 | 8.583566901 | 0 | 0 | 0 | 0 | 0 | 0 | 0 | 1.06E-07 | 1 |
| ZNF75D | XM_024452446 | 0 | 8.853093555 | 0 | 0 | 0 | 0 | 0 | 0 | 7.91E-08 | 1 |
| ZNF770 | NM_014106 | 0 | 9.073822446 | 0 | 0 | 0 | 0 | 0 | 0 | 7.73E-08 | 1 |
| ZNF79 | XM_024447662 | 0 | 0 | 0 | 0 | 6.713888284 | 0 | 0 | 0 | 3.66E-05 | 1 |
| ZNF793 | XM_024451511 | 8.357138871 | 0 | 0 | 0 | 0 | 0 | 0 | 0 | 1.96E-07 | 1 |
| ZNF800 | XM_011515859 | 0 | 0 | 6.559575832 | 0 | 0 | 0 | 0 | 0 | 4.56E-05 | 1 |
| ZNF821 | NM_001201553 | 10.0942935 | 0 | 0 | 0 | 0 | 0 | 0 | 0 | 1.80E-08 | 1 |
| ZNF821 | XM_017023409 | 9.517322653 | 0 | 0 | 0 | 0 | 0 | 0 | 0 | 3.36E-08 | 1 |
| ZNF821 | XM_017023419 | 0 | 9.712905893 | 0 | 0 | 0 | 0 | 0 | 0 | 2.55E-08 | 1 |
| ZNF821 | XM_017023423 | 0 | 8.671801345 | 0 | 0 | 0 | 0 | 0 | 0 | 9.15E-08 | 1 |
| ZNF821 | XM_024450339 | 10.25203635 | 9.145128245 | 0 | 0 | 0 | 11.10694718 | 9.765064406 | 0 | 0.00116894 | 0.02072721 |
| ZNF830 | NM_052857 | 0 | 0 | 0 | 0 | 6.351978551 | 0 | 5.635260462 | 0 | 4.81E-05 | 1 |
| ZNF836 | NM_001102657 | 0 | 0 | 0 | 0 | 6.351978551 | 0 | 0 | 0 | 4.81E-05 | 1 |
| ZNF85 | NM_001256172 | 0 | 9.777165557 | 0 | 0 | 0 | 11.19082201 | 10.14858295 | 10.29647003 | 4.32E-08 | 0.000855759 |
| ZNF85 | XM_011528264 | 0 | 9.749975406 | 0 | 0 | 0 | 0 | 0 | 0 | 2.16E-08 | 1 |
| ZNF91 | XR_001753754 | 11.35941642 | 9.585396421 | 10.36345654 | 10.23604134 | 0 | 0 | 0 | 0 | 0.999953634 | 0.02072721 |
| ZNF91 | XR_001753755 | 0 | 0 | 0 | 0 | 0 | 0 | 6.31926467 | 10.12274908 | 1 | 0.02072721 |
| ZNF91 | XR_001753756 | 0 | 0 | 0 | 8.244627116 | 0 | 0 | 7.032041479 | 0 | 1.67E-05 | 1 |
| ZNRF3 | NM_001206998 | 0 | 0 | 0 | 0 | 0 | 9.9578806 | 9.829916618 | 9.172865883 | 1 | 0.002233122 |
| ZPR1 | NM_003904 | 0 | 0 | 0 | 10.06846137 | 9.481148681 | 0 | 0 | 10.49329345 | 0.016539023 | 0.999932046 |
| ZRSR2 | XM_017029881 | 0 | 9.179501644 | 0 | 0 | 0 | 0 | 0 | 0 | 4.86E-08 | 1 |

| | | | | | | | | | | | |
|---|---|---|---|---|---|---|---|---|---|---|---|
| ZSCAN18 | XM_017027171 | 11.01034435 | 0 | 0 | 0 | 0 | 0 | 0 | 0 | 1.34E-08 | 1 |
| ZSCAN22 | XM_017026747 | 9.434546067 | 0 | 0 | 0 | 0 | 0 | 0 | 0 | 3.90E-08 | 1 |
| ZSCAN9 | NM_006299 | 0 | 8.384152849 | 0 | 0 | 0 | 0 | 0 | 0 | 1.18E-07 | 1 |
| ZSWIM3 | NR_037628 | 0 | 0 | 10.71216955 | 0 | 6.023432489 | 0 | 0 | 0 | 0.020147878 | 4.32E-09 |
| ZUP1 | NM_145062 | 0 | 9.309347676 | 0 | 0 | 0 | 0 | 4.451059805 | 0 | 3.94E-08 | 0.745732931 |
| ZWILCH | XR_001751345 | 9.877828587 | 0 | 0 | 0 | 0 | 0 | 0 | 0 | 2.13E-08 | 1 |
| ZXDC | XR_924169 | 9.843239492 | 0 | 0 | 0 | 0 | 0 | 0 | 0 | 2.27E-08 | 1 |
| ZYG11B | NM_024646 | 10.10406561 | 8.853093555 | 0 | 0 | 0 | 0 | 0 | 0 | 0.00151351 | 1 |
| ZZZ3 | XR_001737104 | 0 | 8.78328601 | 0 | 0 | 0 | 0 | 0 | 0 | 8.40E-08 | 1 |

Supplementary Table S5: Differentially expressed genes (DEGs) and isoforms (DEIs)

| Both DEG and DEI | | |
|---|---|---|
| Gene | DEG | DEI |
| AACS | Y | Y |
| AAR2 | Y | Y |
| ABCA5 | Y | Y |
| ABCA8 | Y | Y |
| ABCB10 | Y | Y |
| ABCC4 | Y | Y |
| ABCC6P2 | Y | Y |
| ABCD3 | Y | Y |
| ABHD11 | Y | Y |
| ABHD12 | Y | Y |
| ABHD14A | Y | Y |
| ABI3 | Y | Y |
| ABRAXAS1 | Y | Y |
| ACAA1 | Y | Y |
| ACAD9 | Y | Y |
| ACADS | Y | Y |
| ACD | Y | Y |
| ACLY | Y | Y |
| ACOT2 | Y | Y |
| ACP5 | Y | Y |
| ACSF2 | Y | Y |
| ACSM3 | Y | Y |
| ACSS1 | Y | Y |
| ACTR1B | Y | Y |
| ACYP2 | Y | Y |
| ADA2 | Y | Y |
| ADAM28 | Y | Y |
| ADAMTS10 | Y | Y |
| ADAMTSL1 | Y | Y |
| ADAMTSL2 | Y | Y |
| ADAT1 | Y | Y |
| ADAT2 | Y | Y |
| ADD2 | Y | Y |
| ADGRL3 | Y | Y |
| ADH1C | Y | Y |
| ADIPOR2 | Y | Y |
| ADORA2B | Y | Y |
| ADPRM | Y | Y |
| ADSSL1 | Y | Y |
| AFMID | Y | Y |

| DEG only | | |
|---|---|---|
| Gene | DEG | DEI |
| AAMDC | Y | N |
| AATF | Y | N |
| ABCC6 | Y | N |
| ABI2 | Y | N |
| ADCY7 | Y | N |
| ADNP2 | Y | N |
| AEBP2 | Y | N |
| AKAP13 | Y | N |
| AKR1B1 | Y | N |
| ALS2 | Y | N |
| ANAPC4 | Y | N |
| ANKRD16 | Y | N |
| APTX | Y | N |
| AQP7 | Y | N |
| ARAP1 | Y | N |
| ARNTL2 | Y | N |
| ASPG | Y | N |
| ATIC | Y | N |
| ATP11B | Y | N |
| ATP6V1F | Y | N |
| B4GALT7 | Y | N |
| BAHCC1 | Y | N |
| BARX2 | Y | N |
| BCL2L1 | Y | N |
| BEX4 | Y | N |
| BMP1 | Y | N |
| BRPF3 | Y | N |
| BST1 | Y | N |
| C16orf87 | Y | N |
| C2 | Y | N |
| C6orf89 | Y | N |
| CAB39L | Y | N |
| CASP10 | Y | N |
| CC2D1A | Y | N |
| CCDC137 | Y | N |
| CCDC146 | Y | N |
| CCDC25 | Y | N |
| CCDC93 | Y | N |
| CCR7 | Y | N |
| CCZ1 | Y | N |

| DEI only | | |
|---|---|---|
| Gene | DEG | DEI |
| A1CF | N | Y |
| AAAS | N | Y |
| AAGAB | N | Y |
| AAMP | N | Y |
| ABCA2 | N | Y |
| ABCB8 | N | Y |
| ABCC1 | N | Y |
| ABCF3 | N | Y |
| ABCG2 | N | Y |
| ABHD13 | N | Y |
| ABHD14B | N | Y |
| ABHD16A | N | Y |
| ABHD17A | N | Y |
| ABHD17B | N | Y |
| ABHD2 | N | Y |
| ABHD3 | N | Y |
| ABHD5 | N | Y |
| ABI1 | N | Y |
| ABLIM1 | N | Y |
| ABR | N | Y |
| ABTB1 | N | Y |
| ACAD8 | N | Y |
| ACADVL | N | Y |
| ACAP3 | N | Y |
| ACIN1 | N | Y |
| ACOT8 | N | Y |
| ACP1 | N | Y |
| ACP2 | N | Y |
| ACSF3 | N | Y |
| ACSL1 | N | Y |
| ACSS2 | N | Y |
| ACTA2 | N | Y |
| ACTL6A | N | Y |
| ACTN4 | N | Y |
| ACTR3 | N | Y |
| ACTR3C | N | Y |
| ACTR6 | N | Y |
| ACVR1 | N | Y |
| ACY1 | N | Y |
| ADA | N | Y |

| | | |
|---|---|---|
| AGAP3 | Y | Y |
| AGFG2 | Y | Y |
| AGO3 | Y | Y |
| AGPAT4 | Y | Y |
| AGTRAP | Y | Y |
| AIM2 | Y | Y |
| AK4 | Y | Y |
| AK6 | Y | Y |
| AKAP11 | Y | Y |
| AKAP8 | Y | Y |
| AKR1B10 | Y | Y |
| ALB | Y | Y |
| ALDH9A1 | Y | Y |
| ALG9 | Y | Y |
| ALK | Y | Y |
| ALKBH5 | Y | Y |
| ALOX15 | Y | Y |
| AMIGO2 | Y | Y |
| AMN | Y | Y |
| AMN1 | Y | Y |
| ANAPC1 | Y | Y |
| ANAPC2 | Y | Y |
| ANGEL1 | Y | Y |
| ANKHD1-EIF4EBP3 | Y | Y |
| ANKRD22 | Y | Y |
| ANKRD27 | Y | Y |
| ANKRD36B | Y | Y |
| ANKRD40 | Y | Y |
| ANKRD52 | Y | Y |
| ANKRD55 | Y | Y |
| ANKS1B | Y | Y |
| ANKZF1 | Y | Y |
| ANO10 | Y | Y |
| ANTXR2 | Y | Y |
| ANXA2R | Y | Y |
| AP1M1 | Y | Y |
| AP2A2 | Y | Y |
| AP3D1 | Y | Y |
| AP3S2 | Y | Y |
| AP4B1 | Y | Y |
| AP4M1 | Y | Y |
| APC | Y | Y |
| APCDD1 | Y | Y |

| | | |
|---|---|---|
| CD79B | Y | N |
| CDC45 | Y | N |
| CDC6 | Y | N |
| CDCA8 | Y | N |
| CENPP | Y | N |
| CLK2 | Y | N |
| CLYBL | Y | N |
| CNEP1R1 | Y | N |
| CNOT2 | Y | N |
| COPS7B | Y | N |
| COQ3 | Y | N |
| COQ7 | Y | N |
| CORO1C | Y | N |
| CRNDE | Y | N |
| CRTC3 | Y | N |
| CRYBB2P1 | Y | N |
| CUL2 | Y | N |
| CWC27 | Y | N |
| CYB5R2 | Y | N |
| CYBB | Y | N |
| DDOST | Y | N |
| DDX11 | Y | N |
| DDX49 | Y | N |
| DGCR5 | Y | N |
| DGKA | Y | N |
| DIEXF | Y | N |
| DKK3 | Y | N |
| DMAP1 | Y | N |
| DNAJA3 | Y | N |
| DNMBP | Y | N |
| DOCK1 | Y | N |
| DPH2 | Y | N |
| DST | Y | N |
| DSTYK | Y | N |
| DUSP1 | Y | N |
| DVL1 | Y | N |
| DXO | Y | N |
| ECI2 | Y | N |
| EEF1G | Y | N |
| EFCAB5 | Y | N |
| EIF1AD | Y | N |
| EIF2AK2 | Y | N |
| ELL2 | Y | N |

| | | |
|---|---|---|
| ADAM15 | N | Y |
| ADAP2 | N | Y |
| ADCK5 | N | Y |
| ADD1 | N | Y |
| ADIPOR1 | N | Y |
| ADK | N | Y |
| ADPRHL2 | N | Y |
| ADRM1 | N | Y |
| ADTRP | N | Y |
| AFDN | N | Y |
| AFF1 | N | Y |
| AFF3 | N | Y |
| AFG1L | N | Y |
| AFG3L2 | N | Y |
| AFTPH | N | Y |
| AGAP1 | N | Y |
| AGBL4 | N | Y |
| AGPAT3 | N | Y |
| AGRN | N | Y |
| AHCY | N | Y |
| AHCYL2 | N | Y |
| AHDC1 | N | Y |
| AHI1 | N | Y |
| AHNAK | N | Y |
| AIDA | N | Y |
| AIF1 | N | Y |
| AIFM1 | N | Y |
| AIG1 | N | Y |
| AIMP2 | N | Y |
| AIP | N | Y |
| AK1 | N | Y |
| AK2 | N | Y |
| AKAP1 | N | Y |
| AKAP8L | N | Y |
| AKAP9 | N | Y |
| AKR1A1 | N | Y |
| AKR7A3 | N | Y |
| AKT1 | N | Y |
| AKT3 | N | Y |
| AKTIP | N | Y |
| ALDH18A1 | N | Y |
| ALDH2 | N | Y |
| ALDOA | N | Y |

| | | | | | | | | |
|---|---|---|---|---|---|---|---|---|
| APOA2 | Y | Y | ELP2 | Y | N | ALG13 | N | Y |
| APOBEC3F | Y | Y | EMX1 | Y | N | ALG3 | N | Y |
| APOL2 | Y | Y | ERCC8 | Y | N | ALG8 | N | Y |
| AQP12A | Y | Y | F12 | Y | N | ALKBH2 | N | Y |
| AQP3 | Y | Y | FAAP24 | Y | N | ALOX5AP | N | Y |
| AR | Y | Y | FAHD2B | Y | N | ALS2CL | N | Y |
| ARAP3 | Y | Y | FAM107B | Y | N | AMD1 | N | Y |
| ARCN1 | Y | Y | FAM120A | Y | N | AMOTL1 | N | Y |
| AREL1 | Y | Y | FAM177A1 | Y | N | AMT | N | Y |
| ARFGEF1 | Y | Y | FAM49A | Y | N | AMZ2 | N | Y |
| ARFIP2 | Y | Y | FAM83H | Y | N | ANAPC11 | N | Y |
| ARHGAP12 | Y | Y | FBXO25 | Y | N | ANAPC13 | N | Y |
| ARHGAP17 | Y | Y | FBXO7 | Y | N | ANAPC15 | N | Y |
| ARHGAP21 | Y | Y | FCN3 | Y | N | ANAPC5 | N | Y |
| ARHGAP27 | Y | Y | FEN1 | Y | N | ANAPC7 | N | Y |
| ARHGAP31 | Y | Y | FIG4 | Y | N | ANGEL2 | N | Y |
| ARHGAP4 | Y | Y | FOXC1 | Y | N | ANGPTL4 | N | Y |
| ARHGAP8 | Y | Y | FPGS | Y | N | ANK3 | N | Y |
| ARHGEF16 | Y | Y | FRMD3 | Y | N | ANKH | N | Y |
| ARHGEF2 | Y | Y | FTL | Y | N | ANKHD1 | N | Y |
| ARHGEF37 | Y | Y | FUOM | Y | N | ANKRD10 | N | Y |
| ARHGEF40 | Y | Y | GATA3-AS1 | Y | N | ANKRD11 | N | Y |
| ARID4B | Y | Y | GBP3 | Y | N | ANKRD13A | N | Y |
| ARID5A | Y | Y | GDPD5 | Y | N | ANKRD17 | N | Y |
| ARL17B | Y | Y | GFM1 | Y | N | ANKRD36 | N | Y |
| ARL2BP | Y | Y | GIGYF2 | Y | N | ANKRD46 | N | Y |
| ARL6 | Y | Y | GLIDR | Y | N | ANXA11 | N | Y |
| ARMC1 | Y | Y | GLIPR1 | Y | N | ANXA7 | N | Y |
| ARMC10 | Y | Y | GOLIM4 | Y | N | AOAH | N | Y |
| ARMC7 | Y | Y | GORASP1 | Y | N | AP1AR | N | Y |
| ARMCX2 | Y | Y | GPATCH4 | Y | N | AP1G2 | N | Y |
| ARMCX5 | Y | Y | GPC3 | Y | N | AP2B1 | N | Y |
| ARPP19 | Y | Y | GPR89B | Y | N | AP2S1 | N | Y |
| ARRDC2 | Y | Y | GRAMD1A | Y | N | AP5M1 | N | Y |
| ARRDC3-AS1 | Y | Y | GTF2H2 | Y | N | APBA3 | N | Y |
| ARSA | Y | Y | GTPBP6 | Y | N | APH1A | N | Y |
| AS3MT | Y | Y | GXYLT2 | Y | N | APLP2 | N | Y |
| ASB3 | Y | Y | H2AFZ | Y | N | APOBEC1 | N | Y |
| ASCC3 | Y | Y | HCAR1 | Y | N | APOC1 | N | Y |
| ASCL2 | Y | Y | HCG18 | Y | N | APOL1 | N | Y |
| ASH1L | Y | Y | HELLS | Y | N | APOL3 | N | Y |
| ASIC2 | Y | Y | HK2 | Y | N | APOL4 | N | Y |
| ASNS | Y | Y | HMGXB3 | Y | N | APP | N | Y |

| | | |
|---|---|---|
| ASNSD1 | Y | Y |
| ATAD1 | Y | Y |
| ATF2 | Y | Y |
| ATF3 | Y | Y |
| ATF7 | Y | Y |
| ATF7IP | Y | Y |
| ATG4A | Y | Y |
| ATG4B | Y | Y |
| ATM | Y | Y |
| ATOH1 | Y | Y |
| ATP13A1 | Y | Y |
| ATP1A1 | Y | Y |
| ATP1A1-AS1 | Y | Y |
| ATP2B1 | Y | Y |
| ATP2B4 | Y | Y |
| ATP2C1 | Y | Y |
| ATPSCKMT | Y | Y |
| ATXN1 | Y | Y |
| ATXN1L | Y | Y |
| ATXN7L2 | Y | Y |
| ATXN7L3 | Y | Y |
| ATXN7L3B | Y | Y |
| AURKB | Y | Y |
| AZGP1 | Y | Y |
| AZIN1-AS1 | Y | Y |
| B3GALT5 | Y | Y |
| B3GAT3 | Y | Y |
| B3GNT2 | Y | Y |
| B3GNT3 | Y | Y |
| B3GNT6 | Y | Y |
| B4GALT1-AS1 | Y | Y |
| B4GALT4 | Y | Y |
| B9D1 | Y | Y |
| B9D2 | Y | Y |
| BAAT | Y | Y |
| BACE2 | Y | Y |
| BAG5 | Y | Y |
| BAMBI | Y | Y |
| BANK1 | Y | Y |
| BASP1 | Y | Y |
| BATF2 | Y | Y |
| BAZ2B | Y | Y |
| BBIP1 | Y | Y |

| | | |
|---|---|---|
| HPS1 | Y | N |
| IFT52 | Y | N |
| IL6ST | Y | N |
| INMT | Y | N |
| IPMK | Y | N |
| ITGB4 | Y | N |
| ITPR2 | Y | N |
| JMJD8 | Y | N |
| KALRN | Y | N |
| KAZALD1 | Y | N |
| KAZN | Y | N |
| KDF1 | Y | N |
| KDM4C | Y | N |
| KDM5A | Y | N |
| KHNYN | Y | N |
| KIFC2 | Y | N |
| KLK10 | Y | N |
| KMT2E | Y | N |
| KRAS | Y | N |
| KYAT3 | Y | N |
| LAMB2 | Y | N |
| LARP1B | Y | N |
| LDHA | Y | N |
| LDLRAP1 | Y | N |
| LEPROTL1 | Y | N |
| LINC01124 | Y | N |
| LIPC | Y | N |
| LOC100128108 | Y | N |
| LOC100996732 | Y | N |
| LOC100996842 | Y | N |
| LOC102723906 | Y | N |
| LOC105369748 | Y | N |
| LOC105370546 | Y | N |
| LOC105373428 | Y | N |
| LOC105377094 | Y | N |
| LOC107987276 | Y | N |
| LOC283710 | Y | N |
| LOC284581 | Y | N |
| LRRC19 | Y | N |
| LZIC | Y | N |
| MANEAL | Y | N |
| MAP3K12 | Y | N |
| MARVELD2 | Y | N |

| | | |
|---|---|---|
| AQP12B | N | Y |
| ARFGAP1 | N | Y |
| ARFGAP2 | N | Y |
| ARFRP1 | N | Y |
| ARHGAP10 | N | Y |
| ARHGAP15 | N | Y |
| ARHGAP24 | N | Y |
| ARHGAP25 | N | Y |
| ARHGAP26 | N | Y |
| ARHGAP45 | N | Y |
| ARHGAP6 | N | Y |
| ARHGEF10L | N | Y |
| ARHGEF7 | N | Y |
| ARHGEF9 | N | Y |
| ARID1B | N | Y |
| ARID2 | N | Y |
| ARIH2 | N | Y |
| ARL1 | N | Y |
| ARL15 | N | Y |
| ARL5A | N | Y |
| ARL6IP4 | N | Y |
| ARL6IP6 | N | Y |
| ARMC8 | N | Y |
| ARMCX3 | N | Y |
| ARNT | N | Y |
| ARNTL | N | Y |
| ARPC2 | N | Y |
| ARPC5L | N | Y |
| ARSE | N | Y |
| ARV1 | N | Y |
| ARVCF | N | Y |
| ASB9 | N | Y |
| ASCC1 | N | Y |
| ASH1L-AS1 | N | Y |
| ASPH | N | Y |
| ASPSCR1 | N | Y |
| ASTN2 | N | Y |
| ASXL1 | N | Y |
| ASXL2 | N | Y |
| ATAD3B | N | Y |
| ATAD5 | N | Y |
| ATE1 | N | Y |
| ATF4 | N | Y |

| | | |
|---|---|---|
| BBOF1 | Y | Y |
| BCAS2 | Y | Y |
| BCL2L14 | Y | Y |
| BCL2L15 | Y | Y |
| BCL2L2 | Y | Y |
| BCL7C | Y | Y |
| BCL9 | Y | Y |
| BCO1 | Y | Y |
| BEST2 | Y | Y |
| BEX3 | Y | Y |
| BIRC5 | Y | Y |
| BIVM | Y | Y |
| BIVM-ERCC5 | Y | Y |
| BMP5 | Y | Y |
| BMP8B | Y | Y |
| BNIP2 | Y | Y |
| BNIP3 | Y | Y |
| BNIP3L | Y | Y |
| BOC | Y | Y |
| BOD1 | Y | Y |
| BORCS6 | Y | Y |
| BORCS8 | Y | Y |
| BPNT1 | Y | Y |
| BRAP | Y | Y |
| BRCC3 | Y | Y |
| BRIP1 | Y | Y |
| BRMS1 | Y | Y |
| BRPF1 | Y | Y |
| BRWD1-AS2 | Y | Y |
| BRWD3 | Y | Y |
| BSDC1 | Y | Y |
| BTBD2 | Y | Y |
| BTD | Y | Y |
| BTNL3 | Y | Y |
| C10orf143 | Y | Y |
| C11orf54 | Y | Y |
| C11orf71 | Y | Y |
| C11orf74 | Y | Y |
| C11orf80 | Y | Y |
| C12orf29 | Y | Y |
| C12orf43 | Y | Y |
| C12orf56 | Y | Y |
| C15orf40 | Y | Y |

| | | |
|---|---|---|
| MAST2 | Y | N |
| MCMBP | Y | N |
| MELK | Y | N |
| MEMO1 | Y | N |
| METTL18 | Y | N |
| METTL4 | Y | N |
| MFAP5 | Y | N |
| MFGE8 | Y | N |
| MIA3 | Y | N |
| MICOS10 | Y | N |
| MICOS13 | Y | N |
| MINDY2 | Y | N |
| MIPOL1 | Y | N |
| MKNK2 | Y | N |
| MLST8 | Y | N |
| MND1 | Y | N |
| MNT | Y | N |
| MRPS33 | Y | N |
| MRRF | Y | N |
| MTIF2 | Y | N |
| MTMR10 | Y | N |
| MZT1 | Y | N |
| NABP1 | Y | N |
| NAF1 | Y | N |
| NDC1 | Y | N |
| NDUFS5 | Y | N |
| NEDD9 | Y | N |
| NKD1 | Y | N |
| NME6 | Y | N |
| NMRK1 | Y | N |
| NPAS3 | Y | N |
| NPRL2 | Y | N |
| NPRL3 | Y | N |
| NRG4 | Y | N |
| NSF | Y | N |
| NTHL1 | Y | N |
| NUDT1 | Y | N |
| NUF2 | Y | N |
| NUP43 | Y | N |
| NUP62CL | Y | N |
| OCLN | Y | N |
| ODF2 | Y | N |
| ODF2L | Y | N |

| | | |
|---|---|---|
| ATF5 | N | Y |
| ATG10 | N | Y |
| ATG101 | N | Y |
| ATG5 | N | Y |
| ATL2 | N | Y |
| ATL3 | N | Y |
| ATP10D | N | Y |
| ATP2A2 | N | Y |
| ATP5ME | N | Y |
| ATP5MF | N | Y |
| ATP5MG | N | Y |
| ATP5PF | N | Y |
| ATP5S | N | Y |
| ATP6V0B | N | Y |
| ATP6V1C1 | N | Y |
| ATP6V1H | N | Y |
| ATP9B | N | Y |
| ATPAF2 | N | Y |
| ATR | N | Y |
| ATRAID | N | Y |
| ATRX | N | Y |
| ATXN2 | N | Y |
| AUH | N | Y |
| AUP1 | N | Y |
| AURKAIP1 | N | Y |
| AVPI1 | N | Y |
| AXIN1 | N | Y |
| AXIN2 | N | Y |
| B3GNT7 | N | Y |
| B3GNT8 | N | Y |
| B4GALNT2 | N | Y |
| B4GALT3 | N | Y |
| BABAM1 | N | Y |
| BABAM2 | N | Y |
| BAG2 | N | Y |
| BAG3 | N | Y |
| BAG6 | N | Y |
| BAIAP2L2 | N | Y |
| BANF1 | N | Y |
| BAP1 | N | Y |
| BARD1 | N | Y |
| BBX | N | Y |
| BCAP29 | N | Y |

| | | |
|---|---|---|
| C15orf41 | Y | Y |
| C1GALT1 | Y | Y |
| C1orf115 | Y | Y |
| C1orf131 | Y | Y |
| C1orf198 | Y | Y |
| C1orf35 | Y | Y |
| C1orf56 | Y | Y |
| C1QL1 | Y | Y |
| C1R | Y | Y |
| C20orf204 | Y | Y |
| C20orf27 | Y | Y |
| C21orf58 | Y | Y |
| C2CD5 | Y | Y |
| C2orf72 | Y | Y |
| C3orf62 | Y | Y |
| C3orf70 | Y | Y |
| C4orf33 | Y | Y |
| C6orf120 | Y | Y |
| C6orf132 | Y | Y |
| C6orf136 | Y | Y |
| C6orf203 | Y | Y |
| C7orf13 | Y | Y |
| C7orf50 | Y | Y |
| C8orf33 | Y | Y |
| C8orf37 | Y | Y |
| C8orf76 | Y | Y |
| C9orf78 | Y | Y |
| CA12 | Y | Y |
| CA2 | Y | Y |
| CA9 | Y | Y |
| CACNA2D1 | Y | Y |
| CALCOCO1 | Y | Y |
| CAMK2G | Y | Y |
| CAMKK2 | Y | Y |
| CAND1 | Y | Y |
| CAPN3 | Y | Y |
| CARD10 | Y | Y |
| CARD16 | Y | Y |
| CARD6 | Y | Y |
| CARS2 | Y | Y |
| CASC4 | Y | Y |
| CASC8 | Y | Y |
| CASC9 | Y | Y |

| | | |
|---|---|---|
| ORC3 | Y | N |
| PACRGL | Y | N |
| PAIP2B | Y | N |
| PARP2 | Y | N |
| PARVB | Y | N |
| PDCD4-AS1 | Y | N |
| PDXDC1 | Y | N |
| PDZD7 | Y | N |
| PHETA2 | Y | N |
| PIK3C2A | Y | N |
| PITRM1 | Y | N |
| PKMYT1 | Y | N |
| PLAT | Y | N |
| PLCE1 | Y | N |
| PLCXD1 | Y | N |
| PLSCR4 | Y | N |
| POLR3C | Y | N |
| PPM1A | Y | N |
| PPP1CA | Y | N |
| PPP1R18 | Y | N |
| PRCP | Y | N |
| PSD4 | Y | N |
| PSMA7 | Y | N |
| PTHLH | Y | N |
| PTPRA | Y | N |
| R3HCC1L | Y | N |
| RABEP1 | Y | N |
| RAC1 | Y | N |
| RAPGEF1 | Y | N |
| RASSF3 | Y | N |
| RASSF8 | Y | N |
| RBAK | Y | N |
| RBM19 | Y | N |
| RENBP | Y | N |
| RFC1 | Y | N |
| RFX7 | Y | N |
| RFXANK | Y | N |
| RHBDL1 | Y | N |
| RMDN2 | Y | N |
| RMDN3 | Y | N |
| RNF2 | Y | N |
| RPL7 | Y | N |
| RPL8 | Y | N |

| | | |
|---|---|---|
| BCAP31 | N | Y |
| BCAR1 | N | Y |
| BCAS1 | N | Y |
| BCAS3 | N | Y |
| BCAT2 | N | Y |
| BCL11A | N | Y |
| BCL2L11 | N | Y |
| BCL3 | N | Y |
| BCL7B | N | Y |
| BDH2 | N | Y |
| BDP1 | N | Y |
| BECN1 | N | Y |
| BEND5 | N | Y |
| BET1L | N | Y |
| BFSP2 | N | Y |
| BHLHE40 | N | Y |
| BICDL2 | N | Y |
| BID | N | Y |
| BIN1 | N | Y |
| BIN3 | N | Y |
| BIRC3 | N | Y |
| BIRC6 | N | Y |
| BLCAP | N | Y |
| BLM | N | Y |
| BLNK | N | Y |
| BLZF1 | N | Y |
| BMI1 | N | Y |
| BMP2K | N | Y |
| BOLA2-SMG1P6 | N | Y |
| BOLA2B | N | Y |
| BOLA3 | N | Y |
| BPTF | N | Y |
| BRAT1 | N | Y |
| BRCA1 | N | Y |
| BRD1 | N | Y |
| BRD8 | N | Y |
| BRD9 | N | Y |
| BSCL2 | N | Y |
| BSG | N | Y |
| BTBD6 | N | Y |
| BTBD9 | N | Y |
| BTF3L4 | N | Y |
| BTN3A1 | N | Y |

| | | |
|---|---|---|
| CASKIN2 | Y | Y |
| CASP3 | Y | Y |
| CASP4 | Y | Y |
| CASP8AP2 | Y | Y |
| CASR | Y | Y |
| CASTOR1 | Y | Y |
| CAV2 | Y | Y |
| CAVIN3 | Y | Y |
| CBR3-AS1 | Y | Y |
| CBX7 | Y | Y |
| CCDC107 | Y | Y |
| CCDC120 | Y | Y |
| CCDC14 | Y | Y |
| CCDC142 | Y | Y |
| CCDC144A | Y | Y |
| CCDC178 | Y | Y |
| CCDC18-AS1 | Y | Y |
| CCDC186 | Y | Y |
| CCDC43 | Y | Y |
| CCDC47 | Y | Y |
| CCDC85B | Y | Y |
| CCDC9 | Y | Y |
| CCDC91 | Y | Y |
| CCL13 | Y | Y |
| CCL19 | Y | Y |
| CCL21 | Y | Y |
| CCL23 | Y | Y |
| CCL24 | Y | Y |
| CCM2 | Y | Y |
| CCN3 | Y | Y |
| CCNB1 | Y | Y |
| CCND2-AS1 | Y | Y |
| CCNO | Y | Y |
| CCNT2 | Y | Y |
| CCPG1 | Y | Y |
| CCSER2 | Y | Y |
| CCT6P1 | Y | Y |
| CCZ1B | Y | Y |
| CCZ1P-OR7E38P | Y | Y |
| CD177 | Y | Y |
| CD27 | Y | Y |
| CD302 | Y | Y |
| CD33 | Y | Y |

| | | |
|---|---|---|
| RPS5 | Y | N |
| RRAGC | Y | N |
| RSF1 | Y | N |
| RSPH3 | Y | N |
| SCAF1 | Y | N |
| SCFD2 | Y | N |
| SEMA4G | Y | N |
| 4-Sep | Y | N |
| 8-Sep | Y | N |
| SERPINF1 | Y | N |
| SIKE1 | Y | N |
| SINHCAF | Y | N |
| SIRT5 | Y | N |
| SKA2 | Y | N |
| SKA3 | Y | N |
| SLC10A3 | Y | N |
| SLC18B1 | Y | N |
| SLC25A10 | Y | N |
| SLC25A33 | Y | N |
| SLC25A35 | Y | N |
| SLC35A2 | Y | N |
| SLC35C1 | Y | N |
| SLC38A3 | Y | N |
| SLC41A3 | Y | N |
| SLC5A1 | Y | N |
| SLC5A6 | Y | N |
| SLC7A1 | Y | N |
| SLC9A4 | Y | N |
| SMAD6 | Y | N |
| SMPD2 | Y | N |
| SNCA | Y | N |
| SNHG12 | Y | N |
| SPAG1 | Y | N |
| SPAG9 | Y | N |
| SPARCL1 | Y | N |
| SPATA13 | Y | N |
| SPATC1L | Y | N |
| SSH3 | Y | N |
| STAMBP | Y | N |
| STT3B | Y | N |
| SUCNR1 | Y | N |
| SUMF1 | Y | N |
| TAOK1 | Y | N |

| | | |
|---|---|---|
| BTNL8 | N | Y |
| BUB3 | N | Y |
| BUD23 | N | Y |
| BZW1 | N | Y |
| BZW2 | N | Y |
| C11orf24 | N | Y |
| C11orf49 | N | Y |
| C11orf68 | N | Y |
| C11orf86 | N | Y |
| C12orf4 | N | Y |
| C12orf49 | N | Y |
| C14orf119 | N | Y |
| C16orf72 | N | Y |
| C19orf12 | N | Y |
| C19orf48 | N | Y |
| C1orf159 | N | Y |
| C1orf43 | N | Y |
| C1orf52 | N | Y |
| C1QTNF3 | N | Y |
| C21orf2 | N | Y |
| C2orf49 | N | Y |
| C3orf38 | N | Y |
| C4orf19 | N | Y |
| C4orf48 | N | Y |
| C5orf51 | N | Y |
| C6orf48 | N | Y |
| C8orf34 | N | Y |
| C9orf16 | N | Y |
| C9orf3 | N | Y |
| CA1 | N | Y |
| CA4 | N | Y |
| CAAP1 | N | Y |
| CADPS | N | Y |
| CALD1 | N | Y |
| CALM2 | N | Y |
| CALML4 | N | Y |
| CAMSAP2 | N | Y |
| CAMTA1 | N | Y |
| CANT1 | N | Y |
| CAPN1 | N | Y |
| CAPN12 | N | Y |
| CAPN2 | N | Y |
| CAPN5 | N | Y |

| | | |
|---|---|---|
| CD37 | Y | Y |
| CD3D | Y | Y |
| CD3E | Y | Y |
| CD3G | Y | Y |
| CD53 | Y | Y |
| CD55 | Y | Y |
| CD79A | Y | Y |
| CD8B | Y | Y |
| CD96 | Y | Y |
| CD99P1 | Y | Y |
| CDCA5 | Y | Y |
| CDCA7L | Y | Y |
| CDH5 | Y | Y |
| CDHR1 | Y | Y |
| CDK1 | Y | Y |
| CDK11A | Y | Y |
| CDK11B | Y | Y |
| CDK7 | Y | Y |
| CDKN1C | Y | Y |
| CDKN2B-AS1 | Y | Y |
| CDKN3 | Y | Y |
| CDYL | Y | Y |
| CEBPZ | Y | Y |
| CECR7 | Y | Y |
| CELF1 | Y | Y |
| CELSR3 | Y | Y |
| CEMIP2 | Y | Y |
| CENPA | Y | Y |
| CENPM | Y | Y |
| CENPN | Y | Y |
| CENPO | Y | Y |
| CEP19 | Y | Y |
| CERKL | Y | Y |
| CERNA2 | Y | Y |
| CETN2 | Y | Y |
| CFAP20 | Y | Y |
| CFAP65 | Y | Y |
| CFD | Y | Y |
| CFTR | Y | Y |
| CH17-125A10.2 | Y | Y |
| CHAF1A | Y | Y |
| CHD5 | Y | Y |
| CHD9 | Y | Y |

| | | |
|---|---|---|
| TAPBPL | Y | N |
| TBC1D31 | Y | N |
| TCAIM | Y | N |
| TEDC1 | Y | N |
| TES | Y | N |
| TESMIN | Y | N |
| TEX30 | Y | N |
| TGDS | Y | N |
| THAP6 | Y | N |
| THBS4 | Y | N |
| THUMPD3-AS1 | Y | N |
| TK2 | Y | N |
| TLDC2 | Y | N |
| TMEM116 | Y | N |
| TMEM129 | Y | N |
| TMEM184A | Y | N |
| TMEM39A | Y | N |
| TMEM69 | Y | N |
| TOLLIP | Y | N |
| TP53I11 | Y | N |
| TPBG | Y | N |
| TPD52L1 | Y | N |
| TSSC4 | Y | N |
| TTC8 | Y | N |
| TUBA1B | Y | N |
| U2AF1L4 | Y | N |
| UBE2D4 | Y | N |
| UIMC1 | Y | N |
| USP30 | Y | N |
| USP8 | Y | N |
| USP9Y | Y | N |
| VANGL1 | Y | N |
| VIPAS39 | Y | N |
| VIPR1 | Y | N |
| VMO1 | Y | N |
| WARS2 | Y | N |
| WASHC2A | Y | N |
| WASHC5 | Y | N |
| WDR73 | Y | N |
| WHAMM | Y | N |
| WRAP73 | Y | N |
| YARS | Y | N |
| YIPF1 | Y | N |

| | | |
|---|---|---|
| CAPNS1 | N | Y |
| CAPRIN1 | N | Y |
| CAPZB | N | Y |
| CARD19 | N | Y |
| CARM1 | N | Y |
| CARS | N | Y |
| CASP1 | N | Y |
| CASP8 | N | Y |
| CAST | N | Y |
| CBLB | N | Y |
| CBLC | N | Y |
| CBR4 | N | Y |
| CBWD1 | N | Y |
| CBWD2 | N | Y |
| CBWD5 | N | Y |
| CBX5 | N | Y |
| CCAR1 | N | Y |
| CCDC12 | N | Y |
| CCDC124 | N | Y |
| CCDC126 | N | Y |
| CCDC159 | N | Y |
| CCDC32 | N | Y |
| CCDC34 | N | Y |
| CCDC51 | N | Y |
| CCDC57 | N | Y |
| CCDC7 | N | Y |
| CCDC88A | N | Y |
| CCL20 | N | Y |
| CCND1 | N | Y |
| CCND3 | N | Y |
| CCNDBP1 | N | Y |
| CCNG2 | N | Y |
| CCNH | N | Y |
| CCNK | N | Y |
| CCNL2 | N | Y |
| CCSER1 | N | Y |
| CCT4 | N | Y |
| CCT7 | N | Y |
| CD151 | N | Y |
| CD164 | N | Y |
| CD1A | N | Y |
| CD226 | N | Y |
| CD274 | N | Y |

| | | |
|---|---|---|
| CHEK2P2 | Y | Y |
| CHFR | Y | Y |
| CHGB | Y | Y |
| CHKB | Y | Y |
| CHMP4C | Y | Y |
| CHMP7 | Y | Y |
| CHODL | Y | Y |
| CHPF | Y | Y |
| CHRAC1 | Y | Y |
| CHRNB1 | Y | Y |
| CHST6 | Y | Y |
| CHST7 | Y | Y |
| CHTF8 | Y | Y |
| CIAO3 | Y | Y |
| CIAPIN1 | Y | Y |
| CIITA | Y | Y |
| CINP | Y | Y |
| CIPC | Y | Y |
| CISD1 | Y | Y |
| CIT | Y | Y |
| CIZ1 | Y | Y |
| CLASRP | Y | Y |
| CLCA1 | Y | Y |
| CLCN3 | Y | Y |
| CLDN10 | Y | Y |
| CLDN12 | Y | Y |
| CLDN5 | Y | Y |
| CLDND1 | Y | Y |
| CLEC18C | Y | Y |
| CLIC3 | Y | Y |
| CLIP2 | Y | Y |
| CLN6 | Y | Y |
| CLSTN2 | Y | Y |
| CLUH | Y | Y |
| CMAHP | Y | Y |
| CMC4 | Y | Y |
| CMKLR1 | Y | Y |
| CMTM3 | Y | Y |
| CMTM4 | Y | Y |
| CMTM6 | Y | Y |
| CNDP2 | Y | Y |
| CNIH3 | Y | Y |
| CNOT8 | Y | Y |

| | | |
|---|---|---|
| YTHDF2 | Y | N |
| ZBTB48 | Y | N |
| ZMYM2 | Y | N |
| ZNF138 | Y | N |
| ZNF282 | Y | N |
| ZNF335 | Y | N |
| ZNF440 | Y | N |
| ZNF451 | Y | N |
| ZNF627 | Y | N |
| ZNF740 | Y | N |
| ZNF766 | Y | N |
| ZSCAN16 | Y | N |

| | | |
|---|---|---|
| CD36 | N | Y |
| CD44 | N | Y |
| CD46 | N | Y |
| CD47 | N | Y |
| CD48 | N | Y |
| CD63 | N | Y |
| CD68 | N | Y |
| CD86 | N | Y |
| CD99 | N | Y |
| CDC16 | N | Y |
| CDC26 | N | Y |
| CDC42BPG | N | Y |
| CDC42SE1 | N | Y |
| CDH1 | N | Y |
| CDH11 | N | Y |
| CDHR5 | N | Y |
| CDK10 | N | Y |
| CDK14 | N | Y |
| CDK19 | N | Y |
| CDK8 | N | Y |
| CDKL3 | N | Y |
| CDKN2A | N | Y |
| CDX2 | N | Y |
| CEACAM1 | N | Y |
| CEACAM5 | N | Y |
| CEBPZOS | N | Y |
| CENPJ | N | Y |
| CENPX | N | Y |
| CEP104 | N | Y |
| CEP164 | N | Y |
| CEP170 | N | Y |
| CEP57 | N | Y |
| CEP70 | N | Y |
| CEP76 | N | Y |
| CEPT1 | N | Y |
| CERS2 | N | Y |
| CERS5 | N | Y |
| CFH | N | Y |
| CFLAR | N | Y |
| CGGBP1 | N | Y |
| CHAC2 | N | Y |
| CHCHD4 | N | Y |
| CHD1 | N | Y |

| | | |
|---|---|---|
| CNPPD1 | Y | Y |
| CNPY3-GNMT | Y | Y |
| CNPY4 | Y | Y |
| COA1 | Y | Y |
| COA6 | Y | Y |
| COL17A1 | Y | Y |
| COL18A1 | Y | Y |
| COL24A1 | Y | Y |
| COL4A2 | Y | Y |
| COL4A5 | Y | Y |
| COMT | Y | Y |
| COPA | Y | Y |
| COPG1 | Y | Y |
| COQ8A | Y | Y |
| CORO1A | Y | Y |
| COX7A1 | Y | Y |
| CPB1 | Y | Y |
| CPEB2 | Y | Y |
| CPEB3 | Y | Y |
| CPLX1 | Y | Y |
| CPQ | Y | Y |
| CPSF1 | Y | Y |
| CPSF3 | Y | Y |
| CR1L | Y | Y |
| CR2 | Y | Y |
| CRAT | Y | Y |
| CRIPAK | Y | Y |
| CRISPLD2 | Y | Y |
| CRLF3 | Y | Y |
| CROCCP2 | Y | Y |
| CRTAP | Y | Y |
| CRY2 | Y | Y |
| CRYBB3 | Y | Y |
| CRYM | Y | Y |
| CS | Y | Y |
| CSK | Y | Y |
| CSMD1 | Y | Y |
| CSNK1D | Y | Y |
| CSNK2A1 | Y | Y |
| CSRNP3 | Y | Y |
| CSRP2 | Y | Y |
| CST7 | Y | Y |
| CSTA | Y | Y |

| | | |
|---|---|---|
| CHD1L | N | Y |
| CHD3 | N | Y |
| CHD4 | N | Y |
| CHEK2 | N | Y |
| CHGA | N | Y |
| CHIC2 | N | Y |
| CHID1 | N | Y |
| CHKA | N | Y |
| CHMP2B | N | Y |
| CHMP3 | N | Y |
| CHTOP | N | Y |
| CHURC1 | N | Y |
| CIAO2B | N | Y |
| CIB1 | N | Y |
| CIC | N | Y |
| CIRBP | N | Y |
| CKLF | N | Y |
| CLASP2 | N | Y |
| CLDN7 | N | Y |
| CLEC16A | N | Y |
| CLEC2D | N | Y |
| CLIC1 | N | Y |
| CLINT1 | N | Y |
| CLIP1 | N | Y |
| CLK3 | N | Y |
| CLMN | N | Y |
| CLPTM1L | N | Y |
| CLTB | N | Y |
| CLTC | N | Y |
| CLTCL1 | N | Y |
| CMC1 | N | Y |
| CNN2 | N | Y |
| CNOT6 | N | Y |
| CNOT9 | N | Y |
| CNPY3 | N | Y |
| CNRIP1 | N | Y |
| COA6-AS1 | N | Y |
| COG6 | N | Y |
| COL16A1 | N | Y |
| COL21A1 | N | Y |
| COL6A2 | N | Y |
| COL9A2 | N | Y |
| COLGALT1 | N | Y |

| | | |
|---|---|---|
| CSTF1 | Y | Y |
| CSTF2 | Y | Y |
| CTB-49A3.2 | Y | Y |
| CTNS | Y | Y |
| CTSV | Y | Y |
| CTSZ | Y | Y |
| CTTNBP2 | Y | Y |
| CTU1 | Y | Y |
| CUEDC2 | Y | Y |
| CUL4A | Y | Y |
| CUL5 | Y | Y |
| CUL7 | Y | Y |
| CUTC | Y | Y |
| CWF19L1 | Y | Y |
| CXCL1 | Y | Y |
| CXCL11 | Y | Y |
| CXCL12 | Y | Y |
| CXCL16 | Y | Y |
| CXCR4 | Y | Y |
| CXCR5 | Y | Y |
| CXorf56 | Y | Y |
| CXorf65 | Y | Y |
| CYB561A3 | Y | Y |
| CYB561D2 | Y | Y |
| CYB5R1 | Y | Y |
| CYB5R4 | Y | Y |
| CYBC1 | Y | Y |
| CYP21A1P | Y | Y |
| CYP27A1 | Y | Y |
| CYP2D6 | Y | Y |
| CYP2S1 | Y | Y |
| CYP3A5 | Y | Y |
| CYP51A1 | Y | Y |
| CYP7B1 | Y | Y |
| CYREN | Y | Y |
| CYTH2 | Y | Y |
| CYTOR | Y | Y |
| CZIB | Y | Y |
| DAGLA | Y | Y |
| DALRD3 | Y | Y |
| DAND5 | Y | Y |
| DAPK1 | Y | Y |
| DAPK2 | Y | Y |

| | | |
|---|---|---|
| COMMD1 | N | Y |
| COMMD10 | N | Y |
| COMMD5 | N | Y |
| COMMD9 | N | Y |
| COMTD1 | N | Y |
| COP1 | N | Y |
| COPB2 | N | Y |
| COPS2 | N | Y |
| COPS3 | N | Y |
| COPZ2 | N | Y |
| COQ6 | N | Y |
| CORO1B | N | Y |
| COX15 | N | Y |
| COX20 | N | Y |
| COX7A2 | N | Y |
| CPM | N | Y |
| CPOX | N | Y |
| CPT1A | N | Y |
| CRACR2B | N | Y |
| CRADD | N | Y |
| CRB3 | N | Y |
| CRBN | N | Y |
| CRCP | N | Y |
| CREBBP | N | Y |
| CREBZF | N | Y |
| CRELD1 | N | Y |
| CRELD2 | N | Y |
| CREM | N | Y |
| CRK | N | Y |
| CRTC1 | N | Y |
| CRYL1 | N | Y |
| CRYZ | N | Y |
| CSF1 | N | Y |
| CSNK1E | N | Y |
| CTBP1 | N | Y |
| CTBP2 | N | Y |
| CTCF | N | Y |
| CTDSP2 | N | Y |
| CTNNA2 | N | Y |
| CTNNAL1 | N | Y |
| CTNNB1 | N | Y |
| CTNNBIP1 | N | Y |
| CTNND1 | N | Y |

| | | |
|---|---|---|
| DCAF15 | Y | Y |
| DCP2 | Y | Y |
| DCTN1 | Y | Y |
| DDA1 | Y | Y |
| DDAH1 | Y | Y |
| DDB1 | Y | Y |
| DDX28 | Y | Y |
| DDX3Y | Y | Y |
| DDX56 | Y | Y |
| DDX58 | Y | Y |
| DEF8 | Y | Y |
| DEFA5 | Y | Y |
| DENND4A | Y | Y |
| DEPP1 | Y | Y |
| DEUP1 | Y | Y |
| DFFA | Y | Y |
| DGCR11 | Y | Y |
| DGKE | Y | Y |
| DHFR | Y | Y |
| DHX16 | Y | Y |
| DHX32 | Y | Y |
| DICER1 | Y | Y |
| DICER1-AS1 | Y | Y |
| DIPK1A | Y | Y |
| DKK4 | Y | Y |
| DLAT | Y | Y |
| DLG1 | Y | Y |
| DMAC2 | Y | Y |
| DMBT1 | Y | Y |
| DMKN | Y | Y |
| DNAH9 | Y | Y |
| DNAJB9 | Y | Y |
| DNAJC21 | Y | Y |
| DNAJC25 | Y | Y |
| DNAJC25-GNG10 | Y | Y |
| DNAJC28 | Y | Y |
| DNAJC9 | Y | Y |
| DNALI1 | Y | Y |
| DNASE2 | Y | Y |
| DNHD1 | Y | Y |
| DNTTIP2 | Y | Y |
| DOCK8 | Y | Y |
| DOCK9 | Y | Y |

| | | |
|---|---|---|
| CTPS1 | N | Y |
| CTSA | N | Y |
| CTTN | N | Y |
| CUL3 | N | Y |
| CUL4B | N | Y |
| CWH43 | N | Y |
| CXCL10 | N | Y |
| CXCL8 | N | Y |
| CXXC1 | N | Y |
| CYB5R3 | N | Y |
| CYBA | N | Y |
| CYC1 | N | Y |
| CYLD | N | Y |
| CYP2C18 | N | Y |
| CYP4F12 | N | Y |
| DACH1 | N | Y |
| DAG1 | N | Y |
| DAP3 | N | Y |
| DAXX | N | Y |
| DAZAP1 | N | Y |
| DAZAP2 | N | Y |
| DBI | N | Y |
| DBNDD2 | N | Y |
| DBNL | N | Y |
| DCAF10 | N | Y |
| DCAF4 | N | Y |
| DCTD | N | Y |
| DCTN5 | N | Y |
| DCUN1D4 | N | Y |
| DCUN1D5 | N | Y |
| DDAH2 | N | Y |
| DDC | N | Y |
| DDHD2 | N | Y |
| DDR1 | N | Y |
| DDX19A | N | Y |
| DDX24 | N | Y |
| DDX31 | N | Y |
| DDX39A | N | Y |
| DDX3X | N | Y |
| DDX46 | N | Y |
| DDX52 | N | Y |
| DEGS2 | N | Y |
| DELE1 | N | Y |

| | | |
|---|---|---|
| DOK1 | Y | Y |
| DOK2 | Y | Y |
| DPAGT1 | Y | Y |
| DPH5 | Y | Y |
| DPP4 | Y | Y |
| DPP8 | Y | Y |
| DPT | Y | Y |
| DPYSL3 | Y | Y |
| DRAM2 | Y | Y |
| DRG2 | Y | Y |
| DROSHA | Y | Y |
| DSC2 | Y | Y |
| DSCC1 | Y | Y |
| DTX3L | Y | Y |
| DTYMK | Y | Y |
| DUOXA2 | Y | Y |
| DUSP11 | Y | Y |
| DUSP26 | Y | Y |
| DUSP6 | Y | Y |
| DVL3 | Y | Y |
| DYNC2H1 | Y | Y |
| DYNC2LI1 | Y | Y |
| DZIP3 | Y | Y |
| EBF4 | Y | Y |
| EBI3 | Y | Y |
| ECM2 | Y | Y |
| EDDM13 | Y | Y |
| EDN2 | Y | Y |
| EDRF1 | Y | Y |
| EEF1AKMT1 | Y | Y |
| EEF1AKMT2 | Y | Y |
| EEF1AKMT4 | Y | Y |
| EEF1B2 | Y | Y |
| EFCAB11 | Y | Y |
| EFEMP1 | Y | Y |
| EFNA3 | Y | Y |
| EFNA5 | Y | Y |
| EFR3A | Y | Y |
| EGLN2 | Y | Y |
| EGLN3 | Y | Y |
| EGR1 | Y | Y |
| EHD1 | Y | Y |
| EID2 | Y | Y |

| | | |
|---|---|---|
| DENND1A | N | Y |
| DENND3 | N | Y |
| DEPDC1B | N | Y |
| DERL2 | N | Y |
| DGAT2 | N | Y |
| DGCR2 | N | Y |
| DGUOK | N | Y |
| DHDDS | N | Y |
| DHODH | N | Y |
| DHPS | N | Y |
| DHRS11 | N | Y |
| DHRS4 | N | Y |
| DHRS7B | N | Y |
| DHX37 | N | Y |
| DHX40 | N | Y |
| DHX8 | N | Y |
| DIABLO | N | Y |
| DIDO1 | N | Y |
| DIP2A | N | Y |
| DIS3 | N | Y |
| DLG3 | N | Y |
| DLG5 | N | Y |
| DLGAP4 | N | Y |
| DLL1 | N | Y |
| DMAC1 | N | Y |
| DMTF1 | N | Y |
| DNAJB1 | N | Y |
| DNAJB4 | N | Y |
| DNAJB6 | N | Y |
| DNAJC19 | N | Y |
| DNAJC2 | N | Y |
| DNAJC4 | N | Y |
| DNAJC7 | N | Y |
| DNASE1 | N | Y |
| DNASE1L1 | N | Y |
| DNPEP | N | Y |
| DNTTIP1 | N | Y |
| DOPEY2 | N | Y |
| DPH3 | N | Y |
| DPP7 | N | Y |
| DPP9 | N | Y |
| DPYD | N | Y |
| DQX1 | N | Y |

| | | |
|---|---|---|
| EIF2AK3 | Y | Y |
| EIF2B1 | Y | Y |
| EIF2D | Y | Y |
| EIF2S2 | Y | Y |
| EIF3J-DT | Y | Y |
| EIF4A1 | Y | Y |
| ELF3-AS1 | Y | Y |
| ELF4 | Y | Y |
| ELMO2 | Y | Y |
| ELMO3 | Y | Y |
| ELMSAN1 | Y | Y |
| ELOVL5 | Y | Y |
| ELP3 | Y | Y |
| ELP6 | Y | Y |
| EMC9 | Y | Y |
| EMCN | Y | Y |
| EML5 | Y | Y |
| EML6 | Y | Y |
| EMP1 | Y | Y |
| ENAH | Y | Y |
| ENTPD4 | Y | Y |
| ENTPD8 | Y | Y |
| EPB41L3 | Y | Y |
| EPHA2 | Y | Y |
| EPHB3 | Y | Y |
| EPN1 | Y | Y |
| EPRS | Y | Y |
| EPS8L1 | Y | Y |
| ERC1 | Y | Y |
| ERICH1 | Y | Y |
| ERICH2 | Y | Y |
| ERVK3-1 | Y | Y |
| ESM1 | Y | Y |
| ESRP1 | Y | Y |
| ETAA1 | Y | Y |
| ETF1 | Y | Y |
| ETFDH | Y | Y |
| EVI2B | Y | Y |
| EVI5L | Y | Y |
| EXD3 | Y | Y |
| EXO5 | Y | Y |
| EXOSC4 | Y | Y |
| EXPH5 | Y | Y |

| | | |
|---|---|---|
| DTL | N | Y |
| DTNA | N | Y |
| DTNB | N | Y |
| DTNBP1 | N | Y |
| DTX2 | N | Y |
| DUS1L | N | Y |
| DUSP16 | N | Y |
| DUSP22 | N | Y |
| DUT | N | Y |
| DYM | N | Y |
| DYNC1LI2 | N | Y |
| DYNLL1 | N | Y |
| DYNLRB1 | N | Y |
| DYRK1A | N | Y |
| DYRK4 | N | Y |
| EBAG9 | N | Y |
| ECE1 | N | Y |
| ECHDC1 | N | Y |
| ECHDC2 | N | Y |
| ECSIT | N | Y |
| EEF1D | N | Y |
| EEF2KMT | N | Y |
| EEFSEC | N | Y |
| EFL1 | N | Y |
| EHBP1L1 | N | Y |
| EHF | N | Y |
| EIF2A | N | Y |
| EIF2AK4 | N | Y |
| EIF2B2 | N | Y |
| EIF2B5 | N | Y |
| EIF3B | N | Y |
| EIF3I | N | Y |
| EIF3L | N | Y |
| EIF4A3 | N | Y |
| EIF4E2 | N | Y |
| EIF4G3 | N | Y |
| EIF5A | N | Y |
| EIF6 | N | Y |
| ELFN1-AS1 | N | Y |
| ELMOD2 | N | Y |
| ELOC | N | Y |
| ELP5 | N | Y |
| EMC2 | N | Y |

| | | |
|---|---|---|
| F2RL1 | Y | Y |
| F8A1 | Y | Y |
| FA2H | Y | Y |
| FABP3 | Y | Y |
| FADS3 | Y | Y |
| FAM102A | Y | Y |
| FAM110C | Y | Y |
| FAM117B | Y | Y |
| FAM120AOS | Y | Y |
| FAM120B | Y | Y |
| FAM129C | Y | Y |
| FAM153C | Y | Y |
| FAM160A2 | Y | Y |
| FAM172A | Y | Y |
| FAM174B | Y | Y |
| FAM185A | Y | Y |
| FAM200A | Y | Y |
| FAM206A | Y | Y |
| FAM20C | Y | Y |
| FAM213A | Y | Y |
| FAM213B | Y | Y |
| FAM214A | Y | Y |
| FAM21EP | Y | Y |
| FAM226A | Y | Y |
| FAM24B | Y | Y |
| FAM30A | Y | Y |
| FAM83D | Y | Y |
| FAM86DP | Y | Y |
| FAM86JP | Y | Y |
| FAM8A1 | Y | Y |
| FAM92A | Y | Y |
| FAN1 | Y | Y |
| FANCD2OS | Y | Y |
| FANCE | Y | Y |
| FANCL | Y | Y |
| FARSB | Y | Y |
| FASTK | Y | Y |
| FASTKD1 | Y | Y |
| FASTKD2 | Y | Y |
| FAT4 | Y | Y |
| FBXL3 | Y | Y |
| FBXL6 | Y | Y |
| FBXO32 | Y | Y |

| | | |
|---|---|---|
| EMD | N | Y |
| EML4 | N | Y |
| ENC1 | N | Y |
| ENGASE | N | Y |
| ENPP2 | N | Y |
| ENSA | N | Y |
| ENTPD1 | N | Y |
| ENTPD5 | N | Y |
| EPB41L2 | N | Y |
| EPHB2 | N | Y |
| EPS15L1 | N | Y |
| EPS8L3 | N | Y |
| ERG | N | Y |
| ERI3 | N | Y |
| ERLIN1 | N | Y |
| ERN2 | N | Y |
| ERO1A | N | Y |
| ERP29 | N | Y |
| ESS2 | N | Y |
| ESYT2 | N | Y |
| ETFA | N | Y |
| ETNK1 | N | Y |
| ETV1 | N | Y |
| ETV6 | N | Y |
| EVA1C | N | Y |
| EVL | N | Y |
| EVPL | N | Y |
| EWSR1 | N | Y |
| EXD2 | N | Y |
| EXOSC10 | N | Y |
| EXOSC2 | N | Y |
| EXOSC8 | N | Y |
| EXT2 | N | Y |
| EXTL2 | N | Y |
| EYA2 | N | Y |
| EYA3 | N | Y |
| EZH2 | N | Y |
| FAF1 | N | Y |
| FAHD2A | N | Y |
| FAIM | N | Y |
| FAM104A | N | Y |
| FAM104B | N | Y |
| FAM114A2 | N | Y |

| | | |
|---|---|---|
| FBXO34 | Y | Y |
| FBXO4 | Y | Y |
| FBXO45 | Y | Y |
| FBXO6 | Y | Y |
| FBXW4 | Y | Y |
| FCER2 | Y | Y |
| FCGBP | Y | Y |
| FCGR1A | Y | Y |
| FCGR2B | Y | Y |
| FCHO1 | Y | Y |
| FCHSD1 | Y | Y |
| FES | Y | Y |
| FFAR4 | Y | Y |
| FGD4 | Y | Y |
| FGF12 | Y | Y |
| FGF18 | Y | Y |
| FGFR1OP2 | Y | Y |
| FGFR4 | Y | Y |
| FGL2 | Y | Y |
| FH | Y | Y |
| FITM1 | Y | Y |
| FKBP1A-SDCBP2 | Y | Y |
| FLAD1 | Y | Y |
| FLI1 | Y | Y |
| FLJ44635 | Y | Y |
| FLOT2 | Y | Y |
| FLVCR2 | Y | Y |
| FLYWCH2 | Y | Y |
| FMNL3 | Y | Y |
| FMO5 | Y | Y |
| FN3KRP | Y | Y |
| FOSL1 | Y | Y |
| FOXA1 | Y | Y |
| FOXA3 | Y | Y |
| FOXF1 | Y | Y |
| FOXJ1 | Y | Y |
| FOXK2 | Y | Y |
| FOXO3B | Y | Y |
| FOXRED1 | Y | Y |
| FPR1 | Y | Y |
| FREM1 | Y | Y |
| FRG1FP | Y | Y |
| FRK | Y | Y |

| | | |
|---|---|---|
| FAM118B | N | Y |
| FAM129B | N | Y |
| FAM133B | N | Y |
| FAM135A | N | Y |
| FAM136A | N | Y |
| FAM156B | N | Y |
| FAM171A1 | N | Y |
| FAM192A | N | Y |
| FAM193A | N | Y |
| FAM208A | N | Y |
| FAM219B | N | Y |
| FAM3C | N | Y |
| FAM3D | N | Y |
| FAM45A | N | Y |
| FAM49B | N | Y |
| FAM83G | N | Y |
| FAM84A | N | Y |
| FAM89B | N | Y |
| FANCA | N | Y |
| FANCC | N | Y |
| FAR2 | N | Y |
| FAS | N | Y |
| FASTKD3 | N | Y |
| FBL | N | Y |
| FBLIM1 | N | Y |
| FBLN5 | N | Y |
| FBP1 | N | Y |
| FBXL12 | N | Y |
| FBXL22 | N | Y |
| FBXO11 | N | Y |
| FBXO21 | N | Y |
| FBXO38 | N | Y |
| FBXO9 | N | Y |
| FBXW2 | N | Y |
| FBXW5 | N | Y |
| FBXW7 | N | Y |
| FCHSD2 | N | Y |
| FDPS | N | Y |
| FER | N | Y |
| FERMT2 | N | Y |
| FETUB | N | Y |
| FGFR1 | N | Y |
| FGFR2 | N | Y |

| | | |
|---|---|---|
| FRMD5 | Y | Y |
| FRY | Y | Y |
| FSD2 | Y | Y |
| FTO | Y | Y |
| FTSJ1 | Y | Y |
| FUT2 | Y | Y |
| FUT8 | Y | Y |
| FXR1 | Y | Y |
| FXYD1 | Y | Y |
| FZD5 | Y | Y |
| FZD8 | Y | Y |
| G6PC3 | Y | Y |
| G6PD | Y | Y |
| GABARAPL1 | Y | Y |
| GABPB1-AS1 | Y | Y |
| GABPB1-IT1 | Y | Y |
| GAL3ST2 | Y | Y |
| GALNS | Y | Y |
| GALNT1 | Y | Y |
| GALNT14 | Y | Y |
| GALNT5 | Y | Y |
| GALT | Y | Y |
| GAPLINC | Y | Y |
| GARS | Y | Y |
| GATAD1 | Y | Y |
| GATD3B | Y | Y |
| GATM | Y | Y |
| GBA2 | Y | Y |
| GBAT2 | Y | Y |
| GBGT1 | Y | Y |
| GBP1 | Y | Y |
| GCC2 | Y | Y |
| GCLM | Y | Y |
| GDPD1 | Y | Y |
| GDPD3 | Y | Y |
| GEMIN2 | Y | Y |
| GEMIN4 | Y | Y |
| GEMIN8 | Y | Y |
| GFER | Y | Y |
| GGA1 | Y | Y |
| GGA3 | Y | Y |
| GGCT | Y | Y |
| GGPS1 | Y | Y |

| | | |
|---|---|---|
| FGFR3 | N | Y |
| FGGY | N | Y |
| FGR | N | Y |
| FHDC1 | N | Y |
| FHIT | N | Y |
| FHL1 | N | Y |
| FHL2 | N | Y |
| FHOD1 | N | Y |
| FIBP | N | Y |
| FIP1L1 | N | Y |
| FKBP2 | N | Y |
| FKBP3 | N | Y |
| FKBP8 | N | Y |
| FLJ37453 | N | Y |
| FLNB | N | Y |
| FLOT1 | N | Y |
| FLT3LG | N | Y |
| FLYWCH1 | N | Y |
| FMO4 | N | Y |
| FN1 | N | Y |
| FNBP1 | N | Y |
| FOCAD | N | Y |
| FOLR1 | N | Y |
| FOPNL | N | Y |
| FOXJ3 | N | Y |
| FOXO1 | N | Y |
| FOXP1 | N | Y |
| FOXRED2 | N | Y |
| FRG1 | N | Y |
| FRMD4B | N | Y |
| FSTL4 | N | Y |
| FUBP1 | N | Y |
| FUBP3 | N | Y |
| FUCA1 | N | Y |
| FUS | N | Y |
| FUT3 | N | Y |
| FXYD3 | N | Y |
| FYN | N | Y |
| FZD6 | N | Y |
| G3BP1 | N | Y |
| GAA | N | Y |
| GAK | N | Y |
| GALE | N | Y |

| | | |
|---|---|---|
| GGT1 | Y | Y |
| GGT6 | Y | Y |
| GGTA1P | Y | Y |
| GIMAP1 | Y | Y |
| GIMAP7 | Y | Y |
| GINS2 | Y | Y |
| GIPC2 | Y | Y |
| GIT1 | Y | Y |
| GJC2 | Y | Y |
| GJC3 | Y | Y |
| GK | Y | Y |
| GLCCI1 | Y | Y |
| GLE1 | Y | Y |
| GLG1 | Y | Y |
| GLI1 | Y | Y |
| GLI4 | Y | Y |
| GLIS2 | Y | Y |
| GLIS3 | Y | Y |
| GLT8D2 | Y | Y |
| GM2A | Y | Y |
| GMFB | Y | Y |
| GMFG | Y | Y |
| GMIP | Y | Y |
| GNA14 | Y | Y |
| GNLY | Y | Y |
| GNPTAB | Y | Y |
| GOLGA1 | Y | Y |
| GOLGA2 | Y | Y |
| GOLGA2P5 | Y | Y |
| GOLGA8B | Y | Y |
| GOLGA8J | Y | Y |
| GOLGA8O | Y | Y |
| GOT1 | Y | Y |
| GP1BA | Y | Y |
| GPALPP1 | Y | Y |
| GPAT4 | Y | Y |
| GPATCH11 | Y | Y |
| GPATCH2 | Y | Y |
| GPLD1 | Y | Y |
| GPR108 | Y | Y |
| GPR183 | Y | Y |
| GPT | Y | Y |
| GPX7 | Y | Y |

| | | |
|---|---|---|
| GALNT3 | N | Y |
| GAPDH | N | Y |
| GAPVD1 | N | Y |
| GAR1 | N | Y |
| GART | N | Y |
| GAS5 | N | Y |
| GATAD2A | N | Y |
| GATD1 | N | Y |
| GDE1 | N | Y |
| GDF15 | N | Y |
| GEM | N | Y |
| GEMIN7 | N | Y |
| GFOD2 | N | Y |
| GGNBP2 | N | Y |
| GID4 | N | Y |
| GIPC1 | N | Y |
| GJA9-MYCBP | N | Y |
| GJB1 | N | Y |
| GLA | N | Y |
| GLCE | N | Y |
| GLIPR2 | N | Y |
| GLRX2 | N | Y |
| GLYR1 | N | Y |
| GMCL1 | N | Y |
| GMNN | N | Y |
| GMPR2 | N | Y |
| GMPS | N | Y |
| GNAS | N | Y |
| GNG10 | N | Y |
| GNPNAT1 | N | Y |
| GOLGA4 | N | Y |
| GOLGB1 | N | Y |
| GOLM1 | N | Y |
| GOLPH3 | N | Y |
| GOSR1 | N | Y |
| GPAT2 | N | Y |
| GPBP1 | N | Y |
| GPBP1L1 | N | Y |
| GPC6 | N | Y |
| GPHN | N | Y |
| GPI | N | Y |
| GPR160 | N | Y |
| GPS1 | N | Y |

| | | |
|---|---|---|
| GRB7 | Y | Y |
| GRID1 | Y | Y |
| GRINA | Y | Y |
| GRIPAP1 | Y | Y |
| GRK6 | Y | Y |
| GRPEL1 | Y | Y |
| GSE1 | Y | Y |
| GSR | Y | Y |
| GSTM2 | Y | Y |
| GSTZ1 | Y | Y |
| GTF2A1 | Y | Y |
| GTF2E2 | Y | Y |
| GTF2H5 | Y | Y |
| GTF2IRD2 | Y | Y |
| GTF3A | Y | Y |
| GTF3C3 | Y | Y |
| GTPBP10 | Y | Y |
| GTPBP2 | Y | Y |
| GUCY2C | Y | Y |
| GUSBP11 | Y | Y |
| GUSBP5 | Y | Y |
| GYG1 | Y | Y |
| GYS1 | Y | Y |
| GZMB | Y | Y |
| HAAO | Y | Y |
| HAPLN1 | Y | Y |
| HARS | Y | Y |
| HAS2 | Y | Y |
| HAUS1 | Y | Y |
| HAUS2 | Y | Y |
| HAUS3 | Y | Y |
| HAUS4 | Y | Y |
| HBEGF | Y | Y |
| HCCS | Y | Y |
| HCP5 | Y | Y |
| HEATR3 | Y | Y |
| HEATR5B | Y | Y |
| HECTD3 | Y | Y |
| HENMT1 | Y | Y |
| HHEX | Y | Y |
| HIF1AN | Y | Y |
| HIKESHI | Y | Y |
| HINFP | Y | Y |

| | | |
|---|---|---|
| GPSM2 | N | Y |
| GPT2 | N | Y |
| GPX1 | N | Y |
| GPX4 | N | Y |
| GRAMD2B | N | Y |
| GRAP2 | N | Y |
| GRB10 | N | Y |
| GRB2 | N | Y |
| GRK2 | N | Y |
| GRK3 | N | Y |
| GRM8 | N | Y |
| GRSF1 | N | Y |
| GRTP1 | N | Y |
| GSDMD | N | Y |
| GSK3B | N | Y |
| GSKIP | N | Y |
| GSN | N | Y |
| GSPT1 | N | Y |
| GSS | N | Y |
| GSTO1 | N | Y |
| GSTT1 | N | Y |
| GTF2F1 | N | Y |
| GTF2H3 | N | Y |
| GTF2IRD1 | N | Y |
| GTF3C2 | N | Y |
| GTF3C5 | N | Y |
| GUK1 | N | Y |
| GUSB | N | Y |
| H2AFJ | N | Y |
| H2AFV | N | Y |
| H2AFY | N | Y |
| H6PD | N | Y |
| HABP4 | N | Y |
| HADHB | N | Y |
| HARBI1 | N | Y |
| HAT1 | N | Y |
| HBP1 | N | Y |
| HBS1L | N | Y |
| HDAC5 | N | Y |
| HDGF | N | Y |
| HDGFL2 | N | Y |
| HDHD2 | N | Y |
| HDHD5-AS1 | N | Y |

| | | |
|---|---|---|
| HIP1R | Y | Y |
| HIPK1 | Y | Y |
| HIPK2 | Y | Y |
| HIST1H1C | Y | Y |
| HIST1H2BH | Y | Y |
| HIST2H2BC | Y | Y |
| HIST3H2A | Y | Y |
| HK1 | Y | Y |
| HLA-DPB1 | Y | Y |
| HLA-G | Y | Y |
| HLF | Y | Y |
| HMCES | Y | Y |
| HMCN1 | Y | Y |
| HMG20B | Y | Y |
| HMGCLL1 | Y | Y |
| HOXA13 | Y | Y |
| HOXA4 | Y | Y |
| HOXB4 | Y | Y |
| HOXD1 | Y | Y |
| HOXD11 | Y | Y |
| HOXD13 | Y | Y |
| HPS6 | Y | Y |
| HR | Y | Y |
| HRASLS | Y | Y |
| HS1BP3 | Y | Y |
| HSD17B7 | Y | Y |
| HSD17B7P2 | Y | Y |
| HSD17B8 | Y | Y |
| HSF1 | Y | Y |
| HSH2D | Y | Y |
| HSPA13 | Y | Y |
| HTD2 | Y | Y |
| HTR7P1 | Y | Y |
| HYAL2 | Y | Y |
| ICAM1 | Y | Y |
| ICAM4 | Y | Y |
| ICMT | Y | Y |
| ID2-AS1 | Y | Y |
| IDH3A | Y | Y |
| IDNK | Y | Y |
| IER2 | Y | Y |
| IER5L | Y | Y |
| IFFO1 | Y | Y |

| | | |
|---|---|---|
| HEBP2 | N | Y |
| HEPACAM2 | N | Y |
| HERPUD2 | N | Y |
| HES4 | N | Y |
| HEYL | N | Y |
| HHLA2 | N | Y |
| HIBCH | N | Y |
| HIGD1A | N | Y |
| HIST1H2BD | N | Y |
| HLA-F | N | Y |
| HLA-F-AS1 | N | Y |
| HMGA1 | N | Y |
| HMGB1 | N | Y |
| HMGCS1 | N | Y |
| HMGCS2 | N | Y |
| HMGN1 | N | Y |
| HMGN3 | N | Y |
| HNF1B | N | Y |
| HNMT | N | Y |
| HNRNPA1 | N | Y |
| HNRNPA3 | N | Y |
| HNRNPC | N | Y |
| HNRNPD | N | Y |
| HNRNPF | N | Y |
| HNRNPH1 | N | Y |
| HNRNPH3 | N | Y |
| HNRNPK | N | Y |
| HNRNPL | N | Y |
| HNRNPM | N | Y |
| HNRNPR | N | Y |
| HNRNPUL1 | N | Y |
| HOMER2 | N | Y |
| HOOK2 | N | Y |
| HOPX | N | Y |
| HOTTIP | N | Y |
| HOXA11-AS | N | Y |
| HOXB-AS3 | N | Y |
| HOXB3 | N | Y |
| HOXB6 | N | Y |
| HOXB8 | N | Y |
| HOXB9 | N | Y |
| HRH1 | N | Y |
| HSD17B11 | N | Y |

| | | |
|---|---|---|
| IFI16 | Y | Y |
| IFIT1 | Y | Y |
| IFIT3 | Y | Y |
| IFNAR2 | Y | Y |
| IFRD1 | Y | Y |
| IFT172 | Y | Y |
| IGF1R | Y | Y |
| IGF2 | Y | Y |
| IGFBP2 | Y | Y |
| IL16 | Y | Y |
| IL1R2 | Y | Y |
| IL20RB | Y | Y |
| IL34 | Y | Y |
| IL4I1 | Y | Y |
| IL4R | Y | Y |
| IL6R | Y | Y |
| IL7 | Y | Y |
| IL7R | Y | Y |
| ILDR1 | Y | Y |
| ILK | Y | Y |
| ILKAP | Y | Y |
| IMMP1L | Y | Y |
| IMPA2 | Y | Y |
| INAVA | Y | Y |
| INCA1 | Y | Y |
| INF2 | Y | Y |
| ING1 | Y | Y |
| INO80B-WBP1 | Y | Y |
| INPP5A | Y | Y |
| INPP5E | Y | Y |
| INSIG2 | Y | Y |
| INTS11 | Y | Y |
| INTS3 | Y | Y |
| INTS5 | Y | Y |
| IPO5 | Y | Y |
| IPO8 | Y | Y |
| IPO9 | Y | Y |
| IQGAP2 | Y | Y |
| IREB2 | Y | Y |
| IRF3 | Y | Y |
| IRF6 | Y | Y |
| ISOC1 | Y | Y |
| ITCH | Y | Y |

| | | |
|---|---|---|
| HSD17B14 | N | Y |
| HSPA1A | N | Y |
| HSPBP1 | N | Y |
| HSPH1 | N | Y |
| HTATIP2 | N | Y |
| HUS1 | N | Y |
| HUWE1 | N | Y |
| HYOU1 | N | Y |
| IAH1 | N | Y |
| ICA1 | N | Y |
| IDH2 | N | Y |
| IDH3B | N | Y |
| IDH3G | N | Y |
| IDI1 | N | Y |
| IDS | N | Y |
| IFI27 | N | Y |
| IFI27L1 | N | Y |
| IFI27L2 | N | Y |
| IFT140 | N | Y |
| IFT80 | N | Y |
| IGBP1 | N | Y |
| IGF2BP2 | N | Y |
| IGSF8 | N | Y |
| IKBKB | N | Y |
| IKZF1 | N | Y |
| IL15RA | N | Y |
| IL17RE | N | Y |
| IL20RA | N | Y |
| IL32 | N | Y |
| ILF3 | N | Y |
| IMMP2L | N | Y |
| IMMT | N | Y |
| IMP4 | N | Y |
| IMPDH1 | N | Y |
| ING3 | N | Y |
| ING4 | N | Y |
| INO80C | N | Y |
| INO80E | N | Y |
| INPP4B | N | Y |
| INPP5D | N | Y |
| INPP5K | N | Y |
| INSC | N | Y |
| INTS10 | N | Y |

| | | |
|---|---|---|
| ITFG2 | Y | Y |
| ITGA1 | Y | Y |
| ITGA3 | Y | Y |
| ITGB3BP | Y | Y |
| ITLN1 | Y | Y |
| ITM2A | Y | Y |
| ITPKA | Y | Y |
| ITPR3 | Y | Y |
| ITPRIP | Y | Y |
| JADE3 | Y | Y |
| JAG1 | Y | Y |
| JMJD4 | Y | Y |
| JMJD6 | Y | Y |
| JMJD7 | Y | Y |
| JMY | Y | Y |
| JRK | Y | Y |
| JUND | Y | Y |
| KANSL2 | Y | Y |
| KANSL3 | Y | Y |
| KAT7 | Y | Y |
| KBTBD3 | Y | Y |
| KCNA6 | Y | Y |
| KCNIP4 | Y | Y |
| KCNK15 | Y | Y |
| KCNK5 | Y | Y |
| KCNMA1 | Y | Y |
| KCNS3 | Y | Y |
| KCTD10 | Y | Y |
| KCTD20 | Y | Y |
| KCTD5 | Y | Y |
| KDELC2 | Y | Y |
| KDM1A | Y | Y |
| KEAP1 | Y | Y |
| KIAA0895L | Y | Y |
| KIAA1143 | Y | Y |
| KIAA1958 | Y | Y |
| KIF12 | Y | Y |
| KIF23 | Y | Y |
| KIF25 | Y | Y |
| KIF2C | Y | Y |
| KLC1 | Y | Y |
| KLF10 | Y | Y |
| KLF3-AS1 | Y | Y |

| | | |
|---|---|---|
| INTS14 | N | Y |
| INTS7 | N | Y |
| INTS9 | N | Y |
| IPO4 | N | Y |
| IPPK | N | Y |
| IQCH | N | Y |
| IRF7 | N | Y |
| IRF8 | N | Y |
| ISG20 | N | Y |
| ITGA6 | N | Y |
| ITGAE | N | Y |
| ITGB1 | N | Y |
| ITGB1BP1 | N | Y |
| IVD | N | Y |
| JAK2 | N | Y |
| JKAMP | N | Y |
| JRKL | N | Y |
| KARS | N | Y |
| KAT5 | N | Y |
| KAT8 | N | Y |
| KATNA1 | N | Y |
| KBTBD2 | N | Y |
| KCNK1 | N | Y |
| KCNN3 | N | Y |
| KCNN4 | N | Y |
| KCTD13 | N | Y |
| KCTD14 | N | Y |
| KDM2A | N | Y |
| KDM4B | N | Y |
| KDM5C | N | Y |
| KDM5D | N | Y |
| KDM6A | N | Y |
| KHDRBS1 | N | Y |
| KHDRBS3 | N | Y |
| KIAA0232 | N | Y |
| KIAA0391 | N | Y |
| KIAA1109 | N | Y |
| KIAA1191 | N | Y |
| KIAA1324 | N | Y |
| KIF13A | N | Y |
| KIF9 | N | Y |
| KIT | N | Y |
| KLC4 | N | Y |

| | | |
|---|---|---|
| KLHDC1 | Y | Y |
| KLHDC9 | Y | Y |
| KLHL21 | Y | Y |
| KLHL36 | Y | Y |
| KLHL42 | Y | Y |
| KLHL7 | Y | Y |
| KLK12 | Y | Y |
| KLK3 | Y | Y |
| KLK6 | Y | Y |
| KMT5A | Y | Y |
| KMT5C | Y | Y |
| KPNA7 | Y | Y |
| KPNB1 | Y | Y |
| KRR1 | Y | Y |
| KRT15 | Y | Y |
| KRT6A | Y | Y |
| L2HGDH | Y | Y |
| LAMA5 | Y | Y |
| LANCL1 | Y | Y |
| LANCL2 | Y | Y |
| LARP4 | Y | Y |
| LAT2 | Y | Y |
| LBR | Y | Y |
| LCA5L | Y | Y |
| LCMT1 | Y | Y |
| LDB1 | Y | Y |
| LDHD | Y | Y |
| LEFTY1 | Y | Y |
| LEMD1 | Y | Y |
| LEMD3 | Y | Y |
| LEO1 | Y | Y |
| LEPROT | Y | Y |
| LGALS9 | Y | Y |
| LGMN | Y | Y |
| LGR4 | Y | Y |
| LGR6 | Y | Y |
| LHFPL2 | Y | Y |
| LHFPL6 | Y | Y |
| LILRA5 | Y | Y |
| LIMA1 | Y | Y |
| LIME1 | Y | Y |
| LIN7A | Y | Y |
| LIN7C | Y | Y |

| | | |
|---|---|---|
| KLF6 | N | Y |
| KLHDC2 | N | Y |
| KLHDC3 | N | Y |
| KLHL2 | N | Y |
| KLHL24 | N | Y |
| KLHL6 | N | Y |
| KLK1 | N | Y |
| KLK11 | N | Y |
| KMT2A | N | Y |
| KMT2C | N | Y |
| KRT10 | N | Y |
| KRTCAP3 | N | Y |
| KTN1 | N | Y |
| KYAT1 | N | Y |
| KYNU | N | Y |
| LAMP2 | N | Y |
| LAMP3 | N | Y |
| LAMTOR3 | N | Y |
| LAMTOR4 | N | Y |
| LARP1 | N | Y |
| LARP4B | N | Y |
| LARP7 | N | Y |
| LATS1 | N | Y |
| LDHB | N | Y |
| LDLR | N | Y |
| LETMD1 | N | Y |
| LFNG | N | Y |
| LGALS3 | N | Y |
| LGALS4 | N | Y |
| LGR5 | N | Y |
| LIAS | N | Y |
| LIMCH1 | N | Y |
| LIMK2 | N | Y |
| LIMS2 | N | Y |
| LINC00543 | N | Y |
| LINC00667 | N | Y |
| LINC00869 | N | Y |
| LINC01123 | N | Y |
| LINC01578 | N | Y |
| LINC02562 | N | Y |
| LITAF | N | Y |
| LLGL2 | N | Y |
| LMBRD2 | N | Y |

| | | |
|---|---|---|
| LINC-PINT | Y | Y |
| LINC00114 | Y | Y |
| LINC00167 | Y | Y |
| LINC00265 | Y | Y |
| LINC00476 | Y | Y |
| LINC00482 | Y | Y |
| LINC00511 | Y | Y |
| LINC00513 | Y | Y |
| LINC00630 | Y | Y |
| LINC00639 | Y | Y |
| LINC00668 | Y | Y |
| LINC00673 | Y | Y |
| LINC00842 | Y | Y |
| LINC00847 | Y | Y |
| LINC00857 | Y | Y |
| LINC01000 | Y | Y |
| LINC01133 | Y | Y |
| LINC01279 | Y | Y |
| LINC01315 | Y | Y |
| LINC01389 | Y | Y |
| LINC01431 | Y | Y |
| LINC01481 | Y | Y |
| LINC01504 | Y | Y |
| LINC01547 | Y | Y |
| LINC01550 | Y | Y |
| LINC01590 | Y | Y |
| LINC01781 | Y | Y |
| LINC01814 | Y | Y |
| LINC01871 | Y | Y |
| LINC01881 | Y | Y |
| LINC02001 | Y | Y |
| LINC02018 | Y | Y |
| LINC02023 | Y | Y |
| LINC02108 | Y | Y |
| LINC02332 | Y | Y |
| LINC02535 | Y | Y |
| LINC02588 | Y | Y |
| LIPH | Y | Y |
| LIPT2 | Y | Y |
| LMNB2 | Y | Y |
| LMOD1 | Y | Y |
| LMTK2 | Y | Y |
| LMTK3 | Y | Y |

| | | |
|---|---|---|
| LMCD1 | N | Y |
| LMF1 | N | Y |
| LMNA | N | Y |
| LMO4 | N | Y |
| LNX1 | N | Y |
| LOC100288798 | N | Y |
| LOC100506922 | N | Y |
| LOC100996717 | N | Y |
| LOC100996763 | N | Y |
| LOC101928069 | N | Y |
| LOC101928881 | N | Y |
| LOC101929243 | N | Y |
| LOC102724715 | N | Y |
| LOC102724985 | N | Y |
| LOC105369315 | N | Y |
| LOC105370969 | N | Y |
| LOC105372629 | N | Y |
| LOC105375566 | N | Y |
| LOC105376567 | N | Y |
| LOC107985200 | N | Y |
| LOC112268277 | N | Y |
| LOC112268389 | N | Y |
| LPCAT3 | N | Y |
| LPP | N | Y |
| LRP6 | N | Y |
| LRPAP1 | N | Y |
| LRRC23 | N | Y |
| LRRC40 | N | Y |
| LRRC6 | N | Y |
| LRRC8A | N | Y |
| LSM1 | N | Y |
| LSM6 | N | Y |
| LSR | N | Y |
| LSS | N | Y |
| LST1 | N | Y |
| LTA4H | N | Y |
| LTBP1 | N | Y |
| LTBP3 | N | Y |
| LTBP4 | N | Y |
| LTF | N | Y |
| LUC7L3 | N | Y |
| LURAP1L | N | Y |
| LYPLA2 | N | Y |

| | | |
|---|---|---|
| LOC100129034 | Y | Y |
| LOC100130298 | Y | Y |
| LOC100132154 | Y | Y |
| LOC100133331 | Y | Y |
| LOC100190986 | Y | Y |
| LOC100233156 | Y | Y |
| LOC100288846 | Y | Y |
| LOC100289230 | Y | Y |
| LOC100289361 | Y | Y |
| LOC100289561 | Y | Y |
| LOC100419583 | Y | Y |
| LOC100505549 | Y | Y |
| LOC100506123 | Y | Y |
| LOC100506472 | Y | Y |
| LOC100506476 | Y | Y |
| LOC100506544 | Y | Y |
| LOC100507507 | Y | Y |
| LOC100996756 | Y | Y |
| LOC101926967 | Y | Y |
| LOC101927069 | Y | Y |
| LOC101927100 | Y | Y |
| LOC101927511 | Y | Y |
| LOC101928076 | Y | Y |
| LOC101928156 | Y | Y |
| LOC101928215 | Y | Y |
| LOC101928323 | Y | Y |
| LOC101928347 | Y | Y |
| LOC101929709 | Y | Y |
| LOC101929796 | Y | Y |
| LOC101930071 | Y | Y |
| LOC102723728 | Y | Y |
| LOC102723878 | Y | Y |
| LOC102724156 | Y | Y |
| LOC102724397 | Y | Y |
| LOC102724720 | Y | Y |
| LOC102724760 | Y | Y |
| LOC103091866 | Y | Y |
| LOC105369347 | Y | Y |
| LOC105369360 | Y | Y |
| LOC105369370 | Y | Y |
| LOC105369656 | Y | Y |
| LOC105369690 | Y | Y |
| LOC105369819 | Y | Y |

| | | |
|---|---|---|
| LYPLAL1 | N | Y |
| LYRM2 | N | Y |
| LYRM4 | N | Y |
| LYSMD2 | N | Y |
| LZTFL1 | N | Y |
| M6PR | N | Y |
| MACF1 | N | Y |
| MAD1L1 | N | Y |
| MADD | N | Y |
| MAFIP | N | Y |
| MAFK | N | Y |
| MAGOHB | N | Y |
| MALL | N | Y |
| MALT1 | N | Y |
| MAN2A1 | N | Y |
| MAN2B2 | N | Y |
| MAP1LC3A | N | Y |
| MAP1S | N | Y |
| MAP2K6 | N | Y |
| MAP3K20 | N | Y |
| MAP4 | N | Y |
| MAP7 | N | Y |
| MAP7D1 | N | Y |
| MAPK10 | N | Y |
| MAPKAP1 | N | Y |
| MAPKAPK5-AS1 | N | Y |
| MAPRE2 | N | Y |
| 5-Mar | N | Y |
| 6-Mar | N | Y |
| MARK2 | N | Y |
| MARK3 | N | Y |
| MATN2 | N | Y |
| MAX | N | Y |
| MAZ | N | Y |
| MBIP | N | Y |
| MBP | N | Y |
| MCAT | N | Y |
| MCM7 | N | Y |
| MCM9 | N | Y |
| MCOLN2 | N | Y |
| MCRIP2 | N | Y |
| MCRS1 | N | Y |
| MCTP1 | N | Y |

| | | |
|---|---|---|
| LOC105370624 | Y | Y |
| LOC105370858 | Y | Y |
| LOC105371257 | Y | Y |
| LOC105371554 | Y | Y |
| LOC105371762 | Y | Y |
| LOC105371763 | Y | Y |
| LOC105371870 | Y | Y |
| LOC105372246 | Y | Y |
| LOC105372401 | Y | Y |
| LOC105372426 | Y | Y |
| LOC105372441 | Y | Y |
| LOC105372480 | Y | Y |
| LOC105372578 | Y | Y |
| LOC105372803 | Y | Y |
| LOC105372962 | Y | Y |
| LOC105373733 | Y | Y |
| LOC105373759 | Y | Y |
| LOC105373785 | Y | Y |
| LOC105374041 | Y | Y |
| LOC105374080 | Y | Y |
| LOC105374148 | Y | Y |
| LOC105376095 | Y | Y |
| LOC105376159 | Y | Y |
| LOC105377763 | Y | Y |
| LOC105378179 | Y | Y |
| LOC105378539 | Y | Y |
| LOC105378720 | Y | Y |
| LOC105378948 | Y | Y |
| LOC105379172 | Y | Y |
| LOC105379280 | Y | Y |
| LOC105379499 | Y | Y |
| LOC105379508 | Y | Y |
| LOC105379524 | Y | Y |
| LOC107984329 | Y | Y |
| LOC107984347 | Y | Y |
| LOC107984813 | Y | Y |
| LOC107984948 | Y | Y |
| LOC107985208 | Y | Y |
| LOC107985216 | Y | Y |
| LOC107985249 | Y | Y |
| LOC107985367 | Y | Y |
| LOC107985404 | Y | Y |
| LOC107985433 | Y | Y |

| | | |
|---|---|---|
| MCU | N | Y |
| MDFI | N | Y |
| MDM2 | N | Y |
| MEA1 | N | Y |
| MEAF6 | N | Y |
| MECR | N | Y |
| MED1 | N | Y |
| MED15 | N | Y |
| MED24 | N | Y |
| MED30 | N | Y |
| MED8 | N | Y |
| MEG3 | N | Y |
| MEN1 | N | Y |
| MERTK | N | Y |
| MESD | N | Y |
| MET | N | Y |
| METTL13 | N | Y |
| METTL17 | N | Y |
| METTL26 | N | Y |
| METTL3 | N | Y |
| METTL5 | N | Y |
| METTL6 | N | Y |
| METTL9 | N | Y |
| MFF | N | Y |
| MFSD1 | N | Y |
| MFSD3 | N | Y |
| MFSD5 | N | Y |
| MFSD9 | N | Y |
| MGA | N | Y |
| MGAT4B | N | Y |
| MGST1 | N | Y |
| MGST2 | N | Y |
| MICAL2 | N | Y |
| MID2 | N | Y |
| MIF4GD | N | Y |
| MIGA1 | N | Y |
| MINDY1 | N | Y |
| MINDY3 | N | Y |
| MINPP1 | N | Y |
| MIS18A | N | Y |
| MISP | N | Y |
| MITF | N | Y |
| MKL1 | N | Y |

| | | |
|---|---|---|
| LOC107985480 | Y | Y |
| LOC107985554 | Y | Y |
| LOC107985567 | Y | Y |
| LOC107985733 | Y | Y |
| LOC107985910 | Y | Y |
| LOC107985917 | Y | Y |
| LOC107985923 | Y | Y |
| LOC107986003 | Y | Y |
| LOC107986298 | Y | Y |
| LOC107986348 | Y | Y |
| LOC107986483 | Y | Y |
| LOC107986854 | Y | Y |
| LOC107986991 | Y | Y |
| LOC107986997 | Y | Y |
| LOC107987043 | Y | Y |
| LOC107987072 | Y | Y |
| LOC107987102 | Y | Y |
| LOC107987135 | Y | Y |
| LOC107987136 | Y | Y |
| LOC107987206 | Y | Y |
| LOC107987251 | Y | Y |
| LOC107987294 | Y | Y |
| LOC107987299 | Y | Y |
| LOC108783654 | Y | Y |
| LOC110384692 | Y | Y |
| LOC112267883 | Y | Y |
| LOC112268035 | Y | Y |
| LOC112268037 | Y | Y |
| LOC112268063 | Y | Y |
| LOC112268088 | Y | Y |
| LOC112268098 | Y | Y |
| LOC112268109 | Y | Y |
| LOC112268124 | Y | Y |
| LOC112268146 | Y | Y |
| LOC112268184 | Y | Y |
| LOC112268313 | Y | Y |
| LOC171391 | Y | Y |
| LOC257396 | Y | Y |
| LOC283788 | Y | Y |
| LOC284454 | Y | Y |
| LOC389602 | Y | Y |
| LOC389831 | Y | Y |
| LOC389906 | Y | Y |

| | | |
|---|---|---|
| MKL2 | N | Y |
| MKRN1 | N | Y |
| MLH1 | N | Y |
| MLLT10 | N | Y |
| MMP19 | N | Y |
| MMP24OS | N | Y |
| MMP28 | N | Y |
| MNAT1 | N | Y |
| MOB1A | N | Y |
| MORC2 | N | Y |
| MORC3 | N | Y |
| MORF4L2 | N | Y |
| MOSPD3 | N | Y |
| MPC1 | N | Y |
| MPDU1 | N | Y |
| MPPE1 | N | Y |
| MRGBP | N | Y |
| MRPL14 | N | Y |
| MRPL21 | N | Y |
| MRPL22 | N | Y |
| MRPL34 | N | Y |
| MRPL42 | N | Y |
| MRPL43 | N | Y |
| MRPL47 | N | Y |
| MRPL49 | N | Y |
| MRPL52 | N | Y |
| MRPL55 | N | Y |
| MRPL57 | N | Y |
| MRPL58 | N | Y |
| MRPL9 | N | Y |
| MRPS11 | N | Y |
| MRPS12 | N | Y |
| MRPS25 | N | Y |
| MRPS35 | N | Y |
| MRPS5 | N | Y |
| MRPS9 | N | Y |
| MRTFA | N | Y |
| MS4A1 | N | Y |
| MSH2 | N | Y |
| MSI2 | N | Y |
| MSLN | N | Y |
| MSMB | N | Y |
| MST1R | N | Y |

| | | |
|---|---|---|
| LOC646626 | Y | Y |
| LOC728743 | Y | Y |
| LOC729737 | Y | Y |
| LOC93622 | Y | Y |
| LONP1 | Y | Y |
| LONP2 | Y | Y |
| LONRF1 | Y | Y |
| LPCAT1 | Y | Y |
| LPIN3 | Y | Y |
| LRCH4 | Y | Y |
| LRFN4 | Y | Y |
| LRIG2 | Y | Y |
| LRP5 | Y | Y |
| LRRC14 | Y | Y |
| LRRC32 | Y | Y |
| LRRC34 | Y | Y |
| LRRC42 | Y | Y |
| LRRC75A | Y | Y |
| LRRFIP2 | Y | Y |
| LSG1 | Y | Y |
| LSM14A | Y | Y |
| LTB | Y | Y |
| LTV1 | Y | Y |
| LYRM1 | Y | Y |
| LYVE1 | Y | Y |
| MAD2L1BP | Y | Y |
| MAF | Y | Y |
| MAFB | Y | Y |
| MAFF | Y | Y |
| MAGED1 | Y | Y |
| MAGED2 | Y | Y |
| MAGEF1 | Y | Y |
| MAGEH1 | Y | Y |
| MAGIX | Y | Y |
| MAIP1 | Y | Y |
| MAMDC4 | Y | Y |
| MAN1B1 | Y | Y |
| MAN2C1 | Y | Y |
| MAOA | Y | Y |
| MAP2K7 | Y | Y |
| MAP3K21 | Y | Y |
| MAP3K4 | Y | Y |
| MAP3K5 | Y | Y |

| | | |
|---|---|---|
| MTA1 | N | Y |
| MTCH1 | N | Y |
| MTCH2 | N | Y |
| MTERF4 | N | Y |
| MTF2 | N | Y |
| MTFR2 | N | Y |
| MTIF3 | N | Y |
| MTMR11 | N | Y |
| MTMR12 | N | Y |
| MTMR14 | N | Y |
| MTMR2 | N | Y |
| MTOR | N | Y |
| MTRF1 | N | Y |
| MTSS1 | N | Y |
| MTX2 | N | Y |
| MUC4 | N | Y |
| MXI1 | N | Y |
| MYB | N | Y |
| MYL9 | N | Y |
| MYLK | N | Y |
| MYO1A | N | Y |
| MYO1D | N | Y |
| MYO6 | N | Y |
| MYOF | N | Y |
| MYRIP | N | Y |
| MZT2A | N | Y |
| N4BP1 | N | Y |
| N4BP2L2 | N | Y |
| N4BP3 | N | Y |
| NAA20 | N | Y |
| NAA38 | N | Y |
| NABP2 | N | Y |
| NACA | N | Y |
| NADK | N | Y |
| NANS | N | Y |
| NAP1L1 | N | Y |
| NAPA | N | Y |
| NAPG | N | Y |
| NAPRT | N | Y |
| NARS2 | N | Y |
| NAXD | N | Y |
| NBL1 | N | Y |
| NCALD | N | Y |

| | | |
|---|---|---|
| MAP3K6 | Y | Y |
| MAP4K4 | Y | Y |
| MAP4K5 | Y | Y |
| MAPK1IP1L | Y | Y |
| MAPKAPK2 | Y | Y |
| MAPRE1 | Y | Y |
| 2-Mar | Y | Y |
| MARF1 | Y | Y |
| MARK1 | Y | Y |
| MARVELD3 | Y | Y |
| MAVS | Y | Y |
| MB | Y | Y |
| MBD6 | Y | Y |
| MBOAT4 | Y | Y |
| MBTPS2 | Y | Y |
| MCCC1 | Y | Y |
| MCEE | Y | Y |
| MCRIP1 | Y | Y |
| MDFIC | Y | Y |
| MDM1 | Y | Y |
| MDP1 | Y | Y |
| ME3 | Y | Y |
| MECOM | Y | Y |
| MED14 | Y | Y |
| MED25 | Y | Y |
| MED27 | Y | Y |
| MED28 | Y | Y |
| MED29 | Y | Y |
| MED6 | Y | Y |
| MEF2D | Y | Y |
| MEGF6 | Y | Y |
| MELTF | Y | Y |
| MEPCE | Y | Y |
| MESP1 | Y | Y |
| METTL1 | Y | Y |
| METTL21A | Y | Y |
| METTL25 | Y | Y |
| METTL2B | Y | Y |
| METTL7B | Y | Y |
| MEX3C | Y | Y |
| MFAP1 | Y | Y |
| MFAP4 | Y | Y |
| MFRP | Y | Y |

| | | |
|---|---|---|
| NCAM2 | N | Y |
| NCAPH | N | Y |
| NCBP3 | N | Y |
| NCKAP5 | N | Y |
| NCOA2 | N | Y |
| NCOA4 | N | Y |
| NCOA7 | N | Y |
| NCOR1 | N | Y |
| NDEL1 | N | Y |
| NDRG2 | N | Y |
| NDUFA10 | N | Y |
| NDUFA11 | N | Y |
| NDUFA2 | N | Y |
| NDUFA3 | N | Y |
| NDUFAB1 | N | Y |
| NDUFAF5 | N | Y |
| NDUFAF7 | N | Y |
| NDUFB3 | N | Y |
| NDUFB6 | N | Y |
| NDUFS1 | N | Y |
| NDUFS2 | N | Y |
| NDUFS4 | N | Y |
| NDUFV2 | N | Y |
| NDUFV3 | N | Y |
| NEAT1 | N | Y |
| NEBL | N | Y |
| NECAP1 | N | Y |
| NEDD4L | N | Y |
| NEK4 | N | Y |
| NEK6 | N | Y |
| NELFCD | N | Y |
| NELL2 | N | Y |
| NEMP2 | N | Y |
| NET1 | N | Y |
| NFIB | N | Y |
| NFIC | N | Y |
| NFS1 | N | Y |
| NFU1 | N | Y |
| NFYC | N | Y |
| NGEF | N | Y |
| NGLY1 | N | Y |
| NHSL1 | N | Y |
| NINJ1 | N | Y |

| | | |
|---|---|---|
| MFSD10 | Y | Y |
| MFSD11 | Y | Y |
| MFSD14A | Y | Y |
| MFSD14B | Y | Y |
| MFSD2A | Y | Y |
| MFSD6 | Y | Y |
| MFSD6L | Y | Y |
| MGAT1 | Y | Y |
| MGAT2 | Y | Y |
| MIA2 | Y | Y |
| MIAT | Y | Y |
| MICALL2 | Y | Y |
| MIPEPP3 | Y | Y |
| MIR200CHG | Y | Y |
| MIS12 | Y | Y |
| MISP3 | Y | Y |
| MITD1 | Y | Y |
| MLLT3 | Y | Y |
| MLPH | Y | Y |
| MMEL1 | Y | Y |
| MMP14 | Y | Y |
| MMP17 | Y | Y |
| MMP2 | Y | Y |
| MMP25 | Y | Y |
| MMUT | Y | Y |
| MOB2 | Y | Y |
| MOB3B | Y | Y |
| MOCS2 | Y | Y |
| MORC4 | Y | Y |
| MORN2 | Y | Y |
| MORN4 | Y | Y |
| MOSPD1 | Y | Y |
| MPEG1 | Y | Y |
| MPG | Y | Y |
| MPHOSPH6 | Y | Y |
| MPND | Y | Y |
| MPP6 | Y | Y |
| MPPED2 | Y | Y |
| MPZL2 | Y | Y |
| MPZL3 | Y | Y |
| MREG | Y | Y |
| MRFAP1L1 | Y | Y |
| MROH3P | Y | Y |

| | | |
|---|---|---|
| NIPA2 | N | Y |
| NISCH | N | Y |
| NIT1 | N | Y |
| NMD3 | N | Y |
| NME3 | N | Y |
| NMI | N | Y |
| NMT1 | N | Y |
| NOB1 | N | Y |
| NOL4 | N | Y |
| NOL8 | N | Y |
| NOSIP | N | Y |
| NOTCH1 | N | Y |
| NOX1 | N | Y |
| NOXA1 | N | Y |
| NPAS2 | N | Y |
| NPIPB4 | N | Y |
| NPTN | N | Y |
| NQO1 | N | Y |
| NR2C1 | N | Y |
| NR2C2 | N | Y |
| NR2F2 | N | Y |
| NRBP1 | N | Y |
| NRP1 | N | Y |
| NSD1 | N | Y |
| NSD2 | N | Y |
| NSDHL | N | Y |
| NSMCE1 | N | Y |
| NSMCE2 | N | Y |
| NSMCE4A | N | Y |
| NSUN2 | N | Y |
| NSUN5 | N | Y |
| NT5C | N | Y |
| NT5C2 | N | Y |
| NT5C3A | N | Y |
| NT5C3B | N | Y |
| NT5DC1 | N | Y |
| NTM | N | Y |
| NTRK3 | N | Y |
| NUCB1 | N | Y |
| NUCB2 | N | Y |
| NUDT16 | N | Y |
| NUDT16L1 | N | Y |
| NUDT4 | N | Y |

| | | |
|---|---|---|
| MRPL10 | Y | Y |
| MRPL20-AS1 | Y | Y |
| MRPL28 | Y | Y |
| MRPL37 | Y | Y |
| MRPL40 | Y | Y |
| MRPL46 | Y | Y |
| MRPS10 | Y | Y |
| MRPS17 | Y | Y |
| MRPS18B | Y | Y |
| MRPS30 | Y | Y |
| MRS2 | Y | Y |
| MRTO4 | Y | Y |
| MS4A7 | Y | Y |
| MSANTD2 | Y | Y |
| MSH5-SAPCD1 | Y | Y |
| MSN | Y | Y |
| MSRB3 | Y | Y |
| MT1A | Y | Y |
| MT1DP | Y | Y |
| MTA3 | Y | Y |
| MTHFD1L | Y | Y |
| MTHFD2 | Y | Y |
| MTHFR | Y | Y |
| MTMR4 | Y | Y |
| MTPN | Y | Y |
| MTR | Y | Y |
| MTRF1L | Y | Y |
| MTRNR2L6 | Y | Y |
| MTURN | Y | Y |
| MUC12 | Y | Y |
| MUC2 | Y | Y |
| MUC5B | Y | Y |
| MUL1 | Y | Y |
| MUSTN1 | Y | Y |
| MXD1 | Y | Y |
| MXD3 | Y | Y |
| MXD4 | Y | Y |
| MYADM | Y | Y |
| MYCBP | Y | Y |
| MYD88 | Y | Y |
| MYL6B | Y | Y |
| MYNN | Y | Y |
| MYO10 | Y | Y |

| | | |
|---|---|---|
| NUFIP2 | N | Y |
| NUMA1 | N | Y |
| NUP210 | N | Y |
| NUP35 | N | Y |
| NUP85 | N | Y |
| NUP88 | N | Y |
| NUPR1 | N | Y |
| NUS1 | N | Y |
| NXF1 | N | Y |
| NXPE2 | N | Y |
| NYNRIN | N | Y |
| OARD1 | N | Y |
| OAS1 | N | Y |
| OAT | N | Y |
| OCEL1 | N | Y |
| OCIAD2 | N | Y |
| OFD1 | N | Y |
| OGDH | N | Y |
| OGFOD3 | N | Y |
| OGT | N | Y |
| OLFM1 | N | Y |
| OLR1 | N | Y |
| ORMDL1 | N | Y |
| OS9 | N | Y |
| OSBPL2 | N | Y |
| OSBPL3 | N | Y |
| OSBPL7 | N | Y |
| OSBPL9 | N | Y |
| OSER1 | N | Y |
| OSTF1 | N | Y |
| OTUB1 | N | Y |
| OTUD5 | N | Y |
| OTUD6B-AS1 | N | Y |
| OXLD1 | N | Y |
| OXNAD1 | N | Y |
| OXR1 | N | Y |
| P2RX4 | N | Y |
| P2RY13 | N | Y |
| P3H1 | N | Y |
| P3H2 | N | Y |
| P4HA2 | N | Y |
| PAAF1 | N | Y |
| PABPC1L | N | Y |

| | | |
|---|---|---|
| MYO1E | Y | Y |
| MYO1G | Y | Y |
| MYO5B | Y | Y |
| MYO7B | Y | Y |
| MYOM1 | Y | Y |
| MYOZ2 | Y | Y |
| MYZAP | Y | Y |
| NAA25 | Y | Y |
| NAB1 | Y | Y |
| NARF | Y | Y |
| NAT10 | Y | Y |
| NAT2 | Y | Y |
| NAT9 | Y | Y |
| NBPF14 | Y | Y |
| NBPF19 | Y | Y |
| NBR1 | Y | Y |
| NCAPG2 | Y | Y |
| NCAPH2 | Y | Y |
| NCBP1 | Y | Y |
| NCKAP1 | Y | Y |
| NCOA1 | Y | Y |
| NCOA5 | Y | Y |
| NDFIP1 | Y | Y |
| NDN | Y | Y |
| NDST1 | Y | Y |
| NDUFAF4 | Y | Y |
| NDUFAF6 | Y | Y |
| NECAP2 | Y | Y |
| NEFL | Y | Y |
| NEIL1 | Y | Y |
| NEK7 | Y | Y |
| NEO1 | Y | Y |
| NEPRO | Y | Y |
| NEURL4 | Y | Y |
| NEXN | Y | Y |
| NFASC | Y | Y |
| NFIX | Y | Y |
| NFKB1 | Y | Y |
| NFKBIB | Y | Y |
| NFX1 | Y | Y |
| NHLRC1 | Y | Y |
| NIPSNAP3A | Y | Y |
| NKAP | Y | Y |

| | | |
|---|---|---|
| PACRG | N | Y |
| PACSIN2 | N | Y |
| PADI2 | N | Y |
| PADI4 | N | Y |
| PAFAH1B2 | N | Y |
| PAG1 | N | Y |
| PAK4 | N | Y |
| PAMR1 | N | Y |
| PAN3 | N | Y |
| PAPOLA | N | Y |
| PARL | N | Y |
| PARN | N | Y |
| PARP6 | N | Y |
| PATJ | N | Y |
| PAWR | N | Y |
| PAX8 | N | Y |
| PAXX | N | Y |
| PBRM1 | N | Y |
| PCCA | N | Y |
| PCDHGC3 | N | Y |
| PCED1A | N | Y |
| PCID2 | N | Y |
| PCIF1 | N | Y |
| PCK1 | N | Y |
| PCM1 | N | Y |
| PCMT1 | N | Y |
| PCNP | N | Y |
| PCNT | N | Y |
| PCTP | N | Y |
| PDCD10 | N | Y |
| PDCD2 | N | Y |
| PDCD2L | N | Y |
| PDE4D | N | Y |
| PDE9A | N | Y |
| PDGFB | N | Y |
| PDGFRA | N | Y |
| PDHA1 | N | Y |
| PDHB | N | Y |
| PDK1 | N | Y |
| PDLIM1 | N | Y |
| PDS5B | N | Y |
| PDXK | N | Y |
| PDZD11 | N | Y |

| | | |
|---|---|---|
| NLN | Y | Y |
| NLRP2 | Y | Y |
| NLRX1 | Y | Y |
| NME4 | Y | Y |
| NOL3 | Y | Y |
| NOMO1 | Y | Y |
| NOMO3 | Y | Y |
| NOP56 | Y | Y |
| NPFFR2 | Y | Y |
| NPIPA3 | Y | Y |
| NPIPA5 | Y | Y |
| NPL | Y | Y |
| NPNT | Y | Y |
| NPPA-AS1 | Y | Y |
| NPPC | Y | Y |
| NPY1R | Y | Y |
| NR1D2 | Y | Y |
| NR1H2 | Y | Y |
| NR1H3 | Y | Y |
| NR1H4 | Y | Y |
| NR2F1 | Y | Y |
| NR2F6 | Y | Y |
| NR3C2 | Y | Y |
| NRAV | Y | Y |
| NRSN2 | Y | Y |
| NSFL1C | Y | Y |
| NSUN5P1 | Y | Y |
| NSUN5P2 | Y | Y |
| NT5E | Y | Y |
| NTAN1 | Y | Y |
| NUB1 | Y | Y |
| NUDT13 | Y | Y |
| NUDT21 | Y | Y |
| NUDT3 | Y | Y |
| NUDT4B | Y | Y |
| NUDT8 | Y | Y |
| NUDT9 | Y | Y |
| NUP54 | Y | Y |
| NUTM2A-AS1 | Y | Y |
| NVL | Y | Y |
| NXPE1 | Y | Y |
| NXPE3 | Y | Y |
| NXPE4 | Y | Y |

| | | |
|---|---|---|
| PDZD3 | N | Y |
| PDZD8 | N | Y |
| PEA15 | N | Y |
| PES1 | N | Y |
| PET100 | N | Y |
| PEX16 | N | Y |
| PEX3 | N | Y |
| PFDN6 | N | Y |
| PFKP | N | Y |
| PGAP2 | N | Y |
| PGAP3 | N | Y |
| PGGT1B | N | Y |
| PGLS | N | Y |
| PGM3 | N | Y |
| PHACTR1 | N | Y |
| PHACTR2 | N | Y |
| PHACTR4 | N | Y |
| PHC3 | N | Y |
| PHF1 | N | Y |
| PHF10 | N | Y |
| PHF14 | N | Y |
| PHLDB1 | N | Y |
| PHTF2 | N | Y |
| PHYH | N | Y |
| PHYKPL | N | Y |
| PIAS1 | N | Y |
| PIAS2 | N | Y |
| PIBF1 | N | Y |
| PICALM | N | Y |
| PID1 | N | Y |
| PIGL | N | Y |
| PIGT | N | Y |
| PIK3C2B | N | Y |
| PIK3C3 | N | Y |
| PIK3CB | N | Y |
| PIN1 | N | Y |
| PIP4K2C | N | Y |
| PISD | N | Y |
| PITPNC1 | N | Y |
| PKIG | N | Y |
| PKM | N | Y |
| PKP4 | N | Y |
| PLA2G10 | N | Y |

| | | |
|---|---|---|
| NXT1 | Y | Y |
| OGA | Y | Y |
| OGN | Y | Y |
| OIP5-AS1 | Y | Y |
| OIT3 | Y | Y |
| OLFM2 | Y | Y |
| OLFM4 | Y | Y |
| OMA1 | Y | Y |
| OPN3 | Y | Y |
| ORAI3 | Y | Y |
| ORC1 | Y | Y |
| OSBPL10 | Y | Y |
| OSCAR | Y | Y |
| OSMR | Y | Y |
| OTOP2 | Y | Y |
| P2RX1 | Y | Y |
| P2RY1 | Y | Y |
| PACS1 | Y | Y |
| PACS2 | Y | Y |
| PAFAH1B1 | Y | Y |
| PAFAH1B3 | Y | Y |
| PAGR1 | Y | Y |
| PAK1 | Y | Y |
| PALD1 | Y | Y |
| PANK3 | Y | Y |
| PAOX | Y | Y |
| PAQR5 | Y | Y |
| PARD3B | Y | Y |
| PARD6A | Y | Y |
| PARP1 | Y | Y |
| PARP10 | Y | Y |
| PARP8 | Y | Y |
| PATL2 | Y | Y |
| PAX5 | Y | Y |
| PBK | Y | Y |
| PC | Y | Y |
| PCDHB13 | Y | Y |
| PCGF1 | Y | Y |
| PCLAF | Y | Y |
| PCNX1 | Y | Y |
| PCSK7 | Y | Y |
| PCYOX1L | Y | Y |
| PDE6A | Y | Y |

| | | |
|---|---|---|
| PLA2G15 | N | Y |
| PLA2G16 | N | Y |
| PLA2G2A | N | Y |
| PLA2G6 | N | Y |
| PLCG1 | N | Y |
| PLD3 | N | Y |
| PLEKHA2 | N | Y |
| PLEKHA7 | N | Y |
| PLEKHB2 | N | Y |
| PLEKHG6 | N | Y |
| PLEKHH1 | N | Y |
| PLEKHM2 | N | Y |
| PLPBP | N | Y |
| PLPP2 | N | Y |
| PLPP5 | N | Y |
| PLSCR1 | N | Y |
| PLTP | N | Y |
| PLXNA3 | N | Y |
| PML | N | Y |
| PMM2 | N | Y |
| PMP22 | N | Y |
| PMS2 | N | Y |
| PNKD | N | Y |
| PNPLA2 | N | Y |
| PNRC2 | N | Y |
| POFUT1 | N | Y |
| POLB | N | Y |
| POLD2 | N | Y |
| POLDIP3 | N | Y |
| POLE | N | Y |
| POLI | N | Y |
| POLR2E | N | Y |
| POLR2F | N | Y |
| POLR3F | N | Y |
| POLR3H | N | Y |
| POLRMT | N | Y |
| POMT1 | N | Y |
| PON2 | N | Y |
| POR | N | Y |
| POT1 | N | Y |
| PPA1 | N | Y |
| PPARG | N | Y |
| PPDPF | N | Y |

| | | |
|---|---|---|
| PDE6B | Y | Y |
| PDE6D | Y | Y |
| PDE8A | Y | Y |
| PDK4 | Y | Y |
| PDLIM5 | Y | Y |
| PDP2 | Y | Y |
| PDPK1 | Y | Y |
| PDS5A | Y | Y |
| PECAM1 | Y | Y |
| PEMT | Y | Y |
| PEPD | Y | Y |
| PER1 | Y | Y |
| PEX10 | Y | Y |
| PEX19 | Y | Y |
| PEX26 | Y | Y |
| PEX5 | Y | Y |
| PEX6 | Y | Y |
| PEX7 | Y | Y |
| PF4 | Y | Y |
| PFKFB2 | Y | Y |
| PFN2 | Y | Y |
| PGBD2 | Y | Y |
| PGF | Y | Y |
| PGM1 | Y | Y |
| PGM2 | Y | Y |
| PGS1 | Y | Y |
| PHC1 | Y | Y |
| PHF12 | Y | Y |
| PHF19 | Y | Y |
| PHF20L1 | Y | Y |
| PHF21A | Y | Y |
| PHKB | Y | Y |
| PHRF1 | Y | Y |
| PI16 | Y | Y |
| PI4KAP1 | Y | Y |
| PIAS3 | Y | Y |
| PIDD1 | Y | Y |
| PIFO | Y | Y |
| PIGBOS1 | Y | Y |
| PIGH | Y | Y |
| PIGQ | Y | Y |
| PIGR | Y | Y |
| PIGV | Y | Y |

| | | |
|---|---|---|
| PPFIA1 | N | Y |
| PPFIA2 | N | Y |
| PPFIBP2 | N | Y |
| PPHLN1 | N | Y |
| PPIE | N | Y |
| PPIL2 | N | Y |
| PPIL3 | N | Y |
| PPL | N | Y |
| PPM1H | N | Y |
| PPP1CC | N | Y |
| PPP1R12C | N | Y |
| PPP1R13B | N | Y |
| PPP1R32 | N | Y |
| PPP1R35 | N | Y |
| PPP1R7 | N | Y |
| PPP2R2A | N | Y |
| PPP2R2D | N | Y |
| PPP2R3C | N | Y |
| PPP4C | N | Y |
| PPP4R1 | N | Y |
| PPP4R3A | N | Y |
| PPP6R2 | N | Y |
| PPP6R3 | N | Y |
| PPT1 | N | Y |
| PRAC2 | N | Y |
| PRAP1 | N | Y |
| PRC1 | N | Y |
| PRDM2 | N | Y |
| PRDX3 | N | Y |
| PRDX5 | N | Y |
| PRELID2 | N | Y |
| PREX1 | N | Y |
| PRKAA1 | N | Y |
| PRKAB1 | N | Y |
| PRKACB | N | Y |
| PRKAG2 | N | Y |
| PRKCD | N | Y |
| PRKCH | N | Y |
| PRKCI | N | Y |
| PRKCSH | N | Y |
| PRKCZ | N | Y |
| PRKRA | N | Y |
| PRMT1 | N | Y |

| | | |
|---|---|---|
| PIGX | Y | Y |
| PIGY-DT | Y | Y |
| PIH1D2 | Y | Y |
| PIK3R2 | Y | Y |
| PIK3R3 | Y | Y |
| PIM2 | Y | Y |
| PINLYP | Y | Y |
| PINX1 | Y | Y |
| PIP4P1 | Y | Y |
| PIP5K1C | Y | Y |
| PIR | Y | Y |
| PITHD1 | Y | Y |
| PITPNM2 | Y | Y |
| PITX1 | Y | Y |
| PKHD1L1 | Y | Y |
| PLAA | Y | Y |
| PLCB4 | Y | Y |
| PLCD4 | Y | Y |
| PLD1 | Y | Y |
| PLEK2 | Y | Y |
| PLEKHA4 | Y | Y |
| PLEKHG5 | Y | Y |
| PLEKHO2 | Y | Y |
| PLIN3 | Y | Y |
| PLK1 | Y | Y |
| PLLP | Y | Y |
| PLOD2 | Y | Y |
| PLPPR2 | Y | Y |
| PLRG1 | Y | Y |
| PLS3 | Y | Y |
| PLVAP | Y | Y |
| PMS2P9 | Y | Y |
| PMVK | Y | Y |
| PNO1 | Y | Y |
| PNPLA4 | Y | Y |
| PNPLA6 | Y | Y |
| PNPLA8 | Y | Y |
| PNPO | Y | Y |
| PODXL2 | Y | Y |
| POGLUT1 | Y | Y |
| POLE3 | Y | Y |
| POLR1B | Y | Y |
| POLR2D | Y | Y |

| | | |
|---|---|---|
| PRMT2 | N | Y |
| PRMT9 | N | Y |
| PROCR | N | Y |
| PRPSAP2 | N | Y |
| PRR15 | N | Y |
| PRRC2A | N | Y |
| PRRG2 | N | Y |
| PRSS16 | N | Y |
| PRSS3 | N | Y |
| PRUNE1 | N | Y |
| PSD3 | N | Y |
| PSIP1 | N | Y |
| PSMA1 | N | Y |
| PSMA3 | N | Y |
| PSMA3-AS1 | N | Y |
| PSMA6 | N | Y |
| PSMB3 | N | Y |
| PSMB5 | N | Y |
| PSMC4 | N | Y |
| PSMC6 | N | Y |
| PSMD1 | N | Y |
| PSMF1 | N | Y |
| PSMG4 | N | Y |
| PSPH | N | Y |
| PSTPIP1 | N | Y |
| PTBP1 | N | Y |
| PTCD2 | N | Y |
| PTGES2 | N | Y |
| PTGES3L | N | Y |
| PTGR1 | N | Y |
| PTGS2 | N | Y |
| PTK2 | N | Y |
| PTK2B | N | Y |
| PTMA | N | Y |
| PTP4A1 | N | Y |
| PTP4A2 | N | Y |
| PTPA | N | Y |
| PTPN11 | N | Y |
| PTPN13 | N | Y |
| PTPN18 | N | Y |
| PTPN2 | N | Y |
| PTPRK | N | Y |
| PTPRN2 | N | Y |

| | | |
|---|---|---|
| POLR2J3 | Y | Y |
| POLR3A | Y | Y |
| POLR3E | Y | Y |
| POMZP3 | Y | Y |
| POSTN | Y | Y |
| POU2AF1 | Y | Y |
| POU2F1 | Y | Y |
| PPARA | Y | Y |
| PPARD | Y | Y |
| PPCDC | Y | Y |
| PPFIA3 | Y | Y |
| PPIG | Y | Y |
| PPIL1 | Y | Y |
| PPIP5K1 | Y | Y |
| PPIP5K2 | Y | Y |
| PPM1B | Y | Y |
| PPM1G | Y | Y |
| PPME1 | Y | Y |
| PPP1R11 | Y | Y |
| PPP1R12B | Y | Y |
| PPP1R27 | Y | Y |
| PPP1R3E | Y | Y |
| PPP1R9B | Y | Y |
| PPP2R3B | Y | Y |
| PPP2R5B | Y | Y |
| PPP4R3B | Y | Y |
| PQBP1 | Y | Y |
| PQLC1 | Y | Y |
| PRADC1 | Y | Y |
| PRDM11 | Y | Y |
| PRICKLE3 | Y | Y |
| PRIM1 | Y | Y |
| PRKAB2 | Y | Y |
| PRKAG1 | Y | Y |
| PRKCA | Y | Y |
| PRKCQ-AS1 | Y | Y |
| PRKRA-AS1 | Y | Y |
| PRLR | Y | Y |
| PRMT3 | Y | Y |
| PROSER2 | Y | Y |
| PRPF19 | Y | Y |
| PRPF40A | Y | Y |
| PRPF40B | Y | Y |

| | | |
|---|---|---|
| PTPRR | N | Y |
| PTTG1IP | N | Y |
| PUF60 | N | Y |
| PUS1 | N | Y |
| PUSL1 | N | Y |
| PVR | N | Y |
| PXN | N | Y |
| PXYLP1 | N | Y |
| PYM1 | N | Y |
| QARS | N | Y |
| QKI | N | Y |
| QPCT | N | Y |
| QPRT | N | Y |
| QRICH1 | N | Y |
| RAB10 | N | Y |
| RAB18 | N | Y |
| RAB24 | N | Y |
| RAB34 | N | Y |
| RAB40B | N | Y |
| RAB40C | N | Y |
| RAB4A | N | Y |
| RAB5A | N | Y |
| RAB5C | N | Y |
| RAB6A | N | Y |
| RAB7A | N | Y |
| RAB8B | N | Y |
| RABGEF1 | N | Y |
| RABL2A | N | Y |
| RAD17 | N | Y |
| RAD23A | N | Y |
| RAD51 | N | Y |
| RAD51C | N | Y |
| RAF1 | N | Y |
| RALB | N | Y |
| RALGPS1 | N | Y |
| RALY | N | Y |
| RALYL | N | Y |
| RAMP1 | N | Y |
| RANBP1 | N | Y |
| RANBP17 | N | Y |
| RANBP3 | N | Y |
| RAP1A | N | Y |
| RAP1GDS1 | N | Y |

| | | |
|---|---|---|
| PRPF6 | Y | Y |
| PRPSAP1 | Y | Y |
| PRR3 | Y | Y |
| PRR34-AS1 | Y | Y |
| PRR7 | Y | Y |
| PRRC1 | Y | Y |
| PRSS12 | Y | Y |
| PRTFDC1 | Y | Y |
| PRXL2C | Y | Y |
| PSAT1 | Y | Y |
| PSKH1 | Y | Y |
| PSMD14 | Y | Y |
| PSMD2 | Y | Y |
| PSME4 | Y | Y |
| PSMG3 | Y | Y |
| PSORS1C1 | Y | Y |
| PTAR1 | Y | Y |
| PTDSS1 | Y | Y |
| PTGDS | Y | Y |
| PTGER3 | Y | Y |
| PTGR2 | Y | Y |
| PTN | Y | Y |
| PTOV1 | Y | Y |
| PTPN23 | Y | Y |
| PTPRCAP | Y | Y |
| PTPRF | Y | Y |
| PTPRH | Y | Y |
| PTPRM | Y | Y |
| PTPRO | Y | Y |
| PTTG1 | Y | Y |
| PUDP | Y | Y |
| PUM1 | Y | Y |
| PUM2 | Y | Y |
| PUM3 | Y | Y |
| PUS10 | Y | Y |
| PUS7L | Y | Y |
| PVALB | Y | Y |
| PWWP2B | Y | Y |
| PXK | Y | Y |
| PXMP2 | Y | Y |
| PXMP4 | Y | Y |
| PYCR1 | Y | Y |
| PYY | Y | Y |

| | | |
|---|---|---|
| RAPGEFL1 | N | Y |
| RARS2 | N | Y |
| RASSF4 | N | Y |
| RBCK1 | N | Y |
| RBM10 | N | Y |
| RBM18 | N | Y |
| RBM26 | N | Y |
| RBM3 | N | Y |
| RBM33 | N | Y |
| RBM34 | N | Y |
| RBM39 | N | Y |
| RBM42 | N | Y |
| RBM47 | N | Y |
| RBM6 | N | Y |
| RBMS1 | N | Y |
| RBPMS | N | Y |
| RCAN1 | N | Y |
| RCE1 | N | Y |
| RCHY1 | N | Y |
| RCOR3 | N | Y |
| RDH11 | N | Y |
| RDH13 | N | Y |
| REEP4 | N | Y |
| RELA | N | Y |
| RELB | N | Y |
| REPS1 | N | Y |
| RERGL | N | Y |
| RETREG3 | N | Y |
| REV1 | N | Y |
| REV3L | N | Y |
| RFC2 | N | Y |
| RFC5 | N | Y |
| RFT1 | N | Y |
| RFWD3 | N | Y |
| RGS13 | N | Y |
| RGS14 | N | Y |
| RHOG | N | Y |
| RHOT1 | N | Y |
| RHOU | N | Y |
| RILP | N | Y |
| RINT1 | N | Y |
| RIOK2 | N | Y |
| RMDN1 | N | Y |

| | | |
|---|---|---|
| RAB17 | Y | Y |
| RAB23 | Y | Y |
| RAB27A | Y | Y |
| RAB30 | Y | Y |
| RAB32 | Y | Y |
| RAB33A | Y | Y |
| RAB35 | Y | Y |
| RAB37 | Y | Y |
| RAB38 | Y | Y |
| RAB3D | Y | Y |
| RAB4B | Y | Y |
| RAB4B-EGLN2 | Y | Y |
| RAB8A | Y | Y |
| RABEP2 | Y | Y |
| RABIF | Y | Y |
| RABL2B | Y | Y |
| RAD1 | Y | Y |
| RAD50 | Y | Y |
| RAD51AP1 | Y | Y |
| RAD54B | Y | Y |
| RALA | Y | Y |
| RALGDS | Y | Y |
| RARRES3 | Y | Y |
| RASA2 | Y | Y |
| RASEF | Y | Y |
| RASSF1 | Y | Y |
| RAVER2 | Y | Y |
| RBBP5 | Y | Y |
| RBBP6 | Y | Y |
| RBBP8 | Y | Y |
| RBM12 | Y | Y |
| RBM17 | Y | Y |
| RBM23 | Y | Y |
| RBM24 | Y | Y |
| RBM27 | Y | Y |
| RBM45 | Y | Y |
| RBM4B | Y | Y |
| RBMXL1 | Y | Y |
| RBP4 | Y | Y |
| RBP5 | Y | Y |
| RC3H2 | Y | Y |
| RCAN3 | Y | Y |
| RCBTB1 | Y | Y |

| | | |
|---|---|---|
| RMND5A | N | Y |
| RNASE1 | N | Y |
| RNASE6 | N | Y |
| RNASEH1 | N | Y |
| RNASET2 | N | Y |
| RND3 | N | Y |
| RNF121 | N | Y |
| RNF126 | N | Y |
| RNF130 | N | Y |
| RNF135 | N | Y |
| RNF139 | N | Y |
| RNF145 | N | Y |
| RNF146 | N | Y |
| RNF149 | N | Y |
| RNF19B | N | Y |
| RNF20 | N | Y |
| RNF216 | N | Y |
| RNF216P1 | N | Y |
| RNF217 | N | Y |
| RNF220 | N | Y |
| RNF38 | N | Y |
| RNF43 | N | Y |
| RNF6 | N | Y |
| RNF7 | N | Y |
| RNF8 | N | Y |
| RNPS1 | N | Y |
| ROGDI | N | Y |
| RPA1 | N | Y |
| RPAIN | N | Y |
| RPE | N | Y |
| RPF2 | N | Y |
| RPL10 | N | Y |
| RPL13A | N | Y |
| RPL14 | N | Y |
| RPL22L1 | N | Y |
| RPL31 | N | Y |
| RPL32 | N | Y |
| RPL34 | N | Y |
| RPL7L1 | N | Y |
| RPL9 | N | Y |
| RPN2 | N | Y |
| RPP14 | N | Y |
| RPRD1A | N | Y |

| | | |
|---|---|---|
| RCC1 | Y | Y |
| RCC2 | Y | Y |
| RCN3 | Y | Y |
| RCSD1 | Y | Y |
| REST | Y | Y |
| RET | Y | Y |
| RETNLB | Y | Y |
| RFC4 | Y | Y |
| RGCC | Y | Y |
| RGPD2 | Y | Y |
| RGS1 | Y | Y |
| RGS19 | Y | Y |
| RHBDD1 | Y | Y |
| RHBDF1 | Y | Y |
| RHBG | Y | Y |
| RHPN1 | Y | Y |
| RHPN2 | Y | Y |
| RILPL2 | Y | Y |
| RIMS3 | Y | Y |
| RIN2 | Y | Y |
| RIOX1 | Y | Y |
| RMC1 | Y | Y |
| RMI1 | Y | Y |
| RMI2 | Y | Y |
| RNA45SN4 | Y | Y |
| RNASEH2A | Y | Y |
| RNF103 | Y | Y |
| RNF111 | Y | Y |
| RNF113A | Y | Y |
| RNF125 | Y | Y |
| RNF13 | Y | Y |
| RNF138 | Y | Y |
| RNF141 | Y | Y |
| RNF152 | Y | Y |
| RNF170 | Y | Y |
| RNF208 | Y | Y |
| RNF219 | Y | Y |
| RNF227 | Y | Y |
| RNF25 | Y | Y |
| RNF40 | Y | Y |
| RNF44 | Y | Y |
| RNGTT | Y | Y |
| RNPEP | Y | Y |

| | | |
|---|---|---|
| RPS19BP1 | N | Y |
| RPS21 | N | Y |
| RPS27 | N | Y |
| RPS3 | N | Y |
| RPS3A | N | Y |
| RPS6KA1 | N | Y |
| RPS6KA3 | N | Y |
| RPS9 | N | Y |
| RRBP1 | N | Y |
| RRS1 | N | Y |
| RSPH1 | N | Y |
| RSPO3 | N | Y |
| RSRC1 | N | Y |
| RSRC2 | N | Y |
| RSRP1 | N | Y |
| RTCA | N | Y |
| RTEL1 | N | Y |
| RTKN | N | Y |
| RTKN2 | N | Y |
| RTL8A | N | Y |
| RTN3 | N | Y |
| RTN4 | N | Y |
| RTN4IP1 | N | Y |
| RUBCNL | N | Y |
| RUNX1 | N | Y |
| RUNX1T1 | N | Y |
| RUVBL1 | N | Y |
| RWDD2A | N | Y |
| RWDD3 | N | Y |
| RXRB | N | Y |
| S100A13 | N | Y |
| S100A16 | N | Y |
| S100A9 | N | Y |
| SAFB | N | Y |
| SAMD12 | N | Y |
| SAMD3 | N | Y |
| SAP18 | N | Y |
| SAP30L | N | Y |
| SARAF | N | Y |
| SART1 | N | Y |
| SASH1 | N | Y |
| SAT2 | N | Y |
| SBDSP1 | N | Y |

| | | |
|---|---|---|
| RNPEPL1 | Y | Y |
| ROBO1 | Y | Y |
| ROBO2 | Y | Y |
| RP9 | Y | Y |
| RPGRIP1L | Y | Y |
| RPL10L | Y | Y |
| RPL23AP7 | Y | Y |
| RPP25 | Y | Y |
| RPP30 | Y | Y |
| RPP38 | Y | Y |
| RPP40 | Y | Y |
| RPS6KL1 | Y | Y |
| RPSAP52 | Y | Y |
| RPSAP58 | Y | Y |
| RPSAP9 | Y | Y |
| RPUSD1 | Y | Y |
| RREB1 | Y | Y |
| RRM2 | Y | Y |
| RRP12 | Y | Y |
| RSAD1 | Y | Y |
| RSAD2 | Y | Y |
| RSBN1 | Y | Y |
| RSPRY1 | Y | Y |
| RTF1 | Y | Y |
| RTL8C | Y | Y |
| RUFY2 | Y | Y |
| RUFY3 | Y | Y |
| RUSC1 | Y | Y |
| RUVBL2 | Y | Y |
| RWDD2B | Y | Y |
| RYK | Y | Y |
| S100A8 | Y | Y |
| SAE1 | Y | Y |
| SAMD13 | Y | Y |
| SAP30BP | Y | Y |
| SAR1B | Y | Y |
| SARS2 | Y | Y |
| SASH3 | Y | Y |
| SAV1 | Y | Y |
| SCAF8 | Y | Y |
| SCAMP1-AS1 | Y | Y |
| SCAMP4 | Y | Y |
| SCAND2P | Y | Y |

| | | |
|---|---|---|
| SBF1 | N | Y |
| SBNO2 | N | Y |
| SCAMP2 | N | Y |
| SCAMP3 | N | Y |
| SCAND1 | N | Y |
| SCEL | N | Y |
| SCIN | N | Y |
| SCNM1 | N | Y |
| SCO1 | N | Y |
| SCYL1 | N | Y |
| SDC1 | N | Y |
| SDCBP | N | Y |
| SDF2 | N | Y |
| SDF4 | N | Y |
| SEC11C | N | Y |
| SEC13 | N | Y |
| SEC16A | N | Y |
| SEC23A | N | Y |
| SEC24A | N | Y |
| SEC62 | N | Y |
| SECISBP2 | N | Y |
| SEH1L | N | Y |
| SELENOS | N | Y |
| SEMA3B | N | Y |
| SENP8 | N | Y |
| SEPHS1 | N | Y |
| 10-Sep | N | Y |
| 2-Sep | N | Y |
| 7-Sep | N | Y |
| 9-Sep | N | Y |
| SERF2 | N | Y |
| SERGEF | N | Y |
| SERINC2 | N | Y |
| SERPINA1 | N | Y |
| SERPINB6 | N | Y |
| SERPINE2 | N | Y |
| SERTAD3 | N | Y |
| SETD2 | N | Y |
| SETD6 | N | Y |
| SETD9 | N | Y |
| SF3B1 | N | Y |
| SFPQ | N | Y |
| SFXN1 | N | Y |

| | | |
|---|---|---|
| SCARB2 | Y | Y |
| SCARF2 | Y | Y |
| SCG5 | Y | Y |
| SCML1 | Y | Y |
| SCNN1B | Y | Y |
| SCNN1G | Y | Y |
| SCX | Y | Y |
| SDC2 | Y | Y |
| SDHAF3 | Y | Y |
| SDHAF4 | Y | Y |
| SDHAP2 | Y | Y |
| SDK2 | Y | Y |
| SDR16C5 | Y | Y |
| SDR39U1 | Y | Y |
| SEC14L1 | Y | Y |
| SEC16B | Y | Y |
| SEC31A | Y | Y |
| SECTM1 | Y | Y |
| SEL1L | Y | Y |
| SELPLG | Y | Y |
| SEMA6A | Y | Y |
| SEMA6D | Y | Y |
| SENP7 | Y | Y |
| SEPT5-GP1BB | Y | Y |
| SERAC1 | Y | Y |
| SERPINB5 | Y | Y |
| SERPINB8 | Y | Y |
| SERPINB9 | Y | Y |
| SERPINI1 | Y | Y |
| SESTD1 | Y | Y |
| SETBP1 | Y | Y |
| SFI1 | Y | Y |
| SFMBT2 | Y | Y |
| SFSWAP | Y | Y |
| SFT2D3 | Y | Y |
| SFXN2 | Y | Y |
| SGK3 | Y | Y |
| SH2D4A | Y | Y |
| SH3BP1 | Y | Y |
| SH3KBP1 | Y | Y |
| SHANK2 | Y | Y |
| SHC2 | Y | Y |
| SHOX | Y | Y |

| | | |
|---|---|---|
| SFXN4 | N | Y |
| SGK2 | N | Y |
| SGPL1 | N | Y |
| SGSM3 | N | Y |
| SH2B1 | N | Y |
| SH3D19 | N | Y |
| SH3GLB1 | N | Y |
| SH3RF2 | N | Y |
| SH3TC1 | N | Y |
| SH3YL1 | N | Y |
| SHC1 | N | Y |
| SHISA5 | N | Y |
| SHKBP1 | N | Y |
| SHMT2 | N | Y |
| SHROOM1 | N | Y |
| SHTN1 | N | Y |
| SIAH1 | N | Y |
| SKAP1 | N | Y |
| SLAMF1 | N | Y |
| SLBP | N | Y |
| SLC16A3 | N | Y |
| SLC16A5 | N | Y |
| SLC16A9 | N | Y |
| SLC17A4 | N | Y |
| SLC20A1 | N | Y |
| SLC20A2 | N | Y |
| SLC22A18 | N | Y |
| SLC22A23 | N | Y |
| SLC25A1 | N | Y |
| SLC25A11 | N | Y |
| SLC25A14 | N | Y |
| SLC25A26 | N | Y |
| SLC25A39 | N | Y |
| SLC25A44 | N | Y |
| SLC26A2 | N | Y |
| SLC26A6 | N | Y |
| SLC29A3 | N | Y |
| SLC2A8 | N | Y |
| SLC30A5 | N | Y |
| SLC30A6 | N | Y |
| SLC35A1 | N | Y |
| SLC35A3 | N | Y |
| SLC35B2 | N | Y |

| | | |
|---|---|---|
| SHPRH | Y | Y |
| SI | Y | Y |
| SIAE | Y | Y |
| SIAH2 | Y | Y |
| SIDT1 | Y | Y |
| SIDT2 | Y | Y |
| SIGLEC10 | Y | Y |
| SIK3 | Y | Y |
| SIL1 | Y | Y |
| SIRT3 | Y | Y |
| SIRT6 | Y | Y |
| SLC15A3 | Y | Y |
| SLC16A7 | Y | Y |
| SLC18A1 | Y | Y |
| SLC25A13 | Y | Y |
| SLC25A34 | Y | Y |
| SLC25A36 | Y | Y |
| SLC25A40 | Y | Y |
| SLC25A45 | Y | Y |
| SLC25A46 | Y | Y |
| SLC27A2 | Y | Y |
| SLC27A6 | Y | Y |
| SLC28A2 | Y | Y |
| SLC29A1 | Y | Y |
| SLC2A1 | Y | Y |
| SLC33A1 | Y | Y |
| SLC35A4 | Y | Y |
| SLC35D2 | Y | Y |
| SLC35E3 | Y | Y |
| SLC35E4 | Y | Y |
| SLC35G1 | Y | Y |
| SLC37A4 | Y | Y |
| SLC39A1 | Y | Y |
| SLC39A13 | Y | Y |
| SLC39A8 | Y | Y |
| SLC41A2 | Y | Y |
| SLC48A1 | Y | Y |
| SLC4A4 | Y | Y |
| SLC6A19 | Y | Y |
| SLC6A8 | Y | Y |
| SLC7A14 | Y | Y |
| SLC7A6OS | Y | Y |
| SLC7A8 | Y | Y |

| | | |
|---|---|---|
| SLC35B3 | N | Y |
| SLC35C2 | N | Y |
| SLC35D1 | N | Y |
| SLC35F5 | N | Y |
| SLC38A2 | N | Y |
| SLC38A9 | N | Y |
| SLC39A11 | N | Y |
| SLC39A14 | N | Y |
| SLC39A4 | N | Y |
| SLC39A5 | N | Y |
| SLC3A2 | N | Y |
| SLC40A1 | N | Y |
| SLC43A1 | N | Y |
| SLC43A2 | N | Y |
| SLC44A5 | N | Y |
| SLC4A2 | N | Y |
| SLC50A1 | N | Y |
| SLC52A2 | N | Y |
| SLC9A3R2 | N | Y |
| SLF2 | N | Y |
| SLIRP | N | Y |
| SLX1B | N | Y |
| SMAD2 | N | Y |
| SMAP2 | N | Y |
| SMARCA4 | N | Y |
| SMDT1 | N | Y |
| SMG6 | N | Y |
| SMIM12 | N | Y |
| SMIM19 | N | Y |
| SMIM22 | N | Y |
| SMIM26 | N | Y |
| SMIM29 | N | Y |
| SMN2 | N | Y |
| SMOC2 | N | Y |
| SMPD1 | N | Y |
| SMPDL3A | N | Y |
| SMTN | N | Y |
| SMURF1 | N | Y |
| SMYD3 | N | Y |
| SNAPIN | N | Y |
| SNHG17 | N | Y |
| SNHG21 | N | Y |
| SNHG29 | N | Y |

| | | |
|---|---|---|
| SLCO1C1 | Y | Y |
| SLCO4A1 | Y | Y |
| SLX1A-SULT1A3 | Y | Y |
| SMARCC1 | Y | Y |
| SMARCC2 | Y | Y |
| SMARCD1 | Y | Y |
| SMARCD2 | Y | Y |
| SMC4 | Y | Y |
| SMG1 | Y | Y |
| SMG1P5 | Y | Y |
| SMG5 | Y | Y |
| SMG7 | Y | Y |
| SMIM11A | Y | Y |
| SMIM27 | Y | Y |
| SMIM3 | Y | Y |
| SMIM35 | Y | Y |
| SMIM8 | Y | Y |
| SMN1 | Y | Y |
| SMURF2 | Y | Y |
| SMYD2 | Y | Y |
| SNAI1 | Y | Y |
| SNAPC1 | Y | Y |
| SNHG10 | Y | Y |
| SNHG15 | Y | Y |
| SNORD3A | Y | Y |
| SNRK | Y | Y |
| SNRNP200 | Y | Y |
| SNRNP25 | Y | Y |
| SNRNP40 | Y | Y |
| SNTG2 | Y | Y |
| SNUPN | Y | Y |
| SNURF | Y | Y |
| SNX32 | Y | Y |
| SNX8 | Y | Y |
| SON | Y | Y |
| SORD | Y | Y |
| SOS2 | Y | Y |
| SOWAHA | Y | Y |
| SOX12 | Y | Y |
| SOX15 | Y | Y |
| SOX4 | Y | Y |
| SOX9 | Y | Y |
| SOX9-AS1 | Y | Y |

| | | |
|---|---|---|
| SNHG6 | N | Y |
| SNHG7 | N | Y |
| SNRNP35 | N | Y |
| SNRPA1 | N | Y |
| SNRPC | N | Y |
| SNW1 | N | Y |
| SNX13 | N | Y |
| SNX15 | N | Y |
| SNX20 | N | Y |
| SNX29 | N | Y |
| SNX3 | N | Y |
| SNX4 | N | Y |
| SNX5 | N | Y |
| SOCS2 | N | Y |
| SOD2 | N | Y |
| SOD3 | N | Y |
| SORBS2 | N | Y |
| SORBS3 | N | Y |
| SOX6 | N | Y |
| SP100 | N | Y |
| SP3 | N | Y |
| SP5 | N | Y |
| SPATA6L | N | Y |
| SPATS2L | N | Y |
| SPC24 | N | Y |
| SPIDR | N | Y |
| SPINDOC | N | Y |
| SPINT1 | N | Y |
| SPINT1-AS1 | N | Y |
| SPIRE2 | N | Y |
| SPNS1 | N | Y |
| SPOCK2 | N | Y |
| SPP1 | N | Y |
| SPTAN1 | N | Y |
| SPTLC1 | N | Y |
| SRBD1 | N | Y |
| SREBF2 | N | Y |
| SREK1 | N | Y |
| SRGN | N | Y |
| SRI | N | Y |
| SRM | N | Y |
| SRPK1 | N | Y |
| SRRM2 | N | Y |

| | | |
|---|---|---|
| SP110 | Y | Y |
| SP2 | Y | Y |
| SPATA21 | Y | Y |
| SPATA7 | Y | Y |
| SPDL1 | Y | Y |
| SPECC1L | Y | Y |
| SPG7 | Y | Y |
| SPHK2 | Y | Y |
| SPINK2 | Y | Y |
| SPINT2 | Y | Y |
| SPN | Y | Y |
| SPOUT1 | Y | Y |
| SPPL3 | Y | Y |
| SPRED1 | Y | Y |
| SPRY4 | Y | Y |
| SPRYD7 | Y | Y |
| SPSB3 | Y | Y |
| SPTBN1 | Y | Y |
| SQLE | Y | Y |
| SRA1 | Y | Y |
| SREK1IP1 | Y | Y |
| SRF | Y | Y |
| SRP68 | Y | Y |
| SRRT | Y | Y |
| SRSF11 | Y | Y |
| SS18L1 | Y | Y |
| SS18L2 | Y | Y |
| SSRP1 | Y | Y |
| ST3GAL4 | Y | Y |
| ST6GALNAC1 | Y | Y |
| STAG3 | Y | Y |
| STAG3L5P-PVRIG2P-PILRB | Y | Y |
| STAM | Y | Y |
| STARD10 | Y | Y |
| STARD7-AS1 | Y | Y |
| STAT1 | Y | Y |
| STBD1 | Y | Y |
| STEAP1 | Y | Y |
| STIM2 | Y | Y |
| STIP1 | Y | Y |
| STK10 | Y | Y |
| STK16 | Y | Y |
| STK17B | Y | Y |

| | | |
|---|---|---|
| SRSF10 | N | Y |
| SRSF2 | N | Y |
| SRSF4 | N | Y |
| SRSF5 | N | Y |
| SRSF6 | N | Y |
| SRSF7 | N | Y |
| SRSF8 | N | Y |
| SS18 | N | Y |
| SSBP1 | N | Y |
| SSBP2 | N | Y |
| SSBP3 | N | Y |
| SSBP4 | N | Y |
| ST3GAL1 | N | Y |
| ST6GALNAC4 | N | Y |
| ST6GALNAC6 | N | Y |
| ST7L | N | Y |
| STAG1 | N | Y |
| STAG2 | N | Y |
| STAMBPL1 | N | Y |
| STARD3 | N | Y |
| STAT5A | N | Y |
| STAT5B | N | Y |
| STAT6 | N | Y |
| STAU2 | N | Y |
| STK11 | N | Y |
| STK17A | N | Y |
| STK19 | N | Y |
| STK24 | N | Y |
| STK25 | N | Y |
| STK32C | N | Y |
| STK35 | N | Y |
| STK38 | N | Y |
| STK38L | N | Y |
| STK39 | N | Y |
| STRAP | N | Y |
| STRBP | N | Y |
| STRN4 | N | Y |
| STT3A | N | Y |
| STX5 | N | Y |
| STX8 | N | Y |
| STXBP6 | N | Y |
| SUFU | N | Y |
| SUGCT | N | Y |

| | | |
|---|---|---|
| STMN1 | Y | Y |
| STMN3 | Y | Y |
| STOM | Y | Y |
| STOX1 | Y | Y |
| STX17-AS1 | Y | Y |
| STX6 | Y | Y |
| SUCLG2 | Y | Y |
| SUCO | Y | Y |
| SULF1 | Y | Y |
| SUN2 | Y | Y |
| SUPT6H | Y | Y |
| SUPT7L | Y | Y |
| SURF4 | Y | Y |
| SVBP | Y | Y |
| SVIL | Y | Y |
| SWI5 | Y | Y |
| SWSAP1 | Y | Y |
| SYMPK | Y | Y |
| SYN2 | Y | Y |
| SYNGR2 | Y | Y |
| SYNJ2 | Y | Y |
| SYNPO | Y | Y |
| SYS1 | Y | Y |
| SYT11 | Y | Y |
| SYT13 | Y | Y |
| TACO1 | Y | Y |
| TADA3 | Y | Y |
| TAF6 | Y | Y |
| TALAM1 | Y | Y |
| TAMM41 | Y | Y |
| TAOK2 | Y | Y |
| TARSL2 | Y | Y |
| TAZ | Y | Y |
| TBC1D22A | Y | Y |
| TBC1D23 | Y | Y |
| TBC1D24 | Y | Y |
| TBCB | Y | Y |
| TBK1 | Y | Y |
| TBL3 | Y | Y |
| TBRG1 | Y | Y |
| TCEA1 | Y | Y |
| TCEAL2 | Y | Y |
| TCEAL7 | Y | Y |

| | | |
|---|---|---|
| SUGP2 | N | Y |
| SUGT1 | N | Y |
| SULT1A1 | N | Y |
| SULT1A2 | N | Y |
| SUOX | N | Y |
| SUPT16H | N | Y |
| SUPT3H | N | Y |
| SUSD3 | N | Y |
| SYF2 | N | Y |
| SYK | N | Y |
| SYNE1 | N | Y |
| SYNPO2 | N | Y |
| SYT7 | N | Y |
| SZRD1 | N | Y |
| SZT2 | N | Y |
| TACC1 | N | Y |
| TACC2 | N | Y |
| TACC3 | N | Y |
| TACSTD2 | N | Y |
| TADA1 | N | Y |
| TAF1D | N | Y |
| TAF6L | N | Y |
| TANK | N | Y |
| TAP2 | N | Y |
| TARBP1 | N | Y |
| TARS2 | N | Y |
| TASP1 | N | Y |
| TATDN1 | N | Y |
| TATDN3 | N | Y |
| TBC1D10B | N | Y |
| TBC1D14 | N | Y |
| TBC1D2 | N | Y |
| TBC1D7 | N | Y |
| TBC1D8 | N | Y |
| TBCA | N | Y |
| TBL1X | N | Y |
| TBL1XR1 | N | Y |
| TBL2 | N | Y |
| TBX10 | N | Y |
| TBX18 | N | Y |
| TBXAS1 | N | Y |
| TCEA3 | N | Y |
| TCEAL1 | N | Y |

| | | |
|---|---|---|
| TCERG1 | Y | Y |
| TCF7L2 | Y | Y |
| TCFL5 | Y | Y |
| TCHP | Y | Y |
| TCIRG1 | Y | Y |
| TCN2 | Y | Y |
| TCTEX1D1 | Y | Y |
| TCTN3 | Y | Y |
| TDG | Y | Y |
| TDRP | Y | Y |
| TEAD3 | Y | Y |
| TEFM | Y | Y |
| TEKT4P2 | Y | Y |
| TENT2 | Y | Y |
| TENT5C | Y | Y |
| TERF2IP | Y | Y |
| TESC | Y | Y |
| TEX261 | Y | Y |
| TFAP4 | Y | Y |
| TFB2M | Y | Y |
| TFCP2 | Y | Y |
| TFEC | Y | Y |
| TFIP11 | Y | Y |
| TFPI | Y | Y |
| TFPI2 | Y | Y |
| TGFB1 | Y | Y |
| TGFB1I1 | Y | Y |
| TGFBI | Y | Y |
| THAP12 | Y | Y |
| THAP4 | Y | Y |
| THBS3 | Y | Y |
| THSD4 | Y | Y |
| TIAF1 | Y | Y |
| TIMM10B | Y | Y |
| TIMM44 | Y | Y |
| TIMM8A | Y | Y |
| TIMM9 | Y | Y |
| TINCR | Y | Y |
| TIPIN | Y | Y |
| TJAP1 | Y | Y |
| TLCD2 | Y | Y |
| TLE2 | Y | Y |
| TLK1 | Y | Y |

| | | |
|---|---|---|
| TCEAL9 | N | Y |
| TCF12 | N | Y |
| TCF3 | N | Y |
| TCOF1 | N | Y |
| TCTEX1D2 | N | Y |
| TDRD3 | N | Y |
| TDRKH | N | Y |
| TEAD1 | N | Y |
| TECR | N | Y |
| TERF1 | N | Y |
| TEX264 | N | Y |
| TFDP2 | N | Y |
| TFEB | N | Y |
| TFF2 | N | Y |
| TGFBR2 | N | Y |
| TGFBR3L | N | Y |
| TGFBRAP1 | N | Y |
| TGIF1 | N | Y |
| THAP1 | N | Y |
| THAP3 | N | Y |
| THBS1 | N | Y |
| THOC7 | N | Y |
| THOP1 | N | Y |
| THRA | N | Y |
| THRAP3 | N | Y |
| THRB | N | Y |
| THSD7A | N | Y |
| THYN1 | N | Y |
| TIAL1 | N | Y |
| TIMM17B | N | Y |
| TIMM50 | N | Y |
| TIMM8B | N | Y |
| TINF2 | N | Y |
| TJP1 | N | Y |
| TJP2 | N | Y |
| TKT | N | Y |
| TLCD1 | N | Y |
| TLE1 | N | Y |
| TM2D1 | N | Y |
| TM2D3 | N | Y |
| TM4SF5 | N | Y |
| TMBIM1 | N | Y |
| TMCO3 | N | Y |

| | | |
|---|---|---|
| TLK2 | Y | Y |
| TLNRD1 | Y | Y |
| TMC6 | Y | Y |
| TMCC1-AS1 | Y | Y |
| TMCC3 | Y | Y |
| TMEM107 | Y | Y |
| TMEM11 | Y | Y |
| TMEM115 | Y | Y |
| TMEM131L | Y | Y |
| TMEM14A | Y | Y |
| TMEM159 | Y | Y |
| TMEM161A | Y | Y |
| TMEM161B-AS1 | Y | Y |
| TMEM163 | Y | Y |
| TMEM167A | Y | Y |
| TMEM171 | Y | Y |
| TMEM175 | Y | Y |
| TMEM185A | Y | Y |
| TMEM189 | Y | Y |
| TMEM191C | Y | Y |
| TMEM199 | Y | Y |
| TMEM200A | Y | Y |
| TMEM204 | Y | Y |
| TMEM236 | Y | Y |
| TMEM237 | Y | Y |
| TMEM246 | Y | Y |
| TMEM250 | Y | Y |
| TMEM251 | Y | Y |
| TMEM253 | Y | Y |
| TMEM26-AS1 | Y | Y |
| TMEM265 | Y | Y |
| TMEM267 | Y | Y |
| TMEM268 | Y | Y |
| TMEM30B | Y | Y |
| TMEM40 | Y | Y |
| TMEM64 | Y | Y |
| TMEM81 | Y | Y |
| TMEM8B | Y | Y |
| TMEM91 | Y | Y |
| TMEM99 | Y | Y |
| TMF1 | Y | Y |
| TMIGD2 | Y | Y |
| TMOD3 | Y | Y |

| | | |
|---|---|---|
| TMED1 | N | Y |
| TMED3 | N | Y |
| TMED4 | N | Y |
| TMEM120A | N | Y |
| TMEM126B | N | Y |
| TMEM131 | N | Y |
| TMEM134 | N | Y |
| TMEM138 | N | Y |
| TMEM139 | N | Y |
| TMEM147 | N | Y |
| TMEM14B | N | Y |
| TMEM150A | N | Y |
| TMEM161B | N | Y |
| TMEM165 | N | Y |
| TMEM173 | N | Y |
| TMEM176A | N | Y |
| TMEM182 | N | Y |
| TMEM183A | N | Y |
| TMEM191B | N | Y |
| TMEM208 | N | Y |
| TMEM216 | N | Y |
| TMEM218 | N | Y |
| TMEM220 | N | Y |
| TMEM230 | N | Y |
| TMEM33 | N | Y |
| TMEM37 | N | Y |
| TMEM38B | N | Y |
| TMEM41B | N | Y |
| TMEM44 | N | Y |
| TMEM50A | N | Y |
| TMEM51 | N | Y |
| TMEM53 | N | Y |
| TMEM54 | N | Y |
| TMEM59 | N | Y |
| TMEM62 | N | Y |
| TMEM63A | N | Y |
| TMEM65 | N | Y |
| TMEM68 | N | Y |
| TMEM80 | N | Y |
| TMEM87A | N | Y |
| TMEM9 | N | Y |
| TMEM94 | N | Y |
| TMEM98 | N | Y |

| | | |
|---|---|---|
| TMX1 | Y | Y |
| TNFAIP8L3 | Y | Y |
| TNFRSF14-AS1 | Y | Y |
| TNFRSF6B | Y | Y |
| TNKS2 | Y | Y |
| TNNT3 | Y | Y |
| TOB2 | Y | Y |
| TOMM34 | Y | Y |
| TOP1 | Y | Y |
| TOP2A | Y | Y |
| TOP2B | Y | Y |
| TOR2A | Y | Y |
| TOX3 | Y | Y |
| TP53I13 | Y | Y |
| TP53I3 | Y | Y |
| TP53INP1 | Y | Y |
| TP53INP2 | Y | Y |
| TPCN1 | Y | Y |
| TPH1 | Y | Y |
| TPSB2 | Y | Y |
| TPTEP1 | Y | Y |
| TPTEP2-CSNK1E | Y | Y |
| TPX2 | Y | Y |
| TRAF2 | Y | Y |
| TRAM1L1 | Y | Y |
| TRANK1 | Y | Y |
| TRAPPC12 | Y | Y |
| TRAPPC2L | Y | Y |
| TRAPPC5 | Y | Y |
| TRAPPC8 | Y | Y |
| TRDMT1 | Y | Y |
| TRIB1 | Y | Y |
| TRIM15 | Y | Y |
| TRIM23 | Y | Y |
| TRIM28 | Y | Y |
| TRIM3 | Y | Y |
| TRIM34 | Y | Y |
| TRIM45 | Y | Y |
| TRIM65 | Y | Y |
| TRIM74 | Y | Y |
| TRMT12 | Y | Y |
| TRMT61A | Y | Y |
| TRMT9B | Y | Y |

| | | |
|---|---|---|
| TMLHE | N | Y |
| TMPO | N | Y |
| TMUB1 | N | Y |
| TMUB2 | N | Y |
| TNFAIP3 | N | Y |
| TNFRSF10B | N | Y |
| TNFRSF1B | N | Y |
| TNFSF10 | N | Y |
| TNIP1 | N | Y |
| TNRC18 | N | Y |
| TNS4 | N | Y |
| TOE1 | N | Y |
| TOM1 | N | Y |
| TOM1L1 | N | Y |
| TOP3A | N | Y |
| TPD52 | N | Y |
| TPD52L2 | N | Y |
| TPGS2 | N | Y |
| TPI1 | N | Y |
| TPM1 | N | Y |
| TPM3 | N | Y |
| TPP2 | N | Y |
| TPRA1 | N | Y |
| TPRKB | N | Y |
| TPST2 | N | Y |
| TRA2A | N | Y |
| TRABD | N | Y |
| TRABD2A | N | Y |
| TRAF3 | N | Y |
| TRAF7 | N | Y |
| TRAK1 | N | Y |
| TRAPPC1 | N | Y |
| TRAPPC3 | N | Y |
| TRAPPC4 | N | Y |
| TRERF1 | N | Y |
| TRIM14 | N | Y |
| TRIM16 | N | Y |
| TRIM29 | N | Y |
| TRIM31 | N | Y |
| TRIP12 | N | Y |
| TRIP4 | N | Y |
| TRIQK | N | Y |
| TRIR | N | Y |

| | | |
|---|---|---|
| TRPM7 | Y | Y |
| TRUB2 | Y | Y |
| TSHZ3 | Y | Y |
| TSPYL4 | Y | Y |
| TSR1 | Y | Y |
| TSTD2 | Y | Y |
| TTC28-AS1 | Y | Y |
| TTC39B | Y | Y |
| TTLL3 | Y | Y |
| TTLL5 | Y | Y |
| TTLL6 | Y | Y |
| TTN-AS1 | Y | Y |
| TTTY15 | Y | Y |
| TUBA1A | Y | Y |
| TUBB2A | Y | Y |
| TUBB6 | Y | Y |
| TUBE1 | Y | Y |
| TUBGCP5 | Y | Y |
| TUSC1 | Y | Y |
| TUT7 | Y | Y |
| TWF2 | Y | Y |
| TWISTNB | Y | Y |
| TXK | Y | Y |
| TXNDC12 | Y | Y |
| TXNDC5 | Y | Y |
| TYMS | Y | Y |
| TYMSOS | Y | Y |
| TYRP1 | Y | Y |
| TYW1 | Y | Y |
| U2AF1L5 | Y | Y |
| UBA2 | Y | Y |
| UBE2E3 | Y | Y |
| UBE2F | Y | Y |
| UBE2G1 | Y | Y |
| UBE2H | Y | Y |
| UBE2M | Y | Y |
| UBE2O | Y | Y |
| UBE2Q2 | Y | Y |
| UBE3D | Y | Y |
| UBE4B | Y | Y |
| UBN2 | Y | Y |
| UBR4 | Y | Y |
| UBXN11 | Y | Y |

| | | |
|---|---|---|
| TRMT10B | N | Y |
| TRMT11 | N | Y |
| TRMT2B | N | Y |
| TRMU | N | Y |
| TRNT1 | N | Y |
| TROAP | N | Y |
| TRPM4 | N | Y |
| TSC22D1 | N | Y |
| TSC22D4 | N | Y |
| TSEN2 | N | Y |
| TSEN54 | N | Y |
| TSFM | N | Y |
| TSHZ2 | N | Y |
| TSPAN1 | N | Y |
| TSPAN12 | N | Y |
| TSPAN15 | N | Y |
| TSPAN33 | N | Y |
| TST | N | Y |
| TTC13 | N | Y |
| TTC22 | N | Y |
| TTC27 | N | Y |
| TTC3 | N | Y |
| TTC33 | N | Y |
| TTC38 | N | Y |
| TTC39A | N | Y |
| TTC7A | N | Y |
| TTLL1 | N | Y |
| TUBB | N | Y |
| TUBD1 | N | Y |
| TUBG1 | N | Y |
| TUSC3 | N | Y |
| TXNDC11 | N | Y |
| TXNIP | N | Y |
| TXNL1 | N | Y |
| TXNRD1 | N | Y |
| TYSND1 | N | Y |
| U2AF1 | N | Y |
| UAP1 | N | Y |
| UBA3 | N | Y |
| UBA5 | N | Y |
| UBAC2 | N | Y |
| UBAP2L | N | Y |
| UBB | N | Y |

| | | |
|---|---|---|
| UBXN2B | Y | Y |
| UBXN7 | Y | Y |
| UCA1 | Y | Y |
| UCHL1 | Y | Y |
| UCHL5 | Y | Y |
| UFL1 | Y | Y |
| UFL1-AS1 | Y | Y |
| UGT1A6 | Y | Y |
| UGT2B17 | Y | Y |
| UGT2B7 | Y | Y |
| UHRF1 | Y | Y |
| UMAD1 | Y | Y |
| UNC119 | Y | Y |
| UNC5B | Y | Y |
| URB1-AS1 | Y | Y |
| USF1 | Y | Y |
| USHBP1 | Y | Y |
| USO1 | Y | Y |
| USP11 | Y | Y |
| USP24 | Y | Y |
| USP3 | Y | Y |
| USP32 | Y | Y |
| USP47 | Y | Y |
| USP5 | Y | Y |
| USP54 | Y | Y |
| USPL1 | Y | Y |
| UTP14C | Y | Y |
| UTP3 | Y | Y |
| UTP4 | Y | Y |
| UTS2 | Y | Y |
| UVRAG | Y | Y |
| VAMP5 | Y | Y |
| VAV1 | Y | Y |
| VAV3 | Y | Y |
| VBP1 | Y | Y |
| VDAC2 | Y | Y |
| VEGFB | Y | Y |
| VIL1 | Y | Y |
| VILL | Y | Y |
| VMAC | Y | Y |
| VPREB3 | Y | Y |
| VPS13A | Y | Y |
| VPS13C | Y | Y |

| | | |
|---|---|---|
| UBE2E1 | N | Y |
| UBE2G2 | N | Y |
| UBE2J2 | N | Y |
| UBE2L3 | N | Y |
| UBE2R2 | N | Y |
| UBE2V1 | N | Y |
| UBE2W | N | Y |
| UBE3B | N | Y |
| UBL4A | N | Y |
| UBL7 | N | Y |
| UBQLN1 | N | Y |
| UBR2 | N | Y |
| UBR3 | N | Y |
| UBR5 | N | Y |
| UBTF | N | Y |
| UBXN1 | N | Y |
| UBXN6 | N | Y |
| UCKL1 | N | Y |
| UCP2 | N | Y |
| UFM1 | N | Y |
| UGDH | N | Y |
| UGGT1 | N | Y |
| UGP2 | N | Y |
| UGT2A3 | N | Y |
| UGT8 | N | Y |
| UHRF1BP1L | N | Y |
| ULK3 | N | Y |
| ULK4 | N | Y |
| UNC45A | N | Y |
| UNK | N | Y |
| UPF2 | N | Y |
| UPF3A | N | Y |
| UPP1 | N | Y |
| UQCC1 | N | Y |
| URGCP | N | Y |
| URM1 | N | Y |
| UROD | N | Y |
| UROS | N | Y |
| USF2 | N | Y |
| USH1C | N | Y |
| USP10 | N | Y |
| USP13 | N | Y |
| USP33 | N | Y |

| | | |
|---|---|---|
| VPS13D | Y | Y |
| VPS16 | Y | Y |
| VPS33A | Y | Y |
| VPS33B | Y | Y |
| VPS37C | Y | Y |
| VPS72 | Y | Y |
| VPS9D1 | Y | Y |
| VSIR | Y | Y |
| WARS | Y | Y |
| WASH7P | Y | Y |
| WASH8P | Y | Y |
| WBP4 | Y | Y |
| WDR11-AS1 | Y | Y |
| WDR18 | Y | Y |
| WDR36 | Y | Y |
| WDR46 | Y | Y |
| WDR47 | Y | Y |
| WDR59 | Y | Y |
| WDR5B | Y | Y |
| WDR61 | Y | Y |
| WDR77 | Y | Y |
| WDR90 | Y | Y |
| WFDC2 | Y | Y |
| WHRN | Y | Y |
| WIPF2 | Y | Y |
| WNK1 | Y | Y |
| WNT2 | Y | Y |
| WWC3 | Y | Y |
| XK | Y | Y |
| XRCC1 | Y | Y |
| XRN1 | Y | Y |
| YAF2 | Y | Y |
| YARS2 | Y | Y |
| YBEY | Y | Y |
| YIPF6 | Y | Y |
| YJU2 | Y | Y |
| YPEL3 | Y | Y |
| YRDC | Y | Y |
| YWHAH | Y | Y |
| ZBED1 | Y | Y |
| ZBED8 | Y | Y |
| ZBTB6 | Y | Y |
| ZBTB7C | Y | Y |

| | | |
|---|---|---|
| USP42 | N | Y |
| USP53 | N | Y |
| UTP18 | N | Y |
| UXS1 | N | Y |
| VAMP7 | N | Y |
| VAMP8 | N | Y |
| VAPA | N | Y |
| VAPB | N | Y |
| VASP | N | Y |
| VDAC1 | N | Y |
| VEZF1 | N | Y |
| VGF | N | Y |
| VKORC1L1 | N | Y |
| VMP1 | N | Y |
| VPS13B | N | Y |
| VPS18 | N | Y |
| VPS29 | N | Y |
| VPS45 | N | Y |
| VPS52 | N | Y |
| VPS53 | N | Y |
| VPS54 | N | Y |
| VRK2 | N | Y |
| VSIG1 | N | Y |
| VSIG2 | N | Y |
| VTI1A | N | Y |
| VWA5A | N | Y |
| WDFY2 | N | Y |
| WDR1 | N | Y |
| WDR13 | N | Y |
| WDR20 | N | Y |
| WDR34 | N | Y |
| WDR41 | N | Y |
| WDR45 | N | Y |
| WDR45B | N | Y |
| WDR74 | N | Y |
| WDYHV1 | N | Y |
| WFDC3 | N | Y |
| WFS1 | N | Y |
| WLS | N | Y |
| WNK2 | N | Y |
| WRAP53 | N | Y |
| WTAP | N | Y |
| WWP2 | N | Y |

| | | |
|---|---|---|
| ZC3H12A | Y | Y |
| ZC3H7A | Y | Y |
| ZC3HC1 | Y | Y |
| ZCCHC17 | Y | Y |
| ZCCHC18 | Y | Y |
| ZCCHC3 | Y | Y |
| ZCCHC7 | Y | Y |
| ZCRB1 | Y | Y |
| ZDHHC11B | Y | Y |
| ZDHHC19 | Y | Y |
| ZDHHC21 | Y | Y |
| ZDHHC3 | Y | Y |
| ZDHHC8 | Y | Y |
| ZDHHC9 | Y | Y |
| ZEB2 | Y | Y |
| ZFP62 | Y | Y |
| ZFP69B | Y | Y |
| ZFP82 | Y | Y |
| ZFPL1 | Y | Y |
| ZFYVE1 | Y | Y |
| ZFYVE28 | Y | Y |
| ZGLP1 | Y | Y |
| ZGPAT | Y | Y |
| ZKSCAN8 | Y | Y |
| ZMIZ1 | Y | Y |
| ZNF101 | Y | Y |
| ZNF107 | Y | Y |
| ZNF121 | Y | Y |
| ZNF134 | Y | Y |
| ZNF169 | Y | Y |
| ZNF184 | Y | Y |
| ZNF20 | Y | Y |
| ZNF217 | Y | Y |
| ZNF219 | Y | Y |
| ZNF251 | Y | Y |
| ZNF26 | Y | Y |
| ZNF263 | Y | Y |
| ZNF264 | Y | Y |
| ZNF296 | Y | Y |
| ZNF326 | Y | Y |
| ZNF330 | Y | Y |
| ZNF362 | Y | Y |
| ZNF366 | Y | Y |

| | | |
|---|---|---|
| WWTR1 | N | Y |
| XKR9 | N | Y |
| XPC | N | Y |
| XRRA1 | N | Y |
| YAP1 | N | Y |
| YBX2 | N | Y |
| YBX3 | N | Y |
| YEATS2 | N | Y |
| YIF1B | N | Y |
| YIPF2 | N | Y |
| YIPF5 | N | Y |
| YME1L1 | N | Y |
| YPEL5 | N | Y |
| YWHAB | N | Y |
| YWHAZ | N | Y |
| ZADH2 | N | Y |
| ZBED5 | N | Y |
| ZBP1 | N | Y |
| ZBTB16 | N | Y |
| ZBTB38 | N | Y |
| ZBTB8OS | N | Y |
| ZC2HC1A | N | Y |
| ZC3H11A | N | Y |
| ZC3H12D | N | Y |
| ZC3H13 | N | Y |
| ZC3H14 | N | Y |
| ZC3H8 | N | Y |
| ZCCHC10 | N | Y |
| ZDHHC1 | N | Y |
| ZDHHC14 | N | Y |
| ZDHHC2 | N | Y |
| ZDHHC20 | N | Y |
| ZDHHC5 | N | Y |
| ZDHHC6 | N | Y |
| ZER1 | N | Y |
| ZFAND1 | N | Y |
| ZFAND2B | N | Y |
| ZFAND5 | N | Y |
| ZFAS1 | N | Y |
| ZFYVE16 | N | Y |
| ZFYVE27 | N | Y |
| ZMIZ2 | N | Y |
| ZMYM3 | N | Y |

| | | |
|---|---|---|
| ZNF37A | Y | Y |
| ZNF408 | Y | Y |
| ZNF410 | Y | Y |
| ZNF414 | Y | Y |
| ZNF439 | Y | Y |
| ZNF44 | Y | Y |
| ZNF444 | Y | Y |
| ZNF445 | Y | Y |
| ZNF454 | Y | Y |
| ZNF468 | Y | Y |
| ZNF496 | Y | Y |
| ZNF503-AS1 | Y | Y |
| ZNF511 | Y | Y |
| ZNF512 | Y | Y |
| ZNF513 | Y | Y |
| ZNF527 | Y | Y |
| ZNF528 | Y | Y |
| ZNF547 | Y | Y |
| ZNF557 | Y | Y |
| ZNF567 | Y | Y |
| ZNF57 | Y | Y |
| ZNF574 | Y | Y |
| ZNF580 | Y | Y |
| ZNF581 | Y | Y |
| ZNF582-AS1 | Y | Y |
| ZNF585A | Y | Y |
| ZNF585B | Y | Y |
| ZNF593 | Y | Y |
| ZNF595 | Y | Y |
| ZNF599 | Y | Y |
| ZNF616 | Y | Y |
| ZNF625 | Y | Y |
| ZNF626 | Y | Y |
| ZNF638 | Y | Y |
| ZNF652 | Y | Y |
| ZNF668 | Y | Y |
| ZNF700 | Y | Y |
| ZNF74 | Y | Y |
| ZNF746 | Y | Y |
| ZNF75D | Y | Y |
| ZNF770 | Y | Y |
| ZNF79 | Y | Y |
| ZNF793 | Y | Y |

| | | |
|---|---|---|
| ZMYM4 | N | Y |
| ZMYND11 | N | Y |
| ZMYND8 | N | Y |
| ZNF207 | N | Y |
| ZNF214 | N | Y |
| ZNF22 | N | Y |
| ZNF250 | N | Y |
| ZNF254 | N | Y |
| ZNF266 | N | Y |
| ZNF271P | N | Y |
| ZNF280D | N | Y |
| ZNF350 | N | Y |
| ZNF358 | N | Y |
| ZNF384 | N | Y |
| ZNF41 | N | Y |
| ZNF431 | N | Y |
| ZNF461 | N | Y |
| ZNF480 | N | Y |
| ZNF485 | N | Y |
| ZNF521 | N | Y |
| ZNF563 | N | Y |
| ZNF639 | N | Y |
| ZNF655 | N | Y |
| ZNF674 | N | Y |
| ZNF69 | N | Y |
| ZNF692 | N | Y |
| ZNF695 | N | Y |
| ZNF7 | N | Y |
| ZNF738 | N | Y |
| ZNF75A | N | Y |
| ZNF800 | N | Y |
| ZNF821 | N | Y |
| ZNF85 | N | Y |
| ZNRF3 | N | Y |
| ZRSR2 | N | Y |
| ZSCAN18 | N | Y |
| ZWILCH | N | Y |
| ZXDC | N | Y |

| | | |
|---|---|---|
| ZNF830 | Y | Y |
| ZNF836 | Y | Y |
| ZNF91 | Y | Y |
| ZPR1 | Y | Y |
| ZSCAN22 | Y | Y |
| ZSCAN9 | Y | Y |
| ZSWIM3 | Y | Y |
| ZUP1 | Y | Y |
| ZYG11B | Y | Y |
| ZZZ3 | Y | Y |

Supplementary Table S6: Total SNVs identified in colon cancer samples

| chr | pos | ref | alt | cosmicID | annotation | gene | mutType | AAanno |
|---|---|---|---|---|---|---|---|---|
| chr1 | 1014228 | G | A | COSV65106391 | exonic | ISG15 | nonsynonymous SNV | ISG15:NM_005101:exon2:c.G248A:p.S83N, |
| chr1 | 109202996 | T | C | COSV64040898 | exonic | KIAA1324 | nonsynonymous SNV | KIAA1324:NM_001284353:exon15:c.T2012C:p.L671P,KIAA1324:NM_020775:exon22:c.T3026C:p.L1009P,KIAA1324:NM_001267048:exon20:c.T2765C:p.L922P, |
| chr1 | 109411043 | C | T | . | exonic | PSMA5 | nonsynonymous SNV | PSMA5:NM_002790:exon7:c.G529A:p.A177T,PSMA5:NM_001199773:exon7:c.G355A:p.A119T,PSMA5:NM_001199774:exon7:c.G355A:p.A119T,PSMA5:NM_001199772:exon6:c.G355A:p.A119T, |
| chr1 | 109757819 | G | A | COSV62609036 | exonic | EPS8L3 | nonsynonymous SNV | EPS8L3:NM_139053:exon10:c.C880T:p.H294Y,EPS8L3:NM_133181:exon10:c.C877T:p.H293Y,EPS8L3:NM_024526:exon10:c.C877T:p.H293Y, |
| chr1 | 11069667 | G | T | . | exonic | EXOSC10 | nonsynonymous SNV | EXOSC10:NM_001001998:exon22:c.C2380A:p.Q794K,EXOSC10:NM_002685:exon21:c.C2305A:p.Q769K, |
| chr1 | 11081110 | C | T | . | exonic | EXOSC10 | stopgain | EXOSC10:NM_001001998:exon11:c.G1409A:p.W470X,EXOSC10:NM_002685:exon11:c.G1409A:p.W470X, |
| chr1 | 112913924 | A | T | COSV63710018 | exonic | SLC16A1 | nonsynonymous SNV | SLC16A1:NM_001166496:exon5:c.T1470A:p.D490E,SLC16A1:NM_003051:exon5:c.T1470A:p.D490E, |
| chr1 | 112914011 | A | T | . | exonic | SLC16A1 | nonsynonymous SNV | SLC16A1:NM_001166496:exon5:c.T1383A:p.S461R,SLC16A1:NM_003051:exon5:c.T1383A:p.S461R, |
| chr1 | 112917721 | C | T | . | exonic | SLC16A1 | nonsynonymous SNV | SLC16A1:NM_001166496:exon4:c.G685A:p.A229T,SLC16A1:NM_003051:exon4:c.G685A:p.A229T, |
| chr1 | 113881979 | C | T | . | exonic | BCL2L15 | nonsynonymous SNV | BCL2L15:NM_001010922:exon3:c.G268A:p.D90N, |
| chr1 | 114405659 | A | G | COSV61821233 | exonic | TRIM33 | nonsynonymous SNV | TRIM33:NM_015906:exon15:c.T2519C:p.I840T,TRIM33:NM_033020:exon15:c.T2519C:p.I840T, |
| chr1 | 114430835 | A | T | . | exonic | TRIM33 | nonsynonymous SNV | TRIM33:NM_015906:exon6:c.T1118A:p.I373N,TRIM33:NM_033020:exon6:c.T1118A:p.I373N, |
| chr1 | 114510877 | A | G | COSV61824840 | exonic | TRIM33 | nonsynonymous SNV | TRIM33:NM_015906:exon1:c.T200C:p.V67A,TRIM33:NM_033020:exon1:c.T200C:p.V67A, |
| chr1 | 114718174 | C | T | . | exonic | CSDE1 | nonsynonymous SNV | CSDE1:NM_001130523:exon20:c.G2437A:p.D813N,CSDE1:NM_001242892:exon19:c.G2392A:p.D798N,CSDE1:NM_001007553:exon20:c.G2392A:p.D798N,CSDE1:NM_001242893:exon18:c.G2299A:p.D767N,CSDE1:NM_001242891:exon21:c.G2530A:p.D844N,CSDE1:NM_007158:exon19:c.G2299A:p.D767N, |
| chr1 | 114718665 | C | T | . | exonic | CSDE1 | nonsynonymous SNV | CSDE1:NM_001130523:exon19:c.G2342A:p.S781N,CSDE1:NM_001242892:exon18:c.G2297A:p.S766N,CSDE1:NM_001007553:exon19:c.G2297A:p.S766N,CSDE1:NM_001242893:exon17:c.G2204A:p.S735N,CSDE1:NM_001242891:exon20:c.G2435A:p.S812N,CSDE1:NM_007158:exon18:c.G2204A:p.S735N, |
| chr1 | 114718716 | C | T | COSV54743242 | exonic | CSDE1 | nonsynonymous SNV | CSDE1:NM_001130523:exon19:c.G2291A:p.R764Q,CSDE1:NM_001242892:exon18:c.G2246A:p.R749Q,CSDE1:NM_001007553:exon19:c.G2246A:p.R749Q,CSDE1:NM_001242893:exon17:c.G2153A:p.R718Q,CSDE1:NM_001242891:exon20:c.G2384A:p.R795Q,CSDE1:NM_007158:exon18:c.G2153A:p.R718Q, |
| chr1 | 114723883 | C | T | . | exonic | CSDE1 | nonsynonymous SNV | CSDE1:NM_001130523:exon16:c.G1918A:p.G640S,CSDE1:NM_001242892:exon15:c.G1873A:p.G625S,CSDE1:NM_001007553:exon16:c.G1873A:p.G625S,CSDE1:NM_001242893:exon14:c.G1780A:p.G594S,CSDE1:NM_001242891:exon17:c.G2011A:p.G671S,CSDE1:NM_007158:exon15:c.G1780A:p.G594S, |
| chr1 | 114723979 | C | T | . | exonic | CSDE1 | nonsynonymous SNV | CSDE1:NM_001130523:exon16:c.G1822A:p.D608N,CSDE1:NM_001242892:exon15:c.G1777A:p.D593N,CSDE1:NM_001007553:exon16:c.G1777A:p.D593N,CSDE1:NM_001242893:exon14:c.G1684A:p.D562N,CSDE1:NM_001242891:exon17:c.G1915A:p.D639N,CSDE1:NM_007158:exon15:c.G1684A:p.D562N, |
| chr1 | 114726284 | C | T | . | exonic | CSDE1 | nonsynonymous SNV | CSDE1:NM_001130523:exon14:c.G1612A:p.V538M,CSDE1:NM_001242892:exon13:c.G1567A:p.V523M,CSDE1:NM_001007553:exon14:c.G1567A:p.V523M,CSDE1:NM_001242893:exon12:c.G1474A:p.V492M,CSDE1:NM_001242891:exon15:c.G1705A:p.V569M,CSDE1:NM_007158:exon13:c.G1474A:p.V492M, |
| chr1 | 114730417 | C | T | . | exonic | CSDE1 | nonsynonymous SNV | CSDE1:NM_001130523:exon12:c.G1242A:p.M414I,CSDE1:NM_001242892:exon11:c.G1197A:p.M399I,CSDE1:NM_001007553:exon12:c.G1197A:p.M399I,CSDE1:NM_001242893:exon10:c.G1104A:p.M368I,CSDE1:NM_001242891:exon13:c.G1335A:p.M445I,CSDE1:NM_007158:exon11:c.G1104A:p.M368I, |
| chr1 | 114730647 | C | T | . | exonic | CSDE1 | nonsynonymous SNV | CSDE1:NM_001130523:exon11:c.G1097A:p.G366D,CSDE1:NM_001242892:exon10:c.G1052A:p.G351D,CSDE1:NM_001007553:exon11:c.G1052A:p.G351D,CSDE1:NM_001242893:exon9:c.G959A:p.G320D,CSDE1:NM_001242891:exon12:c.G1190A:p.G397D,CSDE1:NM_007158:exon10:c.G959A:p.G320D, |
| chr1 | 114732651 | C | T | . | exonic | CSDE1 | nonsynonymous SNV | CSDE1:NM_001130523:exon10:c.G1048A:p.E350K,CSDE1:NM_001242892:exon9:c.G1003A:p.E335K,CSDE1:NM_001007553:exon10:c.G1003A:p.E335K,CSDE1:NM_001242893:exon8:c.G910A:p.E304K,CSDE1:NM_001242891:exon11:c.G1141A:p.E381K,CSDE1:NM_007158:exon9:c.G910A:p.E304K, |
| chr1 | 114732707 | A | G | . | exonic | CSDE1 | nonsynonymous SNV | CSDE1:NM_001130523:exon10:c.T992C:p.V331A,CSDE1:NM_001242892:exon9:c.T947C:p.V316A,CSDE1:NM_001007553:exon10:c.T947C:p.V316A,CSDE1:NM_001242893:exon8:c.T854C:p.V285A,CSDE1:NM_001242891:exon11:c.T1085C:p.V362A,CSDE1:NM_007158:exon9:c.T854C:p.V285A, |
| chr1 | 114736763 | A | T | . | exonic | CSDE1 | nonsynonymous SNV | CSDE1:NM_001130523:exon6:c.T540A:p.N180K,CSDE1:NM_001242892:exon5:c.T495A:p.N165K,CSDE1:NM_001007553:exon6:c.T495A:p.N165K,CSDE1:NM_001242893:exon4:c.T402A:p.N134K,CSDE1:NM_001242891:exon7:c.T633A:p.N211K,CSDE1:NM_007158:exon5:c.T402A:p.N134K, |
| chr1 | 116373516 | G | A | . | exonic | ATP1A1 | nonsynonymous SNV | ATP1A1:NM_000701:exon1:c.G5A:p.G2E, |
| chr1 | 116387304 | G | A | COSV55192926 | exonic | ATP1A1 | nonsynonymous SNV | ATP1A1:NM_001160233:exon4:c.G200A:p.R67H,ATP1A1:NM_001160234:exon4:c.G107A:p.R36H,ATP1A1:NM_000701:exon4:c.G200A:p.R67H, |
| chr1 | 116387385 | G | A | . | exonic | ATP1A1 | nonsynonymous SNV | ATP1A1:NM_001160233:exon4:c.G281A:p.R94Q,ATP1A1:NM_001160234:exon4:c.G188A:p.R63Q,ATP1A1:NM_000701:exon4:c.G281A:p.R94Q, |
| chr1 | 116387395 | T | A | . | exonic | ATP1A1 | nonsynonymous SNV | ATP1A1:NM_001160233:exon4:c.T291A:p.F97L,ATP1A1:NM_001160234:exon4:c.T198A:p.F66L,ATP1A1:NM_000701:exon4:c.T291A:p.F97L, |
| chr1 | 116388756 | C | G | . | exonic | ATP1A1 | nonsynonymous SNV | ATP1A1:NM_001160233:exon6:c.C620G:p.S207C,ATP1A1:NM_001160234:exon6:c.C527G:p.S176C,ATP1A1:NM_000701:exon6:c.C620G:p.S207C, |
| chr1 | 116389529 | C | G | . | exonic | ATP1A1 | nonsynonymous SNV | ATP1A1:NM_001160233:exon8:c.C845G:p.T282S,ATP1A1:NM_001160234:exon8:c.C752G:p.T251S,ATP1A1:NM_000701:exon8:c.C845G:p.T282S, |
| chr1 | 116389583 | T | G | . | exonic | ATP1A1 | nonsynonymous SNV | ATP1A1:NM_001160233:exon8:c.T899G:p.V300G,ATP1A1:NM_001160234:exon8:c.T806G:p.V269G,ATP1A1:NM_000701:exon8:c.T899G:p.V300G, |
| chr1 | 116389696 | G | A | . | exonic | ATP1A1 | nonsynonymous SNV | ATP1A1:NM_001160233:exon8:c.G1012A:p.A338T,ATP1A1:NM_001160234:exon8:c.G919A:p.A307T,ATP1A1:NM_000701:exon8:c.G1012A:p.A338T, |
| chr1 | 116390839 | T | C | . | exonic | ATP1A1 | nonsynonymous SNV | ATP1A1:NM_001160233:exon10:c.T1280C:p.L427P,ATP1A1:NM_001160234:exon10:c.T1187C:p.L396P,ATP1A1:NM_000701:exon10:c.T1280C:p.L427P, |
| chr1 | 116392920 | G | A | . | exonic | ATP1A1 | nonsynonymous SNV | ATP1A1:NM_001160233:exon11:c.G1399A:p.V467M,ATP1A1:NM_001160234:exon11:c.G1306A:p.V436M,ATP1A1:NM_000701:exon11:c.G1399A:p.V467M, |
| chr1 | 116395163 | G | A | . | exonic | ATP1A1 | nonsynonymous SNV | ATP1A1:NM_001160233:exon13:c.G1714A:p.D572N,ATP1A1:NM_001160234:exon13:c.G1621A:p.D541N,ATP1A1:NM_000701:exon13:c.G1714A:p.D572N, |
| chr1 | 116395187 | A | G | COSV55190446 | exonic | ATP1A1 | nonsynonymous SNV | ATP1A1:NM_001160233:exon13:c.A1738G:p.I580V,ATP1A1:NM_001160234:exon13:c.A1645G:p.I549V,ATP1A1:NM_000701:exon13:c.A1738G:p.I580V, |
| chr1 | 116397893 | C | A | . | exonic | ATP1A1 | nonsynonymous SNV | ATP1A1:NM_001160233:exon15:c.C1979A:p.A660D,ATP1A1:NM_001160234:exon15:c.C1886A:p.A629D,ATP1A1:NM_000701:exon15:c.C1979A:p.A660D, |
| chr1 | 116397918 | T | G | . | exonic | ATP1A1 | nonsynonymous SNV | ATP1A1:NM_001160233:exon15:c.T2004G:p.S668R,ATP1A1:NM_001160234:exon15:c.T1911G:p.S637R,ATP1A1:NM_000701:exon15:c.T2004G:p.S668R, |
| chr1 | 116398748 | T | A | . | exonic | ATP1A1 | nonsynonymous SNV | ATP1A1:NM_001160233:exon16:c.T2252A:p.L751Q,ATP1A1:NM_001160234:exon16:c.T2159A:p.L720Q,ATP1A1:NM_000701:exon16:c.T2252A:p.L751Q, |
| chr1 | 116398784 | A | G | . | exonic | ATP1A1 | nonsynonymous SNV | ATP1A1:NM_001160233:exon16:c.A2288G:p.E763G,ATP1A1:NM_001160234:exon16:c.A2195G:p.E732G,ATP1A1:NM_000701:exon16:c.A2288G:p.E763G, |
| chr1 | 116399073 | G | A | . | exonic | ATP1A1 | nonsynonymous SNV | ATP1A1:NM_001160233:exon17:c.G2437A:p.G813S,ATP1A1:NM_001160234:exon17:c.G2344A:p.G782S,ATP1A1:NM_000701:exon17:c.G2437A:p.G813S, |
| chr1 | 116399079 | G | A | . | exonic | ATP1A1 | nonsynonymous SNV | ATP1A1:NM_001160233:exon17:c.G2443A:p.D815N,ATP1A1:NM_001160234:exon17:c.G2350A:p.D784N,ATP1A1:NM_000701:exon17:c.G2443A:p.D815N, |
| chr1 | 116399444 | G | A | . | exonic | ATP1A1 | nonsynonymous SNV | ATP1A1:NM_001160233:exon18:c.G2473A:p.E825K,ATP1A1:NM_001160234:exon18:c.G2380A:p.E794K,ATP1A1:NM_000701:exon18:c.G2473A:p.E825K, |
| chr1 | 116399523 | G | A | . | exonic | ATP1A1 | nonsynonymous SNV | ATP1A1:NM_001160233:exon18:c.G2552A:p.S851N,ATP1A1:NM_001160234:exon18:c.G2459A:p.S820N,ATP1A1:NM_000701:exon18:c.G2552A:p.S851N, |
| chr1 | 116400918 | G | A | . | exonic | ATP1A1 | nonsynonymous SNV | ATP1A1:NM_001160233:exon19:c.G2630A:p.G877D,ATP1A1:NM_001160234:exon19:c.G2537A:p.G846D,ATP1A1:NM_000701:exon19:c.G2630A:p.G877D, |
| chr1 | 116401151 | G | A | COSV55193103 | exonic | ATP1A1 | nonsynonymous SNV | ATP1A1:NM_001160233:exon20:c.G2740A:p.V914M,ATP1A1:NM_001160234:exon20:c.G2647A:p.V883M,ATP1A1:NM_000701:exon20:c.G2740A:p.V914M, |
| chr1 | 116401165 | C | A | . | exonic | ATP1A1 | stopgain | ATP1A1:NM_001160233:exon20:c.C2754A:p.C918X,ATP1A1:NM_001160234:exon20:c.C2661A:p.C887X,ATP1A1:NM_000701:exon20:c.C2754A:p.C918X, |
| chr1 | 116401181 | G | A | . | exonic | ATP1A1 | nonsynonymous SNV | ATP1A1:NM_001160233:exon20:c.G2770A:p.V924I,ATP1A1:NM_001160234:exon20:c.G2677A:p.V893I,ATP1A1:NM_000701:exon20:c.G2770A:p.V924I, |
| chr1 | 116401185 | G | A | . | exonic | ATP1A1 | nonsynonymous SNV | ATP1A1:NM_001160233:exon20:c.G2774A:p.S925N,ATP1A1:NM_001160234:exon20:c.G2681A:p.S894N,ATP1A1:NM_000701:exon20:c.G2774A:p.S925N, |
| chr1 | 116401585 | G | A | . | exonic | ATP1A1 | nonsynonymous SNV | ATP1A1:NM_001160233:exon21:c.G2881A:p.E961K,ATP1A1:NM_001160234:exon21:c.G2788A:p.E930K,ATP1A1:NM_000701:exon21:c.G2881A:p.E961K, |
| chr1 | 116401644 | G | A | . | exonic | ATP1A1 | nonsynonymous SNV | ATP1A1:NM_001160233:exon21:c.G2940A:p.M980I,ATP1A1:NM_001160234:exon21:c.G2847A:p.M949I,ATP1A1:NM_000701:exon21:c.G2940A:p.M980I, |
| chr1 | 116403975 | G | A | COSV55190853 | exonic | ATP1A1 | nonsynonymous SNV | ATP1A1:NM_001160233:exon22:c.G3043A:p.G1015S,ATP1A1:NM_001160234:exon22:c.G2950A:p.G984S,ATP1A1:NM_000701:exon22:c.G3043A:p.G1015S, |
| chr1 | 13816508 | A | G | COSV52443068 | exonic | PRDM2 | nonsynonymous SNV | PRDM2:NM_001135610:exon7:c.A593G:p.Q198R, |
| chr1 | 146156641 | G | A | COSV61245135 | exonic | NOTCH2NL | nonsynonymous SNV | NOTCH2NL:NM_203458:exon4:c.C473T:p.T158I, |
| chr1 | 147268843 | C | G | COSV63614007 | exonic | CHD1L | nonsynonymous SNV | CHD1L:NM_004284:exon10:c.C1050G:p.H350Q,CHD1L:NM_001256338:exon4:c.C438G:p.H146Q,CHD1L:NM_024568:exon10:c.C711G:p.H237Q,CHD1L:NM_001256336:exon9:c.C750G:p.H250Q,CHD1L:NM_001256337:exon5:c.C207G:p.H69Q, |
| chr1 | 147286485 | A | G | . | exonic | CHD1L | nonsynonymous SNV | CHD1L:NM_004284:exon18:c.A2206G:p.I736V,CHD1L:NM_001256338:exon12:c.A1594G:p.I532V,CHD1L:NM_024568:exon18:c.A1867G:p.I623V,CHD1L:NM_001256336:exon17:c.A1906G:p.I636V,CHD1L:NM_001256337:exon13:c.A1363G:p.I455V, |
| chr1 | 147295468 | T | G | . | exonic | CHD1L | nonsynonymous SNV | CHD1L:NM_004284:exon23:c.T2653G:p.S885A,CHD1L:NM_001256338:exon17:c.T2041G:p.S681A,CHD1L:NM_024568:exon23:c.T2314G:p.S772A,CHD1L:NM_001256336:exon22:c.T2353G:p.S785A,CHD1L:NM_001256337:exon18:c.T1810G:p.S604A, |
| chr1 | 150308112 | T | C | . | exonic | MRPS21 | nonsynonymous SNV | MRPS21:NM_018997:exon2:c.T148C:p.C50R,MRPS21:NM_031901:exon3:c.T148C:p.C50R, |
| chr1 | 150755063 | G | A | COSV64566034 | exonic | CTSS | nonsynonymous SNV | CTSS:NM_004079:exon4:c.C337T:p.R113W, |
| chr1 | 150764744 | A | G | COSV64566328 | exonic | CTSS | nonsynonymous SNV | CTSS:NM_001199739:exon2:c.T20C:p.V7A,CTSS:NM_004079:exon2:c.T20C:p.V7A, |
| chr1 | 150968149 | T | G | . | exonic | CERS2 | nonsynonymous SNV | CERS2:NM_022075:exon4:c.A344C:p.E115A,CERS2:NM_181746:exon4:c.A344C:p.E115A, |

| | | | | | | | | |
|---|---|---|---|---|---|---|---|---|
| chr1 | 150998101 | G | T | . | exonic | FAM63A | nonsynonymous SNV | FAM63A:NM_001163259:exon8:c.C869A:p.T290K,FAM63A:NM_001163260:exon8:c.C728A:p.T243K,FAM63A:NM_001040217:exon8:c.C728A:p.T243K,FAM63A:NM_018379:exon9:c.C1154A:p.T385K,FAM63A:NM_001163258:exon9:c.C1298A:p.T433K, |
| chr1 | 151401549 | T | C | COSV51852732 | exonic | PSMB4 | nonsynonymous SNV | PSMB4:NM_002796:exon6:c.T701C:p.I234T, |
| chr1 | 151760851 | T | C | COSV58618941 | exonic | MRPL9 | nonsynonymous SNV | MRPL9:NM_001300733:exon5:c.A535G:p.I179V,MRPL9:NM_031420:exon6:c.A637G:p.I213V, |
| chr1 | 151760859 | T | G | COSV58618735 | exonic | MRPL9 | nonsynonymous SNV | MRPL9:NM_001300733:exon5:c.A527C:p.E176A,MRPL9:NM_031420:exon6:c.A629C:p.E210A, |
| chr1 | 151763100 | T | C | COSV58619003 | exonic | MRPL9 | nonsynonymous SNV | MRPL9:NM_001300733:exon2:c.A200G:p.E67G,MRPL9:NM_031420:exon2:c.A200G:p.E67G, |
| chr1 | 151909409 | A | C | COSV64295606 | exonic | THEM4 | nonsynonymous SNV | THEM4:NM_053055:exon1:c.T50G:p.L17R, |
| chr1 | 153534727 | G | A | . | exonic | S100A6 | nonsynonymous SNV | S100A6:NM_014624:exon3:c.C242T:p.A81V, |
| chr1 | 153534737 | C | T | . | exonic | S100A6 | nonsynonymous SNV | S100A6:NM_014624:exon3:c.G232A:p.G78R, |
| chr1 | 153534809 | C | T | . | exonic | S100A6 | nonsynonymous SNV | S100A6:NM_014624:exon3:c.G160A:p.A54T, |
| chr1 | 153543769 | C | T | . | exonic | S100A4 | nonsynonymous SNV | S100A4:NM_019554:exon4:c.G296A:p.R99K,S100A4:NM_002961:exon3:c.G296A:p.R99K, |
| chr1 | 153543870 | G | T | . | exonic | S100A4 | nonsynonymous SNV | S100A4:NM_019554:exon4:c.C195A:p.N65K,S100A4:NM_002961:exon3:c.C195A:p.N65K, |
| chr1 | 153544773 | C | T | . | exonic | S100A4 | nonsynonymous SNV | S100A4:NM_002961:exon2:c.G22A:p.A8T,S100A4:NM_019554:exon3:c.G22A:p.A8T, |
| chr1 | 153561551 | C | T | . | exonic | S100A2 | nonsynonymous SNV | S100A2:NM_005978:exon3:c.G182A:p.S61N, |
| chr1 | 153618984 | C | T | . | exonic | S100A13 | nonsynonymous SNV | S100A13:NM_001024213:exon3:c.G208A:p.E70K,S100A13:NM_005979:exon4:c.G208A:p.E70K,S100A13:NM_001024210:exon5:c.G208A:p.E70K,S100A13:NM_001024211:exon3:c.G208A:p.E70K,S100A13:NM_001024212:exon3:c.G208A:p.E70K, |
| chr1 | 153626427 | C | T | . | exonic | S100A13 | nonsynonymous SNV | S100A13:NM_001024211:exon2:c.G46A:p.V16M,S100A13:NM_001024213:exon2:c.G46A:p.V16M,S100A13:NM_005979:exon3:c.G46A:p.V16M,S100A13:NM_001024212:exon2:c.G46A:p.V16M,S100A13:NM_001024210:exon4:c.G46A:p.V16M, |
| chr1 | 154170403 | C | T | . | exonic | TPM3 | nonsynonymous SNV | TPM3:NM_152263:exon8:c.G772A:p.E258K,TPM3:NM_001278190:exon6:c.G598A:p.E200K,TPM3:NM_001043352:exon7:c.G661A:p.E221K,TPM3:NM_001043351:exon7:c.G661A:p.E221K,TPM3:NM_001043353:exon7:c.G661A:p.E221K,TPM3:NM_001278191:exon7:c.G391A:p.E131K,TPM3:NM_001278188:exon6:c.G463A:p.E155K,TPM3:NM_001278189:exon7:c.G661A:p.E221K,TPM3:NM_153649:exon7:c.G661A:p.E221K, |
| chr1 | 154170448 | C | T | COSV55136361 | exonic | TPM3 | nonsynonymous SNV | TPM3:NM_152263:exon8:c.G727A:p.A243T,TPM3:NM_001278190:exon6:c.G553A:p.A185T,TPM3:NM_001043352:exon7:c.G616A:p.A206T,TPM3:NM_001043351:exon7:c.G616A:p.A206T,TPM3:NM_001043353:exon7:c.G616A:p.A206T,TPM3:NM_001278191:exon7:c.G346A:p.A116T,TPM3:NM_001278188:exon6:c.G418A:p.A140T,TPM3:NM_001278189:exon7:c.G616A:p.A206T,TPM3:NM_153649:exon7:c.G616A:p.A206T, |
| chr1 | 154172915 | C | T | . | exonic | TPM3 | nonsynonymous SNV | TPM3:NM_152263:exon5:c.G559A:p.A187T,TPM3:NM_001278190:exon4:c.G448A:p.A150T,TPM3:NM_001043352:exon4:c.G448A:p.A150T,TPM3:NM_001043351:exon4:c.G448A:p.A150T,TPM3:NM_001043353:exon4:c.G448A:p.A150T,TPM3:NM_001278191:exon4:c.G178A:p.A60T,TPM3:NM_001278188:exon3:c.G250A:p.A84T,TPM3:NM_001278189:exon4:c.G448A:p.A150T,TPM3:NM_153649:exon4:c.G448A:p.A150T, |
| chr1 | 154172926 | C | T | COSV55137972 | exonic | TPM3 | nonsynonymous SNV | TPM3:NM_152263:exon5:c.G548A:p.R183Q,TPM3:NM_001278190:exon4:c.G437A:p.R146Q,TPM3:NM_001043352:exon4:c.G437A:p.R146Q,TPM3:NM_001043351:exon4:c.G437A:p.R146Q,TPM3:NM_001043353:exon4:c.G437A:p.R146Q,TPM3:NM_001278191:exon4:c.G167A:p.R56Q,TPM3:NM_001278188:exon3:c.G239A:p.R80Q,TPM3:NM_001278189:exon4:c.G437A:p.R146Q,TPM3:NM_153649:exon4:c.G437A:p.R146Q, |
| chr1 | 154173103 | G | T | . | exonic | TPM3 | nonsynonymous SNV | TPM3:NM_152263:exon4:c.C476A:p.A159E,TPM3:NM_001278190:exon3:c.C365A:p.A122E,TPM3:NM_001043352:exon3:c.C365A:p.A122E,TPM3:NM_001043351:exon3:c.C365A:p.A122E,TPM3:NM_001043353:exon3:c.C365A:p.A122E,TPM3:NM_001278191:exon3:c.C95A:p.A32E,TPM3:NM_001278188:exon2:c.C167A:p.A56E,TPM3:NM_001278189:exon3:c.C365A:p.A122E,TPM3:NM_153649:exon3:c.C365A:p.A122E, |
| chr1 | 154173152 | C | T | . | exonic | TPM3 | nonsynonymous SNV | TPM3:NM_152263:exon4:c.G427A:p.E143K,TPM3:NM_001278190:exon3:c.G316A:p.E106K,TPM3:NM_001043352:exon3:c.G316A:p.E106K,TPM3:NM_001043351:exon3:c.G316A:p.E106K,TPM3:NM_001043353:exon3:c.G316A:p.E106K,TPM3:NM_001278191:exon3:c.G46A:p.E16K,TPM3:NM_001278188:exon2:c.G118A:p.E40K,TPM3:NM_001278189:exon3:c.G316A:p.E106K,TPM3:NM_153649:exon3:c.G316A:p.E106K, |
| chr1 | 154173163 | T | C | . | exonic | TPM3 | nonsynonymous SNV | TPM3:NM_152263:exon4:c.A416G:p.E139G,TPM3:NM_001278190:exon3:c.A305G:p.E102G,TPM3:NM_001043352:exon3:c.A305G:p.E102G,TPM3:NM_001043351:exon3:c.A305G:p.E102G,TPM3:NM_001043353:exon3:c.A305G:p.E102G,TPM3:NM_001278191:exon3:c.A35G:p.E12G,TPM3:NM_001278188:exon2:c.A107G:p.E36G,TPM3:NM_001278189:exon3:c.A305G:p.E102G,TPM3:NM_153649:exon3:c.A305G:p.E102G, |
| chr1 | 154173184 | T | C | . | exonic | TPM3 | nonsynonymous SNV | TPM3:NM_152263:exon4:c.A395G:p.E132G,TPM3:NM_001278190:exon3:c.A284G:p.E95G,TPM3:NM_001043352:exon3:c.A284G:p.E95G,TPM3:NM_001043351:exon3:c.A284G:p.E95G,TPM3:NM_001043353:exon3:c.A284G:p.E95G,TPM3:NM_001278191:exon3:c.A14G:p.E5G,TPM3:NM_001278188:exon2:c.A86G:p.E29G,TPM3:NM_001278189:exon3:c.A284G:p.E95G,TPM3:NM_153649:exon3:c.A284G:p.E95G, |
| chr1 | 154176119 | C | T | . | exonic | TPM3 | nonsynonymous SNV | TPM3:NM_152263:exon3:c.G373A:p.E125K,TPM3:NM_001278190:exon2:c.G262A:p.E88K,TPM3:NM_001043352:exon2:c.G262A:p.E88K,TPM3:NM_001043351:exon2:c.G262A:p.E88K,TPM3:NM_001043353:exon2:c.G262A:p.E88K,TPM3:NM_001278189:exon2:c.G262A:p.E88K,TPM3:NM_153649:exon2:c.G262A:p.E88K, |
| chr1 | 154183010 | C | T | . | exonic | TPM3 | nonsynonymous SNV | TPM3:NM_001278190:exon1:c.G110A:p.G37E,TPM3:NM_001043352:exon1:c.G110A:p.G37E,TPM3:NM_001043351:exon1:c.G110A:p.G37E,TPM3:NM_001043353:exon1:c.G110A:p.G37E,TPM3:NM_001278188:exon1:c.G46A:p.E16K,TPM3:NM_001278189:exon1:c.G110A:p.G37E,TPM3:NM_153649:exon1:c.G110A:p.G37E, |
| chr1 | 154183040 | C | T | . | exonic | TPM3 | nonsynonymous SNV | TPM3:NM_001278190:exon1:c.G80A:p.R27Q,TPM3:NM_001043352:exon1:c.G80A:p.R27Q,TPM3:NM_001043351:exon1:c.G80A:p.R27Q,TPM3:NM_001043353:exon1:c.G80A:p.R27Q,TPM3:NM_001278188:exon1:c.G16A:p.E6K,TPM3:NM_001278189:exon1:c.G80A:p.R27Q,TPM3:NM_153649:exon1:c.G80A:p.R27Q, |
| chr1 | 1544935 | A | G | . | exonic | SSU72 | nonsynonymous SNV | SSU72:NM_014188:exon3:c.T292C:p.F98L, |
| chr1 | 155054466 | A | C | . | exonic | ADAM15 | nonsynonymous SNV | ADAM15:NM_003815:exon6:c.A572C:p.K191T,ADAM15:NM_001261465:exon6:c.A572C:p.K191T,ADAM15:NM_207197:exon6:c.A572C:p.K191T,ADAM15:NM_001261466:exon6:c.A524C:p.K175T,ADAM15:NM_207191:exon6:c.A572C:p.K191T,ADAM15:NM_001261464:exon6:c.A602C:p.K201T,ADAM15:NM_207195:exon6:c.A572C:p.K191T,ADAM15:NM_207196:exon6:c.A572C:p.K191T,ADAM15:NM_207194:exon6:c.A572C:p.K191T, |
| chr1 | 155058123 | C | A | . | exonic | ADAM15 | nonsynonymous SNV | ADAM15:NM_003815:exon14:c.C1689A:p.N563K,ADAM15:NM_001261465:exon14:c.C1689A:p.N563K,ADAM15:NM_207197:exon14:c.C1689A:p.N563K,ADAM15:NM_001261466:exon14:c.C1641A:p.N547K,ADAM15:NM_207191:exon14:c.C1689A:p.N563K,ADAM15:NM_001261464:exon14:c.C1719A:p.N573K,ADAM15:NM_207195:exon14:c.C1689A:p.N563K,ADAM15:NM_207196:exon14:c.C1689A:p.N563K,ADAM15:NM_207194:exon14:c.C1689A:p.N563K, |
| chr1 | 155061442 | C | T | COSV55128941 | exonic | ADAM15 | nonsynonymous SNV | ADAM15:NM_207197:exon20:c.C2305T:p.P769S,ADAM15:NM_207196:exon20:c.C2302T:p.P768S,ADAM15:NM_207194:exon20:c.C2305T:p.P769S, |
| chr1 | 155133751 | A | T | COSV57596245 | exonic | EFNA1 | nonsynonymous SNV | EFNA1:NM_004428:exon4:c.A476T:p.D159V,EFNA1:NM_182685:exon3:c.A410T:p.D137V, |
| chr1 | 155320440 | T | C | COSV52740311 | exonic | FDPS | nonsynonymous SNV | FDPS:NM_001242824:exon10:c.T893C:p.V298A,FDPS:NM_001135822:exon10:c.T893C:p.V298A,FDPS:NM_001242825:exon9:c.T578C:p.V193A,FDPS:NM_002004:exon11:c.T1091C:p.V364A,FDPS:NM_001135821:exon11:c.T1091C:p.V364A, |
| chr1 | 156213100 | G | A | . | exonic | PMF1 | nonsynonymous SNV | PMF1:NM_007221:exon1:c.G85A:p.G29S,PMF1-BGLAP:NM_001199662:exon1:c.G85A:p.G29S,PMF1:NM_001199654:exon1:c.G85A:p.G29S,PMF1:NM_001199653:exon1:c.G85A:p.G29S,PMF1-BGLAP:NM_001199664:exon1:c.G85A:p.G29S,PMF1-BGLAP:NM_001199661:exon1:c.G85A:p.G29S,PMF1-BGLAP:NM_001199663:exon1:c.G85A:p.G29S, |
| chr1 | 156320865 | T | C | . | exonic | CCT3 | nonsynonymous SNV | CCT3:NM_001008800:exon5:c.A469G:p.I157V,CCT3:NM_005998:exon7:c.A583G:p.I195V, |
| chr1 | 156591859 | G | C | COSV63992221 | exonic | APOA1BP | nonsynonymous SNV | APOA1BP:NM_144772:exon1:c.G55C:p.V19L, |
| chr1 | 156768177 | C | T | . | exonic | PRCC | nonsynonymous SNV | PRCC:NM_005973:exon1:c.C406T:p.P136S, |
| chr1 | 156786792 | G | A | . | exonic | PRCC | nonsynonymous SNV | PRCC:NM_005973:exon3:c.G701A:p.G234D, |
| chr1 | 156794687 | G | A | . | exonic | PRCC | nonsynonymous SNV | PRCC:NM_005973:exon5:c.G1202A:p.R401K, |
| chr1 | 15770439 | C | T | COSV60030004 | exonic | FBLIM1 | nonsynonymous SNV | FBLIM1:NM_001024215:exon5:c.C572T:p.S191F,FBLIM1:NM_001024216:exon2:c.C281T:p.S94F,FBLIM1:NM_017556:exon6:c.C572T:p.S191F, |
| chr1 | 15928597 | C | A | . | exonic | SPEN | nonsynonymous SNV | SPEN:NM_015001:exon11:c.C2357A:p.T786K, |
| chr1 | 15929512 | T | C | COSV65359297 | exonic | SPEN | nonsynonymous SNV | SPEN:NM_015001:exon11:c.T3272C:p.L1091P, |
| chr1 | 159953321 | T | G | . | exonic | SLAMF9 | nonsynonymous SNV | SLAMF9:NM_001146172:exon2:c.A379C:p.I127L,SLAMF9:NM_033438:exon2:c.A379C:p.I127L, |
| chr1 | 160610759 | G | T | COSV52502577 | exonic | SLAMF1 | nonsynonymous SNV | SLAMF1:NM_003037:exon7:c.C997A:p.P333T, |
| chr1 | 160619820 | C | T | COSV52500314 | exonic | SLAMF1 | nonsynonymous SNV | SLAMF1:NM_003037:exon5:c.G820A:p.E274K, |
| chr1 | 160646913 | G | T | COSV52498809 | exonic | SLAMF1 | nonsynonymous SNV | SLAMF1:NM_003037:exon1:c.C33A:p.F11L, |

| | | | | | | | | |
|---|---|---|---|---|---|---|---|---|
| chr1 | 160880690 | C | T | . | exonic | ITLN1 | nonsynonymous SNV | ITLN1:NM_017625:exon6:c.G583A:p.G195R, |
| chr1 | 160882018 | C | T | . | exonic | ITLN1 | nonsynonymous SNV | ITLN1:NM_017625:exon4:c.G344A:p.G115D, |
| chr1 | 160882036 | A | T | COSV58275427 | exonic | ITLN1 | nonsynonymous SNV | ITLN1:NM_017625:exon4:c.T326A:p.V109D, |
| chr1 | 160882072 | C | T | . | exonic | ITLN1 | nonsynonymous SNV | ITLN1:NM_017625:exon4:c.G290A:p.G97D, |
| chr1 | 160883469 | C | T | . | exonic | ITLN1 | nonsynonymous SNV | ITLN1:NM_017625:exon3:c.G116A:p.R39K, |
| chr1 | 161202443 | C | A | COSV63487223 | exonic | NDUFS2 | nonsynonymous SNV | NDUFS2:NM_001166159:exon1:c.C58A:p.P20T,NDUFS2:NM_004550:exon2:c.C58A:p.P20T, |
| chr1 | 161781951 | A | G | COSV63406746 | exonic | ATF6 | nonsynonymous SNV | ATF6:NM_007348:exon3:c.A199G:p.M67V, |
| chr1 | 16206003 | G | A | COSV54619075 | exonic | ARHGEF19 | nonsynonymous SNV | ARHGEF19:NM_153213:exon8:c.C1379T:p.P460L, |
| chr1 | 164559883 | G | A | COSV61532892 | exonic | PBX1 | nonsynonymous SNV | PBX1:NM_002585:exon1:c.G61A:p.G21S,PBX1:NM_001204961:exon1:c.G61A:p.G21S,PBX1:NM_001204963:exon1:c.G61A:p.G21S, |
| chr1 | 165890345 | T | C | . | exonic | UCK2 | nonsynonymous SNV | UCK2:NM_012474:exon2:c.T241C:p.F81L, |
| chr1 | 165907728 | G | A | . | exonic | UCK2 | nonsynonymous SNV | UCK2:NM_012474:exon7:c.G691A:p.G231R, |
| chr1 | 169529737 | T | C | COSV63121507 | exonic | F5 | nonsynonymous SNV | F5:NM_000130:exon16:c.A5290G:p.M1764V, |
| chr1 | 17024071 | C | T | COSV64964891 | exonic | SDHB | nonsynonymous SNV | SDHB:NM_003000:exon6:c.G544A:p.G182R, |
| chr1 | 17027766 | C | T | . | exonic | SDHB | nonsynonymous SNV | SDHB:NM_003000:exon5:c.G523A:p.E175K, |
| chr1 | 17028671 | C | T | . | exonic | SDHB | nonsynonymous SNV | SDHB:NM_003000:exon4:c.G352A:p.D118N, |
| chr1 | 17028733 | A | G | . | exonic | SDHB | nonsynonymous SNV | SDHB:NM_003000:exon4:c.T290C:p.I97T, |
| chr1 | 17044821 | C | T | . | exonic | SDHB | stopgain | SDHB:NM_003000:exon2:c.G140A:p.W47X, |
| chr1 | 17044848 | C | T | COSV64965099 | exonic | SDHB | nonsynonymous SNV | SDHB:NM_003000:exon2:c.G113A:p.R38H, |
| chr1 | 17044852 | G | A | . | exonic | SDHB | nonsynonymous SNV | SDHB:NM_003000:exon2:c.C109T:p.P37S, |
| chr1 | 17054007 | C | T | . | exonic | SDHB | nonsynonymous SNV | SDHB:NM_003000:exon1:c.G13A:p.V5I, |
| chr1 | 1719406 | G | A | COSV62266388 | exonic | CDK11A | nonsynonymous SNV | CDK11A:NM_033529:exon4:c.C277T:p.R93W,CDK11A:NM_001313896:exon4:c.C277T:p.R93W,CDK11A:NM_001313982:exon4:c.C277T:p.R93W,CDK11A:NM_024011:exon4:c.C277T:p.R93W, |
| chr1 | 172441827 | G | A | COSV51129915 | exonic | PIGC | nonsynonymous SNV | PIGC:NM_002642:exon2:c.C796T:p.P266S,PIGC:NM_153747:exon2:c.C796T:p.P266S, |
| chr1 | 17342014 | G | A | COSV99080965 | exonic | PADI4 | nonsynonymous SNV | PADI4:NM_012387:exon7:c.G724A:p.G242S, |
| chr1 | 175987103 | C | T | COSV58165189 | exonic | RFWD2 | nonsynonymous SNV | RFWD2:NM_001001740:exon17:c.G1901A:p.G634E,RFWD2:NM_001286644:exon16:c.G1253A:p.G418E,RFWD2:NM_022457:exon18:c.G1973A:p.G658E, |
| chr1 | 175988327 | A | G | . | exonic | RFWD2 | nonsynonymous SNV | RFWD2:NM_001286644:exon15:c.T1213C:p.F405L,RFWD2:NM_001001740:exon16:c.T1861C:p.F621L,RFWD2:NM_022457:exon17:c.T1933C:p.F645L, |
| chr1 | 176027641 | C | T | . | exonic | RFWD2 | nonsynonymous SNV | RFWD2:NM_001286644:exon13:c.G940A:p.A314T,RFWD2:NM_001001740:exon14:c.G1588A:p.A530T,RFWD2:NM_022457:exon15:c.G1660A:p.A554T, |
| chr1 | 176043243 | C | T | . | exonic | RFWD2 | nonsynonymous SNV | RFWD2:NM_001286644:exon12:c.G835A:p.V279I,RFWD2:NM_001001740:exon13:c.G1483A:p.V495I,RFWD2:NM_022457:exon14:c.G1555A:p.V519I, |
| chr1 | 176085784 | C | T | . | exonic | RFWD2 | nonsynonymous SNV | RFWD2:NM_001286644:exon8:c.G413A:p.R138H,RFWD2:NM_001001740:exon9:c.G1061A:p.R354H,RFWD2:NM_022457:exon10:c.G1133A:p.R378H, |
| chr1 | 176184637 | A | G | . | exonic | RFWD2 | nonsynonymous SNV | RFWD2:NM_001001740:exon2:c.T463C:p.F155L,RFWD2:NM_022457:exon2:c.T463C:p.F155L, |
| chr1 | 17697195 | A | G | COSV51429640 | exonic | ARHGEF10L | nonsynonymous SNV | ARHGEF10L:NM_001011722:exon27:c.A3538G:p.I1180V,ARHGEF10L:NM_018125:exon29:c.A3655G:p.I1219V, |
| chr1 | 183215555 | C | T | . | exonic | LAMC2 | nonsynonymous SNV | LAMC2:NM_018891:exon3:c.C371T:p.T124M,LAMC2:NM_005562:exon3:c.C371T:p.T124M, |
| chr1 | 183228542 | C | A | . | exonic | LAMC2 | nonsynonymous SNV | LAMC2:NM_018891:exon11:c.C1637A:p.A546D,LAMC2:NM_005562:exon11:c.C1637A:p.A546D, |
| chr1 | 183235696 | C | G | . | exonic | LAMC2 | nonsynonymous SNV | LAMC2:NM_018891:exon16:c.C2422G:p.P808A,LAMC2:NM_005562:exon16:c.C2422G:p.P808A, |
| chr1 | 184590781 | G | A | . | exonic | C1ORF21 | nonsynonymous SNV | C1orf21:NM_030806:exon4:c.G232A:p.E78K, |
| chr1 | 186393987 | C | G | COSV55215663 | exonic | C1ORF27 | nonsynonymous SNV | C1orf27:NM_001164246:exon8:c.C656G:p.S219C,C1orf27:NM_017847:exon9:c.C752G:p.S251C, |
| chr1 | 19282760 | T | C | . | exonic | AKR7A3 | nonsynonymous SNV | AKR7A3:NM_012067:exon7:c.A967G:p.T323A, |
| chr1 | 19284747 | T | C | . | exonic | AKR7A3 | nonsynonymous SNV | AKR7A3:NM_012067:exon5:c.A643G:p.N215D, |
| chr1 | 193125191 | C | T | . | exonic | CDC73 | nonsynonymous SNV | CDC73:NM_024529:exon2:c.C211T:p.H71Y, |
| chr1 | 193233008 | T | G | . | exonic | CDC73 | nonsynonymous SNV | CDC73:NM_024529:exon14:c.T1170G:p.D390E, |
| chr1 | 193233054 | C | A | . | exonic | CDC73 | nonsynonymous SNV | CDC73:NM_024529:exon14:c.C1216A:p.Q406K, |
| chr1 | 193236318 | T | C | . | exonic | CDC73 | nonsynonymous SNV | CDC73:NM_024529:exon15:c.T1379C:p.L460P, |
| chr1 | 19339555 | G | A | . | exonic | CAPZB | nonsynonymous SNV | CAPZB:NM_004930:exon9:c.C794T:p.A265V,CAPZB:NM_001282162:exon9:c.C881T:p.A294V, |
| chr1 | 19339568 | C | T | . | exonic | CAPZB | nonsynonymous SNV | CAPZB:NM_004930:exon9:c.G781A:p.D261N,CAPZB:NM_001282162:exon9:c.G868A:p.D290N, |
| chr1 | 19356654 | C | T | . | exonic | CAPZB | nonsynonymous SNV | CAPZB:NM_004930:exon6:c.G569A:p.G190E,CAPZB:NM_001282162:exon6:c.G656A:p.G219E,CAPZB:NM_001206541:exon7:c.G647A:p.G216E,CAPZB:NM_001206540:exon6:c.G569A:p.G190E, |
| chr1 | 19356670 | C | G | . | exonic | CAPZB | nonsynonymous SNV | CAPZB:NM_004930:exon6:c.G553C:p.G185R,CAPZB:NM_001282162:exon6:c.G640C:p.G214R,CAPZB:NM_001206541:exon7:c.G631C:p.G211R,CAPZB:NM_001206540:exon6:c.G553C:p.G185R, |
| chr1 | 19357478 | C | T | . | exonic | CAPZB | nonsynonymous SNV | CAPZB:NM_004930:exon5:c.G415A:p.D139N,CAPZB:NM_001282162:exon5:c.G502A:p.D168N,CAPZB:NM_001206541:exon6:c.G493A:p.D165N,CAPZB:NM_001206540:exon5:c.G415A:p.D139N, |
| chr1 | 19385557 | C | T | . | exonic | CAPZB | nonsynonymous SNV | CAPZB:NM_004930:exon3:c.G163A:p.V55M,CAPZB:NM_001282162:exon3:c.G250A:p.V84M,CAPZB:NM_001313932:exon3:c.G163A:p.V55M,CAPZB:NM_001206541:exon4:c.G241A:p.V81M,CAPZB:NM_001206540:exon3:c.G163A:p.V55M, |
| chr1 | 201386394 | T | C | COSV66212460 | exonic | LAD1 | nonsynonymous SNV | LAD1:NM_005558:exon3:c.A967G:p.K323E, |
| chr1 | 202011975 | G | A | . | exonic | ELF3 | nonsynonymous SNV | ELF3:NM_001114309:exon3:c.G182A:p.G61E,ELF3:NM_004433:exon3:c.G182A:p.G61E, |
| chr1 | 202012026 | A | G | . | exonic | ELF3 | nonsynonymous SNV | ELF3:NM_001114309:exon3:c.A233G:p.Y78C,ELF3:NM_004433:exon3:c.A233G:p.Y78C, |
| chr1 | 202013945 | T | C | . | exonic | ELF3 | nonsynonymous SNV | ELF3:NM_001114309:exon8:c.T922C:p.S308P,ELF3:NM_004433:exon8:c.T922C:p.S308P, |
| chr1 | 202013948 | G | A | . | exonic | ELF3 | nonsynonymous SNV | ELF3:NM_001114309:exon8:c.G925A:p.E309K,ELF3:NM_004433:exon8:c.G925A:p.E309K, |
| chr1 | 202015319 | A | T | . | exonic | ELF3 | nonsynonymous SNV | ELF3:NM_001114309:exon9:c.A1112T:p.N371I,ELF3:NM_004433:exon9:c.A1112T:p.N371I, |
| chr1 | 20589208 | A | C | COSV66751560 | exonic | CDA | nonsynonymous SNV | CDA:NM_001785:exon1:c.A79C:p.K27Q, |
| chr1 | 20661380 | G | C | COSV56117446 | exonic | DDOST | nonsynonymous SNV | DDOST:NM_005216:exon1:c.C22G:p.R8G, |
| chr1 | 206935771 | C | T | COSV62903928 | exonic | PIGR | nonsynonymous SNV | PIGR:NM_002644:exon5:c.G1093A:p.G365S, |
| chr1 | 207506782 | G | A | . | exonic | CR1 | nonsynonymous SNV | CR1:NM_000651:exon3:c.G370A:p.G124R,CR1:NM_000573:exon3:c.G370A:p.G124R, |
| chr1 | 207506806 | A | C | . | exonic | CR1 | nonsynonymous SNV | CR1:NM_000651:exon3:c.A394C:p.T132P,CR1:NM_000573:exon3:c.A394C:p.T132P, |
| chr1 | 207757633 | G | A | . | exonic | CD46 | nonsynonymous SNV | CD46:NM_172350:exon3:c.G380A:p.C127Y,CD46:NM_153826:exon3:c.G380A:p.C127Y,CD46:NM_172353:exon3:c.G380A:p.C127Y,CD46:NM_172359:exon3:c.G380A:p.C127Y,CD46:NM_002389:exon3:c.G380A:p.C127Y,CD46:NM_172361:exon3:c.G380A:p.C127Y,CD46:NM_172351:exon3:c.G380A:p.C127Y,CD46:NM_172352:exon3:c.G380A:p.C127Y, |

| | | | | | | | | |
|---|---|---|---|---|---|---|---|---|
| chr1 | 207761386 | G | A | . | exonic | CD46 | nonsynonymous SNV | CD46:NM_172350:exon5:c.G613A:p.E205K,CD46:NM_153826:exon5:c.G613A:p.E205K,CD46:NM_172353:exon5:c.G613A:p.E205K,CD46:NM_172359:exon5:c.G613A:p.E205K,CD46:NM_002389:exon5:c.G613A:p.E205K,CD46:NM_172361:exon5:c.G613A:p.E205K,CD46:NM_172351:exon5:c.G613A:p.E205K,CD46:NM_172352:exon5:c.G613A:p.E205K, |
| chr1 | 208038879 | C | T | . | exonic | PLXNA2 | nonsynonymous SNV | PLXNA2:NM_025179:exon25:c.G4606A:p.V1536M, |
| chr1 | 212100297 | C | T | . | exonic | DTL | nonsynonymous SNV | DTL:NM_016448:exon14:c.C1307T:p.A436V,DTL:NM_001286230:exon13:c.C1181T:p.A394V,DTL:NM_001286229:exon12:c.C494T:p.A165V, |
| chr1 | 212101071 | A | C | . | exonic | DTL | nonsynonymous SNV | DTL:NM_016448:exon14:c.A2081C:p.K694T,DTL:NM_001286230:exon13:c.A1955C:p.K652T,DTL:NM_001286229:exon12:c.A1268C:p.K423T, |
| chr1 | 212738494 | T | C | COSV65328273 | exonic | NSL1 | nonsynonymous SNV | NSL1:NM_001297736:exon4:c.A637G:p.M213V,NSL1:NM_015471:exon6:c.A760G:p.M254V, |
| chr1 | 214318943 | G | A | COSV65291854 | exonic | SMYD2 | nonsynonymous SNV | SMYD2:NM_020197:exon5:c.G494A:p.G165E, |
| chr1 | 214663767 | C | G | . | exonic | CENPF | nonsynonymous SNV | CENPF:NM_016343:exon20:c.C9318G:p.N3106K, |
| chr1 | 219210563 | A | G | . | exonic | LYPLAL1 | nonsynonymous SNV | LYPLAL1:NM_138794:exon4:c.A393G:p.I131M,LYPLAL1:NM_001300769:exon3:c.A240G:p.I80M,LYPLAL1:NM_001300771:exon4:c.A279G:p.I93M,LYPLAL1:NM_001300770:exon4:c.A345G:p.I115M,LYPLAL1:NM_001300772:exon4:c.A279G:p.I93M, |
| chr1 | 2193733 | G | A | COSV66031527 | exonic | FAAP20 | nonsynonymous SNV | FAAP20:NM_001282673:exon3:c.C376T:p.P126S,FAAP20:NM_001282670:exon7:c.C685T:p.P229S,FAAP20:NM_001256946:exon3:c.C376T:p.P126S,FAAP20:NM_182533:exon3:c.C376T:p.P126S, |
| chr1 | 2194700 | C | G | COSV66031548 | exonic | FAAP20 | nonsynonymous SNV | FAAP20:NM_001282673:exon1:c.G50C:p.R17P,FAAP20:NM_001256946:exon1:c.G50C:p.R17P,FAAP20:NM_001256947:exon1:c.G50C:p.R17P,FAAP20:NM_182533:exon1:c.G50C:p.R17P,FAAP20:NM_001256945:exon1:c.G50C:p.R17P, |
| chr1 | 223545289 | G | A | COSV64814534 | exonic | CAPN8 | nonsynonymous SNV | CAPN8:NM_001143962:exon17:c.C1775T:p.T592M, |
| chr1 | 225828739 | G | A | . | exonic | EPHX1 | nonsynonymous SNV | EPHX1:NM_001291163:exon2:c.G10A:p.E4K,EPHX1:NM_000120:exon2:c.G10A:p.E4K,EPHX1:NM_001136018:exon2:c.G10A:p.E4K, |
| chr1 | 225828850 | G | A | . | exonic | EPHX1 | nonsynonymous SNV | EPHX1:NM_001291163:exon2:c.G121A:p.A41T,EPHX1:NM_000120:exon2:c.G121A:p.A41T,EPHX1:NM_001136018:exon2:c.G121A:p.A41T, |
| chr1 | 225831932 | T | C | COSV55298811 | exonic | EPHX1 | nonsynonymous SNV | EPHX1:NM_001291163:exon3:c.T337C:p.Y113H,EPHX1:NM_000120:exon3:c.T337C:p.Y113H,EPHX1:NM_001136018:exon3:c.T337C:p.Y113H, |
| chr1 | 225838705 | A | G | COSV55299484 | exonic | EPHX1 | nonsynonymous SNV | EPHX1:NM_001291163:exon4:c.A416G:p.H139R,EPHX1:NM_000120:exon4:c.A416G:p.H139R,EPHX1:NM_001136018:exon4:c.A416G:p.H139R, |
| chr1 | 225886863 | T | G | COSV55282479 | exonic | LEFTY1 | nonsynonymous SNV | LEFTY1:NM_020997:exon4:c.A965C:p.D322A, |
| chr1 | 228735798 | T | C | . | exonic | RHOU | nonsynonymous SNV | RHOU:NM_021205:exon1:c.T56C:p.V19A, |
| chr1 | 22913752 | G | A | COSV65863875 | exonic | EPHB2 | nonsynonymous SNV | EPHB2:NM_001309193:exon17:c.G3050A:p.G1017E, |
| chr1 | 22913770 | G | A | . | exonic | EPHB2 | nonsynonymous SNV | EPHB2:NM_001309193:exon17:c.G3068A:p.G1023E, |
| chr1 | 230978719 | C | G | COSV59617127 | exonic | TTC13 | nonsynonymous SNV | TTC13:NM_024525:exon1:c.G112C:p.G38R,TTC13:NM_001122835:exon1:c.G112C:p.G38R, |
| chr1 | 231272345 | A | G | COSV64152997 | exonic | GNPAT | nonsynonymous SNV | GNPAT:NM_001316350:exon10:c.A1373G:p.D458G,GNPAT:NM_014236:exon11:c.A1556G:p.D519G, |
| chr1 | 23387335 | C | T | . | exonic | TCEA3 | nonsynonymous SNV | TCEA3:NM_003196:exon9:c.G904A:p.G302S, |
| chr1 | 23387378 | C | T | . | exonic | TCEA3 | nonsynonymous SNV | TCEA3:NM_003196:exon9:c.G861A:p.M287I, |
| chr1 | 23393917 | C | T | . | exonic | TCEA3 | nonsynonymous SNV | TCEA3:NM_003196:exon8:c.G781A:p.A261T, |
| chr1 | 234609263 | A | G | COSV64014804 | exonic | IRF2BP2 | nonsynonymous SNV | IRF2BP2:NM_001077397:exon1:c.T232C:p.S78P,IRF2BP2:NM_182972:exon1:c.T232C:p.S78P, |
| chr1 | 23559007 | T | C | . | exonic | ID3 | nonsynonymous SNV | ID3:NM_002167:exon2:c.A313G:p.T105A, |
| chr1 | 236537507 | T | A | COSV53023494 | exonic | LGALS8 | nonsynonymous SNV | LGALS8:NM_201545:exon4:c.T56A:p.F19Y,LGALS8:NM_201543:exon4:c.T56A:p.F19Y,LGALS8:NM_006499:exon4:c.T56A:p.F19Y,LGALS8:NM_201544:exon3:c.T56A:p.F19Y, |
| chr1 | 236537557 | C | T | COSV53023516 | exonic | LGALS8 | nonsynonymous SNV | LGALS8:NM_201545:exon4:c.C106T:p.R36C,LGALS8:NM_201543:exon4:c.C106T:p.R36C,LGALS8:NM_006499:exon4:c.C106T:p.R36C,LGALS8:NM_201544:exon3:c.C106T:p.R36C, |
| chr1 | 236538910 | A | G | COSV53023904 | exonic | LGALS8 | nonsynonymous SNV | LGALS8:NM_201545:exon5:c.A166G:p.M56V,LGALS8:NM_201543:exon5:c.A166G:p.M56V,LGALS8:NM_006499:exon5:c.A166G:p.M56V,LGALS8:NM_201544:exon4:c.A166G:p.M56V, |
| chr1 | 236543562 | G | C | COSV53025054 | exonic | LGALS8 | nonsynonymous SNV | LGALS8:NM_201545:exon10:c.G678C:p.R226S,LGALS8:NM_201543:exon9:c.G552C:p.R184S,LGALS8:NM_006499:exon10:c.G678C:p.R226S,LGALS8:NM_201544:exon8:c.G552C:p.R184S, |
| chr1 | 236554626 | T | C | COSV63979833 | exonic | HEATR1 | nonsynonymous SNV | HEATR1:NM_018072:exon42:c.A6050G:p.E2017G, |
| chr1 | 236555320 | T | C | COSV63979840 | exonic | HEATR1 | nonsynonymous SNV | HEATR1:NM_018072:exon41:c.A5899G:p.N1967D, |
| chr1 | 236555893 | A | G | COSV63979843 | exonic | HEATR1 | nonsynonymous SNV | HEATR1:NM_018072:exon39:c.T5561C:p.V1854A, |
| chr1 | 236558360 | T | C | COSV63981419 | exonic | HEATR1 | nonsynonymous SNV | HEATR1:NM_018072:exon36:c.A5081G:p.N1694S, |
| chr1 | 23868258 | G | C | COSV65692258 | exonic | FUCA1 | nonsynonymous SNV | FUCA1:NM_000147:exon1:c.C29G:p.P10R, |
| chr1 | 23868283 | G | A | COSV65692243 | exonic | FUCA1 | nonsynonymous SNV | FUCA1:NM_000147:exon1:c.C4T:p.R2W, |
| chr1 | 243191489 | C | T | COSV60505743 | exonic | CEP170 | nonsynonymous SNV | CEP170:NM_001042404:exon8:c.G637A:p.G213S,CEP170:NM_001042405:exon8:c.G637A:p.G213S,CEP170:NM_014812:exon8:c.G637A:p.G213S, |
| chr1 | 245858566 | C | T | . | exonic | SMYD3 | nonsynonymous SNV | SMYD3:NM_022743:exon10:c.G829A:p.D277N,SMYD3:NM_001167740:exon10:c.G1006A:p.D336N, |
| chr1 | 245858646 | C | T | . | exonic | SMYD3 | nonsynonymous SNV | SMYD3:NM_022743:exon10:c.G749A:p.C250Y,SMYD3:NM_001167740:exon10:c.G926A:p.C309Y, |
| chr1 | 246327223 | T | A | . | exonic | SMYD3 | nonsynonymous SNV | SMYD3:NM_022743:exon5:c.A332T:p.D111V,SMYD3:NM_001167740:exon5:c.A509T:p.D170V, |
| chr1 | 246327298 | C | T | . | exonic | SMYD3 | nonsynonymous SNV | SMYD3:NM_022743:exon5:c.G257A:p.R86K,SMYD3:NM_001167740:exon5:c.G434A:p.R145K, |
| chr1 | 246335387 | C | T | . | exonic | SMYD3 | nonsynonymous SNV | SMYD3:NM_022743:exon3:c.G139A:p.G47S,SMYD3:NM_001167740:exon3:c.G316A:p.G106S, |
| chr1 | 246335402 | A | G | . | exonic | SMYD3 | nonsynonymous SNV | SMYD3:NM_022743:exon3:c.T124C:p.S42P,SMYD3:NM_001167740:exon3:c.T301C:p.S101P, |
| chr1 | 246507179 | C | G | COSV66438600 | exonic | SMYD3 | nonsynonymous SNV | SMYD3:NM_001167740:exon1:c.G39C:p.K13N, |
| chr1 | 246988255 | A | G | COSV60584996 | exonic | ZNF695 | nonsynonymous SNV | ZNF695:NM_020394:exon4:c.T260C:p.V87A,ZNF695:NM_001204221:exon4:c.T260C:p.V87A, |
| chr1 | 246999356 | C | T | . | exonic | ZNF695 | nonsynonymous SNV | ZNF695:NM_020394:exon3:c.G251A:p.R84K,ZNF695:NM_001204221:exon3:c.G251A:p.R84K, |
| chr1 | 25243590 | T | C | COSV54562947 | exonic | RSRP1 | nonsynonymous SNV | RSRP1:NM_020317:exon4:c.A716G:p.E239G, |
| chr1 | 2556714 | A | G | COSV63186428 | exonic | TNFRSF14 | nonsynonymous SNV | TNFRSF14:NM_003820:exon1:c.A50G:p.K17R,TNFRSF14:NM_001297605:exon1:c.A50G:p.K17R, |
| chr1 | 26284400 | A | C | COSV54624386 | exonic | UBXN11 | nonsynonymous SNV | UBXN11:NM_001077262:exon7:c.T575G:p.L192R,UBXN11:NM_183008:exon12:c.T935G:p.L312R,UBXN11:NM_145345:exon11:c.T836G:p.L279R, |
| chr1 | 26320235 | A | G | COSV57485050 | exonic | CD52 | nonsynonymous SNV | CD52:NM_001803:exon2:c.A119G:p.N40S, |
| chr1 | 26320239 | A | G | COSV57485055 | exonic | CD52 | nonsynonymous SNV | CD52:NM_001803:exon2:c.A123G:p.I41M, |
| chr1 | 26460136 | G | A | COSV52575895 | exonic | DHDDS | nonsynonymous SNV | DHDDS:NM_001243565:exon7:c.G640A:p.V214M,DHDDS:NM_205861:exon8:c.G757A:p.V253M,DHDDS:NM_001243564:exon7:c.G655A:p.V219M,DHDDS:NM_024887:exon8:c.G757A:p.V253M, |
| chr1 | 26863381 | G | A | . | exonic | SFN | nonsynonymous SNV | SFN:NM_006142:exon1:c.G169A:p.A57T, |
| chr1 | 26863411 | C | A | . | exonic | SFN | nonsynonymous SNV | SFN:NM_006142:exon1:c.C199A:p.Q67K, |
| chr1 | 28473868 | C | A | COSV65783948 | exonic | PHACTR4 | nonsynonymous SNV | PHACTR4:NM_001048183:exon7:c.C1138A:p.L380I,PHACTR4:NM_023923:exon6:c.C1168A:p.L390I, |
| chr1 | 29148829 | C | T | COSV65694750 | exonic | SRSF4 | nonsynonymous SNV | SRSF4:NM_005626:exon6:c.G1066A:p.G356S, |
| chr1 | 29148882 | C | G | COSV65694996 | exonic | SRSF4 | nonsynonymous SNV | SRSF4:NM_005626:exon6:c.G1013C:p.G338A, |
| chr1 | 29149136 | T | G | COSV65694019 | exonic | SRSF4 | nonsynonymous SNV | SRSF4:NM_005626:exon6:c.A759C:p.E253D, |
| chr1 | 29216125 | A | G | COSV55285635 | exonic | MECR | nonsynonymous SNV | MECR:NM_001024732:exon3:c.T58C:p.F20L,MECR:NM_016011:exon3:c.T286C:p.F96L, |
| chr1 | 31369473 | T | C | COSV60001252 | exonic | FABP3 | nonsynonymous SNV | FABP3:NM_004102:exon2:c.A158G:p.K53R, |
| chr1 | 31413273 | C | G | . | exonic | SERINC2 | nonsynonymous SNV | SERINC2:NM_178865:exon1:c.C8G:p.A3G, |
| chr1 | 31425370 | G | A | . | exonic | SERINC2 | nonsynonymous SNV | SERINC2:NM_001199039:exon5:c.G268A:p.V90M,SERINC2:NM_178865:exon4:c.G433A:p.V145M,SERINC2:NM_001199037:exon4:c.G445A:p.V149M,SERINC2:NM_018565:exon4:c.G445A:p.V149M,SERINC2:NM_001199038:exon5:c.G460A:p.V154M, |
| chr1 | 31699894 | G | T | COSV54702454 | exonic | COL16A1 | nonsynonymous SNV | COL16A1:NM_001856:exon4:c.C185A:p.T62K, |
| chr1 | 35184478 | G | T | . | exonic | SFPQ | nonsynonymous SNV | SFPQ:NM_005066:exon10:c.C2102A:p.P701Q, |
| chr1 | 35184506 | C | T | . | exonic | SFPQ | nonsynonymous SNV | SFPQ:NM_005066:exon10:c.G2074A:p.G692S, |

| | | | | | | | | |
|---|---|---|---|---|---|---|---|---|
| chr1 | 35184578 | C | T | . | exonic | SFPQ | nonsynonymous SNV | SFPQ:NM_005066:exon10:c.G2002A:p.G668R, |
| chr1 | 35187006 | C | T | . | exonic | SFPQ | nonsynonymous SNV | SFPQ:NM_005066:exon9:c.G1981A:p.D661N, |
| chr1 | 35187060 | C | T | . | exonic | SFPQ | nonsynonymous SNV | SFPQ:NM_005066:exon9:c.G1927A:p.E643K, |
| chr1 | 35187987 | C | T | . | exonic | SFPQ | nonsynonymous SNV | SFPQ:NM_005066:exon7:c.G1801A:p.G601S, |
| chr1 | 35188008 | C | T | . | exonic | SFPQ | nonsynonymous SNV | SFPQ:NM_005066:exon7:c.G1780A:p.E594K, |
| chr1 | 35189057 | C | T | COSV61758334 | exonic | SFPQ | nonsynonymous SNV | SFPQ:NM_005066:exon6:c.G1643A:p.R548H, |
| chr1 | 35189312 | A | G | . | exonic | SFPQ | nonsynonymous SNV | SFPQ:NM_005066:exon5:c.T1486C:p.S496P, |
| chr1 | 35190817 | C | T | . | exonic | SFPQ | nonsynonymous SNV | SFPQ:NM_005066:exon3:c.G1196A:p.R399K, |
| chr1 | 38886599 | G | T | COSV56707149 | exonic | RHBDL2 | nonsynonymous SNV | RHBDL2:NM_001304746:exon8:c.C1057A:p.L353M,RHBDL2:NM_017821:exon8:c.C817A:p.L273M, |
| chr1 | 39028819 | G | A | . | exonic | NDUFS5 | nonsynonymous SNV | NDUFS5:NM_004552:exon2:c.G95A:p.R32Q,NDUFS5:NM_001184979:exon2:c.G95A:p.R32Q, |
| chr1 | 39034471 | T | C | . | exonic | NDUFS5 | nonsynonymous SNV | NDUFS5:NM_004552:exon3:c.T296C:p.I99T,NDUFS5:NM_001184979:exon3:c.T296C:p.I99T, |
| chr1 | 39632656 | T | C | COSV65721876 | exonic | HEYL | nonsynonymous SNV | HEYL:NM_014571:exon2:c.A140G:p.H47R, |
| chr1 | 40067594 | T | G | COSV61222692 | exonic | CAP1 | nonsynonymous SNV | CAP1:NM_006367:exon8:c.T685G:p.C229G,CAP1:NM_001105530:exon8:c.T685G:p.C229G, |
| chr1 | 40067615 | T | G | COSV61222695 | exonic | CAP1 | nonsynonymous SNV | CAP1:NM_006367:exon8:c.T706G:p.C236G,CAP1:NM_001105530:exon8:c.T706G:p.C236G, |
| chr1 | 40067643 | T | G | COSV61222699 | exonic | CAP1 | nonsynonymous SNV | CAP1:NM_006367:exon8:c.T734G:p.I245S,CAP1:NM_001105530:exon8:c.T734G:p.I245S, |
| chr1 | 40067648 | T | G | COSV61222703 | exonic | CAP1 | nonsynonymous SNV | CAP1:NM_006367:exon8:c.T739G:p.C247G,CAP1:NM_001105530:exon8:c.T739G:p.C247G, |
| chr1 | 40067654 | T | G | COSV61222705 | exonic | CAP1 | nonsynonymous SNV | CAP1:NM_006367:exon8:c.T745G:p.Y249D,CAP1:NM_001105530:exon8:c.T745G:p.Y249D, |
| chr1 | 40067675 | T | G | COSV61222709 | exonic | CAP1 | nonsynonymous SNV | CAP1:NM_006367:exon8:c.T766G:p.S256A,CAP1:NM_001105530:exon8:c.T766G:p.S256A, |
| chr1 | 40749673 | G | A | . | exonic | NFYC | nonsynonymous SNV | NFYC:NM_001142588:exon4:c.G278A:p.R93H,NFYC:NM_014223:exon4:c.G278A:p.R93H,NFYC:NM_001142590:exon9:c.G278A:p.R93H,NFYC:NM_001308115:exon4:c.G278A:p.R93H,NFYC:NM_001142587:exon4:c.G278A:p.R93H,NFYC:NM_001308114:exon3:c.G278A:p.R93H, |
| chr1 | 42164693 | G | A | COSV63691329 | exonic | GUCA2A | nonsynonymous SNV | GUCA2A:NM_033553:exon1:c.C20T:p.S7F, |
| chr1 | 45584193 | T | C | . | exonic | NASP | nonsynonymous SNV | NASP:NM_002482:exon1:c.T47C:p.V16A,NASP:NM_001195193:exon1:c.T47C:p.V16A,NASP:NM_152298:exon1:c.T47C:p.V16A, |
| chr1 | 45606501 | G | A | . | exonic | NASP | nonsynonymous SNV | NASP:NM_002482:exon5:c.G319A:p.G107R,NASP:NM_001195193:exon3:c.G127A:p.G43R,NASP:NM_152298:exon5:c.G319A:p.G107R, |
| chr1 | 45606519 | G | A | . | exonic | NASP | nonsynonymous SNV | NASP:NM_002482:exon5:c.G337A:p.V113M,NASP:NM_001195193:exon3:c.G145A:p.V49M,NASP:NM_152298:exon5:c.G337A:p.V113M, |
| chr1 | 45607380 | G | A | . | exonic | NASP | nonsynonymous SNV | NASP:NM_002482:exon6:c.G469A:p.A157T,NASP:NM_001195193:exon4:c.G277A:p.A93T, |
| chr1 | 45607440 | G | A | . | exonic | NASP | nonsynonymous SNV | NASP:NM_002482:exon6:c.G529A:p.E177K,NASP:NM_001195193:exon4:c.G337A:p.E113K, |
| chr1 | 45613204 | G | A | . | exonic | NASP | nonsynonymous SNV | NASP:NM_002482:exon7:c.G1462A:p.D488N,NASP:NM_001195193:exon5:c.G1270A:p.D424N,NASP:NM_152298:exon6:c.G445A:p.D149N, |
| chr1 | 45614157 | A | T | . | exonic | NASP | nonsynonymous SNV | NASP:NM_002482:exon8:c.A1568T:p.D523V,NASP:NM_001195193:exon6:c.A1376T:p.D459V,NASP:NM_152298:exon7:c.A551T:p.D184V, |
| chr1 | 45615123 | G | A | . | exonic | NASP | nonsynonymous SNV | NASP:NM_002482:exon10:c.G1777A:p.A593T,NASP:NM_001195193:exon8:c.G1585A:p.A529T,NASP:NM_152298:exon9:c.G760A:p.A254T, |
| chr1 | 46189486 | T | C | . | exonic | POMGNT1 | nonsynonymous SNV | POMGNT1:NM_001290129:exon20:c.A1801G:p.M601V,POMGNT1:NM_001243766:exon21:c.A1867G:p.M623V,POMGNT1:NM_001290130:exon20:c.A1438G:p.M480V,POMGNT1:NM_017739:exon21:c.A1867G:p.M623V, |
| chr1 | 46309111 | A | G | COSV61175886 | exonic | UQCRH | nonsynonymous SNV | UQCRH:NM_006004:exon2:c.A65G:p.E22G, |
| chr1 | 46405089 | C | A | COSV54544102 | exonic | FAAH | nonsynonymous SNV | FAAH:NM_001441:exon3:c.C385A:p.P129T, |
| chr1 | 46668139 | T | C | COSV61304776 | exonic | ATPAF1 | nonsynonymous SNV | ATPAF1:NM_022745:exon1:c.A253G:p.S85G,ATPAF1:NM_001042546:exon1:c.A253G:p.S85G, |
| chr1 | 47333967 | G | C | COSV64101982 | exonic | CMPK1 | nonsynonymous SNV | CMPK1:NM_001136140:exon1:c.G22C:p.G8R,CMPK1:NM_016308:exon1:c.G22C:p.G8R, |
| chr1 | 51361249 | T | C | COSV65541248 | exonic | EPS15 | nonsynonymous SNV | EPS15:NM_001159969:exon12:c.A1524G:p.I508M,EPS15:NM_001981:exon24:c.A2466G:p.I822M, |
| chr1 | 51772111 | G | A | . | exonic | OSBPL9 | nonsynonymous SNV | OSBPL9:NM_148906:exon12:c.G929A:p.G310E,OSBPL9:NM_148909:exon13:c.G1010A:p.G337E,OSBPL9:NM_148904:exon9:c.G446A:p.G149E,OSBPL9:NM_024586:exon13:c.G980A:p.G327E,OSBPL9:NM_148907:exon9:c.G650A:p.G217E,OSBPL9:NM_148908:exon12:c.G941A:p.G314E,OSBPL9:NM_148905:exon8:c.G446A:p.G149E, |
| chr1 | 51787439 | G | A | . | exonic | OSBPL9 | nonsynonymous SNV | OSBPL9:NM_024586:exon23:c.G2087A:p.R696K,OSBPL9:NM_148906:exon22:c.G2036A:p.R679K,OSBPL9:NM_148907:exon19:c.G1757A:p.R586K,OSBPL9:NM_148908:exon22:c.G2048A:p.R683K,OSBPL9:NM_148909:exon23:c.G2117A:p.R706K,OSBPL9:NM_148904:exon19:c.G1553A:p.R518K,OSBPL9:NM_148905:exon18:c.G1553A:p.R518K, |
| chr1 | 51787742 | G | T | . | exonic | OSBPL9 | nonsynonymous SNV | OSBPL9:NM_024586:exon24:c.G2164T:p.V722F,OSBPL9:NM_148906:exon23:c.G2113T:p.V705F,OSBPL9:NM_148907:exon20:c.G1834T:p.V612F,OSBPL9:NM_148908:exon23:c.G2125T:p.V709F,OSBPL9:NM_148909:exon24:c.G2194T:p.V732F,OSBPL9:NM_148904:exon20:c.G1630T:p.V544F,OSBPL9:NM_148905:exon19:c.G1630T:p.V544F, |
| chr1 | 54216247 | G | C | . | exonic | MRPL37 | nonsynonymous SNV | MRPL37:NM_016491:exon6:c.G1097C:p.C366S, |
| chr1 | 59321678 | T | G | COSV58028591 | exonic | FGGY | nonsynonymous SNV | FGGY:NM_001244714:exon2:c.T129G:p.N43K,FGGY:NM_001113411:exon2:c.T129G:p.N43K,FGGY:NM_018291:exon2:c.T129G:p.N43K, |
| chr1 | 6186721 | C | T | . | exonic | RPL22 | nonsynonymous SNV | RPL22:NM_000983:exon4:c.G338A:p.R113H, |
| chr1 | 6186728 | C | T | . | exonic | RPL22 | nonsynonymous SNV | RPL22:NM_000983:exon4:c.G331A:p.E111K, |
| chr1 | 6186749 | C | T | . | exonic | RPL22 | nonsynonymous SNV | RPL22:NM_000983:exon4:c.G310A:p.A104T, |
| chr1 | 6186799 | G | A | . | exonic | RPL22 | nonsynonymous SNV | RPL22:NM_000983:exon4:c.C260T:p.T87I, |
| chr1 | 6192958 | C | T | COSV52379371 | exonic | RPL22 | nonsynonymous SNV | RPL22:NM_000983:exon3:c.G214A:p.V72M, |
| chr1 | 6192997 | C | T | . | exonic | RPL22 | nonsynonymous SNV | RPL22:NM_000983:exon3:c.G175A:p.G59R, |
| chr1 | 6197667 | C | T | . | exonic | RPL22 | nonsynonymous SNV | RPL22:NM_000983:exon2:c.G102A:p.M34I, |
| chr1 | 6197681 | C | T | . | exonic | RPL22 | nonsynonymous SNV | RPL22:NM_000983:exon2:c.G88A:p.E30K, |
| chr1 | 61990342 | G | A | COSV56249041 | exonic | INADL | nonsynonymous SNV | INADL:NM_176877:exon28:c.G3845A:p.R1282H, |
| chr1 | 62051011 | G | C | COSV56248179 | exonic | INADL | nonsynonymous SNV | INADL:NM_176877:exon31:c.G4078C:p.V1360L, |
| chr1 | 6281268 | G | C | . | exonic | ACOT7 | nonsynonymous SNV | ACOT7:NM_181864:exon8:c.C878G:p.S293W,ACOT7:NM_181865:exon8:c.C788G:p.S263W,ACOT7:NM_007274:exon8:c.C848G:p.S283W,ACOT7:NM_181866:exon8:c.C725G:p.S242W, |
| chr1 | 6327385 | C | T | . | exonic | ACOT7 | nonsynonymous SNV | ACOT7:NM_181864:exon5:c.G569A:p.G190D,ACOT7:NM_181865:exon5:c.G479A:p.G160D,ACOT7:NM_007274:exon5:c.G539A:p.G180D,ACOT7:NM_181866:exon5:c.G416A:p.G139D, |
| chr1 | 6327392 | C | G | . | exonic | ACOT7 | nonsynonymous SNV | ACOT7:NM_181864:exon5:c.G562C:p.E188Q,ACOT7:NM_181865:exon5:c.G472C:p.E158Q,ACOT7:NM_007274:exon5:c.G532C:p.E178Q,ACOT7:NM_181866:exon5:c.G409C:p.E137Q, |
| chr1 | 6589168 | T | G | COSV66553504 | exonic | ZBTB48 | nonsynonymous SNV | ZBTB48:NM_001278647:exon11:c.T2023G:p.S675A,ZBTB48:NM_005341:exon11:c.T2023G:p.S675A,ZBTB48:NM_001278648:exon11:c.T2023G:p.S675A, |
| chr1 | 66801073 | T | A | COSV58788106 | exonic | INSL5 | nonsynonymous SNV | INSL5:NM_005478:exon1:c.A149T:p.Q50L, |
| chr1 | 67705475 | A | T | . | exonic | GNG12 | nonsynonymous SNV | GNG12:NM_018841:exon4:c.T195A:p.D65E, |

| | | | | | | | | |
|---|---|---|---|---|---|---|---|---|
| chr1 | 68137903 | C | T | COSV52037496 | exonic | WLS | nonsynonymous SNV | WLS:NM_001002292:exon11:c.G1387A:p.V463I,WLS:NM_024911:exon11:c.G1393A:p.V465I,WLS:NM_001193334:exon10:c.G1120A:p.V374I, |
| chr1 | 6820206 | G | A | . | exonic | CAMTA1 | nonsynonymous SNV | CAMTA1:NM_001242701:exon2:c.G71A:p.G24E,CAMTA1:NM_001195563:exon2:c.G71A:p.G24E,CAMTA1:NM_015215:exon2:c.G71A:p.G24E, |
| chr1 | 6820250 | G | A | . | exonic | CAMTA1 | nonsynonymous SNV | CAMTA1:NM_001242701:exon2:c.G115A:p.D39N,CAMTA1:NM_001195563:exon2:c.G115A:p.D39N,CAMTA1:NM_015215:exon2:c.G115A:p.D39N, |
| chr1 | 6887657 | C | T | COSV57917420 | exonic | CAMTA1 | nonsynonymous SNV | CAMTA1:NM_001242701:exon4:c.C238T:p.L80F, |
| chr1 | 75733609 | T | C | . | exonic | ACADM | nonsynonymous SNV | ACADM:NM_001286043:exon6:c.T467C:p.I156T,ACADM:NM_001286042:exon4:c.T260C:p.I87T,ACADM:NM_001127328:exon5:c.T380C:p.I127T,ACADM:NM_000016:exon5:c.T368C:p.I123T, |
| chr1 | 75740038 | C | A | . | exonic | ACADM | nonsynonymous SNV | ACADM:NM_001286043:exon8:c.C626A:p.A209E,ACADM:NM_001286042:exon6:c.C419A:p.A140E,ACADM:NM_001127328:exon7:c.C539A:p.A180E,ACADM:NM_000016:exon7:c.C527A:p.A176E, |
| chr1 | 75749432 | G | A | . | exonic | ACADM | nonsynonymous SNV | ACADM:NM_001286043:exon10:c.G821A:p.G274D,ACADM:NM_001286042:exon8:c.G614A:p.G205D,ACADM:NM_001286044:exon6:c.G155A:p.G52D,ACADM:NM_001127328:exon9:c.G734A:p.G245D,ACADM:NM_000016:exon9:c.G722A:p.G241D, |
| chr1 | 75749507 | A | G | COSV57714736 | exonic | ACADM | nonsynonymous SNV | ACADM:NM_001286043:exon10:c.A896G:p.D299G,ACADM:NM_001286042:exon8:c.A689G:p.D230G,ACADM:NM_001286044:exon6:c.A230G:p.D77G,ACADM:NM_001127328:exon9:c.A809G:p.D270G,ACADM:NM_000016:exon9:c.A797G:p.D266G, |
| chr1 | 75749527 | G | A | . | exonic | ACADM | nonsynonymous SNV | ACADM:NM_001286042:exon8:c.G709A:p.A237T,ACADM:NM_001286043:exon10:c.G916A:p.A306T,ACADM:NM_001286044:exon6:c.G250A:p.A84T,ACADM:NM_001127328:exon9:c.G829A:p.A277T,ACADM:NM_000016:exon9:c.G817A:p.A273T, |
| chr1 | 75761165 | T | C | . | exonic | ACADM | nonsynonymous SNV | ACADM:NM_001286043:exon12:c.T1088C:p.V363A,ACADM:NM_001286042:exon10:c.T881C:p.V294A,ACADM:NM_001286044:exon8:c.T422C:p.V141A,ACADM:NM_001127328:exon11:c.T1001C:p.V334A,ACADM:NM_000016:exon11:c.T989C:p.V330A, |
| chr1 | 75761222 | G | A | . | exonic | ACADM | nonsynonymous SNV | ACADM:NM_001286043:exon12:c.G1145A:p.R382Q,ACADM:NM_001286042:exon10:c.G938A:p.R313Q,ACADM:NM_001286044:exon8:c.G479A:p.R160Q,ACADM:NM_001127328:exon11:c.G1058A:p.R353Q,ACADM:NM_000016:exon11:c.G1046A:p.R349Q, |
| chr1 | 75762731 | G | A | . | exonic | ACADM | nonsynonymous SNV | ACADM:NM_001286043:exon13:c.G1333A:p.A445T,ACADM:NM_001286042:exon11:c.G1126A:p.A376T,ACADM:NM_001286044:exon9:c.G667A:p.A223T,ACADM:NM_001127328:exon12:c.G1246A:p.A416T,ACADM:NM_000016:exon12:c.G1234A:p.A412T, |
| chr1 | 77949235 | C | T | . | exonic | FUBP1 | nonsynonymous SNV | FUBP1:NM_003902:exon19:c.G1846A:p.A616T,FUBP1:NM_001303433:exon20:c.G1909A:p.A637T, |
| chr1 | 77960348 | G | T | . | exonic | FUBP1 | nonsynonymous SNV | FUBP1:NM_003902:exon15:c.C1492A:p.P498T,FUBP1:NM_001303433:exon16:c.C1555A:p.P519T, |
| chr1 | 77960356 | C | T | . | exonic | FUBP1 | nonsynonymous SNV | FUBP1:NM_003902:exon15:c.G1484A:p.G495D,FUBP1:NM_001303433:exon16:c.G1547A:p.G516D, |
| chr1 | 77960401 | G | T | . | exonic | FUBP1 | nonsynonymous SNV | FUBP1:NM_003902:exon15:c.C1439A:p.P480Q,FUBP1:NM_001303433:exon16:c.C1502A:p.P501Q, |
| chr1 | 77964263 | A | G | . | exonic | FUBP1 | nonsynonymous SNV | FUBP1:NM_003902:exon11:c.T931C:p.F311L,FUBP1:NM_001303433:exon12:c.T994C:p.F332L, |
| chr1 | 77964325 | A | T | . | exonic | FUBP1 | nonsynonymous SNV | FUBP1:NM_003902:exon11:c.T869A:p.V290E,FUBP1:NM_001303433:exon12:c.T932A:p.V311E, |
| chr1 | 77964356 | C | T | . | exonic | FUBP1 | nonsynonymous SNV | FUBP1:NM_003902:exon11:c.G838A:p.V280I,FUBP1:NM_001303433:exon12:c.G901A:p.V301I, |
| chr1 | 77967057 | A | G | . | exonic | FUBP1 | nonsynonymous SNV | FUBP1:NM_003902:exon5:c.T335C:p.V112A,FUBP1:NM_001303433:exon6:c.T398C:p.V133A, |
| chr1 | 78016135 | C | A | . | exonic | DNAJB4 | nonsynonymous SNV | DNAJB4:NM_001317100:exon4:c.C782A:p.P261Q,DNAJB4:NM_001317102:exon3:c.C557A:p.P186Q,DNAJB4:NM_007034:exon3:c.C902A:p.P301Q,DNAJB4:NM_001317099:exon4:c.C902A:p.P301Q,DNAJB4:NM_001317101:exon3:c.C557A:p.P186Q, |
| chr1 | 84479306 | A | G | COSV65724027 | exonic | RPF1 | nonsynonymous SNV | RPF1:NM_025065:exon1:c.A25G:p.S9G, |
| chr1 | 86473447 | C | T | . | exonic | CLCA1 | nonsynonymous SNV | CLCA1:NM_001285:exon2:c.C193T:p.L65F, |
| chr1 | 86486641 | A | G | COSV52324766 | exonic | CLCA1 | nonsynonymous SNV | CLCA1:NM_001285:exon7:c.A1070G:p.N357S, |
| chr1 | 86493490 | T | C | COSV52326279 | exonic | CLCA1 | nonsynonymous SNV | CLCA1:NM_001285:exon10:c.T1571C:p.M524T, |
| chr1 | 8862903 | A | G | . | exonic | ENO1 | nonsynonymous SNV | ENO1:NM_001201483:exon8:c.T940C:p.Y314H,ENO1:NM_001428:exon11:c.T1219C:p.Y407H, |
| chr1 | 8862909 | C | T | . | exonic | ENO1 | nonsynonymous SNV | ENO1:NM_001201483:exon8:c.G934A:p.A312T,ENO1:NM_001428:exon11:c.G1213A:p.A405T, |
| chr1 | 8862933 | C | G | . | exonic | ENO1 | nonsynonymous SNV | ENO1:NM_001201483:exon8:c.G910C:p.A304P,ENO1:NM_001428:exon11:c.G1189C:p.A397P, |
| chr1 | 8863258 | C | T | . | exonic | ENO1 | nonsynonymous SNV | ENO1:NM_001201483:exon7:c.G874A:p.V292I,ENO1:NM_001428:exon10:c.G1153A:p.V385I, |
| chr1 | 8863288 | C | T | . | exonic | ENO1 | nonsynonymous SNV | ENO1:NM_001201483:exon7:c.G844A:p.E282K,ENO1:NM_001428:exon10:c.G1123A:p.E375K, |
| chr1 | 8863894 | T | C | COSV52304857 | exonic | ENO1 | nonsynonymous SNV | ENO1:NM_001201483:exon6:c.A785G:p.Q262R,ENO1:NM_001428:exon9:c.A1064G:p.Q355R, |
| chr1 | 8863964 | C | T | . | exonic | ENO1 | nonsynonymous SNV | ENO1:NM_001201483:exon6:c.G715A:p.V239M,ENO1:NM_001428:exon9:c.G994A:p.V332M, |
| chr1 | 8863973 | C | T | . | exonic | ENO1 | nonsynonymous SNV | ENO1:NM_001201483:exon6:c.G706A:p.A236T,ENO1:NM_001428:exon9:c.G985A:p.A329T, |
| chr1 | 8864024 | C | T | . | exonic | ENO1 | nonsynonymous SNV | ENO1:NM_001201483:exon6:c.G655A:p.G219R,ENO1:NM_001428:exon9:c.G934A:p.G312R, |
| chr1 | 8864029 | C | T | . | exonic | ENO1 | nonsynonymous SNV | ENO1:NM_001201483:exon6:c.G650A:p.S217N,ENO1:NM_001428:exon9:c.G929A:p.S310N, |
| chr1 | 8864090 | C | T | . | exonic | ENO1 | nonsynonymous SNV | ENO1:NM_001201483:exon6:c.G589A:p.V197M,ENO1:NM_001428:exon9:c.G868A:p.V290M, |
| chr1 | 8865320 | G | T | . | exonic | ENO1 | nonsynonymous SNV | ENO1:NM_001201483:exon5:c.C551A:p.A184D,ENO1:NM_001428:exon8:c.C830A:p.A277D, |
| chr1 | 8865347 | C | T | . | exonic | ENO1 | nonsynonymous SNV | ENO1:NM_001201483:exon5:c.G524A:p.S175N,ENO1:NM_001428:exon8:c.G803A:p.S268N, |
| chr1 | 8865392 | C | T | . | exonic | ENO1 | nonsynonymous SNV | ENO1:NM_001201483:exon5:c.G479A:p.R160K,ENO1:NM_001428:exon8:c.G758A:p.R253K, |
| chr1 | 8865405 | A | G | . | exonic | ENO1 | nonsynonymous SNV | ENO1:NM_001201483:exon5:c.T466C:p.S156P,ENO1:NM_001428:exon8:c.T745C:p.S249P, |
| chr1 | 8865414 | C | T | . | exonic | ENO1 | nonsynonymous SNV | ENO1:NM_001201483:exon5:c.G457A:p.V153I,ENO1:NM_001428:exon8:c.G736A:p.V246I, |
| chr1 | 8865418 | C | T | . | exonic | ENO1 | nonsynonymous SNV | ENO1:NM_001201483:exon5:c.G453A:p.M151I,ENO1:NM_001428:exon8:c.G732A:p.M244I, |
| chr1 | 8866420 | C | T | . | exonic | ENO1 | nonsynonymous SNV | ENO1:NM_001201483:exon4:c.G247A:p.A83T,ENO1:NM_001428:exon7:c.G526A:p.A176T, |
| chr1 | 8866429 | C | T | . | exonic | ENO1 | nonsynonymous SNV | ENO1:NM_001201483:exon4:c.G238A:p.V80I,ENO1:NM_001428:exon7:c.G517A:p.V173I, |
| chr1 | 8866441 | T | C | . | exonic | ENO1 | nonsynonymous SNV | ENO1:NM_001201483:exon4:c.A226G:p.M76V,ENO1:NM_001428:exon7:c.A505G:p.M169V, |
| chr1 | 8866480 | C | T | . | exonic | ENO1 | nonsynonymous SNV | ENO1:NM_001201483:exon4:c.G187A:p.G63S,ENO1:NM_001428:exon7:c.G466A:p.G156S, |
| chr1 | 8867154 | T | C | . | exonic | ENO1 | nonsynonymous SNV | ENO1:NM_001201483:exon3:c.A128G:p.D43G,ENO1:NM_001428:exon6:c.A407G:p.D136G, |
| chr1 | 8867158 | C | T | . | exonic | ENO1 | nonsynonymous SNV | ENO1:NM_001201483:exon3:c.G124A:p.A42T,ENO1:NM_001428:exon6:c.G403A:p.A135T, |
| chr1 | 8868006 | C | T | . | exonic | ENO1 | nonsynonymous SNV | ENO1:NM_001201483:exon2:c.G13A:p.D5N,ENO1:NM_001428:exon5:c.G292A:p.D98N, |
| chr1 | 8868012 | C | T | . | exonic | ENO1 | nonsynonymous SNV | ENO1:NM_001201483:exon2:c.G7A:p.E3K,ENO1:NM_001428:exon5:c.G286A:p.E96K, |
| chr1 | 8868041 | T | C | . | exonic | ENO1 | nonsynonymous SNV | ENO1:NM_001428:exon5:c.A257G:p.E86G, |
| chr1 | 8868048 | C | T | COSV52303828 | exonic | ENO1 | nonsynonymous SNV | ENO1:NM_001428:exon5:c.G250A:p.V84I, |

| | | | | | | | | |
|---|---|---|---|---|---|---|---|---|
| chr1 | 8870510 | C | T | . | exonic | ENO1 | nonsynonymous SNV | ENO1:NM_001428:exon4:c.G182A:p.G61D, |
| chr1 | 8871898 | C | T | . | exonic | ENO1 | nonsynonymous SNV | ENO1:NM_001428:exon3:c.G174A:p.M58I, |
| chr1 | 8871903 | A | G | . | exonic | ENO1 | nonsynonymous SNV | ENO1:NM_001428:exon3:c.T169C:p.Y57H, |
| chr1 | 8871921 | C | T | . | exonic | ENO1 | nonsynonymous SNV | ENO1:NM_001428:exon3:c.G151A:p.D51N, |
| chr1 | 8871948 | C | T | . | exonic | ENO1 | nonsynonymous SNV | ENO1:NM_001428:exon3:c.G124A:p.G42S, |
| chr1 | 8874829 | G | T | . | exonic | ENO1 | stopgain | ENO1:NM_001428:exon2:c.C80A:p.S27X, |
| chr1 | 89114258 | T | C | COSV65068341 | exonic | GBP2 | nonsynonymous SNV | GBP2:NM_004120:exon7:c.A907G:p.S303G, |
| chr1 | 89117007 | G | C | COSV65068867 | exonic | GBP2 | nonsynonymous SNV | GBP2:NM_004120:exon6:c.C853G:p.P285A, |
| chr1 | 8949347 | C | T | COSV66262569 | exonic | CA6 | nonsynonymous SNV | CA6:NM_001215:exon2:c.C164T:p.T55M,CA6:NM_001270500:exon2:c.C164T:p.T55M, |
| chr1 | 8949385 | A | C | COSV66261772 | exonic | CA6 | nonsynonymous SNV | CA6:NM_001215:exon2:c.A202C:p.M68L,CA6:NM_001270500:exon2:c.A202C:p.M68L, |
| chr1 | 8957145 | A | G | . | exonic | CA6 | nonsynonymous SNV | CA6:NM_001270501:exon2:c.A88G:p.S30G,CA6:NM_001215:exon3:c.A268G:p.S90G,CA6:NM_001270500:exon3:c.A268G:p.S90G, |
| chr1 | 92833389 | G | A | . | exonic | RPL5 | nonsynonymous SNV | RPL5:NM_000969:exon2:c.G4A:p.G2R, |
| chr1 | 92833396 | T | C | . | exonic | RPL5 | nonsynonymous SNV | RPL5:NM_000969:exon2:c.T11C:p.V4A, |
| chr1 | 92833401 | G | A | . | exonic | RPL5 | nonsynonymous SNV | RPL5:NM_000969:exon2:c.G16A:p.V6I, |
| chr1 | 92833404 | G | A | . | exonic | RPL5 | nonsynonymous SNV | RPL5:NM_000969:exon2:c.G19A:p.V7I, |
| chr1 | 92833433 | C | A | COSV59873416 | exonic | RPL5 | stopgain | RPL5:NM_000969:exon2:c.C48A:p.Y16X, |
| chr1 | 92833446 | A | G | . | exonic | RPL5 | nonsynonymous SNV | RPL5:NM_000969:exon2:c.A61G:p.R21G, |
| chr1 | 92833556 | G | A | . | exonic | RPL5 | nonsynonymous SNV | RPL5:NM_000969:exon3:c.G85A:p.D29N, |
| chr1 | 92833632 | G | A | . | exonic | RPL5 | nonsynonymous SNV | RPL5:NM_000969:exon3:c.G161A:p.R54H, |
| chr1 | 92833646 | G | A | . | exonic | RPL5 | nonsynonymous SNV | RPL5:NM_000969:exon3:c.G175A:p.D59N, |
| chr1 | 92833652 | A | T | . | exonic | RPL5 | nonsynonymous SNV | RPL5:NM_000969:exon3:c.A181T:p.I61F, |
| chr1 | 92834818 | G | A | COSV59873410 | exonic | RPL5 | nonsynonymous SNV | RPL5:NM_000969:exon4:c.G229A:p.A77T, |
| chr1 | 92834833 | G | A | COSV59872943 | exonic | RPL5 | nonsynonymous SNV | RPL5:NM_000969:exon4:c.G244A:p.E82K, |
| chr1 | 92834849 | G | A | . | exonic | RPL5 | nonsynonymous SNV | RPL5:NM_000969:exon4:c.G260A:p.G87D, |
| chr1 | 92834879 | C | T | . | exonic | RPL5 | nonsynonymous SNV | RPL5:NM_000969:exon4:c.C290T:p.A97V, |
| chr1 | 92834909 | G | A | . | exonic | RPL5 | nonsynonymous SNV | RPL5:NM_000969:exon4:c.G320A:p.R107H, |
| chr1 | 92836226 | G | A | . | exonic | RPL5 | nonsynonymous SNV | RPL5:NM_000969:exon5:c.G361A:p.G121S, |
| chr1 | 92836244 | G | A | . | exonic | RPL5 | nonsynonymous SNV | RPL5:NM_000969:exon5:c.G379A:p.G127S, |
| chr1 | 92836259 | G | A | . | exonic | RPL5 | nonsynonymous SNV | RPL5:NM_000969:exon5:c.G394A:p.V132M, |
| chr1 | 92836299 | A | T | . | exonic | RPL5 | nonsynonymous SNV | RPL5:NM_000969:exon5:c.A434T:p.Y145F, |
| chr1 | 92836347 | G | A | . | exonic | RPL5 | nonsynonymous SNV | RPL5:NM_000969:exon5:c.G482A:p.G161D, |
| chr1 | 92837488 | G | A | . | exonic | RPL5 | nonsynonymous SNV | RPL5:NM_000969:exon6:c.G560A:p.S187N, |
| chr1 | 92837515 | G | A | COSV59874073 | exonic | RPL5 | nonsynonymous SNV | RPL5:NM_000969:exon6:c.G587A:p.R196Q, |
| chr1 | 92837565 | G | A | . | exonic | RPL5 | nonsynonymous SNV | RPL5:NM_000969:exon6:c.G637A:p.E213K, |
| chr1 | 92837619 | G | A | COSV59872895 | exonic | RPL5 | nonsynonymous SNV | RPL5:NM_000969:exon6:c.G691A:p.V231I, |
| chr1 | 92840600 | T | C | . | exonic | RPL5 | nonsynonymous SNV | RPL5:NM_000969:exon7:c.T755C:p.V252A, |
| chr1 | 92840627 | T | A | . | exonic | RPL5 | nonsynonymous SNV | RPL5:NM_000969:exon7:c.T782A:p.V261D, |
| chr1 | 92841810 | T | C | . | exonic | RPL5 | nonsynonymous SNV | RPL5:NM_000969:exon8:c.T839C:p.V280A, |
| chr1 | 95064901 | C | T | . | exonic | ALG14 | nonsynonymous SNV | ALG14:NM_001305242:exon2:c.G253A:p.E85K,ALG14:NM_144988:exon2:c.G253A:p.E85K, |
| chr1 | 95072819 | C | T | . | exonic | ALG14 | nonsynonymous SNV | ALG14:NM_001305242:exon1:c.G80A:p.R27H,ALG14:NM_144988:exon1:c.G80A:p.R27H, |
| chr1 | 95244265 | T | C | . | exonic | RWDD3 | nonsynonymous SNV | RWDD3:NM_001278247:exon3:c.T95C:p.V32A,RWDD3:NM_015485:exon2:c.T140C:p.V47A,RWDD3:NM_001199682:exon2:c.T140C:p.V47A,RWDD3:NM_001128142:exon2:c.T140C:p.V47A,RWDD3:NM_001278248:exon3:c.T95C:p.V32A, |
| chr1 | 953279 | T | C | . | exonic | NOC2L | nonsynonymous SNV | NOC2L:NM_015658:exon9:c.A898G:p.I300V, |
| chr1 | 97515839 | T | C | COSV64593269 | exonic | DPYD | nonsynonymous SNV | DPYD:NM_000110:exon13:c.A1627G:p.I543V, |
| chr1 | 999842 | C | A | COSV59241795 | exonic | HES4 | nonsynonymous SNV | HES4:NM_001142467:exon1:c.G132T:p.R44S, |
| chr10 | 102157495 | C | A | . | exonic | NOLC1 | nonsynonymous SNV | NOLC1:NM_001284388:exon4:c.C381A:p.S127R,NOLC1:NM_001284389:exon4:c.C384A:p.S128R,NOLC1:NM_004741:exon4:c.C381A:p.S127R, |
| chr10 | 102159994 | G | A | . | exonic | NOLC1 | nonsynonymous SNV | NOLC1:NM_001284388:exon8:c.G988A:p.E330K,NOLC1:NM_001284389:exon8:c.G961A:p.E321K,NOLC1:NM_004741:exon8:c.G958A:p.E320K, |
| chr10 | 102160718 | T | C | . | exonic | NOLC1 | nonsynonymous SNV | NOLC1:NM_001284388:exon10:c.T1396C:p.S466P,NOLC1:NM_001284389:exon10:c.T1369C:p.S457P,NOLC1:NM_004741:exon10:c.T1366C:p.S456P, |
| chr10 | 102160941 | A | G | . | exonic | NOLC1 | nonsynonymous SNV | NOLC1:NM_001284388:exon10:c.A1619G:p.E540G,NOLC1:NM_001284389:exon10:c.A1592G:p.E531G,NOLC1:NM_004741:exon10:c.A1589G:p.E530G, |
| chr10 | 102161057 | G | A | . | exonic | NOLC1 | nonsynonymous SNV | NOLC1:NM_001284388:exon10:c.G1735A:p.E579K,NOLC1:NM_001284389:exon10:c.G1708A:p.E570K,NOLC1:NM_004741:exon10:c.G1705A:p.E569K, |
| chr10 | 102813206 | T | C | COSV64009277 | exonic | WBP1L | nonsynonymous SNV | WBP1L:NM_017787:exon4:c.T904C:p.S302P,WBP1L:NM_001083913:exon4:c.T967C:p.S323P, |
| chr10 | 102878966 | T | C | COSV54916951 | exonic | AS3MT | nonsynonymous SNV | AS3MT:NM_020682:exon9:c.T860C:p.M287T, |
| chr10 | 103174952 | T | C | . | exonic | NT5C2 | nonsynonymous SNV | NT5C2:NM_001134373:exon2:c.A7G:p.T3A,NT5C2:NM_012229:exon3:c.A7G:p.T3A, |
| chr10 | 103882691 | G | A | COSV56543963 | exonic | OBFC1 | nonsynonymous SNV | OBFC1:NM_024928:exon10:c.C1100T:p.A367V, |
| chr10 | 103897558 | G | C | . | exonic | OBFC1 | nonsynonymous SNV | OBFC1:NM_024928:exon7:c.C743G:p.S248C, |
| chr10 | 103900068 | T | C | . | exonic | OBFC1 | nonsynonymous SNV | OBFC1:NM_024928:exon5:c.A451G:p.T151A, |
| chr10 | 104263031 | C | A | COSV63846494 | exonic | GSTO1 | nonsynonymous SNV | GSTO1:NM_001191003:exon4:c.C335A:p.A112D,GSTO1:NM_004832:exon4:c.C419A:p.A140D, |
| chr10 | 104279427 | A | G | COSV58537765 | exonic | GSTO2 | nonsynonymous SNV | GSTO2:NM_183239:exon5:c.A424G:p.N142D,GSTO2:NM_001191014:exon3:c.A340G:p.N114D, |
| chr10 | 110295351 | A | T | . | exonic | SMNDC1 | stopgain | SMNDC1:NM_005871:exon5:c.T456A:p.Y152X, |
| chr10 | 110507064 | G | A | . | exonic | DUSP5 | nonsynonymous SNV | DUSP5:NM_004419:exon3:c.G658A:p.A220T, |
| chr10 | 110507065 | C | T | . | exonic | DUSP5 | nonsynonymous SNV | DUSP5:NM_004419:exon3:c.C659T:p.A220V, |
| chr10 | 110881295 | A | G | . | exonic | PDCD4 | nonsynonymous SNV | PDCD4:NM_145341:exon4:c.A73G:p.I25V,PDCD4:NM_014456:exon3:c.A106G:p.I36V,PDCD4:NM_001199492:exon3:c.A64G:p.I22V, |
| chr10 | 113729393 | C | G | COSV61881355 | exonic | CASP7 | nonsynonymous SNV | CASP7:NM_001267056:exon7:c.C765G:p.D255E,CASP7:NM_001267058:exon6:c.C690G:p.D230E,CASP7:NM_033338:exon8:c.C864G:p.D288E,CASP7:NM_001227:exon7:c.C765G:p.D255E,CASP7:NM_001267057:exon7:c.C1020G:p.D340E,CASP7:NM_033339:exon8:c.C765G:p.D255E,CASP7:NM_033340:exon8:c.C731G:p.T244S, |
| chr10 | 113729408 | G | A | COSV61881361 | exonic | CASP7 | nonsynonymous SNV | CASP7:NM_033340:exon8:c.G746A:p.R249K, |
| chr10 | 114162733 | C | T | COSV65159404 | exonic | CCDC186 | nonsynonymous SNV | CCDC186:NM_018017:exon2:c.G536A:p.R179Q, |
| chr10 | 114163015 | G | A | COSV65159408 | exonic | CCDC186 | nonsynonymous SNV | CCDC186:NM_018017:exon2:c.C254T:p.T85I, |
| chr10 | 11747383 | G | A | . | exonic | ECHDC3 | nonsynonymous SNV | ECHDC3:NM_024693:exon2:c.G205A:p.A69T, |
| chr10 | 120355012 | G | T | . | exonic | RPL21 | nonsynonymous SNV | RPL21:NM_000982:exon1:c.C172A:p.H58N, |
| chr10 | 121911030 | A | T | . | exonic | ATE1 | nonsynonymous SNV | ATE1:NM_001288736:exon5:c.T438A:p.S146R,ATE1:NM_001288734:exon4:c.T114A:p.S38R,ATE1:NM_001001976:exon5:c.T459A:p.S153R,ATE1:NM_007041:exon5:c.T459A:p.S153R,ATE1:NM_001288735:exon3:c.T171A:p.S57R, |
| chr10 | 122211015 | T | C | COSV53281976 | exonic | TACC2 | nonsynonymous SNV | TACC2:NM_206861:exon6:c.T1028C:p.V343A,TACC2:NM_206860:exon3:c.T824C:p.V275A,TACC2:NM_001291876:exon8:c.T6455C:p.V2152A,TACC2:NM_001291878:exon3:c.T824C:p.V275A,TACC2:NM_006997:exon3:c.T824C:p.V275A,TACC2:NM_001291877:exon8:c.T6602C:p.V2201A,TACC2:NM_206862:exon9:c.T6590C:p.V2197A, |

| | | | | | | | | |
|---|---|---|---|---|---|---|---|---|
| chr10 | 122211054 | C | T | COSV53280580 | exonic | TACC2 | nonsynonymous SNV | TACC2:NM_206861:exon6:c.C1067T:p.A356V,TACC2:NM_206860:exon3:c.C863T:p.A288V,TACC2:NM_001291876:exon8:c.C6494T:p.A2165V,TACC2:NM_001291878:exon3:c.C863T:p.A288V,TACC2:NM_006997:exon3:c.C863T:p.A288V,TACC2:NM_001291877:exon8:c.C6641T:p.A2214V,TACC2:NM_206862:exon9:c.C6629T:p.A2210V, |
| chr10 | 122211207 | T | A | COSV53280628 | exonic | TACC2 | nonsynonymous SNV | TACC2:NM_206861:exon6:c.T1220A:p.L407H,TACC2:NM_206860:exon3:c.T1016A:p.L339H,TACC2:NM_001291876:exon8:c.T6647A:p.L2216H,TACC2:NM_001291878:exon3:c.T1016A:p.L339H,TACC2:NM_006997:exon3:c.T1016A:p.L339H,TACC2:NM_001291877:exon8:c.T6794A:p.L2265H,TACC2:NM_206862:exon9:c.T6782A:p.L2261H, |
| chr10 | 122237461 | G | A | COSV53279154 | exonic | TACC2 | nonsynonymous SNV | TACC2:NM_206861:exon14:c.G2632A:p.A878T,TACC2:NM_206860:exon10:c.G2338A:p.A780T,TACC2:NM_001291876:exon14:c.G7828A:p.A2610T,TACC2:NM_001291879:exon11:c.G1075A:p.A359T,TACC2:NM_001291878:exon11:c.G2374A:p.A792T,TACC2:NM_006997:exon11:c.G2428A:p.A810T,TACC2:NM_001291877:exon14:c.G7975A:p.A2659T,TACC2:NM_206862:exon17:c.G8194A:p.A2732T, |
| chr10 | 122249609 | G | C | COSV53282365 | exonic | TACC2 | nonsynonymous SNV | TACC2:NM_206861:exon19:c.G3164C:p.S1055T,TACC2:NM_206860:exon15:c.G2870C:p.S957T,TACC2:NM_001291876:exon19:c.G8360C:p.S2787T,TACC2:NM_001291879:exon16:c.G1607C:p.S536T,TACC2:NM_001291878:exon16:c.G2906C:p.S969T,TACC2:NM_006997:exon16:c.G2960C:p.S987T,TACC2:NM_001291877:exon19:c.G8507C:p.S2836T,TACC2:NM_206862:exon22:c.G8726C:p.S2909T, |
| chr10 | 127451398 | G | A | COSV54751969 | exonic | DOCK1 | nonsynonymous SNV | DOCK1:NM_001290223:exon52:c.G5632A:p.A1878T,DOCK1:NM_001380:exon52:c.G5569A:p.A1857T, |
| chr10 | 129536247 | G | A | . | exonic | MGMT | nonsynonymous SNV | MGMT:NM_002412:exon2:c.G88A:p.G30R, |
| chr10 | 129536255 | G | A | . | exonic | MGMT | nonsynonymous SNV | MGMT:NM_002412:exon2:c.G96A:p.M32I, |
| chr10 | 129708019 | C | T | COSV60031127 | exonic | MGMT | nonsynonymous SNV | MGMT:NM_002412:exon3:c.C343T:p.L115F, |
| chr10 | 129766942 | C | A | . | exonic | MGMT | nonsynonymous SNV | MGMT:NM_002412:exon5:c.C662A:p.A221D, |
| chr10 | 133352286 | A | G | . | exonic | PRAP1 | nonsynonymous SNV | PRAP1:NM_001145201:exon5:c.A275G:p.H92R,PRAP1:NM_145202:exon5:c.A302G:p.H101R, |
| chr10 | 133370622 | G | A | . | exonic | ECHS1 | nonsynonymous SNV | ECHS1:NM_004092:exon2:c.C224T:p.T75I, |
| chr10 | 133373302 | A | G | COSV63906597 | exonic | ECHS1 | nonsynonymous SNV | ECHS1:NM_004092:exon1:c.T32C:p.V11A, |
| chr10 | 16593489 | C | T | . | exonic | RSU1 | nonsynonymous SNV | RSU1:NM_152724:exon8:c.G580A:p.G194S,RSU1:NM_012425:exon9:c.G739A:p.G247S, |
| chr10 | 16764391 | C | T | . | exonic | RSU1 | nonsynonymous SNV | RSU1:NM_152724:exon3:c.G121A:p.G41S,RSU1:NM_012425:exon4:c.G280A:p.G94S, |
| chr10 | 16764442 | C | T | . | exonic | RSU1 | nonsynonymous SNV | RSU1:NM_152724:exon3:c.G70A:p.E24K,RSU1:NM_012425:exon4:c.G229A:p.E77K, |
| chr10 | 16816985 | C | T | . | exonic | RSU1 | nonsynonymous SNV | RSU1:NM_012425:exon2:c.G97A:p.V33I, |
| chr10 | 17229690 | G | A | . | exonic | VIM | nonsynonymous SNV | VIM:NM_003380:exon2:c.G268A:p.D90N, |
| chr10 | 17229886 | G | A | . | exonic | VIM | nonsynonymous SNV | VIM:NM_003380:exon2:c.G464A:p.R155Q, |
| chr10 | 17229898 | G | A | . | exonic | VIM | nonsynonymous SNV | VIM:NM_003380:exon2:c.G476A:p.R159Q, |
| chr10 | 17233627 | G | A | . | exonic | VIM | nonsynonymous SNV | VIM:NM_003380:exon4:c.G665A:p.R222H, |
| chr10 | 17233635 | G | A | . | exonic | VIM | nonsynonymous SNV | VIM:NM_003380:exon4:c.G673A:p.E225K, |
| chr10 | 17233677 | G | A | COSV56405834 | exonic | VIM | nonsynonymous SNV | VIM:NM_003380:exon4:c.G715A:p.E239K, |
| chr10 | 17234699 | G | A | . | exonic | VIM | nonsynonymous SNV | VIM:NM_003380:exon6:c.G889A:p.D297N, |
| chr10 | 17234705 | T | C | . | exonic | VIM | nonsynonymous SNV | VIM:NM_003380:exon6:c.T895C:p.S299P, |
| chr10 | 17234714 | G | A | . | exonic | VIM | nonsynonymous SNV | VIM:NM_003380:exon6:c.G904A:p.A302T, |
| chr10 | 17234736 | T | C | . | exonic | VIM | nonsynonymous SNV | VIM:NM_003380:exon6:c.T926C:p.L309P, |
| chr10 | 17234739 | G | A | COSV56406122 | exonic | VIM | nonsynonymous SNV | VIM:NM_003380:exon6:c.G929A:p.R310H, |
| chr10 | 17234769 | G | A | COSV56405803 | exonic | VIM | nonsynonymous SNV | VIM:NM_003380:exon6:c.G959A:p.R320Q, |
| chr10 | 17234793 | G | A | . | exonic | VIM | nonsynonymous SNV | VIM:NM_003380:exon6:c.G983A:p.C328Y, |
| chr10 | 17235223 | G | A | . | exonic | VIM | nonsynonymous SNV | VIM:NM_003380:exon7:c.G1063A:p.A355T, |
| chr10 | 17235344 | T | C | . | exonic | VIM | nonsynonymous SNV | VIM:NM_003380:exon7:c.T1184C:p.I395T, |
| chr10 | 17235352 | G | A | . | exonic | VIM | nonsynonymous SNV | VIM:NM_003380:exon7:c.G1192A:p.A398T, |
| chr10 | 17235362 | G | A | . | exonic | VIM | nonsynonymous SNV | VIM:NM_003380:exon7:c.G1202A:p.R401K, |
| chr10 | 17235368 | T | C | . | exonic | VIM | nonsynonymous SNV | VIM:NM_003380:exon7:c.T1208C:p.L403P, |
| chr10 | 17594310 | G | A | COSV63517260 | exonic | HACD1 | nonsynonymous SNV | HACD1:NM_014241:exon6:c.C679T:p.H227Y, |
| chr10 | 17617132 | C | A | COSV58892476 | exonic | HACD1 | nonsynonymous SNV | HACD1:NM_014241:exon1:c.G208T:p.V70F, |
| chr10 | 17617150 | C | T | COSV58892488 | exonic | HACD1 | nonsynonymous SNV | HACD1:NM_014241:exon1:c.G190A:p.E64K, |
| chr10 | 20817617 | G | T | . | exonic | NEBL | nonsynonymous SNV | NEBL:NM_006393:exon21:c.C2131A:p.Q711K, |
| chr10 | 20840818 | A | T | . | exonic | NEBL | nonsynonymous SNV | NEBL:NM_006393:exon13:c.T1259A:p.I420K, |
| chr10 | 20840822 | C | T | . | exonic | NEBL | nonsynonymous SNV | NEBL:NM_006393:exon13:c.G1255A:p.E419K, |
| chr10 | 22316509 | C | A | . | exonic | COMMD3 | nonsynonymous SNV | COMMD3:NM_012071:exon1:c.C92A:p.A31E,COMMD3-BMI1:NM_001204062:exon1:c.C92A:p.A31E, |
| chr10 | 28617756 | G | A | . | exonic | WAC | nonsynonymous SNV | WAC:NM_016628:exon13:c.G1846A:p.E616K,WAC:NM_100264:exon13:c.G1711A:p.E571K,WAC:NM_100486:exon12:c.G1537A:p.E513K, |
| chr10 | 30849888 | G | A | COSV59195695 | exonic | ZNF438 | nonsynonymous SNV | ZNF438:NM_001143771:exon7:c.C487T:p.P163S,ZNF438:NM_182755:exon7:c.C517T:p.P173S,ZNF438:NM_001143767:exon5:c.C517T:p.P173S,ZNF438:NM_001143769:exon7:c.C370T:p.P124S,ZNF438:NM_001143766:exon7:c.C517T:p.P173S,ZNF438:NM_001143770:exon8:c.C487T:p.P163S,ZNF438:NM_001143768:exon6:c.C517T:p.P173S, |
| chr10 | 31520330 | T | A | . | exonic | ZEB1 | nonsynonymous SNV | ZEB1:NM_001174093:exon6:c.T935A:p.I312K,ZEB1:NM_001174095:exon6:c.T794A:p.I265K,ZEB1:NM_001128128:exon7:c.T947A:p.I316K,ZEB1:NM_030751:exon7:c.T995A:p.I332K,ZEB1:NM_001174096:exon7:c.T998A:p.I333K,ZEB1:NM_001174094:exon7:c.T944A:p.I315K, |
| chr10 | 31520332 | G | A | . | exonic | ZEB1 | nonsynonymous SNV | ZEB1:NM_001174093:exon6:c.G937A:p.E313K,ZEB1:NM_001174095:exon6:c.G796A:p.E266K,ZEB1:NM_001128128:exon7:c.G949A:p.E317K,ZEB1:NM_030751:exon7:c.G997A:p.E333K,ZEB1:NM_001174096:exon7:c.G1000A:p.E334K,ZEB1:NM_001174094:exon7:c.G946A:p.E316K, |
| chr10 | 31520803 | C | A | . | exonic | ZEB1 | nonsynonymous SNV | ZEB1:NM_001174093:exon6:c.C1408A:p.Q470K,ZEB1:NM_001174095:exon6:c.C1267A:p.Q423K,ZEB1:NM_030751:exon7:c.C1468A:p.Q490K,ZEB1:NM_001174094:exon7:c.C1417A:p.Q473K,ZEB1:NM_001128128:exon7:c.C1420A:p.Q474K,ZEB1:NM_001174096:exon7:c.C1471A:p.Q491K, |
| chr10 | 31520809 | T | A | . | exonic | ZEB1 | nonsynonymous SNV | ZEB1:NM_001174093:exon6:c.T1414A:p.L472I,ZEB1:NM_001174095:exon6:c.T1273A:p.L425I,ZEB1:NM_001128128:exon7:c.T1426A:p.L476I,ZEB1:NM_030751:exon7:c.T1474A:p.L492I,ZEB1:NM_001174096:exon7:c.T1477A:p.L493I,ZEB1:NM_001174094:exon7:c.T1423A:p.L475I, |
| chr10 | 31524006 | C | A | . | exonic | ZEB1 | nonsynonymous SNV | ZEB1:NM_001174093:exon7:c.C2615A:p.P872H,ZEB1:NM_001174095:exon7:c.C2474A:p.P825H,ZEB1:NM_030751:exon8:c.C2675A:p.P892H,ZEB1:NM_001174094:exon8:c.C2624A:p.P875H,ZEB1:NM_001128128:exon8:c.C2627A:p.P876H,ZEB1:NM_001174096:exon8:c.C2678A:p.P893H, |
| chr10 | 32686041 | A | G | . | exonic | CCDC7 | nonsynonymous SNV | CCDC7:NM_024688:exon4:c.A115G:p.I39V, |
| chr10 | 33263768 | A | G | . | exonic | NRP1 | nonsynonymous SNV | NRP1:NM_001024628:exon4:c.T536C:p.V179A,NRP1:NM_001244973:exon4:c.T536C:p.V179A,NRP1:NM_001024629:exon4:c.T536C:p.V179A,NRP1:NM_003873:exon4:c.T536C:p.V179A,NRP1:NM_001244972:exon4:c.T536C:p.V179A, |
| chr10 | 34317194 | C | T | . | exonic | PARD3 | nonsynonymous SNV | PARD3:NM_001184794:exon19:c.G2849A:p.G950E,PARD3:NM_001184791:exon17:c.G2762A:p.G921E,PARD3:NM_001184786:exon19:c.G2939A:p.G980E,PARD3:NM_001184792:exon20:c.G2978A:p.G993E,PARD3:NM_019619:exon20:c.G2987A:p.G996E,PARD3:NM_001184785:exon20:c.G2978A:p.G993E,PARD3:NM_001184788:exon19:c.G2849A:p.G950E,PARD3:NM_001184793:exon20:c.G2885A:p.G962E,PARD3:NM_001184787:exon20:c.G2987A:p.G996E,PARD3:NM_001184789:exon19:c.G2849A:p.G950E,PARD3:NM_001184790:exon18:c.G2717A:p.G906E, |
| chr10 | 46009424 | C | T | . | exonic | NCOA4 | nonsynonymous SNV | NCOA4:NM_001145261:exon10:c.G1874A:p.R625Q,NCOA4:NM_001145260:exon10:c.G1874A:p.R625Q,NCOA4:NM_005437:exon9:c.G1826A:p.R609Q,NCOA4:NM_001145263:exon9:c.G1826A:p.R609Q,NCOA4:NM_001145262:exon9:c.G1826A:p.R609Q, |
| chr10 | 46009493 | C | T | . | exonic | NCOA4 | nonsynonymous SNV | NCOA4:NM_001145261:exon10:c.G1805A:p.G602D,NCOA4:NM_001145260:exon10:c.G1805A:p.G602D,NCOA4:NM_005437:exon9:c.G1757A:p.G586D,NCOA4:NM_001145263:exon9:c.G1757A:p.G586D,NCOA4:NM_001145262:exon9:c.G1757A:p.G586D, |
| chr10 | 46010293 | C | T | . | exonic | NCOA4 | nonsynonymous SNV | NCOA4:NM_001145261:exon9:c.G1676A:p.G559E,NCOA4:NM_001145260:exon9:c.G1676A:p.G559E,NCOA4:NM_005437:exon8:c.G1628A:p.G543E,NCOA4:NM_001145263:exon8:c.G1628A:p.G543E,NCOA4:NM_001145262:exon8:c.G1628A:p.G543E, |
| chr10 | 46010365 | C | T | . | exonic | NCOA4 | nonsynonymous SNV | NCOA4:NM_001145261:exon9:c.G1604A:p.G535D,NCOA4:NM_001145260:exon9:c.G1604A:p.G535D,NCOA4:NM_005437:exon8:c.G1556A:p.G519D,NCOA4:NM_001145263:exon8:c.G1556A:p.G519D,NCOA4:NM_001145262:exon8:c.G1556A:p.G519D, |
| chr10 | 46010369 | C | T | . | exonic | NCOA4 | nonsynonymous SNV | NCOA4:NM_001145261:exon9:c.G1600A:p.A534T,NCOA4:NM_001145260:exon9:c.G1600A:p.A534T,NCOA4:NM_005437:exon8:c.G1552A:p.A518T,NCOA4:NM_001145263:exon8:c.G1552A:p.A518T,NCOA4:NM_001145262:exon8:c.G1552A:p.A518T, |

| | | | | | | | | |
|---|---|---|---|---|---|---|---|---|
| chr10 | 46010595 | C | T | . | exonic | NCOA4 | nonsynonymous SNV | NCOA4:NM_001145261:exon9:c.G1374A:p.M458I,NCOA4:NM_001145260:exon9:c.G1374A:p.M458I,NCOA4:NM_005437:exon8:c.G1326A:p.M442I,NCOA4:NM_001145263:exon8:c.G1326A:p.M442I,NCOA4:NM_001145262:exon8:c.G1326A:p.M442I, |
| chr10 | 46010599 | C | T | . | exonic | NCOA4 | nonsynonymous SNV | NCOA4:NM_001145261:exon9:c.G1370A:p.G457E,NCOA4:NM_001145260:exon9:c.G1370A:p.G457E,NCOA4:NM_005437:exon8:c.G1322A:p.G441E,NCOA4:NM_001145263:exon8:c.G1322A:p.G441E,NCOA4:NM_001145262:exon8:c.G1322A:p.G441E, |
| chr10 | 46010618 | C | T | . | exonic | NCOA4 | nonsynonymous SNV | NCOA4:NM_001145261:exon9:c.G1351A:p.E451K,NCOA4:NM_001145260:exon9:c.G1351A:p.E451K,NCOA4:NM_005437:exon8:c.G1303A:p.E435K,NCOA4:NM_001145263:exon8:c.G1303A:p.E435K,NCOA4:NM_001145262:exon8:c.G1303A:p.E435K, |
| chr10 | 46010714 | C | T | . | exonic | NCOA4 | nonsynonymous SNV | NCOA4:NM_001145261:exon9:c.G1255A:p.V419M,NCOA4:NM_001145260:exon9:c.G1255A:p.V419M,NCOA4:NM_005437:exon8:c.G1207A:p.V403M,NCOA4:NM_001145263:exon8:c.G1207A:p.V403M,NCOA4:NM_001145262:exon8:c.G1207A:p.V403M, |
| chr10 | 46010750 | C | T | . | exonic | NCOA4 | nonsynonymous SNV | NCOA4:NM_001145263:exon8:c.G1171A:p.V391I,NCOA4:NM_001145262:exon8:c.G1171A:p.V391I,NCOA4:NM_001145261:exon9:c.G1219A:p.V407I,NCOA4:NM_001145260:exon9:c.G1219A:p.V407I,NCOA4:NM_005437:exon8:c.G1171A:p.V391I, |
| chr10 | 46010818 | C | T | . | exonic | NCOA4 | nonsynonymous SNV | NCOA4:NM_001145261:exon9:c.G1151A:p.C384Y,NCOA4:NM_001145260:exon9:c.G1151A:p.C384Y,NCOA4:NM_005437:exon8:c.G1103A:p.C368Y,NCOA4:NM_001145263:exon8:c.G1103A:p.C368Y,NCOA4:NM_001145262:exon8:c.G1103A:p.C368Y, |
| chr10 | 46010921 | A | G | . | exonic | NCOA4 | nonsynonymous SNV | NCOA4:NM_001145261:exon9:c.T1048C:p.S350P,NCOA4:NM_001145260:exon9:c.T1048C:p.S350P,NCOA4:NM_005437:exon8:c.T1000C:p.S334P,NCOA4:NM_001145263:exon8:c.T1000C:p.S334P,NCOA4:NM_001145262:exon8:c.T1000C:p.S334P, |
| chr10 | 46010959 | C | T | . | exonic | NCOA4 | nonsynonymous SNV | NCOA4:NM_001145261:exon9:c.G1010A:p.S337N,NCOA4:NM_001145260:exon9:c.G1010A:p.S337N,NCOA4:NM_005437:exon8:c.G962A:p.S321N,NCOA4:NM_001145263:exon8:c.G962A:p.S321N,NCOA4:NM_001145262:exon8:c.G962A:p.S321N, |
| chr10 | 46013593 | C | T | . | exonic | NCOA4 | nonsynonymous SNV | NCOA4:NM_001145261:exon7:c.G575A:p.G192E,NCOA4:NM_001145260:exon7:c.G575A:p.G192E,NCOA4:NM_005437:exon6:c.G527A:p.G176E,NCOA4:NM_001145263:exon6:c.G527A:p.G176E,NCOA4:NM_001145262:exon6:c.G527A:p.G176E, |
| chr10 | 46013603 | C | T | . | exonic | NCOA4 | nonsynonymous SNV | NCOA4:NM_001145261:exon7:c.G565A:p.A189T,NCOA4:NM_001145260:exon7:c.G565A:p.A189T,NCOA4:NM_005437:exon6:c.G517A:p.A173T,NCOA4:NM_001145263:exon6:c.G517A:p.A173T,NCOA4:NM_001145262:exon6:c.G517A:p.A173T, |
| chr10 | 5094459 | C | G | COSV65910253 | exonic | AKR1C3 | nonsynonymous SNV | AKR1C3:NM_003739:exon1:c.C15G:p.H5Q,AKR1C3:NM_001253909:exon1:c.C15G:p.H5Q, |
| chr10 | 59792934 | G | T | COSV54054295 | exonic | CCDC6 | nonsynonymous SNV | CCDC6:NM_005436:exon9:c.C1408A:p.P470T, |
| chr10 | 62198353 | T | C | . | exonic | RTKN2 | nonsynonymous SNV | RTKN2:NM_145307:exon12:c.A1385G:p.H462R, |
| chr10 | 68757297 | G | A | . | exonic | CCAR1 | nonsynonymous SNV | CCAR1:NM_001282959:exon14:c.G1795A:p.E599K,CCAR1:NM_001282960:exon14:c.G1795A:p.E599K,CCAR1:NM_018237:exon15:c.G1840A:p.E614K, |
| chr10 | 68771261 | G | A | . | exonic | CCAR1 | nonsynonymous SNV | CCAR1:NM_001282959:exon17:c.G2309A:p.R770H,CCAR1:NM_001282960:exon17:c.G2309A:p.R770H,CCAR1:NM_018237:exon18:c.G2354A:p.R785H, |
| chr10 | 68771301 | C | A | . | exonic | CCAR1 | nonsynonymous SNV | CCAR1:NM_001282959:exon17:c.C2349A:p.D783E,CCAR1:NM_001282960:exon17:c.C2349A:p.D783E,CCAR1:NM_018237:exon18:c.C2394A:p.D798E, |
| chr10 | 68771325 | C | A | COSV56268323 | exonic | CCAR1 | nonsynonymous SNV | CCAR1:NM_001282959:exon17:c.C2373A:p.S791R,CCAR1:NM_001282960:exon17:c.C2373A:p.S791R,CCAR1:NM_018237:exon18:c.C2418A:p.S806R, |
| chr10 | 68771417 | A | G | . | exonic | CCAR1 | nonsynonymous SNV | CCAR1:NM_001282959:exon17:c.A2465G:p.D822G,CCAR1:NM_001282960:exon17:c.A2465G:p.D822G,CCAR1:NM_018237:exon18:c.A2510G:p.D837G, |
| chr10 | 68989028 | G | A | COSV62555322 | exonic | KIAA1279 | nonsynonymous SNV | KIAA1279:NM_015634:exon1:c.G196A:p.G66S, |
| chr10 | 69097096 | G | A | COSV54345640 | exonic | SRGN | nonsynonymous SNV | SRGN:NM_002727:exon2:c.G92A:p.R31Q, |
| chr10 | 71322806 | A | G | . | exonic | SLC29A3 | nonsynonymous SNV | SLC29A3:NM_001174098:exon2:c.A52G:p.R18G,SLC29A3:NM_018344:exon2:c.A52G:p.R18G, |
| chr10 | 71351651 | C | T | COSV64385485 | exonic | SLC29A3 | nonsynonymous SNV | SLC29A3:NM_001174098:exon4:c.C473T:p.S158F,SLC29A3:NM_018344:exon4:c.C473T:p.S158F, |
| chr10 | 71356185 | G | A | . | exonic | SLC29A3 | nonsynonymous SNV | SLC29A3:NM_001174098:exon5:c.G715A:p.V239I,SLC29A3:NM_018344:exon5:c.G715A:p.V239I, |
| chr10 | 71362156 | A | G | COSV64386549 | exonic | SLC29A3 | nonsynonymous SNV | SLC29A3:NM_018344:exon6:c.A976G:p.I326V, |
| chr10 | 71760875 | A | C | COSV56463939 | exonic | C10ORF54 | nonsynonymous SNV | C10orf54:NM_022153:exon3:c.T561G:p.D187E, |
| chr10 | 71818628 | C | T | COSV56473009 | exonic | PSAP | nonsynonymous SNV | PSAP:NM_002778:exon13:c.G1528A:p.A510T,PSAP:NM_001042465:exon14:c.G1537A:p.A513T,PSAP:NM_001042466:exon13:c.G1534A:p.A512T, |
| chr10 | 71818665 | C | T | . | exonic | PSAP | stopgain | PSAP:NM_001042465:exon14:c.G1500A:p.W500X,PSAP:NM_002778:exon13:c.G1491A:p.W497X,PSAP:NM_001042466:exon13:c.G1497A:p.W499X, |
| chr10 | 71818711 | C | T | . | exonic | PSAP | nonsynonymous SNV | PSAP:NM_002778:exon13:c.G1445A:p.C482Y,PSAP:NM_001042465:exon14:c.G1454A:p.C485Y,PSAP:NM_001042466:exon13:c.G1451A:p.C484Y, |
| chr10 | 71819039 | C | T | COSV56459891 | exonic | PSAP | nonsynonymous SNV | PSAP:NM_002778:exon12:c.G1423A:p.V475M,PSAP:NM_001042465:exon13:c.G1432A:p.V478M,PSAP:NM_001042466:exon12:c.G1429A:p.V477M, |
| chr10 | 71819072 | C | T | . | exonic | PSAP | nonsynonymous SNV | PSAP:NM_002778:exon12:c.G1390A:p.E464K,PSAP:NM_001042465:exon13:c.G1399A:p.E467K,PSAP:NM_001042466:exon12:c.G1396A:p.E466K, |
| chr10 | 71819553 | C | T | . | exonic | PSAP | nonsynonymous SNV | PSAP:NM_001042465:exon12:c.G1271A:p.R424H,PSAP:NM_002778:exon11:c.G1262A:p.R421H,PSAP:NM_001042466:exon11:c.G1268A:p.R423H, |
| chr10 | 71819580 | C | T | . | exonic | PSAP | nonsynonymous SNV | PSAP:NM_002778:exon11:c.G1235A:p.C412Y,PSAP:NM_001042465:exon12:c.G1244A:p.C415Y,PSAP:NM_001042466:exon11:c.G1241A:p.C414Y, |
| chr10 | 71819592 | A | G | . | exonic | PSAP | nonsynonymous SNV | PSAP:NM_002778:exon11:c.T1223C:p.F408S,PSAP:NM_001042465:exon12:c.T1232C:p.F411S,PSAP:NM_001042466:exon11:c.T1229C:p.F410S, |
| chr10 | 71819726 | G | A | . | exonic | PSAP | nonsynonymous SNV | PSAP:NM_002778:exon10:c.C1180T:p.P394S,PSAP:NM_001042465:exon11:c.C1189T:p.P397S,PSAP:NM_001042466:exon10:c.C1186T:p.P396S, |
| chr10 | 71819731 | C | T | . | exonic | PSAP | nonsynonymous SNV | PSAP:NM_002778:exon10:c.G1175A:p.R392Q,PSAP:NM_001042465:exon11:c.G1184A:p.R395Q,PSAP:NM_001042466:exon10:c.G1181A:p.R394Q, |
| chr10 | 71819758 | C | T | . | exonic | PSAP | nonsynonymous SNV | PSAP:NM_002778:exon10:c.G1148A:p.S383N,PSAP:NM_001042465:exon11:c.G1157A:p.S386N,PSAP:NM_001042466:exon10:c.G1154A:p.S385N, |
| chr10 | 71819781 | C | A | . | exonic | PSAP | nonsynonymous SNV | PSAP:NM_002778:exon10:c.G1125T:p.E375D,PSAP:NM_001042465:exon11:c.G1134T:p.E378D,PSAP:NM_001042466:exon10:c.G1131T:p.E377D, |
| chr10 | 71819836 | C | T | . | exonic | PSAP | nonsynonymous SNV | PSAP:NM_002778:exon10:c.G1070A:p.C357Y,PSAP:NM_001042465:exon11:c.G1079A:p.C360Y,PSAP:NM_001042466:exon10:c.G1076A:p.C359Y, |
| chr10 | 71820329 | C | A | . | exonic | PSAP | stopgain | PSAP:NM_002778:exon9:c.G916T:p.E306X,PSAP:NM_001042465:exon10:c.G925T:p.E309X,PSAP:NM_001042466:exon9:c.G922T:p.E308X, |
| chr10 | 71821902 | C | T | . | exonic | PSAP | nonsynonymous SNV | PSAP:NM_001042465:exon9:c.G892A:p.A298T,PSAP:NM_002778:exon8:c.G883A:p.A295T,PSAP:NM_001042466:exon8:c.G889A:p.A297T, |
| chr10 | 71821938 | C | T | . | exonic | PSAP | nonsynonymous SNV | PSAP:NM_002778:exon8:c.G847A:p.V283I,PSAP:NM_001042465:exon9:c.G856A:p.V286I,PSAP:NM_001042466:exon8:c.G853A:p.V285I, |
| chr10 | 71821991 | C | T | . | exonic | PSAP | nonsynonymous SNV | PSAP:NM_002778:exon8:c.G794A:p.C265Y,PSAP:NM_001042465:exon9:c.G803A:p.C268Y,PSAP:NM_001042466:exon8:c.G800A:p.C267Y, |
| chr10 | 71828936 | C | T | COSV67551161 | exonic | PSAP | nonsynonymous SNV | PSAP:NM_001042465:exon5:c.G517A:p.A173T,PSAP:NM_002778:exon5:c.G517A:p.A173T,PSAP:NM_001042466:exon5:c.G517A:p.A173T, |
| chr10 | 71829073 | C | T | . | exonic | PSAP | nonsynonymous SNV | PSAP:NM_002778:exon5:c.G380A:p.R127H,PSAP:NM_001042465:exon5:c.G380A:p.R127H,PSAP:NM_001042466:exon5:c.G380A:p.R127H, |
| chr10 | 71831854 | C | A | . | exonic | PSAP | nonsynonymous SNV | PSAP:NM_002778:exon3:c.G241T:p.A81S,PSAP:NM_001042465:exon3:c.G241T:p.A81S,PSAP:NM_001042466:exon3:c.G241T:p.A81S, |
| chr10 | 71831887 | C | T | . | exonic | PSAP | nonsynonymous SNV | PSAP:NM_002778:exon3:c.G208A:p.V70I,PSAP:NM_001042465:exon3:c.G208A:p.V70I,PSAP:NM_001042466:exon3:c.G208A:p.V70I, |
| chr10 | 71834421 | C | T | . | exonic | PSAP | nonsynonymous SNV | PSAP:NM_001042465:exon2:c.G125A:p.C42Y,PSAP:NM_002778:exon2:c.G125A:p.C42Y,PSAP:NM_001042466:exon2:c.G125A:p.C42Y, |
| chr10 | 74394142 | G | A | . | exonic | ADK | stopgain | ADK:NM_001202450:exon5:c.G275A:p.W92X,ADK:NM_006721:exon5:c.G275A:p.W92X,ADK:NM_001202449:exon4:c.G170A:p.W57X,ADK:NM_001123:exon5:c.G224A:p.W75X, |
| chr10 | 74394190 | G | A | . | exonic | ADK | nonsynonymous SNV | ADK:NM_001202450:exon5:c.G323A:p.G108E,ADK:NM_006721:exon5:c.G323A:p.G108E,ADK:NM_001202449:exon4:c.G218A:p.G73E,ADK:NM_001123:exon5:c.G272A:p.G91E, |
| chr10 | 74394258 | G | A | . | exonic | ADK | nonsynonymous SNV | ADK:NM_001202450:exon5:c.G391A:p.E131K,ADK:NM_006721:exon5:c.G391A:p.E131K,ADK:NM_001202449:exon4:c.G286A:p.E96K,ADK:NM_001123:exon5:c.G340A:p.E114K, |
| chr10 | 74398526 | G | A | . | exonic | ADK | nonsynonymous SNV | ADK:NM_001202450:exon6:c.G502A:p.D168N,ADK:NM_006721:exon6:c.G502A:p.D168N,ADK:NM_001202449:exon5:c.G397A:p.D133N,ADK:NM_001123:exon6:c.G451A:p.D151N, |
| chr10 | 74398546 | G | A | . | exonic | ADK | nonsynonymous SNV | ADK:NM_001202450:exon6:c.G522A:p.M174I,ADK:NM_006721:exon6:c.G522A:p.M174I,ADK:NM_001202449:exon5:c.G417A:p.M139I,ADK:NM_001123:exon6:c.G471A:p.M157I, |
| chr10 | 74525410 | T | C | . | exonic | ADK | nonsynonymous SNV | ADK:NM_006721:exon7:c.T710C:p.L237P,ADK:NM_001202449:exon6:c.T605C:p.L202P,ADK:NM_001123:exon7:c.T659C:p.L220P, |
| chr10 | 74589285 | G | A | . | exonic | ADK | nonsynonymous SNV | ADK:NM_001202450:exon7:c.G559A:p.A187T,ADK:NM_006721:exon8:c.G730A:p.A244T,ADK:NM_001202449:exon7:c.G625A:p.A209T,ADK:NM_001123:exon8:c.G679A:p.A227T, |
| chr10 | 74600440 | G | A | . | exonic | ADK | nonsynonymous SNV | ADK:NM_001202450:exon8:c.G653A:p.R218K,ADK:NM_006721:exon9:c.G824A:p.R275K,ADK:NM_001202449:exon8:c.G719A:p.R240K,ADK:NM_001123:exon9:c.G773A:p.R258K, |
| chr10 | 74600493 | G | A | . | exonic | ADK | nonsynonymous SNV | ADK:NM_001202450:exon8:c.G706A:p.E236K,ADK:NM_006721:exon9:c.G877A:p.E293K,ADK:NM_001202449:exon8:c.G772A:p.E258K,ADK:NM_001123:exon9:c.G826A:p.E276K, |
| chr10 | 74670220 | C | A | . | exonic | ADK | nonsynonymous SNV | ADK:NM_001202450:exon9:c.C744A:p.D248E,ADK:NM_006721:exon10:c.C915A:p.D305E,ADK:NM_001202449:exon9:c.C810A:p.D270E,ADK:NM_001123:exon10:c.C864A:p.D288E, |

| | | | | | | | | |
|---|---|---|---|---|---|---|---|---|
| chr10 | 74670224 | A | G | . | exonic | ADK | nonsynonymous SNV | ADK:NM_001202450:exon9:c.A748G:p.K250E,ADK:NM_006721:exon10:c.A919G:p.K307E,ADK:NM_001202449:exon9:c.A814G:p.K272E,ADK:NM_001123:exon10:c.A868G:p.K290E, |
| chr10 | 74708410 | G | A | . | exonic | ADK | nonsynonymous SNV | ADK:NM_001202450:exon10:c.G883A:p.G295S,ADK:NM_006721:exon11:c.G1054A:p.G352S,ADK:NM_001202449:exon10:c.G949A:p.G317S,ADK:NM_001123:exon11:c.G1003A:p.G335S, |
| chr10 | 78037198 | G | A | . | exonic | RPS24 | nonsynonymous SNV | RPS24:NM_001142283:exon4:c.G284A:p.G95D,RPS24:NM_001142285:exon4:c.G284A:p.G95D,RPS24:NM_001142284:exon4:c.G284A:p.G95D,RPS24:NM_001026:exon4:c.G284A:p.G95D,RPS24:NM_033022:exon4:c.G284A:p.G95D,RPS24:NM_001142282:exon4:c.G284A:p.G95D, |
| chr10 | 86876022 | C | A | COSV64409048 | exonic | BMPR1A | nonsynonymous SNV | BMPR1A:NM_004329:exon3:c.C4A:p.P2T, |
| chr10 | 86919290 | A | C | . | exonic | BMPR1A | nonsynonymous SNV | BMPR1A:NM_004329:exon10:c.A987C:p.R329S, |
| chr10 | 87513141 | T | C | . | exonic | MINPP1 | nonsynonymous SNV | MINPP1:NM_001178118:exon3:c.T250C:p.F84L,MINPP1:NM_004897:exon3:c.T853C:p.F285L, |
| chr10 | 87864144 | G | C | . | exonic | PTEN | nonsynonymous SNV | PTEN:NM_001304717:exon2:c.G194C:p.C65S, |
| chr10 | 87933135 | G | A | COSV64288974 | exonic | PTEN | nonsynonymous SNV | PTEN:NM_000314:exon5:c.G376A:p.A126T,PTEN:NM_001304717:exon6:c.G895A:p.A299T, |
| chr10 | 87933148 | G | A | COSV64288376 | exonic | PTEN | nonsynonymous SNV | PTEN:NM_000314:exon5:c.G389A:p.R130Q,PTEN:NM_001304717:exon6:c.G908A:p.R303Q, |
| chr10 | 87933153 | G | A | COSV64288639 | exonic | PTEN | nonsynonymous SNV | PTEN:NM_000314:exon5:c.G394A:p.G132S,PTEN:NM_001304717:exon6:c.G913A:p.G305S, |
| chr10 | 87933225 | G | A | COSV64288799 | exonic | PTEN | nonsynonymous SNV | PTEN:NM_000314:exon5:c.G466A:p.G156R,PTEN:NM_001304717:exon6:c.G985A:p.G329R, |
| chr10 | 87952143 | G | A | COSV64288961 | exonic | PTEN | nonsynonymous SNV | PTEN:NM_000314:exon6:c.G518A:p.R173H,PTEN:NM_001304717:exon7:c.G1037A:p.R346H, |
| chr10 | 87957924 | G | A | . | exonic | PTEN | nonsynonymous SNV | PTEN:NM_000314:exon7:c.G706A:p.D236N,PTEN:NM_001304717:exon8:c.G1225A:p.D409N,PTEN:NM_001304718:exon7:c.G115A:p.D39N, |
| chr10 | 87957972 | G | A | COSV64302293 | exonic | PTEN | nonsynonymous SNV | PTEN:NM_000314:exon7:c.G754A:p.D252N,PTEN:NM_001304717:exon8:c.G1273A:p.D425N,PTEN:NM_001304718:exon7:c.G163A:p.D55N, |
| chr10 | 87958004 | C | A | . | exonic | PTEN | nonsynonymous SNV | PTEN:NM_000314:exon7:c.C786A:p.N262K,PTEN:NM_001304717:exon8:c.C1305A:p.N435K,PTEN:NM_001304718:exon7:c.C195A:p.N65K, |
| chr10 | 87960961 | T | A | . | exonic | PTEN | nonsynonymous SNV | PTEN:NM_000314:exon8:c.T869A:p.V290E,PTEN:NM_001304717:exon9:c.T1388A:p.V463E,PTEN:NM_001304718:exon8:c.T278A:p.V93E, |
| chr10 | 87960969 | G | A | . | exonic | PTEN | nonsynonymous SNV | PTEN:NM_000314:exon8:c.G877A:p.G293R,PTEN:NM_001304717:exon9:c.G1396A:p.G466R,PTEN:NM_001304718:exon8:c.G286A:p.G96R, |
| chr10 | 87965287 | G | A | . | exonic | PTEN | nonsynonymous SNV | PTEN:NM_000314:exon9:c.G1027A:p.V343M,PTEN:NM_001304717:exon10:c.G1546A:p.V516M,PTEN:NM_001304718:exon9:c.G436A:p.V146M, |
| chr10 | 87965335 | G | A | . | exonic | PTEN | nonsynonymous SNV | PTEN:NM_000314:exon9:c.G1075A:p.A359T,PTEN:NM_001304717:exon10:c.G1594A:p.A532T,PTEN:NM_001304718:exon9:c.G484A:p.A162T, |
| chr10 | 87965468 | T | C | . | exonic | PTEN | nonsynonymous SNV | PTEN:NM_000314:exon9:c.T1208C:p.V403A,PTEN:NM_001304717:exon10:c.T1727C:p.V576A,PTEN:NM_001304718:exon9:c.T617C:p.V206A, |
| chr10 | 88583080 | C | G | COSV59290594 | exonic | RNLS | nonsynonymous SNV | RNLS:NM_018363:exon1:c.G111C:p.E37D,RNLS:NM_001031709:exon1:c.G111C:p.E37D, |
| chr10 | 89008889 | G | A | COSV58239617 | exonic | FAS | nonsynonymous SNV | FAS:NM_152871:exon4:c.G335A:p.G112D,FAS:NM_152872:exon4:c.G335A:p.G112D,FAS:NM_000043:exon4:c.G335A:p.G112D, |
| chr10 | 89012029 | G | A | . | exonic | FAS | nonsynonymous SNV | FAS:NM_152871:exon6:c.G536A:p.R179K,FAS:NM_152872:exon7:c.G599A:p.R200K,FAS:NM_000043:exon7:c.G599A:p.R200K, |
| chr10 | 89014124 | G | A | COSV58239581 | exonic | FAS | nonsynonymous SNV | FAS:NM_152871:exon8:c.G619A:p.D207N,FAS:NM_000043:exon9:c.G682A:p.D228N, |
| chr10 | 89014285 | G | A | COSV58240270 | exonic | FAS | stopgain | FAS:NM_152871:exon8:c.G780A:p.W260X,FAS:NM_000043:exon9:c.G843A:p.W281X, |
| chr10 | 89014343 | G | A | . | exonic | FAS | nonsynonymous SNV | FAS:NM_152871:exon8:c.G838A:p.A280T,FAS:NM_000043:exon9:c.G901A:p.A301T, |
| chr10 | 89709168 | C | G | COSV53307358 | exonic | KIF20B | nonsynonymous SNV | KIF20B:NM_016195:exon3:c.C149G:p.A50G,KIF20B:NM_001284259:exon3:c.C149G:p.A50G, |
| chr10 | 89719454 | G | T | COSV53306008 | exonic | KIF20B | nonsynonymous SNV | KIF20B:NM_016195:exon13:c.G1470T:p.E490D,KIF20B:NM_001284259:exon13:c.G1470T:p.E490D, |
| chr10 | 89727892 | A | T | COSV53305837 | exonic | KIF20B | nonsynonymous SNV | KIF20B:NM_016195:exon17:c.A2147T:p.N716I,KIF20B:NM_001284259:exon17:c.A2267T:p.N756I, |
| chr10 | 89729222 | A | T | COSV53307436 | exonic | KIF20B | nonsynonymous SNV | KIF20B:NM_016195:exon18:c.A2246T:p.H749L,KIF20B:NM_001284259:exon18:c.A2366T:p.H789L, |
| chr10 | 89738204 | A | G | . | exonic | KIF20B | nonsynonymous SNV | KIF20B:NM_016195:exon20:c.A3243G:p.I1081M,KIF20B:NM_001284259:exon20:c.A3363G:p.I1121M, |
| chr10 | 89738370 | T | C | COSV53308149 | exonic | KIF20B | nonsynonymous SNV | KIF20B:NM_016195:exon20:c.T3409C:p.C1137R,KIF20B:NM_001284259:exon20:c.T3529C:p.C1177R, |
| chr10 | 89754613 | T | A | . | exonic | KIF20B | nonsynonymous SNV | KIF20B:NM_016195:exon26:c.T4323A:p.N1441K,KIF20B:NM_001284259:exon26:c.T4443A:p.N1481K, |
| chr10 | 89762791 | A | G | . | exonic | KIF20B | nonsynonymous SNV | KIF20B:NM_016195:exon29:c.A4825G:p.K1609E,KIF20B:NM_001284259:exon29:c.A4945G:p.K1649E, |
| chr10 | 93519749 | A | T | COSV65184775 | exonic | CEP55 | nonsynonymous SNV | CEP55:NM_001127182:exon8:c.A1133T:p.H378L,CEP55:NM_018131:exon8:c.A1133T:p.H378L, |
| chr10 | 93700060 | C | T | COSV63272927 | exonic | FRA10AC1 | nonsynonymous SNV | FRA10AC1:NM_145246:exon2:c.G47A:p.R16H, |
| chr10 | 94694947 | G | A | . | exonic | CYP2C18 | nonsynonymous SNV | CYP2C18:NM_000772:exon4:c.G512A:p.G171D,CYP2C18:NM_001128925:exon4:c.G512A:p.G171D, |
| chr10 | 94941985 | T | G | . | exonic | CYP2C9 | nonsynonymous SNV | CYP2C9:NM_000771:exon2:c.T296G:p.I99S, |
| chr10 | 94942017 | T | C | . | exonic | CYP2C9 | nonsynonymous SNV | CYP2C9:NM_000771:exon2:c.T328C:p.F110L, |
| chr10 | 94942197 | G | C | . | exonic | CYP2C9 | nonsynonymous SNV | CYP2C9:NM_000771:exon3:c.G337C:p.V113L, |
| chr10 | 95263873 | T | C | COSV61474172 | exonic | PDLIM1 | nonsynonymous SNV | PDLIM1:NM_020992:exon4:c.A524G:p.N175S, |
| chr10 | 97744873 | G | T | COSV61747241 | exonic | ZFYVE27 | nonsynonymous SNV | ZFYVE27:NM_001174121:exon2:c.G119T:p.G40V,ZFYVE27:NM_001174119:exon3:c.G317T:p.G106V,ZFYVE27:NM_001002262:exon4:c.G413T:p.G138V,ZFYVE27:NM_144588:exon4:c.G413T:p.G138V,ZFYVE27:NM_001002261:exon3:c.G413T:p.G138V, |
| chr11 | 108688967 | G | A | . | exonic | DDX10 | nonsynonymous SNV | DDX10:NM_004398:exon7:c.G880A:p.V294I, |
| chr11 | 108719836 | G | A | . | exonic | DDX10 | nonsynonymous SNV | DDX10:NM_004398:exon12:c.G1450A:p.D484N, |
| chr11 | 108838501 | C | A | . | exonic | DDX10 | nonsynonymous SNV | DDX10:NM_004398:exon14:c.C2021A:p.A674E, |
| chr11 | 108841436 | A | G | . | exonic | DDX10 | nonsynonymous SNV | DDX10:NM_004398:exon15:c.A2207G:p.D736G, |
| chr11 | 112194711 | T | C | . | exonic | BCO2 | nonsynonymous SNV | BCO2:NM_001037290:exon5:c.T590C:p.L197P,BCO2:NM_001256400:exon4:c.T377C:p.L126P,BCO2:NM_001256397:exon5:c.T590C:p.L197P,BCO2:NM_031938:exon5:c.T692C:p.L231P, |
| chr11 | 112217776 | A | C | . | exonic | BCO2 | nonsynonymous SNV | BCO2:NM_001256398:exon10:c.A1423C:p.I475L,BCO2:NM_001037290:exon12:c.A1540C:p.I514L,BCO2:NM_001256400:exon11:c.A1327C:p.I443L,BCO2:NM_001256397:exon12:c.A1522C:p.I508L,BCO2:NM_031938:exon12:c.A1642C:p.I548L, |
| chr11 | 114698723 | A | G | COSV99080990 | exonic | NXPE2 | nonsynonymous SNV | NXPE2:NM_182495:exon3:c.A811G:p.T271A, |
| chr11 | 114705904 | A | T | COSV64940629 | exonic | NXPE2 | nonsynonymous SNV | NXPE2:NM_182495:exon5:c.A1052T:p.N351I, |
| chr11 | 116784884 | G | A | . | exonic | ZPR1 | nonsynonymous SNV | ZPR1:NM_003904:exon8:c.C791T:p.A264V,ZPR1:NM_001317086:exon7:c.C629T:p.A210V, |
| chr11 | 117351876 | G | A | COSV54039014 | exonic | CEP164 | nonsynonymous SNV | CEP164:NM_014956:exon5:c.G281A:p.S94N,CEP164:NM_001271933:exon4:c.G281A:p.S94N, |
| chr11 | 117351975 | C | A | . | exonic | CEP164 | nonsynonymous SNV | CEP164:NM_014956:exon5:c.C380A:p.P127H,CEP164:NM_001271933:exon4:c.C380A:p.P127H, |
| chr11 | 119018832 | G | A | . | exonic | TRAPPC4 | nonsynonymous SNV | TRAPPC4:NM_016146:exon1:c.G37A:p.G13S, |
| chr11 | 119019178 | G | A | . | exonic | TRAPPC4 | nonsynonymous SNV | TRAPPC4:NM_016146:exon2:c.G211A:p.V71M, |
| chr11 | 119019301 | G | A | . | exonic | TRAPPC4 | nonsynonymous SNV | TRAPPC4:NM_016146:exon2:c.G334A:p.A112T, |
| chr11 | 119021783 | G | A | . | exonic | TRAPPC4 | nonsynonymous SNV | TRAPPC4:NM_016146:exon4:c.G478A:p.D160N, |
| chr11 | 119023373 | G | A | . | exonic | TRAPPC4 | nonsynonymous SNV | TRAPPC4:NM_016146:exon5:c.G634A:p.A212T, |
| chr11 | 119188695 | G | A | COSV52702606 | exonic | PDZD3 | nonsynonymous SNV | PDZD3:NM_001168468:exon9:c.G1115A:p.R372Q,PDZD3:NM_024791:exon9:c.G1073A:p.R358Q, |

| | | | | | | | | |
|---|---|---|---|---|---|---|---|---|
| chr11 | 119206433 | A | C | . | exonic | CBL | nonsynonymous SNV | CBL:NM_005188:exon1:c.A16C:p.K6Q, |
| chr11 | 120228949 | A | G | COSV61109223 | exonic | OAF | nonsynonymous SNV | OAF:NM_178507:exon4:c.A629G:p.H210R, |
| chr11 | 120228970 | G | A | COSV61109375 | exonic | OAF | nonsynonymous SNV | OAF:NM_178507:exon4:c.G650A:p.R217H, |
| chr11 | 120305040 | A | G | COSV52793130 | exonic | POU2F3 | nonsynonymous SNV | POU2F3:NM_014352:exon7:c.A455G:p.H152R,POU2F3:NM_001244682:exon7:c.A461G:p.H154R, |
| chr11 | 121307243 | G | A | COSV99074056 | exonic | SC5D | nonsynonymous SNV | SC5D:NM_006918:exon5:c.G631A:p.G211S,SC5D:NM_001024956:exon5:c.G631A:p.G211S, |
| chr11 | 12477925 | G | A | . | exonic | PARVA | nonsynonymous SNV | PARVA:NM_018222:exon4:c.G496A:p.G166R, |
| chr11 | 12477926 | G | A | . | exonic | PARVA | nonsynonymous SNV | PARVA:NM_018222:exon4:c.G497A:p.G166E, |
| chr11 | 125102312 | A | G | COSV54444751 | exonic | TMEM218 | nonsynonymous SNV | TMEM218:NM_001258246:exon3:c.T35C:p.L12S,TMEM218:NM_001258243:exon4:c.T35C:p.L12S,TMEM218:NM_001258239:exon3:c.T35C:p.L12S,TMEM218:NM_001258242:exon3:c.T35C:p.L12S, |
| chr11 | 125655300 | A | G | . | exonic | CHEK1 | nonsynonymous SNV | CHEK1:NM_001114122:exon13:c.A1411G:p.I471V,CHEK1:NM_001244846:exon12:c.A1309G:p.I437V,CHEK1:NM_001274:exon13:c.A1411G:p.I471V,CHEK1:NM_001114121:exon13:c.A1411G:p.I471V, |
| chr11 | 18245480 | C | T | . | exonic | SAA2 | nonsynonymous SNV | SAA2:NM_030754:exon4:c.G266A:p.R89H, |
| chr11 | 18269312 | C | T | COSV62945873 | exonic | SAA1 | nonsynonymous SNV | SAA1:NM_199161:exon3:c.C209T:p.A70V,SAA1:NM_001178006:exon4:c.C209T:p.A70V,SAA1:NM_000331:exon3:c.C209T:p.A70V, |
| chr11 | 18269327 | T | C | COSV62945890 | exonic | SAA1 | nonsynonymous SNV | SAA1:NM_199161:exon3:c.T224C:p.V75A,SAA1:NM_001178006:exon4:c.T224C:p.V75A,SAA1:NM_000331:exon3:c.T224C:p.V75A, |
| chr11 | 20383067 | T | G | COSV58178278 | exonic | HTATIP2 | nonsynonymous SNV | HTATIP2:NM_006410:exon6:c.T591G:p.S197R,HTATIP2:NM_001098522:exon5:c.T591G:p.S197R,HTATIP2:NM_001098520:exon6:c.T693G:p.S231R,HTATIP2:NM_001098521:exon6:c.T591G:p.S197R, |
| chr11 | 237087 | A | G | COSV61500926 | exonic | PSMD13 | nonsynonymous SNV | PSMD13:NM_175932:exon1:c.A38G:p.N13S,PSMD13:NM_002817:exon1:c.A38G:p.N13S, |
| chr11 | 2903380 | G | A | COSV56539474 | exonic | SLC22A18 | nonsynonymous SNV | SLC22A18:NM_002555:exon2:c.G35A:p.R12Q,SLC22A18:NM_183233:exon2:c.G35A:p.R12Q,SLC22A18:NM_001315501:exon2:c.G290A:p.R97Q,SLC22A18:NM_001315502:exon2:c.G35A:p.R12Q, |
| chr11 | 2951282 | C | T | . | exonic | NAP1L4 | nonsynonymous SNV | NAP1L4:NM_005969:exon14:c.G1099A:p.D367N, |
| chr11 | 2969848 | C | A | . | exonic | NAP1L4 | nonsynonymous SNV | NAP1L4:NM_005969:exon7:c.G489T:p.W163C, |
| chr11 | 3001233 | C | T | COSV53461688 | exonic | CARS | nonsynonymous SNV | CARS:NM_001751:exon22:c.G2128A:p.D710N,CARS:NM_001194997:exon23:c.G2385A:p.M795I,CARS:NM_139273:exon22:c.G2136A:p.M712I,CARS:NM_001014437:exon23:c.G2377A:p.D793N, |
| chr11 | 3006927 | G | A | COSV53454323 | exonic | CARS | nonsynonymous SNV | CARS:NM_001751:exon18:c.C1852T:p.R618W,CARS:NM_001194997:exon19:c.C2101T:p.R701W,CARS:NM_139273:exon18:c.C1852T:p.R618W,CARS:NM_001014437:exon19:c.C2101T:p.R701W, |
| chr11 | 3015801 | C | T | . | exonic | CARS | nonsynonymous SNV | CARS:NM_001751:exon16:c.G1717A:p.G573R,CARS:NM_139273:exon16:c.G1717A:p.G573R,CARS:NM_001014437:exon17:c.G1966A:p.G656R,CARS:NM_001194997:exon17:c.G1966A:p.G656R, |
| chr11 | 3017206 | C | T | . | exonic | CARS | nonsynonymous SNV | CARS:NM_001751:exon15:c.G1568A:p.S523N,CARS:NM_001194997:exon16:c.G1817A:p.S606N,CARS:NM_139273:exon15:c.G1568A:p.S523N,CARS:NM_001014437:exon16:c.G1817A:p.S606N, |
| chr11 | 3017237 | T | G | . | exonic | CARS | nonsynonymous SNV | CARS:NM_001751:exon15:c.A1537C:p.T513P,CARS:NM_001194997:exon16:c.A1786C:p.T596P,CARS:NM_139273:exon15:c.A1537C:p.T513P,CARS:NM_001014437:exon16:c.A1786C:p.T596P, |
| chr11 | 3018455 | A | G | . | exonic | CARS | nonsynonymous SNV | CARS:NM_001751:exon13:c.T1333C:p.S445P,CARS:NM_001194997:exon14:c.T1582C:p.S528P,CARS:NM_139273:exon13:c.T1333C:p.S445P,CARS:NM_001014437:exon14:c.T1582C:p.S528P, |
| chr11 | 3026739 | G | A | . | exonic | CARS | nonsynonymous SNV | CARS:NM_001751:exon9:c.C841T:p.H281Y,CARS:NM_139273:exon9:c.C841T:p.H281Y,CARS:NM_001014437:exon10:c.C1090T:p.H364Y,CARS:NM_001194997:exon10:c.C1090T:p.H364Y, |
| chr11 | 3038130 | C | T | COSV53457881 | exonic | CARS | nonsynonymous SNV | CARS:NM_001751:exon6:c.G472A:p.A158T,CARS:NM_001194997:exon7:c.G721A:p.A241T,CARS:NM_139273:exon6:c.G472A:p.A158T,CARS:NM_001014437:exon7:c.G721A:p.A241T, |
| chr11 | 3039870 | A | G | . | exonic | CARS | nonsynonymous SNV | CARS:NM_001751:exon4:c.T268C:p.F90L,CARS:NM_001194997:exon5:c.T517C:p.F173L,CARS:NM_139273:exon4:c.T268C:p.F90L,CARS:NM_001014437:exon5:c.T517C:p.F173L, |
| chr11 | 3042169 | T | A | . | exonic | CARS | nonsynonymous SNV | CARS:NM_001751:exon2:c.A113T:p.N38I,CARS:NM_001194997:exon3:c.A362T:p.N121I,CARS:NM_139273:exon2:c.A113T:p.N38I,CARS:NM_001014437:exon3:c.A362T:p.N121I, |
| chr11 | 3042256 | C | T | . | exonic | CARS | nonsynonymous SNV | CARS:NM_001751:exon2:c.G26A:p.G9D,CARS:NM_001194997:exon3:c.G275A:p.S92N,CARS:NM_139273:exon2:c.G26A:p.G9D,CARS:NM_001014437:exon3:c.G275A:p.S92N, |
| chr11 | 308290 | T | C | COSV67714640 | exonic | IFITM2 | nonsynonymous SNV | IFITM2:NM_006435:exon1:c.T98C:p.V33A, |
| chr11 | 308314 | T | C | COSV67714346 | exonic | IFITM2 | nonsynonymous SNV | IFITM2:NM_006435:exon1:c.T122C:p.M41T, |
| chr11 | 308361 | G | A | . | exonic | IFITM2 | nonsynonymous SNV | IFITM2:NM_006435:exon1:c.G169A:p.V57M, |
| chr11 | 308393 | G | A | . | exonic | IFITM2 | nonsynonymous SNV | IFITM2:NM_006435:exon1:c.G201A:p.M67I, |
| chr11 | 308433 | G | A | . | exonic | IFITM2 | nonsynonymous SNV | IFITM2:NM_006435:exon1:c.G241A:p.V81M, |
| chr11 | 309055 | G | A | . | exonic | IFITM2 | nonsynonymous SNV | IFITM2:NM_006435:exon2:c.G289A:p.A97T, |
| chr11 | 309090 | G | A | . | exonic | IFITM2 | stopgain | IFITM2:NM_006435:exon2:c.G324A:p.W108X, |
| chr11 | 309127 | A | G | COSV60249689 | exonic | IFITM2 | nonsynonymous SNV | IFITM2:NM_006435:exon2:c.A361G:p.I121V, |
| chr11 | 309149 | T | C | . | exonic | IFITM2 | nonsynonymous SNV | IFITM2:NM_006435:exon2:c.T383C:p.V128A, |
| chr11 | 314207 | C | G | COSV60248916 | exonic | IFITM1 | nonsynonymous SNV | IFITM1:NM_003641:exon1:c.C37G:p.P13A, |
| chr11 | 33755902 | C | T | . | exonic | FBXO3 | nonsynonymous SNV | FBXO3:NM_033406:exon5:c.G547A:p.G183R,FBXO3:NM_012175:exon5:c.G547A:p.G183R, |
| chr11 | 34882722 | C | T | . | exonic | APIP | nonsynonymous SNV | APIP:NM_015957:exon7:c.G724A:p.V242I, |
| chr11 | 34883353 | C | T | . | exonic | APIP | nonsynonymous SNV | APIP:NM_015957:exon6:c.G613A:p.E205K, |
| chr11 | 34888839 | C | T | . | exonic | APIP | nonsynonymous SNV | APIP:NM_015957:exon4:c.G238A:p.E80K, |
| chr11 | 34888850 | C | T | COSV53505953 | exonic | APIP | nonsynonymous SNV | APIP:NM_015957:exon4:c.G227A:p.C76Y, |
| chr11 | 34895110 | C | T | COSV53506741 | exonic | APIP | nonsynonymous SNV | APIP:NM_015957:exon2:c.G58A:p.D20N, |

| | | | | | | | | |
|---|---|---|---|---|---|---|---|---|
| chr11 | 34916253 | C | T | . | exonic | APIP | nonsynonymous SNV | APIP:NM_015957:exon1:c.G32A:p.C11Y, |
| chr11 | 34916266 | G | A | COSV53507442 | exonic | APIP | nonsynonymous SNV | APIP:NM_015957:exon1:c.C19T:p.R7W, |
| chr11 | 34916722 | C | T | . | exonic | PDHX | nonsynonymous SNV | PDHX:NM_001166158:exon1:c.C67T:p.R23C,PDHX:NM_003477:exon1:c.C67T:p.R23C, |
| chr11 | 34931445 | A | G | . | exonic | PDHX | nonsynonymous SNV | PDHX:NM_001135024:exon2:c.A157G:p.M53V,PDHX:NM_001166158:exon2:c.A202G:p.M68V,PDHX:NM_003477:exon2:c.A202G:p.M68V, |
| chr11 | 34931464 | T | C | . | exonic | PDHX | nonsynonymous SNV | PDHX:NM_001135024:exon2:c.T176C:p.V59A,PDHX:NM_001166158:exon2:c.T221C:p.V74A,PDHX:NM_003477:exon2:c.T221C:p.V74A, |
| chr11 | 34947556 | G | A | . | exonic | PDHX | nonsynonymous SNV | PDHX:NM_001135024:exon3:c.G247A:p.A83T,PDHX:NM_001166158:exon3:c.G292A:p.A98T,PDHX:NM_003477:exon3:c.G292A:p.A98T, |
| chr11 | 34947565 | A | G | COSV57166122 | exonic | PDHX | nonsynonymous SNV | PDHX:NM_001135024:exon3:c.A256G:p.T86A,PDHX:NM_001166158:exon3:c.A301G:p.T101A,PDHX:NM_003477:exon3:c.A301G:p.T101A, |
| chr11 | 34957583 | G | A | COSV57166828 | exonic | PDHX | nonsynonymous SNV | PDHX:NM_001135024:exon4:c.G497A:p.R166Q,PDHX:NM_003477:exon4:c.G542A:p.R181Q, |
| chr11 | 34966693 | G | A | . | exonic | PDHX | nonsynonymous SNV | PDHX:NM_001135024:exon6:c.G650A:p.R217K,PDHX:NM_003477:exon6:c.G695A:p.R232K, |
| chr11 | 34970184 | G | A | . | exonic | PDHX | nonsynonymous SNV | PDHX:NM_001135024:exon7:c.G817A:p.A273T,PDHX:NM_003477:exon7:c.G862A:p.A288T, |
| chr11 | 34970235 | G | A | . | exonic | PDHX | nonsynonymous SNV | PDHX:NM_001135024:exon7:c.G868A:p.A290T,PDHX:NM_003477:exon7:c.G913A:p.A305T, |
| chr11 | 34984607 | G | A | . | exonic | PDHX | nonsynonymous SNV | PDHX:NM_001135024:exon9:c.G1016A:p.G339D,PDHX:NM_001166158:exon4:c.G380A:p.G127D,PDHX:NM_003477:exon9:c.G1061A:p.G354D, |
| chr11 | 34984655 | A | G | . | exonic | PDHX | nonsynonymous SNV | PDHX:NM_001135024:exon9:c.A1064G:p.D355G,PDHX:NM_001166158:exon4:c.A428G:p.D143G,PDHX:NM_003477:exon9:c.A1109G:p.D370G, |
| chr11 | 34984708 | G | A | . | exonic | PDHX | nonsynonymous SNV | PDHX:NM_001135024:exon9:c.G1117A:p.E373K,PDHX:NM_001166158:exon4:c.G481A:p.E161K,PDHX:NM_003477:exon9:c.G1162A:p.E388K, |
| chr11 | 34995036 | G | A | . | exonic | PDHX | nonsynonymous SNV | PDHX:NM_001135024:exon11:c.G1325A:p.G442E,PDHX:NM_001166158:exon6:c.G689A:p.G230E,PDHX:NM_003477:exon11:c.G1370A:p.G457E, |
| chr11 | 34995093 | G | A | COSV57170013 | exonic | PDHX | nonsynonymous SNV | PDHX:NM_001135024:exon11:c.G1382A:p.R461Q,PDHX:NM_001166158:exon6:c.G746A:p.R249Q,PDHX:NM_003477:exon11:c.G1427A:p.R476Q, |
| chr11 | 35180332 | G | A | . | exonic | CD44 | nonsynonymous SNV | CD44:NM_001001389:exon3:c.G292A:p.A98T,CD44:NM_001202556:exon3:c.G292A:p.A98T,CD44:NM_001202555:exon3:c.G292A:p.A98T,CD44:NM_001001390:exon3:c.G292A:p.A98T,CD44:NM_000610:exon3:c.G292A:p.A98T,CD44:NM_001202557:exon3:c.G292A:p.A98T,CD44:NM_001001391:exon3:c.G292A:p.A98T, |
| chr11 | 35186843 | G | A | . | exonic | CD44 | nonsynonymous SNV | CD44:NM_001001389:exon4:c.G379A:p.E127K,CD44:NM_001202556:exon4:c.G379A:p.E127K,CD44:NM_001202555:exon4:c.G379A:p.E127K,CD44:NM_001001390:exon4:c.G379A:p.E127K,CD44:NM_000610:exon4:c.G379A:p.E127K,CD44:NM_001202557:exon4:c.G379A:p.E127K,CD44:NM_001001391:exon4:c.G379A:p.E127K, |
| chr11 | 35189910 | G | A | . | exonic | CD44 | nonsynonymous SNV | CD44:NM_001001389:exon5:c.G512A:p.S171N,CD44:NM_000610:exon5:c.G512A:p.S171N,CD44:NM_001202556:exon5:c.G512A:p.S171N,CD44:NM_001202555:exon5:c.G512A:p.S171N,CD44:NM_001202557:exon5:c.G512A:p.S171N,CD44:NM_001001390:exon5:c.G512A:p.S171N,CD44:NM_001001391:exon5:c.G512A:p.S171N, |
| chr11 | 35189981 | T | C | . | exonic | CD44 | nonsynonymous SNV | CD44:NM_001001389:exon5:c.T583C:p.F195L,CD44:NM_000610:exon5:c.T583C:p.F195L,CD44:NM_001202556:exon5:c.T583C:p.F195L,CD44:NM_001202555:exon5:c.T583C:p.F195L,CD44:NM_001202557:exon5:c.T583C:p.F195L,CD44:NM_001001390:exon5:c.T583C:p.F195L,CD44:NM_001001391:exon5:c.T583C:p.F195L, |
| chr11 | 35204596 | G | A | . | exonic | CD44 | nonsynonymous SNV | CD44:NM_001001389:exon9:c.G1109A:p.R370H,CD44:NM_000610:exon10:c.G1238A:p.R413H, |
| chr11 | 35204608 | A | G | COSV53530507 | exonic | CD44 | nonsynonymous SNV | CD44:NM_001001389:exon9:c.A1121G:p.K374R,CD44:NM_000610:exon10:c.A1250G:p.K417R, |
| chr11 | 35208126 | T | C | . | exonic | CD44 | nonsynonymous SNV | CD44:NM_001001389:exon11:c.T1307C:p.I436T,CD44:NM_000610:exon12:c.T1436C:p.I479T,CD44:NM_001001390:exon6:c.T689C:p.I230T, |
| chr11 | 35208180 | G | A | . | exonic | CD44 | nonsynonymous SNV | CD44:NM_001001389:exon11:c.G1361A:p.R454K,CD44:NM_000610:exon12:c.G1490A:p.R497K,CD44:NM_001001390:exon6:c.G743A:p.R248K, |
| chr11 | 35221716 | G | A | . | exonic | CD44 | nonsynonymous SNV | CD44:NM_001001389:exon16:c.G1879A:p.V627I,CD44:NM_001202556:exon7:c.G802A:p.V268I,CD44:NM_001202555:exon9:c.G1069A:p.V357I,CD44:NM_001001390:exon11:c.G1261A:p.V421I,CD44:NM_001001392:exon3:c.G245A:p.S82N,CD44:NM_000610:exon17:c.G2008A:p.V670I,CD44:NM_001202557:exon8:c.G865A:p.V289I,CD44:NM_001001391:exon8:c.G865A:p.V289I, |
| chr11 | 35221726 | G | A | COSV53530853 | exonic | CD44 | nonsynonymous SNV | CD44:NM_001001389:exon16:c.G1889A:p.R630Q,CD44:NM_000610:exon17:c.G2018A:p.R673Q,CD44:NM_001202556:exon7:c.G812A:p.R271Q,CD44:NM_001202555:exon9:c.G1079A:p.R360Q,CD44:NM_001202557:exon8:c.G875A:p.R292Q,CD44:NM_001001390:exon11:c.G1271A:p.R424Q,CD44:NM_001001391:exon8:c.G875A:p.R292Q, |
| chr11 | 35229141 | G | A | . | exonic | CD44 | nonsynonymous SNV | CD44:NM_001001392:exon4:c.G274A:p.E92K, |
| chr11 | 35229180 | G | A | . | exonic | CD44 | nonsynonymous SNV | CD44:NM_001001392:exon4:c.G313A:p.G105R, |
| chr11 | 35229232 | G | A | . | exonic | CD44 | nonsynonymous SNV | CD44:NM_001001389:exon17:c.G1999A:p.V667M,CD44:NM_000610:exon18:c.G2128A:p.V710M,CD44:NM_001202556:exon8:c.G922A:p.V308M,CD44:NM_001202555:exon10:c.G1189A:p.V397M,CD44:NM_001001390:exon12:c.G1381A:p.V461M,CD44:NM_001001392:exon4:c.G365A:p.G122D,CD44:NM_001001391:exon9:c.G985A:p.V329M, |
| chr11 | 35434513 | T | C | COSV53510283 | exonic | PAMR1 | nonsynonymous SNV | PAMR1:NM_015430:exon11:c.A1676G:p.Q559R,PAMR1:NM_001282675:exon12:c.A1505G:p.Q502R,PAMR1:NM_001001991:exon10:c.A1625G:p.Q542R,PAMR1:NM_001282676:exon8:c.A1292G:p.Q431R, |
| chr11 | 35436106 | G | A | COSV53512813 | exonic | PAMR1 | nonsynonymous SNV | PAMR1:NM_015430:exon10:c.C1181T:p.A394V,PAMR1:NM_001282675:exon11:c.C1010T:p.A337V,PAMR1:NM_001001991:exon9:c.C1130T:p.A377V,PAMR1:NM_001282676:exon7:c.C797T:p.A266V, |
| chr11 | 3723330 | C | T | . | exonic | NUP98 | nonsynonymous SNV | NUP98:NM_016320:exon16:c.G1973A:p.G658E,NUP98:NM_139132:exon16:c.G1973A:p.G658E,NUP98:NM_005387:exon16:c.G2024A:p.G675E,NUP98:NM_139131:exon16:c.G1973A:p.G658E, |
| chr11 | 406483 | T | C | . | exonic | SIGIRR | nonsynonymous SNV | SIGIRR:NM_021805:exon9:c.A935G:p.Q312R,SIGIRR:NM_001135053:exon9:c.A935G:p.Q312R,SIGIRR:NM_001135054:exon9:c.A935G:p.Q312R, |
| chr11 | 43831003 | G | A | . | exonic | HSD17B12 | nonsynonymous SNV | HSD17B12:NM_016142:exon7:c.G529A:p.V177M, |
| chr11 | 43855148 | C | T | COSV53496916 | exonic | HSD17B12 | nonsynonymous SNV | HSD17B12:NM_016142:exon11:c.C839T:p.S280L, |
| chr11 | 43901546 | C | T | COSV57023500 | exonic | ALKBH3 | nonsynonymous SNV | ALKBH3:NM_139178:exon8:c.C490T:p.R164C, |
| chr11 | 43919052 | C | G | COSV57023880 | exonic | ALKBH3 | nonsynonymous SNV | ALKBH3:NM_139178:exon9:c.C684G:p.D228E, |

| | | | | | | | | |
|---|---|---|---|---|---|---|---|---|
| chr11 | 44126945 | G | A | . | exonic | EXT2 | nonsynonymous SNV | EXT2:NM_000401:exon6:c.G1168A:p.D390N,EXT2:NM_207122:exon6:c.G1069A:p.D357N,EXT2:NM_001178083:exon6:c.G1069A:p.D357N, |
| chr11 | 44234166 | G | A | . | exonic | EXT2 | nonsynonymous SNV | EXT2:NM_207122:exon12:c.G1858A:p.V620I,EXT2:NM_000401:exon12:c.G1957A:p.V653I,EXT2:NM_001178083:exon13:c.G1888A:p.V630I, |
| chr11 | 44236293 | G | A | . | exonic | EXT2 | nonsynonymous SNV | EXT2:NM_207122:exon13:c.G1936A:p.V646I,EXT2:NM_000401:exon13:c.G2035A:p.V679I,EXT2:NM_001178083:exon14:c.G1966A:p.V656I, |
| chr11 | 45915716 | C | T | COSV53800685 | exonic | PEX16 | nonsynonymous SNV | PEX16:NM_004813:exon4:c.G346A:p.V116I,PEX16:NM_057174:exon4:c.G346A:p.V116I, |
| chr11 | 46936748 | G | T | COSV53566094 | exonic | C11ORF49 | nonsynonymous SNV | C11orf49:NM_001278222:exon1:c.G19T:p.G7C, |
| chr11 | 47166860 | C | T | COSV53563682 | exonic | ARFGAP2 | nonsynonymous SNV | ARFGAP2:NM_032389:exon13:c.G1232A:p.S411N,ARFGAP2:NM_001242832:exon12:c.G1148A:p.S383N, |
| chr11 | 47618877 | G | C | COSV56766406 | exonic | MTCH2 | nonsynonymous SNV | MTCH2:NM_001317232:exon12:c.C841G:p.P281A,MTCH2:NM_001317233:exon12:c.C424G:p.P142A,MTCH2:NM_014342:exon13:c.C868G:p.P290A, |
| chr11 | 562421 | G | A | COSV60346289 | exonic | RASSF7 | nonsynonymous SNV | RASSF7:NM_003475:exon3:c.G467A:p.R156Q,RASSF7:NM_001143994:exon3:c.G467A:p.R156Q,RASSF7:NM_001143993:exon3:c.G467A:p.R156Q, |
| chr11 | 57614516 | G | A | COSV53542407 | exonic | SERPING1 | nonsynonymous SNV | SERPING1:NM_001032295:exon7:c.G1438A:p.V480M,SERPING1:NM_000062:exon8:c.G1438A:p.V480M, |
| chr11 | 57712658 | G | A | . | exonic | TMX2 | nonsynonymous SNV | TMX2:NM_015959:exon1:c.G40A:p.V14M,TMX2:NM_001144012:exon1:c.G40A:p.V14M, |
| chr11 | 57737930 | A | G | . | exonic | TMX2 | nonsynonymous SNV | TMX2:NM_015959:exon3:c.A268G:p.I90V, |
| chr11 | 57738368 | T | C | . | exonic | TMX2 | nonsynonymous SNV | TMX2:NM_015959:exon4:c.T379C:p.C127R,TMX2:NM_001144012:exon3:c.T265C:p.C89R, |
| chr11 | 57738418 | T | A | . | exonic | TMX2 | nonsynonymous SNV | TMX2:NM_015959:exon4:c.T429A:p.D143E,TMX2:NM_001144012:exon3:c.T315A:p.D105E, |
| chr11 | 57738428 | G | A | . | exonic | TMX2 | nonsynonymous SNV | TMX2:NM_015959:exon4:c.G439A:p.D147N,TMX2:NM_001144012:exon3:c.G325A:p.D109N, |
| chr11 | 60497529 | C | T | COSV50014135 | exonic | MS4A12 | stopgain | MS4A12:NM_001164470:exon2:c.C211T:p.Q71X,MS4A12:NM_017716:exon2:c.C211T:p.Q71X, |
| chr11 | 61398269 | G | C | COSV58426369 | exonic;splicing | TMEM216 | nonsynonymous SNV | TMEM216:NM_001173991:exon5:c.G440C:p.R147T,TMEM216:NM_016499:exon5:c.G257C:p.R86T, |
| chr11 | 61430154 | T | C | . | exonic | SDHAF2 | nonsynonymous SNV | SDHAF2:NM_017841:exon1:c.T8C:p.V3A, |
| chr11 | 61437683 | G | A | . | exonic | SDHAF2 | nonsynonymous SNV | SDHAF2:NM_017841:exon2:c.G95A:p.R32K, |
| chr11 | 61437704 | G | A | . | exonic | SDHAF2 | nonsynonymous SNV | SDHAF2:NM_017841:exon2:c.G116A:p.S39N, |
| chr11 | 61437753 | G | A | . | exonic | SDHAF2 | stopgain | SDHAF2:NM_017841:exon2:c.G165A:p.W55X, |
| chr11 | 61437794 | G | A | . | exonic | SDHAF2 | nonsynonymous SNV | SDHAF2:NM_017841:exon2:c.G206A:p.R69H, |
| chr11 | 61438059 | G | A | . | exonic | SDHAF2 | nonsynonymous SNV | SDHAF2:NM_017841:exon3:c.G316A:p.D106N, |
| chr11 | 61445981 | G | A | COSV57099212 | exonic | SDHAF2 | nonsynonymous SNV | SDHAF2:NM_017841:exon4:c.G411A:p.M137I, |
| chr11 | 61446067 | G | A | . | exonic | SDHAF2 | nonsynonymous SNV | SDHAF2:NM_017841:exon4:c.G497A:p.R166H, |
| chr11 | 61795807 | G | A | . | exonic | FEN1 | nonsynonymous SNV | FEN1:NM_004111:exon2:c.G446A:p.G149D, |
| chr11 | 61795849 | G | A | . | exonic | FEN1 | nonsynonymous SNV | FEN1:NM_004111:exon2:c.G488A:p.C163Y, |
| chr11 | 617967 | C | G | . | exonic | CDHR5 | nonsynonymous SNV | CDHR5:NM_031264:exon13:c.G1523C:p.C508S,CDHR5:NM_021924:exon14:c.G2105C:p.C702S,CDHR5:NM_001171968:exon14:c.G2087C:p.C696S, |
| chr11 | 62672097 | G | A | COSV53656335 | exonic | UQCC3 | nonsynonymous SNV | UQCC3:NM_001085372:exon2:c.G265A:p.G89S, |
| chr11 | 62807528 | C | T | . | exonic | STX5 | nonsynonymous SNV | STX5:NM_003164:exon11:c.G1009A:p.V337I, |
| chr11 | 62824182 | C | T | . | exonic | STX5 | nonsynonymous SNV | STX5:NM_001244666:exon10:c.G892A:p.E298K,STX5:NM_003164:exon10:c.G892A:p.E298K, |
| chr11 | 62825500 | C | T | . | exonic | STX5 | nonsynonymous SNV | STX5:NM_001244666:exon6:c.G463A:p.D155N,STX5:NM_003164:exon6:c.G463A:p.D155N, |
| chr11 | 62831205 | A | C | . | exonic | STX5 | nonsynonymous SNV | STX5:NM_001244666:exon2:c.T39G:p.D13E,STX5:NM_003164:exon2:c.T39G:p.D13E, |
| chr11 | 6319476 | A | G | COSV58268956 | exonic | PRKCDBP | nonsynonymous SNV | PRKCDBP:NM_145040:exon2:c.T473C:p.L158P, |
| chr11 | 6320454 | C | G | . | exonic | PRKCDBP | nonsynonymous SNV | PRKCDBP:NM_145040:exon1:c.G23C:p.R8P, |
| chr11 | 63558439 | C | T | . | exonic | HRASLS2 | nonsynonymous SNV | HRASLS2:NM_017878:exon3:c.G340A:p.E114K, |
| chr11 | 63558592 | C | T | COSV55374981 | exonic | HRASLS2 | nonsynonymous SNV | HRASLS2:NM_017878:exon3:c.G187A:p.E63K, |
| chr11 | 64225815 | C | T | . | exonic | TRPT1 | nonsynonymous SNV | TRPT1:NM_001160392:exon2:c.G46A:p.G16S,TRPT1:NM_001033678:exon2:c.G46A:p.G16S,TRPT1:NM_001160390:exon2:c.G46A:p.G16S,TRPT1:NM_001160389:exon2:c.G46A:p.G16S,TRPT1:NM_001160393:exon1:c.G46A:p.G16S, |
| chr11 | 64229857 | A | G | . | exonic | NUDT22 | nonsynonymous SNV | NUDT22:NM_001271831:exon5:c.A680G:p.Q227R,NUDT22:NM_001128613:exon5:c.A680G:p.Q227R,NUDT22:NM_001128612:exon6:c.A779G:p.Q260R,NUDT22:NM_032344:exon6:c.A779G:p.Q260R, |
| chr11 | 64229866 | T | C | . | exonic | NUDT22 | nonsynonymous SNV | NUDT22:NM_001271831:exon5:c.T689C:p.L230P,NUDT22:NM_001128613:exon5:c.T689C:p.L230P,NUDT22:NM_001128612:exon6:c.T788C:p.L263P,NUDT22:NM_032344:exon6:c.T788C:p.L263P, |
| chr11 | 64318313 | A | G | COSV50005053 | exonic | PRDX5 | nonsynonymous SNV | PRDX5:NM_012094:exon1:c.A98G:p.Y33C,PRDX5:NM_181652:exon1:c.A98G:p.Y33C,PRDX5:NM_181651:exon1:c.A98G:p.Y33C, |
| chr11 | 64804546 | T | C | COSV53639974 | exonic | MEN1 | nonsynonymous SNV | MEN1:NM_130801:exon10:c.A1636G:p.T546A,MEN1:NM_130800:exon10:c.A1636G:p.T546A,MEN1:NM_130804:exon11:c.A1636G:p.T546A,MEN1:NM_130803:exon10:c.A1636G:p.T546A,MEN1:NM_000244:exon10:c.A1636G:p.T546A,MEN1:NM_130799:exon10:c.A1621G:p.T541A,MEN1:NM_130802:exon10:c.A1636G:p.T546A, |
| chr11 | 65580165 | G | C | . | exonic | EHBP1L1 | nonsynonymous SNV | EHBP1L1:NM_001099409:exon5:c.G397C:p.V133L, |
| chr11 | 65581592 | G | A | COSV58572498 | exonic | EHBP1L1 | nonsynonymous SNV | EHBP1L1:NM_001099409:exon9:c.G920A:p.R307Q, |
| chr11 | 65868582 | T | C | . | exonic | EFEMP2 | nonsynonymous SNV | EFEMP2:NM_016938:exon8:c.A775G:p.I259V, |
| chr11 | 65917754 | T | C | . | exonic | C11ORF68 | nonsynonymous SNV | C11orf68:NM_031450:exon2:c.A584G:p.Q195R,C11orf68:NM_001135635:exon2:c.A587G:p.Q196R, |
| chr11 | 67399035 | C | T | COSV56702423 | exonic | PPP1CA | nonsynonymous SNV | PPP1CA:NM_002708:exon5:c.G652A:p.E218K,PPP1CA:NM_001008709:exon5:c.G685A:p.E229K,PPP1CA:NM_206873:exon4:c.G520A:p.E174K, |
| chr11 | 67399044 | C | T | . | exonic | PPP1CA | nonsynonymous SNV | PPP1CA:NM_002708:exon5:c.G643A:p.G215S,PPP1CA:NM_001008709:exon5:c.G676A:p.G226S,PPP1CA:NM_206873:exon4:c.G511A:p.G171S, |
| chr11 | 67400704 | C | T | . | exonic | PPP1CA | nonsynonymous SNV | PPP1CA:NM_002708:exon3:c.G403A:p.G135S,PPP1CA:NM_001008709:exon3:c.G436A:p.G146S,PPP1CA:NM_206873:exon2:c.G271A:p.G91S, |
| chr11 | 67400908 | C | T | . | exonic | PPP1CA | nonsynonymous SNV | PPP1CA:NM_002708:exon3:c.G199A:p.G67S,PPP1CA:NM_001008709:exon3:c.G232A:p.G78S,PPP1CA:NM_206873:exon2:c.G67A:p.G23S, |
| chr11 | 67401755 | C | T | . | exonic | PPP1CA | nonsynonymous SNV | PPP1CA:NM_002708:exon1:c.G28A:p.D10N,PPP1CA:NM_001008709:exon1:c.G28A:p.D10N,PPP1CA:NM_206873:exon1:c.G28A:p.D10N, |
| chr11 | 67490352 | C | A | COSV54161477 | exonic | AIP | nonsynonymous SNV | AIP:NM_001302960:exon5:c.C682A:p.Q228K,AIP:NM_003977:exon5:c.C682A:p.Q228K,AIP:NM_001302959:exon5:c.C505A:p.Q169K, |
| chr11 | 67585218 | A | G | COSV66992376 | exonic | GSTP1 | nonsynonymous SNV | GSTP1:NM_000852:exon5:c.A313G:p.I105V, |
| chr11 | 67608465 | C | A | . | exonic | NDUFV1 | nonsynonymous SNV | NDUFV1:NM_001166102:exon2:c.C115A:p.R39S,NDUFV1:NM_007103:exon2:c.C142A:p.R48S, |
| chr11 | 68032155 | C | T | COSV50288755 | exonic | NDUFS8 | nonsynonymous SNV | NDUFS8:NM_002496:exon2:c.C4T:p.R2C, |
| chr11 | 68775395 | G | A | COSV55757794 | exonic | CPT1A | nonsynonymous SNV | CPT1A:NM_001031847:exon13:c.C1496T:p.A499V,CPT1A:NM_001876:exon13:c.C1496T:p.A499V, |
| chr11 | 68794860 | C | T | . | exonic | CPT1A | nonsynonymous SNV | CPT1A:NM_001031847:exon8:c.G823A:p.A275T,CPT1A:NM_001876:exon8:c.G823A:p.A275T, |
| chr11 | 68812482 | G | C | . | exonic | CPT1A | nonsynonymous SNV | CPT1A:NM_001031847:exon3:c.C236G:p.P79R,CPT1A:NM_001876:exon3:c.C236G:p.P79R, |
| chr11 | 7000929 | A | G | . | exonic | ZNF214 | nonsynonymous SNV | ZNF214:NM_013249:exon3:c.T754C:p.C252R, |
| chr11 | 7001129 | A | C | . | exonic | ZNF214 | nonsynonymous SNV | ZNF214:NM_013249:exon3:c.T554G:p.I185R, |
| chr11 | 7001300 | A | T | COSV53481866 | exonic | ZNF214 | nonsynonymous SNV | ZNF214:NM_013249:exon3:c.T383A:p.L128H, |
| chr11 | 7001486 | T | C | COSV53480686 | exonic | ZNF214 | nonsynonymous SNV | ZNF214:NM_013249:exon3:c.A197G:p.Y66C, |
| chr11 | 70272383 | G | A | COSV54131640 | exonic | PPFIA1 | nonsynonymous SNV | PPFIA1:NM_177423:exon2:c.G211A:p.V71I,PPFIA1:NM_003626:exon2:c.G211A:p.V71I, |
| chr11 | 70343834 | G | A | . | exonic | PPFIA1 | nonsynonymous SNV | PPFIA1:NM_177423:exon15:c.G1873A:p.A625T,PPFIA1:NM_003626:exon15:c.G1873A:p.A625T, |
| chr11 | 72192177 | G | A | . | exonic | FOLR1 | nonsynonymous SNV | FOLR1:NM_016729:exon1:c.G4A:p.A2T,FOLR1:NM_000802:exon2:c.G4A:p.A2T,FOLR1:NM_016725:exon2:c.G4A:p.A2T,FOLR1:NM_016724:exon3:c.G4A:p.A2T, |
| chr11 | 72195272 | G | A | . | exonic | FOLR1 | nonsynonymous SNV | FOLR1:NM_016729:exon2:c.G170A:p.C57Y,FOLR1:NM_000802:exon3:c.G170A:p.C57Y,FOLR1:NM_016725:exon3:c.G170A:p.C57Y,FOLR1:NM_016724:exon4:c.G170A:p.C57Y, |

| | | | | | | | | |
|---|---|---|---|---|---|---|---|---|
| chr11 | 72195293 | C | T | . | exonic | FOLR1 | nonsynonymous SNV | FOLR1:NM_016729:exon2:c.C191T:p.A64V,FOLR1:NM_000802:exon3:c.C191T:p.A64V,FOLR1:NM_016725:exon3:c.C191T:p.A64V,FOLR1:NM_016724:exon4:c.C191T:p.A64V, |
| chr11 | 72195331 | G | A | COSV56616313 | exonic | FOLR1 | nonsynonymous SNV | FOLR1:NM_016729:exon2:c.G229A:p.D77N,FOLR1:NM_000802:exon3:c.G229A:p.D77N,FOLR1:NM_016725:exon3:c.G229A:p.D77N,FOLR1:NM_016724:exon4:c.G229A:p.D77N, |
| chr11 | 72195694 | G | A | . | exonic | FOLR1 | nonsynonymous SNV | FOLR1:NM_016729:exon3:c.G440A:p.R147H,FOLR1:NM_000802:exon4:c.G440A:p.R147H,FOLR1:NM_016725:exon4:c.G440A:p.R147H,FOLR1:NM_016724:exon5:c.G440A:p.R147H, |
| chr11 | 72196097 | G | A | . | exonic | FOLR1 | nonsynonymous SNV | FOLR1:NM_016729:exon4:c.G694A:p.A232T,FOLR1:NM_000802:exon5:c.G694A:p.A232T,FOLR1:NM_016725:exon5:c.G694A:p.A232T,FOLR1:NM_016724:exon6:c.G694A:p.A232T, |
| chr11 | 73716312 | C | T | . | exonic | RAB6A | nonsynonymous SNV | RAB6A:NM_198896:exon5:c.G340A:p.E114K,RAB6A:NM_001243719:exon5:c.G241A:p.E81K,RAB6A:NM_002869:exon5:c.G340A:p.E114K, |
| chr11 | 73808313 | G | T | COSV99063341 | exonic | MRPL48 | nonsynonymous SNV | MRPL48:NM_016055:exon3:c.G75T:p.R25S, |
| chr11 | 73844966 | G | A | . | exonic | MRPL48 | nonsynonymous SNV | MRPL48:NM_016055:exon5:c.G361A:p.V121I, |
| chr11 | 73859960 | G | A | . | exonic | MRPL48 | nonsynonymous SNV | MRPL48:NM_016055:exon6:c.G425A:p.S142N, |
| chr11 | 73859999 | G | A | COSV58638902 | exonic | MRPL48 | nonsynonymous SNV | MRPL48:NM_016055:exon6:c.G464A:p.R155Q, |
| chr11 | 73863211 | G | A | . | exonic | MRPL48 | nonsynonymous SNV | MRPL48:NM_016055:exon7:c.G514A:p.E172K, |
| chr11 | 73863245 | G | A | . | exonic | MRPL48 | nonsynonymous SNV | MRPL48:NM_016055:exon7:c.G548A:p.R183K, |
| chr11 | 73864333 | G | A | . | exonic | MRPL48 | nonsynonymous SNV | MRPL48:NM_016055:exon8:c.G602A:p.R201Q, |
| chr11 | 75404052 | A | T | . | exonic | RPS3 | nonsynonymous SNV | RPS3:NM_001256802:exon5:c.A383T:p.E128V,RPS3:NM_001260506:exon5:c.A431T:p.E144V,RPS3:NM_001005:exon5:c.A383T:p.E128V,RPS3:NM_001260507:exon4:c.A5T:p.E2V, |
| chr11 | 75404779 | G | A | . | exonic | RPS3 | nonsynonymous SNV | RPS3:NM_001256802:exon6:c.G646A:p.E216K,RPS3:NM_001260506:exon6:c.G694A:p.E232K,RPS3:NM_001005:exon6:c.G646A:p.E216K,RPS3:NM_001260507:exon5:c.G268A:p.E90K, |
| chr11 | 75441198 | C | T | COSV61126831 | exonic | GDPD5 | nonsynonymous SNV | GDPD5:NM_030792:exon14:c.G1438A:p.A480T, |
| chr11 | 77683795 | C | T | . | exonic | RSF1 | nonsynonymous SNV | RSF1:NM_016578:exon11:c.G2980A:p.E994K, |
| chr11 | 77701806 | A | G | COSV57839549 | exonic | RSF1 | nonsynonymous SNV | RSF1:NM_016578:exon6:c.T1423C:p.S475P, |
| chr11 | 77702301 | C | T | COSV57836647 | exonic | RSF1 | nonsynonymous SNV | RSF1:NM_016578:exon6:c.G928A:p.E310K, |
| chr11 | 77702322 | C | T | . | exonic | RSF1 | nonsynonymous SNV | RSF1:NM_016578:exon6:c.G907A:p.E303K, |
| chr11 | 77702323 | A | T | . | exonic | RSF1 | nonsynonymous SNV | RSF1:NM_016578:exon6:c.T906A:p.N302K, |
| chr11 | 77702448 | C | T | . | exonic | RSF1 | nonsynonymous SNV | RSF1:NM_016578:exon6:c.G781A:p.D261N, |
| chr11 | 77702471 | A | T | . | exonic | RSF1 | nonsynonymous SNV | RSF1:NM_016578:exon6:c.T758A:p.M253K, |
| chr11 | 78079609 | G | C | COSV55247515 | exonic | NDUFC2 | nonsynonymous SNV | NDUFC2-KCTD14:NM_001203261:exon1:c.C136G:p.L46V,NDUFC2:NM_004549:exon1:c.C136G:p.L46V,NDUFC2-KCTD14:NM_001203262:exon1:c.C136G:p.L46V,NDUFC2:NM_001204054:exon1:c.C136G:p.L46V,NDUFC2:NM_001204055:exon1:c.C136G:p.L46V,NDUFC2-KCTD14:NM_001203260:exon1:c.C136G:p.L46V, |
| chr11 | 78559613 | C | A | . | exonic | NARS2 | nonsynonymous SNV | NARS2:NM_024678:exon5:c.G520T:p.G174C, |
| chr11 | 78568744 | T | G | COSV55250015 | exonic | NARS2 | nonsynonymous SNV | NARS2:NM_024678:exon3:c.A260C:p.N87T, |
| chr11 | 7987468 | C | T | COSV59141570 | exonic | EIF3F | nonsynonymous SNV | EIF3F:NM_003754:exon1:c.C116T:p.P39L, |
| chr11 | 810326 | G | A | COSV58252909 | exonic | RPLP2 | nonsynonymous SNV | RPLP2:NM_001004:exon2:c.G92A:p.G31D, |
| chr11 | 824789 | T | C | COSV60743886 | exonic | PNPLA2 | nonsynonymous SNV | PNPLA2:NM_020376:exon10:c.T1442C:p.L481P, |
| chr11 | 85628665 | G | A | . | exonic | TMEM126B | nonsynonymous SNV | TMEM126B:NM_018480:exon1:c.G58A:p.V20M, |
| chr11 | 85631741 | G | A | . | exonic | TMEM126B | nonsynonymous SNV | TMEM126B:NM_001256546:exon3:c.G46A:p.A16T,TMEM126B:NM_001193538:exon3:c.G46A:p.A16T,TMEM126B:NM_001193537:exon3:c.G76A:p.A26T,TMEM126B:NM_018480:exon2:c.G136A:p.A46T, |
| chr11 | 85631762 | G | A | . | exonic | TMEM126B | nonsynonymous SNV | TMEM126B:NM_001256546:exon3:c.G67A:p.V23I,TMEM126B:NM_001193538:exon3:c.G67A:p.V23I,TMEM126B:NM_001193537:exon3:c.G97A:p.V33I,TMEM126B:NM_018480:exon2:c.G157A:p.V53I, |
| chr11 | 85634117 | G | A | . | exonic | TMEM126B | nonsynonymous SNV | TMEM126B:NM_001256546:exon4:c.G145A:p.G49R,TMEM126B:NM_001193538:exon4:c.G145A:p.G49R,TMEM126B:NM_001256547:exon2:c.G97A:p.G33R,TMEM126B:NM_001193537:exon4:c.G175A:p.G59R,TMEM126B:NM_018480:exon3:c.G235A:p.G79R, |
| chr11 | 870446 | G | A | COSV60260399 | exonic | CHID1 | nonsynonymous SNV | CHID1:NM_023947:exon11:c.C1013T:p.A338V,CHID1:NM_001142675:exon11:c.C1013T:p.A338V,CHID1:NM_001142674:exon12:c.C1013T:p.A338V,CHID1:NM_001142676:exon12:c.C1088T:p.A363V,CHID1:NM_001142677:exon10:c.C920T:p.A307V, |
| chr11 | 87302361 | T | A | . | exonic | TMEM135 | nonsynonymous SNV | TMEM135:NM_022918:exon8:c.T617A:p.V206E,TMEM135:NM_001168724:exon7:c.T551A:p.V184E, |
| chr11 | 87302396 | G | A | COSV59699111 | exonic | TMEM135 | nonsynonymous SNV | TMEM135:NM_022918:exon8:c.G652A:p.G218R,TMEM135:NM_001168724:exon7:c.G586A:p.G196R, |
| chr11 | 88312415 | A | G | COSV57060752 | exonic | CTSC | nonsynonymous SNV | CTSC:NM_001814:exon3:c.T458C:p.I153T, |
| chr11 | 8953249 | C | T | COSV58427716 | exonic | TMEM9B | nonsynonymous SNV | TMEM9B:NM_001286095:exon4:c.G173A:p.R58H,TMEM9B:NM_020644:exon4:c.G395A:p.R132H,TMEM9B:NM_001286094:exon3:c.G173A:p.R58H, |
| chr11 | 8962128 | C | T | . | exonic | TMEM9B | nonsynonymous SNV | TMEM9B:NM_020644:exon2:c.G161A:p.G54E, |
| chr11 | 90202418 | G | T | . | exonic | CHORDC1 | nonsynonymous SNV | CHORDC1:NM_012124:exon11:c.C986A:p.A329D,CHORDC1:NM_001144073:exon10:c.C929A:p.A310D, |
| chr11 | 95923947 | T | G | COSV60579185 | exonic | MTMR2 | nonsynonymous SNV | MTMR2:NM_016156:exon1:c.A8C:p.K3T, |
| chr11 | 96092591 | G | C | . | exonic | MAML2 | nonsynonymous SNV | MAML2:NM_032427:exon2:c.C1440G:p.I480M, |
| chr12 | 100624303 | C | A | . | exonic | GAS2L3 | nonsynonymous SNV | GAS2L3:NM_001303130:exon9:c.C1498A:p.P500T,GAS2L3:NM_001303131:exon10:c.C1186A:p.P396T,GAS2L3:NM_174942:exon10:c.C1498A:p.P500T, |
| chr12 | 101714567 | T | C | COSV57533603 | exonic | CHPT1 | nonsynonymous SNV | CHPT1:NM_020244:exon3:c.T485C:p.F162S, |
| chr12 | 102074401 | C | T | . | exonic | NUP37 | nonsynonymous SNV | NUP37:NM_024057:exon9:c.G934A:p.V312I, |
| chr12 | 102074454 | C | T | . | exonic | NUP37 | nonsynonymous SNV | NUP37:NM_024057:exon9:c.G881A:p.G294D, |
| chr12 | 102075091 | C | T | . | exonic | NUP37 | stopgain | NUP37:NM_024057:exon8:c.G777A:p.W259X, |
| chr12 | 102077404 | C | T | COSV51837132 | exonic | NUP37 | nonsynonymous SNV | NUP37:NM_024057:exon6:c.G640A:p.A214T, |
| chr12 | 102099169 | C | A | . | exonic | NUP37 | nonsynonymous SNV | NUP37:NM_024057:exon4:c.G386T:p.G129V, |
| chr12 | 102118430 | C | T | . | exonic | NUP37 | nonsynonymous SNV | NUP37:NM_024057:exon1:c.G89A:p.G30E, |
| chr12 | 10418366 | C | G | . | exonic | KLRC3 | nonsynonymous SNV | KLRC3:NM_002261:exon4:c.G464C:p.C155S,KLRC3:NM_007333:exon4:c.G464C:p.C155S, |
| chr12 | 10418492 | T | G | COSV67249807 | exonic | KLRC3 | nonsynonymous SNV | KLRC3:NM_002261:exon4:c.A338C:p.H113P,KLRC3:NM_007333:exon4:c.A338C:p.H113P, |
| chr12 | 10419059 | C | T | COSV67249816 | exonic | KLRC3 | nonsynonymous SNV | KLRC3:NM_002261:exon3:c.G316A:p.A106T,KLRC3:NM_007333:exon3:c.G316A:p.A106T, |
| chr12 | 10420495 | C | G | COSV67249876 | exonic | KLRC3 | nonsynonymous SNV | KLRC3:NM_002261:exon1:c.G56C:p.W19S,KLRC3:NM_007333:exon1:c.G56C:p.W19S, |
| chr12 | 10420496 | A | G | COSV67249820 | exonic | KLRC3 | nonsynonymous SNV | KLRC3:NM_002261:exon1:c.T55C:p.W19R,KLRC3:NM_007333:exon1:c.T55C:p.W19R, |
| chr12 | 104757724 | A | T | . | exonic | CHST11 | nonsynonymous SNV | CHST11:NM_018413:exon3:c.A980T:p.Q327L,CHST11:NM_001173982:exon3:c.A965T:p.Q322L, |
| chr12 | 108562641 | T | G | COSV57206465 | exonic | ISCU | nonsynonymous SNV | ISCU:NM_001301141:exon1:c.T19G:p.F7V,ISCU:NM_001301140:exon1:c.T19G:p.F7V,ISCU:NM_213595:exon1:c.T19G:p.F7V, |
| chr12 | 108562642 | T | G | COSV57206471 | exonic | ISCU | nonsynonymous SNV | ISCU:NM_001301141:exon1:c.T20G:p.F7C,ISCU:NM_001301140:exon1:c.T20G:p.F7C,ISCU:NM_213595:exon1:c.T20G:p.F7C, |
| chr12 | 108562657 | C | T | COSV57206596 | exonic | ISCU | nonsynonymous SNV | ISCU:NM_001301141:exon1:c.C35T:p.A12V,ISCU:NM_001301140:exon1:c.C35T:p.A12V,ISCU:NM_213595:exon1:c.C35T:p.A12V, |
| chr12 | 109099210 | G | A | . | exonic | UNG | nonsynonymous SNV | UNG:NM_080911:exon3:c.G361A:p.E121K,UNG:NM_003362:exon2:c.G334A:p.E112K, |
| chr12 | 109103552 | G | A | . | exonic | UNG | nonsynonymous SNV | UNG:NM_080911:exon6:c.G742A:p.G248S,UNG:NM_003362:exon5:c.G715A:p.G239S, |
| chr12 | 109557065 | A | T | COSV57169652 | exonic | MMAB | nonsynonymous SNV | MMAB:NM_052845:exon9:c.T716A:p.M239K, |
| chr12 | 109573425 | C | T | COSV57169511 | exonic | MMAB | nonsynonymous SNV | MMAB:NM_052845:exon1:c.G56A:p.R19H, |
| chr12 | 110496121 | C | T | . | exonic | VPS29 | nonsynonymous SNV | VPS29:NM_057180:exon3:c.G98A:p.G33E,VPS29:NM_001282151:exon2:c.G86A:p.G29E,VPS29:NM_001282150:exon4:c.G182A:p.G61E,VPS29:NM_016226:exon2:c.G86A:p.G29E, |

| | | | | | | | | |
|---|---|---|---|---|---|---|---|---|
| chr12 | 112450442 | T | A | . | exonic | PTPN11 | nonsynonymous SNV | PTPN11:NM_002834:exon3:c.T262A:p.L88I,PTPN11:NM_080601:exon3:c.T262A:p.L88I, |
| chr12 | 112453244 | G | A | . | exonic | PTPN11 | nonsynonymous SNV | PTPN11:NM_002834:exon4:c.G382A:p.E128K,PTPN11:NM_080601:exon4:c.G382A:p.E128K, |
| chr12 | 112453250 | G | A | . | exonic | PTPN11 | nonsynonymous SNV | PTPN11:NM_002834:exon4:c.G388A:p.G130R,PTPN11:NM_080601:exon4:c.G388A:p.G130R, |
| chr12 | 112477993 | C | A | . | exonic | PTPN11 | nonsynonymous SNV | PTPN11:NM_080601:exon9:c.C1070A:p.T357K,PTPN11:NM_002834:exon9:c.C1070A:p.T357K, |
| chr12 | 112911065 | G | A | COSV52534422 | exonic | OAS1 | nonsynonymous SNV | OAS1:NM_016816:exon3:c.G484A:p.G162S,OAS1:NM_002534:exon3:c.G484A:p.G162S,OAS1:NM_001032409:exon3:c.G484A:p.G162S, |
| chr12 | 112919637 | G | A | COSV52534024 | exonic | OAS1 | nonsynonymous SNV | OAS1:NM_001032409:exon6:c.G1189A:p.G397R, |
| chr12 | 116723171 | T | C | COSV54339873 | exonic | C12ORF49 | nonsynonymous SNV | C12orf49:NM_024738:exon2:c.A164G:p.Q55R, |
| chr12 | 11752469 | C | T | . | exonic | ETV6 | nonsynonymous SNV | ETV6:NM_001987:exon2:c.C53T:p.T18I, |
| chr12 | 11884577 | G | A | COSV56765875 | exonic | ETV6 | nonsynonymous SNV | ETV6:NM_001987:exon6:c.G1142A:p.G381E, |
| chr12 | 120224174 | T | C | . | exonic | PXN | nonsynonymous SNV | PXN:NM_002859:exon2:c.A217G:p.S73G,PXN:NM_001080855:exon2:c.A217G:p.S73G,PXN:NM_001243756:exon2:c.A217G:p.S73G, |
| chr12 | 121888906 | T | C | COSV55838997 | exonic | PSMD9 | nonsynonymous SNV | PSMD9:NM_002813:exon1:c.T50C:p.V17A,PSMD9:NM_001261400:exon1:c.T50C:p.V17A, |
| chr12 | 121894772 | G | A | . | exonic | PSMD9 | nonsynonymous SNV | PSMD9:NM_002813:exon2:c.G172A:p.D58N, |
| chr12 | 121899757 | G | A | . | exonic | PSMD9 | nonsynonymous SNV | PSMD9:NM_002813:exon3:c.G365A:p.R122H, |
| chr12 | 121903040 | G | A | . | exonic | PSMD9 | nonsynonymous SNV | PSMD9:NM_002813:exon4:c.G488A:p.G163D,PSMD9:NM_001261400:exon2:c.G173A:p.G58D, |
| chr12 | 121915874 | G | A | . | exonic | PSMD9 | nonsynonymous SNV | PSMD9:NM_002813:exon5:c.G574A:p.V192M,PSMD9:NM_001261400:exon3:c.G259A:p.V87M, |
| chr12 | 122133442 | A | G | COSV59834494 | exonic | MLXIP | nonsynonymous SNV | MLXIP:NM_014938:exon9:c.A1187G:p.E396G, |
| chr12 | 122327956 | G | C | COSV56798569 | exonic | CLIP1 | nonsynonymous SNV | CLIP1:NM_002956:exon16:c.C3207G:p.D1069E,CLIP1:NM_001247997:exon17:c.C3240G:p.D1080E,CLIP1:NM_198240:exon15:c.C3102G:p.D1034E, |
| chr12 | 122328296 | C | T | . | exonic | CLIP1 | nonsynonymous SNV | CLIP1:NM_002956:exon15:c.G2965A:p.E989K,CLIP1:NM_001247997:exon16:c.G2998A:p.E1000K,CLIP1:NM_198240:exon14:c.G2860A:p.E954K, |
| chr12 | 122508364 | C | T | . | exonic | RSRC2 | nonsynonymous SNV | RSRC2:NM_023012:exon8:c.G889A:p.A297T, |
| chr12 | 122518905 | C | T | . | exonic | RSRC2 | nonsynonymous SNV | RSRC2:NM_023012:exon4:c.G332A:p.R111K, |
| chr12 | 122521416 | C | T | COSV59206104 | exonic | RSRC2 | nonsynonymous SNV | RSRC2:NM_023012:exon3:c.G176A:p.R59K, |
| chr12 | 122521423 | C | T | . | exonic | RSRC2 | nonsynonymous SNV | RSRC2:NM_023012:exon3:c.G169A:p.E57K, |
| chr12 | 122522239 | T | C | . | exonic | RSRC2 | nonsynonymous SNV | RSRC2:NM_023012:exon2:c.A67G:p.K23E, |
| chr12 | 124786500 | C | A | . | exonic | SCARB1 | nonsynonymous SNV | SCARB1:NM_001082959:exon11:c.G1258T:p.G420W,SCARB1:NM_005505:exon11:c.G1258T:p.G420W, |
| chr12 | 124807895 | A | T | . | exonic | SCARB1 | nonsynonymous SNV | SCARB1:NM_001082959:exon7:c.T875A:p.V292E,SCARB1:NM_005505:exon7:c.T875A:p.V292E, |
| chr12 | 124814300 | C | T | COSV55544722 | exonic | SCARB1 | nonsynonymous SNV | SCARB1:NM_001082959:exon4:c.G532A:p.E178K,SCARB1:NM_005505:exon4:c.G532A:p.E178K, |
| chr12 | 124863632 | A | C | . | exonic | SCARB1 | nonsynonymous SNV | SCARB1:NM_001082959:exon1:c.T89G:p.V30G,SCARB1:NM_005505:exon1:c.T89G:p.V30G, |
| chr12 | 124863717 | C | T | COSV55547637 | exonic | SCARB1 | nonsynonymous SNV | SCARB1:NM_001082959:exon1:c.G4A:p.G2S,SCARB1:NM_005505:exon1:c.G4A:p.G2S, |
| chr12 | 124950034 | C | T | COSV58132345 | exonic | DHX37 | nonsynonymous SNV | DHX37:NM_032656:exon25:c.G3242A:p.R1081Q, |
| chr12 | 124953970 | T | C | COSV58134449 | exonic | DHX37 | nonsynonymous SNV | DHX37:NM_032656:exon20:c.A2605G:p.S869G, |
| chr12 | 132152094 | G | A | . | exonic | NOC4L | nonsynonymous SNV | NOC4L:NM_024078:exon14:c.G1328A:p.R443H, |
| chr12 | 132730003 | C | T | COSV61708335 | exonic | ANKLE2 | nonsynonymous SNV | ANKLE2:NM_015114:exon11:c.G2159A:p.R720H, |
| chr12 | 132754951 | G | A | COSV61708344 | exonic | ANKLE2 | nonsynonymous SNV | ANKLE2:NM_015114:exon2:c.C364T:p.H122Y, |
| chr12 | 1444631 | A | G | COSV61692803 | exonic | ERC1 | nonsynonymous SNV | ERC1:NM_001301248:exon17:c.A3094G:p.T1032A,ERC1:NM_178039:exon17:c.A3010G:p.T1004A,ERC1:NM_178040:exon18:c.A3094G:p.T1032A, |
| chr12 | 14503834 | G | C | COSV53694489 | exonic | PLBD1 | nonsynonymous SNV | PLBD1:NM_024829:exon11:c.C1600G:p.P534A, |
| chr12 | 14511316 | A | G | COSV53692200 | exonic | PLBD1 | nonsynonymous SNV | PLBD1:NM_024829:exon8:c.T1130C:p.V377A, |
| chr12 | 14535710 | C | T | . | exonic | PLBD1 | nonsynonymous SNV | PLBD1:NM_024829:exon6:c.G793A:p.V265I, |
| chr12 | 14567620 | G | T | COSV53693941 | exonic | PLBD1 | nonsynonymous SNV | PLBD1:NM_024829:exon1:c.C77A:p.P26Q, |
| chr12 | 14882147 | T | C | COSV57453935 | exonic | MGP | nonsynonymous SNV | MGP:NM_000900:exon4:c.A304G:p.T102A,MGP:NM_001190839:exon5:c.A379G:p.T127A, |
| chr12 | 20858546 | T | G | COSV53937987 | exonic | SLCO1B3 | nonsynonymous SNV | SLCO1B3:NM_019844:exon5:c.T334G:p.S112A, |
| chr12 | 21635571 | C | T | . | exonic | LDHB | nonsynonymous SNV | LDHB:NM_002300:exon8:c.G976A:p.D326N,LDHB:NM_001174097:exon8:c.G976A:p.D326N, |
| chr12 | 21635572 | C | T | . | exonic | LDHB | stopgain | LDHB:NM_002300:exon8:c.G975A:p.W325X,LDHB:NM_001174097:exon8:c.G975A:p.W325X, |
| chr12 | 21635648 | C | T | . | exonic | LDHB | nonsynonymous SNV | LDHB:NM_002300:exon8:c.G899A:p.G300E,LDHB:NM_001174097:exon8:c.G899A:p.G300E, |
| chr12 | 21635651 | C | T | . | exonic | LDHB | nonsynonymous SNV | LDHB:NM_002300:exon8:c.G896A:p.R299Q,LDHB:NM_001174097:exon8:c.G896A:p.R299Q, |
| chr12 | 21635700 | C | T | . | exonic | LDHB | nonsynonymous SNV | LDHB:NM_002300:exon8:c.G847A:p.G283S,LDHB:NM_001174097:exon8:c.G847A:p.G283S, |
| chr12 | 21637154 | C | T | COSV63364527 | exonic | LDHB | nonsynonymous SNV | LDHB:NM_002300:exon7:c.G754A:p.A252T,LDHB:NM_001174097:exon7:c.G754A:p.A252T, |
| chr12 | 21638389 | T | C | . | exonic | LDHB | nonsynonymous SNV | LDHB:NM_002300:exon6:c.A677G:p.E226G,LDHB:NM_001174097:exon6:c.A677G:p.E226G, |
| chr12 | 21638402 | C | T | . | exonic | LDHB | nonsynonymous SNV | LDHB:NM_002300:exon6:c.G664A:p.D222N,LDHB:NM_001174097:exon6:c.G664A:p.D222N, |
| chr12 | 21638407 | C | T | . | exonic | LDHB | nonsynonymous SNV | LDHB:NM_002300:exon6:c.G659A:p.G220E,LDHB:NM_001174097:exon6:c.G659A:p.G220E, |
| chr12 | 21638426 | C | T | . | exonic | LDHB | nonsynonymous SNV | LDHB:NM_002300:exon6:c.G640A:p.E214K,LDHB:NM_001174097:exon6:c.G640A:p.E214K, |
| chr12 | 21638460 | C | T | . | exonic | LDHB | stopgain | LDHB:NM_002300:exon6:c.G606A:p.W202X,LDHB:NM_001174097:exon6:c.G606A:p.W202X, |
| chr12 | 21641972 | C | T | . | exonic | LDHB | nonsynonymous SNV | LDHB:NM_002300:exon5:c.G575A:p.G192E,LDHB:NM_001174097:exon5:c.G575A:p.G192E, |
| chr12 | 21641973 | C | T | . | exonic | LDHB | nonsynonymous SNV | LDHB:NM_002300:exon5:c.G574A:p.G192R,LDHB:NM_001174097:exon5:c.G574A:p.G192R, |
| chr12 | 21641979 | T | A | . | exonic | LDHB | nonsynonymous SNV | LDHB:NM_002300:exon5:c.A568T:p.I190F,LDHB:NM_001174097:exon5:c.A568T:p.I190F, |
| chr12 | 21641980 | C | T | . | exonic | LDHB | stopgain | LDHB:NM_002300:exon5:c.G567A:p.W189X,LDHB:NM_001174097:exon5:c.G567A:p.W189X, |
| chr12 | 21642032 | C | T | . | exonic | LDHB | nonsynonymous SNV | LDHB:NM_002300:exon5:c.G515A:p.R172H,LDHB:NM_001174097:exon5:c.G515A:p.R172H, |
| chr12 | 21642066 | C | T | COSV63365537 | exonic | LDHB | nonsynonymous SNV | LDHB:NM_002300:exon5:c.G481A:p.G161R,LDHB:NM_001174097:exon5:c.G481A:p.G161R, |
| chr12 | 21643947 | C | T | . | exonic | LDHB | nonsynonymous SNV | LDHB:NM_002300:exon4:c.G409A:p.V137I,LDHB:NM_001174097:exon4:c.G409A:p.V137I, |
| chr12 | 21644036 | C | T | . | exonic | LDHB | nonsynonymous SNV | LDHB:NM_002300:exon4:c.G320A:p.R107Q,LDHB:NM_001174097:exon4:c.G320A:p.R107Q, |
| chr12 | 21644046 | C | T | . | exonic | LDHB | nonsynonymous SNV | LDHB:NM_002300:exon4:c.G310A:p.G104R,LDHB:NM_001174097:exon4:c.G310A:p.G104R, |
| chr12 | 21644057 | C | T | . | exonic | LDHB | nonsynonymous SNV | LDHB:NM_002300:exon4:c.G299A:p.R100H,LDHB:NM_001174097:exon4:c.G299A:p.R100H, |
| chr12 | 21644108 | T | C | . | exonic | LDHB | nonsynonymous SNV | LDHB:NM_002300:exon4:c.A248G:p.D83G,LDHB:NM_001174097:exon4:c.A248G:p.D83G, |
| chr12 | 21646992 | C | T | . | exonic | LDHB | nonsynonymous SNV | LDHB:NM_002300:exon3:c.G154A:p.V52M,LDHB:NM_001174097:exon3:c.G154A:p.V52M, |
| chr12 | 21647010 | C | T | . | exonic | LDHB | nonsynonymous SNV | LDHB:NM_002300:exon3:c.G136A:p.A46T,LDHB:NM_001174097:exon3:c.G136A:p.A46T, |
| chr12 | 21654556 | C | T | . | exonic | LDHB | nonsynonymous SNV | LDHB:NM_002300:exon2:c.G116A:p.S39N,LDHB:NM_001174097:exon2:c.G116A:p.S39N, |
| chr12 | 21654570 | C | T | . | exonic | LDHB | nonsynonymous SNV | LDHB:NM_002300:exon2:c.G102A:p.M34I,LDHB:NM_001174097:exon2:c.G102A:p.M34I, |
| chr12 | 21654669 | C | T | . | exonic | LDHB | nonsynonymous SNV | LDHB:NM_002300:exon2:c.G3A:p.M1I,LDHB:NM_001174097:exon2:c.G3A:p.M1I, |
| chr12 | 22055223 | G | A | . | exonic | CMAS | nonsynonymous SNV | CMAS:NM_018686:exon2:c.G335A:p.S112N, |
| chr12 | 22061376 | G | C | . | exonic | CMAS | nonsynonymous SNV | CMAS:NM_018686:exon6:c.G884C:p.G295A, |
| chr12 | 22062404 | G | A | . | exonic | CMAS | nonsynonymous SNV | CMAS:NM_018686:exon7:c.G1084A:p.G362S, |

| | | | | | | | | |
|---|---|---|---|---|---|---|---|---|
| chr12 | 25245350 | C | A | COSV55497419 | exonic | KRAS | nonsynonymous SNV | KRAS:NM_004985:exon2:c.G35T:p.G12V,KRAS:NM_033360:exon2:c.G35T:p.G12V, |
| chr12 | 26398925 | G | T | . | exonic | ITPR2 | nonsynonymous SNV | ITPR2:NM_002223:exon54:c.C7647A:p.S2549R, |
| chr12 | 27696863 | A | G | . | exonic | REP15 | nonsynonymous SNV | REP15:NM_001029874:exon1:c.A301G:p.N101D, |
| chr12 | 285482 | T | C | COSV66996660 | exonic | KDM5A | nonsynonymous SNV | KDM5A:NM_001042603:exon28:c.A5047G:p.M1683V, |
| chr12 | 292872 | C | T | . | exonic | KDM5A | nonsynonymous SNV | KDM5A:NM_001042603:exon27:c.G4753A:p.E1585K, |
| chr12 | 292888 | G | T | . | exonic | KDM5A | nonsynonymous SNV | KDM5A:NM_001042603:exon27:c.C4737A:p.S1579R, |
| chr12 | 307975 | C | T | . | exonic | KDM5A | nonsynonymous SNV | KDM5A:NM_001042603:exon23:c.G3409A:p.E1137K, |
| chr12 | 309877 | C | T | COSV66991870 | exonic | KDM5A | nonsynonymous SNV | KDM5A:NM_001042603:exon22:c.G3304A:p.E1102K, |
| chr12 | 309883 | C | T | COSV66991284 | exonic | KDM5A | nonsynonymous SNV | KDM5A:NM_001042603:exon22:c.G3298A:p.E1100K, |
| chr12 | 309891 | A | T | . | exonic | KDM5A | nonsynonymous SNV | KDM5A:NM_001042603:exon22:c.T3290A:p.I1097K, |
| chr12 | 309922 | T | A | . | exonic | KDM5A | stopgain | KDM5A:NM_001042603:exon22:c.A3259T:p.K1087X, |
| chr12 | 318409 | A | G | COSV66991468 | exonic | KDM5A | nonsynonymous SNV | KDM5A:NM_001042603:exon19:c.T2594C:p.M865T, |
| chr12 | 3678979 | C | T | COSV52913838 | exonic | CRACR2A | nonsynonymous SNV | CRACR2A:NM_032680:exon6:c.G460A:p.E154K,CRACR2A:NM_001144958:exon6:c.G460A:p.E154K, |
| chr12 | 3679056 | G | A | COSV52919219 | exonic | CRACR2A | nonsynonymous SNV | CRACR2A:NM_032680:exon6:c.C383T:p.A128V,CRACR2A:NM_001144958:exon6:c.C383T:p.A128V, |
| chr12 | 438517 | T | C | COSV53475389 | exonic | CCDC77 | nonsynonymous SNV | CCDC77:NM_032358:exon10:c.T1004C:p.M335T,CCDC77:NM_001130146:exon9:c.T908C:p.M303T,CCDC77:NM_001130147:exon9:c.T908C:p.M303T,CCDC77:NM_001130148:exon8:c.T908C:p.M303T, |
| chr12 | 48843970 | T | A | . | exonic | DDX23 | nonsynonymous SNV | DDX23:NM_004818:exon3:c.A290T:p.H97L, |
| chr12 | 50819655 | G | A | . | exonic | ATF1 | nonsynonymous SNV | ATF1:NM_005171:exon7:c.G692A:p.R231H, |
| chr12 | 51049161 | G | A | . | exonic | LETMD1 | nonsynonymous SNV | LETMD1:NM_015416:exon2:c.G250A:p.V84I,LETMD1:NM_001300765:exon2:c.G250A:p.V84I,LETMD1:NM_001243689:exon2:c.G250A:p.V84I, |
| chr12 | 53299748 | A | G | . | exonic | C12ORF10 | nonsynonymous SNV | C12orf10:NM_021640:exon1:c.A11G:p.Q4R, |
| chr12 | 53299749 | A | C | . | exonic | C12ORF10 | nonsynonymous SNV | C12orf10:NM_021640:exon1:c.A12C:p.Q4H, |
| chr12 | 53307064 | C | T | . | exonic | C12ORF10 | nonsynonymous SNV | C12orf10:NM_021640:exon7:c.C1046T:p.T349I, |
| chr12 | 53307076 | A | G | COSV52934545 | exonic | C12ORF10 | nonsynonymous SNV | C12orf10:NM_021640:exon7:c.A1058G:p.H353R, |
| chr12 | 53454850 | G | A | . | exonic | PCBP2 | nonsynonymous SNV | PCBP2:NM_005016:exon2:c.G50A:p.R17Q,PCBP2:NM_001128911:exon2:c.G50A:p.R17Q,PCBP2:NM_001128914:exon2:c.G50A:p.R17Q,PCBP2:NM_001098620:exon2:c.G50A:p.R17Q,PCBP2:NM_031989:exon2:c.G50A:p.R17Q,PCBP2:NM_001128913:exon2:c.G50A:p.R17Q,PCBP2:NM_001128912:exon2:c.G50A:p.R17Q, |
| chr12 | 53455957 | G | A | . | exonic | PCBP2 | nonsynonymous SNV | PCBP2:NM_005016:exon5:c.G199A:p.A67T,PCBP2:NM_001128911:exon5:c.G199A:p.A67T,PCBP2:NM_001128914:exon5:c.G199A:p.A67T,PCBP2:NM_001098620:exon5:c.G199A:p.A67T,PCBP2:NM_031989:exon5:c.G199A:p.A67T,PCBP2:NM_001128913:exon5:c.G199A:p.A67T,PCBP2:NM_001128912:exon5:c.G199A:p.A67T, |
| chr12 | 53455987 | G | A | . | exonic | PCBP2 | nonsynonymous SNV | PCBP2:NM_005016:exon5:c.G229A:p.D77N,PCBP2:NM_001128911:exon5:c.G229A:p.D77N,PCBP2:NM_001128914:exon5:c.G229A:p.D77N,PCBP2:NM_001098620:exon5:c.G229A:p.D77N,PCBP2:NM_031989:exon5:c.G229A:p.D77N,PCBP2:NM_001128913:exon5:c.G229A:p.D77N,PCBP2:NM_001128912:exon5:c.G229A:p.D77N, |
| chr12 | 53459377 | G | A | . | exonic | PCBP2 | nonsynonymous SNV | PCBP2:NM_005016:exon6:c.G349A:p.G117R,PCBP2:NM_001128911:exon6:c.G349A:p.G117R,PCBP2:NM_001128914:exon6:c.G349A:p.G117R,PCBP2:NM_001098620:exon6:c.G349A:p.G117R,PCBP2:NM_031989:exon6:c.G349A:p.G117R,PCBP2:NM_001128913:exon6:c.G349A:p.G117R,PCBP2:NM_001128912:exon6:c.G349A:p.G117R, |
| chr12 | 53461030 | G | A | . | exonic | PCBP2 | nonsynonymous SNV | PCBP2:NM_005016:exon7:c.G391A:p.V131I,PCBP2:NM_001128911:exon7:c.G391A:p.V131I,PCBP2:NM_001128914:exon7:c.G391A:p.V131I,PCBP2:NM_001098620:exon7:c.G391A:p.V131I,PCBP2:NM_031989:exon7:c.G391A:p.V131I,PCBP2:NM_001128913:exon7:c.G391A:p.V131I,PCBP2:NM_001128912:exon7:c.G391A:p.V131I, |
| chr12 | 53467242 | G | A | . | exonic | PCBP2 | nonsynonymous SNV | PCBP2:NM_005016:exon11:c.G748A:p.A250T,PCBP2:NM_001128911:exon11:c.G748A:p.A250T,PCBP2:NM_001128914:exon10:c.G643A:p.A215T,PCBP2:NM_001098620:exon10:c.G643A:p.A215T,PCBP2:NM_031989:exon11:c.G736A:p.A246T,PCBP2:NM_001128913:exon10:c.G655A:p.A219T,PCBP2:NM_001128912:exon11:c.G736A:p.A246T, |
| chr12 | 53468825 | C | A | . | exonic | PCBP2 | nonsynonymous SNV | PCBP2:NM_005016:exon13:c.C887A:p.P296Q,PCBP2:NM_001128911:exon13:c.C884A:p.P295Q,PCBP2:NM_001128914:exon11:c.C743A:p.P248Q,PCBP2:NM_001098620:exon12:c.C782A:p.P261Q,PCBP2:NM_031989:exon13:c.C875A:p.P292Q,PCBP2:NM_001128913:exon12:c.C794A:p.P265Q,PCBP2:NM_001128912:exon13:c.C872A:p.P291Q, |
| chr12 | 53471656 | G | A | . | exonic | PCBP2 | nonsynonymous SNV | PCBP2:NM_005016:exon14:c.G913A:p.G305R,PCBP2:NM_001128911:exon14:c.G910A:p.G304R,PCBP2:NM_001128914:exon12:c.G769A:p.G257R,PCBP2:NM_001098620:exon13:c.G808A:p.G270R,PCBP2:NM_031989:exon14:c.G901A:p.G301R,PCBP2:NM_001128913:exon13:c.G820A:p.G274R,PCBP2:NM_001128912:exon14:c.G898A:p.G300R, |
| chr12 | 55789103 | C | T | . | exonic | SARNP | nonsynonymous SNV | SARNP:NM_033082:exon9:c.G473A:p.G158D, |
| chr12 | 55800868 | C | T | . | exonic | SARNP | nonsynonymous SNV | SARNP:NM_033082:exon3:c.G169A:p.G57R, |
| chr12 | 55842818 | G | A | . | exonic | MMP19 | stopgain | MMP19:NM_001272101:exon1:c.C13T:p.Q5X,MMP19:NM_002429:exon1:c.C13T:p.Q5X, |
| chr12 | 56004279 | G | A | . | exonic | SUOX | nonsynonymous SNV | SUOX:NM_001032386:exon5:c.G890A:p.R297Q,SUOX:NM_001032387:exon4:c.G890A:p.R297Q,SUOX:NM_000456:exon6:c.G890A:p.R297Q, |
| chr12 | 56083835 | G | A | . | exonic | ERBB3 | nonsynonymous SNV | ERBB3:NM_001982:exon2:c.G167A:p.C56Y,ERBB3:NM_001005915:exon2:c.G167A:p.C56Y, |
| chr12 | 56086599 | A | T | . | exonic | ERBB3 | nonsynonymous SNV | ERBB3:NM_001982:exon4:c.A490T:p.R164W, |
| chr12 | 56159625 | G | A | . | exonic | MYL6 | nonsynonymous SNV | MYL6:NM_079423:exon3:c.G70A:p.D24N,MYL6:NM_021019:exon3:c.G70A:p.D24N, |
| chr12 | 56267173 | C | A | . | exonic | COQ10A | nonsynonymous SNV | COQ10A:NM_144576:exon1:c.C55A:p.R19S, |
| chr12 | 56666218 | C | T | . | exonic | PTGES3 | nonsynonymous SNV | PTGES3:NM_001282603:exon5:c.G325A:p.D109N,PTGES3:NM_006601:exon6:c.G424A:p.D142N,PTGES3:NM_001282602:exon5:c.G334A:p.D112N,PTGES3:NM_001282604:exon7:c.G436A:p.D146N, |
| chr12 | 56671771 | C | T | . | exonic | PTGES3 | nonsynonymous SNV | PTGES3:NM_006601:exon4:c.G263A:p.R88K,PTGES3:NM_001282602:exon4:c.G263A:p.R88K,PTGES3:NM_001282601:exon4:c.G263A:p.R88K,PTGES3:NM_001282605:exon4:c.G263A:p.R88K,PTGES3:NM_001282604:exon5:c.G275A:p.R92K, |
| chr12 | 56672997 | C | T | . | exonic | PTGES3 | nonsynonymous SNV | PTGES3:NM_001282603:exon2:c.G71A:p.S24N,PTGES3:NM_006601:exon2:c.G71A:p.S24N,PTGES3:NM_001282602:exon2:c.G71A:p.S24N,PTGES3:NM_001282601:exon2:c.G71A:p.S24N,PTGES3:NM_001282605:exon2:c.G71A:p.S24N,PTGES3:NM_001282604:exon3:c.G83A:p.S28N, |
| chr12 | 56712541 | C | T | . | exonic | NACA | nonsynonymous SNV | NACA:NM_001113203:exon11:c.G2775A:p.M925I,NACA:NM_001113202:exon8:c.G645A:p.M215I, |
| chr12 | 56712827 | C | T | . | exonic | NACA | nonsynonymous SNV | NACA:NM_001113203:exon10:c.G2722A:p.A908T,NACA:NM_001113202:exon7:c.G592A:p.A198T, |
| chr12 | 56712838 | T | C | . | exonic | NACA | nonsynonymous SNV | NACA:NM_001113203:exon10:c.A2711G:p.K904R,NACA:NM_001113202:exon7:c.A581G:p.K194R, |
| chr12 | 56712844 | C | T | . | exonic | NACA | nonsynonymous SNV | NACA:NM_001113203:exon10:c.G2705A:p.R902K,NACA:NM_001113202:exon7:c.G575A:p.R192K, |
| chr12 | 56712902 | C | T | COSV63269367 | exonic | NACA | nonsynonymous SNV | NACA:NM_001113203:exon10:c.G2647A:p.E883K,NACA:NM_001113202:exon7:c.G517A:p.E173K, |
| chr12 | 56713177 | G | A | . | exonic | NACA | nonsynonymous SNV | NACA:NM_001113203:exon9:c.C2525T:p.S842F,NACA:NM_001113202:exon6:c.C395T:p.S132F, |
| chr12 | 56713551 | A | G | . | exonic | NACA | nonsynonymous SNV | NACA:NM_001113203:exon8:c.T2497C:p.F833L,NACA:NM_001113202:exon5:c.T367C:p.F123L, |
| chr12 | 56713572 | C | T | . | exonic | NACA | nonsynonymous SNV | NACA:NM_001113203:exon8:c.G2476A:p.A826T,NACA:NM_001113202:exon5:c.G346A:p.A116T, |
| chr12 | 56713638 | C | A | . | exonic | NACA | nonsynonymous SNV | NACA:NM_001113203:exon8:c.G2410T:p.V804F,NACA:NM_001113202:exon5:c.G280T:p.V94F, |
| chr12 | 56714379 | C | T | . | exonic | NACA | nonsynonymous SNV | NACA:NM_001113203:exon7:c.G2347A:p.E783K,NACA:NM_001113202:exon4:c.G217A:p.E73K, |
| chr12 | 56714670 | C | T | . | exonic | NACA | nonsynonymous SNV | NACA:NM_001113203:exon6:c.G2218A:p.D740N,NACA:NM_001113202:exon3:c.G88A:p.D30N, |

| | | | | | | | | |
|---|---|---|---|---|---|---|---|---|
| chr12 | 56724452 | C | T | . | exonic | NACA | nonsynonymous SNV | NACA:NM_001113203:exon2:c.G70A:p.A24T,NACA:NM_001113202:exon2:c.G70A:p.G24R, |
| chr12 | 56724496 | A | G | . | exonic | NACA | nonsynonymous SNV | NACA:NM_001113203:exon2:c.T26C:p.V9A,NACA:NM_001113202:exon2:c.T26C:p.V9A, |
| chr12 | 57516963 | C | T | . | exonic | DDIT3 | nonsynonymous SNV | DDIT3:NM_001195056:exon3:c.G425A:p.R142H,DDIT3:NM_001195053:exon4:c.G425A:p.R142H,DDIT3:NM_001195057:exon4:c.G356A:p.R119H,DDIT3:NM_001195055:exon4:c.G425A:p.R142H,DDIT3:NM_001195054:exon4:c.G425A:p.R142H,DDIT3:NM_004083:exon4:c.G356A:p.R119H, |
| chr12 | 57517102 | C | T | . | exonic | DDIT3 | nonsynonymous SNV | DDIT3:NM_001195056:exon3:c.G286A:p.V96I,DDIT3:NM_001195053:exon4:c.G286A:p.V96I,DDIT3:NM_001195057:exon4:c.G217A:p.V73I,DDIT3:NM_001195055:exon4:c.G286A:p.V96I,DDIT3:NM_001195054:exon4:c.G286A:p.V96I,DDIT3:NM_004083:exon4:c.G217A:p.V73I, |
| chr12 | 57517133 | C | T | . | exonic | DDIT3 | stopgain | DDIT3:NM_001195056:exon3:c.G255A:p.W85X,DDIT3:NM_001195053:exon4:c.G255A:p.W85X,DDIT3:NM_001195057:exon4:c.G186A:p.W62X,DDIT3:NM_001195055:exon4:c.G255A:p.W85X,DDIT3:NM_001195054:exon4:c.G255A:p.W85X,DDIT3:NM_004083:exon4:c.G186A:p.W62X, |
| chr12 | 57786199 | G | A | . | exonic | TSFM | nonsynonymous SNV | TSFM:NM_005726:exon3:c.G268A:p.G90S,TSFM:NM_001172697:exon3:c.G268A:p.G90S,TSFM:NM_001172696:exon3:c.G268A:p.G90S,TSFM:NM_001172695:exon3:c.G268A:p.G90S, |
| chr12 | 57796188 | G | A | . | exonic | TSFM | nonsynonymous SNV | TSFM:NM_005726:exon6:c.G583A:p.E195K,TSFM:NM_001172696:exon7:c.G646A:p.E216K, |
| chr12 | 57941843 | A | T | COSV55686289 | exonic | XRCC6BP1 | nonsynonymous SNV | XRCC6BP1:NM_033276:exon1:c.A142T:p.S48C, |
| chr12 | 57956855 | T | C | . | exonic | XRCC6BP1 | nonsynonymous SNV | XRCC6BP1:NM_033276:exon6:c.T706C:p.F236L, |
| chr12 | 6312329 | G | A | COSV50607477 | exonic | PLEKHG6 | nonsynonymous SNV | PLEKHG6:NM_018173:exon2:c.G103A:p.A35T,PLEKHG6:NM_001144856:exon2:c.G103A:p.A35T, |
| chr12 | 6333094 | C | T | . | exonic | TNFRSF1A | nonsynonymous SNV | TNFRSF1A:NM_001065:exon5:c.G526A:p.E176K, |
| chr12 | 6333108 | C | A | . | exonic | TNFRSF1A | nonsynonymous SNV | TNFRSF1A:NM_001065:exon5:c.G512T:p.G171V, |
| chr12 | 6333477 | C | T | COSV50610888 | exonic | TNFRSF1A | nonsynonymous SNV | TNFRSF1A:NM_001065:exon4:c.G362A:p.R121Q, |
| chr12 | 6347896 | T | C | COSV57434232 | exonic | SCNN1A | nonsynonymous SNV | SCNN1A:NM_001159576:exon12:c.A2164G:p.T722A,SCNN1A:NM_001038:exon13:c.A1987G:p.T663A,SCNN1A:NM_001159575:exon13:c.A2056G:p.T686A, |
| chr12 | 6534851 | G | A | . | exonic | GAPDH | nonsynonymous SNV | GAPDH:NM_001289745:exon2:c.G19A:p.G7R,GAPDH:NM_001289746:exon1:c.G19A:p.G7R,GAPDH:NM_002046:exon2:c.G19A:p.G7R, |
| chr12 | 6536591 | A | T | . | exonic | GAPDH | nonsynonymous SNV | GAPDH:NM_001256799:exon2:c.A1T:p.M1L,GAPDH:NM_001289745:exon3:c.A127T:p.M43L,GAPDH:NM_001289746:exon2:c.A127T:p.M43L,GAPDH:NM_002046:exon3:c.A127T:p.M43L, |
| chr12 | 6536756 | G | A | . | exonic | GAPDH | nonsynonymous SNV | GAPDH:NM_001256799:exon3:c.G76A:p.V26I,GAPDH:NM_001289745:exon4:c.G202A:p.V68I,GAPDH:NM_001289746:exon3:c.G202A:p.V68I,GAPDH:NM_002046:exon4:c.G202A:p.V68I, |
| chr12 | 6536957 | G | A | COSV57521174 | exonic | GAPDH | nonsynonymous SNV | GAPDH:NM_001256799:exon4:c.G148A:p.A50T,GAPDH:NM_001289745:exon5:c.G274A:p.A92T,GAPDH:NM_001289746:exon4:c.G274A:p.A92T,GAPDH:NM_002046:exon5:c.G274A:p.A92T, |
| chr12 | 6536969 | G | A | . | exonic | GAPDH | nonsynonymous SNV | GAPDH:NM_001256799:exon4:c.G160A:p.V54M,GAPDH:NM_001289745:exon5:c.G286A:p.V96M,GAPDH:NM_001289746:exon4:c.G286A:p.V96M,GAPDH:NM_002046:exon5:c.G286A:p.V96M, |
| chr12 | 6537101 | G | A | . | exonic | GAPDH | nonsynonymous SNV | GAPDH:NM_001256799:exon5:c.G202A:p.A68T,GAPDH:NM_001289745:exon6:c.G328A:p.A110T,GAPDH:NM_001289746:exon5:c.G328A:p.A110T,GAPDH:NM_002046:exon6:c.G328A:p.A110T, |
| chr12 | 6537116 | G | A | COSV57520634 | exonic | GAPDH | nonsynonymous SNV | GAPDH:NM_001256799:exon5:c.G217A:p.G73R,GAPDH:NM_001289745:exon6:c.G343A:p.G115R,GAPDH:NM_001289746:exon5:c.G343A:p.G115R,GAPDH:NM_002046:exon6:c.G343A:p.G115R, |
| chr12 | 6537140 | G | A | . | exonic | GAPDH | nonsynonymous SNV | GAPDH:NM_001256799:exon5:c.G241A:p.A81T,GAPDH:NM_001289745:exon6:c.G367A:p.A123T,GAPDH:NM_001289746:exon5:c.G367A:p.A123T,GAPDH:NM_002046:exon6:c.G367A:p.A123T, |
| chr12 | 6537158 | C | A | . | exonic | GAPDH | nonsynonymous SNV | GAPDH:NM_001256799:exon5:c.C259A:p.P87T,GAPDH:NM_001289745:exon6:c.C385A:p.P129T,GAPDH:NM_001289746:exon5:c.C385A:p.P129T,GAPDH:NM_002046:exon6:c.C385A:p.P129T, |
| chr12 | 6537167 | G | A | . | exonic | GAPDH | nonsynonymous SNV | GAPDH:NM_001256799:exon5:c.G268A:p.V90I,GAPDH:NM_001289745:exon6:c.G394A:p.V132I,GAPDH:NM_001289746:exon5:c.G394A:p.V132I,GAPDH:NM_002046:exon6:c.G394A:p.V132I, |
| chr12 | 6537216 | G | A | . | exonic | GAPDH | nonsynonymous SNV | GAPDH:NM_001256799:exon5:c.G317A:p.S106N,GAPDH:NM_001289745:exon6:c.G443A:p.S148N,GAPDH:NM_001289746:exon5:c.G443A:p.S148N,GAPDH:NM_002046:exon6:c.G443A:p.S148N, |
| chr12 | 6537361 | G | A | . | exonic | GAPDH | nonsynonymous SNV | GAPDH:NM_001256799:exon6:c.G370A:p.D124N,GAPDH:NM_001289745:exon7:c.G496A:p.D166N,GAPDH:NM_001289746:exon6:c.G496A:p.D166N,GAPDH:NM_002046:exon7:c.G496A:p.D166N, |
| chr12 | 6537660 | G | A | . | exonic | GAPDH | nonsynonymous SNV | GAPDH:NM_001256799:exon7:c.G476A:p.G159E,GAPDH:NM_001289745:exon8:c.G602A:p.G201E,GAPDH:NM_001289746:exon7:c.G602A:p.G201E,GAPDH:NM_002046:exon8:c.G602A:p.G201E, |
| chr12 | 6537734 | G | A | . | exonic | GAPDH | nonsynonymous SNV | GAPDH:NM_001256799:exon7:c.G550A:p.G184R,GAPDH:NM_001289745:exon8:c.G676A:p.G226R,GAPDH:NM_001289746:exon7:c.G676A:p.G226R,GAPDH:NM_002046:exon8:c.G676A:p.G226R, |
| chr12 | 6537820 | A | T | . | exonic | GAPDH | nonsynonymous SNV | GAPDH:NM_001256799:exon7:c.A636T:p.K212N,GAPDH:NM_001289745:exon8:c.A762T:p.K254N,GAPDH:NM_001289746:exon7:c.A762T:p.K254N,GAPDH:NM_002046:exon8:c.A762T:p.K254N, |
| chr12 | 6537821 | T | A | . | exonic | GAPDH | nonsynonymous SNV | GAPDH:NM_001256799:exon7:c.T637A:p.Y213N,GAPDH:NM_001289745:exon8:c.T763A:p.Y255N,GAPDH:NM_001289746:exon7:c.T763A:p.Y255N,GAPDH:NM_002046:exon8:c.T763A:p.Y255N, |
| chr12 | 6537872 | G | A | . | exonic | GAPDH | nonsynonymous SNV | GAPDH:NM_001256799:exon7:c.G688A:p.G230S,GAPDH:NM_001289745:exon8:c.G814A:p.G272S,GAPDH:NM_001289746:exon7:c.G814A:p.G272S,GAPDH:NM_002046:exon8:c.G814A:p.G272S, |
| chr12 | 6537873 | G | A | . | exonic | GAPDH | nonsynonymous SNV | GAPDH:NM_001256799:exon7:c.G689A:p.G230D,GAPDH:NM_001289745:exon8:c.G815A:p.G272D,GAPDH:NM_001289746:exon7:c.G815A:p.G272D,GAPDH:NM_002046:exon8:c.G815A:p.G272D, |
| chr12 | 6537882 | G | A | . | exonic | GAPDH | nonsynonymous SNV | GAPDH:NM_001256799:exon7:c.G698A:p.G233D,GAPDH:NM_001289745:exon8:c.G824A:p.G275D,GAPDH:NM_001289746:exon7:c.G824A:p.G275D,GAPDH:NM_002046:exon8:c.G824A:p.G275D, |
| chr12 | 66152320 | A | G | . | exonic | TMBIM4 | nonsynonymous SNV | TMBIM4:NM_001282606:exon4:c.T404C:p.I135T,TMBIM4:NM_016056:exon3:c.T263C:p.I88T,TMBIM4:NM_001282610:exon3:c.T170C:p.I57T,TMBIM4:NM_001282609:exon3:c.T263C:p.I88T, |
| chr12 | 68726526 | T | C | . | exonic | NUP107 | nonsynonymous SNV | NUP107:NM_020401:exon19:c.T1604C:p.L535P, |
| chr12 | 68733472 | G | A | . | exonic | NUP107 | nonsynonymous SNV | NUP107:NM_020401:exon24:c.G2122A:p.A708T, |
| chr12 | 68741848 | G | A | . | exonic | NUP107 | nonsynonymous SNV | NUP107:NM_020401:exon27:c.G2538A:p.M846I, |
| chr12 | 68867041 | C | T | . | exonic | CPM | nonsynonymous SNV | CPM:NM_001874:exon7:c.G795A:p.M265I,CPM:NM_198320:exon7:c.G795A:p.M265I,CPM:NM_001005502:exon7:c.G795A:p.M265I, |
| chr12 | 6912921 | T | A | COSV50387059 | exonic | LRRC23 | nonsynonymous SNV | LRRC23:NM_001135217:exon7:c.T950A:p.V317E,LRRC23:NM_201650:exon7:c.T950A:p.V317E, |
| chr12 | 6913959 | A | G | COSV50386845 | exonic | LRRC23 | nonsynonymous SNV | LRRC23:NM_006992:exon7:c.A827G:p.E276G, |
| chr12 | 69251270 | G | A | . | exonic | CPSF6 | nonsynonymous SNV | CPSF6:NM_007007:exon2:c.G202A:p.A68T,CPSF6:NM_001300947:exon2:c.G202A:p.A68T, |
| chr12 | 69253133 | G | A | . | exonic | CPSF6 | nonsynonymous SNV | CPSF6:NM_007007:exon3:c.G353A:p.R118Q,CPSF6:NM_001300947:exon3:c.G353A:p.R118Q, |
| chr12 | 69257786 | G | A | . | exonic | CPSF6 | nonsynonymous SNV | CPSF6:NM_001300947:exon5:c.G575A:p.S192N,CPSF6:NM_007007:exon5:c.G575A:p.S192N, |
| chr12 | 69257824 | G | A | . | exonic | CPSF6 | nonsynonymous SNV | CPSF6:NM_001300947:exon5:c.G613A:p.G205S,CPSF6:NM_007007:exon5:c.G613A:p.G205S, |
| chr12 | 69257828 | G | T | . | exonic | CPSF6 | nonsynonymous SNV | CPSF6:NM_007007:exon5:c.G617T:p.R206L,CPSF6:NM_001300947:exon5:c.G617T:p.R206L, |
| chr12 | 69257831 | T | C | . | exonic | CPSF6 | nonsynonymous SNV | CPSF6:NM_001300947:exon5:c.T620C:p.F207S,CPSF6:NM_007007:exon5:c.T620C:p.F207S, |
| chr12 | 69258629 | G | A | . | exonic | CPSF6 | nonsynonymous SNV | CPSF6:NM_007007:exon6:c.G734A:p.G245D,CPSF6:NM_001300947:exon7:c.G845A:p.G282D, |
| chr12 | 69258638 | G | A | . | exonic | CPSF6 | nonsynonymous SNV | CPSF6:NM_007007:exon6:c.G743A:p.G248D,CPSF6:NM_001300947:exon7:c.G854A:p.G285D, |
| chr12 | 69258950 | C | A | . | exonic | CPSF6 | nonsynonymous SNV | CPSF6:NM_007007:exon6:c.C1055A:p.P352Q,CPSF6:NM_001300947:exon7:c.C1166A:p.P389Q, |
| chr12 | 69259520 | G | A | . | exonic | CPSF6 | nonsynonymous SNV | CPSF6:NM_001300947:exon8:c.G1403A:p.R468K,CPSF6:NM_007007:exon7:c.G1292A:p.R431K, |
| chr12 | 69259543 | G | A | . | exonic | CPSF6 | nonsynonymous SNV | CPSF6:NM_001300947:exon8:c.G1426A:p.G476S,CPSF6:NM_007007:exon7:c.G1315A:p.G439S, |
| chr12 | 69262411 | G | A | COSV57012371 | exonic | CPSF6 | nonsynonymous SNV | CPSF6:NM_001300947:exon10:c.G1619A:p.R540Q,CPSF6:NM_007007:exon9:c.G1508A:p.R503Q, |

| | | | | | | | | |
|---|---|---|---|---|---|---|---|---|
| chr12 | 69262413 | G | A | . | exonic | CPSF6 | nonsynonymous SNV | CPSF6:NM_001300947:exon10:c.G1621A:p.E541K,CPSF6:NM_007007:exon9:c.G1510A:p.E504K, |
| chr12 | 69262446 | G | A | . | exonic | CPSF6 | nonsynonymous SNV | CPSF6:NM_007007:exon9:c.G1543A:p.D515N,CPSF6:NM_001300947:exon10:c.G1654A:p.D552N, |
| chr12 | 69262455 | G | A | COSV57012330 | exonic | CPSF6 | nonsynonymous SNV | CPSF6:NM_001300947:exon10:c.G1663A:p.D555N,CPSF6:NM_007007:exon9:c.G1552A:p.D518N, |
| chr12 | 69262458 | G | A | COSV57013366 | exonic | CPSF6 | nonsynonymous SNV | CPSF6:NM_001300947:exon10:c.G1666A:p.D556N,CPSF6:NM_007007:exon9:c.G1555A:p.D519N, |
| chr12 | 69262500 | G | A | . | exonic | CPSF6 | nonsynonymous SNV | CPSF6:NM_007007:exon9:c.G1597A:p.D533N,CPSF6:NM_001300947:exon10:c.G1708A:p.D570N, |
| chr12 | 69262555 | G | A | . | exonic | CPSF6 | nonsynonymous SNV | CPSF6:NM_001300947:exon10:c.G1763A:p.R588H,CPSF6:NM_007007:exon9:c.G1652A:p.R551H, |
| chr12 | 69348466 | G | A | . | exonic | LYZ | nonsynonymous SNV | LYZ:NM_000239:exon1:c.G58A:p.V20I, |
| chr12 | 69348472 | G | A | . | exonic | LYZ | nonsynonymous SNV | LYZ:NM_000239:exon1:c.G64A:p.E22K, |
| chr12 | 69348476 | G | A | . | exonic | LYZ | nonsynonymous SNV | LYZ:NM_000239:exon1:c.G68A:p.R23K, |
| chr12 | 69348538 | G | A | . | exonic | LYZ | nonsynonymous SNV | LYZ:NM_000239:exon1:c.G130A:p.A44T, |
| chr12 | 69350109 | G | A | . | exonic | LYZ | stopgain | LYZ:NM_000239:exon2:c.G138A:p.W46X, |
| chr12 | 69350147 | G | A | . | exonic | LYZ | nonsynonymous SNV | LYZ:NM_000239:exon2:c.G176A:p.R59Q, |
| chr12 | 69350195 | T | C | . | exonic | LYZ | nonsynonymous SNV | LYZ:NM_000239:exon2:c.T224C:p.F75S, |
| chr12 | 69350242 | G | A | . | exonic | LYZ | nonsynonymous SNV | LYZ:NM_000239:exon2:c.G271A:p.A91T, |
| chr12 | 69353162 | G | A | . | exonic | LYZ | stopgain | LYZ:NM_000239:exon4:c.G390A:p.W130X, |
| chr12 | 69362831 | G | A | . | exonic | YEATS4 | nonsynonymous SNV | YEATS4:NM_001300950:exon2:c.G95A:p.R32Q,YEATS4:NM_006530:exon2:c.G95A:p.R32Q, |
| chr12 | 6951727 | G | A | . | exonic | PTPN6 | nonsynonymous SNV | PTPN6:NM_080549:exon2:c.G127A:p.V43I,PTPN6:NM_080548:exon2:c.G133A:p.V45I,PTPN6:NM_002831:exon2:c.G127A:p.V43I, |
| chr12 | 6956514 | G | A | . | exonic | PTPN6 | stopgain | PTPN6:NM_080549:exon9:c.G1020A:p.W340X,PTPN6:NM_080548:exon9:c.G1026A:p.W342X,PTPN6:NM_002831:exon9:c.G1020A:p.W340X, |
| chr12 | 69655049 | T | C | COSV58299766 | exonic | BEST3 | nonsynonymous SNV | BEST3:NM_032735:exon10:c.A1865G:p.E622G,BEST3:NM_001282613:exon8:c.A1547G:p.E516G,BEST3:NM_152439:exon6:c.A1226G:p.E409G, |
| chr12 | 70701175 | C | T | . | exonic | PTPRR | nonsynonymous SNV | PTPRR:NM_001207016:exon4:c.G538A:p.V180I,PTPRR:NM_001207015:exon6:c.G820A:p.V274I,PTPRR:NM_130846:exon3:c.G421A:p.V141I,PTPRR:NM_002849:exon7:c.G1156A:p.V386I, |
| chr12 | 70745884 | T | C | COSV51736040 | exonic | PTPRR | nonsynonymous SNV | PTPRR:NM_001207016:exon3:c.A323G:p.K108R,PTPRR:NM_001207015:exon5:c.A605G:p.K202R,PTPRR:NM_130846:exon2:c.A206G:p.K69R,PTPRR:NM_002849:exon6:c.A941G:p.K314R, |
| chr12 | 70745915 | C | T | . | exonic | PTPRR | nonsynonymous SNV | PTPRR:NM_001207016:exon3:c.G292A:p.D98N,PTPRR:NM_001207015:exon5:c.G574A:p.D192N,PTPRR:NM_130846:exon2:c.G175A:p.D59N,PTPRR:NM_002849:exon6:c.G910A:p.D304N, |
| chr12 | 70745958 | C | T | . | exonic | PTPRR | nonsynonymous SNV | PTPRR:NM_001207016:exon3:c.G249A:p.M83I,PTPRR:NM_001207015:exon5:c.G531A:p.M177I,PTPRR:NM_130846:exon2:c.G132A:p.M44I,PTPRR:NM_002849:exon6:c.G867A:p.M289I, |
| chr12 | 71129354 | A | C | COSV56081147 | exonic | TSPAN8 | nonsynonymous SNV | TSPAN8:NM_004616:exon8:c.T637G:p.S213A, |
| chr12 | 71139754 | C | G | COSV56077446 | exonic | TSPAN8 | nonsynonymous SNV | TSPAN8:NM_004616:exon4:c.G218C:p.G73A, |
| chr12 | 7652640 | C | G | COSV57548535 | exonic | APOBEC1 | nonsynonymous SNV | APOBEC1:NM_005889:exon2:c.G105C:p.M35I,APOBEC1:NM_001304566:exon4:c.G240C:p.M80I,APOBEC1:NM_001644:exon3:c.G240C:p.M80I, |
| chr12 | 88046899 | G | C | COSV58354252 | exonic | C12ORF29 | nonsynonymous SNV | C12orf29:NM_001009894:exon6:c.G712C:p.V238L, |
| chr12 | 93678803 | G | A | COSV60554501 | exonic | CRADD | nonsynonymous SNV | CRADD:NM_003805:exon2:c.G29A:p.R10H, |
| chr12 | 93678886 | G | A | . | exonic | CRADD | nonsynonymous SNV | CRADD:NM_003805:exon2:c.G112A:p.E38K, |
| chr13 | 110641568 | T | G | . | exonic | CARS2 | nonsynonymous SNV | CARS2:NM_024537:exon15:c.A1664C:p.Q555P, |
| chr13 | 114247004 | G | A | . | exonic | CDC16 | nonsynonymous SNV | CDC16:NM_001078645:exon11:c.G971A:p.S324N,CDC16:NM_003903:exon11:c.G971A:p.S324N, |
| chr13 | 114286386 | G | A | . | exonic | UPF3A | nonsynonymous SNV | UPF3A:NM_023011:exon4:c.G506A:p.G169E, |
| chr13 | 114286524 | G | A | . | exonic | UPF3A | nonsynonymous SNV | UPF3A:NM_023011:exon5:c.G526A:p.E176K,UPF3A:NM_080687:exon4:c.G427A:p.E143K, |
| chr13 | 114286552 | G | A | COSV60896282 | exonic | UPF3A | nonsynonymous SNV | UPF3A:NM_023011:exon5:c.G554A:p.C185Y,UPF3A:NM_080687:exon4:c.G455A:p.C152Y, |
| chr13 | 114291534 | T | A | . | exonic | UPF3A | stopgain | UPF3A:NM_023011:exon6:c.T677A:p.L226X,UPF3A:NM_080687:exon5:c.T578A:p.L193X, |
| chr13 | 114291734 | G | A | COSV60894561 | exonic | UPF3A | nonsynonymous SNV | UPF3A:NM_023011:exon7:c.G788A:p.C263Y,UPF3A:NM_080687:exon6:c.G689A:p.C230Y, |
| chr13 | 114298886 | C | A | . | exonic | UPF3A | nonsynonymous SNV | UPF3A:NM_023011:exon8:c.C893A:p.P298Q,UPF3A:NM_080687:exon7:c.C794A:p.P265Q, |
| chr13 | 21382981 | G | A | COSV99059267 | exonic | ZDHHC20 | nonsynonymous SNV | ZDHHC20:NM_153251:exon10:c.C883T:p.R295C,ZDHHC20:NM_001286638:exon9:c.C694T:p.R232C, |
| chr13 | 21493305 | C | T | . | exonic | MICU2 | nonsynonymous SNV | MICU2:NM_152726:exon12:c.G1249A:p.E417K, |
| chr13 | 21495228 | C | T | . | exonic | MICU2 | nonsynonymous SNV | MICU2:NM_152726:exon11:c.G1133A:p.G378D, |
| chr13 | 21502979 | C | T | . | exonic | MICU2 | nonsynonymous SNV | MICU2:NM_152726:exon9:c.G880A:p.E294K, |
| chr13 | 21510035 | G | T | . | exonic | MICU2 | nonsynonymous SNV | MICU2:NM_152726:exon8:c.C730A:p.Q244K, |
| chr13 | 21521269 | C | T | . | exonic | MICU2 | nonsynonymous SNV | MICU2:NM_152726:exon6:c.G573A:p.M191I, |
| chr13 | 21603944 | C | T | . | exonic | MICU2 | nonsynonymous SNV | MICU2:NM_152726:exon1:c.G205A:p.A69T, |
| chr13 | 23836316 | G | A | . | exonic | MIPEP | nonsynonymous SNV | MIPEP:NM_005932:exon14:c.C1577T:p.P526L, |
| chr13 | 23836320 | C | A | . | exonic | MIPEP | nonsynonymous SNV | MIPEP:NM_005932:exon14:c.G1573T:p.V525F, |
| chr13 | 23836331 | T | C | . | exonic | MIPEP | nonsynonymous SNV | MIPEP:NM_005932:exon14:c.A1562G:p.D521G, |

| | | | | | | | | |
|---|---|---|---|---|---|---|---|---|
| chr13 | 23837633 | T | C | . | exonic | MIPEP | nonsynonymous SNV | MIPEP:NM_005932:exon13:c.A1462G:p.S488G, |
| chr13 | 23841429 | C | T | . | exonic | MIPEP | nonsynonymous SNV | MIPEP:NM_005932:exon11:c.G1166A:p.G389D, |
| chr13 | 27434894 | G | C | COSV63528570 | exonic | GTF3A | nonsynonymous SNV | GTF3A:NM_002097:exon7:c.G733C:p.V245L, |
| chr13 | 27435783 | G | C | COSV63528713 | exonic | MTIF3 | nonsynonymous SNV | MTIF3:NM_001166261:exon5:c.C729G:p.F243L,MTIF3:NM_152912:exon5:c.C729G:p.F243L,MTIF3:NM_001166262:exon5:c.C729G:p.F243L,MTIF3:NM_001166263:exon6:c.C729G:p.F243L, |
| chr13 | 27440036 | T | C | . | exonic | MTIF3 | nonsynonymous SNV | MTIF3:NM_001166261:exon3:c.A413G:p.Q138R,MTIF3:NM_001166262:exon3:c.A413G:p.Q138R,MTIF3:NM_001166263:exon4:c.A413G:p.Q138R,MTIF3:NM_152912:exon3:c.A413G:p.Q138R, |
| chr13 | 27665862 | A | G | . | exonic | POLR1D | nonsynonymous SNV | POLR1D:NM_152705:exon3:c.A278G:p.H93R,POLR1D:NM_001206559:exon3:c.A194G:p.H65R, |
| chr13 | 27963180 | G | A | COSV66829935 | exonic | CDX2 | nonsynonymous SNV | CDX2:NM_001265:exon3:c.C877T:p.P293S, |
| chr13 | 31155639 | T | G | COSV99062651 | exonic | HSPH1 | nonsynonymous SNV | HSPH1:NM_001286505:exon3:c.A187C:p.N63H,HSPH1:NM_001286503:exon3:c.A181C:p.N61H,HSPH1:NM_006644:exon3:c.A181C:p.N61H,HSPH1:NM_001286504:exon3:c.A187C:p.N63H, |
| chr13 | 32332343 | A | C | COSV66448377 | exonic | BRCA2 | nonsynonymous SNV | BRCA2:NM_000059:exon10:c.A865C:p.N289H, |
| chr13 | 32332521 | T | A | . | exonic | BRCA2 | nonsynonymous SNV | BRCA2:NM_000059:exon10:c.T1043A:p.V348E, |
| chr13 | 32332592 | A | C | COSV66448817 | exonic | BRCA2 | nonsynonymous SNV | BRCA2:NM_000059:exon10:c.A1114C:p.N372H, |
| chr13 | 32333238 | C | G | . | exonic | BRCA2 | nonsynonymous SNV | BRCA2:NM_000059:exon10:c.C1760G:p.T587R, |
| chr13 | 32333278 | T | A | . | exonic | BRCA2 | stopgain | BRCA2:NM_000059:exon10:c.T1800A:p.Y600X, |
| chr13 | 32337303 | C | A | . | exonic | BRCA2 | nonsynonymous SNV | BRCA2:NM_000059:exon11:c.C2948A:p.P983Q, |
| chr13 | 32770436 | G | A | . | exonic | PDS5B | nonsynonymous SNV | PDS5B:NM_015032:exon32:c.G3940A:p.G1314R, |
| chr13 | 37005941 | C | A | COSV99050029 | exonic | EXOSC8 | nonsynonymous SNV | EXOSC8:NM_181503:exon6:c.C260A:p.P87H, |
| chr13 | 39655754 | G | A | COSV62995992 | exonic | COG6 | nonsynonymous SNV | COG6:NM_020751:exon1:c.G28A:p.A10T,COG6:NM_001145079:exon1:c.G28A:p.A10T, |
| chr13 | 39655820 | T | A | COSV62995068 | exonic | COG6 | nonsynonymous SNV | COG6:NM_020751:exon1:c.T94A:p.C32S,COG6:NM_001145079:exon1:c.T94A:p.C32S, |
| chr13 | 39694708 | T | G | . | exonic | COG6 | nonsynonymous SNV | COG6:NM_020751:exon12:c.T1149G:p.F383L,COG6:NM_001145079:exon12:c.T1149G:p.F383L, |
| chr13 | 40941150 | T | A | COSV53497091 | exonic | ELF1 | nonsynonymous SNV | ELF1:NM_001145353:exon7:c.A955T:p.T319S,ELF1:NM_172373:exon8:c.A1027T:p.T343S, |
| chr13 | 40958916 | T | C | COSV53497107 | exonic | ELF1 | nonsynonymous SNV | ELF1:NM_001145353:exon2:c.A173G:p.N58S,ELF1:NM_172373:exon3:c.A173G:p.N58S, |
| chr13 | 45338712 | C | T | . | exonic | TPT1 | nonsynonymous SNV | TPT1:NM_003295:exon5:c.G464A:p.G155D,TPT1:NM_001286272:exon5:c.G464A:p.G155D,TPT1:NM_001286273:exon4:c.G362A:p.G121D, |
| chr13 | 45338721 | C | T | . | exonic | TPT1 | nonsynonymous SNV | TPT1:NM_003295:exon5:c.G455A:p.R152H,TPT1:NM_001286272:exon5:c.G455A:p.R152H,TPT1:NM_001286273:exon4:c.G353A:p.R118H, |
| chr13 | 45338737 | C | T | . | exonic | TPT1 | nonsynonymous SNV | TPT1:NM_003295:exon5:c.G439A:p.A147T,TPT1:NM_001286272:exon5:c.G439A:p.A147T,TPT1:NM_001286273:exon4:c.G337A:p.A113T, |
| chr13 | 45339510 | A | G | . | exonic | TPT1 | nonsynonymous SNV | TPT1:NM_003295:exon4:c.T386C:p.F129S,TPT1:NM_001286272:exon4:c.T386C:p.F129S,TPT1:NM_001286273:exon3:c.T284C:p.F95S, |
| chr13 | 45339541 | C | T | . | exonic | TPT1 | nonsynonymous SNV | TPT1:NM_003295:exon4:c.G355A:p.A119T,TPT1:NM_001286272:exon4:c.G355A:p.A119T,TPT1:NM_001286273:exon3:c.G253A:p.A85T, |
| chr13 | 45339594 | C | T | . | exonic | TPT1 | nonsynonymous SNV | TPT1:NM_003295:exon4:c.G302A:p.G101E,TPT1:NM_001286272:exon4:c.G302A:p.G101E,TPT1:NM_001286273:exon3:c.G200A:p.G67E, |
| chr13 | 45340076 | C | T | . | exonic | TPT1 | nonsynonymous SNV | TPT1:NM_003295:exon3:c.G211A:p.D71N,TPT1:NM_001286272:exon3:c.G211A:p.D71N,TPT1:NM_001286273:exon2:c.G109A:p.D37N, |
| chr13 | 45340082 | C | T | . | exonic | TPT1 | nonsynonymous SNV | TPT1:NM_003295:exon3:c.G205A:p.G69S,TPT1:NM_001286272:exon3:c.G205A:p.G69S,TPT1:NM_001286273:exon2:c.G103A:p.G35S, |
| chr13 | 45340100 | C | T | . | exonic | TPT1 | nonsynonymous SNV | TPT1:NM_003295:exon3:c.G187A:p.E63K,TPT1:NM_001286272:exon3:c.G187A:p.E63K,TPT1:NM_001286273:exon2:c.G85A:p.E29K, |
| chr13 | 45340174 | C | T | . | exonic | TPT1 | nonsynonymous SNV | TPT1:NM_003295:exon3:c.G113A:p.R38K,TPT1:NM_001286272:exon3:c.G113A:p.R38K,TPT1:NM_001286273:exon2:c.G11A:p.R4K, |
| chr13 | 45341067 | C | T | COSV58539175 | exonic | TPT1 | nonsynonymous SNV | TPT1:NM_003295:exon1:c.G3A:p.M1I,TPT1:NM_001286272:exon1:c.G3A:p.M1I, |
| chr13 | 45969538 | G | T | . | exonic | ZC3H13 | nonsynonymous SNV | ZC3H13:NM_015070:exon14:c.C3006A:p.S1002R, |
| chr13 | 47973332 | A | T | . | exonic | SUCLA2 | nonsynonymous SNV | SUCLA2:NM_003850:exon5:c.T595A:p.S199T, |
| chr13 | 48077198 | C | T | . | exonic | MED4 | nonsynonymous SNV | MED4:NM_001270629:exon7:c.G616A:p.E206K,MED4:NM_014166:exon7:c.G754A:p.E252K, |
| chr13 | 48079895 | C | T | . | exonic | MED4 | nonsynonymous SNV | MED4:NM_001270629:exon6:c.G451A:p.G151S,MED4:NM_014166:exon6:c.G589A:p.G197S, |
| chr13 | 48079921 | C | T | . | exonic | MED4 | nonsynonymous SNV | MED4:NM_001270629:exon6:c.G425A:p.G142D,MED4:NM_014166:exon6:c.G563A:p.G188D, |
| chr13 | 48079975 | C | T | . | exonic | MED4 | nonsynonymous SNV | MED4:NM_014166:exon6:c.G509A:p.G170E,MED4:NM_001270629:exon6:c.G371A:p.G124E, |
| chr13 | 48081668 | C | T | . | exonic | MED4 | nonsynonymous SNV | MED4:NM_014166:exon5:c.G485A:p.C162Y,MED4:NM_001270629:exon5:c.G347A:p.C116Y, |
| chr13 | 48083379 | G | T | . | exonic | MED4 | nonsynonymous SNV | MED4:NM_001270629:exon4:c.C275A:p.A92E,MED4:NM_014166:exon4:c.C413A:p.A138E, |
| chr13 | 48083380 | C | T | . | exonic | MED4 | nonsynonymous SNV | MED4:NM_001270629:exon4:c.G274A:p.A92T,MED4:NM_014166:exon4:c.G412A:p.A138T, |
| chr13 | 48086358 | A | G | . | exonic | MED4 | nonsynonymous SNV | MED4:NM_001270629:exon3:c.T149C:p.L50S,MED4:NM_014166:exon3:c.T287C:p.L96S, |
| chr13 | 48086366 | C | T | . | exonic | MED4 | nonsynonymous SNV | MED4:NM_001270629:exon3:c.G141A:p.M47I,MED4:NM_014166:exon3:c.G279A:p.M93I, |
| chr13 | 48094997 | C | T | . | exonic | MED4 | nonsynonymous SNV | MED4:NM_014166:exon1:c.G82A:p.E28K, |
| chr13 | 48362896 | T | A | . | exonic | RB1 | nonsynonymous SNV | RB1:NM_000321:exon8:c.T800A:p.L267Q, |
| chr13 | 48367538 | T | A | . | exonic | RB1 | nonsynonymous SNV | RB1:NM_000321:exon10:c.T984A:p.N328K, |
| chr13 | 48480024 | C | A | . | exonic | RB1 | nonsynonymous SNV | RB1:NM_000321:exon27:c.C2740A:p.Q914K, |
| chr13 | 50020311 | G | C | . | exonic | KCNRG | nonsynonymous SNV | KCNRG:NM_173605:exon2:c.G676C:p.V226L, |
| chr13 | 52461790 | A | G | . | exonic | CKAP2 | nonsynonymous SNV | CKAP2:NM_001286686:exon4:c.A820G:p.I274V,CKAP2:NM_018204:exon4:c.A964G:p.I322V,CKAP2:NM_001286687:exon4:c.A964G:p.I322V,CKAP2:NM_001098525:exon4:c.A967G:p.I323V, |
| chr13 | 72727590 | T | C | COSV64695018 | exonic | MZT1 | nonsynonymous SNV | MZT1:NM_001071775:exon1:c.A13G:p.S5G, |
| chr13 | 72795504 | A | G | COSV58323477 | exonic | PIBF1 | nonsynonymous SNV | PIBF1:NM_006346:exon4:c.A499G:p.I167V, |
| chr13 | 72821865 | G | A | . | exonic | PIBF1 | nonsynonymous SNV | PIBF1:NM_006346:exon6:c.G689A:p.R230Q, |
| chr13 | 72893887 | G | A | . | exonic | PIBF1 | nonsynonymous SNV | PIBF1:NM_006346:exon11:c.G1426A:p.E476K, |
| chr13 | 75849112 | T | C | COSV58530213 | exonic | LMO7 | nonsynonymous SNV | LMO7:NM_001306080:exon27:c.T4184C:p.M1395T,LMO7:NM_015842:exon24:c.T3485C:p.M1162T,LMO7:NM_005358:exon26:c.T3338C:p.M1113T, |
| chr13 | 77613943 | A | G | COSV62993129 | exonic | SCEL | nonsynonymous SNV | SCEL:NM_003843:exon23:c.A1379G:p.K460R,SCEL:NM_001160706:exon22:c.A1313G:p.K438R,SCEL:NM_144777:exon24:c.A1439G:p.K480R, |
| chr14 | 102081731 | C | T | . | exonic | HSP90AA1 | nonsynonymous SNV | HSP90AA1:NM_005348:exon11:c.G2180A:p.R727H,HSP90AA1:NM_001017963:exon12:c.G2546A:p.R849H, |
| chr14 | 102083272 | G | T | . | exonic | HSP90AA1 | nonsynonymous SNV | HSP90AA1:NM_005348:exon9:c.C1517A:p.A506D,HSP90AA1:NM_001017963:exon10:c.C1883A:p.A628D, |
| chr14 | 102083681 | C | T | . | exonic | HSP90AA1 | nonsynonymous SNV | HSP90AA1:NM_005348:exon8:c.G1351A:p.E451K,HSP90AA1:NM_001017963:exon9:c.G1717A:p.E573K, |
| chr14 | 102083841 | A | T | . | exonic | HSP90AA1 | nonsynonymous SNV | HSP90AA1:NM_005348:exon7:c.T1290A:p.D430E,HSP90AA1:NM_001017963:exon8:c.T1656A:p.D552E, |
| chr14 | 102083930 | C | T | . | exonic | HSP90AA1 | nonsynonymous SNV | HSP90AA1:NM_005348:exon7:c.G1201A:p.E401K,HSP90AA1:NM_001017963:exon8:c.G1567A:p.E523K, |
| chr14 | 102083972 | C | T | . | exonic | HSP90AA1 | nonsynonymous SNV | HSP90AA1:NM_005348:exon7:c.G1159A:p.G387R,HSP90AA1:NM_001017963:exon8:c.G1525A:p.G509R, |
| chr14 | 102084408 | C | T | . | exonic | HSP90AA1 | nonsynonymous SNV | HSP90AA1:NM_005348:exon6:c.G1138A:p.E380K,HSP90AA1:NM_001017963:exon7:c.G1504A:p.E502K, |
| chr14 | 102084449 | C | T | . | exonic | HSP90AA1 | nonsynonymous SNV | HSP90AA1:NM_005348:exon6:c.G1097A:p.R366H,HSP90AA1:NM_001017963:exon7:c.G1463A:p.R488H, |
| chr14 | 102084938 | C | T | . | exonic | HSP90AA1 | nonsynonymous SNV | HSP90AA1:NM_005348:exon5:c.G724A:p.E242K,HSP90AA1:NM_001017963:exon6:c.G1090A:p.E364K, |
| chr14 | 102085785 | C | T | . | exonic | HSP90AA1 | nonsynonymous SNV | HSP90AA1:NM_005348:exon3:c.G502A:p.G168R,HSP90AA1:NM_001017963:exon4:c.G868A:p.G290R, |
| chr14 | 102086242 | C | T | . | exonic | HSP90AA1 | nonsynonymous SNV | HSP90AA1:NM_005348:exon2:c.G137A:p.R46K,HSP90AA1:NM_001017963:exon3:c.G503A:p.R168K, |
| chr14 | 102102030 | T | A | COSV58233514 | exonic | HSP90AA1 | nonsynonymous SNV | HSP90AA1:NM_001017963:exon2:c.A211T:p.M71L, |
| chr14 | 102348705 | C | T | COSV53736670 | exonic | CINP | nonsynonymous SNV | CINP:NM_032630:exon5:c.G491A:p.R164H, |
| chr14 | 102359436 | G | C | COSV53736637 | exonic | CINP | nonsynonymous SNV | CINP:NM_032630:exon2:c.C159G:p.N53K, |
| chr14 | 102875712 | T | C | COSV61023607 | exonic | TRAF3 | nonsynonymous SNV | TRAF3:NM_001199427:exon4:c.T386C:p.M129T,TRAF3:NM_145725:exon5:c.T386C:p.M129T,TRAF3:NM_145726:exon5:c.T386C:p.M129T,TRAF3:NM_003300:exon4:c.T386C:p.M129T, |
| chr14 | 103915097 | C | T | . | exonic | C14ORF2 | nonsynonymous SNV | C14orf2:NM_001127393:exon3:c.G144A:p.M48I,C14orf2:NM_004894:exon2:c.G93A:p.M31I, |
| chr14 | 105488368 | C | T | COSV57635229 | exonic | CRIP1 | nonsynonymous SNV | CRIP1:NM_001311:exon4:c.C173T:p.A58V, |

| | | | | | | | | |
|---|---|---|---|---|---|---|---|---|
| chr14 | 20456995 | T | G | COSV51657755 | exonic | APEX1 | nonsynonymous SNV | APEX1:NM_080648:exon5:c.T444G:p.D148E,APEX1:NM_080649:exon5:c.T444G:p.D148E,APEX1:NM_001244249:exon5:c.T444G:p.D148E,APEX1:NM_001641:exon5:c.T444G:p.D148E, |
| chr14 | 20699417 | A | T | COSV59011747 | exonic | RNASE4 | nonsynonymous SNV | RNASE4:NM_001282193:exon2:c.A46T:p.T16S,RNASE4:NM_002937:exon2:c.A46T:p.T16S,RNASE4:NM_001282192:exon3:c.A46T:p.T16S,RNASE4:NM_194431:exon2:c.A46T:p.T16S, |
| chr14 | 22767315 | C | T | . | exonic | OXA1L | nonsynonymous SNV | OXA1L:NM_005015:exon2:c.C311T:p.A104V, |
| chr14 | 22830077 | T | C | . | exonic | MRPL52 | nonsynonymous SNV | MRPL52:NM_181307:exon2:c.T56C:p.V19A,MRPL52:NM_180982:exon2:c.T53C:p.V18A,MRPL52:NM_178336:exon2:c.T56C:p.V19A, |
| chr14 | 23953798 | G | A | COSV57481616 | exonic | DHRS4 | nonsynonymous SNV | DHRS4:NM_001282990:exon1:c.G10A:p.A4T,DHRS4:NM_001282989:exon1:c.G10A:p.A4T,DHRS4:NM_021004:exon1:c.G10A:p.A4T,DHRS4:NM_001282988:exon1:c.G10A:p.A4T,DHRS4:NM_001282987:exon1:c.G10A:p.A4T,DHRS4:NM_001282991:exon1:c.G10A:p.A4T, |
| chr14 | 23988953 | G | C | COSV58729325 | exonic | DHRS4L2 | nonsynonymous SNV | DHRS4L2:NM_198083:exon1:c.G6C:p.Q2H, |
| chr14 | 24145152 | T | G | . | exonic | PSME2 | nonsynonymous SNV | PSME2:NM_002818:exon6:c.A266C:p.H89P, |
| chr14 | 24210671 | C | T | COSV55302396 | exonic | CHMP4A | nonsynonymous SNV | CHMP4A:NM_014169:exon4:c.G586A:p.G196R, |
| chr14 | 24267755 | T | C | COSV52861986 | exonic | RABGGTA | nonsynonymous SNV | RABGGTA:NM_004581:exon13:c.A1258G:p.T420A,RABGGTA:NM_182836:exon14:c.A1258G:p.T420A, |
| chr14 | 24267920 | C | T | . | exonic | RABGGTA | nonsynonymous SNV | RABGGTA:NM_004581:exon12:c.G1186A:p.D396N,RABGGTA:NM_182836:exon13:c.G1186A:p.D396N, |
| chr14 | 24336257 | G | T | COSV53475639 | exonic | RIPK3 | nonsynonymous SNV | RIPK3:NM_006871:exon10:c.C1475A:p.P492Q, |
| chr14 | 24440156 | T | C | COSV52162406 | exonic | SDR39U1 | nonsynonymous SNV | SDR39U1:NM_001290292:exon5:c.A563G:p.Q188R,SDR39U1:NM_020195:exon6:c.A809G:p.Q270R,SDR39U1:NM_001308075:exon5:c.A563G:p.Q188R,SDR39U1:NM_001290293:exon5:c.A485G:p.Q162R, |
| chr14 | 24440269 | C | G | COSV52161794 | exonic | SDR39U1 | nonsynonymous SNV | SDR39U1:NM_001290292:exon5:c.G450C:p.L150F,SDR39U1:NM_020195:exon6:c.G696C:p.L232F,SDR39U1:NM_001308075:exon5:c.G450C:p.L150F,SDR39U1:NM_001290293:exon5:c.G372C:p.L124F, |
| chr14 | 24441767 | T | G | COSV52162442 | exonic | SDR39U1 | nonsynonymous SNV | SDR39U1:NM_020195:exon4:c.A235C:p.I79L, |
| chr14 | 34800273 | C | T | . | exonic | BAZ1A | nonsynonymous SNV | BAZ1A:NM_182648:exon9:c.G1079A:p.R360K,BAZ1A:NM_013448:exon9:c.G1079A:p.R360K, |
| chr14 | 34800320 | A | T | . | exonic | BAZ1A | nonsynonymous SNV | BAZ1A:NM_182648:exon9:c.T1032A:p.D344E,BAZ1A:NM_013448:exon9:c.T1032A:p.D344E, |
| chr14 | 34800325 | C | T | . | exonic | BAZ1A | nonsynonymous SNV | BAZ1A:NM_182648:exon9:c.G1027A:p.E343K,BAZ1A:NM_013448:exon9:c.G1027A:p.E343K, |
| chr14 | 34801128 | A | T | . | exonic | BAZ1A | nonsynonymous SNV | BAZ1A:NM_182648:exon8:c.T927A:p.D309E,BAZ1A:NM_013448:exon8:c.T927A:p.D309E, |
| chr14 | 35053359 | T | C | . | exonic | FAM177A1 | nonsynonymous SNV | FAM177A1:NM_001289022:exon3:c.T178C:p.F60L,FAM177A1:NM_001079519:exon4:c.T178C:p.F60L,FAM177A1:NM_173607:exon2:c.T247C:p.F83L, |
| chr14 | 35053368 | G | A | . | exonic | FAM177A1 | nonsynonymous SNV | FAM177A1:NM_001289022:exon3:c.G187A:p.G63S,FAM177A1:NM_001079519:exon4:c.G187A:p.G63S,FAM177A1:NM_173607:exon2:c.G256A:p.G86S, |
| chr14 | 35053431 | G | A | . | exonic | FAM177A1 | nonsynonymous SNV | FAM177A1:NM_001289022:exon3:c.G250A:p.V84I,FAM177A1:NM_001079519:exon4:c.G250A:p.V84I,FAM177A1:NM_173607:exon2:c.G319A:p.V107I, |
| chr14 | 35078972 | G | A | . | exonic | FAM177A1 | nonsynonymous SNV | FAM177A1:NM_001289022:exon5:c.G383A:p.S128N,FAM177A1:NM_001079519:exon6:c.G383A:p.S128N,FAM177A1:NM_173607:exon4:c.G452A:p.S151N, |
| chr14 | 35266761 | A | G | COSV51617748 | exonic | KIAA0391 | nonsynonymous SNV | KIAA0391:NM_001256680:exon5:c.A194G:p.N65S,KIAA0391:NM_014672:exon6:c.A1310G:p.N437S,KIAA0391:NM_001256678:exon5:c.A1262G:p.N421S,KIAA0391:NM_001256681:exon4:c.A194G:p.N65S,KIAA0391:NM_001256679:exon6:c.A1025G:p.N342S, |
| chr14 | 36320523 | T | G | COSV59253593 | exonic | MBIP | nonsynonymous SNV | MBIP:NM_001144891:exon1:c.A66C:p.R22S,MBIP:NM_016586:exon1:c.A66C:p.R22S,MBIP:NM_001308110:exon1:c.A66C:p.R22S, |
| chr14 | 36320569 | A | T | . | exonic | MBIP | nonsynonymous SNV | MBIP:NM_001144891:exon1:c.T20A:p.L7H,MBIP:NM_016586:exon1:c.T20A:p.L7H,MBIP:NM_001308110:exon1:c.T20A:p.L7H, |
| chr14 | 50312098 | A | C | COSV55413053 | exonic | L2HGDH | nonsynonymous SNV | L2HGDH:NM_024884:exon1:c.T53G:p.L18R, |
| chr14 | 50321495 | C | T | COSV55412785 | exonic | ATP5S | nonsynonymous SNV | ATP5S:NM_015684:exon2:c.C53T:p.P18L,ATP5S:NM_001003805:exon2:c.C53T:p.P18L,ATP5S:NM_001003803:exon2:c.C53T:p.P18L, |
| chr14 | 53950804 | A | G | COSV55416014 | exonic | BMP4 | nonsynonymous SNV | BMP4:NM_130850:exon4:c.T455C:p.V152A,BMP4:NM_130851:exon4:c.T455C:p.V152A,BMP4:NM_001202:exon4:c.T455C:p.V152A, |
| chr14 | 55138217 | C | A | . | exonic | LGALS3 | nonsynonymous SNV | LGALS3:NM_001177388:exon3:c.C191A:p.P64H,LGALS3:NM_002306:exon3:c.C191A:p.P64H, |
| chr14 | 55138318 | A | C | . | exonic | LGALS3 | nonsynonymous SNV | LGALS3:NM_001177388:exon3:c.A292C:p.T98P,LGALS3:NM_002306:exon3:c.A292C:p.T98P, |
| chr14 | 61720399 | G | A | COSV60188236 | exonic | HIF1A | nonsynonymous SNV | HIF1A:NM_001530:exon2:c.G53A:p.R18Q,HIF1A:NM_181054:exon2:c.G53A:p.R18Q,HIF1A:NM_001243084:exon2:c.G125A:p.R42Q, |
| chr14 | 65078029 | C | T | COSV52414939 | exonic | MAX | nonsynonymous SNV | MAX:NM_145113:exon4:c.G179A:p.R60Q,MAX:NM_145112:exon3:c.G152A:p.R51Q,MAX:NM_002382:exon4:c.G179A:p.R60Q, |
| chr14 | 65078036 | C | T | . | exonic | MAX | nonsynonymous SNV | MAX:NM_145113:exon4:c.G172A:p.A58T,MAX:NM_145112:exon3:c.G145A:p.A49T,MAX:NM_002382:exon4:c.G172A:p.A58T, |
| chr14 | 65093716 | C | T | . | exonic | MAX | nonsynonymous SNV | MAX:NM_001271069:exon2:c.G136A:p.G46R,MAX:NM_145114:exon3:c.G163A:p.G55R,MAX:NM_001271068:exon2:c.G136A:p.G46R,MAX:NM_002382:exon3:c.G163A:p.G55R,MAX:NM_145113:exon3:c.G163A:p.G55R,MAX:NM_145112:exon2:c.G136A:p.G46R,MAX:NM_197957:exon3:c.G163A:p.G55R, |
| chr14 | 65093767 | G | T | . | exonic | MAX | nonsynonymous SNV | MAX:NM_001271069:exon2:c.C85A:p.H29N,MAX:NM_145114:exon3:c.C112A:p.H38N,MAX:NM_001271068:exon2:c.C85A:p.H29N,MAX:NM_002382:exon3:c.C112A:p.H38N,MAX:NM_145113:exon3:c.C112A:p.H38N,MAX:NM_145112:exon2:c.C85A:p.H29N,MAX:NM_197957:exon3:c.C112A:p.H38N, |
| chr14 | 66924264 | A | G | . | exonic | GPHN | nonsynonymous SNV | GPHN:NM_020806:exon8:c.A800G:p.N267S, |
| chr14 | 68874936 | T | A | . | exonic | ACTN1 | nonsynonymous SNV | ACTN1:NM_001130005:exon21:c.A2587T:p.T863S,ACTN1:NM_001130004:exon22:c.A2668T:p.T890S,ACTN1:NM_001102:exon21:c.A2602T:p.T868S, |
| chr14 | 68885575 | C | T | . | exonic | ACTN1 | nonsynonymous SNV | ACTN1:NM_001130005:exon12:c.G1235A:p.G412D,ACTN1:NM_001130004:exon12:c.G1235A:p.G412D,ACTN1:NM_001102:exon12:c.G1235A:p.G412D, |
| chr14 | 68902530 | C | T | . | exonic | ACTN1 | nonsynonymous SNV | ACTN1:NM_001130005:exon8:c.G709A:p.A237T,ACTN1:NM_001130004:exon8:c.G709A:p.A237T,ACTN1:NM_001102:exon8:c.G709A:p.A237T, |
| chr14 | 69237836 | G | T | . | exonic | EXD2 | nonsynonymous SNV | EXD2:NM_001193361:exon8:c.G1554T:p.Q518H,EXD2:NM_001193363:exon8:c.G1554T:p.Q518H,EXD2:NM_001193362:exon9:c.G1554T:p.Q518H,EXD2:NM_001193360:exon9:c.G1554T:p.Q518H,EXD2:NM_018199:exon7:c.G1179T:p.Q393H, |
| chr14 | 73105860 | G | A | COSV56201910 | exonic | RBM25 | nonsynonymous SNV | RBM25:NM_021239:exon11:c.G1156A:p.E386K, |
| chr14 | 73105954 | G | A | COSV56202337 | exonic | RBM25 | nonsynonymous SNV | RBM25:NM_021239:exon11:c.G1250A:p.R417Q, |
| chr14 | 73105988 | C | A | . | exonic | RBM25 | nonsynonymous SNV | RBM25:NM_021239:exon11:c.C1284A:p.D428E, |
| chr14 | 73106253 | G | A | . | exonic | RBM25 | nonsynonymous SNV | RBM25:NM_021239:exon12:c.G1435A:p.A479T, |
| chr14 | 73110885 | T | A | . | exonic | RBM25 | nonsynonymous SNV | RBM25:NM_021239:exon15:c.T1747A:p.S583T, |
| chr14 | 73110909 | C | A | . | exonic | RBM25 | nonsynonymous SNV | RBM25:NM_021239:exon15:c.C1771A:p.Q591K, |
| chr14 | 77723227 | C | T | . | exonic | SNW1 | nonsynonymous SNV | SNW1:NM_012245:exon11:c.G1084A:p.E362K, |
| chr14 | 77731069 | C | T | . | exonic | SNW1 | nonsynonymous SNV | SNW1:NM_012245:exon10:c.G952A:p.E318K, |
| chr14 | 91802271 | T | G | . | exonic | TC2N | nonsynonymous SNV | TC2N:NM_001289134:exon4:c.A452C:p.K151T,TC2N:NM_001128595:exon4:c.A452C:p.K151T,TC2N:NM_152332:exon4:c.A452C:p.K151T,TC2N:NM_001128596:exon4:c.A452C:p.K151T, |
| chr14 | 92732735 | C | T | COSV58403407 | exonic | LGMN | nonsynonymous SNV | LGMN:NM_001008530:exon3:c.G52A:p.V18I,LGMN:NM_005606:exon2:c.G52A:p.V18I, |
| chr14 | 94378506 | T | G | COSV63345434 | exonic | SERPINA1 | nonsynonymous SNV | SERPINA1:NM_001002235:exon5:c.A1200C:p.E400D,SERPINA1:NM_001127704:exon7:c.A1200C:p.E400D,SERPINA1:NM_001127703:exon7:c.A1200C:p.E400D,SERPINA1:NM_001127700:exon5:c.A1200C:p.E400D,SERPINA1:NM_000295:exon5:c.A1200C:p.E400D,SERPINA1:NM_001127706:exon6:c.A1200C:p.E400D,SERPINA1:NM_001127705:exon7:c.A1200C:p.E400D,SERPINA1:NM_001002236:exon7:c.A1200C:p.E400D,SERPINA1:NM_001127702:exon6:c.A1200C:p.E400D,SERPINA1:NM_001127707:exon6:c.A1200C:p.E400D,SERPINA1:NM_001127701:exon7:c.A1200C:p.E400D, |
| chr14 | 94378613 | C | T | . | exonic | SERPINA1 | nonsynonymous SNV | SERPINA1:NM_001002235:exon5:c.G1093A:p.D365N,SERPINA1:NM_001127704:exon7:c.G1093A:p.D365N,SERPINA1:NM_001127703:exon7:c.G1093A:p.D365N,SERPINA1:NM_001127700:exon5:c.G1093A:p.D365N,SERPINA1:NM_000295:exon5:c.G1093A:p.D365N,SERPINA1:NM_001127706:exon6:c.G1093A:p.D365N,SERPINA1:NM_001127705:exon7:c.G1093A:p.D365N,SERPINA1:NM_001002236:exon7:c.G1093A:p.D365N,SERPINA1:NM_001127702:exon6:c.G1093A:p.D365N,SERPINA1:NM_001127707:exon6:c.G1093A:p.D365N,SERPINA1:NM_001127701:exon7:c.G1093A:p.D365N, |
| chr14 | 94379484 | C | T | . | exonic | SERPINA1 | nonsynonymous SNV | SERPINA1:NM_001002235:exon4:c.G1045A:p.A349T,SERPINA1:NM_001127704:exon6:c.G1045A:p.A349T,SERPINA1:NM_001127703:exon6:c.G1045A:p.A349T,SERPINA1:NM_001127700:exon4:c.G1045A:p.A349T,SERPINA1:NM_000295:exon4:c.G1045A:p.A349T,SERPINA1:NM_001127706:exon5:c.G1045A:p.A349T,SERPINA1:NM_001127705:exon6:c.G1045A:p.A349T,SERPINA1:NM_001002236:exon6:c.G1045A:p.A349T,SERPINA1:NM_001127702:exon5:c.G1045A:p.A349T,SERPINA1:NM_001127707:exon5:c.G1045A:p.A349T,SERPINA1:NM_001127701:exon6:c.G1045A:p.A349T, |

| | | | | | | | | |
|---|---|---|---|---|---|---|---|---|
| chr14 | 94379511 | C | T | . | exonic | SERPINA1 | nonsynonymous SNV | SERPINA1:NM_001002235:exon4:c.G1018A:p.A340T,SERPINA1:NM_001127704:exon6:c.G1018A:p.A340T,SERPINA1:NM_001127703:exon6:c.G1018A:p.A340T,SERPINA1:NM_001127700:exon4:c.G1018A:p.A340T,SERPINA1:NM_000295:exon4:c.G1018A:p.A340T,SERPINA1:NM_001127706:exon5:c.G1018A:p.A340T,SERPINA1:NM_001127705:exon6:c.G1018A:p.A340T,SERPINA1:NM_001002236:exon6:c.G1018A:p.A340T,SERPINA1:NM_001127702:exon5:c.G1018A:p.A340T,SERPINA1:NM_001127707:exon5:c.G1018A:p.A340T,SERPINA1:NM_001127701:exon6:c.G1018A:p.A340T, |
| chr14 | 94379513 | C | T | COSV63346165 | exonic | SERPINA1 | nonsynonymous SNV | SERPINA1:NM_001002235:exon4:c.G1016A:p.G339E,SERPINA1:NM_001127704:exon6:c.G1016A:p.G339E,SERPINA1:NM_001127703:exon6:c.G1016A:p.G339E,SERPINA1:NM_001127700:exon4:c.G1016A:p.G339E,SERPINA1:NM_000295:exon4:c.G1016A:p.G339E,SERPINA1:NM_001127706:exon5:c.G1016A:p.G339E,SERPINA1:NM_001127705:exon6:c.G1016A:p.G339E,SERPINA1:NM_001002236:exon6:c.G1016A:p.G339E,SERPINA1:NM_001127702:exon5:c.G1016A:p.G339E,SERPINA1:NM_001127707:exon5:c.G1016A:p.G339E,SERPINA1:NM_001127701:exon6:c.G1016A:p.G339E, |
| chr14 | 94379547 | C | T | . | exonic | SERPINA1 | nonsynonymous SNV | SERPINA1:NM_001002235:exon4:c.G982A:p.G328S,SERPINA1:NM_001127704:exon6:c.G982A:p.G328S,SERPINA1:NM_001127703:exon6:c.G982A:p.G328S,SERPINA1:NM_001127700:exon4:c.G982A:p.G328S,SERPINA1:NM_000295:exon4:c.G982A:p.G328S,SERPINA1:NM_001127706:exon5:c.G982A:p.G328S,SERPINA1:NM_001127705:exon6:c.G982A:p.G328S,SERPINA1:NM_001002236:exon6:c.G982A:p.G328S,SERPINA1:NM_001127702:exon5:c.G982A:p.G328S,SERPINA1:NM_001127707:exon5:c.G982A:p.G328S,SERPINA1:NM_001127701:exon6:c.G982A:p.G328S, |
| chr14 | 94380871 | C | T | . | exonic | SERPINA1 | nonsynonymous SNV | SERPINA1:NM_001002235:exon3:c.G917A:p.R306K,SERPINA1:NM_001127704:exon5:c.G917A:p.R306K,SERPINA1:NM_001127703:exon5:c.G917A:p.R306K,SERPINA1:NM_001127700:exon3:c.G917A:p.R306K,SERPINA1:NM_000295:exon3:c.G917A:p.R306K,SERPINA1:NM_001127706:exon4:c.G917A:p.R306K,SERPINA1:NM_001127705:exon5:c.G917A:p.R306K,SERPINA1:NM_001002236:exon5:c.G917A:p.R306K,SERPINA1:NM_001127702:exon4:c.G917A:p.R306K,SERPINA1:NM_001127707:exon4:c.G917A:p.R306K,SERPINA1:NM_001127701:exon5:c.G917A:p.R306K, |
| chr14 | 94382691 | C | T | COSV63346090 | exonic | SERPINA1 | nonsynonymous SNV | SERPINA1:NM_001002235:exon2:c.G547A:p.D183N,SERPINA1:NM_001127704:exon4:c.G547A:p.D183N,SERPINA1:NM_001127703:exon4:c.G547A:p.D183N,SERPINA1:NM_001127700:exon2:c.G547A:p.D183N,SERPINA1:NM_000295:exon2:c.G547A:p.D183N,SERPINA1:NM_001127706:exon3:c.G547A:p.D183N,SERPINA1:NM_001127705:exon4:c.G547A:p.D183N,SERPINA1:NM_001002236:exon4:c.G547A:p.D183N,SERPINA1:NM_001127702:exon3:c.G547A:p.D183N,SERPINA1:NM_001127707:exon3:c.G547A:p.D183N,SERPINA1:NM_001127701:exon4:c.G547A:p.D183N, |
| chr14 | 94382766 | C | T | . | exonic | SERPINA1 | nonsynonymous SNV | SERPINA1:NM_001002235:exon2:c.G472A:p.V158I,SERPINA1:NM_001127704:exon4:c.G472A:p.V158I,SERPINA1:NM_001127703:exon4:c.G472A:p.V158I,SERPINA1:NM_001127700:exon2:c.G472A:p.V158I,SERPINA1:NM_000295:exon2:c.G472A:p.V158I,SERPINA1:NM_001127706:exon3:c.G472A:p.V158I,SERPINA1:NM_001127705:exon4:c.G472A:p.V158I,SERPINA1:NM_001002236:exon4:c.G472A:p.V158I,SERPINA1:NM_001127702:exon3:c.G472A:p.V158I,SERPINA1:NM_001127707:exon3:c.G472A:p.V158I,SERPINA1:NM_001127701:exon4:c.G472A:p.V158I, |
| chr14 | 94382864 | C | T | COSV63345368 | exonic | SERPINA1 | nonsynonymous SNV | SERPINA1:NM_001002235:exon2:c.G374A:p.R125H,SERPINA1:NM_001127704:exon4:c.G374A:p.R125H,SERPINA1:NM_001127703:exon4:c.G374A:p.R125H,SERPINA1:NM_001127700:exon2:c.G374A:p.R125H,SERPINA1:NM_000295:exon2:c.G374A:p.R125H,SERPINA1:NM_001127706:exon3:c.G374A:p.R125H,SERPINA1:NM_001127705:exon4:c.G374A:p.R125H,SERPINA1:NM_001002236:exon4:c.G374A:p.R125H,SERPINA1:NM_001127702:exon3:c.G374A:p.R125H,SERPINA1:NM_001127707:exon3:c.G374A:p.R125H,SERPINA1:NM_001127701:exon4:c.G374A:p.R125H, |
| chr14 | 94383218 | C | T | . | exonic | SERPINA1 | stopgain | SERPINA1:NM_001002235:exon2:c.G20A:p.W7X,SERPINA1:NM_001127704:exon4:c.G20A:p.W7X,SERPINA1:NM_001127703:exon4:c.G20A:p.W7X,SERPINA1:NM_001127700:exon2:c.G20A:p.W7X,SERPINA1:NM_000295:exon2:c.G20A:p.W7X,SERPINA1:NM_001127706:exon3:c.G20A:p.W7X,SERPINA1:NM_001127705:exon4:c.G20A:p.W7X,SERPINA1:NM_001002236:exon4:c.G20A:p.W7X,SERPINA1:NM_001127702:exon3:c.G20A:p.W7X,SERPINA1:NM_001127707:exon3:c.G20A:p.W7X,SERPINA1:NM_001127701:exon4:c.G20A:p.W7X, |
| chr14 | 95129545 | C | T | . | exonic | DICER1 | nonsynonymous SNV | DICER1:NM_030621:exon8:c.G661A:p.E221K,DICER1:NM_001271282:exon6:c.G661A:p.E221K,DICER1:NM_001291628:exon6:c.G661A:p.E221K,DICER1:NM_177438:exon6:c.G661A:p.E221K,DICER1:NM_001195573:exon5:c.G661A:p.E221K, |
| chr14 | 95196631 | A | T | . | exonic | CLMN | nonsynonymous SNV | CLMN:NM_024734:exon10:c.T2575A:p.S859T, |
| chr14 | 95203149 | C | A | . | exonic | CLMN | nonsynonymous SNV | CLMN:NM_024734:exon9:c.G2200T:p.V734F, |
| chr14 | 95203707 | C | T | . | exonic | CLMN | nonsynonymous SNV | CLMN:NM_024734:exon9:c.G1642A:p.E548K, |
| chr14 | 95544087 | G | A | COSV58763541 | exonic | GLRX5 | nonsynonymous SNV | GLRX5:NM_016417:exon2:c.G436A:p.A146T, |
| chr14 | 96837806 | G | A | . | exonic | VRK1 | nonsynonymous SNV | VRK1:NM_003384:exon3:c.G205A:p.V69I, |
| chr14 | 96847336 | T | A | . | exonic | VRK1 | nonsynonymous SNV | VRK1:NM_003384:exon5:c.T366A:p.N122K, |
| chr14 | 96855270 | G | A | . | exonic | VRK1 | nonsynonymous SNV | VRK1:NM_003384:exon8:c.G623A:p.G208E, |
| chr15 | 101652345 | A | C | COSV60109095 | exonic | TM2D3 | nonsynonymous SNV | TM2D3:NM_001307960:exon1:c.T17G:p.L6R,TM2D3:NM_001308026:exon1:c.T17G:p.L6R,TM2D3:NM_078474:exon1:c.T17G:p.L6R,TM2D3:NM_025141:exon1:c.T17G:p.L6R, |
| chr15 | 40036374 | G | C | COSV51115776 | exonic | SRP14 | nonsynonymous SNV | SRP14:NM_003134:exon5:c.C370G:p.P124A,SRP14:NM_001309434:exon6:c.C226G:p.P76A, |
| chr15 | 40183821 | T | A | . | exonic | BUB1B | nonsynonymous SNV | BUB1B:NM_001211:exon6:c.T689A:p.L230Q, |
| chr15 | 40183828 | C | A | . | exonic | BUB1B | nonsynonymous SNV | BUB1B:NM_001211:exon6:c.C696A:p.S232R, |
| chr15 | 40185630 | G | A | COSV55009718 | exonic | BUB1B | nonsynonymous SNV | BUB1B:NM_001211:exon8:c.G1046A:p.R349Q, |
| chr15 | 40199659 | C | A | . | exonic | BUB1B | nonsynonymous SNV | BUB1B:NM_001211:exon10:c.C1333A:p.Q445K, |
| chr15 | 40199687 | A | G | . | exonic | BUB1B | nonsynonymous SNV | BUB1B:NM_001211:exon10:c.A1361G:p.K454R, |
| chr15 | 40199710 | G | A | COSV55013795 | exonic | BUB1B | nonsynonymous SNV | BUB1B:NM_001211:exon10:c.G1384A:p.E462K, |
| chr15 | 40356131 | G | A | . | exonic | PHGR1 | nonsynonymous SNV | PHGR1:NM_001145643:exon3:c.G77A:p.G26D, |
| chr15 | 40563011 | T | A | COSV63634548 | exonic | C15ORF57 | nonsynonymous SNV | C15orf57:NM_001080792:exon2:c.A5T:p.K2I,C15orf57:NM_001080791:exon2:c.A32T:p.K11I,C15orf57:NM_052849:exon2:c.A5T:p.K2I,C15orf57:NM_001289132:exon2:c.A5T:p.K2I, |
| chr15 | 40606445 | G | C | COSV61150583 | exonic | CASC5 | nonsynonymous SNV | CASC5:NM_144508:exon4:c.G128C:p.R43T,CASC5:NM_170589:exon4:c.G128C:p.R43T, |
| chr15 | 40611486 | A | G | COSV61154720 | exonic | CASC5 | nonsynonymous SNV | CASC5:NM_144508:exon7:c.A259G:p.T87A,CASC5:NM_170589:exon8:c.A337G:p.T113A, |
| chr15 | 40621642 | G | T | COSV61152671 | exonic | CASC5 | nonsynonymous SNV | CASC5:NM_144508:exon10:c.G1378T:p.A460S,CASC5:NM_170589:exon11:c.G1456T:p.A486S, |
| chr15 | 40621979 | T | C | COSV61151299 | exonic | CASC5 | nonsynonymous SNV | CASC5:NM_144508:exon10:c.T1715C:p.M572T,CASC5:NM_170589:exon11:c.T1793C:p.M598T, |
| chr15 | 40622992 | A | G | COSV61150613 | exonic | CASC5 | nonsynonymous SNV | CASC5:NM_144508:exon10:c.A2728G:p.R910G,CASC5:NM_170589:exon11:c.A2806G:p.R936G, |
| chr15 | 40813728 | T | G | . | exonic | ZFYVE19 | nonsynonymous SNV | ZFYVE19:NM_001077268:exon9:c.T1126G:p.S376A,ZFYVE19:NM_032850:exon10:c.T1096G:p.S366A,ZFYVE19:NM_001258421:exon9:c.T601G:p.S201A,ZFYVE19:NM_001258420:exon8:c.T922G:p.S308A, |
| chr15 | 43475576 | G | C | COSV55496034 | exonic | TP53BP1 | nonsynonymous SNV | TP53BP1:NM_005657:exon9:c.C1059G:p.D353E,TP53BP1:NM_001141980:exon9:c.C1074G:p.D358E,TP53BP1:NM_001141979:exon9:c.C1074G:p.D358E, |
| chr15 | 43599215 | G | A | . | exonic | CKMT1B | nonsynonymous SNV | CKMT1B:NM_020990:exon10:c.G1196A:p.R399H, |
| chr15 | 44711577 | G | A | . | exonic | B2M | nonsynonymous SNV | B2M:NM_004048:exon1:c.G31A:p.A11T, |
| chr15 | 44711595 | G | A | . | exonic | B2M | nonsynonymous SNV | B2M:NM_004048:exon1:c.G49A:p.G17S, |
| chr15 | 44715450 | G | A | . | exonic | B2M | nonsynonymous SNV | B2M:NM_004048:exon2:c.G95A:p.R32H, |
| chr15 | 44715489 | G | A | . | exonic | B2M | nonsynonymous SNV | B2M:NM_004048:exon2:c.G134A:p.C45Y, |
| chr15 | 44715585 | G | A | . | exonic | B2M | nonsynonymous SNV | B2M:NM_004048:exon2:c.G230A:p.S77N, |
| chr15 | 44715595 | G | A | . | exonic | B2M | stopgain | B2M:NM_004048:exon2:c.G240A:p.W80X, |
| chr15 | 44716332 | G | A | COSV62564948 | exonic | B2M | nonsynonymous SNV | B2M:NM_004048:exon3:c.G350A:p.R117Q, |
| chr15 | 45068982 | A | T | COSV51042839 | exonic | SORD | nonsynonymous SNV | SORD:NM_003104:exon7:c.A716T:p.Q239L, |
| chr15 | 45072336 | A | C | COSV51042844 | exonic | SORD | nonsynonymous SNV | SORD:NM_003104:exon8:c.A806C:p.N269T, |
| chr15 | 45116216 | C | G | . | exonic | DUOXA2 | nonsynonymous SNV | DUOXA2:NM_207581:exon3:c.C298G:p.R100G, |
| chr15 | 45676237 | T | C | COSV52940775 | exonic | SQRDL | nonsynonymous SNV | SQRDL:NM_021199:exon6:c.T791C:p.I264T,SQRDL:NM_001271213:exon7:c.T791C:p.I264T, |
| chr15 | 50449422 | G | A | . | exonic | USP8 | nonsynonymous SNV | USP8:NM_001128610:exon4:c.G272A:p.G91E,USP8:NM_005154:exon4:c.G272A:p.G91E, |
| chr15 | 50459029 | T | C | . | exonic | USP8 | nonsynonymous SNV | USP8:NM_001283049:exon3:c.T134C:p.L45P,USP8:NM_001128610:exon5:c.T365C:p.L122P,USP8:NM_005154:exon5:c.T365C:p.L122P, |
| chr15 | 50459034 | G | A | . | exonic | USP8 | nonsynonymous SNV | USP8:NM_001283049:exon3:c.G139A:p.E47K,USP8:NM_001128610:exon5:c.G370A:p.E124K,USP8:NM_005154:exon5:c.G370A:p.E124K, |
| chr15 | 50471637 | A | G | . | exonic | USP8 | nonsynonymous SNV | USP8:NM_001283049:exon6:c.A460G:p.T154A,USP8:NM_001128610:exon8:c.A691G:p.T231A,USP8:NM_005154:exon8:c.A691G:p.T231A, |
| chr15 | 50476913 | T | C | . | exonic | USP8 | nonsynonymous SNV | USP8:NM_001283049:exon7:c.T683C:p.L228P,USP8:NM_001128610:exon9:c.T914C:p.L305P,USP8:NM_005154:exon9:c.T914C:p.L305P, |

| | | | | | | | | |
|---|---|---|---|---|---|---|---|---|
| chr15 | 50476967 | G | A | COSV56168389 | exonic | USP8 | nonsynonymous SNV | USP8:NM_001283049:exon7:c.G737A:p.R246H,USP8:NM_001128610:exon9:c.G968A:p.R323H,USP8:NM_005154:exon9:c.G968A:p.R323H, |
| chr15 | 50477326 | G | A | . | exonic | USP8 | nonsynonymous SNV | USP8:NM_001283049:exon8:c.G814A:p.A272T,USP8:NM_001128610:exon10:c.G1045A:p.A349T,USP8:NM_005154:exon10:c.G1045A:p.A349T, |
| chr15 | 50481625 | C | A | . | exonic | USP8 | nonsynonymous SNV | USP8:NM_001283049:exon9:c.C1132A:p.L378I,USP8:NM_001128610:exon11:c.C1363A:p.L455I,USP8:NM_005154:exon11:c.C1363A:p.L455I, |
| chr15 | 50481664 | G | A | COSV56164169 | exonic | USP8 | nonsynonymous SNV | USP8:NM_001283049:exon9:c.G1171A:p.E391K,USP8:NM_001128610:exon11:c.G1402A:p.E468K,USP8:NM_005154:exon11:c.G1402A:p.E468K, |
| chr15 | 50481694 | C | A | . | exonic | USP8 | nonsynonymous SNV | USP8:NM_001283049:exon9:c.C1201A:p.Q401K,USP8:NM_001128610:exon11:c.C1432A:p.Q478K,USP8:NM_005154:exon11:c.C1432A:p.Q478K, |
| chr15 | 50481757 | C | A | . | exonic | USP8 | nonsynonymous SNV | USP8:NM_001283049:exon9:c.C1264A:p.Q422K,USP8:NM_001128610:exon11:c.C1495A:p.Q499K,USP8:NM_005154:exon11:c.C1495A:p.Q499K, |
| chr15 | 50481908 | C | A | . | exonic | USP8 | nonsynonymous SNV | USP8:NM_001128610:exon11:c.C1646A:p.T549K,USP8:NM_005154:exon11:c.C1646A:p.T549K,USP8:NM_001283049:exon9:c.C1415A:p.T472K, |
| chr15 | 50490506 | A | G | . | exonic | USP8 | nonsynonymous SNV | USP8:NM_001283049:exon11:c.A1897G:p.T633A,USP8:NM_001128610:exon14:c.A2215G:p.T739A,USP8:NM_005154:exon14:c.A2215G:p.T739A, |
| chr15 | 55360395 | C | T | . | exonic | CCPG1 | nonsynonymous SNV | CCPG1:NM_004748:exon8:c.G1378A:p.G460R,CCPG1:NM_001204450:exon8:c.G1378A:p.G460R,CCPG1:NM_020739:exon8:c.G1378A:p.G460R, |
| chr15 | 55360686 | G | T | . | exonic | CCPG1 | nonsynonymous SNV | CCPG1:NM_004748:exon8:c.C1087A:p.Q363K,CCPG1:NM_001204450:exon8:c.C1087A:p.Q363K,CCPG1:NM_020739:exon8:c.C1087A:p.Q363K, |
| chr15 | 55377003 | A | G | . | exonic | CCPG1 | nonsynonymous SNV | CCPG1:NM_001204451:exon5:c.T400C:p.F134L,CCPG1:NM_004748:exon5:c.T400C:p.F134L,CCPG1:NM_001204450:exon5:c.T400C:p.F134L,CCPG1:NM_020739:exon5:c.T400C:p.F134L, |
| chr15 | 55377045 | G | T | . | exonic | CCPG1 | nonsynonymous SNV | CCPG1:NM_001204451:exon5:c.C358A:p.Q120K,CCPG1:NM_020739:exon5:c.C358A:p.Q120K,CCPG1:NM_004748:exon5:c.C358A:p.Q120K,CCPG1:NM_001204450:exon5:c.C358A:p.Q120K, |
| chr15 | 58545811 | A | G | COSV54420926 | exonic | LIPC | nonsynonymous SNV | LIPC:NM_000236:exon5:c.A644G:p.N215S, |
| chr15 | 58560880 | C | A | . | exonic | LIPC | nonsynonymous SNV | LIPC:NM_000236:exon7:c.C1068A:p.F356L, |
| chr15 | 63059612 | G | A | . | exonic | TPM1 | nonsynonymous SNV | TPM1:NM_001301289:exon3:c.G316A:p.E106K,TPM1:NM_000366:exon4:c.G424A:p.E142K,TPM1:NM_001018004:exon4:c.G424A:p.E142K,TPM1:NM_001018020:exon4:c.G424A:p.E142K,TPM1:NM_001018008:exon3:c.G316A:p.E106K,TPM1:NM_001301244:exon4:c.G424A:p.E142K,TPM1:NM_001018007:exon4:c.G424A:p.E142K,TPM1:NM_001018006:exon4:c.G424A:p.E142K,TPM1:NM_001018005:exon4:c.G424A:p.E142K, |
| chr15 | 63059660 | G | A | . | exonic | TPM1 | nonsynonymous SNV | TPM1:NM_001301289:exon3:c.G364A:p.A122T,TPM1:NM_001018020:exon4:c.G472A:p.A158T,TPM1:NM_001018008:exon3:c.G364A:p.A122T,TPM1:NM_001301244:exon4:c.G472A:p.A158T,TPM1:NM_000366:exon4:c.G472A:p.A158T,TPM1:NM_001018007:exon4:c.G472A:p.A158T,TPM1:NM_001018006:exon4:c.G472A:p.A158T,TPM1:NM_001018005:exon4:c.G472A:p.A158T,TPM1:NM_001018004:exon4:c.G472A:p.A158T, |
| chr15 | 63060876 | G | A | . | exonic | TPM1 | nonsynonymous SNV | TPM1:NM_001301289:exon4:c.G392A:p.R131H,TPM1:NM_000366:exon5:c.G500A:p.R167H,TPM1:NM_001018004:exon5:c.G500A:p.R167H,TPM1:NM_001018020:exon5:c.G500A:p.R167H,TPM1:NM_001018008:exon4:c.G392A:p.R131H,TPM1:NM_001301244:exon5:c.G500A:p.R167H,TPM1:NM_001018007:exon5:c.G500A:p.R167H,TPM1:NM_001018006:exon5:c.G500A:p.R167H,TPM1:NM_001018005:exon5:c.G500A:p.R167H, |
| chr15 | 63060909 | G | A | . | exonic | TPM1 | nonsynonymous SNV | TPM1:NM_001301289:exon4:c.G425A:p.R142H,TPM1:NM_001018020:exon5:c.G533A:p.R178H,TPM1:NM_001018008:exon4:c.G425A:p.R142H,TPM1:NM_001301244:exon5:c.G533A:p.R178H,TPM1:NM_000366:exon5:c.G533A:p.R178H,TPM1:NM_001018007:exon5:c.G533A:p.R178H,TPM1:NM_001018006:exon5:c.G533A:p.R178H,TPM1:NM_001018005:exon5:c.G533A:p.R178H,TPM1:NM_001018004:exon5:c.G533A:p.R178H, |
| chr15 | 63061262 | G | A | . | exonic | TPM1 | nonsynonymous SNV | TPM1:NM_000366:exon6:c.G628A:p.A210T,TPM1:NM_001018020:exon6:c.G628A:p.A210T,TPM1:NM_001018006:exon6:c.G628A:p.A210T, |
| chr15 | 63071104 | G | A | . | exonic | TPM1 | nonsynonymous SNV | TPM1:NM_001301289:exon8:c.G679A:p.A227T,TPM1:NM_001018020:exon9:c.G787A:p.A263T,TPM1:NM_001018007:exon9:c.G787A:p.A263T,TPM1:NM_001018006:exon9:c.G787A:p.A263T,TPM1:NM_001018004:exon9:c.G787A:p.A263T, |
| chr15 | 65119810 | C | T | . | exonic | PDCD7 | nonsynonymous SNV | PDCD7:NM_005707:exon3:c.G1154A:p.R385K, |
| chr15 | 65119820 | C | T | . | exonic | PDCD7 | nonsynonymous SNV | PDCD7:NM_005707:exon3:c.G1144A:p.E382K, |
| chr15 | 65593489 | A | G | . | exonic | VWA9 | nonsynonymous SNV | VWA9:NM_001136043:exon8:c.T688C:p.F230L,VWA9:NM_001207058:exon8:c.T871C:p.F291L,VWA9:NM_001207059:exon7:c.T754C:p.F252L, |
| chr15 | 66503424 | C | T | . | exonic | RPL4 | nonsynonymous SNV | RPL4:NM_000968:exon2:c.G109A:p.V37M, |
| chr15 | 67236036 | T | G | COSV56015936 | exonic | AAGAB | nonsynonymous SNV | AAGAB:NM_001271885:exon4:c.A67C:p.I23L,AAGAB:NM_001271886:exon4:c.A67C:p.I23L,AAGAB:NM_024666:exon4:c.A394C:p.I132L, |
| chr15 | 67236426 | C | T | . | exonic | AAGAB | nonsynonymous SNV | AAGAB:NM_001271886:exon3:c.G16A:p.D6N,AAGAB:NM_024666:exon3:c.G343A:p.D115N,AAGAB:NM_001271885:exon3:c.G16A:p.D6N, |
| chr15 | 67236650 | C | A | . | exonic | AAGAB | nonsynonymous SNV | AAGAB:NM_024666:exon2:c.G244T:p.V82F, |
| chr15 | 67475813 | G | C | . | exonic | IQCH | nonsynonymous SNV | IQCH:NM_001031715:exon18:c.G2794C:p.V932L,IQCH:NM_001284347:exon13:c.G1765C:p.V589L,IQCH:NM_001284348:exon15:c.G1777C:p.V593L, |
| chr15 | 68176524 | T | C | . | exonic | PIAS1 | nonsynonymous SNV | PIAS1:NM_016166:exon11:c.T1351C:p.S451P, |
| chr15 | 68176535 | C | A | . | exonic | PIAS1 | nonsynonymous SNV | PIAS1:NM_016166:exon11:c.C1362A:p.N454K, |
| chr15 | 68199564 | C | T | . | exonic | CALML4 | nonsynonymous SNV | CALML4:NM_001286694:exon2:c.G53A:p.R18Q,CALML4:NM_033429:exon3:c.G281A:p.R94Q, |
| chr15 | 70667983 | T | C | . | exonic | UACA | nonsynonymous SNV | UACA:NM_001008224:exon16:c.A2662G:p.K888E,UACA:NM_018003:exon16:c.A2701G:p.K901E, |
| chr15 | 70671053 | T | C | . | exonic | UACA | nonsynonymous SNV | UACA:NM_001008224:exon15:c.A1168G:p.M390V,UACA:NM_018003:exon15:c.A1207G:p.M403V, |
| chr15 | 70678124 | T | C | . | exonic | UACA | nonsynonymous SNV | UACA:NM_001008224:exon11:c.A935G:p.N312S,UACA:NM_018003:exon11:c.A974G:p.N325S, |
| chr15 | 72140242 | A | G | . | exonic | SENP8 | nonsynonymous SNV | SENP8:NM_001166340:exon2:c.A619G:p.T207A,SENP8:NM_001172111:exon3:c.A619G:p.T207A,SENP8:NM_001172110:exon3:c.A619G:p.T207A,SENP8:NM_001172109:exon3:c.A619G:p.T207A,SENP8:NM_145204:exon2:c.A619G:p.T207A, |
| chr15 | 72202609 | C | A | . | exonic | PKM | nonsynonymous SNV | PKM:NM_001206797:exon8:c.G930T:p.E310D,PKM:NM_001206798:exon9:c.G1107T:p.E369D,PKM:NM_002654:exon9:c.G1152T:p.E384D, |
| chr15 | 72202620 | T | C | . | exonic | PKM | nonsynonymous SNV | PKM:NM_001206797:exon8:c.A919G:p.I307V,PKM:NM_001206798:exon9:c.A1096G:p.I366V,PKM:NM_002654:exon9:c.A1141G:p.I381V, |
| chr15 | 72206841 | C | T | . | exonic | PKM | nonsynonymous SNV | PKM:NM_001206799:exon8:c.G1042A:p.A348T,PKM:NM_182471:exon8:c.G1027A:p.A343T,PKM:NM_001316318:exon8:c.G1132A:p.A378T,PKM:NM_182470:exon8:c.G1027A:p.A343T,PKM:NM_001206797:exon7:c.G805A:p.A269T,PKM:NM_001206796:exon9:c.G1249A:p.A417T,PKM:NM_001206798:exon8:c.G982A:p.A328T,PKM:NM_002654:exon8:c.G1027A:p.A343T, |
| chr15 | 72207129 | G | A | . | exonic | PKM | stopgain | PKM:NM_001206799:exon7:c.C1000T:p.Q334X,PKM:NM_182471:exon7:c.C985T:p.Q329X,PKM:NM_001316318:exon7:c.C1090T:p.Q364X,PKM:NM_182470:exon7:c.C985T:p.Q329X,PKM:NM_001206797:exon6:c.C763T:p.Q255X,PKM:NM_001206796:exon8:c.C1207T:p.Q403X,PKM:NM_001206798:exon7:c.C940T:p.Q314X,PKM:NM_002654:exon7:c.C985T:p.Q329X, |
| chr15 | 72208630 | C | T | . | exonic | PKM | nonsynonymous SNV | PKM:NM_001206799:exon6:c.G842A:p.G281E,PKM:NM_182471:exon6:c.G827A:p.G276E,PKM:NM_001316318:exon6:c.G932A:p.G311E,PKM:NM_182470:exon6:c.G827A:p.G276E,PKM:NM_001206797:exon5:c.G605A:p.G202E,PKM:NM_001206796:exon7:c.G1049A:p.G350E,PKM:NM_001206798:exon6:c.G782A:p.G261E,PKM:NM_002654:exon6:c.G827A:p.G276E, |
| chr15 | 72208736 | A | G | . | exonic | PKM | nonsynonymous SNV | PKM:NM_001206799:exon6:c.T736C:p.F246L,PKM:NM_182471:exon6:c.T721C:p.F241L,PKM:NM_001316318:exon6:c.T826C:p.F276L,PKM:NM_182470:exon6:c.T721C:p.F241L,PKM:NM_001206797:exon5:c.T499C:p.F167L,PKM:NM_001206796:exon7:c.T943C:p.F315L,PKM:NM_001206798:exon6:c.T676C:p.F226L,PKM:NM_002654:exon6:c.T721C:p.F241L, |

| | | | | | | | | |
|---|---|---|---|---|---|---|---|---|
| chr15 | 72209741 | T | A | . | exonic | PKM | nonsynonymous SNV | PKM:NM_001206799:exon5:c.A512T:p.K171M,PKM:NM_182471:exon5:c.A497T:p.K166M,PKM:NM_001316318:exon5:c.A602T:p.K201M,PKM:NM_182470:exon5:c.A497T:p.K166M,PKM:NM_001206797:exon4:c.A275T:p.K92M,PKM:NM_001206796:exon6:c.A719T:p.K240M,PKM:NM_001206798:exon5:c.A452T:p.K151M,PKM:NM_002654:exon5:c.A497T:p.K166M, |
| chr15 | 72209745 | A | T | . | exonic | PKM | nonsynonymous SNV | PKM:NM_001206799:exon5:c.T508A:p.C170S,PKM:NM_182471:exon5:c.T493A:p.C165S,PKM:NM_001316318:exon5:c.T598A:p.C200S,PKM:NM_182470:exon5:c.T493A:p.C165S,PKM:NM_001206797:exon4:c.T271A:p.C91S,PKM:NM_001206796:exon6:c.T715A:p.C239S,PKM:NM_001206798:exon5:c.T448A:p.C150S,PKM:NM_002654:exon5:c.T493A:p.C165S, |
| chr15 | 72209855 | C | A | COSV99069963 | exonic | PKM | nonsynonymous SNV | PKM:NM_001206799:exon5:c.G398T:p.G133V,PKM:NM_182471:exon5:c.G383T:p.G128V,PKM:NM_001316318:exon5:c.G488T:p.G163V,PKM:NM_182470:exon5:c.G383T:p.G128V,PKM:NM_001206796:exon6:c.G605T:p.G202V,PKM:NM_001206798:exon5:c.G338T:p.G113V,PKM:NM_002654:exon5:c.G383T:p.G128V, |
| chr15 | 72209855 | C | T | . | exonic | PKM | nonsynonymous SNV | PKM:NM_001206799:exon5:c.G398A:p.G133D,PKM:NM_001206798:exon5:c.G338A:p.G113D,PKM:NM_002654:exon5:c.G383A:p.G128D,PKM:NM_182471:exon5:c.G383A:p.G128D,PKM:NM_001316318:exon5:c.G488A:p.G163D,PKM:NM_182470:exon5:c.G383A:p.G128D,PKM:NM_001206796:exon6:c.G605A:p.G202D, |
| chr15 | 72209856 | C | A | . | exonic | PKM | nonsynonymous SNV | PKM:NM_001206799:exon5:c.G397T:p.G133C,PKM:NM_182471:exon5:c.G382T:p.G128C,PKM:NM_001316318:exon5:c.G487T:p.G163C,PKM:NM_182470:exon5:c.G382T:p.G128C,PKM:NM_001206796:exon6:c.G604T:p.G202C,PKM:NM_001206798:exon5:c.G337T:p.G113C,PKM:NM_002654:exon5:c.G382T:p.G128C, |
| chr15 | 72210433 | A | G | . | exonic | PKM | nonsynonymous SNV | PKM:NM_001206799:exon4:c.T307C:p.F103L,PKM:NM_182471:exon4:c.T292C:p.F98L,PKM:NM_001316318:exon4:c.T397C:p.F133L,PKM:NM_182470:exon4:c.T292C:p.F98L,PKM:NM_001206796:exon5:c.T514C:p.F172L,PKM:NM_001206798:exon4:c.T247C:p.F83L,PKM:NM_002654:exon4:c.T292C:p.F98L, |
| chr15 | 72210439 | C | T | . | exonic | PKM | nonsynonymous SNV | PKM:NM_001206799:exon4:c.G301A:p.E101K,PKM:NM_182471:exon4:c.G286A:p.E96K,PKM:NM_001316318:exon4:c.G391A:p.E131K,PKM:NM_182470:exon4:c.G286A:p.E96K,PKM:NM_001206796:exon5:c.G508A:p.E170K,PKM:NM_001206798:exon4:c.G241A:p.E81K,PKM:NM_002654:exon4:c.G286A:p.E96K, |
| chr15 | 72210450 | C | T | . | exonic | PKM | nonsynonymous SNV | PKM:NM_001206799:exon4:c.G290A:p.R97H,PKM:NM_182471:exon4:c.G275A:p.R92H,PKM:NM_001316318:exon4:c.G380A:p.R127H,PKM:NM_182470:exon4:c.G275A:p.R92H,PKM:NM_001206796:exon5:c.G497A:p.R166H,PKM:NM_001206798:exon4:c.G230A:p.R77H,PKM:NM_002654:exon4:c.G275A:p.R92H, |
| chr15 | 72217417 | T | C | . | exonic | PKM | nonsynonymous SNV | PKM:NM_001206799:exon3:c.A253G:p.T85A,PKM:NM_002654:exon3:c.A238G:p.T80A,PKM:NM_182471:exon3:c.A238G:p.T80A,PKM:NM_001316318:exon3:c.A343G:p.T115A,PKM:NM_182470:exon3:c.A238G:p.T80A,PKM:NM_001206796:exon4:c.A460G:p.T154A, |
| chr15 | 72217468 | C | T | . | exonic | PKM | nonsynonymous SNV | PKM:NM_001206799:exon3:c.G202A:p.E68K,PKM:NM_182471:exon3:c.G187A:p.E63K,PKM:NM_001316318:exon3:c.G292A:p.E98K,PKM:NM_182470:exon3:c.G187A:p.E63K,PKM:NM_001206796:exon4:c.G409A:p.E137K,PKM:NM_002654:exon3:c.G187A:p.E63K, |
| chr15 | 72217486 | A | G | . | exonic | PKM | nonsynonymous SNV | PKM:NM_001206799:exon3:c.T184C:p.S62P,PKM:NM_182471:exon3:c.T169C:p.S57P,PKM:NM_001316318:exon3:c.T274C:p.S92P,PKM:NM_182470:exon3:c.T169C:p.S57P,PKM:NM_001206797:exon3:c.T169C:p.S57P,PKM:NM_001206796:exon4:c.T391C:p.S131P,PKM:NM_002654:exon3:c.T169C:p.S57P, |
| chr15 | 72218954 | G | C | . | exonic | PKM | nonsynonymous SNV | PKM:NM_001206799:exon2:c.C159G:p.I53M,PKM:NM_182471:exon2:c.C144G:p.I48M,PKM:NM_001316318:exon2:c.C249G:p.I83M,PKM:NM_182470:exon2:c.C144G:p.I48M,PKM:NM_001206797:exon2:c.C144G:p.I48M,PKM:NM_001206796:exon3:c.C366G:p.I122M,PKM:NM_001206798:exon3:c.C191G:p.S64C,PKM:NM_002654:exon2:c.C144G:p.I48M, |
| chr15 | 72219068 | A | T | . | exonic | PKM | nonsynonymous SNV | PKM:NM_001206798:exon3:c.T77A:p.L26Q, |
| chr15 | 72346551 | T | C | . | exonic | HEXA | nonsynonymous SNV | HEXA:NM_000520:exon11:c.A1306G:p.I436V, |
| chr15 | 74035775 | A | G | COSV51443885 | exonic | PML | nonsynonymous SNV | PML:NM_033250:exon7:c.A2170G:p.S724G,PML:NM_033239:exon8:c.A2314G:p.S772G, |
| chr15 | 74035800 | G | T | COSV51443888 | exonic | PML | nonsynonymous SNV | PML:NM_033250:exon7:c.G2195T:p.G732V,PML:NM_033239:exon8:c.G2339T:p.G780V, |
| chr15 | 76286421 | G | A | COSV51211583 | exonic | ETFA | nonsynonymous SNV | ETFA:NM_001127716:exon5:c.C365T:p.T122I,ETFA:NM_000126:exon6:c.C512T:p.T171I, |
| chr15 | 77046911 | C | T | . | exonic | TSPAN3 | nonsynonymous SNV | TSPAN3:NM_001168412:exon6:c.G494A:p.C165Y,TSPAN3:NM_005724:exon7:c.G686A:p.C229Y,TSPAN3:NM_198902:exon6:c.G611A:p.C204Y, |
| chr15 | 77054189 | C | T | . | exonic | TSPAN3 | nonsynonymous SNV | TSPAN3:NM_001168412:exon3:c.G229A:p.V77I,TSPAN3:NM_005724:exon4:c.G421A:p.V141I,TSPAN3:NM_198902:exon3:c.G346A:p.V116I, |
| chr15 | 77056087 | C | T | . | exonic | TSPAN3 | nonsynonymous SNV | TSPAN3:NM_005724:exon2:c.G232A:p.E78K,TSPAN3:NM_198902:exon2:c.G232A:p.E78K, |
| chr15 | 77056128 | A | G | . | exonic | TSPAN3 | nonsynonymous SNV | TSPAN3:NM_005724:exon2:c.T191C:p.L64P,TSPAN3:NM_198902:exon2:c.T191C:p.L64P, |
| chr15 | 77056183 | C | T | . | exonic | TSPAN3 | nonsynonymous SNV | TSPAN3:NM_005724:exon2:c.G136A:p.E46K,TSPAN3:NM_198902:exon2:c.G136A:p.E46K, |
| chr15 | 78944905 | C | G | COSV54984032 | exonic | CTSH | nonsynonymous SNV | CTSH:NM_004390:exon1:c.G77C:p.C26S, |
| chr15 | 79899001 | G | A | COSV51912919 | exonic | ST20 | nonsynonymous SNV | ST20:NM_001100879:exon3:c.C170T:p.P57L,ST20:NM_001199757:exon3:c.C170T:p.P57L,ST20:NM_001100880:exon3:c.C170T:p.P57L, |
| chr15 | 79970875 | C | T | COSV51212886 | exonic | BCL2A1 | nonsynonymous SNV | BCL2A1:NM_001114735:exon1:c.G245A:p.G82D,BCL2A1:NM_004049:exon1:c.G245A:p.G82D, |
| chr15 | 79971003 | A | C | COSV51213210 | exonic | BCL2A1 | nonsynonymous SNV | BCL2A1:NM_001114735:exon1:c.T117G:p.N39K,BCL2A1:NM_004049:exon1:c.T117G:p.N39K, |
| chr15 | 79971064 | C | T | COSV51213224 | exonic | BCL2A1 | nonsynonymous SNV | BCL2A1:NM_001114735:exon1:c.G56A:p.C19Y,BCL2A1:NM_004049:exon1:c.G56A:p.C19Y, |
| chr15 | 82540031 | G | T | . | exonic | RPS17 | stopgain | RPS17:NM_001021:exon2:c.C105A:p.C35X, |
| chr15 | 83011535 | A | G | COSV55602266 | exonic | C15ORF40 | nonsynonymous SNV | C15orf40:NM_001160114:exon1:c.T73C:p.C25R,C15orf40:NM_001160115:exon1:c.T73C:p.C25R,C15orf40:NM_144597:exon1:c.T73C:p.C25R,C15orf40:NM_001160116:exon1:c.T73C:p.C25R,C15orf40:NM_001160113:exon1:c.T73C:p.C25R, |
| chr15 | 83011577 | G | A | COSV55602272 | exonic | C15ORF40 | nonsynonymous SNV | C15orf40:NM_001160114:exon1:c.C31T:p.L11F,C15orf40:NM_001160115:exon1:c.C31T:p.L11F,C15orf40:NM_144597:exon1:c.C31T:p.L11F,C15orf40:NM_001160116:exon1:c.C31T:p.L11F,C15orf40:NM_001160113:exon1:c.C31T:p.L11F, |
| chr15 | 84670026 | C | T | COSV51612248 | exonic | SEC11A | nonsynonymous SNV | SEC11A:NM_001271918:exon7:c.G455A:p.R152H,SEC11A:NM_001271922:exon5:c.G475A:p.V159M,SEC11A:NM_014300:exon6:c.G533A:p.R178H,SEC11A:NM_001271921:exon5:c.G383A:p.R128H,SEC11A:NM_001271919:exon5:c.G413A:p.R138H, |
| chr15 | 84670033 | C | T | . | exonic | SEC11A | stopgain | SEC11A:NM_001271922:exon5:c.G468A:p.W156X, |
| chr15 | 84670047 | C | A | . | exonic | SEC11A | nonsynonymous SNV | SEC11A:NM_001271918:exon7:c.G434T:p.G145V,SEC11A:NM_001271922:exon5:c.G454T:p.V152F,SEC11A:NM_014300:exon6:c.G512T:p.G171V,SEC11A:NM_001271921:exon5:c.G362T:p.G121V,SEC11A:NM_001271919:exon5:c.G392T:p.G131V, |
| chr15 | 84680779 | T | A | . | exonic | SEC11A | nonsynonymous SNV | SEC11A:NM_001271918:exon5:c.A287T:p.D96V,SEC11A:NM_001271922:exon4:c.A365T:p.D122V,SEC11A:NM_014300:exon4:c.A365T:p.D122V,SEC11A:NM_001271921:exon3:c.A215T:p.D72V,SEC11A:NM_001271920:exon4:c.A365T:p.D122V, |
| chr15 | 84716037 | C | T | . | exonic | SEC11A | nonsynonymous SNV | SEC11A:NM_001271922:exon1:c.G39A:p.M13I,SEC11A:NM_014300:exon1:c.G39A:p.M13I,SEC11A:NM_001271921:exon1:c.G39A:p.M13I,SEC11A:NM_001271920:exon1:c.G39A:p.M13I,SEC11A:NM_001271919:exon1:c.G39A:p.M13I, |
| chr15 | 85543936 | C | G | COSV99057187 | exonic | AKAP13 | nonsynonymous SNV | AKAP13:NM_006738:exon5:c.C643G:p.L215V,AKAP13:NM_007200:exon5:c.C643G:p.L215V, |
| chr15 | 85579423 | T | C | COSV63448142 | exonic | AKAP13 | nonsynonymous SNV | AKAP13:NM_006738:exon7:c.T1355C:p.M452T,AKAP13:NM_007200:exon7:c.T1355C:p.M452T, |
| chr15 | 85579548 | T | C | COSV63448149 | exonic | AKAP13 | nonsynonymous SNV | AKAP13:NM_006738:exon7:c.T1480C:p.W494R,AKAP13:NM_007200:exon7:c.T1480C:p.W494R, |
| chr15 | 85579788 | C | T | COSV63448171 | exonic | AKAP13 | nonsynonymous SNV | AKAP13:NM_006738:exon7:c.C1720T:p.R574C,AKAP13:NM_007200:exon7:c.C1720T:p.R574C, |
| chr15 | 85579939 | G | T | COSV63448180 | exonic | AKAP13 | nonsynonymous SNV | AKAP13:NM_006738:exon7:c.G1871T:p.G624V,AKAP13:NM_007200:exon7:c.G1871T:p.G624V, |
| chr15 | 85580602 | T | C | COSV63448191 | exonic | AKAP13 | nonsynonymous SNV | AKAP13:NM_006738:exon7:c.T2534C:p.V845A,AKAP13:NM_007200:exon7:c.T2534C:p.V845A, |
| chr15 | 85580757 | G | A | COSV63448207 | exonic | AKAP13 | nonsynonymous SNV | AKAP13:NM_006738:exon7:c.G2689A:p.V897M,AKAP13:NM_007200:exon7:c.G2689A:p.V897M, |
| chr15 | 85581252 | C | G | COSV63448224 | exonic | AKAP13 | nonsynonymous SNV | AKAP13:NM_006738:exon7:c.C3184G:p.P1062A,AKAP13:NM_007200:exon7:c.C3184G:p.P1062A, |
| chr15 | 85581324 | G | A | COSV63448234 | exonic | AKAP13 | nonsynonymous SNV | AKAP13:NM_006738:exon7:c.G3256A:p.D1086N,AKAP13:NM_007200:exon7:c.G3256A:p.D1086N, |
| chr15 | 85581385 | A | G | COSV63450063 | exonic | AKAP13 | nonsynonymous SNV | AKAP13:NM_006738:exon7:c.A3317G:p.E1106G,AKAP13:NM_007200:exon7:c.A3317G:p.E1106G, |
| chr15 | 85581715 | T | C | COSV63447944 | exonic | AKAP13 | nonsynonymous SNV | AKAP13:NM_006738:exon7:c.T3647C:p.M1216T,AKAP13:NM_007200:exon7:c.T3647C:p.M1216T, |
| chr15 | 85684869 | G | A | . | exonic | AKAP13 | nonsynonymous SNV | AKAP13:NM_006738:exon16:c.G5297A:p.S1766N,AKAP13:NM_007200:exon16:c.G5285A:p.S1762N,AKAP13:NM_001270546:exon8:c.G1151A:p.S384N, |
| chr15 | 85735078 | G | A | COSV63448267 | exonic | AKAP13 | nonsynonymous SNV | AKAP13:NM_006738:exon31:c.G7381A:p.G2461S,AKAP13:NM_007200:exon31:c.G7369A:p.G2457S,AKAP13:NM_001270546:exon23:c.G3232A:p.G1078S, |
| chr15 | 89792557 | C | T | COSV55591836 | exonic | ANPEP | nonsynonymous SNV | ANPEP:NM_001150:exon17:c.G2255A:p.S752N, |
| chr15 | 89801121 | T | C | COSV55589989 | exonic | ANPEP | nonsynonymous SNV | ANPEP:NM_001150:exon12:c.A1809G:p.I603M, |
| chr15 | 90360079 | A | G | . | exonic | ZNF774 | nonsynonymous SNV | ZNF774:NM_001004309:exon4:c.A248G:p.N83S, |
| chr15 | 90494752 | C | A | . | exonic | IQGAP1 | nonsynonymous SNV | IQGAP1:NM_003870:exon36:c.C4668A:p.S1556R, |
| chr15 | 90974173 | C | T | . | exonic | PRC1 | nonsynonymous SNV | PRC1:NM_001267580:exon10:c.G1301A:p.R434Q,PRC1:NM_199413:exon11:c.G1424A:p.R475Q,PRC1:NM_003981:exon11:c.G1424A:p.R475Q, |
| chr15 | 90974696 | C | T | . | exonic | PRC1 | stopgain | PRC1:NM_199413:exon10:c.G1239A:p.W413X,PRC1:NM_003981:exon10:c.G1239A:p.W413X,PRC1:NM_001267580:exon9:c.G1116A:p.W372X, |
| chr15 | 90981611 | G | T | . | exonic | PRC1 | nonsynonymous SNV | PRC1:NM_001267580:exon4:c.C437A:p.A146E,PRC1:NM_199413:exon5:c.C560A:p.A187E,PRC1:NM_003981:exon5:c.C560A:p.A187E, |

| | | | | | | | | |
|---|---|---|---|---|---|---|---|---|
| chr15 | 92939625 | G | A | . | exonic | CHD2 | nonsynonymous SNV | CHD2:NM_001271:exon7:c.G599A:p.G200E,CHD2:NM_001042572:exon7:c.G599A:p.G200E, |
| chr15 | 92943036 | C | A | . | exonic | CHD2 | nonsynonymous SNV | CHD2:NM_001271:exon9:c.C1020A:p.N340K,CHD2:NM_001042572:exon9:c.C1020A:p.N340K, |
| chr16 | 10744056 | C | G | COSV51593232 | exonic | NUBP1 | nonsynonymous SNV | NUBP1:NM_002484:exon2:c.C115G:p.P39A,NUBP1:NM_001278506:exon2:c.C115G:p.P39A, |
| chr16 | 10757974 | C | T | . | exonic | NUBP1 | nonsynonymous SNV | NUBP1:NM_002484:exon7:c.C553T:p.R185W,NUBP1:NM_001278506:exon6:c.C520T:p.R174W, |
| chr16 | 11679806 | C | G | . | exonic | TXNDC11 | nonsynonymous SNV | TXNDC11:NM_015914:exon12:c.G2266C:p.V756L,TXNDC11:NM_001303447:exon13:c.G2347C:p.V783L, |
| chr16 | 11839806 | C | A | . | exonic | RSL1D1 | nonsynonymous SNV | RSL1D1:NM_015659:exon8:c.G1035T:p.K345N, |
| chr16 | 11915422 | A | G | COSV70452514 | exonic | GSPT1 | nonsynonymous SNV | GSPT1:NM_002094:exon1:c.T299C:p.V100A,GSPT1:NM_001130006:exon1:c.T299C:p.V100A, |
| chr16 | 11966306 | A | G | . | exonic | TNFRSF17 | nonsynonymous SNV | TNFRSF17:NM_001192:exon2:c.A242G:p.N81S, |
| chr16 | 12051934 | G | C | . | exonic | SNX29 | nonsynonymous SNV | SNX29:NM_032167:exon8:c.G836C:p.G279A, |
| chr16 | 1452856 | C | T | COSV51970417 | exonic | CLCN7 | nonsynonymous SNV | CLCN7:NM_001114331:exon14:c.G1180A:p.V394M,CLCN7:NM_001287:exon15:c.G1252A:p.V418M, |
| chr16 | 15038105 | A | G | COSV55074188 | exonic | NTAN1 | nonsynonymous SNV | NTAN1:NM_001270767:exon8:c.T544C:p.S182P,NTAN1:NM_001270766:exon9:c.T544C:p.S182P,NTAN1:NM_173474:exon10:c.T859C:p.S287P, |
| chr16 | 15038117 | G | T | COSV55074343 | exonic | NTAN1 | nonsynonymous SNV | NTAN1:NM_001270767:exon8:c.C532A:p.H178N,NTAN1:NM_001270766:exon9:c.C532A:p.H178N,NTAN1:NM_173474:exon10:c.C847A:p.H283N, |
| chr16 | 15727006 | C | T | COSV55543853 | exonic | MYH11 | nonsynonymous SNV | MYH11:NM_022844:exon28:c.G3700A:p.A1234T,MYH11:NM_001040114:exon29:c.G3721A:p.A1241T,MYH11:NM_001040113:exon29:c.G3721A:p.A1241T,MYH11:NM_002474:exon28:c.G3700A:p.A1234T, |
| chr16 | 1788639 | T | C | . | exonic | NUBP2 | nonsynonymous SNV | NUBP2:NM_001284502:exon6:c.T295C:p.W99R, |
| chr16 | 1788835 | C | G | . | exonic | NUBP2 | nonsynonymous SNV | NUBP2:NM_001284502:exon6:c.C491G:p.P164R, |
| chr16 | 18789111 | C | T | . | exonic | RPS15A | nonsynonymous SNV | RPS15A:NM_001030009:exon2:c.G3A:p.M1I,RPS15A:NM_001019:exon2:c.G3A:p.M1I, |
| chr16 | 19764081 | A | C | . | exonic | IQCK | nonsynonymous SNV | IQCK:NM_153208:exon7:c.A574C:p.I192L,IQCK:NM_001305122:exon7:c.A574C:p.I192L,IQCK:NM_001305121:exon6:c.A310C:p.I104L, |
| chr16 | 2020199 | G | C | COSV61585942 | exonic | NPW | nonsynonymous SNV | NPW:NM_001099456:exon1:c.G298C:p.E100Q, |
| chr16 | 2020567 | A | C | . | exonic | NPW | nonsynonymous SNV | NPW:NM_001099456:exon2:c.A446C:p.D149A, |
| chr16 | 2047780 | A | G | COSV54593304 | exonic | NTHL1 | nonsynonymous SNV | NTHL1:NM_002528:exon1:c.T68C:p.L23P, |
| chr16 | 206123 | C | T | . | exonic | LUC7L | nonsynonymous SNV | LUC7L:NM_201412:exon5:c.G391A:p.E131K,LUC7L:NM_018032:exon5:c.G391A:p.E131K, |
| chr16 | 206124 | A | T | . | exonic | LUC7L | nonsynonymous SNV | LUC7L:NM_201412:exon5:c.T390A:p.N130K,LUC7L:NM_018032:exon5:c.T390A:p.N130K, |
| chr16 | 206128 | A | T | . | exonic | LUC7L | stopgain | LUC7L:NM_201412:exon5:c.T386A:p.L129X,LUC7L:NM_018032:exon5:c.T386A:p.L129X, |
| chr16 | 21179532 | A | C | . | exonic | TMEM159 | nonsynonymous SNV | TMEM159:NM_001301775:exon6:c.A534C:p.E178D,TMEM159:NM_020422:exon5:c.A462C:p.E154D,TMEM159:NM_001301769:exon6:c.A462C:p.E154D,TMEM159:NM_001301771:exon5:c.A462C:p.E154D, |
| chr16 | 21834992 | C | T | . | exonic | NPIPB4 | nonsynonymous SNV | NPIPB4:NM_001310148:exon8:c.G3395A:p.R1132Q, |
| chr16 | 25228484 | G | C | COSV54844539 | exonic | AQP8 | nonsynonymous SNV | AQP8:NM_001169:exon6:c.G778C:p.A260P, |
| chr16 | 27235296 | T | C | COSV63863773 | exonic | NSMCE1 | nonsynonymous SNV | NSMCE1:NM_145080:exon3:c.A140G:p.N47S, |
| chr16 | 27257458 | G | C | COSV63864235 | exonic | NSMCE1 | nonsynonymous SNV | NSMCE1:NM_145080:exon2:c.C113G:p.T38R, |
| chr16 | 28477622 | T | C | COSV61107434 | exonic | CLN3 | nonsynonymous SNV | CLN3:NM_001286105:exon13:c.A911G:p.H304R,CLN3:NM_001286110:exon14:c.A1049G:p.H350R,CLN3:NM_001042432:exon16:c.A1211G:p.H404R,CLN3:NM_001286109:exon13:c.A977G:p.H326R,CLN3:NM_000086:exon15:c.A1211G:p.H404R,CLN3:NM_001286104:exon15:c.A1139G:p.H380R, |
| chr16 | 28606164 | C | T | . | exonic | SULT1A1 | nonsynonymous SNV | SULT1A1:NM_001055:exon7:c.G667A:p.V223M,SULT1A1:NM_177530:exon7:c.G667A:p.V223M,SULT1A1:NM_177529:exon7:c.G667A:p.V223M,SULT1A1:NM_177536:exon5:c.G433A:p.V145M,SULT1A1:NM_177534:exon6:c.G667A:p.V223M, |
| chr16 | 29695123 | C | T | COSV54880380 | exonic | QPRT | nonsynonymous SNV | QPRT:NM_014298:exon2:c.C473T:p.A158V, |
| chr16 | 29697029 | A | G | COSV54880251 | exonic | QPRT | nonsynonymous SNV | QPRT:NM_014298:exon3:c.A583G:p.T195A, |
| chr16 | 29779543 | G | A | . | exonic | ZG16 | nonsynonymous SNV | ZG16:NM_152338:exon3:c.G94A:p.G32S, |
| chr16 | 29780400 | G | C | COSV68783707 | exonic | ZG16 | nonsynonymous SNV | ZG16:NM_152338:exon4:c.G485C:p.S162T, |
| chr16 | 30067327 | G | A | . | exonic | ALDOA | nonsynonymous SNV | ALDOA:NM_184043:exon2:c.G73A:p.A25T,ALDOA:NM_184041:exon2:c.G73A:p.A25T,ALDOA:NM_001127617:exon2:c.G73A:p.A25T,ALDOA:NM_000034:exon7:c.G73A:p.A25T,ALDOA:NM_001243177:exon3:c.G235A:p.A79T, |
| chr16 | 30067485 | G | A | . | exonic | ALDOA | nonsynonymous SNV | ALDOA:NM_001127617:exon3:c.G148A:p.E50K,ALDOA:NM_000034:exon8:c.G148A:p.E50K,ALDOA:NM_184043:exon3:c.G148A:p.E50K,ALDOA:NM_184041:exon3:c.G148A:p.E50K,ALDOA:NM_001243177:exon4:c.G310A:p.E104K, |
| chr16 | 30067509 | T | C | . | exonic | ALDOA | nonsynonymous SNV | ALDOA:NM_001127617:exon3:c.T172C:p.F58L,ALDOA:NM_000034:exon8:c.T172C:p.F58L,ALDOA:NM_184043:exon3:c.T172C:p.F58L,ALDOA:NM_184041:exon3:c.T172C:p.F58L,ALDOA:NM_001243177:exon4:c.T334C:p.F112L, |
| chr16 | 30067561 | G | A | . | exonic | ALDOA | nonsynonymous SNV | ALDOA:NM_001127617:exon3:c.G224A:p.G75E,ALDOA:NM_000034:exon8:c.G224A:p.G75E,ALDOA:NM_184043:exon3:c.G224A:p.G75E,ALDOA:NM_184041:exon3:c.G224A:p.G75E,ALDOA:NM_001243177:exon4:c.G386A:p.G129E, |
| chr16 | 30067612 | G | A | . | exonic | ALDOA | nonsynonymous SNV | ALDOA:NM_001127617:exon3:c.G275A:p.R92H,ALDOA:NM_000034:exon8:c.G275A:p.R92H,ALDOA:NM_184043:exon3:c.G275A:p.R92H,ALDOA:NM_184041:exon3:c.G275A:p.R92H,ALDOA:NM_001243177:exon4:c.G437A:p.R146H, |
| chr16 | 30067630 | T | C | . | exonic | ALDOA | nonsynonymous SNV | ALDOA:NM_001127617:exon3:c.T293C:p.I98T,ALDOA:NM_000034:exon8:c.T293C:p.I98T,ALDOA:NM_184043:exon3:c.T293C:p.I98T,ALDOA:NM_184041:exon3:c.T293C:p.I98T,ALDOA:NM_001243177:exon4:c.T455C:p.I152T, |
| chr16 | 30067642 | G | A | . | exonic | ALDOA | nonsynonymous SNV | ALDOA:NM_001127617:exon3:c.G305A:p.G102D,ALDOA:NM_000034:exon8:c.G305A:p.G102D,ALDOA:NM_184043:exon3:c.G305A:p.G102D,ALDOA:NM_184041:exon3:c.G305A:p.G102D,ALDOA:NM_001243177:exon4:c.G467A:p.G156D, |
| chr16 | 30067654 | G | A | . | exonic | ALDOA | nonsynonymous SNV | ALDOA:NM_001127617:exon3:c.G317A:p.G106D,ALDOA:NM_000034:exon8:c.G317A:p.G106D,ALDOA:NM_184043:exon3:c.G317A:p.G106D,ALDOA:NM_184041:exon3:c.G317A:p.G106D,ALDOA:NM_001243177:exon4:c.G479A:p.G160D, |
| chr16 | 30068823 | G | A | COSV57632467 | exonic | ALDOA | nonsynonymous SNV | ALDOA:NM_001127617:exon5:c.G385A:p.D129N,ALDOA:NM_000034:exon10:c.G385A:p.D129N,ALDOA:NM_184043:exon5:c.G385A:p.D129N,ALDOA:NM_184041:exon5:c.G385A:p.D129N,ALDOA:NM_001243177:exon6:c.G547A:p.D183N, |
| chr16 | 30068827 | G | A | . | exonic | ALDOA | nonsynonymous SNV | ALDOA:NM_001127617:exon5:c.G389A:p.G130E,ALDOA:NM_000034:exon10:c.G389A:p.G130E,ALDOA:NM_184043:exon5:c.G389A:p.G130E,ALDOA:NM_184041:exon5:c.G389A:p.G130E,ALDOA:NM_001243177:exon6:c.G551A:p.G184E, |
| chr16 | 30068839 | G | A | . | exonic | ALDOA | nonsynonymous SNV | ALDOA:NM_001127617:exon5:c.G401A:p.R134H,ALDOA:NM_000034:exon10:c.G401A:p.R134H,ALDOA:NM_184043:exon5:c.G401A:p.R134H,ALDOA:NM_184041:exon5:c.G401A:p.R134H,ALDOA:NM_001243177:exon6:c.G563A:p.R188H, |
| chr16 | 30068901 | G | A | . | exonic | ALDOA | nonsynonymous SNV | ALDOA:NM_001127617:exon5:c.G463A:p.G155R,ALDOA:NM_000034:exon10:c.G463A:p.G155R,ALDOA:NM_184043:exon5:c.G463A:p.G155R,ALDOA:NM_184041:exon5:c.G463A:p.G155R,ALDOA:NM_001243177:exon6:c.G625A:p.G209R, |
| chr16 | 30068904 | G | A | . | exonic | ALDOA | nonsynonymous SNV | ALDOA:NM_001127617:exon5:c.G466A:p.E156K,ALDOA:NM_000034:exon10:c.G466A:p.E156K,ALDOA:NM_184043:exon5:c.G466A:p.E156K,ALDOA:NM_184041:exon5:c.G466A:p.E156K,ALDOA:NM_001243177:exon6:c.G628A:p.E210K, |
| chr16 | 30068926 | C | A | . | exonic | ALDOA | nonsynonymous SNV | ALDOA:NM_001127617:exon5:c.C488A:p.A163D,ALDOA:NM_000034:exon10:c.C488A:p.A163D,ALDOA:NM_184043:exon5:c.C488A:p.A163D,ALDOA:NM_184041:exon5:c.C488A:p.A163D,ALDOA:NM_001243177:exon6:c.C650A:p.A217D, |
| chr16 | 30068929 | T | C | . | exonic | ALDOA | nonsynonymous SNV | ALDOA:NM_001127617:exon5:c.T491C:p.I164T,ALDOA:NM_000034:exon10:c.T491C:p.I164T,ALDOA:NM_184043:exon5:c.T491C:p.I164T,ALDOA:NM_184041:exon5:c.T491C:p.I164T,ALDOA:NM_001243177:exon6:c.T653C:p.I218T, |
| chr16 | 30068933 | G | A | . | exonic | ALDOA | nonsynonymous SNV | ALDOA:NM_001127617:exon5:c.G495A:p.M165I,ALDOA:NM_000034:exon10:c.G495A:p.M165I,ALDOA:NM_184043:exon5:c.G495A:p.M165I,ALDOA:NM_184041:exon5:c.G495A:p.M165I,ALDOA:NM_001243177:exon6:c.G657A:p.M219I, |
| chr16 | 30068935 | A | G | . | exonic | ALDOA | nonsynonymous SNV | ALDOA:NM_001127617:exon5:c.A497G:p.E166G,ALDOA:NM_000034:exon10:c.A497G:p.E166G,ALDOA:NM_184043:exon5:c.A497G:p.E166G,ALDOA:NM_184041:exon5:c.A497G:p.E166G,ALDOA:NM_001243177:exon6:c.A659G:p.E220G, |
| chr16 | 30068956 | G | A | . | exonic | ALDOA | nonsynonymous SNV | ALDOA:NM_001127617:exon5:c.G518A:p.R173H,ALDOA:NM_000034:exon10:c.G518A:p.R173H,ALDOA:NM_184043:exon5:c.G518A:p.R173H,ALDOA:NM_184041:exon5:c.G518A:p.R173H,ALDOA:NM_001243177:exon6:c.G680A:p.R227H, |

| | | | | | | | | |
|---|---|---|---|---|---|---|---|---|
| chr16 | 30069349 | G | A | COSV57632343 | exonic | ALDOA | nonsynonymous SNV | ALDOA:NM_001127617:exon6:c.G584A:p.G195E,ALDOA:NM_000034:exon11:c.G584A:p.G195E,ALDOA:NM_184043:exon6:c.G584A:p.G195E,ALDOA:NM_184041:exon6:c.G584A:p.G195E,ALDOA:NM_001243177:exon7:c.G746A:p.G249E, |
| chr16 | 30069505 | G | A | . | exonic | ALDOA | nonsynonymous SNV | ALDOA:NM_001127617:exon7:c.G631A:p.A211T,ALDOA:NM_000034:exon12:c.G631A:p.A211T,ALDOA:NM_184043:exon7:c.G631A:p.A211T,ALDOA:NM_184041:exon7:c.G631A:p.A211T,ALDOA:NM_001243177:exon8:c.G793A:p.A265T, |
| chr16 | 30069664 | G | A | COSV57632104 | exonic | ALDOA | nonsynonymous SNV | ALDOA:NM_001127617:exon7:c.G790A:p.A264T,ALDOA:NM_000034:exon12:c.G790A:p.A264T,ALDOA:NM_184043:exon7:c.G790A:p.A264T,ALDOA:NM_184041:exon7:c.G790A:p.A264T,ALDOA:NM_001243177:exon8:c.G952A:p.A318T, |
| chr16 | 30069881 | T | C | . | exonic | ALDOA | nonsynonymous SNV | ALDOA:NM_001127617:exon8:c.T851C:p.L284P,ALDOA:NM_000034:exon13:c.T851C:p.L284P,ALDOA:NM_184043:exon8:c.T851C:p.L284P,ALDOA:NM_184041:exon8:c.T851C:p.L284P,ALDOA:NM_001243177:exon9:c.T1013C:p.L338P, |
| chr16 | 30069977 | G | A | . | exonic | ALDOA | nonsynonymous SNV | ALDOA:NM_001127617:exon8:c.G947A:p.G316E,ALDOA:NM_000034:exon13:c.G947A:p.G316E,ALDOA:NM_184043:exon8:c.G947A:p.G316E,ALDOA:NM_184041:exon8:c.G947A:p.G316E,ALDOA:NM_001243177:exon9:c.G1109A:p.G370E, |
| chr16 | 30070117 | G | A | . | exonic | ALDOA | nonsynonymous SNV | ALDOA:NM_001127617:exon9:c.G1000A:p.A334T,ALDOA:NM_000034:exon14:c.G1000A:p.A334T,ALDOA:NM_184043:exon9:c.G1000A:p.A334T,ALDOA:NM_184041:exon9:c.G1000A:p.A334T,ALDOA:NM_001243177:exon10:c.G1162A:p.A388T, |
| chr16 | 30070156 | G | A | . | exonic | ALDOA | nonsynonymous SNV | ALDOA:NM_001127617:exon9:c.G1039A:p.G347S,ALDOA:NM_000034:exon14:c.G1039A:p.G347S,ALDOA:NM_184043:exon9:c.G1039A:p.G347S,ALDOA:NM_184041:exon9:c.G1039A:p.G347S,ALDOA:NM_001243177:exon10:c.G1201A:p.G401S, |
| chr16 | 31182545 | G | A | . | exonic | FUS | nonsynonymous SNV | FUS:NM_004960:exon3:c.G71A:p.G24D,FUS:NM_001170634:exon3:c.G71A:p.G24D,FUS:NM_001170937:exon3:c.G71A:p.G24D, |
| chr16 | 31183893 | G | A | . | exonic | FUS | nonsynonymous SNV | FUS:NM_001170634:exon4:c.G223A:p.G75S,FUS:NM_001170937:exon4:c.G226A:p.G76S,FUS:NM_004960:exon4:c.G226A:p.G76S, |
| chr16 | 31183906 | G | A | . | exonic | FUS | nonsynonymous SNV | FUS:NM_004960:exon4:c.G239A:p.G80D,FUS:NM_001170634:exon4:c.G236A:p.G79D,FUS:NM_001170937:exon4:c.G239A:p.G80D, |
| chr16 | 31184352 | A | G | . | exonic | FUS | nonsynonymous SNV | FUS:NM_004960:exon5:c.A479G:p.Q160R,FUS:NM_001170634:exon5:c.A476G:p.Q159R,FUS:NM_001170937:exon5:c.A479G:p.Q160R, |
| chr16 | 31184983 | A | T | . | exonic | FUS | nonsynonymous SNV | FUS:NM_004960:exon6:c.A568T:p.S190C,FUS:NM_001170634:exon6:c.A565T:p.S189C,FUS:NM_001170937:exon6:c.A556T:p.S186C, |
| chr16 | 31185078 | T | G | . | exonic | FUS | nonsynonymous SNV | FUS:NM_004960:exon6:c.T663G:p.S221R,FUS:NM_001170634:exon6:c.T660G:p.S220R,FUS:NM_001170937:exon6:c.T651G:p.S217R, |
| chr16 | 31189720 | G | A | . | exonic | FUS | nonsynonymous SNV | FUS:NM_001170634:exon10:c.G989A:p.G330D,FUS:NM_001170937:exon10:c.G980A:p.G327D,FUS:NM_004960:exon10:c.G992A:p.G331D, |
| chr16 | 31190821 | G | A | . | exonic | FUS | nonsynonymous SNV | FUS:NM_001170634:exon13:c.G1369A:p.G457R,FUS:NM_001170937:exon13:c.G1360A:p.G454R,FUS:NM_004960:exon13:c.G1372A:p.G458R, |
| chr16 | 31191419 | G | A | . | exonic | FUS | nonsynonymous SNV | FUS:NM_004960:exon15:c.G1562A:p.R521H,FUS:NM_001170634:exon15:c.G1559A:p.R520H,FUS:NM_001170937:exon15:c.G1550A:p.R517H, |
| chr16 | 314606 | T | C | . | exonic | AXIN1 | nonsynonymous SNV | AXIN1:NM_003502:exon3:c.A956G:p.N319S,AXIN1:NM_181050:exon3:c.A956G:p.N319S, |
| chr16 | 3484892 | G | A | COSV62654782 | exonic | NAA60 | nonsynonymous SNV | NAA60:NM_001317096:exon5:c.G539A:p.R180Q, |
| chr16 | 3657065 | C | T | . | exonic | DNASE1 | nonsynonymous SNV | DNASE1:NM_005223:exon6:c.C503T:p.A168V, |
| chr16 | 3657746 | G | A | COSV55912793 | exonic | DNASE1 | nonsynonymous SNV | DNASE1:NM_005223:exon8:c.G731A:p.R244Q, |
| chr16 | 3658169 | C | T | COSV55914199 | exonic | TRAP1 | nonsynonymous SNV | TRAP1:NM_016292:exon18:c.G2075A:p.R692H,TRAP1:NM_001272049:exon17:c.G1916A:p.R639H, |
| chr16 | 3674464 | G | C | COSV55915372 | exonic | TRAP1 | nonsynonymous SNV | TRAP1:NM_016292:exon9:c.C919G:p.R307G,TRAP1:NM_001272049:exon8:c.C760G:p.R254G, |
| chr16 | 3676074 | G | A | . | exonic | TRAP1 | nonsynonymous SNV | TRAP1:NM_016292:exon7:c.C776T:p.S259F,TRAP1:NM_001272049:exon6:c.C617T:p.S206F, |
| chr16 | 3717472 | G | A | COSV55917281 | exonic | TRAP1 | nonsynonymous SNV | TRAP1:NM_016292:exon1:c.C37T:p.R13C,TRAP1:NM_001272049:exon1:c.C37T:p.R13C, |
| chr16 | 4426088 | T | C | COSV52163376 | exonic | DNAJA3 | nonsynonymous SNV | DNAJA3:NM_001286516:exon1:c.T97C:p.Y33H, |
| chr16 | 50326301 | C | T | . | exonic | BRD7 | nonsynonymous SNV | BRD7:NM_013263:exon10:c.G1178A:p.R393K,BRD7:NM_001173984:exon10:c.G1178A:p.R393K, |
| chr16 | 56626904 | G | A | COSV57842018 | exonic | MT1E | nonsynonymous SNV | MT1E:NM_175617:exon3:c.G119A:p.G40D, |
| chr16 | 56633370 | C | A | . | exonic | MT1M | nonsynonymous SNV | MT1M:NM_176870:exon2:c.C59A:p.T20K, |
| chr16 | 56939326 | A | G | . | exonic | HERPUD1 | nonsynonymous SNV | HERPUD1:NM_014685:exon5:c.A521G:p.Q174R,HERPUD1:NM_001272103:exon5:c.A521G:p.Q174R,HERPUD1:NM_001010989:exon5:c.A518G:p.Q173R, |
| chr16 | 58709550 | A | C | COSV55331033 | exonic | GOT2 | nonsynonymous SNV | GOT2:NM_001286220:exon8:c.T908G:p.V303G,GOT2:NM_002080:exon9:c.T1037G:p.V346G, |
| chr16 | 64982184 | A | C | . | exonic | CDH11 | nonsynonymous SNV | CDH11:NM_001308392:exon8:c.T1117G:p.S373A,CDH11:NM_001797:exon8:c.T1117G:p.S373A, |
| chr16 | 64991815 | G | A | COSV51767367 | exonic | CDH11 | nonsynonymous SNV | CDH11:NM_001308392:exon6:c.C764T:p.T255M,CDH11:NM_001797:exon6:c.C764T:p.T255M, |
| chr16 | 66813637 | C | T | . | exonic | NAE1 | nonsynonymous SNV | NAE1:NM_003905:exon13:c.G961A:p.G321R,NAE1:NM_001286500:exon13:c.G970A:p.G324R,NAE1:NM_001018159:exon14:c.G943A:p.G315R,NAE1:NM_001018160:exon12:c.G694A:p.G232R, |
| chr16 | 66813809 | C | T | . | exonic | NAE1 | nonsynonymous SNV | NAE1:NM_003905:exon12:c.G878A:p.R293H,NAE1:NM_001286500:exon12:c.G887A:p.R296H,NAE1:NM_001018159:exon13:c.G860A:p.R287H,NAE1:NM_001018160:exon11:c.G611A:p.R204H, |
| chr16 | 66816657 | T | C | . | exonic | NAE1 | nonsynonymous SNV | NAE1:NM_003905:exon11:c.A764G:p.E255G,NAE1:NM_001286500:exon11:c.A773G:p.E258G,NAE1:NM_001018159:exon12:c.A746G:p.E249G,NAE1:NM_001018160:exon10:c.A497G:p.E166G, |
| chr16 | 66817004 | A | T | . | exonic | NAE1 | nonsynonymous SNV | NAE1:NM_003905:exon10:c.T709A:p.Y237N,NAE1:NM_001286500:exon10:c.T718A:p.Y240N,NAE1:NM_001018159:exon11:c.T691A:p.Y231N,NAE1:NM_001018160:exon9:c.T442A:p.Y148N, |
| chr16 | 66826722 | C | T | . | exonic | NAE1 | nonsynonymous SNV | NAE1:NM_003905:exon2:c.G112A:p.A38T,NAE1:NM_001286500:exon2:c.G112A:p.A38T,NAE1:NM_001018159:exon3:c.G94A:p.A32T, |
| chr16 | 66826755 | C | T | COSV51977029 | exonic | NAE1 | nonsynonymous SNV | NAE1:NM_003905:exon2:c.G79A:p.A27T,NAE1:NM_001286500:exon2:c.G79A:p.A27T,NAE1:NM_001018159:exon3:c.G61A:p.A21T, |
| chr16 | 66938138 | G | A | . | exonic | CES2 | nonsynonymous SNV | CES2:NM_198061:exon2:c.G370A:p.G124R,CES2:NM_003869:exon2:c.G370A:p.G124R, |
| chr16 | 67199363 | A | C | COSV62605808 | exonic | ELMO3 | nonsynonymous SNV | ELMO3:NM_024712:exon1:c.A196C:p.K66Q, |
| chr16 | 67611443 | C | A | COSV50477534 | exonic | CTCF | nonsynonymous SNV | CTCF:NM_006565:exon3:c.C611A:p.T204K, |
| chr16 | 67611452 | C | G | . | exonic | CTCF | nonsynonymous SNV | CTCF:NM_006565:exon3:c.C620G:p.T207S, |
| chr16 | 67611595 | A | G | . | exonic | CTCF | nonsynonymous SNV | CTCF:NM_006565:exon3:c.A763G:p.T255A, |
| chr16 | 67657765 | A | G | COSV54667315 | exonic | ACD | nonsynonymous SNV | ACD:NM_001082487:exon11:c.T1505C:p.V502A,ACD:NM_001082486:exon11:c.T1553C:p.V518A,ACD:NM_022914:exon11:c.T1544C:p.V515A, |
| chr16 | 685282 | C | T | . | exonic | WDR24 | nonsynonymous SNV | WDR24:NM_032259:exon7:c.G1994A:p.R665H, |
| chr16 | 68738336 | C | A | . | exonic | CDH1 | nonsynonymous SNV | CDH1:NM_001317184:exon2:c.C88A:p.P30T,CDH1:NM_004360:exon2:c.C88A:p.P30T, |
| chr16 | 68813430 | G | A | COSV55743232 | exonic | CDH1 | nonsynonymous SNV | CDH1:NM_004360:exon9:c.G1255A:p.D419N, |
| chr16 | 69330540 | A | G | COSV55340449 | exonic | PDF | nonsynonymous SNV | PDF:NM_022341:exon1:c.T31C:p.W11R, |
| chr16 | 69714966 | G | A | COSV57731384 | exonic | NQO1 | nonsynonymous SNV | NQO1:NM_001025433:exon4:c.C415T:p.R139W,NQO1:NM_000903:exon4:c.C415T:p.R139W, |
| chr16 | 70514394 | G | A | . | exonic | COG4 | nonsynonymous SNV | COG4:NM_001195139:exon4:c.C485T:p.T162I,COG4:NM_015386:exon4:c.C485T:p.T162I, |
| chr16 | 70564003 | G | A | . | exonic | SF3B3 | nonsynonymous SNV | SF3B3:NM_012426:exon18:c.G2416A:p.A806T, |
| chr16 | 71626407 | G | A | COSV51704980 | exonic | MARVELD3 | nonsynonymous SNV | MARVELD3:NM_052858:exon1:c.G178A:p.E60K,MARVELD3:NM_001271329:exon1:c.G178A:p.E60K,MARVELD3:NM_001017967:exon1:c.G178A:p.E60K, |
| chr16 | 74630845 | T | C | COSV63097488 | exonic | RFWD3 | nonsynonymous SNV | RFWD3:NM_018124:exon10:c.A1690G:p.I564V, |
| chr16 | 74661181 | G | T | . | exonic | RFWD3 | nonsynonymous SNV | RFWD3:NM_018124:exon2:c.C269A:p.T90N, |
| chr16 | 766977 | G | A | COSV53502132 | exonic | MSLN | nonsynonymous SNV | MSLN:NM_001177355:exon15:c.G1466A:p.G489E,MSLN:NM_005823:exon15:c.G1466A:p.G489E,MSLN:NM_013404:exon14:c.G1490A:p.G497E, |
| chr16 | 81017360 | A | T | . | exonic | CENPN | nonsynonymous SNV | CENPN:NM_001100625:exon4:c.A252T:p.E84D,CENPN:NM_001100624:exon4:c.A252T:p.E84D,CENPN:NM_018455:exon4:c.A252T:p.E84D,CENPN:NM_001270474:exon4:c.A252T:p.E84D, |
| chr16 | 81096217 | G | A | COSV59601803 | exonic | GCSH | nonsynonymous SNV | GCSH:NM_004483:exon1:c.C62T:p.S21L, |

| | | | | | | | | |
|---|---|---|---|---|---|---|---|---|
| chr16 | 82090913 | A | G | COSV52275659 | exonic | HSD17B2 | nonsynonymous SNV | HSD17B2:NM_002153:exon4:c.A676G:p.M226V, |
| chr16 | 84745079 | A | G | COSV54746826 | exonic | USP10 | nonsynonymous SNV | USP10:NM_001272075:exon5:c.A610G:p.M204V,USP10:NM_005153:exon4:c.A598G:p.M200V, |
| chr16 | 84745088 | T | C | COSV54751736 | exonic | USP10 | nonsynonymous SNV | USP10:NM_001272075:exon5:c.T619C:p.S207P,USP10:NM_005153:exon4:c.T607C:p.S203P, |
| chr16 | 84745091 | G | C | COSV54751750 | exonic | USP10 | nonsynonymous SNV | USP10:NM_001272075:exon5:c.G622C:p.V208L,USP10:NM_005153:exon4:c.G610C:p.V204L, |
| chr16 | 84764218 | A | T | . | exonic | USP10 | nonsynonymous SNV | USP10:NM_001272075:exon11:c.A1799T:p.D600V,USP10:NM_005153:exon10:c.A1787T:p.D596V, |
| chr16 | 85801212 | G | A | COSV53665976 | exonic | COX4I1 | nonsynonymous SNV | COX4I1:NM_001861:exon2:c.G7A:p.A3T, |
| chr16 | 88643420 | A | G | COSV54918013 | exonic | CYBA | nonsynonymous SNV | CYBA:NM_000101:exon6:c.T521C:p.V174A, |
| chr16 | 88646828 | A | G | COSV55368224 | exonic | CYBA | nonsynonymous SNV | CYBA:NM_000101:exon4:c.T214C:p.Y72H, |
| chr16 | 89100686 | T | C | COSV58076417 | exonic | ACSF3 | nonsynonymous SNV | ACSF3:NM_001127214:exon2:c.T5C:p.L2P,ACSF3:NM_001243279:exon3:c.T5C:p.L2P,ACSF3:NM_174917:exon3:c.T5C:p.L2P, |
| chr16 | 89114475 | G | A | COSV58077067 | exonic | ACSF3 | nonsynonymous SNV | ACSF3:NM_001127214:exon5:c.G1114A:p.V372M,ACSF3:NM_001243279:exon6:c.G1114A:p.V372M,ACSF3:NM_174917:exon6:c.G1114A:p.V372M,ACSF3:NM_001284316:exon4:c.G319A:p.V107M , |
| chr16 | 89283630 | G | A | COSV56354485 | exonic | ANKRD11 | nonsynonymous SNV | ANKRD11:NM_001256183:exon9:c.C2912T:p.A971V,ANKRD11:NM_001256182:exon10:c.C2912T:p.A971V,ANKRD11:NM_013275:exon9:c.C2912T:p.A971V, |
| chr16 | 89561665 | G | A | COSV51948976 | exonic | RPL13 | nonsynonymous SNV | RPL13:NM_001243130:exon4:c.G277A:p.A93T,RPL13:NM_033251:exon3:c.G334A:p.A112T,RPL13:NM_000977:exon4:c.G334A:p.A112T, |
| chr16 | 89637957 | G | C | COSV55359253 | exonic | DPEP1 | nonsynonymous SNV | DPEP1:NM_004413:exon10:c.G1051C:p.E351Q,DPEP1:NM_001128141:exon10:c.G1051C:p.E351Q, |
| chr16 | 89646633 | C | T | . | exonic | CHMP1A | nonsynonymous SNV | CHMP1A:NM_001083314:exon5:c.G443A:p.R148Q,CHMP1A:NM_002768:exon6:c.G463A:p.E155K, |
| chr16 | 89647248 | C | T | . | exonic | CHMP1A | nonsynonymous SNV | CHMP1A:NM_001083314:exon4:c.G316A:p.G106R,CHMP1A:NM_002768:exon5:c.G336A:p.M112I, |
| chr16 | 89649365 | C | T | . | exonic | CHMP1A | nonsynonymous SNV | CHMP1A:NM_001083314:exon3:c.G218A:p.C73Y,CHMP1A:NM_002768:exon4:c.G238A:p.V80M, |
| chr17 | 1345458 | C | T | COSV51985336 | exonic | YWHAE | nonsynonymous SNV | YWHAE:NM_006761:exon6:c.G757A:p.E253K, |
| chr17 | 1354316 | C | T | COSV51984710 | exonic | YWHAE | nonsynonymous SNV | YWHAE:NM_006761:exon5:c.G610A:p.A204T, |
| chr17 | 1354342 | G | A | . | exonic | YWHAE | nonsynonymous SNV | YWHAE:NM_006761:exon5:c.C584T:p.A195V, |
| chr17 | 1361101 | C | T | . | exonic | YWHAE | nonsynonymous SNV | YWHAE:NM_006761:exon4:c.G569A:p.R190H, |
| chr17 | 1361143 | T | A | . | exonic | YWHAE | nonsynonymous SNV | YWHAE:NM_006761:exon4:c.A527T:p.N176I, |
| chr17 | 1361156 | C | T | . | exonic | YWHAE | nonsynonymous SNV | YWHAE:NM_006761:exon4:c.G514A:p.G172S, |
| chr17 | 1361161 | C | T | . | exonic | YWHAE | nonsynonymous SNV | YWHAE:NM_006761:exon4:c.G509A:p.R170H, |
| chr17 | 1361927 | C | T | . | exonic | YWHAE | nonsynonymous SNV | YWHAE:NM_006761:exon3:c.G346A:p.E116K, |
| chr17 | 1361947 | A | G | . | exonic | YWHAE | nonsynonymous SNV | YWHAE:NM_006761:exon3:c.T326C:p.I109T, |
| chr17 | 1364864 | G | T | . | exonic | YWHAE | nonsynonymous SNV | YWHAE:NM_006761:exon2:c.C259A:p.Q87K, |
| chr17 | 1364880 | C | T | . | exonic | YWHAE | nonsynonymous SNV | YWHAE:NM_006761:exon2:c.G243A:p.M81I, |
| chr17 | 1364911 | T | C | . | exonic | YWHAE | nonsynonymous SNV | YWHAE:NM_006761:exon2:c.A212G:p.E71G, |
| chr17 | 1364924 | C | T | . | exonic | YWHAE | nonsynonymous SNV | YWHAE:NM_006761:exon2:c.G199A:p.E67K, |
| chr17 | 1364941 | C | T | . | exonic | YWHAE | nonsynonymous SNV | YWHAE:NM_006761:exon2:c.G182A:p.R61K, |
| chr17 | 1364943 | C | T | . | exonic | YWHAE | stopgain | YWHAE:NM_006761:exon2:c.G180A:p.W60X, |
| chr17 | 1400108 | C | T | . | exonic | YWHAE | nonsynonymous SNV | YWHAE:NM_006761:exon1:c.G3A:p.M1I, |
| chr17 | 1496164 | C | A | . | exonic | INPP5K | nonsynonymous SNV | INPP5K:NM_130766:exon12:c.G958T:p.V320F,INPP5K:NM_016532:exon11:c.G1186T:p.V396F,INPP5K:NM_001135642:exon13:c.G958T:p.V320F, |
| chr17 | 16121231 | G | T | . | exonic | NCOR1 | nonsynonymous SNV | NCOR1:NM_001190440:exon15:c.C1673A:p.A558E,NCOR1:NM_001190438:exon13:c.C1346A:p.A449E,NCOR1:NM_006311:exon16:c.C1673A:p.A558E, |
| chr17 | 16126110 | C | T | . | exonic | NCOR1 | nonsynonymous SNV | NCOR1:NM_001190440:exon14:c.G1606A:p.E536K,NCOR1:NM_001190438:exon12:c.G1279A:p.E427K,NCOR1:NM_006311:exon15:c.G1606A:p.E536K, |
| chr17 | 16126142 | G | T | . | exonic | NCOR1 | nonsynonymous SNV | NCOR1:NM_001190440:exon14:c.C1574A:p.T525K,NCOR1:NM_001190438:exon12:c.C1247A:p.T416K,NCOR1:NM_006311:exon15:c.C1574A:p.T525K, |
| chr17 | 16126190 | G | T | . | exonic | NCOR1 | stopgain | NCOR1:NM_001190440:exon14:c.C1526A:p.S509X,NCOR1:NM_001190438:exon12:c.C1199A:p.S400X,NCOR1:NM_006311:exon15:c.C1526A:p.S509X, |
| chr17 | 16126196 | C | T | . | exonic | NCOR1 | nonsynonymous SNV | NCOR1:NM_001190440:exon14:c.G1520A:p.R507Q,NCOR1:NM_001190438:exon12:c.G1193A:p.R398Q,NCOR1:NM_006311:exon15:c.G1520A:p.R507Q, |
| chr17 | 16137334 | C | T | . | exonic | NCOR1 | nonsynonymous SNV | NCOR1:NM_001190440:exon13:c.G1486A:p.G496R,NCOR1:NM_001190438:exon11:c.G1159A:p.G387R,NCOR1:NM_006311:exon14:c.G1486A:p.G496R, |

| | | | | | | | | |
|---|---|---|---|---|---|---|---|---|
| chr17 | 16138178 | C | T | . | exonic | NCOR1 | nonsynonymous SNV | NCOR1:NM_001190440:exon12:c.G1387A:p.A463T,NCOR1:NM_001190438:exon10:c.G1060A:p.A354T,NCOR1:NM_006311:exon13:c.G1387A:p.A463T, |
| chr17 | 16139095 | C | T | . | exonic | NCOR1 | nonsynonymous SNV | NCOR1:NM_001190440:exon11:c.G1265A:p.G422E,NCOR1:NM_001190438:exon9:c.G938A:p.G313E,NCOR1:NM_006311:exon12:c.G1265A:p.G422E, |
| chr17 | 16139119 | C | T | COSV52000356 | exonic | NCOR1 | nonsynonymous SNV | NCOR1:NM_001190440:exon11:c.G1241A:p.R414Q,NCOR1:NM_001190438:exon9:c.G914A:p.R305Q,NCOR1:NM_006311:exon12:c.G1241A:p.R414Q, |
| chr17 | 16143692 | C | T | . | exonic | NCOR1 | nonsynonymous SNV | NCOR1:NM_001190440:exon10:c.G1087A:p.G363R,NCOR1:NM_001190438:exon8:c.G760A:p.G254R,NCOR1:NM_006311:exon11:c.G1087A:p.G363R, |
| chr17 | 16146392 | G | T | . | exonic | NCOR1 | nonsynonymous SNV | NCOR1:NM_001190440:exon9:c.C1066A:p.Q356K,NCOR1:NM_001190438:exon7:c.C739A:p.Q247K,NCOR1:NM_006311:exon10:c.C1066A:p.Q356K, |
| chr17 | 16146482 | C | T | . | exonic | NCOR1 | nonsynonymous SNV | NCOR1:NM_001190440:exon9:c.G976A:p.E326K,NCOR1:NM_001190438:exon7:c.G649A:p.E217K,NCOR1:NM_006311:exon10:c.G976A:p.E326K, |
| chr17 | 16146509 | C | T | . | exonic | NCOR1 | nonsynonymous SNV | NCOR1:NM_001190440:exon9:c.G949A:p.A317T,NCOR1:NM_001190438:exon7:c.G622A:p.A208T,NCOR1:NM_006311:exon10:c.G949A:p.A317T, |
| chr17 | 16149456 | G | T | . | exonic | NCOR1 | nonsynonymous SNV | NCOR1:NM_001190440:exon8:c.C904A:p.Q302K,NCOR1:NM_001190438:exon6:c.C577A:p.Q193K,NCOR1:NM_006311:exon9:c.C904A:p.Q302K, |
| chr17 | 16151983 | G | A | . | exonic | NCOR1 | nonsynonymous SNV | NCOR1:NM_001190440:exon7:c.C805T:p.P269S,NCOR1:NM_001190438:exon5:c.C478T:p.P160S,NCOR1:NM_006311:exon8:c.C805T:p.P269S, |
| chr17 | 16151992 | A | G | . | exonic | NCOR1 | nonsynonymous SNV | NCOR1:NM_001190440:exon7:c.T796C:p.Y266H,NCOR1:NM_001190438:exon5:c.T469C:p.Y157H,NCOR1:NM_006311:exon8:c.T796C:p.Y266H, |
| chr17 | 16158825 | C | T | . | exonic | NCOR1 | nonsynonymous SNV | NCOR1:NM_001190440:exon5:c.G667A:p.V223M,NCOR1:NM_001190438:exon3:c.G340A:p.V114M,NCOR1:NM_006311:exon6:c.G667A:p.V223M, |
| chr17 | 16165062 | C | T | . | exonic | NCOR1 | nonsynonymous SNV | NCOR1:NM_001190440:exon4:c.G535A:p.E179K,NCOR1:NM_001190438:exon2:c.G208A:p.E70K,NCOR1:NM_006311:exon5:c.G535A:p.E179K, |

| | | | | | | | | |
|---|---|---|---|---|---|---|---|---|
| chr17 | 16165149 | C | T | COSV51959167 | exonic | NCOR1 | nonsynonymous SNV | NCOR1:NM_001190440:exon4:c.G448A:p.G150R,NCOR1:NM_001190438:exon2:c.G121A:p.G41R,NCOR1:NM_006311:exon5:c.G448A:p.G150R, |
| chr17 | 16194514 | C | T | COSV99063219 | exonic | NCOR1 | nonsynonymous SNV | NCOR1:NM_001190440:exon1:c.G56A:p.R19H,NCOR1:NM_001190438:exon1:c.G56A:p.R19H,NCOR1:NM_006311:exon2:c.G56A:p.R19H, |
| chr17 | 16194524 | C | T | . | exonic | NCOR1 | nonsynonymous SNV | NCOR1:NM_001190440:exon1:c.G46A:p.E16K,NCOR1:NM_001190438:exon1:c.G46A:p.E16K,NCOR1:NM_006311:exon2:c.G46A:p.E16K, |
| chr17 | 16382134 | G | A | . | exonic | UBB | nonsynonymous SNV | UBB:NM_001281719:exon2:c.G227A:p.G76D,UBB:NM_001281717:exon2:c.G227A:p.G76D,UBB:NM_001281716:exon2:c.G227A:p.G76D,UBB:NM_001281718:exon2:c.G227A:p.G76D,UBB:NM_001281720:exon2:c.G227A:p.G76D,UBB:NM_018955:exon2:c.G227A:p.G76D, |
| chr17 | 17270778 | C | T | . | exonic | COPS3 | nonsynonymous SNV | COPS3:NM_003653:exon4:c.G328A:p.A110T,COPS3:NM_001316355:exon4:c.G328A:p.A110T,COPS3:NM_001199125:exon4:c.G268A:p.A90T,COPS3:NM_001316356:exon3:c.G127A:p.A43T, |
| chr17 | 17506246 | C | T | COSV55131323 | exonic | PEMT | nonsynonymous SNV | PEMT:NM_007169:exon6:c.G523A:p.V175M,PEMT:NM_001267552:exon7:c.G665A:p.S222N,PEMT:NM_148173:exon6:c.G523A:p.V175M,PEMT:NM_148172:exon6:c.G634A:p.V212M,PEMT:NM_001267551:exon6:c.G568A:p.V190M, |
| chr17 | 17522317 | C | T | COSV55131208 | exonic | PEMT | nonsynonymous SNV | PEMT:NM_007169:exon3:c.G172A:p.V58I,PEMT:NM_001267552:exon3:c.G283A:p.V95I,PEMT:NM_148173:exon3:c.G172A:p.V58I,PEMT:NM_148172:exon3:c.G283A:p.V95I,PEMT:NM_001267551:exon3:c.G217A:p.V73I, |
| chr17 | 1769982 | C | T | COSV54616828 | exonic | SERPINF1 | nonsynonymous SNV | SERPINF1:NM_002615:exon3:c.C215T:p.T72M, |
| chr17 | 18245220 | C | T | COSV57501487 | exonic | FLII | nonsynonymous SNV | FLII:NM_001256264:exon30:c.G3695A:p.R1232H,FLII:NM_002018:exon30:c.G3728A:p.R1243H,FLII:NM_001256265:exon29:c.G3563A:p.R1188H, |
| chr17 | 27631278 | G | A | COSV56349194 | exonic | LGALS9 | nonsynonymous SNV | LGALS9:NM_009587:exon1:c.G13A:p.G5S,LGALS9:NM_002308:exon1:c.G13A:p.G5S, |
| chr17 | 27643607 | G | A | COSV56348080 | exonic | LGALS9 | nonsynonymous SNV | LGALS9:NM_009587:exon5:c.G527A:p.G176E, |
| chr17 | 27643616 | A | T | COSV56348908 | exonic | LGALS9 | nonsynonymous SNV | LGALS9:NM_009587:exon5:c.A536T:p.Q179L, |
| chr17 | 29993230 | T | A | . | exonic | EFCAB5 | nonsynonymous SNV | EFCAB5:NM_198529:exon5:c.T833A:p.I278K,EFCAB5:NM_001145053:exon5:c.T665A:p.I222K, |
| chr17 | 30971286 | C | G | COSV60433295 | exonic | RNF135 | nonsynonymous SNV | RNF135:NM_197939:exon1:c.C213G:p.H71Q,RNF135:NM_001184992:exon1:c.C213G:p.H71Q,RNF135:NM_032322:exon1:c.C213G:p.H71Q, |
| chr17 | 30971395 | T | C | COSV60433303 | exonic | RNF135 | nonsynonymous SNV | RNF135:NM_197939:exon1:c.T322C:p.S108P,RNF135:NM_001184992:exon1:c.T322C:p.S108P,RNF135:NM_032322:exon1:c.T322C:p.S108P, |
| chr17 | 31252972 | T | A | . | exonic | NF1 | nonsynonymous SNV | NF1:NM_001042492:exon31:c.T4145A:p.V1382E, |
| chr17 | 31856838 | T | C | . | exonic | COPRS | nonsynonymous SNV | COPRS:NM_018405:exon2:c.A127G:p.S43G, |
| chr17 | 31947652 | T | A | . | exonic | SUZ12 | nonsynonymous SNV | SUZ12:NM_015355:exon4:c.T422A:p.V141E, |
| chr17 | 31993275 | C | A | . | exonic | SUZ12 | nonsynonymous SNV | SUZ12:NM_015355:exon11:c.C1235A:p.T412K, |
| chr17 | 31993308 | G | A | COSV59500263 | exonic | SUZ12 | nonsynonymous SNV | SUZ12:NM_015355:exon11:c.G1268A:p.R423Q, |
| chr17 | 32454518 | G | A | . | exonic | PSMD11 | nonsynonymous SNV | PSMD11:NM_001270482:exon3:c.G217A:p.V73I,PSMD11:NM_002815:exon3:c.G217A:p.V73I, |
| chr17 | 32454557 | G | A | . | exonic | PSMD11 | nonsynonymous SNV | PSMD11:NM_001270482:exon3:c.G256A:p.A86T,PSMD11:NM_002815:exon3:c.G256A:p.A86T, |
| chr17 | 32454612 | G | A | . | exonic | PSMD11 | nonsynonymous SNV | PSMD11:NM_001270482:exon3:c.G311A:p.G104E,PSMD11:NM_002815:exon3:c.G311A:p.G104E, |
| chr17 | 32464049 | G | A | COSV55580427 | exonic | PSMD11 | nonsynonymous SNV | PSMD11:NM_001270482:exon4:c.G319A:p.V107I,PSMD11:NM_002815:exon4:c.G319A:p.V107I, |
| chr17 | 32469044 | T | C | . | exonic | PSMD11 | nonsynonymous SNV | PSMD11:NM_001270482:exon6:c.T494C:p.L165P,PSMD11:NM_002815:exon6:c.T494C:p.L165P, |
| chr17 | 32469085 | G | A | . | exonic | PSMD11 | nonsynonymous SNV | PSMD11:NM_001270482:exon6:c.G535A:p.A179T,PSMD11:NM_002815:exon6:c.G535A:p.A179T, |
| chr17 | 32469110 | G | A | COSV55577078 | exonic | PSMD11 | nonsynonymous SNV | PSMD11:NM_001270482:exon6:c.G560A:p.R187Q,PSMD11:NM_002815:exon6:c.G560A:p.R187Q, |
| chr17 | 32473815 | G | A | . | exonic | PSMD11 | nonsynonymous SNV | PSMD11:NM_001270482:exon7:c.G658A:p.A220T,PSMD11:NM_002815:exon7:c.G658A:p.A220T, |
| chr17 | 32473831 | G | A | . | exonic | PSMD11 | stopgain | PSMD11:NM_001270482:exon7:c.G674A:p.W225X,PSMD11:NM_002815:exon7:c.G674A:p.W225X, |
| chr17 | 35717225 | A | G | . | exonic | AP2B1 | nonsynonymous SNV | AP2B1:NM_001282:exon20:c.A2615G:p.N872S,AP2B1:NM_001030006:exon21:c.A2657G:p.N886S, |
| chr17 | 35824128 | G | A | . | exonic | TAF15 | nonsynonymous SNV | TAF15:NM_139215:exon7:c.G535A:p.G179R,TAF15:NM_003487:exon7:c.G526A:p.G176R, |
| chr17 | 35844650 | A | G | . | exonic | TAF15 | nonsynonymous SNV | TAF15:NM_139215:exon15:c.A1351G:p.S451G,TAF15:NM_003487:exon15:c.A1342G:p.S448G, |
| chr17 | 35844989 | G | A | . | exonic | TAF15 | nonsynonymous SNV | TAF15:NM_139215:exon15:c.G1690A:p.G564R,TAF15:NM_003487:exon15:c.G1681A:p.G561R, |
| chr17 | 36001422 | G | A | . | exonic | CCL15 | nonsynonymous SNV | CCL15:NM_032965:exon1:c.C71T:p.T24I, |
| chr17 | 37621247 | C | T | . | exonic | DDX52 | nonsynonymous SNV | DDX52:NM_007010:exon11:c.G1381A:p.G461R,DDX52:NM_001291476:exon12:c.G1057A:p.G353R, |
| chr17 | 37624364 | C | T | . | exonic | DDX52 | nonsynonymous SNV | DDX52:NM_001291476:exon10:c.G883A:p.V295M,DDX52:NM_007010:exon9:c.G1207A:p.V403M, |
| chr17 | 37626060 | A | G | . | exonic | DDX52 | nonsynonymous SNV | DDX52:NM_001291476:exon9:c.T647C:p.F216S,DDX52:NM_007010:exon8:c.T971C:p.F324S, |
| chr17 | 38322567 | G | T | . | exonic | MRPL45 | nonsynonymous SNV | MRPL45:NM_001278279:exon7:c.G743T:p.G248V,MRPL45:NM_032351:exon8:c.G893T:p.G298V, |
| chr17 | 38733306 | C | T | . | exonic | CISD3 | nonsynonymous SNV | CISD3:NM_001136498:exon4:c.C235T:p.R79C, |
| chr17 | 38733414 | A | G | . | exonic | CISD3 | nonsynonymous SNV | CISD3:NM_001136498:exon4:c.A343G:p.S115G, |
| chr17 | 38753202 | G | A | . | exonic | PSMB3 | nonsynonymous SNV | PSMB3:NM_002795:exon2:c.G56A:p.C19Y, |
| chr17 | 38753246 | A | T | . | exonic | PSMB3 | nonsynonymous SNV | PSMB3:NM_002795:exon2:c.A100T:p.M34L, |
| chr17 | 38755965 | G | A | . | exonic | PSMB3 | nonsynonymous SNV | PSMB3:NM_002795:exon3:c.G271A:p.V91M, |
| chr17 | 38755990 | G | A | . | exonic | PSMB3 | nonsynonymous SNV | PSMB3:NM_002795:exon3:c.G296A:p.R99Q, |
| chr17 | 38760534 | G | A | . | exonic | PSMB3 | nonsynonymous SNV | PSMB3:NM_002795:exon4:c.G400A:p.D134N, |
| chr17 | 38762466 | G | A | . | exonic | PSMB3 | nonsynonymous SNV | PSMB3:NM_002795:exon5:c.G530A:p.R177Q, |
| chr17 | 38764142 | G | A | . | exonic | PSMB3 | nonsynonymous SNV | PSMB3:NM_002795:exon6:c.G593A:p.R198K, |
| chr17 | 38810562 | T | C | . | exonic | CWC25 | nonsynonymous SNV | CWC25:NM_017748:exon5:c.A532G:p.K178E, |
| chr17 | 38810564 | T | A | . | exonic | CWC25 | nonsynonymous SNV | CWC25:NM_017748:exon5:c.A530T:p.K177M, |
| chr17 | 38812814 | A | T | . | exonic | CWC25 | nonsynonymous SNV | CWC25:NM_017748:exon4:c.T479A:p.M160K, |
| chr17 | 38812820 | A | T | . | exonic | CWC25 | nonsynonymous SNV | CWC25:NM_017748:exon4:c.T473A:p.V158E, |
| chr17 | 39471045 | G | A | COSV71003036 | exonic | CDK12 | nonsynonymous SNV | CDK12:NM_015083:exon2:c.G1213A:p.E405K,CDK12:NM_016507:exon2:c.G1213A:p.E405K, |
| chr17 | 39630026 | T | C | . | exonic | PPP1R1B | nonsynonymous SNV | PPP1R1B:NM_181505:exon4:c.T112C:p.Y38H,PPP1R1B:NM_001242464:exon4:c.T112C:p.Y38H,PPP1R1B:NM_032192:exon4:c.T220C:p.Y74H, |
| chr17 | 39633969 | G | A | . | exonic | PPP1R1B | nonsynonymous SNV | PPP1R1B:NM_181505:exon5:c.G220A:p.E74K,PPP1R1B:NM_001242464:exon5:c.G220A:p.E74K,PPP1R1B:NM_032192:exon5:c.G328A:p.E110K, |
| chr17 | 39634038 | G | A | . | exonic | PPP1R1B | nonsynonymous SNV | PPP1R1B:NM_181505:exon5:c.G289A:p.E97K,PPP1R1B:NM_001242464:exon5:c.G289A:p.E97K,PPP1R1B:NM_032192:exon5:c.G397A:p.E133K, |
| chr17 | 39634069 | A | T | . | exonic | PPP1R1B | nonsynonymous SNV | PPP1R1B:NM_181505:exon5:c.A320T:p.K107M,PPP1R1B:NM_001242464:exon5:c.A320T:p.K107M,PPP1R1B:NM_032192:exon5:c.A428T:p.K143M, |

| | | | | | | | | |
|---|---|---|---|---|---|---|---|---|
| chr17 | 39635609 | G | A | . | exonic | PPP1R1B | nonsynonymous SNV | PPP1R1B:NM_181505:exon6:c.G340A:p.G114R,PPP1R1B:NM_001242464:exon6:c.G340A:p.G114R,PPP1R1B:NM_032192:exon6:c.G448A:p.G150R, |
| chr17 | 39635675 | T | C | . | exonic | PPP1R1B | nonsynonymous SNV | PPP1R1B:NM_181505:exon6:c.T406C:p.S136P,PPP1R1B:NM_001242464:exon6:c.T406C:p.S136P,PPP1R1B:NM_032192:exon6:c.T514C:p.S172P, |
| chr17 | 39635678 | G | A | . | exonic | PPP1R1B | nonsynonymous SNV | PPP1R1B:NM_181505:exon6:c.G409A:p.E137K,PPP1R1B:NM_001242464:exon6:c.G409A:p.E137K,PPP1R1B:NM_032192:exon6:c.G517A:p.E173K, |
| chr17 | 39635691 | G | A | . | exonic | PPP1R1B | nonsynonymous SNV | PPP1R1B:NM_181505:exon6:c.G422A:p.G141D,PPP1R1B:NM_001242464:exon6:c.G422A:p.G141D,PPP1R1B:NM_032192:exon6:c.G530A:p.G177D, |
| chr17 | 39657827 | G | A | COSV60419443 | exonic | STARD3 | nonsynonymous SNV | STARD3:NM_001165937:exon4:c.G350A:p.R117Q,STARD3:NM_006804:exon4:c.G350A:p.R117Q,STARD3:NM_001165938:exon4:c.G350A:p.R117Q, |
| chr17 | 39684607 | G | C | . | exonic | PGAP3 | nonsynonymous SNV | PGAP3:NM_001291728:exon3:c.C422G:p.A141G,PGAP3:NM_001291730:exon3:c.C422G:p.A141G,PGAP3:NM_033419:exon3:c.C422G:p.A141G,PGAP3:NM_001291733:exon3:c.C422G:p.A141G,<br>PGAP3:NM_001291732:exon3:c.C422G:p.A141G, |
| chr17 | 39727739 | G | T | . | exonic | ERBB2 | nonsynonymous SNV | ERBB2:NM_001289936:exon31:c.G3418T:p.G1140C,ERBB2:NM_004448:exon27:c.G3463T:p.G1155C,ERBB2:NM_001005862:exon30:c.G3373T:p.G1125C, |
| chr17 | 39727784 | C | G | COSV54064104 | exonic | ERBB2 | nonsynonymous SNV | ERBB2:NM_001289936:exon31:c.C3463G:p.P1155A,ERBB2:NM_004448:exon27:c.C3508G:p.P1170A,ERBB2:NM_001005862:exon30:c.C3418G:p.P1140A, |
| chr17 | 39727835 | T | C | . | exonic | ERBB2 | nonsynonymous SNV | ERBB2:NM_001289936:exon31:c.T3514C:p.F1172L,ERBB2:NM_004448:exon27:c.T3559C:p.F1187L,ERBB2:NM_001005862:exon30:c.T3469C:p.F1157L, |
| chr17 | 39742290 | T | C | . | exonic | GRB7 | nonsynonymous SNV | GRB7:NM_001242442:exon2:c.T58C:p.C20R, |
| chr17 | 39905943 | G | A | COSV58779652 | exonic | GSDMB | nonsynonymous SNV | GSDMB:NM_018530:exon6:c.C865T:p.P289S,GSDMB:NM_001165959:exon8:c.C904T:p.P302S,GSDMB:NM_001042471:exon8:c.C892T:p.P298S,GSDMB:NM_001165958:exon9:c.C931T:p.P311S, |
| chr17 | 39905964 | C | T | COSV58779510 | exonic | GSDMB | nonsynonymous SNV | GSDMB:NM_018530:exon6:c.G844A:p.G282R,GSDMB:NM_001165959:exon8:c.G883A:p.G295R,GSDMB:NM_001042471:exon8:c.G871A:p.G291R,GSDMB:NM_001165958:exon9:c.G910A:p.G304<br>R, |
| chr17 | 40300899 | G | A | COSV52929970 | exonic | CDC6 | nonsynonymous SNV | CDC6:NM_001254:exon10:c.G1321A:p.V441I, |
| chr17 | 40301973 | G | A | . | exonic | CDC6 | nonsynonymous SNV | CDC6:NM_001254:exon12:c.G1655A:p.G552E, |
| chr17 | 40356122 | G | A | COSV54190350 | exonic | RARA | nonsynonymous SNV | RARA:NM_001145301:exon9:c.G1285A:p.G429S,RARA:NM_001024809:exon8:c.G1270A:p.G424S,RARA:NM_001145302:exon7:c.G994A:p.G332S,RARA:NM_000964:exon9:c.G1285A:p.G429S, |
| chr17 | 40484492 | C | T | COSV54184711 | exonic | TNS4 | nonsynonymous SNV | TNS4:NM_032865:exon6:c.G1493A:p.S498N, |
| chr17 | 40488873 | A | G | COSV54183743 | exonic | TNS4 | nonsynonymous SNV | TNS4:NM_032865:exon3:c.T536C:p.L179P, |
| chr17 | 40628800 | A | T | . | exonic | SMARCE1 | nonsynonymous SNV | SMARCE1:NM_003079:exon11:c.T1221A:p.D407E, |
| chr17 | 40630756 | C | T | . | exonic | SMARCE1 | nonsynonymous SNV | SMARCE1:NM_003079:exon10:c.G985A:p.G329S, |
| chr17 | 40630845 | C | T | . | exonic | SMARCE1 | nonsynonymous SNV | SMARCE1:NM_003079:exon10:c.G896A:p.R299K, |
| chr17 | 40631682 | T | A | . | exonic | SMARCE1 | nonsynonymous SNV | SMARCE1:NM_003079:exon9:c.A726T:p.E242D, |
| chr17 | 40636474 | C | T | . | exonic | SMARCE1 | nonsynonymous SNV | SMARCE1:NM_003079:exon6:c.G290A:p.G97D, |
| chr17 | 40636522 | C | T | . | exonic | SMARCE1 | stopgain | SMARCE1:NM_003079:exon6:c.G242A:p.W81X, |
| chr17 | 41523789 | C | T | . | exonic | KRT19 | nonsynonymous SNV | KRT19:NM_002276:exon6:c.G1157A:p.G386E, |
| chr17 | 41523886 | C | T | . | exonic | KRT19 | nonsynonymous SNV | KRT19:NM_002276:exon6:c.G1060A:p.A354T, |
| chr17 | 41523949 | C | T | . | exonic | KRT19 | nonsynonymous SNV | KRT19:NM_002276:exon6:c.G997A:p.A333T, |
| chr17 | 41523955 | A | G | . | exonic | KRT19 | nonsynonymous SNV | KRT19:NM_002276:exon6:c.T991C:p.F331L, |
| chr17 | 41524199 | C | T | . | exonic | KRT19 | nonsynonymous SNV | KRT19:NM_002276:exon5:c.G892A:p.D298N, |
| chr17 | 41524424 | C | T | . | exonic | KRT19 | nonsynonymous SNV | KRT19:NM_002276:exon4:c.G777A:p.M259I, |
| chr17 | 41524450 | C | T | . | exonic | KRT19 | nonsynonymous SNV | KRT19:NM_002276:exon4:c.G751A:p.D251N, |
| chr17 | 41524452 | C | T | . | exonic | KRT19 | nonsynonymous SNV | KRT19:NM_002276:exon4:c.G749A:p.S250N, |
| chr17 | 41524848 | C | T | . | exonic | KRT19 | nonsynonymous SNV | KRT19:NM_002276:exon3:c.G655A:p.E219K, |
| chr17 | 41524917 | C | T | . | exonic | KRT19 | nonsynonymous SNV | KRT19:NM_002276:exon3:c.G586A:p.A196T, |
| chr17 | 41527932 | C | T | COSV64239348 | exonic | KRT19 | nonsynonymous SNV | KRT19:NM_002276:exon1:c.G316A:p.G106S, |
| chr17 | 41527977 | G | C | . | exonic | KRT19 | nonsynonymous SNV | KRT19:NM_002276:exon1:c.C271G:p.L91V, |
| chr17 | 41527982 | T | G | . | exonic | KRT19 | nonsynonymous SNV | KRT19:NM_002276:exon1:c.A266C:p.D89A, |
| chr17 | 41527997 | A | T | . | exonic | KRT19 | nonsynonymous SNV | KRT19:NM_002276:exon1:c.T251A:p.M84K, |
| chr17 | 41528069 | G | C | COSV64237813 | exonic | KRT19 | nonsynonymous SNV | KRT19:NM_002276:exon1:c.C179G:p.A60G, |
| chr17 | 41528117 | C | A | . | exonic | KRT19 | nonsynonymous SNV | KRT19:NM_002276:exon1:c.G131T:p.G44V, |
| chr17 | 41528163 | C | T | . | exonic | KRT19 | nonsynonymous SNV | KRT19:NM_002276:exon1:c.G85A:p.V29I, |
| chr17 | 41755775 | G | A | . | exonic | JUP | nonsynonymous SNV | JUP:NM_021991:exon14:c.C2207T:p.P736L,JUP:NM_002230:exon14:c.C2207T:p.P736L, |
| chr17 | 41755893 | T | A | COSV60277352 | exonic | JUP | nonsynonymous SNV | JUP:NM_021991:exon14:c.A2089T:p.M697L,JUP:NM_002230:exon14:c.A2089T:p.M697L, |
| chr17 | 41827556 | G | C | . | exonic | NT5C3B | nonsynonymous SNV | NT5C3B:NM_052935:exon8:c.C638G:p.S213C, |
| chr17 | 41827568 | G | A | COSV54056904 | exonic | NT5C3B | nonsynonymous SNV | NT5C3B:NM_052935:exon8:c.C626T:p.A209V, |
| chr17 | 41909521 | T | G | COSV61250479 | exonic | ACLY | nonsynonymous SNV | ACLY:NM_001096:exon5:c.A525C:p.E175D,ACLY:NM_001303275:exon5:c.A687C:p.E229D,ACLY:NM_198830:exon5:c.A525C:p.E175D,ACLY:NM_001303274:exon5:c.A687C:p.E229D, |
| chr17 | 42017404 | C | T | . | exonic | DNAJC7 | nonsynonymous SNV | DNAJC7:NM_003315:exon1:c.G13A:p.A5T, |
| chr17 | 42324726 | C | T | . | exonic | STAT3 | nonsynonymous SNV | STAT3:NM_139276:exon17:c.G1585A:p.A529T,STAT3:NM_003150:exon17:c.G1585A:p.A529T,STAT3:NM_213662:exon17:c.G1585A:p.A529T, |
| chr17 | 42666681 | A | G | . | exonic | TUBG2 | nonsynonymous SNV | TUBG2:NM_016437:exon11:c.A1237G:p.M413V, |
| chr17 | 42957178 | C | T | . | exonic | AARSD1 | nonsynonymous SNV | AARSD1:NM_001261434:exon4:c.G349A:p.A117T,PTGES3L-AARSD1:NM_001136042:exon9:c.G871A:p.A291T,PTGES3L-AARSD1:NM_025267:exon9:c.G688A:p.A230T, |
| chr17 | 42961251 | C | T | . | exonic | AARSD1 | nonsynonymous SNV | AARSD1:NM_001261434:exon3:c.G272A:p.S91N,PTGES3L-AARSD1:NM_001136042:exon8:c.G794A:p.S265N,PTGES3L-AARSD1:NM_025267:exon8:c.G611A:p.S204N, |
| chr17 | 42964161 | C | T | . | exonic | AARSD1 | nonsynonymous SNV | AARSD1:NM_001261434:exon2:c.G116A:p.S39N,PTGES3L-AARSD1:NM_001136042:exon7:c.G638A:p.S213N,PTGES3L-AARSD1:NM_025267:exon7:c.G455A:p.S152N, |
| chr17 | 42964180 | C | T | . | exonic | AARSD1 | nonsynonymous SNV | AARSD1:NM_001261434:exon2:c.G97A:p.G33S,PTGES3L-AARSD1:NM_001136042:exon7:c.G619A:p.G207S,PTGES3L-AARSD1:NM_025267:exon7:c.G436A:p.G146S, |
| chr17 | 43013476 | A | G | . | exonic | IFI35 | nonsynonymous SNV | IFI35:NM_005533:exon5:c.A382G:p.M128V, |
| chr17 | 43071077 | T | C | COSV58783381 | exonic | BRCA1 | nonsynonymous SNV | BRCA1:NM_007300:exon16:c.A4900G:p.S1634G,BRCA1:NM_007298:exon14:c.A1525G:p.S509G,BRCA1:NM_007294:exon15:c.A4837G:p.S1613G,BRCA1:NM_007297:exon14:c.A4696G:p.S1566G,<br>BRCA1:NM_007299:exon15:c.A1525G:p.S509G, |
| chr17 | 43492697 | A | G | . | exonic | DHX8 | nonsynonymous SNV | DHX8:NM_001302623:exon6:c.A520G:p.K174E,DHX8:NM_004941:exon6:c.A520G:p.K174E, |
| chr17 | 43517240 | G | A | . | exonic | DHX8 | nonsynonymous SNV | DHX8:NM_001302623:exon18:c.G2717A:p.R906H,DHX8:NM_004941:exon18:c.G2717A:p.R906H, |
| chr17 | 43529209 | T | G | . | exonic | ETV4 | nonsynonymous SNV | ETV4:NM_001079675:exon12:c.A1156C:p.N386H,ETV4:NM_001261438:exon11:c.A1039C:p.N347H,ETV4:NM_001261437:exon11:c.A1039C:p.N347H,ETV4:NM_001986:exon12:c.A1156C:p.N386H,<br>ETV4:NM_001261439:exon4:c.A325C:p.N109H, |
| chr17 | 43529220 | C | G | . | exonic | ETV4 | nonsynonymous SNV | ETV4:NM_001079675:exon12:c.G1145C:p.G382A,ETV4:NM_001261438:exon11:c.G1028C:p.G343A,ETV4:NM_001261437:exon11:c.G1028C:p.G343A,ETV4:NM_001986:exon12:c.G1145C:p.G382<br>A,ETV4:NM_001261439:exon4:c.G314C:p.G105A, |
| chr17 | 43529223 | C | G | . | exonic | ETV4 | nonsynonymous SNV | ETV4:NM_001079675:exon12:c.G1142C:p.W381S,ETV4:NM_001261438:exon11:c.G1025C:p.W342S,ETV4:NM_001261437:exon11:c.G1025C:p.W342S,ETV4:NM_001986:exon12:c.G1142C:p.W3<br>81S,ETV4:NM_001261439:exon4:c.G311C:p.W104S, |
| chr17 | 43953163 | G | C | COSV63970394 | exonic | PYY | nonsynonymous SNV | PYY:NM_004160:exon6:c.C215G:p.T72R, |
| chr17 | 43953375 | G | C | COSV63969765 | exonic | PYY | nonsynonymous SNV | PYY:NM_004160:exon5:c.C109G:p.R37G, |
| chr17 | 44207343 | C | T | . | exonic | UBTF | nonsynonymous SNV | UBTF:NM_001076683:exon20:c.G2083A:p.D695N,UBTF:NM_014233:exon21:c.G2194A:p.D732N,UBTF:NM_001076684:exon20:c.G2083A:p.D695N, |
| chr17 | 4559728 | G | A | . | exonic | GGT6 | nonsynonymous SNV | GGT6:NM_001122890:exon2:c.C173T:p.A58V,GGT6:NM_001288704:exon2:c.C173T:p.A58V,GGT6:NM_001288703:exon2:c.C173T:p.A58V,GGT6:NM_001288702:exon2:c.C173T:p.A58V,GGT6:N<br>M_153338:exon2:c.C173T:p.A58V, |
| chr17 | 47827054 | C | T | COSV51621557 | exonic | MRPL10 | nonsynonymous SNV | MRPL10:NM_145255:exon3:c.G373A:p.V125I,MRPL10:NM_148887:exon4:c.G403A:p.V135I, |

| | | | | | | | | |
|---|---|---|---|---|---|---|---|---|
| chr17 | 47975224 | G | A | . | exonic | CDK5RAP3 | nonsynonymous SNV | CDK5RAP3:NM_176096:exon6:c.G400A:p.A134T,CDK5RAP3:NM_001278217:exon5:c.G139A:p.A47T,CDK5RAP3:NM_001278197:exon6:c.G475A:p.A159T, |
| chr17 | 47975970 | G | A | COSV58114545 | exonic | CDK5RAP3 | nonsynonymous SNV | CDK5RAP3:NM_001278198:exon6:c.G80A:p.R27Q,CDK5RAP3:NM_176096:exon8:c.G755A:p.R252Q,CDK5RAP3:NM_001278217:exon7:c.G494A:p.R165Q,CDK5RAP3:NM_001278216:exon6:c.G80A:p.R27Q,CDK5RAP3:NM_001278197:exon8:c.G830A:p.R277Q, |
| chr17 | 47981705 | T | C | COSV58114512 | exonic | CDK5RAP3 | nonsynonymous SNV | CDK5RAP3:NM_001278198:exon13:c.T922C:p.S308P, |
| chr17 | 48184809 | C | T | COSV61146844 | exonic | SKAP1 | nonsynonymous SNV | SKAP1:NM_001075099:exon7:c.G481A:p.G161S,SKAP1:NM_003726:exon7:c.G481A:p.G161S, |
| chr17 | 48849253 | G | A | COSV51951123 | exonic | CALCOCO2 | nonsynonymous SNV | CALCOCO2:NM_001261390:exon6:c.G491A:p.G164E,CALCOCO2:NM_005831:exon5:c.G419A:p.G140E,CALCOCO2:NM_001261395:exon4:c.G203A:p.G68E,CALCOCO2:NM_001261391:exon6:c.G482A:p.G161E, |
| chr17 | 48862296 | C | G | COSV51953043 | exonic | CALCOCO2 | nonsynonymous SNV | CALCOCO2:NM_001261393:exon11:c.C1039G:p.P347A,CALCOCO2:NM_001261390:exon13:c.C1237G:p.P413A,CALCOCO2:NM_005831:exon12:c.C1165G:p.P389A,CALCOCO2:NM_001261395:exon11:c.C949G:p.P317A,CALCOCO2:NM_001261391:exon13:c.C1228G:p.P410A, |
| chr17 | 49600418 | G | A | . | exonic | SPOP | nonsynonymous SNV | SPOP:NM_001007226:exon12:c.C1085T:p.P362L,SPOP:NM_001007227:exon11:c.C1085T:p.P362L,SPOP:NM_001007228:exon10:c.C1085T:p.P362L,SPOP:NM_001007229:exon11:c.C1085T:p.P362L,SPOP:NM_001007230:exon12:c.C1085T:p.P362L,SPOP:NM_003563:exon11:c.C1085T:p.P362L, |
| chr17 | 49600458 | C | T | . | exonic | SPOP | nonsynonymous SNV | SPOP:NM_001007226:exon12:c.G1045A:p.V349M,SPOP:NM_001007227:exon11:c.G1045A:p.V349M,SPOP:NM_001007228:exon10:c.G1045A:p.V349M,SPOP:NM_001007229:exon11:c.G1045A:p.V349M,SPOP:NM_001007230:exon12:c.G1045A:p.V349M,SPOP:NM_003563:exon11:c.G1045A:p.V349M, |
| chr17 | 49601941 | C | T | COSV61652823 | exonic | SPOP | nonsynonymous SNV | SPOP:NM_001007226:exon11:c.G904A:p.A302T,SPOP:NM_001007227:exon10:c.G904A:p.A302T,SPOP:NM_001007228:exon9:c.G904A:p.A302T,SPOP:NM_001007229:exon10:c.G904A:p.A302T,SPOP:NM_001007230:exon11:c.G904A:p.A302T,SPOP:NM_003563:exon10:c.G904A:p.A302T, |
| chr17 | 4960274 | C | T | . | exonic | SPAG7 | nonsynonymous SNV | SPAG7:NM_004890:exon4:c.G287A:p.G96E, |
| chr17 | 4960513 | C | T | . | exonic | SPAG7 | nonsynonymous SNV | SPAG7:NM_004890:exon3:c.G188A:p.S63N, |
| chr17 | 49607368 | C | T | . | exonic | SPOP | nonsynonymous SNV | SPOP:NM_001007226:exon10:c.G719A:p.R240Q,SPOP:NM_001007227:exon9:c.G719A:p.R240Q,SPOP:NM_001007228:exon8:c.G719A:p.R240Q,SPOP:NM_001007229:exon9:c.G719A:p.R240Q,SPOP:NM_001007230:exon10:c.G719A:p.R240Q,SPOP:NM_003563:exon9:c.G719A:p.R240Q, |
| chr17 | 49619069 | C | T | . | exonic | SPOP | stopgain | SPOP:NM_001007226:exon7:c.G392A:p.W131X,SPOP:NM_001007227:exon6:c.G392A:p.W131X,SPOP:NM_001007228:exon5:c.G392A:p.W131X,SPOP:NM_001007229:exon6:c.G392A:p.W131X,SPOP:NM_001007230:exon7:c.G392A:p.W131X,SPOP:NM_003563:exon6:c.G392A:p.W131X, |
| chr17 | 49619261 | C | T | . | exonic | SPOP | nonsynonymous SNV | SPOP:NM_001007226:exon6:c.G325A:p.A109T,SPOP:NM_001007227:exon5:c.G325A:p.A109T,SPOP:NM_001007228:exon4:c.G325A:p.A109T,SPOP:NM_001007229:exon5:c.G325A:p.A109T,SPOP:NM_001007230:exon6:c.G325A:p.A109T,SPOP:NM_003563:exon5:c.G325A:p.A109T, |
| chr17 | 49619333 | A | G | . | exonic | SPOP | nonsynonymous SNV | SPOP:NM_001007227:exon5:c.T253C:p.S85P,SPOP:NM_001007228:exon4:c.T253C:p.S85P,SPOP:NM_001007226:exon6:c.T253C:p.S85P,SPOP:NM_001007229:exon5:c.T253C:p.S85P,SPOP:NM_001007230:exon6:c.T253C:p.S85P,SPOP:NM_003563:exon5:c.T253C:p.S85P, |
| chr17 | 49619362 | C | T | . | exonic | SPOP | nonsynonymous SNV | SPOP:NM_001007226:exon6:c.G224A:p.G75E,SPOP:NM_001007227:exon5:c.G224A:p.G75E,SPOP:NM_001007228:exon4:c.G224A:p.G75E,SPOP:NM_001007229:exon5:c.G224A:p.G75E,SPOP:NM_001007230:exon6:c.G224A:p.G75E,SPOP:NM_003563:exon5:c.G224A:p.G75E, |
| chr17 | 49622001 | C | T | . | exonic | SPOP | nonsynonymous SNV | SPOP:NM_001007226:exon5:c.G145A:p.G49S,SPOP:NM_001007227:exon4:c.G145A:p.G49S,SPOP:NM_001007228:exon3:c.G145A:p.G49S,SPOP:NM_001007229:exon4:c.G145A:p.G49S,SPOP:NM_001007230:exon5:c.G145A:p.G49S,SPOP:NM_003563:exon4:c.G145A:p.G49S, |
| chr17 | 49622007 | C | T | COSV61653952 | exonic | SPOP | nonsynonymous SNV | SPOP:NM_001007226:exon5:c.G139A:p.E47K,SPOP:NM_001007227:exon4:c.G139A:p.E47K,SPOP:NM_001007228:exon3:c.G139A:p.E47K,SPOP:NM_001007229:exon4:c.G139A:p.E47K,SPOP:NM_001007230:exon5:c.G139A:p.E47K,SPOP:NM_003563:exon4:c.G139A:p.E47K, |
| chr17 | 49703193 | C | T | . | exonic | SLC35B1 | nonsynonymous SNV | SLC35B1:NM_001278784:exon7:c.G556A:p.G186S,SLC35B1:NM_005827:exon7:c.G868A:p.G290S, |
| chr17 | 49703231 | A | G | . | exonic | SLC35B1 | nonsynonymous SNV | SLC35B1:NM_005827:exon7:c.T830C:p.I277T,SLC35B1:NM_001278784:exon7:c.T518C:p.I173T, |
| chr17 | 49703250 | C | T | . | exonic | SLC35B1 | nonsynonymous SNV | SLC35B1:NM_001278784:exon7:c.G499A:p.E167K,SLC35B1:NM_005827:exon7:c.G811A:p.E271K, |
| chr17 | 49703253 | C | T | . | exonic | SLC35B1 | nonsynonymous SNV | SLC35B1:NM_005827:exon7:c.G808A:p.A270T,SLC35B1:NM_001278784:exon7:c.G496A:p.A166T, |
| chr17 | 49705198 | C | T | . | exonic | SLC35B1 | nonsynonymous SNV | SLC35B1:NM_005827:exon5:c.G565A:p.A189T,SLC35B1:NM_001278784:exon5:c.G253A:p.A85T, |
| chr17 | 49706301 | C | T | . | exonic | SLC35B1 | nonsynonymous SNV | SLC35B1:NM_001278784:exon3:c.G41A:p.R14H,SLC35B1:NM_005827:exon3:c.G353A:p.R118H, |
| chr17 | 49706974 | C | T | . | exonic | SLC35B1 | nonsynonymous SNV | SLC35B1:NM_001278784:exon2:c.G23A:p.C8Y,SLC35B1:NM_005827:exon2:c.G310A:p.A104T, |
| chr17 | 50360095 | C | G | COSV50017604 | exonic | XYLT2 | nonsynonymous SNV | XYLT2:NM_022167:exon11:c.C2402G:p.T801R, |
| chr17 | 50385154 | C | T | . | exonic | LRRC59 | nonsynonymous SNV | LRRC59:NM_018509:exon6:c.G640A:p.E214K, |
| chr17 | 5335325 | T | A | . | exonic | RABEP1 | stopgain | RABEP1:NM_001083585:exon4:c.T509A:p.L170X,RABEP1:NM_001291581:exon3:c.T380A:p.L127X,RABEP1:NM_001291582:exon4:c.T509A:p.L170X,RABEP1:NM_004703:exon4:c.T509A:p.L170X, |
| chr17 | 5335332 | T | A | COSV52584305 | exonic | RABEP1 | nonsynonymous SNV | RABEP1:NM_001083585:exon4:c.T516A:p.N172K,RABEP1:NM_001291581:exon3:c.T387A:p.N129K,RABEP1:NM_001291582:exon4:c.T516A:p.N172K,RABEP1:NM_004703:exon4:c.T516A:p.N172K, |
| chr17 | 5422825 | C | G | COSV56705547 | exonic | RPAIN | nonsynonymous SNV | RPAIN:NM_001160266:exon3:c.C309G:p.N103K,RPAIN:NM_001160244:exon3:c.C309G:p.N103K,RPAIN:NM_001033002:exon3:c.C309G:p.N103K,RPAIN:NM_001160246:exon3:c.C309G:p.N103K,RPAIN:NM_001160243:exon3:c.C309G:p.N103K, |
| chr17 | 5521757 | T | C | COSV52564879 | exonic | NLRP1 | nonsynonymous SNV | NLRP1:NM_033006:exon12:c.A3460G:p.M1154V,NLRP1:NM_033004:exon13:c.A3550G:p.M1184V,NLRP1:NM_014922:exon13:c.A3550G:p.M1184V,NLRP1:NM_033007:exon12:c.A3460G:p.M1154V,NLRP1:NM_001033053:exon13:c.A3562G:p.M1188V, |
| chr17 | 55751132 | A | C | . | exonic | PCTP | nonsynonymous SNV | PCTP:NM_021213:exon1:c.A29C:p.E10A, |
| chr17 | 57849294 | C | T | . | exonic | MRPS23 | nonsynonymous SNV | MRPS23:NM_016070:exon2:c.G161A:p.G54D, |
| chr17 | 57849295 | C | T | . | exonic | MRPS23 | nonsynonymous SNV | MRPS23:NM_016070:exon2:c.G160A:p.G54S, |
| chr17 | 57849358 | C | T | . | exonic | MRPS23 | nonsynonymous SNV | MRPS23:NM_016070:exon2:c.G97A:p.D33N, |
| chr17 | 57849986 | C | T | . | exonic | MRPS23 | nonsynonymous SNV | MRPS23:NM_016070:exon1:c.G25A:p.V9I, |
| chr17 | 58357515 | C | G | . | exonic | RNF43 | nonsynonymous SNV | RNF43:NM_017763:exon9:c.G2261C:p.G754A,RNF43:NM_001305545:exon8:c.G1880C:p.G627A,RNF43:NM_001305544:exon9:c.G2261C:p.G754A, |
| chr17 | 58357536 | C | T | . | exonic | RNF43 | stopgain | RNF43:NM_017763:exon9:c.G2240A:p.W747X,RNF43:NM_001305545:exon8:c.G1859A:p.W620X,RNF43:NM_001305544:exon9:c.G2240A:p.W747X, |
| chr17 | 58357719 | G | C | . | exonic | RNF43 | nonsynonymous SNV | RNF43:NM_017763:exon9:c.C2057G:p.P686R,RNF43:NM_001305545:exon8:c.C1676G:p.P559R,RNF43:NM_001305544:exon9:c.C2057G:p.P686R, |
| chr17 | 58358524 | G | T | COSV68456749 | exonic | RNF43 | nonsynonymous SNV | RNF43:NM_017763:exon9:c.C1252A:p.L418M,RNF43:NM_001305545:exon8:c.C871A:p.L291M,RNF43:NM_001305544:exon9:c.C1252A:p.L418M, |
| chr17 | 58358748 | C | T | . | exonic | RNF43 | nonsynonymous SNV | RNF43:NM_017763:exon9:c.G1028A:p.R343H,RNF43:NM_001305545:exon8:c.G647A:p.R216H,RNF43:NM_001305544:exon9:c.G1028A:p.R343H, |
| chr17 | 58360940 | G | A | COSV68457515 | exonic | RNF43 | nonsynonymous SNV | RNF43:NM_017763:exon7:c.C692T:p.P231L,RNF43:NM_001305545:exon6:c.C311T:p.P104L,RNF43:NM_001305544:exon7:c.C692T:p.P231L, |
| chr17 | 58363317 | T | G | . | exonic | RNF43 | nonsynonymous SNV | RNF43:NM_017763:exon5:c.A540C:p.Q180H,RNF43:NM_001305545:exon4:c.A159C:p.Q53H,RNF43:NM_001305544:exon5:c.A540C:p.Q180H, |
| chr17 | 58415439 | T | C | COSV60415571 | exonic | RNF43 | nonsynonymous SNV | RNF43:NM_017763:exon2:c.A139G:p.I47V,RNF43:NM_001305544:exon2:c.A139G:p.I47V, |
| chr17 | 59599379 | G | A | . | exonic | DHX40 | nonsynonymous SNV | DHX40:NM_001166301:exon13:c.G1471A:p.A491T,DHX40:NM_024612:exon14:c.G1702A:p.A568T, |
| chr17 | 59682649 | T | G | . | exonic | CLTC | nonsynonymous SNV | CLTC:NM_001288653:exon23:c.T3633G:p.D1211E,CLTC:NM_004859:exon23:c.T3621G:p.D1207E, |
| chr17 | 59697747 | C | T | . | exonic | PTRH2 | nonsynonymous SNV | PTRH2:NM_016077:exon2:c.G232A:p.G78R,PTRH2:NM_001015509:exon3:c.G235A:p.G79R, |
| chr17 | 59886176 | A | G | COSV57872241 | exonic | TUBD1 | nonsynonymous SNV | TUBD1:NM_001193609:exon3:c.T227C:p.M76T,TUBD1:NM_001193611:exon3:c.T227C:p.M76T,TUBD1:NM_016261:exon3:c.T227C:p.M76T,TUBD1:NM_001193610:exon3:c.T227C:p.M76T, |
| chr17 | 60709264 | A | G | . | exonic | BCAS3 | nonsynonymous SNV | BCAS3:NM_001099432:exon5:c.A260G:p.N87S,BCAS3:NM_017679:exon5:c.A260G:p.N87S, |

| | | | | | | | | |
|---|---|---|---|---|---|---|---|---|
| chr17 | 61368392 | C | T | COSV66773169 | exonic | BCAS3 | nonsynonymous SNV | BCAS3:NM_001099432:exon24:c.C2536T:p.R846W,BCAS3:NM_017679:exon23:c.C2491T:p.R831W, |
| chr17 | 623667 | G | T | COSV52126287 | exonic | VPS53 | nonsynonymous SNV | VPS53:NM_018289:exon10:c.C895A:p.L299I,VPS53:NM_001128159:exon11:c.C982A:p.L328I, |
| chr17 | 64002851 | C | T | . | exonic | ICAM2 | nonsynonymous SNV | ICAM2:NM_001099786:exon6:c.G724A:p.V242I,ICAM2:NM_001099787:exon5:c.G724A:p.V242I,ICAM2:NM_001099788:exon6:c.G724A:p.V242I,ICAM2:NM_001099789:exon5:c.G724A:p.V242I,ICAM2:NM_000873:exon4:c.G724A:p.V242I, |
| chr17 | 64499944 | C | T | . | exonic | DDX5 | nonsynonymous SNV | DDX5:NM_004396:exon13:c.G1824A:p.M608I, |
| chr17 | 64500188 | C | T | . | exonic | DDX5 | nonsynonymous SNV | DDX5:NM_004396:exon13:c.G1580A:p.G527E, |
| chr17 | 64502025 | C | T | . | exonic | DDX5 | nonsynonymous SNV | DDX5:NM_004396:exon11:c.G1201A:p.A401T, |
| chr17 | 64502200 | C | T | . | exonic | DDX5 | nonsynonymous SNV | DDX5:NM_004396:exon10:c.G1118A:p.G373D, |
| chr17 | 64502216 | C | T | . | exonic | DDX5 | nonsynonymous SNV | DDX5:NM_004396:exon10:c.G1102A:p.A368T, |
| chr17 | 64502220 | C | T | . | exonic | DDX5 | stopgain | DDX5:NM_004396:exon10:c.G1098A:p.W366X, |
| chr17 | 64503063 | C | T | . | exonic | DDX5 | stopgain | DDX5:NM_004396:exon8:c.G846A:p.W282X, |
| chr17 | 64504015 | C | T | . | exonic | DDX5 | nonsynonymous SNV | DDX5:NM_004396:exon4:c.G409A:p.V137M, |
| chr17 | 64504033 | C | T | . | exonic | DDX5 | nonsynonymous SNV | DDX5:NM_004396:exon4:c.G391A:p.G131R, |
| chr17 | 65558473 | G | A | COSV61057354 | exonic | AXIN2 | nonsynonymous SNV | AXIN2:NM_004655:exon2:c.C148T:p.P50S, |
| chr17 | 6589787 | T | C | . | exonic | KIAA0753 | nonsynonymous SNV | KIAA0753:NM_014804:exon18:c.A2778G:p.I926M, |
| chr17 | 67348597 | G | T | . | exonic | PSMD12 | nonsynonymous SNV | PSMD12:NM_174871:exon4:c.C403A:p.Q135K,PSMD12:NM_001316341:exon7:c.C286A:p.Q96K,PSMD12:NM_002816:exon5:c.C463A:p.Q155K, |
| chr17 | 67724070 | G | A | . | exonic | NOL11 | nonsynonymous SNV | NOL11:NM_015462:exon6:c.G541A:p.V181M, |
| chr17 | 68250275 | A | G | COSV63082480 | exonic | AMZ2 | nonsynonymous SNV | AMZ2:NM_001033574:exon2:c.A88G:p.N30D,AMZ2:NM_001033571:exon3:c.A88G:p.N30D,AMZ2:NM_001289054:exon3:c.A88G:p.N30D,AMZ2:NM_016627:exon2:c.A88G:p.N30D,AMZ2:NM_001033569:exon3:c.A88G:p.N30D,AMZ2:NM_001033570:exon3:c.A88G:p.N30D,AMZ2:NM_001289056:exon3:c.A88G:p.N30D,AMZ2:NM_001033572:exon3:c.A88G:p.N30D, |
| chr17 | 68452981 | G | A | COSV50929487 | exonic | WIPI1 | nonsynonymous SNV | WIPI1:NM_017983:exon2:c.C92T:p.T31I, |
| chr17 | 7260420 | A | G | COSV63038719 | exonic | CLDN7 | nonsynonymous SNV | CLDN7:NM_001185022:exon5:c.T590C:p.V197A,CLDN7:NM_001307:exon4:c.T590C:p.V197A, |
| chr17 | 72947851 | T | C | . | exonic | SLC39A11 | nonsynonymous SNV | SLC39A11:NM_139177:exon5:c.A331G:p.T111A,SLC39A11:NM_001159770:exon5:c.A331G:p.T111A, |
| chr17 | 73209715 | A | G | COSV55428466 | exonic | FAM104A | nonsynonymous SNV | FAM104A:NM_001289411:exon3:c.T293C:p.I98T,FAM104A:NM_001289410:exon2:c.T275C:p.I92T, |
| chr17 | 73212717 | A | T | . | exonic | FAM104A | nonsynonymous SNV | FAM104A:NM_001098832:exon3:c.T345A:p.S115R,FAM104A:NM_001289412:exon3:c.T144A:p.S48R, |
| chr17 | 75225432 | G | A | . | exonic | NUP85 | nonsynonymous SNV | NUP85:NM_001303276:exon8:c.G685A:p.A229T,NUP85:NM_024844:exon9:c.G823A:p.A275T, |
| chr17 | 75261905 | C | T | COSV55452937 | exonic | MRPS7 | nonsynonymous SNV | MRPS7:NM_015971:exon1:c.C5T:p.A2V, |
| chr17 | 75556104 | G | A | COSV51344491 | exonic | LLGL2 | nonsynonymous SNV | LLGL2:NM_001015002:exon3:c.G134A:p.R45H,LLGL2:NM_001031803:exon3:c.G134A:p.R45H,LLGL2:NM_004524:exon3:c.G134A:p.R45H, |
| chr17 | 760232 | C | A | . | exonic | GLOD4 | nonsynonymous SNV | GLOD4:NM_016080:exon9:c.G838T:p.A280S, |
| chr17 | 7676071 | G | A | COSV52702138 | exonic | TP53 | stopgain | TP53:NM_001126118:exon3:c.C181T:p.Q61X,TP53:NM_001276696:exon4:c.C181T:p.Q61X,TP53:NM_001126113:exon4:c.C298T:p.Q100X,TP53:NM_001126112:exon4:c.C298T:p.Q100X,TP53:NM_001276695:exon4:c.C181T:p.Q61X,TP53:NM_001276761:exon4:c.C181T:p.Q61X,TP53:NM_001126114:exon4:c.C298T:p.Q100X,TP53:NM_000546:exon4:c.C298T:p.Q100X,TP53:NM_001276760:exon4:c.C181T:p.Q61X, |
| chr17 | 7676154 | G | C | COSV52666208 | exonic | TP53 | nonsynonymous SNV | TP53:NM_001126118:exon3:c.C98G:p.P33R,TP53:NM_001276696:exon4:c.C98G:p.P33R,TP53:NM_001126113:exon4:c.C215G:p.P72R,TP53:NM_001126112:exon4:c.C215G:p.P72R,TP53:NM_001276695:exon4:c.C98G:p.P33R,TP53:NM_001276761:exon4:c.C98G:p.P33R,TP53:NM_001126114:exon4:c.C215G:p.P72R,TP53:NM_000546:exon4:c.C215G:p.P72R,TP53:NM_001276760:exon4:c.C98G:p.P33R, |
| chr17 | 7688850 | C | G | COSV52834815 | exonic | WRAP53 | nonsynonymous SNV | WRAP53:NM_001143990:exon2:c.C202G:p.R68G,WRAP53:NM_001143991:exon2:c.C202G:p.R68G,WRAP53:NM_001143992:exon2:c.C202G:p.R68G,WRAP53:NM_018081:exon1:c.C202G:p.R68G, |
| chr17 | 7703404 | C | G | COSV56832921 | exonic | WRAP53 | nonsynonymous SNV | WRAP53:NM_001143990:exon11:c.C1565G:p.A522G,WRAP53:NM_001143991:exon11:c.C1565G:p.A522G,WRAP53:NM_001143992:exon11:c.C1565G:p.A522G,WRAP53:NM_018081:exon10:c.C1565G:p.A522G, |
| chr17 | 776971 | C | T | . | exonic | GLOD4 | stopgain | GLOD4:NM_016080:exon3:c.G158A:p.W53X, |
| chr17 | 78223510 | G | A | . | exonic | BIRC5 | nonsynonymous SNV | BIRC5:NM_001168:exon4:c.G385A:p.E129K,BIRC5:NM_001012271:exon5:c.G454A:p.E152K, |
| chr17 | 81246424 | G | C | COSV55844005 | exonic | SLC38A10 | nonsynonymous SNV | SLC38A10:NM_001037984:exon16:c.C2492G:p.A831G, |
| chr17 | 81510694 | A | G | . | exonic | ACTG1 | nonsynonymous SNV | ACTG1:NM_001199954:exon6:c.T1124C:p.F375S,ACTG1:NM_001614:exon6:c.T1124C:p.F375S, |
| chr17 | 81510703 | C | T | . | exonic | ACTG1 | nonsynonymous SNV | ACTG1:NM_001199954:exon6:c.G1115A:p.R372H,ACTG1:NM_001614:exon6:c.G1115A:p.R372H, |
| chr17 | 81510763 | A | G | COSV59510406 | exonic | ACTG1 | nonsynonymous SNV | ACTG1:NM_001199954:exon6:c.T1055C:p.F352S,ACTG1:NM_001614:exon6:c.T1055C:p.F352S, |
| chr17 | 81510790 | C | T | COSV59509690 | exonic | ACTG1 | nonsynonymous SNV | ACTG1:NM_001199954:exon6:c.G1028A:p.G343D,ACTG1:NM_001614:exon6:c.G1028A:p.G343D, |
| chr17 | 81510814 | C | T | . | exonic | ACTG1 | nonsynonymous SNV | ACTG1:NM_001199954:exon6:c.G1004A:p.R335H,ACTG1:NM_001614:exon6:c.G1004A:p.R335H, |
| chr17 | 81510976 | C | T | . | exonic | ACTG1 | nonsynonymous SNV | ACTG1:NM_001199954:exon5:c.G935A:p.R312K,ACTG1:NM_001614:exon5:c.G935A:p.R312K, |
| chr17 | 81511058 | A | G | . | exonic | ACTG1 | nonsynonymous SNV | ACTG1:NM_001199954:exon5:c.T853C:p.C285R,ACTG1:NM_001614:exon5:c.T853C:p.C285R, |
| chr17 | 81511094 | C | T | COSV59510324 | exonic | ACTG1 | nonsynonymous SNV | ACTG1:NM_001199954:exon5:c.G817A:p.G273S,ACTG1:NM_001614:exon5:c.G817A:p.G273S, |
| chr17 | 81511238 | C | T | . | exonic | ACTG1 | nonsynonymous SNV | ACTG1:NM_001199954:exon4:c.G752A:p.G251D,ACTG1:NM_001614:exon4:c.G752A:p.G251D, |
| chr17 | 81511256 | C | T | . | exonic | ACTG1 | nonsynonymous SNV | ACTG1:NM_001199954:exon4:c.G734A:p.G245D,ACTG1:NM_001614:exon4:c.G734A:p.G245D, |
| chr17 | 81511299 | C | T | . | exonic | ACTG1 | nonsynonymous SNV | ACTG1:NM_001199954:exon4:c.G691A:p.A231T,ACTG1:NM_001614:exon4:c.G691A:p.A231T, |
| chr17 | 81511343 | A | G | . | exonic | ACTG1 | nonsynonymous SNV | ACTG1:NM_001199954:exon4:c.T647C:p.L216P,ACTG1:NM_001614:exon4:c.T647C:p.L216P, |
| chr17 | 81511361 | C | T | . | exonic | ACTG1 | nonsynonymous SNV | ACTG1:NM_001199954:exon4:c.G629A:p.R210H,ACTG1:NM_001614:exon4:c.G629A:p.R210H, |
| chr17 | 81511373 | C | T | . | exonic | ACTG1 | nonsynonymous SNV | ACTG1:NM_001199954:exon4:c.G617A:p.R206Q,ACTG1:NM_001614:exon4:c.G617A:p.R206Q, |
| chr17 | 81511400 | C | T | . | exonic | ACTG1 | nonsynonymous SNV | ACTG1:NM_001199954:exon4:c.G590A:p.G197D,ACTG1:NM_001614:exon4:c.G590A:p.G197D, |
| chr17 | 81511407 | C | T | . | exonic | ACTG1 | nonsynonymous SNV | ACTG1:NM_001199954:exon4:c.G583A:p.E195K,ACTG1:NM_001614:exon4:c.G583A:p.E195K, |
| chr17 | 81511428 | A | G | . | exonic | ACTG1 | nonsynonymous SNV | ACTG1:NM_001199954:exon4:c.T562C:p.Y188H,ACTG1:NM_001614:exon4:c.T562C:p.Y188H, |
| chr17 | 81511455 | C | T | . | exonic | ACTG1 | nonsynonymous SNV | ACTG1:NM_001199954:exon4:c.G535A:p.D179N,ACTG1:NM_001614:exon4:c.G535A:p.D179N, |
| chr17 | 81511523 | C | T | COSV59509822 | exonic | ACTG1 | nonsynonymous SNV | ACTG1:NM_001199954:exon4:c.G467A:p.G156E,ACTG1:NM_001614:exon4:c.G467A:p.G156E, |
| chr17 | 81511571 | A | G | . | exonic | ACTG1 | nonsynonymous SNV | ACTG1:NM_001199954:exon4:c.T419C:p.L140P,ACTG1:NM_001614:exon4:c.T419C:p.L140P, |
| chr17 | 81511594 | C | T | . | exonic | ACTG1 | nonsynonymous SNV | ACTG1:NM_001199954:exon4:c.G396A:p.M132I,ACTG1:NM_001614:exon4:c.G396A:p.M132I, |
| chr17 | 81511926 | C | T | . | exonic | ACTG1 | nonsynonymous SNV | ACTG1:NM_001199954:exon3:c.G340A:p.A114T,ACTG1:NM_001614:exon3:c.G340A:p.A114T, |
| chr17 | 81511934 | T | G | . | exonic | ACTG1 | nonsynonymous SNV | ACTG1:NM_001199954:exon3:c.A332C:p.N111T,ACTG1:NM_001614:exon3:c.A332C:p.N111T, |
| chr17 | 81511943 | G | A | . | exonic | ACTG1 | nonsynonymous SNV | ACTG1:NM_001199954:exon3:c.C323T:p.A108V,ACTG1:NM_001614:exon3:c.C323T:p.A108V, |
| chr17 | 81511944 | C | T | . | exonic | ACTG1 | nonsynonymous SNV | ACTG1:NM_001199954:exon3:c.G322A:p.A108T,ACTG1:NM_001614:exon3:c.G322A:p.A108T, |
| chr17 | 81511980 | C | T | . | exonic | ACTG1 | nonsynonymous SNV | ACTG1:NM_001199954:exon3:c.G286A:p.V96M,ACTG1:NM_001614:exon3:c.G286A:p.V96M, |
| chr17 | 81512025 | C | T | . | exonic | ACTG1 | nonsynonymous SNV | ACTG1:NM_001199954:exon3:c.G241A:p.D81N,ACTG1:NM_001614:exon3:c.G241A:p.D81N, |
| chr17 | 81512045 | C | T | . | exonic | ACTG1 | nonsynonymous SNV | ACTG1:NM_001199954:exon3:c.G221A:p.G74D,ACTG1:NM_001614:exon3:c.G221A:p.G74D, |
| chr17 | 81512088 | T | A | . | exonic | ACTG1 | nonsynonymous SNV | ACTG1:NM_001199954:exon3:c.A178T:p.S60C,ACTG1:NM_001614:exon3:c.A178T:p.S60C, |
| chr17 | 81512130 | C | T | COSV59509545 | exonic | ACTG1 | nonsynonymous SNV | ACTG1:NM_001199954:exon3:c.G136A:p.G46S,ACTG1:NM_001614:exon3:c.G136A:p.G46S, |
| chr17 | 81512285 | C | T | . | exonic | ACTG1 | nonsynonymous SNV | ACTG1:NM_001199954:exon2:c.G70A:p.D24N,ACTG1:NM_001614:exon2:c.G70A:p.D24N, |
| chr17 | 81512315 | A | G | . | exonic | ACTG1 | nonsynonymous SNV | ACTG1:NM_001199954:exon2:c.T40C:p.S14P,ACTG1:NM_001614:exon2:c.T40C:p.S14P, |
| chr17 | 81512336 | C | T | . | exonic | ACTG1 | nonsynonymous SNV | ACTG1:NM_001199954:exon2:c.G19A:p.A7T,ACTG1:NM_001614:exon2:c.G19A:p.A7T, |
| chr17 | 81512339 | C | T | . | exonic | ACTG1 | nonsynonymous SNV | ACTG1:NM_001199954:exon2:c.G16A:p.A6T,ACTG1:NM_001614:exon2:c.G16A:p.A6T, |

| | | | | | | | | |
|---|---|---|---|---|---|---|---|---|
| chr17 | 81704684 | T | C | COSV61270949 | exonic | MRPL12 | nonsynonymous SNV | MRPL12:NM_002949:exon3:c.T313C:p.S105P, |
| chr17 | 81846461 | A | G | . | exonic | P4HB | nonsynonymous SNV | P4HB:NM_000918:exon7:c.T1024C:p.F342L, |
| chr17 | 81846512 | T | A | . | exonic | P4HB | nonsynonymous SNV | P4HB:NM_000918:exon7:c.A973T:p.T325S, |
| chr17 | 81847265 | A | T | . | exonic | P4HB | nonsynonymous SNV | P4HB:NM_000918:exon5:c.T707A:p.L236H, |
| chr17 | 81847323 | C | T | COSV58939761 | exonic | P4HB | nonsynonymous SNV | P4HB:NM_000918:exon5:c.G649A:p.E217K, |
| chr17 | 81859334 | C | T | . | exonic | P4HB | nonsynonymous SNV | P4HB:NM_000918:exon2:c.G199A:p.A67T, |
| chr17 | 81860397 | G | C | . | exonic | P4HB | nonsynonymous SNV | P4HB:NM_000918:exon1:c.C75G:p.H25Q, |
| chr17 | 81868822 | A | C | . | exonic | ARHGDIA | nonsynonymous SNV | ARHGDIA:NM_001301242:exon7:c.T557G:p.M186R, |
| chr17 | 81869013 | T | A | . | exonic | ARHGDIA | nonsynonymous SNV | ARHGDIA:NM_001301241:exon6:c.A478T:p.T160S,ARHGDIA:NM_004309:exon6:c.A478T:p.T160S,ARHGDIA:NM_001301243:exon5:c.A613T:p.T205S,ARHGDIA:NM_001301240:exon6:c.A478T:p.T160S,ARHGDIA:NM_001185077:exon6:c.A478T:p.T160S, |
| chr17 | 81996668 | T | A | COSV60727771 | exonic | ASPSCR1 | nonsynonymous SNV | ASPSCR1:NM_001251888:exon7:c.T755A:p.L252Q,ASPSCR1:NM_024083:exon7:c.T755A:p.L252Q, |
| chr17 | 8205013 | A | G | COSV60243948 | exonic | AURKB | nonsynonymous SNV | AURKB:NM_001313955:exon7:c.T389C:p.M130T,AURKB:NM_001313950:exon9:c.T893C:p.M298T,AURKB:NM_001313954:exon9:c.T437C:p.M146T,AURKB:NM_001313951:exon9:c.T770C:p.M257T,AURKB:NM_001256834:exon8:c.T770C:p.M257T,AURKB:NM_001313953:exon8:c.T797C:p.M266T,AURKB:NM_004217:exon9:c.T893C:p.M298T,AURKB:NM_001284526:exon9:c.T896C:p.M299T,AURKB:NM_001313952:exon8:c.T773C:p.M258T, |
| chr17 | 82061312 | G | T | . | exonic | DUS1L | nonsynonymous SNV | DUS1L:NM_022156:exon8:c.C739A:p.P247T, |
| chr18 | 12274105 | G | T | COSV57599921 | exonic | CIDEA | nonsynonymous SNV | CIDEA:NM_001279:exon4:c.G343T:p.V115F, |
| chr18 | 20979942 | A | T | . | exonic | ROCK1 | nonsynonymous SNV | ROCK1:NM_005406:exon22:c.T2622A:p.N874K, |
| chr18 | 20992868 | C | T | . | exonic | ROCK1 | nonsynonymous SNV | ROCK1:NM_005406:exon17:c.G1955A:p.R652K, |
| chr18 | 3457608 | C | T | . | exonic | TGIF1 | nonsynonymous SNV | TGIF1:NM_173208:exon4:c.C487T:p.P163S,TGIF1:NM_170695:exon3:c.C874T:p.P292S,TGIF1:NM_173209:exon3:c.C427T:p.P143S,TGIF1:NM_001278684:exon4:c.C487T:p.P163S,TGIF1:NM_173210:exon3:c.C427T:p.P143S,TGIF1:NM_001278686:exon4:c.C427T:p.P143S,TGIF1:NM_173207:exon3:c.C529T:p.P177S,TGIF1:NM_173211:exon3:c.C427T:p.P143S,TGIF1:NM_003244:exon3:c.C487T:p.P163S,TGIF1:NM_174886:exon4:c.C427T:p.P143S,TGIF1:NM_001278682:exon3:c.C496T:p.P166S, |
| chr18 | 34890327 | G | A | COSV51991943 | exonic | DTNA | nonsynonymous SNV | DTNA:NM_001198938:exon20:c.G2062A:p.E688K, |
| chr18 | 47896582 | C | T | COSV50993158 | exonic | SMAD2 | nonsynonymous SNV | SMAD2:NM_005901:exon2:c.G175A:p.D59N,SMAD2:NM_001003652:exon2:c.G175A:p.D59N,SMAD2:NM_001135937:exon2:c.G175A:p.D59N, |
| chr18 | 47896587 | C | T | COSV50994318 | exonic | SMAD2 | nonsynonymous SNV | SMAD2:NM_005901:exon2:c.G170A:p.R57Q,SMAD2:NM_001003652:exon2:c.G170A:p.R57Q,SMAD2:NM_001135937:exon2:c.G170A:p.R57Q, |
| chr18 | 47896638 | C | T | . | exonic | SMAD2 | stopgain | SMAD2:NM_005901:exon2:c.G119A:p.W40X,SMAD2:NM_001003652:exon2:c.G119A:p.W40X,SMAD2:NM_001135937:exon2:c.G119A:p.W40X, |
| chr18 | 56603034 | C | T | . | exonic | TXNL1 | nonsynonymous SNV | TXNL1:NM_004786:exon8:c.G863A:p.S288N, |
| chr18 | 56611070 | C | T | . | exonic | TXNL1 | nonsynonymous SNV | TXNL1:NM_004786:exon7:c.G763A:p.E255K, |
| chr18 | 56618086 | C | T | . | exonic | TXNL1 | nonsynonymous SNV | TXNL1:NM_004786:exon4:c.G410A:p.C137Y, |
| chr18 | 56624460 | C | T | . | exonic | TXNL1 | nonsynonymous SNV | TXNL1:NM_004786:exon3:c.G197A:p.G66E, |
| chr18 | 56624461 | C | T | . | exonic | TXNL1 | nonsynonymous SNV | TXNL1:NM_004786:exon3:c.G196A:p.G66R, |
| chr18 | 76880129 | C | T | . | exonic | ZNF236 | nonsynonymous SNV | ZNF236:NM_007345:exon8:c.C995T:p.T332M,ZNF236:NM_001306089:exon8:c.C1001T:p.T334M, |
| chr18 | 79307094 | G | A | COSV56948748 | exonic | ATP9B | nonsynonymous SNV | ATP9B:NM_198531:exon15:c.G1633A:p.V545M,ATP9B:NM_001306085:exon15:c.G1633A:p.V545M, |
| chr18 | 79337360 | A | T | . | exonic | ATP9B | nonsynonymous SNV | ATP9B:NM_198531:exon19:c.A2194T:p.M732L,ATP9B:NM_001306085:exon19:c.A2194T:p.M732L, |
| chr18 | 79968135 | G | T | . | exonic | HSBP1L1 | nonsynonymous SNV | HSBP1L1:NM_001136180:exon3:c.G165T:p.K55N, |
| chr18 | 9117869 | T | C | COSV59186794 | exonic | NDUFV2 | nonsynonymous SNV | NDUFV2:NM_021074:exon2:c.T86C:p.V29A, |
| chr19 | 1010397 | G | C | . | exonic | TMEM259 | nonsynonymous SNV | TMEM259:NM_001033026:exon11:c.C1816G:p.P606A, |
| chr19 | 10252575 | A | C | . | exonic | MRPL4 | nonsynonymous SNV | MRPL4:NM_146387:exon4:c.A149C:p.K50T,MRPL4:NM_146388:exon3:c.A149C:p.K50T,MRPL4:NM_015956:exon3:c.A149C:p.K50T, |
| chr19 | 10631494 | A | G | COSV59835474 | exonic | SLC44A2 | nonsynonymous SNV | SLC44A2:NM_020428:exon7:c.A461G:p.Q154R,SLC44A2:NM_001145056:exon7:c.A455G:p.Q152R, |
| chr19 | 10638257 | G | A | COSV59837549 | exonic | SLC44A2 | nonsynonymous SNV | SLC44A2:NM_001145056:exon20:c.G1865A:p.R622H,SLC44A2:NM_020428:exon20:c.G1871A:p.R624H, |
| chr19 | 10775793 | T | A | . | exonic | DNM2 | nonsynonymous SNV | DNM2:NM_001190716:exon4:c.T476A:p.M159K,DNM2:NM_001005360:exon4:c.T476A:p.M159K,DNM2:NM_004945:exon4:c.T476A:p.M159K,DNM2:NM_001005361:exon4:c.T476A:p.M159K,DNM2:NM_001005362:exon4:c.T476A:p.M159K, |
| chr19 | 10775873 | G | A | . | exonic | DNM2 | nonsynonymous SNV | DNM2:NM_001190716:exon4:c.G556A:p.A186T,DNM2:NM_001005360:exon4:c.G556A:p.A186T,DNM2:NM_004945:exon4:c.G556A:p.A186T,DNM2:NM_001005361:exon4:c.G556A:p.A186T,DNM2:NM_001005362:exon4:c.G556A:p.A186T, |
| chr19 | 10777166 | G | A | . | exonic | DNM2 | nonsynonymous SNV | DNM2:NM_001190716:exon5:c.G638A:p.G213D,DNM2:NM_001005360:exon5:c.G638A:p.G213D,DNM2:NM_004945:exon5:c.G638A:p.G213D,DNM2:NM_001005361:exon5:c.G638A:p.G213D,DNM2:NM_001005362:exon5:c.G638A:p.G213D, |
| chr19 | 10783113 | T | C | COSV58971997 | exonic | DNM2 | nonsynonymous SNV | DNM2:NM_001190716:exon6:c.T842C:p.L281P,DNM2:NM_001005360:exon6:c.T842C:p.L281P,DNM2:NM_004945:exon6:c.T842C:p.L281P,DNM2:NM_001005361:exon6:c.T842C:p.L281P,DNM2:NM_001005362:exon6:c.T842C:p.L281P, |
| chr19 | 10783116 | A | C | . | exonic | DNM2 | nonsynonymous SNV | DNM2:NM_004945:exon6:c.A845C:p.N282T,DNM2:NM_001005361:exon6:c.A845C:p.N282T,DNM2:NM_001190716:exon6:c.A845C:p.N282T,DNM2:NM_001005360:exon6:c.A845C:p.N282T,DNM2:NM_001005362:exon6:c.A845C:p.N282T, |
| chr19 | 10795384 | G | A | . | exonic | DNM2 | nonsynonymous SNV | DNM2:NM_001190716:exon9:c.G1141A:p.E381K,DNM2:NM_001005360:exon9:c.G1141A:p.E381K,DNM2:NM_004945:exon9:c.G1141A:p.E381K,DNM2:NM_001005361:exon9:c.G1141A:p.E381K,DNM2:NM_001005362:exon9:c.G1141A:p.E381K, |
| chr19 | 10796146 | G | A | . | exonic | DNM2 | nonsynonymous SNV | DNM2:NM_001190716:exon10:c.G1282A:p.V428I,DNM2:NM_001005360:exon10:c.G1282A:p.V428I,DNM2:NM_004945:exon10:c.G1282A:p.V428I, |
| chr19 | 10812271 | G | A | COSV58957808 | exonic | DNM2 | nonsynonymous SNV | DNM2:NM_001190716:exon15:c.G1565A:p.R522H,DNM2:NM_001005360:exon15:c.G1565A:p.R522H,DNM2:NM_004945:exon14:c.G1553A:p.R518H,DNM2:NM_001005361:exon15:c.G1565A:p.R522H,DNM2:NM_001005362:exon14:c.G1553A:p.R518H, |
| chr19 | 10812274 | G | A | . | exonic | DNM2 | nonsynonymous SNV | DNM2:NM_001190716:exon15:c.G1568A:p.R523K,DNM2:NM_001005360:exon15:c.G1568A:p.R523K,DNM2:NM_004945:exon14:c.G1556A:p.R519K,DNM2:NM_001005361:exon15:c.G1568A:p.R523K,DNM2:NM_001005362:exon14:c.G1556A:p.R519K, |
| chr19 | 10820026 | G | A | . | exonic | DNM2 | nonsynonymous SNV | DNM2:NM_004945:exon15:c.G1706A:p.R569H,DNM2:NM_001005361:exon16:c.G1718A:p.R573H,DNM2:NM_001190716:exon16:c.G1718A:p.R573H,DNM2:NM_001005360:exon16:c.G1718A:p.R573H,DNM2:NM_001005362:exon15:c.G1706A:p.R569H, |
| chr19 | 10829148 | G | A | COSV53033495 | exonic | DNM2 | nonsynonymous SNV | DNM2:NM_001190716:exon19:c.G2171A:p.R724H,DNM2:NM_001005360:exon19:c.G2171A:p.R724H,DNM2:NM_004945:exon18:c.G2159A:p.R720H,DNM2:NM_001005361:exon19:c.G2171A:p.R724H,DNM2:NM_001005362:exon18:c.G2159A:p.R720H, |
| chr19 | 1094005 | G | A | . | exonic | POLR2E | nonsynonymous SNV | POLR2E:NM_002695:exon2:c.C131T:p.S44F, |
| chr19 | 11003383 | G | A | . | exonic | SMARCA4 | nonsynonymous SNV | SMARCA4:NM_001128844:exon14:c.G1987A:p.E663K,SMARCA4:NM_001128846:exon12:c.G1987A:p.E663K,SMARCA4:NM_001128847:exon12:c.G1987A:p.E663K,SMARCA4:NM_003072:exon13:c.G1987A:p.E663K,SMARCA4:NM_001128849:exon13:c.G1987A:p.E663K,SMARCA4:NM_001128848:exon12:c.G1987A:p.E663K,SMARCA4:NM_001128845:exon12:c.G1987A:p.E663K, |
| chr19 | 11059834 | G | A | . | exonic | SMARCA4 | nonsynonymous SNV | SMARCA4:NM_001128846:exon31:c.G4624A:p.E1542K,SMARCA4:NM_001128847:exon31:c.G4618A:p.E1540K,SMARCA4:NM_003072:exon33:c.G4717A:p.E1573K,SMARCA4:NM_001128848:exon31:c.G4615A:p.E1539K,SMARCA4:NM_001128844:exon34:c.G4717A:p.E1573K,SMARCA4:NM_001128849:exon34:c.G4813A:p.E1605K,SMARCA4:NM_001128845:exon31:c.G4627A:p.E1543K, |
| chr19 | 11111624 | G | A | COSV52943212 | exonic | LDLR | nonsynonymous SNV | LDLR:NM_001195799:exon7:c.G1048A:p.A350T,LDLR:NM_000527:exon8:c.G1171A:p.A391T,LDLR:NM_001195798:exon8:c.G1171A:p.A391T,LDLR:NM_001195803:exon7:c.G790A:p.A264T,LDLR:NM_001195800:exon6:c.G667A:p.A223T, |
| chr19 | 11214289 | C | T | COSV53914202 | exonic | DOCK6 | nonsynonymous SNV | DOCK6:NM_020812:exon34:c.G4324A:p.A1442T, |
| chr19 | 1223095 | T | C | . | exonic | STK11 | nonsynonymous SNV | STK11:NM_000455:exon8:c.T1031C:p.L344P, |
| chr19 | 12431839 | T | G | COSV56890258 | exonic | ZNF443 | nonsynonymous SNV | ZNF443:NM_005815:exon4:c.A333C:p.K111N, |
| chr19 | 12431840 | T | A | COSV56890270 | exonic | ZNF443 | nonsynonymous SNV | ZNF443:NM_005815:exon4:c.A332T:p.K111I, |

| | | | | | | | | |
|---|---|---|---|---|---|---|---|---|
| chr19 | 12432437 | A | C | COSV56890302 | exonic | ZNF443 | nonsynonymous SNV | ZNF443:NM_005815:exon3:c.T131G:p.V44G, |
| chr19 | 12899706 | A | G | COSV53365821 | exonic | GCDH | nonsynonymous SNV | GCDH:NM_013976:exon12:c.A1250G:p.Q417R, |
| chr19 | 12924908 | T | C | . | exonic | FARSA | nonsynonymous SNV | FARSA:NM_004461:exon9:c.A1022G:p.Q341R, |
| chr19 | 12930628 | G | A | . | exonic | FARSA | nonsynonymous SNV | FARSA:NM_004461:exon2:c.C269T:p.A90V, |
| chr19 | 12930703 | G | A | . | exonic | FARSA | nonsynonymous SNV | FARSA:NM_004461:exon2:c.C194T:p.A65V, |
| chr19 | 12954294 | G | T | . | exonic | GADD45GIP1 | nonsynonymous SNV | GADD45GIP1:NM_052850:exon2:c.C583A:p.Q195K, |
| chr19 | 12956978 | G | C | COSV57534227 | exonic | GADD45GIP1 | nonsynonymous SNV | GADD45GIP1:NM_052850:exon1:c.C235G:p.P79A, |
| chr19 | 1360576 | G | A | COSV60900886 | exonic | MUM1 | nonsynonymous SNV | MUM1:NM_032853:exon5:c.G658A:p.G220R, |
| chr19 | 13774670 | A | G | COSV50004416 | exonic | C19ORF53 | nonsynonymous SNV | C19orf53:NM_014047:exon2:c.A116G:p.K39R, |
| chr19 | 1388539 | C | T | COSV52039612 | exonic | NDUFS7 | nonsynonymous SNV | NDUFS7:NM_024407:exon3:c.C68T:p.P23L, |
| chr19 | 15256162 | G | T | . | exonic | BRD4 | nonsynonymous SNV | BRD4:NM_014299:exon9:c.C1653A:p.H551Q,BRD4:NM_058243:exon9:c.C1653A:p.H551Q, |
| chr19 | 15697074 | A | G | COSV61175313 | exonic | CYP4F12 | nonsynonymous SNV | CYP4F12:NM_023944:exon13:c.A1564G:p.S522G, |
| chr19 | 16654066 | A | G | . | exonic | SMIM7 | nonsynonymous SNV | SMIM7:NM_001300925:exon4:c.T181C:p.W61R,SMIM7:NM_024104:exon4:c.T181C:p.W61R, |
| chr19 | 16660086 | C | T | . | exonic | SMIM7 | nonsynonymous SNV | SMIM7:NM_001300925:exon1:c.G25A:p.G9R,SMIM7:NM_024104:exon1:c.G25A:p.G9R, |
| chr19 | 17060075 | C | T | COSV53725996 | exonic | HAUS8 | nonsynonymous SNV | HAUS8:NM_033417:exon5:c.G247A:p.G83R,HAUS8:NM_001011699:exon5:c.G244A:p.G82R, |
| chr19 | 17219251 | T | C | . | exonic | USE1 | nonsynonymous SNV | USE1:NM_018467:exon7:c.T461C:p.L154S, |
| chr19 | 17424663 | G | A | COSV57679130 | exonic | MVB12A | nonsynonymous SNV | MVB12A:NM_001304547:exon9:c.G469A:p.D157N,MVB12A:NM_138401:exon8:c.G745A:p.D249N, |
| chr19 | 17511745 | C | A | . | exonic | PGLS | nonsynonymous SNV | PGLS:NM_012088:exon1:c.C73A:p.L25M, |
| chr19 | 17726616 | C | G | COSV60721781 | exonic | MAP1S | nonsynonymous SNV | MAP1S:NM_001308363:exon5:c.C1154G:p.S385C,MAP1S:NM_018174:exon5:c.C1232G:p.S411C, |
| chr19 | 18175134 | G | A | COSV55847770 | exonic | IFI30 | nonsynonymous SNV | IFI30:NM_006332:exon2:c.G227A:p.R76Q, |
| chr19 | 18386214 | G | C | COSV53250533 | exonic | GDF15 | nonsynonymous SNV | GDF15:NM_004864:exon1:c.G25C:p.V9L, |
| chr19 | 18386331 | T | A | COSV53250556 | exonic | GDF15 | nonsynonymous SNV | GDF15:NM_004864:exon1:c.T142A:p.S48T, |
| chr19 | 18451628 | C | T | COSV53211465 | exonic | ELL | nonsynonymous SNV | ELL:NM_006532:exon7:c.G890A:p.S297N, |
| chr19 | 18537611 | C | T | . | exonic | FKBP8 | nonsynonymous SNV | FKBP8:NM_001308373:exon6:c.G932A:p.R311H,FKBP8:NM_012181:exon6:c.G935A:p.R312H, |
| chr19 | 18568569 | C | G | COSV55889000 | exonic | KXD1 | nonsynonymous SNV | KXD1:NM_001171948:exon6:c.C469G:p.P157A,KXD1:NM_001171949:exon5:c.C469G:p.P157A,KXD1:NM_024069:exon5:c.C469G:p.P157A, |
| chr19 | 19120007 | T | C | COSV50777255 | exonic | TMEM161A | nonsynonymous SNV | TMEM161A:NM_017814:exon12:c.A1363G:p.I455V,TMEM161A:NM_001256766:exon10:c.A1054G:p.I352V, |
| chr19 | 19891759 | T | C | . | exonic | ZNF253 | nonsynonymous SNV | ZNF253:NM_021047:exon4:c.T512C:p.L171P, |
| chr19 | 19891817 | T | A | . | exonic | ZNF253 | nonsynonymous SNV | ZNF253:NM_021047:exon4:c.T570A:p.H190Q, |
| chr19 | 19891985 | T | A | . | exonic | ZNF253 | nonsynonymous SNV | ZNF253:NM_021047:exon4:c.T738A:p.H246Q, |
| chr19 | 19892444 | T | A | COSV63036584 | exonic | ZNF253 | nonsynonymous SNV | ZNF253:NM_021047:exon4:c.T1197A:p.D399E, |
| chr19 | 19892457 | A | G | COSV63036591 | exonic | ZNF253 | nonsynonymous SNV | ZNF253:NM_021047:exon4:c.A1210G:p.T404A, |
| chr19 | 19892464 | C | A | . | exonic | ZNF253 | nonsynonymous SNV | ZNF253:NM_021047:exon4:c.C1217A:p.T406N, |
| chr19 | 19892468 | G | C | . | exonic | ZNF253 | nonsynonymous SNV | ZNF253:NM_021047:exon4:c.G1221C:p.W407C, |
| chr19 | 2040173 | G | A | . | exonic | MKNK2 | nonsynonymous SNV | MKNK2:NM_199054:exon13:c.C1115T:p.A372V,MKNK2:NM_017572:exon13:c.C1115T:p.A372V, |
| chr19 | 2050824 | G | T | COSV51731205 | exonic | MKNK2 | nonsynonymous SNV | MKNK2:NM_199054:exon2:c.C28A:p.Q10K,MKNK2:NM_017572:exon2:c.C28A:p.Q10K, |
| chr19 | 2109158 | T | C | COSV61427281 | exonic | AP3D1 | nonsynonymous SNV | AP3D1:NM_001261826:exon30:c.A3400G:p.I1134V,AP3D1:NM_003938:exon28:c.A3214G:p.I1072V, |
| chr19 | 23359488 | A | G | COSV56084671 | exonic | ZNF91 | nonsynonymous SNV | ZNF91:NM_003430:exon4:c.T3491C:p.L1164P,ZNF91:NM_001300951:exon3:c.T3395C:p.L1132P, |
| chr19 | 23359489 | G | C | COSV56084686 | exonic | ZNF91 | nonsynonymous SNV | ZNF91:NM_003430:exon4:c.C3490G:p.L1164V,ZNF91:NM_001300951:exon3:c.C3394G:p.L1132V, |
| chr19 | 23360293 | T | C | COSV56080094 | exonic | ZNF91 | nonsynonymous SNV | ZNF91:NM_003430:exon4:c.A2686G:p.T896A,ZNF91:NM_001300951:exon3:c.A2590G:p.T864A, |
| chr19 | 23360692 | T | A | . | exonic | ZNF91 | nonsynonymous SNV | ZNF91:NM_003430:exon4:c.A2287T:p.I763F,ZNF91:NM_001300951:exon3:c.A2191T:p.I731F, |
| chr19 | 23361049 | C | T | COSV56085308 | exonic | ZNF91 | nonsynonymous SNV | ZNF91:NM_003430:exon4:c.G1930A:p.A644T,ZNF91:NM_001300951:exon3:c.G1834A:p.A612T, |
| chr19 | 23361823 | C | T | COSV56079882 | exonic | ZNF91 | nonsynonymous SNV | ZNF91:NM_003430:exon4:c.G1156A:p.A386T,ZNF91:NM_001300951:exon3:c.G1060A:p.A354T, |
| chr19 | 23362043 | A | T | . | exonic | ZNF91 | nonsynonymous SNV | ZNF91:NM_003430:exon4:c.T936A:p.H312Q,ZNF91:NM_001300951:exon3:c.T840A:p.H280Q, |
| chr19 | 23362045 | G | T | . | exonic | ZNF91 | nonsynonymous SNV | ZNF91:NM_003430:exon4:c.C934A:p.H312N,ZNF91:NM_001300951:exon3:c.C838A:p.H280N, |
| chr19 | 23362645 | A | G | COSV56080539 | exonic | ZNF91 | nonsynonymous SNV | ZNF91:NM_003430:exon4:c.T334C:p.Y112H,ZNF91:NM_001300951:exon3:c.T238C:p.Y80H, |
| chr19 | 23362714 | G | C | . | exonic | ZNF91 | nonsynonymous SNV | ZNF91:NM_003430:exon4:c.C265G:p.H89D,ZNF91:NM_001300951:exon3:c.C169G:p.H57D, |
| chr19 | 23374700 | G | A | COSV56080556 | exonic | ZNF91 | nonsynonymous SNV | ZNF91:NM_003430:exon2:c.C95T:p.T32I,ZNF91:NM_001300951:exon2:c.C95T:p.T32I, |
| chr19 | 282753 | G | A | COSV54120817 | exonic | PLPP2 | nonsynonymous SNV | PLPP2:NM_177526:exon4:c.C371T:p.A124V,PLPP2:NM_003712:exon4:c.C539T:p.A180V,PLPP2:NM_177543:exon4:c.C602T:p.A201V, |
| chr19 | 29213103 | A | C | COSV59184287 | exonic | UQCRFS1 | nonsynonymous SNV | UQCRFS1:NM_006003:exon1:c.T16G:p.S6A, |
| chr19 | 30009243 | G | A | . | exonic | URI1 | nonsynonymous SNV | URI1:NM_003796:exon8:c.G925A:p.D309N,URI1:NM_001252641:exon8:c.G871A:p.D291N, |
| chr19 | 30011135 | T | A | . | exonic | URI1 | nonsynonymous SNV | URI1:NM_003796:exon9:c.T1077A:p.S359R,URI1:NM_001252641:exon9:c.T1023A:p.S341R, |
| chr19 | 33109858 | T | C | COSV50111027 | exonic | GPATCH1 | nonsynonymous SNV | GPATCH1:NM_018025:exon11:c.T1427C:p.L476P, |
| chr19 | 33114394 | A | G | COSV50114971 | exonic | GPATCH1 | nonsynonymous SNV | GPATCH1:NM_018025:exon15:c.A2171G:p.H724R, |
| chr19 | 33114406 | T | C | COSV50114723 | exonic | GPATCH1 | nonsynonymous SNV | GPATCH1:NM_018025:exon15:c.T2183C:p.L728S, |
| chr19 | 33125143 | C | A | . | exonic | GPATCH1 | nonsynonymous SNV | GPATCH1:NM_018025:exon18:c.C2560A:p.H854N, |
| chr19 | 33125171 | C | T | . | exonic | GPATCH1 | nonsynonymous SNV | GPATCH1:NM_018025:exon18:c.C2588T:p.A863V, |
| chr19 | 33126632 | T | A | COSV50118646 | exonic | GPATCH1 | nonsynonymous SNV | GPATCH1:NM_018025:exon19:c.T2664A:p.N888K, |
| chr19 | 34215626 | A | G | COSV70885886 | exonic | LSM14A | nonsynonymous SNV | LSM14A:NM_001114093:exon6:c.A746G:p.N249S,LSM14A:NM_015578:exon6:c.A746G:p.N249S, |
| chr19 | 35157462 | T | G | COSV54343702 | exonic | FXYD5 | nonsynonymous SNV | FXYD5:NM_001164605:exon3:c.T103G:p.S35A,FXYD5:NM_144779:exon3:c.T103G:p.S35A,FXYD5:NM_014164:exon3:c.T103G:p.S35A, |
| chr19 | 35169605 | G | A | . | exonic | FXYD5 | nonsynonymous SNV | FXYD5:NM_001164605:exon9:c.G527A:p.R176H,FXYD5:NM_144779:exon9:c.G527A:p.R176H,FXYD5:NM_014164:exon9:c.G527A:p.R176H, |
| chr19 | 35500540 | T | G | . | exonic | DMKN | nonsynonymous SNV | DMKN:NM_001308383:exon7:c.A368C:p.D123A,DMKN:NM_001126056:exon14:c.A1247C:p.D416A,DMKN:NM_001035516:exon2:c.A122C:p.D41A,DMKN:NM_001308380:exon8:c.A419C:p.D140A,DM<br>KN:NM_001126059:exon8:c.A419C:p.D140A,DMKN:NM_001190347:exon12:c.A1157C:p.D386A,DMKN:NM_033317:exon12:c.A1280C:p.D427A, |
| chr19 | 35995543 | G | C | COSV66677829 | exonic | SDHAF1 | nonsynonymous SNV | SDHAF1:NM_001042631:exon1:c.G269C:p.C90S, |
| chr19 | 36104534 | A | C | COSV54335267 | exonic | WDR62 | nonsynonymous SNV | WDR62:NM_001083961:exon31:c.A4170C:p.L1390F,WDR62:NM_173636:exon31:c.A4155C:p.L1385F, |
| chr19 | 36656520 | C | T | COSV64453504 | exonic | ZNF461 | nonsynonymous SNV | ZNF461:NM_001297623:exon4:c.G160A:p.V54I,ZNF461:NM_153257:exon4:c.G160A:p.V54I, |
| chr19 | 3750617 | T | C | . | exonic | TJP3 | nonsynonymous SNV | TJP3:NM_001267561:exon21:c.T2720C:p.M907T,TJP3:NM_001267560:exon21:c.T2693C:p.M898T, |
| chr19 | 38624111 | G | A | . | exonic | EIF3K | nonsynonymous SNV | EIF3K:NM_001300992:exon3:c.G193A:p.V65I,EIF3K:NM_013234:exon3:c.G193A:p.V65I, |
| chr19 | 38624192 | G | A | . | exonic | EIF3K | nonsynonymous SNV | EIF3K:NM_001300992:exon3:c.G274A:p.A92T,EIF3K:NM_013234:exon3:c.G274A:p.A92T,EIF3K:NM_001308393:exon3:c.G13A:p.A5T, |
| chr19 | 38626047 | G | A | COSV50263701 | exonic | EIF3K | nonsynonymous SNV | EIF3K:NM_001300992:exon4:c.G299A:p.R100Q,EIF3K:NM_013234:exon4:c.G299A:p.R100Q,EIF3K:NM_001308393:exon4:c.G38A:p.R13Q, |
| chr19 | 38632643 | G | A | . | exonic | EIF3K | nonsynonymous SNV | EIF3K:NM_013234:exon6:c.G464A:p.R155H,EIF3K:NM_001308393:exon6:c.G203A:p.R68H, |
| chr19 | 38636906 | G | A | . | exonic | EIF3K | nonsynonymous SNV | EIF3K:NM_001300992:exon7:c.G565A:p.A189T,EIF3K:NM_013234:exon8:c.G643A:p.A215T,EIF3K:NM_001308393:exon8:c.G382A:p.A128T, |
| chr19 | 38710325 | T | C | . | exonic | ACTN4 | nonsynonymous SNV | ACTN4:NM_004924:exon8:c.T802C:p.F268L, |
| chr19 | 38816463 | C | T | COSV55489256 | exonic | ECH1 | nonsynonymous SNV | ECH1:NM_001398:exon7:c.G649A:p.G217R, |
| chr19 | 38831447 | T | G | COSV55489637 | exonic | ECH1 | nonsynonymous SNV | ECH1:NM_001398:exon2:c.A122C:p.E41A, |
| chr19 | 39435675 | C | T | . | exonic | RPS16 | nonsynonymous SNV | RPS16:NM_001020:exon2:c.G82A:p.G28S, |
| chr19 | 3977950 | C | T | . | exonic | EEF2 | nonsynonymous SNV | EEF2:NM_001961:exon12:c.G1936A:p.A646T, |
| chr19 | 3977970 | T | C | . | exonic | EEF2 | nonsynonymous SNV | EEF2:NM_001961:exon12:c.A1916G:p.Y639C, |
| chr19 | 3977991 | G | A | . | exonic | EEF2 | nonsynonymous SNV | EEF2:NM_001961:exon12:c.C1895T:p.A632V, |

| | | | | | | | | |
|---|---|---|---|---|---|---|---|---|
| chr19 | 3981373 | T | A | . | exonic | EEF2 | nonsynonymous SNV | EEF2:NM_001961:exon7:c.A977T:p.E326V, |
| chr19 | 3982000 | C | T | . | exonic | EEF2 | nonsynonymous SNV | EEF2:NM_001961:exon6:c.G844A:p.G282R, |
| chr19 | 3982321 | T | A | . | exonic | EEF2 | nonsynonymous SNV | EEF2:NM_001961:exon5:c.A716T:p.K239M, |
| chr19 | 3982354 | A | G | . | exonic | EEF2 | nonsynonymous SNV | EEF2:NM_001961:exon5:c.T683C:p.F228S, |
| chr19 | 3982839 | C | T | . | exonic | EEF2 | nonsynonymous SNV | EEF2:NM_001961:exon4:c.G580A:p.E194K, |
| chr19 | 3982868 | T | A | . | exonic | EEF2 | nonsynonymous SNV | EEF2:NM_001961:exon4:c.A551T:p.N184I, |
| chr19 | 3982948 | C | T | . | exonic | EEF2 | nonsynonymous SNV | EEF2:NM_001961:exon4:c.G471A:p.M157I, |
| chr19 | 3982961 | G | T | . | exonic | EEF2 | nonsynonymous SNV | EEF2:NM_001961:exon4:c.C458A:p.P153H, |
| chr19 | 3982995 | C | T | . | exonic | EEF2 | nonsynonymous SNV | EEF2:NM_001961:exon4:c.G424A:p.V142M, |
| chr19 | 3983207 | G | C | . | exonic | EEF2 | nonsynonymous SNV | EEF2:NM_001961:exon3:c.C303G:p.N101K, |
| chr19 | 3984149 | T | G | . | exonic | EEF2 | nonsynonymous SNV | EEF2:NM_001961:exon2:c.A205C:p.T69P, |
| chr19 | 3984161 | C | T | . | exonic | EEF2 | nonsynonymous SNV | EEF2:NM_001961:exon2:c.G193A:p.E65K, |
| chr19 | 3984187 | A | T | . | exonic | EEF2 | nonsynonymous SNV | EEF2:NM_001961:exon2:c.T167A:p.F56Y, |
| chr19 | 3984241 | G | A | . | exonic | EEF2 | nonsynonymous SNV | EEF2:NM_001961:exon2:c.C113T:p.S38F, |
| chr19 | 3984263 | C | T | . | exonic | EEF2 | nonsynonymous SNV | EEF2:NM_001961:exon2:c.G91A:p.G31S, |
| chr19 | 39873276 | C | T | . | exonic | FCGBP | nonsynonymous SNV | FCGBP:NM_003890:exon23:c.G11123A:p.R3708H, |
| chr19 | 4123845 | C | T | . | exonic | MAP2K2 | nonsynonymous SNV | MAP2K2:NM_030662:exon1:c.G31A:p.A11T, |
| chr19 | 41397315 | G | A | COSV54195409 | exonic | EXOSC5 | nonsynonymous SNV | EXOSC5:NM_020158:exon1:c.C14T:p.T5M, |
| chr19 | 41686928 | T | A | . | exonic | CEACAM7 | nonsynonymous SNV | CEACAM7:NM_001291485:exon2:c.A358T:p.I120F,CEACAM7:NM_006890:exon2:c.A358T:p.I120F, |
| chr19 | 41761981 | T | G | . | exonic | CEACAM6 | nonsynonymous SNV | CEACAM6:NM_002483:exon4:c.T716G:p.V239G, |
| chr19 | 4333714 | C | T | . | exonic | STAP2 | nonsynonymous SNV | STAP2:NM_017720:exon3:c.G277A:p.D93N,STAP2:NM_001013841:exon3:c.G277A:p.D93N, |
| chr19 | 43353721 | G | C | COSV65121323 | exonic | CD177 | nonsynonymous SNV | CD177:NM_020406:exon1:c.G7C:p.A3P, |
| chr19 | 43356040 | T | G | COSV65120741 | exonic | CD177 | nonsynonymous SNV | CD177:NM_020406:exon5:c.T551G:p.V184G, |
| chr19 | 43356099 | G | A | COSV65120743 | exonic | CD177 | nonsynonymous SNV | CD177:NM_020406:exon5:c.G610A:p.D204N, |
| chr19 | 43356103 | T | G | COSV65120745 | exonic | CD177 | nonsynonymous SNV | CD177:NM_020406:exon5:c.T614G:p.M205R, |
| chr19 | 43361540 | G | A | COSV65120749 | exonic | CD177 | nonsynonymous SNV | CD177:NM_020406:exon8:c.G1042A:p.A348T, |
| chr19 | 43652320 | T | C | COSV55389007 | exonic | PLAUR | nonsynonymous SNV | PLAUR:NM_001005376:exon6:c.A659G:p.K220R,PLAUR:NM_001005377:exon5:c.A524G:p.K175R,PLAUR:NM_002659:exon6:c.A659G:p.K220R, |
| chr19 | 44905910 | C | G | COSV52986063 | exonic | APOE | nonsynonymous SNV | APOE:NM_001302688:exon1:c.C42G:p.N14K, |
| chr19 | 44908684 | T | C | COSV52985231 | exonic | APOE | nonsynonymous SNV | APOE:NM_000041:exon4:c.T388C:p.C130R,APOE:NM_001302691:exon4:c.T388C:p.C130R,APOE:NM_001302689:exon4:c.T388C:p.C130R,APOE:NM_001302690:exon4:c.T388C:p.C130R,APOE:NM_001302688:exon4:c.T466C:p.C156R, |
| chr19 | 45351661 | T | G | COSV67266431 | exonic | ERCC2 | nonsynonymous SNV | ERCC2:NM_000400:exon23:c.A2251C:p.K751Q, |
| chr19 | 45355704 | C | G | . | exonic | ERCC2 | nonsynonymous SNV | ERCC2:NM_000400:exon16:c.G1504C:p.V502L, |
| chr19 | 45364001 | C | T | COSV55538614 | exonic | ERCC2 | nonsynonymous SNV | ERCC2:NM_000400:exon10:c.G934A:p.D312N,ERCC2:NM_001130867:exon9:c.G862A:p.D288N, |
| chr19 | 45507774 | G | C | . | exonic | VASP | nonsynonymous SNV | VASP:NM_003370:exon1:c.G3C:p.M1I, |
| chr19 | 4670301 | C | G | COSV53560730 | exonic | MYDGF | nonsynonymous SNV | MYDGF:NM_019107:exon1:c.G34C:p.G12R, |
| chr19 | 46775602 | C | G | COSV69467128 | exonic | SLC1A5 | nonsynonymous SNV | SLC1A5:NM_001145145:exon7:c.G928C:p.V310L,SLC1A5:NM_005628:exon8:c.G1534C:p.V512L,SLC1A5:NM_001145144:exon8:c.G850C:p.V284L, |
| chr19 | 46787917 | G | C | COSV69467084 | exonic | SLC1A5 | nonsynonymous SNV | SLC1A5:NM_005628:exon1:c.C49G:p.P17A, |
| chr19 | 47046196 | C | T | COSV53409969 | exonic | TMEM160 | nonsynonymous SNV | TMEM160:NM_017854:exon3:c.G358A:p.G120S, |
| chr19 | 47755460 | A | G | COSV55881187 | exonic | GLTSCR2 | nonsynonymous SNV | GLTSCR2:NM_015710:exon9:c.A1166G:p.Q389R, |
| chr19 | 4847701 | A | G | . | exonic | PLIN3 | nonsynonymous SNV | PLIN3:NM_005817:exon6:c.T824C:p.V275A,PLIN3:NM_001164194:exon6:c.T788C:p.V263A,PLIN3:NM_001164189:exon6:c.T824C:p.V275A, |
| chr19 | 4859925 | T | C | COSV55735432 | exonic | PLIN3 | nonsynonymous SNV | PLIN3:NM_005817:exon3:c.A166G:p.I56V,PLIN3:NM_001164194:exon3:c.A166G:p.I56V,PLIN3:NM_001164189:exon3:c.A166G:p.I56V, |
| chr19 | 48835841 | T | C | COSV54361404 | exonic | HSD17B14 | nonsynonymous SNV | HSD17B14:NM_016246:exon2:c.A91G:p.N31D, |
| chr19 | 48956250 | G | A | . | exonic | BAX | nonsynonymous SNV | BAX:NM_001291428:exon4:c.G286A:p.A96T,BAX:NM_001291431:exon2:c.G52A:p.A18T,BAX:NM_138763:exon3:c.G139A:p.A47T,BAX:NM_001291430:exon2:c.G52A:p.A18T,BAX:NM_001291429:exon3:c.G175A:p.A59T,BAX:NM_138764:exon4:c.G286A:p.A96T,BAX:NM_004324:exon4:c.G286A:p.A96T,BAX:NM_138761:exon4:c.G286A:p.A96T, |
| chr19 | 48961655 | A | T | . | exonic | BAX | nonsynonymous SNV | BAX:NM_001291428:exon6:c.A647T:p.D216V, |
| chr19 | 49107866 | G | A | . | exonic | SNRNP70 | nonsynonymous SNV | SNRNP70:NM_001301069:exon10:c.G710A:p.R237Q,SNRNP70:NM_003089:exon10:c.G737A:p.R246Q, |
| chr19 | 49446395 | G | A | COSV51807265 | exonic | PIH1D1 | nonsynonymous SNV | PIH1D1:NM_017916:exon9:c.C860T:p.P287L, |
| chr19 | 49447041 | C | T | COSV51807272 | exonic | PIH1D1 | nonsynonymous SNV | PIH1D1:NM_017916:exon7:c.G670A:p.V224I, |
| chr19 | 49451546 | C | T | COSV51806224 | exonic | PIH1D1 | nonsynonymous SNV | PIH1D1:NM_017916:exon1:c.G29A:p.G10E, |
| chr19 | 49451550 | T | G | COSV51806239 | exonic | PIH1D1 | nonsynonymous SNV | PIH1D1:NM_017916:exon1:c.A25C:p.M9L, |
| chr19 | 49880379 | T | C | COSV55578602 | exonic | TBC1D17 | nonsynonymous SNV | TBC1D17:NM_001168222:exon3:c.T197C:p.L66P,TBC1D17:NM_024682:exon4:c.T296C:p.L99P, |
| chr19 | 50820217 | C | G | . | exonic | KLK1 | nonsynonymous SNV | KLK1:NM_002257:exon3:c.G433C:p.E145Q, |
| chr19 | 51015980 | A | G | COSV59399939 | exonic | KLK10 | nonsynonymous SNV | KLK10:NM_001077500:exon4:c.T446C:p.L149P,KLK10:NM_002776:exon4:c.T446C:p.L149P,KLK10:NM_145888:exon4:c.T446C:p.L149P, |
| chr19 | 51017231 | A | C | COSV59397518 | exonic | KLK10 | nonsynonymous SNV | KLK10:NM_001077500:exon3:c.T148G:p.S50A,KLK10:NM_002776:exon3:c.T148G:p.S50A,KLK10:NM_145888:exon3:c.T148G:p.S50A, |
| chr19 | 51024714 | C | T | COSV99031948 | exonic | KLK11 | nonsynonymous SNV | KLK11:NM_144947:exon3:c.G217A:p.E73K,KLK11:NM_001136032:exon3:c.G121A:p.E41K,KLK11:NM_001167605:exon3:c.G121A:p.E41K,KLK11:NM_006853:exon3:c.G121A:p.E41K, |
| chr19 | 51347036 | G | A | COSV58535703 | exonic | ETFB | nonsynonymous SNV | ETFB:NM_001014763:exon4:c.C734T:p.T245M,ETFB:NM_001985:exon5:c.C461T:p.T154M, |
| chr19 | 51965078 | C | T | . | exonic | ZNF350 | nonsynonymous SNV | ZNF350:NM_021632:exon5:c.G1375A:p.G459R, |
| chr19 | 51965857 | G | T | . | exonic | ZNF350 | nonsynonymous SNV | ZNF350:NM_021632:exon5:c.C596A:p.T199K, |
| chr19 | 52211311 | G | A | . | exonic | PPP2R1A | nonsynonymous SNV | PPP2R1A:NM_014225:exon4:c.G322A:p.A108T, |
| chr19 | 52212718 | C | A | . | exonic | PPP2R1A | nonsynonymous SNV | PPP2R1A:NM_014225:exon5:c.C536A:p.P179H, |
| chr19 | 52212721 | T | A | . | exonic | PPP2R1A | nonsynonymous SNV | PPP2R1A:NM_014225:exon5:c.T539A:p.M180K, |
| chr19 | 52213078 | G | A | . | exonic | PPP2R1A | nonsynonymous SNV | PPP2R1A:NM_014225:exon6:c.G775A:p.V259I, |
| chr19 | 52216031 | G | A | . | exonic | PPP2R1A | nonsynonymous SNV | PPP2R1A:NM_014225:exon8:c.G950A:p.C317Y, |
| chr19 | 52219704 | G | A | . | exonic | PPP2R1A | nonsynonymous SNV | PPP2R1A:NM_014225:exon10:c.G1142A:p.R381Q, |
| chr19 | 52219820 | C | T | . | exonic | PPP2R1A | nonsynonymous SNV | PPP2R1A:NM_014225:exon10:c.C1258T:p.R420W, |
| chr19 | 52384174 | A | C | COSV69906033 | exonic | ZNF880 | nonsynonymous SNV | ZNF880:NM_001145434:exon4:c.A594C:p.R198S, |
| chr19 | 52384184 | A | C | COSV69905703 | exonic | ZNF880 | nonsynonymous SNV | ZNF880:NM_001145434:exon4:c.A604C:p.N202H, |
| chr19 | 52384818 | G | T | COSV69905689 | exonic | ZNF880 | nonsynonymous SNV | ZNF880:NM_001145434:exon4:c.G1238T:p.C413F, |
| chr19 | 52384992 | A | G | COSV69906039 | exonic | ZNF880 | nonsynonymous SNV | ZNF880:NM_001145434:exon4:c.A1412G:p.K471R, |
| chr19 | 54160472 | C | G | . | exonic | TMC4 | nonsynonymous SNV | TMC4:NM_144686:exon14:c.G2047C:p.E683Q,TMC4:NM_001145303:exon14:c.G2065C:p.E689Q, |
| chr19 | 54173068 | T | C | . | exonic | TMC4 | nonsynonymous SNV | TMC4:NM_144686:exon1:c.A50G:p.E17G,TMC4:NM_001145303:exon1:c.A50G:p.E17G, |
| chr19 | 55384220 | G | A | COSV59804634 | exonic | TMEM238 | nonsynonymous SNV | TMEM238:NM_001190764:exon1:c.C40T:p.P14S, |
| chr19 | 5692181 | C | T | COSV56798284 | exonic | LONP1 | nonsynonymous SNV | LONP1:NM_001276480:exon18:c.G2143A:p.V715I,LONP1:NM_001276479:exon19:c.G2539A:p.V847I,LONP1:NM_004793:exon18:c.G2731A:p.V911I, |
| chr19 | 6737470 | T | C | COSV51184367 | exonic | GPR108 | nonsynonymous SNV | GPR108:NM_001080452:exon1:c.A107G:p.Q36R, |
| chr19 | 7293887 | G | C | . | exonic | INSR | nonsynonymous SNV | INSR:NM_000208:exon1:c.C5G:p.A2G,INSR:NM_001079817:exon1:c.C5G:p.A2G, |
| chr19 | 7647391 | A | G | COSV55401668 | exonic | STXBP2 | nonsynonymous SNV | STXBP2:NM_006949:exon18:c.A1576G:p.I526V,STXBP2:NM_001272034:exon18:c.A1609G:p.I537V,STXBP2:NM_001127396:exon18:c.A1567G:p.I523V, |
| chr19 | 7682407 | T | G | COSV58039587 | exonic | TRAPPC5 | nonsynonymous SNV | TRAPPC5:NM_001042461:exon2:c.T154G:p.S52A,TRAPPC5:NM_174894:exon2:c.T154G:p.S52A,TRAPPC5:NM_001042462:exon2:c.T154G:p.S52A, |

| | | | | | | | | |
|---|---|---|---|---|---|---|---|---|
| chr19 | 8371280 | C | T | COSV56844530 | exonic | ANGPTL4 | nonsynonymous SNV | ANGPTL4:NM_001039667:exon5:c.C683T:p.T228M,ANGPTL4:NM_139314:exon6:c.C797T:p.T266M, |
| chr19 | 990281 | G | A | COSV52120984 | exonic | WDR18 | nonsynonymous SNV | WDR18:NM_024100:exon4:c.G514A:p.A172T, |
| chr2 | 100974842 | A | G | COSV59555566 | exonic | NPAS2 | nonsynonymous SNV | NPAS2:NM_002518:exon13:c.A1180G:p.T394A, |
| chr2 | 10122793 | T | G | COSV58815817 | exonic | RRM2 | nonsynonymous SNV | RRM2:NM_001165931:exon1:c.T175G:p.S59A, |
| chr2 | 101281950 | G | C | . | exonic | RNF149 | nonsynonymous SNV | RNF149:NM_173647:exon6:c.C1068G:p.D356E, |
| chr2 | 101308564 | T | C | COSV54863782 | exonic | RNF149 | nonsynonymous SNV | RNF149:NM_173647:exon1:c.A25G:p.S9G, |
| chr2 | 102814945 | T | C | COSV68425595 | exonic | TMEM182 | nonsynonymous SNV | TMEM182:NM_144632:exon5:c.T667C:p.W223R, |
| chr2 | 10790777 | T | C | COSV55349252 | exonic | PDIA6 | nonsynonymous SNV | PDIA6:NM_001282705:exon8:c.A785G:p.K262R,PDIA6:NM_001282707:exon8:c.A632G:p.K211R,PDIA6:NM_001282706:exon8:c.A656G:p.K219R,PDIA6:NM_005742:exon7:c.A641G:p.K214R,PDIA6:NM_001282704:exon9:c.A797G:p.K266R, |
| chr2 | 111929411 | G | A | COSV54933511 | exonic | MERTK | nonsynonymous SNV | MERTK:NM_006343:exon2:c.G353A:p.S118N, |
| chr2 | 111994351 | G | A | COSV54924715 | exonic | MERTK | nonsynonymous SNV | MERTK:NM_006343:exon9:c.G1397A:p.R466K, |
| chr2 | 111997424 | A | G | COSV54924734 | exonic | MERTK | nonsynonymous SNV | MERTK:NM_006343:exon10:c.A1552G:p.I518V, |
| chr2 | 112585994 | C | T | . | exonic | CHCHD5 | nonsynonymous SNV | CHCHD5:NM_032309:exon2:c.C23T:p.T8I, |
| chr2 | 119372265 | A | G | COSV61055619 | exonic | DBI | nonsynonymous SNV | DBI:NM_020548:exon5:c.A262G:p.M88V,DBI:NM_001282633:exon5:c.A262G:p.M88V,DBI:NM_001282634:exon5:c.A262G:p.M88V,DBI:NM_001178041:exon5:c.A337G:p.M113V,DBI:NM_001079863:exon4:c.A214G:p.M72V,DBI:NM_001282636:exon4:c.A139G:p.M47V,DBI:NM_001178017:exon4:c.A394G:p.M132V,DBI:NM_001282635:exon5:c.A262G:p.M88V,DBI:NM_001178043:exon4:c.A241G:p.M81V,DBI:NM_001079862:exon4:c.A211G:p.M71V,DBI:NM_001178042:exon5:c.A262G:p.M88V, |
| chr2 | 119680947 | A | G | . | exonic | TMEM177 | nonsynonymous SNV | TMEM177:NM_001105198:exon2:c.A94G:p.I32V,TMEM177:NM_030577:exon2:c.A94G:p.I32V,TMEM177:NM_001105199:exon2:c.A94G:p.I32V, |
| chr2 | 127770816 | T | C | . | exonic | WDR33 | nonsynonymous SNV | WDR33:NM_001006622:exon2:c.A166G:p.T56A,WDR33:NM_001006623:exon2:c.A166G:p.T56A,WDR33:NM_018383:exon2:c.A166G:p.T56A, |
| chr2 | 132782163 | C | T | . | exonic | NCKAP5 | nonsynonymous SNV | NCKAP5:NM_207363:exon14:c.G4648A:p.E1550K, |
| chr2 | 132783032 | G | T | COSV58445700 | exonic | NCKAP5 | nonsynonymous SNV | NCKAP5:NM_207363:exon14:c.C3779A:p.P1260Q, |
| chr2 | 132783534 | T | A | COSV58444336 | exonic | NCKAP5 | nonsynonymous SNV | NCKAP5:NM_207363:exon14:c.A3277T:p.N1093Y, |
| chr2 | 132783881 | A | G | COSV58434578 | exonic | NCKAP5 | nonsynonymous SNV | NCKAP5:NM_207363:exon14:c.T2930C:p.I977T, |
| chr2 | 132963761 | C | T | . | exonic | NCKAP5 | nonsynonymous SNV | NCKAP5:NM_207363:exon8:c.G538A:p.E180K,NCKAP5:NM_207481:exon8:c.G538A:p.E180K, |
| chr2 | 135135696 | G | A | . | exonic | RAB3GAP1 | nonsynonymous SNV | RAB3GAP1:NM_001172435:exon17:c.G1687A:p.E563K,RAB3GAP1:NM_012233:exon17:c.G1687A:p.E563K, |
| chr2 | 135135700 | C | A | . | exonic | RAB3GAP1 | nonsynonymous SNV | RAB3GAP1:NM_001172435:exon17:c.C1691A:p.T564K,RAB3GAP1:NM_012233:exon17:c.C1691A:p.T564K, |
| chr2 | 135135704 | T | A | . | exonic | RAB3GAP1 | nonsynonymous SNV | RAB3GAP1:NM_001172435:exon17:c.T1695A:p.D565E,RAB3GAP1:NM_012233:exon17:c.T1695A:p.D565E, |
| chr2 | 135135802 | A | G | COSV51482881 | exonic | RAB3GAP1 | nonsynonymous SNV | RAB3GAP1:NM_001172435:exon17:c.A1793G:p.N598S,RAB3GAP1:NM_012233:exon17:c.A1793G:p.N598S, |
| chr2 | 136114924 | C | T | . | exonic | CXCR4 | nonsynonymous SNV | CXCR4:NM_003467:exon2:c.G1004A:p.G335D,CXCR4:NM_001008540:exon1:c.G1016A:p.G339D, |
| chr2 | 136114991 | C | T | . | exonic | CXCR4 | nonsynonymous SNV | CXCR4:NM_003467:exon2:c.G937A:p.A313T,CXCR4:NM_001008540:exon1:c.G949A:p.A317T, |
| chr2 | 136115155 | C | T | . | exonic | CXCR4 | nonsynonymous SNV | CXCR4:NM_003467:exon2:c.G773A:p.G258E,CXCR4:NM_001008540:exon1:c.G785A:p.G262E, |
| chr2 | 136115189 | C | T | . | exonic | CXCR4 | nonsynonymous SNV | CXCR4:NM_003467:exon2:c.G739A:p.A247T,CXCR4:NM_001008540:exon1:c.G751A:p.A251T, |
| chr2 | 136115646 | C | T | . | exonic | CXCR4 | stopgain | CXCR4:NM_003467:exon2:c.G282A:p.W94X,CXCR4:NM_001008540:exon1:c.G294A:p.W98X, |
| chr2 | 136115708 | C | T | . | exonic | CXCR4 | nonsynonymous SNV | CXCR4:NM_003467:exon2:c.G220A:p.D74N,CXCR4:NM_001008540:exon1:c.G232A:p.D78N, |
| chr2 | 143040620 | A | G | COSV51580314 | exonic | KYNU | nonsynonymous SNV | KYNU:NM_001199241:exon14:c.A1234G:p.K412E,KYNU:NM_003937:exon13:c.A1234G:p.K412E, |
| chr2 | 151282902 | G | A | COSV54638715 | exonic | NMI | nonsynonymous SNV | NMI:NM_004688:exon2:c.C47T:p.S16L, |
| chr2 | 157738519 | G | T | . | exonic | ACVR1 | nonsynonymous SNV | ACVR1:NM_001105:exon10:c.C1316A:p.P439Q,ACVR1:NM_001111067:exon10:c.C1316A:p.P439Q, |
| chr2 | 162267541 | C | T | COSV55125663 | exonic | IFIH1 | nonsynonymous SNV | IFIH1:NM_022168:exon15:c.G2836A:p.A946T, |
| chr2 | 162272314 | T | C | COSV55127127 | exonic | IFIH1 | nonsynonymous SNV | IFIH1:NM_022168:exon13:c.A2528G:p.H843R, |
| chr2 | 171976274 | T | C | . | exonic | HAT1 | nonsynonymous SNV | HAT1:NM_003642:exon9:c.T941C:p.V314A, |
| chr2 | 177230913 | C | T | . | exonic | NFE2L2 | nonsynonymous SNV | NFE2L2:NM_001145413:exon5:c.G1621A:p.V541I,NFE2L2:NM_001313901:exon5:c.G1642A:p.V548I,NFE2L2:NM_001313902:exon4:c.G1600A:p.V534I,NFE2L2:NM_001313900:exon5:c.G1642A:p.V548I,NFE2L2:NM_001313904:exon5:c.G1390A:p.V464I,NFE2L2:NM_001313903:exon5:c.G1471A:p.V491I,NFE2L2:NM_006164:exon5:c.G1690A:p.V564I,NFE2L2:NM_001145412:exon5:c.G1642A:p.V548I, |
| chr2 | 177231000 | C | T | . | exonic | NFE2L2 | nonsynonymous SNV | NFE2L2:NM_001145413:exon5:c.G1534A:p.E512K,NFE2L2:NM_001313901:exon5:c.G1555A:p.E519K,NFE2L2:NM_001313902:exon4:c.G1513A:p.E505K,NFE2L2:NM_001313900:exon5:c.G1555A:p.E519K,NFE2L2:NM_001313904:exon5:c.G1303A:p.E435K,NFE2L2:NM_001313903:exon5:c.G1384A:p.E462K,NFE2L2:NM_006164:exon5:c.G1603A:p.E535K,NFE2L2:NM_001145412:exon5:c.G1555A:p.E519K, |
| chr2 | 177231062 | C | T | . | exonic | NFE2L2 | nonsynonymous SNV | NFE2L2:NM_001145413:exon5:c.G1472A:p.C491Y,NFE2L2:NM_001313901:exon5:c.G1493A:p.C498Y,NFE2L2:NM_001313902:exon4:c.G1451A:p.C484Y,NFE2L2:NM_001313900:exon5:c.G1493A:p.C498Y,NFE2L2:NM_001313904:exon5:c.G1241A:p.C414Y,NFE2L2:NM_001313903:exon5:c.G1322A:p.C441Y,NFE2L2:NM_006164:exon5:c.G1541A:p.C514Y,NFE2L2:NM_001145412:exon5:c.G1493A:p.C498Y, |
| chr2 | 177231064 | A | T | . | exonic | NFE2L2 | nonsynonymous SNV | NFE2L2:NM_001145413:exon5:c.T1470A:p.N490K,NFE2L2:NM_001313901:exon5:c.T1491A:p.N497K,NFE2L2:NM_001313902:exon4:c.T1449A:p.N483K,NFE2L2:NM_001313900:exon5:c.T1491A:p.N497K,NFE2L2:NM_001313904:exon5:c.T1239A:p.N413K,NFE2L2:NM_001313903:exon5:c.T1320A:p.N440K,NFE2L2:NM_006164:exon5:c.T1539A:p.N513K,NFE2L2:NM_001145412:exon5:c.T1491A:p.N497K, |
| chr2 | 177231095 | C | T | . | exonic | NFE2L2 | nonsynonymous SNV | NFE2L2:NM_001145413:exon5:c.G1439A:p.R480K,NFE2L2:NM_001313901:exon5:c.G1460A:p.R487K,NFE2L2:NM_001313902:exon4:c.G1418A:p.R473K,NFE2L2:NM_001313900:exon5:c.G1460A:p.R487K,NFE2L2:NM_001313904:exon5:c.G1208A:p.R403K,NFE2L2:NM_001313903:exon5:c.G1289A:p.R430K,NFE2L2:NM_006164:exon5:c.G1508A:p.R503K,NFE2L2:NM_001145412:exon5:c.G1460A:p.R487K, |
| chr2 | 177231336 | C | T | . | exonic | NFE2L2 | nonsynonymous SNV | NFE2L2:NM_001145413:exon5:c.G1198A:p.E400K,NFE2L2:NM_001313901:exon5:c.G1219A:p.E407K,NFE2L2:NM_001313902:exon4:c.G1177A:p.E393K,NFE2L2:NM_001313900:exon5:c.G1219A:p.E407K,NFE2L2:NM_001313904:exon5:c.G967A:p.E323K,NFE2L2:NM_001313903:exon5:c.G1048A:p.E350K,NFE2L2:NM_006164:exon5:c.G1267A:p.E423K,NFE2L2:NM_001145412:exon5:c.G1219A:p.E407K, |
| chr2 | 177231740 | G | A | . | exonic | NFE2L2 | nonsynonymous SNV | NFE2L2:NM_001145413:exon5:c.C794T:p.A265V,NFE2L2:NM_001313901:exon5:c.C815T:p.A272V,NFE2L2:NM_001313902:exon4:c.C773T:p.A258V,NFE2L2:NM_001313900:exon5:c.C815T:p.A272V,NFE2L2:NM_001313904:exon5:c.C563T:p.A188V,NFE2L2:NM_001313903:exon5:c.C644T:p.A215V,NFE2L2:NM_006164:exon5:c.C863T:p.A288V,NFE2L2:NM_001145412:exon5:c.C815T:p.A272V, |
| chr2 | 177231758 | C | T | . | exonic | NFE2L2 | nonsynonymous SNV | NFE2L2:NM_001145413:exon5:c.G776A:p.G259D,NFE2L2:NM_001313901:exon5:c.G797A:p.G266D,NFE2L2:NM_001313902:exon4:c.G755A:p.G252D,NFE2L2:NM_001313900:exon5:c.G797A:p.G266D,NFE2L2:NM_001313904:exon5:c.G545A:p.G182D,NFE2L2:NM_001313903:exon5:c.G626A:p.G209D,NFE2L2:NM_006164:exon5:c.G845A:p.G282D,NFE2L2:NM_001145412:exon5:c.G797A:p.G266D, |
| chr2 | 177231891 | C | T | . | exonic | NFE2L2 | nonsynonymous SNV | NFE2L2:NM_001313902:exon4:c.G622A:p.E208K,NFE2L2:NM_001313900:exon5:c.G664A:p.E222K,NFE2L2:NM_001313904:exon5:c.G412A:p.E138K,NFE2L2:NM_001145412:exon5:c.G664A:p.E222K,NFE2L2:NM_001145413:exon5:c.G643A:p.E215K,NFE2L2:NM_001313901:exon5:c.G664A:p.E222K,NFE2L2:NM_001313903:exon5:c.G493A:p.E165K,NFE2L2:NM_006164:exon5:c.G712A:p.E238K, |
| chr2 | 177231951 | C | T | . | exonic | NFE2L2 | nonsynonymous SNV | NFE2L2:NM_001145413:exon5:c.G583A:p.A195T,NFE2L2:NM_001313901:exon5:c.G604A:p.A202T,NFE2L2:NM_001313902:exon4:c.G562A:p.A188T,NFE2L2:NM_001313900:exon5:c.G604A:p.A202T,NFE2L2:NM_001313904:exon5:c.G352A:p.A118T,NFE2L2:NM_001313903:exon5:c.G433A:p.A145T,NFE2L2:NM_006164:exon5:c.G652A:p.A218T,NFE2L2:NM_001145412:exon5:c.G604A:p.A202T, |
| chr2 | 177232008 | A | G | . | exonic | NFE2L2 | nonsynonymous SNV | NFE2L2:NM_001313902:exon4:c.T505C:p.C169R,NFE2L2:NM_001313900:exon5:c.T547C:p.C183R,NFE2L2:NM_001313904:exon5:c.T295C:p.C99R,NFE2L2:NM_001145412:exon5:c.T547C:p.C183R,NFE2L2:NM_001145413:exon5:c.T526C:p.C176R,NFE2L2:NM_001313901:exon5:c.T547C:p.C183R,NFE2L2:NM_001313903:exon5:c.T376C:p.C126R,NFE2L2:NM_006164:exon5:c.T595C:p.C199R, |
| chr2 | 177232454 | C | T | . | exonic | NFE2L2 | nonsynonymous SNV | NFE2L2:NM_001145413:exon4:c.G463A:p.D155N,NFE2L2:NM_001313901:exon4:c.G484A:p.D162N,NFE2L2:NM_001313902:exon3:c.G442A:p.D148N,NFE2L2:NM_001313900:exon4:c.G484A:p.D162N,NFE2L2:NM_001313904:exon4:c.G232A:p.D78N,NFE2L2:NM_001313903:exon4:c.G313A:p.D105N,NFE2L2:NM_006164:exon4:c.G532A:p.D178N,NFE2L2:NM_001145412:exon4:c.G484A:p.D162N, |

| | | | | | | | | |
|---|---|---|---|---|---|---|---|---|
| chr2 | 177234073 | C | T | . | exonic | NFE2L2 | nonsynonymous SNV | NFE2L2:NM_001313902:exon2:c.G244A:p.E82K,NFE2L2:NM_001313900:exon2:c.G196A:p.E66K,NFE2L2:NM_001145412:exon2:c.G196A:p.E66K,NFE2L2:NM_001145413:exon2:c.G196A:p.E66K,NFE2L2:NM_001313901:exon2:c.G196A:p.E66K,NFE2L2:NM_006164:exon2:c.G244A:p.E82K, |
| chr2 | 177234109 | A | G | . | exonic | NFE2L2 | nonsynonymous SNV | NFE2L2:NM_001145413:exon2:c.T160C:p.F54L,NFE2L2:NM_001313901:exon2:c.T160C:p.F54L,NFE2L2:NM_001313902:exon2:c.T208C:p.F70L,NFE2L2:NM_001313900:exon2:c.T160C:p.F54L,NFE2L2:NM_006164:exon2:c.T208C:p.F70L,NFE2L2:NM_001145412:exon2:c.T160C:p.F54L, |
| chr2 | 177513866 | G | A | . | exonic | AGPS | nonsynonymous SNV | AGPS:NM_003659:exon17:c.G1655A:p.C552Y, |
| chr2 | 17932356 | A | G | COSV58406825 | exonic | KCNS3 | nonsynonymous SNV | KCNS3:NM_002252:exon3:c.A1348G:p.T450A,KCNS3:NM_001282428:exon3:c.A1348G:p.T450A, |
| chr2 | 186501387 | G | A | . | exonic | ZC3H15 | nonsynonymous SNV | ZC3H15:NM_018471:exon4:c.G404A:p.S135N, |
| chr2 | 186504041 | C | T | . | exonic | ZC3H15 | nonsynonymous SNV | ZC3H15:NM_018471:exon6:c.C544T:p.H182Y, |
| chr2 | 186504053 | G | A | . | exonic | ZC3H15 | nonsynonymous SNV | ZC3H15:NM_018471:exon6:c.G556A:p.A186T, |
| chr2 | 189010695 | T | G | . | exonic | COL3A1 | nonsynonymous SNV | COL3A1:NM_000090:exon50:c.T4059G:p.H1353Q, |
| chr2 | 190200027 | C | T | COSV61409827 | exonic | C2ORF88 | nonsynonymous SNV | C2orf88:NM_001042520:exon2:c.C167T:p.T56I,C2orf88:NM_001042521:exon3:c.C167T:p.T56I,C2orf88:NM_001042519:exon2:c.C167T:p.T56I,C2orf88:NM_032321:exon2:c.C167T:p.T56I, |
| chr2 | 190296896 | T | C | COSV62891843 | exonic | HIBCH | nonsynonymous SNV | HIBCH:NM_198047:exon3:c.A136G:p.T46A,HIBCH:NM_014362:exon3:c.A136G:p.T46A, |
| chr2 | 190319749 | A | G | COSV62891757 | exonic | HIBCH | nonsynonymous SNV | HIBCH:NM_198047:exon1:c.T2C:p.M1T,HIBCH:NM_014362:exon1:c.T2C:p.M1T, |
| chr2 | 190534652 | C | G | COSV58108655 | exonic | NEMP2 | nonsynonymous SNV | NEMP2:NM_001142645:exon1:c.G4C:p.G2R, |
| chr2 | 197398025 | C | T | COSV59231028 | exonic | SF3B1 | nonsynonymous SNV | SF3B1:NM_012433:exon22:c.G3226A:p.A1076T, |
| chr2 | 197398088 | A | G | . | exonic | SF3B1 | nonsynonymous SNV | SF3B1:NM_012433:exon22:c.T3163C:p.W1055R, |
| chr2 | 197405149 | G | T | . | exonic | SF3B1 | nonsynonymous SNV | SF3B1:NM_012433:exon11:c.C1466A:p.P489Q, |
| chr2 | 197408047 | C | T | COSV59230816 | exonic | SF3B1 | nonsynonymous SNV | SF3B1:NM_012433:exon9:c.G1190A:p.R397H, |
| chr2 | 197408525 | C | T | . | exonic | SF3B1 | nonsynonymous SNV | SF3B1:NM_012433:exon8:c.G961A:p.D321N, |
| chr2 | 197409869 | C | T | . | exonic | SF3B1 | nonsynonymous SNV | SF3B1:NM_012433:exon7:c.G805A:p.G269R, |
| chr2 | 197460008 | G | A | . | exonic | COQ10B | nonsynonymous SNV | COQ10B:NM_025147:exon2:c.G181A:p.A61T, |
| chr2 | 197487063 | C | T | . | exonic | HSPD1 | nonsynonymous SNV | HSPD1:NM_002156:exon12:c.G1705A:p.G569R,HSPD1:NM_199440:exon12:c.G1705A:p.G569R, |
| chr2 | 197487080 | C | G | COSV61445925 | exonic | HSPD1 | nonsynonymous SNV | HSPD1:NM_002156:exon12:c.G1688C:p.G563A,HSPD1:NM_199440:exon12:c.G1688C:p.G563A, |
| chr2 | 197487083 | C | T | . | exonic | HSPD1 | nonsynonymous SNV | HSPD1:NM_002156:exon12:c.G1685A:p.G562D,HSPD1:NM_199440:exon12:c.G1685A:p.G562D, |
| chr2 | 197487094 | C | T | . | exonic | HSPD1 | nonsynonymous SNV | HSPD1:NM_002156:exon12:c.G1674A:p.M558I,HSPD1:NM_199440:exon12:c.G1674A:p.M558I, |
| chr2 | 197487141 | C | T | . | exonic | HSPD1 | nonsynonymous SNV | HSPD1:NM_002156:exon12:c.G1627A:p.E543K,HSPD1:NM_199440:exon12:c.G1627A:p.E543K, |
| chr2 | 197487167 | C | T | . | exonic | HSPD1 | nonsynonymous SNV | HSPD1:NM_002156:exon12:c.G1601A:p.G534D,HSPD1:NM_199440:exon12:c.G1601A:p.G534D, |
| chr2 | 197487195 | C | T | . | exonic | HSPD1 | nonsynonymous SNV | HSPD1:NM_002156:exon12:c.G1573A:p.V525M,HSPD1:NM_199440:exon12:c.G1573A:p.V525M, |
| chr2 | 197487895 | A | G | . | exonic | HSPD1 | nonsynonymous SNV | HSPD1:NM_002156:exon11:c.T1532C:p.V511A,HSPD1:NM_199440:exon11:c.T1532C:p.V511A, |
| chr2 | 197487973 | A | G | . | exonic | HSPD1 | nonsynonymous SNV | HSPD1:NM_002156:exon11:c.T1454C:p.V485A,HSPD1:NM_199440:exon11:c.T1454C:p.V485A, |
| chr2 | 197488001 | C | T | . | exonic | HSPD1 | nonsynonymous SNV | HSPD1:NM_002156:exon11:c.G1426A:p.A476T,HSPD1:NM_199440:exon11:c.G1426A:p.A476T, |
| chr2 | 197488332 | C | T | . | exonic | HSPD1 | nonsynonymous SNV | HSPD1:NM_002156:exon10:c.G1375A:p.E459K,HSPD1:NM_199440:exon10:c.G1375A:p.E459K, |
| chr2 | 197488374 | G | A | . | exonic | HSPD1 | nonsynonymous SNV | HSPD1:NM_002156:exon10:c.C1333T:p.L445F,HSPD1:NM_199440:exon10:c.C1333T:p.L445F, |
| chr2 | 197488391 | C | T | . | exonic | HSPD1 | nonsynonymous SNV | HSPD1:NM_002156:exon10:c.G1316A:p.G439E,HSPD1:NM_199440:exon10:c.G1316A:p.G439E, |
| chr2 | 197488416 | C | T | . | exonic | HSPD1 | nonsynonymous SNV | HSPD1:NM_002156:exon10:c.G1291A:p.A431T,HSPD1:NM_199440:exon10:c.G1291A:p.A431T, |
| chr2 | 197488473 | C | T | . | exonic | HSPD1 | nonsynonymous SNV | HSPD1:NM_002156:exon10:c.G1234A:p.V412I,HSPD1:NM_199440:exon10:c.G1234A:p.V412I, |
| chr2 | 197488484 | C | T | . | exonic | HSPD1 | nonsynonymous SNV | HSPD1:NM_002156:exon10:c.G1223A:p.G408E,HSPD1:NM_199440:exon10:c.G1223A:p.G408E, |
| chr2 | 197489109 | G | T | COSV61444464 | exonic | HSPD1 | nonsynonymous SNV | HSPD1:NM_002156:exon9:c.C1108A:p.R370S,HSPD1:NM_199440:exon9:c.C1108A:p.R370S, |
| chr2 | 197489115 | C | T | . | exonic | HSPD1 | nonsynonymous SNV | HSPD1:NM_002156:exon9:c.G1102A:p.E368K,HSPD1:NM_199440:exon9:c.G1102A:p.E368K, |
| chr2 | 197489132 | C | T | . | exonic | HSPD1 | nonsynonymous SNV | HSPD1:NM_002156:exon9:c.G1085A:p.G362D,HSPD1:NM_199440:exon9:c.G1085A:p.G362D, |
| chr2 | 197489145 | A | T | . | exonic | HSPD1 | nonsynonymous SNV | HSPD1:NM_002156:exon9:c.T1072A:p.L358I,HSPD1:NM_199440:exon9:c.T1072A:p.L358I, |
| chr2 | 197489178 | C | T | . | exonic | HSPD1 | nonsynonymous SNV | HSPD1:NM_002156:exon9:c.G1039A:p.E347K,HSPD1:NM_199440:exon9:c.G1039A:p.E347K, |
| chr2 | 197489231 | C | T | . | exonic | HSPD1 | nonsynonymous SNV | HSPD1:NM_002156:exon9:c.G986A:p.G329E,HSPD1:NM_199440:exon9:c.G986A:p.G329E, |
| chr2 | 197489241 | C | T | . | exonic | HSPD1 | nonsynonymous SNV | HSPD1:NM_002156:exon9:c.G976A:p.G326R,HSPD1:NM_199440:exon9:c.G976A:p.G326R, |
| chr2 | 197490277 | C | A | . | exonic | HSPD1 | nonsynonymous SNV | HSPD1:NM_002156:exon8:c.G889T:p.V297F,HSPD1:NM_199440:exon8:c.G889T:p.V297F, |
| chr2 | 197493385 | G | T | . | exonic | HSPD1 | nonsynonymous SNV | HSPD1:NM_002156:exon7:c.C808A:p.P270T,HSPD1:NM_199440:exon7:c.C808A:p.P270T, |
| chr2 | 197493390 | C | T | COSV61443847 | exonic | HSPD1 | nonsynonymous SNV | HSPD1:NM_002156:exon7:c.G803A:p.R268H,HSPD1:NM_199440:exon7:c.G803A:p.R268H, |
| chr2 | 197493415 | C | T | . | exonic | HSPD1 | nonsynonymous SNV | HSPD1:NM_002156:exon7:c.G778A:p.A260T,HSPD1:NM_199440:exon7:c.G778A:p.A260T, |
| chr2 | 197494716 | C | T | . | exonic | HSPD1 | nonsynonymous SNV | HSPD1:NM_002156:exon5:c.G547A:p.G183S,HSPD1:NM_199440:exon5:c.G547A:p.G183S, |
| chr2 | 197494731 | C | T | . | exonic | HSPD1 | nonsynonymous SNV | HSPD1:NM_002156:exon5:c.G532A:p.G178R,HSPD1:NM_199440:exon5:c.G532A:p.G178R, |
| chr2 | 197495302 | T | A | . | exonic | HSPD1 | nonsynonymous SNV | HSPD1:NM_002156:exon4:c.A502T:p.I168F,HSPD1:NM_199440:exon4:c.A502T:p.I168F, |
| chr2 | 197497155 | C | T | . | exonic | HSPD1 | nonsynonymous SNV | HSPD1:NM_002156:exon3:c.G412A:p.V138M,HSPD1:NM_199440:exon3:c.G412A:p.V138M, |
| chr2 | 197497182 | C | T | COSV52099976 | exonic | HSPD1 | nonsynonymous SNV | HSPD1:NM_002156:exon3:c.G385A:p.E129K,HSPD1:NM_199440:exon3:c.G385A:p.E129K, |
| chr2 | 197497188 | C | T | . | exonic | HSPD1 | nonsynonymous SNV | HSPD1:NM_002156:exon3:c.G379A:p.G127S,HSPD1:NM_199440:exon3:c.G379A:p.G127S, |
| chr2 | 197497242 | C | T | . | exonic | HSPD1 | nonsynonymous SNV | HSPD1:NM_002156:exon3:c.G325A:p.A109T,HSPD1:NM_199440:exon3:c.G325A:p.A109T, |
| chr2 | 197497287 | C | T | . | exonic | HSPD1 | nonsynonymous SNV | HSPD1:NM_002156:exon3:c.G280A:p.G94R,HSPD1:NM_199440:exon3:c.G280A:p.G94R, |
| chr2 | 197497334 | A | G | . | exonic | HSPD1 | nonsynonymous SNV | HSPD1:NM_002156:exon3:c.T233C:p.V78A,HSPD1:NM_199440:exon3:c.T233C:p.V78A, |
| chr2 | 197497392 | C | T | . | exonic | HSPD1 | nonsynonymous SNV | HSPD1:NM_002156:exon3:c.G175A:p.G59R,HSPD1:NM_199440:exon3:c.G175A:p.G59R, |
| chr2 | 197498682 | C | T | COSV52098272 | exonic | HSPD1 | nonsynonymous SNV | HSPD1:NM_002156:exon2:c.G167A:p.G56E,HSPD1:NM_199440:exon2:c.G167A:p.G56E, |
| chr2 | 197498823 | C | T | . | exonic | HSPD1 | nonsynonymous SNV | HSPD1:NM_002156:exon2:c.G26A:p.R9H,HSPD1:NM_199440:exon2:c.G26A:p.R9H, |
| chr2 | 201129878 | G | A | . | exonic | CFLAR | nonsynonymous SNV | CFLAR:NM_001308042:exon1:c.G13A:p.V5I,CFLAR:NM_001202516:exon2:c.G13A:p.V5I,CFLAR:NM_001127184:exon2:c.G13A:p.V5I,CFLAR:NM_001308043:exon2:c.G13A:p.V5I,CFLAR:NM_001127183:exon2:c.G13A:p.V5I,CFLAR:NM_003879:exon2:c.G13A:p.V5I, |
| chr2 | 201140386 | G | A | . | exonic | CFLAR | nonsynonymous SNV | CFLAR:NM_001308042:exon4:c.G553A:p.V185I,CFLAR:NM_001202516:exon5:c.G553A:p.V185I,CFLAR:NM_001127184:exon5:c.G553A:p.V185I,CFLAR:NM_001202519:exon4:c.G265A:p.V89I,CFLAR:NM_001202517:exon4:c.G265A:p.V89I,CFLAR:NM_001202518:exon3:c.G265A:p.V89I,CFLAR:NM_001127183:exon5:c.G553A:p.V185I,CFLAR:NM_003879:exon5:c.G553A:p.V185I, |
| chr2 | 201258272 | A | G | COSV51848843 | exonic | CASP8 | nonsynonymous SNV | CASP8:NM_001080125:exon1:c.A41G:p.K14R, |
| chr2 | 201258379 | G | A | . | exonic | CASP8 | nonsynonymous SNV | CASP8:NM_001080125:exon1:c.G148A:p.A50T, |
| chr2 | 201272772 | C | A | . | exonic | CASP8 | nonsynonymous SNV | CASP8:NM_033358:exon5:c.C546A:p.S182R,CASP8:NM_001080124:exon5:c.C546A:p.S182R,CASP8:NM_001228:exon6:c.C642A:p.S214R,CASP8:NM_033355:exon5:c.C546A:p.S182R,CASP8:NM_001080125:exon4:c.C723A:p.S241R,CASP8:NM_033356:exon4:c.C546A:p.S182R, |
| chr2 | 20204033 | A | T | . | exonic | SDC1 | nonsynonymous SNV | SDC1:NM_002997:exon3:c.T407A:p.L136Q,SDC1:NM_001006946:exon4:c.T407A:p.L136Q, |
| chr2 | 205104496 | G | A | . | exonic | PARD3B | nonsynonymous SNV | PARD3B:NM_001302769:exon5:c.G575A:p.R192K,PARD3B:NM_205863:exon5:c.G575A:p.R192K,PARD3B:NM_057177:exon5:c.G575A:p.R192K,PARD3B:NM_152526:exon5:c.G575A:p.R192K, |
| chr2 | 213317301 | G | C | . | exonic | SPAG16 | nonsynonymous SNV | SPAG16:NM_024532:exon5:c.G481C:p.E161Q,SPAG16:NM_001025436:exon5:c.G481C:p.E161Q, |
| chr2 | 213317306 | T | A | . | exonic | SPAG16 | nonsynonymous SNV | SPAG16:NM_024532:exon5:c.T486A:p.N162K,SPAG16:NM_001025436:exon5:c.T486A:p.N162K, |
| chr2 | 214767531 | C | T | COSV53610001 | exonic | BARD1 | nonsynonymous SNV | BARD1:NM_001282543:exon5:c.G1462A:p.V488M,BARD1:NM_000465:exon6:c.G1519A:p.V507M, |
| chr2 | 214780740 | C | G | COSV53612030 | exonic | BARD1 | nonsynonymous SNV | BARD1:NM_001282543:exon3:c.G1077C:p.R359S,BARD1:NM_000465:exon4:c.G1134C:p.R378S, |

| | | | | | | | | |
|---|---|---|---|---|---|---|---|---|
| chr2 | 214809500 | G | A | COSV53608734 | exonic | BARD1 | nonsynonymous SNV | BARD1:NM_001282543:exon1:c.C70T:p.P24S,BARD1:NM_001282545:exon1:c.C70T:p.P24S,BARD1:NM_001282549:exon1:c.C70T:p.P24S,BARD1:NM_001282548:exon1:c.C70T:p.P24S,BARD1:NM_000465:exon1:c.C70T:p.P24S, |
| chr2 | 215370366 | C | T | . | exonic | FN1 | nonsynonymous SNV | FN1:NM_001306130:exon40:c.G6151A:p.V2051I,FN1:NM_212474:exon39:c.G5878A:p.V1960I,FN1:NM_212476:exon39:c.G6238A:p.V2080I,FN1:NM_001306129:exon42:c.G6688A:p.V2230I,FN1:NM_001306131:exon40:c.G6145A:p.V2049I,FN1:NM_001306132:exon40:c.G6070A:p.V2024I,FN1:NM_002026:exon41:c.G6415A:p.V2139I,FN1:NM_212478:exon41:c.G6340A:p.V2114I,FN1:NM_212482:exon41:c.G6781A:p.V2261I, |
| chr2 | 215370392 | C | T | . | exonic | FN1 | nonsynonymous SNV | FN1:NM_001306130:exon40:c.G6125A:p.G2042D,FN1:NM_212474:exon39:c.G5852A:p.G1951D,FN1:NM_212476:exon39:c.G6212A:p.G2071D,FN1:NM_001306129:exon42:c.G6662A:p.G2221D,FN1:NM_001306131:exon40:c.G6119A:p.G2040D,FN1:NM_001306132:exon40:c.G6044A:p.G2015D,FN1:NM_002026:exon41:c.G6389A:p.G2130D,FN1:NM_212478:exon41:c.G6314A:p.G2105D,FN1:NM_212482:exon41:c.G6755A:p.G2252D, |
| chr2 | 215407182 | A | T | . | exonic | FN1 | nonsynonymous SNV | FN1:NM_001306130:exon18:c.T2658A:p.N886K,FN1:NM_212474:exon18:c.T2658A:p.N886K,FN1:NM_212476:exon18:c.T2658A:p.N886K,FN1:NM_001306129:exon18:c.T2658A:p.N886K,FN1:NM_001306131:exon18:c.T2658A:p.N886K,FN1:NM_001306132:exon18:c.T2658A:p.N886K,FN1:NM_002026:exon18:c.T2658A:p.N886K,FN1:NM_212478:exon18:c.T2658A:p.N886K,FN1:NM_212482:exon18:c.T2658A:p.N886K, |
| chr2 | 215408177 | T | G | . | exonic | FN1 | nonsynonymous SNV | FN1:NM_001306130:exon17:c.A2449C:p.T817P,FN1:NM_212474:exon17:c.A2449C:p.T817P,FN1:NM_212476:exon17:c.A2449C:p.T817P,FN1:NM_001306129:exon17:c.A2449C:p.T817P,FN1:NM_001306131:exon17:c.A2449C:p.T817P,FN1:NM_001306132:exon17:c.A2449C:p.T817P,FN1:NM_002026:exon17:c.A2449C:p.T817P,FN1:NM_212478:exon17:c.A2449C:p.T817P,FN1:NM_212482:exon17:c.A2449C:p.T817P, |
| chr2 | 215435759 | T | A | COSV60547598 | exonic | FN1 | nonsynonymous SNV | FN1:NM_001306130:exon1:c.A44T:p.Q15L,FN1:NM_212474:exon1:c.A44T:p.Q15L,FN1:NM_212476:exon1:c.A44T:p.Q15L,FN1:NM_001306129:exon1:c.A44T:p.Q15L,FN1:NM_054034:exon1:c.A44T:p.Q15L,FN1:NM_001306131:exon1:c.A44T:p.Q15L,FN1:NM_001306132:exon1:c.A44T:p.Q15L,FN1:NM_002026:exon1:c.A44T:p.Q15L,FN1:NM_212478:exon1:c.A44T:p.Q15L,FN1:NM_212482:exon1:c.A44T:p.Q15L, |
| chr2 | 219172671 | A | G | . | exonic | CNPPD1 | nonsynonymous SNV | CNPPD1:NM_015680:exon8:c.T1148C:p.L383P, |
| chr2 | 219172944 | A | G | COSV55405643 | exonic | CNPPD1 | nonsynonymous SNV | CNPPD1:NM_015680:exon8:c.T875C:p.L292P, |
| chr2 | 219173034 | A | G | COSV55407502 | exonic | CNPPD1 | nonsynonymous SNV | CNPPD1:NM_015680:exon8:c.T785C:p.I262T, |
| chr2 | 222571888 | C | T | . | exonic | FARSB | nonsynonymous SNV | FARSB:NM_005687:exon17:c.G1753A:p.V585I, |
| chr2 | 224506893 | C | T | . | exonic | CUL3 | nonsynonymous SNV | CUL3:NM_003590:exon7:c.G994A:p.E332K,CUL3:NM_001257198:exon7:c.G1012A:p.E338K,CUL3:NM_001257197:exon6:c.G796A:p.E266K, |
| chr2 | 224511367 | A | T | . | exonic | CUL3 | nonsynonymous SNV | CUL3:NM_001257198:exon6:c.T888A:p.N296K,CUL3:NM_001257197:exon5:c.T672A:p.N224K,CUL3:NM_003590:exon6:c.T870A:p.N290K, |
| chr2 | 224511533 | T | C | . | exonic | CUL3 | nonsynonymous SNV | CUL3:NM_001257198:exon6:c.A722G:p.K241R,CUL3:NM_001257197:exon5:c.A506G:p.K169R,CUL3:NM_003590:exon6:c.A704G:p.K235R, |
| chr2 | 227379189 | G | A | COSV58818572 | exonic | TM4SF20 | nonsynonymous SNV | TM4SF20:NM_024795:exon1:c.C80T:p.A27V, |
| chr2 | 230469048 | A | G | COSV50977261 | exonic | SP100 | nonsynonymous SNV | SP100:NM_001206704:exon15:c.A1192G:p.M398V,SP100:NM_001206701:exon14:c.A1297G:p.M433V,SP100:NM_003113:exon14:c.A1297G:p.M433V,SP100:NM_001080391:exon14:c.A1297G:p.M433V,SP100:NM_001206703:exon13:c.A1213G:p.M405V,SP100:NM_001206702:exon14:c.A1297G:p.M433V, |
| chr2 | 230873453 | G | A | . | exonic | ITM2C | nonsynonymous SNV | ITM2C:NM_001287241:exon3:c.G157A:p.G53S,ITM2C:NM_001012516:exon2:c.G157A:p.G53S,ITM2C:NM_030926:exon2:c.G157A:p.G53S, |
| chr2 | 231455235 | C | T | . | exonic | NCL | nonsynonymous SNV | NCL:NM_005381:exon14:c.G2089A:p.G697R, |
| chr2 | 231455491 | G | A | . | exonic | NCL | nonsynonymous SNV | NCL:NM_005381:exon13:c.C1966T:p.R656C, |
| chr2 | 231455503 | C | T | . | exonic | NCL | nonsynonymous SNV | NCL:NM_005381:exon13:c.G1954A:p.G652S, |
| chr2 | 231458365 | G | T | . | exonic | NCL | nonsynonymous SNV | NCL:NM_005381:exon8:c.C1190A:p.A397D, |
| chr2 | 231461664 | C | A | . | exonic | NCL | nonsynonymous SNV | NCL:NM_005381:exon3:c.G489T:p.E163D, |
| chr2 | 231461667 | G | T | . | exonic | NCL | nonsynonymous SNV | NCL:NM_005381:exon3:c.C486A:p.D162E, |
| chr2 | 231461670 | C | A | . | exonic | NCL | nonsynonymous SNV | NCL:NM_005381:exon3:c.G483T:p.E161D, |
| chr2 | 231461674 | T | A | . | exonic | NCL | nonsynonymous SNV | NCL:NM_005381:exon3:c.A479T:p.D160V, |
| chr2 | 232970265 | A | G | COSV50914443 | exonic | NGEF | nonsynonymous SNV | NGEF:NM_019850:exon3:c.T332C:p.M111T, |
| chr2 | 233204655 | C | T | COSV64037273 | exonic | INPP5D | nonsynonymous SNV | INPP5D:NM_001017915:exon26:c.C3505T:p.H1169Y,INPP5D:NM_005541:exon26:c.C3502T:p.H1168Y, |
| chr2 | 236049178 | G | A | COSV58326802 | exonic | AGAP1 | nonsynonymous SNV | AGAP1:NM_001037131:exon16:c.G2011A:p.V671I,AGAP1:NM_014914:exon15:c.G1852A:p.V618I, |
| chr2 | 237525791 | C | T | COSV52820483 | exonic | MLPH | nonsynonymous SNV | MLPH:NM_001281473:exon6:c.C746T:p.T249I,MLPH:NM_024101:exon7:c.C866T:p.T289I,MLPH:NM_001042467:exon7:c.C866T:p.T289I, |
| chr2 | 237540364 | T | C | COSV52819570 | exonic | MLPH | nonsynonymous SNV | MLPH:NM_001281473:exon8:c.T917C:p.V306A,MLPH:NM_024101:exon10:c.T1121C:p.V374A,MLPH:NM_001281474:exon8:c.T692C:p.V231A,MLPH:NM_001042467:exon9:c.T1037C:p.V346A, |
| chr2 | 237540465 | G | T | COSV52821423 | exonic | MLPH | nonsynonymous SNV | MLPH:NM_001281473:exon8:c.G1018T:p.A340S,MLPH:NM_024101:exon10:c.G1222T:p.A408S,MLPH:NM_001281474:exon8:c.G793T:p.A265S,MLPH:NM_001042467:exon9:c.G1138T:p.A380S, |
| chr2 | 240005279 | T | C | . | exonic | NDUFA10 | nonsynonymous SNV | NDUFA10:NM_004544:exon8:c.A821G:p.E274G, |
| chr2 | 240011617 | C | T | . | exonic | NDUFA10 | nonsynonymous SNV | NDUFA10:NM_004544:exon6:c.G749A:p.S250N, |
| chr2 | 240018561 | C | T | COSV53152766 | exonic | NDUFA10 | nonsynonymous SNV | NDUFA10:NM_004544:exon4:c.G539A:p.R180Q, |
| chr2 | 240018606 | C | T | . | exonic | NDUFA10 | nonsynonymous SNV | NDUFA10:NM_004544:exon4:c.G494A:p.S165N, |
| chr2 | 240021286 | T | C | . | exonic | NDUFA10 | nonsynonymous SNV | NDUFA10:NM_004544:exon3:c.A371G:p.N124S, |
| chr2 | 240021344 | C | T | . | exonic | NDUFA10 | nonsynonymous SNV | NDUFA10:NM_004544:exon3:c.G313A:p.A105T, |
| chr2 | 240022294 | C | T | . | exonic | NDUFA10 | stopgain | NDUFA10:NM_004544:exon2:c.G122A:p.W41X, |
| chr2 | 240025229 | C | T | . | exonic | NDUFA10 | nonsynonymous SNV | NDUFA10:NM_004544:exon1:c.G73A:p.V25M, |
| chr2 | 240025297 | G | C | COSV53155269 | exonic | NDUFA10 | nonsynonymous SNV | NDUFA10:NM_004544:exon1:c.C5G:p.A2G, |
| chr2 | 24039190 | G | A | COSV54591813 | exonic | C2ORF44 | nonsynonymous SNV | C2orf44:NM_001142319:exon2:c.C305T:p.T102M,C2orf44:NM_025203:exon2:c.C305T:p.T102M, |
| chr2 | 240598657 | A | G | . | exonic | CAPN10 | nonsynonymous SNV | CAPN10:NM_023085:exon10:c.A1531G:p.I511V,CAPN10:NM_023083:exon12:c.A1996G:p.I666V, |
| chr2 | 240630275 | C | T | COSV60576792 | exonic | GPR35 | nonsynonymous SNV | GPR35:NM_001195382:exon6:c.C416T:p.T139M,GPR35:NM_005301:exon2:c.C323T:p.T108M,GPR35:NM_001195381:exon6:c.C416T:p.T139M, |
| chr2 | 240630832 | A | C | COSV60576322 | exonic | GPR35 | nonsynonymous SNV | GPR35:NM_001195382:exon6:c.A973C:p.S325R,GPR35:NM_005301:exon2:c.A880C:p.S294R,GPR35:NM_001195381:exon6:c.A973C:p.S325R, |
| chr2 | 241264600 | T | C | . | exonic | HDLBP | nonsynonymous SNV | HDLBP:NM_203346:exon4:c.A82G:p.T28A,HDLBP:NM_001243900:exon5:c.A190G:p.T64A,HDLBP:NM_005336:exon4:c.A82G:p.T28A, |
| chr2 | 241633796 | T | C | COSV67190580 | exonic | THAP4 | nonsynonymous SNV | THAP4:NM_015963:exon2:c.A361G:p.S121G, |
| chr2 | 26191351 | C | T | . | exonic | HADHA | nonsynonymous SNV | HADHA:NM_000182:exon20:c.G2191A:p.D731N, |
| chr2 | 26191519 | C | T | . | exonic | HADHA | nonsynonymous SNV | HADHA:NM_000182:exon19:c.G2110A:p.A704T, |
| chr2 | 26191554 | C | T | . | exonic | HADHA | nonsynonymous SNV | HADHA:NM_000182:exon19:c.G2075A:p.G692E, |
| chr2 | 26191602 | C | T | COSV66106823 | exonic | HADHA | nonsynonymous SNV | HADHA:NM_000182:exon19:c.G2027A:p.R676H, |
| chr2 | 26192404 | A | G | . | exonic | HADHA | nonsynonymous SNV | HADHA:NM_000182:exon18:c.T1906C:p.F636L, |
| chr2 | 26193613 | C | T | . | exonic | HADHA | nonsynonymous SNV | HADHA:NM_000182:exon17:c.G1849A:p.E617K, |
| chr2 | 26195221 | C | T | . | exonic | HADHA | nonsynonymous SNV | HADHA:NM_000182:exon15:c.G1491A:p.M497I, |
| chr2 | 26201151 | C | T | . | exonic | HADHA | nonsynonymous SNV | HADHA:NM_000182:exon13:c.G1390A:p.A464T, |
| chr2 | 26201212 | C | T | . | exonic | HADHA | nonsynonymous SNV | HADHA:NM_000182:exon13:c.G1329A:p.M443I, |
| chr2 | 26201217 | C | T | COSV66106966 | exonic | HADHA | nonsynonymous SNV | HADHA:NM_000182:exon13:c.G1324A:p.D442N, |
| chr2 | 26201231 | C | A | . | exonic | HADHA | nonsynonymous SNV | HADHA:NM_000182:exon13:c.G1310T:p.G437V, |
| chr2 | 26201249 | C | T | . | exonic | HADHA | nonsynonymous SNV | HADHA:NM_000182:exon13:c.G1292A:p.G431E, |
| chr2 | 26201280 | C | T | . | exonic | HADHA | nonsynonymous SNV | HADHA:NM_000182:exon13:c.G1261A:p.E421K, |
| chr2 | 26204083 | C | T | . | exonic | HADHA | nonsynonymous SNV | HADHA:NM_000182:exon12:c.G1199A:p.G400E, |
| chr2 | 26204141 | C | T | . | exonic | HADHA | nonsynonymous SNV | HADHA:NM_000182:exon12:c.G1141A:p.V381M, |
| chr2 | 26214512 | G | C | . | exonic | HADHA | stopgain | HADHA:NM_000182:exon9:c.C849G:p.Y283X, |
| chr2 | 26232192 | C | T | . | exonic | HADHA | nonsynonymous SNV | HADHA:NM_000182:exon6:c.G541A:p.G181R, |
| chr2 | 26232237 | T | A | . | exonic | HADHA | stopgain | HADHA:NM_000182:exon6:c.A496T:p.K166X, |
| chr2 | 27212382 | A | G | COSV60160028 | exonic | ATRAID | nonsynonymous SNV | ATRAID:NM_080592:exon1:c.A179G:p.D60G,ATRAID:NM_001170795:exon1:c.A14G:p.D5G, |
| chr2 | 27216883 | G | T | . | exonic | ATRAID | nonsynonymous SNV | ATRAID:NM_080592:exon7:c.G790T:p.A264S,ATRAID:NM_001170795:exon7:c.G625T:p.A209S,ATRAID:NM_016085:exon7:c.G451T:p.A151S, |
| chr2 | 272203 | C | T | COSV55243186 | exonic | ACP1 | nonsynonymous SNV | ACP1:NM_001040649:exon3:c.C284T:p.T95I, |

| | | | | | | | | |
|---|---|---|---|---|---|---|---|---|
| chr2 | 27342108 | G | C | . | exonic | GTF3C2 | nonsynonymous SNV | GTF3C2:NM_001521:exon5:c.C695G:p.A232G,GTF3C2:NM_001035521:exon4:c.C695G:p.A232G, |
| chr2 | 27342117 | T | A | . | exonic | GTF3C2 | nonsynonymous SNV | GTF3C2:NM_001521:exon5:c.A686T:p.Q229L,GTF3C2:NM_001035521:exon4:c.A686T:p.Q229L, |
| chr2 | 27342124 | T | C | . | exonic | GTF3C2 | nonsynonymous SNV | GTF3C2:NM_001521:exon5:c.A679G:p.I227V,GTF3C2:NM_001035521:exon4:c.A679G:p.I227V, |
| chr2 | 27367171 | G | C | COSV52007864 | exonic | EIF2B4 | nonsynonymous SNV | EIF2B4:NM_172195:exon9:c.C976G:p.R326G,EIF2B4:NM_001034116:exon10:c.C916G:p.R306G,EIF2B4:NM_015636:exon10:c.C913G:p.R305G, |
| chr2 | 27439799 | G | A | . | exonic | NRBP1 | nonsynonymous SNV | NRBP1:NM_013392:exon11:c.G937A:p.A313T, |
| chr2 | 277003 | A | G | COSV55243196 | exonic | ACP1 | nonsynonymous SNV | ACP1:NM_004300:exon5:c.A317G:p.Q106R,ACP1:NM_007099:exon5:c.A317G:p.Q106R, |
| chr2 | 28851255 | C | T | . | exonic | TRMT61B | nonsynonymous SNV | TRMT61B:NM_017910:exon5:c.G1129A:p.E377K, |
| chr2 | 30158683 | G | A | . | exonic | YPEL5 | nonsynonymous SNV | YPEL5:NM_001127400:exon4:c.G206A:p.R69Q,YPEL5:NM_001127401:exon5:c.G206A:p.R69Q,YPEL5:NM_016061:exon3:c.G206A:p.R69Q,YPEL5:NM_001127399:exon4:c.G206A:p.R69Q, |
| chr2 | 30158784 | C | T | COSV54395171 | exonic | YPEL5 | nonsynonymous SNV | YPEL5:NM_001127400:exon4:c.C307T:p.R103C,YPEL5:NM_001127401:exon5:c.C307T:p.R103C,YPEL5:NM_016061:exon3:c.C307T:p.R103C,YPEL5:NM_001127399:exon4:c.C307T:p.R103C, |
| chr2 | 30158785 | G | A | . | exonic | YPEL5 | nonsynonymous SNV | YPEL5:NM_001127400:exon4:c.G308A:p.R103H,YPEL5:NM_001127401:exon5:c.G308A:p.R103H,YPEL5:NM_016061:exon3:c.G308A:p.R103H,YPEL5:NM_001127399:exon4:c.G308A:p.R103H, |
| chr2 | 32476232 | C | A | . | exonic | BIRC6 | nonsynonymous SNV | BIRC6:NM_016252:exon34:c.C6740A:p.A2247E, |
| chr2 | 32543343 | G | A | . | exonic | BIRC6 | nonsynonymous SNV | BIRC6:NM_016252:exon62:c.G12394A:p.A4132T, |
| chr2 | 32736791 | A | G | COSV58653535 | exonic | TTC27 | nonsynonymous SNV | TTC27:NM_017735:exon12:c.A1427G:p.Y476C,TTC27:NM_001193509:exon12:c.A1277G:p.Y426C, |
| chr2 | 33360729 | T | C | . | exonic | LTBP1 | nonsynonymous SNV | LTBP1:NM_000627:exon23:c.T3155C:p.V1052A,LTBP1:NM_001166264:exon22:c.T3029C:p.V1010A,LTBP1:NM_001166266:exon22:c.T2870C:p.V957A,LTBP1:NM_001166265:exon23:c.T2996C:p.V999A,LTBP1:NM_206943:exon27:c.T4133C:p.V1378A, |
| chr2 | 38564205 | A | G | . | exonic | HNRNPLL | nonsynonymous SNV | HNRNPLL:NM_138394:exon13:c.T1606C:p.F536L,HNRNPLL:NM_001142650:exon14:c.T1591C:p.F531L, |
| chr2 | 38602506 | T | C | . | exonic | HNRNPLL | nonsynonymous SNV | HNRNPLL:NM_138394:exon1:c.A121G:p.N41D,HNRNPLL:NM_001142650:exon2:c.A106G:p.N36D, |
| chr2 | 42282878 | A | G | COSV59293461 | exonic | EML4 | nonsynonymous SNV | EML4:NM_001145076:exon7:c.A673G:p.K225E,EML4:NM_019063:exon8:c.A847G:p.K283E, |
| chr2 | 42288248 | A | G | COSV59293527 | exonic | EML4 | nonsynonymous SNV | EML4:NM_001145076:exon10:c.A970G:p.I324V,EML4:NM_019063:exon11:c.A1144G:p.I382V, |
| chr2 | 42288297 | A | G | COSV59293537 | exonic | EML4 | nonsynonymous SNV | EML4:NM_001145076:exon10:c.A1019G:p.K340R,EML4:NM_019063:exon11:c.A1193G:p.K398R, |
| chr2 | 44275948 | G | A | . | exonic | SLC3A1 | nonsynonymous SNV | SLC3A1:NM_000341:exon1:c.G413A:p.G138E, |
| chr2 | 44286045 | G | A | . | exonic | SLC3A1 | nonsynonymous SNV | SLC3A1:NM_000341:exon4:c.G779A:p.G260E, |
| chr2 | 44301117 | G | A | COSV53224754 | exonic | SLC3A1 | nonsynonymous SNV | SLC3A1:NM_000341:exon6:c.G1126A:p.G376S, |
| chr2 | 44312694 | G | A | . | exonic | SLC3A1 | nonsynonymous SNV | SLC3A1:NM_000341:exon8:c.G1441A:p.G481R, |
| chr2 | 44313895 | G | A | . | exonic | SLC3A1 | nonsynonymous SNV | SLC3A1:NM_000341:exon9:c.G1561A:p.E521K, |
| chr2 | 44320435 | G | A | COSV53215689 | exonic | SLC3A1 | nonsynonymous SNV | SLC3A1:NM_000341:exon10:c.G1854A:p.M618I, |
| chr2 | 46909059 | C | T | . | exonic | MCFD2 | nonsynonymous SNV | MCFD2:NM_001171507:exon2:c.G113A:p.S38N,MCFD2:NM_001171506:exon3:c.G113A:p.S38N,MCFD2:NM_139279:exon2:c.G113A:p.S38N,MCFD2:NM_001171508:exon2:c.G113A:p.S38N, |
| chr2 | 47373493 | T | C | . | exonic | EPCAM | nonsynonymous SNV | EPCAM:NM_002354:exon2:c.T107C:p.V36A, |
| chr2 | 47373859 | G | A | . | exonic | EPCAM | nonsynonymous SNV | EPCAM:NM_002354:exon3:c.G236A:p.G79E, |
| chr2 | 47373862 | G | A | . | exonic | EPCAM | nonsynonymous SNV | EPCAM:NM_002354:exon3:c.G239A:p.R80K, |
| chr2 | 47373967 | T | C | COSV55392922 | exonic | EPCAM | nonsynonymous SNV | EPCAM:NM_002354:exon3:c.T344C:p.M115T, |
| chr2 | 47374036 | G | A | . | exonic | EPCAM | nonsynonymous SNV | EPCAM:NM_002354:exon3:c.G413A:p.R138Q, |
| chr2 | 47375236 | G | A | . | exonic | EPCAM | stopgain | EPCAM:NM_002354:exon4:c.G428A:p.W143X, |
| chr2 | 47375268 | G | A | . | exonic | EPCAM | nonsynonymous SNV | EPCAM:NM_002354:exon4:c.G460A:p.E154K, |
| chr2 | 47379025 | G | A | COSV55394795 | exonic | EPCAM | nonsynonymous SNV | EPCAM:NM_002354:exon6:c.G628A:p.A210T, |
| chr2 | 47379776 | G | A | . | exonic | EPCAM | nonsynonymous SNV | EPCAM:NM_002354:exon7:c.G665A:p.G222D, |
| chr2 | 47379778 | G | A | . | exonic | EPCAM | nonsynonymous SNV | EPCAM:NM_002354:exon7:c.G667A:p.E223K, |
| chr2 | 47379823 | G | A | . | exonic | EPCAM | nonsynonymous SNV | EPCAM:NM_002354:exon7:c.G712A:p.E238K, |
| chr2 | 47379844 | G | A | . | exonic | EPCAM | nonsynonymous SNV | EPCAM:NM_002354:exon7:c.G733A:p.G245S, |
| chr2 | 47379847 | C | A | . | exonic | EPCAM | nonsynonymous SNV | EPCAM:NM_002354:exon7:c.C736A:p.Q246K, |
| chr2 | 47379871 | G | A | . | exonic | EPCAM | nonsynonymous SNV | EPCAM:NM_002354:exon7:c.G760A:p.E254K, |
| chr2 | 47379883 | G | A | . | exonic | EPCAM | nonsynonymous SNV | EPCAM:NM_002354:exon7:c.G772A:p.E258K, |
| chr2 | 47379961 | G | A | . | exonic | EPCAM | nonsynonymous SNV | EPCAM:NM_002354:exon7:c.G850A:p.V284I, |
| chr2 | 47379964 | G | A | . | exonic | EPCAM | nonsynonymous SNV | EPCAM:NM_002354:exon7:c.G853A:p.V285M, |
| chr2 | 47386599 | G | A | . | exonic | EPCAM | nonsynonymous SNV | EPCAM:NM_002354:exon9:c.G931A:p.E311K, |
| chr2 | 47445602 | G | A | COSV51883444 | exonic | MSH2 | nonsynonymous SNV | MSH2:NM_000251:exon8:c.G1331A:p.R444H,MSH2:NM_001258281:exon9:c.G1133A:p.R378H, |
| chr2 | 47463098 | T | A | . | exonic | MSH2 | nonsynonymous SNV | MSH2:NM_000251:exon9:c.T1454A:p.M485K,MSH2:NM_001258281:exon10:c.T1256A:p.M419K, |
| chr2 | 47471001 | T | A | COSV51879859 | exonic | MSH2 | nonsynonymous SNV | MSH2:NM_000251:exon11:c.T1698A:p.N566K,MSH2:NM_001258281:exon12:c.T1500A:p.N500K, |
| chr2 | 47475195 | G | A | . | exonic | MSH2 | nonsynonymous SNV | MSH2:NM_000251:exon12:c.G1930A:p.V644I,MSH2:NM_001258281:exon13:c.G1732A:p.V578I, |
| chr2 | 47475201 | G | A | . | exonic | MSH2 | nonsynonymous SNV | MSH2:NM_000251:exon12:c.G1936A:p.D646N,MSH2:NM_001258281:exon13:c.G1738A:p.D580N, |
| chr2 | 47475249 | C | A | . | exonic | MSH2 | nonsynonymous SNV | MSH2:NM_000251:exon12:c.C1984A:p.Q662K,MSH2:NM_001258281:exon13:c.C1786A:p.Q596K, |
| chr2 | 47478343 | G | A | . | exonic | MSH2 | nonsynonymous SNV | MSH2:NM_000251:exon14:c.G2282A:p.G761E,MSH2:NM_001258281:exon15:c.G2084A:p.G695E, |
| chr2 | 47478394 | G | A | . | exonic | MSH2 | nonsynonymous SNV | MSH2:NM_000251:exon14:c.G2333A:p.C778Y,MSH2:NM_001258281:exon15:c.G2135A:p.C712Y, |
| chr2 | 47478426 | G | A | . | exonic | MSH2 | nonsynonymous SNV | MSH2:NM_000251:exon14:c.G2365A:p.A789T,MSH2:NM_001258281:exon15:c.G2167A:p.A723T, |
| chr2 | 47819056 | A | T | . | exonic | FBXO11 | nonsynonymous SNV | FBXO11:NM_025133:exon15:c.T1568A:p.I523K,FBXO11:NM_001190274:exon15:c.T1820A:p.I607K, |
| chr2 | 55235552 | G | A | . | exonic | RPS27A | nonsynonymous SNV | RPS27A:NM_002954:exon6:c.G446A:p.C149Y,RPS27A:NM_001135592:exon6:c.G446A:p.C149Y,RPS27A:NM_001177413:exon5:c.G446A:p.C149Y, |
| chr2 | 60546080 | C | T | . | exonic | BCL11A | nonsynonymous SNV | BCL11A:NM_138559:exon2:c.G276A:p.M92I,BCL11A:NM_018014:exon2:c.G276A:p.M92I,BCL11A:NM_022893:exon2:c.G276A:p.M92I, |
| chr2 | 60546181 | C | T | . | exonic | BCL11A | nonsynonymous SNV | BCL11A:NM_138559:exon2:c.G175A:p.D59N,BCL11A:NM_018014:exon2:c.G175A:p.D59N,BCL11A:NM_022893:exon2:c.G175A:p.D59N, |
| chr2 | 61869495 | C | T | . | exonic | CCT4 | nonsynonymous SNV | CCT4:NM_006430:exon13:c.G1550A:p.S517N,CCT4:NM_001256721:exon12:c.G1460A:p.S487N, |
| chr2 | 61869505 | C | T | . | exonic | CCT4 | nonsynonymous SNV | CCT4:NM_006430:exon13:c.G1540A:p.V514I,CCT4:NM_001256721:exon12:c.G1450A:p.V484I, |
| chr2 | 61872165 | C | T | . | exonic | CCT4 | nonsynonymous SNV | CCT4:NM_006430:exon12:c.G1408A:p.G470S,CCT4:NM_001256721:exon11:c.G1318A:p.G440S, |
| chr2 | 61872230 | G | T | . | exonic | CCT4 | nonsynonymous SNV | CCT4:NM_006430:exon12:c.C1343A:p.S448Y,CCT4:NM_001256721:exon11:c.C1253A:p.S418Y, |
| chr2 | 61872269 | C | T | . | exonic | CCT4 | nonsynonymous SNV | CCT4:NM_006430:exon12:c.G1304A:p.R435Q,CCT4:NM_001256721:exon11:c.G1214A:p.R405Q, |
| chr2 | 61872571 | G | T | . | exonic | CCT4 | nonsynonymous SNV | CCT4:NM_006430:exon11:c.C1143A:p.S381R,CCT4:NM_001256721:exon10:c.C1053A:p.S351R, |
| chr2 | 61873274 | C | T | . | exonic | CCT4 | nonsynonymous SNV | CCT4:NM_006430:exon9:c.G937A:p.A313T,CCT4:NM_001256721:exon8:c.G847A:p.A283T, |
| chr2 | 61876191 | C | T | . | exonic | CCT4 | nonsynonymous SNV | CCT4:NM_006430:exon8:c.G821A:p.R274Q,CCT4:NM_001256721:exon7:c.G731A:p.R244Q, |
| chr2 | 61876204 | C | T | . | exonic | CCT4 | nonsynonymous SNV | CCT4:NM_006430:exon8:c.G808A:p.A270T,CCT4:NM_001256721:exon7:c.G718A:p.A240T, |
| chr2 | 61877015 | C | T | COSV66702218 | exonic | CCT4 | nonsynonymous SNV | CCT4:NM_006430:exon7:c.G682A:p.V228I,CCT4:NM_001256721:exon6:c.G592A:p.V198I, |
| chr2 | 61877439 | C | T | . | exonic | CCT4 | nonsynonymous SNV | CCT4:NM_006430:exon6:c.G598A:p.A200T,CCT4:NM_001256721:exon5:c.G508A:p.A170T, |
| chr2 | 61885033 | C | T | . | exonic | CCT4 | nonsynonymous SNV | CCT4:NM_006430:exon2:c.G167A:p.G56E,CCT4:NM_001256721:exon2:c.G167A:p.G56E, |
| chr2 | 68822809 | T | C | COSV54900269 | exonic | ARHGAP25 | nonsynonymous SNV | ARHGAP25:NM_001166276:exon9:c.T1649C:p.M550T,ARHGAP25:NM_014882:exon9:c.T1646C:p.M549T,ARHGAP25:NM_001007231:exon10:c.T1670C:p.M557T,ARHGAP25:NM_001166277:exon9:c.T1550C:p.M517T, |
| chr2 | 69326220 | C | T | . | exonic | GFPT1 | nonsynonymous SNV | GFPT1:NM_002056:exon19:c.G2015A:p.R672Q,GFPT1:NM_001244710:exon20:c.G2069A:p.R690Q, |
| chr2 | 69326979 | C | T | . | exonic | GFPT1 | nonsynonymous SNV | GFPT1:NM_002056:exon18:c.G1936A:p.G646S,GFPT1:NM_001244710:exon19:c.G1990A:p.G664S, |
| chr2 | 69327048 | C | T | . | exonic | GFPT1 | nonsynonymous SNV | GFPT1:NM_002056:exon18:c.G1867A:p.E623K,GFPT1:NM_001244710:exon19:c.G1921A:p.E641K, |
| chr2 | 69327071 | C | T | COSV61947419 | exonic | GFPT1 | nonsynonymous SNV | GFPT1:NM_002056:exon18:c.G1844A:p.R615Q,GFPT1:NM_001244710:exon19:c.G1898A:p.R633Q, |
| chr2 | 69329332 | C | T | . | exonic | GFPT1 | nonsynonymous SNV | GFPT1:NM_002056:exon16:c.G1636A:p.G546S,GFPT1:NM_001244710:exon17:c.G1690A:p.G564S, |

| | | | | | | | | |
|---|---|---|---|---|---|---|---|---|
| chr2 | 69329410 | C | T | . | exonic | GFPT1 | nonsynonymous SNV | GFPT1:NM_002056:exon16:c.G1558A:p.V520I,GFPT1:NM_001244710:exon17:c.G1612A:p.V538I, |
| chr2 | 69329735 | A | G | . | exonic | GFPT1 | nonsynonymous SNV | GFPT1:NM_002056:exon15:c.T1492C:p.S498P,GFPT1:NM_001244710:exon16:c.T1546C:p.S516P, |
| chr2 | 69329740 | C | T | . | exonic | GFPT1 | nonsynonymous SNV | GFPT1:NM_002056:exon15:c.G1487A:p.R496Q,GFPT1:NM_001244710:exon16:c.G1541A:p.R514Q, |
| chr2 | 69329750 | A | G | . | exonic | GFPT1 | nonsynonymous SNV | GFPT1:NM_002056:exon15:c.T1477C:p.C493R,GFPT1:NM_001244710:exon16:c.T1531C:p.C511R, |
| chr2 | 69329779 | A | G | . | exonic | GFPT1 | nonsynonymous SNV | GFPT1:NM_002056:exon15:c.T1448C:p.V483A,GFPT1:NM_001244710:exon16:c.T1502C:p.V501A, |
| chr2 | 69337926 | C | T | . | exonic | GFPT1 | nonsynonymous SNV | GFPT1:NM_002056:exon14:c.G1400A:p.G467D,GFPT1:NM_001244710:exon15:c.G1454A:p.G485D, |
| chr2 | 69338463 | A | G | . | exonic | GFPT1 | nonsynonymous SNV | GFPT1:NM_002056:exon13:c.T1252C:p.F418L,GFPT1:NM_001244710:exon14:c.T1306C:p.F436L, |
| chr2 | 69338561 | C | T | . | exonic | GFPT1 | nonsynonymous SNV | GFPT1:NM_002056:exon13:c.G1154A:p.R385H,GFPT1:NM_001244710:exon14:c.G1208A:p.R403H, |
| chr2 | 69348193 | C | T | . | exonic | GFPT1 | nonsynonymous SNV | GFPT1:NM_002056:exon10:c.G933A:p.M311I,GFPT1:NM_001244710:exon11:c.G987A:p.M329I, |
| chr2 | 69348266 | C | T | . | exonic | GFPT1 | nonsynonymous SNV | GFPT1:NM_002056:exon10:c.G860A:p.G287E,GFPT1:NM_001244710:exon11:c.G914A:p.G305E, |
| chr2 | 69350094 | A | G | . | exonic | GFPT1 | nonsynonymous SNV | GFPT1:NM_002056:exon9:c.T775C:p.F259L,GFPT1:NM_001244710:exon10:c.T829C:p.F277L, |
| chr2 | 69358436 | C | T | . | exonic | GFPT1 | nonsynonymous SNV | GFPT1:NM_002056:exon6:c.G436A:p.E146K,GFPT1:NM_001244710:exon6:c.G436A:p.E146K, |
| chr2 | 69431994 | A | T | COSV58006358 | exonic | NFU1 | nonsynonymous SNV | NFU1:NM_015700:exon2:c.T2A:p.M1K,NFU1:NM_001002755:exon2:c.T74A:p.M25K, |
| chr2 | 69804572 | G | A | . | exonic | ANXA4 | nonsynonymous SNV | ANXA4:NM_001153:exon4:c.G137A:p.R46H, |
| chr2 | 69804622 | G | A | . | exonic | ANXA4 | nonsynonymous SNV | ANXA4:NM_001153:exon4:c.G187A:p.G63S, |
| chr2 | 69806452 | C | T | COSV67848858 | exonic | ANXA4 | nonsynonymous SNV | ANXA4:NM_001153:exon5:c.C260T:p.T87M, |
| chr2 | 69807913 | G | A | . | exonic | ANXA4 | nonsynonymous SNV | ANXA4:NM_001153:exon6:c.G314A:p.G105D, |
| chr2 | 69807924 | G | A | . | exonic | ANXA4 | nonsynonymous SNV | ANXA4:NM_001153:exon6:c.G325A:p.G109S, |
| chr2 | 69807952 | G | A | COSV67848884 | exonic | ANXA4 | nonsynonymous SNV | ANXA4:NM_001153:exon6:c.G353A:p.R118Q, |
| chr2 | 69816107 | T | C | . | exonic | ANXA4 | nonsynonymous SNV | ANXA4:NM_001153:exon9:c.T541C:p.Y181H, |
| chr2 | 69818664 | G | A | . | exonic | ANXA4 | nonsynonymous SNV | ANXA4:NM_001153:exon10:c.G694A:p.G232S, |
| chr2 | 69820699 | G | A | . | exonic | ANXA4 | nonsynonymous SNV | ANXA4:NM_001153:exon12:c.G784A:p.G262S, |
| chr2 | 69825510 | G | A | COSV57003814 | exonic | ANXA4 | nonsynonymous SNV | ANXA4:NM_001153:exon13:c.G961A:p.D321N, |
| chr2 | 74958729 | G | T | COSV52050939 | exonic | POLE4 | nonsynonymous SNV | POLE4:NM_019896:exon1:c.G50T:p.G17V, |
| chr2 | 84528925 | G | A | . | exonic | DNAH6 | nonsynonymous SNV | DNAH6:NM_001370:exon4:c.G421A:p.V141M, |
| chr2 | 84823998 | G | A | COSV52874802 | exonic | TRABD2A | nonsynonymous SNV | TRABD2A:NM_001080824:exon5:c.C1142T:p.P381L,TRABD2A:NM_001277053:exon6:c.C1289T:p.P430L, |
| chr2 | 85394936 | T | C | . | exonic | CAPG | nonsynonymous SNV | CAPG:NM_001747:exon10:c.A1004G:p.H335R,CAPG:NM_001256140:exon10:c.A959G:p.H320R,CAPG:NM_001256139:exon10:c.A1004G:p.H335R, |
| chr2 | 85401860 | C | T | COSV55709070 | exonic | CAPG | nonsynonymous SNV | CAPG:NM_001747:exon3:c.G121A:p.V41I,CAPG:NM_001256140:exon3:c.G121A:p.V41I,CAPG:NM_001256139:exon3:c.G121A:p.V41I, |
| chr2 | 85597128 | T | C | COSV57817795 | exonic | RNF181 | nonsynonymous SNV | RNF181:NM_016494:exon4:c.T352C:p.Y118H, |
| chr2 | 86173701 | G | A | COSV54485553 | exonic | IMMT | nonsynonymous SNV | IMMT:NM_006839:exon4:c.C370T:p.P124S,IMMT:NM_001100170:exon4:c.C370T:p.P124S,IMMT:NM_001100169:exon4:c.C370T:p.P124S, |
| chr2 | 88124547 | T | C | COSV55559009 | exonic | FABP1 | nonsynonymous SNV | FABP1:NM_001443:exon3:c.A280G:p.T94A, |
| chr2 | 88126337 | C | T | . | exonic | FABP1 | nonsynonymous SNV | FABP1:NM_001443:exon2:c.G79A:p.E27K, |
| chr2 | 88127981 | C | T | . | exonic | FABP1 | nonsynonymous SNV | FABP1:NM_001443:exon1:c.G37A:p.E13K, |
| chr2 | 99439044 | A | G | COSV51482947 | exonic | REV1 | nonsynonymous SNV | REV1:NM_016316:exon6:c.T770C:p.F257S,REV1:NM_001037872:exon6:c.T770C:p.F257S, |
| chr2 | 99442408 | C | T | COSV51481954 | exonic | REV1 | nonsynonymous SNV | REV1:NM_016316:exon5:c.G412A:p.V138M,REV1:NM_001037872:exon5:c.G412A:p.V138M, |
| chr20 | 1118880 | T | G | COSV55672896 | exonic | PSMF1 | nonsynonymous SNV | PSMF1:NM_178578:exon2:c.T107G:p.F36C,PSMF1:NM_006814:exon1:c.T107G:p.F36C, |
| chr20 | 1445748 | C | T | COSV53793742 | exonic | NSFL1C | nonsynonymous SNV | NSFL1C:NM_016143:exon8:c.G868A:p.D290N,NSFL1C:NM_018839:exon7:c.G775A:p.D259N, |
| chr20 | 17624595 | A | T | . | exonic | RRBP1 | nonsynonymous SNV | RRBP1:NM_004587:exon13:c.T1829A:p.L610H,RRBP1:NM_001042576:exon14:c.T1829A:p.L610H, |
| chr20 | 17969902 | A | T | COSV66624187 | exonic | MGME1 | nonsynonymous SNV | MGME1:NM_052865:exon2:c.A43T:p.S15C,MGME1:NM_001310339:exon2:c.A43T:p.S15C,MGME1:NM_001310338:exon2:c.A43T:p.S15C, |
| chr20 | 17990053 | G | A | . | exonic | MGME1 | nonsynonymous SNV | MGME1:NM_052865:exon5:c.G979A:p.E327K,MGME1:NM_001310339:exon3:c.G626A:p.R209K,MGME1:NM_001310338:exon6:c.G1024A:p.E342K, |
| chr20 | 17990061 | G | A | . | exonic | MGME1 | nonsynonymous SNV | MGME1:NM_001310339:exon3:c.G634A:p.G212R, |
| chr20 | 17990067 | G | A | . | exonic | MGME1 | nonsynonymous SNV | MGME1:NM_001310339:exon3:c.G640A:p.E214K, |
| chr20 | 18162473 | T | G | . | exonic | CSRP2BP | nonsynonymous SNV | CSRP2BP:NM_020536:exon6:c.T1199G:p.V400G, |
| chr20 | 18524991 | A | T | . | exonic | SEC23B | nonsynonymous SNV | SEC23B:NM_001172745:exon6:c.A660T:p.Q220H,SEC23B:NM_032985:exon6:c.A660T:p.Q220H,SEC23B:NM_001172746:exon6:c.A606T:p.Q202H,SEC23B:NM_006363:exon6:c.A660T:p.Q220H,SEC23B:NM_032986:exon6:c.A660T:p.Q220H, |
| chr20 | 18532728 | C | T | COSV52719453 | exonic | SEC23B | nonsynonymous SNV | SEC23B:NM_001172745:exon11:c.C1298T:p.P433L,SEC23B:NM_032985:exon11:c.C1298T:p.P433L,SEC23B:NM_001172746:exon11:c.C1244T:p.P415L,SEC23B:NM_006363:exon11:c.C1298T:p.P433L,SEC23B:NM_032986:exon11:c.C1298T:p.P433L, |
| chr20 | 18535653 | G | A | . | exonic | SEC23B | nonsynonymous SNV | SEC23B:NM_001172745:exon12:c.G1315A:p.E439K,SEC23B:NM_032985:exon12:c.G1315A:p.E439K,SEC23B:NM_006363:exon12:c.G1315A:p.E439K,SEC23B:NM_001172746:exon12:c.G1261A:p.E421K,SEC23B:NM_032986:exon12:c.G1315A:p.E439K, |
| chr20 | 18542358 | C | G | COSV52719183 | exonic | SEC23B | nonsynonymous SNV | SEC23B:NM_001172745:exon13:c.C1467G:p.H489Q,SEC23B:NM_032985:exon13:c.C1467G:p.H489Q,SEC23B:NM_001172746:exon13:c.C1413G:p.H471Q,SEC23B:NM_006363:exon13:c.C1467G:p.H489Q,SEC23B:NM_032986:exon13:c.C1467G:p.H489Q, |
| chr20 | 18744172 | G | A | COSV66281766 | exonic | DTD1 | nonsynonymous SNV | DTD1:NM_080820:exon5:c.G550A:p.E184K, |
| chr20 | 23637790 | C | T | COSV65361855 | exonic | CST3 | nonsynonymous SNV | CST3:NM_001288614:exon1:c.G73A:p.A25T,CST3:NM_000099:exon1:c.G73A:p.A25T, |
| chr20 | 25214873 | A | G | . | exonic | ENTPD6 | nonsynonymous SNV | ENTPD6:NM_001247:exon6:c.A604G:p.K202E,ENTPD6:NM_001114089:exon5:c.A553G:p.K185E, |
| chr20 | 25278370 | G | T | COSV53816254 | exonic | PYGB | nonsynonymous SNV | PYGB:NM_002862:exon8:c.G907T:p.A303S, |
| chr20 | 25281024 | C | T | . | exonic | PYGB | nonsynonymous SNV | PYGB:NM_002862:exon11:c.C1315T:p.R439W, |
| chr20 | 25282133 | G | A | COSV53816266 | exonic | PYGB | nonsynonymous SNV | PYGB:NM_002862:exon12:c.G1504A:p.D502N, |
| chr20 | 31766583 | A | G | . | exonic | TPX2 | nonsynonymous SNV | TPX2:NM_012112:exon5:c.A257G:p.E86G, |
| chr20 | 31783887 | G | A | . | exonic | TPX2 | nonsynonymous SNV | TPX2:NM_012112:exon12:c.G1379A:p.R460K, |
| chr20 | 31793914 | C | A | . | exonic | TPX2 | nonsynonymous SNV | TPX2:NM_012112:exon14:c.C1576A:p.Q526K, |
| chr20 | 31798403 | C | A | . | exonic | TPX2 | nonsynonymous SNV | TPX2:NM_012112:exon17:c.C1984A:p.L662M, |
| chr20 | 31946540 | C | T | COSV52431464 | exonic | PDRG1 | nonsynonymous SNV | PDRG1:NM_030815:exon4:c.G275A:p.R92Q, |
| chr20 | 32165386 | G | A | . | exonic | TM9SF4 | nonsynonymous SNV | TM9SF4:NM_014742:exon18:c.G1871A:p.G624D, |
| chr20 | 32432923 | G | A | . | exonic | ASXL1 | nonsynonymous SNV | ASXL1:NM_015338:exon10:c.G1023A:p.M341I, |

| | | | | | | | | |
|---|---|---|---|---|---|---|---|---|
| chr20 | 32432924 | G | A | . | exonic | ASXL1 | nonsynonymous SNV | ASXL1:NM_015338:exon10:c.G1024A:p.E342K, |
| chr20 | 32432949 | G | A | . | exonic | ASXL1 | stopgain | ASXL1:NM_015338:exon10:c.G1049A:p.W350X, |
| chr20 | 34928883 | G | T | . | exonic | GSS | nonsynonymous SNV | GSS:NM_000178:exon13:c.C1370A:p.A457E, |
| chr20 | 34928896 | C | T | . | exonic | GSS | nonsynonymous SNV | GSS:NM_000178:exon13:c.G1357A:p.A453T, |
| chr20 | 34928917 | C | T | . | exonic | GSS | nonsynonymous SNV | GSS:NM_000178:exon13:c.G1336A:p.G446R, |
| chr20 | 34928920 | C | T | . | exonic | GSS | nonsynonymous SNV | GSS:NM_000178:exon13:c.G1333A:p.V445M, |
| chr20 | 34929476 | C | T | . | exonic | GSS | nonsynonymous SNV | GSS:NM_000178:exon12:c.G1226A:p.C409Y, |
| chr20 | 34931396 | T | C | . | exonic | GSS | nonsynonymous SNV | GSS:NM_000178:exon11:c.A1051G:p.I351V, |
| chr20 | 34931405 | C | T | . | exonic | GSS | nonsynonymous SNV | GSS:NM_000178:exon11:c.G1042A:p.D348N, |
| chr20 | 34932127 | C | T | . | exonic | GSS | nonsynonymous SNV | GSS:NM_000178:exon10:c.G841A:p.E281K, |
| chr20 | 34935583 | C | T | . | exonic | GSS | nonsynonymous SNV | GSS:NM_000178:exon9:c.G827A:p.S276N, |
| chr20 | 34936982 | A | G | . | exonic | GSS | nonsynonymous SNV | GSS:NM_000178:exon7:c.T650C:p.I217T, |
| chr20 | 34941717 | G | T | . | exonic | GSS | nonsynonymous SNV | GSS:NM_000178:exon6:c.C604A:p.P202T, |
| chr20 | 34941732 | C | T | . | exonic | GSS | nonsynonymous SNV | GSS:NM_000178:exon6:c.G589A:p.E197K, |
| chr20 | 34942579 | C | T | . | exonic | GSS | nonsynonymous SNV | GSS:NM_000178:exon5:c.G400A:p.A134T, |
| chr20 | 34942969 | C | T | . | exonic | GSS | nonsynonymous SNV | GSS:NM_000178:exon4:c.G313A:p.D105N, |
| chr20 | 34951791 | T | A | . | exonic | GSS | nonsynonymous SNV | GSS:NM_000178:exon2:c.A62T:p.Q21L, |
| chr20 | 34951800 | A | G | . | exonic | GSS | nonsynonymous SNV | GSS:NM_000178:exon2:c.T53C:p.L18P, |
| chr20 | 35384111 | C | T | . | exonic | UQCC1 | nonsynonymous SNV | UQCC1:NM_199487:exon3:c.G152A:p.R51Q,UQCC1:NM_018244:exon3:c.G152A:p.R51Q, |
| chr20 | 35631574 | T | C | COSV58290412 | exonic | CPNE1 | nonsynonymous SNV | CPNE1:NM_152927:exon9:c.A632G:p.Q211R,CPNE1:NM_003915:exon8:c.A647G:p.Q216R,CPNE1:NM_152925:exon8:c.A632G:p.Q211R,CPNE1:NM_152926:exon8:c.A632G:p.Q211R,CPNE1:NM_152928:exon9:c.A632G:p.Q211R,CPNE1:NM_001198863:exon8:c.A632G:p.Q211R, |
| chr20 | 37223968 | C | T | . | exonic | RPN2 | nonsynonymous SNV | RPN2:NM_002951:exon10:c.C1183T:p.R395W,RPN2:NM_001135771:exon9:c.C1087T:p.R363W, |
| chr20 | 37316338 | G | A | . | exonic | MANBAL | nonsynonymous SNV | MANBAL:NM_022077:exon4:c.G181A:p.E61K,MANBAL:NM_001003897:exon3:c.G181A:p.E61K, |
| chr20 | 41168825 | T | C | COSV54816568 | exonic | PLCG1 | nonsynonymous SNV | PLCG1:NM_002660:exon21:c.T2438C:p.I813T,PLCG1:NM_182811:exon21:c.T2438C:p.I813T, |
| chr20 | 45174972 | C | T | COSV54783425 | exonic | PI3 | nonsynonymous SNV | PI3:NM_002638:exon1:c.C50T:p.T17M, |
| chr20 | 45175881 | A | C | COSV54782855 | exonic | PI3 | nonsynonymous SNV | PI3:NM_002638:exon2:c.A100C:p.T34P, |
| chr20 | 45409934 | A | G | COSV61919052 | exonic | DBNDD2 | nonsynonymous SNV | DBNDD2:NM_001048221:exon4:c.A280G:p.M94V,DBNDD2:NM_001197140:exon4:c.A280G:p.M94V,DBNDD2:NM_001048223:exon4:c.A280G:p.M94V,DBNDD2:NM_018478:exon4:c.A574G:p.M192V,DBNDD2:NM_001197139:exon4:c.A280G:p.M94V,DBNDD2:NM_001048225:exon3:c.A586G:p.M196V, |
| chr20 | 50153622 | T | C | COSV59009885 | exonic | TMEM189 | nonsynonymous SNV | TMEM189:NM_199129:exon1:c.A16G:p.N6D,TMEM189-UBE2V1:NM_199203:exon1:c.A16G:p.N6D,TMEM189:NM_001162505:exon1:c.A16G:p.N6D, |
| chr20 | 50936179 | C | T | . | exonic | DPM1 | nonsynonymous SNV | DPM1:NM_001317035:exon9:c.G728A:p.R243Q,DPM1:NM_003859:exon8:c.G647A:p.R216Q,DPM1:NM_001317034:exon9:c.G752A:p.R251Q,DPM1:NM_001317036:exon7:c.G578A:p.R193Q, |
| chr20 | 50936254 | C | T | . | exonic | DPM1 | nonsynonymous SNV | DPM1:NM_001317035:exon9:c.G653A:p.R218Q,DPM1:NM_003859:exon8:c.G572A:p.R191Q,DPM1:NM_001317034:exon9:c.G677A:p.R226Q,DPM1:NM_001317036:exon7:c.G503A:p.R168Q, |
| chr20 | 50940920 | A | T | . | exonic | DPM1 | nonsynonymous SNV | DPM1:NM_001317035:exon8:c.T589A:p.F197I,DPM1:NM_003859:exon7:c.T508A:p.F170I,DPM1:NM_001317034:exon8:c.T613A:p.F205I, |
| chr20 | 50942116 | C | T | . | exonic | DPM1 | nonsynonymous SNV | DPM1:NM_001317035:exon6:c.G409A:p.E137K,DPM1:NM_003859:exon6:c.G409A:p.E137K,DPM1:NM_001317034:exon6:c.G409A:p.E137K,DPM1:NM_001317036:exon6:c.G409A:p.E137K, |
| chr20 | 50945887 | C | T | . | exonic | DPM1 | nonsynonymous SNV | DPM1:NM_001317034:exon4:c.G332A:p.G111E,DPM1:NM_001317036:exon4:c.G332A:p.G111E,DPM1:NM_001317035:exon4:c.G332A:p.G111E,DPM1:NM_003859:exon4:c.G332A:p.G111E, |
| chr20 | 50958492 | C | T | . | exonic | DPM1 | nonsynonymous SNV | DPM1:NM_001317034:exon1:c.G32A:p.R11H,DPM1:NM_001317036:exon1:c.G32A:p.R11H,DPM1:NM_001317035:exon1:c.G32A:p.R11H,DPM1:NM_003859:exon1:c.G32A:p.R11H, |
| chr20 | 52152840 | C | T | COSV53799554 | exonic | ZFP64 | nonsynonymous SNV | ZFP64:NM_199426:exon6:c.G1346A:p.S449N,ZFP64:NM_022088:exon5:c.G1190A:p.S397N,ZFP64:NM_018197:exon6:c.G1352A:p.S451N, |
| chr20 | 54214409 | G | A | . | exonic | PFDN4 | nonsynonymous SNV | PFDN4:NM_002623:exon2:c.G83A:p.R28Q, |
| chr20 | 54219064 | T | C | . | exonic | PFDN4 | nonsynonymous SNV | PFDN4:NM_002623:exon4:c.T319C:p.S107P, |
| chr20 | 54219119 | G | A | . | exonic | PFDN4 | nonsynonymous SNV | PFDN4:NM_002623:exon4:c.G374A:p.S125N, |
| chr20 | 56513348 | A | G | COSV50130089 | exonic | RTFDC1 | nonsynonymous SNV | RTFDC1:NM_001283035:exon7:c.A601G:p.M201V,RTFDC1:NM_016407:exon6:c.A511G:p.M171V,RTFDC1:NM_001283037:exon6:c.A511G:p.M171V,RTFDC1:NM_001283036:exon6:c.A511G:p.M171V, |
| chr20 | 57354069 | G | A | . | exonic | RAE1 | nonsynonymous SNV | RAE1:NM_001015885:exon2:c.G31A:p.G11R,RAE1:NM_003610:exon2:c.G31A:p.G11R, |
| chr20 | 57365417 | G | A | . | exonic | RAE1 | nonsynonymous SNV | RAE1:NM_001015885:exon5:c.G350A:p.S117N,RAE1:NM_003610:exon5:c.G350A:p.S117N, |
| chr20 | 57366847 | C | A | . | exonic | RAE1 | nonsynonymous SNV | RAE1:NM_001015885:exon6:c.C416A:p.P139Q,RAE1:NM_003610:exon6:c.C416A:p.P139Q, |
| chr20 | 57366861 | G | A | . | exonic | RAE1 | nonsynonymous SNV | RAE1:NM_001015885:exon6:c.G430A:p.V144M,RAE1:NM_003610:exon6:c.G430A:p.V144M, |
| chr20 | 57373682 | T | C | . | exonic | RAE1 | nonsynonymous SNV | RAE1:NM_001015885:exon10:c.T769C:p.F257L,RAE1:NM_003610:exon10:c.T769C:p.F257L, |
| chr20 | 57373704 | G | A | . | exonic | RAE1 | nonsynonymous SNV | RAE1:NM_001015885:exon10:c.G791A:p.G264E,RAE1:NM_003610:exon10:c.G791A:p.G264E, |
| chr20 | 57374734 | T | C | . | exonic | RAE1 | nonsynonymous SNV | RAE1:NM_001015885:exon11:c.T953C:p.I318T,RAE1:NM_003610:exon11:c.T953C:p.I318T, |
| chr20 | 57562839 | G | C | COSV60128476 | exonic | PCK1 | nonsynonymous SNV | PCK1:NM_002591:exon4:c.G550C:p.V184L, |
| chr20 | 58898946 | G | A | . | exonic | GNAS | nonsynonymous SNV | GNAS:NM_001309842:exon3:c.G218A:p.G73D,GNAS:NM_080425:exon3:c.G2147A:p.G716D,GNAS:NM_001309840:exon3:c.G41A:p.G14D,GNAS:NM_001309861:exon3:c.G41A:p.G14D,GNAS:NM_000516:exon3:c.G218A:p.G73D,GNAS:NM_001077488:exon3:c.G218A:p.G73D, |
| chr20 | 58903571 | A | C | . | exonic | GNAS | nonsynonymous SNV | GNAS:NM_080426:exon3:c.A256C:p.K86Q,GNAS:NM_080425:exon4:c.A2227C:p.K743Q,GNAS:NM_001309840:exon4:c.A121C:p.K41Q,GNAS:NM_001077489:exon3:c.A253C:p.K85Q,GNAS:NM_001309861:exon4:c.A121C:p.K41Q,GNAS:NM_000516:exon4:c.A298C:p.K100Q,GNAS:NM_001077488:exon4:c.A301C:p.K101Q, |
| chr20 | 58903682 | C | A | . | exonic | GNAS | nonsynonymous SNV | GNAS:NM_080426:exon4:c.C281A:p.A94D,GNAS:NM_080425:exon5:c.C2252A:p.A751D,GNAS:NM_001309840:exon5:c.C146A:p.A49D,GNAS:NM_001077489:exon4:c.C278A:p.A93D,GNAS:NM_001309861:exon5:c.C146A:p.A49D,GNAS:NM_000516:exon5:c.C323A:p.A108D,GNAS:NM_001077488:exon5:c.C326A:p.A109D, |
| chr20 | 58903684 | G | A | COSV55672684 | exonic | GNAS | nonsynonymous SNV | GNAS:NM_080426:exon4:c.G283A:p.A95T,GNAS:NM_080425:exon5:c.G2254A:p.A752T,GNAS:NM_001309840:exon5:c.G148A:p.A50T,GNAS:NM_001077489:exon4:c.G280A:p.A94T,GNAS:NM_001309861:exon5:c.G148A:p.A50T,GNAS:NM_000516:exon5:c.G325A:p.A109T,GNAS:NM_001077488:exon5:c.G328A:p.A110T, |
| chr20 | 58903691 | G | A | COSV99039588 | exonic | GNAS | nonsynonymous SNV | GNAS:NM_080426:exon4:c.G290A:p.S97N,GNAS:NM_080425:exon5:c.G2261A:p.S754N,GNAS:NM_001309840:exon5:c.G155A:p.S52N,GNAS:NM_001077489:exon4:c.G287A:p.S96N,GNAS:NM_001309861:exon5:c.G155A:p.S52N,GNAS:NM_000516:exon5:c.G332A:p.S111N,GNAS:NM_001077488:exon5:c.G335A:p.S112N, |

| | | | | | | | | |
|---|---|---|---|---|---|---|---|---|
| chr20 | 58903717 | G | A | . | exonic | GNAS | nonsynonymous SNV | GNAS:NM_080426:exon4:c.G316A:p.A106T,GNAS:NM_080425:exon5:c.G2287A:p.A763T,GNAS:NM_001309840:exon5:c.G181A:p.A61T,GNAS:NM_001077489:exon4:c.G313A:p.A105T,GNAS:NM_001309861:exon5:c.G181A:p.A61T,GNAS:NM_000516:exon5:c.G358A:p.A120T,GNAS:NM_001077488:exon5:c.G361A:p.A121T, |
| chr20 | 58903726 | G | A | . | exonic | GNAS | nonsynonymous SNV | GNAS:NM_080426:exon4:c.G325A:p.E109K,GNAS:NM_080425:exon5:c.G2296A:p.E766K,GNAS:NM_001309840:exon5:c.G190A:p.E64K,GNAS:NM_001077489:exon4:c.G322A:p.E108K,GNAS:NM_001309861:exon5:c.G190A:p.E64K,GNAS:NM_000516:exon5:c.G367A:p.E123K,GNAS:NM_001077488:exon5:c.G370A:p.E124K, |
| chr20 | 58903768 | G | A | . | exonic | GNAS | nonsynonymous SNV | GNAS:NM_080426:exon4:c.G367A:p.V123M,GNAS:NM_080425:exon5:c.G2338A:p.V780M,GNAS:NM_001309840:exon5:c.G232A:p.V78M,GNAS:NM_001077489:exon4:c.G364A:p.V122M,GNAS:NM_001309861:exon5:c.G232A:p.V78M,GNAS:NM_000516:exon5:c.G409A:p.V137M,GNAS:NM_001077488:exon5:c.G412A:p.V138M, |
| chr20 | 58905383 | G | A | . | exonic | GNAS | nonsynonymous SNV | GNAS:NM_080426:exon5:c.G391A:p.E131K,GNAS:NM_080425:exon6:c.G2362A:p.E788K,GNAS:NM_001309840:exon6:c.G256A:p.E86K,GNAS:NM_001077489:exon5:c.G388A:p.E130K,GNAS:NM_001309861:exon6:c.G256A:p.E86K,GNAS:NM_000516:exon6:c.G433A:p.E145K,GNAS:NM_001077488:exon6:c.G436A:p.E146K, |
| chr20 | 58905398 | G | A | . | exonic | GNAS | nonsynonymous SNV | GNAS:NM_080426:exon5:c.G406A:p.A136T,GNAS:NM_080425:exon6:c.G2377A:p.A793T,GNAS:NM_001309840:exon6:c.G271A:p.A91T,GNAS:NM_001077489:exon5:c.G403A:p.A135T,GNAS:NM_001309861:exon6:c.G271A:p.A91T,GNAS:NM_000516:exon6:c.G448A:p.A150T,GNAS:NM_001077488:exon6:c.G451A:p.A151T, |
| chr20 | 58905399 | C | T | COSV55688594 | exonic | GNAS | nonsynonymous SNV | GNAS:NM_001309840:exon6:c.C272T:p.A91V,GNAS:NM_000516:exon6:c.C449T:p.A150V,GNAS:NM_080426:exon5:c.C407T:p.A136V,GNAS:NM_080425:exon6:c.C2378T:p.A793V,GNAS:NM_001077489:exon5:c.C404T:p.A135V,GNAS:NM_001309861:exon6:c.C272T:p.A91V,GNAS:NM_001077488:exon6:c.C452T:p.A151V, |
| chr20 | 58905411 | G | A | . | exonic | GNAS | stopgain | GNAS:NM_080426:exon5:c.G419A:p.W140X,GNAS:NM_080425:exon6:c.G2390A:p.W797X,GNAS:NM_001309840:exon6:c.G284A:p.W95X,GNAS:NM_001077489:exon5:c.G416A:p.W139X,GNAS:NM_001309861:exon6:c.G284A:p.W95X,GNAS:NM_000516:exon6:c.G461A:p.W154X,GNAS:NM_001077488:exon6:c.G464A:p.W155X, |
| chr20 | 58905425 | G | A | . | exonic | GNAS | nonsynonymous SNV | GNAS:NM_080426:exon5:c.G433A:p.V145M,GNAS:NM_080425:exon6:c.G2404A:p.V802M,GNAS:NM_001309840:exon6:c.G298A:p.V100M,GNAS:NM_001077489:exon5:c.G430A:p.V144M,GNAS:NM_001309861:exon6:c.G298A:p.V100M,GNAS:NM_000516:exon6:c.G475A:p.V159M,GNAS:NM_001077488:exon6:c.G478A:p.V160M, |
| chr20 | 58905455 | T | C | . | exonic | GNAS | nonsynonymous SNV | GNAS:NM_080426:exon5:c.T463C:p.Y155H,GNAS:NM_080425:exon6:c.T2434C:p.Y812H,GNAS:NM_001309840:exon6:c.T328C:p.Y110H,GNAS:NM_001077489:exon5:c.T460C:p.Y154H,GNAS:NM_001309861:exon6:c.T328C:p.Y110H,GNAS:NM_000516:exon6:c.T505C:p.Y169H,GNAS:NM_001077488:exon6:c.T508C:p.Y170H, |
| chr20 | 58905471 | G | A | . | exonic | GNAS | nonsynonymous SNV | GNAS:NM_080426:exon5:c.G479A:p.C160Y,GNAS:NM_080425:exon6:c.G2450A:p.C817Y,GNAS:NM_001309840:exon6:c.G344A:p.C115Y,GNAS:NM_001077489:exon5:c.G476A:p.C159Y,GNAS:NM_001309861:exon6:c.G344A:p.C115Y,GNAS:NM_000516:exon6:c.G521A:p.C174Y,GNAS:NM_001077488:exon6:c.G524A:p.C175Y, |
| chr20 | 58905479 | T | C | . | exonic | GNAS | nonsynonymous SNV | GNAS:NM_080426:exon5:c.T487C:p.Y163H,GNAS:NM_080425:exon6:c.T2458C:p.Y820H,GNAS:NM_001309840:exon6:c.T352C:p.Y118H,GNAS:NM_001077489:exon5:c.T484C:p.Y162H,GNAS:NM_001309861:exon6:c.T352C:p.Y118H,GNAS:NM_000516:exon6:c.T529C:p.Y177H,GNAS:NM_001077488:exon6:c.T532C:p.Y178H, |
| chr20 | 58909211 | G | A | . | exonic | GNAS | nonsynonymous SNV | GNAS:NM_080426:exon6:c.G538A:p.D180N,GNAS:NM_080425:exon7:c.G2509A:p.D837N,GNAS:NM_001309840:exon7:c.G403A:p.D135N,GNAS:NM_001077489:exon6:c.G535A:p.D179N,GNAS:NM_001309861:exon7:c.G403A:p.D135N,GNAS:NM_000516:exon7:c.G580A:p.D194N,GNAS:NM_001077488:exon7:c.G583A:p.D195N, |
| chr20 | 58909380 | G | A | . | exonic | GNAS | nonsynonymous SNV | GNAS:NM_080426:exon7:c.G574A:p.G192R,GNAS:NM_080425:exon8:c.G2545A:p.G849R,GNAS:NM_001309840:exon8:c.G439A:p.G147R,GNAS:NM_001077489:exon7:c.G571A:p.G191R,GNAS:NM_001309861:exon8:c.G439A:p.G147R,GNAS:NM_000516:exon8:c.G616A:p.G206R,GNAS:NM_001077488:exon8:c.G619A:p.G207R, |
| chr20 | 58909398 | T | C | COSV55672955 | exonic | GNAS | nonsynonymous SNV | GNAS:NM_001309840:exon8:c.T457C:p.F153L,GNAS:NM_000516:exon8:c.T634C:p.F212L,GNAS:NM_080426:exon7:c.T592C:p.F198L,GNAS:NM_080425:exon8:c.T2563C:p.F855L,GNAS:NM_001077489:exon7:c.T589C:p.F197L,GNAS:NM_001309861:exon8:c.T457C:p.F153L,GNAS:NM_001077488:exon8:c.T637C:p.F213L, |
| chr20 | 58909410 | A | C | . | exonic | GNAS | nonsynonymous SNV | GNAS:NM_080426:exon7:c.A604C:p.K202Q,GNAS:NM_080425:exon8:c.A2575C:p.K859Q,GNAS:NM_001309840:exon8:c.A469C:p.K157Q,GNAS:NM_001077489:exon7:c.A601C:p.K201Q,GNAS:NM_001309861:exon8:c.A469C:p.K157Q,GNAS:NM_000516:exon8:c.A646C:p.K216Q,GNAS:NM_001077488:exon8:c.A649C:p.K217Q, |
| chr20 | 58909417 | A | G | . | exonic | GNAS | nonsynonymous SNV | GNAS:NM_080426:exon7:c.A611G:p.N204S,GNAS:NM_080425:exon8:c.A2582G:p.N861S,GNAS:NM_001309840:exon8:c.A476G:p.N159S,GNAS:NM_001077489:exon7:c.A608G:p.N203S,GNAS:NM_001309861:exon8:c.A476G:p.N159S,GNAS:NM_000516:exon8:c.A653G:p.N218S,GNAS:NM_001077488:exon8:c.A656G:p.N219S, |
| chr20 | 58909556 | G | A | COSV55674033 | exonic | GNAS | nonsynonymous SNV | GNAS:NM_080426:exon8:c.G653A:p.R218H,GNAS:NM_080425:exon9:c.G2624A:p.R875H,GNAS:NM_001309840:exon9:c.G518A:p.R173H,GNAS:NM_001077489:exon8:c.G650A:p.R217H,GNAS:NM_001309861:exon9:c.G518A:p.R173H,GNAS:NM_000516:exon9:c.G695A:p.R232H,GNAS:NM_001077488:exon9:c.G698A:p.R233H, |
| chr20 | 58909686 | G | A | . | exonic | GNAS | nonsynonymous SNV | GNAS:NM_080426:exon9:c.G679A:p.V227M,GNAS:NM_080425:exon10:c.G2650A:p.V884M,GNAS:NM_001309840:exon10:c.G544A:p.V182M,GNAS:NM_001077489:exon9:c.G676A:p.V226M,GNAS:NM_001309861:exon10:c.G544A:p.V182M,GNAS:NM_000516:exon10:c.G721A:p.V241M,GNAS:NM_001077488:exon10:c.G724A:p.V242M, |
| chr20 | 58909717 | G | A | . | exonic | GNAS | nonsynonymous SNV | GNAS:NM_001309840:exon10:c.G575A:p.S192N,GNAS:NM_000516:exon10:c.G752A:p.S251N,GNAS:NM_080426:exon9:c.G710A:p.S237N,GNAS:NM_080425:exon10:c.G2681A:p.S894N,GNAS:NM_001077489:exon9:c.G707A:p.S236N,GNAS:NM_001309861:exon10:c.G575A:p.S192N,GNAS:NM_001077488:exon10:c.G755A:p.S252N, |

| | | | | | | | | |
|---|---|---|---|---|---|---|---|---|
| chr20 | 58909721 | C | A | . | exonic | GNAS | nonsynonymous SNV | GNAS:NM_080426:exon9:c.C714A:p.S238R,GNAS:NM_080425:exon10:c.C2685A:p.S895R,GNAS:NM_001309840:exon10:c.C579A:p.S193R,GNAS:NM_001077489:exon9:c.C711A:p.S237R,GNAS:NM_001309861:exon10:c.C579A:p.S193R,GNAS:NM_000516:exon10:c.C756A:p.S252R,GNAS:NM_001077488:exon10:c.C759A:p.S253R, |
| chr20 | 58909753 | C | A | . | exonic | GNAS | nonsynonymous SNV | GNAS:NM_080426:exon9:c.C746A:p.T249N,GNAS:NM_080425:exon10:c.C2717A:p.T906N,GNAS:NM_001309840:exon10:c.C611A:p.T204N,GNAS:NM_001077489:exon9:c.C743A:p.T248N,GNAS:NM_001309861:exon10:c.C611A:p.T204N,GNAS:NM_000516:exon10:c.C788A:p.T263N,GNAS:NM_001077488:exon10:c.C791A:p.T264N, |
| chr20 | 58909759 | G | A | COSV55672773 | exonic | GNAS | nonsynonymous SNV | GNAS:NM_080426:exon9:c.G752A:p.R251H,GNAS:NM_080425:exon10:c.G2723A:p.R908H,GNAS:NM_001309840:exon10:c.G617A:p.R206H,GNAS:NM_001077489:exon9:c.G749A:p.R250H,GNAS:NM_001309861:exon10:c.G617A:p.R206H,GNAS:NM_000516:exon10:c.G794A:p.R265H,GNAS:NM_001077488:exon10:c.G797A:p.R266H, |
| chr20 | 58909795 | G | A | . | exonic | GNAS | stopgain | GNAS:NM_080426:exon9:c.G788A:p.W263X,GNAS:NM_080425:exon10:c.G2759A:p.W920X,GNAS:NM_001309840:exon10:c.G653A:p.W218X,GNAS:NM_001077489:exon9:c.G785A:p.W262X,GNAS:NM_001309861:exon10:c.G653A:p.W218X,GNAS:NM_000516:exon10:c.G830A:p.W277X,GNAS:NM_001077488:exon10:c.G833A:p.W278X, |
| chr20 | 58909952 | T | C | . | exonic | GNAS | nonsynonymous SNV | GNAS:NM_001309840:exon11:c.T664C:p.W222R,GNAS:NM_000516:exon11:c.T841C:p.W281R,GNAS:NM_080426:exon10:c.T799C:p.W267R,GNAS:NM_080425:exon11:c.T2770C:p.W924R,GNAS:NM_001077489:exon10:c.T796C:p.W266R,GNAS:NM_001309861:exon11:c.T664C:p.W222R,GNAS:NM_001077488:exon11:c.T844C:p.W282R, |
| chr20 | 58910021 | G | A | . | exonic | GNAS | nonsynonymous SNV | GNAS:NM_080426:exon10:c.G868A:p.G290R,GNAS:NM_080425:exon11:c.G2839A:p.G947R,GNAS:NM_001309840:exon11:c.G733A:p.G245R,GNAS:NM_001077489:exon10:c.G865A:p.G289R,GNAS:NM_001309861:exon11:c.G733A:p.G245R,GNAS:NM_000516:exon11:c.G910A:p.G304R,GNAS:NM_001077488:exon11:c.G913A:p.G305R, |
| chr20 | 58910022 | G | A | . | exonic | GNAS | nonsynonymous SNV | GNAS:NM_080426:exon10:c.G869A:p.G290E,GNAS:NM_080425:exon11:c.G2840A:p.G947E,GNAS:NM_001309840:exon11:c.G734A:p.G245E,GNAS:NM_001077489:exon10:c.G866A:p.G289E,GNAS:NM_001309861:exon11:c.G734A:p.G245E,GNAS:NM_000516:exon11:c.G911A:p.G304E,GNAS:NM_001077488:exon11:c.G914A:p.G305E, |
| chr20 | 58910039 | G | A | . | exonic | GNAS | nonsynonymous SNV | GNAS:NM_080426:exon10:c.G886A:p.D296N,GNAS:NM_080425:exon11:c.G2857A:p.D953N,GNAS:NM_001309840:exon11:c.G751A:p.D251N,GNAS:NM_001077489:exon10:c.G883A:p.D295N,GNAS:NM_001309861:exon11:c.G751A:p.D251N,GNAS:NM_000516:exon11:c.G928A:p.D310N,GNAS:NM_001077488:exon11:c.G931A:p.D311N, |
| chr20 | 58910045 | T | C | . | exonic | GNAS | nonsynonymous SNV | GNAS:NM_080426:exon10:c.T892C:p.F298L,GNAS:NM_080425:exon11:c.T2863C:p.F955L,GNAS:NM_001309840:exon11:c.T757C:p.F253L,GNAS:NM_001077489:exon10:c.T889C:p.F297L,GNAS:NM_001309861:exon11:c.T757C:p.F253L,GNAS:NM_000516:exon11:c.T934C:p.F312L,GNAS:NM_001077488:exon11:c.T937C:p.F313L, |
| chr20 | 58910061 | G | A | COSV55679114 | exonic | GNAS | nonsynonymous SNV | GNAS:NM_080426:exon10:c.G908A:p.R303H,GNAS:NM_080425:exon11:c.G2879A:p.R960H,GNAS:NM_001309840:exon11:c.G773A:p.R258H,GNAS:NM_001077489:exon10:c.G905A:p.R302H,GNAS:NM_001309861:exon11:c.G773A:p.R258H,GNAS:NM_000516:exon11:c.G950A:p.R317H,GNAS:NM_001077488:exon11:c.G953A:p.R318H, |
| chr20 | 58910348 | G | A | COSV55690777 | exonic | GNAS | nonsynonymous SNV | GNAS:NM_080426:exon11:c.G943A:p.G315R,GNAS:NM_080425:exon12:c.G2914A:p.G972R,GNAS:NM_001309840:exon12:c.G808A:p.G270R,GNAS:NM_001077489:exon11:c.G940A:p.G314R,GNAS:NM_001309861:exon12:c.G808A:p.G270R,GNAS:NM_000516:exon12:c.G985A:p.G329R,GNAS:NM_001077488:exon12:c.G988A:p.G330R, |
| chr20 | 58910364 | T | C | . | exonic | GNAS | nonsynonymous SNV | GNAS:NM_080426:exon11:c.T959C:p.V320A,GNAS:NM_080425:exon12:c.T2930C:p.V977A,GNAS:NM_001309840:exon12:c.T824C:p.V275A,GNAS:NM_001077489:exon11:c.T956C:p.V319A,GNAS:NM_001309861:exon12:c.T824C:p.V275A,GNAS:NM_000516:exon12:c.T1001C:p.V334A,GNAS:NM_001077488:exon12:c.T1004C:p.V335A, |
| chr20 | 58910373 | C | A | . | exonic | GNAS | nonsynonymous SNV | GNAS:NM_080426:exon11:c.C968A:p.A323D,GNAS:NM_080425:exon12:c.C2939A:p.A980D,GNAS:NM_001309840:exon12:c.C833A:p.A278D,GNAS:NM_001077489:exon11:c.C965A:p.A322D,GNAS:NM_001309861:exon12:c.C833A:p.A278D,GNAS:NM_000516:exon12:c.C1010A:p.A337D,GNAS:NM_001077488:exon12:c.C1013A:p.A338D, |
| chr20 | 58910385 | T | C | . | exonic | GNAS | nonsynonymous SNV | GNAS:NM_001309840:exon12:c.T845C:p.I282T,GNAS:NM_000516:exon12:c.T1022C:p.I341T,GNAS:NM_080426:exon11:c.T980C:p.I327T,GNAS:NM_080425:exon12:c.T2951C:p.I984T,GNAS:NM_001077489:exon11:c.T977C:p.I326T,GNAS:NM_001309861:exon12:c.T845C:p.I282T,GNAS:NM_001077488:exon12:c.T1025C:p.I342T, |
| chr20 | 58910684 | G | A | . | exonic | GNAS | nonsynonymous SNV | GNAS:NM_080426:exon12:c.G998A:p.R333K,GNAS:NM_080425:exon13:c.G2969A:p.R990K,GNAS:NM_001309840:exon13:c.G863A:p.R288K,GNAS:NM_001077489:exon12:c.G995A:p.R332K,GNAS:NM_001309861:exon13:c.G863A:p.R288K,GNAS:NM_000516:exon13:c.G1040A:p.R347K,GNAS:NM_001077488:exon13:c.G1043A:p.R348K, |
| chr20 | 58910699 | G | A | . | exonic | GNAS | nonsynonymous SNV | GNAS:NM_080426:exon12:c.G1013A:p.S338N,GNAS:NM_080425:exon13:c.G2984A:p.S995N,GNAS:NM_001309840:exon13:c.G878A:p.S293N,GNAS:NM_001077489:exon12:c.G1010A:p.S337N,GNAS:NM_001309861:exon13:c.G878A:p.S293N,GNAS:NM_000516:exon13:c.G1055A:p.S352N,GNAS:NM_001077488:exon13:c.G1058A:p.S353N, |
| chr20 | 58910704 | G | A | COSV55681349 | exonic | GNAS | nonsynonymous SNV | GNAS:NM_080426:exon12:c.G1018A:p.D340N,GNAS:NM_080425:exon13:c.G2989A:p.D997N,GNAS:NM_001309840:exon13:c.G883A:p.D295N,GNAS:NM_001077489:exon12:c.G1015A:p.D339N,GNAS:NM_001309861:exon13:c.G883A:p.D295N,GNAS:NM_000516:exon13:c.G1060A:p.D354N,GNAS:NM_001077488:exon13:c.G1063A:p.D355N, |
| chr20 | 58910738 | G | A | . | exonic | GNAS | nonsynonymous SNV | GNAS:NM_080426:exon12:c.G1052A:p.C351Y,GNAS:NM_080425:exon13:c.G3023A:p.C1008Y,GNAS:NM_001309840:exon13:c.G917A:p.C306Y,GNAS:NM_001077489:exon12:c.G1049A:p.C350Y,GNAS:NM_001309861:exon13:c.G917A:p.C306Y,GNAS:NM_000516:exon13:c.G1094A:p.C365Y,GNAS:NM_001077488:exon13:c.G1097A:p.C366Y, |
| chr20 | 58910746 | G | A | . | exonic | GNAS | nonsynonymous SNV | GNAS:NM_080426:exon12:c.G1060A:p.D354N,GNAS:NM_080425:exon13:c.G3031A:p.D1011N,GNAS:NM_001309840:exon13:c.G925A:p.D309N,GNAS:NM_001077489:exon12:c.G1057A:p.D353N,GNAS:NM_001309861:exon13:c.G925A:p.D309N,GNAS:NM_000516:exon13:c.G1102A:p.D368N,GNAS:NM_001077488:exon13:c.G1105A:p.D369N, |

| | | | | | | | | |
|---|---|---|---|---|---|---|---|---|
| chr20 | 58910762 | G | A | . | exonic | GNAS | nonsynonymous SNV | GNAS:NM_080426:exon12:c.G1076A:p.R359H,GNAS:NM_080425:exon13:c.G3047A:p.R1016H,GNAS:NM_001309840:exon13:c.G941A:p.R314H,GNAS:NM_001077489:exon12:c.G1073A:p.R358H,GNAS:NM_001309861:exon13:c.G941A:p.R314H,GNAS:NM_000516:exon13:c.G1118A:p.R373H,GNAS:NM_001077488:exon13:c.G1121A:p.R374H, |
| chr20 | 58910809 | C | A | . | exonic | GNAS | nonsynonymous SNV | GNAS:NM_001309840:exon13:c.C988A:p.R330S,GNAS:NM_000516:exon13:c.C1165A:p.R389S,GNAS:NM_080426:exon12:c.C1123A:p.R375S,GNAS:NM_080425:exon13:c.C3094A:p.R1032S,GNAS:NM_001077489:exon12:c.C1120A:p.R374S,GNAS:NM_001309861:exon13:c.C988A:p.R330S,GNAS:NM_001077488:exon13:c.C1168A:p.R390S, |
| chr20 | 5922421 | T | A | COSV66759908 | exonic | CHGB | nonsynonymous SNV | CHGB:NM_001819:exon4:c.T277A:p.S93T, |
| chr20 | 5922677 | G | A | . | exonic | CHGB | nonsynonymous SNV | CHGB:NM_001819:exon4:c.G533A:p.R178Q, |
| chr20 | 5922871 | A | G | . | exonic | CHGB | nonsynonymous SNV | CHGB:NM_001819:exon4:c.A727G:p.T243A, |
| chr20 | 5923202 | C | G | . | exonic | CHGB | nonsynonymous SNV | CHGB:NM_001819:exon4:c.C1058G:p.A353G, |
| chr20 | 5923394 | G | A | COSV66759649 | exonic | CHGB | nonsynonymous SNV | CHGB:NM_001819:exon4:c.G1250A:p.R417H, |
| chr20 | 6006304 | G | A | COSV66651511 | exonic | CRLS1 | nonsynonymous SNV | CRLS1:NM_019095:exon1:c.G58A:p.A20T, |
| chr20 | 6031288 | C | T | . | exonic | CRLS1 | nonsynonymous SNV | CRLS1:NM_001127458:exon4:c.C281T:p.P94L,CRLS1:NM_019095:exon4:c.C578T:p.P193L, |
| chr20 | 62307691 | C | A | . | exonic | ADRM1 | nonsynonymous SNV | ADRM1:NM_001281437:exon6:c.C602A:p.T201N,ADRM1:NM_001281438:exon6:c.C602A:p.T201N,ADRM1:NM_175573:exon7:c.C719A:p.T240N,ADRM1:NM_007002:exon7:c.C719A:p.T240N, |
| chr20 | 62817114 | G | A | COSV51702900 | exonic | COL9A3 | nonsynonymous SNV | COL9A3:NM_001853:exon1:c.G50A:p.G17E, |
| chr20 | 62819980 | C | T | COSV59651862 | exonic | COL9A3 | nonsynonymous SNV | COL9A3:NM_001853:exon5:c.C307T:p.R103W, |
| chr20 | 63708763 | C | G | COSV60832599 | exonic | ZGPAT | nonsynonymous SNV | ZGPAT:NM_001195653:exon2:c.C183G:p.S61R,ZGPAT:NM_181485:exon2:c.C183G:p.S61R,ZGPAT:NM_032527:exon2:c.C183G:p.S61R,ZGPAT:NM_001083113:exon2:c.C183G:p.S61R,ZGPAT:NM_001195654:exon2:c.C183G:p.S61R, |
| chr20 | 63742354 | G | T | COSV55507914 | exonic | SLC2A4RG | nonsynonymous SNV | SLC2A4RG:NM_020062:exon6:c.G699T:p.E233D, |
| chr21 | 25606638 | A | G | . | exonic | MRPL39 | nonsynonymous SNV | MRPL39:NM_080794:exon2:c.T91C:p.S31P,MRPL39:NM_017446:exon2:c.T91C:p.S31P, |
| chr21 | 33268483 | A | G | COSV51614715 | exonic | IL10RB | nonsynonymous SNV | IL10RB:NM_000628:exon2:c.A139G:p.K47E, |
| chr21 | 33343393 | G | C | COSV54251383 | exonic | IFNAR1 | nonsynonymous SNV | IFNAR1:NM_000629:exon4:c.G502C:p.V168L, |
| chr21 | 33543109 | A | T | . | exonic | SON | nonsynonymous SNV | SON:NM_001291411:exon1:c.A17T:p.E6V,SON:NM_001291412:exon1:c.A17T:p.E6V,SON:NM_032195:exon1:c.A17T:p.E6V,SON:NM_138927:exon1:c.A17T:p.E6V, |
| chr21 | 33543145 | G | A | . | exonic | SON | nonsynonymous SNV | SON:NM_001291412:exon1:c.G53A:p.R18Q,SON:NM_138927:exon1:c.G53A:p.R18Q,SON:NM_001291411:exon1:c.G53A:p.R18Q,SON:NM_032195:exon1:c.G53A:p.R18Q, |
| chr21 | 33543159 | G | A | . | exonic | SON | nonsynonymous SNV | SON:NM_001291412:exon1:c.G67A:p.E23K,SON:NM_138927:exon1:c.G67A:p.E23K,SON:NM_001291411:exon1:c.G67A:p.E23K,SON:NM_032195:exon1:c.G67A:p.E23K, |
| chr21 | 33546280 | G | A | . | exonic | SON | nonsynonymous SNV | SON:NM_001291411:exon2:c.G145A:p.A49T,SON:NM_001291412:exon2:c.G145A:p.A49T,SON:NM_032195:exon2:c.G145A:p.A49T,SON:NM_138927:exon2:c.G145A:p.A49T, |
| chr21 | 33549563 | A | G | . | exonic | SON | nonsynonymous SNV | SON:NM_001291411:exon3:c.A332G:p.H111R,SON:NM_032195:exon3:c.A332G:p.H111R,SON:NM_138927:exon3:c.A332G:p.H111R, |
| chr21 | 33549579 | C | A | . | exonic | SON | nonsynonymous SNV | SON:NM_001291411:exon3:c.C348A:p.N116K,SON:NM_032195:exon3:c.C348A:p.N116K,SON:NM_138927:exon3:c.C348A:p.N116K, |
| chr21 | 33549584 | A | G | . | exonic | SON | nonsynonymous SNV | SON:NM_001291411:exon3:c.A353G:p.K118R,SON:NM_032195:exon3:c.A353G:p.K118R,SON:NM_138927:exon3:c.A353G:p.K118R, |
| chr21 | 33549616 | T | A | COSV51654122 | exonic | SON | nonsynonymous SNV | SON:NM_001291411:exon3:c.T385A:p.Y129N,SON:NM_032195:exon3:c.T385A:p.Y129N,SON:NM_138927:exon3:c.T385A:p.Y129N, |
| chr21 | 33549618 | T | A | . | exonic | SON | stopgain | SON:NM_001291411:exon3:c.T387A:p.Y129X,SON:NM_032195:exon3:c.T387A:p.Y129X,SON:NM_138927:exon3:c.T387A:p.Y129X, |
| chr21 | 33552836 | C | T | . | exonic | SON | nonsynonymous SNV | SON:NM_001291411:exon3:c.C3605T:p.S1202L,SON:NM_032195:exon3:c.C3605T:p.S1202L,SON:NM_138927:exon3:c.C3605T:p.S1202L, |
| chr21 | 33553954 | C | T | COSV51652492 | exonic | SON | nonsynonymous SNV | SON:NM_001291411:exon3:c.C4723T:p.R1575C,SON:NM_032195:exon3:c.C4723T:p.R1575C,SON:NM_138927:exon3:c.C4723T:p.R1575C, |
| chr21 | 38806118 | A | G | . | exonic | ETS2 | nonsynonymous SNV | ETS2:NM_001256295:exon2:c.A418G:p.R140G, |
| chr21 | 38806119 | G | C | . | exonic | ETS2 | nonsynonymous SNV | ETS2:NM_001256295:exon2:c.G419C:p.R140T, |
| chr21 | 39298534 | G | C | COSV58122425 | exonic | BRWD1 | nonsynonymous SNV | BRWD1:NM_033656:exon5:c.C247G:p.Q83E,BRWD1:NM_001007246:exon5:c.C247G:p.Q83E,BRWD1:NM_018963:exon5:c.C247G:p.Q83E, |
| chr21 | 39390705 | G | A | . | exonic | WRB | nonsynonymous SNV | WRB:NM_004627:exon2:c.G110A:p.R37K,WRB:NM_001146218:exon2:c.G8A:p.R3K, |
| chr21 | 39406256 | C | T | COSV55748125 | exonic | LCA5L | nonsynonymous SNV | LCA5L:NM_152505:exon10:c.G1639A:p.G547S, |
| chr21 | 41480570 | C | T | COSV59820912 | exonic | TMPRSS2 | nonsynonymous SNV | TMPRSS2:NM_001135099:exon6:c.G589A:p.V197M,TMPRSS2:NM_005656:exon6:c.G478A:p.V160M, |
| chr21 | 41489560 | A | G | . | exonic | TMPRSS2 | nonsynonymous SNV | TMPRSS2:NM_001135099:exon4:c.T383C:p.L128P,TMPRSS2:NM_005656:exon4:c.T272C:p.L91P, |
| chr21 | 41507982 | C | A | COSV59827803 | exonic | TMPRSS2 | nonsynonymous SNV | TMPRSS2:NM_001135099:exon1:c.G23T:p.G8V, |
| chr21 | 44851537 | C | T | COSV58365573 | exonic | PTTG1IP | nonsynonymous SNV | PTTG1IP:NM_001286822:exon3:c.G259A:p.A87T, |
| chr21 | 46003595 | C | T | COSV62611834 | exonic | COL6A1 | nonsynonymous SNV | COL6A1:NM_001848:exon35:c.C2669T:p.S890L, |
| chr22 | 17159155 | G | A | COSV50084409 | exonic | CECR5 | nonsynonymous SNV | CECR5:NM_033070:exon1:c.C97T:p.R33C, |
| chr22 | 19134359 | G | A | COSV52817839 | exonic | DGCR14 | nonsynonymous SNV | DGCR14:NM_022719:exon10:c.C1268T:p.A423V, |
| chr22 | 19963748 | G | A | COSV52889256 | exonic | COMT | nonsynonymous SNV | COMT:NM_001135162:exon4:c.G472A:p.V158M,COMT:NM_000754:exon4:c.G472A:p.V158M,COMT:NM_007310:exon2:c.G322A:p.V108M,COMT:NM_001135161:exon4:c.G472A:p.V158M, |
| chr22 | 19968635 | T | C | . | exonic | COMT | nonsynonymous SNV | COMT:NM_001135161:exon6:c.T715C:p.F239L,COMT:NM_001135162:exon6:c.T715C:p.F239L,COMT:NM_000754:exon6:c.T715C:p.F239L,COMT:NM_007310:exon4:c.T565C:p.F189L, |
| chr22 | 21000681 | C | G | COSV53145905 | exonic | THAP7 | nonsynonymous SNV | THAP7:NM_030573:exon3:c.G343C:p.A115P,THAP7:NM_001008695:exon4:c.G343C:p.A115P, |
| chr22 | 22893773 | T | A | . | exonic | IGLL5 | nonsynonymous SNV | IGLL5:NM_001178126:exon2:c.T280A:p.S94T, |
| chr22 | 22893794 | A | G | . | exonic | IGLL5 | nonsynonymous SNV | IGLL5:NM_001178126:exon2:c.A301G:p.T101A, |
| chr22 | 22895680 | A | G | COSV66535329 | exonic | IGLL5 | nonsynonymous SNV | IGLL5:NM_001178126:exon3:c.A631G:p.T211A,IGLL5:NM_001256296:exon2:c.A406G:p.T136A, |
| chr22 | 23285182 | A | G | COSV59932309 | exonic | BCR | nonsynonymous SNV | BCR:NM_021574:exon10:c.A2387G:p.N796S,BCR:NM_004327:exon10:c.A2387G:p.N796S, |
| chr22 | 23779191 | C | T | COSV53155825 | exonic | MMP11 | nonsynonymous SNV | MMP11:NM_005940:exon2:c.C113T:p.A38V, |
| chr22 | 23791772 | G | A | . | exonic | SMARCB1 | nonsynonymous SNV | SMARCB1:NM_003073:exon2:c.G110A:p.R37H,SMARCB1:NM_001007468:exon2:c.G110A:p.R37H, |
| chr22 | 23803341 | C | A | . | exonic | SMARCB1 | nonsynonymous SNV | SMARCB1:NM_003073:exon5:c.C547A:p.P183T,SMARCB1:NM_001007468:exon5:c.C520A:p.P174T, |
| chr22 | 28695719 | A | G | . | exonic | CHEK2 | nonsynonymous SNV | CHEK2:NM_001005735:exon12:c.T1379C:p.L460P,CHEK2:NM_001257387:exon12:c.T587C:p.L196P,CHEK2:NM_007194:exon11:c.T1250C:p.L417P,CHEK2:NM_145862:exon10:c.T1163C:p.L388P, |
| chr22 | 29282401 | G | A | . | exonic | EWSR1 | nonsynonymous SNV | EWSR1:NM_005243:exon6:c.G425A:p.G142E,EWSR1:NM_013986:exon7:c.G443A:p.G148E,EWSR1:NM_001163285:exon6:c.G425A:p.G142E,EWSR1:NM_001163287:exon6:c.G425A:p.G142E, |
| chr22 | 29282472 | G | A | . | exonic | EWSR1 | nonsynonymous SNV | EWSR1:NM_005243:exon6:c.G496A:p.G166R,EWSR1:NM_001163287:exon6:c.G496A:p.G166R,EWSR1:NM_013986:exon7:c.G514A:p.G172R,EWSR1:NM_001163285:exon6:c.G496A:p.G166R, |
| chr22 | 29299271 | T | C | . | exonic | EWSR1 | nonsynonymous SNV | EWSR1:NM_005243:exon15:c.T1618C:p.C540R,EWSR1:NM_013986:exon16:c.T1633C:p.C545R,EWSR1:NM_001163285:exon15:c.T1615C:p.C539R,EWSR1:NM_001163286:exon14:c.T1450C:p.C484R, |
| chr22 | 36191656 | T | C | COSV60654226 | exonic | APOL4 | nonsynonymous SNV | APOL4:NM_030643:exon6:c.A466G:p.M156V,APOL4:NM_145660:exon5:c.A475G:p.M159V, |
| chr22 | 36202003 | C | G | COSV60495655 | exonic | APOL4 | nonsynonymous SNV | APOL4:NM_145660:exon2:c.G34C:p.V12L, |
| chr22 | 36202012 | T | C | COSV60495932 | exonic | APOL4 | nonsynonymous SNV | APOL4:NM_145660:exon2:c.A25G:p.I9V, |
| chr22 | 36202035 | A | T | COSV60495853 | exonic | APOL4 | nonsynonymous SNV | APOL4:NM_145660:exon2:c.T2A:p.M1K, |

| | | | | | | | | |
|---|---|---|---|---|---|---|---|---|
| chr22 | 36257404 | C | T | . | exonic | APOL1 | nonsynonymous SNV | APOL1:NM_145343:exon5:c.C232T:p.P78S,APOL1:NM_003661:exon4:c.C184T:p.P62S,APOL1:NM_001136540:exon4:c.C184T:p.P62S,APOL1:NM_001136541:exon3:c.C130T:p.P44S, |
| chr22 | 36265284 | G | A | COSV59868177 | exonic | APOL1 | nonsynonymous SNV | APOL1:NM_001136541:exon5:c.G394A:p.E132K,APOL1:NM_145343:exon7:c.G496A:p.E166K,APOL1:NM_003661:exon6:c.G448A:p.E150K,APOL1:NM_001136540:exon6:c.G448A:p.E150K, |
| chr22 | 36265520 | G | A | COSV59868421 | exonic | APOL1 | nonsynonymous SNV | APOL1:NM_001136541:exon5:c.G630A:p.M210I,APOL1:NM_145343:exon7:c.G732A:p.M244I,APOL1:NM_003661:exon6:c.G684A:p.M228I,APOL1:NM_001136540:exon6:c.G684A:p.M228I, |
| chr22 | 36265600 | G | A | COSV59868427 | exonic | APOL1 | nonsynonymous SNV | APOL1:NM_001136541:exon5:c.G710A:p.R237K,APOL1:NM_145343:exon7:c.G812A:p.R271K,APOL1:NM_003661:exon6:c.G764A:p.R255K,APOL1:NM_001136540:exon6:c.G764A:p.R255K, |
| chr22 | 36284418 | C | A | . | exonic | MYH9 | nonsynonymous SNV | MYH9:NM_002473:exon39:c.G5577T:p.E1859D, |
| chr22 | 36285664 | G | C | . | exonic | MYH9 | nonsynonymous SNV | MYH9:NM_002473:exon37:c.C5268G:p.N1756K, |
| chr22 | 36288308 | T | C | COSV53381533 | exonic | MYH9 | nonsynonymous SNV | MYH9:NM_002473:exon34:c.A4876G:p.I1626V, |
| chr22 | 36293365 | G | C | . | exonic | MYH9 | nonsynonymous SNV | MYH9:NM_002473:exon30:c.C4059G:p.H1353Q, |
| chr22 | 36294155 | C | A | . | exonic | MYH9 | nonsynonymous SNV | MYH9:NM_002473:exon28:c.G3774T:p.Q1258H, |
| chr22 | 36298982 | C | T | . | exonic | MYH9 | nonsynonymous SNV | MYH9:NM_002473:exon24:c.G3037A:p.E1013K, |
| chr22 | 36300967 | C | T | . | exonic | MYH9 | nonsynonymous SNV | MYH9:NM_002473:exon22:c.G2722A:p.A908T, |
| chr22 | 36322464 | C | T | COSV53385150 | exonic | MYH9 | nonsynonymous SNV | MYH9:NM_002473:exon6:c.G670A:p.A224T, |
| chr22 | 36349142 | A | T | . | exonic | MYH9 | nonsynonymous SNV | MYH9:NM_002473:exon2:c.T95A:p.V32E, |
| chr22 | 36467860 | C | T | . | exonic | TXN2 | nonsynonymous SNV | TXN2:NM_012473:exon4:c.G445A:p.V149M, |
| chr22 | 36467871 | A | G | . | exonic | TXN2 | nonsynonymous SNV | TXN2:NM_012473:exon4:c.T434C:p.V145A, |
| chr22 | 36467911 | C | T | . | exonic | TXN2 | nonsynonymous SNV | TXN2:NM_012473:exon4:c.G394A:p.A132T, |
| chr22 | 36476789 | C | T | . | exonic | TXN2 | nonsynonymous SNV | TXN2:NM_012473:exon3:c.G331A:p.V111M, |
| chr22 | 36476848 | C | T | COSV53398335 | exonic | TXN2 | nonsynonymous SNV | TXN2:NM_012473:exon3:c.G272A:p.G91E, |
| chr22 | 36504224 | T | C | COSV53400124 | exonic | FOXRED2 | nonsynonymous SNV | FOXRED2:NM_001102371:exon4:c.A923G:p.N308S,FOXRED2:NM_024955:exon4:c.A923G:p.N308S, |
| chr22 | 36511605 | C | T | . | exonic | EIF3D | nonsynonymous SNV | EIF3D:NM_003753:exon14:c.G1531A:p.D511N, |
| chr22 | 36511670 | C | T | . | exonic | EIF3D | nonsynonymous SNV | EIF3D:NM_003753:exon14:c.G1466A:p.G489D, |
| chr22 | 36511677 | C | T | . | exonic | EIF3D | nonsynonymous SNV | EIF3D:NM_003753:exon14:c.G1459A:p.A487T, |
| chr22 | 36511683 | C | T | . | exonic | EIF3D | nonsynonymous SNV | EIF3D:NM_003753:exon14:c.G1453A:p.E485K, |
| chr22 | 36516525 | C | T | . | exonic | EIF3D | nonsynonymous SNV | EIF3D:NM_003753:exon12:c.G1159A:p.E387K, |
| chr22 | 36516528 | C | T | . | exonic | EIF3D | nonsynonymous SNV | EIF3D:NM_003753:exon12:c.G1156A:p.G386R, |
| chr22 | 36516546 | C | T | . | exonic | EIF3D | nonsynonymous SNV | EIF3D:NM_003753:exon12:c.G1138A:p.V380I, |
| chr22 | 36516762 | G | T | . | exonic | EIF3D | nonsynonymous SNV | EIF3D:NM_003753:exon11:c.C1019A:p.P340Q, |
| chr22 | 36518817 | C | T | COSV53403638 | exonic | EIF3D | nonsynonymous SNV | EIF3D:NM_003753:exon9:c.G805A:p.V269I, |
| chr22 | 36518843 | C | T | . | exonic | EIF3D | nonsynonymous SNV | EIF3D:NM_003753:exon9:c.G779A:p.R260H, |
| chr22 | 36523226 | A | G | . | exonic | EIF3D | nonsynonymous SNV | EIF3D:NM_003753:exon6:c.T448C:p.W150R, |
| chr22 | 36523895 | C | T | . | exonic | EIF3D | nonsynonymous SNV | EIF3D:NM_003753:exon5:c.G392A:p.R131K, |
| chr22 | 36523899 | C | T | COSV53403626 | exonic | EIF3D | nonsynonymous SNV | EIF3D:NM_003753:exon5:c.G388A:p.E130K, |
| chr22 | 36523952 | C | T | . | exonic | EIF3D | nonsynonymous SNV | EIF3D:NM_003753:exon5:c.G335A:p.R112Q, |
| chr22 | 36526073 | C | T | COSV53403959 | exonic | EIF3D | nonsynonymous SNV | EIF3D:NM_003753:exon2:c.G49A:p.G17S, |
| chr22 | 37808082 | C | T | COSV50649782 | exonic | GCAT | nonsynonymous SNV | GCAT:NM_001171690:exon1:c.C115T:p.R39C,GCAT:NM_014291:exon1:c.C115T:p.R39C, |
| chr22 | 38089533 | A | G | COSV60201964 | exonic | BAIAP2L2 | nonsynonymous SNV | BAIAP2L2:NM_025045:exon8:c.T754C:p.C252R, |
| chr22 | 39521510 | A | C | COSV57478471 | exonic | ATF4 | nonsynonymous SNV | ATF4:NM_182810:exon2:c.A65C:p.Q22P,ATF4:NM_001675:exon1:c.A65C:p.Q22P, |
| chr22 | 40021816 | A | G | COSV60999568 | exonic | FAM83F | nonsynonymous SNV | FAM83F:NM_138435:exon4:c.A1306G:p.R436G, |
| chr22 | 40156115 | G | A | COSV57292828 | exonic | TNRC6B | nonsynonymous SNV | TNRC6B:NM_001024843:exon4:c.G46A:p.V16M, |
| chr22 | 40827186 | C | T | . | exonic | ST13 | nonsynonymous SNV | ST13:NM_003932:exon11:c.G891A:p.M297I,ST13:NM_001278589:exon11:c.G861A:p.M287I, |
| chr22 | 40967827 | G | A | COSV53430854 | exonic | RBX1 | nonsynonymous SNV | RBX1:NM_014248:exon4:c.G257A:p.R86H, |
| chr22 | 40967872 | G | A | . | exonic | RBX1 | stopgain | RBX1:NM_014248:exon4:c.G302A:p.W101X, |
| chr22 | 41173676 | C | A | . | exonic | EP300 | nonsynonymous SNV | EP300:NM_001429:exon29:c.C4671A:p.S1557R, |
| chr22 | 41532697 | A | G | . | exonic | POLR3H | nonsynonymous SNV | POLR3H:NM_001282884:exon4:c.T257C:p.I86T,POLR3H:NM_001282885:exon4:c.T257C:p.I86T,POLR3H:NM_138338:exon4:c.T257C:p.I86T,POLR3H:NM_001018050:exon3:c.T257C:p.I86T, |
| chr22 | 41675009 | A | G | . | exonic | SNU13 | nonsynonymous SNV | SNU13:NM_001003796:exon3:c.T311C:p.V104A,SNU13:NM_005008:exon4:c.T311C:p.V104A, |
| chr22 | 41675142 | C | T | . | exonic | SNU13 | nonsynonymous SNV | SNU13:NM_001003796:exon3:c.G178A:p.A60T,SNU13:NM_005008:exon4:c.G178A:p.A60T, |
| chr22 | 41675186 | G | T | . | exonic | SNU13 | nonsynonymous SNV | SNU13:NM_001003796:exon3:c.C134A:p.T45N,SNU13:NM_005008:exon4:c.C134A:p.T45N, |
| chr22 | 42090719 | G | A | COSV68911946 | exonic | NDUFA6 | nonsynonymous SNV | NDUFA6:NM_002490:exon1:c.C104T:p.A35V, |
| chr22 | 42516130 | G | T | COSV59899209 | exonic | RRP7A | nonsynonymous SNV | RRP7A:NM_015703:exon3:c.C223A:p.L75M, |
| chr22 | 42912031 | C | T | . | exonic | PACSIN2 | nonsynonymous SNV | PACSIN2:NM_007229:exon2:c.G50A:p.S17N,PACSIN2:NM_001184971:exon2:c.G50A:p.S17N,PACSIN2:NM_001184970:exon2:c.G50A:p.S17N, |
| chr22 | 43162920 | A | G | COSV53355454 | exonic | TSPO | nonsynonymous SNV | TSPO:NM_000714:exon4:c.A439G:p.T147A,TSPO:NM_001256530:exon4:c.A439G:p.T147A,TSPO:NM_001256531:exon4:c.A439G:p.T147A, |
| chr22 | 43162966 | G | A | COSV53354339 | exonic | TSPO | nonsynonymous SNV | TSPO:NM_000714:exon4:c.G485A:p.R162H,TSPO:NM_001256530:exon4:c.G485A:p.R162H,TSPO:NM_001256531:exon4:c.G485A:p.R162H, |
| chr22 | 43983958 | A | G | . | exonic | SAMM50 | nonsynonymous SNV | SAMM50:NM_015380:exon12:c.A1033G:p.I345V, |
| chr22 | 44132959 | G | A | . | exonic | PARVB | nonsynonymous SNV | PARVB:NM_013327:exon6:c.G583A:p.A195T,PARVB:NM_001243386:exon6:c.G427A:p.A143T,PARVB:NM_001003828:exon7:c.G682A:p.A228T,PARVB:NM_001243385:exon6:c.G472A:p.A158T, |
| chr22 | 45527947 | A | G | . | exonic | FBLN1 | nonsynonymous SNV | FBLN1:NM_001996:exon4:c.A422G:p.Q141R,FBLN1:NM_006485:exon4:c.A422G:p.Q141R,FBLN1:NM_006486:exon4:c.A422G:p.Q141R,FBLN1:NM_006487:exon4:c.A422G:p.Q141R, |
| chr22 | 49922662 | G | A | . | exonic | CRELD2 | nonsynonymous SNV | CRELD2:NM_001284317:exon6:c.G643A:p.G215S,CRELD2:NM_001284318:exon6:c.G643A:p.G215S,CRELD2:NM_024324:exon6:c.G643A:p.G215S,CRELD2:NM_001135101:exon7:c.G790A:p.G264S, |
| chr22 | 50525807 | G | A | COSV52688025 | exonic | TYMP | nonsynonymous SNV | TYMP:NM_001257989:exon10:c.C1427T:p.S476L,TYMP:NM_001953:exon10:c.C1412T:p.S471L,TYMP:NM_001113755:exon10:c.C1412T:p.S471L,TYMP:NM_001257988:exon10:c.C1412T:p.S471L,TYMP:NM_001113756:exon9:c.C1412T:p.S471L, |
| chr22 | 50525826 | C | T | COSV52687812 | exonic | TYMP | nonsynonymous SNV | TYMP:NM_001257989:exon10:c.G1408A:p.A470T,TYMP:NM_001953:exon10:c.G1393A:p.A465T,TYMP:NM_001257988:exon10:c.G1393A:p.A465T,TYMP:NM_001113755:exon10:c.G1393A:p.A465T,TYMP:NM_001113756:exon9:c.G1393A:p.A465T, |
| chr3 | 10064848 | C | T | COSV55033303 | exonic | FANCD2 | nonsynonymous SNV | FANCD2:NM_033084:exon23:c.C2141T:p.P714L,FANCD2:NM_001018115:exon23:c.C2141T:p.P714L, |
| chr3 | 100713788 | G | A | . | exonic | TFG | nonsynonymous SNV | TFG:NM_006070:exon2:c.G103A:p.E35K,TFG:NM_001195479:exon2:c.G103A:p.E35K,TFG:NM_001195478:exon2:c.G103A:p.E35K,TFG:NM_001007565:exon2:c.G103A:p.E35K, |
| chr3 | 100720041 | T | C | . | exonic | TFG | nonsynonymous SNV | TFG:NM_006070:exon3:c.T251C:p.L84P,TFG:NM_001195479:exon3:c.T251C:p.L84P,TFG:NM_001195478:exon3:c.T251C:p.L84P,TFG:NM_001007565:exon3:c.T251C:p.L84P, |
| chr3 | 100728747 | G | A | . | exonic | TFG | nonsynonymous SNV | TFG:NM_006070:exon4:c.G304A:p.V102M,TFG:NM_001195479:exon4:c.G304A:p.V102M,TFG:NM_001195478:exon4:c.G304A:p.V102M,TFG:NM_001007565:exon4:c.G304A:p.V102M, |
| chr3 | 10072973 | G | A | COSV55034331 | exonic | FANCD2 | nonsynonymous SNV | FANCD2:NM_033084:exon27:c.G2597A:p.G866E,FANCD2:NM_001018115:exon27:c.G2597A:p.G866E, |
| chr3 | 100732552 | G | A | . | exonic | TFG | nonsynonymous SNV | TFG:NM_006070:exon5:c.G460A:p.G154R,TFG:NM_001195479:exon5:c.G460A:p.G154R,TFG:NM_001195478:exon5:c.G460A:p.G154R,TFG:NM_001007565:exon5:c.G460A:p.G154R, |
| chr3 | 100732553 | G | A | COSV53748082 | exonic | TFG | nonsynonymous SNV | TFG:NM_006070:exon5:c.G461A:p.G154E,TFG:NM_001195479:exon5:c.G461A:p.G154E,TFG:NM_001195478:exon5:c.G461A:p.G154E,TFG:NM_001007565:exon5:c.G461A:p.G154E, |
| chr3 | 100732579 | G | A | . | exonic | TFG | nonsynonymous SNV | TFG:NM_006070:exon5:c.G487A:p.A163T,TFG:NM_001195479:exon5:c.G487A:p.A163T,TFG:NM_001195478:exon5:c.G487A:p.A163T,TFG:NM_001007565:exon5:c.G487A:p.A163T, |
| chr3 | 100732618 | G | A | . | exonic | TFG | nonsynonymous SNV | TFG:NM_006070:exon5:c.G526A:p.E176K,TFG:NM_001195479:exon5:c.G526A:p.E176K,TFG:NM_001195478:exon5:c.G526A:p.E176K,TFG:NM_001007565:exon5:c.G526A:p.E176K, |
| chr3 | 100736585 | G | A | . | exonic | TFG | nonsynonymous SNV | TFG:NM_006070:exon6:c.G590A:p.S197N,TFG:NM_001195479:exon6:c.G590A:p.S197N,TFG:NM_001195478:exon6:c.G590A:p.S197N,TFG:NM_001007565:exon6:c.G590A:p.S197N, |

| | | | | | | | | |
|---|---|---|---|---|---|---|---|---|
| chr3 | 100744893 | C | T | . | exonic | TFG | nonsynonymous SNV | TFG:NM_006070:exon7:c.C782T:p.P261L,TFG:NM_001195479:exon7:c.C770T:p.P257L,TFG:NM_001195478:exon7:c.C782T:p.P261L,TFG:NM_001007565:exon7:c.C782T:p.P261L, |
| chr3 | 100748170 | G | A | . | exonic | TFG | nonsynonymous SNV | TFG:NM_006070:exon8:c.G842A:p.G281E,TFG:NM_001195479:exon8:c.G830A:p.G277E,TFG:NM_001195478:exon8:c.G842A:p.G281E,TFG:NM_001007565:exon8:c.G842A:p.G281E, |
| chr3 | 100748190 | T | C | . | exonic | TFG | nonsynonymous SNV | TFG:NM_001195479:exon8:c.T850C:p.F284L,TFG:NM_001195478:exon8:c.T862C:p.F288L,TFG:NM_001007565:exon8:c.T862C:p.F288L,TFG:NM_006070:exon8:c.T862C:p.F288L, |
| chr3 | 100748407 | G | A | . | exonic | TFG | nonsynonymous SNV | TFG:NM_006070:exon8:c.G1079A:p.R360K,TFG:NM_001195479:exon8:c.G1067A:p.R356K,TFG:NM_001195478:exon8:c.G1079A:p.R360K,TFG:NM_001007565:exon8:c.G1079A:p.R360K, |
| chr3 | 100748434 | G | A | . | exonic | TFG | nonsynonymous SNV | TFG:NM_006070:exon8:c.G1106A:p.S369N,TFG:NM_001195479:exon8:c.G1094A:p.S365N,TFG:NM_001195478:exon8:c.G1106A:p.S369N,TFG:NM_001007565:exon8:c.G1106A:p.S369N, |
| chr3 | 10301280 | C | T | . | exonic | SEC13 | nonsynonymous SNV | SEC13:NM_183352:exon9:c.G950A:p.G317D,SEC13:NM_030673:exon10:c.G959A:p.G320D,SEC13:NM_001136232:exon9:c.G908A:p.G303D,SEC13:NM_001136026:exon10:c.G1088A:p.G363D, |
| chr3 | 10301340 | C | T | . | exonic | SEC13 | stopgain | SEC13:NM_183352:exon9:c.G890A:p.W297X,SEC13:NM_030673:exon10:c.G899A:p.W300X,SEC13:NM_001136232:exon9:c.G848A:p.W283X,SEC13:NM_001136026:exon10:c.G1028A:p.W343X, |
| chr3 | 10301364 | C | T | . | exonic | SEC13 | stopgain | SEC13:NM_183352:exon9:c.G866A:p.W289X,SEC13:NM_030673:exon10:c.G875A:p.W292X,SEC13:NM_001136232:exon9:c.G824A:p.W275X,SEC13:NM_001136026:exon10:c.G1004A:p.W335X, |
| chr3 | 10304123 | C | T | . | exonic | SEC13 | stopgain | SEC13:NM_183352:exon8:c.G758A:p.W253X,SEC13:NM_030673:exon9:c.G767A:p.W256X,SEC13:NM_001136232:exon8:c.G716A:p.W239X,SEC13:NM_001136026:exon9:c.G896A:p.W299X, |
| chr3 | 10304165 | C | T | COSV61602678 | exonic | SEC13 | nonsynonymous SNV | SEC13:NM_183352:exon8:c.G716A:p.R239H,SEC13:NM_030673:exon9:c.G725A:p.R242H,SEC13:NM_001136232:exon8:c.G674A:p.R225H,SEC13:NM_001136026:exon9:c.G854A:p.R285H, |
| chr3 | 10312033 | C | T | . | exonic | SEC13 | nonsynonymous SNV | SEC13:NM_183352:exon5:c.G382A:p.A128T,SEC13:NM_030673:exon6:c.G391A:p.A131T,SEC13:NM_001136232:exon5:c.G340A:p.A114T,SEC13:NM_001278946:exon5:c.G382A:p.A128T,SEC13:NM_001136026:exon6:c.G520A:p.A174T, |
| chr3 | 10312082 | C | T | . | exonic | SEC13 | stopgain | SEC13:NM_183352:exon5:c.G333A:p.W111X,SEC13:NM_030673:exon6:c.G342A:p.W114X,SEC13:NM_001136232:exon5:c.G291A:p.W97X,SEC13:NM_001278946:exon5:c.G333A:p.W111X,SEC13:NM_001136026:exon6:c.G471A:p.W157X, |
| chr3 | 10312608 | C | T | . | exonic | SEC13 | nonsynonymous SNV | SEC13:NM_030673:exon5:c.G296A:p.S99N,SEC13:NM_001278946:exon4:c.G287A:p.S96N,SEC13:NM_001136026:exon5:c.G425A:p.S142N,SEC13:NM_183352:exon4:c.G287A:p.S96N,SEC13:NM_001136232:exon4:c.G245A:p.S82N, |
| chr3 | 105659148 | C | T | . | exonic | CBLB | nonsynonymous SNV | CBLB:NM_170662:exon19:c.G2771A:p.G924E, |
| chr3 | 107716655 | G | A | . | exonic | BBX | nonsynonymous SNV | BBX:NM_020235:exon5:c.G211A:p.E71K,BBX:NM_001142568:exon5:c.G211A:p.E71K,BBX:NM_001276286:exon5:c.G211A:p.E71K, |
| chr3 | 107716707 | T | C | . | exonic | BBX | nonsynonymous SNV | BBX:NM_020235:exon5:c.T263C:p.L88P,BBX:NM_001142568:exon5:c.T263C:p.L88P,BBX:NM_001276286:exon5:c.T263C:p.L88P, |
| chr3 | 107716728 | G | A | COSV57881501 | exonic | BBX | nonsynonymous SNV | BBX:NM_020235:exon5:c.G284A:p.R95H,BBX:NM_001142568:exon5:c.G284A:p.R95H,BBX:NM_001276286:exon5:c.G284A:p.R95H, |
| chr3 | 107772704 | T | A | . | exonic | BBX | nonsynonymous SNV | BBX:NM_020235:exon11:c.T983A:p.I328N,BBX:NM_001142568:exon11:c.T983A:p.I328N, |
| chr3 | 107772736 | G | A | . | exonic | BBX | nonsynonymous SNV | BBX:NM_020235:exon11:c.G1015A:p.E339K,BBX:NM_001142568:exon11:c.G1015A:p.E339K, |
| chr3 | 107773061 | C | A | . | exonic | BBX | nonsynonymous SNV | BBX:NM_020235:exon11:c.C1340A:p.P447Q,BBX:NM_001142568:exon11:c.C1340A:p.P447Q, |
| chr3 | 107773304 | A | G | . | exonic | BBX | nonsynonymous SNV | BBX:NM_020235:exon11:c.A1583G:p.K528R,BBX:NM_001142568:exon11:c.A1583G:p.K528R, |
| chr3 | 107773313 | T | A | . | exonic | BBX | nonsynonymous SNV | BBX:NM_020235:exon11:c.T1592A:p.V531E,BBX:NM_001142568:exon11:c.T1592A:p.V531E, |
| chr3 | 108353451 | T | C | COSV63315693 | exonic | HHLA2 | nonsynonymous SNV | HHLA2:NM_001282556:exon4:c.T89C:p.I30T,HHLA2:NM_001282557:exon4:c.T89C:p.I30T,HHLA2:NM_001282558:exon5:c.T89C:p.I30T,HHLA2:NM_007072:exon4:c.T89C:p.I30T, |
| chr3 | 108362430 | C | A | . | exonic | HHLA2 | nonsynonymous SNV | HHLA2:NM_001282556:exon7:c.C1092A:p.S364R,HHLA2:NM_001282559:exon8:c.C900A:p.S300R,HHLA2:NM_001282557:exon7:c.C1092A:p.S364R,HHLA2:NM_001282558:exon8:c.C1092A:p.S364R,HHLA2:NM_007072:exon7:c.C1092A:p.S364R, |
| chr3 | 108553699 | G | T | . | exonic | KIAA1524 | nonsynonymous SNV | KIAA1524:NM_020890:exon19:c.C2356A:p.Q786K, |
| chr3 | 108757127 | G | A | COSV55465500 | exonic | RETNLB | nonsynonymous SNV | RETNLB:NM_032579:exon1:c.C59T:p.P20L, |
| chr3 | 112102900 | A | G | COSV53495730 | exonic | C3ORF52 | nonsynonymous SNV | C3orf52:NM_024616:exon3:c.A331G:p.I111V,C3orf52:NM_001171747:exon3:c.A331G:p.I111V, |
| chr3 | 112109576 | G | A | . | exonic | C3ORF52 | nonsynonymous SNV | C3orf52:NM_024616:exon4:c.G430A:p.G144S, |
| chr3 | 11358589 | G | A | . | exonic | ATG7 | nonsynonymous SNV | ATG7:NM_001136031:exon13:c.G1456A:p.V486I,ATG7:NM_001144912:exon12:c.G1339A:p.V447I,ATG7:NM_006395:exon13:c.G1456A:p.V486I, |
| chr3 | 120670469 | T | A | COSV52199452 | exonic | HGD | nonsynonymous SNV | HGD:NM_000187:exon4:c.A240T:p.Q80H, |
| chr3 | 122106350 | G | A | COSV52608918 | exonic | CD86 | nonsynonymous SNV | CD86:NM_176892:exon4:c.G535A:p.V179I,CD86:NM_175862:exon4:c.G553A:p.V185I,CD86:NM_006889:exon4:c.G535A:p.V179I,CD86:NM_001206924:exon3:c.G217A:p.V73I,CD86:NM_001206925:exon3:c.G307A:p.V103I, |
| chr3 | 122407300 | G | A | . | exonic | FAM162A | nonsynonymous SNV | FAM162A:NM_014367:exon4:c.G283A:p.A95T, |
| chr3 | 122409790 | C | T | COSV51657361 | exonic | FAM162A | nonsynonymous SNV | FAM162A:NM_014367:exon5:c.C424T:p.R142C, |
| chr3 | 12351626 | C | G | COSV55140710 | exonic | PPARG | nonsynonymous SNV | PPARG:NM_015869:exon1:c.C34G:p.P12A, |
| chr3 | 12379708 | G | A | . | exonic | PPARG | nonsynonymous SNV | PPARG:NM_005037:exon2:c.G3A:p.M1I,PPARG:NM_138712:exon3:c.G3A:p.M1I,PPARG:NM_015869:exon2:c.G87A:p.M29I,PPARG:NM_138711:exon3:c.G3A:p.M1I, |
| chr3 | 12379892 | G | A | . | exonic | PPARG | nonsynonymous SNV | PPARG:NM_005037:exon2:c.G187A:p.A63T,PPARG:NM_138712:exon3:c.G187A:p.A63T,PPARG:NM_015869:exon2:c.G271A:p.A91T,PPARG:NM_138711:exon3:c.G187A:p.A63T, |

| | | | | | | | | |
|---|---|---|---|---|---|---|---|---|
| chr3 | 12381324 | G | A | . | exonic | PPARG | nonsynonymous SNV | PPARG:NM_005037:exon3:c.G229A:p.A77T,PPARG:NM_138712:exon4:c.G229A:p.A77T,PPARG:NM_015869:exon3:c.G313A:p.A105T,PPARG:NM_138711:exon4:c.G229A:p.A77T, |
| chr3 | 12381402 | T | C | . | exonic | PPARG | nonsynonymous SNV | PPARG:NM_005037:exon3:c.T307C:p.S103P,PPARG:NM_138712:exon4:c.T307C:p.S103P,PPARG:NM_015869:exon3:c.T391C:p.S131P,PPARG:NM_138711:exon4:c.T307C:p.S103P, |
| chr3 | 12381423 | G | A | COSV55145297 | exonic | PPARG | nonsynonymous SNV | PPARG:NM_005037:exon3:c.G328A:p.E110K,PPARG:NM_138712:exon4:c.G328A:p.E110K,PPARG:NM_015869:exon3:c.G412A:p.E138K,PPARG:NM_138711:exon4:c.G328A:p.E110K, |
| chr3 | 12381441 | G | A | . | exonic | PPARG | nonsynonymous SNV | PPARG:NM_005037:exon3:c.G346A:p.D116N,PPARG:NM_138712:exon4:c.G346A:p.D116N,PPARG:NM_015869:exon3:c.G430A:p.D144N,PPARG:NM_138711:exon4:c.G346A:p.D116N, |
| chr3 | 12392624 | G | A | COSV55140645 | exonic | PPARG | nonsynonymous SNV | PPARG:NM_005037:exon4:c.G407A:p.R136Q,PPARG:NM_138712:exon5:c.G407A:p.R136Q,PPARG:NM_015869:exon4:c.G491A:p.R164Q,PPARG:NM_138711:exon5:c.G407A:p.R136Q, |
| chr3 | 12392691 | T | A | . | exonic | PPARG | nonsynonymous SNV | PPARG:NM_005037:exon4:c.T474A:p.S158R,PPARG:NM_138712:exon5:c.T474A:p.S158R,PPARG:NM_015869:exon4:c.T558A:p.S186R,PPARG:NM_138711:exon5:c.T474A:p.S158R, |
| chr3 | 12392697 | T | A | . | exonic | PPARG | nonsynonymous SNV | PPARG:NM_005037:exon4:c.T480A:p.N160K,PPARG:NM_138712:exon5:c.T480A:p.N160K,PPARG:NM_015869:exon4:c.T564A:p.N188K,PPARG:NM_138711:exon5:c.T480A:p.N160K, |
| chr3 | 12392731 | G | A | . | exonic | PPARG | nonsynonymous SNV | PPARG:NM_005037:exon4:c.G514A:p.A172T,PPARG:NM_138712:exon5:c.G514A:p.A172T,PPARG:NM_015869:exon4:c.G598A:p.A200T,PPARG:NM_138711:exon5:c.G514A:p.A172T, |
| chr3 | 12405911 | G | A | . | exonic | PPARG | nonsynonymous SNV | PPARG:NM_015869:exon5:c.G649A:p.E217K,PPARG:NM_138711:exon6:c.G565A:p.E189K,PPARG:NM_005037:exon5:c.G565A:p.E189K,PPARG:NM_138712:exon6:c.G565A:p.E189K, |
| chr3 | 12405971 | G | A | . | exonic | PPARG | nonsynonymous SNV | PPARG:NM_005037:exon5:c.G625A:p.A209T,PPARG:NM_138712:exon6:c.G625A:p.A209T,PPARG:NM_015869:exon5:c.G709A:p.A237T,PPARG:NM_138711:exon6:c.G625A:p.A209T, |
| chr3 | 12405984 | C | A | . | exonic | PPARG | nonsynonymous SNV | PPARG:NM_005037:exon5:c.C638A:p.A213D,PPARG:NM_138712:exon6:c.C638A:p.A213D,PPARG:NM_015869:exon5:c.C722A:p.A241D,PPARG:NM_138711:exon6:c.C638A:p.A213D, |
| chr3 | 12406037 | G | A | . | exonic | PPARG | nonsynonymous SNV | PPARG:NM_005037:exon5:c.G691A:p.A231T,PPARG:NM_138712:exon6:c.G691A:p.A231T,PPARG:NM_015869:exon5:c.G775A:p.A259T,PPARG:NM_138711:exon6:c.G691A:p.A231T, |
| chr3 | 12416831 | G | A | . | exonic | PPARG | nonsynonymous SNV | PPARG:NM_005037:exon6:c.G863A:p.R288H,PPARG:NM_138712:exon7:c.G863A:p.R288H,PPARG:NM_015869:exon6:c.G947A:p.R316H,PPARG:NM_138711:exon7:c.G863A:p.R288H, |
| chr3 | 12433907 | G | T | . | exonic | PPARG | nonsynonymous SNV | PPARG:NM_005037:exon7:c.G1196T:p.G399V,PPARG:NM_138712:exon8:c.G1196T:p.G399V,PPARG:NM_015869:exon7:c.G1280T:p.G427V,PPARG:NM_138711:exon8:c.G1196T:p.G399V, |
| chr3 | 12434072 | A | G | . | exonic | PPARG | nonsynonymous SNV | PPARG:NM_005037:exon7:c.A1361G:p.Q454R,PPARG:NM_138712:exon8:c.A1361G:p.Q454R,PPARG:NM_015869:exon7:c.A1445G:p.Q482R,PPARG:NM_138711:exon8:c.A1361G:p.Q454R, |
| chr3 | 124859365 | C | G | . | exonic | ITGB5 | nonsynonymous SNV | ITGB5:NM_002213:exon3:c.G238C:p.E80Q, |
| chr3 | 124908177 | T | G | COSV60698847 | exonic | MUC13 | nonsynonymous SNV | MUC13:NM_033049:exon11:c.A1509C:p.R503S, |
| chr3 | 12585743 | C | T | . | exonic | RAF1 | nonsynonymous SNV | RAF1:NM_002880:exon14:c.G1474A:p.V492I, |
| chr3 | 12600412 | C | T | . | exonic | RAF1 | nonsynonymous SNV | RAF1:NM_002880:exon8:c.G838A:p.A280T, |
| chr3 | 126107160 | T | C | COSV56418368 | exonic | ALDH1L1 | nonsynonymous SNV | ALDH1L1:NM_001270365:exon19:c.A2131G:p.I711V,ALDH1L1:NM_012190:exon21:c.A2434G:p.I812V,ALDH1L1:NM_001270364:exon21:c.A2464G:p.I822V, |
| chr3 | 126107216 | T | C | COSV56412084 | exonic | ALDH1L1 | nonsynonymous SNV | ALDH1L1:NM_001270365:exon19:c.A2075G:p.D692G,ALDH1L1:NM_012190:exon21:c.A2378G:p.D793G,ALDH1L1:NM_001270364:exon21:c.A2408G:p.D803G, |
| chr3 | 126153541 | A | G | . | exonic | ALDH1L1 | nonsynonymous SNV | ALDH1L1:NM_001270365:exon5:c.T458C:p.L153P,ALDH1L1:NM_012190:exon7:c.T761C:p.L254P,ALDH1L1:NM_001270364:exon7:c.T791C:p.L264P, |
| chr3 | 12769256 | G | C | COSV53145197 | exonic | TMEM40 | nonsynonymous SNV | TMEM40:NM_001284406:exon1:c.C35G:p.P12R, |
| chr3 | 12769284 | C | T | COSV53145207 | exonic | TMEM40 | nonsynonymous SNV | TMEM40:NM_001284406:exon1:c.G7A:p.E3K, |
| chr3 | 12839508 | C | T | . | exonic | RPL32 | nonsynonymous SNV | RPL32:NM_001007073:exon3:c.G119A:p.G40D,RPL32:NM_000994:exon3:c.G119A:p.G40D,RPL32:NM_001007074:exon2:c.G119A:p.G40D, |
| chr3 | 129170475 | C | T | . | exonic | CNBP | nonsynonymous SNV | CNBP:NM_001127195:exon5:c.G494A:p.C165Y,CNBP:NM_001127193:exon5:c.G515A:p.C172Y,CNBP:NM_001127196:exon5:c.G491A:p.C164Y,CNBP:NM_001127192:exon5:c.G518A:p.C173Y,CNBP:NM_003418:exon5:c.G512A:p.C171Y,CNBP:NM_001127194:exon5:c.G497A:p.C166Y, |
| chr3 | 129170479 | C | T | . | exonic | CNBP | nonsynonymous SNV | CNBP:NM_001127195:exon5:c.G490A:p.E164K,CNBP:NM_001127193:exon5:c.G511A:p.E171K,CNBP:NM_001127196:exon5:c.G487A:p.E163K,CNBP:NM_001127192:exon5:c.G514A:p.E172K,CNBP:NM_003418:exon5:c.G508A:p.E170K,CNBP:NM_001127194:exon5:c.G493A:p.E165K, |
| chr3 | 129170485 | C | T | . | exonic | CNBP | nonsynonymous SNV | CNBP:NM_001127195:exon5:c.G484A:p.A162T,CNBP:NM_001127193:exon5:c.G505A:p.A169T,CNBP:NM_001127196:exon5:c.G481A:p.A161T,CNBP:NM_001127192:exon5:c.G508A:p.A170T,CNBP:NM_003418:exon5:c.G502A:p.A168T,CNBP:NM_001127194:exon5:c.G487A:p.A163T, |
| chr3 | 129170535 | C | T | . | exonic | CNBP | nonsynonymous SNV | CNBP:NM_001127195:exon5:c.G434A:p.S145N,CNBP:NM_001127193:exon5:c.G455A:p.S152N,CNBP:NM_001127196:exon5:c.G431A:p.S144N,CNBP:NM_001127192:exon5:c.G458A:p.S153N,CNBP:NM_003418:exon5:c.G452A:p.S151N,CNBP:NM_001127194:exon5:c.G437A:p.S146N, |
| chr3 | 129171245 | C | T | . | exonic | CNBP | nonsynonymous SNV | CNBP:NM_001127195:exon4:c.G232A:p.A78T,CNBP:NM_001127193:exon4:c.G253A:p.A85T,CNBP:NM_001127196:exon4:c.G229A:p.A77T,CNBP:NM_001127192:exon4:c.G256A:p.A86T,CNBP:NM_003418:exon4:c.G250A:p.A84T,CNBP:NM_001127194:exon4:c.G235A:p.A79T, |
| chr3 | 129171262 | C | T | . | exonic | CNBP | nonsynonymous SNV | CNBP:NM_001127195:exon4:c.G215A:p.G72D,CNBP:NM_001127193:exon4:c.G236A:p.G79D,CNBP:NM_001127196:exon4:c.G212A:p.G71D,CNBP:NM_001127192:exon4:c.G239A:p.G80D,CNBP:NM_003418:exon4:c.G233A:p.G78D,CNBP:NM_001127194:exon4:c.G218A:p.G73D, |
| chr3 | 129171265 | C | T | . | exonic | CNBP | nonsynonymous SNV | CNBP:NM_001127195:exon4:c.G212A:p.C71Y,CNBP:NM_001127193:exon4:c.G233A:p.C78Y,CNBP:NM_001127196:exon4:c.G209A:p.C70Y,CNBP:NM_001127192:exon4:c.G236A:p.C79Y,CNBP:NM_003418:exon4:c.G230A:p.C77Y,CNBP:NM_001127194:exon4:c.G215A:p.C72Y, |
| chr3 | 129171488 | C | T | . | exonic | CNBP | nonsynonymous SNV | CNBP:NM_001127195:exon3:c.G154A:p.E52K,CNBP:NM_001127193:exon3:c.G175A:p.E59K,CNBP:NM_001127196:exon3:c.G154A:p.E52K,CNBP:NM_001127192:exon3:c.G175A:p.E59K,CNBP:NM_003418:exon3:c.G175A:p.E59K,CNBP:NM_001127194:exon3:c.G154A:p.E52K, |
| chr3 | 129171671 | C | T | . | exonic | CNBP | nonsynonymous SNV | CNBP:NM_001127195:exon2:c.G87A:p.M29I,CNBP:NM_001127193:exon2:c.G87A:p.M29I,CNBP:NM_001127196:exon2:c.G87A:p.M29I,CNBP:NM_001127192:exon2:c.G87A:p.M29I,CNBP:NM_003418:exon2:c.G87A:p.M29I,CNBP:NM_001127194:exon2:c.G87A:p.M29I, |
| chr3 | 129171694 | C | T | . | exonic | CNBP | nonsynonymous SNV | CNBP:NM_001127195:exon2:c.G64A:p.G22S,CNBP:NM_001127193:exon2:c.G64A:p.G22S,CNBP:NM_001127196:exon2:c.G64A:p.G22S,CNBP:NM_001127194:exon2:c.G64A:p.G22S,CNBP:NM_001127192:exon2:c.G64A:p.G22S,CNBP:NM_003418:exon2:c.G64A:p.G22S, |
| chr3 | 133623144 | T | C | COSV53439574 | exonic | TOPBP1 | nonsynonymous SNV | TOPBP1:NM_007027:exon19:c.A3125G:p.N1042S, |
| chr3 | 133649518 | T | G | COSV53439449 | exonic | TOPBP1 | nonsynonymous SNV | TOPBP1:NM_007027:exon10:c.A1369C:p.K457Q, |
| chr3 | 13571436 | A | G | COSV55481423 | exonic | FBLN2 | nonsynonymous SNV | FBLN2:NM_001165035:exon2:c.A1081G:p.S361G,FBLN2:NM_001998:exon2:c.A1081G:p.S361G,FBLN2:NM_001004019:exon2:c.A1081G:p.S361G, |
| chr3 | 13629036 | A | G | COSV55488788 | exonic | FBLN2 | nonsynonymous SNV | FBLN2:NM_001165035:exon12:c.A2701G:p.T901A,FBLN2:NM_001998:exon11:c.A2560G:p.T854A,FBLN2:NM_001004019:exon12:c.A2701G:p.T901A, |
| chr3 | 136328837 | C | T | . | exonic | PCCB | nonsynonymous SNV | PCCB:NM_001178014:exon15:c.C1538T:p.P513L,PCCB:NM_000532:exon14:c.C1478T:p.P493L, |
| chr3 | 136542144 | C | T | . | exonic | STAG1 | nonsynonymous SNV | STAG1:NM_005862:exon6:c.G446A:p.R149K, |
| chr3 | 136604433 | C | A | . | exonic | STAG1 | nonsynonymous SNV | STAG1:NM_005862:exon4:c.G173T:p.S58I, |
| chr3 | 136623187 | C | T | . | exonic | STAG1 | nonsynonymous SNV | STAG1:NM_005862:exon3:c.G91A:p.V31I, |
| chr3 | 136928170 | G | A | . | exonic | NCK1 | nonsynonymous SNV | NCK1:NM_006153:exon2:c.G169A:p.E57K,NCK1:NM_001291999:exon2:c.G169A:p.E57K, |
| chr3 | 136945895 | C | T | COSV56637558 | exonic | NCK1 | nonsynonymous SNV | NCK1:NM_001190796:exon2:c.C347T:p.A116V,NCK1:NM_006153:exon3:c.C539T:p.A180V,NCK1:NM_001291999:exon3:c.C539T:p.A180V, |
| chr3 | 138295865 | G | C | . | exonic | ARMC8 | nonsynonymous SNV | ARMC8:NM_015396:exon23:c.G1953C:p.K651N,ARMC8:NM_001282342:exon20:c.G1776C:p.K592N,ARMC8:NM_001267041:exon21:c.G1902C:p.K634N,ARMC8:NM_001267042:exon21:c.G1794C:p.K598N, |
| chr3 | 138570379 | C | T | . | exonic | CEP70 | nonsynonymous SNV | CEP70:NM_001288966:exon6:c.G404A:p.S135N,CEP70:NM_001288964:exon6:c.G350A:p.S117N,CEP70:NM_024491:exon6:c.G404A:p.S135N,CEP70:NM_001288965:exon5:c.G344A:p.S115N, |

| | | | | | | | | |
|---|---|---|---|---|---|---|---|---|
| chr3 | 139361084 | C | T | . | exonic | COPB2 | nonsynonymous SNV | COPB2:NM_004766:exon17:c.G2207A:p.G736D, |
| chr3 | 139361198 | C | T | COSV60349232 | exonic | COPB2 | nonsynonymous SNV | COPB2:NM_004766:exon17:c.G2093A:p.G698D, |
| chr3 | 139361225 | C | T | . | exonic | COPB2 | nonsynonymous SNV | COPB2:NM_004766:exon17:c.G2066A:p.C689Y, |
| chr3 | 139367091 | A | G | . | exonic | COPB2 | nonsynonymous SNV | COPB2:NM_004766:exon14:c.T1600C:p.F534L, |
| chr3 | 139369332 | C | T | . | exonic | COPB2 | nonsynonymous SNV | COPB2:NM_004766:exon12:c.G1330A:p.V444I, |
| chr3 | 139369344 | C | T | . | exonic | COPB2 | nonsynonymous SNV | COPB2:NM_004766:exon12:c.G1318A:p.G440R, |
| chr3 | 139369498 | A | T | . | exonic | COPB2 | nonsynonymous SNV | COPB2:NM_004766:exon11:c.T1252A:p.F418I, |
| chr3 | 139373715 | C | T | . | exonic | COPB2 | nonsynonymous SNV | COPB2:NM_004766:exon8:c.G845A:p.G282E, |
| chr3 | 139375512 | T | A | . | exonic | COPB2 | nonsynonymous SNV | COPB2:NM_004766:exon6:c.A607T:p.I203F, |
| chr3 | 141442720 | G | A | . | exonic | ZBTB38 | nonsynonymous SNV | ZBTB38:NM_001080412:exon8:c.G332A:p.G111E, |
| chr3 | 141442835 | T | A | . | exonic | ZBTB38 | nonsynonymous SNV | ZBTB38:NM_001080412:exon8:c.T447A:p.N149K, |
| chr3 | 141444232 | C | T | . | exonic | ZBTB38 | nonsynonymous SNV | ZBTB38:NM_001080412:exon8:c.C1844T:p.T615M, |
| chr3 | 14145949 | G | T | COSV53207797 | exonic | XPC | nonsynonymous SNV | XPC:NM_004628:exon16:c.C2815A:p.Q939K, |
| chr3 | 14158387 | G | A | COSV53203534 | exonic | XPC | nonsynonymous SNV | XPC:NM_004628:exon9:c.C1496T:p.A499V, |
| chr3 | 14158408 | C | T | COSV53203815 | exonic | XPC | nonsynonymous SNV | XPC:NM_004628:exon9:c.G1475A:p.R492H, |
| chr3 | 14158440 | C | A | COSV53206797 | exonic | XPC | nonsynonymous SNV | XPC:NM_004628:exon9:c.G1443T:p.K481N, |
| chr3 | 14170484 | G | T | . | exonic | XPC | nonsynonymous SNV | XPC:NM_004628:exon3:c.C366A:p.S122R, |
| chr3 | 141974094 | C | T | . | exonic | TFDP2 | stopgain | TFDP2:NM_001178140:exon5:c.G533A:p.W178X,TFDP2:NM_001178139:exon8:c.G617A:p.W206X,TFDP2:NM_006286:exon6:c.G437A:p.W146X,TFDP2:NM_001178138:exon9:c.G437A:p.W146X,TFDP2:NM_001178142:exon3:c.G209A:p.W70X,TFDP2:NM_001178141:exon5:c.G326A:p.W109X, |
| chr3 | 141974124 | A | T | . | exonic | TFDP2 | nonsynonymous SNV | TFDP2:NM_001178140:exon5:c.T503A:p.I168K,TFDP2:NM_001178139:exon8:c.T587A:p.I196K,TFDP2:NM_006286:exon6:c.T407A:p.I136K,TFDP2:NM_001178138:exon9:c.T407A:p.I136K,TFDP2:NM_001178142:exon3:c.T179A:p.I60K,TFDP2:NM_001178141:exon5:c.T296A:p.I99K, |
| chr3 | 142536136 | A | T | . | exonic | ATR | nonsynonymous SNV | ATR:NM_001184:exon20:c.T3791A:p.I1264K, |
| chr3 | 142562770 | A | G | COSV63383325 | exonic | ATR | nonsynonymous SNV | ATR:NM_001184:exon4:c.T632C:p.M211T, |
| chr3 | 151328520 | G | A | COSV56380424 | exonic | P2RY13 | nonsynonymous SNV | P2RY13:NM_176894:exon2:c.C536T:p.T179M, |
| chr3 | 158596932 | T | A | . | exonic | MLF1 | stopgain | MLF1:NM_001195434:exon5:c.T236A:p.L79X,MLF1:NM_001130156:exon3:c.T191A:p.L64X,MLF1:NM_001195433:exon4:c.T191A:p.L64X,MLF1:NM_001195432:exon5:c.T359A:p.L120X,MLF1:NM_022443:exon3:c.T266A:p.L89X,MLF1:NM_001130157:exon4:c.T191A:p.L64X, |
| chr3 | 158602914 | C | A | COSV63035321 | exonic | MLF1 | nonsynonymous SNV | MLF1:NM_001195434:exon8:c.C646A:p.P216T,MLF1:NM_001130156:exon6:c.C601A:p.P201T,MLF1:NM_001195433:exon6:c.C472A:p.P158T,MLF1:NM_001195432:exon8:c.C769A:p.P257T,MLF1:NM_022443:exon6:c.C676A:p.P226T,MLF1:NM_001130157:exon7:c.C601A:p.P201T, |
| chr3 | 158649111 | G | A | COSV51833379 | exonic | GFM1 | nonsynonymous SNV | GFM1:NM_001308164:exon5:c.G643A:p.V215I,GFM1:NM_024996:exon5:c.G643A:p.V215I,GFM1:NM_001308166:exon5:c.G643A:p.V215I, |
| chr3 | 158670991 | T | C | COSV51834864 | exonic | LXN | nonsynonymous SNV | LXN:NM_020169:exon2:c.A158G:p.H53R, |
| chr3 | 158710887 | C | T | . | exonic | RARRES1 | nonsynonymous SNV | RARRES1:NM_002888:exon3:c.G386A:p.R129Q,RARRES1:NM_206963:exon3:c.G386A:p.R129Q, |
| chr3 | 160279273 | T | A | COSV58451332 | exonic | IFT80 | nonsynonymous SNV | IFT80:NM_020800:exon16:c.A1756T:p.T586S,IFT80:NM_001190242:exon15:c.A1345T:p.T449S,IFT80:NM_001190241:exon17:c.A1345T:p.T449S, |
| chr3 | 168010802 | C | T | . | exonic | GOLIM4 | nonsynonymous SNV | GOLIM4:NM_001308155:exon14:c.G1798A:p.E600K,GOLIM4:NM_014498:exon15:c.G1882A:p.E628K, |
| chr3 | 168024587 | C | T | . | exonic | GOLIM4 | nonsynonymous SNV | GOLIM4:NM_001308155:exon13:c.G1715A:p.G572E,GOLIM4:NM_014498:exon14:c.G1799A:p.G600E, |
| chr3 | 168029234 | C | T | COSV58329953 | exonic | GOLIM4 | nonsynonymous SNV | GOLIM4:NM_001308155:exon10:c.G1418A:p.G473E,GOLIM4:NM_014498:exon11:c.G1502A:p.G501E, |
| chr3 | 169090018 | C | T | . | exonic | MECOM | nonsynonymous SNV | MECOM:NM_001105078:exon14:c.G2819A:p.C940Y,MECOM:NM_001163999:exon13:c.G2795A:p.C932Y,MECOM:NM_001105077:exon15:c.G3014A:p.C1005Y,MECOM:NM_004991:exon15:c.G3383A:p.C1128Y,MECOM:NM_001164000:exon13:c.G2792A:p.C931Y,MECOM:NM_001205194:exon14:c.G2819A:p.C940Y,MECOM:NM_005241:exon14:c.G2819A:p.C940Y, |
| chr3 | 169090127 | C | T | COSV52972789 | exonic | MECOM | nonsynonymous SNV | MECOM:NM_001105078:exon14:c.G2710A:p.E904K,MECOM:NM_001163999:exon13:c.G2686A:p.E896K,MECOM:NM_001105077:exon15:c.G2905A:p.E969K,MECOM:NM_004991:exon15:c.G3274A:p.E1092K,MECOM:NM_001164000:exon13:c.G2683A:p.E895K,MECOM:NM_001205194:exon14:c.G2710A:p.E904K,MECOM:NM_005241:exon14:c.G2710A:p.E904K, |
| chr3 | 169115970 | A | T | . | exonic | MECOM | nonsynonymous SNV | MECOM:NM_001105078:exon7:c.T1338A:p.N446K,MECOM:NM_001163999:exon7:c.T1341A:p.N447K,MECOM:NM_001105077:exon8:c.T1533A:p.N511K,MECOM:NM_004991:exon8:c.T1902A:p.N634K,MECOM:NM_001164000:exon7:c.T1338A:p.N446K,MECOM:NM_001205194:exon7:c.T1338A:p.N446K,MECOM:NM_005241:exon7:c.T1338A:p.N446K, |
| chr3 | 169116038 | A | T | . | exonic | MECOM | nonsynonymous SNV | MECOM:NM_001105078:exon7:c.T1270A:p.F424I,MECOM:NM_001163999:exon7:c.T1273A:p.F425I,MECOM:NM_001105077:exon8:c.T1465A:p.F489I,MECOM:NM_004991:exon8:c.T1834A:p.F612I,MECOM:NM_001164000:exon7:c.T1270A:p.F424I,MECOM:NM_001205194:exon7:c.T1270A:p.F424I,MECOM:NM_005241:exon7:c.T1270A:p.F424I, |
| chr3 | 169116044 | C | T | . | exonic | MECOM | nonsynonymous SNV | MECOM:NM_001105078:exon7:c.G1264A:p.E422K,MECOM:NM_001163999:exon7:c.G1267A:p.E423K,MECOM:NM_001105077:exon8:c.G1459A:p.E487K,MECOM:NM_004991:exon8:c.G1828A:p.E610K,MECOM:NM_001164000:exon7:c.G1264A:p.E422K,MECOM:NM_001205194:exon7:c.G1264A:p.E422K,MECOM:NM_005241:exon7:c.G1264A:p.E422K, |
| chr3 | 169116050 | C | T | COSV52964849 | exonic | MECOM | nonsynonymous SNV | MECOM:NM_001105078:exon7:c.G1258A:p.D420N,MECOM:NM_004991:exon8:c.G1822A:p.D608N,MECOM:NM_001164000:exon7:c.G1258A:p.D420N,MECOM:NM_005241:exon7:c.G1258A:p.D420N,MECOM:NM_001163999:exon7:c.G1261A:p.D421N,MECOM:NM_001105077:exon8:c.G1453A:p.D485N,MECOM:NM_001205194:exon7:c.G1258A:p.D420N, |
| chr3 | 169122594 | C | T | . | exonic | MECOM | nonsynonymous SNV | MECOM:NM_001105078:exon5:c.G400A:p.E134K,MECOM:NM_001163999:exon5:c.G400A:p.E134K,MECOM:NM_001105077:exon6:c.G592A:p.E198K,MECOM:NM_004991:exon6:c.G964A:p.E322K,MECOM:NM_001164000:exon5:c.G400A:p.E134K,MECOM:NM_001205194:exon5:c.G400A:p.E134K,MECOM:NM_005241:exon5:c.G400A:p.E134K, |
| chr3 | 169381204 | G | A | COSV72109887 | exonic | MECOM | nonsynonymous SNV | MECOM:NM_004991:exon2:c.C358T:p.P120S, |
| chr3 | 169796797 | A | T | COSV57186258 | exonic | LRRC34 | nonsynonymous SNV | LRRC34:NM_001172780:exon8:c.T856A:p.L286I,LRRC34:NM_001172779:exon8:c.T856A:p.L286I,LRRC34:NM_153353:exon7:c.T760A:p.L254I, |
| chr3 | 169800667 | T | C | COSV57186839 | exonic | LRRC34 | nonsynonymous SNV | LRRC34:NM_001172780:exon7:c.A745G:p.S249G,LRRC34:NM_001172779:exon7:c.A745G:p.S249G, |
| chr3 | 172506539 | C | T | . | exonic | TNFSF10 | nonsynonymous SNV | TNFSF10:NM_003810:exon5:c.G799A:p.D267N, |
| chr3 | 172506729 | G | T | . | exonic | TNFSF10 | nonsynonymous SNV | TNFSF10:NM_003810:exon5:c.C609A:p.D203E, |
| chr3 | 172506761 | G | T | . | exonic | TNFSF10 | nonsynonymous SNV | TNFSF10:NM_003810:exon5:c.C577A:p.Q193K, |
| chr3 | 172509318 | T | C | . | exonic | TNFSF10 | nonsynonymous SNV | TNFSF10:NM_003810:exon4:c.A317G:p.K106R, |
| chr3 | 172710992 | T | C | COSV56436612 | exonic | NCEH1 | nonsynonymous SNV | NCEH1:NM_001146276:exon1:c.A89G:p.Q30R,NCEH1:NM_020792:exon1:c.A89G:p.Q30R, |
| chr3 | 177046143 | A | G | . | exonic | TBL1XR1 | nonsynonymous SNV | TBL1XR1:NM_024665:exon10:c.T911C:p.F304S, |
| chr3 | 177053882 | C | T | . | exonic | TBL1XR1 | nonsynonymous SNV | TBL1XR1:NM_024665:exon4:c.G95A:p.S32N, |
| chr3 | 183135882 | T | C | COSV55616290 | exonic | LAMP3 | nonsynonymous SNV | LAMP3:NM_014398:exon5:c.A952G:p.I318V, |
| chr3 | 183840614 | C | G | COSV57700167 | exonic | PARL | nonsynonymous SNV | PARL:NM_018622:exon7:c.G784C:p.V262L,PARL:NM_001037639:exon6:c.G634C:p.V212L, |
| chr3 | 184178897 | G | C | . | exonic | AP2M1 | nonsynonymous SNV | AP2M1:NM_001025205:exon3:c.G115C:p.A39P, |
| chr3 | 184178961 | G | A | . | exonic | AP2M1 | nonsynonymous SNV | AP2M1:NM_001025205:exon3:c.G179A:p.R60Q, |
| chr3 | 184179005 | G | A | . | exonic | AP2M1 | nonsynonymous SNV | AP2M1:NM_001025205:exon3:c.G223A:p.A75T, |
| chr3 | 184180238 | G | A | . | exonic | AP2M1 | nonsynonymous SNV | AP2M1:NM_001025205:exon4:c.G410A:p.G137D, |
| chr3 | 184181091 | T | A | . | exonic | AP2M1 | nonsynonymous SNV | AP2M1:NM_001025205:exon6:c.T566A:p.V189E, |
| chr3 | 184181167 | G | A | . | exonic | AP2M1 | nonsynonymous SNV | AP2M1:NM_001025205:exon6:c.G642A:p.M214I, |
| chr3 | 184181977 | G | A | . | exonic | AP2M1 | nonsynonymous SNV | AP2M1:NM_001025205:exon8:c.G887A:p.R296H, |
| chr3 | 184182033 | G | A | . | exonic | AP2M1 | nonsynonymous SNV | AP2M1:NM_001025205:exon8:c.G943A:p.A315T, |

| | | | | | | | | |
|---|---|---|---|---|---|---|---|---|
| chr3 | 184182235 | G | C | . | exonic | AP2M1 | nonsynonymous SNV | AP2M1:NM_001025205:exon9:c.G1042C:p.A348P, |
| chr3 | 184182770 | G | A | . | exonic | AP2M1 | nonsynonymous SNV | AP2M1:NM_001025205:exon10:c.G1069A:p.A357T, |
| chr3 | 184182797 | G | A | . | exonic | AP2M1 | nonsynonymous SNV | AP2M1:NM_001025205:exon10:c.G1096A:p.A366T, |
| chr3 | 184182806 | G | A | . | exonic | AP2M1 | nonsynonymous SNV | AP2M1:NM_001025205:exon10:c.G1105A:p.E369K, |
| chr3 | 184182866 | G | A | . | exonic | AP2M1 | nonsynonymous SNV | AP2M1:NM_001025205:exon10:c.G1165A:p.E389K, |
| chr3 | 184183530 | G | A | . | exonic | AP2M1 | nonsynonymous SNV | AP2M1:NM_001025205:exon11:c.G1216A:p.E406K, |
| chr3 | 184183557 | G | A | . | exonic | AP2M1 | nonsynonymous SNV | AP2M1:NM_001025205:exon11:c.G1243A:p.D415N, |
| chr3 | 184711115 | T | G | COSV58633357 | exonic | MAGEF1 | nonsynonymous SNV | MAGEF1:NM_022149:exon1:c.A707C:p.E236A, |
| chr3 | 184982638 | C | T | COSV54982221 | exonic | VPS8 | nonsynonymous SNV | VPS8:NM_001009921:exon40:c.C3493T:p.H1165Y,VPS8:NM_015303:exon40:c.C3487T:p.H1163Y, |
| chr3 | 185048513 | T | C | COSV54997868 | exonic | VPS8 | nonsynonymous SNV | VPS8:NM_001009921:exon46:c.T4091C:p.I1364T,VPS8:NM_015303:exon46:c.T4085C:p.I1362T, |
| chr3 | 186048763 | T | G | . | exonic | ETV5 | nonsynonymous SNV | ETV5:NM_004454:exon13:c.A1409C:p.E470A, |
| chr3 | 195725009 | A | G | COSV57846887 | exonic | MUC20 | nonsynonymous SNV | MUC20:NM_020790:exon2:c.A406G:p.R136G,MUC20:NM_001291833:exon2:c.A406G:p.R136G,MUC20:NM_001282506:exon2:c.A406G:p.R136G,MUC20:NM_152673:exon2:c.A406G:p.R136G, |
| chr3 | 195725010 | G | A | COSV57846894 | exonic | MUC20 | nonsynonymous SNV | MUC20:NM_020790:exon2:c.G407A:p.R136K,MUC20:NM_001291833:exon2:c.G407A:p.R136K,MUC20:NM_001282506:exon2:c.G407A:p.R136K,MUC20:NM_152673:exon2:c.G407A:p.R136K, |
| chr3 | 195725146 | C | G | . | exonic | MUC20 | nonsynonymous SNV | MUC20:NM_001282506:exon2:c.C543G:p.D181E, |
| chr3 | 195725147 | A | G | COSV57850489 | exonic | MUC20 | nonsynonymous SNV | MUC20:NM_001282506:exon2:c.A544G:p.S182G, |
| chr3 | 195725928 | C | T | COSV57846901 | exonic | MUC20 | nonsynonymous SNV | MUC20:NM_020790:exon4:c.C869T:p.T290I,MUC20:NM_001291833:exon4:c.C812T:p.T271I,MUC20:NM_001282506:exon2:c.C1325T:p.T442I,MUC20:NM_152673:exon4:c.C812T:p.T271I, |
| chr3 | 195726080 | G | C | COSV57846804 | exonic | MUC20 | nonsynonymous SNV | MUC20:NM_020790:exon4:c.G1021C:p.D341H,MUC20:NM_001291833:exon4:c.G964C:p.D322H,MUC20:NM_001282506:exon2:c.G1477C:p.D493H,MUC20:NM_152673:exon4:c.G964C:p.D322H, |
| chr3 | 195762138 | C | A | COSV57763148 | exonic | MUC4 | nonsynonymous SNV | MUC4:NM_138297:exon12:c.G1600T:p.A534S,MUC4:NM_018406:exon14:c.G14461T:p.A4821S,MUC4:NM_004532:exon13:c.G1753T:p.A585S, |
| chr3 | 195769045 | G | C | COSV57780809 | exonic | MUC4 | nonsynonymous SNV | MUC4:NM_138297:exon5:c.C645G:p.N215K,MUC4:NM_018406:exon7:c.C13506G:p.N4502K,MUC4:NM_004532:exon6:c.C798G:p.N266K, |
| chr3 | 195791459 | C | G | . | exonic | MUC4 | nonsynonymous SNV | MUC4:NM_018406:exon2:c.G121C:p.A41P, |
| chr3 | 196247477 | C | T | . | exonic | PCYT1A | nonsynonymous SNV | PCYT1A:NM_005017:exon6:c.G376A:p.V126M,PCYT1A:NM_001312673:exon5:c.G376A:p.V126M, |
| chr3 | 32997076 | T | C | . | exonic | GLB1 | nonsynonymous SNV | GLB1:NM_001317040:exon17:c.A2147G:p.K716R,GLB1:NM_001135602:exon13:c.A1610G:p.K537R,GLB1:NM_000404:exon16:c.A2003G:p.K668R,GLB1:NM_001079811:exon16:c.A1913G:p.K638R, |
| chr3 | 32997085 | G | T | . | exonic | GLB1 | nonsynonymous SNV | GLB1:NM_001317040:exon17:c.C2138A:p.P713H,GLB1:NM_000404:exon16:c.C1994A:p.P665H,GLB1:NM_001135602:exon13:c.C1601A:p.P534H,GLB1:NM_001079811:exon16:c.C1904A:p.P635H, |
| chr3 | 32997107 | C | T | . | exonic | GLB1 | nonsynonymous SNV | GLB1:NM_001317040:exon17:c.G2116A:p.E706K,GLB1:NM_000404:exon16:c.G1972A:p.E658K,GLB1:NM_001135602:exon13:c.G1579A:p.E527K,GLB1:NM_001079811:exon16:c.G1882A:p.E628K, |
| chr3 | 33014174 | C | T | . | exonic | GLB1 | stopgain | GLB1:NM_001317040:exon16:c.G1760A:p.W587X,GLB1:NM_000404:exon15:c.G1616A:p.W539X,GLB1:NM_001135602:exon12:c.G1223A:p.W408X,GLB1:NM_001079811:exon15:c.G1526A:p.W509X, |
| chr3 | 33014196 | T | C | . | exonic | GLB1 | nonsynonymous SNV | GLB1:NM_001317040:exon16:c.A1738G:p.S580G,GLB1:NM_000404:exon15:c.A1594G:p.S532G,GLB1:NM_001135602:exon12:c.A1201G:p.S401G,GLB1:NM_001079811:exon15:c.A1504G:p.S502G, |
| chr3 | 33014229 | A | G | . | exonic | GLB1 | nonsynonymous SNV | GLB1:NM_001317040:exon16:c.T1705C:p.C569R,GLB1:NM_000404:exon15:c.T1561C:p.C521R,GLB1:NM_001135602:exon12:c.T1168C:p.C390R,GLB1:NM_001079811:exon15:c.T1471C:p.C491R, |
| chr3 | 33051938 | C | T | COSV56561196 | exonic | GLB1 | nonsynonymous SNV | GLB1:NM_001317040:exon9:c.G1003A:p.E335K,GLB1:NM_000404:exon8:c.G859A:p.E287K,GLB1:NM_001135602:exon5:c.G466A:p.E156K,GLB1:NM_001079811:exon8:c.G769A:p.E257K, |
| chr3 | 33065527 | C | T | . | exonic | GLB1 | nonsynonymous SNV | GLB1:NM_001317040:exon6:c.G632A:p.G211E,GLB1:NM_000404:exon5:c.G488A:p.G163E,GLB1:NM_001079811:exon5:c.G398A:p.G133E, |
| chr3 | 33065549 | C | T | . | exonic | GLB1 | nonsynonymous SNV | GLB1:NM_001317040:exon6:c.G610A:p.A204T,GLB1:NM_000404:exon5:c.G466A:p.A156T,GLB1:NM_001135602:exon3:c.G254A:p.G85D,GLB1:NM_001079811:exon5:c.G376A:p.A126T, |
| chr3 | 33097057 | G | A | COSV56561124 | exonic | GLB1 | nonsynonymous SNV | GLB1:NM_001317040:exon1:c.C29T:p.P10L,GLB1:NM_000404:exon1:c.C29T:p.P10L,GLB1:NM_001135602:exon1:c.C29T:p.P10L, |
| chr3 | 37012077 | A | G | COSV51613800 | exonic | MLH1 | nonsynonymous SNV | MLH1:NM_000249:exon8:c.A655G:p.I219V,MLH1:NM_001258271:exon8:c.A655G:p.I219V,MLH1:NM_001167617:exon8:c.A361G:p.I121V, |
| chr3 | 37012096 | G | A | . | exonic | MLH1 | nonsynonymous SNV | MLH1:NM_000249:exon8:c.G674A:p.S225N,MLH1:NM_001167617:exon8:c.G380A:p.S127N,MLH1:NM_001258271:exon8:c.G674A:p.S225N, |
| chr3 | 37020420 | G | A | . | exonic | MLH1 | nonsynonymous SNV | MLH1:NM_000249:exon11:c.G995A:p.S332N,MLH1:NM_001167618:exon11:c.G272A:p.S91N,MLH1:NM_001258271:exon11:c.G995A:p.S332N,MLH1:NM_001167619:exon10:c.G272A:p.S91N,MLH1:NM_001258273:exon10:c.G272A:p.S91N,MLH1:NM_001258274:exon12:c.G272A:p.S91N,MLH1:NM_001167617:exon11:c.G701A:p.S234N, |
| chr3 | 37025998 | G | A | . | exonic | MLH1 | nonsynonymous SNV | MLH1:NM_000249:exon12:c.G1400A:p.S467N,MLH1:NM_001167618:exon12:c.G677A:p.S226N,MLH1:NM_001258271:exon12:c.G1400A:p.S467N,MLH1:NM_001167619:exon11:c.G677A:p.S226N,MLH1:NM_001258273:exon11:c.G677A:p.S226N,MLH1:NM_001258274:exon13:c.G677A:p.S226N,MLH1:NM_001167617:exon12:c.G1106A:p.S369N, |
| chr3 | 37028795 | G | A | . | exonic | MLH1 | nonsynonymous SNV | MLH1:NM_000249:exon13:c.G1421A:p.R474Q,MLH1:NM_001167618:exon13:c.G698A:p.R233Q,MLH1:NM_001258271:exon13:c.G1421A:p.R474Q,MLH1:NM_001167619:exon12:c.G698A:p.R233Q,MLH1:NM_001258273:exon12:c.G698A:p.R233Q,MLH1:NM_001258274:exon14:c.G698A:p.R233Q,MLH1:NM_001167617:exon13:c.G1127A:p.R376Q, |
| chr3 | 37040225 | G | A | . | exonic | MLH1 | nonsynonymous SNV | MLH1:NM_000249:exon14:c.G1598A:p.C533Y,MLH1:NM_001167618:exon14:c.G875A:p.C292Y,MLH1:NM_001258271:exon14:c.G1598A:p.C533Y,MLH1:NM_001167619:exon13:c.G875A:p.C292Y,MLH1:NM_001258273:exon13:c.G875A:p.C292Y,MLH1:NM_001258274:exon15:c.G875A:p.C292Y,MLH1:NM_001167617:exon14:c.G1304A:p.C435Y, |
| chr3 | 37048550 | G | A | . | exonic | MLH1 | nonsynonymous SNV | MLH1:NM_000249:exon17:c.G1930A:p.D644N,MLH1:NM_001167617:exon17:c.G1636A:p.D546N,MLH1:NM_001167618:exon17:c.G1207A:p.D403N,MLH1:NM_001167619:exon16:c.G1207A:p.D403N,MLH1:NM_001258273:exon16:c.G1207A:p.D403N,MLH1:NM_001258274:exon18:c.G1207A:p.D403N, |
| chr3 | 37048939 | C | A | . | exonic | MLH1 | nonsynonymous SNV | MLH1:NM_000249:exon18:c.C2025A:p.S675R,MLH1:NM_001167618:exon18:c.C1302A:p.S434R,MLH1:NM_001167619:exon17:c.C1302A:p.S434R,MLH1:NM_001258273:exon17:c.C1302A:p.S434R,MLH1:NM_001258274:exon19:c.C1302A:p.S434R,MLH1:NM_001167617:exon18:c.C1731A:p.S577R, |
| chr3 | 37302221 | G | A | . | exonic | GOLGA4 | nonsynonymous SNV | GOLGA4:NM_002078:exon10:c.G1123A:p.E375K,GOLGA4:NM_001172713:exon11:c.G1189A:p.E397K, |
| chr3 | 37315518 | G | A | . | exonic | GOLGA4 | nonsynonymous SNV | GOLGA4:NM_002078:exon11:c.G1333A:p.E445K,GOLGA4:NM_001172713:exon12:c.G1399A:p.E467K, |
| chr3 | 37323683 | T | A | . | exonic | GOLGA4 | nonsynonymous SNV | GOLGA4:NM_002078:exon14:c.T1797A:p.N599K,GOLGA4:NM_001172713:exon15:c.T1863A:p.N621K, |
| chr3 | 37323837 | C | A | . | exonic | GOLGA4 | nonsynonymous SNV | GOLGA4:NM_002078:exon14:c.C1951A:p.Q651K,GOLGA4:NM_001172713:exon15:c.C2017A:p.Q673K, |

| | | | | | | | | |
|---|---|---|---|---|---|---|---|---|
| chr3 | 37324258 | G | A | . | exonic | GOLGA4 | nonsynonymous SNV | GOLGA4:NM_002078:exon14:c.G2372A:p.G791E,GOLGA4:NM_001172713:exon15:c.G2438A:p.G813E, |
| chr3 | 37324327 | C | A | . | exonic | GOLGA4 | nonsynonymous SNV | GOLGA4:NM_002078:exon14:c.C2441A:p.T814K,GOLGA4:NM_001172713:exon15:c.C2507A:p.T836K, |
| chr3 | 37324512 | G | A | . | exonic | GOLGA4 | nonsynonymous SNV | GOLGA4:NM_002078:exon14:c.G2626A:p.E876K,GOLGA4:NM_001172713:exon15:c.G2692A:p.E898K, |
| chr3 | 37324641 | G | A | COSV62710454 | exonic | GOLGA4 | nonsynonymous SNV | GOLGA4:NM_002078:exon14:c.G2755A:p.G919R,GOLGA4:NM_001172713:exon15:c.G2821A:p.G941R, |
| chr3 | 37324792 | C | A | . | exonic | GOLGA4 | nonsynonymous SNV | GOLGA4:NM_002078:exon14:c.C2906A:p.T969K,GOLGA4:NM_001172713:exon15:c.C2972A:p.T991K, |
| chr3 | 37324794 | T | A | . | exonic | GOLGA4 | nonsynonymous SNV | GOLGA4:NM_002078:exon14:c.T2908A:p.L970I,GOLGA4:NM_001172713:exon15:c.T2974A:p.L992I, |
| chr3 | 37324968 | C | A | COSV62709296 | exonic | GOLGA4 | nonsynonymous SNV | GOLGA4:NM_002078:exon14:c.C3082A:p.Q1028K,GOLGA4:NM_001172713:exon15:c.C3148A:p.Q1050K, |
| chr3 | 37325148 | G | A | . | exonic | GOLGA4 | nonsynonymous SNV | GOLGA4:NM_002078:exon14:c.G3262A:p.E1088K,GOLGA4:NM_001172713:exon15:c.G3328A:p.E1110K, |
| chr3 | 37327264 | T | A | COSV62710975 | exonic | GOLGA4 | nonsynonymous SNV | GOLGA4:NM_002078:exon14:c.T5378A:p.M1793K,GOLGA4:NM_001172713:exon15:c.T5444A:p.M1815K, |
| chr3 | 38126634 | A | C | . | exonic | ACAA1 | nonsynonymous SNV | ACAA1:NM_001607:exon8:c.T693G:p.H231Q,ACAA1:NM_001130410:exon7:c.T594G:p.H198Q, |
| chr3 | 41225688 | G | A | . | exonic | CTNNB1 | nonsynonymous SNV | CTNNB1:NM_001904:exon6:c.G763A:p.A255T,CTNNB1:NM_001098210:exon6:c.G763A:p.A255T,CTNNB1:NM_001098209:exon6:c.G763A:p.A255T, |
| chr3 | 41227257 | G | A | . | exonic | CTNNB1 | nonsynonymous SNV | CTNNB1:NM_001904:exon7:c.G986A:p.R329K,CTNNB1:NM_001098210:exon7:c.G986A:p.R329K,CTNNB1:NM_001098209:exon7:c.G986A:p.R329K, |
| chr3 | 41233679 | G | A | . | exonic | CTNNB1 | nonsynonymous SNV | CTNNB1:NM_001904:exon9:c.G1336A:p.A446T,CTNNB1:NM_001098210:exon9:c.G1336A:p.A446T,CTNNB1:NM_001098209:exon9:c.G1336A:p.A446T, |
| chr3 | 41234145 | G | A | . | exonic | CTNNB1 | nonsynonymous SNV | CTNNB1:NM_001904:exon10:c.G1531A:p.V511I,CTNNB1:NM_001098210:exon10:c.G1531A:p.V511I,CTNNB1:NM_001098209:exon10:c.G1531A:p.V511I, |
| chr3 | 41234260 | G | A | . | exonic | CTNNB1 | nonsynonymous SNV | CTNNB1:NM_001904:exon10:c.G1646A:p.R549H,CTNNB1:NM_001098210:exon10:c.G1646A:p.R549H,CTNNB1:NM_001098209:exon10:c.G1646A:p.R549H, |
| chr3 | 41235730 | G | A | . | exonic | CTNNB1 | nonsynonymous SNV | CTNNB1:NM_001904:exon11:c.G1690A:p.V564I,CTNNB1:NM_001098210:exon11:c.G1690A:p.V564I,CTNNB1:NM_001098209:exon11:c.G1690A:p.V564I, |
| chr3 | 41235790 | G | A | . | exonic | CTNNB1 | nonsynonymous SNV | CTNNB1:NM_001098210:exon11:c.G1750A:p.V584I,CTNNB1:NM_001098209:exon11:c.G1750A:p.V584I,CTNNB1:NM_001904:exon11:c.G1750A:p.V584I, |
| chr3 | 41235800 | G | A | COSV62694859 | exonic | CTNNB1 | nonsynonymous SNV | CTNNB1:NM_001904:exon11:c.G1760A:p.R587Q,CTNNB1:NM_001098210:exon11:c.G1760A:p.R587Q,CTNNB1:NM_001098209:exon11:c.G1760A:p.R587Q, |
| chr3 | 41235835 | T | A | . | exonic | CTNNB1 | nonsynonymous SNV | CTNNB1:NM_001098210:exon11:c.T1795A:p.F599I,CTNNB1:NM_001098209:exon11:c.T1795A:p.F599I,CTNNB1:NM_001904:exon11:c.T1795A:p.F599I, |
| chr3 | 41236415 | G | A | COSV62712530 | exonic | CTNNB1 | nonsynonymous SNV | CTNNB1:NM_001904:exon12:c.G1870A:p.D624N,CTNNB1:NM_001098210:exon12:c.G1870A:p.D624N,CTNNB1:NM_001098209:exon12:c.G1870A:p.D624N, |
| chr3 | 41236455 | C | T | . | exonic | CTNNB1 | nonsynonymous SNV | CTNNB1:NM_001098210:exon12:c.C1910T:p.T637I,CTNNB1:NM_001098209:exon12:c.C1910T:p.T637I,CTNNB1:NM_001904:exon12:c.C1910T:p.T637I, |
| chr3 | 41236618 | T | G | . | exonic | CTNNB1 | nonsynonymous SNV | CTNNB1:NM_001904:exon13:c.T1985G:p.M662R,CTNNB1:NM_001098210:exon13:c.T1985G:p.M662R,CTNNB1:NM_001098209:exon13:c.T1985G:p.M662R, |
| chr3 | 41238041 | G | A | . | exonic | CTNNB1 | nonsynonymous SNV | CTNNB1:NM_001098210:exon14:c.G2102A:p.G701D,CTNNB1:NM_001098209:exon14:c.G2102A:p.G701D,CTNNB1:NM_001904:exon14:c.G2102A:p.G701D, |
| chr3 | 41239160 | G | A | . | exonic | CTNNB1 | nonsynonymous SNV | CTNNB1:NM_001904:exon15:c.G2164A:p.G722S,CTNNB1:NM_001098210:exon15:c.G2164A:p.G722S,CTNNB1:NM_001098209:exon15:c.G2164A:p.G722S, |
| chr3 | 41239218 | G | A | . | exonic | CTNNB1 | nonsynonymous SNV | CTNNB1:NM_001098210:exon15:c.G2222A:p.G741D,CTNNB1:NM_001098209:exon15:c.G2222A:p.G741D,CTNNB1:NM_001904:exon15:c.G2222A:p.G741D, |
| chr3 | 41239247 | G | A | COSV62706846 | exonic | CTNNB1 | nonsynonymous SNV | CTNNB1:NM_001904:exon15:c.G2251A:p.D751N,CTNNB1:NM_001098210:exon15:c.G2251A:p.D751N,CTNNB1:NM_001098209:exon15:c.G2251A:p.D751N, |
| chr3 | 41239286 | G | A | . | exonic | CTNNB1 | nonsynonymous SNV | CTNNB1:NM_001098210:exon15:c.G2290A:p.D764N,CTNNB1:NM_001098209:exon15:c.G2290A:p.D764N,CTNNB1:NM_001904:exon15:c.G2290A:p.D764N, |
| chr3 | 41800224 | A | C | COSV57206179 | exonic | ULK4 | nonsynonymous SNV | ULK4:NM_017886:exon20:c.T1918G:p.S640A, |
| chr3 | 43561311 | C | T | . | exonic | ANO10 | nonsynonymous SNV | ANO10:NM_001204833:exon7:c.G1052A:p.R351Q,ANO10:NM_018075:exon9:c.G1385A:p.R462Q,ANO10:NM_001204832:exon8:c.G1187A:p.R396Q,ANO10:NM_001204831:exon9:c.G1385A:p.R462Q,ANO10:NM_001204834:exon8:c.G815A:p.R272Q, |
| chr3 | 45516118 | C | A | . | exonic | LARS2 | nonsynonymous SNV | LARS2:NM_015340:exon17:c.C1886A:p.T629K, |
| chr3 | 46408491 | T | C | . | exonic | CCRL2 | nonsynonymous SNV | CCRL2:NM_001130910:exon2:c.T448C:p.F150L,CCRL2:NM_003965:exon2:c.T412C:p.F138L, |
| chr3 | 46408579 | T | A | COSV62193340 | exonic | CCRL2 | nonsynonymous SNV | CCRL2:NM_003965:exon2:c.T500A:p.F167Y,CCRL2:NM_001130910:exon2:c.T536A:p.F179Y, |
| chr3 | 46408581 | G | A | COSV62194015 | exonic | CCRL2 | nonsynonymous SNV | CCRL2:NM_003965:exon2:c.G502A:p.V168M,CCRL2:NM_001130910:exon2:c.G538A:p.V180M, |
| chr3 | 46408864 | C | T | . | exonic | CCRL2 | nonsynonymous SNV | CCRL2:NM_003965:exon2:c.C785T:p.T262I,CCRL2:NM_001130910:exon2:c.C821T:p.T274I, |
| chr3 | 46439467 | C | G | COSV51606384 | exonic | LTF | nonsynonymous SNV | LTF:NM_001199149:exon15:c.G1605C:p.E535D,LTF:NM_002343:exon15:c.G1737C:p.E579D, |
| chr3 | 47062226 | C | T | COSV57435899 | exonic | SETD2 | nonsynonymous SNV | SETD2:NM_014159:exon14:c.G6230A:p.R2077Q, |
| chr3 | 47083895 | G | A | COSV57428846 | exonic | SETD2 | nonsynonymous SNV | SETD2:NM_014159:exon12:c.C5885T:p.P1962L, |
| chr3 | 47418189 | C | T | COSV55542934 | exonic | SCAP | nonsynonymous SNV | SCAP:NM_012235:exon16:c.G2392A:p.V798I, |
| chr3 | 47914934 | C | T | COSV53135994 | exonic | MAP4 | nonsynonymous SNV | MAP4:NM_002375:exon8:c.G1882A:p.V628I,MAP4:NM_001134364:exon8:c.G1882A:p.V628I, |
| chr3 | 47916041 | G | T | . | exonic | MAP4 | nonsynonymous SNV | MAP4:NM_002375:exon7:c.C1786A:p.Q596K,MAP4:NM_001134364:exon7:c.C1786A:p.Q596K, |
| chr3 | 47916547 | G | T | COSV64236753 | exonic | MAP4 | nonsynonymous SNV | MAP4:NM_002375:exon7:c.C1280A:p.S427Y,MAP4:NM_001134364:exon7:c.C1280A:p.S427Y, |
| chr3 | 47916670 | A | T | . | exonic | MAP4 | nonsynonymous SNV | MAP4:NM_002375:exon7:c.T1157A:p.I386K,MAP4:NM_001134364:exon7:c.T1157A:p.I386K, |

| | | | | | | | | |
|---|---|---|---|---|---|---|---|---|
| chr3 | 47916694 | T | A | . | exonic | MAP4 | nonsynonymous SNV | MAP4:NM_002375:exon7:c.A1133T:p.K378I,MAP4:NM_001134364:exon7:c.A1133T:p.K378I, |
| chr3 | 47916700 | T | C | . | exonic | MAP4 | nonsynonymous SNV | MAP4:NM_002375:exon7:c.A1127G:p.N376S,MAP4:NM_001134364:exon7:c.A1127G:p.N376S, |
| chr3 | 47916709 | G | T | . | exonic | MAP4 | nonsynonymous SNV | MAP4:NM_002375:exon7:c.C1118A:p.P373H,MAP4:NM_001134364:exon7:c.C1118A:p.P373H, |
| chr3 | 47916712 | G | T | . | exonic | MAP4 | nonsynonymous SNV | MAP4:NM_002375:exon7:c.C1115A:p.P372Q,MAP4:NM_001134364:exon7:c.C1115A:p.P372Q, |
| chr3 | 49030471 | T | C | COSV58708022 | exonic | QRICH1 | nonsynonymous SNV | QRICH1:NM_017730:exon11:c.A2312G:p.N771S,QRICH1:NM_198880:exon10:c.A2312G:p.N771S, |
| chr3 | 49360262 | C | T | . | exonic | RHOA | nonsynonymous SNV | RHOA:NM_001313945:exon5:c.G286A:p.A96T,RHOA:NM_001664:exon5:c.G529A:p.A177T,RHOA:NM_001313946:exon3:c.G277A:p.A93T,RHOA:NM_001313941:exon6:c.G529A:p.A177T,RHOA:NM_001313944:exon5:c.G469A:p.A157T, |
| chr3 | 49360264 | C | T | . | exonic | RHOA | nonsynonymous SNV | RHOA:NM_001313945:exon5:c.G284A:p.R95K,RHOA:NM_001664:exon5:c.G527A:p.R176K,RHOA:NM_001313946:exon3:c.G275A:p.R92K,RHOA:NM_001313941:exon6:c.G527A:p.R176K,RHOA:NM_001313944:exon5:c.G467A:p.R156K, |
| chr3 | 49360277 | C | T | COSV69041753 | exonic | RHOA | nonsynonymous SNV | RHOA:NM_001313945:exon5:c.G271A:p.E91K,RHOA:NM_001664:exon5:c.G514A:p.E172K,RHOA:NM_001313946:exon3:c.G262A:p.E88K,RHOA:NM_001313941:exon6:c.G514A:p.E172K,RHOA:NM_001313944:exon5:c.G454A:p.E152K, |
| chr3 | 49360350 | C | A | . | exonic | RHOA | nonsynonymous SNV | RHOA:NM_001313945:exon5:c.G198T:p.M66I,RHOA:NM_001313946:exon3:c.G189T:p.M63I,RHOA:NM_001313941:exon6:c.G441T:p.M147I,RHOA:NM_001664:exon5:c.G441T:p.M147I,RHOA:NM_001313944:exon5:c.G381T:p.M127I, |
| chr3 | 49362534 | C | T | . | exonic | RHOA | nonsynonymous SNV | RHOA:NM_001313945:exon4:c.G127A:p.D43N,RHOA:NM_001664:exon4:c.G370A:p.D124N,RHOA:NM_001313941:exon5:c.G370A:p.D124N,RHOA:NM_001313947:exon3:c.G249A:p.M83I,RHOA:NM_001313944:exon4:c.G310A:p.D104N,RHOA:NM_001313943:exon4:c.G370A:p.D124N, |
| chr3 | 49362608 | C | T | . | exonic | RHOA | stopgain | RHOA:NM_001313945:exon4:c.G53A:p.W18X,RHOA:NM_001664:exon4:c.G296A:p.W99X,RHOA:NM_001313941:exon5:c.G296A:p.W99X,RHOA:NM_001313944:exon4:c.G236A:p.W79X,RHOA:NM_001313943:exon4:c.G296A:p.W99X, |
| chr3 | 49368531 | C | T | . | exonic | RHOA | stopgain | RHOA:NM_001664:exon3:c.G174A:p.W58X,RHOA:NM_001313941:exon4:c.G174A:p.W58X,RHOA:NM_001313943:exon3:c.G174A:p.W58X, |
| chr3 | 49510575 | C | G | . | exonic | DAG1 | nonsynonymous SNV | DAG1:NM_001177637:exon3:c.C41G:p.S14W,DAG1:NM_004393:exon2:c.C41G:p.S14W,DAG1:NM_001177636:exon4:c.C41G:p.S14W,DAG1:NM_001177642:exon3:c.C41G:p.S14W,DAG1:NM_001177635:exon3:c.C41G:p.S14W,DAG1:NM_001177634:exon5:c.C41G:p.S14W,DAG1:NM_001177641:exon3:c.C41G:p.S14W,DAG1:NM_001177644:exon2:c.C41G:p.S14W,DAG1:NM_001177638:exon3:c.C41G:p.S14W,DAG1:NM_001177639:exon2:c.C41G:p.S14W,DAG1:NM_001177643:exon2:c.C41G:p.S14W,DAG1:NM_001177640:exon3:c.C41G:p.S14W,DAG1:NM_001165928:exon5:c.C41G:p.S14W, |
| chr3 | 49722606 | T | C | . | exonic | GMPPB | nonsynonymous SNV | GMPPB:NM_021971:exon5:c.A551G:p.Q184R,GMPPB:NM_013334:exon5:c.A551G:p.Q184R, |
| chr3 | 5172157 | A | G | . | exonic | ARL8B | nonsynonymous SNV | ARL8B:NM_018184:exon3:c.A212G:p.D71G, |
| chr3 | 52476576 | G | A | COSV61900229 | exonic | NISCH | nonsynonymous SNV | NISCH:NM_007184:exon8:c.G895A:p.V299I,NISCH:NM_001276293:exon8:c.G895A:p.V299I,NISCH:NM_001276294:exon8:c.G895A:p.V299I, |
| chr3 | 52603593 | C | T | . | exonic | PBRM1 | nonsynonymous SNV | PBRM1:NM_018313:exon18:c.G2707A:p.E903K, |
| chr3 | 52634665 | G | T | . | exonic | PBRM1 | stopgain | PBRM1:NM_018313:exon12:c.C1238A:p.S413X, |
| chr3 | 52687262 | G | A | . | exonic | GNL3 | nonsynonymous SNV | GNL3:NM_206826:exon3:c.G53A:p.R18Q,GNL3:NM_014366:exon3:c.G89A:p.R30Q,GNL3:NM_206825:exon3:c.G53A:p.R18Q, |
| chr3 | 52687289 | G | A | COSV66920847 | exonic | GNL3 | nonsynonymous SNV | GNL3:NM_206826:exon3:c.G80A:p.R27Q,GNL3:NM_014366:exon3:c.G116A:p.R39Q,GNL3:NM_206825:exon3:c.G80A:p.R27Q, |
| chr3 | 52687536 | G | A | . | exonic | GNL3 | nonsynonymous SNV | GNL3:NM_206826:exon4:c.G209A:p.R70K,GNL3:NM_014366:exon4:c.G245A:p.R82K,GNL3:NM_206825:exon4:c.G209A:p.R70K, |
| chr3 | 52688178 | C | T | . | exonic | GNL3 | stopgain | GNL3:NM_206826:exon5:c.C358T:p.Q120X,GNL3:NM_014366:exon5:c.C394T:p.Q132X,GNL3:NM_206825:exon5:c.C358T:p.Q120X, |
| chr3 | 52689185 | G | A | . | exonic | GNL3 | nonsynonymous SNV | GNL3:NM_206826:exon6:c.G484A:p.V162I,GNL3:NM_014366:exon6:c.G520A:p.V174I,GNL3:NM_206825:exon6:c.G484A:p.V162I, |
| chr3 | 52690686 | T | A | . | exonic | GNL3 | nonsynonymous SNV | GNL3:NM_206826:exon7:c.T600A:p.D200E,GNL3:NM_014366:exon7:c.T636A:p.D212E,GNL3:NM_206825:exon7:c.T600A:p.D200E, |
| chr3 | 52691045 | G | A | . | exonic | GNL3 | nonsynonymous SNV | GNL3:NM_206826:exon8:c.G719A:p.S240N,GNL3:NM_014366:exon8:c.G755A:p.S252N,GNL3:NM_206825:exon8:c.G719A:p.S240N, |
| chr3 | 52691059 | G | A | . | exonic | GNL3 | nonsynonymous SNV | GNL3:NM_206826:exon8:c.G733A:p.V245I,GNL3:NM_014366:exon8:c.G769A:p.V257I,GNL3:NM_206825:exon8:c.G733A:p.V245I, |
| chr3 | 52693241 | G | A | COSV56476624 | exonic | GNL3 | nonsynonymous SNV | GNL3:NM_206826:exon11:c.G1063A:p.V355M,GNL3:NM_014366:exon11:c.G1099A:p.V367M,GNL3:NM_206825:exon11:c.G1063A:p.V355M, |
| chr3 | 52693275 | G | A | . | exonic | GNL3 | nonsynonymous SNV | GNL3:NM_206826:exon11:c.G1097A:p.G366D,GNL3:NM_014366:exon11:c.G1133A:p.G378D,GNL3:NM_206825:exon11:c.G1097A:p.G366D, |
| chr3 | 52693315 | G | A | . | exonic | GNL3 | stopgain | GNL3:NM_206826:exon11:c.G1137A:p.W379X,GNL3:NM_014366:exon11:c.G1173A:p.W391X,GNL3:NM_206825:exon11:c.G1137A:p.W379X, |
| chr3 | 52693518 | A | G | . | exonic | GNL3 | nonsynonymous SNV | GNL3:NM_206826:exon12:c.A1262G:p.E421G,GNL3:NM_014366:exon12:c.A1298G:p.E433G,GNL3:NM_206825:exon12:c.A1262G:p.E421G, |
| chr3 | 52693700 | G | A | . | exonic | GNL3 | nonsynonymous SNV | GNL3:NM_206826:exon13:c.G1357A:p.E453K,GNL3:NM_014366:exon13:c.G1393A:p.E465K,GNL3:NM_206825:exon13:c.G1357A:p.E453K, |
| chr3 | 52693728 | C | A | . | exonic | GNL3 | nonsynonymous SNV | GNL3:NM_206826:exon13:c.C1385A:p.P462Q,GNL3:NM_014366:exon13:c.C1421A:p.P474Q,GNL3:NM_206825:exon13:c.C1385A:p.P462Q, |
| chr3 | 52693734 | G | A | COSV56477800 | exonic | GNL3 | nonsynonymous SNV | GNL3:NM_206826:exon13:c.G1391A:p.R464Q,GNL3:NM_014366:exon13:c.G1427A:p.R476Q,GNL3:NM_206825:exon13:c.G1391A:p.R464Q, |
| chr3 | 52694061 | G | A | . | exonic | GNL3 | nonsynonymous SNV | GNL3:NM_206826:exon14:c.G1489A:p.A497T,GNL3:NM_014366:exon14:c.G1525A:p.A509T,GNL3:NM_206825:exon14:c.G1489A:p.A497T, |
| chr3 | 52694079 | G | A | . | exonic | GNL3 | nonsynonymous SNV | GNL3:NM_206826:exon14:c.G1507A:p.A503T,GNL3:NM_014366:exon14:c.G1543A:p.A515T,GNL3:NM_206825:exon14:c.G1507A:p.A503T, |
| chr3 | 52694211 | C | T | . | exonic | GNL3 | nonsynonymous SNV | GNL3:NM_206826:exon15:c.C1550T:p.S517F,GNL3:NM_014366:exon15:c.C1586T:p.S529F,GNL3:NM_206825:exon15:c.C1550T:p.S517F, |
| chr3 | 52706166 | C | G | COSV51776812 | exonic | SPCS1 | nonsynonymous SNV | SPCS1:NM_014041:exon1:c.C121G:p.P41A, |
| chr3 | 52763618 | G | C | COSV51778826 | exonic | NEK4 | nonsynonymous SNV | NEK4:NM_003157:exon5:c.C673G:p.P225A,NEK4:NM_001193533:exon4:c.C406G:p.P136A, |
| chr3 | 53184922 | T | A | . | exonic | PRKCD | nonsynonymous SNV | PRKCD:NM_212539:exon9:c.T836A:p.L279H,PRKCD:NM_006254:exon10:c.T836A:p.L279H,PRKCD:NM_001316327:exon10:c.T836A:p.L279H, |
| chr3 | 53226803 | C | T | COSV56246245 | exonic | TKT | nonsynonymous SNV | TKT:NM_001135055:exon13:c.G1649A:p.R550H,TKT:NM_001064:exon13:c.G1649A:p.R550H,TKT:NM_001258028:exon14:c.G1673A:p.R558H, |
| chr3 | 57556446 | C | T | COSV60818263 | exonic | PDE12 | nonsynonymous SNV | PDE12:NM_177966:exon1:c.C67T:p.R23W, |
| chr3 | 66369532 | C | T | COSV60914960 | exonic | SLC25A26 | nonsynonymous SNV | SLC25A26:NM_001164796:exon7:c.C359T:p.T120M,SLC25A26:NM_173471:exon9:c.C623T:p.T208M, |
| chr3 | 67375857 | T | C | COSV56206071 | exonic | SUCLG2 | nonsynonymous SNV | SUCLG2:NM_003848:exon11:c.A1186G:p.T396A, |
| chr3 | 69180910 | G | A | COSV68330003 | exonic | FRMD4B | nonsynonymous SNV | FRMD4B:NM_015123:exon21:c.C2840T:p.S947L, |
| chr3 | 69196303 | T | C | . | exonic | FRMD4B | nonsynonymous SNV | FRMD4B:NM_015123:exon14:c.A1186G:p.T396A, |
| chr3 | 70972576 | C | T | COSV59547066 | exonic | FOXP1 | nonsynonymous SNV | FOXP1:NM_001244814:exon14:c.G1631A:p.R544Q,FOXP1:NM_032682:exon18:c.G1631A:p.R544Q,FOXP1:NM_001244813:exon12:c.G1331A:p.R444Q,FOXP1:NM_001244812:exon13:c.G1403A:p.R468Q,FOXP1:NM_001244815:exon13:c.G1637A:p.R546Q,FOXP1:NM_001244808:exon18:c.G1628A:p.R543Q,FOXP1:NM_001244816:exon17:c.G1631A:p.R544Q, |
| chr3 | 70988068 | C | T | . | exonic | FOXP1 | nonsynonymous SNV | FOXP1:NM_001244814:exon10:c.G1072A:p.D358N,FOXP1:NM_001244810:exon14:c.G1072A:p.D358N,FOXP1:NM_032682:exon14:c.G1072A:p.D358N,FOXP1:NM_001244813:exon8:c.G772A:p.D258N,FOXP1:NM_001244812:exon9:c.G844A:p.D282N,FOXP1:NM_001244815:exon9:c.G1078A:p.D360N,FOXP1:NM_001244808:exon14:c.G1072A:p.D358N,FOXP1:NM_001244816:exon13:c.G1072A:p.D358N, |
| chr3 | 77477557 | G | A | . | exonic | ROBO2 | nonsynonymous SNV | ROBO2:NM_001290039:exon3:c.G532A:p.E178K,ROBO2:NM_001290040:exon3:c.G532A:p.E178K,ROBO2:NM_002942:exon3:c.G532A:p.E178K, |
| chr3 | 9684666 | G | C | . | exonic | MTMR14 | nonsynonymous SNV | MTMR14:NM_001077525:exon11:c.G1046C:p.W349S,MTMR14:NM_022485:exon11:c.G1046C:p.W349S,MTMR14:NM_001077526:exon11:c.G1046C:p.W349S, |

| | | | | | | | | |
|---|---|---|---|---|---|---|---|---|
| chr3 | 9690083 | C | A | . | exonic | MTMR14 | stopgain | MTMR14:NM_001077525:exon17:c.C1553A:p.S518X,MTMR14:NM_001077526:exon17:c.C1553A:p.S518X, |
| chr3 | 9701861 | G | A | . | exonic | MTMR14 | nonsynonymous SNV | MTMR14:NM_001077525:exon19:c.G1841A:p.S614N,MTMR14:NM_022485:exon17:c.G1505A:p.S502N,MTMR14:NM_001077526:exon18:c.G1685A:p.S562N, |
| chr3 | 9914595 | G | A | . | exonic | IL17RE | nonsynonymous SNV | IL17RE:NM_001193380:exon14:c.G1417A:p.G473R, |
| chr3 | 9934475 | A | G | . | exonic | CRELD1 | nonsynonymous SNV | CRELD1:NM_001031717:exon2:c.A37G:p.M13V,CRELD1:NM_001077415:exon2:c.A37G:p.M13V,CRELD1:NM_015513:exon1:c.A37G:p.M13V, |
| chr3 | 9964092 | C | T | . | exonic | EMC3 | nonsynonymous SNV | EMC3:NM_018447:exon8:c.G763A:p.E255K, |
| chr3 | 9969746 | C | T | . | exonic | EMC3 | nonsynonymous SNV | EMC3:NM_018447:exon7:c.G630A:p.M210I, |
| chr3 | 9970612 | T | A | . | exonic | EMC3 | nonsynonymous SNV | EMC3:NM_018447:exon6:c.A544T:p.I182F, |
| chr3 | 9970648 | A | G | . | exonic | EMC3 | nonsynonymous SNV | EMC3:NM_018447:exon6:c.T508C:p.S170P, |
| chr3 | 9970651 | C | T | . | exonic | EMC3 | nonsynonymous SNV | EMC3:NM_018447:exon6:c.G505A:p.A169T, |
| chr3 | 9974393 | A | G | . | exonic | EMC3 | nonsynonymous SNV | EMC3:NM_018447:exon4:c.T403C:p.F135L, |
| chr3 | 9974458 | C | T | . | exonic | EMC3 | nonsynonymous SNV | EMC3:NM_018447:exon4:c.G338A:p.G113E, |
| chr3 | 9974462 | T | C | . | exonic | EMC3 | nonsynonymous SNV | EMC3:NM_018447:exon4:c.A334G:p.K112E, |
| chr3 | 9986541 | C | T | . | exonic | EMC3 | nonsynonymous SNV | EMC3:NM_018447:exon1:c.G121A:p.D41N, |
| chr3 | 9986637 | C | T | COSV55277193 | exonic | EMC3 | nonsynonymous SNV | EMC3:NM_018447:exon1:c.G25A:p.D9N, |
| chr3 | 9986654 | C | T | . | exonic | EMC3 | nonsynonymous SNV | EMC3:NM_018447:exon1:c.G8A:p.G3E, |
| chr4 | 10097716 | T | C | COSV53331677 | exonic | WDR1 | nonsynonymous SNV | WDR1:NM_017491:exon5:c.A553G:p.I185V, |
| chr4 | 103092639 | T | C | COSV56477889 | exonic | BDH2 | nonsynonymous SNV | BDH2:NM_020139:exon4:c.A209G:p.N70S, |
| chr4 | 105235006 | G | A | COSV54398666 | exonic | TET2 | nonsynonymous SNV | TET2:NM_001127208:exon3:c.G1064A:p.G355D,TET2:NM_017628:exon3:c.G1064A:p.G355D, |
| chr4 | 105235030 | C | T | COSV54399299 | exonic | TET2 | nonsynonymous SNV | TET2:NM_001127208:exon3:c.C1088T:p.P363L,TET2:NM_017628:exon3:c.C1088T:p.P363L, |
| chr4 | 105236187 | C | A | . | exonic | TET2 | nonsynonymous SNV | TET2:NM_001127208:exon3:c.C2245A:p.Q749K,TET2:NM_017628:exon3:c.C2245A:p.Q749K, |
| chr4 | 105396272 | C | G | COSV58995765 | exonic | PPA2 | nonsynonymous SNV | PPA2:NM_176866:exon5:c.G540C:p.K180N,PPA2:NM_006903:exon8:c.G759C:p.K253N,PPA2:NM_176869:exon9:c.G846C:p.K282N,PPA2:NM_176867:exon3:c.G348C:p.K116N, |
| chr4 | 108009883 | T | C | . | exonic | HADH | nonsynonymous SNV | HADH:NM_005327:exon2:c.T257C:p.L86P,HADH:NM_001184705:exon2:c.T257C:p.L86P, |
| chr4 | 108622597 | G | A | . | exonic | RPL34 | nonsynonymous SNV | RPL34:NM_033625:exon4:c.G248A:p.C83Y,RPL34:NM_000995:exon4:c.G248A:p.C83Y, |
| chr4 | 112646387 | T | A | . | exonic | LARP7 | nonsynonymous SNV | LARP7:NM_016648:exon3:c.T239A:p.M80K,LARP7:NM_001267039:exon5:c.T260A:p.M87K,LARP7:NM_015454:exon3:c.T239A:p.M80K, |
| chr4 | 112647299 | T | A | . | exonic | LARP7 | nonsynonymous SNV | LARP7:NM_016648:exon7:c.T747A:p.S249R,LARP7:NM_001267039:exon9:c.T768A:p.S256R,LARP7:NM_015454:exon7:c.T747A:p.S249R, |
| chr4 | 112647304 | T | A | . | exonic | LARP7 | nonsynonymous SNV | LARP7:NM_016648:exon7:c.T752A:p.M251K,LARP7:NM_001267039:exon9:c.T773A:p.M258K,LARP7:NM_015454:exon7:c.T752A:p.M251K, |
| chr4 | 112647305 | G | A | . | exonic | LARP7 | nonsynonymous SNV | LARP7:NM_016648:exon7:c.G753A:p.M251I,LARP7:NM_001267039:exon9:c.G774A:p.M258I,LARP7:NM_015454:exon7:c.G753A:p.M251I, |
| chr4 | 112647363 | C | A | COSV60520772 | exonic | LARP7 | nonsynonymous SNV | LARP7:NM_016648:exon7:c.C811A:p.Q271K,LARP7:NM_001267039:exon9:c.C832A:p.Q278K,LARP7:NM_015454:exon7:c.C811A:p.Q271K, |
| chr4 | 112647751 | G | C | . | exonic | LARP7 | nonsynonymous SNV | LARP7:NM_016648:exon8:c.G1059C:p.L353F,LARP7:NM_001267039:exon10:c.G1080C:p.L360F,LARP7:NM_015454:exon8:c.G1059C:p.L353F, |
| chr4 | 112647767 | C | A | . | exonic | LARP7 | nonsynonymous SNV | LARP7:NM_016648:exon8:c.C1075A:p.H359N,LARP7:NM_015454:exon8:c.C1075A:p.H359N,LARP7:NM_001267039:exon10:c.C1096A:p.H366N, |
| chr4 | 119320747 | T | C | COSV56788103 | exonic | FABP2 | nonsynonymous SNV | FABP2:NM_000134:exon2:c.A163G:p.T55A, |
| chr4 | 121816828 | G | A | . | exonic | EXOSC9 | nonsynonymous SNV | EXOSC9:NM_005033:exon12:c.G1292A:p.R431K,EXOSC9:NM_001034194:exon13:c.G1343A:p.R448K, |
| chr4 | 121821062 | T | C | . | exonic | CCNA2 | nonsynonymous SNV | CCNA2:NM_001237:exon3:c.A487G:p.I163V, |
| chr4 | 122917603 | A | G | COSV58645888 | exonic | NUDT6 | nonsynonymous SNV | NUDT6:NM_007083:exon2:c.T340C:p.C114R, |
| chr4 | 1336971 | T | C | COSV53262916 | exonic | MAEA | nonsynonymous SNV | MAEA:NM_001297431:exon5:c.T676C:p.W226R, |
| chr4 | 142402981 | A | T | . | exonic | INPP4B | nonsynonymous SNV | INPP4B:NM_003866:exon8:c.T329A:p.I110K,INPP4B:NM_001101669:exon7:c.T329A:p.I110K, |
| chr4 | 150277928 | G | A | COSV63949643 | exonic | LRBA | nonsynonymous SNV | LRBA:NM_006726:exon57:c.C8426T:p.S2809L,LRBA:NM_001199282:exon56:c.C8390T:p.S2797L, |
| chr4 | 150285975 | C | T | COSV63966388 | exonic | LRBA | nonsynonymous SNV | LRBA:NM_006726:exon55:c.G8110A:p.A2704T,LRBA:NM_001199282:exon54:c.G8074A:p.A2692T, |
| chr4 | 151099696 | A | G | . | exonic | RPS3A | nonsynonymous SNV | RPS3A:NM_001267699:exon1:c.A44G:p.K15R,RPS3A:NM_001006:exon1:c.A44G:p.K15R, |
| chr4 | 151104209 | A | G | . | exonic | RPS3A | nonsynonymous SNV | RPS3A:NM_001006:exon5:c.A596G:p.K199R, |
| chr4 | 15688610 | C | A | . | exonic | FAM200B | nonsynonymous SNV | FAM200B:NM_001145191:exon2:c.C1633A:p.H545N, |
| chr4 | 163514113 | A | C | COSV62187111 | exonic | TMA16 | nonsynonymous SNV | TMA16:NM_018352:exon4:c.A194C:p.Q65P, |
| chr4 | 163519429 | T | C | COSV56808231 | exonic | TMA16 | nonsynonymous SNV | TMA16:NM_018352:exon7:c.T527C:p.I176T, |
| chr4 | 169007691 | G | T | . | exonic | CBR4 | nonsynonymous SNV | CBR4:NM_032783:exon2:c.C208A:p.L70M, |
| chr4 | 169742084 | C | T | . | exonic | C4ORF27 | nonsynonymous SNV | C4orf27:NM_017867:exon5:c.G521A:p.R174K, |
| chr4 | 1728226 | G | A | COSV57603699 | exonic | TACC3 | nonsynonymous SNV | TACC3:NM_006342:exon4:c.G824A:p.C275Y, |
| chr4 | 1728261 | G | A | COSV57603712 | exonic | TACC3 | nonsynonymous SNV | TACC3:NM_006342:exon4:c.G859A:p.G287S, |
| chr4 | 1731251 | G | A | COSV57605388 | exonic | TACC3 | nonsynonymous SNV | TACC3:NM_006342:exon6:c.G1541A:p.G514E, |
| chr4 | 1740865 | T | A | . | exonic | TACC3 | nonsynonymous SNV | TACC3:NM_006342:exon13:c.T2102A:p.I701N, |
| chr4 | 184388978 | A | T | . | exonic | IRF2 | nonsynonymous SNV | IRF2:NM_002199:exon9:c.T830A:p.M277K, |
| chr4 | 184388982 | C | T | COSV66929604 | exonic | IRF2 | nonsynonymous SNV | IRF2:NM_002199:exon9:c.G826A:p.G276S, |
| chr4 | 184388984 | G | T | . | exonic | IRF2 | nonsynonymous SNV | IRF2:NM_002199:exon9:c.C824A:p.P275H, |
| chr4 | 184762452 | G | C | . | exonic | ACSL1 | nonsynonymous SNV | ACSL1:NM_001286710:exon17:c.C1593G:p.D531E,ACSL1:NM_001286712:exon18:c.C1080G:p.D360E,ACSL1:NM_001286711:exon16:c.C1491G:p.D497E,ACSL1:NM_001995:exon17:c.C1593G:p.D531E,ACSL1:NM_001286708:exon17:c.C1593G:p.D531E, |
| chr4 | 185190485 | T | C | COSV57111528 | exonic | CFAP97 | nonsynonymous SNV | CFAP97:NM_020827:exon2:c.A712G:p.T238A,CFAP97:NM_001292033:exon2:c.A712G:p.T238A, |
| chr4 | 185638960 | C | T | . | exonic | SORBS2 | nonsynonymous SNV | SORBS2:NM_001145671:exon12:c.G1018A:p.E340K,SORBS2:NM_001145673:exon12:c.G1297A:p.E433K, |
| chr4 | 186598052 | C | G | . | exonic | FAT1 | nonsynonymous SNV | FAT1:NM_005245:exon23:c.G12177C:p.K4059N, |
| chr4 | 186603866 | A | C | COSV71673371 | exonic | FAT1 | nonsynonymous SNV | FAT1:NM_005245:exon19:c.T10660G:p.S3554A, |
| chr4 | 186621601 | T | C | COSV71672102 | exonic | FAT1 | nonsynonymous SNV | FAT1:NM_005245:exon10:c.A4985G:p.N1662S, |
| chr4 | 186636739 | T | C | COSV71672989 | exonic | FAT1 | nonsynonymous SNV | FAT1:NM_005245:exon5:c.A3818G:p.H1273R, |
| chr4 | 186663440 | C | T | . | exonic | FAT1 | nonsynonymous SNV | FAT1:NM_005245:exon3:c.G3439A:p.E1147K, |
| chr4 | 186706638 | T | C | COSV71672014 | exonic | FAT1 | nonsynonymous SNV | FAT1:NM_005245:exon2:c.A3190G:p.R1064G, |
| chr4 | 186707244 | C | G | COSV71672016 | exonic | FAT1 | nonsynonymous SNV | FAT1:NM_005245:exon2:c.G2584C:p.V862L, |
| chr4 | 186707986 | G | C | COSV71672018 | exonic | FAT1 | nonsynonymous SNV | FAT1:NM_005245:exon2:c.C1842G:p.F614L, |
| chr4 | 186708384 | C | T | COSV71672959 | exonic | FAT1 | nonsynonymous SNV | FAT1:NM_005245:exon2:c.G1444A:p.V482I, |
| chr4 | 186708616 | A | C | COSV71672025 | exonic | FAT1 | nonsynonymous SNV | FAT1:NM_005245:exon2:c.T1212G:p.S404R, |
| chr4 | 186709443 | C | G | COSV71676532 | exonic | FAT1 | nonsynonymous SNV | FAT1:NM_005245:exon2:c.G385C:p.V129L, |

| | | | | | | | | |
|---|---|---|---|---|---|---|---|---|
| chr4 | 1935170 | C | A | . | exonic | WHSC1 | nonsynonymous SNV | WHSC1:NM_133335:exon7:c.C1582A:p.H528N,WHSC1:NM_007331:exon9:c.C1582A:p.H528N,WHSC1:NM_133330:exon9:c.C1582A:p.H528N,WHSC1:NM_001042424:exon7:c.C1582A:p.H528N,WHSC1:NM_133331:exon8:c.C1582A:p.H528N,WHSC1:NM_133334:exon6:c.C1582A:p.H528N, |
| chr4 | 2042401 | C | T | . | exonic | C4ORF48 | nonsynonymous SNV | C4orf48:NM_001168243:exon2:c.C149T:p.P50L,C4orf48:NM_001141936:exon2:c.C50T:p.P17L, |
| chr4 | 24542978 | C | T | . | exonic | DHX15 | nonsynonymous SNV | DHX15:NM_001358:exon7:c.G1297A:p.V433M, |
| chr4 | 24554823 | C | T | . | exonic | DHX15 | nonsynonymous SNV | DHX15:NM_001358:exon5:c.G982A:p.E328K, |
| chr4 | 24799693 | G | A | COSV66125434 | exonic | SOD3 | nonsynonymous SNV | SOD3:NM_003102:exon2:c.G172A:p.A58T, |
| chr4 | 40425993 | T | C | COSV55931179 | exonic | RBM47 | nonsynonymous SNV | RBM47:NM_019027:exon5:c.A1486G:p.M496V,RBM47:NM_001098634:exon7:c.A1693G:p.M565V, |
| chr4 | 41646183 | G | A | . | exonic | LIMCH1 | nonsynonymous SNV | LIMCH1:NM_001289122:exon10:c.G1123A:p.E375K,LIMCH1:NM_001112719:exon6:c.G697A:p.E233K,LIMCH1:NM_001112720:exon5:c.G661A:p.E221K,LIMCH1:NM_014988:exon11:c.G1159A:p.E387K,LIMCH1:NM_001289124:exon10:c.G646A:p.E216K,LIMCH1:NM_001112717:exon11:c.G1159A:p.E387K,LIMCH1:NM_001112718:exon11:c.G1159A:p.E387K, |
| chr4 | 41646750 | G | A | COSV99047572 | exonic | LIMCH1 | nonsynonymous SNV | LIMCH1:NM_001289122:exon11:c.G1486A:p.A496T,LIMCH1:NM_001112719:exon7:c.G1060A:p.A354T,LIMCH1:NM_001112720:exon6:c.G1024A:p.A342T,LIMCH1:NM_014988:exon12:c.G1522A:p.A508T,LIMCH1:NM_001289124:exon11:c.G1009A:p.A337T,LIMCH1:NM_001112717:exon12:c.G1522A:p.A508T,LIMCH1:NM_001112718:exon12:c.G1522A:p.A508T, |
| chr4 | 41671587 | T | C | COSV58284968 | exonic | LIMCH1 | nonsynonymous SNV | LIMCH1:NM_001289122:exon17:c.T2315C:p.M772T,LIMCH1:NM_001112719:exon12:c.T1814C:p.M605T,LIMCH1:NM_001112720:exon11:c.T1778C:p.M593T,LIMCH1:NM_014988:exon17:c.T2276C:p.M759T,LIMCH1:NM_001289124:exon17:c.T1838C:p.M613T,LIMCH1:NM_001112717:exon17:c.T2276C:p.M759T,LIMCH1:NM_001112718:exon17:c.T2276C:p.M759T, |
| chr4 | 42001654 | A | G | . | exonic | SLC30A9 | nonsynonymous SNV | SLC30A9:NM_006345:exon2:c.A148G:p.M50V, |
| chr4 | 42018125 | A | G | . | exonic | SLC30A9 | nonsynonymous SNV | SLC30A9:NM_006345:exon3:c.A289G:p.T97A, |
| chr4 | 4274597 | C | T | . | exonic | LYAR | nonsynonymous SNV | LYAR:NM_001145725:exon7:c.G602A:p.R201K,LYAR:NM_017816:exon7:c.G602A:p.R201K, |
| chr4 | 4274607 | T | C | . | exonic | LYAR | nonsynonymous SNV | LYAR:NM_001145725:exon7:c.A592G:p.K198E,LYAR:NM_017816:exon7:c.A592G:p.K198E, |
| chr4 | 4274649 | C | T | . | exonic | LYAR | nonsynonymous SNV | LYAR:NM_001145725:exon7:c.G550A:p.V184M,LYAR:NM_017816:exon7:c.G550A:p.V184M, |
| chr4 | 442278 | C | A | . | exonic | ZNF721 | nonsynonymous SNV | ZNF721:NM_133474:exon3:c.G2189T:p.R730L, |
| chr4 | 442869 | A | T | COSV59089336 | exonic | ZNF721 | nonsynonymous SNV | ZNF721:NM_133474:exon3:c.T1598A:p.V533E, |
| chr4 | 443038 | G | C | . | exonic | ZNF721 | nonsynonymous SNV | ZNF721:NM_133474:exon3:c.C1429G:p.Q477E, |
| chr4 | 443692 | A | T | . | exonic | ZNF721 | nonsynonymous SNV | ZNF721:NM_133474:exon3:c.T775A:p.S259T, |
| chr4 | 444042 | C | T | COSV59092947 | exonic | ZNF721 | nonsynonymous SNV | ZNF721:NM_133474:exon3:c.G425A:p.R142H, |
| chr4 | 48842657 | G | A | COSV51902955 | exonic | OCIAD1 | nonsynonymous SNV | OCIAD1:NM_001079840:exon4:c.G161A:p.S54N,OCIAD1:NM_001079841:exon4:c.G161A:p.S54N,OCIAD1:NM_001079842:exon4:c.G161A:p.S54N,OCIAD1:NM_001079839:exon4:c.G161A:p.S54N,OCIAD1:NM_001168254:exon4:c.G176A:p.S59N,OCIAD1:NM_017830:exon4:c.G161A:p.S54N, |
| chr4 | 48850049 | G | A | . | exonic | OCIAD1 | nonsynonymous SNV | OCIAD1:NM_001079839:exon6:c.G344A:p.R115Q,OCIAD1:NM_001079840:exon6:c.G344A:p.R115Q,OCIAD1:NM_001079841:exon6:c.G344A:p.R115Q,OCIAD1:NM_001079842:exon6:c.G344A:p.R115Q,OCIAD1:NM_001168254:exon6:c.G359A:p.R120Q,OCIAD1:NM_017830:exon6:c.G344A:p.R115Q, |
| chr4 | 48851847 | G | A | . | exonic | OCIAD1 | nonsynonymous SNV | OCIAD1:NM_001079839:exon7:c.G419A:p.G140D,OCIAD1:NM_001079840:exon7:c.G419A:p.G140D,OCIAD1:NM_001079841:exon7:c.G419A:p.G140D,OCIAD1:NM_001079842:exon7:c.G419A:p.G140D,OCIAD1:NM_001168254:exon7:c.G434A:p.G145D,OCIAD1:NM_017830:exon7:c.G419A:p.G140D, |
| chr4 | 48851876 | G | A | . | exonic | OCIAD1 | nonsynonymous SNV | OCIAD1:NM_001079839:exon7:c.G448A:p.A150T,OCIAD1:NM_001079840:exon7:c.G448A:p.A150T,OCIAD1:NM_001079841:exon7:c.G448A:p.A150T,OCIAD1:NM_001079842:exon7:c.G448A:p.A150T,OCIAD1:NM_001168254:exon7:c.G463A:p.A155T,OCIAD1:NM_017830:exon7:c.G448A:p.A150T, |
| chr4 | 48851942 | G | A | . | exonic | OCIAD1 | nonsynonymous SNV | OCIAD1:NM_001079839:exon7:c.G514A:p.A172T,OCIAD1:NM_001079840:exon7:c.G514A:p.A172T,OCIAD1:NM_001079841:exon7:c.G514A:p.A172T,OCIAD1:NM_001079842:exon7:c.G514A:p.A172T,OCIAD1:NM_001168254:exon7:c.G529A:p.A177T,OCIAD1:NM_017830:exon7:c.G514A:p.A172T, |
| chr4 | 48860745 | G | A | . | exonic | OCIAD1 | nonsynonymous SNV | OCIAD1:NM_001079839:exon9:c.G721A:p.D241N,OCIAD1:NM_001079840:exon8:c.G568A:p.D190N,OCIAD1:NM_001079842:exon8:c.G568A:p.D190N,OCIAD1:NM_001168254:exon9:c.G736A:p.D246N,OCIAD1:NM_017830:exon9:c.G721A:p.D241N, |
| chr4 | 48986433 | C | A | COSV56939573 | exonic | CWH43 | nonsynonymous SNV | CWH43:NM_025087:exon1:c.C4A:p.P2T, |
| chr4 | 49061855 | C | A | COSV56936745 | exonic | CWH43 | nonsynonymous SNV | CWH43:NM_001286791:exon16:c.C1984A:p.H662N,CWH43:NM_025087:exon16:c.C2065A:p.H689N, |
| chr4 | 53453034 | G | A | . | exonic | FIP1L1 | nonsynonymous SNV | FIP1L1:NM_030917:exon16:c.G1400A:p.R467K,FIP1L1:NM_001134938:exon13:c.G1178A:p.R393K,FIP1L1:NM_001134937:exon16:c.G1382A:p.R461K, |
| chr4 | 53458720 | G | A | . | exonic | FIP1L1 | nonsynonymous SNV | FIP1L1:NM_030917:exon17:c.G1567A:p.E523K,FIP1L1:NM_001134938:exon14:c.G1345A:p.E449K,FIP1L1:NM_001134937:exon17:c.G1549A:p.E517K, |
| chr4 | 53459366 | T | A | . | exonic | FIP1L1 | nonsynonymous SNV | FIP1L1:NM_030917:exon18:c.T1702A:p.S568T,FIP1L1:NM_001134938:exon15:c.T1480A:p.S494T,FIP1L1:NM_001134937:exon18:c.T1684A:p.S562T, |
| chr4 | 53459377 | C | A | . | exonic | FIP1L1 | nonsynonymous SNV | FIP1L1:NM_030917:exon18:c.C1713A:p.S571R,FIP1L1:NM_001134938:exon15:c.C1491A:p.S497R,FIP1L1:NM_001134937:exon18:c.C1695A:p.S565R, |
| chr4 | 56441726 | G | A | . | exonic | PAICS | nonsynonymous SNV | PAICS:NM_001079525:exon2:c.G80A:p.S27N,PAICS:NM_006452:exon3:c.G80A:p.S27N,PAICS:NM_001079524:exon2:c.G80A:p.S27N, |
| chr4 | 56441810 | C | A | . | exonic | PAICS | nonsynonymous SNV | PAICS:NM_001079524:exon2:c.C164A:p.A55E,PAICS:NM_001079525:exon2:c.C164A:p.A55E,PAICS:NM_006452:exon3:c.C164A:p.A55E, |
| chr4 | 56446800 | C | T | . | exonic | PAICS | nonsynonymous SNV | PAICS:NM_001079524:exon3:c.C320T:p.S107F,PAICS:NM_001079525:exon4:c.C341T:p.S114F,PAICS:NM_006452:exon4:c.C320T:p.S107F, |
| chr4 | 56448443 | G | A | . | exonic | PAICS | stopgain | PAICS:NM_001079525:exon5:c.G440A:p.W147X,PAICS:NM_006452:exon5:c.G419A:p.W140X,PAICS:NM_001079524:exon4:c.G419A:p.W140X, |
| chr4 | 56448739 | A | C | . | exonic | PAICS | nonsynonymous SNV | PAICS:NM_001079525:exon6:c.A624C:p.K208N,PAICS:NM_006452:exon6:c.A603C:p.K201N,PAICS:NM_001079524:exon5:c.A603C:p.K201N, |
| chr4 | 56448764 | G | A | . | exonic | PAICS | nonsynonymous SNV | PAICS:NM_001079525:exon6:c.G649A:p.D217N,PAICS:NM_006452:exon6:c.G628A:p.D210N,PAICS:NM_001079524:exon5:c.G628A:p.D210N, |
| chr4 | 56450664 | G | A | . | exonic | PAICS | nonsynonymous SNV | PAICS:NM_001079525:exon7:c.G754A:p.V252I,PAICS:NM_006452:exon7:c.G733A:p.V245I,PAICS:NM_001079524:exon6:c.G733A:p.V245I, |
| chr4 | 56451918 | G | A | . | exonic | PAICS | nonsynonymous SNV | PAICS:NM_001079524:exon7:c.G818A:p.G273D,PAICS:NM_001079525:exon8:c.G839A:p.G280D,PAICS:NM_006452:exon8:c.G818A:p.G273D, |
| chr4 | 56650730 | G | A | COSV58499621 | exonic | HOPX | nonsynonymous SNV | HOPX:NM_001145460:exon4:c.C281T:p.P94L, |
| chr4 | 57023510 | G | A | . | exonic | POLR2B | nonsynonymous SNV | POLR2B:NM_001303269:exon20:c.G2675A:p.S892N,POLR2B:NM_000938:exon19:c.G2696A:p.S899N,POLR2B:NM_001303268:exon18:c.G2471A:p.S824N, |
| chr4 | 57023518 | G | A | . | exonic | POLR2B | nonsynonymous SNV | POLR2B:NM_001303269:exon20:c.G2683A:p.G895S,POLR2B:NM_000938:exon19:c.G2704A:p.G902S,POLR2B:NM_001303268:exon18:c.G2479A:p.G827S, |
| chr4 | 57110321 | G | A | COSV55277369 | exonic | IGFBP7 | nonsynonymous SNV | IGFBP7:NM_001553:exon1:c.C31T:p.L11F,IGFBP7:NM_001253835:exon1:c.C31T:p.L11F, |
| chr4 | 6301627 | G | A | COSV56987772 | exonic | WFS1 | nonsynonymous SNV | WFS1:NM_001145853:exon8:c.G1832A:p.R611H,WFS1:NM_006005:exon8:c.G1832A:p.R611H, |
| chr4 | 70839545 | A | G | . | exonic | GRSF1 | nonsynonymous SNV | GRSF1:NM_002092:exon1:c.T283C:p.S95P, |
| chr4 | 76132681 | T | C | . | exonic | NUP54 | nonsynonymous SNV | NUP54:NM_017426:exon6:c.A749G:p.N250S,NUP54:NM_001278603:exon5:c.A605G:p.N202S, |
| chr4 | 78597384 | G | A | . | exonic | ANXA3 | nonsynonymous SNV | ANXA3:NM_005139:exon10:c.G700A:p.G234R, |
| chr4 | 87482427 | A | T | . | exonic | SPARCL1 | nonsynonymous SNV | SPARCL1:NM_001291977:exon6:c.T1290A:p.N430K,SPARCL1:NM_001291976:exon9:c.T1290A:p.N430K,SPARCL1:NM_001128310:exon9:c.T1665A:p.N555K,SPARCL1:NM_004684:exon8:c.T1665A:p.N555K, |
| chr4 | 87491654 | T | C | COSV56803313 | exonic | SPARCL1 | nonsynonymous SNV | SPARCL1:NM_001291977:exon3:c.A880G:p.T294A,SPARCL1:NM_001291976:exon6:c.A880G:p.T294A,SPARCL1:NM_001128310:exon6:c.A1255G:p.T419A,SPARCL1:NM_004684:exon5:c.A1255G:p.T419A, |
| chr4 | 87494149 | G | C | . | exonic | SPARCL1 | nonsynonymous SNV | SPARCL1:NM_001291977:exon2:c.C276G:p.N92K,SPARCL1:NM_001291976:exon5:c.C276G:p.N92K,SPARCL1:NM_001128310:exon5:c.C651G:p.N217K,SPARCL1:NM_004684:exon4:c.C651G:p.N217K, |
| chr4 | 87494337 | C | T | . | exonic | SPARCL1 | nonsynonymous SNV | SPARCL1:NM_001291977:exon2:c.G88A:p.E30K,SPARCL1:NM_001291976:exon5:c.G88A:p.E30K,SPARCL1:NM_001128310:exon5:c.G463A:p.E155K,SPARCL1:NM_004684:exon4:c.G463A:p.E155K, |
| chr4 | 87494484 | G | C | COSV56802379 | exonic | SPARCL1 | nonsynonymous SNV | SPARCL1:NM_001128310:exon5:c.C316G:p.H106D,SPARCL1:NM_004684:exon4:c.C316G:p.H106D, |
| chr4 | 87495034 | C | T | . | exonic | SPARCL1 | nonsynonymous SNV | SPARCL1:NM_001128310:exon4:c.G148A:p.E50K,SPARCL1:NM_004684:exon3:c.G148A:p.E50K, |
| chr4 | 87495036 | G | T | COSV56802387 | exonic | SPARCL1 | nonsynonymous SNV | SPARCL1:NM_001128310:exon4:c.C146A:p.A49D,SPARCL1:NM_004684:exon3:c.C146A:p.A49D, |

| | | | | | | | | |
|---|---|---|---|---|---|---|---|---|
| chr4 | 90468303 | T | A | . | exonic | CCSER1 | nonsynonymous SNV | CCSER1:NM_207491:exon5:c.T1673A:p.M558K,CCSER1:NM_001145065:exon5:c.T1673A:p.M558K, |
| chr4 | 98379034 | G | A | . | exonic | RAP1GDS1 | nonsynonymous SNV | RAP1GDS1:NM_021159:exon5:c.G382A:p.V128I,RAP1GDS1:NM_001100427:exon5:c.G379A:p.V127I,RAP1GDS1:NM_001100426:exon5:c.G382A:p.V128I, |
| chr4 | 99318162 | T | C | COSV59295640 | exonic | ADH1B | nonsynonymous SNV | ADH1B:NM_000668:exon3:c.A143G:p.H48R,ADH1B:NM_001286650:exon4:c.A23G:p.H8R, |
| chr4 | 99339632 | T | C | COSV72624794 | exonic | ADH1C | nonsynonymous SNV | ADH1C:NM_000669:exon8:c.A1048G:p.I350V, |
| chr4 | 99342808 | C | T | COSV72463409 | exonic | ADH1C | nonsynonymous SNV | ADH1C:NM_000669:exon6:c.G815A:p.R272Q, |
| chr5 | 10256060 | A | T | . | exonic | CCT5 | nonsynonymous SNV | CCT5:NM_001306155:exon3:c.A158T:p.E53V,CCT5:NM_001306153:exon4:c.A374T:p.E125V,CCT5:NM_012073:exon4:c.A437T:p.E146V,CCT5:NM_001306156:exon4:c.A323T:p.E108V,CCT5:NM_001306154:exon3:c.A272T:p.E91V, |
| chr5 | 103276233 | G | A | . | exonic | C5ORF30 | nonsynonymous SNV | C5orf30:NM_001316969:exon3:c.G314A:p.R105H,C5orf30:NM_001316968:exon3:c.G314A:p.R105H,C5orf30:NM_033211:exon3:c.G314A:p.R105H, |
| chr5 | 10680885 | G | A | COSV50143482 | exonic | DAP | nonsynonymous SNV | DAP:NM_001291963:exon3:c.C437T:p.S146F, |
| chr5 | 115870087 | G | A | . | exonic | AP3S1 | nonsynonymous SNV | AP3S1:NM_001284:exon3:c.G232A:p.D78N, |
| chr5 | 115902914 | G | A | . | exonic | AP3S1 | nonsynonymous SNV | AP3S1:NM_001284:exon5:c.G375A:p.M125I, |
| chr5 | 116092684 | T | G | COSV57234444 | exonic | COMMD10 | nonsynonymous SNV | COMMD10:NM_016144:exon4:c.T383G:p.I128S,COMMD10:NM_001308080:exon4:c.T341G:p.I114S, |
| chr5 | 116134084 | G | A | . | exonic | COMMD10 | stopgain | COMMD10:NM_016144:exon5:c.G416A:p.W139X,COMMD10:NM_001308080:exon5:c.G374A:p.W125X, |
| chr5 | 116134143 | G | A | . | exonic | COMMD10 | nonsynonymous SNV | COMMD10:NM_016144:exon5:c.G475A:p.V159M,COMMD10:NM_001308080:exon5:c.G433A:p.V145M, |
| chr5 | 123023945 | T | C | COSV60574680 | exonic | PPIC | nonsynonymous SNV | PPIC:NM_000943:exon5:c.A569G:p.N190S, |
| chr5 | 132209275 | C | T | . | exonic | P4HA2 | nonsynonymous SNV | P4HA2:NM_001017974:exon7:c.G766A:p.E256K,P4HA2:NM_001142599:exon8:c.G766A:p.E256K,P4HA2:NM_004199:exon7:c.G766A:p.E256K,P4HA2:NM_001142598:exon8:c.G766A:p.E256K,P4HA2:NM_001017973:exon7:c.G766A:p.E256K, |
| chr5 | 132214053 | C | A | . | exonic | P4HA2 | nonsynonymous SNV | P4HA2:NM_001017974:exon5:c.G332T:p.G111V,P4HA2:NM_001142599:exon6:c.G332T:p.G111V,P4HA2:NM_004199:exon5:c.G332T:p.G111V,P4HA2:NM_001142598:exon6:c.G332T:p.G111V,P4HA2:NM_001017973:exon5:c.G332T:p.G111V, |
| chr5 | 132218548 | T | G | . | exonic | P4HA2 | nonsynonymous SNV | P4HA2:NM_001017974:exon2:c.A79C:p.I27L,P4HA2:NM_001142599:exon3:c.A79C:p.I27L,P4HA2:NM_004199:exon2:c.A79C:p.I27L,P4HA2:NM_001142598:exon3:c.A79C:p.I27L,P4HA2:NM_001017973:exon2:c.A79C:p.I27L, |
| chr5 | 132747766 | G | C | . | exonic | CCNI2 | nonsynonymous SNV | CCNI2:NM_001287252:exon1:c.G271C:p.A91P,CCNI2:NM_001287253:exon1:c.G271C:p.A91P,CCNI2:NM_001039780:exon1:c.G271C:p.A91P, |
| chr5 | 133225776 | C | A | COSV54773531 | exonic | FSTL4 | nonsynonymous SNV | FSTL4:NM_015082:exon9:c.G1059T:p.E353D, |
| chr5 | 135335037 | A | G | . | exonic | H2AFY | nonsynonymous SNV | H2AFY:NM_004893:exon9:c.T1055C:p.L352P,H2AFY:NM_001040158:exon9:c.T1055C:p.L352P,H2AFY:NM_138610:exon9:c.T1058C:p.L353P,H2AFY:NM_138609:exon9:c.T1049C:p.L350P, |
| chr5 | 135335098 | C | T | . | exonic | H2AFY | nonsynonymous SNV | H2AFY:NM_004893:exon9:c.G994A:p.A332T,H2AFY:NM_001040158:exon9:c.G994A:p.A332T,H2AFY:NM_138610:exon9:c.G997A:p.A333T,H2AFY:NM_138609:exon9:c.G988A:p.A330T, |
| chr5 | 135360597 | C | T | . | exonic | H2AFY | nonsynonymous SNV | H2AFY:NM_004893:exon5:c.G485A:p.G162D,H2AFY:NM_001040158:exon5:c.G485A:p.G162D,H2AFY:NM_138610:exon5:c.G488A:p.G163D,H2AFY:NM_138609:exon5:c.G488A:p.G163D, |
| chr5 | 135388952 | A | G | . | exonic | H2AFY | nonsynonymous SNV | H2AFY:NM_004893:exon2:c.T142C:p.Y48H,H2AFY:NM_001040158:exon2:c.T142C:p.Y48H,H2AFY:NM_138610:exon2:c.T142C:p.Y48H,H2AFY:NM_138609:exon2:c.T142C:p.Y48H, |
| chr5 | 139478334 | T | C | . | exonic | TMEM173 | nonsynonymous SNV | TMEM173:NM_198282:exon6:c.A695G:p.H232R,TMEM173:NM_001301738:exon5:c.A695G:p.H232R, |
| chr5 | 140557175 | G | T | COSV60365268 | exonic | SRA1 | stopgain | SRA1:NM_001253764:exon1:c.C278A:p.S93X, |
| chr5 | 140668682 | T | C | . | exonic | WDR55 | nonsynonymous SNV | WDR55:NM_017706:exon4:c.T451C:p.C151R, |
| chr5 | 14581949 | C | G | COSV57036647 | exonic | FAM105A | nonsynonymous SNV | FAM105A:NM_019018:exon1:c.C55G:p.P19A, |
| chr5 | 146013801 | C | T | COSV63039708 | exonic | SH3RF2 | nonsynonymous SNV | SH3RF2:NM_152550:exon5:c.C799T:p.R267C, |
| chr5 | 146060084 | T | C | COSV63040514 | exonic | SH3RF2 | nonsynonymous SNV | SH3RF2:NM_152550:exon9:c.T1774C:p.W592R, |
| chr5 | 146062640 | G | C | COSV63040479 | exonic | SH3RF2 | nonsynonymous SNV | SH3RF2:NM_152550:exon10:c.G2129C:p.G710A, |
| chr5 | 14610200 | C | G | COSV57035619 | exonic | FAM105A | nonsynonymous SNV | FAM105A:NM_019018:exon8:c.C957G:p.F319L, |
| chr5 | 146120433 | C | T | COSV50972264 | exonic | LARS | nonsynonymous SNV | LARS:NM_020117:exon31:c.G3263A:p.R1088K, |
| chr5 | 146129073 | T | C | COSV50973199 | exonic | LARS | nonsynonymous SNV | LARS:NM_020117:exon26:c.A2674G:p.N892D, |
| chr5 | 147906491 | G | A | COSV59153993 | exonic | C5ORF46 | nonsynonymous SNV | C5orf46:NM_206966:exon1:c.C11T:p.S4L, |
| chr5 | 150396669 | C | T | COSV60347011 | exonic | TCOF1 | nonsynonymous SNV | TCOF1:NM_000356:exon23:c.C3938T:p.A1313V,TCOF1:NM_001135243:exon24:c.C4169T:p.A1390V,TCOF1:NM_001195141:exon23:c.C4055T:p.A1352V,TCOF1:NM_001135244:exon23:c.C4058T:p.A1353V,TCOF1:NM_001135245:exon23:c.C3941T:p.A1314V, |
| chr5 | 150404692 | C | T | . | exonic | CD74 | nonsynonymous SNV | CD74:NM_004355:exon6:c.G613A:p.A205T,CD74:NM_001025159:exon6:c.G613A:p.A205T, |
| chr5 | 150405089 | C | T | . | exonic | CD74 | stopgain | CD74:NM_004355:exon5:c.G533A:p.W178X,CD74:NM_001025159:exon5:c.G533A:p.W178X, |
| chr5 | 150406271 | C | T | . | exonic | CD74 | nonsynonymous SNV | CD74:NM_004355:exon4:c.G429A:p.M143I,CD74:NM_001025158:exon4:c.G429A:p.M143I,CD74:NM_001025159:exon4:c.G429A:p.M143I, |
| chr5 | 150406277 | A | T | . | exonic | CD74 | nonsynonymous SNV | CD74:NM_004355:exon4:c.T423A:p.H141Q,CD74:NM_001025159:exon4:c.T423A:p.H141Q,CD74:NM_001025158:exon4:c.T423A:p.H141Q, |
| chr5 | 150406296 | C | T | . | exonic | CD74 | nonsynonymous SNV | CD74:NM_004355:exon4:c.G404A:p.G135D,CD74:NM_001025158:exon4:c.G404A:p.G135D,CD74:NM_001025159:exon4:c.G404A:p.G135D, |
| chr5 | 150406931 | C | T | . | exonic | CD74 | nonsynonymous SNV | CD74:NM_004355:exon3:c.G328A:p.A110T,CD74:NM_001025158:exon3:c.G328A:p.A110T,CD74:NM_001025159:exon3:c.G328A:p.A110T, |
| chr5 | 150406957 | G | T | . | exonic | CD74 | nonsynonymous SNV | CD74:NM_004355:exon3:c.C302A:p.P101H,CD74:NM_001025158:exon3:c.C302A:p.P101H,CD74:NM_001025159:exon3:c.C302A:p.P101H, |
| chr5 | 150407199 | G | A | . | exonic | CD74 | nonsynonymous SNV | CD74:NM_004355:exon2:c.C251T:p.T84I,CD74:NM_001025158:exon2:c.C251T:p.T84I,CD74:NM_001025159:exon2:c.C251T:p.T84I, |
| chr5 | 150407308 | C | T | . | exonic | CD74 | nonsynonymous SNV | CD74:NM_004355:exon2:c.G142A:p.A48T,CD74:NM_001025158:exon2:c.G142A:p.A48T,CD74:NM_001025159:exon2:c.G142A:p.A48T, |
| chr5 | 150412635 | C | G | . | exonic | CD74 | nonsynonymous SNV | CD74:NM_004355:exon1:c.G115C:p.A39P,CD74:NM_001025158:exon1:c.G115C:p.A39P,CD74:NM_001025159:exon1:c.G115C:p.A39P, |
| chr5 | 150412686 | C | T | COSV50532067 | exonic | CD74 | nonsynonymous SNV | CD74:NM_004355:exon1:c.G64A:p.D22N,CD74:NM_001025158:exon1:c.G64A:p.D22N,CD74:NM_001025159:exon1:c.G64A:p.D22N, |
| chr5 | 150412727 | C | T | . | exonic | CD74 | nonsynonymous SNV | CD74:NM_004355:exon1:c.G23A:p.S8N,CD74:NM_001025159:exon1:c.G23A:p.S8N,CD74:NM_001025158:exon1:c.G23A:p.S8N, |
| chr5 | 154002288 | C | T | COSV61078046 | exonic | FAM114A2 | nonsynonymous SNV | FAM114A2:NM_018691:exon11:c.G1219A:p.E407K, |

| | | | | | | | | |
|---|---|---|---|---|---|---|---|---|
| chr5 | 154002906 | C | T | . | exonic | FAM114A2 | nonsynonymous SNV | FAM114A2:NM_018691:exon10:c.G1057A:p.E353K, |
| chr5 | 154002910 | A | T | . | exonic | FAM114A2 | nonsynonymous SNV | FAM114A2:NM_018691:exon10:c.T1053A:p.N351K, |
| chr5 | 154033830 | C | T | COSV61077353 | exonic | FAM114A2 | nonsynonymous SNV | FAM114A2:NM_018691:exon4:c.G364A:p.G122S, |
| chr5 | 160232266 | C | T | COSV67437401 | exonic | FABP6 | nonsynonymous SNV | FABP6:NM_001130958:exon5:c.C383T:p.T128M,FABP6:NM_001445:exon2:c.C236T:p.T79M,FABP6:NM_001040442:exon4:c.C383T:p.T128M, |
| chr5 | 16477692 | C | T | . | exonic | FAM134B | nonsynonymous SNV | FAM134B:NM_019000:exon6:c.G547A:p.G183R,FAM134B:NM_001034850:exon8:c.G970A:p.G324R, |
| chr5 | 171392799 | G | A | . | exonic | NPM1 | nonsynonymous SNV | NPM1:NM_001037738:exon5:c.G442A:p.G148S,NPM1:NM_199185:exon5:c.G442A:p.G148S,NPM1:NM_002520:exon5:c.G442A:p.G148S, |
| chr5 | 171400846 | G | A | . | exonic | NPM1 | nonsynonymous SNV | NPM1:NM_001037738:exon8:c.G590A:p.R197Q,NPM1:NM_002520:exon8:c.G590A:p.R197Q, |
| chr5 | 171405303 | G | A | . | exonic | NPM1 | nonsynonymous SNV | NPM1:NM_001037738:exon9:c.G671A:p.G224E,NPM1:NM_199185:exon8:c.G584A:p.G195E,NPM1:NM_002520:exon9:c.G671A:p.G224E, |
| chr5 | 171405389 | G | A | . | exonic | NPM1 | nonsynonymous SNV | NPM1:NM_001037738:exon9:c.G757A:p.A253T,NPM1:NM_199185:exon8:c.G670A:p.A224T,NPM1:NM_002520:exon9:c.G757A:p.A253T, |
| chr5 | 171407742 | G | A | . | exonic | NPM1 | nonsynonymous SNV | NPM1:NM_199185:exon9:c.G727A:p.V243M,NPM1:NM_002520:exon10:c.G814A:p.V272M, |
| chr5 | 171407758 | G | A | COSV51564036 | exonic | NPM1 | nonsynonymous SNV | NPM1:NM_199185:exon9:c.G743A:p.R248Q,NPM1:NM_002520:exon10:c.G830A:p.R277Q, |
| chr5 | 171407762 | G | A | . | exonic | NPM1 | nonsynonymous SNV | NPM1:NM_199185:exon9:c.G747A:p.M249I,NPM1:NM_002520:exon10:c.G834A:p.M278I, |
| chr5 | 172914743 | G | A | . | exonic | ERGIC1 | nonsynonymous SNV | ERGIC1:NM_001031711:exon5:c.G280A:p.G94S, |
| chr5 | 174729165 | T | C | COSV53327173 | exonic | MSX2 | nonsynonymous SNV | MSX2:NM_002449:exon2:c.T386C:p.M129T, |
| chr5 | 177592261 | C | G | . | exonic | TMED9 | nonsynonymous SNV | TMED9:NM_017510:exon1:c.C47G:p.T16S, |
| chr5 | 179617063 | C | T | . | exonic | HNRNPH1 | nonsynonymous SNV | HNRNPH1:NM_001257293:exon10:c.G1105A:p.G369S,HNRNPH1:NM_005520:exon10:c.G1105A:p.G369S, |
| chr5 | 179822979 | C | T | . | exonic | SQSTM1 | nonsynonymous SNV | SQSTM1:NM_003900:exon2:c.C227T:p.A76V, |
| chr5 | 179823007 | G | A | . | exonic | SQSTM1 | nonsynonymous SNV | SQSTM1:NM_001142299:exon3:c.G3A:p.M1I,SQSTM1:NM_003900:exon2:c.G255A:p.M85I,SQSTM1:NM_001142298:exon3:c.G3A:p.M1I, |
| chr5 | 179823014 | T | C | COSV62434739 | exonic | SQSTM1 | nonsynonymous SNV | SQSTM1:NM_001142299:exon3:c.T10C:p.S4P,SQSTM1:NM_003900:exon2:c.T262C:p.S88P,SQSTM1:NM_001142298:exon3:c.T10C:p.S4P, |
| chr5 | 179824055 | G | A | COSV62435663 | exonic | SQSTM1 | nonsynonymous SNV | SQSTM1:NM_001142299:exon4:c.G247A:p.A83T,SQSTM1:NM_003900:exon3:c.G499A:p.A167T,SQSTM1:NM_001142298:exon4:c.G247A:p.A83T, |
| chr5 | 179824203 | C | T | . | exonic | SQSTM1 | nonsynonymous SNV | SQSTM1:NM_003900:exon4:c.C553T:p.L185F,SQSTM1:NM_001142299:exon5:c.C301T:p.L101F,SQSTM1:NM_001142298:exon5:c.C301T:p.L101F, |
| chr5 | 179824227 | T | A | . | exonic | SQSTM1 | nonsynonymous SNV | SQSTM1:NM_003900:exon4:c.T577A:p.F193I,SQSTM1:NM_001142299:exon5:c.T325A:p.F109I,SQSTM1:NM_001142298:exon5:c.T325A:p.F109I, |
| chr5 | 179824227 | T | C | . | exonic | SQSTM1 | nonsynonymous SNV | SQSTM1:NM_001142299:exon5:c.T325C:p.F109L,SQSTM1:NM_003900:exon4:c.T577C:p.F193L,SQSTM1:NM_001142298:exon5:c.T325C:p.F109L, |
| chr5 | 179824230 | G | A | . | exonic | SQSTM1 | nonsynonymous SNV | SQSTM1:NM_001142299:exon5:c.G328A:p.G110R,SQSTM1:NM_003900:exon4:c.G580A:p.G194R,SQSTM1:NM_001142298:exon5:c.G328A:p.G110R, |
| chr5 | 179824244 | G | A | . | exonic | SQSTM1 | stopgain | SQSTM1:NM_001142299:exon5:c.G342A:p.W114X,SQSTM1:NM_003900:exon4:c.G594A:p.W198X,SQSTM1:NM_001142298:exon5:c.G342A:p.W114X, |
| chr5 | 179824245 | G | A | . | exonic | SQSTM1 | nonsynonymous SNV | SQSTM1:NM_001142299:exon5:c.G343A:p.E115K,SQSTM1:NM_003900:exon4:c.G595A:p.E199K,SQSTM1:NM_001142298:exon5:c.G343A:p.E115K, |
| chr5 | 179825191 | T | C | . | exonic | SQSTM1 | nonsynonymous SNV | SQSTM1:NM_001142299:exon6:c.T467C:p.V156A,SQSTM1:NM_003900:exon5:c.T719C:p.V240A,SQSTM1:NM_001142298:exon6:c.T467C:p.V156A, |
| chr5 | 179833077 | G | A | . | exonic | SQSTM1 | nonsynonymous SNV | SQSTM1:NM_003900:exon6:c.G800A:p.R267H,SQSTM1:NM_001142299:exon7:c.G548A:p.R183H,SQSTM1:NM_001142298:exon7:c.G548A:p.R183H, |
| chr5 | 179833099 | G | C | COSV52975046 | exonic | SQSTM1 | nonsynonymous SNV | SQSTM1:NM_001142299:exon7:c.G570C:p.E190D,SQSTM1:NM_003900:exon6:c.G822C:p.E274D,SQSTM1:NM_001142298:exon7:c.G570C:p.E190D, |
| chr5 | 179833169 | G | A | . | exonic | SQSTM1 | nonsynonymous SNV | SQSTM1:NM_001142299:exon7:c.G640A:p.G214R,SQSTM1:NM_003900:exon6:c.G892A:p.G298R,SQSTM1:NM_001142298:exon7:c.G640A:p.G214R, |
| chr5 | 179833722 | C | A | . | exonic | SQSTM1 | nonsynonymous SNV | SQSTM1:NM_001142299:exon8:c.C853A:p.P285T,SQSTM1:NM_003900:exon7:c.C1105A:p.P369T,SQSTM1:NM_001142298:exon8:c.C853A:p.P285T, |
| chr5 | 179836463 | T | C | . | exonic | SQSTM1 | nonsynonymous SNV | SQSTM1:NM_001142299:exon9:c.T941C:p.L314P,SQSTM1:NM_003900:exon8:c.T1193C:p.L398P,SQSTM1:NM_001142298:exon9:c.T941C:p.L314P, |
| chr5 | 179836477 | T | C | . | exonic | SQSTM1 | nonsynonymous SNV | SQSTM1:NM_001142299:exon9:c.T955C:p.S319P,SQSTM1:NM_003900:exon8:c.T1207C:p.S403P,SQSTM1:NM_001142298:exon9:c.T955C:p.S319P, |
| chr5 | 179836492 | G | A | . | exonic | SQSTM1 | nonsynonymous SNV | SQSTM1:NM_001142299:exon9:c.G970A:p.D324N,SQSTM1:NM_003900:exon8:c.G1222A:p.D408N,SQSTM1:NM_001142298:exon9:c.G970A:p.D324N, |
| chr5 | 179836502 | G | A | . | exonic | SQSTM1 | nonsynonymous SNV | SQSTM1:NM_001142299:exon9:c.G980A:p.G327D,SQSTM1:NM_003900:exon8:c.G1232A:p.G411D,SQSTM1:NM_001142298:exon9:c.G980A:p.G327D, |
| chr5 | 179836555 | G | A | . | exonic | SQSTM1 | nonsynonymous SNV | SQSTM1:NM_001142299:exon9:c.G1033A:p.D345N,SQSTM1:NM_003900:exon8:c.G1285A:p.D429N,SQSTM1:NM_001142298:exon9:c.G1033A:p.D345N, |
| chr5 | 179837731 | T | C | COSV52973228 | exonic | C5ORF45 | nonsynonymous SNV | C5orf45:NM_016175:exon7:c.A692G:p.Q231R,C5orf45:NM_001017987:exon5:c.A527G:p.Q176R, |
| chr5 | 180911368 | A | G | COSV51457265 | exonic | BTNL8 | nonsynonymous SNV | BTNL8:NM_024850:exon3:c.A427G:p.T143A,BTNL8:NM_001159709:exon3:c.A52G:p.T18A,BTNL8:NM_001040462:exon3:c.A427G:p.T143A,BTNL8:NM_001159707:exon2:c.A79G:p.T27A,BTNL8:NM_001159708:exon3:c.A427G:p.T143A, |
| chr5 | 180949972 | C | G | COSV51457334 | exonic | BTNL8 | nonsynonymous SNV | BTNL8:NM_001159710:exon7:c.C379G:p.H127D,BTNL8:NM_001159709:exon8:c.C556G:p.H186D,BTNL8:NM_001040462:exon8:c.C931G:p.H311D,BTNL8:NM_001159707:exon7:c.C583G:p.H195D, |
| chr5 | 180949974 | T | G | COSV51460539 | exonic | BTNL8 | nonsynonymous SNV | BTNL8:NM_001159710:exon7:c.T381G:p.H127Q,BTNL8:NM_001159709:exon8:c.T558G:p.H186Q,BTNL8:NM_001040462:exon8:c.T933G:p.H311Q,BTNL8:NM_001159707:exon7:c.T585G:p.H195Q, |
| chr5 | 181237624 | C | T | . | exonic | GNB2L1 | stopgain | GNB2L1:NM_006098:exon7:c.G873A:p.W291X, |
| chr5 | 181237656 | C | T | . | exonic | GNB2L1 | nonsynonymous SNV | GNB2L1:NM_006098:exon7:c.G841A:p.A281T, |
| chr5 | 181237669 | A | C | . | exonic | GNB2L1 | nonsynonymous SNV | GNB2L1:NM_006098:exon7:c.T828G:p.S276R, |
| chr5 | 181238118 | C | G | . | exonic | GNB2L1 | nonsynonymous SNV | GNB2L1:NM_006098:exon6:c.G758C:p.G253A, |
| chr5 | 181239134 | C | T | . | exonic | GNB2L1 | nonsynonymous SNV | GNB2L1:NM_006098:exon5:c.G569A:p.G190D, |
| chr5 | 181239158 | C | T | . | exonic | GNB2L1 | nonsynonymous SNV | GNB2L1:NM_006098:exon5:c.G545A:p.C182Y, |
| chr5 | 181241551 | A | G | . | exonic | GNB2L1 | nonsynonymous SNV | GNB2L1:NM_006098:exon3:c.T370C:p.S124P, |
| chr5 | 181242185 | C | T | . | exonic | GNB2L1 | stopgain | GNB2L1:NM_006098:exon2:c.G270A:p.W90X, |
| chr5 | 181242262 | A | G | . | exonic | GNB2L1 | nonsynonymous SNV | GNB2L1:NM_006098:exon2:c.T193C:p.F65L, |
| chr5 | 181242273 | C | T | . | exonic | GNB2L1 | nonsynonymous SNV | GNB2L1:NM_006098:exon2:c.G182A:p.G61D, |
| chr5 | 181243763 | C | T | . | exonic | GNB2L1 | nonsynonymous SNV | GNB2L1:NM_006098:exon1:c.G38A:p.G13D, |
| chr5 | 181243776 | C | T | . | exonic | GNB2L1 | nonsynonymous SNV | GNB2L1:NM_006098:exon1:c.G25A:p.G9S, |
| chr5 | 205457 | C | T | . | exonic | CCDC127 | nonsynonymous SNV | CCDC127:NM_145265:exon3:c.G623A:p.G208D, |
| chr5 | 311335 | A | G | . | exonic | PDCD6 | nonsynonymous SNV | PDCD6:NM_001267557:exon3:c.A206G:p.K69R,PDCD6:NM_001267558:exon6:c.A200G:p.K67R,PDCD6:NM_013232:exon5:c.A410G:p.K137R,PDCD6:NM_001267556:exon5:c.A404G:p.K135R, |
| chr5 | 32388508 | T | C | . | exonic | ZFR | nonsynonymous SNV | ZFR:NM_016107:exon13:c.A2309G:p.N770S, |
| chr5 | 32400161 | A | G | COSV54053200 | exonic | ZFR | nonsynonymous SNV | ZFR:NM_016107:exon9:c.T1559C:p.I520T, |
| chr5 | 32403241 | C | T | . | exonic | ZFR | nonsynonymous SNV | ZFR:NM_016107:exon8:c.G1381A:p.V461I, |
| chr5 | 32406858 | A | T | . | exonic | ZFR | nonsynonymous SNV | ZFR:NM_016107:exon6:c.T948A:p.N316K, |
| chr5 | 33989413 | C | T | COSV56871911 | exonic | AMACR | nonsynonymous SNV | AMACR:NM_001167595:exon5:c.G829A:p.E277K,AMACR:NM_014324:exon5:c.G829A:p.E277K, |
| chr5 | 33998663 | C | A | COSV59460034 | exonic | AMACR | nonsynonymous SNV | AMACR:NM_001167595:exon4:c.G717T:p.Q239H,AMACR:NM_203382:exon3:c.G556T:p.V186F,AMACR:NM_014324:exon4:c.G717T:p.Q239H, |
| chr5 | 33998778 | A | G | COSV59461114 | exonic | AMACR | nonsynonymous SNV | AMACR:NM_001167595:exon4:c.T602C:p.L201S,AMACR:NM_014324:exon4:c.T602C:p.L201S, |
| chr5 | 34004602 | C | T | COSV59461134 | exonic | AMACR | nonsynonymous SNV | AMACR:NM_001167595:exon3:c.G524A:p.G175D,AMACR:NM_014324:exon3:c.G524A:p.G175D, |
| chr5 | 34007995 | C | T | COSV59461909 | exonic | AMACR | nonsynonymous SNV | AMACR:NM_001167595:exon1:c.G25A:p.V9M,AMACR:NM_203382:exon1:c.G25A:p.V9M,AMACR:NM_014324:exon1:c.G25A:p.V9M, |
| chr5 | 36985201 | A | G | COSV56947629 | exonic | NIPBL | nonsynonymous SNV | NIPBL:NM_133433:exon10:c.A2021G:p.N674S,NIPBL:NM_015384:exon10:c.A2021G:p.N674S, |
| chr5 | 37381631 | G | A | . | exonic | WDR70 | nonsynonymous SNV | WDR70:NM_018034:exon3:c.G121A:p.A41T, |
| chr5 | 37605228 | G | A | . | exonic | WDR70 | nonsynonymous SNV | WDR70:NM_018034:exon10:c.G1082A:p.R361Q, |
| chr5 | 37752562 | C | A | . | exonic | WDR70 | nonsynonymous SNV | WDR70:NM_018034:exon18:c.C1954A:p.R652S, |

| | | | | | | | | |
|---|---|---|---|---|---|---|---|---|
| chr5 | 55425334 | C | T | COSV53308023 | exonic | PLPP1 | nonsynonymous SNV | PLPP1:NM_003711:exon6:c.G727A:p.A243T,PLPP1:NM_176895:exon6:c.G730A:p.A244T, |
| chr5 | 55467989 | C | T | . | exonic | PLPP1 | nonsynonymous SNV | PLPP1:NM_003711:exon3:c.G371A:p.G124D,PLPP1:NM_176895:exon3:c.G374A:p.G125D, |
| chr5 | 55475402 | C | T | COSV53306302 | exonic | PLPP1 | nonsynonymous SNV | PLPP1:NM_003711:exon2:c.G107A:p.R36Q, |
| chr5 | 55664845 | C | G | COSV59422666 | exonic | SLC38A9 | nonsynonymous SNV | SLC38A9:NM_001282429:exon7:c.G173C:p.S58T,SLC38A9:NM_001258286:exon6:c.G356C:p.S119T,SLC38A9:NM_173514:exon8:c.G545C:p.S182T,SLC38A9:NM_001258287:exon7:c.G464C:p.S155T, |
| chr5 | 55956103 | G | C | COSV61140369 | exonic | IL6ST | nonsynonymous SNV | IL6ST:NM_001190981:exon10:c.C1189G:p.L397V,IL6ST:NM_002184:exon10:c.C1189G:p.L397V, |
| chr5 | 65563377 | G | A | . | exonic | PPWD1 | nonsynonymous SNV | PPWD1:NM_015342:exon1:c.G67A:p.E23K, |
| chr5 | 65572006 | G | A | . | exonic | PPWD1 | nonsynonymous SNV | PPWD1:NM_001278929:exon4:c.G221A:p.R74Q,PPWD1:NM_001278927:exon5:c.G269A:p.R90Q,PPWD1:NM_001278926:exon6:c.G599A:p.R200Q,PPWD1:NM_015342:exon5:c.G689A:p.R230Q, |
| chr5 | 6599951 | G | T | . | exonic | NSUN2 | nonsynonymous SNV | NSUN2:NM_001193455:exon18:c.C2174A:p.P725Q,NSUN2:NM_017755:exon19:c.C2279A:p.P760Q, |
| chr5 | 68292320 | G | A | COSV57127731 | exonic | PIK3R1 | nonsynonymous SNV | PIK3R1:NM_181523:exon8:c.G978A:p.M326I,PIK3R1:NM_181524:exon2:c.G78A:p.M26I,PIK3R1:NM_181504:exon2:c.G168A:p.M56I, |
| chr5 | 69400113 | T | G | COSV51456873 | exonic | RAD17 | nonsynonymous SNV | RAD17:NM_133344:exon15:c.T1637G:p.L546R,RAD17:NM_133342:exon16:c.T1637G:p.L546R,RAD17:NM_133339:exon14:c.T1670G:p.L557R,RAD17:NM_002873:exon15:c.T1637G:p.L546R,RAD17:NM_133340:exon14:c.T1142G:p.L381R,RAD17:NM_133338:exon17:c.T1637G:p.L546R,RAD17:NM_133341:exon14:c.T1379G:p.L460R,RAD17:NM_001278622:exon16:c.T1637G:p.L546R,RAD17:NM_133343:exon16:c.T1637G:p.L546R, |
| chr5 | 69414161 | G | A | . | exonic | RAD17 | nonsynonymous SNV | RAD17:NM_133344:exon17:c.G1882A:p.E628K,RAD17:NM_133342:exon18:c.G1882A:p.E628K,RAD17:NM_133339:exon16:c.G1915A:p.E639K,RAD17:NM_002873:exon17:c.G1882A:p.E628K,RAD17:NM_133340:exon16:c.G1387A:p.E463K,RAD17:NM_133338:exon19:c.G1882A:p.E628K,RAD17:NM_133341:exon16:c.G1624A:p.E542K,RAD17:NM_001278622:exon18:c.G1882A:p.E628K,RAD17:NM_133343:exon18:c.G1882A:p.E628K, |
| chr5 | 71455991 | T | G | COSV62429550 | exonic | BDP1 | nonsynonymous SNV | BDP1:NM_018429:exon1:c.T114G:p.D38E, |
| chr5 | 71502714 | A | G | . | exonic | BDP1 | nonsynonymous SNV | BDP1:NM_018429:exon15:c.A2164G:p.K722E, |
| chr5 | 71504711 | G | A | COSV62429660 | exonic | BDP1 | nonsynonymous SNV | BDP1:NM_018429:exon16:c.G2332A:p.V778M, |
| chr5 | 71510630 | G | A | COSV62429841 | exonic | BDP1 | nonsynonymous SNV | BDP1:NM_018429:exon17:c.G3538A:p.G1180S, |
| chr5 | 71510822 | T | A | COSV62430362 | exonic | BDP1 | nonsynonymous SNV | BDP1:NM_018429:exon17:c.T3730A:p.F1244I, |
| chr5 | 74685445 | T | C | COSV54668597 | exonic | HEXB | nonsynonymous SNV | HEXB:NM_000521:exon1:c.T185C:p.L62S, |
| chr5 | 74689390 | A | G | COSV54666892 | exonic | HEXB | nonsynonymous SNV | HEXB:NM_000521:exon2:c.A362G:p.K121R, |
| chr5 | 74697056 | A | G | COSV54667253 | exonic | HEXB | nonsynonymous SNV | HEXB:NM_000521:exon5:c.A619G:p.I207V, |
| chr5 | 78116234 | C | T | COSV54878379 | exonic | AP3B1 | nonsynonymous SNV | AP3B1:NM_003664:exon18:c.G1969A:p.A657T,AP3B1:NM_001271769:exon18:c.G1822A:p.A608T, |
| chr5 | 78129204 | A | T | COSV54884128 | exonic | AP3B1 | nonsynonymous SNV | AP3B1:NM_001271769:exon16:c.T1607A:p.V536E,AP3B1:NM_003664:exon16:c.T1754A:p.V585E, |
| chr5 | 7867450 | C | T | . | exonic | FASTKD3 | nonsynonymous SNV | FASTKD3:NM_024091:exon2:c.G634A:p.G212S, |
| chr5 | 7867917 | T | C | . | exonic | FASTKD3 | nonsynonymous SNV | FASTKD3:NM_024091:exon2:c.A167G:p.K56R, |
| chr5 | 82253397 | C | T | COSV56424284 | exonic | ATG10 | nonsynonymous SNV | ATG10:NM_031482:exon7:c.C635T:p.T212M,ATG10:NM_001131028:exon8:c.C635T:p.T212M, |
| chr5 | 82253421 | C | A | . | exonic | ATG10 | nonsynonymous SNV | ATG10:NM_031482:exon7:c.C659A:p.P220H,ATG10:NM_001131028:exon8:c.C659A:p.P220H, |
| chr5 | 82276126 | C | T | . | exonic | RPS23 | nonsynonymous SNV | RPS23:NM_001025:exon4:c.G415A:p.E139K, |
| chr5 | 87399457 | A | G | COSV56924372 | exonic | CCNH | nonsynonymous SNV | CCNH:NM_001239:exon7:c.T809C:p.V270A,CCNH:NM_001199189:exon7:c.T650C:p.V217A, |
| chr5 | 891703 | C | T | . | exonic | BRD9 | nonsynonymous SNV | BRD9:NM_001009877:exon2:c.G50A:p.R17K, |
| chr5 | 891709 | C | T | . | exonic | BRD9 | nonsynonymous SNV | BRD9:NM_001009877:exon2:c.G44A:p.R15K, |
| chr5 | 891718 | G | T | . | exonic | BRD9 | nonsynonymous SNV | BRD9:NM_001009877:exon2:c.C35A:p.T12K,BRD9:NM_023924:exon2:c.C189A:p.H63Q, |
| chr6 | 105278161 | C | T | COSV64870385 | exonic | PREP | nonsynonymous SNV | PREP:NM_002726:exon15:c.G2116A:p.V706I, |
| chr6 | 106640356 | C | T | . | exonic | QRSL1 | nonsynonymous SNV | QRSL1:NM_018292:exon2:c.C32T:p.A11V, |
| chr6 | 106665854 | G | A | . | exonic | QRSL1 | nonsynonymous SNV | QRSL1:NM_018292:exon11:c.G1439A:p.R480H, |
| chr6 | 106971734 | G | A | . | exonic | CD24 | nonsynonymous SNV | CD24:NM_001291737:exon3:c.C170T:p.A57V,CD24:NM_013230:exon3:c.C170T:p.A57V,CD24:NM_001291738:exon3:c.C170T:p.A57V,CD24:NM_001291739:exon2:c.C317T:p.A106V, |
| chr6 | 107883003 | G | T | . | exonic | SEC63 | nonsynonymous SNV | SEC63:NM_007214:exon17:c.C1818A:p.N606K, |
| chr6 | 110456925 | T | C | COSV57930020 | exonic | SLC22A16 | nonsynonymous SNV | SLC22A16:NM_033125:exon2:c.A146G:p.H49R, |
| chr6 | 122801319 | C | T | COSV63755382 | exonic | SMPDL3A | nonsynonymous SNV | SMPDL3A:NM_006714:exon4:c.C481T:p.P161S,SMPDL3A:NM_001286138:exon3:c.C88T:p.P30S, |
| chr6 | 122803797 | A | C | . | exonic | SMPDL3A | nonsynonymous SNV | SMPDL3A:NM_006714:exon5:c.A702C:p.E234D,SMPDL3A:NM_001286138:exon4:c.A309C:p.E103D, |
| chr6 | 124963311 | T | C | COSV71376720 | exonic | RNF217 | nonsynonymous SNV | RNF217:NM_001286398:exon1:c.T767C:p.V256A, |
| chr6 | 125957084 | G | C | COSV57649843 | exonic | HINT3 | nonsynonymous SNV | HINT3:NM_138571:exon1:c.G107C:p.G36A, |
| chr6 | 125999566 | G | A | . | exonic | TRMT11 | nonsynonymous SNV | TRMT11:NM_001031712:exon7:c.G632A:p.G211E, |
| chr6 | 127289985 | C | T | . | exonic | ECHDC1 | nonsynonymous SNV | ECHDC1:NM_001002030:exon6:c.G790A:p.G264S,ECHDC1:NM_001139510:exon6:c.G808A:p.G270S,ECHDC1:NM_001105544:exon5:c.G565A:p.G189S, |
| chr6 | 127289994 | C | T | . | exonic | ECHDC1 | nonsynonymous SNV | ECHDC1:NM_001139510:exon6:c.G799A:p.V267I,ECHDC1:NM_001105544:exon5:c.G556A:p.V186I,ECHDC1:NM_001002030:exon6:c.G781A:p.V261I, |
| chr6 | 127290035 | A | G | . | exonic | ECHDC1 | nonsynonymous SNV | ECHDC1:NM_001002030:exon6:c.T740C:p.I247T,ECHDC1:NM_001139510:exon6:c.T758C:p.I253T,ECHDC1:NM_001105544:exon5:c.T515C:p.I172T, |
| chr6 | 127316482 | C | T | . | exonic | ECHDC1 | nonsynonymous SNV | ECHDC1:NM_001002030:exon4:c.G384A:p.M128I,ECHDC1:NM_001139510:exon4:c.G402A:p.M134I,ECHDC1:NM_001105544:exon3:c.G159A:p.M53I, |
| chr6 | 127327004 | C | T | . | exonic | ECHDC1 | nonsynonymous SNV | ECHDC1:NM_001002030:exon3:c.G361A:p.E121K,ECHDC1:NM_001139510:exon3:c.G379A:p.E127K,ECHDC1:NM_001105544:exon2:c.G136A:p.E46K,ECHDC1:NM_018479:exon3:c.G361A:p.E121K,ECHDC1:NM_001105545:exon2:c.G136A:p.E46K, |
| chr6 | 127327012 | C | T | . | exonic | ECHDC1 | nonsynonymous SNV | ECHDC1:NM_001002030:exon3:c.G353A:p.G118E,ECHDC1:NM_001139510:exon3:c.G371A:p.G124E,ECHDC1:NM_001105544:exon2:c.G128A:p.G43E,ECHDC1:NM_018479:exon3:c.G353A:p.G118E,ECHDC1:NM_001105545:exon2:c.G128A:p.G43E, |
| chr6 | 127327037 | C | T | . | exonic | ECHDC1 | nonsynonymous SNV | ECHDC1:NM_001002030:exon3:c.G328A:p.D110N,ECHDC1:NM_001139510:exon3:c.G346A:p.D116N,ECHDC1:NM_001105544:exon2:c.G103A:p.D35N,ECHDC1:NM_018479:exon3:c.G328A:p.D110N,ECHDC1:NM_001105545:exon2:c.G103A:p.D35N, |
| chr6 | 127327072 | A | G | . | exonic | ECHDC1 | nonsynonymous SNV | ECHDC1:NM_001139510:exon3:c.T311C:p.V104A,ECHDC1:NM_001105544:exon2:c.T68C:p.V23A,ECHDC1:NM_018479:exon3:c.T293C:p.V98A,ECHDC1:NM_001002030:exon3:c.T293C:p.V98A,ECHDC1:NM_001105545:exon2:c.T68C:p.V23A, |

| | | | | | | | | |
|---|---|---|---|---|---|---|---|---|
| chr6 | 127996951 | G | T | . | exonic | PTPRK | nonsynonymous SNV | PTPRK:NM_002844:exon17:c.C2717A:p.A906D,PTPRK:NM_001135648:exon17:c.C2717A:p.A906D,PTPRK:NM_001291984:exon17:c.C2714A:p.A905D,PTPRK:NM_001291981:exon19:c.C2765A:p.A922D, |
| chr6 | 128240094 | C | T | COSV63888698 | exonic | PTPRK | nonsynonymous SNV | PTPRK:NM_002844:exon5:c.G634A:p.A212T,PTPRK:NM_001291983:exon4:c.G247A:p.A83T,PTPRK:NM_001135648:exon5:c.G634A:p.A212T,PTPRK:NM_001291984:exon5:c.G634A:p.A212T,PTPRK:NM_001291982:exon5:c.G634A:p.A212T,PTPRK:NM_001291981:exon5:c.G634A:p.A212T, |
| chr6 | 129599245 | A | T | . | exonic | ARHGAP18 | nonsynonymous SNV | ARHGAP18:NM_033515:exon12:c.T1684A:p.Y562N, |
| chr6 | 129599297 | A | T | . | exonic | ARHGAP18 | nonsynonymous SNV | ARHGAP18:NM_033515:exon12:c.T1632A:p.D544E, |
| chr6 | 129599299 | C | T | . | exonic | ARHGAP18 | nonsynonymous SNV | ARHGAP18:NM_033515:exon12:c.G1630A:p.D544N, |
| chr6 | 129599306 | A | T | . | exonic | ARHGAP18 | nonsynonymous SNV | ARHGAP18:NM_033515:exon12:c.T1623A:p.H541Q, |
| chr6 | 129607973 | C | T | . | exonic | ARHGAP18 | nonsynonymous SNV | ARHGAP18:NM_033515:exon9:c.G1202A:p.S401N, |
| chr6 | 129618698 | A | G | . | exonic | ARHGAP18 | nonsynonymous SNV | ARHGAP18:NM_033515:exon6:c.T941C:p.I314T, |
| chr6 | 129710070 | T | C | COSV63763931 | exonic | ARHGAP18 | nonsynonymous SNV | ARHGAP18:NM_033515:exon1:c.A67G:p.T23A, |
| chr6 | 130184126 | G | C | . | exonic | SAMD3 | nonsynonymous SNV | SAMD3:NM_001017373:exon7:c.C631G:p.L211V,SAMD3:NM_001258275:exon9:c.C631G:p.L211V,SAMD3:NM_001277185:exon6:c.C703G:p.L235V, |
| chr6 | 130869910 | C | A | . | exonic | EPB41L2 | stopgain | EPB41L2:NM_001431:exon15:c.G2260T:p.G754X,EPB41L2:NM_001199388:exon13:c.G2050T:p.G684X, |
| chr6 | 131950812 | G | C | COSV63466445 | exonic | CTGF | nonsynonymous SNV | CTGF:NM_001901:exon2:c.C247G:p.H83D, |
| chr6 | 132815015 | G | T | . | exonic | RPS12 | stopgain | RPS12:NM_001016:exon3:c.G58T:p.E20X, |
| chr6 | 135195839 | G | A | . | exonic | MYB | nonsynonymous SNV | MYB:NM_001161656:exon9:c.G1031A:p.C344Y,MYB:NM_001130172:exon9:c.G1031A:p.C344Y,MYB:NM_001161660:exon8:c.G935A:p.C312Y,MYB:NM_001161658:exon9:c.G1040A:p.C347Y,MYB:NM_005375:exon9:c.G1040A:p.C347Y,MYB:NM_001161659:exon9:c.G1040A:p.C347Y,MYB:NM_001130173:exon9:c.G1040A:p.C347Y, |
| chr6 | 136361034 | G | A | COSV63420650 | exonic | MAP7 | nonsynonymous SNV | MAP7:NM_001198617:exon10:c.C1390T:p.R464W,MAP7:NM_001198619:exon12:c.C1234T:p.R412W,MAP7:NM_003980:exon12:c.C1672T:p.R558W,MAP7:NM_001198611:exon11:c.C1627T:p.R543W,MAP7:NM_001198615:exon12:c.C1627T:p.R543W,MAP7:NM_001198616:exon11:c.C1561T:p.R521W,MAP7:NM_001198609:exon12:c.C1762T:p.R588W,MAP7:NM_001198618:exon12:c.C1234T:p.R412W,MAP7:NM_001198608:exon12:c.C1738T:p.R580W,MAP7:NM_001198614:exon13:c.C1738T:p.R580W, |
| chr6 | 137002076 | G | A | COSV57358376 | exonic | IL20RA | nonsynonymous SNV | IL20RA:NM_014432:exon7:c.C1144T:p.L382F,IL20RA:NM_001278724:exon6:c.C745T:p.L249F,IL20RA:NM_001278723:exon6:c.C811T:p.L271F,IL20RA:NM_001278722:exon7:c.C997T:p.L333F, |
| chr6 | 137004710 | C | T | . | exonic | IL20RA | nonsynonymous SNV | IL20RA:NM_014432:exon6:c.G775A:p.V259I,IL20RA:NM_001278724:exon5:c.G376A:p.V126I,IL20RA:NM_001278723:exon5:c.G442A:p.V148I,IL20RA:NM_001278722:exon6:c.G628A:p.V210I, |
| chr6 | 138092196 | G | C | . | exonic | PERP | nonsynonymous SNV | PERP:NM_022121:exon3:c.C428G:p.P143R, |
| chr6 | 138442036 | A | T | . | exonic | NHSL1 | nonsynonymous SNV | NHSL1:NM_001144060:exon5:c.T611A:p.V204D, |
| chr6 | 143136899 | G | A | . | exonic | AIG1 | nonsynonymous SNV | AIG1:NM_001286588:exon2:c.G206A:p.G69E,AIG1:NM_016108:exon2:c.G206A:p.G69E,AIG1:NM_001286589:exon2:c.G206A:p.G69E,AIG1:NM_001286587:exon3:c.G176A:p.G59E, |
| chr6 | 143165112 | G | A | . | exonic | AIG1 | nonsynonymous SNV | AIG1:NM_001286588:exon3:c.G328A:p.A110T,AIG1:NM_016108:exon3:c.G328A:p.A110T,AIG1:NM_001286589:exon3:c.G328A:p.A110T,AIG1:NM_001286587:exon4:c.G298A:p.A100T, |
| chr6 | 143339644 | G | A | . | exonic | AIG1 | nonsynonymous SNV | AIG1:NM_016108:exon6:c.G685A:p.E229K,AIG1:NM_001286587:exon7:c.G655A:p.E219K, |
| chr6 | 143503967 | G | T | COSV50019718 | exonic | FUCA2 | nonsynonymous SNV | FUCA2:NM_032020:exon3:c.C698A:p.A233E, |
| chr6 | 149595175 | C | T | . | exonic | KATNA1 | nonsynonymous SNV | KATNA1:NM_007044:exon11:c.G1337A:p.R446Q, |
| chr6 | 150905656 | G | A | COSV66209095 | exonic | MTHFD1L | nonsynonymous SNV | MTHFD1L:NM_015440:exon8:c.G787A:p.E263K,MTHFD1L:NM_001242767:exon8:c.G790A:p.E264K,MTHFD1L:NM_001242768:exon8:c.G592A:p.E198K, |
| chr6 | 151573370 | T | C | COSV53338703 | exonic | CCDC170 | nonsynonymous SNV | CCDC170:NM_025059:exon6:c.T971C:p.F324S, |
| chr6 | 151615542 | G | A | COSV53337542 | exonic | CCDC170 | nonsynonymous SNV | CCDC170:NM_025059:exon10:c.G1810A:p.V604I, |
| chr6 | 151618046 | G | A | COSV53344090 | exonic | CCDC170 | nonsynonymous SNV | CCDC170:NM_025059:exon11:c.G2047A:p.V683I, |
| chr6 | 152225839 | A | T | . | exonic | SYNE1 | nonsynonymous SNV | SYNE1:NM_033071:exon115:c.T21020A:p.V7007E,SYNE1:NM_182961:exon116:c.T21233A:p.V7078E, |
| chr6 | 152225849 | C | T | . | exonic | SYNE1 | nonsynonymous SNV | SYNE1:NM_033071:exon115:c.G21010A:p.E7004K,SYNE1:NM_182961:exon116:c.G21223A:p.E7075K, |
| chr6 | 152975406 | G | C | COSV57670920 | exonic | FBXO5 | nonsynonymous SNV | FBXO5:NM_012177:exon2:c.C319G:p.Q107E,FBXO5:NM_001142522:exon2:c.C181G:p.Q61E, |
| chr6 | 155257877 | C | T | . | exonic | TFB1M | nonsynonymous SNV | TFB1M:NM_016020:exon7:c.G1000A:p.E334K, |
| chr6 | 155257885 | C | T | . | exonic | TFB1M | nonsynonymous SNV | TFB1M:NM_016020:exon7:c.G992A:p.S331N, |
| chr6 | 155257894 | C | T | . | exonic | TFB1M | nonsynonymous SNV | TFB1M:NM_016020:exon7:c.G983A:p.R328Q, |
| chr6 | 155257964 | C | T | . | exonic | TFB1M | nonsynonymous SNV | TFB1M:NM_016020:exon7:c.G913A:p.D305N, |
| chr6 | 155257971 | G | T | . | exonic | TFB1M | nonsynonymous SNV | TFB1M:NM_016020:exon7:c.C906A:p.S302R, |
| chr6 | 155296958 | C | T | . | exonic | TFB1M | nonsynonymous SNV | TFB1M:NM_016020:exon4:c.G541A:p.A181T, |
| chr6 | 155311216 | T | C | . | exonic | TFB1M | nonsynonymous SNV | TFB1M:NM_016020:exon2:c.A257G:p.K86R, |
| chr6 | 156779221 | G | A | . | exonic | ARID1B | nonsynonymous SNV | ARID1B:NM_020732:exon1:c.G1292A:p.R431Q,ARID1B:NM_017519:exon1:c.G1292A:p.R431Q, |
| chr6 | 158767403 | A | G | . | exonic | EZR | nonsynonymous SNV | EZR:NM_003379:exon12:c.T1454C:p.V485A,EZR:NM_001111077:exon13:c.T1454C:p.V485A, |
| chr6 | 158769941 | A | G | . | exonic | EZR | nonsynonymous SNV | EZR:NM_001111077:exon11:c.T1094C:p.L365P,EZR:NM_003379:exon10:c.T1094C:p.L365P, |
| chr6 | 158770806 | G | A | COSV61456427 | exonic | EZR | nonsynonymous SNV | EZR:NM_001111077:exon10:c.C1048T:p.R350W,EZR:NM_003379:exon9:c.C1048T:p.R350W, |
| chr6 | 158771347 | C | T | . | exonic | EZR | nonsynonymous SNV | EZR:NM_001111077:exon9:c.G856A:p.G286S,EZR:NM_003379:exon8:c.G856A:p.G286S, |
| chr6 | 158784683 | C | T | . | exonic | EZR | nonsynonymous SNV | EZR:NM_001111077:exon6:c.G512A:p.R171Q,EZR:NM_003379:exon5:c.G512A:p.R171Q, |
| chr6 | 158785351 | T | A | . | exonic | EZR | nonsynonymous SNV | EZR:NM_003379:exon4:c.A425T:p.H142L,EZR:NM_001111077:exon5:c.A425T:p.H142L, |
| chr6 | 158785397 | A | G | . | exonic | EZR | nonsynonymous SNV | EZR:NM_001111077:exon5:c.T379C:p.S127P,EZR:NM_003379:exon4:c.T379C:p.S127P, |
| chr6 | 158785400 | C | T | . | exonic | EZR | nonsynonymous SNV | EZR:NM_001111077:exon5:c.G376A:p.G126R,EZR:NM_003379:exon4:c.G376A:p.G126R, |
| chr6 | 158785426 | C | T | . | exonic | EZR | nonsynonymous SNV | EZR:NM_001111077:exon5:c.G350A:p.C117Y,EZR:NM_003379:exon4:c.G350A:p.C117Y, |
| chr6 | 158785450 | C | T | . | exonic | EZR | nonsynonymous SNV | EZR:NM_001111077:exon5:c.G326A:p.G109E,EZR:NM_003379:exon4:c.G326A:p.G109E, |
| chr6 | 158785459 | A | G | . | exonic | EZR | nonsynonymous SNV | EZR:NM_001111077:exon5:c.T317C:p.V106A,EZR:NM_003379:exon4:c.T317C:p.V106A, |
| chr6 | 158785517 | C | T | . | exonic | EZR | nonsynonymous SNV | EZR:NM_001111077:exon5:c.G259A:p.E87K,EZR:NM_003379:exon4:c.G259A:p.E87K, |
| chr6 | 158787181 | C | T | . | exonic | EZR | nonsynonymous SNV | EZR:NM_001111077:exon4:c.G119A:p.R40Q,EZR:NM_003379:exon3:c.G119A:p.R40Q, |
| chr6 | 158789365 | C | T | . | exonic | EZR | nonsynonymous SNV | EZR:NM_001111077:exon3:c.G19A:p.V7I,EZR:NM_003379:exon2:c.G19A:p.V7I, |
| chr6 | 159692840 | A | G | COSV61622991 | exonic | SOD2 | nonsynonymous SNV | SOD2:NM_001024465:exon2:c.T47C:p.V16A,SOD2:NM_001024466:exon2:c.T47C:p.V16A,SOD2:NM_000636:exon2:c.T47C:p.V16A, |
| chr6 | 159775311 | A | G | COSV58460357 | exonic | ACAT2 | nonsynonymous SNV | ACAT2:NM_001303253:exon5:c.A719G:p.K240R,ACAT2:NM_005891:exon5:c.A632G:p.K211R, |
| chr6 | 159790604 | G | A | COSV58459001 | exonic | MRPL18 | nonsynonymous SNV | MRPL18:NM_014161:exon1:c.G17A:p.R6Q, |
| chr6 | 160073377 | A | G | . | exonic | IGF2R | nonsynonymous SNV | IGF2R:NM_000876:exon34:c.A4855G:p.R1619G, |
| chr6 | 1624213 | C | T | . | exonic | GMDS | nonsynonymous SNV | GMDS:NM_001500:exon11:c.G1075A:p.V359M,GMDS:NM_001253846:exon11:c.G985A:p.V329M, |
| chr6 | 166326158 | T | C | COSV62277048 | exonic | SFT2D1 | nonsynonymous SNV | SFT2D1:NM_145169:exon5:c.A325G:p.I109V, |
| chr6 | 166929653 | G | A | COSV50351546 | exonic | RNASET2 | nonsynonymous SNV | RNASET2:NM_003730:exon9:c.C706T:p.R236W, |
| chr6 | 167305948 | G | A | COSV57807281 | exonic | UNC93A | nonsynonymous SNV | UNC93A:NM_001143947:exon5:c.G748A:p.V250I,UNC93A:NM_018974:exon6:c.G874A:p.V292I, |
| chr6 | 167315286 | T | C | COSV57806804 | exonic | UNC93A | nonsynonymous SNV | UNC93A:NM_001143947:exon7:c.T1082C:p.M361T,UNC93A:NM_018974:exon8:c.T1208C:p.M403T, |
| chr6 | 167315287 | G | A | COSV57807036 | exonic | UNC93A | nonsynonymous SNV | UNC93A:NM_001143947:exon7:c.G1083A:p.M361I,UNC93A:NM_018974:exon8:c.G1209A:p.M403I, |
| chr6 | 167315303 | G | A | COSV57807274 | exonic | UNC93A | nonsynonymous SNV | UNC93A:NM_001143947:exon7:c.G1099A:p.V367I,UNC93A:NM_018974:exon8:c.G1225A:p.V409I, |
| chr6 | 167827136 | T | G | . | exonic | MLLT4 | nonsynonymous SNV | MLLT4:NM_001040000:exon1:c.T4G:p.S2A,MLLT4:NM_001207008:exon1:c.T4G:p.S2A, |
| chr6 | 167870392 | G | A | . | exonic | MLLT4 | nonsynonymous SNV | MLLT4:NM_001291964:exon3:c.G185A:p.R62K,MLLT4:NM_001040000:exon3:c.G308A:p.R103K,MLLT4:NM_001207008:exon3:c.G308A:p.R103K, |
| chr6 | 167870421 | G | A | . | exonic | MLLT4 | nonsynonymous SNV | MLLT4:NM_001207008:exon3:c.G337A:p.V113I,MLLT4:NM_001291964:exon3:c.G214A:p.V72I,MLLT4:NM_001040000:exon3:c.G337A:p.V113I, |

| | | | | | | | | |
|---|---|---|---|---|---|---|---|---|
| chr6 | 167875447 | G | A | . | exonic | MLLT4 | nonsynonymous SNV | MLLT4:NM_001207008:exon5:c.G688A:p.E230K,MLLT4:NM_001291964:exon5:c.G568A:p.E190K,MLLT4:NM_001040000:exon5:c.G691A:p.E231K, |
| chr6 | 168029668 | A | T | COSV63026416 | exonic | KIF25 | nonsynonymous SNV | KIF25:NM_005355:exon2:c.A83T:p.K28M,KIF25:NM_030615:exon2:c.A83T:p.K28M, |
| chr6 | 170553212 | G | C | COSV51529993 | exonic | PSMB1 | nonsynonymous SNV | PSMB1:NM_002793:exon1:c.C31G:p.P11A, |
| chr6 | 18130729 | C | T | COSV59428760 | exonic | TPMT | nonsynonymous SNV | TPMT:NM_000367:exon9:c.G677A:p.R226Q, |
| chr6 | 18139681 | T | A | . | exonic | TPMT | nonsynonymous SNV | TPMT:NM_000367:exon5:c.A403T:p.I135F, |
| chr6 | 18143697 | C | T | . | exonic | TPMT | nonsynonymous SNV | TPMT:NM_000367:exon4:c.G265A:p.V89M, |
| chr6 | 18149028 | C | T | . | exonic | TPMT | nonsynonymous SNV | TPMT:NM_000367:exon2:c.G100A:p.V34M, |
| chr6 | 18249708 | A | T | . | exonic | DEK | nonsynonymous SNV | DEK:NM_003472:exon7:c.T705A:p.D235E,DEK:NM_001134709:exon6:c.T603A:p.D201E, |
| chr6 | 18249748 | C | T | . | exonic | DEK | nonsynonymous SNV | DEK:NM_003472:exon7:c.G665A:p.C222Y,DEK:NM_001134709:exon6:c.G563A:p.C188Y, |
| chr6 | 18256392 | G | T | . | exonic | DEK | nonsynonymous SNV | DEK:NM_003472:exon5:c.C421A:p.Q141K,DEK:NM_001134709:exon4:c.C319A:p.Q107K, |
| chr6 | 18256394 | A | G | . | exonic | DEK | nonsynonymous SNV | DEK:NM_003472:exon5:c.T419C:p.V140A,DEK:NM_001134709:exon4:c.T317C:p.V106A, |
| chr6 | 18258385 | C | T | . | exonic | DEK | nonsynonymous SNV | DEK:NM_003472:exon3:c.G166A:p.G56S, |
| chr6 | 18263865 | C | G | COSV55236589 | exonic | DEK | nonsynonymous SNV | DEK:NM_003472:exon2:c.G123C:p.E41D,DEK:NM_001134709:exon2:c.G123C:p.E41D, |
| chr6 | 1960930 | C | T | . | exonic | GMDS | nonsynonymous SNV | GMDS:NM_001500:exon5:c.G382A:p.D128N,GMDS:NM_001253846:exon5:c.G292A:p.D98N, |
| chr6 | 2117471 | C | T | . | exonic | GMDS | nonsynonymous SNV | GMDS:NM_001500:exon3:c.G233A:p.G78E,GMDS:NM_001253846:exon3:c.G143A:p.G48E, |
| chr6 | 24403227 | C | T | COSV51233989 | exonic | MRS2 | nonsynonymous SNV | MRS2:NM_001286264:exon1:c.C181T:p.R61C,MRS2:NM_020662:exon1:c.C181T:p.R61C,MRS2:NM_001286266:exon1:c.C181T:p.R61C,MRS2:NM_001286265:exon1:c.C181T:p.R61C, |
| chr6 | 28923304 | T | C | . | exonic | TRIM27 | nonsynonymous SNV | TRIM27:NM_006510:exon1:c.A329G:p.D110G, |
| chr6 | 2955568 | T | C | COSV59565275 | exonic | SERPINB6 | nonsynonymous SNV | SERPINB6:NM_001271823:exon3:c.A325G:p.M109V,SERPINB6:NM_001271825:exon3:c.A268G:p.M90V,SERPINB6:NM_001271822:exon3:c.A310G:p.M104V,SERPINB6:NM_001271824:exon3:c.A268G:p.M90V,SERPINB6:NM_001297699:exon3:c.A268G:p.M90V,SERPINB6:NM_001195291:exon4:c.A280G:p.M94V,SERPINB6:NM_001297700:exon3:c.A268G:p.M90V,SERPINB6:NM_004568:exon3:c.A268G:p.M90V, |
| chr6 | 29555899 | C | G | . | exonic | UBD | nonsynonymous SNV | UBD:NM_006398:exon2:c.G479C:p.C160S, |
| chr6 | 29556175 | A | G | COSV63542234 | exonic | UBD | nonsynonymous SNV | UBD:NM_006398:exon2:c.T203C:p.I68T, |
| chr6 | 29725234 | C | T | . | exonic | HLA-F | nonsynonymous SNV | HLA-F:NM_018950:exon4:c.C814T:p.P272S,HLA-F:NM_001098479:exon4:c.C814T:p.P272S, |
| chr6 | 29942581 | C | G | COSV65136471 | exonic | HLA-A | nonsynonymous SNV | HLA-A:NM_001242758:exon1:c.C28G:p.L10V,HLA-A:NM_002116:exon1:c.C28G:p.L10V, |
| chr6 | 29942781 | T | C | COSV65138565 | exonic | HLA-A | nonsynonymous SNV | HLA-A:NM_001242758:exon2:c.T98C:p.F33S,HLA-A:NM_002116:exon2:c.T98C:p.F33S, |
| chr6 | 29942886 | G | A | COSV65139535 | exonic | HLA-A | nonsynonymous SNV | HLA-A:NM_001242758:exon2:c.G203A:p.R68K,HLA-A:NM_002116:exon2:c.G203A:p.R68K, |
| chr6 | 29942939 | C | G | COSV65136733 | exonic | HLA-A | nonsynonymous SNV | HLA-A:NM_001242758:exon2:c.C256G:p.Q86E,HLA-A:NM_002116:exon2:c.C256G:p.Q86E, |
| chr6 | 29942948 | C | G | COSV65136740 | exonic | HLA-A | nonsynonymous SNV | HLA-A:NM_001242758:exon2:c.C265G:p.R89G,HLA-A:NM_002116:exon2:c.C265G:p.R89G, |
| chr6 | 29942953 | T | A | COSV65137266 | exonic | HLA-A | nonsynonymous SNV | HLA-A:NM_001242758:exon2:c.T270A:p.N90K,HLA-A:NM_002116:exon2:c.T270A:p.N90K, |
| chr6 | 29942954 | G | A | . | exonic | HLA-A | nonsynonymous SNV | HLA-A:NM_001242758:exon2:c.G271A:p.V91M,HLA-A:NM_002116:exon2:c.G271A:p.V91M, |
| chr6 | 29942965 | G | C | COSV65140297 | exonic | HLA-A | nonsynonymous SNV | HLA-A:NM_001242758:exon2:c.G282C:p.Q94H,HLA-A:NM_002116:exon2:c.G282C:p.Q94H, |
| chr6 | 29942982 | T | A | COSV65136627 | exonic | HLA-A | nonsynonymous SNV | HLA-A:NM_001242758:exon2:c.T299A:p.V100E,HLA-A:NM_002116:exon2:c.T299A:p.V100E, |
| chr6 | 29942984 | G | A | COSV65136637 | exonic | HLA-A | nonsynonymous SNV | HLA-A:NM_001242758:exon2:c.G301A:p.D101N,HLA-A:NM_002116:exon2:c.G301A:p.D101N, |
| chr6 | 29942990 | G | C | COSV65141826 | exonic | HLA-A | nonsynonymous SNV | HLA-A:NM_001242758:exon2:c.G307C:p.G103R,HLA-A:NM_002116:exon2:c.G307C:p.G103R, |
| chr6 | 29942994 | C | T | COSV65136647 | exonic | HLA-A | nonsynonymous SNV | HLA-A:NM_001242758:exon2:c.C311T:p.T104I,HLA-A:NM_002116:exon2:c.C311T:p.T104I, |
| chr6 | 29942996 | C | G | COSV65136655 | exonic | HLA-A | nonsynonymous SNV | HLA-A:NM_001242758:exon2:c.C313G:p.L105V,HLA-A:NM_002116:exon2:c.C313G:p.L105V, |
| chr6 | 29942997 | T | C | COSV65136680 | exonic | HLA-A | nonsynonymous SNV | HLA-A:NM_001242758:exon2:c.T314C:p.L105P,HLA-A:NM_002116:exon2:c.T314C:p.L105P, |
| chr6 | 29943000 | G | T | COSV65136690 | exonic | HLA-A | nonsynonymous SNV | HLA-A:NM_001242758:exon2:c.G317T:p.R106L,HLA-A:NM_002116:exon2:c.G317T:p.R106L, |
| chr6 | 29943002 | G | C | COSV65141194 | exonic | HLA-A | nonsynonymous SNV | HLA-A:NM_001242758:exon2:c.G319C:p.G107R,HLA-A:NM_002116:exon2:c.G319C:p.G107R, |
| chr6 | 29943024 | C | A | COSV65140734 | exonic | HLA-A | nonsynonymous SNV | HLA-A:NM_001242758:exon2:c.C341A:p.A114D,HLA-A:NM_002116:exon2:c.C341A:p.A114D, |
| chr6 | 29943279 | A | G | COSV65137506 | exonic | HLA-A | nonsynonymous SNV | HLA-A:NM_001242758:exon3:c.A355G:p.I119V,HLA-A:NM_002116:exon3:c.A355G:p.I119V, |
| chr6 | 29943287 | A | G | COSV65136818 | exonic | HLA-A | nonsynonymous SNV | HLA-A:NM_001242758:exon3:c.A363G:p.I121M,HLA-A:NM_002116:exon3:c.A363G:p.I121M, |
| chr6 | 29943292 | A | T | COSV65145084 | exonic | HLA-A | nonsynonymous SNV | HLA-A:NM_001242758:exon3:c.A368T:p.Y123F,HLA-A:NM_002116:exon3:c.A368T:p.Y123F, |
| chr6 | 29943309 | T | C | COSV65136992 | exonic | HLA-A | nonsynonymous SNV | HLA-A:NM_001242758:exon3:c.T385C:p.S129P,HLA-A:NM_002116:exon3:c.T385C:p.S129P, |
| chr6 | 29943337 | G | A | COSV65137272 | exonic | HLA-A | nonsynonymous SNV | HLA-A:NM_001242758:exon3:c.G413A:p.R138Q,HLA-A:NM_002116:exon3:c.G413A:p.R138Q, |
| chr6 | 29943342 | G | T | COSV65143116 | exonic | HLA-A | nonsynonymous SNV | HLA-A:NM_001242758:exon3:c.G418T:p.D140Y,HLA-A:NM_002116:exon3:c.G418T:p.D140Y, |
| chr6 | 29943377 | C | A | COSV65137229 | exonic | HLA-A | nonsynonymous SNV | HLA-A:NM_001242758:exon3:c.C453A:p.N151K,HLA-A:NM_002116:exon3:c.C453A:p.N151K, |
| chr6 | 29943445 | C | T | COSV65137144 | exonic | HLA-A | nonsynonymous SNV | HLA-A:NM_001242758:exon3:c.C521T:p.A174V,HLA-A:NM_002116:exon3:c.C521T:p.A174V, |
| chr6 | 29943451 | A | T | COSV65136508 | exonic | HLA-A | nonsynonymous SNV | HLA-A:NM_001242758:exon3:c.A527T:p.E176V,HLA-A:NM_002116:exon3:c.A527T:p.E176V, |
| chr6 | 29943463 | T | A | COSV65136709 | exonic | HLA-A | stopgain | HLA-A:NM_001242758:exon3:c.T539A:p.L180X,HLA-A:NM_002116:exon3:c.T539A:p.L180X, |
| chr6 | 29943463 | T | G | COSV65140034 | exonic | HLA-A | nonsynonymous SNV | HLA-A:NM_001242758:exon3:c.T539G:p.L180W,HLA-A:NM_002116:exon3:c.T539G:p.L180W, |
| chr6 | 29943469 | C | T | COSV65137164 | exonic | HLA-A | nonsynonymous SNV | HLA-A:NM_001242758:exon3:c.C545T:p.A182V,HLA-A:NM_002116:exon3:c.C545T:p.A182V, |
| chr6 | 29943479 | T | G | COSV65143681 | exonic | HLA-A | nonsynonymous SNV | HLA-A:NM_001242758:exon3:c.T555G:p.D185E,HLA-A:NM_002116:exon3:c.T555G:p.D185E, |
| chr6 | 29943483 | A | C | COSV65137172 | exonic | HLA-A | nonsynonymous SNV | HLA-A:NM_001242758:exon3:c.A559C:p.T187P,HLA-A:NM_002116:exon3:c.A559C:p.T187P, |
| chr6 | 29943484 | C | G | COSV65137185 | exonic | HLA-A | nonsynonymous SNV | HLA-A:NM_001242758:exon3:c.C560G:p.T187R,HLA-A:NM_002116:exon3:c.C560G:p.T187R, |
| chr6 | 29943494 | G | C | COSV65141379 | exonic | HLA-A | nonsynonymous SNV | HLA-A:NM_001242758:exon3:c.G570C:p.E190D,HLA-A:NM_002116:exon3:c.G570C:p.E190D, |
| chr6 | 29943495 | T | G | COSV65137199 | exonic | HLA-A | nonsynonymous SNV | HLA-A:NM_001242758:exon3:c.T571G:p.W191G,HLA-A:NM_002116:exon3:c.T571G:p.W191G, |
| chr6 | 29944193 | G | A | COSV65137221 | exonic | HLA-A | nonsynonymous SNV | HLA-A:NM_001242758:exon4:c.G691A:p.G231S,HLA-A:NM_002116:exon4:c.G691A:p.G231S, |
| chr6 | 29944310 | G | T | COSV65136926 | exonic | HLA-A | nonsynonymous SNV | HLA-A:NM_001242758:exon4:c.G808T:p.A270S,HLA-A:NM_002116:exon4:c.G808T:p.A270S, |
| chr6 | 29944331 | G | C | COSV65136521 | exonic | HLA-A | nonsynonymous SNV | HLA-A:NM_001242758:exon4:c.G829C:p.E277Q,HLA-A:NM_002116:exon4:c.G829C:p.E277Q, |
| chr6 | 29944503 | T | C | COSV65136543 | exonic | HLA-A | nonsynonymous SNV | HLA-A:NM_001242758:exon5:c.T899C:p.L300P,HLA-A:NM_002116:exon5:c.T899C:p.L300P, |
| chr6 | 29944520 | A | G | COSV65137836 | exonic | HLA-A | nonsynonymous SNV | HLA-A:NM_001242758:exon5:c.A916G:p.I306V,HLA-A:NM_002116:exon5:c.A916G:p.I306V, |
| chr6 | 29944556 | C | T | COSV65136547 | exonic | HLA-A | nonsynonymous SNV | HLA-A:NM_001242758:exon5:c.C952T:p.L318F,HLA-A:NM_002116:exon5:c.C952T:p.L318F, |
| chr6 | 29944565 | G | A | COSV65141873 | exonic | HLA-A | nonsynonymous SNV | HLA-A:NM_001242758:exon5:c.G961A:p.V321M,HLA-A:NM_002116:exon5:c.G961A:p.V321M, |
| chr6 | 29944568 | A | T | COSV65136823 | exonic | HLA-A | nonsynonymous SNV | HLA-A:NM_001242758:exon5:c.A964T:p.I322F,HLA-A:NM_002116:exon5:c.A964T:p.I322F, |
| chr6 | 29944571 | A | G | COSV65138386 | exonic | HLA-A | nonsynonymous SNV | HLA-A:NM_001242758:exon5:c.A967G:p.T323A,HLA-A:NM_002116:exon5:c.A967G:p.T323A, |
| chr6 | 29944609 | G | C | COSV65137133 | exonic | HLA-A | nonsynonymous SNV | HLA-A:NM_001242758:exon5:c.G1005C:p.K335N,HLA-A:NM_002116:exon5:c.G1005C:p.K335N, |
| chr6 | 29945079 | A | T | COSV65136576 | exonic | HLA-A | nonsynonymous SNV | HLA-A:NM_001242758:exon6:c.A1033T:p.T345S,HLA-A:NM_002116:exon6:c.A1033T:p.T345S, |
| chr6 | 30062243 | G | A | . | exonic | ZNRD1 | nonsynonymous SNV | ZNRD1:NM_014596:exon4:c.G266A:p.R89Q,ZNRD1:NM_001278785:exon4:c.G266A:p.R89Q,ZNRD1:NM_001278786:exon4:c.G266A:p.R89Q,ZNRD1:NM_170783:exon3:c.G266A:p.R89Q, |
| chr6 | 30062299 | G | A | . | exonic | ZNRD1 | nonsynonymous SNV | ZNRD1:NM_014596:exon4:c.G322A:p.G108R,ZNRD1:NM_001278785:exon4:c.G322A:p.G108R,ZNRD1:NM_001278786:exon4:c.G322A:p.G108R,ZNRD1:NM_170783:exon3:c.G322A:p.G108R, |
| chr6 | 30062330 | G | A | . | exonic | ZNRD1 | nonsynonymous SNV | ZNRD1:NM_014596:exon4:c.G353A:p.C118Y,ZNRD1:NM_001278785:exon4:c.G353A:p.C118Y,ZNRD1:NM_001278786:exon4:c.G353A:p.C118Y,ZNRD1:NM_170783:exon3:c.G353A:p.C118Y, |
| chr6 | 3010156 | C | T | COSV57643512 | exonic | NQO2 | nonsynonymous SNV | NQO2:NM_001290222:exon3:c.C139T:p.L47F,NQO2:NM_000904:exon3:c.C139T:p.L47F,NQO2:NM_001290221:exon6:c.C139T:p.L47F, |
| chr6 | 30103553 | C | T | COSV65060120 | exonic | TRIM31 | nonsynonymous SNV | TRIM31:NM_007028:exon9:c.G1261A:p.E421K, |
| chr6 | 30345181 | C | T | . | exonic | RPP21 | nonsynonymous SNV | RPP21:NM_001199120:exon1:c.C10T:p.P4S,RPP21:NM_024839:exon1:c.C10T:p.P4S,RPP21:NM_001199121:exon1:c.C10T:p.P4S, |
| chr6 | 30490287 | G | A | COSV64928040 | exonic | HLA-E | nonsynonymous SNV | HLA-E:NM_005516:exon3:c.G382A:p.G128R, |
| chr6 | 30744028 | C | G | . | exonic | IER3 | nonsynonymous SNV | IER3:NM_003897:exon2:c.G379C:p.A127P, |
| chr6 | 31161839 | A | G | . | exonic | TCF19 | nonsynonymous SNV | TCF19:NM_001077511:exon3:c.A631G:p.M211V,TCF19:NM_007109:exon3:c.A631G:p.M211V, |
| chr6 | 31269347 | T | C | COSV66117127 | exonic | HLA-C | nonsynonymous SNV | HLA-C:NM_002117:exon7:c.A1087G:p.T363A,HLA-C:NM_001243042:exon7:c.A1087G:p.T363A, |
| chr6 | 31269385 | C | G | COSV66115057 | exonic | HLA-C | nonsynonymous SNV | HLA-C:NM_002117:exon7:c.G1049C:p.C350S,HLA-C:NM_001243042:exon7:c.G1049C:p.C350S, |
| chr6 | 31269990 | T | C | COSV66110395 | exonic | HLA-C | nonsynonymous SNV | HLA-C:NM_002117:exon5:c.A991G:p.M331V,HLA-C:NM_001243042:exon5:c.A991G:p.M331V, |

| | | | | | | | | |
|---|---|---|---|---|---|---|---|---|
| chr6 | 31269992 | G | A | COSV66110576 | exonic | HLA-C | nonsynonymous SNV | HLA-C:NM_002117:exon5:c.C989T:p.A330V,HLA-C:NM_001243042:exon5:c.C989T:p.A330V, |
| chr6 | 31269996 | T | C | COSV66110404 | exonic | HLA-C | nonsynonymous SNV | HLA-C:NM_002117:exon5:c.A985G:p.T329A,HLA-C:NM_001243042:exon5:c.A985G:p.T329A, |
| chr6 | 31269999 | C | T | COSV66112417 | exonic | HLA-C | nonsynonymous SNV | HLA-C:NM_002117:exon5:c.G982A:p.V328I,HLA-C:NM_001243042:exon5:c.G982A:p.V328I, |
| chr6 | 31270002 | C | T | COSV66110360 | exonic | HLA-C | nonsynonymous SNV | HLA-C:NM_002117:exon5:c.G979A:p.V327M,HLA-C:NM_001243042:exon5:c.G979A:p.V327M, |
| chr6 | 31270025 | A | G | COSV66111096 | exonic | HLA-C | nonsynonymous SNV | HLA-C:NM_002117:exon5:c.T956C:p.V319A,HLA-C:NM_001243042:exon5:c.T956C:p.V319A, |
| chr6 | 31270056 | T | C | COSV66114266 | exonic | HLA-C | nonsynonymous SNV | HLA-C:NM_002117:exon5:c.A925G:p.M309V,HLA-C:NM_001243042:exon5:c.A925G:p.M309V, |
| chr6 | 31270085 | T | C | COSV66111578 | exonic | HLA-C | nonsynonymous SNV | HLA-C:NM_002117:exon5:c.A896G:p.E299G,HLA-C:NM_001243042:exon5:c.A896G:p.E299G, |
| chr6 | 31270210 | C | T | COSV66110876 | exonic | HLA-C | nonsynonymous SNV | HLA-C:NM_002117:exon4:c.G895A:p.E299K,HLA-C:NM_001243042:exon4:c.G895A:p.E299K, |
| chr6 | 31270214 | G | T | COSV66110582 | exonic | HLA-C | nonsynonymous SNV | HLA-C:NM_002117:exon4:c.C891A:p.S297R,HLA-C:NM_001243042:exon4:c.C891A:p.S297R, |
| chr6 | 31270233 | T | G | COSV66110591 | exonic | HLA-C | nonsynonymous SNV | HLA-C:NM_002117:exon4:c.A872C:p.Q291P,HLA-C:NM_001243042:exon4:c.A872C:p.Q291P, |
| chr6 | 31270252 | T | C | COSV66110598 | exonic | HLA-C | nonsynonymous SNV | HLA-C:NM_002117:exon4:c.A853G:p.M285V,HLA-C:NM_001243042:exon4:c.A853G:p.M285V, |
| chr6 | 31270276 | G | C | COSV66110239 | exonic | HLA-C | nonsynonymous SNV | HLA-C:NM_002117:exon4:c.C829G:p.Q277E,HLA-C:NM_001243042:exon4:c.C829G:p.Q277E, |
| chr6 | 31270293 | A | C | . | exonic | HLA-C | nonsynonymous SNV | HLA-C:NM_002117:exon4:c.T812G:p.V271G,HLA-C:NM_001243042:exon4:c.T812G:p.V271G, |
| chr6 | 31270359 | G | T | . | exonic | HLA-C | nonsynonymous SNV | HLA-C:NM_002117:exon4:c.C746A:p.T249N,HLA-C:NM_001243042:exon4:c.C746A:p.T249N, |
| chr6 | 31270378 | G | A | COSV66117576 | exonic | HLA-C | nonsynonymous SNV | HLA-C:NM_002117:exon4:c.C727T:p.R243W,HLA-C:NM_001243042:exon4:c.C727T:p.R243W, |
| chr6 | 31270482 | G | T | COSV66109964 | exonic | HLA-C | nonsynonymous SNV | HLA-C:NM_002117:exon4:c.C623A:p.P208H,HLA-C:NM_001243042:exon4:c.C623A:p.P208H, |
| chr6 | 31271091 | C | T | COSV66110610 | exonic | HLA-C | nonsynonymous SNV | HLA-C:NM_002117:exon3:c.G601A:p.E201K,HLA-C:NM_001243042:exon3:c.G601A:p.E201K, |
| chr6 | 31271103 | C | T | COSV66112160 | exonic | HLA-C | nonsynonymous SNV | HLA-C:NM_002117:exon3:c.G589A:p.E197K,HLA-C:NM_001243042:exon3:c.G589A:p.E197K, |
| chr6 | 31271109 | A | G | COSV66110011 | exonic | HLA-C | nonsynonymous SNV | HLA-C:NM_002117:exon3:c.T583C:p.Y195H,HLA-C:NM_001243042:exon3:c.T583C:p.Y195H, |
| chr6 | 31271120 | C | G | COSV66111341 | exonic | HLA-C | nonsynonymous SNV | HLA-C:NM_002117:exon3:c.G572C:p.W191S,HLA-C:NM_001243042:exon3:c.G572C:p.W191S, |
| chr6 | 31271132 | G | A | COSV66110017 | exonic | HLA-C | nonsynonymous SNV | HLA-C:NM_002117:exon3:c.C560T:p.T187M,HLA-C:NM_001243042:exon3:c.C560T:p.T187M, |
| chr6 | 31271133 | T | G | COSV66110024 | exonic | HLA-C | nonsynonymous SNV | HLA-C:NM_002117:exon3:c.A559C:p.T187P,HLA-C:NM_001243042:exon3:c.A559C:p.T187P, |
| chr6 | 31271148 | C | T | COSV66113730 | exonic | HLA-C | nonsynonymous SNV | HLA-C:NM_002117:exon3:c.G544A:p.A182T,HLA-C:NM_001243042:exon3:c.G544A:p.A182T, |
| chr6 | 31271153 | A | C | COSV66109994 | exonic | HLA-C | nonsynonymous SNV | HLA-C:NM_002117:exon3:c.T539G:p.L180R,HLA-C:NM_001243042:exon3:c.T539G:p.L180R, |
| chr6 | 31271165 | G | T | COSV66110458 | exonic | HLA-C | nonsynonymous SNV | HLA-C:NM_002117:exon3:c.C527A:p.A176E,HLA-C:NM_001243042:exon3:c.C527A:p.A176E, |
| chr6 | 31271180 | A | C | COSV66110087 | exonic | HLA-C | nonsynonymous SNV | HLA-C:NM_002117:exon3:c.T512G:p.L171W,HLA-C:NM_001243042:exon3:c.T512G:p.L171W, |
| chr6 | 31271193 | T | A | COSV66110720 | exonic | HLA-C | nonsynonymous SNV | HLA-C:NM_002117:exon3:c.A499T:p.T167S,HLA-C:NM_001243042:exon3:c.A499T:p.T167S, |
| chr6 | 31271207 | G | T | COSV66110642 | exonic | HLA-C | nonsynonymous SNV | HLA-C:NM_002117:exon3:c.C485A:p.T162N,HLA-C:NM_001243042:exon3:c.C485A:p.T162N, |
| chr6 | 31271229 | G | T | COSV66110004 | exonic | HLA-C | nonsynonymous SNV | HLA-C:NM_002117:exon3:c.C463A:p.R155S,HLA-C:NM_001243042:exon3:c.C463A:p.R155S, |
| chr6 | 31271273 | G | A | COSV66110477 | exonic | HLA-C | nonsynonymous SNV | HLA-C:NM_002117:exon3:c.C419T:p.S140F,HLA-C:NM_001243042:exon3:c.C419T:p.S140F, |
| chr6 | 31271274 | A | C | COSV66109986 | exonic | HLA-C | nonsynonymous SNV | HLA-C:NM_002117:exon3:c.T418G:p.S140A,HLA-C:NM_001243042:exon3:c.T418G:p.S140A, |
| chr6 | 31271280 | C | T | COSV66110483 | exonic | HLA-C | nonsynonymous SNV | HLA-C:NM_002117:exon3:c.G412A:p.D138N,HLA-C:NM_001243042:exon3:c.G412A:p.D138N, |
| chr6 | 31271283 | A | G | COSV66110655 | exonic | HLA-C | nonsynonymous SNV | HLA-C:NM_002117:exon3:c.T409C:p.Y137H,HLA-C:NM_001243042:exon3:c.T409C:p.Y137H, |
| chr6 | 31271292 | G | T | . | exonic | HLA-C | nonsynonymous SNV | HLA-C:NM_002117:exon3:c.C400A:p.L134I,HLA-C:NM_001243042:exon3:c.C400A:p.L134I, |
| chr6 | 31271313 | G | C | COSV66114851 | exonic | HLA-C | nonsynonymous SNV | HLA-C:NM_002117:exon3:c.C379G:p.L127V,HLA-C:NM_001243042:exon3:c.C379G:p.L127V, |
| chr6 | 31271324 | G | T | COSV66110489 | exonic | HLA-C | nonsynonymous SNV | HLA-C:NM_002117:exon3:c.C368A:p.S123Y,HLA-C:NM_001243042:exon3:c.C368A:p.S123Y, |
| chr6 | 31271331 | T | A | COSV66110137 | exonic | HLA-C | nonsynonymous SNV | HLA-C:NM_002117:exon3:c.A361T:p.R121W,HLA-C:NM_001243042:exon3:c.A361T:p.R121W, |
| chr6 | 31271336 | A | C | . | exonic | HLA-C | nonsynonymous SNV | HLA-C:NM_002117:exon3:c.T356G:p.L119R,HLA-C:NM_001243042:exon3:c.T356G:p.L119R, |
| chr6 | 31271337 | G | A | . | exonic | HLA-C | nonsynonymous SNV | HLA-C:NM_002117:exon3:c.C355T:p.L119F,HLA-C:NM_001243042:exon3:c.C355T:p.L119F, |
| chr6 | 31271337 | G | T | COSV66110498 | exonic | HLA-C | nonsynonymous SNV | HLA-C:NM_002117:exon3:c.C355A:p.L119I,HLA-C:NM_001243042:exon3:c.C355A:p.L119I, |
| chr6 | 31271339 | G | A | COSV66110997 | exonic | HLA-C | nonsynonymous SNV | HLA-C:NM_002117:exon3:c.C353T:p.T118I,HLA-C:NM_001243042:exon3:c.C353T:p.T118I, |
| chr6 | 31271601 | T | G | COSV66110096 | exonic | HLA-C | nonsynonymous SNV | HLA-C:NM_002117:exon2:c.A341C:p.D114A,HLA-C:NM_001243042:exon2:c.A341C:p.D114A, |
| chr6 | 31271625 | C | A | COSV66111615 | exonic | HLA-C | nonsynonymous SNV | HLA-C:NM_002117:exon2:c.G317T:p.R106L,HLA-C:NM_001243042:exon2:c.G317T:p.R106L, |
| chr6 | 31271630 | G | T | COSV66115598 | exonic | HLA-C | nonsynonymous SNV | HLA-C:NM_002117:exon2:c.C312A:p.N104K,HLA-C:NM_001243042:exon2:c.C312A:p.N104K, |
| chr6 | 31271640 | C | T | COSV66115617 | exonic | HLA-C | nonsynonymous SNV | HLA-C:NM_002117:exon2:c.G302A:p.S101N,HLA-C:NM_001243042:exon2:c.G302A:p.S101N, |
| chr6 | 31271643 | A | T | . | exonic | HLA-C | nonsynonymous SNV | HLA-C:NM_002117:exon2:c.T299A:p.V100E,HLA-C:NM_001243042:exon2:c.T299A:p.V100E, |
| chr6 | 31271653 | C | T | COSV66110668 | exonic | HLA-C | nonsynonymous SNV | HLA-C:NM_002117:exon2:c.G289A:p.A97T,HLA-C:NM_001243042:exon2:c.G289A:p.A97T, |
| chr6 | 31271659 | C | T | . | exonic | HLA-C | nonsynonymous SNV | HLA-C:NM_002117:exon2:c.G283A:p.A95T,HLA-C:NM_001243042:exon2:c.G283A:p.A95T, |
| chr6 | 31271670 | T | G | COSV66109953 | exonic | HLA-C | nonsynonymous SNV | HLA-C:NM_002117:exon2:c.A272C:p.Y91S,HLA-C:NM_001243042:exon2:c.A272C:p.Y91S, |
| chr6 | 31271672 | C | G | COSV66110203 | exonic | HLA-C | nonsynonymous SNV | HLA-C:NM_002117:exon2:c.G270C:p.K90N,HLA-C:NM_001243042:exon2:c.G270C:p.K90N, |
| chr6 | 31271674 | T | G | . | exonic | HLA-C | nonsynonymous SNV | HLA-C:NM_002117:exon2:c.A268C:p.K90Q,HLA-C:NM_001243042:exon2:c.A268C:p.K90Q, |
| chr6 | 31271701 | G | T | . | exonic | HLA-C | nonsynonymous SNV | HLA-C:NM_002117:exon2:c.C241A:p.P81T,HLA-C:NM_001243042:exon2:c.C241A:p.P81T, |
| chr6 | 31271724 | G | T | COSV66115644 | exonic | HLA-C | nonsynonymous SNV | HLA-C:NM_002117:exon2:c.C218A:p.A73E,HLA-C:NM_001243042:exon2:c.C218A:p.A73E, |
| chr6 | 31271749 | C | T | . | exonic | HLA-C | nonsynonymous SNV | HLA-C:NM_002117:exon2:c.G193A:p.A65T,HLA-C:NM_001243042:exon2:c.G193A:p.A65T, |
| chr6 | 31271766 | C | T | COSV66117347 | exonic | HLA-C | nonsynonymous SNV | HLA-C:NM_002117:exon2:c.G176A:p.R59Q,HLA-C:NM_001243042:exon2:c.G176A:p.R59Q, |
| chr6 | 31271800 | A | C | COSV66110102 | exonic | HLA-C | nonsynonymous SNV | HLA-C:NM_002117:exon2:c.T142G:p.S48A,HLA-C:NM_001243042:exon2:c.T142G:p.S48A, |
| chr6 | 31271808 | C | T | COSV66110108 | exonic | HLA-C | nonsynonymous SNV | HLA-C:NM_002117:exon2:c.G134A:p.R45H,HLA-C:NM_001243042:exon2:c.G134A:p.R45H, |
| chr6 | 31271830 | G | A | COSV66111027 | exonic | HLA-C | nonsynonymous SNV | HLA-C:NM_002117:exon2:c.C112T:p.R38W,HLA-C:NM_001243042:exon2:c.C112T:p.R38W, |
| chr6 | 31271836 | C | T | COSV66110844 | exonic | HLA-C | nonsynonymous SNV | HLA-C:NM_002117:exon2:c.G106A:p.V36M,HLA-C:NM_001243042:exon2:c.G106A:p.V36M, |
| chr6 | 31271839 | C | A | COSV66109978 | exonic | HLA-C | nonsynonymous SNV | HLA-C:NM_002117:exon2:c.G103T:p.A35S,HLA-C:NM_001243042:exon2:c.G103T:p.A35S, |
| chr6 | 31271844 | T | G | COSV66111042 | exonic | HLA-C | nonsynonymous SNV | HLA-C:NM_002117:exon2:c.A98C:p.D33A,HLA-C:NM_001243042:exon2:c.A98C:p.D33A, |
| chr6 | 31271845 | C | A | COSV66119680 | exonic | HLA-C | nonsynonymous SNV | HLA-C:NM_002117:exon2:c.G97T:p.D33Y,HLA-C:NM_001243042:exon2:c.G97T:p.D33Y, |
| chr6 | 31271999 | A | C | . | exonic | HLA-C | nonsynonymous SNV | HLA-C:NM_002117:exon1:c.T73G:p.C25G,HLA-C:NM_001243042:exon1:c.T73G:p.C25G, |
| chr6 | 31272025 | C | G | COSV66117644 | exonic | HLA-C | nonsynonymous SNV | HLA-C:NM_002117:exon1:c.G47C:p.G16A,HLA-C:NM_001243042:exon1:c.G47C:p.G16A, |
| chr6 | 31272028 | C | G | . | exonic | HLA-C | nonsynonymous SNV | HLA-C:NM_002117:exon1:c.G44C:p.G15A,HLA-C:NM_001243042:exon1:c.G44C:p.G15A, |
| chr6 | 31272044 | G | T | COSV66117654 | exonic | HLA-C | nonsynonymous SNV | HLA-C:NM_002117:exon1:c.C28A:p.L10I,HLA-C:NM_001243042:exon1:c.C28A:p.L10I, |
| chr6 | 31272050 | C | T | COSV66117662 | exonic | HLA-C | nonsynonymous SNV | HLA-C:NM_002117:exon1:c.G22A:p.A8T,HLA-C:NM_001243042:exon1:c.G22A:p.A8T, |
| chr6 | 3127301 | G | A | . | exonic | BPHL | nonsynonymous SNV | BPHL:NM_001302777:exon4:c.G220A:p.V74M,BPHL:NM_004332:exon3:c.G271A:p.V91M, |
| chr6 | 31354526 | C | G | . | exonic | HLA-B | nonsynonymous SNV | HLA-B:NM_005514:exon7:c.G1046C:p.C349S, |
| chr6 | 31354662 | C | T | . | exonic | HLA-B | nonsynonymous SNV | HLA-B:NM_005514:exon6:c.G1016A:p.G339E, |
| chr6 | 31355134 | C | T | COSV69520203 | exonic | HLA-B | nonsynonymous SNV | HLA-B:NM_005514:exon5:c.G985A:p.A329T, |
| chr6 | 31355140 | C | T | . | exonic | HLA-B | nonsynonymous SNV | HLA-B:NM_005514:exon5:c.G979A:p.V327I, |
| chr6 | 31355203 | C | T | . | exonic | HLA-B | nonsynonymous SNV | HLA-B:NM_005514:exon5:c.G916A:p.V306I, |
| chr6 | 31355407 | C | T | . | exonic | HLA-B | nonsynonymous SNV | HLA-B:NM_005514:exon4:c.G805A:p.A269T, |
| chr6 | 31355526 | G | T | . | exonic | HLA-B | nonsynonymous SNV | HLA-B:NM_005514:exon4:c.C686A:p.A229D, |
| chr6 | 31355527 | C | G | . | exonic | HLA-B | nonsynonymous SNV | HLA-B:NM_005514:exon4:c.G685C:p.A229P, |
| chr6 | 31355532 | C | T | . | exonic | HLA-B | nonsynonymous SNV | HLA-B:NM_005514:exon4:c.G680A:p.C227Y, |
| chr6 | 31355544 | G | A | COSV69521058 | exonic | HLA-B | nonsynonymous SNV | HLA-B:NM_005514:exon4:c.C668T:p.A223V, |
| chr6 | 31355592 | T | G | . | exonic | HLA-B | nonsynonymous SNV | HLA-B:NM_005514:exon4:c.A620C:p.D207A, |
| chr6 | 31356176 | C | G | COSV69520285 | exonic | HLA-B | nonsynonymous SNV | HLA-B:NM_005514:exon3:c.G610C:p.E204Q, |

| | | | | | | | | |
|---|---|---|---|---|---|---|---|---|
| chr6 | 31356181 | T | G | COSV69520287 | exonic | HLA-B | nonsynonymous SNV | HLA-B:NM_005514:exon3:c.A605C:p.K202T, |
| chr6 | 31356183 | G | C | COSV69520290 | exonic | HLA-B | nonsynonymous SNV | HLA-B:NM_005514:exon3:c.C603G:p.D201E, |
| chr6 | 31356197 | C | T | COSV69521691 | exonic | HLA-B | nonsynonymous SNV | HLA-B:NM_005514:exon3:c.G589A:p.E197K, |
| chr6 | 31356203 | A | G | COSV69520683 | exonic | HLA-B | nonsynonymous SNV | HLA-B:NM_005514:exon3:c.T583C:p.Y195H, |
| chr6 | 31356214 | C | G | COSV69520483 | exonic | HLA-B | nonsynonymous SNV | HLA-B:NM_005514:exon3:c.G572C:p.W191S, |
| chr6 | 31356217 | T | A | . | exonic | HLA-B | nonsynonymous SNV | HLA-B:NM_005514:exon3:c.A569T:p.E190V, |
| chr6 | 31356222 | G | C | . | exonic | HLA-B | nonsynonymous SNV | HLA-B:NM_005514:exon3:c.C564G:p.C188W, |
| chr6 | 31356224 | A | T | . | exonic | HLA-B | nonsynonymous SNV | HLA-B:NM_005514:exon3:c.T562A:p.C188S, |
| chr6 | 31356226 | T | G | COSV69520451 | exonic | HLA-B | nonsynonymous SNV | HLA-B:NM_005514:exon3:c.A560C:p.E187A, |
| chr6 | 31356227 | C | T | COSV69520455 | exonic | HLA-B | nonsynonymous SNV | HLA-B:NM_005514:exon3:c.G559A:p.E187K, |
| chr6 | 31356242 | C | T | . | exonic | HLA-B | nonsynonymous SNV | HLA-B:NM_005514:exon3:c.G544A:p.A182T, |
| chr6 | 31356247 | C | A | COSV69520219 | exonic | HLA-B | nonsynonymous SNV | HLA-B:NM_005514:exon3:c.G539T:p.R180L, |
| chr6 | 31356259 | T | A | COSV69520258 | exonic | HLA-B | nonsynonymous SNV | HLA-B:NM_005514:exon3:c.A527T:p.E176V, |
| chr6 | 31356274 | C | A | COSV69520985 | exonic | HLA-B | nonsynonymous SNV | HLA-B:NM_005514:exon3:c.G512T:p.W171L, |
| chr6 | 31356323 | G | T | COSV69520301 | exonic | HLA-B | nonsynonymous SNV | HLA-B:NM_005514:exon3:c.C463A:p.R155S, |
| chr6 | 31356366 | G | T | COSV69520643 | exonic | HLA-B | stopgain | HLA-B:NM_005514:exon3:c.C420A:p.Y140X, |
| chr6 | 31356367 | T | A | COSV69520463 | exonic | HLA-B | nonsynonymous SNV | HLA-B:NM_005514:exon3:c.A419T:p.Y140F, |
| chr6 | 31356368 | A | C | COSV69521076 | exonic | HLA-B | nonsynonymous SNV | HLA-B:NM_005514:exon3:c.T418G:p.Y140D, |
| chr6 | 31356374 | C | T | COSV69520768 | exonic | HLA-B | nonsynonymous SNV | HLA-B:NM_005514:exon3:c.G412A:p.D138N, |
| chr6 | 31356377 | G | A | COSV69520464 | exonic | HLA-B | nonsynonymous SNV | HLA-B:NM_005514:exon3:c.C409T:p.H137Y, |
| chr6 | 31356391 | C | A | . | exonic | HLA-B | nonsynonymous SNV | HLA-B:NM_005514:exon3:c.G395T:p.R132L, |
| chr6 | 31356392 | G | A | . | exonic | HLA-B | nonsynonymous SNV | HLA-B:NM_005514:exon3:c.C394T:p.R132C, |
| chr6 | 31356407 | C | G | COSV69520491 | exonic | HLA-B | nonsynonymous SNV | HLA-B:NM_005514:exon3:c.G379C:p.V127L, |
| chr6 | 31356423 | G | C | COSV69520372 | exonic | HLA-B | nonsynonymous SNV | HLA-B:NM_005514:exon3:c.C363G:p.S121R, |
| chr6 | 31356425 | T | A | . | exonic | HLA-B | nonsynonymous SNV | HLA-B:NM_005514:exon3:c.A361T:p.S121C, |
| chr6 | 31356430 | A | C | COSV69520377 | exonic | HLA-B | nonsynonymous SNV | HLA-B:NM_005514:exon3:c.T356G:p.L119R, |
| chr6 | 31356431 | G | A | COSV69520378 | exonic | HLA-B | nonsynonymous SNV | HLA-B:NM_005514:exon3:c.C355T:p.L119F, |
| chr6 | 31356431 | G | T | COSV69520495 | exonic | HLA-B | nonsynonymous SNV | HLA-B:NM_005514:exon3:c.C355A:p.L119I, |
| chr6 | 31356433 | G | A | COSV69520255 | exonic | HLA-B | nonsynonymous SNV | HLA-B:NM_005514:exon3:c.C353T:p.T118I, |
| chr6 | 31356704 | G | C | . | exonic | HLA-B | stopgain | HLA-B:NM_005514:exon2:c.C327G:p.Y109X, |
| chr6 | 31356711 | C | T | . | exonic | HLA-B | nonsynonymous SNV | HLA-B:NM_005514:exon2:c.G320A:p.G107D, |
| chr6 | 31356712 | C | G | COSV69520745 | exonic | HLA-B | nonsynonymous SNV | HLA-B:NM_005514:exon2:c.G319C:p.G107R, |
| chr6 | 31356714 | C | A | COSV69520695 | exonic | HLA-B | nonsynonymous SNV | HLA-B:NM_005514:exon2:c.G317T:p.R106L, |
| chr6 | 31356717 | A | G | COSV69520426 | exonic | HLA-B | nonsynonymous SNV | HLA-B:NM_005514:exon2:c.T314C:p.L105P, |
| chr6 | 31356718 | G | C | COSV69520618 | exonic | HLA-B | nonsynonymous SNV | HLA-B:NM_005514:exon2:c.C313G:p.L105V, |
| chr6 | 31356720 | T | A | COSV69520629 | exonic | HLA-B | nonsynonymous SNV | HLA-B:NM_005514:exon2:c.A311T:p.N104I, |
| chr6 | 31356729 | C | T | COSV69520964 | exonic | HLA-B | nonsynonymous SNV | HLA-B:NM_005514:exon2:c.G302A:p.S101N, |
| chr6 | 31356735 | C | A | . | exonic | HLA-B | nonsynonymous SNV | HLA-B:NM_005514:exon2:c.G296T:p.R99L, |
| chr6 | 31356739 | C | A | COSV69520380 | exonic | HLA-B | nonsynonymous SNV | HLA-B:NM_005514:exon2:c.G292T:p.D98Y, |
| chr6 | 31356748 | C | T | COSV69520384 | exonic | HLA-B | nonsynonymous SNV | HLA-B:NM_005514:exon2:c.G283A:p.A95T, |
| chr6 | 31356749 | C | G | COSV69520387 | exonic | HLA-B | nonsynonymous SNV | HLA-B:NM_005514:exon2:c.G282C:p.Q94H, |
| chr6 | 31356751 | G | T | COSV69520389 | exonic | HLA-B | nonsynonymous SNV | HLA-B:NM_005514:exon2:c.C280A:p.Q94K, |
| chr6 | 31356754 | C | T | COSV69520472 | exonic | HLA-B | nonsynonymous SNV | HLA-B:NM_005514:exon2:c.G277A:p.A93T, |
| chr6 | 31356770 | G | C | COSV69520395 | exonic | HLA-B | nonsynonymous SNV | HLA-B:NM_005514:exon2:c.C261G:p.N87K, |
| chr6 | 31356772 | T | C | COSV69520398 | exonic | HLA-B | nonsynonymous SNV | HLA-B:NM_005514:exon2:c.A259G:p.N87D, |
| chr6 | 31356826 | C | T | COSV69520466 | exonic | HLA-B | nonsynonymous SNV | HLA-B:NM_005514:exon2:c.G205A:p.E69K, |
| chr6 | 31356838 | C | T | COSV69520400 | exonic | HLA-B | nonsynonymous SNV | HLA-B:NM_005514:exon2:c.G193A:p.A65T, |
| chr6 | 31356864 | T | A | COSV69520244 | exonic | HLA-B | nonsynonymous SNV | HLA-B:NM_005514:exon2:c.A167T:p.Q56L, |
| chr6 | 31356889 | A | T | COSV69520355 | exonic | HLA-B | nonsynonymous SNV | HLA-B:NM_005514:exon2:c.T142A:p.S48T, |
| chr6 | 31356909 | C | T | . | exonic | HLA-B | nonsynonymous SNV | HLA-B:NM_005514:exon2:c.G122A:p.R41H, |
| chr6 | 31356910 | G | T | . | exonic | HLA-B | nonsynonymous SNV | HLA-B:NM_005514:exon2:c.C121A:p.R41S, |
| chr6 | 31356922 | A | T | . | exonic | HLA-B | nonsynonymous SNV | HLA-B:NM_005514:exon2:c.T109A:p.S37T, |
| chr6 | 31356925 | C | T | COSV69520570 | exonic | HLA-B | nonsynonymous SNV | HLA-B:NM_005514:exon2:c.G106A:p.V36M, |
| chr6 | 31356928 | A | C | COSV69520756 | exonic | HLA-B | nonsynonymous SNV | HLA-B:NM_005514:exon2:c.T103G:p.S35A, |
| chr6 | 31356934 | A | G | COSV69520552 | exonic | HLA-B | nonsynonymous SNV | HLA-B:NM_005514:exon2:c.T97C:p.Y33H, |
| chr6 | 31357095 | T | G | . | exonic | HLA-B | nonsynonymous SNV | HLA-B:NM_005514:exon1:c.A64C:p.T22P, |
| chr6 | 31357110 | G | C | COSV69520883 | exonic | HLA-B | nonsynonymous SNV | HLA-B:NM_005514:exon1:c.C49G:p.L17V, |
| chr6 | 31357115 | G | C | COSV69520887 | exonic | HLA-B | nonsynonymous SNV | HLA-B:NM_005514:exon1:c.C44G:p.A15G, |
| chr6 | 31357118 | G | C | COSV69520786 | exonic | HLA-B | nonsynonymous SNV | HLA-B:NM_005514:exon1:c.C41G:p.S14W, |
| chr6 | 31357134 | C | G | COSV69520789 | exonic | HLA-B | nonsynonymous SNV | HLA-B:NM_005514:exon1:c.G25C:p.V9L, |
| chr6 | 31357148 | A | G | COSV69520322 | exonic | HLA-B | nonsynonymous SNV | HLA-B:NM_005514:exon1:c.T11C:p.M4T, |
| chr6 | 31357154 | A | C | COSV69520325 | exonic | HLA-B | nonsynonymous SNV | HLA-B:NM_005514:exon1:c.T5G:p.L2R, |
| chr6 | 31870664 | T | C | COSV57667713 | exonic | SLC44A4 | nonsynonymous SNV | SLC44A4:NM_001178045:exon11:c.A748G:p.M250V,SLC44A4:NM_001178044:exon10:c.A850G:p.M284V,SLC44A4:NM_025257:exon11:c.A976G:p.M326V, |
| chr6 | 31871532 | C | T | COSV57666266 | exonic | SLC44A4 | nonsynonymous SNV | SLC44A4:NM_001178045:exon8:c.G331A:p.V111I,SLC44A4:NM_001178044:exon7:c.G433A:p.V145I,SLC44A4:NM_025257:exon8:c.G559A:p.V187I, |
| chr6 | 31874821 | T | A | . | exonic | SLC44A4 | nonsynonymous SNV | SLC44A4:NM_001178045:exon6:c.A140T:p.D47V,SLC44A4:NM_025257:exon6:c.A368T:p.D123V, |
| chr6 | 31896770 | G | T | . | exonic | EHMT2 | nonsynonymous SNV | EHMT2:NM_001289413:exon2:c.C335A:p.T112N,EHMT2:NM_025256:exon3:c.C164A:p.T55N,EHMT2:NM_006709:exon3:c.C164A:p.T55N, |
| chr6 | 31946402 | C | T | COSV54959347 | exonic | CFB | nonsynonymous SNV | CFB:NM_001710:exon2:c.C94T:p.R32W, |
| chr6 | 31947752 | G | A | . | exonic | CFB | nonsynonymous SNV | CFB:NM_001710:exon5:c.G669A:p.M223I, |
| chr6 | 31948465 | G | A | . | exonic | CFB | nonsynonymous SNV | CFB:NM_001710:exon7:c.G989A:p.S330N, |
| chr6 | 31949243 | G | A | . | exonic | CFB | nonsynonymous SNV | CFB:NM_001710:exon9:c.G1169A:p.G390E, |
| chr6 | 31951473 | G | A | . | exonic | CFB | nonsynonymous SNV | CFB:NM_001710:exon16:c.G2089A:p.G697S, |
| chr6 | 31952020 | G | T | . | exonic | CFB | nonsynonymous SNV | CFB:NM_001710:exon18:c.G2285T:p.G762V, |
| chr6 | 32443869 | T | G | COSV66622530 | exonic | HLA-DRA | nonsynonymous SNV | HLA-DRA:NM_019111:exon4:c.T724G:p.L242V, |
| chr6 | 32580249 | G | C | COSV63511877 | exonic | HLA-DRB1 | nonsynonymous SNV | HLA-DRB1:NM_002124:exon5:c.C785G:p.T262R, |
| chr6 | 32581675 | C | G | COSV63511369 | exonic | HLA-DRB1 | nonsynonymous SNV | HLA-DRB1:NM_002124:exon3:c.G534C:p.Q178H, |
| chr6 | 32581698 | T | C | COSV63511375 | exonic | HLA-DRB1 | nonsynonymous SNV | HLA-DRB1:NM_002124:exon3:c.A511G:p.M171V, |
| chr6 | 32581704 | C | T | COSV63511807 | exonic | HLA-DRB1 | nonsynonymous SNV | HLA-DRB1:NM_002124:exon3:c.G505A:p.A169T, |
| chr6 | 32581724 | A | C | COSV63513549 | exonic | HLA-DRB1 | nonsynonymous SNV | HLA-DRB1:NM_002124:exon3:c.T485G:p.L162R, |
| chr6 | 32581834 | T | A | COSV63511918 | exonic | HLA-DRB1 | nonsynonymous SNV | HLA-DRB1:NM_002124:exon3:c.A375T:p.Q125H, |
| chr6 | 32581836 | G | A | . | exonic | HLA-DRB1 | stopgain | HLA-DRB1:NM_002124:exon3:c.C373T:p.Q125X, |

| | | | | | | | | |
|---|---|---|---|---|---|---|---|---|
| chr6 | 32584151 | G | A | COSV63512099 | exonic | HLA-DRB1 | nonsynonymous SNV | HLA-DRB1:NM_002124:exon2:c.C328T:p.H110Y, |
| chr6 | 32584181 | C | T | COSV63515717 | exonic | HLA-DRB1 | nonsynonymous SNV | HLA-DRB1:NM_002124:exon2:c.G298A:p.A100T, |
| chr6 | 32584193 | T | G | COSV63511955 | exonic | HLA-DRB1 | nonsynonymous SNV | HLA-DRB1:NM_002124:exon2:c.A286C:p.I96L, |
| chr6 | 32584219 | G | T | COSV63512333 | exonic | HLA-DRB1 | nonsynonymous SNV | HLA-DRB1:NM_002124:exon2:c.C260A:p.A87D, |
| chr6 | 32584252 | A | T | COSV63512106 | exonic | HLA-DRB1 | nonsynonymous SNV | HLA-DRB1:NM_002124:exon2:c.T227A:p.F76Y, |
| chr6 | 32584282 | G | T | COSV63511637 | exonic | HLA-DRB1 | nonsynonymous SNV | HLA-DRB1:NM_002124:exon2:c.C197A:p.S66Y, |
| chr6 | 32584301 | A | T | COSV63512990 | exonic | HLA-DRB1 | nonsynonymous SNV | HLA-DRB1:NM_002124:exon2:c.T178A:p.F60I, |
| chr6 | 32584353 | C | A | COSV63517279 | exonic;splicing | HLA-DRB1 | nonsynonymous SNV | HLA-DRB1:NM_002124:exon2:c.G126T:p.R42S, |
| chr6 | 32584354 | C | T | . | exonic;splicing | HLA-DRB1 | nonsynonymous SNV | HLA-DRB1:NM_002124:exon2:c.G125A:p.R42K, |
| chr6 | 32584355 | T | A | COSV63517292 | exonic | HLA-DRB1 | nonsynonymous SNV | HLA-DRB1:NM_002124:exon2:c.A124T:p.R42W, |
| chr6 | 32584357 | T | G | COSV63515882 | exonic | HLA-DRB1 | nonsynonymous SNV | HLA-DRB1:NM_002124:exon2:c.A122C:p.K41T, |
| chr6 | 32584360 | G | A | COSV63512628 | exonic | HLA-DRB1 | nonsynonymous SNV | HLA-DRB1:NM_002124:exon2:c.C119T:p.P40L, |
| chr6 | 32584361 | G | A | . | exonic | HLA-DRB1 | nonsynonymous SNV | HLA-DRB1:NM_002124:exon2:c.C118T:p.P40S, |
| chr6 | 32584361 | G | C | . | exonic | HLA-DRB1 | nonsynonymous SNV | HLA-DRB1:NM_002124:exon2:c.C118G:p.P40A, |
| chr6 | 32584362 | C | G | COSV63515888 | exonic | HLA-DRB1 | nonsynonymous SNV | HLA-DRB1:NM_002124:exon2:c.G117C:p.Q39H, |
| chr6 | 32584364 | G | A | COSV63515896 | exonic | HLA-DRB1 | stopgain | HLA-DRB1:NM_002124:exon2:c.C115T:p.Q39X, |
| chr6 | 32584366 | C | T | COSV63511380 | exonic | HLA-DRB1 | stopgain | HLA-DRB1:NM_002124:exon2:c.G113A:p.W38X, |
| chr6 | 32584367 | A | C | COSV63511385 | exonic | HLA-DRB1 | nonsynonymous SNV | HLA-DRB1:NM_002124:exon2:c.T112G:p.W38G, |
| chr6 | 32589658 | A | C | COSV63512004 | exonic | HLA-DRB1 | nonsynonymous SNV | HLA-DRB1:NM_002124:exon1:c.T85G:p.S29A, |
| chr6 | 32589702 | G | A | COSV63511399 | exonic | HLA-DRB1 | nonsynonymous SNV | HLA-DRB1:NM_002124:exon1:c.C41T:p.A14V, |
| chr6 | 32589706 | T | C | COSV63512025 | exonic | HLA-DRB1 | nonsynonymous SNV | HLA-DRB1:NM_002124:exon1:c.A37G:p.T13A, |
| chr6 | 32589729 | T | C | COSV63511656 | exonic | HLA-DRB1 | nonsynonymous SNV | HLA-DRB1:NM_002124:exon1:c.A14G:p.K5R, |
| chr6 | 32637480 | C | A | COSV58241001 | exonic | HLA-DQA1 | nonsynonymous SNV | HLA-DQA1:NM_002122:exon1:c.C22A:p.L8M, |
| chr6 | 32641328 | G | A | COSV58237735 | exonic | HLA-DQA1 | nonsynonymous SNV | HLA-DQA1:NM_002122:exon2:c.G101A:p.C34Y, |
| chr6 | 32641349 | T | C | COSV58239937 | exonic | HLA-DQA1 | nonsynonymous SNV | HLA-DQA1:NM_002122:exon2:c.T122C:p.F41S, |
| chr6 | 32641396 | C | G | COSV58238975 | exonic | HLA-DQA1 | nonsynonymous SNV | HLA-DQA1:NM_002122:exon2:c.C169G:p.Q57E, |
| chr6 | 32641430 | C | T | COSV58237057 | exonic | HLA-DQA1 | nonsynonymous SNV | HLA-DQA1:NM_002122:exon2:c.C203T:p.A68V, |
| chr6 | 32641439 | G | T | COSV58237192 | exonic | HLA-DQA1 | nonsynonymous SNV | HLA-DQA1:NM_002122:exon2:c.G212T:p.W71L, |
| chr6 | 32641445 | A | T | COSV58237205 | exonic | HLA-DQA1 | nonsynonymous SNV | HLA-DQA1:NM_002122:exon2:c.A218T:p.E73V, |
| chr6 | 32641459 | G | A | COSV58237229 | exonic | HLA-DQA1 | nonsynonymous SNV | HLA-DQA1:NM_002122:exon2:c.G232A:p.G78R, |
| chr6 | 32641477 | G | T | COSV58237257 | exonic | HLA-DQA1 | nonsynonymous SNV | HLA-DQA1:NM_002122:exon2:c.G250T:p.G84C, |
| chr6 | 32641478 | G | T | COSV58237262 | exonic | HLA-DQA1 | nonsynonymous SNV | HLA-DQA1:NM_002122:exon2:c.G251T:p.G84V, |
| chr6 | 32641487 | G | C | COSV58238124 | exonic | HLA-DQA1 | nonsynonymous SNV | HLA-DQA1:NM_002122:exon2:c.G260C:p.R87T, |
| chr6 | 32641494 | G | C | COSV58238131 | exonic | HLA-DQA1 | nonsynonymous SNV | HLA-DQA1:NM_002122:exon2:c.G267C:p.M89I, |
| chr6 | 32641501 | G | C | COSV58238145 | exonic | HLA-DQA1 | nonsynonymous SNV | HLA-DQA1:NM_002122:exon2:c.G274C:p.A92P, |
| chr6 | 32641502 | C | T | COSV58238153 | exonic | HLA-DQA1 | nonsynonymous SNV | HLA-DQA1:NM_002122:exon2:c.C275T:p.A92V, |
| chr6 | 32641535 | A | C | COSV58237540 | exonic | HLA-DQA1 | nonsynonymous SNV | HLA-DQA1:NM_002122:exon2:c.A308C:p.Y103S, |
| chr6 | 32642096 | G | C | COSV58237000 | exonic | HLA-DQA1 | nonsynonymous SNV | HLA-DQA1:NM_002122:exon3:c.G456C:p.Q152H, |
| chr6 | 32642232 | C | G | COSV58237862 | exonic | HLA-DQA1 | nonsynonymous SNV | HLA-DQA1:NM_002122:exon3:c.C592G:p.Q198E, |
| chr6 | 3264253 | T | C | . | exonic | PSMG4 | nonsynonymous SNV | PSMG4:NM_001128592:exon3:c.T295C:p.W99R, |
| chr6 | 3264268 | G | C | . | exonic | PSMG4 | nonsynonymous SNV | PSMG4:NM_001128592:exon3:c.G310C:p.G104R, |
| chr6 | 32642684 | A | G | COSV58237233 | exonic | HLA-DQA1 | nonsynonymous SNV | HLA-DQA1:NM_002122:exon4:c.A688G:p.M230V, |
| chr6 | 3264292 | C | T | COSV61476587 | exonic | PSMG4 | nonsynonymous SNV | PSMG4:NM_001128592:exon3:c.C334T:p.L112F, |
| chr6 | 3264293 | T | C | COSV61476414 | exonic | PSMG4 | nonsynonymous SNV | PSMG4:NM_001128592:exon3:c.T335C:p.L112P, |
| chr6 | 32661352 | T | C | COSV66570628 | exonic | HLA-DQB1 | nonsynonymous SNV | HLA-DQB1:NM_001243961:exon4:c.A767G:p.Q256R,HLA-DQB1:NM_002123:exon4:c.A767G:p.Q256R, |
| chr6 | 32661360 | T | G | COSV66569175 | exonic | HLA-DQB1 | nonsynonymous SNV | HLA-DQB1:NM_001243961:exon4:c.A759C:p.Q253H,HLA-DQB1:NM_002123:exon4:c.A759C:p.Q253H, |
| chr6 | 32661364 | C | T | COSV66569234 | exonic | HLA-DQB1 | nonsynonymous SNV | HLA-DQB1:NM_001243961:exon4:c.G755A:p.R252H,HLA-DQB1:NM_002123:exon4:c.G755A:p.R252H, |
| chr6 | 32661416 | C | T | COSV66569072 | exonic | HLA-DQB1 | nonsynonymous SNV | HLA-DQB1:NM_001243961:exon4:c.G703A:p.V235I,HLA-DQB1:NM_002123:exon4:c.G703A:p.V235I, |
| chr6 | 32662158 | C | G | COSV66569169 | exonic | HLA-DQB1 | nonsynonymous SNV | HLA-DQB1:NM_001243961:exon3:c.G470C:p.G157A,HLA-DQB1:NM_002123:exon3:c.G470C:p.G157A, |
| chr6 | 32662159 | C | T | COSV66571389 | exonic | HLA-DQB1 | nonsynonymous SNV | HLA-DQB1:NM_001243961:exon3:c.G469A:p.G157S,HLA-DQB1:NM_002123:exon3:c.G469A:p.G157S, |
| chr6 | 32662186 | C | T | COSV66571396 | exonic | HLA-DQB1 | nonsynonymous SNV | HLA-DQB1:NM_001243961:exon3:c.G442A:p.V148I,HLA-DQB1:NM_002123:exon3:c.G442A:p.V148I, |
| chr6 | 32664812 | A | G | COSV66572445 | exonic | HLA-DQB1 | nonsynonymous SNV | HLA-DQB1:NM_001243961:exon2:c.T365C:p.I122T,HLA-DQB1:NM_002123:exon2:c.T365C:p.I122T, |
| chr6 | 32664815 | C | G | COSV66570777 | exonic | HLA-DQB1 | nonsynonymous SNV | HLA-DQB1:NM_001243961:exon2:c.G362C:p.G121A,HLA-DQB1:NM_002123:exon2:c.G362C:p.G121A, |
| chr6 | 32664816 | C | T | COSV66570784 | exonic | HLA-DQB1 | nonsynonymous SNV | HLA-DQB1:NM_001243961:exon2:c.G361A:p.G121R,HLA-DQB1:NM_002123:exon2:c.G361A:p.G121R, |
| chr6 | 32664821 | A | T | COSV66571213 | exonic | HLA-DQB1 | nonsynonymous SNV | HLA-DQB1:NM_001243961:exon2:c.T356A:p.F119Y,HLA-DQB1:NM_002123:exon2:c.T356A:p.F119Y, |
| chr6 | 32664822 | A | G | COSV66571708 | exonic | HLA-DQB1 | nonsynonymous SNV | HLA-DQB1:NM_001243961:exon2:c.T355C:p.F119L,HLA-DQB1:NM_002123:exon2:c.T355C:p.F119L, |
| chr6 | 32664824 | G | T | COSV66570235 | exonic | HLA-DQB1 | nonsynonymous SNV | HLA-DQB1:NM_001243961:exon2:c.C353A:p.A118E,HLA-DQB1:NM_002123:exon2:c.C353A:p.A118E, |
| chr6 | 32664828 | C | A | COSV66569295 | exonic | HLA-DQB1 | nonsynonymous SNV | HLA-DQB1:NM_001243961:exon2:c.G349T:p.V117L,HLA-DQB1:NM_002123:exon2:c.G349T:p.V117L, |
| chr6 | 32664831 | C | G | COSV66569005 | exonic | HLA-DQB1 | nonsynonymous SNV | HLA-DQB1:NM_001243961:exon2:c.G346C:p.E116Q,HLA-DQB1:NM_002123:exon2:c.G346C:p.E116Q, |
| chr6 | 32664851 | G | C | COSV66569250 | exonic | HLA-DQB1 | nonsynonymous SNV | HLA-DQB1:NM_001243961:exon2:c.C326G:p.T109R,HLA-DQB1:NM_002123:exon2:c.C326G:p.T109R, |
| chr6 | 32664858 | A | C | COSV66571226 | exonic | HLA-DQB1 | nonsynonymous SNV | HLA-DQB1:NM_001243961:exon2:c.T319G:p.L107V,HLA-DQB1:NM_002123:exon2:c.T319G:p.L107V, |
| chr6 | 32664860 | T | G | COSV66569728 | exonic | HLA-DQB1 | nonsynonymous SNV | HLA-DQB1:NM_001243961:exon2:c.A317C:p.E106A,HLA-DQB1:NM_002123:exon2:c.A317C:p.E106A, |
| chr6 | 32664861 | C | A | . | exonic | HLA-DQB1 | stopgain | HLA-DQB1:NM_001243961:exon2:c.G316T:p.E106X,HLA-DQB1:NM_002123:exon2:c.G316T:p.E106X, |
| chr6 | 32664870 | T | C | COSV66574305 | exonic | HLA-DQB1 | nonsynonymous SNV | HLA-DQB1:NM_001243961:exon2:c.A307G:p.T103A,HLA-DQB1:NM_002123:exon2:c.A307G:p.T103A, |
| chr6 | 32664873 | C | T | COSV66569694 | exonic | HLA-DQB1 | nonsynonymous SNV | HLA-DQB1:NM_001243961:exon2:c.G304A:p.G102R,HLA-DQB1:NM_002123:exon2:c.G304A:p.G102R, |
| chr6 | 32664911 | T | A | COSV66574314 | exonic | HLA-DQB1 | nonsynonymous SNV | HLA-DQB1:NM_001243961:exon2:c.A266T:p.D89V,HLA-DQB1:NM_002123:exon2:c.A266T:p.D89V, |
| chr6 | 32664911 | T | G | COSV66569310 | exonic | HLA-DQB1 | nonsynonymous SNV | HLA-DQB1:NM_001243961:exon2:c.A266C:p.D89A,HLA-DQB1:NM_002123:exon2:c.A266C:p.D89A, |
| chr6 | 32664923 | T | A | COSV66569317 | exonic | HLA-DQB1 | nonsynonymous SNV | HLA-DQB1:NM_001243961:exon2:c.A254T:p.Q85L,HLA-DQB1:NM_002123:exon2:c.A254T:p.Q85L, |
| chr6 | 32664968 | G | A | COSV66571252 | exonic | HLA-DQB1 | nonsynonymous SNV | HLA-DQB1:NM_001243961:exon2:c.C209T:p.A70V,HLA-DQB1:NM_002123:exon2:c.C209T:p.A70V, |
| chr6 | 32664993 | A | G | COSV66574347 | exonic | HLA-DQB1 | nonsynonymous SNV | HLA-DQB1:NM_001243961:exon2:c.T184C:p.Y62H,HLA-DQB1:NM_002123:exon2:c.T184C:p.Y62H, |
| chr6 | 32664993 | A | T | COSV66569205 | exonic | HLA-DQB1 | nonsynonymous SNV | HLA-DQB1:NM_001243961:exon2:c.T184A:p.Y62N,HLA-DQB1:NM_002123:exon2:c.T184A:p.Y62N, |
| chr6 | 32665004 | A | C | COSV66569625 | exonic | HLA-DQB1 | nonsynonymous SNV | HLA-DQB1:NM_001243961:exon2:c.T173G:p.L58R,HLA-DQB1:NM_002123:exon2:c.T173G:p.L58R, |
| chr6 | 32665005 | G | C | COSV66569632 | exonic | HLA-DQB1 | nonsynonymous SNV | HLA-DQB1:NM_001243961:exon2:c.C172G:p.L58V,HLA-DQB1:NM_002123:exon2:c.C172G:p.L58V, |
| chr6 | 32665041 | T | G | COSV66570981 | exonic | HLA-DQB1 | nonsynonymous SNV | HLA-DQB1:NM_001243961:exon2:c.A136C:p.M46L,HLA-DQB1:NM_002123:exon2:c.A136C:p.M46L, |
| chr6 | 32665055 | A | T | COSV66570989 | exonic | HLA-DQB1 | nonsynonymous SNV | HLA-DQB1:NM_001243961:exon2:c.T122A:p.F41Y,HLA-DQB1:NM_002123:exon2:c.T122A:p.F41Y, |
| chr6 | 32666523 | G | C | COSV66569489 | exonic | HLA-DQB1 | nonsynonymous SNV | HLA-DQB1:NM_001243961:exon1:c.C85G:p.L29V,HLA-DQB1:NM_002123:exon1:c.C85G:p.L29V, |
| chr6 | 32666525 | A | G | COSV66570860 | exonic | HLA-DQB1 | nonsynonymous SNV | HLA-DQB1:NM_001243961:exon1:c.T83C:p.L28P,HLA-DQB1:NM_002123:exon1:c.T83C:p.L28P, |
| chr6 | 32666529 | A | T | COSV66569215 | exonic | HLA-DQB1 | nonsynonymous SNV | HLA-DQB1:NM_001243961:exon1:c.T79A:p.S27T,HLA-DQB1:NM_002123:exon1:c.T79A:p.S27T, |

| | | | | | | | | |
|---|---|---|---|---|---|---|---|---|
| chr6 | 32666536 | C | G | COSV66571583 | exonic | HLA-DQB1 | nonsynonymous SNV | HLA-DQB1:NM_001243961:exon1:c.G72C:p.M24I,HLA-DQB1:NM_002123:exon1:c.G72C:p.M24I, |
| chr6 | 32666564 | A | G | COSV66571272 | exonic | HLA-DQB1 | nonsynonymous SNV | HLA-DQB1:NM_001243961:exon1:c.T44C:p.V15A,HLA-DQB1:NM_002123:exon1:c.T44C:p.V15A, |
| chr6 | 32666573 | T | C | COSV66569687 | exonic | HLA-DQB1 | nonsynonymous SNV | HLA-DQB1:NM_001243961:exon1:c.A35G:p.D12G,HLA-DQB1:NM_002123:exon1:c.A35G:p.D12G, |
| chr6 | 32666592 | C | A | . | exonic | HLA-DQB1 | nonsynonymous SNV | HLA-DQB1:NM_001243961:exon1:c.G16T:p.A6S,HLA-DQB1:NM_002123:exon1:c.G16T:p.A6S, |
| chr6 | 32847198 | T | C | COSV62754137 | exonic | TAP1 | nonsynonymous SNV | TAP1:NM_000593:exon10:c.A2090G:p.D697G,TAP1:NM_001292022:exon10:c.A1307G:p.D436G, |
| chr6 | 32857313 | G | A | COSV62754598 | exonic | PSMB9 | nonsynonymous SNV | PSMB9:NM_002800:exon3:c.G179A:p.R60H, |
| chr6 | 32976408 | G | A | . | exonic | BRD2 | nonsynonymous SNV | BRD2:NM_001113182:exon6:c.G769A:p.A257T,BRD2:NM_005104:exon6:c.G769A:p.A257T,BRD2:NM_001291986:exon7:c.G409A:p.A137T,BRD2:NM_001199456:exon5:c.G628A:p.A210T,BRD2:NM_001199455:exon5:c.G769A:p.A257T, |
| chr6 | 32976800 | G | A | COSV66374303 | exonic | BRD2 | nonsynonymous SNV | BRD2:NM_001113182:exon7:c.G1064A:p.G355D,BRD2:NM_005104:exon7:c.G1064A:p.G355D,BRD2:NM_001291986:exon8:c.G704A:p.G235D,BRD2:NM_001199456:exon6:c.G923A:p.G308D,BRD2:NM_001199455:exon6:c.G1064A:p.G355D, |
| chr6 | 32976832 | G | A | . | exonic | BRD2 | nonsynonymous SNV | BRD2:NM_001113182:exon7:c.G1096A:p.A366T,BRD2:NM_005104:exon7:c.G1096A:p.A366T,BRD2:NM_001291986:exon8:c.G736A:p.A246T,BRD2:NM_001199456:exon6:c.G955A:p.A319T,BRD2:NM_001199455:exon6:c.G1096A:p.A366T, |
| chr6 | 32977847 | G | A | . | exonic | BRD2 | nonsynonymous SNV | BRD2:NM_001113182:exon9:c.G1420A:p.A474T,BRD2:NM_005104:exon9:c.G1420A:p.A474T,BRD2:NM_001291986:exon10:c.G1060A:p.A354T,BRD2:NM_001199456:exon8:c.G1279A:p.A427T,BRD2:NM_001199455:exon8:c.G1420A:p.A474T, |
| chr6 | 33068658 | T | G | COSV70088544 | exonic | HLA-DPA1 | nonsynonymous SNV | HLA-DPA1:NM_001242525:exon5:c.A775C:p.T259P,HLA-DPA1:NM_033554:exon4:c.A775C:p.T259P,HLA-DPA1:NM_001242524:exon5:c.A775C:p.T259P, |
| chr6 | 33069076 | A | C | COSV70089042 | exonic | HLA-DPA1 | nonsynonymous SNV | HLA-DPA1:NM_001242525:exon4:c.T571G:p.F191V,HLA-DPA1:NM_033554:exon3:c.T571G:p.F191V,HLA-DPA1:NM_001242524:exon4:c.T571G:p.F191V, |
| chr6 | 33069174 | A | G | COSV70089044 | exonic | HLA-DPA1 | nonsynonymous SNV | HLA-DPA1:NM_001242525:exon4:c.T473C:p.L158P,HLA-DPA1:NM_033554:exon3:c.T473C:p.L158P,HLA-DPA1:NM_001242524:exon4:c.T473C:p.L158P, |
| chr6 | 33069222 | T | C | . | exonic | HLA-DPA1 | nonsynonymous SNV | HLA-DPA1:NM_001242525:exon4:c.A425G:p.K142R,HLA-DPA1:NM_033554:exon3:c.A425G:p.K142R,HLA-DPA1:NM_001242524:exon4:c.A425G:p.K142R, |
| chr6 | 33069647 | T | C | COSV70088423 | exonic | HLA-DPA1 | nonsynonymous SNV | HLA-DPA1:NM_001242525:exon3:c.A340G:p.T114A,HLA-DPA1:NM_033554:exon2:c.A340G:p.T114A,HLA-DPA1:NM_001242524:exon3:c.A340G:p.T114A, |
| chr6 | 33069745 | T | C | COSV70089057 | exonic | HLA-DPA1 | nonsynonymous SNV | HLA-DPA1:NM_001242525:exon3:c.A242G:p.Q81R,HLA-DPA1:NM_033554:exon2:c.A242G:p.Q81R,HLA-DPA1:NM_001242524:exon3:c.A242G:p.Q81R, |
| chr6 | 33069802 | A | T | . | exonic | HLA-DPA1 | nonsynonymous SNV | HLA-DPA1:NM_001242525:exon3:c.T185A:p.M62K,HLA-DPA1:NM_033554:exon2:c.T185A:p.M62K,HLA-DPA1:NM_001242524:exon3:c.T185A:p.M62K, |
| chr6 | 33069803 | T | G | . | exonic | HLA-DPA1 | nonsynonymous SNV | HLA-DPA1:NM_001242525:exon3:c.A184C:p.M62L,HLA-DPA1:NM_033554:exon2:c.A184C:p.M62L,HLA-DPA1:NM_001242524:exon3:c.A184C:p.M62L, |
| chr6 | 33080680 | C | G | COSV69604293 | exonic | HLA-DPB1 | nonsynonymous SNV | HLA-DPB1:NM_002121:exon2:c.C109G:p.L37V, |
| chr6 | 33080683 | T | C | COSV69602618 | exonic | HLA-DPB1 | nonsynonymous SNV | HLA-DPB1:NM_002121:exon2:c.T112C:p.F38L, |
| chr6 | 33080684 | T | A | COSV69604303 | exonic | HLA-DPB1 | nonsynonymous SNV | HLA-DPB1:NM_002121:exon2:c.T113A:p.F38Y, |
| chr6 | 33080689 | G | T | COSV69602627 | exonic | HLA-DPB1 | stopgain | HLA-DPB1:NM_002121:exon2:c.G118T:p.G40X, |
| chr6 | 33080690 | G | T | COSV69602634 | exonic | HLA-DPB1 | nonsynonymous SNV | HLA-DPB1:NM_002121:exon2:c.G119T:p.G40V, |
| chr6 | 33080761 | T | C | COSV69604555 | exonic | HLA-DPB1 | nonsynonymous SNV | HLA-DPB1:NM_002121:exon2:c.T190C:p.F64L, |
| chr6 | 33080765 | C | T | COSV69603920 | exonic | HLA-DPB1 | nonsynonymous SNV | HLA-DPB1:NM_002121:exon2:c.C194T:p.A65V, |
| chr6 | 33080822 | C | A | COSV69602760 | exonic | HLA-DPB1 | nonsynonymous SNV | HLA-DPB1:NM_002121:exon2:c.C251A:p.A84D, |
| chr6 | 33080825 | C | A | COSV69602767 | exonic | HLA-DPB1 | nonsynonymous SNV | HLA-DPB1:NM_002121:exon2:c.C254A:p.A85E, |
| chr6 | 33080829 | G | C | COSV69602641 | exonic | HLA-DPB1 | nonsynonymous SNV | HLA-DPB1:NM_002121:exon2:c.G258C:p.E86D, |
| chr6 | 33080851 | A | C | COSV69602396 | exonic | HLA-DPB1 | nonsynonymous SNV | HLA-DPB1:NM_002121:exon2:c.A280C:p.I94L, |
| chr6 | 33080863 | A | G | COSV69602323 | exonic | HLA-DPB1 | nonsynonymous SNV | HLA-DPB1:NM_002121:exon2:c.A292G:p.K98E, |
| chr6 | 33080884 | A | G | COSV69602522 | exonic | HLA-DPB1 | nonsynonymous SNV | HLA-DPB1:NM_002121:exon2:c.A313G:p.M105V, |
| chr6 | 33080886 | G | A | COSV69602532 | exonic | HLA-DPB1 | nonsynonymous SNV | HLA-DPB1:NM_002121:exon2:c.G315A:p.M105I, |
| chr6 | 33080909 | G | A | COSV69602658 | exonic | HLA-DPB1 | nonsynonymous SNV | HLA-DPB1:NM_002121:exon2:c.G338A:p.G113D, |
| chr6 | 33080912 | G | A | COSV69602664 | exonic | HLA-DPB1 | nonsynonymous SNV | HLA-DPB1:NM_002121:exon2:c.G341A:p.G114E, |
| chr6 | 33080914 | C | G | COSV69602509 | exonic | HLA-DPB1 | nonsynonymous SNV | HLA-DPB1:NM_002121:exon2:c.C343G:p.P115A, |
| chr6 | 33080917 | A | G | COSV69602675 | exonic | HLA-DPB1 | nonsynonymous SNV | HLA-DPB1:NM_002121:exon2:c.A346G:p.M116V, |
| chr6 | 33084959 | G | A | COSV69602680 | exonic | HLA-DPB1 | nonsynonymous SNV | HLA-DPB1:NM_002121:exon3:c.G374A:p.R125K, |
| chr6 | 33085181 | C | T | . | exonic | HLA-DPB1 | nonsynonymous SNV | HLA-DPB1:NM_002121:exon3:c.C596T:p.T199I, |
| chr6 | 33085832 | G | A | COSV69602711 | exonic | HLA-DPB1 | nonsynonymous SNV | HLA-DPB1:NM_002121:exon4:c.G700A:p.V234M, |
| chr6 | 34246545 | C | G | COSV58988465 | exonic | C6ORF1 | nonsynonymous SNV | C6orf1:NM_001287398:exon4:c.G389C:p.G130A,C6orf1:NM_001008703:exon5:c.G449C:p.G150A,C6orf1:NM_001287397:exon4:c.G389C:p.G130A,C6orf1:NM_001008704:exon5:c.G449C:p.G150A,C6orf1:NM_178508:exon5:c.G449C:p.G150A,C6orf1:NM_001287396:exon5:c.G449C:p.G150A, |
| chr6 | 36319535 | G | C | COSV71325706 | exonic | C6ORF222 | nonsynonymous SNV | C6orf222:NM_001010903:exon11:c.C1744G:p.L582V, |
| chr6 | 3727577 | C | G | COSV66661258 | exonic | PXDC1 | nonsynonymous SNV | PXDC1:NM_183373:exon4:c.G552C:p.Q184H, |
| chr6 | 41936044 | A | C | COSV57827647 | exonic | CCND3 | nonsynonymous SNV | CCND3:NM_001760:exon5:c.T775G:p.S259A,CCND3:NM_001136017:exon5:c.T532G:p.S178A,CCND3:NM_001287434:exon5:c.T187G:p.S63A,CCND3:NM_001136126:exon4:c.T187G:p.S63A,CCND3:NM_001136125:exon4:c.T559G:p.S187A,CCND3:NM_001287427:exon5:c.T625G:p.S209A, |
| chr6 | 42745880 | A | G | . | exonic | TBCC | nonsynonymous SNV | TBCC:NM_003192:exon1:c.T194C:p.V65A, |
| chr6 | 42938646 | G | T | COSV65709724 | exonic | CNPY3 | nonsynonymous SNV | CNPY3:NM_006586:exon6:c.G692T:p.S231I, |
| chr6 | 42964462 | G | T | COSV55103186 | exonic | PEX6 | nonsynonymous SNV | PEX6:NM_000287:exon17:c.C2816A:p.P939Q,PEX6:NM_001316313:exon17:c.C2552A:p.P851Q, |
| chr6 | 43777496 | G | A | COSV57880232 | exonic | VEGFA | nonsynonymous SNV | VEGFA:NM_001025370:exon3:c.G686A:p.R229H,VEGFA:NM_001033756:exon3:c.G686A:p.R229H,VEGFA:NM_001171622:exon3:c.G686A:p.R229H,VEGFA:NM_001025366:exon3:c.G686A:p.R229H,VEGFA:NM_001171630:exon3:c.G146A:p.R49H,VEGFA:NM_001171626:exon3:c.G146A:p.R49H,VEGFA:NM_001171625:exon3:c.G146A:p.R49H,VEGFA:NM_001204385:exon3:c.G686A:p.R229H,VEGFA:NM_001171623:exon3:c.G146A:p.R49H,VEGFA:NM_001171624:exon3:c.G146A:p.R49H,VEGFA:NM_001025368:exon3:c.G686A:p.R229H,VEGFA:NM_001171628:exon3:c.G146A:p.R49H,VEGFA:NM_001204384:exon3:c.G146A:p.R49H,VEGFA:NM_001025367:exon3:c.G686A:p.R229H,VEGFA:NM_001171627:exon3:c.G146A:p.R49H,VEGFA:NM_001171629:exon3:c.G146A:p.R49H,VEGFA:NM_001287044:exon3:c.G62A:p.R21H,VEGFA:NM_003376:exon3:c.G686A:p.R229H,VEGFA:NM_001025369:exon3:c.G686A:p.R229H, |
| chr6 | 44229463 | G | A | . | exonic | SLC29A1 | nonsynonymous SNV | SLC29A1:NM_001304462:exon4:c.G340A:p.A114T,SLC29A1:NM_001304463:exon3:c.G229A:p.A77T,SLC29A1:NM_001078177:exon4:c.G103A:p.A35T,SLC29A1:NM_001304466:exon3:c.G178A:p.A60T,SLC29A1:NM_001078175:exon4:c.G103A:p.A35T,SLC29A1:NM_001304465:exon3:c.G181A:p.A61T, |
| chr6 | 44230616 | G | A | . | exonic | SLC29A1 | nonsynonymous SNV | SLC29A1:NM_001304462:exon8:c.G875A:p.C292Y,SLC29A1:NM_001304463:exon7:c.G764A:p.C255Y,SLC29A1:NM_001078177:exon8:c.G638A:p.C213Y,SLC29A1:NM_001304466:exon7:c.G713A:p.C238Y,SLC29A1:NM_001078175:exon8:c.G638A:p.C213Y,SLC29A1:NM_001304465:exon7:c.G716A:p.C239Y, |
| chr6 | 44232847 | C | T | . | exonic | SLC29A1 | nonsynonymous SNV | SLC29A1:NM_001304462:exon13:c.C1337T:p.A446V,SLC29A1:NM_001304463:exon12:c.C1226T:p.A409V,SLC29A1:NM_001078177:exon13:c.C1100T:p.A367V,SLC29A1:NM_001304466:exon12:c.C1175T:p.A392V,SLC29A1:NM_001078175:exon13:c.C1100T:p.A367V,SLC29A1:NM_001304465:exon12:c.C1178T:p.A393V, |
| chr6 | 44251773 | C | A | . | exonic | HSP90AB1 | nonsynonymous SNV | HSP90AB1:NM_001271971:exon9:c.C1207A:p.L403M,HSP90AB1:NM_001271972:exon9:c.C1321A:p.L441M,HSP90AB1:NM_001271970:exon9:c.C1351A:p.L451M,HSP90AB1:NM_001271969:exon9:c.C1351A:p.L451M,HSP90AB1:NM_007355:exon9:c.C1351A:p.L451M, |
| chr6 | 46140015 | C | G | COSV58088141 | exonic | ENPP4 | nonsynonymous SNV | ENPP4:NM_014936:exon2:c.C432G:p.H144Q, |
| chr6 | 46839072 | C | T | COSV57919061 | exonic | MEP1A | nonsynonymous SNV | MEP1A:NM_005588:exon14:c.C2177T:p.T726M, |
| chr6 | 54321349 | T | C | COSV52510662 | exonic | TINAG | nonsynonymous SNV | TINAG:NM_014464:exon3:c.T472C:p.S158P, |
| chr6 | 69697573 | A | T | COSV65296211 | exonic | LMBRD1 | nonsynonymous SNV | LMBRD1:NM_018368:exon14:c.T1407A:p.D469E, |
| chr6 | 70836999 | C | T | COSV57640107 | exonic | SMAP1 | nonsynonymous SNV | SMAP1:NM_001044305:exon7:c.C635T:p.A212V,SMAP1:NM_001281440:exon7:c.C605T:p.A202V,SMAP1:NM_001281439:exon6:c.C554T:p.A185V,SMAP1:NM_021940:exon6:c.C554T:p.A185V, |
| chr6 | 7393172 | G | A | . | exonic | RIOK1 | nonsynonymous SNV | RIOK1:NM_031480:exon2:c.G145A:p.E49K, |
| chr6 | 7393257 | G | A | . | exonic | RIOK1 | nonsynonymous SNV | RIOK1:NM_031480:exon2:c.G230A:p.G77E, |
| chr6 | 7395101 | G | A | . | exonic | RIOK1 | nonsynonymous SNV | RIOK1:NM_031480:exon3:c.G325A:p.A109T, |
| chr6 | 7402828 | G | A | COSV53572602 | exonic | RIOK1 | nonsynonymous SNV | RIOK1:NM_031480:exon8:c.G698A:p.G233D, |
| chr6 | 7404970 | G | A | . | exonic | RIOK1 | nonsynonymous SNV | RIOK1:NM_153005:exon4:c.G322A:p.D108N,RIOK1:NM_031480:exon11:c.G1045A:p.D349N, |
| chr6 | 7405015 | G | A | . | exonic | RIOK1 | nonsynonymous SNV | RIOK1:NM_153005:exon4:c.G367A:p.V123I,RIOK1:NM_031480:exon11:c.G1090A:p.V364I, |
| chr6 | 7411389 | G | A | . | exonic | RIOK1 | nonsynonymous SNV | RIOK1:NM_153005:exon7:c.G604A:p.D202N,RIOK1:NM_031480:exon14:c.G1327A:p.D443N, |
| chr6 | 75634715 | C | T | . | exonic | SENP6 | nonsynonymous SNV | SENP6:NM_015571:exon5:c.C362T:p.T121M,SENP6:NM_001304792:exon5:c.C362T:p.T121M,SENP6:NM_001100409:exon5:c.C362T:p.T121M, |

| | | | | | | | | |
|---|---|---|---|---|---|---|---|---|
| chr6 | 7565494 | A | T | COSV65792675 | exonic | DSP | nonsynonymous SNV | DSP:NM_001008844:exon7:c.A913T:p.I305F,DSP:NM_004415:exon7:c.A913T:p.I305F, |
| chr6 | 75715572 | A | G | . | exonic | SENP6 | nonsynonymous SNV | SENP6:NM_015571:exon24:c.A3317G:p.Y1106C,SENP6:NM_001100409:exon23:c.A3296G:p.Y1099C, |
| chr6 | 7581403 | G | A | COSV65791527 | exonic | DSP | nonsynonymous SNV | DSP:NM_004415:exon23:c.G5213A:p.R1738Q, |
| chr6 | 75841286 | G | A | . | exonic | MYO6 | nonsynonymous SNV | MYO6:NM_001300899:exon9:c.G724A:p.E242K,MYO6:NM_004999:exon9:c.G724A:p.E242K, |
| chr6 | 75867077 | G | A | . | exonic | MYO6 | nonsynonymous SNV | MYO6:NM_001300899:exon18:c.G1916A:p.S639N,MYO6:NM_004999:exon18:c.G1916A:p.S639N, |
| chr6 | 75890204 | G | A | COSV64120693 | exonic | MYO6 | nonsynonymous SNV | MYO6:NM_001300899:exon26:c.G2806A:p.E936K,MYO6:NM_004999:exon26:c.G2806A:p.E936K, |
| chr6 | 85489515 | A | G | . | exonic | NT5E | nonsynonymous SNV | NT5E:NM_001204813:exon6:c.A1126G:p.T376A,NT5E:NM_002526:exon6:c.A1126G:p.T376A, |
| chr6 | 99545173 | C | T | . | exonic | CCNC | nonsynonymous SNV | CCNC:NM_005190:exon11:c.G736A:p.E246K,CCNC:NM_001013399:exon11:c.G481A:p.E161K, |
| chr6 | 99546409 | C | T | . | exonic | CCNC | nonsynonymous SNV | CCNC:NM_005190:exon10:c.G664A:p.V222M,CCNC:NM_001013399:exon10:c.G409A:p.V137M, |
| chr6 | 99546415 | G | A | . | exonic | CCNC | nonsynonymous SNV | CCNC:NM_005190:exon10:c.C658T:p.L220F,CCNC:NM_001013399:exon10:c.C403T:p.L135F, |
| chr7 | 100105324 | G | A | COSV57996467 | exonic | AP4M1 | nonsynonymous SNV | AP4M1:NM_004722:exon10:c.G812A:p.R271H, |
| chr7 | 101916133 | G | A | COSV52908115 | exonic | CUX1 | nonsynonymous SNV | CUX1:NM_181552:exon2:c.G49A:p.A17T,CUX1:NM_181500:exon2:c.G82A:p.A28T,CUX1:NM_001202545:exon2:c.G82A:p.A28T,CUX1:NM_001202544:exon2:c.G82A:p.A28T,CUX1:NM_001202543:exon2:c.G82A:p.A28T,CUX1:NM_001913:exon2:c.G82A:p.A28T, |
| chr7 | 101916152 | G | A | . | exonic | CUX1 | nonsynonymous SNV | CUX1:NM_181552:exon2:c.G68A:p.R23Q,CUX1:NM_181500:exon2:c.G101A:p.R34Q,CUX1:NM_001202545:exon2:c.G101A:p.R34Q,CUX1:NM_001202544:exon2:c.G101A:p.R34Q,CUX1:NM_001202543:exon2:c.G101A:p.R34Q,CUX1:NM_001913:exon2:c.G101A:p.R34Q, |
| chr7 | 102070358 | G | A | . | exonic | CUX1 | nonsynonymous SNV | CUX1:NM_181552:exon4:c.G209A:p.R70K,CUX1:NM_001202546:exon3:c.G131A:p.R44K,CUX1:NM_181500:exon4:c.G242A:p.R81K,CUX1:NM_001202545:exon4:c.G242A:p.R81K,CUX1:NM_001202544:exon3:c.G194A:p.R65K,CUX1:NM_001202543:exon4:c.G242A:p.R81K,CUX1:NM_001913:exon4:c.G242A:p.R81K, |
| chr7 | 102274250 | G | A | COSV52922844 | exonic | CUX1 | nonsynonymous SNV | CUX1:NM_001202546:exon15:c.G1273A:p.A425T,CUX1:NM_181500:exon16:c.G1384A:p.A462T,CUX1:NM_001202545:exon15:c.G1252A:p.A418T,CUX1:NM_001202544:exon15:c.G1342A:p.A448T,CUX1:NM_001913:exon16:c.G1390A:p.A464T, |
| chr7 | 102278012 | G | A | . | exonic | CUX1 | nonsynonymous SNV | CUX1:NM_001202546:exon17:c.G1510A:p.D504N,CUX1:NM_001202545:exon17:c.G1489A:p.D497N,CUX1:NM_001913:exon18:c.G1627A:p.D543N,CUX1:NM_181500:exon18:c.G1621A:p.D541N,CUX1:NM_001202544:exon17:c.G1579A:p.D527N, |
| chr7 | 102278021 | A | C | . | exonic | CUX1 | nonsynonymous SNV | CUX1:NM_001202546:exon17:c.A1519C:p.K507Q,CUX1:NM_181500:exon18:c.A1630C:p.K544Q,CUX1:NM_001202545:exon17:c.A1498C:p.K500Q,CUX1:NM_001202544:exon17:c.A1588C:p.K530Q,CUX1:NM_001913:exon18:c.A1636C:p.K546Q, |
| chr7 | 102283050 | A | G | . | exonic | CUX1 | nonsynonymous SNV | CUX1:NM_001202546:exon22:c.A1880G:p.E627G,CUX1:NM_181500:exon23:c.A1991G:p.E664G,CUX1:NM_001202545:exon22:c.A1859G:p.E620G,CUX1:NM_001202544:exon22:c.A1949G:p.E650G,CUX1:NM_001913:exon23:c.A1997G:p.E666G, |
| chr7 | 102283059 | A | G | . | exonic | CUX1 | nonsynonymous SNV | CUX1:NM_001202546:exon22:c.A1889G:p.N630S,CUX1:NM_181500:exon23:c.A2000G:p.N667S,CUX1:NM_001202545:exon22:c.A1868G:p.N623S,CUX1:NM_001202544:exon22:c.A1958G:p.N653S,CUX1:NM_001913:exon23:c.A2006G:p.N669S, |
| chr7 | 102668971 | C | A | COSV61015656 | exonic | POLR2J2 | nonsynonymous SNV | POLR2J2:NM_032959:exon2:c.G84T:p.K28N, |
| chr7 | 105062265 | G | T | . | exonic | KMT2E | nonsynonymous SNV | KMT2E:NM_182931:exon4:c.G173T:p.G58V,KMT2E:NM_018682:exon3:c.G173T:p.G58V, |
| chr7 | 105063364 | G | A | . | exonic | KMT2E | nonsynonymous SNV | KMT2E:NM_018682:exon4:c.G200A:p.G67D,KMT2E:NM_182931:exon5:c.G200A:p.G67D, |
| chr7 | 105063420 | A | G | . | exonic | KMT2E | nonsynonymous SNV | KMT2E:NM_018682:exon4:c.A256G:p.K86E,KMT2E:NM_182931:exon5:c.A256G:p.K86E, |
| chr7 | 105063492 | G | A | . | exonic | KMT2E | nonsynonymous SNV | KMT2E:NM_018682:exon4:c.G328A:p.E110K,KMT2E:NM_182931:exon5:c.G328A:p.E110K, |
| chr7 | 105063532 | G | A | . | exonic | KMT2E | nonsynonymous SNV | KMT2E:NM_018682:exon4:c.G368A:p.C123Y,KMT2E:NM_182931:exon5:c.G368A:p.C123Y, |
| chr7 | 105066747 | G | A | . | exonic | KMT2E | nonsynonymous SNV | KMT2E:NM_018682:exon5:c.G437A:p.C146Y,KMT2E:NM_182931:exon6:c.G437A:p.C146Y, |
| chr7 | 105073642 | T | G | . | exonic | KMT2E | nonsynonymous SNV | KMT2E:NM_018682:exon6:c.T521G:p.V174G,KMT2E:NM_182931:exon7:c.T521G:p.V174G, |
| chr7 | 105076981 | G | A | . | exonic | KMT2E | nonsynonymous SNV | KMT2E:NM_182931:exon10:c.G787A:p.D263N,KMT2E:NM_018682:exon9:c.G787A:p.D263N, |
| chr7 | 105077127 | T | A | . | exonic | KMT2E | nonsynonymous SNV | KMT2E:NM_018682:exon9:c.T933A:p.N311K,KMT2E:NM_182931:exon10:c.T933A:p.N311K, |
| chr7 | 105077304 | G | A | . | exonic | KMT2E | nonsynonymous SNV | KMT2E:NM_018682:exon10:c.G1001A:p.S334N,KMT2E:NM_182931:exon11:c.G1001A:p.S334N, |
| chr7 | 105090032 | G | A | . | exonic | KMT2E | nonsynonymous SNV | KMT2E:NM_182931:exon14:c.G1382A:p.C461Y,KMT2E:NM_018682:exon13:c.G1382A:p.C461Y, |
| chr7 | 105090106 | G | A | . | exonic | KMT2E | nonsynonymous SNV | KMT2E:NM_182931:exon14:c.G1456A:p.E486K,KMT2E:NM_018682:exon13:c.G1456A:p.E486K, |
| chr7 | 105090113 | G | A | . | exonic | KMT2E | nonsynonymous SNV | KMT2E:NM_018682:exon13:c.G1463A:p.R488K,KMT2E:NM_182931:exon14:c.G1463A:p.R488K, |
| chr7 | 105090116 | G | A | COSV57576667 | exonic | KMT2E | nonsynonymous SNV | KMT2E:NM_018682:exon13:c.G1466A:p.R489Q,KMT2E:NM_182931:exon14:c.G1466A:p.R489Q, |
| chr7 | 105090133 | G | A | . | exonic | KMT2E | nonsynonymous SNV | KMT2E:NM_018682:exon13:c.G1483A:p.D495N,KMT2E:NM_182931:exon14:c.G1483A:p.D495N, |
| chr7 | 105091303 | A | T | . | exonic | KMT2E | stopgain | KMT2E:NM_018682:exon14:c.A1711T:p.K571X,KMT2E:NM_182931:exon15:c.A1711T:p.K571X, |
| chr7 | 105143206 | A | T | . | exonic | SRPK2 | nonsynonymous SNV | SRPK2:NM_182691:exon9:c.T905A:p.I302K,SRPK2:NM_182692:exon10:c.T938A:p.I313K,SRPK2:NM_001278273:exon11:c.T905A:p.I302K, |
| chr7 | 105143209 | A | C | . | exonic | SRPK2 | nonsynonymous SNV | SRPK2:NM_182691:exon9:c.T902G:p.I301R,SRPK2:NM_182692:exon10:c.T935G:p.I312R,SRPK2:NM_001278273:exon11:c.T902G:p.I301R, |
| chr7 | 111487298 | A | G | . | exonic | IMMP2L | nonsynonymous SNV | IMMP2L:NM_001244606:exon4:c.T179C:p.L60P,IMMP2L:NM_032549:exon3:c.T179C:p.L60P, |
| chr7 | 116526621 | G | A | . | exonic | CAV1 | nonsynonymous SNV | CAV1:NM_001172895:exon2:c.G34A:p.D12N,CAV1:NM_001172896:exon1:c.G34A:p.D12N,CAV1:NM_001172897:exon2:c.G34A:p.D12N,CAV1:NM_001753:exon2:c.G127A:p.D43N, |
| chr7 | 116558997 | G | A | COSV61954750 | exonic | CAV1 | nonsynonymous SNV | CAV1:NM_001172895:exon3:c.G154A:p.G52S,CAV1:NM_001172896:exon2:c.G154A:p.G52S,CAV1:NM_001172897:exon3:c.G154A:p.G52S,CAV1:NM_001753:exon3:c.G247A:p.G83S, |
| chr7 | 116559052 | G | A | . | exonic | CAV1 | nonsynonymous SNV | CAV1:NM_001753:exon3:c.G302A:p.R101H,CAV1:NM_001172895:exon3:c.G209A:p.R70H,CAV1:NM_001172896:exon2:c.G209A:p.R70H,CAV1:NM_001172897:exon3:c.G209A:p.R70H, |
| chr7 | 116763143 | T | C | . | exonic | MET | nonsynonymous SNV | MET:NM_000245:exon11:c.T2458C:p.F820L,MET:NM_001127500:exon11:c.T2512C:p.F838L, |
| chr7 | 116771529 | G | A | . | exonic | MET | nonsynonymous SNV | MET:NM_000245:exon13:c.G2762A:p.G921E,MET:NM_001127500:exon13:c.G2816A:p.G939E, |
| chr7 | 116904177 | G | A | . | exonic | CAPZA2 | nonsynonymous SNV | CAPZA2:NM_006136:exon5:c.G220A:p.V74I, |
| chr7 | 116904195 | G | A | . | exonic | CAPZA2 | nonsynonymous SNV | CAPZA2:NM_006136:exon5:c.G238A:p.G80S, |
| chr7 | 116912087 | G | A | . | exonic | CAPZA2 | nonsynonymous SNV | CAPZA2:NM_006136:exon8:c.G604A:p.G202S, |
| chr7 | 116917796 | G | A | . | exonic | CAPZA2 | nonsynonymous SNV | CAPZA2:NM_006136:exon10:c.G790A:p.V264I, |
| chr7 | 117320651 | A | T | . | exonic | WNT2 | nonsynonymous SNV | WNT2:NM_003391:exon2:c.T226A:p.C76S, |
| chr7 | 124851918 | C | A | . | exonic | POT1 | nonsynonymous SNV | POT1:NM_001042594:exon10:c.G510T:p.Q170H,POT1:NM_015450:exon11:c.G903T:p.Q301H, |
| chr7 | 127686613 | G | A | . | exonic | SND1 | nonsynonymous SNV | SND1:NM_014390:exon2:c.G79A:p.V27I, |
| chr7 | 127686719 | G | A | . | exonic | SND1 | nonsynonymous SNV | SND1:NM_014390:exon2:c.G185A:p.R62Q, |
| chr7 | 127721296 | G | A | . | exonic | SND1 | nonsynonymous SNV | SND1:NM_014390:exon10:c.G1048A:p.V350I, |
| chr7 | 127807503 | G | A | COSV61241417 | exonic | SND1 | nonsynonymous SNV | SND1:NM_014390:exon11:c.G1172A:p.R391H, |
| chr7 | 127844384 | T | C | . | exonic | SND1 | nonsynonymous SNV | SND1:NM_014390:exon12:c.T1303C:p.F435L, |
| chr7 | 127844417 | G | A | . | exonic | SND1 | nonsynonymous SNV | SND1:NM_014390:exon12:c.G1336A:p.G446R, |
| chr7 | 127887963 | G | A | . | exonic | SND1 | nonsynonymous SNV | SND1:NM_014390:exon13:c.G1405A:p.D469N, |
| chr7 | 127888002 | G | A | COSV61246863 | exonic | SND1 | nonsynonymous SNV | SND1:NM_014390:exon13:c.G1444A:p.A482T, |
| chr7 | 128074541 | T | C | . | exonic | SND1 | nonsynonymous SNV | SND1:NM_014390:exon17:c.T1819C:p.F607L, |
| chr7 | 128081366 | G | A | . | exonic | SND1 | nonsynonymous SNV | SND1:NM_014390:exon18:c.G1975A:p.A659T, |
| chr7 | 128081388 | T | C | . | exonic | SND1 | nonsynonymous SNV | SND1:NM_014390:exon18:c.T1997C:p.V666A, |
| chr7 | 128081405 | G | A | . | exonic | SND1 | nonsynonymous SNV | SND1:NM_014390:exon18:c.G2014A:p.V672M, |

| | | | | | | | | |
|---|---|---|---|---|---|---|---|---|
| chr7 | 128084805 | G | A | . | exonic | SND1 | nonsynonymous SNV | SND1:NM_014390:exon19:c.G2192A:p.R731H, |
| chr7 | 128084841 | G | A | . | exonic | SND1 | nonsynonymous SNV | SND1:NM_014390:exon19:c.G2228A:p.G743E, |
| chr7 | 128086980 | G | A | . | exonic | SND1 | nonsynonymous SNV | SND1:NM_014390:exon21:c.G2347A:p.A783T, |
| chr7 | 128087004 | G | A | . | exonic | SND1 | nonsynonymous SNV | SND1:NM_014390:exon21:c.G2371A:p.A791T, |
| chr7 | 128089571 | G | A | . | exonic | SND1 | nonsynonymous SNV | SND1:NM_014390:exon22:c.G2501A:p.S834N, |
| chr7 | 128089572 | T | G | . | exonic | SND1 | nonsynonymous SNV | SND1:NM_014390:exon22:c.T2502G:p.S834R, |
| chr7 | 130554653 | T | A | . | exonic | COPG2 | nonsynonymous SNV | COPG2:NM_012133:exon14:c.A1296T:p.K432N,COPG2:NM_001290033:exon14:c.A1296T:p.K432N, |
| chr7 | 134005051 | G | A | . | exonic | EXOC4 | nonsynonymous SNV | EXOC4:NM_021807:exon16:c.G2488A:p.A830T, |
| chr7 | 134933194 | C | A | . | exonic | CALD1 | nonsynonymous SNV | CALD1:NM_033157:exon5:c.C425A:p.A142E,CALD1:NM_033139:exon3:c.C407A:p.A136E,CALD1:NM_004342:exon5:c.C425A:p.A142E,CALD1:NM_033140:exon3:c.C407A:p.A136E,CALD1:NM_033138:exon5:c.C425A:p.A142E, |
| chr7 | 134933201 | T | A | . | exonic | CALD1 | nonsynonymous SNV | CALD1:NM_033157:exon5:c.T432A:p.N144K,CALD1:NM_033139:exon3:c.T414A:p.N138K,CALD1:NM_004342:exon5:c.T432A:p.N144K,CALD1:NM_033140:exon3:c.T414A:p.N138K,CALD1:NM_033138:exon5:c.T432A:p.N144K, |
| chr7 | 134933313 | G | A | . | exonic | CALD1 | nonsynonymous SNV | CALD1:NM_033157:exon5:c.G544A:p.E182K,CALD1:NM_033139:exon3:c.G526A:p.E176K,CALD1:NM_004342:exon5:c.G544A:p.E182K,CALD1:NM_033140:exon3:c.G526A:p.E176K,CALD1:NM_033138:exon5:c.G544A:p.E182K, |
| chr7 | 137915944 | C | T | . | exonic | CREB3L2 | nonsynonymous SNV | CREB3L2:NM_001253775:exon3:c.G388A:p.V130I,CREB3L2:NM_194071:exon3:c.G388A:p.V130I, |
| chr7 | 138627377 | A | C | COSV55991488 | exonic | SVOPL | nonsynonymous SNV | SVOPL:NM_001139456:exon11:c.T1154G:p.F385C,SVOPL:NM_174959:exon8:c.T698G:p.F233C, |
| chr7 | 139055291 | G | C | COSV54295377 | exonic | ZC3HAV1 | nonsynonymous SNV | ZC3HAV1:NM_020119:exon10:c.C2101G:p.Q701E, |
| chr7 | 139341406 | T | G | COSV52545769 | exonic | C7ORF55 | nonsynonymous SNV | C7orf55-LUC7L2:NM_001244584:exon1:c.T22G:p.S8A,C7orf55:NM_197964:exon1:c.T22G:p.S8A, |
| chr7 | 13935774 | C | T | COSV54145283 | exonic | ETV1 | nonsynonymous SNV | ETV1:NM_004956:exon8:c.G488A:p.R163Q,ETV1:NM_001163151:exon3:c.G314A:p.R105Q,ETV1:NM_001163150:exon4:c.G368A:p.R123Q,ETV1:NM_001163148:exon7:c.G434A:p.R145Q,ETV1:NM_001163147:exon7:c.G488A:p.R163Q,ETV1:NM_001163149:exon6:c.G434A:p.R145Q, |
| chr7 | 13939184 | T | C | COSV54136900 | exonic | ETV1 | nonsynonymous SNV | ETV1:NM_001163152:exon3:c.A178G:p.S60G,ETV1:NM_004956:exon7:c.A298G:p.S100G,ETV1:NM_001163151:exon2:c.A124G:p.S42G,ETV1:NM_001163150:exon3:c.A178G:p.S60G,ETV1:NM_001163148:exon6:c.A244G:p.S82G,ETV1:NM_001163147:exon6:c.A298G:p.S100G,ETV1:NM_001163149:exon5:c.A244G:p.S82G, |
| chr7 | 140098970 | A | T | . | exonic | KDM7A | nonsynonymous SNV | KDM7A:NM_030647:exon14:c.T1827A:p.D609E, |
| chr7 | 140098979 | A | T | . | exonic | KDM7A | nonsynonymous SNV | KDM7A:NM_030647:exon14:c.T1818A:p.N606K, |
| chr7 | 140459051 | C | G | COSV55436481 | exonic | MKRN1 | nonsynonymous SNV | MKRN1:NM_001291663:exon6:c.G535C:p.V179L,MKRN1:NM_013446:exon4:c.G727C:p.V243L,MKRN1:NM_001145125:exon4:c.G727C:p.V243L, |
| chr7 | 140753336 | A | T | COSV56056643 | exonic | BRAF | nonsynonymous SNV | BRAF:NM_004333:exon15:c.T1799A:p.V600E, |
| chr7 | 141010482 | C | T | . | exonic | MRPS33 | stopgain | MRPS33:NM_016071:exon2:c.G152A:p.W51X,MRPS33:NM_053035:exon2:c.G152A:p.W51X, |
| chr7 | 141010541 | C | T | . | exonic | MRPS33 | nonsynonymous SNV | MRPS33:NM_016071:exon2:c.G93A:p.M31I,MRPS33:NM_053035:exon2:c.G93A:p.M31I, |
| chr7 | 142771005 | C | T | . | exonic | PRSS2 | nonsynonymous SNV | PRSS2:NM_001303414:exon1:c.C23T:p.T8I,PRSS2:NM_002770:exon1:c.C23T:p.T8I, |
| chr7 | 143388875 | C | T | . | exonic | ZYX | nonsynonymous SNV | ZYX:NM_003461:exon8:c.C1423T:p.R475C,ZYX:NM_001010972:exon8:c.C1423T:p.R475C, |
| chr7 | 148757106 | C | T | COSV57390008 | exonic | CUL1 | nonsynonymous SNV | CUL1:NM_003592:exon4:c.C439T:p.R147C, |
| chr7 | 148757125 | G | A | . | exonic | CUL1 | nonsynonymous SNV | CUL1:NM_003592:exon4:c.G458A:p.R153Q, |
| chr7 | 148828812 | C | G | COSV57449162 | exonic | EZH2 | nonsynonymous SNV | EZH2:NM_001203249:exon6:c.G526C:p.D176H,EZH2:NM_004456:exon6:c.G553C:p.D185H,EZH2:NM_001203247:exon6:c.G553C:p.D185H,EZH2:NM_001203248:exon6:c.G526C:p.D176H,EZH2:NM_152998:exon5:c.G436C:p.D146H, |
| chr7 | 150371282 | T | C | COSV68292042 | exonic | REPIN1 | nonsynonymous SNV | REPIN1:NM_014374:exon1:c.T41C:p.L14P,REPIN1:NM_001099695:exon3:c.T212C:p.L71P,REPIN1:NM_013400:exon4:c.T41C:p.L14P,REPIN1:NM_001099696:exon2:c.T41C:p.L14P, |
| chr7 | 150371516 | G | A | COSV68291804 | exonic | REPIN1 | nonsynonymous SNV | REPIN1:NM_014374:exon1:c.G275A:p.R92H,REPIN1:NM_001099695:exon3:c.G446A:p.R149H,REPIN1:NM_013400:exon4:c.G275A:p.R92H,REPIN1:NM_001099696:exon2:c.G275A:p.R92H, |
| chr7 | 150371531 | C | T | COSV68291808 | exonic | REPIN1 | nonsynonymous SNV | REPIN1:NM_014374:exon1:c.C290T:p.A97V,REPIN1:NM_001099695:exon3:c.C461T:p.A154V,REPIN1:NM_013400:exon4:c.C290T:p.A97V,REPIN1:NM_001099696:exon2:c.C290T:p.A97V, |
| chr7 | 150792139 | C | T | . | exonic | TMEM176B | nonsynonymous SNV | TMEM176B:NM_014020:exon6:c.G637A:p.V213I,TMEM176B:NM_001101312:exon6:c.G637A:p.V213I,TMEM176B:NM_001101311:exon6:c.G637A:p.V213I,TMEM176B:NM_001101314:exon5:c.G526A:p.V176I, |
| chr7 | 150793105 | A | G | . | exonic | TMEM176B | nonsynonymous SNV | TMEM176B:NM_014020:exon5:c.T583C:p.Y195H,TMEM176B:NM_001101312:exon5:c.T583C:p.Y195H,TMEM176B:NM_001101311:exon5:c.T583C:p.Y195H,TMEM176B:NM_001101314:exon4:c.T472C:p.Y158H, |
| chr7 | 150793191 | C | T | COSV58441166 | exonic | TMEM176B | nonsynonymous SNV | TMEM176B:NM_014020:exon5:c.G497A:p.R166H,TMEM176B:NM_001101312:exon5:c.G497A:p.R166H,TMEM176B:NM_001101311:exon5:c.G497A:p.R166H,TMEM176B:NM_001101314:exon4:c.G386A:p.R129H, |
| chr7 | 150793200 | A | C | . | exonic | TMEM176B | nonsynonymous SNV | TMEM176B:NM_014020:exon5:c.T488G:p.V163G,TMEM176B:NM_001101312:exon5:c.T488G:p.V163G,TMEM176B:NM_001101311:exon5:c.T488G:p.V163G,TMEM176B:NM_001101314:exon4:c.T377G:p.V126G, |
| chr7 | 150793219 | A | G | . | exonic | TMEM176B | nonsynonymous SNV | TMEM176B:NM_014020:exon5:c.T469C:p.F157L,TMEM176B:NM_001101312:exon5:c.T469C:p.F157L,TMEM176B:NM_001101311:exon5:c.T469C:p.F157L,TMEM176B:NM_001101314:exon4:c.T358C:p.F120L, |
| chr7 | 150793249 | C | T | COSV58439227 | exonic | TMEM176B | nonsynonymous SNV | TMEM176B:NM_014020:exon5:c.G439A:p.V147M,TMEM176B:NM_001101312:exon5:c.G439A:p.V147M,TMEM176B:NM_001101311:exon5:c.G439A:p.V147M,TMEM176B:NM_001101314:exon4:c.G328A:p.V110M, |
| chr7 | 150793260 | A | G | . | exonic | TMEM176B | nonsynonymous SNV | TMEM176B:NM_014020:exon5:c.T428C:p.V143A,TMEM176B:NM_001101312:exon5:c.T428C:p.V143A,TMEM176B:NM_001101311:exon5:c.T428C:p.V143A,TMEM176B:NM_001101314:exon4:c.T317C:p.V106A, |
| chr7 | 150793288 | C | T | COSV58438904 | exonic | TMEM176B | nonsynonymous SNV | TMEM176B:NM_014020:exon5:c.G400A:p.A134T,TMEM176B:NM_001101312:exon5:c.G400A:p.A134T,TMEM176B:NM_001101311:exon5:c.G400A:p.A134T,TMEM176B:NM_001101314:exon4:c.G289A:p.A97T, |
| chr7 | 150793302 | C | T | . | exonic | TMEM176B | nonsynonymous SNV | TMEM176B:NM_014020:exon5:c.G386A:p.S129N,TMEM176B:NM_001101312:exon5:c.G386A:p.S129N,TMEM176B:NM_001101311:exon5:c.G386A:p.S129N,TMEM176B:NM_001101314:exon4:c.G275A:p.S92N, |
| chr7 | 150793578 | G | T | . | exonic | TMEM176B | nonsynonymous SNV | TMEM176B:NM_014020:exon4:c.C338A:p.A113D,TMEM176B:NM_001101312:exon4:c.C338A:p.A113D,TMEM176B:NM_001101311:exon4:c.C338A:p.A113D,TMEM176B:NM_001101314:exon3:c.C227A:p.A76D, |
| chr7 | 150793587 | C | T | . | exonic | TMEM176B | nonsynonymous SNV | TMEM176B:NM_014020:exon4:c.G329A:p.G110E,TMEM176B:NM_001101312:exon4:c.G329A:p.G110E,TMEM176B:NM_001101311:exon4:c.G329A:p.G110E,TMEM176B:NM_001101314:exon3:c.G218A:p.G73E, |
| chr7 | 150793996 | T | G | COSV58439821 | exonic | TMEM176B | nonsynonymous SNV | TMEM176B:NM_014020:exon3:c.A280C:p.S94R,TMEM176B:NM_001101312:exon3:c.A280C:p.S94R,TMEM176B:NM_001101311:exon3:c.A280C:p.S94R, |
| chr7 | 150794034 | A | G | . | exonic | TMEM176B | nonsynonymous SNV | TMEM176B:NM_014020:exon3:c.T242C:p.L81P,TMEM176B:NM_001101312:exon3:c.T242C:p.L81P,TMEM176B:NM_001101311:exon3:c.T242C:p.L81P, |
| chr7 | 150796412 | G | C | . | exonic | TMEM176B | nonsynonymous SNV | TMEM176B:NM_014020:exon2:c.C158G:p.T53S,TMEM176B:NM_001101312:exon2:c.C158G:p.T53S,TMEM176B:NM_001101311:exon2:c.C158G:p.T53S,TMEM176B:NM_001101314:exon2:c.C158G:p.T53S, |
| chr7 | 150796518 | A | G | . | exonic | TMEM176B | nonsynonymous SNV | TMEM176B:NM_014020:exon2:c.T52C:p.S18P,TMEM176B:NM_001101312:exon2:c.T52C:p.S18P,TMEM176B:NM_001101311:exon2:c.T52C:p.S18P,TMEM176B:NM_001101314:exon2:c.T52C:p.S18P, |
| chr7 | 150803641 | A | G | COSV50246101 | exonic | TMEM176A | nonsynonymous SNV | TMEM176A:NM_018487:exon5:c.A364G:p.T122A, |
| chr7 | 150804428 | A | G | COSV50241405 | exonic | TMEM176A | nonsynonymous SNV | TMEM176A:NM_018487:exon6:c.A622G:p.T208A, |
| chr7 | 150858967 | C | G | COSV62866708 | exonic | AOC1 | nonsynonymous SNV | AOC1:NM_001091:exon3:c.C1775G:p.T592R,AOC1:NM_001272072:exon3:c.C1775G:p.T592R, |
| chr7 | 150860577 | C | G | COSV62867576 | exonic | AOC1 | nonsynonymous SNV | AOC1:NM_001091:exon4:c.C1933G:p.H645D,AOC1:NM_001272072:exon4:c.C1990G:p.H664D, |
| chr7 | 1547026 | T | C | COSV52472075 | exonic | TMEM184A | nonsynonymous SNV | TMEM184A:NM_001097620:exon9:c.A1168G:p.S390G, |
| chr7 | 155964808 | T | C | . | exonic | LOC389602 | nonsynonymous SNV | LOC389602:NM_001291913:exon2:c.T803C:p.V268A, |
| chr7 | 1936767 | C | T | COSV56229834 | exonic | MAD1L1 | nonsynonymous SNV | MAD1L1:NM_001013837:exon17:c.G1727A:p.R576H,MAD1L1:NM_001304523:exon16:c.G1727A:p.R576H,MAD1L1:NM_001304524:exon15:c.G1451A:p.R484H,MAD1L1:NM_001013836:exon17:c.G1727A:p.R576H,MAD1L1:NM_001304525:exon2:c.G95A:p.R32H,MAD1L1:NM_003550:exon17:c.G1727A:p.R576H, |

| | | | | | | | | |
|---|---|---|---|---|---|---|---|---|
| chr7 | 1936821 | C | T | COSV56229694 | exonic | MAD1L1 | nonsynonymous SNV | MAD1L1:NM_001013837:exon17:c.G1673A:p.R558H,MAD1L1:NM_001304523:exon16:c.G1673A:p.R558H,MAD1L1:NM_001304524:exon15:c.G1397A:p.R466H,MAD1L1:NM_001013836:exon17:c.G1673A:p.R558H,MAD1L1:NM_001304525:exon2:c.G41A:p.R14H,MAD1L1:NM_003550:exon17:c.G1673A:p.R558H, |
| chr7 | 23505784 | C | T | COSV51719159 | exonic | TRA2A | nonsynonymous SNV | TRA2A:NM_001282757:exon8:c.G497A:p.R166Q,TRA2A:NM_001282758:exon8:c.G497A:p.R166Q,TRA2A:NM_013293:exon7:c.G800A:p.R267Q,TRA2A:NM_001282759:exon9:c.G494A:p.R165Q, |
| chr7 | 23506241 | C | T | . | exonic | TRA2A | nonsynonymous SNV | TRA2A:NM_001282757:exon7:c.G364A:p.G122S,TRA2A:NM_001282758:exon7:c.G364A:p.G122S,TRA2A:NM_013293:exon6:c.G667A:p.G223S,TRA2A:NM_001282759:exon8:c.G364A:p.G122S, |
| chr7 | 23507429 | C | T | . | exonic | TRA2A | nonsynonymous SNV | TRA2A:NM_001282757:exon6:c.G329A:p.R110K,TRA2A:NM_001282758:exon6:c.G329A:p.R110K,TRA2A:NM_013293:exon5:c.G632A:p.R211K,TRA2A:NM_001282759:exon7:c.G329A:p.R110K, |
| chr7 | 23781461 | G | A | . | exonic | STK31 | nonsynonymous SNV | STK31:NM_032944:exon16:c.G1939A:p.E647K,STK31:NM_031414:exon16:c.G2008A:p.E670K,STK31:NM_001260504:exon16:c.G1939A:p.E647K,STK31:NM_001260505:exon16:c.G2008A:p.E670K, |
| chr7 | 26192560 | C | T | . | exonic | HNRNPA2B1 | nonsynonymous SNV | HNRNPA2B1:NM_002137:exon10:c.G982A:p.G328S,HNRNPA2B1:NM_031243:exon11:c.G1018A:p.G340S, |
| chr7 | 26193286 | C | T | . | exonic | HNRNPA2B1 | nonsynonymous SNV | HNRNPA2B1:NM_002137:exon9:c.G929A:p.G310D,HNRNPA2B1:NM_031243:exon10:c.G965A:p.G322D, |
| chr7 | 26193298 | C | T | . | exonic | HNRNPA2B1 | nonsynonymous SNV | HNRNPA2B1:NM_002137:exon9:c.G917A:p.S306N,HNRNPA2B1:NM_031243:exon10:c.G953A:p.S318N, |
| chr7 | 26193611 | C | T | . | exonic | HNRNPA2B1 | nonsynonymous SNV | HNRNPA2B1:NM_002137:exon8:c.G805A:p.G269R,HNRNPA2B1:NM_031243:exon9:c.G841A:p.G281R, |
| chr7 | 26193623 | C | T | . | exonic | HNRNPA2B1 | nonsynonymous SNV | HNRNPA2B1:NM_002137:exon8:c.G793A:p.G265S,HNRNPA2B1:NM_031243:exon9:c.G829A:p.G277S, |
| chr7 | 26193629 | C | T | . | exonic | HNRNPA2B1 | nonsynonymous SNV | HNRNPA2B1:NM_002137:exon8:c.G787A:p.G263R,HNRNPA2B1:NM_031243:exon9:c.G823A:p.G275R, |
| chr7 | 26193649 | C | A | . | exonic | HNRNPA2B1 | nonsynonymous SNV | HNRNPA2B1:NM_002137:exon8:c.G767T:p.G256V,HNRNPA2B1:NM_031243:exon9:c.G803T:p.G268V, |
| chr7 | 26193670 | C | T | . | exonic | HNRNPA2B1 | nonsynonymous SNV | HNRNPA2B1:NM_002137:exon8:c.G746A:p.G249D,HNRNPA2B1:NM_031243:exon9:c.G782A:p.G261D, |
| chr7 | 26193680 | C | T | . | exonic | HNRNPA2B1 | nonsynonymous SNV | HNRNPA2B1:NM_002137:exon8:c.G736A:p.G246S,HNRNPA2B1:NM_031243:exon9:c.G772A:p.G258S, |
| chr7 | 26193683 | C | T | . | exonic | HNRNPA2B1 | nonsynonymous SNV | HNRNPA2B1:NM_002137:exon8:c.G733A:p.G245R,HNRNPA2B1:NM_031243:exon9:c.G769A:p.G257R, |
| chr7 | 26196437 | C | T | . | exonic | HNRNPA2B1 | nonsynonymous SNV | HNRNPA2B1:NM_002137:exon6:c.G622A:p.G208R,HNRNPA2B1:NM_031243:exon7:c.G658A:p.G220R, |
| chr7 | 26196476 | A | G | . | exonic | HNRNPA2B1 | nonsynonymous SNV | HNRNPA2B1:NM_002137:exon6:c.T583C:p.F195L,HNRNPA2B1:NM_031243:exon7:c.T619C:p.F207L, |
| chr7 | 26196837 | C | T | . | exonic | HNRNPA2B1 | nonsynonymous SNV | HNRNPA2B1:NM_002137:exon4:c.G445A:p.D149N,HNRNPA2B1:NM_031243:exon5:c.G481A:p.D161N, |
| chr7 | 26196849 | A | G | . | exonic | HNRNPA2B1 | nonsynonymous SNV | HNRNPA2B1:NM_002137:exon4:c.T433C:p.F145L,HNRNPA2B1:NM_031243:exon5:c.T469C:p.F157L, |
| chr7 | 26196870 | C | T | . | exonic | HNRNPA2B1 | nonsynonymous SNV | HNRNPA2B1:NM_002137:exon4:c.G412A:p.G138R,HNRNPA2B1:NM_031243:exon5:c.G448A:p.G150R, |
| chr7 | 26196896 | A | G | . | exonic | HNRNPA2B1 | nonsynonymous SNV | HNRNPA2B1:NM_002137:exon4:c.T386C:p.I129T,HNRNPA2B1:NM_031243:exon5:c.T422C:p.I141T, |
| chr7 | 26196903 | C | T | . | exonic | HNRNPA2B1 | nonsynonymous SNV | HNRNPA2B1:NM_002137:exon4:c.G379A:p.D127N,HNRNPA2B1:NM_031243:exon5:c.G415A:p.D139N, |

| | | | | | | | | |
|---|---|---|---|---|---|---|---|---|
| chr7 | 26197017 | C | T | . | exonic | HNRNPA2B1 | nonsynonymous SNV | HNRNPA2B1:NM_002137:exon4:c.G265A:p.E89K,HNRNPA2B1:NM_031243:exon5:c.G301A:p.E101K, |
| chr7 | 26197626 | C | T | . | exonic | HNRNPA2B1 | nonsynonymous SNV | HNRNPA2B1:NM_002137:exon2:c.G113A:p.C38Y,HNRNPA2B1:NM_031243:exon3:c.G149A:p.C50Y, |
| chr7 | 27538413 | G | T | . | exonic | HIBADH | nonsynonymous SNV | HIBADH:NM_152740:exon6:c.C623A:p.A208E, |
| chr7 | 27543035 | C | T | . | exonic | HIBADH | nonsynonymous SNV | HIBADH:NM_152740:exon5:c.G550A:p.A184T, |
| chr7 | 27632370 | T | C | . | exonic | HIBADH | nonsynonymous SNV | HIBADH:NM_152740:exon3:c.A328G:p.I110V, |
| chr7 | 27649567 | G | T | . | exonic | HIBADH | nonsynonymous SNV | HIBADH:NM_152740:exon2:c.C158A:p.P53Q, |
| chr7 | 27662785 | C | T | . | exonic | HIBADH | nonsynonymous SNV | HIBADH:NM_152740:exon1:c.G4A:p.A2T, |
| chr7 | 27758116 | G | A | . | exonic | TAX1BP1 | nonsynonymous SNV | TAX1BP1:NM_006024:exon3:c.G248A:p.C83Y,TAX1BP1:NM_001079864:exon3:c.G248A:p.C83Y,TAX1BP1:NM_001206901:exon3:c.G248A:p.C83Y, |
| chr7 | 27769748 | G | A | . | exonic | TAX1BP1 | nonsynonymous SNV | TAX1BP1:NM_006024:exon5:c.G526A:p.A176T,TAX1BP1:NM_001206901:exon5:c.G526A:p.A176T,TAX1BP1:NM_001206902:exon4:c.G55A:p.A19T,TAX1BP1:NM_001079864:exon5:c.G526A:p.A176T, |
| chr7 | 27769826 | G | A | . | exonic | TAX1BP1 | nonsynonymous SNV | TAX1BP1:NM_006024:exon5:c.G604A:p.E202K,TAX1BP1:NM_001079864:exon5:c.G604A:p.E202K,TAX1BP1:NM_001206901:exon5:c.G604A:p.E202K,TAX1BP1:NM_001206902:exon4:c.G133A:p.E45K, |
| chr7 | 27785163 | G | A | . | exonic | TAX1BP1 | nonsynonymous SNV | TAX1BP1:NM_006024:exon6:c.G613A:p.G205S,TAX1BP1:NM_001206901:exon6:c.G613A:p.G205S,TAX1BP1:NM_001206902:exon5:c.G142A:p.G48S,TAX1BP1:NM_001079864:exon6:c.G613A:p.G205S, |
| chr7 | 27787476 | T | A | . | exonic | TAX1BP1 | stopgain | TAX1BP1:NM_006024:exon8:c.T911A:p.L304X,TAX1BP1:NM_001206901:exon8:c.T911A:p.L304X,TAX1BP1:NM_001206902:exon7:c.T440A:p.L147X,TAX1BP1:NM_001079864:exon8:c.T911A:p.L304X, |
| chr7 | 27792093 | G | A | . | exonic | TAX1BP1 | nonsynonymous SNV | TAX1BP1:NM_006024:exon9:c.G1126A:p.E376K,TAX1BP1:NM_001079864:exon9:c.G1126A:p.E376K,TAX1BP1:NM_001206901:exon9:c.G1126A:p.E376K,TAX1BP1:NM_001206902:exon8:c.G655A:p.E219K, |
| chr7 | 27792114 | G | A | COSV55291444 | exonic | TAX1BP1 | nonsynonymous SNV | TAX1BP1:NM_006024:exon9:c.G1147A:p.V383I,TAX1BP1:NM_001206901:exon9:c.G1147A:p.V383I,TAX1BP1:NM_001206902:exon8:c.G676A:p.V226I,TAX1BP1:NM_001079864:exon9:c.G1147A:p.V383I, |
| chr7 | 27792131 | G | A | . | exonic | TAX1BP1 | nonsynonymous SNV | TAX1BP1:NM_006024:exon9:c.G1164A:p.M388I,TAX1BP1:NM_001079864:exon9:c.G1164A:p.M388I,TAX1BP1:NM_001206901:exon9:c.G1164A:p.M388I,TAX1BP1:NM_001206902:exon8:c.G693A:p.M231I, |
| chr7 | 27792161 | C | A | . | exonic | TAX1BP1 | nonsynonymous SNV | TAX1BP1:NM_006024:exon9:c.C1194A:p.N398K,TAX1BP1:NM_001206901:exon9:c.C1194A:p.N398K,TAX1BP1:NM_001206902:exon8:c.C723A:p.N241K,TAX1BP1:NM_001079864:exon9:c.C1194A:p.N398K, |
| chr7 | 27792169 | T | A | . | exonic | TAX1BP1 | nonsynonymous SNV | TAX1BP1:NM_006024:exon9:c.T1202A:p.V401E,TAX1BP1:NM_001206901:exon9:c.T1202A:p.V401E,TAX1BP1:NM_001206902:exon8:c.T731A:p.V244E,TAX1BP1:NM_001079864:exon9:c.T1202A:p.V401E, |
| chr7 | 27793144 | C | A | . | exonic | TAX1BP1 | nonsynonymous SNV | TAX1BP1:NM_006024:exon10:c.C1342A:p.H448N,TAX1BP1:NM_001206901:exon10:c.C1342A:p.H448N,TAX1BP1:NM_001206902:exon9:c.C871A:p.H291N,TAX1BP1:NM_001079864:exon10:c.C1342A:p.H448N, |
| chr7 | 27796162 | G | A | . | exonic | TAX1BP1 | nonsynonymous SNV | TAX1BP1:NM_006024:exon12:c.G1581A:p.M527I,TAX1BP1:NM_001079864:exon12:c.G1581A:p.M527I,TAX1BP1:NM_001206901:exon12:c.G1581A:p.M527I,TAX1BP1:NM_001206902:exon11:c.G1110A:p.M370I, |
| chr7 | 27800028 | G | A | . | exonic | TAX1BP1 | nonsynonymous SNV | TAX1BP1:NM_006024:exon13:c.G1702A:p.E568K,TAX1BP1:NM_001206901:exon13:c.G1702A:p.E568K,TAX1BP1:NM_001206902:exon12:c.G1231A:p.E411K,TAX1BP1:NM_001079864:exon13:c.G1702A:p.E568K, |
| chr7 | 27817021 | G | A | . | exonic | TAX1BP1 | nonsynonymous SNV | TAX1BP1:NM_006024:exon15:c.G2068A:p.D690N,TAX1BP1:NM_001079864:exon15:c.G1942A:p.D648N,TAX1BP1:NM_001206901:exon15:c.G1942A:p.D648N,TAX1BP1:NM_001206902:exon14:c.G1471A:p.D491N, |
| chr7 | 27817035 | C | A | . | exonic | TAX1BP1 | nonsynonymous SNV | TAX1BP1:NM_006024:exon15:c.C2082A:p.S694R,TAX1BP1:NM_001079864:exon15:c.C1956A:p.S652R,TAX1BP1:NM_001206901:exon15:c.C1956A:p.S652R,TAX1BP1:NM_001206902:exon14:c.C1485A:p.S495R, |
| chr7 | 27827804 | T | C | . | exonic | TAX1BP1 | nonsynonymous SNV | TAX1BP1:NM_006024:exon16:c.T2152C:p.F718L,TAX1BP1:NM_001206901:exon16:c.T2026C:p.F676L,TAX1BP1:NM_001206902:exon15:c.T1555C:p.F519L,TAX1BP1:NM_001079864:exon16:c.T2026C:p.F676L, |
| chr7 | 27828701 | G | A | . | exonic | TAX1BP1 | nonsynonymous SNV | TAX1BP1:NM_006024:exon17:c.G2242A:p.E748K,TAX1BP1:NM_001206901:exon17:c.G2116A:p.E706K,TAX1BP1:NM_001206902:exon16:c.G1645A:p.E549K,TAX1BP1:NM_001079864:exon17:c.G2116A:p.E706K, |
| chr7 | 27828706 | T | G | . | exonic | TAX1BP1 | nonsynonymous SNV | TAX1BP1:NM_006024:exon17:c.T2247G:p.H749Q,TAX1BP1:NM_001079864:exon17:c.T2121G:p.H707Q,TAX1BP1:NM_001206901:exon17:c.T2121G:p.H707Q,TAX1BP1:NM_001206902:exon16:c.T1650G:p.H550Q, |
| chr7 | 27828720 | G | A | . | exonic | TAX1BP1 | stopgain | TAX1BP1:NM_006024:exon17:c.G2261A:p.W754X,TAX1BP1:NM_001206901:exon17:c.G2135A:p.W712X,TAX1BP1:NM_001206902:exon16:c.G1664A:p.W555X,TAX1BP1:NM_001079864:exon17:c.G2135A:p.W712X, |
| chr7 | 29566672 | C | T | COSV60501727 | exonic | PRR15 | nonsynonymous SNV | PRR15:NM_175887:exon2:c.C343T:p.P115S, |
| chr7 | 29566700 | C | A | COSV60501902 | exonic | PRR15 | nonsynonymous SNV | PRR15:NM_175887:exon2:c.C371A:p.P124H, |
| chr7 | 30595045 | C | G | COSV66827983 | exonic | GARS | nonsynonymous SNV | GARS:NM_002047:exon1:c.C124G:p.P42A, |
| chr7 | 35670232 | C | T | COSV60943904 | exonic | HERPUD2 | nonsynonymous SNV | HERPUD2:NM_022373:exon4:c.G322A:p.A108T, |
| chr7 | 36513048 | G | A | COSV51735862 | exonic | AOAH | nonsynonymous SNV | AOAH:NM_001177506:exon22:c.C2051T:p.P684L, |
| chr7 | 36616429 | G | C | COSV51741603 | exonic | AOAH | nonsynonymous SNV | AOAH:NM_001637:exon11:c.C797G:p.A266G,AOAH:NM_001177506:exon11:c.C797G:p.A266G,AOAH:NM_001177507:exon10:c.C701G:p.A234G, |
| chr7 | 36632067 | C | A | . | exonic | AOAH | nonsynonymous SNV | AOAH:NM_001637:exon6:c.G490T:p.V164F,AOAH:NM_001177506:exon6:c.G490T:p.V164F,AOAH:NM_001177507:exon5:c.G394T:p.V132F, |
| chr7 | 36724067 | C | T | COSV51752564 | exonic | AOAH | nonsynonymous SNV | AOAH:NM_001637:exon1:c.G82A:p.D28N,AOAH:NM_001177506:exon1:c.G82A:p.D28N,AOAH:NM_001177507:exon1:c.G82A:p.D28N, |
| chr7 | 39570578 | A | G | COSV56232709 | exonic | YAE1D1 | nonsynonymous SNV | YAE1D1:NM_001282446:exon2:c.A202G:p.K68E,YAE1D1:NM_020192:exon2:c.A202G:p.K68E, |
| chr7 | 43624681 | A | G | COSV99069595 | exonic | STK17A | nonsynonymous SNV | STK17A:NM_004760:exon7:c.A1084G:p.K362E, |
| chr7 | 43866914 | A | G | COSV58182079 | exonic | MRPS24 | nonsynonymous SNV | MRPS24:NM_032014:exon4:c.T289C:p.W97R, |
| chr7 | 44114181 | A | G | . | exonic | AEBP1 | nonsynonymous SNV | AEBP1:NM_001129:exon21:c.A3397G:p.K1133E, |
| chr7 | 55161562 | G | A | COSV51770369 | exonic | EGFR | nonsynonymous SNV | EGFR:NM_201282:exon13:c.G1562A:p.R521K,EGFR:NM_005228:exon13:c.G1562A:p.R521K,EGFR:NM_201284:exon13:c.G1562A:p.R521K, |
| chr7 | 5527828 | A | G | . | exonic | ACTB | nonsynonymous SNV | ACTB:NM_001101:exon6:c.T1048C:p.S350P, |
| chr7 | 5527852 | C | T | COSV59321582 | exonic | ACTB | nonsynonymous SNV | ACTB:NM_001101:exon6:c.G1024A:p.G342S, |
| chr7 | 5528020 | C | T | . | exonic | ACTB | nonsynonymous SNV | ACTB:NM_001101:exon5:c.G968A:p.S323N, |
| chr7 | 5528105 | C | T | . | exonic | ACTB | nonsynonymous SNV | ACTB:NM_001101:exon5:c.G883A:p.A295T, |
| chr7 | 5529382 | C | T | COSV59317454 | exonic | ACTB | nonsynonymous SNV | ACTB:NM_001101:exon3:c.G142A:p.G48S, |
| chr7 | 5529548 | C | T | . | exonic | ACTB | nonsynonymous SNV | ACTB:NM_001101:exon2:c.G110A:p.R37H, |
| chr7 | 56011772 | A | G | . | exonic | PSPH | nonsynonymous SNV | PSPH:NM_004577:exon8:c.T668C:p.L223P, |
| chr7 | 56021132 | T | A | COSV51908933 | exonic | PSPH | nonsynonymous SNV | PSPH:NM_004577:exon4:c.A81T:p.R27S, |
| chr7 | 56021209 | C | T | COSV51908733 | exonic | PSPH | nonsynonymous SNV | PSPH:NM_004577:exon4:c.G4A:p.V2I, |
| chr7 | 56068567 | C | A | . | exonic | SUMF2 | nonsynonymous SNV | SUMF2:NM_001042469:exon2:c.C210A:p.D70E,SUMF2:NM_001130069:exon2:c.C210A:p.D70E,SUMF2:NM_015411:exon2:c.C210A:p.D70E,SUMF2:NM_001042470:exon2:c.C210A:p.D70E, |
| chr7 | 5725380 | C | T | . | exonic | RNF216 | nonsynonymous SNV | RNF216:NM_207111:exon8:c.G1448A:p.R483K,RNF216:NM_207116:exon8:c.G1277A:p.R426K, |
| chr7 | 5905213 | G | C | COSV58073639 | exonic | CCZ1 | nonsynonymous SNV | CCZ1:NM_015622:exon7:c.G642C:p.E214D, |

| | | | | | | | | |
|---|---|---|---|---|---|---|---|---|
| chr7 | 5919854 | G | A | COSV58073320 | exonic | CCZ1 | nonsynonymous SNV | CCZ1:NM_015622:exon12:c.G994A:p.V332I, |
| chr7 | 5919961 | G | C | COSV99073124 | exonic | CCZ1 | nonsynonymous SNV | CCZ1:NM_015622:exon12:c.G1101C:p.M367I, |
| chr7 | 5986771 | C | T | . | exonic | PMS2 | nonsynonymous SNV | PMS2:NM_000535:exon11:c.G1994A:p.R665K, |
| chr7 | 5986899 | C | T | COSV56221681 | exonic | PMS2 | nonsynonymous SNV | PMS2:NM_000535:exon11:c.G1866A:p.M622I, |
| chr7 | 5987144 | T | C | COSV56223170 | exonic | PMS2 | nonsynonymous SNV | PMS2:NM_000535:exon11:c.A1621G:p.K541E, |
| chr7 | 5987357 | G | A | COSV56220401 | exonic | PMS2 | nonsynonymous SNV | PMS2:NM_000535:exon11:c.C1408T:p.P470S, |
| chr7 | 6009457 | G | A | . | exonic | AIMP2 | nonsynonymous SNV | AIMP2:NM_006303:exon1:c.G94A:p.G32S, |
| chr7 | 6015183 | G | A | . | exonic | AIMP2 | nonsynonymous SNV | AIMP2:NM_006303:exon2:c.G173A:p.R58H, |
| chr7 | 6015275 | G | A | . | exonic | AIMP2 | nonsynonymous SNV | AIMP2:NM_006303:exon2:c.G265A:p.V89I, |
| chr7 | 6023474 | G | A | . | exonic | AIMP2 | nonsynonymous SNV | AIMP2:NM_006303:exon4:c.G746A:p.S249N, |
| chr7 | 6023498 | G | A | . | exonic | AIMP2 | nonsynonymous SNV | AIMP2:NM_006303:exon4:c.G770A:p.R257H, |
| chr7 | 6023578 | G | A | . | exonic | AIMP2 | nonsynonymous SNV | AIMP2:NM_006303:exon4:c.G850A:p.V284I, |
| chr7 | 6036192 | A | T | COSV52237423 | exonic | ANKRD61 | nonsynonymous SNV | ANKRD61:NM_001271700:exon3:c.A1063T:p.M355L, |
| chr7 | 6054578 | G | T | . | exonic | EIF2AK1 | nonsynonymous SNV | EIF2AK1:NM_001134335:exon2:c.C245A:p.P82Q,EIF2AK1:NM_014413:exon2:c.C245A:p.P82Q, |
| chr7 | 6154643 | T | C | COSV60359814 | exonic | USP42 | nonsynonymous SNV | USP42:NM_032172:exon15:c.T3089C:p.L1030P, |
| chr7 | 66988489 | A | G | COSV55886995 | exonic | SBDS | nonsynonymous SNV | SBDS:NM_016038:exon5:c.T635C:p.I212T, |
| chr7 | 76048186 | C | T | COSV50388126 | exonic | MDH2 | nonsynonymous SNV | MDH2:NM_001282403:exon1:c.C26T:p.A9V,MDH2:NM_005918:exon1:c.C26T:p.A9V, |
| chr7 | 77922810 | G | A | . | exonic | PHTF2 | nonsynonymous SNV | PHTF2:NM_001127359:exon9:c.G1049A:p.R350H,PHTF2:NM_001127360:exon9:c.G1037A:p.R346H, |
| chr7 | 87171048 | G | C | COSV58683589 | exonic | DMTF1 | nonsynonymous SNV | DMTF1:NM_021145:exon7:c.G286C:p.A96P,DMTF1:NM_001142327:exon5:c.G286C:p.A96P,DMTF1:NM_001142326:exon4:c.G22C:p.A8P, |
| chr7 | 900544 | C | T | COSV56214400 | exonic | ADAP1 | nonsynonymous SNV | ADAP1:NM_001284309:exon7:c.G505A:p.G169S,ADAP1:NM_006869:exon7:c.G721A:p.G241S,ADAP1:NM_001284308:exon7:c.G754A:p.G252S,ADAP1:NM_001284311:exon5:c.G436A:p.G146S,ADAP1:NM_001284310:exon6:c.G505A:p.G169S, |
| chr7 | 92076897 | C | A | . | exonic | AKAP9 | nonsynonymous SNV | AKAP9:NM_005751:exon29:c.C6655A:p.L2219M,AKAP9:NM_147185:exon29:c.C6631A:p.L2211M, |
| chr7 | 92079584 | A | G | . | exonic | AKAP9 | nonsynonymous SNV | AKAP9:NM_005751:exon31:c.A7451G:p.K2484R,AKAP9:NM_147185:exon31:c.A7427G:p.K2476R, |
| chr7 | 92085597 | C | T | . | exonic | AKAP9 | nonsynonymous SNV | AKAP9:NM_005751:exon36:c.C8935T:p.P2979S,AKAP9:NM_147185:exon36:c.C8911T:p.P2971S, |
| chr7 | 92510995 | A | C | . | exonic | PEX1 | nonsynonymous SNV | PEX1:NM_000466:exon8:c.T1536G:p.D512E,PEX1:NM_001282678:exon8:c.T912G:p.D304E,PEX1:NM_001282677:exon8:c.T1536G:p.D512E, |
| chr7 | 93195855 | A | G | . | exonic | HEPACAM2 | nonsynonymous SNV | HEPACAM2:NM_001288810:exon6:c.T1187C:p.L396P, |
| chr7 | 93208659 | C | T | . | exonic | HEPACAM2 | nonsynonymous SNV | HEPACAM2:NM_001288810:exon3:c.G897A:p.M299I,HEPACAM2:NM_198151:exon3:c.G897A:p.M299I,HEPACAM2:NM_001288804:exon5:c.G1002A:p.M334I,HEPACAM2:NM_001039372:exon4:c.G933A:p.M311I, |
| chr7 | 93215607 | C | T | . | exonic | HEPACAM2 | nonsynonymous SNV | HEPACAM2:NM_001288810:exon2:c.G473A:p.C158Y,HEPACAM2:NM_198151:exon2:c.G473A:p.C158Y,HEPACAM2:NM_001288804:exon4:c.G578A:p.C193Y,HEPACAM2:NM_001039372:exon3:c.G509A:p.C170Y, |
| chr7 | 97117880 | T | C | . | exonic | SDHAF3 | nonsynonymous SNV | SDHAF3:NM_020186:exon1:c.T157C:p.F53L, |
| chr7 | 99923585 | G | A | . | exonic | GJC3 | nonsynonymous SNV | GJC3:NM_181538:exon2:c.C800T:p.A267V, |
| chr8 | 100704315 | C | T | . | exonic | PABPC1 | nonsynonymous SNV | PABPC1:NM_002568:exon14:c.G1894A:p.G632S, |
| chr8 | 100704992 | C | T | COSV59397438 | exonic | PABPC1 | nonsynonymous SNV | PABPC1:NM_002568:exon13:c.G1752A:p.M584I, |
| chr8 | 100704996 | C | T | COSV59400112 | exonic | PABPC1 | nonsynonymous SNV | PABPC1:NM_002568:exon13:c.G1748A:p.G583D, |
| chr8 | 100705009 | C | T | COSV59397447 | exonic | PABPC1 | nonsynonymous SNV | PABPC1:NM_002568:exon13:c.G1735A:p.G579S, |
| chr8 | 100705025 | C | T | COSV59400560 | exonic | PABPC1 | nonsynonymous SNV | PABPC1:NM_002568:exon13:c.G1719A:p.M573I, |
| chr8 | 100705050 | C | T | . | exonic | PABPC1 | nonsynonymous SNV | PABPC1:NM_002568:exon13:c.G1694A:p.R565Q, |
| chr8 | 100705051 | G | A | COSV59397476 | exonic | PABPC1 | nonsynonymous SNV | PABPC1:NM_002568:exon13:c.C1693T:p.R565W, |
| chr8 | 100705654 | C | T | . | exonic | PABPC1 | nonsynonymous SNV | PABPC1:NM_002568:exon12:c.G1622A:p.G541D, |
| chr8 | 100706700 | C | T | . | exonic | PABPC1 | nonsynonymous SNV | PABPC1:NM_002568:exon11:c.G1553A:p.R518H, |
| chr8 | 100706713 | C | T | . | exonic | PABPC1 | nonsynonymous SNV | PABPC1:NM_002568:exon11:c.G1540A:p.A514T, |
| chr8 | 100706740 | C | T | COSV59397513 | exonic | PABPC1 | nonsynonymous SNV | PABPC1:NM_002568:exon11:c.G1513A:p.V505I, |
| chr8 | 100706910 | C | T | COSV59397547 | exonic | PABPC1 | nonsynonymous SNV | PABPC1:NM_002568:exon10:c.G1424A:p.R475Q, |
| chr8 | 100706929 | C | T | . | exonic | PABPC1 | nonsynonymous SNV | PABPC1:NM_002568:exon10:c.G1405A:p.A469T, |
| chr8 | 100706947 | A | G | . | exonic | PABPC1 | nonsynonymous SNV | PABPC1:NM_002568:exon10:c.T1387C:p.F463L, |
| chr8 | 100709162 | C | T | . | exonic | PABPC1 | nonsynonymous SNV | PABPC1:NM_002568:exon9:c.G1307A:p.R436H, |
| chr8 | 100709490 | G | A | . | exonic | PABPC1 | nonsynonymous SNV | PABPC1:NM_002568:exon8:c.C1214T:p.P405L, |
| chr8 | 100709497 | C | T | . | exonic | PABPC1 | nonsynonymous SNV | PABPC1:NM_002568:exon8:c.G1207A:p.A403T, |
| chr8 | 100709573 | G | T | . | exonic | PABPC1 | nonsynonymous SNV | PABPC1:NM_002568:exon8:c.C1131A:p.H377Q, |
| chr8 | 100709578 | C | T | . | exonic | PABPC1 | nonsynonymous SNV | PABPC1:NM_002568:exon8:c.G1126A:p.A376T, |
| chr8 | 100709587 | C | T | . | exonic | PABPC1 | nonsynonymous SNV | PABPC1:NM_002568:exon8:c.G1117A:p.E373K, |
| chr8 | 100709641 | C | T | . | exonic | PABPC1 | nonsynonymous SNV | PABPC1:NM_002568:exon8:c.G1063A:p.G355S, |
| chr8 | 100709667 | G | T | . | exonic | PABPC1 | nonsynonymous SNV | PABPC1:NM_002568:exon8:c.C1037A:p.A346D, |
| chr8 | 100709668 | C | T | . | exonic | PABPC1 | nonsynonymous SNV | PABPC1:NM_002568:exon8:c.G1036A:p.A346T, |
| chr8 | 100709674 | C | T | . | exonic | PABPC1 | nonsynonymous SNV | PABPC1:NM_002568:exon8:c.G1030A:p.E344K, |
| chr8 | 100709708 | G | T | . | exonic | PABPC1 | nonsynonymous SNV | PABPC1:NM_002568:exon8:c.C996A:p.S332R, |
| chr8 | 100709709 | C | T | . | exonic | PABPC1 | nonsynonymous SNV | PABPC1:NM_002568:exon8:c.G995A:p.S332N, |
| chr8 | 100712384 | A | G | . | exonic | PABPC1 | nonsynonymous SNV | PABPC1:NM_002568:exon7:c.T950C:p.F317S, |
| chr8 | 100712402 | C | T | COSV59399520 | exonic | PABPC1 | nonsynonymous SNV | PABPC1:NM_002568:exon7:c.G932A:p.R311Q, |
| chr8 | 100712405 | A | G | . | exonic | PABPC1 | nonsynonymous SNV | PABPC1:NM_002568:exon7:c.T929C:p.L310P, |
| chr8 | 100712765 | C | T | COSV59400273 | exonic | PABPC1 | nonsynonymous SNV | PABPC1:NM_002568:exon6:c.G763A:p.E255K, |
| chr8 | 100712771 | C | T | . | exonic | PABPC1 | nonsynonymous SNV | PABPC1:NM_002568:exon6:c.G757A:p.G253R, |
| chr8 | 100713101 | C | T | . | exonic | PABPC1 | nonsynonymous SNV | PABPC1:NM_002568:exon5:c.G724A:p.E242K, |
| chr8 | 100713145 | C | T | . | exonic | PABPC1 | nonsynonymous SNV | PABPC1:NM_002568:exon5:c.G680A:p.S227N, |
| chr8 | 100713148 | T | C | . | exonic | PABPC1 | nonsynonymous SNV | PABPC1:NM_002568:exon5:c.A677G:p.E226G, |
| chr8 | 100715467 | T | C | . | exonic | PABPC1 | nonsynonymous SNV | PABPC1:NM_002568:exon4:c.A638G:p.K213R, |

| | | | | | | | | |
|---|---|---|---|---|---|---|---|---|
| chr8 | 100715488 | C | T | COSV59397665 | exonic | PABPC1 | nonsynonymous SNV | PABPC1:NM_002568:exon4:c.G617A:p.R206H, |
| chr8 | 100715492 | C | T | . | exonic | PABPC1 | nonsynonymous SNV | PABPC1:NM_002568:exon4:c.G613A:p.E205K, |
| chr8 | 100715549 | T | C | . | exonic | PABPC1 | nonsynonymous SNV | PABPC1:NM_002568:exon4:c.A556G:p.R186G, |
| chr8 | 100715564 | C | T | COSV59397693 | exonic | PABPC1 | nonsynonymous SNV | PABPC1:NM_002568:exon4:c.G541A:p.A181T, |
| chr8 | 100715573 | C | T | COSV59397705 | exonic | PABPC1 | nonsynonymous SNV | PABPC1:NM_002568:exon4:c.G532A:p.E178K, |
| chr8 | 100717798 | C | T | . | exonic | PABPC1 | nonsynonymous SNV | PABPC1:NM_002568:exon3:c.G478A:p.G160R, |
| chr8 | 100717822 | C | T | COSV59399238 | exonic | PABPC1 | nonsynonymous SNV | PABPC1:NM_002568:exon3:c.G454A:p.E152K, |
| chr8 | 100717827 | G | A | . | exonic | PABPC1 | nonsynonymous SNV | PABPC1:NM_002568:exon3:c.C449T:p.A150V, |
| chr8 | 100717831 | C | T | . | exonic | PABPC1 | nonsynonymous SNV | PABPC1:NM_002568:exon3:c.G445A:p.E149K, |
| chr8 | 100717885 | C | A | . | exonic | PABPC1 | nonsynonymous SNV | PABPC1:NM_002568:exon3:c.G391T:p.V131F, |
| chr8 | 100721459 | A | G | . | exonic | PABPC1 | nonsynonymous SNV | PABPC1:NM_002568:exon1:c.T125C:p.V42A, |
| chr8 | 101692841 | G | T | . | exonic | NCALD | nonsynonymous SNV | NCALD:NM_001040624:exon7:c.C434A:p.P145Q,NCALD:NM_001040626:exon6:c.C434A:p.P145Q,NCALD:NM_001040630:exon5:c.C434A:p.P145Q,NCALD:NM_001040627:exon5:c.C434A:p.P145Q,NCALD:NM_001040628:exon7:c.C434A:p.P145Q,NCALD:NM_032041:exon3:c.C434A:p.P145Q,NCALD:NM_001040625:exon6:c.C434A:p.P145Q,NCALD:NM_001040629:exon6:c.C434A:p.P145Q, |
| chr8 | 102305099 | G | T | . | exonic | UBR5 | nonsynonymous SNV | UBR5:NM_001282873:exon21:c.C2813A:p.T938N,UBR5:NM_015902:exon21:c.C2813A:p.T938N, |
| chr8 | 102828585 | G | T | . | exonic | AZIN1 | nonsynonymous SNV | AZIN1:NM_001301668:exon12:c.C1081A:p.L361I,AZIN1:NM_015878:exon13:c.C1329A:p.S443R,AZIN1:NM_148174:exon12:c.C1329A:p.S443R, |
| chr8 | 103404817 | C | T | . | exonic | SLC25A32 | nonsynonymous SNV | SLC25A32:NM_030780:exon3:c.G350A:p.R117H, |
| chr8 | 103420317 | A | G | COSV52571616 | exonic | DCAF13 | nonsynonymous SNV | DCAF13:NM_015420:exon2:c.A580G:p.I194V, |
| chr8 | 106710628 | C | T | . | exonic | OXR1 | nonsynonymous SNV | OXR1:NM_018002:exon10:c.C1631T:p.A544V,OXR1:NM_001198533:exon10:c.C1631T:p.A544V,OXR1:NM_181354:exon8:c.C1610T:p.A537V,OXR1:NM_001198532:exon9:c.C1634T:p.A545V, |
| chr8 | 106713899 | G | A | . | exonic | OXR1 | nonsynonymous SNV | OXR1:NM_018002:exon11:c.G1870A:p.E624K,OXR1:NM_001198533:exon11:c.G1870A:p.E624K,OXR1:NM_181354:exon9:c.G1849A:p.E617K,OXR1:NM_001198532:exon10:c.G1873A:p.E625K, |
| chr8 | 109336161 | G | A | . | exonic | ENY2 | nonsynonymous SNV | ENY2:NM_001193557:exon2:c.G25A:p.A9T,ENY2:NM_020189:exon2:c.G40A:p.A14T, |
| chr8 | 109339385 | G | A | . | exonic | ENY2 | nonsynonymous SNV | ENY2:NM_001193557:exon3:c.G134A:p.C45Y,ENY2:NM_020189:exon3:c.G149A:p.C50Y, |
| chr8 | 109340507 | G | A | . | exonic | ENY2 | nonsynonymous SNV | ENY2:NM_001193557:exon4:c.G158A:p.G53E,ENY2:NM_020189:exon4:c.G173A:p.G58E, |
| chr8 | 109340542 | G | A | . | exonic | ENY2 | nonsynonymous SNV | ENY2:NM_001193557:exon4:c.G193A:p.E65K,ENY2:NM_020189:exon4:c.G208A:p.E70K, |
| chr8 | 109343474 | G | A | . | exonic | ENY2 | nonsynonymous SNV | ENY2:NM_001193557:exon5:c.G284A:p.S95N,ENY2:NM_020189:exon5:c.G299A:p.S100N, |
| chr8 | 116645041 | A | G | . | exonic | EIF3H | nonsynonymous SNV | EIF3H:NM_003756:exon8:c.T1024C:p.F342L, |
| chr8 | 116645044 | G | A | . | exonic | EIF3H | nonsynonymous SNV | EIF3H:NM_003756:exon8:c.C1021T:p.L341F, |
| chr8 | 116645049 | C | T | . | exonic | EIF3H | nonsynonymous SNV | EIF3H:NM_003756:exon8:c.G1016A:p.G339D, |
| chr8 | 116645050 | C | T | . | exonic | EIF3H | nonsynonymous SNV | EIF3H:NM_003756:exon8:c.G1015A:p.G339S, |
| chr8 | 116648862 | C | T | . | exonic | EIF3H | nonsynonymous SNV | EIF3H:NM_003756:exon6:c.G772A:p.V258I, |
| chr8 | 116648888 | C | T | . | exonic | EIF3H | nonsynonymous SNV | EIF3H:NM_003756:exon6:c.G746A:p.R249K, |
| chr8 | 116655916 | C | T | . | exonic | EIF3H | stopgain | EIF3H:NM_003756:exon5:c.G647A:p.W216X, |
| chr8 | 116655926 | C | T | . | exonic | EIF3H | nonsynonymous SNV | EIF3H:NM_003756:exon5:c.G637A:p.V213I, |
| chr8 | 116655961 | A | G | . | exonic | EIF3H | nonsynonymous SNV | EIF3H:NM_003756:exon5:c.T602C:p.V201A, |
| chr8 | 116655994 | G | T | . | exonic | EIF3H | nonsynonymous SNV | EIF3H:NM_003756:exon5:c.C569A:p.A190E, |
| chr8 | 116657230 | T | A | . | exonic | EIF3H | nonsynonymous SNV | EIF3H:NM_003756:exon4:c.A542T:p.D181V, |
| chr8 | 116657234 | T | C | . | exonic | EIF3H | nonsynonymous SNV | EIF3H:NM_003756:exon4:c.A538G:p.K180E, |
| chr8 | 116657250 | C | T | . | exonic | EIF3H | nonsynonymous SNV | EIF3H:NM_003756:exon4:c.G522A:p.M174I, |
| chr8 | 116658850 | G | T | . | exonic | EIF3H | stopgain | EIF3H:NM_003756:exon3:c.C420A:p.Y140X, |
| chr8 | 116658911 | T | C | . | exonic | EIF3H | nonsynonymous SNV | EIF3H:NM_003756:exon3:c.A359G:p.Y120C, |
| chr8 | 116658918 | C | T | . | exonic | EIF3H | nonsynonymous SNV | EIF3H:NM_003756:exon3:c.G352A:p.G118S, |
| chr8 | 116658921 | C | T | COSV52664467 | exonic | EIF3H | nonsynonymous SNV | EIF3H:NM_003756:exon3:c.G349A:p.V117M, |
| chr8 | 116726108 | A | G | . | exonic | EIF3H | nonsynonymous SNV | EIF3H:NM_003756:exon2:c.T197C:p.V66A, |
| chr8 | 116755725 | C | T | . | exonic | EIF3H | nonsynonymous SNV | EIF3H:NM_003756:exon1:c.G73A:p.G25S, |
| chr8 | 116755743 | C | T | . | exonic | EIF3H | nonsynonymous SNV | EIF3H:NM_003756:exon1:c.G55A:p.A19T, |
| chr8 | 116755778 | C | T | . | exonic | EIF3H | nonsynonymous SNV | EIF3H:NM_003756:exon1:c.G20A:p.G7D, |
| chr8 | 116755797 | T | A | . | exonic | EIF3H | nonsynonymous SNV | EIF3H:NM_003756:exon1:c.A1T:p.M1L, |
| chr8 | 116847613 | C | T | COSV52063702 | exonic | RAD21 | nonsynonymous SNV | RAD21:NM_006265:exon14:c.G1783A:p.A595T, |
| chr8 | 116849011 | C | T | . | exonic | RAD21 | nonsynonymous SNV | RAD21:NM_006265:exon13:c.G1639A:p.G547S, |
| chr8 | 116849014 | C | T | . | exonic | RAD21 | nonsynonymous SNV | RAD21:NM_006265:exon13:c.G1636A:p.G546R, |
| chr8 | 116850670 | G | T | COSV52056756 | exonic | RAD21 | nonsynonymous SNV | RAD21:NM_006265:exon12:c.C1568A:p.P523Q, |
| chr8 | 116851977 | C | T | . | exonic | RAD21 | nonsynonymous SNV | RAD21:NM_006265:exon11:c.G1441A:p.G481R, |
| chr8 | 116852014 | C | T | . | exonic | RAD21 | nonsynonymous SNV | RAD21:NM_006265:exon11:c.G1404A:p.M468I, |
| chr8 | 116852549 | C | T | . | exonic | RAD21 | nonsynonymous SNV | RAD21:NM_006265:exon10:c.G1321A:p.D441N, |
| chr8 | 116856280 | G | T | . | exonic | RAD21 | nonsynonymous SNV | RAD21:NM_006265:exon8:c.C823A:p.P275T, |
| chr8 | 116856742 | C | T | COSV52059248 | exonic | RAD21 | nonsynonymous SNV | RAD21:NM_006265:exon7:c.G718A:p.G240S, |
| chr8 | 116857282 | C | T | COSV52061325 | exonic | RAD21 | nonsynonymous SNV | RAD21:NM_006265:exon6:c.G673A:p.D225N, |
| chr8 | 116857288 | C | T | . | exonic | RAD21 | nonsynonymous SNV | RAD21:NM_006265:exon6:c.G667A:p.G223R, |
| chr8 | 116857401 | C | T | . | exonic | RAD21 | nonsynonymous SNV | RAD21:NM_006265:exon6:c.G554A:p.S185N, |
| chr8 | 116861896 | C | T | . | exonic | RAD21 | nonsynonymous SNV | RAD21:NM_006265:exon4:c.G319A:p.A107T, |
| chr8 | 116863235 | C | T | . | exonic | RAD21 | nonsynonymous SNV | RAD21:NM_006265:exon3:c.G169A:p.G57R, |
| chr8 | 117799731 | C | T | . | exonic | EXT1 | nonsynonymous SNV | EXT1:NM_000127:exon11:c.G2222A:p.R741Q, |
| chr8 | 117804842 | C | T | . | exonic | EXT1 | nonsynonymous SNV | EXT1:NM_000127:exon10:c.G1935A:p.M645I, |
| chr8 | 117812961 | C | T | . | exonic | EXT1 | nonsynonymous SNV | EXT1:NM_000127:exon8:c.G1633A:p.V545I, |
| chr8 | 117818467 | C | T | . | exonic | EXT1 | nonsynonymous SNV | EXT1:NM_000127:exon7:c.G1600A:p.V534M, |
| chr8 | 117822534 | C | T | . | exonic | EXT1 | nonsynonymous SNV | EXT1:NM_000127:exon5:c.G1348A:p.G450R, |
| chr8 | 11802853 | T | C | COSV55043146 | exonic | FDFT1 | nonsynonymous SNV | FDFT1:NM_001287756:exon1:c.T7C:p.W3R, |
| chr8 | 11802926 | G | A | . | exonic | FDFT1 | nonsynonymous SNV | FDFT1:NM_004462:exon1:c.G94A:p.D32N,FDFT1:NM_001287743:exon2:c.G94A:p.D32N,FDFT1:NM_001287742:exon3:c.G94A:p.D32N, |
| chr8 | 11808849 | G | A | . | exonic | FDFT1 | nonsynonymous SNV | FDFT1:NM_004462:exon2:c.G155A:p.R52H,FDFT1:NM_001287743:exon3:c.G155A:p.R52H,FDFT1:NM_001287750:exon1:c.G332A:p.R111H,FDFT1:NM_001287742:exon4:c.G155A:p.R52H, |
| chr8 | 11808860 | G | A | . | exonic | FDFT1 | nonsynonymous SNV | FDFT1:NM_004462:exon2:c.G166A:p.A56T,FDFT1:NM_001287743:exon3:c.G166A:p.A56T,FDFT1:NM_001287750:exon1:c.G343A:p.A115T,FDFT1:NM_001287742:exon4:c.G166A:p.A56T, |
| chr8 | 11809701 | G | A | . | exonic | FDFT1 | nonsynonymous SNV | FDFT1:NM_004462:exon3:c.G232A:p.A78T,FDFT1:NM_001287743:exon4:c.G232A:p.A78T,FDFT1:NM_001287750:exon2:c.G409A:p.A137T,FDFT1:NM_001287742:exon5:c.G232A:p.A78T,FDFT1:NM_001287747:exon3:c.G40A:p.A14T,FDFT1:NM_001287745:exon4:c.G40A:p.A14T,FDFT1:NM_001287744:exon3:c.G40A:p.A14T,FDFT1:NM_001287749:exon3:c.G40A:p.A14T,FDFT1:NM_001287748:exon3:c.G40A:p.A14T, |

| | | | | | | | | |
|---|---|---|---|---|---|---|---|---|
| chr8 | 11809707 | G | A | . | exonic | FDFT1 | nonsynonymous SNV | FDFT1:NM_004462:exon3:c.G238A:p.D80N,FDFT1:NM_001287743:exon4:c.G238A:p.D80N,FDFT1:NM_001287750:exon2:c.G415A:p.D139N,FDFT1:NM_001287742:exon5:c.G238A:p.D80N,FDFT1:NM_001287747:exon3:c.G46A:p.D16N,FDFT1:NM_001287745:exon4:c.G46A:p.D16N,FDFT1:NM_001287744:exon3:c.G46A:p.D16N,FDFT1:NM_001287749:exon3:c.G46A:p.D16N,FDFT1:NM_001287748:exon3:c.G46A:p.D16N, |
| chr8 | 11809722 | G | A | . | exonic | FDFT1 | nonsynonymous SNV | FDFT1:NM_004462:exon3:c.G253A:p.D85N,FDFT1:NM_001287743:exon4:c.G253A:p.D85N,FDFT1:NM_001287750:exon2:c.G430A:p.D144N,FDFT1:NM_001287742:exon5:c.G253A:p.D85N,FDFT1:NM_001287747:exon3:c.G61A:p.D21N,FDFT1:NM_001287745:exon4:c.G61A:p.D21N,FDFT1:NM_001287744:exon3:c.G61A:p.D21N,FDFT1:NM_001287749:exon3:c.G61A:p.D21N,FDFT1:NM_001287748:exon3:c.G61A:p.D21N, |
| chr8 | 11809789 | C | G | . | exonic | FDFT1 | nonsynonymous SNV | FDFT1:NM_004462:exon3:c.C320G:p.P107R,FDFT1:NM_001287751:exon2:c.C65G:p.P22R,FDFT1:NM_001287743:exon4:c.C320G:p.P107R,FDFT1:NM_001287750:exon2:c.C497G:p.P166R,FDFT1:NM_001287742:exon5:c.C320G:p.P107R,FDFT1:NM_001287747:exon3:c.C128G:p.P43R,FDFT1:NM_001287745:exon4:c.C128G:p.P43R,FDFT1:NM_001287744:exon3:c.C128G:p.P43R,FDFT1:NM_001287749:exon3:c.C128G:p.P43R,FDFT1:NM_001287748:exon3:c.C128G:p.P43R, |
| chr8 | 11821774 | G | A | . | exonic | FDFT1 | nonsynonymous SNV | FDFT1:NM_004462:exon4:c.G406A:p.A136T,FDFT1:NM_001287751:exon3:c.G151A:p.A51T,FDFT1:NM_001287743:exon5:c.G406A:p.A136T,FDFT1:NM_001287750:exon3:c.G583A:p.A195T,FDFT1:NM_001287742:exon6:c.G406A:p.A136T,FDFT1:NM_001287747:exon4:c.G214A:p.A72T,FDFT1:NM_001287745:exon5:c.G214A:p.A72T,FDFT1:NM_001287744:exon4:c.G214A:p.A72T,FDFT1:NM_001287749:exon4:c.G214A:p.A72T,FDFT1:NM_001287748:exon4:c.G214A:p.A72T, |
| chr8 | 11821811 | G | A | COSV55042913 | exonic | FDFT1 | nonsynonymous SNV | FDFT1:NM_004462:exon4:c.G443A:p.R148Q,FDFT1:NM_001287751:exon3:c.G188A:p.R63Q,FDFT1:NM_001287743:exon5:c.G443A:p.R148Q,FDFT1:NM_001287750:exon3:c.G620A:p.R207Q,FDFT1:NM_001287742:exon6:c.G443A:p.R148Q,FDFT1:NM_001287747:exon4:c.G251A:p.R84Q,FDFT1:NM_001287745:exon5:c.G251A:p.R84Q,FDFT1:NM_001287744:exon4:c.G251A:p.R84Q,FDFT1:NM_001287749:exon4:c.G251A:p.R84Q,FDFT1:NM_001287748:exon4:c.G251A:p.R84Q, |
| chr8 | 11821861 | G | A | . | exonic | FDFT1 | nonsynonymous SNV | FDFT1:NM_004462:exon4:c.G493A:p.E165K,FDFT1:NM_001287751:exon3:c.G238A:p.E80K,FDFT1:NM_001287743:exon5:c.G493A:p.E165K,FDFT1:NM_001287750:exon3:c.G670A:p.E224K,FDFT1:NM_001287742:exon6:c.G493A:p.E165K,FDFT1:NM_001287747:exon4:c.G301A:p.E101K,FDFT1:NM_001287745:exon5:c.G301A:p.E101K,FDFT1:NM_001287744:exon4:c.G301A:p.E101K,FDFT1:NM_001287749:exon4:c.G301A:p.E101K,FDFT1:NM_001287748:exon4:c.G301A:p.E101K, |
| chr8 | 11830275 | A | T | . | exonic | FDFT1 | nonsynonymous SNV | FDFT1:NM_004462:exon6:c.A734T:p.D245V,FDFT1:NM_001287751:exon5:c.A479T:p.D160V,FDFT1:NM_001287743:exon7:c.A734T:p.D245V,FDFT1:NM_001287750:exon5:c.A911T:p.D304V,FDFT1:NM_001287742:exon8:c.A734T:p.D245V,FDFT1:NM_001287747:exon6:c.A542T:p.D181V,FDFT1:NM_001287745:exon7:c.A542T:p.D181V,FDFT1:NM_001287744:exon6:c.A542T:p.D181V,FDFT1:NM_001287756:exon3:c.A233T:p.D78V,FDFT1:NM_001287749:exon6:c.A542T:p.D181V,FDFT1:NM_001287748:exon6:c.A542T:p.D181V, |
| chr8 | 11830314 | G | A | . | exonic | FDFT1 | nonsynonymous SNV | FDFT1:NM_004462:exon6:c.G773A:p.C258Y,FDFT1:NM_001287751:exon5:c.G518A:p.C173Y,FDFT1:NM_001287743:exon7:c.G773A:p.C258Y,FDFT1:NM_001287750:exon5:c.G950A:p.C317Y,FDFT1:NM_001287742:exon8:c.G773A:p.C258Y,FDFT1:NM_001287747:exon6:c.G581A:p.C194Y,FDFT1:NM_001287745:exon7:c.G581A:p.C194Y,FDFT1:NM_001287744:exon6:c.G581A:p.C194Y,FDFT1:NM_001287756:exon3:c.G272A:p.C91Y,FDFT1:NM_001287749:exon6:c.G581A:p.C194Y,FDFT1:NM_001287748:exon6:c.G581A:p.C194Y, |
| chr8 | 11830358 | G | A | . | exonic | FDFT1 | nonsynonymous SNV | FDFT1:NM_004462:exon6:c.G817A:p.V273I,FDFT1:NM_001287751:exon5:c.G562A:p.V188I,FDFT1:NM_001287743:exon7:c.G817A:p.V273I,FDFT1:NM_001287750:exon5:c.G994A:p.V332I,FDFT1:NM_001287742:exon8:c.G817A:p.V273I,FDFT1:NM_001287747:exon6:c.G625A:p.V209I,FDFT1:NM_001287745:exon7:c.G625A:p.V209I,FDFT1:NM_001287744:exon6:c.G625A:p.V209I,FDFT1:NM_001287756:exon3:c.G316A:p.V106I,FDFT1:NM_001287749:exon6:c.G625A:p.V209I,FDFT1:NM_001287748:exon6:c.G625A:p.V209I, |
| chr8 | 11831620 | G | A | . | exonic | FDFT1 | nonsynonymous SNV | FDFT1:NM_004462:exon7:c.G982A:p.A328T,FDFT1:NM_001287750:exon6:c.G1159A:p.A387T,FDFT1:NM_001287742:exon9:c.G982A:p.A328T,FDFT1:NM_001287744:exon7:c.G790A:p.A264T,FDFT1:NM_001287756:exon4:c.G481A:p.A161T,FDFT1:NM_001287749:exon7:c.G790A:p.A264T,FDFT1:NM_001287751:exon6:c.G727A:p.A243T,FDFT1:NM_001287743:exon8:c.G982A:p.A328T,FDFT1:NM_001287747:exon7:c.G790A:p.A264T,FDFT1:NM_001287745:exon8:c.G790A:p.A264T,FDFT1:NM_001287748:exon7:c.G790A:p.A264T, |
| chr8 | 11838530 | T | C | . | exonic | FDFT1 | nonsynonymous SNV | FDFT1:NM_004462:exon8:c.T1175C:p.L392P,FDFT1:NM_001287751:exon7:c.T920C:p.L307P,FDFT1:NM_001287743:exon9:c.T1175C:p.L392P,FDFT1:NM_001287750:exon7:c.T1352C:p.L451P,FDFT1:NM_001287742:exon10:c.T1175C:p.L392P,FDFT1:NM_001287747:exon8:c.T983C:p.L328P,FDFT1:NM_001287745:exon9:c.T983C:p.L328P,FDFT1:NM_001287744:exon8:c.T983C:p.L328P,FDFT1:NM_001287756:exon5:c.T674C:p.L225P,FDFT1:NM_001287749:exon8:c.T983C:p.L328P,FDFT1:NM_001287748:exon8:c.T983C:p.L328P, |
| chr8 | 11838584 | A | T | . | exonic | FDFT1 | nonsynonymous SNV | FDFT1:NM_004462:exon8:c.A1229T:p.D410V,FDFT1:NM_001287751:exon7:c.A974T:p.D325V,FDFT1:NM_001287743:exon9:c.A1229T:p.D410V,FDFT1:NM_001287750:exon7:c.A1406T:p.D469V,FDFT1:NM_001287742:exon10:c.A1229T:p.D410V,FDFT1:NM_001287747:exon8:c.A1037T:p.D346V,FDFT1:NM_001287745:exon9:c.A1037T:p.D346V,FDFT1:NM_001287744:exon8:c.A1037T:p.D346V,FDFT1:NM_001287756:exon5:c.A728T:p.D243V,FDFT1:NM_001287749:exon8:c.A1037T:p.D346V,FDFT1:NM_001287748:exon8:c.A1037T:p.D346V, |
| chr8 | 11852665 | T | C | . | exonic | CTSB | nonsynonymous SNV | CTSB:NM_001908:exon3:c.A157G:p.S53G,CTSB:NM_147783:exon4:c.A157G:p.S53G,CTSB:NM_147781:exon4:c.A157G:p.S53G,CTSB:NM_147780:exon5:c.A157G:p.S53G,CTSB:NM_147782:exon4:c.A157G:p.S53G, |
| chr8 | 11853379 | G | C | COSV61540513 | exonic | CTSB | nonsynonymous SNV | CTSB:NM_001908:exon2:c.C76G:p.L26V,CTSB:NM_147783:exon3:c.C76G:p.L26V,CTSB:NM_147781:exon3:c.C76G:p.L26V,CTSB:NM_147780:exon4:c.C76G:p.L26V,CTSB:NM_147782:exon3:c.C76G:p.L26V, |
| chr8 | 119583783 | A | G | . | exonic | ENPP2 | nonsynonymous SNV | ENPP2:NM_001040092:exon17:c.T1477C:p.S493P,ENPP2:NM_006209:exon18:c.T1633C:p.S545P,ENPP2:NM_001130863:exon17:c.T1477C:p.S493P, |
| chr8 | 119834948 | T | C | COSV58086368 | exonic | DSCC1 | nonsynonymous SNV | DSCC1:NM_024094:exon9:c.A1127G:p.H376R, |
| chr8 | 123427934 | A | G | COSV54873851 | exonic | WDYHV1 | nonsynonymous SNV | WDYHV1:NM_018024:exon2:c.A94G:p.I32V, |
| chr8 | 123436496 | A | G | COSV54874032 | exonic | WDYHV1 | nonsynonymous SNV | WDYHV1:NM_001283024:exon4:c.A98G:p.N33S,WDYHV1:NM_001283027:exon5:c.A74G:p.N25S,WDYHV1:NM_018024:exon4:c.A278G:p.N93S, |
| chr8 | 123436564 | T | A | COSV54873861 | exonic | WDYHV1 | nonsynonymous SNV | WDYHV1:NM_001283024:exon4:c.T166A:p.F56I,WDYHV1:NM_001283027:exon5:c.T142A:p.F48I,WDYHV1:NM_018024:exon4:c.T346A:p.F116I, |
| chr8 | 123437226 | C | T | COSV54874614 | exonic | WDYHV1 | nonsynonymous SNV | WDYHV1:NM_001283024:exon5:c.C220T:p.R74C,WDYHV1:NM_001283027:exon6:c.C196T:p.R66C,WDYHV1:NM_018024:exon5:c.C400T:p.R134C, |
| chr8 | 123698489 | C | T | COSV51624677 | exonic | ANXA13 | nonsynonymous SNV | ANXA13:NM_001003954:exon5:c.G380A:p.R127H,ANXA13:NM_004306:exon4:c.G257A:p.R86H, |
| chr8 | 124539205 | G | A | . | exonic | NDUFB9 | nonsynonymous SNV | NDUFB9:NM_001311168:exon1:c.G19A:p.G7R,NDUFB9:NM_005005:exon1:c.G19A:p.G7R, |
| chr8 | 124539232 | G | A | . | exonic | NDUFB9 | nonsynonymous SNV | NDUFB9:NM_001311168:exon1:c.G46A:p.V16M,NDUFB9:NM_005005:exon1:c.G46A:p.V16M, |
| chr8 | 124539242 | T | C | . | exonic | NDUFB9 | nonsynonymous SNV | NDUFB9:NM_001311168:exon1:c.T56C:p.L19P,NDUFB9:NM_005005:exon1:c.T56C:p.L19P, |
| chr8 | 124539253 | G | A | . | exonic | NDUFB9 | nonsynonymous SNV | NDUFB9:NM_001311168:exon1:c.G67A:p.A23T,NDUFB9:NM_005005:exon1:c.G67A:p.A23T, |
| chr8 | 124539280 | G | A | . | exonic | NDUFB9 | nonsynonymous SNV | NDUFB9:NM_001311168:exon1:c.G94A:p.V32I,NDUFB9:NM_005005:exon1:c.G94A:p.V32I, |
| chr8 | 124543098 | G | A | . | exonic | NDUFB9 | nonsynonymous SNV | NDUFB9:NM_005005:exon2:c.G113A:p.R38Q, |
| chr8 | 124543106 | G | A | . | exonic | NDUFB9 | nonsynonymous SNV | NDUFB9:NM_005005:exon2:c.G121A:p.A41T, |
| chr8 | 124543156 | G | A | . | exonic | NDUFB9 | nonsynonymous SNV | NDUFB9:NM_001278646:exon2:c.G42A:p.M14I,NDUFB9:NM_001278645:exon2:c.G3A:p.M1I,NDUFB9:NM_001311168:exon2:c.G138A:p.M46I,NDUFB9:NM_005005:exon2:c.G171A:p.M57I, |
| chr8 | 124543163 | G | A | . | exonic | NDUFB9 | nonsynonymous SNV | NDUFB9:NM_001278646:exon2:c.G49A:p.A17T,NDUFB9:NM_001311168:exon2:c.G145A:p.A49T,NDUFB9:NM_001278645:exon2:c.G10A:p.A4T,NDUFB9:NM_005005:exon2:c.G178A:p.A60T, |
| chr8 | 124543173 | T | C | . | exonic | NDUFB9 | nonsynonymous SNV | NDUFB9:NM_001278646:exon2:c.T59C:p.L20P,NDUFB9:NM_001278645:exon2:c.T20C:p.L7P,NDUFB9:NM_001311168:exon2:c.T155C:p.L52P,NDUFB9:NM_005005:exon2:c.T188C:p.L63P, |
| chr8 | 124543206 | G | A | . | exonic | NDUFB9 | nonsynonymous SNV | NDUFB9:NM_001278646:exon2:c.G92A:p.R31H,NDUFB9:NM_001278645:exon2:c.G53A:p.R18H,NDUFB9:NM_001311168:exon2:c.G188A:p.R63H,NDUFB9:NM_005005:exon2:c.G221A:p.R74H, |
| chr8 | 124543248 | G | A | . | exonic | NDUFB9 | nonsynonymous SNV | NDUFB9:NM_001278646:exon2:c.G134A:p.G45D,NDUFB9:NM_001311168:exon2:c.G230A:p.G77D,NDUFB9:NM_001278645:exon2:c.G95A:p.G32D,NDUFB9:NM_005005:exon2:c.G263A:p.G88D, |
| chr8 | 124543268 | G | A | . | exonic | NDUFB9 | nonsynonymous SNV | NDUFB9:NM_001278646:exon2:c.G154A:p.D52N,NDUFB9:NM_001278645:exon2:c.G115A:p.D39N,NDUFB9:NM_001311168:exon2:c.G250A:p.D84N,NDUFB9:NM_005005:exon2:c.G283A:p.D95N, |
| chr8 | 124547042 | G | A | . | exonic | NDUFB9 | nonsynonymous SNV | NDUFB9:NM_001278646:exon3:c.G208A:p.A70T,NDUFB9:NM_001278645:exon3:c.G169A:p.A57T,NDUFB9:NM_001311168:exon3:c.G304A:p.A102T,NDUFB9:NM_005005:exon3:c.G337A:p.A113T, |
| chr8 | 124547080 | G | A | . | exonic | NDUFB9 | stopgain | NDUFB9:NM_001278646:exon3:c.G246A:p.W82X,NDUFB9:NM_001278645:exon3:c.G207A:p.W69X,NDUFB9:NM_001311168:exon3:c.G342A:p.W114X,NDUFB9:NM_005005:exon3:c.G375A:p.W125X, |
| chr8 | 124547096 | G | A | . | exonic | NDUFB9 | nonsynonymous SNV | NDUFB9:NM_001278646:exon3:c.G262A:p.E88K,NDUFB9:NM_001278645:exon3:c.G223A:p.E75K,NDUFB9:NM_001311168:exon3:c.G358A:p.E120K,NDUFB9:NM_005005:exon3:c.G391A:p.E131K, |
| chr8 | 124547109 | G | A | . | exonic | NDUFB9 | nonsynonymous SNV | NDUFB9:NM_001278646:exon3:c.G275A:p.R92Q,NDUFB9:NM_001278645:exon3:c.G236A:p.R79Q,NDUFB9:NM_001311168:exon3:c.G371A:p.R124Q,NDUFB9:NM_005005:exon3:c.G404A:p.R135Q, |
| chr8 | 124549776 | G | A | . | exonic | NDUFB9 | nonsynonymous SNV | NDUFB9:NM_001278646:exon4:c.G295A:p.E99K,NDUFB9:NM_001278645:exon4:c.G256A:p.E86K,NDUFB9:NM_001311168:exon4:c.G391A:p.E131K,NDUFB9:NM_005005:exon4:c.G424A:p.E142K, |
| chr8 | 124549788 | C | T | . | exonic | NDUFB9 | nonsynonymous SNV | NDUFB9:NM_001278646:exon4:c.C307T:p.P103S,NDUFB9:NM_001278645:exon4:c.C268T:p.P90S,NDUFB9:NM_001311168:exon4:c.C403T:p.P135S,NDUFB9:NM_005005:exon4:c.C436T:p.P146S, |

| | | | | | | | | |
|---|---|---|---|---|---|---|---|---|
| chr8 | 124549792 | G | A | . | exonic | NDUFB9 | nonsynonymous SNV | NDUFB9:NM_001278646:exon4:c.G311A:p.G104D,NDUFB9:NM_001278645:exon4:c.G272A:p.G91D,NDUFB9:NM_001311168:exon4:c.G407A:p.G136D,NDUFB9:NM_005005:exon4:c.G440A:p.G147D, |
| chr8 | 124549809 | G | A | . | exonic | NDUFB9 | nonsynonymous SNV | NDUFB9:NM_001278646:exon4:c.G328A:p.A110T,NDUFB9:NM_001278645:exon4:c.G289A:p.A97T,NDUFB9:NM_001311168:exon4:c.G424A:p.A142T,NDUFB9:NM_005005:exon4:c.G457A:p.A153T, |
| chr8 | 124549834 | G | A | . | exonic | NDUFB9 | nonsynonymous SNV | NDUFB9:NM_001278646:exon4:c.G353A:p.G118D,NDUFB9:NM_001278645:exon4:c.G314A:p.G105D,NDUFB9:NM_001311168:exon4:c.G449A:p.G150D,NDUFB9:NM_005005:exon4:c.G482A:p.G161D, |
| chr8 | 124549842 | C | T | . | exonic | NDUFB9 | nonsynonymous SNV | NDUFB9:NM_001278646:exon4:c.C361T:p.P121S,NDUFB9:NM_001278645:exon4:c.C322T:p.P108S,NDUFB9:NM_001311168:exon4:c.C457T:p.P153S,NDUFB9:NM_005005:exon4:c.C490T:p.P164S, |
| chr8 | 124549856 | G | A | . | exonic | NDUFB9 | stopgain | NDUFB9:NM_001278646:exon4:c.G375A:p.W125X,NDUFB9:NM_001278645:exon4:c.G336A:p.W112X,NDUFB9:NM_001311168:exon4:c.G471A:p.W157X,NDUFB9:NM_005005:exon4:c.G504A:p.W168X, |
| chr8 | 124549861 | T | C | . | exonic | NDUFB9 | nonsynonymous SNV | NDUFB9:NM_001278646:exon4:c.T380C:p.I127T,NDUFB9:NM_001278645:exon4:c.T341C:p.I114T,NDUFB9:NM_001311168:exon4:c.T476C:p.I159T,NDUFB9:NM_005005:exon4:c.T509C:p.I170T, |
| chr8 | 124549882 | G | A | COSV52674143 | exonic | NDUFB9 | nonsynonymous SNV | NDUFB9:NM_001278646:exon4:c.G401A:p.R134Q,NDUFB9:NM_001278645:exon4:c.G362A:p.R121Q,NDUFB9:NM_001311168:exon4:c.G497A:p.R166Q,NDUFB9:NM_005005:exon4:c.G530A:p.R177Q, |
| chr8 | 127740569 | G | A | . | exonic | MYC | nonsynonymous SNV | MYC:NM_002467:exon3:c.G976A:p.A326T, |
| chr8 | 132572310 | A | G | COSV51539954 | exonic | LRRC6 | nonsynonymous SNV | LRRC6:NM_012472:exon12:c.T1397C:p.I466T, |
| chr8 | 132625413 | G | A | COSV51539603 | exonic | LRRC6 | nonsynonymous SNV | LRRC6:NM_012472:exon6:c.C695T:p.T232I, |
| chr8 | 133254568 | C | T | . | exonic | NDRG1 | nonsynonymous SNV | NDRG1:NM_001258433:exon8:c.G322A:p.D108N,NDRG1:NM_001135242:exon9:c.G565A:p.D189N,NDRG1:NM_001258432:exon7:c.G367A:p.D123N,NDRG1:NM_006096:exon9:c.G565A:p.D189N, |
| chr8 | 133256785 | C | T | . | exonic | NDRG1 | nonsynonymous SNV | NDRG1:NM_001258433:exon7:c.G286A:p.A96T,NDRG1:NM_001135242:exon8:c.G529A:p.A177T,NDRG1:NM_001258432:exon6:c.G331A:p.A111T,NDRG1:NM_006096:exon8:c.G529A:p.A177T, |
| chr8 | 133258368 | C | T | . | exonic | NDRG1 | nonsynonymous SNV | NDRG1:NM_001258433:exon6:c.G205A:p.A69T,NDRG1:NM_001135242:exon7:c.G448A:p.A150T,NDRG1:NM_001258432:exon5:c.G250A:p.A84T,NDRG1:NM_006096:exon7:c.G448A:p.A150T, |
| chr8 | 133284281 | C | T | . | exonic | NDRG1 | nonsynonymous SNV | NDRG1:NM_001135242:exon2:c.G31A:p.A11T,NDRG1:NM_006096:exon2:c.G31A:p.A11T, |
| chr8 | 143276211 | G | A | COSV60681816 | exonic | GLI4 | nonsynonymous SNV | GLI4:NM_138465:exon4:c.G538A:p.A180T, |
| chr8 | 144095929 | A | G | COSV59644891 | exonic | CYC1 | nonsynonymous SNV | CYC1:NM_001916:exon2:c.A226G:p.M76V, |
| chr8 | 144095930 | T | C | . | exonic | CYC1 | nonsynonymous SNV | CYC1:NM_001916:exon2:c.T227C:p.M76T, |
| chr8 | 144106920 | G | A | . | exonic | MAF1 | nonsynonymous SNV | MAF1:NM_032272:exon7:c.G706A:p.G236R, |
| chr8 | 144355665 | C | T | . | exonic | FBXL6 | nonsynonymous SNV | FBXL6:NM_024555:exon9:c.G1468A:p.G490S,FBXL6:NM_012162:exon9:c.G1486A:p.G496S, |
| chr8 | 144379425 | C | A | . | exonic | ADCK5 | nonsynonymous SNV | ADCK5:NM_174922:exon2:c.C51A:p.S17R, |
| chr8 | 144414297 | C | G | . | exonic | SLC39A4 | nonsynonymous SNV | SLC39A4:NM_017767:exon5:c.G1039C:p.V347L,SLC39A4:NM_130849:exon6:c.G1114C:p.V372L, |
| chr8 | 144414342 | T | C | COSV52777858 | exonic | SLC39A4 | nonsynonymous SNV | SLC39A4:NM_017767:exon5:c.A994G:p.T332A,SLC39A4:NM_130849:exon6:c.A1069G:p.T357A, |
| chr8 | 144415811 | A | G | . | exonic | SLC39A4 | nonsynonymous SNV | SLC39A4:NM_017767:exon1:c.T398C:p.M133T,SLC39A4:NM_130849:exon2:c.T473C:p.M158T, |
| chr8 | 144415944 | C | T | COSV52777498 | exonic | SLC39A4 | nonsynonymous SNV | SLC39A4:NM_017767:exon1:c.G265A:p.A89T,SLC39A4:NM_130849:exon2:c.G340A:p.A114T, |
| chr8 | 144416618 | C | T | COSV52777818 | exonic | SLC39A4 | nonsynonymous SNV | SLC39A4:NM_130849:exon1:c.G172A:p.A58T, |
| chr8 | 144517130 | A | G | COSV52881402 | exonic | RECQL4 | nonsynonymous SNV | RECQL4:NM_004260:exon4:c.T274C:p.S92P, |
| chr8 | 144851323 | C | T | COSV59310992 | exonic | COMMD5 | nonsynonymous SNV | COMMD5:NM_001081003:exon2:c.G16A:p.A6T,COMMD5:NM_001287237:exon2:c.G16A:p.A6T,COMMD5:NM_014066:exon2:c.G16A:p.A6T,COMMD5:NM_001081004:exon2:c.G16A:p.A6T, |
| chr8 | 18009676 | A | T | COSV59586229 | exonic | PCM1 | nonsynonymous SNV | PCM1:NM_001315508:exon30:c.A4930T:p.I1644L,PCM1:NM_006197:exon31:c.A5092T:p.I1698L,PCM1:NM_001315507:exon31:c.A5068T:p.I1690L, |
| chr8 | 18061425 | A | G | . | exonic | ASAH1 | nonsynonymous SNV | ASAH1:NM_001127505:exon10:c.T719C:p.V240A,ASAH1:NM_177924:exon10:c.T737C:p.V246A,ASAH1:NM_004315:exon10:c.T785C:p.V262A, |
| chr8 | 18069818 | T | C | COSV50501249 | exonic | ASAH1 | nonsynonymous SNV | ASAH1:NM_177924:exon4:c.A277G:p.I93V,ASAH1:NM_004315:exon4:c.A325G:p.I109V, |
| chr8 | 18071302 | C | T | COSV50502085 | exonic | ASAH1 | nonsynonymous SNV | ASAH1:NM_001127505:exon4:c.G283A:p.V95M,ASAH1:NM_177924:exon3:c.G214A:p.V72M,ASAH1:NM_004315:exon3:c.G262A:p.V88M, |
| chr8 | 23288582 | A | G | COSV56120569 | exonic | R3HCC1 | nonsynonymous SNV | R3HCC1:NM_001136108:exon2:c.A59G:p.H20R,R3HCC1:NM_001301650:exon2:c.A59G:p.H20R, |
| chr8 | 23290051 | G | A | COSV56120064 | exonic;splicing | R3HCC1 | nonsynonymous SNV | R3HCC1:NM_001136108:exon4:c.G434A:p.R145K, |
| chr8 | 23291427 | G | A | COSV56119757 | exonic | R3HCC1 | nonsynonymous SNV | R3HCC1:NM_001136108:exon5:c.G919A:p.V307M,R3HCC1:NM_001301650:exon6:c.G793A:p.V265M, |
| chr8 | 23293365 | T | G | COSV56119293 | exonic | R3HCC1 | nonsynonymous SNV | R3HCC1:NM_001136108:exon6:c.T1088G:p.L363R,R3HCC1:NM_001301650:exon7:c.T962G:p.L321R, |
| chr8 | 26354556 | T | C | . | exonic | PPP2R2A | nonsynonymous SNV | PPP2R2A:NM_002717:exon4:c.T269C:p.L90S,PPP2R2A:NM_001177591:exon4:c.T299C:p.L100S, |
| chr8 | 26370277 | G | A | . | exonic | PPP2R2A | nonsynonymous SNV | PPP2R2A:NM_002717:exon10:c.G1208A:p.R403Q,PPP2R2A:NM_001177591:exon10:c.G1238A:p.R413Q, |
| chr8 | 27500988 | A | G | COSV66844518 | exonic | EPHX2 | nonsynonymous SNV | EPHX2:NM_001256482:exon2:c.A5G:p.K2R,EPHX2:NM_001256484:exon2:c.A5G:p.K2R,EPHX2:NM_001979:exon2:c.A164G:p.K55R, |
| chr8 | 27516348 | G | A | COSV66844144 | exonic | EPHX2 | nonsynonymous SNV | EPHX2:NM_001256482:exon8:c.G701A:p.R234Q,EPHX2:NM_001256484:exon8:c.G701A:p.R234Q,EPHX2:NM_001979:exon8:c.G860A:p.R287Q,EPHX2:NM_001256483:exon7:c.G662A:p.R221Q, |
| chr8 | 31081188 | G | A | COSV53296756 | exonic | WRN | nonsynonymous SNV | WRN:NM_000553:exon9:c.G1161A:p.M387I, |
| chr8 | 31141764 | G | T | COSV53297644 | exonic | WRN | nonsynonymous SNV | WRN:NM_000553:exon26:c.G3222T:p.L1074F, |
| chr8 | 31167138 | T | C | COSV53296936 | exonic | WRN | nonsynonymous SNV | WRN:NM_000553:exon34:c.T4099C:p.C1367R, |
| chr8 | 32595840 | G | A | COSV55177787 | exonic | NRG1 | nonsynonymous SNV | NRG1:NM_001160004:exon2:c.G113A:p.R38Q,NRG1:NM_001160005:exon2:c.G113A:p.R38Q,NRG1:NM_001160002:exon2:c.G113A:p.R38Q,NRG1:NM_013962:exon2:c.G758A:p.R253Q,NRG1:NM_013957:exon2:c.G113A:p.R38Q,NRG1:NM_001160007:exon2:c.G113A:p.R38Q,NRG1:NM_013964:exon2:c.G113A:p.R38Q,NRG1:NM_013958:exon2:c.G113A:p.R38Q,NRG1:NM_004495:exon2:c.G113A:p.R38Q,NRG1:NM_013960:exon2:c.G113A:p.R38Q,NRG1:NM_001160008:exon2:c.G113A:p.R38Q,NRG1:NM_013956:exon2:c.G113A:p.R38Q, |
| chr8 | 32756465 | T | C | . | exonic | NRG1 | nonsynonymous SNV | NRG1:NM_001160004:exon9:c.T857C:p.M286T,NRG1:NM_013957:exon9:c.T857C:p.M286T,NRG1:NM_013964:exon9:c.T866C:p.M289T,NRG1:NM_001159996:exon5:c.T404C:p.M135T,NRG1:NM_013960:exon9:c.T866C:p.M289T,NRG1:NM_001160008:exon9:c.T857C:p.M286T,NRG1:NM_013956:exon10:c.T881C:p.M294T, |
| chr8 | 38288542 | A | G | . | exonic | WHSC1L1 | nonsynonymous SNV | WHSC1L1:NM_023034:exon19:c.T3446C:p.I1149T, |
| chr8 | 38289464 | G | T | . | exonic | WHSC1L1 | nonsynonymous SNV | WHSC1L1:NM_023034:exon18:c.C3160A:p.Q1054K, |
| chr8 | 40153560 | G | A | . | exonic | C8ORF4 | nonsynonymous SNV | C8orf4:NM_020130:exon1:c.G28A:p.V10I, |
| chr8 | 413192 | G | A | COSV52338688 | exonic | FBXO25 | nonsynonymous SNV | FBXO25:NM_183421:exon2:c.G113A:p.R38H,FBXO25:NM_183420:exon2:c.G113A:p.R38H, |
| chr8 | 42357203 | G | A | . | exonic | POLB | nonsynonymous SNV | POLB:NM_002690:exon8:c.G457A:p.E153K, |
| chr8 | 42973323 | C | A | . | exonic | HOOK3 | stopgain | HOOK3:NM_032410:exon12:c.C1157A:p.S386X, |

| | | | | | | | | |
|---|---|---|---|---|---|---|---|---|
| chr8 | 47595875 | G | A | . | exonic | SPIDR | nonsynonymous SNV | SPIDR:NM_001282916:exon8:c.G952A:p.V318I,SPIDR:NM_001282919:exon9:c.G982A:p.V328I,SPIDR:NM_001080394:exon9:c.G1162A:p.V388I, |
| chr8 | 47599013 | G | A | COSV52368528 | exonic | SPIDR | nonsynonymous SNV | SPIDR:NM_001282916:exon9:c.G1151A:p.R384H,SPIDR:NM_001282919:exon10:c.G1181A:p.R394H,SPIDR:NM_001080394:exon10:c.G1361A:p.R454H, |
| chr8 | 47599142 | G | A | . | exonic | SPIDR | nonsynonymous SNV | SPIDR:NM_001282916:exon9:c.G1280A:p.S427N,SPIDR:NM_001282919:exon10:c.G1310A:p.S437N,SPIDR:NM_001080394:exon10:c.G1490A:p.S497N, |
| chr8 | 47712692 | G | A | . | exonic | SPIDR | nonsynonymous SNV | SPIDR:NM_001282916:exon14:c.G1798A:p.E600K,SPIDR:NM_001282919:exon15:c.G1828A:p.E610K,SPIDR:NM_001080394:exon15:c.G2008A:p.E670K, |
| chr8 | 47712815 | G | A | COSV52369911 | exonic | SPIDR | nonsynonymous SNV | SPIDR:NM_001282916:exon14:c.G1921A:p.A641T,SPIDR:NM_001282919:exon15:c.G1951A:p.A651T,SPIDR:NM_001080394:exon15:c.G2131A:p.A711T, |
| chr8 | 47713593 | G | A | COSV52370111 | exonic | SPIDR | nonsynonymous SNV | SPIDR:NM_001282916:exon15:c.G2083A:p.D695N,SPIDR:NM_001282919:exon16:c.G2113A:p.D705N,SPIDR:NM_001080394:exon16:c.G2293A:p.D765N, |
| chr8 | 47729031 | G | A | . | exonic | SPIDR | nonsynonymous SNV | SPIDR:NM_001282916:exon17:c.G2324A:p.C775Y,SPIDR:NM_001282919:exon18:c.G2354A:p.C785Y,SPIDR:NM_001080394:exon18:c.G2534A:p.C845Y, |
| chr8 | 47735424 | G | A | . | exonic | SPIDR | nonsynonymous SNV | SPIDR:NM_001282916:exon19:c.G2512A:p.A838T,SPIDR:NM_001282919:exon19:c.G2543A:p.C848Y,SPIDR:NM_001080394:exon20:c.G2722A:p.A908T, |
| chr8 | 51820489 | G | T | . | exonic | PCMTD1 | nonsynonymous SNV | PCMTD1:NM_001286783:exon6:c.C408A:p.N136K,PCMTD1:NM_052937:exon6:c.C936A:p.N312K,PCMTD1:NM_001286782:exon4:c.C708A:p.N236K, |
| chr8 | 51820490 | T | A | COSV62119484 | exonic | PCMTD1 | nonsynonymous SNV | PCMTD1:NM_001286783:exon6:c.A407T:p.N136I,PCMTD1:NM_052937:exon6:c.A935T:p.N312I,PCMTD1:NM_001286782:exon4:c.A707T:p.N236I, |
| chr8 | 51820592 | C | T | . | exonic | PCMTD1 | nonsynonymous SNV | PCMTD1:NM_001286783:exon6:c.G305A:p.R102K,PCMTD1:NM_052937:exon6:c.G833A:p.R278K,PCMTD1:NM_001286782:exon4:c.G605A:p.R202K, |
| chr8 | 51820598 | C | T | COSV62124907 | exonic | PCMTD1 | nonsynonymous SNV | PCMTD1:NM_001286783:exon6:c.G299A:p.R100K,PCMTD1:NM_052937:exon6:c.G827A:p.R276K,PCMTD1:NM_001286782:exon4:c.G599A:p.R200K, |
| chr8 | 51820604 | G | T | . | exonic | PCMTD1 | nonsynonymous SNV | PCMTD1:NM_001286783:exon6:c.C293A:p.P98H,PCMTD1:NM_052937:exon6:c.C821A:p.P274H,PCMTD1:NM_001286782:exon4:c.C593A:p.P198H, |
| chr8 | 51820605 | G | T | . | exonic | PCMTD1 | nonsynonymous SNV | PCMTD1:NM_001286783:exon6:c.C292A:p.P98T,PCMTD1:NM_052937:exon6:c.C820A:p.P274T,PCMTD1:NM_001286782:exon4:c.C592A:p.P198T, |
| chr8 | 53743645 | C | T | . | exonic | ATP6V1H | nonsynonymous SNV | ATP6V1H:NM_015941:exon13:c.G1323A:p.M441I,ATP6V1H:NM_213620:exon13:c.G1323A:p.M441I,ATP6V1H:NM_213619:exon12:c.G1269A:p.M423I, |
| chr8 | 53769661 | C | T | . | exonic | ATP6V1H | nonsynonymous SNV | ATP6V1H:NM_015941:exon11:c.G1132A:p.E378K,ATP6V1H:NM_213620:exon11:c.G1132A:p.E378K,ATP6V1H:NM_213619:exon10:c.G1078A:p.E360K, |
| chr8 | 53795687 | A | G | . | exonic | ATP6V1H | nonsynonymous SNV | ATP6V1H:NM_015941:exon9:c.T830C:p.V277A,ATP6V1H:NM_213620:exon9:c.T830C:p.V277A,ATP6V1H:NM_213619:exon8:c.T776C:p.V259A, |
| chr8 | 53795762 | A | G | . | exonic | ATP6V1H | nonsynonymous SNV | ATP6V1H:NM_015941:exon9:c.T755C:p.F252S,ATP6V1H:NM_213620:exon9:c.T755C:p.F252S,ATP6V1H:NM_213619:exon8:c.T701C:p.F234S, |
| chr8 | 53795801 | A | G | . | exonic | ATP6V1H | nonsynonymous SNV | ATP6V1H:NM_015941:exon9:c.T716C:p.L239P,ATP6V1H:NM_213620:exon9:c.T716C:p.L239P,ATP6V1H:NM_213619:exon8:c.T662C:p.L221P, |
| chr8 | 53801883 | A | G | . | exonic | ATP6V1H | nonsynonymous SNV | ATP6V1H:NM_015941:exon8:c.T593C:p.V198A,ATP6V1H:NM_213620:exon8:c.T593C:p.V198A,ATP6V1H:NM_213619:exon7:c.T539C:p.V180A, |
| chr8 | 53817517 | C | T | COSV62220815 | exonic | ATP6V1H | nonsynonymous SNV | ATP6V1H:NM_015941:exon5:c.G320A:p.R107H,ATP6V1H:NM_213620:exon5:c.G320A:p.R107H,ATP6V1H:NM_213619:exon5:c.G320A:p.R107H, |
| chr8 | 60741446 | G | A | . | exonic | CHD7 | nonsynonymous SNV | CHD7:NM_001316690:exon1:c.G14A:p.G5E,CHD7:NM_017780:exon2:c.G14A:p.G5E, |
| chr8 | 60742412 | G | T | . | exonic | CHD7 | nonsynonymous SNV | CHD7:NM_001316690:exon1:c.G980T:p.G327V,CHD7:NM_017780:exon2:c.G980T:p.G327V, |
| chr8 | 60742756 | G | A | . | exonic | CHD7 | nonsynonymous SNV | CHD7:NM_001316690:exon1:c.G1324A:p.A442T,CHD7:NM_017780:exon2:c.G1324A:p.A442T, |
| chr8 | 60742862 | C | A | . | exonic | CHD7 | nonsynonymous SNV | CHD7:NM_001316690:exon1:c.C1430A:p.P477Q,CHD7:NM_017780:exon2:c.C1430A:p.P477Q, |
| chr8 | 60781138 | A | G | . | exonic | CHD7 | nonsynonymous SNV | CHD7:NM_017780:exon3:c.A1804G:p.K602E, |
| chr8 | 60781153 | G | A | . | exonic | CHD7 | nonsynonymous SNV | CHD7:NM_017780:exon3:c.G1819A:p.V607I, |
| chr8 | 60781220 | T | A | . | exonic | CHD7 | nonsynonymous SNV | CHD7:NM_017780:exon3:c.T1886A:p.L629Q, |
| chr8 | 60781249 | G | C | . | exonic | CHD7 | nonsynonymous SNV | CHD7:NM_017780:exon3:c.G1915C:p.E639Q, |
| chr8 | 60781252 | G | A | COSV71113211 | exonic | CHD7 | nonsynonymous SNV | CHD7:NM_017780:exon3:c.G1918A:p.E640K, |
| chr8 | 60781258 | A | G | . | exonic | CHD7 | nonsynonymous SNV | CHD7:NM_017780:exon3:c.A1924G:p.K642E, |
| chr8 | 60781279 | G | A | . | exonic | CHD7 | nonsynonymous SNV | CHD7:NM_017780:exon3:c.G1945A:p.A649T, |
| chr8 | 60781280 | C | T | . | exonic | CHD7 | nonsynonymous SNV | CHD7:NM_017780:exon3:c.C1946T:p.A649V, |
| chr8 | 63026205 | G | A | COSV52649428 | exonic | GGH | nonsynonymous SNV | GGH:NM_003878:exon5:c.C452T:p.T151I, |
| chr8 | 63038678 | C | T | COSV52649507 | exonic | GGH | nonsynonymous SNV | GGH:NM_003878:exon1:c.G91A:p.A31T, |

| | | | | | | | | |
|---|---|---|---|---|---|---|---|---|
| chr8 | 63038753 | A | G | COSV52650077 | exonic | GGH | nonsynonymous SNV | GGH:NM_003878:exon1:c.T16C:p.C6R, |
| chr8 | 70707000 | G | T | . | exonic | XKR9 | nonsynonymous SNV | XKR9:NM_001287259:exon5:c.G340T:p.V114L,XKR9:NM_001011720:exon4:c.G340T:p.V114L,XKR9:NM_001287260:exon4:c.G340T:p.V114L, |
| chr8 | 73291102 | C | T | . | exonic | RPL7 | nonsynonymous SNV | RPL7:NM_000971:exon6:c.G689A:p.G230D, |
| chr8 | 73291818 | G | A | . | exonic | RPL7 | nonsynonymous SNV | RPL7:NM_000971:exon4:c.C383T:p.A128V, |
| chr8 | 73292711 | C | T | . | exonic | RPL7 | nonsynonymous SNV | RPL7:NM_000971:exon2:c.G101A:p.R34K, |
| chr8 | 73292717 | C | T | . | exonic | RPL7 | nonsynonymous SNV | RPL7:NM_000971:exon2:c.G95A:p.R32H, |
| chr8 | 76983629 | A | G | . | exonic | PEX2 | nonsynonymous SNV | PEX2:NM_001172086:exon5:c.T550C:p.C184R,PEX2:NM_001079867:exon3:c.T550C:p.C184R,PEX2:NM_001172087:exon3:c.T550C:p.C184R,PEX2:NM_000318:exon4:c.T550C:p.C184R, |
| chr8 | 79641420 | C | T | . | exonic | STMN2 | nonsynonymous SNV | STMN2:NM_007029:exon3:c.C158T:p.A53V,STMN2:NM_001199214:exon3:c.C158T:p.A53V, |
| chr8 | 79654941 | C | T | . | exonic | STMN2 | nonsynonymous SNV | STMN2:NM_007029:exon4:c.C359T:p.A120V,STMN2:NM_001199214:exon4:c.C359T:p.A120V, |
| chr8 | 85328643 | C | T | . | exonic | CA1 | nonsynonymous SNV | CA1:NM_001164830:exon7:c.G703A:p.E235K,CA1:NM_001291967:exon7:c.G505A:p.E169K,CA1:NM_001128830:exon9:c.G703A:p.E235K,CA1:NM_001128831:exon8:c.G703A:p.E235K,CA1:NM_001291968:exon6:c.G364A:p.E122K,CA1:NM_001738:exon9:c.G703A:p.E235K,CA1:NM_001128829:exon9:c.G703A:p.E235K, |
| chr8 | 85328663 | C | T | . | exonic | CA1 | nonsynonymous SNV | CA1:NM_001164830:exon7:c.G683A:p.R228H,CA1:NM_001291967:exon7:c.G485A:p.R162H,CA1:NM_001128830:exon9:c.G683A:p.R228H,CA1:NM_001128831:exon8:c.G683A:p.R228H,CA1:NM_001291968:exon6:c.G344A:p.R115H,CA1:NM_001738:exon9:c.G683A:p.R228H,CA1:NM_001128829:exon9:c.G683A:p.R228H, |
| chr8 | 85329729 | C | T | . | exonic | CA1 | stopgain | CA1:NM_001164830:exon6:c.G629A:p.W210X,CA1:NM_001291967:exon6:c.G431A:p.W144X,CA1:NM_001128830:exon8:c.G629A:p.W210X,CA1:NM_001128831:exon7:c.G629A:p.W210X,CA1:NM_001291968:exon5:c.G290A:p.W97X,CA1:NM_001738:exon8:c.G629A:p.W210X,CA1:NM_001128829:exon8:c.G629A:p.W210X, |
| chr8 | 85329736 | C | T | . | exonic | CA1 | nonsynonymous SNV | CA1:NM_001164830:exon6:c.G622A:p.V208I,CA1:NM_001291967:exon6:c.G424A:p.V142I,CA1:NM_001128830:exon8:c.G622A:p.V208I,CA1:NM_001128831:exon7:c.G622A:p.V208I,CA1:NM_001291968:exon5:c.G283A:p.V95I,CA1:NM_001738:exon8:c.G622A:p.V208I,CA1:NM_001128829:exon8:c.G622A:p.V208I, |
| chr8 | 85329768 | C | T | . | exonic | CA1 | nonsynonymous SNV | CA1:NM_001164830:exon6:c.G590A:p.G197D,CA1:NM_001291967:exon6:c.G392A:p.G131D,CA1:NM_001128830:exon8:c.G590A:p.G197D,CA1:NM_001128831:exon7:c.G590A:p.G197D,CA1:NM_001291968:exon5:c.G251A:p.G84D,CA1:NM_001738:exon8:c.G590A:p.G197D,CA1:NM_001128829:exon8:c.G590A:p.G197D, |
| chr8 | 85329802 | G | A | COSV56279080 | exonic | CA1 | nonsynonymous SNV | CA1:NM_001164830:exon6:c.C556T:p.L186F,CA1:NM_001291967:exon6:c.C358T:p.L120F,CA1:NM_001128830:exon8:c.C556T:p.L186F,CA1:NM_001128831:exon7:c.C556T:p.L186F,CA1:NM_001291968:exon5:c.C217T:p.L73F,CA1:NM_001738:exon8:c.C556T:p.L186F,CA1:NM_001128829:exon8:c.C556T:p.L186F, |
| chr8 | 85332531 | G | T | . | exonic | CA1 | nonsynonymous SNV | CA1:NM_001164830:exon5:c.C472A:p.L158M,CA1:NM_001291967:exon5:c.C274A:p.L92M,CA1:NM_001128830:exon7:c.C472A:p.L158M,CA1:NM_001128831:exon6:c.C472A:p.L158M,CA1:NM_001291968:exon4:c.C133A:p.L45M,CA1:NM_001738:exon7:c.C472A:p.L158M,CA1:NM_001128829:exon7:c.C472A:p.L158M, |
| chr8 | 85333535 | A | G | . | exonic | CA1 | nonsynonymous SNV | CA1:NM_001164830:exon4:c.T440C:p.V147A,CA1:NM_001291967:exon4:c.T242C:p.V81A,CA1:NM_001128830:exon6:c.T440C:p.V147A,CA1:NM_001128831:exon5:c.T440C:p.V147A,CA1:NM_001291968:exon3:c.T101C:p.V34A,CA1:NM_001738:exon6:c.T440C:p.V147A,CA1:NM_001128829:exon6:c.T440C:p.V147A, |
| chr8 | 85336964 | C | T | . | exonic | CA1 | nonsynonymous SNV | CA1:NM_001164830:exon3:c.G335A:p.G112E,CA1:NM_001291967:exon3:c.G137A:p.G46E,CA1:NM_001128830:exon5:c.G335A:p.G112E,CA1:NM_001128831:exon4:c.G335A:p.G112E,CA1:NM_001738:exon5:c.G335A:p.G112E,CA1:NM_001128829:exon5:c.G335A:p.G112E, |
| chr8 | 85337019 | A | G | . | exonic | CA1 | nonsynonymous SNV | CA1:NM_001164830:exon3:c.T280C:p.F94L,CA1:NM_001291967:exon3:c.T82C:p.F28L,CA1:NM_001128830:exon5:c.T280C:p.F94L,CA1:NM_001128831:exon4:c.T280C:p.F94L,CA1:NM_001738:exon5:c.T280C:p.F94L,CA1:NM_001128829:exon5:c.T280C:p.F94L, |
| chr8 | 85337036 | C | T | . | exonic | CA1 | nonsynonymous SNV | CA1:NM_001164830:exon3:c.G263A:p.S88N,CA1:NM_001291967:exon3:c.G65A:p.S22N,CA1:NM_001128830:exon5:c.G263A:p.S88N,CA1:NM_001128831:exon4:c.G263A:p.S88N,CA1:NM_001738:exon5:c.G263A:p.S88N,CA1:NM_001128829:exon5:c.G263A:p.S88N, |
| chr8 | 85338277 | A | T | . | exonic | CA1 | nonsynonymous SNV | CA1:NM_001164830:exon2:c.T210A:p.N70K,CA1:NM_001128830:exon4:c.T210A:p.N70K,CA1:NM_001128831:exon3:c.T210A:p.N70K,CA1:NM_001738:exon4:c.T210A:p.N70K,CA1:NM_001128829:exon4:c.T210A:p.N70K, |
| chr8 | 85338318 | C | T | . | exonic | CA1 | nonsynonymous SNV | CA1:NM_001164830:exon2:c.G169A:p.A57T,CA1:NM_001128830:exon4:c.G169A:p.A57T,CA1:NM_001128831:exon3:c.G169A:p.A57T,CA1:NM_001738:exon4:c.G169A:p.A57T,CA1:NM_001128829:exon4:c.G169A:p.A57T, |
| chr8 | 85338375 | C | T | . | exonic | CA1 | nonsynonymous SNV | CA1:NM_001164830:exon2:c.G112A:p.E38K,CA1:NM_001128830:exon4:c.G112A:p.E38K,CA1:NM_001128831:exon3:c.G112A:p.E38K,CA1:NM_001738:exon4:c.G112A:p.E38K,CA1:NM_001128829:exon4:c.G112A:p.E38K, |
| chr8 | 85338380 | G | T | . | exonic | CA1 | nonsynonymous SNV | CA1:NM_001164830:exon2:c.C107A:p.T36N,CA1:NM_001128830:exon4:c.C107A:p.T36N,CA1:NM_001128831:exon3:c.C107A:p.T36N,CA1:NM_001738:exon4:c.C107A:p.T36N,CA1:NM_001128829:exon4:c.C107A:p.T36N, |
| chr8 | 85338411 | C | T | COSV56279142 | exonic | CA1 | nonsynonymous SNV | CA1:NM_001164830:exon2:c.G76A:p.G26R,CA1:NM_001128830:exon4:c.G76A:p.G26R,CA1:NM_001128831:exon3:c.G76A:p.G26R,CA1:NM_001738:exon4:c.G76A:p.G26R,CA1:NM_001128829:exon4:c.G76A:p.G26R, |
| chr8 | 85338417 | C | T | . | exonic | CA1 | nonsynonymous SNV | CA1:NM_001164830:exon2:c.G70A:p.A24T,CA1:NM_001128830:exon4:c.G70A:p.A24T,CA1:NM_001128831:exon3:c.G70A:p.A24T,CA1:NM_001738:exon4:c.G70A:p.A24T,CA1:NM_001128829:exon4:c.G70A:p.A24T, |
| chr8 | 85338423 | G | A | . | exonic | CA1 | nonsynonymous SNV | CA1:NM_001164830:exon2:c.C64T:p.P22S,CA1:NM_001128830:exon4:c.C64T:p.P22S,CA1:NM_001128831:exon3:c.C64T:p.P22S,CA1:NM_001738:exon4:c.C64T:p.P22S,CA1:NM_001128829:exon4:c.C64T:p.P22S, |
| chr8 | 86507086 | T | G | COSV52206582 | exonic | RMDN1 | nonsynonymous SNV | RMDN1:NM_016033:exon2:c.A156C:p.K52N,RMDN1:NM_001286719:exon2:c.A156C:p.K52N,RMDN1:NM_001286707:exon2:c.A156C:p.K52N, |
| chr8 | 89769874 | G | A | . | exonic | RIPK2 | nonsynonymous SNV | RIPK2:NM_003821:exon4:c.G586A:p.E196K, |
| chr8 | 89914658 | A | C | . | exonic | OSGIN2 | nonsynonymous SNV | OSGIN2:NM_001126111:exon4:c.A440C:p.Y147S,OSGIN2:NM_004337:exon4:c.A308C:p.Y103S, |
| chr8 | 89946258 | G | T | . | exonic | NBN | nonsynonymous SNV | NBN:NM_002485:exon13:c.C1952A:p.P651Q,NBN:NM_001024688:exon14:c.C1706A:p.P569Q, |
| chr8 | 89953445 | A | T | . | exonic | NBN | nonsynonymous SNV | NBN:NM_002485:exon11:c.T1644A:p.N548K,NBN:NM_001024688:exon12:c.T1398A:p.N466K, |
| chr8 | 89953453 | T | C | . | exonic | NBN | nonsynonymous SNV | NBN:NM_002485:exon11:c.A1636G:p.R546G,NBN:NM_001024688:exon12:c.A1390G:p.R464G, |
| chr8 | 89953586 | C | T | . | exonic | NBN | stopgain | NBN:NM_002485:exon11:c.G1503A:p.W501X,NBN:NM_001024688:exon12:c.G1257A:p.W419X, |
| chr8 | 89953587 | C | T | . | exonic | NBN | stopgain | NBN:NM_002485:exon11:c.G1502A:p.W501X,NBN:NM_001024688:exon12:c.G1256A:p.W419X, |
| chr8 | 89953641 | A | T | . | exonic | NBN | nonsynonymous SNV | NBN:NM_002485:exon11:c.T1448A:p.I483K,NBN:NM_001024688:exon12:c.T1202A:p.I401K, |
| chr8 | 89955360 | A | T | . | exonic | NBN | nonsynonymous SNV | NBN:NM_002485:exon10:c.T1320A:p.N440K,NBN:NM_001024688:exon11:c.T1074A:p.N358K, |
| chr8 | 89955460 | C | T | . | exonic | NBN | nonsynonymous SNV | NBN:NM_002485:exon10:c.G1220A:p.C407Y,NBN:NM_001024688:exon11:c.G974A:p.C325Y, |
| chr8 | 89958756 | C | T | . | exonic | NBN | nonsynonymous SNV | NBN:NM_002485:exon9:c.G1093A:p.A365T,NBN:NM_001024688:exon10:c.G847A:p.A283T, |
| chr8 | 89970467 | C | T | . | exonic | NBN | nonsynonymous SNV | NBN:NM_002485:exon7:c.G793A:p.A265T,NBN:NM_001024688:exon8:c.G547A:p.A183T, |
| chr8 | 89970473 | A | G | COSV55378087 | exonic | NBN | nonsynonymous SNV | NBN:NM_002485:exon7:c.T787C:p.F263L,NBN:NM_001024688:exon8:c.T541C:p.F181L, |
| chr8 | 89970527 | C | T | . | exonic | NBN | nonsynonymous SNV | NBN:NM_002485:exon7:c.G733A:p.G245R,NBN:NM_001024688:exon8:c.G487A:p.G163R, |
| chr8 | 89971226 | C | T | . | exonic | NBN | nonsynonymous SNV | NBN:NM_001024688:exon7:c.G403A:p.E135K,NBN:NM_002485:exon6:c.G649A:p.E217K, |

| | | | | | | | | |
|---|---|---|---|---|---|---|---|---|
| chr8 | 89971255 | C | T | COSV55371813 | exonic | NBN | nonsynonymous SNV | NBN:NM_001024688:exon7:c.G374A:p.S125N,NBN:NM_002485:exon6:c.G620A:p.S207N, |
| chr8 | 89978251 | C | G | COSV55371261 | exonic | NBN | nonsynonymous SNV | NBN:NM_002485:exon5:c.G553C:p.E185Q,NBN:NM_001024688:exon6:c.G307C:p.E103Q, |
| chr8 | 89978302 | C | T | . | exonic | NBN | nonsynonymous SNV | NBN:NM_001024688:exon6:c.G256A:p.G86R,NBN:NM_002485:exon5:c.G502A:p.G168R, |
| chr8 | 94128313 | C | T | COSV50327363 | exonic | CDH17 | nonsynonymous SNV | CDH17:NM_001144663:exon18:c.G2426A:p.R809H,CDH17:NM_004063:exon18:c.G2426A:p.R809H, |
| chr8 | 94128329 | C | T | . | exonic | CDH17 | nonsynonymous SNV | CDH17:NM_001144663:exon18:c.G2410A:p.A804T,CDH17:NM_004063:exon18:c.G2410A:p.A804T, |
| chr8 | 94130682 | C | T | . | exonic | CDH17 | nonsynonymous SNV | CDH17:NM_001144663:exon17:c.G2342A:p.G781E,CDH17:NM_004063:exon17:c.G2342A:p.G781E, |
| chr8 | 94130926 | C | T | COSV50345009 | exonic | CDH17 | nonsynonymous SNV | CDH17:NM_001144663:exon16:c.G2234A:p.R745H,CDH17:NM_004063:exon16:c.G2234A:p.R745H, |
| chr8 | 94130944 | T | G | COSV50337541 | exonic | CDH17 | nonsynonymous SNV | CDH17:NM_001144663:exon16:c.A2216C:p.E739A,CDH17:NM_004063:exon16:c.A2216C:p.E739A, |
| chr8 | 94130958 | C | G | COSV50336268 | exonic | CDH17 | nonsynonymous SNV | CDH17:NM_001144663:exon16:c.G2202C:p.E734D,CDH17:NM_004063:exon16:c.G2202C:p.E734D, |
| chr8 | 94145972 | C | T | . | exonic | CDH17 | nonsynonymous SNV | CDH17:NM_001144663:exon15:c.G2123A:p.G708D,CDH17:NM_004063:exon15:c.G2123A:p.G708D, |
| chr8 | 94148791 | C | T | . | exonic | CDH17 | nonsynonymous SNV | CDH17:NM_001144663:exon14:c.G1880A:p.R627K,CDH17:NM_004063:exon14:c.G1880A:p.R627K, |
| chr8 | 94151940 | A | G | . | exonic | CDH17 | nonsynonymous SNV | CDH17:NM_001144663:exon13:c.T1724C:p.V575A,CDH17:NM_004063:exon13:c.T1724C:p.V575A, |
| chr8 | 94151980 | C | T | . | exonic | CDH17 | nonsynonymous SNV | CDH17:NM_001144663:exon13:c.G1684A:p.E562K,CDH17:NM_004063:exon13:c.G1684A:p.E562K, |
| chr8 | 94165842 | C | T | . | exonic | CDH17 | nonsynonymous SNV | CDH17:NM_001144663:exon10:c.G1201A:p.G401R,CDH17:NM_004063:exon10:c.G1201A:p.G401R, |
| chr8 | 94170444 | C | T | . | exonic | CDH17 | nonsynonymous SNV | CDH17:NM_001144663:exon9:c.G1019A:p.C340Y,CDH17:NM_004063:exon9:c.G1019A:p.C340Y, |
| chr8 | 94176622 | T | C | COSV50326476 | exonic | CDH17 | nonsynonymous SNV | CDH17:NM_001144663:exon5:c.A343G:p.K115E,CDH17:NM_004063:exon5:c.A343G:p.K115E, |
| chr8 | 94189252 | C | T | . | exonic | CDH17 | nonsynonymous SNV | CDH17:NM_001144663:exon3:c.G85A:p.G29R,CDH17:NM_004063:exon3:c.G85A:p.G29R, |
| chr8 | 94467452 | G | C | COSV52594351 | exonic | RAD54B | nonsynonymous SNV | RAD54B:NM_012415:exon2:c.C88G:p.L30V,RAD54B:NM_001205262:exon2:c.C88G:p.L30V, |
| chr8 | 94527128 | A | T | . | exonic | KIAA1429 | nonsynonymous SNV | KIAA1429:NM_015496:exon8:c.T1116A:p.D372E,KIAA1429:NM_183009:exon8:c.T1116A:p.D372E, |
| chr8 | 94529205 | C | T | . | exonic | KIAA1429 | nonsynonymous SNV | KIAA1429:NM_015496:exon7:c.G745A:p.E249K,KIAA1429:NM_183009:exon7:c.G745A:p.E249K, |
| chr8 | 94529220 | G | T | . | exonic | KIAA1429 | nonsynonymous SNV | KIAA1429:NM_015496:exon7:c.C730A:p.Q244K,KIAA1429:NM_183009:exon7:c.C730A:p.Q244K, |
| chr8 | 94529253 | G | T | . | exonic | KIAA1429 | nonsynonymous SNV | KIAA1429:NM_015496:exon7:c.C697A:p.Q233K,KIAA1429:NM_183009:exon7:c.C697A:p.Q233K, |
| chr8 | 96231068 | C | T | . | exonic | UQCRB | stopgain | UQCRB:NM_001199975:exon5:c.G227A:p.W76X,UQCRB:NM_006294:exon4:c.G323A:p.W108X, |
| chr8 | 96231075 | C | T | . | exonic | UQCRB | nonsynonymous SNV | UQCRB:NM_001199975:exon5:c.G220A:p.E74K,UQCRB:NM_006294:exon4:c.G316A:p.E106K, |
| chr8 | 96231113 | G | A | COSV54629990 | exonic | UQCRB | nonsynonymous SNV | UQCRB:NM_001254752:exon5:c.C415T:p.R139C,UQCRB:NM_001199975:exon5:c.C182T:p.P61L,UQCRB:NM_006294:exon4:c.C278T:p.P93L, |
| chr8 | 96231860 | C | T | . | exonic | UQCRB | nonsynonymous SNV | UQCRB:NM_001254752:exon3:c.G172A:p.D58N,UQCRB:NM_001199975:exon4:c.G76A:p.D26N,UQCRB:NM_006294:exon3:c.G172A:p.D58N, |
| chr8 | 96233194 | C | T | . | exonic | UQCRB | nonsynonymous SNV | UQCRB:NM_001254752:exon2:c.G53A:p.R18Q,UQCRB:NM_006294:exon2:c.G53A:p.R18Q, |
| chr8 | 96233201 | C | T | . | exonic | UQCRB | nonsynonymous SNV | UQCRB:NM_001254752:exon2:c.G46A:p.G16S,UQCRB:NM_006294:exon2:c.G46A:p.G16S, |
| chr8 | 96235512 | C | T | . | exonic | UQCRB | nonsynonymous SNV | UQCRB:NM_001254752:exon1:c.G19A:p.V7I,UQCRB:NM_006294:exon1:c.G19A:p.V7I, |
| chr8 | 96235527 | C | T | COSV54629256 | exonic | UQCRB | nonsynonymous SNV | UQCRB:NM_001254752:exon1:c.G4A:p.A2T,UQCRB:NM_006294:exon1:c.G4A:p.A2T, |
| chr8 | 97776065 | G | A | . | exonic | LAPTM4B | nonsynonymous SNV | LAPTM4B:NM_018407:exon1:c.G329A:p.C110Y, |
| chr8 | 97805395 | G | A | . | exonic | LAPTM4B | nonsynonymous SNV | LAPTM4B:NM_018407:exon2:c.G415A:p.A139T, |
| chr8 | 97851439 | G | A | . | exonic | LAPTM4B | nonsynonymous SNV | LAPTM4B:NM_018407:exon7:c.G919A:p.A307T, |
| chr8 | 98042660 | C | T | . | exonic | RPL30 | nonsynonymous SNV | RPL30:NM_000989:exon4:c.G283A:p.A95T, |
| chr8 | 98044965 | C | T | . | exonic | RPL30 | nonsynonymous SNV | RPL30:NM_000989:exon3:c.G145A:p.A49T, |
| chr9 | 100006522 | C | T | . | exonic | ERP44 | nonsynonymous SNV | ERP44:NM_015051:exon10:c.G1000A:p.D334N, |
| chr9 | 100007623 | C | T | . | exonic | ERP44 | nonsynonymous SNV | ERP44:NM_015051:exon9:c.G829A:p.E277K, |
| chr9 | 100020637 | C | T | . | exonic | ERP44 | nonsynonymous SNV | ERP44:NM_015051:exon6:c.G566A:p.C189Y, |
| chr9 | 100052486 | C | T | . | exonic | ERP44 | nonsynonymous SNV | ERP44:NM_015051:exon4:c.G217A:p.D73N, |
| chr9 | 104752933 | G | A | COSV66113413 | exonic | NIPSNAP3A | nonsynonymous SNV | NIPSNAP3A:NM_015469:exon3:c.G299A:p.R100Q, |
| chr9 | 111597344 | C | A | COSV53028155 | exonic | PTGR1 | nonsynonymous SNV | PTGR1:NM_012212:exon2:c.G79T:p.A27S,PTGR1:NM_001146108:exon2:c.G79T:p.A27S,PTGR1:NM_001146109:exon2:c.G79T:p.A27S, |
| chr9 | 112830816 | T | A | . | exonic | SNX30 | nonsynonymous SNV | SNX30:NM_001012994:exon4:c.T551A:p.L184Q, |
| chr9 | 113188426 | A | C | COSV53035634 | exonic | FKBP15 | nonsynonymous SNV | FKBP15:NM_015258:exon13:c.T1239G:p.H413Q, |
| chr9 | 113362390 | G | A | COSV65242448 | exonic | BSPRY | nonsynonymous SNV | BSPRY:NM_017688:exon4:c.G553A:p.E185K, |
| chr9 | 113370054 | C | T | COSV53057170 | exonic | BSPRY | nonsynonymous SNV | BSPRY:NM_017688:exon6:c.C1121T:p.T374I, |
| chr9 | 113373918 | C | T | COSV53056665 | exonic | HDHD3 | nonsynonymous SNV | HDHD3:NM_001304509:exon3:c.G437A:p.G146E,HDHD3:NM_001304510:exon3:c.G437A:p.G146E,HDHD3:NM_001304511:exon2:c.G437A:p.G146E,HDHD3:NM_031219:exon2:c.G437A:p.G146E, |
| chr9 | 114323246 | G | A | COSV52277829 | exonic | ORM1 | nonsynonymous SNV | ORM1:NM_000607:exon1:c.G113A:p.R38Q, |
| chr9 | 115030304 | C | G | COSV60785451 | exonic | TNC | nonsynonymous SNV | TNC:NM_002160:exon24:c.G6022C:p.E2008Q, |
| chr9 | 121326630 | C | G | . | exonic | GSN | nonsynonymous SNV | GSN:NM_000177:exon12:c.C1688G:p.T563S,GSN:NM_001127665:exon14:c.C1535G:p.T512S,GSN:NM_001127667:exon14:c.C1568G:p.T523S,GSN:NM_001127664:exon14:c.C1535G:p.T512S,GSN:NM_001127666:exon14:c.C1568G:p.T523S,GSN:NM_001127663:exon14:c.C1643G:p.T548S,GSN:NM_198252:exon13:c.C1535G:p.T512S,GSN:NM_001258030:exon12:c.C1559G:p.T520S,GSN:NM_001127662:exon12:c.C1535G:p.T512S,GSN:NM_001258029:exon12:c.C1586G:p.T529S, |
| chr9 | 121327414 | C | T | . | exonic | GSN | nonsynonymous SNV | GSN:NM_000177:exon13:c.C1847T:p.T616M,GSN:NM_001127665:exon15:c.C1694T:p.T565M,GSN:NM_001127667:exon15:c.C1727T:p.T576M,GSN:NM_001127664:exon15:c.C1694T:p.T565M,GSN:NM_001127666:exon15:c.C1727T:p.T576M,GSN:NM_001127663:exon15:c.C1802T:p.T601M,GSN:NM_198252:exon14:c.C1694T:p.T565M,GSN:NM_001258030:exon13:c.C1718T:p.T573M,GSN:NM_001127662:exon13:c.C1694T:p.T565M,GSN:NM_001258029:exon13:c.C1745T:p.T582M, |
| chr9 | 121341245 | A | G | . | exonic | STOM | nonsynonymous SNV | STOM:NM_004099:exon7:c.T824C:p.M275T,STOM:NM_198194:exon3:c.T329C:p.M110T, |
| chr9 | 124414882 | A | G | COSV52330411 | exonic | PSMB7 | nonsynonymous SNV | PSMB7:NM_002799:exon2:c.T116C:p.V39A, |
| chr9 | 12775890 | A | G | COSV59983395 | exonic | LURAP1L | nonsynonymous SNV | LURAP1L:NM_203403:exon1:c.A175G:p.S59G, |
| chr9 | 127935750 | G | C | COSV58709021 | exonic | DPM2 | nonsynonymous SNV | DPM2:NM_003863:exon4:c.C227G:p.T76S, |
| chr9 | 128149550 | G | A | . | exonic | LCN2 | nonsynonymous SNV | LCN2:NM_005564:exon1:c.G25A:p.G9S, |
| chr9 | 128149589 | G | A | . | exonic | LCN2 | nonsynonymous SNV | LCN2:NM_005564:exon1:c.G64A:p.D22N, |
| chr9 | 128323094 | G | C | . | exonic | COQ4 | nonsynonymous SNV | COQ4:NM_001305942:exon2:c.G149C:p.G50A,COQ4:NM_016035:exon2:c.G149C:p.G50A, |
| chr9 | 128721470 | T | A | COSV52579129 | exonic | ZDHHC12 | nonsynonymous SNV | ZDHHC12:NM_032799:exon5:c.A515T:p.Q172L, |
| chr9 | 130634691 | G | A | . | exonic | FUBP3 | nonsynonymous SNV | FUBP3:NM_003934:exon17:c.G1535A:p.S512N, |
| chr9 | 130695494 | G | A | . | exonic | EXOSC2 | nonsynonymous SNV | EXOSC2:NM_014285:exon2:c.G125A:p.G42D,EXOSC2:NM_001282709:exon2:c.G125A:p.G42D,EXOSC2:NM_001282708:exon2:c.G125A:p.G42D, |
| chr9 | 132595027 | T | C | COSV55036973 | exonic | DDX31 | nonsynonymous SNV | DDX31:NM_022779:exon20:c.A2395G:p.I799V, |
| chr9 | 132662629 | C | T | COSV60137948 | exonic | DDX31 | nonsynonymous SNV | DDX31:NM_022779:exon2:c.G457A:p.E153K,DDX31:NM_138620:exon2:c.G457A:p.E153K, |
| chr9 | 132906741 | A | T | . | exonic | TSC1 | nonsynonymous SNV | TSC1:NM_001162426:exon14:c.T1425A:p.D475E,TSC1:NM_000368:exon14:c.T1428A:p.D476E,TSC1:NM_001162427:exon13:c.T1275A:p.D425E, |
| chr9 | 132911517 | A | G | COSV53763263 | exonic | TSC1 | nonsynonymous SNV | TSC1:NM_001162426:exon10:c.T965C:p.M322T,TSC1:NM_000368:exon10:c.T965C:p.M322T,TSC1:NM_001162427:exon9:c.T812C:p.M271T, |
| chr9 | 133071462 | T | G | . | exonic | CEL | nonsynonymous SNV | CEL:NM_001807:exon11:c.T1969G:p.S657A, |
| chr9 | 133071468 | G | C | . | exonic | CEL | nonsynonymous SNV | CEL:NM_001807:exon11:c.G1975C:p.A659P, |
| chr9 | 133071561 | T | G | . | exonic | CEL | nonsynonymous SNV | CEL:NM_001807:exon11:c.T2068G:p.S690A, |

| | | | | | | | | |
|---|---|---|---|---|---|---|---|---|
| chr9 | 133071567 | G | C | COSV60827582 | exonic | CEL | nonsynonymous SNV | CEL:NM_001807:exon11:c.G2074C:p.A692P, |
| chr9 | 133360384 | G | A | . | exonic | SURF2 | nonsynonymous SNV | SURF2:NM_017503:exon5:c.G637A:p.G213S,SURF2:NM_001278928:exon5:c.G637A:p.G213S, |
| chr9 | 133361131 | C | T | COSV99058110 | exonic | SURF2 | nonsynonymous SNV | SURF2:NM_017503:exon6:c.C763T:p.P255S,SURF2:NM_001278928:exon6:c.C760T:p.P254S, |
| chr9 | 133778872 | T | C | COSV64080812 | exonic | VAV2 | nonsynonymous SNV | VAV2:NM_003371:exon20:c.A1750G:p.M584V,VAV2:NM_001134398:exon22:c.A1780G:p.M594V, |
| chr9 | 134040214 | G | T | . | exonic | BRD3 | nonsynonymous SNV | BRD3:NM_007371:exon9:c.C1463A:p.P488Q, |
| chr9 | 134041792 | G | A | . | exonic | BRD3 | nonsynonymous SNV | BRD3:NM_007371:exon8:c.C1375T:p.R459W, |
| chr9 | 136837879 | G | C | . | exonic | RABL6 | nonsynonymous SNV | RABL6:NM_024718:exon10:c.G1144C:p.E382Q,RABL6:NM_001173988:exon10:c.G1147C:p.E383Q, |
| chr9 | 137114298 | G | C | COSV65370924 | exonic | DPP7 | nonsynonymous SNV | DPP7:NM_013379:exon3:c.C266G:p.A89G, |
| chr9 | 15474113 | C | T | . | exonic | PSIP1 | nonsynonymous SNV | PSIP1:NM_001128217:exon9:c.G754A:p.E252K,PSIP1:NM_021144:exon8:c.G754A:p.E252K,PSIP1:NM_033222:exon9:c.G754A:p.E252K, |
| chr9 | 15474188 | G | T | . | exonic | PSIP1 | nonsynonymous SNV | PSIP1:NM_001128217:exon9:c.C679A:p.P227T,PSIP1:NM_021144:exon8:c.C679A:p.P227T,PSIP1:NM_033222:exon9:c.C679A:p.P227T, |
| chr9 | 15474200 | C | T | . | exonic | PSIP1 | nonsynonymous SNV | PSIP1:NM_001128217:exon9:c.G667A:p.E223K,PSIP1:NM_021144:exon8:c.G667A:p.E223K,PSIP1:NM_033222:exon9:c.G667A:p.E223K, |
| chr9 | 15478498 | G | T | . | exonic | PSIP1 | nonsynonymous SNV | PSIP1:NM_001128217:exon8:c.C608A:p.P203H,PSIP1:NM_021144:exon7:c.C608A:p.P203H,PSIP1:NM_033222:exon8:c.C608A:p.P203H, |
| chr9 | 19376615 | C | T | . | exonic | RPS6 | nonsynonymous SNV | RPS6:NM_001010:exon5:c.G533A:p.R178H, |
| chr9 | 19376631 | C | T | . | exonic | RPS6 | nonsynonymous SNV | RPS6:NM_001010:exon5:c.G517A:p.A173T, |
| chr9 | 19378379 | A | T | . | exonic | RPS6 | stopgain | RPS6:NM_001010:exon4:c.T485A:p.L162X, |
| chr9 | 19378403 | C | T | . | exonic | RPS6 | nonsynonymous SNV | RPS6:NM_001010:exon4:c.G461A:p.R154H, |
| chr9 | 19378764 | C | T | . | exonic | RPS6 | nonsynonymous SNV | RPS6:NM_001010:exon3:c.G293A:p.R98H, |
| chr9 | 19378803 | C | T | . | exonic | RPS6 | nonsynonymous SNV | RPS6:NM_001010:exon3:c.G254A:p.R85K, |
| chr9 | 19378899 | C | T | . | exonic | RPS6 | nonsynonymous SNV | RPS6:NM_001010:exon3:c.G158A:p.S53N, |
| chr9 | 19379522 | C | T | . | exonic | RPS6 | nonsynonymous SNV | RPS6:NM_001010:exon2:c.G103A:p.E35K, |
| chr9 | 19379567 | C | T | . | exonic | RPS6 | nonsynonymous SNV | RPS6:NM_001010:exon2:c.G58A:p.D20N, |
| chr9 | 20988427 | A | G | . | exonic | FOCAD | nonsynonymous SNV | FOCAD:NM_017794:exon43:c.A5002G:p.K1668E, |
| chr9 | 21815499 | G | A | . | exonic | MTAP | nonsynonymous SNV | MTAP:NM_002451:exon2:c.G100A:p.V34M, |
| chr9 | 21816759 | G | A | COSV66477132 | exonic | MTAP | nonsynonymous SNV | MTAP:NM_002451:exon3:c.G166A:p.V56I, |
| chr9 | 21854631 | G | A | . | exonic | MTAP | nonsynonymous SNV | MTAP:NM_002451:exon6:c.G451A:p.V151I, |
| chr9 | 21854668 | G | A | . | exonic | MTAP | nonsynonymous SNV | MTAP:NM_002451:exon6:c.G488A:p.C163Y, |
| chr9 | 286491 | G | A | COSV66634213 | exonic | DOCK8 | nonsynonymous SNV | DOCK8:NM_203447:exon3:c.G187A:p.D63N, |
| chr9 | 286593 | C | A | COSV66633951 | exonic | DOCK8 | nonsynonymous SNV | DOCK8:NM_203447:exon3:c.C289A:p.P97T,DOCK8:NM_001190458:exon2:c.C85A:p.P29T,DOCK8:NM_001193536:exon2:c.C85A:p.P29T, |
| chr9 | 33264542 | C | G | COSV56303169 | exonic | BAG1 | nonsynonymous SNV | BAG1:NM_004323:exon1:c.G133C:p.G45R, |
| chr9 | 33798019 | A | G | COSV61553087 | exonic | PRSS3 | nonsynonymous SNV | PRSS3:NM_001197098:exon3:c.A370G:p.T124A,PRSS3:NM_007343:exon3:c.A562G:p.T188A,PRSS3:NM_001197097:exon4:c.A433G:p.T145A,PRSS3:NM_002771:exon3:c.A391G:p.T131A, |
| chr9 | 35660993 | A | G | COSV58397237 | exonic | CCDC107 | nonsynonymous SNV | CCDC107:NM_001195200:exon6:c.A577G:p.I193V,CCDC107:NM_174923:exon5:c.A658G:p.I220V, |
| chr9 | 35674056 | G | A | COSV65675780 | exonic | CA9 | nonsynonymous SNV | CA9:NM_001216:exon1:c.G97A:p.V33M, |
| chr9 | 37432117 | C | T | . | exonic | GRHPR | nonsynonymous SNV | GRHPR:NM_012203:exon8:c.C844T:p.L282F, |
| chr9 | 4860259 | G | A | . | exonic | RCL1 | nonsynonymous SNV | RCL1:NM_005772:exon9:c.G1106A:p.S369N,RCL1:NM_001286699:exon7:c.G632A:p.S211N,RCL1:NM_001286700:exon8:c.G632A:p.S211N,RCL1:NM_001286701:exon5:c.G548A:p.S183N, |
| chr9 | 65682821 | G | T | COSV66214404 | exonic | CBWD5 | nonsynonymous SNV | CBWD5:NM_001024916:exon4:c.G394T:p.D132Y, |
| chr9 | 65690975 | A | G | . | exonic | CBWD5 | nonsynonymous SNV | CBWD5:NM_001024916:exon5:c.A439G:p.T147A, |
| chr9 | 69228107 | C | A | . | exonic | TJP2 | nonsynonymous SNV | TJP2:NM_001170630:exon9:c.C1446A:p.D482E,TJP2:NM_001170416:exon9:c.C1539A:p.D513E,TJP2:NM_001170415:exon9:c.C1458A:p.D486E,TJP2:NM_201629:exon9:c.C1446A:p.D482E,TJP2:NM_004817:exon9:c.C1446A:p.D482E,TJP2:NM_001170414:exon10:c.C1377A:p.D459E, |
| chr9 | 710992 | C | T | . | exonic | KANK1 | nonsynonymous SNV | KANK1:NM_001256876:exon7:c.C226T:p.P76S,KANK1:NM_001256877:exon4:c.C226T:p.P76S,KANK1:NM_015158:exon3:c.C226T:p.P76S, |
| chr9 | 712060 | G | C | COSV63211042 | exonic | KANK1 | nonsynonymous SNV | KANK1:NM_001256876:exon7:c.G1294C:p.E432Q,KANK1:NM_001256877:exon4:c.G1294C:p.E432Q,KANK1:NM_015158:exon3:c.G1294C:p.E432Q,KANK1:NM_153186:exon2:c.G820C:p.E274Q, |
| chr9 | 7799653 | G | T | COSV64065665 | exonic | TMEM261 | nonsynonymous SNV | TMEM261:NM_033428:exon1:c.C82A:p.P28T, |
| chr9 | 92032377 | C | T | . | exonic | SPTLC1 | nonsynonymous SNV | SPTLC1:NM_001281303:exon15:c.G1478A:p.S493N, |
| chr9 | 92032395 | C | T | . | exonic | SPTLC1 | stopgain | SPTLC1:NM_001281303:exon15:c.G1460A:p.W487X, |
| chr9 | 92038307 | C | T | . | exonic | SPTLC1 | nonsynonymous SNV | SPTLC1:NM_001281303:exon13:c.G1195A:p.E399K,SPTLC1:NM_006415:exon13:c.G1195A:p.E399K, |
| chr9 | 92046024 | C | T | . | exonic | SPTLC1 | nonsynonymous SNV | SPTLC1:NM_001281303:exon12:c.G1111A:p.G371R,SPTLC1:NM_006415:exon12:c.G1111A:p.G371R, |
| chr9 | 92046045 | C | T | . | exonic | SPTLC1 | nonsynonymous SNV | SPTLC1:NM_001281303:exon12:c.G1090A:p.A364T,SPTLC1:NM_006415:exon12:c.G1090A:p.A364T, |
| chr9 | 92047238 | C | T | . | exonic | SPTLC1 | nonsynonymous SNV | SPTLC1:NM_001281303:exon11:c.G1015A:p.A339T,SPTLC1:NM_006415:exon11:c.G1015A:p.A339T, |
| chr9 | 92047244 | A | G | . | exonic | SPTLC1 | nonsynonymous SNV | SPTLC1:NM_001281303:exon11:c.T1009C:p.F337L,SPTLC1:NM_006415:exon11:c.T1009C:p.F337L, |
| chr9 | 92047258 | C | T | . | exonic | SPTLC1 | nonsynonymous SNV | SPTLC1:NM_001281303:exon11:c.G995A:p.G332D,SPTLC1:NM_006415:exon11:c.G995A:p.G332D, |
| chr9 | 92047627 | C | T | . | exonic | SPTLC1 | nonsynonymous SNV | SPTLC1:NM_001281303:exon10:c.G970A:p.V324I,SPTLC1:NM_006415:exon10:c.G970A:p.V324I, |
| chr9 | 92047653 | C | T | . | exonic | SPTLC1 | nonsynonymous SNV | SPTLC1:NM_001281303:exon10:c.G944A:p.G315E,SPTLC1:NM_006415:exon10:c.G944A:p.G315E, |
| chr9 | 92050010 | C | T | . | exonic | SPTLC1 | nonsynonymous SNV | SPTLC1:NM_001281303:exon9:c.G838A:p.G280R,SPTLC1:NM_006415:exon9:c.G838A:p.G280R, |
| chr9 | 92055430 | C | T | . | exonic | SPTLC1 | nonsynonymous SNV | SPTLC1:NM_001281303:exon8:c.G755A:p.G252E,SPTLC1:NM_006415:exon8:c.G755A:p.G252E, |
| chr9 | 92067982 | C | T | . | exonic | SPTLC1 | nonsynonymous SNV | SPTLC1:NM_001281303:exon6:c.G544A:p.G182R,SPTLC1:NM_006415:exon6:c.G544A:p.G182R, |
| chr9 | 92080051 | C | T | . | exonic | SPTLC1 | nonsynonymous SNV | SPTLC1:NM_001281303:exon5:c.G392A:p.G131E,SPTLC1:NM_006415:exon5:c.G392A:p.G131E,SPTLC1:NM_178324:exon5:c.G392A:p.G131E, |
| chr9 | 92108741 | C | T | . | exonic | SPTLC1 | nonsynonymous SNV | SPTLC1:NM_001281303:exon3:c.G259A:p.G87S,SPTLC1:NM_006415:exon3:c.G259A:p.G87S,SPTLC1:NM_178324:exon3:c.G259A:p.G87S, |
| chr9 | 92649479 | G | A | . | exonic | IPPK | nonsynonymous SNV | IPPK:NM_022755:exon5:c.C388T:p.R130W, |
| chr9 | 93077974 | A | G | COSV64937050 | exonic | SUSD3 | nonsynonymous SNV | SUSD3:NM_001287007:exon2:c.A217G:p.K73E,SUSD3:NM_001287008:exon2:c.A217G:p.K73E,SUSD3:NM_001287005:exon4:c.A367G:p.K123E,SUSD3:NM_001287006:exon3:c.A406G:p.K136E,SUSD3:NM_145006:exon3:c.A406G:p.K136E, |
| chr9 | 93113099 | G | C | . | exonic | CARD19 | nonsynonymous SNV | CARD19:NM_032310:exon6:c.G544C:p.G182R, |
| chr9 | 95005558 | G | A | . | exonic | C9ORF3 | nonsynonymous SNV | C9orf3:NM_032823:exon11:c.G1760A:p.G587E,C9orf3:NM_001193329:exon12:c.G2057A:p.G686E, |
| chr9 | 95150025 | T | A | . | exonic | FANCC | nonsynonymous SNV | FANCC:NM_001243743:exon7:c.A584T:p.D195V,FANCC:NM_000136:exon7:c.A584T:p.D195V,FANCC:NM_001243744:exon7:c.A584T:p.D195V, |
| chr9 | 95449256 | C | T | . | exonic | PTCH1 | nonsynonymous SNV | PTCH1:NM_001083605:exon22:c.G3164A:p.R1055H,PTCH1:NM_001083602:exon22:c.G3419A:p.R1140H,PTCH1:NM_001083604:exon22:c.G3164A:p.R1055H,PTCH1:NM_001083606:exon22:c.G3164A:p.R1055H,PTCH1:NM_000264:exon22:c.G3617A:p.R1206H,PTCH1:NM_001083603:exon22:c.G3614A:p.R1205H,PTCH1:NM_001083607:exon22:c.G3164A:p.R1055H, |
| chr9 | 96465465 | G | A | . | exonic | HABP4 | nonsynonymous SNV | HABP4:NM_014282:exon3:c.G641A:p.R214Q, |
| chr9 | 98060853 | G | C | COSV52962972 | exonic | NANS | nonsynonymous SNV | NANS:NM_018946:exon2:c.G204C:p.E68D, |
| chr9 | 99928791 | G | A | . | exonic | STX17 | nonsynonymous SNV | STX17:NM_017919:exon3:c.G137A:p.R46K, |
| chr9 | 99951114 | G | A | . | exonic | STX17 | nonsynonymous SNV | STX17:NM_017919:exon4:c.G244A:p.D82N, |
| chr9 | 99968585 | T | A | . | exonic | STX17 | stopgain | STX17:NM_017919:exon8:c.T821A:p.L274X, |
| chr9 | 99985025 | C | T | . | exonic | ERP44 | nonsynonymous SNV | ERP44:NM_015051:exon11:c.G1061A:p.G354E, |

| Isoform | 1M__WT | 1M__MUT | 1N__WT | 1N__MUT | 1T__WT | 1T__MUT | 2L__WT | 2L__MUT | 2T__WT | 2T__MUT |
|---|---|---|---|---|---|---|---|---|---|---|
| NM_005101 | 0 | 2 | 3 | 2 | 3 | 4 | 0 | 0 | 0 | 0 |
| NM_020775:XM_017001883 | 0:00 | 0:00 | 0:00 | 0:00 | 0:00 | 0:00 | 0:00 | 0:00 | 0:00 | 1:01 |
| NM_002790:NM_001199773:NM_001199774:NM_001199772 | 276:5:21:0 | 0:0:0:0 | 273:11:25:0 | 0:0:0:0 | 239:2:11:1 | 0:0:0:0 | 2056:64:115:6 | 2:0:0:0 | 574:16:33:2 | 0:0:0:0 |
| NM_133181:XM_011542134:NM_024526:XR_001737407 | 0:0:0:0 | 0:0:0:0 | 0:0:0:0 | 0:0:0:0 | 1:0:0:0 | 0:0:0:0 | 0:0:0:0 | 1:1:0:0 | 0:0:0:0 | 1:0:1:0 |
| XM_005263475:NM_001001998:XR_002956819:XR_002956821:XM_017001497:NM_002685:XR_002956822 | 5:30:0:0:0:0:0 | 0:0:0:0:0:0:0 | 0:12:0:1:6:0:0 | 0:0:1:0:0:0:0 | 5:52:4:0:4:3:0 | 0:0:0:0:0:0:0 | 7:98:1:0:31:0:1 | 0:0:2:0:0:0:0 | 8:77:3:0:3:2:0 | 0:0:0:0:0:0:0 |
| XM_005263475:NM_001001998:XR_002956822:XR_002956819:XM_017001497:NM_002685:XR_002956820 | 5:30:1:0:0:0:0 | 0:0:0:0:0:0:0 | 0:11:0:15:0:0:0 | 0:0:0:0:0:0:0 | 5:49:1:4:1:3:0 | 0:0:0:0:1:0:0 | 4:91:1:68:5:2:0 | 0:2:1:0:0:0:0 | 8:75:0:7:1:2:0 | 0:0:0:0:0:0:0 |
| NM_003051:NM_001166496 | 0:00 | 9:00 | 0:00 | 18:00 | 0:00 | 20:00 | 53:00:00 | 73:00:00 | 9:00 | 10:01 |
| NM_003051:NM_001166496 | 9:00 | 1:00 | 16:00 | 4:00 | 20:00 | 0:00 | 120:00:00 | 26:01:00 | 18:01 | 1:00 |
| NM_003051:NM_001166496 | 9:00 | 0:00 | 14:00 | 0:00 | 19:00 | 0:00 | 116:01:00 | 2:00 | 18:01 | 0:00 |
| NM_001010922 | 0 | 0 | 0 | 0 | 0 | 1 | 0 | 0 | 0 | 0 |
| NM_015906:XM_005270936:XM_011541568:XM_024447584:NM_033020:XM_005270937 | 0:0:0:0:0:0 | 3:1:1:1:1:0 | 0:0:0:0:0:0 | 5:1:3:1:12:1 | 0:0:0:0:0:0 | 7:0:0:3:3:3 | 0:0:0:0:0:0 | 26:5:8:14:35:13 | 0:0:0:0:0:0 | 2:1:0:1:2:2 |
| NM_015906:XM_011541568:NM_033020:XM_017001454:XM_005270936:XM_005270937:XM_024447584 | 3:1:1:2:0:0:0 | 0:0:0:0:0:0:0 | 0:1:6:2:1:0:0 | 0:0:0:1:0:0:0 | 6:0:3:1:0:1:0 | 0:0:0:3:0:0:0 | 11:1:15:1:3:1:0 | 0:0:0:1:0:0:0 | 1:0:0:1:1:0:0 | 0:0:0:0:0:0:0 |
| NM_015906:XM_017001454:XM_011541568:NM_033020:XM_005270936:XM_005270937:XM_024447584 | 1:0:1:1:0:0:0 | 1:1:0:0:0:0:0 | 1:5:0:2:0:0:0 | 0:0:0:0:0:0:0 | 1:2:0:1:0:0:0 | 3:2:0:2:0:0:0 | 2:0:1:1:0:0:0 | 4:2:0:1:0:0:0 | 1:1:0:0:0:0:0 | 0:0:0:0:1:0:0 |
| NM_007158:NM_001242893:NM_001007553:NM_001242892 | 324:3:32:3 | 0:0:0:0 | 350:1:3:6 | 2:0:0:0 | 241:1:26:6 | 1:0:0:0 | 3526:1:64:46 | 3:1:0:0 | 419:0:35:7 | 1:0:0:0 |
| NM_007158:NM_001242893:NM_001007553:NM_001242892 | 324:3:32:3 | 0:0:0:0 | 351:1:3:6 | 1:0:0:0 | 242:1:26:6 | 0:0:0:0 | 3544:2:64:46 | 9:0:0:0 | 418:0:35:7 | 0:0:0:0 |
| NM_007158:NM_001242893:NM_001007553:NM_001242892 | 324:3:32:3 | 0:0:0:0 | 350:1:3:6 | 1:0:0:0 | 241:1:26:6 | 1:0:0:0 | 3525:2:65:46 | 11:0:0:0 | 419:0:35:7 | 0:0:0:0 |
| NM_007158:NM_001242893:NM_001007553:NM_001242892 | 319:3:32:3 | 1:0:0:0 | 332:1:3:6 | 2:0:0:0 | 244:1:26:6 | 0:0:0:0 | 3327:2:67:46 | 6:0:0:0 | 414:0:35:7 | 1:0:0:0 |
| NM_007158:NM_001242893:NM_001007553:NM_001242892 | 320:3:32:3 | 1:0:0:0 | 325:1:3:6 | 1:0:0:0 | 242:1:26:6 | 0:0:0:0 | 3286:2:67:46 | 5:0:0:0 | 416:0:35:7 | 0:0:0:0 |
| NM_007158:NM_001242893:NM_001007553:NM_001242892 | 321:3:32:3 | 0:0:0:0 | 313:1:3:6 | 0:0:0:0 | 241:1:26:6 | 0:0:0:0 | 2959:2:71:47 | 8:0:0:0 | 411:0:35:7 | 0:0:0:0 |
| NM_007158:NM_001007553:NM_001242893:NM_001242892 | 318:30:3:3 | 1:1:0:0 | 222:3:1:6 | 1:0:0:0 | 237:25:1:6 | 0:0:0:0 | 2067:72:2:47 | 5:0:0:0 | 396:35:0:7 | 0:0:0:0 |
| NM_007158:NM_001242893:NM_001007553:NM_001242892 | 307:3:31:3 | 0:0:0:0 | 183:1:3:5 | 1:0:0:1 | 232:1:25:6 | 1:0:0:0 | 1688:2:72:48 | 10:0:0:0 | 379:0:35:7 | 0:0:0:0 |
| NM_007158:NM_001242893:NM_001007553:NM_001242892 | 307:3:32:3 | 0:0:0:0 | 176:1:3:6 | 0:0:0:0 | 233:1:26:6 | 0:0:0:0 | 1590:2:72:48 | 4:0:0:0 | 371:0:35:7 | 3:0:0:0 |
| NM_007158:NM_001242893:NM_001007553:NM_001242892 | 304:3:32:3 | 0:0:0:0 | 163:1:3:6 | 0:0:0:0 | 228:1:26:6 | 0:0:0:0 | 1477:2:72:48 | 4:0:0:0 | 369:0:35:7 | 0:0:0:0 |
| NM_007158:NM_001242893:NM_001007553:NM_001242892 | 281:3:32:3 | 1:0:0:0 | 78:1:4:6 | 0:0:0:0 | 211:1:26:6 | 0:0:0:0 | 812:3:76:48 | 8:0:0:0 | 310:0:35:7 | 0:0:0:0 |
| NM_001160234:XM_017001361:XR_002956654:NM_000701 | 41:5:35:117 | 2:0:0:0 | 49:26:69:0 | 0:0:0:0 | 77:13:79:118 | 0:0:0:2 | 88:51:102:44 | 0:1:0:0 | 44:8:21:143 | 0:0:0:0 |
| NM_001160233:XM_017001361:XR_002956654:NM_000701:NM_001160234 | 2:9:31:124:54 | 0:0:0:0:0 | 0:19:29:0:33 | 0:0:0:0:0 | 0:17:76:128:76 | 0:0:0:0:0 | 1:30:90:47:73 | 0:1:0:0:0 | 1:13:29:152:76 | 0:0:0:0:0 |
| NM_001160233:XM_017001361:XR_002956654:NM_000701:NM_001160234 | 2:11:32:124:56 | 0:0:0:0:0 | 0:15:30:0:32 | 0:2:0:0:1 | 0:23:76:128:75 | 0:0:0:0:0 | 1:26:90:47:71 | 0:1:0:0:0 | 1:14:27:152:86 | 0:0:1:0:1 |
| NM_001160233:XM_017001361:XR_002956654:NM_000701:NM_001160234 | 2:11:32:124:56 | 0:0:0:0:0 | 0:15:30:0:32 | 0:2:0:0:1 | 0:23:75:128:74 | 0:0:0:0:0 | 1:25:91:47:69 | 0:1:0:0:1 | 1:15:28:152:86 | 0:0:0:0:1 |
| NM_001160234:XR_002956654:NM_001160233:NM_000701:XM_017001361 | 48:20:2:124:9 | 1:1:0:0:0 | 20:23:0:0:7 | 1:1:0:0:1 | 82:53:0:128:18 | 1:0:0:0:0 | 66:71:0:47:24 | 1:0:1:0:1 | 91:25:1:152:24 | 0:0:0:0:0 |
| NM_001160233:NM_000701:XR_002956654:NM_001160234:XM_017001361 | 2:124:18:55:11 | 0:0:0:0:0 | 0:0:24:32:15 | 0:0:0:1:1 | 0:128:34:90:29 | 0:0:1:0:0 | 0:47:67:92:33 | 0:0:3:1:3 | 1:152:22:115:32 | 0:0:0:0:0 |
| NM_000701:NM_001160233:XR_002956654:NM_001160234:XM_017001361 | 123:2:18:55:11 | 1:0:0:0:0 | 0:0:24:35:16 | 0:0:0:1:2 | 128:0:35:95:31 | 0:0:1:0:0 | 47:0:68:93:42 | 0:0:1:2:1 | 152:1:20:118:33 | 0:0:0:0:1 |
| NM_000701:NM_001160233:XR_002956654:NM_001160234:XM_017001361 | 123:2:12:58:12 | 1:0:0:0:0 | 0:0:24:46:27 | 0:0:0:1:0 | 127:0:32:101:33 | 1:0:1:0:0 | 47:0:64:111:52 | 0:0:1:0:1 | 152:1:19:120:34 | 0:0:0:0:2 |
| NM_001160234:NM_001160233:NM_000701:XR_002956654:XM_017001361 | 69:2:124:12:11 | 1:0:0:0:0 | 76:0:0:21:52 | 1:0:0:0:1 | 116:0:128:28:36 | 0:0:0:0:0 | 167:0:47:66:92 | 2:0:0:0:0 | 133:1:152:17:42 | 0:0:0:0:0 |
| XR_002956654:NM_000701:NM_001160233:NM_001160234:XM_017001361 | 10:123:2:72:11 | 1:1:0:0:0 | 20:0:0:87:62 | 0:0:0:2:0 | 26:128:0:121:38 | 0:0:0:0:0 | 66:47:0:195:120 | 1:0:0:1:1 | 16:151:1:137:45 | 0:1:0:0:0 |
| NM_001160234:NM_001160233:NM_000701:XR_002956654:XM_017001361 | 81:2:124:9:18 | 1:0:0:0:0 | 126:0:0:23:112 | 0:0:0:0:0 | 160:0:128:23:52 | 0:0:0:0:0 | 297:0:47:66:246 | 2:0:0:0:2 | 157:1:150:15:57 | 0:0:2:0:0 |
| NM_001160233:NM_000701:XR_002956654:NM_001160234:XM_017001361 | 2:124:9:85:18 | 0:0:0:0:0 | 0:0:23:128:117 | 0:0:0:0:1 | 0:128:22:161:52 | 0:0:0:0:0 | 0:47:64:308:251 | 0:0:0:0:2 | 1:152:15:159:58 | 0:0:0:1:1 |
| NM_001160233:NM_000701:NM_001160234:XR_002956654:XM_017001361 | 2:124:106:7:21 | 0:0:0:0:0 | 0:0:159:18:145 | 0:0:1:0:2 | 0:128:184:14:61 | 0:0:1:0:0 | 0:47:400:41:341 | 0:0:1:0:1 | 1:152:190:13:61 | 0:0:0:0:1 |
| NM_001160234:NM_001160233:NM_000701:XR_002956654:XM_017001361 | 109:2:124:7:21 | 1:0:0:0:0 | 163:0:0:17:147 | 0:0:0:0:2 | 191:0:128:14:63 | 0:0:0:0:0 | 410:0:47:39:357 | 3:0:0:0:1 | 194:1:152:13:64 | 0:0:0:0:0 |
| NM_001160234:NM_001160233:XM_017001361:NM_000701:XR_002956654 | 139:2:27:124:7 | 2:0:0:0:0 | 213:0:185:0:15 | 1:0:1:0:0 | 248:0:79:128:12 | 1:0:0:0:0 | 522:0:490:47:27 | 3:0:2:0:1 | 230:1:75:152:10 | 0:0:0:0:0 |
| NM_001160233:XM_017001361:NM_000701:NM_001160234:XR_002956654 | 2:27:124:142:6 | 0:0:0:0:0 | 0:188:0:220:13 | 0:0:0:0:0 | 0:81:128:250:10 | 0:0:0:1:0 | 0:505:47:532:24 | 0:1:0:2:1 | 1:76:152:235:9 | 0:0:0:0:0 |
| NM_001160234:NM_001160233:XM_017001361:NM_000701:XR_002956654 | 148:2:35:124:6 | 1:0:0:0:0 | 239:0:212:0:10 | 0:0:1:0:0 | 290:0:92:127:12 | 0:0:0:1:0 | 578:0:585:47:18 | 1:0:1:0:1 | 250:1:85:152:10 | 0:0:0:0:0 |
| NM_001160233:XM_017001361:NM_000701:NM_001160234:XR_002956654 | 2:35:124:149:6 | 0:0:0:0:0 | 0:212:0:239:10 | 0:1:0:0:0 | 0:92:128:290:12 | 0:0:0:0:0 | 0:586:47:578:17 | 0:0:0:1:0 | 1:85:152:250:10 | 0:0:0:0:0 |
| NM_001160234:XM_017001361:NM_001160233:NM_000701:XR_002956654 | 153:34:2:124:6 | 1:1:0:0:0 | 245:217:0:0:13 | 0:1:0:0:0 | 297:92:0:128:14 | 1:0:0:0:0 | 592:603:0:47:22 | 1:2:0:0:0 | 262:89:1:152:10 | 0:0:0:0:0 |
| NM_001160233:XM_017001361:NM_001160234:NM_000701:XR_002956654 | 2:37:162:124:4 | 0:0:0:0:0 | 0:225:261:0:11 | 0:1:0:0:0 | 0:95:315:128:14 | 0:1:0:0:0 | 0:639:613:47:20 | 0:0:1:0:0 | 1:94:273:152:11 | 0:0:0:0:1 |
| NM_000701:NM_001160234:NM_001160233:XM_017001361:XR_002956654 | 123:174:2:39:6 | 1:1:0:0:0 | 0:279:0:238:12 | 0:1:0:0:0 | 128:332:0:100:16 | 0:0:0:0:0 | 47:640:0:661:23 | 0:0:0:3:0 | 152:290:1:100:15 | 0:1:0:0:0 |
| NM_001160233:XM_017001361:XR_002956654:NM_001160234:NM_000701 | 2:39:16:187:124 | 0:0:0:0:0 | 0:250:32:315:0 | 0:1:1:0:0 | 0:110:48:354:128 | 0:0:1:1:0 | 0:689:83:678:47 | 0:2:1:1:0 | 1:105:22:312:151 | 0:1:0:0:1 |
| NM_001160233:XM_017001361:XR_002956654:NM_001160234:NM_000701 | 2:39:16:188:124 | 0:0:0:0:0 | 0:251:35:316:0 | 0:0:0:1:0 | 0:110:53:353:128 | 0:0:0:0:0 | 0:690:85:681:47 | 0:2:1:2:0 | 1:105:23:312:151 | 0:1:0:1:1 |
| NM_001160234:NM_001160233:XM_017001361:XR_002956654:NM_000701 | 188:2:39:18:124 | 1:0:0:0:0 | 319:0:252:39:0 | 0:0:0:0:0 | 358:0:110:59:128 | 0:0:0:0:0 | 683:0:688:98:47 | 0:0:3:1:0 | 312:1:105:27:152 | 1:0:1:0:0 |
| NM_001160233:XM_017001361:XR_002956654:NM_001160234:NM_000701 | 2:39:18:189:124 | 0:0:0:0:0 | 0:252:40:320:0 | 0:0:0:1:0 | 0:110:61:360:128 | 0:0:0:1:0 | 0:690:100:690:47 | 0:1:1:1:0 | 1:105:26:313:152 | 0:1:1:0:0 |
| NM_001160233:XM_017001361:XR_002956654:NM_001160234:NM_000701 | 2:36:28:194:124 | 0:0:0:0:0 | 0:246:64:367:0 | 0:1:0:0:0 | 0:108:83:384:128 | 0:0:0:0:0 | 0:684:151:726:47 | 0:1:3:3:0 | 1:104:33:331:152 | 0:0:0:0:0 |
| NM_001160233:XM_017001361:XR_002956654:NM_001160234:NM_000701 | 2:36:33:197:124 | 0:0:0:0:0 | 0:245:74:382:0 | 0:1:0:0:0 | 0:105:87:390:127 | 0:0:1:1:1 | 0:680:170:734:47 | 0:1:1:1:0 | 1:103:36:335:152 | 0:0:0:1:0 |
| XR_002956654:NM_001160234:NM_001160233:XM_017001361:NM_000701 | 35:220:2:36:124 | 1:1:0:0:0 | 130:496:0:244:0 | 0:1:0:0:0 | 99:424:0:104:128 | 0:0:0:0:0 | 201:769:0:678:46 | 0:2:0:1:0 | 47:354:1:102:152 | 0:2:0:0:0 |
| XM_017002264:XM_017002263:NM_001135610:XM_017002256:NM_012231:XR_002957562 | 0:0:0:0:0:0 | 4:3:0:0:0:0 | 0:0:0:0:0:0 | 2:0:0:0:0:0 | 0:0:0:0:0:0 | 3:0:1:0:0:0 | 0:0:0:0:0:0 | 12:3:0:1:0:0 | 0:0:0:0:0:0 | 7:1:2:0:0:0 |
| NM_203458 | 1 | 0 | 2 | 0 | 4 | 0 | 1 | 1 | 0 | 1 |
| NM_001348451:NM_004284:NM_024568:NM_001348462:XR_001737550:NR_145682:NR_046070:NM_001348452:NR_145695 | 1:2:1:0:0:0:0:0:0 | 0:0:0:0:0:0:0:0:0 | 0:0:0:0:0:0:0:0:0 | 0:0:0:0:0:0:0:0:0 | 0:4:0:0:0:0:0:0:0 | 0:0:0:0:0:0:0:0:0 | 0:3:3:0:2:1:0:0:0 | 0:7:5:1:1:0:0:0:0 | 0:4:1:0:0:0:0:0:0 | 0:4:1:0:0:0:0:0:0 |
| NM_001348451:NM_004284:NM_024568:XM_024451051:XR_001737550:NR_145682:NM_001348462:NR_145695:NR_046070:NM_001348452 | 1:1:1:0:0:0:0:0:0:0 | 0:0:0:0:0:0:0:0:0:0 | 0:0:0:2:0:0:0:0:0:0 | 0:0:0:0:0:0:0:0:0:0 | 0:4:0:0:1:0:0:0:0:0 | 0:0:0:0:0:0:0:0:0:0 | 0:5:7:5:5:1:1:3:0:0 | 0:0:0:0:0:0:0:0:0:0 | 0:7:2:1:0:0:0:0:0:0 | 0:0:0:0:0:0:0:0:0:0 |
| NM_004284:NM_001348451:NM_024568:XM_024451051:NM_001348462:XR_001737550:NR_145695:NM_001348452 | 0:1:1:0:0:0:0:0 | 1:0:0:0:0:0:0:0 | 0:0:0:0:0:0:0:0 | 0:0:0:1:0:0:0:0 | 2:0:0:0:0:0:0:0 | 2:0:0:0:0:0:0:0 | 5:0:4:0:1:3:1:0 | 0:0:0:0:0:0:0:0 | 6:0:2:0:0:0:0:0 | 0:0:0:0:0:0:0:0 |
| NM_018997:NM_031901 | 0:00 | 2:03 | 0:00 | 0:01 | 0:00 | 0:00 | 0:00 | 1:06 | 0:00 | 3:04 |
| NM_004079 | 0 | 1 | 2 | 5 | 2 | 7 | 1 | 1 | 0 | 1 |
| NM_004079 | 0 | 0 | 7 | 0 | 9 | 0 | 2 | 0 | 1 | 0 |
| NM_022075:NM_181746 | 0:00 | 0:00 | 0:00 | 0:00 | 0:00 | 0:00 | 1:00 | 2:01 | 0:00 | 0:00 |

| | | | | | | | | | | |
|---|---|---|---|---|---|---|---|---|---|---|
| XM_017001774:NM_018379:XM_024448340:XM_017001771 | 0:0:0:0 | 0:0:0:0 | 0:0:0:0 | 0:0:0:0 | 0:0:0:0 | 1:0:0:0 | 0:0:0:0 | 0:1:0:0 | 0:0:0:0 | 0:0:0:0 |
| NM_002796 | 1 | 0 | 3 | 0 | 3 | 0 | 1 | 0 | 4 | 0 |
| NM_031420:NR_125331:XR_921910 | 1:00:00 | 0:00:00 | 0:01:00 | 0:00:00 | 0:00:00 | 0:00:00 | 4:00:00 | 0:00:00 | 3:00:00 | 0:00:00 |
| NM_031420:NR_125331:XR_921910 | 0:00:00 | 1:00:00 | 0:00:00 | 0:01:00 | 0:00:00 | 0:00:00 | 0:00:00 | 4:00:00 | 0:00:00 | 3:00:00 |
| NM_031420:XR_921910 | 1:00 | 0:00 | 0:00 | 0:00 | 0:00 | 0:00 | 3:00 | 1:00 | 0:00 | 3:00 |
| XM_011509142:NM_053055 | 0:00 | 0:00 | 0:00 | 0:00 | 0:00 | 0:00 | 0:00 | 0:00 | 0:00 | 0:00 |
| NM_014624:XM_017002033 | 779:40:00 | 0:00 | 2329:199 | 4:02 | 753:34:00 | 2:00 | 4343:165 | 2:00 | 1207:64 | 1:01 |
| NM_014624:XM_017002033 | 779:40:00 | 0:00 | 2330:200 | 4:01 | 751:34:00 | 4:00 | 4343:165 | 2:00 | 1207:65 | 1:00 |
| NM_014624:XM_017002033 | 780:40:00 | 1:00 | 2351:197 | 2:02 | 757:34:00 | 0:00 | 4343:165 | 7:00 | 1208:64 | 1:01 |
| NM_002961:NM_019554 | 379:26:00 | 1:00 | 490:48:00 | 0:00 | 210:23:00 | 1:01 | 486:05:00 | 0:00 | 196:21:00 | 0:00 |
| NM_002961:NM_019554 | 381:26:00 | 0:00 | 491:47:00 | 2:00 | 213:24:00 | 0:00 | 491:04:00 | 0:00 | 196:21:00 | 0:00 |
| NM_002961:NM_019554 | 379:22:00 | 1:00 | 493:44:00 | 0:00 | 212:22:00 | 1:01 | 488:04:00 | 4:00 | 196:21:00 | 0:00 |
| NM_005978 | 0 | 0 | 1 | 0 | 0 | 0 | 1 | 0 | 2 | 0 |
| XM_017002034:XM_017002036:NM_005979:NM_001024211:NM_001024212:NM_001024213:XM_017002035:XM_011509863:XM_011509864:NM_001024210:XM_005245434:XM_011509862 | 59:3:126:29:15:11:4:6:1:1:0:0 | 0:0:0:0:0:0:0:0:0:0:0:0 | 92:5:123:69:10:4:1:3:1:8:1:1 | 0:0:0:0:0:0:0:0:0:0:0 | 86:5:173:154:16:7:0:1:0:7:0:3 | 0:0:0:0:0:0:0:0:0:0:0 | 118:5:141:116:36:11:17:17:1:3:0:3 | 0:0:0:0:0:1:0:0:0:0:0 | 41:4:66:47:7:5:0:1:0:0:0:1 | 0:0:0:0:0:0:0:0:0:0 |
| XM_017002034:XM_017002036:NM_005979:NM_001024211:NM_001024212:NM_001024213:XM_017002035:XM_011509863:XM_011509864:NM_001024210:XM_005245434:XM_011509862 | 59:3:126:29:15:11:4:6:1:1:0:0 | 0:0:0:0:0:0:0:0:0:0:0:0 | 92:6:123:71:10:4:1:3:1:8:1:1 | 0:0:0:0:0:0:0:0:0:0:0 | 85:4:170:154:16:7:0:1:0:7:0:3 | 0:0:2:0:0:0:0:0:0:0:0 | 118:5:141:116:36:11:17:17:1:3:0:3 | 0:0:0:0:0:0:0:0:0:0:0 | 41:4:65:46:7:5:0:1:0:0:0:1 | 0:0:0:1:0:0:0:0:0:0 |
| NM_153649:NR_103461:NM_001043351:NM_001278189:NM_001349679:NM_001278188:NM_001043352:NM_001278190:NM_152263:NM_001278191:NM_001043353 | 590:9:2:7:1:2:3:0:0:0:0 | 0:0:0:0:0:0:0:0:0:0:0 | 909:18:2:5:0:4:3:1:1:0:0 | 0:1:0:0:0:0:0:0:0:0:0 | 968:8:4:4:0:5:7:0:0:0:0 | 0:0:0:0:0:0:0:0:0:1:0 | 2460:56:6:71:3:16:43:2:0:6:0 | 4:0:0:0:0:0:0:0:0:0:0 | 1097:7:7:13:0:11:41:0:0:0:0 | 0:0:0:0:0:0:0:0:0:0 |
| NM_153649:NR_103461:NM_001043351:NM_001278189:NM_001349679:NM_001278188:NM_001043352:NM_001278190:NM_152263:NM_001278191:NM_001043353 | 591:8:2:7:1:2:3:0:0:0:0 | 0:0:0:0:0:0:0:0:0:0:0 | 908:17:2:5:0:4:3:1:1:0:0 | 2:1:0:0:0:0:0:0:0:0:0 | 968:7:4:4:0:5:7:0:0:1:0 | 0:0:0:0:0:0:0:0:0:0:0 | 2467:32:6:71:1:16:43:2:0:6:0 | 0:2:0:0:0:0:0:0:0:0:0 | 1096:7:7:13:0:11:41:0:0:0:0 | 1:0:0:0:0:0:0:0:0:0 |
| NM_153649:NR_103461:NM_001043351:NM_001278189:NM_001349679:NM_001278188:NM_001043352:NM_001278190:NM_152263:NM_001278191 | 582:7:2:7:1:2:3:0:0:0 | 0:0:0:0:0:0:0:0:0:0 | 864:27:2:5:0:4:3:1:1:0 | 1:0:0:0:0:0:0:0:0:0 | 949:7:4:4:0:5:7:0:0:0 | 0:0:0:0:0:0:0:0:0:0 | 2358:102:6:67:0:16:42:2:0:6 | 6:0:0:0:0:0:0:0:0:0 | 1083:8:7:13:0:11:41:0:0:0 | 0:0:0:0:0:0:0:0:0:0 |
| NM_153649:NR_103461:NM_001043351:NM_001278189:NM_001349679:NM_001278188:NM_001043352:NM_001278190:NM_152263:NM_001278191 | 581:7:2:7:1:2:3:0:0:0 | 1:0:0:0:0:0:0:0:0:0 | 863:29:2:5:0:4:3:1:1:0 | 2:0:0:0:0:0:0:0:0:0 | 949:7:4:4:0:5:7:0:0:0 | 0:0:0:0:0:0:0:0:0:0 | 2361:102:6:67:0:16:42:2:0:6 | 1:0:0:0:0:0:0:0:0:0 | 1083:8:7:13:0:11:41:0:0:0 | 0:0:0:0:0:0:0:0:0:0 |
| NM_153649:NR_103461:NM_001043351:NM_001278189:NM_001349679:NM_001278188:NM_001043352:NM_001278190:NM_152263:NM_001278191 | 575:7:2:7:1:2:3:0:0:0 | 0:0:0:0:0:0:0:0:0:0 | 846:31:2:5:0:4:3:1:1:0 | 0:0:0:0:0:0:0:0:0:0 | 943:9:4:4:0:5:7:0:0:0 | 0:0:0:0:0:0:0:0:0:0 | 2328:103:6:67:0:16:42:2:0:5 | 2:0:0:0:0:0:0:0:0:0 | 1073:8:7:13:0:11:40:0:0:0 | 1:0:0:0:0:0:0:0:0:0 |
| NM_153649:NR_103461:NM_001043351:NM_001278189:NM_001349679:NM_001278188:NM_001043352:NM_001278190:NM_152263:NM_001278191 | 575:8:2:7:1:2:3:0:0:0 | 0:0:0:0:0:0:0:0:0:0 | 836:31:2:4:0:4:3:1:1:0 | 0:0:0:0:0:0:0:0:0:0 | 936:10:4:4:0:5:7:0:0:0 | 2:0:0:0:0:0:0:0:0:0 | 2306:102:6:65:0:16:42:2:0:5 | 4:0:0:1:0:0:0:0:0:0 | 1068:9:7:13:0:11:40:0:0:0 | 1:0:0:0:0:0:0:0:0:0 |
| NM_153649:NR_103461:NM_001043351:NM_001278189:NM_001349679:NM_001278188:NM_001043352:NM_001278190:NM_152263:NM_001278191 | 574:8:2:7:1:2:2:0:0:0 | 0:0:0:0:0:0:0:0:0:0 | 828:31:2:4:0:4:3:1:1:0 | 0:0:0:0:0:0:0:0:0:0 | 934:10:4:4:0:5:7:0:0:0 | 2:0:0:0:0:0:0:0:0:0 | 2286:102:6:65:0:16:42:2:0:5 | 0:0:0:0:0:0:0:0:0:0 | 1064:9:7:13:0:11:40:0:0:0 | 0:0:0:0:0:0:0:0:0:0 |
| NM_153649:NR_103461:NM_001043351:NM_001278189:NM_001349679:NM_001278188:NM_001043352:NM_001278190:NM_152263:NM_001278191 | 573:8:2:7:1:2:2:0:0:0 | 0:0:0:0:0:0:0:0:0:0 | 824:30:2:4:0:4:3:1:1:0 | 2:0:0:0:0:0:0:0:0:0 | 933:10:4:4:0:5:7:0:0:0 | 0:0:0:0:0:0:0:0:0:0 | 2277:100:6:64:0:16:42:2:0:5 | 1:0:0:0:0:0:0:0:0:0 | 1060:9:7:13:0:11:40:0:0:0 | 1:0:0:0:0:0:0:0:0:0 |
| NM_153649:NR_103461:NM_001043351:NM_001278189:NM_001349679:NM_001278188:NM_001043352:NM_001278190:NM_001278191 | 569:8:2:7:1:2:2:0:0 | 1:0:0:0:0:0:0:0:0 | 816:32:2:4:0:1:3:1:0 | 2:0:0:0:0:0:0:0:0 | 933:10:4:4:0:1:6:0:0 | 0:0:0:0:0:0:0:0:0 | 2256:101:6:63:0:9:41:2:3 | 0:0:0:0:0:0:0:0:0 | 1054:9:7:11:0:40:0:0 | 1:0:0:0:0:0:0:0:0 |
| NM_153649:NM_001043352:NR_103461:NM_001043351:NM_001278189:NM_001349679:NM_001278188:NM_001278191:NM_001278190 | 565:2:9:2:7:1:0:0:0 | 1:1:0:0:0:0:0:0:0 | 793:3:31:2:5:0:2:0:0 | 2:0:0:0:0:0:0:0:0 | 920:6:10:4:4:0:3:0:0 | 0:0:0:0:0:0:0:0:0 | 2188:36:96:6:60:0:8:4:2 | 4:0:0:0:0:0:0:0:0 | 1041:37:9:7:12:0:10:0:0 | 2:0:0:0:0:0:0:0:0 |
| NM_153649:NR_103461:NM_001043351:NM_001278189:NM_001349679:NM_001043352:NM_001278188:NM_001278191:NM_001278190 | 566:9:2:7:1:3:0:0:0 | 0:0:0:0:0:0:0:0:0 | 792:31:2:5:0:3:3:0:0 | 0:0:0:0:0:0:0:0:0 | 916:10:4:4:0:6:3:0:0 | 2:0:0:0:0:0:0:0:0 | 2181:95:6:60:0:36:10:4:2 | 4:0:0:0:0:0:0:0:0 | 1040:8:7:12:0:37:10:0:0 | 0:0:0:0:0:0:0:0:0 |
| NM_014188 | 57 | 0 | 73 | 0 | 221 | 0 | 77 | 2 | 60 | 0 |
| NR_048579:NM_207196:NM_207197:NM_003815:NM_001261464:NM_207194 | 0:0:0:0:0:0 | 0:0:0:0:0:0 | 0:0:0:0:0:0 | 2:0:0:0:0:0 | 0:0:0:0:0:0 | 0:1:0:0:0:0 | 0:0:0:0:0:0 | 0:0:1:0:0:0 | 0:0:0:0:0:0 | 0:0:0:0:0:0 |
| NM_207196:NM_003815:NM_207197:NM_207195:NM_207191:NR_048579:NM_207194:NM_001261464 | 0:0:0:0:0:0:0:0 | 0:0:0:0:0:0:0:0 | 1:0:0:0:0:0:0:0 | 0:0:0:0:0:0:0:0 | 1:1:0:0:0:0:0:0 | 0:0:0:0:0:0:0:0 | 0:0:1:1:0:0:0:0 | 0:0:0:0:0:0:0:0 | 0:0:0:0:0:0:0:0 | 0:0:0:0:0:0:0:0 |
| NM_207196:NM_207197:NM_207194 | 0:00:00 | 0:00:00 | 1:00:00 | 0:00:00 | 1:00:00 | 0:00:00 | 0:00:00 | 0:03:00 | 0:00:00 | 0:00:00 |
| NM_004428:NM_182685 | 1:00 | 0:00 | 1:00 | 0:00 | 0:00 | 0:00 | 8:00 | 1:00 | 0:00 | 0:00 |
| XM_005244963:NM_001135822:NM_001242824:NM_002004 | 0:0:0:0 | 0:0:0:0 | 2:1:0:0 | 0:0:0:0 | 1:1:1:0 | 0:0:0:0 | 0:0:0:0 | 0:0:0:0 | 0:1:0:1 | 1:3:2:0 |
| NM_007221:NM_001199662 | 0:00 | 1:00 | 0:00 | 1:00 | 2:00 | 0:00 | 0:00 | 0:00 | 0:01 | 0:00 |
| NM_005998 | 2 | 0 | 1 | 0 | 1 | 0 | 5 | 0 | 3 | 0 |
| NM_144772:XM_017000319 | 0:00 | 0:00 | 0:00 | 0:00 | 0:00 | 2:00 | 0:00 | 3:00 | 0:00 | 0:00 |
| NM_005973:XR_921868:XM_005245314:XM_005245313 | 10:0:1:0 | 1:1:0:0 | 3:1:0:0 | 3:0:1:0 | 15:0:1:0 | 5:3:0:0 | 50:6:3:2 | 1:0:0:0 | 23:0:0:0 | 0:0:0:0 |
| NM_005973:XM_005245314:XR_921868:XM_005245313 | 12:1:0:0 | 0:0:0:0 | 6:1:0:0 | 0:0:0:0 | 25:1:0:0 | 0:0:0:0 | 56:3:6:2 | 2:0:0:0 | 27:0:0:0 | 0:0:0:0 |
| NM_005973:XM_005245314:XR_921868:XM_005245313 | 12:1:0:0 | 0:0:0:0 | 6:1:0:0 | 0:0:0:0 | 25:1:2:0 | 0:0:0:0 | 59:3:4:2 | 2:0:0:0 | 28:0:0:1 | 0:0:0:0 |
| XM_017001525:XM_006710704:XM_024447751:XM_017001526:XM_017001519:NM_001350151:XM_006710705:XM_011541617:NM_001024216:XM_011541616:NM_017556 | 0:0:0:0:0:0:0:0:0:0:0 | 0:0:0:0:0:0:0:0:0:0:0 | 0:1:2:0:0:0:0:0:0:0:0 | 1:0:0:0:0:0:0:0:0:0:0 | 4:0:1:1:1:1:0:0:0:0:0 | 0:0:0:0:0:0:0:0:0:0:0 | 0:0:0:0:0:0:0:0:0:0:0 | 2:0:1:1:0:0:1:0:0:0:0 | 0:0:0:0:0:0:0:0:0:0:0 | 1:0:0:0:0:1:0:1:1:0:0 |
| NM_015001 | 8 | 2 | 23 | 2 | 3 | 0 | 43 | 5 | 14 | 1 |
| NM_015001 | 0 | 0 | 3 | 0 | 1 | 0 | 0 | 8 | 2 | 0 |
| NM_001146172:NM_033438 | 0:00 | 1:00 | 0:00 | 0:00 | 0:00 | 0:00 | 1:00 | 0:00 | 0:00 | 0:00 |
| NR_104400:NM_003037:XM_005245456:XM_011509905:NM_001330754:NR_104399 | 1:3:0:1:0:0 | 2:2:1:0:0:0 | 1:4:0:1:0:0 | 0:1:0:0:0:0 | 0:3:0:0:1:0 | 0:1:0:0:0:0 | 0:0:0:0:0:0 | 0:0:0:0:0:0 | 0:1:0:0:0:0 | 0:0:0:0:0:0 |
| NM_003037:XM_011509905:XM_005245456:NM_001330754 | 7:1:1:0 | 1:0:0:0 | 6:1:0:0 | 1:0:0:0 | 5:0:0:1 | 0:0:0:0 | 0:0:0:0 | 1:0:0:0 | 1:0:0:0 | 0:0:0:0 |
| NR_104400:XM_011509905:NM_003037:XM_005245456:NM_001330754:XM_017002131:XM_017002130:NR_104401:NR_104399 | 3:1:8:1:0:0:0:0:0 | 0:0:0:0:0:0:0:0:0 | 1:0:7:0:1:1:0:0:0 | 0:0:0:0:0:0:0:0:0 | 0:0:5:0:1:0:1:1:0 | 0:0:0:0:0:0:0:0:0 | 0:0:1:0:0:0:0:1:0 | 0:0:0:0:0:0:0:0:0 | 0:0:1:0:0:0:0:0:0 | 0:0:0:0:0:0:0:0:0 |

| | | | | | | | | | | |
|---|---|---|---|---|---|---|---|---|---|---|
| NM_017625 | 0 | 0 | 246 | 0 | 1063 | 1 | 2 | 0 | 100 | 0 |
| NM_017625 | 0 | 0 | 245 | 0 | 1045 | 2 | 2 | 0 | 101 | 0 |
| NM_017625 | 0 | 0 | 0 | 245 | 0 | 1047 | 2 | 0 | 56 | 45 |
| NM_017625 | 0 | 0 | 244 | 0 | 1047 | 2 | 2 | 0 | 100 | 0 |
| NM_017625 | 0 | 0 | 242 | 1 | 1039 | 3 | 2 | 0 | 99 | 0 |
| NM_004550:XM_005245209:NM_001166159 | 1:00:00 | 0:00:00 | 0:00:00 | 0:00:00 | 1:00:00 | 1:00:00 | 1:00:00 | 0:00:00 | 2:00:00 | 2:00:00 |
| NM_007348:XM_006711224:XM_011509310:XM_011509308 | 9:1:1:0 | 10:0:0:0 | 10:1:4:0 | 14:1:2:0 | 29:0:0:1 | 11:1:1:0 | 57:5:10:0 | 43:2:8:0 | 18:0:1:0 | 24:1:2:0 |
| XM_011540706 | 0 | 0 | 0 | 0 | 0 | 0 | 0 | 0 | 0 | 0 |
| XM_017001395:NM_002585:XM_005245229:NM_001204961 | 0:0:0:0 | 0:0:0:0 | 1:0:0:0 | 0:0:0:0 | 2:1:0:0 | 0:2:0:0 | 3:0:0:2 | 1:0:1:1 | 0:0:0:1 | 0:0:0:0 |
| NM_012474 | 37 | 0 | 84 | 0 | 72 | 0 | 289 | 0 | 95 | 0 |
| NM_012474 | 34 | 0 | 84 | 0 | 71 | 0 | 287 | 0 | 94 | 0 |
| XM_017000660 | 1 | 0 | 0 | 1 | 0 | 0 | 0 | 0 | 0 | 0 |
| NM_003000 | 269 | 0 | 426 | 1 | 2612 | 4 | 602 | 0 | 349 | 0 |
| NM_003000 | 269 | 0 | 426 | 0 | 2614 | 7 | 601 | 2 | 347 | 0 |
| NM_003000 | 271 | 0 | 425 | 0 | 2632 | 1 | 604 | 0 | 343 | 0 |
| NM_003000 | 272 | 0 | 425 | 0 | 2621 | 2 | 602 | 2 | 342 | 0 |
| NM_003000 | 267 | 0 | 419 | 1 | 2595 | 11 | 597 | 0 | 340 | 0 |
| NM_003000 | 266 | 0 | 419 | 1 | 2585 | 15 | 596 | 0 | 339 | 0 |
| NM_003000 | 266 | 0 | 418 | 1 | 2589 | 10 | 596 | 0 | 339 | 0 |
| NM_003000 | 261 | 0 | 414 | 0 | 2526 | 17 | 591 | 0 | 340 | 0 |
| NM_033529:NR_132740 | 0:00 | 0:00 | 0:00 | 0:00 | 0:00 | 1:00 | 0:00 | 0:01 | 0:00 | 0:01 |
| NM_153747 | 0 | 0 | 0 | 0 | 0 | 0 | 1 | 0 | 1 | 0 |
| NM_012387 | 1 | 0 | 0 | 0 | 0 | 2 | 0 | 0 | 0 | 0 |
| NM_022457:XM_017002069:XM_005245447:XM_005245448:NM_001001740:XM_011509881:XM_017002076:XM_006711487:XM_017002075:XM_017002066:NM_001286644:XM_017002079:XM_017002061:XM_017002078:XM_017002065:XM_017002067:XM_017002060:XM_017002080:XM_017002062 | 16:2:13:4:1:1:1:1:0:0:0:0:0:0:0:0:0:0:0 | 0:0:0:0:0:0:0:0:0:0:0:0:0:0:0:0:0:0:0 | 20:1:10:6:1:0:21:1:1:1:0:0:0:0:0:0:0:0:0 | 0:0:0:0:0:0:0:0:0:0:0:0:0:0:0:0:0:0:0 | 21:0:7:17:3:2:2:0:0:0:1:2:1:0:0:0:0:0:0 | 0:0:0:0:0:0:0:0:0:0:0:0:0:0:0:0:0:0:0 | 96:1:18:33:10:1:104:7:1:0:6:2:2:1:2:1:0:0:0 | 0:0:0:0:0:0:0:0:0:0:0:0:0:0:0:0:0:0:0 | 39:5:15:15:8:0:3:1:0:0:3:1:1:0:0:0:0:0:0 | 0:0:0:0:0:0:0:0:0:0:0:0:0:0:0:0:0:0:0 |
| NM_022457:XM_017002069:XM_005245447:XM_005245448:NM_001001740:XM_011509881:XM_017002076:XM_006711487:XM_017002075:XM_017002066:NM_001286644:XM_017002079:XM_017002061:XM_017002078:XM_017002065:XM_017002067:XM_017002060:XM_017002080:XM_017002062 | 16:2:13:4:1:1:1:1:0:0:0:0:0:0:0:0:0:0:0 | 0:0:0:0:0:0:0:0:0:0:0:0:0:0:0:0:0:0:0 | 20:1:10:6:1:0:21:1:1:1:0:0:0:0:0:0:0:0:0 | 0:0:0:0:0:0:0:0:0:0:0:0:0:0:0:0:0:0:0 | 21:0:7:17:3:2:2:0:0:0:1:2:1:0:0:0:0:0:0 | 0:0:0:0:0:0:0:0:0:0:0:0:0:0:0:0:0:0:0 | 96:1:18:33:10:1:101:7:1:0:6:2:2:1:2:1:0:0:0 | 0:0:0:0:0:0:0:1:0:0:0:0:0:0:0:0:0:0:0 | 39:5:15:15:8:0:3:1:0:0:3:1:1:0:0:0:0:0:0 | 0:0:0:0:0:0:0:0:0:0:0:0:0:0:0:0:0:0:0 |
| NM_022457:XM_017002069:XM_005245447:XM_005245448:NM_001001740:XM_011509881:XM_017002076:XM_006711487:XM_017002080:XM_017002075:XM_017002066:NM_001286644:XM_017002079:XM_017002061:XM_017002078:XM_017002065:XM_017002067:XM_017002060:XM_017002062 | 16:2:13:4:1:1:1:1:1:0:0:0:0:0:0:0:0:0:0 | 0:0:0:0:0:0:0:0:0:0:0:0:0:0:0:0:0:0:0 | 20:1:10:6:1:0:20:1:0:1:1:0:0:0:0:0:0:0:0 | 0:0:0:0:0:0:0:0:0:0:0:0:0:0:0:0:0:0:0 | 21:0:7:17:3:2:2:0:0:0:0:1:2:1:0:0:0:0:0 | 0:0:0:0:0:0:0:0:0:0:0:0:0:0:0:0:0:0:0 | 93:1:18:32:10:1:93:7:0:1:1:6:2:1:1:2:1:0:0 | 2:0:0:1:0:0:0:0:0:0:0:0:0:0:0:0:0:0:0 | 39:5:15:15:8:0:2:1:2:0:0:3:1:1:0:0:0:0:0 | 0:0:0:0:0:0:0:0:0:0:0:0:0:0:0:0:0:0:0 |
| NM_022457:XM_017002069:XM_005245447:XM_005245448:NM_001001740:XM_011509881:XM_017002076:XM_006711487:XM_017002080:XM_017002075:XM_017002066:NM_001286644:XM_017002079:XM_017002061:XM_017002078:XM_017002065:XM_017002067:XM_017002060:XM_017002062 | 16:2:13:4:1:1:1:1:1:0:0:0:0:0:0:0:0:0:0 | 0:0:0:0:0:0:0:0:0:0:0:0:0:0:0:0:0:0:0 | 19:1:10:6:1:0:20:1:1:1:1:0:0:0:0:0:0:0:0 | 0:0:0:0:0:0:0:0:0:0:0:0:0:0:0:0:0:0:0 | 21:0:7:17:3:2:2:0:0:0:0:1:2:1:0:0:0:0:0 | 0:0:0:0:0:0:0:0:0:0:0:0:0:0:0:0:0:0:0 | 93:1:17:33:10:1:89:7:0:1:1:6:2:2:1:2:1:0:0 | 1:0:1:0:0:0:0:0:0:0:0:0:0:0:0:0:0:0:0 | 39:5:14:15:8:0:2:1:2:0:0:3:1:1:0:0:0:0:0 | 0:0:1:0:0:0:0:0:0:0:0:0:0:0:0:0:0:0:0 |
| NM_022457:XM_017002069:XM_005245447:XM_005245448:NM_001001740:XM_011509881:XM_017002076:XM_006711487:XM_017002075:XM_017002066:NM_001286644:XM_017002079:XM_017002061:XM_017002078:XM_017002065:XM_017002067:XM_017002063:XM_017002080:XM_017002060:XM_017002062 | 16:2:13:4:1:1:1:1:0:0:0:0:0:0:0:0:0:0:0:0 | 0:0:0:0:0:0:0:0:0:0:0:0:0:0:0:0:0:0:0:0 | 18:1:10:6:1:0:13:1:1:1:0:0:0:0:0:0:0:0:0:0 | 0:0:0:0:0:0:0:0:0:0:0:0:0:0:0:0:0:0:0:0 | 20:0:7:17:3:2:2:0:0:0:1:2:1:0:0:0:0:0:0:0 | 0:0:0:0:0:0:0:0:0:0:0:0:0:0:0:0:0:0:0:0 | 86:1:18:33:10:1:59:6:1:1:6:2:3:1:2:1:1:0:0:0 | 1:0:0:0:0:0:0:1:0:0:0:0:0:0:0:0:0:0:0:0 | 38:5:15:15:8:0:2:1:0:0:3:1:1:0:0:0:0:1:0:0 | 0:0:0:0:0:0:0:0:0:0:0:0:0:0:0:0:0:0:0:0 |
| NM_022457:XM_017002069:XM_005245447:XM_005245448:NM_001001740:XM_011509881:XM_017002076:XM_006711487:XM_017002075:XM_017002066:NM_001286644:XM_017002079:XM_017002061:XM_017002078:XM_017002065:XM_017002067:XM_017002063:XM_017002060:XM_017002062 | 15:2:13:4:1:1:1:1:0:0:0:0:0:0:0:0:0:0:0 | 0:0:0:0:0:0:0:0:0:0:0:0:0:0:0:0:0:0:0 | 10:1:10:4:0:0:1:1:1:1:0:0:0:0:0:0:0:0:0 | 0:0:0:0:0:0:0:0:0:0:0:0:0:0:0:0:0:0:0 | 20:0:7:16:3:2:1:0:0:0:1:2:1:0:0:0:0:0:0 | 0:0:0:1:0:0:0:0:0:0:0:0:0:0:0:0:0:0:0 | 63:1:17:19:9:0:3:6:1:1:5:1:3:1:2:1:1:0:0 | 2:0:0:0:0:0:0:0:0:0:1:0:0:0:0:0:0:0:0 | 34:5:15:15:7:0:2:1:0:0:3:1:1:0:0:0:0:0:0 | 0:0:0:0:0:0:0:0:0:0:0:0:0:0:0:0:0:0:0 |
| NR_137288:XR_002956989:XR_001737277:XM_017001622:XR_946690:XR_946686:XM_024448062:NM_001328124:NM_001319838:XM_017001620 | 2:3:1:1:0:0:0:0:0:0 | 1:2:0:0:0:0:0:0:0:0 | 2:2:0:0:0:1:2:0:0:0 | 2:2:0:1:1:0:0:0:0:0 | 3:4:0:0:0:0:0:1:0:0 | 2:2:0:0:0:0:1:1:1:0 | 0:0:0:0:0:0:0:0:0:0 | 3:9:0:0:0:0:1:0:0:0 | 0:0:0:0:0:0:0:0:0:0 | 3:2:1:0:0:1:2:0:0:1 |
| NM_018891:XM_017001273:NM_005562 | 1:01:00 | 0:00:00 | 1:00:00 | 0:00:00 | 1:01:00 | 0:00:00 | 0:00:00 | 0:00:00 | 0:01:00 | 0:00:00 |
| NM_018891:XM_017001273:NM_005562 | 2:01:00 | 0:00:00 | 0:00:02 | 0:00:00 | 1:00:02 | 0:00:00 | 0:00:00 | 1:00:00 | 1:00:00 | 1:00:01 |
| NM_018891:NM_005562:XM_017001273 | 0:00:00 | 2:00:00 | 0:02:00 | 0:00:00 | 1:00:00 | 0:02:00 | 2:01:00 | 0:00:00 | 2:01:01 | 0:00:00 |
| NM_030806 | 98 | 0 | 208 | 1 | 175 | 0 | 544 | 0 | 89 | 0 |
| NM_017847:XM_011509661 | 0:00 | 0:00 | 0:00 | 0:00 | 0:00 | 0:00 | 0:00 | 0:00 | 0:00 | 0:00 |
| NM_012067:XR_001737055 | 0:00 | 0:00 | 1:00 | 1:00 | 0:00 | 0:00 | 0:00 | 0:00 | 1:00 | 1:00 |
| NM_012067:XR_001737055 | 0:00 | 0:00 | 1:00 | 1:00 | 0:00 | 0:00 | 0:00 | 0:00 | 1:00 | 1:00 |
| NM_024529:XM_006711537 | 13:00 | 9:00 | 26:00:00 | 0:00 | 28:00:00 | 0:00 | 159:07:00 | 0:00 | 37:01:00 | 0:00 |
| NM_024529 | 22 | 0 | 24 | 1 | 27 | 0 | 174 | 1 | 34 | 0 |
| NM_024529 | 21 | 0 | 8 | 8 | 23 | 3 | 113 | 30 | 29 | 5 |
| NM_024529 | 20 | 0 | 5 | 0 | 21 | 0 | 91 | 1 | 28 | 0 |
| NM_004930:XM_024450097:NM_001206540:XM_011542228:NM_001282162:NM_001206541:XM_017002428 | 329:17:2:0:0:0:0 | 3:0:0:0:0:0:0 | 450:62:12:1:4:0:0 | 6:0:0:0:0:0:0 | 871:35:14:0:6:1:0 | 6:0:2:0:0:0:0 | 727:63:1:0:2:0:0 | 1:0:0:0:0:0:0 | 576:41:0:0:1:1:0 | 3:0:0:0:0:0:0 |
| NM_004930:XM_024450097:NM_001206540:XM_011542228:NM_001282162:NM_001206541:XM_017002428 | 332:17:2:0:0:0:0 | 1:0:0:0:0:0:0 | 454:63:12:1:4:0:0 | 2:0:0:0:0:0:0 | 877:35:16:0:6:1:0 | 0:0:0:0:0:0:0 | 728:64:1:0:2:0:0 | 1:0:0:0:0:0:0 | 579:41:0:0:1:1:0 | 0:0:0:0:0:0:0 |
| NM_004930:XM_024450097:NM_001206540:XM_017002428:XM_011542228:NM_001282162:NM_001206541 | 336:15:2:1:0:0:0 | 0:0:0:0:0:0:0 | 464:56:12:0:1:4:0 | 0:0:0:0:0:0:0 | 885:32:16:1:0:6:1 | 2:1:0:0:0:0:0 | 742:56:1:0:0:2:0 | 1:0:0:0:0:0:0 | 578:34:0:0:0:1:1 | 2:0:0:0:0:0:0 |
| NM_004930:XM_024450097:NM_001206540:XM_017002428:XM_011542228:NM_001282162:NM_001206541 | 335:15:2:1:0:0:0 | 1:0:0:0:0:0:0 | 464:55:12:0:1:4:0 | 0:1:0:0:0:0:0 | 888:33:16:1:0:6:1 | 0:0:0:0:0:0:0 | 743:55:1:0:0:2:0 | 0:0:0:0:0:0:0 | 580:34:0:0:0:1:1 | 0:0:0:0:0:0:0 |
| NM_004930:XM_024450097:NM_001206540:XM_017002428:XM_011542228:NM_001282162:NM_001206541 | 335:11:2:1:0:0:0 | 1:0:0:0:0:0:0 | 464:35:12:1:1:4:0 | 2:1:0:0:0:0:0 | 888:34:16:2:0:6:1 | 0:0:0:0:0:0:0 | 740:37:1:2:0:2:0 | 1:0:0:0:0:0:0 | 582:27:0:0:0:1:1 | 0:0:0:0:0:0:0 |
| NM_004930:XM_024450097:NM_001206540:XM_017002428:NM_001313932:XM_011542228:NM_001282162:NM_001206541:XM_017002430 | 335:6:2:2:1:0:0:0:0 | 0:0:0:0:0:0:0:0:0 | 466:6:12:2:13:1:4:0:0 | 0:0:0:0:0:0:0:0:0 | 885:13:16:2:23:0:6:1:0 | 1:0:0:0:0:0:0:0:0 | 741:11:1:2:13:0:2:0:0 | 0:0:0:0:0:0:0:0:0 | 581:6:0:1:5:0:1:1:0 | 0:0:0:0:0:0:0:0:0 |
| NM_005558 | 0 | 0 | 0 | 0 | 1 | 1 | 0 | 0 | 1 | 3 |
| NM_004433:XM_005244942:NM_001114309 | 314:06:01 | 1:00:00 | 627:15:10 | 1:00:00 | 888:13:12 | 1:00:00 | 1909:46:22 | 6:00:00 | 608:06:02 | 0:00:00 |
| NM_004433:XM_005244942:NM_001114309 | 313:06:01 | 0:00:00 | 625:14:10 | 1:00:00 | 894:12:12 | 1:00:00 | 1909:46:22 | 8:00:00 | 609:06:02 | 1:00:00 |
| NM_004433:XM_005244942:NM_001114309 | 330:02:01 | 0:00:00 | 794:08:10 | 4:00:00 | 1013:04:12 | 1:00:00 | 2241:24:22 | 4:00:00 | 674:02:02 | 0:00:00 |
| NM_004433:XM_005244942:NM_001114309 | 331:02:01 | 0:00:00 | 797:08:10 | 3:00:00 | 1012:04:12 | 2:00:00 | 2241:24:22 | 7:00:00 | 673:02:02 | 1:00:00 |
| NM_004433:NM_001114309 | 201:01:00 | 0:00 | 504:04:00 | 4:00 | 560:05:00 | 3:00 | 1140:09:00 | 3:00 | 411:02:00 | 0:00 |
| NM_001785 | 0 | 0 | 1 | 0 | 2 | 0 | 0 | 4 | 1 | 1 |
| NM_005216 | 0 | 0 | 0 | 0 | 0 | 3 | 0 | 2 | 0 | 6 |
| NM_002644:XM_011509629 | 0:00 | 0:00 | 1:00 | 2:02 | 3:02 | 2:01 | 0:00 | 0:00 | 0:00 | 0:00 |
| XM_024453287 | 4 | 0 | 2 | 0 | 0 | 0 | 2 | 0 | 1 | 0 |
| XM_024453287 | 4 | 0 | 2 | 0 | 0 | 0 | 2 | 0 | 1 | 0 |
| NM_172353:NM_172351:NM_153826:NM_172352:NM_172359:XR_002956622:NM_002389:NM_172361:XR_002956621:XM_011509563 | 183:40:122:38:1:3:1:1:0:0 | 2:0:0:0:0:0:0:0:0:0 | 0:0:4:0:0:1:0:0:0:0 | 0:0:0:0:0:0:0:0:0:0 | 6:10:76:2:2:2:0:0:2:0 | 0:0:0:0:0:0:0:0:0:0 | 151:37:41:109:0:13:0:3:2:0 | 0:0:0:0:0:0:0:0:0:0 | 77:22:39:33:0:6:0:4:3:0 | 0:0:0:1:0:0:0:0:0:0 |

| | | | | | | | | | | |
|---|---|---|---|---|---|---|---|---|---|---|
| NM_153826:NM_172351:NM_172353:NM_172352:NM_172359:XR_002956622:NM_002389:NM_172361:XR_002956621:XM_011509563 | 120:40:186:41:1:3:2:1:0:0 | 2:0:0:0:0:0:0:0:0:0 | 7:0:0:0:0:1:0:0:0:0 | 0:0:0:0:0:0:0:0:0:0 | 78:11:6:2:2:2:0:0:2:0 | 0:0:0:0:0:0:0:0:0:0 | 50:41:177:127:0:11:0:4:2:0 | 0:0:0:0:0:0:0:0:0:0 | 45:24:81:39:0:6:0:4:3:0 | 0:0:0:0:0:0:0:0:0:0 |
| NM_025179 | 0 | 0 | 0 | 0 | 0 | 2 | 1 | 0 | 0 | 0 |
| NM_016448:NM_001286230:NM_001286229 | 0:00:00 | 4:00:00 | 0:00:00 | 0:00:00 | 0:00:00 | 2:00:00 | 0:00:00 | 9:00:00 | 0:00:00 | 11:00:00 |
| NM_016448:NM_001286230:NM_001286229 | 0:00:00 | 5:00:00 | 0:00:00 | 0:00:00 | 0:00:00 | 2:00:00 | 0:00:00 | 11:00:00 | 0:00:00 | 15:00:00 |
| NM_015471 | 0 | 0 | 0 | 0 | 0 | 0 | 0 | 0 | 0 | 0 |
| NM_020197 | 0 | 0 | 0 | 0 | 0 | 0 | 0 | 0 | 0 | 2 |
| XM_017000086 | 0 | 0 | 0 | 0 | 0 | 1 | 0 | 0 | 0 | 0 |
| XR_002959345:NM_001300769:XR_001736968 | 0:00:00 | 0:00:00 | 0:00:00 | 0:00:00 | 0:00:00 | 1:01:00 | 0:00:00 | 1:00:00 | 0:00:00 | 0:00:00 |
| NM_182533:NM_001282670:NM_001256947 | 0:00:00 | 0:00:00 | 1:00:00 | 0:00:00 | 2:00:00 | 0:00:00 | 1:00:00 | 0:00:00 | 4:01:00 | 0:00:00 |
| NM_182533:NM_001256947 | 0:00 | 0:00 | 0:00 | 1:00 | 0:00 | 2:00 | 0:00 | 1:00 | 0:00 | 4:00 |
| XM_017001266 | 0 | 0 | 0 | 0 | 0 | 0 | 0 | 0 | 0 | 1 |
| NM_001136018:NM_001291163 | 24:01:00 | 0:00 | 5:00 | 0:00 | 25:01:00 | 0:01 | 125:09:00 | 0:00 | 7:01 | 0:00 |
| NM_001136018:NM_001291163 | 24:00:00 | 0:00 | 5:00 | 0:00 | 25:01:00 | 0:01 | 122:05:00 | 2:00 | 7:01 | 0:00 |
| NM_001136018:NM_001291163 | 18:01 | 6:00 | 2:01 | 3:00 | 13:02 | 12:01 | 124:08:00 | 0:00 | 7:01 | 0:00 |
| NM_001136018:NM_001291163 | 6:00 | 18:01 | 3:00 | 2:01 | 12:01 | 13:02 | 0:00 | 124:04:00 | 0:00 | 7:00 |
| NM_020997 | 0 | 0 | 0 | 0 | 0 | 1 | 0 | 0 | 1 | 0 |
| NM_021205:NR_037962 | 0:00 | 0:00 | 1:00 | 1:00 | 5:00 | 4:01 | 2:00 | 0:00 | 1:00 | 0:00 |
| XM_024453895:NM_017449:XM_006710442:NM_001309193:NM_004442 | 3:0:0:0:0 | 0:0:0:0:0 | 1:0:0:0:0 | 0:1:0:0:0 | 0:2:0:0:0 | 0:2:1:0:0 | 16:6:1:0:0 | 0:1:1:0:0 | 5:4:1:0:0 | 1:1:0:0:0 |
| XM_024453895:NM_017449:XM_006710442:NM_004442 | 2:0:0:0 | 1:0:0:0 | 1:0:0:0 | 0:1:0:0 | 0:1:0:0 | 0:3:1:0 | 12:1:1:0 | 3:5:1:0 | 5:0:1:0 | 1:5:0:0 |
| XM_006711815:XM_005273262:NM_024525:NM_001122835:XM_005273260 | 0:0:0:0:0 | 1:1:0:0:0 | 0:0:0:0:0 | 0:0:0:0:0 | 0:0:0:0:0 | 0:0:1:0:0 | 0:0:3:0:0 | 0:0:0:0:0 | 0:0:2:0:0 | 0:0:0:0:0 |
| XM_011544304:NM_014236 | 0:00 | 0:00 | 0:00 | 0:00 | 0:00 | 0:00 | 0:00 | 0:00 | 0:00 | 0:00 |
| NM_003196:XM_017002200:XM_024449399:XM_017002201:XM_017002204:XM_006710864:XM_011542053 | 16:1:1:0:0:0:0 | 0:0:0:0:0:0:0 | 129:3:7:1:0:0:0 | 0:0:0:0:0:0:0 | 403:9:33:0:4:1:0 | 0:0:0:0:0:0:0 | 365:13:23:0:2:1:1 | 0:0:0:0:0:0:0 | 80:1:0:0:0:0:0 | 2:0:0:0:0:0:0 |
| NM_003196:XM_017002200:XM_024449399:XM_017002201:XM_017002204:XM_006710864:XM_011542053 | 16:1:1:0:0:0:0 | 0:0:0:0:0:0:0 | 129:3:7:1:0:0:0 | 0:0:0:0:0:0:0 | 402:9:33:0:4:1:0 | 1:0:0:0:0:0:0 | 363:12:26:0:2:1:1 | 2:0:0:0:0:0:0 | 82:1:0:0:0:0:0 | 0:0:0:0:0:0:0 |
| NM_003196:XM_024449399:XM_017002201:XM_017002200:XM_017002204:XM_006710864:XM_011542053 | 16:1:0:0:0:0:0 | 0:0:0:0:0:0:0 | 129:6:1:3:0:0:0 | 1:0:0:0:0:0:0 | 399:25:0:8:2:1:0 | 2:0:0:0:0:0:0 | 363:27:0:9:1:1:1 | 0:0:0:1:0:0:0 | 82:0:0:1:0:0:0 | 0:0:0:0:0:0:0 |
| NM_182972:NM_001077397 | 0:00 | 2:00 | 0:00 | 1:00 | 0:00 | 0:00 | 0:00 | 1:02 | 0:00 | 0:02 |
| NM_002167 | 2 | 3 | 7 | 4 | 3 | 2 | 4 | 0 | 3 | 1 |
| NM_201544:NM_201545:NM_006499 | 0:00:00 | 1:00:00 | 0:00:00 | 0:00:00 | 0:00:00 | 0:00:00 | 0:00:00 | 0:00:00 | 0:00:00 | 0:00:00 |
| NM_201544:NM_201545:NM_006499 | 0:00:00 | 1:00:00 | 0:00:00 | 0:00:00 | 0:00:00 | 0:00:00 | 0:00:00 | 1:00:00 | 0:00:00 | 0:00:00 |
| NM_201544:NM_201545:NM_006499 | 0:00:00 | 1:00:00 | 0:00:00 | 0:00:00 | 0:00:00 | 0:00:00 | 0:00:00 | 1:00:00 | 0:00:00 | 0:00:00 |
| NM_201544:NM_006499 | 0:00 | 1:00 | 0:00 | 0:00 | 0:00 | 0:00 | 0:00 | 1:00 | 0:00 | 0:00 |
| NM_018072 | 0 | 1 | 0 | 3 | 0 | 1 | 0 | 36 | 0 | 7 |
| NM_018072 | 0 | 1 | 0 | 2 | 0 | 1 | 0 | 32 | 0 | 8 |
| NM_018072 | 0 | 1 | 0 | 1 | 0 | 1 | 0 | 26 | 0 | 7 |
| NM_018072 | 0 | 0 | 0 | 0 | 0 | 0 | 0 | 13 | 0 | 3 |
| XM_017000905:NM_000147 | 0:00 | 0:00 | 0:00 | 0:00 | 0:00 | 1:01 | 0:00 | 0:01 | 0:00 | 0:00 |
| XM_017000905:NM_000147 | 0:00 | 0:00 | 0:00 | 0:00 | 0:00 | 1:01 | 0:01 | 0:00 | 0:00 | 0:00 |
| XM_011544342:XM_017002943:XM_011544337:XM_017002942 | 0:0:0:0 | 2:1:0:0 | 0:0:0:0 | 0:0:1:1 | 0:0:0:0 | 0:0:0:0 | 0:0:0:0 | 2:0:0:3 | 0:0:0:0 | 0:0:0:0 |
| NM_001167740:XM_011544253:NM_022743:XM_024449136:XM_024449137:XM_024449166:XM_024449141:XM_011544258:XM_011544257:XM_024449162:XM_011544254:XM_011544261:XM_024449149:XM_024449148:XM_011544259 | 174:20:2:1:6:7:0:0:0:0:0:0:0:0:0 | 1:0:0:0:0:0:0:0:0:0:0:0:0:0:0 | 33:5:1:0:2:2:1:3:0:0:0:0:0:0:0 | 0:0:0:0:0:0:0:0:0:0:0:0:0:0:0 | 96:7:0:0:4:3:0:4:2:1:0:0:0:0:0 | 0:0:0:0:0:0:0:0:0:0:0:0:0:0:0 | 395:53:2:1:14:11:0:3:1:4:1:1:1:0:0 | 0:0:0:0:0:0:0:0:0:0:0:0:0:0:0 | 131:16:1:1:4:3:0:2:1:1:1:0:0:0:0 | 0:0:0:0:0:0:0:0:0:0:0:0:0:0:0 |
| NM_001167740:XM_011544253:NM_022743:XM_024449136:XM_024449137:XM_024449166:XM_024449141:XM_011544258:XM_011544257:XM_024449162:XM_011544254:XM_011544261:XM_024449149:XM_024449148:XM_011544259 | 175:20:2:1:6:7:0:0:0:0:0:0:0:0:0 | 0:0:0:0:0:0:0:0:0:0:0:0:0:0:0 | 33:5:1:0:2:2:1:3:0:0:0:0:0:0:0 | 0:0:0:0:0:0:0:0:0:0:0:0:0:0:0 | 96:7:0:0:4:2:0:4:2:1:0:0:0:0:0 | 0:0:0:0:0:0:0:0:0:0:0:0:0:0:0 | 396:53:2:1:14:11:0:3:1:4:1:1:1:0:0 | 0:0:0:0:0:0:0:0:0:0:0:0:0:0:0 | 129:16:1:1:4:2:0:2:1:1:1:0:0:0:0 | 2:0:0:0:0:0:0:0:0:0:0:0:0:0:0 |
| XM_011544253:NM_001167740:XM_024449137:XM_024449166:XM_024449136:NM_022743:XM_011544258:XM_024449149:XM_011544257:XM_011544261:XM_011544254 | 21:176:3:2:1:2:0:0:0:0:0 | 0:0:0:0:0:0:0:0:0:0:0 | 9:34:3:0:0:1:1:0:0:0:0 | 0:0:0:0:0:0:0:0:0:0:0 | 11:97:4:0:0:0:2:0:0:0:0 | 0:0:0:0:0:0:0:0:0:0:0 | 58:399:14:5:1:0:1:1:1:1:0 | 0:0:0:0:0:0:0:0:0:0:0 | 17:133:5:0:1:0:2:0:0:0:1 | 0:0:0:0:0:0:0:0:0:0:0 |
| XM_011544253:NM_001167740:XM_024449137:XM_024449166:XM_024449136:NM_022743:XM_011544258:XM_024449149:XM_011544257:XM_011544261:XM_011544254 | 22:176:3:2:1:2:0:0:0:0:0 | 0:0:0:0:0:0:0:0:0:0:0 | 9:33:3:0:0:1:1:0:0:0:0 | 0:1:0:0:0:0:0:0:0:0:0 | 11:97:4:0:0:0:2:0:0:0:0 | 0:0:0:0:0:0:0:0:0:0:0 | 58:397:14:5:1:0:1:1:1:1:0 | 2:2:0:0:0:0:0:0:0:0:0 | 17:133:5:0:1:0:2:0:0:0:1 | 0:0:0:0:0:0:0:0:0:0:0 |
| XM_011544253:NM_001167740:XM_024449137:XM_024449166:XM_024449136:NM_022743:XM_011544258:XM_024449149:XM_011544257:XM_011544261:XM_011544254 | 24:176:2:2:1:1:0:0:0:0:0 | 0:0:0:0:0:0:0:0:0:0:0 | 9:34:3:0:0:1:1:0:0:0:0 | 0:0:0:0:0:0:0:0:0:0:0 | 11:97:3:0:0:0:2:0:0:0:0 | 0:0:0:0:0:0:0:0:0:0:0 | 65:397:8:5:1:0:3:1:1:1:0 | 0:2:0:0:0:0:0:0:0:0:0 | 18:133:3:0:1:0:2:0:0:0:1 | 0:0:0:0:0:0:0:0:0:0:0 |
| XM_011544253:NM_001167740:XM_024449137:XM_024449166:XM_024449136:NM_022743:XM_011544258:XM_024449149:XM_011544257:XM_011544261:XM_011544254 | 24:176:2:2:1:1:0:0:0:0:0 | 0:0:0:0:0:0:0:0:0:0:0 | 9:34:3:0:0:1:1:0:0:0:0 | 0:0:0:0:0:0:0:0:0:0:0 | 11:97:3:0:0:0:2:0:0:0:0 | 0:0:0:0:0:0:0:0:0:0:0 | 64:398:8:5:1:0:3:1:1:1:0 | 1:1:0:0:0:0:0:0:0:0:0 | 18:133:3:0:1:0:2:0:0:0:1 | 0:0:0:0:0:0:0:0:0:0:0 |
| XM_011544253:NM_001167740:XM_024449137:XM_024449166:NM_022743:XM_024449136:XM_011544258:XM_024449149:XM_011544261:XM_011544254:XM_024449162 | 7:68:1:1:0:1:0:0:0:0:0 | 17:108:2:1:1:0:0:0:0:0:0 | 5:16:0:0:1:0:1:0:0:0:0 | 4:16:0:0:0:0:0:0:0:0:0 | 8:43:2:0:0:0:0:0:0:0:0 | 3:54:0:0:0:0:1:0:0:0:0 | 0:0:0:0:0:0:0:0:0:0:0 | 62:393:4:6:0:1:3:1:1:0:0 | 0:0:0:0:0:0:0:0:0:0:0 | 18:131:0:0:0:1:2:0:0:1:0 |
| NM_020394:NR_037892 | 0:00 | 0:00 | 1:01 | 0:00 | 0:01 | 0:00 | 0:03 | 0:00 | 0:01 | 0:02 |
| NM_020394:NR_037892 | 0:00 | 0:00 | 0:00 | 1:01 | 0:00 | 0:01 | 0:00 | 0:03 | 0:00 | 0:02 |
| NR_135780:NM_020317:XR_946709:NR_135789:XM_011541797:NR_135783:NR_135782 | 0:0:0:0:0:0:0 | 1:0:0:0:0:0:0 | 0:0:0:0:0:0:0 | 0:0:0:0:0:0:0 | 0:0:0:0:0:0:0 | 0:0:0:0:0:0:0 | 0:0:0:0:0:0:0 | 0:1:1:0:0:0:0 | 0:0:0:0:0:0:0 | 0:0:0:1:0:0:0 |
| NM_003820:XR_426638:XR_001737513:XM_017002720:XR_001737511:XR_001737512 | 0:1:0:0:0:0 | 3:2:0:0:0:0 | 4:2:2:0:0:0 | 4:1:1:0:0:0 | 1:0:2:1:0:0 | 0:0:1:0:0:0 | 7:2:3:0:2:0 | 0:0:0:0:0:0 | 5:1:2:0:0:0 | 0:0:0:0:0:0 |
| NM_183008:NM_145345 | 0:00 | 0:00 | 0:00 | 0:00 | 0:00 | 2:00 | 0:00 | 0:00 | 0:00 | 1:00 |
| NM_001803 | 0 | 3 | 0 | 1 | 0 | 2 | 0 | 1 | 0 | 0 |
| NM_001803 | 0 | 3 | 0 | 1 | 0 | 2 | 0 | 1 | 0 | 0 |
| NM_205861:NM_001243564:NM_024887:NM_001319959:NM_001243565 | 2:0:1:0:0 | 6:1:1:0:0 | 8:1:4:0:0 | 12:0:0:0:0 | 27:4:7:0:0 | 15:3:9:0:0 | 22:4:6:3:0 | 0:0:0:0:0 | 7:3:1:1:0 | 0:0:0:0:0 |
| NM_006142 | 114 | 1 | 289 | 1 | 437 | 1 | 291 | 1 | 190 | 0 |
| NM_006142 | 112 | 1 | 292 | 1 | 440 | 2 | 294 | 0 | 190 | 0 |
| NM_023923:NM_001350161:NM_001350158:NM_001048183:NM_001350160 | 1:1:0:0:0 | 1:0:0:0:0 | 1:0:0:0:0 | 3:0:0:0:0 | 1:0:0:4:0 | 1:0:2:1:0 | 0:0:0:0:0 | 7:0:0:0:0 | 1:0:0:1:1 | 0:2:0:1:0 |
| XM_011541951:NM_005626 | 0:00 | 0:00 | 0:00 | 0:00 | 1:00 | 0:00 | 0:00 | 0:00 | 0:00 | 0:00 |
| XM_011541951:NM_005626 | 0:00 | 0:00 | 0:00 | 0:00 | 1:00 | 0:00 | 0:00 | 0:00 | 0:00 | 0:00 |
| NM_005626:XM_011541951 | 0:00 | 0:00 | 0:00 | 0:00 | 0:00 | 0:00 | 0:00 | 0:00 | 0:00 | 0:00 |
| NM_016011 | 0 | 0 | 0 | 0 | 0 | 0 | 0 | 0 | 0 | 0 |
| NM_004102 | 0 | 0 | 0 | 0 | 1 | 0 | 0 | 0 | 0 | 0 |
| NM_178865:NM_001199039 | 32:00:00 | 0:00 | 103:03:00 | 0:00 | 249:03:00 | 2:01 | 97:02:00 | 0:00 | 65:00:00 | 0:00 |
| NM_178865:NM_001199039:NM_001199037:NM_018565 | 32:0:0:0 | 0:0:0:0 | 110:3:0:0 | 2:0:0:0 | 264:6:1:2 | 2:0:0:0 | 115:6:0:3 | 1:0:0:0 | 71:1:0:0 | 0:0:0:0 |
| XR_946546 | 0 | 1 | 0 | 1 | 0 | 0 | 0 | 0 | 0 | 0 |
| XM_017002054:XM_011541950 | 228:05:00 | 0:00 | 213:04:00 | 2:00 | 221:02:00 | 1:00 | 1115:07:00 | 5:00 | 358:01:00 | 0:00 |
| XM_017002054:XM_011541950 | 228:05:00 | 0:00 | 213:04:00 | 3:00 | 222:02:00 | 1:00 | 1117:07:00 | 3:00 | 358:01:00 | 0:00 |

| | | | | | | | | | | |
|---|---|---|---|---|---|---|---|---|---|---|
| XM_017002054:XM_011541950 | 229:05:00 | 1:00 | 219:04:00 | 0:00 | 224:02:00 | 1:00 | 1122:07:00 | 0:00 | 357:01:00 | 0:00 |
| NR_136702:XM_005271115:XM_017002054:XM_011541950:NR_136703 | 30:19:230:5:0 | 0:0:0:0:0 | 57:19:220:5:0 | 0:0:0:0:0 | 18:7:224:2:0 | 0:0:0:0:0 | 94:50:1116:7:1 | 0:1:2:0:0 | 24:15:357:3:0 | 0:0:0:0:0 |
| NR_136702:XM_005271115:XM_017002054:XM_011541950:NR_136703 | 30:19:231:5:0 | 0:0:0:0:0 | 57:19:221:5:0 | 0:0:0:0:0 | 18:7:224:2:0 | 0:0:0:0:0 | 94:51:1122:7:1 | 0:0:1:0:0 | 24:15:357:3:0 | 0:0:0:0:0 |
| NR_136702:XM_005271115:XM_017002054:XM_011541950:NR_136703 | 30:19:227:5:0 | 0:0:0:0:0 | 53:16:215:5:0 | 0:0:3:0:0 | 18:7:222:2:0 | 0:0:0:0:0 | 94:50:1102:7:1 | 0:0:3:0:0 | 24:15:354:3:0 | 0:0:0:0:0 |
| NR_136702:XM_005271115:XM_017002054:XM_011541950:NR_136703 | 30:19:227:5:0 | 0:0:0:0:0 | 52:16:215:5:0 | 0:0:15:0:0 | 18:7:222:1:0 | 0:0:0:0:0 | 92:49:1094:7:1 | 1:0:8:0:0 | 23:15:352:3:0 | 1:0:1:0:0 |
| NR_136702:XM_005271115:XM_017002054:XM_011541950:NR_136703 | 30:19:225:5:0 | 0:0:0:0:0 | 51:16:258:5:0 | 0:0:2:0:0 | 17:7:221:1:0 | 0:0:0:0:0 | 92:49:1104:7:1 | 0:0:2:0:0 | 24:15:349:3:0 | 0:0:0:0:0 |
| NR_136702:XM_017002054:XM_005271115:XM_011541950:NR_136703 | 29:215:18:4:0 | 1:2:0:0:0 | 51:248:14:5:0 | 0:1:0:0:0 | 16:214:7:1:0 | 0:0:0:0:0 | 91:1059:48:6:1 | 0:4:0:0:0 | 23:334:15:3:0 | 0:0:0:0:0 |
| NR_136702:XM_005271115:XM_017002054:XM_011541950:NR_136703 | 29:18:212:4:0 | 0:0:0:0:0 | 47:10:184:3:0 | 0:0:0:0:0 | 14:6:192:1:0 | 0:0:1:0:0 | 84:41:912:6:1 | 0:0:3:0:0 | 20:15:313:3:0 | 0:0:0:0:0 |
| NM_017821 | 0 | 0 | 0 | 0 | 0 | 0 | 0 | 0 | 0 | 0 |
| NM_001184979:NM_004552 | 310:88 | 0:00 | 525:225 | 1:00 | 298:114 | 0:00 | 1170:1673 | 1:03 | 388:538 | 0:02 |
| NM_001184979:NM_004552 | 308:87 | 0:00 | 517:222 | 0:02 | 296:114 | 0:01 | 1156:1660 | 2:00 | 384:539 | 1:00 |
| XM_005270745 | 0 | 0 | 0 | 0 | 0 | 0 | 0 | 0 | 0 | 0 |
| NM_001350478:NM_006367:NM_001350477:NM_001105530 | 0:0:0:0 | 0:0:0:0 | 0:0:0:0 | 0:0:0:0 | 0:0:0:0 | 1:0:0:0 | 0:0:0:0 | 0:0:0:0 | 0:0:0:0 | 0:2:1:0 |
| NM_001350478:NM_006367:NM_001350477:NM_001105530 | 0:0:0:0 | 0:0:0:0 | 0:0:0:0 | 0:0:0:0 | 0:0:0:0 | 1:0:0:0 | 0:0:0:0 | 0:0:0:0 | 0:0:0:0 | 0:2:1:0 |
| NM_001350478:NM_006367:NM_001350477:NM_001105530 | 0:0:0:0 | 0:0:0:0 | 0:0:0:0 | 0:0:0:0 | 0:0:0:0 | 1:0:0:0 | 0:0:0:0 | 0:0:0:0 | 0:0:0:0 | 0:2:1:0 |
| NM_001350478:NM_006367:NM_001350477:NM_001105530 | 0:0:0:0 | 0:0:0:0 | 0:0:0:0 | 0:0:0:0 | 0:0:0:0 | 1:0:0:0 | 0:0:0:0 | 0:0:0:0 | 0:0:0:0 | 0:2:1:0 |
| NM_001350478:NM_006367:NM_001350477:NM_001105530 | 0:0:0:0 | 0:0:0:0 | 0:0:0:0 | 0:0:0:0 | 0:0:0:0 | 1:0:0:0 | 0:0:0:0 | 0:0:0:0 | 0:0:0:0 | 0:2:1:0 |
| NM_001350478:NM_006367:NM_001350477:NM_001105530 | 0:0:0:0 | 0:0:0:0 | 0:0:0:0 | 0:0:0:0 | 0:0:0:0 | 1:0:0:0 | 0:0:0:0 | 0:0:0:0 | 0:0:0:0 | 0:2:1:0 |
| NM_014223:NM_001142587:XM_006710661:XM_024447357:NM_001142588:XM_017001365:XM_011541517:XM_017001368:XM_017001369:NM_001308115:XM_017001364:XM_024447356:NM_001142590:XM_024447355:XM_017001367 | 20:1:2:0:0:0:0:0:0:0:0:0:0:0:0 | 0:0:0:0:0:0:0:0:0:0:0:0:0:0:0 | 24:0:5:2:1:1:1:1:0:0:0:0:0:0:0 | 0:0:0:0:0:0:0:0:0:0:0:0:0:0:0 | 45:2:8:1:1:1:0:1:1:2:1:0:0:0:0 | 0:0:0:0:0:0:0:0:0:0:0:0:0:0:0 | 188:8:22:16:14:2:1:6:4:4:3:1:2:0:0 | 2:0:1:0:0:0:0:0:0:0:0:0:0:0:0 | 42:1:2:0:0:0:0:0:1:1:1:0:0:1:0 | 0:0:0:0:0:0:0:0:0:0:0:0:0:0:0 |
| NM_033553 | 0 | 0 | 0 | 34 | 0 | 60 | 0 | 0 | 1 | 1 |
| NM_152298:NM_002482:XM_005270889:XM_017001356:XM_005270888:XM_011541509:NM_001195193 | 29:7:11:2:1:0:0 | 0:0:0:0:0:0:0 | 32:2:5:34:0:2:0 | 0:0:0:0:0:0:0 | 39:2:3:2:0:0:0 | 0:0:0:0:0:0:0 | 173:60:27:97:8:8:1 | 0:0:0:1:0:0:0 | 111:14:8:13:3:1:0 | 0:0:0:0:0:0:0 |
| NM_152298:NM_002482:XM_005270889:XM_017001356:XM_005270888:XM_011541509:NM_001195193 | 29:7:11:2:1:0:0 | 0:0:0:0:0:0:0 | 32:2:5:36:0:1:0 | 0:0:0:0:0:0:0 | 39:2:3:2:0:0:0 | 0:0:0:0:0:0:0 | 172:60:27:98:7:11:1 | 0:0:0:1:1:0:0 | 112:15:6:14:2:2:0 | 0:0:0:0:0:0:0 |
| NM_152298:NM_002482:XM_005270889:XM_017001356:XM_005270888:XM_011541509:NM_001195193 | 29:7:11:2:1:0:0 | 0:0:0:0:0:0:0 | 32:2:5:36:0:1:0 | 0:0:0:0:0:0:0 | 39:2:3:2:0:0:0 | 0:0:0:0:0:0:0 | 172:60:28:98:8:9:1 | 0:0:0:1:0:1:0 | 112:15:6:14:2:2:0 | 0:0:0:0:0:0:0 |
| NM_002482:XM_005270888:XM_011541509:NM_001195193 | 5:1:0:0 | 0:0:0:0 | 2:0:0:0 | 0:0:0:0 | 2:0:0:0 | 0:0:0:0 | 46:8:13:1 | 2:0:0:0 | 15:2:1:0 | 0:0:0:0 |
| NM_002482:XM_005270888:XM_011541509:NM_001195193 | 5:1:0:0 | 0:0:0:0 | 1:0:0:0 | 1:0:0:0 | 2:0:0:0 | 0:0:0:0 | 44:9:11:1 | 1:0:0:0 | 14:2:0:0 | 0:0:0:0 |
| NM_152298:NM_002482:XM_005270889:XM_005270888:XM_017001356:XM_011541509:NM_001195193 | 28:5:11:1:0:0:0 | 0:0:0:0:0:0:0 | 28:1:5:0:22:1:0 | 0:0:0:0:2:0:0 | 39:2:3:0:1:0:0 | 0:0:0:0:0:0:0 | 161:29:28:17:75:15:1 | 3:2:1:0:3:3:0 | 109:11:6:3:11:0:0 | 0:0:0:0:0:0:0 |
| NM_152298:NM_002482:XM_005270889:XM_005270888:XM_017001356:XM_011541509:NM_001195193 | 27:5:11:1:0:0:0 | 0:0:0:0:0:0:0 | 28:1:6:0:22:1:0 | 0:0:0:0:0:0:0 | 39:2:3:0:1:0:0 | 0:0:0:0:0:0:0 | 160:29:28:17:75:15:1 | 1:0:0:0:0:0:0 | 108:11:6:3:12:0:0 | 0:0:0:0:0:0:0 |
| NM_152298:NM_002482:XM_005270889:XM_005270888:XM_017001356:XM_011541509:NM_001195193 | 27:5:12:1:0:0:0 | 0:0:0:0:0:0:0 | 28:1:7:0:22:1:0 | 0:0:0:0:0:0:0 | 38:2:3:0:0:0:0 | 1:0:0:0:0:0:0 | 160:29:32:17:69:15:1 | 0:0:1:0:1:0:0 | 108:11:7:3:11:0:0 | 0:0:0:0:0:0:0 |
| NM_017739 | 0 | 0 | 0 | 0 | 0 | 0 | 0 | 2 | 0 | 1 |
| NM_006004:NM_001297565 | 9:00 | 0:00 | 7:00 | 0:00 | 13:00 | 0:00 | 12:00 | 8:02 | 10:00 | 8:00 |
| NM_001441 | 0 | 0 | 0 | 0 | 1 | 0 | 1 | 1 | 0 | 0 |
| NM_022745 | 0 | 0 | 0 | 0 | 0 | 1 | 0 | 1 | 0 | 2 |
| NM_016308:NR_046394 | 0:00 | 0:00 | 1:00 | 0:00 | 4:00 | 0:00 | 0:00 | 0:00 | 0:00 | 0:00 |
| XM_017000618:XM_017000620:NM_001981:XM_017000617:XM_017000615:XM_017000619:XM_017000616:XM_005270618 | 1:28:12:0:1:0:0:0 | 1:17:5:1:0:0:0:0 | 0:8:0:0:0:0:0:0 | 0:10:0:0:0:0:0:0 | 2:32:9:0:0:2:0:0 | 2:15:6:0:0:1:0:0 | 0:1:5:1:0:0:0:0 | 0:119:4:0:0:2:0:0 | 0:1:0:0:0:0:0:0 | 3:78:4:0:0:1:0:0 |
| NM_024586:NM_148908:NR_036662:NM_148904:NM_001330580:NM_148907:XM_006710320:NM_148906:XM_011540604:XM_017000224:XM_006710324:XM_006710323:XM_006710318:NM_148909:XM_006710325:XM_011540599:XM_011540600:XR_001736955:XR_001736954:XM_017000221:NM_001350210:XM_017000218:NM_001350208:XM_006710326 | 12:5:5:10:7:2:2:0:0:0:0:0:0:0:0:0:0:0:0:0:0:0:0:0 | 0:0:0:0:0:0:0:0:0:0:0:0:0:0:0:0:0:0:0:0:0:0:0:0 | 3:1:2:1:3:1:4:2:2:8:0:0:0:0:0:0:0:0:0:0:0:0:0:0 | 0:0:0:0:0:0:0:0:0:0:0:0:0:0:0:0:0:0:0:0:0:0:0:0 | 21:1:2:1:9:1:5:2:1:2:1:1:0:0:0:0:0:0:0:0:0:0:0:0 | 0:0:0:0:0:0:0:0:0:0:0:0:0:0:0:0:0:0:0:0:0:0:0:0 | 16:8:6:16:27:2:16:0:20:84:3:2:2:2:1:1:2:3:1:9:0:0:0:0 | 0:0:0:0:0:0:0:0:0:0:0:0:0:0:0:0:0:0:0:0:0:0:0:0 | 15:1:2:11:20:1:4:1:0:1:4:1:0:0:0:0:0:0:0:0:3:0:0:0 | 0:0:0:0:0:0:0:0:1:0:0:0:0:0:0:0:0:0:0:0:0:0:0:0 |
| NM_024586:NM_148908:NR_036662:NM_148904:NM_001330580:NM_148907:XM_006710320:XM_017000224:XR_001736955:NM_148906:XM_011540604:XM_006710324:XM_006710323:XR_001736954:XM_006710318:NM_148909:XM_006710325:XM_011540599:XM_011540600:XM_017000221:NM_001350210:XM_017000218:NM_001350208:XM_006710326 | 12:5:5:10:7:2:2:3:2:0:0:0:0:0:0:0:0:0:0:0:0:0:0:0 | 0:0:0:0:0:0:0:0:0:0:0:0:0:0:0:0:0:0:0:0:0:0:0:0 | 3:1:2:1:3:1:4:11:6:2:2:0:0:0:0:0:0:0:0:0:0:0:0:0 | 0:0:0:0:0:0:0:0:0:0:0:0:0:0:0:0:0:0:0:0:0:0:0:0 | 21:1:2:1:9:1:5:3:1:2:1:1:1:0:0:0:0:0:0:0:0:0:0:0 | 0:0:0:0:0:0:0:0:0:0:0:0:0:0:0:0:0:0:0:0:0:0:0:0 | 15:7:11:16:26:2:15:125:63:0:20:3:2:13:2:2:1:1:2:9:0:0:0:0 | 0:0:0:0:1:0:0:0:0:0:0:0:0:1:0:0:0:0:0:0:0:0:0:0 | 15:1:2:11:19:1:4:3:1:1:0:4:1:0:0:0:0:0:0:0:3:0:0:0 | 0:0:0:0:0:0:0:0:0:0:0:0:0:0:0:0:0:0:0:0:0:0:0:0 |
| NM_024586:NM_148908:NR_036662:NM_148904:NM_001330580:NM_148907:XM_006710320:XM_017000224:XR_001736955:NM_148906:XM_011540604:XM_006710324:XM_006710323:XM_006710318:NM_148909:XM_006710325:XM_011540599:XM_011540600:XR_001736954:XM_017000221:NM_001350210:XM_017000218:NM_001350208:XM_006710326 | 12:5:5:10:7:2:2:3:2:0:0:0:0:0:0:0:0:0:0:0:0:0:0:0 | 0:0:0:0:0:0:0:0:0:0:0:0:0:0:0:0:0:0:0:0:0:0:0:0 | 3:1:2:1:3:1:4:11:5:1:2:0:0:0:0:0:0:0:0:0:0:0:0:0 | 0:0:0:0:0:0:0:0:0:0:0:0:0:0:0:0:0:0:0:0:0:0:0:0 | 21:1:2:1:9:1:5:3:0:2:1:1:1:0:0:0:0:0:0:0:0:0:0:0 | 0:0:0:0:0:0:0:0:0:0:0:0:0:0:0:0:0:0:0:0:0:0:0:0 | 14:7:11:16:27:2:15:123:61:0:20:3:2:2:2:1:1:2:14:9:0:0:0:0 | 0:0:0:0:0:0:0:2:0:0:0:0:0:0:0:0:0:0:0:0:0:0:0:0 | 15:1:2:11:19:1:4:3:1:1:0:4:1:0:0:0:0:0:0:0:3:0:0:0 | 0:0:0:0:0:0:0:0:0:0:0:0:0:0:0:0:0:0:0:0:0:0:0:0 |
| NM_016491 | 0 | 0 | 0 | 0 | 0 | 2 | 0 | 0 | 1 | 0 |
| XM_017001646:XM_024448207:XR_001737287:NM_018291:XM_024448229:XM_017001645:XM_017001652:XM_017001679:NM_001350793:NM_001350794:XM_017001656:XM_017001657:XM_011541730:NM_001244714:XR_001737285:NM_001113411:NR_103473:XM_017001649:XM_017001673:XM_024448176:NM_001350799:NM_001350797:XM_024448220:XM_017001655 | 0:1:0:1:0:0:0:0:0:0:0:0:0:0:0:0:0:0:0:0:0:0:0:0 | 1:1:1:0:0:0:0:0:0:0:0:0:0:0:0:0:0:0:0:0:0:0:0:0 | 0:0:0:1:0:1:0:0:0:0:0:0:0:0:0:0:0:0:0:0:0:0:0:0 | 0:0:0:2:1:1:1:1:0:0:0:0:0:0:0:0:0:0:0:0:0:0:0:0 | 1:2:0:3:0:2:1:0:1:0:0:0:0:0:0:0:0:0:0:0:0:0:0:0 | 1:0:0:5:0:2:0:1:0:0:0:0:0:0:0:0:0:0:0:0:0:0:0:0 | 5:0:0:30:2:6:6:0:0:1:0:0:0:0:1:2:0:1:0:1:2:1:1:0 | 7:1:3:35:0:7:11:0:0:7:1:1:1:1:1:2:1:1:1:0:0:0:0:0 | 0:1:1:2:0:0:0:0:0:0:0:0:0:0:0:0:0:0:0:0:0:0:0:0 | 1:2:0:6:0:3:2:0:0:0:0:0:0:0:0:0:0:1:0:0:0:0:0:0 |
| NM_000983 | 6298 | 8 | 5908 | 30 | 4823 | 15 | 25351 | 62 | 9162 | 12 |
| NM_000983 | 6295 | 16 | 5916 | 31 | 4828 | 11 | 25352 | 69 | 9168 | 10 |
| NM_000983 | 6313 | 4 | 5983 | 8 | 4846 | 4 | 25454 | 20 | 9173 | 11 |
| NM_000983 | 6540 | 4 | 6771 | 11 | 4956 | 3 | 26601 | 15 | 9473 | 2 |
| NM_000983 | 6588 | 3 | 7050 | 7 | 4990 | 2 | 27237 | 23 | 9530 | 4 |
| NM_000983 | 6588 | 2 | 7048 | 5 | 4979 | 11 | 27217 | 33 | 9516 | 12 |
| NM_000983 | 6578 | 3 | 7043 | 2 | 4962 | 2 | 27192 | 12 | 9508 | 4 |
| NM_000983 | 6565 | 6 | 7024 | 7 | 4955 | 5 | 27163 | 24 | 9499 | 6 |
| NM_176877:XM_024448614:XM_011540465:XR_001736900:XM_017000001:XR_002957157:XM_011540469:NM_001350145 | 1:1:0:0:0:0:0:0 | 0:0:0:0:0:0:0:0 | 1:0:0:0:0:0:0:0 | 0:0:0:0:0:0:0:0 | 1:1:1:0:0:0:0:0 | 0:0:0:0:0:0:0:0 | 3:0:2:0:0:0:1:1 | 5:0:0:2:1:1:0:0 | 0:0:0:0:0:0:0:0 | 0:0:2:0:0:0:0:0 |
| XM_024448614:NM_176877:XR_002957157:XM_011540465:NM_001350145:XR_001736900:XM_017000001:XM_011540469:XM_016999999 | 0:2:1:0:0:0:0:0:0 | 1:0:0:0:0:0:0:0:0 | 0:0:0:0:0:0:0:0:0 | 0:1:0:0:0:0:0:0:0 | 1:1:0:0:0:0:0:0:0 | 0:0:0:1:1:0:0:0:0 | 0:4:0:3:2:0:0:1:0 | 0:6:0:0:0:1:1:0:0 | 0:0:0:0:0:0:0:0:0 | 0:0:0:2:0:0:0:0:0 |
| NM_007274:NM_181866:NM_181865:NM_181864 | 45:28:8:0 | 0:0:0:0 | 20:52:33:3 | 0:1:1:0 | 57:101:76:4 | 0:1:0:0 | 32:40:9:2 | 0:1:0:0 | 114:48:13:0 | 0:1:0:0 |
| NM_007274:NM_181865:NM_181866:NM_181864 | 48:8:13:0 | 1:0:0:0 | 20:34:37:3 | 0:0:1:0 | 57:76:66:4 | 0:0:0:0 | 35:9:10:2 | 0:0:0:0 | 116:13:25:0 | 1:0:1:0 |
| NM_007274:NM_181865:NM_181866:NM_181864 | 49:8:11:0 | 0:0:0:0 | 20:34:36:3 | 0:0:2:0 | 57:76:66:4 | 0:0:0:0 | 35:9:10:2 | 0:0:0:0 | 117:13:26:0 | 0:0:0:0 |
| NM_005341:NM_001278648 | 0:00 | 0:00 | 0:00 | 0:00 | 0:00 | 0:00 | 0:00 | 0:00 | 1:00 | 0:00 |
| NM_005478 | 0 | 0 | 0 | 0 | 0 | 0 | 0 | 0 | 0 | 0 |
| NM_018841:XM_024448405:XM_017001809 | 22:01:00 | 0:00:00 | 60:06:02 | 0:00:00 | 377:33:04 | 2:00:00 | 205:09:02 | 0:00:00 | 31:02:01 | 0:00:00 |

| | | | | | | | | | | |
|---|---|---|---|---|---|---|---|---|---|---|
| XM_011542191:XM_011542192:NM_001002292:NM_024911:XM_017002390:NM_001193334 | 16:7:4:32:6:0 | 0:0:0:0:0:0 | 11:5:5:13:27:0 | 0:0:0:0:0:0 | 6:2:0:24:7:0 | 0:0:0:0:0:0 | 30:13:3:24:71:1 | 22:10:6:19:58:1 | 11:0:1:8:1:1 | 2:2:0:5:6:0 |
| NM_001195563:XR_001737066:XR_001737065:NR_146203:XM_017000780:NM_015215:XM_017000781:NM_001242701:NR_038934:XR_001737061:XR_001737063:NR_146204:XR_001737062 | 165:2:22:1:3:0:0:0:0:0:0:0 | 0:0:0:0:0:0:0:0:0:0:0:0 | 423:11:56:1:2:1:1:3:1:0:0:0:0 | 1:0:0:0:0:0:0:0:0:0:0:0:0 | 670:5:141:0:0:0:1:4:1:1:1:0:0 | 0:0:0:0:0:0:0:0:0:0:0:0 | 529:10:86:1:0:0:0:2:0:0:0:0:0 | 0:0:0:0:0:0:0:0:0:0:0:0 | 354:4:62:1:1:0:0:4:0:0:0:0:0 | 0:0:0:0:0:0:0:0:0:0:0:0 |
| NM_001195563:XR_001737066:XR_001737065:NR_146203:XM_017000780:NM_015215:XM_017000781:NM_001242701:NR_038934:XR_001737061:XR_001737063:NR_146204:XR_001737062 | 165:2:22:1:3:0:0:0:0:0:0:0 | 0:0:0:0:0:0:0:0:0:0:0:0 | 424:11:56:1:2:1:1:3:1:0:0:0:0 | 0:0:0:0:0:0:0:0:0:0:0:0 | 670:5:141:0:0:0:1:4:1:1:1:0:0 | 1:0:0:0:0:0:0:0:0:0:0:0 | 528:9:86:1:0:0:0:2:0:0:0:0:0 | 1:1:0:0:0:0:0:0:0:0:0:0 | 354:4:62:1:1:0:0:4:0:0:0:0:0 | 0:0:0:0:0:0:0:0:0:0:0:0 |
| XR_001737065:NR_038934:NM_001242701:XR_001737066:NR_146202:NR_146203:XR_001737061:XR_001737063:XR_001737062 | 9:5:1:2:1:0:0:0:0 | 13:2:2:0:0:0:0:0:0 | 26:17:3:0:1:0:0:0:0 | 30:16:0:2:0:1:0:0:0 | 66:39:2:2:2:0:1:1:0 | 82:37:3:2:1:0:0:0:0 | 87:39:5:4:1:0:0:0:0 | 0:0:0:0:0:0:0:0:0 | 64:25:8:1:1:1:0:0:0 | 0:0:0:0:0:0:0:0:0 |
| NM_000016:NM_001286042:NM_001127328:NM_001286044:NM_001286043 | 23:3:1:0:0 | 0:0:0:0:0 | 83:12:1:0:0 | 0:0:0:0:0 | 70:13:1:0:0 | 0:0:0:0:0 | 587:66:30:3:10 | 1:1:0:0:0 | 99:2:0:2:0 | 0:0:0:0:0 |
| NM_001286043:NM_000016:NM_001286042:NM_001127328:NM_001286044 | 1:23:3:1:1 | 0:0:0:0:0 | 2:72:7:1:0 | 0:2:0:0:0 | 0:67:12:1:0 | 0:0:0:0:0 | 24:536:59:28:11 | 0:1:0:0:0 | 0:93:2:0:3 | 0:1:0:0:0 |
| NM_001286043:NM_000016:NM_001286042:NM_001127328:NM_001286044 | 1:22:3:1:1 | 0:0:0:0:0 | 2:61:5:1:4 | 0:0:0:0:0 | 0:63:12:1:3 | 0:0:0:0:0 | 23:505:58:25:40 | 0:1:0:0:1 | 0:88:2:0:2 | 0:0:0:0:0 |
| NM_001286043:NM_000016:NM_001286042:NM_001127328:NM_001286044 | 0:5:1:0:0 | 1:17:2:1:1 | 1:30:4:0:3 | 1:31:1:1:1 | 0:32:2:0:3 | 0:31:10:1:0 | 23:506:58:25:45 | 0:0:0:0:0 | 0:87:2:0:2 | 0:0:0:0:0 |
| NM_001286043:NM_000016:NM_001286042:NM_001127328:NM_001286044 | 1:22:3:1:1 | 0:0:0:0:0 | 2:61:5:1:4 | 0:0:0:0:0 | 0:63:12:1:3 | 0:0:0:0:0 | 23:503:58:25:47 | 0:1:0:0:0 | 0:87:2:0:2 | 0:0:0:0:0 |
| NM_001286043:NM_000016:NM_001286042:NM_001127328:NM_001286044 | 1:22:3:1:1 | 0:0:0:0:0 | 2:61:5:1:5 | 0:0:0:0:0 | 0:63:12:1:3 | 0:0:0:0:0 | 23:498:58:25:54 | 0:2:0:0:0 | 0:87:2:0:3 | 0:0:0:0:0 |
| NM_001286043:NM_000016:NM_001286042:NM_001127328:NM_001286044 | 1:22:3:1:1 | 0:0:0:0:0 | 2:61:5:1:5 | 0:0:0:0:0 | 0:63:12:1:3 | 0:0:0:0:0 | 23:499:58:25:56 | 0:2:0:0:0 | 0:87:2:0:3 | 0:0:0:0:0 |
| NM_000016:NM_001286043:NM_001286042:NM_001127328:NM_001286044 | 21:1:3:1:1 | 1:0:0:0:0 | 61:2:5:1:4 | 0:0:0:0:0 | 63:0:12:1:4 | 0:0:0:0:0 | 493:22:57:24:64 | 3:0:0:0:0 | 87:0:2:0:3 | 0:0:0:0:0 |
| XR_001737516:NR_146539:XR_001737521:XR_001737520:NR_146540:XM_024450628:XR_001737524:XM_017002744:NM_003902:NM_001303433:XM_017002742:XR_946791:XR_001737522:XM_011542393:XR_946790:XR_001737517 | 1:6:1:1:2:1:4:69:150:5:3:10:1:0:0:0 | 0:0:0:0:0:0:0:0:0:0:0:0:0:0:0:0 | 0:0:0:0:1:0:2:50:24:0:4:17:1:1:0:0 | 0:0:0:0:0:0:0:0:0:0:0:0:0:0:0:0 | 0:1:1:0:0:0:1:78:134:1:1:21:3:1:0:0 | 0:0:0:0:0:0:0:0:0:0:0:0:0:0:0:0 | 0:9:0:13:14:9:13:259:242:2:13:292:3:0:3:1 | 0:0:0:0:1:0:0:0:0:0:0:0:0:0:0:0 | 0:7:0:0:0:1:7:85:129:1:3:13:3:0:0:0 | 0:0:0:0:0:0:0:0:0:0:0:0:0:0:0:0 |
| XR_001737516:NR_146539:XR_001737520:NR_146540:XM_024450628:XR_001737524:XM_017002744:NM_003902:NM_001303433:XM_017002742:XR_946791:XM_011542393:XR_001737522:XR_946790:XR_001737517 | 1:6:1:2:1:3:69:149:5:3:10:0:0:0:0 | 0:0:0:0:0:0:0:0:0:0:0:0:0:0:0 | 0:0:0:1:0:3:50:24:0:4:11:1:0:0:0 | 0:0:0:0:0:0:0:0:0:0:0:0:0:0:0 | 0:1:0:0:0:0:78:132:1:1:16:1:3:0:0 | 0:0:0:0:0:0:0:0:0:1:0:0:0:0:0 | 0:9:12:12:8:13:260:240:2:13:285:0:5:3:1 | 0:0:0:0:0:0:0:0:0:3:0:0:0:0:0 | 0:7:0:0:0:5:85:129:1:3:12:0:2:0:0 | 0:0:0:0:0:0:0:0:0:0:0:0:0:0:0 |
| XR_001737516:NR_146539:XR_001737520:NR_146540:XM_024450628:XR_001737524:XM_017002744:NM_003902:NM_001303433:XM_017002742:XR_946791:XM_011542393:XR_001737522:XR_946790:XR_001737517 | 1:6:1:2:1:3:69:149:5:3:10:0:0:0:0 | 0:0:0:0:0:0:0:0:0:0:0:0:0:0:0 | 0:0:0:1:0:3:50:24:0:4:11:1:0:0:0 | 0:0:0:0:0:0:0:0:0:0:0:0:0:0:0 | 0:1:0:0:0:0:77:132:1:1:17:1:3:0:0 | 0:0:0:0:0:0:0:1:0:0:0:0:0:0:0 | 0:9:12:12:8:13:260:240:2:13:287:0:5:3:1 | 0:0:0:0:0:0:0:0:0:1:0:0:0:0:0 | 0:7:0:0:0:5:85:129:1:3:12:0:2:0:0 | 0:0:0:0:0:0:0:0:0:0:0:0:0:0:0 |
| XR_001737516:NR_146539:XR_001737520:NR_146540:XM_024450628:XR_001737524:XM_017002744:NM_003902:NM_001303433:XM_017002742:XR_946791:XM_011542393:XR_001737522:XR_946790:XR_001737517 | 1:6:1:2:1:3:69:149:5:3:8:0:0:0:0 | 0:0:0:0:0:0:0:0:0:0:0:0:0:0:0 | 0:0:0:1:0:3:50:24:0:4:11:1:0:0:0 | 0:0:0:0:0:0:0:0:0:0:0:0:0:0:0 | 0:1:0:0:0:0:78:131:1:1:17:1:3:0:0 | 0:0:0:0:0:0:0:1:0:0:0:0:0:0:0 | 0:9:11:12:8:14:260:240:2:13:287:0:5:3:1 | 0:0:0:0:0:0:0:0:0:1:0:0:0:0:0 | 0:7:0:0:0:4:85:128:1:3:11:0:2:0:0 | 0:0:0:0:0:0:0:0:0:0:0:0:0:0:0 |
| NR_146539:XR_001737520:NR_146540:XM_024450628:XR_001737524:XM_017002744:NM_003902:NM_001303433:XM_017002742:XR_946791:XR_001737522:XM_011542393:XR_946790:XR_001737517 | 6:1:2:1:11:69:145:5:3:6:1:0:0:0 | 0:0:0:0:0:0:0:0:0:0:0:0:0:0 | 0:0:5:0:11:50:22:0:4:6:3:1:0:0 | 0:0:0:0:0:0:0:0:0:0:0:0:0:0 | 2:0:0:0:12:78:127:1:1:11:5:1:0:0 | 0:0:0:0:0:0:0:0:0:0:0:0:0:0 | 13:11:31:3:63:259:231:2:13:187:18:0:2:1 | 0:0:0:0:2:1:0:0:0:1:0:0:0:0 | 7:0:0:0:12:85:125:1:3:7:4:0:0:0 | 0:0:0:0:0:0:0:0:0:0:0:0:0:0 |
| XR_001737524:NR_146539:XR_001737520:NR_146540:XM_024450628:XM_017002744:NM_003902:NM_001303433:XM_017002742:XR_946791:XR_001737522:XM_011542393:XR_946790:XR_001737517 | 27:6:1:6:1:69:145:5:3:7:2:0:0:0 | 4:0:0:0:0:0:0:0:0:0:0:0:0:0 | 19:0:0:9:0:50:22:0:4:4:9:1:0:0 | 0:0:1:0:0:0:0:0:0:0:0:0:0:0 | 24:2:0:3:0:78:127:1:1:9:9:1:0:0 | 3:0:0:0:0:0:0:0:0:0:2:0:0:0 | 117:22:13:50:3:260:232:2:13:172:26:0:2:1 | 7:0:0:2:0:0:0:0:0:1:4:0:0:0 | 31:8:1:5:0:85:124:1:3:7:5:0:0:0 | 3:0:0:1:0:0:0:0:0:0:1:0:0:0 |
| NM_003902:NR_146539:XR_001737520:NR_146540:XM_024450628:XR_001737524:XM_017002744:NM_001303433:XM_017002742:XR_946791:XR_001737522:XM_011542393:XR_946790:XR_001737517 | 143:6:1:6:1:31:69:5:3:7:2:0:0:0 | 2:0:0:0:0:0:0:0:0:0:0:0:0:0 | 22:0:1:9:0:20:50:0:4:4:9:1:0:0 | 0:0:0:0:0:0:0:0:0:0:0:0:0:0 | 127:2:0:3:0:27:78:1:1:9:11:1:0:0 | 0:0:0:0:0:0:0:0:0:0:0:0:0:0 | 230:22:13:52:3:124:260:2:13:163:29:0:2:1 | 1:0:0:0:0:0:0:0:0:1:0:0:0:0 | 124:8:1:6:0:34:85:1:3:6:6:0:0:0 | 0:0:0:0:0:0:0:0:0:0:0:0:0:0 |
| NR_146539:XR_001737520:NR_146540:XM_024450628:XM_017002744:XR_001737524:NM_003902:NM_001303433:XM_017002742:XR_946791:XR_001737522:XM_011542393:XR_001737517 | 6:3:11:1:69:37:146:5:3:5:3:0:0 | 0:0:0:0:0:0:0:0:0:0:0:0:0 | 1:1:10:0:50:28:22:0:4:1:14:1:0 | 0:0:0:0:0:0:0:0:0:0:0:0:0 | 2:2:8:0:78:35:127:1:1:3:12:1:0 | 0:0:0:0:0:0:0:0:0:0:0:0:0 | 24:13:59:3:258:143:226:2:13:38:35:0:1 | 0:0:0:0:2:0:0:0:0:0:0:0:0 | 8:1:9:0:85:36:124:1:3:6:9:0:0 | 0:0:0:0:0:0:0:0:0:0:0:0:0 |
| NM_007034:NM_001317102:NM_001317101:NM_001317100:NM_001317103 | 27:6:1:0:0 | 1:0:0:0:0 | 31:2:0:1:0 | 0:0:0:0:0 | 46:10:1:0:0 | 0:0:0:0:0 | 81:25:7:1:0 | 1:2:0:0:0 | 7:7:1:0:0 | 0:0:1:0:0 |
| NM_025065 | 0 | 0 | 1 | 0 | 0 | 0 | 0 | 1 | 0 | 0 |
| NM_001285 | 0 | 0 | 0 | 1 | 3 | 6 | 0 | 0 | 0 | 0 |
| NM_001285 | 0 | 0 | 2 | 0 | 6 | 3 | 0 | 0 | 0 | 0 |
| NM_001285 | 0 | 0 | 0 | 2 | 0 | 11 | 0 | 0 | 0 | 0 |
| NM_001428:NM_001201483 | 2354:01:00 | 0:00 | 1505:01:00 | 1:00 | 1581:00:00 | 2:00 | 5157:02:00 | 2:00 | 4334:01:00 | 1:00 |
| NM_001428:NM_001201483 | 2350:01:00 | 4:00 | 1506:01:00 | 1:00 | 1583:00:00 | 1:00 | 5151:02:00 | 8:00 | 4333:01:00 | 1:00 |
| NM_001428:NM_001201483 | 2352:01:00 | 1:00 | 1488:01:00 | 4:00 | 1582:00:00 | 0:00 | 5135:02:00 | 3:00 | 4331:01:00 | 1:00 |
| NM_001428:NM_001201483 | 2338:01:00 | 1:00 | 1454:01:00 | 4:00 | 1568:00:00 | 3:00 | 5080:02:00 | 3:00 | 4291:01:00 | 7:00 |
| NM_001428:NM_001201483 | 2330:01:00 | 2:00 | 1421:01:00 | 5:00 | 1564:00:00 | 2:00 | 5044:02:00 | 6:00 | 4276:01:00 | 5:00 |
| NM_001428:NM_001201483 | 2304:01:00 | 2:00 | 1366:01:00 | 2:00 | 1545:00:00 | 0:00 | 4993:02:00 | 0:00 | 4223:01:00 | 2:00 |
| NM_001428:NM_001201483 | 2271:01:00 | 4:00 | 1318:01:00 | 3:00 | 1520:00:00 | 1:00 | 4928:02:00 | 6:00 | 4191:01:00 | 6:00 |
| NM_001428:NM_001201483 | 2268:01:00 | 4:00 | 1314:01:00 | 6:00 | 1514:00:00 | 3:00 | 4926:02:00 | 8:00 | 4182:01:00 | 12:00 |
| NM_001428:NM_001201483 | 2250:01:00 | 2:00 | 1279:01:00 | 5:00 | 1499:00:00 | 3:00 | 4893:02:00 | 5:00 | 4141:01:00 | 2:00 |
| NM_001428:NM_001201483 | 2248:01:00 | 2:00 | 1271:01:00 | 5:00 | 1495:00:00 | 2:00 | 4888:02:00 | 8:00 | 4134:01:00 | 3:00 |
| NM_001428:NM_001201483 | 2229:01:00 | 4:00 | 1234:01:00 | 4:00 | 1480:00:00 | 1:00 | 4851:02:00 | 5:00 | 4088:01:00 | 4:00 |
| NM_001428:NM_001201483 | 2241:01:00 | 2:00 | 1229:01:00 | 3:00 | 1472:00:00 | 1:00 | 4855:02:00 | 2:00 | 4064:01:00 | 6:00 |
| NM_001428:NM_001201483 | 2224:01:00 | 3:00 | 1223:01:00 | 0:00 | 1460:00:00 | 3:00 | 4804:02:00 | 0:00 | 4050:01:00 | 7:00 |
| NM_001428:NM_001201483 | 2215:01:00 | 3:00 | 1196:01:00 | 3:00 | 1447:00:00 | 0:00 | 4747:02:00 | 8:00 | 4033:01:00 | 4:00 |
| NM_001428:NM_001201483 | 2212:01:00 | 3:00 | 1195:01:00 | 3:00 | 1444:00:00 | 0:00 | 4740:02:00 | 6:00 | 4029:01:00 | 4:00 |
| NM_001428:NM_001201483 | 2208:01:00 | 1:00 | 1193:01:00 | 3:00 | 1442:00:00 | 1:00 | 4731:02:00 | 10:00 | 4023:01:00 | 4:00 |
| NM_001428:NM_001201483 | 2210:01:00 | 0:00 | 1195:01:00 | 2:00 | 1443:00:00 | 0:00 | 4722:02:00 | 8:00 | 4017:01:00 | 8:00 |
| NM_001428:NM_001201483 | 2211:01:00 | 6:00 | 1190:01:00 | 3:00 | 1463:00:00 | 2:00 | 4709:02:00 | 12:00 | 3952:01:00 | 6:00 |
| NM_001428:NM_001201483 | 2226:01:00 | 0:00 | 1195:01:00 | 2:00 | 1464:00:00 | 2:00 | 4710:02:00 | 11:00 | 3949:01:00 | 7:00 |
| NM_001428:NM_001201483 | 2224:01:00 | 1:00 | 1191:01:00 | 2:00 | 1461:00:00 | 0:00 | 4705:02:00 | 10:00 | 3942:01:00 | 6:00 |
| NM_001428:NM_001201483 | 2214:01:00 | 3:00 | 1170:01:00 | 4:00 | 1443:00:00 | 1:00 | 4653:02:00 | 12:00 | 3917:01:00 | 4:00 |
| NM_001428:NM_001201483 | 2203:01:00 | 3:00 | 1159:01:00 | 1:00 | 1434:00:00 | 0:00 | 4630:02:00 | 4:00 | 3895:01:00 | 1:00 |
| NM_001428:NM_001201483 | 2197:01:00 | 4:00 | 1154:01:00 | 2:00 | 1434:00:00 | 1:00 | 4623:02:00 | 8:00 | 3886:01:00 | 8:00 |
| NM_001428:NM_001201483 | 2197:01:00 | 1:00 | 1146:01:00 | 2:00 | 1441:00:00 | 0:00 | 4577:02:00 | 1:00 | 3868:01:00 | 5:00 |
| NM_001428:NM_001201483 | 2199:01:00 | 5:00 | 1150:01:00 | 1:00 | 1443:00:00 | 1:00 | 4578:02:00 | 5:00 | 3869:01:00 | 6:00 |
| NM_001428:NM_001201483 | 2214:01:00 | 2:00 | 1151:01:00 | 1:00 | 1446:00:00 | 1:00 | 4571:02:00 | 7:00 | 3878:01:00 | 5:00 |
| NM_001428:NM_001201483 | 2214:01:00 | 0:00 | 1150:01:00 | 0:00 | 1444:00:00 | 1:00 | 4562:02:00 | 7:00 | 3873:01:00 | 4:00 |

| | | | | | | | | | | |
|---|---|---|---|---|---|---|---|---|---|---|
| NM_001428:NM_001201483 | 2220:01:00 | 2:00 | 1139:01:00 | 0:00 | 1455:00:00 | 2:00 | 4567:02:00 | 4:00 | 3861:01:00 | 6:00 |
| NM_001428:NM_001201483:NM_001353346 | 2215:01:00 | 4:00:00 | 1135:00:00 | 0:00:00 | 1455:00:00 | 1:00:00 | 4554:01:00 | 4:00:00 | 3859:00:00 | 2:00:00 |
| NM_001428:NM_001201483:NM_001353346 | 2220:01:00 | 0:00:00 | 1136:00:00 | 0:00:00 | 1454:00:00 | 3:00:00 | 4558:01:00 | 3:00:00 | 3859:00:00 | 2:00:00 |
| NM_001428:NM_001201483:NM_001353346 | 2216:01:00 | 3:00:00 | 1133:00:00 | 3:00:00 | 1457:00:00 | 1:00:00 | 4547:01:00 | 7:00:00 | 3859:00:00 | 3:00:00 |
| NM_001428:NM_001201483:NM_001353346 | 2207:01:00 | 6:00:00 | 1133:00:00 | 1:00:00 | 1455:00:00 | 1:00:00 | 4536:01:00 | 7:00:00 | 3855:00:00 | 4:00:00 |
| NM_001428:NM_001201483:NM_001353346 | 2198:01:00 | 1:00:00 | 1120:00:00 | 2:00:00 | 1448:00:00 | 0:00:00 | 4500:02:00 | 4:00:00 | 3839:00:00 | 3:00:00 |
| NM_004120 | 0 | 0 | 0 | 0 | 0 | 2 | 0 | 1 | 0 | 0 |
| NM_004120 | 0 | 0 | 0 | 0 | 0 | 2 | 0 | 1 | 0 | 0 |
| NM_001215:XM_011542084:XM_011542083 | 2:00:00 | 2:01:01 | 0:00:00 | 0:00:00 | 0:00:00 | 0:00:00 | 0:00:00 | 0:00:00 | 0:00:00 | 0:00:00 |
| NM_001215:XM_011542084:XM_011542083 | 2:00:00 | 2:01:01 | 0:00:00 | 0:00:00 | 0:00:00 | 0:00:00 | 0:00:00 | 0:00:00 | 0:00:00 | 0:00:00 |
| NM_001215:XM_011542084:XM_011542083 | 2:01:01 | 2:00:00 | 0:00:00 | 0:00:00 | 0:00:00 | 0:00:00 | 0:00:00 | 0:00:00 | 0:00:00 | 0:00:00 |
| NM_000969:NR_146333 | 4404:17:00 | 9:00 | 2288:13:00 | 6:00 | 1614:04:00 | 10:00 | 24487:38 | 24:00:00 | 4808:18:00 | 8:00 |
| NM_000969:NR_146333 | 4404:17:00 | 9:00 | 2290:13:00 | 5:00 | 1618:04:00 | 8:00 | 24504:37 | 16:01 | 4810:18:00 | 8:00 |
| NM_000969:NR_146333 | 4403:17:00 | 13:00 | 2290:13:00 | 6:00 | 1617:04:00 | 9:00 | 24502:37 | 27:01:00 | 4813:18:00 | 7:00 |
| NM_000969:NR_146333 | 4401:17:00 | 15:00 | 2291:13:00 | 7:00 | 1618:04:00 | 11:00 | 24505:38 | 30:00:00 | 4816:18:00 | 7:00 |
| NM_000969:NR_146333 | 4424:17:00 | 7:00 | 2298:13:00 | 5:00 | 1631:04:00 | 6:00 | 24580:37 | 9:00 | 4836:18:00 | 3:00 |
| NM_000969:NR_146333 | 4432:17:00 | 4:00 | 2302:13:00 | 2:00 | 1633:04:00 | 5:00 | 24600:37 | 6:00 | 4841:18:00 | 2:00 |
| NM_000969:NR_146333 | 4460:17:00 | 9:00 | 2322:11:00 | 3:01 | 1650:04:00 | 3:00 | 25248:37 | 19:00 | 4897:18:00 | 2:00 |
| NM_000969:NR_146333 | 4526:18:00 | 8:00 | 2354:11:00 | 1:00 | 1675:04:00 | 2:00 | 25450:37 | 22:00 | 4932:18:00 | 9:00 |
| NM_000969:NR_146333 | 4537:18:00 | 4:00 | 2359:10:00 | 1:01 | 1680:04:00 | 0:00 | 25478:37 | 13:00 | 4948:18:00 | 6:00 |
| NM_000969:NR_146333 | 4536:18:00 | 4:00 | 2360:10:00 | 0:01 | 1678:04:00 | 2:00 | 25479:36 | 10:00 | 4952:18:00 | 1:00 |
| NM_000969:NR_146333 | 4675:14:00 | 4:00 | 2469:03:00 | 7:00 | 1734:01:00 | 2:00 | 26089:22 | 23:00 | 5023:09:00 | 2:01 |
| NM_000969:NR_146333 | 4687:12:00 | 2:00 | 2479:03:00 | 3:00 | 1736:01:00 | 2:00 | 26124:18 | 24:01:00 | 5027:08:00 | 3:00 |
| NM_000969:NR_146333 | 4707:09:00 | 3:00 | 2507:01:00 | 4:00 | 1744:01:00 | 2:00 | 26191:14 | 24:01:00 | 5038:06:00 | 7:00 |
| NM_000969:NR_146333 | 4731:01:00 | 1:00 | 2531:00:00 | 0:00 | 1747:00:00 | 1:00 | 26279:6 | 9:01 | 5054:00:00 | 4:00 |
| NM_000969:NR_146333 | 4733:01:00 | 2:00 | 2533:00:00 | 2:00 | 1747:00:00 | 1:00 | 26293:3 | 20:00 | 5063:00:00 | 3:00 |
| NM_000969:NR_146333 | 4751:00:00 | 2:01 | 2551:00:00 | 1:00 | 1755:00:00 | 1:00 | 26374:4 | 26:04:00 | 5083:00:00 | 3:00 |
| NM_000969:NR_146333 | 4736:07:00 | 11:00 | 2547:05:00 | 4:00 | 1753:02:00 | 1:00 | 26364:43 | 30:03:00 | 5084:05:00 | 2:00 |
| NM_000969:NR_146333 | 4741:12:00 | 1:00 | 2548:10:00 | 1:00 | 1755:06:00 | 0:00 | 26369:92 | 21:01 | 5079:07:00 | 4:00 |
| NM_000969:NR_146333 | 4737:28:00 | 3:00 | 2542:18:00 | 3:01 | 1753:09:00 | 0:00 | 26351:152 | 12:06 | 5075:12:00 | 0:01 |
| NM_000969:NR_146333 | 4721:46:00 | 5:01 | 2540:38:00 | 2:00 | 1748:11:00 | 0:00 | 26313:230 | 24:04:00 | 5066:26:00 | 0:00 |
| NM_000969:NR_146333 | 4692:82 | 5:01 | 2534:103 | 1:01 | 1739:27:00 | 2:00 | 26243:414 | 32:03:00 | 5048:63 | 4:03 |
| NM_000969:NR_146333 | 4670:94 | 3:02 | 2525:127 | 2:00 | 1732:35:00 | 0:00 | 26206:466 | 17:01 | 5032:77 | 11:01 |
| NM_000969:NR_146333 | 4666:103 | 5:01 | 2525:167 | 2:01 | 1725:41:00 | 2:00 | 26175:549 | 33:02:00 | 5036:94 | 5:00 |
| NM_000969:NR_146333 | 4655:120 | 3:00 | 2519:232 | 2:00 | 1718:51:00 | 3:00 | 26108:660 | 32:01:00 | 5023:106 | 8:00 |
| NM_000969:NR_146333 | 4628:127 | 3:01 | 2513:330 | 1:01 | 1715:60 | 1:00 | 26019:678 | 23:00 | 5023:116 | 5:00 |
| NM_000969:NR_146333 | 4226:118 | 304:02:00 | 2274:320 | 144:04:00 | 1559:57:00 | 123:02:00 | 23298:648 | 1775:09:00 | 4526:114 | 377:01:00 |
| NM_000969:NR_146333 | 4196:108 | 3:00 | 1441:254 | 2:00 | 1552:56:00 | 1:00 | 15757:404 | 26:02:00 | 4501:110 | 2:00 |
| XR_001737025:XR_001737024:NM_144988:XM_011540897:XR_946568:XM_005270582:NM_001305242 | 5:4:151:1:2:2:0 | 0:0:0:0:0:0:0 | 1:2:163:1:2:2:0 | 0:0:0:0:0:0:0 | 5:3:190:1:0:2:0 | 0:0:0:0:0:0:0 | 12:13:509:3:9:6:0 | 0:0:2:0:0:0:0 | 4:10:199:3:2:4:0 | 0:0:0:0:0:0:0 |
| XR_001737025:XR_001737024:NM_144988:NR_131032:XM_011540897:XR_946568:XM_005270582:NM_001305242 | 5:4:150:3:1:2:2:0 | 0:0:0:0:0:0:0:0 | 1:2:160:0:1:2:2:0 | 0:0:0:0:0:0:0:0 | 5:3:187:1:1:0:2:0 | 0:0:1:0:0:0:0:0 | 12:12:507:0:3:9:6:0 | 0:0:0:0:0:0:0:0 | 4:9:195:1:3:2:4:0 | 0:0:0:0:0:0:0:0 |
| NM_001278248:NM_015485:NR_103484 | 0:00:00 | 0:00:00 | 0:00:00 | 1:00:00 | 0:00:00 | 0:00:00 | 0:00:00 | 0:01:00 | 0:00:00 | 0:00:00 |
| NM_015658 | 0 | 0 | 0 | 0 | 0 | 0 | 0 | 0 | 0 | 3 |
| NM_000110:XM_017000510:XM_017000507:XM_006710397 | 0:0:0:0 | 2:2:0:0 | 0:0:0:0 | 0:1:1:1 | 0:0:0:0 | 2:2:0:1 | 0:0:0:0 | 0:0:0:0 | 0:0:0:0 | 0:0:0:0 |
| XM_005244771:NM_001142467 | 0:00 | 0:00 | 0:00 | 0:00 | 0:00 | 1:00 | 0:00 | 0:00 | 0:01 | 0:01 |
| NM_001284389:NM_004741:XM_005270273:NM_001284388 | 19:17:11:5 | 0:0:0:0 | 5:2:1:0 | 0:0:0:0 | 6:4:2:1 | 0:0:0:0 | 100:137:46:12 | 0:0:2:0 | 26:44:10:4 | 0:0:0:0 |
| NM_001284389:NM_004741:XM_005270273:NM_001284388 | 15:16:7:5 | 0:0:0:0 | 3:1:1:0 | 0:0:0:0 | 4:1:4:0 | 0:0:0:0 | 68:110:37:10 | 1:1:0:0 | 20:29:14:3 | 0:1:0:0 |
| NM_004741:XM_005270273:NM_001284389:NM_001284388 | 7:3:6:1 | 12:7:7:4 | 2:0:0:0 | 0:1:3:0 | 1:4:2:0 | 1:1:2:0 | 33:31:25:4 | 68:29:35:3 | 18:9:10:2 | 13:9:9:1 |
| NM_001284389:NM_004741:XM_005270273:NM_001284388 | 13:19:12:5 | 0:0:0:0 | 3:2:1:0 | 0:0:0:0 | 4:3:5:0 | 0:0:0:0 | 35:69:39:3 | 27:33:33:4 | 10:19:9:1 | 9:17:9:2 |
| NM_004741:NM_001284389:XM_005270273:NM_001284388 | 19:12:13:5 | 2:1:0:0 | 2:2:1:0 | 0:1:0:0 | 3:4:4:0 | 0:0:0:0 | 91:51:75:7 | 11:9:2:0 | 35:17:18:3 | 0:1:0:0 |
| XM_011539913:NM_017787:XM_017016360:NM_001083913 | 0:1:0:0 | 1:0:0:0 | 0:0:0:0 | 0:0:0:0 | 0:0:0:1 | 0:1:1:0 | 0:0:0:0 | 0:0:1:0 | 0:0:0:0 | 0:0:0:0 |
| NM_020682 | 0 | 0 | 0 | 0 | 0 | 0 | 7 | 0 | 2 | 0 |
| NM_001351171:NM_001351183:NM_001351172:NM_001134373:NM_012229:NM_001351169:XM_024447902:NM_001351181:NM_001351175:NM_001351182:XM_024447904:NM_001351185:NM_001351170:NM_001351190:NM_001351194:XM_024447911:XM_024447905:XM_024447907:XM_024447908:XM_024447910:XM_011539537:XM_017015966:NM_001351189:NM_001351197:NM_001351187:XM_024447901:NM_001351178:XM_024447909:NM_001351180:NM_001351176:NM_001351177:XM_006717723 | 2:1:2:5:3:4:0:2:1:1:1:0:0:0:0:0:0:0:0:0:0:0:0:0:0:0:0:0:0:0:0:0:0:0 | 1:2:1:4:1:1:1:0:0:0:0:0:0:0:0:0:0:0:0:0:0:0:0:0:0:0:0:0:0:0:0:0 | 0:0:0:0:2:1:0:0:0:0:0:0:1:1:0:1:1:1:1:1:0:0:0:0:0:0:0:0:0:0:0:0 | 0:0:0:5:0:0:1:0:0:0:0:1:1:1:1:0:0:0:0:0:0:0:0:0:0:0:0:0:0:0:0:0 | 1:1:2:7:3:5:0:0:1:0:1:0:1:0:0:0:0:1:0:1:1:0:2:1:1:0:0:0:0:0:0:0 | 0:1:3:2:2:6:0:0:0:0:0:0:2:0:0:0:1:0:0:0:1:1:0:0:0:0:0:0:0:0:0:0 | 1:1:6:4:6:4:2:1:3:0:0:0:6:1:0:2:0:0:0:10:1:0:0:0:0:1:1:6:0:0:0:0 | 0:0:0:0:0:0:0:0:0:0:0:0:0:0:0:0:0:0:0:0:0:0:0:0:0:0:0:0:0:0:0:0:0:0:0 | 4:1:5:14:3:4:0:0:0:0:1:0:5:0:0:2:0:0:0:1:0:0:0:0:0:0:0:0:1:1:0:0 | 0:0:0:0:0:0:0:0:0:0:0:0:0:0:0:0:0:0:0:0:0:0:0:0:0:0:0:0:0:0:0:0 |
| NM_024928 | 3 | 2 | 5 | 4 | 2 | 2 | 40 | 0 | 7 | 0 |
| NM_024928 | 0 | 5 | 0 | 8 | 0 | 4 | 0 | 40 | 0 | 7 |
| NM_024928 | 0 | 5 | 0 | 8 | 0 | 4 | 0 | 40 | 0 | 7 |
| NM_004832 | 1 | 1 | 0 | 0 | 2 | 2 | 4 | 0 | 3 | 0 |
| XM_011539270:XM_006717624 | 0:00 | 0:00 | 0:00 | 0:00 | 0:00 | 0:00 | 1:00 | 0:00 | 0:00 | 0:00 |
| NM_005871:XM_005269382 | 32:01:00 | 0:00 | 15:02 | 1:01 | 15:01 | 0:00 | 132:08:00 | 5:00 | 25:04:00 | 1:00 |
| NM_004419 | 0 | 0 | 0 | 0 | 0 | 2 | 0 | 0 | 0 | 0 |
| NM_004419 | 0 | 0 | 0 | 0 | 0 | 2 | 0 | 0 | 0 | 0 |
| NM_014456:NM_145341:NM_001199492 | 0:00:00 | 0:00:00 | 0:00:00 | 3:02:00 | 0:00:00 | 9:01:01 | 0:00:00 | 4:01:01 | 0:00:00 | 1:00:00 |
| NM_001227:XM_011540260 | 1:00 | 0:00 | 1:00 | 0:00 | 1:02 | 3:00 | 0:00 | 0:00 | 0:00 | 0:00 |
| NM_001227:XM_011540260 | 1:00 | 0:00 | 1:00 | 0:00 | 1:02 | 3:00 | 0:00 | 0:00 | 0:00 | 0:00 |
| XM_011539916:NM_018017:NR_135815:NM_153249 | 0:0:0:0 | 0:0:0:0 | 1:0:0:0 | 0:0:0:0 | 0:0:0:0 | 0:0:0:0 | 0:0:1:0 | 0:3:0:0 | 0:0:0:0 | 0:0:0:0 |
| XM_011539916:NM_018017:NR_135815:NM_153249 | 0:0:0:0 | 0:0:0:0 | 1:0:0:0 | 0:0:0:0 | 0:0:0:0 | 0:0:0:0 | 0:0:1:0 | 0:3:0:0 | 0:0:0:0 | 0:0:0:0 |
| NM_024693 | 0 | 0 | 0 | 0 | 0 | 0 | 0 | 2 | 0 | 0 |
| NM_000982 | 151 | 0 | 32 | 1 | 47 | 0 | 174 | 2 | 241 | 0 |
| NM_007041:XM_017015598:XM_017015599:NM_001288736:NM_001001976:XM_017015604:XM_005269459:XM_005269458:NR_110006 | 4:2:0:0:0:0:0:0:0 | 0:0:0:0:0:0:0:0:0 | 1:3:8:1:1:1:0:0:0 | 1:1:2:0:0:0:0:0:0 | 1:2:1:1:2:0:1:0:0 | 0:0:0:0:0:0:0:0:0 | 4:11:9:1:3:1:0:2:0 | 2:1:4:0:0:0:0:0:0 | 2:2:0:0:0:0:1:0:0 | 0:0:1:0:0:0:0:0:0 |
| NM_001291877:XM_024447768:XM_011539134:XM_006717553:XM_011539133:NM_001291879:XM_017015495:XM_006717554 | 0:0:0:0:0:0:0:0 | 1:1:1:0:0:0:0:0 | 0:0:0:0:0:0:0:0 | 0:0:0:1:0:0:0:0 | 1:0:0:0:0:1:0:0 | 0:0:0:0:1:0:0:0 | 1:1:1:0:2:0:5:0 | 0:0:0:0:0:0:0:0 | 1:0:0:0:0:0:1:0 | 0:0:0:0:0:0:0:0 |

| | | | | | | | | | | |
|---|---|---|---|---|---|---|---|---|---|---|
| NM_001291877:XM_024447768:XM_011539134:XM_006717553:XM_011539133:NM_001291879:XM_017015495:XM_006717554 | 0:0:0:0:0:0:0:0 | 1:1:1:0:0:0:0:0 | 0:0:0:0:0:0:0:0 | 0:0:0:1:0:0:0:0 | 1:0:0:0:0:1:0:0 | 0:0:0:0:1:0:0:0 | 1:1:1:0:2:0:5:0 | 0:0:0:0:0:0:0:0 | 1:0:0:0:0:0:1:0 | 0:0:0:0:0:0:0:0 |
| NM_001291877:XM_024447768:XM_011539134:XM_006717553:XM_017015495:XM_011539133:NM_001291879:XM_006717554 | 0:0:0:0:0:0:0:0 | 1:1:1:0:0:0:0:0 | 0:0:0:0:0:0:0:0 | 0:0:0:1:1:0:0:0 | 1:0:0:0:0:0:1:0 | 0:0:0:0:0:1:0:0 | 1:1:1:0:5:2:0:0 | 0:0:0:0:0:0:0:0 | 1:0:0:0:1:0:0:0 | 0:0:0:0:0:0:0:0 |
| NM_001291877:XM_011539134:NM_001291879:XM_006717553:XM_017015495:XM_024447768:XM_011539133:NM_206860:XM_006717554:XM_024447762:XM_006717552:NM_206862 | 1:1:2:0:0:0:0:0:0:0:0:0 | 0:0:0:0:0:0:0:0:0:0:0:0 | 0:0:2:1:7:1:0:0:0:0:0:0 | 0:0:0:0:0:0:0:0:0:0:0:0 | 1:0:5:0:5:0:1:1:0:0:0:0 | 0:0:0:0:0:0:0:0:0:0:0:0 | 0:1:2:0:15:0:1:0:1:1:1:0 | 1:0:4:0:26:0:1:0:1:0:0:0 | 1:0:0:0:1:0:0:0:0:0:0:0 | 0:0:2:0:7:0:0:0:0:0:0:0 |
| NM_001291877:XM_024447768:XM_011539134:NM_001291879:XM_017015495:XM_011539133:NM_206860:NM_206862:XM_006717552:XM_006717554:XM_017015499:XM_006717553 | 1:1:1:2:0:0:0:0:0:0:0:0 | 0:0:0:0:0:0:0:0:0:0:0:0 | 0:0:0:2:5:0:0:0:0:0:0:0 | 0:0:0:0:0:0:0:0:0:0:0:0 | 1:0:0:5:5:1:1:1:0:0:0:0 | 0:0:0:0:0:0:0:0:0:0:0:0 | 1:2:0:4:23:0:0:0:0:1:1:0 | 0:2:1:2:17:1:0:0:1:1:0:0 | 0:0:0:2:6:0:0:0:0:0:0:0 | 1:0:0:0:0:0:0:0:0:0:0:0 |
| XM_024447859:NM_001380 | 1:02 | 0:00 | 2:00 | 0:02 | 1:01 | 0:00 | 0:00 | 0:08 | 0:00 | 0:02 |
| NM_002412:XM_017016275:XM_005252682 | 137:02:00 | 0:00:00 | 165:02:03 | 0:00:00 | 401:04:06 | 2:00:02 | 17:01:00 | 0:00:00 | 18:01:00 | 0:00:00 |
| NM_002412:XM_017016275:XM_005252682 | 137:02:00 | 0:00:00 | 165:02:03 | 0:00:00 | 402:04:06 | 1:00:02 | 17:01:00 | 0:00:00 | 18:01:00 | 0:00:00 |
| NM_002412:XM_017016275:XM_005252682 | 52:04:00 | 82:02:01 | 69:01:03 | 92:00:05 | 193:07:10 | 204:06:05 | 17:00:00 | 0:00:00 | 18:01:00 | 0:00:00 |
| NM_002412:XM_017016275:XM_005252682 | 134:10:01 | 0:00:00 | 161:03:07 | 1:00:00 | 390:15:13 | 2:01:00 | 17:00:00 | 0:00:00 | 18:01:00 | 0:00:00 |
| NM_145202:NM_001145201 | 0:00 | 3:00 | 0:00 | 10:01 | 0:00 | 21:01 | 0:00 | 9:00 | 0:00 | 7:00 |
| NM_004092 | 0 | 1 | 0 | 1 | 0 | 10 | 0 | 8 | 0 | 7 |
| NM_004092 | 0 | 1 | 0 | 1 | 0 | 10 | 0 | 8 | 0 | 6 |
| NM_152724:NM_012425:XM_005252552 | 46:243:14 | 0:00:00 | 74:302:18 | 0:00:00 | 125:716:29 | 0:02:00 | 431:1571:15 | 2:02:00 | 94:440:1 | 0:01:00 |
| NM_012425:NM_152724:XM_005252552 | 242:42:13 | 2:00:00 | 303:50:20 | 1:00:00 | 720:100:30 | 2:00:00 | 1588:348:24 | 1:00:00 | 441:74:1 | 0:00:00 |
| NM_012425:NM_152724:XM_005252552 | 244:42:13 | 0:00:00 | 304:49:20 | 0:00:00 | 722:96:31 | 0:00:00 | 1587:337:23 | 2:01:00 | 439:73:1 | 1:00:00 |
| NM_012425:NM_152724:XM_005252552 | 243:03:13 | 1:00:00 | 303:02:18 | 0:00:00 | 720:03:28 | 2:00:00 | 1586:21:20 | 2:00:00 | 441:06:01 | 0:00:00 |
| XM_006717500:NM_003380 | 415:19:00 | 1:00 | 351:08:00 | 2:00 | 191:06:00 | 0:00 | 399:05:00 | 3:00 | 210:11:00 | 0:00 |
| XM_006717500:NM_003380 | 431:20:00 | 0:00 | 384:08:00 | 3:00 | 210:07:00 | 1:01 | 437:06:00 | 2:00 | 224:12:00 | 0:00 |
| XM_006717500:NM_003380 | 429:20:00 | 3:00 | 387:08:00 | 1:00 | 212:07:00 | 1:01 | 438:06:00 | 2:00 | 226:12:00 | 0:00 |
| XM_006717500:NM_003380 | 467:22:00 | 3:00 | 426:09:00 | 1:00 | 230:08:00 | 0:00 | 478:08:00 | 1:00 | 238:12:00 | 1:00 |
| XM_006717500:NM_003380 | 468:22:00 | 4:00 | 427:09:00 | 1:00 | 231:08:00 | 0:00 | 481:08:00 | 1:00 | 239:12:00 | 1:00 |
| XM_006717500:NM_003380 | 475:22:00 | 1:00 | 434:09:00 | 1:00 | 233:08:00 | 1:00 | 486:08:00 | 0:00 | 241:12:00 | 0:00 |
| XM_006717500:NM_003380 | 489:25:00 | 1:00 | 468:09:00 | 0:00 | 248:08:00 | 1:00 | 520:10:00 | 2:00 | 251:12:00 | 0:00 |
| XM_006717500:NM_003380 | 490:25:00 | 1:00 | 468:09:00 | 0:00 | 249:08:00 | 0:00 | 522:10:00 | 2:00 | 251:12:00 | 0:00 |
| XM_006717500:NM_003380 | 494:25:00 | 1:00 | 466:09:00 | 1:00 | 249:08:00 | 0:00 | 525:10:00 | 0:00 | 251:12:00 | 0:00 |
| XM_006717500:NM_003380 | 495:25:00 | 1:00 | 468:09:00 | 0:00 | 248:08:00 | 1:00 | 530:10:00 | 2:00 | 251:12:00 | 0:00 |
| XM_006717500:NM_003380 | 494:25:00 | 2:00 | 470:09:00 | 1:00 | 250:08:00 | 0:00 | 533:10:00 | 0:00 | 252:12:00 | 0:00 |
| XM_006717500:NM_003380 | 499:25:00 | 1:00 | 477:09:00 | 0:00 | 255:08:00 | 0:00 | 548:12:00 | 0:00 | 258:12:00 | 0:00 |
| XM_006717500:NM_003380 | 501:25:00 | 0:00 | 482:09:00 | 0:00 | 256:08:00 | 0:00 | 551:12:00 | 0:00 | 259:12:00 | 0:00 |
| XM_006717500:NM_003380 | 503:26:00 | 0:00 | 501:09:00 | 0:00 | 261:09:00 | 0:00 | 564:12:00 | 0:00 | 267:12:00 | 0:00 |
| XM_006717500:NM_003380 | 511:26:00 | 0:00 | 528:09:00 | 2:00 | 266:09:00 | 0:00 | 593:12:00 | 0:00 | 277:12:00 | 0:00 |
| XM_006717500:NM_003380 | 511:26:00 | 0:00 | 532:09:00 | 0:00 | 266:09:00 | 0:00 | 593:12:00 | 1:00 | 276:13:00 | 1:00 |
| XM_006717500:NM_003380 | 511:26:00 | 1:00 | 533:09:00 | 0:00 | 267:09:00 | 0:00 | 597:12:00 | 0:00 | 277:15:00 | 0:00 |
| XM_006717500:NM_003380 | 511:26:00 | 0:00 | 533:09:00 | 0:00 | 267:09:00 | 0:00 | 595:12:00 | 3:00 | 277:15:00 | 0:00 |
| NM_014241:XR_428651 | 1:00 | 0:00 | 3:00 | 0:00 | 0:00 | 0:00 | 4:00 | 0:00 | 0:00 | 0:00 |
| NM_014241:XR_428651 | 0:00 | 1:00 | 0:00 | 3:01 | 0:00 | 0:00 | 0:00 | 3:03 | 0:00 | 0:00 |
| NM_014241:XR_428651 | 0:00 | 1:00 | 0:00 | 3:01 | 0:00 | 0:00 | 0:00 | 3:03 | 0:00 | 0:00 |
| XR_001746996:XR_001746995:XR_242691 | 0:00:00 | 0:00:00 | 0:00:00 | 0:00:00 | 1:00:00 | 0:00:00 | 0:00:01 | 0:01:01 | 0:00:00 | 0:00:00 |
| XM_005252342:XR_001746996:XR_001746995:XR_242691 | 0:0:0:0 | 1:0:0:0 | 0:0:0:0 | 0:1:0:0 | 0:1:0:0 | 0:0:0:0 | 0:7:1:7 | 0:8:0:0 | 0:0:0:0 | 0:0:0:0 |
| XM_005252342:XR_001746996:XR_001746995:XR_242691 | 0:0:0:0 | 1:0:0:0 | 0:0:0:0 | 0:1:0:0 | 0:1:0:0 | 0:0:0:0 | 0:7:1:7 | 0:9:0:0 | 0:0:0:0 | 0:0:0:0 |
| NM_012071 | 1 | 0 | 0 | 0 | 4 | 0 | 0 | 2 | 2 | 0 |
| XM_017016318:XM_024448036:XM_017016317:NM_100264:NM_016628:NM_100486:XM_017016315:XR_001747110:XR_930491 | 1:3:1:2:5:1:0:0:0 | 0:0:0:0:0:0:0:0:0 | 2:3:0:1:1:0:2:2:1 | 0:0:0:0:0:0:0:0:0 | 2:1:0:1:5:3:0:0:0 | 0:0:0:0:0:0:0:0:0 | 7:21:1:1:6:1:3:5:9 | 0:0:1:0:0:1:0:0:0 | 0:3:0:0:3:4:0:0:1 | 0:0:0:0:0:0:0:0:0 |
| XM_024447875:XM_006717399:NM_182755 | 0:00:00 | 0:00:00 | 0:00:00 | 1:00:00 | 0:00:00 | 0:00:00 | 0:00:00 | 0:00:00 | 0:00:00 | 0:00:00 |
| NM_001323647:NM_001174096:NM_030751:NM_001174093:NM_001323655:XM_017016597:NM_001323649:NM_001174095:NM_001323653 | 0:3:2:1:0:0:0:0:0 | 1:0:0:0:0:0:0:0:0 | 1:11:3:0:1:3:0:0:0 | 0:1:0:0:0:0:0:0:0 | 1:9:3:0:0:1:0:0:0 | 1:1:0:0:0:0:1:0:0 | 0:0:1:0:0:0:1:0:0 | 0:0:0:0:0:0:0:0:0 | 0:2:0:0:0:0:0:0:0 | 0:1:0:0:0:0:0:0:0 |
| NM_001323647:NM_001174096:NM_030751:NM_001174093:NM_001323655:XM_017016597:NM_001323649:NM_001174095:NM_001323653 | 0:3:2:1:0:0:0:0:0 | 1:0:0:0:0:0:0:0:0 | 1:11:3:0:1:3:0:0:0 | 0:1:0:0:0:0:0:0:0 | 1:9:3:0:0:1:0:0:0 | 1:1:0:0:0:0:1:0:0 | 0:0:1:0:0:0:1:0:0 | 0:0:0:0:0:0:0:0:0 | 0:2:0:0:0:0:0:0:0 | 0:1:0:0:0:0:0:0:0 |
| NM_001174096:NM_030751:NM_001174093:NM_001323647:NM_001323655:XM_017016597:NM_001323649:NM_001174095:NM_001323653 | 3:2:1:0:0:0:0:0:0 | 0:0:0:0:0:0:0:0:0 | 10:3:0:1:1:5:0:0:0 | 1:0:0:0:0:0:0:0:0 | 9:3:0:1:0:1:0:0:0 | 0:0:0:0:0:0:0:0:0 | 0:1:0:0:0:2:1:0:0 | 0:0:0:0:0:0:0:0:0 | 2:0:0:0:0:0:0:0:0 | 0:0:0:0:0:0:0:0:0 |
| NM_001174096:NM_001174093:NM_030751:NM_001323655:NM_001323647:XM_017016597:NM_001174095:NM_001323653 | 2:1:0:0:0:0:0:0 | 1:0:0:0:0:0:0:0 | 4:0:1:0:1:5:0:0 | 6:0:2:1:0:0:0:0 | 3:0:1:0:0:1:0:0 | 3:0:2:0:1:0:0:0 | 0:0:0:0:0:2:0:0 | 0:0:1:0:0:0:0:0 | 0:0:0:0:0:0:0:0 | 1:0:0:0:0:0:0:0 |
| NM_001174093:NM_001174096:XM_017016597:NM_030751 | 1:2:4:0 | 0:0:0:0 | 0:0:8:0 | 0:0:1:0 | 0:2:3:1 | 0:1:0:0 | 0:0:3:0 | 0:0:0:0 | 0:0:0:0 | 0:0:0:0 |
| XM_017016651:NM_145023:XM_006717506:XM_011519678:NR_109827:XM_024448168:XM_011519686:XM_024448169:XM_024448170:NM_001026383:NR_109826 | 0:0:0:0:0:0:0:0:0:0:0 | 0:0:0:0:0:0:0:0:0:0:0 | 0:0:0:0:1:1:0:0:0:0:0 | 1:1:2:1:0:0:0:0:0:0:0 | 0:0:0:0:0:0:0:0:0:0:0 | 0:0:0:0:0:1:0:0:0:0:0 | 0:0:0:0:0:0:0:0:0:0:0 | 0:0:0:0:1:0:0:1:1:0:0 | 0:0:0:0:0:0:0:0:0:0:0 | 0:0:0:0:0:0:0:0:0:0:0 |
| NM_001024628:NM_001244973:NM_003873:XM_006717526:NM_001024629 | 0:0:0:0:0 | 3:1:1:0:0 | 0:0:0:0:0 | 0:0:1:0:0 | 0:0:0:0:0 | 1:0:1:0:0 | 0:0:0:0:0 | 2:1:1:0:0 | 0:0:0:0:0 | 5:0:4:1:0 |
| NM_019619:NM_001184786:NM_001184791:NM_001184792:NM_001184789:NM_001184790:NM_001184793:NM_001184785:NM_001184794:NM_001184787 | 1:1:0:0:0:0:0:0:0:0 | 1:0:0:0:0:0:0:0:0:0 | 2:1:1:2:0:0:0:0:0:0 | 2:0:0:0:0:0:0:0:0:0 | 0:1:0:2:1:1:1:0:0:0 | 1:0:0:0:0:0:0:0:0:0 | 6:8:1:10:0:0:1:1:4:0 | 1:0:0:1:0:0:0:0:0:0 | 0:3:0:2:0:0:0:0:1:0 | 0:0:0:0:0:0:0:0:0:0 |
| NM_001145263:NM_005437:NM_001145261:NM_001145260:NM_001145262 | 351:186:1:1:0 | 1:0:0:0:0 | 19:494:0:0:0 | 0:1:0:0:0 | 805:940:6:3:3 | 3:2:0:0:0 | 235:1228:4:2:1 | 0:2:0:0:0 | 274:332:2:0:1 | 0:0:0:0:0 |
| NM_005437:NM_001145263:NM_001145261:NM_001145260:NM_001145262 | 184:352:1:1:0 | 0:0:0:0:0 | 476:19:0:0:0 | 0:0:0:0:0 | 930:808:6:3:3 | 2:0:0:0:0 | 1168:236:4:2:1 | 2:0:0:0:0 | 329:274:2:0:1 | 0:0:0:0:0 |
| NM_001145263:NM_005437:NM_001145261:NM_001145260:NM_001145262 | 348:172:1:1:0 | 3:0:0:0:0 | 18:397:0:0:0 | 0:4:0:0:0 | 801:878:6:3:3 | 9:3:0:0:0 | 236:995:4:2:1 | 0:6:0:0:0 | 270:316:2:0:1 | 2:1:0:0:0 |
| NM_005437:NM_001145263:NM_001145261:NM_001145260:NM_001145262 | 170:353:1:1:0 | 0:0:0:0:0 | 365:18:0:0:0 | 5:0:0:0:0 | 874:808:6:3:3 | 1:2:0:0:0 | 911:237:4:2:1 | 6:0:0:0:0 | 316:273:2:0:1 | 0:0:0:0:0 |
| NM_005437:NM_001145263:NM_001145261:NM_001145260:NM_001145262 | 170:353:1:1:0 | 0:0:0:0:0 | 364:17:0:0:0 | 3:1:0:0:0 | 871:809:6:3:3 | 1:1:0:0:0 | 904:237:4:2:1 | 4:0:0:0:0 | 316:273:2:0:1 | 0:0:0:0:0 |

| | | | | | | | | | | |
|---|---|---|---|---|---|---|---|---|---|---|
| NM_001145263:NM_005437:NM_001145261:NM_001145260:NM_001145262 | 358:169:1:1:0 | 1:0:0:0:0 | 32:248:1:0:0 | 15:8:0:0:0 | 820:828:6:3:3 | 1:1:0:0:0 | 274:698:3:2:1 | 16:4:1:0:0 | 279:302:2:0:1 | 2:1:0:0:0 |
| NM_001145263:NM_005437:NM_001145261:NM_001145260:NM_001145262 | 359:169:1:1:0 | 1:0:0:0:0 | 33:247:1:0:0 | 17:9:0:0:0 | 818:826:6:3:3 | 11:1:0:0:0 | 273:695:4:2:1 | 20:3:0:0:0 | 279:302:2:0:1 | 5:1:0:0:0 |
| NM_001145263:NM_005437:NM_001145261:NM_001145260:NM_001145262 | 359:167:1:1:0 | 3:0:0:0:0 | 46:240:1:0:0 | 5:3:0:0:0 | 820:816:6:3:3 | 16:6:0:0:0 | 286:684:4:2:1 | 14:1:0:0:0 | 281:301:2:0:1 | 5:1:0:0:0 |
| NM_001145263:NM_005437:NM_001145261:NM_001145260:NM_001145262 | 361:161:1:1:0 | 1:0:0:0:0 | 52:207:1:0:0 | 0:1:0:0:0 | 839:797:6:3:3 | 1:4:0:0:0 | 306:623:4:2:1 | 1:1:0:0:0 | 286:293:2:0:1 | 0:0:0:0:0 |
| NM_005437:NM_001145263:NM_001145261:NM_001145260:NM_001145262 | 160:362:1:1:0 | 0:0:0:0:0 | 199:51:1:0:0 | 1:1:0:0:0 | 789:838:6:3:3 | 2:2:0:0:0 | 600:306:4:2:1 | 4:1:0:0:0 | 290:286:2:0:1 | 0:0:0:0:0 |
| NM_005437:NM_001145263:NM_001145261:NM_001145260:NM_001145262 | 159:362:1:1:0 | 0:0:0:0:0 | 187:52:1:0:0 | 2:0:0:0:0 | 767:839:6:3:3 | 2:1:0:0:0 | 563:305:4:2:1 | 1:2:0:0:0 | 282:285:2:0:1 | 0:1:0:0:0 |
| NM_005437:NM_001145263:NM_001145261:NM_001145260:NM_001145262 | 153:362:1:1:0 | 0:0:0:0:0 | 161:52:1:0:0 | 2:0:0:0:0 | 744:840:6:3:3 | 2:0:0:0:0 | 511:306:4:2:1 | 1:2:0:0:0 | 275:285:2:0:1 | 1:1:0:0:0 |
| NM_001145263:NM_005437:NM_001145261:NM_001145260:NM_001145262 | 361:152:1:1:0 | 1:0:0:0:0 | 52:146:1:0:0 | 0:2:0:0:0 | 840:736:6:3:3 | 0:2:0:0:0 | 307:488:4:2:1 | 1:3:0:0:0 | 286:272:2:0:1 | 0:0:0:0:0 |
| NM_005437:NM_001145263:NM_001145261:NM_001145260:NM_001145262 | 136:362:1:1:0 | 0:0:0:0:0 | 55:59:1:0:0 | 0:0:0:0:0 | 582:846:6:2:3 | 1:3:0:0:0 | 305:322:4:2:1 | 0:3:0:0:0 | 194:288:2:0:1 | 1:0:0:0:0 |
| NM_005437:NM_001145263:NM_001145261:NM_001145260:NM_001145262 | 136:362:1:1:0 | 0:0:0:0:0 | 55:58:1:0:0 | 0:1:0:0:0 | 581:848:5:2:3 | 1:1:1:0:0 | 300:325:4:2:1 | 0:0:0:0:0 | 190:288:2:0:1 | 2:0:0:0:0 |
| NM_003739 | 3 | 1 | 1 | 0 | 1 | 4 | 0 | 3 | 2 | 0 |
| NM_005436 | 17 | 11 | 14 | 9 | 29 | 16 | 84 | 86 | 15 | 32 |
| XM_017015844:NM_145307:XR_001747053 | 0:00:00 | 0:00:00 | 0:00:00 | 0:00:00 | 0:00:00 | 0:00:00 | 0:00:00 | 1:00:00 | 0:00:00 | 0:01:01 |
| NM_018237:NM_001282960:NR_104262:NM_001282959 | 40:2:1:0 | 0:0:0:0 | 63:2:10:0 | 0:0:0:0 | 29:0:6:0 | 0:0:0:0 | 320:15:41:0 | 0:0:0:0 | 34:5:5:1 | 0:0:0:0 |
| NM_018237:NR_104262:NM_001282960:NM_001282959 | 36:1:1:0 | 0:0:0:0 | 47:24:1:0 | 0:0:0:0 | 24:4:0:0 | 0:0:0:0 | 252:44:13:0 | 0:0:0:0 | 26:7:4:1 | 0:0:0:0 |
| NM_018237:NR_104262:NM_001282960:NM_001282959 | 36:1:1:0 | 0:0:0:0 | 40:23:1:0 | 4:0:0:0 | 24:3:0:0 | 0:0:0:0 | 231:43:11:0 | 12:0:1:0 | 25:7:4:1 | 0:0:0:0 |
| NM_018237:NR_104262:NM_001282960:NM_001282959 | 27:1:1:0 | 1:0:0:0 | 26:18:0:0 | 9:5:1:0 | 17:3:0:0 | 1:0:0:0 | 138:31:4:0 | 26:6:3:0 | 16:3:2:1 | 1:0:1:0 |
| NM_018237:NM_001282960:NR_104262:NM_001282959 | 18:0:0:0 | 1:1:0:0 | 16:0:15:0 | 1:0:0:0 | 14:0:3:0 | 0:0:0:0 | 86:1:24:0 | 7:1:1:0 | 8:2:3:0 | 2:0:0:0 |
| NM_015634 | 0 | 0 | 0 | 0 | 0 | 0 | 0 | 0 | 0 | 0 |
| XM_024448066:NM_002727 | 0:00 | 1:00 | 1:00 | 0:00 | 1:00 | 3:01 | 0:00 | 0:00 | 0:00 | 1:00 |
| NM_018344:NR_033414:XM_017016378:NR_033413:XM_017016377:NM_001174098 | 0:0:1:0:0:0 | 2:1:0:0:0:0 | 3:0:0:0:0:0 | 0:0:0:0:0:0 | 3:0:0:0:0:0 | 0:0:1:0:0:0 | 8:0:2:4:0:0 | 5:3:0:4:0:0 | 0:0:0:1:0:0 | 2:0:0:1:0:0 |
| XM_017016378:NM_018344:NR_033414:XM_017016377:NR_033413:NM_001174098 | 0:2:1:0:0:0 | 1:0:0:0:0:0 | 0:0:0:0:0:0 | 0:3:0:0:0:0 | 1:0:0:1:0:0 | 0:3:0:0:0:0 | 0:0:0:0:1:0 | 0:15:0:3:9:0 | 0:0:0:0:0:0 | 0:3:0:0:2:0 |
| NM_018344:NR_033414:XM_017016378:XM_017016377:NR_033413:NM_001174098 | 0:0:0:0:0:0 | 2:1:0:0:0:0 | 0:0:0:0:0:0 | 3:1:0:0:0:0 | 0:0:0:0:0:0 | 3:1:2:1:1:0 | 0:0:0:0:0:0 | 15:4:3:5:12:0 | 0:0:0:0:0:0 | 3:0:0:0:2:0 |
| NM_018344:NR_033414:XM_017016378:XM_017016377:NR_033413:NM_001174098 | 0:0:0:0:0:0 | 2:2:1:0:0:0 | 0:0:0:0:0:0 | 3:1:0:0:0:0 | 0:0:0:0:0:0 | 3:1:1:1:1:0 | 0:0:0:0:0:0 | 15:8:9:5:12:0 | 0:0:0:0:0:0 | 3:0:0:0:2:0 |
| NM_022153 | 0 | 0 | 0 | 0 | 0 | 0 | 0 | 0 | 0 | 0 |
| NM_001042465:NM_002778:NM_001042466 | 92:255:22 | 0:00:00 | 1.10568287 | 0:00:00 | 56:650:5 | 0:00:00 | 246:1135:61 | 0:03:00 | 24:255:3 | 0:00:00 |
| NM_001042465:NM_002778:NM_001042466 | 92:256:24 | 0:00:00 | 1.104293981 | 0:00:00 | 57:650:5 | 0:00:00 | 248:1129:60 | 0:03:01 | 24:255:3 | 0:00:00 |
| NM_002778:NM_001042465:NM_001042466 | 255:92:24 | 2:00:00 | 210:24:12 | 0:00:00 | 648:58:05 | 0:00:00 | 1113:248:62 | 2:01:00 | 254:24:03 | 0:00:00 |
| NM_001042465:NM_002778:NM_001042466 | 92:254:24 | 0:00:00 | 24:209:12 | 0:01:00 | 59:644:5 | 0:02:00 | 251:1104:63 | 1:01:01 | 24:254:3 | 0:00:00 |
| NM_001042465:NM_002778:NM_001042466 | 92:255:24 | 0:00:00 | 24:210:12 | 0:00:00 | 58:643:5 | 1:03:00 | 252:1096:64 | 0:04:00 | 24:255:3 | 0:01:00 |
| NM_001042465:NM_002778:NM_001042466 | 93:256:25 | 0:00:00 | 26:203:12 | 0:01:00 | 59:647:5 | 0:00:00 | 256:1034:65 | 0:07:01 | 24:254:3 | 0:01:00 |
| NM_001042465:NM_002778:NM_001042466 | 94:255:25 | 0:00:00 | 26:199:12 | 0:01:00 | 59:643:5 | 0:02:00 | 258:1024:65 | 0:03:01 | 24:254:3 | 0:01:00 |
| NM_001042465:NM_002778:NM_001042466 | 94:257:25 | 0:00:00 | 26:199:12 | 0:02:00 | 59:648:5 | 0:00:00 | 259:1011:65 | 0:02:01 | 24:254:3 | 0:01:00 |
| NM_001042465:NM_002778:NM_001042466 | 94:255:25 | 0:00:00 | 26:195:12 | 0:00:00 | 59:640:5 | 0:02:00 | 261:976:65 | 0:04:01 | 24:254:3 | 0:00:00 |
| NM_001042465:NM_002778:NM_001042466 | 94:255:25 | 0:00:00 | 26:194:12 | 0:01:00 | 59:641:5 | 0:01:00 | 260:975:65 | 1:05:01 | 24:253:3 | 0:01:00 |
| NM_002778:NM_001042465:NM_001042466 | 254:94:25 | 1:00:00 | 191:26:12 | 0:00:00 | 640:59:05 | 2:00:00 | 969:261:65 | 3:00:01 | 253:24:03 | 0:00:00 |
| NM_002778:NM_001042465:NM_001042466 | 255:95:25 | 1:00:00 | 190:25:12 | 0:01:00 | 646:59:05 | 1:00:00 | 958:262:65 | 4:00:01 | 253:24:03 | 1:00:00 |
| NM_001042466:NM_001042465:NM_002778 | 24:95:254 | 1:00:00 | 0.520196759 | 0:00:01 | 6:60:647 | 0:00:02 | 65:262:938 | 1:00:00 | 0.144583333 | 0:00:01 |
| NM_001042465:NM_002778:NM_001042466 | 95:242:25 | 0:00:00 | 26:171:12 | 0:00:00 | 60:635:6 | 0:00:00 | 262:798:66 | 0:04:00 | 24:244:3 | 0:02:00 |
| NM_001042465:NM_002778:NM_001042466 | 95:240:25 | 0:00:00 | 26:169:12 | 0:00:00 | 60:633:6 | 0:01:00 | 261:772:66 | 1:01:00 | 24:241:3 | 0:02:00 |
| NM_002778:NM_001042465:NM_001042466 | 237:95:25 | 1:00:00 | 166:26:12 | 0:00:00 | 627:60:6 | 0:00:00 | 749:262:66 | 2:00:00 | 240:24:03 | 0:00:00 |
| NM_002778:NM_001042465:NM_001042466 | 235:95:25 | 1:00:00 | 159:26:12 | 0:00:00 | 618:60:6 | 2:00:00 | 713:262:66 | 1:00:00 | 233:24:03 | 1:00:00 |
| NM_001042465:NM_002778:NM_001042466 | 88:219:20 | 0:00:00 | 0.963969907 | 0:01:00 | 52:573:6 | 0:01:00 | 208:524:51 | 1:00:00 | 1.10974537 | 0:00:00 |
| NM_001042465:NM_002778:NM_001042466 | 89:212:19 | 0:00:00 | 0.793125 | 0:01:00 | 51:568:6 | 0:00:00 | 190:462:41 | 0:02:00 | 1.063217593 | 0:02:00 |
| NM_001042465:NM_002778:NM_001042466 | 89:209:20 | 0:00:00 | 0.745208333 | 0:01:00 | 50:564:5 | 0:04:00 | 178:431:41 | 1:01:00 | 1.021539352 | 0:00:00 |
| NM_001042465:NM_002778:NM_001042466 | 90:210:20 | 0:00:00 | 0.828541667 | 0:01:00 | 50:564:4 | 0:01:01 | 176:431:41 | 0:01:00 | 1.021527778 | 0:00:01 |
| NM_002778:NM_001042465:NM_001042466 | 214:89:21 | 1:00:00 | 113:16:06 | 1:00:00 | 563:53:05 | 1:00:00 | 434:175:40 | 1:00:00 | 215:22:01 | 0:00:00 |
| XM_017015704:NM_006721:NM_001202450:XM_017015698:NM_001123:XM_017015702:XM_017015703:XM_017015706:XM_017015705:NM_001202449:XM_017015699 | 1:72:3:2:52:1:0:0:0:0:0 | 0:0:0:0:0:0:0:0:0:0:0 | 0:32:1:0:23:1:0:3:0:0:0 | 0:0:0:0:0:0:1:0:0:0:0 | 0:48:0:0:38:0:0:0:0:0:0 | 0:0:0:0:0:0:0:0:0:0:0 | 3:416:11:19:219:7:3:7:2:2:0 | 0:0:0:0:0:0:2:0:0:0:0 | 1:111:2:0:130:3:1:3:0:1:0 | 0:0:0:0:0:0:0:0:0:0:0 |
| XM_017015704:NM_006721:NM_001202450:XM_017015698:NM_001123:XM_017015702:XM_017015703:XM_017015706:XM_017015705:NM_001202449:XM_017015699 | 1:72:3:2:52:1:1:0:0:0:0 | 0:0:0:0:0:0:0:0:0:0:0 | 0:32:1:0:23:1:1:3:0:0:0 | 0:0:0:0:0:0:0:0:0:0:0 | 0:48:0:0:38:0:0:0:0:0:0 | 0:0:0:0:0:0:0:0:0:0:0 | 3:416:11:19:218:7:7:6:2:2:0 | 0:0:0:0:1:0:0:0:0:0:0 | 1:110:2:0:129:3:3:3:0:1:0 | 0:1:0:0:1:0:0:0:0:0:0 |
| XM_017015704:NM_006721:NM_001202450:XM_017015698:NM_001123:XM_017015702:XM_017015703:XM_017015706:XM_017015705:NM_001202449:XM_017015699 | 1:72:3:2:52:1:3:0:0:0:0 | 0:0:0:0:0:0:0:0:0:0:0 | 0:32:1:0:23:1:1:3:0:0:0 | 0:0:0:0:0:0:0:0:0:0:0 | 0:48:0:0:38:0:0:0:0:0:0 | 0:0:0:0:0:0:0:0:0:0:0 | 3:416:11:19:218:7:11:7:1:2:0 | 0:1:0:0:1:0:0:0:0:0:0 | 1:111:2:0:130:3:5:2:0:1:0 | 0:0:0:0:0:0:0:1:0:0:0 |
| XM_017015704:NM_006721:NM_001202450:XM_017015698:NM_001123:XM_017015702:XM_017015703:XM_017015706:XM_017015705:NM_001202449:XM_017015699 | 1:72:3:2:52:1:5:0:0:0:0 | 0:0:0:0:0:0:0:0:0:0:0 | 0:32:1:0:23:1:2:2:0:0:0 | 0:0:0:0:0:0:0:1:0:0:0 | 0:48:0:0:38:0:0:0:0:0:0 | 0:0:0:0:0:0:0:0:0:0:0 | 3:418:11:19:219:7:19:5:0:2:0 | 0:0:0:0:0:0:1:2:1:0:0 | 1:111:2:0:130:3:9:3:0:1:0 | 0:0:0:0:0:0:0:0:0:0:0 |
| XM_017015704:NM_006721:NM_001202450:XM_017015698:NM_001123:XM_017015702:XM_017015703:XM_017015706:NM_001202449:XM_017015699 | 1:72:3:2:52:1:5:0:0:0 | 0:0:0:0:0:0:0:0:0:0 | 0:32:1:0:23:1:2:2:0:0 | 0:0:0:0:0:0:0:0:0:0 | 0:48:0:0:38:0:0:0:0:0 | 0:0:0:0:0:0:0:0:0:0 | 3:419:11:20:219:7:21:5:2:0 | 0:1:0:0:0:0:1:0:0:0 | 1:111:2:0:129:3:9:3:1:0 | 0:0:0:0:1:0:0:0:0:0 |
| XM_017015704:NM_006721:XM_017015698:NM_001123:XM_017015702:XM_017015703:XM_017015706:NM_001202449:XM_017015699 | 1:72:2:52:1:6:0:0:0 | 0:0:0:0:0:0:0:0:0 | 0:32:0:23:1:5:2:0:0 | 0:0:0:0:0:0:0:0:0 | 0:48:0:38:0:1:0:0:0 | 0:0:0:0:0:0:0:0:0 | 3:418:20:219:7:42:5:2:0 | 0:2:0:0:0:0:0:0:0 | 1:110:0:130:3:11:1:1:0 | 0:1:0:0:0:0:0:0:0 |
| NM_006721:NM_001202450:XM_017015698:NM_001123:XM_017015702:XM_017015703:NM_001202449:XM_017015699 | 72:3:2:52:1:6:0:0 | 0:0:0:0:0:0:0:0 | 32:1:0:23:1:5:0:0 | 0:0:0:0:0:0:0:0 | 48:0:0:38:0:1:0:0 | 0:0:0:0:0:0:0:0 | 421:11:20:219:3:43:2:0 | 0:0:0:0:0:0:0:0 | 111:2:0:130:3:11:1:0 | 0:0:0:0:0:0:0:0 |
| XM_017015704:NM_006721:NM_001202450:XM_017015698:NM_001123:XM_017015703:NM_001202449:XM_017015699:XM_017015702 | 1:72:3:2:52:5:0:0:0 | 0:0:0:0:0:0:0:0:0 | 0:32:1:0:23:6:0:0:0 | 0:0:0:0:0:0:0:0:0 | 0:48:0:0:37:1:0:0:0 | 0:0:0:0:0:0:0:0:0 | 1:421:11:19:219:47:2:0:0 | 0:0:0:0:0:1:0:0:0 | 0:111:2:0:130:12:1:0:0 | 0:0:0:0:0:0:0:0:0 |
| XM_017015704:NM_006721:NM_001202450:XM_017015698:NM_001123:XM_017015703:NM_001202449:XM_017015706:XM_017015699:XM_017015702 | 1:72:3:2:52:5:0:0:0:0 | 0:0:0:0:0:0:0:0:0:0 | 0:32:1:0:23:6:0:0:0:0 | 0:0:0:0:0:0:0:0:0:0 | 0:48:0:0:37:1:0:0:0:0 | 0:0:0:0:0:0:0:0:0:0 | 1:419:11:19:217:48:2:0:0:0 | 0:2:0:0:1:0:0:0:0:0 | 0:111:2:0:130:10:1:1:0:0 | 0:0:0:0:0:0:0:0:0:0 |
| NM_006721:XM_017015704:NM_001202450:XM_017015698:NM_001123:XM_017015702:XM_017015703:NM_001202449:XM_017015705:XM_017015706:XM_017015699 | 71:1:3:1:52:1:5:0:0:0:0 | 1:0:0:0:0:0:0:0:0:0:0 | 32:0:1:0:23:1:6:0:0:0:0 | 0:0:0:0:0:0:0:0:0:0:0 | 48:0:0:0:37:0:1:0:0:0:0 | 0:0:0:0:0:0:0:0:0:0:0 | 419:1:11:1:215:3:47:2:1:0:0 | 2:0:0:0:1:0:0:0:0:0:0 | 111:0:2:0:129:2:11:1:0:1:0 | 0:0:0:0:0:0:0:0:0:0:0 |

| | | | | | | | | | | |
|---|---|---|---|---|---|---|---|---|---|---|
| XM_017015704:NM_006721:NM_001202450:XM_017015698:NM_001123:XM_017015702:XM_017015703:NM_001202449:XM_017015705:XM_017015706:XM_017015699 | 1:72:3:1:52:1:5:0:0:0:0 | 0:0:0:0:0:0:0:0:0:0:0 | 0:32:1:0:23:1:6:0:0:0:0 | 0:0:0:0:0:0:0:0:0:0 | 0:48:0:0:37:0:1:0:0:0:0 | 0:0:0:0:0:0:0:0:0:0:0 | 1:419:11:1:216:3:47:2:1:0:0 | 0:2:0:0:0:0:0:0:0:0:0 | 0:111:2:0:129:2:11:1:0:1:0 | 0:0:0:0:0:0:0:0:0:0 |
| XM_017015704:NM_006721:NM_001202450:XM_017015698:NM_001123:XM_017015702:XM_017015703:NM_001202449:XM_017015705:XM_017015706:XM_017015699 | 1:70:3:1:51:1:4:0:0:0:0 | 0:0:0:0:0:0:0:0:0:0:0 | 0:32:1:0:22:1:6:0:0:0:0 | 0:0:0:0:0:0:0:0:0:0 | 0:48:0:0:37:0:1:0:0:0:0 | 0:0:0:0:0:0:0:0:0:0:0 | 1:415:11:1:214:3:40:2:2:0:0 | 0:0:0:0:1:0:1:0:0:0:0 | 0:111:2:0:126:3:11:1:1:1:0 | 0:0:0:0:0:0:0:0:0:0 |
| NM_001142282:NM_033022:NM_001026 | 13:07:01 | 0:00:00 | 18:04:00 | 0:00:00 | 14:03:00 | 0:00:00 | 39:19:08 | 0:00:00 | 9:06:11 | 0:00:00 |
| NM_004329:XM_011540104 | 5:00 | 3:00 | 1:00 | 0:00 | 3:00 | 4:00 | 4:03 | 3:04 | 4:00 | 4:01 |
| NM_004329:XM_011540104 | 5:01 | 3:00 | 3:00 | 3:00 | 3:00 | 10:00 | 23:07 | 0:00 | 13:01 | 0:00 |
| NM_004897:XM_011540379:XM_006718078:NM_001178117:NM_001178118:XM_017016965 | 57:3:1:0:0:0 | 0:0:0:0:0:0 | 53:13:0:0:0:0 | 0:0:0:0:0:0 | 148:4:1:0:0:0 | 0:0:0:0:0:0 | 352:118:1:1:1:0 | 2:0:0:0:0:0 | 71:5:1:0:0:0 | 1:0:0:0:0:0 |
| NM_001304717:NM_001304718 | 0:00 | 177:01:00 | 0:00 | 187:03:00 | 0:00 | 147:02:00 | 1:00 | 765:08:00 | 0:00 | 149:03:00 |
| NM_001304717:NM_001304718 | 284:02:00 | 0:00 | 328:04:00 | 1:00 | 216:03:00 | 0:00 | 1459:14:00 | 3:00 | 243:04:00 | 1:00 |
| NM_001304717:NM_001304718 | 283:02:00 | 1:00 | 328:04:00 | 2:00 | 216:03:00 | 0:00 | 1463:14:00 | 2:00 | 244:04:00 | 0:00 |
| NM_001304717:NM_001304718 | 283:02:00 | 0:00 | 329:04:00 | 1:00 | 215:03:00 | 1:00 | 1465:14:00 | 2:00 | 244:04:00 | 0:00 |
| NM_001304717:NM_001304718 | 285:02:00 | 0:00 | 338:03:00 | 0:00 | 217:03:00 | 0:00 | 1488:14:00 | 4:00 | 244:04:00 | 0:00 |
| NM_001304717:NM_001304718 | 285:01:00 | 0:00 | 337:06:00 | 0:00 | 217:02:00 | 0:00 | 1470:15:00 | 1:01 | 237:02:00 | 2:00 |
| NM_001304717:NM_001304718 | 286:07:00 | 0:00 | 333:19:00 | 1:00 | 216:07:00 | 1:00 | 1487:56:00 | 3:01 | 240:06:00 | 1:00 |
| NM_001304717:NM_001304718 | 286:08:00 | 0:00 | 334:22:00 | 0:00 | 216:07:00 | 1:00 | 1484:65 | 4:00 | 240:06:00 | 0:00 |
| NM_001304717:NM_001304718 | 286:09:00 | 0:00 | 331:22:00 | 3:00 | 215:08:00 | 2:00 | 1479:69 | 7:00 | 240:06:00 | 0:00 |
| NM_001304717:NM_001304718 | 284:10:00 | 1:00 | 294:26:00 | 14:02 | 209:08:00 | 4:00 | 1463:73 | 6:01 | 233:07:00 | 6:00 |
| NM_001304717:NM_001304718 | 284:10:00 | 1:00 | 296:27:00 | 12:01 | 209:08:00 | 3:00 | 1458:75 | 11:01 | 233:07:00 | 6:00 |
| NM_001304717:NM_001304718 | 280:10:00 | 0:00 | 280:30:00 | 0:01 | 199:07:00 | 0:00 | 1392:81 | 2:00 | 229:06:00 | 0:00 |
| NM_001304717:NM_001304718 | 279:08:00 | 1:00 | 280:30:00 | 0:01 | 198:07:00 | 1:00 | 1392:82 | 2:00 | 228:06:00 | 1:00 |
| NM_001304717:NM_001304718 | 279:08:00 | 0:00 | 273:30:00 | 3:01 | 198:09:00 | 1:00 | 1368:86 | 10:01 | 228:05:00 | 0:00 |
| NM_001031709:XM_005269946:XM_017016385:XM_005269948:XM_005269947:XM_017016384:XR_001747122 | 5:0:0:0:0:0:0 | 6:0:0:0:0:0:0 | 7:0:1:0:0:0:0 | 7:1:0:0:0:0:0 | 16:1:0:1:0:0:0 | 12:0:0:0:0:0:0 | 59:2:0:5:3:1:1 | 0:0:0:0:0:0:0 | 5:0:0:0:0:0:0 | 0:0:0:0:0:0:0 |
| XM_011539766:NM_000043:NM_152871:XM_011539765:NM_152872:XM_011539767:NR_135314:XR_945733:NM_001320619:XR_945732:NR_135313 | 1:11:3:1:0:0:0:0:0:0:0 | 0:0:0:0:0:0:0:0:0:0:0 | 0:88:5:0:6:3:1:0:0:0:0 | 0:1:0:0:0:0:0:0:0:0:0 | 0:181:9:0:3:5:4:1:1:1:0 | 0:1:1:0:0:0:0:0:0:0:0 | 0:27:4:0:3:2:1:0:0:0:0 | 0:0:0:0:0:0:0:0:0:0:0 | 0:13:1:0:1:3:2:0:0:0:0 | 0:0:0:0:0:0:0:0:0:0 |
| XM_011539766:NM_000043:NM_152871:XM_011539765:NM_152872:NR_028036:XM_011539767:NR_028035:NR_135314:XR_945732:XR_945733:NR_028034:NR_135313:NR_135315 | 1:11:3:1:0:0:0:0:0:0:0:0:0:0 | 0:0:0:0:0:0:0:0:0:0:0:0:0:0 | 0:89:5:0:0:1:3:1:1:2:1:0:0:0 | 0:0:0:0:3:0:0:0:0:0:0:0:0:0 | 0:182:10:0:1:0:8:0:4:2:1:1:0:0 | 0:0:0:0:0:0:0:0:0:0:0:0:0:0 | 0:27:4:0:0:0:4:0:1:0:0:0:0:0 | 0:0:0:0:0:0:0:0:0:0:0:0:0:0 | 0:13:1:0:0:0:2:0:2:0:1:0:0:0 | 0:0:0:0:0:0:0:0:0:0:0:0:0 |
| XM_011539766:NM_000043:NM_152871:XM_011539765:NR_028036:XM_011539767:NR_028035:NR_135314:XR_945732:XR_945733:NR_135313:NR_028034:NM_152872:NM_001320619:NR_135315 | 1:11:3:1:0:0:0:0:0:0:0:0:0:0:0 | 0:0:0:0:0:0:0:0:0:0:0:0:0:0:0 | 0:87:4:0:1:3:1:1:2:1:1:0:0:0:0 | 0:2:0:0:0:0:0:0:0:0:0:0:0:0:0 | 0:181:9:0:0:7:0:4:2:4:0:1:1:1:1 | 0:1:0:0:0:0:0:0:0:0:0:0:0:0:0 | 0:27:4:0:0:4:0:1:0:3:0:0:0:0:0 | 0:0:0:0:0:0:0:0:0:0:0:0:0:0:0 | 0:13:1:0:0:2:0:2:0:2:0:0:0:0:0 | 0:0:0:0:0:0:0:0:0:0:0:0:0:0 |
| XM_011539766:NM_000043:NM_152871:XM_011539765:XR_945733:NR_028036:XM_011539767:NR_028035:NR_135314:XR_945732:NR_135313:NR_028034:NM_152872:NM_001320619:NR_135315 | 1:11:3:1:0:0:0:0:0:0:0:0:0:0:0 | 0:0:0:0:0:0:0:0:0:0:0:0:0:0:0 | 0:86:3:0:3:1:3:1:1:2:1:0:0:0:0 | 0:1:1:0:1:0:0:0:0:0:0:0:0:0:0 | 0:179:9:0:6:0:6:0:4:2:0:1:1:1:1 | 0:1:0:0:0:0:0:0:0:0:0:0:0:0:0 | 0:27:4:0:3:0:4:0:1:0:0:0:0:0:0 | 0:0:0:0:0:0:0:0:0:0:0:0:0:0:0 | 0:13:1:0:2:0:2:0:2:0:0:0:0:0:0 | 0:0:0:0:0:0:0:0:0:0:0:0:0:0 |
| XM_011539766:NM_000043:NM_152871:XM_011539765:NR_028036:XM_011539767:NR_028035:NR_135314:XR_945732:XR_945733:NR_135313:NR_028034:NM_152872:NM_001320619:NR_135315 | 1:11:3:1:0:0:0:0:0:0:0:0:0:0:0 | 0:0:0:0:0:0:0:0:0:0:0:0:0:0:0 | 0:84:4:0:1:3:1:1:2:4:1:0:0:0:0 | 0:0:0:0:0:0:0:0:0:0:0:0:0:0:0 | 0:175:8:0:0:5:0:4:2:7:0:1:1:1:1 | 0:2:0:0:0:0:0:0:0:0:0:0:0:0:0 | 0:26:4:0:0:4:0:1:0:3:0:0:0:0:0 | 0:0:0:0:0:0:0:0:0:0:0:0:0:0:0 | 0:13:1:0:0:2:0:2:0:2:0:0:0:0:0 | 0:0:0:0:0:0:0:0:0:0:0:0:0:0 |
| NM_016195:NM_001284259 | 3:00 | 0:00 | 0:00 | 0:00 | 2:00 | 0:00 | 5:00 | 0:00 | 8:00 | 0:00 |
| NM_016195:NM_001284259 | 2:00 | 1:00 | 0:00 | 1:00 | 2:00 | 2:00 | 15:00 | 0:00 | 12:00 | 0:00 |
| NM_016195:NM_001284259 | 3:00 | 0:00 | 3:00 | 0:00 | 4:00 | 0:00 | 18:00 | 0:00 | 13:00 | 0:00 |
| NM_016195:NM_001284259 | 3:00 | 0:00 | 3:00 | 0:00 | 4:00 | 0:00 | 19:00 | 0:00 | 13:00 | 0:00 |
| NM_016195 | 1 | 0 | 3 | 0 | 2 | 0 | 0 | 5 | 6 | 2 |
| NM_016195 | 1 | 0 | 1 | 2 | 1 | 1 | 4 | 0 | 8 | 0 |
| NM_016195 | 1 | 0 | 0 | 0 | 2 | 0 | 0 | 1 | 2 | 0 |
| NM_016195 | 1 | 0 | 0 | 0 | 1 | 1 | 0 | 0 | 3 | 0 |
| XM_017016373 | 0 | 0 | 0 | 0 | 0 | 0 | 0 | 0 | 0 | 0 |
| NM_145246:NM_001347713 | 0:00 | 0:00 | 0:00 | 0:00 | 0:00 | 0:00 | 0:00 | 1:01 | 0:00 | 1:00 |
| NM_000772:NM_001128925 | 0:00 | 0:00 | 37:01:00 | 0:00 | 81:00:00 | 1:00 | 5:00 | 0:00 | 4:00 | 0:00 |
| NM_000771 | 0 | 0 | 0 | 0 | 0 | 2 | 0 | 0 | 0 | 0 |
| NM_000771 | 0 | 0 | 0 | 0 | 0 | 2 | 0 | 0 | 0 | 0 |
| NM_000771 | 0 | 0 | 0 | 0 | 0 | 2 | 0 | 0 | 0 | 0 |
| NM_020992 | 1 | 0 | 2 | 1 | 3 | 3 | 0 | 3 | 0 | 2 |
| XM_017015653:XR_945594:XM_017015654 | 0:00:00 | 0:00:00 | 0:00:00 | 0:00:00 | 0:00:00 | 0:00:00 | 0:00:00 | 0:00:00 | 0:00:00 | 1:00:00 |
| NM_004398 | 22 | 0 | 7 | 0 | 6 | 0 | 143 | 0 | 40 | 0 |
| NM_004398 | 26 | 0 | 16 | 0 | 8 | 0 | 184 | 0 | 43 | 0 |
| NM_004398 | 21 | 2 | 33 | 0 | 9 | 0 | 179 | 10 | 40 | 2 |
| NM_004398 | 16 | 1 | 18 | 2 | 9 | 0 | 123 | 6 | 31 | 3 |
| NM_031938 | 0 | 1 | 0 | 1 | 0 | 1 | 0 | 0 | 0 | 0 |
| NM_031938 | 0 | 1 | 0 | 1 | 0 | 1 | 0 | 1 | 0 | 0 |
| XM_017017209:XM_011542604:XR_001747769:NM_182495 | 1:0:0:0 | 0:0:0:0 | 2:4:2:0 | 0:0:0:0 | 1:1:1:1 | 0:0:0:0 | 0:0:0:0 | 0:0:1:0 | 0:0:2:0 | 0:1:0:0 |
| XM_017017209:XM_011542604:XR_001747769:NM_182495 | 0:0:0:0 | 1:0:0:0 | 1:2:1:0 | 2:2:0:0 | 1:0:1:0 | 0:0:0:0 | 0:0:0:0 | 0:0:0:0 | 0:0:2:0 | 0:1:0:0 |
| NM_003904:XR_001748023 | 0:00 | 0:00 | 0:00 | 0:00 | 0:00 | 0:00 | 0:00 | 0:00 | 1:00 | 0:00 |
| XM_017017365:XR_428971:XM_017017382:XM_017017384:XM_017017367:XM_017017377:XM_006718794:XM_017017381:XM_017017366:XM_017017380:XM_017017379:XM_005271456:XM_017017385:XR_002957133:XM_017017386 | 1:8:1:1:0:0:0:0:0:0:0:0:0:0:0 | 0:0:0:0:0:0:0:0:0:0:0:0:0:0:0 | 0:10:3:0:6:1:0:0:0:0:0:0:0:0:0 | 0:0:0:0:0:0:0:0:0:0:0:0:0:0:0 | 1:9:4:0:3:0:1:1:0:0:0:0:0:0:0 | 0:1:0:0:0:0:0:0:0:0:0:0:0:0:0 | 7:30:6:0:4:1:0:0:2:2:2:1:0:0:0 | 0:0:0:0:0:0:0:0:0:0:0:0:0:0:0 | 2:6:1:0:0:0:0:0:0:0:0:0:0:0:0 | 0:0:0:0:0:0:0:0:0:0:0:0:0:0 |
| XM_017017382:XM_017017384:XR_428971:XM_006718794:XM_017017381:XM_005271456:XM_017017379:XM_017017380:XM_017017367:XM_017017385:XM_017017386 | 1:1:0:0:0:0:0:0:0:0:0 | 0:0:0:0:0:0:0:0:0:0:0 | 0:0:0:0:0:0:0:0:0:0:0 | 0:0:0:0:0:0:0:0:0:0 | 0:0:0:1:1:0:0:0:0:0:0 | 0:0:1:0:0:0:0:0:0:0:0 | 1:0:0:0:0:0:1:2:0:0:0 | 0:0:0:0:0:1:1:0:0:0:0 | 0:0:0:0:0:0:0:0:0:0:0 | 0:0:0:0:0:0:0:0:0:0 |
| NM_016146:NM_001318486:NM_001318492:NM_001318490:NM_001318489:NM_001318488:NM_001318494 | 164:5:3:1:0:0:0 | 0:0:0:0:0:0:0 | 232:9:6:1:1:0:0 | 1:0:0:0:0:0:0 | 472:10:6:6:0:0:0 | 2:0:1:0:0:0:0 | 1248:35:14:21:1:6:6 | 3:0:0:0:0:0:0 | 198:5:5:1:0:0:1 | 0:0:0:0:0:0:0 |
| NM_016146:NM_001318486:NM_001318492:NM_001318490:NM_001318488:NM_001318489 | 168:6:3:1:0:0 | 0:0:0:0:0:0 | 234:10:6:1:0:0 | 0:0:0:0:0:0 | 479:10:7:6:0:0 | 0:0:0:0:0:0 | 1273:35:15:21:6:1 | 1:0:0:0:0:0 | 200:5:5:1:0:0 | 0:0:0:0:0:0 |
| NM_016146:NM_001318486:NM_001318492:NM_001318488 | 173:6:3:0 | 0:0:0:0 | 236:10:6:0 | 0:0:0:0 | 489:11:7:0 | 0:0:0:0 | 1287:33:15:6 | 2:0:0:0 | 201:6:5:0 | 1:0:0:0 |
| NM_016146:NM_001318486:NM_001318492:NM_001318490:NM_001318489:NM_001318494 | 176:6:3:1:0:0 | 0:0:0:0:0:0 | 239:10:5:1:1:0 | 0:0:0:0:0:0 | 494:11:6:6:0:0 | 1:0:0:0:0:0 | 1302:35:13:21:1:6 | 2:0:0:0:0:0 | 205:6:5:1:0:1 | 1:0:0:0:0:0 |
| NM_016146:NM_001318486:NM_001318492:NM_001318490:NM_001318489:NM_001318488:NM_001318494 | 176:6:3:1:0:0:0 | 0:0:0:0:0:0:0 | 255:10:6:1:1:0:0 | 0:0:0:0:0:0:0 | 499:11:7:6:0:0:0 | 0:0:0:0:0:0:0 | 1313:34:15:21:1:6:6 | 3:1:0:0:0:0:0 | 208:6:5:1:0:0:1 | 0:0:0:0:0:0:0 |
| XM_017018334:NM_001168468:XM_017018335:XM_011543005 | 0:0:0:0 | 0:0:0:0 | 3:0:0:0 | 3:0:0:0 | 2:0:0:1 | 3:1:1:0 | 0:0:0:0 | 11:1:0:1 | 0:0:0:0 | 1:0:0:0 |

| | | | | | | | | | | |
|---|---|---|---|---|---|---|---|---|---|---|
| NM_005188 | 0 | 1 | 0 | 3 | 1 | 1 | 1 | 0 | 1 | 0 |
| NM_178507 | 0 | 1 | 0 | 10 | 0 | 9 | 1 | 7 | 1 | 0 |
| NM_178507 | 0 | 1 | 3 | 7 | 6 | 3 | 1 | 7 | 1 | 0 |
| XM_011542739:NM_014352:XM_024448424 | 0:00:00 | 0:00:00 | 0:00:00 | 0:00:00 | 1:00:00 | 0:00:00 | 0:00:00 | 0:01:03 | 0:00:00 | 0:00:00 |
| NM_006918:NM_001024956 | 11:00 | 0:00 | 17:01 | 0:00 | 16:00 | 0:00 | 85:02:00 | 0:00 | 15:00 | 0:00 |
| XM_005253015:NM_018222 | 5:51 | 0:00 | 1:47 | 0:00 | 0.483333333 | 0:00 | 0.755555556 | 0:01 | 0.338888889 | 0:00 |
| XM_005253015:NM_018222 | 5:51 | 0:00 | 1:47 | 0:00 | 0.440972222 | 1:01 | 0.75625 | 0:00 | 0.338888889 | 0:00 |
| NM_001258239:NM_001258238:NM_001258244:XM_024448388:XM_024448384:XM_006718783:NM_001080546:XM_017017324:XM_024448386:NM_001258245:NM_001258242:NR_047588 | 0:0:0:0:0:0:0:0:0:0:0:0:0 | 1:0:0:0:0:0:0:0:0:0:0:0 | 0:0:0:0:0:0:0:0:0:0:0:0 | 0:1:1:1:0:0:0:0:0:0:0:0 | 0:0:0:0:0:0:0:0:0:0:0:0:0 | 0:0:0:0:3:0:0:0:0:0:0:0 | 0:0:0:0:0:0:0:0:0:0:0:0:0 | 1:2:0:0:2:1:3:1:0:0:0:0 | 0:0:0:0:0:0:0:0:0:0:0:0 | 0:0:1:0:1:0:0:0:1:0:0:0 |
| NM_001114122:XM_011542560 | 0:00 | 0:00 | 0:00 | 0:00 | 0:00 | 0:00 | 0:00 | 0:00 | 0:00 | 1:01 |
| NM_030754 | 0 | 2 | 0 | 0 | 0 | 0 | 0 | 0 | 0 | 0 |
| NM_199161 | 1 | 3 | 0 | 0 | 0 | 0 | 0 | 0 | 0 | 0 |
| NM_199161 | 1 | 3 | 0 | 0 | 0 | 0 | 0 | 0 | 0 | 0 |
| NM_001098520:NM_006410:NM_001098521:NM_001098522 | 0:0:0:0 | 2:0:0:0 | 0:0:0:0 | 0:2:1:1 | 0:0:0:0 | 3:2:0:1 | 0:0:0:0 | 0:1:0:0 | 0:0:0:0 | 2:0:1:1 |
| NM_002817:NM_175932 | 0:00 | 1:00 | 0:00 | 1:00 | 0:00 | 0:00 | 0:00 | 1:00 | 0:00 | 1:00 |
| NM_001315502:NM_183233:NM_002555 | 0:00:00 | 1:00:00 | 0:00:00 | 0:00:00 | 0:00:00 | 0:00:00 | 0:00:00 | 1:01:00 | 0:00:00 | 0:00:01 |
| NM_005969 | 60 | 0 | 35 | 0 | 29 | 0 | 248 | 2 | 88 | 0 |
| NM_005969 | 65 | 1 | 47 | 1 | 36 | 0 | 230 | 4 | 98 | 0 |
| NM_001751:XM_006718341:NM_139273:XR_930914:XM_017018391:XM_017018389:NM_001014437:XR_002957203:XR_001747995:NR_036542:NM_001194997:XR_001747996:XR_001747998:XR_930913:XR_001747997:XM_017018392:XM_017018390:XM_024448705:XM_024448703 | 99:28:57:22:8:12:7:3:3:8:7:4:4:3:1:2:0:0:0 | 1:0:0:0:0:0:0:0:0:0:0:0:0:0:0:0:0:0:0 | 29:10:21:18:17:4:3:3:2:22:2:0:5:2:0:11:0:0:0 | 0:0:0:0:0:0:0:0:0:0:0:0:0:0:0:0:0:0:0 | 127:23:66:33:16:20:13:3:0:29:9:7:9:4:1:4:0:0:0 | 0:0:0:0:0:0:0:0:0:0:0:0:0:0:0:0:0:0:0 | 168:30:146:94:152:13:13:18:19:133:5:7:26:15:3:92:4:1:0 | 0:0:1:0:1:0:0:0:0:0:0:0:0:0:0:0:0:0:0 | 116:31:71:24:13:17:17:1:2:22:3:3:8:1:0:5:1:0:0 | 0:0:0:0:0:0:0:0:0:0:0:0:0:0:0:0:0:0:0 |
| XM_006718341:NM_139273:XR_930914:NM_001751:XM_017018391:XM_017018389:NM_001014437:XR_001747995:NR_036542:NM_001194997:XR_001747996:XR_001747998:XR_930913:XR_001747997:XR_002957203:XM_017018392:XM_017018390:XM_024448705:XR_002957204:XM_024448703 | 28:57:22:100:8:12:7:3:8:7:6:4:4:1:1:2:0:0:0:0 | 0:0:0:0:0:0:0:0:0:0:0:0:0:0:0:0:0:0:0:0 | 10:22:13:29:17:6:3:2:15:2:9:5:2:0:0:12:0:0:0:0 | 0:0:0:0:0:0:0:0:0:0:0:0:0:0:0:0:0:0:0:0 | 23:65:28:127:16:22:13:0:25:10:21:7:7:1:2:5:0:0:0:0 | 0:0:1:0:0:0:0:0:0:0:0:0:0:0:0:0:0:0:0:0 | 30:140:86:168:154:17:13:19:124:6:38:23:29:2:11:95:4:1:0:0 | 0:0:0:0:0:0:0:0:0:0:0:0:0:0:0:0:0:0:0:0 | 31:71:19:116:13:17:17:2:21:3:11:8:3:0:0:6:1:0:0:0 | 0:0:1:0:0:0:0:0:0:0:0:0:0:0:0:0:0:0:0:0 |
| XM_006718341:NM_139273:XR_930914:NM_001751:XM_017018391:XM_017018389:NM_001014437:XR_001747995:NR_036542:NM_001194997:XR_930913:XR_001747997:XR_001747998:XR_001747996:XM_017018392:XR_002957203:XM_017018390:XM_024448705:XR_002957204:XM_024448703 | 28:54:19:100:8:11:7:3:6:7:4:1:1:4:2:0:0:0:0:0 | 0:0:0:0:0:0:0:0:0:0:0:0:0:0:0:0:0:0:0:0 | 10:19:8:29:17:6:3:2:12:2:3:0:4:9:12:0:0:0:0:0 | 0:0:0:0:0:0:0:0:0:0:0:0:0:0:0:0:0:0:0:0 | 23:64:21:126:16:22:13:0:20:10:7:1:6:16:5:0:0:0:0:0 | 0:0:1:0:0:0:0:0:0:0:0:0:0:0:0:0:0:0:0:0 | 30:137:68:168:154:16:13:19:116:6:27:2:20:37:95:8:4:1:0:0 | 0:0:0:0:0:0:0:0:0:0:0:0:0:0:0:0:0:0:0:0 | 30:69:13:116:13:17:17:2:17:3:3:0:6:9:6:0:1:0:0:0 | 1:0:0:0:0:0:0:0:0:0:0:0:0:0:0:0:0:0:0:0 |
| XM_006718341:NM_139273:XR_930914:NM_001751:XM_017018391:XM_017018389:NM_001014437:XR_001747995:NR_036542:NM_001194997:XR_001747996:XR_930913:XR_001747997:XR_001747998:XM_017018392:XR_002957203:XM_017018390:XM_024448705:XR_002957204:XM_024448703 | 28:51:14:100:8:12:7:3:5:7:4:4:1:1:2:0:0:0:0:0 | 0:0:0:0:0:0:0:0:0:0:0:0:0:0:0:0:0:0:0:0 | 10:18:5:29:17:6:3:2:11:2:9:4:0:4:12:0:0:0:0:0 | 0:0:0:0:0:0:0:0:0:0:0:0:0:0:0:0:0:0:0:0 | 23:64:17:127:16:22:13:0:19:10:20:8:1:5:5:0:0:0:0:0 | 0:0:0:0:0:0:0:0:0:0:0:0:0:0:0:0:0:0:0:0 | 30:137:45:168:154:17:13:19:109:6:40:31:2:11:94:7:4:1:0:0 | 0:0:0:0:0:0:0:0:1:0:0:0:0:0:1:0:0:0:0:0 | 31:69:8:116:13:17:17:2:15:3:11:3:0:4:6:0:1:0:0:0 | 0:0:0:0:0:0:0:0:0:0:0:0:0:0:0:0:0:0:0:0 |
| XM_006718341:NM_139273:XR_930914:NM_001751:XM_017018391:XM_017018389:NM_001014437:XR_001747995:NR_036542:NM_001194997:XR_001747996:XR_930913:XR_001747997:XR_001747998:XM_017018392:XR_002957203:XM_017018390:XM_024448705:XR_002957204:XM_024448703 | 28:51:14:100:8:12:7:3:5:7:4:4:1:1:2:0:0:0:0:0 | 0:0:0:0:0:0:0:0:0:0:0:0:0:0:0:0:0:0:0:0 | 10:18:4:29:17:6:3:2:11:2:9:4:0:4:12:0:0:0:0:0 | 0:0:0:0:0:0:0:0:0:0:0:0:0:0:0:0:0:0:0:0 | 23:64:17:127:16:22:13:0:19:10:20:8:1:4:5:0:0:0:0:0 | 0:0:0:0:0:0:0:0:0:0:0:0:0:0:0:0:0:0:0:0 | 30:135:41:168:154:17:13:19:107:6:40:30:2:11:95:6:4:1:0:0 | 0:0:1:0:0:0:0:0:1:0:0:0:0:0:0:0:0:0:0:0 | 31:68:7:116:13:17:17:2:14:3:11:3:0:4:6:0:1:0:0:0 | 0:0:0:0:0:0:0:0:0:0:0:0:0:0:0:0:0:0:0:0 |
| XM_006718341:NM_139273:XR_930914:NM_001751:XM_017018391:XM_017018389:NM_001014437:XR_001747995:NR_036542:NM_001194997:XR_001747996:XR_930913:XR_001747997:XM_017018392:XR_001747998:XM_017018390:XM_024448705:XR_002957203:XR_002957204:XM_024448703 | 28:51:6:100:8:12:7:3:6:7:6:4:1:2:0:0:0:0:0:0 | 0:0:0:0:0:0:0:0:0:0:0:0:0:0:0:0:0:0:0:0 | 10:17:2:29:17:6:3:2:9:2:9:4:0:12:3:0:0:0:0:0 | 0:0:0:0:0:0:0:0:0:0:0:0:0:0:0:0:0:0:0:0 | 23:62:9:127:16:24:13:0:16:10:21:8:1:5:0:0:0:0:0:0 | 0:0:0:0:0:0:0:0:0:0:0:0:0:0:0:0:0:0:0:0 | 30:129:14:168:153:17:13:19:99:6:43:33:2:95:3:4:1:2:0:0 | 0:1:0:0:1:0:0:0:0:0:0:0:0:0:0:0:0:0:0:0 | 31:67:2:116:14:17:17:2:12:3:12:3:0:6:3:1:0:0:0:0 | 0:0:0:0:0:0:0:0:0:0:0:0:0:0:0:0:0:0:0:0 |
| XM_006718341:NM_139273:XR_930914:NM_001751:XM_017018391:XM_017018389:NM_001014437:XR_001747995:NR_036542:NM_001194997:XR_001747996:XR_930913:XR_001747997:XM_017018392:XR_001747998:XM_017018390:XM_024448705:XR_002957204:XM_024448703 | 28:52:5:100:5:12:7:3:3:7:8:3:1:1:0:0:0:0:0 | 0:0:0:0:0:0:0:0:0:0:0:0:0:0:0:0:0:0:0 | 10:15:0:29:12:6:3:1:4:2:10:4:0:7:1:0:0:0:0 | 0:0:0:0:0:0:0:0:0:0:0:0:0:0:0:0:0:0:0 | 23:61:11:127:14:24:13:0:9:10:23:6:1:2:0:0:0:0:0 | 0:0:0:0:0:0:0:0:0:0:0:0:0:0:0:0:0:0:0 | 30:110:9:168:92:17:13:15:67:6:45:26:2:56:2:4:1:0:0 | 0:0:0:0:0:0:0:0:0:0:0:0:1:0:0:0:0:0:0 | 31:65:1:116:7:17:17:2:8:3:12:2:0:4:3:1:0:0:0 | 0:0:0:0:0:0:0:0:0:0:0:0:0:0:0:0:0:0:0 |
| XM_006718341:NM_139273:XR_930914:NM_001751:XM_017018389:NM_001014437:XR_001747995:NR_036542:NM_001194997:XM_017018391:XR_001747996:XR_930913:XR_001747997:XR_001747998:XM_017018392:XM_017018390:XR_002957203:XM_024448705:XM_024448703 | 28:51:7:100:12:7:3:2:7:4:8:2:1:1:1:0:0:0:0 | 0:0:0:0:0:0:0:0:0:0:0:0:0:0:0:0:0:0:0 | 10:14:0:29:6:3:1:3:2:8:10:2:0:0:6:0:0:0:0 | 0:0:0:0:0:0:0:1:0:0:0:0:0:0:0:0:0:0:0 | 23:60:11:127:24:13:0:7:10:7:23:5:1:1:1:0:0:0:0 | 0:0:0:0:0:0:0:0:0:0:0:0:0:0:0:0:0:0:0 | 30:94:11:167:18:13:6:39:6:43:47:20:2:3:25:4:1:0:0 | 0:1:0:1:0:0:0:0:0:0:0:0:0:0:1:0:0:0:0 | 31:64:3:116:17:17:2:8:3:7:12:2:0:4:3:1:0:0:0 | 0:0:0:0:0:0:0:0:0:0:0:0:0:0:0:0:0:0:0 |
| NM_139273:XM_006718341:XR_930914:NM_001751:XM_017018389:NM_001014437:XR_001747995:NR_036542:NM_001194997:XM_017018391:XR_001747996:XR_930913:XR_001747997:XR_001747998:XM_017018392:XM_017018390:XR_002957203:XM_024448705:XM_024448703 | 50:28:6:100:12:7:3:2:7:4:8:2:2:1:1:0:0:0:0 | 1:0:0:0:0:0:0:0:0:0:0:0:0:0:0:0:0:0:0 | 13:10:1:29:6:3:1:3:2:3:10:2:0:0:3:0:0:0:0 | 0:0:0:0:0:0:0:0:0:0:0:0:0:0:0:0:0:0:0 | 60:23:12:127:24:13:0:6:10:5:24:4:1:1:0:0:0:0:0 | 0:0:0:0:0:0:0:0:0:0:0:0:0:0:0:0:0:0:0 | 82:30:13:168:18:13:6:16:6:17:47:13:3:1:12:4:2:0:0 | 0:0:0:0:0:0:0:0:0:0:0:0:0:0:0:0:0:0:0 | 64:31:4:116:17:17:2:7:3:3:12:1:0:4:3:1:0:0:0 | 1:0:0:0:0:0:0:0:0:0:0:0:0:0:0:0:0:0:0 |
| NM_139273:XM_006718341:XR_930914:NM_001751:XM_017018389:NM_001014437:XR_001747995:NR_036542:NM_001194997:XM_017018391:XR_001747996:XR_930913:XR_001747997:XR_001747998:XM_017018392:XM_017018390:XR_002957203:XM_024448703 | 51:28:9:100:12:7:3:1:7:3:8:2:1:1:0:0:0:0 | 1:0:0:0:0:0:0:0:0:0:0:0:0:0:0:0:0:0 | 13:9:1:29:6:3:1:3:2:0:10:2:0:0:2:0:0:0 | 0:0:0:0:0:0:0:0:0:0:0:0:0:0:0:0:0:0 | 60:21:15:127:24:13:0:12:10:5:24:2:1:4:0:0:0:0 | 0:0:0:0:0:0:0:0:0:0:0:0:0:0:0:0:0:0 | 80:20:15:166:18:13:5:13:6:5:48:8:3:3:6:2:3:0 | 0:0:1:0:0:0:0:1:0:0:0:0:0:0:0:0:0:0 | 64:28:6:116:17:17:2:3:3:2:12:1:0:5:1:0:0:0 | 0:0:0:0:0:0:0:0:0:0:0:0:0:0:0:0:0:0 |
| NM_001751:XR_001747996:XM_006718341:NM_139273:XR_930914:XM_017018389:NM_001014437:XR_001747995:NR_036542:NM_001194997:XM_017018391:XR_930913:XR_001747997:XR_001747998:XR_002957203:XM_017018392:XM_017018390:XM_024448703 | 98:7:28:53:9:12:7:3:2:7:3:2:1:1:1:0:0:0 | 1:1:0:0:0:0:0:0:0:0:0:0:0:0:0:0:0:0 | 26:10:9:13:1:6:3:1:4:2:0:1:0:0:0:2:0:0 | 1:0:0:0:0:0:0:0:0:0:0:0:0:0:0:0:0:0 | 126:24:21:59:14:24:13:0:13:10:4:2:1:4:0:0:0:0 | 1:0:0:1:0:0:0:0:0:0:0:0:0:0:0:0:0:0 | 157:48:19:81:23:18:13:3:18:6:4:8:3:3:3:6:2:0 | 1:0:0:0:0:0:0:1:0:0:0:1:0:0:0:0:0:0 | 116:12:28:64:7:17:17:2:5:3:2:1:0:5:0:0:0:0 | 0:0:0:0:0:0:0:0:0:0:0:0:0:0:0:0:0:0 |
| NM_006435 | 0 | 1384 | 1 | 547 | 1 | 1044 | 3256 | 1 | 960 | 2 |
| NM_006435 | 962 | 423 | 296 | 253 | 597 | 450 | 3258 | 3 | 963 | 1 |
| NM_006435 | 1391 | 0 | 553 | 1 | 1047 | 6 | 3269 | 3 | 962 | 3 |
| NM_006435 | 1392 | 0 | 555 | 0 | 1053 | 2 | 3273 | 2 | 963 | 3 |
| NM_006435 | 1391 | 0 | 554 | 2 | 1056 | 0 | 3268 | 4 | 964 | 2 |
| NM_006435 | 1388 | 1 | 555 | 0 | 1056 | 4 | 3270 | 5 | 966 | 1 |
| NM_006435 | 1385 | 5 | 552 | 0 | 1060 | 2 | 3273 | 4 | 966 | 0 |
| NM_006435 | 967 | 424 | 297 | 254 | 603 | 458 | 3274 | 1 | 966 | 0 |
| NM_006435 | 1388 | 0 | 548 | 0 | 1058 | 2 | 3272 | 2 | 964 | 1 |
| NM_003641 | 0 | 14 | 0 | 8 | 0 | 10 | 0 | 66 | 0 | 28 |
| NM_012175:XM_011519980:XM_011519981:NM_033406:XM_017017533 | 18:1:6:6:0 | 0:0:0:0:0 | 17:0:1:1:0 | 0:0:0:0:1 | 33:1:6:4:0 | 0:0:0:0:0 | 76:0:15:8:6 | 1:0:1:0:0 | 25:0:4:2:0 | 1:0:0:0:0 |
| NM_015957:XM_017017875:XM_011520154 | 125:06:01 | 0:00:00 | 88:00:00 | 0:00:00 | 264:16:05 | 0:00:00 | 596:11:04 | 2:00:00 | 182:09:02 | 1:00:00 |
| NM_015957:XM_017017875:XM_011520154 | 125:04:01 | 0:00:00 | 87:00:00 | 0:00:00 | 265:16:05 | 0:00:00 | 598:13:04 | 0:00:00 | 184:09:02 | 0:00:00 |
| NM_015957:XM_017017875:XM_011520154 | 138:04:01 | 0:00:00 | 121:03:00 | 0:00:00 | 282:11:05 | 1:00:00 | 790:22:10 | 2:00:00 | 196:08:03 | 1:00:00 |
| NM_015957:XM_017017875:XM_011520154 | 36:03:00 | 102:01:01 | 39:02:00 | 82:01:00 | 100:04:00 | 183:07:05 | 792:21:10 | 0:00:00 | 197:08:03 | 0:00:00 |
| NM_015957:XM_017017875:XM_011520154 | 138:04:01 | 0:00:00 | 122:02:00 | 0:00:00 | 284:06:05 | 0:01:00 | 791:15:09 | 1:01:00 | 196:06:03 | 1:00:00 |

| | | | | | | | | | | |
|---|---|---|---|---|---|---|---|---|---|---|
| NM_015957:XM_017017875:XM_011520154 | 138:04:01 | 0:00:00 | 121:01:00 | 1:00:00 | 283:08:05 | 0:00:00 | 791:16:10 | 0:00:00 | 197:07:03 | 0:00:00 |
| NM_015957:XM_017017875:XM_011520154 | 35:03:00 | 102:01:01 | 39:01:00 | 83:00:00 | 100:04:00 | 183:04:05 | 790:16:10 | 0:00:00 | 197:07:03 | 0:00:00 |
| NM_003477:NM_001166158:XM_011520390:NM_001135024 | 30:2:2:0 | 22:1:2:1 | 25:1:1:0 | 19:1:1:0 | 72:3:3:0 | 83:3:2:0 | 154:11:8:0 | 158:12:13:0 | 46:0:2:0 | 52:0:3:0 |
| NM_003477:NM_001166158:XM_011520390:NM_001135024 | 53:3:4:1 | 0:0:0:0 | 45:2:2:0 | 0:0:0:0 | 158:7:6:0 | 0:0:0:0 | 325:23:22:2 | 0:0:2:0 | 98:1:5:0 | 0:0:0:0 |
| NM_003477:NM_001166158:XM_011520390:NM_001135024 | 53:3:4:1 | 0:0:0:0 | 45:2:2:0 | 0:0:0:0 | 158:7:6:0 | 0:0:0:0 | 325:23:23:2 | 0:0:0:0 | 98:1:5:0 | 0:0:0:0 |
| NM_003477:NM_001166158:XM_011520390:NM_001135024 | 50:3:4:1 | 0:0:0:0 | 42:2:4:0 | 1:0:0:0 | 158:7:7:0 | 0:0:0:0 | 324:23:29:2 | 0:0:0:0 | 97:1:5:0 | 0:0:0:0 |
| NM_003477:NM_001166158:XM_011520390:NM_001135024 | 50:3:4:1 | 0:0:0:0 | 43:2:4:0 | 0:0:1:0 | 158:7:7:0 | 0:0:0:0 | 324:23:29:2 | 0:0:0:0 | 97:1:5:0 | 0:0:0:0 |
| NM_003477:XM_011520390:NM_001135024 | 48:04:01 | 0:00:00 | 36:04:00 | 0:00:00 | 156:04:00 | 0:00:00 | 306:68:2 | 3:00:00 | 92:05:00 | 0:00:00 |
| NM_003477:XM_011520390:NM_001135024 | 47:03:01 | 0:00:00 | 36:05:00 | 0:00:00 | 155:06:00 | 0:00:00 | 307:88:2 | 0:01:00 | 92:05:00 | 0:00:00 |
| NM_003477:XM_011520390:NM_001135024 | 47:04:01 | 0:00:00 | 35:09:00 | 1:00:00 | 154:07:00 | 0:00:00 | 303:123:2 | 1:00:00 | 92:06:00 | 0:00:00 |
| NM_003477:XM_011520390:NM_001135024 | 47:04:01 | 0:00:00 | 35:09:00 | 0:00:00 | 154:07:00 | 0:00:00 | 303:129:2 | 0:00:00 | 92:06:00 | 0:00:00 |
| NM_003477:NM_001166158:XM_011520390:NM_001135024 | 47:3:3:1 | 0:0:0:0 | 35:0:9:0 | 0:0:0:0 | 152:7:7:0 | 2:0:0:0 | 275:23:147:2 | 0:0:0:0 | 91:1:6:0 | 0:0:0:0 |
| NM_003477:NM_001166158:XM_011520390:NM_001135024 | 47:3:3:1 | 0:0:0:0 | 35:0:9:0 | 0:0:0:0 | 154:7:8:0 | 0:0:0:0 | 275:23:156:2 | 1:0:1:0 | 91:1:6:0 | 0:0:0:0 |
| NM_003477:NM_001166158:XM_011520390:NM_001135024 | 47:3:3:1 | 0:0:0:0 | 34:0:9:0 | 1:0:0:0 | 153:7:8:0 | 0:0:0:0 | 275:23:163:2 | 1:0:1:0 | 91:1:7:0 | 0:0:0:0 |
| NM_003477:NM_001166158:XM_011520390:NM_001135024 | 45:3:4:1 | 0:0:0:0 | 33:0:13:0 | 0:0:0:0 | 151:7:9:0 | 0:0:0:0 | 264:22:173:2 | 2:0:0:0 | 85:1:8:0 | 2:0:0:0 |
| NM_003477:NM_001166158:XM_011520390:NM_001135024 | 45:3:4:1 | 0:0:0:0 | 33:0:13:0 | 0:0:0:0 | 151:7:9:0 | 0:0:0:0 | 266:22:172:2 | 0:0:0:0 | 87:1:7:0 | 0:0:0:0 |
| NM_001001390:NM_001202555:XM_005253238:NM_001001389:NM_001001391:XM_017018585:XM_011520488:XM_011520486:XM_011520483:XM_005253240:XM_005253235:XM_017018583:XM_006718388:XM_011520485:NM_001202557:XM_005253239:NM_000610:XM_011520487:XM_011520484:XM_017018584:XM_005253231:NM_001202556:XM_005253232:XM_011520489:XM_011520482 | 85:3:16:3:67:20:2:3:2:4:2:4:1:1:0:0:0:0:0:0:0:0:0:0:0 | 0:0:0:0:0:0:0:0:0:0:0:0:0:0:0:0:0:0:0:0:0:0 | 4:0:1:1:51:6:2:0:0:1:0:1:0:0:2:1:0:0:0:0:0:0:0:0:0:0 | 0:0:0:0:0:0:0:0:0:0:0:0:0:0:0:0:0:0:0:0:0:0:0:0:0 | 28:1:4:4:121:34:5:0:0:1:0:3:0:0:2:1:0:0:0:0:0:0:0:0:0 | 0:0:0:0:1:0:0:0:0:0:0:0:0:0:0:0:0:0:0:0:0:0:0:0:0 | 316:0:162:47:66:201:16:6:6:124:41:31:5:2:1:112:50:8:9:12:46:1:13:10:4 | 0:0:0:0:0:0:0:0:0:0:0:0:0:0:0:0:0:0:0:0:0:0:0:0:0 | 53:0:46:8:43:32:8:5:0:20:5:12:1:0:1:1:5:1:2:2:3:0:0:0:0 | 0:0:0:0:0:0:0:0:0:0:0:0:0:0:0:0:0:0:0:0:0:0:0:0:0 |
| NM_001001390:NM_001202555:XM_005253238:NM_001001389:NM_001001391:XM_017018585:XM_011520488:XM_011520486:XM_011520483:XM_005253240:XM_005253235:XM_017018583:XM_006718388:XM_011520485:NM_001202557:XM_005253239:NM_000610:XM_011520487:XM_017018584:XM_005253231:NM_001202556:XM_005253232:XM_011520484:XM_011520489:XM_011520482 | 84:3:16:3:68:20:2:3:2:4:2:4:1:1:0:0:0:0:0:0:0:0:0:0:0 | 0:0:0:0:0:0:0:0:0:0:0:0:0:0:0:0:0:0:0:0:0:0 | 4:0:1:1:52:5:2:0:0:1:0:2:0:0:2:1:0:0:0:0:0:0:0:0:0:0 | 0:0:0:0:0:0:0:0:0:0:0:0:0:0:0:0:0:0:0:0:0:0:0:0:0 | 28:1:4:4:124:35:5:0:0:1:0:3:0:0:2:1:0:0:0:0:0:0:0:0:0 | 0:0:0:0:0:0:0:0:0:0:0:0:0:0:0:0:0:0:0:0:0:0:0:0:0 | 319:0:163:48:66:194:16:5:6:127:41:31:5:2:1:116:52:8:12:46:1:13:9:10:4 | 0:0:0:0:0:0:0:0:0:0:1:0:0:0:1:0:0:0:0:0:0:0:0:0:0 | 55:0:46:8:43:29:8:5:0:20:5:12:1:0:0:1:5:1:2:3:0:0:2:0:0 | 0:0:0:0:0:0:0:0:0:0:0:0:0:0:0:0:0:0:0:0:0:0:0:0:0 |
| NM_001001390:NM_001202555:XM_005253238:NM_001001389:NM_001001391:XM_017018585:XM_011520488:XM_011520486:XM_011520483:XM_005253240:XM_005253235:XM_017018583:XM_006718388:XM_011520485:NM_001202557:XM_005253239:NM_000610:XM_011520487:XM_017018584:XM_005253231:NM_001202556:XM_005253232:XM_011520489:XM_011520482:XM_011520484 | 84:3:16:3:69:20:2:3:2:4:2:4:1:1:1:0:0:0:0:0:0:0:0:0:0 | 0:0:0:0:0:0:0:0:0:0:0:0:0:0:0:0:0:0:0:0:0:0 | 4:0:1:1:51:5:2:0:0:1:0:2:0:0:2:1:0:0:0:0:0:0:0:0:0:0 | 0:0:0:0:1:0:0:0:0:0:0:0:0:0:0:0:0:0:0:0:0:0:0:0:0 | 28:2:4:4:125:33:5:0:0:1:0:2:0:0:3:1:0:0:0:0:0:0:0:0:0 | 0:0:0:0:0:0:0:0:0:0:0:0:0:0:0:0:0:0:0:0:0:0:0:0:0 | 323:0:165:48:66:186:16:5:6:135:44:33:5:2:1:123:56:8:12:46:1:14:10:4:8 | 0:0:1:0:0:1:0:0:0:0:0:0:0:0:0:0:0:0:0:0:0:0:0:0:0 | 56:0:46:8:44:27:8:5:0:20:5:12:1:0:0:1:5:1:3:3:0:0:0:0:2 | 0:0:0:0:0:0:0:0:0:0:0:0:0:0:0:0:0:0:0:0:0:0:0:0:0 |
| NM_001001390:NM_001202555:XM_005253238:NM_001001389:NM_001001391:XM_017018585:XM_011520488:XM_011520486:XM_011520483:XM_005253240:XM_005253235:XM_017018583:XM_006718388:XM_011520485:NM_001202557:XM_005253239:NM_000610:XM_011520487:XM_017018584:XM_005253231:NM_001202556:XM_005253232:XM_011520489:XM_011520482:XM_011520484 | 84:3:16:3:71:20:2:3:2:4:2:2:1:1:1:0:0:0:0:0:0:0:0:0:0 | 0:0:0:0:0:0:0:0:0:0:0:0:0:0:0:0:0:0:0:0:0:0 | 4:0:1:1:52:5:2:0:0:1:0:1:0:0:3:1:0:0:0:0:0:0:0:0:0:0 | 0:0:0:0:0:0:0:0:0:0:0:0:0:0:0:0:0:0:0:0:0:0:0:0:0 | 28:2:3:4:124:31:5:0:0:1:0:1:0:0:2:1:0:0:0:0:0:0:0:0:0 | 0:0:0:0:0:0:0:0:0:0:0:0:0:0:0:0:0:0:0:0:0:0:0:0:0 | 325:0:167:49:66:184:16:5:6:137:44:28:5:1:2:125:58:8:12:48:1:14:10:4:8 | 1:0:1:0:0:0:0:0:0:0:0:0:0:0:0:0:0:0:0:0:0:0:0:0:0 | 56:0:47:8:44:27:8:5:0:20:5:10:1:0:0:1:5:1:3:3:0:0:0:0:1 | 0:0:0:0:0:0:0:0:0:0:0:0:0:0:0:0:0:0:0:0:0:0:0:0:0 |
| XM_005253238:NM_001001389:XM_011520483:XM_005253235:XM_017018583:XM_006718388:XM_011520486:XM_005253231:XM_011520484:NM_000610:XM_011520487:XM_005253232:XM_011520489:XM_011520482:XM_011520485:XM_017018585 | 16:2:2:2:1:1:1:0:0:0:0:0:0:0:0:0 | 0:0:0:0:0:0:0:0:0:0:0:0:0:0:0:0 | 1:0:0:0:1:0:0:0:0:0:0:0:0:0:0:0 | 0:0:0:0:0:0:0:0:0:0:0:0:0:0:0:0 | 4:0:0:0:1:0:0:0:0:0:0:0:0:0:0:0 | 0:0:0:0:0:0:0:0:0:0:0:0:0:0:0:0 | 171:56:18:48:14:5:0:55:41:59:9:16:9:5:4:2 | 0:0:0:0:0:0:0:1:1:0:0:0:0:0:0:0 | 47:4:0:5:5:1:1:3:2:5:1:0:0:0:0:0 | 0:0:0:0:0:0:0:0:0:0:0:0:0:0:0:0 |
| XM_005253238:XM_005253235:NM_001001389:XM_011520483:XM_011520486:XM_017018583:XM_006718388:NM_000610:XM_011520487:XM_011520484:XM_005253231:XM_005253232:XM_011520489:XM_011520482:XM_011520485:XM_017018585 | 7:1:1:1:0:1:1:0:0:0:0:0:0:0:0:0 | 9:1:1:1:1:0:0:0:0:0:0:0:0:0:0:0 | 0:0:0:0:0:1:0:0:0:0:0:0:0:0:0:0 | 1:0:0:0:0:0:0:0:0:0:0:0:0:0:0:0 | 3:0:0:0:0:1:0:0:0:0:0:0:0:0:0:0 | 1:0:0:0:0:0:0:0:0:0:0:0:0:0:0:0 | 171:48:56:18:0:12:5:59:9:45:56:16:9:5:4:2 | 0:0:0:0:0:2:0:0:0:0:0:0:0:0:0:0 | 47:5:4:0:1:5:1:5:1:2:3:0:0:0:0:0 | 0:0:0:0:0:0:0:0:0:0:0:0:0:0:0:0 |
| NM_001001390:XM_005253238:NM_001001389:XM_011520488:XM_011520486:XM_011520483:XM_005253240:XM_005253235:XM_017018583:XM_006718388:XM_011520485:XM_005253239:NM_000610:XM_011520484:XM_005253231:XM_005253232:XM_011520482:XM_017018585:XM_011520489 | 0:0:0:0:0:0:0:0:0:0:0:0:0:0:0:0:0:0:0 | 85:16:3:2:3:2:4:2:2:1:1:0:0:0:0:0:0:0:0 | 0:0:0:0:0:0:0:0:0:0:0:0:0:0:0:0:0:0:0 | 4:1:0:2:0:0:1:0:1:0:0:1:0:0:0:0:0:0:0 | 0:0:0:0:0:0:0:0:0:0:0:0:0:0:0:0:0:0:0 | 29:3:0:5:0:0:1:0:2:0:0:1:0:0:0:0:0:0:0 | 334:150:55:14:29:16:151:46:20:5:9:132:59:53:56:16:3:1:0 | 0:0:0:0:0:0:0:0:1:0:0:0:0:0:0:0:0:0:0 | 56:39:4:8:5:0:20:5:5:1:0:1:5:2:3:0:0:0:0 | 0:0:0:0:0:0:0:0:0:0:0:0:0:0:0:0:0:0:0 |
| NM_001001390:XM_005253238:NM_001001389:XM_011520488:XM_011520486:XM_011520483:XM_005253240:XM_005253235:XM_017018583:XM_006718388:XM_011520485:XM_005253239:NM_000610:XM_011520484:XM_005253231:XM_005253232:XM_011520482:XM_017018585:XM_011520489 | 85:16:3:2:3:2:4:2:2:1:1:0:0:0:0:0:0:0:0 | 0:0:0:0:0:0:0:0:0:0:0:0:0:0:0:0:0:0:0 | 4:1:0:2:0:0:1:0:1:0:0:1:0:0:0:0:0:0:0 | 0:0:0:0:0:0:0:0:0:0:0:0:0:0:0:0:0:0:0 | 29:3:0:5:0:0:1:0:2:0:0:1:0:0:0:0:0:0:0 | 0:0:0:0:0:0:0:0:0:0:0:0:0:0:0:0:0:0:0 | 329:150:55:14:29:16:151:46:34:5:9:132:59:52:56:16:3:1:0 | 4:0:0:0:0:0:0:0:1:0:0:0:0:0:0:0:0:0:0 | 56:39:4:8:5:0:20:5:5:1:0:1:5:2:3:0:0:0:0 | 0:0:0:0:0:0:0:0:0:0:0:0:0:0:0:0:0:0:0 |
| NM_001001390:NM_001202555:XM_005253238:NM_001001391:NM_001001389:XM_011520488:XM_011520486:XM_011520483:XM_005253240:XM_005253235:XM_017018583:XM_006718388:XM_011520485:XM_017018585:NM_001202556:NM_001202557:XM_005253239:NM_001001392:NM_000610:XM_011520484:XM_005253231:XM_005253232:XM_011520482:XM_017018584:XM_011520489 | 85:4:16:71:3:2:3:2:4:2:4:1:1:2:4:0:0:0:0:0:0:0:0:0:0 | 0:0:0:0:0:0:0:0:0:0:0:0:0:0:0:0:0:0:0:0:0:0 | 4:0:1:52:0:1:0:0:1:0:2:0:0:2:0:2:1:0:0:0:0:0:0:0:0:0 | 0:0:0:0:0:0:0:0:0:0:0:0:0:0:0:0:0:0:0:0:0:0:0:0:0 | 29:2:3:125:0:5:0:0:1:0:4:0:0:5:1:1:1:1:0:0:0:0:0:0:0 | 0:0:0:0:0:0:0:0:0:0:0:0:0:0:0:0:0:0:0:0:0:0:0:0:0 | 329:0:149:66:54:14:28:16:148:46:136:5:9:32:2:0:131:0:57:50:55:16:2:1:0 | 0:0:0:0:0:0:0:0:0:0:0:0:0:0:0:0:0:0:0:0:0:0:0:0:0 | 55:0:39:44:3:8:5:0:20:5:17:1:0:3:0:0:1:1:5:2:3:0:0:0:0 | 0:0:0:0:0:0:0:0:0:0:0:0:0:0:0:0:0:0:0:0:0:0:0:0:0 |
| NM_001001390:NM_001202555:XM_005253238:NM_001001391:NM_001001389:XM_011520488:XM_011520486:XM_011520483:XM_005253240:XM_005253235:XM_017018583:XM_006718388:XM_011520485:XM_017018585:NM_001202556:NM_001202557:XM_005253239:NM_001001392:NM_000610:XM_005253232:XM_005253231:XM_011520484:XM_017018584:XM_011520482:XM_011520489 | 85:4:16:71:3:2:3:2:4:2:4:1:1:2:4:0:0:0:0:0:0:0:0:0:0 | 0:0:0:0:0:0:0:0:0:0:0:0:0:0:0:0:0:0:0:0:0:0 | 4:0:1:52:0:1:0:0:1:0:2:0:0:2:0:2:1:0:0:0:0:0:0:0:0:0 | 0:0:0:0:0:0:0:0:0:0:0:0:0:0:0:0:0:0:0:0:0:0:0:0:0 | 29:2:3:124:0:5:0:0:1:0:4:0:0:5:1:1:1:1:0:0:0:0:0:0:0 | 0:0:0:1:0:0:0:0:0:0:0:0:0:0:0:0:0:0:0:0:0:0:0:0:0 | 155:0:75:46:26:8:11:9:73:19:51:3:5:13:1:0:62:0:26:10:34:31:0:1:0 | 173:0:74:20:28:6:17:7:75:27:85:2:4:19:1:0:69:0:31:6:21:19:1:1:0 | 27:0:16:20:3:4:0:0:8:3:7:1:0:2:0:0:0:1:3:0:2:2:0:0:0 | 28:0:22:24:0:4:5:0:12:2:10:0:0:1:0:0:1:0:2:0:1:0:0:0:0 |
| NM_001001390:NM_001202555:XM_005253238:NM_001001391:NM_001001389:XM_011520488:XM_011520486:XM_011520483:XM_005253240:XM_005253235:XM_017018583:XM_006718388:XM_011520485:XM_017018585:NM_001202556:NM_001202557:XM_005253239:NM_001001392:NM_000610:XM_011520484:XM_005253231:XM_005253232:XM_011520482:XM_017018584:XM_011520489 | 85:4:16:71:3:2:3:2:4:2:4:1:1:2:4:0:0:0:0:0:0:0:0:0:0 | 0:0:0:0:0:0:0:0:0:0:0:0:0:0:0:0:0:0:0:0:0:0 | 4:0:1:51:0:1:0:0:1:0:2:0:0:2:0:1:1:0:0:0:0:0:0:0:0:0 | 0:0:0:0:0:0:0:0:0:0:0:0:0:0:0:0:0:0:0:0:0:0:0:0:0 | 28:2:3:125:0:5:0:0:1:0:4:0:0:6:1:1:1:1:0:0:0:0:0:0:0 | 0:0:0:0:0:0:0:0:0:0:0:0:0:0:0:0:0:0:0:0:0:0:0:0:0 | 329:0:148:65:54:15:28:16:147:46:134:5:9:34:2:0:131:0:57:50:55:16:2:1:0 | 0:0:1:1:0:0:0:0:0:0:1:0:0:0:1:0:0:0:0:0:0:0:0:0:0 | 55:0:39:44:3:8:5:0:20:5:17:1:0:3:1:0:1:1:5:2:3:0:0:0:0 | 0:0:0:0:0:0:0:0:0:0:0:0:0:0:0:0:0:0:0:0:0:0:0:0:0 |
| NM_001001391:NM_001001390:NM_001202555:XM_005253238:NM_001001389:XM_011520488:XM_011520486:XM_011520483:XM_005253240:XM_005253235:XM_017018583:XM_006718388:XM_011520485:XM_017018585:NM_001202556:NM_001202557:XM_005253239:NM_001001392:NM_000610:XM_011520484:XM_005253231:XM_005253232:XM_011520482:XM_017018584:XM_011520489 | 70:85:4:16:3:2:3:2:4:2:4:1:1:3:6:0:0:0:0:0:0:0:0:0:0 | 1:0:0:0:0:0:0:0:0:0:0:0:0:0:0:0:0:0:0:0:0:0 | 51:4:0:1:0:1:0:0:1:0:2:0:0:2:0:1:1:0:0:0:0:0:0:0:0:0 | 0:0:0:0:0:0:0:0:0:0:0:0:0:0:0:0:0:0:0:0:0:0:0:0:0 | 125:28:2:3:0:5:0:0:1:0:4:0:0:6:1:1:1:1:0:0:0:0:0:0:0 | 0:0:0:0:0:0:0:0:0:0:0:0:0:0:0:0:0:0:0:0:0:0:0:0:0 | 66:328:0:149:54:15:28:16:146:46:133:5:9:35:2:0:131:0:57:50:55:16:2:1:0 | 0:1:0:0:0:0:0:0:1:0:0:0:0:0:1:0:0:0:0:0:0:0:0:0:0 | 44:55:0:39:3:8:5:0:20:5:17:1:0:3:1:0:1:1:5:2:3:0:0:0:0 | 0:0:0:0:0:0:0:0:0:0:0:0:0:0:0:0:0:0:0:0:0:0:0:0:0 |
| NM_001001390:NM_001202555:XM_005253238:NM_001001391:NM_001001389:XM_011520488:XM_011520486:XM_011520483:XM_005253240:XM_005253235:XM_017018583:XM_006718388:XM_011520485:XM_017018585:NM_001202556:NM_001202557:XM_005253239:NM_001001392:NM_000610:XM_011520484:XM_005253231:XM_005253232:XM_011520482:XM_017018584:XM_011520489 | 84:4:16:71:3:2:3:2:4:2:4:1:1:3:6:0:0:0:0:0:0:0:0:0:0 | 1:0:0:0:0:0:0:0:0:0:0:0:0:0:0:0:0:0:0:0:0:0 | 4:0:1:51:0:1:0:0:1:0:2:0:0:2:0:1:1:0:0:0:0:0:0:0:0:0 | 0:0:0:0:0:0:0:0:0:0:0:0:0:0:0:0:0:0:0:0:0:0:0:0:0 | 28:2:3:124:0:5:0:0:1:0:4:0:0:7:2:1:1:1:0:0:0:0:0:0:0 | 0:0:0:0:0:0:0:0:0:0:0:0:0:0:0:0:0:0:0:0:0:0:0:0:0 | 329:0:149:66:54:15:28:16:146:46:132:5:9:38:2:0:131:0:57:49:55:16:2:2:0 | 1:0:0:0:0:0:0:0:1:0:0:0:0:0:1:0:0:0:0:0:0:0:0:0:0 | 55:0:38:44:3:8:5:0:20:5:16:1:0:4:1:0:1:1:5:2:3:0:0:0:0 | 0:0:0:0:0:0:0:0:0:0:0:0:0:0:0:0:0:0:0:0:0:0:0:0:0 |
| NM_001001991:NM_001282675:NM_015430 | 0:00:00 | 0:00:00 | 1:01:00 | 0:00:00 | 1:00:00 | 0:00:00 | 0:00:00 | 0:00:00 | 1:00:00 | 0:00:00 |
| NM_001001991:NM_001282675:NM_015430:NM_001282676 | 0:0:0:0 | 0:0:0:0 | 0:1:0:0 | 2:0:0:0 | 1:0:0:0 | 0:0:1:0 | 0:0:0:0 | 0:0:0:0 | 1:0:0:0 | 0:0:0:0 |
| NM_005387:NM_139131:NM_139132 | 7:19:06 | 0:00:00 | 2:19:09 | 0:01:00 | 11:41:08 | 0:00:00 | 0.545578704 | 0:01:00 | 8:34:05 | 0:00:00 |
| NM_021805:NM_001135053:XR_001747933 | 0:00:00 | 0:00:00 | 0:00:00 | 1:00:00 | 0:00:00 | 0:02:00 | 0:00:00 | 3:00:01 | 0:00:00 | 1:00:00 |
| NM_016142:XM_024448573 | 54:00:00 | 0:00 | 6:07 | 0:00 | 75:04:00 | 0:00 | 254:51:00 | 0:01 | 235:02:00 | 1:00 |
| NM_016142:XM_024448573 | 0:00 | 53:02:00 | 0:00 | 6:08 | 0:00 | 75:04:00 | 275:84 | 0:00 | 239:04:00 | 0:00 |
| NM_139178 | 20 | 0 | 33 | 0 | 23 | 0 | 0 | 69 | 0 | 30 |
| NM_139178 | 20 | 0 | 40 | 0 | 27 | 0 | 72 | 0 | 33 | 0 |

| | | | | | | | | | | |
|---|---|---|---|---|---|---|---|---|---|---|
| NM_207122:XM_011519951:NM_000401:XM_024448383:XM_011519950 | 50:13:2:0:0 | 0:0:0:0:0 | 6:48:1:1:0 | 0:1:0:0:0 | 63:26:21:0:1 | 0:0:0:0:0 | 49:128:7:1:0 | 2:1:0:0:0 | 66:24:5:0:1 | 0:0:0:0:0 |
| NM_207122:XM_011519951:NM_000401:XM_011519950 | 53:22:2:0 | 0:0:0:0 | 16:102:1:0 | 0:0:0:0 | 65:50:21:1 | 0:0:0:0 | 60:315:7:0 | 0:0:0:0 | 66:40:5:1 | 0:0:0:0 |
| NM_207122:XM_011519951:NM_000401:XM_011519950 | 54:23:2:0 | 0:0:0:0 | 16:104:1:0 | 0:1:0:0 | 65:51:20:1 | 0:0:0:0 | 59:322:7:0 | 0:1:0:0 | 66:41:5:1 | 0:0:0:0 |
| NM_004813:NM_057174 | 0:00 | 0:00 | 0:00 | 0:00 | 0:00 | 1:00 | 0:00 | 1:00 | 0:00 | 1:00 |
| NM_024113:XM_006718315:XM_017018282:NM_001003676:NM_001003677:XM_011520364:NM_001278222:NM_001003678:NR_103472:XM_017018281:XM_017018276:XM_017018279 | 3:5:0:0:0:0:0:0:0:0:0 | 0:0:0:0:0:0:0:0:0:0 | 5:2:0:2:1:0:0:0:0:0:0:0 | 0:0:1:0:0:0:0:0:0:0:0 | 26:0:1:1:7:1:3:1:0:0:0:0 | 0:0:0:0:0:0:0:0:0:0 | 19:3:1:1:1:0:4:1:1:0:0:0 | 8:1:0:1:1:1:1:0:0:0:0:0 | 3:3:0:1:1:1:3:0:1:0:0:0 | 4:1:0:0:1:0:1:0:0:0:0 |
| NM_032389:XM_005253168:XM_011520405:XM_005253166:XM_017018415:XM_005253167:XM_017018413:NM_001242832 | 0:1:0:0:0:0:0:0 | 4:2:0:0:0:0:0:0 | 1:1:0:0:0:0:0:0 | 0:0:0:0:0:0:0:0 | 4:1:0:0:0:0:0:0 | 1:0:0:0:0:0:0:0 | 8:6:0:0:1:0:0:0 | 16:4:1:1:2:0:0:0 | 1:4:1:0:0:0:1:0 | 2:3:0:0:0:1:0:0 |
| NM_014342:NM_001317232:XM_017017462:XM_006718172 | 0:0:0:0 | 0:0:0:0 | 1:0:0:0 | 0:0:0:0 | 1:1:0:0 | 1:0:0:0 | 0:0:0:0 | 4:0:1:0 | 0:0:0:0 | 5:0:0:1 |
| NM_001143993:NM_003475:NM_001143994 | 0:00:00 | 0:00:00 | 3:00:00 | 0:00:00 | 0:00:00 | 0:00:00 | 0:00:00 | 0:00:00 | 0:00:00 | 3:00:00 |
| NM_000062:NM_001032295 | 1:00 | 0:00 | 0:01 | 0:00 | 0:01 | 0:00 | 0:00 | 0:00 | 0:00 | 0:00 |
| NM_001144012:NM_015959:NM_001347894:NR_037645:NM_001347898:NM_001347893:NM_001347890:NM_001347896:NR_144935:NM_001347895:NM_001347892:NR_144933:NR_144934 | 24:197:3:2:1:1:1:0:0:0:0:0 | 0:0:0:0:0:0:0:0:0:0:0 | 11:149:7:1:1:0:0:2:1:0:0:0:0 | 1:0:0:0:0:0:0:0:0:0:0 | 9:137:3:1:0:0:0:0:0:1:0:0:0 | 0:0:0:0:0:0:0:0:0:0:0 | 148:1101:30:17:6:2:5:0:12:3:3:3:0 | 0:3:1:0:0:0:0:0:0:0:0 | 71:318:4:8:3:1:0:0:1:0:1:0:0 | 0:1:0:0:0:0:0:0:0:0:0:0 |
| NM_015959:NM_001347894:NM_001347898:NM_001347893:NR_037645:NM_001347890:NM_001347896:NM_001347895:NR_144935:NM_001347892:NR_144933:NR_144934 | 198:3:1:1:1:1:0:0:0:0:0:0 | 0:0:0:0:0:0:0:0:0:0:0 | 150:7:1:0:1:0:2:0:0:0:0:0 | 0:0:0:0:0:0:0:0:0:0:0 | 137:3:0:0:1:0:0:1:0:0:0:0 | 1:0:0:0:0:0:0:0:0:0 | 1132:31:6:2:7:5:0:3:11:3:3:0 | 0:0:0:0:0:0:0:0:0:0 | 326:4:3:0:6:0:0:0:1:1:0:0 | 0:0:0:0:0:0:0:0:0:0:0 |
| NM_001144012:NM_015959:NM_001347894:NM_001347898:NM_001347893:NR_037645:NM_001347890:NR_144934:NM_001347896:NR_144935:NM_001347895:NM_001347892:NM_001347891 | 24:199:3:1:1:1:1:1:0:0:0:0:0 | 0:0:0:0:0:0:0:0:0:0:0 | 12:151:7:1:0:2:0:0:2:1:0:0:0 | 0:0:0:0:0:0:0:0:0:0:0 | 9:139:3:0:0:1:0:0:0:0:1:0:0 | 0:0:0:0:0:0:0:0:0:0 | 150:1148:31:6:4:6:5:0:0:8:3:3:0 | 0:2:0:0:0:0:0:0:0:0:0 | 72:329:4:3:1:5:0:0:0:0:0:1:1 | 0:0:0:0:0:0:0:0:0:0:0 |
| NM_001144012:NM_015959:NM_001347894:NM_001347898:NM_001347893:NR_037645:NM_001347890:NR_144934:NM_001347896:NR_144935:NM_001347895:NM_001347892:NR_144933:NM_001347891 | 24:199:3:1:1:1:1:1:0:0:0:0:0:0 | 0:0:0:0:0:0:0:0:0:0:0:0 | 12:152:7:1:0:2:0:0:2:1:0:0:0:0 | 0:0:0:0:0:0:0:0:0:0:0:0 | 8:139:3:0:0:1:0:0:0:0:1:0:0:0 | 0:0:0:0:0:0:0:0:0:0:0 | 150:1156:31:6:4:6:5:0:0:8:3:3:1:0 | 0:0:0:0:0:0:0:0:0:0:0:0 | 72:326:4:3:1:5:0:0:0:0:0:1:0:1 | 0:2:0:0:0:0:0:0:0:0:0:0 |
| NM_001144012:NM_015959:NM_001347894:NM_001347898:NM_001347893:NR_037645:NM_001347890:NR_144934:NM_001347896:NR_144935:NM_001347895:NM_001347892:NR_144933:NM_001347891 | 24:199:3:1:1:1:1:1:0:0:0:0:0:0 | 0:0:0:0:0:0:0:0:0:0:0:0 | 12:152:7:1:0:2:0:0:2:1:0:0:0:0 | 0:0:0:0:0:0:0:0:0:0:0:0 | 8:139:3:0:0:1:0:0:0:0:1:0:0:0 | 0:0:0:0:0:0:0:0:0:0:0 | 149:1154:31:6:4:6:5:0:0:8:3:3:1:0 | 1:2:0:0:0:0:0:0:0:0:0:0 | 72:328:4:3:1:5:0:0:0:0:0:1:0:1 | 0:0:0:0:0:0:0:0:0:0:0:0 |
| XM_011545117:NM_017716:NM_001164470 | 0:00:00 | 0:00:00 | 0:13:00 | 1:04:00 | 6:38:01 | 2:06:00 | 0:00:00 | 0:00:00 | 0:00:00 | 0:00:00 |
| NM_016499:XM_005274039:NM_001173991 | 0:00:00 | 0:00:00 | 0:00:00 | 1:00:00 | 0:00:00 | 0:01:01 | 0:00:00 | 2:00:01 | 0:00:00 | 0:00:00 |
| NM_017841 | 328 | 0 | 418 | 1 | 402 | 2 | 2230 | 2 | 408 | 0 |
| NM_017841 | 300 | 0 | 355 | 1 | 354 | 1 | 2083 | 1 | 366 | 1 |
| NM_017841 | 300 | 0 | 357 | 0 | 354 | 2 | 2080 | 8 | 365 | 1 |
| NM_017841 | 301 | 2 | 361 | 1 | 359 | 2 | 2093 | 2 | 367 | 0 |
| NM_017841 | 304 | 1 | 365 | 0 | 364 | 0 | 2098 | 0 | 371 | 0 |
| NM_017841 | 309 | 0 | 370 | 0 | 373 | 1 | 2120 | 4 | 374 | 0 |
| NM_017841 | 338 | 0 | 449 | 0 | 422 | 1 | 2311 | 4 | 425 | 0 |
| NM_017841 | 331 | 1 | 440 | 0 | 413 | 1 | 2284 | 3 | 416 | 1 |
| NM_004111 | 35 | 0 | 21 | 0 | 75 | 1 | 190 | 0 | 128 | 1 |
| NM_004111 | 35 | 0 | 21 | 0 | 76 | 0 | 192 | 0 | 127 | 0 |
| NM_021924:XM_006718253 | 0:00 | 0:00 | 0:00 | 1:00 | 0:00 | 2:01 | 0:00 | 0:00 | 0:00 | 0:00 |
| NM_001085372 | 2 | 0 | 1 | 2 | 0 | 1 | 0 | 4 | 0 | 7 |
| NM_003164:XM_011545224:NM_001244666:NM_001330294 | 86:3:1:0 | 0:0:0:0 | 69:14:1:0 | 0:0:0:0 | 361:46:2:1 | 2:2:0:0 | 412:76:3:1 | 0:0:0:0 | 112:5:2:0 | 0:0:0:0 |
| NM_003164:XM_011545224:NM_001244666:NM_001330294 | 89:4:1:0 | 0:0:0:0 | 72:14:1:0 | 0:0:0:0 | 373:43:2:1 | 0:0:0:0 | 417:74:2:1 | 1:0:0:0 | 114:4:2:0 | 0:0:0:0 |
| NM_003164:XM_011545224:NM_001244666:NM_001330294 | 88:2:1:0 | 1:0:0:0 | 72:4:1:0 | 0:0:0:0 | 380:20:1:1 | 1:0:0:0 | 422:20:2:1 | 2:0:0:0 | 115:2:2:0 | 0:0:0:0 |
| NM_003164:XM_011545224:NM_001244666 | 88:02:00 | 0:00:00 | 58:06:01 | 0:01:00 | 333:27:02 | 0:00:00 | 321:33:01 | 0:02:00 | 106:00:01 | 0:00:00 |
| NM_145040 | 0 | 0 | 0 | 0 | 0 | 1 | 0 | 0 | 1 | 0 |
| NM_145040 | 0 | 0 | 0 | 0 | 0 | 1 | 0 | 0 | 0 | 1 |
| NM_017878:XM_011545120 | 4:01 | 0:00 | 110:07:00 | 1:00 | 749:25:00 | 2:00 | 4:03 | 0:00 | 0:00 | 0:00 |
| NM_017878:XM_011545120 | 4:01 | 0:00 | 111:06:00 | 0:00 | 746:15:00 | 2:00 | 4:03 | 0:00 | 0:00 | 0:00 |
| NM_001033678:XM_005274345 | 0:00 | 1:00 | 0:00 | 1:00 | 0:00 | 1:01 | 1:00 | 0:00 | 1:00 | 0:00 |
| NM_032344:NM_001128612 | 0:00 | 0:00 | 0:00 | 2:00 | 0:00 | 1:00 | 0:00 | 1:00 | 0:00 | 1:00 |
| NM_032344:NM_001128612 | 0:00 | 0:00 | 0:00 | 2:00 | 0:00 | 1:00 | 0:00 | 1:00 | 0:00 | 1:00 |
| NM_012094:NM_001358511:NM_181651:NM_181652 | 0:1:0:0 | 4:0:0:0 | 0:0:0:0 | 6:0:1:0 | 0:0:0:0 | 19:2:1:1 | 0:0:0:0 | 79:3:2:1 | 0:0:0:0 | 57:0:1:0 |
| NM_130804 | 0 | 0 | 0 | 1 | 0 | 0 | 0 | 2 | 0 | 0 |
| NM_001099409:XM_005273868:XM_011544886:NM_001351087 | 0:0:0:0 | 0:0:0:0 | 0:0:0:0 | 1:0:0:0 | 0:0:0:0 | 2:1:0:0 | 0:0:0:0 | 0:0:1:0 | 0:0:0:0 | 0:0:0:0 |
| NM_001099409:XM_005273868 | 0:00 | 0:00 | 0:00 | 1:00 | 0:00 | 2:01 | 0:00 | 0:00 | 0:00 | 0:00 |
| NM_016938:NR_037718 | 0:00 | 2:00 | 0:00 | 0:00 | 0:00 | 0:01 | 0:00 | 0:00 | 0:00 | 0:00 |
| NM_031450:NM_001135635 | 0:00 | 0:00 | 0:00 | 0:00 | 0:00 | 0:00 | 0:00 | 0:00 | 0:00 | 1:00 |
| NM_002708:NM_001008709:NM_206873 | 147:01:02 | 0:00:00 | 160:00:15 | 2:00:00 | 725:06:26 | 1:00:01 | 662:04:22 | 1:00:00 | 277:01:06 | 0:00:00 |
| NM_002708:NM_001008709:NM_206873 | 147:01:01 | 0:00:00 | 161:00:14 | 1:00:00 | 726:06:25 | 0:00:01 | 661:04:22 | 2:00:00 | 277:01:05 | 0:00:00 |
| NM_002708:NM_206873:NM_001008709 | 150:03:01 | 1:00:00 | 162:09:00 | 1:00:00 | 732:17:06 | 1:00:00 | 667:13:04 | 0:00:00 | 277:04:01 | 0:00:00 |
| NM_002708:NM_206873:NM_001008709 | 152:02:01 | 0:00:00 | 165:01:00 | 0:00:00 | 734:06:06 | 2:00:00 | 675:09:04 | 0:01:00 | 279:03:01 | 1:00:00 |
| NM_002708:NM_206873:NM_001008709 | 147:03:01 | 0:00:00 | 168:06:00 | 0:00:00 | 721:11:06 | 3:00:00 | 660:14:02 | 1:00:00 | 276:04:01 | 0:00:00 |
| NM_003977:XM_024448761 | 0:00 | 0:00 | 0:00 | 0:00 | 0:00 | 1:00 | 0:00 | 1:03 | 0:00 | 0:00 |
| NM_000852 | 6 | 1 | 6 | 0 | 14 | 0 | 25 | 21 | 15 | 24 |
| NM_007103 | 1 | 0 | 2 | 0 | 5 | 0 | 1 | 0 | 6 | 0 |
| NM_002496 | 1 | 0 | 1 | 0 | 1 | 0 | 2 | 0 | 9 | 0 |
| NM_001876:NM_001031847:XM_005273763:XM_017017220 | 0:0:0:0 | 0:0:0:0 | 2:0:0:0 | 0:0:0:0 | 6:2:1:0 | 1:0:0:0 | 0:0:0:1 | 4:0:0:0 | 1:0:0:0 | 0:0:0:0 |
| NM_001876:XM_005273763:NM_001031847:XM_017017220 | 0:0:0:0 | 0:0:0:0 | 2:0:0:0 | 1:0:0:0 | 6:0:2:0 | 5:1:0:0 | 5:0:0:1 | 0:0:0:0 | 1:0:0:0 | 0:0:0:0 |
| NM_001876:NM_001031847:XM_005273762:XM_005273763:XM_017017220 | 0:0:0:0:0 | 0:0:0:0:0 | 7:0:0:0:0 | 0:0:0:0:0 | 16:3:3:1:0 | 1:0:0:0:0 | 15:0:3:0:0 | 0:0:0:0:0 | 2:0:1:0:0 | 0:0:0:0:0 |
| NM_001354833:NM_001354831:XM_017018269:NM_013249 | 0:0:0:0 | 0:0:0:0 | 0:0:0:0 | 1:0:0:0 | 0:0:0:0 | 0:0:0:0 | 0:0:0:0 | 1:1:2:0 | 0:0:0:0 | 0:0:0:1 |
| NM_001354833:NM_001354831:NM_013249:NM_001354832:XM_017018269 | 0:0:0:0:0 | 0:0:0:0:0 | 1:0:0:0:0 | 0:0:0:0:0 | 0:0:0:0:0 | 0:0:0:0:0 | 0:0:0:0:0 | 1:1:0:0:0 | 0:0:0:0:0 | 0:0:1:0:0 |
| NM_001354833:NM_001354831:NM_013249:NM_001354832 | 0:0:0:0 | 0:0:0:0 | 0:0:0:0 | 1:0:0:0 | 0:0:0:0 | 0:0:0:0 | 1:1:0:0 | 0:0:0:0 | 0:0:1:0 | 0:0:0:0 |
| NM_001354833:NM_001354831:NM_013249:NM_001354832 | 0:0:0:0 | 0:0:0:0 | 0:0:0:0 | 1:0:0:0 | 0:0:0:0 | 0:0:0:0 | 1:1:0:0 | 0:0:0:0 | 0:0:1:0 | 0:0:0:0 |
| XM_011545317:NM_177423:XM_017018450:XM_017018449:XM_011545310 | 1:0:0:0:0 | 0:0:0:0:0 | 0:1:1:0:0 | 0:0:0:0:0 | 0:3:0:1:1 | 0:0:0:0:0 | 0:0:0:0:0 | 0:1:1:0:0 | 0:0:0:0:0 | 0:0:0:1:0 |
| XM_011545317:XM_017018448:NM_177423:XM_011545314:XM_011545310:XM_017018449 | 1:1:0:0:0:0 | 0:0:0:0:0:0 | 0:1:2:1:0:0 | 0:0:0:0:0:0 | 3:0:3:0:1:1 | 0:0:0:0:0:0 | 7:1:3:0:0:2 | 0:0:0:0:0:0 | 0:0:0:0:0:0 | 0:0:0:0:0:0 |
| NM_016725:NM_016724:NM_000802:NM_016729 | 14:13:1:37 | 0:0:0:0 | 0:0:0:0 | 0:0:0:0 | 0:1:0:0 | 0:0:0:0 | 141:351:3:1694 | 0:1:0:3 | 0:7:1:6 | 0:0:0:0 |
| NM_016725:NM_016724:NM_000802:NM_016729 | 14:13:1:37 | 0:0:0:0 | 0:0:0:0 | 0:0:0:0 | 0:1:0:0 | 0:0:0:0 | 140:344:3:1653 | 0:1:0:4 | 0:7:1:6 | 0:0:0:0 |

| | | | | | | | | | | |
|---|---|---|---|---|---|---|---|---|---|---|
| NM_016725:NM_016724:NM_000802:NM_016729 | 14:13:1:37 | 0:0:0:0 | 0:0:0:0 | 0:0:0:0 | 0:1:0:0 | 0:0:0:0 | 140:343:3:1657 | 0:2:0:2 | 0:7:1:6 | 0:0:0:0 |
| NM_016725:NM_016724:NM_000802:NM_016729 | 14:13:1:37 | 0:0:0:0 | 0:0:0:0 | 0:0:0:0 | 0:1:0:0 | 0:0:0:0 | 139:345:3:1657 | 1:0:0:5 | 0:7:1:6 | 0:0:0:0 |
| NM_016725:NM_016724:NM_000802:NM_016729 | 14:13:1:39 | 0:0:0:0 | 0:0:0:0 | 0:0:0:0 | 0:1:0:0 | 0:0:0:0 | 138:341:3:1698 | 1:1:0:0 | 0:7:1:6 | 0:0:0:0 |
| NM_016725:NM_016724:NM_000802:NM_016729 | 14:13:1:37 | 0:0:0:0 | 0:0:0:0 | 0:0:0:0 | 0:1:0:0 | 0:0:0:0 | 134:339:3:1690 | 0:0:0:3 | 0:7:1:6 | 0:0:0:0 |
| NM_002869:NM_198896:NM_001243719 | 92:66:0 | 0:00:00 | 16:12:00 | 0:00:00 | 120:138:0 | 2:02:00 | 237:96:2 | 1:00:00 | 156:92:0 | 0:00:00 |
| NM_016055:NM_001318500:NM_001318499:NM_001318498:XM_017017919:NR_134660:NR_134661:NR_134659:NR_134658 | 200:3:4:4:4:1:1:1:0 | 0:0:0:0:0:0:0:0:0 | 120:1:2:0:3:0:1:1:0 | 0:0:0:0:0:0:0:0 | 125:0:4:1:1:1:2:2:0 | 0:0:0:0:0:0:0:0:0 | 1254:13:31:6:33:8:11:12:2 | 6:0:1:0:0:0:0:0:0 | 357:8:8:0:6:2:3:1:0 | 0:0:0:0:0:0:0:0:0 |
| NM_016055:NM_001318500:NR_134660:NM_001318499:NM_001318498:NR_134658:XM_017017919:NR_134661:NR_134659 | 199:1:8:4:4:6:7:5:2 | 0:0:0:0:0:0:0:0:0 | 114:0:6:2:0:5:11:1:2 | 1:0:0:0:0:0:0:0 | 123:0:5:5:1:8:6:2:3 | 1:0:0:0:0:0:0:0:0 | 1242:2:25:35:7:58:65:17:15 | 2:0:0:0:0:0:0:0:0 | 354:0:9:8:1:16:11:9:3 | 0:0:0:0:0:0:0:0:0 |
| NM_016055:NR_134660:NM_001318499:NM_001318498:NR_134658:XM_017017919:NR_134659:NM_001318500:NR_134661 | 196:9:4:4:6:7:2:0:0 | 0:0:0:0:0:0:0:0:0 | 113:6:2:0:5:11:2:0:0 | 0:0:0:0:0:0:0:0 | 123:5:5:1:8:6:3:0:0 | 0:0:0:0:0:0:0:0:0 | 1234:28:34:7:57:72:13:2:0 | 0:0:0:0:0:0:0:0:0 | 351:9:8:1:16:14:3:0:0 | 0:0:0:0:0:0:0:0:0 |
| NM_016055:NR_134660:NM_001318499:NM_001318498:NR_134658:XM_017017919:NR_134659:NM_001318500:NR_134661 | 196:9:4:4:6:7:2:0:0 | 0:0:0:0:0:0:0:0:0 | 113:6:2:0:5:11:2:0:0 | 0:0:0:0:0:0:0:0 | 122:5:5:1:8:6:2:0:0 | 0:0:0:0:0:0:0:0:0 | 1231:28:34:7:57:74:13:2:0 | 2:0:0:0:0:0:0:0:0 | 351:9:8:1:16:14:3:0:0 | 0:0:0:0:0:0:0:0:0 |
| NM_016055:NR_134660:NM_001318499:NM_001318498:NR_134658:NM_001318500:XM_017017919:NR_134661:NR_134659 | 196:9:4:4:6:2:7:5:2 | 0:0:0:0:0:0:0:0:0 | 113:6:2:0:5:1:11:1:2 | 0:0:0:0:0:0:0:0 | 122:5:5:1:8:0:4:2:2 | 0:0:0:0:0:0:0:0:0 | 1223:28:35:7:57:15:76:16:12 | 4:0:0:0:0:0:0:0:0 | 349:9:8:1:16:8:15:9:3 | 0:0:0:0:0:0:0:0:0 |
| NM_016055:NR_134660:NM_001318499:NM_001318498:NR_134658:NM_001318500:XM_017017919:NR_134661:NR_134659 | 196:9:4:4:6:2:5:5:2 | 0:0:0:0:0:0:0:0:0 | 113:6:2:0:5:1:11:1:2 | 0:0:0:0:0:0:0:0 | 122:5:5:1:8:0:4:2:2 | 0:0:0:0:0:0:0:0:0 | 1225:28:35:7:56:15:78:16:12 | 1:0:0:0:1:0:0:0:0 | 349:9:8:1:16:8:15:9:2 | 0:0:0:0:0:0:0:0:0 |
| NM_016055:NR_134660:NM_001318499:NM_001318498:NR_134658:NM_001318500:XM_017017919:NR_134661:NR_134659 | 197:9:4:4:6:2:4:5:2 | 0:0:0:0:0:0:0:0:0 | 113:6:2:0:5:1:11:1:2 | 0:0:0:0:0:0:0:0 | 122:5:5:1:8:0:4:2:2 | 0:0:0:0:0:0:0:0:0 | 1228:28:35:7:58:15:78:16:13 | 3:0:0:0:0:0:0:0:0 | 351:9:8:1:16:8:15:9:2 | 1:0:0:0:0:0:0:0:0 |
| NM_001005:NM_001256802:NM_001260507 | 36:03:03 | 0:00:00 | 16:01:01 | 0:01:00 | 39:01:01 | 0:00:00 | 120:04:07 | 0:01:00 | 101:01:05 | 0:00:00 |
| NM_001005:NM_001256802:NM_001260507:NM_001260506 | 36:4:3:0 | 0:0:0:0 | 16:2:4:0 | 0:0:0:0 | 39:0:1:0 | 0:0:0:0 | 120:5:8:0 | 0:0:0:0 | 101:1:5:1 | 0:0:0:0 |
| NM_030792:XM_011545277 | 0:00 | 0:00 | 0:00 | 0:00 | 0:00 | 0:00 | 0:00 | 3:00 | 0:00 | 0:00 |
| NM_016578:XM_005274051:XM_017017923 | 0:00:00 | 2:00:00 | 0:00:00 | 0:00:00 | 1:00:00 | 1:00:00 | 1:00:01 | 0:01:00 | 1:00:00 | 0:00:00 |
| NM_016578:XM_005274051:XM_017017923 | 1:00:00 | 1:02:00 | 0:02:00 | 0:02:00 | 2:01:00 | 0:00:00 | 0:00:00 | 2:08:06 | 0:00:00 | 1:02:00 |
| XM_005274051:NM_016578:XM_017017923 | 3:02:00 | 4:00:00 | 4:00:00 | 25:01:01 | 4:02:00 | 5:00:00 | 17:02:05 | 39:00:03 | 6:01:01 | 7:01:00 |
| XM_005274051:NM_016578:XM_017017923 | 5:02:00 | 2:00:00 | 10:00:01 | 6:00:00 | 9:02:00 | 3:00:00 | 30:02:07 | 10:00:00 | 12:02:01 | 7:00:00 |
| XM_005274051:NM_016578:XM_017017923 | 5:02:00 | 4:00:00 | 10:00:01 | 17:01:00 | 5:02:00 | 7:00:00 | 28:02:07 | 28:00:01 | 11:02:01 | 8:00:00 |
| NM_016578:XM_005274051:XM_017017923 | 2:15:00 | 0:00:00 | 0.090983796 | 0:01:00 | 2:21:00 | 0:00:00 | 0.265451389 | 0:01:00 | 2:25:01 | 0:00:00 |
| NM_016578:XM_005274051:XM_017017923 | 2:15:00 | 0:00:00 | 0.09099537 | 0:01:00 | 2:21:00 | 0:00:00 | 0.266157407 | 0:02:00 | 2:25:01 | 0:01:00 |
| NM_004549:NM_001204055 | 0:00 | 2:00 | 1:00 | 0:00 | 3:00 | 4:00 | 11:00 | 0:00 | 7:00 | 0:00 |
| NM_024678:NM_001243251:XM_017018303:XR_001747964:XR_001747966:XM_017018304:XM_011545253:XR_001747965:XR_001747963 | 1:3:0:0:0:0:0:0:0 | 2:0:0:0:0:0:0:0:0 | 2:1:0:0:0:0:0:0:0 | 0:0:0:0:0:0:0:0 | 1:3:1:1:0:0:0:0:0 | 0:0:0:0:0:0:0:0:0 | 20:19:5:0:7:2:0:0:0 | 0:0:0:0:0:0:0:0:0 | 5:7:1:0:1:1:1:0:0 | 0:0:0:0:0:0:0:0:0 |
| NM_001243251:NM_024678:XM_017018303:XR_001747964:XR_001747966:XM_017018304:XM_011545253:XR_001747965:XR_001747963 | 0:3:0:0:0:0:0:0:0 | 3:0:0:0:0:0:0:0:0 | 0:2:0:0:0:0:0:0:0 | 1:0:0:0:0:0:0:0 | 0:0:0:1:0:0:0:0:0 | 3:1:1:0:0:0:0:0:0 | 0:0:0:0:0:0:0:0:0 | 19:16:6:0:8:1:0:0:0 | 0:0:0:0:0:0:0:0:0 | 7:5:1:0:1:1:1:0:0 |
| NM_003754 | 1 | 1 | 0 | 2 | 1 | 3 | 10 | 0 | 10 | 0 |
| NM_001004 | 26 | 0 | 18 | 0 | 13 | 0 | 79 | 0 | 60 | 0 |
| XM_024448618:NM_020376:XM_017018028 | 0:00:00 | 1:00:00 | 0:00:00 | 0:01:01 | 0:00:00 | 0:00:00 | 0:00:00 | 0:01:04 | 0:00:00 | 0:00:00 |
| NM_018480:NM_001193538:NM_001256547:NM_001350394:XM_011545165:NR_146645:NM_001350396:NM_001193537:NR_146646:NM_001350395:NM_001256546 | 24:42:3:1:1:1:1:0:0:0:0 | 0:0:0:0:0:0:0:0:0:0:0 | 52:35:1:1:6:0:2:1:0:0:0 | 0:0:0:0:0:0:0:0:0:0 | 34:28:4:0:1:0:0:1:1:0:0 | 0:0:0:0:0:0:0:0:0:0:0 | 257:157:7:2:62:3:2:98:1:2:15 | 0:0:0:0:0:0:0:0:0:0 | 91:41:2:6:12:0:0:15:1:0:3 | 0:0:0:0:0:0:0:0:0:0:0 |
| NM_018480:NM_001193538:XM_011545165:NM_001350394:NM_001350396:NM_001193537:NR_146646:NM_001350395:NM_001256546:NR_146645 | 24:42:2:1:1:0:0:0:0:0 | 0:0:0:0:0:0:0:0:0:0 | 53:36:8:1:2:1:0:0:0:0 | 0:0:0:0:0:0:0:0:0 | 34:28:3:0:0:1:1:0:0:0 | 0:0:0:0:0:0:0:0:0:0 | 257:157:67:2:2:100:1:2:15:2 | 0:0:1:0:0:0:0:0:0:0 | 91:41:14:6:0:16:1:0:4:0 | 0:0:0:0:0:0:0:0:0:0 |
| NM_018480:NM_001193538:XM_011545165:NM_001350394:NM_001350396:NM_001193537:NR_146646:NM_001350395:NM_001256546:NR_146645 | 24:42:2:1:1:0:0:0:0:0 | 0:0:0:0:0:0:0:0:0:0 | 53:36:8:1:2:1:0:0:0:0 | 0:0:0:0:0:0:0:0:0 | 34:28:4:0:0:1:1:0:0:0 | 0:0:0:0:0:0:0:0:0:0 | 255:157:71:2:2:100:1:2:15:2 | 2:0:0:0:0:0:0:0:0:0 | 91:41:14:6:0:16:1:0:4:0 | 0:0:0:0:0:0:0:0:0:0 |
| NM_018480:NM_001193538:NM_001256547:XM_011545165:NM_001350394:NM_001350396:NM_001193537:NR_146646:NM_001256546:NR_146645:NM_001350395 | 24:42:3:3:1:1:0:0:0:0:0 | 0:0:0:0:0:0:0:0:0:0:0 | 53:36:1:8:1:2:1:0:0:0:0 | 0:0:0:0:0:0:0:0:0:0 | 34:28:4:4:0:0:1:1:0:0:0 | 0:0:0:0:0:0:0:0:0:0:0 | 257:157:7:73:2:2:100:2:15:2:0 | 0:0:0:0:0:0:0:0:0:0 | 91:41:2:16:6:0:16:1:4:0:0 | 0:0:0:0:0:0:0:0:0:0:0 |
| XM_024448663:XM_005253073:NM_023947:NM_001142674:XM_017018166:NM_001142676 | 1:1:0:0:0:0 | 0:0:0:0:0:0 | 0:0:0:0:0:0 | 0:0:0:0:0:0 | 0:0:0:0:0:0 | 0:0:0:0:0:0 | 0:0:0:0:0:0 | 0:0:0:0:0:0 | 0:0:1:0:0:0 | 0:0:0:0:0:0 |
| NM_022918:NM_001168724:XM_017018141:XM_017018142:NR_033149 | 22:3:2:1:0 | 0:0:0:0:0 | 5:2:0:0:0 | 0:0:0:0:0 | 39:2:2:8:0 | 1:0:0:0:0 | 19:3:1:4:0 | 2:0:0:0:0 | 18:0:2:2:0 | 0:0:0:0:0 |
| NM_022918:NM_001168724:XM_017018141:XM_017018142:NR_033149 | 0:0:0:0:0 | 22:3:2:1:0 | 0:0:0:0:0 | 5:2:0:0:0 | 0:0:0:0:0 | 40:2:2:8:0 | 11:1:0:3:0 | 7:2:1:2:0 | 8:0:0:2:0 | 10:0:2:0:0 |
| NM_001814 | 0 | 1 | 0 | 0 | 0 | 1 | 0 | 0 | 0 | 1 |
| NM_020644:NM_001286095:NM_001286094 | 102:15:08 | 0:00:00 | 273:12:09 | 0:00:00 | 761:35:23 | 2:00:00 | 430:53:23 | 1:00:00 | 147:13:07 | 0:00:00 |
| NM_020644:NM_001286095 | 102:15:00 | 0:00 | 266:12:00 | 1:00 | 757:35:00 | 1:00 | 419:53:00 | 1:00 | 146:13:00 | 1:00 |
| NM_012124:XR_001747832 | 0:00 | 1:00 | 0:00 | 0:00 | 0:00 | 0:00 | 0:00 | 0:00 | 0:00 | 0:00 |
| NM_016156:NM_201281:NM_201278:XM_005274375:NM_001243571:XM_005274374 | 5:2:2:1:0:0 | 0:0:0:0:0:0 | 0:0:1:0:0:0 | 0:0:0:0:0:0 | 1:2:0:1:0:0 | 0:0:0:0:0:0 | 2:2:0:1:1:0 | 1:3:1:0:0:0 | 0:1:0:0:0:1 | 1:1:0:3:0:0 |
| XM_011543025:XM_011543024:NM_032427 | 1:00:00 | 0:00:00 | 0:01:00 | 1:00:00 | 0:00:00 | 0:00:00 | 1:01:01 | 2:01:00 | 0:00:00 | 0:00:01 |
| XM_017019206:NM_001303130:NM_174942:XM_005268804:XM_011538220 | 0:0:0:0:0 | 0:0:0:0:0 | 4:1:1:0:0 | 0:0:0:0:0 | 0:0:0:3:0 | 0:0:0:0:0 | 5:3:6:0:0 | 0:0:0:0:0 | 2:4:0:0:1 | 0:0:0:0:0 |
| NM_020244:XR_245946 | 1:00 | 0:00 | 1:00 | 0:00 | 0:00 | 0:00 | 2:00 | 1:01 | 0:00 | 0:00 |
| NM_024057:XR_944712 | 44:03:00 | 0:00 | 28:00:00 | 0:00 | 27:01:00 | 0:00 | 418:06:00 | 2:00 | 145:00:00 | 0:00 |
| NM_024057:XR_944712 | 44:03:00 | 0:00 | 28:00:00 | 0:00 | 27:01:00 | 0:00 | 418:06:00 | 2:00 | 145:00:00 | 0:00 |
| NM_024057:XR_944712 | 44:02:00 | 0:00 | 29:01:00 | 0:00 | 27:00:00 | 0:00 | 421:05:00 | 0:00 | 145:00:00 | 0:00 |
| NM_024057:XR_944712 | 44:02:00 | 0:00 | 29:01:00 | 0:00 | 27:00:00 | 0:00 | 422:05:00 | 0:00 | 147:00:00 | 0:00 |
| NM_024057:XR_944712 | 44:01:00 | 0:00 | 29:00:00 | 0:00 | 27:00:00 | 0:00 | 419:07:00 | 1:00 | 146:00:00 | 1:00 |
| NM_024057:XR_944712 | 44:01:00 | 0:00 | 28:01:00 | 0:00 | 27:00:00 | 0:00 | 416:10:00 | 0:00 | 143:00:00 | 0:00 |
| NM_002261 | 0 | 0 | 0 | 1 | 0 | 0 | 0 | 5 | 0 | 0 |
| NM_002261 | 0 | 0 | 0 | 0 | 0 | 0 | 0 | 5 | 0 | 0 |
| NM_002261 | 0 | 0 | 0 | 0 | 0 | 0 | 0 | 5 | 0 | 0 |
| NM_002261 | 0 | 0 | 0 | 0 | 0 | 0 | 1 | 1 | 0 | 0 |
| NM_002261 | 0 | 0 | 0 | 0 | 0 | 0 | 0 | 2 | 0 | 0 |
| NM_018413:XM_017019369:NM_001173982 | 3:00:00 | 1:01:00 | 1:00:07 | 0:00:00 | 5:00:01 | 0:00:00 | 13:04:10 | 0:00:00 | 0:00:02 | 0:00:00 |
| NM_213595:NM_001301141:XM_011538101 | 0:00:00 | 4:00:00 | 0:00:00 | 2:00:00 | 0:00:00 | 5:00:00 | 0:00:00 | 8:01:01 | 0:00:00 | 8:00:00 |
| NM_213595:NM_001301141:XM_011538101 | 0:00:00 | 4:00:00 | 0:00:00 | 2:00:00 | 0:00:00 | 5:00:00 | 0:00:00 | 8:01:01 | 0:00:00 | 8:00:00 |
| NM_213595:NM_001301141:XM_011538101 | 0:00:00 | 4:00:00 | 0:00:00 | 2:00:00 | 0:00:00 | 5:00:00 | 0:00:00 | 8:01:01 | 0:00:00 | 8:00:00 |
| NM_080911:NM_003362 | 28:34:00 | 0:00 | 3:37 | 0:00 | 35:68 | 0:00 | 116:189 | 1:00 | 54:54:00 | 0:00 |
| NM_080911:NM_003362 | 33:34:00 | 0:00 | 9:37 | 0:00 | 42:68 | 0:00 | 151:187 | 1:01 | 63:54:00 | 0:00 |
| NM_052845:NR_038118 | 0:00 | 2:00 | 0:00 | 0:00 | 0:00 | 0:00 | 0:01 | 0:00 | 0:00 | 0:00 |
| NM_052845:NR_038118 | 0:00 | 2:00 | 0:00 | 0:00 | 0:00 | 0:00 | 0:01 | 0:00 | 0:00 | 0:00 |
| NR_104101:XM_017019459:XM_006719459:NM_016226:NM_057180:NR_104099 | 0:0:0:0:0:0 | 0:0:0:0:0:0 | 0:0:0:0:0:0 | 0:0:0:0:0:0 | 0:1:1:0:0:0 | 2:0:0:0:0:0 | 0:0:0:4:3:0 | 0:0:0:0:0:0 | 0:0:0:0:3:1 | 0:0:0:0:0:0 |

| | | | | | | | | | | |
|---|---|---|---|---|---|---|---|---|---|---|
| NM_002834:NM_080601:XM_017019722:NM_001330437 | 2:19:1:1 | 0:0:0:0 | 3:25:1:1 | 0:0:0:0 | 1:37:4:2 | 0:5:0:0 | 7:187:10:0 | 0:1:0:0 | 1:47:4:0 | 0:1:0:0 |
| NM_080601:XM_017019722:NM_002834:NM_001330437:XM_011538613 | 17:0:2:1:0 | 2:1:0:0:0 | 23:0:3:1:0 | 1:0:0:0:0 | 32:3:1:2:0 | 3:0:0:0:0 | 171:12:7:0:0 | 16:0:0:0:0 | 42:5:1:0:0 | 4:0:0:0:0 |
| NM_080601:XM_017019722:NM_002834:NM_001330437:XM_011538613 | 17:0:2:1:0 | 2:1:0:0:0 | 23:0:3:1:0 | 1:0:0:0:0 | 33:3:1:2:0 | 3:0:0:0:0 | 164:11:7:0:0 | 22:1:0:0:0 | 43:5:1:0:0 | 3:0:0:0:0 |
| NM_002834:NM_080601:NM_001330437:XM_017019722:XM_011538613 | 2:2:1:0:0 | 0:0:0:0:0 | 3:0:1:2:0 | 0:1:0:0:0 | 1:0:2:0:0 | 0:0:0:1:0 | 7:1:0:4:0 | 0:1:0:0:0 | 1:1:0:0:1 | 0:1:0:0:0 |
| NM_001032409:NM_002534:NM_016816:XM_006719434 | 0:0:0:0 | 0:0:0:0 | 0:0:0:0 | 1:1:0:0 | 0:0:0:0 | 1:0:1:0 | 0:0:3:0 | 0:0:0:0 | 0:0:1:0 | 0:0:0:0 |
| NM_001032409:XM_011538413:XM_017019362:NM_016816:XM_006719434 | 0:0:0:0:0 | 0:0:0:0:0 | 0:0:0:0:0 | 1:1:1:0:0 | 0:0:0:0:0 | 1:0:0:1:0 | 0:0:1:3:0 | 0:0:0:0:0 | 0:0:0:1:0 | 0:0:0:0:0 |
| NM_024738:XM_024449189 | 1:00 | 0:00 | 3:01 | 0:00 | 9:00 | 0:00 | 8:00 | 0:00 | 2:00 | 0:00 |
| XM_011520607:NM_001987:XM_017018990:XM_011520608:XM_017018991:XM_011520611 | 8:5:0:0:0:0 | 0:0:0:0:0:0 | 3:10:2:1:0:0 | 0:0:0:0:0:0 | 13:14:2:0:0:0 | 0:0:0:0:0:0 | 10:22:3:1:9:0 | 0:0:0:0:0:0 | 3:6:2:0:4:2 | 0:0:0:0:0:0 |
| XM_011520607:NM_001987:XM_017018991:XM_011520608:XM_011520611:XM_017018990:XM_011520612 | 6:4:1:0:0:0:0 | 0:0:0:0:0:0:0 | 3:7:9:1:2:0:0 | 0:0:0:0:0:0:0 | 10:14:5:0:2:1:1 | 0:0:0:0:0:0:0 | 7:21:23:2:1:0:0 | 0:0:0:0:0:0:0 | 3:5:7:0:1:0:0 | 0:0:0:0:1:0:0 |
| XM_024449110:XM_017019743:NM_002859:XM_006719532 | 0:0:0:0 | 0:0:0:0 | 0:0:0:0 | 0:0:0:0 | 0:0:0:0 | 0:0:0:0 | 0:0:0:0 | 1:0:0:0 | 0:0:0:0 | 0:1:0:0 |
| NM_002813:NM_001261400:NR_048555 | 0:00:00 | 87:01:01 | 0:00:00 | 100:02:01 | 0:00:00 | 247:02:03 | 0:00:00 | 381:03:12 | 0:00:00 | 161:03:00 |
| NM_002813:NR_048555:NM_001261400 | 89:01:00 | 0:00:00 | 105:01:00 | 0:00:00 | 254:01:00 | 1:00:00 | 390:12:00 | 0:00:00 | 166:00:00 | 0:00:00 |
| NM_002813:NM_001261400:NR_048555 | 94:01:00 | 0:00:00 | 105:00:00 | 0:00:00 | 263:00:00 | 1:00:00 | 395:00:01 | 3:00:00 | 176:00:00 | 0:00:00 |
| NM_002813:NM_001261400:NR_048555 | 96:01:01 | 0:00:00 | 108:02:02 | 0:00:00 | 268:02:01 | 0:00:00 | 400:03:14 | 1:00:00 | 175:03:00 | 0:00:00 |
| NM_002813:NM_001261400:NR_048555 | 96:01:02 | 0:00:00 | 108:02:07 | 0:00:00 | 268:02:07 | 0:00:00 | 398:03:19 | 2:00:00 | 175:03:01 | 0:00:00 |
| XR_944509:XM_006719294 | 0:00 | 0:00 | 0:00 | 0:00 | 0:00 | 0:00 | 2:00 | 3:00 | 0:00 | 0:00 |
| XM_006719552:XM_017019788:XM_017019792:XM_017019790:NM_198240:XM_017019786 | 1:1:0:0:0:0 | 0:0:0:0:0:0 | 0:0:0:2:0:0 | 0:0:1:1:0:0 | 0:0:0:3:0:0 | 0:0:0:2:0:0 | 0:0:0:0:0:0 | 0:0:0:6:1:0 | 0:0:0:0:0:0 | 0:1:0:4:0:0 |
| XM_017019790:XM_006719552:XM_017019788:XM_017019786:XM_017019792:NM_198240:XM_005253593 | 1:1:1:0:0:0:0 | 1:0:0:0:0:0:0 | 3:0:0:0:1:0:0 | 2:0:0:1:2:0:0 | 6:1:0:0:0:0:0 | 1:0:3:0:0:0:0 | 11:0:0:0:1:1:0 | 13:0:0:0:0:0:0 | 4:0:1:0:0:0:0 | 0:0:0:0:0:0:0 |
| NM_023012:XM_005253601:XR_001748845:XM_017019833:NR_036434:XR_002957370:XR_001748844:XM_017019830:XM_011538688:NR_036435:XM_005253604:XR_001748848:XM_017019832:XM_017019831:XM_011538689:XR_001748846:NR_036436:XR_001748847:XM_005253602 | 38:42:2:7:1:2:4:2:1:1:1:0:0:0:0:0:0:0:0 | 0:0:0:0:0:0:0:0:0:0:0:0:0:0:0:0 | 10:24:1:6:5:4:7:14:20:4:1:2:11:1:4:0:0:0:0 | 0:0:0:0:0:0:0:0:0:0:0:0:0:0:0:0:0:0:0 | 28:47:1:3:4:1:1:0:1:3:4:0:0:0:1:1:0:0:0 | 0:0:0:0:0:0:0:0:0:0:0:0:0:0:0:0:0:0:0 | 137:139:11:21:3:26:25:102:97:11:1:10:21:0:1:2:2:5:0 | 0:0:0:0:0:0:0:1:0:0:0:1:0:0:0:0:0:0:0 | 55:45:5:4:3:1:0:1:2:7:0:1:4:1:2:0:0:0:1 | 1:0:0:0:0:0:0:0:0:0:0:0:0:0:0:0:0:0:0 |
| NM_023012:XM_005253601:XR_001748845:XM_017019833:NR_036434:XR_002957370:XR_001748844:XM_017019830:XM_011538688:NR_036435:XM_005253604:XM_017019831:XM_017019832:XM_011538689:XR_001748846:NR_036436:XR_001748847:XR_001748848:XM_005253602 | 38:42:3:3:2:2:4:2:1:1:1:0:0:0:0:0:0:0:0 | 0:0:0:0:0:0:0:0:0:0:0:0:0:0:0:0 | 10:24:1:3:5:0:4:15:20:4:1:1:5:4:0:0:0:0:0 | 0:0:0:0:0:0:0:0:0:0:0:0:0:0:0:0:0:0:0 | 28:47:2:1:4:1:1:1:1:3:4:0:0:1:1:0:0:0:0 | 0:0:0:0:0:0:0:0:0:0:0:0:0:0:0:0:0:0:0 | 137:138:10:10:3:9:20:113:98:11:1:0:8:2:2:1:3:2:0 | 0:1:1:0:0:1:0:0:0:0:0:0:0:0:0:0:0:0:0 | 56:45:4:2:3:2:0:3:2:7:0:1:2:3:0:0:0:0:1 | 0:0:0:0:0:0:0:0:0:0:0:0:0:0:0:0:0:0:0 |
| NM_023012:XM_005253601:XR_001748845:XM_017019833:NR_036434:XR_002957370:XM_017019830:XR_001748844:XM_011538688:NR_036435:XM_005253604:XM_017019831:XM_017019832:XM_011538689:XR_001748846:NR_036436:XR_001748847:XR_001748848:XM_005253602 | 37:42:3:2:2:2:2:2:1:1:1:0:0:0:0:0:0:0:0 | 0:0:0:0:0:0:0:0:0:0:0:0:0:0:0:0 | 10:23:1:2:5:0:15:4:19:4:1:1:4:1:0:0:0:0:0 | 0:1:0:0:0:0:0:0:0:0:0:0:0:0:0:0:0:0:0 | 28:47:2:0:4:2:1:1:1:3:4:0:0:1:1:0:0:0:0 | 0:0:0:0:0:0:0:0:0:0:0:0:0:0:0:0:0:0:0 | 137:137:7:5:2:9:113:17:98:8:1:0:2:2:2:1:2:2:0 | 0:1:0:0:0:0:0:0:0:0:0:0:0:0:0:0:0:0:0 | 55:45:4:2:3:1:3:0:2:6:0:0:0:2:0:0:0:0:1 | 1:0:0:0:0:0:0:0:0:0:0:0:1:0:0:0:0:0:0 |
| NM_023012:XM_005253601:XR_001748845:XM_017019833:NR_036434:XR_002957370:XM_017019830:XR_001748844:XM_011538688:NR_036435:XM_005253604:XM_017019831:XM_017019832:XM_011538689:XR_001748846:NR_036436:XR_001748847:XR_001748848:XM_005253602 | 37:42:3:2:2:2:2:2:1:1:1:0:0:0:0:0:0:0:0 | 0:0:0:0:0:0:0:0:0:0:0:0:0:0:0:0 | 10:24:1:2:5:0:15:4:19:4:1:1:4:1:0:0:0:0:0 | 0:0:0:0:0:0:0:0:0:0:0:0:0:0:0:0:0:0:0 | 28:47:2:0:4:2:1:1:1:3:4:0:0:1:1:0:0:0:0 | 0:0:0:0:0:0:0:0:0:0:0:0:0:0:0:0:0:0:0 | 137:139:7:5:2:8:113:17:96:8:1:0:2:2:2:1:2:2:0 | 0:0:0:0:1:0:0:1:0:0:0:0:0:0:0:0:0:0:0 | 56:45:4:2:3:1:3:0:2:6:0:1:0:2:0:0:0:0:1 | 0:0:0:0:0:0:0:0:0:0:0:0:0:0:0:0:0:0:0 |
| NM_023012:XM_005253601:XR_001748845:XM_017019833:NR_036434:XR_002957370:XM_017019830:XR_001748844:XM_011538688:NR_036435:XM_005253604:XM_017019831:XM_017019832:XM_011538689:NR_036436:XR_001748847:XR_001748848:XR_001748846:XM_005253602 | 37:42:3:2:2:1:2:2:1:1:1:0:0:0:0:0:0:0:0 | 0:0:0:0:0:0:0:0:0:0:0:0:0:0:0:0 | 10:24:0:0:5:0:15:4:19:4:1:1:3:1:0:0:0:0:0 | 0:0:0:0:0:0:0:0:0:0:0:0:0:0:0:0:0:0:0 | 26:47:2:0:4:1:1:1:1:3:4:0:0:0:0:0:0:0:0 | 1:0:0:0:0:0:0:0:0:0:0:0:0:0:0:0:0:0:0 | 136:136:7:6:2:10:112:17:95:8:1:0:2:2:1:2:2:2:0 | 0:0:0:0:0:0:0:0:0:0:0:0:0:0:0:0:0:0:0 | 56:45:4:2:3:1:3:0:2:4:0:1:0:2:0:0:0:0:1 | 0:0:0:0:0:0:0:0:0:0:0:0:0:0:0:0:0:0:0 |
| NM_005505:NM_001082959 | 67:04:00 | 0:00 | 122:12:00 | 1:00 | 211:14:00 | 0:00 | 341:11:00 | 0:00 | 93:01:00 | 0:00 |
| NM_005505:NM_001082959 | 68:04:00 | 0:00 | 102:09:00 | 1:00 | 199:13:00 | 0:00 | 268:10:00 | 1:00 | 89:00:00 | 0:00 |
| NM_005505:NM_001082959 | 66:04:00 | 0:00 | 88:08:00 | 0:00 | 186:13:00 | 1:00 | 195:07:00 | 2:00 | 81:00:00 | 0:00 |
| NM_005505:NM_001082959 | 46:03:00 | 17:01 | 52:04:00 | 31:02:00 | 95:09:00 | 83:03:00 | 136:04:00 | 0:00 | 77:00:00 | 0:00 |
| NM_005505:NM_001082959 | 62:04:00 | 0:00 | 77:06:00 | 0:00 | 171:11:00 | 0:00 | 122:02:00 | 2:00 | 72:00:00 | 0:00 |
| NM_032656:XR_001748820 | 0:00 | 1:00 | 0:00 | 0:01 | 0:00 | 1:00 | 0:00 | 0:00 | 1:00 | 0:00 |
| XR_001748820:NM_032656:XM_005253590 | 0:00:00 | 0:00:00 | 0:00:00 | 1:00:00 | 0:00:00 | 0:01:00 | 0:00:01 | 0:00:00 | 0:01:00 | 0:00:00 |
| NM_024078:XR_944714:XM_017019960 | 4:01:01 | 0:00:00 | 2:00:00 | 0:00:00 | 13:01:01 | 0:00:00 | 34:01:06 | 2:00:00 | 20:02:01 | 0:00:00 |
| NM_015114:XR_001748638 | 0:00 | 0:00 | 0:00 | 0:00 | 0:00 | 0:00 | 0:00 | 0:00 | 0:00 | 0:00 |
| NM_015114 | 0 | 0 | 0 | 0 | 0 | 0 | 0 | 0 | 0 | 0 |
| XM_017019067:XM_017019068:XM_017019075:XR_001748637 | 0:0:0:0 | 0:0:0:0 | 1:1:0:0 | 1:0:0:0 | 0:0:1:0 | 0:0:0:0 | 0:0:2:0 | 0:0:2:0 | 0:0:0:0 | 0:0:0:0 |
| NM_024829 | 0 | 0 | 0 | 0 | 0 | 0 | 0 | 3 | 0 | 1 |
| NM_024829 | 0 | 0 | 0 | 0 | 0 | 0 | 2 | 2 | 1 | 0 |
| NM_024829 | 0 | 0 | 0 | 0 | 0 | 0 | 0 | 3 | 0 | 1 |
| NM_024829 | 0 | 0 | 0 | 0 | 0 | 0 | 3 | 0 | 1 | 0 |
| NM_000900 | 13 | 0 | 1 | 0 | 3 | 0 | 4 | 0 | 5 | 0 |
| NM_001349920 | 0 | 0 | 0 | 0 | 0 | 0 | 0 | 0 | 0 | 0 |
| NM_002300:NM_001174097 | 214:00:00 | 1:00 | 277:00:00 | 0:00 | 378:00:00 | 0:00 | 7247:07:00 | 3:00 | 1753:01:00 | 1:00 |
| NM_002300:NM_001174097 | 215:00:00 | 0:00 | 276:00:00 | 1:00 | 378:00:00 | 0:00 | 7242:07:00 | 8:00 | 1754:01:00 | 1:00 |
| NM_002300:NM_001174097 | 215:00:00 | 0:00 | 276:00:00 | 1:00 | 377:00:00 | 0:00 | 7258:07:00 | 9:00 | 1761:01:00 | 0:00 |
| NM_002300:NM_001174097 | 216:00:00 | 0:00 | 275:00:00 | 3:00 | 377:00:00 | 0:00 | 7257:07:00 | 9:00 | 1760:01:00 | 2:00 |
| NM_002300:NM_001174097 | 216:00:00 | 0:00 | 278:00:00 | 0:00 | 377:00:00 | 0:00 | 7269:07:00 | 0:00 | 1762:01:00 | 0:00 |
| NM_002300:XM_006719074:NM_001174097 | 217:01:00 | 0:00:00 | 274:00:00 | 1:00:00 | 375:00:00 | 0:00:00 | 7252:13:07 | 8:00:00 | 1763:03:01 | 1:00:00 |
| NM_002300:XM_006719074:NM_001174097 | 217:02:00 | 0:00:00 | 270:00:00 | 1:00:00 | 371:00:00 | 0:00:00 | 7220:20:07 | 8:00:00 | 1755:04:01 | 0:00:00 |
| NM_002300:XM_006719074:NM_001174097 | 216:02:00 | 1:00:00 | 270:00:00 | 0:00:00 | 371:00:00 | 0:00:00 | 7207:21:07 | 9:00:00 | 1754:04:01 | 1:00:00 |
| NM_002300:XM_006719074:NM_001174097 | 217:02:00 | 0:00:00 | 267:00:00 | 3:00:00 | 371:00:00 | 0:00:00 | 7205:21:07 | 9:00:00 | 1753:04:01 | 1:00:00 |
| NM_002300:XM_006719074:NM_001174097 | 216:02:00 | 0:00:00 | 268:00:00 | 0:00:00 | 371:00:00 | 0:00:00 | 7184:22:07 | 15:00:00 | 1750:04:01 | 0:00:00 |
| NM_002300:XM_006719074:NM_001174097 | 216:02:00 | 0:00:00 | 266:00:00 | 0:00:00 | 368:00:00 | 0:00:00 | 7173:23:07 | 7:00:00 | 1744:05:01 | 1:00:00 |
| NM_002300:XM_006719074:NM_001174097 | 214:02:00 | 0:00:00 | 264:00:00 | 0:00:00 | 365:00:00 | 1:00:00 | 7158:24:07 | 9:00:00 | 1737:05:01 | 2:00:00 |
| NM_002300:XM_006719074:NM_001174097 | 213:02:00 | 1:00:00 | 263:00:00 | 1:00:00 | 366:00:00 | 0:00:00 | 7157:24:07 | 10:00:00 | 1737:05:01 | 2:00:00 |
| NM_002300:XM_006719074:NM_001174097 | 214:02:00 | 0:00:00 | 263:00:00 | 1:00:00 | 365:00:00 | 1:00:00 | 7153:25:07 | 11:00:00 | 1737:05:01 | 1:00:00 |
| NM_002300:XM_006719074:NM_001174097 | 214:02:00 | 0:00:00 | 264:00:00 | 0:00:00 | 365:00:00 | 1:00:00 | 7153:25:07 | 11:00:00 | 1735:05:01 | 3:00:00 |
| NM_002300:XM_006719074:NM_001174097 | 214:01:00 | 0:00:00 | 262:00:00 | 0:00:00 | 364:00:00 | 0:00:00 | 7129:26:07 | 9:00:00 | 1726:05:01 | 4:00:00 |
| NM_002300:XM_006719074:NM_001174097 | 213:01:00 | 0:00:00 | 260:00:00 | 0:00:00 | 360:00:00 | 1:00:00 | 7123:23:07 | 7:00:00 | 1725:04:01 | 1:00:00 |
| NM_002300:XM_006719074:NM_001174097 | 211:01:00 | 2:00:00 | 259:00:00 | 0:00:00 | 357:00:00 | 0:00:00 | 7055:15:07 | 9:00:00 | 1708:03:01 | 4:00:00 |
| NM_002300:NM_001174097:XM_006719074 | 212:00:00 | 0:00:00 | 257:00:00 | 0:00:00 | 353:00:00 | 0:00:00 | 7013:07:06 | 11:00:00 | 1700:01:03 | 0:00:00 |
| NM_002300:NM_001174097:XM_006719074 | 212:00:00 | 0:00:00 | 257:00:00 | 0:00:00 | 355:00:00 | 0:00:00 | 7007:07:05 | 12:00:00 | 1698:01:02 | 4:00:00 |
| NM_002300:NM_001174097:XM_006719074 | 212:00:00 | 0:00:00 | 256:00:00 | 0:00:00 | 355:00:00 | 0:00:00 | 6987:07:04 | 19:00:00 | 1701:01:01 | 0:00:01 |
| NM_002300:NM_001174097 | 212:00:00 | 0:00 | 255:00:00 | 0:00 | 355:00:00 | 0:00 | 6969:07:00 | 3:00 | 1694:01:00 | 0:00 |
| NM_002300:XM_006719074:NM_001174097 | 212:00:00 | 0:00:00 | 253:01:00 | 0:00:00 | 351:01:00 | 0:00:00 | 6923:06:07 | 7:00:00 | 1679:00:01 | 3:00:00 |
| NM_002300:XM_006719074:NM_001174097 | 211:00:00 | 0:00:00 | 252:01:00 | 1:00:00 | 351:01:00 | 0:00:00 | 6917:06:07 | 6:00:00 | 1679:00:01 | 1:00:00 |
| NM_002300:XM_006719074:NM_001174097 | 210:00:00 | 0:00:00 | 252:01:00 | 0:00:00 | 349:01:00 | 0:00:00 | 6891:06:07 | 9:00:00 | 1672:00:01 | 2:00:00 |
| NM_002300:XM_006719074:NM_001174097 | 209:00:00 | 1:00:00 | 251:01:00 | 0:00:00 | 349:01:00 | 0:00:00 | 6884:06:07 | 12:00:00 | 1674:00:01 | 0:00:00 |
| NM_002300:XM_006719074:NM_001174097 | 205:00:00 | 1:00:00 | 248:01:00 | 0:00:00 | 344:01:00 | 1:00:00 | 6844:06:07 | 13:00:00 | 1654:00:01 | 1:00:00 |
| NM_018686:NR_135117 | 30:00:00 | 0:00 | 89:03:00 | 0:00 | 65:00:00 | 0:00 | 188:05:00 | 1:00 | 59:03:00 | 0:00 |
| NM_018686 | 29 | 0 | 87 | 0 | 66 | 2 | 193 | 1 | 61 | 0 |
| NM_018686:NR_135117 | 28:00:00 | 0:00 | 73:01:00 | 9:00 | 63:01:00 | 0:00 | 182:02:00 | 2:00 | 55:00:00 | 4:00 |

| | | | | | | | | | | |
|---|---|---|---|---|---|---|---|---|---|---|
| NM_004985:NM_033360 | 1:02 | 0:00 | 17:19 | 0:00 | 7:08 | 0:00 | 6:09 | 20:05 | 3:06 | 0:01 |
| XM_017019266:XM_017019267 | 1:10 | 0:00 | 0:15 | 0:03 | 0:02 | 0:00 | 1:16 | 0:00 | 0:07 | 0:00 |
| NM_001029874 | 0 | 0 | 0 | 0 | 0 | 2 | 0 | 0 | 0 | 0 |
| NM_001042603 | 5 | 0 | 3 | 0 | 2 | 0 | 8 | 9 | 2 | 4 |
| NM_001042603 | 5 | 2 | 6 | 2 | 6 | 1 | 16 | 4 | 7 | 0 |
| NM_001042603 | 7 | 0 | 19 | 7 | 6 | 0 | 28 | 2 | 7 | 0 |
| NM_001042603 | 9 | 0 | 17 | 0 | 11 | 0 | 59 | 2 | 6 | 0 |
| NM_001042603 | 8 | 1 | 21 | 15 | 15 | 2 | 78 | 39 | 6 | 6 |
| NM_001042603 | 9 | 0 | 32 | 6 | 13 | 2 | 104 | 10 | 11 | 4 |
| NM_001042603 | 7 | 4 | 21 | 5 | 15 | 9 | 63 | 44 | 6 | 10 |
| NM_001042603 | 11 | 0 | 43 | 2 | 25 | 0 | 159 | 0 | 20 | 1 |
| NM_001042603 | 5 | 3 | 15 | 16 | 10 | 8 | 133 | 0 | 20 | 0 |
| XM_006719021:NM_032680:XM_011521036 | 1:00:00 | 0:00:00 | 1:00:00 | 1:00:00 | 0:00:00 | 1:01:00 | 2:00:00 | 0:00:00 | 0:00:00 | 0:00:00 |
| XM_006719021:NM_032680:XM_011521036 | 1:00:00 | 0:00:00 | 1:00:00 | 1:00:00 | 0:00:00 | 1:01:00 | 2:00:00 | 0:00:00 | 0:00:00 | 0:00:00 |
| NM_032358:NM_001130146:NM_001130147 | 3:01:00 | 0:00:00 | 2:00:00 | 0:00:00 | 0:01:00 | 0:00:00 | 0:00:00 | 16:08:01 | 0:00:00 | 4:02:01 |
| NM_004818 | 12 | 0 | 14 | 0 | 35 | 0 | 63 | 2 | 14 | 0 |
| XM_017019336:NM_005171:XM_011538388:XM_017019333:XM_017019335:XM_017019334:XM_011538387:XM_011538386:XM_017019331 | 2:11:1:1:1:1:0:0:0 | 0:0:0:0:0:0:0:0:0 | 5:16:0:2:1:1:1:0:0 | 0:0:0:0:0:0:0:0:0 | 5:17:1:3:0:0:2:3:0 | 0:0:0:0:0:0:0:0:0 | 37:80:2:4:6:22:12:4:1 | 0:5:0:0:0:0:0:0:0 | 3:22:0:0:0:2:2:2:0 | 0:0:0:0:0:0:0:0:0 |
| NM_015416:NR_045020:NR_147120:XM_006719335:NM_001351313 | 0:0:0:0:0 | 0:0:0:0:0 | 1:0:0:0:0 | 0:0:0:0:0 | 1:1:0:0:0 | 0:0:0:0:0 | 0:0:0:0:0 | 0:0:1:2:0 | 0:0:0:0:0 | 0:0:0:0:1 |
| NM_021640 | 0 | 0 | 0 | 0 | 0 | 0 | 0 | 0 | 0 | 0 |
| NM_021640 | 0 | 0 | 0 | 0 | 0 | 0 | 0 | 0 | 0 | 0 |
| NM_021640 | 0 | 0 | 0 | 0 | 0 | 0 | 0 | 0 | 0 | 0 |
| NM_021640 | 0 | 0 | 0 | 0 | 0 | 0 | 0 | 0 | 0 | 0 |
| NM_005016:NM_031989:NM_001128913:NM_001098620:NM_001128912:NM_001128911:NM_001128914 | 87:130:55:98:20:13:46 | 0:0:0:0:0:0:0 | 83:95:34:47:10:4:56 | 0:0:0:0:0:0:0 | 425:460:178:196:65:29:317 | 0:0:0:0:1:0:0 | 712:1139:409:722:216:140:214 | 0:1:0:0:0:0:0 | 190:264:127:151:49:31:44 | 0:0:0:1:0:0:0 |
| NM_031989:NM_005016:NM_001128913:NM_001098620:NM_001128912:NM_001128911:NM_001128914 | 133:89:56:102:21:14:48 | 2:0:0:0:0:0:0 | 102:85:35:48:10:4:60 | 0:0:0:0:0:0:0 | 474:442:187:204:70:30:334 | 1:0:2:0:0:0:0 | 1167:721:420:745:227:142:215 | 1:1:3:2:0:0:0 | 270:192:131:155:53:32:45 | 1:0:2:0:0:0:0 |
| NM_031989:NM_005016:NM_001128913:NM_001098620:NM_001128912:NM_001128911:NM_001128914 | 133:89:56:103:21:14:49 | 1:0:0:0:0:0:0 | 104:85:35:49:10:4:60 | 0:0:0:0:0:0:0 | 474:445:191:207:73:31:340 | 2:2:0:0:0:0:0 | 1170:726:423:748:228:142:217 | 1:0:1:0:1:1:0 | 271:193:132:155:54:33:46 | 0:0:1:0:0:0:0 |
| NM_001098620:NM_005016:NM_031989:NM_001128913:NM_001128912:NM_001128911:NM_001128914 | 101:90:135:55:21:14:49 | 1:0:0:0:0:0:0 | 51:86:104:34:10:5:64 | 0:0:1:0:0:0:0 | 214:447:487:196:74:32:354 | 0:1:0:1:0:0:1 | 751:730:1179:419:240:145:221 | 0:0:2:0:1:1:0 | 155:188:274:131:53:33:48 | 0:1:1:0:0:0:0 |
| NM_001128913:NM_001098620:NM_005016:NM_031989:NM_001128912:NM_001128911:NM_001128914 | 55:106:92:138:21:14:51 | 1:1:0:0:0:0:0 | 35:53:88:109:10:6:64 | 0:0:0:0:0:0:0 | 200:219:454:490:76:34:367 | 1:0:0:1:0:0:2 | 431:766:740:1212:248:154:232 | 0:1:0:2:0:0:1 | 135:157:193:277:55:33:49 | 2:0:0:0:1:0:0 |
| NM_005016:NM_031989:NM_001128913:NM_001098620:NM_001128912:NM_001128911:NM_001128914 | 95:142:57:108:25:16:58 | 0:0:0:0:0:0:0 | 94:113:41:57:10:7:75 | 0:0:0:0:0:0:0 | 458:498:209:226:78:35:410 | 2:1:0:0:0:0:0 | 742:1266:442:789:267:159:263 | 2:0:0:0:1:0:0 | 195:287:137:160:61:33:51 | 0:0:0:0:0:0:0 |
| NM_005016:NM_031989:NM_001128913:NM_001098620:NM_001128912:NM_001128911:NM_001128914 | 95:141:57:108:26:16:58 | 0:0:0:0:0:0:0 | 94:115:41:56:11:7:77 | 0:0:0:0:0:0:0 | 458:508:208:225:80:35:410 | 0:0:0:0:0:0:0 | 741:1288:438:787:271:158:260 | 0:2:0:0:0:0:1 | 194:291:136:160:61:33:51 | 0:0:0:0:0:0:0 |
| NM_005016:NM_031989:NM_001128913:NM_001098620:NM_001128912:NM_001128911:NM_001128914 | 95:141:57:108:26:15:58 | 0:0:0:0:0:0:0 | 92:117:41:56:11:7:76 | 1:0:0:0:0:0:0 | 457:510:208:225:80:34:410 | 1:1:0:0:0:1:0 | 739:1294:435:785:271:158:260 | 2:0:2:2:0:0:1 | 194:292:135:160:61:33:51 | 0:0:0:0:0:0:0 |
| NM_033082:NR_026723:NR_026722 | 277:95:6 | 2:00:00 | 247:52:04 | 0:00:00 | 594:125:8 | 0:00:00 | 901:550:20 | 2:01:00 | 394:115:3 | 1:00:00 |
| NR_026723:NM_033082 | 99:276 | 0:00 | 60:245 | 0:00 | 113:588 | 0:00 | 554:898 | 1:02 | 115:392 | 0:01 |
| NM_002429:XM_011538359:XR_944553:XM_017019309 | 1:1:0:0 | 1:0:0:0 | 0:0:0:0 | 0:0:0:0 | 0:0:0:0 | 1:0:0:0 | 0:0:0:0 | 0:0:0:0 | 0:0:0:0 | 0:0:0:0 |
| NM_001032386:NM_000456:NM_001032387:XM_024449167:XM_017019905:XM_017019907 | 1:2:1:0:0:0 | 0:0:0:0:0:0 | 2:1:2:2:0:0 | 0:0:0:0:0:0 | 11:5:2:1:1:2 | 0:0:0:0:0:0 | 8:22:14:17:2:9 | 1:0:1:0:0:0 | 2:2:4:0:0:5 | 0:0:0:0:0:0 |
| NM_001982:NM_001005915 | 5:06 | 0:00 | 12:36 | 1:00 | 0.888888889 | 1:00 | 45:40:00 | 0:02 | 12:16 | 0:00 |
| NM_001982 | 2 | 0 | 4 | 1 | 16 | 0 | 24 | 2 | 8 | 0 |
| NM_021019:NM_079423 | 17:11 | 0:00 | 7:33 | 0:00 | 0.158333333 | 0:02 | 12:29 | 0:00 | 10:37 | 0:00 |
| NM_144576 | 1 | 2 | 3 | 0 | 2 | 0 | 16 | 0 | 6 | 0 |
| NM_006601:NM_001282602:XM_005268576:XM_006719199:NR_104219:NM_001282604:XM_011537774:XM_011537773:XM_017018716:NM_001282601 | 254:9:8:1:0:0:0:0:0:0 | 0:0:0:0:0:0:0:0:0:0 | 182:6:4:0:1:0:0:0:0:0 | 1:0:0:0:0:0:0:0:0:0 | 758:16:17:5:1:1:0:0:0:0 | 2:0:0:0:0:0:0:0:0:0 | 1220:51:64:4:2:9:1:1:1:0 | 4:0:0:0:0:0:0:0:0:0 | 520:9:17:6:0:3:0:0:0:0 | 0:0:0:0:0:0:0:0:0:0 |
| NM_006601:NM_001282601:NM_001282602:XM_005268576:XM_006719199:XM_011537773:NM_001282605:NR_104219:NM_001282604:XM_011537774:XM_017018716 | 254:5:9:7:1:3:0:0:0:0:0 | 0:0:0:0:0:0:0:0:0:0:0 | 183:6:6:3:0:1:1:1:0:0:0 | 0:0:0:0:0:18:0:0:0:0:0 | 761:7:16:14:1:2:4:1:1:0:0 | 0:0:0:0:0:1:0:0:0:0:0 | 1221:17:51:56:2:12:9:2:9:2:1 | 6:0:0:0:0:29:0:0:0:0:0 | 519:7:9:12:1:3:3:0:5:0:0 | 1:0:0:1:0:0:0:0:0:0:0 |
| NM_006601:NM_001282601:NM_001282602:XM_005268576:XM_006719199:XM_011537773:NM_001282605:NR_104219:NM_001282604:XM_011537774 | 255:5:9:3:1:10:0:0:0:0 | 0:0:0:0:0:0:0:0:0:0 | 183:6:6:2:0:33:1:1:0:0 | 1:0:0:0:0:0:0:0:0:0 | 760:7:16:7:1:15:4:1:1:0 | 1:0:0:1:0:0:0:0:0:0 | 1227:17:51:23:1:105:8:2:9:2 | 1:0:0:2:0:0:0:0:0:0 | 519:7:9:6:1:13:3:0:5:0 | 1:0:0:0:0:0:0:0:0:0 |
| NM_001113202:NM_001320194:NM_005594:NM_001320193:NM_001113201:NM_001113203 | 3031:16:193:112:3:0 | 4:1:0:0:0:0 | 1453:12:80:62:1:0 | 1:0:0:0:0:0 | 1448:10:106:73:1:1 | 1:6:1:0:0:0 | 18048:90:777:931:7:2 | 15:3:2:0:0:0 | 3983:16:254:185:2:0 | 2:1:1:0:0:0 |
| NM_001113202:NM_005594:NM_001320193:NM_001320194:NM_001113201:NM_001113203 | 3033:193:112:25:3:0 | 4:0:0:0:0:0 | 1451:80:62:20:1:0 | 4:0:0:0:0:0 | 1443:107:73:15:1:1 | 6:0:0:3:0:0 | 18055:781:933:134:7:2 | 34:0:0:9:0:0 | 3984:254:185:20:2:0 | 3:1:0:1:0:0 |
| NM_001113202:NM_005594:NM_001320193:NM_001320194:NM_001113201:NM_001113203 | 3031:193:112:27:3:0 | 2:0:0:0:0:0 | 1454:80:62:20:1:0 | 0:0:0:0:0:0 | 1447:106:73:18:1:1 | 0:0:0:1:0:0 | 18060:779:933:151:7:2 | 10:0:0:3:0:0 | 3982:255:185:23:2:0 | 0:0:0:0:0:0 |
| NM_005594:NM_001113202:NM_001320194:NM_001320193:NM_001113201:NM_001113203 | 192:3035:26:112:3:0 | 1:2:1:0:0:0 | 80:1453:20:62:1:0 | 0:2:0:0:0:0 | 107:1447:15:73:1:1 | 0:2:4:0:0:0 | 781:18076:152:933:7:2 | 0:11:8:0:0:0 | 255:3980:24:185:2:0 | 0:5:0:0:0:0 |
| NM_001113202:NM_005594:NM_001320193:NM_001320194:NM_001113201:NM_001113203 | 3027:193:112:33:3:0 | 10:0:0:0:0:0 | 1451:80:62:24:1:0 | 4:0:0:0:0:0 | 1445:107:73:21:1:1 | 4:0:0:0:0:0 | 18051:777:931:185:7:2 | 31:4:2:0:0:0 | 3979:257:185:30:2:0 | 6:0:0:0:0:0 |
| NM_001113202:NM_005594:NM_001320193:NM_001320194:NM_001113201:NM_001113203 | 3033:189:112:53:3:0 | 4:0:0:0:0:0 | 1455:81:63:43:1:0 | 0:0:0:0:0:0 | 1449:106:73:27:1:1 | 0:0:0:0:0:0 | 18080:781:934:291:7:2 | 1:1:1:0:0:0 | 3986:257:185:56:2:0 | 0:0:0:0:0:0 |
| NM_001113202:NM_005594:NM_001320193:NM_001320194:NM_001113201:NM_001113203 | 3035:188:112:57:3:0 | 3:1:0:0:0:0 | 1455:81:63:46:1:0 | 0:0:0:0:0:0 | 1449:106:73:27:1:1 | 0:0:0:0:0:0 | 18078:783:934:315:7:2 | 3:1:1:2:0:0 | 3986:256:185:58:2:0 | 0:0:0:1:0:0 |
| NM_001113202:NM_005594:NM_001320193:NM_001320194:NM_001113201:NM_001113203 | 3033:189:112:61:3:0 | 4:0:0:0:0:0 | 1454:81:63:48:1:0 | 1:0:0:0:0:0 | 1449:106:73:27:1:1 | 0:0:0:0:0:0 | 18064:782:932:323:7:2 | 17:2:3:5:0:0 | 3983:256:185:58:2:0 | 3:0:0:2:0:0 |
| NM_001113202:NM_005594:NM_001320194:NM_001320193:NM_001113201:NM_001113203 | 3035:186:75:113:3:0 | 3:1:1:0:0:0 | 1453:80:51:63:1:0 | 2:0:0:0:0:0 | 1448:106:29:73:1:1 | 1:0:0:0:0:0 | 18065:781:361:934:7:2 | 16:0:0:1:0:0 | 3980:251:74:185:2:0 | 6:2:0:0:0:0 |
| NM_001113202:NM_001320193:NM_001320194:NM_005594:NM_001113201:NM_001113203 | 3035:112:86:187:3:0 | 2:1:3:0:0:0 | 1453:63:59:81:1:0 | 2:0:11:1:0:0 | 1447:73:31:105:1:1 | 2:0:0:1:0:0 | 18054:935:384:781:7:2 | 26:1:22:3:0:0 | 3978:186:84:253:2:0 | 8:0:3:0:0:0 |
| NM_005594:NM_001113202:NM_001320193:NM_001320194:NM_001113201:NM_001113203 | 185:3033:113:92:3:0 | 1:3:0:0:0:0 | 79:1454:63:71:1:0 | 2:1:0:1:0:0 | 103:1446:73:31:1:1 | 1:3:0:0:0:0 | 773:18077:937:414:7:2 | 3:5:0:0:0:0 | 249:3986:186:88:2:0 | 1:0:0:0:0:0 |

| | | | | | | | | | | |
|---|---|---|---|---|---|---|---|---|---|---|
| NM_001113202:NM_001320194:NM_005594:NM_001320193:NM_001113201 | 3032:91:177:113:3 | 4:1:0:0:0 | 1454:72:72:63:1 | 1:0:0:0:0 | 1448:31:98:73:1 | 1:0:0:0:0 | 18066:412:742:936:7 | 17:2:2:1:0 | 3985:88:237:186:2 | 1:0:0:0:0 |
| NM_001113202:NM_005594:NM_001320193:NM_001320194:NM_001113201 | 3034:162:113:92:3 | 2:0:0:0:0 | 1455:62:63:72:1 | 0:2:0:0:0 | 1447:94:73:31:1 | 2:0:0:0:0 | 18075:711:937:414:7 | 10:7:0:0:0 | 3985:227:186:88:2 | 1:5:0:0:0 |
| NM_004083:NM_001195057:NM_001195056:NM_001195053:NM_001195055:NM_001195054 | 159:19:19:2:0:0 | 0:0:0:0:0:0 | 140:25:38:3:1:0 | 0:0:0:0:0:0 | 511:76:117:5:2:0 | 2:0:0:0:0:0 | 1527:223:80:4:11:1 | 3:1:0:0:0:0 | 205:39:10:1:1:0 | 0:0:0:0:0:0 |
| NM_004083:NM_001195057:NM_001195056:NM_001195053:NM_001195055:NM_001195054 | 157:19:19:2:0:0 | 0:0:0:0:0:0 | 137:25:36:3:1:0 | 0:0:0:0:0:0 | 511:76:117:5:2:0 | 4:0:0:0:0:0 | 1517:228:81:4:11:1 | 1:0:0:0:0:0 | 203:39:10:1:1:0 | 0:0:0:0:0:0 |
| NM_004083:NM_001195057:NM_001195056:NM_001195053:NM_001195055:NM_001195054 | 156:19:19:2:0:0 | 0:0:0:0:0:0 | 137:25:36:3:1:0 | 0:0:0:0:0:0 | 513:76:117:5:2:0 | 2:0:0:0:0:0 | 1515:228:81:4:11:1 | 0:0:0:0:0:0 | 203:39:10:1:1:0 | 0:0:0:0:0:0 |
| NM_005726:NM_001172695:NM_001172696:NM_001172697 | 51:2:2:0 | 1:0:0:0 | 65:3:0:0 | 0:0:0:0 | 45:0:0:1 | 0:0:0:0 | 481:7:6:0 | 2:0:0:0 | 100:2:0:0 | 0:0:0:0 |
| NM_001172695:NM_005726:NM_001172696 | 2:54:02 | 0:00:00 | 0.16875 | 0:00:00 | 0:47:00 | 0:00:00 | 0.675763889 | 0:01:00 | 0.194444444 | 0:01:00 |
| NM_033276:NM_001320408 | 1:00 | 2:00 | 7:00 | 4:00 | 4:00 | 6:00 | 0:00 | 23:00 | 0:00 | 7:00 |
| NM_033276:XM_011538976:XM_017020206:NM_001320408 | 3:0:0:0 | 0:0:0:0 | 11:0:0:0 | 0:0:0:0 | 8:0:0:0 | 2:0:0:0 | 22:1:1:0 | 0:0:0:0 | 7:0:2:0 | 0:0:0:0 |
| XR_931514:NM_018173:XM_011520967:XM_017019555:XM_006718985:NM_001144856 | 0:0:0:0:0:0 | 0:0:0:0:0:0 | 4:1:1:0:0:0 | 0:0:0:0:0:0 | 6:0:1:1:0:0 | 0:0:0:0:0:0 | 4:1:0:1:0:0 | 0:0:0:0:0:0 | 1:0:0:0:0:0 | 0:0:0:0:0:0 |
| NM_001065:NR_144351:NM_001346092:NM_001346091 | 51:1:5:2 | 0:0:0:0 | 157:5:6:17 | 0:0:0:0 | 491:4:10:17 | 1:0:0:1 | 239:9:23:40 | 1:1:0:0 | 59:1:4:0 | 0:0:0:0 |
| NM_001065:NR_144351:NM_001346092:NM_001346091 | 51:1:5:2 | 0:0:0:0 | 157:5:6:17 | 0:0:0:0 | 489:4:10:17 | 4:0:0:0 | 238:9:23:35 | 3:1:0:2 | 59:1:4:0 | 0:0:0:0 |
| NM_001065:NR_144351:NM_001346092:NM_001346091 | 51:1:5:2 | 0:0:0:0 | 156:6:6:9 | 1:1:0:1 | 492:11:10:12 | 0:0:0:0 | 241:12:23:23 | 0:0:0:0 | 59:1:4:0 | 0:0:0:0 |
| NM_001159576:NM_001038 | 0:00 | 1:00 | 0:00 | 0:00 | 0:00 | 0:01 | 0:00 | 0:00 | 0:00 | 1:00 |
| NM_002046:NR_152150:NM_001289746:NM_001289745:NM_001357943 | 2571:8:15:0:0 | 2:0:0:0:0 | 2516:18:17:1:0 | 5:0:0:0:0 | 3416:15:23:6:0 | 5:0:0:0:0 | 24249:95:139:12:3 | 14:1:0:0:0 | 10005:48:56:8:1 | 9:0:0:0:0 |
| NM_002046:NR_152150:NM_001289746:NM_001289745:NM_001357943:NM_001256799 | 2604:12:15:0:0:0 | 0:0:0:0:0:0 | 2564:19:17:1:0:0 | 3:1:0:0:0:0 | 3476:16:23:6:0:0 | 3:0:0:0:0:0 | 24509:118:139:12:3:3 | 16:0:0:0:0:0 | 10114:56:55:8:0:2 | 5:0:0:0:0:0 |
| NM_002046:NR_152150:NM_001289746:NM_001289745:NM_001357943:NM_001256799 | 2626:13:15:0:0:0 | 0:0:0:0:0:0 | 2600:18:17:1:0:0 | 4:0:0:0:0:0 | 3521:17:23:6:0:0 | 5:0:0:0:0:0 | 24681:116:139:12:3:3 | 41:0:0:0:0:0 | 10224:57:55:8:1:2 | 11:1:0:0:0:0 |
| NM_002046:NR_152150:NM_001289746:NM_001289745:NM_001357943:NM_001256799 | 2625:8:15:0:0:0 | 4:0:0:0:0:0 | 2618:9:17:1:0:0 | 4:0:0:0:0:0 | 3539:10:23:6:0:0 | 2:0:0:0:0:0 | 24694:74:138:12:4:3 | 36:2:0:0:0:0 | 10267:34:55:8:1:2 | 15:0:0:0:0:0 |
| NM_002046:NR_152150:NM_001289746:NM_001289745:NM_001357943:NM_001256799 | 2630:8:15:0:0:0 | 2:0:0:0:0:0 | 2620:9:17:1:0:0 | 6:0:0:0:0:0 | 3532:9:23:6:0:0 | 8:0:0:0:0:0 | 24687:72:138:12:4:3 | 36:0:0:0:0:0 | 10273:34:55:8:1:2 | 14:0:0:0:0:0 |
| NM_002046:NR_152150:NM_001289746:NM_001289745:NM_001357943:NM_001256799 | 2638:7:15:0:0:0 | 4:0:0:0:0:0 | 2643:5:17:1:0:0 | 8:0:0:0:0:0 | 3558:7:23:6:0:0 | 2:0:0:0:0:0 | 24771:47:137:12:4:3 | 43:1:0:0:0:0 | 10299:24:55:8:1:2 | 15:0:0:0:0:0 |
| NM_002046:NR_152150:NM_001289746:NM_001289745:NM_001357943:NM_001256799 | 2640:7:15:0:0:0 | 3:0:0:0:0:0 | 2653:5:17:1:0:0 | 8:0:0:0:0:0 | 3566:7:23:6:0:0 | 5:0:0:0:0:0 | 24818:47:136:12:4:3 | 36:1:1:0:0:0 | 10323:24:55:8:1:2 | 6:0:0:0:0:0 |
| NM_002046:NR_152150:NM_001289746:NM_001289745:NM_001357943:NM_001256799 | 2647:7:15:0:0:0 | 4:0:0:0:0:0 | 2678:3:17:1:0:0 | 9:0:0:0:0:0 | 3572:6:23:6:0:0 | 11:0:0:0:0:0 | 24871:43:137:12:4:3 | 55:0:0:0:0:0 | 10338:18:55:8:1:2 | 21:1:0:0:0:0 |
| NM_002046:NR_152150:NM_001289746:NM_001289745:NM_001357943:NM_001256799 | 2663:6:15:0:0:0 | 2:0:0:0:0:0 | 2699:3:17:1:0:0 | 7:0:0:0:0:0 | 3592:6:23:6:0:0 | 2:0:0:0:0:0 | 25043:38:137:12:4:3 | 27:0:0:0:0:0 | 10404:18:55:8:1:2 | 4:0:0:0:0:0 |
| NM_002046:NR_152150:NM_001289746:NM_001289745:NM_001357943:NM_001256799 | 2672:6:15:0:0:0 | 2:0:0:0:0:0 | 2712:3:17:1:0:0 | 6:0:0:0:0:0 | 3603:6:23:6:0:0 | 6:0:0:0:0:0 | 25119:34:137:12:4:3 | 39:0:0:0:0:0 | 10409:16:55:8:1:2 | 15:1:0:0:0:0 |
| NM_002046:NM_001289746:NM_001289745:NR_152150:NM_001357943:NM_001256799 | 2688:15:0:0:0:0 | 2:0:0:0:0:0 | 2742:17:1:0:0:0 | 5:0:0:0:0:0 | 3629:23:6:0:0:0 | 2:0:0:0:0:0 | 25233:137:12:7:4:3 | 34:0:0:1:0:0 | 10455:55:8:3:1:2 | 8:0:0:1:0:0 |
| NM_002046:NM_001289746:NM_001289745:NR_152150:NM_001357943:NM_001256799 | 2684:15:0:0:0:0 | 5:0:0:0:0:0 | 2759:17:1:0:0:0 | 3:0:0:0:0:0 | 3646:23:6:0:0:0 | 4:0:0:0:0:0 | 25224:137:10:2:4:3 | 34:0:0:2:0:0 | 10468:55:8:1:1:2 | 10:0:0:0:0:0 |
| NM_002046:NM_001289746:NM_001289745:NR_152150:NM_001357943:NM_001256799 | 2701:14:0:0:0:0 | 6:0:0:0:0:0 | 2827:17:1:0:0:0 | 9:0:0:0:0:0 | 3675:23:6:0:0:0 | 4:0:0:0:0:0 | 25479:135:10:0:4:3 | 49:0:0:1:0:0 | 10569:55:8:0:1:2 | 13:0:0:0:0:0 |
| NM_002046:NM_001289746:NM_001289745:NR_152150:NM_001357943:NM_001256799 | 2732:14:0:0:0:0 | 2:0:0:0:0:0 | 2864:17:1:0:0:0 | 11:0:0:0:0:0 | 3678:23:6:0:0:0 | 3:0:0:0:0:0 | 25617:135:10:0:4:2 | 36:0:0:1:0:0 | 10618:55:8:0:1:2 | 10:0:0:0:0:0 |
| NM_002046:NM_001289746:NM_001289745:NR_152150:NM_001357943:NM_001256799 | 2737:14:0:0:0:0 | 3:0:0:0:0:0 | 2926:17:1:1:0:0 | 22:0:0:0:0:0 | 3702:23:6:0:0:0 | 3:0:0:0:0:0 | 25904:136:11:1:4:2 | 72:0:0:0:0:0 | 10685:55:8:0:1:2 | 7:0:0:0:0:0 |
| NM_002046:NM_001289746:NM_001289745:NR_152150:NM_001357943:NM_001256799 | 2735:14:0:0:0:0 | 5:0:0:0:0:0 | 2928:17:1:1:0:0 | 23:0:0:0:0:0 | 3702:23:6:0:0:0 | 5:0:0:0:0:0 | 25922:136:11:1:4:2 | 59:0:0:0:0:0 | 10684:55:8:0:1:2 | 6:0:0:0:0:0 |
| NM_002046:NM_001289746:NM_001289745:NR_152150:NM_001357943:NM_001256799 | 2733:14:0:0:0:0 | 2:0:0:0:0:0 | 2961:17:1:1:0:0 | 10:0:0:0:0:0 | 3679:22:5:0:0:0 | 7:0:1:0:0:0 | 25862:135:11:2:4:2 | 52:1:0:0:0:0 | 10673:55:8:1:1:2 | 15:0:0:0:0:0 |
| NM_002046:NM_001289746:NM_001289745:NR_152150:NM_001357943:NM_001256799 | 2734:14:0:0:0:0 | 1:0:0:0:0:0 | 2965:17:1:1:0:0 | 7:0:0:0:0:0 | 3680:22:6:0:0:0 | 6:0:0:0:0:0 | 25881:136:11:2:4:2 | 36:0:0:0:0:0 | 10675:55:8:1:1:2 | 14:0:0:0:0:0 |
| NM_002046:NM_001289746:NM_001289745:NR_152150:NM_001357943:NM_001256799 | 2734:14:0:0:0:0 | 2:0:0:0:0:0 | 2967:17:1:1:0:0 | 10:0:0:0:0:0 | 3683:21:6:0:0:0 | 6:1:0:0:0:0 | 25883:136:11:2:4:2 | 41:0:0:0:0:0 | 10682:55:8:1:1:2 | 10:0:0:0:0:0 |
| NM_016056:NM_001282609 | 1:00 | 0:00 | 1:00 | 0:00 | 3:00 | 0:00 | 0:00 | 2:00 | 1:00 | 1:01 |
| NM_020401:XM_005269037:NM_001330192 | 88:06:00 | 0:00:00 | 48:12:00 | 0:00:00 | 87:03:00 | 0:00:00 | 382:146:0 | 0:00:00 | 179:05:01 | 2:00:00 |
| NM_020401:XM_005269037:NM_001330192 | 88:12:00 | 0:00:00 | 45:35:00 | 0:00:00 | 86:06:00 | 0:00:00 | 368:272:0 | 1:00:00 | 177:17:01 | 0:00:00 |
| NM_020401:XM_005269037:NM_001330192 | 83:11:00 | 0:00:00 | 3:21:00 | 0:00:00 | 80:09:00 | 0:00:00 | 124:94:0 | 0:00:00 | 167:17:00 | 1:00:01 |
| NM_198320:XR_001748581:XR_001748577:XR_001748578:NM_001005502:XR_001748580:XR_001748583:XR_001748579:XR_001748584:XR_001748582 | 3:0:0:0:0:0:0:0:0:0 | 0:0:0:0:0:0:0:0:0:0 | 27:2:2:10:2:1:1:1:0:0 | 0:0:0:0:0:0:0:0:0:0 | 91:1:6:19:2:4:0:0:1:1 | 2:0:0:0:0:0:0:0:0:0 | 6:1:0:0:0:0:0:0:0:0 | 0:0:0:0:0:0:0:0:0:0 | 2:0:0:2:0:0:0:0:0:1 | 0:0:0:0:0:0:0:0:0:0 |
| NM_201650 | 3 | 0 | 1 | 0 | 3 | 0 | 20 | 0 | 2 | 0 |
| NM_006992:NM_201650 | 1:05 | 0:00 | 0:02 | 0:00 | 0:03 | 0:00 | 5:23 | 0:00 | 2:02 | 0:00 |
| NM_007007:XM_005268590:NM_001300947:XM_005268588 | 111:31:11:1 | 1:0:0:0 | 65:27:6:2 | 0:0:0:0 | 182:50:17:9 | 0:0:0:1 | 339:78:140:44 | 0:0:0:0 | 168:42:35:11 | 0:0:0:0 |
| NM_007007:XM_005268590:NM_001300947:XM_005268588 | 111:31:11:1 | 0:0:0:0 | 70:27:6:2 | 0:0:0:0 | 184:50:17:10 | 0:0:0:0 | 384:85:148:54 | 0:0:0:1 | 170:43:36:11 | 0:0:0:0 |
| NM_007007:XM_005268590:NM_001300947:XM_005268588 | 113:28:11:1 | 0:0:0:0 | 70:28:7:2 | 0:0:0:0 | 184:49:17:10 | 0:0:0:0 | 428:97:168:70 | 1:0:0:0 | 173:43:38:11 | 1:1:0:0 |
| NM_007007:XM_005268590:NM_001300947:XM_005268588 | 113:28:11:1 | 0:0:0:0 | 72:27:7:2 | 1:0:0:0 | 178:49:18:10 | 2:0:0:0 | 425:96:173:70 | 1:1:1:0 | 172:43:38:11 | 0:0:0:0 |
| NM_007007:XM_005268590:NM_001300947:XM_005268588 | 113:28:11:1 | 0:0:0:0 | 72:27:7:2 | 1:0:0:0 | 180:49:18:10 | 0:0:0:0 | 426:99:175:70 | 1:0:0:0 | 172:42:38:11 | 0:1:0:0 |
| NM_007007:XM_005268590:NM_001300947:XM_005268588 | 113:28:11:1 | 0:0:0:0 | 72:27:7:2 | 1:0:0:0 | 180:49:18:10 | 0:0:0:0 | 426:98:174:70 | 1:0:1:0 | 172:43:38:11 | 0:0:0:0 |
| NM_007007:XM_005268590:NM_001300947:XM_005268588 | 114:28:11:1 | 0:0:0:0 | 77:28:9:3 | 1:0:0:0 | 181:48:18:10 | 0:0:0:1 | 467:96:191:74 | 5:1:0:0 | 173:43:39:11 | 0:0:0:0 |
| NM_007007:XM_005268590:NM_001300947:XM_005268588 | 113:29:11:1 | 0:0:0:0 | 77:27:9:3 | 1:0:0:0 | 181:48:18:11 | 0:0:0:0 | 470:98:191:73 | 4:1:0:1 | 173:43:39:11 | 1:0:0:0 |
| NM_007007:XM_005268590:NM_001300947:XM_005268588 | 125:34:11:1 | 0:0:0:0 | 90:31:9:3 | 0:0:0:0 | 200:59:18:10 | 0:0:0:0 | 573:131:191:71 | 0:0:0:1 | 200:49:39:11 | 1:0:0:0 |
| NM_007007:XM_005268590:NM_001300947:XM_005268588 | 135:35:11:1 | 0:0:0:0 | 97:38:9:3 | 1:0:0:0 | 215:62:18:11 | 0:0:0:0 | 611:145:190:68 | 0:1:1:0 | 207:50:39:11 | 1:0:0:0 |
| NM_007007:XM_005268590:NM_001300947:XM_005268588 | 134:35:11:1 | 1:0:0:0 | 98:37:9:3 | 0:1:0:0 | 216:62:18:11 | 0:0:0:0 | 610:146:189:67 | 2:0:2:1 | 207:50:39:11 | 1:0:0:0 |
| NM_007007:XM_005268590:NM_001300947:XM_005268588 | 137:36:11:1 | 0:0:0:0 | 104:40:9:3 | 0:1:0:0 | 219:69:18:11 | 0:0:0:0 | 636:156:191:68 | 1:1:0:0 | 213:55:39:11 | 1:0:0:0 |

| | | | | | | | | | | |
|---|---|---|---|---|---|---|---|---|---|---|
| NM_007007:XM_005268590:NM_001300947:XM_005268588 | 137:36:11:1 | 0:0:0:0 | 103:41:9:3 | 1:0:0:0 | 219:69:18:11 | 0:0:0:0 | 636:158:190:68 | 1:0:1:0 | 213:55:39:11 | 1:0:0:0 |
| NM_007007:XM_005268590:NM_001300947:XM_005268588 | 137:37:11:1 | 0:0:0:0 | 103:42:9:3 | 1:0:0:0 | 218:69:18:11 | 0:0:0:0 | 636:161:191:68 | 0:0:0:0 | 214:55:39:11 | 0:0:0:0 |
| NM_007007:XM_005268590:NM_001300947:XM_005268588 | 136:37:11:1 | 0:0:0:0 | 104:47:9:3 | 0:0:0:0 | 218:69:18:11 | 0:0:0:0 | 636:163:190:68 | 0:0:0:0 | 213:54:39:11 | 1:1:0:0 |
| NM_007007:XM_005268590:NM_001300947:XM_005268588 | 136:37:11:1 | 0:0:0:0 | 104:47:9:3 | 0:0:0:0 | 216:70:18:11 | 2:0:0:0 | 636:163:189:68 | 0:0:1:0 | 212:55:39:11 | 2:0:0:0 |
| NM_007007:XM_005268590:NM_001300947:XM_005268588 | 136:37:11:1 | 0:0:0:0 | 104:48:9:3 | 0:0:0:0 | 218:72:18:11 | 0:0:0:0 | 637:170:190:68 | 0:0:0:0 | 213:57:39:11 | 0:0:0:0 |
| NM_007007:XM_005268590:NM_001300947:XM_005268588 | 135:37:11:1 | 0:0:0:0 | 103:48:9:3 | 1:1:0:0 | 218:73:18:11 | 0:0:0:0 | 634:173:191:68 | 2:0:0:0 | 212:58:39:11 | 1:0:0:0 |
| NM_000239 | 725 | 2 | 413 | 2 | 412 | 1 | 694 | 2 | 195 | 1 |
| NM_000239 | 723 | 4 | 411 | 4 | 411 | 2 | 695 | 2 | 197 | 0 |
| NM_000239 | 728 | 1 | 413 | 2 | 413 | 0 | 697 | 1 | 197 | 0 |
| NM_000239 | 749 | 0 | 416 | 0 | 415 | 1 | 701 | 0 | 201 | 0 |
| NM_000239 | 750 | 1 | 417 | 0 | 416 | 0 | 700 | 4 | 202 | 0 |
| NM_000239 | 755 | 0 | 419 | 0 | 417 | 0 | 705 | 1 | 206 | 0 |
| NM_000239 | 757 | 0 | 419 | 0 | 417 | 1 | 707 | 0 | 206 | 0 |
| NM_000239 | 759 | 0 | 420 | 0 | 420 | 0 | 709 | 1 | 207 | 0 |
| NM_000239 | 763 | 0 | 428 | 1 | 424 | 1 | 720 | 2 | 209 | 0 |
| NM_006530:XR_944742:NM_001300950:XR_001748876 | 58:1:3:0 | 1:0:0:0 | 30:2:0:0 | 0:0:0:0 | 26:0:0:0 | 0:0:0:0 | 403:20:23:2 | 0:0:0:0 | 91:1:3:1 | 1:0:0:0 |
| NM_080548:XM_006718994:NM_002831:XM_024449106:XM_011520988:NM_080549 | 12:2:10:1:0:0 | 0:0:0:0:0:0 | 23:2:14:2:2:0 | 0:0:0:0:0:0 | 45:1:19:0:2:1 | 0:0:0:0:0:0 | 121:16:14:4:11:0 | 1:1:0:0:0:0 | 20:0:6:0:1:0 | 0:0:0:0:0:0 |
| NM_080548:XM_006718994:NM_002831:XM_024449106:NM_080549:XM_011520988 | 20:2:10:0:0:0 | 0:0:0:0:0:0 | 42:3:14:3:3:0 | 0:0:0:0:0:0 | 63:3:19:3:2:1 | 0:0:0:0:0:0 | 164:36:14:8:1:6 | 1:1:0:0:0:0 | 24:4:6:0:0:1 | 0:0:0:0:0:0 |
| NM_152439 | 0 | 0 | 0 | 0 | 0 | 0 | 0 | 3 | 0 | 0 |
| NM_001207016:NM_130846:XM_011538615:NM_002849:XR_001748831:NR_073474 | 1:1:1:0:0:0 | 2:2:0:0:0:0 | 16:48:9:0:5:0 | 16:38:14:2:2:0 | 52:145:6:0:14:3 | 38:144:2:0:10:0 | 0:2:1:0:1:0 | 0:0:0:0:0:0 | 0:1:0:0:0:0 | 0:0:0:0:0:0 |
| NM_130846:NM_001207016:XM_011538615:NR_073474:NM_002849:XR_001748831 | 2:1:1:1:0:0 | 6:2:0:0:0:0 | 51:16:4:0:0:4 | 43:16:20:5:2:2 | 151:52:7:5:0:13 | 176:38:23:7:0:11 | 2:0:2:1:0:1 | 0:0:0:0:0:0 | 1:0:0:0:0:0 | 0:0:0:0:0:0 |
| NM_130846:NM_001207016:XM_011538615:NR_073474:NM_002849:XR_001748831 | 8:3:1:1:0:0 | 0:0:0:0:0:0 | 94:32:24:6:2:6 | 0:0:0:0:0:0 | 325:90:30:12:0:24 | 2:0:0:0:0:0 | 2:0:2:1:0:1 | 0:0:0:0:0:0 | 1:0:0:0:0:0 | 0:0:0:0:0:0 |
| NM_130846:NM_001207016:XM_011538615:NR_073474:NM_002849:XR_001748831 | 8:3:1:1:0:0 | 0:0:0:0:0:0 | 89:32:23:5:2:6 | 0:0:0:1:0:0 | 325:90:28:12:0:23 | 1:0:0:0:0:1 | 2:0:2:1:0:1 | 0:0:0:0:0:0 | 1:0:0:0:0:0 | 0:0:0:0:0:0 |
| NM_004616:XM_006719583 | 1:00 | 2:00 | 1:00 | 0:00 | 4:00 | 6:00 | 9:00 | 0:00 | 12:01 | 0:00 |
| NM_004616:XM_006719583 | 1:00 | 2:00 | 0:00 | 0:00 | 3:00 | 5:00 | 9:00 | 0:00 | 11:01 | 0:00 |
| NM_005889 | 0 | 1 | 0 | 0 | 0 | 1 | 0 | 0 | 0 | 0 |
| NM_001009894 | 0 | 0 | 0 | 0 | 0 | 1 | 0 | 0 | 2 | 0 |
| NM_003805:XM_024449252:XM_017020145:XR_001748908:XR_001748910:XR_001748909:NM_001320101:NM_001320100:XM_017020144:NM_001320099 | 38:4:0:0:0:0:0:0:0:0 | 0:0:0:0:0:0:0:0:0:0 | 118:1:3:1:0:0:0:0:0:0 | 1:0:0:0:0:0:0:0:0:0 | 336:10:2:0:3:1:1:0:0:0 | 3:0:0:0:0:0:0:0:0:0 | 259:10:3:0:0:2:1:1:1:3 | 1:0:0:0:0:0:0:0:0:0 | 64:1:0:0:0:0:0:0:0:0 | 0:0:0:0:0:0:0:0:0:0 |
| NM_003805:XM_024449252:XM_017020145:XR_001748908:NM_001330126:XR_001748910:XR_001748909:NM_001320101:NM_001320100:XM_017020144:NM_001320099 | 39:4:0:0:0:0:0:0:0:0:0 | 0:0:0:0:0:0:0:0:0:0:0 | 119:1:3:1:1:0:0:0:0:0:0 | 0:0:0:0:0:0:0:0:0:0:0 | 343:10:2:0:0:3:1:1:0:0:0 | 0:0:0:0:0:0:0:0:0:0:0 | 261:11:3:0:0:0:2:1:1:1:3 | 3:0:0:0:0:0:0:0:0:0:0 | 64:1:0:0:0:0:0:0:0:0:0 | 0:0:0:0:0:0:0:0:0:0:0 |
| XR_243051:NM_024537 | 1:00 | 0:00 | 0:00 | 0:00 | 1:00 | 0:00 | 0:00 | 0:00 | 0:00 | 1:02 |
| NM_001078645:NM_003903:XM_017020833:NM_001330101:XM_011534853:NM_001330104:NM_001318517:XM_005266206:NM_001318518:XR_001749713:XR_941424:XR_245358:XR_001749714:XM_017020831:XM_011534851:NM_001330105:XR_245360:XM_017020830 | 66:24:8:6:2:3:3:1:1:1:2:1:1:0:0:0:0:0 | 0:0:0:0:0:0:0:0:0:0:0:0:0:0:0:0:0:0 | 14:17:13:2:2:1:0:1:0:0:1:1:1:1:1:1:1:0 | 0:0:0:0:0:0:0:0:0:0:0:0:0:0:0:0:0:0 | 11:7:3:3:2:2:0:1:0:0:0:0:0:2:0:1:0:0 | 0:0:0:0:0:0:0:0:0:0:0:0:0:0:0:0:0:0 | 71:49:84:7:11:42:2:5:1:0:0:1:3:1:2:11:3:0 | 0:0:0:0:0:0:0:0:0:0:0:0:0:0:0:0:0:0 | 28:16:11:5:1:10:1:5:2:0:0:0:0:0:1:5:2:0 | 0:0:0:0:0:0:0:0:0:0:0:0:0:0:0:0:0:0 |
| NM_001353652:NM_001353647:NM_023011:NR_148483:XM_006719991:NM_001353644:XR_001749644:NM_001353650:XR_941422:NR_148487:XM_017020709:NR_148492:NR_148491:NR_148494:XM_011534848:XM_011534847:XM_024449400:XR_001749647:NR_148482:NR_148486:NR_148493:NM_001353645:XR_001749649:XM_011534845:XM_024449401:XM_017020711:NM_001353651:NM_080687:NR_148485:NR_148484 | 4:13:43:3:5:1:33:7:2:58:4:7:1:1:8:6:3:0:0:0:0:0:0:0:0:0:0:0:0:0 | 1:1:0:0:0:0:0:0:0:0:0:0:0:0:0:0:0:0:0:0:0:0:0:0:0:0:0:0:0:0 | 0:1:27:1:5:0:27:0:0:21:30:2:1:0:11:1:3:0:1:1:1:0:0:0:0:0:0:0:0:0 | 0:0:0:0:0:0:0:0:0:0:0:0:0:0:0:1:0:0:0:0:0:0:0:0:0:0:0:0:0:0 | 1:3:33:0:1:0:11:1:1:15:2:0:0:0:13:2:0:1:0:0:0:2:3:2:0:0:0:0:0:0 | 0:0:0:0:0:0:0:0:0:0:0:0:0:0:0:0:0:0:0:0:0:0:0:0:0:0:0:0:0:0 | 5:9:81:3:15:0:40:19:5:68:3:4:2:0:38:35:3:0:1:0:0:4:2:2:0:1:1:0:0:0 | 0:0:0:0:0:0:0:0:0:0:0:0:0:0:0:0:0:0:0:0:0:0:0:1:0:0:0:0:0:0 | 4:4:39:0:0:0:13:1:3:1:8:6:0:0:3:15:5:0:0:0:0:0:0:0:1:0:0:1:0:0:0 | 0:0:0:0:0:0:0:1:0:0:0:0:0:0:0:0:0:0:0:0:0:0:0:0:0:0:0:0:0:0 |
| XR_001749647:NM_080687:NM_023011:NR_148483:XM_006719991:NM_001353652:NM_001353644:XM_017020711:NR_148485:XR_001749644:NM_001353650:XR_941422:NR_148487:NR_148489:NM_001353646:NR_148496:NM_001353647:XM_017020709:XM_005266202:NR_148492:NR_148491:NM_001353651:NR_148490:NR_148494:XM_024449401:XM_011534848:XM_011534847:XM_024449400:NR_148482:NR_148486:NR_148493:NM_001353645:XR_001749649:XM_011534845:NM_001353649:NR_148484:NR_148488 | 37:7:43:8:5:5:1:23:1:33:7:2:58:44:16:2:1:4:4:14:7:1:2:1:2:1:9:6:3:0:0:0:0:0:0:0:0 | 1:0:0:0:0:0:0:0:0:0:0:0:0:0:0:0:0:0:0:0:0:0:0:0:0:0:0:0:0:0:0:0:0:0:0:0:0 | 23:2:27:4:5:0:0:11:0:27:0:0:21:4:5:4:2:1:30:3:2:1:0:1:4:2:13:1:3:1:1:1:0:0:0:0:0 | 1:0:0:0:0:0:0:0:0:0:0:0:0:0:0:0:0:0:0:0:0:0:0:0:0:0:0:0:0:0:0:0:0:0:0:0:0 | 19:9:33:2:1:1:0:8:0:11:1:2:15:25:7:1:3:2:2:0:0:0:0:2:0:14:4:0:0:0:1:2:3:2:0:0:0 | 0:0:0:0:0:0:0:0:0:0:0:0:0:0:0:0:0:0:0:0:0:0:0:0:0:0:0:0:0:0:0:0:0:0:0:0:0 | 125:30:80:12:15:5:1:168:0:40:19:6:68:1:52:24:12:9:3:26:4:2:10:2:17:7:47:40:3:1:2:1:4:2:2:2:1:0 | 0:0:1:0:0:0:0:0:0:0:0:0:0:0:0:0:0:0:0:0:0:0:0:1:0:0:0:0:0:0:0:0:0:0:0:0:0 | 39:10:39:3:0:4:0:31:2:14:1:5:18:32:6:3:4:7:15:0:0:9:1:5:3:19:5:0:0:1:0:0:0:1:1:0:1 | 0:0:0:0:0:0:0:0:0:0:0:0:0:0:0:0:0:0:0:0:0:0:0:0:0:0:0:0:0:0:0:0:0:0:0:0:0 |
| NM_080687:NM_023011:NR_148483:XM_006719991:NM_001353652:XR_001749647:NM_001353644:XM_017020711:NR_148485:XR_001749644:NM_001353650:XR_941422:NR_148487:NR_148489:NM_001353646:NR_148496:NM_001353647:XM_017020709:XM_005266202:NR_148492:NR_148491:NM_001353651:NR_148490:NR_148494:XM_024449401:XM_011534848:XM_011534847:XM_024449400:NR_148482:NR_148486:NR_148493:NM_001353645:XR_001749649:XM_011534845:NM_001353649:NR_148484:NR_148488 | 7:43:8:5:5:38:1:23:1:33:7:1:58:44:16:2:1:4:4:14:7:1:2:1:2:1:9:6:3:0:0:0:0:0:0:0:0 | 0:0:0:0:0:0:0:0:0:0:0:0:0:0:0:0:0:0:0:0:0:0:0:0:0:0:0:0:0:0:0:0:0:0:0:0:0 | 2:27:4:5:0:25:0:11:0:26:0:0:20:4:5:4:2:1:29:3:2:1:0:1:4:2:13:1:3:1:1:1:0:0:0:0:0 | 0:0:0:0:0:0:0:0:1:0:0:0:0:0:0:1:0:0:0:0:0:0:0:0:0:0:0:0:0:0:0:0:0:0:0:0:0 | 9:33:2:1:1:17:0:8:0:11:1:2:15:25:7:1:3:2:2:0:0:0:0:2:1:14:4:0:0:0:1:2:3:2:0:0:0 | 0:0:0:0:0:2:0:0:0:0:0:0:0:0:0:0:0:0:0:0:0:0:0:0:0:0:0:0:0:0:0:0:0:0:0:0:0 | 30:81:12:15:5:126:1:167:0:40:19:6:68:1:52:24:12:9:3:26:4:2:10:2:17:7:47:40:3:1:2:1:4:2:2:2:1:0 | 0:0:0:0:0:1:0:1:0:0:0:0:0:0:0:0:0:0:0:0:0:0:0:1:0:0:0:0:0:0:0:0:0:0:0:0:0 | 10:39:3:0:4:40:0:31:2:14:1:5:18:32:6:3:4:7:15:0:0:9:1:5:3:19:5:0:0:1:0:0:0:1:1:0:1 | 0:0:0:0:0:0:0:0:0:0:0:0:0:0:0:0:0:0:0:0:0:0:0:0:0:0:0:0:0:0:0:0:0:0:0:0:0 |
| NM_023011:XR_001749644:NR_148487:NR_148489:XM_017020711:XR_001749647:XM_017020709:NM_080687:XM_006719991:NM_001353644:NR_148485:NM_001353650:NM_001353646:NM_001353647:XM_005266202:NR_148494:XM_024449401:XM_011534848:XM_011534847:XM_024449400:XM_011534845:NM_001353645:XR_941422:NR_148484:NM_001353649 | 37:28:56:41:22:35:3:7:5:1:1:7:16:13:14:2:1:6:6:3:0:0:0:0:0 | 6:4:2:2:1:2:1:0:0:0:0:0:0:0:0:0:0:0:0:0:0:0:0:0:0 | 18:23:16:36:10:21:24:2:5:0:0:0:4:1:3:0:4:10:1:1:0:0:0:0:0 | 8:4:5:9:1:4:4:0:0:0:0:0:0:0:0:0:0:1:0:1:0:0:0:0:0 | 27:11:11:24:7:19:2:9:1:0:0:1:7:3:1:0:1:1:2:1:0:1:1:1:0:0 | 6:0:1:1:0:0:0:0:0:0:0:0:0:0:0:0:0:0:0:1:0:1:0:0:0:0 | 72:34:60:112:129:106:3:30:11:0:0:12:24:9:21:1:6:26:29:2:2:4:1:0:2 | 8:6:7:37:18:14:0:1:4:1:0:6:0:0:4:0:3:14:7:0:0:0:0:1:0 | 36:13:18:29:27:39:5:10:0:0:2:1:6:4:14:5:2:14:5:0:1:0:1:0:1 | 0:1:0:2:1:0:1:0:0:0:0:0:0:0:0:0:1:2:0:0:0:0:0:0:0 |
| NM_023011:NR_148487:XM_017020711:NR_148489:NM_001353650:XM_006719991:XR_001749644:XR_001749647:XM_024449401:XM_011534848:XM_011534847:NM_001353644:NR_148485:NM_001353646:NM_001353647:XM_017020709:NR_148491:NR_148492:XM_005266202:NM_080687:NR_148490:NR_148494:XM_024449400:NR_148482:NR_148496:NR_148486:NR_148493:XR_941422:NM_001353645:XR_001749649:NM_001353649:XM_011534845 | 19:27:10:21:2:2:10:21:0:1:2:1:1:16:14:2:1:3:7:6:1:2:1:0:0:0:0:0:0:0:0:0 | 5:5:1:2:1:2:3:3:1:1:3:0:0:0:0:0:0:0:0:0:0:0:0:0:0:0:0:0:0:0:0:0 | 7:5:5:16:0:4:9:5:2:3:0:0:0:4:1:5:1:0:2:1:1:2:1:1:1:1:1:1:0:0:0:0 | 0:1:0:1:0:0:0:1:0:0:0:0:0:0:1:0:0:0:0:0:0:0:0:0:0:0:0:0:0:0:0:0 | 12:8:5:10:0:1:5:13:1:7:1:0:0:7:3:0:0:0:0:8:0:2:0:0:0:0:1:1:2:1:0:0 | 2:0:0:2:0:0:0:2:0:1:0:0:0:0:0:1:0:0:0:0:0:0:0:0:0:0:0:0:0:0:0:0 | 36:24:47:41:3:5:12:39:4:14:14:0:0:24:9:2:1:0:11:27:1:4:2:0:4:2:0:9:4:1:2:0 | 2:0:6:2:0:1:2:3:0:2:1:0:0:0:0:0:0:0:1:0:1:1:0:0:0:0:0:0:0:0:0:0 | 21:8:20:12:1:0:4:17:1:7:3:0:0:6:4:1:0:0:8:10:0:5:0:0:1:1:0:3:0:0:1:0 | 1:2:1:3:0:0:0:4:0:0:0:0:0:0:0:0:0:0:2:0:0:0:0:0:0:0:0:0:0:0:0:0 |
| NM_023011:NM_001353647:NM_001353646:NR_148490:XM_017020709:NR_148492:NM_080687:NR_148494:XM_017020711:XM_011534848:XM_024449400:XM_005266202:XM_024449401:XM_006719991:NR_148486:XR_941422:NM_001353645:XR_001749649:NM_001353649:XM_011534847:NR_148484:NR_148488 | 9:10:9:0:1:1:5:2:1:1:1:0:0:0:0:0:0:0:0:0:0:0 | 2:2:2:1:0:0:0:0:0:0:0:0:0:0:0:0:0:0:0:0:0:0 | 1:0:4:0:1:0:0:0:0:1:0:1:0:1:1:1:0:0:0:0:0:0 | 0:0:0:0:0:0:0:0:0:1:2:0:0:0:0:0:0:0:0:0:0:0 | 6:2:4:0:0:0:6:0:0:1:0:0:1:1:0:1:1:1:0:0:0:0 | 0:1:0:0:0:0:2:1:0:1:0:0:0:0:0:0:0:0:0:0:0:0 | 12:8:19:0:0:0:19:2:9:6:2:3:2:1:2:8:3:0:2:2:1:0 | 3:0:2:0:0:0:2:0:2:0:0:1:2:0:0:0:0:0:0:0:0:0 | 7:3:4:0:0:0:7:5:2:3:0:2:1:0:1:3:0:0:1:0:0:1 | 1:0:1:0:0:0:1:0:1:0:0:2:0:0:0:0:0:0:0:0:0:0 |
| NM_153251:XM_017020527:XM_006719797:NR_104487:NR_104486:XR_429215:NM_001286638:NM_001330059:XR_001749530:XR_941546:XM_006719796 | 4:1:2:0:0:0:0:0:0:0:0 | 0:0:0:0:0:0:0:0:0:0:0 | 0:1:1:1:0:0:0:0:0:0:0 | 0:0:0:0:0:0:0:0:0:0:0 | 1:0:1:0:0:0:0:0:0:0:0 | 0:0:0:0:0:0:0:0:0:0:0 | 0:0:5:3:0:1:0:0:0:0:0 | 1:0:0:0:1:0:0:0:0:0:0 | 0:0:1:1:0:0:1:1:0:0:0 | 0:0:0:0:0:0:0:0:0:0:0 |
| NM_152726:XR_429214:XR_001749501:XM_017020433 | 182:6:2:3 | 1:0:0:0 | 163:3:1:17 | 3:0:0:0 | 143:4:0:5 | 0:0:0:0 | 561:9:5:67 | 1:0:0:0 | 213:3:0:8 | 1:0:0:0 |
| NM_152726:XR_429214:XR_001749501:XM_017020433 | 184:4:2:4 | 0:0:0:0 | 166:1:1:18 | 0:0:0:0 | 143:0:0:5 | 0:0:0:0 | 563:5:5:68 | 0:0:0:0 | 212:1:0:9 | 2:0:0:0 |
| NM_152726:XR_429214:XR_001749501:XM_017020433 | 184:10:7:2 | 1:0:0:0 | 166:7:1:13 | 0:0:0:0 | 142:2:1:3 | 0:0:0:0 | 564:19:5:64 | 0:0:0:0 | 214:11:0:8 | 0:0:0:0 |
| NM_152726:XR_429214:XR_001749501:XM_017020433 | 185:12:7:1 | 0:0:0:0 | 165:12:1:11 | 1:7:0:0 | 142:3:1:3 | 0:0:0:0 | 563:27:5:50 | 0:4:0:0 | 214:13:0:7 | 0:0:0:0 |
| NM_152726:XR_429214:XR_001749501:XM_017020433 | 185:12:7:1 | 0:0:0:0 | 165:19:1:4 | 0:0:0:0 | 142:4:1:2 | 0:0:0:0 | 561:34:4:38 | 0:0:0:0 | 214:13:0:6 | 0:0:0:0 |
| NM_152726:XR_429214:XR_001749501:XM_017020433 | 184:12:7:0 | 0:0:0:0 | 159:16:1:1 | 0:0:0:0 | 138:4:1:2 | 0:0:0:0 | 528:30:2:9 | 1:1:0:0 | 207:13:0:2 | 0:0:0:0 |
| NM_005932:XM_011535097:XM_011535098 | 75:03:00 | 1:00:00 | 36:03:00 | 0:00:00 | 95:02:03 | 0:00:00 | 527:15:06 | 1:00:01 | 123:04:00 | 1:00:00 |
| NM_005932:XM_011535097:XM_011535098 | 76:03:00 | 0:00:00 | 36:03:00 | 0:00:00 | 95:02:03 | 0:00:00 | 527:15:07 | 1:00:00 | 124:04:00 | 0:00:00 |
| NM_005932:XM_011535097:XM_011535098 | 76:03:00 | 0:00:00 | 36:03:00 | 0:00:00 | 95:01:03 | 0:00:00 | 527:13:07 | 0:00:00 | 124:04:00 | 0:00:00 |

| | | | | | | | | | | |
|---|---|---|---|---|---|---|---|---|---|---|
| NM_005932:XM_011535097:XM_011535098 | 0:00:00 | 75:01:00 | 0:00:00 | 35:01:00 | 0:00:00 | 92:01:03 | 2:00:00 | 516:07:07 | 0:00:00 | 123:02:00 |
| NM_005932:XM_011535097:XM_011535098 | 69:01:00 | 0:00:00 | 33:01:00 | 0:00:00 | 85:01:05 | 0:00:00 | 445:08:10 | 2:00:00 | 116:02:02 | 0:00:00 |
| NM_002097 | 1 | 1 | 0 | 0 | 1 | 1 | 5 | 0 | 3 | 0 |
| XM_024449331:NM_152912:NM_001166261:NM_001166262:NM_001166263:XM_006719772:XM_011534959 | 0:0:0:0:0:0:0 | 1:0:0:0:0:0:0 | 0:0:0:0:0:0:0 | 0:0:0:0:0:0:0 | 0:0:0:0:0:0:0 | 0:2:1:0:0:0:0 | 0:0:1:0:1:0:0 | 0:1:0:1:0:0:0 | 0:3:0:0:0:0:0 | 0:0:0:0:0:0:0 |
| XM_024449331:NM_152912:NM_001166261:NM_001166262:NM_001166263:XM_006719772:XM_011534959 | 1:0:0:0:0:0:0 | 0:0:0:0:0:0:0 | 0:0:0:0:0:0:0 | 0:0:0:0:0:0:0 | 0:2:1:0:0:0:0 | 0:0:0:0:0:0:0 | 0:1:2:1:1:0:0 | 0:0:0:0:0:0:0 | 0:3:0:0:0:0:0 | 0:0:0:0:0:0:0 |
| NM_152705 | 2 | 0 | 0 | 0 | 0 | 0 | 0 | 0 | 0 | 2 |
| NM_001265:XM_011534879:NM_001354700 | 0:00:00 | 0:00:00 | 0:01:00 | 1:00:00 | 0:00:00 | 1:00:00 | 0:00:00 | 1:01:01 | 0:00:00 | 0:00:00 |
| NM_001286504:NM_006644:NM_001286503:XM_011534887:XM_017020364:XM_017020363:XM_005266236:XM_017020362 | 3:15:1:2:1:0:0:0 | 0:0:0:0:0:0:0:0 | 0:2:0:0:0:0:0:0 | 0:0:0:0:0:0:0:0 | 1:0:0:1:0:0:0:0 | 0:0:0:0:0:0:0:0 | 1:16:0:3:0:2:1:1 | 1:9:0:3:0:1:0:0 | 8:4:0:2:0:0:0:0 | 1:3:0:0:1:1:0:0 |
| NM_000059 | 7 | 1 | 4 | 4 | 0 | 1 | 46 | 0 | 20 | 0 |
| NM_000059 | 8 | 0 | 7 | 0 | 1 | 0 | 31 | 8 | 17 | 4 |
| NM_000059 | 8 | 0 | 7 | 0 | 1 | 0 | 23 | 8 | 13 | 5 |
| NM_000059 | 2 | 4 | 1 | 3 | 0 | 1 | 11 | 13 | 8 | 5 |
| NM_000059 | 2 | 0 | 0 | 1 | 0 | 0 | 11 | 1 | 7 | 1 |
| NM_000059 | 0 | 0 | 0 | 0 | 0 | 0 | 2 | 0 | 3 | 2 |
| XM_017020450:XM_017020453:XM_017020452:XM_017020449 | 1:0:0:0 | 0:0:0:0 | 0:0:0:0 | 0:0:0:0 | 0:0:0:0 | 0:0:0:0 | 0:1:1:1 | 0:0:0:0 | 0:0:0:0 | 0:0:1:0 |
| NM_181503:XR_941480 | 0:00 | 4:00 | 0:00 | 0:00 | 1:00 | 0:00 | 1:00 | 0:00 | 7:01 | 0:00 |
| NM_020751:NR_026745:XM_011535168:NM_001145079 | 51:2:1:0 | 0:0:0:0 | 3:1:1:1 | 0:0:0:0 | 7:0:0:0 | 0:0:0:0 | 14:0:0:1 | 17:2:2:1 | 3:0:0:0 | 8:2:0:0 |
| NM_020751:NR_026745:XM_011535168:NM_001145079 | 51:2:1:0 | 0:0:0:0 | 3:1:1:1 | 0:0:0:0 | 8:0:0:0 | 0:0:0:0 | 15:0:0:1 | 19:2:2:1 | 3:0:0:0 | 10:2:0:0 |
| NM_020751:NR_026745:XM_011535168:NM_001145079 | 49:3:1:0 | 3:7:0:0 | 3:3:1:1 | 0:2:1:0 | 6:2:0:0 | 4:4:0:0 | 36:59:5:2 | 0:0:0:0 | 15:15:1:0 | 0:0:0:0 |
| NM_172373 | 0 | 21 | 0 | 1 | 0 | 4 | 0 | 13 | 0 | 10 |
| NM_172373:NM_001145353 | 0:00 | 44:00:00 | 0:00 | 45:00:00 | 0:00 | 13:00 | 0:00 | 187:01:00 | 0:00 | 32:00:00 |
| NM_003295:NM_001286273:NM_001286272 | 9527:66:14 | 14:00:00 | 4671:25:06 | 6:00:00 | 2663:18:08 | 0:00:00 | 49153:459:69 | 41:00:00 | 7409:53:15 | 10:01:00 |
| NM_003295:NM_001286273:NM_001286272 | 9543:66:14 | 9:00:00 | 4670:25:06 | 7:00:00 | 2661:18:08 | 3:00:00 | 49168:459:69 | 49:00:00 | 7414:54:15 | 8:00:00 |
| NM_003295:NM_001286273:NM_001286272 | 9541:66:14 | 18:00:00 | 4645:25:06 | 8:00:00 | 2663:18:08 | 1:00:00 | 49166:459:69 | 47:00:00 | 7407:54:15 | 6:00:00 |
| NM_003295:NM_001286273:NM_001286272 | 9588:66:14 | 5:00:00 | 4571:25:06 | 1:00:00 | 2662:18:08 | 1:00:00 | 48876:461:69 | 19:00:00 | 7393:53:15 | 0:02:00 |
| NM_003295:NM_001286273:NM_001286272 | 9593:66:14 | 16:00:00 | 4567:26:06 | 6:00:00 | 2659:19:08 | 2:00:00 | 48878:460:69 | 62:01:00 | 7393:54:15 | 4:01:00 |
| NM_003295:NM_001286273:NM_001286272 | 9606:67:14 | 13:00:00 | 4553:26:06 | 11:00:00 | 2657:19:08 | 2:00:00 | 48903:461:67 | 55:00:01 | 7371:54:15 | 13:01:00 |
| NM_003295:NM_001286273:NM_001286272 | 9554:67:14 | 13:00:00 | 4485:26:06 | 9:00:00 | 2638:19:08 | 3:00:00 | 48700:461:66 | 55:00:01 | 7347:55:15 | 5:00:00 |
| NM_003295:NM_001286273:NM_001286272 | 9548:67:14 | 16:00:00 | 4477:26:06 | 5:00:00 | 2641:19:08 | 2:00:00 | 48669:461:66 | 41:00:00 | 7341:55:15 | 7:00:00 |
| NM_003295:NM_001286273:NM_001286272 | 9536:67:14 | 19:00:00 | 4465:26:05 | 10:00:00 | 2635:19:08 | 8:00:00 | 48504:461:65 | 121:00:00 | 7312:54:15 | 26:01:00 |
| NM_003295:NM_001286273:NM_001286272 | 9460:67:13 | 8:00:00 | 4432:26:05 | 7:00:00 | 2617:19:08 | 4:00:00 | 47991:459:65 | 68:00:00 | 7288:54:15 | 6:00:00 |
| NM_003295:NM_001286273:NM_001286272 | 9150:67:13 | 12:00:00 | 4300:26:05 | 13:00:00 | 2531:19:08 | 6:00:00 | 47226:460:65 | 42:00:00 | 7086:51:15 | 10:00:00 |
| XM_024449333:XM_017020480:NM_015070:NM_001330567 | 0:0:0:0 | 1:0:0:0 | 0:0:0:0 | 1:0:0:0 | 1:0:0:0 | 0:0:0:0 | 3:0:0:0 | 6:1:0:0 | 1:0:0:0 | 0:0:1:0 |
| NM_003850 | 0 | 0 | 0 | 0 | 0 | 0 | 0 | 1 | 4 | 0 |
| NM_014166:NM_001270629:XM_017020549 | 569:177:17 | 12:04:00 | 97:50:03 | 99:44:03 | 171:66:8 | 7:02:00 | 1127:356:67 | 668:223:18 | 438:157:11 | 8:03:00 |
| NM_014166:NM_001270629:XM_017020549 | 600:190:20 | 0:00:00 | 202:103:7 | 1:00:00 | 184:71:8 | 1:02:00 | 1847:604:74 | 1:00:00 | 473:186:12 | 0:00:00 |
| NM_001270629:NM_014166:XM_017020549 | 189:600:19 | 1:00:00 | 103:203:7 | 0:00:00 | 73:185:7 | 0:00:01 | 602:1847:73 | 2:01:00 | 185:473:12 | 1:00:00 |
| NM_014166:NM_001270629:XM_017020549 | 601:190:19 | 0:00:00 | 203:103:6 | 1:00:00 | 186:73:8 | 0:00:00 | 1848:605:72 | 1:00:00 | 473:186:12 | 1:00:00 |
| NM_014166:NM_001270629:XM_017020549 | 601:190:19 | 1:00:00 | 203:103:6 | 1:01:00 | 188:73:8 | 0:00:00 | 1848:603:71 | 2:02:00 | 474:186:13 | 0:00:00 |
| NM_014166:NM_001270629:XM_017020549 | 604:190:15 | 11:00:00 | 207:106:4 | 1:01:00 | 188:73:6 | 6:00:00 | 1863:607:62 | 37:10:00 | 476:186:10 | 20:03:00 |
| NM_014166:NM_001270629:XM_017020549 | 609:190:15 | 8:00:00 | 208:107:4 | 0:00:00 | 192:73:6 | 2:00:00 | 1880:614:62 | 24:04:00 | 483:187:10 | 15:02:00 |
| NM_014166:NM_001270629:XM_017020549 | 636:190:14 | 12:00:00 | 260:108:3 | 20:00:00 | 202:72:6 | 3:00:00 | 2106:625:54 | 92:01:00 | 516:190:9 | 7:00:00 |
| NM_014166:NM_001270629:XM_017020549 | 646:190:14 | 8:00:00 | 274:108:3 | 8:00:00 | 205:73:6 | 0:00:00 | 2204:626:54 | 28:01:00 | 523:190:9 | 4:00:00 |
| NM_014166:XM_017020549 | 647:04:00 | 0:00 | 282:00:00 | 2:00 | 202:00:00 | 1:00 | 2293:10:00 | 3:00 | 526:02:00 | 0:00 |
| NM_000321:XM_011535171 | 21:00 | 2:00 | 9:01 | 1:00 | 5:01 | 0:00 | 19:02 | 5:01 | 13:00 | 0:00 |
| NM_000321:XM_011535171 | 17:02 | 3:00 | 8:00 | 0:00 | 5:02 | 1:00 | 13:04 | 0:00 | 13:00 | 0:00 |
| NM_000321:XM_011535171 | 34:32:00 | 0:00 | 0:02 | 0:00 | 12:10 | 0:00 | 9:24 | 0:02 | 27:16:00 | 0:00 |
| NM_199464 | 2 | 0 | 2 | 0 | 0 | 0 | 7 | 0 | 2 | 0 |
| NM_001098525:XM_011535043 | 0:00 | 0:00 | 0:00 | 0:00 | 0:00 | 0:00 | 0:00 | 0:00 | 0:00 | 1:01 |
| NM_001071775 | 0 | 0 | 0 | 0 | 0 | 2 | 0 | 0 | 0 | 5 |
| NM_006346:XM_017020350:XM_011534885:XM_017020354:NR_146206:NM_001349655:XR_001749460:XR_002957449:XM_017020351:XR_001749457:XR_001749459:XM_024449314:NR_146205:XR_941461:XM_017020352 | 16:1:1:0:0:0:0:0:0:0:0:0:0:0:0 | 1:0:0:0:0:0:0:0:0:0:0:0:0:0:0 | 3:0:0:0:0:0:0:0:0:0:0:0:0:0:0 | 2:0:0:0:0:0:0:0:0:0:0:0:0:0:0 | 2:0:0:0:0:0:0:0:0:0:0:0:0:0:0 | 2:0:0:0:0:0:0:0:0:0:0:0:0:0:0 | 8:7:0:0:0:0:0:0:0:1:1:1:1:0:0 | 17:5:1:3:1:1:1:2:0:0:0:0:0:0:0 | 2:1:0:0:0:0:0:0:0:0:0:0:0:0:0 | 13:1:0:3:0:1:0:1:0:0:0:0:0:0:0 |
| NM_006346:XM_017020350:XM_011534885:XM_017020351:XM_017020354:XR_001749457:NR_146206:NM_001349655:XR_001749460:XR_002957449:XR_001749459:XM_024449314:XM_011534886:XR_001749458:NR_146205:XR_941461:XM_017020352 | 17:2:1:0:0:0:0:0:0:0:0:0:0:0:0:0:0 | 0:0:0:0:0:0:0:0:0:0:0:0:0:0:0:0:0 | 5:1:0:0:0:0:0:0:0:0:0:0:0:0:0:0:0 | 0:0:0:0:0:0:0:0:0:0:0:0:0:0:0:0:0 | 4:0:0:0:0:0:0:0:0:0:0:0:0:0:0:0:0 | 0:0:0:0:0:0:0:0:0:0:0:0:0:0:0:0:0 | 23:19:1:0:2:1:1:1:1:5:2:1:1:1:0:0:0 | 2:0:0:1:1:0:0:0:0:0:0:0:0:0:0:0:0 | 15:2:0:0:3:0:0:1:0:1:0:0:0:0:0:0:0 | 0:0:0:0:0:0:0:0:0:0:0:0:0:0:0:0:0 |
| NM_006346:XM_017020350:XM_011534885:XM_017020354:XM_017020351:XR_001749457:NM_001349655:XR_002957449:XR_001749459:XM_024449314:XR_001749458:XM_017020352:NR_146206:NR_146205:XR_941461 | 17:2:1:0:0:0:0:0:0:0:0:0:0:0:0 | 0:0:0:0:0:0:0:0:0:0:0:0:0:0:0 | 5:3:0:2:0:0:0:0:0:0:0:0:0:0:0 | 0:0:0:0:0:0:0:0:0:0:0:0:0:0:0 | 4:0:0:0:0:0:0:0:0:0:0:0:0:0:0 | 0:0:0:0:0:0:0:0:0:0:0:0:0:0:0 | 24:23:3:10:1:1:1:4:3:1:1:1:0:0:0 | 0:2:0:0:0:0:0:0:0:0:0:0:0:0:0 | 15:2:0:4:0:0:1:1:0:0:0:0:0:0:0 | 0:0:0:0:0:0:0:0:0:0:0:0:0:0:0 |
| XM_024449363 | 0 | 0 | 0 | 0 | 0 | 0 | 0 | 0 | 0 | 1 |
| XM_011535289:NM_144777:NM_003843:XM_011535288:XM_011535290:XM_006719885 | 0:0:0:0:0:0 | 0:0:0:0:0:0 | 1:0:0:0:0:0 | 0:0:0:0:0:0 | 0:0:0:0:0:0 | 0:0:0:0:0:0 | 0:0:0:0:0:0 | 0:0:0:0:0:0 | 0:0:0:0:0:0 | 0:0:0:0:0:0 |
| NM_005348:XM_011536718 | 59:20:00 | 0:00 | 0.516666667 | 0:00 | 36:12:00 | 0:00 | 86:442 | 0:00 | 63:32:00 | 0:00 |
| NM_005348:XM_011536718:NM_001017963 | 133:30:00 | 0:00:00 | 67:97:0 | 0:00:00 | 96:25:00 | 0:00:00 | 308:448:0 | 1:02:00 | 134:43:00 | 0:00:00 |
| NM_005348:XM_011536718:NM_001017963 | 134:33:00 | 0:00:00 | 67:97:0 | 0:00:00 | 96:25:00 | 0:00:00 | 306:411:0 | 1:02:00 | 136:41:00 | 0:00:00 |
| NM_005348:XM_011536718:NM_001017963 | 134:33:00 | 0:00:00 | 67:92:0 | 0:00:00 | 96:25:00 | 0:00:00 | 308:383:0 | 1:01:00 | 137:41:00 | 0:00:00 |
| NM_005348:XM_011536718:NM_001017963 | 134:34:00 | 0:00:00 | 67:87:0 | 0:00:00 | 97:25:00 | 0:00:00 | 312:332:0 | 1:01:00 | 138:40:00 | 0:00:00 |
| NM_005348:XM_011536718:NM_001017963 | 134:32:00 | 0:00:00 | 67:80:0 | 0:01:00 | 97:24:00 | 0:00:00 | 312:312:0 | 1:03:00 | 137:40:00 | 1:00:00 |
| NM_005348:XM_011536718:NM_001017963 | 134:30:00 | 0:00:00 | 67:77:0 | 0:00:00 | 97:25:00 | 0:00:00 | 313:296:0 | 0:02:00 | 138:40:00 | 0:00:00 |
| NM_005348:XM_011536718:NM_001017963 | 133:31:00 | 1:00:00 | 68:73:0 | 0:00:00 | 97:25:00 | 0:00:00 | 312:276:0 | 1:03:00 | 137:40:00 | 1:00:00 |
| NM_005348:XM_011536718:NM_001017963 | 138:23:00 | 0:00:00 | 76:46:00 | 0:01:00 | 105:21:00 | 0:00:00 | 384:159:0 | 5:01:00 | 145:31:00 | 0:00:00 |
| NM_005348:XM_011536718:NM_001017963 | 139:09:00 | 0:00:00 | 75:31:00 | 0:00:00 | 104:13:00 | 1:00:00 | 388:98:0 | 2:00:00 | 145:16:00 | 0:00:00 |
| NM_005348:XM_011536718:NM_001017963 | 140:01:00 | 0:00:00 | 75:03:00 | 0:00:00 | 105:01:00 | 0:00:00 | 387:15:00 | 3:00:00 | 145:06:00 | 0:00:00 |
| XM_011536718:NM_001017963 | 0:00 | 0:00 | 0:00 | 0:00 | 0:00 | 0:00 | 0:00 | 1:00 | 0:00 | 0:00 |
| NM_032630 | 0 | 0 | 0 | 0 | 2 | 0 | 1 | 0 | 2 | 0 |
| NM_032630 | 0 | 0 | 0 | 0 | 2 | 0 | 1 | 0 | 2 | 0 |
| NM_145725:XM_017021618:NM_003300:XM_011537117:NM_145726:XM_017021619:XM_017021620 | 1:0:0:0:0:0:0 | 2:1:1:0:0:0:0 | 5:0:1:0:0:0:0 | 1:1:0:1:0:0:0 | 7:0:3:0:0:0:0 | 3:2:1:0:1:0:0 | 0:0:0:0:0:0:0 | 8:2:4:0:0:0:0 | 1:0:0:0:0:0:0 | 3:1:0:0:0:0:0 |
| NM_004894:NM_001127393 | 177:00:00 | 0:00 | 1135:00:00 | 1:00 | 322:07:00 | 0:00 | 2797:16:00 | 1:00 | 340:03:00 | 0:00 |
| NM_001311 | 1 | 0 | 6 | 0 | 6 | 0 | 1 | 0 | 3 | 4 |

| | | | | | | | | | | |
|---|---|---|---|---|---|---|---|---|---|---|
| NM_080648:NM_001641:NM_080649 | 0:00:00 | 0:00:00 | 0:00:00 | 0:00:00 | 0:00:00 | 0:00:00 | 0:00:00 | 2:02:01 | 0:00:00 | 0:00:00 |
| NM_194431:NM_002937:NM_001282192 | 0:00:00 | 0:00:00 | 0:00:00 | 1:00:00 | 0:04:00 | 0:02:00 | 0:00:01 | 0:00:00 | 0:01:00 | 0:00:00 |
| NM_005015 | 0 | 0 | 0 | 0 | 0 | 0 | 0 | 0 | 0 | 0 |
| NM_180982:NM_178336:NM_181307:NM_181304:NM_181306:NM_181305:XM_005267328:XM_005267325 | 0:0:0:0:0:0:0:0 | 1:0:0:0:0:0:0:0 | 0:0:0:0:0:0:0:0 | 0:2:0:0:0:0:0:0 | 0:0:0:0:0:0:0:0 | 0:0:1:1:0:0:0:0 | 0:0:0:0:0:0:0:0 | 1:3:0:0:1:0:0:0 | 0:0:0:0:0:0:0:0 | 0:1:0:0:1:1:0:0 |
| NM_001282991:XM_006720005:NM_021004:NM_001282989:XM_011536366 | 1:0:0:0:0 | 0:0:0:0:0 | 0:0:0:0:0 | 0:0:0:0:0 | 0:1:1:1:0 | 0:0:0:0:0 | 0:0:1:0:1 | 0:0:0:0:0 | 0:0:0:0:0 | 0:0:0:0:0 |
| NM_198083 | 0 | 0 | 0 | 0 | 0 | 1 | 0 | 0 | 0 | 0 |
| NM_002818:XM_006720213 | 0:00 | 7:00 | 0:00 | 5:00 | 0:00 | 9:02 | 0:00 | 4:00 | 0:00 | 5:00 |
| NM_014169 | 2 | 0 | 1 | 0 | 1 | 1 | 0 | 1 | 0 | 1 |
| NM_182836:NM_004581 | 2:00 | 16:01 | 14:00 | 10:01 | 31:03:00 | 32:03:00 | 67:03:00 | 0:00 | 20:00 | 5:00 |
| NM_182836:NM_004581 | 18:01 | 0:00 | 22:01 | 0:00 | 62:06:00 | 0:00 | 66:03:00 | 0:00 | 24:00:00 | 0:00 |
| NM_006871 | 0 | 0 | 0 | 1 | 0 | 2 | 0 | 0 | 0 | 0 |
| NM_020195 | 0 | 0 | 0 | 0 | 0 | 1 | 0 | 0 | 0 | 2 |
| NM_020195 | 0 | 0 | 0 | 0 | 0 | 1 | 0 | 0 | 0 | 2 |
| NM_020195 | 0 | 0 | 0 | 0 | 0 | 1 | 0 | 0 | 0 | 2 |
| XR_943381:XM_011536374:XM_024449460:NM_182648 | 9:1:0:0 | 2:0:0:0 | 40:2:0:1 | 8:0:3:1 | 24:0:1:1 | 2:0:0:0 | 41:1:4:5 | 14:0:0:1 | 21:0:3:0 | 0:0:0:0 |
| XR_943381:XM_011536374:XM_024449460:NM_182648 | 11:1:1:0 | 3:0:0:0 | 59:2:3:2 | 47:0:5:2 | 36:0:2:1 | 5:0:0:1 | 63:1:5:6 | 46:0:9:1 | 25:1:3:1 | 5:0:1:1 |
| XM_024449460:XR_943381:XM_011536374:NM_182648 | 1:13:1:0 | 1:8:0:0 | 3:65:2:4 | 1:21:0:1 | 2:40:0:2 | 0:18:3:1 | 5:69:1:6 | 8:39:1:1 | 4:27:1:2 | 2:13:0:1 |
| XR_943381:XM_011536374:XM_024449460:NM_182648 | 21:1:2:0 | 0:0:0:0 | 159:4:8:6 | 1:0:0:0 | 66:3:3:3 | 1:0:0:0 | 176:3:17:8 | 4:0:0:0 | 48:1:9:4 | 0:0:0:0 |
| NM_001289022:NM_173607:NM_001079519 | 47:63:0 | 0:00:00 | 149:283:1 | 0:01:00 | 67:206:0 | 0:00:00 | 399:352:1 | 1:02:00 | 70:83:0 | 0:00:00 |
| NM_001289022:NM_173607:NM_001079519 | 48:63:0 | 0:00:00 | 149:284:1 | 0:00:00 | 67:206:0 | 0:00:00 | 399:354:1 | 1:01:00 | 71:83:0 | 0:00:00 |
| NM_001289022:NM_173607:NM_001079519 | 47:63:0 | 0:00:00 | 149:289:1 | 0:01:00 | 66:206:0 | 0:01:00 | 399:358:1 | 0:00:00 | 70:84:0 | 0:00:00 |
| NM_173607:NM_001289022:NM_001079519 | 66:46:00 | 0:00:00 | 297:150:1 | 0:00:00 | 207:67:0 | 0:00:00 | 366:398:1 | 0:01:00 | 86:71:0 | 0:00:00 |
| NM_014672:XM_005268237:NM_001256680:NM_001256681:XM_017021836:XM_011537410:NM_001256678 | 2:1:7:1:2:0:0 | 0:0:0:0:0:0:0 | 0:6:5:1:4:11:0 | 0:0:0:0:0:0:0 | 2:1:6:4:4:6:1 | 0:0:0:0:0:0:0 | 8:0:21:3:4:18:0 | 0:0:0:0:0:0:0 | 5:5:12:2:1:3:2 | 0:0:0:0:0:1:0 |
| XR_001750371:NM_016586:XM_005267756 | 0:00:00 | 0:00:00 | 0:00:00 | 1:00:00 | 0:00:00 | 0:00:00 | 0:01:00 | 0:00:00 | 0:00:00 | 0:00:00 |
| XR_001750371:NM_016586:XM_005267756 | 0:00:00 | 0:00:00 | 0:00:00 | 1:00:00 | 0:00:00 | 0:00:00 | 0:01:00 | 0:00:00 | 0:00:00 | 0:00:00 |
| XM_005268075:NM_024884:XM_017021657:XM_011537166:XM_017021658:XM_017021656 | 2:0:0:0:0:0 | 0:0:0:0:0:0 | 0:0:0:0:0:0 | 0:1:0:0:0:0 | 3:0:1:0:0:0 | 0:0:0:0:0:0 | 3:2:0:0:0:0 | 0:0:0:0:0:0 | 3:0:0:0:0:0 | 0:0:0:0:0:0 |
| XM_011536652:XM_011536657:XM_024449540:XM_011536655 | 2:0:0:0 | 0:0:0:0 | 0:0:0:0 | 0:0:0:0 | 0:0:0:0 | 0:0:0:0 | 0:0:0:0 | 0:0:0:0 | 0:0:0:0 | 0:0:0:0 |
| NM_001202:NM_130850:NM_130851 | 0:00:00 | 0:00:00 | 0:00:00 | 0:00:00 | 0:00:00 | 0:00:00 | 0:00:00 | 0:00:00 | 0:00:00 | 0:00:00 |
| NM_002306:NM_001357678:NR_003225 | 7:00:00 | 0:00:00 | 65:01:01 | 0:00:00 | 125:02:04 | 0:00:00 | 0:00:00 | 17:00:00 | 13:00:00 | 27:01:00 |
| NM_002306:NR_003225:NM_001357678 | 7:01:00 | 0:00:00 | 65:03:01 | 0:00:00 | 125:08:02 | 0:00:00 | 0:00:00 | 17:00:00 | 13:00:00 | 27:00:01 |
| NM_001530:NM_181054 | 6:02 | 0:00 | 1:00 | 0:00 | 3:03 | 0:01 | 1:02 | 0:00 | 2:03 | 0:02 |
| NM_145112:NM_002382:XM_017021312:NR_073137:NM_001320415:XR_002957553:XR_001750327:XR_943452:NM_145113:XR_943451:XM_017021313:XR_943450:XR_001750326 | 93:45:10:3:3:2:3:1:1:0:0:0:0 | 0:0:0:0:0:0:0:0:0:0:0:0:0 | 204:128:2:9:5:0:3:6:7:1:0:0:0 | 0:0:0:0:0:0:0:0:0:0:0:0:0 | 361:233:6:13:12:1:4:4:6:1:3:0:0 | 0:0:0:0:0:0:0:0:0:0:0:0:0 | 164:184:2:16:13:3:6:8:8:5:1:3:6 | 0:0:0:1:0:0:0:0:0:0:0:0:0 | 85:85:3:4:4:1:2:1:1:2:0:3:2 | 0:0:0:0:0:0:0:0:0:0:0:0:0 |
| NM_145112:NM_002382:XM_017021312:NR_073137:NM_001320415:XR_002957553:XR_001750327:XR_943452:NM_145113:XR_943451:XM_017021313:XR_943450:XR_001750326 | 93:45:10:3:3:2:3:1:1:0:0:0:0 | 0:0:0:0:0:0:0:0:0:0:0:0:0 | 203:127:2:8:5:0:3:6:7:1:0:0:0 | 1:1:0:1:0:0:0:0:0:0:0:0:0 | 361:233:6:12:12:1:4:4:6:1:3:0:0 | 0:1:0:1:0:0:0:0:0:0:0:0:0 | 164:184:2:15:13:3:5:8:8:5:1:3:6 | 0:0:0:1:0:0:0:0:0:0:0:0:0 | 85:85:3:4:4:1:2:1:1:2:0:3:2 | 0:0:0:0:0:0:0:0:0:0:0:0:0 |
| NM_145112:NM_002382:XM_017021312:NR_073137:NM_001320415:XR_002957553:XR_001750327:XR_943452:NM_145113:NM_001271068:NM_145114:XR_943451:XM_017021313:NR_073138:XR_943450:XR_001750326:NM_197957:NM_001271069 | 93:45:10:3:3:2:3:3:1:19:12:0:0:0:0:0:0:0 | 0:0:0:0:0:0:0:0:0:0:0:0:0:0:0:0:0:0 | 204:128:2:5:5:0:3:8:7:21:12:1:0:0:0:0:0:0 | 0:0:0:0:0:0:0:0:0:0:0:0:0:0:0:0:0:0 | 361:234:6:13:13:1:4:4:6:35:30:2:3:1:0:0:0:0 | 0:0:0:0:0:0:0:0:0:0:0:0:0:0:0:0:0:0 | 164:184:2:14:13:3:5:10:8:60:42:5:1:1:3:6:1:0 | 0:0:0:0:0:0:0:0:0:0:0:0:0:0:0:0:0:0 | 85:85:3:5:5:1:2:2:1:12:16:2:0:0:3:2:0:0 | 0:0:0:0:0:0:0:0:0:0:0:0:0:0:0:0:0:0 |
| NM_145112:NM_002382:XM_017021312:NR_073137:NM_001320415:XR_002957553:XR_001750327:XR_943452:NM_145113:NM_001271068:NM_145114:XR_943451:XM_017021313:NR_073138:XR_943450:XR_001750326:NM_197957:NM_001271069 | 93:45:10:2:2:2:3:3:1:19:12:0:0:0:0:0:0:0 | 0:0:0:0:0:0:0:0:0:0:0:0:0:0:0:0:0:0 | 204:128:2:4:5:0:3:9:7:21:12:1:0:0:0:0:0:0 | 0:0:0:0:0:0:0:0:0:0:0:0:0:0:0:0:0:0 | 361:234:6:10:13:1:4:4:6:35:30:2:3:1:0:0:0:0 | 0:0:0:1:0:0:0:0:0:0:0:0:0:0:0:0:0:0 | 164:184:2:10:13:3:5:11:8:60:42:6:1:1:3:6:1:0 | 0:0:0:0:0:0:0:0:0:0:0:0:0:0:0:0:0:0 | 85:85:3:4:5:1:2:2:1:12:16:2:0:0:3:2:0:0 | 0:0:0:0:0:0:0:0:0:0:0:0:0:0:0:0:0:0 |
| NM_020806:XM_017020923:XM_017020926:XM_011536345 | 3:1:0:0 | 0:0:0:0 | 3:2:0:0 | 0:1:0:0 | 3:0:0:0 | 3:1:0:0 | 9:8:1:0 | 0:0:0:0 | 2:3:0:1 | 0:0:0:0 |
| NM_001130005:NM_001102:XM_011537267:XM_017021728:XM_011537265:XM_017021723:NM_001130004:XM_017021727:XM_017021726 | 4:10:2:0:0:0:0:0:0 | 1:1:0:0:0:0:0:0:0 | 1:1:4:1:0:0:0:0:0 | 1:1:7:2:0:0:0:0:0 | 3:3:3:2:0:0:0:0:0 | 5:3:9:1:0:0:0:0:0 | 2:21:61:9:1:1:2:0:0 | 0:0:0:0:0:0:0:0:0 | 1:22:8:0:0:1:0:0:0 | 0:0:0:0:0:0:0:0:0 |
| NM_001102:NM_001130005:XM_017021728:XM_011537267:XM_017021723:XM_011537265:XM_017021727:NM_001130004:XM_017021726 | 14:5:2:0:0:0:0:0:0 | 0:0:0:0:0:0:0:0:0 | 2:1:0:2:1:0:0:0:0 | 0:0:0:0:0:0:0:0:0 | 6:9:2:5:4:0:0:0:0 | 0:0:0:0:0:0:0:0:0 | 23:2:13:14:2:1:0:0:0 | 0:1:0:0:0:0:0:0:0 | 23:2:0:2:1:0:0:0:0 | 0:0:0:0:0:0:0:0:0 |
| NM_001102:NM_001130005:XM_017021728:XM_017021723:XM_011537267:XM_017021727:NM_001130004 | 14:5:2:0:0:0:0 | 0:0:0:0:0:0:0 | 2:2:0:1:0:0:0 | 0:0:0:0:0:0:0 | 4:9:0:4:2:0:0 | 0:0:0:0:0:0:0 | 21:3:9:2:4:1:0 | 0:0:0:0:0:0:0 | 23:2:0:1:1:0:0 | 0:0:0:0:0:0:0 |
| NM_001193360:NM_001193363:XM_017021421:NM_001193361:NM_018199:XM_011536908 | 0:0:0:0:0:0 | 2:1:1:0:0:0 | 0:0:1:0:0:0 | 0:0:3:0:0:0 | 3:1:0:1:0:0 | 4:0:1:1:0:0 | 0:0:0:1:0:0 | 2:3:19:2:1:0 | 0:0:2:0:0:0 | 2:0:1:0:0:0 |
| XR_943501:NM_021239:XM_011537044:XM_024449680 | 30:7:2:2 | 1:0:0:0 | 261:9:8:4 | 9:0:0:0 | 48:11:5:0 | 1:0:0:0 | 182:25:10:5 | 3:0:0:0 | 42:7:1:0 | 0:0:0:0 |
| NM_021239:XM_011537044:XR_943501:XM_024449680 | 7:2:31:2 | 0:0:0:0 | 8:7:265:4 | 1:0:5:0 | 10:5:47:0 | 1:0:0:0 | 25:10:183:5 | 0:0:1:0 | 7:1:40:0 | 0:0:1:0 |
| XR_943501:NM_021239:XM_011537044:XM_024449680 | 27:7:2:2 | 3:0:0:0 | 245:5:5:4 | 21:3:2:0 | 46:10:4:0 | 2:0:0:0 | 170:21:8:6 | 11:4:0:0 | 38:7:1:0 | 1:0:0:0 |
| NM_021239:XR_943501:XM_011537044:XM_024449680 | 5:16:2:3 | 1:3:0:0 | 3:173:4:2 | 0:7:0:0 | 5:32:2:0 | 1:2:0:0 | 10:104:5:6 | 0:5:1:0 | 7:27:1:0 | 0:2:0:0 |
| NM_021239:XM_024449680:XR_943501:XM_011537044 | 3:3:5:1 | 0:0:0:0 | 0:4:26:2 | 0:0:5:1 | 4:0:15:1 | 0:0:1:0 | 3:7:27:4 | 1:0:1:0 | 1:0:4:0 | 0:0:0:0 |
| XR_943501:XM_024449680:NM_021239:XM_011537044 | 0:3:2:1 | 5:0:0:0 | 8:4:0:1 | 17:0:0:0 | 4:0:4:1 | 10:0:0:0 | 6:7:1:3 | 19:0:2:0 | 3:0:0:0 | 1:0:1:0 |
| NM_012245:NM_001318844 | 25:00:00 | 2:00 | 36:00:00 | 0:00 | 30:01:00 | 0:00 | 112:04:00 | 2:01 | 42:00:00 | 0:00 |
| XM_005267413:NM_012245:NM_001318844 | 7:27:00 | 3:00:00 | 4:35:00 | 13:00:00 | 7:30:01 | 2:00:00 | 0.828518519 | 19:00:00 | 4:42:00 | 3:00:00 |
| NM_001128595:NM_001289134:NM_152332 | 0:00:00 | 0:00:00 | 0:00:00 | 0:00:00 | 0:00:00 | 0:00:00 | 0:00:00 | 1:01:00 | 0:00:00 | 0:00:00 |
| XM_011536970:NM_005606:NM_001008530:XM_017021465 | 0:0:0:0 | 0:0:0:0 | 0:0:0:0 | 1:0:0:0 | 0:0:0:0 | 0:2:1:0 | 0:0:0:0 | 0:0:0:0 | 0:0:0:1 | 0:0:0:0 |
| NM_001002235:NM_001127707:NM_000295:NM_001127706:XM_017021370:NM_001002236:NM_001127702:NM_001127705:NM_001127703:NM_001127700:NM_001127704:NM_001127701 | 24:46:18:5:4:1:1:0:0:0:0:0 | 0:0:0:0:0:0:0:0:0:0:0:0 | 67:7:1:9:24:0:2:1:2:1:0:0 | 0:0:0:0:0:0:0:0:0:0:0:0 | 659:58:2:45:86:6:2:2:10:15:5:2 | 0:0:0:0:0:0:0:0:0:0:0:0 | 42:73:17:5:27:2:1:0:0:0:1:0 | 0:0:0:0:0:0:0:0:0:0:0:0 | 35:74:25:4:18:0:1:0:1:2:0:0 | 0:0:0:0:0:0:0:0:0:0:0:0 |
| NM_001002235:NM_001127707:NM_000295:NM_001127706:XM_017021370:NM_001002236:NM_001127702:NM_001127705:NM_001127703:NM_001127700:NM_001127704:NM_001127701 | 24:46:18:5:4:1:1:0:0:0:0:0 | 0:0:0:0:0:0:0:0:0:0:0:0 | 66:5:1:9:24:0:2:1:2:1:0:0 | 0:1:0:0:0:0:0:0:0:0:0:0 | 652:58:2:45:91:6:2:2:10:15:4:2 | 2:0:0:0:0:0:0:0:0:0:0:0 | 42:73:17:5:28:2:1:0:0:1:1:0 | 0:0:0:0:0:0:0:0:0:0:0:0 | 35:73:25:4:19:0:1:0:1:2:0:0 | 0:0:0:0:0:0:0:0:0:0:0:0 |
| NM_001002235:NM_001127707:NM_000295:NM_001127706:XM_017021370:NM_001002236:NM_001127702:NM_001127705:NM_001127703:NM_001127700:NM_001127704:NM_001127701 | 24:46:18:5:4:1:1:0:0:0:0:0 | 0:0:0:0:0:0:0:0:0:0:0:0 | 66:5:1:9:22:0:2:1:2:1:0:0 | 0:0:0:0:0:0:0:0:0:0:0:0 | 650:58:2:45:88:6:2:2:9:15:4:2 | 2:0:0:0:1:0:0:0:1:0:0:0 | 42:73:16:5:28:2:1:0:0:1:1:0 | 0:0:1:0:0:0:0:0:0:0:0:0 | 35:73:25:4:19:0:1:0:1:2:0:0 | 0:0:0:0:0:0:0:0:0:0:0:0 |

| | | | | | | | | | | |
|---|---|---|---|---|---|---|---|---|---|---|
| NM_001002235:NM_001127707:NM_000295:NM_001127706:XM_017021370:NM_001002236:NM_001127702:NM_001127705:NM_001127703:NM_001127700:NM_001127704:NM_001127701 | 24:46:18:5:4:1:1:0:0:0:0 | 0:0:0:0:0:0:0:0:0:0:0 | 65:5:1:9:20:0:2:1:2:1:0:0 | 1:0:0:0:0:0:0:0:0:0:0 | 651:57:2:45:89:6:2:2:10:14:4:2 | 1:0:0:0:0:0:0:0:0:1:0:0 | 42:73:17:5:28:2:1:0:0:1:1:0 | 0:0:0:0:0:0:0:0:0:0:0 | 35:73:25:4:19:0:1:0:1:2:0:0 | 0:0:0:0:0:0:0:0:0:0:0 |
| NM_001127707:NM_001002235:NM_000295:NM_001127706:XM_017021370:NM_001002236:NM_001127702:NM_001127705:NM_001127703:NM_001127700:NM_001127704:NM_001127701 | 45:24:18:5:4:1:1:0:0:0:0 | 1:0:0:0:0:0:0:0:0:0:0 | 5:66:1:9:20:0:2:1:2:1:0:0 | 0:0:0:0:0:0:0:0:0:0:0 | 57:650:2:45:89:6:2:2:10:15:4:2 | 0:2:0:0:0:0:0:0:0:0:0 | 73:42:17:5:28:2:1:0:0:1:1:0 | 0:0:0:0:0:0:0:0:0:0:0 | 73:35:25:4:19:0:1:0:1:2:0:0 | 0:0:0:0:0:0:0:0:0:0:0 |
| NM_001002235:NM_001127707:NM_000295:NM_001127706:XM_017021370:NM_001002236:NM_001127702:NM_001127705:NM_001127703:NM_001127700:NM_001127704:NM_001127701 | 24:46:18:5:4:1:1:0:0:0:0 | 0:0:0:0:0:0:0:0:0:0:0 | 66:5:1:9:19:0:2:1:2:1:0:0 | 0:0:0:0:0:0:0:0:0:0:0 | 652:57:2:45:87:6:2:2:10:15:4:2 | 1:0:0:0:1:0:0:0:0:0:0 | 42:73:17:5:27:2:1:0:0:1:1:0 | 0:0:0:0:0:0:0:0:0:0:0 | 35:73:25:4:18:0:1:0:1:2:0:0 | 0:0:0:0:0:0:0:0:0:0:0 |
| NM_001002235:NM_001127707:NM_000295:NM_001127706:XM_017021370:NM_001002236:NM_001127702:NM_001127705:NM_001127703:NM_001127700:NM_001127704:NM_001127701 | 24:45:18:5:4:1:1:0:0:0:0 | 0:0:0:0:0:0:0:0:0:0:0 | 66:5:1:9:15:0:2:1:2:1:0:0 | 0:0:0:0:0:0:0:0:0:0:0 | 651:55:2:45:74:6:2:2:10:14:4:2 | 1:0:0:0:0:0:0:0:0:1:0:0 | 42:73:17:5:24:2:1:0:0:1:1:0 | 0:0:0:0:0:0:0:0:0:0:0 | 35:73:25:4:13:0:0:0:1:2:0:0 | 0:0:0:0:0:0:1:0:0:0:0 |
| NM_001002235:NM_001127707:NM_000295:NM_001127706:XM_017021370:NM_001002236:NM_001127702:NM_001127705:NM_001127703:NM_001127700:NM_001127704:NM_001127701 | 24:45:19:5:3:1:1:0:0:0:0 | 0:0:0:0:0:0:0:0:0:0:0 | 67:4:1:9:11:0:2:1:2:1:0:0 | 0:0:0:0:0:0:0:0:0:0:0 | 653:50:2:45:56:5:2:2:10:15:4:2 | 2:0:0:0:1:1:0:0:0:0:0 | 42:72:18:5:15:2:1:0:0:1:1:0 | 0:1:0:0:0:0:0:0:0:0:0 | 35:73:25:4:6:0:1:0:1:2:0:0 | 0:0:0:0:0:0:0:0:0:0:0 |
| NM_001002235:NM_001127707:NM_000295:NM_001127706:XM_017021370:NM_001002236:NM_001127702:NM_001127705:NM_001127703:NM_001127700:NM_001127704:NM_001127701 | 24:45:19:5:2:1:1:0:0:0:0 | 0:0:0:0:0:0:0:0:0:0:0 | 67:4:1:9:11:0:2:1:2:1:0:0 | 0:0:0:0:0:0:0:0:0:0:0 | 654:48:2:45:47:6:2:2:10:15:4:2 | 1:0:0:0:0:0:0:0:0:0:0 | 42:72:18:5:13:2:1:0:0:1:1:0 | 0:1:0:0:0:0:0:0:0:0:0 | 34:73:25:4:6:0:1:0:1:2:0:0 | 1:0:0:0:0:0:0:0:0:0:0 |
| NM_001002235:NM_001127707:NM_000295:NM_001127706:NM_001002236:NM_001127702:XM_017021370:NM_001127705:NM_001127703:NM_001127700:NM_001127704:NM_001127701 | 24:45:19:5:1:1:1:0:0:0:0 | 0:0:0:0:0:0:0:0:0:0:0 | 67:4:1:9:0:2:8:1:2:1:0:0 | 0:0:0:0:0:0:0:0:0:0:0 | 655:48:2:45:6:2:40:2:10:15:4:2 | 0:0:0:0:0:0:1:0:0:0:0 | 42:72:18:5:2:1:10:0:0:1:1:0 | 0:1:0:0:0:0:0:0:0:0:0 | 35:73:25:4:0:1:4:0:1:2:0:0 | 0:0:0:0:0:0:0:0:0:0:0 |
| NM_001002235:NM_001127707:NM_000295:NM_001127706:XM_017021370:NM_001002236:NM_001127702:NM_001127705:NM_001127703:NM_001127700:NM_001127704:NM_001127701 | 24:44:19:5:1:1:1:0:0:0:0 | 0:0:0:0:0:0:0:0:0:0:0 | 67:3:1:9:3:0:2:1:2:1:0:0 | 0:0:0:0:0:0:0:0:0:0:0 | 655:45:2:45:8:6:2:2:10:15:4:2 | 3:0:0:0:0:0:0:0:0:0:0 | 42:72:18:5:7:2:1:0:0:1:1:0 | 0:0:0:0:0:0:0:0:0:0:0 | 35:73:25:4:1:0:1:0:1:2:0:0 | 0:0:0:0:0:0:0:0:0:0:0 |
| XM_017021121:XM_011536599:NM_177438:XM_017021123:XM_011536601:XM_011536600:XM_011536602 | 1:2:0:0:0:0:0 | 0:0:0:0:0:0:0 | 0:4:6:2:2:1:0 | 1:0:0:0:0:0:0 | 0:0:2:1:0:0:0 | 0:0:0:0:0:0:0 | 2:1:5:1:1:0:0 | 0:1:0:0:0:0:0 | 1:1:2:0:1:0:0 | 0:0:0:0:0:0:0 |
| NM_024734:XR_429332:XM_017021647:XR_001750558:XM_017021646:XR_429330 | 1:1:0:0:0:0 | 0:0:0:0:0:0 | 20:206:0:0:0:0 | 0:10:0:0:0:0 | 14:28:1:0:0:0 | 0:3:0:0:0:0 | 4:41:0:1:0:0 | 0:3:0:0:0:0 | 4:10:0:0:0:0 | 0:1:0:0:0:0 |
| NM_024734:XR_429332:XM_017021647:XM_017021646:XR_001750558 | 1:1:0:0:0 | 0:0:0:0:0 | 18:183:0:0:0 | 0:0:0:0:0 | 14:33:1:1:0 | 0:0:0:0:0 | 2:54:0:0:1 | 0:0:0:0:0 | 4:10:0:0:0 | 0:0:0:0:0 |
| NM_024734:XR_429332:XM_017021647:XM_017021646 | 1:1:0:0 | 0:0:0:0 | 16:99:0:0 | 0:0:0:0 | 14:27:1:1 | 0:0:0:0 | 2:39:0:0 | 0:0:0:0 | 4:9:0:0 | 0:0:0:0 |
| NM_016417 | 0 | 0 | 1 | 1 | 3 | 0 | 1 | 0 | 0 | 0 |
| NM_003384:XM_006720247:XM_017021624:XR_001750539:XM_017021625:XM_017021626 | 24:1:0:0:0:0 | 0:0:0:0:0:0 | 12:8:1:3:1:0 | 0:0:0:0:0:0 | 48:0:2:1:0:0 | 0:0:0:0:0:0 | 89:46:7:6:1:4 | 0:0:0:0:0:0 | 83:2:4:2:1:0 | 0:0:0:0:0:0 |
| NM_003384:XM_006720247:XR_001750539:XM_017021624:XM_017021625:XM_017021626 | 24:1:0:0:0:0 | 0:0:0:0:0:0 | 12:8:0:1:2:0 | 0:0:3:0:0:0 | 48:0:1:2:0:0 | 0:0:0:0:0:0 | 85:46:2:7:1:4 | 3:0:4:0:0:0 | 82:2:1:4:1:0 | 0:0:0:0:0:0 |
| NM_003384:XM_006720247:XM_017021624:XM_017021625:XM_017021626:XR_001750539 | 24:1:0:0:0:0 | 0:0:0:0:0:0 | 12:8:1:2:0:0 | 0:0:0:0:0:0 | 48:0:2:0:0:0 | 0:0:0:0:0:0 | 84:46:8:4:4:2 | 1:0:0:0:0:0 | 81:2:4:1:0:0 | 0:0:0:0:0:0 |
| NM_025141:NM_078474 | 0:01 | 1:00 | 0:00 | 0:00 | 0:00 | 1:00 | 0:00 | 1:02 | 0:00 | 2:00 |
| NM_003134 | 0 | 10 | 0 | 20 | 0 | 42 | 0 | 62 | 0 | 52 |
| NM_001211 | 11 | 0 | 0 | 2 | 7 | 0 | 33 | 0 | 39 | 0 |
| NM_001211 | 11 | 0 | 0 | 2 | 7 | 0 | 32 | 1 | 39 | 0 |
| NM_001211 | 0 | 10 | 0 | 1 | 2 | 6 | 36 | 0 | 46 | 0 |
| NM_001211 | 10 | 0 | 1 | 0 | 6 | 0 | 32 | 3 | 47 | 0 |
| NM_001211 | 9 | 0 | 1 | 0 | 6 | 0 | 27 | 0 | 45 | 0 |
| NM_001211 | 9 | 0 | 1 | 0 | 6 | 0 | 26 | 1 | 44 | 1 |
| NM_001145643 | 5 | 0 | 98 | 2 | 139 | 2 | 29 | 0 | 37 | 0 |
| NM_052849:NM_001080792 | 0:00 | 0:00 | 0:00 | 0:00 | 0:00 | 1:00 | 0:01 | 0:00 | 0:00 | 0:00 |
| NM_144508:XM_017022432 | 0:00 | 4:00 | 0:00 | 7:00 | 0:00 | 3:01 | 20:05 | 0:00 | 11:03 | 0:00 |
| NM_144508:XM_017022432:NM_170589 | 0:00:00 | 4:00:00 | 0:00:00 | 5:00:00 | 0:00:00 | 3:00:00 | 19:01:00 | 0:00:00 | 11:01:00 | 0:00:00 |
| NM_144508:XM_017022432 | 0:00 | 3:00 | 0:00 | 0:02 | 0:00 | 0:01 | 6:06 | 0:00 | 5:00 | 0:00 |
| NM_144508:XM_017022432 | 2:00 | 0:00 | 0:02 | 0:00 | 0:01 | 0:00 | 0:05 | 0:00 | 3:00 | 0:00 |
| NM_144508:XM_017022432 | 0:00 | 1:00 | 0:00 | 0:02 | 0:00 | 0:00 | 0:06 | 0:00 | 2:00 | 0:00 |
| NM_001077268:NM_001258420 | 0:00 | 0:00 | 0:00 | 0:00 | 0:00 | 0:00 | 0:00 | 1:01 | 0:00 | 0:00 |
| NM_001355001:XR_001751382:XR_931899 | 1:01:00 | 0:00:00 | 0:00:00 | 0:00:00 | 0:00:01 | 0:00:00 | 5:01:09 | 0:00:00 | 0:00:00 | 0:00:00 |
| NM_020990:XM_017021902:XM_011521197 | 0:00:00 | 0:00:00 | 3:01:00 | 0:00:00 | 3:01:00 | 0:00:00 | 0:00:00 | 1:00:00 | 3:00:00 | 2:00:00 |
| NM_004048:XR_002957658:XM_005254549 | 13000:137:22 | 7:00:00 | 44586:2859:43 | 51:05:00 | 136792:1344:127 | 102:06:00 | 42988:238:47 | 30:00:00 | 17778:163:15 | 18:00:00 |
| NM_004048:XR_002957658:XM_005254549 | 13009:138:22 | 6:00:00 | 44649:2870:43 | 20:01:00 | 136969:1352:127 | 54:04:00 | 43013:240:47 | 15:01:00 | 17799:165:15 | 10:00:00 |
| NM_004048:XR_002957658:XM_005254549 | 13126:139:22 | 23:01:01 | 44947:2887:42 | 110:19:00 | 137989:1368:129 | 347:15:00 | 43057:239:47 | 69:03:00 | 17913:164:16 | 30:00:00 |
| NM_004048:XR_002957658:XM_005254549 | 26446:282:48 | 30:00:00 | 90456:5764:84 | 172:28:00 | 278032:2724:258 | 604:28:00 | 86286:476:94 | 122:06:00 | 36048:328:32 | 50:00:00 |
| XR_002957658:NM_004048:XM_005254549 | 132:13357:24 | 5:11:00 | 2528:45918:42 | 76:82:1 | 1287:140956:132 | 0.756944444 | 213:43356:47 | 6:52:00 | 155:18194:17 | 2:24:00 |
| NM_004048:XR_002957658:XM_005254549 | 13378:130:24 | 10:05:00 | 46004:2392:42 | 67:73:1 | 141172:1261:132 | 224:17:00 | 43383:207:47 | 45:08:00 | 18212:149:17 | 19:03:00 |
| NM_004048:XR_002957658:XM_005254549 | 26998:44:48 | 42:02:00 | 92722:176:102 | 108:08:00 | 284706:378:308 | 412:16:02 | 87102:66:114 | 98:00:00 | 36674:48:38 | 38:00:00 |
| NM_003104 | 0 | 0 | 0 | 0 | 0 | 0 | 0 | 0 | 0 | 0 |
| NM_003104 | 0 | 0 | 0 | 0 | 0 | 0 | 0 | 0 | 0 | 0 |
| NM_207581 | 0 | 0 | 0 | 0 | 0 | 0 | 0 | 0 | 0 | 1 |
| NM_021199 | 0 | 0 | 0 | 0 | 4 | 3 | 1 | 0 | 4 | 0 |
| NM_001128610:XM_017022720:XM_017022718:XM_017022719:XM_017022722:XM_006720762:XM_024450098:XM_011522193:NM_005154:XM_017022721 | 4:12:57:20:2:3:1:1:0:0 | 0:0:0:0:0:0:0:0:0:0 | 15:68:193:87:17:22:15:0:1:5 | 0:0:1:0:0:0:0:0:0:0 | 1:21:82:34:3:5:1:0:1:2 | 0:0:0:0:0:0:0:0:0:0 | 37:193:668:359:23:81:31:2:10:18 | 0:0:0:1:1:0:0:0:0:0 | 4:31:74:42:2:9:2:0:5:4 | 0:0:0:0:0:0:0:0:0:0 |
| NM_001128610:XM_017022720:XM_017022718:NM_001283049:XM_017022719:XM_017022722:XM_006720762:XM_011522193:XM_024450098:NM_005154:XM_017022721 | 4:12:57:2:20:2:3:1:1:0:0 | 0:0:0:0:0:0:0:0:0:0:0 | 15:66:194:0:87:19:22:0:15:1:5 | 0:0:0:0:0:0:0:0:1:0:0 | 1:22:81:0:34:3:5:0:2:1:2 | 0:0:1:0:0:0:0:0:0:0:0 | 36:192:666:0:360:33:80:2:32:10:18 | 1:0:2:0:0:0:1:0:1:0:0 | 4:31:74:0:40:5:9:0:1:5:4 | 0:0:0:0:2:0:0:0:0:0 |
| NM_001128610:XM_017022720:XM_017022718:NM_001283049:XM_017022719:XM_017022722:XM_006720762:XM_011522193:XM_024450098:NM_005154:XM_017022721 | 4:12:57:2:20:2:3:1:1:0:0 | 0:0:0:0:0:0:0:0:0:0:0 | 15:66:194:0:87:19:22:0:15:1:5 | 0:0:0:0:0:0:0:0:1:0:0 | 1:22:82:0:34:3:5:0:2:1:2 | 0:0:0:0:0:0:0:0:0:0:0 | 36:190:667:0:355:33:81:2:33:10:18 | 1:1:1:0:5:0:0:0:1:0:0 | 4:31:73:0:41:5:9:0:1:5:4 | 0:0:1:0:1:0:0:0:0:0 |
| NM_001128610:XM_017022720:XM_017022718:NM_001283049:XM_017022719:XM_017022722:XM_006720762:XM_024450098:XM_011522193:NM_005154:XM_017022721 | 4:12:52:2:19:2:3:7:1:0:0 | 0:0:0:0:0:0:0:0:0:0:0 | 14:58:184:0:81:17:21:36:0:1:4 | 0:0:0:0:0:0:0:0:0:0:0 | 1:24:78:0:32:3:5:4:0:1:2 | 0:0:0:0:0:0:0:0:0:0:0 | 34:183:634:0:348:31:77:75:2:9:17 | 0:0:2:0:0:0:0:1:0:0:0 | 4:26:69:0:33:4:7:9:0:5:3 | 0:0:0:0:0:0:0:0:0:0 |
| NM_001128610:XM_017022720:XM_017022718:NM_001283049:XM_017022719:XM_017022722:XM_006720762:XM_024450098:XM_011522193:NM_005154:XM_017022721 | 4:11:51:2:19:2:3:7:1:0:0 | 0:0:0:0:0:0:0:0:0:0:0 | 14:57:181:0:81:17:21:61:0:1:4 | 0:0:1:0:0:0:0:1:0:0:0 | 1:23:79:0:32:4:5:7:0:1:2 | 0:0:0:0:0:0:0:0:0:0:0 | 33:179:634:0:347:30:77:103:2:9:17 | 0:0:1:0:0:0:0:0:0:0:0 | 3:30:68:0:34:4:7:11:0:5:3 | 0:0:0:0:0:0:0:0:0:0 |

| | | | | | | | | | | |
|---|---|---|---|---|---|---|---|---|---|---|
| NM_001128610:XM_017022720:XM_017022718:NM_001283049:XM_017022719:XM_017022722:XM_006720762:XM_024450098:XM_011522193:NM_005154:XM_017022721 | 4:10:52:2:19:2:3:8:1:0:0 | 0:0:0:0:0:0:0:0:0:0:0 | 14:58:182:0:80:17:21:70:0:1:4 | 0:0:0:0:0:0:0:0:0:0 | 1:23:79:0:32:4:5:9:0:1:2 | 0:0:0:0:0:0:0:0:0:0 | 33:180:634:0:345:30:77:111:2:9:17 | 0:0:1:0:2:0:0:0:0:0:0 | 3:30:68:0:34:4:7:11:0:5:3 | 0:0:0:0:0:0:0:0:0:0 |
| NM_001128610:XM_017022720:XM_017022718:NM_001283049:XM_017022719:XM_017022722:XM_006720762:XM_024450098:XM_011522193:NM_005154:XM_017022721 | 4:10:52:2:19:2:3:6:1:0:0 | 0:0:0:0:0:0:0:0:0:0:0 | 14:57:181:0:80:17:21:85:0:1:4 | 0:0:0:0:0:0:0:0:0:0 | 1:22:79:0:32:4:5:11:0:1:2 | 0:0:0:0:0:0:0:0:0:0 | 33:180:633:0:348:30:77:126:2:9:17 | 0:1:2:0:0:0:0:0:0:0:0 | 3:30:68:0:35:4:7:12:0:5:3 | 0:0:0:0:0:0:0:0:0:0 |
| NM_001128610:XM_017022720:XM_017022718:NM_001283049:XM_017022719:XM_017022722:XM_006720762:XM_024450098:XM_011522193:NM_005154:XM_017022721 | 4:10:50:2:19:2:3:8:1:0:0 | 0:0:0:0:0:0:0:0:0:0:0 | 14:56:180:0:79:17:21:108:0:1:4 | 0:0:0:0:0:0:0:0:0:0 | 1:22:77:0:32:4:5:11:0:1:2 | 0:0:2:0:0:0:0:0:0:0 | 33:183:623:0:342:29:76:144:2:9:17 | 0:1:1:0:0:0:0:1:0:0:0 | 3:30:68:0:35:4:7:16:0:5:3 | 0:0:0:0:0:0:0:0:0:0 |
| NM_001128610:XM_017022720:XM_017022718:NM_001283049:XM_017022719:XM_017022722:XM_006720762:XM_024450098:XM_011522193:NM_005154:XM_017022721 | 8:20:98:4:38:4:6:14:2:0:0 | 0:0:0:0:0:0:0:0:0:0:0 | 28:114:360:0:158:34:42:214:0:2:8 | 0:0:0:0:0:0:0:0:0:0 | 2:44:154:0:64:8:10:20:0:2:4 | 0:0:4:0:0:0:0:2:0:0:0 | 66:368:1242:0:684:56:152:286:4:18:34 | 0:0:6:0:0:2:0:0:0:0:0 | 6:60:136:0:70:8:14:30:0:10:6 | 0:0:0:0:0:0:0:2:0:0:0 |
| XM_017022718:XM_017022720:XM_017022719:XM_017022722:XM_024450098:NM_001128610:NM_001283049:XM_006720762:XM_011522193:XM_017022721:NM_005154 | 43:8:17:0:5:4:2:3:1:0:0 | 6:1:1:1:2:0:0:0:0:0:0 | 163:55:75:15:98:11:0:20:0:3:1 | 15:2:4:2:7:3:0:1:0:1:0 | 72:20:25:3:9:1:0:5:0:2:1 | 7:2:7:1:1:0:0:0:0:0:0 | 602:183:332:29:138:32:0:75:2:14:9 | 20:1:8:0:4:1:0:1:0:3:0 | 64:27:30:4:13:3:0:5:0:3:5 | 3:2:4:0:3:0:0:1:0:0:0 |
| XM_017022718:XM_017022720:NM_001128610:NM_001283049:XM_017022719:XM_006720762:XM_024450098:XM_011522193:NM_005154:XM_017022721:XM_017022722 | 72:12:8:2:32:6:10:2:0:0:0 | 6:4:0:0:0:0:0:0:0:0:0 | 276:94:20:0:132:34:168:0:0:4:24 | 26:8:2:0:14:2:14:0:2:2:6 | 118:36:2:0:46:10:18:0:2:2:6 | 8:0:0:0:2:0:0:0:0:2:0 | 1086:324:60:0:610:126:232:4:16:26:54 | 56:18:0:0:24:16:12:0:2:2:2 | 106:52:4:0:52:10:24:0:6:6:8 | 8:2:2:0:4:0:0:0:2:0:0 |
| XM_017022719:XM_017022718:NM_001128610:XM_011522193:XM_024450098:XM_017022722:XM_017022720:NM_005154:XM_006720762 | 1:2:1:1:2:0:0:0:0 | 0:0:0:0:0:0:0:0:0 | 0:2:0:0:3:1:3:0:0 | 0:0:0:0:1:0:0:0:0 | 3:1:1:0:2:1:6:0:0 | 2:1:0:0:0:0:0:1:0 | 0:6:1:2:11:1:11:1:2 | 7:4:0:0:1:0:1:0:0 | 0:1:0:0:5:2:5:1:0 | 0:0:0:0:0:0:0:1:0:0 |
| XM_017022719:XM_017022718:NM_001128610:XM_011522193:XM_024450098:XM_017022720:XM_017022722:NM_005154:XM_006720762 | 1:1:1:1:3:0:0:0:0 | 0:0:0:0:0:0:0:0:0 | 0:1:0:0:4:1:0:0:0 | 0:0:0:0:0:1:0:0:0 | 2:1:0:0:4:6:1:0:0 | 0:0:0:0:0:0:0:0:0 | 0:1:0:1:31:10:1:1:2 | 0:0:0:0:0:0:0:0:0 | 0:0:0:0:7:5:2:1:0 | 0:0:0:0:0:0:0:0:0:0 |
| NM_020739:NM_001204450:NM_004748 | 58:08:00 | 2:00:00 | 148:00:00 | 30:00:02 | 106:26:04 | 6:00:00 | 290:06:22 | 60:00:12 | 54:02:00 | 2:00:00 |
| NM_020739:NM_001204450:NM_004748 | 72:08:00 | 6:00:00 | 296:00:06 | 16:00:02 | 150:26:04 | 12:00:00 | 552:06:54 | 40:00:00 | 62:02:02 | 4:00:02 |
| NM_020739:NM_001204450:NM_004748:NM_001204451 | 40:4:0:0 | 1:0:0:0 | 164:0:4:0 | 0:0:0:0 | 91:13:2:1 | 0:0:0:0 | 290:3:29:0 | 2:0:1:0 | 37:1:2:0 | 0:0:0:0 |
| NM_001204450:NM_020739:NM_004748:NM_001204451 | 4:41:0:0 | 0:0:0:0 | 0:163:4:0 | 0:6:0:0 | 13:91:2:1 | 0:2:0:0 | 3:290:30:0 | 0:4:3:0 | 1:37:2:0 | 0:2:0:0 |
| XM_006720502:XM_005254374 | 0:00 | 0:00 | 0:00 | 2:00 | 0:00 | 2:01 | 0:00 | 0:00 | 0:00 | 0:00 |
| XM_006720502:XM_005254374:XM_024449917 | 0:00:00 | 0:00:00 | 0:00:00 | 2:00:00 | 0:00:00 | 2:01:00 | 0:00:00 | 0:00:00 | 0:00:00 | 0:00:00 |
| NM_001018020:NM_001018004:NM_001018006:NM_001018007:XM_005254639:XM_017022537:XM_017022534:NM_001330344:NM_001301289:XM_005254650:XM_017022538:XM_024450040:XM_017022539:XR_002957675:XM_017022535:NM_001301244:XM_005254646:XM_006720667:XM_017022540:NM_001330351:NM_000366:NM_001018005 | 13:45:25:26:2:1:1:49:6:1:1:1:0:0:0:0:0:0:0:0:0:0 | 0:0:0:0:0:0:0:0:0:0:0:0:0:0:0:0:0:0:0:0:0:0 | 8:5:72:68:1:1:0:160:4:0:0:4:1:6:2:1:1:0:0:0:0:0 | 0:0:0:0:0:0:0:0:0:0:0:1:0:0:0:0:0:0:0:0:0:0 | 11:6:142:102:2:0:0:192:3:0:1:2:1:3:1:0:0:1:1:0:0:0 | 0:0:0:0:0:1:0:0:0:0:0:0:0:0:0:0:0:0:0:0:0:0 | 11:17:18:12:0:0:0:464:0:1:0:1:0:7:0:0:0:0:0:3:0:0 | 0:0:0:0:0:0:0:1:0:0:0:0:0:0:0:0:0:0:0:0:0:0 | 6:3:45:8:0:0:0:138:2:0:2:0:1:2:0:0:0:0:0:0:0:0 | 0:0:0:0:0:0:0:0:0:0:0:0:0:0:0:0:0:0:0:0:0:0 |
| NM_001018020:NM_001018004:NM_001018006:NM_001018007:XM_005254639:XM_017022537:XM_017022534:NM_001330344:NM_001301289:XM_005254650:XM_017022538:XM_024450040:XR_002957675:XM_017022535:XM_017022539:NM_001301244:XM_005254646:XM_006720667:XM_017022540:NM_001330351:NM_000366:NM_001018005 | 13:45:25:26:2:1:1:49:6:1:1:1:0:0:0:0:0:0:0:0:0:0 | 0:0:0:0:0:0:0:0:0:0:0:0:0:0:0:0:0:0:0:0:0:0 | 8:5:72:66:1:1:0:161:4:0:0:5:6:2:1:1:1:0:0:0:0:0 | 0:0:0:2:0:0:0:0:0:0:0:0:0:0:0:0:0:0:0:0:0:0 | 11:6:142:102:2:0:0:192:3:0:1:1:3:1:1:0:0:1:1:0:0:0 | 0:0:0:0:0:0:0:0:0:0:0:0:0:0:0:0:0:0:0:0:0:0 | 11:17:18:12:0:0:0:469:0:1:1:1:7:0:0:0:0:0:1:0:0 | 0:0:0:0:0:0:0:0:0:0:0:0:0:0:0:0:0:0:0:0:0:0 | 6:3:45:8:0:0:0:138:2:0:2:0:2:0:0:0:0:0:0:0:1:0 | 0:0:0:0:0:0:0:0:0:0:0:0:0:0:0:0:0:0:0:0:0:0 |
| NM_001018020:NM_001018004:NM_001018006:NM_001018007:XM_005254639:XM_017022537:XM_017022534:NM_001330344:NM_001301289:XM_005254650:XM_017022538:XM_024450040:XR_002957675:XM_017022535:XM_017022539:NM_001301244:XM_005254646:XM_006720667:XM_017022540:NM_001330351:NM_000366:NM_001018005 | 13:45:25:26:2:1:1:53:6:1:1:1:0:0:0:0:0:0:0:0:0:0 | 0:0:0:0:0:0:0:0:0:0:0:0:0:0:0:0:0:0:0:0:0:0 | 8:5:72:69:1:1:0:166:4:0:0:5:5:2:1:1:1:0:0:0:0:0 | 0:0:0:0:0:0:0:0:0:0:0:0:0:0:0:0:0:0:0:0:0:0 | 11:6:142:102:2:0:0:191:3:0:1:1:3:1:1:0:0:1:1:0:0:0 | 0:0:0:0:0:0:0:1:0:0:0:0:0:0:0:0:0:0:0:0:0:0 | 11:17:18:12:0:0:0:470:0:1:1:1:7:0:0:0:0:0:1:0:0 | 0:0:0:0:0:0:0:0:0:0:0:0:0:0:0:0:0:0:0:0:0:0 | 6:3:45:8:0:0:0:139:2:0:2:0:2:0:0:0:0:0:0:0:1:0 | 0:0:0:0:0:0:0:0:0:0:0:0:0:0:0:0:0:0:0:0:0:0 |
| NM_001018020:NM_001018004:NM_001018006:NM_001018007:XM_005254639:XM_017022537:XM_017022534:NM_001330344:NM_001301289:XM_005254650:XM_017022538:XM_024450040:XR_002957675:XM_017022535:XM_017022539:NM_001301244:XM_005254646:XM_006720667:XM_017022540:NM_001330351:NM_000366:NM_001018005 | 13:45:25:26:2:1:1:53:6:1:1:1:0:0:0:0:0:0:0:0:0:0 | 0:0:0:0:0:0:0:0:0:0:0:0:0:0:0:0:0:0:0:0:0:0 | 8:5:72:69:1:1:0:168:4:0:0:5:5:2:1:1:1:0:0:0:0:0 | 0:0:0:0:0:0:0:0:0:0:0:0:0:0:0:0:0:0:0:0:0:0 | 11:6:142:101:2:0:0:191:3:0:1:1:3:1:1:0:0:1:1:0:0:0 | 0:0:0:1:0:0:0:1:0:0:0:0:0:0:0:0:0:0:0:0:0:0 | 11:17:18:12:0:0:0:471:0:1:1:1:7:0:0:0:0:0:1:0:0 | 0:0:0:0:0:0:0:1:0:0:0:0:0:0:0:0:0:0:0:0:0:0 | 6:3:45:8:0:0:0:141:2:0:2:0:1:0:0:0:0:0:0:0:1:0 | 0:0:0:0:0:0:0:0:0:0:0:0:0:0:0:0:0:0:0:0:0:0 |
| NM_001018020:NM_001018006:NM_001330344:XM_005254639:XM_017022537:XM_005254650:XM_017022538:XM_024450040:XR_002957675:XM_017022539:XM_005254646:NM_001018007:XM_006720667:NM_000366:NM_001330351 | 13:25:55:2:1:1:1:1:0:0:0:0:0:0:0 | 0:0:0:0:0:0:0:0:0:0:0:0:0:0:0 | 8:71:172:1:1:0:0:5:5:1:1:0:0:0:0 | 0:1:0:0:0:0:0:0:0:0:0:0:0:0:0 | 11:142:199:2:0:0:1:1:2:1:0:2:1:0:0 | 0:0:0:0:0:0:0:0:0:0:0:0:0:0:0 | 11:18:482:0:0:1:1:1:7:0:0:2:0:0:0 | 0:0:0:0:0:0:0:0:0:0:0:0:0:0:0 | 6:45:143:0:0:0:2:0:1:1:0:1:0:1:0 | 0:0:0:0:0:0:0:0:0:0:0:0:0:0:0 |
| NM_001018004:NM_001018006:NM_001018020:NM_001018007:NM_001330344:NM_001301289:XR_002957675:XM_017022537:XM_024450040:XM_017022539:NM_001330351 | 43:24:13:26:57:6:1:0:0:0:0 | 1:1:0:0:0:0:0:0:0:0:0 | 5:72:8:70:174:4:11:1:2:1:0 | 0:0:0:0:0:0:0:0:0:0 | 6:142:11:102:205:3:7:0:0:0:0 | 0:0:0:0:0:0:0:0:0:0 | 17:18:11:13:485:0:12:0:0:0:1 | 0:0:0:0:1:0:0:0:0:0:0 | 3:45:6:8:147:2:2:0:0:0:0 | 0:0:0:0:0:0:0:0:0:0 |
| NM_005707 | 8 | 1 | 35 | 2 | 6 | 2 | 110 | 7 | 23 | 3 |
| NM_005707 | 18 | 2 | 78 | 6 | 12 | 4 | 222 | 20 | 48 | 4 |
| NR_045106:NR_045104:NM_001207058:NM_001136043:NR_045105:NR_045901:XM_006720711:NM_001207059:XM_011522101 | 2:9:1:2:1:0:0:0:0 | 0:0:0:0:0:0:0:0:0 | 3:10:0:5:2:0:0:0:0 | 0:0:0:0:0:0:0:0:0 | 4:22:1:2:0:2:1:0:0 | 0:0:0:0:0:0:0:0:0 | 33:116:3:30:7:10:2:7:1 | 0:0:0:0:0:0:0:0:0 | 6:35:1:2:3:2:0:1:0 | 0:0:0:0:0:0:0:0:0 |
| NM_000968 | 14 | 0 | 7 | 0 | 13 | 0 | 42 | 0 | 46 | 1 |
| NM_024666:NM_001271886:XM_011522020:XM_024450052:XM_024450053:NM_001271885 | 8:0:4:0:0:0 | 80:6:6:2:0:0 | 54:4:8:0:0:0 | 84:8:24:0:0:0 | 158:12:14:0:0:0 | 112:8:14:0:0:0 | 0:0:2:0:0:0 | 352:14:100:4:2:0 | 0:0:0:0:0:0 | 318:14:18:0:0:2 |
| NM_024666:NM_001271886:XM_011522020:XM_024450052:XM_024450053:NM_001271885 | 44:3:4:1:0:0 | 0:0:0:0:0:0 | 69:6:15:0:0:0 | 0:0:0:0:0:0 | 135:10:11:0:0:0 | 0:0:0:0:0:0 | 176:7:46:2:1:0 | 0:0:0:0:0:0 | 159:7:8:0:0:1 | 0:0:0:0:0:0 |
| NM_024666:NM_001271886:XM_011522020:XM_024450052:XM_024450053:NM_001271885 | 43:3:4:1:0:0 | 1:0:0:0:0:0 | 69:6:10:0:0:0 | 0:0:0:0:0:0 | 134:10:10:0:0:0 | 1:0:0:0:0:0 | 176:7:36:2:1:0 | 0:0:0:0:0:0 | 159:7:8:0:0:1 | 0:0:0:0:0:0 |
| NM_001322475:NM_001284348:NR_147832:NM_001284347 | 0:0:0:0 | 0:0:0:0 | 0:0:0:0 | 0:0:0:0 | 0:0:0:0 | 2:0:0:0 | 0:0:0:0 | 4:2:0:0 | 0:0:0:0 | 0:0:0:0 |
| NM_016166:XM_024450094:XM_017022689:XM_011522127:XM_017022691:XM_017022690:XM_011522126 | 32:2:0:0:0:0:0 | 0:0:0:0:0:0:0 | 42:3:2:1:7:0:0 | 0:0:0:0:0:0:0 | 44:5:0:1:2:0:0 | 1:1:0:0:0:0:0 | 154:37:0:0:41:0:0 | 2:0:0:0:0:0:0 | 34:1:0:0:2:0:0 | 0:0:0:0:0:0:0 |
| NM_016166:XM_024450094:XM_017022691:XM_017022689:XM_011522127:XM_017022690:XM_011522126 | 64:4:0:0:0:0:0 | 0:0:0:0:0:0:0 | 76:6:12:4:2:0:0 | 0:0:2:0:0:0:0 | 84:6:4:0:2:0:0 | 4:4:0:0:0:0:0 | 290:76:82:0:0:0:0 | 10:0:0:0:0:0:0 | 60:2:4:0:0:0:0 | 2:0:0:0:0:0:0 |
| NM_033429:NM_001286694 | 0:00 | 0:00 | 3:00 | 1:00 | 4:02 | 4:00 | 3:00 | 0:00 | 4:01 | 0:00 |
| XM_017022395:NM_018003:NM_001008224:XM_017022394:XM_011521752:XM_011521754 | 1:9:1:2:0:0 | 0:0:0:0:0:0 | 4:0:0:0:0:0 | 1:0:0:0:0:0 | 1:9:0:3:1:0 | 0:0:0:0:0:0 | 9:6:0:0:0:0 | 0:0:0:1:0:0 | 2:5:0:2:0:0 | 0:0:0:0:0:0 |
| NM_001008224:XM_017022394:NM_018003:XM_017022395:XM_011521754 | 2:1:21:1:1 | 1:2:5:0:0 | 1:4:9:4:0 | 1:3:5:4:0 | 0:3:12:1:0 | 0:2:8:1:0 | 1:1:50:15:0 | 0:0:0:0:0 | 2:3:16:2:0 | 0:0:0:0:0 |
| NM_018003:NM_001008224:XM_017022394:XM_011521754:XM_017022395 | 27:3:3:1:0 | 0:0:0:0:0 | 16:2:7:0:7 | 0:0:0:0:0 | 21:0:7:0:2 | 0:0:0:0:0 | 52:1:2:0:14 | 0:0:0:0:0 | 17:2:3:0:2 | 0:0:0:0:0 |
| NM_145204:XM_005254157:NM_001172111:XM_011521217:XM_011521215 | 0:0:0:0:0 | 4:0:0:0:0 | 0:0:0:0:0 | 6:0:0:0:0 | 0:0:0:0:0 | 26:0:0:0:0 | 0:0:0:0:0 | 84:4:2:2:0 | 0:0:0:0:0 | 18:0:0:0:0 |
| XR_001751314:NM_002654:XM_011521670:XM_005254445:NM_001206797:NM_182471:XM_006720570:NM_001206798 | 4:253:32:1:0:0:0:0 | 1:1:0:0:0:0:0:0 | 3:185:53:1:3:1:0:0 | 0:0:0:0:0:0:0:0 | 2:176:16:0:0:0:0:0 | 0:2:0:0:0:0:0:0 | 19:1059:350:3:10:3:1:0 | 4:7:0:0:0:0:0:0 | 4:695:79:3:2:1:1:0 | 1:2:0:0:0:0:0:0 |
| NM_002654:XM_011521670:XM_005254445:XR_001751314:NM_001206797:NM_182471:XM_006720570:NM_001206798 | 251:32:1:5:0:0:0:0 | 2:0:0:0:0:0:0:0 | 183:52:1:1:3:1:0:0 | 0:0:0:2:0:0:0:0 | 177:16:0:1:0:0:0:0 | 1:0:0:0:0:0:0:0 | 1054:349:3:15:10:3:1:0 | 7:0:0:3:0:0:0:0 | 695:79:3:4:2:1:1:0 | 0:0:0:1:0:0:0:0 |
| NM_002654:XR_001751314:XM_011521670:XM_005254445:NM_182471:XM_017022313:NM_001206797:NM_182470:XM_006720570:NM_001206798 | 236:7:21:1:14:1:0:0:0:0 | 0:0:0:0:0:0:0:0:0:0 | 164:8:43:1:6:0:3:1:0:0 | 0:0:0:0:0:0:0:0:0:0 | 165:3:12:0:7:0:1:0:0:0 | 0:0:0:0:0:0:0:0:0:0 | 976:45:279:3:12:2:11:0:1:0 | 0:2:3:0:0:0:0:0:0:0 | 655:10:61:3:13:4:2:1:1:0 | 0:1:0:0:0:0:0:0:0:0 |
| XR_001751314:NM_002654:XM_011521670:XM_005254445:NM_182471:XM_017022313:NM_001206797:NM_182470:XM_006720570:NM_001206798 | 6:231:19:1:14:1:0:0:0:0 | 2:0:0:0:0:0:0:0:0:0 | 9:157:40:1:5:0:3:1:0:0 | 0:0:0:0:0:0:0:0:0:0 | 4:162:11:0:7:0:1:0:0:0 | 0:0:0:0:0:0:0:0:0:0 | 64:972:269:3:13:2:11:0:1:0 | 1:0:0:0:0:0:0:0:0:0 | 10:642:55:3:13:4:2:1:1:0 | 1:0:0:0:0:0:0:0:0:0 |
| NM_002654:NM_001206797:XR_001751314:NM_182471:XM_011521670:XM_005254445:XM_017022313:NM_182470:XM_006720570:NM_001206798 | 452:2:36:30:34:2:2:0:0:0 | 0:0:0:0:0:0:0:0:0:0 | 306:6:32:10:64:2:2:2:0:0 | 0:0:0:0:0:0:0:0:0:0 | 316:2:18:12:16:0:0:0:0:0 | 0:0:0:0:0:0:0:0:0:0 | 1874:22:246:26:452:6:0:0:2:0 | 0:0:6:0:0:0:0:0:0:0 | 1244:4:60:28:106:6:8:2:2:0 | 2:0:2:0:0:0:0:0:0:0 |
| NM_002654:NM_001206797:XR_001751314:NM_182471:XM_011521670:XM_005254445:XM_017022313:NM_182470:XM_006720570:NM_001206798 | 222:1:30:16:12:1:1:0:0:0 | 1:0:0:0:0:0:0:0:0:0 | 149:3:32:5:30:1:1:1:0:0 | 0:0:0:0:0:0:0:0:0:0 | 157:1:17:6:6:0:0:0:0:0 | 0:0:0:0:0:0:0:0:0:0 | 930:12:175:19:200:3:2:0:1:0 | 4:0:1:0:3:0:0:0:0:0 | 614:2:51:16:47:3:3:1:1:0 | 0:0:0:0:0:0:0:0:0:0 |

| | | | | | | | | | | |
|---|---|---|---|---|---|---|---|---|---|---|
| NM_002654:XR_001751314:NM_001206797:NM_182471:XM_011521670:XM_005254445:XM_017022313:NM_182470:XM_006720570:NM_001206799:NM_001206798 | 213:52:1:16:9:1:0:0:0:0:0 | 0:0:0:0:0:0:0:0:0:0:0 | 141:43:3:8:20:1:1:1:0:0:0 | 0:0:0:0:1:0:0:0:0:0:0 | 149:31:1:7:6:0:0:0:0:0:0 | 0:0:0:0:0:0:0:0:0:0 | 897:265:12:31:153:3:2:0:1:0:0 | 3:4:0:0:3:0:0:0:0:0:0 | 599:74:2:23:38:3:3:1:1:0:0 | 1:3:0:0:0:0:0:0:0:0:0 |
| NM_002654:XR_001751314:NM_001206797:NM_182471:XM_011521670:XM_005254445:XM_017022313:NM_182470:XM_006720570:NM_001206799:NM_001206798 | 215:52:1:16:9:1:0:0:0:0:0 | 0:0:0:0:0:0:0:0:0:0:0 | 141:45:3:8:20:1:1:1:0:0:0 | 1:0:0:0:1:0:0:0:0:0:0 | 148:31:1:8:6:0:0:0:0:0:0 | 1:0:0:0:0:0:0:0:0:0 | 921:282:12:32:157:3:2:0:1:0:0 | 2:2:0:0:0:0:0:0:0:0:0 | 607:80:2:23:36:3:3:1:1:0:0 | 0:1:0:0:1:0:0:0:0:0:0 |
| NM_002654:XR_001751314:NM_182471:NM_001206797:XM_011521670:XM_005254445:XM_017022313:NM_182470:XM_006720570:NM_001206798:NM_001206799 | 214:60:21:1:6:1:0:0:0:0:0 | 0:0:0:0:0:0:0:0:0:0:0 | 144:49:8:1:16:1:1:1:0:0:0 | 0:0:0:1:1:0:0:0:0:0:0 | 149:34:8:0:5:0:0:0:0:0:0 | 0:0:0:0:0:0:0:0:0:0 | 912:342:49:2:134:3:3:0:1:0:0 | 2:0:0:7:0:0:0:0:0:0:0 | 607:103:26:1:30:3:4:1:1:0:0 | 0:0:0:1:0:0:0:0:0:0:0 |
| NM_002654:XR_001751314:NM_182471:NM_001206797:XM_011521670:XM_005254445:XM_017022313:NM_182470:XM_006720570:NM_001206798:NM_001206799 | 214:60:21:1:6:1:0:0:0:0:0 | 0:0:0:0:0:0:0:0:0:0:0 | 144:49:8:1:16:1:1:1:0:0:0 | 0:0:0:1:1:0:0:0:0:0:0 | 149:34:8:0:5:0:0:0:0:0:0 | 0:0:0:0:0:0:0:0:0:0 | 912:342:49:2:134:3:3:0:1:0:0 | 2:0:0:7:0:0:0:0:0:0:0 | 607:103:26:1:30:3:4:1:1:0:0 | 0:0:0:1:0:0:0:0:0:0:0 |
| NM_002654:XR_001751314:NM_182471:NM_001206797:XM_011521670:XM_005254445:XM_017022313:NM_182470:XM_006720570:NM_001206798:NM_001206799 | 213:60:21:1:6:1:0:0:0:0:0 | 1:0:0:0:0:0:0:0:0:0:0 | 144:49:8:1:16:1:1:1:0:0:0 | 0:0:0:1:1:0:0:0:0:0:0 | 149:34:8:0:5:0:0:0:0:0:0 | 0:0:0:0:0:0:0:0:0:0 | 913:342:49:2:134:3:2:0:1:0:0 | 1:0:0:7:1:0:1:0:0:0:0 | 607:103:26:1:30:3:4:1:1:0:0 | 0:0:0:1:0:0:0:0:0:0:0 |
| NM_002654:XR_001751314:NM_182471:XM_011521670:XM_005254445:XM_017022313:NM_182470:XM_006720570:NM_001206798:NM_001206799 | 213:68:21:8:1:1:0:0:0:0 | 0:0:0:0:0:0:0:0:0:0 | 143:56:12:17:1:1:1:0:0:0 | 1:0:0:0:0:0:0:0:0:0 | 150:40:10:5:0:0:0:0:0:0 | 0:0:0:0:0:0:0:0:0:0 | 894:387:61:111:3:3:0:1:0:0 | 1:0:0:0:0:0:0:0:0:0 | 604:117:28:24:3:4:1:1:0:0 | 0:0:0:1:0:0:0:0:0:0 |
| NM_002654:XR_001751314:NM_182471:XM_011521670:XM_005254445:XM_017022313:NM_182470:XM_006720570:NM_001206798:NM_001206799 | 211:68:21:8:1:1:0:0:0:0 | 2:0:0:0:0:0:0:0:0:0 | 144:56:12:19:1:1:1:0:0:0 | 0:0:0:0:0:0:0:0:0:0 | 150:39:10:5:0:0:0:0:0:0 | 0:1:0:0:0:0:0:0:0:0 | 892:388:61:110:3:3:0:1:0:0 | 3:0:0:0:0:0:0:0:0:0 | 602:117:28:25:3:4:1:1:0:0 | 2:0:0:0:0:0:0:0:0:0 |
| NM_002654:XR_001751314:NM_182471:XM_011521670:XM_005254445:XM_017022313:NM_182470:XM_006720570:NM_001206798:NM_001206799 | 213:66:21:8:1:1:0:0:0:0 | 0:0:0:0:0:0:0:0:0:0 | 145:56:12:17:1:1:1:0:0:0 | 0:0:0:0:0:0:0:0:0:0 | 149:40:10:5:0:0:0:0:0:0 | 0:0:0:0:0:0:0:0:0:0 | 892:386:60:107:3:3:0:1:0:0 | 2:0:0:2:0:0:0:0:0:0 | 602:116:29:24:3:4:1:1:0:0 | 0:0:0:1:0:0:0:0:0:0 |
| NM_002654:NM_182471:XR_001751314:XM_011521670:XM_005254445:XM_017022313:NM_182470:XM_006720570:NM_001206797:NM_001206799 | 216:21:64:9:1:1:0:0:0:0 | 0:0:0:0:0:0:0:0:0:0 | 148:12:56:17:1:1:1:0:0:0 | 0:0:0:0:0:0:0:0:0:0 | 149:11:40:5:0:0:0:0:0:0 | 0:0:0:0:0:0:0:0:0:0 | 890:64:392:105:3:2:0:1:1:0 | 0:0:0:0:0:0:0:0:0:0 | 599:30:117:26:3:3:1:1:0:0 | 0:0:0:0:0:0:0:0:0:0 |
| NM_002654:NM_182471:XR_001751314:XM_011521670:XM_005254445:XM_017022313:NM_001206797:NM_182470:XM_006720570:NM_001206799 | 217:24:66:11:1:1:0:0:0:0 | 0:0:0:0:0:0:0:0:0:0 | 147:14:57:17:1:1:1:1:0:0 | 1:0:0:0:0:0:0:0:0:0 | 150:11:41:5:0:0:1:0:0:0 | 0:0:0:0:0:0:0:0:0:0 | 900:74:403:97:3:3:2:0:1:0 | 3:0:0:1:0:0:0:0:0:0 | 607:33:120:30:3:3:0:1:1:0 | 1:0:0:0:0:0:0:0:0:0 |
| NM_002654:NM_182471:XR_001751314:XM_011521670:XM_005254445:XM_017022313:NM_001206797:NM_182470:XM_006720570:NM_001206799 | 216:24:66:12:1:1:0:0:0:0 | 0:0:0:0:0:0:0:0:0:0 | 148:14:56:16:1:1:1:1:0:0 | 0:0:0:0:0:0:0:0:0:0 | 150:12:41:4:0:0:1:0:0:0 | 0:0:0:0:0:0:0:0:0:0 | 899:74:405:98:3:3:2:0:1:0 | 2:0:0:1:0:0:0:0:0:0 | 608:33:119:32:3:3:0:1:1:0 | 0:0:0:0:0:0:0:0:0:0 |
| NM_002654:NM_182471:XR_001751314:XM_011521670:XM_005254445:XM_017022313:NM_001206797:NM_182470:XM_006720570:NM_001206798:NM_001206799 | 215:28:66:12:1:1:0:0:0:0:0 | 0:0:0:0:0:0:0:0:0:0:0 | 149:18:58:20:1:1:3:1:0:0:0 | 0:0:1:0:0:0:0:0:0:0:0 | 150:12:41:4:0:0:1:0:0:0:0 | 0:0:0:0:0:0:0:0:0:0 | 895:101:426:98:3:4:11:0:1:0:0 | 1:0:2:0:0:0:0:0:0:0:0 | 612:42:126:32:3:3:1:1:1:0:0 | 0:0:2:0:0:0:0:0:0:0:0 |
| NM_002654:NM_182471:NM_001206797:XR_001751314:XM_005254445:XM_011521670:XM_017022313:NM_182470:XM_006720570:NM_001206798:NM_001206799 | 214:38:1:77:1:11:2:0:0:0:0 | 0:0:0:0:0:0:0:0:0:0:0 | 149:26:3:72:1:20:1:1:0:0:0 | 0:0:0:0:0:1:0:0:0:0:0 | 153:18:1:41:0:4:0:0:0:0:0 | 1:0:0:0:0:0:0:0:0:0 | 905:152:12:463:3:85:4:0:1:0:0 | 0:1:0:1:0:2:0:0:0:0 | 621:67:2:138:3:25:3:1:1:0:0 | 0:0:0:0:0:0:0:0:0:0:0 |
| NM_001318825:NM_000520 | 0:00 | 0:00 | 0:00 | 0:00 | 0:00 | 0:00 | 0:00 | 2:02 | 0:00 | 0:02 |
| NM_033250:NM_033240:NM_033244:NM_033239 | 0:0:0:0 | 0:0:0:0 | 0:0:0:0 | 0:0:0:0 | 0:0:0:0 | 2:2:0:0 | 0:0:0:0 | 16:6:2:0 | 0:0:0:0 | 2:0:0:0 |
| NM_033240:NM_033250:NM_033244:NM_033239 | 0:0:0:0 | 0:0:0:0 | 0:0:0:0 | 0:0:0:0 | 0:2:0:0 | 2:0:0:0 | 4:8:0:0 | 2:8:2:0 | 0:2:0:0 | 2:0:0:0 |
| NM_000126:NM_001127716:XR_931766 | 0:00:00 | 0:00:00 | 4:00:00 | 0:00:00 | 2:00:00 | 2:01:00 | 4:01:00 | 0:00:00 | 6:06:01 | 0:00:00 |
| NM_005724:XM_017021857:NM_198902 | 162:02:00 | 0:00:00 | 456:19:00 | 0:00:00 | 690:18:00 | 0:00:00 | 473:20:01 | 0:00:00 | 203:06:00 | 0:00:00 |
| NM_005724:XM_017021857:NM_198902 | 162:03:00 | 0:00:00 | 461:10:00 | 0:00:00 | 693:12:00 | 1:00:00 | 473:12:01 | 1:00:00 | 204:00:00 | 0:00:00 |
| NM_005724:XM_017021857:NM_198902 | 162:00:00 | 0:00:00 | 460:06:00 | 0:00:00 | 695:04:00 | 0:00:00 | 474:04:01 | 0:00:00 | 203:00:00 | 1:00:00 |
| NM_005724:XM_017021857:NM_198902 | 324:00:00 | 0:00:00 | 916:10:00 | 6:00:00 | 1388:02:00 | 4:00:00 | 940:04:02 | 8:02:00 | 402:00:00 | 6:00:00 |
| NM_005724:XM_017021857:NM_198902 | 324:00:00 | 0:00:00 | 920:04:00 | 0:00:00 | 1392:02:00 | 0:00:00 | 936:02:02 | 4:00:00 | 408:00:00 | 0:00:00 |
| NM_004390:XM_024449855 | 0:00 | 2:00 | 0:00 | 0:00 | 0:00 | 0:00 | 0:00 | 0:00 | 0:00 | 0:00 |
| NR_037653:NR_037652 | 0:00 | 0:00 | 0:00 | 1:01 | 0:00 | 0:01 | 0:00 | 0:00 | 0:00 | 0:00 |
| NM_004049:NM_001114735 | 14:00 | 4:04 | 18:04 | 16:10 | 8:00 | 4:04 | 20:04 | 22:08 | 2:00 | 4:04 |
| NM_004049:NM_001114735 | 14:00 | 4:04 | 18:02 | 14:10 | 8:00 | 4:04 | 20:04 | 22:08 | 2:00 | 4:00 |
| NM_004049:NM_001114735 | 14:00 | 4:04 | 18:02 | 14:10 | 8:00 | 4:04 | 20:04 | 18:08 | 2:00 | 4:00 |
| NM_001021:NR_111943:NR_111944 | 25:01:00 | 0:00:00 | 17:01:00 | 0:00:00 | 24:01:00 | 0:00:00 | 106:01:01 | 0:00:00 | 83:04:00 | 1:01:00 |
| NM_001160114:NR_027650 | 0:00 | 0:00 | 0:00 | 0:00 | 0:00 | 0:00 | 0:00 | 0:00 | 1:00 | 0:00 |
| NM_001160114:NR_027650 | 0:00 | 0:00 | 0:00 | 0:00 | 0:00 | 0:00 | 0:00 | 0:00 | 1:00 | 0:00 |
| NM_014300:NM_001271920:NM_001271919:NM_001271921:NM_001271918:NR_073518:NM_001271922 | 213:2:9:4:5:1:0 | 0:0:0:0:0:0:0 | 233:0:9:2:12:3:0 | 2:0:0:0:0:0:0 | 223:2:10:1:19:1:0 | 0:0:0:0:0:0:0 | 1184:0:22:9:50:4:2 | 1:0:0:0:0:0:0 | 363:0:9:7:28:0:3 | 0:0:0:0:0:0:0 |
| NM_014300:NM_001271920:NM_001271919:NM_001271921:NM_001271918:NR_073518:NM_001271922 | 213:2:9:4:5:1:0 | 0:0:0:0:0:0:0 | 234:0:9:2:12:2:0 | 1:0:0:0:0:0:0 | 223:2:10:1:19:1:0 | 0:0:0:0:0:0:0 | 1184:0:21:9:49:4:2 | 1:0:1:0:1:0:0 | 363:0:9:7:28:0:3 | 0:0:0:0:0:0:0 |
| NM_014300:NM_001271920:NM_001271919:NM_001271921:NM_001271918:NR_073518:NM_001271922 | 214:2:9:4:5:1:0 | 0:0:0:0:0:0:0 | 235:0:9:2:12:2:0 | 0:0:0:0:0:0:0 | 223:2:10:1:19:1:0 | 0:0:0:0:0:0:0 | 1183:0:22:9:50:4:2 | 1:0:0:0:0:0:0 | 363:0:9:7:28:0:3 | 0:0:0:0:0:0:0 |
| NM_014300:NM_001271920:NM_001271921:NM_001271918:NR_073518:NM_001271922 | 214:2:4:5:2:0 | 0:0:0:0:0:0 | 235:0:2:12:1:0 | 0:0:0:0:0:0 | 223:2:1:19:0:0 | 0:0:0:0:0:0 | 1182:0:10:50:7:2 | 2:0:0:0:0:0 | 362:0:7:28:0:3 | 0:0:0:0:0:0 |
| NM_001271921:NM_014300:NM_001271920:NM_001271919:NM_001271918:NR_073518:NM_001271922 | 3:207:2:9:6:1:0 | 1:0:0:0:0:0:0 | 2:231:0:8:11:5:0 | 0:0:0:0:0:0:0 | 1:216:2:9:18:1:0 | 0:0:0:0:0:0:0 | 10:1170:0:23:47:17:2 | 0:1:0:0:0:0:0 | 7:356:0:9:27:1:3 | 0:0:0:0:0:0:0 |
| NM_007200 | 2 | 0 | 4 | 0 | 5 | 0 | 6 | 0 | 0 | 0 |
| NM_007200 | 0 | 2 | 0 | 0 | 0 | 2 | 3 | 0 | 0 | 0 |
| NM_007200 | 0 | 2 | 0 | 0 | 0 | 2 | 2 | 0 | 0 | 0 |
| NM_007200 | 0 | 1 | 0 | 0 | 0 | 2 | 1 | 0 | 0 | 0 |
| NM_007200 | 1 | 0 | 0 | 0 | 1 | 2 | 1 | 0 | 0 | 0 |
| NM_007200:NM_001270546 | 0:00 | 2:00 | 0:00 | 5:01 | 0:00 | 3:00 | 4:00 | 0:00 | 1:00 | 0:00 |
| NM_007200:NM_001270546 | 0:00 | 2:00 | 0:00 | 5:00 | 0:00 | 4:00 | 4:00 | 0:00 | 1:00 | 0:00 |
| NM_007200:NM_001270546:NM_006738 | 0:00:00 | 2:00:00 | 0:00:00 | 8:01:01 | 0:00:00 | 6:00:00 | 5:00:00 | 0:00:00 | 1:00:00 | 0:00:00 |
| NM_007200:NM_001270546:NM_006738 | 0:00:00 | 2:00:00 | 0:00:00 | 9:01:01 | 0:00:00 | 6:00:00 | 5:00:00 | 0:00:00 | 1:00:00 | 0:00:00 |
| NM_007200:NM_001270546:NM_006738 | 0:00:00 | 2:00:00 | 4:00:01 | 6:01:00 | 5:00:00 | 1:00:00 | 6:00:00 | 0:00:00 | 1:00:00 | 0:00:00 |
| NM_007200:NM_001270546:NM_006738 | 0:00:00 | 2:00:00 | 0:00:00 | 12:02:01 | 0:00:00 | 8:00:00 | 9:00:00 | 0:00:00 | 1:00:00 | 0:00:00 |
| NM_001270546:NM_007200:NM_006738 | 2:08:00 | 4:00:00 | 32:54:04 | 0:04:02 | 10:12:00 | 2:06:00 | 4:50:04 | 2:06:02 | 2:06:02 | 0:00:00 |
| NM_007200:NM_001270546 | 2:00 | 0:00 | 1:00 | 3:00 | 3:00 | 4:01 | 3:00 | 0:00 | 0:00 | 0:00 |
| NM_001150:XM_011521473 | 1:00 | 0:00 | 24:25:00 | 26:39:00 | 84:51:00 | 101:55:00 | 5:04 | 0:00 | 1:01 | 0:00 |
| NM_001150:XM_011521473 | 0:00 | 1:00 | 12:25 | 17:16 | 75:42:00 | 56:34:00 | 2:02 | 0:00 | 1:00 | 0:00 |
| NM_001004309:XR_001751249:XM_017022127:XM_017022128:XM_017022129 | 0:0:0:0:0 | 0:0:0:0:0 | 2:0:0:0:0 | 0:0:0:0:0 | 2:0:0:0:0 | 0:0:0:0:0 | 2:0:0:2:0 | 8:2:2:0:0 | 2:0:0:0:0 | 0:2:0:0:0 |
| XR_001751409:NM_003870 | 2:02 | 0:00 | 2:00 | 0:00 | 8:08 | 0:02 | 14:08 | 0:02 | 16:04 | 2:02 |
| XM_011522187:XM_006720760:XM_005254987:XM_006720759:XM_011522188:XM_017022716:NM_199413:NM_001267580:NM_003981:XM_017022717:XM_017022713:XM_017022715:XM_011522191 | 1:4:4:1:1:0:0:0:0:0:0:0:0 | 0:0:0:0:0:0:0:0:0:0:0:0:0 | 0:0:2:0:0:2:0:0:0:0:0:0:0 | 0:0:0:0:0:0:0:0:0:0:0:0:0 | 0:1:3:0:1:0:2:0:0:0:0:0:0 | 0:0:0:0:0:0:0:0:0:0:0:0:0 | 0:4:10:1:2:6:12:10:3:18:1:4:1 | 0:0:0:0:0:0:1:1:0:0:0:0:0 | 0:2:12:2:1:2:17:5:8:2:3:1:0 | 0:0:0:0:0:0:0:0:0:0:0:0:0 |
| XM_011522187:XM_006720760:XM_005254987:XM_006720759:XM_011522188:XM_017022716:NM_199413:XM_017022717:NM_001267580:NM_003981:XM_017022713:XM_011522191:XM_017022715 | 1:3:4:1:1:0:0:0:0:0:0:0:0 | 0:0:0:0:0:0:0:0:0:0:0:0:0 | 0:0:2:0:0:2:0:0:0:0:0:0:0 | 0:0:0:0:0:0:0:0:0:0:0:0:0 | 0:1:3:0:1:0:2:0:0:0:0:0:0 | 0:0:0:0:0:0:0:0:0:0:0:0:0 | 0:4:10:1:2:6:14:13:10:3:1:1:1 | 0:0:0:0:0:0:0:1:0:0:0:0:0 | 0:2:12:2:0:1:17:2:5:7:3:0:1 | 0:0:0:0:0:0:0:0:0:0:0:0:0 |
| XM_011522187:XM_006720760:XM_005254987:XM_006720759:NM_199413:XM_011522188:NM_001267580:NM_003981:XM_017022717:XM_017022713:XM_011522191 | 0:0:0:0:0:0:0:0:0:0:0 | 2:6:8:2:0:0:0:0:0:0:0 | 0:0:0:0:0:0:0:0:0:0:0 | 0:0:4:0:0:0:0:0:0:0:0 | 0:0:0:0:0:0:0:0:0:0:0 | 0:0:6:0:4:2:0:0:0:0:0 | 0:0:0:0:0:0:0:0:0:0:0 | 0:8:20:2:34:2:10:6:6:2:2 | 0:0:0:0:0:0:0:0:0:0:0 | 0:2:24:4:34:0:6:12:0:6:0 |

| | | | | | | | | | | |
|---|---|---|---|---|---|---|---|---|---|---|
| NM_001271:NM_001042572 | 16:20 | 0:00 | 24:04:00 | 10:00 | 12:20 | 2:00 | 72:40:00 | 30:00:00 | 18:18 | 2:00 |
| NM_001271:NM_001042572 | 4:10 | 1:00 | 0:03 | 3:00 | 5:10 | 0:00 | 11:20 | 5:00 | 6:10 | 1:00 |
| NM_002484:NM_001323595:NM_001278506:XM_017023252 | 15:2:0:0 | 0:0:0:0 | 12:2:0:0 | 0:0:0:0 | 11:0:2:0 | 0:0:0:0 | 30:2:2:1 | 26:2:0:0 | 10:0:0:0 | 10:0:0:0 |
| NM_001323595:NM_002484:NM_001278506:XM_017023252:NM_001323596 | 1:12:0:0:0 | 1:3:0:0:0 | 2:12:0:0:0 | 0:0:0:0:0 | 0:13:2:0:0 | 0:0:0:0:0 | 6:58:2:1:0 | 0:0:0:0:0 | 0:20:0:0:0 | 0:0:0:0:0 |
| XM_017023268:NM_015914:NR_136672 | 0:00:00 | 0:00:00 | 0:00:00 | 1:00:00 | 0:00:00 | 1:00:00 | 0:00:00 | 0:01:01 | 0:00:00 | 0:00:00 |
| NM_015659 | 0 | 0 | 0 | 0 | 0 | 0 | 1 | 3 | 0 | 0 |
| NM_002094:NM_001130006 | 0:00 | 0:00 | 0:00 | 0:00 | 0:00 | 0:00 | 0:00 | 0:00 | 0:00 | 1:00 |
| NM_001192 | 0 | 9 | 0 | 21 | 0 | 46 | 0 | 0 | 0 | 2 |
| NM_032167:XR_001752024:XM_017023877:XM_017023874:XM_011522744 | 0:0:0:0:0 | 0:0:0:0:0 | 1:1:0:0:0 | 0:0:0:0:0 | 0:0:1:1:0 | 0:0:0:0:0 | 1:0:1:0:0 | 0:0:1:0:0 | 0:0:0:0:0 | 0:1:0:0:0 |
| NM_001287:XM_011522354 | 1:00 | 0:00 | 0:00 | 0:00 | 1:05 | 0:01 | 5:03 | 2:02 | 0:00 | 0:00 |
| NM_173474:NM_001270766 | 0:00 | 0:00 | 0:00 | 0:00 | 1:00 | 0:00 | 0:00 | 0:00 | 1:01 | 0:00 |
| NM_173474:NM_001270766 | 0:00 | 0:00 | 0:00 | 0:00 | 1:00 | 0:00 | 0:00 | 0:00 | 1:01 | 0:00 |
| NM_002474:XM_017023250:NM_022844 | 0:01:00 | 1:00:00 | 0:02:00 | 0:00:00 | 1:03:02 | 6:01:03 | 1:00:00 | 0:00:00 | 0:01:00 | 0:00:00 |
| XM_024450135:NM_012225:XM_011522338 | 0:00:00 | 0:00:00 | 0:00:00 | 1:00:00 | 0:00:00 | 0:00:00 | 0:00:00 | 2:00:00 | 0:00:00 | 1:00:00 |
| XM_024450135:NM_012225:XM_011522338 | 0:00:00 | 0:00:00 | 0:00:00 | 0:00:00 | 0:00:00 | 0:00:00 | 0:00:00 | 2:00:00 | 0:00:00 | 1:00:00 |
| NM_001030009:NM_001019 | 48:00:00 | 1:00 | 14:00 | 0:00 | 36:02:00 | 0:00 | 154:18:00 | 0:01 | 114:09:00 | 1:00 |
| NM_001305122 | 0 | 0 | 0 | 0 | 0 | 0 | 0 | 0 | 0 | 0 |
| NM_001099456 | 0 | 0 | 0 | 0 | 0 | 0 | 0 | 0 | 0 | 0 |
| NM_001099456 | 0 | 0 | 0 | 0 | 0 | 0 | 0 | 0 | 0 | 0 |
| NM_001318193:NM_002528 | 1:07 | 0:00 | 0:02 | 0:00 | 0:20 | 0:00 | 3:34 | 0:02 | 0:23 | 0:00 |
| XM_017023440:XM_017023438:XM_017023437:NM_001320226:NM_001330420:NM_201412:XM_005255429:XM_005255427:XM_011522561:NM_018032:XM_024450344 | 6:3:6:2:1:3:1:0:0:0:0 | 0:0:0:0:0:0:0:0:0:0:0 | 10:15:21:0:2:8:3:4:1:3:1 | 1:0:5:0:0:0:0:0:0:0:0 | 11:10:19:1:4:21:2:3:1:5:0 | 0:0:0:0:0:0:0:0:0:0:0 | 19:28:14:6:7:20:14:2:2:5:0 | 1:0:1:0:0:2:0:1:0:0:1 | 7:6:6:2:6:12:2:4:1:0:1 | 0:0:1:0:0:1:0:0:0:0:0 |
| XM_017023440:XM_017023438:XM_017023437:NM_001320226:NM_001330420:NM_201412:XM_005255429:XM_005255427:XM_011522561:NM_018032:XM_024450344 | 6:3:6:2:1:3:1:0:0:0:0 | 0:0:0:0:0:0:0:0:0:0:0 | 10:15:21:0:2:8:3:4:1:3:1 | 1:0:6:0:0:0:0:0:0:0:0 | 11:10:19:1:4:21:2:3:1:5:0 | 0:0:0:0:0:0:0:0:0:0:0 | 19:27:14:6:7:20:14:2:2:5:0 | 1:1:1:0:0:2:0:1:0:0:1 | 7:6:6:2:6:12:2:4:1:0:1 | 0:0:1:0:0:1:0:0:0:0:0 |
| XM_017023440:XM_017023438:XM_017023437:NM_001320226:NM_001330420:NM_201412:XM_005255429:XM_005255427:XM_011522561:NM_018032:XM_024450344 | 6:3:6:2:1:3:1:0:0:0:0 | 0:0:0:0:0:0:0:0:0:0:0 | 10:15:22:0:2:8:3:4:1:3:1 | 1:0:5:0:0:0:0:0:0:0:0 | 11:10:19:1:4:21:2:3:1:5:0 | 0:0:0:0:0:0:0:0:0:0:0 | 19:27:14:6:7:21:14:3:2:5:1 | 1:0:1:0:0:1:0:0:0:0:0 | 7:6:6:2:6:12:2:4:1:0:1 | 0:0:1:0:0:1:0:0:0:0:0 |
| NM_020422:NM_001301771 | 0:00 | 0:00 | 0:00 | 0:00 | 0:00 | 2:00 | 0:00 | 0:00 | 1:00 | 0:00 |
| XM_024450289 | 0 | 0 | 0 | 0 | 0 | 0 | 0 | 0 | 0 | 0 |
| XM_011545822:XM_011545823:NM_001169 | 0:00:00 | 0:00:00 | 9:00:00 | 2:00:00 | 3:00:03 | 8:01:02 | 0:00:00 | 0:00:00 | 0:00:00 | 0:00:00 |
| NM_145080:XM_006721023:XM_017023042 | 0:00:00 | 0:00:00 | 1:00:00 | 0:00:00 | 1:00:00 | 0:00:00 | 1:00:00 | 0:00:00 | 2:01:00 | 0:00:00 |
| NM_145080:XM_006721023 | 0:00 | 0:00 | 1:00 | 0:00 | 1:00 | 0:00 | 1:00 | 0:00 | 1:00 | 1:01 |
| NM_001042432:NM_001286110:NM_000086 | 1:00:00 | 0:00:00 | 0:01:00 | 0:00:00 | 1:01:01 | 0:00:00 | 1:00:00 | 1:00:00 | 0:00:00 | 2:00:00 |
| XM_017023612:XM_024450411:NM_001055 | 0:00:00 | 0:00:00 | 0:00:00 | 1:00:00 | 0:00:00 | 4:01:00 | 0:00:00 | 0:00:00 | 0:00:00 | 0:00:00 |
| NM_014298:XM_017023101 | 0:00 | 0:00 | 0:00 | 0:00 | 1:00 | 0:00 | 3:00 | 1:00 | 1:00 | 0:02 |
| NM_014298:NM_001318249:XM_005255223 | 0:00:00 | 0:00:00 | 0:00:00 | 0:00:00 | 0:00:00 | 1:00:00 | 0:00:00 | 5:01:01 | 0:00:00 | 1:00:00 |
| NM_152338:XM_011545921 | 0:00 | 0:00 | 0:00 | 37:00:00 | 0:00 | 146:01:00 | 0:00 | 0:00 | 0:00 | 6:00 |
| NM_152338:XM_011545921 | 0:00 | 0:00 | 0:00 | 39:00:00 | 0:00 | 151:01:00 | 0:00 | 0:00 | 0:00 | 6:00 |
| NM_184041:NM_184043:NM_001127617:NM_001243177 | 1776:29:3:3 | 1:0:0:0 | 1622:20:1:2 | 1:1:0:0 | 3393:36:3:6 | 4:0:0:0 | 4646:74:9:19 | 4:0:0:0 | 2371:33:6:6 | 1:0:0:0 |
| NM_184041:NM_184043:NM_001127617:NM_001243177 | 1775:32:3:3 | 0:0:0:0 | 1618:21:1:2 | 1:2:0:0 | 3390:44:3:6 | 2:0:0:0 | 4635:84:9:19 | 9:0:0:0 | 2369:38:6:6 | 2:0:0:0 |
| NM_184041:NM_184043:NM_001127617:NM_001243177 | 1770:33:3:3 | 2:1:0:0 | 1614:21:1:2 | 0:2:0:0 | 3387:46:3:6 | 1:0:0:0 | 4636:82:9:19 | 1:0:0:0 | 2368:38:6:6 | 2:0:0:0 |
| NM_184041:NM_184043:NM_001127617:NM_001243177 | 1769:42:3:3 | 1:0:0:0 | 1609:23:1:2 | 2:1:0:0 | 3379:52:3:6 | 5:0:0:0 | 4625:83:9:19 | 4:2:0:0 | 2365:40:6:6 | 3:0:0:0 |
| NM_184041:NM_184043:NM_001127617:NM_001243177 | 1763:48:3:3 | 1:0:0:0 | 1586:20:1:2 | 4:1:0:0 | 3361:48:3:6 | 4:0:0:0 | 4585:76:9:19 | 5:0:0:0 | 2361:46:6:6 | 4:0:0:0 |
| NM_184041:NM_184043:NM_001127617:NM_001243177 | 1761:49:3:3 | 1:0:0:0 | 1588:21:1:2 | 1:0:0:0 | 3359:54:3:6 | 1:0:0:0 | 4585:77:9:19 | 2:0:0:0 | 2364:46:6:6 | 0:0:0:0 |
| NM_184041:NM_184043:NM_001127617:NM_001243177 | 1758:50:3:3 | 2:0:0:0 | 1583:22:1:2 | 4:0:0:0 | 3350:54:3:6 | 6:0:0:0 | 4579:76:9:19 | 3:0:0:0 | 2362:50:6:6 | 1:0:0:0 |
| NM_184041:NM_184043:NM_001127617:NM_001243177 | 1758:50:3:3 | 2:0:0:0 | 1582:22:1:2 | 0:0:0:0 | 3351:54:3:6 | 2:0:0:0 | 4575:76:9:19 | 5:0:0:0 | 2360:50:6:6 | 2:0:0:0 |
| NM_184041:NM_184043:NM_001127617:NM_001243177 | 1755:57:3:3 | 0:0:0:0 | 1573:23:1:2 | 1:0:0:0 | 3344:72:3:6 | 4:0:0:0 | 4560:91:9:18 | 5:0:0:0 | 2358:62:6:6 | 1:0:0:0 |
| NM_184041:NM_184043:NM_001127617:NM_001243177 | 1755:58:3:3 | 0:0:0:0 | 1573:24:1:2 | 1:0:0:0 | 3344:72:3:6 | 4:0:0:0 | 4558:91:9:18 | 7:0:0:0 | 2359:63:6:6 | 0:0:0:0 |
| NM_184041:NM_184043:NM_001127617:NM_001243177 | 1753:59:3:3 | 2:0:0:0 | 1571:24:1:2 | 3:0:0:0 | 3339:73:3:6 | 9:0:0:0 | 4558:92:9:18 | 5:0:0:0 | 2358:63:6:6 | 1:0:0:0 |
| NM_184041:NM_184043:NM_001127617:NM_001243177 | 1751:64:3:3 | 2:0:0:0 | 1572:25:1:2 | 1:0:0:0 | 3344:84:3:6 | 4:0:0:0 | 4560:100:9:17 | 3:0:0:1 | 2358:66:6:6 | 1:1:0:0 |
| NM_184041:NM_184043:NM_001127617:NM_001243177 | 1750:63:3:3 | 3:1:0:0 | 1572:25:1:2 | 0:0:0:0 | 3344:84:3:6 | 4:0:0:0 | 4562:100:9:18 | 1:0:0:0 | 2358:66:6:6 | 1:1:0:0 |
| NM_184043:NM_184041:NM_001127617:NM_001243177 | 66:1753:3:3 | 1:0:0:0 | 26:1571:1:2 | 0:0:0:0 | 86:3344:3:6 | 1:3:0:0 | 101:4560:9:18 | 0:1:0:0 | 69:2356:6:6 | 1:0:0:0 |
| NM_184041:NM_184043:NM_001127617:NM_001243177 | 1752:68:3:3 | 1:0:0:0 | 1570:26:1:2 | 1:0:0:0 | 3346:87:3:6 | 2:4:0:0 | 4559:101:9:18 | 1:4:0:0 | 2357:70:6:6 | 1:1:0:0 |
| NM_184041:NM_184043:NM_001127617:NM_001243177 | 1749:69:3:3 | 1:0:0:0 | 1569:27:1:2 | 1:0:0:0 | 3342:91:3:6 | 3:0:0:0 | 4557:105:9:18 | 3:0:0:0 | 2357:73:6:6 | 1:1:0:0 |
| NM_184041:NM_184043:NM_001127617:NM_001243177 | 1747:69:3:3 | 3:0:0:0 | 1568:27:1:2 | 1:0:0:0 | 3334:91:3:6 | 5:0:0:0 | 4551:105:9:18 | 2:0:0:0 | 2357:72:6:6 | 1:2:0:0 |
| NM_184041:NM_184043:NM_001127617:NM_001243177 | 1751:69:3:3 | 1:0:0:0 | 1568:27:1:2 | 1:0:0:0 | 3336:97:3:6 | 6:1:0:0 | 4552:113:9:18 | 4:0:0:0 | 2357:75:6:6 | 2:1:0:0 |

| | | | | | | | | | | |
|---|---|---|---|---|---|---|---|---|---|---|
| NM_184041:NM_184043:NM_001127617:NM_001243177 | 1750:79:3:3 | 1:0:0:0 | 1560:28:1:2 | 2:0:0:0 | 3333:103:3:5 | 5:1:0:1 | 4540:125:9:18 | 5:1:0:0 | 2354:81:6:6 | 2:1:0:0 |
| NM_184041:NM_184043:NM_001127617:NM_001243177 | 1745:81:3:3 | 3:2:0:0 | 1561:30:1:2 | 2:0:0:0 | 3331:113:3:6 | 4:0:0:0 | 4531:131:9:18 | 5:2:0:0 | 2350:92:6:6 | 0:0:0:0 |
| NM_184043:NM_184041:NM_001127617:NM_001243177 | 89:1725:3:3 | 2:0:0:0 | 42:1520:1:2 | 0:1:0:0 | 132:3304:3:6 | 1:4:0:0 | 146:4491:9:18 | 1:4:0:0 | 121:2341:6:6 | 0:2:0:0 |
| NM_184041:NM_184043:NM_001127617:NM_001243177 | 1726:96:3:3 | 1:1:0:0 | 1508:54:1:2 | 4:0:0:0 | 3308:153:3:6 | 3:0:0:0 | 4484:173:9:18 | 1:0:0:0 | 2341:137:6:6 | 1:0:0:0 |
| NM_184041:NM_184043:NM_001127617:NM_001243177 | 1729:101:3:3 | 3:0:0:0 | 1504:73:1:2 | 9:0:0:0 | 3303:160:3:6 | 7:1:0:0 | 4475:183:9:18 | 7:2:0:0 | 2342:146:6:6 | 0:0:0:0 |
| NM_184041:NM_184043:NM_001127617:NM_001243177 | 1727:105:3:3 | 4:0:0:0 | 1513:81:1:2 | 4:0:0:0 | 3304:170:3:6 | 9:0:0:0 | 4478:189:9:18 | 3:0:0:0 | 2341:150:6:6 | 2:1:0:0 |
| NM_184041:NM_184043:NM_001127617:NM_001243177 | 1726:108:3:3 | 1:0:0:0 | 1511:95:1:2 | 6:0:0:0 | 3304:172:3:6 | 4:0:0:0 | 4476:190:9:18 | 4:0:0:0 | 2339:156:6:6 | 3:1:0:0 |
| NR_028388:NM_004960:NM_001170634:XM_024450221:XM_011545782:XM_005255233:XM_011545781:NM_001170937 | 8:27:14:6:3:1:0:0 | 1:0:0:0:0:0:0:0 | 16:36:17:12:0:0:0:0 | 0:0:0:0:0:0:0:0 | 4:61:25:4:0:0:1:0 | 0:0:0:0:0:0:0:0 | 63:198:148:40:1:2:7:0 | 0:2:0:0:0:0:0:0 | 5:53:28:8:2:0:0:0 | 0:0:0:0:0:0:0:0 |
| NM_004960:NM_001170634:NR_028388:XM_024450221:XM_011545782:XM_005255233:XM_011545781:NM_001170937 | 26:14:10:6:3:1:0:0 | 0:0:0:0:0:0:0:0 | 37:17:17:11:0:0:0:0 | 0:0:0:0:0:0:0:0 | 60:25:4:5:0:0:1:0 | 0:0:0:0:0:0:0:0 | 202:150:65:38:1:4:7:0 | 3:0:0:1:0:0:0:0 | 53:28:5:9:2:0:0:0 | 0:0:0:0:0:0:0:0 |
| NM_004960:NM_001170634:NR_028388:XM_024450221:XM_011545782:XM_005255233:XM_011545781:NM_001170937 | 26:14:10:6:3:1:0:0 | 0:0:0:0:0:0:0:0 | 37:17:17:11:0:0:0:0 | 0:0:0:0:0:0:0:0 | 60:25:4:6:0:0:1:0 | 0:0:0:0:0:0:0:0 | 202:150:64:37:1:4:7:0 | 3:0:1:1:0:0:0:0 | 52:28:5:9:2:0:0:0 | 1:0:0:0:0:0:0:0 |
| NM_004960:NM_001170634:NR_028388:XM_024450221:XM_011545782:XM_005255233:XM_011545781:NM_001170937 | 29:14:10:5:3:1:0:0 | 0:0:0:0:0:0:0:0 | 41:17:17:9:0:1:0:0 | 0:0:0:0:0:0:0:0 | 61:25:4:7:0:0:0:0 | 0:0:0:0:0:0:0:0 | 222:149:65:33:1:4:3:0 | 0:0:0:3:0:0:0:0 | 55:28:5:9:2:1:0:0 | 0:0:0:0:0:0:0:0 |
| NM_001170634:NM_004960:NR_028388:XM_024450221:XM_011545782:XM_005255233:NM_001170937:XM_011545781 | 13:29:9:2:1:1:0:0 | 1:0:0:0:0:0:0:0 | 16:41:15:5:0:1:0:0 | 0:0:0:3:0:0:0:0 | 25:61:3:4:0:0:0:0 | 0:0:0:0:0:0:0:0 | 145:223:52:20:0:4:0:0 | 1:0:2:3:0:0:0:0 | 26:55:4:7:0:1:0:0 | 1:0:0:0:0:0:0:0 |
| NM_001170634:NM_004960:XM_024450221:NR_028388:XM_005255233:XM_011545781:XM_011545782:NM_001170937 | 12:29:3:7:1:0:0:0 | 2:0:0:0:0:0:0:0 | 16:41:3:8:1:0:0:0 | 1:0:1:0:0:0:0:0 | 25:60:2:1:0:0:0:0 | 0:0:0:1:0:0:0:0 | 131:223:13:23:3:5:0:0 | 2:0:2:7:0:0:0:0 | 24:55:3:1:1:0:0:0 | 2:0:0:0:0:0:0:0 |
| NM_004960:NM_001170634:XM_024450221:XM_011545782:NR_028388:XM_005255233 | 29:12:9:3:5:0 | 0:0:0:0:0:0 | 16:6:24:1:0:1 | 0:0:0:0:0:0 | 60:24:11:0:0:0 | 0:0:0:0:0:0 | 122:59:93:2:3:0 | 0:1:0:0:0:0 | 54:24:17:3:1:1 | 0:0:0:0:0:0 |
| NM_004960:NM_001170634:XM_024450221:XM_011545782:NR_028388:XM_005255233:XM_011545781 | 28:12:11:3:6:0:0 | 0:0:0:0:0:0:0 | 16:6:25:1:3:1:0 | 0:0:0:0:0:0:0 | 60:23:11:0:2:0:1 | 0:1:0:0:0:0:0 | 119:65:91:2:11:0:0 | 0:0:0:0:0:0:0 | 54:23:17:3:2:0:0 | 0:0:0:0:0:0:0 |
| NM_004960:NM_001170634:XM_024450221:XM_011545782:NR_028388:XM_005255233:XM_011545781 | 29:12:10:3:8:0:0 | 0:0:0:0:0:0:0 | 16:6:25:1:6:1:0 | 0:0:0:0:0:0:0 | 60:24:10:0:2:0:1 | 0:0:0:0:0:0:0 | 123:70:90:2:28:1:0 | 1:1:0:0:0:0:0 | 54:26:17:3:4:1:0 | 0:0:0:0:0:0:0 |
| XR_001751996:NM_181050:NR_134879:XM_017023748:NM_003502:XM_017023746 | 0:0:0:0:0:0 | 1:0:0:0:0:0 | 2:1:0:0:0:0 | 0:0:0:0:0:0 | 1:0:1:1:0:0 | 0:0:0:0:0:0 | 6:0:2:0:1:3 | 1:0:0:0:0:0 | 2:0:1:0:0:0 | 0:0:0:0:0:0 |
| NM_024845:NM_001083600:NM_001317098 | 0:00:00 | 0:00:00 | 0:00:00 | 0:00:00 | 1:00:00 | 0:00:00 | 0:00:00 | 0:00:00 | 0:00:00 | 0:00:00 |
| XM_011522393:NM_005223:XM_024450171:XM_024450170:XR_002957783:XM_017022993:NM_001351825:XM_017023001 | 1:2:1:0:0:0:0:0 | 3:0:0:0:0:0:0:0 | 1:0:0:0:0:0:0:0 | 0:0:0:1:0:0:0:0 | 0:0:0:0:0:0:0:0 | 1:0:0:0:0:0:0:0 | 5:13:1:3:2:1:1:0 | 0:0:0:0:0:0:0:0 | 0:0:0:0:1:0:0:0 | 0:0:0:0:0:0:0:0 |
| XM_011522393:NM_005223:XM_024450171:XM_024450170:XR_002957783:XM_017022993:XR_002957785:XM_024450172:NM_001351825:XM_017023001 | 1:2:1:0:0:0:0:0:0:0 | 3:0:0:0:0:0:0:0:0:0 | 1:0:0:0:0:0:0:0:0:0 | 0:0:0:1:0:0:0:0:0:0 | 0:0:0:0:0:0:0:0:0:0 | 1:0:0:0:0:0:0:0:0:0 | 3:11:1:1:1:0:1:0:0:0 | 2:2:0:1:1:1:0:0:0:0 | 0:0:0:0:1:0:0:1:0:0 | 0:0:0:0:0:0:0:0:0:0 |
| NM_016292:XM_011522345:NM_001272049 | 3:01:00 | 0:00:00 | 3:01:00 | 0:00:00 | 6:01:01 | 0:00:00 | 21:06:02 | 23:05:02 | 8:00:01 | 7:00:01 |
| NM_016292:XM_011522345:NM_001272049 | 0:00:00 | 2:00:00 | 1:00:00 | 2:00:00 | 1:00:00 | 3:01:00 | 15:01:02 | 17:03:01 | 6:00:00 | 4:00:00 |
| NM_016292:XM_011522345:NM_001272049 | 2:00:00 | 0:00:00 | 3:00:00 | 0:00:00 | 4:01:00 | 0:00:00 | 15:01:01 | 17:03:01 | 6:00:00 | 4:00:00 |
| NM_016292:XM_011522345:NM_001272049 | 3:00:00 | 0:00:00 | 2:01:00 | 0:00:00 | 5:00:01 | 2:00:00 | 20:00:01 | 21:00:02 | 8:00:01 | 7:00:00 |
| NM_001135110 | 0 | 0 | 0 | 0 | 0 | 0 | 0 | 2 | 0 | 1 |
| XM_011523047:NM_001173984:NM_013263:XM_017023181:XM_011523050:XM_011523046:XR_933290:XR_001751896:XR_933291 | 1:5:23:3:2:0:0:0:0 | 0:0:0:0:0:0:0:0:0 | 0:9:6:0:1:0:0:0:0 | 0:0:0:0:0:0:0:0:0 | 1:9:16:1:1:2:0:0:0 | 0:0:0:0:0:0:0:0:0 | 23:36:113:35:12:15:1:0:0 | 0:0:3:1:0:0:0:0:0 | 1:9:27:6:0:0:0:0:0 | 0:0:0:0:0:0:0:0:0 |
| NM_175617 | 0 | 0 | 27 | 0 | 18 | 0 | 1 | 3 | 1 | 0 |
| NM_176870 | 0 | 0 | 3 | 8 | 2 | 2 | 0 | 0 | 0 | 0 |
| NM_014685:NM_001010989 | 0:00 | 0:00 | 0:00 | 2:00 | 1:00 | 1:00 | 4:00 | 0:00 | 1:01 | 0:00 |
| NM_002080 | 0 | 0 | 0 | 0 | 0 | 0 | 0 | 0 | 0 | 0 |
| NM_001797:XM_024450134:NM_001308392 | 0:00:00 | 3:01:00 | 0:00:00 | 0:03:00 | 0:00:00 | 0:01:00 | 0:00:00 | 0:02:00 | 0:00:00 | 0:00:00 |
| XM_024450134:NM_001797 | 0:02 | 1:01 | 0:00 | 0:00 | 1:00 | 0:00 | 1:00 | 1:00 | 0:00 | 0:00 |
| NM_003905:NM_001018159:NM_001018160:XM_005256215:NM_001286500 | 111:16:4:3:0 | 2:0:0:0:0 | 85:2:1:3:1 | 0:0:0:0:0 | 138:6:3:3:0 | 0:0:0:1:0 | 628:109:48:52:1 | 1:0:0:0:0 | 215:20:11:4:0 | 1:0:0:0:0 |
| NM_001018159:NM_003905:NM_001018160:XM_005256215:NM_001286500 | 16:113:4:3:0 | 0:0:0:0:0 | 2:85:1:3:1 | 0:0:0:0:0 | 6:138:3:4:0 | 0:0:0:0:0 | 109:629:49:48:1 | 0:3:0:0:0 | 21:217:11:3:0 | 0:0:0:0:0 |
| NM_001018159:NM_003905:NM_001018160:XM_005256215:NM_001286500 | 16:115:4:5:0 | 0:0:0:0:0 | 2:89:1:3:1 | 0:0:0:0:0 | 6:140:3:3:0 | 0:0:0:0:0 | 109:640:49:44:1 | 0:1:0:0:0 | 21:219:11:3:0 | 0:0:0:0:0 |
| NM_003905:NM_001018159:NM_001018160:XM_005256215:NM_001286500 | 118:16:4:4:0 | 1:0:0:0:0 | 98:2:1:4:1 | 2:0:0:0:0 | 145:6:3:3:0 | 0:0:0:0:0 | 682:113:54:44:1 | 22:1:3:2:0 | 225:22:12:3:0 | 4:0:0:0:0 |
| NM_001018159:NM_003905:XM_005256215:NM_001286500 | 16:122:2:0 | 0:0:0:0 | 4:108:1:1 | 0:0:0:0 | 7:146:1:0 | 0:0:0:0 | 121:751:23:1 | 0:0:0:0 | 22:229:3:0 | 0:0:0:0 |
| NM_001018159:NM_003905:XM_005256215:NM_001286500 | 16:120:2:0 | 0:0:0:0 | 4:108:1:1 | 0:0:0:0 | 7:145:1:0 | 0:1:0:0 | 120:744:23:1 | 1:1:0:0 | 22:228:3:0 | 0:0:0:0 |
| NM_003869:NM_198061 | 34:04:00 | 0:00 | 57:31:00 | 0:00 | 327:69 | 1:00 | 12:08 | 0:00 | 28:04:00 | 0:00 |
| NM_024712 | 0 | 0 | 0 | 0 | 0 | 1 | 0 | 0 | 0 | 0 |
| XM_005255775:NM_006565:XM_017022868 | 2:11:01 | 1:00:00 | 6:03:00 | 6:01:00 | 12:07:01 | 2:01:00 | 25:09:05 | 16:02:00 | 1:08:02 | 2:00:00 |
| NM_006565:XM_005255775:XM_017022868 | 11:02:01 | 0:00:00 | 3:05:00 | 1:07:00 | 7:10:01 | 0:01:00 | 9:21:05 | 1:16:00 | 8:01:02 | 0:00:00 |
| XM_005255775:NM_006565:XM_017022868 | 1:10:01 | 1:00:00 | 4:03:00 | 1:00:00 | 8:08:01 | 2:00:00 | 19:10:05 | 0:00:00 | 1:09:02 | 0:00:00 |
| XM_005256115:NM_001082486:XR_429728:NM_022914 | 0:0:0:0 | 0:0:0:0 | 0:0:0:0 | 0:0:0:0 | 0:0:0:0 | 0:0:0:0 | 1:0:0:0 | 0:0:0:0 | 0:0:1:0 | 0:1:0:0 |
| NM_032259:XM_011522699 | 0:00 | 0:00 | 1:01 | 0:00 | 0:01 | 0:00 | 1:02 | 0:00 | 1:00 | 0:00 |
| NM_004360:NM_001317186 | 19:00 | 1:00 | 23:00 | 0:00 | 82:00:00 | 0:00 | 30:01:00 | 34:00:00 | 5:00 | 6:00 |
| NM_004360:NM_001317186 | 33:00:00 | 0:00 | 21:00 | 0:00 | 75:01:00 | 2:00 | 42:01:00 | 0:00 | 20:01 | 0:00 |
| NM_022341 | 0 | 0 | 0 | 0 | 0 | 0 | 0 | 0 | 0 | 2 |
| NM_000903:NM_001025433 | 3:00 | 0:00 | 0:00 | 2:00 | 2:00 | 1:00 | 18:00 | 0:00 | 11:00 | 0:00 |
| XM_024450224:NM_015386:XR_933267:NM_001195139 | 0:0:0:0 | 3:10:0:0 | 0:0:0:0 | 0:9:0:0 | 0:0:0:0 | 2:15:0:0 | 0:0:0:0 | 5:29:7:0 | 0:0:0:0 | 5:25:4:0 |
| NM_012426 | 15 | 0 | 6 | 0 | 6 | 0 | 62 | 0 | 13 | 0 |
| NM_001017967:NM_052858 | 0:00 | 0:00 | 0:00 | 0:00 | 1:00 | 0:00 | 0:00 | 0:00 | 0:00 | 0:00 |
| NM_018124:XM_017023391 | 0:00 | 0:00 | 1:00 | 0:00 | 1:00 | 2:00 | 1:02 | 0:00 | 3:00 | 0:00 |
| XM_017023391:NM_018124:XM_011523191:XM_006721228 | 0:0:0:0 | 1:0:0:0 | 0:0:0:0 | 0:1:0:0 | 0:0:0:0 | 0:3:0:0 | 0:1:0:0 | 0:0:0:0 | 0:0:0:0 | 0:3:1:0 |
| NM_005823:NM_013404 | 0:00 | 0:00 | 0:00 | 0:00 | 0:00 | 0:00 | 3:00 | 0:00 | 0:00 | 0:00 |
| NM_018455:XM_017023456 | 0:00 | 0:00 | 0:00 | 0:00 | 0:00 | 0:00 | 0:00 | 0:00 | 0:00 | 1:01 |
| NM_004483 | 0 | 0 | 0 | 0 | 0 | 0 | 0 | 2 | 0 | 5 |

| | | | | | | | | | | |
|---|---|---|---|---|---|---|---|---|---|---|
| NM_002153:XR_001751898 | 0:00 | 0:00 | 0:00 | 2:01 | 4:00 | 3:00 | 0:00 | 0:00 | 0:00 | 0:00 |
| NM_005153:XM_017023869:XM_006721332:NM_001272075:XM_017023864 | 13:1:0:0:0 | 0:0:0:0:0 | 1:1:0:0:0 | 0:0:0:0:0 | 12:0:1:0:0 | 0:0:0:0:0 | 8:4:0:0:0 | 17:5:0:1:1 | 20:3:1:1:0 | 16:0:0:0:0 |
| NM_005153:XM_017023869:XM_006721332:NM_001272075:XM_017023864 | 0:0:0:0:0 | 13:1:0:0:0 | 0:0:0:0:0 | 1:2:0:0:0 | 0:0:0:0:0 | 12:0:1:0:0 | 25:9:0:1:1 | 0:0:0:0:0 | 36:3:1:1:0 | 0:0:0:0:0 |
| NM_005153:XM_017023869:XM_006721332:NM_001272075:XM_017023864 | 0:0:0:0:0 | 13:1:0:0:0 | 0:0:0:0:0 | 1:2:0:0:0 | 0:0:0:0:0 | 12:0:1:0:0 | 25:10:0:1:1 | 0:0:0:0:0 | 36:3:1:1:0 | 0:0:0:0:0 |
| NM_005153:NR_073577:XM_017023869:NR_073578:XM_006721332:NM_001272075:XM_017023864 | 13:3:2:0:0:0:0 | 0:0:0:0:0:0:0 | 4:1:7:2:0:0:0 | 0:0:0:0:0:0:0 | 16:1:5:0:1:0:0 | 0:0:0:0:0:0:0 | 30:9:91:17:0:1:1 | 1:0:0:0:0:0:0 | 45:12:8:1:1:1:0 | 0:0:0:0:0:0:0 |
| NM_001861:XM_024450156:NM_001318797:NM_001318794:NM_001318786 | 7:0:0:0:0 | 5:1:0:0:0 | 10:0:0:0:0 | 11:1:0:0:0 | 17:1:0:0:0 | 23:0:0:0:0 | 26:1:0:0:0 | 0:0:0:0:0 | 42:1:0:0:0 | 0:0:0:0:0 |
| NM_000101 | 2 | 1 | 3 | 2 | 6 | 4 | 5 | 2 | 5 | 5 |
| NM_000101 | 0 | 3 | 0 | 4 | 0 | 10 | 2 | 5 | 5 | 5 |
| XM_011522943:NM_001127214:NM_174917:NM_001243279:XR_933238 | 0:0:0:0:0 | 0:0:0:0:0 | 0:0:0:0:0 | 0:0:0:0:0 | 0:0:0:0:0 | 0:0:0:0:0 | 0:0:0:0:0 | 1:2:2:0:0 | 0:0:0:0:0 | 0:0:0:1:1 |
| XM_024450187:NM_174917:XR_933240:NR_147928:XM_024450186:NM_001127214:XM_017023019:NM_001243279:XR_933238 | 0:0:0:0:0:0:0:0:0 | 2:0:0:0:0:0:0:0:0 | 0:0:0:0:0:0:0:0:0 | 4:2:1:0:0:0:0:0:0 | 0:0:0:0:0:0:0:0:0 | 5:0:0:1:1:0:0:0:0 | 0:0:0:0:0:0:0:0:0 | 3:5:1:0:0:2:1:0:0 | 0:0:0:0:0:0:0:0:0 | 2:0:0:0:0:0:1:1:1 |
| XM_024450244:XM_011523057 | 1:00 | 0:00 | 0:00 | 0:00 | 0:00 | 0:00 | 0:00 | 2:00 | 0:00 | 0:01 |
| NM_000977:NM_033251 | 25:04:00 | 16:00 | 15:00 | 15:02 | 10:02 | 14:03 | 142:18:00 | 0:00 | 97:09:00 | 0:00 |
| XM_017023009:XM_005256285:XM_024450173:XM_011522925:NM_004413 | 0:0:0:0:0 | 1:0:0:0:0 | 0:0:0:0:0 | 0:0:0:0:0 | 0:0:0:0:0 | 0:0:0:0:0 | 0:1:3:0:1 | 0:4:3:3:3 | 0:0:0:1:0 | 1:2:0:0:1 |
| NR_046418:NM_002768:NM_001083314 | 22:31:02 | 1:00:00 | 44:56:01 | 0:00:00 | 67:144:11 | 1:00:01 | 90:114:5 | 0:01:00 | 13:58:01 | 0:00:00 |
| NR_046418:NM_002768:NM_001083314:XM_011523098 | 29:32:2:0 | 0:0:0:0 | 56:58:1:0 | 0:0:0:0 | 88:148:11:0 | 1:0:1:0 | 122:122:5:1 | 0:0:0:0 | 17:59:1:0 | 0:0:0:0 |
| NR_046418:NM_002768:NM_001083314 | 30:32:02 | 0:00:00 | 70:55:01 | 0:00:00 | 111:144:12 | 0:00:00 | 156:114:4 | 1:00:01 | 21:57:01 | 0:00:00 |
| NR_024058:NM_006761:XM_017025005:XM_005256784 | 162:1720:5:0 | 1:4:0:0 | 210:1323:6:0 | 1:4:0:0 | 573:6002:25:0 | 3:30:2:0 | 1019:7198:72:1 | 3:30:9:0 | 202:2375:6:1 | 1:6:1:0 |
| NM_006761:NR_024058:XM_017025005:XM_005256784 | 1728:151:3:1 | 0:0:0:0 | 1328:195:0:4 | 1:0:0:0 | 6035:559:12:16 | 9:2:0:0 | 7244:964:12:16 | 6:6:0:0 | 2386:196:5:3 | 1:0:0:0 |
| NM_006761:NR_024058:XM_017025005:XM_005256784 | 1727:151:3:1 | 0:0:0:0 | 1329:193:0:4 | 0:0:0:0 | 6042:553:12:17 | 3:0:0:0 | 7246:955:12:17 | 4:2:0:0 | 2386:195:5:4 | 1:0:0:0 |
| NM_006761:NR_024058:XM_017025005:XM_005256784 | 1727:151:3:1 | 1:0:0:0 | 1328:192:1:6 | 2:0:0:0 | 6034:544:20:19 | 14:1:0:0 | 7241:945:15:20 | 7:0:0:0 | 2384:196:6:4 | 3:1:0:0 |
| NM_006761:NR_024058:XM_017025005:XM_005256784 | 1727:151:3:1 | 1:0:0:0 | 1330:184:1:5 | 0:0:0:1 | 6038:533:23:27 | 12:0:0:1 | 7241:923:15:19 | 8:0:1:1 | 2383:196:7:5 | 3:0:0:0 |
| NM_006761:NR_024058:XM_017025005:XM_005256784 | 1725:149:3:1 | 3:2:0:0 | 1330:183:1:6 | 0:0:0:0 | 6045:532:23:31 | 6:0:0:0 | 7240:910:16:20 | 9:2:0:0 | 2387:193:7:5 | 0:1:0:0 |
| NM_006761:NR_024058:XM_017025005:XM_005256784 | 1723:150:3:1 | 5:1:0:0 | 1330:180:1:5 | 0:3:0:1 | 6041:532:23:32 | 10:0:0:0 | 7245:908:17:19 | 4:1:0:1 | 2386:193:7:5 | 1:1:0:0 |
| NM_006761:NR_024058:XM_017025005:XM_005256784 | 1729:136:8:8 | 2:0:0:0 | 1328:147:2:12 | 2:0:0:0 | 6054:491:32:65 | 11:1:0:1 | 7244:733:20:25 | 10:0:0:0 | 2386:177:6:7 | 2:0:0:0 |
| NM_006761:NR_024058:XM_005256784:XM_017025005 | 1731:139:8:7 | 0:0:0:0 | 1329:150:12:2 | 1:0:1:0 | 6059:499:66:31 | 6:0:0:0 | 7252:723:25:19 | 2:1:0:0 | 2388:177:7:6 | 0:0:0:0 |
| NM_006761:NR_024058:XM_005256784:XM_017025005 | 1731:143:11:7 | 0:0:0:0 | 1330:146:16:2 | 2:0:18:0 | 6064:499:77:31 | 3:0:3:0 | 7255:675:28:17 | 1:2:2:0 | 2387:174:10:6 | 0:1:3:0 |
| NM_006761:XM_005256784:NR_024058:XM_017025005 | 1730:11:143:7 | 1:3:0:0 | 1330:28:145:2 | 5:13:0:1 | 6056:81:495:31 | 12:3:3:0 | 7249:29:675:18 | 11:5:1:1 | 2385:13:175:5 | 4:1:0:0 |
| NM_006761:NR_024058:XM_005256784:XM_017025005 | 1748:141:56:12 | 1:1:0:0 | 1455:142:505:45 | 2:1:2:0 | 6099:491:255:48 | 2:2:0:0 | 7440:660:357:53 | 4:2:0:0 | 2435:173:84:9 | 0:0:0:0 |
| NM_006761:XM_017025005:XM_005256784:NR_024058 | 1744:9:51:140 | 6:2:3:0 | 1449:42:481:142 | 4:3:21:1 | 6088:46:238:489 | 12:4:14:3 | 7423:50:312:652 | 22:7:40:6 | 2429:8:79:172 | 3:0:2:0 |
| NR_024058:NM_006761:XM_005256784:XM_017025005 | 137:1750:56:11 | 1:0:0:0 | 141:1453:507:45 | 1:5:1:0 | 488:6092:258:51 | 4:10:3:1 | 652:7432:356:55 | 3:13:3:1 | 171:2435:85:9 | 0:0:0:0 |
| NM_006761:NR_024058:XM_005256784:XM_017025005 | 1748:139:56:11 | 2:0:0:0 | 1456:141:506:44 | 2:1:2:1 | 6092:488:260:52 | 9:3:1:0 | 7442:651:356:56 | 3:4:2:0 | 2435:171:85:9 | 0:0:0:0 |
| NM_006761:NR_024058:XM_005256784:XM_017025005 | 1746:135:58:8 | 2:0:0:0 | 1461:131:549:54 | 3:0:0:0 | 6067:453:253:38 | 17:1:0:0 | 7484:588:471:62 | 7:2:0:1 | 2429:158:86:9 | 1:1:0:0 |
| NM_130766:NM_016532:XM_024450802:XM_005256683:XM_011523934:NM_001135642:XM_017024756:XM_005256686 | 2:0:0:0:0:0:0:0 | 0:0:0:0:0:0:0:0 | 0:6:0:0:0:0:0:0 | 0:0:0:0:0:0:0:0 | 2:7:1:2:2:0:0:0 | 0:1:1:0:0:0:0:0 | 2:5:1:0:1:1:1:0 | 0:0:0:0:0:0:0:0 | 0:3:1:0:0:0:0:0 | 0:1:0:0:0:0:0:0 |
| XM_017025419:XM_017025409:NM_006311:XM_024451043:XM_024451042:XM_005256871:XM_011524086:XM_017025420:XM_024451041:XM_017025411:XM_017025413:XM_005256868:XM_006721602:XM_017025403:XM_006721601:XM_005256866:XM_017025404:NM_001190438:XM_005256872:XM_006721603 | 1:1:1:0:0:0:0:0:0:0:0:0:0:0:0:0:0:0:0:0 | 0:0:0:0:0:0:0:0:0:0:0:0:0:0:0:0:0:0:0:0 | 0:0:2:0:1:0:3:1:0:0:0:0:0:0:0:0:0:0:0:0 | 0:0:0:2:2:1:0:0:0:0:0:0:0:0:0:0:0:0:0:0 | 0:0:0:0:0:0:1:0:1:0:0:0:0:0:0:0:0:0:0:0 | 0:0:0:0:0:0:0:0:0:0:0:0:0:0:0:0:0:0:0:0 | 0:0:1:3:0:0:0:0:0:0:0:1:1:0:0:0:0:0:0:0 | 0:0:0:0:0:0:0:0:1:1:1:0:0:0:0:0:0:0:0:0 | 0:0:0:1:0:1:0:0:0:0:0:0:0:1:0:0:0:0:0:0 | 0:0:0:0:0:0:1:0:0:0:0:0:0:0:0:0:0:0:0:0 |
| XM_011524086:XM_017025411:XM_017025409:XM_017025396:NM_006311:XM_017025420:XM_005256866:XM_006721602:XM_017025419:XM_005256871:XM_017025403:XM_005256868:XM_024451043:XM_024451042:XM_017025404:XM_017025405:XM_017025407:XM_006721601:XM_024451041:XM_017025406:NM_001190438:XM_017025413:XM_017025416:XM_017025397:XM_017025400:XM_017025398:XM_017025402:XM_005256872:XM_006721603 | 1:0:2:0:1:1:0:1:1:1:1:0:0:0:0:0:0:0:0:0:0:0:0:0:0:0:0:0:0 | 3:1:3:1:2:1:1:1:0:0:0:0:0:0:0:0:0:0:0:0:0:0:0:0:0:0:0:0:0 | 12:0:5:1:6:3:1:2:0:13:1:3:11:5:0:0:0:1:2:3:1:0:0:0:0:0:0:0:0 | 20:0:17:1:12:7:1:3:0:32:12:11:13:9:2:2:1:3:1:0:0:0:0:0:0:0:0:0:0 | 6:0:1:1:2:2:1:1:0:6:4:1:1:2:0:0:0:0:2:0:0:0:0:0:0:0:0:0:0 | 5:0:3:1:4:0:2:0:0:6:2:0:1:2:0:0:0:0:0:0:0:1:0:0:0:0:0:0:0 | 5:1:5:0:7:8:4:6:0:8:6:9:4:5:0:0:0:4:4:0:0:1:0:2:0:1:0:0:0 | 15:1:16:0:25:17:6:13:0:20:11:24:11:6:2:1:1:9:2:0:0:1:5:1:1:0:0:0:0 | 2:0:1:0:2:0:1:0:0:3:1:0:2:1:0:0:0:0:0:0:0:0:0:1:0:0:0:1:0:0 | 1:0:1:0:0:0:0:0:0:5:1:4:0:0:0:0:0:1:0:0:0:0:0:0:1:0:0:0:0 |
| XM_017025403:XM_011524086:XM_006721601:NM_006311:XM_005256868:XM_005256871:XM_017025420:XM_017025409:XM_024451041:XM_006721602:XM_024451042:XM_017025419:XM_017025397:XM_005256866:XM_017025411:XM_017025396:XM_024451043:XM_017025406:NM_001190438:XM_017025404:XM_017025405:XM_017025407:XM_017025416:XM_017025413:XM_017025399:XM_017025412:XM_017025398:XM_017025400:XM_017025401:XM_017025402:XM_011524083:XM_005256872:XM_006721603 | 4:6:1:11:4:6:6:11:0:6:1:1:1:4:1:1:0:0:0:0:0:0:0:0:0:0:0:0:0:0:0:0:0 | 4:10:6:6:4:7:2:2:1:2:1:0:0:0:0:0:0:0:0:0:0:0:0:0:0:0:0:0:0:0:0:0:0 | 17:40:4:24:17:62:14:25:5:6:15:0:0:4:0:2:29:4:2:3:2:1:0:0:0:0:0:0:0:0:0:0:0 | 2:1:0:1:2:4:0:3:0:0:0:0:0:0:0:0:0:0:0:0:0:0:0:0:0:0:0:0:0:0:0:0:0 | 13:26:1:15:6:58:12:12:3:9:10:0:0:4:1:2:7:0:0:0:0:1:0:1:0:0:0:0:0:0:0:0:0 | 1:24:0:19:7:32:8:7:0:0:9:0:0:3:0:0:3:0:1:0:0:0:1:0:0:0:0:0:0:0:0:0:0 | 26:36:22:59:69:49:40:39:11:29:19:0:3:20:2:0:20:1:1:6:1:1:7:2:0:2:1:2:1:0:0:0:0 | 4:9:2:14:10:24:9:5:2:12:6:0:1:8:0:0:3:0:0:0:0:0:1:0:1:0:0:0:0:0:0:0:0 | 6:17:1:15:10:16:8:10:1:3:5:0:1:2:0:0:3:0:0:0:0:0:2:0:0:0:0:2:0:1:0:0:0 | 3:11:4:3:3:10:6:5:0:3:2:0:0:5:0:2:4:0:0:0:0:0:0:0:0:0:0:0:0:0:1:0:0 |
| XM_017025419:XM_017025409:NM_006311:XM_006721602:XM_011524086:XM_017025420:XM_017025397:XM_005256871:XM_017025403:XM_005256866:XM_017025411:XM_017025396:XM_005256868:XM_006721601:XM_024451042:XM_024451041:XM_024451043:XM_017025406:NM_001190438:XM_017025404:XM_017025405:XM_017025407:XM_017025416:XM_011524083:XM_017025402:XM_017025413:XM_017025399:XM_017025412:XM_017025398:XM_017025400:XM_017025401:XM_006721603:XM_005256872 | 1:13:24:9:21:8:2:18:10:6:1:1:13:8:3:2:1:0:0:0:0:0:0:0:0:0:0:0:0:0:0:0:0 | 0:0:0:0:0:0:0:0:0:0:0:0:0:0:0:0:0:0:0:0:0:0:0:0:0:0:0:0:0:0:0:0:0 | 0:48:49:11:83:30:0:123:40:8:0:3:31:6:34:11:48:5:3:4:3:1:2:1:1:1:1:0:0:0:0:0:0 | 0:0:0:0:0:0:0:0:1:0:0:0:0:0:0:0:0:0:0:0:0:0:0:0:0:0:0:0:0:0:0:0:0 | 0:22:41:11:61:24:0:110:25:7:1:2:17:2:23:4:13:1:1:0:0:1:2:0:0:1:0:0:0:0:0:0:0 | 0:0:0:0:0:0:0:0:0:0:0:0:0:0:0:0:0:0:0:0:0:0:0:0:0:0:0:0:0:0:0:0:0 | 0:81:116:73:95:79:7:151:59:40:3:0:144:44:46:24:42:1:1:13:1:2:12:1:1:3:3:4:1:2:4:0:0 | 0:0:1:0:0:0:0:0:1:0:0:0:0:0:0:0:0:0:0:0:0:0:0:0:0:0:0:0:0:0:0:0:0 | 0:23:21:7:31:17:1:37:13:7:0:2:20:6:8:2:8:0:0:0:0:2:0:2:0:0:0:0:2:1:0:0 | 0:0:0:0:0:0:0:0:0:0:0:0:0:0:0:0:0:0:0:0:0:0:0:0:0:0:0:0:0:0:0:0:0 |
| XM_017025419:XM_017025409:NM_006311:XM_006721602:XM_011524086:XM_017025420:XM_017025397:XM_005256871:XM_017025403:XM_005256866:XM_017025411:XM_017025396:XM_005256868:XM_006721601:XM_024451042:XM_024451041:XM_024451043:XM_017025406:NM_001190438:XM_017025404:XM_017025405:XM_017025407:XM_017025416:XM_011524083:XM_017025402:XM_017025413:XM_017025399:XM_017025412:XM_017025398:XM_017025400:XM_017025401:XM_006721603:XM_005256872 | 1:13:24:9:21:8:2:18:10:6:1:1:13:8:3:2:1:0:0:0:0:0:0:0:0:0:0:0:0:0:0:0:0 | 0:0:0:0:0:0:0:0:0:0:0:0:0:0:0:0:0:0:0:0:0:0:0:0:0:0:0:0:0:0:0:0:0 | 0:48:49:11:83:30:0:124:40:8:0:3:31:6:34:11:47:5:3:4:3:1:2:1:1:1:1:0:0:0:0:0:0 | 0:0:0:0:0:0:0:0:0:0:0:0:0:0:0:0:1:0:0:0:0:0:0:0:0:0:0:0:0:0:0:0:0 | 0:22:41:11:60:24:0:110:25:7:1:2:17:2:23:4:13:1:1:0:0:1:2:0:0:1:0:0:0:0:0:0:0 | 0:0:0:0:1:0:0:0:0:0:0:0:0:0:0:0:0:0:0:0:0:0:0:0:0:0:0:0:0:0:0:0:0 | 0:81:117:73:95:79:7:151:60:40:3:0:144:44:46:24:42:1:1:13:1:2:12:1:1:3:3:4:1:2:4:0:0 | 0:0:0:0:0:0:0:0:0:0:0:0:0:0:0:0:0:0:0:0:0:0:0:0:0:0:0:0:0:0:0:0:0 | 0:23:21:7:30:17:1:36:13:7:0:2:20:6:8:2:8:0:0:0:0:2:0:2:0:0:0:0:2:1:0:0 | 0:0:0:0:1:0:0:1:0:0:0:0:0:0:0:0:0:0:0:0:0:0:0:0:0:0:0:0:0:0:0:0:0 |
| XM_017025419:XM_017025409:NM_006311:XM_006721602:XM_011524086:XM_017025420:XM_017025397:XM_005256871:XM_017025403:XM_005256866:XM_017025411:XM_017025396:XM_005256868:XM_006721601:XM_024451042:XM_024451041:XM_024451043:XM_017025406:NM_001190438:XM_017025404:XM_017025405:XM_017025407:XM_017025416:XM_011524083:XM_017025402:XM_017025413:XM_017025399:XM_017025412:XM_017025398:XM_017025400:XM_017025401:XM_005256872 | 1:14:24:9:21:8:2:18:10:6:1:1:13:8:3:2:1:0:0:0:0:0:0:0:0:0:0:0:0:0:0:0 | 0:0:0:0:0:0:0:0:0:0:0:0:0:0:0:0:0:0:0:0:0:0:0:0:0:0:0:0:0:0:0:0 | 0:48:49:11:83:30:0:124:40:8:0:3:31:6:35:11:50:5:3:4:3:1:2:1:1:1:1:0:0:0:0:0 | 0:0:0:0:0:0:0:0:0:0:0:0:0:0:0:0:0:0:0:0:0:0:0:0:0:0:0:0:0:0:0:0 | 0:21:41:11:61:24:0:110:25:7:1:2:17:2:22:5:13:1:1:0:0:1:2:0:0:1:0:0:0:0:0:0 | 0:1:1:0:0:0:0:0:0:0:0:0:0:0:0:0:0:0:0:0:0:0:0:0:0:0:0:0:0:0:0:0 | 0:83:118:74:97:79:7:151:60:40:3:0:143:44:46:24:48:1:1:13:1:2:12:1:1:3:3:4:1:2:4:0 | 0:0:0:0:0:0:0:0:0:0:0:1:0:0:0:0:0:0:0:0:0:0:0:0:0:0:0:0:0:0:0:0 | 0:23:21:7:33:17:1:37:13:7:0:2:20:6:9:2:8:0:0:0:0:2:0:2:0:0:0:0:2:1:0 | 0:0:0:0:0:0:0:0:0:0:0:0:0:0:0:0:0:0:0:0:0:0:0:0:0:0:0:0:0:0:0:0 |

| | | | | | | | | | | |
|---|---|---|---|---|---|---|---|---|---|---|
| XM_017025419:XM_017025409:NM_006311:XM_006721602:XM_011524086:XM_017025420:XM_017025397:XM_005256871:XM_017025403:XM_005256866:XM_017025411:XM_017025396:XM_005256868:XM_006721601:XM_024451042:XM_024451041:XM_024451043:XM_017025404:XM_017025406:NM_001190438:XM_017025405:XM_017025407:XM_017025416:XM_011524083:XM_017025402:XM_017025413:XM_017025399:XM_017025412:XM_017025398:XM_017025400:XM_017025401:XM_005256872 | 1:14:25:11:21:8:2:18:10:6:1:1:13:8:3:2:1:0:0:0:0:0:0:0:0:0:0:0:0:0:0:0 | 0:0:0:0:0:0:0:0:0:0:0:0:0:0:0:0:0:0:0:0:0:0:0:0:0:0:0:0:0:0:0:0 | 0:49:50:14:84:31:0:124:40:8:0:3:31:6:36:11:56:3:5:3:3:1:2:1:1:1:1:0:0:0:0:0 | 0:0:0:0:0:0:0:0:0:0:0:0:0:0:0:0:0:1:0:0:0:0:0:0:0:0:0:0:0:0:0:0 | 0:22:45:11:71:26:0:110:25:8:1:2:17:2:22:6:17:0:1:1:0:1:1:0:0:1:0:0:0:0:0:0 | 0:0:0:0:0:0:0:0:0:0:0:0:0:0:0:0:0:0:0:0:0:0:1:0:0:0:0:0:0:0:0:0 | 0:91:131:76:100:83:7:151:60:43:4:0:144:44:47:25:60:13:1:1:1:2:12:1:1:3:3:4:1:2:4:0 | 0:0:0:0:0:0:0:0:0:0:0:0:0:0:0:0:0:0:0:0:0:0:0:0:0:0:0:0:0:0:0:0 | 0:24:22:7:36:18:1:37:13:9:0:2:20:6:9:2:10:0:0:0:0:0:2:0:2:0:0:0:0:2:1:0 | 0:0:0:0:0:0:0:0:0:0:0:0:0:0:0:0:0:0:0:0:0:0:0:0:0:0:0:0:0:0:0:0 |
| XM_017025419:XM_017025409:NM_006311:XM_006721602:XM_011524086:XM_017025420:XM_017025397:XM_005256871:XM_017025403:XM_005256866:XM_017025411:XM_017025396:XM_005256868:XM_006721601:XM_024451042:XM_024451041:XM_024451043:XM_017025406:NM_001190438:XM_017025404:XM_017025405:XM_017025407:XM_017025416:XM_011524083:XM_017025402:XM_017025413:XM_017025399:XM_017025412:XM_017025398:XM_017025400:XM_017025401:XM_005256872 | 1:14:25:11:21:8:2:18:10:6:1:1:13:8:3:2:1:0:0:0:0:0:0:0:0:0:0:0:0:0:0:0 | 0:0:0:0:0:0:0:0:0:0:0:0:0:0:0:0:0:0:0:0:0:0:0:0:0:0:0:0:0:0:0:0 | 0:50:50:14:86:31:0:123:39:8:0:3:31:6:36:11:57:5:3:4:3:1:2:1:1:1:1:0:0:0:0:0 | 0:0:0:0:0:0:0:1:1:0:0:0:0:0:0:0:0:0:0:0:0:0:0:0:0:0:0:0:0:0:0:0 | 0:22:45:11:73:26:0:110:25:8:1:2:17:2:21:7:20:1:1:0:0:1:2:0:0:1:0:0:0:0:0:0 | 0:0:0:0:0:0:0:0:0:0:0:0:0:0:0:0:0:0:0:0:0:0:0:0:0:0:0:0:0:0:0:0 | 0:91:131:76:100:83:7:151:60:43:4:0:144:44:47:25:61:1:1:13:1:2:12:1:1:3:3:4:1:2:4:0 | 0:0:0:0:0:0:0:0:0:0:0:0:0:0:0:0:1:0:0:0:0:0:0:0:0:0:0:0:0:0:0:0 | 0:25:22:7:36:19:1:37:13:9:0:2:20:6:8:2:10:0:0:0:0:0:2:0:2:0:0:0:0:2:1:0 | 0:0:0:0:0:0:0:0:0:0:0:0:0:0:0:0:0:0:0:0:0:0:0:0:0:0:0:0:0:0:0:0 |
| XM_017025419:XM_017025409:NM_006311:XM_006721602:XM_011524086:XM_017025420:XM_017025397:XM_005256871:XM_017025403:XM_005256866:XM_017025411:XM_017025396:XM_005256868:XM_006721601:XM_024451042:XM_024451041:XM_024451043:XM_017025406:NM_001190438:XM_017025404:XM_017025405:XM_017025407:XM_017025416:XM_011524083:XM_017025402:XM_017025413:XM_017025399:XM_017025412:XM_017025398:XM_017025400:XM_017025401:XM_005256872 | 1:14:25:11:21:8:2:18:10:6:1:1:13:8:3:2:1:0:0:0:0:0:0:0:0:0:0:0:0:0:0:0 | 0:0:0:0:0:0:0:0:0:0:0:0:0:0:0:0:0:0:0:0:0:0:0:0:0:0:0:0:0:0:0:0 | 0:50:50:14:86:31:0:124:40:8:0:3:31:6:36:11:57:5:3:4:3:1:2:1:1:1:1:0:0:0:0:0 | 0:0:0:0:0:0:0:0:0:0:0:0:0:0:0:0:0:0:0:0:0:0:0:0:0:0:0:0:0:0:0:0 | 0:22:45:11:73:26:0:110:25:8:1:2:17:2:21:7:20:1:1:0:0:1:2:0:0:1:0:0:0:0:0:0 | 0:0:0:0:0:0:0:0:0:0:0:0:0:0:0:0:0:0:0:0:0:0:0:0:0:0:0:0:0:0:0:0 | 0:91:131:75:100:83:7:151:60:43:4:0:144:44:47:26:63:1:1:13:1:2:12:1:1:3:3:4:1:2:4:0 | 0:0:1:0:0:0:0:0:0:0:0:0:0:0:0:0:1:0:0:0:0:0:0:0:0:0:0:0:0:0:0:0 | 0:25:22:7:36:19:1:37:13:9:0:2:20:6:8:2:10:0:0:0:0:0:2:0:2:0:0:0:0:2:1:0 | 0:0:0:0:0:0:0:0:0:0:0:0:0:0:0:0:0:0:0:0:0:0:0:0:0:0:0:0:0:0:0:0 |
| XM_017025419:XM_017025409:NM_006311:XM_006721602:XM_011524086:XM_017025420:XM_017025397:XM_005256871:XM_017025403:XM_005256866:XM_017025411:XM_017025396:XM_005256868:XM_006721601:XM_024451042:XM_024451041:XM_024451043:XM_017025406:NM_001190438:XM_017025404:XM_017025405:XM_017025407:XM_017025416:XM_011524083:XM_017025402:XM_017025413:XM_017025399:XM_017025412:XM_017025398:XM_017025400:XM_017025401:XM_005256872 | 1:14:25:11:21:8:2:18:10:6:1:1:13:8:3:2:1:0:0:0:0:0:0:0:0:0:0:0:0:0:0:0 | 0:0:0:0:0:0:0:0:0:0:0:0:0:0:0:0:0:0:0:0:0:0:0:0:0:0:0:0:0:0:0:0 | 0:50:50:14:84:31:0:124:40:8:0:3:31:6:36:12:57:5:3:4:3:1:2:1:1:1:1:0:0:0:0:0 | 0:0:0:0:1:0:0:0:0:0:0:0:0:0:0:0:0:0:0:0:0:0:0:0:0:0:0:0:0:0:0:0 | 0:22:45:11:73:26:0:109:25:8:1:2:17:2:21:6:17:1:1:0:0:1:2:0:0:1:0:0:0:0:0:0 | 0:0:0:0:0:0:0:1:0:0:0:0:0:0:0:0:1:0:0:0:0:0:0:0:0:0:0:0:0:0:0:0 | 0:91:131:76:100:83:7:150:60:44:4:0:144:44:47:28:60:1:1:13:1:2:13:1:1:3:3:4:1:2:4:0 | 0:0:0:0:0:0:0:1:0:0:0:0:0:0:0:0:0:0:0:0:0:0:0:0:0:0:0:0:0:0:0:0 | 0:25:22:7:36:19:1:37:13:9:0:2:20:6:8:2:10:0:0:0:0:0:2:0:2:0:0:0:0:2:1:0 | 0:0:0:0:0:0:0:0:0:0:0:0:0:0:0:0:0:0:0:0:0:0:0:0:0:0:0:0:0:0:0:0 |
| NM_006311:XM_011524086:XM_017025409:XM_017025419:XM_006721602:XM_017025420:XM_017025397:XM_005256871:XM_017025403:XM_005256866:XM_017025411:XM_017025396:XM_005256868:XM_006721601:XM_024451042:XM_024451041:XM_024451043:XM_017025406:NM_001190438:XM_017025404:XM_017025405:XM_017025407:XM_017025416:XM_011524083:XM_017025402:XM_017025413:XM_017025399:XM_017025401:XM_017025412:XM_017025398:XM_017025400:XM_005256872 | 25:21:14:1:11:8:2:18:10:6:1:1:13:8:3:2:2:0:0:0:0:0:0:0:0:0:0:0:0:0:0:0 | 1:4:2:0:0:0:0:0:0:0:0:0:0:0:0:0:0:0:0:0:0:0:0:0:0:0:0:0:0:0:0:0 | 50:87:50:0:14:32:0:124:40:8:0:3:31:6:36:12:56:5:3:4:3:1:2:1:1:1:1:0:0:0:0:0 | 6:5:4:0:1:3:0:0:0:1:0:0:0:0:0:2:11:0:0:0:0:0:0:0:0:0:0:0:0:0:0:0 | 45:73:22:0:11:26:0:110:25:8:1:2:17:2:21:6:18:1:1:0:0:1:2:0:0:1:0:0:0:0:0:0 | 3:6:2:0:0:0:0:0:0:1:0:0:0:0:0:1:4:0:0:0:0:0:0:0:0:0:0:0:0:0:0:0 | 132:100:91:0:76:83:7:151:60:45:4:0:144:44:47:28:61:1:1:13:1:2:13:1:1:3:3:4:4:1:2:0 | 5:7:9:0:4:3:0:0:0:4:0:0:0:1:0:4:2:0:0:0:0:0:3:0:0:0:0:1:0:0:0:0 | 23:37:25:0:7:19:1:37:13:9:0:2:20:6:8:2:10:0:0:0:0:0:2:0:2:0:0:1:0:0:2:0 | 5:3:4:0:1:4:0:0:0:1:0:0:0:1:0:0:1:0:0:0:0:0:0:0:0:0:0:1:0:0:0:0 |
| XM_011524086:NM_006311:XM_017025419:XM_017025409:XM_006721602:XM_017025420:XM_017025397:XM_005256871:XM_017025403:XM_005256866:XM_017025411:XM_017025396:XM_005256868:XM_006721601:XM_024451042:XM_024451041:XM_024451043:XM_017025400:XM_017025406:NM_001190438:XM_017025404:XM_017025405:XM_017025407:XM_017025416:XM_011524083:XM_017025402:XM_017025413:XM_017025399:XM_017025412:XM_017025398:XM_017025401:XM_005256872 | 25:27:1:16:11:8:2:18:10:6:1:1:13:8:3:2:3:1:0:0:0:0:0:0:0:0:0:0:0:0:0:0 | 1:1:0:0:0:0:0:0:0:0:0:0:0:0:0:0:0:0:0:0:0:0:0:0:0:0:0:0:0:0:0:0 | 95:63:0:60:15:36:0:124:40:9:0:3:31:6:35:19:72:0:5:3:4:3:1:3:1:1:1:1:0:0:0:0 | 0:0:0:0:0:0:0:0:0:0:0:0:0:0:0:0:1:0:0:0:0:0:0:0:0:0:0:0:0:0:0:0 | 83:52:0:24:11:27:0:110:25:9:1:2:17:2:21:7:22:0:1:1:0:0:1:2:0:0:1:0:0:0:0:0 | 2:1:0:1:0:2:0:0:0:0:0:0:0:0:0:0:0:0:0:0:0:0:0:0:0:0:0:0:0:0:0:0 | 115:143:0:107:86:87:7:151:60:52:4:0:144:45:47:33:64:2:1:1:13:1:2:16:1:1:3:3:3:1:5:0 | 1:2:0:1:0:0:0:0:0:1:0:0:0:1:0:0:1:0:0:0:0:0:0:0:0:0:0:0:1:0:0:0 | 41:32:0:29:10:24:1:37:13:11:0:2:20:7:8:2:11:2:0:0:0:0:0:3:0:2:0:0:0:0:2:0 | 0:0:0:0:0:0:0:0:0:0:0:0:0:0:0:0:1:0:0:0:0:0:0:0:0:0:0:0:0:0:0:0 |
| XM_017025419:XM_017025409:NM_006311:XM_006721602:XM_011524086:XM_017025420:XM_017025397:XM_005256871:XM_017025403:XM_005256866:XM_017025411:XM_017025396:XM_005256868:XM_006721601:XM_024451042:XM_024451041:XM_024451043:XM_017025400:XM_017025406:NM_001190438:XM_017025404:XM_017025405:XM_017025407:XM_017025416:XM_011524083:XM_017025402:XM_017025413:XM_017025399:XM_017025412:XM_017025398:XM_017025401:XM_005256872 | 1:16:28:11:26:8:2:18:10:6:1:1:13:8:3:2:3:1:0:0:0:0:0:0:0:0:0:0:0:0:0:0 | 0:0:0:0:0:0:0:0:0:0:0:0:0:0:0:0:0:0:0:0:0:0:0:0:0:0:0:0:0:0:0:0 | 0:60:63:15:95:36:0:124:40:9:0:3:31:6:35:19:73:0:5:3:4:3:1:3:1:1:1:1:0:0:0:0 | 0:0:0:0:0:0:0:0:0:0:0:0:0:0:0:0:0:0:0:0:0:0:0:0:0:0:0:0:0:0:0:0 | 0:24:53:11:85:29:0:110:25:9:1:2:17:2:20:7:21:0:1:1:0:0:1:2:0:0:1:0:0:0:0:0 | 0:1:0:0:0:0:0:0:0:0:0:0:0:0:0:0:0:0:0:0:0:0:0:0:0:0:0:0:0:0:0:0 | 0:108:145:86:116:87:7:151:60:53:4:0:143:46:47:33:64:2:1:1:13:1:2:16:1:1:3:3:4:1:5:0 | 0:0:0:0:0:0:0:0:0:0:0:1:0:0:0:0:0:0:0:0:0:0:0:0:0:0:0:0:0:0:0:0 | 0:29:32:10:41:24:1:37:13:11:0:2:20:7:8:3:12:2:0:0:0:0:0:3:0:2:0:0:0:0:2:0 | 0:0:0:0:0:0:0:0:0:0:0:0:0:0:0:0:0:0:0:0:0:0:0:0:0:0:0:0:0:0:0:0 |
| XM_017025409:NM_006311:XM_017025419:XM_006721602:XM_011524086:XM_017025420:XM_017025397:XM_005256871:XM_017025403:XM_005256866:XM_017025411:XM_017025396:XM_005256868:XM_006721601:XM_024451042:XM_024451041:XM_024451043:XM_017025400:XM_017025416:XM_017025406:NM_001190438:XM_017025404:XM_017025405:XM_017025407:XM_011524083:XM_017025402:XM_017025413:XM_017025399:XM_017025401:XM_017025412:XM_017025398:XM_005256872 | 16:29:1:11:26:8:2:18:10:6:1:1:13:8:3:2:3:1:0:0:0:0:0:0:0:0:0:0:0:0:0:0 | 2:2:0:0:0:0:0:0:0:0:0:0:0:0:0:0:0:0:0:0:0:0:0:0:0:0:0:0:0:0:0:0 | 61:63:0:15:95:36:0:124:40:9:0:3:31:6:35:19:73:0:3:5:3:4:3:1:1:1:1:1:0:0:0:0 | 6:11:0:2:15:1:0:0:0:0:0:0:0:0:0:5:12:1:1:0:0:0:0:0:0:0:0:0:0:0:0:0 | 25:53:0:11:87:30:0:110:25:9:1:2:17:2:20:8:23:0:2:1:1:0:0:1:0:0:1:0:0:0:0:0 | 2:6:0:0:9:2:0:0:0:1:0:0:0:0:0:0:0:0:0:0:0:0:0:0:0:0:0:0:0:0:0:0 | 109:146:0:86:117:87:7:151:60:55:4:0:144:46:46:33:61:2:16:1:1:13:1:2:1:2:3:3:5:4:1:0 | 10:15:0:0:11:5:0:0:0:3:0:0:0:0:0:1:0:0:0:0:0:0:0:0:0:0:0:0:1:0:0:0 | 32:32:0:10:41:24:1:37:13:11:0:2:20:7:8:3:12:2:3:0:0:0:0:0:0:2:0:0:2:0:0:0 | 3:2:0:4:2:0:0:0:0:2:1:0:0:0:0:1:1:0:0:0:0:0:0:0:0:0:0:0:0:0:0:0 |
| XM_017025419:XM_017025409:NM_006311:XM_006721602:XM_011524086:XM_017025420:XM_017025397:XM_005256871:XM_017025403:XM_005256866:XM_017025411:XM_017025396:XM_005256868:XM_006721601:XM_024451042:XM_024451041:XM_024451043:XM_017025400:XM_017025406:NM_001190438:XM_017025404:XM_017025405:XM_017025407:XM_017025416:XM_011524083:XM_017025402:XM_017025413:XM_017025399:XM_017025412:XM_017025398:XM_017025401:XM_005256872 | 1:21:33:11:26:9:2:18:10:6:1:1:13:8:4:3:3:1:0:0:0:0:0:0:0:0:0:0:0:0:0:0 | 0:0:0:0:0:0:0:0:0:0:0:0:0:0:0:0:0:0:0:0:0:0:0:0:0:0:0:0:0:0:0:0 | 0:72:77:20:114:42:0:124:40:10:1:3:31:6:33:28:92:1:5:3:4:3:1:5:1:1:1:1:0:0:0:0 | 0:0:0:0:0:0:0:0:0:0:0:0:0:0:0:0:0:0:0:0:0:0:0:0:0:0:0:0:0:0:0:0 | 0:29:72:11:105:34:0:110:25:10:1:2:17:2:21:8:26:0:1:1:0:0:1:2:0:0:1:0:0:0:0:0 | 0:0:0:0:0:0:0:0:0:0:0:0:0:0:0:0:0:0:0:0:0:0:0:0:0:0:0:0:0:0:0:0 | 0:134:175:96:149:107:7:151:60:64:4:0:144:46:41:41:56:5:1:1:13:1:2:22:1:2:3:3:4:1:6:0 | 0:0:0:0:0:0:0:0:0:0:0:0:0:0:0:0:0:0:0:0:0:0:0:0:0:0:0:0:0:0:0:0 | 0:36:36:14:45:24:1:37:13:16:1:2:20:7:9:4:14:2:0:0:0:0:0:3:0:2:0:0:0:0:2:0 | 0:0:0:0:0:0:0:0:0:0:0:0:0:0:0:0:0:0:0:0:0:0:0:0:0:0:0:0:0:0:0:0 |
| XM_017025419:XM_017025409:NM_006311:XM_006721602:XM_011524086:XM_017025420:XM_017025397:XM_005256871:XM_017025403:XM_005256866:XM_017025411:XM_017025396:XM_005256868:XM_006721601:XM_024451042:XM_024451041:XM_024451043:XM_017025400:XM_017025406:NM_001190438:XM_017025404:XM_017025405:XM_017025407:XM_017025416:XM_011524083:XM_017025402:XM_017025413:XM_017025399:XM_017025412:XM_017025398:XM_017025401:XM_005256872 | 1:21:33:11:26:9:2:18:10:6:1:1:13:8:4:3:3:1:0:0:0:0:0:0:0:0:0:0:0:0:0:0 | 0:0:0:0:0:0:0:0:0:0:0:0:0:0:0:0:0:0:0:0:0:0:0:0:0:0:0:0:0:0:0:0 | 0:72:77:20:114:42:0:124:40:10:1:3:31:6:32:28:90:1:5:3:4:3:1:5:1:1:1:1:0:0:0:0 | 0:0:0:0:0:0:0:0:0:0:0:0:0:0:0:0:2:0:0:0:0:0:0:0:0:0:0:0:0:0:0:0 | 0:29:72:11:105:34:0:110:25:10:1:2:17:2:21:8:26:0:1:1:0:0:1:2:0:0:1:0:0:0:0:0 | 0:0:0:0:0:0:0:0:0:0:0:0:0:0:0:0:0:0:0:0:0:0:0:0:0:0:0:0:0:0:0:0 | 0:134:175:96:149:107:7:151:60:64:4:0:144:46:41:41:56:5:1:1:13:1:2:22:1:2:3:3:4:1:6:0 | 0:0:0:0:0:0:0:0:0:0:0:0:0:0:0:0:0:0:0:0:0:0:0:0:0:0:0:0:0:0:0:0 | 0:36:36:14:45:24:1:37:13:16:1:2:20:7:9:4:14:2:0:0:0:0:0:3:0:2:0:0:0:0:2:0 | 0:0:0:0:0:0:0:0:0:0:0:0:0:0:0:0:0:0:0:0:0:0:0:0:0:0:0:0:0:0:0:0 |
| XM_017025419:XM_017025409:NM_006311:XM_006721602:XM_011524086:XM_017025420:XM_017025397:XM_005256871:XM_017025403:XM_005256866:XM_017025411:XM_017025396:XM_005256868:XM_006721601:XM_024451042:XM_024451041:XM_024451043:XM_017025400:XM_017025406:NM_001190438:XM_017025404:XM_017025405:XM_017025407:XM_017025416:XM_011524083:XM_017025402:XM_017025413:XM_017025399:XM_017025412:XM_017025398:XM_017025401:XM_005256872 | 1:22:33:11:26:9:2:18:10:6:1:1:13:8:4:3:1:1:0:0:0:0:0:0:0:0:0:0:0:0:0:0 | 0:0:0:0:0:0:0:0:0:0:0:0:0:0:0:0:0:0:0:0:0:0:0:0:0:0:0:0:0:0:0:0 | 0:72:79:20:114:41:0:124:40:10:1:3:31:6:27:26:90:1:5:3:4:3:1:5:1:1:1:1:0:0:0:0 | 0:0:0:0:0:0:0:0:0:0:0:0:0:0:0:0:0:0:0:0:0:0:0:0:0:0:0:0:0:0:0:0 | 0:30:71:11:105:34:0:110:25:10:1:2:17:2:18:6:24:0:1:1:0:0:1:2:0:0:1:0:0:0:0:0 | 0:0:1:0:0:0:0:0:0:0:0:0:0:0:1:0:0:0:0:0:0:0:0:0:0:0:0:0:0:0:0:0 | 0:134:176:96:148:105:7:151:60:64:4:0:144:46:37:37:53:5:1:1:13:1:2:22:1:2:3:3:4:1:6:0 | 0:0:0:0:1:0:0:0:0:0:0:0:0:0:0:0:0:0:0:0:0:0:0:0:0:0:0:0:0:0:0:0 | 0:36:36:14:45:26:1:37:13:16:1:2:20:7:8:4:14:2:0:0:0:0:0:3:0:2:0:0:0:0:2:0 | 0:0:0:0:0:0:0:0:0:0:0:0:0:0:0:0:0:0:0:0:0:0:0:0:0:0:0:0:0:0:0:0 |
| XM_017025419:XM_017025409:NM_006311:XM_006721602:XM_011524086:XM_017025420:XM_017025397:XM_005256871:XM_017025403:XM_005256866:XM_017025396:XM_005256868:XM_006721601:XM_024451042:XM_024451041:XM_024451043:XM_017025400:XM_017025416:XM_017025406:NM_001190438:XM_017025404:XM_017025405:XM_017025407:XM_011524083:XM_017025402:XM_017025413:XM_017025399:XM_017025411:XM_017025412:XM_017025398:XM_017025401:XM_005256872 | 1:23:35:11:26:12:2:18:10:6:1:13:8:4:3:2:1:1:0:0:0:0:0:0:0:0:0:0:0:0:0:0 | 0:0:0:0:0:0:0:0:0:0:0:0:0:0:0:0:0:0:0:0:0:0:0:0:0:0:0:0:0:0:0:0 | 0:75:82:20:113:54:0:123:40:10:3:31:6:23:21:82:1:5:5:3:4:3:1:1:1:1:1:1:0:0:0:0 | 0:0:0:0:1:0:0:1:0:0:0:0:0:0:0:0:0:0:0:0:0:0:0:0:0:0:0:0:0:0:0:0 | 0:31:72:11:105:37:0:110:25:10:2:17:2:16:6:20:1:2:1:1:0:0:1:0:0:1:0:1:0:0:0:0 | 0:1:0:0:0:1:0:0:0:0:0:0:0:0:0:0:0:0:0:0:0:0:0:0:0:0:0:0:0:0:0:0 | 0:152:184:96:149:116:7:151:60:64:0:143:46:34:32:51:6:23:1:1:13:1:2:1:2:3:3:4:4:1:6:0 | 0:2:2:0:0:1:0:0:0:0:0:1:0:0:0:1:0:0:0:0:0:0:0:0:0:0:0:0:0:0:0:0 | 0:35:37:14:45:28:1:37:13:16:2:20:7:7:4:12:2:4:0:0:0:0:0:0:2:0:0:1:0:0:2:0 | 0:1:0:0:0:0:0:0:0:0:0:0:0:0:0:0:0:0:0:0:0:0:0:0:0:0:0:0:0:0:0:0 |

| | | | | | | | | | | |
|---|---|---|---|---|---|---|---|---|---|---|
| XM_017025419:XM_017025409:NM_006311:XM_006721602:XM_011524086:XM_017025420:XM_017025397:XM_005256871:XM_017025403:XM_005256866:XM_017025396:XM_005256868:XM_006721601:XM_024451042:XM_024451041:XM_024451043:XM_017025400:XM_017025416:XM_017025406:NM_001190438:XM_017025404:XM_017025405:XM_017025407:XM_011524083:XM_017025402:XM_017025413:XM_017025399:XM_017025411:XM_017025412:XM_017025398:XM_017025401:XM_005256872 | 1:23:35:11:26:12:2:18:10:6:1:13:8:4:3:2:1:1:0:0:0:0:0:0:0:0:0:0:0:0:0:0 | 0:0:0:0:0:0:0:0:0:0:0:0:0:0:0:0:0:0:0:0:0:0:0:0:0:0:0:0:0:0 | 0:75:82:20:114:53:0:124:40:10:3:31:6:19:17:64:1:5:5:3:4:3:1:1:1:1:1:1:0:0:0:0 | 0:0:0:0:0:0:0:0:0:0:0:0:1:0:3:0:0:0:0:0:0:0:0:0:0:0:0:0:0:0 | 0:32:72:11:105:38:0:110:25:10:2:17:2:14:5:19:1:2:1:1:0:0:1:0:0:1:0:1:0:0:0:0 | 0:0:0:0:0:0:0:0:0:0:0:0:0:0:0:0:0:0:0:0:0:0:0:0:0:0:0:0:0:0:0:0 | 0:154:186:96:149:114:7:151:60:64:0:144:46:29:28:45:6:23:1:1:13:1:2:1:2:3:3:4:4:1:6:0 | 0:0:0:0:0:1:0:0:0:0:0:0:0:0:1:0:0:0:0:0:0:0:0:0:0:0:0:0:0 | 0:36:37:14:45:28:1:37:13:16:2:20:7:7:4:11:2:4:0:0:0:0:0:0:2:0:0:1:0:0:2:0 | 0:0:0:0:0:0:0:0:0:0:0:0:0:0:0:0:0:0:0:0:0:0:0:0:0:0:0:0:0:0 |
| XM_011524086:XM_017025409:NM_006311:XM_006721602:XM_017025420:XM_017025397:XM_005256871:XM_017025403:XM_005256866:XM_017025411:XM_017025396:XM_005256868:XM_006721601:XM_024451041:XM_024451042:XM_024451043:XM_017025400:XM_017025416:XM_017025406:NM_001190438:XM_017025404:XM_017025405:XM_017025407:XM_011524083:XM_017025402:XM_017025413:XM_017025399:XM_017025412:XM_017025398:XM_017025401:XM_005256872 | 25:23:35:11:11:2:18:10:6:1:1:13:8:2:2:2:1:2:0:0:0:0:0:0:0:0:0:0:0:0:0 | 1:0:0:0:0:0:0:0:0:0:0:0:0:0:0:0:0:0:0:0:0:0:0:0:0:0:0:0:0 | 114:75:82:20:46:0:124:40:10:1:3:31:6:9:6:23:1:5:5:3:4:3:1:1:1:1:1:1:0:0:0:0 | 0:0:0:0:0:0:0:0:0:0:0:0:0:0:1:0:0:0:0:0:0:0:0:0:0:0:0:0:0 | 105:33:73:11:40:0:109:25:10:1:2:17:2:3:10:11:1:2:1:1:0:0:1:0:0:1:0:0:0:0:0 | 0:0:0:0:0:0:1:0:0:0:0:0:0:0:0:0:0:0:0:0:0:0:0:0:0:0:0:0:0:0:0 | 149:155:188:96:110:7:150:60:64:4:0:144:46:15:16:25:6:23:1:1:13:1:2:1:2:3:3:4:1:6:0 | 0:0:0:0:0:0:1:0:0:0:0:0:0:0:0:0:0:0:0:0:0:0:0:0:0:0:0:0:0 | 45:36:37:14:26:1:37:13:16:1:2:20:7:1:4:5:2:4:0:0:0:0:0:0:2:0:0:0:0:2:0 | 0:0:0:0:0:0:0:0:0:0:0:0:0:0:0:0:0:0:0:0:0:0:0:0:0:0:0:0:0:0 |
| XM_017025409:NM_006311:XM_006721602:XM_011524086:XM_017025420:XM_017025397:XM_005256871:XM_017025403:XM_005256866:XM_017025411:XM_017025396:XM_005256868:XM_006721601:XM_024451041:XM_024451042:XM_024451043:XM_017025400:XM_017025416:XM_017025406:NM_001190438:XM_017025404:XM_017025405:XM_017025407:XM_011524083:XM_017025402:XM_017025413:XM_017025399:XM_017025412:XM_017025398:XM_017025401:XM_005256872 | 23:35:11:26:11:2:18:10:6:1:1:13:8:2:2:2:1:2:0:0:0:0:0:0:0:0:0:0:0:0:0 | 0:0:0:0:0:0:0:0:0:0:0:0:0:0:0:0:0:0:0:0:0:0:0:0:0:0:0:0:0 | 75:82:20:114:46:0:124:40:10:1:3:31:6:9:6:23:1:5:5:3:4:3:1:1:1:1:1:1:0:0:0:0 | 0:0:0:0:0:0:0:0:0:0:0:0:0:0:1:0:0:0:0:0:0:0:0:0:0:0:0:0:0 | 33:73:11:105:40:0:110:25:10:1:2:17:2:3:10:10:1:2:1:1:0:0:1:0:0:1:0:0:0:0:0 | 0:0:0:0:0:0:0:0:0:0:0:0:0:0:0:0:0:0:0:0:0:0:0:0:0:0:0:0:0:0:0 | 155:188:95:149:109:7:151:60:64:4:0:144:46:14:15:24:6:23:1:1:13:1:2:1:2:3:3:4:1:6:0 | 0:0:1:0:1:0:0:0:0:0:0:0:0:0:0:0:0:0:0:0:0:0:0:0:0:0:0:0:0 | 36:37:14:45:26:1:37:13:16:1:2:20:7:1:4:5:2:4:0:0:0:0:0:0:2:0:0:0:0:2:0 | 0:0:0:0:0:0:0:0:0:0:0:0:0:0:0:0:0:0:0:0:0:0:0:0:0:0:0:0:0:0 |
| NM_018955:NM_001281719 | 6:00 | 0:00 | 9:00 | 0:00 | 14:00 | 1:00 | 8:00 | 1:00 | 8:00 | 1:00 |
| NM_003653:NM_001316356:XM_005256837:XM_005256840:XM_005256842:NM_001316355:NM_001316354:NM_001199125:XM_017025246 | 36:4:4:3:1:1:0:0:0 | 0:0:0:0:0:0:0:0:0 | 45:1:9:0:0:0:0:0:0 | 0:0:0:0:0:0:0:0:0 | 23:3:4:0:0:0:0:0:0 | 0:0:0:0:0:0:0:0:0 | 140:4:9:1:0:6:2:1:0 | 0:0:0:0:0:0:0:0:0 | 53:2:5:2:1:3:1:0:0 | 0:0:0:0:0:0:0:0:0 |
| NM_148172:NM_001267551 | 0:00 | 0:00 | 0:00 | 0:00 | 1:00 | 0:00 | 0:00 | 0:00 | 1:00 | 0:01 |
| NM_148172:NM_001267551 | 0:00 | 0:00 | 0:00 | 0:00 | 1:00 | 0:00 | 0:00 | 0:00 | 1:00 | 0:01 |
| NM_002615:NM_001329904 | 0:00 | 0:00 | 0:00 | 0:00 | 0:00 | 0:00 | 0:00 | 0:00 | 1:00 | 0:00 |
| XM_005256558 | 0 | 0 | 1 | 0 | 0 | 0 | 1 | 0 | 0 | 0 |
| NM_009587:XM_006721893:XM_017024623 | 0:00:00 | 0:00:00 | 0:00:00 | 0:00:00 | 1:01:00 | 0:00:00 | 0:00:00 | 0:00:00 | 0:00:00 | 1:00:00 |
| NM_009587:XM_006721893:XM_017024623 | 0:00:00 | 0:00:00 | 0:00:00 | 0:00:00 | 1:01:00 | 0:00:00 | 0:00:00 | 0:00:00 | 0:00:00 | 1:00:00 |
| NM_009587:XM_006721893:XM_017024623 | 0:00:00 | 0:00:00 | 0:00:00 | 0:00:00 | 0:00:00 | 1:01:00 | 0:00:00 | 0:00:00 | 1:00:00 | 0:00:00 |
| XM_017024591:NM_001145053:NR_026738 | 0:00:00 | 0:00:00 | 0:00:00 | 0:00:00 | 0:00:00 | 1:01:00 | 0:00:00 | 0:00:00 | 0:00:00 | 0:00:00 |
| NM_032322:XM_024451000:NM_197939:XM_024451001:NM_001184992:XR_002958077:XR_002958078 | 2:1:2:0:0:0:0 | 0:0:0:0:0:0:0 | 15:0:2:1:4:1:1 | 0:0:0:0:0:0:0 | 28:0:4:0:5:1:0 | 0:0:0:0:0:0:0 | 43:0:9:0:6:1:1 | 0:0:1:1:0:0:0 | 8:1:1:0:3:0:0 | 0:0:0:0:0:0:0 |
| XM_024451000:NM_197939:NM_032322:XM_024451001:NM_001184992:XR_002958077:XR_002958078 | 1:2:1:0:0:0:0 | 0:0:0:0:0:0:0 | 0:3:15:2:4:1:1 | 0:0:0:0:0:0:0 | 0:4:31:0:5:2:0 | 0:0:0:0:0:0:0 | 0:10:47:0:7:1:1 | 0:1:0:2:0:0:0 | 1:1:8:0:3:0:0 | 0:0:0:0:0:0:0 |
| NM_001042492 | 0 | 0 | 7 | 16 | 0 | 2 | 13 | 10 | 0 | 1 |
| NM_018405 | 0 | 0 | 1 | 0 | 0 | 0 | 0 | 2 | 0 | 3 |
| NM_015355:XR_001752462:XM_006721794:XM_017024409 | 22:1:0:0 | 0:0:0:0 | 34:3:1:0 | 0:0:0:0 | 34:1:3:0 | 0:0:0:0 | 78:7:4:0 | 1:1:0:0 | 22:0:1:0 | 0:0:0:0 |
| NM_015355:NM_001321207:XR_001752462:XM_006721794:XM_017024409 | 16:2:4:1:0 | 3:1:0:0:0 | 31:4:3:5:3 | 2:0:0:0:1 | 21:2:6:9:2 | 5:0:0:0:0 | 58:13:6:11:2 | 8:0:1:1:0 | 16:2:4:5:1 | 1:0:0:0:0 |
| NM_015355:NM_001321207:XR_001752462:XM_006721794:XM_017024409 | 13:2:4:1:0 | 1:0:0:0:0 | 9:1:2:0:3 | 11:2:1:4:0 | 15:1:6:7:2 | 1:1:0:1:0 | 17:4:7:10:2 | 22:5:1:1:0 | 12:2:4:5:1 | 1:0:0:0:0 |
| NM_002815:NM_001270482 | 127:07:00 | 0:00 | 195:14:00 | 0:00 | 381:15:00 | 0:00 | 451:20:00 | 1:00 | 208:11:00 | 0:00 |
| NM_002815:NM_001270482 | 127:07:00 | 0:00 | 194:14:00 | 1:00 | 384:15:00 | 0:00 | 452:19:00 | 0:00 | 209:11:00 | 0:00 |
| NM_002815:NM_001270482 | 127:07:00 | 0:00 | 194:14:00 | 1:00 | 384:15:00 | 0:00 | 453:20:00 | 0:00 | 209:11:00 | 0:00 |
| NM_002815:NM_001270482 | 127:07:00 | 1:00 | 196:14:00 | 0:00 | 384:15:00 | 0:00 | 454:20:00 | 0:00 | 209:11:00 | 0:00 |
| NM_002815:NM_001270482 | 128:07:00 | 0:00 | 197:14:00 | 1:00 | 384:15:00 | 1:00 | 455:19:00 | 1:01 | 209:11:00 | 0:00 |
| NM_002815:NM_001270482 | 126:07:00 | 2:00 | 197:14:00 | 1:00 | 384:15:00 | 0:00 | 456:20:00 | 1:00 | 209:11:00 | 0:00 |
| NM_002815:NM_001270482 | 127:07:00 | 0:00 | 200:14:00 | 0:00 | 384:15:00 | 1:00 | 458:20:00 | 2:00 | 209:11:00 | 0:00 |
| NM_002815:NM_001270482 | 127:07:00 | 0:00 | 202:14:00 | 0:00 | 389:15:00 | 0:00 | 460:21:00 | 0:00 | 209:11:00 | 0:00 |
| NM_002815:NM_001270482 | 126:07:00 | 0:00 | 201:14:00 | 0:00 | 389:15:00 | 0:00 | 457:21:00 | 3:00 | 210:11:00 | 0:00 |
| NM_001282:NM_001030006:XM_017024285:XM_017024286:XM_017024284:XM_011524454:XM_011524451:XM_011524449:XM_005257937 | 2:1:1:1:0:0:0:0:0 | 0:0:0:0:0:0:0:0:0 | 0:0:3:4:0:0:0:0:0 | 0:0:0:0:0:0:0:0:0 | 1:0:0:0:2:2:0:0:0 | 0:0:0:0:0:0:0:0:0 | 0:1:2:5:1:0:1:0:0 | 0:0:0:0:0:0:0:0:0 | 2:3:1:1:3:1:0:1:0 | 0:0:0:0:0:0:0:0:0 |
| NM_139215:NM_003487:XM_024450959 | 48:10:01 | 0:00:00 | 72:16:02 | 0:00:00 | 36:11:00 | 0:00:00 | 175:32:02 | 0:00:00 | 35:04:00 | 0:00:00 |
| NM_003487:NM_139215:XM_024450959 | 7:40:01 | 0:00:00 | 12:56:05 | 1:00:00 | 7:27:00 | 0:00:00 | 0.707789352 | 0:02:00 | 2:26:01 | 0:01:00 |
| NM_003487:NM_139215:XM_024450959 | 9:44:02 | 0:00:00 | 0.541759259 | 1:00:00 | 7:29:02 | 0:00:00 | 0.965439815 | 0:03:00 | 2:28:02 | 0:01:00 |
| NM_032965:NR_027922 | 0:00 | 0:00 | 6:00 | 0:00 | 6:03 | 0:00 | 2:01 | 0:00 | 1:00 | 2:00 |
| NM_007010:XM_011524233:XM_011524232:NM_001291476 | 5:4:12:0 | 0:0:0:0 | 16:5:9:1 | 0:0:0:0 | 14:2:11:0 | 0:0:0:0 | 33:21:34:0 | 0:1:0:0 | 10:2:13:0 | 0:0:0:0 |
| NM_007010:XM_011524233:XM_011524232:NM_001291476 | 5:3:12:0 | 0:0:0:0 | 16:2:9:1 | 0:0:0:0 | 14:3:12:0 | 0:0:0:0 | 0:1:0:0 | 33:20:34:0 | 1:0:2:0 | 9:2:11:0 |
| NM_007010:XM_011524233:XM_011524232:NM_001291476 | 5:3:12:0 | 0:0:0:0 | 16:2:9:1 | 0:0:0:0 | 14:2:12:0 | 0:0:0:0 | 33:20:35:0 | 0:0:0:0 | 10:2:13:0 | 0:0:0:0 |
| NM_032351 | 0 | 0 | 0 | 0 | 1 | 0 | 1 | 0 | 1 | 0 |
| NM_001136498 | 34 | 0 | 79 | 0 | 199 | 0 | 263 | 0 | 101 | 0 |
| NM_001136498 | 34 | 0 | 79 | 0 | 199 | 0 | 261 | 2 | 101 | 0 |
| NM_002795:NR_104195:NR_104194 | 652:02:00 | 2:00:00 | 676:05:00 | 2:00:00 | 1955:05:00 | 7:00:00 | 3821:04:00 | 6:00:00 | 1161:01:00 | 1:00:00 |
| NM_002795:NR_104195:NR_104194 | 659:02:00 | 0:00:00 | 684:05:00 | 1:00:00 | 1976:05:00 | 3:00:00 | 3847:04:00 | 6:00:00 | 1172:01:00 | 2:00:00 |
| NM_002795:NR_104195:NR_104194 | 671:00:00 | 1:00:00 | 715:01:00 | 1:00:00 | 2012:01:00 | 3:00:00 | 3913:03:00 | 7:00:00 | 1187:01:00 | 1:00:00 |
| NM_002795:NR_104195:NR_104194 | 673:00:00 | 0:00:00 | 715:00:00 | 0:00:00 | 2013:00:00 | 3:00:00 | 3917:00:00 | 3:01:00 | 1187:00:00 | 1:00:00 |
| NM_002795:NR_104195 | 673:00:00 | 1:00 | 727:00:00 | 1:00 | 2019:00:00 | 3:00 | 3929:00:00 | 4:00 | 1190:00:00 | 2:00 |
| NM_002795:NR_104194:NR_104195 | 668:00:00 | 0:00:00 | 723:00:00 | 2:00:00 | 1996:00:00 | 5:00:00 | 3900:00:00 | 3:00:00 | 1184:00:00 | 3:00:00 |
| NM_002795:NR_104194:NR_104195 | 662:00:00 | 0:00:00 | 716:00:00 | 0:00:00 | 1985:00:00 | 5:00:00 | 3875:00:00 | 3:00:00 | 1178:00:00 | 1:00:00 |
| XR_001752548:NR_073428:NM_017748 | 33:06:14 | 1:01:01 | 147:08:57 | 0:00:01 | 189:12:108 | 12:01:04 | 309:23:102 | 5:00:01 | 50:03:29 | 3:00:01 |
| NR_073428:NM_017748:XR_001752548 | 7:15:34 | 0:00:00 | 0.333622685 | 0:00:00 | 13:112:201 | 0:00:00 | 23:102:311 | 0:00:02 | 3:30:52 | 0:00:01 |
| NM_017748:XR_001752548:NR_073428 | 18:40:07 | 4:08:00 | 81:190:8 | 5:07:00 | 130:291:18 | 15:42:00 | 127:415:28 | 8:27:00 | 37:77:5 | 2:16:00 |
| NM_017748:XR_001752548:NR_073428 | 20:43:07 | 2:05:00 | 85:196:8 | 1:07:00 | 135:308:18 | 10:25:00 | 132:431:28 | 3:13:00 | 38:86:5 | 1:08:00 |
| XM_024450801:XM_017024753 | 0:00 | 0:00 | 3:02 | 1:02 | 0:00 | 2:00 | 4:00 | 1:01 | 1:00 | 0:00 |
| XM_017025217:NM_032192:NM_181505:XM_017025216:NM_001242464 | 9:38:31:3:9 | 1:0:0:0:0 | 35:49:79:9:13 | 1:0:0:0:0 | 40:157:192:14:28 | 0:0:1:0:0 | 129:568:423:66:115 | 1:0:1:1:0 | 8:49:87:9:21 | 1:0:0:0:0 |
| NM_181505:NM_032192:XM_017025217:XM_017025216:NM_001242464 | 37:38:11:3:9 | 1:0:0:0:0 | 88:49:47:9:13 | 1:0:0:0:0 | 210:157:57:14:28 | 1:0:0:0:0 | 481:572:167:67:114 | 3:1:1:0:1 | 95:49:18:9:21 | 0:0:0:0:0 |
| NM_032192:NM_181505:XM_017025217:XM_017025216:NM_001242464 | 35:41:15:2:8 | 0:0:0:0:0 | 41:82:37:6:12 | 6:10:11:2:0 | 156:238:64:12:28 | 0:0:2:1:0 | 541:477:187:62:111 | 20:10:1:2:1 | 47:94:18:8:21 | 1:2:0:0:0 |
| NM_032192:NM_181505:XM_017025217:XM_017025216:NM_001242464 | 34:41:15:2:8 | 0:0:0:0:0 | 41:77:37:6:12 | 0:5:0:0:0 | 156:239:65:12:28 | 0:2:1:0:0 | 538:476:188:61:110 | 0:15:6:0:0 | 48:95:18:8:21 | 0:0:0:0:0 |

| | | | | | | | | | | |
|---|---|---|---|---|---|---|---|---|---|---|
| NM_032192:NM_181505:XM_017025217:XM_017025216:NM_001242464 | 34:41:15:2:8 | 0:0:0:0:0 | 41:79:38:6:12 | 0:0:0:0:0 | 155:246:67:12:28 | 1:0:0:0:0 | 533:490:195:61:110 | 0:1:0:0:0 | 48:95:18:8:21 | 0:0:0:0:0 |
| NM_032192:NM_181505:XM_017025217:XM_017025216:NM_001242464 | 34:40:14:2:8 | 0:0:0:0:0 | 40:78:40:6:12 | 0:0:0:0:0 | 156:245:67:12:28 | 0:0:0:0:0 | 529:487:204:58:109 | 0:1:0:1:1 | 48:96:18:8:21 | 0:0:0:0:0 |
| NM_032192:NM_181505:XM_017025217:XM_017025216:NM_001242464 | 34:40:14:2:8 | 0:0:0:0:0 | 40:78:40:6:12 | 0:0:0:0:0 | 155:243:67:12:28 | 1:2:0:0:0 | 527:487:203:59:109 | 2:1:1:0:1 | 48:96:18:8:21 | 0:0:0:0:0 |
| NM_032192:NM_181505:XM_017025217:XM_017025216:NM_001242464 | 34:40:14:2:8 | 0:0:0:0:0 | 40:78:41:6:12 | 0:0:0:0:0 | 156:243:67:12:28 | 0:1:0:0:0 | 528:485:203:59:109 | 1:3:2:0:1 | 48:96:18:8:21 | 0:0:0:0:0 |
| XM_024450549:NM_006804 | 0:00 | 0:00 | 0:00 | 0:00 | 0:00 | 0:00 | 0:00 | 1:01 | 0:00 | 0:01 |
| NM_033419:NM_001291730:XM_011525480:XR_002958086:XM_011525481:NM_001291733:XR_934601 | 9:1:0:0:0:0:0 | 0:0:0:0:0:0:0 | 13:1:4:1:1:0:0 | 0:0:0:0:0:0:0 | 43:0:1:1:0:0:0 | 0:0:0:0:0:0:0 | 10:1:1:2:0:1:0 | 9:0:1:2:1:0:0 | 3:0:0:0:0:0:0 | 2:0:0:0:0:0:0 |
| XM_024450643:NM_004448:NR_110535:NM_001289937:NM_001289938 | 20:1:8:0:0 | 0:0:0:0:0 | 97:0:22:0:0 | 1:0:0:0:0 | 115:2:50:0:0 | 0:0:0:0:0 | 284:2:57:1:0 | 4:0:0:0:0 | 32:1:19:0:0 | 0:0:0:0:0 |
| XM_024450643:NM_004448:NR_110535:NM_001289937:NM_001289938 | 21:1:8:0:0 | 0:0:0:0:0 | 60:0:10:0:0 | 40:0:12:0:0 | 68:1:29:0:0 | 50:3:22:0:0 | 0:0:0:0:0 | 300:2:59:1:0 | 0:0:0:0:0 | 32:2:19:0:0 |
| XM_024450643:NM_004448:NR_110535:NM_001289937:NM_001289938 | 21:1:8:0:0 | 0:0:0:0:0 | 100:0:23:0:0 | 1:0:0:0:0 | 119:4:51:0:0 | 0:0:0:0:0 | 309:2:60:1:0 | 4:0:1:0:0 | 33:2:20:0:0 | 0:0:0:0:0 |
| NM_005310:NM_001030002:NM_001330207:XM_024450729:NM_001242443 | 0:0:0:0:0 | 5:2:0:0:0 | 0:0:0:0:0 | 2:0:1:0:0 | 0:0:0:0:0 | 7:2:0:1:0 | 0:0:0:0:0 | 56:31:4:11:4 | 0:0:0:0:0 | 11:6:1:2:0 |
| NM_001165959:NM_001165958:NM_001042471:NM_018530 | 4:2:2:0 | 0:0:0:0 | 7:33:12:0 | 9:10:23:1 | 10:32:13:1 | 3:6:20:1 | 6:28:11:2 | 6:2:13:1 | 2:15:13:0 | 0:1:5:0 |
| NM_001165959:NM_001165958:NM_001042471:NM_018530 | 4:2:2:0 | 0:0:0:0 | 7:33:12:0 | 9:10:23:1 | 10:32:13:1 | 3:6:20:1 | 6:28:11:2 | 6:2:13:1 | 2:15:13:0 | 0:1:5:0 |
| NM_001254:XM_011525542 | 10:00 | 0:00 | 1:00 | 0:00 | 3:01 | 0:00 | 17:01 | 13:01 | 11:00 | 5:01 |
| NM_001254:XM_011525542 | 10:00 | 0:00 | 1:00 | 0:00 | 3:01 | 0:00 | 24:03:00 | 0:00 | 14:00 | 1:01 |
| XM_011525095:NM_001145301:NM_001024809:XM_017024920:NM_000964:NM_001145302:XM_005257553 | 2:7:10:2:0:0:0 | 1:0:0:0:0:0:0 | 1:1:14:9:1:0:0 | 0:0:0:1:0:0:0 | 1:5:36:9:2:0:0 | 0:0:0:0:0:0:0 | 7:7:52:29:6:1:0 | 0:0:2:1:0:0:0 | 0:3:7:3:2:0:0 | 0:0:0:1:0:0:0 |
| NM_032865:XM_005257744:XM_017025236:XR_001752669 | 1:0:0:0 | 0:0:0:0 | 0:0:0:0 | 0:0:0:0 | 0:3:0:0 | 0:0:0:0 | 3:9:2:0 | 0:0:0:0 | 0:4:0:0 | 0:0:0:0 |
| XR_001752669:XM_005257744:NM_032865 | 0:00:00 | 0:00:00 | 0:00:00 | 0:00:00 | 1:00:00 | 0:00:00 | 0:02:00 | 0:00:00 | 0:00:00 | 0:00:00 |
| NM_003079 | 77 | 0 | 74 | 4 | 93 | 0 | 700 | 28 | 155 | 0 |
| NM_003079 | 81 | 1 | 78 | 4 | 90 | 0 | 755 | 11 | 158 | 3 |
| NM_003079 | 85 | 0 | 82 | 0 | 91 | 0 | 771 | 4 | 164 | 1 |
| NM_003079 | 85 | 0 | 82 | 0 | 90 | 0 | 766 | 3 | 165 | 0 |
| NM_003079 | 83 | 0 | 77 | 0 | 87 | 0 | 722 | 3 | 161 | 0 |
| NM_003079 | 83 | 0 | 77 | 0 | 87 | 0 | 723 | 0 | 159 | 0 |
| NM_002276 | 1320 | 1 | 5794 | 15 | 4933 | 7 | 3899 | 6 | 2308 | 2 |
| NM_002276 | 1317 | 0 | 5693 | 30 | 4899 | 5 | 3864 | 5 | 2286 | 5 |
| NM_002276 | 1308 | 0 | 5554 | 11 | 4879 | 8 | 3818 | 3 | 2270 | 1 |
| NM_002276 | 1304 | 4 | 5520 | 41 | 4882 | 6 | 3816 | 2 | 2271 | 0 |
| NM_002276 | 1297 | 1 | 5257 | 16 | 4819 | 7 | 3748 | 4 | 2227 | 3 |
| NM_002276 | 1256 | 2 | 4938 | 13 | 4656 | 9 | 3625 | 8 | 2160 | 5 |
| NM_002276 | 1253 | 2 | 4893 | 13 | 4645 | 7 | 3611 | 2 | 2151 | 1 |
| NM_002276 | 1253 | 1 | 4883 | 14 | 4641 | 7 | 3605 | 5 | 2146 | 2 |
| NM_002276 | 1217 | 2 | 4582 | 2 | 4483 | 4 | 3470 | 3 | 2084 | 0 |
| NM_002276 | 1206 | 3 | 4490 | 4 | 4426 | 5 | 3436 | 7 | 2079 | 1 |
| NM_002276 | 1143 | 3 | 4072 | 12 | 4234 | 5 | 3253 | 12 | 1999 | 0 |
| NM_002276 | 1141 | 0 | 4015 | 24 | 4217 | 5 | 3228 | 11 | 1987 | 0 |
| NM_002276 | 1147 | 0 | 4042 | 16 | 4230 | 5 | 3237 | 13 | 1996 | 3 |
| NM_002276 | 1162 | 5 | 4108 | 32 | 4274 | 10 | 3270 | 20 | 2011 | 3 |
| NM_002276 | 3 | 1174 | 10 | 4121 | 5 | 4288 | 10 | 3285 | 0 | 2027 |
| NM_002276 | 1162 | 5 | 4074 | 29 | 4282 | 13 | 3254 | 17 | 2021 | 1 |
| NM_002276 | 1164 | 2 | 4059 | 34 | 4263 | 13 | 3251 | 15 | 2018 | 3 |
| XM_017024590:NM_021991:NM_002230 | 0:00:00 | 0:00:00 | 0:00:00 | 0:00:00 | 0:01:00 | 2:00:00 | 1:00:00 | 0:00:00 | 0:00:00 | 0:00:00 |
| XM_017024590:NM_021991:NM_002230 | 0:00:00 | 0:00:00 | 0:00:00 | 0:00:00 | 0:00:00 | 2:01:00 | 0:00:00 | 1:00:00 | 0:00:00 | 0:00:00 |
| NM_052935:NR_033465 | 0:00 | 0:00 | 0:00 | 0:00 | 0:00 | 0:00 | 0:00 | 1:01 | 0:00 | 0:00 |
| NM_052935:NR_033465 | 0:00 | 0:00 | 0:00 | 0:00 | 0:00 | 0:00 | 0:00 | 1:01 | 0:00 | 0:00 |
| NM_001096:XM_005257395:NM_198830:NM_001303274 | 2:1:0:0 | 0:0:0:0 | 0:0:0:0 | 0:0:0:0 | 2:0:1:0 | 0:0:0:0 | 1:1:0:0 | 0:0:0:0 | 1:2:0:1 | 0:0:0:0 |
| NM_003315:XR_001752602:XM_017024994:XM_011525167:XR_001752603:NR_029431 | 100:3:4:1:2:0 | 1:0:0:0:0:0 | 134:3:4:1:0:0 | 0:0:0:0:0:0 | 97:3:4:2:0:1 | 0:0:0:0:0:0 | 459:14:16:11:4:0 | 1:0:0:0:0:0 | 122:1:6:4:2:1 | 0:0:0:0:0:0 |
| NM_139276:NM_003150:XM_024450896:NM_213662:XM_017024973 | 5:0:0:0:0 | 0:0:0:0:0 | 0:2:1:0:0 | 0:0:0:0:0 | 5:2:5:1:1 | 0:0:0:0:0 | 13:10:10:3:1 | 0:0:0:0:0 | 4:4:0:0:1 | 0:0:0:0:0 |
| XM_024450703:NM_016437 | 0:00 | 0:00 | 0:00 | 0:00 | 0:00 | 0:00 | 1:00 | 1:00 | 0:00 | 0:00 |
| NM_001261434 | 100 | 0 | 48 | 0 | 200 | 1 | 685 | 2 | 189 | 0 |
| NM_001261434 | 102 | 0 | 49 | 0 | 201 | 0 | 686 | 2 | 190 | 0 |
| NM_001261434 | 102 | 0 | 49 | 0 | 200 | 0 | 685 | 1 | 190 | 0 |
| NM_001261434 | 102 | 0 | 49 | 0 | 199 | 0 | 685 | 1 | 190 | 0 |
| NM_001330230:NM_005533 | 0:00 | 1:00 | 0:00 | 0:00 | 0:00 | 0:01 | 0:00 | 0:01 | 0:00 | 0:00 |
| NM_007298:NM_007299:NR_027676 | 0:00:00 | 1:00:00 | 0:00:00 | 0:00:00 | 0:00:01 | 0:01:01 | 0:00:00 | 0:00:01 | 0:00:02 | 0:00:00 |
| NM_004941:NR_136228:NM_001322219:NM_001322217:NM_001322221:NR_136226 | 0:1:0:0:0:0 | 1:2:0:0:0:0 | 0:2:0:0:0:0 | 1:1:0:0:0:0 | 0:7:0:0:0:0 | 0:3:0:0:0:0 | 2:10:1:0:0:0 | 0:0:0:0:0:0 | 0:1:0:0:0:0 | 0:0:0:0:0:0 |
| NR_136228:NM_001322218:NM_004941:NM_001322219:NM_001322220:NR_136226 | 2:0:0:0:0:0 | 0:0:0:0:0:0 | 1:1:1:0:0:0 | 0:0:0:0:0:0 | 0:0:2:0:0:0 | 0:0:0:0:0:0 | 11:0:10:1:0:0 | 1:0:0:0:0:0 | 1:0:4:0:1:0 | 0:0:0:0:0:0 |
| NM_001261438:NM_001986:XM_024450644:NM_001079675:NM_001261439:NM_001261437 | 5:17:3:7:0:0 | 0:0:0:0:0:0 | 0:0:0:0:1:0 | 0:0:0:0:0:0 | 0:1:0:0:0:0 | 0:0:0:0:0:0 | 87:345:94:33:9:6 | 0:2:2:0:0:0 | 24:44:6:17:2:1 | 0:1:1:0:0:0 |
| NM_001261438:NM_001986:XM_024450644:NM_001079675:NM_001261439:NM_001261437 | 5:17:3:7:0:0 | 0:0:0:0:0:0 | 0:0:0:0:1:0 | 0:0:0:0:0:0 | 0:1:0:0:0:0 | 0:0:0:0:0:0 | 87:338:90:33:9:6 | 0:2:0:0:0:0 | 24:44:6:17:2:1 | 0:0:0:0:0:0 |
| NM_001261438:NM_001986:XM_024450644:NM_001079675:NM_001261439:NM_001261437 | 5:17:3:7:0:0 | 0:0:0:0:0:0 | 0:0:0:0:1:0 | 0:0:0:0:0:0 | 0:1:0:0:0:0 | 0:0:0:0:0:0 | 87:340:89:33:9:6 | 0:2:0:0:0:0 | 24:43:6:17:2:1 | 0:1:0:0:0:0 |
| NM_004160 | 0 | 0 | 0 | 0 | 1 | 1 | 0 | 0 | 0 | 0 |
| NM_004160 | 0 | 0 | 0 | 0 | 0 | 2 | 0 | 0 | 0 | 0 |
| NM_001076684:XM_017025004:NR_045058:XM_006722061:NM_001076683:XM_006722059:XM_017025003:XM_024450907:XM_017025000:NM_014233:XM_017024997:XM_017025001:XM_017024996:XM_017024995 | 3:4:1:1:1:0:0:0:0:0:0:0:0:0 | 0:0:0:0:0:0:0:0:0:0:0:0:0:0 | 0:19:0:1:0:1:2:2:1:0:0:0:0:0 | 0:1:0:0:0:0:0:0:0:0:0:0:0:0 | 13:16:0:1:4:2:2:1:0:4:0:0:0:0 | 0:0:0:0:0:0:0:0:0:0:0:0:0:0 | 2:92:1:2:4:3:2:3:0:0:2:1:0:0 | 0:0:0:0:0:0:0:0:0:0:0:0:0:0 | 3:9:1:0:1:1:8:0:0:2:0:0:0:0 | 0:0:0:0:0:0:0:0:0:0:0:0:0:0 |
| NM_153338 | 0 | 0 | 0 | 2 | 0 | 0 | 0 | 0 | 0 | 2 |
| NM_145255 | 0 | 0 | 0 | 0 | 1 | 0 | 0 | 0 | 2 | 0 |

| | | | | | | | | | | |
|---|---|---|---|---|---|---|---|---|---|---|
| XM_017025166:NM_176096:NM_001278216:XM_017025164:XM_017025165:XM_011525297:NM_001278217:NM_001278198 | 1:15:6:0:0:0:0:0 | 1:0:0:0:0:0:0:0 | 9:17:3:1:3:1:0:0 | 0:0:0:0:0:0:0:0 | 1:17:8:0:1:1:1:1 | 0:0:0:0:0:0:0:0 | 28:130:13:0:13:2:1:2 | 0:0:0:0:0:0:1:0 | 7:26:6:1:0:1:1:0 | 0:2:0:0:0:0:0:0 |
| NM_176096:NM_001278216:XM_017025165:XM_017025166:XM_011525297:NM_001278217:NM_001278198:XM_017025164 | 14:7:0:0:0:0:0:0 | 0:0:0:0:0:0:0:0 | 16:6:3:6:1:0:0:0 | 0:0:0:0:0:0:0:0 | 16:8:1:0:1:1:0:0 | 0:0:0:0:0:0:0:0 | 116:19:12:32:2:0:2:0 | 3:0:0:0:0:0:0:0 | 28:6:0:4:0:1:0:1 | 0:0:0:0:0:0:0:0 |
| NM_176096:XM_017025166:NM_001278216:NM_001278198:XM_017025165:NM_001278217:XM_011525297:XM_017025164 | 2:1:0:0:0:0:0:0 | 14:3:7:1:0:0:0:0 | 8:4:3:0:1:1:1:0 | 9:6:3:1:2:0:0:0 | 8:0:7:1:1:1:1:0 | 9:1:1:0:0:0:0:2 | 76:35:10:1:10:2:2:0 | 52:16:9:1:3:0:0:0 | 19:10:3:0:0:1:0:0 | 10:2:3:0:0:0:0:1 |
| NM_003726:NM_001075099:XM_024451012:XM_017025260:XM_005257755:XM_017025258:XM_017025259 | 0:0:0:0:0:0:0 | 38:22:4:1:1:0:0 | 2:0:1:0:2:0:0 | 0:3:0:0:2:0:0 | 2:2:1:0:0:1:1 | 2:1:0:0:0:0:0 | 66:26:4:0:9:0:0 | 79:23:2:0:10:1:2 | 8:4:0:0:0:0:0 | 14:5:0:0:2:1:0 |
| NM_005831 | 0 | 0 | 0 | 0 | 5 | 0 | 0 | 2 | 1 | 0 |
| NM_005831 | 0 | 0 | 0 | 0 | 2 | 3 | 1 | 0 | 1 | 0 |
| NM_003563:NM_001007229:NM_001007228:NM_001007230:NM_001007227:NM_001007226:XM_024450995 | 14:17:149:4:14:7:12 | 0:0:0:0:0:0:0 | 75:43:171:8:23:4:46 | 0:0:1:1:0:0:0 | 49:133:539:20:66:15:69 | 0:1:2:0:0:0:1 | 359:86:847:67:232:51:247 | 2:1:3:0:0:0:2 | 13:12:137:12:29:11:16 | 0:0:0:0:0:0:0 |
| NM_003563:NM_001007229:NM_001007228:NM_001007230:NM_001007227:NM_001007226:XM_024450995 | 14:17:149:4:14:7:12 | 0:0:0:0:0:0:0 | 74:43:171:9:23:4:44 | 0:0:0:0:0:0:0 | 46:134:537:20:65:15:70 | 0:0:3:0:0:0:0 | 351:87:849:67:232:51:245 | 2:0:1:0:0:0:0 | 13:12:137:12:29:11:16 | 0:0:0:0:0:0:0 |
| NM_003563:NM_001007229:NM_001007228:NM_001007230:NM_001007227:NM_001007226:XM_024450995 | 12:17:151:4:14:7:12 | 0:0:0:0:0:0:0 | 70:43:170:9:23:4:42 | 0:0:0:0:0:0:0 | 42:132:542:20:66:15:70 | 0:0:1:0:0:0:0 | 313:87:857:67:234:52:235 | 1:0:0:0:0:0:1 | 12:12:137:12:29:10:16 | 1:0:0:0:0:1:0 |
| NM_004890:XM_006721600 | 139:01:00 | 0:00 | 158:01:00 | 0:00 | 1059:24:00 | 4:00 | 758:07:00 | 0:00 | 143:01:00 | 0:00 |
| NM_004890:XM_006721600 | 140:01:00 | 0:00 | 161:01:00 | 0:00 | 1064:15:00 | 2:00 | 762:07:00 | 0:00 | 142:01:00 | 0:00 |
| NM_001007228:NM_003563:NM_001007229:NM_001007230:NM_001007227:NM_001007226:XM_024450995 | 151:10:17:4:14:7:13 | 1:0:0:0:0:0:0 | 174:63:45:10:23:4:39 | 0:1:0:0:0:0:0 | 541:40:132:20:67:15:67 | 4:0:0:0:0:0:0 | 866:273:88:67:234:52:211 | 0:0:0:0:0:0:1 | 138:12:12:12:29:11:15 | 0:0:0:0:0:0:0 |
| NM_001007229:NM_001007228:NM_003563:NM_001007230:NM_001007227:NM_001007226:XM_024450995 | 17:152:8:4:14:7:7 | 0:0:0:0:0:0:0 | 45:177:40:10:23:4:26 | 0:0:0:0:0:0:0 | 135:548:36:20:67:15:50 | 0:2:0:0:0:0:0 | 90:872:181:67:241:51:132 | 0:0:2:0:0:1:0 | 12:139:11:13:30:11:16 | 0:0:0:0:0:0:0 |
| NM_001007229:NM_001007228:NM_003563:NM_001007230:NM_001007227:NM_001007226:XM_024450995 | 17:152:9:4:14:7:6 | 0:0:0:0:0:0:0 | 45:177:37:10:23:4:25 | 0:0:0:0:0:0:0 | 135:549:36:20:67:15:50 | 0:0:0:0:0:0:0 | 90:873:158:68:241:52:114 | 0:0:0:0:0:0:1 | 12:138:11:13:30:11:15 | 0:1:0:0:0:0:0 |
| NM_003563:NM_001007229:NM_001007228:NM_001007230:NM_001007227:NM_001007226:XM_024450995 | 10:17:152:4:14:7:6 | 0:0:0:0:0:0:0 | 36:45:177:10:23:4:20 | 0:0:1:0:0:0:0 | 36:135:549:20:67:15:45 | 0:0:1:0:0:0:0 | 130:90:874:68:241:52:103 | 0:0:0:0:0:0:1 | 11:12:139:13:30:11:14 | 0:0:0:0:0:0:0 |
| NM_003563:NM_001007229:NM_001007228:NM_001007230:NM_001007227:NM_001007226:XM_024450995 | 10:17:152:4:14:7:6 | 0:0:0:0:0:0:0 | 34:45:175:10:23:4:20 | 0:0:2:0:0:0:0 | 35:135:550:20:67:15:43 | 1:0:0:0:0:0:0 | 122:90:875:68:241:53:96 | 0:0:0:0:0:0:0 | 10:12:139:13:30:11:14 | 0:0:0:0:0:0:0 |
| NM_001007227:NM_003563:NM_001007229:NM_001007228:NM_001007230:NM_001007226:XM_024450995 | 13:9:17:152:4:7:5 | 1:0:0:0:0:0:0 | 23:34:45:178:10:4:16 | 0:0:0:0:0:0:0 | 68:37:134:549:20:15:41 | 0:1:1:1:0:0:0 | 242:91:90:877:69:53:75 | 0:1:0:1:0:0:0 | 30:10:12:139:13:11:13 | 0:0:0:0:0:0:0 |
| NM_003563:NM_001007229:NM_001007228:NM_001007230:NM_001007227:NM_001007226:XM_024450995 | 9:17:152:4:14:7:5 | 0:0:0:0:0:0:0 | 34:45:178:10:23:4:16 | 0:0:0:0:0:0:0 | 38:134:548:20:68:15:40 | 0:1:2:0:0:0:0 | 92:90:876:69:241:53:74 | 0:0:2:0:1:0:0 | 9:12:138:13:30:11:13 | 1:0:1:0:0:0:0 |
| NM_005827:XM_011524180:NM_001278784:XM_006721632:XM_011524179:XM_005256929 | 113:11:42:4:2:0 | 0:0:0:0:0:0 | 91:36:37:7:4:0 | 0:0:0:0:0:0 | 189:22:56:10:8:1 | 0:0:1:0:0:0 | 566:109:356:48:96:5 | 3:0:0:0:0:0 | 140:14:56:9:27:3 | 0:0:0:0:0:0 |
| NM_005827:XM_011524180:NM_001278784:XM_006721632:XM_011524179:XM_005256929 | 112:11:42:4:2:0 | 1:0:0:0:0:0 | 91:36:37:7:4:0 | 0:0:0:0:0:0 | 189:21:57:10:8:1 | 0:0:0:0:0:0 | 569:107:356:48:96:5 | 0:0:0:0:0:0 | 140:14:56:9:27:3 | 0:0:0:0:0:0 |
| NM_005827:XM_011524180:NM_001278784:XM_006721632:XM_011524179:XM_005256929 | 113:11:43:4:2:0 | 0:0:0:0:0:0 | 90:36:37:7:4:0 | 1:0:0:0:0:0 | 188:21:57:10:8:1 | 1:0:0:0:0:0 | 567:106:355:48:96:5 | 2:0:1:0:0:0 | 140:14:56:9:27:3 | 0:0:0:0:0:0 |
| NM_005827:XM_011524180:NM_001278784:XM_006721632:XM_011524179:XM_005256929 | 113:11:43:4:2:0 | 0:0:0:0:0:0 | 91:36:37:7:4:0 | 0:0:0:0:0:0 | 189:20:57:10:8:1 | 0:1:0:0:0:0 | 569:106:356:48:96:5 | 0:0:0:0:0:0 | 140:14:56:9:27:3 | 0:0:0:0:0:0 |
| NM_005827:XM_011524180:NM_001278784:XM_006721632:XM_011524179:XM_005256929 | 113:5:43:5:2:0 | 0:0:0:0:0:0 | 92:30:36:8:4:0 | 0:0:1:0:0:0 | 189:12:57:10:8:1 | 1:0:0:0:0:0 | 568:85:356:52:96:5 | 2:0:1:0:0:0 | 141:11:56:10:27:3 | 0:0:0:0:0:0 |
| NM_005827:NM_001278784:XM_011524180:XM_006721632:XM_011524179:XM_005256929 | 102:35:3:5:2:0 | 12:9:0:0:0:0 | 54:18:5:6:2:0 | 40:21:4:2:2:0 | 118:42:3:5:4:1 | 72:16:2:5:4:0 | 301:192:16:30:46:2 | 270:165:14:22:50:3 | 81:27:6:6:21:1 | 60:30:0:4:7:2 |
| NM_005827:XM_011524180:NM_001278784:XM_006721632:XM_005256929 | 114:4:44:2:0 | 0:0:0:0:0 | 93:9:39:6:0 | 1:0:0:0:0 | 190:8:57:7:1 | 0:0:1:0:0 | 571:52:356:39:5 | 1:0:1:0:0 | 141:9:57:8:3 | 0:0:0:0:0 |
| XM_005257572:NR_110010 | 0:00 | 0:00 | 1:00 | 0:00 | 0:00 | 0:00 | 0:00 | 1:01 | 0:00 | 0:00 |
| NM_018509 | 31 | 0 | 10 | 1 | 70 | 1 | 96 | 1 | 71 | 0 |
| NM_004703:XM_011524060:XM_017025295:NM_001291582:NM_001083585:NM_001291581 | 2:9:0:0:0:0 | 1:2:0:0:0:0 | 3:12:1:4:0:0 | 1:3:1:0:0:0 | 2:3:1:0:0:0 | 0:0:0:0:0:0 | 9:20:0:35:1:1 | 2:13:0:0:0:0 | 2:3:0:5:0:0 | 0:1:0:0:0:0 |
| NM_004703:XM_011524060:NM_001291582:XM_017025295:NM_001083585:NM_001291581 | 2:9:0:0:0:0 | 1:1:0:0:0:0 | 4:12:4:1:0:0 | 0:1:0:0:0:0 | 2:3:0:1:0:0 | 0:0:0:0:0:0 | 8:19:35:0:1:1 | 1:5:0:0:0:0 | 2:3:5:0:0:0 | 0:1:0:0:0:0 |
| NM_001033002:NM_001160246:NR_027683:NM_001160266:NM_001160244:XR_001752663 | 0:0:0:0:0:0 | 0:0:0:0:0:0 | 0:0:0:0:0:0 | 0:0:0:0:0:0 | 1:0:0:0:0:0 | 0:0:0:0:0:0 | 1:1:0:0:0:0 | 0:0:0:0:0:0 | 1:0:2:1:0:0 | 0:0:0:0:0:0 |
| NM_001033053 | 0 | 0 | 0 | 0 | 0 | 0 | 0 | 0 | 0 | 0 |
| NM_021213:NM_001330377:NM_001330378:XR_001752577 | 11:0:0:0 | 11:1:0:0 | 6:0:1:0 | 5:0:0:0 | 10:0:0:0 | 9:0:0:0 | 21:1:0:0 | 20:0:0:2 | 9:0:0:0 | 9:1:0:0 |
| NM_016070 | 173 | 0 | 91 | 0 | 136 | 0 | 996 | 2 | 250 | 2 |
| NM_016070 | 173 | 0 | 91 | 0 | 136 | 0 | 998 | 0 | 252 | 0 |
| NM_016070 | 172 | 0 | 87 | 1 | 134 | 1 | 992 | 1 | 252 | 0 |
| NM_016070 | 170 | 0 | 88 | 0 | 135 | 0 | 974 | 2 | 252 | 0 |
| NM_017763:XM_017024800:NM_001305544:XM_011524955:XM_011524956 | 25:0:0:0:0 | 0:0:0:0:0 | 1:1:0:0:0 | 0:0:0:0:0 | 2:0:0:0:0 | 0:0:0:0:0 | 94:1:0:0:2 | 81:2:1:1:0 | 10:1:0:0:0 | 12:1:0:0:0 |
| NM_017763:XM_017024800:XM_011524955:NM_001305544:XM_011524956 | 24:0:0:0:0 | 0:0:0:0:0 | 0:1:0:0:0 | 0:0:0:0:0 | 2:0:0:0:0 | 0:0:0:0:0 | 162:4:0:1:2 | 2:1:1:0:0 | 22:2:0:0:0 | 0:0:0:0:0 |
| NM_017763:XM_017024800:XM_011524955:XM_011524956 | 23:0:0:0 | 0:0:0:0 | 0:1:0:0 | 0:0:0:0 | 2:0:0:0 | 0:0:0:0 | 60:3:0:2 | 51:3:1:0 | 10:1:0:0 | 12:1:0:0 |
| NM_017763:XM_017024800:XM_011524956:XM_011524955 | 9:0:0:0 | 9:0:0:0 | 0:1:0:0 | 0:0:0:0 | 1:0:0:0 | 0:0:0:0 | 11:4:1:0 | 17:4:1:1 | 5:1:0:0 | 2:1:0:0 |
| NM_017763:XM_017024800:XM_011524956:NM_001305544:XM_011524955 | 8:0:0:0:0 | 9:0:0:0:0 | 0:1:0:0:0 | 0:0:0:0:0 | 1:1:0:0:0 | 0:0:0:0:0 | 25:7:1:2:1 | 2:1:0:0:0 | 7:2:0:0:0 | 0:0:0:0:0 |
| NM_017763:XM_017024800:XM_011524956:NM_001305544:XM_011524955 | 9:0:0:0:0 | 4:0:0:0:0 | 0:0:0:0:0 | 0:0:0:0:0 | 0:0:0:0:0 | 1:1:0:0:0 | 15:10:1:4:1 | 0:0:0:0:0 | 4:1:0:0:0 | 0:0:0:0:0 |
| NM_017763:XM_011524955:XM_017024800:XM_011524956:NM_001305544 | 13:1:1:0:0 | 0:0:0:0:0 | 0:0:0:0:0 | 0:0:0:0:0 | 1:0:1:0:0 | 0:0:0:0:0 | 22:2:15:2:5 | 0:0:0:2:0 | 4:0:1:0:0 | 0:0:0:0:0 |
| NM_017763:XM_017024800:XM_011524955:NM_001305544 | 2:1:2:1 | 9:3:0:0 | 1:0:0:0 | 0:0:0:1 | 1:1:0:0 | 1:0:0:0 | 32:77:5:17 | 32:80:11:6 | 2:0:0:0 | 1:2:0:0 |
| NM_024612:XM_011525253:XM_017025081:XM_017025083:XM_006722083:XM_017025080:XM_011525254:XM_017025082 | 11:2:1:1:0:0:0:0 | 0:0:0:0:0:0:0:0 | 4:5:0:4:0:0:0:0 | 0:0:0:0:0:0:0:0 | 10:3:0:0:1:0:0:0 | 0:0:0:0:0:0:0:0 | 20:15:0:56:0:2:0:0 | 0:0:0:0:0:0:0:0 | 4:4:1:2:0:0:0:0 | 0:0:0:0:0:0:0:0 |
| NM_004859 | 48 | 0 | 11 | 0 | 53 | 0 | 59 | 0 | 46 | 0 |
| XM_011524887:NM_016077:NM_001015509 | 0.257013889 | 2:00:00 | 5:56:01 | 0:00:00 | 3:56:01 | 0:00:00 | 27:343:56 | 0:02:01 | 0.321597222 | 0:00:00 |
| XM_005257425:NM_001193609:NM_016261:XM_024450784:NM_001193612:XM_024450783:XM_017024720:XM_017024716 | 3:1:0:0:0:0:0:0 | 0:0:0:0:0:0:0:0 | 2:1:3:1:0:0:0:0 | 0:0:0:0:0:0:0:0 | 0:1:1:0:1:0:0:0 | 0:0:0:0:0:0:0:0 | 5:2:13:1:1:1:2:2 | 0:0:0:0:0:0:0:0 | 0:0:4:0:0:0:0:0 | 0:0:0:0:0:0:0:0 |
| NM_001320470:NM_017679:NM_001330414:NM_001353145 | 0:0:0:0 | 1:0:0:0 | 0:0:0:0 | 0:0:0:0 | 0:0:0:0 | 0:5:1:0 | 0:0:0:0 | 1:5:0:1 | 0:0:0:0 | 0:0:0:0 |

| | | | | | | | | | | |
|---|---|---|---|---|---|---|---|---|---|---|
| NM_001320470:NM_001330414:NM_017679:XM_017024787:XM_017024788 | 1:0:0:0:0 | 0:0:0:0:0 | 2:0:0:0:0 | 0:0:0:0:0 | 0:1:4:0:0 | 0:0:0:0:0 | 1:0:1:0:0 | 0:0:3:0:0 | 1:0:0:0:0 | 0:0:0:0:0 |
| NM_018289:XR_001752553 | 0:00 | 0:00 | 0:00 | 0:00 | 0:00 | 0:00 | 2:00 | 0:00 | 0:00 | 0:00 |
| NM_001099789:NM_000873:NM_001099787:NM_001099788:NM_001099786 | 26:16:2:1:0 | 1:0:0:0:0 | 25:9:2:4:0 | 0:0:0:0:0 | 32:2:3:1:0 | 2:0:0:0:0 | 67:8:8:2:0 | 0:0:0:0:0 | 15:6:2:1:0 | 0:0:0:0:0 |
| NM_004396:NM_001320596:NM_001320595:NM_001320597 | 475:1:6:0 | 0:0:0:0 | 512:0:6:4 | 4:0:0:0 | 303:0:5:1 | 1:0:0:0 | 1565:7:27:1 | 4:0:0:0 | 363:3:3:0 | 1:0:0:0 |
| NM_004396:NM_001320596:NM_001320595:NM_001320597 | 463:1:6:0 | 0:0:0:0 | 452:0:6:4 | 1:0:0:0 | 298:0:5:1 | 0:0:0:0 | 1524:5:27:1 | 1:0:0:0 | 353:3:3:0 | 0:0:0:0 |
| NM_001320595:NM_004396:NM_001320596:NM_001320597 | 5:431:1:0 | 1:0:0:0 | 6:355:0:4 | 0:2:0:0 | 5:287:0:1 | 0:0:0:0 | 27:1376:3:1 | 0:1:0:0 | 3:328:2:0 | 0:0:0:0 |
| NM_004396:NM_001320596:NM_001320595:NM_001320597 | 429:1:6:0 | 0:0:0:0 | 354:0:6:4 | 1:0:0:0 | 287:0:5:1 | 0:0:0:0 | 1363:3:27:1 | 0:0:0:0 | 324:2:3:0 | 0:0:0:0 |
| NM_004396:NM_001320596:NM_001320595:NM_001320597 | 425:1:6:0 | 0:0:0:0 | 353:0:6:4 | 1:0:0:0 | 287:0:5:1 | 0:0:0:0 | 1356:3:27:1 | 3:0:0:0 | 322:2:3:0 | 1:0:0:0 |
| NM_004396:NM_001320596:NM_001320595:NM_001320597 | 425:1:6:0 | 0:0:0:0 | 353:0:6:4 | 1:0:0:0 | 286:0:5:1 | 0:0:0:0 | 1356:3:27:1 | 3:0:0:0 | 323:2:3:0 | 0:0:0:0 |
| NM_004396:NM_001320596:NM_001320595:NM_001320597 | 415:1:6:0 | 1:0:0:0 | 339:0:6:4 | 0:0:0:0 | 275:0:5:1 | 2:0:0:0 | 1306:2:27:1 | 1:0:0:0 | 315:2:3:0 | 1:0:0:0 |
| NM_004396:NM_001320596:NM_001320595:NM_001320597 | 403:1:6:0 | 1:0:0:0 | 305:0:6:4 | 1:0:0:0 | 267:0:5:1 | 2:0:0:0 | 1111:2:27:1 | 5:0:0:0 | 297:2:3:0 | 0:0:0:0 |
| NM_004396:NM_001320596:NM_001320595:NM_001320597 | 402:1:6:0 | 1:0:0:0 | 306:0:6:4 | 0:0:0:0 | 269:0:5:1 | 0:0:0:0 | 1101:2:27:1 | 8:0:0:0 | 297:1:3:0 | 0:0:0:0 |
| XM_017025192:XM_011525320:XM_017025193:XM_011525319:NM_004655 | 0:0:0:0:0 | 2:1:0:0:0 | 0:0:0:0:0 | 0:0:1:0:0 | 0:0:0:0:0 | 0:0:0:1:0 | 0:0:0:0:0 | 1:0:0:3:4 | 0:0:0:0:0 | 0:0:0:0:0 |
| XR_934126:NR_147088:NM_014804:XR_001752707 | 0:0:0:0 | 0:0:0:0 | 1:0:0:0 | 0:0:0:0 | 0:0:0:0 | 0:0:0:0 | 0:0:0:0 | 0:0:0:0 | 0:1:0:0 | 0:0:0:0 |
| NM_002816:XR_001752571:XM_024450843:XM_024450842 | 21:2:0:0 | 0:0:0:0 | 7:2:0:0 | 0:1:0:0 | 13:2:1:0 | 0:1:0:0 | 46:17:2:0 | 1:0:0:0 | 13:6:0:0 | 0:1:0:0 |
| NM_015462 | 29 | 0 | 19 | 0 | 24 | 0 | 174 | 2 | 37 | 0 |
| NM_001033570:NR_144442:NM_001033571:NM_001346478 | 1:0:0:0 | 0:0:0:0 | 0:1:0:0 | 0:0:0:0 | 0:0:0:0 | 0:0:0:0 | 0:0:1:1 | 0:0:0:0 | 0:0:0:0 | 0:0:0:0 |
| NM_017983 | 0 | 1 | 0 | 0 | 0 | 1 | 0 | 0 | 0 | 0 |
| NM_001185022:NM_001307:NM_001185023 | 1:00:00 | 0:00:00 | 2:12:00 | 0:00:00 | 15:18:00 | 0:00:00 | 0:06:00 | 0:00:00 | 1:08:00 | 0:00:00 |
| NM_139177:NM_001159770:XM_006721757:NM_001352692:XM_017024342:XM_011524496:XM_017024339:XM_017024330:NM_001352693:XM_017024332:XM_011524494:XM_011524495:XM_006721756:XM_017024337:XM_017024335:XM_006721759:XM_017024340:XM_017024341:XM_011524498:XM_006721758:XM_006721755 | 0:0:0:0:0:0:0:0:0:0:0:0:0:0:0:0:0:0:0:0:0:0:0:0 | 22:2:1:0:0:0:0:0:0:0:0:0:0:0:0:0:0:0:0:0:0 | 0:0:0:0:0:0:0:0:0:0:0:0:0:0:0:0:0:0:0:0:0 | 14:0:0:1:2:1:1:0:0:0:0:0:0:0:0:0:0:0:0:0:0 | 0:0:0:0:0:0:0:0:0:0:0:0:0:0:0:0:0:0:0:0:0 | 38:7:0:2:4:0:1:1:2:2:1:1:3:1:0:0:0:0:0:0:0 | 0:0:0:0:0:0:0:0:0:0:0:0:0:0:0:0:0:0:0:0:0 | 21:7:1:2:3:0:0:0:2:5:0:1:5:2:1:3:1:1:2:0:0 | 0:0:0:0:0:0:0:0:0:0:0:0:0:0:0:0:0:0:0:0:0 | 12:0:0:2:0:1:0:0:1:0:0:0:1:0:1:0:0:0:0:0:0 |
| NM_001098832:NM_032837:NM_001289410:XM_024451008 | 2:2:0:0 | 1:3:1:0 | 0:2:0:0 | 1:0:0:0 | 4:9:0:1 | 4:15:0:0 | 8:9:0:0 | 0:9:0:0 | 1:2:0:0 | 2:0:0:0 |
| NM_001098832:XM_024451008 | 4:00 | 0:00 | 1:00 | 0:00 | 8:01 | 2:00 | 9:00 | 1:00 | 3:00 | 0:00 |
| NM_024844:XM_017025110:XR_934552:NM_001303276:XM_017025108:XM_011525267:XR_001752628:XR_001752629:XM_024450951:XR_001752627:XR_002958072 | 29:1:2:0:0:0:0:0:0:0:0 | 0:0:0:0:0:0:0:0:0:0:0 | 14:1:1:1:1:1:1:0:0:0:0 | 0:0:0:0:0:0:0:0:0:0:0 | 52:1:1:2:0:0:2:1:0:0:0 | 0:0:0:0:0:0:0:0:0:0:0 | 37:3:1:2:0:1:2:0:0:0:0 | 44:2:5:0:1:1:2:0:1:0:0 | 11:0:0:0:0:0:1:0:0:0:0 | 20:0:1:0:0:0:1:1:0:0:0 |
| NM_015971 | 0 | 1 | 0 | 1 | 0 | 3 | 0 | 2 | 0 | 3 |
| XR_001752508:XR_002958000:NM_001015002 | 0:00:00 | 0:00:00 | 0:00:00 | 0:00:00 | 0:00:00 | 0:00:00 | 0:00:00 | 1:01:00 | 0:00:00 | 0:00:00 |
| NM_016080:XM_024450778 | 74:02:00 | 1:00 | 111:03:00 | 0:00 | 229:10:00 | 0:00 | 372:27:00 | 4:00 | 80:05:00 | 15:00 |
| NM_001276761:NM_001276760:NM_001126118:NM_001276696 | 3:5:0:0 | 0:0:0:0 | 2:8:0:0 | 0:0:0:0 | 7:33:1:0 | 0:1:0:0 | 1:3:0:0 | 2:2:0:0 | 4:8:0:0 | 0:1:0:0 |
| NM_001276761:NM_001276760:NM_001126118:NM_001276696 | 1:3:0:0 | 1:2:0:0 | 1:2:0:0 | 0:6:0:0 | 1:13:0:0 | 4:21:1:0 | 0:0:0:0 | 2:5:0:0 | 0:0:0:0 | 4:9:0:0 |
| NM_001143992:NM_018081:XR_001752551 | 1:00:00 | 0:00:00 | 3:01:01 | 0:00:00 | 2:02:00 | 0:00:00 | 8:01:00 | 0:00:00 | 4:02:00 | 0:00:00 |
| NM_001143992:NM_018081:XR_001752551:XM_024450824:XM_011523952 | 1:0:0:0:0 | 0:0:0:0:0 | 2:2:1:1:0 | 0:0:0:0:0 | 2:2:0:0:0 | 0:0:0:0:0 | 8:3:1:0:0 | 0:0:0:0:0 | 4:2:0:0:0 | 0:0:0:0:0 |
| NM_016080:XM_024450778 | 82:00:00 | 0:00 | 130:00:00 | 1:00 | 236:04:00 | 0:00 | 405:12:00 | 2:00 | 97:02:00 | 0:00 |
| NM_001168 | 0 | 0 | 0 | 0 | 0 | 0 | 0 | 0 | 0 | 1 |
| XM_011524290:XM_011524289 | 0:00 | 0:00 | 0:00 | 0:00 | 0:01 | 1:00 | 0:00 | 0:02 | 0:00 | 0:00 |
| NM_001614:NR_037688:NM_001199954 | 7765:116:31 | 10:00:00 | 9003:255:45 | 24:04:00 | 10652:168:56 | 22:05:00 | 22060:560:143 | 32:11:00 | 9309:154:65 | 11:01:00 |
| NM_001614:NR_037688:NM_001199954 | 7749:116:31 | 12:00:00 | 8981:253:45 | 31:04:00 | 10630:168:56 | 26:03:00 | 21996:565:143 | 45:10:00 | 9289:155:65 | 16:00:00 |
| NM_001614:NR_037688:NM_001199954 | 7575:118:30 | 26:00:00 | 8616:261:45 | 52:00:00 | 10310:171:56 | 37:03:00 | 21526:604:145 | 63:05:00 | 9062:156:65 | 30:00:00 |
| NM_001614:NR_037688:NM_001199954 | 7468:115:30 | 37:01:00 | 8427:254:45 | 73:02:00 | 10170:168:56 | 47:03:00 | 21237:599:145 | 90:06:00 | 8958:154:65 | 35:00:00 |
| NM_001614:NR_037688:NM_001199954 | 7405:121:30 | 32:00:00 | 8305:261:45 | 68:01:00 | 10074:173:56 | 47:04:00 | 20992:619:145 | 95:04:00 | 8883:153:65 | 33:00:00 |
| NM_001614:NR_037688:NM_001199954 | 7215:119:30 | 37:02:00 | 7786:271:46 | 76:02:00 | 9675:171:56 | 43:03:00 | 20426:665:145 | 81:05:01 | 8519:158:65 | 25:00:00 |
| NM_001614:NR_037688:NM_001199954 | 7073:124:30 | 33:00:00 | 7441:273:46 | 75:03:00 | 9409:171:56 | 49:02:00 | 19959:709:146 | 117:06:00 | 8272:158:65 | 26:00:00 |
| NM_001614:NR_037688:NM_001199954 | 6983:125:30 | 28:01:00 | 7283:271:46 | 82:05:00 | 9253:162:56 | 45:01:00 | 19741:720:145 | 93:05:01 | 8182:155:65 | 28:00:00 |
| NM_001614:NR_037688:NM_001199954 | 6864:125:31 | 29:00:00 | 7007:269:46 | 72:06:00 | 9045:166:56 | 35:01:00 | 19327:730:146 | 73:06:00 | 8046:156:65 | 19:00:00 |
| NM_001614:NR_037688:NM_001199954 | 6810:125:31 | 28:01:00 | 6820:270:46 | 73:03:00 | 8914:166:56 | 41:01:00 | 19113:726:146 | 88:03:00 | 7937:153:65 | 24:01:00 |
| NM_001614:NR_037688:NM_001199954 | 6728:123:31 | 28:01:00 | 6542:269:46 | 103:04:00 | 8775:160:56 | 47:05:00 | 18837:730:146 | 95:06:00 | 7818:156:65 | 24:00:00 |
| NM_001614:NR_037688:NM_001199954 | 6639:118:31 | 32:02:00 | 6413:259:46 | 116:05:00 | 8635:157:56 | 57:02:00 | 18546:734:146 | 124:05:00 | 7735:153:65 | 19:00:00 |
| NM_001614:NR_037688:NM_001199954 | 6572:112:31 | 40:04:00 | 6351:252:46 | 102:06:00 | 8552:157:56 | 59:01:00 | 18367:732:146 | 128:03:00 | 7690:151:65 | 26:00:00 |
| NM_001614:NR_037688:NM_001199954 | 6567:113:31 | 34:03:00 | 6324:250:46 | 112:06:00 | 8547:157:56 | 51:02:00 | 18328:732:146 | 132:06:00 | 7673:151:65 | 27:00:00 |
| NM_001614:NR_037688:NM_001199954 | 6518:114:31 | 34:03:00 | 6253:246:46 | 102:05:00 | 8502:156:56 | 38:02:00 | 18251:726:146 | 95:06:00 | 7625:151:65 | 23:00:00 |
| NM_001614:NR_037688:NM_001199954 | 6525:116:31 | 28:01:00 | 6253:248:46 | 98:05:00 | 8495:155:56 | 52:02:00 | 18269:729:146 | 100:04:00 | 7629:150:65 | 22:00:00 |
| NM_001614:NR_037688:NM_001199954 | 6488:116:31 | 37:02:00 | 6201:244:46 | 113:06:00 | 8433:153:56 | 66:04:00 | 18176:726:146 | 133:08:00 | 7604:150:66 | 23:00:00 |
| NM_001614:NR_037688:NM_001199954 | 6428:116:31 | 40:02:00 | 6152:235:46 | 106:03:00 | 8376:152:56 | 64:03:00 | 18013:715:146 | 147:03:00 | 7534:147:66 | 31:01:00 |
| NM_001614:NR_037688:NM_001199954 | 6318:116:31 | 45:02:00 | 5924:219:46 | 121:03:00 | 8177:152:56 | 64:01:00 | 17566:690:146 | 143:05:00 | 7362:150:66 | 31:00:00 |
| NM_001614:NR_037688:NM_001199954 | 6255:120:31 | 41:03:00 | 5828:210:46 | 129:08:00 | 8060:154:56 | 62:01:00 | 17281:671:146 | 168:05:00 | 7267:146:66 | 30:00:00 |
| NM_001614:NR_037688:NM_001199954 | 6232:124:31 | 37:03:00 | 5796:209:46 | 113:05:00 | 8034:153:56 | 57:02:00 | 17178:666:146 | 130:04:00 | 7237:141:66 | 28:01:00 |
| NR_037688:NM_001614:NM_001199954 | 125:6094:31 | 2:24:00 | 203:5697:46 | 4:56:00 | 149:7852:56 | 1:35:00 | 643:16786:146 | 0.222222222 | 136:7116:66 | 0:12:00 |
| NM_001614:NR_037688:NM_001199954 | 6083:127:31 | 25:00:00 | 5664:205:46 | 73:02:00 | 7821:147:56 | 39:00:00 | 16734:641:146 | 93:04:00 | 7085:135:66 | 15:00:00 |
| NM_001614:NR_037688:NM_001199954 | 6056:124:31 | 41:01:00 | 5639:205:46 | 77:05:00 | 7786:144:56 | 52:01:00 | 16600:638:146 | 111:04:00 | 7050:133:66 | 24:00:00 |
| NM_001614:NR_037688:NM_001199954 | 6048:125:31 | 47:00:00 | 5621:206:46 | 89:05:00 | 7775:144:56 | 61:01:00 | 16569:634:145 | 130:08:01 | 7043:133:66 | 32:00:00 |
| NM_001614:NR_037688:NM_001199954 | 5991:124:31 | 41:00:00 | 5564:209:47 | 86:03:00 | 7715:144:56 | 60:00:00 | 16379:627:145 | 142:06:00 | 6979:135:66 | 37:00:00 |
| NM_001614:NR_037688:NM_001199954 | 5979:115:31 | 42:01:00 | 5566:218:47 | 94:03:00 | 7725:140:56 | 70:01:00 | 16372:628:147 | 157:03:00 | 6979:138:66 | 29:00:00 |
| NR_037688:NM_001614:NM_001199954 | 113:5948:31 | 1:38:00 | 223:5553:47 | 0.147222222 | 142:7707:56 | 0:46:00 | 627:16296:147 | 0.222222222 | 139:6941:65 | 0:33:01 |
| NM_001614:NR_037688:NM_001199954 | 5947:117:31 | 43:00:00 | 5575:232:47 | 87:03:00 | 7711:143:56 | 62:00:00 | 16322:620:146 | 129:01:01 | 6940:136:66 | 33:02:00 |
| NM_001614:NR_037688:NM_001199954 | 5890:115:31 | 31:01:00 | 5509:227:47 | 82:02:00 | 7646:139:56 | 42:00:00 | 16131:611:146 | 108:03:01 | 6868:133:66 | 29:00:00 |
| NM_001614:NR_037688:NM_001199954 | 5862:102:31 | 30:00:00 | 5525:223:47 | 43:02:00 | 7610:128:56 | 27:00:00 | 16022:598:146 | 74:00:01 | 6836:127:66 | 16:00:00 |
| NM_001614:NR_037688:NM_001199954 | 5842:100:31 | 31:00:00 | 5491:217:47 | 64:02:00 | 7574:123:56 | 28:02:00 | 15932:586:145 | 82:00:01 | 6797:125:66 | 19:00:00 |
| NM_001614:NR_037688:NM_001199954 | 5812:98:31 | 36:00:00 | 5476:209:47 | 56:02:00 | 7548:124:56 | 29:00:00 | 15877:579:146 | 66:01:01 | 6766:127:66 | 25:01:00 |
| NM_001614:NR_037688:NM_001199954 | 5813:96:31 | 33:02:00 | 5473:209:47 | 58:02:00 | 7550:123:56 | 25:01:00 | 15883:578:145 | 60:02:02 | 6770:128:66 | 20:00:00 |

| | | | | | | | | | | |
|---|---|---|---|---|---|---|---|---|---|---|
| NM_002949 | 0 | 0 | 1 | 2 | 1 | 1 | 2 | 0 | 2 | 0 |
| NM_000918:XR_002958014:XM_024450777 | 339:16:01 | 0:00:00 | 266:31:03 | 2:00:00 | 218:14:00 | 0:00:00 | 1694:209:10 | 3:01:00 | 347:16:01 | 0:00:00 |
| NM_000918:XR_002958014:XM_024450777 | 332:15:01 | 0:00:00 | 258:32:03 | 2:00:00 | 214:14:00 | 0:00:00 | 1650:246:9 | 4:00:00 | 344:23:01 | 2:00:00 |
| NM_000918:XR_002958014:XM_024450777 | 315:35:01 | 0:00:00 | 238:40:01 | 1:00:00 | 202:23:00 | 1:00:00 | 1494:390:11 | 3:02:01 | 329:41:01 | 1:00:00 |
| NM_000918:XR_002958014:XM_024450777 | 317:37:01 | 0:00:00 | 234:40:01 | 0:00:00 | 201:23:00 | 0:00:00 | 1474:409:12 | 4:02:01 | 330:46:01 | 0:00:00 |
| NM_000918:XR_002958014:XM_024450777 | 316:40:00 | 0:00:00 | 213:43:02 | 1:00:00 | 191:27:00 | 0:00:00 | 1171:448:18 | 3:00:00 | 308:50:01 | 0:00:00 |
| NM_000918:XR_002958014:XM_024450777 | 315:42:00 | 0:00:00 | 216:45:01 | 1:00:00 | 194:27:00 | 0:00:00 | 1124:449:16 | 2:02:00 | 308:51:01 | 0:00:00 |
| NM_004309:XM_011523574:NR_125441:NM_001185078:NM_001185077:NM_001301242:NM_001301243:NM_001301241 | 47:2:0:0:0:0:0:0 | 0:0:0:0:0:0:0:0 | 36:4:0:1:0:0:0:0 | 0:2:2:0:0:0:0:0 | 95:4:4:1:1:1:0:0 | 0:2:1:0:0:0:0:0 | 124:12:1:1:1:2:0:0 | 2:11:5:0:0:0:0:0 | 52:1:4:0:0:0:1:0 | 0:2:1:0:0:0:0:0 |
| NM_004309:XM_011523574:NR_125441:NM_001301243:NM_001185077:NM_001301240:NM_001301241 | 53:1:0:0:0:0:0 | 0:0:0:0:0:0:0 | 47:3:0:1:0:0:0 | 0:1:1:0:0:0:0 | 104:2:4:0:1:3:0 | 0:1:1:0:0:0:0 | 149:10:4:0:3:2:1 | 2:7:3:0:0:0:0 | 62:0:1:1:0:1:0 | 0:0:0:0:0:0:0 |
| XR_001752618:NM_024083:XM_017025044 | 0:00:00 | 1:01:00 | 0:00:00 | 0:00:00 | 0:00:00 | 0:00:00 | 0:00:00 | 0:00:00 | 0:00:00 | 0:00:00 |
| NM_004217:XM_017025311:NM_001256834 | 0:00:00 | 0:00:00 | 0:00:00 | 0:00:00 | 0:00:00 | 0:00:00 | 0:00:00 | 0:00:00 | 0:00:00 | 1:00:00 |
| NM_022156:XR_933938:XR_933939:XM_005256393 | 0:0:0:0 | 0:0:0:0 | 0:0:0:0 | 0:0:0:0 | 0:0:0:0 | 0:0:0:0 | 0:0:0:0 | 0:0:0:0 | 1:0:0:0 | 2:0:0:0 |
| NM_001279:NR_134607 | 6:00 | 10:00 | 0:00 | 2:00 | 2:00 | 5:01 | 0:00 | 0:00 | 0:00 | 0:00 |
| NM_005406 | 1 | 5 | 5 | 1 | 2 | 1 | 5 | 6 | 1 | 3 |
| NM_005406 | 12 | 5 | 27 | 2 | 8 | 2 | 39 | 4 | 9 | 2 |
| NM_174886:XM_017025959:NM_003244:NM_001278686:XM_017025958:NM_001278682 | 5:1:0:0:0:0 | 0:0:0:0:0:0 | 0:0:0:0:0:0 | 0:0:0:0:0:0 | 8:0:2:0:0:0 | 0:0:0:0:0:0 | 18:2:1:1:0:0 | 0:0:0:0:0:0 | 4:1:1:0:1:0 | 0:0:0:0:0:0 |
| XM_024451096:NM_001198938:XM_017025591:XR_001753161:XR_001753159:XR_001753155 | 0:0:0:0:0:0 | 1:0:0:0:0:0 | 0:0:0:0:0:0 | 0:2:1:1:0:0 | 0:0:0:0:0:0 | 0:0:0:0:0:0 | 0:0:0:0:0:0 | 0:0:0:0:0:0 | 0:0:0:0:1:0 | 0:0:0:0:0:0 |
| XM_024451173:XM_005258259:NM_005901:NM_001003652:XM_006722451:XM_017025749:XM_017025746:XM_017025750:XM_017025745:XM_011525985:NM_001135937:XM_017025747 | 6:5:22:5:2:1:0:0:0:0:0:0 | 0:0:0:0:0:0:0:0:0:0:0:0 | 3:8:7:11:0:1:2:1:0:0:0:0 | 1:0:0:0:0:0:0:0:0:0:0:0 | 19:18:43:6:0:2:2:0:2:1:0:0 | 0:0:0:0:0:0:0:0:0:0:0:0 | 17:41:72:17:0:1:3:1:0:1:0:0 | 0:0:0:0:0:0:1:0:0:0:0:0 | 10:14:37:10:1:1:0:0:1:0:0:0 | 0:0:0:0:0:0:0:0:0:0:0:0 |
| XM_024451173:XM_005258259:NM_005901:NM_001003652:XM_006722451:XM_017025749:XM_017025746:XM_017025750:XM_017025745:XM_011525985:NM_001135937:XM_017025747 | 6:5:22:5:2:1:0:0:0:0:0:0 | 0:0:0:0:0:0:0:0:0:0:0:0 | 3:8:7:11:0:1:2:1:0:0:0:0 | 1:0:0:0:0:0:0:0:0:0:0:0 | 19:18:43:6:0:2:2:0:2:1:0:0 | 0:0:0:0:0:0:0:0:0:0:0:0 | 17:41:71:17:0:1:4:1:0:1:0:0 | 0:0:1:0:0:0:0:0:0:0:0:0 | 10:14:37:10:1:1:0:0:1:0:0:0 | 0:0:0:0:0:0:0:0:0:0:0:0 |
| XM_024451173:XM_005258259:NM_005901:NM_001003652:XM_006722451:XM_017025749:XM_017025746:XM_017025750:XM_017025745:XM_011525985:NM_001135937:XM_017025747 | 6:5:22:5:2:1:0:0:0:0:0:0 | 0:0:0:0:0:0:0:0:0:0:0:0 | 3:8:7:11:0:1:2:1:0:0:0:0 | 0:0:0:0:0:1:0:0:0:0:0:0 | 18:18:43:6:0:2:2:0:2:1:0:0 | 0:0:0:0:0:1:0:0:0:0:0:0 | 12:41:72:17:0:1:3:1:0:1:0:0 | 0:0:0:0:0:0:1:0:0:0:0:0 | 9:14:37:10:1:2:0:0:1:0:0:0 | 0:0:0:0:0:0:0:0:0:0:0:0 |
| NR_024546:NM_004786:XM_024451290:XM_024451289 | 6:265:1:0 | 0:0:0:0 | 10:260:1:0 | 0:0:0:0 | 28:1130:1:1 | 0:2:0:0 | 22:1086:2:0 | 0:2:0:0 | 5:278:1:0 | 0:0:0:0 |
| NR_024546:NM_004786:XM_024451290:XM_024451289 | 6:265:2:0 | 0:0:0:0 | 10:261:2:0 | 0:2:1:0 | 28:1132:7:1 | 0:2:1:0 | 22:1093:10:0 | 0:1:1:0 | 5:281:2:1 | 0:0:0:0 |
| NR_024546:NM_004786:XM_024451290:XM_024451289 | 5:265:4:0 | 0:0:0:0 | 6:262:13:0 | 0:1:0:0 | 16:1135:27:1 | 0:0:1:0 | 16:1092:24:0 | 0:2:0:0 | 3:281:4:1 | 0:0:0:0 |
| NR_024546:NM_004786:XM_024451290:XM_024451289 | 3:265:1:0 | 0:0:0:0 | 2:263:11:0 | 0:0:0:0 | 5:1134:24:1 | 0:1:0:0 | 3:1094:32:0 | 0:0:0:0 | 3:280:4:1 | 0:1:0:0 |
| NR_024546:NM_004786:XM_024451290:XM_024451289 | 3:265:1:0 | 0:0:0:0 | 2:263:11:0 | 0:0:0:0 | 5:1134:24:1 | 0:1:0:0 | 3:1092:31:0 | 0:2:0:0 | 3:280:4:1 | 0:1:0:0 |
| XM_011526169:XM_011526165 | 2:00 | 0:00 | 0:00 | 0:00 | 1:00 | 0:00 | 2:01 | 0:00 | 0:00 | 0:00 |
| XM_017025728:XR_001753195:XR_002958170:NM_198531:XM_017025730:XM_024451169:XR_002958169:XM_017025742:XM_024451171:XM_024451172 | 1:0:0:0:0:0:0:0:0:0 | 0:0:0:0:0:0:0:0:0:0 | 0:0:0:0:0:0:0:0:0:0 | 0:1:1:0:0:0:0:0:0:0 | 0:0:0:0:1:0:0:0:0:0 | 0:0:0:1:0:0:0:0:0:0 | 0:1:0:0:0:0:0:0:0:0 | 0:0:0:0:0:0:0:0:0:0 | 0:0:0:0:0:1:0:0:0:0 | 0:0:0:0:0:0:0:0:0:0 |
| XM_017025728:XR_002958170:NM_198531:XM_017025730:XR_001753195:XM_024451169:XM_024451171:XM_024451172 | 1:0:0:0:0:0:0:0 | 0:0:0:0:0:0:0:0 | 0:1:0:0:0:0:0:0 | 0:0:0:0:0:0:0:0 | 0:0:1:1:0:0:0:0 | 0:0:0:0:0:0:0:0 | 0:0:1:0:1:0:0:0 | 0:0:0:0:0:0:0:0 | 0:0:0:0:0:1:0:0 | 0:0:0:0:0:0:0:0 |
| NM_001136180 | 0 | 3 | 0 | 1 | 0 | 2 | 0 | 3 | 0 | 0 |
| NM_021074 | 1 | 1 | 2 | 1 | 4 | 4 | 0 | 3 | 0 | 7 |
| XM_024451773:NM_001033026 | 0:00 | 0:00 | 1:00 | 0:00 | 0:00 | 0:00 | 0:00 | 2:01 | 0:00 | 0:00 |
| NM_146387:NM_015956:XM_011528045:NM_146388 | 1:0:0:0 | 3:0:0:0 | 1:0:0:0 | 2:1:1:0 | 7:0:0:2 | 7:0:2:1 | 8:1:3:1 | 0:0:0:0 | 10:0:3:2 | 0:0:0:0 |
| NM_001145056:NM_020428:XM_005259999 | 0:00:00 | 11:26:01 | 0:00:00 | 1:08:01 | 0:00:00 | 0.163888889 | 0:00:00 | 2:24:00 | 4:13:00 | 2:29:00 |
| NM_001145056:NM_020428:XM_005259999 | 11:32:01 | 0:00:00 | 0.090983796 | 0:01:00 | 0.208333333 | 0:00:00 | 0.145868056 | 0:00:00 | 0.291666667 | 0:00:00 |
| NM_001005360:NM_001190716:NM_001005361:NM_004945:NM_001005362 | 51:16:13:12:0 | 0:0:0:0:0 | 1:10:4:55:6 | 0:0:1:2:0 | 155:64:29:378:42 | 0:0:0:0:0 | 49:43:2:39:2 | 0:0:0:0:0 | 57:34:6:27:4 | 0:0:1:0:0 |
| NM_001005360:NM_001190716:NM_001005361:NM_004945:NM_001005362 | 52:16:13:12:0 | 0:0:0:0:0 | 2:14:4:59:6 | 0:0:0:1:0 | 162:70:25:391:42 | 1:0:2:0:0 | 50:44:1:43:2 | 0:0:1:0:0 | 59:35:6:28:4 | 0:0:0:0:0 |
| NM_001005360:NM_001190716:NM_001005361:NM_004945:NM_001005362 | 53:16:13:12:0 | 0:0:0:0:0 | 2:14:4:64:7 | 0:0:0:0:0 | 164:70:29:401:47 | 1:1:0:0:0 | 50:46:2:42:2 | 0:0:0:0:0 | 59:36:6:28:4 | 0:0:0:0:0 |
| NM_001005360:NM_001190716:NM_001005361:NM_004945:NM_001005362 | 56:16:13:11:0 | 0:0:0:0:0 | 5:21:5:75:7 | 0:0:0:1:0 | 176:84:32:424:48 | 0:1:1:0:0 | 52:62:4:33:3 | 0:0:0:0:0 | 64:41:6:26:4 | 0:0:0:0:0 |
| NM_004945:NM_001005360:NM_001190716:NM_001005361:NM_001005362 | 10:56:16:13:0 | 1:0:0:0:0 | 74:5:21:5:7 | 2:0:0:0:0 | 423:176:85:33:48 | 1:0:0:0:0 | 31:52:62:4:3 | 1:0:0:0:0 | 26:63:41:6:4 | 0:1:0:0:0 |
| NM_001005360:NM_001190716:NM_001005361:NM_004945:NM_001005362 | 60:18:13:9:0 | 0:0:0:0:0 | 15:42:9:88:7 | 0:0:0:0:0 | 204:94:26:440:55 | 0:0:1:1:0 | 71:79:5:14:3 | 0:0:0:0:0 | 78:47:7:21:6 | 0:0:0:0:0 |
| NM_001005360:NM_001190716:NM_004945:NM_001005361:NM_001005362 | 61:21:9:1:0 | 0:0:0:0:0 | 25:48:108:1:0 | 0:1:1:0:0 | 215:98:446:0:2 | 0:0:3:0:0 | 78:91:10:1:0 | 0:0:1:0:0 | 84:49:20:0:0 | 0:0:0:0:0 |
| NM_001005360:NM_001190716:NM_001005361:NM_004945:NM_001005362 | 62:20:18:7:0 | 0:0:0:0:0 | 27:64:29:146:11 | 0:0:0:1:0 | 217:101:54:496:58 | 1:0:0:2:0 | 81:114:22:8:2 | 0:0:0:1:0 | 84:53:13:20:6 | 0:0:0:0:0 |
| NM_001005360:NM_001190716:NM_001005361:NM_004945:NM_001005362 | 62:20:18:7:0 | 0:0:0:0:0 | 27:64:29:148:11 | 0:0:0:0:0 | 218:102:54:496:58 | 0:0:0:2:0 | 80:114:22:10:2 | 1:0:0:0:0 | 84:53:13:20:6 | 0:0:0:0:0 |
| NM_001005360:NM_001190716:NM_001005361:NM_004945:NM_001005362 | 62:22:18:10:0 | 0:0:0:0:0 | 27:66:29:161:8 | 0:0:0:2:0 | 218:99:54:514:57 | 0:0:0:3:0 | 81:110:22:23:1 | 0:1:0:0:1 | 84:49:13:22:6 | 0:0:0:0:0 |
| NM_001005360:NM_001190716:NM_001005361:NM_004945:NM_001005362 | 62:22:18:13:0 | 0:0:0:0:0 | 27:71:27:235:7 | 0:0:1:2:0 | 218:90:54:541:53 | 0:1:0:2:0 | 81:90:20:79:1 | 0:0:0:1:0 | 84:45:13:27:6 | 0:0:0:0:0 |
| NM_002695 | 0 | 3 | 0 | 1 | 0 | 3 | 0 | 2 | 0 | 2 |
| XM_006722845:XM_024451667:XM_024451664:XM_024451665 | 0:2:1:0 | 1:0:0:0 | 0:6:0:0 | 0:5:0:0 | 0:4:0:1 | 0:1:0:0 | 0:10:0:0 | 0:2:0:0 | 0:2:0:0 | 0:0:0:0 |
| XM_024451667:XM_024451664:XM_024451661:XM_024451665:XM_024451660:NM_003072:XM_024451658:XM_024451662:NM_001128848:XM_024451659:NM_001128846:XM_024451663 | 5:2:1:1:0:0:0:0:0:0:0:0 | 0:0:0:0:0:0:0:0:0:0:0:0 | 10:5:0:1:1:0:0:0:0:0:0:0 | 0:0:0:0:0:0:0:0:0:0:0:0 | 7:1:2:3:1:4:1:1:0:0:0:0 | 0:0:0:0:0:0:0:0:0:0:0:0 | 27:5:3:4:1:0:0:1:1:1:1:0 | 0:0:0:0:0:0:0:0:0:0:0:0 | 10:0:0:0:0:1:0:0:0:0:1:0 | 0:0:0:0:0:0:0:0:0:0:0:0 |
| NM_000527:NM_001195799:NM_001195800:XR_001753686:XR_001753685:XM_011528010:NM_001195798 | 1:1:1:0:0:0:0 | 0:0:0:0:0:0:0 | 0:0:0:2:1:1:0 | 0:0:0:0:0:0:0 | 2:0:1:1:0:0:1 | 0:0:0:0:0:0:0 | 0:0:0:1:0:0:0 | 0:0:0:0:0:0:0 | 0:0:0:0:1:0:0 | 0:0:0:0:0:0:0 |
| XM_011528152 | 1 | 0 | 1 | 0 | 2 | 0 | 0 | 2 | 0 | 0 |
| NM_000455 | 5 | 2 | 4 | 4 | 6 | 6 | 22 | 0 | 13 | 0 |
| NM_005815 | 0 | 2 | 0 | 0 | 0 | 0 | 0 | 0 | 0 | 0 |
| NM_005815 | 0 | 2 | 0 | 0 | 0 | 0 | 0 | 0 | 0 | 0 |

| | | | | | | | | | | |
|---|---|---|---|---|---|---|---|---|---|---|
| NM_005815 | 0 | 2 | 0 | 0 | 0 | 0 | 0 | 0 | 0 | 0 |
| NM_000159:NR_102316 | 0:00 | 0:00 | 0:00 | 0:00 | 0:00 | 0:00 | 0:00 | 0:00 | 0:00 | 0:00 |
| NM_004461 | 38 | 0 | 26 | 0 | 95 | 0 | 167 | 5 | 38 | 28 |
| NM_004461 | 30 | 0 | 16 | 0 | 71 | 0 | 117 | 1 | 30 | 23 |
| NM_004461 | 10 | 20 | 12 | 4 | 37 | 34 | 115 | 0 | 52 | 0 |
| NM_052850 | 9 | 0 | 7 | 0 | 13 | 0 | 11 | 0 | 6 | 0 |
| NM_052850 | 9 | 0 | 4 | 0 | 13 | 0 | 10 | 0 | 6 | 0 |
| NM_032853:XM_017027401:XM_011528382:XM_011528383 | 0:0:0:0 | 2:2:0:0 | 0:0:0:0 | 2:1:1:0 | 0:0:0:0 | 0:2:0:0 | 4:0:1:0 | 0:1:0:0 | 1:0:0:0 | 1:0:0:0 |
| NM_014047 | 2 | 1 | 1 | 2 | 0 | 4 | 3 | 0 | 0 | 1 |
| NM_024407 | 0 | 3 | 0 | 2 | 0 | 7 | 0 | 0 | 2 | 0 |
| XM_011527856:NM_001330384 | 1:00 | 0:00 | 0:00 | 1:00 | 1:00 | 0:02 | 0:00 | 1:00 | 0:00 | 0:00 |
| XM_011528208:NR_117085:XM_017027172:NM_023944 | 0:0:0:0 | 0:0:0:0 | 0:0:0:0 | 0:0:0:0 | 0:0:0:0 | 1:1:1:0 | 0:0:0:0 | 1:0:0:0 | 0:0:0:0 | 0:0:0:0 |
| XR_244086:XR_244084:NM_024104:XR_244083:XR_244085:XR_001753759:NM_001300925:XR_001753761 | 8:6:146:23:6:0:0:0 | 0:0:0:0:0:0:0:0 | 10:3:185:10:11:1:0:0 | 0:0:1:0:0:0:0:0 | 11:7:444:27:18:2:0:0 | 0:0:0:0:0:0:0:0 | 17:23:390:95:46:19:1:0 | 0:0:0:0:0:0:0:0 | 7:11:135:31:10:3:0:0 | 0:0:0:0:0:0:0:0 |
| XR_244086:XR_244084:NM_024104:XR_001753761:XR_001753759:XR_244083:XR_001753760:XR_244085:NM_001300925 | 10:6:148:4:8:21:4:6:0 | 0:0:0:0:0:0:0:0:0 | 11:3:172:1:17:10:8:11:0 | 0:0:1:0:0:0:0:0:0 | 12:8:447:0:50:27:10:18:0 | 0:0:1:0:1:0:0:0:0 | 20:27:396:17:45:98:35:46:1 | 0:0:0:0:0:0:0:0:0 | 9:12:141:3:13:31:11:10:0 | 0:0:0:0:0:0:0:0:0 |
| NM_033417:NM_001011699:XM_005260154 | 1:01:00 | 6:01:01 | 14:01:01 | 4:01:01 | 43:08:03 | 26:06:00 | 25:08:00 | 0:00:00 | 20:04:02 | 0:00:00 |
| NM_018467 | 0 | 0 | 0 | 0 | 0 | 1 | 0 | 1 | 0 | 2 |
| NM_138401:NM_001304547 | 1:00 | 0:00 | 0:00 | 0:00 | 0:00 | 0:00 | 2:01 | 0:00 | 0:00 | 0:00 |
| NM_012088 | 0 | 0 | 1 | 0 | 1 | 0 | 1 | 0 | 0 | 0 |
| XM_017026931:XM_017026930 | 0:00 | 0:00 | 0:00 | 0:00 | 0:00 | 0:00 | 0:00 | 1:00 | 0:01 | 0:00 |
| NM_006332 | 1 | 0 | 2 | 2 | 0 | 3 | 1 | 2 | 0 | 1 |
| XM_024451789:NM_004864 | 2:00 | 0:00 | 0:00 | 0:00 | 3:00 | 0:00 | 0:00 | 1:00 | 0:00 | 3:01 |
| XM_024451789:NM_004864 | 2:00 | 0:00 | 0:00 | 0:00 | 3:00 | 0:00 | 0:00 | 1:00 | 0:00 | 3:01 |
| NM_006532:XM_017027335:XR_001753769:XM_017027336 | 0:0:0:0 | 5:0:0:0 | 0:0:0:0 | 0:0:0:0 | 4:0:0:0 | 5:0:0:0 | 5:1:0:0 | 0:0:0:0 | 4:0:0:0 | 0:0:0:0 |
| NM_012181:NM_001308373 | 21:05 | 0:00 | 19:04 | 0:00 | 74:24:00 | 0:01 | 60:32:00 | 0:00 | 12:13 | 0:00 |
| XM_006722883:NM_024069:XM_011528273:NM_001171948 | 0:0:0:0 | 0:0:0:0 | 1:0:0:0 | 0:0:0:0 | 0:0:0:0 | 0:0:0:0 | 0:0:0:0 | 0:0:0:0 | 0:0:0:0 | 1:0:0:0 |
| NM_017814:XM_017026916 | 0:00 | 0:00 | 0:00 | 0:00 | 0:00 | 0:00 | 0:00 | 0:00 | 1:01 | 0:00 |
| NM_021047:NM_001331134:NM_001331133 | 4:01:01 | 0:00:00 | 19:00:03 | 0:00:01 | 18:02:03 | 0:00:00 | 39:01:02 | 1:00:00 | 13:01:00 | 0:00:00 |
| NM_021047:NM_001331134:NM_001331133 | 4:01:01 | 0:00:00 | 16:00:02 | 0:00:00 | 16:02:03 | 0:00:00 | 36:01:02 | 0:00:00 | 13:01:00 | 0:00:00 |
| NM_021047:NM_001331134:NM_001331133 | 4:01:01 | 0:00:00 | 14:00:02 | 2:00:00 | 16:01:03 | 0:00:00 | 35:01:02 | 1:00:00 | 14:01:00 | 0:00:00 |
| NM_021047:NM_001331134:NM_001331133 | 4:01:01 | 0:00:00 | 9:00:02 | 0:00:00 | 12:01:03 | 0:00:00 | 45:01:01 | 0:00:00 | 13:01:00 | 0:00:00 |
| NM_021047:NM_001331134:NM_001331133 | 4:01:01 | 0:00:00 | 9:00:02 | 0:00:00 | 12:01:03 | 0:00:00 | 45:01:01 | 0:00:00 | 13:01:00 | 0:00:00 |
| NM_021047:NM_001331134:NM_001331133 | 4:01:01 | 0:00:00 | 9:00:02 | 0:00:00 | 12:01:03 | 0:00:00 | 46:01:01 | 0:00:00 | 13:01:00 | 0:00:00 |
| NM_021047:NM_001331134:NM_001331133 | 4:01:01 | 0:00:00 | 9:00:02 | 0:00:00 | 12:01:03 | 0:00:00 | 47:01:01 | 0:00:00 | 13:01:00 | 0:00:00 |
| NM_017572:NM_199054:XM_024451471 | 3:03:00 | 0:00:00 | 6:01:01 | 1:00:00 | 24:12:06 | 0:00:00 | 5:00:03 | 0:00:00 | 7:02:01 | 0:00:00 |
| NM_199054:NM_017572:XM_024451471 | 4:03:00 | 0:00:00 | 1:02:00 | 1:05:01 | 11:10:01 | 8:13:03 | 5:05:01 | 0:00:00 | 3:07:00 | 0:00:00 |
| NM_003938:XM_017027422:NM_001261826 | 0:00:00 | 0:00:00 | 0:00:00 | 0:00:00 | 0:00:00 | 0:00:00 | 0:00:00 | 0:00:00 | 1:00:00 | 0:00:00 |
| XR_002958354:NM_003430:XR_001753756 | 0:00:00 | 0:00:00 | 5:00:00 | 3:00:00 | 2:02:00 | 5:01:00 | 0:00:00 | 3:00:00 | 0:00:00 | 0:01:01 |
| XR_002958354:NM_003430:XR_001753756 | 0:00:00 | 0:00:00 | 5:00:00 | 3:00:00 | 2:02:00 | 5:01:00 | 0:00:00 | 3:00:00 | 0:00:00 | 0:01:01 |
| XR_002958354:NM_003430 | 0:00 | 0:00 | 12:00 | 0:00 | 8:02 | 0:00 | 0:00 | 8:00 | 0:00 | 1:01 |
| XR_002958354:NM_003430 | 0:00 | 0:00 | 5:00 | 0:00 | 7:01 | 0:00 | 8:00 | 0:00 | 2:00 | 0:00 |
| XR_002958354:NM_003430:XM_011528266:XM_011528265 | 4:0:0:0 | 0:0:0:0 | 7:0:0:0 | 0:0:0:0 | 7:1:1:0 | 1:0:0:0 | 14:0:0:1 | 1:0:0:0 | 3:0:0:0 | 0:0:0:0 |
| XR_002958354:XM_011528265:XM_011528266:XM_017027236:NM_003430 | 1:0:0:0:0 | 2:0:0:0:0 | 1:0:0:0:0 | 5:1:0:0:0 | 2:0:0:0:0 | 1:0:1:0:0 | 12:1:0:1:0 | 1:0:0:0:0 | 3:0:0:0:0 | 0:0:0:0:0 |
| XR_002958354:XM_011528266:XM_011528265:XM_017027236 | 5:1:1:0 | 1:0:0:0 | 12:0:1:0 | 6:0:1:0 | 6:1:0:0 | 2:0:0:0 | 23:0:3:2 | 13:0:1:1 | 3:0:2:0 | 0:0:0:0 |
| XR_002958354:XM_011528266:XM_011528265:XM_017027236 | 5:1:1:0 | 1:0:0:0 | 15:0:1:0 | 3:0:1:0 | 7:1:0:0 | 1:0:0:0 | 27:0:4:3 | 9:0:0:0 | 3:0:2:0 | 0:0:0:0 |
| XR_002958354:XM_011528266:XM_011528265:XM_017027236:NM_003430 | 10:1:1:1:0 | 0:0:0:0:0 | 47:2:2:0:0 | 5:0:0:0:0 | 20:1:0:0:0 | 2:0:0:0:0 | 1:0:0:0:0 | 61:0:8:4:0 | 8:2:0:0:0 | 6:0:2:0:0 |
| XR_002958354:XM_011528266:XM_011528265:XM_017027236:NM_003430 | 10:1:1:1:0 | 0:0:0:0:0 | 55:2:2:0:0 | 0:0:0:0:0 | 22:1:0:0:0 | 0:0:0:0:0 | 61:0:8:4:0 | 1:0:0:0:0 | 6:0:2:0:0 | 8:2:0:0:0 |
| XR_002958354:XM_011528266:XM_011528265:XR_001753754:XM_024451693:XR_001753755:NM_003430 | 12:1:1:0:0:0:0 | 0:0:0:0:0:0:0 | 60:3:2:1:0:0:0 | 0:0:0:0:0:0:0 | 25:1:0:1:0:0:0 | 0:0:0:0:0:0:0 | 1:0:0:0:1:0:0 | 59:0:8:0:2:0:0 | 9:3:0:0:1:0:0 | 6:0:2:1:0:0:0 |
| XM_011528396:NM_003712 | 0:00 | 0:00 | 1:00 | 1:00 | 0:01 | 0:01 | 0:00 | 0:00 | 0:02 | 0:00 |
| NM_006003 | 0 | 0 | 0 | 2 | 0 | 8 | 0 | 2 | 0 | 8 |
| NR_045557:NM_003796:XM_005259363:XM_024451751:XM_005259362:NM_001252641:XM_011527435 | 8:20:1:2:0:0:0 | 0:0:0:0:0:0:0 | 5:17:4:21:0:0:0 | 0:0:0:0:0:0:0 | 12:22:3:2:2:0:0 | 0:0:0:0:0:0:0 | 32:50:33:45:2:0:0 | 0:1:0:1:0:0:0 | 17:23:6:1:0:1:0 | 0:0:0:0:0:0:0 |
| NR_045557:NM_003796:XM_005259363:XM_024451751:XM_005259362:NM_001252641:XM_011527435 | 8:19:1:2:0:0:0 | 0:0:0:0:0:0:0 | 3:15:4:21:0:0:0 | 0:0:0:0:0:0:0 | 12:20:2:2:2:0:0 | 0:2:0:0:0:0:0 | 26:43:32:52:2:0:0 | 3:4:2:0:0:0:0 | 16:21:5:2:0:1:0 | 0:1:0:0:0:0:0 |
| NM_018025:NR_135270:XM_006723255 | 8:01:00 | 0:00:00 | 9:00:01 | 0:00:00 | 11:00:00 | 0:00:00 | 28:01:02 | 0:00:00 | 6:00:01 | 3:01:00 |
| NM_018025:NR_135270:XM_006723255 | 11:01:00 | 0:00:00 | 15:00:00 | 0:00:00 | 13:00:00 | 0:00:00 | 39:01:02 | 0:00:00 | 7:00:01 | 3:01:00 |
| NM_018025:NR_135270:XM_006723255 | 11:01:00 | 0:00:00 | 15:00:00 | 0:00:00 | 13:00:00 | 0:00:00 | 0:00:00 | 39:01:02 | 0:00:00 | 10:01:01 |
| NM_018025:NR_135270:XM_006723255 | 10:01:00 | 1:00:00 | 15:01:01 | 1:00:00 | 12:08:00 | 0:00:00 | 35:02:03 | 0:00:00 | 10:01:01 | 0:00:00 |
| NM_018025:NR_135270:XM_006723255 | 4:00:00 | 6:01:00 | 7:01:01 | 8:01:00 | 3:05:00 | 9:04:00 | 34:02:03 | 1:00:00 | 10:01:01 | 0:00:00 |
| NM_018025:NR_135270:XM_006723255 | 7:01:00 | 0:00:00 | 6:01:00 | 0:00:00 | 7:08:00 | 1:00:00 | 9:01:00 | 2:00:00 | 7:01:01 | 1:00:00 |
| NM_015578 | 0 | 0 | 0 | 0 | 1 | 0 | 0 | 0 | 1 | 0 |
| NM_014164:NM_144779 | 0:00 | 2:00 | 0:00 | 1:00 | 0:00 | 0:00 | 2:00 | 0:00 | 5:00 | 0:00 |
| NM_014164:NR_028406:NM_144779 | 0:00:00 | 2:01:00 | 0:00:00 | 1:00:00 | 0:00:00 | 0:00:00 | 2:00:00 | 0:00:00 | 4:01:00 | 1:00:00 |
| NM_001352329:XM_024451779:NM_001352330:NM_001308383 | 0:0:0:0 | 0:0:0:0 | 0:0:0:0 | 0:0:0:0 | 0:0:0:0 | 1:0:0:0 | 0:0:0:0 | 0:0:0:0 | 0:0:0:0 | 0:0:0:0 |
| NM_001042631 | 0 | 0 | 0 | 1 | 0 | 1 | 0 | 0 | 0 | 1 |
| XM_017026665:XM_011526844 | 0:00 | 0:00 | 0:00 | 0:00 | 0:00 | 0:00 | 0:00 | 0:00 | 0:00 | 0:00 |
| NM_001297623:NM_001322825:NM_153257:NM_001322826:NM_001322821:NM_001322827 | 1:0:0:0:0:0 | 0:0:0:0:0:0 | 0:1:1:0:0:0 | 1:0:0:0:0:0 | 1:0:2:0:0:0 | 0:0:2:1:0:0 | 1:0:5:0:2:1 | 0:0:0:0:0:0 | 2:0:1:0:0:0 | 0:0:0:0:0:0 |
| NM_001267560:NM_001267561 | 0:00 | 0:00 | 0:00 | 1:00 | 0:00 | 0:00 | 0:00 | 0:00 | 0:00 | 0:00 |
| NM_013234:NM_001308393:NM_001300992 | 1033:68:10 | 1:00:00 | 1168:73:7 | 1:00:00 | 1006:45:05 | 2:00:00 | 2410:175:50 | 3:01:01 | 1396:64:18 | 1:00:00 |
| NM_013234:NM_001308393:NM_001300992 | 1033:68:10 | 1:00:00 | 1183:70:7 | 0:00:00 | 1009:45:05 | 3:00:00 | 2419:175:51 | 2:01:00 | 1401:64:18 | 2:00:00 |
| NM_013234:NM_001308393:NM_001300992 | 1035:68:10 | 2:00:00 | 1200:70:7 | 1:00:00 | 1017:45:05 | 2:00:00 | 2431:173:51 | 0:00:00 | 1406:64:18 | 2:00:00 |
| NM_013234:NM_001308393 | 1044:68 | 2:00 | 1215:70 | 2:00 | 1015:45:00 | 1:00 | 2427:171 | 1:00 | 1412:64 | 0:00 |
| NM_013234:NM_001308393:NM_001300992 | 1016:68:9 | 1:00:00 | 1170:69:6 | 1:00:01 | 990:43:04 | 1:00:00 | 2385:168:52 | 5:01:00 | 1385:62:18 | 1:00:00 |
| NM_004924:XM_017027331:XM_006723406 | 3:04:01 | 0:00:00 | 0:03:00 | 0:00:00 | 6:06:00 | 0:00:00 | 6:02:01 | 0:00:00 | 4:02:00 | 0:00:00 |
| NM_001398:XM_024451409 | 1:00 | 1:00 | 2:00 | 1:00 | 2:00 | 4:00 | 0:00 | 5:00 | 0:00 | 4:00 |
| NM_001398:XM_017026448:XM_024451408:XM_024451409 | 1:0:0:0 | 1:0:0:0 | 0:0:0:0 | 2:0:0:0 | 4:0:0:0 | 2:1:0:0 | 5:0:0:0 | 0:0:0:0 | 4:0:0:0 | 0:0:1:0 |
| NM_001020:NM_001321111 | 15:01 | 0:00 | 14:01 | 0:00 | 14:02 | 0:00 | 28:03:00 | 0:00 | 50:00:00 | 0:00 |
| NM_001961 | 1510 | 4 | 796 | 4 | 666 | 4 | 3107 | 25 | 1518 | 1 |
| NM_001961 | 1484 | 7 | 778 | 8 | 658 | 4 | 3046 | 27 | 1486 | 2 |
| NM_001961 | 1477 | 1 | 770 | 8 | 651 | 4 | 2991 | 25 | 1478 | 3 |

| | | | | | | | | | | |
|---|---|---|---|---|---|---|---|---|---|---|
| NM_001961 | 1085 | 4 | 407 | 3 | 473 | 4 | 1303 | 10 | 1020 | 4 |
| NM_001961 | 1075 | 4 | 410 | 3 | 480 | 0 | 1323 | 7 | 1033 | 2 |
| NM_001961 | 1133 | 3 | 423 | 6 | 489 | 3 | 1431 | 21 | 1070 | 3 |
| NM_001961 | 1175 | 2 | 444 | 4 | 507 | 3 | 1539 | 15 | 1124 | 2 |
| NM_001961 | 1205 | 2 | 440 | 6 | 517 | 1 | 1574 | 8 | 1136 | 5 |
| NM_001961 | 1214 | 0 | 442 | 3 | 520 | 2 | 1579 | 14 | 1147 | 2 |
| NM_001961 | 1228 | 2 | 440 | 4 | 536 | 0 | 1632 | 6 | 1171 | 2 |
| NM_001961 | 1232 | 3 | 446 | 3 | 535 | 1 | 1644 | 12 | 1176 | 4 |
| NM_001961 | 1224 | 3 | 434 | 3 | 530 | 3 | 1635 | 11 | 1172 | 1 |
| NM_001961 | 1282 | 7 | 446 | 4 | 537 | 4 | 1698 | 6 | 1191 | 5 |
| NM_001961 | 1367 | 3 | 468 | 1 | 568 | 1 | 1866 | 6 | 1268 | 5 |
| NM_001961 | 1374 | 0 | 470 | 3 | 572 | 1 | 1875 | 4 | 1273 | 2 |
| NM_001961 | 1396 | 0 | 480 | 1 | 584 | 2 | 1918 | 7 | 1291 | 3 |
| NM_001961 | 1403 | 1 | 483 | 2 | 591 | 3 | 1942 | 7 | 1293 | 2 |
| NM_001961 | 1395 | 1 | 482 | 3 | 590 | 0 | 1931 | 6 | 1283 | 4 |
| NM_003890 | 0 | 0 | 0 | 0 | 0 | 2 | 0 | 0 | 2 | 0 |
| NM_030662:XM_006722799:XM_017026989:XM_017026991 | 34:1:1:3 | 0:0:0:0 | 57:1:1:1 | 0:0:0:0 | 79:1:0:4 | 0:0:0:0 | 82:3:3:5 | 1:0:0:0 | 28:1:2:2 | 0:0:0:0 |
| NM_020158 | 2 | 3 | 0 | 0 | 1 | 0 | 0 | 3 | 0 | 2 |
| NM_001291485:NM_006890 | 0:00 | 0:00 | 0:00 | 8:02 | 0:00 | 21:05 | 0:00 | 0:00 | 0:00 | 0:00 |
| XM_011526990:NM_002483 | 0:00 | 0:00 | 0:00 | 0:00 | 0:01 | 1:01 | 0:00 | 0:00 | 0:00 | 0:06 |
| NM_001013841:XM_011528123:NM_017720 | 0:00:00 | 1:00:00 | 0:00:00 | 5:01:00 | 0:00:00 | 2:00:01 | 0:00:00 | 0:00:00 | 0:00:01 | 0:00:00 |
| XM_017027022:NM_020406 | 0:00 | 0:00 | 2:03 | 0:00 | 2:11 | 0:00 | 0:00 | 0:00 | 0:00 | 0:00 |
| XM_017027022:NM_020406 | 0:00 | 0:00 | 0:00 | 1:05 | 0:00 | 2:11 | 0:00 | 0:00 | 0:00 | 0:00 |
| XM_017027022:NM_020406 | 0:00 | 0:00 | 0:00 | 1:05 | 0:00 | 2:11 | 0:00 | 0:00 | 0:00 | 0:00 |
| XM_017027022:NM_020406 | 0:00 | 0:00 | 0:00 | 1:05 | 0:00 | 2:11 | 0:00 | 0:00 | 0:00 | 0:00 |
| NM_020406 | 0 | 0 | 0 | 5 | 0 | 12 | 0 | 0 | 0 | 0 |
| NM_002659:NM_001005376:NM_001005377:XM_017026872:XM_017026873 | 0:0:0:0:0 | 0:0:0:0:0 | 2:0:0:0:0 | 3:0:0:0:0 | 0:1:0:0:0 | 1:0:0:0:0 | 0:0:0:0:0 | 0:0:0:0:0 | 0:0:0:0:0 | 0:0:0:0:0 |
| NM_001302688:NM_001302691 | 0:01 | 1:00 | 0:00 | 0:00 | 0:00 | 0:00 | 3:00 | 2:00 | 2:00 | 0:00 |
| NM_000041:NM_001302688:NM_001302691 | 9:01:01 | 0:00:00 | 7:00:01 | 0:00:00 | 8:00:01 | 0:00:00 | 31:01:02 | 39:01:05 | 1:00:01 | 3:00:00 |
| XR_001753633:XR_001753634:XM_011526611:XM_017026467 | 3:0:0:0 | 1:0:0:0 | 3:1:0:0 | 2:0:0:0 | 4:1:0:0 | 5:1:1:0 | 39:4:0:3 | 1:0:0:0 | 9:0:0:0 | 5:0:0:0 |
| XR_001753633:XM_011526611:XR_001753634:XM_017026467 | 4:0:0:0 | 0:0:0:0 | 5:0:0:0 | 0:0:0:0 | 9:1:1:0 | 0:0:0:0 | 39:0:1:2 | 0:0:0:0 | 14:0:0:0 | 0:0:0:0 |
| XR_001753633:NM_001130867:XM_011526611:XR_001753634:XM_017026467 | 3:0:0:0:0 | 1:1:0:0:0 | 2:1:0:0:0 | 2:0:0:0:0 | 4:0:0:0:0 | 4:0:1:1:0 | 30:3:0:1:3 | 0:0:0:0:0 | 9:2:0:0:0 | 5:0:0:0:0 |
| XM_017027200:NM_003370:XM_005259199 | 0:00:00 | 0:00:00 | 1:01:00 | 0:00:00 | 0:00:00 | 0:00:00 | 0:00:00 | 0:00:00 | 0:00:00 | 0:00:00 |
| NM_019107 | 0 | 0 | 2 | 0 | 0 | 0 | 3 | 0 | 4 | 0 |
| NM_005628:NM_001145144:NM_001145145 | 58:01:00 | 0:00:00 | 23:01:00 | 0:01:00 | 26:01:00 | 0:00:00 | 0:00:00 | 255:25:00 | 22:03:00 | 29:04:00 |
| NM_005628 | 34 | 0 | 2 | 0 | 10 | 0 | 33 | 1 | 22 | 1 |
| NM_017854 | 2 | 0 | 1 | 0 | 1 | 0 | 0 | 0 | 1 | 0 |
| NM_015710 | 0 | 1 | 0 | 2 | 0 | 2 | 0 | 1 | 0 | 5 |
| NM_005817 | 0 | 0 | 0 | 0 | 1 | 0 | 0 | 0 | 0 | 0 |
| NM_005817 | 0 | 0 | 0 | 0 | 0 | 2 | 0 | 0 | 0 | 0 |
| XM_005258970:XM_005258969:NM_016246:XR_243938 | 1:0:0:0 | 0:0:0:0 | 0:0:0:0 | 0:0:0:0 | 1:1:2:1 | 0:0:0:0 | 4:1:2:1 | 0:0:0:0 | 0:0:0:0 | 0:0:0:0 |
| NM_138761:NR_027882:XM_017027077:NM_001291428:NM_138763:NM_138764:NM_001291431:NM_004324:NM_001291429 | 68:1:8:4:3:0:0:0:0 | 0:0:0:0:0:0:0:0:0 | 131:2:29:16:0:1:1:2:0 | 0:0:0:0:0:0:0:0:0 | 345:1:37:17:1:1:1:2:0 | 0:0:1:0:0:0:0:0:0 | 264:1:72:20:2:2:4:2:0 | 1:0:1:0:0:0:0:0:0 | 117:0:14:8:2:0:1:0:0 | 0:0:0:0:0:0:0:0:0 |
| NM_138761:NR_027882:XM_017027077:NM_001291428:NM_138763:NM_138764:NM_001291431:NM_004324:NM_001291429 | 68:1:8:4:3:0:0:0:0 | 0:0:0:0:0:0:0:0:0 | 130:3:37:17:1:1:1:2:0 | 1:0:0:0:0:0:0:0:0 | 343:3:37:18:1:0:1:2:0 | 0:0:0:0:0:0:0:0:0 | 264:2:85:20:3:0:4:2:0 | 1:0:0:0:0:0:0:0:0 | 117:0:14:8:3:0:1:0:0 | 0:0:0:0:0:0:0:0:0 |
| NM_003089:XM_011527241:NM_001301069 | 4:01:02 | 0:00:00 | 6:05:06 | 0:00:00 | 9:02:03 | 0:00:00 | 48:05:10 | 0:00:00 | 5:03:03 | 0:00:00 |
| NM_017916:XM_024451570 | 0:00 | 1:00 | 0:00 | 0:00 | 1:00 | 0:00 | 2:00 | 0:00 | 0:00 | 0:00 |
| NM_017916:XM_024451570 | 0:00 | 1:00 | 0:00 | 0:00 | 1:00 | 0:00 | 2:00 | 0:00 | 0:00 | 0:00 |
| NM_017916:XM_024451570 | 0:00 | 1:00 | 0:00 | 0:00 | 1:00 | 0:00 | 2:00 | 0:00 | 0:00 | 0:00 |
| NM_017916:XM_024451570 | 0:00 | 1:00 | 0:00 | 0:00 | 1:00 | 0:00 | 2:00 | 0:00 | 0:00 | 0:00 |
| NM_024682:XM_011527317:NM_001168222 | 1:00:00 | 0:00:00 | 2:01:00 | 0:00:00 | 1:00:01 | 0:00:00 | 0:00:00 | 3:00:02 | 0:00:00 | 1:00:00 |
| XM_011526942:NM_002257 | 0:00 | 0:00 | 0:00 | 2:00 | 0:00 | 0:03 | 0:00 | 0:00 | 2:01 | 0:00 |
| XM_005259061:NM_002776:XM_017026993 | 0:00:00 | 0:00:00 | 0:00:00 | 0:00:00 | 0:00:00 | 0:00:00 | 0:00:00 | 0:00:00 | 0:00:00 | 2:00:00 |
| XM_005259061:NM_002776:XM_017026993 | 0:00:00 | 0:00:00 | 0:00:00 | 0:00:00 | 0:00:00 | 0:00:00 | 0:00:00 | 0:00:00 | 0:00:00 | 2:00:00 |
| NM_001136032:NM_144947:NM_006853:XM_011526373 | 0:0:0:0 | 0:0:0:0 | 0:0:0:0 | 0:0:0:0 | 1:1:0:0 | 0:0:0:0 | 2:3:0:0 | 0:0:0:0 | 1:1:0:0 | 1:4:0:0 |
| NM_001985:XM_024451418 | 1:00 | 0:00 | 0:00 | 0:00 | 1:00 | 0:00 | 0:00 | 1:00 | 0:00 | 4:00 |
| XM_017027099:NM_021632:XM_017027103:XM_017027102:XM_017027100:XM_017027101:XM_017027095:XM_011527196:XM_017027098:XM_017027094:XM_024451635:XM_024451633 | 0:1:1:0:0:0:0:0:0:0:0:0 | 1:1:0:0:0:0:0:0:0:0:0:0 | 0:3:0:0:1:0:0:0:0:0:0:0 | 0:3:0:1:0:0:0:0:0:0:0:0 | 0:5:0:1:0:0:0:0:0:0:0:0 | 0:3:0:0:0:0:0:0:0:0:0:0 | 0:16:2:4:3:9:1:1:3:1:7:1 | 0:0:0:0:0:0:0:0:0:0:0:0 | 0:4:0:1:0:0:1:0:0:0:0:2 | 0:0:0:0:0:0:0:0:0:0:0:0 |
| NM_021632:XM_017027099:XM_017027103:XM_017027100:XM_017027102:XM_017027101:XM_017027095:XM_011527196:XM_017027098:XM_017027094:XM_024451635:XM_024451633 | 2:1:1:0:0:0:0:0:0:0:0:0 | 0:0:0:0:0:0:0:0:0:0:0:0 | 6:0:0:1:1:0:0:0:0:0:0:0 | 0:0:0:0:0:0:0:0:0:0:0:0 | 9:0:0:0:1:0:0:0:0:0:0:0 | 0:0:0:0:0:0:0:0:0:0:0:0 | 15:0:2:3:4:11:1:1:3:2:6:1 | 2:0:0:0:0:0:0:0:0:0:0:0 | 4:0:0:0:1:1:1:0:0:0:0:2 | 0:0:0:0:0:0:0:0:0:0:0:0 |
| NM_014225:NR_033500 | 210:01:00 | 1:00 | 48:00:00 | 0:00 | 298:00:00 | 0:00 | 420:02:00 | 0:00 | 215:00:00 | 0:01 |
| NM_014225:NR_033500 | 210:01:00 | 0:00 | 46:00:00 | 0:00 | 295:01:00 | 0:00 | 413:04:00 | 1:00 | 215:00:00 | 0:00 |
| NM_014225:NR_033500 | 210:01:00 | 0:00 | 46:00:00 | 0:00 | 295:01:00 | 0:00 | 413:04:00 | 1:00 | 214:00:00 | 1:00 |
| NM_014225:NR_033500 | 209:01:00 | 0:00 | 45:00:00 | 0:00 | 287:02:00 | 1:00 | 401:04:00 | 2:00 | 214:00:00 | 0:00 |
| NM_014225:NR_033500 | 208:01:00 | 0:00 | 45:00:00 | 0:00 | 285:02:00 | 1:00 | 402:04:00 | 0:00 | 213:01:00 | 0:00 |
| NM_014225:NR_033500 | 204:01:00 | 0:00 | 45:00:00 | 0:00 | 277:03:00 | 0:00 | 390:04:00 | 2:00 | 211:01:00 | 0:00 |
| NM_014225:NR_033500 | 204:01:00 | 0:00 | 45:00:00 | 0:00 | 281:03:00 | 0:00 | 387:06:00 | 0:00 | 211:01:00 | 0:00 |
| XR_001753688:NM_001145434 | 0:00 | 0:00 | 0:00 | 0:00 | 1:00 | 0:00 | 0:00 | 0:00 | 0:00 | 0:00 |
| XR_001753688:NM_001145434 | 0:00 | 0:00 | 0:00 | 0:00 | 0:00 | 1:00 | 0:00 | 0:00 | 0:00 | 0:00 |
| XR_001753688:NM_001145434:XM_017026812 | 0:00:00 | 0:00:00 | 0:00:00 | 0:00:00 | 0:00:00 | 0:00:00 | 0:00:00 | 0:00:00 | 0:00:00 | 0:00:00 |
| XR_001753688:NM_001145434 | 0:00 | 0:00 | 0:00 | 0:00 | 0:00 | 0:00 | 0:00 | 0:00 | 0:00 | 0:00 |
| NM_144686:XM_011526486 | 1:00 | 0:00 | 0:01 | 1:02 | 0:00 | 2:00 | 0:00 | 1:00 | 0:00 | 0:00 |
| NM_144686 | 1 | 0 | 0 | 0 | 0 | 2 | 0 | 0 | 0 | 0 |
| NM_001190764 | 0 | 1 | 0 | 1 | 0 | 5 | 0 | 3 | 7 | 2 |
| NR_076392 | 0 | 0 | 0 | 0 | 0 | 0 | 0 | 0 | 0 | 1 |
| XM_024451618:NM_001080452 | 0:00 | 0:00 | 0:00 | 0:00 | 0:00 | 1:00 | 0:00 | 0:00 | 0:00 | 0:00 |
| XM_011527989 | 0 | 0 | 0 | 0 | 0 | 2 | 0 | 0 | 0 | 0 |
| NM_006949:NR_073560 | 0:00 | 0:00 | 0:00 | 0:00 | 2:00 | 2:00 | 3:01 | 0:00 | 1:00 | 0:00 |
| NM_001042462:NM_001042461 | 0:00 | 0:00 | 0:00 | 0:00 | 0:00 | 3:00 | 2:00 | 0:00 | 1:00 | 0:00 |

| | | | | | | | | | | |
|---|---|---|---|---|---|---|---|---|---|---|
| NM_139314:NM_001039667:XM_005272484 | 0:00:00 | 0:00:00 | 0:00:00 | 1:00:00 | 0:00:00 | 0:00:00 | 0:00:00 | 0:00:00 | 0:00:00 | 0:01:00 |
| NM_024100 | 0 | 0 | 0 | 0 | 0 | 1 | 0 | 0 | 0 | 1 |
| XM_005263959:XM_011511242:XM_005263960:NM_002518:XM_017004216:XM_005263961:XM_017004215:XM_017004217:XM_017004214 | 0:0:1:1:0:0:0:0:0 | 1:1:0:0:0:0:0:0:0 | 1:0:0:0:0:0:0:0:0 | 0:0:0:0:0:0:0:0:0 | 1:2:0:1:1:1:0:0:0 | 1:0:0:2:0:0:0:0:0 | 5:0:0:0:0:0:1:3:0 | 7:1:0:0:2:0:1:1:0 | 0:2:0:0:0:0:0:1:0 | 1:0:0:0:0:0:0:0:0 |
| NM_001165931 | 0 | 0 | 0 | 0 | 0 | 0 | 0 | 0 | 0 | 2 |
| NM_173647:XR_922908:XR_244888 | 0:00:00 | 1:00:00 | 0:00:00 | 1:00:00 | 0:00:00 | 0:00:00 | 0:00:00 | 0:00:00 | 0:00:00 | 2:00:00 |
| NM_173647:XM_005263920:XR_922908 | 0:00:00 | 1:00:00 | 0:00:00 | 1:00:00 | 0:00:00 | 0:00:00 | 0:01:00 | 0:00:00 | 2:00:00 | 0:00:00 |
| NM_001321345:NM_144632:NM_001321346:NM_001321344:NM_001321343 | 0:0:0:0:0 | 1:0:0:0:0 | 0:0:0:0:0 | 1:1:0:0:0 | 0:0:0:0:0 | 0:1:1:0:0 | 0:0:0:0:0 | 15:5:7:2:0 | 0:0:0:0:0 | 2:3:1:2:0 |
| XM_011510308:NM_005742 | 0:00 | 0:00 | 1:00 | 0:00 | 0:00 | 0:00 | 3:01 | 1:01 | 0:01 | 0:04 |
| NM_006343 | 0 | 0 | 0 | 0 | 2 | 0 | 0 | 1 | 0 | 0 |
| NM_006343 | 0 | 0 | 0 | 0 | 3 | 0 | 0 | 2 | 0 | 0 |
| NM_006343 | 0 | 0 | 0 | 0 | 3 | 0 | 0 | 2 | 0 | 0 |
| NM_032309:NM_001304353 | 0:00 | 0:00 | 2:00 | 0:00 | 0:00 | 0:00 | 0:00 | 0:00 | 2:00 | 0:00 |
| NM_001079862:NM_001178042:NM_001079863:NM_001282634:NM_001178017:NM_001282636:NM_020548 | 2:0:0:0:0:0:0 | 0:0:0:0:0:0:0 | 6:0:0:0:0:0:0 | 0:0:0:0:0:0:0 | 12:1:0:0:0:0:0 | 0:0:0:0:0:0:0 | 17:0:1:0:0:0:0 | 0:0:0:0:0:0:0 | 17:0:0:0:0:0:0 | 0:0:0:0:0:0:0 |
| NM_030577:NM_001105199 | 1:00 | 0:00 | 0:00 | 0:00 | 0:00 | 0:00 | 0:00 | 0:00 | 0:00 | 0:01 |
| NM_001006622:XM_017004436 | 1:00 | 0:00 | 0:00 | 0:00 | 2:00 | 0:00 | 0:00 | 0:00 | 0:00 | 0:00 |
| XM_017003978:XM_017003977 | 0:00 | 0:00 | 1:00 | 0:00 | 0:00 | 0:02 | 3:00 | 1:00 | 1:00 | 0:00 |
| XM_017003977:XM_017003978 | 1:00 | 0:00 | 0:01 | 0:02 | 1:00 | 1:00 | 0:06 | 0:00 | 0:01 | 0:00 |
| XM_017003977:XM_017003978 | 0:00 | 1:00 | 0:01 | 0:01 | 1:00 | 1:00 | 0:00 | 0:04 | 0:00 | 0:01 |
| XM_017003977:XM_017003978 | 0:00 | 1:00 | 0:00 | 0:01 | 0:00 | 2:00 | 0:00 | 0:02 | 0:00 | 0:01 |
| XM_011511101:XM_017003977:NM_207481:XM_017003978:XM_017003980:XM_011511099 | 0:1:0:0:0:0 | 1:0:0:0:0:0 | 3:0:1:1:1:0 | 1:0:0:0:0:0 | 7:2:0:0:0:0 | 3:0:0:0:1:0 | 6:0:0:0:0:0 | 0:0:0:0:0:0 | 1:0:0:0:0:0 | 0:0:0:0:0:0 |
| NM_012233:XM_011510825:XR_001738674:XM_011510823 | 7:1:3:0 | 0:0:0:0 | 3:1:4:0 | 0:0:1:0 | 7:0:10:0 | 0:0:1:0 | 24:4:23:0 | 0:0:3:0 | 4:0:1:0 | 0:0:2:0 |
| NM_012233:XM_011510825:XR_001738674:XM_011510823 | 7:1:3:0 | 0:0:0:0 | 3:1:3:0 | 0:0:2:0 | 7:0:6:0 | 0:0:4:0 | 24:4:20:0 | 0:0:7:0 | 4:0:1:0 | 0:0:2:0 |
| NM_012233:XM_011510825:XR_001738674:XM_011510823 | 7:1:3:0 | 0:0:0:0 | 3:1:3:0 | 0:0:2:0 | 7:0:6:0 | 0:0:4:0 | 24:4:18:0 | 0:0:9:0 | 4:0:1:0 | 0:0:2:0 |
| XM_011510825:XR_001738674:NM_012233:XM_011510823 | 0:1:3:0 | 1:1:4:0 | 0:3:0:0 | 1:0:3:0 | 0:3:4:0 | 0:0:3:0 | 5:14:26:0 | 0:1:0:0 | 0:1:5:0 | 0:0:0:0 |
| NM_003467:NM_001008540 | 281:00:00 | 0:00 | 244:00:00 | 0:00 | 124:00:00 | 1:00 | 157:00:00 | 0:00 | 47:00:00 | 0:00 |
| NM_003467:NM_001008540 | 280:00:00 | 0:00 | 243:00:00 | 1:00 | 125:00:00 | 0:00 | 157:00:00 | 0:00 | 47:00:00 | 0:00 |
| NM_003467:NM_001008540 | 279:00:00 | 0:00 | 245:00:00 | 1:00 | 124:00:00 | 1:00 | 158:00:00 | 0:00 | 47:00:00 | 0:00 |
| NM_003467:NM_001008540 | 277:00:00 | 1:00 | 243:00:00 | 2:00 | 125:00:00 | 0:00 | 158:00:00 | 0:00 | 47:00:00 | 0:00 |
| NM_003467:NM_001008540 | 269:00:00 | 1:00 | 241:00:00 | 0:00 | 125:00:00 | 0:00 | 154:00:00 | 0:00 | 48:00:00 | 0:00 |
| NM_003467:NM_001008540 | 270:00:00 | 0:00 | 241:00:00 | 0:00 | 124:00:00 | 1:00 | 152:00:00 | 0:00 | 48:00:00 | 0:00 |
| NM_003937:XM_024453208:NM_001199241:XR_923058 | 13:1:0:0 | 0:0:0:0 | 2:0:0:0 | 0:0:0:0 | 6:1:0:0 | 0:0:0:0 | 6:1:1:0 | 5:0:0:0 | 0:0:0:0 | 1:0:0:0 |
| NM_004688 | 2 | 0 | 0 | 0 | 1 | 0 | 0 | 0 | 0 | 0 |
| XM_011512108:NM_001111067:NM_001347663:NM_001347665:NM_001347664:NM_001347666:XM_006712825:NM_001347667 | 3:15:3:3:2:0:0:0 | 0:0:0:0:0:0:0:0 | 13:6:0:0:0:1:0:0 | 0:0:0:0:0:0:0:0 | 0:11:2:0:1:1:1:0 | 0:0:0:0:0:0:0:0 | 51:42:1:7:2:3:0:0 | 1:1:0:0:0:0:0:0 | 2:5:2:1:1:1:0:0 | 0:0:0:0:0:0:0:0 |
| NM_022168 | 0 | 0 | 0 | 0 | 0 | 0 | 0 | 0 | 0 | 0 |
| NM_022168 | 0 | 0 | 0 | 0 | 0 | 0 | 0 | 0 | 0 | 0 |
| NM_003642:NR_027862 | 0:00 | 0:00 | 0:00 | 0:00 | 0:00 | 1:00 | 0:00 | 1:00 | 0:00 | 0:00 |
| NM_006164:NM_001145412:NM_001313901:NM_001313904:NM_001313900 | 178:54:1:2:0 | 0:0:0:0:0 | 67:6:0:0:0 | 0:0:0:0:0 | 332:39:0:0:0 | 2:0:0:0:0 | 764:198:0:1:0 | 5:0:0:0:0 | 247:33:2:1:1 | 0:0:0:0:0 |
| NM_006164:NM_001145412:NM_001313901:NM_001313904:NM_001313900:NM_001313902:NM_001313903 | 192:54:1:2:0:0:0 | 1:0:0:0:0:0:0 | 265:24:0:1:0:0:0 | 3:0:0:0:0:0:0 | 363:42:0:0:0:0:0 | 1:0:0:0:0:0:0 | 2122:392:1:1:0:0:0 | 10:3:0:0:0:0:0 | 270:34:2:1:1:0:0 | 1:1:0:0:0:0:0 |
| NM_006164:NM_001145412:NM_001313901:NM_001313904:NM_001313900:NM_001313903:NM_001313902 | 213:62:1:2:0:0:0 | 1:0:0:0:0:0:0 | 346:34:0:1:0:0:0 | 8:3:0:0:0:0:0 | 394:44:0:0:0:0:0 | 6:0:0:0:0:0:0 | 2442:497:2:1:1:0:0 | 84:22:0:0:0:0:0 | 293:35:2:1:1:0:0 | 0:1:0:0:0:0:0 |
| NM_006164:NM_001145412:NM_001313901:NM_001313904:NM_001313900:NM_001313903:NM_001313902 | 213:62:1:2:0:0:0 | 0:0:0:0:0:0:0 | 354:37:0:1:0:0:0 | 0:0:0:0:0:0:0 | 398:44:0:0:0:0:0 | 3:0:0:0:0:0:0 | 2520:519:2:1:1:0:0 | 11:1:0:0:0:0:0 | 293:35:2:1:1:0:0 | 1:1:0:0:0:0:0 |
| NM_006164:NM_001145412:NM_001313901:NM_001313904:NM_001313900:NM_001313903:NM_001313902 | 215:62:1:2:0:0:0 | 0:0:0:0:0:0:0 | 356:36:0:1:0:0:0 | 1:1:0:0:0:0:0 | 401:44:0:0:0:0:0 | 0:0:0:0:0:0:0 | 2550:520:2:1:1:0:0 | 8:1:0:0:0:0:0 | 296:36:2:1:1:0:0 | 0:0:0:0:0:0:0 |
| NM_006164:NM_001145412:NM_001313901:NM_001313904:NM_001313900:NM_001313903:NM_001313902 | 218:62:1:2:0:0:0 | 0:0:0:0:0:0:0 | 367:37:0:1:0:0:0 | 0:1:0:0:0:0:0 | 407:45:1:0:0:0:0 | 3:0:0:0:0:0:0 | 2582:531:2:0:1:0:0 | 2:0:0:1:0:0:0 | 297:37:2:1:2:0:0 | 0:0:0:0:0:0:0 |
| NM_006164:NM_001145412:NM_001313901:NM_001313904:NM_001313900:NM_001313903:NM_001313902 | 211:62:1:2:0:0:0 | 1:0:0:0:0:0:0 | 353:38:0:1:0:0:0 | 0:0:0:0:0:0:0 | 408:45:1:0:0:0:0 | 0:0:0:0:0:0:0 | 2428:530:2:1:1:0:0 | 4:2:0:0:0:0:0 | 291:37:2:1:2:0:0 | 0:0:0:0:0:0:0 |
| NM_006164:NM_001145412:NM_001313901:NM_001313904:NM_001313900:NM_001313903:NM_001313902 | 211:62:1:2:0:0:0 | 1:0:0:0:0:0:0 | 350:38:0:1:0:0:0 | 2:0:0:0:0:0:0 | 406:45:1:0:0:0:0 | 0:0:0:0:0:0:0 | 2399:531:2:1:1:0:0 | 8:2:0:0:0:0:0 | 290:37:2:1:2:0:0 | 0:0:0:0:0:0:0 |
| NM_006164:NM_001145412:NM_001313901:NM_001313904:NM_001313900:NM_001313903:NM_001313902 | 210:62:1:2:0:0:0 | 1:0:0:0:0:0:0 | 345:39:0:1:0:0:0 | 3:0:0:0:0:0:0 | 404:46:1:0:0:0:0 | 1:0:0:0:0:0:0 | 2262:532:2:1:1:0:0 | 12:1:0:0:0:0:0 | 290:37:2:1:2:0:0 | 1:0:0:0:0:0:0 |
| NM_006164:NM_001145412:NM_001313901:NM_001313904:NM_001313900:NM_001313903:NM_001313902 | 209:62:1:2:0:0:0 | 0:0:0:0:0:0:0 | 342:40:0:1:0:0:0 | 0:0:0:0:0:0:0 | 404:46:1:0:0:0:0 | 0:0:0:0:0:0:0 | 2189:533:2:1:1:0:0 | 7:1:0:0:0:0:0 | 291:37:2:1:2:0:0 | 0:0:0:0:0:0:0 |
| NM_006164:NM_001145412:NM_001313901:NM_001313904:NM_001313900:NM_001313903:NM_001313902 | 209:62:1:2:0:0:0 | 0:0:0:0:0:0:0 | 338:40:0:1:0:0:0 | 1:0:0:0:0:0:0 | 404:46:1:0:0:0:0 | 0:0:0:0:0:0:0 | 2102:534:2:1:1:0:0 | 2:0:0:0:0:0:0 | 291:37:2:1:2:0:0 | 0:0:0:0:0:0:0 |
| NM_006164:NM_001145412:NM_001313901:NM_001313904:NM_001313900:NM_001313903:NM_001313902 | 209:62:1:2:0:0:0 | 0:0:0:0:0:0:0 | 333:40:0:1:0:0:0 | 0:0:0:0:0:0:0 | 403:46:1:0:0:0:0 | 0:0:0:0:0:0:0 | 2026:537:2:1:1:0:0 | 7:0:0:0:0:0:0 | 290:37:2:1:2:0:0 | 0:0:0:0:0:0:0 |

| | | | | | | | | | | |
|---|---|---|---|---|---|---|---|---|---|---|
| NM_006164:NM_001145412:NM_001313901:NM_001313900 | 200:62:1:0 | 1:0:0:0 | 286:40:0:0 | 0:0:0:0 | 397:46:1:0 | 0:0:0:0 | 1619:537:2:1 | 8:1:0:0 | 283:37:2:2 | 1:0:0:0 |
| NM_006164:NM_001145412:NM_001313901:NM_001313904:NM_001313900:NM_001313902 | 200:62:1:2:0:0 | 1:0:0:0:0:0 | 283:40:0:1:0:0 | 0:0:0:0:0:0 | 394:46:1:0:0:0 | 2:0:0:0:0:0 | 1550:538:2:1:1:0 | 7:0:0:0:0:0 | 284:36:2:1:2:0 | 0:1:0:0:0:0 |
| NM_003659:XR_001739007:XM_011512042:XM_011512043:XM_011512041 | 2:3:1:0:0 | 0:0:0:0:0 | 0:6:0:5:0 | 0:0:0:0:0 | 3:0:0:1:1 | 0:0:0:0:0 | 12:9:1:4:0 | 0:0:0:0:0 | 8:2:2:2:0 | 0:0:0:0:0 |
| NM_002252 | 0 | 0 | 0 | 0 | 0 | 2 | 0 | 0 | 0 | 1 |
| NM_018471:XM_017004502 | 44:02:00 | 0:00 | 108:02:00 | 0:00 | 44:03:00 | 0:02 | 487:13:00 | 0:00 | 100:03:00 | 0:00 |
| NM_018471:XM_017004502 | 44:01:00 | 0:00 | 108:01:00 | 0:00 | 43:02:00 | 0:00 | 455:10:00 | 0:00 | 100:01:00 | 0:00 |
| NM_018471:XM_017004502 | 44:01:00 | 0:00 | 108:01:00 | 0:00 | 43:02:00 | 0:00 | 452:10:00 | 3:00 | 100:01:00 | 0:00 |
| NM_000090 | 0 | 2 | 0 | 1 | 0 | 0 | 0 | 0 | 0 | 0 |
| NM_001042519 | 0 | 0 | 0 | 1 | 2 | 3 | 0 | 0 | 0 | 0 |
| XM_011510954:NM_014362:NM_198047:XR_922903:XM_011510953 | 1:0:0:0:0 | 0:0:0:0:0 | 0:1:2:1:0 | 0:3:0:0:0 | 0:1:0:0:0 | 0:0:1:0:0 | 0:9:0:0:0 | 0:5:0:1:0 | 0:2:2:0:0 | 0:2:0:0:0 |
| XM_011510954:NM_014362:NM_198047:XR_922903:XM_011510953 | 1:0:0:0:0 | 0:0:0:0:0 | 0:4:2:1:0 | 0:0:0:0:0 | 0:1:1:0:0 | 0:0:0:0:0 | 0:9:0:0:0 | 0:5:0:1:0 | 0:2:2:0:0 | 0:2:0:0:0 |
| NM_001142645 | 1 | 0 | 0 | 0 | 0 | 0 | 0 | 0 | 0 | 0 |
| NM_012433:XR_001738680 | 44:03:00 | 0:00 | 48:10:00 | 0:00 | 46:03:00 | 0:00 | 238:39:00 | 1:01 | 57:00:00 | 0:00 |
| NM_012433:XR_001738680 | 44:03:00 | 0:00 | 48:09:00 | 0:00 | 46:03:00 | 0:00 | 237:31:00 | 2:00 | 57:00:00 | 0:00 |
| NM_012433:XR_001738680:NM_001308824:XR_241302:XR_001738681 | 36:31:1:2:0 | 0:0:0:0:0 | 6:69:0:1:0 | 0:1:0:0:0 | 37:25:0:3:0 | 0:0:0:0:0 | 48:175:1:5:1 | 0:3:0:0:0 | 35:44:1:3:2 | 0:0:0:0:0 |
| NM_012433:XR_001738680:NM_001308824:XR_241302:XR_001738681 | 28:31:1:2:0 | 0:0:0:0:0 | 5:70:0:2:0 | 0:0:0:0:0 | 21:26:0:3:0 | 0:0:0:0:0 | 33:176:1:8:1 | 0:4:0:0:0 | 15:44:2:3:2 | 1:0:0:0:0 |
| XR_001738680:NM_012433:NM_001308824:XR_241302:XR_001738681 | 30:17:1:2:0 | 1:0:0:0:0 | 84:6:0:5:0 | 0:0:0:0:0 | 24:15:0:4:0 | 0:0:0:0:0 | 227:32:1:12:1 | 0:0:0:0:0 | 44:8:2:3:2 | 0:0:0:0:0 |
| NM_012433:XR_001738680:NM_001308824:XR_241302:XR_001738681 | 13:30:1:2:0 | 0:0:0:0:0 | 6:81:1:6:0 | 0:0:0:0:0 | 11:24:0:4:0 | 0:0:0:0:0 | 31:231:1:11:1 | 0:0:0:0:0 | 7:43:2:3:2 | 0:0:0:0:0 |
| NM_025147:NM_001320819 | 18:00 | 0:00 | 39:00:00 | 0:00 | 51:01:00 | 0:00 | 114:03:00 | 0:00 | 26:00:00 | 0:00 |
| NM_002156:NM_199440 | 1616:124 | 1:00 | 140:83 | 0:00 | 816:110 | 0:00 | 5429:3493 | 10:10 | 1922:191 | 3:00 |
| NM_002156:NM_199440 | 1614:125 | 3:00 | 140:83 | 0:00 | 813:110 | 3:00 | 5429:3492 | 10:10 | 1924:191 | 1:00 |
| NM_002156:NM_199440 | 1612:125 | 4:00 | 140:83 | 0:02 | 815:110 | 1:00 | 5432:3497 | 17:09 | 1922:191 | 3:00 |
| NM_002156:NM_199440 | 1617:125 | 1:00 | 140:82 | 1:03 | 815:110 | 1:00 | 5430:3493 | 24:10:00 | 1921:190 | 4:00 |
| NM_002156:NM_199440 | 1617:123 | 3:00 | 141:86 | 0:01 | 817:109 | 0:00 | 5464:3495 | 5:06 | 1924:190 | 4:00 |
| NM_002156:NM_199440 | 1619:123 | 1:00 | 141:85 | 0:02 | 817:108 | 0:00 | 5459:3489 | 10:10 | 1926:190 | 2:01 |
| NM_002156:NM_199440 | 1619:118 | 1:00 | 141:86 | 0:01 | 816:108 | 1:00 | 5460:3490 | 8:04 | 1926:192 | 3:00 |
| NM_002156:NM_199440 | 1624:117 | 2:00 | 161:87 | 0:01 | 824:109 | 1:01 | 5691:3498 | 7:05 | 1948:194 | 1:01 |
| NM_002156:NM_199440 | 1626:116 | 0:00 | 161:88 | 0:00 | 825:112 | 0:00 | 5696:3461 | 3:06 | 1945:198 | 2:01 |
| NM_002156:NM_199440 | 1625:117 | 1:00 | 160:88 | 1:00 | 825:113 | 0:00 | 5694:3440 | 14:09 | 1948:197 | 1:00 |
| NM_002156:NM_199440 | 1682:117 | 3:00 | 170:86 | 0:00 | 878:114 | 1:00 | 6212:3407 | 10:04 | 2025:205 | 3:00 |
| NM_002156:NM_199440 | 1683:116 | 2:00 | 169:84 | 0:00 | 879:112 | 0:00 | 6217:3376 | 5:09 | 2026:202 | 2:00 |
| NM_002156:NM_199440 | 1685:116 | 0:00 | 169:84 | 0:00 | 877:113 | 2:00 | 6221:3367 | 4:05 | 2026:202 | 2:00 |
| NM_002156:NM_199440 | 1685:117 | 1:00 | 169:82 | 1:00 | 877:113 | 2:00 | 6213:3339 | 11:06 | 2026:201 | 2:00 |
| NM_002156:NM_199440 | 1707:120 | 3:00 | 179:89 | 0:00 | 893:109 | 1:01 | 6395:3318 | 10:04 | 2085:194 | 2:01 |
| NM_002156:NM_199440 | 1711:119 | 0:00 | 179:89 | 0:00 | 892:109 | 1:01 | 6395:3309 | 10:08 | 2084:193 | 3:00 |
| NM_002156:NM_199440 | 1943:121 | 0:00 | 355:106 | 14:05 | 986:124 | 0:00 | 10874:3715 | 217:08:00 | 2312:199 | 10:01 |
| NM_002156:NM_199440 | 1940:121 | 4:00 | 356:107 | 23:06 | 986:124 | 2:00 | 10881:3708 | 460:23:00 | 2317:198 | 6:02 |
| NM_002156:NM_199440 | 1944:120 | 52:03:00 | 356:107 | 30:09:00 | 985:122 | 25:01:00 | 10899:3686 | 844:65 | 2318:198 | 41:03:00 |
| NM_002156:NM_199440 | 1985:123 | 11:01 | 374:114 | 13:02 | 1005:124 | 7:00 | 11353:3724 | 399:20:00 | 2347:196 | 12:02 |
| NM_002156:NM_199440 | 1996:122 | 3:00 | 387:116 | 1:01 | 1011:125 | 4:00 | 11737:3734 | 34:06:00 | 2364:195 | 2:01 |
| NM_002156:NM_199440 | 1997:122 | 3:00 | 389:117 | 0:00 | 1013:122 | 2:00 | 11758:3696 | 15:14 | 2367:195 | 1:00 |
| NM_002156:NM_199440 | 1998:122 | 2:00 | 388:117 | 1:00 | 1013:122 | 2:00 | 11761:3686 | 13:10 | 2365:195 | 3:00 |
| NM_002156:NM_199440 | 2001:119 | 1:00 | 390:114 | 0:00 | 1015:121 | 1:01 | 11775:3589 | 3:05 | 2368:194 | 1:00 |
| NM_199440:NM_002156 | 111:2002 | 1:00 | 94:390 | 1:00 | 109:1017 | 0:01 | 3397:11777 | 11:00 | 175:2368 | 0:01 |
| NM_002156:NM_199440 | 1999:111 | 3:00 | 390:93 | 0:02 | 1017:109 | 1:00 | 11773:3392 | 4:08 | 2367:173 | 2:01 |
| NM_002156:NM_199440 | 2000:109 | 2:00 | 390:86 | 0:01 | 1016:107 | 3:00 | 11768:3311 | 8:07 | 2367:172 | 2:00 |
| NM_002156:NM_199440 | 2005:75 | 0:00 | 392:41:00 | 0:00 | 1019:72 | 0:00 | 11850:2163 | 11:07 | 2365:123 | 5:01 |
| NM_199440:NM_002156 | 73:2005 | 1:00 | 38:392 | 2:00 | 72:1018 | 0:01 | 2105:11846 | 14:15 | 119:2369 | 0:01 |
| NM_002156:NM_199440 | 2005:71 | 0:00 | 392:36:00 | 0:00 | 1020:73 | 0:00 | 11871:1990 | 3:01 | 2372:115 | 0:00 |
| NM_002156:NM_199440 | 2000:67 | 4:00 | 391:30:00 | 0:00 | 1019:67 | 1:01 | 11858:1615 | 6:05 | 2368:105 | 2:00 |
| NM_002156:NM_199440 | 2005:67 | 0:00 | 391:28:00 | 1:00 | 1019:65 | 1:00 | 11866:1484 | 12:10 | 2370:97 | 2:01 |
| NM_002156:NM_199440 | 2001:65 | 4:00 | 392:26:00 | 0:01 | 1019:64 | 1:00 | 11869:1450 | 9:08 | 2370:98 | 2:00 |
| NM_002156:NM_199440 | 2002:62 | 3:01 | 390:24:00 | 2:00 | 1018:59:00 | 2:00 | 11867:1285 | 17:09 | 2369:90 | 3:00 |
| NM_002156:NM_199440 | 2005:59:00 | 3:01 | 392:20:00 | 0:01 | 1019:57:00 | 1:00 | 11873:1118 | 17:16 | 2371:83 | 1:00 |
| NM_002156:NM_199440 | 2009:58:00 | 1:00 | 393:20:00 | 0:00 | 1020:51:00 | 0:00 | 11890:929 | 10:19 | 2373:79 | 1:00 |
| NM_002156:NM_199440 | 2010:48:00 | 0:00 | 393:17:00 | 0:00 | 1018:46:00 | 2:00 | 11896:696 | 12:07 | 2373:68 | 1:00 |
| NM_002156:NM_199440 | 2009:45:00 | 1:00 | 392:15:00 | 1:00 | 1018:42:00 | 2:00 | 11899:635 | 9:01 | 2371:63 | 2:00 |
| NM_002156:NM_199440 | 2009:18:00 | 1:02 | 391:09:00 | 2:01 | 1019:23:00 | 1:01 | 11901:191 | 8:29 | 2373:25:00 | 2:04 |
| NM_001308043:NM_003879:NM_001127184:NR_147251:XR_001739017:XR_001739013:NM_001351590:XR_002959354:NM_001351592:NM_001308042:NM_001351591:NM_001127183:XM_017005190:XM_017005191:NR_147245:NR_147246:XM_017005195:XR_001739016:XR_001739018:XM_017005193:NR_147250:NM_001351594:NR_147248:NM_001202516:NR_147253 | 2:9:51:2:3:1:1:1:2:1:7:2:1:1:0:0:0:0:0:0:0:0:0:0:0 | 0:0:0:0:0:0:0:0:0:0:0:0:0:0:0:0:0:0:0:0:0:0:0:0:0 | 1:12:58:2:18:3:0:4:2:0:10:1:0:4:10:1:2:2:1:1:0:0:0:0:0 | 0:0:0:0:0:0:0:0:0:0:0:0:0:1:0:0:0:0:0:0:0:0:0:0:0 | 0:17:53:2:17:2:0:0:1:0:4:1:0:1:1:2:1:2:1:0:0:0:0:0:0 | 0:0:0:0:0:0:0:0:0:0:0:0:0:0:0:0:0:0:0:0:0:0:0:0:0 | 3:10:360:14:11:10:9:24:17:0:18:0:0:3:6:1:13:4:3:5:1:2:1:0:0 | 0:0:0:0:0:0:0:1:0:0:0:0:0:0:0:0:0:0:0:0:0:0:0:0:0 | 1:4:83:1:6:2:7:5:4:0:7:0:0:3:1:2:3:0:0:0:0:2:0:0:0 | 0:0:0:0:0:0:0:0:0:0:0:0:0:0:0:0:0:0:0:0:0:0:0:0:0 |
| NM_003879:NM_001127184:XR_001739013:NR_147251:NM_001351590:NM_001351592:NM_001308042:NM_001351591:NM_001127183:XM_017005190:XM_017005191:NR_147253:NR_147255:NR_147245:NR_147246:XR_002959354:XR_001739016:XM_017005193:XR_001739018:XM_017005195:NM_001351594:NR_147248:NM_001202516 | 9:51:2:1:1:3:1:7:2:1:1:1:1:0:0:0:0:0:0:0:0:0:0:0 | 0:0:0:0:0:0:0:0:0:0:0:0:0:0:0:0:0:0:0:0:0:0:0 | 12:58:3:1:0:4:0:10:1:0:4:0:0:10:1:1:2:1:1:0:0:0:0 | 0:0:0:0:0:0:0:0:0:0:0:0:0:0:0:0:0:0:0:0:0:0:0 | 17:52:2:2:0:1:0:4:1:0:1:0:2:1:1:0:3:0:0:0:0:0:0 | 0:1:0:0:0:0:0:0:0:0:0:0:0:0:0:0:0:0:0:0:0:0:0 | 10:360:10:13:9:21:0:18:0:0:5:1:4:6:1:1:6:7:2:10:2:3:0 | 0:0:0:0:0:0:0:0:0:0:0:0:0:0:0:0:0:0:0:0:0:0:0 | 4:83:2:1:7:4:0:7:0:0:3:0:1:1:1:0:0:1:0:2:2:0:0 | 0:0:0:0:0:0:0:0:0:0:0:0:0:0:0:0:0:0:0:0:0:0:0 |
| NM_001080125:NR_111983:NM_033356:XM_005246885 | 1:0:0:0 | 0:0:0:0 | 1:2:0:0 | 0:0:0:0 | 2:0:0:0 | 0:0:0:0 | 0:1:3:0 | 0:2:1:0 | 0:0:0:0 | 0:0:0:0 |
| NM_001080125:NM_033356:NR_111983:XM_011511969:NM_033355:XM_005246885:XM_005246895:XM_005246892:XR_923035 | 1:1:0:0:0:0:0:0:0:0 | 0:0:0:0:0:0:0:0:0 | 2:0:2:0:0:0:0:0:0:0 | 0:0:0:0:0:0:0:0:0 | 2:0:0:0:0:0:0:0:0:0 | 0:0:0:0:0:0:0:0:0:0 | 1:4:3:1:1:1:1:1:0:0 | 0:0:0:0:0:0:0:0:0:0 | 0:1:3:0:0:1:0:0:0 | 0:0:0:0:0:0:0:0:0 |
| NM_033355:NM_001080124:NM_001080125:NM_033356:XM_006712790:XM_006712789:XM_005246887:XM_005246888:XR_001738971:NR_111983:XM_005246889:XM_005246892:XM_005246895:XM_011511969:XM_005246885:NM_033358:XM_005246894:XR_923035 | 3:4:1:1:1:0:0:0:0:0:0:0:0:0:0:0:0:0 | 0:0:0:0:0:0:0:0:0:0:0:0:0:0:0:0:0:0 | 10:5:2:4:0:2:3:1:1:1:1:2:1:1:0:0:0:0 | 0:0:0:0:0:0:0:0:0:0:0:0:0:0:0:0:0:0 | 5:14:3:4:1:3:4:0:0:3:0:2:0:0:0:0:0:0 | 0:0:0:0:0:0:0:0:0:0:0:0:0:0:0:0:0:0 | 7:3:3:10:4:6:2:2:0:5:7:4:2:1:1:0:0:0 | 0:0:0:0:0:0:0:0:0:0:0:0:0:0:0:0:0:0 | 10:2:5:5:2:3:4:0:0:5:2:0:1:1:1:1:1:0 | 0:0:0:0:0:0:0:0:0:0:0:0:0:0:0:0:0:0 |
| NM_002997:XM_005262622 | 0:00 | 0:00 | 0:00 | 0:00 | 0:00 | 2:00 | 0:00 | 0:00 | 0:00 | 0:01 |
| NM_205863:XM_017003290 | 0:00 | 0:00 | 0:00 | 0:00 | 0:00 | 1:00 | 0:00 | 0:00 | 0:00 | 0:00 |
| NM_001025436:XM_017004899:NM_024532:XM_011511837:XM_017004896:XM_011511814 | 9:2:0:0:0:0 | 0:0:0:0:0:0 | 51:7:0:0:0:0 | 0:2:1:0:0:0 | 45:3:1:0:0:0 | 0:0:0:0:0:0 | 267:23:5:2:0:0 | 0:6:2:0:0:0 | 58:1:1:0:0:0 | 0:0:0:0:0:0 |
| NM_001025436:XM_017004899:NM_024532:XM_011511837:XM_017004896:XM_011511814 | 9:2:0:0:0:0 | 0:0:0:0:0:0 | 51:3:0:0:0:0 | 0:6:1:0:0:0 | 45:2:1:0:0:0 | 0:1:0:0:0:0 | 267:14:1:2:0:0 | 0:15:6:0:0:0 | 58:1:1:0:0:0 | 0:0:0:0:0:0 |
| NM_000465:XR_002959322:XM_017004613:XM_017004614:NR_104212 | 0:0:0:0:0 | 0:0:0:0:0 | 1:1:0:0:0 | 0:0:0:0:0 | 0:0:0:0:0 | 0:0:0:0:0 | 0:0:0:0:0 | 0:0:0:0:0 | 2:0:1:1:0 | 0:0:0:0:0 |
| NM_000465:XR_002959322:XM_017004613:XM_017004614:NR_104212 | 0:0:0:0:0 | 0:0:0:0:0 | 0:1:0:0:0 | 1:0:0:0:0 | 0:0:0:0:0 | 0:0:0:0:0 | 0:0:0:0:0 | 0:0:0:0:0 | 0:0:1:1:0 | 2:0:0:0:0 |

| | | | | | | | | | | |
|---|---|---|---|---|---|---|---|---|---|---|
| NM_000465:NR_104212 | 0:00 | 0:00 | 1:00 | 0:00 | 0:00 | 0:00 | 0:00 | 0:00 | 2:00 | 0:00 |
| NM_212482:NM_212474:XM_017003692:NM_212476:NM_002026:NM_001306132:NM_212478:NM_001306131:NM_001306130 | 0:0:0:0:0:0:0:0:0 | 4:9:3:1:4:1:3:1:0 | 0:0:0:0:0:0:0:0:0 | 0:0:0:0:0:0:0:0:0 | 0:0:0:0:0:0:0:0:0 | 1:0:0:0:0:0:0:0:0 | 0:0:0:0:0:0:0:0:0 | 2:1:0:0:0:0:0:0:0 | 0:0:0:0:0:0:0:0:0 | 0:0:0:0:1:0:1:0:0 |
| NM_212482:NM_212474:XM_017003692:NM_212476:NM_002026:NM_001306132:NM_212478:NM_001306131:NM_001306130 | 4:9:3:1:4:1:3:1:0 | 0:0:0:0:0:0:0:0:0 | 0:0:0:0:0:0:0:0:0 | 0:0:0:0:0:0:0:0:0 | 1:0:0:0:0:0:0:0:0 | 0:0:0:0:0:0:0:0:0 | 1:1:0:0:0:0:0:0:0 | 0:0:0:0:0:0:0:0:0 | 0:0:0:0:1:0:1:0:0 | 0:0:0:0:0:0:0:0:0 |
| NM_212482:NM_212474:NM_212478:NM_212476 | 1:2:2:1 | 6:0:0:0 | 0:0:0:0 | 0:0:0:0 | 0:0:0:0 | 1:0:0:0 | 0:0:0:0 | 2:0:0:0 | 0:0:0:0 | 0:0:0:0 |
| NM_212474:NM_212478:NM_212476:NM_212482 | 0:0:0:0 | 2:2:1:7 | 0:0:0:0 | 0:0:0:0 | 0:0:0:0 | 0:0:0:1 | 0:0:0:0 | 0:0:0:2 | 0:0:0:0 | 0:0:0:0 |
| NM_002026:NM_212482:NM_212478:NM_212474:NM_054034 | 2:1:0:0:0 | 1:2:0:0:0 | 0:0:0:0:0 | 0:0:0:0:0 | 0:1:0:0:0 | 0:0:0:0:0 | 0:0:0:0:0 | 0:0:0:0:0 | 0:0:0:0:0 | 0:0:0:0:0 |
| NM_015680 | 0 | 1 | 0 | 0 | 0 | 1 | 0 | 0 | 0 | 0 |
| NM_015680 | 0 | 1 | 0 | 0 | 0 | 1 | 0 | 0 | 0 | 0 |
| NM_015680 | 0 | 1 | 0 | 0 | 0 | 1 | 0 | 0 | 0 | 0 |
| NM_005687:XM_024452492 | 0:00 | 0:00 | 0:00 | 0:00 | 0:00 | 0:00 | 0:00 | 0:00 | 0:00 | 1:00 |
| NM_003590:XM_011511996:XM_011511995:XM_011511994:NM_001257197 | 3:0:0:0:0 | 0:0:0:0:0 | 7:16:3:0:0 | 0:0:0:0:0 | 18:2:0:1:0 | 0:0:0:0:0 | 27:42:8:0:0 | 0:0:0:0:0 | 11:2:2:0:2 | 0:0:0:0:0 |
| NM_003590:XM_011511996:XM_011511995:XM_011511994:NM_001257197 | 3:0:0:0:0 | 1:0:0:0:0 | 7:16:3:0:0 | 0:0:0:0:0 | 18:2:0:1:0 | 1:0:0:0:0 | 27:37:8:0:0 | 5:0:0:0:0 | 11:2:2:0:2 | 1:0:0:0:0 |
| NM_003590:XM_011511996:XM_011511995:XM_011511994:NM_001257197 | 4:0:0:0:0 | 0:0:0:0:0 | 12:11:3:0:0 | 2:0:0:0:0 | 22:2:0:1:0 | 0:0:0:0:0 | 57:28:8:0:1 | 1:0:0:0:0 | 18:2:2:0:2 | 0:0:0:0:0 |
| NM_024795 | 1 | 0 | 1 | 0 | 0 | 2 | 1 | 0 | 0 | 0 |
| NM_001206702:NM_001206704 | 0:00 | 1:01 | 0:00 | 0:00 | 0:00 | 1:00 | 0:00 | 0:00 | 2:00 | 0:00 |
| NM_030926 | 0 | 0 | 3 | 3 | 7 | 4 | 2 | 0 | 2 | 0 |
| NM_005381 | 100 | 0 | 71 | 0 | 50 | 0 | 765 | 2 | 132 | 3 |
| NM_005381 | 91 | 1 | 60 | 1 | 44 | 0 | 715 | 3 | 122 | 0 |
| NM_005381 | 91 | 1 | 58 | 1 | 44 | 0 | 713 | 2 | 122 | 0 |
| NM_005381 | 78 | 0 | 42 | 0 | 35 | 0 | 555 | 2 | 108 | 0 |
| NM_005381 | 74 | 0 | 30 | 0 | 33 | 0 | 321 | 2 | 102 | 0 |
| NM_005381 | 74 | 0 | 30 | 0 | 33 | 0 | 318 | 3 | 101 | 1 |
| NM_005381 | 74 | 0 | 30 | 0 | 33 | 0 | 319 | 1 | 102 | 0 |
| NM_005381 | 74 | 0 | 31 | 0 | 34 | 0 | 320 | 1 | 102 | 0 |
| NM_019850:XM_011510923 | 0:00 | 0:00 | 0:00 | 0:00 | 0:00 | 1:00 | 0:00 | 1:00 | 0:00 | 1:00 |
| NM_001017915:NM_005541:XM_017004004 | 0:00:01 | 1:05:00 | 0:00:01 | 0:00:00 | 0:00:00 | 0:00:00 | 0:01:02 | 0:00:00 | 0:01:00 | 0:00:00 |
| XM_024452672:XM_011510549 | 0:00 | 0:00 | 0:00 | 1:00 | 0:00 | 0:01 | 0:00 | 0:02 | 0:00 | 1:00 |
| NM_001042467:XM_006712740:NM_001281473 | 0:00:00 | 0:00:00 | 0:00:00 | 0:00:00 | 0:00:00 | 0:00:00 | 0:00:00 | 0:00:00 | 1:00:00 | 0:00:00 |
| NM_001042467:XM_006712740:NM_001281473 | 0:00:00 | 0:00:00 | 0:00:00 | 0:00:00 | 0:00:00 | 0:00:00 | 0:00:00 | 0:00:00 | 1:00:00 | 0:00:00 |
| NM_001042467:XM_006712740:NM_001281473 | 0:00:00 | 0:00:00 | 0:00:00 | 0:00:00 | 0:00:00 | 0:00:00 | 0:00:00 | 0:00:00 | 1:00:00 | 0:00:00 |
| NM_004544:NR_136157:NM_001322019:XM_011511228:XR_001738750:NR_136158 | 139:7:1:1:0:0 | 0:0:0:0:0:0 | 202:12:0:6:1:0 | 0:0:0:0:0:0 | 249:3:0:2:0:0 | 1:0:0:0:0:0 | 959:33:1:14:7:1 | 1:0:0:1:0:0 | 287:11:1:5:0:0 | 0:0:0:0:0:0 |
| NM_004544:NR_136157:NM_001322019:XM_011511228:XR_001738750:NR_136158 | 139:2:1:1:0:0 | 0:0:0:0:0:0 | 202:8:0:6:1:0 | 0:0:0:0:0:0 | 250:2:0:2:0:0 | 0:0:0:0:0:0 | 960:17:1:15:9:1 | 0:0:0:0:0:0 | 287:7:1:5:0:0 | 0:0:0:0:0:0 |
| NM_004544:NM_001322019:XM_011511228:XR_001738750:NR_136157:NR_136158 | 135:1:0:0:0:0 | 0:0:0:0:0:0 | 198:0:4:2:0:0 | 0:0:0:0:0:0 | 242:0:2:3:1:0 | 0:0:0:0:0:0 | 941:1:12:13:7:2 | 3:0:0:0:0:0 | 283:1:4:0:2:0 | 0:0:0:0:0:0 |
| NM_004544:NM_001322019:XM_011511228:XR_001738750:NR_136157:NR_136158 | 135:1:0:0:0:0 | 0:0:0:0:0:0 | 197:0:4:2:0:0 | 0:0:0:0:0:0 | 241:0:2:3:1:0 | 1:0:0:0:0:0 | 942:1:12:15:7:2 | 1:0:0:0:0:0 | 281:1:4:0:2:0 | 1:0:0:0:0:0 |
| NM_004544:NR_136157:NM_001322019:XR_001738750:XM_011511228:NR_136158 | 136:1:1:1:0:0 | 0:0:0:0:0:0 | 195:0:0:3:2:0 | 0:0:0:0:0:0 | 242:1:0:3:2:0 | 1:0:0:0:0:0 | 930:7:1:15:10:2 | 2:0:0:0:0:0 | 282:2:1:0:4:0 | 0:0:0:0:0:0 |
| NM_004544:NR_136157:NM_001322019:XR_001738750:XM_011511228:NR_136158 | 134:1:1:1:0:0 | 1:0:0:0:0:0 | 191:0:0:5:2:0 | 3:0:0:0:0:0 | 241:1:0:3:2:0 | 0:0:0:1:0:0 | 927:7:1:15:10:2 | 2:0:0:0:0:0 | 282:2:1:0:4:0 | 0:0:0:0:0:0 |
| NM_004544:NR_136157:NM_001322019:XM_011511228:XR_001738750:NR_136158 | 133:1:1:0:0:0 | 0:0:0:0:0:0 | 191:0:0:2:5:0 | 0:0:0:0:0:0 | 240:1:0:2:3:0 | 0:0:0:0:0:0 | 912:9:1:10:10:2 | 2:0:0:0:0:0 | 273:2:1:2:0:0 | 0:0:0:0:0:0 |
| NM_004544:NR_136157:NM_001322019:XM_011511228:XR_001738750:NR_136158 | 130:1:1:0:0:0 | 2:0:0:0:0:0 | 187:0:0:2:4:0 | 0:0:0:0:0:0 | 238:1:0:2:3:0 | 0:0:0:0:0:0 | 902:9:1:10:9:2 | 0:0:0:0:0:0 | 269:2:1:2:0:0 | 1:0:0:0:0:0 |
| NM_004544:NR_136157:NM_001322019:XM_011511228:XR_001738750:NR_136158 | 51:1:1:0:0:0 | 81:0:0:0:0:0 | 104:0:0:0:4:0 | 84:0:0:2:0:0 | 138:1:0:1:2:0 | 102:0:0:1:1:0 | 904:9:1:10:9:2 | 0:0:0:0:0:0 | 271:2:1:2:0:0 | 0:0:0:0:0:0 |
| NM_025203:XM_017005029 | 0:00 | 2:00 | 0:02 | 0:00 | 0:00 | 0:00 | 0:01 | 0:00 | 0:00 | 0:00 |
| NM_023085:NM_023083 | 0:00 | 0:00 | 0:00 | 0:00 | 0:00 | 0:00 | 0:00 | 0:00 | 0:00 | 0:00 |
| NM_001195381:NM_001195382:NM_005301 | 0:06:00 | 1:02:00 | 0:07:00 | 1:09:00 | 1:05:00 | 5:02:00 | 0.583576389 | 0:00:00 | 4:12:03 | 0:00:00 |
| NM_001195382:NM_001195381:NM_005301 | 8:01:00 | 0:00:00 | 16:01:00 | 0:00:00 | 7:06:00 | 0:00:00 | 30:08:14 | 29:05:18 | 7:02:03 | 5:02:00 |
| XM_017003942:XM_024452833:NM_005336 | 0:00:00 | 0:00:00 | 0:00:00 | 0:00:00 | 0:00:00 | 0:00:00 | 1:00:00 | 0:00:00 | 0:00:00 | 0:00:00 |
| NM_015963 | 0 | 0 | 0 | 0 | 1 | 0 | 0 | 0 | 0 | 0 |
| NM_000182 | 129 | 0 | 199 | 0 | 194 | 1 | 675 | 0 | 179 | 0 |
| NM_000182 | 128 | 0 | 199 | 0 | 194 | 1 | 670 | 0 | 177 | 0 |
| NM_000182 | 128 | 0 | 198 | 0 | 195 | 0 | 665 | 0 | 177 | 0 |
| NM_000182 | 126 | 1 | 196 | 1 | 196 | 0 | 657 | 2 | 177 | 0 |
| NM_000182 | 127 | 0 | 196 | 0 | 195 | 0 | 654 | 0 | 175 | 0 |
| NM_000182 | 127 | 0 | 196 | 0 | 192 | 0 | 650 | 3 | 176 | 0 |
| NM_000182 | 124 | 0 | 186 | 0 | 190 | 0 | 603 | 2 | 175 | 0 |
| NM_000182 | 125 | 0 | 178 | 0 | 189 | 0 | 586 | 0 | 173 | 0 |
| NM_000182 | 126 | 0 | 175 | 2 | 188 | 0 | 580 | 1 | 173 | 0 |
| NM_000182 | 126 | 0 | 176 | 1 | 188 | 0 | 578 | 3 | 173 | 0 |
| NM_000182 | 126 | 0 | 177 | 0 | 188 | 0 | 575 | 2 | 173 | 0 |
| NM_000182 | 124 | 2 | 175 | 0 | 188 | 0 | 569 | 2 | 173 | 0 |
| NM_000182 | 125 | 1 | 171 | 1 | 187 | 1 | 567 | 1 | 172 | 2 |
| NM_000182 | 127 | 0 | 169 | 0 | 189 | 0 | 556 | 1 | 172 | 1 |
| NM_000182 | 125 | 1 | 164 | 0 | 186 | 0 | 545 | 2 | 171 | 0 |
| NM_000182 | 142 | 3 | 178 | 2 | 203 | 2 | 530 | 7 | 197 | 1 |
| NM_000182 | 140 | 0 | 169 | 0 | 198 | 0 | 443 | 0 | 190 | 1 |
| NM_000182 | 140 | 0 | 164 | 0 | 197 | 0 | 431 | 3 | 189 | 0 |
| NM_016085 | 0 | 0 | 0 | 2 | 0 | 0 | 0 | 2 | 0 | 0 |
| NM_016085 | 0 | 0 | 2 | 0 | 0 | 0 | 2 | 0 | 0 | 0 |
| NM_007099:NR_024080 | 1:00 | 1:00 | 1:00 | 0:01 | 0:00 | 0:00 | 8:00 | 0:00 | 4:00 | 2:00 |

| | | | | | | | | | | |
|---|---|---|---|---|---|---|---|---|---|---|
| NM_001521:NM_001035521:XM_005264272:NM_001318909 | 5:9:2:0 | 3:2:0:0 | 5:2:2:0 | 3:0:1:0 | 19:18:4:1 | 8:4:5:1 | 21:32:20:0 | 10:7:15:2 | 6:11:2:0 | 3:1:2:0 |
| NM_001521:NM_001035521:XM_005264272:NM_001318909 | 6:10:2:0 | 3:1:0:0 | 7:2:2:0 | 1:0:1:0 | 22:20:3:1 | 5:2:5:1 | 27:34:33:2 | 4:5:7:0 | 8:11:2:0 | 2:1:2:0 |
| NM_001521:NM_001035521:XM_005264272:NM_001318909 | 9:11:2:0 | 1:0:0:0 | 8:2:3:0 | 0:0:0:0 | 26:22:8:1 | 1:0:0:1 | 30:38:40:2 | 2:1:0:0 | 10:12:4:0 | 0:0:0:0 |
| NM_001034116:NM_015636 | 1:00 | 0:00 | 0:00 | 0:00 | 0:01 | 0:00 | 0:01 | 0:00 | 1:00 | 0:00 |
| NM_001321357:NM_013392:NM_001321358:NM_001321363:NM_001321362 | 40:11:1:0:0 | 0:0:0:0:0 | 46:18:0:0:0 | 0:0:0:0:0 | 70:14:0:2:0 | 0:0:0:0:0 | 166:57:0:0:0 | 1:1:0:0:0 | 39:2:0:0:1 | 0:0:0:0:0 |
| NM_007099:NM_004300:NR_024080 | 1:03:00 | 1:00:00 | 1:01:00 | 0:00:01 | 0:00:00 | 0:00:00 | 8:00:00 | 0:03:00 | 4:00:00 | 2:02:00 |
| NM_017910:XM_005264393:XR_244945:XM_017004403:XR_939691 | 19:5:1:1:0 | 0:0:0:0:0 | 13:10:0:1:0 | 0:0:0:0:0 | 17:2:0:3:0 | 0:0:0:0:0 | 77:80:7:13:6 | 0:0:0:1:0 | 22:5:0:5:0 | 1:0:0:0:0 |
| NM_016061:NM_001127400:XM_017004319:XM_017004320:XM_017004318:NM_001127401:XM_017004322:NM_001127399:XM_017004321 | 87:4:4:2:2:1:4:0:0 | 1:0:0:0:0:0:0:0:0 | 135:4:2:1:0:1:8:2:0 | 2:0:0:0:0:0:0:0:0 | 127:5:2:1:0:0:2:3:0 | 0:0:0:0:0:0:0:0:0 | 374:31:15:11:3:2:23:20:3 | 1:1:0:0:0:0:0:0:0 | 60:3:1:2:0:2:1:1:0 | 0:0:0:0:0:0:0:0:0 |
| NM_016061:NM_001127400:XM_017004319:XM_017004320:XM_017004318:NM_001127401:XM_017004322:NM_001127399:XM_017004321 | 88:4:4:2:2:1:4:0:0 | 0:0:0:0:0:0:0:0:0 | 138:4:2:1:0:1:8:2:0 | 0:0:0:0:0:0:0:0:0 | 127:5:2:1:0:0:2:3:0 | 0:0:0:0:0:0:0:0:0 | 378:32:15:11:3:2:28:20:3 | 1:0:0:0:0:0:0:0:0 | 60:3:1:2:0:2:1:1:0 | 0:0:0:0:0:0:0:0:0 |
| NM_016061:NM_001127400:XM_017004319:XM_017004320:XM_017004318:NM_001127401:XM_017004322:NM_001127399:XM_017004321 | 87:4:4:2:2:1:4:0:0 | 1:0:0:0:0:0:0:0:0 | 137:4:2:1:0:1:8:2:0 | 1:0:0:0:0:0:0:0:0 | 127:5:2:1:0:0:2:3:0 | 0:0:0:0:0:0:0:0:0 | 377:31:15:11:3:2:28:20:3 | 2:1:0:0:0:0:0:0:0 | 60:3:1:2:0:2:1:1:0 | 0:0:0:0:0:0:0:0:0 |
| XM_017004560:XR_244950:XM_005264450:NM_016252:XM_005264453:XM_017004557 | 1:0:0:0:0:0 | 0:0:0:0:0:0 | 1:6:1:0:0:0 | 0:0:0:0:0:0 | 0:0:0:0:0:0 | 0:0:0:0:0:0 | 1:10:1:1:3:0 | 0:0:0:0:0:0 | 0:0:1:0:0:1 | 0:0:0:0:0:0 |
| XM_017004558:XR_244950:XM_017004559:XM_011533003:NM_016252 | 2:0:0:0:0 | 4:0:0:0:0 | 0:2:0:0:0 | 1:0:0:0:0 | 5:0:1:1:0 | 1:1:0:0:0 | 9:1:3:0:0 | 0:0:0:0:0 | 9:1:0:0:1 | 0:0:0:0:0 |
| NM_017735:XR_002959314:NM_001193509:XM_011532958 | 17:0:0:0 | 0:0:0:0 | 12:2:0:0 | 0:0:0:0 | 11:1:0:0 | 0:0:0:0 | 78:5:1:2 | 67:3:7:3 | 7:0:0:0 | 5:0:0:1 |
| NM_206943:NM_001166265:NM_001166266:XM_011532859 | 0:0:0:0 | 0:0:0:0 | 0:0:0:0 | 1:0:0:0 | 0:0:0:0 | 2:1:0:0 | 0:0:0:0 | 0:0:1:0 | 0:0:0:0 | 1:0:0:0 |
| NM_138394:NM_001142650:XM_005264639:XM_005264640:XM_011533165 | 22:2:1:0:0 | 0:0:0:0:0 | 36:6:4:1:0 | 0:0:0:0:0 | 24:7:2:0:0 | 0:0:0:0:0 | 172:34:5:10:0 | 0:1:0:0:0 | 46:5:0:1:0 | 0:0:0:0:0 |
| NM_138394:NM_001142650:XM_005264639:XM_011533165:XM_011533166:XM_005264640 | 15:2:1:1:1:0 | 0:0:0:0:0:0 | 27:6:3:0:0:1 | 0:0:0:0:0:0 | 13:7:3:0:0:0 | 0:0:0:0:0:0 | 73:35:2:0:4:6 | 2:0:0:0:1:0 | 33:5:0:0:1:1 | 0:0:0:0:0:0 |
| NM_019063:XR_001738710:XR_001738709:XM_006711992 | 0:0:0:0 | 6:0:0:0 | 0:0:0:0 | 2:0:0:0 | 0:0:0:0 | 2:0:0:0 | 0:0:0:0 | 9:0:0:0 | 0:0:0:0 | 2:0:0:0 |
| NM_019063:XR_001738710:XR_001738709:XM_006711992 | 0:0:0:0 | 6:0:0:0 | 0:0:0:0 | 2:0:0:0 | 0:0:0:0 | 2:0:0:0 | 0:0:0:0 | 13:0:0:0 | 0:0:0:0 | 3:0:0:0 |
| NM_019063:XR_001738710:XR_001738709:XM_006711992 | 3:0:0:0 | 3:0:0:0 | 0:0:0:0 | 2:0:0:0 | 0:0:0:0 | 2:0:0:0 | 0:0:0:0 | 13:0:0:0 | 0:0:0:0 | 3:0:0:0 |
| XM_011533047:NM_000341 | 0:00 | 0:00 | 0.431944444 | 0:00 | 0.7125 | 0:02 | 15:45 | 0:00 | 2:05 | 0:00 |
| XM_011533047:NM_000341 | 0:00 | 0:00 | 0.647916667 | 0:00 | 0.759027778 | 0:02 | 0.796527778 | 0:00 | 1:07 | 0:00 |
| XM_011533047:NM_000341 | 0:00 | 0:00 | 0.694444444 | 0:00 | 0.853472222 | 0:02 | 0.847916667 | 0:00 | 1:08 | 0:00 |
| NM_000341:XM_011533047 | 0:00 | 0:00 | 110:14:00 | 1:00 | 213:17:00 | 2:00 | 87:20:00 | 0:00 | 9:01 | 0:00 |
| NM_000341:XM_011533047 | 0:00 | 0:00 | 111:12:00 | 2:00 | 216:14:00 | 1:00 | 87:20:00 | 0:00 | 9:01 | 0:00 |
| XM_011533047:NM_000341 | 0:00 | 0:00 | 0:00 | 0.329166667 | 0:00 | 0.484027778 | 5:50 | 4:37 | 0:04 | 1:05 |
| NM_139279:NM_001171506 | 59:00:00 | 0:00 | 16:01 | 0:00 | 59:03:00 | 0:00 | 271:03:00 | 2:00 | 60:02:00 | 0:00 |
| NM_002354 | 1707 | 2 | 5288 | 3 | 4003 | 1 | 13927 | 6 | 3400 | 3 |
| NM_002354 | 1717 | 3 | 5330 | 6 | 4020 | 1 | 14036 | 7 | 3431 | 3 |
| NM_002354 | 1720 | 0 | 5330 | 6 | 4018 | 2 | 14031 | 13 | 3432 | 3 |
| NM_002354 | 1729 | 1 | 5398 | 3 | 4044 | 3 | 6961 | 7239 | 1737 | 1726 |
| NM_002354 | 1731 | 1 | 5414 | 3 | 4061 | 2 | 14252 | 11 | 3472 | 3 |
| NM_002354 | 1735 | 3 | 5430 | 2 | 4069 | 2 | 14291 | 11 | 3489 | 1 |
| NM_002354 | 1730 | 10 | 5456 | 12 | 4078 | 7 | 14340 | 33 | 3497 | 12 |
| NM_002354 | 1758 | 1 | 5651 | 5 | 4143 | 4 | 14581 | 12 | 3558 | 5 |
| NM_002354 | 1761 | 1 | 5682 | 12 | 4172 | 3 | 14623 | 20 | 3573 | 1 |
| NM_002354 | 1760 | 2 | 5685 | 9 | 4170 | 5 | 14630 | 14 | 3569 | 5 |
| NM_002354 | 1762 | 3 | 5724 | 11 | 4182 | 5 | 14690 | 10 | 3584 | 4 |
| NM_002354 | 1768 | 0 | 5751 | 9 | 4190 | 1 | 14717 | 12 | 3586 | 3 |
| NM_002354 | 1767 | 1 | 5755 | 7 | 4189 | 2 | 14715 | 20 | 3589 | 4 |
| NM_002354 | 1772 | 1 | 5813 | 11 | 4198 | 7 | 14749 | 31 | 3593 | 8 |
| NM_002354 | 1774 | 1 | 5818 | 13 | 4204 | 1 | 14773 | 16 | 3604 | 4 |
| NM_002354 | 1781 | 2 | 5902 | 5 | 4215 | 2 | 14852 | 14 | 3623 | 5 |
| NM_002354 | 1782 | 1 | 5898 | 9 | 4214 | 3 | 14850 | 14 | 3623 | 5 |
| NM_002354 | 1775 | 2 | 5891 | 6 | 4194 | 6 | 14815 | 22 | 3615 | 0 |
| NM_000251:XR_939685:NM_001258281:XM_011532867:XM_005264332:XR_001738747 | 49:5:6:0:0:0 | 0:0:0:0:0:0 | 4:10:15:0:0:0 | 0:0:0:0:0:0 | 49:5:14:0:0:0 | 0:0:0:0:0:0 | 51:64:102:1:1:0 | 0:0:0:0:0:0 | 93:10:35:0:1:0 | 0:0:0:0:0:0 |
| NM_000251:XR_939685:NM_001258281:XM_011532867:XM_005264332:XR_001738747 | 49:3:6:0:0:0 | 0:0:0:0:0:0 | 4:10:18:0:0:0 | 0:0:0:0:0:0 | 49:4:14:0:0:0 | 0:0:0:0:0:0 | 49:64:119:1:1:0 | 2:1:0:0:0:0 | 93:7:36:0:1:0 | 0:0:0:0:0:0 |
| NM_000251:XR_939685:NM_001258281:XM_011532867:XM_005264332:XR_001738747 | 45:3:6:0:0:0 | 4:0:0:0:0:0 | 2:7:23:0:0:0 | 2:1:0:0:0:0 | 48:5:14:0:0:0 | 1:1:0:0:0:0 | 41:49:184:1:2:0 | 7:9:0:0:0:0 | 91:5:36:0:1:0 | 2:4:0:0:0:0 |
| NM_000251:XR_939685:NM_001258281:XM_011532867:XM_005264332:XR_001738747 | 45:3:6:0:0:0 | 0:0:0:0:0:0 | 1:6:26:0:0:0 | 0:0:0:0:0:0 | 47:4:15:0:0:0 | 0:0:0:0:0:0 | 41:47:216:1:3:0 | 0:0:0:0:0:0 | 91:5:37:0:1:0 | 0:0:0:0:0:0 |
| NM_000251:XR_939685:NM_001258281:XM_011532867:XM_005264332:XR_001738747 | 45:3:6:0:0:0 | 0:0:0:0:0:0 | 1:6:26:0:0:0 | 0:0:0:0:0:0 | 47:4:15:0:0:0 | 0:0:0:0:0:0 | 41:48:216:1:3:0 | 0:0:1:0:0:0 | 91:5:37:0:1:0 | 0:0:0:0:0:0 |
| NM_000251:XR_939685:NM_001258281:XM_011532867:XM_005264332:XR_001738747 | 45:3:6:0:0:0 | 0:0:0:0:0:0 | 1:0:26:0:0:0 | 0:6:0:0:0:0 | 47:4:15:0:0:0 | 0:0:0:0:0:0 | 39:10:225:1:3:0 | 2:37:0:0:0:0 | 91:5:37:0:1:0 | 0:0:0:0:0:0 |
| NM_000251:XR_939685:NM_001258281:XM_011532867:XM_005264332:XR_001738747 | 45:4:7:0:0:0 | 0:0:0:0:0:0 | 1:0:27:0:0:0 | 0:0:0:0:0:0 | 47:3:16:0:0:0 | 0:0:0:0:0:0 | 39:10:237:1:4:0 | 0:0:0:0:0:0 | 91:5:37:0:1:0 | 0:0:0:0:0:0 |
| NM_000251:XR_939685:NM_001258281:XM_011532867:XM_005264332:XR_001738747 | 45:4:7:0:0:0 | 0:0:0:0:0:0 | 1:0:28:0:0:0 | 0:0:0:0:0:0 | 47:3:16:0:0:0 | 0:0:0:0:0:0 | 39:10:241:1:4:0 | 0:0:0:0:0:0 | 91:5:37:0:1:0 | 0:0:0:0:0:0 |
| NM_000251:XR_939685:NM_001258281:XM_011532867:XM_005264332:XR_001738747 | 45:4:8:0:0:0 | 0:0:0:0:0:0 | 1:0:26:0:0:0 | 0:0:2:0:0:0 | 47:3:16:0:0:0 | 0:0:0:0:0:0 | 39:11:240:1:4:0 | 0:0:1:0:0:0 | 90:5:37:0:1:0 | 1:0:0:0:0:0 |
| XM_005264572:XM_017005017:NM_001190274:NM_025133 | 0:2:3:0 | 1:0:0:0 | 0:5:0:0 | 1:2:0:0 | 0:1:1:0 | 0:0:0:0 | 3:17:1:5 | 2:1:1:0 | 0:0:1:6 | 0:0:1:0 |
| NM_002954:NM_001177413:NM_001135592 | 38:02:00 | 0:00:00 | 27:02:00 | 0:00:00 | 39:01:00 | 0:00:00 | 131:00:00 | 0:00:00 | 97:06:00 | 0:00:00 |
| NM_138559:XM_011532910:XM_017004335:NM_022893:XM_024452963:XM_017004333 | 8:1:0:0:0:0 | 0:0:0:0:0:0 | 7:0:0:0:0:0 | 0:0:0:0:0:0 | 8:0:0:0:0:0 | 0:0:0:0:0:0 | 130:4:6:1:1:1 | 6:0:0:0:0:0 | 50:1:2:0:1:0 | 0:0:0:0:0:0 |
| NM_138559:XM_011532910:XM_017004335:NM_022893:XM_024452963:XM_017004333 | 8:1:0:0:0:0 | 0:0:0:0:0:0 | 7:0:0:0:0:0 | 0:0:0:0:0:0 | 8:0:0:0:0:0 | 0:0:0:0:0:0 | 134:4:6:1:1:1 | 2:0:0:0:0:0 | 49:1:2:0:1:0 | 0:0:0:0:0:0 |
| NM_006430:NM_001256721 | 156:01:00 | 0:00 | 58:00:00 | 1:00 | 57:01:00 | 0:00 | 1101:00:00 | 2:00 | 264:01:00 | 0:00 |
| NM_006430:NM_001256721 | 156:01:00 | 0:00 | 60:00:00 | 0:00 | 56:01:00 | 1:00 | 1101:00:00 | 2:00 | 264:01:00 | 0:00 |
| NM_006430:NM_001256721 | 158:01:00 | 1:00 | 60:00:00 | 0:00 | 57:01:00 | 0:00 | 1093:00:00 | 3:00 | 267:01:00 | 1:00 |
| NM_006430:NM_001256721 | 159:01:00 | 0:00 | 60:00:00 | 0:00 | 57:01:00 | 0:00 | 1096:00:00 | 2:00 | 268:01:00 | 0:00 |
| NM_006430:NM_001256721 | 159:01:00 | 0:00 | 60:00:00 | 0:00 | 57:01:00 | 0:00 | 1092:00:00 | 3:00 | 267:01:00 | 0:00 |
| NM_006430:NM_001256721 | 158:01:00 | 0:00 | 57:00:00 | 1:00 | 58:01:00 | 0:00 | 1084:00:00 | 2:00 | 264:01:00 | 0:00 |
| NM_006430:NM_001256721 | 157:01:00 | 0:00 | 57:00:00 | 0:00 | 57:01:00 | 0:00 | 1079:00:00 | 2:00 | 264:01:00 | 0:00 |
| NM_006430:NM_001256721 | 157:01:00 | 0:00 | 58:00:00 | 0:00 | 56:01:00 | 0:00 | 1065:00:00 | 3:00 | 267:01:00 | 0:00 |
| NM_006430:NM_001256721 | 157:01:00 | 0:00 | 58:00:00 | 0:00 | 56:01:00 | 0:00 | 1067:00:00 | 0:00 | 267:01:00 | 0:00 |
| NM_006430:NM_001256721 | 153:01:00 | 1:00 | 58:00:00 | 0:00 | 56:01:00 | 0:00 | 1037:00:00 | 2:00 | 265:01:00 | 0:00 |
| NM_006430:NM_001256721 | 153:01:00 | 0:00 | 57:00:00 | 0:00 | 56:01:00 | 0:00 | 1022:00:00 | 2:00 | 264:01:00 | 0:00 |
| NM_006430:NM_001256721 | 149:01:00 | 0:00 | 49:00:00 | 0:00 | 54:01:00 | 0:00 | 934:00:00 | 0:00 | 254:01:00 | 2:00 |
| NM_001007231:NM_014882:XM_017005427:NM_001166276:NM_001166277:XM_017005423:XM_011533207 | 0:0:0:0:0:0:0 | 0:0:0:0:0:0:0 | 0:0:1:0:0:0:0 | 2:1:0:0:0:0:0 | 1:0:1:1:0:0:0 | 0:1:0:2:0:0:0 | 0:0:0:0:0:0:0 | 0:0:1:0:0:0:0 | 0:0:0:0:0:0:0 | 0:0:0:0:0:0:0 |
| XM_017003801:NM_002056:XM_017003802:NM_001244710 | 3:64:8:0 | 0:0:0:0 | 6:18:96:0 | 0:0:0:0 | 8:68:17:0 | 0:0:0:0 | 43:220:666:0 | 0:0:2:0 | 10:225:82:0 | 0:0:1:0 |
| XM_017003801:NM_002056:XM_017003802:NM_001244710 | 3:64:8:0 | 0:0:0:0 | 6:18:97:0 | 0:0:0:0 | 8:68:17:0 | 0:0:0:0 | 42:219:660:0 | 0:0:1:0 | 10:225:83:0 | 0:0:0:0 |
| XM_017003801:NM_002056:XM_017003802:NM_001244710 | 3:64:8:0 | 0:0:0:0 | 6:18:97:0 | 0:0:0:0 | 8:68:16:0 | 0:0:0:0 | 36:221:661:0 | 0:0:1:0 | 10:225:83:0 | 0:0:0:0 |
| XM_017003801:NM_002056:XM_017003802:NM_001244710 | 3:64:8:0 | 0:0:0:0 | 6:18:97:0 | 0:0:1:0 | 7:68:17:0 | 0:0:0:0 | 34:221:660:0 | 0:0:2:0 | 10:225:83:0 | 0:0:0:0 |
| NM_002056:XM_017003802:XM_017003801:NM_001244710 | 64:8:1:0 | 0:0:0:0 | 19:93:5:0 | 0:0:0:0 | 68:17:7:0 | 0:0:0:0 | 226:624:29:0 | 0:0:0:0 | 227:82:10:0 | 0:0:0:0 |

| | | | | | | | | | | |
|---|---|---|---|---|---|---|---|---|---|---|
| NM_002056:XM_017003802:XM_017003801:NM_001244710 | 64:8:1:0 | 0:0:0:0 | 19:91:5:0 | 0:0:0:0 | 67:17:7:0 | 1:0:0:0 | 226:607:27:0 | 0:0:0:0 | 226:80:10:0 | 0:0:0:0 |
| NM_002056:XM_017003802:XM_017003801:NM_001244710 | 64:8:1:0 | 0:0:0:0 | 19:89:4:0 | 0:0:0:0 | 68:17:7:0 | 0:0:0:0 | 226:591:25:0 | 0:0:0:0 | 227:80:9:0 | 0:0:0:0 |
| NM_002056:XM_017003802:XM_017003801:NM_001244710 | 64:8:1:0 | 0:0:0:0 | 19:89:4:0 | 0:0:0:0 | 68:17:7:0 | 0:0:0:0 | 226:586:24:0 | 0:2:0:0 | 227:80:9:0 | 0:0:0:0 |
| NM_002056:XM_017003802:XM_017003801:NM_001244710 | 64:8:1:0 | 0:0:0:0 | 19:89:4:0 | 0:0:0:0 | 68:17:6:0 | 0:0:0:0 | 226:581:23:0 | 0:2:0:0 | 227:80:9:0 | 0:0:0:0 |
| NM_002056:XM_017003802:XM_017003801:NM_001244710 | 64:8:1:0 | 0:0:0:0 | 19:88:4:0 | 0:0:0:0 | 68:17:6:0 | 0:0:0:0 | 226:574:24:0 | 0:2:0:0 | 228:80:9:0 | 0:0:0:0 |
| NM_002056:XM_017003802:XM_017003801:NM_001244710 | 64:8:1:0 | 0:0:0:0 | 19:86:4:0 | 0:0:0:0 | 68:17:6:0 | 0:0:0:0 | 224:554:21:0 | 0:1:0:0 | 227:79:8:0 | 1:0:0:0 |
| NM_002056:XM_017003802:XM_017003801:NM_001244710 | 64:8:1:0 | 0:0:0:0 | 19:82:2:0 | 0:1:0:0 | 68:15:3:0 | 0:0:0:0 | 223:484:20:0 | 1:4:0:0 | 228:64:7:0 | 0:0:0:0 |
| NM_002056:XM_017003802:XM_017003801:NM_001244710 | 64:7:1:0 | 0:0:0:0 | 19:74:2:0 | 0:0:0:0 | 68:15:1:0 | 0:0:0:0 | 224:398:12:0 | 0:4:0:0 | 228:58:6:0 | 0:0:0:0 |
| NM_002056:XM_017003802:XM_017003801:NM_001244710 | 64:5:1:0 | 0:0:0:0 | 19:57:0:0 | 0:0:0:0 | 68:11:0:0 | 0:0:0:0 | 226:286:6:0 | 0:0:0:0 | 228:54:3:0 | 0:0:0:0 |
| NM_002056:XM_017003802:XM_017003801:NM_001244710 | 64:4:1:0 | 0:0:0:0 | 19:50:0:0 | 0:0:0:0 | 68:11:0:0 | 0:0:0:0 | 226:252:1:0 | 0:2:0:0 | 227:50:1:0 | 1:0:0:0 |
| NM_002056:XM_017003802:XM_017003801:NM_001244710 | 64:4:1:0 | 0:0:0:0 | 19:45:0:0 | 0:0:0:0 | 68:11:0:0 | 0:0:0:0 | 226:211:1:0 | 0:0:0:0 | 228:45:1:0 | 0:0:0:0 |
| NM_002056:XM_017003802:NM_001244710 | 64:03:00 | 0:00:00 | 17:17:00 | 0:00:00 | 68:07:00 | 0:00:00 | 221:82:0 | 1:00:00 | 226:31:00 | 2:00:00 |
| NR_045631:NM_001002755 | 0:00 | 0:00 | 0:00 | 1:00 | 0:00 | 0:02 | 0:00 | 0:03 | 0:00 | 0:00 |
| XM_024452835:NM_001153:NM_001320702:NM_001320700:XR_001738734:XM_017003944:NM_001320698 | 7:497:2:20:7:1:0 | 0:0:0:0:0:0:0 | 19:1148:7:21:22:1:3 | 0:3:0:0:0:0:0 | 28:1638:5:35:14:1:12 | 0:3:1:0:0:0:0 | 36:2688:4:101:41:3:7 | 0:1:1:0:0:0:0 | 11:1192:4:64:11:2:2 | 0:1:0:0:0:0:0 |
| XM_024452835:NM_001153:NM_001320702:NM_001320700:XR_001738734:XM_017003944:NM_001320698 | 7:497:2:20:6:1:0 | 0:0:0:0:0:0:0 | 19:1150:7:21:21:1:3 | 0:2:0:0:0:0:0 | 28:1641:5:32:13:1:12 | 0:0:0:0:1:0:0 | 36:2688:4:98:39:3:7 | 0:1:0:0:0:0:0 | 11:1189:4:64:11:2:2 | 0:4:0:0:0:0:0 |
| NM_001153:XM_024452835:NM_001320702:XR_001738734:XM_017003944:NM_001320700:NM_001320698 | 495:7:2:6:1:1:0 | 2:0:0:0:0:0:0 | 1152:19:7:13:1:3:3 | 0:0:0:0:0:0:0 | 1641:28:5:11:1:8:12 | 0:0:0:0:0:0:0 | 2688:36:6:33:3:8:7 | 1:0:0:1:0:0:0 | 1193:11:4:11:2:1:2 | 0:0:0:0:0:0:0 |
| XM_024452835:NM_001153:NM_001320702:NM_001320700:XM_017003944:XR_001738734:NM_001320698 | 7:497:2:19:1:5:0 | 0:0:0:0:0:0:0 | 19:1150:7:18:1:10:3 | 0:1:0:0:0:0:0 | 28:1640:5:27:1:10:12 | 0:1:0:0:0:0:0 | 36:2687:6:93:3:28:7 | 0:1:0:0:0:0:0 | 11:1192:4:64:2:10:2 | 0:1:0:0:0:0:0 |
| NM_001153:XM_024452835:NM_001320702:NM_001320700:XM_017003944:XR_001738734:NM_001320698 | 496:7:2:19:1:5:0 | 1:0:0:0:0:0:0 | 1152:19:7:18:1:9:3 | 0:0:0:0:0:0:0 | 1638:28:6:31:1:9:12 | 3:0:0:0:0:0:0 | 2686:36:6:93:3:25:7 | 3:0:0:1:0:0:0 | 1193:11:4:64:2:10:2 | 0:0:0:0:0:0:0 |
| NM_001153:XM_024452835:NM_001320702:NM_001320700:XM_017003944:XR_001738734:NM_001320698 | 494:7:2:19:1:5:0 | 1:0:0:0:0:0:0 | 1149:19:7:18:1:7:3 | 1:0:0:0:0:0:0 | 1635:28:6:33:1:8:12 | 2:0:0:0:0:0:0 | 2677:36:6:95:3:22:7 | 1:0:0:0:0:0:0 | 1193:11:5:64:2:10:2 | 0:0:0:0:0:0:0 |
| NM_001153:XM_024452835:NM_001320702:NM_001320700:XR_001738734:XM_017003944:NM_001320698 | 496:7:2:20:6:1:0 | 1:0:0:0:0:0:0 | 1149:19:9:21:16:1:3 | 0:0:0:0:0:0:0 | 1639:28:11:35:11:1:12 | 1:0:0:0:0:0:0 | 2685:36:11:99:40:3:7 | 1:0:0:0:1:0:0 | 1189:11:5:65:12:2:2 | 1:0:0:0:0:0:0 |
| XM_024452835:NM_001153:NM_001320702:NM_001320700:XR_001738734:XM_017003944:NM_001320698 | 7:496:2:19:8:1:0 | 0:0:0:0:0:0:0 | 18:1139:9:21:28:1:3 | 0:2:0:0:0:0:0 | 28:1634:12:35:21:1:12 | 0:2:0:0:0:0:0 | 36:2668:13:98:76:3:7 | 0:0:0:0:0:0:0 | 11:1187:5:64:18:2:2 | 0:2:0:0:0:0:0 |
| XM_024452835:NM_001153:NM_001320702:NM_001320700:XR_001738734:XM_017003944:NM_001320698 | 7:493:2:19:9:1:0 | 0:0:0:0:0:0:0 | 18:1139:9:21:33:1:3 | 0:0:0:0:0:0:0 | 28:1631:11:35:25:1:12 | 0:3:0:0:0:0:0 | 36:2655:14:98:88:3:7 | 0:3:0:0:1:0:0 | 11:1185:6:63:16:2:2 | 0:1:0:1:0:0:0 |
| XM_024452835:NM_001153:NM_001320702:NM_001320700:XR_001738734:XM_017003944:NM_001320698 | 7:495:2:19:12:1:0 | 0:0:0:0:0:0:0 | 18:1134:12:21:34:1:3 | 0:0:0:0:0:0:0 | 28:1628:12:34:34:1:12 | 0:7:0:0:0:0:0 | 36:2645:19:98:104:3:7 | 0:6:1:0:0:0:0 | 11:1183:6:64:18:2:2 | 0:1:0:0:0:0:0 |
| NM_019896 | 0 | 0 | 3 | 0 | 0 | 0 | 3 | 4 | 1 | 4 |
| XM_017003524 | 0 | 0 | 0 | 0 | 0 | 0 | 0 | 0 | 0 | 0 |
| NM_001307978:NM_001277053:NM_001080824:XM_011532504 | 1:0:0:0 | 0:0:0:0 | 0:1:0:0 | 0:0:0:0 | 0:1:1:0 | 0:0:0:0 | 0:0:0:0 | 0:1:0:0 | 0:0:0:0 | 0:0:0:2 |
| NM_001747:XM_011533123:NM_001256139 | 0:00:00 | 1:01:00 | 0:00:00 | 1:00:00 | 0:00:00 | 1:01:00 | 0:00:00 | 0:00:03 | 1:00:00 | 0:00:00 |
| NM_001747:XM_011533123:NM_001256139 | 1:00:00 | 0:00:00 | 1:00:00 | 0:00:00 | 1:01:00 | 0:00:00 | 0:00:00 | 0:00:03 | 1:00:00 | 0:00:00 |
| NM_016494 | 0 | 0 | 2 | 0 | 3 | 0 | 3 | 3 | 4 | 3 |
| XM_005264110:NM_001100170:NM_001100169 | 0:00:00 | 0:00:00 | 0:00:00 | 0:00:00 | 0:00:00 | 1:01:00 | 0:00:01 | 0:00:00 | 0:00:00 | 0:00:00 |
| NM_001443 | 0 | 2 | 131 | 155 | 180 | 218 | 8 | 1 | 21 | 13 |
| NM_001443 | 2 | 0 | 279 | 1 | 394 | 2 | 9 | 0 | 34 | 0 |
| NM_001443 | 2 | 0 | 276 | 1 | 391 | 3 | 9 | 0 | 34 | 0 |
| NM_001321459:NR_135653:NR_135650:XM_017004313:NR_135651:XM_024452960:NM_001321454:XM_017004311 | 1:0:0:0:0:0:0:0 | 0:0:0:0:0:0:0:0 | 0:2:1:0:0:0:0:0 | 0:0:0:0:0:0:0:0 | 0:1:0:1:0:0:0:0 | 0:0:0:0:0:0:0:0 | 0:4:0:0:0:0:1:0 | 0:2:1:0:1:4:0:0 | 0:0:0:0:0:0:1:0 | 0:1:2:0:1:1:0:0 |
| NM_001321459:NR_135653:NR_135650:NR_135651:XM_024452960:NM_001321454:XM_017004311 | 1:1:0:0:0:0:0 | 0:0:0:0:0:0:0 | 0:3:1:0:0:0:0 | 0:0:0:0:0:0:0 | 0:1:0:0:0:0:0 | 0:0:0:0:0:0:0 | 0:4:0:0:0:1:0 | 0:2:1:1:4:0:0 | 0:0:0:0:0:1:0 | 0:1:2:1:1:0:0 |
| NM_006814 | 0 | 0 | 0 | 0 | 0 | 0 | 0 | 2 | 0 | 1 |
| XM_017027958:NM_016143 | 0:00 | 0:00 | 0:00 | 0:00 | 0:00 | 0:00 | 0:00 | 1:00 | 0:00 | 0:01 |
| NM_004587 | 0 | 0 | 0 | 1 | 0 | 0 | 0 | 1 | 0 | 1 |
| XM_017028128:NM_052865:XM_017028127:XM_006723663:XM_011529394:NM_001310338:XM_005260870 | 1:13:2:0:0:0:0 | 0:0:0:0:0:0:0 | 2:12:1:1:1:0:0 | 0:0:0:0:0:0:0 | 3:14:2:0:0:2:0 | 0:0:0:0:0:0:0 | 7:70:4:1:0:1:0 | 0:0:0:0:0:0:0 | 3:36:0:0:0:2:0 | 1:3:0:0:0:0:0 |
| NM_052865:XM_017028128:XM_017028127:NM_001310338:XM_024452017:XM_006723663:XM_011529394 | 23:8:2:2:1:0:0 | 1:0:0:0:0:0:0 | 21:13:0:1:1:0:1 | 0:1:0:0:0:1:0 | 33:7:2:2:0:0:0 | 0:0:0:0:0:0:0 | 94:48:4:1:5:1:2 | 0:0:0:0:0:0:0 | 45:8:0:2:2:0:0 | 0:0:0:0:0:0:0 |
| XM_017028128:NM_052865:XM_017028127:NM_001310338:XM_024452017:XM_006723663:XM_011529394 | 8:23:2:2:1:0:0 | 0:0:0:0:0:0:0 | 13:21:0:1:1:0:1 | 0:0:0:0:0:1:0 | 7:33:2:2:0:0:0 | 0:0:0:0:0:0:0 | 46:90:4:1:5:1:2 | 1:3:0:0:0:0:0 | 8:45:0:2:2:0:0 | 0:0:0:0:0:0:0 |
| XM_024452017:XM_017028128:NM_052865:XM_017028127:NM_001310338:XM_006723663:XM_011529394 | 0:8:24:2:2:0:0 | 1:0:0:0:0:0:0 | 1:14:21:0:1:1:1 | 0:0:0:0:0:0:0 | 0:7:33:2:2:0:0 | 0:0:0:0:0:0:0 | 5:47:90:4:1:1:2 | 0:0:2:0:0:0:0 | 1:6:44:0:2:0:0 | 1:2:1:0:0:0:0 |
| NM_020536:NR_028402 | 0:00 | 5:01 | 0:00 | 2:03 | 0:00 | 12:03 | 0:00 | 9:18 | 0:00 | 3:01 |
| NM_006363:NM_032985:XM_017027593:NM_032986:NM_001172745 | 12:14:2:0:0 | 0:0:0:0:0 | 2:6:6:1:0 | 0:0:0:0:0 | 14:9:5:0:0 | 0:0:0:0:0 | 19:28:46:6:0 | 0:1:1:1:0 | 19:42:5:2:2 | 0:0:0:0:0 |
| NM_006363:NM_032985:XM_017027593:NM_032986:NM_001172745 | 12:14:3:0:0 | 0:0:0:0:0 | 2:5:11:1:0 | 0:0:0:0:0 | 14:9:4:0:0 | 0:0:0:0:0 | 18:27:99:10:0 | 0:0:0:0:0 | 19:42:8:2:2 | 0:0:0:0:0 |
| NM_006363:NM_032985:XM_017027593:NM_032986:NM_001172745 | 12:14:2:0:0 | 0:0:0:0:0 | 2:5:11:1:0 | 0:0:0:0:0 | 14:9:4:0:0 | 0:0:0:0:0 | 18:27:99:10:0 | 0:0:0:0:0 | 18:41:8:2:2 | 0:1:0:0:0 |
| NM_006363:NM_032985:XM_017027593:NM_032986:NM_001172745 | 1:2:0:0:0 | 11:12:3:0:0 | 2:3:5:1:0 | 0:2:8:0:0 | 6:3:2:0:0 | 8:6:2:0:0 | 18:25:107:11:0 | 0:0:0:0:0 | 19:42:8:2:2 | 0:0:0:0:0 |
| NM_080820:XM_005260872:NM_001318043 | 17:03:00 | 0:00:00 | 29:02:00 | 0:00:00 | 39:03:00 | 0:00:00 | 92:11:01 | 1:00:00 | 82:03:00 | 0:00:00 |
| NM_000099:NM_001288614 | 6:00 | 0:00 | 15:00 | 0:00 | 44:01:00 | 0:00 | 11:00 | 0:00 | 15:00 | 0:00 |
| XM_024452020:NM_001322393:XM_024452021:NM_001322383:NM_001247:NM_001322398:NM_001114089 | 0:0:0:0:0:0:0 | 0:0:0:0:0:0:0 | 0:0:0:0:0:0:0 | 0:0:0:0:0:0:0 | 0:0:0:0:0:0:0 | 0:0:0:0:0:0:0 | 0:0:0:0:0:0:0 | 1:1:0:0:0:0:0 | 0:0:0:0:0:0:0 | 0:0:1:1:0:0:0 |
| NM_002862:XR_002958496 | 7:00 | 0:00 | 1:01 | 0:00 | 9:00 | 0:00 | 0:00 | 23:21 | 0:00 | 1:01 |
| NM_002862:XR_002958496 | 8:00 | 0:00 | 1:00 | 0:00 | 11:00 | 0:00 | 32:01:00 | 0:00 | 4:00 | 0:00 |
| NM_002862:XR_002958496 | 6:00 | 0:00 | 1:00 | 0:00 | 14:00 | 0:00 | 0:00 | 34:01:00 | 0:00 | 6:00 |
| NM_012112:XM_011528700:XM_011528699:XM_011528697 | 11:4:1:0 | 0:0:0:0 | 0:3:0:0 | 0:0:0:0 | 3:5:1:0 | 0:0:0:0 | 33:60:1:11 | 0:0:0:0 | 56:28:2:3 | 0:0:0:0 |
| NM_012112:XM_011528699:XM_011528700:XM_011528697 | 11:1:1:0 | 0:0:0:0 | 0:1:1:0 | 0:0:0:0 | 3:1:2:0 | 0:0:0:0 | 70:50:37:5 | 0:0:0:0 | 56:7:11:3 | 0:0:0:0 |
| NM_012112:XM_011528699:XM_011528700:XM_011528697 | 11:1:2:0 | 0:0:0:0 | 0:1:1:0 | 0:0:0:0 | 3:1:2:0 | 0:0:0:0 | 79:56:36:5 | 0:1:0:0 | 56:7:10:2 | 0:0:0:0 |
| NM_012112:XM_011528699:XM_011528700:XM_011528697 | 11:1:0:0 | 0:0:0:0 | 0:1:1:0 | 0:0:0:0 | 3:1:0:0 | 0:0:0:0 | 83:69:15:4 | 0:0:0:0 | 55:7:8:2 | 0:0:0:0 |
| NM_030815 | 66 | 0 | 39 | 2 | 33 | 1 | 695 | 9 | 125 | 1 |
| NM_014742:XM_017028160:XM_017028161:XM_017028157:XM_017028154:XM_017028156:XM_017028159:XM_017028158 | 15:1:2:1:0:0:0:0 | 0:0:0:0:0:0:0:0 | 3:2:0:0:1:1:0:0 | 0:0:0:0:0:0:0:0 | 12:1:0:1:0:0:1:0 | 1:0:0:0:0:0:0:0 | 103:35:10:13:6:1:1:0 | 1:1:1:0:0:0:0:0 | 33:5:1:2:1:0:0:0 | 0:0:0:0:0:0:0:0 |
| XM_017027706:XM_006723727:XM_006723728:NM_015338:XM_011528648:XM_017027705:XM_006723733:XM_017027704:XM_006723730 | 1:0:0:0:0:0:0:0:0 | 0:0:0:0:0:0:0:0:0 | 0:0:0:0:0:0:0:0:0 | 0:0:0:0:0:0:0:0:0 | 0:0:0:0:0:0:0:0:0 | 1:0:0:0:0:0:0:0:0 | 8:0:0:7:1:1:1:0:0 | 2:1:2:0:0:0:0:0:0 | 0:0:0:0:0:0:0:0:0 | 2:0:0:1:0:0:0:0:0 |

| | | | | | | | | | | |
|---|---|---|---|---|---|---|---|---|---|---|
| XM_017027706:XM_006723727:NM_015338:XM_006723728:XM_006723733:XM_011528648:XM_017027705:XM_017027704:XM_006723730 | 1:0:0:0:0:0:0:0:0 | 0:0:0:0:0:0:0:0:0 | 0:0:0:0:0:0:0:0:0 | 0:0:0:0:0:0:0:0:0 | 0:0:0:0:0:0:0:0:0 | 1:0:0:0:0:0:0:0:0 | 7:0:4:0:0:1:1:0:0 | 3:1:3:2:1:0:0:0:0 | 0:0:0:0:0:0:0:0:0 | 2:0:1:0:0:0:0:0:0 |
| NM_015338:XM_017027706:XM_006723728:XM_011528648:XM_017027705:XM_017027704:XM_006723730 | 0:0:0:0:0:0:0 | 0:0:0:0:0:0:0 | 0:0:0:0:0:0:0 | 0:0:0:0:0:0:0 | 0:0:0:0:0:0:0 | 0:0:0:0:0:0:0 | 0:3:0:0:1:0:0 | 6:1:1:1:0:0:0 | 0:0:0:0:0:0:0 | 0:0:0:0:0:0:0 |
| NM_000178:NM_001322495:NM_001322494 | 105:26:00 | 0:00:00 | 75:41:00 | 0:00:00 | 239:51:00 | 0:00:00 | 571:181:0 | 2:00:00 | 269:42:01 | 0:00:00 |
| NM_000178:NM_001322495:NM_001322494 | 105:26:00 | 0:00:00 | 75:41:00 | 0:00:00 | 239:52:00 | 0:00:00 | 572:182:0 | 0:00:00 | 268:42:01 | 1:00:00 |
| NM_000178:NM_001322495:NM_001322494 | 105:26:00 | 0:00:00 | 75:41:00 | 0:00:00 | 239:52:00 | 0:00:00 | 572:181:0 | 0:01:00 | 267:42:01 | 2:00:00 |
| NM_000178:NM_001322495:NM_001322494 | 105:26:00 | 0:00:00 | 75:41:00 | 0:00:00 | 239:52:00 | 0:00:00 | 572:182:0 | 0:00:00 | 266:42:01 | 3:00:00 |
| NM_000178:NM_001322495:NM_001322494 | 104:22:00 | 0:00:00 | 76:39:00 | 1:00:00 | 238:50:00 | 0:00:00 | 565:172:0 | 0:01:00 | 255:41:01 | 0:00:00 |
| NM_000178:NM_001322495:NM_001322494 | 105:22:00 | 0:00:00 | 78:36:00 | 0:00:00 | 239:45:00 | 0:01:00 | 570:151:1 | 0:00:00 | 269:36:01 | 0:00:00 |
| NM_000178:NM_001322495:NM_001322494 | 105:19:00 | 0:00:00 | 78:35:00 | 0:00:00 | 239:44:00 | 0:00:00 | 570:150:1 | 0:00:00 | 266:33:01 | 2:00:00 |
| NM_000178:NM_001322495:NM_001322494 | 105:18:00 | 0:00:00 | 78:26:00 | 0:00:00 | 240:33:00 | 0:00:00 | 577:123:1 | 1:00:00 | 267:30:01 | 2:00:00 |
| NM_000178:NM_001322495:NM_001322494 | 104:18:00 | 1:00:00 | 78:27:00 | 0:00:00 | 240:33:00 | 0:00:00 | 579:121:1 | 1:00:00 | 269:30:01 | 0:00:00 |
| NM_000178:NM_001322495:NM_001322494 | 105:17:00 | 1:00:00 | 78:27:00 | 5:00:00 | 243:33:00 | 0:00:00 | 580:114:1 | 12:00:00 | 271:31:01 | 0:00:00 |
| NM_000178:NM_001322495:NM_001322494 | 106:17:00 | 0:00:00 | 83:25:00 | 0:00:00 | 243:33:00 | 0:00:00 | 592:105:1 | 2:00:00 | 271:28:01 | 0:00:00 |
| NM_000178:NM_001322495:NM_001322494 | 104:17:00 | 2:00:00 | 83:25:00 | 0:00:00 | 242:33:00 | 1:00:00 | 594:105:1 | 0:00:00 | 269:28:01 | 2:00:00 |
| NM_000178:NM_001322495:NM_001322494 | 105:15:00 | 1:00:00 | 83:21:00 | 0:00:00 | 244:35:00 | 0:00:00 | 595:87:1 | 1:01:00 | 272:26:01 | 1:00:00 |
| NM_000178:NM_001322495:NM_001322494 | 108:19:00 | 0:00:00 | 83:19:00 | 0:00:00 | 243:31:00 | 1:00:00 | 599:84:1 | 2:00:00 | 273:27:01 | 0:00:00 |
| NM_000178:NM_001322495:NM_001322494 | 108:19:00 | 0:00:00 | 86:29:00 | 0:00:00 | 247:41:00 | 0:01:00 | 619:116:1 | 0:01:00 | 276:35:01 | 0:00:00 |
| NM_001322495:NM_000178:NM_001322494 | 0.825 | 1:00:00 | 29:86:0 | 0:00:00 | 42:247:0 | 0:00:00 | 117:619:1 | 0:00:00 | 35:276:1 | 0:00:00 |
| NM_018244:XM_011528881:XM_011528884:XM_011528880:XM_011528877:NM_001184977:XM_011528882:XM_011528879:NM_199487 | 0:0:0:0:0:0:0:0:0 | 9:1:1:0:0:0:0:0:0 | 0:0:0:0:0:0:0:0:0 | 13:0:0:2:1:0:0:0:0 | 0:0:0:0:0:0:0:0:0 | 25:0:0:0:0:0:0:0:0 | 27:0:0:2:1:0:0:0:1 | 131:1:2:12:3:1:3:1:2 | 17:0:0:2:1:0:0:0:0 | 20:0:0:2:1:0:0:0:1 |
| NM_001198863:NR_037188:NM_152926:NM_152925:NM_152928 | 0:0:0:0:0 | 0:0:0:0:0 | 0:0:0:0:0 | 0:0:0:0:0 | 0:0:0:0:0 | 0:0:0:0:0 | 2:1:1:0:0 | 0:0:0:0:0 | 0:0:0:0:0 | 0:0:0:0:0 |
| NM_002951:XM_006723852:NM_001324302:NM_001324305 | 2:0:0:0 | 0:0:0:0 | 2:0:0:0 | 0:0:0:0 | 2:0:0:0 | 0:0:0:0 | 1:0:0:0 | 2:1:0:0 | 2:0:0:0 | 1:0:3:0 |
| NM_022077:NM_001003897:XM_017028007:XM_017028006 | 47:80:3:18 | 0:0:0:0 | 89:69:5:10 | 0:0:0:0 | 141:170:6:26 | 0:0:0:0 | 717:797:60:300 | 0:1:0:1 | 101:120:11:27 | 0:0:0:0 |
| XM_011528865:XR_001754295:NM_002660:XR_001754296:NM_182811 | 0:0:0:0:0 | 0:0:0:0:0 | 1:0:0:0:0 | 0:0:0:0:0 | 0:0:0:0:0 | 0:1:1:0:0 | 1:0:2:0:0 | 0:0:0:0:0 | 0:0:0:1:0 | 0:0:0:0:0 |
| NM_002638 | 0 | 0 | 5 | 0 | 8 | 0 | 0 | 0 | 5 | 0 |
| NM_002638 | 0 | 0 | 5 | 0 | 8 | 0 | 0 | 0 | 5 | 0 |
| NM_001048223:NM_001048221:NM_001197140 | 0:00:00 | 1:00:00 | 0:00:00 | 0:00:00 | 0:00:00 | 1:01:00 | 3:00:00 | 1:00:00 | 1:00:00 | 0:00:00 |
| NM_199129 | 0 | 0 | 0 | 1 | 0 | 0 | 0 | 0 | 0 | 1 |
| NM_003859:NM_001317035:NM_001317034:NM_001317036:XR_002958551:XR_002958550:NR_133648 | 56:9:6:1:1:2:0 | 0:0:0:0:0:0:0 | 70:2:2:0:1:1:1 | 0:0:0:0:0:0:0 | 54:6:1:3:0:3:0 | 0:0:0:0:0:0:0 | 920:140:54:40:4:18:13 | 2:1:0:0:0:0:0 | 145:13:7:12:0:2:2 | 0:0:0:0:0:0:0 |
| NM_003859:NM_001317035:NM_001317034:NM_001317036:XR_002958551:XR_002958550:NR_133648 | 56:9:6:2:1:2:0 | 0:0:0:0:0:0:0 | 70:2:2:0:1:1:1 | 0:0:0:0:0:0:0 | 55:6:1:3:0:3:0 | 0:0:0:0:0:0:0 | 920:142:54:40:2:18:13 | 3:0:0:0:0:0:0 | 143:13:7:12:0:2:2 | 2:0:0:0:0:0:0 |
| NM_003859:NM_001317035:NM_001317034:XR_002958551:XR_002958550:NR_133648 | 56:9:6:1:1:0 | 0:0:0:0:0:0 | 73:2:2:0:1:1 | 0:0:0:0:0:0 | 55:6:1:0:3:0 | 0:0:0:0:0:0 | 929:146:55:0:16:10 | 2:0:0:0:0:0 | 146:14:7:0:2:2 | 0:0:0:0:0:0 |
| NM_003859:NM_001317035:NM_001317034:NM_001317036:XR_002958550:NR_133648:XR_002958551 | 56:9:6:2:1:0:0 | 0:0:0:0:0:0:0 | 72:2:2:0:7:1:0 | 0:0:0:0:0:0:0 | 55:6:1:3:1:0:0 | 0:0:0:0:0:0:0 | 926:144:56:40:22:12:4 | 2:1:0:0:0:0:0 | 146:13:7:12:0:2:0 | 0:1:0:0:0:0:0 |
| NM_003859:NM_001317035:NM_001317034:NM_001317036:XR_002958551:XR_002958550:NR_133648 | 56:9:6:2:1:2:0 | 0:0:0:0:0:0:0 | 72:2:2:0:0:7:1 | 0:0:0:0:0:0:0 | 55:6:1:3:0:4:0 | 0:0:0:0:0:0:0 | 927:143:53:40:7:31:13 | 0:0:1:0:0:0:0 | 146:14:7:11:0:3:2 | 0:0:0:0:0:0:0 |
| NM_003859:NM_001317035:NM_001317034:NM_001317036:XR_002958551:XR_002958550:NR_133648 | 56:9:6:1:1:1:0 | 0:0:0:0:0:0:0 | 72:2:2:0:0:7:1 | 0:0:0:0:0:0:0 | 53:6:1:3:0:4:0 | 0:0:0:0:0:0:0 | 919:141:54:40:13:29:13 | 2:0:0:0:0:0:0 | 145:14:7:11:0:2:2 | 0:0:0:0:0:0:0 |
| NM_199426:NM_018197 | 0:00 | 0:00 | 0:00 | 0:00 | 0:00 | 0:00 | 0:00 | 0:00 | 0:00 | 0:00 |
| NM_002623:XM_017027879 | 77:02:00 | 0:00 | 138:06:00 | 0:00 | 46:01:00 | 0:00 | 3211:38:00 | 5:00 | 243:06:00 | 2:00 |
| NM_002623:XM_017027879 | 78:04:00 | 0:00 | 135:10:00 | 1:00 | 46:01:00 | 0:00 | 3176:52:00 | 2:00 | 246:06:00 | 0:00 |
| NM_002623:XM_017027879 | 78:04:00 | 0:00 | 136:10:00 | 0:00 | 45:01:00 | 0:00 | 3135:51:00 | 5:00 | 245:05:00 | 0:00 |
| NM_016407:XM_017027872 | 0:00 | 1:00 | 0:00 | 0:01 | 0:00 | 2:00 | 0:00 | 1:01 | 0:00 | 5:00 |
| NM_003610:XM_005260582:XM_017028108:XM_011529087:XM_011529089:XM_011529088 | 77:10:4:0:0:0 | 0:0:0:0:0:0 | 35:1:3:1:0:0 | 0:0:0:0:0:0 | 30:0:1:0:0:0 | 0:0:0:0:0:0 | 707:101:42:1:26:1 | 0:0:0:0:1:0 | 174:14:12:0:0:0 | 0:0:0:0:1:0 |
| NM_003610:XM_005260582:XM_017028108:XM_011529087:XM_011529089:XM_011529088 | 77:10:6:0:0:0 | 0:0:0:0:0:0 | 35:1:3:1:2:0 | 0:0:0:0:0:0 | 30:0:1:0:0:0 | 0:0:0:0:0:0 | 705:101:47:1:51:1 | 2:0:0:0:0:0 | 175:14:12:0:5:0 | 0:0:0:0:0:0 |
| NM_003610:XM_005260582:XM_017028108:XM_011529087:XM_011529089:XM_011529088 | 77:10:6:0:0:0 | 0:0:0:0:0:0 | 35:1:3:1:2:0 | 0:0:0:0:0:0 | 30:0:1:0:0:0 | 0:0:0:0:0:0 | 706:101:46:1:53:1 | 1:0:0:0:1:0 | 175:14:11:0:6:0 | 0:0:0:0:0:0 |
| NM_003610:XM_005260582:XM_017028108:XM_011529087:XM_011529089:XM_011529088 | 77:10:6:0:0:0 | 0:0:0:0:0:0 | 35:1:3:1:2:0 | 0:0:0:0:0:0 | 30:0:1:0:0:0 | 0:0:0:0:0:0 | 706:101:45:1:54:1 | 0:0:0:0:0:0 | 174:14:11:0:5:0 | 1:0:0:0:1:0 |
| NM_003610:XM_005260582:XM_017028108:XM_011529089:XM_011529087:XM_011529088 | 77:10:5:1:0:0 | 0:0:0:0:0:0 | 35:1:2:2:1:0 | 0:0:0:0:0:0 | 30:0:2:0:0:0 | 0:0:0:0:0:0 | 705:100:45:75:1:1 | 2:1:0:0:0:0 | 175:14:11:6:0:0 | 0:0:0:0:0:0 |
| NM_003610:XM_005260582:XM_017028108:XM_011529089:XM_011529087:XM_011529088 | 77:10:5:2:0:0 | 0:0:0:0:0:0 | 35:1:2:2:1:0 | 0:0:0:0:0:0 | 30:0:2:0:0:0 | 0:0:0:0:0:0 | 706:101:45:76:1:1 | 1:0:0:0:0:0 | 175:14:11:6:0:0 | 0:0:0:0:0:0 |
| NM_003610:XM_005260582:XM_017028108:XM_011529089:XM_011529088:XM_011529087 | 77:10:3:2:0:0 | 0:0:0:0:0:0 | 35:1:1:2:0:0 | 0:0:0:0:0:0 | 29:0:1:0:0:0 | 0:0:0:0:0:0 | 705:101:34:82:1:0 | 1:0:0:0:0:0 | 175:14:11:7:0:0 | 0:0:0:0:0:0 |
| NM_002591:XM_024451888 | 0:00 | 0:00 | 0:00 | 0:00 | 0:00 | 2:00 | 0:00 | 0:00 | 0:00 | 0:00 |
| NM_001309840:NM_001077488:NM_000516:XM_024451874:NM_080426:NM_080425:XM_024451875:XR_002958471:XM_024451873:NM_001309883:XM_017027812:NR_132273:XM_024451872:NM_016592:NM_001077489:NM_001309842:NM_001309861:NR_132272 | 12:9:87:2:1:2:3:1:0:0:0:0:0:0:0:0:0:0 | 0:0:0:0:0:0:0:0:0:0:0:0:0:0:0:0:0:0 | 14:15:67:0:1:2:5:2:7:0:0:0:0:0:0:0:0:0 | 0:0:0:0:0:0:0:0:0:0:0:0:0:0:0:0:0:0 | 9:23:111:1:1:1:8:2:2:1:2:1:0:0:0:0:0:0 | 0:0:0:0:0:0:0:0:0:0:0:0:0:0:0:0:0:0 | 67:58:492:3:4:12:37:15:48:1:1:0:4:2:1:1:1:0 | 0:0:1:0:0:0:0:0:0:0:0:0:0:0:0:0:0:0 | 20:24:175:1:1:3:11:5:0:0:0:0:1:0:0:1:0 | 0:0:0:0:0:0:0:0:0:0:0:0:0:0:0:0:0:0 |
| NM_000516:NM_080426:NM_001077489:XM_017027817:NM_001309840:NM_001077488:XM_017027818:XM_017027813:XM_024451874:XR_002958471:NM_080425:XM_024451875:XM_017027819:XM_024451873:XM_017027815:XM_017027812:XM_024451872:NM_016592:XM_017027820:XM_017027814:NM_001309861 | 86:266:105:40:12:9:10:17:2:9:2:3:7:0:0:0:0:0:0:0:0 | 1:1:0:0:0:0:0:0:0:0:0:0:0:0:0:0:0:0:0:0:0 | 67:211:73:33:14:15:3:10:0:5:2:8:4:7:0:0:0:0:0:0:0 | 0:0:0:0:0:0:0:0:1:0:0:0:0:0:0:0:0:0:0:0:0 | 111:256:101:42:9:23:3:10:1:5:1:8:7:2:2:2:0:0:0:0:0 | 0:0:0:0:0:0:0:0:0:0:0:0:0:0:0:0:0:0:0:0:0 | 493:1710:922:244:67:58:142:123:3:119:14:46:79:48:10:4:4:3:2:5:1 | 0:7:0:0:0:0:0:0:24:0:0:0:0:0:0:0:0:0:0:0:0 | 175:281:156:32:20:24:5:29:1:29:3:11:11:0:0:1:0:1:0:0:1 | 0:1:0:0:0:0:0:0:0:0:0:0:0:0:0:0:0:0:0:0:0 |
| XM_017027818:XR_002958471:NM_080426:NM_001077489:XM_017027817:NM_001309840:NM_001077488:XM_017027813:NM_000516:XM_024451874:NM_080425:XM_024451875:XM_017027819:XM_024451873:XM_017027815:XM_017027812:XM_024451872:NM_016592:XM_017027820:XM_017027814:NM_001309861 | 9:16:267:105:40:12:9:17:87:2:2:3:7:0:0:0:0:0:0:0:0 | 1:1:1:0:0:0:0:0:0:0:0:0:0:0:0:0:0:0:0:0:0 | 3:10:211:73:33:14:15:10:66:0:2:8:4:7:0:0:0:0:0:0:0 | 0:0:0:0:0:0:0:0:0:0:0:0:0:0:0:0:0:0:0:0:0 | 3:10:257:101:42:9:23:10:110:1:1:8:7:2:2:2:0:0:0:0:0 | 0:0:0:0:0:0:0:0:0:0:0:0:0:0:0:0:0:0:0:0:0 | 142:145:1711:922:244:67:57:123:491:3:14:46:79:47:10:4:4:3:2:5:1 | 0:24:6:0:0:0:0:0:0:0:0:0:0:0:0:0:0:0:0:0:0 | 5:31:284:156:32:20:24:29:173:1:3:11:11:0:0:1:0:1:0:0:1 | 0:1:0:0:0:0:0:0:0:0:0:0:0:0:0:0:0:0:0:0:0 |
| XR_002958471:NM_001077489:XM_017027817:NM_001309840:NM_001077488:NM_080426:XM_017027818:XM_017027813:NM_000516:XM_024451874:NM_080425:XM_024451875:XM_017027819:XM_024451873:XM_017027815:XM_017027812:XM_024451872:NM_016592:XM_017027820:XM_017027814:NM_001309861 | 16:105:40:12:9:268:10:17:87:2:2:3:7:0:0:0:0:0:0:0:0 | 1:0:0:0:0:0:0:0:0:0:0:0:0:0:0:0:0:0:0:0:0 | 10:73:33:14:15:211:3:10:66:0:2:8:4:7:0:0:0:0:0:0:0 | 0:0:0:0:0:0:0:0:0:0:0:0:0:0:0:0:0:0:0:0:0 | 10:101:42:9:23:256:3:10:110:1:1:8:7:2:2:2:0:0:0:0:0 | 0:0:0:0:0:1:0:0:0:0:0:0:0:0:0:0:0:0:0:0:0 | 152:920:244:67:57:1707:142:123:491:3:14:46:79:47:10:4:4:3:2:5:1 | 19:2:0:0:0:10:0:0:0:0:0:0:0:0:0:0:0:0:0:0:0 | 31:155:32:20:24:284:5:29:172:1:3:11:11:0:0:1:0:1:0:0:1 | 1:1:0:0:0:0:0:0:1:0:0:0:0:0:0:0:0:0:0:0:0 |
| NM_080426:NM_001077489:XM_017027817:NM_001309840:NM_001077488:XM_017027818:XM_017027813:NM_000516:XR_002958471:XM_024451874:NM_080425:XM_024451875:XM_017027819:XM_024451873:XM_017027815:XM_017027812:XM_024451872:NM_016592:XM_017027820:XM_017027814:NM_001309861 | 267:105:40:12:9:10:17:87:18:2:2:3:7:0:0:0:0:0:0:0:0 | 1:0:0:0:0:0:0:0:0:0:0:0:0:0:0:0:0:0:0:0:0 | 210:73:33:14:15:3:10:65:11:0:2:8:4:7:0:0:0:0:0:0:0 | 1:0:0:0:0:0:0:0:0:0:0:0:0:0:0:0:0:0:0:0:0 | 257:101:42:9:23:3:10:110:11:1:1:8:7:2:2:2:0:0:0:0:0 | 0:0:0:0:0:0:0:0:0:0:0:0:0:0:0:0:0:0:0:0:0 | 1712:923:243:67:57:142:123:491:158:3:14:46:79:47:10:4:4:3:2:5:1 | 8:0:1:0:0:0:0:0:22:0:0:0:0:0:0:0:0:0:0:0:0 | 285:156:32:20:24:5:29:173:34:1:3:11:11:0:0:1:0:1:0:0:1 | 0:0:0:0:0:0:0:0:0:0:0:0:0:0:0:0:0:0:0:0:0 |

| | | | | | | | | | | |
|---|---|---|---|---|---|---|---|---|---|---|
| NM_080426:NM_001077489:XM_017027817:NM_001309840:NM_001077488:XM_017027818:XM_017027813:NM_000516:XR_002958471:XM_024451874:NM_080425:XM_024451875:XM_017027819:XM_024451873:XM_017027815:XM_017027812:XM_024451872:NM_016592:XM_017027820:XM_017027814:NM_001309861 | 266:105:40:12:9:10:17:87:21:2:2:3:7:0:0:0:0:0:0:0:0 | 3:0:0:0:0:0:0:0:0:0:0:0:0:0:0:0:0:0:0:0:0 | 209:73:33:14:15:3:10:65:16:0:2:8:4:7:0:0:0:0:0:0:0 | 1:0:0:0:0:0:0:0:0:0:0:0:0:0:0:0:0:0:0:0:0 | 255:101:42:9:23:3:10:110:14:1:1:8:7:2:2:2:0:0:0:0:0 | 2:0:0:0:0:0:0:0:0:0:0:0:0:0:0:0:0:0:0:0:0 | 1721:923:244:67:57:142:123:490:199:3:14:46:79:47:10:4:4:3:2:5:1 | 6:0:0:0:0:0:0:1:15:0:0:0:0:0:0:0:0:0:0:0:0 | 283:156:32:20:24:4:29:173:39:1:3:11:11:0:0:1:0:1:0:0:1 | 1:0:0:0:0:1:0:0:0:0:0:0:0:0:0:0:0:0:0:0:0 |
| NM_080426:NM_001077489:XM_017027817:NM_001309840:NM_001077488:XM_017027818:XM_017027813:NM_000516:XR_002958471:XM_024451874:NM_080425:XM_024451875:XM_017027819:XM_024451873:XM_017027815:XM_017027812:XM_024451872:NM_016592:XM_017027820:XM_017027814:NM_001309861 | 268:105:40:12:9:10:17:87:23:2:2:3:7:0:0:0:0:0:0:0:0 | 1:0:0:0:0:0:0:0:0:0:0:0:0:0:0:0:0:0:0:0:0 | 210:73:33:14:15:2:10:65:18:0:2:8:4:7:0:0:0:0:0:0:0 | 0:0:0:0:0:1:0:0:0:0:0:0:0:0:0:0:0:0:0:0:0 | 256:101:42:9:23:3:10:110:17:1:1:8:7:2:2:2:0:0:0:0:0 | 1:0:0:0:0:0:0:0:0:0:0:0:0:0:0:0:0:0:0:0:0 | 1722:923:244:66:57:140:123:491:217:3:14:46:79:47:10:4:4:3:2:5:1 | 5:0:0:1:0:1:0:0:10:0:0:0:0:0:0:0:0:0:0:0:0 | 285:156:32:20:24:5:29:173:42:1:3:11:11:0:0:1:0:1:0:0:1 | 0:0:0:0:0:0:0:0:0:0:0:0:0:0:0:0:0:0:0:0:0 |
| NM_001077489:XM_017027817:NM_001309840:NM_001077488:XM_017027818:NM_080426:XM_017027813:NM_000516:XR_002958471:XM_024451874:NM_080425:XM_024451875:XM_017027819:XM_024451873:XM_017027815:XM_017027812:XM_024451872:NM_016592:XM_017027820:XM_017027814:NM_001309861 | 105:40:12:9:10:272:17:87:28:2:2:3:7:0:0:0:0:0:0:0:0 | 0:0:0:0:0:0:0:0:0:0:0:0:0:0:0:0:0:0:0:0:0 | 73:33:14:15:3:211:10:65:26:0:2:8:4:7:0:0:0:0:0:0:0 | 0:0:0:0:0:0:0:0:0:0:0:0:0:0:0:0:0:0:0:0:0 | 100:42:9:23:3:260:10:109:21:1:1:8:7:2:2:2:0:0:0:0:0 | 1:0:0:0:0:0:0:1:0:0:0:0:0:0:0:0:0:0:0:0:0 | 919:244:67:57:141:1729:122:490:272:3:14:46:79:47:10:4:4:3:2:5:1 | 3:0:0:0:0:1:1:0:1:0:0:0:0:0:0:0:0:0:0:0:0 | 156:32:20:24:5:288:29:173:53:1:3:11:11:0:0:1:0:1:0:0:1 | 0:0:0:0:0:0:0:0:0:0:0:0:0:0:0:0:0:0:0:0:0 |
| NM_001077489:XM_017027817:NM_001309840:NM_001077488:XM_017027818:NM_080426:XM_017027813:NM_000516:XR_002958471:XM_024451874:NM_080425:XM_024451875:XM_017027819:XM_024451873:XM_017027815:XM_017027812:XM_024451872:NM_016592:XM_017027820:XM_017027814:NM_001309861 | 105:40:12:9:10:273:17:87:31:2:2:3:7:0:0:0:0:0:0:0:0 | 0:0:0:0:0:0:0:0:0:0:0:0:0:0:0:0:0:0:0:0:0 | 73:33:14:15:3:210:10:65:31:0:2:8:4:7:0:0:0:0:0:0:0 | 0:0:0:0:0:1:0:0:2:0:0:0:0:0:0:0:0:0:0:0:0 | 100:42:9:23:3:260:10:110:24:1:1:8:7:2:2:2:0:0:0:0:0 | 1:0:0:0:0:0:0:0:0:0:0:0:0:0:0:0:0:0:0:0:0 | 921:244:67:57:141:1732:123:490:307:3:14:46:78:47:10:4:4:3:2:5:1 | 1:0:0:0:0:2:0:0:7:0:0:0:1:0:0:0:0:0:0:0:0 | 156:32:20:24:5:290:29:172:60:1:3:11:11:0:0:1:0:1:0:0:1 | 0:0:0:0:0:1:0:1:0:0:0:0:0:0:0:0:0:0:0:0:0 |
| XR_002958471:NM_001077489:XM_017027817:NM_001309840:NM_001077488:XM_017027818:NM_080426:XM_017027813:NM_000516:XM_024451874:NM_080425:XM_024451875:XM_017027819:XM_024451873:XM_017027815:XM_017027812:XM_024451872:NM_016592:XM_017027820:XM_017027814:NM_001309861 | 32:105:40:12:9:10:274:17:87:2:2:3:7:0:0:0:0:0:0:0:0 | 1:0:0:0:0:0:0:0:0:0:0:0:0:0:0:0:0:0:0:0:0 | 36:73:33:14:15:3:213:10:65:0:2:8:4:7:0:0:0:0:0:0:0 | 1:0:0:0:0:0:0:0:0:0:0:0:0:0:0:0:0:0:0:0:0 | 26:101:42:9:23:3:261:10:110:1:1:8:7:2:2:2:0:0:0:0:0 | 0:0:0:0:0:0:0:0:0:0:0:0:0:0:0:0:0:0:0:0:0 | 335:922:244:67:57:141:1739:123:490:3:14:46:79:47:10:4:4:3:2:5:1 | 4:0:0:0:0:0:1:0:0:0:0:0:0:0:0:0:0:0:0:0:0 | 63:156:32:20:24:5:292:28:172:1:3:10:11:0:0:1:0:1:0:0:1 | 0:0:0:0:0:0:0:1:1:0:0:1:0:0:0:0:0:0:0:0:0 |
| XR_002958471:NM_001077489:XM_017027817:NM_001309840:NM_001077488:XM_017027818:NM_080426:XM_017027813:NM_000516:XM_024451874:NM_080425:XM_024451875:XM_017027819:XM_024451873:XM_017027815:XM_017027812:XM_024451872:NM_016592:XM_017027820:XM_017027814:NM_001309861 | 32:105:40:12:9:10:274:17:87:2:2:3:7:0:0:0:0:0:0:0:0 | 1:0:0:0:0:0:0:0:0:0:0:0:0:0:0:0:0:0:0:0:0 | 35:73:33:14:15:3:213:10:65:0:2:8:4:7:0:0:0:0:0:0:0 | 2:0:0:0:0:0:0:0:0:0:0:0:0:0:0:0:0:0:0:0:0 | 26:101:42:9:23:3:260:10:110:1:1:8:7:2:2:2:0:0:0:0:0 | 0:0:0:0:0:0:0:1:0:0:0:0:0:0:0:0:0:0:0:0:0 | 334:921:244:67:57:141:1738:123:490:3:14:46:79:47:10:4:4:3:2:5:1 | 7:0:0:0:0:0:3:0:0:0:0:0:0:0:0:0:0:0:0:0:0 | 62:156:32:20:24:5:291:29:173:1:3:11:11:0:0:1:0:1:0:0:1 | 1:0:0:0:0:0:2:0:0:0:0:0:0:0:0:0:0:0:0:0:0 |
| NM_000516:NM_001077489:XM_017027817:NM_001309840:NM_001077488:XM_017027818:NM_080426:XM_017027813:XR_002958471:XM_024451874:NM_080425:XM_024451875:XM_017027819:XM_024451873:XM_017027815:XM_017027812:XM_024451872:NM_016592:XM_017027820:XM_017027814:NM_001309861 | 86:105:40:12:9:10:274:17:36:2:2:3:7:0:0:0:0:0:0:0:0 | 1:0:0:0:0:0:0:0:0:0:0:0:0:0:0:0:0:0:0:0:0 | 65:72:33:14:15:3:213:10:41:0:2:8:4:7:0:0:0:0:0:0:0 | 0:1:0:0:0:0:1:0:1:0:0:0:0:0:0:0:0:0:0:0:0 | 110:101:42:9:23:3:262:10:28:1:1:8:7:2:2:2:0:0:0:0:0 | 0:0:0:0:0:0:0:0:0:0:0:0:0:0:0:0:0:0:0:0:0 | 490:921:244:67:56:141:1749:123:360:3:14:46:79:47:10:4:4:3:2:5:1 | 0:1:0:0:0:0:0:0:4:0:0:0:0:0:0:0:0:0:0:0:0 | 173:156:32:20:24:5:293:29:67:1:3:11:11:0:0:1:0:1:0:0:1 | 0:0:0:0:0:0:0:0:0:0:0:0:0:0:0:0:0:0:0:0:0 |
| NM_001077489:XM_017027817:NM_001309840:NM_001077488:XM_017027818:NM_080426:XM_017027813:NM_000516:XR_002958471:XM_024451874:NM_080425:XM_024451875:XM_017027819:XM_024451873:XM_017027815:XM_017027812:XM_024451872:NM_016592:XM_017027820:XM_017027814:NM_001309861 | 104:40:12:9:10:276:17:87:38:2:2:3:7:0:0:0:0:0:0:0:0 | 1:0:0:0:0:0:0:0:0:0:0:0:0:0:0:0:0:0:0:0:0 | 73:33:14:15:3:215:10:65:42:0:2:8:4:7:0:0:0:0:0:0:0 | 0:0:0:0:0:0:0:0:2:0:0:0:0:0:0:0:0:0:0:0:0 | 101:42:9:23:3:262:10:110:30:1:1:8:7:2:2:2:0:0:0:0:0 | 0:0:0:0:0:0:0:0:0:0:0:0:0:0:0:0:0:0:0:0:0 | 921:244:67:56:140:1752:123:489:387:3:14:46:79:47:10:4:4:3:2:5:1 | 1:0:0:0:1:2:0:0:3:0:0:0:0:0:0:0:0:0:0:0:0 | 156:32:20:24:5:293:29:172:73:1:3:11:11:0:0:1:0:1:0:0:1 | 0:0:0:0:0:0:0:0:0:0:0:0:0:0:0:0:0:0:0:0:0 |
| NM_080426:NM_001077489:XM_017027817:NM_001309840:NM_001077488:XM_017027818:XM_017027813:NM_000516:XR_002958471:XM_024451874:NM_080425:XM_024451875:XM_017027819:XM_024451873:XM_017027815:XM_017027812:XM_024451872:NM_016592:XM_017027820:XM_017027814:NM_001309861 | 277:105:40:12:9:10:17:87:44:2:2:3:7:0:0:0:0:0:0:0:0 | 1:0:0:0:0:0:0:0:0:0:0:0:0:0:0:0:0:0:0:0:0 | 216:73:33:14:15:3:10:65:52:0:2:8:4:7:0:0:0:0:0:0:0 | 0:0:0:0:0:0:0:0:1:0:0:0:0:0:0:0:0:0:0:0:0 | 263:101:42:9:23:3:10:109:33:1:1:8:7:2:2:2:0:0:0:0:0 | 0:0:0:0:0:0:0:1:0:0:0:0:0:0:0:0:0:0:0:0:0 | 1756:921:244:67:56:141:123:489:429:3:14:46:79:47:10:4:4:3:2:5:1 | 4:1:0:0:0:0:0:0:2:0:0:0:0:0:0:0:0:0:0:0:0 | 294:156:32:20:24:5:29:171:80:1:3:11:11:0:0:1:0:1:0:0:1 | 0:0:0:0:0:0:0:1:1:0:0:0:0:0:0:0:0:0:0:0:0 |
| NM_080426:NM_001077489:XM_017027817:NM_001309840:NM_001077488:XM_017027818:XM_017027813:NM_000516:XR_002958471:XM_024451874:NM_080425:XM_024451875:XM_017027819:XM_024451873:XM_017027815:XM_017027812:XM_024451872:NM_016592:XM_017027820:XM_017027814:NM_001309861 | 277:105:40:12:9:10:17:87:44:2:2:3:7:0:0:0:0:0:0:0:0 | 1:0:0:0:0:0:0:0:0:0:0:0:0:0:0:0:0:0:0:0:0 | 216:73:33:14:15:3:10:65:53:0:2:8:4:6:0:0:0:0:0:0:0 | 0:0:0:0:0:0:0:0:0:0:0:0:0:0:0:0:0:0:0:0:0 | 263:101:42:9:23:3:10:109:34:1:1:8:7:2:2:2:0:0:0:0:0 | 0:0:0:0:0:0:0:0:0:0:0:0:0:0:0:0:0:0:0:0:0 | 1758:922:244:66:56:141:123:489:441:3:14:46:78:47:10:4:4:3:2:5:1 | 2:0:0:1:0:0:0:0:0:0:0:0:0:0:0:0:0:0:0:0:0 | 294:156:32:20:24:5:29:172:83:1:3:11:10:0:0:1:0:1:0:0:1 | 0:0:0:0:0:0:0:0:0:0:0:0:0:0:0:0:0:0:0:0:0 |
| NM_001077489:XM_017027817:NM_001309840:NM_001077488:XM_017027818:NM_080426:XM_017027813:NM_000516:XR_002958471:XM_024451874:NM_080425:XM_024451875:XM_017027819:XM_024451873:XM_017027815:XM_017027812:XM_024451872:NM_016592:XM_017027820:XM_017027814:NM_001309861 | 105:40:12:9:10:276:17:87:44:2:2:3:7:0:0:0:0:0:0:0:0 | 0:0:0:0:0:0:0:0:0:0:0:0:0:0:0:0:0:0:0:0:0 | 73:33:14:15:3:216:10:65:53:0:2:8:3:6:0:0:0:0:0:0:0 | 0:0:0:0:0:0:0:0:0:0:0:0:0:0:0:0:0:0:0:0:0 | 101:42:9:23:3:263:10:110:34:1:1:8:7:2:2:2:0:0:0:0:0 | 0:0:0:0:0:0:0:0:0:0:0:0:0:0:0:0:0:0:0:0:0 | 922:244:66:56:141:1759:123:488:441:3:14:46:78:47:10:4:4:3:2:5:1 | 0:0:1:0:0:1:0:1:0:0:0:0:0:0:0:0:0:0:0:0:0 | 156:32:20:24:5:294:29:172:83:1:3:11:10:0:0:1:0:1:0:0:1 | 0:0:0:0:0:0:0:0:0:0:0:0:0:0:0:0:0:0:0:0:0 |
| NM_001077489:XM_017027817:NM_001309840:NM_001077488:NM_080426:XM_017027818:XM_017027813:NM_000516:XR_002958471:XM_024451874:NM_080425:XM_024451875:XM_017027819:XM_024451873:XM_017027815:XM_017027812:XM_024451872:NM_016592:XM_017027820:XM_017027814:NM_001309861 | 105:40:12:9:277:10:16:87:56:2:2:3:7:0:0:0:0:0:0:0:0 | 0:0:0:0:0:0:0:0:0:0:0:0:0:0:0:0:0:0:0:0:0 | 73:33:14:15:219:3:10:65:61:0:2:8:3:6:0:0:0:0:0:0:0 | 0:0:0:0:0:0:0:0:0:0:0:0:0:0:0:0:0:0:0:0:0 | 101:41:9:23:264:3:10:110:37:1:1:8:7:2:2:2:0:0:0:0:0 | 0:1:0:0:0:0:0:0:0:0:0:0:0:0:0:0:0:0:0:0:0 | 919:243:67:56:1777:141:121:488:539:3:14:46:77:47:10:4:4:3:2:5:1 | 1:0:0:0:5:0:0:0:3:0:0:0:0:0:0:0:0:0:0:0:0 | 156:32:20:24:300:5:29:172:92:1:3:11:10:0:0:1:0:1:0:0:1 | 0:0:0:0:0:0:0:0:0:0:0:0:0:0:0:0:0:0:0:0:0 |
| NM_000516:XR_002958471:NM_001077489:XM_017027817:NM_001309840:NM_001077488:NM_080426:XM_017027818:XM_017027813:XM_024451874:NM_080425:XM_024451875:XM_017027819:XM_024451873:XM_017027815:XM_017027812:XM_024451872:NM_016592:XM_017027820:XM_017027814:NM_001309861 | 86:61:105:40:12:9:277:10:16:2:2:3:7:0:0:0:0:0:0:0:0 | 1:2:0:0:0:0:0:0:0:0:0:0:0:0:0:0:0:0:0:0:0 | 65:74:73:33:14:15:219:3:10:0:2:8:3:6:0:0:0:0:0:0:0 | 0:1:0:0:0:0:0:0:0:0:0:0:0:0:0:0:0:0:0:0:0 | 110:41:101:42:9:23:265:3:10:1:1:8:7:2:2:2:0:0:0:0:0 | 0:0:0:0:0:0:0:0:0:0:0:0:0:0:0:0:0:0:0:0:0 | 487:599:918:243:67:56:1789:141:118:3:14:46:77:47:10:4:4:3:2:5:1 | 0:20:2:0:0:0:8:0:1:0:0:0:0:0:0:0:0:0:0:0:0 | 171:103:156:32:20:24:300:5:29:1:3:11:10:0:0:1:0:1:0:0:1 | 1:2:0:0:0:0:1:0:0:0:0:0:0:0:0:0:0:0:0:0:0 |
| XR_002958471:NM_001077489:XM_017027817:NM_001309840:NM_001077488:NM_080426:XM_017027818:XM_017027813:NM_000516:XM_024451874:NM_080425:XM_024451875:XM_017027819:XM_024451873:XM_017027815:XM_017027812:XM_024451872:NM_016592:XM_017027820:XM_017027814:NM_001309861 | 66:105:40:12:9:278:10:16:87:2:2:3:7:0:0:0:0:0:0:0:0 | 1:0:0:0:0:0:0:0:0:0:0:0:0:0:0:0:0:0:0:0:0 | 79:73:33:14:15:219:3:10:65:0:2:8:3:6:0:0:0:0:0:0:0 | 1:0:0:0:0:0:0:0:0:0:0:0:0:0:0:0:0:0:0:0:0 | 42:101:42:9:23:265:3:10:110:1:1:8:7:2:2:2:0:0:0:0:0 | 0:0:0:0:0:0:0:0:0:0:0:0:0:0:0:0:0:0:0:0:0 | 622:919:243:67:56:1794:141:119:487:3:14:46:76:47:10:4:4:3:2:5:1 | 20:0:0:0:0:6:0:0:0:0:0:0:0:0:0:0:0:0:0:0:0 | 107:155:32:20:24:299:5:29:172:1:3:11:10:0:0:1:0:1:0:0:1 | 0:1:0:0:0:1:0:0:0:0:0:0:0:0:0:0:0:0:0:0:0 |
| XR_002958471:NM_001077489:XM_017027817:NM_001309840:NM_001077488:NM_080426:XM_017027818:XM_017027813:NM_000516:XM_024451874:NM_080425:XM_024451875:XM_017027819:XM_024451873:XM_017027815:XM_017027812:XM_024451872:NM_016592:XM_017027820:XM_017027814:NM_001309861 | 66:105:40:12:9:278:10:15:87:2:2:3:7:0:0:0:0:0:0:0:0 | 1:0:0:0:0:0:0:0:0:0:0:0:0:0:0:0:0:0:0:0:0 | 81:73:33:14:15:219:3:10:65:0:2:8:3:6:0:0:0:0:0:0:0 | 1:0:0:0:0:0:0:0:0:0:0:0:0:0:0:0:0:0:0:0:0 | 42:101:42:9:23:266:3:10:110:1:1:8:7:2:2:2:0:0:0:0:0 | 0:0:0:0:0:0:0:0:0:0:0:0:0:0:0:0:0:0:0:0:0 | 629:918:243:67:56:1796:139:118:487:3:14:46:76:47:10:4:4:3:2:5:1 | 26:0:0:0:0:6:0:1:0:0:0:0:0:0:0:0:0:0:0:0:0 | 108:156:32:20:24:300:5:29:172:1:3:10:10:0:0:1:0:1:0:0:1 | 0:0:0:0:0:1:0:0:0:0:0:0:0:0:0:0:0:0:0:0:0 |
| XR_002958471:NM_001077489:XM_017027817:NM_001309840:NM_001077488:NM_080426:XM_017027818:XM_017027813:NM_000516:XM_024451874:NM_080425:XM_024451875:XM_017027819:XM_024451873:XM_017027815:XM_017027812:XM_024451872:NM_016592:XM_017027820:XM_017027814:NM_001309861 | 66:105:40:12:9:278:10:15:87:2:2:3:7:0:0:0:0:0:0:0:0 | 1:0:0:0:0:0:0:0:0:0:0:0:0:0:0:0:0:0:0:0:0 | 81:73:33:14:15:219:3:10:65:0:2:8:3:6:0:0:0:0:0:0:0 | 1:0:0:0:0:0:0:0:0:0:0:0:0:0:0:0:0:0:0:0:0 | 42:101:42:9:23:266:3:10:110:1:1:8:7:2:2:2:0:0:0:0:0 | 0:0:0:0:0:0:0:0:0:0:0:0:0:0:0:0:0:0:0:0:0 | 634:919:243:67:56:1798:141:118:487:3:14:46:76:47:10:4:4:3:2:5:1 | 21:0:0:0:0:5:0:0:0:0:0:0:0:0:0:0:0:0:0:0:0 | 107:156:32:20:24:300:5:29:172:1:3:11:10:0:0:1:0:1:0:0:1 | 1:0:0:0:0:1:0:0:0:0:0:0:0:0:0:0:0:0:0:0:0 |
| XR_002958471:NM_001077489:XM_017027817:NM_001309840:NM_001077488:NM_080426:XM_017027818:XM_017027813:NM_000516:XM_024451874:NM_080425:XM_024451875:XM_017027819:XM_024451873:XM_017027815:XM_017027812:XM_024451872:NM_016592:XM_017027820:XM_017027814:NM_001309861 | 73:105:40:12:9:281:10:15:87:2:2:3:7:0:0:0:0:0:0:0:0 | 1:0:0:0:0:0:0:0:0:0:0:0:0:0:0:0:0:0:0:0:0 | 87:72:33:14:15:228:3:10:65:0:2:8:3:6:0:0:0:0:0:0:0 | 3:0:0:0:0:1:0:0:0:0:0:0:0:0:0:0:0:0:0:0:0 | 46:101:42:9:22:268:3:10:110:1:1:8:7:2:2:2:0:0:0:0:0 | 0:0:0:0:1:1:0:0:0:0:0:0:0:0:0:0:0:0:0:0:0 | 729:919:243:67:56:1812:141:118:487:3:14:45:76:47:10:4:4:3:2:5:1 | 19:0:0:0:0:8:0:0:0:0:0:1:0:0:0:0:0:0:0:0:0 | 119:156:32:20:24:307:5:29:171:1:3:11:10:0:0:1:0:1:0:0:1 | 1:0:0:0:0:3:0:0:1:0:0:0:0:0:0:0:0:0:0:0:0 |
| NM_080426:XR_002958471:NM_001077489:XM_017027817:NM_001309840:NM_001077488:XM_017027818:XM_017027813:NM_000516:XM_024451874:NM_080425:XM_024451875:XM_017027819:XM_024451873:XM_017027815:XM_017027812:XM_024451872:NM_016592:XM_017027820:XM_017027814:NM_001309861 | 281:78:105:40:12:9:10:14:87:2:2:3:7:0:0:0:0:0:0:0:0 | 2:1:0:0:0:0:0:0:0:0:0:0:0:0:0:0:0:0:0:0:0 | 230:92:72:33:14:15:3:10:65:0:2:8:3:6:0:0:0:0:0:0:0 | 0:1:0:0:0:0:0:0:0:0:0:0:0:0:0:0:0:0:0:0:0 | 268:48:100:42:9:22:3:10:110:1:1:8:7:2:2:2:0:0:0:0:0 | 2:1:0:0:0:0:0:0:0:0:0:0:0:0:0:0:0:0:0:0:0 | 1818:782:919:242:65:56:140:117:487:3:14:46:76:47:10:4:4:3:2:5:1 | 5:19:0:0:0:0:1:0:0:0:0:0:0:0:0:0:0:0:0:0:0 | 307:127:156:32:20:24:5:29:171:1:3:11:10:0:0:1:0:1:0:0:1 | 3:1:0:0:0:0:0:0:1:0:0:0:0:0:0:0:0:0:0:0:0 |
| NM_080426:XR_002958471:NM_001077489:XM_017027817:NM_001309840:NM_001077488:XM_017027818:XM_017027813:NM_000516:XM_024451874:XM_024451875:XM_017027819:NM_080425:XM_024451873:XM_017027815:XM_017027812:XM_024451872:NM_016592:XM_017027820:XM_017027814:NM_001309861 | 282:80:105:40:12:9:10:14:87:2:3:7:1:0:0:0:0:0:0:0:0 | 1:1:0:0:0:0:0:0:0:0:0:0:0:0:0:0:0:0:0:0:0 | 234:98:71:33:14:15:3:10:65:0:8:3:2:6:0:0:0:0:0:0:0 | 1:1:0:0:0:0:0:0:0:0:0:0:0:0:0:0:0:0:0:0:0 | 272:53:100:42:9:22:3:10:110:1:8:7:1:2:2:2:0:0:0:0:0 | 0:1:0:0:0:0:0:0:0:0:0:0:0:0:0:0:0:0:0:0:0 | 1821:803:918:242:64:56:141:117:487:3:46:76:13:47:10:4:4:3:2:5:1 | 8:28:0:0:0:0:0:0:0:0:0:0:1:0:0:0:0:0:0:0:0 | 313:135:156:32:20:24:5:29:172:1:11:10:3:0:0:1:0:1:0:0:1 | 2:1:0:0:0:0:0:0:0:0:0:0:0:0:0:0:0:0:0:0:0 |

| | | | | | | | | | | |
|---|---|---|---|---|---|---|---|---|---|---|
| NM_080426:XR_002958471:NM_001077489:XM_017027817:NM_001309840:NM_001077488:XM_017027818:XM_017027813:NM_000516:XM_024451874:NM_080425:XM_024451875:XM_017027819:XM_024451873:XM_017027815:XM_017027812:XM_024451872:NM_016592:XM_017027820:XM_017027814:NM_001309861 | 282:81:105:40:12:9:10:14:87:2:2:3:7:0:0:0:0:0:0:0:0 | 1:1:0:0:0:0:0:0:0:0:0:0:0:0:0:0:0:0:0 | 235:97:71:33:14:15:3:10:65:0:2:8:3:6:0:0:0:0:0:0:0 | 0:4:0:0:0:0:0:0:0:0:0:0:0:0:0:0:0:0:0 | 271:54:100:42:9:22:3:10:110:1:1:8:7:2:2:2:0:0:0:0:0 | 1:2:0:0:0:0:0:0:0:0:0:0:0:0:0:0:0:0:0 | 1824:816:918:242:64:56:141:116:487:3:14:46:76:47:10:4:4:3:2:5:1 | 6:25:0:0:0:0:0:1:0:0:0:0:0:0:0:0:0:0:0 | 314:136:156:32:20:24:5:29:172:1:3:11:10:0:0:1:0:1:0:0:1 | 2:2:0:0:0:0:0:0:0:0:0:0:0:0:0:0:0:0:0 |
| NM_001077489:XM_017027817:NM_001309840:NM_001077488:NM_080426:XM_017027818:XM_017027813:NM_000516:XR_002958471:XM_024451874:NM_080425:XM_024451875:XM_017027819:XM_024451873:XM_017027815:XM_017027812:XM_024451872:NM_016592:XM_017027820:XM_017027814:NM_001309861 | 105:40:12:9:285:10:14:87:86:2:2:3:7:0:0:0:0:0:0:0:0 | 0:0:0:0:0:0:0:0:0:0:0:0:0:0:0:0:0:0:0 | 70:33:14:15:236:3:10:65:109:0:1:8:2:6:0:0:0:0:0:0:0 | 1:0:0:0:0:0:0:0:2:0:1:0:0:0:0:0:0:0:0:0 | 100:42:9:22:276:3:10:110:57:1:0:8:7:2:2:2:0:0:0:0:0 | 0:0:0:0:0:0:0:0:2:0:0:0:0:0:0:0:0:0:0 | 916:242:64:56:1839:141:117:486:891:3:14:46:75:47:10:4:4:3:2:5:1 | 1:0:0:0:6:0:0:0:18:0:0:0:0:0:0:0:0:0:0:0 | 155:32:20:24:320:5:29:172:145:1:3:11:10:0:0:1:0:1:0:0:1 | 0:0:0:0:2:0:0:0:0:0:0:0:0:0:0:0:0:0:0 |
| NM_001077489:XM_017027817:NM_001309840:NM_001077488:NM_080426:XM_017027818:XM_017027813:NM_000516:XR_002958471:XM_024451874:NM_080425:XM_024451875:XM_017027819:XM_024451873:XM_017027815:XM_017027812:XM_024451872:NM_016592:XM_017027820:XM_017027814:NM_001309861 | 105:40:12:9:285:10:14:87:87:2:2:3:7:0:0:0:0:0:0:0:0 | 0:0:0:0:0:0:0:0:0:0:0:0:0:0:0:0:0:0:0 | 71:33:14:15:236:3:10:65:111:0:2:8:2:6:0:0:0:0:0:0:0 | 0:0:0:0:0:0:0:1:0:0:0:0:0:0:0:0:0:0:0 | 100:42:9:22:276:3:10:110:58:1:0:7:7:2:2:2:0:0:0:0:0 | 0:0:0:0:0:0:0:0:3:0:0:1:0:0:0:0:0:0:0:0 | 917:241:64:56:1847:140:117:486:903:3:13:46:75:47:10:4:4:3:2:5:1 | 0:1:0:0:1:1:0:0:16:0:1:0:0:0:0:0:0:0:0:0 | 155:32:20:24:321:5:29:172:145:1:3:11:10:0:0:1:0:1:0:0:1 | 0:0:0:0:1:0:0:0:0:0:0:0:0:0:0:0:0:0:0 |
| XR_002958471:NM_001077489:XM_017027817:NM_001309840:NM_001077488:NM_080426:XM_017027818:XM_017027813:NM_000516:XM_024451874:NM_080425:XM_024451875:XM_017027819:XM_024451873:XM_017027815:XM_017027812:XM_024451872:NM_016592:XM_017027820:XM_017027814:NM_001309861 | 86:105:40:12:9:286:10:14:87:2:2:2:7:0:0:0:0:0:0:0:0 | 2:0:0:0:0:0:0:0:0:0:0:0:0:0:0:0:0:0:0 | 116:71:33:14:15:238:3:10:65:0:2:8:2:6:0:0:0:0:0:0:0 | 1:0:0:0:0:0:0:0:0:0:0:0:0:0:0:0:0:0:0 | 62:100:42:9:22:278:3:10:108:1:0:8:7:2:2:2:0:0:0:0:0 | 0:0:0:0:0:0:0:0:1:0:0:0:0:0:0:0:0:0:0:0 | 977:916:242:64:56:1855:141:117:486:3:14:46:75:47:10:4:4:3:2:5:1 | 2:0:0:0:0:3:0:0:0:0:0:0:0:0:0:0:0:0:0 | 144:156:32:20:24:325:5:27:172:1:3:11:10:0:0:1:0:1:0:0:1 | 0:0:0:0:0:0:0:1:0:0:0:0:0:0:0:0:0:0:0 |
| XR_002958471:NM_001077489:XM_017027817:NM_001309840:NM_001077488:NM_080426:XM_017027818:XM_017027813:NM_000516:XM_024451874:NM_080425:XM_024451875:XM_017027819:XM_024451873:XM_017027815:XM_017027812:XM_024451872:NM_016592:XM_017027820:XM_017027814:NM_001309861 | 91:105:40:12:9:286:10:14:87:2:2:2:7:0:0:0:0:0:0:0:0 | 1:0:0:0:0:0:0:0:0:0:0:0:0:0:0:0:0:0:0 | 121:71:33:14:15:239:3:10:65:0:2:8:2:6:0:0:0:0:0:0:0 | 0:0:0:0:0:0:0:0:0:0:0:0:0:0:0:0:0:0:0 | 63:100:42:9:22:278:3:10:108:1:0:8:7:2:2:2:0:0:0:0:0 | 1:0:0:0:0:0:0:0:0:0:0:0:0:0:0:0:0:0:0 | 996:915:242:64:56:1861:141:117:486:3:14:46:75:47:10:4:4:3:2:5:1 | 7:1:0:1:0:2:0:0:0:0:0:0:0:0:0:0:0:0:0 | 145:156:32:20:24:326:5:28:172:1:3:11:10:0:0:1:0:1:0:0:1 | 0:0:0:0:0:0:0:0:0:0:0:0:0:0:0:0:0:0:0 |
| NM_001077489:XM_017027817:NM_001309840:NM_001077488:NM_080426:XM_017027818:XM_017027813:NM_000516:XR_002958471:XM_024451874:NM_080425:XM_024451875:XM_017027819:XM_024451873:XM_017027815:XM_017027812:XM_024451872:NM_016592:XM_017027820:XM_017027814:NM_001309861 | 105:40:12:9:286:10:14:87:94:2:2:2:7:0:0:0:0:0:0:0:0 | 0:0:0:0:0:0:0:0:0:0:0:0:0:0:0:0:0:0:0 | 70:33:14:15:239:3:10:65:140:0:2:8:2:6:0:0:0:0:0:0:0 | 0:0:0:0:2:0:0:0:0:0:0:0:0:0:0:0:0:0:0 | 98:42:9:22:281:3:10:107:71:1:0:7:7:2:2:2:0:0:0:0:0 | 1:0:0:0:0:0:0:1:0:0:0:0:0:0:0:0:0:0:0 | 913:242:65:56:1877:141:116:486:1081:3:14:45:75:46:10:4:4:3:2:5:1 | 1:0:0:0:1:0:1:0:7:0:0:0:0:1:0:0:0:0:0:0:0 | 155:32:20:24:327:5:27:170:156:1:3:11:10:0:0:1:0:1:0:0:1 | 0:0:0:0:0:0:0:1:0:0:0:0:0:0:0:0:0:0:0 |
| NM_080426:XR_002958471:NM_001077489:XM_017027817:NM_001309840:NM_001077488:XM_017027818:XM_017027813:NM_000516:XM_024451874:NM_080425:XM_024451875:XM_017027819:XM_024451873:XM_017027815:XM_017027812:XM_024451872:NM_016592:XM_017027820:XM_017027814:NM_001309861 | 285:93:105:40:12:9:10:14:87:2:2:2:7:0:0:0:0:0:0:0:0 | 1:1:0:0:0:0:0:0:0:0:0:0:0:0:0:0:0:0:0 | 241:138:70:33:14:15:3:10:65:0:2:8:2:6:0:0:0:0:0:0:0 | 0:1:0:0:0:0:0:0:0:0:0:0:0:0:0:0:0:0:0 | 281:70:100:42:9:22:3:10:107:1:0:7:7:2:2:2:0:0:0:0:0 | 0:0:0:0:0:0:0:0:1:0:0:0:0:0:0:0:0:0:0:0 | 1876:1085:913:242:65:56:141:116:485:3:14:45:75:47:10:4:4:3:2:5:1 | 2:3:2:0:0:0:0:1:1:0:0:0:0:0:0:0:0:0:0:0 | 327:157:155:32:20:24:5:27:171:1:3:11:10:0:0:1:0:1:0:0:1 | 0:0:0:0:0:0:0:0:0:0:0:0:0:0:0:0:0:0:0 |
| NM_001077489:XM_017027817:NM_001309840:NM_001077488:NM_080426:XM_017027818:XM_017027813:NM_000516:XR_002958471:XM_024451874:NM_080425:XM_024451875:XM_017027819:XM_024451873:XM_017027815:XM_017027812:XM_024451872:NM_016592:XM_017027820:XM_017027814:NM_001309861 | 105:40:12:9:286:10:14:87:96:2:2:2:7:0:0:0:0:0:0:0:0 | 0:0:0:0:0:0:0:0:0:0:0:0:0:0:0:0:0:0:0 | 70:33:14:15:243:3:10:65:145:0:2:8:2:6:0:0:0:0:0:0:0 | 0:0:0:0:0:0:0:1:0:0:0:0:0:0:0:0:0:0:0 | 99:42:9:22:281:3:10:107:72:1:0:7:7:2:2:2:0:0:0:0:0 | 1:0:0:0:0:0:0:0:0:0:0:0:0:0:0:0:0:0:0 | 912:242:65:56:1879:141:116:485:1105:3:14:45:75:47:10:4:4:3:2:5:1 | 2:0:0:0:2:0:1:0:5:0:0:0:0:0:0:0:0:0:0:0:0 | 155:32:20:24:327:5:26:171:157:1:3:11:10:0:0:1:0:1:0:0:1 | 0:0:0:0:0:0:0:0:0:0:0:0:0:0:0:0:0:0:0 |
| NM_001077489:XM_017027817:NM_001309840:NM_001077488:NM_080426:XM_017027818:XM_017027813:NM_000516:XR_002958471:XM_024451874:NM_080425:XM_024451875:XM_017027819:XM_024451873:XM_017027815:XM_017027812:XM_024451872:NM_016592:XM_017027820:XM_017027814:NM_001309861 | 105:40:12:9:286:10:14:87:96:2:2:2:7:0:0:0:0:0:0:0:0 | 0:0:0:0:0:0:0:0:0:0:0:0:0:0:0:0:0:0:0 | 69:33:14:15:243:3:9:65:144:0:2:8:2:6:0:0:0:0:0:0:0 | 1:0:0:0:0:0:1:0:2:0:0:0:0:0:0:0:0:0:0:0 | 100:42:9:22:280:3:10:107:72:1:0:7:7:2:2:2:0:0:0:0:0 | 0:0:0:0:1:0:0:0:0:0:0:0:0:0:0:0:0:0:0 | 910:241:65:56:1879:140:116:484:1104:3:14:45:74:47:10:4:4:3:2:5:1 | 3:1:0:0:0:0:1:1:8:0:0:0:1:0:0:0:0:0:0:0:0 | 155:32:20:24:326:5:26:171:157:1:3:11:10:0:0:1:0:1:0:0:1 | 0:0:0:0:1:0:0:0:0:0:0:0:0:0:0:0:0:0:0 |
| NM_000516:NM_001077489:XM_017027817:NM_001309840:NM_001077488:NM_080426:XM_017027818:XM_017027813:XR_002958471:XM_024451874:NM_080425:XM_024451875:XM_017027819:XM_024451873:XM_017027815:XM_017027812:XM_024451872:NM_016592:XM_017027820:XM_017027814:NM_001309861 | 86:104:40:12:9:286:10:14:97:2:2:2:7:0:0:0:0:0:0:0:0 | 1:1:0:0:0:0:0:0:0:0:0:0:0:0:0:0:0:0:0 | 65:69:33:14:15:243:3:9:146:0:2:8:2:6:0:0:0:0:0:0:0 | 0:1:0:0:0:0:0:1:1:0:0:0:0:0:0:0:0:0:0:0 | 107:100:42:9:22:282:3:9:72:1:0:7:7:2:2:2:0:0:0:0:0 | 0:0:0:0:0:0:0:1:0:0:0:0:0:0:0:0:0:0:0 | 485:912:242:65:56:1876:141:116:1123:3:14:44:75:47:10:4:4:3:2:5:1 | 0:1:0:0:0:3:0:1:7:0:0:0:0:0:0:0:0:0:0:0 | 171:155:32:20:24:327:5:26:157:1:3:11:10:0:0:1:0:1:0:0:1 | 0:0:0:0:0:0:0:0:0:0:0:0:0:0:0:0:0:0:0 |
| XR_002958471:NM_001077489:XM_017027817:NM_001309840:NM_001077488:NM_080426:XM_017027818:XM_017027813:NM_000516:XM_024451874:NM_080425:XM_024451875:XM_017027819:XM_024451873:XM_017027815:XM_017027812:XM_024451872:NM_016592:XM_017027820:XM_017027814:NM_001309861 | 96:105:40:12:9:287:10:13:87:2:2:2:7:0:0:0:0:0:0:0:0 | 2:0:0:0:0:0:0:0:0:0:0:0:0:0:0:0:0:0:0 | 152:70:33:14:15:243:3:9:65:0:2:8:2:6:0:0:0:0:0:0:0 | 0:0:0:0:0:2:0:0:0:0:0:0:0:0:0:0:0:0:0 | 75:100:41:9:22:283:3:10:106:1:0:7:7:2:2:2:0:0:0:0:0 | 1:0:1:0:0:0:0:0:1:0:0:0:0:0:0:0:0:0:0:0 | 1142:913:242:65:56:1885:141:116:485:3:14:44:75:47:10:4:4:3:2:5:1 | 8:0:0:0:0:2:0:0:0:0:0:0:0:0:0:0:0:0:0 | 163:155:32:20:24:329:5:26:170:1:3:11:10:0:0:1:0:1:0:0:1 | 0:0:0:0:0:0:0:1:0:0:0:0:0:0:0:0:0:0:0 |
| XR_002958471:NM_001077489:XM_017027817:NM_001309840:NM_001077488:NM_080426:XM_017027818:XM_017027813:NM_000516:XM_024451874:NM_080425:XM_024451875:XM_017027819:XM_024451873:XM_017027815:XM_017027812:XM_024451872:NM_016592:XM_017027820:XM_017027814:NM_001309861 | 98:105:40:12:9:287:10:13:87:2:2:2:7:0:0:0:0:0:0:0:0 | 1:0:0:0:0:0:0:0:0:0:0:0:0:0:0:0:0:0:0 | 156:70:33:14:15:245:3:9:65:0:2:8:2:6:0:0:0:0:0:0:0 | 1:0:0:0:0:1:0:0:0:0:0:0:0:0:0:0:0:0:0 | 76:100:42:9:22:284:3:10:107:1:0:7:7:2:2:2:0:0:0:0:0 | 0:0:0:0:0:0:0:0:0:0:0:0:0:0:0:0:0:0:0 | 1155:913:240:65:55:1890:141:116:485:3:14:44:75:47:10:4:4:3:2:5:1 | 7:0:1:0:0:3:0:0:0:0:0:0:0:0:0:0:0:0:0 | 161:155:32:20:24:330:5:25:171:1:3:11:10:0:0:1:0:1:0:0:1 | 0:0:0:0:0:0:0:0:0:0:0:0:0:0:0:0:0:0:0 |
| NM_001077489:XM_017027817:NM_001309840:NM_001077488:NM_080426:XM_017027818:XM_017027813:NM_000516:XR_002958471:XM_024451874:NM_080425:XM_024451875:XM_017027819:XM_024451873:XM_017027815:XM_017027812:XM_024451872:NM_016592:XM_017027820:XM_017027814:NM_001309861 | 105:40:12:9:286:10:13:87:100:2:2:2:7:0:0:0:0:0:0:0:0 | 0:0:0:0:0:0:0:0:0:0:0:0:0:0:0:0:0:0:0 | 70:33:14:15:246:3:9:65:156:0:2:8:2:6:0:0:0:0:0:0:0 | 0:0:0:0:0:0:0:1:0:0:0:0:0:0:0:0:0:0:0 | 100:42:9:22:284:3:10:107:76:1:0:7:7:2:2:2:0:0:0:0:0 | 0:0:0:0:0:0:0:0:0:0:0:0:0:0:0:0:0:0:0 | 913:240:65:54:1891:141:116:484:1165:3:14:44:75:47:10:4:4:3:2:5:1 | 0:1:0:1:3:0:0:0:5:0:0:0:0:0:0:0:0:0:0:0:0 | 155:32:20:24:332:5:25:171:160:1:3:11:10:0:0:1:0:1:0:0:1 | 0:0:0:0:0:0:0:0:0:0:0:0:0:0:0:0:0:0:0 |
| NM_080426:NM_001077489:XM_017027817:NM_001309840:NM_001077488:XM_017027818:XM_017027813:NM_000516:XR_002958471:XM_024451874:NM_080425:XM_024451875:XM_017027819:XM_024451873:XM_017027815:XM_017027812:XM_024451872:NM_016592:XM_017027820:XM_017027814:NM_001309861 | 287:105:40:12:9:10:13:87:101:2:2:2:7:0:0:0:0:0:0:0:0 | 1:0:0:0:0:0:0:0:0:0:0:0:0:0:0:0:0:0:0 | 251:70:33:14:15:3:9:65:159:0:2:8:2:6:0:0:0:0:0:0:0 | 0:0:0:0:0:0:0:0:0:0:0:0:0:0:0:0:0:0:0 | 285:100:42:9:22:3:10:107:78:1:0:7:7:2:2:2:0:0:0:0:0 | 0:0:0:0:0:0:0:0:0:0:0:0:0:0:0:0:0:0:0 | 1898:913:241:65:55:141:116:484:1195:3:14:44:75:47:10:4:4:3:2:5:1 | 1:0:0:0:0:0:0:0:2:0:0:0:0:0:0:0:0:0:0:0:0 | 333:155:32:20:24:5:25:171:160:1:3:11:10:0:0:1:0:1:0:0:1 | 0:0:0:0:0:0:0:0:0:0:0:0:0:0:0:0:0:0:0 |
| NM_080426:NM_001077489:XM_017027817:NM_001309840:NM_001077488:XM_017027818:XM_017027813:NM_000516:XR_002958471:XM_024451874:NM_080425:XM_024451875:XM_017027819:XM_024451873:XM_017027815:XM_017027812:XM_024451872:NM_016592:XM_017027820:XM_017027814:NM_001309861 | 290:105:40:12:9:10:13:87:104:2:2:2:7:0:0:0:0:0:0:0:0 | 1:0:0:0:0:0:0:0:0:0:0:0:0:0:0:0:0:0:0 | 252:70:33:14:15:3:9:65:161:0:2:8:2:6:0:0:0:0:0:0:0 | 0:0:0:0:0:0:0:1:0:0:0:0:0:0:0:0:0:0:0 | 283:100:42:9:23:3:10:107:78:1:0:7:7:2:2:2:0:0:0:0:0 | 2:0:0:0:0:0:0:0:0:0:0:0:0:0:0:0:0:0:0 | 1909:913:241:65:55:141:116:483:1221:3:14:44:75:47:10:4:4:3:2:5:1 | 2:0:0:0:0:0:0:0:4:0:0:0:0:0:0:0:0:0:0:0:0 | 333:155:32:20:24:5:25:171:156:1:3:11:10:0:0:1:0:1:0:0:1 | 0:0:0:0:0:0:0:0:0:0:0:0:0:0:0:0:0:0:0 |
| NM_000516:NM_080426:NM_001077489:XM_017027817:NM_001309840:NM_001077488:XM_017027818:XM_017027813:XR_002958471:XM_024451874:NM_080425:XM_024451875:XM_017027819:XM_024451873:XM_017027815:XM_017027812:XM_024451872:NM_016592:XM_017027820:XM_017027814:NM_001309861 | 86:291:105:40:12:9:10:13:105:2:2:2:7:0:0:0:0:0:0:0:0 | 1:1:0:0:0:0:0:0:0:0:0:0:0:0:0:0:0:0:0 | 65:251:70:33:14:15:3:9:168:0:2:8:2:6:0:0:0:0:0:0:0 | 0:1:0:0:0:0:0:0:0:0:0:0:0:0:0:0:0:0:0 | 107:285:100:42:9:23:3:10:78:1:0:7:7:2:2:2:0:0:0:0:0 | 0:0:0:0:0:0:0:0:0:0:0:0:0:0:0:0:0:0:0 | 484:1913:913:241:65:55:141:116:1236:3:14:44:75:47:10:4:4:3:2:5:1 | 0:4:0:0:0:0:0:0:4:0:0:0:0:0:0:0:0:0:0:0:0 | 171:334:155:32:20:24:5:25:157:1:3:11:10:0:0:1:0:1:0:0:1 | 0:0:0:0:0:0:0:0:0:0:0:0:0:0:0:0:0:0:0 |
| NM_000516:NM_001077489:XM_017027817:NM_001309840:NM_001077488:NM_080426:XM_017027818:XM_017027813:XR_002958471:XM_024451874:NM_080425:XM_024451875:XM_017027819:XM_024451873:XM_017027815:XM_017027812:XM_024451872:NM_016592:XM_017027820:XM_017027814:NM_001309861 | 86:105:40:12:9:292:10:13:104:2:2:2:7:0:0:0:0:0:0:0:0 | 1:0:0:0:0:0:0:0:0:0:0:0:0:0:0:0:0:0:0 | 65:70:33:14:15:254:3:8:167:0:2:8:2:6:0:0:0:0:0:0:0 | 0:0:0:0:0:0:0:2:0:0:0:0:0:0:0:0:0:0:0 | 107:100:42:9:23:284:3:10:78:1:0:7:7:2:2:2:0:0:0:0:0 | 0:0:0:0:0:1:0:0:0:0:0:0:0:0:0:0:0:0:0 | 484:912:240:65:55:1918:141:116:1249:3:14:44:75:47:10:4:4:3:2:5:1 | 0:1:1:0:0:2:0:0:2:0:0:0:0:0:0:0:0:0:0:0:0 | 171:155:32:20:24:334:5:25:157:1:3:11:10:0:0:1:0:1:0:0:1 | 0:0:0:0:0:0:0:0:0:0:0:0:0:0:0:0:0:0:0 |
| NM_001077489:XM_017027817:NM_001309840:NM_001077488:NM_080426:XM_017027818:XM_017027813:NM_000516:XR_002958471:XM_024451874:NM_080425:XM_024451875:XM_017027819:XM_024451873:XM_017027815:XM_017027812:XM_024451872:NM_016592:XM_017027820:XM_017027814:NM_001309861 | 105:40:12:9:295:10:13:87:105:2:2:2:7:0:0:0:0:0:0:0:0 | 0:0:0:0:0:0:0:0:0:0:0:0:0:0:0:0:0:0:0 | 70:33:14:15:257:3:8:65:172:0:2:8:2:6:0:0:0:0:0:0:0 | 0:0:0:0:1:0:0:0:1:0:0:0:0:0:0:0:0:0:0:0 | 100:42:9:23:285:3:10:107:78:1:0:7:7:2:2:2:0:0:0:0:0 | 0:0:0:0:1:0:0:0:0:0:0:0:0:0:0:0:0:0:0 | 913:241:65:55:1920:141:116:484:1270:3:14:44:75:47:10:4:4:3:2:5:1 | 1:0:0:0:2:0:0:0:5:0:0:0:0:0:0:0:0:0:0:0:0 | 155:32:20:24:337:5:25:170:159:1:3:11:10:0:0:1:0:1:0:0:1 | 0:0:0:0:1:0:0:1:1:0:0:0:0:0:0:0:0:0:0 |
| NM_001077489:XM_017027817:NM_001309840:NM_001077488:NM_080426:XM_017027818:XM_017027813:NM_000516:XR_002958471:XM_024451874:NM_080425:XM_024451875:XM_017027819:XM_024451873:XM_017027815:XM_017027812:XM_024451872:NM_016592:XM_017027820:XM_017027814:NM_001309861 | 105:40:12:9:295:10:13:87:104:2:2:2:7:0:0:0:0:0:0:0:0 | 0:0:0:0:0:0:0:0:0:0:0:0:0:0:0:0:0:0:0 | 70:33:14:15:258:3:8:65:173:0:2:8:2:6:0:0:0:0:0:0:0 | 0:0:0:0:0:0:0:1:0:0:0:0:0:0:0:0:0:0:0 | 100:42:9:22:285:3:10:106:77:1:0:7:7:2:2:2:0:0:0:0:0 | 0:0:0:1:1:0:0:1:1:0:0:0:0:0:0:0:0:0:0:0 | 914:241:65:55:1920:141:116:484:1271:3:14:44:75:47:10:4:4:3:2:5:1 | 0:0:0:0:3:0:0:0:4:0:0:0:0:0:0:0:0:0:0:0:0 | 155:32:20:24:337:5:25:171:158:1:3:11:10:0:0:1:0:1:0:0:1 | 0:0:0:0:1:0:0:0:1:0:0:0:0:0:0:0:0:0:0 |

| | | | | | | | | | | |
|---|---|---|---|---|---|---|---|---|---|---|
| NM_001077489:XM_017027817:NM_001309840:NM_001077488:NM_080426:XM_017027818:XM_017027813:NM_000516:XR_002958471:XM_024451874:NM_080425:XM_024451875:XM_017027819:XM_024451873:XM_017027815:XM_017027812:XM_024451872:NM_016592:XM_017027820:XM_017027814:NM_001309861 | 105:40:12:9:295:10:13:87:104:2:2:2:7:0:0:0:0:0:0:0:0:0 | 0:0:0:0:0:0:0:0:0:0:0:0:0:0:0:0:0:0:0:0:0 | 70:33:14:15:258:3:8:65:172:0:2:8:2:6:0:0:0:0:0:0:0 | 0:0:0:0:0:0:0:0:1:0:0:0:0:0:0:0:0:0:0:0:0 | 100:42:9:23:286:3:10:107:78:1:0:7:7:2:2:2:0:0:0:0:0 | 0:0:0:0:0:0:0:0:0:0:0:0:0:0:0:0:0:0:0:0:0 | 914:240:65:55:1922:141:116:484:1274:3:14:44:74:47:10:4:4:3:2:5:1 | 0:1:0:0:2:0:0:0:5:0:0:0:0:0:0:0:0:0:0:0:0 | 155:32:20:24:338:5:25:170:164:1:3:11:10:0:0:1:0:1:0:0:1 | 0:0:0:0:0:0:0:0:1:0:0:0:0:0:0:0:0:0:0:0:0 |
| NM_001077489:XM_017027817:NM_001309840:NM_001077488:NM_080426:XM_017027818:XM_017027813:NM_000516:XR_002958471:XM_024451874:NM_080425:XM_024451875:XM_017027819:XM_024451873:XM_017027815:XM_017027812:XM_024451872:NM_016592:XM_017027820:XM_017027814:NM_001309861 | 105:40:12:9:296:10:12:87:103:2:2:2:7:0:0:0:0:0:0:0:0:0 | 0:0:0:0:0:0:0:0:0:0:0:0:0:0:0:0:0:0:0:0:0 | 70:33:14:15:258:3:8:65:183:0:2:8:2:6:0:0:0:0:0:0:0 | 0:0:0:0:0:0:0:0:0:0:0:0:0:0:0:0:0:0:0:0:0 | 100:42:9:23:285:3:9:107:76:1:0:7:7:2:2:2:0:0:0:0:0 | 0:0:0:0:1:0:0:0:0:0:0:0:0:0:0:0:0:0:0:0:0 | 914:242:65:55:1925:141:116:483:1294:3:14:44:74:47:10:4:4:3:2:5:1 | 0:0:0:0:1:0:0:0:3:0:0:0:0:0:0:0:0:0:0:0:0 | 155:32:20:24:338:5:25:171:163:1:3:11:10:0:0:1:0:1:0:0:1 | 0:0:0:0:0:0:0:0:0:0:0:0:0:0:0:0:0:0:0:0:0 |
| NM_001819 | 0 | 0 | 1 | 0 | 1 | 1 | 0 | 0 | 0 | 0 |
| NM_001819 | 0 | 0 | 1 | 0 | 0 | 1 | 0 | 0 | 0 | 0 |
| NM_001819 | 0 | 0 | 0 | 1 | 0 | 2 | 0 | 0 | 0 | 0 |
| NM_001819 | 0 | 0 | 1 | 0 | 2 | 1 | 0 | 0 | 0 | 0 |
| NM_001819 | 0 | 0 | 1 | 0 | 2 | 1 | 0 | 0 | 0 | 0 |
| NM_019095:XR_001754298 | 0:00 | 1:00 | 0:00 | 0:00 | 0:00 | 0:00 | 0:00 | 0:00 | 0:00 | 1:00 |
| NM_019095:XM_005260738 | 0:00 | 1:00 | 0:00 | 0:00 | 0:00 | 0:00 | 0:00 | 0:00 | 0:00 | 1:00 |
| NM_007002:NM_175573:NM_001281438:XM_024451818 | 0:0:0:0 | 0:0:0:0 | 0:0:0:0 | 0:0:0:0 | 0:0:0:0 | 0:0:0:0 | 1:0:0:0 | 2:1:0:0 | 1:0:0:0 | 1:0:0:0 |
| NM_001853 | 0 | 0 | 0 | 0 | 0 | 0 | 0 | 2 | 0 | 0 |
| NM_001853 | 0 | 0 | 0 | 0 | 0 | 0 | 0 | 2 | 0 | 0 |
| NM_181485 | 0 | 0 | 0 | 0 | 0 | 3 | 0 | 1 | 0 | 1 |
| NM_020062 | 0 | 0 | 0 | 3 | 0 | 1 | 0 | 2 | 0 | 0 |
| NM_017446 | 0 | 0 | 0 | 0 | 0 | 0 | 0 | 2 | 0 | 0 |
| NM_000628 | 0 | 0 | 0 | 1 | 2 | 0 | 0 | 1 | 0 | 0 |
| XM_005260964:NM_000629 | 1:07 | 0:00 | 0:03 | 0:00 | 0:03 | 0:00 | 7:11 | 5:13 | 0:00 | 0:03 |
| NR_103797:NM_001291411:NM_001291412:NM_032195 | 340:1:0:0 | 2:0:0:0 | 1377:0:0:0 | 8:0:0:0 | 319:0:0:0 | 1:0:0:0 | 3876:0:3:1 | 5:0:0:0 | 580:0:0:0 | 0:0:0:0 |
| NR_103797:NM_001291411:NM_001291412:NM_032195 | 342:1:0:0 | 0:0:0:0 | 1399:0:0:0 | 2:0:0:0 | 319:0:0:0 | 1:0:0:0 | 3873:0:3:1 | 8:0:0:0 | 580:0:0:0 | 1:0:0:0 |
| NR_103797:NM_001291411:NM_001291412:NM_032195 | 342:1:0:0 | 0:0:0:0 | 1400:0:0:0 | 1:0:0:0 | 320:0:0:0 | 0:0:0:0 | 3874:0:3:1 | 9:0:0:0 | 581:0:0:0 | 0:0:0:0 |
| NR_103797:NM_001291411:NM_001291412:NM_032195 | 340:1:0:0 | 2:0:0:0 | 1408:0:0:0 | 2:0:0:0 | 320:0:0:0 | 1:0:0:0 | 3896:0:3:1 | 0:0:0:0 | 580:0:0:0 | 2:0:0:0 |
| NR_103797:NM_001291411 | 338:01:00 | 0:00 | 1372:00:00 | 2:00 | 310:00:00 | 2:00 | 3672:00:00 | 18:00 | 573:00:00 | 0:00 |
| NR_103797:NM_001291411 | 258:01:00 | 16:00 | 1198:00:00 | 159:00:00 | 248:00:00 | 18:00 | 3074:00:00 | 362:00:00 | 417:00:00 | 28:00:00 |
| NR_103797:NM_001291411 | 320:01:00 | 3:00 | 1335:00:00 | 2:00 | 305:00:00 | 1:00 | 3318:00:00 | 21:00 | 559:00:00 | 1:00 |
| NR_103797:NM_001291411 | 0:01 | 13:00 | 5:00 | 909:00:00 | 0:00 | 25:00:00 | 17:00 | 1320:00:00 | 1:00 | 31:00:00 |
| NR_103797:NM_001291411 | 0:01 | 5:00 | 4:00 | 916:00:00 | 0:00 | 8:00 | 18:00 | 1324:00:00 | 1:00 | 3:00 |
| NR_103797 | 0 | 2 | 0 | 3 | 0 | 0 | 0 | 6 | 0 | 5 |
| NR_103797:NM_138927 | 0:00 | 2:01 | 0:00 | 5:00 | 0:00 | 0:00 | 6:00 | 8:00 | 3:00 | 2:00 |
| NM_005239 | 0 | 0 | 1 | 0 | 0 | 0 | 2 | 2 | 0 | 0 |
| NM_005239 | 0 | 0 | 1 | 0 | 0 | 0 | 2 | 2 | 0 | 0 |
| NM_001007246 | 0 | 0 | 0 | 0 | 0 | 2 | 0 | 0 | 0 | 0 |
| NM_004627:NR_146615:NR_146616:NR_146614:NM_001146218 | 94:2:2:0:0 | 0:0:0:0:0 | 40:1:2:2:0 | 0:0:0:0:0 | 54:0:2:1:0 | 0:0:0:0:0 | 239:3:6:3:1 | 2:0:0:0:0 | 68:1:2:0:0 | 0:0:0:0:0 |
| XM_024452056 | 0 | 0 | 0 | 0 | 0 | 0 | 3 | 2 | 0 | 1 |
| NM_005656:NM_001135099 | 21:05 | 28:05:00 | 55:04:00 | 52:02:00 | 256:17:00 | 294:19:00 | 70:07:00 | 71:08:00 | 44:05:00 | 33:08:00 |
| NM_005656:NM_001135099 | 54:10:00 | 0:00 | 106:06:00 | 0:00 | 542:38:00 | 2:00 | 151:19:00 | 0:00 | 76:13:00 | 0:00 |
| NM_005656:NM_001135099 | 1:06 | 1:05 | 1:03 | 0:02 | 4:18 | 2:18 | 4:17 | 0:00 | 2:13 | 0:00 |
| NM_004339:NR_104597 | 0:00 | 0:00 | 0:00 | 0:00 | 1:00 | 2:00 | 0:00 | 0:00 | 0:00 | 0:01 |
| NM_001848 | 0 | 0 | 0 | 1 | 0 | 0 | 0 | 0 | 1 | 0 |
| NM_033070:XR_002958690 | 2:00 | 0:00 | 2:00 | 0:00 | 0:00 | 0:00 | 0:00 | 0:00 | 0:00 | 0:00 |
| NM_022719:XM_017028958:XM_006724330:XR_002958714 | 1:0:0:0 | 1:0:0:0 | 1:1:0:0 | 0:0:0:0 | 3:2:1:0 | 2:0:0:0 | 6:1:0:0 | 0:0:0:0 | 3:2:0:0 | 0:0:0:0 |
| NM_000754:NM_001135162:NM_001135161 | 0:00:00 | 99:01:00 | 0:00:00 | 11:00:00 | 0:00:00 | 90:01:00 | 0:00:00 | 136:01:00 | 0:00:00 | 54:01:00 |
| NM_000754:NM_001135162:NM_001135161 | 99:01:00 | 0:00:00 | 10:00:00 | 0:00:00 | 89:01:00 | 0:00:00 | 130:01:00 | 1:00:00 | 54:01:00 | 0:00:00 |
| NM_030573:NM_001008695 | 4:03 | 6:01 | 4:02 | 5:02 | 10:08 | 8:05 | 0:00 | 37:09:00 | 0:00 | 4:04 |
| NM_001178126 | 4 | 0 | 4 | 2 | 10 | 5 | 0 | 0 | 1 | 0 |
| NM_001178126 | 4 | 0 | 7 | 1 | 18 | 2 | 0 | 0 | 1 | 0 |
| NM_001178126:NM_001256296 | 1:00 | 4:00 | 4:00 | 5:00 | 11:00 | 11:01 | 0:00 | 0:00 | 3:01 | 0:00 |
| NM_004327:NM_021574 | 0:00 | 3:00 | 0:00 | 2:00 | 0:00 | 2:00 | 0:00 | 16:01 | 0:00 | 5:00 |
| NM_005940:NR_133013 | 0:00 | 0:00 | 0:00 | 0:00 | 0:00 | 0:00 | 0:00 | 0:00 | 0:00 | 0:00 |
| NM_001007468:NM_003073:NM_001317946 | 70:35:03 | 0:00:00 | 52:31:01 | 0:00:00 | 129:73:6 | 0:00:00 | 222:122:7 | 0:00:00 | 54:50:00 | 0:00:00 |
| NM_001007468:NM_003073:NM_001317946 | 70:37:02 | 0:00:00 | 53:56:00 | 0:00:00 | 130:98:7 | 0:00:00 | 225:150:4 | 0:00:00 | 54:53:00 | 0:00:00 |
| NM_007194:XR_937805:NM_145862:XM_017028560:NM_001005735:XM_011529839:NM_001349956:XM_024452148:NM_001257387:XM_024452149:XM_017028561:XM_011529842 | 12:2:0:0:0:0:0:0:0:0:0:0 | 0:0:0:0:0:0:0:0:0:0:0:0 | 6:0:1:1:0:0:0:0:0:0:0:0 | 0:0:0:0:0:0:0:0:0:0:0:0 | 13:1:0:0:1:0:0:0:0:0:0:0 | 0:0:0:0:0:0:0:0:0:0:0:0 | 50:0:2:0:0:4:1:5:0:0:0:0 | 0:0:0:0:0:0:0:0:0:0:0:0 | 14:3:0:0:2:0:0:0:0:0:0:0 | 0:0:0:0:0:0:0:0:0:0:0:0 |
| NM_005243:NM_001163285:XM_017028648:XM_017028645:XM_017028647:XM_017028660:XM_017028665:NM_001163287:XR_002958676:XM_011529998:XM_017028652:NM_013986:XM_011529997:XM_024452180:XM_017028661:XM_011530000:XM_017028644:XM_017028666:XM_011530002:XM_017028664:XM_024452181:XM_017028651:XM_017028656:XM_017028653:XM_017028657:XM_017028659:XM_005261389:XM_017028646:XM_011529996:XM_011529995:XM_017028662:XM_017028650:XM_017028649 | 48:64:1:5:3:3:6:2:8:3:1:1:0:0:0:0:0:0:0:0:0:0:0:0:0:0:0:0:0:0:0:0:0 | 1:0:0:0:0:0:0:0:0:0:0:0:0:0:0:0:0:0:0:0:0:0:0:0:0:0:0:0:0:0:0:0:0 | 19:26:2:1:1:0:5:0:10:2:1:0:3:1:2:1:1:1:0:0:0:0:0:0:0:0:0:0:0:0:0:0:0 | 0:0:0:0:0:0:0:0:0:0:0:0:0:0:0:0:0:0:0:0:0:0:0:0:0:0:0:0:0:0:0:0:0 | 67:55:2:3:1:1:6:2:5:8:2:0:4:1:1:4:4:1:1:1:2:1:0:0:0:0:0:0:0:0:0:0:0 | 0:0:0:0:0:0:0:0:0:0:0:0:0:0:0:0:0:0:0:0:0:0:0:0:0:0:0:0:0:0:0:0:0 | 161:215:13:3:3:13:14:6:31:17:1:0:27:0:12:3:6:7:0:2:2:3:9:3:3:1:2:1:1:1:0:0:0 | 0:2:0:0:0:0:0:0:0:0:0:0:0:0:0:0:0:0:0:0:0:0:0:0:0:0:0:0:0:0:0:0:0 | 46:88:1:3:1:0:4:2:4:3:1:0:2:0:2:1:0:3:0:0:1:2:0:1:0:0:0:0:0:1:0:0:0 | 1:0:0:0:0:0:0:0:0:0:0:0:0:0:0:0:0:0:0:0:0:0:0:0:0:0:0:0:0:0:0:0:0 |
| NM_001163285:NM_005243:XM_017028648:XM_017028645:XM_017028647:XM_017028660:XM_017028665:NM_001163287:XR_002958676:XM_011529998:XM_017028652:NM_013986:XM_011529997:XM_024452180:XM_017028661:XM_011530000:XM_017028644:XM_017028666:XM_011530002:XM_017028664:XM_024452181:XM_017028651:XM_017028656:XM_017028653:XM_017028657:XM_005261389:XM_017028646:XM_011529996:XM_011529995:XM_017028662:XM_017028650:XM_017028649 | 64:49:1:5:3:3:6:2:5:3:1:1:0:0:0:0:0:0:0:0:0:0:0:0:0:0:0:0:0:0:0:0 | 0:0:0:0:0:0:0:0:0:0:0:0:0:0:0:0:0:0:0:0:0:0:0:0:0:0:0:0:0:0:0:0 | 26:19:2:1:1:0:5:0:9:2:1:0:4:1:3:1:1:1:0:0:0:0:0:0:0:0:0:0:0:0:0:0 | 0:0:0:0:0:0:0:0:0:0:0:0:0:0:0:0:0:0:0:0:0:0:0:0:0:0:0:0:0:0:0:0 | 55:67:2:3:1:1:6:2:5:8:2:0:4:1:1:4:4:1:1:1:2:1:0:0:0:0:0:0:0:0:0:0 | 0:0:0:0:0:0:0:0:0:0:0:0:0:0:0:0:0:0:0:0:0:0:0:0:0:0:0:0:0:0:0:0 | 217:161:13:3:3:13:14:6:28:17:1:0:32:0:18:3:6:7:0:2:2:3:9:3:4:2:1:1:1:0:0:0 | 0:0:0:0:0:0:0:0:0:0:0:0:0:0:0:0:0:0:0:0:0:0:0:0:0:0:0:0:0:0:0:0 | 88:47:1:3:1:0:4:2:3:3:1:0:2:0:2:1:0:3:0:0:1:2:0:1:0:0:0:0:1:0:0:0 | 0:0:0:0:0:0:0:0:0:0:0:0:0:0:0:0:0:0:0:0:0:0:0:0:0:0:0:0:0:0:0:0 |
| NM_001163285:NM_005243:XM_017028648:XM_017028666:XM_017028645:XM_017028659:XM_017028647:XM_017028665:NM_001163286:XR_002958676:XM_011529998:NM_013986:XM_017028657:XM_017028661:XM_011529997:XM_024452180:XM_017028644:XM_011530002:XM_017028664:XM_011530000:XM_024452181:XM_017028660:XM_017028651:XM_017028646:XM_017028662:XM_017028663:XM_017028653:XM_017028655:XM_005261389:XM_011529996:XM_011529995:XM_017028652:XM_017028656:XM_017028650:XM_017028658:XM_017028649 | 64:49:1:2:2:2:3:6:1:8:3:1:1:1:0:0:0:0:0:0:0:0:0:0:0:0:0:0:0:0:0:0:0:0:0:0 | 0:0:0:0:0:0:0:0:0:0:0:0:0:0:0:0:0:0:0:0:0:0:0:0:0:0:0:0:0:0:0:0:0:0:0:0 | 26:19:1:4:0:1:1:5:2:4:2:0:0:3:4:1:1:0:0:0:0:0:0:0:0:0:0:0:0:0:0:0:0:0:0:0 | 0:0:0:0:0:0:0:0:0:0:0:0:0:0:0:0:0:0:0:0:0:0:0:0:0:0:0:0:0:0:0:0:0:0:0:0 | 54:67:2:3:2:0:1:6:0:5:8:2:0:1:6:1:4:1:1:3:2:1:1:1:0:0:0:0:0:0:0:0:0:0:0:0 | 1:0:0:0:0:0:0:0:0:0:0:0:0:0:0:0:0:0:0:0:0:0:0:0:0:0:0:0:0:0:0:0:0:0:0:0 | 213:159:5:29:2:10:3:14:6:29:17:2:1:36:52:1:6:0:2:3:2:1:3:1:1:1:4:1:2:1:1:0:0:0:0:0 | 0:0:0:0:0:0:0:0:0:0:1:0:0:0:0:0:0:0:0:0:0:0:0:0:0:0:0:0:0:0:0:0:0:0:0:0 | 88:46:1:6:2:1:1:4:2:4:3:1:0:2:3:0:0:0:0:1:1:0:2:0:0:0:1:0:0:0:1:1:0:0:0:0 | 0:1:0:0:0:0:0:0:0:0:0:0:0:0:0:0:0:1:0:0:0:0:0:0:0:0:0:0:0:0:0:0:0:0:0:0 |
| NM_145660:NM_030643 | 2:00 | 0:00 | 1:00 | 0:00 | 0:00 | 0:00 | 0:00 | 5:00 | 0:00 | 0:01 |
| NM_145660 | 0 | 2 | 0 | 1 | 0 | 0 | 0 | 5 | 0 | 0 |
| NM_145660 | 0 | 2 | 0 | 1 | 0 | 0 | 0 | 5 | 0 | 0 |
| NM_145660 | 0 | 2 | 0 | 1 | 0 | 0 | 0 | 5 | 0 | 0 |

| | | | | | | | | | | |
|---|---|---|---|---|---|---|---|---|---|---|
| NM_003661:XM_011530478:NM_145343:NM_001136541:NM_001136540 | 8:1:0:0:0 | 0:0:0:0:0 | 20:0:3:1:0 | 0:0:0:0:0 | 11:0:1:0:0 | 0:0:0:0:0 | 20:0:0:0:1 | 0:0:0:0:0 | 3:0:0:1:1 | 0:0:0:0:0 |
| NM_003661:NM_145343:NM_001136540:NM_001136541:XM_011530478 | 4:0:0:0:0 | 2:0:0:0:0 | 10:0:0:0:0 | 9:1:0:0:0 | 6:0:0:0:0 | 5:0:0:0:0 | 0:0:0:0:0 | 20:0:1:0:0 | 0:0:0:0:0 | 3:0:1:1:0 |
| NM_003661:NM_145343:NM_001136540:XM_011530478:NM_001136541 | 4:0:0:0:0 | 2:0:0:0:0 | 10:0:0:0:0 | 9:1:0:0:0 | 6:0:0:0:0 | 5:0:0:0:0 | 0:0:0:0:0 | 20:0:1:2:0 | 0:0:0:0:0 | 3:0:1:0:1 |
| NM_003661:NM_145343:XM_011530478:NM_001136540:NM_001136541 | 4:0:0:0:0 | 2:0:0:0:0 | 9:0:0:0:0 | 9:1:1:0:0 | 5:0:0:0:0 | 5:0:0:0:0 | 0:0:0:0:0 | 18:0:2:1:0 | 0:0:0:0:0 | 3:0:0:1:1 |
| NM_002473 | 43 | 0 | 66 | 0 | 67 | 0 | 91 | 1 | 55 | 0 |
| NM_002473 | 34 | 0 | 30 | 0 | 45 | 0 | 64 | 1 | 33 | 0 |
| NM_002473 | 22 | 0 | 9 | 0 | 31 | 0 | 14 | 21 | 13 | 10 |
| NM_002473 | 14 | 0 | 18 | 0 | 17 | 1 | 26 | 0 | 8 | 0 |
| NM_002473 | 13 | 0 | 13 | 0 | 11 | 0 | 18 | 0 | 4 | 0 |
| NM_002473 | 3 | 0 | 2 | 0 | 8 | 2 | 11 | 1 | 1 | 0 |
| NM_002473 | 1 | 0 | 4 | 0 | 23 | 0 | 16 | 1 | 3 | 0 |
| NM_002473 | 20 | 0 | 27 | 0 | 70 | 0 | 63 | 1 | 22 | 0 |
| NM_002473 | 38 | 0 | 76 | 0 | 117 | 1 | 124 | 1 | 46 | 0 |
| XM_005261508:XM_011530111:XM_006724226:NM_012473 | 534:34:61:66 | 2:0:0:0 | 669:52:127:44 | 0:0:0:0 | 1877:123:260:211 | 5:1:0:0 | 2499:127:420:192 | 4:0:0:0 | 582:44:85:87 | 1:0:0:0 |
| NM_012473:XM_011530111:XM_005261508:XM_006724226 | 65:32:536:62 | 1:1:1:0 | 44:48:669:127 | 0:1:0:0 | 211:121:1884:259 | 0:1:1:1 | 192:121:2511:421 | 0:0:1:0 | 87:43:583:86 | 0:0:0:0 |
| XM_005261508:XM_011530111:XM_006724226:NM_012473 | 537:32:62:64 | 1:0:0:0 | 672:47:128:44 | 1:0:0:0 | 1886:115:264:211 | 5:1:1:0 | 2516:109:422:193 | 1:2:0:0 | 583:39:86:87 | 0:0:0:0 |
| XM_005261508:XM_006724226:XM_011530111:NM_012473 | 536:59:2:62 | 3:0:0:0 | 679:103:4:42 | 0:0:0:0 | 1893:247:8:210 | 4:1:0:0 | 2528:401:13:190 | 5:1:1:0 | 585:82:3:86 | 1:0:0:0 |
| XM_005261508:XM_006724226:XM_011530111:NM_012473 | 539:52:2:63 | 0:0:0:0 | 683:88:5:41 | 1:0:0:0 | 1899:231:12:208 | 1:1:0:0 | 2533:376:16:189 | 7:0:0:0 | 586:77:3:86 | 1:0:0:0 |
| NM_024955 | 0 | 0 | 0 | 0 | 0 | 0 | 0 | 0 | 0 | 0 |
| NM_003753 | 694 | 0 | 274 | 1 | 994 | 0 | 2020 | 0 | 567 | 0 |
| NM_003753 | 692 | 2 | 274 | 0 | 994 | 1 | 2016 | 2 | 568 | 0 |
| NM_003753 | 696 | 0 | 274 | 0 | 995 | 0 | 2016 | 3 | 568 | 0 |
| NM_003753 | 696 | 0 | 273 | 1 | 995 | 0 | 2016 | 2 | 568 | 0 |
| NM_003753 | 695 | 1 | 253 | 0 | 975 | 1 | 1993 | 2 | 562 | 2 |
| NM_003753 | 691 | 2 | 254 | 0 | 971 | 5 | 1993 | 1 | 562 | 2 |
| NM_003753 | 692 | 1 | 253 | 0 | 971 | 2 | 1985 | 2 | 562 | 1 |
| NM_003753 | 688 | 1 | 240 | 0 | 957 | 2 | 1952 | 2 | 554 | 0 |
| NM_003753 | 678 | 2 | 225 | 1 | 927 | 2 | 1901 | 2 | 544 | 1 |
| NM_003753 | 678 | 0 | 221 | 0 | 923 | 3 | 1886 | 2 | 540 | 0 |
| NM_003753 | 660 | 3 | 206 | 0 | 882 | 1 | 1795 | 2 | 526 | 0 |
| NM_003753 | 661 | 0 | 206 | 0 | 885 | 2 | 1780 | 1 | 524 | 0 |
| NM_003753 | 661 | 0 | 205 | 1 | 883 | 4 | 1778 | 3 | 524 | 0 |
| NM_003753 | 658 | 0 | 205 | 0 | 883 | 2 | 1777 | 3 | 523 | 0 |
| NM_003753 | 646 | 0 | 205 | 0 | 873 | 3 | 1742 | 0 | 520 | 0 |
| NM_014291 | 0 | 0 | 0 | 0 | 0 | 0 | 0 | 2 | 0 | 0 |
| XM_005261751 | 0 | 0 | 0 | 0 | 0 | 1 | 0 | 0 | 0 | 0 |
| XM_017028807:NM_001675:NM_182810 | 2:00:00 | 0:00:00 | 1:00:00 | 1:01:00 | 0:00:00 | 1:00:01 | 1:00:00 | 0:00:00 | 9:00:00 | 0:00:00 |
| NM_138435 | 0 | 0 | 1 | 1 | 1 | 1 | 1 | 3 | 1 | 0 |
| NM_001024843:NM_015088 | 18:00 | 9:00 | 88:00:00 | 88:00:00 | 40:01:00 | 31:00:00 | 236:03:00 | 0:00 | 17:00 | 0:00 |
| NM_003932 | 3 | 2 | 0 | 1 | 0 | 1 | 5 | 0 | 0 | 0 |
| NM_014248 | 340 | 0 | 586 | 0 | 540 | 1 | 1368 | 0 | 434 | 0 |
| NM_014248 | 340 | 0 | 582 | 1 | 539 | 0 | 1366 | 1 | 431 | 2 |
| NM_001429 | 0 | 0 | 0 | 0 | 1 | 0 | 0 | 2 | 0 | 0 |
| NM_001018050:NM_138338:NM_001282885:NM_001018052:XR_244356 | 16:9:1:1:3 | 0:0:0:0:0 | 2:6:0:0:0 | 0:0:0:0:0 | 7:4:1:0:1 | 0:0:0:0:0 | 7:8:2:0:1 | 29:21:3:0:4 | 5:3:0:0:1 | 8:2:1:0:5 |
| NM_001003796:NM_005008 | 608:162 | 1:00 | 388:184 | 2:00 | 518:303 | 0:00 | 2577:1552 | 2:02 | 558:198 | 0:00 |
| NM_001003796:NM_005008 | 609:160 | 0:00 | 395:184 | 0:00 | 518:298 | 1:01 | 2587:1554 | 1:02 | 561:198 | 1:00 |
| NM_001003796:NM_005008 | 609:159 | 0:00 | 395:184 | 0:00 | 519:298 | 0:00 | 2587:1550 | 1:00 | 562:195 | 0:00 |
| NM_002490 | 0 | 3 | 0 | 6 | 0 | 5 | 4 | 0 | 1 | 0 |
| NM_015703 | 0 | 0 | 0 | 0 | 2 | 1 | 3 | 0 | 0 | 0 |
| NM_007229:NM_001184970:NM_001349974:NM_001349971:NM_001184971:NM_001349969 | 2:10:0:0:0:0 | 0:0:0:0:0:0 | 9:4:1:0:0:0 | 0:0:0:0:0:0 | 23:100:1:1:3:0 | 0:1:0:0:0:0 | 9:28:0:0:1:1 | 0:0:0:0:0:0 | 5:7:0:0:0:0 | 0:0:0:0:0:0 |
| NM_000714:NR_046308:NM_001256531:NM_001256530 | 1:0:0:0 | 2:1:0:0 | 0:0:1:0 | 0:0:0:0 | 6:0:0:0 | 7:0:0:0 | 0:0:0:0 | 7:1:0:1 | 0:0:0:0 | 9:0:0:0 |
| NM_000714:NR_046308:NM_001256531:NM_001256530 | 1:0:0:0 | 2:1:0:0 | 0:0:1:0 | 0:0:0:0 | 6:0:0:0 | 7:0:0:0 | 3:0:0:1 | 4:1:0:0 | 6:0:0:0 | 3:0:0:0 |
| NM_015380 | 0 | 0 | 0 | 0 | 0 | 1 | 0 | 3 | 0 | 1 |
| NM_013327 | 0 | 0 | 0 | 0 | 0 | 0 | 1 | 0 | 2 | 2 |
| NM_006486:NM_006487 | 0:00 | 4:00 | 0:00 | 1:00 | 0:00 | 17:00 | 0:00 | 0:00 | 0:00 | 1:00 |
| NM_024324:NR_104295 | 1:00 | 0:00 | 0:00 | 0:00 | 2:00 | 0:00 | 1:00 | 0:00 | 1:00 | 0:00 |
| NM_001953:NM_001257989:NM_001113755 | 1:00:00 | 0:00:00 | 2:00:00 | 0:00:00 | 1:00:00 | 0:00:00 | 0:01:00 | 0:00:00 | 1:00:00 | 0:00:00 |
| NM_001953:NM_001257989:NM_001113755 | 1:00:00 | 0:00:00 | 0:00:00 | 2:00:00 | 0:00:00 | 1:00:00 | 0:01:00 | 0:00:00 | 1:00:00 | 0:00:00 |
| NM_033084:NM_001319984 | 0:00 | 0:00 | 6:00 | 0:00 | 1:01 | 0:00 | 15:01 | 7:01 | 10:01 | 3:00 |
| NM_001007565:NM_006070:XM_005247066:NM_001195479:NM_001195478:XM_017005527:XM_011512334:XM_006713472 | 15:244:2:96:6:2:2:0 | 0:0:0:0:0:0:0:0 | 8:233:4:116:1:1:5:0 | 0:1:0:0:0:0:0:0 | 19:458:8:168:3:5:3:2 | 0:0:0:0:0:0:0:0 | 46:1840:27:643:24:20:41:10 | 0:2:0:1:0:0:0:0 | 17:500:5:145:5:3:6:2 | 0:1:0:0:0:0:0:0 |
| NM_001007565:NM_006070:XM_005247066:NM_001195479:NM_001195478:XM_017005527:XM_011512334:XM_006713472 | 15:246:2:99:6:2:3:0 | 0:0:0:0:0:0:0:0 | 8:236:4:117:1:4:9:0 | 0:1:0:1:0:0:0:0 | 19:461:8:170:3:7:7:2 | 0:0:0:0:0:0:0:0 | 46:1866:27:651:24:38:111:10 | 0:0:0:1:0:0:0:0 | 17:508:5:146:5:4:8:2 | 0:0:0:0:0:0:0:0 |
| NM_001007565:NM_006070:XM_005247066:NM_001195479:NM_001195478:XM_017005527:XM_011512334:XM_006713472 | 15:247:2:99:6:3:3:0 | 0:0:0:0:0:0:0:0 | 8:239:4:119:1:4:9:0 | 0:0:0:0:0:0:0:0 | 19:462:8:170:3:9:7:2 | 0:0:0:0:0:0:0:0 | 46:1876:27:657:24:46:125:10 | 0:1:0:0:0:2:0:0 | 17:507:5:146:5:5:8:2 | 0:1:0:0:0:0:0:0 |
| NM_033084:NM_001319984 | 0:00 | 0:00 | 1:00 | 3:00 | 1:01 | 0:00 | 5:02 | 9:00 | 9:01 | 2:01 |
| NM_001007565:NM_006070:XM_005247066:NM_001195479:NM_001195478:XM_017005527:XM_011512334:XM_006713472 | 15:248:2:98:6:3:4:0 | 0:0:0:0:0:0:0:0 | 8:242:4:119:1:6:12:0 | 0:0:0:0:0:0:0:0 | 19:464:8:173:3:8:8:2 | 0:1:0:0:0:0:0:0 | 46:1890:27:667:24:63:153:10 | 0:1:0:1:0:0:1:0 | 17:511:5:146:5:4:8:2 | 0:0:0:0:0:0:0:0 |
| NM_006070:NM_001007565:XM_005247066:NM_001195479:NM_001195478:XM_017005527:XM_011512334:XM_006713472 | 247:15:2:99:6:3:4:0 | 1:0:0:0:0:0:0:0 | 242:8:4:119:1:6:12:0 | 0:0:0:0:0:0:0:0 | 465:19:8:173:3:9:8:2 | 0:0:0:0:0:0:0:0 | 1889:45:27:668:24:63:154:10 | 3:1:0:1:0:0:0:0 | 508:17:5:146:5:4:8:2 | 3:0:0:0:0:0:0:0 |
| NM_006070:NM_001007565:XM_005247066:NM_001195479:NM_001195478:XM_017005527:XM_011512334:XM_006713472 | 248:15:2:99:6:3:4:0 | 1:0:0:0:0:0:0:0 | 242:8:4:119:1:6:13:0 | 0:0:0:0:0:0:0:0 | 466:19:8:172:3:10:9:2 | 1:0:0:1:0:0:0:0 | 1895:46:27:669:24:65:160:10 | 1:0:0:1:0:0:0:0 | 510:17:5:147:5:5:9:2 | 1:0:0:0:0:0:0:0 |
| NM_001007565:NM_006070:XM_005247066:NM_001195479:NM_001195478:XM_017005527:XM_011512334:XM_006713472 | 14:249:2:99:6:3:4:0 | 1:1:0:0:0:0:0:0 | 8:245:4:120:1:8:15:0 | 0:0:0:0:0:0:0:0 | 19:468:8:174:3:11:9:2 | 0:0:0:0:0:0:0:0 | 46:1896:27:671:24:66:171:10 | 0:5:0:1:0:0:0:0 | 17:510:5:147:5:6:10:2 | 0:1:0:1:0:0:0:0 |
| NM_001007565:NM_006070:XM_005247066:NM_001195479:NM_001195478:XM_017005527:XM_011512334:XM_006713472 | 15:250:2:100:6:3:4:0 | 0:0:0:0:0:0:0:0 | 8:249:4:119:1:8:16:0 | 0:0:0:1:0:0:0:0 | 19:472:8:174:3:11:10:2 | 0:0:0:0:0:0:0:0 | 46:1902:27:674:24:66:183:10 | 0:0:0:1:0:1:3:0 | 17:511:5:148:5:5:11:2 | 0:0:0:0:0:0:0:0 |

| | | | | | | | | | | |
|---|---|---|---|---|---|---|---|---|---|---|
| NM_006070:NM_001007565:XM_005247066:NM_001195479:NM_001195478:XM_017005527:XM_011512334:XM_006713472 | 250:15:2:100:6:3:3:0 | 1:0:0:0:0:0:0:0 | 242:8:4:121:1:8:17:0 | 1:0:0:0:0:0:0:0 | 474:19:8:174:3:11:11:2 | 0:0:0:0:0:0:0:0 | 1866:46:27:670:24:74:188:10 | 5:0:0:0:0:0:0:0 | 512:17:5:149:5:7:13:2 | 0:0:0:0:0:0:0:0 |
| NM_006070:NM_001007565:XM_005247066:NM_001195479:NM_001195478:XM_017005527:XM_011512334:XM_006713472 | 250:15:2:100:6:4:2:0 | 1:0:0:0:0:0:0:0 | 245:8:4:122:1:10:16:0 | 0:0:0:0:0:0:0:0 | 472:19:8:173:3:11:11:2 | 1:0:0:0:0:0:0:0 | 1896:46:27:670:24:87:185:9 | 0:0:0:2:1:0:0:0 | 512:17:5:149:5:7:13:2 | 0:0:0:0:0:0:0:0 |
| NM_001007565:NM_006070:XM_005247066:NM_001195479:NM_001195478:XM_017005527:XM_011512334:XM_006713472 | 15:251:2:100:6:4:2:0 | 0:0:0:0:0:0:0:0 | 8:245:4:122:1:10:15:0 | 0:0:0:0:0:0:0:0 | 19:474:8:173:3:11:10:2 | 0:0:0:0:0:0:0:0 | 46:1887:27:673:24:91:184:9 | 0:4:0:0:1:0:0:0 | 17:512:5:149:5:7:13:2 | 0:0:0:0:0:0:0:0 |
| NM_006070:NM_001007565:XM_005247066:NM_001195479:NM_001195478:XM_017005527:XM_011512334:XM_006713472 | 250:15:2:100:6:6:2:0 | 1:0:0:0:0:0:0:0 | 244:8:4:121:1:14:15:0 | 1:0:0:0:0:0:0:0 | 476:18:8:173:3:13:7:2 | 0:1:0:0:0:0:1:0 | 1895:46:27:670:24:110:167:9 | 3:0:0:0:1:0:0:0 | 511:17:5:149:5:9:11:2 | 1:0:0:0:0:0:1:0 |
| NM_001007565:NM_006070:XM_005247066:NM_001195479:NM_001195478:XM_017005527:XM_011512334:XM_006713472 | 15:251:2:100:6:7:2:0 | 0:0:0:0:0:0:0:0 | 8:244:4:121:1:14:12:0 | 0:1:0:0:0:0:1:0 | 19:476:8:173:3:15:8:2 | 0:0:0:0:0:0:0:0 | 46:1895:27:670:24:112:164:9 | 0:3:0:0:1:0:0:0 | 17:512:5:149:5:9:11:2 | 0:0:0:0:0:0:1:0 |
| NM_183352:XM_024453704:NM_001136026:NM_030673:XM_024453703:XM_017007021:NM_001136232:XM_024453702:XM_005265379:XM_024453701:XM_024453705 | 335:49:1:4:2:5:0:0:0:0:0 | 1:0:0:0:0:0:0:0:0:0:0 | 185:38:0:0:0:1:1:0:0:0:0 | 0:0:0:0:0:0:0:0:0:0:0 | 626:112:0:5:5:2:0:2:1:0:0 | 0:0:0:1:0:0:0:0:0:0:0 | 1729:291:2:23:9:12:17:3:3:1:0 | 2:0:0:0:0:0:0:0:0:0:0 | 434:48:1:12:2:0:0:1:1:0:0 | 0:0:0:0:0:0:0:0:0:0:0 |
| NM_183352:XM_024453704:NM_001136026:NM_030673:XM_024453703:XM_017007021:NM_001136232:XM_024453702:XM_005265379:XM_024453701:XM_024453705 | 334:49:1:4:2:5:0:0:0:0:0 | 2:0:0:0:0:0:0:0:0:0:0 | 185:38:0:0:0:1:1:0:0:0:0 | 0:0:0:0:0:0:0:0:0:0:0 | 625:115:0:6:5:2:0:2:1:0:0 | 1:0:0:0:0:0:0:0:0:0:0 | 1729:298:2:23:9:10:17:3:3:1:0 | 4:0:0:0:0:0:0:0:0:0:0 | 433:48:1:12:2:0:0:1:1:0:0 | 1:0:0:0:0:0:0:0:0:0:0 |
| NM_183352:XM_024453704:NM_001136026:NM_030673:XM_024453703:XM_017007021:NM_001136232:XM_024453702:XM_005265379:XM_024453701:XM_024453705 | 336:49:1:4:2:5:0:0:0:0:0 | 0:0:0:0:0:0:0:0:0:0:0 | 184:38:0:0:0:1:1:0:0:0:0 | 1:0:0:0:0:0:0:0:0:0:0 | 625:115:0:6:5:1:0:2:1:0:0 | 1:0:0:0:0:0:0:0:0:0:0 | 1730:297:2:23:9:8:17:3:3:1:0 | 2:0:0:0:0:0:0:0:0:0:0 | 434:48:1:12:1:0:0:1:1:0:0 | 0:0:0:0:1:0:0:0:0:0:0 |
| NM_183352:XM_017007021:XM_024453704:NM_001136026:XM_024453705:NM_030673:XM_024453703:XM_017007020:NM_001136232:XM_024453702:XM_005265379:XM_024453701 | 335:61:49:1:7:4:2:2:0:0:0:0 | 1:0:0:0:0:0:0:0:0:0:0:0 | 185:29:37:0:4:0:0:0:1:0:0:0 | 0:0:0:0:0:0:0:0:0:0:0:0 | 626:115:114:0:12:6:5:0:0:2:1:0 | 0:0:0:0:0:0:0:0:0:0:0:0 | 1733:152:292:2:19:24:9:2:17:3:3:1 | 2:0:0:0:0:0:0:0:0:0:0:0 | 433:78:48:1:12:12:2:1:0:1:1:0 | 1:0:0:0:0:0:0:0:0:0:0:0 |
| NM_183352:XM_017007021:XM_024453704:NM_001136026:XM_024453705:NM_030673:XM_024453703:XM_017007020:NM_001136232:XM_024453702:XM_005265379:XM_024453701 | 336:61:49:1:7:4:2:2:0:0:0:0 | 0:0:0:0:0:0:0:0:0:0:0:0 | 185:29:37:0:4:0:0:0:1:0:0:0 | 0:0:0:0:0:0:0:0:0:0:0:0 | 624:114:112:0:13:6:5:0:0:2:1:0 | 2:0:0:0:0:0:0:0:0:0:0:0 | 1734:154:288:2:21:24:9:2:17:3:3:1 | 1:0:0:0:0:0:0:0:0:0:0:0 | 434:79:46:1:14:12:2:1:0:1:1:0 | 0:0:0:0:0:0:0:0:0:0:0:0 |
| NM_183352:XM_017007021:XM_024453704:NM_001136026:XM_024453705:NM_030673:XM_024453703:XM_017007020:NM_001136232:XM_024453702:XM_005265379:XM_024453701:NM_001278946 | 336:68:42:1:8:4:2:2:0:0:0:0:0 | 0:0:0:0:0:0:0:0:0:0:0:0:0 | 184:29:27:0:9:0:0:0:1:0:0:0:0 | 0:1:0:0:0:0:0:0:0:0:0:0:0 | 625:119:90:0:18:6:5:0:0:2:1:0:0 | 1:1:1:0:0:0:0:0:0:0:0:0:0 | 1731:169:221:2:27:24:9:2:17:3:3:1:0 | 2:0:2:0:0:0:0:0:0:0:0:0:0 | 433:82:37:1:11:12:2:1:0:1:1:0:0 | 1:0:0:0:0:0:0:0:0:0:0:0:0 |
| NM_183352:XM_017007021:XM_024453704:NM_001136026:XM_024453705:NM_030673:XM_024453703:XM_017007020:NM_001136232:XM_024453702:XM_005265379:XM_024453701:NM_001278946 | 336:69:38:1:9:4:2:2:0:0:0:0:0 | 0:0:0:0:0:0:0:0:0:0:0:0:0 | 185:30:24:0:7:0:0:0:1:0:0:0:0 | 0:0:0:0:0:0:0:0:0:0:0:0:0 | 622:120:86:0:15:6:5:0:0:2:1:0:0 | 4:0:0:0:0:0:0:0:0:0:0:0:0 | 1734:170:201:2:26:24:9:2:17:3:3:1:0 | 1:0:7:0:0:0:0:0:0:0:0:0:0 | 433:83:35:1:12:12:2:1:0:1:1:0:0 | 0:0:0:0:0:0:0:0:0:0:0:0:0 |
| NM_183352:XM_017007021:XM_024453704:NM_001136026:XM_024453705:NM_030673:XM_024453703:XM_017007020:NM_001136232:XM_024453702:XM_005265379:XM_024453701:NM_001278946 | 336:69:37:1:8:4:2:2:0:0:0:0:0 | 0:0:0:0:0:0:0:0:0:0:0:0:0 | 185:30:24:0:8:0:0:0:1:0:0:0:0 | 0:0:0:0:0:0:0:0:0:0:0:0:0 | 626:120:82:0:13:6:5:0:0:2:1:0:0 | 0:0:0:0:0:0:0:0:0:0:0:0:0 | 1735:169:202:2:23:24:9:2:17:3:3:1:0 | 0:2:4:0:0:0:0:0:0:0:0:0:0 | 433:82:31:1:12:12:2:1:0:1:1:0:0 | 0:1:0:0:0:0:0:0:0:0:0:0:0 |
| XM_017007395:NM_170662:XM_017007399:NM_001321791:XM_017007400:XR_001740339:NR_135809:NM_001321788:XR_001740338:XM_017007396:NR_135812:NR_135811:NM_001321813 | 1:2:1:1:0:0:0:0:0:0:0:0:0 | 0:0:0:0:0:0:0:0:0:0:0:0:0 | 0:0:2:0:6:1:0:0:0:0:0:0:0 | 0:0:0:0:0:0:0:0:0:0:0:0:0 | 2:1:1:0:1:0:1:2:1:0:0:0:0 | 0:0:0:0:0:0:0:0:0:0:0:0:0 | 0:0:35:0:14:5:1:0:0:1:1:0:0 | 0:0:0:0:0:0:0:0:0:0:0:0:0 | 6:0:2:1:0:0:1:2:0:2:0:0:0 | 1:0:0:0:0:0:0:0:0:0:0:0:0 |
| NM_020235:XM_024453659:XM_011513001:XM_024453646:NM_001142568:XM_024453648:XM_024453647:XM_024453667:XM_024453650:XM_005247642:XM_005247643:XM_024453653:XM_024453661:XM_017006882:XM_024453666:XM_024453655:XM_024453660:XM_024453651:NM_001276286 | 30:1:1:1:1:1:1:1:1:0:0:0:0:0:0:0:0:0:0 | 0:0:0:0:0:0:0:0:0:0:0:0:0:0:0:0:0:0:0 | 44:4:1:2:0:2:1:0:2:3:1:1:2:0:0:0:0:0:0 | 0:0:0:0:0:0:0:0:0:0:0:0:0:0:0:0:0:0:0 | 15:1:0:0:2:2:1:0:0:0:1:0:0:0:0:0:0:0:0 | 0:0:0:0:0:0:0:0:0:0:0:0:0:0:0:0:0:0:0 | 243:15:1:7:0:10:19:1:2:9:1:0:5:10:4:4:1:0:0 | 1:0:0:1:0:0:0:0:0:0:0:0:0:0:0:0:0:0:0 | 24:1:0:1:1:2:5:0:0:2:0:0:1:0:0:0:0:0:0 | 0:0:0:0:0:0:0:0:0:0:0:0:0:0:0:0:0:0:0 |
| NM_020235:XM_024453659:XM_011513001:XM_024453646:NM_001142568:XM_024453648:XM_024453647:XM_024453667:XM_024453650:XM_005247642:XM_005247643:XM_024453653:XM_024453661:XM_017006882:XM_024453666:XM_024453655:XM_024453660:XM_024453651:NM_001276286 | 30:1:1:1:1:1:1:1:1:0:0:0:0:0:0:0:0:0:0 | 0:0:0:0:0:0:0:0:0:0:0:0:0:0:0:0:0:0:0 | 44:4:1:2:0:2:1:0:2:3:1:1:2:0:0:0:0:0:0 | 0:0:0:0:0:0:0:0:0:0:0:0:0:0:0:0:0:0:0 | 15:1:0:0:2:2:1:0:0:0:1:0:0:0:0:0:0:0:0 | 0:0:0:0:0:0:0:0:0:0:0:0:0:0:0:0:0:0:0 | 244:15:1:8:0:10:19:2:2:9:1:0:11:10:2:4:1:0:0 | 0:0:0:0:0:0:0:0:0:0:0:0:0:0:0:0:0:0:0 | 24:1:0:1:1:2:5:0:0:2:0:0:1:0:0:0:0:0:0 | 0:0:0:0:0:0:0:0:0:0:0:0:0:0:0:0:0:0:0 |
| NM_020235:XM_024453659:XM_011513001:XM_024453646:NM_001142568:XM_024453648:XM_024453647:XM_024453667:XM_024453650:XM_005247642:XM_005247643:XM_024453653:XM_024453661:XM_017006882:XM_024453666:XM_024453655:XM_024453660:XM_024453651:NM_001276286 | 30:1:1:1:1:1:1:1:1:0:0:0:0:0:0:0:0:0:0 | 0:0:0:0:0:0:0:0:0:0:0:0:0:0:0:0:0:0:0 | 44:4:1:2:0:2:1:0:2:3:1:1:2:0:0:0:0:0:0 | 0:0:0:0:0:0:0:0:0:0:0:0:0:0:0:0:0:0:0 | 14:1:0:0:2:2:1:0:0:0:1:0:0:0:0:0:0:0:0 | 1:0:0:0:0:0:0:0:0:0:0:0:0:0:0:0:0:0:0 | 244:15:1:8:0:10:19:2:2:9:1:0:11:10:2:4:1:0:0 | 0:0:0:0:0:0:0:0:0:0:0:0:0:0:0:0:0:0:0 | 24:1:0:1:1:2:5:0:0:2:0:0:0:0:0:0:0:0:0 | 0:0:0:0:0:0:0:0:0:0:0:1:0:0:0:0:0:0:0 |
| NM_020235:XM_024453659:XM_011513001:XM_024453646:NM_001142568:XM_024453648:XM_024453647:XM_024453653:XM_024453650:XM_005247642:XM_005247643:XM_024453661:XM_017006882:XM_024453655:XM_024453660:XM_024453644:XM_024453666:XM_024453651:XM_024453656:XM_024453667 | 28:1:1:1:1:1:1:1:0:0:0:0:0:0:0:0:0:0:0:0 | 0:0:0:0:0:0:0:0:0:0:0:0:0:0:0:0:0:0:0:0 | 44:4:1:2:0:2:1:0:2:3:1:19:0:0:0:0:0:0:0:0 | 0:0:0:0:0:0:0:0:1:0:0:0:0:0:0:0:0:0:0:0 | 14:1:0:0:2:2:1:0:0:0:0:0:0:0:0:0:0:0:0:0 | 0:0:0:0:0:0:0:0:0:0:0:0:0:0:0:0:0:0:0:0 | 226:14:1:8:0:10:19:0:2:8:1:27:10:3:1:1:0:0:0:0 | 10:0:0:0:0:0:0:0:0:0:0:0:0:0:0:0:0:0:0 | 23:1:0:1:1:2:3:0:0:2:0:4:0:0:0:0:0:0:0:0 | 0:0:0:0:0:0:2:0:0:0:0:0:0:0:0:0:0:0:0:0 |
| NM_020235:XM_024453648:XM_024453659:XM_024453646:NM_001142568:XM_024453647:XM_005247643:XM_024453653:XM_024453661:XM_024453650:XM_005247642:XM_011513001:XM_017006882:XM_024453655:XM_024453660:XM_024453644:XM_024453666:XM_024453651:XM_024453656:XM_024453667 | 17:0:1:1:1:1:1:0:0:0:0:0:0:0:0:0:0:0:0:0 | 10:1:0:0:0:0:0:0:0:0:0:0:0:0:0:0:0:0:0 | 31:2:4:1:0:1:0:0:15:2:2:1:0:0:0:0:0:0:0:0 | 12:0:0:1:0:0:1:1:4:0:0:0:0:0:0:0:0:0:0 | 10:1:1:0:2:0:0:0:0:0:0:0:0:0:0:0:0:0:0:0 | 3:1:0:0:0:1:0:0:0:0:0:0:0:0:0:0:0:0:0 | 163:8:11:8:0:14:1:0:24:2:7:1:5:3:1:1:0:0:0:0 | 54:2:4:0:0:2:0:0:0:0:0:0:3:0:0:0:0:0:0 | 15:1:0:1:1:2:0:0:0:2:0:0:0:0:0:0:0:0:0:0 | 4:1:1:0:0:1:0:0:4:0:0:0:0:0:0:0:0:0:0 |
| XM_024453646:XM_024453659:NM_020235:NM_001142568:XM_024453647:XM_024453661:XM_024453648:XM_005247642:XM_011513001:XM_024453656:XM_024453655:XM_017006882:XM_024453650:XM_024453660:XM_024453644:XM_024453666:XM_024453651:XM_005247643:XM_024453653 | 0:1:12:1:1:0:0:0:0:0:0:0:0:0:0:0:0:0:0 | 1:0:0:0:0:0:0:0:0:0:0:0:0:0:0:0:0:0:0 | 1:4:20:0:1:11:1:1:1:1:0:0:0:0:0:0:0:0:0 | 0:0:3:0:0:1:0:0:0:0:0:0:0:0:0:0:0:0:0 | 0:1:6:2:0:0:0:0:0:0:0:0:0:0:0:0:0:0:0 | 0:0:0:0:0:0:0:0:0:0:0:0:0:0:0:0:0:0:0 | 6:7:89:0:9:17:4:6:1:0:5:3:1:1:1:0:0:0:0 | 0:0:13:0:1:0:1:0:0:0:1:0:0:0:0:0:0:0:0 | 1:0:13:1:2:0:1:0:0:0:0:0:0:0:0:0:0:0:0 | 0:0:0:0:0:0:0:0:0:0:0:0:0:0:0:0:0:0:0 |
| NM_001142568:NM_020235:XM_005247642:XM_024453659:XM_024453656:XM_024453655:XM_024453646:XM_017006882:XM_024453648:XM_024453647:XM_024453661:XM_024453644:XM_024453660:XM_024453658 | 1:0:0:0:0:0:0:0:0:0:0:0:0:0 | 0:0:0:0:0:0:0:0:0:0:0:0:0:0 | 0:2:1:1:1:0:0:0:0:0:0:0:0:0 | 0:2:0:0:0:0:0:0:0:0:0:0:0:0 | 2:0:0:0:0:0:0:0:0:0:0:0:0:0 | 0:0:0:0:0:0:0:0:0:0:0:0:0:0 | 0:6:0:1:0:2:2:1:2:1:1:2:0:0 | 0:2:0:0:0:3:0:0:0:0:0:0:0:0 | 1:3:0:0:0:0:0:0:0:0:0:0:0:0 | 0:0:0:0:0:0:0:0:0:0:0:0:0:0 |
| NM_001142568:NM_020235:XM_005247642:XM_024453659:XM_024453656:XM_017006882:XM_024453655:XM_024453648:XM_024453646:XM_024453661:XM_024453647:XM_024453644:XM_024453660:XM_024453658 | 1:0:0:0:0:0:0:0:0:0:0:0:0:0 | 0:0:0:0:0:0:0:0:0:0:0:0:0:0 | 0:0:0:1:1:0:0:0:0:0:0:0:0:0 | 0:4:1:0:0:0:0:0:0:0:0:0:0:0 | 2:0:0:0:0:0:0:0:0:0:0:0:0:0 | 0:0:0:0:0:0:0:0:0:0:0:0:0:0 | 0:1:0:0:0:0:2:1:1:0:1:2:0:0 | 0:7:0:0:0:1:3:1:1:1:0:0:0:0 | 1:1:0:0:0:0:0:0:0:0:0:0:0:0 | 0:2:0:0:0:0:0:0:0:0:0:0:0:0 |
| NM_001282556:XM_005247080:XM_011512367:XM_011512364:XM_011512362:NM_001282558:NM_007072:NM_001282557:XM_024453326:XM_011512363 | 0:0:0:0:0:0:0:0:0:0 | 1:1:0:0:0:0:0:0:0:0 | 32:41:5:1:1:4:6:1:0:0 | 33:23:6:3:2:2:1:0:0:0 | 34:52:5:1:2:1:2:0:0:0 | 34:55:4:1:1:0:0:0:0:0 | 0:0:0:0:0:0:0:0:0:0 | 13:19:1:0:0:3:1:1:1:0 | 0:0:0:0:0:0:0:0:0:0 | 6:9:0:0:0:0:0:0:1:0 |
| NM_001282556:XM_005247080:NM_007072:NM_001282558:XM_011512364:XM_011512362:NM_001282557:XM_024453326:XM_011512363 | 1:1:0:0:0:0:0:0:0 | 0:0:0:0:0:0:0:0:0 | 65:64:44:7:3:2:1:2:0 | 0:0:0:0:0:0:0:0:0 | 68:107:5:1:1:3:0:1:0 | 0:0:0:0:0:0:0:0:0 | 13:19:7:3:0:0:1:1:0 | 0:0:0:0:0:0:0:0:0 | 6:9:0:0:0:0:0:1:0 | 0:0:0:0:0:0:0:0:0 |
| NM_020890:XM_011513057:XM_006713717:XM_011513056 | 0:0:0:0 | 1:0:0:0 | 0:0:0:0 | 0:0:0:0 | 1:0:0:0 | 0:0:0:0 | 2:1:1:5 | 0:1:0:0 | 3:1:0:1 | 2:0:0:0 |
| NM_032579 | 0 | 0 | 0 | 0 | 0 | 1 | 0 | 0 | 0 | 0 |
| NM_024616 | 0 | 0 | 0 | 0 | 0 | 2 | 0 | 0 | 0 | 0 |
| NM_024616 | 0 | 0 | 0 | 0 | 0 | 2 | 0 | 0 | 0 | 0 |
| NM_006395:NM_001349238:NM_001349237:XR_001739980:XM_024453315:NM_001349235:NM_001349233:XR_940363:XM_024453312:XM_011533284:XM_011533278:NM_001349236:NM_001136031:NM_001349232:XM_011533286:NM_001349234:XM_006712931:XM_006712932:XM_024453314:XM_017005542:XM_017005547:XM_017005550:XM_011533283:XM_017005543:XM_017005551 | 20:1:3:1:4:1:2:1:0:0:0:0:0:0:0:0:0:0:0:0:0:0:0:0:0 | 0:0:0:0:0:0:0:0:0:0:0:0:0:0:0:0:0:0:0:0:0:0:0:0:0 | 10:0:2:1:3:0:1:0:1:1:1:1:0:0:0:0:0:0:0:0:0:0:0:0:0 | 0:0:0:0:0:0:0:0:0:0:0:0:0:0:0:0:0:0:0:0:0:0:0:0:0 | 48:0:2:0:5:2:2:0:0:1:0:0:4:1:1:0:0:0:0:0:0:0:0:0:0 | 0:0:0:0:0:0:0:0:0:0:0:0:0:0:0:0:0:0:0:0:0:0:0:0:0 | 93:0:9:4:28:0:0:5:0:0:0:1:1:2:2:2:2:1:2:2:1:1:1:0:0 | 0:0:0:0:0:0:0:0:0:0:0:0:0:0:0:0:0:0:0:0:0:0:0:0:0 | 32:0:4:0:9:0:1:1:0:0:0:0:0:0:0:0:0:0:0:0:0:0:0:0:0 | 0:0:0:0:0:0:0:0:0:0:0:0:0:0:0:0:0:0:0:0:0:0:0:0:0 |
| NM_000187:XM_011512746:XM_005247413:XM_005247414:XM_005247412:XM_017006277 | 0:0:0:0:0:0 | 2:2:1:1:0:0 | 0:0:0:0:0:0 | 18:1:8:1:2:3 | 0:0:0:0:0:0 | 23:4:8:1:2:3 | 0:0:0:0:0:0 | 11:11:17:9:3:2 | 0:0:0:0:0:0 | 10:1:3:1:1:4 |
| NM_175862:NM_006889:NM_176892 | 4:00:00 | 14:00:00 | 2:01:00 | 2:01:00 | 4:00:00 | 5:00:00 | 0:00:00 | 9:01:00 | 0:00:00 | 0:00:00 |
| NM_014367 | 1 | 2 | 2 | 0 | 11 | 0 | 6 | 0 | 7 | 0 |
| NM_014367 | 2 | 1 | 0 | 3 | 4 | 6 | 1 | 5 | 7 | 0 |
| NM_015869:NM_138712 | 4:00 | 0:00 | 2:00 | 0:00 | 2:00 | 0:00 | 15:01 | 0:00 | 0:00 | 0:00 |
| NM_005037:NM_138712:NM_001330615:NM_015869:NM_138711:NM_001354667:NM_001354666:NM_001354669:NM_001354670:NM_001354668 | 17:31:1:5:18:2:0:0:0:0 | 0:0:0:0:0:0:0:0:0:0 | 59:105:1:2:631:30:1:7:3:0 | 0:0:0:0:0:0:0:0:0:0 | 53:140:1:5:817:50:0:0:0:0 | 0:0:0:0:2:0:0:0:0:0 | 274:486:2:17:1580:67:1:10:4:1 | 0:0:0:1:1:0:0:0:0:0 | 31:56:2:1:181:12:0:0:0:0 | 0:0:0:0:1:0:0:0:0:0 |
| NM_005037:NM_138712:NM_001330615:NM_015869:NM_138711:NM_001354667:NM_001354666:NM_001354669:NM_001354670:NM_001354668 | 17:30:1:5:18:2:0:0:0:0 | 0:0:0:0:0:0:0:0:0:0 | 59:105:1:2:628:30:1:7:3:0 | 0:0:0:0:1:0:0:0:0:0 | 53:138:1:5:818:50:0:0:0:0 | 0:0:0:0:1:0:0:0:0:0 | 274:483:2:17:1576:67:1:10:4:1 | 0:0:0:0:5:0:0:0:0:0 | 31:57:2:1:182:12:0:0:0:0 | 0:0:0:0:0:0:0:0:0:0 |

| | | | | | | | | | | |
|---|---|---|---|---|---|---|---|---|---|---|
| NM_005037:NM_138712:NM_001330615:NM_015869:NM_138711:NM_001354667:NM_001354666:NM_001354669:NM_001354670:NM_001354668 | 17:30:1:5:18:2:0:0:0:0 | 0:0:0:0:0:0:0:0:0:0 | 59:105:1:2:626:30:1:7:3:0 | 0:0:0:0:2:0:0:0:0 | 53:139:1:5:819:50:0:0:0:0 | 0:0:0:0:0:0:0:0:0 | 274:482:2:17:1580:67:1:10:4:1 | 0:0:0:0:1:0:0:0:0 | 31:57:3:1:182:12:0:0:0:0 | 0:0:0:0:0:0:0:0:0 |
| NM_005037:NM_138712:NM_001330615:NM_015869:NM_138711:NM_001354667:NM_001354666:NM_001354669:NM_001354670:NM_001354668 | 17:30:1:5:18:2:0:0:0:0 | 0:0:0:0:0:0:0:0:0:0 | 59:105:1:2:626:30:1:7:3:0 | 0:0:0:0:2:0:0:0:0 | 53:140:1:5:816:50:0:0:0:0 | 0:0:0:0:0:0:0:0:0 | 274:482:2:17:1576:67:1:10:3:1 | 0:0:0:0:2:0:0:0:0 | 31:57:3:1:182:12:0:0:0:0 | 0:0:0:0:0:0:0:0:0 |
| NM_005037:NM_138712:NM_001330615:NM_015869:NM_138711:NM_001354667:NM_001354666:NM_001354669:NM_001354670:NM_001354668 | 17:30:1:5:18:2:0:0:0:0 | 0:0:0:0:0:0:0:0:0:0 | 59:105:1:2:628:30:1:7:3:0 | 0:0:0:0:0:0:0:0:0 | 53:140:1:5:817:50:0:0:0:0 | 0:0:0:0:0:0:0:0:0 | 274:481:2:17:1578:67:1:10:3:1 | 0:1:0:0:1:0:0:0:0 | 31:57:3:1:182:12:0:0:0:0 | 0:0:0:0:0:0:0:0:0 |
| NM_138712:NM_005037:NM_001330615:NM_015869:NM_138711:NM_001354667:NM_001354666:NM_001354669:NM_001354670:NM_001354668 | 29:17:1:5:18:2:0:0:0:0 | 1:0:0:0:0:0:0:0:0:0 | 105:59:1:2:628:30:1:7:3:0 | 0:0:0:0:0:0:0:0:0 | 140:53:1:5:815:50:0:0:0:0 | 0:0:0:0:2:0:0:0:0 | 482:273:2:17:1577:67:1:10:3:1 | 0:1:0:0:2:0:0:0:0 | 57:31:3:1:182:12:0:0:0:0 | 0:0:0:0:0:0:0:0:0 |
| NM_005037:NM_138712:NM_001330615:NM_015869:NM_138711:NM_001354667:NM_001354666:NM_001354669:NM_001354670:NM_001354668 | 17:30:1:5:18:2:0:0:0:0 | 0:0:0:0:0:0:0:0:0:0 | 59:105:1:2:627:30:1:7:3:0 | 0:0:0:0:0:0:0:0:0 | 53:141:1:5:814:50:0:0:0:0 | 0:0:0:0:1:0:0:0:0 | 273:484:2:17:1574:67:1:10:3:1 | 1:0:0:0:1:0:0:0:0 | 30:57:3:1:182:12:0:0:0:0 | 0:0:0:0:0:0:0:0:0 |
| NM_138712:NM_005037:NM_001330615:NM_015869:NM_138711:NM_001354667:NM_001354666:NM_001354669:NM_001354670:NM_001354668 | 29:16:1:5:18:2:0:0:0:0 | 1:0:0:0:0:0:0:0:0:0 | 98:55:1:2:536:30:1:7:3:0 | 7:4:0:0:83:0:0:0:0 | 141:53:1:5:807:50:0:0:0:0 | 1:0:0:0:6:0:0:0:0 | 460:268:2:17:1448:64:1:10:3:1 | 23:6:0:0:110:3:0:0:0:0 | 57:30:3:1:182:11:0:0:0:0 | 0:0:0:0:0:1:0:0:0 |
| NM_005037:NM_138712:NM_001330615:NM_015869:NM_138711:NM_001354667:NM_001354666:NM_001354669:NM_001354670:NM_001354668 | 16:29:1:5:18:2:0:0:0:0 | 1:1:0:0:0:0:0:0:0:0 | 55:98:1:2:536:30:1:7:3:0 | 4:7:0:0:87:0:0:0:0 | 53:141:1:5:807:50:0:0:0:0 | 0:1:0:0:8:0:0:0:0 | 268:459:2:17:1448:64:1:10:3:1 | 6:25:0:0:115:3:0:0:0:0 | 30:57:3:1:182:11:0:0:0:0 | 0:0:0:0:0:1:0:0:0 |
| NM_005037:NM_138712:NM_001330615:NM_015869:NM_138711:NM_001354667:NM_001354666:NM_001354669:NM_001354670:NM_001354668 | 16:29:1:5:18:2:0:0:0:0 | 0:0:0:0:0:0:0:0:0:0 | 55:98:1:2:535:30:1:7:3:0 | 0:0:0:0:0:0:0:0:0 | 53:141:1:5:805:50:0:0:0:0 | 0:0:0:0:2:0:0:0:0 | 268:456:2:17:1447:64:1:10:3:1 | 0:1:0:0:1:0:0:0:0 | 29:56:3:1:182:11:0:0:0:0 | 1:0:0:0:0:0:0:0:0 |
| NM_005037:NM_138712:NM_001330615:NM_015869:NM_138711:NM_001354667:NM_001354666:NM_001354670:NM_001354668 | 16:29:1:5:18:2:0:0:0 | 0:0:0:0:0:0:0:0:0 | 55:99:1:2:531:29:1:3:0 | 0:0:0:0:1:0:0:0 | 53:141:1:5:803:50:0:0 | 0:0:0:0:0:0:0:0:0 | 266:453:2:17:1440:63:1:3:1 | 0:1:0:0:1:0:0:0:0 | 30:55:3:1:176:11:0:0:0 | 0:0:0:0:0:0:0:0 |
| NM_005037:NM_138712:NM_001330615:NM_015869:NM_138711:NM_001354667:NM_001354666:NM_001354670:NM_001354668 | 16:29:1:5:18:2:0:0:0 | 0:0:0:0:0:0:0:0:0 | 55:99:1:2:530:29:1:3:0 | 0:0:0:0:0:0:0:0 | 52:143:1:5:802:50:0:0 | 1:1:0:0:1:0:0:0:0 | 265:459:2:17:1438:63:1:3:1 | 0:0:0:0:2:0:0:0:0 | 29:56:3:1:176:11:0:0:0 | 1:0:0:0:0:0:0:0 |
| NM_005037:NM_138712:NM_001330615:NM_015869:NM_138711:NM_001354667:NM_001354666:NM_001354670:NM_001354668 | 16:30:1:5:18:2:0:0:0 | 0:0:0:0:0:0:0:0:0 | 55:99:1:2:530:29:1:3:0 | 0:0:0:0:1:0:0:0 | 53:145:1:5:802:50:0:0 | 0:0:0:0:0:0:0:0:0 | 265:459:2:17:1438:63:1:3:1 | 0:1:0:0:2:0:0:0:0 | 30:56:3:1:176:11:0:0:0 | 0:0:0:0:0:0:0:0 |
| NM_005037:NM_138712:NM_001330615:NM_015869:NM_138711:NM_001354667:NM_001354666:NM_001354670:NM_001354668:NM_001354669 | 16:30:1:5:18:2:0:0:0:0 | 0:0:0:0:0:0:0:0:0:0 | 55:99:1:2:530:29:1:3:0:0 | 0:0:0:0:0:0:0:0:0 | 53:143:1:5:798:50:0:0:0:0 | 0:1:0:0:1:0:0:0:0 | 265:459:2:17:1433:63:1:3:1:0 | 0:0:0:0:2:0:0:0:0 | 30:56:3:1:175:11:0:0:0:0 | 0:0:0:0:0:0:0:0:0 |
| NM_005037:NM_138712:NM_015869:NM_138711:NM_001354667:NM_001354666:NM_001354669 | 16:30:5:18:2:0:0 | 0:0:0:0:0:0:0 | 55:98:2:528:28:1:7 | 0:0:0:0:0:0:0 | 52:145:5:794:50:0:1 | 0:0:0:0:0:0:0 | 264:457:17:1423:62:1:11 | 1:1:0:0:0:0:0 | 30:56:1:174:11:0:0 | 0:0:0:0:0:0:0 |
| NM_005037:NM_138712:NM_001330615:NM_015869:NM_138711:NM_001354667:NM_001354666:NM_001354669 | 16:31:1:5:18:2:0:0 | 0:0:0:0:0:0:0:0 | 54:99:2:2:522:27:1:7 | 0:1:0:0:3:0:0:0 | 52:149:1:5:796:50:0:1 | 0:0:0:0:3:0:0:0 | 267:462:2:17:1429:61:1:11 | 0:2:0:0:7:0:0:0 | 30:56:3:1:179:11:0:0 | 0:1:0:0:1:0:0:0 |
| NM_005037:NM_138712:NM_001330615:NM_015869:NM_138711:NM_001354667:NM_001354666:NM_001354669 | 16:31:1:5:18:2:0:0 | 0:0:0:0:0:0:0:0 | 54:99:2:2:525:27:1:7 | 0:0:0:0:0:0:0:0 | 52:149:1:5:799:50:0:1 | 0:0:0:0:0:0:0:0 | 267:464:2:17:1438:61:1:11 | 0:1:0:0:1:0:0:0 | 30:57:4:1:180:11:0:0 | 0:0:0:0:0:0:0:0 |
| XM_017006355:NM_002213:XM_017006353:NM_001354764 | 0:2:1:0 | 1:2:0:0 | 0:0:0:0 | 0:1:0:0 | 0:2:0:0 | 0:1:2:0 | 0:3:1:1 | 0:0:0:0 | 0:2:0:0 | 0:0:0:0 |
| NM_033049 | 0 | 0 | 3 | 2 | 1 | 1 | 0 | 1 | 1 | 1 |
| NM_002880:NM_001354693:XM_011533974:NM_001354692:XM_017006966:NR_148940:NM_001354694:NM_001354695:NM_001354690:NR_148941:NM_001354689 | 42:4:8:1:8:1:0:0:0:0:0 | 0:0:0:0:0:0:0:0:0:0:0 | 14:1:18:0:65:0:1:0:0:0:0 | 0:0:1:0:1:0:0:0:0:0:0 | 93:11:23:3:24:1:0:1:0:0:0 | 0:0:0:0:0:0:0:0:0:0:0 | 94:10:79:2:286:1:0:1:1:0:0 | 0:0:0:0:1:0:0:0:0:0:0 | 77:10:13:2:17:0:1:0:1:1:0 | 0:0:0:0:0:0:0:0:0:0:0 |
| NM_002880:NM_001354693:XM_011533974:NM_001354692:XM_017006966:NR_148940:XR_001740227:NM_001354694:NR_148942:NM_001354689:NM_001354695:NM_001354690:NR_148941 | 39:4:8:1:2:1:1:0:0:0:0:0:0 | 0:0:0:0:0:0:0:0:0:0:0:0:0 | 12:1:22:0:21:0:0:1:3:1:0:0:0 | 0:0:0:0:0:0:0:0:0:0:0:0:0 | 94:11:24:3:10:0:0:0:1:0:1:0:0 | 0:0:0:0:0:0:0:0:0:0:0:0:0 | 88:10:84:2:44:1:0:0:3:1:1:1:0 | 0:0:0:0:0:0:0:0:0:0:0:0:0 | 73:10:14:2:7:0:0:1:0:0:0:1:1 | 1:0:0:0:0:0:0:0:0:0:0:0:0 |
| XM_024453325:XM_011512355:XM_017005614:NM_012190:NR_072979 | 0:0:0:0:0 | 0:0:0:0:0 | 0:0:0:0:0 | 0:0:0:0:0 | 3:2:1:1:0 | 0:0:0:0:0 | 37:0:3:12:2 | 0:0:0:0:0 | 5:1:1:2:1 | 0:0:0:0:0 |
| XM_024453325:XM_011512355:XM_017005614:NM_012190:NR_072979 | 0:0:0:0:0 | 0:0:0:0:0 | 0:0:0:0:0 | 0:0:0:0:0 | 1:0:0:0:0 | 2:2:1:1:0 | 14:0:3:4:1 | 22:0:0:8:1 | 1:1:0:0:1 | 4:0:1:2:0 |
| XM_024453325:XM_011512355:NM_012190:NR_072979:XM_017005614:XM_006713481 | 0:0:0:0:0:0 | 0:0:0:0:0:0 | 0:0:0:0:0:0 | 0:0:0:0:0:0 | 1:2:1:0:0:0 | 0:0:0:0:0:0 | 7:0:6:1:5:0 | 0:0:0:0:0:0 | 2:1:2:0:0:0 | 0:0:0:0:0:0 |
| NM_001284408:NM_001284406 | 0:00 | 0:00 | 0:00 | 0:00 | 0:00 | 0:00 | 1:00 | 2:01 | 0:00 | 0:00 |
| NM_001284408:NM_001284406 | 0:00 | 0:00 | 0:00 | 0:00 | 0:00 | 0:00 | 1:00 | 2:01 | 0:00 | 0:00 |
| NM_000994:NM_001007074:NM_001007073 | 64:00:00 | 0:00:00 | 23:01:00 | 0:00:00 | 42:00:00 | 0:00:00 | 136:02:01 | 2:00:00 | 125:02:00 | 0:00:00 |
| NM_001127195:NM_001127196:NM_003418:NM_001127193:NM_001127194:NM_001127192 | 129:440:460:157:31:5 | 1:1:0:0:0:0 | 119:279:239:76:23:4 | 0:0:0:0:0:0 | 684:1730:1487:575:111:40 | 1:7:5:2:1:0 | 716:2105:3105:1003:195:65 | 1:3:4:0:0:0 | 192:509:617:178:40:9 | 0:0:1:0:0:0 |
| NM_001127195:NM_003418:NM_001127193:NM_001127196:NM_001127194:NM_001127192 | 128:460:157:441:31:5 | 1:0:0:0:0:0 | 119:238:75:278:23:4 | 0:0:1:1:0:0 | 683:1485:574:1733:112:40 | 3:6:2:5:0:0 | 716:3105:1001:2103:195:65 | 1:3:2:5:0:0 | 192:616:178:509:40:9 | 0:1:0:1:0:0 |
| NM_003418:NM_001127193:NM_001127196:NM_001127194:NM_001127195:NM_001127192 | 459:157:441:31:130:5 | 0:0:0:0:0:0 | 238:76:279:23:119:4 | 0:0:0:0:0:0 | 1484:575:1736:112:685:40 | 5:2:3:0:1:0 | 3101:1003:2105:195:717:65 | 5:0:3:0:0:0 | 616:177:510:39:192:9 | 1:1:0:1:0:0 |
| NM_003418:NM_001127193:NM_001127196:NM_001127194:NM_001127195:NM_001127192 | 456:157:442:31:131:5 | 0:0:0:0:0:0 | 235:76:278:23:119:4 | 0:0:2:0:0:0 | 1475:575:1737:113:686:41 | 4:4:4:0:2:0 | 3081:1005:2103:195:717:65 | 8:0:5:0:0:0 | 611:179:510:40:192:9 | 4:0:0:0:0:0 |
| NM_003418:NM_001127193:NM_001127196:NM_001127194:NM_001127195:NM_001127192 | 446:157:444:31:131:5 | 0:0:0:0:0:0 | 224:76:281:23:120:4 | 0:0:0:0:0:0 | 1423:580:1745:114:684:41 | 3:2:4:0:3:0 | 3019:1007:2112:195:722:65 | 5:0:1:0:1:0 | 608:179:511:40:193:9 | 0:0:0:0:0:0 |
| NM_001127196:NM_001127193:NM_003418:NM_001127194:NM_001127195:NM_001127192 | 442:156:445:31:131:5 | 2:1:0:0:0:0 | 280:76:224:23:119:4 | 1:0:0:0:1:0 | 1745:581:1420:115:688:41 | 4:1:4:0:1:0 | 2109:1005:3016:195:723:64 | 3:2:3:0:1:1 | 510:179:606:40:193:9 | 1:0:1:0:0:0 |
| NM_003418:NM_001127193:NM_001127196:NM_001127194:NM_001127195:NM_001127192 | 445:157:444:31:131:5 | 0:0:0:0:0:0 | 224:76:281:23:120:4 | 0:0:0:0:0:0 | 1423:581:1743:114:689:41 | 1:1:6:1:0:0 | 3010:1004:2110:195:724:65 | 8:3:2:0:0:0 | 605:179:511:40:193:9 | 1:0:0:0:0:0 |
| NM_003418:NM_001127193:NM_001127196:NM_001127194:NM_001127195:NM_001127192 | 447:157:445:31:130:5 | 0:0:0:0:0:0 | 222:75:282:23:120:4 | 0:1:0:0:0:0 | 1419:581:1751:115:682:41 | 3:0:3:0:1:0 | 3006:1007:2117:194:720:65 | 1:0:2:0:1:0 | 605:179:513:39:191:9 | 0:0:0:0:2:0 |
| NM_001127196:NM_003418:NM_001127193:NM_001127194:NM_001127195:NM_001127192 | 442:446:157:31:130:5 | 2:0:0:0:0:0 | 282:219:76:23:118:4 | 0:2:0:0:0:0 | 1748:1403:580:113:680:41 | 4:4:0:0:0:0 | 2118:2971:1001:189:714:65 | 1:3:3:0:1:0 | 512:603:179:39:191:9 | 1:0:0:0:1:0 |
| NM_003418:NM_001127193:NM_001127196:NM_001127194:NM_001127195:NM_001127192 | 446:157:444:31:129:5 | 0:0:0:0:0:0 | 220:76:282:23:118:4 | 0:0:0:0:0:0 | 1401:578:1746:113:679:40 | 5:1:4:0:0:0 | 2964:1003:2115:189:713:65 | 6:0:3:0:1:0 | 603:179:513:39:191:9 | 0:0:0:0:0:0 |
| NM_007027:XM_017005637:XM_017005636:XR_001739988 | 1:1:0:0 | 0:0:0:0 | 0:1:0:0 | 0:0:0:0 | 0:0:0:0 | 0:0:1:0 | 0:1:0:0 | 0:8:0:2 | 0:4:0:0 | 1:3:0:0 |
| XM_017005636:XM_017005637 | 0:00 | 0:00 | 0:00 | 0:00 | 0:00 | 1:00 | 0:00 | 0:00 | 0:00 | 0:01 |
| NM_001004019:NM_001165035:XM_006713026:NM_001998 | 1:0:0:0 | 0:0:0:0 | 0:0:0:0 | 0:0:0:0 | 0:2:0:0 | 0:0:0:0 | 0:0:0:0 | 0:0:0:0 | 0:0:0:0 | 0:0:0:0 |
| NM_001004019:NM_001998:XM_006713026:NM_001165035 | 0:1:0:1 | 1:1:1:0 | 0:0:0:0 | 0:0:0:0 | 0:3:0:1 | 0:2:0:4 | 0:0:0:0 | 0:8:3:0 | 0:0:0:0 | 0:0:0:0 |
| NM_000532:NM_001178014:XM_011512873 | 17:02:01 | 0:00:00 | 23:05:02 | 0:00:00 | 43:07:02 | 0:00:00 | 90:16:03 | 0:00:00 | 52:07:00 | 0:00:00 |
| NM_005862:XR_001739978:XM_017005523:XM_011512329:XM_017005524:XM_011512331 | 3:2:0:0:0:0 | 0:0:0:0:0:0 | 1:8:1:0:0:0 | 0:3:0:0:0:0 | 1:21:0:0:0:0 | 0:2:0:0:0:0 | 6:39:0:1:0:0 | 2:1:0:0:0:0 | 1:11:0:0:0:0 | 0:1:0:0:0:0 |
| NM_005862:XR_001739978:XM_017005523:XM_011512329:XM_017005525:XM_011512331:XM_017005524 | 3:2:1:0:0:0:0 | 0:0:0:0:0:0:0 | 1:12:2:0:0:0:0 | 0:0:0:0:0:0:0 | 1:26:1:8:0:0:0 | 0:0:0:0:0:0:0 | 8:39:2:27:0:2:0 | 0:0:0:0:1:0:0 | 1:14:1:1:0:0:0 | 0:0:0:0:0:0:0 |
| XR_001739978:NM_005862:XM_017005523:XM_011512331:XM_011512329:XM_017005525 | 4:3:1:0:0:0 | 1:0:0:0:0:0 | 27:1:2:2:0:0 | 6:0:0:1:0:0 | 38:1:1:1:7:0 | 0:0:0:0:0:0 | 68:8:2:2:26:1 | 12:0:0:2:0:1 | 19:1:1:1:1:0 | 2:0:0:0:0:0 |
| NM_006153:XM_024453538:NM_001291999 | 33:02:02 | 0:00:00 | 19:08:03 | 0:01:00 | 49:06:05 | 0:00:00 | 160:28:25 | 1:00:01 | 28:08:06 | 2:00:00 |
| NM_006153:NM_001291999:XM_024453538 | 33:02:02 | 0:00:00 | 17:02:07 | 0:00:00 | 48:05:03 | 0:00:00 | 82:13:11 | 71:11:10 | 13:03:00 | 13:03:00 |
| XM_017006095:XM_024453441:NM_001267042:XR_001740087:XM_006713566:NM_015396:XM_017006098:NM_001267041:XM_017006099:XR_002959506:XM_006713565:XR_002959505:XM_024453442:XM_017006097:XM_011512632:XM_017006100 | 1:1:1:1:0:0:0:0:0:0:0:0:0:0:0:0 | 0:0:0:0:0:0:0:0:0:0:0:0:0:0:0:0 | 1:4:0:3:1:1:1:1:1:3:0:0:0:0:0:0 | 0:0:0:0:0:0:0:0:0:0:0:0:0:0:0:0 | 0:0:0:2:0:5:0:1:0:0:4:0:0:0:0:0 | 0:0:0:0:0:0:0:0:0:0:0:0:0:0:0:0 | 1:17:0:11:1:3:0:0:0:5:3:0:3:0:0:0 | 0:20:1:12:1:2:0:2:0:4:0:1:6:0:0:0 | 1:0:0:2:1:5:0:0:0:0:1:1:0:0:0:0 | 0:0:0:2:0:1:0:0:0:0:0:0:0:0:0:0 |
| NM_001320598:NM_024491:XM_017007271:NM_001288964:XM_017007283:NM_001288965:XM_017007275:NM_001320600:XM_024453781:XM_017007278 | 1:0:0:0:0:0:0:0:0:0 | 0:0:0:0:0:0:0:0:0:0 | 0:0:0:1:0:0:0:0:0:0 | 0:1:1:0:0:0:0:0:0:0 | 0:3:0:0:0:1:0:0:0:0 | 0:2:0:0:1:0:0:0:0:0 | 2:6:0:0:0:0:0:0:1:0 | 0:6:0:0:0:1:1:1:0:0 | 1:1:0:0:0:0:0:0:0:0 | 1:2:0:0:0:0:0:0:0:0 |

| | | | | | | | | | | |
|---|---|---|---|---|---|---|---|---|---|---|
| NM_004766:XM_024453831:NR_023350 | 86:13:25 | 0:00:00 | 9:40:37 | 0:00:00 | 92:09:19 | 0:00:00 | 94:394:275 | 1:01:01 | 93:15:34 | 0:00:00 |
| NM_004766:XM_024453831:NR_023350 | 86:13:25 | 0:00:00 | 9:39:37 | 0:00:00 | 92:09:19 | 0:00:00 | 94:385:274 | 0:00:00 | 91:15:34 | 2:00:00 |
| NM_004766:XM_024453831:NR_023350 | 86:13:25 | 0:00:00 | 9:39:37 | 0:00:00 | 92:08:19 | 0:00:00 | 94:377:274 | 0:01:00 | 94:14:33 | 0:00:00 |
| NM_004766:XM_024453831:NR_023350 | 85:10:25 | 0:00:00 | 11:31:36 | 0:00:00 | 92:04:17 | 0:00:00 | 97:300:259 | 0:00:00 | 94:09:33 | 0:00:00 |
| NM_004766:XM_024453831:NR_023350 | 87:09:24 | 0:00:00 | 11:21:35 | 0:00:00 | 92:03:18 | 0:00:00 | 98:177:224 | 0:02:00 | 95:06:33 | 0:00:00 |
| NM_004766:XM_024453831:NR_023350 | 87:09:24 | 0:00:00 | 11:19:35 | 0:01:00 | 91:03:18 | 1:00:00 | 98:171:221 | 0:02:02 | 95:06:33 | 0:00:00 |
| NM_004766:XM_024453831:NR_023350 | 90:09:24 | 12:00:00 | 14:16:35 | 4:01:00 | 94:02:18 | 5:00:01 | 107:130:204 | 19:02:01 | 97:06:33 | 7:00:00 |
| NM_004766:XM_024453831:NR_023350 | 104:03:22 | 0:00:00 | 19:00:26 | 0:00:00 | 98:01:17 | 2:00:00 | 139:4:149 | 1:00:00 | 103:05:25 | 1:00:00 |
| NM_004766:XM_024453831:NR_023350:XM_017007513 | 105:4:19:0 | 0:0:0:0 | 19:1:19:1 | 0:0:0:0 | 100:3:16:0 | 1:0:0:0 | 144:6:93:0 | 0:0:0:0 | 105:3:22:0 | 0:0:0:0 |
| XM_017006058:XM_005247261:XM_024453431:NM_001080412:XM_024453433:XM_005247257:XM_017006050:XM_017006057:XM_024453434:NM_001350099:XM_005247258:NM_001350100 | 3:1:0:0:0:0:0:0:0:0:0:0 | 0:0:0:0:0:0:0:0:0:0:0 | 1:7:5:2:4:1:1:1:7:0:0:0 | 0:0:0:0:0:0:0:0:0:0:0 | 0:1:5:0:0:0:0:0:0:0:0:0 | 0:0:0:0:0:0:0:0:0:0:0 | 7:14:38:0:38:0:3:3:7:1:1:1 | 1:0:6:0:1:0:0:0:2:0:0:0 | 4:2:12:0:8:0:0:0:3:0:0:0 | 0:0:0:0:0:0:0:0:1:0:0:0 |
| XM_017006058:XM_005247261:XM_024453431:XM_017006057:XM_024453434:XM_024453433:XM_017006050:NM_001350099:NM_001350100:XM_005247258 | 2:1:0:0:0:0:0:0:0:0 | 0:0:0:0:0:0:0:0:0:0 | 0:2:1:1:7:0:0:0:0:0 | 0:0:0:0:0:0:0:0:0:0 | 0:1:2:0:0:0:0:0:0:0 | 0:0:2:0:0:0:0:0:0:0 | 4:5:15:0:11:16:2:1:1:0 | 1:0:0:0:0:1:0:0:0:0 | 4:0:4:0:5:4:0:0:0:0 | 0:1:5:0:0:3:0:0:0:0 |
| XM_017006058:XM_005247261:XM_024453431:XM_017006057:XM_024453434:XM_024453433 | 1:1:0:0:0:0 | 0:0:0:0:0:0 | 0:2:1:1:9:0 | 0:0:0:0:0:0 | 0:1:1:0:0:0 | 0:0:0:0:0:0 | 3:2:0:0:41:2 | 0:0:0:0:0:0 | 1:0:1:0:6:3 | 0:0:0:0:0:0 |
| NM_004628:XR_002959581:XR_002959580 | 0:00:00 | 10:04:00 | 0:00:00 | 9:02:00 | 0:00:00 | 13:01:00 | 13:11:00 | 18:12:00 | 5:02:00 | 4:02:00 |
| NM_004628:XR_002959581 | 3:00 | 4:00 | 1:00 | 4:00 | 5:00 | 6:00 | 6:01 | 7:01 | 4:01 | 4:00 |
| NM_004628:XR_002959581 | 7:00 | 0:00 | 5:00 | 0:00 | 11:00 | 0:00 | 6:01 | 7:01 | 4:01 | 4:00 |
| NM_004628:XR_002959581 | 4:00 | 3:00 | 3:00 | 1:00 | 6:00 | 5:00 | 12:03 | 0:00 | 8:01 | 0:00 |
| XR_002959581:NM_004628:NR_148951:NR_148950:NM_001354729 | 8:8:1:0:0 | 2:0:0:0:0 | 20:0:0:0:0 | 2:0:0:0:0 | 12:10:1:1:0 | 1:0:0:0:0 | 71:6:12:2:0 | 12:0:1:0:0 | 9:8:0:0:0 | 2:0:1:0:0 |
| XM_017007097:XM_011513107:XM_017007100:XM_011513106:XM_005247731:NM_001178138:NM_001178139:NM_006286:XM_011513111:XM_024453729:XM_017007091:XM_024453730:XM_024453726:XM_011513109:XM_024453728:XM_017007098:XM_024453725:XM_011513102:XM_011513101:NM_001178141:XM_017007102:XM_024453727 | 1:0:1:0:1:1:2:6:1:1:0:0:0:0:0:0:0:0:0:0:0:0 | 1:1:1:1:0:0:0:0:0:0:0:0:0:0:0:0:0:0:0:0:0:0 | 0:0:0:0:2:0:0:2:0:2:1:0:0:0:0:0:0:0:0:0:0:0 | 2:0:4:0:0:2:0:0:0:3:1:0:0:0:0:0:0:0:0:0:0:0 | 0:0:0:0:0:0:3:1:0:1:0:1:0:0:0:0:0:0:0:0:0:0 | 1:2:0:0:0:1:0:0:0:0:0:0:0:0:0:0:0:0:0:0:0:0 | 9:0:4:1:4:3:25:59:0:34:12:4:0:1:1:2:7:2:1:2:1:0 | 7:13:20:2:0:5:1:1:0:0:2:5:0:1:0:0:0:0:0:0:0:0 | 2:0:0:1:2:0:9:7:0:2:1:0:0:0:0:0:0:0:0:0:0:0 | 1:0:4:0:0:0:1:0:0:1:1:0:0:0:0:1:0:0:0:0:0:0 |
| XM_017007097:XM_017007100:XM_005247731:NM_001178138:NM_001178139:NM_006286:XM_011513111:XM_024453729:XM_011513107:XM_011513106:XM_017007091:XM_024453730:XM_011513109:XM_024453728:XM_017007098:XM_024453725:XM_011513102:XM_011513101:NM_001178141:XM_017007102:XM_024453726:XM_024453727 | 2:2:1:1:2:6:1:1:1:1:0:0:0:0:0:0:0:0:0:0:0:0 | 1:1:0:0:0:0:0:0:0:0:0:0:0:0:0:0:0:0:0:0:0:0 | 2:4:2:2:0:2:0:5:0:0:2:0:0:0:0:0:0:0:0:0:0:0 | 0:0:0:0:0:0:0:0:0:0:0:0:0:0:0:0:0:0:0:0:0:0 | 1:0:0:1:3:1:0:1:2:0:0:1:0:0:0:0:0:0:0:0:0:0 | 1:0:0:0:0:0:0:0:0:0:0:0:0:0:0:0:0:0:0:0:0:0 | 15:27:4:8:26:70:0:36:13:3:17:4:1:1:3:7:2:1:2:1:1:0 | 1:0:0:0:0:0:0:0:0:0:0:0:0:0:0:0:0:0:0:0:0:0 | 3:5:2:0:10:7:0:3:0:1:2:0:0:0:1:0:0:0:0:0:0:0 | 0:2:0:0:0:0:0:0:0:0:0:0:0:0:0:0:0:0:0:0:0:0 |
| XR_001740179:XR_924148 | 0:00 | 0:00 | 0:00 | 0:00 | 0:00 | 0:00 | 2:00 | 0:00 | 1:00 | 0:00 |
| XR_924148:NM_001184:XM_011512925:NM_001354579 | 1:0:0:0 | 0:0:0:0 | 0:0:0:0 | 1:1:0:0 | 2:0:0:0 | 1:0:0:0 | 2:0:0:0 | 3:0:0:0 | 1:0:0:0 | 2:0:0:0 |
| NM_176894 | 3 | 1 | 1 | 3 | 4 | 2 | 0 | 1 | 0 | 0 |
| NM_001130156:NM_001130157:NM_022443 | 14:03:19 | 0:00:00 | 2:01:09 | 0:00:00 | 4:00:05 | 0:00:00 | 64:9:112 | 1:00:01 | 11:02:10 | 0:00:00 |
| NM_001130156:NM_022443:NM_001130157 | 5:07:03 | 9:12:00 | 0:03:01 | 2:06:00 | 4:02:00 | 0:03:00 | 0:00:00 | 63:112:9 | 0:00:00 | 11:10:02 |
| NM_024996 | 0 | 0 | 0 | 0 | 0 | 2 | 0 | 0 | 0 | 0 |
| NM_020169 | 0 | 0 | 0 | 1 | 0 | 0 | 0 | 0 | 0 | 2 |
| XM_005247686:NM_206963:NM_002888:XR_001740228 | 8:27:2:0 | 0:0:0:0 | 23:94:7:0 | 0:0:0:0 | 26:103:5:0 | 0:0:0:0 | 21:101:9:0 | 0:0:0:0 | 8:33:6:0 | 0:0:0:0 |
| NM_020800:NM_001190241 | 0:00 | 0:00 | 1:00 | 0:00 | 1:02 | 0:00 | 0:01 | 0:00 | 2:01 | 0:00 |
| XM_024453455:XM_005247364:NM_014498:XM_017006183:XM_005247365:NM_001308155 | 9:1:8:5:1:1 | 0:0:0:0:0:0 | 42:3:14:6:0:0 | 11:2:5:2:0:0 | 13:2:10:4:1:1 | 0:0:0:0:0:0 | 45:16:36:12:5:6 | 2:0:3:2:0:3 | 9:1:8:1:1:1 | 0:0:1:0:0:0 |
| XM_024453455:XM_005247364:NM_014498:XM_017006183:XM_005247365:NM_001308155 | 11:1:9:5:1:1 | 0:0:0:0:0:0 | 64:6:21:9:0:0 | 0:0:0:0:0:0 | 12:2:11:4:1:1 | 0:0:0:0:0:0 | 46:17:45:14:5:10 | 0:0:0:0:0:0 | 9:1:8:1:1:1 | 0:0:1:0:0:0 |
| XM_005247364:NM_014498:XM_024453455:XM_017006183:XM_005247365:NM_001308155 | 1:9:5:5:1:1 | 0:0:0:0:0:0 | 6:21:29:8:0:0 | 0:0:0:0:0:0 | 2:11:7:4:1:1 | 0:0:0:0:0:0 | 17:46:29:13:5:10 | 0:0:0:0:0:0 | 1:9:6:1:1:1 | 0:0:0:0:0:0 |
| NM_005241:XM_017005874:XM_017005877:XM_005247225:XM_005247214:XM_005247224:NM_001164000:XM_011512546:XM_005247213:NM_001105078:NM_004991 | 1:2:2:2:1:0:0:0:0:0:0 | 0:0:0:0:0:0:0:0:0:0:0 | 0:22:11:1:0:4:0:0:0:0:0 | 0:0:0:1:0:0:0:0:0:0:0 | 0:11:16:9:0:5:2:8:1:2:0 | 0:0:0:0:0:0:0:0:0:0:0 | 1:63:41:22:0:8:1:3:0:0:0 | 0:0:1:0:0:0:0:0:0:0:0 | 0:20:25:11:0:4:0:11:0:1:1 | 0:0:0:0:0:0:0:0:0:0:0 |
| NM_005241:XM_017005874:XM_017005877:XM_005247225:XM_005247214:XM_005247213:XM_005247224:NM_001164000:XM_011512546:NM_001105078:NM_004991 | 1:2:2:2:1:1:0:0:0:0:0 | 0:0:0:0:0:0:0:0:0:0:0 | 0:22:9:3:0:0:3:0:0:0:0 | 0:0:0:0:0:0:1:0:0:0:0 | 0:11:16:9:0:1:5:2:9:2:0 | 0:0:0:0:0:0:0:0:0:0:0 | 0:63:37:22:0:0:8:1:4:0:0 | 0:0:2:0:0:0:0:0:0:0:0 | 0:20:24:11:0:0:4:0:12:1:1 | 0:0:0:0:0:0:0:0:0:0:0 |
| NM_005241:XM_005247214:XM_005247213:XM_011512546:XM_017005874:NM_001164000:NM_001105078:NM_004991 | 1:1:1:0:0:0:0:0 | 0:0:0:0:0:0:0:0 | 0:0:0:3:7:0:0:0 | 0:0:0:1:0:0:0:0 | 0:0:1:11:5:2:2:0 | 0:0:0:1:0:0:0:0 | 0:0:0:10:9:2:0:0 | 0:0:0:1:1:0:0:0 | 0:0:0:13:12:0:1:1 | 0:0:0:2:0:0:0:0 |
| XM_011512546:NM_005241:XM_005247214:XM_005247213:NM_001164000:NM_001105078:XM_017005874:XM_005247221:NM_004991 | 0:1:1:1:0:0:0:0:0 | 1:0:0:0:0:0:0:0:0 | 5:0:0:0:0:0:6:0:0 | 7:0:1:0:1:1:1:0:0 | 12:0:0:1:2:2:5:0:0 | 3:0:0:0:0:0:0:0:0 | 14:0:0:0:2:0:7:0:0 | 48:0:1:0:1:0:1:1:0 | 17:0:0:0:0:1:11:0:1 | 7:0:0:0:0:0:0:0:0 |
| NM_005241:XM_005247214:XM_005247213:XM_011512546:NM_001164000:NM_001105078:XM_017005874:XM_005247221:NM_004991:XM_017005877 | 1:1:1:1:0:0:0:0:0:0 | 0:0:0:0:0:0:0:0:0:0 | 0:0:0:7:0:0:6:0:0:0 | 0:1:0:5:2:1:1:0:0:0 | 0:0:1:12:2:2:5:0:0:0 | 0:0:0:3:0:0:0:0:0:0 | 0:0:0:22:2:0:7:0:0:0 | 0:1:0:40:1:0:1:1:0:0 | 0:0:0:17:0:1:11:0:1:0 | 0:0:0:7:0:0:0:0:0:0 |
| NM_005241:XM_005247214:XM_005247213:XM_011512546:NM_001164000:XM_017005874:NM_001105078:XM_005247221:NM_004991 | 1:1:1:1:0:0:0:0:0 | 0:0:0:0:0:0:0:0:0 | 0:1:0:7:1:7:1:0:0 | 0:0:0:6:1:0:0:0:0 | 0:0:1:13:2:5:2:0:0 | 0:0:0:2:0:0:0:0:0 | 0:1:0:46:2:7:0:1:0 | 0:0:0:23:1:1:0:0:0 | 0:0:0:19:0:11:1:0:1 | 0:0:0:6:0:0:0:0:0 |
| XM_005247225:XM_005247214:XM_017005877:XM_005247213:XM_011512546:XM_005247224:XM_017005874:NM_001164000:NM_001105078:XM_005247221:NM_004991 | 2:1:1:1:1:0:0:0:0:0:0 | 0:0:0:0:0:0:0:0:0:0:0 | 3:1:5:0:17:4:2:2:1:0:0 | 0:0:0:0:0:0:0:0:0:0:0 | 7:0:7:1:15:5:1:2:2:0:0 | 0:0:0:0:0:0:0:0:0:0:0 | 21:1:5:0:78:8:3:3:0:1:0 | 0:0:0:0:0:0:0:0:0:0:0 | 11:0:3:0:25:4:3:0:1:0:1 | 0:0:0:0:0:0:0:0:0:0:0 |
| XM_005247225:XM_005247213:XM_017005877:XM_005247214:NM_005241:NM_004991:XM_005247224:XM_017005874 | 1:1:1:1:0:0:0:0 | 0:0:0:0:0:0:0:0 | 0:0:2:1:1:0:0:0 | 0:0:0:0:0:0:0:0 | 1:1:2:0:0:0:0:0 | 0:0:0:0:0:0:0:0 | 0:0:0:1:0:0:0:0 | 0:0:0:0:0:0:0:0 | 1:0:4:0:0:1:0:0 | 0:0:0:0:0:0:0:0 |
| NM_001172780:NM_001172779:NM_153353 | 1:00:00 | 0:00:00 | 0:00:00 | 0:00:00 | 0:00:00 | 0:00:00 | 1:00:00 | 0:00:00 | 0:00:00 | 0:00:00 |
| NM_001172780:NM_001172779 | 1:00 | 0:00 | 0:00 | 0:00 | 0:00 | 0:00 | 1:00 | 0:00 | 0:00 | 0:00 |
| NM_003810:NR_033994:NM_001190942 | 53:05:00 | 0:00:00 | 623:43:01 | 2:01:00 | 422:13:00 | 2:00:00 | 293:22:00 | 1:01:00 | 90:04:00 | 0:00:00 |
| NR_033994:NM_003810:NM_001190942 | 5:57:00 | 1:00:00 | 37:641:1 | 0:39:00 | 0.849305556 | 0:05:00 | 0.959722222 | 0:15:00 | 0.277083333 | 0:02:00 |
| NM_003810:NR_033994:NM_001190942:NM_001190943 | 59:6:0:0 | 0:0:0:0 | 713:38:1:0 | 13:0:0:0 | 464:12:0:0 | 5:0:0:0 | 347:18:0:0 | 4:0:0:0 | 102:5:0:0 | 1:0:0:0 |
| NM_003810:NR_033994:NM_001190943 | 57:06:00 | 2:00:00 | 743:27:00 | 0:00:00 | 473:10:00 | 0:00:00 | 357:09:00 | 0:00:00 | 103:03:00 | 0:00:00 |
| NM_020792:NM_001146276 | 0:00 | 0:00 | 0:00 | 0:00 | 0:00 | 3:00 | 0:00 | 0:00 | 0:00 | 2:00 |
| XM_024453752:NM_001321194:XM_005247775:NM_024665:NM_001321195:XM_006713746:XM_011513142:XM_024453751:XM_011513141:XM_011513143:NM_001321193:XM_017007185 | 8:4:7:20:1:1:0:0:0:0:0:0 | 0:0:0:0:0:0:0:0:0:0:0:0 | 16:0:1:8:1:1:2:1:1:0:0:0 | 0:0:0:0:0:0:0:0:0:0:0:0 | 23:2:4:27:1:0:0:0:0:2:1:0 | 0:0:0:0:0:0:0:0:0:0:0:0 | 80:9:13:42:8:0:1:0:0:0:6:1 | 0:0:0:0:0:0:0:0:0:0:0:0 | 16:5:8:27:1:0:2:0:1:1:2:1 | 0:0:0:0:0:0:0:0:0:0:0:0 |
| XM_024453752:NM_001321194:XM_005247775:NM_024665:NM_001321195:XM_006713746:XM_011513142:XM_024453751:XM_011513141:XM_017007185:XM_011513143:NM_001321193 | 3:4:8:25:2:1:0:0:0:0:0:0 | 0:0:0:0:0:0:0:0:0:0:0:0 | 10:2:5:25:4:1:3:1:1:1:0:0 | 0:0:0:0:0:0:0:0:0:0:0:0 | 8:7:6:45:2:2:1:0:0:1:2:3 | 0:0:0:0:0:0:0:0:0:0:0:0 | 22:16:38:109:14:0:6:1:0:2:0:10 | 0:0:0:1:0:0:0:0:0:0:0:0 | 9:6:11:42:2:0:2:0:1:1:1:3 | 0:0:0:0:0:0:0:0:0:0:0:0 |
| NM_014398:XM_005247360:XM_024453453 | 0:00:00 | 2:01:00 | 0:00:00 | 1:01:00 | 0:00:00 | 0:00:00 | 3:02:02 | 0:02:01 | 1:00:00 | 0:00:00 |
| NM_018622:XM_024453633:NM_001324436:XM_017006803:NM_001037639 | 0:0:0:0:0 | 0:0:0:0:0 | 1:0:0:0:0 | 0:0:0:0:0 | 0:0:0:0:0 | 0:0:0:0:0 | 0:0:0:0:0 | 0:1:1:0:0 | 0:0:0:0:0 | 0:0:0:0:0 |
| NM_004068:NM_001025205 | 244:57:00 | 1:00 | 142:50:00 | 1:00 | 200:83 | 0:00 | 1057:207 | 2:01 | 307:73 | 2:00 |
| NM_004068:NM_001025205 | 246:57:00 | 0:00 | 142:50:00 | 0:00 | 201:84 | 0:00 | 1063:213 | 1:00 | 309:74 | 0:00 |
| NM_004068:NM_001025205 | 244:57:00 | 1:00 | 143:50:00 | 0:00 | 202:84 | 0:00 | 1070:215 | 0:00 | 310:73 | 0:00 |
| NM_004068:NM_001025205 | 250:60 | 0:00 | 150:50:00 | 1:00 | 204:87 | 0:00 | 1111:224 | 2:00 | 316:76 | 1:00 |
| NM_004068:NM_001025205 | 253:60 | 2:00 | 155:50:00 | 3:00 | 208:87 | 0:00 | 1144:223 | 2:00 | 318:76 | 0:00 |
| NM_004068:NM_001025205 | 259:59:00 | 0:00 | 164:50:00 | 0:00 | 210:87 | 1:00 | 1159:222 | 5:01 | 321:76 | 0:00 |
| NM_001025205:NM_004068 | 57:262 | 1:00 | 50:168 | 0:02 | 87:219 | 0:00 | 221:1196 | 1:01 | 76:330 | 0:00 |
| NM_004068:NM_001025205 | 263:58:00 | 1:00 | 167:50:00 | 1:00 | 220:87 | 0:00 | 1197:222 | 1:00 | 330:75 | 0:01 |

| | | | | | | | | | | |
|---|---|---|---|---|---|---|---|---|---|---|
| NM_004068:NM_001025205 | 269:58:00 | 0:00 | 177:50:00 | 0:00 | 232:86 | 0:01 | 1210:221 | 1:00 | 340:76 | 0:00 |
| NM_004068:NM_001025205 | 269:58:00 | 1:00 | 177:50:00 | 0:00 | 231:87 | 1:00 | 1217:220 | 2:00 | 340:76 | 1:00 |
| NM_004068:NM_001025205 | 271:58:00 | 0:00 | 178:50:00 | 0:00 | 234:87 | 0:00 | 1223:219 | 2:00 | 342:76 | 0:00 |
| NM_004068:NM_001025205 | 271:58:00 | 0:00 | 178:50:00 | 0:00 | 234:86 | 0:01 | 1224:219 | 1:00 | 342:76 | 0:00 |
| NM_004068:NM_001025205 | 273:58:00 | 0:00 | 178:50:00 | 0:00 | 234:87 | 0:00 | 1224:219 | 0:00 | 345:76 | 0:00 |
| NM_004068:NM_001025205 | 274:58:00 | 1:00 | 179:50:00 | 1:00 | 234:87 | 1:00 | 1240:219 | 3:00 | 349:76 | 0:00 |
| NM_004068:NM_001025205 | 276:58:00 | 0:00 | 185:50:00 | 1:00 | 238:87 | 0:00 | 1250:219 | 1:00 | 351:76 | 0:00 |
| NM_022149 | 0 | 1 | 0 | 0 | 0 | 1 | 0 | 0 | 0 | 0 |
| XM_024453430:NM_015303:NR_146113 | 1:00:00 | 0:00:00 | 0:00:00 | 0:00:00 | 2:02:00 | 0:02:00 | 2:02:01 | 0:00:00 | 1:00:01 | 0:00:00 |
| XM_024453430:NM_015303:NR_146113 | 1:01:00 | 0:00:00 | 4:00:02 | 0:00:00 | 6:04:04 | 0:00:00 | 1:01:01 | 2:01:00 | 1:00:00 | 0:00:01 |
| NM_004454 | 4 | 0 | 1 | 0 | 0 | 0 | 4 | 0 | 1 | 0 |
| NM_152673 | 0 | 0 | 0 | 1 | 0 | 2 | 0 | 0 | 0 | 0 |
| NM_152673 | 0 | 0 | 0 | 1 | 0 | 2 | 0 | 0 | 0 | 0 |
| NM_152673 | 0 | 0 | 0 | 0 | 2 | 0 | 0 | 0 | 0 | 0 |
| NM_152673 | 0 | 0 | 0 | 0 | 0 | 2 | 0 | 0 | 0 | 0 |
| NM_152673:NM_001282506 | 0:00 | 0:00 | 0:00 | 2:00 | 0:00 | 2:00 | 0:01 | 0:00 | 0:00 | 0:00 |
| NM_152673:NM_001282506 | 0:00 | 0:00 | 0:00 | 2:00 | 0:00 | 2:00 | 0:01 | 0:00 | 0:00 | 0:00 |
| NM_138297 | 0 | 0 | 0 | 0 | 0 | 5 | 0 | 0 | 0 | 0 |
| NM_138297 | 0 | 0 | 0 | 0 | 0 | 1 | 0 | 0 | 0 | 0 |
| NM_001322468:NM_018406 | 0:00 | 0:00 | 0:00 | 2:00 | 0:00 | 1:00 | 0:00 | 0:00 | 0:01 | 0:00 |
| NM_001312673:NM_005017 | 32:03:00 | 0:00 | 26:06:00 | 0:00 | 99:26:00 | 0:00 | 46:12:00 | 0:00 | 30:02:00 | 0:00 |
| NM_001079811:NM_000404:NM_001317040:NM_001135602 | 10:37:7:0 | 0:0:0:0 | 8:43:19:0 | 0:0:1:0 | 18:68:12:0 | 0:0:0:0 | 38:157:250:2 | 0:0:0:0 | 25:70:16:0 | 0:0:0:0 |
| NM_001079811:NM_000404:NM_001317040:NM_001135602 | 10:37:7:0 | 0:0:0:0 | 8:43:20:0 | 0:0:0:0 | 18:68:11:0 | 0:0:1:0 | 38:157:247:2 | 0:0:3:0 | 25:70:16:0 | 0:0:0:0 |
| NM_001079811:NM_000404:NM_001317040:NM_001135602 | 10:37:7:0 | 0:0:0:0 | 8:43:20:0 | 0:0:0:0 | 18:68:11:0 | 0:0:0:0 | 38:157:246:2 | 0:1:4:0 | 25:70:16:0 | 0:0:0:0 |
| NM_001079811:NM_000404:NM_001317040:NM_001135602 | 9:35:7:0 | 0:0:0:0 | 8:39:22:0 | 0:0:0:0 | 17:65:8:0 | 0:0:0:0 | 35:151:237:2 | 0:1:1:0 | 25:69:17:0 | 0:0:0:0 |
| NM_001079811:NM_000404:NM_001317040:NM_001135602 | 2:15:3:0 | 7:19:4:0 | 4:15:13:0 | 4:24:9:0 | 7:26:6:0 | 11:37:2:0 | 35:151:235:2 | 0:0:0:0 | 25:69:17:0 | 0:0:0:0 |
| NM_001079811:NM_000404:NM_001317040:NM_001135602 | 0:0:0:0 | 9:34:7:0 | 0:0:0:0 | 8:39:20:0 | 0:0:0:0 | 18:63:7:0 | 0:0:0:0 | 36:148:236:2 | 0:0:0:0 | 25:69:17:0 |
| NM_001079811:NM_000404:NM_001317040:NM_001135602 | 10:38:5:0 | 0:0:0:0 | 8:43:13:0 | 0:0:0:0 | 19:68:5:0 | 0:0:0:0 | 40:162:156:2 | 1:1:1:0 | 26:70:16:0 | 0:0:0:0 |
| NM_001079811:NM_000404:NM_001317040:NM_001135602 | 11:39:6:0 | 0:0:0:0 | 8:44:10:0 | 0:0:1:0 | 19:72:5:0 | 0:0:0:0 | 44:170:105:2 | 0:0:0:0 | 26:71:14:0 | 0:0:0:0 |
| NM_001079811:NM_000404:NM_001317040:NM_001135602 | 11:39:6:0 | 0:0:0:0 | 8:44:11:0 | 0:0:0:0 | 19:72:5:0 | 0:0:0:0 | 44:167:100:2 | 0:3:2:0 | 26:71:14:0 | 0:0:0:0 |
| NM_000404:NM_001317040:NM_001135602 | 0:00:00 | 40:01:00 | 0:00:00 | 43:01:00 | 0:00:00 | 72:01:00 | 78:18:01 | 93:11:00 | 27:06:00 | 43:03:00 |
| NM_000249:NM_001354628:NM_001258273:NM_001354617:NM_001354619:NM_001354618:NM_001167617:NM_001354629:NM_001354622:XM_017006450:NM_001354627:NM_001354630:NM_001167618:NM_001167619:NM_001354620:NM_001354616:NM_001354624:XM_005265161:NM_001258271:NM_001354621:NM_001258274:NM_001354623 | 17:2:1:1:1:7:0:3:1:1:1:0:0:0:0:0:0:0:0:0:0:0 | 9:2:1:1:1:3:1:1:0:0:0:0:0:0:0:0:0:0:0:0:0:0 | 8:1:4:1:0:2:0:2:0:0:0:1:1:1:0:0:0:0:0:0:0:0 | 9:4:3:0:0:2:0:0:0:0:0:0:0:0:0:0:0:0:0:0:0:0 | 19:2:1:0:2:2:0:0:1:0:0:0:0:0:0:0:0:0:0:0:0 | 13:0:1:1:0:3:0:0:0:0:1:0:0:1:0:0:0:0:0:0:0:0 | 36:9:7:0:9:23:3:10:1:0:4:7:0:0:1:0:1:0:0:0:0:0 | 20:11:14:1:14:8:3:13:0:0:2:4:0:0:1:1:1:0:0:0:0:0 | 29:3:1:0:1:1:0:1:0:0:0:2:0:1:0:0:1:0:0:0:0:0 | 18:3:0:0:6:3:2:0:0:0:0:1:1:1:0:0:0:0:0:0:0:0 |
| NM_000249:NM_001354628:NM_001354629:NM_001354622:NM_001354617:XM_017006450:NM_001258273:NM_001354618:NM_001354619:NM_001167617:NM_001354627:NM_001354630:NM_001167618:NM_001167619:NM_001354620:NM_001354616:NM_001354624:XM_005265161:NM_001258271:NM_001354621:NM_001258274:NM_001354623 | 26:4:4:1:2:1:2:10:2:1:1:0:0:0:0:0:0:0:0:0:0:0:0 | 0:0:0:0:0:0:0:0:0:0:0:0:0:0:0:0:0:0:0:0 | 17:5:2:0:1:0:7:4:0:0:0:1:1:1:0:0:0:0:0:0:0:0 | 0:0:0:0:0:0:0:0:0:0:0:0:0:0:0:0:0:0:0:0:0:0 | 32:2:0:1:1:0:2:5:2:0:1:0:0:1:0:0:0:0:0:0:0:0:0 | 0:0:0:0:0:0:0:0:0:0:0:0:0:0:0:0:0:0 | 56:20:23:1:1:0:21:31:23:6:6:11:0:0:2:1:2:0:0:0:0:0 | 0:0:0:0:0:0:0:0:0:0:0:0:0:0:0:0:0:0 | 47:6:1:0:0:0:1:4:7:2:0:3:1:2:0:0:1:0:0:0:0:0 | 0:0:0:0:0:0:0:0:0:0:0:0:0:0:0:0:0:0:0:0 |
| NM_000249:NM_001354628:NM_001354629:NM_001354622:NM_001354617:XM_017006450:NM_001258273:NM_001354618:NM_001354619:NM_001167617:NM_001354630:NM_001167618:NM_001167619:NM_001354620:NM_001354616:NM_001354623:XM_005265161:NM_001258271:NM_001354621:NM_001258274 | 26:4:4:1:2:1:2:10:2:1:0:0:0:0:0:0:0:0:0:0:0:0 | 0:0:0:0:0:0:0:0:0:0:0:0:0:0:0:0:0 | 17:5:5:0:1:0:6:4:0:0:1:1:1:0:0:0:0:0:0 | 0:0:0:0:0:0:0:0:0:0:0:0:0:0:0:0:0:0:0:0 | 32:2:0:1:1:0:3:5:2:0:0:0:1:0:0:0:0:0:0:0:0 | 0:0:0:0:0:0:0:0:0:0:0:0:0:0:0:0 | 55:22:43:1:1:8:37:30:23:6:11:0:0:2:1:0:0:0:0 | 1:0:0:0:0:0:1:0:0:0:0:0:0:0:0:0 | 47:6:1:0:0:0:2:4:7:2:3:1:2:0:0:0:0:0:0:0 | 0:0:0:0:0:0:0:0:0:0:0:0:0:0:0:0:0:0:0 |
| NM_000249:NM_001354628:NM_001354629:NM_001354622:NM_001354617:XM_017006450:NM_001258273:NM_001354618:NM_001354619:NM_001167617:NM_001354627:NM_001354630:NM_001167618:NM_001167619:NM_001354620:NM_001354616:NM_001354624:NM_001354623:XM_005265161:NM_001258271:NM_001354621:NM_001258274 | 26:4:4:1:2:1:2:10:2:1:1:0:0:0:0:0:0:0:0:0:0:0:0 | 0:0:0:0:0:0:0:0:0:0:0:0:0:0:0:0:0:0:0:0 | 17:5:5:0:1:2:6:4:0:0:0:1:1:1:0:0:0:0:0:0:0:0 | 0:0:0:0:0:0:0:0:0:0:0:0:0:0:0:0:0:0:0:0:0:0 | 32:3:0:1:1:0:7:5:2:0:1:0:0:1:0:0:0:0:0:0:0:0:0 | 0:0:0:0:0:0:0:0:0:0:0:0:0:0:0:0:0:0 | 56:22:43:1:1:29:46:31:22:4:6:9:0:0:2:1:2:0:0:0:0:0 | 0:0:0:0:0:0:1:0:1:0:0:0:0:0:0:0:0:0:0:0:0 | 47:6:1:0:0:1:4:4:6:2:0:3:1:2:0:0:1:0:0:0:0:0 | 0:0:0:0:0:0:0:0:0:0:0:0:0:0:0:0:0:0:0:0 |
| NM_000249:NM_001354628:NM_001354629:NM_001354622:NM_001354617:XM_017006450:NM_001258273:NM_001354618:NM_001354619:NM_001167617:NM_001354627:NM_001354630:NM_001167618:NM_001167619:NM_001354620:NM_001354616:NM_001354624:NM_001354623:XM_005265161:NM_001258271:NM_001354621:NM_001258274 | 26:4:4:1:2:2:2:10:2:1:1:0:0:0:0:0:0:0:0:0:0:0:0 | 0:0:0:0:0:0:0:0:0:0:0:0:0:0:0:0:0:0:0:0 | 17:5:5:0:1:2:7:4:0:0:0:1:1:1:0:0:0:0:0:0:0:0 | 0:0:0:0:0:0:0:0:0:0:0:0:0:0:0:0:0:0:0:0:0:0 | 32:3:0:1:1:0:7:5:2:0:1:0:0:1:0:0:0:0:0:0:0:0:0 | 0:0:0:0:0:0:0:0:0:0:0:0:0:0:0:0:0:0 | 54:24:43:1:1:31:47:30:23:4:6:7:0:0:2:1:2:0:0:0:0:0 | 2:0:0:0:0:0:0:0:0:0:0:0:0:0:0:0:0:0 | 47:6:1:0:0:1:4:4:6:2:0:3:1:2:0:0:1:0:0:0:0:0 | 0:0:0:0:0:0:0:0:0:0:0:0:0:0:0:0:0:0:0:0 |
| NM_000249:NM_001354628:NM_001354629:NM_001354622:NM_001354617:XM_017006450:NM_001258273:NM_001354618:NM_001354619:NM_001167617:NM_001354627:NM_001167618:NM_001167619:NM_001354620:NM_001354630:NM_001354616:NM_001354624:NM_001354623:XM_005265161:NM_001258271:NM_001354621:NM_001258274 | 26:4:4:1:2:3:4:10:2:1:1:0:0:0:0:0:0:0:0:0:0:0:0 | 0:0:0:0:0:0:0:0:0:0:0:0:0:0:0:0:0:0:0:0 | 17:4:5:0:1:2:8:4:0:0:0:1:1:0:0:0:0:0:0:0:0:0 | 0:0:0:0:0:0:0:0:0:0:0:0:0:0:0:0:0:0:0:0:0:0 | 32:3:0:1:1:2:7:5:2:0:1:0:1:0:0:0:0:0:0:0:0:0:0 | 0:0:0:0:0:0:0:0:0:0:0:0:0:0:0:0:0:0 | 56:24:43:1:1:38:49:30:23:5:6:0:0:2:4:1:2:0:0:0:0:0 | 0:0:0:0:1:1:0:0:0:0:0:0:0:0:0:0:0:0 | 47:7:1:0:0:2:5:4:7:2:0:1:2:0:3:0:1:0:0:0:0:0 | 0:0:0:0:0:0:0:0:0:0:0:0:0:0:0:0:0:0:0:0 |
| NM_000249:NM_001354629:NM_001354622:NM_001354617:XM_017006450:NM_001258273:NM_001354618:NM_001354619:NM_001167617:NM_001354627:NM_001167618:NM_001167619:NM_001354620:NM_001354630:NM_001354616:NM_001354624:NM_001354623:XM_005265161:NM_001354621:NM_001258274 | 26:4:1:2:3:4:10:2:1:1:0:0:0:0:0:0:0:0:0:0:0:0 | 0:0:0:0:0:0:0:0:0:0:0:0:0:0:0:0:0 | 17:5:0:1:5:10:4:0:0:0:1:1:0:0:0:0:0:0:0:0 | 0:0:0:0:0:0:0:0:0:0:0:0:0:0:0:0:0:0:0:0 | 32:0:1:1:4:7:4:2:0:1:0:1:0:0:0:0:0:0:0:0 | 0:0:0:0:0:0:0:0:0:0:0:0:0:0:0:0 | 55:42:1:1:49:54:25:22:4:5:0:0:2:2:1:1:0:0:0:0 | 1:0:0:0:0:0:0:0:0:0:0:0:0:0:0:0 | 47:1:0:0:3:6:4:7:2:0:0:2:0:2:0:1:0:0:0:0 | 0:0:0:0:0:0:0:0:0:0:0:0:0:0:0:0:0:0:0 |
| NM_000249:NM_001354628:NM_001354629:NM_001354622:NM_001354617:XM_017006450:NM_001258273:NM_001354618:NM_001354619:NM_001167617:NM_001354627:NM_001167618:NM_001167619:NM_001354620:NM_001354630:NM_001354616:NM_001354624:NM_001354623:XM_005265161:NM_001354621:NM_001258274 | 26:1:4:1:2:3:4:10:2:1:1:0:0:0:0:0:0:0:0:0:0:0:0 | 0:0:0:0:0:0:0:0:0:0:0:0:0:0:0:0:0:0 | 17:3:5:0:1:5:10:4:0:0:0:1:1:0:0:0:0:0:0:0:0 | 0:0:0:0:0:0:0:0:0:0:0:0:0:0:0:0:0:0:0:0 | 32:0:0:1:1:4:7:4:2:0:1:0:1:0:0:0:0:0:0:0:0:0 | 0:0:0:0:0:0:0:0:0:0:0:0:0:0:0:0 | 56:18:42:1:1:47:54:25:21:4:5:0:0:2:2:1:2:0:0:0:0:0 | 0:0:0:0:1:0:0:1:0:0:0:0:0:0:0:0:0 | 47:5:1:0:0:3:6:4:7:2:0:0:2:0:2:0:1:0:0:0:0 | 0:0:0:0:0:0:0:0:0:0:0:0:0:0:0:0:0:0:0 |
| XM_005265074:NM_001172713:NM_002078:XM_024453457:XM_006713110:XM_017006187:XM_024453458:XM_005265075:XM_024453456:XM_017006188:XM_017006186:XM_011533598:XM_017006184 | 9:3:5:3:3:4:3:0:0:0:0:0:0 | 1:0:0:0:0:0:0:0:0:0:0:0:0 | 19:6:3:11:1:2:27:2:0:0:0:0:0 | 0:0:0:0:0:0:1:0:0:0:0:0:0 | 14:4:3:1:0:3:2:0:2:0:0:0:0 | 0:0:0:0:0:0:0:0:0:0:0:0:0 | 58:19:18:22:4:13:62:5:1:1:1:1:0 | 0:0:0:0:0:0:0:0:0:0:0:0:0 | 24:3:2:6:0:3:6:1:0:0:0:0:0 | 0:0:0:0:0:0:0:0:0:0:0:0:0 |
| NM_001172713:NM_002078:XM_024453457:XM_006713110:XM_005265074:XM_017006187:XM_024453458:XM_005265075:XM_024453456:XM_017006188:XM_017006186:XM_011533598:XM_017006184 | 3:5:3:3:10:4:4:0:0:0:0:0:0 | 0:0:0:0:0:0:0:0:0:0:0:0:0 | 6:2:11:1:19:2:40:2:0:0:0:0:0 | 0:0:0:0:0:0:1:0:0:0:0:0:0 | 4:2:1:0:14:3:2:0:2:0:0:0:0 | 0:1:0:0:0:0:0:0:0:0:0:0:0 | 18:18:20:4:54:13:89:5:1:1:1:1:0 | 0:0:1:0:1:0:0:0:0:0:0:0:0 | 3:2:6:0:23:3:7:1:0:0:0:1:0 | 0:0:0:0:0:0:1:0:0:0:0:0:0 |
| XM_005265074:NM_001172713:XM_024453457:XM_006713110:NM_002078:XM_017006187:XM_024453458:XM_011533598:XM_005265075:XM_024453456:XM_017006188:XM_017006186:XM_017006184 | 7:1:3:3:4:4:5:2:0:0:0:0:0 | 2:0:0:0:0:0:0:0:0:0:0:0:0 | 15:5:10:1:0:2:44:1:1:0:0:0:0 | 0:0:0:0:0:0:0:0:0:0:0:0:0 | 11:4:1:0:2:1:1:1:0:1:0:0:0 | 2:0:0:0:1:0:0:0:0:1:0:0:0 | 41:15:19:4:12:9:108:2:3:1:1:1:0 | 6:0:1:0:2:0:1:0:0:0:0:0:0 | 18:2:3:0:2:3:9:0:1:0:0:0:0 | 3:0:0:0:0:0:1:0:0:0:0:0:0 |
| NM_001172713:XM_024453457:XM_005265074:XM_006713110:NM_002078:XM_017006187:XM_024453458:XM_011533598:XM_005265075:XM_024453456:XM_017006188:XM_017006186:XM_017006184 | 1:3:5:2:3:4:5:2:0:0:0:0:0 | 0:0:0:0:0:0:0:0:0:0:0:0:0 | 4:10:11:0:0:1:43:1:0:0:0:0:0 | 0:0:1:1:0:1:1:0:0:0:0:0:0 | 3:0:10:0:2:0:1:1:0:0:0:0:0 | 1:0:0:0:0:0:0:0:0:0:0:0:0 | 10:18:31:3:12:6:104:1:1:1:1:1:0 | 2:1:7:1:0:2:2:0:1:0:0:0:0 | 2:3:16:0:1:3:9:0:1:0:0:0:0 | 0:0:0:0:0:0:0:0:0:0:0:0:0 |

| | | | | | | | | | | |
|---|---|---|---|---|---|---|---|---|---|---|
| NM_001172713:XM_024453457:XM_005265074:XM_006713110:NM_002078:XM_017006187:XM_024453458:XM_011533598:XM_024453456:XM_005265075:XM_017006188:XM_017006184 | 1:3:5:2:3:4:4:2:0:0:0:0 | 0:0:0:0:0:0:0:0:0:0:0:0 | 0:9:5:0:0:0:35:1:0:0:0:0 | 0:0:0:0:0:0:0:0:0:0:0:0 | 2:0:7:0:2:0:2:1:0:0:0:0 | 0:0:0:0:0:0:0:0:0:0:0:0 | 4:14:10:1:3:3:102:1:1:1:0:0 | 0:0:1:0:1:0:0:0:0:0:0:0 | 1:3:12:0:1:3:10:0:0:1:0:0 | 0:0:0:0:0:0:0:0:0:0:0:0 |
| NM_001172713:XM_024453457:XM_005265074:XM_006713110:NM_002078:XM_017006187:XM_024453458:XM_011533598:XM_024453456:XM_005265075:XM_017006188:XM_017006184 | 1:3:5:2:3:4:4:2:0:0:0:0 | 0:0:0:0:0:0:0:0:0:0:0:0 | 0:9:5:0:0:0:36:1:0:0:0:0 | 0:0:0:0:0:0:0:0:0:0:0:0 | 2:0:7:0:2:0:2:1:0:0:0:0 | 0:0:0:0:0:0:0:0:0:0:0:0 | 4:14:11:1:4:3:102:1:1:1:0:0 | 0:0:0:0:0:0:0:0:0:0:0:0 | 1:3:12:0:1:3:10:0:0:1:0:0 | 0:0:0:0:0:0:0:0:0:0:0:0 |
| XM_011533598:NM_001172713:XM_024453457:XM_005265074:XM_006713110:NM_002078:XM_017006187:XM_024453458:XM_024453456:XM_005265075:XM_017006188:XM_017006184 | 1:1:3:5:2:3:4:4:0:0:0:0 | 1:0:0:0:0:0:0:0:0:0:0:0 | 1:0:8:4:0:0:0:33:0:0:0:0 | 0:0:1:1:0:0:0:2:0:0:0:0 | 1:2:0:7:0:2:0:2:0:0:0:0 | 0:0:0:0:0:0:0:0:0:0:0:0 | 1:3:13:10:1:4:3:95:1:1:0:0 | 0:1:1:1:0:0:0:3:0:0:0:0 | 0:1:3:12:0:1:3:8:0:1:0:0 | 0:0:0:0:0:0:0:1:0:0:0:0 |
| XM_006713110:XM_024453458:NM_001172713:XM_024453457:XM_005265074:NM_002078:XM_017006187:XM_011533598:XM_024453456:XM_005265075:XM_017006188:XM_017006184 | 1:3:1:2:5:3:4:1:0:0:0:0 | 1:1:0:0:0:0:0:0:0:0:0:0 | 0:25:0:6:2:0:0:1:0:0:0:0 | 0:3:0:1:2:0:0:0:0:0:0:0 | 0:2:1:0:7:0:0:1:0:0:0:0 | 0:0:0:0:0:0:0:0:0:0:0:0 | 1:83:3:10:9:4:3:1:1:1:0:0 | 0:5:0:1:0:0:0:0:0:0:0:0 | 0:7:1:3:9:1:3:0:0:1:0:0 | 0:0:0:0:1:0:0:0:0:0:0:0 |
| XM_024453458:XM_005265074:XM_017006187:XM_024453457:XM_011533598:XM_006713110:NM_001172713:XM_005265075:XM_017006188:NM_002078 | 1:3:2:1:1:0:0:0:0:0 | 1:0:0:0:0:0:0:0:0:0 | 6:0:0:2:0:0:0:0:0:0 | 3:0:0:1:0:0:0:0:0:0 | 1:1:0:0:1:0:0:0:0:0 | 0:0:0:0:0:0:0:0:0:0 | 19:1:1:1:1:0:1:0:0:0 | 6:0:0:2:0:1:0:0:0:0 | 2:2:2:1:0:0:0:1:0:0 | 2:0:0:0:0:0:0:0:0:0 |
| XM_024453458:XM_005265074:XM_017006187:XM_024453457:XM_011533598:XM_006713110:NM_001172713:XM_005265075:XM_017006188:NM_002078 | 1:3:2:1:1:0:0:0:0:0 | 1:0:0:0:0:0:0:0:0:0 | 6:0:0:0:0:0:0:0:0:0 | 3:0:0:3:0:0:0:0:0:0 | 1:1:0:0:1:0:0:0:0:0 | 0:0:0:0:0:0:0:0:0:0 | 18:1:1:1:1:0:1:0:0:0 | 7:0:0:2:0:1:0:0:0:0 | 2:2:2:1:0:0:0:1:0:0 | 2:0:0:0:0:0:0:0:0:0 |
| XM_005265074:XM_017006187:XM_024453457:XM_011533598:XM_024453458:NM_001172713:XM_005265075:XM_017006188:NM_002078 | 0:0:1:1:0:0:0:0:0 | 3:2:0:0:0:0:0:0:0 | 0:0:0:0:4:0:0:0:0 | 0:0:0:0:2:0:0:0:0 | 1:0:0:0:1:0:0:0:0 | 0:0:0:1:0:0:0:0:0 | 0:0:0:0:5:0:0:0:0 | 1:1:1:0:7:1:0:0:0 | 0:2:1:0:0:0:0:0:0 | 1:0:0:0:2:0:1:0:0 |
| XM_005265074:XM_024453457:XM_011533598:XM_024453458:XM_017006187:XM_017006188:NM_002078 | 1:1:1:0:0:0:0 | 0:0:0:0:0:0:0 | 0:0:0:1:0:0:0 | 0:0:0:0:0:0:0 | 1:0:0:1:0:0:0 | 0:0:0:0:0:0:0 | 0:0:0:7:0:0:0 | 0:0:0:1:0:0:0 | 0:1:0:1:0:0:0 | 0:0:0:0:2:0:0 |
| XM_011533598:XM_024453458 | 1:00 | 0:00 | 0:00 | 0:01 | 0:00 | 0:00 | 0:01 | 0:02 | 0:00 | 0:00 |
| XM_006713123:NR_024024:NM_001607:XM_011533650 | 1:0:0:0 | 0:0:0:0 | 0:0:0:0 | 0:0:0:0 | 0:2:1:0 | 0:0:0:0 | 1:1:1:0 | 0:0:0:0 | 0:0:1:1 | 0:0:0:0 |
| NM_001904:XM_024453360:XM_024453359:NM_001098210:XM_024453356:NM_001098209:XM_024453357:XM_006712985:XM_017005738:XM_024453358:NM_001330729 | 49:18:37:119:5:25:7:2:1:3:0 | 1:2:1:0:0:0:0:0:0:0:0 | 0:12:1:0:0:0:2:0:0:0:0 | 0:0:0:0:0:0:0:0:0:0:0 | 20:14:36:28:6:20:16:2:0:0:0 | 0:0:0:0:0:0:0:0:0:0:0 | 5:78:32:38:3:13:33:3:3:5:1 | 0:0:0:0:0:0:0:0:0:0:0 | 12:57:30:98:4:30:41:3:4:11:0 | 0:0:0:0:0:0:0:0:0:0:0 |
| NM_001098210:XM_024453356:NM_001904:NM_001098209:XM_024453360:XM_024453357:XM_006712985:XM_024453359:XM_017005738:XM_024453358:NM_001330729 | 119:5:50:25:25:8:2:40:1:3:0 | 0:0:0:0:0:0:0:0:0:0:0 | 0:0:0:0:18:5:0:4:0:0:0 | 0:0:0:0:0:0:0:0:0:0:0 | 28:6:21:20:14:17:2:47:0:0:0 | 0:0:0:0:0:0:0:0:0:0:0 | 38:3:5:13:124:55:3:40:3:5:1 | 0:0:0:0:0:0:0:0:0:0:0 | 98:4:12:29:65:48:3:37:4:11:1 | 0:0:0:1:1:0:0:0:0:0:0 |
| NM_001098210:XM_024453356:NM_001904:NM_001098209:XM_024453360:XM_024453357:XM_024453359:XM_006712985:XM_017005738:XM_024453358:NM_001330729 | 119:5:50:25:39:10:49:1:1:3:0 | 0:0:0:0:0:0:0:0:0:0:0 | 0:0:0:0:27:17:11:0:0:0:1 | 0:0:0:0:0:0:0:0:0:0:0 | 28:6:21:20:17:20:55:2:0:0:0 | 0:0:0:0:0:0:0:0:0:0:0 | 38:3:5:13:235:102:68:0:3:5:1 | 0:0:0:0:3:0:0:2:0:0:0 | 98:4:12:30:84:56:49:3:4:11:1 | 0:0:0:0:0:0:0:1:0:0:0:0 |
| XM_024453357:NM_001098210:XM_024453356:NM_001904:NM_001098209:XM_024453360:XM_024453359:XM_006712985:XM_017005738:XM_024453358:NM_001330729 | 10:119:5:50:25:46:58:1:1:3:0 | 1:0:0:0:0:0:0:0:0:0:0 | 26:0:0:0:0:33:18:0:0:0:1 | 0:0:0:0:0:1:0:0:0:0:0 | 21:28:6:21:20:16:64:2:0:0:0 | 0:0:0:0:0:0:0:0:0:0:0 | 157:38:3:5:13:325:98:1:3:5:1 | 0:0:0:0:0:0:0:0:0:0:0 | 63:98:4:12:30:101:53:2:4:11:1 | 0:0:0:0:0:0:2:0:0:0:0 |
| NM_001098210:XM_024453356:NM_001904:NM_001098209:XM_024453360:XM_024453357:XM_024453359:XM_017005738:XM_024453358:NM_001330729:XM_006712985 | 119:5:50:25:47:13:61:1:3:0:0 | 0:0:0:0:0:0:0:0:0:0:0 | 0:0:0:0:41:32:25:0:0:1:0 | 0:0:0:0:0:0:0:0:0:0:0 | 28:6:21:20:19:22:68:0:0:0:2 | 0:0:0:0:0:0:0:0:0:0:0 | 38:3:5:13:377:178:118:3:5:1:0 | 0:0:0:0:0:0:0:0:0:0:0 | 98:4:12:30:106:65:59:4:11:1:2 | 0:0:0:0:0:0:0:0:0:0:0 |
| NM_001098210:XM_024453356:NM_001904:NM_001098209:XM_024453360:XM_024453357:XM_024453359:XM_017005738:XM_024453358:NM_001330729:XM_006712985 | 119:5:50:25:52:14:63:1:3:0:0 | 0:0:0:0:0:0:0:0:0:0:0 | 0:0:0:0:41:32:28:0:0:1:0 | 0:0:0:0:0:0:0:0:0:0:0 | 28:6:21:20:20:23:71:0:0:0:2 | 0:0:0:0:0:0:0:0:0:0:0 | 39:3:5:13:395:185:125:3:5:1:0 | 0:0:0:0:1:1:1:0:0:0:0 | 98:4:12:30:113:69:64:4:11:1:2 | 0:0:0:0:0:0:0:0:0:0:0 |
| NM_001098210:XM_024453356:NM_001904:NM_001098209:XM_024453360:XM_024453357:XM_024453359:XM_006712985:XM_017005738:XM_024453358:NM_001330729 | 119:5:50:25:53:15:64:1:1:3:0 | 0:0:0:0:0:0:0:0:0:0:0 | 0:0:0:0:42:36:33:0:0:0:1 | 0:0:0:0:0:1:0:0:0:0:0 | 28:6:21:20:22:23:72:2:0:0:1 | 0:0:0:0:0:0:0:0:0:0:0 | 39:3:5:13:416:203:139:0:3:5:1 | 0:0:0:0:4:1:0:0:0:0:0 | 98:4:12:30:117:74:67:2:4:11:1 | 0:0:0:0:0:0:0:0:0:0:0 |
| XM_024453360:NM_001098210:XM_024453356:NM_001904:NM_001098209:XM_024453357:XM_024453359:XM_006712985:XM_017005738:XM_024453358:NM_001330729 | 52:119:5:50:25:15:64:1:1:3:0 | 1:0:0:0:0:0:0:0:0:0:0 | 42:0:0:0:0:38:35:0:0:0:1 | 0:0:0:0:0:0:0:0:0:0:0 | 23:28:6:21:20:23:72:2:0:0:1 | 0:0:0:0:0:0:0:0:0:0:0 | 421:39:3:5:13:207:141:0:3:5:1 | 2:0:0:0:0:2:0:0:0:0:0 | 118:98:4:12:30:74:71:2:4:11:1 | 0:0:0:0:0:0:0:0:0:0:0 |
| NM_001098210:XM_024453356:NM_001904:NM_001098209:XM_024453360:XM_024453357:XM_024453359:XM_006712985:XM_017005738:XM_024453358:NM_001330729 | 119:5:50:25:54:15:64:1:1:3:0 | 0:0:0:0:0:0:0:0:0:0:0 | 0:0:0:0:42:39:38:0:0:0:1 | 0:0:0:0:0:0:0:0:0:0:0 | 28:6:21:20:23:23:72:2:0:0:1 | 0:0:0:0:0:0:0:0:0:0:0 | 39:3:5:13:432:215:150:0:3:5:1 | 0:0:0:0:1:0:0:0:0:0:0 | 98:4:12:30:120:76:71:2:4:11:1 | 0:0:0:0:0:0:0:0:0:0:0 |
| NM_001098210:XM_024453356:NM_001904:NM_001098209:XM_024453360:XM_024453357:XM_024453359:XM_006712985:XM_017005738:XM_024453358:NM_001330729 | 119:5:50:25:54:20:66:1:1:3:0 | 0:0:0:0:0:0:0:0:0:0:0 | 0:0:0:0:44:50:56:0:0:0:1 | 0:0:0:0:0:1:0:0:0:0:0 | 28:6:21:20:24:26:75:2:0:0:1 | 0:0:0:0:0:0:0:0:0:0:0 | 39:3:5:13:455:248:171:0:3:5:2 | 0:0:0:0:0:2:0:0:0:0:0 | 98:4:12:30:125:79:73:2:4:11:1 | 0:0:0:0:0:0:0:0:0:0:0 |
| NM_001098210:XM_024453356:NM_001904:NM_001098209:XM_024453360:XM_024453357:XM_006712985:XM_024453359:XM_017005738:XM_024453358:NM_001330729 | 119:5:50:25:54:20:2:69:1:3:0 | 0:0:0:0:0:0:0:0:0:0:0 | 0:0:0:0:47:51:0:64:0:0:1 | 0:0:0:0:0:0:0:0:0:0:0 | 28:6:21:20:24:26:2:76:0:0:1 | 0:0:0:0:0:0:0:0:0:0:0 | 39:3:5:13:460:254:0:177:3:5:2 | 0:0:0:0:0:1:0:1:0:0:0 | 98:4:12:30:127:79:2:76:4:11:1 | 0:0:0:0:0:0:0:0:0:0:0 |
| NM_001098210:XM_024453356:NM_001904:NM_001098209:XM_024453360:XM_024453357:XM_006712985:XM_024453359:XM_017005738:XM_024453358:NM_001330729 | 119:5:50:25:56:23:2:71:1:3:0 | 0:0:0:0:0:0:0:0:0:0:0 | 0:0:0:0:53:57:0:71:0:0:1 | 0:0:0:0:0:0:0:0:0:0:0 | 28:6:21:20:23:28:2:78:0:0:1 | 0:0:0:0:0:0:0:1:0:0:0 | 39:3:5:13:482:282:0:202:3:5:2 | 0:0:0:0:3:0:0:0:0:0:0 | 98:4:12:30:135:86:2:77:4:11:1 | 0:0:0:0:0:0:0:0:0:0:0 |
| NM_001098210:XM_024453356:NM_001904:NM_001098209:XM_024453360:XM_024453357:XM_006712985:XM_024453359:XM_017005738:XM_024453358:NM_001330729 | 119:5:50:25:59:26:2:79:1:3:0 | 0:0:0:0:0:0:0:0:0:0:0 | 0:0:0:0:58:61:0:83:0:0:1 | 0:0:0:0:0:0:0:1:0:0:0 | 28:6:21:20:27:29:0:81:0:0:2 | 0:0:0:0:0:0:0:0:0:0:0 | 39:3:5:13:487:317:0:240:3:5:3 | 0:0:0:0:0:1:0:0:0:0:0 | 98:4:12:30:149:92:1:86:4:11:2 | 0:0:0:0:0:0:0:0:0:0:0 |
| NM_001098209:NM_001098210:XM_024453356:NM_001904:XM_024453360:XM_024453357:XM_006712985:XM_024453359:XM_017005738:XM_024453358:NM_001330729 | 23:119:5:50:62:27:2:80:1:3:0 | 2:0:0:0:0:0:0:0:0:0:0 | 0:0:0:0:62:64:0:94:0:0:1 | 0:0:0:0:0:0:0:0:0:0:0 | 20:28:6:21:28:29:0:87:0:0:2 | 0:0:0:0:0:0:0:0:0:0:0 | 13:39:3:5:488:329:0:255:3:5:4 | 0:0:0:0:2:0:0:0:0:0:0 | 30:98:4:14:154:94:1:92:4:11:2 | 0:0:0:0:0:0:0:0:0:0:0 |
| XM_024453359:NM_001098210:XM_024453356:NM_001904:NM_001098209:XM_024453360:XM_024453357:XM_006712985:XM_017005738:XM_024453358:NM_001330729 | 81:119:5:50:25:63:29:2:1:3:0 | 1:0:0:0:0:0:0:0:0:0:0 | 101:0:0:0:0:61:67:0:0:0:1 | 0:0:0:0:0:0:0:0:0:0:0 | 89:29:6:21:20:31:31:0:0:0:2 | 0:0:0:0:0:0:0:0:0:0:0 | 270:39:3:5:13:492:343:0:3:5:6 | 0:0:0:0:0:0:0:0:0:0:0 | 97:98:4:14:30:160:98:1:4:11:3 | 0:0:0:0:0:0:0:0:0:0:0 |
| NM_001098210:XM_024453356:NM_001904:NM_001098209:XM_024453360:XM_024453357:XM_006712985:XM_024453359:XM_017005738:XM_024453358:NM_001330729 | 119:5:50:25:63:30:2:82:1:3:0 | 0:0:0:0:0:0:0:0:0:0:0 | 0:0:0:0:61:67:1:105:0:0:1 | 0:0:0:0:0:0:0:0:0:0:0 | 29:6:21:20:31:31:0:89:0:0:2 | 0:0:0:0:0:0:0:0:0:0:0 | 39:3:5:13:490:344:0:281:3:5:6 | 0:0:0:0:1:1:0:0:0:0:0 | 98:4:14:30:159:100:1:97:4:11:3 | 0:0:0:0:0:0:0:0:0:0:0 |
| NM_001098210:XM_024453356:NM_001904:NM_001098209:XM_024453360:XM_024453357:XM_006712985:XM_024453359:XM_017005738:XM_024453358:NM_001330729 | 119:5:50:25:62:30:2:87:1:3:0 | 0:0:0:0:0:0:0:0:0:0:0 | 0:0:0:0:63:68:1:109:0:0:2 | 0:0:0:0:0:0:0:0:0:0:0 | 29:6:21:20:31:31:1:92:0:0:2 | 0:0:0:0:0:0:0:0:0:0:0 | 39:3:5:13:490:352:0:294:3:5:6 | 0:0:0:0:0:0:0:2:0:0:0 | 98:4:14:30:162:102:1:104:4:11:3 | 0:0:0:0:0:0:0:0:0:0:0 |
| XM_011533880:XM_024453613 | 0:00 | 0:00 | 0:00 | 0:00 | 0:00 | 0:00 | 0:00 | 0:00 | 0:00 | 0:00 |
| NM_018075 | 0 | 0 | 0 | 0 | 0 | 2 | 0 | 0 | 0 | 1 |
| NM_015340:XM_017006042 | 1:01 | 0:00 | 0:00 | 0:01 | 0:00 | 0:01 | 0:01 | 0:00 | 0:00 | 0:02 |
| NM_003965:XM_017007436:XM_011534209:XM_011534208:NM_001130910 | 13:1:0:0:0 | 0:0:0:0:0 | 33:3:1:3:2 | 0:0:0:0:0 | 53:5:0:4:5 | 0:0:0:0:0 | 140:27:5:15:2 | 1:0:0:0:0 | 23:4:3:2:1 | 0:0:0:0:0 |
| NM_003965:XM_017007436:XM_011534209:XM_011534208:NM_001130910 | 13:1:0:0:0 | 0:0:0:0:0 | 33:3:1:3:2 | 0:0:0:0:0 | 53:5:0:4:5 | 0:0:0:0:0 | 70:15:3:8:1 | 71:14:2:7:1 | 7:5:1:2:1 | 16:1:2:0:0 |
| NM_003965:XM_017007436:XM_011534209:XM_011534208:NM_001130910 | 13:1:0:0:0 | 0:0:0:0:0 | 33:3:1:3:2 | 0:0:0:0:0 | 53:5:0:4:5 | 0:0:0:0:0 | 71:14:2:7:1 | 70:15:3:8:1 | 16:1:2:0:0 | 7:5:1:2:1 |
| NM_003965:XM_017007436:XM_011534209:XM_011534208:NM_001130910 | 13:1:0:0:0 | 0:0:0:0:0 | 33:3:1:3:2 | 0:0:0:0:0 | 52:5:0:4:5 | 0:1:0:0:0 | 138:33:5:14:2 | 2:1:0:0:0 | 23:4:3:2:1 | 0:0:0:0:0 |
| NM_002343:NM_001199149:NM_001321122 | 0:00:00 | 2:00:00 | 7:01:00 | 6:00:00 | 0:00:01 | 0:00:00 | 2:00:00 | 0:00:00 | 1:00:00 | 0:00:00 |
| XR_002959510:NR_146158:NM_014159:XR_002959515:XR_002959514:XM_011533634:XR_002959512:XR_002959517:XR_002959511:XM_024453488:XM_024453487 | 2:2:3:0:0:0:0:0:0:0:0 | 1:0:0:0:0:0:0:0:0:0:0 | 1:2:0:2:0:1:0:0:0:0:0 | 11:0:0:1:1:0:0:0:0:0:0 | 2:0:9:0:0:0:1:0:0:0:0 | 2:0:0:0:0:0:0:0:0:0:0 | 4:11:14:3:0:0:0:0:1:2:0 | 8:1:0:2:0:0:0:1:0:0:0 | 0:3:6:0:0:0:0:0:0:0:0 | 0:0:0:0:0:0:0:0:0:0:0 |
| NM_014159:XR_002959510:XM_011533634:XR_002959511:XR_002959514:XR_002959515:XR_002959516:NR_146158:XR_002959512:XR_002959517:XR_002959513:XM_024453487:XM_024453488 | 2:2:0:0:0:0:0:0:0:0:0:0:0 | 1:2:0:0:0:0:0:0:0:0:0:0:0 | 0:8:0:0:3:1:2:0:0:0:0:0:0 | 0:4:1:1:1:0:0:0:0:0:0:0:0 | 5:3:0:0:1:0:0:0:1:0:0:0:0 | 1:3:0:1:1:0:1:1:0:0:0:0:0 | 6:8:0:5:3:1:3:2:0:0:0:0:0 | 8:5:0:2:3:1:1:1:0:1:0:0:0 | 4:1:0:0:0:0:0:0:0:0:1:0:0 | 3:1:0:0:0:0:1:0:0:0:0:0:0 |
| XM_017005921:XM_005264967:XM_017005918:NM_012235:NM_001320044 | 0:1:0:0:0 | 2:1:0:0:0 | 1:0:0:0:0 | 5:0:0:0:0 | 2:5:0:0:0 | 1:6:0:0:0 | 11:1:1:0:0 | 9:1:1:0:0 | 2:0:0:0:0 | 1:2:2:1:1 |
| NM_002375:XM_024453530:XR_002959534:XM_017006412:XM_017006402:XR_002959526:XR_002959525:XR_002959533:XM_017006410:XR_002959532:XR_002959527:XR_002959524:NM_001134364:XR_002959528:XM_024453528:XM_017006416 | 1:3:1:1:1:1:1:1:1:0:0:0:0:0:0:0 | 0:0:0:0:0:0:0:0:0:0:0:0:0:0:0:0 | 0:0:0:0:0:1:0:1:2:1:0:0:0:0:0:0 | 0:0:0:0:0:0:0:0:0:0:0:0:0:0:0:0 | 0:3:0:1:0:0:0:2:0:3:0:0:0:0:0:0 | 0:0:0:0:0:0:0:0:0:0:0:0:0:0:0:0 | 0:1:8:0:0:0:0:1:0:5:0:0:0:0:0:0 | 0:2:4:0:1:2:0:0:1:3:2:1:0:0:0:0 | 0:0:0:0:0:1:0:0:1:3:0:0:0:0:0:0 | 0:1:2:2:0:0:0:0:0:4:0:0:1:0:0:0 |
| XR_002959533:NM_002375:XM_024453530:XR_002959534:XM_017006412:XM_017006402:XR_002959526:XR_002959525:XR_002959532:XM_017006410:XR_002959527:XR_002959524:XR_002959530:XR_002959528:XM_017006416 | 2:1:1:1:1:1:1:1:1:3:0:0:0:0:0:0 | 1:0:0:0:0:0:0:0:0:0:0:0:0:0:0 | 3:0:0:1:0:0:1:0:7:2:0:0:0:0:0 | 0:0:0:0:0:0:0:1:0:0:0:0:0:0:0 | 2:0:3:1:1:0:0:0:8:0:0:0:0:0:0 | 0:0:0:1:0:0:0:0:0:0:0:0:0:0:0 | 2:0:3:15:0:1:2:0:43:0:2:2:1:0:0 | 0:0:0:1:0:0:0:0:5:0:0:0:0:0:0 | 1:0:1:2:1:0:1:0:8:1:0:0:0:0:0 | 0:0:0:0:0:0:0:0:0:0:1:0:0:0:0:0 |
| NM_002375:XR_002959534:XM_017006412:XM_017006402:XR_002959526:XR_002959525:XR_002959533:XR_002959532:XM_024453530:XR_002959527:XR_002959524:XR_002959530:XR_002959528:XM_017006410 | 1:1:1:1:1:1:1:3:6:0:0:0:0:0 | 0:0:0:0:0:0:0:0:0:0:0:0:0:0 | 0:2:0:0:1:0:3:14:0:0:0:0:0:0 | 0:0:0:0:0:0:0:0:0:0:0:0:0:0 | 0:3:1:0:0:0:2:10:2:0:0:0:0:0 | 0:0:0:0:0:0:0:0:0:0:0:0:0:0 | 0:7:0:0:0:0:1:37:0:0:1:1:0:0 | 0:3:0:1:2:0:1:29:0:2:2:0:0:0 | 0:0:0:0:1:0:1:6:0:0:0:0:0:0 | 0:1:1:0:0:0:1:4:1:0:0:0:0:0 |
| NM_002375:XR_002959534:XM_017006412:XM_017006402:XR_002959526:XR_002959525:XR_002959533:XR_002959532:XR_002959524:XM_017006414:XM_024453530:XR_002959530:XR_002959527:XR_002959528 | 1:1:1:1:1:1:1:3:6:0:0:0:0:0 | 0:0:0:0:0:0:0:0:0:0:0:0:0:0 | 0:2:0:0:1:0:3:14:0:1:0:0:0:0 | 0:3:0:0:0:0:0:28:2:0:0:0:0:0 | 0:3:0:0:0:0:2:12:0:0:2:2:0:0 | 0:0:0:0:0:0:0:0:0:0:0:0:0:0 | 0:8:0:1:2:0:3:76:3:0:0:1:2:0 | 0:7:0:0:0:0:3:100:0:0:0:0:0:0 | 0:1:1:0:1:0:2:12:0:0:0:0:0:0 | 0:0:0:0:0:0:0:0:0:0:0:0:0:0 |

| | | | | | | | | | | |
|---|---|---|---|---|---|---|---|---|---|---|
| NM_002375:XR_002959534:XM_017006412:XM_017006402:XR_002959526:XR_002959525:XR_002959533:XR_002959532:XM_017006410:XR_002959524:XM_017006414:XR_002959529:XM_024453530:XR_002959530:XR_002959527:XR_002959528 | 1:3:1:1:1:1:6:34:1:1:0:0:0:0:0:0 | 0:0:0:0:0:0:0:0:0:0:0:0:0:0:0:0 | 0:5:0:0:1:0:4:58:0:3:1:1:0:0:0:0 | 0:0:0:0:0:0:0:0:0:0:0:0:0:0:0:0 | 0:6:0:0:0:0:3:78:0:3:0:0:2:5:0:0 | 0:0:1:0:0:0:0:0:0:0:0:0:1:0:0:0 | 0:22:0:1:2:0:12:306:0:11:0:0:0:2:2:0 | 0:1:0:0:0:0:0:0:0:0:0:0:0:0:0:0 | 0:2:1:0:1:0:4:39:0:1:0:0:0:0:0:0 | 0:0:0:0:0:0:0:1:0:0:0:0:0:0:0:0 |
| NM_002375:XR_002959534:XM_017006412:XM_017006402:XR_002959526:XR_002959525:XR_002959533:XR_002959532:XM_017006410:XR_002959524:XM_017006414:XR_002959529:XM_024453530:XR_002959530:XR_002959527:XR_002959528 | 1:3:1:1:1:1:6:34:1:1:0:0:0:0:0:0 | 0:0:0:0:0:0:0:0:0:0:0:0:0:0:0:0 | 0:6:0:0:1:0:4:50:0:3:1:1:0:0:0:0 | 0:0:0:0:0:0:0:8:0:0:0:0:0:0:0:0 | 0:6:1:0:0:0:3:79:0:3:0:0:3:5:0:0 | 0:0:0:0:0:0:0:0:0:0:0:0:0:0:0:0 | 0:23:0:1:2:0:11:274:0:10:0:0:0:2:2:0 | 0:1:0:0:0:0:1:42:0:1:0:0:0:0:0:0 | 0:2:1:0:1:0:4:40:0:1:0:0:0:0:0:0 | 0:0:0:0:0:0:0:0:0:0:0:0:0:0:0:0 |
| XR_002959532:XM_017006410:NM_002375:XR_002959534:XM_017006412:XM_017006402:XR_002959526:XR_002959525:XR_002959533:XR_002959524:XM_017006414:XR_002959529:XR_002959530:XM_024453530:XR_002959527:XR_002959528 | 28:0:1:3:1:1:1:1:6:1:0:0:0:0:0:0 | 5:1:0:0:0:0:0:0:0:0:0:0:0:0:0:0 | 57:0:0:6:0:0:1:0:4:3:1:1:0:0:0:0 | 1:0:0:0:0:0:0:0:0:0:0:0:0:0:0:0 | 62:0:0:4:1:0:0:0:3:2:0:0:4:3:0:0 | 17:0:0:2:0:0:0:0:0:1:0:0:1:0:0:0 | 300:0:0:24:0:1:2:0:12:11:0:0:2:0:2:0 | 19:0:0:0:0:0:0:0:0:0:0:0:0:0:0:0 | 34:0:0:1:1:0:1:0:3:0:0:0:0:0:0:0 | 6:0:0:1:0:0:0:0:1:1:0:0:0:0:0:0 |
| NM_002375:XR_002959534:XM_017006412:XM_017006402:XR_002959526:XR_002959525:XR_002959533:XR_002959532:XM_017006410:XR_002959524:XM_017006414:XR_002959529:XM_024453530:XR_002959530:XR_002959527:XR_002959528 | 1:3:1:1:1:1:6:34:1:1:0:0:0:0:0:0 | 0:0:0:0:0:0:0:0:0:0:0:0:0:0:0:0 | 0:6:0:0:1:0:4:58:0:3:1:1:0:0:0:0 | 0:0:0:0:0:0:0:1:0:0:0:0:0:0:0:0 | 0:6:1:0:0:0:3:79:0:3:0:0:3:5:0:0 | 0:0:0:0:0:0:0:0:0:0:0:0:0:0:0:0 | 0:25:0:1:2:0:12:318:0:11:0:0:0:2:2:0 | 0:0:0:0:0:0:0:1:0:0:0:0:0:0:0:0 | 0:2:1:0:1:0:4:41:0:1:0:0:0:0:0:0 | 0:0:0:0:0:0:0:0:0:0:0:0:0:0:0:0 |
| NM_198880:NM_017730:XM_011533863:NM_001320584:NM_001320581 | 8:1:4:1:0 | 0:0:0:0:0 | 7:0:14:0:0 | 0:0:0:0:0 | 12:3:9:0:0 | 0:0:0:0:0 | 12:0:24:0:0 | 6:0:22:1:0 | 4:0:6:0:0 | 12:0:3:0:0 |
| NM_001664:NM_001313943:NM_001313947:NM_001313944:NM_001313946:NM_001313941:NM_001313945 | 3222:21:8:3:2:1:0 | 0:0:0:0:0:0:0 | 3548:7:4:4:5:2:0 | 4:0:0:0:0:0:0 | 8868:29:9:7:16:2:0 | 18:0:0:0:0:0:0 | 15023:155:18:16:20:15:2 | 17:0:0:0:0:0:0 | 4444:27:4:2:9:5:0 | 2:0:0:0:0:0:0 |
| NM_001664:NM_001313943:NM_001313947:NM_001313944:NM_001313946:NM_001313941:NM_001313945 | 3219:21:8:3:2:1:0 | 3:0:0:0:0:0:0 | 3545:6:4:4:5:2:0 | 7:1:0:0:0:0:0 | 8869:29:9:7:16:2:0 | 17:0:0:0:0:0:0 | 15016:155:18:16:20:15:2 | 23:0:0:0:0:0:0 | 4444:27:4:2:9:5:0 | 4:0:0:0:0:0:0 |
| NM_001664:NM_001313943:NM_001313947:NM_001313944:NM_001313946:NM_001313941:NM_001313945 | 3216:21:8:3:2:1:0 | 4:0:0:0:0:0:0 | 3544:7:4:4:5:2:0 | 8:0:0:0:0:0:0 | 8860:29:9:7:16:2:0 | 22:0:0:0:0:0:0 | 15012:155:18:16:20:15:2 | 23:0:0:0:0:0:0 | 4440:27:4:2:9:5:0 | 4:0:0:0:0:0:0 |
| NM_001664:NM_001313943:NM_001313947:NM_001313944:NM_001313946:NM_001313941:NM_001313945 | 3219:21:8:3:2:1:0 | 2:0:0:0:0:0:0 | 3539:7:4:4:5:2:0 | 5:0:0:0:0:0:0 | 8863:28:9:7:16:1:0 | 13:1:0:0:0:1:0 | 14971:155:18:16:20:15:2 | 22:0:0:0:0:0:0 | 4438:27:4:2:9:5:0 | 10:0:0:0:0:0:0 |
| NM_001664:NM_001313943:NM_001313947:NM_001313944:NM_001313941:NM_001313946:NM_001313945 | 3206:21:8:3:1:0:0 | 3:0:0:0:0:0:0 | 3542:7:4:4:2:0:0 | 4:0:0:0:0:0:0 | 8838:29:9:7:2:0:0 | 12:0:0:0:0:0:0 | 14944:153:18:16:15:1:2 | 14:0:0:0:0:0:0 | 4440:26:4:2:5:1:0 | 4:0:0:0:0:0:0 |
| NM_001664:NM_001313943:NM_001313947:NM_001313944:NM_001313941:NM_001313945 | 3208:21:8:3:1:0 | 1:0:0:0:0:0 | 3526:7:4:4:2:0 | 3:0:0:0:0:0 | 8833:29:9:7:2:0 | 5:0:0:0:0:0 | 14906:153:17:16:15:2 | 13:0:0:0:0:0 | 4433:26:4:2:5:0 | 6:0:0:0:0:0 |
| NM_001664:NM_001313943:NM_001313944:NM_001313941:NM_001313945:NM_001313947 | 3194:21:1:1:0:0 | 0:0:0:0:0:0 | 3484:7:0:2:0:0 | 7:0:0:0:0:0 | 8787:29:1:2:0:0 | 8:0:0:0:0:0 | 14754:151:0:15:1:1 | 13:0:0:0:0:0 | 4420:26:1:5:0:0 | 3:0:0:0:0:0 |
| NM_001177639:NM_001177636:NM_004393:NM_001177638 | 0:0:0:0 | 1:0:0:0 | 0:0:0:0 | 0:1:0:0 | 0:0:0:0 | 0:0:2:0 | 0:0:0:0 | 1:0:1:0 | 0:0:0:0 | 0:0:0:0 |
| NM_021971 | 0 | 0 | 0 | 1 | 0 | 0 | 0 | 2 | 0 | 1 |
| NM_018184 | 49 | 0 | 20 | 0 | 106 | 0 | 118 | 2 | 48 | 0 |
| XM_005264839:NM_007184:NM_001276293:NM_001276294:XM_006712955 | 0:0:0:0:0 | 1:1:0:0:0 | 0:0:0:0:0 | 0:1:1:2:0 | 0:0:0:0:0 | 0:1:1:6:0 | 0:0:0:0:0 | 2:1:0:14:0 | 0:0:0:0:0 | 0:1:0:1:0 |
| XM_017006750:NM_181042:XM_017006748:XM_017006764:XR_001740194:XM_017006759:XM_017006749:XM_017006745:XM_005265280:XM_024453619:XM_017006732:NM_018313 | 5:2:1:0:0:0:0:0:0:0:0:0:0 | 0:0:0:0:0:0:0:0:0:0:0:0 | 3:0:0:18:4:1:2:0:0:0:0:0 | 0:0:0:1:0:0:0:0:0:0:0:0 | 3:2:0:2:1:0:0:1:0:0:0:0 | 0:0:0:0:0:0:0:0:0:0:0:0 | 2:5:1:56:17:1:1:1:1:1:0:0 | 0:1:0:3:2:0:0:0:0:0:0:0 | 9:2:2:4:0:0:1:1:3:0:0:0 | 0:0:0:0:0:0:0:0:0:0:0:0 |
| XM_017006750:NM_181042:XM_017006748:XM_017006749:XM_017006764:XM_017006744:XM_005265282:XM_024453619:XR_001740194:XM_017006747:XM_017006745:XM_005265280:XM_017006738:XM_017006758:XM_017006746:NM_018313:XM_005265279:XM_017006732:XM_017006734:XM_017006736 | 10:2:4:1:1:1:1:1:1:0:0:0:0:0:0:0:0:0:0:0 | 0:0:0:0:0:0:0:0:0:0:0:0:0:0:0:0:0:0:0:0 | 7:0:1:2:19:0:3:0:2:1:0:0:0:0:0:0:0:0:0:0 | 0:0:0:0:0:0:0:0:0:0:0:0:0:0:0:0:0:0:0:0 | 9:2:0:0:4:0:2:0:1:1:1:1:0:0:0:0:0:0:0:0 | 0:0:0:0:0:0:0:0:0:0:0:0:0:0:0:0:0:0:0:0 | 15:6:3:3:58:1:10:3:12:2:1:1:1:1:2:1:0:0:0:0 | 0:0:0:0:0:0:1:0:0:0:0:0:0:0:0:0:0:0:0:0 | 24:2:4:3:6:0:3:0:0:0:1:4:0:1:1:0:0:0:0:0 | 0:0:0:0:0:0:0:0:0:0:0:0:0:0:0:0:0:0:0:0 |
| NM_206826:NM_014366:NM_206825 | 0.561168981 | 0:00:00 | 0.33681713 | 0:00:00 | 0.208344907 | 0:00:00 | 110:1863:29 | 1:03:00 | 35:346:10 | 0:00:00 |
| NM_206826:NM_014366:NM_206825 | 0.217384259 | 0.343784722 | 5:37:01 | 2:28:00 | 4:28:01 | 1:32:00 | 120:1865:29 | 1:00:00 | 36:345:10 | 0:01:00 |
| NM_206826:NM_014366:NM_206825 | 0.602835648 | 0:00:00 | 0.33681713 | 0:00:00 | 0.250011574 | 0:00:00 | 151:1864:29 | 1:01:00 | 41:345:10 | 0:01:00 |
| NM_206826:NM_014366:NM_206825 | 0.602835648 | 0:00:00 | 0.46181713 | 0:00:00 | 0.291678241 | 0:00:00 | 185:1861:29 | 0:00:00 | 45:346:10 | 0:00:00 |
| NM_206826:NM_014366:NM_206825 | 0.727835648 | 0:00:00 | 0.46181713 | 0:00:00 | 0.250011574 | 0:00:00 | 236:1858:29 | 0:03:00 | 50:345:10 | 1:00:00 |
| NM_014366:NM_206825:NM_206826 | 203:04:15 | 4:01:00 | 64:01:13 | 0:00:00 | 59:01:05 | 1:00:00 | 1831:27:288 | 24:02:01 | 338:10:54 | 7:00:00 |
| NM_206826:NM_014366:NM_206825 | 0.766018519 | 0:00:00 | 0.710428241 | 0:00:00 | 5:59:01 | 0:00:00 | 309:1825:27 | 1:04:00 | 56:339:10 | 0:00:00 |
| NM_206826:NM_014366:NM_206825 | 0.766018519 | 0:00:00 | 0.710428241 | 0:00:00 | 4:59:01 | 0:00:00 | 312:1827:26 | 0:02:01 | 56:339:10 | 0:00:00 |
| NM_206826:NM_014366:NM_206825 | 0.297928241 | 0.925034722 | 9:37:01 | 8:26:00 | 3:27:01 | 2:31:00 | 384:1809:27 | 1:00:00 | 61:337:10 | 0:00:00 |
| NM_206826:NM_014366:NM_206825 | 25:201:4 | 0:00:00 | 0.751400463 | 0:01:00 | 6:58:01 | 0:00:00 | 390:1808:27 | 0:02:00 | 61:336:10 | 0:01:00 |
| NM_206826:NM_014366:NM_206825 | 26:202:4 | 0:00:00 | 0.752094907 | 0:00:00 | 6:58:01 | 0:00:00 | 390:1809:27 | 0:03:00 | 60:337:10 | 0:00:00 |
| NM_014366:NM_206826:NM_206825 | 198:23:04 | 1:00:00 | 62:17:01 | 0:00:00 | 57:08:01 | 0:00:00 | 1808:391:27 | 3:01:00 | 336:62:10 | 0:00:00 |
| NM_014366:NM_206826:NM_206825 | 194:23:04 | 6:00:00 | 63:18:01 | 0:00:00 | 55:08:01 | 3:00:00 | 1769:391:26 | 29:05:00 | 332:59:10 | 5:01:00 |
| NM_014366:NM_206826:NM_206825 | 192:22:04 | 2:01:00 | 53:17:01 | 9:01:00 | 55:08:01 | 0:00:00 | 1659:381:25 | 79:07:01 | 327:61:10 | 5:00:00 |
| NM_014366:NM_206826:NM_206825 | 192:22:04 | 2:01:00 | 51:18:00 | 0:00:00 | 55:08:01 | 0:00:00 | 1624:374:24 | 32:02:01 | 318:60:9 | 14:01:01 |
| NM_206826:NM_014366:NM_206825 | 1.00837963 | 0:00:00 | 15:48:00 | 0:00:00 | 7:54:01 | 0:01:00 | 367:1592:24 | 0:03:00 | 58:312:9 | 0:00:00 |
| NM_206826:NM_014366:NM_206825 | 1.00837963 | 0:00:00 | 15:48:00 | 0:00:00 | 7:55:01 | 0:00:00 | 365:1594:24 | 1:01:00 | 57:312:9 | 0:00:00 |
| NM_206826:NM_014366:NM_206825 | 0.966712963 | 0:00:00 | 15:48:00 | 0:00:00 | 7:55:01 | 0:00:00 | 362:1594:24 | 2:01:00 | 57:312:9 | 0:00:00 |
| NM_014041 | 3 | 4 | 2 | 1 | 3 | 4 | 15 | 0 | 19 | 0 |
| NM_003157:XM_017007085:NM_001348412:XM_017007086 | 0:0:1:0 | 4:1:1:0 | 1:0:0:0 | 1:0:0:1 | 3:0:2:0 | 1:1:0:0 | 5:0:0:3 | 0:0:0:0 | 5:0:2:0 | 0:0:0:0 |
| NM_212539:NM_006254:XR_002959550:NM_001354678:NM_001354680:NM_001354679 | 17:4:1:0:0:0 | 0:0:0:0:0:0 | 41:8:2:1:0:0 | 0:0:0:0:0:0 | 64:22:1:1:1:0 | 0:0:0:0:0:0 | 64:7:2:0:3:7 | 1:0:0:0:0:0 | 19:2:0:0:1:2 | 1:0:0:0:0:0 |
| NM_001064:XM_011534055 | 3:01 | 0:00 | 0:02 | 0:00 | 2:01 | 0:00 | 2:01 | 0:00 | 2:00 | 0:00 |
| NM_177966:NM_001322176 | 0:00 | 0:00 | 0:00 | 1:00 | 0:00 | 0:00 | 0:00 | 3:00 | 0:00 | 0:00 |
| XM_017005674:NM_001350992:NM_001350991:NM_173471:XR_001739996:NM_001164796 | 0:0:0:0:0:0 | 0:0:0:0:0:0 | 0:0:0:0:0:0 | 0:0:0:0:0:0 | 1:0:0:0:0:0 | 0:0:0:0:0:0 | 0:0:0:0:0:0 | 0:0:0:0:0:0 | 0:2:0:0:0:0 | 0:0:0:0:0:0 |
| NM_003848 | 0 | 0 | 0 | 2 | 0 | 0 | 0 | 0 | 2 | 0 |
| XM_017005994:XM_005264723:NM_015123:XM_017005997 | 2:0:0:0 | 5:0:0:0 | 5:0:0:0 | 1:0:0:0 | 7:1:0:0 | 3:0:0:0 | 0:0:0:0 | 6:1:1:0 | 0:0:0:0 | 1:0:0:0 |
| XM_017005994:XM_005264723:NM_015123 | 0:00:00 | 0:00:00 | 0:00:00 | 0:00:00 | 3:01:00 | 0:00:00 | 0:00:00 | 0:01:01 | 0:00:00 | 0:00:00 |
| NM_001244815:NM_001244813:XM_017006165:XM_006713102:NM_001349344:XR_001740102:XM_005264736:NM_001244808:NM_001349337:XM_017006167:NM_001244814:NM_001244812:NM_001349341:NR_146143:NR_146142:NM_001349338:NM_001349342:XM_011533585:NM_001349343:NM_001349340:XM_006713103:NM_001244816:NM_032682 | 2:6:2:1:1:1:1:0:0:0:0:0:0:0:0:0:0:0:0:0:0:0:0 | 0:0:0:0:0:0:0:0:0:0:0:0:0:0:0:0:0:0:0:0:0:0:0 | 17:32:31:3:3:0:1:10:4:3:1:1:3:1:1:0:0:0:0:0:0:0:0 | 0:0:0:0:0:0:0:0:0:0:0:0:0:0:0:0:0:0:0:0:0:0:0 | 12:9:5:0:1:0:2:2:1:2:0:0:1:0:0:4:1:1:0:0:0:0:0 | 0:0:0:0:1:0:0:0:0:0:0:0:0:0:0:0:0:0:0:0:0:0:0 | 11:15:26:3:0:0:4:12:2:0:0:3:3:0:1:4:0:0:1:1:1:0:0 | 0:0:0:0:0:0:0:0:0:0:0:0:0:0:0:0:0:0:0:0:0:0:0 | 4:6:9:0:4:0:1:2:1:0:0:0:1:0:0:1:1:1:0:0:0:0:0 | 0:0:0:0:0:0:0:0:0:0:0:0:0:0:0:0:0:0:0:0:0:0:0 |
| NM_001349341:NM_001244815:NM_001244813:XM_017006165:XM_006713102:NM_001349344:XM_005264736:NM_001244808:NM_001349337:XM_017006167:NM_001244814:NM_001244812:NR_146143:NR_146142:XR_001740102:NM_001349338:NM_001349342:XM_011533585:XM_006713103:NM_001349343:NM_001244810:NM_001349340:NM_001244816 | 1:2:7:2:1:1:1:0:0:0:0:0:0:0:0:0:0:0:0:0:0:0:0 | 0:0:0:0:0:0:0:0:0:0:0:0:0:0:0:0:0:0:0:0:0:0:0 | 3:19:32:28:3:3:1:10:5:2:1:1:1:1:5:0:0:0:0:0:0:0:0 | 0:0:0:0:0:0:0:0:0:0:0:0:0:0:0:0:0:0:0:0:0:0:0 | 1:12:10:6:0:2:2:2:1:3:0:0:1:0:1:5:1:1:1:0:0:0:0 | 0:0:0:0:0:0:0:0:0:0:0:0:0:0:0:0:0:0:0:0:0:0:0 | 3:12:15:19:3:0:4:12:2:0:0:3:0:1:0:4:0:0:1:1:1:1:0 | 0:0:0:0:0:0:0:0:0:0:0:0:0:0:0:0:0:0:0:0:0:0:0 | 1:5:7:5:1:4:1:2:1:0:0:0:0:0:1:3:1:1:0:0:0:0:0 | 0:0:0:0:0:0:0:0:0:0:0:0:0:0:0:0:0:0:0:0:0:0:0 |
| XM_017007006 | 0 | 0 | 0 | 0 | 0 | 0 | 0 | 0 | 0 | 1 |
| NM_001077526:NM_022485:NM_001077525:XR_245152:XM_017007039:XM_024453709:XM_024453710:XM_017007034:XM_017007043:XM_017007038:XM_005265400:XM_017007036:XM_017007042:XM_017007045:XR_001740231:XM_017007035:XM_017007044:XM_011534012:XR_001740233 | 14:8:9:2:1:1:0:0:0:0:0:0:0:0:0:0:0:0:0 | 0:0:0:0:0:0:0:0:0:0:0:0:0:0:0:0:0:0:0 | 9:5:7:4:1:3:5:3:1:0:0:0:0:0:0:0:0:0:0 | 0:0:0:0:0:0:0:0:0:0:0:0:0:0:0:0:0:0:0 | 12:5:14:1:1:3:1:0:0:3:1:1:0:0:0:0:0:0:0 | 0:0:0:0:0:0:0:0:0:0:0:0:0:0:0:0:0:0:0 | 49:51:41:16:12:10:11:4:1:5:2:1:4:1:1:1:3:0:0 | 0:0:0:1:0:0:1:0:0:0:0:0:0:0:0:0:0:0:0 | 12:10:17:0:1:0:2:0:0:2:0:0:1:0:0:0:1:0 | 0:0:0:0:0:0:0:0:0:0:0:0:0:0:0:0:0:0:0 |

| | | | | | | | | | | |
|---|---|---|---|---|---|---|---|---|---|---|
| NM_001077526:NM_001077525:XR_245152:XM_024453709:XM_024453710:XM_017007034:XM_017007043:NM_022485:XM_017007038:XM_005265400:XM_017007036:XR_001740231:XM_017007035:XM_017007044:XM_017007045:XM_011534012 | 14:9:2:1:0:0:0:0:0:0:0:0:0:0:0:0 | 0:0:0:0:0:0:0:0:0:0:0:0:0:0:0:0 | 9:7:1:6:7:3:1:0:0:0:0:0:0:0:0:0 | 0:0:0:0:0:0:0:0:0:0:0:0:0:0:0:0 | 12:14:0:4:1:0:0:1:3:1:1:0:0:0:0:0 | 0:0:0:1:0:0:0:0:0:0:0:0:0:0:0:0 | 49:43:2:17:20:4:1:1:5:2:1:1:1:3:0:0 | 0:0:0:0:0:0:0:0:0:0:0:0:0:0:0:0 | 12:17:0:0:2:0:0:0:2:0:0:0:0:0:1:1 | 0:0:0:0:0:0:0:0:0:0:0:0:0:0:0:0 |
| NM_001077526:NM_022485:NM_001077525:XR_245152:XM_017007039:XM_024453709:XM_024453710:XM_017007034:XM_017007043:XM_017007038:XM_005265400:XM_017007036:XM_017007042:XR_001740231:XM_017007035:XM_017007044:XM_017007045:XM_011534012 | 14:8:9:2:1:1:0:0:0:0:0:0:0:0:0:0:0:0 | 0:0:0:0:0:0:0:0:0:0:0:0:0:0:0:0:0:0 | 9:5:7:4:1:6:8:3:1:0:0:0:0:0:0:0:0:0 | 0:0:0:0:0:1:0:0:0:0:0:0:0:0:0:0:0:0 | 12:6:14:0:1:5:1:0:0:3:1:1:0:0:0:0:0:0 | 0:0:0:0:0:0:0:0:0:0:0:0:0:0:0:0:0:0 | 48:54:43:13:12:17:19:4:1:5:2:1:7:1:1:3:0:0 | 0:0:0:0:0:0:0:0:0:0:0:0:0:0:0:0:0:0 | 12:10:17:0:1:0:2:0:0:2:0:0:0:0:0:0:1:1 | 0:0:0:0:0:0:0:0:0:0:0:0:0:0:0:0:0:0 |
| XM_006712975:NM_153480:XM_011533362:NR_104198:NM_001193380:NM_153481:XM_011533361:XM_006712976:NM_153483:XR_940382:XM_011533363:XR_940380 | 0:0:0:0:0:0:0:0:0:0:0:0 | 1:12:1:1:1:3:1:1:0:0:0:0 | 0:0:0:0:0:0:0:0:0:0:0:0 | 0:12:0:1:8:1:15:5:2:1:0:0 | 0:0:0:0:0:0:0:0:0:0:0:0 | 4:70:3:6:6:10:16:20:6:2:2:0 | 0:0:0:0:0:0:0:0:0:0:0:0 | 3:40:1:5:11:14:107:14:10:2:0:1 | 0:0:0:0:0:0:0:0:0:0:0:0 | 1:14:0:0:2:1:5:1:2:0:0:0 |
| NM_015513:NM_001077415 | 0:00 | 0:00 | 0:00 | 0:00 | 0:00 | 0:00 | 0:00 | 1:00 | 0:00 | 0:00 |
| NM_018447:XM_005265321 | 400:01:00 | 2:00 | 466:02:00 | 4:00 | 1106:09:00 | 12:00 | 2041:01:00 | 36:00:00 | 487:03:00 | 4:00 |
| NM_018447:XM_005265321 | 405:09:00 | 0:00 | 472:08:00 | 1:00 | 1120:33:00 | 1:00 | 2088:40:00 | 2:00 | 492:26:00 | 1:01 |
| NM_018447:XM_005265321 | 405:09:00 | 0:00 | 468:10:00 | 1:00 | 1119:42:00 | 2:00 | 2082:44:00 | 2:00 | 490:29:00 | 0:00 |
| NM_018447:XM_005265321 | 404:09:00 | 1:00 | 465:11:00 | 0:00 | 1117:43:00 | 1:00 | 2073:46:00 | 4:00 | 487:31:00 | 0:00 |
| NM_018447:XM_005265321 | 403:09:00 | 1:00 | 464:11:00 | 1:00 | 1115:43:00 | 3:00 | 2075:46:00 | 2:00 | 487:31:00 | 0:00 |
| NM_018447:XM_005265321 | 396:10:00 | 0:00 | 443:10:00 | 3:00 | 1100:46:00 | 2:02 | 2042:53:00 | 4:00 | 480:33:00 | 1:00 |
| NM_018447:XM_005265321 | 393:12:00 | 2:00 | 438:10:00 | 0:00 | 1088:46:00 | 0:00 | 2028:56:00 | 2:00 | 479:33:00 | 1:00 |
| NM_018447:XM_005265321 | 395:12:00 | 0:00 | 437:10:00 | 0:00 | 1087:45:00 | 1:01 | 2029:56:00 | 0:00 | 479:31:00 | 1:01 |
| NM_018447:XM_005265321 | 379:16:00 | 1:00 | 409:10:00 | 3:00 | 1052:46:00 | 2:00 | 1980:63 | 1:00 | 469:28:00 | 1:00 |
| NM_018447:XM_005265321 | 376:16:00 | 0:00 | 400:10:00 | 3:00 | 1045:44:00 | 0:00 | 1954:56:00 | 4:00 | 464:28:00 | 0:00 |
| NM_018447:XM_005265321 | 376:15:00 | 0:00 | 398:10:00 | 4:00 | 1040:44:00 | 1:00 | 1952:54:00 | 1:00 | 461:28:00 | 2:00 |
| NM_017491 | 0 | 0 | 0 | 0 | 0 | 0 | 0 | 0 | 0 | 2 |
| NM_020139 | 0 | 0 | 0 | 0 | 0 | 1 | 1 | 1 | 0 | 1 |
| XM_005263082:XR_938747:NM_001127208:XM_024454102 | 1:0:0:0 | 0:0:0:0 | 4:21:0:0 | 0:0:0:0 | 7:6:1:1 | 0:0:0:0 | 1:21:0:1 | 0:0:0:0 | 0:0:0:0 | 0:0:0:0 |
| XM_005263082:XR_938747:NM_001127208:XM_024454102 | 1:0:0:0 | 0:0:0:0 | 4:21:0:0 | 0:0:0:0 | 7:6:1:1 | 0:0:0:0 | 1:21:0:1 | 0:0:0:0 | 0:0:0:0 | 0:0:0:0 |
| XM_005263082:XR_938747:XM_024454102:NM_001127208 | 0:0:0:0 | 1:0:0:0 | 0:4:0:0 | 3:18:0:0 | 1:1:0:0 | 5:5:1:0 | 0:2:0:0 | 0:11:1:0 | 0:0:0:0 | 0:0:0:0 |
| NM_176869 | 2 | 0 | 0 | 2 | 2 | 1 | 2 | 0 | 0 | 3 |
| NM_005327 | 0 | 0 | 0 | 1 | 0 | 2 | 0 | 5 | 0 | 1 |
| NM_001319236:NM_000995:NM_033625 | 39:00:00 | 0:00:00 | 33:01:01 | 0:00:00 | 28:01:02 | 0:00:00 | 140:00:03 | 0:00:00 | 83:02:01 | 2:00:00 |
| NM_016648:XM_024454083:NM_001267039:NM_015454:XM_024454089:XM_024454086:XM_024454087:XM_024454084 | 6:3:1:14:0:0:0:0 | 0:0:0:0:0:0:0:0 | 13:1:0:21:1:5:1:0 | 0:0:0:1:0:0:0:0 | 18:0:0:12:1:4:0:0 | 0:0:0:0:0:0:0:0 | 65:4:1:98:5:11:0:0 | 0:1:0:0:0:0:0:0 | 8:0:0:14:4:2:1:0 | 0:0:0:0:0:0:0:0 |
| NM_016648:XM_024454083:NM_015454:XM_024454089:XM_024454086:XM_024454087:NM_001267039:XM_024454084 | 6:1:9:0:0:0:0:0 | 0:0:0:0:0:0:0:0 | 9:0:11:1:0:0:0:0 | 0:0:1:0:0:0:0:0 | 18:0:9:0:1:0:0:0 | 0:0:0:0:0:0:0:0 | 60:2:63:2:0:0:0:0 | 1:1:1:0:1:0:0:0 | 8:0:7:1:0:1:0:0 | 0:0:0:0:0:0:0:0 |
| NM_015454:NM_016648:XM_024454083:XM_024454089:XM_024454086:XM_024454087:NM_001267039:XM_024454084 | 8:6:1:0:0:0:0:0 | 1:0:0:0:0:0:0:0 | 11:8:0:1:0:0:0:0 | 1:1:0:0:0:0:0:0 | 9:18:0:0:1:0:0:0 | 0:0:0:0:0:0:0:0 | 60:60:2:2:0:0:0:0 | 4:1:1:0:1:0:0:0 | 7:8:0:1:0:1:0:0 | 0:0:0:1:0:0:0:0 |
| NM_015454:NM_016648:XM_024454083:XM_024454089:XM_024454086:XM_024454087:NM_001267039:XM_024454084 | 8:6:1:0:0:0:0:0 | 1:0:0:0:0:0:0:0 | 12:8:0:1:0:0:0:0 | 0:1:0:0:0:0:0:0 | 9:18:0:0:1:0:0:0 | 0:0:0:0:0:0:0:0 | 61:60:2:2:0:0:0:0 | 3:1:1:0:1:0:0:0 | 7:8:0:1:0:1:0:0 | 0:0:0:1:0:0:0:0 |
| NM_016648:XM_024454083:NM_015454:XM_024454086:XM_024454089:XM_024454087:NM_001267039:XM_024454084 | 6:1:8:0:0:0:0:0 | 0:0:0:0:0:0:0:0 | 9:0:11:0:0:0:0:0 | 0:0:0:0:0:0:0:0 | 18:0:9:1:0:0:0:0 | 0:0:0:0:0:0:0:0 | 60:2:58:0:2:0:0:0 | 0:0:1:0:0:0:0:0 | 8:0:7:0:1:1:0:0 | 0:0:0:0:0:0:0:0 |
| NM_016648:XM_024454083:NM_015454:XM_024454089:XM_024454086:XM_024454087:NM_001267039:XM_024454084 | 6:1:8:0:0:0:0:0 | 0:0:0:0:0:0:0:0 | 8:0:10:1:0:0:0:0 | 0:0:0:0:0:0:0:0 | 18:0:7:0:1:0:0:0 | 0:0:2:0:0:0:0:0 | 57:1:51:2:0:0:0:0 | 0:0:1:0:0:0:0:0 | 8:0:6:1:0:1:0:0 | 0:0:0:0:0:0:0:0 |
| NM_016648:NM_015454:XM_024454089:XM_024454086:XM_024454083:XM_024454087:NM_001267039:XM_024454084 | 5:7:0:0:0:0:0:0 | 1:0:0:0:0:0:0:0 | 6:8:1:0:0:0:0:0 | 0:1:0:0:0:0:0:0 | 15:6:0:1:0:0:0:0 | 2:3:0:0:0:0:0:0 | 40:42:2:0:1:0:0:0 | 3:3:0:0:0:0:0:0 | 8:5:1:0:0:1:0:0 | 0:1:0:0:0:0:0:0 |
| NM_000134 | 0 | 0 | 0 | 0 | 0 | 2 | 0 | 1 | 0 | 1 |
| NM_005033:NM_001034194:XR_001741241 | 14:01:01 | 2:00:00 | 18:00:01 | 6:00:00 | 11:02:00 | 1:00:00 | 77:03:02 | 19:01:02 | 28:00:00 | 4:00:00 |
| NM_001237 | 0 | 16 | 0 | 2 | 0 | 10 | 31 | 15 | 39 | 12 |
| NM_007083 | 0 | 0 | 0 | 1 | 0 | 0 | 0 | 0 | 0 | 0 |
| NM_001017405:XM_017007637:XR_924910 | 1:00:00 | 0:00:00 | 0:00:00 | 0:00:00 | 1:01:00 | 0:00:00 | 0:01:01 | 0:00:00 | 0:00:00 | 0:00:00 |
| XR_001741343:XM_024454275:NM_003866:XM_017008794:XM_017008796:XM_024454273 | 1:0:0:0:0:0 | 0:0:0:0:0:0 | 0:3:1:1:0:0 | 0:0:0:0:0:0 | 0:0:3:0:2:1 | 0:0:0:0:0:0 | 4:13:1:2:0:5 | 1:0:0:0:0:0 | 0:0:1:0:0:2 | 0:0:0:0:0:0 |
| NM_006726:NM_001199282:XM_017008874:XM_011532434:XM_017008873 | 2:1:0:0:0 | 0:0:0:0:0 | 4:0:4:1:0 | 0:0:0:0:0 | 5:0:3:2:0 | 0:0:0:0:0 | 21:0:25:3:0 | 0:0:0:0:0 | 1:0:4:0:0 | 0:0:0:0:0 |
| NM_006726:NM_001199282:XM_017008874:XM_011532434:XM_017008873 | 2:1:0:0:0 | 0:0:0:0:0 | 2:0:4:1:0 | 1:0:0:0:0 | 5:0:3:2:0 | 0:0:0:0:0 | 19:0:21:3:0 | 0:0:0:0:0 | 1:0:4:0:0 | 0:0:0:0:0 |
| NM_001006:NM_001267699 | 61:00:00 | 0:00 | 22:00 | 0:00 | 59:00:00 | 0:00 | 137:00:00 | 2:00 | 112:01:00 | 0:00 |
| NM_001006:NM_001267699 | 64:00:00 | 0:00 | 23:00 | 0:00 | 62:00:00 | 0:00 | 145:00:00 | 0:00 | 116:00:00 | 0:00 |
| XM_024454011 | 0 | 0 | 0 | 1 | 0 | 0 | 0 | 0 | 0 | 0 |
| NM_018352 | 0 | 0 | 0 | 0 | 0 | 0 | 0 | 0 | 0 | 0 |
| NM_018352 | 0 | 0 | 0 | 0 | 0 | 0 | 0 | 0 | 0 | 0 |
| XM_005263316:NM_032783 | 0:00 | 0:00 | 0:00 | 0:00 | 0:00 | 1:01 | 0:00 | 0:00 | 0:00 | 0:00 |
| NM_017867 | 0 | 0 | 0 | 0 | 0 | 0 | 0 | 0 | 0 | 1 |
| NM_006342:XM_017007653:XM_005247929:XM_011513386 | 5:1:0:0 | 0:0:0:0 | 3:2:1:0 | 0:0:0:0 | 5:3:0:2 | 0:0:0:0 | 13:5:1:1 | 0:0:0:0 | 13:2:1:0 | 0:0:0:0 |
| NM_006342:XM_017007653:XM_005247929:XM_011513386 | 5:1:0:0 | 0:0:0:0 | 3:2:1:0 | 0:0:0:0 | 5:3:0:2 | 0:0:0:0 | 13:7:1:1 | 0:0:0:0 | 13:3:1:0 | 0:0:0:0 |
| NM_006342:XM_017007653:XM_005247929:XM_011513386:XM_005247930 | 5:1:0:0:0 | 0:0:0:0:0 | 3:7:1:0:0 | 0:0:0:0:0 | 6:7:0:0:0 | 0:0:0:0:0 | 15:19:1:1:0 | 0:0:0:0:0 | 15:8:1:0:1 | 0:0:0:0:0 |
| NM_006342:XM_017007653:XM_005247929:XM_005247930:XM_011513386 | 5:1:0:0:0 | 0:0:0:0:0 | 4:7:1:1:0 | 0:0:0:0:0 | 7:11:0:0:2 | 0:0:0:0:0 | 19:29:1:0:3 | 0:0:0:0:0 | 19:10:1:1:0 | 1:0:0:0:0 |
| NM_002199:XM_024454037:XM_024454036:XM_024454039:XM_024454034:XM_024454038 | 44:3:2:2:0:0 | 0:0:0:0:0:0 | 48:3:4:3:3:0 | 1:0:0:0:0:0 | 193:8:15:14:3:1 | 0:0:0:2:0:0 | 195:9:28:43:1:1 | 0:0:0:1:0:0 | 35:3:3:1:1:0 | 0:0:0:0:0:0 |
| NM_002199:XM_024454037:XM_024454036:XM_024454039:XM_024454034:XM_024454038 | 44:3:2:2:0:0 | 0:0:0:0:0:0 | 49:3:4:3:3:0 | 0:0:0:0:0:0 | 192:8:15:13:3:1 | 0:0:0:1:0:0 | 194:9:28:43:1:1 | 1:0:0:0:0:0 | 35:3:3:1:1:0 | 0:0:0:0:0:0 |
| NM_002199:XM_024454037:XM_024454036:XM_024454039:XM_024454034:XM_024454038 | 44:3:2:2:0:0 | 0:0:0:0:0:0 | 49:3:4:3:3:0 | 0:0:0:0:0:0 | 192:8:15:14:3:1 | 0:0:0:0:0:0 | 195:9:28:43:1:1 | 0:0:0:0:0:0 | 35:3:3:1:1:0 | 0:0:0:0:0:0 |
| NM_001995:NM_001286710:XM_017007888:XM_005262828:XM_005262829:NM_001286711:XM_017007889 | 13:2:1:0:0:0:0 | 0:0:0:0:0:0:0 | 2:0:0:0:0:0:0 | 0:0:0:0:0:0:0 | 8:0:0:2:1:0:0 | 0:0:0:0:0:0:0 | 13:7:0:0:0:0:0 | 0:0:0:0:0:0:0 | 2:0:0:1:0:1:0 | 0:0:0:0:0:0:0 |
| XM_024454160:XM_017008483:NM_020827 | 1:01:00 | 0:00:00 | 1:00:00 | 0:00:00 | 0:00:00 | 0:00:00 | 0:00:00 | 15:03:01 | 0:00:00 | 3:00:00 |
| XM_017008774:XM_005263308 | 0:00 | 0:00 | 0:00 | 0:00 | 0:00 | 0:00 | 2:01 | 0:00 | 0:00 | 0:00 |
| XM_006714139 | 0 | 0 | 0 | 0 | 0 | 1 | 0 | 12 | 0 | 0 |
| XM_006714139 | 0 | 0 | 0 | 1 | 0 | 1 | 2 | 4 | 1 | 1 |
| XM_006714139 | 1 | 0 | 0 | 0 | 1 | 0 | 11 | 13 | 2 | 0 |
| XM_006714139 | 0 | 1 | 0 | 0 | 0 | 1 | 13 | 6 | 1 | 1 |
| XM_006714139 | 1 | 0 | 1 | 0 | 2 | 0 | 23 | 1 | 3 | 0 |
| XM_006714139 | 0 | 1 | 1 | 0 | 1 | 0 | 0 | 25 | 0 | 2 |
| XM_006714139 | 0 | 1 | 0 | 0 | 1 | 0 | 0 | 16 | 0 | 3 |
| XM_006714139 | 0 | 0 | 0 | 0 | 1 | 0 | 1 | 3 | 0 | 1 |
| XM_006714139 | 0 | 0 | 0 | 0 | 1 | 0 | 1 | 4 | 0 | 1 |
| XM_006714139 | 0 | 0 | 1 | 0 | 1 | 0 | 0 | 9 | 0 | 2 |
| XM_006714139 | 0 | 0 | 0 | 2 | 0 | 2 | 14 | 4 | 1 | 0 |

| | | | | | | | | | | |
|---|---|---|---|---|---|---|---|---|---|---|
| XM_011513557:XM_005248005:NM_007331:XM_006713914:NM_001042424:XM_005248001:XM_005248002 | 0:1:1:1:0:0:0 | 1:0:0:0:0:0:0 | 0:0:0:1:0:0:0 | 11:1:0:0:4:0:0 | 0:0:0:0:0:0:0 | 0:0:0:1:0:0:0 | 2:1:0:9:0:0:0 | 5:4:0:0:2:0:0 | 1:0:1:3:0:0:1 | 0:0:0:0:1:1:0 |
| NM_001168243:NM_001141936:XM_011513469 | 0:00:00 | 2:00:00 | 0:00:00 | 4:00:00 | 1:00:00 | 2:00:00 | 0:00:00 | 1:02:00 | 0:00:00 | 6:00:02 |
| NM_001358:XR_001741152 | 67:16:00 | 0:00 | 52:18:00 | 0:00 | 122:35:00 | 0:00 | 406:101 | 3:00 | 106:21:00 | 0:00 |
| NM_001358:XR_001741152 | 67:16:00 | 0:00 | 41:20:00 | 0:00 | 118:37:00 | 1:00 | 295:94 | 1:00 | 103:23:00 | 2:00 |
| NM_003102:XR_427488 | 0:00 | 0:00 | 1:00 | 0:00 | 0:00 | 0:00 | 1:00 | 0:00 | 0:01 | 0:00 |
| NM_001098634:XM_005248103 | 0:02 | 1:00 | 0:00 | 0:00 | 0:00 | 2:03 | 0:00 | 0:00 | 1:01 | 0:00 |
| XM_017007900:XM_024453938:XM_024453937:NM_014988:NM_001330790:NM_001330792:NM_001289124:XM_011513646:NM_001112718:NM_001289122 | 1:1:1:0:0:0:0:0:0:0 | 1:1:0:0:0:0:0:0:0:0 | 4:0:0:0:1:0:0:0:0:0 | 6:0:0:1:0:0:0:0:0:0 | 0:0:1:0:0:0:0:0:0:0 | 0:0:0:0:0:0:0:0:0:0 | 2:0:7:0:0:2:1:1:0:0 | 0:0:10:0:0:1:0:0:0:0 | 0:0:0:0:0:1:0:0:0:0 | 0:0:0:0:0:1:0:0:0:0 |
| XM_024453938:XM_024453937:NM_001330790:NM_001289124:XM_011513646:NM_001330792:NM_001112718:XM_006713996:NM_001289122:XM_017007900:NM_001330982 | 0:0:0:0:0:0:0:0:0:0:0 | 1:1:0:0:0:0:0:0:0:0:0 | 0:0:0:0:0:0:0:0:0:0:0 | 0:0:1:0:0:0:0:0:0:0:0 | 0:0:0:0:0:0:0:0:0:0:0 | 0:0:0:0:0:0:0:0:0:0:0 | 0:1:0:1:1:0:0:0:0:0:0 | 0:0:0:0:0:0:0:0:0:0:0 | 0:0:0:0:0:1:0:0:0:0:0 | 0:0:0:0:0:0:0:0:0:0:0 |
| NM_001330790:XM_006713996:NM_001289124:XM_011513646:XM_024453938:NM_001330792:NM_001112718:NM_001289122:NM_001330982 | 0:0:0:0:0:0:0:0:0 | 0:0:0:0:0:0:0:0:0 | 1:1:0:0:0:0:0:0:0 | 0:0:0:0:0:0:0:0:0 | 0:0:0:0:0:0:0:0:0 | 0:0:0:0:0:0:0:0:0 | 0:0:0:0:0:0:0:0:0 | 0:2:2:1:1:0:0:0:0 | 0:0:0:0:0:0:0:0:0 | 0:0:0:0:0:1:0:0:0 |
| NM_006345:XR_001741095:XM_017007654 | 0:00:00 | 1:00:00 | 0:00:00 | 0:01:00 | 0:00:00 | 1:00:00 | 0:00:00 | 0:00:01 | 0:00:00 | 0:00:00 |
| NM_006345:XR_001741095:XM_017007654 | 0:00:00 | 1:00:00 | 0:00:00 | 0:01:00 | 0:00:00 | 1:00:00 | 0:00:00 | 0:00:01 | 0:00:00 | 0:00:00 |
| NM_017816:XM_011513506:XM_011513505:NM_001145725 | 44:2:0:0 | 23:2:0:0 | 7:2:0:0 | 19:3:1:0 | 28:3:0:0 | 21:0:0:0 | 226:22:2:0 | 268:9:0:1 | 75:3:1:0 | 43:3:0:1 |
| NM_017816:XM_011513506:XM_011513505:NM_001145725 | 67:4:0:0 | 0:0:0:0 | 28:5:1:0 | 0:0:0:0 | 51:3:0:0 | 0:0:0:0 | 514:31:2:1 | 1:0:0:0 | 119:6:2:1 | 0:0:0:0 |
| NM_017816:XM_011513506:XM_011513505:NM_001145725 | 72:5:0:0 | 2:0:0:0 | 28:5:1:0 | 0:0:0:0 | 57:3:0:0 | 0:0:0:0 | 570:34:2:1 | 8:0:0:0 | 132:7:2:1 | 1:0:0:0 |
| NM_133474 | 2 | 0 | 4 | 1 | 1 | 0 | 15 | 0 | 3 | 0 |
| NM_133474 | 2 | 0 | 3 | 0 | 1 | 0 | 14 | 2 | 2 | 0 |
| NM_133474 | 1 | 0 | 3 | 0 | 1 | 0 | 14 | 3 | 1 | 0 |
| NM_133474 | 2 | 0 | 1 | 0 | 0 | 0 | 23 | 1 | 0 | 0 |
| NM_133474 | 0 | 3 | 1 | 1 | 1 | 2 | 41 | 0 | 1 | 0 |
| NM_017830:XM_024454108:XM_024454106:XM_024454105:NM_001079839:XR_001741265:XM_024454107:XR_002959740:NM_001168254:XR_001741266 | 140:22:3:1:10:1:2:1:0:0 | 1:0:0:0:0:0:0:0:0:0 | 257:41:7:2:52:2:4:2:4:0 | 0:0:0:0:0:0:0:0:0:0 | 188:29:9:5:40:1:5:0:1:0 | 0:0:0:0:0:0:0:0:0:0 | 1827:174:22:38:121:11:11:5:14:13 | 0:1:0:0:0:0:0:0:0:0 | 200:17:1:6:13:0:1:1:3:0 | 0:0:0:0:0:0:0:0:0:0 |
| XM_024454108:NM_017830:XM_024454106:XM_024454105:NM_001079839:XR_001741265:XM_024454107:XR_002959740:NM_001168254:XR_001741266 | 26:141:3:1:10:1:2:1:0:0 | 0:0:0:0:0:0:0:0:0:0 | 61:258:7:2:52:2:4:2:4:0 | 0:1:0:0:0:0:0:0:0:0 | 34:187:9:5:40:0:5:0:1:0 | 0:1:0:0:0:0:0:0:0:0 | 235:1830:22:38:121:8:11:5:14:13 | 0:3:0:0:0:0:0:0:0:0 | 21:199:1:6:13:0:1:0:3:0 | 0:1:0:0:0:0:0:0:0:0 |
| XM_024454108:NM_017830:XM_024454106:XM_024454105:NM_001079839:XM_024454107:NM_001168254:XR_001741265:XR_002959740:XR_001741266 | 28:141:3:1:10:2:0:0:0:0 | 0:0:0:0:0:0:0:0:0:0 | 70:257:7:2:52:4:4:2:2:0 | 0:2:0:0:0:0:0:0:0:0 | 34:189:8:5:40:5:1:0:0:0 | 0:0:1:0:0:0:0:0:0:0 | 258:1836:22:38:121:11:14:8:5:13 | 1:0:0:0:0:0:0:0:0:0 | 23:200:1:6:13:1:3:0:0:0 | 0:0:0:0:0:0:0:0:0:0 |
| XM_024454108:NM_017830:XM_024454106:XM_024454105:NM_001079839:XM_024454107:NM_001168254:XR_001741265:XR_002959740:XR_001741266 | 28:141:3:1:10:2:0:0:0:0 | 0:0:0:0:0:0:0:0:0:0 | 75:259:7:2:52:4:4:2:2:0 | 0:0:0:0:0:0:0:0:0:0 | 34:190:9:5:40:5:1:0:1:0 | 0:0:0:0:0:0:0:0:0:0 | 264:1837:21:38:121:11:14:8:4:13 | 2:2:1:0:0:0:0:0:0:0 | 22:199:1:6:13:1:3:0:0:0 | 1:1:0:0:0:0:0:0:0:0 |
| XM_024454108:NM_017830:XM_024454106:XM_024454105:NM_001079839:XM_024454107:NM_001168254:XR_001741265:XR_002959740:XR_001741266 | 28:141:3:1:10:2:0:0:0:0 | 0:0:0:0:0:0:0:0:0:0 | 81:260:7:2:51:4:4:2:0:0 | 0:0:0:0:1:0:0:0:0:0 | 34:190:9:5:40:5:1:0:1:0 | 0:0:0:0:0:0:0:0:0:0 | 279:1839:22:38:120:11:14:7:3:13 | 1:2:0:0:0:0:0:0:0:0 | 28:200:1:6:13:1:3:0:0:0 | 0:0:0:0:0:0:0:0:0:0 |
| XM_024454108:NM_017830:NM_001079839:XM_024454106:XM_024454107:XR_002959740:NM_001168254:XM_024454105:XR_001741265 | 16:90:8:2:2:3:0:0:0 | 0:0:0:0:0:0:0:0:0 | 91:131:23:0:2:14:3:1:1 | 0:1:0:0:0:0:0:0:0 | 24:145:33:5:5:4:1:5:0 | 0:0:0:0:0:0:0:0:0 | 252:892:67:13:5:39:11:18:1 | 0:2:0:0:0:2:0:0:0 | 17:142:10:0:1:5:3:2:0 | 0:0:0:0:0:0:0:0:0 |
| NM_025087:XM_011513758:XM_011513755 | 0:00:00 | 0:00:00 | 0:00:00 | 25:03:01 | 0:00:00 | 29:02:01 | 0:01:00 | 0:00:00 | 2:00:00 | 0:00:00 |
| NM_025087:XM_011513758:XM_011513755:XM_011513757:XM_011513759 | 0:0:0:0:0 | 0:0:0:0:0 | 0:0:0:0:0 | 45:3:1:0:0 | 0:0:0:0:0 | 31:2:1:0:0 | 2:1:0:1:1 | 0:0:0:0:0 | 2:0:0:0:0 | 0:0:0:0:0 |
| NM_030917:XM_017008666:XM_005265781:XM_005265774:XM_017008665:XM_005265769:XM_005265782:XM_017008669:NM_001134938:XM_005265773:XM_017008662:XM_017008664:XM_017008663:XM_017008667:XM_017008668:NM_001134937:XM_005265779:XM_017008671:XM_017008670:XM_005265778 | 1:8:3:8:3:3:7:9:6:4:9:2:2:1:3:0:0:0:0:0 | 1:1:0:0:0:0:0:0:0:0:0:0:0:0:0:0:0:0:0:0 | 4:17:6:10:6:8:6:11:5:7:6:4:3:3:6:4:0:0:0:0 | 0:0:0:0:0:0:0:0:0:0:0:0:0:0:0:0:0:0:0:0 | 7:17:6:12:1:3:6:6:5:8:0:9:1:0:3:4:1:1:0:0 | 0:0:0:0:0:0:0:0:0:0:0:0:0:0:0:0:0:0:0:0 | 62:33:15:40:37:29:36:38:31:20:43:28:19:13:64:14:0:1:1:1 | 0:0:0:0:0:0:0:0:0:1:0:0:0:0:1:0:0:0:0:0 | 4:12:1:9:2:2:8:2:5:6:3:4:2:2:8:2:1:0:0:0 | 0:0:0:0:0:0:0:0:0:0:0:0:0:0:0:0:0:0:0:0 |
| XM_017008662:XM_017008669:XM_005265781:XM_005265774:NM_030917:XM_017008666:XM_017008665:XM_005265769:XM_005265782:XM_005265778:NM_001134938:XM_005265773:XM_017008664:XM_017008663:XM_005265779:XM_017008667:XM_017008668:XM_017008671:NM_001134937:XM_017008670 | 7:7:3:8:2:9:3:3:7:1:6:4:2:2:1:1:3:1:0:0 | 2:1:0:0:0:0:0:0:0:0:0:0:0:0:0:0:0:0:0:0 | 4:9:6:9:3:17:4:7:6:0:4:7:3:4:0:3:6:1:4:1 | 1:3:0:1:1:0:2:0:0:0:0:0:1:0:0:0:0:0:0:0 | 0:6:6:11:6:16:1:3:4:0:5:7:9:1:1:0:3:1:3:0 | 0:0:0:1:1:1:0:0:2:0:0:1:0:0:0:0:0:0:1:0 | 39:36:15:39:56:32:34:28:32:2:29:19:28:16:0:12:62:1:16:1 | 2:2:0:0:6:0:3:1:2:0:2:0:0:3:0:1:1:0:0:0 | 3:3:1:9:4:12:3:2:8:0:5:6:4:2:1:2:8:0:2:0 | 0:0:0:0:0:0:0:0:0:0:0:1:0:0:0:0:0:0:0:0 |
| NM_030917:XM_005265774:XM_017008666:XM_017008665:XM_005265769:XM_005265782:XM_005265778:XM_005265773:XM_017008662:XM_017008664:XM_017008669:NM_001134938:XM_017008663:XM_005265779:XM_005265781:XM_017008668:XM_017008671:NM_001134937:XM_017008670:XM_017008667 | 1:7:4:2:1:3:1:4:5:2:4:3:2:1:1:1:1:0:0:0 | 1:0:0:0:0:0:0:0:0:0:0:0:0:0:0:0:0:0:0:0 | 1:6:8:3:6:3:0:3:1:2:8:2:1:0:2:5:0:2:1:3 | 0:0:1:0:0:1:0:0:2:0:0:0:0:0:0:0:0:0:0:0 | 3:7:13:0:1:1:0:5:0:5:5:2:1:0:3:3:1:2:0:0 | 0:0:0:0:0:0:0:0:0:0:0:0:0:0:0:0:0:0:0:0 | 23:19:18:13:13:19:0:12:18:10:16:15:11:0:8:28:1:10:0:4 | 3:2:1:0:1:0:0:1:2:2:0:0:0:0:3:0:0:0:0:0 | 3:7:7:3:2:6:0:5:3:2:2:4:1:0:0:2:0:2:0:2 | 0:0:0:0:0:0:0:0:1:0:0:0:0:0:0:0:0:0:0:0 |
| XM_005265769:NM_030917:XM_017008666:NM_001134938:XM_005265781:XM_017008665:XM_005265782:XM_005265774:XM_017008662:XM_017008667:XM_017008668:XM_017008669:XM_005265778:XM_005265773:XM_017008664:XM_017008663:XM_005265779:XM_017008671:NM_001134937:XM_017008670 | 1:1:4:3:1:2:2:7:5:0:1:4:1:4:2:2:1:1:0:0 | 2:1:4:3:1:1:3:1:1:1:1:1:0:0:0:0:0:0:0:0 | 5:0:6:1:2:2:2:6:1:2:3:8:0:3:2:1:0:0:2:1 | 2:3:4:1:0:2:4:2:2:1:2:0:0:1:0:2:0:1:0:0 | 1:2:12:2:3:0:1:7:0:0:3:4:0:5:5:1:0:1:2:0 | 2:4:2:3:3:1:1:4:0:0:0:0:0:2:4:0:1:0:1:0 | 7:17:13:14:6:8:13:12:13:1:18:10:0:8:6:7:0:1:9:0 | 12:16:9:5:3:10:11:16:10:4:20:7:1:6:8:6:0:0:4:1 | 2:2:7:4:0:3:6:7:2:2:2:2:0:5:2:1:0:0:2:0 | 0:1:5:1:1:0:1:2:0:0:5:1:0:1:1:1:1:1:0:0:0 |
| NM_006452:NM_001079524:XM_011534366:NM_001079525 | 28:88:3:0 | 0:0:0:0 | 6:19:1:0 | 0:0:0:0 | 11:38:5:0 | 0:0:0:0 | 267:673:34:0 | 2:1:0:0 | 65:145:9:0 | 0:0:0:0 |
| NM_006452:NM_001079524:XM_011534366:NM_001079525 | 28:88:3:0 | 0:0:0:0 | 6:19:1:0 | 0:0:0:0 | 11:38:5:0 | 0:0:0:0 | 290:673:37:0 | 1:1:0:0 | 66:145:9:0 | 0:0:0:0 |
| NM_006452:NM_001079524:XM_011534366:NM_001079525 | 30:88:4:0 | 0:0:0:0 | 6:19:1:0 | 0:0:0:0 | 13:38:6:0 | 0:0:0:0 | 311:673:44:0 | 1:0:0:0 | 69:144:9:0 | 0:1:0:0 |
| NM_006452:NM_001079524:XM_011534366:NM_001079525 | 31:88:4:0 | 0:0:0:0 | 6:19:1:0 | 0:0:0:0 | 13:38:6:0 | 0:0:0:0 | 329:671:45:0 | 1:1:1:0 | 70:144:9:0 | 1:1:0:0 |
| NM_006452:NM_001079524:XM_011534366:NM_001079525 | 35:88:3:0 | 0:0:0:0 | 7:19:1:0 | 0:0:0:0 | 15:38:5:0 | 0:0:0:0 | 366:672:49:0 | 0:0:0:0 | 76:144:10:0 | 0:0:0:0 |
| NM_006452:NM_001079524:XM_011534366:NM_001079525 | 35:88:3:0 | 0:0:0:0 | 7:19:1:0 | 0:0:0:0 | 15:38:5:0 | 0:0:0:0 | 370:670:48:0 | 1:2:0:0 | 77:143:10:0 | 0:1:0:0 |
| NM_001079524:NM_006452:XM_011534366:NM_001079525 | 87:39:3:0 | 1:0:0:0 | 19:7:2:0 | 0:0:0:0 | 38:16:5:0 | 0:0:0:0 | 669:388:46:0 | 1:0:1:0 | 144:80:11:0 | 0:0:0:0 |
| NM_006452:NM_001079524:XM_011534366:NM_001079525 | 39:88:3:0 | 0:0:0:0 | 7:19:2:0 | 0:0:0:0 | 16:38:4:0 | 0:0:0:0 | 406:670:44:0 | 0:0:0:0 | 81:143:10:0 | 0:0:0:0 |
| NM_001145460 | 0 | 0 | 0 | 0 | 0 | 0 | 0 | 0 | 0 | 0 |
| NM_000938:NM_001303269 | 6:11 | 0:00 | 0:47 | 0:01 | 6:30 | 0:00 | 0.227083333 | 0:02 | 10:37 | 0:00 |
| NM_000938:NM_001303269 | 6:11 | 0:00 | 0:46 | 0:02 | 6:30 | 0:00 | 0.229861111 | 0:01 | 10:37 | 0:00 |
| NM_001553:NM_001253835 | 7:01 | 0:00 | 3:00 | 0:00 | 2:00 | 0:00 | 0:00 | 0:00 | 4:00 | 3:00 |
| XM_017008586 | 1 | 0 | 0 | 0 | 0 | 0 | 0 | 2 | 0 | 0 |
| NM_002092:XM_011531897 | 0:00 | 0:00 | 0:00 | 0:00 | 0:00 | 0:00 | 0:00 | 2:00 | 0:00 | 1:00 |
| NM_017426 | 0 | 0 | 0 | 0 | 0 | 0 | 0 | 0 | 0 | 0 |
| NM_005139 | 0 | 0 | 0 | 0 | 3 | 2 | 1 | 0 | 3 | 0 |
| NM_004684:NM_001128310 | 36:00:00 | 0:00 | 46:00:00 | 0:00 | 48:01:00 | 1:00 | 29:00:00 | 0:00 | 45:00:00 | 1:00 |
| NM_004684:NM_001128310 | 37:00:00 | 0:00 | 43:00:00 | 0:00 | 50:01:00 | 0:00 | 25:00:00 | 0:00 | 43:00:00 | 0:00 |
| NM_004684:NM_001128310 | 44:01:00 | 3:00 | 79:02:00 | 4:00 | 58:02:00 | 2:00 | 42:00:00 | 4:00 | 54:00:00 | 3:01 |
| NM_004684:NM_001128310:NM_001291976 | 56:01:00 | 0:00:00 | 112:02:00 | 5:00:00 | 65:02:00 | 4:01:00 | 50:00:00 | 6:00:00 | 62:01:00 | 1:01:00 |
| NM_004684:NM_001128310:NM_001291976 | 54:01:00 | 0:00:00 | 117:02:00 | 0:00:00 | 70:03:00 | 0:00:00 | 32:00:00 | 26:00:00 | 30:02:00 | 33:00:00 |
| NM_004684:NM_001128310:NM_001291976 | 54:01:00 | 0:00:00 | 129:02:00 | 0:00:00 | 71:03:00 | 0:00:00 | 59:00:00 | 1:00:00 | 65:02:00 | 0:00:00 |
| NM_004684:NM_001128310:NM_001291976 | 54:01:00 | 0:00:00 | 129:03:00 | 0:00:00 | 71:03:00 | 0:00:00 | 33:00:00 | 27:00:00 | 30:02:00 | 35:00:00 |

| | | | | | | | | | | |
|---|---|---|---|---|---|---|---|---|---|---|
| XM_017008199:XM_017008197 | 1:00 | 0:00 | 2:00 | 0:00 | 0:00 | 0:00 | 1:03 | 0:00 | 0:00 | 0:00 |
| NM_001100427:NM_021159:XM_024454167:NM_001100426:XR_002959742:XM_024454164:XM_024454175:XM_024454168:XM_024454176:XM_024454174:NM_001100430:XM_024454172:XM_024454173 | 23:1:1:7:2:1:1:1:0:0:0:0:0 | 0:0:0:0:0:0:0:0:0:0:0:0 | 8:1:1:4:0:3:0:2:0:0:0:0 | 0:0:0:0:0:0:0:0:0:0:0 | 23:1:0:7:1:1:0:1:0:0:0:0:0 | 0:0:0:0:0:0:0:0:0:0:0:0 | 103:3:3:29:22:39:1:20:7:4:0:0:0 | 0:0:0:0:0:0:0:1:0:0:0:0 | 36:1:0:14:2:8:0:2:1:0:1:0:0 | 0:0:0:0:0:0:0:0:0:0:0 |
| NM_000668 | 0 | 2 | 0 | 0 | 0 | 0 | 0 | 0 | 0 | 0 |
| NM_000669:NR_133005 | 0:00 | 0:00 | 0:00 | 3:00 | 0:00 | 12:01 | 0:00 | 0:00 | 0:00 | 1:00 |
| NM_000669 | 0 | 0 | 0 | 3 | 0 | 12 | 0 | 0 | 0 | 1 |
| NM_012073 | 0 | 0 | 0 | 0 | 1 | 0 | 5 | 0 | 4 | 0 |
| NM_001316968:NM_033211:NM_001316969 | 0:00:00 | 1:00:00 | 0:01:00 | 0:02:01 | 0:04:00 | 0:01:00 | 0:00:01 | 0:00:00 | 0:00:01 | 0:00:00 |
| NM_004394 | 37 | 0 | 24 | 0 | 33 | 0 | 210 | 0 | 75 | 0 |
| NM_001284:NM_001002924:NM_001318094:NM_001318093:XM_017009024 | 104:1:3:0:0 | 0:0:0:0:0 | 120:1:0:1:0 | 2:0:0:0:0 | 85:1:1:1:1 | 0:0:0:0:0 | 298:14:1:1:1 | 1:0:0:0:0 | 108:3:2:0:0 | 0:0:0:0:0 |
| NM_001284:NM_001002924:NM_001318090:NM_001318091:NM_001318093:XM_017009024:XM_017009025:NM_001318094 | 104:1:1:2:0:0:0:0 | 0:0:0:0:0:0:0:0 | 122:1:1:0:0:0:0:0 | 0:0:0:0:0:0:0:0 | 85:1:1:1:1:1:0:0 | 0:0:0:0:0:0:0:0 | 299:14:3:3:1:0:0:0 | 0:0:0:0:0:0:0:0 | 108:3:2:0:0:0:0:0 | 1:0:0:0:0:0:0:0 |
| NM_016144:NR_146220:NM_001308080 | 110:01:01 | 0:00:00 | 54:01:00 | 0:00:00 | 72:00:00 | 0:00:00 | 151:00:01 | 8:00:00 | 79:01:00 | 72:00:00 |
| NM_016144:NR_146220:NM_001308080 | 110:01:01 | 0:00:00 | 54:01:00 | 0:00:00 | 72:00:00 | 0:00:00 | 159:00:01 | 0:00:00 | 151:01:00 | 0:00:00 |
| NM_016144:NR_146220:NM_001308080 | 110:01:01 | 0:00:00 | 53:01:00 | 1:00:00 | 72:00:00 | 0:00:00 | 159:00:01 | 0:00:00 | 151:01:00 | 0:00:00 |
| NM_000943 | 0 | 0 | 0 | 1 | 0 | 2 | 1 | 0 | 1 | 0 |
| XM_024446256:NM_001017974:NM_001142598:NM_001142599:XM_005272117:XM_005272118:XM_024446255:NM_004199:XM_006714728:XM_005272120 | 0:46:21:5:2:2:1:1:1:0 | 2:0:0:0:0:0:0:0:0:0 | 4:20:11:5:0:1:0:0:2:0 | 0:0:0:0:0:0:0:0:0:0 | 1:39:31:12:6:2:0:0:1:1 | 1:1:0:0:0:0:0:0:0:0 | 6:65:39:6:2:0:0:4:3:0 | 0:0:0:0:0:0:0:0:0:0 | 0:20:13:3:0:1:0:0:0:0 | 0:0:0:0:0:0:0:0:0:0 |
| NM_001017974:NM_001142598:NM_001142599:XM_005272117:XM_005272118:XM_024446255:XM_006714728:XM_024446256:XM_005272120:NM_004199 | 46:20:5:2:2:1:1:3:0:0 | 0:0:0:0:0:0:0:0:0:0 | 20:12:6:0:1:0:2:5:0:0 | 0:0:0:0:0:0:0:0:0:0 | 42:30:12:5:2:0:0:3:1:0 | 0:2:0:1:0:0:0:0:0:0 | 64:40:6:2:0:0:0:9:0:3 | 2:0:0:0:0:0:0:0:0:0 | 19:12:3:0:1:0:0:2:0:0 | 1:0:0:0:0:0:0:0:0:0 |
| NM_004199:NM_001017974:NM_001142598:NM_001142599:XM_005272117:XM_005272118:XM_024446255:XM_024446256:XM_006714728:XM_005272120 | 0:46:22:5:2:2:1:4:1:0 | 1:0:0:0:0:0:0:0:0:0 | 0:20:13:6:0:1:0:5:1:0 | 0:0:0:0:0:0:0:0:0:0 | 0:42:32:12:6:2:0:4:0:1 | 0:0:0:0:0:0:0:0:0:0 | 1:66:40:6:2:0:0:8:0:0 | 0:0:0:0:0:0:0:0:0:0 | 0:20:13:3:0:1:0:1:0:0 | 0:0:0:0:0:0:0:0:0:0 |
| NM_001287252:NM_001039780 | 0:00 | 0:00 | 0:00 | 0:00 | 0:00 | 0:00 | 0:00 | 0:00 | 0:00 | 0:00 |
| XM_011543287:XM_017009251:XM_011543283 | 0:00:00 | 0:00:00 | 0:00:00 | 0:00:00 | 0:00:00 | 0:00:00 | 1:00:00 | 0:00:00 | 1:00:00 | 0:03:00 |
| NM_001040158:XR_948311:NM_004893:NM_138610:XM_011543729:XR_001742353:XR_948306:XM_011543730:XM_005272132:XM_011543732:XM_011543731:NM_138609:XR_001742358:XR_948309:XR_948310:XR_948307:XR_948308 | 163:6:22:118:8:1:2:11:1:0:0:0:0:0:0:0:0 | 2:1:0:0:0:0:0:0:0:0:0:0:0:0:0:0:0 | 76:2:11:54:2:1:2:16:0:2:0:0:0:0:0:0:0 | 0:0:0:0:0:0:0:0:0:0:0:0:0:0:0:0:0 | 60:2:7:50:2:2:2:9:0:0:1:1:0:0:0:0:0 | 0:0:0:0:0:0:0:0:0:0:0:0:0:0:0:0:0 | 347:19:39:243:14:11:2:53:0:6:0:0:1:0:0:0:0 | 1:0:0:2:0:0:0:0:0:0:0:0:0:0:0:0:0 | 213:2:29:135:14:1:1:17:1:3:1:2:0:0:0:0:0 | 0:0:1:1:0:0:0:0:0:0:0:0:0:0:0:0:0 |
| NM_001040158:NM_004893:NM_138610:XM_011543729:XR_001742353:XR_948311:XR_948306:XM_011543730:XM_005272132:XM_011543732:XM_011543731:NM_138609:XR_001742358:XR_948309:XR_948310:XR_948307:XR_948308 | 165:22:118:8:1:7:2:11:1:0:0:0:0:0:0:0:0 | 0:0:0:0:0:0:0:0:0:0:0:0:0:0:0:0:0 | 76:11:53:2:1:2:2:16:0:2:0:0:0:0:0:0:0 | 0:0:1:0:0:0:0:0:0:0:0:0:0:0:0:0:0 | 60:7:50:2:2:2:2:9:0:0:1:1:0:0:0:0:0 | 0:0:0:0:0:0:0:0:0:0:0:0:0:0:0:0:0 | 348:39:246:15:10:16:2:54:0:6:0:0:1:0:0:0:0 | 0:0:0:0:0:1:0:0:0:0:0:0:0:0:0:0:0 | 211:30:136:14:1:2:1:17:1:3:1:2:0:0:0:0:0 | 2:0:0:0:0:0:0:0:0:0:0:0:0:0:0:0:0 |
| XM_011543730:NM_001040158:NM_004893:NM_138610:XM_011543729:XR_001742353:XR_948306:XM_005272132:XR_948311:XR_948308:XM_011543732:XM_011543731:NM_138609:XR_001742358:XR_948309:XR_948310:XR_948307:XM_011543734:XM_011543735 | 4:165:11:117:8:1:4:1:1:1:0:0:0:0:0:0:0:0:0 | 1:0:0:0:0:0:0:0:0:0:0:0:0:0:0:0:0:0:0 | 10:76:5:53:2:1:1:0:1:0:2:0:0:0:0:0:0:0:0 | 0:0:1:0:0:0:0:0:0:0:0:0:0:0:0:0:0:0:0 | 5:60:6:50:2:2:3:0:0:1:0:1:1:0:0:0:0:0:0 | 0:0:0:0:0:0:0:0:0:0:0:0:0:0:0:0:0:0:0 | 32:348:25:246:15:17:4:0:2:2:6:0:0:1:2:1:0:0:0 | 0:0:0:0:0:0:0:0:0:0:0:0:0:0:0:0:0:0:0 | 11:213:15:136:14:3:2:1:0:0:3:1:2:0:0:0:0:0:0 | 0:0:0:0:0:0:0:0:0:0:0:0:0:0:0:0:0:0:0 |
| NM_138610:NM_001040158:NM_004893:XR_001742353:XR_948306:XM_011543729:XR_948311:XM_005272132:XM_011543730:XR_948308:XM_011543732:XM_011543731:NM_138609:XR_001742358:XR_948309:XR_948310:XR_948307:XM_011543734:XM_011543735 | 115:164:7:1:3:4:4:1:2:1:0:0:0:0:0:0:0:0:0 | 1:1:0:0:0:0:0:0:0:0:0:0:0:0:0:0:0:0:0 | 53:76:4:1:1:1:2:0:5:0:1:0:0:0:0:0:0:0:0 | 0:0:0:0:0:0:0:0:0:0:0:0:0:0:0:0:0:0:0 | 48:60:3:2:2:1:0:0:2:0:0:1:1:0:0:0:0:0:0 | 0:0:0:0:0:0:0:0:0:0:0:0:0:0:0:0:0:0:0 | 242:348:18:14:4:6:2:0:21:2:5:0:0:1:1:1:0:0:0 | 0:0:0:0:0:0:0:0:0:0:0:0:0:0:0:0:0:0:0 | 134:213:6:3:0:2:1:1:5:0:3:1:2:0:0:0:0:0:0 | 0:0:0:0:0:0:0:0:0:0:0:0:0:0:0:0:0:0:0 |
| NM_198282 | 0 | 0 | 0 | 0 | 0 | 0 | 0 | 0 | 0 | 1 |
| NR_045587:NM_001035235 | 0:00 | 0:00 | 0:00 | 1:00 | 0:00 | 0:00 | 0:00 | 0:00 | 0:03 | 0:01 |
| NM_017706 | 0 | 0 | 0 | 0 | 0 | 0 | 0 | 0 | 0 | 0 |
| NM_019018 | 4 | 0 | 1 | 0 | 1 | 0 | 3 | 2 | 1 | 2 |
| XM_011537569:NM_152550:XM_017009136:XM_011537567:XM_006714757:XM_005268380:XM_011537568 | 0:0:0:0:0:0:0 | 1:1:0:0:0:0:0 | 0:3:2:0:0:0:0 | 1:1:0:0:0:0:0 | 4:3:0:0:0:0:0 | 3:3:1:1:0:0:0 | 0:0:0:0:0:0:0 | 12:18:2:0:1:0:0 | 0:0:0:0:0:0:0 | 2:5:1:1:0:2:1 |
| NM_152550:XM_011537569:XM_017009136:XM_011537568:XM_011537567:XM_006714757:XM_005268380 | 0:0:0:0:0:0:0 | 1:0:0:0:0:0:0 | 0:0:0:0:0:0:0 | 5:1:1:1:0:0:0 | 0:0:0:0:0:0:0 | 7:5:1:1:1:0:0 | 0:0:0:0:0:0:0 | 27:11:1:0:0:1:0 | 0:0:0:0:0:0:0 | 6:2:1:1:1:0:2 |
| NM_152550:XM_011537569:XM_011537568:XM_017009136:XM_011537567:XM_006714757:XM_005268380 | 0:0:0:0:0:0:0 | 1:0:0:0:0:0:0 | 3:0:0:1:0:0:0 | 2:2:1:0:0:0:0 | 4:4:1:0:0:0:0 | 3:3:0:1:1:0:0 | 0:0:0:0:0:0:0 | 28:12:0:2:0:1:0 | 0:0:0:0:0:0:0 | 5:2:1:1:1:0:2 |
| NM_019018:XM_011514056 | 4:01 | 0:00 | 1:00 | 0:00 | 1:00 | 0:00 | 2:00 | 3:00 | 2:00 | 1:00 |
| XM_011537656 | 1 | 0 | 0 | 0 | 1 | 0 | 0 | 0 | 1 | 0 |
| XM_011537656 | 1 | 0 | 0 | 0 | 0 | 0 | 0 | 0 | 1 | 0 |
| XM_017009459 | 1 | 1 | 0 | 0 | 0 | 0 | 0 | 0 | 0 | 0 |
| XM_017009794 | 0 | 0 | 0 | 0 | 0 | 0 | 0 | 0 | 0 | 0 |
| NM_004355:NM_001025159:XM_017010089:NM_001025158:XM_017010090:XR_002956198 | 6062:1270:12:3:41:29 | 6:0:0:0:0:0 | 3287:328:4:2:29:14 | 1:0:0:0:0:1 | 7469:1019:10:3:64:27 | 6:1:0:0:0:1 | 6548:934:6:2:43:17 | 4:0:0:0:0:2 | 1152:108:0:1:3:0 | 1:0:0:0:0:0 |
| NM_004355:NM_001025159:XR_002956198:XM_017010089:XM_017010090:NM_001025158 | 6013:1257:66:12:40:0 | 10:1:1:0:0:0 | 3210:320:26:3:29:0 | 7:2:0:0:0:0 | 7398:1010:46:10:64:0 | 9:0:0:0:0:0 | 6396:903:46:5:43:3 | 8:2:0:0:0:0 | 1140:106:6:0:3:0 | 0:0:0:0:0:0 |
| NM_004355:NM_001025159:XM_017010089:XM_017010090:XR_002956198:NM_001025158 | 5932:1233:12:40:92:3 | 4:2:0:0:0:0 | 3114:307:3:27:34:6 | 5:0:0:1:0:0 | 7234:981:8:64:62:5 | 7:3:0:0:0:0 | 6233:870:5:43:57:9 | 9:3:0:0:0:0 | 1122:101:0:3:6:1 | 0:0:0:0:0:0 |
| NM_004355:XR_002956198:NM_001025159:NM_001025158:XM_017010089:XM_017010090 | 5939:92:1233:4:12:40 | 4:1:3:1:0:0 | 3118:34:308:6:3:28 | 1:0:0:0:0:0 | 7240:65:986:7:8:64 | 8:0:0:0:0:0 | 6246:58:874:10:5:43 | 1:2:2:1:0:0 | 1123:6:101:1:0:3 | 0:0:0:0:0:0 |
| NM_004355:NM_001025159:XM_017010089:XM_017010090:XR_002956198:NM_001025158 | 5931:1232:11:40:99:5 | 6:1:0:0:0:0 | 3098:304:3:28:35:7 | 3:0:0:0:0:0 | 7212:979:8:64:70:7 | 20:1:0:0:0:0 | 6212:868:5:43:65:11 | 9:2:0:0:0:0 | 1121:101:0:3:8:2 | 2:0:0:0:0:0 |
| NM_004355:NM_001025159:XM_017010089:NM_001025158:XM_017010090:XR_002956198 | 5816:1211:10:6:35:101 | 7:3:0:0:0:0 | 2970:291:3:8:25:40 | 9:0:0:0:0:0 | 7021:949:9:8:61:71 | 11:1:0:0:0:0 | 5934:828:5:12:41:68 | 12:0:0:0:0:0 | 1092:99:0:2:3:7 | 1:1:0:0:0:0 |
| NM_001025159:NM_004355:XM_017010089:NM_001025158:XM_017010090:XR_002956198 | 1208:5807:10:6:34:101 | 4:2:0:0:0:0 | 287:2958:3:8:22:41 | 0:6:0:0:0:0 | 947:7007:9:8:58:72 | 0:7:0:0:0:0 | 823:5899:5:12:41:68 | 1:16:0:0:0:0 | 100:1090:0:2:3:7 | 0:1:0:0:0:0 |
| NM_004355:XR_002956198:NM_001025159:XM_017010089:NM_001025158:XM_017010090 | 5876:104:1217:10:6:33 | 2:2:2:0:0:0 | 2967:41:293:3:8:21 | 11:3:0:0:0:0 | 7052:78:954:9:8:56 | 6:0:2:0:0:0 | 5981:72:834:5:12:39 | 11:1:1:0:0:0 | 1096:8:100:0:2:3 | 0:0:0:0:0:0 |
| NM_004355:NM_001025159:XM_017010089:NM_001025158:XM_017010090:XR_002956198 | 6000:1223:8:6:28:127 | 12:2:1:0:0:0 | 2980:291:3:8:18:55 | 21:2:0:0:0:0 | 7140:960:8:8:50:110 | 29:5:0:0:1:1 | 6117:850:5:11:39:175 | 18:4:0:0:0:0 | 1115:102:0:2:3:14 | 2:0:0:0:0:0 |
| NM_004355:NM_001025159:XM_017010089:NM_001025158:XM_017010090:XR_002956198 | 5991:1219:8:6:28:119 | 6:2:0:0:0:0 | 2968:290:3:7:18:60 | 12:1:0:0:0:0 | 7118:958:8:8:47:107 | 8:3:0:0:0:0 | 6083:847:5:11:39:172 | 12:0:0:0:0:1 | 1113:103:0:2:4:14 | 1:0:0:0:0:0 |
| NM_004355:NM_001025159:XR_002956198:XM_017010089:NM_001025158:XM_017010090 | 6024:1221:120:8:6:29 | 9:3:1:0:0:0 | 2988:293:58:2:7:17 | 4:0:0:0:0:0 | 7140:956:108:7:8:45 | 15:2:1:0:0:1 | 6156:851:193:5:12:38 | 5:1:0:0:0:0 | 1115:103:14:0:2:4 | 2:0:0:0:0:0 |
| NM_004355:NM_001025159:XM_017010089:NM_001025158:XM_017010090:XR_002956198 | 6014:1216:7:6:27:116 | 8:0:0:0:0:0 | 2978:290:2:7:17:58 | 6:0:0:0:0:0 | 7109:951:7:8:46:104 | 19:3:0:0:0:1 | 6136:847:4:12:37:192 | 9:1:0:0:0:0 | 1111:101:0:2:4:15 | 2:0:0:0:0:0 |
| NM_018691:NM_001317993:XM_005268360:XM_011537542:XR_001741975:XM_024454339:XM_005268361:NM_001317994:XM_017008967:XM_011537540 | 8:1:1:0:0:3:0:0:0:0 | 9:2:2:1:1:1:0:0:0:0 | 5:0:2:0:0:3:0:0:0:0 | 3:0:1:1:0:6:1:0:0:0 | 13:0:1:0:0:1:0:0:0:0 | 12:1:3:0:1:5:0:0:0:0 | 80:2:6:0:11:74:1:3:0:0 | 0:0:0:0:0:0:0:0:0:0 | 19:0:9:2:1:9:2:1:1:0 | 0:0:0:0:0:0:0:0:0:0 |

| | | | | | | | | | | |
|---|---|---|---|---|---|---|---|---|---|---|
| NM_018691:NM_001317993:XM_005268360:XM_011537542:XM_024454339:XR_001741975:XM_005268361:NM_001317994:XM_017008967:XM_011537540 | 17:3:3:1:5:1:0:0:0:0 | 0:0:0:0:0:0:0:0:0:0 | 9:0:3:1:9:1:1:0:0:0 | 1:0:0:0:0:0:0:0:0 | 26:1:4:0:6:1:0:0:0:0 | 1:0:0:0:0:0:0:0:0:0 | 81:2:6:0:75:4:1:3:0:0 | 6:0:0:0:1:0:0:0:0:0 | 20:0:9:2:9:0:2:1:1:0 | 1:0:0:0:0:0:0:0:0:0 |
| NM_018691:NM_001317993:XM_005268360:XM_011537542:XM_024454339:XR_001741975:XM_005268361:NM_001317994:XM_017008967:XM_011537540 | 17:3:3:1:5:1:0:0:0:0 | 0:0:0:0:0:0:0:0:0:0 | 9:0:3:1:9:1:1:0:0:0 | 1:0:0:0:0:0:0:0:0 | 26:1:4:0:6:1:0:0:0:0 | 1:0:0:0:0:0:0:0:0:0 | 81:2:6:0:75:4:1:3:0:0 | 5:0:0:0:1:0:0:0:0:0 | 19:0:9:2:9:0:2:1:1:0 | 2:0:0:0:0:0:0:0:0:0 |
| NM_018691:NM_001317993:XM_005268360:XM_011537542:XM_024454339:XR_001741975:XM_005268361:NM_001317994:XM_017008967:XM_011537540 | 0:0:0:0:0:0:0:0:0:0 | 17:3:3:1:1:1:0:0:0:0 | 0:0:0:0:0:0:0:0:0:0 | 10:0:3:1:5:1:1:0:0:0 | 0:0:0:0:0:0:0:0:0:0 | 27:1:4:0:3:1:0:0:0:0 | 51:0:3:0:15:3:1:2:0:0 | 36:2:4:0:5:1:0:1:0:0 | 12:0:1:0:2:0:2:0:1:0 | 9:0:8:2:2:0:0:1:0:0 |
| NM_001445 | 0 | 0 | 0 | 0 | 0 | 0 | 0 | 0 | 0 | 0 |
| NM_019000:XM_024446118:XM_011514054 | 3:00:00 | 0:00:00 | 90:16:04 | 0:00:00 | 100:02:04 | 0:00:00 | 66:13:03 | 2:00:00 | 8:00:00 | 0:00:00 |
| NM_199185:NM_002520:NM_001355007:NM_001037738:NM_001355010:NM_001355009:NR_149149:NM_001355006 | 51:1804:5:0:0:0:0:0 | 1:0:0:0:0:0:0:0 | 26:536:6:0:0:0:0:0 | 0:0:0:0:0:0:0:0 | 18:477:3:0:0:0:0:0 | 0:1:0:0:0:0:0:0 | 306:9109:17:16:2:0:0:0 | 1:5:0:0:0:0:0:0 | 65:2076:0:2:0:0:0:0 | 0:2:0:0:0:0:0:0 |
| NM_002520:NM_001355007:NR_149149:NM_001037738:NM_199185:NM_001355006 | 1811:5:7:0:0:0 | 3:0:0:0:0:0 | 537:6:7:0:0:0 | 1:0:0:0:0:0 | 478:3:4:0:0:0 | 2:0:0:0:0:0 | 9149:17:38:16:0:0 | 10:0:0:0:0:0 | 2078:0:13:2:0:0 | 4:0:0:0:0:0 |
| NM_002520:NM_199185:NM_001355007:NR_149149:NM_001037738:NM_001355010:NM_001355009:NM_001355006 | 1803:39:5:13:0:0:0:0 | 2:0:0:0:0:0:0:0 | 537:15:6:15:0:0:0:0 | 0:0:0:1:0:0:0:0 | 474:12:3:7:0:0:0:0 | 1:0:0:0:0:0:0:0 | 9129:195:17:78:16:2:0:0 | 6:0:0:0:0:0:0:0 | 2077:57:0:24:3:0:0:0 | 3:0:0:0:0:0:0:0 |
| NM_002520:NM_199185:NM_001355007:NR_149149:NM_001037738:NM_001355010:NM_001355009:NM_001355006 | 1794:39:5:16:0:0:0:0 | 2:0:0:0:0:0:0:0 | 530:15:6:20:0:0:0:0 | 1:0:0:0:0:0:0:0 | 474:12:3:10:0:0:0:0 | 1:0:0:0:0:0:0:0 | 9010:192:16:98:16:2:0:0 | 12:1:0:0:0:0:0:0 | 2075:56:0:29:3:0:0:0 | 3:0:0:0:0:0:0:0 |
| NM_002520:NM_199185:NM_001355007:NR_149149:NM_001355010:NM_001037738:NM_001355006 | 1786:39:5:17:0:0:0 | 3:0:0:0:0:0:0 | 530:15:6:29:0:0:0 | 0:0:0:0:0:0:0 | 473:12:3:11:0:0:0 | 1:0:0:0:0:0:0 | 8998:193:16:136:2:0:0 | 9:0:0:0:0:0:0 | 2074:56:0:36:0:0:0 | 0:0:0:0:0:0:0 |
| NM_002520:NM_199185:NM_001355007:NR_149149:NM_001355010:NM_001037738:NM_001355006 | 1786:39:5:18:0:0:0 | 2:0:0:0:0:0:0 | 529:15:6:30:0:0:0 | 1:0:0:0:0:0:0 | 474:12:3:11:0:0:0 | 0:0:0:0:0:0:0 | 8995:192:16:138:2:0:0 | 6:1:0:1:0:0:0 | 2073:56:0:36:0:0:0 | 0:0:0:0:0:0:0 |
| NM_002520:NM_199185:NM_001355007:NR_149149:NM_001355010:NM_001037738:NM_001355006 | 1785:39:5:18:0:0:0 | 3:0:0:0:0:0:0 | 530:15:6:30:0:0:0 | 0:0:0:0:0:0:0 | 471:12:3:11:0:0:0 | 2:0:0:0:0:0:0 | 8992:193:16:139:2:0:0 | 7:0:0:0:0:0:0 | 2071:56:0:36:0:0:0 | 2:0:0:0:0:0:0 |
| NM_001031711:XR_001742159:XM_011534597:XM_017009655:XM_024446135 | 39:2:4:1:0 | 0:0:0:0:0 | 9:0:2:0:0 | 0:0:0:0:0 | 68:3:2:0:0 | 0:0:0:0:0 | 112:9:12:0:1 | 1:0:0:0:0 | 46:2:1:0:1 | 1:0:1:0:0 |
| NM_002449 | 0 | 0 | 0 | 0 | 0 | 0 | 0 | 2 | 0 | 0 |
| NM_017510 | 2 | 0 | 1 | 0 | 3 | 0 | 3 | 0 | 7 | 0 |
| NM_001257293:NM_005520 | 25:44:00 | 0:00 | 6:10 | 0:00 | 10:14 | 0:00 | 46:38:00 | 1:00 | 26:24:00 | 1:01 |
| NM_003900:NM_001142298:NM_001142299 | 253:01:08 | 1:00:00 | 214:07:11 | 0:00:00 | 1086:06:31 | 2:00:00 | 1807:18:62 | 2:00:00 | 379:04:23 | 0:00:00 |
| NM_001142298:NM_001142299:NM_003900 | 0.050150463 | 0:00:00 | 0.301793981 | 0:00:00 | 0.284097222 | 0:00:02 | 18:62:1808 | 0:00:01 | 0.187037037 | 0:00:00 |
| NM_003900:NM_001142298:NM_001142299 | 252:01:08 | 1:00:00 | 215:07:11 | 0:00:00 | 1088:06:31 | 0:00:00 | 1809:18:62 | 1:00:00 | 380:04:23 | 0:00:00 |
| NM_001142298:NM_001142299:NM_003900 | 0.050138889 | 0:00:00 | 0.301006944 | 0:00:00 | 0.284027778 | 0:00:00 | 18:62:1775 | 0:00:00 | 0.187013889 | 0:00:00 |
| NM_001142298:NM_001142299:NM_003900 | 0.050127315 | 0:00:00 | 0.30099537 | 0:00:00 | 0.284016204 | 0:00:01 | 18:62:1775 | 0:00:01 | 0.187013889 | 0:00:00 |
| NM_001142298:NM_001142299:NM_003900 | 0.050127315 | 0:00:00 | 0.301006944 | 0:00:01 | 0.284027778 | 0:00:00 | 18:62:1774 | 0:00:02 | 0.187013889 | 0:00:01 |
| NM_001142298:NM_001142299:NM_003900 | 0.050127315 | 0:00:00 | 0.301006944 | 0:00:01 | 0.284027778 | 0:00:00 | 18:62:1774 | 0:00:02 | 0.187013889 | 0:00:01 |
| NM_001142298:NM_001142299:NM_003900 | 0.050127315 | 0:00:00 | 0.301018519 | 0:00:00 | 0.284016204 | 0:00:01 | 18:62:1775 | 0:00:01 | 0.187025463 | 0:00:00 |
| NM_001142298:NM_001142299:NM_003900 | 0.050127315 | 0:00:00 | 0.301006944 | 0:00:00 | 0.284027778 | 0:00:00 | 18:62:1776 | 0:00:01 | 0.187025463 | 0:00:00 |
| NM_001142298:NM_001142299:NM_003900 | 0.050127315 | 0:00:00 | 0.301006944 | 0:00:00 | 0.284027778 | 0:00:00 | 18:62:1772 | 0:00:05 | 0.186319444 | 0:01:01 |
| NM_001142298:NM_001142299:NM_003900 | 0.050127315 | 0:00:00 | 0.301030093 | 0:00:00 | 0.28400463 | 0:00:01 | 18:61:1768 | 0:00:03 | 0.187048611 | 0:00:00 |
| NM_003900:NM_001142298:NM_001142299 | 250:01:08 | 1:00:00 | 210:07:10 | 0:00:00 | 1076:06:31 | 0:00:00 | 1766:18:61 | 2:00:00 | 381:03:23 | 0:00:00 |
| NM_001142298:NM_001142299:NM_003900 | 0.004895833 | 0.045231481 | 0.21380787 | 0.087222222 | 0.22412037 | 0.059837963 | 18:61:1763 | 0:00:04 | 0.145358796 | 0:00:00 |
| NM_003900:NM_001142298:NM_001142299 | 251:01:08 | 1:00:00 | 210:07:10 | 0:00:00 | 1064:06:31 | 1:00:00 | 1747:18:61 | 1:00:00 | 377:03:23 | 0:00:00 |
| NM_001142298:NM_001142299:NM_003900 | 0.050081019 | 0:00:00 | 0.299594907 | 0:00:02 | 0.283541667 | 0:00:02 | 18:61:1679 | 0:00:03 | 0.144618056 | 0:00:00 |
| NM_001142298:NM_001142299:NM_003900 | 0.050115741 | 0:00:00 | 0.301018519 | 0:00:00 | 0.282361111 | 0:00:01 | 18:60:1714 | 0:00:02 | 0.186990741 | 0:00:00 |
| NM_001142298:NM_001142299:NM_003900 | 0.050127315 | 0:00:00 | 0.300347222 | 0:00:00 | 0.283900463 | 0:00:00 | 18:61:1747 | 0:00:04 | 0.187048611 | 0:00:00 |
| NM_001142298:NM_001142299:NM_003900 | 0.050127315 | 0:00:00 | 0.300347222 | 0:00:00 | 0.283912037 | 0:00:02 | 18:61:1759 | 0:00:00 | 0.187048611 | 0:00:00 |
| NM_001142298:NM_001142299:NM_003900 | 0.050138889 | 0:00:00 | 0.300335648 | 0:00:00 | 0.283912037 | 0:00:02 | 18:61:1759 | 0:00:01 | 0.187048611 | 0:00:00 |
| NM_003900:NM_001142298:NM_001142299 | 251:01:08 | 1:00:00 | 208:07:09 | 0:00:00 | 1074:06:31 | 1:00:00 | 1762:18:61 | 1:00:00 | 380:04:23 | 1:00:00 |
| NM_016175 | 0 | 0 | 0 | 0 | 0 | 1 | 0 | 0 | 0 | 0 |
| NM_024850:XM_011534649:XM_011534653:NM_001040462 | 0:0:0:0 | 0:0:0:0 | 1:0:0:0 | 0:0:0:0 | 0:0:0:0 | 0:0:0:0 | 0:0:0:0 | 0:0:0:0 | 0:0:0:0 | 0:0:0:0 |
| XM_011534653:XM_011534649:NM_001040462 | 0:00:00 | 0:00:00 | 0:00:00 | 1:00:00 | 0:00:00 | 0:00:00 | 0:00:00 | 0:00:00 | 0:00:00 | 0:00:00 |
| XM_011534653:XM_011534649:NM_001040462 | 0:00:00 | 0:00:00 | 0:00:00 | 1:00:00 | 0:00:00 | 0:00:00 | 0:00:00 | 0:00:00 | 0:00:00 | 0:00:00 |
| NM_006098 | 19894 | 37 | 5449 | 9 | 11840 | 34 | 80376 | 118 | 14449 | 24 |
| NM_006098 | 19958 | 40 | 5461 | 11 | 11872 | 26 | 80504 | 150 | 14473 | 23 |
| NM_006098 | 19992 | 28 | 5470 | 10 | 11892 | 15 | 80626 | 94 | 14494 | 12 |
| NM_006098 | 20102 | 55 | 5443 | 16 | 11936 | 21 | 80900 | 190 | 14526 | 33 |
| NM_006098 | 20210 | 26 | 5202 | 5 | 11892 | 20 | 80638 | 111 | 14499 | 15 |
| NM_006098 | 20238 | 23 | 5197 | 3 | 11889 | 20 | 80655 | 71 | 14503 | 17 |
| NM_006098 | 20258 | 19 | 5034 | 8 | 11854 | 5 | 79567 | 43 | 14447 | 3 |
| NM_006098 | 20013 | 16 | 4932 | 7 | 11707 | 14 | 77879 | 95 | 14284 | 19 |
| NM_006098 | 20003 | 25 | 4939 | 8 | 11707 | 13 | 78209 | 120 | 14310 | 14 |
| NM_006098 | 19970 | 22 | 4939 | 4 | 11689 | 21 | 78124 | 101 | 14296 | 17 |
| NM_006098 | 19756 | 28 | 4927 | 10 | 11633 | 27 | 78291 | 83 | 14176 | 12 |
| NM_006098 | 19726 | 27 | 4917 | 11 | 11625 | 22 | 78240 | 70 | 14142 | 21 |
| NM_145265 | 38 | 0 | 13 | 1 | 52 | 0 | 173 | 2 | 22 | 0 |
| NM_001267558:NM_001267556:NM_013232:NM_001267557 | 3:10:47:0 | 0:0:0:0 | 2:11:67:1 | 0:0:0:0 | 8:38:138:0 | 0:0:1:0 | 8:59:211:1 | 0:0:1:0 | 4:13:77:0 | 0:0:0:0 |
| NM_016107:NR_144318 | 3:00 | 0:00 | 1:00 | 0:00 | 12:04 | 0:02 | 3:02 | 0:01 | 4:01 | 0:00 |
| NM_016107:NR_144318 | 1:00 | 2:00 | 0:04 | 1:02 | 5:11 | 3:22 | 0:00 | 2:20 | 0:00 | 1:01 |
| NM_016107:NR_144318 | 0:00 | 2:00 | 0:00 | 1:06 | 0:00 | 6:34 | 0:00 | 2:19 | 0:00 | 1:01 |
| NR_144318:NM_016107 | 9:00 | 2:00 | 15:00 | 0:00 | 201:02:00 | 9:00 | 66:00:00 | 5:00 | 11:00 | 0:00 |
| NM_014324:NM_203382:NM_001167595 | 0:00:00 | 18:02:00 | 0:00:00 | 29:01:01 | 0:00:00 | 82:04:02 | 0:00:00 | 279:10:08 | 0:00:00 | 33:00:00 |
| NM_014324:NM_203382:NM_001167595 | 18:02:00 | 0:00:00 | 29:01:01 | 0:00:00 | 84:04:02 | 0:00:00 | 135:00:03 | 146:10:05 | 15:00:00 | 20:00:00 |
| NM_014324:NM_203382:NM_001167595 | 0:00:00 | 18:02:00 | 0:00:00 | 29:01:01 | 1:00:00 | 83:04:02 | 0:00:00 | 282:10:08 | 0:00:00 | 35:00:00 |
| NM_014324:NM_001167595 | 11:00 | 7:00 | 7:01 | 22:00 | 32:01:00 | 52:01:00 | 144:05:00 | 137:03:00 | 20:00 | 15:00 |
| NM_014324:NM_203382:NM_001167595 | 3:01:00 | 3:00:00 | 6:00:00 | 14:00:00 | 25:04:01 | 25:00:01 | 84:06:01 | 64:00:02 | 7:00:00 | 7:00:00 |
| XM_017009329:XM_017009330:XM_006714468 | 3:01:00 | 0:00:00 | 5:05:00 | 0:00:00 | 1:00:00 | 0:00:00 | 11:06:00 | 8:08:01 | 0:00:00 | 1:00:00 |
| NM_018034:XM_011514061:NM_001345998:NM_001345999 | 11:0:0:0 | 0:0:0:0 | 12:1:1:0 | 1:0:0:0 | 16:0:0:0 | 0:0:0:0 | 61:1:3:2 | 0:0:0:0 | 13:0:1:0 | 0:0:0:0 |
| NM_018034:XM_011514061:NM_001345998:NM_001345999 | 12:0:0:0 | 0:0:0:0 | 13:1:1:0 | 0:0:0:0 | 16:0:0:0 | 0:0:0:0 | 64:7:3:1 | 0:0:0:0 | 14:0:1:0 | 0:0:0:0 |
| NM_018034:XM_011514061:NM_001345998:NM_001345999 | 12:1:0:0 | 0:0:0:0 | 13:3:0:0 | 0:0:0:0 | 16:0:0:0 | 0:0:0:0 | 63:7:1:1 | 0:0:0:0 | 12:0:1:0 | 0:0:0:0 |

| | | | | | | | | | | |
|---|---|---|---|---|---|---|---|---|---|---|
| NM_176895:NM_003711:XM_006714724:NR_103485 | 15:41:3:1 | 0:0:0:0 | 38:246:23:5 | 0:0:0:0 | 125:945:78:19 | 0:2:0:1 | 17:96:7:2 | 0:0:0:0 | 1:16:2:1 | 0:0:0:0 |
| NM_176895:NM_003711:NR_103485:XM_006714724 | 15:42:1:1 | 0:0:0:0 | 38:248:6:17 | 0:1:0:0 | 125:947:23:55 | 1:1:0:0 | 18:95:2:3 | 0:1:0:0 | 1:16:1:1 | 0:0:0:0 |
| NM_003711:NR_103485:XM_006714724 | 42:01:00 | 0:00:00 | 250:05:00 | 0:00:00 | 948:19:04 | 2:00:00 | 96:02:00 | 0:00:00 | 16:01:00 | 0:00:00 |
| NM_173514:NM_001349384:XM_017009082:XR_948238:XM_011543174:NM_001349382:NM_001349385:XM_024454369:XM_017009077:NR_146165:XM_011543176:XM_017009080:XR_948237:NM_001258286:NM_001349383 | 0:0:0:0:0:0:0:0:0:0:0:0:0:0 | 1:1:0:0:0:0:0:0:0:0:0:0:0:0 | 0:0:0:0:0:0:0:0:0:0:0:0:0:0 | 0:0:1:1:1:0:0:0:0:0:0:0:0:0 | 0:0:0:0:0:0:0:0:0:0:0:0:0:0 | 1:2:0:0:0:2:1:1:2:0:0:0:0:0:0 | 5:1:0:1:0:2:0:0:0:2:0:0:0:0 | 0:0:0:0:0:0:0:0:0:0:0:0:0:0 | 3:1:0:0:0:0:0:0:0:0:0:0:1:0 | 2:1:0:0:0:0:1:0:1:0:1:1:1:0:0 |
| NM_002184:NM_001190981 | 1:00 | 1:00 | 3:00 | 2:00 | 2:00 | 1:00 | 1:00 | 0:00 | 1:00 | 0:00 |
| NM_015342:XM_006714583:NM_001278926:XM_017009286 | 28:1:1:0 | 3:0:0:0 | 23:1:0:0 | 5:2:0:0 | 20:1:1:0 | 4:0:0:0 | 58:3:0:0 | 14:1:0:0 | 40:2:0:0 | 16:0:0:0 |
| NM_015342:XM_006714583:NM_001278926:NM_001278929:XM_017009286 | 34:9:1:1:0 | 0:0:0:0:0 | 28:7:0:0:0 | 0:0:0:0:0 | 23:4:1:0:0 | 0:0:0:0:0 | 86:21:0:1:1 | 0:0:0:0:0 | 56:7:0:0:0 | 0:0:0:0:0 |
| NM_017755:NR_037947:NM_001193455 | 11:00:00 | 0:00:00 | 5:06:00 | 0:00:00 | 4:01:00 | 0:00:00 | 13:28:00 | 17:27:00 | 13:03:01 | 11:00:00 |
| XM_005248542:NM_181523:XM_017009586:XM_017009585 | 4:4:1:1 | 2:1:0:0 | 1:3:1:0 | 2:1:0:0 | 0:1:0:0 | 0:2:0:0 | 0:1:1:0 | 2:4:3:2 | 3:5:0:1 | 1:1:1:0 |
| NM_133343:NM_133338:NM_133344:NM_133342:NM_001278622:NM_002873:XM_017009681:NM_133340:NM_133341 | 3:1:4:16:5:1:1:0:0 | 0:0:0:0:0:0:0:0:0 | 0:0:4:24:0:0:30:1:0 | 0:0:0:0:0:0:0:0:0 | 2:0:5:13:2:0:2:7:0 | 0:0:0:0:0:0:0:0:0 | 0:0:0:2:0:0:0:0:0 | 0:2:16:84:1:2:120:7:2 | 2:0:2:11:1:0:6:3:3 | 1:1:3:15:0:0:2:3:0 |
| NM_133343:NM_133338:NM_133344:NM_133342:NM_001278622:NM_002873:XM_017009681:NM_133340:NM_133341 | 3:1:4:16:5:1:1:0:0 | 0:0:0:0:0:0:0:0:0 | 0:0:4:25:0:0:30:1:0 | 0:0:0:0:0:0:1:0:0 | 2:0:5:13:2:0:2:7:0 | 0:0:1:0:0:0:0:0:0 | 0:2:16:87:1:2:126:7:1 | 0:0:0:0:0:0:0:0:0 | 3:1:5:26:1:0:8:6:3 | 0:0:0:0:0:0:0:0:0 |
| XM_017009634:NM_018429 | 5:00 | 0:00 | 3:01 | 0:00 | 4:00 | 0:00 | 0:00 | 6:00 | 0:00 | 2:00 |
| XM_017009634:NM_018429 | 6:00 | 0:00 | 4:01 | 0:00 | 4:00 | 0:00 | 8:00 | 0:00 | 1:00 | 2:00 |
| XM_017009634:NM_018429 | 6:00 | 0:00 | 2:01 | 0:00 | 4:00 | 0:00 | 0:00 | 9:00 | 0:00 | 2:00 |
| XM_017009634 | 2 | 0 | 3 | 0 | 1 | 0 | 0 | 2 | 0 | 1 |
| XM_017009634 | 1 | 0 | 3 | 0 | 1 | 0 | 0 | 2 | 0 | 1 |
| NM_000521 | 0 | 2 | 0 | 1 | 0 | 4 | 0 | 1 | 0 | 2 |
| NM_000521 | 2 | 0 | 1 | 0 | 4 | 0 | 1 | 0 | 2 | 0 |
| NM_000521 | 2 | 0 | 1 | 0 | 4 | 0 | 1 | 0 | 2 | 0 |
| NM_003664:XM_005248618:XM_017010001 | 11:01:00 | 0:00:00 | 20:00:03 | 0:00:00 | 11:00:01 | 0:00:00 | 23:00:07 | 0:00:00 | 22:01:01 | 0:00:00 |
| NM_003664:XM_005248618:XM_017010001:XM_005248619 | 0:0:0:0 | 10:1:0:0 | 0:0:0:0 | 14:0:2:0 | 0:0:0:0 | 11:0:1:0 | 0:0:0:0 | 12:0:6:0 | 0:0:0:0 | 21:1:1:1 |
| NM_024091:XR_427664:XM_006714498:XR_001742253 | 3:1:0:0 | 0:0:0:0 | 3:1:10:0 | 0:0:0:0 | 2:1:8:1 | 0:0:0:0 | 16:6:15:0 | 0:0:2:0 | 5:1:1:1 | 0:0:0:0 |
| NM_024091:XR_427664:XM_006714498:XR_001742253 | 0:0:0:0 | 3:1:0:0 | 0:0:0:0 | 3:1:10:0 | 0:0:0:0 | 2:1:8:1 | 7:1:5:0 | 6:2:5:0 | 1:1:0:1 | 3:0:1:0 |
| NM_001131028:XM_017009944:NM_031482:XM_011543660 | 0:1:0:0 | 1:0:0:0 | 0:0:1:1 | 2:1:2:0 | 0:0:0:0 | 0:0:0:0 | 3:0:7:1 | 0:0:0:0 | 2:0:1:0 | 0:0:0:0 |
| NM_001131028:XM_017009944:NM_031482:XM_011543660 | 0:0:0:0 | 1:1:0:0 | 0:0:0:0 | 2:1:3:1 | 0:0:0:0 | 0:0:0:0 | 3:0:7:1 | 0:0:0:0 | 2:0:1:0 | 0:0:0:0 |
| NM_001025 | 64 | 0 | 33 | 1 | 34 | 0 | 113 | 0 | 92 | 0 |
| NM_001239:XR_001742327:XM_005248627:NM_001199189 | 25:1:2:0 | 0:0:0:0 | 33:0:0:1 | 0:0:0:0 | 26:1:0:0 | 0:0:0:0 | 2:0:0:0 | 142:1:0:5 | 38:0:0:0 | 37:0:0:0 |
| NM_023924:NR_134296:XR_002956175:XM_024446192:XM_024446199:XM_024446198:XR_002956174:NM_001317951:XM_024446193:XM_024446200:XM_006714494:XM_024446196:NM_001009877:XM_024446197 | 3:3:5:0:0:0:0:0:0:0:0:0:0:0 | 0:0:0:0:0:0:0:0:0:0:0:0:0:0 | 3:0:15:1:1:0:0:0:0:0:0:0:0:0 | 0:0:0:0:0:0:0:0:0:0:0:0:0:0 | 9:0:14:1:5:3:1:0:0:0:0:0:0:0 | 0:0:0:0:0:0:0:0:0:0:0:0:0:0 | 3:1:18:3:2:2:1:2:1:1:0:0:0:0 | 0:0:1:0:0:0:0:0:0:0:0:0:0:0 | 5:0:4:1:0:0:1:0:3:1:1:0:0:0 | 0:0:0:0:0:0:0:0:0:0:0:0:0:0 |
| XR_002956175:NM_023924:NR_134296:XM_024446192:XM_024446199:XM_024446198:XR_002956174:NM_001317951:XM_024446193:XM_024446200:XM_006714494:XM_024446196:NM_001009877:XM_024446197 | 2:3:3:0:0:0:0:0:0:0:0:0:0:0 | 3:0:0:0:0:0:0:0:0:0:0:0:0:0 | 19:3:0:1:1:0:0:0:0:0:0:0:0:0 | 0:0:0:0:0:0:0:0:0:0:0:0:0:0 | 10:9:0:1:5:3:1:0:0:0:0:0:0:0 | 4:0:0:0:0:0:0:0:0:0:0:0:0:0 | 21:3:1:3:2:2:1:2:1:1:0:0:0:0 | 2:0:0:0:0:0:0:0:0:0:0:0:0:0 | 4:5:0:1:0:0:1:0:3:1:1:0:0:0 | 0:0:0:0:0:0:0:0:0:0:0:0:0:0 |
| NM_023924:NR_134296:XR_002956175:XM_024446192:XM_024446199:XM_024446198:XR_002956174:NM_001317951:XM_024446193:XM_024446200:XM_006714494:XM_024446196:NM_001009877:XM_024446197 | 3:3:5:0:0:0:0:0:0:0:0:0:0:0 | 0:0:0:0:0:0:0:0:0:0:0:0:0:0 | 3:0:19:1:1:0:0:0:0:0:0:0:0:0 | 0:0:0:0:0:0:0:0:0:0:0:0:0:0 | 9:0:14:1:5:3:1:0:0:0:0:0:0:0 | 0:0:0:0:0:0:0:0:0:0:0:0:0:0 | 3:1:22:3:2:2:1:2:1:1:0:0:0:0 | 0:0:2:0:0:0:0:0:0:0:0:0:0:0 | 5:0:6:1:0:0:1:0:3:1:1:0:0:0 | 0:0:0:0:0:0:0:0:0:0:0:0:0:0 |
| NM_002726:XM_011535925:XM_005267044 | 4:00:00 | 16:00:00 | 19:00:00 | 13:00:00 | 17:00:00 | 16:00:00 | 123:01:00 | 0:00:00 | 54:00:00 | 0:00:00 |
| NM_018292:XM_011535924 | 36:00:00 | 0:00 | 34:02:00 | 0:00 | 29:02:00 | 0:00 | 142:16:00 | 92:06:00 | 19:02 | 25:01:00 |
| NM_018292:XM_011535924 | 29:04:00 | 0:00 | 18:05 | 0:00 | 27:03:00 | 0:00 | 132:55:00 | 1:02 | 35:04:00 | 0:00 |
| XM_024446293 | 1 | 3 | 7 | 8 | 7 | 5 | 4 | 2 | 4 | 5 |
| NM_007214:XM_017010218 | 1:00 | 0:00 | 0:04 | 1:00 | 1:01 | 0:00 | 8:04 | 3:01 | 1:00 | 2:00 |
| NM_033125 | 0 | 0 | 1 | 0 | 0 | 0 | 0 | 0 | 0 | 0 |
| XM_005266797:NM_006714:XM_005266799 | 0:00:00 | 0:00:00 | 0:00:00 | 1:00:00 | 0:00:00 | 0:06:00 | 0:02:01 | 0:00:00 | 0:01:00 | 0:00:00 |
| XM_005266797:NM_006714:XM_005266799:XM_017010204:XM_005266801 | 0:0:0:0:0 | 0:0:0:0:0 | 1:0:0:0:0 | 0:0:0:0:0 | 0:6:0:0:0 | 0:0:0:0:0 | 0:2:0:0:0 | 0:0:1:0:0 | 0:1:0:0:0 | 0:0:0:0:0 |
| XM_011535495:XR_942305:NM_001286398:XM_011535494 | 0:0:0:0 | 1:0:0:0 | 0:0:0:0 | 0:0:0:0 | 0:0:0:0 | 1:2:0:0 | 2:0:0:0 | 0:1:0:0 | 0:0:0:0 | 0:0:0:0 |
| NM_138571 | 0 | 1 | 0 | 1 | 0 | 0 | 0 | 19 | 0 | 2 |
| XR_002956294:NM_001031712:NM_001350582:NR_146799:NM_001350580:NR_146802:NM_001350581:NR_146796:NR_146798:NM_001350583:NR_146800:NM_001350586:NR_146801:NM_001350584:NM_001350588:XM_017011168:NR_146804:NM_001350589:NM_001350592:NM_001350591:NR_146795:NM_001350585:NM_001350596:NM_001350597:NR_146803:NM_001350594:XM_024446516 | 3:59:1:5:1:1:2:2:1:1:2:1:3:1:1:1:0:0:0:0:0:0:0:0:0:0:0 | 2:0:0:0:0:0:0:0:0:0:0:0:0:0:0:0:0:0:0:0:0:0:0:0:0:0:0 | 5:23:1:6:2:0:1:1:2:0:1:0:2:0:0:1:0:0:0:0:0:0:0:0:0:0:0 | 0:0:0:0:0:0:0:0:0:0:0:0:0:0:0:0:0:0:0:0:0:0:0:0:0:0:0 | 8:76:0:10:1:1:3:4:1:0:1:0:5:2:2:2:1:1:1:1:0:0:0:0:0:0:0 | 0:0:0:0:0:0:0:0:0:0:0:0:0:0:0:0:0:1:0:0:0:0:0:0:0:0:0 | 4:235:0:43:0:0:10:6:5:0:0:1:0:18:0:42:1:0:0:0:2:1:0:0:0:0:0 | 1:0:0:0:0:0:0:0:0:0:0:0:0:0:0:0:0:0:0:0:0:0:0:0:0:0:0 | 2:69:0:7:1:1:7:5:3:0:0:1:0:0:0:3:0:0:2:0:0:1:1:0:0:0:0 | 1:0:0:1:0:0:0:0:0:0:0:0:0:0:0:0:0:0:0:0:0:0:0:0:0:0:0 |
| XM_017011056:NM_001105544:NM_001002030:NM_018479:XM_005267047:XM_011535942:XM_005267049:XM_024446490:NM_001139510:XM_005267048:XM_005267050:NM_001105545 | 1:2:80:21:6:1:2:3:5:4:1:1 | 0:0:0:0:0:0:0:0:0:0:0:0 | 7:0:113:14:5:2:0:5:7:3:0:2 | 0:0:1:0:0:0:0:0:0:0:0:0 | 0:2:114:14:4:1:1:1:8:2:0:1 | 0:0:2:0:0:0:0:0:0:0:0:0 | 36:22:472:219:49:36:7:40:63:23:7:5 | 0:0:2:1:0:0:0:0:0:0:0:0 | 1:7:160:35:8:6:0:4:10:3:4:0 | 0:0:0:0:0:0:0:0:0:0:0:0 |
| XM_017011056:NM_001105544:NM_001002030:NM_018479:XM_005267047:XM_011535942:XM_005267049:XM_024446490:NM_001139510:XM_005267048:XM_005267050:NM_001105545 | 1:2:80:21:6:1:2:3:5:4:1:1 | 0:0:0:0:0:0:0:0:0:0:0:0 | 7:0:113:14:5:2:0:5:7:3:0:2 | 0:0:1:0:0:0:0:0:0:0:0:0 | 0:2:116:14:4:1:1:1:8:2:0:1 | 0:0:0:0:0:0:0:0:0:0:0:0 | 36:22:474:219:49:36:7:40:63:23:7:5 | 0:0:0:0:0:0:0:0:0:0:0:0 | 1:7:160:35:8:6:0:4:10:3:4:0 | 0:0:0:0:0:0:0:0:0:0:0:0 |
| XM_017011056:NM_001105544:NM_001002030:NM_018479:XM_005267047:XM_011535942:XM_005267049:XM_024446490:NM_001139510:XM_005267048:XM_005267050:NM_001105545 | 1:2:80:21:6:1:2:3:5:4:1:1 | 0:0:0:0:0:0:0:0:0:0:0:0 | 7:0:114:14:5:2:0:5:7:3:0:2 | 0:0:0:0:0:0:0:0:0:0:0:0 | 0:2:116:14:4:1:1:0:8:2:0:1 | 0:0:0:0:0:0:0:1:0:0:0:0 | 36:22:473:217:48:36:7:34:63:23:7:5 | 1:0:1:1:1:0:0:0:0:0:0:0 | 1:7:160:34:8:6:0:4:10:3:4:0 | 0:0:0:0:0:0:0:0:0:0:0:0 |
| XM_017011056:NM_001105544:NM_001002030:XM_005267047:XM_011535942:NM_001139510:XM_005267048:XM_024446490:NM_018479 | 2:2:80:6:2:5:5:0:0 | 0:0:0:0:0:0:0:0:0 | 7:0:114:5:3:7:3:0:0 | 0:0:0:0:0:0:0:0:0 | 1:2:119:4:1:8:2:0:0 | 0:0:0:0:0:0:0:0:0 | 40:22:477:50:37:60:25:1:0 | 0:0:1:0:0:1:0:0:0 | 1:7:161:8:6:10:4:0:0 | 0:0:0:0:0:0:0:0:0 |
| XM_017011056:NM_001105544:NM_001002030:NM_018479:XM_005267047:XM_011535942:XM_005267049:NM_001139510:XM_024446490:XM_005267048:XM_005267050:NM_001105545 | 2:2:80:21:6:2:2:5:1:5:1:1 | 0:0:0:0:0:0:0:0:0:0:0:0 | 4:0:116:14:5:3:0:7:0:3:0:2 | 0:0:0:1:0:0:0:0:0:0:0:0 | 1:1:119:13:4:1:1:8:0:2:0:1 | 0:0:0:0:0:0:0:0:0:0:0:0 | 32:21:479:209:50:38:5:59:15:25:7:5 | 0:0:0:0:0:0:2:0:0:0:0:0 | 1:7:161:34:8:6:0:10:0:4:4:0 | 0:0:0:0:0:0:0:0:0:0:0:0 |
| XM_017011056:NM_001105544:NM_001002030:NM_018479:XM_005267047:XM_011535942:XM_005267049:NM_001139510:XM_024446490:XM_005267048:XM_005267050:NM_001105545 | 2:2:80:21:6:2:2:5:1:5:1:1 | 0:0:0:0:0:0:0:0:0:0:0:0 | 4:0:116:14:5:3:0:7:0:3:0:2 | 0:0:0:1:0:0:0:0:0:0:0:0 | 1:1:118:13:4:1:1:8:0:2:0:1 | 0:0:1:0:0:0:0:0:0:0:0:0 | 32:21:478:208:50:38:7:59:15:25:7:5 | 0:0:1:1:0:0:0:0:0:0:0:0 | 1:7:161:34:8:6:0:9:0:4:4:0 | 0:0:0:0:0:0:0:0:1:0:0:0 |
| XM_017011056:NM_001105544:NM_001002030:NM_018479:XM_005267047:XM_011535942:XM_005267049:NM_001139510:XM_024446490:XM_005267048:XM_005267050:NM_001105545 | 2:2:80:21:6:2:2:5:1:5:1:1 | 0:0:0:0:0:0:0:0:0:0:0:0 | 4:0:116:14:5:3:0:7:0:3:0:2 | 0:0:0:1:0:0:0:0:0:0:0:0 | 1:1:119:13:4:1:1:8:0:2:0:1 | 0:0:0:0:0:0:0:0:0:0:0:0 | 30:20:479:209:50:38:7:58:14:25:7:5 | 1:0:0:0:0:0:0:0:0:0:0:0 | 1:7:161:34:8:6:0:10:0:4:4:0 | 0:0:0:0:0:0:0:0:0:0:0:0 |
| XM_017011056:NM_001105544:NM_001002030:NM_018479:XM_005267047:XM_011535942:XM_005267049:NM_001139510:XM_024446490:XM_005267048:XM_005267050:NM_001105545 | 2:2:80:21:6:2:2:5:1:5:1:1 | 0:0:0:0:0:0:0:0:0:0:0:0 | 4:0:116:14:5:2:0:7:1:3:0:2 | 0:0:0:1:0:0:0:0:0:0:0:0 | 1:1:119:13:4:1:1:8:0:2:0:1 | 0:0:0:0:0:0:0:0:0:0:0:0 | 23:20:478:209:50:38:7:58:14:25:7:5 | 0:0:1:0:0:0:0:0:0:0:0:0 | 1:7:161:34:8:6:0:10:0:4:4:0 | 0:0:0:0:0:0:0:0:0:0:0:0 |

| | | | | | | | | | | |
|---|---|---|---|---|---|---|---|---|---|---|
| XM_017011148:NM_002844:XM_006715537:XM_017011146:XM_017011145:XM_017011150 | 4:4:1:0:0:0 | 0:0:0:0:0:0 | 1:5:1:8:0:0 | 0:1:0:2:1:0 | 7:9:0:0:0:1 | 0:0:0:0:0:0 | 18:7:2:24:0:0 | 0:2:0:6:0:0 | 7:6:0:0:0:0 | 0:0:0:0:0:0 |
| XM_006715537:XM_011536021:XM_017011148:NM_002844:XM_017011146:XM_017011145:XM_011536016:NM_001291982:XM_011536014:XM_011536020:XM_017011147:XM_017011150:XM_011536015:NM_001291983 | 1:7:1:0:0:0:0:0:0:0:0:0:0:0 | 0:0:0:0:0:0:0:0:0:0:0:0:0:0 | 1:36:7:8:10:2:6:0:0:0:0:0:0:0 | 0:0:0:0:0:0:0:0:0:0:0:0:0:0 | 0:38:3:1:0:0:0:1:0:0:0:0:0:0 | 0:0:0:0:0:0:0:0:0:0:0:0:0:0 | 2:126:34:7:22:3:9:0:1:2:2:1:0:0 | 0:1:0:0:0:0:0:0:0:0:0:0:0:0 | 0:18:1:3:5:0:1:0:0:0:1:0:0:0 | 0:0:0:0:0:0:0:0:0:0:0:0:0:0 |
| XM_005267213:XM_017011494:NM_033515:XM_005267212 | 3:0:25:4 | 5:1:0:0 | 0:0:0:2 | 2:0:0:0 | 1:0:8:1 | 8:0:0:0 | 5:1:29:74 | 24:2:0:0 | 1:0:16:4 | 4:0:0:0 |
| XM_005267213:NM_033515:XM_005267212:XM_017011494 | 9:25:4:1 | 3:0:0:0 | 2:0:2:0 | 4:0:0:0 | 10:8:1:0 | 0:0:0:0 | 29:29:73:4 | 24:0:0:2 | 5:16:4:0 | 0:0:0:0 |
| XM_005267213:NM_033515:XM_005267212:XM_017011494 | 9:25:4:1 | 3:0:0:0 | 2:0:2:0 | 4:0:0:0 | 10:8:1:0 | 0:0:0:0 | 31:29:73:4 | 22:0:0:2 | 5:16:4:0 | 0:0:0:0 |
| XM_005267213:NM_033515:XM_005267212:XM_017011494 | 9:25:4:1 | 3:0:0:0 | 3:0:2:0 | 3:0:0:0 | 10:8:1:0 | 0:0:0:0 | 34:29:73:4 | 18:0:0:2 | 5:16:4:0 | 0:0:0:0 |
| NM_033515:XM_005267213:XM_005267212:XM_017011494 | 25:30:3:3 | 0:0:0:0 | 0:15:1:4 | 0:0:0:0 | 8:21:1:1 | 0:0:0:0 | 29:250:51:39 | 0:2:1:0 | 16:27:4:1 | 0:0:0:0 |
| NM_033515:XM_005267213:XM_005267212:XM_017011494 | 25:32:2:3 | 0:0:0:0 | 0:15:0:2 | 0:1:0:0 | 8:23:1:1 | 0:0:0:0 | 29:261:35:40 | 0:10:0:2 | 16:27:4:1 | 0:1:0:0 |
| NM_033515:XM_005267213:XM_005267212:XM_017011494 | 0:0:0:0 | 24:31:0:0 | 0:0:0:0 | 0:17:0:0 | 0:0:0:0 | 7:22:1:0 | 0:0:0:0 | 29:274:0:7 | 0:1:0:0 | 14:27:0:0 |
| NM_001017373:XM_017010307:XR_001743174:XM_024446336:NM_001258275:XM_024446334:XM_024446335:XM_024446337 | 1:1:0:0:0:0:0:0 | 0:0:0:0:0:0:0:0 | 0:0:1:0:0:0:0:0 | 0:0:0:0:0:0:0:0 | 0:0:0:1:0:0:0:0 | 0:0:0:0:0:0:0:0 | 0:0:0:1:0:0:0:0 | 0:0:0:0:0:0:0:0 | 0:0:0:0:1:0:0:0 | 0:0:0:0:0:0:0:0 |
| XR_001743216:NM_001350320:NM_001350306:NM_001350305:NM_001350304:NM_001431:XM_017010356:XM_011535525:NM_001350310:NM_001252660:XM_024446349:XM_017010364:NM_001350308:NM_001199388:NM_001350301:XR_001743215:XR_001743213:XR_001743217:XM_011535528 | 1:7:5:2:5:3:1:1:1:0:0:0:0:0:0:0:0:0:0 | 1:0:0:0:0:0:0:0:0:0:0:0:0:0:0:0:0:0:0 | 0:5:0:0:0:1:0:1:0:1:2:0:0:0:0:0:0:0:0 | 0:0:0:0:0:1:0:0:0:0:0:0:0:0:0:0:0:0:0 | 0:18:4:1:6:3:0:1:0:0:2:1:1:0:0:0:0:0:0 | 0:0:0:0:0:0:0:0:0:0:0:0:0:0:0:0:0:0:0 | 0:13:3:2:1:2:0:0:2:0:0:0:0:1:0:0:0:0:0 | 1:1:0:0:0:0:0:0:0:0:0:0:0:0:0:0:0:0:0 | 2:44:7:2:4:4:0:1:1:1:1:0:0:1:0:0:0:0:0 | 0:0:0:0:0:0:0:0:0:0:0:0:0:0:0:0:0:0:0 |
| NM_001901 | 0 | 3 | 0 | 0 | 0 | 1 | 0 | 1 | 0 | 0 |
| NM_001016 | 49 | 0 | 23 | 0 | 40 | 0 | 175 | 2 | 97 | 0 |
| NM_005375:NR_134958:NM_001130172:NR_134960:NM_001130173:XR_942444:NM_001161656:NR_134965:NR_134963:NR_134962:NR_134959:NR_134964:NM_001161660 | 54:1:9:4:2:1:1:1:0:0:0:0:0 | 0:0:0:0:0:0:0:0:0:0:0:0:0 | 7:0:1:0:0:1:0:0:3:1:0:0:0 | 0:0:0:0:0:0:0:0:0:0:0:0:0 | 50:0:12:5:1:0:0:1:0:2:1:0:0 | 0:0:0:0:0:0:0:0:0:0:0:0:0 | 69:1:21:9:5:2:3:3:11:23:0:4:0 | 2:0:0:0:0:0:0:0:0:0:0:0:0 | 106:5:22:3:7:0:3:1:0:4:0:0:0 | 0:0:0:0:0:0:0:0:0:0:0:0:0 |
| XM_011536246:XM_017011471 | 0:00 | 1:00 | 0:00 | 0:00 | 0:00 | 0:00 | 0:00 | 0:00 | 0:00 | 0:00 |
| NM_014432:XM_017010954:NM_001278723:XM_017010955:NM_001278722:XM_011535904:NM_001278724 | 2:0:0:0:0:0:0 | 1:0:0:0:0:0:0 | 0:0:0:0:0:0:0 | 0:0:0:0:0:0:0 | 0:0:1:0:0:0:0 | 1:1:1:0:0:0:0 | 19:0:7:4:2:3:0 | 0:0:0:0:0:0:0 | 6:0:4:1:1:0:0 | 0:0:0:0:0:0:0 |
| NM_014432:NM_001278723:XM_017010954:NM_001278722:XM_011535904:XM_017010955:NM_001278724 | 0:0:0:0:0:0:0 | 3:0:0:0:0:0:0 | 0:0:0:0:0:0:0 | 0:0:0:0:0:0:0 | 0:0:0:0:0:0:0 | 1:2:1:0:0:0:0 | 0:0:0:0:0:0:0 | 20:7:0:2:3:0:0 | 0:0:0:0:0:0:0 | 6:4:0:1:0:1:0 |
| NM_022121:XM_024446520 | 0:00 | 4:00 | 0:00 | 2:01 | 0:00 | 7:02 | 0:00 | 5:01 | 0:00 | 11:00 |
| NM_001144060:XM_017011089:XM_024446497 | 0:00:00 | 1:00:00 | 1:00:00 | 3:01:00 | 1:00:02 | 3:01:00 | 1:00:00 | 2:00:00 | 0:00:00 | 0:00:00 |
| NM_001286587:NM_016108 | 0.131944444 | 0:00 | 0:59 | 0:00 | 0.054166667 | 0:00 | 0.780555556 | 0:01 | 0.111111111 | 0:00 |
| NM_001286587:NM_016108 | 0.175694444 | 0:00 | 0:59 | 0:00 | 0.054166667 | 0:00 | 0.789583333 | 0:01 | 0.118055556 | 0:00 |
| NM_001286587:NM_016108 | 0.134722222 | 0:00 | 0:50 | 0:00 | 0.054861111 | 0:00 | 0.679166667 | 0:00 | 0.114583333 | 0:00 |
| NM_032020 | 0 | 0 | 1 | 0 | 1 | 0 | 0 | 2 | 1 | 1 |
| NM_007044:XM_005266805:XM_017010210:XM_017010208:XM_024446317:XM_017010207 | 18:2:1:0:0:0 | 0:0:0:0:0:0 | 14:1:0:1:0:0 | 0:0:0:0:0:0 | 14:0:0:0:0:0 | 0:0:0:0:0:0 | 91:6:10:1:2:0 | 0:0:0:0:0:0 | 28:3:2:0:0:1 | 0:0:0:0:0:0 |
| NM_015440:NM_001242767:NM_001242768:XM_024446395:XR_002956274:XM_017010703:NM_001350493:NR_146720 | 6:4:4:2:0:0:0:0 | 0:0:0:0:0:0:0:0 | 0:0:0:0:0:0:0:0 | 0:0:0:0:0:0:0:0 | 1:0:0:0:0:0:0:0 | 0:0:0:0:0:0:0:0 | 5:5:3:1:0:0:0:0 | 0:0:0:0:0:0:0:0 | 4:3:0:1:1:1:1:0 | 0:0:0:0:0:0:0:0 |
| NM_025059:XM_011536147:XM_011536148 | 1:01:00 | 2:00:00 | 0:00:00 | 0:00:00 | 0:00:00 | 0:00:00 | 0:00:00 | 0:00:01 | 0:00:00 | 0:00:00 |
| XM_011536147:NM_025059:XM_011536148 | 1:03:08 | 0:00:00 | 0:00:11 | 0:00:00 | 0:00:04 | 0:00:00 | 0.001643519 | 0.043263889 | 0:00:16 | 0:00:11 |
| XM_011536147:NM_025059:XM_011536148 | 0:02:06 | 1:01:02 | 0:00:01 | 0:00:09 | 0:00:00 | 0:00:04 | 0.00150463 | 0.043043981 | 0:00:15 | 0:00:09 |
| XM_017010615:XM_017010619 | 0:00 | 1:00 | 0:00 | 0:00 | 0:00 | 0:02 | 0:00 | 0:00 | 0:00 | 0:00 |
| XM_017010615:XM_017010619 | 0:00 | 1:00 | 0:00 | 0:00 | 0:00 | 0:02 | 0:00 | 0:00 | 0:00 | 0:00 |
| NM_001142522:NM_012177 | 0:00 | 0:00 | 0:00 | 0:00 | 0:00 | 0:00 | 0:00 | 7:01 | 0:00 | 1:00 |
| NM_016020:XR_001743453:NM_001350502:XR_942460:XM_017010917 | 48:5:2:2:0 | 5:0:0:0:0 | 26:4:1:2:6 | 0:0:0:0:0 | 51:10:2:0:4 | 17:0:1:1:0 | 252:12:6:4:33 | 16:0:0:0:1 | 66:10:2:1:5 | 5:1:0:0:1 |
| NM_016020:XR_001743453:NM_001350502:XR_942460:XM_017010917 | 63:6:2:2:0 | 8:0:0:0:0 | 30:4:1:3:6 | 2:0:0:0:0 | 61:11:2:0:5 | 14:0:1:1:1 | 285:13:6:4:33 | 19:1:0:0:1 | 82:11:2:1:5 | 9:1:0:0:2 |
| NM_016020:XR_001743453:NM_001350502:XR_942460:XM_017010917 | 68:6:2:2:0 | 5:0:0:0:0 | 31:4:1:3:5 | 0:0:0:0:1 | 66:11:3:0:6 | 9:0:0:1:0 | 284:14:6:4:32 | 16:0:0:0:3 | 88:11:2:1:7 | 3:1:0:0:0 |
| NM_016020:NM_001350502:XR_942460:XM_017010917:XR_001743453 | 73:2:2:0:0 | 0:0:0:0:0 | 32:1:3:5:0 | 0:0:0:0:0 | 75:3:1:6:0 | 1:0:0:0:0 | 310:6:5:38:0 | 0:0:0:0:0 | 91:2:1:7:0 | 0:0:0:0:0 |
| NM_016020:NM_001350502:XR_942460:XM_017010917:XR_001743453 | 73:2:2:1:0 | 0:0:0:0:0 | 32:1:3:5:0 | 0:0:0:0:0 | 76:3:1:6:0 | 0:0:0:0:0 | 310:6:5:38:0 | 0:0:0:0:0 | 91:2:1:6:0 | 0:0:0:0:0 |
| NM_016020:XR_001743453:NM_001350502:XR_942460:XM_017010917 | 73:6:2:2:1 | 0:0:0:0:0 | 32:5:1:0:3 | 0:0:0:0:0 | 76:11:3:2:4 | 0:0:0:0:0 | 309:14:6:5:31 | 1:1:0:0:0 | 91:11:2:1:4 | 0:1:0:0:0 |
| NM_016020:XR_001743453:NM_001350502:XR_942460:XM_017010917 | 73:6:2:2:1 | 0:0:0:0:0 | 30:3:1:0:2 | 2:1:0:0:0 | 76:10:3:3:4 | 0:0:0:0:0 | 309:14:6:8:19 | 0:0:0:0:1 | 91:12:2:1:3 | 0:0:0:0:0 |
| NM_001346813:XR_002956289:NM_017519:XM_017011107:XM_017011106:XM_017011103 | 0:1:0:0:0:0 | 1:0:0:0:0:0 | 0:1:1:0:0:0 | 0:1:2:0:0:0 | 0:0:1:0:0:0 | 0:2:2:0:0:0 | 0:2:11:1:1:1 | 0:0:0:0:0:0 | 0:0:1:0:0:2 | 0:0:0:0:0:0 |
| NM_001111077:NM_003379:XM_011536110 | 113:72:9 | 0:00:00 | 95:209:133 | 1:01:00 | 586:696:83 | 2:02:01 | 223:228:231 | 0:00:00 | 267:129:19 | 1:00:00 |
| NM_001111077:NM_003379:XM_011536110 | 121:80:10 | 0:00:00 | 102:209:84 | 0:01:04 | 603:702:50 | 1:00:01 | 240:217:131 | 0:00:02 | 273:123:10 | 0:00:00 |
| NM_001111077:NM_003379:XM_011536110 | 122:81:10 | 1:00:00 | 110:219:82 | 0:01:01 | 621:725:51 | 3:01:01 | 261:221:126 | 2:03:03 | 279:126:13 | 1:00:00 |
| NM_001111077:NM_003379:XM_011536110 | 153:98:12 | 1:00:00 | 250:355:81 | 0:01:00 | 821:896:69 | 2:05:00 | 525:309:102 | 1:00:00 | 337:143:11 | 1:01:00 |
| NM_003379:NM_001111077:XM_011536110 | 106:176:4 | 1:00:00 | 455:351:39 | 0:00:00 | 958:860:44 | 0:01:01 | 344:633:33 | 0:02:00 | 148:345:8 | 0:00:00 |
| NM_001111077:NM_003379:XM_011536110 | 176:107:4 | 0:00:00 | 352:446:30 | 0:00:00 | 865:953:43 | 1:00:02 | 644:336:27 | 0:00:00 | 346:147:10 | 0:00:00 |
| NM_001111077:NM_003379:XM_011536110 | 176:106:4 | 0:00:00 | 354:438:32 | 0:01:00 | 870:950:45 | 0:01:01 | 648:326:26 | 1:03:00 | 347:149:10 | 0:00:00 |
| NM_001111077:NM_003379:XM_011536110 | 176:106:4 | 0:00:00 | 353:439:32 | 0:01:00 | 868:948:45 | 2:01:01 | 648:327:26 | 1:02:00 | 344:149:10 | 3:00:00 |
| NM_001111077:NM_003379:XM_011536110 | 176:106:4 | 0:00:00 | 355:437:33 | 0:01:00 | 870:947:46 | 1:01:01 | 649:320:26 | 2:02:00 | 347:148:10 | 0:01:00 |
| NM_001111077:NM_003379:XM_011536110 | 176:106:4 | 0:00:00 | 355:431:33 | 0:02:01 | 870:946:46 | 1:01:01 | 649:321:27 | 3:01:00 | 346:149:10 | 1:00:00 |
| XM_011536110:NM_001111077:NM_003379 | 3:175:105 | 1:01:00 | 33:356:430 | 1:00:01 | 45:870:947 | 1:01:00 | 26:651:317 | 0:01:02 | 10:347:149 | 0:00:00 |
| NM_001111077:NM_003379:XM_011536110 | 176:105:4 | 0:00:00 | 355:407:33 | 0:02:00 | 874:951:49 | 1:02:01 | 653:306:28 | 1:02:00 | 347:148:11 | 0:01:00 |
| NM_001111077:NM_003379:XM_011536110 | 176:99:6 | 0:00:00 | 356:376:29 | 0:01:00 | 874:903:52 | 2:03:00 | 657:259:32 | 1:01:00 | 348:143:14 | 0:00:00 |
| NM_001111077:NM_003379:XM_011536110 | 176:100:6 | 0:00:00 | 358:359:30 | 0:00:00 | 877:876:50 | 0:02:00 | 656:235:36 | 2:00:00 | 348:138:16 | 0:01:00 |
| NM_001024465:NM_000636 | 1:01 | 0:00 | 0:00 | 0:00 | 0:00 | 0:00 | 2:01 | 0:00 | 0:00 | 0:00 |
| NM_005891 | 0 | 0 | 1 | 0 | 4 | 0 | 0 | 0 | 0 | 1 |
| NM_014161:NM_001318817 | 1:00 | 0:00 | 1:01 | 0:00 | 0:00 | 0:00 | 1:00 | 1:00 | 2:00 | 1:01 |
| NM_000876 | 0 | 1 | 0 | 0 | 0 | 0 | 0 | 2 | 0 | 0 |
| NM_001500:XM_011514500:XR_001743349:XM_017010752:XM_011514507:XR_001743350 | 36:2:1:2:0:0 | 0:0:0:0:0:0 | 65:4:1:6:0:0 | 0:0:0:0:0:0 | 68:5:4:10:0:0 | 0:0:1:0:0:0 | 125:4:3:19:1:0 | 0:0:3:0:0:0 | 132:2:2:8:0:0 | 1:0:1:1:0:0 |
| NM_145169:NR_130112 | 3:01 | 0:00 | 1:00 | 0:00 | 4:01 | 0:00 | 2:00 | 0:00 | 5:01 | 0:00 |
| NM_003730:XM_024446576:XM_017011397:XM_017011399 | 1:1:0:0 | 0:0:0:0 | 1:1:0:0 | 0:0:0:0 | 3:0:1:0 | 0:0:0:0 | 9:0:0:0 | 0:0:0:0 | 3:0:0:1 | 0:0:0:0 |
| XM_017010958:XM_011535905:NM_001143947:NM_018974:XM_011535906:XM_011535908:XM_011535907 | 2:6:3:1:0:0:0 | 0:0:0:0:0:0:0 | 0:0:0:0:0:0:0 | 0:0:0:0:0:0:0 | 0:1:0:0:0:0:0 | 0:0:0:0:0:0:0 | 0:0:0:0:0:0:0 | 0:4:0:0:0:0:0 | 0:0:0:0:0:0:0 | 0:3:0:1:0:0:0 |
| XM_017010958:XM_011535905:NM_001143947:NM_018974:XM_011535906:XM_011535907 | 3:6:3:1:0:0 | 0:0:0:0:0:0 | 0:0:0:0:0:0 | 0:0:0:0:0:0 | 0:1:0:0:0:0 | 0:0:0:0:0:0 | 0:0:0:0:0:0 | 0:7:0:0:0:0 | 0:0:0:0:0:0 | 0:4:0:1:0:0 |
| XM_017010958:XM_011535905:NM_001143947:NM_018974:XM_011535906:XM_011535907 | 3:6:3:1:0:0 | 0:0:0:0:0:0 | 0:0:0:0:0:0 | 0:0:0:0:0:0 | 0:1:0:0:0:0 | 0:0:0:0:0:0 | 0:0:0:0:0:0 | 0:7:0:0:0:0 | 0:0:0:0:0:0 | 0:4:0:1:0:0 |
| XM_017010958:XM_011535905:NM_001143947:NM_018974:XM_011535906:XM_011535907 | 3:6:3:1:0:0 | 0:0:0:0:0:0 | 0:0:0:0:0:0 | 0:0:0:0:0:0 | 0:1:0:0:0:0 | 0:0:0:0:0:0 | 0:7:0:0:0:0 | 0:0:0:0:0:0 | 0:4:0:1:0:0 | 0:0:0:0:0:0 |
| XR_001743427:XM_017010885:XM_017010880:XM_017010884:XM_017010887:XM_017010886 | 21:7:10:0:0:0 | 1:0:0:0:0:0 | 51:22:29:0:0:0 | 2:0:0:0:0:0 | 22:15:8:1:0:0 | 1:0:0:0:0:0 | 132:49:62:0:0:0 | 9:4:2:0:0:0 | 39:10:20:0:0:0 | 0:0:0:0:0:0 |
| XR_001743427:XM_017010880:XM_017010885:NM_001291964:XM_017010884:XM_017010887:XM_017010886 | 25:12:7:1:0:0:0 | 0:0:0:0:0:0:0 | 76:49:37:3:2:0:0 | 0:0:0:0:0:0:0 | 27:11:16:1:1:0:0 | 0:0:0:0:0:0:0 | 202:105:90:44:0:0:0 | 0:0:0:0:0:0:0 | 47:23:13:1:0:0:0 | 0:0:0:0:0:0:0 |
| XR_001743427:XM_017010880:XM_017010885:NM_001291964:XM_017010884:XM_017010887:XM_017010886 | 25:12:7:1:0:0:0 | 0:0:0:0:0:0:0 | 76:49:37:3:2:0:0 | 0:0:0:0:0:0:0 | 27:11:16:1:1:0:0 | 0:0:0:0:0:0:0 | 202:105:89:44:0:0:0 | 0:0:2:0:0:0:0 | 47:23:13:1:0:0:0 | 0:0:0:0:0:0:0 |

| | | | | | | | | | | |
|---|---|---|---|---|---|---|---|---|---|---|
| XM_017010884:XM_017010880:XR_001743427:XM_017010885 | 1:0:0:0 | 0:0:0:0 | 1:0:4:2 | 0:2:2:1 | 1:1:1:1 | 0:0:0:0 | 1:0:4:3 | 0:1:2:0 | 0:1:2:1 | 0:0:1:0 |
| NM_030615:XM_011535803:XM_011535802 | 0:00:00 | 0:00:00 | 0:00:00 | 0:00:00 | 0:00:00 | 0:00:00 | 0:02:02 | 1:01:00 | 0:00:00 | 0:00:00 |
| NM_002793 | 1 | 1 | 2 | 1 | 2 | 0 | 2 | 2 | 5 | 3 |
| NM_000367:NM_001346818:NM_001346817 | 34:02:02 | 0:00:00 | 106:05:13 | 0:00:00 | 101:02:05 | 0:00:00 | 279:10:31 | 1:00:00 | 107:02:13 | 1:00:00 |
| NM_000367:NM_001346818:NM_001346817 | 34:03:02 | 0:00:00 | 106:06:11 | 0:00:01 | 101:02:05 | 0:00:00 | 280:15:25 | 0:00:00 | 108:02:13 | 0:00:00 |
| NM_000367:NM_001346818:NM_001346817 | 34:03:02 | 0:00:00 | 107:11:07 | 0:00:00 | 103:02:05 | 0:00:00 | 280:20:19 | 0:00:00 | 106:02:10 | 2:00:00 |
| NM_000367:NM_001346818:NM_001346817 | 35:04:02 | 0:00:00 | 107:14:05 | 0:00:00 | 103:02:03 | 0:00:00 | 279:21:09 | 1:00:00 | 108:02:09 | 0:00:00 |
| XM_024446544:XR_001743635:NM_003472:NM_001134709:XM_017011273 | 13:2:71:1:20 | 1:4:1:0:0 | 29:6:23:0:39 | 0:1:0:0:0 | 11:4:43:0:15 | 1:2:0:0:0 | 107:36:150:2:170 | 3:11:3:0:0 | 54:6:110:1:45 | 4:8:2:0:0 |
| NM_001134709:NM_003472:XM_024446544:XR_001743635:XM_017011273 | 1:72:14:6:20 | 0:0:0:0:0 | 0:23:29:7:39 | 0:0:0:0:0 | 0:43:11:6:15 | 0:0:0:0:0 | 1:153:106:51:168 | 0:0:0:0:1 | 1:112:58:14:44 | 0:0:0:0:1 |
| NM_003472:XM_024446544:XR_001743635:XM_017011273:NM_001134709 | 73:12:7:20:0 | 0:0:0:0:0 | 23:20:12:39:0 | 0:1:0:0:0 | 42:9:9:15:0 | 0:0:0:0:0 | 154:86:63:167:1 | 1:0:0:2:0 | 115:57:17:45:0 | 0:1:0:0:0 |
| NM_003472:XM_024446544:XR_001743635:XM_017011273:NM_001134709 | 73:12:7:20:0 | 0:0:0:0:0 | 23:20:12:39:0 | 0:1:0:0:0 | 42:9:9:15:0 | 0:0:0:0:0 | 76:40:30:90:0 | 79:46:32:79:1 | 54:32:11:21:0 | 61:26:6:24:0 |
| NM_003472:XM_024446544:XR_001743635:XM_017011273:NM_001134709 | 73:11:12:20:0 | 0:0:0:0:0 | 26:12:15:39:0 | 0:1:0:0:0 | 43:8:8:15:0 | 0:0:0:0:0 | 161:69:81:169:1 | 1:0:1:0:0 | 115:55:21:45:0 | 1:0:0:0:0 |
| NM_003472:XM_024446544:XR_001743635:XM_017011273:NM_001134709 | 72:11:15:20:0 | 1:0:0:0:0 | 27:13:19:39:0 | 0:0:0:0:0 | 43:11:12:15:0 | 0:0:0:0:0 | 163:69:89:168:1 | 1:0:0:1:0 | 116:56:24:45:0 | 0:0:0:0:0 |
| NM_001500:XM_011514500:XR_001743349:XM_011514502:XM_017010752:XR_001743350:XM_011514507 | 36:2:3:0:0:0:0 | 0:0:0:0:0:0:0 | 64:4:1:2:0:0:0 | 0:0:0:0:0:0:0 | 68:5:0:0:2:1:0 | 0:0:0:0:0:0:0 | 126:4:3:3:2:1:1 | 0:0:0:0:0:0:0 | 133:2:2:0:2:0:0 | 0:0:0:0:0:0:0 |
| NM_001500:XM_011514500:XR_001743349:XM_017010752:XM_011514502:XR_001743350:XM_011514507 | 36:1:3:1:0:0:0 | 0:0:0:0:0:0:0 | 63:1:1:1:2:0:0 | 0:0:0:0:0:0:0 | 68:2:0:1:0:1:0 | 0:0:0:0:0:0:0 | 126:2:3:1:3:1:1 | 0:0:0:0:0:0:0 | 133:1:2:0:0:1:0 | 0:0:0:0:0:0:0 |
| NM_020662:XR_001743533:NM_001286265 | 0:00:00 | 0:00:00 | 0:00:00 | 0:00:00 | 0:00:00 | 0:00:00 | 0:01:00 | 1:00:00 | 0:00:00 | 0:00:00 |
| NM_006510 | 23 | 0 | 12 | 0 | 40 | 1 | 143 | 2 | 32 | 0 |
| NM_004568:XM_011514674:NM_001271824:XM_024446465:NM_001297699:NM_001271825:NM_001195291:NM_001271822:XM_024446463 | 0:0:0:0:0:0:0:0:0 | 3:1:0:0:0:0:0:0:0 | 0:0:0:0:0:0:0:0:0 | 1:0:1:1:0:0:0:0:0 | 0:0:0:0:0:0:0:0:0 | 1:0:1:0:1:0:0:0:0 | 1:0:0:0:1:0:0:0:0 | 0:0:0:0:0:0:0:0:0 | 2:0:0:0:0:1:0:0:0 | 1:0:0:0:0:0:0:0:0 |
| NM_006398 | 0 | 2 | 0 | 6 | 0 | 0 | 1 | 2 | 0 | 0 |
| NM_006398 | 0 | 2 | 5 | 2 | 0 | 0 | 3 | 0 | 0 | 0 |
| NM_018950:XM_017010815:XR_001743374 | 0:00:00 | 0:00:00 | 0:00:00 | 1:00:00 | 0:00:00 | 3:01:00 | 0:00:00 | 0:01:00 | 0:00:00 | 1:00:00 |
| NM_002116:NM_001242758 | 0:00 | 1:00 | 0:00 | 14:00 | 0:00 | 15:00 | 0:07 | 0:00 | 0:04 | 0:00 |
| NM_002116:NM_001242758 | 0:00 | 1:00 | 0:00 | 14:00 | 0:00 | 17:00 | 0:07 | 0:00 | 0:04 | 0:00 |
| NM_002116:NM_001242758 | 1:00 | 0:00 | 14:00 | 0:00 | 16:00 | 0:00 | 0:00 | 0:07 | 0:00 | 0:04 |
| NM_002116:NM_001242758 | 0:00 | 1:00 | 0:00 | 14:00 | 0:00 | 16:00 | 0:07 | 0:00 | 0:05 | 0:00 |
| NM_002116:NM_001242758 | 0:00 | 1:00 | 0:00 | 14:00 | 0:00 | 17:00 | 0:07 | 0:00 | 0:05 | 0:00 |
| NM_002116:NM_001242758 | 0:00 | 1:00 | 0:00 | 14:00 | 0:00 | 17:00 | 0:07 | 0:00 | 0:05 | 0:00 |
| NM_002116:NM_001242758 | 1:00 | 0:00 | 14:00 | 0:00 | 17:00 | 0:00 | 0:00 | 0:07 | 0:00 | 0:05 |
| NM_002116:NM_001242758 | 0:00 | 1:00 | 0:00 | 14:00 | 0:00 | 17:00 | 0:00 | 0:07 | 0:00 | 0:05 |
| NM_002116:NM_001242758 | 0:00 | 1:00 | 0:00 | 14:00 | 0:00 | 17:00 | 0:00 | 0:07 | 0:00 | 0:06 |
| NM_002116:NM_001242758 | 0:00 | 1:00 | 0:00 | 14:00 | 0:00 | 17:00 | 0:00 | 0:07 | 0:00 | 0:06 |
| NM_002116:NM_001242758 | 0:00 | 1:00 | 0:00 | 14:00 | 0:00 | 17:00 | 0:07 | 0:00 | 0:06 | 0:00 |
| NM_002116:NM_001242758 | 0:00 | 1:00 | 0:00 | 14:00 | 0:00 | 17:00 | 0:07 | 0:00 | 0:06 | 0:00 |
| NM_002116:NM_001242758 | 0:00 | 1:00 | 0:00 | 14:00 | 0:00 | 17:00 | 0:07 | 0:00 | 0:06 | 0:00 |
| NM_002116:NM_001242758 | 0:00 | 1:00 | 0:00 | 14:00 | 0:00 | 17:00 | 0:07 | 0:00 | 0:06 | 0:00 |
| NM_002116:NM_001242758 | 0:00 | 1:00 | 0:00 | 14:00 | 0:00 | 17:00 | 0:07 | 0:00 | 0:06 | 0:00 |
| NM_002116:NM_001242758 | 0:00 | 1:00 | 0:00 | 14:00 | 0:00 | 17:00 | 0:07 | 0:00 | 0:06 | 0:00 |
| NM_002116:NM_001242758 | 1:00 | 0:00 | 14:00 | 0:00 | 17:00 | 0:00 | 0:00 | 0:07 | 0:00 | 0:06 |
| NM_002116:NM_001242758 | 0:00 | 1:00 | 0:00 | 15:00 | 0:00 | 18:00 | 0:07 | 0:00 | 0:06 | 0:00 |
| NM_002116:NM_001242758 | 0:00 | 1:00 | 0:00 | 15:00 | 0:00 | 18:00 | 0:07 | 0:00 | 0:06 | 0:00 |
| NM_002116:NM_001242758 | 0:00 | 1:00 | 0:00 | 15:00 | 0:00 | 18:00 | 0:07 | 0:00 | 0:06 | 0:00 |
| NM_002116:NM_001242758 | 1:00 | 0:00 | 15:00 | 0:00 | 18:00 | 0:00 | 0:00 | 0:07 | 0:00 | 0:06 |
| NM_002116:NM_001242758 | 0:00 | 1:00 | 0:00 | 15:00 | 0:00 | 18:00 | 0:07 | 0:00 | 0:06 | 0:00 |
| NM_002116:NM_001242758 | 0:00 | 1:00 | 0:00 | 15:00 | 0:00 | 18:00 | 0:07 | 0:00 | 0:06 | 0:00 |
| NM_002116:NM_001242758 | 0:00 | 1:00 | 0:00 | 15:00 | 0:00 | 18:00 | 0:07 | 0:00 | 0:06 | 0:00 |
| NM_002116:NM_001242758 | 1:00 | 0:00 | 15:00 | 0:00 | 18:00 | 0:00 | 0:00 | 0:07 | 0:00 | 0:07 |
| NM_002116:NM_001242758 | 0:00 | 1:00 | 0:00 | 15:00 | 0:00 | 18:00 | 0:00 | 0:07 | 0:00 | 0:07 |
| NM_002116:NM_001242758 | 0:00 | 1:00 | 0:00 | 15:00 | 0:00 | 19:00 | 0:00 | 0:07 | 0:00 | 0:07 |
| NM_002116:NM_001242758 | 0:00 | 1:00 | 0:00 | 15:00 | 0:00 | 19:00 | 0:00 | 0:07 | 0:00 | 0:07 |
| NM_002116:NM_001242758 | 1:00 | 0:00 | 15:00 | 0:00 | 19:00 | 0:00 | 0:00 | 0:07 | 0:00 | 0:07 |
| NM_002116:NM_001242758 | 0:00 | 1:00 | 0:00 | 15:00 | 0:00 | 19:00 | 0:00 | 0:07 | 0:00 | 0:08 |
| NM_002116:NM_001242758 | 1:00 | 0:00 | 15:00 | 0:00 | 19:00 | 0:00 | 0:00 | 0:07 | 0:00 | 0:08 |
| NM_002116:NM_001242758 | 1:00 | 0:00 | 15:00 | 0:00 | 19:00 | 0:00 | 0:00 | 0:07 | 0:00 | 0:08 |
| NM_002116:NM_001242758 | 0:00 | 1:00 | 0:00 | 15:00 | 0:00 | 19:00 | 0:00 | 0:07 | 0:00 | 0:08 |
| NM_002116:NM_001242758 | 0:00 | 1:00 | 0:00 | 15:00 | 0:00 | 19:00 | 0:00 | 0:07 | 0:00 | 0:08 |
| NM_002116:NM_001242758 | 1:00 | 0:00 | 15:00 | 0:00 | 19:01 | 0:00 | 0:05 | 0:01 | 0:08 | 0:00 |
| NM_002116:NM_001242758 | 1:00 | 0:00 | 12:00 | 0:00 | 18:01 | 1:00 | 0:07 | 0:00 | 0:09 | 0:00 |
| NM_002116:NM_001242758 | 1:00 | 0:00 | 12:00 | 0:00 | 18:01 | 1:00 | 0:06 | 0:01 | 0:09 | 0:00 |
| NM_002116:NM_001242758 | 0:00 | 1:00 | 0:00 | 12:00 | 0:00 | 19:01 | 0:07 | 0:00 | 0:09 | 0:00 |
| NM_002116:NM_001242758 | 0:00 | 1:00 | 0:00 | 13:00 | 1:00 | 18:01 | 0:07 | 0:00 | 0:09 | 0:00 |
| NM_002116:NM_001242758 | 1:00 | 0:00 | 13:00 | 0:00 | 18:00 | 1:00 | 0:07 | 0:00 | 0:10 | 0:00 |
| NM_002116:NM_001242758 | 1:00 | 0:00 | 13:00 | 0:00 | 18:00 | 1:00 | 0:07 | 0:00 | 0:10 | 0:00 |
| NM_002116:NM_001242758 | 1:00 | 0:00 | 13:00 | 0:00 | 18:00 | 1:00 | 0:07 | 0:00 | 0:10 | 0:00 |
| NM_002116:NM_001242758 | 1:00 | 0:00 | 13:00 | 0:00 | 18:00 | 1:00 | 0:07 | 0:00 | 0:10 | 0:00 |
| NM_002116:NM_001242758 | 0:00 | 1:00 | 0:00 | 13:00 | 0:00 | 18:00 | 0:07 | 0:00 | 0:10 | 0:00 |
| NM_002116:NM_001242758 | 0:00 | 1:00 | 0:00 | 12:00 | 0:00 | 17:00 | 0:06 | 0:00 | 0:10 | 0:00 |
| NM_170783:NM_014596:NM_001278785:NM_001278786:NR_103864 | 109:40:6:3:0 | 1:0:0:0:0 | 117:39:8:3:6 | 0:0:0:0:0 | 271:56:15:7:5 | 0:0:0:0:0 | 661:430:87:19:12 | 2:0:0:0:0 | 172:66:3:3:1 | 0:0:0:0:0 |
| NM_170783:NM_014596:NM_001278785:NM_001278786:NR_103864 | 109:40:6:3:0 | 1:0:0:0:0 | 117:39:8:3:6 | 0:0:0:0:0 | 268:56:15:7:5 | 2:0:0:0:0 | 662:430:87:19:12 | 0:0:0:0:0 | 172:66:3:3:1 | 0:0:0:0:0 |
| NM_014596:NM_170783:NM_001278785:NM_001278786:NR_103864 | 40:109:6:3:0 | 0:0:0:0:0 | 39:116:8:3:6 | 0:0:0:0:0 | 56:269:15:7:5 | 0:1:0:0:0 | 428:661:87:19:12 | 2:1:0:0:0 | 65:172:3:3:1 | 1:0:0:0:0 |
| NM_000904:NM_001290221 | 0:00 | 1:00 | 0:00 | 0:00 | 0:00 | 0:00 | 0:00 | 0:00 | 0:00 | 0:00 |
| NM_007028:NR_134870:NR_134871:XM_011514264 | 0:0:0:0 | 0:0:0:0 | 1:0:0:0 | 0:0:0:0 | 0:1:2:0 | 0:0:0:0 | 1:0:0:0 | 2:0:0:0 | 1:0:0:0 | 0:0:0:0 |
| NM_024839:NM_001199120 | 2:00 | 0:00 | 1:00 | 0:00 | 4:00 | 0:00 | 2:01 | 0:00 | 3:00 | 0:00 |
| NM_005516:XM_017010809:XM_017010807 | 2:01:00 | 0:00:00 | 0:00:00 | 0:00:00 | 5:00:01 | 0:00:00 | 0:00:00 | 2:00:02 | 0:00:00 | 0:00:00 |
| NM_003897 | 0 | 1 | 0 | 3 | 0 | 3 | 1 | 8 | 0 | 2 |
| NM_001318908:NM_007109 | 0:00 | 0:00 | 0:00 | 0:00 | 0:00 | 0:00 | 0:00 | 0:00 | 0:00 | 1:01 |
| NM_002117:NM_001243042 | 0:07 | 61:03:00 | 0:37 | 203:10:00 | 0:26 | 214:04:00 | 0:00 | 316:07:00 | 0:00 | 91:01:00 |
| NM_002117:NM_001243042 | 0:07 | 60:03:00 | 2:37 | 199:10:00 | 1:25 | 213:04:00 | 0:00 | 312:07:00 | 0:00 | 91:01:00 |
| NM_002117:NM_001243042 | 0:07 | 60:03:00 | 1:36 | 199:10:00 | 0:25 | 206:04:00 | 0:00 | 308:07:00 | 0:00 | 91:01:00 |

| | | | | | | | | | | |
|---|---|---|---|---|---|---|---|---|---|---|
| NM_002117:NM_001243042 | 0:07 | 60:03:00 | 1:36 | 199:10:00 | 0:25 | 206:04:00 | 0:00 | 308:07:00 | 0:00 | 91:01:00 |
| NM_002117:NM_001243042 | 0:07 | 60:03:00 | 1:36 | 197:10:00 | 0:25 | 206:04:00 | 0:00 | 307:07:00 | 0:00 | 91:01:00 |
| NM_002117:NM_001243042 | 0:07 | 60:03:00 | 1:36 | 198:10:00 | 0:25 | 206:04:00 | 13:00 | 295:07:00 | 6:01 | 85:00:00 |
| NM_002117:NM_001243042 | 60:10:00 | 0:00 | 198:46:00 | 1:00 | 206:29:00 | 0:00 | 308:07:00 | 0:00 | 91:01:00 | 0:00 |
| NM_002117:NM_001243042 | 0:07 | 60:03:00 | 1:36 | 197:10:00 | 0:25 | 203:04:00 | 1:00 | 303:07:00 | 0:00 | 90:01:00 |
| NM_002117:NM_001243042 | 0:07 | 60:03:00 | 1:35 | 192:10:00 | 0:25 | 202:04:00 | 1:00 | 297:07:00 | 0:00 | 90:01:00 |
| NM_002117:NM_001243042 | 0:07 | 60:03:00 | 2:33 | 189:10:00 | 1:24 | 199:04:00 | 297:07:00 | 0:00 | 91:01:00 | 0:00 |
| NM_002117:NM_001243042 | 60:10:00 | 0:00 | 189:43:00 | 1:00 | 198:27:00 | 1:00 | 297:07:00 | 0:00 | 93:01:00 | 0:00 |
| NM_002117:NM_001243042 | 0:07 | 60:03:00 | 1:33 | 189:10:00 | 0:23 | 199:04:00 | 0:00 | 296:07:00 | 0:00 | 93:01:00 |
| NM_002117:NM_001243042 | 0:07 | 60:03:00 | 1:33 | 190:10:00 | 0:23 | 198:04:00 | 0:00 | 297:07:00 | 0:00 | 93:01:00 |
| NM_002117:NM_001243042 | 0:07 | 60:03:00 | 3:33 | 188:10:00 | 0:23 | 197:04:00 | 0:00 | 295:07:00 | 0:00 | 91:01:00 |
| NM_002117:NM_001243042 | 0:07 | 60:03:00 | 1:33 | 189:10:00 | 0:23 | 195:04:00 | 0:00 | 292:07:00 | 0:00 | 91:01:00 |
| NM_002117:NM_001243042 | 60:10:00 | 0:00 | 186:42:00 | 2:01 | 194:27:00 | 0:00 | 292:07:00 | 0:00 | 89:01:00 | 0:00 |
| NM_002117:NM_001243042 | 58:10:00 | 2:00 | 185:43:00 | 3:01 | 190:28:00 | 3:00 | 294:06:00 | 1:01 | 86:01:00 | 1:00 |
| NM_002117:NM_001243042 | 59:10:00 | 1:00 | 184:45:00 | 1:00 | 183:28:00 | 2:00 | 279:07:00 | 10:00 | 78:00:00 | 5:01 |
| NM_002117:NM_001243042 | 1:07 | 54:03:00 | 5:36 | 165:10:00 | 3:26 | 178:04:00 | 0:00 | 284:08:00 | 1:01 | 83:01:00 |
| NM_002117:NM_001243042 | 1:07 | 55:03:00 | 5:40 | 164:10:00 | 2:27 | 179:04:00 | 281:10:00 | 1:00 | 82:01:00 | 0:00 |
| NM_002117:NM_001243042 | 56:10:00 | 0:00 | 169:51:00 | 0:00 | 179:31:00 | 1:00 | 272:10:00 | 9:00 | 76:00:00 | 6:01 |
| NM_002117:NM_001243042 | 54:10:00 | 1:00 | 166:47:00 | 2:03 | 177:28:00 | 3:03 | 281:10:00 | 0:00 | 82:01:00 | 0:00 |
| NM_002117:NM_001243042 | 54:10:00 | 0:00 | 166:50:00 | 0:00 | 178:31:00 | 0:00 | 278:10:00 | 0:00 | 81:01:00 | 0:00 |
| NM_002117:NM_001243042 | 54:10:00 | 0:00 | 161:45:00 | 4:06 | 178:28:00 | 0:03 | 269:10:00 | 9:00 | 75:00:00 | 6:01 |
| NM_002117:NM_001243042 | 54:10:00 | 0:00 | 160:45:00 | 5:06 | 178:28:00 | 0:03 | 269:10:00 | 9:00 | 75:00:00 | 6:01 |
| NM_002117:NM_001243042 | 54:10:00 | 0:00 | 167:51:00 | 0:00 | 178:30:00 | 0:00 | 273:08:00 | 1:01 | 79:01:00 | 0:00 |
| NM_002117:NM_001243042 | 1:07 | 53:03:00 | 6:41 | 161:10:00 | 2:26 | 176:04:00 | 11:01 | 261:08:00 | 6:01 | 73:00:00 |
| NM_002117:NM_001243042 | 0:07 | 54:03:00 | 0:31 | 166:20:00 | 0:20 | 177:10:00 | 0:00 | 264:09:00 | 0:00 | 79:01:00 |
| NM_002117:NM_001243042 | 0:07 | 54:03:00 | 0:31 | 167:20:00 | 0:20 | 177:10:00 | 0:00 | 257:09:00 | 0:00 | 79:01:00 |
| NM_002117:NM_001243042 | 54:11:00 | 0:00 | 168:51:00 | 0:00 | 175:30:00 | 0:00 | 253:09:00 | 2:00 | 79:01:00 | 0:00 |
| NM_002117:NM_001243042 | 1:08 | 53:03:00 | 6:40 | 160:10:00 | 2:26 | 173:04:00 | 249:09:00 | 1:00 | 79:01:00 | 0:00 |
| NM_002117:NM_001243042 | 53:10:00 | 1:01 | 159:40:00 | 5:10 | 171:24:00 | 2:06 | 249:08:00 | 1:01 | 78:01:00 | 0:00 |
| NM_002117:NM_001243042 | 0:07 | 54:04:00 | 0:29 | 162:20:00 | 0:20 | 171:10:00 | 237:08:00 | 10:01 | 71:00:00 | 6:01 |
| NM_002117:NM_001243042 | 54:11:00 | 0:00 | 162:49:00 | 0:00 | 171:30:00 | 0:00 | 247:09:00 | 1:00 | 77:01:00 | 0:00 |
| NM_002117:NM_001243042 | 0:07 | 53:04:00 | 0:29 | 161:20:00 | 0:20 | 170:10:00 | 245:08:00 | 2:01 | 78:01:00 | 0:00 |
| NM_002117:NM_001243042 | 53:11:00 | 0:00 | 160:49:00 | 1:00 | 170:30:00 | 0:00 | 247:08:00 | 0:01 | 78:01:00 | 0:00 |
| NM_002117:NM_001243042 | 53:11:00 | 0:00 | 157:49:00 | 1:00 | 171:30:00 | 0:00 | 247:09:00 | 1:00 | 78:01:00 | 0:00 |
| NM_002117:NM_001243042 | 51:11:00 | 1:00 | 156:47:00 | 1:02 | 166:28:00 | 2:02 | 241:08:00 | 10:01 | 72:00:00 | 6:01 |
| NM_002117:NM_001243042 | 0:00 | 51:11:00 | 1:00 | 155:49:00 | 0:00 | 167:30:00 | 1:00 | 250:09:00 | 0:00 | 78:01:00 |
| NM_002117:NM_001243042 | 51:11:00 | 0:00 | 154:46:00 | 1:03 | 165:27:00 | 2:03 | 9:01 | 242:08:00 | 6:01 | 72:00:00 |
| NM_001243042:NM_002117 | 10:51 | 1:00 | 39:154 | 7:01 | 24:167 | 6:00 | 0.548611111 | 0:01 | 0.095833333 | 0:00 |
| NM_001243042:NM_002117 | 10:51 | 1:00 | 39:154 | 7:01 | 24:167 | 5:00 | 0.542361111 | 0:10 | 0.05 | 1:06 |
| NM_001243042:NM_002117 | 10:51 | 1:00 | 39:154 | 7:01 | 24:167 | 5:00 | 0.542361111 | 0:10 | 0.05 | 1:06 |
| NM_002117:NM_001243042 | 51:11:00 | 0:00 | 154:46:00 | 1:00 | 167:28:00 | 0:01 | 241:09:00 | 10:00 | 72:00:00 | 6:01 |
| NM_002117:NM_001243042 | 0:06 | 52:03:00 | 0:28 | 153:10:00 | 3:20 | 168:04:00 | 1:00 | 254:03:00 | 0:00 | 80:00:00 |
| NM_002117:NM_001243042 | 52:09:00 | 0:00 | 157:30:00 | 0:06 | 173:22:00 | 0:02 | 260:03:00 | 0:00 | 81:00:00 | 0:00 |
| NM_002117:NM_001243042 | 52:09:00 | 0:00 | 157:36:00 | 0:00 | 173:22:00 | 0:00 | 261:02:00 | 0:00 | 81:00:00 | 0:00 |
| NM_002117:NM_001243042 | 51:09:00 | 1:00 | 157:29:00 | 0:07 | 175:21:00 | 1:01 | 261:02:00 | 0:00 | 80:00:00 | 0:00 |
| NM_002117:NM_001243042 | 52:09:00 | 0:00 | 157:29:00 | 0:07 | 175:20:00 | 1:02 | 261:02:00 | 0:00 | 80:00:00 | 0:00 |
| NM_002117:NM_001243042 | 1:06 | 52:03:00 | 1:29 | 160:07:00 | 0:19 | 177:02:00 | 252:00:00 | 9:00 | 75:00:00 | 6:00 |
| NM_002117:NM_001243042 | 53:08:00 | 0:00 | 161:30:00 | 0:06 | 178:20:00 | 0:01 | 261:00:00 | 0:00 | 81:00:00 | 0:00 |
| NM_002117:NM_001243042 | 53:08:00 | 0:00 | 161:30:00 | 0:04 | 178:20:00 | 0:01 | 262:00:00 | 0:00 | 81:00:00 | 0:00 |
| NM_001243042:NM_002117 | 2:54 | 6:00 | 0.195833333 | 33:00:00 | 0.123611111 | 21:00 | 0.181944444 | 0:00 | 0.05625 | 0:00 |
| NM_002117:NM_001243042 | 54:09:00 | 1:00 | 163:32:00 | 0:02 | 180:21:00 | 0:00 | 264:00:00 | 2:00 | 81:00:00 | 0:00 |
| NM_002117:NM_001243042 | 56:09:00 | 0:00 | 162:33:00 | 2:01 | 184:19:00 | 1:00 | 269:00:00 | 0:00 | 80:00:00 | 0:00 |
| NM_002117:NM_001243042 | 56:09:00 | 0:00 | 161:30:00 | 3:01 | 183:19:00 | 1:00 | 266:00:00 | 2:00 | 79:00:00 | 0:00 |
| NM_002117:NM_001243042 | 55:09:00 | 0:00 | 161:31:00 | 2:00 | 182:19:00 | 2:00 | 267:00:00 | 0:00 | 79:00:00 | 0:00 |
| NM_002117:NM_001243042 | 0:07 | 55:02:00 | 3:27 | 161:04:00 | 0:19 | 183:00:00 | 266:00:00 | 0:00 | 79:00:00 | 0:00 |
| NM_002117:NM_001243042 | 0:07 | 55:02:00 | 1:27 | 162:04:00 | 0:19 | 182:00:00 | 0:00 | 265:00:00 | 0:00 | 79:00:00 |
| NM_002117:NM_001243042 | 55:09:00 | 0:00 | 160:30:00 | 3:01 | 180:19:00 | 2:00 | 255:00:00 | 9:00 | 74:00:00 | 6:00 |
| NM_002117:NM_001243042 | 54:09:00 | 0:00 | 157:30:00 | 5:00 | 180:20:00 | 1:00 | 262:00:00 | 1:00 | 80:00:00 | 0:00 |
| NM_002117:NM_001243042 | 54:09:00 | 0:00 | 159:30:00 | 2:00 | 179:20:00 | 2:00 | 262:00:00 | 0:00 | 80:00:00 | 0:00 |
| NM_002117:NM_001243042 | 55:09:00 | 0:00 | 157:30:00 | 4:00 | 180:20:00 | 1:00 | 264:01:00 | 1:00 | 80:00:00 | 0:00 |
| NM_002117:NM_001243042 | 55:09:00 | 0:00 | 157:30:00 | 4:00 | 180:20:00 | 1:00 | 263:01:00 | 2:00 | 80:00:00 | 0:00 |
| NM_002117:NM_001243042 | 0:07 | 55:02:00 | 3:27 | 158:03:00 | 1:20 | 180:00:00 | 1:00 | 264:01:00 | 0:00 | 80:00:00 |
| NM_002117:NM_001243042 | 55:09:00 | 0:00 | 156:30:00 | 3:00 | 179:20:00 | 1:00 | 252:01:00 | 9:00 | 73:00:00 | 6:00 |
| NM_002117:NM_001243042 | 0:07 | 55:03:00 | 2:27 | 156:03:00 | 0:20 | 181:00:00 | 0:00 | 266:01:00 | 0:00 | 79:00:00 |
| NM_002117:NM_001243042 | 55:10:00 | 0:00 | 157:30:00 | 1:00 | 180:20:00 | 1:00 | 266:01:00 | 0:00 | 79:00:00 | 0:00 |
| NM_002117:NM_001243042 | 0:07 | 55:03:00 | 2:27 | 156:03:00 | 0:20 | 181:00:00 | 0:00 | 267:01:00 | 0:00 | 80:00:00 |
| NM_002117:NM_001243042 | 0:07 | 55:03:00 | 3:27 | 155:03:00 | 1:20 | 180:00:00 | 0:00 | 268:01:00 | 0:00 | 80:00:00 |
| NM_004332:NR_026649:NM_001302777:XR_002956299:XR_002956297:XR_002956298:NR_026648 | 34:1:4:1:0:0:0 | 0:0:0:0:0:0:0 | 28:2:3:0:2:0:0 | 0:0:0:0:0:0:0 | 212:3:8:4:4:2:0 | 0:0:0:0:0:0:0 | 172:8:6:5:5:3:1 | 2:0:0:0:0:0:0 | 28:1:0:0:1:0:0 | 0:0:0:0:0:0:0 |
| NM_005514 | 1 | 1123 | 14 | 4810 | 2 | 3668 | 452 | 2477 | 137 | 893 |
| NM_005514 | 1117 | 1 | 4770 | 18 | 3656 | 12 | 2912 | 7 | 1023 | 4 |
| NM_005514 | 687 | 428 | 2243 | 2481 | 1751 | 1894 | 2906 | 3 | 1024 | 0 |
| NM_005514 | 1114 | 1 | 4714 | 6 | 3643 | 2 | 2908 | 3 | 1024 | 0 |
| NM_005514 | 680 | 418 | 2173 | 2431 | 1721 | 1872 | 2871 | 5 | 1014 | 0 |
| NM_005514 | 1061 | 2 | 4335 | 14 | 3477 | 4 | 2781 | 7 | 990 | 0 |
| NM_005514 | 1028 | 2 | 4114 | 19 | 3377 | 7 | 2694 | 5 | 955 | 2 |
| NM_005514 | 1025 | 5 | 4109 | 24 | 3374 | 10 | 2691 | 7 | 955 | 2 |
| NM_005514 | 1027 | 2 | 4117 | 13 | 3378 | 5 | 2686 | 10 | 956 | 1 |
| NM_005514 | 1032 | 3 | 4125 | 16 | 3382 | 7 | 2695 | 9 | 961 | 1 |
| NM_005514 | 1019 | 1 | 4023 | 28 | 3334 | 14 | 2648 | 20 | 942 | 1 |
| NM_005514 | 0 | 989 | 1 | 3931 | 0 | 3290 | 383 | 2212 | 119 | 807 |

| | | | | | | | | | | |
|---|---|---|---|---|---|---|---|---|---|---|
| NM_005514 | 0 | 989 | 1 | 3930 | 0 | 3289 | 384 | 2211 | 119 | 807 |
| NM_005514 | 1 | 990 | 3 | 3929 | 0 | 3290 | 382 | 2213 | 119 | 807 |
| NM_005514 | 994 | 0 | 3938 | 4 | 3293 | 0 | 2590 | 3 | 926 | 1 |
| NM_005514 | 388 | 602 | 2108 | 1827 | 1734 | 1556 | 2591 | 3 | 925 | 1 |
| NM_005514 | 990 | 1 | 3916 | 4 | 3274 | 4 | 2556 | 7 | 921 | 0 |
| NM_005514 | 989 | 2 | 3902 | 11 | 3274 | 4 | 2553 | 6 | 921 | 1 |
| NM_005514 | 982 | 8 | 3880 | 25 | 3262 | 10 | 2548 | 7 | 920 | 1 |
| NM_005514 | 980 | 7 | 3874 | 26 | 3257 | 12 | 2545 | 5 | 918 | 3 |
| NM_005514 | 5 | 982 | 20 | 3871 | 11 | 3255 | 370 | 2174 | 121 | 800 |
| NM_005514 | 1 | 981 | 7 | 3871 | 5 | 3256 | 373 | 2171 | 120 | 801 |
| NM_005514 | 974 | 3 | 3868 | 9 | 3243 | 4 | 360 | 2162 | 116 | 801 |
| NM_005514 | 2 | 971 | 12 | 3849 | 6 | 3238 | 6 | 2517 | 1 | 917 |
| NM_005514 | 963 | 4 | 3818 | 8 | 3218 | 7 | 0 | 2512 | 0 | 913 |
| NM_005514 | 961 | 0 | 3817 | 4 | 3214 | 3 | 2149 | 362 | 794 | 116 |
| NM_005514 | 0 | 952 | 11 | 3746 | 1 | 3180 | 354 | 2122 | 117 | 788 |
| NM_005514 | 577 | 372 | 1722 | 2002 | 1486 | 1668 | 2471 | 4 | 901 | 0 |
| NM_005514 | 1 | 948 | 7 | 3721 | 7 | 3154 | 356 | 2119 | 118 | 783 |
| NM_005514 | 948 | 1 | 3716 | 13 | 3155 | 7 | 2469 | 6 | 899 | 3 |
| NM_005514 | 1 | 948 | 7 | 3696 | 2 | 3148 | 1 | 2469 | 3 | 895 |
| NM_005514 | 0 | 949 | 4 | 3695 | 3 | 3145 | 2463 | 7 | 897 | 0 |
| NM_005514 | 943 | 1 | 3679 | 11 | 3140 | 4 | 2456 | 5 | 895 | 0 |
| NM_005514 | 941 | 3 | 3683 | 6 | 3137 | 7 | 2456 | 5 | 894 | 1 |
| NM_005514 | 571 | 370 | 1693 | 1976 | 1480 | 1659 | 2456 | 6 | 892 | 0 |
| NM_005514 | 0 | 940 | 3 | 3649 | 2 | 3126 | 2 | 2456 | 0 | 888 |
| NM_005514 | 370 | 570 | 1965 | 1687 | 1648 | 1481 | 2451 | 6 | 888 | 0 |
| NM_005514 | 571 | 369 | 1680 | 1965 | 1474 | 1649 | 2444 | 7 | 887 | 0 |
| NM_005514 | 570 | 370 | 1681 | 1963 | 1470 | 1653 | 2444 | 6 | 887 | 0 |
| NM_005514 | 570 | 370 | 1681 | 1963 | 1470 | 1653 | 2444 | 6 | 887 | 0 |
| NM_005514 | 939 | 1 | 3639 | 3 | 3115 | 6 | 2445 | 4 | 887 | 0 |
| NM_005514 | 958 | 0 | 3725 | 11 | 3145 | 12 | 2480 | 4 | 900 | 3 |
| NM_005514 | 962 | 1 | 3741 | 7 | 3156 | 7 | 2480 | 3 | 898 | 0 |
| NM_005514 | 583 | 379 | 1731 | 2014 | 1503 | 1659 | 371 | 2111 | 116 | 782 |
| NM_005514 | 583 | 379 | 1734 | 2010 | 1499 | 1658 | 368 | 2111 | 117 | 782 |
| NM_005514 | 582 | 379 | 1719 | 2011 | 1495 | 1657 | 366 | 2111 | 116 | 781 |
| NM_005514 | 584 | 378 | 1728 | 2010 | 1498 | 1658 | 369 | 2108 | 116 | 781 |
| NM_005514 | 583 | 382 | 1732 | 2016 | 1501 | 1669 | 366 | 2113 | 116 | 784 |
| NM_005514 | 580 | 385 | 1734 | 2036 | 1499 | 1678 | 371 | 2118 | 117 | 786 |
| NM_005514 | 962 | 4 | 3779 | 10 | 3178 | 9 | 2496 | 6 | 903 | 0 |
| NM_005514 | 586 | 380 | 1752 | 2048 | 1509 | 1680 | 7 | 2503 | 1 | 905 |
| NM_005514 | 2 | 966 | 12 | 3789 | 6 | 3177 | 6 | 2507 | 0 | 906 |
| NM_005514 | 3 | 967 | 16 | 3790 | 5 | 3180 | 7 | 2508 | 0 | 906 |
| NM_005514 | 1 | 969 | 16 | 3809 | 10 | 3184 | 11 | 2506 | 0 | 907 |
| NM_005514 | 1 | 971 | 9 | 3826 | 4 | 3196 | 7 | 2522 | 1 | 907 |
| NM_005514 | 590 | 383 | 1795 | 2060 | 1534 | 1688 | 2172 | 376 | 795 | 118 |
| NM_005514 | 592 | 382 | 1794 | 2060 | 1535 | 1688 | 2171 | 376 | 794 | 118 |
| NM_005514 | 579 | 374 | 1763 | 2023 | 1530 | 1663 | 2160 | 373 | 791 | 114 |
| NM_005514 | 577 | 376 | 1758 | 2028 | 1526 | 1662 | 2159 | 371 | 788 | 115 |
| NM_005514 | 576 | 375 | 1753 | 2011 | 1522 | 1655 | 2158 | 369 | 786 | 115 |
| NM_005514 | 572 | 372 | 1743 | 2012 | 1516 | 1650 | 2148 | 372 | 780 | 114 |
| NM_005514 | 943 | 2 | 3729 | 13 | 3136 | 11 | 2507 | 8 | 893 | 0 |
| NM_005514 | 939 | 4 | 3722 | 15 | 3137 | 7 | 2507 | 6 | 893 | 0 |
| NM_005514 | 940 | 4 | 3715 | 13 | 3134 | 9 | 2500 | 8 | 890 | 2 |
| NM_005514 | 573 | 371 | 1741 | 1986 | 1512 | 1631 | 2138 | 370 | 781 | 111 |
| NM_005514 | 1 | 944 | 13 | 3732 | 1 | 3145 | 2135 | 374 | 778 | 113 |
| NM_005514 | 572 | 375 | 1747 | 2008 | 1513 | 1639 | 2137 | 372 | 778 | 113 |
| NM_005514 | 944 | 2 | 3761 | 13 | 3124 | 13 | 2494 | 8 | 884 | 2 |
| NM_005514 | 941 | 5 | 3783 | 11 | 3133 | 5 | 2505 | 10 | 883 | 4 |
| NM_005514 | 942 | 1 | 3776 | 14 | 3127 | 11 | 2500 | 15 | 882 | 4 |
| NM_005514 | 942 | 1 | 3776 | 18 | 3124 | 14 | 2505 | 9 | 884 | 1 |
| NM_005514 | 939 | 3 | 3779 | 13 | 3123 | 15 | 2504 | 9 | 883 | 1 |
| NM_005514 | 574 | 369 | 1763 | 2031 | 1506 | 1638 | 2135 | 380 | 771 | 117 |
| NM_005514 | 573 | 369 | 1759 | 2034 | 1504 | 1634 | 2126 | 392 | 771 | 116 |
| NM_025257 | 0 | 0 | 0 | 4 | 0 | 1 | 0 | 0 | 0 | 0 |
| NM_025257 | 0 | 0 | 0 | 3 | 0 | 1 | 0 | 0 | 0 | 0 |
| NM_025257 | 0 | 0 | 2 | 0 | 0 | 0 | 0 | 0 | 0 | 0 |
| NM_025256:XM_017010202:XM_006714976:NM_006709 | 1:1:0:0 | 0:0:0:0 | 0:0:0:0 | 0:0:0:0 | 0:0:0:0 | 0:0:0:0 | 0:0:0:0 | 5:2:0:0 | 0:0:0:0 | 1:2:1:0 |
| NM_001710 | 19 | 0 | 30 | 1 | 82 | 0 | 197 | 1 | 46 | 0 |
| NM_001710 | 22 | 0 | 40 | 0 | 86 | 0 | 260 | 2 | 51 | 0 |
| NM_001710 | 23 | 0 | 41 | 0 | 90 | 0 | 305 | 2 | 59 | 0 |
| NM_001710 | 24 | 0 | 52 | 0 | 96 | 0 | 327 | 0 | 62 | 0 |
| NM_001710 | 32 | 1 | 102 | 0 | 135 | 0 | 460 | 2 | 95 | 0 |
| NM_001710 | 32 | 0 | 105 | 0 | 131 | 2 | 463 | 2 | 95 | 0 |
| NM_019111 | 0 | 14 | 0 | 5 | 0 | 6 | 2 | 0 | 2 | 0 |
| NM_001243965:NM_002124:XM_024452554:XM_024452553 | 0:1:0:0 | 1:0:0:0 | 0:1:0:0 | 0:0:2:0 | 0:1:0:0 | 0:0:1:1 | 0:0:0:0 | 0:0:2:0 | 0:0:0:0 | 0:0:1:0 |
| NM_001243965:NM_002124:XM_024452554:XM_024452553 | 0:1:0:0 | 1:0:0:0 | 0:1:0:0 | 0:0:2:0 | 0:1:0:0 | 0:0:1:1 | 0:0:0:0 | 0:0:2:0 | 0:0:0:0 | 0:0:1:0 |
| NM_001243965:NM_002124:XM_024452554:XM_024452553 | 0:0:0:0 | 1:1:0:0 | 0:0:0:0 | 0:1:2:0 | 0:0:0:0 | 0:1:1:1 | 0:0:0:0 | 0:0:2:0 | 0:0:0:0 | 0:0:1:0 |
| NM_001243965:NM_002124:XM_024452554:XM_024452553 | 0:1:0:0 | 1:0:0:0 | 0:1:0:0 | 0:0:2:0 | 0:1:0:0 | 0:0:1:1 | 0:0:0:0 | 0:0:2:0 | 0:0:0:0 | 0:0:1:0 |
| NM_002124:XM_024452554:XM_024452553 | 0:00:00 | 1:00:00 | 0:00:00 | 1:02:00 | 0:00:00 | 1:01:01 | 0:00:00 | 0:02:00 | 0:00:00 | 0:01:00 |
| NM_002124:XM_024452554:XM_024452553 | 0:00:00 | 1:00:00 | 0:00:00 | 1:02:00 | 0:00:00 | 1:01:01 | 0:00:00 | 0:01:00 | 0:00:00 | 0:00:00 |
| NM_002124:XM_024452554:XM_024452553 | 0:00:00 | 1:00:00 | 0:02:00 | 1:00:00 | 0:01:01 | 1:00:00 | 0:01:00 | 0:00:00 | 0:00:00 | 0:00:00 |

| | | | | | | | | | | |
|---|---|---|---|---|---|---|---|---|---|---|
| NM_002124:XM_024452554:XM_024452553 | 1:00:00 | 0:00:00 | 1:02:00 | 0:00:00 | 1:00:01 | 0:01:00 | 0:01:00 | 0:00:00 | 0:00:00 | 0:00:00 |
| NM_002124:XM_024452554:XM_024452553 | 0:00:00 | 1:00:00 | 0:02:00 | 1:00:00 | 0:00:01 | 1:00:00 | 0:01:00 | 0:00:00 | 0:00:00 | 0:00:00 |
| NM_002124:XM_024452554:XM_024452553 | 0:00:00 | 1:00:00 | 0:00:00 | 1:02:00 | 0:00:00 | 1:00:01 | 0:01:00 | 0:00:00 | 0:00:00 | 0:00:00 |
| NM_002124:XM_024452554:XM_024452553 | 1:00:00 | 0:00:00 | 1:00:00 | 0:02:00 | 1:00:00 | 0:00:01 | 0:01:00 | 0:00:00 | 0:00:00 | 0:00:00 |
| NM_002124:XM_024452554:XM_024452553 | 0:00:00 | 1:00:00 | 0:02:00 | 1:00:00 | 0:00:01 | 1:00:00 | 0:01:00 | 0:00:00 | 0:00:00 | 0:00:00 |
| NM_002124:XM_024452554:XM_024452553 | 1:00:00 | 0:00:00 | 1:00:00 | 0:02:00 | 1:00:00 | 0:00:01 | 0:00:00 | 0:01:00 | 0:00:00 | 0:00:00 |
| NM_002124:XM_024452554:XM_024452553 | 0:00:00 | 1:00:00 | 0:02:00 | 1:00:00 | 0:00:01 | 1:00:00 | 0:01:00 | 0:00:00 | 0:00:00 | 0:00:00 |
| NM_002124:XM_024452554:XM_024452553 | 0:00:00 | 1:00:00 | 0:00:00 | 1:02:00 | 0:00:00 | 1:00:01 | 0:00:00 | 0:01:00 | 0:00:00 | 0:00:00 |
| NM_002124:XM_024452554:XM_024452553 | 0:00:00 | 1:00:00 | 0:00:00 | 1:02:00 | 0:00:00 | 1:00:01 | 0:00:00 | 0:01:00 | 0:00:00 | 0:00:00 |
| NM_002124:XM_024452554:XM_024452553 | 0:00:00 | 1:00:00 | 0:00:00 | 1:02:00 | 0:00:00 | 1:00:01 | 0:00:00 | 0:01:00 | 0:00:00 | 0:00:00 |
| NM_002124:XM_024452554:XM_024452553 | 1:00:00 | 0:00:00 | 1:00:00 | 0:02:00 | 1:00:00 | 0:00:01 | 0:00:00 | 0:01:00 | 0:00:00 | 0:00:00 |
| NM_002124:XM_024452554:XM_024452553 | 0:00:00 | 1:00:00 | 0:02:00 | 1:00:00 | 0:00:01 | 1:00:00 | 0:01:00 | 0:00:00 | 0:00:00 | 0:00:00 |
| NM_002124:XM_024452554:XM_024452553 | 1:00:00 | 0:00:00 | 1:00:00 | 0:02:00 | 1:00:00 | 0:00:01 | 0:00:00 | 0:01:00 | 0:00:00 | 0:00:00 |
| NM_002124:XM_024452554:XM_024452553 | 1:00:00 | 0:00:00 | 1:00:00 | 0:02:00 | 1:00:00 | 0:00:01 | 0:00:00 | 0:01:00 | 0:00:00 | 0:00:00 |
| NM_002124:XM_024452554:XM_024452553 | 1:00:00 | 0:00:00 | 1:00:00 | 0:02:00 | 1:00:00 | 0:00:01 | 0:00:00 | 0:01:00 | 0:00:00 | 0:00:00 |
| NM_002124:XM_024452554:XM_024452553 | 1:00:00 | 0:00:00 | 1:00:00 | 0:02:00 | 1:00:00 | 0:00:01 | 0:00:00 | 0:01:00 | 0:00:00 | 0:00:00 |
| NM_002124:XM_024452554:XM_024452553 | 1:00:00 | 0:00:00 | 1:00:00 | 0:02:00 | 1:00:00 | 0:00:01 | 0:00:00 | 0:01:00 | 0:00:00 | 0:00:00 |
| NM_002124:XM_024452554:XM_024452553 | 1:00:00 | 0:00:00 | 1:00:00 | 0:02:00 | 1:00:00 | 0:00:01 | 0:00:00 | 0:01:00 | 0:00:00 | 0:00:00 |
| NM_002124:XM_024452554 | 0:00 | 1:00 | 0:00 | 1:02 | 0:00 | 1:00 | 0:00 | 0:01 | 0:00 | 0:00 |
| NM_002124:XM_024452554 | 1:00 | 0:00 | 1:00 | 0:02 | 1:00 | 0:00 | 0:00 | 0:01 | 0:00 | 0:00 |
| NM_002124:XM_024452554 | 1:00 | 0:00 | 1:00 | 0:02 | 1:00 | 0:00 | 0:00 | 0:01 | 0:00 | 0:00 |
| NM_002124:XM_024452554 | 1:00 | 0:00 | 1:00 | 0:02 | 1:00 | 0:00 | 0:00 | 0:01 | 0:00 | 0:00 |
| NM_002122:XM_006715079 | 2:00 | 1:00 | 0:00 | 0:00 | 0:00 | 1:00 | 0:01 | 0:00 | 0:00 | 0:00 |
| NM_002122:XM_006715079 | 2:00 | 1:00 | 0:00 | 0:00 | 0:00 | 1:00 | 0:01 | 0:00 | 0:00 | 0:00 |
| NM_002122:XM_006715079 | 2:00 | 1:00 | 0:00 | 0:00 | 0:00 | 1:00 | 0:01 | 0:00 | 0:00 | 0:00 |
| NM_002122:XM_006715079 | 1:00 | 2:00 | 0:00 | 0:00 | 1:00 | 0:00 | 0:01 | 0:00 | 0:00 | 0:00 |
| NM_002122:XM_006715079 | 2:00 | 1:00 | 0:00 | 0:00 | 0:00 | 1:00 | 0:01 | 0:00 | 0:00 | 0:00 |
| NM_002122:XM_006715079 | 2:00 | 1:00 | 0:00 | 0:00 | 0:00 | 1:00 | 0:01 | 0:00 | 0:00 | 0:00 |
| NM_002122:XM_006715079 | 2:00 | 1:00 | 0:00 | 0:00 | 0:00 | 1:00 | 0:01 | 0:00 | 0:00 | 0:00 |
| NM_002122:XM_006715079 | 2:00 | 1:00 | 0:00 | 0:00 | 0:00 | 1:00 | 0:01 | 0:00 | 0:00 | 0:00 |
| NM_002122:XM_006715079 | 2:00 | 1:00 | 0:00 | 0:00 | 0:00 | 1:00 | 0:01 | 0:00 | 0:00 | 0:00 |
| NM_002122:XM_006715079 | 2:00 | 1:00 | 0:00 | 0:00 | 0:00 | 1:00 | 0:01 | 0:00 | 0:00 | 0:00 |
| NM_002122:XM_006715079 | 2:00 | 1:00 | 0:00 | 0:00 | 0:00 | 1:00 | 0:01 | 0:00 | 0:00 | 0:00 |
| NM_002122:XM_006715079 | 2:00 | 1:00 | 0:00 | 0:00 | 0:00 | 1:00 | 0:01 | 0:00 | 0:00 | 0:00 |
| NM_002122:XM_006715079 | 2:00 | 1:00 | 0:00 | 0:00 | 0:00 | 1:00 | 0:01 | 0:00 | 0:00 | 0:00 |
| NM_002122:XM_006715079 | 2:00 | 1:00 | 0:00 | 0:00 | 0:00 | 1:00 | 0:01 | 0:00 | 0:00 | 0:00 |
| NM_002122:XM_006715079 | 2:00 | 1:00 | 0:00 | 0:00 | 0:00 | 1:00 | 0:01 | 0:00 | 0:00 | 0:00 |
| NM_002122:XM_006715079 | 2:00 | 1:00 | 0:00 | 0:00 | 0:00 | 1:00 | 0:00 | 0:01 | 0:00 | 0:00 |
| NM_002122:XM_006715079 | 2:00 | 1:00 | 0:00 | 0:00 | 0:00 | 1:00 | 0:01 | 0:00 | 0:00 | 0:00 |
| NM_001128592 | 0 | 0 | 0 | 0 | 0 | 0 | 0 | 2 | 0 | 0 |
| NM_001128592 | 0 | 0 | 0 | 0 | 0 | 0 | 0 | 2 | 0 | 0 |
| NM_002122 | 0 | 3 | 0 | 0 | 0 | 1 | 0 | 0 | 0 | 0 |
| NM_001128592 | 0 | 0 | 0 | 0 | 0 | 0 | 0 | 2 | 0 | 0 |
| NM_001128592 | 0 | 0 | 0 | 0 | 0 | 0 | 0 | 2 | 0 | 0 |
| NM_002123:NM_001243962 | 0:01 | 1:00 | 0:00 | 2:00 | 0:00 | 2:00 | 5:00 | 0:00 | 1:00 | 0:00 |
| NM_001243962:NM_002123 | 0:01 | 1:00 | 0:02 | 0:00 | 0:02 | 0:00 | 0:05 | 0:00 | 0:01 | 0:00 |
| NM_001243962:NM_002123 | 0:01 | 1:00 | 0:02 | 0:00 | 0:02 | 0:00 | 0:05 | 0:00 | 0:01 | 0:00 |
| NM_001243962:NM_002123 | 0:01 | 1:00 | 0:02 | 0:00 | 0:02 | 1:00 | 0:05 | 0:00 | 0:01 | 0:00 |
| NM_001243962:NM_002123 | 0:01 | 1:00 | 0:02 | 0:00 | 0:02 | 0:00 | 0:03 | 0:00 | 0:01 | 0:00 |
| NM_002123:NM_001243962 | 0:01 | 1:00 | 0:00 | 2:00 | 0:00 | 2:00 | 3:00 | 0:00 | 1:00 | 0:00 |
| NM_002123:NM_001243962 | 0:01 | 1:00 | 0:00 | 2:00 | 0:00 | 2:00 | 3:00 | 0:00 | 1:00 | 0:00 |
| NM_001243962:NM_002123 | 0:01 | 1:00 | 0:02 | 0:00 | 0:02 | 0:00 | 0:03 | 0:00 | 0:01 | 0:00 |
| NM_001243962:NM_002123 | 0:01 | 1:00 | 0:02 | 0:00 | 0:02 | 0:00 | 0:03 | 0:00 | 0:01 | 0:00 |
| NM_001243962:NM_002123 | 0:01 | 1:00 | 0:02 | 0:00 | 0:02 | 0:00 | 0:03 | 0:00 | 0:01 | 0:00 |
| NM_002123:NM_001243962 | 0:01 | 1:00 | 0:00 | 2:00 | 0:00 | 2:00 | 3:00 | 0:00 | 1:00 | 0:00 |
| NM_001243962:NM_002123 | 0:01 | 1:00 | 0:02 | 0:00 | 0:02 | 0:00 | 0:03 | 0:00 | 0:01 | 0:00 |
| NM_001243962:NM_002123 | 0:01 | 1:00 | 0:02 | 0:00 | 0:02 | 0:00 | 0:03 | 0:00 | 0:01 | 0:00 |
| NM_001243962:NM_002123 | 0:01 | 1:00 | 0:02 | 0:00 | 0:02 | 0:00 | 0:03 | 0:00 | 0:01 | 0:00 |
| NM_001243962:NM_002123 | 0:01 | 1:00 | 0:02 | 0:00 | 0:02 | 0:00 | 0:03 | 0:00 | 0:01 | 0:00 |
| NM_002123:NM_001243962 | 0:01 | 1:00 | 0:00 | 2:00 | 0:00 | 2:00 | 3:00 | 0:00 | 1:00 | 0:00 |
| NM_002123:NM_001243962 | 0:01 | 1:00 | 0:00 | 2:00 | 0:00 | 2:00 | 3:00 | 0:00 | 1:00 | 0:00 |
| NM_002123:NM_001243962 | 0:01 | 1:00 | 0:00 | 2:00 | 0:00 | 2:00 | 3:00 | 0:00 | 1:00 | 0:00 |
| NM_002123:NM_001243962 | 0:01 | 1:00 | 0:00 | 2:00 | 0:00 | 2:00 | 3:00 | 0:00 | 1:00 | 0:00 |
| NM_002123:NM_001243962 | 0:01 | 1:00 | 0:00 | 2:00 | 0:00 | 2:00 | 3:00 | 0:00 | 1:00 | 0:00 |
| NM_001243962:NM_002123 | 0:01 | 1:00 | 0:02 | 0:00 | 0:02 | 0:00 | 0:03 | 0:00 | 0:01 | 0:00 |
| NM_002123:NM_001243962 | 0:01 | 1:00 | 0:00 | 2:00 | 0:00 | 2:00 | 3:00 | 0:00 | 1:00 | 0:00 |
| NM_002123:NM_001243962 | 0:01 | 1:00 | 0:00 | 2:00 | 0:00 | 2:00 | 3:00 | 0:00 | 1:00 | 0:00 |
| NM_001243962:NM_002123 | 0:01 | 1:00 | 0:02 | 0:00 | 0:02 | 0:00 | 0:03 | 0:00 | 0:01 | 0:00 |
| NM_002123:NM_001243962 | 0:01 | 1:00 | 0:00 | 2:00 | 0:00 | 2:00 | 3:00 | 0:00 | 1:00 | 0:00 |
| NM_002123:NM_001243962 | 0:01 | 1:00 | 0:00 | 2:00 | 0:00 | 2:00 | 0:00 | 3:00 | 0:00 | 1:00 |
| NM_002123:NM_001243962 | 0:01 | 1:00 | 0:00 | 2:00 | 0:00 | 2:00 | 0:00 | 3:00 | 0:00 | 1:00 |
| NM_002123:NM_001243962 | 0:00 | 1:01 | 0:00 | 2:00 | 0:00 | 2:00 | 3:00 | 0:00 | 1:00 | 0:00 |
| NM_002123:NM_001243962 | 0:00 | 1:01 | 0:00 | 2:00 | 0:00 | 2:00 | 3:00 | 0:00 | 1:00 | 0:00 |
| NM_002123:NM_001243962 | 0:01 | 1:00 | 0:00 | 2:00 | 0:00 | 2:00 | 3:00 | 0:00 | 1:00 | 0:00 |
| NM_002123:NM_001243962 | 0:00 | 1:01 | 0:00 | 2:00 | 0:00 | 2:00 | 0:00 | 3:00 | 0:00 | 1:00 |
| NM_001243962:NM_002123 | 0:01 | 1:00 | 0:02 | 0:00 | 0:02 | 0:00 | 0:03 | 0:00 | 0:01 | 0:00 |
| NM_002123:NM_001243962 | 0:00 | 1:01 | 0:00 | 2:00 | 0:00 | 2:00 | 3:00 | 0:00 | 1:00 | 0:00 |
| NM_001243962:NM_002123 | 0:01 | 1:00 | 0:02 | 0:00 | 0:02 | 0:00 | 0:03 | 0:00 | 0:01 | 0:00 |

| | | | | | | | | | | |
|---|---|---|---|---|---|---|---|---|---|---|
| NM_002123:NM_001243962 | 0:01 | 1:00 | 0:00 | 2:00 | 0:00 | 2:00 | 3:00 | 0:00 | 1:00 | 0:00 |
| NM_001243962:NM_002123 | 0:01 | 1:00 | 0:02 | 0:00 | 0:02 | 0:00 | 0:03 | 0:00 | 0:01 | 0:00 |
| NM_001243962:NM_002123 | 0:01 | 1:00 | 0:02 | 0:00 | 0:02 | 0:00 | 0:03 | 0:00 | 0:01 | 0:00 |
| NM_002123:NM_001243962 | 0:01 | 1:00 | 0:00 | 2:00 | 0:00 | 2:00 | 3:00 | 0:00 | 1:00 | 0:00 |
| NM_001292022 | 0 | 1 | 0 | 1 | 0 | 0 | 1 | 2 | 0 | 0 |
| NM_002800 | 0 | 3 | 0 | 0 | 2 | 3 | 4 | 0 | 3 | 0 |
| NM_001291986:NM_001199456:NM_005104:NM_001113182:XR_001743557:XR_241911 | 2:2:9:1:16:2 | 0:0:0:0:0:0 | 7:0:10:1:70:3 | 0:0:0:0:1:0 | 9:0:9:0:19:0 | 0:0:0:0:0:0 | 61:3:26:0:381:15 | 0:0:0:0:0:1 | 9:0:9:1:28:2 | 0:0:0:0:1:0 |
| NM_001291986:NM_001199456:NM_005104:NM_001113182:XR_001743557:XR_241911:NM_001199455 | 2:2:8:1:16:1:0 | 0:0:0:0:0:0:0 | 7:0:13:1:79:4:0 | 0:0:0:0:0:0:1 | 9:0:9:0:20:0:0 | 0:0:0:0:0:0:0 | 62:3:40:0:416:13:0 | 0:0:0:0:0:0:0 | 9:0:10:1:29:1:0 | 0:0:0:0:0:0:0 |
| NM_001291986:NM_001199456:NM_005104:NM_001113182:XR_001743557:XR_241911:NM_001199455 | 2:2:8:1:16:1:0 | 0:0:0:0:0:0:0 | 7:0:13:1:79:4:0 | 0:0:0:0:0:0:1 | 9:0:8:0:20:0:0 | 0:0:1:0:0:0:0 | 62:3:40:0:416:12:0 | 0:0:0:0:0:0:0 | 9:0:10:1:29:1:0 | 0:0:0:0:0:0:0 |
| NM_001291986:NM_001199456:NM_005104:NM_001113182:XR_001743557:XR_241911 | 2:2:10:1:17:1 | 0:0:0:0:0:0 | 7:0:13:1:81:6 | 0:0:0:0:0:0 | 9:0:9:0:20:1 | 0:0:0:0:0:0 | 61:3:49:0:425:10 | 0:0:0:0:0:0 | 9:0:10:1:29:0 | 0:0:0:0:0:0 |
| NM_033554 | 5 | 0 | 2 | 0 | 4 | 0 | 1 | 0 | 0 | 0 |
| NM_033554 | 5 | 0 | 2 | 0 | 4 | 0 | 0 | 0 | 0 | 0 |
| NM_033554 | 4 | 0 | 2 | 0 | 4 | 0 | 0 | 0 | 0 | 0 |
| NM_033554 | 5 | 0 | 2 | 0 | 4 | 0 | 0 | 0 | 0 | 0 |
| NM_033554 | 5 | 0 | 2 | 0 | 4 | 0 | 0 | 0 | 0 | 0 |
| NM_033554 | 5 | 0 | 2 | 0 | 4 | 0 | 0 | 0 | 0 | 0 |
| NM_033554 | 5 | 0 | 2 | 0 | 4 | 0 | 0 | 0 | 0 | 0 |
| NM_033554 | 5 | 0 | 2 | 0 | 4 | 0 | 0 | 0 | 0 | 0 |
| NM_002121 | 4 | 0 | 2 | 0 | 0 | 0 | 2 | 0 | 0 | 0 |
| NM_002121 | 4 | 0 | 2 | 0 | 0 | 0 | 2 | 0 | 0 | 0 |
| NM_002121 | 4 | 0 | 2 | 0 | 0 | 0 | 2 | 0 | 0 | 0 |
| NM_002121 | 4 | 0 | 2 | 0 | 0 | 0 | 2 | 0 | 0 | 0 |
| NM_002121 | 4 | 0 | 2 | 0 | 0 | 0 | 2 | 0 | 0 | 0 |
| NM_002121 | 3 | 1 | 2 | 0 | 0 | 0 | 0 | 2 | 0 | 0 |
| NM_002121 | 0 | 4 | 0 | 2 | 0 | 0 | 0 | 2 | 0 | 0 |
| NM_002121 | 1 | 3 | 0 | 2 | 0 | 0 | 0 | 2 | 0 | 0 |
| NM_002121 | 1 | 3 | 0 | 2 | 0 | 0 | 2 | 0 | 0 | 0 |
| NM_002121 | 4 | 0 | 2 | 0 | 0 | 0 | 2 | 0 | 0 | 0 |
| NM_002121 | 3 | 1 | 2 | 0 | 0 | 0 | 2 | 0 | 0 | 0 |
| NM_002121 | 1 | 3 | 0 | 2 | 0 | 0 | 2 | 0 | 0 | 0 |
| NM_002121 | 4 | 0 | 2 | 0 | 0 | 0 | 2 | 0 | 0 | 0 |
| NM_002121 | 4 | 0 | 2 | 0 | 0 | 0 | 2 | 0 | 0 | 0 |
| NM_002121 | 3 | 1 | 2 | 0 | 0 | 0 | 0 | 2 | 0 | 0 |
| NM_002121 | 4 | 0 | 2 | 0 | 0 | 0 | 0 | 2 | 0 | 0 |
| NM_002121 | 4 | 0 | 2 | 0 | 0 | 0 | 0 | 2 | 0 | 0 |
| NM_002121 | 4 | 0 | 2 | 0 | 0 | 0 | 0 | 2 | 0 | 0 |
| NM_002121 | 2 | 1 | 2 | 0 | 0 | 0 | 0 | 2 | 0 | 0 |
| NM_002121 | 3 | 1 | 3 | 0 | 0 | 0 | 0 | 2 | 0 | 0 |
| NM_002121 | 3 | 1 | 3 | 0 | 0 | 0 | 0 | 2 | 0 | 0 |
| NM_001008703:NM_178508 | 0:00 | 0:00 | 0:00 | 1:01 | 0:00 | 0:00 | 0:00 | 2:00 | 0:00 | 1:00 |
| NM_001010903:XM_011514598 | 0:00 | 0:00 | 0:00 | 0:00 | 2:00 | 0:00 | 0:00 | 4:02 | 0:00 | 1:01 |
| NM_183373 | 0 | 0 | 0 | 0 | 0 | 0 | 0 | 2 | 0 | 0 |
| NM_001760:NM_001136017:XM_011514972:NM_001287427:XM_011514971:NM_001136126:NM_001136125 | 1:0:0:0:0:0:0 | 3:1:0:0:0:0:0 | 1:1:0:1:1:0:0 | 7:0:2:0:0:0:0 | 17:1:2:0:0:0:1 | 20:1:1:0:0:4:0 | 0:0:0:0:0:0:0 | 43:12:4:1:1:9:1 | 0:0:0:0:0:0:0 | 24:0:0:0:0:0:0 |
| NM_003192 | 0 | 0 | 0 | 2 | 0 | 1 | 0 | 0 | 0 | 2 |
| NM_006586:NM_001318845:NR_134888:NR_134882 | 0:0:0:0 | 1:0:0:0 | 0:0:0:0 | 1:0:0:0 | 0:0:0:0 | 0:1:1:0 | 0:0:0:0 | 2:0:1:0 | 0:0:0:0 | 2:0:0:0 |
| XR_001743466 | 0 | 0 | 0 | 0 | 0 | 0 | 0 | 2 | 0 | 0 |
| NM_003376:NM_001171628:NM_001287044:NM_001317010 | 14:0:0:0 | 0:0:0:0 | 23:1:0:0 | 1:0:0:0 | 24:0:0:0 | 0:0:0:0 | 46:2:5:0 | 0:2:0:0 | 5:1:1:0 | 0:0:0:0 |
| XM_011514341:NM_001304466:XM_005248878:NM_001304465:NM_001078175:XM_005248880:XM_005248876:XM_024446348:XM_005248879:NM_001078177 | 3:18:1:1:4:1:1:1:0:0 | 0:0:0:0:0:0:0:0:0:0 | 0:1:0:1:1:0:0:0:0:0 | 0:0:0:0:0:0:0:0:0:0 | 2:11:1:0:1:0:0:0:0:0 | 0:0:0:0:0:0:0:0:0:0 | 24:70:3:4:31:6:4:13:1:0 | 0:0:0:0:1:0:0:0:0:0 | 3:19:2:1:14:2:1:2:1:0 | 0:0:0:0:0:0:0:0:0:0 |
| XM_011514341:NM_001304466:NM_001304465:NM_001078175:XM_005248876:XM_005248878:XM_024446348:XM_005248880:XM_005248879:NM_001078177 | 2:18:1:4:1:0:0:0:0:0 | 0:0:0:0:0:0:0:0:0:0 | 0:1:1:1:0:0:0:0:0:0 | 0:0:0:0:0:0:0:0:0:0 | 1:10:0:1:0:1:0:0:0:0 | 0:0:0:0:0:0:0:0:0:0 | 63:71:4:31:3:3:26:6:1:0 | 0:0:0:0:0:0:0:0:0:0 | 2:19:1:14:1:2:2:2:1:0 | 0:0:0:0:0:0:0:0:0:0 |
| XM_011514341:NM_001304466:NM_001304465:NM_001078175:XM_005248876:XM_024446348:XM_005248878:XM_005248880:XM_005248879:NM_001078177 | 2:18:1:4:1:0:0:0:0:0 | 0:0:0:0:0:0:0:0:0:0 | 0:1:1:1:0:1:0:0:0:0 | 0:0:0:0:0:0:0:0:0:0 | 1:10:0:1:0:0:1:0:0:0 | 0:0:0:0:0:0:0:0:0:0 | 96:67:4:30:2:40:3:3:1:0 | 0:1:0:0:0:0:0:0:0:0 | 3:19:1:14:1:5:2:2:1:0 | 0:0:0:0:0:0:0:0:0:0 |
| NM_007355 | 3 | 0 | 0 | 0 | 1 | 0 | 7 | 0 | 6 | 0 |
| NM_014936 | 0 | 1 | 0 | 5 | 0 | 7 | 0 | 8 | 0 | 4 |
| XM_011514628:NM_005588:XM_011514629 | 0:00:00 | 0:00:00 | 0:00:00 | 2:00:00 | 1:01:00 | 1:00:00 | 0:00:00 | 0:00:01 | 0:00:00 | 0:00:00 |
| NM_014464:XM_006715062:XR_001743344:XM_017010746 | 0:0:0:0 | 0:0:0:0 | 0:0:0:0 | 1:0:0:0 | 0:0:0:0 | 0:1:1:0 | 0:0:0:0 | 0:0:0:1 | 0:0:0:0 | 2:0:0:0 |
| NM_018368 | 0 | 1 | 0 | 0 | 0 | 3 | 0 | 0 | 0 | 0 |
| NM_021940:XM_017011170 | 0:00 | 1:01 | 0:00 | 0:00 | 0:00 | 0:00 | 0:00 | 0:00 | 1:00 | 0:00 |
| NM_031480:NM_001348194:XM_011514933 | 62:03:02 | 0:00:00 | 49:02:01 | 0:00:00 | 44:07:01 | 0:00:00 | 305:38:10 | 2:00:00 | 67:10:03 | 0:00:00 |
| NM_031480:NM_001348194:XM_011514933 | 62:03:02 | 0:00:00 | 49:02:01 | 0:00:00 | 44:07:01 | 0:00:00 | 305:39:13 | 0:00:01 | 67:10:03 | 0:00:00 |
| NM_031480:NM_001348194:XM_011514933 | 62:03:02 | 0:00:00 | 49:02:02 | 0:00:00 | 43:07:01 | 1:00:00 | 305:41:18 | 2:00:00 | 67:10:04 | 0:00:00 |
| NM_031480:NM_001348194:XM_011514933 | 61:03:02 | 0:00:00 | 49:02:11 | 0:00:00 | 45:07:01 | 0:00:00 | 310:41:58 | 2:00:01 | 66:09:08 | 1:00:00 |
| NM_031480:NM_001348194:XM_011514933 | 61:03:02 | 0:00:00 | 48:02:14 | 0:00:00 | 45:07:03 | 0:00:00 | 305:40:82 | 0:00:00 | 67:09:10 | 0:00:00 |
| NM_031480:NM_001348194:XM_011514933 | 61:03:02 | 0:00:00 | 48:02:16 | 0:00:00 | 45:07:03 | 0:00:00 | 305:40:82 | 1:00:01 | 68:09:10 | 0:00:00 |
| NM_031480:NM_001348194:XM_011514933 | 61:03:06 | 0:00:00 | 48:02:35 | 0:00:01 | 45:07:08 | 0:00:00 | 302:40:105 | 4:00:00 | 68:09:14 | 0:00:00 |
| NM_001100409:NM_015571:XM_017010719:XR_241847:XM_017010725:XM_006715457:XM_017010721:XR_001743330:XM_017010724:XM_024446400 | 1:2:1:0:0:0:0:0:0:0 | 0:0:0:0:0:0:0:0:0:0 | 0:0:0:1:0:0:0:0:0:0 | 0:0:0:0:0:0:0:0:0:0 | 1:3:0:0:1:0:0:0:0:0 | 0:0:0:0:0:0:0:0:0:0 | 1:0:0:0:0:0:0:1:0:0 | 0:0:1:0:0:1:1:0:0:0 | 0:0:3:0:0:0:0:0:0:0 | 0:0:0:0:0:0:0:0:0:0 |

| | | | | | | | | | | |
|---|---|---|---|---|---|---|---|---|---|---|
| NM_004415:NM_001008844 | 1:00 | 0:00 | 5:00 | 0:00 | 4:01 | 0:00 | 14:00 | 0:00 | 6:00 | 0:00 |
| NM_001100409:XM_017010719:XM_024446400:XR_241847:NM_015571:XM_017010724 | 1:1:2:0:0:0 | 0:0:0:0:0:0 | 0:0:8:1:0:0 | 0:0:0:0:0:0 | 1:0:3:0:1:0 | 0:0:0:0:0:0 | 0:0:8:3:0:0 | 0:0:5:1:0:0 | 0:0:2:1:0:0 | 0:0:4:1:0:0 |
| NM_004415 | 1 | 0 | 3 | 0 | 1 | 0 | 5 | 3 | 0 | 0 |
| NM_001300899:XM_024446447:XM_005248722 | 0:00:00 | 0:00:00 | 0:00:00 | 0:00:00 | 0:00:00 | 1:00:00 | 1:02:00 | 0:00:00 | 0:02:00 | 0:00:00 |
| XM_024446447:NM_001300899:XM_005248722 | 0:00:00 | 0:00:00 | 1:00:00 | 0:00:00 | 0:00:00 | 0:00:00 | 1:01:00 | 1:00:00 | 2:00:00 | 0:00:00 |
| XM_024446447:XM_017010899:XM_005248722 | 0:00:00 | 0:00:00 | 1:00:00 | 0:00:00 | 0:00:00 | 0:00:00 | 1:01:00 | 0:00:00 | 1:00:00 | 1:00:00 |
| NM_002526 | 0 | 0 | 0 | 0 | 0 | 0 | 0 | 0 | 0 | 0 |
| NM_005190:XR_001743714:XM_017011436 | 115:02:00 | 0:00:00 | 89:02:00 | 0:00:00 | 266:04:00 | 1:00:00 | 715:14:00 | 6:00:00 | 250:02:00 | 1:00:00 |
| NM_005190:XR_001743714:XM_017011436 | 115:02:00 | 0:00:00 | 89:02:00 | 0:00:00 | 267:04:00 | 0:00:00 | 718:12:00 | 0:00:00 | 251:02:00 | 0:00:00 |
| NM_005190:XR_001743714:XM_017011436 | 115:02:00 | 0:00:00 | 89:02:00 | 0:00:00 | 266:04:00 | 1:00:00 | 715:12:00 | 3:00:00 | 251:02:00 | 0:00:00 |
| NM_004722:XM_024446996:XM_024446995 | 0:00:00 | 0:00:00 | 0:00:00 | 0:00:00 | 0:00:00 | 0:00:00 | 0:00:00 | 0:00:00 | 1:00:00 | 0:00:00 |
| NM_181500:NM_001202546:NM_001913:XM_017011760:XM_011515825:XM_024446668:XM_006715855:NM_181552:XM_005250154:NM_001202545:NM_001202543:NM_001202544 | 42:2:104:40:5:4:7:1:0:0:0:0 | 0:0:0:0:0:0:0:0:0:0:0 | 25:0:22:85:5:2:2:1:0:0:0:0 | 1:0:0:0:0:0:0:0:0:0:0 | 21:0:58:30:7:0:1:0:2:1:1:0 | 0:0:0:0:0:0:0:0:0:0:0 | 121:1:148:325:40:9:29:0:1:2:3:4 | 0:0:2:0:1:0:0:0:0:0:0 | 31:0:82:36:8:0:7:0:2:0:0:0 | 0:0:0:0:0:0:0:0:0:0:0 |
| NM_181500:NM_001202546:NM_001913:XM_017011760:XM_011515825:XM_024446668:XM_006715855:NM_181552:XM_005250154:NM_001202545:NM_001202543:NM_001202544 | 42:2:104:40:5:4:7:1:1:0:0:0 | 0:0:0:0:0:0:0:0:0:0:0 | 25:0:22:85:5:5:2:1:0:0:0:0 | 1:0:0:0:0:0:0:0:0:0:0 | 21:0:59:30:7:0:1:0:2:1:1:0 | 0:0:0:0:0:0:0:0:0:0:0 | 120:1:151:323:41:10:29:0:2:2:3:4 | 1:0:0:2:0:0:0:0:0:0:0 | 31:0:81:36:8:0:8:0:2:0:0:0 | 0:0:1:0:0:0:0:0:0:0:0 |
| NM_181500:NM_001202546:NM_001913:XM_017011760:XM_011515825:XM_024446668:XM_006715855:NM_181552:XM_005250154:NM_001202543:NM_001202544:NM_001202545 | 40:2:105:40:5:4:6:1:2:0:0:0 | 0:0:0:0:0:0:0:0:0:0:0 | 21:0:22:83:5:5:3:1:1:0:0:0 | 2:0:0:0:0:0:0:0:0:0:0 | 21:0:60:30:7:0:1:0:4:1:0:0 | 0:0:0:0:0:0:0:0:0:0:0 | 117:3:151:320:42:15:27:0:9:3:4:2 | 0:0:0:3:0:0:0:0:0:0:0 | 31:1:82:36:8:1:8:0:2:0:0:0 | 1:0:0:0:0:0:0:0:0:0:0 |
| NM_001913:NM_181500:XM_006715855:XM_005250154:NM_001202546:XM_017011760:NM_001202545:NM_001202544:XM_024446668:XM_011515825 | 51:16:4:3:2:0:0:0:0:0 | 53:13:8:2:0:0:0:0:0 | 9:7:12:4:0:0:0:0:0:0 | 9:6:12:5:0:1:0:0:0:0 | 24:7:1:5:0:0:0:0:0:0 | 34:5:0:3:0:0:1:0:0:0 | 78:59:77:30:0:0:1:0:0:0 | 71:44:85:23:0:0:0:1:1:0 | 37:16:5:1:0:0:0:0:0:0 | 42:10:6:3:0:0:0:0:0:0 |
| NM_001202546:NM_001913:NM_181500:XM_006715855:XM_005250154:NM_001202545:XM_011515825:NM_001202544:XM_017011760:XM_024446668 | 2:102:29:14:5:0:0:0:0:0 | 0:0:0:0:0:0:0:0:0:0 | 0:16:20:34:9:0:0:0:0:0 | 0:0:0:0:0:0:0:0:0:0 | 0:57:14:2:7:1:0:0:0:0 | 0:0:0:0:0:0:0:0:0:0 | 0:140:158:285:47:1:1:1:1:1 | 0:0:0:1:0:0:0:0:0:0 | 0:79:29:11:4:0:0:0:0:0 | 0:0:0:0:0:0:0:0:0:0 |
| NM_001202546:NM_001913:NM_181500:XM_006715855:XM_005250154:NM_001202545:XM_011515825:NM_001202544:XM_017011760:XM_024446668 | 2:101:29:14:5:0:0:0:0:0 | 0:0:0:0:0:0:0:0:0:0 | 0:16:20:34:9:0:0:0:0:0 | 0:0:0:0:0:0:0:0:0:0 | 0:57:14:2:7:1:0:0:0:0 | 0:0:0:0:0:0:0:0:0:0 | 0:65:78:149:22:0:1:1:1:1 | 0:74:80:140:25:1:0:0:0:0 | 0:41:13:6:3:0:0:0:0:0 | 0:38:16:5:1:0:0:0:0:0 |
| XM_006715855:NM_181500:NM_001202546:NM_001913:XM_005250154:XM_011515825:NM_001202545:XM_017011760:NM_001202544:XM_024446668 | 14:33:2:96:5:0:0:0:0:0 | 3:1:0:0:0:0:0:0:0:0 | 43:30:0:15:9:1:0:0:0:0 | 0:0:0:0:0:0:0:0:0:0 | 7:13:0:55:6:0:1:0:0:0 | 0:1:0:0:0:0:0:0:0:0 | 427:192:0:119:35:6:1:1:1:1 | 5:5:0:0:0:0:0:0:0:0 | 17:33:0:75:4:0:0:1:0:0 | 0:0:0:0:0:0:0:0:0:0 |
| NM_001202546:NM_001913:NM_181500:XM_006715855:XM_005250154:XM_011515825:NM_001202545:XM_017011760:NM_001202544:XM_024446668 | 2:96:34:17:5:0:0:0:0:0 | 0:0:0:0:0:0:0:0:0:0 | 0:13:28:43:9:1:0:0:0:0 | 0:0:2:0:0:0:0:0:0:0 | 0:55:13:7:6:0:1:0:0:0 | 0:0:1:0:0:0:0:0:0:0 | 0:119:193:423:35:6:1:1:1:1 | 0:0:5:6:0:0:0:0:0:0 | 0:74:33:17:4:0:0:1:0:0 | 0:0:0:0:0:0:0:0:0:0 |
| NM_032959 | 0 | 0 | 0 | 0 | 0 | 0 | 0 | 0 | 0 | 0 |
| XR_001744836:XM_017012435:NM_182931:XM_011516400:NM_018682:XM_024446837 | 95:3:23:4:1:0 | 1:0:0:0:0:0 | 408:6:129:8:1:9 | 1:0:0:0:0:0 | 196:3:65:7:1:3 | 0:0:2:0:0:0 | 798:13:248:13:4:9 | 3:0:2:0:0:0 | 76:3:15:0:0:1 | 0:0:0:0:0:0 |
| XM_017012435:XR_001744836:NM_182931:XM_011516400:NM_018682:XM_024446837 | 3:96:23:4:1:0 | 0:0:0:0:0:0 | 6:421:129:8:1:9 | 0:1:0:0:0:0 | 3:197:66:7:1:3 | 0:0:1:0:0:0 | 13:809:250:13:4:9 | 0:4:0:0:0:0 | 3:76:15:0:0:1 | 0:0:0:0:0:0 |
| XM_017012435:XR_001744836:NM_182931:XM_011516400:NM_018682:XM_024446837 | 3:95:23:4:1:0 | 0:0:0:0:0:0 | 6:419:126:8:1:9 | 0:1:0:0:0:0 | 3:196:66:7:1:3 | 0:1:1:0:0:0 | 14:811:247:13:4:8 | 0:0:0:0:0:0 | 3:76:15:0:0:1 | 0:0:0:0:0:0 |
| XM_017012435:XR_001744836:NM_182931:XM_011516400:NM_018682:XM_024446837 | 1:96:23:4:1:0 | 0:0:0:0:0:0 | 5:431:128:8:1:8 | 0:1:0:0:0:0 | 2:199:64:7:1:3 | 0:0:1:0:0:0 | 14:825:248:13:4:9 | 0:0:0:0:0:0 | 3:78:15:0:0:1 | 0:0:0:0:0:0 |
| XM_017012435:XR_001744836:NM_182931:XM_011516400:NM_018682:XM_024446837 | 1:96:23:4:1:0 | 0:0:0:0:0:0 | 5:436:128:8:1:8 | 0:0:0:0:0:0 | 2:198:65:7:1:3 | 0:0:0:0:0:0 | 14:828:246:13:4:9 | 0:1:1:0:0:0 | 3:79:15:0:0:1 | 0:0:0:0:0:0 |
| XM_017012435:XR_001744836:NM_182931:XM_011516400:NM_018682:XM_024446837 | 2:97:23:4:1:0 | 0:0:0:0:0:0 | 19:449:129:8:1:9 | 0:0:0:0:0:0 | 2:198:64:7:1:3 | 0:0:0:0:0:0 | 17:837:246:13:4:9 | 0:1:1:0:0:0 | 4:80:15:0:0:1 | 0:0:0:0:0:0 |
| XM_017012435:XR_001744836:NM_182931:XM_011516400:NM_018682:XM_024446837 | 2:95:23:4:1:0 | 0:0:0:0:0:0 | 19:449:126:8:1:8 | 0:0:0:0:0:0 | 2:199:64:7:1:3 | 0:0:0:0:0:0 | 17:843:247:12:4:9 | 0:1:0:1:0:0 | 4:80:15:0:0:1 | 0:0:0:0:0:0 |
| XM_017012435:XR_001744836:NM_182931:XM_011516400:NM_018682:XM_024446837 | 2:72:20:3:0:0 | 0:0:0:0:0:0 | 7:235:59:6:1:5 | 0:0:0:0:0:0 | 1:140:42:5:0:3 | 0:0:0:0:0:0 | 14:609:163:10:3:7 | 0:0:0:0:0:0 | 4:60:14:0:0:1 | 0:1:0:0:0:0 |
| NM_182931:XM_017012435:XR_001744836:XM_011516400:NM_018682:XM_024446837 | 19:2:68:3:0:0 | 1:0:0:0:0:0 | 59:7:227:6:1:5 | 0:0:2:0:0:0 | 41:1:138:5:0:3 | 0:0:1:0:0:0 | 161:14:600:10:3:7 | 2:0:1:0:0:0 | 14:4:57:0:0:1 | 0:0:3:0:0:0 |
| XM_017012435:XR_001744836:NM_182931:XM_011516400:NM_018682:XM_024446837 | 2:58:18:2:0:0 | 0:0:0:0:0:0 | 6:220:58:6:1:4 | 0:0:0:0:0:0 | 1:125:34:4:0:2 | 0:1:0:0:0:0 | 11:576:150:10:3:7 | 0:1:0:0:0:0 | 3:57:13:0:0:1 | 0:0:0:0:0:0 |
| XM_017012435:XR_001744836:NM_182931:XM_011516400:NM_018682:XM_024446837 | 2:59:18:2:0:0 | 0:0:0:0:0:0 | 6:202:51:3:1:4 | 0:0:0:0:0:0 | 1:124:34:4:0:2 | 0:0:0:0:0:0 | 10:486:128:9:3:5 | 0:0:0:0:0:0 | 3:55:13:0:0:1 | 0:0:0:0:0:0 |
| XM_017012435:XR_001744836:NM_182931:XM_011516400:NM_018682:XM_024446837 | 2:59:18:2:0:0 | 0:0:0:0:0:0 | 6:198:51:3:1:4 | 0:1:0:0:0:0 | 1:123:33:4:0:2 | 0:0:0:0:0:0 | 10:486:127:9:3:5 | 0:0:1:0:0:0 | 3:53:13:0:0:1 | 0:2:0:0:0:0 |
| XR_001744836:XM_017012435:NM_182931:XM_011516400:NM_018682:XM_024446837 | 58:2:18:2:0:0 | 1:0:0:0:0:0 | 194:5:50:3:1:4 | 3:0:0:0:0:0 | 123:1:32:4:0:2 | 0:0:0:0:0:0 | 472:10:123:9:3:5 | 6:0:3:0:0:0 | 52:3:13:0:0:1 | 1:0:0:0:0:0 |
| XR_001744836:NM_182931:XM_011516400:XM_017012435:NM_018682:XM_024446837 | 53:17:1:2:0:0 | 5:1:1:0:0:0 | 141:38:2:4:1:4 | 25:6:0:0:0:0 | 119:30:4:1:0:2 | 4:2:0:0:0:0 | 370:91:8:9:2:3 | 59:17:0:1:1:0 | 48:11:0:3:0:1 | 4:2:0:0:0:0 |
| NM_182931:XR_001744836:XM_011516400:XM_017012435:NM_018682:XM_024446837 | 15:41:1:1:0:0 | 3:18:1:1:0:0 | 45:174:3:4:1:4 | 5:21:0:2:0:0 | 21:93:2:1:0:1 | 12:28:2:0:0:1 | 101:401:8:6:3:3 | 22:71:1:4:0:2 | 9:39:0:2:0:1 | 4:13:0:1:0:0 |
| XM_017012435:XR_001744836:NM_182931:XM_024446837:NM_018682:XM_011516400 | 1:26:10:0:0:0 | 0:0:0:0:0:0 | 1:42:13:1:1:1 | 0:2:2:1:0:0 | 1:52:16:0:0:1 | 0:0:0:0:0:0 | 1:111:30:0:0:2 | 0:12:1:0:0:1 | 1:22:4:1:0:0 | 0:0:0:0:0:0 |
| NM_182692:NM_001350743:XM_024446894:XR_001744862:XR_001744863:NM_001350746:XM_024446893 | 1:0:0:0:0:0:0 | 0:0:0:0:0:0:0 | 0:0:0:0:0:0:0 | 0:0:0:0:0:0:0 | 0:0:0:0:0:0:0 | 0:1:0:0:0:0:0 | 1:0:1:1:1:0:0 | 3:0:0:0:0:0:0 | 2:0:0:0:0:0:0 | 1:0:0:0:0:0:0 |
| NM_182692:NM_001350743:XR_001744863:XM_024446894:XR_001744862:NM_001350746:XM_024446893 | 1:0:0:0:0:0:0 | 0:0:0:0:0:0:0 | 0:0:0:0:0:0:0 | 0:0:0:0:0:0:0 | 0:0:0:0:0:0:0 | 2:1:1:0:0:0:0 | 1:0:1:1:1:0:0 | 3:0:0:0:0:0:0 | 2:0:0:0:0:0:0 | 1:0:0:0:0:0:0 |
| NM_001350961:XM_024446956:NM_001350959:NM_001244606:NM_001350963:NM_032549:NM_001350960:XM_024446962:XM_011516609:XM_017012704:XM_017012701:XM_017012702 | 11:8:5:81:10:7:0:0:0:0:0:0 | 0:0:0:0:0:0:0:0:0:0:0:0 | 1:0:0:25:1:0:0:0:0:0:0:0 | 0:0:0:0:0:0:0:0:0:0:0:0 | 8:1:4:69:1:9:3:0:0:0:0:0 | 0:0:0:0:0:0:0:0:0:0:0:0 | 95:37:21:355:25:43:5:3:19:1:0:0 | 0:0:0:0:0:1:0:0:0:0:0:0 | 6:8:0:57:2:7:0:0:0:0:1:0 | 0:0:0:1:0:0:0:0:0:0:0:0 |
| NM_001172896:NM_001753:NM_001172895:NM_001172897 | 68:51:2:8 | 1:0:0:0 | 105:148:10:12 | 0:0:0:0 | 42:59:1:8 | 0:0:0:0 | 13:21:0:1 | 0:0:0:0 | 25:47:1:3 | 0:0:0:0 |
| NM_001753:NM_001172895:NM_001172896:NM_001172897 | 51:2:69:8 | 0:0:0:0 | 152:10:104:12 | 0:0:1:0 | 59:1:42:8 | 0:0:0:0 | 21:0:13:1 | 0:0:0:0 | 47:1:25:3 | 0:0:0:0 |
| NM_001753:NM_001172895:NM_001172896:NM_001172897 | 51:2:69:8 | 0:0:0:0 | 154:10:103:12 | 0:0:2:0 | 59:1:42:8 | 0:0:0:0 | 22:0:13:1 | 0:0:0:0 | 48:1:25:3 | 0:0:0:0 |
| XM_006715990:NM_001324401:XM_011516223:NM_001324402:NM_000245:NM_001127500:XR_001744772 | 1:3:1:1:0:0:0 | 0:0:0:0:0:0:0 | 0:2:3:0:0:0:0 | 0:0:0:0:0:0:0 | 0:1:1:0:0:0:0 | 0:1:0:0:0:0:0 | 29:25:57:0:6:1:1 | 0:0:3:0:0:0:0 | 1:8:3:1:2:0:0 | 0:0:0:0:0:0:0 |
| XM_006715990:NM_001324402:XR_001744772:XM_011516223:NM_000245 | 1:2:1:0:0 | 0:0:0:0:0 | 0:0:0:3:0 | 0:0:0:0:0 | 0:0:0:1:0 | 0:0:0:0:0 | 10:0:5:46:4 | 1:0:1:4:2 | 0:0:1:3:2 | 0:0:0:0:0 |
| NM_006136 | 888 | 2 | 819 | 1 | 1016 | 0 | 2883 | 5 | 783 | 1 |
| NM_006136 | 889 | 2 | 822 | 0 | 1016 | 0 | 2890 | 6 | 783 | 1 |
| NM_006136 | 898 | 1 | 859 | 1 | 1028 | 0 | 3065 | 2 | 794 | 1 |
| NM_006136 | 898 | 2 | 856 | 0 | 1020 | 1 | 3041 | 2 | 786 | 3 |
| NM_003391:NR_024047 | 25:01:00 | 0:00 | 0:00 | 0:00 | 1:00 | 0:00 | 0:00 | 0:00 | 4:00 | 0:00 |
| NM_015450:NM_001042594:NR_003102:NR_003104 | 0:0:0:0 | 0:0:0:0 | 0:0:0:0 | 0:0:0:0 | 2:1:0:0 | 0:0:0:0 | 4:1:0:0 | 0:0:0:0 | 3:1:0:0 | 0:0:0:0 |
| NM_014390:XM_017011987 | 328:12:00 | 1:00 | 10:03 | 0:00 | 135:07:00 | 1:00 | 288:65 | 0:01 | 232:12:00 | 2:00 |
| NM_014390:XM_017011987 | 331:12:00 | 2:00 | 9:04 | 2:00 | 142:07:00 | 2:00 | 293:75 | 18:00 | 251:13:00 | 3:00 |
| NM_014390:XM_017011987 | 385:13:00 | 2:00 | 34:05:00 | 0:00 | 190:09:00 | 1:00 | 605:77 | 3:00 | 329:14:00 | 0:00 |
| XM_017011987:NM_014390 | 0.738194444 | 1:01 | 4:42 | 0:00 | 0.514583333 | 0:01 | 71:689 | 2:02 | 0.736805556 | 0:00 |
| NM_014390:XM_017011987 | 420:07:00 | 1:03 | 59:03:00 | 1:00 | 209:09:00 | 2:01 | 810:53:00 | 4:01 | 348:08:00 | 1:00 |
| XM_017011987:NM_014390 | 0.543055556 | 4:00 | 0.16875 | 0:00 | 0.521527778 | 1:00 | 53:827 | 1:02 | 0.532638889 | 1:01 |
| NM_014390:XM_017011987 | 434:06:00 | 1:00 | 69:03:00 | 0:00 | 212:08:00 | 2:00 | 926:50:00 | 4:00 | 358:07:00 | 1:00 |
| NM_014390:XM_017011987 | 437:06:00 | 0:00 | 72:02:00 | 0:00 | 217:08:00 | 0:00 | 955:50:00 | 2:00 | 363:07:00 | 0:00 |
| NM_014390:XM_017011987 | 450:00:00 | 1:00 | 112:00:00 | 2:00 | 248:02:00 | 0:00 | 1430:02:00 | 16:02 | 386:02:00 | 1:00 |
| NM_014390:XM_017011987 | 449:02:00 | 3:00 | 120:00:00 | 0:00 | 249:02:00 | 1:00 | 1585:04:00 | 14:00 | 398:03:00 | 1:00 |
| XM_017011987:NM_014390 | 0.397222222 | 1:02 | 0.084027778 | 0:00 | 0.259027778 | 0:00 | 1.283333333 | 0:13 | 0.402083333 | 0:01 |
| NM_014390:XM_017011987 | 454:03:00 | 2:00 | 121:00:00 | 1:00 | 253:02:00 | 0:00 | 1634:04:00 | 5:00 | 401:03:00 | 0:00 |

| | | | | | | | | | | |
|---|---|---|---|---|---|---|---|---|---|---|
| NM_014390:XM_017011987 | 454:03:00 | 1:00 | 128:00:00 | 0:00 | 255:01:00 | 0:00 | 1771:04:00 | 10:01 | 402:03:00 | 0:00 |
| NM_014390:XM_017011987 | 454:03:00 | 1:00 | 130:00:00 | 0:00 | 256:01:00 | 0:00 | 1803:05:00 | 1:00 | 400:03:00 | 2:00 |
| NM_014390:XM_017011987 | 452:02:00 | 0:00 | 130:00:00 | 0:00 | 255:02:00 | 0:00 | 1890:05:00 | 5:00 | 406:03:00 | 0:00 |
| NM_014390:XM_017011987 | 450:02:00 | 2:00 | 130:00:00 | 0:00 | 256:02:00 | 0:00 | 1906:05:00 | 2:00 | 406:03:00 | 0:00 |
| NM_014390:XM_017011987 | 451:02:00 | 2:00 | 128:00:00 | 1:00 | 258:02:00 | 0:00 | 1949:05:00 | 3:00 | 403:03:00 | 2:00 |
| NM_014390:XM_017011987 | 453:02:00 | 1:00 | 129:00:00 | 0:00 | 258:02:00 | 0:00 | 1952:05:00 | 7:00 | 405:03:00 | 0:00 |
| NM_012133 | 7 | 0 | 3 | 0 | 4 | 0 | 15 | 0 | 4 | 0 |
| NM_021807:XR_001744845:XM_017012494 | 18:01:00 | 0:00:00 | 6:01:01 | 0:00:00 | 11:01:01 | 0:00:00 | 40:02:13 | 0:00:00 | 27:02:01 | 0:00:00 |
| XR_927542:NM_033140:XM_017012654:XR_927537:XR_001744879:XR_002956490 | 14:1:1:0:0:0 | 1:0:0:0:0:0 | 45:7:0:5:0:0 | 1:0:0:0:0:0 | 50:10:0:4:1:0 | 2:0:0:0:0:0 | 3:0:0:0:0:0 | 0:1:0:0:0:0 | 13:2:0:2:0:0 | 0:0:0:0:0:0 |
| XR_927542:NM_033140:XM_017012654:XR_927537:XR_001744879:XR_002956490 | 13:1:1:0:0:0 | 2:0:0:0:0:0 | 30:6:0:5:0:0 | 16:1:0:0:0:0 | 42:9:0:4:1:0 | 10:1:0:0:0:0 | 3:0:0:0:0:0 | 0:1:0:0:0:0 | 12:2:0:2:0:0 | 2:0:0:0:0:0 |
| NM_033140:XR_927542:XM_017012654:XR_927537:XR_001744879 | 1:8:1:0:0 | 0:0:0:0:0 | 5:13:0:5:0 | 0:0:0:0:0 | 6:22:0:4:1 | 0:2:0:0:0 | 0:1:0:0:0 | 0:0:0:0:0 | 1:7:0:2:0 | 1:3:0:0:0 |
| NM_194071:NM_001253775 | 0:00 | 0:00 | 1:00 | 0:00 | 0:04 | 0:00 | 0:01 | 0:00 | 1:00 | 0:00 |
| XM_024446655:NM_001331192 | 0:00 | 1:00 | 0:00 | 1:00 | 0:00 | 1:01 | 0:00 | 1:00 | 0:00 | 0:00 |
| NM_020119 | 0 | 0 | 1 | 0 | 1 | 2 | 3 | 0 | 1 | 0 |
| NM_197964 | 1 | 0 | 2 | 0 | 0 | 0 | 0 | 1 | 0 | 2 |
| XM_005249635:XM_024446679:XM_011515168:NM_001163150:NM_004956:NR_120445:XM_011515167:XM_011515170:NM_001163147 | 1:1:1:0:0:0:0:0:0 | 0:0:0:0:0:0:0:0:0 | 1:0:0:1:0:0:0:0:0 | 0:0:0:0:0:0:0:0:0 | 1:2:0:1:0:0:0:0:0 | 0:0:0:0:0:0:0:0:0 | 0:0:0:0:0:0:0:0:0 | 0:0:0:0:0:0:0:0:0 | 2:0:0:0:1:0:0:0:0 | 0:0:0:0:0:0:0:0:0 |
| XM_005249635:XM_011515168:XM_024446679:NM_001163150:NM_004956:NR_120445:XM_011515167:XM_011515170:NM_001163147 | 1:1:0:0:0:0:0:0:0 | 0:0:0:0:0:0:0:0:0 | 1:0:1:1:0:0:0:0:0 | 0:0:0:0:0:0:0:0:0 | 1:0:1:1:0:0:0:0:0 | 0:0:0:0:0:0:0:0:0 | 0:0:0:0:0:0:0:0:0 | 0:0:0:0:0:0:0:0:0 | 1:0:0:0:1:0:0:0:0 | 1:0:0:0:0:0:0:0:0 |
| NM_030647:XM_011516587 | 1:02 | 1:00 | 3:00 | 4:00 | 2:01 | 2:00 | 40:05:00 | 9:00 | 3:01 | 0:00 |
| XM_011516587:NM_030647 | 2:02 | 0:00 | 0:03 | 0:04 | 1:02 | 0:02 | 5:44 | 0:04 | 1:03 | 0:00 |
| NM_013446:XM_011515998:NM_001145125:NM_001291663:XM_011515997 | 0:1:0:0:0 | 24:6:8:3:0 | 0:0:0:0:0 | 13:21:16:3:0 | 0:0:0:0:0 | 59:20:37:4:3 | 76:43:82:10:1 | 0:0:0:0:0 | 41:8:14:2:0 | 0:0:0:0:0 |
| XM_017012558:XM_017012559:NM_004333:NM_001354609:NR_148928:XR_001744857:XR_001744858 | 2:2:8:0:0:0:0 | 0:0:0:0:0:0:0 | 0:0:9:1:1:0:0 | 0:0:0:0:0:0:0 | 0:2:8:4:0:1:0 | 0:0:0:0:0:0:0 | 3:9:96:1:1:0:1 | 0:0:0:0:0:0:0 | 0:0:19:1:0:0:0 | 0:0:0:0:0:0:0 |
| NM_053035:NM_016071 | 228:13:00 | 0:00 | 308:28:00 | 0:00 | 159:09:00 | 0:00 | 1569:145 | 0:00 | 255:05:00 | 3:00 |
| NM_053035:NM_016071 | 226:13:00 | 0:00 | 307:28:00 | 0:00 | 158:09:00 | 0:00 | 1563:146 | 3:00 | 257:05:00 | 1:00 |
| NM_002770 | 0 | 0 | 0 | 0 | 0 | 0 | 0 | 0 | 0 | 0 |
| XM_017012587 | 0 | 0 | 1 | 0 | 0 | 0 | 0 | 1 | 1 | 2 |
| NM_003592 | 6 | 0 | 6 | 0 | 4 | 0 | 7 | 7 | 7 | 3 |
| NM_003592 | 6 | 0 | 6 | 0 | 4 | 0 | 10 | 4 | 9 | 1 |
| NM_001203247:XM_005249962:XM_024446680:XM_011515884:XR_002956414:NM_004456:XM_017011818:XM_017011821:XM_011515894:XR_002956413:XM_017011819:XM_011515897:NM_152998:XM_011515887:XM_017011820:XM_011515889:XM_011515898:NM_001203248 | 3:1:1:1:1:1:0:0:0:0:0:0:0:0:0:0:0:0 | 0:0:0:0:0:0:0:0:0:0:0:0:0:0:0:0:0:0 | 4:0:1:0:1:0:1:1:1:0:0:0:0:0:0:0:0:0 | 0:0:0:0:0:0:0:0:0:0:0:0:0:0:0:0:0:0 | 11:0:1:0:0:0:0:1:0:1:0:0:0:0:0:0:0:0 | 0:0:0:0:0:0:0:0:0:0:0:0:0:0:0:0:0:0 | 2:0:1:0:1:3:0:2:0:0:1:0:1:0:0:0:0:0 | 4:0:1:0:0:0:0:1:0:1:1:1:0:0:0:0:0:0 | 9:0:0:0:0:2:0:2:0:0:0:0:0:0:1:0:0:0 | 4:0:0:0:0:4:0:1:0:2:0:0:0:1:0:0:0:0 |
| XM_006715947:NM_001099696 | 0:00 | 0:00 | 0:00 | 0:00 | 0:00 | 0:00 | 0:00 | 0:00 | 0:00 | 1:00 |
| XM_006715947:NM_001099696 | 0:00 | 0:00 | 0:00 | 0:00 | 0:00 | 0:00 | 0:00 | 0:00 | 0:00 | 1:00 |
| XM_006715947:NM_001099696 | 0:00 | 0:00 | 0:00 | 0:00 | 0:00 | 0:00 | 0:00 | 0:00 | 0:00 | 1:00 |
| NM_001101312:NM_014020:NM_001101311:NM_001101314:XM_024446733:XM_024446734:XM_006715933 | 1097:23:88:16:10:52:3 | 1:0:0:0:0:0:0 | 1420:31:63:18:11:86:4 | 2:0:0:0:0:0:0 | 3026:93:136:30:19:141:3 | 3:0:0:0:0:0:0 | 5632:100:1062:107:174:350:27 | 3:0:3:0:0:1:0 | 919:23:92:28:15:40:7 | 0:0:1:0:0:0:0 |
| NM_001101312:NM_014020:NM_001101311:NM_001101314:XM_024446733:XM_024446734:XM_006715933 | 1100:22:89:16:10:50:3 | 1:0:0:0:0:0:0 | 1425:29:63:18:11:85:4 | 1:0:0:0:0:2:0 | 3032:88:136:30:19:135:3 | 2:0:0:0:0:0:0 | 5641:99:1066:108:174:336:27 | 7:0:0:0:0:1:0 | 922:23:93:28:15:40:7 | 0:0:0:0:0:0:0 |
| NM_014020:NM_001101312:NM_001101311:NM_001101314:XM_024446733:XM_024446734:XM_006715933 | 20:1105:89:16:10:50:3 | 0:0:0:0:0:0:0 | 29:1433:63:18:11:77:4 | 0:0:0:0:0:1:0 | 84:3038:136:30:19:126:3 | 0:7:0:0:0:0:0 | 94:5670:1071:108:179:323:27 | 0:10:0:0:0:0:0 | 21:922:94:28:15:39:7 | 0:1:0:0:0:0:0 |
| NM_001101312:XM_024446734:NM_014020:NM_001101311:NM_001101314:XM_024446733:XM_006715933 | 1104:48:20:89:16:10:3 | 1:1:0:0:0:0:0 | 1434:76:29:63:18:11:4 | 0:1:0:0:0:0:0 | 3044:126:85:136:30:18:3 | 1:0:0:0:0:1:0 | 5675:320:94:1070:108:179:27 | 5:1:0:0:0:0:0 | 922:38:21:94:28:15:7 | 1:0:0:0:0:0:0 |
| XM_024446734:NM_014020:NM_001101312:NM_001101311:NM_001101314:XM_024446733:XM_006715933 | 44:20:1105:89:16:10:3 | 3:0:0:0:0:0:0 | 76:29:1433:63:19:11:4 | 0:0:0:0:0:0:0 | 122:85:3041:136:31:19:3 | 1:0:5:0:0:0:0 | 318:93:5675:1071:108:179:28 | 0:1:5:0:0:0:0 | 38:21:924:94:28:15:7 | 0:0:1:0:0:0:0 |
| NM_014020:NM_001101312:NM_001101311:NM_001101314:XM_024446733:XM_024446734:XM_006715933 | 20:1105:89:16:10:46:3 | 0:0:0:0:0:0:0 | 29:1432:63:19:11:72:4 | 0:1:0:0:0:0:0 | 85:3044:137:31:19:120:3 | 0:2:0:0:0:0:0 | 94:5675:1073:108:179:304:28 | 0:5:0:0:0:0:0 | 21:924:94:28:15:34:7 | 0:1:0:0:0:0:0 |
| NM_014020:NM_001101312:NM_001101311:NM_001101314:XM_024446733:XM_024446734:XM_006715933 | 20:1106:89:16:10:46:3 | 0:0:0:0:0:0:0 | 29:1433:63:19:11:69:4 | 0:0:0:0:0:0:0 | 85:3044:137:31:19:116:3 | 0:3:0:0:0:1:0 | 92:5679:1073:108:179:296:28 | 0:4:1:0:0:1:0 | 21:925:94:28:15:33:7 | 0:0:0:0:0:0:0 |
| NM_001101312:NM_001101311:NM_014020:XM_024446733:NM_001101314:XM_024446734:XM_006715933 | 568:52:9:6:9:26:1 | 538:37:10:4:7:18:2 | 805:41:15:7:10:42:1 | 628:22:11:4:8:19:3 | 1660:67:39:11:17:69:1 | 1388:70:45:8:14:43:2 | 2837:564:41:41:52:124:12 | 2852:510:49:139:55:149:16 | 474:57:9:4:17:13:4 | 452:37:10:11:11:19:3 |
| NM_014020:NM_001101312:NM_001101311:NM_001101314:XM_024446733:XM_024446734:XM_006715933 | 19:1106:89:16:10:44:3 | 0:0:0:0:0:0:0 | 25:1434:63:18:11:59:4 | 0:1:0:0:0:0:0 | 84:3043:137:31:19:109:3 | 0:5:0:0:0:0:0 | 90:5687:1074:107:180:262:28 | 0:2:1:0:0:0:0 | 19:926:94:28:15:32:7 | 0:0:0:0:0:0:0 |
| NM_014020:NM_001101312:NM_001101311:NM_001101314:XM_024446733:XM_024446734:XM_006715933 | 18:1108:89:15:10:43:2 | 0:0:0:0:0:0:0 | 24:1437:63:15:11:51:3 | 0:0:0:0:0:0:0 | 78:3053:137:26:19:92:3 | 0:2:0:0:0:0:0 | 84:5699:1076:101:180:203:24 | 0:2:0:0:0:0:0 | 16:928:94:27:15:26:7 | 0:0:0:0:0:0:0 |
| NM_001101312:NM_014020:NM_001101311:XM_024446733:NM_001101314:XM_024446734:XM_006715933 | 1107:17:89:10:12:43:2 | 2:0:0:0:0:0:0 | 1437:24:63:11:14:50:3 | 2:0:0:0:0:0:0 | 3051:77:137:19:19:86:3 | 4:0:0:0:2:0:0 | 5695:81:1074:180:87:196:17 | 6:0:2:0:6:0:0 | 927:16:93:15:25:26:7 | 1:0:1:0:1:0:0 |
| NM_014020:NM_001101312:NM_001101311:XM_024446733:XM_024446734:NM_001101314:XM_006715933 | 1:0:0:0:1:0:0 | 17:1113:89:10:37:0:0 | 1:0:0:0:0:0:0 | 22:1446:63:11:40:3:3 | 1:1:0:0:1:0:0 | 73:3059:137:19:66:1:1 | 2:4:0:0:0:0:0 | 71:5714:1078:180:161:9:1 | 0:0:0:0:0:0:0 | 14:930:95:15:21:2:0 |
| XM_024446734:NM_014020:NM_001101312:NM_001101311:XM_024446733:NM_001101314:XM_006715933 | 37:24:1117:89:10:1:0 | 1:0:0:0:0:0:0 | 39:24:1455:62:11:1:1 | 0:0:1:1:0:0:0 | 73:80:3065:138:19:2:1 | 1:0:2:0:0:0:0 | 169:86:5742:1081:182:6:2 | 0:1:2:0:0:0:0 | 23:17:932:96:15:1:1 | 0:0:0:0:0:0:0 |
| XM_006715933:NM_014020:NM_001101312:NM_001101311:NM_001101314:XM_024446733:XM_024446734 | 3:29:1119:90:19:10:38 | 1:0:0:0:0:0:0 | 3:27:1467:63:24:11:39 | 0:1:0:0:0:0:0 | 2:86:3078:138:37:19:82 | 0:0:1:0:1:0:1 | 31:90:5785:1094:126:183:177 | 1:3:1:1:0:0:11 | 7:20:932:96:34:15:23 | 0:1:0:0:0:0:1 |
| NM_001101312:NM_014020:NM_001101311:NM_001101314:XM_024446733:XM_024446734:XM_006715933 | 1115:33:90:21:10:23:4 | 4:0:0:0:0:0:0 | 1465:36:63:28:11:42:4 | 2:1:0:0:0:1:0 | 3078:85:138:40:19:79:2 | 1:5:0:0:0:3:0 | 5780:106:1095:136:183:179:40 | 6:0:1:0:0:1:0 | 931:24:96:34:15:21:7 | 1:0:0:0:0:0:0 |
| XM_011516377:XM_011516376:NM_018487:XM_017012393:XM_011516378 | 0:0:1:0:0 | 1:3:0:0:0 | 0:0:1:0:0 | 0:0:0:0:0 | 1:2:2:2:0 | 0:0:0:1:0 | 0:4:0:1:0 | 0:1:3:0:0 | 0:1:1:0:0 | 0:0:1:0:0 |
| XM_011516377:XM_011516376:NM_018487:XM_017012393 | 0:0:1:0 | 1:3:0:0 | 0:0:1:0 | 0:0:0:0 | 1:2:2:2 | 0:1:0:1 | 0:5:3:1 | 0:0:0:0 | 0:1:2:0 | 0:0:0:0 |
| XM_017011947:NM_001091 | 0:00 | 0:00 | 0:00 | 2:00 | 0:00 | 1:01 | 0:00 | 1:00 | 0:00 | 0:00 |
| XM_017011947:NM_001091:NM_001272072 | 0:00:00 | 0:00:00 | 2:00:00 | 0:00:00 | 1:01:00 | 0:00:00 | 1:00:00 | 0:00:00 | 0:00:00 | 0:00:00 |
| NM_001097620 | 0 | 0 | 0 | 0 | 0 | 2 | 0 | 0 | 0 | 0 |
| NR_144629 | 0 | 0 | 0 | 0 | 0 | 0 | 0 | 1 | 0 | 0 |
| NM_003550:NM_001013837:NM_001304525:NM_001013836 | 0:0:0:0 | 0:0:0:0 | 0:0:0:0 | 0:0:0:0 | 0:0:0:0 | 0:0:0:0 | 3:1:1:1 | 0:0:0:0 | 0:0:1:0 | 0:0:0:0 |

| | | | | | | | | | | |
|---|---|---|---|---|---|---|---|---|---|---|
| NM_003550:NM_001013837:NM_001304525:NM_001013836 | 0:0:0:0 | 0:0:0:0 | 0:0:0:0 | 0:0:0:0 | 0:0:0:0 | 0:0:0:0 | 2:0:0:0 | 1:1:1:1 | 0:0:1:0 | 0:0:0:0 |
| NM_013293:NM_001282758:NM_001282757:XM_017012090:NM_001282759 | 49:3:3:1:0 | 0:0:0:0:0 | 59:4:0:20:0 | 0:0:0:0:0 | 86:1:2:4:2 | 0:0:0:0:0 | 142:7:5:37:3 | 0:0:0:0:0 | 57:2:1:5:0 | 0:0:0:0:0 |
| NM_013293:NM_001282758:NM_001282757:XM_017012090:NM_001282759 | 49:3:3:1:0 | 0:0:0:0:0 | 60:4:0:17:0 | 0:0:0:0:0 | 89:1:2:3:2 | 0:0:0:1:0 | 158:7:5:35:3 | 0:0:0:0:0 | 57:2:1:6:0 | 0:0:0:0:0 |
| NM_013293:NM_001282758:NM_001282757:XM_017012090:NM_001282759 | 49:4:3:1:0 | 0:0:0:0:0 | 60:4:0:17:0 | 0:0:0:0:0 | 89:1:2:4:2 | 0:0:0:0:0 | 163:7:5:31:3 | 0:0:0:0:0 | 57:2:1:6:0 | 0:0:0:0:0 |
| XM_006715758:NM_031414:XM_011515452:XM_011515450:XM_011515449:NM_001260505 | 0:0:0:0:0:0 | 0:0:0:0:0:0 | 0:0:0:0:0:0 | 0:0:0:0:0:0 | 0:0:0:0:0:0 | 0:0:0:0:0:0 | 0:0:0:0:0:0 | 0:0:0:0:0:0 | 0:0:0:0:0:0 | 0:0:0:0:0:0 |
| XR_001744669:XR_001744671:XM_017012110:NM_031243:XR_001744664:XR_001744673:XR_242076:XR_002956425:XM_005249729:NM_002137:XR_242077:XR_001744674:XR_428077:XR_001744672:XR_001744675:XR_001744667:XM_017012109:XR_001744668:XR_002956424:XR_001744670:XR_001744666 | 411:36:16:16:82:6:6:13:1:19:1:1:3:0:0:0:0:0:0:0:0 | 1:0:0:0:0:0:0:0:0:0:0:0:0:0:0:0:0:0:0:0:0 | 297:22:20:2:39:0:4:14:3:17:1:1:5:2:0:0:0:0:0:0:0:0 | 0:0:0:0:0:0:0:0:0:0:0:0:0:0:0:0:0:0:0:0:0 | 294:22:14:5:51:3:3:16:2:26:1:3:5:1:1:0:0:0:0:0:0 | 0:0:0:0:0:0:0:0:0:0:0:1:0:0:0:0:0:0:0:0:0 | 1618:143:9:17:191:31:27:66:1:123:5:19:12:4:2:1:1:1:2:0:0 | 0:1:0:0:0:0:0:0:0:0:0:1:0:0:0:0:0:0:0:0:0 | 626:25:2:4:100:2:7:33:1:34:3:5:4:1:1:0:1:0:0:0:0 | 0:0:0:0:0:0:0:0:0:0:0:0:0:0:0:0:0:0:0:0:0 |
| XR_001744671:XM_017012110:NM_031243:XR_001744664:XR_001744673:XR_242076:XR_002956425:XM_005249729:NM_002137:XR_242077:XR_001744669:XR_001744674:XR_428077:XR_001744672:XR_001744675:XR_001744667:XM_017012109:XR_001744668:XR_002956424:XR_001744670:XR_001744666 | 36:16:16:82:6:6:9:1:20:1:412:2:4:0:0:0:0:0:0:0:0 | 0:0:0:0:0:0:0:0:0:0:0:0:0:0:0:0:0:0:0:0:0 | 22:20:2:40:0:4:13:3:17:1:296:1:5:2:0:0:0:0:0:0:0:0 | 0:0:0:0:0:0:0:0:1:0:0:0:0:0:0:0:0:0:0:0:0 | 22:14:5:51:3:2:15:2:27:1:294:4:5:1:1:0:0:0:0:0:0 | 0:0:0:0:0:0:0:0:0:0:0:0:0:0:0:0:0:0:0:0:0 | 144:9:17:191:31:26:62:1:122:5:1618:24:18:4:2:1:1:1:2:0:0 | 0:0:0:0:0:0:0:0:0:0:3:0:0:0:0:0:0:0:0:0:0 | 25:2:4:99:2:7:28:1:34:3:624:7:6:1:1:0:1:0:0:0:0 | 0:0:0:1:0:0:0:0:0:0:2:1:0:0:0:0:0:0:0:0:0 |
| XR_001744671:XM_017012110:NM_031243:XR_001744664:XR_001744673:XR_242076:XR_002956425:XM_005249729:NM_002137:XR_242077:XR_001744669:XR_001744674:XR_428077:XR_001744672:XR_001744675:XR_001744667:XM_017012109:XR_001744668:XR_002956424:XR_001744670:XR_001744666 | 36:16:16:82:6:6:8:1:20:1:412:2:4:0:0:0:0:0:0:0:0 | 0:0:0:0:0:0:0:0:0:0:0:0:0:0:0:0:0:0:0:0:0 | 22:20:2:40:0:4:13:3:17:1:296:1:5:2:0:0:0:0:0:0:0:0 | 0:0:0:0:0:0:0:0:1:0:0:0:0:0:0:0:0:0:0:0:0 | 22:14:5:51:3:2:15:2:27:1:294:5:5:1:1:0:0:0:0:0:0 | 0:0:0:0:0:0:0:0:0:0:0:0:0:0:0:0:0:0:0:0:0 | 144:9:17:191:31:26:62:1:122:5:1617:24:17:4:2:1:1:1:2:0:0 | 0:0:0:0:0:0:0:0:0:1:3:1:0:0:0:0:0:0:0:0:0 | 25:2:4:100:2:7:28:1:34:3:626:7:6:1:1:0:1:0:0:0:0 | 0:0:0:0:0:0:0:0:0:0:1:0:0:0:0:0:0:0:0:0:0 |
| XR_001744669:XR_001744671:XM_017012110:NM_031243:XR_001744673:XR_242076:XR_002956425:XM_005249729:NM_002137:XR_242077:XR_001744664:XR_001744674:XR_428077:XR_001744672:XR_001744667:XM_017012109:XR_001744668:XR_001744675:XR_001744670:XR_002956424:XR_001744666 | 410:36:16:15:6:6:6:1:20:1:81:3:8:0:0:0:0:0:0:0:0 | 1:0:0:0:0:0:0:0:0:0:0:0:0:0:0:0:0:0:0:0:0 | 295:22:20:2:0:4:6:3:18:1:40:3:9:2:0:0:0:0:0:0:0:0 | 1:0:0:0:0:0:0:0:1:0:0:0:0:0:0:0:0:0:0:0:0 | 291:22:14:5:3:2:8:2:27:1:51:8:5:1:0:0:0:0:0:0:0 | 1:0:0:0:0:0:0:0:0:0:0:0:0:0:0:0:0:0:0:0:0 | 1613:144:9:16:31:26:29:1:123:4:191:35:23:4:1:1:1:1:0:0:0 | 0:0:0:0:0:0:0:0:1:0:0:3:0:0:0:0:0:0:0:0:0 | 623:25:2:4:2:7:16:1:34:3:100:12:5:1:0:1:0:1:0:0:0 | 0:0:0:0:0:0:0:0:0:0:0:0:0:0:0:0:0:0:0:0:0 |
| XR_001744671:XM_017012110:NM_031243:XR_001744673:XR_242076:XR_002956425:XM_005249729:NM_002137:XR_242077:XR_001744669:XR_001744664:XR_001744674:XR_428077:XR_001744672:XR_001744667:XM_017012109:XR_001744668:XR_001744675:XR_001744670:XR_002956424:XR_001744666 | 36:16:15:6:6:5:1:20:1:411:81:3:8:0:0:0:0:0:0:0:0 | 0:0:0:0:0:0:0:0:0:0:0:0:0:0:0:0:0:0:0:0:0 | 22:20:2:0:4:6:3:19:1:296:40:3:9:2:0:0:0:0:0:0:0 | 0:0:0:0:0:0:0:0:0:0:0:0:0:0:0:0:0:0:0:0:0 | 22:14:5:3:2:8:2:27:1:291:51:8:5:1:0:0:0:0:0:0:0 | 0:0:0:0:0:0:0:0:0:0:0:0:0:0:0:0:0:0:0:0:0 | 144:9:16:31:26:29:1:125:4:1610:191:34:23:4:1:1:1:1:0:0:0 | 0:0:0:0:0:0:0:0:2:0:2:0:0:0:0:0:0:0:0:0:0 | 25:2:4:2:6:15:1:33:3:622:100:11:6:1:0:1:0:0:0:0:0 | 0:0:0:0:0:0:0:0:0:0:1:0:0:0:0:0:0:0:0:0:0 |
| XR_001744671:XM_017012110:NM_031243:XR_001744673:XR_242076:XR_002956425:XM_005249729:NM_002137:XR_242077:XR_001744669:XR_001744664:XR_001744674:XR_428077:XR_001744672:XR_001744667:XM_017012109:XR_001744668:XR_001744675:XR_001744670:XR_002956424:XR_001744666 | 36:16:15:6:6:5:1:20:1:411:81:2:8:0:0:0:0:0:0:0:0 | 0:0:0:0:0:0:0:0:0:0:0:0:0:0:0:0:0:0:0:0:0 | 22:20:2:0:4:6:3:18:1:295:40:3:9:2:0:0:0:0:0:0:0 | 0:0:0:0:0:0:0:1:0:1:0:3:0:0:0:0:0:0:0:0:0 | 22:14:5:3:2:7:2:27:1:291:51:8:5:1:0:0:0:0:0:0:0 | 0:0:0:0:0:0:0:0:0:0:0:0:0:0:0:0:0:0:0:0:0 | 144:9:15:31:26:28:1:124:4:1612:191:34:23:4:1:1:1:1:0:0:0 | 0:0:1:0:0:0:0:1:0:0:0:3:0:0:0:0:0:0:0:0:0 | 25:2:4:2:6:15:1:33:3:621:100:11:6:1:0:1:0:0:0:0:0 | 0:0:0:0:0:0:0:0:0:0:1:0:0:0:0:0:0:0:0:0:0 |
| NM_031243:XR_428077:XR_001744671:XM_017012110:XR_001744673:XR_242076:XR_002956425:XM_005249729:XR_242077:XR_001744669:NM_002137:XR_001744664:XR_001744674:XR_001744672:XR_001744667:XM_017012109:XR_001744668:XR_001744670:XR_001744675:XR_001744666 | 14:9:36:16:6:6:5:1:1:410:19:81:2:0:0:0:0:0:0 | 1:1:0:0:0:0:0:0:0:0:0:0:0:0:0:0:0:0:0:0 | 2:9:22:20:0:4:5:3:1:294:18:40:4:2:0:0:0:0:0:0:0 | 0:0:0:0:0:0:0:0:0:0:1:0:2:0:0:0:0:0:0:0 | 5:5:22:14:3:2:7:2:1:289:27:51:8:1:0:0:0:0:0:0 | 0:1:0:0:0:0:0:0:0:1:0:0:2:0:0:0:0:0:0:0 | 16:26:143:9:31:26:20:1:4:1608:125:191:34:4:1:1:1:0:0:0 | 0:1:1:0:0:0:0:0:0:2:2:0:7:0:0:0:0:0:0:0 | 4:6:25:2:2:6:11:1:3:616:33:99:11:1:0:1:0:0:0:0 | 0:0:0:0:0:0:0:0:0:3:1:0:4:0:0:0:0:0:0:0 |
| XR_001744671:XM_017012110:NM_031243:XR_001744673:XR_242076:XR_002956425:XM_005249729:XR_242077:XR_001744669:NM_002137:XR_001744664:XR_001744674:XR_428077:XR_001744672:XR_001744667:XM_017012109:XR_001744668:XR_001744670:XR_001744666 | 36:16:15:6:6:5:1:1:410:19:81:2:10:0:0:0:0:0:0 | 0:0:0:0:0:0:0:0:0:0:0:0:0:0:0:0:0:0 | 22:20:2:0:4:5:3:1:292:18:40:7:9:2:0:0:0:0:0 | 0:0:0:0:0:0:0:0:1:1:0:0:0:0:0:0:0:0:0 | 22:14:5:3:2:6:2:1:290:27:51:12:7:1:0:0:0:0:0 | 0:0:0:0:0:0:0:0:0:0:0:0:0:0:0:0:0 | 144:9:15:31:26:14:1:4:1607:127:191:45:25:4:1:1:1:0:0 | 0:0:0:0:0:0:0:0:1:0:0:1:0:0:0:0:0:0:0 | 25:2:4:2:6:10:1:3:618:34:99:17:6:1:0:1:0:0:0 | 0:0:0:0:0:0:0:0:0:0:2:0:0:0:0:0:0:0:0 |
| XR_001744671:XM_017012110:NM_031243:XR_001744673:XR_242076:XR_002956425:XM_005249729:XR_242077:XR_001744669:NM_002137:XR_001744664:XR_001744674:XR_428077:XR_001744672:XR_001744667:XM_017012109:XR_001744668:XR_002956424:XR_001744670:XR_001744666 | 36:16:15:6:6:4:1:1:410:19:81:2:10:0:0:0:0:0:0:0 | 0:0:0:0:0:0:0:0:0:0:0:0:0:0:0:0:0:0:0:0 | 22:20:2:0:4:5:3:1:293:19:40:7:9:2:0:0:0:0:0:0 | 0:0:0:0:0:0:0:0:0:0:0:0:0:0:0:0:0:0:0:0 | 21:14:5:3:2:6:2:1:290:27:51:12:7:1:0:0:0:0:0:0 | 1:0:0:0:0:0:0:0:0:0:0:0:0:0:0:0:0:0:0 | 144:9:15:31:25:13:1:4:1607:126:191:47:25:4:1:1:1:1:0:0 | 0:0:0:0:0:0:0:0:1:1:0:0:0:0:0:0:0:0:0 | 25:2:4:2:6:10:1:3:620:35:99:19:6:1:0:1:0:0:0 | 0:0:0:0:0:0:0:0:0:0:0:0:0:0:0:0:0:0:0:0 |
| XR_001744669:XR_001744671:XM_017012110:NM_031243:XR_001744673:XR_242076:XR_002956425:XM_005249729:XR_242077:NM_002137:XR_001744664:XR_001744674:XR_428077:XR_001744672:XR_001744667:XM_017012109:XR_001744668:XR_002956424:XR_001744670:XR_001744666 | 409:36:16:15:6:6:4:1:1:19:81:2:10:0:0:0:0:0:0:0:0 | 1:0:0:0:0:0:0:0:0:0:0:0:0:0:0:0:0:0:0:0 | 293:22:20:2:0:4:5:3:1:19:40:7:9:2:0:0:0:0:0:0:0 | 0:0:0:0:0:0:0:0:0:0:0:0:0:0:0:0:0:0:0:0 | 290:22:14:5:3:2:6:2:1:27:51:12:7:1:0:0:0:0:0:0 | 0:0:0:0:0:0:0:0:0:0:0:0:0:0:0:0:0:0:0 | 1605:144:9:15:31:25:13:1:4:127:191:47:25:4:1:1:1:1:0:0 | 3:0:0:0:0:0:0:0:0:0:0:0:0:0:0:0:0:0:0 | 619:25:2:4:2:6:10:1:3:35:99:19:6:1:0:1:0:0:0 | 1:0:0:0:0:0:0:0:0:0:0:0:0:0:0:0:0:0:0:0 |
| XR_002956425:XR_001744671:XM_017012110:NM_031243:XR_001744673:XR_242076:XM_005249729:NM_002137:XR_242077:XR_001744669:XR_001744664:XR_001744674:XR_428077:XR_001744672:XR_001744675:XR_001744667:XM_017012109:XR_001744668:XR_002956424:XR_001744670:XR_001744666 | 8:36:16:15:6:6:1:20:1:408:81:3:11:0:0:0:0:0:0:0:0 | 1:0:0:0:0:0:0:0:0:0:0:0:0:0:0:0:0:0:0:0:0 | 14:22:20:2:0:4:3:19:1:292:40:7:9:2:0:0:0:0:0:0:0:0 | 0:0:0:0:0:0:0:1:0:0:0:0:0:0:0:0:0:0:0:0:0 | 13:22:14:5:3:2:2:27:0:284:51:12:7:1:1:0:0:0:0:0:0 | 0:0:0:0:0:0:0:0:0:0:0:0:0:0:0:0:0:0:0:0:0 | 64:144:9:14:31:25:1:130:4:1598:191:54:27:4:2:1:1:1:2:0:0 | 2:0:0:0:0:0:0:0:3:0:0:1:0:0:0:0:0:0:0:0:0 | 27:25:2:4:2:6:1:35:3:617:100:20:13:1:1:0:1:0:0:0:0 | 0:0:0:0:0:0:0:0:0:0:0:0:0:0:0:0:0:0:0:0:0 |
| XR_002956425:XR_001744671:XM_017012110:NM_031243:XR_001744673:XR_242076:XM_005249729:NM_002137:XR_242077:XR_001744669:XR_001744664:XR_001744674:XR_428077:XR_001744672:XR_001744675:XR_001744667:XM_017012109:XR_001744668:XR_002956424:XR_001744670:XR_001744666 | 8:36:15:15:6:6:1:20:1:408:81:3:11:0:0:0:0:0:0:0:0 | 1:0:0:0:0:0:0:0:0:0:0:0:0:0:0:0:0:0:0:0:0 | 14:22:20:2:0:4:3:19:1:292:40:7:9:2:0:0:0:0:0:0:0:0 | 0:0:0:0:0:0:0:1:0:0:0:0:0:0:0:0:0:0:0:0:0 | 13:22:14:5:3:2:2:27:0:282:51:12:7:1:1:0:0:0:0:0:0 | 0:0:0:0:0:0:0:0:0:0:0:0:0:0:0:0:0:0:0:0:0 | 67:144:9:14:31:25:1:130:4:1598:191:54:27:4:2:1:1:1:2:0:0 | 0:0:0:0:0:0:0:0:1:0:0:1:0:0:0:0:0:0:0:0:0 | 28:25:2:3:2:6:1:35:3:616:99:20:13:1:1:0:1:0:0:0:0 | 0:0:0:0:0:0:0:0:0:1:1:0:0:0:0:0:0:0:0:0:0 |
| XR_001744671:XM_017012110:NM_031243:XR_001744673:XR_242076:XR_002956425:XM_005249729:NM_002137:XR_242077:XR_001744669:XR_001744664:XR_001744674:XR_428077:XR_001744672:XR_001744675:XR_001744667:XM_017012109:XR_001744668:XR_002956424:XR_001744670:XR_001744666 | 36:16:15:6:6:11:1:20:1:406:81:3:9:0:0:0:0:0:0:0:0 | 0:0:0:0:0:0:0:0:0:0:0:0:0:0:0:0:0:0:0:0:0 | 22:20:2:0:4:12:3:20:1:284:40:7:10:2:0:0:0:0:0:0:0:0 | 0:0:0:0:0:0:0:0:0:0:0:0:0:0:0:0:0:0:0:0:0 | 22:14:5:3:2:13:2:28:0:280:51:12:8:1:1:0:0:0:0:0:0 | 0:0:0:0:0:0:0:0:0:0:0:0:0:0:0:0:0:0:0:0:0 | 144:9:14:31:23:76:1:130:3:1576:191:54:28:4:2:1:1:1:2:0:0 | 0:0:0:0:0:0:0:0:1:0:0:0:0:0:0:0:0:0:0:0:0 | 25:2:3:2:4:35:1:35:2:607:100:20:12:1:1:0:1:0:0:0:0 | 0:0:0:0:0:0:0:0:0:0:0:0:0:0:0:0:0:0:0:0:0 |
| XR_001744671:XM_017012110:NM_031243:XR_001744673:XR_242076:XR_002956425:XM_005249729:NM_002137:XR_242077:XR_001744669:XR_001744664:XR_001744674:XR_428077:XR_001744672:XR_001744675:XR_001744667:XM_017012109:XR_001744668:XR_002956424:XR_001744670:XR_001744666 | 36:16:15:6:6:11:1:20:1:406:81:3:9:0:0:0:0:0:0:0:0 | 0:0:0:0:0:0:0:0:0:0:0:0:0:0:0:0:0:0:0:0:0 | 22:20:2:0:4:12:3:20:1:283:39:7:10:2:0:0:0:0:0:0:0:0 | 0:0:0:0:0:0:0:0:1:0:0:0:0:0:0:0:0:0:0:0:0 | 22:13:5:3:2:13:2:28:0:278:51:12:8:1:1:0:0:0:0:0:0 | 0:1:0:0:0:0:0:0:0:0:0:0:0:0:0:0:0:0:0:0:0 | 143:9:14:31:22:76:1:129:3:1576:191:54:29:4:2:1:1:1:2:0:0 | 1:0:0:0:1:0:0:1:0:0:0:0:0:0:0:0:0:0:0:0:0 | 25:2:3:2:4:35:1:35:2:605:99:20:12:1:1:0:1:0:0:0:0 | 0:0:0:0:0:0:0:0:0:1:0:0:0:0:0:0:0:0:0:0:0 |
| XR_001744669:XR_001744671:XM_017012110:NM_031243:XR_001744673:XR_242076:XR_002956425:XM_005249729:NM_002137:XR_242077:XR_001744664:XR_001744674:XR_428077:XR_001744672:XR_001744675:XR_001744667:XM_017012109:XR_001744668:XR_002956424:XR_001744670:XR_001744666 | 403:36:16:15:6:6:11:1:20:1:81:3:9:0:0:0:0:0:0:0:0 | 2:0:0:0:0:0:0:0:0:0:0:0:0:0:0:0:0:0:0:0:0 | 281:22:20:2:0:4:12:3:20:1:40:7:10:2:0:0:0:0:0:0:0:0 | 1:0:0:0:0:0:0:0:0:0:0:0:0:0:0:0:0:0:0:0:0 | 277:22:13:5:3:2:13:2:28:0:51:12:8:1:1:0:0:0:0:0:0 | 0:0:0:0:0:0:0:0:0:0:0:0:0:0:0:0:0:0:0:0:0 | 1566:144:9:14:31:22:76:1:130:3:191:54:26:4:2:1:1:1:2:0:0 | 3:0:0:0:0:0:0:0:0:0:0:0:0:0:0:0:0:0:0:0:0 | 604:25:2:3:2:4:35:1:35:2:100:20:12:1:1:0:1:0:0:0:0 | 1:0:0:0:0:0:0:0:0:0:0:0:0:0:0:0:0:0:0:0:0 |
| XR_001744671:XM_017012110:NM_031243:XR_001744673:XR_242076:XR_002956425:XM_005249729:NM_002137:XR_242077:XR_001744669:XR_001744664:XR_001744674:XR_428077:XR_001744672:XR_001744675:XR_001744667:XM_017012109:XR_001744668:XR_002956424:XR_001744670:XR_001744666 | 36:15:15:6:6:11:1:20:1:402:81:3:8:0:0:0:0:0:0:0:0 | 0:0:0:0:0:0:0:0:0:0:0:0:0:0:0:0:0:0:0:0:0 | 22:20:2:0:3:12:3:20:1:280:40:7:8:2:0:0:0:0:0:0:0:0 | 0:0:0:0:0:0:0:0:0:0:0:0:0:0:0:0:0:0:0:0:0 | 22:13:5:3:2:13:2:28:0:276:51:12:7:1:1:0:0:0:0:0:0 | 0:0:0:0:0:0:0:0:0:0:0:0:0:0:0:0:0:0:0:0:0 | 144:9:14:31:21:74:1:130:3:1556:191:54:26:4:2:1:1:1:2:0:0 | 0:0:0:0:1:0:0:0:0:1:0:0:0:0:0:0:0:0:0:0:0 | 25:2:3:2:4:35:1:34:2:602:100:20:13:1:1:0:1:0:0:0:0 | 0:0:0:0:0:1:0:1:0:0:0:0:0:0:0:0:0:0:0:0:0 |
| XR_001744671:XM_017012110:NM_031243:XR_001744673:XR_242076:XR_002956425:XM_005249729:NM_002137:XR_242077:XR_001744669:XR_001744664:XR_001744674:XR_428077:XR_001744672:XR_001744675:XR_001744667:XM_017012109:XR_001744668:XR_002956424:XR_001744670:XR_001744666 | 36:15:15:6:6:11:1:20:1:402:81:3:8:0:0:0:0:0:0:0:0 | 0:0:0:0:0:0:0:0:0:0:0:0:0:0:0:0:0:0:0:0:0 | 22:20:2:0:3:12:3:20:1:279:40:7:8:2:0:0:0:0:0:0:0:0 | 0:0:0:0:0:0:0:0:1:0:0:0:0:0:0:0:0:0:0:0:0 | 22:13:5:3:2:13:2:28:0:276:51:12:7:1:1:0:0:0:0:0:0 | 0:0:0:0:0:0:0:0:0:0:0:0:0:0:0:0:0:0:0:0:0 | 144:9:14:31:21:74:1:130:3:1555:191:54:26:4:2:1:1:1:2:0:0 | 0:0:0:0:1:0:0:0:0:1:0:0:0:0:0:0:0:0:0:0:0 | 25:2:3:2:4:35:1:35:2:600:100:20:13:1:1:0:1:0:0:0:0 | 0:0:0:0:0:1:0:0:0:2:0:0:0:0:0:0:0:0:0:0:0 |

| | | | | | | | | | | |
|---|---|---|---|---|---|---|---|---|---|---|
| XR_001744671:XR_001744669:XM_017012110:NM_031243:XR_001744673:XR_242076:XR_002956425:XM_005249729:NM_002137:XR_242077:XR_001744664:XR_001744674:XR_428077:XR_001744672:XR_001744675:XR_001744667:XM_017012109:XR_002956424:XR_001744670:XR_001744666 | 35:397:14:15:6:5:11:1:20:1:81:3:5:0:0:0:0:0:0:0 | 1:1:0:0:0:0:0:0:0:0:0:0:0:0:0:0:0:0:0:0 | 22:263:20:2:0:3:11:2:20:1:40:7:9:2:0:0:0:0:0:0 | 0:2:0:0:0:0:0:1:0:0:0:0:0:0:0:0:0:0:0:0 | 22:273:12:5:3:1:14:2:28:0:51:11:5:1:1:0:0:0:0:0 | 0:0:0:0:0:0:0:0:0:0:1:0:0:0:0:0:0:0:0:0 | 144:1520:9:14:31:19:77:1:130:3:191:54:20:4:2:1:1:2:0:0 | 0:3:0:0:0:0:0:0:0:0:0:0:0:0:0:0:0:0:0:0 | 25:586:2:3:2:3:37:1:35:2:99:20:12:1:1:0:1:0:0:0 | 0:2:0:0:0:0:1:0:0:0:1:0:0:0:0:0:0:0:0:0 |
| XR_001744664:XR_001744669:XR_001744671:XM_017012110:NM_031243:XR_001744673:XR_242076:XR_002956425:XM_005249729:NM_002137:XR_242077:XR_001744674:XR_428077:XR_001744672:XR_001744675:XR_001744667:XM_017012109:XR_002956424:XR_001744670:XR_001744666 | 80:395:36:13:14:6:5:11:1:20:1:3:4:0:0:0:0:0:0:0 | 1:1:0:0:0:0:0:0:0:0:0:0:0:0:0:0:0:0:0:0 | 40:258:22:19:2:0:3:12:3:20:1:7:5:2:0:0:0:0:0:0 | 0:1:0:0:0:0:0:0:0:0:0:0:0:0:0:0:0:0:0:0 | 51:268:22:11:3:3:1:14:2:28:0:12:5:1:1:0:0:0:0:0 | 0:0:0:0:0:0:0:0:0:0:0:0:0:0:0:0:0:0:0:0 | 190:1479:144:9:13:31:14:75:1:130:2:54:16:4:2:1:1:2:0:0 | 1:2:0:0:0:0:0:0:0:0:0:0:0:0:0:0:0:0:0:0 | 99:566:25:2:3:2:3:38:1:35:2:20:8:1:1:0:1:0:0:0 | 0:0:0:0:0:0:0:0:0:0:0:0:0:0:0:0:0:0:0:0 |
| NM_152740:XM_017011719 | 114:08:00 | 0:00 | 77:09:00 | 0:00 | 156:10:00 | 0:00 | 448:57:00 | 3:00 | 90:03:00 | 0:00 |
| NM_152740:XM_017011719 | 114:07:00 | 0:00 | 77:08:00 | 0:00 | 157:08:00 | 0:00 | 485:55:00 | 3:00 | 92:03:00 | 2:00 |
| NM_152740:XM_017011719 | 114:04:00 | 0:00 | 77:03:00 | 0:00 | 157:05:00 | 0:00 | 486:20:00 | 2:00 | 94:00:00 | 0:00 |
| NM_152740:XM_017011719 | 112:01:00 | 2:00 | 76:03:00 | 0:00 | 155:03:00 | 1:00 | 473:06:00 | 2:00 | 93:00:00 | 0:00 |
| NM_152740:XM_017011719 | 110:01:00 | 0:00 | 72:02:00 | 0:00 | 150:04:00 | 1:00 | 441:05:00 | 0:00 | 90:00:00 | 0:00 |
| NM_006024:NM_001079864:NM_001206902:NM_001206901 | 45:102:1:4 | 0:0:0:0 | 58:44:0:15 | 0:0:0:0 | 66:92:1:2 | 0:0:0:0 | 122:145:4:44 | 0:1:0:0 | 45:65:1:6 | 0:0:0:0 |
| NM_001079864:NM_006024:NM_001206902:NM_001206901 | 103:45:1:4 | 0:0:0:0 | 44:55:4:19 | 0:1:0:0 | 92:66:0:6 | 0:0:0:0 | 146:133:5:60 | 0:1:1:1 | 65:45:4:8 | 0:0:0:0 |
| NM_001079864:NM_006024:NM_001206902:NM_001206901 | 103:46:1:4 | 0:0:0:0 | 44:57:6:19 | 0:0:0:0 | 92:66:1:6 | 0:0:0:0 | 146:135:12:61 | 0:0:0:0 | 65:45:6:8 | 0:0:0:0 |
| NM_001079864:NM_006024:NM_001206902:NM_001206901 | 103:45:1:4 | 0:0:0:0 | 44:57:6:19 | 0:0:1:0 | 92:66:1:6 | 0:0:0:0 | 145:135:12:61 | 1:0:1:0 | 65:46:6:8 | 0:0:0:0 |
| NM_001079864:NM_006024:NM_001206902:NM_001206901 | 103:45:5:4 | 0:0:0:0 | 44:62:24:19 | 0:0:0:0 | 91:68:4:6 | 1:0:0:0 | 146:154:67:61 | 0:1:2:0 | 65:45:9:8 | 0:0:0:0 |
| NM_001079864:NM_006024:NM_001206902:NM_001206901 | 103:46:5:4 | 0:0:0:0 | 44:70:39:19 | 0:0:1:0 | 92:69:6:6 | 0:0:0:0 | 146:157:119:61 | 0:0:0:0 | 65:45:11:8 | 0:0:0:0 |
| NM_001079864:NM_006024:NM_001206902:NM_001206901 | 103:46:5:4 | 0:0:0:0 | 44:70:40:19 | 0:0:1:0 | 92:69:6:6 | 0:0:0:0 | 145:160:122:61 | 1:0:0:0 | 65:45:13:8 | 0:0:0:0 |
| NM_001079864:NM_006024:NM_001206902:NM_001206901 | 103:46:5:4 | 0:0:0:0 | 44:70:40:19 | 0:0:1:0 | 92:68:7:6 | 0:1:0:0 | 146:160:124:61 | 0:1:1:0 | 65:45:14:8 | 0:0:0:0 |
| NM_001079864:NM_006024:NM_001206902:NM_001206901 | 103:46:6:4 | 0:0:0:0 | 44:68:41:19 | 0:2:0:0 | 92:66:7:6 | 0:4:0:0 | 146:160:126:61 | 0:3:1:0 | 65:44:14:8 | 0:1:0:0 |
| NM_001079864:NM_006024:NM_001206902:NM_001206901 | 103:46:6:4 | 0:0:0:0 | 44:67:42:19 | 0:3:0:0 | 92:66:7:6 | 0:3:0:0 | 146:158:126:61 | 0:5:1:0 | 65:43:14:8 | 0:2:0:0 |
| NM_001079864:NM_006024:NM_001206902:NM_001206901 | 103:43:9:4 | 0:0:0:0 | 44:56:48:19 | 0:0:0:0 | 92:67:9:6 | 0:0:0:0 | 146:147:155:61 | 0:1:0:0 | 65:33:16:8 | 0:0:0:0 |
| NM_001079864:NM_006024:NM_001206902:NM_001206901 | 102:40:11:4 | 1:0:0:0 | 44:57:57:19 | 0:0:0:0 | 92:59:11:6 | 0:0:0:0 | 146:145:201:61 | 0:1:0:0 | 65:30:20:8 | 0:0:0:0 |
| NM_006024:NM_001079864:NM_001206902:NM_001206901 | 30:103:12:4 | 8:0:0:0 | 57:44:62:19 | 1:0:0:0 | 52:91:14:6 | 7:1:0:0 | 136:146:219:61 | 6:0:2:0 | 27:64:23:8 | 1:1:0:0 |
| NM_001079864:NM_006024:NM_001206902:NM_001206901 | 103:29:12:4 | 0:0:0:0 | 44:58:66:19 | 0:0:0:0 | 92:48:15:6 | 0:0:0:0 | 145:148:245:61 | 0:1:0:0 | 64:27:26:8 | 0:0:0:0 |
| NM_001079864:NM_006024:NM_001206902:NM_001206901 | 102:29:12:4 | 1:0:0:0 | 44:58:66:19 | 0:0:0:0 | 92:48:15:6 | 0:0:0:0 | 144:149:243:61 | 1:0:1:0 | 64:27:26:8 | 0:0:0:0 |
| NM_001079864:NM_006024:NM_001206902:NM_001206901 | 102:30:12:4 | 0:0:0:0 | 43:60:65:18 | 0:0:0:0 | 90:49:15:6 | 1:0:0:0 | 143:146:242:59 | 0:1:1:0 | 64:29:26:8 | 0:0:0:0 |
| NM_001079864:NM_006024:NM_001206902:NM_001206901 | 101:30:12:4 | 1:0:0:0 | 43:60:66:18 | 0:0:0:0 | 91:48:15:6 | 0:0:0:0 | 143:150:240:60 | 0:1:2:0 | 64:29:26:8 | 0:0:0:0 |
| NM_001079864:NM_006024:NM_001206902:NM_001206901 | 102:30:12:4 | 0:0:0:0 | 43:60:66:18 | 0:0:0:0 | 91:48:15:5 | 0:0:0:1 | 143:151:241:60 | 0:0:1:0 | 64:29:25:8 | 0:0:0:0 |
| NM_001079864:NM_006024:NM_001206902:NM_001206901 | 102:30:12:4 | 0:0:0:0 | 43:60:66:18 | 0:0:0:0 | 90:48:15:6 | 1:0:0:0 | 143:151:241:59 | 0:0:1:1 | 64:29:24:8 | 0:0:0:0 |
| NM_175887:XM_011515199:NM_001329996 | 0:00:00 | 0:00:00 | 1:01:00 | 0:00:00 | 0:00:00 | 0:00:00 | 0:00:00 | 3:01:00 | 0:00:00 | 2:01:00 |
| NM_175887:XM_011515199:NM_001329996 | 0:00:00 | 0:00:00 | 1:01:00 | 0:00:00 | 0:00:00 | 0:00:00 | 3:01:00 | 0:00:00 | 2:01:00 | 0:00:00 |
| NM_002047:NM_001316772 | 0:00 | 0:00 | 0:00 | 0:00 | 0:00 | 0:00 | 0:00 | 1:00 | 0:00 | 1:02 |
| XM_017012506:NM_022373:XR_926944:XR_001744846 | 3:1:0:0 | 2:3:0:0 | 1:0:1:0 | 2:3:0:0 | 2:1:0:0 | 3:2:0:0 | 13:4:0:0 | 17:7:0:1 | 2:2:0:0 | 0:0:0:0 |
| XM_017012102:NM_001637:XM_011515336:XM_017012106:XM_011515335:XM_011515340:XM_011515341:NM_001177507 | 2:51:1:0:0:0:0:0 | 0:0:0:0:0:0:0:0 | 0:13:1:1:0:0:0:0 | 0:0:0:0:0:0:0:0 | 0:18:0:0:1:0:0:0 | 0:0:0:0:0:0:0:0 | 6:24:0:0:1:1:0:0 | 0:0:0:0:0:0:0:0 | 0:10:0:0:0:0:0:0 | 0:0:0:0:0:0:0:0 |
| XM_017012102:NM_001637:XM_011515336:XM_011515339:XM_011515340:XM_017012106:XM_011515341:XM_011515335:XM_017012104:XM_011515338:NM_001177507 | 2:52:2:1:16:7:1:0:0:0:0 | 0:0:0:0:0:0:0:0:0:0:0 | 0:14:2:1:7:6:0:0:0:0:0 | 0:0:0:0:0:0:0:0:0:0:0 | 0:19:1:1:7:3:1:1:1:0:0 | 0:0:0:0:0:0:0:0:0:0:0 | 0:6:0:2:30:27:1:0:3:0:0 | 3:20:1:7:71:67:2:1:15:1:0 | 0:4:0:0:6:5:0:0:0:0:0 | 0:6:0:2:23:15:1:0:2:0:0 |
| NM_001637:XM_017012102:XM_011515336:XM_011515339:XM_011515340:XM_017012106:XM_011515341:XM_011515335:XM_017012104:XM_011515338:NM_001177507 | 49:2:2:1:16:9:1:0:0:0:0 | 1:0:0:0:0:0:0:0:0:0:0 | 12:0:1:1:7:6:0:0:0:0:0 | 0:0:0:0:0:0:0:0:0:0:0 | 17:0:1:1:7:3:1:1:1:0:0 | 1:0:0:0:0:0:0:0:0:0:0 | 23:3:1:9:101:91:3:1:16:1:0 | 0:0:0:0:0:2:0:0:0:0:0 | 10:0:0:2:29:20:1:0:2:0:0 | 0:0:0:0:0:0:0:0:0:0:0 |
| NM_001637:XM_017012106:XM_011515335:XM_011515340:XM_011515341:NM_001177507 | 0:0:0:0:0:0 | 6:1:0:0:0:0 | 0:0:0:0:0:0 | 2:0:0:0:0:0 | 0:0:0:0:0:0 | 6:0:1:0:0:0 | 0:0:0:0:0:0 | 2:0:0:1:0:0 | 0:0:0:0:0:0 | 1:0:0:0:0:0 |
| NM_020192:NM_001282446 | 0:00 | 0:00 | 0:00 | 0:00 | 0:00 | 0:00 | 3:00 | 0:00 | 0:00 | 0:00 |
| NM_004760:XM_017012792 | 0:00 | 0:00 | 0:00 | 3:00 | 0:00 | 2:00 | 0:00 | 0:00 | 0:00 | 0:01 |
| NM_032014 | 3 | 0 | 1 | 0 | 3 | 0 | 8 | 0 | 7 | 0 |
| XM_011515162:NM_001129 | 0:00 | 0:00 | 1:00 | 0:00 | 0:00 | 0:00 | 0:00 | 0:00 | 0:00 | 0:00 |
| NM_201284:NM_005228 | 0:00 | 0:00 | 0:01 | 1:00 | 0:01 | 0:00 | 0:00 | 0:01 | 0:00 | 0:00 |
| NM_001101 | 87 | 0 | 177 | 2 | 134 | 0 | 459 | 0 | 250 | 1 |
| NM_001101 | 85 | 0 | 175 | 3 | 133 | 0 | 446 | 3 | 247 | 2 |
| NM_001101 | 82 | 0 | 166 | 2 | 132 | 0 | 443 | 2 | 242 | 1 |
| NM_001101 | 80 | 1 | 159 | 1 | 131 | 0 | 442 | 1 | 240 | 0 |
| NM_001101 | 73 | 1 | 120 | 0 | 122 | 1 | 358 | 2 | 212 | 1 |
| NM_001101 | 73 | 1 | 115 | 0 | 122 | 1 | 353 | 3 | 211 | 1 |
| XR_002956470 | 0 | 0 | 0 | 0 | 0 | 0 | 0 | 1 | 0 | 1 |
| XR_002956470 | 0 | 0 | 0 | 0 | 0 | 0 | 0 | 1 | 0 | 1 |
| XR_002956470 | 0 | 0 | 0 | 0 | 0 | 0 | 0 | 1 | 0 | 1 |
| NM_015411 | 0 | 1 | 0 | 0 | 0 | 1 | 0 | 1 | 0 | 1 |
| XM_017012363:NM_207116:XM_024446805:NM_207111:XM_017012364:XM_024446807:XM_024446806 | 0:1:1:0:0:0:0 | 1:0:0:0:0:0:0 | 0:3:0:1:0:1:0 | 0:8:0:1:3:0:0 | 0:2:1:0:0:0:0 | 0:0:0:0:0:0:0 | 1:1:3:3:0:7:0 | 0:4:0:3:0:0:3 | 0:1:2:0:0:0:0 | 0:0:0:0:0:0:0 |
| NM_015622 | 1 | 0 | 0 | 0 | 1 | 1 | 0 | 0 | 1 | 0 |

| | | | | | | | | | | |
|---|---|---|---|---|---|---|---|---|---|---|
| NM_015622 | 1 | 0 | 0 | 0 | 1 | 1 | 0 | 0 | 1 | 0 |
| NM_015622 | 1 | 0 | 0 | 0 | 2 | 0 | 0 | 0 | 0 | 1 |
| NM_001322005:NM_000535:NM_001322006:NM_001322015:XM_024446800:NM_001322004:NM_001322003:NR_136154:NM_001322008:NM_001322007:XM_017012342:NM_001322014:NM_001322012:NM_001322011 | 1:6:0:0:0:0:0:0:0:0:0:0:0:0:0 | 0:0:0:0:0:0:0:0:0:0:0:0:0:0:0 | 0:0:0:1:1:0:0:0:0:0:0:0:0:0:0 | 0:0:1:0:0:0:0:0:0:0:0:0:0:0:0 | 1:6:0:1:0:0:1:0:0:0:0:0:0:0:0 | 0:0:0:0:0:1:0:0:0:0:0:0:0:0:0 | 5:10:0:13:14:0:0:4:0:0:4:2:0:0 | 0:0:0:0:0:0:0:5:1:1:0:0:0:0 | 0:7:0:0:0:0:2:0:0:0:0:0:0:0 | 0:0:0:0:0:0:0:0:0:0:0:0:0:0:0 |
| NM_001322005:NM_000535:NM_001322015:XM_024446800:NM_001322006:NM_001322003:NM_001322004:NR_136154:NM_001322014:XM_017012342:NM_001322008:NM_001322007:NM_001322012:NM_001322011 | 1:6:0:0:0:0:0:0:0:0:0:0:0:0:0 | 0:0:0:0:0:0:0:0:0:0:0:0:0:0:0 | 0:0:1:1:1:0:0:0:0:0:0:0:0:0:0 | 0:0:0:0:0:0:0:0:0:0:0:0:0:0:0 | 1:6:1:1:0:1:1:0:0:0:0:0:0:0:0 | 0:0:0:0:0:0:0:0:0:0:0:0:0:0:0 | 3:7:5:6:0:0:0:6:1:3:0:1:0:0 | 2:3:8:7:0:0:0:4:1:1:1:0:0:0 | 0:4:0:0:0:2:0:0:0:0:0:0:0:0 | 0:3:0:0:0:0:0:0:0:0:0:0:0:0:0 |
| NM_001322005:NM_000535:NM_001322015:XM_024446800:NM_001322006:NR_136154:NM_001322003:NM_001322004:XM_017012342:NM_001322014:NM_001322007:NM_001322012:NM_001322008:NM_001322011 | 0:0:0:0:0:0:0:0:0:0:0:0:0:0:0 | 1:6:0:0:0:0:0:0:0:0:0:0:0:0:0 | 0:0:0:0:0:0:0:0:0:0:0:0:0:0:0 | 0:0:1:1:1:1:1:1:0:0:0:0:0:0:0 | 0:0:0:0:0:0:0:0:0:0:0:0:0:0:0 | 1:6:1:1:0:0:1:1:0:0:0:0:0:0:0 | 0:0:0:0:0:0:0:0:0:0:0:0:0:1:0 | 5:10:13:10:0:22:0:0:4:2:1:1:0:0 | 0:0:0:0:0:0:0:0:0:0:0:0:0:0 | 0:7:0:0:0:1:3:0:1:0:0:0:0:0 |
| NM_001322005:NM_000535:NM_001322015:XM_024446800:NM_001322006:NR_136154:NM_001322003:NM_001322004:XM_017012342:NM_001322014:NM_001322007:NM_001322008:NM_001322012:NM_001322011 | 0:0:0:0:0:0:0:0:0:0:0:0:0:0:0 | 1:6:0:0:0:0:0:0:0:0:0:0:0:0:0 | 0:0:0:0:0:0:0:0:0:0:0:0:0:0:0 | 0:0:1:1:1:1:1:1:0:0:0:0:0:0:0 | 0:0:0:0:0:0:0:0:0:0:0:0:0:0:0 | 1:6:1:1:0:0:1:1:0:0:0:0:0:0:0 | 2:3:8:5:0:9:0:0:1:1:0:1:1:0 | 3:7:5:4:0:13:0:0:3:1:1:0:0:0 | 0:3:0:0:0:0:1:0:0:0:0:0:0:0 | 0:4:0:0:0:1:2:0:1:0:0:0:0:0 |
| NM_006303:NM_001326610:NM_001326609:XM_005249847:NM_001326607 | 99:1:1:0:0 | 0:0:0:0:0 | 82:0:0:0:0 | 0:0:0:0:0 | 130:0:0:0:0 | 0:0:0:0:0 | 473:1:4:1:1 | 1:0:0:0:0 | 211:0:1:0:0 | 3:0:0:0:0 |
| NM_006303:NM_001326606:NM_001326610:NM_001326609:XM_005249847:NM_001326607 | 99:8:1:1:0:0 | 0:0:0:0:0:0 | 82:5:1:0:2:0 | 0:0:0:0:0:0 | 132:10:0:1:1:0 | 1:0:0:0:0:0 | 474:63:1:7:3:1 | 2:0:0:0:0:0 | 215:27:0:2:2:0 | 0:0:0:0:0:0 |
| NM_006303:NM_001326606:NM_001326610:NM_001326609:XM_005249847 | 99:8:1:1:0 | 0:0:0:0:0 | 81:5:1:0:2 | 0:0:0:0:0 | 133:10:0:1:1 | 0:0:0:0:0 | 476:63:1:7:3 | 0:0:0:0:0 | 215:27:0:2:2 | 0:0:0:0:0 |
| NM_006303:NM_001326606:NM_001326610:NM_001326609:XM_005249847:NM_001326607 | 98:7:1:1:0:0 | 0:0:0:0:0:0 | 81:5:1:0:2:0 | 0:0:0:0:0:0 | 132:10:0:1:1:0 | 1:0:0:0:0:0 | 471:63:1:7:3:1 | 2:0:0:0:0:0 | 214:27:0:2:2:0 | 0:0:0:0:0:0 |
| NM_006303:NM_001326606:NM_001326610:NM_001326609:XM_005249847:NM_001326607 | 98:7:1:1:0:0 | 0:0:0:0:0:0 | 81:5:1:0:2:0 | 0:0:0:0:0:0 | 132:10:0:1:1:0 | 0:0:0:0:0:0 | 471:63:1:7:3:1 | 0:0:0:0:0:0 | 214:27:0:2:2:0 | 0:0:0:0:0:0 |
| NM_006303:NM_001326606:NM_001326610:NM_001326609:XM_005249847:NM_001326607 | 98:8:1:1:0:0 | 0:0:0:0:0:0 | 81:5:1:0:2:0 | 0:0:0:0:0:0 | 131:10:0:1:1:0 | 1:0:0:0:0:0 | 471:63:1:7:3:1 | 0:0:0:0:0:0 | 213:27:0:2:2:0 | 1:0:0:0:0:0 |
| NM_001271700 | 0 | 0 | 0 | 0 | 0 | 0 | 0 | 0 | 0 | 0 |
| NM_014413:NM_001134335:XM_024446718 | 18:05:00 | 0:00:00 | 14:00:01 | 0:00:00 | 25:03:01 | 0:00:00 | 71:07:04 | 1:00:00 | 26:02:01 | 0:00:00 |
| NM_032172:XR_002956494 | 0:00 | 0:00 | 0:00 | 0:00 | 0:00 | 0:00 | 1:02 | 0:00 | 0:00 | 0:00 |
| NM_016038 | 1 | 0 | 1 | 0 | 1 | 0 | 2 | 2 | 0 | 0 |
| NM_005918:NR_104165:NM_001282404 | 1:00:00 | 0:00:00 | 0:00:00 | 0:00:00 | 2:00:00 | 2:00:00 | 9:00:00 | 0:00:00 | 10:00:00 | 0:00:00 |
| NM_001127360 | 0 | 0 | 0 | 0 | 0 | 1 | 0 | 2 | 0 | 0 |
| NR_024550:NM_001142327:XR_001744895:XM_017012877:XM_017012868:XM_024447023:XM_017012875:XM_017012874:NR_024549:XM_017012878:XM_017012870:XM_024447024:XM_017012871:XM_024447021:XR_002956500:XM_024447016:XM_024447014:XM_024447022:XM_024447025:XM_017012869:XM_024447020:XM_017012873:XM_024447026 | 2:4:1:1:1:1:2:0:0:0:0:0:0:0:0:0:0:0:0:0:0:0:0 | 0:0:0:0:0:0:0:0:0:0:0:0:0:0:0:0:0:0:0:0:0:0:0 | 1:0:3:3:2:0:0:3:2:1:1:1:2:1:2:0:0:0:0:0:0:0:0 | 0:0:0:0:0:0:0:0:0:0:0:0:0:0:0:0:0:0:0:0:0:0:0 | 3:4:1:0:0:0:1:1:1:0:0:1:0:0:1:1:1:1:0:0:0:0:0 | 0:0:0:0:0:0:0:0:0:0:0:0:0:0:0:0:0:0:0:0:0:0:0 | 1:3:13:10:0:0:0:3:2:0:1:1:2:0:3:0:0:0:4:1:1:1:0 | 0:0:0:0:0:0:0:0:0:0:0:0:0:0:0:0:0:0:0:0:0:0:0 | 0:2:0:1:1:0:0:1:0:0:0:1:0:0:0:0:0:2:1:1:0:0:0 | 0:0:0:0:0:0:0:0:0:0:0:0:0:0:0:0:0:0:0:0:0:0:0 |
| NM_001284309:NM_001284308:NM_006869:NM_001284311 | 0:0:0:0 | 16:1:0:0 | 0:0:0:0 | 16:0:10:1 | 0:0:0:0 | 56:0:17:0 | 0:0:0:0 | 53:0:19:0 | 0:0:0:0 | 6:0:4:0 |
| XM_017011654:XM_017011650:XM_017011649:XM_024446631:XM_017011653:XM_017011660:NM_147185:XM_017011658 | 3:1:2:1:0:0:0:0 | 0:0:0:0:0:0:0:0 | 7:0:15:24:21:24:6:1 | 0:0:0:1:4:1:0:0 | 3:0:4:7:21:24:2:0 | 0:0:0:0:1:0:0:0 | 7:0:2:11:2:8:0:0 | 0:0:0:1:0:0:0:0 | 1:0:1:6:1:1:0:0 | 0:0:0:0:0:0:0:0 |
| XM_017011654:XM_017011649:NM_147185:XM_017011653:XM_017011660:XM_024446631 | 1:1:0:0:0:0 | 0:0:0:0:0:0 | 4:6:1:1:7:14 | 0:0:0:0:0:0 | 0:1:0:7:8:6 | 0:0:0:0:0:0 | 0:0:0:1:2:4 | 2:0:0:0:1:3 | 0:0:0:0:0:3 | 0:0:0:1:0:2 |
| XM_024446631:XM_017011660 | 0:00 | 1:01 | 0:00 | 2:00 | 0:00 | 3:01 | 0:00 | 1:00 | 0:00 | 0:00 |
| NM_001282678:NM_001282677:XR_242246 | 1:00:00 | 0:00:00 | 0:00:00 | 1:00:00 | 1:00:00 | 0:01:00 | 0:01:00 | 0:01:00 | 0:00:00 | 0:00:01 |
| NM_001039372:NM_001288810:XM_011516001:NM_198151:NM_001288804 | 0:0:0:0:0 | 0:0:0:0:0 | 56:2:33:23:0 | 1:0:0:0:0 | 258:2:47:95:2 | 3:0:0:1:0 | 2:0:2:0:0 | 0:0:0:0:0 | 6:0:1:0:0 | 0:0:0:0:0 |
| NM_001039372:NM_001288810:XM_011516001:NM_198151:NM_001288804 | 0:0:0:0:0 | 0:0:0:0:0 | 61:2:21:25:0 | 0:0:0:0:0 | 261:2:42:95:2 | 1:0:0:1:0 | 2:0:2:0:0 | 0:0:0:0:0 | 6:0:1:0:0 | 0:0:0:0:0 |
| NM_001039372:NM_001288810:XM_011516001:NM_198151:NM_001288804 | 0:0:0:0:0 | 0:0:0:0:0 | 61:2:9:25:0 | 0:0:0:0:0 | 259:2:23:95:2 | 3:0:0:0:0 | 2:0:2:0:0 | 0:0:0:0:0 | 6:0:1:0:0 | 0:0:0:0:0 |
| NM_020186 | 0 | 0 | 1 | 0 | 1 | 0 | 0 | 0 | 0 | 2 |
| NM_181538 | 0 | 0 | 1 | 0 | 0 | 0 | 20 | 13 | 0 | 0 |
| NM_002568:XM_005250861 | 4164:129 | 4:02 | 1269:30:00 | 4:00 | 968:34:00 | 0:01 | 21216:637 | 12:08 | 1982:36:00 | 2:00 |
| NM_002568:XM_005250861 | 4143:141 | 2:00 | 1220:39:00 | 2:00 | 960:38:00 | 1:00 | 20965:813 | 18:01 | 1963:43:00 | 1:00 |
| NM_002568:XM_005250861 | 4135:141 | 8:00 | 1217:38:00 | 2:00 | 959:37:00 | 2:00 | 20958:808 | 15:05 | 1962:43:00 | 2:00 |
| NM_002568:XM_005250861 | 4137:144 | 3:00 | 1212:38:00 | 2:00 | 961:36:00 | 0:00 | 20915:807 | 23:04 | 1962:43:00 | 2:00 |
| NM_002568:XM_005250861 | 4132:143 | 5:00 | 1209:38:00 | 4:00 | 957:36:00 | 3:00 | 20892:812 | 28:02:00 | 1957:43:00 | 4:00 |
| NM_002568:XM_005250861 | 4128:142 | 5:00 | 1191:35:00 | 2:00 | 958:36:00 | 1:00 | 20852:798 | 26:03:00 | 1951:42:00 | 4:00 |
| NM_002568:XM_005250861 | 4130:142 | 3:00 | 1191:35:00 | 2:00 | 959:36:00 | 0:00 | 20862:801 | 16:00 | 1954:42:00 | 1:00 |
| NM_002568:XM_005250861 | 4108:138 | 6:00 | 1152:38:00 | 0:00 | 951:37:00 | 0:00 | 20697:845 | 26:02:00 | 1937:40:00 | 0:00 |
| NM_002568:XM_005250861 | 4075:120 | 4:00 | 1103:38:00 | 1:01 | 943:33:00 | 0:01 | 20446:671 | 32:03:00 | 1922:37:00 | 2:00 |
| NM_002568:XM_005250861 | 4070:116 | 5:03 | 1094:38:00 | 3:00 | 941:33:00 | 2:01 | 20407:643 | 29:07:00 | 1920:38:00 | 2:00 |
| NM_002568:XM_005250861 | 4038:107 | 9:00 | 1076:37:00 | 3:00 | 936:34:00 | 0:01 | 20247:583 | 52:07:00 | 1912:36:00 | 1:00 |
| NM_002568:XM_005250861 | 3967:116 | 10:04 | 1026:38:00 | 5:01 | 909:32:00 | 3:00 | 19856:604 | 61:25:00 | 1869:32:00 | 2:00 |
| NM_002568:XM_005250861 | 3949:119 | 9:00 | 1003:39:00 | 4:00 | 899:31:00 | 1:00 | 19740:630 | 53:01:00 | 1846:31:00 | 2:00 |
| NM_002568:XM_005250861 | 3923:121 | 10:00 | 984:38:00 | 7:01 | 890:32:00 | 1:00 | 19616:623 | 49:04:00 | 1823:30:00 | 6:00 |
| NM_002568:XM_005250861 | 3857:135 | 4:00 | 959:40:00 | 1:00 | 869:32:00 | 0:00 | 19183:752 | 19:03 | 1793:29:00 | 3:00 |
| NM_002568:XM_005250861 | 3775:129 | 11:00 | 933:42:00 | 3:00 | 836:27:00 | 2:01 | 18643:740 | 45:06:00 | 1751:27:00 | 4:00 |
| NM_002568:XM_005250861 | 3773:131 | 9:01 | 927:42:00 | 6:00 | 836:27:00 | 0:00 | 18623:742 | 35:06:00 | 1751:28:00 | 3:00 |
| NM_002568:XM_005250861 | 3712:122 | 17:01 | 907:40:00 | 1:00 | 822:24:00 | 2:00 | 17924:781 | 103:03:00 | 1731:32:00 | 2:00 |
| NM_002568:XM_005250861 | 3708:122 | 16:01 | 904:40:00 | 2:00 | 822:24:00 | 1:00 | 17876:779 | 82:04:00 | 1731:32:00 | 0:00 |
| NM_002568:XM_005250861 | 3701:123 | 15:00 | 898:40:00 | 3:00 | 821:25:00 | 1:00 | 17833:785 | 58:05:00 | 1728:32:00 | 1:00 |
| NM_002568:XM_005250861 | 3669:124 | 14:01 | 885:39:00 | 1:00 | 813:25:00 | 1:00 | 17450:812 | 64:03:00 | 1709:35:00 | 1:00 |
| NM_002568:XM_005250861 | 3643:129 | 16:00 | 878:40:00 | 2:00 | 806:24:00 | 1:00 | 17242:835 | 73:04:00 | 1693:36:00 | 4:00 |
| NM_002568:XM_005250861 | 3640:127 | 19:02 | 879:41:00 | 1:00 | 805:24:00 | 2:00 | 17244:841 | 76:05:00 | 1693:37:00 | 4:00 |
| NM_002568:XM_005250861 | 3647:129 | 10:01 | 874:41:00 | 5:00 | 802:25:00 | 4:00 | 17195:841 | 77:05:00 | 1694:39:00 | 1:00 |
| NM_002568:XM_005250861 | 3623:130 | 10:00 | 859:42:00 | 2:00 | 799:26:00 | 0:00 | 16899:842 | 76:02:00 | 1684:42:00 | 2:00 |
| NM_002568:XM_005250861 | 3615:128 | 18:02 | 854:42:00 | 6:00 | 799:26:00 | 0:00 | 16872:838 | 79:02:00 | 1684:42:00 | 2:00 |
| NM_002568:XM_005250861 | 3569:138 | 5:02 | 847:41:00 | 1:00 | 787:31:00 | 1:00 | 16544:890 | 19:02 | 1670:45:00 | 1:00 |
| NM_002568:XM_005250861 | 3554:140 | 6:01 | 840:41:00 | 3:00 | 785:33:00 | 1:01 | 16412:884 | 33:05:00 | 1668:46:00 | 0:00 |
| NM_002568:XM_005250861 | 3551:139 | 8:02 | 836:41:00 | 6:00 | 784:34:00 | 2:00 | 16396:884 | 32:05:00 | 1666:46:00 | 2:00 |
| NM_002568:XM_005250861 | 3365:138 | 3:00 | 738:36:00 | 0:00 | 729:35:00 | 2:00 | 14557:863 | 51:02:00 | 1593:48:00 | 0:00 |
| NM_002568:XM_005250861 | 3356:137 | 7:00 | 736:36:00 | 2:00 | 727:35:00 | 2:00 | 14500:860 | 60:03:00 | 1589:47:00 | 3:00 |
| NM_002568:XM_005250861 | 3340:142 | 5:00 | 722:35:00 | 2:00 | 721:36:00 | 2:00 | 14286:867 | 15:03 | 1584:50:00 | 3:00 |
| NM_002568:XM_005250861 | 3331:136 | 3:01 | 712:33:00 | 5:00 | 720:36:00 | 0:01 | 13984:855 | 34:03:00 | 1581:50:00 | 1:00 |
| NM_002568:XM_005250861 | 3327:138 | 1:00 | 715:33:00 | 2:00 | 719:35:00 | 0:01 | 13965:856 | 17:01 | 1579:50:00 | 2:00 |
| NM_002568:XM_005250861 | 3221:131 | 1:00 | 694:31:00 | 0:00 | 684:35:00 | 0:00 | 13204:814 | 8:01 | 1526:48:00 | 0:00 |

| | | | | | | | | | | |
|---|---|---|---|---|---|---|---|---|---|---|
| NM_002568:XM_005250861 | 3297:134 | 4:00 | 707:32:00 | 1:00 | 707:35:00 | 2:00 | 13513:841 | 15:02 | 1556:49:00 | 2:00 |
| NM_002568:XM_005250861 | 3297:134 | 3:00 | 706:32:00 | 1:00 | 708:35:00 | 1:00 | 13485:840 | 20:02 | 1556:48:00 | 2:00 |
| NM_002568:XM_005250861 | 3258:131 | 7:00 | 690:31:00 | 2:00 | 698:35:00 | 1:00 | 12956:831 | 45:02:00 | 1546:51:00 | 2:00 |
| NM_002568:XM_005250861 | 3252:131 | 3:00 | 685:30:00 | 0:01 | 698:34:00 | 0:00 | 12817:824 | 40:02:00 | 1542:51:00 | 3:00 |
| NM_002568:XM_005250861 | 3253:132 | 2:00 | 682:30:00 | 2:00 | 697:33:00 | 0:00 | 12757:821 | 24:00:00 | 1544:51:00 | 0:00 |
| NM_002568:XM_005250861 | 3215:130 | 2:00 | 658:30:00 | 2:00 | 692:32:00 | 0:00 | 12225:811 | 10:03 | 1534:50:00 | 2:00 |
| NM_002568:XM_005250861 | 3199:133 | 4:00 | 648:30:00 | 0:00 | 685:32:00 | 1:00 | 11992:801 | 30:05:00 | 1528:48:00 | 1:01 |
| NM_002568:XM_005250861 | 3201:133 | 1:00 | 647:30:00 | 0:00 | 686:32:00 | 0:00 | 11941:800 | 30:03:00 | 1528:48:00 | 0:00 |
| NM_002568:XM_005250861 | 3197:133 | 1:00 | 643:30:00 | 0:00 | 684:32:00 | 1:00 | 11895:799 | 23:00 | 1528:48:00 | 0:00 |
| NM_002568:XM_005250861 | 3161:129 | 8:01 | 616:29:00 | 4:01 | 678:32:00 | 1:00 | 11316:774 | 50:03:00 | 1513:45:00 | 1:01 |
| NM_002568:XM_005250861 | 2982:123 | 18:00 | 478:22:00 | 3:00 | 617:26:00 | 2:00 | 7955:593 | 55:04:00 | 1449:40:00 | 3:00 |
| NM_001040627:NM_032041:NM_001040625:XM_024447297:XM_024447299:NM_001040629:XM_024447304:XM_024447301:XM_024447296:NM_001040624:XM_024447303:XM_024447302:NM_001040628:XM_011517332:NM_001040626:XM_024447298 | 2:2:1:1:0:0:0:0:0:0:0:0:0:0:0:0 | 0:0:0:0:0:0:0:0:0:0:0:0:0:0:0:0 | 0:1:1:1:1:1:0:0:0:0:0:0:0:0:0:0 | 0:0:0:0:0:0:0:0:0:0:0:0:0:0:0:0 | 3:4:0:1:0:0:2:1:0:0:0:0:0:0:0:0 | 0:0:0:0:0:0:0:0:0:0:0:0:0:0:0:0 | 0:1:9:7:2:0:2:0:2:2:2:1:1:1:3:0 | 0:0:1:0:0:0:1:0:0:0:0:0:0:0:0:0 | 0:3:1:5:0:0:1:0:1:0:2:0:0:1:3:1 | 0:0:1:0:0:0:0:0:0:0:0:0:0:0:0:0 |
| NM_015902:XR_001745541:XM_024447179 | 1:00:00 | 1:00:00 | 0:01:00 | 0:02:00 | 0:00:00 | 0:00:00 | 0:01:01 | 1:07:04 | 0:00:00 | 0:00:00 |
| XM_017013566:NM_148174 | 0:00 | 0:00 | 0:00 | 0:00 | 0:00 | 0:00 | 0:00 | 0:00 | 1:00 | 0:00 |
| NM_030780:NR_102338:NR_102337 | 5:01:00 | 13:00:00 | 1:00:00 | 3:00:00 | 1:00:00 | 3:00:00 | 36:15:01 | 0:00:00 | 12:03:02 | 0:00:00 |
| NM_015420 | 2 | 3 | 0 | 0 | 1 | 0 | 6 | 0 | 2 | 0 |
| XM_017013594:NM_181354:XM_017013589:XM_017013595:XM_017013593:NM_001198533:XM_017013600:XM_017013598:XR_001745548:XM_017013590:XM_017013599:XM_006716595:XR_001745547 | 17:37:10:1:2:7:7:1:0:0:0:0:0 | 0:0:0:0:0:0:0:0:0:0:0:0:0 | 1:0:3:0:3:0:3:1:2:0:0:0:0 | 0:0:0:0:0:0:0:0:0:0:0:0:0 | 3:11:2:0:1:3:3:3:0:0:0:0:0 | 0:0:0:0:0:0:0:0:0:0:0:0:0 | 12:21:10:0:21:6:27:13:4:1:2:0:0 | 0:0:0:0:0:0:0:0:0:0:0:0:0 | 1:8:3:0:1:1:1:2:0:0:0:0:0 | 0:0:0:0:0:0:0:0:0:0:0:0:0 |
| XM_017013594:NM_181354:XM_017013589:XM_017013595:XM_017013593:NM_001198533:XM_017013600:XM_017013598:XR_001745548:XM_017013590:XM_017013599:XM_006716595:XR_001745547 | 17:39:10:1:2:8:9:2:0:0:0:0:0 | 0:0:0:0:0:0:0:0:0:0:0:0:0 | 1:0:3:0:3:0:4:3:2:0:0:0:0 | 0:0:0:0:0:0:0:0:0:0:0:0:0 | 3:12:2:0:1:3:4:3:0:0:0:0:0 | 0:0:0:0:0:0:0:0:0:0:0:0:0 | 12:24:9:0:21:9:43:24:1:1:2:0:0 | 0:0:0:0:0:0:1:1:0:0:0:0:0 | 1:8:3:0:1:2:2:3:0:0:0:0:0 | 0:0:0:0:0:0:0:0:0:0:0:0:0 |
| NM_001193557:NM_020189:NR_036472:NR_036471 | 54:343:3:19 | 0:0:0:0 | 70:167:3:9 | 0:0:0:0 | 50:120:2:16 | 0:1:0:0 | 1149:1622:28:135 | 0:2:0:0 | 82:256:4:28 | 0:1:1:0 |
| NM_001193557:NM_020189:NR_036472:NR_036471 | 54:346:3:19 | 0:0:0:0 | 70:169:3:9 | 0:0:0:0 | 50:122:2:16 | 0:0:0:0 | 1148:1649:27:137 | 0:0:0:0 | 82:260:5:28 | 0:0:0:0 |
| NM_001193557:NM_020189:NR_036472:NR_036471 | 54:346:4:19 | 0:0:0:0 | 69:169:4:9 | 1:1:0:0 | 50:122:3:16 | 1:0:0:0 | 1148:1645:27:137 | 0:2:0:0 | 82:259:6:28 | 0:1:0:0 |
| NM_001193557:NM_020189:NR_036472:NR_036471 | 54:343:5:19 | 0:0:0:0 | 70:168:5:9 | 0:1:0:0 | 51:122:3:15 | 0:0:0:0 | 1145:1639:31:137 | 2:7:0:0 | 82:258:6:28 | 0:2:0:0 |
| NM_001193557:NM_020189:NR_036472:NR_036471 | 54:343:5:19 | 0:0:0:0 | 70:166:5:9 | 0:1:0:0 | 49:121:3:15 | 0:0:0:0 | 1142:1637:35:137 | 0:1:0:0 | 82:260:6:28 | 0:0:0:0 |
| NM_003756 | 2589 | 1 | 543 | 0 | 542 | 3 | 14026 | 5 | 1493 | 2 |
| NM_003756 | 2589 | 1 | 543 | 0 | 545 | 1 | 14028 | 5 | 1495 | 1 |
| NM_003756 | 2590 | 2 | 542 | 1 | 547 | 0 | 14014 | 20 | 1495 | 2 |
| NM_003756 | 2591 | 1 | 543 | 0 | 545 | 2 | 14023 | 11 | 1494 | 3 |
| NM_003756 | 2611 | 3 | 545 | 0 | 550 | 3 | 14103 | 17 | 1517 | 3 |
| NM_003756 | 2616 | 3 | 545 | 1 | 557 | 0 | 14105 | 15 | 1521 | 1 |
| NM_003756 | 2618 | 2 | 544 | 2 | 557 | 1 | 14078 | 11 | 1519 | 0 |
| NM_003756 | 2615 | 4 | 546 | 0 | 558 | 0 | 14069 | 15 | 1518 | 1 |
| NM_003756 | 2616 | 1 | 545 | 0 | 556 | 2 | 14064 | 4 | 1516 | 1 |
| NM_003756 | 2615 | 0 | 543 | 0 | 556 | 0 | 14042 | 6 | 1513 | 1 |
| NM_003756 | 2616 | 1 | 547 | 3 | 558 | 0 | 14210 | 12 | 1512 | 2 |
| NM_003756 | 2617 | 0 | 549 | 1 | 558 | 0 | 14216 | 5 | 1513 | 1 |
| NM_003756 | 2612 | 5 | 550 | 0 | 559 | 0 | 14205 | 10 | 1510 | 3 |
| NM_003756 | 2609 | 0 | 549 | 0 | 559 | 0 | 14163 | 5 | 1500 | 0 |
| NM_003756 | 2600 | 1 | 550 | 0 | 557 | 0 | 14119 | 4 | 1493 | 0 |
| NM_003756 | 2596 | 3 | 549 | 1 | 557 | 0 | 14113 | 9 | 1490 | 3 |
| NM_003756 | 2596 | 3 | 550 | 0 | 556 | 1 | 14103 | 17 | 1492 | 1 |
| NM_003756 | 2580 | 2 | 550 | 0 | 547 | 0 | 14029 | 9 | 1492 | 1 |
| NM_003756 | 2541 | 5 | 544 | 1 | 537 | 2 | 13838 | 29 | 1473 | 1 |
| NM_003756 | 2538 | 3 | 544 | 0 | 539 | 1 | 13835 | 21 | 1470 | 2 |
| NM_003756 | 2526 | 8 | 540 | 2 | 536 | 2 | 13799 | 34 | 1462 | 6 |
| NM_003756 | 2522 | 7 | 541 | 1 | 535 | 1 | 13819 | 27 | 1461 | 2 |
| NM_006265 | 121 | 0 | 29 | 0 | 49 | 0 | 385 | 3 | 84 | 1 |
| NM_006265 | 121 | 1 | 24 | 0 | 49 | 0 | 363 | 0 | 87 | 0 |
| NM_006265 | 121 | 1 | 24 | 0 | 49 | 0 | 361 | 2 | 87 | 0 |
| NM_006265 | 162 | 0 | 73 | 1 | 68 | 1 | 721 | 30 | 118 | 2 |
| NM_006265 | 163 | 1 | 70 | 0 | 69 | 0 | 733 | 4 | 117 | 1 |
| NM_006265 | 164 | 0 | 69 | 0 | 69 | 0 | 726 | 2 | 117 | 1 |
| NM_006265 | 164 | 0 | 68 | 0 | 69 | 0 | 701 | 0 | 116 | 0 |
| NM_006265 | 163 | 0 | 68 | 1 | 68 | 0 | 672 | 1 | 114 | 0 |
| NM_006265 | 157 | 1 | 65 | 0 | 65 | 0 | 656 | 3 | 109 | 0 |
| NM_006265 | 160 | 0 | 66 | 2 | 68 | 1 | 652 | 6 | 108 | 1 |
| NM_006265 | 157 | 3 | 66 | 2 | 66 | 2 | 654 | 4 | 107 | 2 |
| NM_006265 | 158 | 0 | 66 | 0 | 69 | 0 | 647 | 2 | 107 | 0 |
| NM_006265 | 152 | 0 | 61 | 0 | 70 | 0 | 630 | 3 | 101 | 0 |
| NM_006265 | 144 | 0 | 54 | 2 | 63 | 0 | 604 | 1 | 85 | 0 |
| NM_000127 | 56 | 0 | 171 | 1 | 115 | 0 | 285 | 3 | 45 | 0 |
| NM_000127 | 55 | 0 | 168 | 0 | 114 | 1 | 288 | 2 | 45 | 0 |
| NM_000127 | 49 | 0 | 144 | 0 | 100 | 0 | 257 | 0 | 43 | 0 |
| NM_000127 | 48 | 0 | 143 | 0 | 97 | 0 | 255 | 0 | 44 | 0 |
| NM_000127 | 45 | 0 | 123 | 0 | 81 | 0 | 212 | 0 | 42 | 0 |
| NM_001287743:NM_004462:NM_001287756:NM_001287745:NM_001287742:NM_001287751 | 0:0:0:0:0:0 | 5:461:4:3:22:0 | 0:0:0:0:0:0 | 0:646:3:4:38:0 | 0:1:0:0:0:0 | 4:2393:10:9:120:0 | 29:964:2:4:49:0 | 0:2:0:0:0:0 | 5:664:2:2:26:0 | 0:0:0:0:0:0 |
| NM_001287743:NM_004462:NM_001287756:NM_001287745:NM_001287742:NM_001287744:NM_001287751 | 5:460:4:3:22:0:0 | 0:0:0:0:0:0:0 | 0:647:3:4:38:0:0 | 0:1:0:0:0:0:0 | 4:2402:10:9:120:1:0 | 0:2:0:0:0:0:0 | 29:962:1:4:49:0:0 | 0:0:0:0:0:0:0 | 5:663:1:2:25:0:0 | 0:0:0:0:0:0:0 |
| NM_004462:NM_001287743:NM_001287756:NM_001287742:NM_001287745:NM_001287744:NM_001287750 | 461:5:4:22:1:0:0 | 1:0:0:0:0:0:0 | 649:0:3:38:0:0:0 | 1:0:0:0:0:0:0 | 2406:4:10:120:1:1:0 | 5:0:0:0:0:0:0 | 971:29:1:48:2:0:0 | 2:0:0:1:0:0:0 | 665:5:1:26:0:0:1 | 2:0:0:0:0:0:0 |
| NM_001287743:NM_004462:NM_001287756:NM_001287742:NM_001287745:NM_001287744:NM_001287750 | 5:462:4:22:1:0:0 | 0:0:0:0:0:0:0 | 0:650:3:38:0:0:0 | 0:0:0:0:0:0:0 | 4:2410:10:120:1:1:0 | 0:3:0:0:0:0:0 | 29:972:1:48:2:0:0 | 0:2:0:1:0:0:0 | 5:667:1:26:0:0:1 | 0:0:0:0:0:0:0 |
| NM_001287742:NM_004462:NM_001287743:NM_001287756:NM_001287745:NM_001287744:NM_001287750:NM_001287751 | 21:461:5:4:1:0:0:0 | 1:1:0:0:0:0:0:0 | 37:649:0:3:0:0:0:0 | 1:4:0:0:0:0:0:0 | 120:2409:4:10:1:1:0:0 | 0:5:0:0:0:0:0:0 | 48:981:29:1:2:0:0:0 | 0:0:0:0:0:0:0:0 | 26:672:5:1:0:0:1:0 | 0:1:0:0:0:0:0:0 |

| | | | | | | | | | | |
|---|---|---|---|---|---|---|---|---|---|---|
| NM_004462:NM_001287743:NM_001287756:NM_001287742:NM_001287745:NM_001287744:NM_001287750:NM_001287751 | 461:5:4:22:1:0:0:0 | 1:0:0:0:0:0:0:0 | 650:0:3:38:0:0:0:0 | 3:0:0:0:0:0:0:0 | 2413:4:10:120:1:1:0:0 | 3:0:0:0:0:0:0:0 | 981:29:1:48:2:0:0:0 | 0:0:0:0:0:0:0:0 | 672:5:1:26:0:0:1:0 | 1:0:0:0:0:0:0:0 |
| NM_001287743:NM_004462:NM_001287756:NM_001287742:NM_001287745:NM_001287744:NM_001287750:NM_001287751 | 5:462:4:22:1:0:0:0 | 0:0:0:0:0:0:0:0 | 0:654:3:38:0:0:0:0 | 0:1:0:0:0:0:0:0 | 4:2413:10:120:1:1:0:0 | 0:3:0:0:0:0:0:0 | 29:978:1:48:2:0:0:0 | 0:0:0:0:0:0:0:0 | 5:673:1:26:0:0:1:0 | 0:0:0:0:0:0:0:0 |
| NM_001287743:NM_004462:NM_001287756:NM_001287742:NM_001287745:NM_001287744:NM_001287750:NM_001287751 | 5:464:4:22:1:0:0:0 | 0:0:0:0:0:0:0:0 | 0:652:3:38:0:0:0:0 | 0:3:0:0:0:0:0:0 | 4:2410:10:120:1:1:0:0 | 0:0:0:0:0:0:0:0 | 29:977:1:48:2:0:0:0 | 0:2:0:0:0:0:0:0 | 5:673:1:26:0:0:1:0 | 0:0:0:0:0:0:0:0 |
| NM_004462:NM_001287743:NM_001287756:NM_001287742:NM_001287745:NM_001287744:NM_001287750:NM_001287751 | 460:5:4:22:1:0:0:0 | 1:0:0:0:0:0:0:0 | 659:0:3:38:0:0:0:0 | 3:0:0:0:0:0:0:0 | 2407:4:10:118:1:1:0:0 | 3:0:0:1:0:0:0:0 | 981:27:1:48:2:0:0:0 | 0:1:0:0:0:0:0:0 | 674:5:1:26:0:0:1:0 | 2:0:0:0:0:0:0:0 |
| NM_004462:NM_001287743:NM_001287756:NM_001287742:NM_001287745:NM_001287744:NM_001287750:NM_001287751 | 460:5:4:22:1:0:0:0 | 1:0:0:0:0:0:0:0 | 660:0:3:38:0:0:0:0 | 0:0:0:0:0:0:0:0 | 2395:4:10:118:1:1:0:0 | 13:0:0:0:0:0:0:0 | 981:28:1:48:2:0:0:0 | 2:0:0:0:0:0:0:0 | 677:5:1:26:0:0:1:0 | 0:0:0:0:0:0:0:0 |
| NM_001287743:NM_004462:NM_001287756:NM_001287742:NM_001287745:NM_001287744:NM_001287750:NM_001287751 | 3:433:4:22:1:0:0:0 | 0:0:0:0:0:0:0:0 | 0:619:3:37:0:0:0:0 | 0:2:0:0:0:0:0:0 | 4:2232:10:112:1:1:0:0 | 0:3:0:0:0:0:0:0 | 26:941:1:44:2:0:0:0 | 0:4:0:0:0:0:0:0 | 5:669:1:26:0:0:1:0 | 0:0:0:0:0:0:0:0 |
| NM_001287743:NM_004462:NM_001287756:NM_001287742:NM_001287744:NM_001287745:NM_001287750:NM_001287751 | 3:427:4:22:0:0:0:0 | 0:0:0:0:0:0:0:0 | 0:615:3:36:0:0:0:0 | 0:0:0:0:0:0:0:0 | 4:2204:10:111:1:1:0:0 | 0:3:0:0:0:0:0:0 | 26:925:2:44:0:2:0:0 | 0:1:0:0:0:0:0:0 | 5:658:2:26:0:0:1:0 | 0:0:0:0:0:0:0:0 |
| NM_001287743:NM_004462:NM_001287756:NM_001287742:NM_001287744:NM_001287745:NM_001287750:NM_001287751 | 3:427:4:22:0:0:0:0 | 0:0:0:0:0:0:0:0 | 0:615:3:36:0:0:0:0 | 0:0:0:0:0:0:0:0 | 4:2208:10:111:1:1:0:0 | 0:2:0:0:0:0:0:0 | 26:930:2:43:0:2:0:0 | 0:0:0:1:0:0:0:0 | 5:655:2:26:0:0:1:0 | 0:2:0:0:0:0:0:0 |
| NM_001287743:NM_004462:NM_001287756:NM_001287742:NM_001287744:NM_001287745:NM_001287750:NM_001287751 | 3:427:4:22:0:0:0:0 | 0:0:0:0:0:0:0:0 | 0:614:3:36:0:0:0:0 | 0:0:0:0:0:0:0:0 | 4:2209:10:111:1:1:0:0 | 0:2:0:0:0:0:0:0 | 25:928:2:44:0:2:0:0 | 1:2:0:0:0:0:0:0 | 5:658:2:26:0:0:1:0 | 0:0:0:0:0:0:0:0 |
| NM_001287743:NM_004462:NM_001287756:NM_001287742:NM_001287745:NM_001287744:NM_001287750:NM_001287751 | 3:441:4:22:1:0:0:0 | 0:0:0:0:0:0:0:0 | 0:627:3:37:0:0:0:0 | 0:2:0:0:0:0:0:0 | 4:2262:10:111:1:1:0:0 | 0:4:0:0:0:0:0:0 | 27:953:2:43:2:0:0:0 | 0:0:0:0:0:0:0:0 | 5:676:2:26:0:0:1:0 | 0:1:0:0:0:0:0:0 |
| NM_004462:NM_001287743:NM_001287756:NM_001287742:NM_001287745:NM_001287744:NM_001287750:NM_001287751 | 436:3:4:22:1:0:0:0 | 1:0:0:0:0:0:0:0 | 639:0:3:36:0:0:0:0 | 1:0:0:0:0:0:0:0 | 2269:4:10:111:1:1:0:0 | 7:0:0:0:0:0:0:0 | 955:27:2:41:2:0:0:0 | 2:0:0:0:0:0:0:0 | 671:5:2:26:0:0:1:0 | 1:0:0:0:0:0:0:0 |
| NM_001287743:NM_004462:NM_001287756:NM_001287742:NM_001287745:NM_001287744:NM_001287750:NM_001287751 | 5:464:4:20:1:0:0:0 | 0:0:0:0:0:0:0:0 | 0:675:3:37:0:0:0:0 | 0:2:0:0:0:0:0:0 | 4:2451:10:116:1:1:0:0 | 0:4:0:0:0:0:0:0 | 28:997:2:47:2:0:0:0 | 0:1:0:0:0:0:0:0 | 5:672:2:26:0:0:1:0 | 0:0:0:0:0:0:0:0 |
| NM_001908:XM_024447071:NM_147782:NM_147780:NM_147783:XM_017013100:XM_017013097:XM_017013099:NM_001317237 | 20:2:2:1:0:0:0:0:0 | 8:1:1:0:0:0:0:0:0 | 9:2:1:0:0:0:0:0:0 | 10:0:2:0:0:0:0:0:0 | 11:1:1:0:0:1:1:0:0 | 10:1:2:0:1:0:0:0:0 | 111:21:12:0:0:0:0:1:0 | 0:0:0:0:0:0:0:0:0 | 27:6:3:0:0:0:0:0:0 | 0:0:0:0:0:0:0:0:0 |
| NM_001908:NM_147782:XM_024447071:NM_147780:XM_017013100:NM_147783:XM_017013097:XM_017013099:NM_001317237 | 0:0:0:0:0:0:0:0:0 | 28:3:5:1:0:0:0:0:0 | 0:0:0:0:0:0:0:0:0 | 19:3:2:0:0:0:0:0:0 | 0:0:0:0:0:0:0:0:0 | 21:3:2:0:1:1:1:0:0 | 111:12:28:0:0:0:0:1:0 | 0:0:0:0:0:0:0:0:0 | 27:3:6:0:0:0:0:0:0 | 0:0:0:0:0:0:0:0:0 |
| NM_001040092:XM_017013574:XM_017013575:NR_045555:NM_001130863:XM_017013572:XM_024447182:XM_006716587:NM_006209:XM_017013573:XM_024447181 | 14:2:10:1:2:2:0:0:0:0:0 | 3:1:1:0:0:0:0:0:0:0:0 | 1:2:0:0:0:0:2:1:0:0:0 | 0:0:0:0:0:0:0:0:0:0:0 | 0:0:3:0:0:0:0:0:0:0:0 | 3:0:1:0:0:0:0:0:1:0:0 | 0:0:0:0:0:0:0:0:0:0:0 | 2:2:2:0:0:0:2:1:0:0:0 | 0:0:0:0:0:0:0:0:0:0:0 | 3:2:1:0:1:0:2:1:0:0:0 |
| NM_024094:XM_005251065 | 0:00 | 0:00 | 0:00 | 0:00 | 0:00 | 0:00 | 0:00 | 0:00 | 0:00 | 1:00 |
| NM_001283024:NM_001283027:NM_018024:XM_017013603:XR_002956636:XM_011517151 | 15:6:19:1:1:0 | 0:0:0:0:0:0 | 3:0:6:0:0:1 | 0:0:0:0:0:0 | 1:5:1:1:0:0 | 0:0:0:0:0:0 | 37:29:46:1:0:15 | 0:0:0:0:0:0 | 5:6:11:0:0:2 | 0:0:0:0:0:0 |
| NM_001283024:NM_001283027:NM_018024:XM_017013603:XR_002956636:XM_011517151 | 15:6:19:1:1:0 | 0:0:0:0:0:0 | 3:0:6:0:0:1 | 0:0:0:0:0:0 | 1:5:1:1:0:0 | 0:0:0:0:0:0 | 37:29:46:1:0:14 | 0:0:0:0:0:0 | 5:6:11:0:0:2 | 0:0:0:0:0:0 |
| NM_001283027:NM_001283024:NM_018024:XM_017013603:XR_002956636:XM_011517151 | 5:15:19:1:1:0 | 1:0:0:0:0:0 | 0:3:6:0:0:1 | 0:0:0:0:0:0 | 5:1:1:1:0:0 | 0:0:0:0:0:0 | 29:37:45:1:0:15 | 0:0:0:0:0:0 | 6:5:11:0:0:2 | 0:0:0:0:0:0 |
| NM_001283024:NM_001283027:NM_018024:XM_017013603:XR_002956636:XM_011517151 | 15:6:19:1:1:0 | 0:0:0:0:0:0 | 3:0:6:0:0:1 | 0:0:0:0:0:0 | 1:5:1:1:0:0 | 0:0:0:0:0:0 | 37:29:45:1:0:18 | 0:0:1:0:0:0 | 5:6:11:0:0:3 | 0:0:0:0:0:0 |
| NM_001003954:NM_004306 | 0:00 | 0:00 | 0:00 | 0:00 | 0:00 | 0:00 | 0:00 | 0:00 | 0:00 | 0:00 |
| NM_005005:NM_001311168:NM_001278646:NM_001278645 | 4752:12:45:1 | 16:0:0:0 | 4098:31:34:3 | 10:0:0:0 | 6666:58:42:4 | 33:0:0:0 | 32693:119:325:9 | 35:0:0:0 | 3507:14:24:1 | 4:0:0:0 |
| NM_005005:NM_001311168:NM_001278646:NM_001278645 | 4772:12:45:1 | 11:0:0:0 | 4122:32:35:3 | 8:0:0:0 | 6682:58:42:4 | 30:0:0:0 | 32708:119:323:9 | 48:0:1:0 | 3522:13:24:1 | 3:1:0:0 |
| NM_005005:NM_001311168:NM_001278646:NM_001278645 | 4778:12:45:1 | 13:0:0:0 | 4126:32:35:3 | 13:0:0:0 | 6705:58:42:4 | 18:0:0:0 | 32723:119:323:9 | 47:0:0:0 | 3524:14:24:1 | 5:0:0:0 |
| NM_005005:NM_001311168:NM_001278646:NM_001278645 | 4795:12:45:1 | 2:0:0:0 | 4144:32:35:3 | 4:0:0:0 | 6717:58:42:4 | 13:0:0:0 | 32774:119:324:9 | 25:0:0:0 | 3533:14:24:1 | 1:0:0:0 |
| NM_005005:NM_001311168:NM_001278646:NM_001278645 | 4803:13:16:1 | 1:0:0:0 | 4168:33:22:3 | 2:0:0:0 | 6744:58:24:4 | 2:0:0:0 | 32779:120:130:9 | 25:0:1:0 | 3540:14:9:1 | 3:0:0:0 |
| NM_005005:NM_001278646:NM_001278645 | 4824:45:00 | 7:00:00 | 4197:35:00 | 8:00:00 | 6767:42:00 | 35:00:00 | 32842:324:0 | 43:00:00 | 3551:24:00 | 4:00:00 |
| NM_005005:NM_001278646:NM_001278645 | 4832:45:00 | 3:00:00 | 4200:35:00 | 9:00:00 | 6793:41:00 | 21:01:00 | 32870:322:0 | 46:01:00 | 3557:24:00 | 4:00:00 |
| NM_005005:NM_001311168:NM_001278646:NM_001278645 | 4858:13:45:1 | 5:0:0:0 | 4246:33:35:3 | 7:0:0:0 | 6853:58:42:4 | 9:0:0:0 | 32985:120:323:9 | 34:0:0:0 | 3573:14:24:1 | 5:0:0:0 |
| NM_005005:NM_001311168:NM_001278646:NM_001278645 | 4859:13:45:1 | 4:0:0:0 | 4248:33:35:3 | 10:0:0:0 | 6860:58:42:4 | 11:0:0:0 | 32986:120:323:9 | 48:0:0:0 | 3574:14:24:1 | 4:0:0:0 |
| NM_005005:NM_001311168:NM_001278646:NM_001278645 | 4863:13:45:1 | 1:0:0:0 | 4260:33:35:3 | 6:0:0:0 | 6865:58:42:4 | 8:0:0:0 | 33024:120:323:9 | 19:0:0:0 | 3576:14:24:1 | 3:0:0:0 |
| NM_005005:NM_001311168:NM_001278646:NM_001278645 | 4876:13:46:1 | 3:0:0:0 | 4276:33:35:3 | 2:0:0:0 | 6880:58:42:4 | 9:0:0:0 | 33096:120:321:9 | 36:0:2:0 | 3587:14:24:1 | 4:0:0:0 |
| NM_005005:NM_001311168:NM_001278646:NM_001278645 | 4880:13:46:1 | 3:0:0:0 | 4302:33:34:3 | 5:0:1:0 | 6895:58:41:4 | 5:0:1:0 | 33145:120:323:9 | 28:0:0:0 | 3604:14:24:1 | 1:0:0:0 |
| NM_005005:NM_001311168:NM_001278646:NM_001278645 | 4879:13:46:1 | 1:0:0:0 | 4293:32:35:3 | 6:0:0:0 | 6890:58:42:4 | 10:0:0:0 | 33134:120:323:9 | 27:0:0:0 | 3603:14:24:1 | 2:0:0:0 |
| NM_005005:NM_001311168:NM_001278646:NM_001278645 | 4862:13:46:1 | 5:0:0:0 | 4273:32:34:3 | 5:0:0:0 | 6878:58:42:4 | 16:0:0:0 | 33062:119:322:9 | 30:1:0:0 | 3586:14:24:1 | 3:0:0:0 |
| NM_005005:NM_001311168:NM_001278646:NM_001278645 | 4854:13:46:1 | 4:0:0:0 | 4260:31:33:3 | 7:0:1:0 | 6867:58:42:4 | 13:0:0:0 | 33004:118:322:9 | 38:1:0:0 | 3585:14:24:1 | 4:0:0:0 |
| NM_005005:NM_001311168:NM_001278646:NM_001278645 | 4842:13:46:1 | 9:0:0:0 | 4250:30:33:2 | 15:1:1:1 | 6852:58:42:4 | 18:0:0:0 | 32913:119:322:9 | 108:0:0:0 | 3579:14:24:1 | 8:0:0:0 |
| NM_005005:NM_001278646:NM_001311168:NM_001278645 | 4845:45:13:1 | 4:1:0:0 | 4249:34:31:3 | 10:0:0:0 | 6852:42:58:4 | 7:0:0:0 | 32952:322:119:9 | 48:0:0:0 | 3582:24:14:1 | 3:0:0:0 |
| NM_005005:NM_001311168:NM_001278646:NM_001278645 | 4832:13:45:1 | 7:0:0:0 | 4240:30:35:3 | 7:0:0:0 | 6841:57:41:4 | 11:0:1:0 | 32949:119:322:9 | 41:0:0:0 | 3583:14:24:1 | 7:0:0:0 |
| NM_005005:NM_001311168:NM_001278646:NM_001278645 | 4832:13:45:1 | 1:0:0:0 | 4242:30:35:3 | 3:0:0:0 | 6840:57:42:4 | 4:0:0:0 | 32970:118:322:9 | 10:0:0:0 | 3589:14:24:1 | 0:0:0:0 |

| | | | | | | | | | | |
|---|---|---|---|---|---|---|---|---|---|---|
| NM_005005:NM_001311168:NM_001278646:NM_001278645 | 4827:13:45:1 | 6:0:0:0 | 4241:30:35:3 | 2:0:0:0 | 6826:57:42:4 | 17:0:0:0 | 32942:118:322:9 | 34:0:0:0 | 3584:14:24:1 | 5:0:0:0 |
| NM_005005:NM_001311168:NM_001278646:NM_001278645 | 4819:13:45:1 | 9:0:0:0 | 4239:30:35:3 | 5:0:0:0 | 6820:56:42:4 | 12:0:0:0 | 32930:117:322:9 | 32:0:0:0 | 3585:14:24:1 | 2:0:0:0 |
| NM_005005:NM_001311168:NM_001278646:NM_001278645 | 4815:13:45:1 | 13:0:0:0 | 4225:30:35:3 | 13:0:0:0 | 6801:56:42:4 | 19:0:0:0 | 32887:117:319:9 | 41:0:3:0 | 3582:14:23:1 | 3:0:1:0 |
| NM_005005:NM_001311168:NM_001278646:NM_001278645 | 4811:13:45:1 | 8:0:0:0 | 4230:30:35:3 | 5:0:0:0 | 6796:56:42:4 | 19:0:0:0 | 32887:117:322:9 | 28:0:0:0 | 3582:14:24:1 | 2:0:0:0 |
| NM_005005:NM_001311168:NM_001278646:NM_001278645 | 4806:13:45:1 | 8:0:0:0 | 4218:30:35:3 | 11:0:0:0 | 6774:56:42:4 | 31:0:0:0 | 32863:116:322:9 | 42:1:0:0 | 3580:14:24:1 | 4:0:0:0 |
| NM_005005:NM_001311168:NM_001278646:NM_001278645 | 4801:13:45:1 | 7:0:0:0 | 4220:30:35:3 | 8:0:0:0 | 6784:56:42:4 | 20:0:0:0 | 32879:117:322:9 | 23:0:0:0 | 3583:14:24:1 | 0:0:0:0 |
| NM_005005:NM_001311168:NM_001278646:NM_001278645 | 4780:13:45:1 | 14:0:0:0 | 4202:30:35:3 | 10:0:0:0 | 6740:56:42:4 | 40:0:0:0 | 32788:117:322:9 | 41:0:0:0 | 3574:14:24:1 | 5:0:0:0 |
| NM_002467:NM_001354870 | 19:00 | 0:00 | 6:00 | 0:00 | 14:07 | 0:00 | 125:17:00 | 0:00 | 24:02:00 | 1:00 |
| NM_012472:NR_073525:XM_017013297:XM_011516950:XM_017013298:NR_135913:NM_001321962:XM_017013296:NR_135907:NR_135906:NM_001321961:NR_135909 | 0:1:0:0:0:0:0:0:0:0:0:0 | 1:0:0:0:0:0:0:0:0:0:0 | 0:0:0:0:0:0:0:0:0:0:0 | 0:0:1:0:0:0:0:0:0:0:0 | 2:1:0:0:0:0:0:0:0:0:0 | 3:0:0:1:0:0:0:0:0:0:0 | 6:5:1:0:3:1:1:2:0:0:0:0 | 0:0:0:0:0:0:0:0:0:0:0 | 3:2:0:0:0:0:0:0:2:0:0:0 | 0:0:0:0:0:0:0:0:0:0:0 |
| NM_012472:NR_073525:XM_017013296:XM_017013297:NR_135907:NR_135909:NM_001321962:NR_135908:XM_011516950:NR_135913:NR_135906:NM_001321961:XM_017013298:NM_001321963 | 0:2:0:0:0:0:0:0:0:0:0:0:0:0 | 1:1:1:0:0:0:0:0:0:0:0:0:0:0 | 0:1:0:0:0:0:0:2:0:0:0:0:0:0 | 0:3:0:1:3:1:1:0:0:0:0:0:0:0 | 3:8:0:0:0:0:0:0:0:0:0:0:0:0 | 3:8:0:0:0:0:0:0:1:0:0:0:0:0 | 5:13:0:2:0:0:1:0:0:0:0:0:0:0 | 3:14:16:0:0:0:8:0:0:4:0:0:0:0 | 3:2:0:0:2:0:0:0:0:0:0:0:0:0 | 0:2:1:0:0:0:0:0:0:0:0:0:0:0 |
| NM_006096:XM_011516792:XM_011516791 | 52:01:00 | 0:00:00 | 43:04:00 | 0:00:00 | 82:00:00 | 0:00:00 | 54:03:00 | 0:00:00 | 13:00:00 | 0:00:00 |
| NM_006096:XM_011516792:XM_011516791 | 52:01:00 | 0:00:00 | 43:04:00 | 0:00:00 | 80:00:00 | 2:00:00 | 52:03:00 | 0:00:00 | 13:00:00 | 0:00:00 |
| NM_006096:XM_011516792:XM_011516791 | 52:01:00 | 0:00:00 | 33:03:06 | 1:01:00 | 81:00:00 | 0:00:00 | 48:03:04 | 0:00:00 | 13:00:00 | 0:00:00 |
| NM_006096:XM_011516791:XM_011516792 | 51:01:03 | 0:00:00 | 33:08:06 | 0:00:00 | 89:03:00 | 0:00:00 | 12:00:05 | 36:08:10 | 5:00:00 | 7:00:00 |
| NM_138465 | 0 | 1 | 0 | 0 | 0 | 0 | 0 | 1 | 0 | 0 |
| NM_001916:XM_024447072:XM_017013102 | 0:00:00 | 5:01:00 | 0:00:00 | 1:00:00 | 0:00:00 | 4:01:01 | 0:00:00 | 12:02:00 | 0:00:00 | 6:00:00 |
| NM_001916:XM_024447072:XM_017013102 | 5:01:00 | 0:00:00 | 1:00:00 | 0:00:00 | 4:01:01 | 0:00:00 | 12:02:00 | 0:00:00 | 2:00:00 | 4:00:00 |
| NM_032272 | 0 | 0 | 1 | 0 | 0 | 0 | 0 | 2 | 0 | 0 |
| NM_024555:NM_012162 | 0:00 | 1:00 | 0:00 | 0:00 | 0:00 | 0:01 | 0:00 | 4:01 | 0:00 | 1:00 |
| XM_011516910:XR_928305:XM_006716527:NM_174922 | 0:0:0:0 | 0:0:0:0 | 0:0:0:0 | 1:0:0:0 | 0:0:0:0 | 0:0:0:0 | 0:0:0:0 | 0:0:0:0 | 0:0:0:0 | 0:0:0:0 |
| NM_130849 | 0 | 0 | 0 | 0 | 0 | 0 | 0 | 10 | 0 | 0 |
| NM_130849 | 0 | 0 | 0 | 0 | 0 | 0 | 0 | 10 | 0 | 0 |
| NM_130849 | 0 | 0 | 0 | 0 | 0 | 0 | 0 | 6 | 0 | 0 |
| NM_130849 | 0 | 0 | 0 | 0 | 0 | 0 | 0 | 6 | 0 | 0 |
| NM_130849 | 0 | 0 | 0 | 0 | 0 | 0 | 0 | 6 | 0 | 0 |
| XM_017014001 | 0 | 0 | 0 | 0 | 0 | 0 | 0 | 0 | 0 | 0 |
| NM_014066:NM_001081003:NM_001081004 | 0:00:00 | 2:00:00 | 0:00:00 | 0:00:00 | 0:00:00 | 0:00:00 | 0:00:00 | 0:01:01 | 0:00:00 | 0:00:00 |
| XM_024447170:XM_017013496:XM_024447168:XM_024447167:XM_017013493:XM_006716340:XM_024447171:NM_001352641 | 1:0:0:0:0:0:0:0 | 0:0:0:0:0:0:0:0 | 0:1:1:1:0:0:0:0 | 0:0:0:0:0:0:0:0 | 0:3:0:0:1:1:0:0 | 0:0:0:0:0:0:0:0 | 0:0:0:0:0:0:1:0 | 0:1:5:1:0:0:8:1 | 0:0:0:0:0:0:0:0 | 0:0:0:0:0:2:0:0 |
| NM_177924 | 0 | 1 | 0 | 0 | 0 | 0 | 0 | 0 | 0 | 0 |
| NM_177924 | 0 | 1 | 1 | 0 | 0 | 0 | 0 | 0 | 0 | 0 |
| NM_177924 | 0 | 1 | 1 | 0 | 0 | 0 | 0 | 0 | 0 | 0 |
| NM_001136108 | 0 | 0 | 0 | 0 | 0 | 2 | 0 | 0 | 0 | 0 |
| NM_001136108 | 0 | 0 | 0 | 0 | 0 | 2 | 0 | 0 | 0 | 0 |
| NM_001136108 | 0 | 0 | 0 | 0 | 0 | 2 | 0 | 0 | 0 | 0 |
| NM_001136108 | 0 | 0 | 0 | 0 | 0 | 2 | 0 | 0 | 0 | 0 |
| XR_002956638:NM_002717:XM_011544567:XM_005273559:XM_024447187:XM_011544568:XM_005273560:NM_001177591:XM_011544566:XM_024447186:XM_011544569:XM_011544565:XR_949430 | 0:18:2:1:1:1:0:0:0:0:0:0:0 | 2:0:0:0:0:0:0:0:0:0:0:0:0 | 3:22:0:0:1:0:1:1:1:1:0:0:0 | 1:2:0:0:0:0:0:0:0:0:0:0:0 | 0:12:1:0:0:0:0:1:0:0:0:0:0 | 0:0:0:0:0:0:0:0:0:0:0:0:0 | 4:37:4:1:4:0:0:0:1:3:1:0:0 | 2:2:1:0:0:0:1:0:0:0:0:0:0 | 0:18:0:2:0:0:0:0:1:0:0:0:0 | 2:1:0:0:0:0:0:0:0:0:0:0:0 |
| NM_002717:XM_011544567:XM_005273559:XM_024447187:XM_011544568:NM_001177591:XM_011544566:XM_024447186:XR_949430:XM_011544569:XM_005273560:XM_011544565 | 19:2:1:1:1:0:0:0:0:0:0:0 | 0:0:0:0:0:0:0:0:0:0:0:0 | 19:0:0:3:0:1:1:1:1:0:0:0 | 0:0:0:0:0:0:0:0:0:0:0:0 | 11:1:0:1:0:1:0:0:0:0:0:0 | 1:0:0:1:0:0:0:0:0:0:0:0 | 30:3:0:18:0:0:1:3:2:0:0:0 | 2:0:0:0:0:0:0:0:0:1:0:0 | 19:0:2:2:0:0:1:0:0:0:0:0 | 0:0:0:0:0:0:0:0:0:0:0:0 |
| NM_001979:XR_001745491:XM_017013199:XM_017013200 | 0:0:0:0 | 23:1:0:0 | 16:2:1:2 | 25:5:2:2 | 52:2:1:2 | 54:3:3:0 | 48:13:3:0 | 0:0:0:0 | 8:0:1:0 | 0:0:0:0 |
| NM_001979:XR_001745491:XM_017013199 | 22:01:00 | 0:00:00 | 25:04:01 | 15:00:01 | 56:02:02 | 52:00:02 | 59:03:03 | 0:00:00 | 8:00:01 | 0:00:00 |
| XR_949472:XM_011544639 | 0:00 | 2:00 | 0:00 | 4:00 | 2:02 | 3:01 | 5:01 | 0:00 | 1:00 | 0:00 |
| XM_024447265:XR_949472:XM_011544639 | 1:00:00 | 0:00:00 | 4:01:00 | 0:00:00 | 0:00:01 | 0:01:02 | 8:01:00 | 0:00:00 | 1:00:00 | 0:01:00 |
| XM_024447265:XM_011544639 | 1:00 | 0:00 | 6:00 | 0:00 | 0:02 | 0:00 | 0:00 | 11:00 | 0:00 | 2:00 |
| NM_013958:NM_004495:NM_013956:NM_013964:XM_024447143:NM_001160008:NM_013960 | 0:0:0:0:0:0:0 | 1:0:0:0:0:0:0 | 0:1:0:3:1:1:0 | 1:1:1:0:0:0:0 | 0:0:0:1:2:1:0 | 0:0:0:0:0:0:0 | 0:0:0:0:0:0:0 | 0:0:0:0:0:0:0 | 0:2:1:0:0:0:0 | 1:4:0:0:0:0:0 |
| NM_001160008:NM_013964:NM_013956:NM_013960:XM_017013366:XM_017013370 | 0:0:0:0:0:0 | 0:0:0:0:0:0 | 0:0:0:0:0:0 | 1:1:0:0:0:0 | 0:0:0:0:0:0 | 1:0:0:0:0:0 | 1:0:0:0:0:0 | 0:0:0:0:0:0 | 0:0:1:1:0:0 | 0:0:0:0:0:0 |
| NM_023034 | 0 | 0 | 10 | 0 | 2 | 0 | 6 | 0 | 1 | 0 |
| NM_023034 | 0 | 0 | 9 | 1 | 1 | 0 | 5 | 3 | 1 | 0 |
| NM_020130 | 0 | 5 | 0 | 4 | 0 | 5 | 0 | 0 | 0 | 3 |
| NM_183420:NM_012173 | 0:00 | 0:00 | 0:00 | 1:00 | 0:00 | 1:00 | 0:00 | 0:00 | 0:00 | 0:00 |
| XM_005273539:NM_002690:XM_005273537:XM_005273536:XM_005273538:XM_017013584:XM_005273535:XM_006716353:XR_001745544:XM_005273540:XR_428311:XM_017013583 | 41:37:4:3:1:2:0:0:0:0:0:0 | 0:0:0:0:0:0:0:0:0:0:0:0 | 6:4:2:0:1:0:0:0:0:0:0:0 | 0:0:0:0:0:0:0:0:0:0:0:0 | 5:6:2:0:0:0:0:0:0:0:0:0 | 0:0:0:0:0:0:0:0:0:0:0:0 | 99:210:30:5:9:4:32:1:2:0:0:0 | 0:0:0:0:0:0:0:0:0:0:0:0 | 10:31:6:1:1:1:7:0:2:0:0:0 | 0:0:0:0:0:0:0:0:0:0:0:0 |
| NM_032410 | 8 | 3 | 7 | 3 | 2 | 0 | 36 | 3 | 10 | 0 |

| | | | | | | | | | | |
|---|---|---|---|---|---|---|---|---|---|---|
| NM_001080394:XR_001745508:NR_148205:XM_017013269:NM_001352956:NM_001352940:NM_001352934:NM_001352933:NM_001352931:NM_001352958:XM_017013271:NM_001352950:NR_104581:NM_001352942:NM_001352937:XM_017013279:NM_001352951:NR_148203:NM_001282916:XM_017013281:NM_001352943:NM_001352935:NM_001352949:NM_001352961:NM_001352932:XM_017013268:XM_017013273:NM_001352955:NM_001352944:NM_001352936:NM_001352939:XM_017013270:NM_001352957:NM_001352948:XM_011517497:NM_001352945:XM_024447117:XM_024447116:NM_001352941:XM_017013274:XM_017013280:XM_017013289 | 21:3:2:5:3:2:1:2:2:2:1:0:0:0:0:0:0:0:0:0:0:0:0:0:0:0:0:0:0:0:0:0:0:0:0:0:0:0:0:0:0:0:0:0:0:0:0:0 | 0:0:0:0:0:0:0:0:0:0:0:0:0:0:0:0:0:0:0:0:0:0:0:0:0:0:0:0:0:0:0:0:0:0:0:0:0:0:0:0:0:0 | 1:0:2:0:4:0:0:0:0:1:0:1:1:1:1:1:1:0:0:0:0:0:0:0:0:0:0:0:0:0:0:0:0:0:0:0:0:0:0:0:0:0:0 | 0:0:0:0:0:0:0:0:0:0:0:0:0:0:0:0:0:0:0:0:0:0:0:0:0:0:0:0:0:0:0:0:0:0:0:0:0:0:0:0:0:0:0:0:0:0:0:0:0 | 18:1:4:2:1:1:1:1:4:1:1:0:0:0:0:0:1:2:1:1:1:1:1:1:1:1:0:0:0:0:0:0:0:0:0:0:0:0:0:0:0:0:0:0 | 0:0:0:0:0:0:0:0:0:0:0:0:0:0:0:0:0:0:0:0:0:0:0:0:0:0:0:0:0:0:0:0:0:0:0:0:0:0:0:0:0:0:0:0:0:0:0:0:0:0:0 | 24:13:14:6:26:1:1:1:1:14:4:2:0:13:0:2:0:4:2:0:1:0:0:15:5:3:4:2:2:1:1:1:4:1:1:1:2:1:0:0:0:0 | 0:0:0:0:0:0:0:0:0:0:0:0:0:0:0:0:0:0:0:0:0:0:0:0:0:0:0:0:0:0:0:0:0:0:0:0:0:0:0:0:0:0:0:0:0:0:0:0:0:0:0 | 10:2:2:2:1:0:0:1:1:1:0:1:0:4:0:0:1:0:0:0:0:0:0:5:2:0:1:0:0:0:0:1:0:0:0:0:1:1:1:0:0:0 | 0:0:0:0:0:0:0:0:0:0:0:0:0:0:0:0:0:0:0:0:0:0:0:0:0:0:0:0:0:0:0:0:0:0:0:0:0:0:0:0:0:0:0:0:0:0:0:0:0:0:0:0 |
| NM_001080394:XR_001745508:NR_148205:XM_017013269:NM_001352956:NM_001352940:NM_001352934:NM_001352933:NM_001352931:NM_001352958:XM_017013271:NM_001352961:NM_001352950:NR_104581:NM_001352942:NM_001352937:XM_017013279:NM_001352951:NM_001352957:NR_148203:NM_001282916:XM_017013281:NM_001352943:NM_001352935:NM_001352949:NM_001352932:XM_017013268:XM_017013273:NM_001352955:NM_001352944:NM_001352936:NM_001352939:XM_017013270:NM_001352948:XM_011517497:NM_001352945:XM_024447117:XM_024447116:NR_148204:NM_001352941:XM_017013274:XM_017013280:XM_017013289 | 21:3:2:5:3:2:1:2:2:6:1:1:0:0:0:0:0:0:0:0:0:0:0:0:0:0:0:0:0:0:0:0:0:0:0:0:0:0:0:0:0:0:0:0:0:0:0:0:0 | 0:0:0:0:0:0:0:0:0:0:0:0:0:0:0:0:0:0:0:0:0:0:0:0:0:0:0:0:0:0:0:0:0:0:0:0:0:0:0:0:0:0:0 | 1:0:2:0:4:0:0:0:0:4:0:0:1:1:1:1:1:1:1:1:0:0:0:0:0:0:0:0:0:0:0:0:0:0:0:0:0:0:0:0:0:0:0:0:0:0:0:0 | 0:0:0:0:0:0:0:0:0:1:0:0:0:0:0:0:0:0:0:0:0:0:0:0:0:0:0:0:0:0:0:0:0:0:0:0:0:0:0:0:0:0:0 | 18:2:4:2:1:1:1:1:4:2:1:1:0:0:0:0:0:1:0:2:1:1:1:1:1:1:1:0:0:0:0:0:0:0:0:0:0:0:0:0:0:0:0:0:0 | 0:0:0:0:0:0:0:0:0:2:0:0:0:0:0:0:0:0:0:0:0:0:0:0:0:0:0:0:0:0:0:0:0:0:0:0:0:0:0:0:0:0:0:0:0:0:0:0:0:0:0 | 24:16:15:6:26:1:1:1:1:37:4:18:2:0:12:0:2:1:5:4:4:0:1:0:0:5:3:4:2:2:1:1:1:1:2:1:5:1:1:0:0:0:0 | 0:0:0:0:0:0:0:0:0:2:0:1:0:0:1:0:0:1:0:0:0:0:0:0:0:0:0:0:0:0:0:0:0:0:0:0:0:0:0:0:0:0:0:0:0:0:0:0:0:0:0 | 10:2:2:2:1:0:0:1:1:2:0:5:1:0:4:0:0:1:0:0:0:0:0:0:0:2:0:1:0:0:0:0:1:0:0:0:1:1:0:1:0:0:0 | 0:0:0:0:0:0:0:0:0:0:0:0:0:0:0:0:0:0:0:0:0:0:0:0:0:0:0:0:0:0:0:0:0:0:0:0:0:0:0:0:0:0:0:0:0:0:0:0:0:0:0:0 |
| NM_001080394:XR_001745508:NR_148205:XM_017013269:NM_001352956:NM_001352940:NM_001352934:NM_001352933:NM_001352958:NM_001352931:XM_017013271:NM_001352961:NM_001352950:NR_104581:NM_001352942:NM_001352937:XM_017013279:NM_001352951:NM_001352957:NR_148203:NM_001282916:XM_017013281:NM_001352943:NM_001352935:NM_001352949:NM_001352932:XM_017013268:XM_017013273:NM_001352955:NM_001352944:NM_001352936:NM_001352939:XM_017013270:NM_001352948:XM_011517497:NM_001352945:XM_024447117:XM_024447116:NR_148204:NM_001352941:XM_017013274:XM_017013280:XM_017013289 | 21:3:3:5:3:2:1:2:13:2:1:1:0:0:0:0:0:0:0:0:0:0:0:0:0:0:0:0:0:0:0:0:0:0:0:0:0:0:0:0:0:0:0:0:0:0:0:0:0:0 | 0:0:0:0:0:0:0:0:0:0:0:0:0:0:0:0:0:0:0:0:0:0:0:0:0:0:0:0:0:0:0:0:0:0:0:0:0:0:0:0:0:0:0 | 1:0:2:0:4:0:0:0:6:0:0:0:1:1:1:1:1:1:1:0:0:0:0:0:0:0:0:0:0:0:0:0:0:0:0:0:0:0:0:0:0:0:0:0:0:0 | 0:0:0:0:0:0:0:0:0:0:0:0:0:0:0:0:0:0:0:0:0:0:0:0:0:0:0:0:0:0:0:0:0:0:0:0:0:0:0:0:0:0:0 | 18:2:4:2:1:1:1:1:4:4:1:1:0:0:0:0:0:1:0:2:1:1:1:1:1:1:1:0:0:0:0:0:0:0:0:0:0:0:0:0:0:0:0:0:0 | 0:0:0:0:0:0:0:0:0:0:0:0:0:0:0:0:0:0:0:0:0:0:0:0:0:0:0:0:0:0:0:0:0:0:0:0:0:0:0:0:0:0 | 24:18:15:10:26:1:1:1:57:1:4:22:2:0:13:0:2:3:5:4:4:0:1:0:0:5:3:4:2:2:1:1:1:1:2:1:5:1:1:0:0:0:0 | 0:1:0:0:0:0:0:0:2:0:0:1:0:0:0:0:0:0:0:0:0:0:0:0:0:0:0:0:0:0:0:0:0:0:0:0:0:0:0:0:0:0 | 10:4:3:2:1:0:0:1:2:1:0:6:1:0:4:0:0:1:0:0:0:0:0:0:0:2:0:1:0:0:0:0:1:0:0:0:1:1:0:1:0:0:0 | 0:0:0:0:0:0:0:0:0:1:0:0:0:0:0:0:0:0:0:0:0:0:0:0:0:0:0:0:0:0:0:0:0:0:0:0:0:0:0:0:0:0:0:0:0:0:0:0:0:0:0:0 |
| NM_001080394:NM_001282916:XR_001745508:XM_017013269:NM_001352956:NM_001352940:NM_001352934:NM_001352933:NM_001352958:NR_148205:NM_001352931:NM_001352961:XM_024447117:NM_001352950:NR_104581:NM_001352942:NM_001352937:XM_017013279:NM_001352951:NM_001352957:NR_148203:XM_017013281:NM_001352943:NM_001352935:NM_001352949:NM_001352932:XM_017013268:XM_017013273:NM_001352955:NM_001352939:XM_017013271:NM_001352948:XM_011517497:NM_001352945:XM_024447116:NR_148204:NR_148202:NM_001352941:XM_017013280 | 21:1:3:3:3:2:1:2:15:2:2:1:1:0:0:0:0:0:0:0:0:0:0:0:0:0:0:0:0:0:0:0:0:0:0:0:0:0:0:0:0:0:0:0:0 | 0:0:0:0:0:0:0:0:0:0:0:0:0:0:0:0:0:0:0:0:0:0:0:0:0:0:0:0:0:0:0:0:0:0:0:0:0:0:0:0 | 1:0:0:0:4:0:0:0:5:1:0:0:2:1:1:1:1:1:2:1:0:0:0:0:0:0:0:0:0:0:0:0:0:0:0:0:0:0:0:0:0 | 0:0:0:0:0:0:0:0:0:0:0:0:1:0:0:0:0:0:0:0:0:0:0:0:0:0:0:0:0:0:0:0:0:0:0:0:0:0:0:0:0:0:0:0:0:0 | 18:1:1:1:1:1:1:1:1:5:1:4:1:0:0:0:0:0:0:2:0:2:1:1:1:1:1:1:0:0:0:0:0:0:0:0:0:0:0:0:0:0 | 0:0:0:0:0:0:0:0:0:0:0:0:0:0:0:0:0:0:0:0:0:0:0:0:0:0:0:0:0:0:0:0:0:0:0:0:0:0:0 | 24:4:15:8:25:1:1:0:67:3:1:27:77:2:1:13:0:2:4:8:0:0:1:0:0:3:2:4:2:1:1:1:3:1:1:3:1:0:0 | 0:0:0:0:1:0:0:1:0:0:0:0:1:0:0:0:0:0:0:0:0:0:0:0:0:0:0:0:0:0:0:0:0:0:0:0:0:0:0 | 10:0:5:0:1:0:0:1:3:1:1:6:0:1:0:4:0:0:1:0:0:0:0:0:0:2:0:1:0:0:0:0:0:0:1:0:0:1:0 | 0:0:0:0:0:0:0:0:0:0:0:0:0:0:0:0:0:0:0:0:0:0:0:0:0:0:0:0:0:0:0:0:0:0:0:0:0:0:0:0:0:0:0:0:0:0:0:0:0 |
| NM_001080394:NM_001282916:XR_001745508:XM_017013269:NM_001352956:NM_001352940:NM_001352934:NM_001352933:NM_001352958:NR_148205:NM_001352931:NM_001352961:XM_024447117:NM_001352950:NR_104581:NM_001352942:NM_001352937:XM_017013279:NM_001352951:NM_001352957:NR_148203:XM_017013281:NM_001352943:NM_001352935:NM_001352949:NM_001352932:XM_017013268:XM_017013273:NM_001352955:NM_001352939:XM_017013271:NM_001352948:XM_011517497:NM_001352945:XM_024447116:NR_148204:NR_148202:NM_001352941:XM_017013280 | 21:1:3:4:3:2:1:2:15:2:2:1:1:0:0:0:0:0:0:0:0:0:0:0:0:0:0:0:0:0:0:0:0:0:0:0:0:0:0:0:0:0:0:0:0 | 0:0:0:0:0:0:0:0:0:0:0:0:0:0:0:0:0:0:0:0:0:0:0:0:0:0:0:0:0:0:0:0:0:0:0:0:0:0:0:0 | 1:0:0:0:4:0:0:0:5:1:0:0:3:1:1:1:1:1:2:1:0:0:0:0:0:0:0:0:0:0:0:0:0:0:0:0:0:0:0:0:0 | 0:0:0:0:0:0:0:0:0:0:0:0:0:0:0:0:0:0:0:0:0:0:0:0:0:0:0:0:0:0:0:0:0:0:0:0:0:0:0:0:0:0:0:0:0:0 | 18:1:1:1:1:1:1:1:1:5:1:4:1:1:0:0:0:0:0:2:0:1:1:1:1:1:1:1:0:0:0:0:0:0:0:0:0:0:0:0:0:0 | 0:0:0:0:0:0:0:0:0:0:0:0:0:0:0:0:0:0:0:0:0:0:0:0:0:0:0:0:0:0:0:0:0:0:0:0:0:0:0 | 24:4:15:8:26:1:1:1:67:4:1:29:89:2:1:13:0:1:4:8:0:0:1:0:0:3:3:4:2:1:1:1:3:1:1:4:1:0:0 | 0:0:0:0:0:0:0:0:0:0:0:0:0:0:0:0:0:1:0:0:0:0:0:0:0:0:0:0:0:0:0:0:0:0:0:0:0:0:0 | 10:0:4:0:1:0:0:1:3:1:1:6:1:1:0:4:0:0:1:0:0:0:0:0:0:2:0:1:0:0:0:0:0:0:1:0:0:1:0 | 0:0:0:0:0:0:0:0:0:0:0:0:0:0:0:0:0:0:0:0:0:0:0:0:0:0:0:0:0:0:0:0:0:0:0:0:0:0:0:0:0:0:0:0:0:0:0:0:0 |
| NM_001080394:NM_001282916:XR_001745508:XM_017013269:NM_001352956:NM_001352940:NM_001352934:NM_001352933:NM_001352958:NM_001352931:XM_017013271:NM_001352961:NR_148205:XM_024447117:NM_001352950:NR_104581:NM_001352942:NM_001352937:XM_017013279:NM_001352951:NM_001352957:NR_148203:XM_017013281:NM_001352943:NM_001352935:NM_001352949:NM_001352932:XM_017013273:NM_001352955:XM_017013268:NM_001352939:NM_001352948:XM_011517497:NM_001352945:XM_024447116:NR_148204:NR_148202:NM_001352941:XM_017013274 | 21:1:3:4:3:2:1:2:15:2:1:1:1:2:0:0:0:0:0:0:0:0:0:0:0:0:0:0:0:0:0:0:0:0:0:0:0:0:0:0:0:0:0:0:0 | 0:0:0:0:0:0:0:0:0:0:0:0:0:0:0:0:0:0:0:0:0:0:0:0:0:0:0:0:0:0:0:0:0:0:0:0:0:0:0:0 | 1:0:2:0:4:0:0:0:5:0:0:0:0:4:1:1:1:1:1:3:1:0:0:0:0:0:0:0:0:0:0:0:0:0:0:0:0:0:0:0:0 | 0:0:0:0:0:0:0:0:0:0:0:0:0:0:0:0:0:0:0:0:0:0:0:0:0:0:0:0:0:0:0:0:0:0:0:0:0:0:0:0:0:0:0:0:0:0 | 17:1:3:1:1:1:1:1:1:5:4:0:1:1:1:0:0:0:0:0:2:0:1:1:1:1:1:1:0:0:0:0:0:0:0:0:0:0:0:0:0:0 | 1:0:0:0:0:0:0:0:0:0:0:0:0:0:0:0:0:0:0:0:0:0:0:0:0:0:0:0:0:0:0:0:0:0:0:0:0:0:0 | 24:4:33:8:26:1:1:1:66:1:1:30:3:103:2:1:13:0:2:4:9:0:0:1:0:0:3:4:2:3:1:1:5:1:1:4:1:0:0 | 0:0:0:0:0:0:0:0:1:0:0:0:0:2:0:0:0:0:0:0:0:0:0:0:0:0:0:0:0:0:0:0:0:0:0:0:0:0:0 | 10:0:5:0:1:0:0:1:3:1:0:6:1:2:1:0:4:0:0:1:0:0:0:0:0:0:2:1:0:0:0:0:0:0:1:0:0:1:0 | 0:0:0:0:0:0:0:0:0:0:0:0:0:0:0:0:0:0:0:0:0:0:0:0:0:0:0:0:0:0:0:0:0:0:0:0:0:0:0:0:0:0:0:0:0:0:0:0:0 |
| NM_001080394:NM_001282916:XM_017013269:NM_001352956:NM_001352940:NM_001352933:XR_001745508:NM_001352958:NM_001352931:XM_017013271:XM_024447117:NR_148205:NR_104581:NM_001352942:XM_017013279:NM_001352951:NR_148203:NM_001352935:NM_001352949:NM_001352932:XM_017013268:XM_017013273:NM_001352955:NM_001352939:NM_001352948:XM_011517497:NR_148204:NR_148202:NM_001352941:XM_017013274:NM_001352945:XM_024447116:XM_017013289 | 21:1:4:3:2:2:2:15:2:1:2:0:0:0:0:0:0:0:0:0:0:0:0:0:0:0:0:0:0:0:0:0:0:0:0:0:0 | 0:0:0:0:0:0:0:0:0:0:0:0:0:0:0:0:0:0:0:0:0:0:0:0:0:0:0:0:0:0:0:0:0:0:0:0:0:0:0:0:0 | 1:0:0:4:0:0:2:5:0:0:4:1:1:1:1:3:0:0:0:0:0:0:0:0:0:0:0:0:0:0:0:0:0:0:0:0 | 0:0:0:0:0:0:0:0:0:0:0:0:0:0:0:0:0:0:0:0:0:0:0:0:0:0:0:0:0:0:0:0:0:0:0:0:0:0:0:0 | 18:1:0:1:1:1:4:5:4:0:1:0:0:0:0:2:2:1:1:1:1:0:0:0:0:0:0:0:0:0:0:0:0:0:0 | 0:0:0:0:0:0:0:0:0:0:0:0:0:0:0:0:0:0:0:0:0:0:0:0:0:0:0:0:0:0:0:0:0:0:0:0:0:0:0:0:0:0 | 24:4:7:26:1:1:30:67:1:1:113:4:1:13:0:4:0:0:0:3:1:4:2:1:1:5:4:1:0:0:0:0:0 | 0:0:0:0:0:0:0:0:0:0:0:0:0:0:0:0:0:0:0:0:0:0:0:0:0:0:0:0:0:0:0:0:0:0:0:0:0:0:0:0:0:0 | 10:0:0:1:0:1:5:3:1:0:4:1:0:4:0:1:0:0:0:2:0:1:0:0:0:0:0:0:1:0:0:0:0 | 0:0:0:0:0:0:0:0:0:0:0:0:0:0:0:0:0:0:0:0:0:0:0:0:0:0:0:0:0:0:0:0:0:0:0:0:0:0:0 |
| XM_024447117:NM_001080394:NM_001282916:XR_001745508:XM_017013269:NM_001352956:NM_001352940:NM_001352934:NM_001352933:NM_001352958:NM_001352931:XM_017013271:NM_001352961:NM_001352950:NR_148205:NR_104581:NM_001352942:NM_001352937:XM_017013279:NM_001352951:NM_001352957:NR_148203:XM_017013281:NM_001352943:NM_001352935:NM_001352949:NM_001352932:XM_017013268:XM_017013273:NM_001352955:NM_001352939:NM_001352948:XM_011517497:NM_001352945:XM_024447116:NR_148204:NR_148202:XM_017013289:NM_001352941:XM_017013274 | 2:21:1:4:4:3:2:1:2:15:2:1:1:0:0:0:0:0:0:0:0:0:0:0:0:0:0:0:0:0:0:0:0:0:0:0:0:0:0:0:0:0:0:0:0:0:0:0 | 1:0:0:0:0:0:0:0:0:0:0:0:0:0:0:0:0:0:0:0:0:0:0:0:0:0:0:0:0:0:0:0:0:0:0:0:0:0:0:0:0:0:0:0:0:0:0:0 | 5:1:0:1:0:4:0:0:0:5:0:0:0:1:1:1:1:1:1:3:1:0:0:0:0:0:0:0:0:0:0:0:0:0:0:0:0:0:0:0:0:0:0 | 0:0:0:0:0:0:0:0:0:0:0:0:0:0:0:0:0:0:0:0:0:0:0:0:0:0:0:0:0:0:0:0:0:0:0:0:0:0:0:0 | 1:18:1:4:1:1:1:1:1:5:4:0:1:0:1:0:0:0:0:2:0:2:1:1:1:1:1:1:0:0:0:0:0:0:0:0:0:0:0:0:0 | 0:0:0:0:0:0:0:0:0:0:0:0:0:0:0:0:0:0:0:0:0:0:0:0:0:0:0:0:0:0:0:0:0:0:0:0:0:0:0:0 | 121:24:4:38:8:26:1:1:1:66:1:1:30:1:4:1:13:0:2:4:9:0:0:1:0:0:3:3:4:2:1:1:5:1:1:4:1:1:0:0 | 0:0:0:0:0:0:0:0:0:0:0:0:0:0:0:0:0:0:0:0:0:0:0:0:0:0:0:0:0:0:0:0:0:0:0:0:0:0:0:0 | 4:10:0:5:0:1:0:0:1:3:1:0:6:1:1:0:4:0:0:1:0:0:0:0:0:0:2:0:1:0:0:0:0:0:1:0:0:0:1:0 | 0:0:0:0:0:0:0:0:0:0:0:0:0:0:0:0:0:0:0:0:0:0:0:0:0:0:0:0:0:0:0:0:0:0:0:0:0:0:0:0:0:0:0:0:0:0:0:0:0:0 |
| NM_052937:NM_001286783:XR_001745466 | 55:03:04 | 7:00:00 | 20:01:00 | 6:00:00 | 44:01:02 | 4:00:00 | 276:00:09 | 58:00:00 | 21:00:02 | 0:00:00 |
| NM_052937:NM_001286783:XR_001745466 | 62:03:04 | 1:00:00 | 27:01:00 | 0:00:00 | 46:01:02 | 2:00:00 | 138:00:02 | 210:00:07 | 10:00:00 | 12:00:02 |
| NM_052937:NM_001286783:XR_001745466 | 78:04:03 | 11:00:00 | 30:01:00 | 0:00:00 | 65:02:01 | 18:02:01 | 403:00:08 | 41:00:00 | 24:00:01 | 1:00:00 |
| NM_052937:XR_001745466:NM_001286783 | 78:02:04 | 11:01:00 | 30:00:01 | 3:00:00 | 67:02:03 | 17:00:01 | 431:09:00 | 31:01:00 | 23:01:00 | 2:00:00 |
| NM_052937:NM_001286783:XR_001745466 | 86:04:02 | 4:00:00 | 33:01:00 | 3:00:00 | 79:04:02 | 6:00:00 | 478:00:10 | 10:00:00 | 24:00:01 | 1:00:00 |
| NM_001286783:NM_052937:XR_001745466 | 0.229189815 | 0:00:00 | 1:35:00 | 0:01:00 | 0.225717593 | 0:00:00 | 0.337615741 | 0:02:00 | 0:25:01 | 0:00:00 |
| NM_015941:NM_213620:XM_011517542:NM_213619:XM_006716455 | 109:11:9:4:0 | 1:0:0:0:0 | 39:6:7:1:0 | 0:0:0:0:0 | 28:1:4:0:0 | 0:0:0:0:0 | 826:223:116:19:9 | 3:0:0:0:0 | 56:6:7:0:1 | 0:0:0:0:0 |
| NM_015941:XM_011517542:NM_213620:NM_213619:XM_006716455 | 111:10:11:4:0 | 0:0:0:0:0 | 43:7:6:1:0 | 0:0:0:0:0 | 29:5:1:0:0 | 0:0:0:0:0 | 856:116:226:19:9 | 1:1:0:0:0 | 56:7:6:0:1 | 0:0:0:0:0 |
| NM_015941:XM_011517542:NM_213620:NM_213619:XM_006716455 | 111:4:14:4:0 | 0:0:0:0:0 | 43:3:6:1:0 | 0:0:0:0:0 | 29:4:1:0:0 | 0:0:0:0:0 | 860:69:229:19:11 | 2:0:0:0:0 | 56:3:6:0:1 | 0:0:0:0:0 |
| NM_015941:XM_011517542:NM_213620:NM_213619:XM_006716455 | 111:4:14:4:0 | 0:0:0:0:0 | 43:2:6:1:0 | 0:0:0:0:0 | 29:4:1:0:0 | 0:0:0:0:0 | 861:55:229:19:11 | 2:1:0:0:0 | 56:2:6:0:1 | 0:0:0:0:0 |
| NM_015941:XM_011517542:NM_213620:NM_213619:XM_006716455 | 111:3:14:4:0 | 0:0:0:0:0 | 43:1:6:1:0 | 0:0:0:0:0 | 29:4:1:0:0 | 0:0:0:0:0 | 865:45:229:18:10 | 0:2:0:1:1 | 56:1:6:0:1 | 0:0:0:0:0 |
| NM_015941:NM_213620:NM_213619:XM_011517542:XM_006716455 | 110:13:4:1:0 | 1:1:0:0:0 | 43:6:1:1:0 | 0:0:0:0:0 | 29:1:0:3:0 | 0:0:0:0:0 | 867:228:17:13:11 | 0:0:0:1:0 | 56:6:0:0:1 | 0:0:0:0:0 |
| NM_015941:NM_213620:NM_213619:XM_011517542:XM_006716455 | 111:10:4:0:0 | 0:0:0:0:0 | 43:5:1:1:0 | 0:0:0:0:0 | 29:1:0:2:0 | 0:0:0:0:0 | 866:158:19:8:9 | 1:3:0:0:0 | 56:6:0:0:1 | 0:0:0:0:0 |
| XM_017013613:XM_011517554:XM_017013612 | 134:24:00 | 0:00:00 | 77:10:01 | 0:00:00 | 66:09:00 | 0:00:00 | 654:62:8 | 4:00:01 | 92:09:03 | 4:00:00 |
| XM_017013613:XM_011517554:XM_017013612 | 157:22:00 | 0:00:00 | 99:09:01 | 1:00:00 | 75:09:00 | 2:00:00 | 952:59:08 | 6:00:01 | 110:09:03 | 0:00:00 |
| XM_017013613:XM_011517554:XM_017013612 | 174:23:00 | 0:00:00 | 135:10:01 | 0:00:00 | 81:09:00 | 1:00:00 | 1085:58:09 | 3:01:00 | 114:09:03 | 0:00:00 |
| XM_017013613:XM_011517554:XM_017013612 | 180:23:00 | 0:00:00 | 159:09:01 | 0:01:00 | 87:09:00 | 0:00:00 | 1112:60:9 | 5:00:00 | 119:09:03 | 0:00:00 |
| XM_017013613:XM_011517554:XM_017013612 | 182:23:00 | 1:00:00 | 195:09:01 | 0:00:00 | 87:09:00 | 0:00:00 | 1045:56:08 | 9:00:00 | 116:09:03 | 0:00:00 |
| XM_017013613:XM_011517554:XM_017013612 | 76:09:00 | 1:01:00 | 104:04:00 | 63:05:00 | 35:03:00 | 1:00:00 | 454:18:01 | 228:16:02 | 30:05:00 | 1:00:00 |
| XM_017013613:XM_011517554:XM_017013612 | 76:09:00 | 1:00:00 | 103:02:00 | 0:00:00 | 35:03:00 | 0:00:00 | 430:18:00 | 5:00:00 | 30:05:00 | 0:00:00 |
| XM_017013613:XM_011517554:XM_017013612 | 66:09:00 | 8:00:00 | 79:02:00 | 23:00:00 | 31:03:00 | 2:00:00 | 321:13:00 | 105:04:00 | 23:05:00 | 2:00:00 |
| XM_017013613:XM_011517554:XM_017013612 | 36:01:00 | 38:08:00 | 97:01:00 | 5:01:00 | 8:01:00 | 25:02:00 | 339:14:00 | 87:03:00 | 11:04:00 | 15:01:00 |
| XM_017013613:XM_011517554:XM_017013612 | 76:09:00 | 1:00:00 | 102:02:00 | 0:00:00 | 35:03:00 | 0:00:00 | 427:18:00 | 4:00:00 | 30:05:00 | 0:00:00 |
| XM_017013613:XM_011517554 | 31:01:00 | 5:02 | 20:00 | 5:00 | 8:01 | 0:00 | 82:05:00 | 22:02 | 10:04 | 1:00 |
| XM_017013613:XM_011517554 | 30:01:00 | 5:02 | 22:00 | 1:00 | 8:01 | 0:00 | 86:06:00 | 17:01 | 9:03 | 2:01 |
| NM_003878:XM_011517623 | 4:00 | 0:00 | 2:00 | 0:00 | 3:00 | 0:00 | 13:01 | 0:00 | 10:00 | 0:00 |
| NM_003878:XM_011517623 | 3:00 | 1:00 | 2:00 | 0:00 | 2:00 | 1:00 | 0:00 | 11:01 | 5:00 | 4:00 |

| | | | | | | | | | | |
|---|---|---|---|---|---|---|---|---|---|---|
| NM_003878:XM_011517623 | 3:00 | 1:00 | 2:00 | 0:00 | 2:00 | 1:00 | 0:00 | 11:01 | 5:00 | 4:00 |
| XM_011517526:NM_001287260:XM_017013404:XM_011517525:XM_017013405:NM_001011720:NM_001287258:XM_006716447:XM_011517524:XM_011517527 | 0:0:0:0:0:0:0:0:0:0 | 0:0:0:0:0:0:0:0:0:0 | 2:3:0:0:0:0:0:0:0:0 | 0:0:0:0:0:0:0:0:0:0 | 0:0:1:0:0:0:0:0:0:0 | 0:0:0:0:0:0:0:0:0:0 | 3:9:4:10:2:24:4:2:0:0 | 0:3:1:3:1:1:1:1:0:0 | 0:1:0:0:0:1:0:1:0:0 | 0:0:0:1:1:0:0:0:0:0 |
| NM_000971 | 213 | 2 | 14 | 0 | 50 | 0 | 367 | 0 | 119 | 0 |
| NM_000971 | 216 | 0 | 14 | 0 | 51 | 0 | 371 | 2 | 120 | 0 |
| NM_000971 | 217 | 0 | 15 | 0 | 51 | 0 | 375 | 0 | 121 | 0 |
| NM_000971 | 217 | 0 | 15 | 0 | 51 | 0 | 375 | 0 | 121 | 0 |
| NM_001172086:NM_001172087:NM_001079867 | 0:00:00 | 1:00:00 | 0:00:00 | 1:00:00 | 0:00:00 | 0:00:00 | 1:01:00 | 0:00:00 | 2:00:00 | 0:00:00 |
| NM_007029:XM_005251142 | 9:00 | 0:00 | 65:01:00 | 0:00 | 52:01:00 | 0:00 | 1:00 | 0:00 | 1:00 | 0:00 |
| NM_007029:XM_005251142 | 8:00 | 1:00 | 63:01:00 | 1:00 | 51:01:00 | 0:00 | 1:00 | 0:00 | 1:00 | 0:00 |
| NM_001164830:NM_001738:NM_001291967:NM_001291968:NM_001128830 | 0:0:0:0:0 | 0:0:0:0:0 | 6940:611:76:2:0 | 7:0:0:0:0 | 14257:667:166:1:1 | 21:2:0:0:0 | 4:1:0:0:0 | 0:0:0:0:0 | 10:0:0:0:0 | 0:0:0:0:0 |
| NM_001164830:NM_001738:NM_001291967:NM_001291968:NM_001128830 | 0:0:0:0:0 | 0:0:0:0:0 | 6939:612:76:2:0 | 7:0:0:0:0 | 14253:674:166:1:1 | 26:4:0:0:0 | 4:1:0:0:0 | 0:0:0:0:0 | 10:0:0:0:0 | 0:0:0:0:0 |
| NM_001164830:NM_001738:NM_001291967:NM_001291968:NM_001128830 | 0:0:0:0:0 | 0:0:0:0:0 | 6951:630:76:2:0 | 6:2:0:0:0 | 14264:694:167:1:1 | 27:0:0:0:0 | 4:1:0:0:0 | 0:0:0:0:0 | 10:0:0:0:0 | 0:0:0:0:0 |
| NM_001164830:NM_001738:NM_001291967:NM_001291968:NM_001128830 | 0:0:0:0:0 | 0:0:0:0:0 | 6954:633:76:2:0 | 3:1:0:0:0 | 14274:701:166:1:1 | 22:0:1:0:0 | 4:1:0:0:0 | 0:0:0:0:0 | 10:0:0:0:0 | 0:0:0:0:0 |
| NM_001164830:NM_001738:NM_001291967:NM_001291968:NM_001128830 | 0:0:0:0:0 | 0:0:0:0:0 | 6952:642:77:2:0 | 7:2:0:0:0 | 14267:708:167:1:1 | 33:1:0:0:0 | 4:1:0:0:0 | 0:0:0:0:0 | 10:0:0:0:0 | 0:0:0:0:0 |
| NM_001164830:NM_001738:NM_001291967:NM_001291968:NM_001128830 | 0:0:0:0:0 | 0:0:0:0:0 | 6955:635:79:2:0 | 4:5:0:0:0 | 14292:717:167:1:1 | 11:0:0:0:0 | 4:1:0:0:0 | 0:0:0:0:0 | 10:0:0:0:0 | 0:0:0:0:0 |
| NM_001164830:NM_001738:NM_001291967:NM_001291968:NM_001128830 | 0:0:0:0:0 | 0:0:0:0:0 | 6974:613:79:2:0 | 3:0:0:0:0 | 14314:712:167:1:1 | 6:1:0:0:0 | 4:1:0:0:0 | 0:0:0:0:0 | 10:0:0:0:0 | 0:0:0:0:0 |
| NM_001164830:NM_001738:NM_001291967:NM_001291968:NM_001128830 | 0:0:0:0:0 | 0:0:0:0:0 | 6985:610:79:2:0 | 4:1:0:0:0 | 14322:724:171:1:1 | 9:2:0:0:0 | 4:1:0:0:0 | 0:0:0:0:0 | 10:0:0:0:0 | 0:0:0:0:0 |
| NM_001164830:NM_001738:NM_001291967:NM_001128830 | 0:0:0:0 | 0:0:0:0 | 6994:541:80:0 | 9:2:0:0 | 14334:666:171:1 | 14:0:1:0 | 4:1:0:0 | 0:0:0:0 | 10:0:0:0 | 0:0:0:0 |
| NM_001164830:NM_001738:NM_001291967:NM_001128830 | 0:0:0:0 | 0:0:0:0 | 7010:440:79:0 | 2:6:1:0 | 14356:567:172:1 | 2:2:0:0 | 4:1:0:0 | 0:0:0:0 | 10:0:0:0 | 0:0:0:0 |
| NM_001164830:NM_001738:NM_001291967:NM_001128830 | 0:0:0:0 | 0:0:0:0 | 7011:408:79:0 | 6:1:1:0 | 14351:544:171:1 | 17:1:0:0 | 4:1:0:0 | 0:0:0:0 | 10:0:0:0 | 0:0:0:0 |
| NM_001164830:NM_001738:NM_001291967:NM_001128830 | 0:0:0:0 | 0:0:0:0 | 7034:337:0:0 | 8:0:0:0 | 14382:484:7:1 | 8:3:0:0 | 4:1:0:0 | 0:0:0:0 | 10:0:0:0 | 0:0:0:0 |
| NM_001164830:NM_001738:NM_001291967:NM_001128830 | 0:0:0:0 | 0:0:0:0 | 7048:287:0:0 | 6:6:0:0 | 14376:425:7:1 | 16:3:0:0 | 4:1:0:0 | 0:0:0:0 | 10:0:0:0 | 0:0:0:0 |
| NM_001164830:NM_001738:NM_001291967:NM_001128830 | 0:0:0:0 | 0:0:0:0 | 7044:213:0:0 | 12:7:0:0 | 14366:327:7:1 | 28:12:0:0 | 4:0:0:0 | 0:0:0:0 | 10:0:0:0 | 0:0:0:0 |
| NM_001164830:NM_001738:NM_001291967:NM_001128830 | 0:0:0:0 | 0:0:0:0 | 7054:202:0:0 | 2:11:0:0 | 14392:314:7:1 | 2:19:0:0 | 4:0:0:0 | 0:0:0:0 | 10:0:0:0 | 0:0:0:0 |
| NM_001164830:NM_001738:NM_001291967:NM_001128830 | 0:0:0:0 | 0:0:0:0 | 7050:171:0:0 | 7:5:0:0 | 14374:263:7:1 | 20:16:0:0 | 4:0:0:0 | 0:0:0:0 | 10:0:0:0 | 0:0:0:0 |
| NM_001164830:NM_001738:NM_001291967:NM_001128830 | 0:0:0:0 | 0:0:0:0 | 7051:163:0:0 | 6:8:0:0 | 14383:257:7:1 | 11:18:0:0 | 4:0:0:0 | 0:0:0:0 | 10:0:0:0 | 0:0:0:0 |
| NM_001164830:NM_001738:NM_001291967:NM_001128830 | 0:0:0:0 | 0:0:0:0 | 7055:159:0:0 | 2:5:0:0 | 14389:249:7:1 | 5:15:0:0 | 4:0:0:0 | 0:0:0:0 | 10:0:0:0 | 0:0:0:0 |
| NM_001286719:NM_016033:XM_017013518 | 0:00:00 | 0:00:00 | 0:00:00 | 0:00:00 | 0:00:00 | 1:00:00 | 0:00:00 | 0:00:00 | 0:01:00 | 0:00:00 |
| NM_003821:XM_005251092:XM_011517357 | 60:31:06 | 0:00:00 | 7:03:00 | 0:00:00 | 8:01:00 | 0:00:00 | 172:64:8 | 0:00:00 | 34:10:02 | 0:00:00 |
| NM_001126111 | 0 | 0 | 0 | 0 | 0 | 0 | 0 | 0 | 0 | 0 |
| XM_024447165:XM_024447164:NM_002485:NM_001024688:XM_024447163:XM_017013462:XM_011517046:XM_011517045 | 17:2:15:0:0:0:0:0 | 0:0:0:0:0:0:0:0 | 3:1:4:2:3:0:0:0 | 0:0:0:0:0:0:0:0 | 6:3:7:0:0:0:0:0 | 0:0:0:0:0:0:0:0 | 57:27:55:8:9:6:1:2 | 1:1:0:0:0:0:0:0 | 13:0:16:10:0:0:0:0 | 0:0:0:0:0:0:0:0 |
| NM_002485:XM_024447165:XM_024447164:NM_001024688:XM_024447163:XM_011517045:XM_017013462:XM_011517046:XM_017013460 | 21:16:2:2:0:0:0:0:0 | 6:0:0:0:0:0:0:0:0 | 7:4:1:4:3:0:0:0:0 | 5:1:0:3:0:0:0:0:0 | 8:4:3:0:0:0:0:0:0 | 2:0:0:0:0:0:0:0:0 | 94:55:30:20:10:3:6:1:0 | 63:10:4:6:0:1:2:0:0 | 25:11:0:14:0:0:1:1:0 | 3:0:0:1:1:0:0:0:0 |
| XM_024447165:XM_024447164:NM_002485:NM_001024688:XM_024447163:XM_017013462:XM_011517046:XM_011517045:XM_017013460 | 16:2:34:2:0:0:0:0:0 | 0:0:0:0:0:0:0:0:0 | 5:1:12:8:3:0:0:0:0 | 0:0:0:0:0:0:0:0:0 | 4:4:12:0:0:0:0:0:0 | 0:0:0:0:0:0:0:0:0 | 67:33:162:30:10:7:1:4:0 | 0:1:2:0:0:2:0:0:0 | 12:1:33:17:1:1:1:0:0 | 0:0:0:0:0:0:0:0:0 |
| XM_024447165:XM_024447164:NM_002485:NM_001024688:XM_024447163:XM_017013462:XM_011517046:XM_011517045:XM_017013460 | 14:2:36:2:0:0:0:0:0 | 0:0:0:0:0:0:0:0:0 | 6:1:13:7:3:0:0:0:0 | 0:0:1:1:0:0:0:0:0 | 4:4:12:0:0:0:0:0:0 | 0:0:0:0:0:0:0:0:0 | 70:36:173:31:10:8:1:4:0 | 0:0:2:0:1:1:0:0:0 | 8:1:32:17:1:1:1:0:0 | 0:0:0:0:0:0:0:0:0 |
| XM_024447165:XM_024447164:NM_002485:NM_001024688:XM_024447163:XM_017013462:XM_011517046:XM_011517045:XM_017013460 | 14:2:36:2:0:0:0:0:0 | 0:0:0:0:0:0:0:0:0 | 6:1:14:8:3:0:0:0:0 | 0:0:0:0:0:0:0:0:0 | 4:4:12:0:0:0:0:0:0 | 0:0:0:0:0:0:0:0:0 | 70:36:173:30:11:8:1:4:0 | 0:0:2:1:0:1:0:0:0 | 7:1:32:17:1:1:1:0:0 | 0:0:0:0:0:0:0:0:0 |
| XM_024447165:XM_024447164:NM_002485:NM_001024688:XM_011517046:XM_024447163:XM_017013462:XM_011517045:XM_017013460 | 11:2:36:2:0:0:0:0:0 | 0:0:0:0:0:0:0:0:0 | 6:1:14:8:0:3:0:0:0 | 0:0:1:0:1:0:0:0:0 | 4:4:12:0:0:0:0:0:0 | 0:0:0:0:0:0:0:0:0 | 69:37:175:32:1:11:9:4:0 | 0:0:3:0:0:0:0:0:0 | 7:1:32:17:1:1:1:0:0 | 0:0:0:0:0:0:0:0:0 |
| NM_001024688:XM_011517046:XM_024447165:XM_024447164:NM_002485:XM_024447163:XM_017013462:XM_011517045:XM_017013460 | 5:11:10:2:38:0:0:0:0 | 1:1:1:0:0:0:0:0:0 | 13:16:11:10:50:5:0:0:0 | 0:0:0:0:0:0:0:0:0 | 0:5:3:4:11:0:0:0:0 | 0:1:0:0:0:0:0:0:0 | 82:94:89:50:349:16:16:6:0 | 0:1:0:0:0:0:1:0:0 | 20:13:7:2:32:1:1:0:0 | 0:2:0:0:0:0:0:0:0 |
| XM_024447165:XM_024447164:NM_002485:NM_001024688:XM_011517046:XM_024447163:XM_017013462:XM_011517045:XM_017013460 | 9:2:38:6:12:0:0:0:0 | 0:0:0:0:0:0:0:0:0 | 13:10:50:13:17:5:0:0:0 | 0:0:0:0:0:0:0:0:0 | 3:4:11:0:6:0:0:0:0 | 0:0:0:0:0:0:0:0:0 | 86:55:347:82:97:16:17:6:0 | 0:0:2:0:1:0:0:0:0 | 7:2:32:20:16:1:1:0:0 | 0:0:0:0:0:0:0:0:0 |
| XM_024447165:XM_024447164:NM_002485:NM_001024688:XM_011517046:XM_024447163:XM_017013462:XM_011517045:XM_017013460 | 8:2:38:6:12:0:0:0:0 | 0:0:0:0:0:0:0:0:0 | 14:10:50:13:17:5:0:0:0 | 0:0:0:0:0:0:0:0:0 | 2:4:11:0:6:0:0:0:0 | 0:0:0:0:0:0:0:0:0 | 86:57:349:81:98:16:17:5:0 | 0:0:0:0:0:0:0:0:0 | 7:2:32:18:16:1:1:0:0 | 0:0:0:2:0:0:0:0:0 |
| XM_024447164:XM_024447165:NM_002485:NM_001024688:XM_011517046:XM_024447163:XM_017013462:XM_011517045:XM_017013460 | 2:4:38:6:12:0:0:0:0 | 0:0:0:0:0:0:0:0:0 | 10:11:50:13:17:5:0:0:0 | 0:0:0:1:2:0:0:0:0 | 4:1:11:0:6:0:0:0:0 | 0:0:0:0:0:0:0:0:0 | 57:63:348:81:99:17:17:6:0 | 0:1:0:2:1:0:0:0:0 | 2:3:32:20:17:1:1:0:0 | 0:0:0:0:0:0:0:0:0 |
| XM_024447164:XM_024447165:NM_002485:NM_001024688:XM_011517046:XM_024447163:XM_017013462:XM_011517045:XM_017013460 | 2:4:38:6:12:0:0:0:0 | 0:0:0:0:0:0:0:0:0 | 10:11:50:13:17:5:0:0:0 | 0:0:0:1:3:0:0:0:0 | 5:1:11:0:6:0:0:0:0 | 0:0:0:0:0:0:0:0:0 | 57:63:348:83:99:17:17:6:0 | 0:1:0:0:2:0:0:0:0 | 2:3:32:20:17:1:1:0:0 | 0:0:0:0:0:0:0:0:0 |
| XM_024447164:XM_024447165:NM_002485:NM_001024688:XM_011517046:XM_024447163:XM_017013462:XM_011517045:XM_017013460 | 2:3:38:6:12:0:0:0:0 | 0:0:0:0:0:0:0:0:0 | 10:8:49:14:21:5:0:0:0 | 0:1:1:0:0:0:0:0:0 | 5:1:11:0:6:0:0:0:0 | 0:0:0:0:0:0:0:0:0 | 58:57:348:82:102:17:17:6:0 | 0:0:0:1:1:0:0:0:0 | 2:3:32:20:17:1:1:0:0 | 0:0:0:0:0:0:0:0:0 |
| XM_024447164:XM_024447165:NM_002485:NM_001024688:XM_011517046:XM_024447163:XM_017013462:XM_011517045:XM_017013460 | 2:2:38:6:12:1:0:0:0 | 0:0:0:0:0:0:0:0:0 | 10:5:50:14:21:5:0:0:0 | 0:0:0:0:0:0:0:0:0 | 6:1:11:1:6:0:0:0:0 | 0:0:0:0:0:0:0:0:0 | 57:44:348:83:103:17:17:6:0 | 1:0:0:0:1:0:0:0:0 | 2:2:32:20:17:1:1:0:0 | 0:0:0:0:0:0:0:0:0 |

| | | | | | | | | | | |
|---|---|---|---|---|---|---|---|---|---|---|
| XM_024447164:XM_024447165:NM_002485:NM_001024688:XM_011517046:XM_024447163:XM_017013462:XM_011517045:XM_017013460 | 2:2:38:6:12:1:0:0:0 | 0:0:0:0:0:0:0:0:0 | 10:5:50:14:21:5:0:0:0 | 0:0:0:0:0:0:0:0 | 6:1:11:1:6:0:0:0:0 | 0:0:0:0:0:0:0:0:0 | 58:39:348:83:104:17:17:6:0 | 0:1:0:0:0:0:0:0:0 | 2:1:32:20:17:1:1:0:0 | 0:0:0:0:0:0:0:0:0 |
| XM_024447164:XM_024447165:NM_002485:NM_001024688:XM_011517046:XM_024447163:XM_011517045:XM_017013462:XM_017013460 | 2:2:38:6:12:1:0:0:0 | 0:0:0:0:0:0:0:0:0 | 10:4:50:14:21:5:0:0:0 | 0:0:0:0:0:0:0:0 | 6:1:11:1:6:0:0:0:0 | 0:0:0:0:0:0:0:0:0 | 0:0:0:0:0:0:0:0:0 | 59:18:348:83:104:17:6:0:0 | 0:0:0:0:0:0:0:0:0 | 2:0:32:20:17:1:0:0:0 |
| XM_024447164:NM_002485:NM_001024688:XM_011517046:XM_024447163:XM_024447165:XM_011517045:XM_017013462:XM_017013460 | 2:38:6:12:1:0:0:0:0 | 0:0:0:0:0:0:0:0:0 | 10:50:14:21:5:1:0:0:0 | 0:0:0:0:0:0:0:0 | 6:11:1:6:0:0:0:0:0 | 0:0:0:0:0:0:0:0:0 | 59:347:83:105:17:9:6:0:0 | 0:0:0:0:0:0:0:0:0 | 2:32:20:17:1:0:0:0:0 | 0:0:0:0:0:0:0:0:0 |
| NM_004063:XM_011516790 | 576:17:00 | 2:00 | 540:55:00 | 3:00 | 712:27:00 | 1:00 | 2649:372 | 8:01 | 379:22:00 | 0:00 |
| NM_004063:XM_011516790 | 577:17:00 | 1:00 | 542:54:00 | 0:00 | 711:26:00 | 2:00 | 2655:338 | 6:02 | 379:22:00 | 0:00 |
| NM_004063:XM_011516790 | 575:14:00 | 2:00 | 540:38:00 | 2:00 | 714:22:00 | 1:00 | 2667:234 | 5:03 | 377:18:00 | 0:00 |
| NM_004063:XM_011516790 | 577:12:00 | 2:00 | 544:23:00 | 0:00 | 715:17:00 | 2:00 | 2688:140 | 9:02 | 376:17:00 | 2:00 |
| NM_004063:XM_011516790 | 110:02:00 | 469:10:00 | 293:13:00 | 252:10:00 | 359:06:00 | 358:10:00 | 470:11:00 | 2228:105 | 145:06:00 | 233:11:00 |
| NM_004063:XM_011516790 | 0:00 | 579:12:00 | 0:01 | 545:22:00 | 0:00 | 717:16:00 | 468:11:00 | 2233:85 | 144:06:00 | 234:10:00 |
| NM_004063:XM_011516790 | 579:02:00 | 0:00 | 539:04:00 | 1:00 | 718:00:00 | 0:00 | 2643:31:00 | 8:01 | 381:02:00 | 1:00 |
| NM_004063:XM_011516790 | 575:13:00 | 2:00 | 461:18:00 | 0:00 | 710:15:00 | 0:00 | 2197:44:00 | 2:00 | 372:14:00 | 1:00 |
| XM_011516790:NM_004063 | 0.927777778 | 1:01 | 1.0375 | 0:00 | 1.1875 | 0:01 | 42:1890 | 1:04 | 0.829861111 | 0:01 |
| NM_004063:XM_011516790 | 553:13:00 | 1:00 | 403:18:00 | 0:00 | 684:17:00 | 1:00 | 1827:42:00 | 6:02 | 352:13:00 | 0:00 |
| NM_004063:XM_011516790 | 507:11:00 | 2:00 | 148:11:00 | 0:00 | 612:15:00 | 1:00 | 859:35:00 | 3:00 | 302:12:00 | 0:00 |
| NM_004063:XM_011516790 | 491:11:00 | 1:00 | 83:05:00 | 0:00 | 583:15:00 | 1:00 | 618:24:00 | 0:00 | 272:12:00 | 0:00 |
| NM_004063:XM_011516790 | 426:10:00 | 0:00 | 13:02 | 0:00 | 461:12:00 | 0:00 | 0:00 | 278:13:00 | 0:00 | 193:09:00 |
| NM_004063:XM_011516790 | 403:11:00 | 0:00 | 9:02 | 0:00 | 416:12:00 | 2:00 | 213:07:00 | 0:00 | 151:08:00 | 0:00 |
| NM_012415:NM_001205262 | 3:00 | 0:00 | 0:00 | 0:00 | 2:00 | 0:00 | 7:00 | 3:00 | 7:00 | 3:00 |
| NM_183009:NM_015496:XM_017013308 | 4:01:03 | 0:00:00 | 1:00:00 | 1:00:01 | 6:00:01 | 0:00:00 | 18:00:02 | 7:00:00 | 8:00:00 | 0:00:00 |
| NM_183009:NM_015496:XM_017013308 | 14:01:03 | 8:00:00 | 16:00:05 | 33:00:00 | 15:00:01 | 5:00:00 | 116:00:04 | 193:00:02 | 20:00:00 | 8:00:00 |
| NM_183009:NM_015496:XM_017013308 | 20:01:03 | 2:00:00 | 42:00:05 | 7:00:00 | 19:00:01 | 1:00:00 | 269:00:05 | 40:00:01 | 26:00:00 | 2:00:00 |
| NM_183009:NM_015496:XM_017013308 | 23:01:03 | 0:00:00 | 49:00:06 | 0:00:00 | 20:00:01 | 0:00:00 | 321:00:05 | 0:00:01 | 28:00:00 | 0:00:00 |
| NM_006294:NR_045639:NM_001254752:NM_001199975 | 1164:4:10:2 | 14:1:0:0 | 1606:1:2:0 | 102:0:0:0 | 620:3:2:1 | 4:0:0:0 | 13171:8:104:13 | 398:0:3:0 | 1135:3:10:0 | 9:0:1:0 |
| NM_006294:NM_001254752:NR_045639:NM_001199975 | 1170:10:5:2 | 35:0:0:0 | 1616:2:1:0 | 1303:3:5:1 | 621:2:3:1 | 9:0:0:0 | 13238:104:8:13 | 11600:102:63:1 | 1140:11:3:0 | 18:0:0:0 |
| NM_006294:NR_045639:NM_001254752:NM_001199975 | 1202:4:10:2 | 1:1:0:0 | 3027:6:5:1 | 1:0:0:0 | 630:3:2:1 | 0:0:0:0 | 25272:72:209:14 | 2:0:0:0 | 1158:2:11:0 | 0:0:0:0 |
| NM_006294:NM_001254752:NR_045639:NM_001199975 | 1204:10:11:2 | 2:0:0:0 | 3039:5:53:1 | 2:0:0:0 | 631:2:10:1 | 0:0:0:0 | 25306:211:177:14 | 23:0:0:0 | 1153:11:11:0 | 5:0:0:0 |
| NM_006294:NM_001254752:NR_045639:NM_001199975 | 1200:10:11:2 | 2:0:0:0 | 3022:5:53:1 | 6:0:0:0 | 625:2:10:1 | 2:0:0:0 | 25259:210:180:14 | 28:0:0:0 | 1152:11:10:0 | 0:0:1:0 |
| NM_006294:NM_001254752:NR_045639:NM_001199975 | 1201:10:11:2 | 1:0:0:0 | 3025:5:53:1 | 3:0:0:0 | 625:2:10:1 | 2:0:0:0 | 25274:209:180:14 | 13:0:0:0 | 1151:11:11:0 | 1:0:0:0 |
| NM_006294:NM_001254752:NR_045639:NM_001199975 | 1190:10:11:2 | 2:0:0:0 | 3003:5:53:0 | 6:0:0:0 | 615:2:10:1 | 1:0:0:0 | 25174:207:179:13 | 17:1:0:0 | 1141:10:11:0 | 0:0:0:0 |
| NM_006294:NM_001254752:NR_045639:NM_001199975 | 1190:10:11:2 | 3:0:0:0 | 3011:5:53:0 | 5:0:0:0 | 618:2:10:1 | 0:0:0:0 | 25206:208:180:12 | 20:0:0:0 | 1141:10:11:0 | 1:0:0:0 |
| NM_018407 | 820 | 2 | 93 | 0 | 165 | 0 | 4039 | 2 | 481 | 0 |
| NM_018407 | 825 | 1 | 97 | 0 | 167 | 0 | 4170 | 16 | 487 | 1 |
| NM_018407 | 878 | 2 | 111 | 0 | 182 | 0 | 4971 | 3 | 526 | 0 |
| NM_000989 | 100 | 0 | 17 | 0 | 30 | 0 | 214 | 0 | 85 | 0 |
| NM_000989 | 99 | 0 | 16 | 1 | 27 | 3 | 215 | 0 | 85 | 0 |
| NM_015051 | 183 | 0 | 153 | 2 | 139 | 0 | 925 | 2 | 217 | 0 |
| NM_015051 | 181 | 0 | 153 | 0 | 138 | 0 | 919 | 5 | 217 | 2 |
| NM_015051 | 184 | 0 | 155 | 0 | 138 | 0 | 908 | 2 | 218 | 1 |
| NM_015051 | 181 | 0 | 147 | 0 | 135 | 0 | 889 | 2 | 218 | 0 |
| NM_015469:NM_001329570 | 0:00 | 0:00 | 0:00 | 2:00 | 0:00 | 2:00 | 0:00 | 0:00 | 1:00 | 2:00 |
| NM_001146108:NM_012212:XR_001746250 | 0:00:00 | 0:00:00 | 0:00:00 | 0:00:00 | 1:00:00 | 1:01:00 | 0:00:00 | 1:00:00 | 0:00:00 | 0:00:00 |
| XM_011518691:NM_001012994:XM_024447544 | 0:00:00 | 0:00:00 | 5:01:00 | 0:00:00 | 2:00:00 | 0:00:00 | 13:01:10 | 1:00:00 | 1:01:00 | 0:00:00 |
| XM_006717019:NM_015258 | 0:00 | 1:01 | 0:00 | 0:00 | 0:00 | 0:00 | 2:00 | 0:00 | 0:01 | 0:00 |
| NM_017688:NM_001317944:XM_006717149:NM_001317943 | 23:2:3:0 | 0:0:0:0 | 16:3:0:0 | 0:0:0:0 | 41:3:2:2 | 0:0:0:0 | 69:3:4:0 | 1:0:0:0 | 26:0:3:0 | 0:0:0:0 |
| NM_001317944:NM_017688:XM_006717149:NM_001317943 | 0:9:2:0 | 2:12:1:0 | 3:9:2:0 | 0:3:1:0 | 1:22:2:0 | 2:11:1:2 | 3:60:17:0 | 0:0:0:0 | 0:21:5:0 | 0:0:0:0 |
| NM_001304509:NM_001304510:NM_031219 | 0:00:00 | 0:00:00 | 1:00:00 | 1:01:00 | 0:01:00 | 3:00:00 | 1:00:01 | 0:01:01 | 0:00:00 | 0:00:00 |
| NM_000607 | 0 | 0 | 0 | 0 | 0 | 0 | 0 | 0 | 0 | 0 |
| XM_024447530:NM_002160 | 0:00 | 0:00 | 0:00 | 0:00 | 0:00 | 0:00 | 0:00 | 0:00 | 0:00 | 3:00 |
| NM_198252:XM_024447524:NM_000177:NM_001353060:XR_001746283 | 1:0:0:0:0 | 0:0:0:0:0 | 0:3:0:0:0 | 0:0:0:0:0 | 2:3:2:0:0 | 0:0:0:0:0 | 0:0:0:0:0 | 0:0:0:0:0 | 1:1:0:0:0 | 0:0:0:0:0 |
| NM_198252:XM_024447524:XR_001746283:NM_000177:NM_001353060 | 1:0:0:0:0 | 0:0:0:0:0 | 0:3:1:0:0 | 0:0:0:0:0 | 2:3:0:2:0 | 0:0:0:0:0 | 0:0:0:0:0 | 0:0:0:0:0 | 1:1:0:0:0 | 0:0:0:0:0 |
| NM_004099 | 0 | 0 | 0 | 0 | 0 | 0 | 0 | 0 | 1 | 0 |
| NM_002799 | 0 | 0 | 1 | 1 | 4 | 0 | 4 | 4 | 3 | 1 |
| NM_203403:XM_005251443 | 0:00 | 0:00 | 0:00 | 1:00 | 0:00 | 1:00 | 0:00 | 0:01 | 0:00 | 0:00 |
| NM_003863 | 0 | 0 | 0 | 0 | 0 | 0 | 0 | 0 | 0 | 2 |
| NM_005564 | 339 | 2 | 741 | 4 | 825 | 2 | 4141 | 4 | 1006 | 1 |
| NM_005564 | 342 | 0 | 750 | 1 | 831 | 1 | 4160 | 7 | 1014 | 0 |
| NM_016035:NM_001305942 | 0:00 | 0:00 | 0:00 | 0:00 | 0:00 | 1:00 | 0:00 | 1:00 | 0:00 | 0:01 |
| NM_032799 | 0 | 1 | 0 | 0 | 0 | 0 | 0 | 0 | 0 | 1 |
| XM_011519174:NM_003934:XM_011519173:XR_001746411:XM_017015272:XM_017015270:XR_001746413 | 1:18:1:1:1:0:0 | 0:0:0:0:0:0:0 | 0:0:0:1:3:0:0 | 0:0:0:0:0:0:0 | 0:12:0:0:2:1:0 | 0:0:0:0:0:0:0 | 1:15:0:7:7:0:1 | 0:2:0:0:0:0:0 | 1:9:0:2:1:0:0 | 0:0:0:0:0:0:0 |
| NM_014285:XM_005272176:NM_001282709:NM_001282708:XR_001746262:XM_017014558:NR_104230:XM_006717023 | 25:1:2:1:1:0:0:0 | 0:0:0:0:0:0:0:0 | 7:0:0:0:0:0:0:0 | 0:0:0:0:0:0:0:0 | 8:0:0:0:1:0:0:0 | 0:0:0:0:0:0:0:0 | 211:4:12:10:7:16:3:0 | 2:0:0:0:0:0:0:0 | 48:0:3:3:2:1:0:0 | 1:0:0:0:0:0:0:0 |
| XM_005272207:NM_022779:XR_929836:XR_246600:XM_011518921:XM_006717236:NM_001322340:XM_011518922 | 0:1:1:0:0:0:0:0 | 1:0:0:0:0:0:0:0 | 0:0:0:0:0:0:0:0 | 1:0:0:0:0:0:0:0 | 0:2:0:1:0:0:0:0 | 0:2:0:1:0:0:0:0 | 0:0:0:0:0:0:0:0 | 27:4:0:2:2:1:1:0 | 0:0:0:0:0:0:0:0 | 4:6:0:0:0:0:0:0 |
| NM_022779:XR_929836:NM_001322344:XM_011518921:XR_246600:XM_006717236:XM_005272207:NM_001322340:NM_001322341 | 1:4:1:0:0:0:0:0:0 | 0:0:0:0:0:0:0:0:0 | 0:0:2:0:0:0:0:0:0 | 0:0:0:0:0:0:0:0:0 | 4:0:0:1:0:0:0:0:0 | 0:0:0:0:0:0:0:0:0 | 4:4:5:1:1:1:4:1:0 | 0:0:0:0:0:0:0:0:0 | 6:0:1:0:0:0:0:0:0 | 0:0:0:0:0:0:0:0:0 |
| XM_005272211:NM_001162426:NM_001162427:XM_017015100:NM_000368:XM_017015101:XM_017015098 | 1:0:0:0:0:0:0 | 0:0:0:0:0:0:0 | 0:0:0:0:0:0:0 | 0:0:0:0:0:0:0 | 1:0:0:0:0:0:0 | 0:0:0:0:0:0:0 | 0:1:0:2:1:2:0 | 0:1:1:0:0:0:0 | 0:0:0:0:1:0:0 | 0:0:0:0:0:0:0 |
| XM_005272211:XM_006717271:NM_001162426:NM_001162427:XM_017015098:NM_000368:XM_017015100:XM_017015101:XM_017015097:XM_011518979 | 2:1:2:0:0:0:0:0:0:0 | 0:0:0:0:0:0:0:0:0:0 | 0:0:0:0:0:0:0:0:0:0 | 0:0:0:0:0:0:0:0:0:0 | 1:0:0:1:1:2:0:0:0:0 | 0:0:0:0:0:0:0:0:0:0 | 0:0:9:2:2:2:5:1:0:0 | 0:0:0:0:0:0:0:0:0:0 | 0:0:0:1:1:5:0:0:0:0 | 0:0:0:0:0:0:0:0:0:0 |
| NM_001807 | 1 | 0 | 0 | 0 | 0 | 0 | 8 | 2 | 1 | 0 |
| NM_001807 | 1 | 0 | 0 | 0 | 0 | 0 | 8 | 2 | 1 | 0 |
| NM_001807 | 1 | 0 | 0 | 0 | 0 | 0 | 1 | 2 | 0 | 1 |

| | | | | | | | | | | |
|---|---|---|---|---|---|---|---|---|---|---|
| NM_001807 | 1 | 0 | 0 | 0 | 0 | 0 | 1 | 2 | 0 | 1 |
| NM_017503 | 1 | 2 | 0 | 0 | 0 | 0 | 0 | 0 | 0 | 0 |
| NM_017503 | 3 | 0 | 0 | 0 | 0 | 0 | 0 | 0 | 0 | 0 |
| XM_017015112 | 0 | 0 | 0 | 0 | 0 | 0 | 0 | 2 | 0 | 0 |
| XM_017015165:NM_007371:XM_017015166:XM_006717291 | 1:1:0:0 | 0:0:0:0 | 0:2:0:0 | 0:3:0:0 | 0:3:1:0 | 0:0:0:0 | 3:12:2:1 | 0:1:1:0 | 0:1:0:0 | 0:0:0:0 |
| XM_017015165:NM_007371:XM_017015166:XM_006717291 | 1:1:0:0 | 0:0:0:0 | 0:5:0:0 | 0:0:0:0 | 0:3:1:0 | 0:0:0:0 | 3:5:2:1 | 0:8:1:0 | 0:1:0:0 | 0:0:0:0 |
| NM_024718 | 0 | 0 | 0 | 1 | 0 | 0 | 0 | 2 | 0 | 0 |
| NM_013379:XM_017014652:XM_024447525:XM_011518599 | 0:0:0:0 | 0:0:0:0 | 0:0:0:0 | 0:0:0:0 | 0:0:0:0 | 0:0:0:0 | 1:1:0:0 | 1:0:0:0 | 0:0:0:0 | 0:0:0:0 |
| NM_001317898:XM_024447402:NM_033222:NM_001128217:XM_024447400:XM_024447399:NM_001317900:NM_021144:XM_024447401 | 7:5:10:8:3:0:0:0:0 | 5:3:1:0:0:0:0:0:0 | 24:32:8:5:4:0:0:0:0 | 7:7:1:0:0:0:0:0:0 | 1:1:9:5:4:0:0:0:0 | 5:4:0:0:0:0:0:0:0 | 138:95:26:13:11:3:1:0:0 | 42:25:0:0:0:0:0:0:0 | 16:15:7:8:14:1:0:1:0 | 7:8:0:0:0:0:0:0:0 |
| NM_033222:NM_001128217:XM_024447400:NM_001317898:XM_024447402:XM_024447399:NM_001317900:NM_021144:XM_024447401 | 11:8:3:13:8:0:0:0:0 | 0:0:0:0:0:0:0:0:0 | 9:5:4:31:41:0:0:0:0 | 0:0:0:0:3:0:0:0:0 | 9:5:4:7:5:0:0:0:0 | 0:0:0:0:0:0:0:0:0 | 27:13:11:190:132:3:1:0:0 | 0:0:0:8:1:0:0:0:0 | 7:8:14:26:26:1:0:1:0 | 0:0:0:0:0:0:0:0:0 |
| NM_001317898:XM_024447402:NM_033222:NM_001128217:XM_024447400:XM_024447399:NM_001317900:NM_021144:XM_024447401 | 13:8:11:8:3:0:0:0:0 | 1:1:0:0:0:0:0:0:0 | 31:42:9:5:4:0:0:0:0 | 0:3:0:0:0:0:0:0:0 | 7:5:9:5:4:0:0:0:0 | 1:1:0:0:0:0:0:0:0 | 193:135:27:13:11:3:1:0:0 | 5:2:0:0:0:0:0:0:0 | 26:26:7:8:14:1:0:1:0 | 2:0:0:0:0:0:0:0:0 |
| NM_033222:NM_001128217:XM_024447400:NM_001317898:XM_024447402:XM_024447399:NM_001317900:NM_021144:XM_024447401 | 11:8:3:14:10:0:0:0:0 | 0:0:0:0:0:0:0:0:0 | 9:5:4:31:48:0:0:0:0 | 0:0:0:0:0:0:0:0:0 | 9:5:4:8:6:0:0:0:0 | 0:0:0:0:2:0:0:0:0 | 27:13:11:204:140:3:1:0:0 | 0:0:0:3:2:0:0:0:0 | 7:8:14:29:26:1:0:1:0 | 0:0:0:0:0:0:0:0:0 |
| NM_001010 | 8058 | 10 | 2851 | 2 | 3044 | 4 | 31677 | 26 | 7262 | 10 |
| NM_001010 | 8067 | 10 | 2848 | 4 | 3045 | 3 | 31674 | 38 | 7270 | 8 |
| NM_001010 | 8092 | 5 | 2829 | 1 | 3058 | 1 | 31750 | 11 | 7299 | 3 |
| NM_001010 | 8119 | 2 | 2813 | 5 | 3060 | 4 | 31752 | 42 | 7296 | 9 |
| NM_001010 | 8109 | 4 | 2756 | 28 | 3044 | 1 | 31609 | 216 | 7289 | 6 |
| NM_001010 | 8063 | 8 | 2771 | 8 | 3032 | 3 | 31717 | 46 | 7275 | 7 |
| NM_001010 | 7971 | 16 | 2722 | 8 | 3009 | 3 | 31494 | 28 | 7192 | 11 |
| NM_001010 | 7945 | 11 | 2712 | 8 | 2992 | 2 | 31415 | 36 | 7164 | 5 |
| NM_001010 | 7894 | 17 | 2698 | 10 | 2971 | 8 | 31336 | 40 | 7137 | 11 |
| XM_024447586:NM_017794 | 0:00 | 0:00 | 1:00 | 0:00 | 0:00 | 0:00 | 0:00 | 0:00 | 0:01 | 0:00 |
| NM_002451 | 76 | 1 | 44 | 0 | 31 | 0 | 426 | 1 | 118 | 0 |
| NM_002451 | 0 | 75 | 0 | 41 | 0 | 32 | 434 | 1 | 118 | 0 |
| NM_002451 | 77 | 1 | 43 | 0 | 32 | 0 | 442 | 2 | 118 | 0 |
| NM_002451 | 78 | 0 | 43 | 0 | 32 | 0 | 447 | 0 | 118 | 0 |
| XM_011518045:NM_001193536:XM_011518047:XM_017015174:XM_011518046:NM_203447:XM_011518048 | 0:1:0:0:0:0:0 | 1:0:0:0:0:0:0 | 0:5:0:1:1:0:0 | 0:1:1:0:0:0:0 | 0:0:0:0:0:0:0 | 0:1:0:0:0:0:0 | 0:0:0:0:0:1:0 | 0:0:0:0:1:6:0 | 0:0:0:1:0:0:0 | 0:0:0:0:0:0:0 |
| NM_001193536:XM_011518045:XM_011518046:XM_011518047:NM_203447:XM_017015174:XM_011518048 | 0:1:0:0:0:0:0 | 1:0:0:0:0:0:0 | 1:0:0:1:0:0:0 | 5:0:1:0:0:0:0 | 1:0:0:0:0:0:0 | 0:0:0:0:0:0:0 | 0:0:1:0:7:0:0 | 0:0:0:0:0:0:0 | 0:0:0:0:0:1:0 | 0:0:0:0:0:0:0 |
| NM_004323 | 0 | 11 | 0 | 8 | 1 | 24 | 1 | 36 | 0 | 14 |
| NM_007343:NM_001197098:XM_011517965 | 0:00:00 | 2:00:00 | 0:00:00 | 1:00:00 | 0:00:00 | 10:01:00 | 0:00:00 | 2:01:00 | 0:00:00 | 0:00:00 |
| XM_005251403:NM_174923:NM_001195200 | 0:00:00 | 0:00:00 | 0:00:00 | 0:00:00 | 0:00:00 | 2:01:00 | 0:00:00 | 0:00:00 | 0:00:00 | 0:01:00 |
| NM_001216 | 0 | 0 | 0 | 0 | 0 | 0 | 0 | 0 | 0 | 3 |
| NM_012203 | 2 | 0 | 0 | 0 | 1 | 0 | 1 | 3 | 1 | 1 |
| NM_005772:XM_006716715:XM_017014176:NM_001286699:NM_001286700 | 27:1:0:0:0 | 0:0:0:0:0 | 8:2:0:0:0 | 0:0:0:0:0 | 26:1:0:0:0 | 0:0:0:0:0 | 229:29:8:1:2 | 0:0:0:0:0 | 35:3:2:1:0 | 0:0:0:0:0 |
| NM_001330668:XM_005272751:XR_428504 | 0:00:00 | 1:00:00 | 0:00:00 | 0:00:00 | 0:00:00 | 0:00:00 | 0:00:01 | 1:01:00 | 0:00:00 | 1:01:00 |
| NM_001330668:XM_005272751:XR_428504 | 0:00:00 | 1:00:00 | 0:00:00 | 0:00:00 | 0:00:00 | 0:00:00 | 0:00:01 | 1:01:00 | 0:00:00 | 1:01:00 |
| XM_011519209:XM_011519208:XM_011519206 | 0:00:00 | 1:00:00 | 0:00:00 | 0:00:00 | 0:00:00 | 0:01:00 | 0:00:00 | 0:00:00 | 0:00:00 | 2:00:00 |
| NM_001256876:NM_153186:XM_017014522:XR_002956771:NR_148869:NM_001354333:NM_015158:XM_024447465:NM_001354344 | 1:0:0:0:0:0:0:0:0 | 1:1:0:0:0:0:0:0:0 | 0:0:0:0:0:0:0:0:0 | 0:0:0:0:0:0:0:0:0 | 0:0:0:0:0:0:0:0:0 | 0:0:1:0:0:0:0:0:0 | 0:2:1:7:5:1:2:1:0 | 0:0:0:0:0:0:0:0:0 | 0:0:0:1:0:0:0:0:1 | 0:0:0:0:0:0:0:0:0 |
| NM_001256876:NM_153186:NM_015158:XM_017014522:XR_002956771 | 0:0:0:0:0 | 2:3:0:0:0 | 0:0:0:0:0 | 0:0:0:0:0 | 0:0:0:0:0 | 0:0:0:0:0 | 0:0:1:0:1 | 0:0:1:1:0 | 0:0:0:0:0 | 0:0:0:0:0 |
| NM_033428:NM_001318058 | 1:00 | 0:00 | 0:00 | 0:00 | 1:00 | 0:00 | 2:00 | 5:00 | 4:00 | 2:01 |
| NM_006415:XM_024447379:NM_001281303:XM_011518138 | 141:1:0:0 | 0:0:0:0 | 53:0:0:0 | 0:0:0:0 | 198:0:0:0 | 0:0:0:0 | 232:1:0:0 | 0:0:0:0 | 209:3:0:0 | 1:0:0:0 |
| NM_006415:XM_024447379:NM_001281303:XM_011518138 | 141:1:0:0 | 0:0:0:0 | 53:0:0:0 | 0:0:0:0 | 198:0:0:0 | 0:0:0:0 | 231:1:0:0 | 1:0:0:0 | 210:3:0:0 | 0:0:0:0 |
| NM_006415:XM_024447379:NM_001281303:XM_011518138 | 141:1:0:0 | 0:0:0:0 | 54:0:0:0 | 0:0:0:0 | 198:0:0:0 | 0:0:0:0 | 235:1:0:0 | 0:0:0:0 | 210:3:0:0 | 0:0:0:0 |
| NM_006415:XM_024447379:NM_001281303:XM_011518138 | 140:1:0:0 | 0:0:0:0 | 54:0:0:0 | 0:0:0:0 | 197:0:0:0 | 1:0:0:0 | 235:1:0:0 | 0:0:0:0 | 210:3:0:0 | 0:0:0:0 |
| NM_006415:XM_024447379:NM_001281303:XM_011518138 | 140:1:0:0 | 0:0:0:0 | 54:0:0:0 | 0:0:0:0 | 195:0:0:0 | 3:0:0:0 | 235:1:0:0 | 0:0:0:0 | 210:3:0:0 | 0:0:0:0 |
| NM_006415:XM_024447379:XM_011518138:NM_001281303 | 140:1:0:0 | 0:0:0:0 | 54:0:0:0 | 0:0:0:0 | 198:0:0:0 | 0:0:0:0 | 234:1:1:0 | 1:0:0:0 | 210:3:0:0 | 0:0:0:0 |
| NM_006415:XM_024447379:XM_011518138:NM_001281303 | 140:1:0:0 | 0:0:0:0 | 54:0:0:0 | 0:0:0:0 | 198:0:0:0 | 0:0:0:0 | 234:1:1:0 | 1:0:0:0 | 210:3:0:0 | 0:0:0:0 |
| NM_006415:XM_024447379:XM_011518138:NM_001281303 | 140:1:0:0 | 0:0:0:0 | 54:0:0:0 | 0:0:0:0 | 197:0:0:0 | 1:0:0:0 | 233:1:1:0 | 2:0:0:0 | 210:3:0:0 | 0:0:0:0 |
| NM_006415:XM_024447379:XM_011518138:NM_001281303 | 140:1:0:0 | 0:0:0:0 | 54:0:0:0 | 0:0:0:0 | 198:0:0:0 | 0:0:0:0 | 235:1:1:0 | 0:0:0:0 | 210:3:0:0 | 0:0:0:0 |
| NM_006415:XM_024447379:XM_011518138:NM_001281303 | 140:1:0:0 | 0:0:0:0 | 53:0:0:0 | 1:0:0:0 | 198:0:0:0 | 0:0:0:0 | 236:1:1:0 | 0:0:0:0 | 209:3:0:0 | 1:0:0:0 |
| NM_006415:XM_024447379:XM_011518138:NM_001281303 | 140:1:0:0 | 0:0:0:0 | 54:0:0:0 | 0:0:0:0 | 198:0:0:0 | 0:0:0:0 | 237:1:1:0 | 0:0:0:0 | 210:3:0:0 | 0:0:0:0 |
| NM_006415:XM_024447379:XM_011518138:NM_001281303 | 140:1:0:0 | 1:0:0:0 | 54:0:0:0 | 0:0:0:0 | 197:0:0:0 | 1:0:0:0 | 237:1:1:0 | 0:0:0:0 | 210:3:0:0 | 0:0:0:0 |
| NM_006415:XM_024447379:XM_011518138:NM_001281303 | 141:1:0:0 | 0:0:0:0 | 53:0:0:0 | 0:0:0:0 | 197:0:0:0 | 0:0:0:0 | 234:1:3:0 | 1:0:0:0 | 210:3:0:0 | 0:0:0:0 |
| NM_006415:XM_024447379:XM_011518138:NM_178324:NM_001281303 | 141:1:0:0:0 | 0:0:0:0:0 | 53:0:0:0:0 | 0:0:0:0:0 | 197:0:0:0:0 | 0:0:0:0:0 | 234:1:3:0:0 | 0:0:0:0:0 | 209:3:0:0:0 | 1:0:0:0:0 |
| NM_006415:XM_024447379:XM_011518138:NM_178324:NM_001281303 | 141:1:0:0:0 | 0:0:0:0:0 | 53:0:0:0:0 | 0:0:0:0:0 | 195:0:0:0:0 | 2:0:0:0:0 | 233:1:4:1:0 | 0:0:0:0:0 | 209:3:0:0:0 | 0:0:0:0:0 |
| NM_022755:XR_001746365:XM_017015041 | 5:01:00 | 0:00:00 | 2:00:00 | 0:00:00 | 10:01:00 | 0:00:00 | 1:00:00 | 5:00:00 | 4:00:00 | 0:00:00 |
| NM_145006:XM_011518359:XM_011518361:XM_011518358:XM_017014449:NM_001287005:XM_017014450:XM_017014448 | 0:0:0:0:0:0:0:0 | 4:1:1:1:1:1:1:0 | 0:0:0:0:0:0:0:0 | 0:0:0:0:0:0:0:0 | 0:0:0:0:0:0:0:0 | 0:0:0:2:2:0:0:0 | 1:0:1:4:2:0:0:1 | 0:0:0:0:0:0:0:0 | 3:0:0:1:1:0:0:0 | 0:0:0:0:0:0:0:0 |
| NR_134466:NM_001318011:NR_134468:NM_032310:XR_929848:NR_134465:NR_134467:NM_001318010 | 3:0:1:2:1:0:0:0 | 6:1:1:6:0:0:0:0 | 3:1:2:8:0:1:0:0 | 5:2:0:4:0:1:0:0 | 12:2:5:15:1:0:1:0 | 11:1:0:15:2:1:1:0 | 16:5:1:50:3:0:7:2 | 0:0:0:0:0:0:0:0 | 15:1:0:17:2:0:1:1 | 0:0:0:0:0:0:0:0 |
| XM_017015236:XR_001746404:XR_929852:XR_001746403:XM_017015235:XR_929854:NM_032823:XM_017015237:XR_001746402:XR_929856:XM_006717306:XM_005252284:XM_017015234 | 10:11:1:1:0:0:0:0:0:0:0:0:0 | 0:0:0:0:0:0:0:0:0:0:0:0:0 | 9:14:0:0:0:0:0:0:0:0:0:0:0 | 0:0:0:0:0:0:0:0:0:0:0:0:0 | 24:22:0:0:1:1:0:0:0:0:0:0:0 | 0:0:0:0:0:0:0:0:0:0:0:0:0 | 103:109:18:10:3:1:4:1:11:13:0:0:0 | 0:0:0:0:0:0:0:0:0:0:0:0:0 | 10:10:1:1:0:0:0:0:0:3:1:0:0 | 0:0:0:0:0:0:0:0:0:0:0:0:0 |
| XM_024447451:NM_000136:NM_001243744:XM_011518366:XM_006717002:XM_017014453:NM_001243743:XM_017014452:XM_006717004 | 2:1:2:1:0:0:0:0:0 | 0:0:0:0:0:0:0:0:0 | 1:0:0:2:1:0:0:0:0 | 0:0:0:0:0:0:0:0:0 | 1:2:1:1:3:1:2:0:0 | 0:0:0:0:0:0:0:0:0 | 2:2:9:13:2:1:0:1:0 | 0:0:0:0:0:0:0:0:0 | 1:6:3:4:1:0:0:0:0 | 0:0:0:0:0:0:0:0:0 |
| NR_149061 | 0 | 0 | 1 | 0 | 0 | 0 | 0 | 2 | 0 | 0 |
| NM_014282:XM_017014484 | 0:00 | 0:00 | 0:00 | 0:00 | 0:00 | 0:00 | 0:00 | 2:00 | 0:00 | 0:00 |
| NM_018946 | 1 | 0 | 1 | 0 | 1 | 0 | 0 | 0 | 7 | 0 |
| NM_017919:XM_024447593:XR_001746332:XM_011518821:XM_011518820 | 13:2:3:0:0 | 0:0:0:0:0 | 46:4:8:4:2 | 0:0:0:0:0 | 18:3:0:1:0 | 0:0:0:0:0 | 132:28:30:8:4 | 1:0:1:0:0 | 14:1:5:0:1 | 0:0:0:0:0 |
| NM_017919:XM_024447593:XR_001746332:XM_011518821:XM_011518820 | 13:2:2:0:0 | 0:0:0:0:0 | 45:4:6:4:2 | 1:0:1:0:0 | 18:4:0:1:0 | 0:0:0:0:0 | 133:28:27:8:4 | 0:0:0:0:0 | 14:1:5:0:1 | 0:0:0:0:0 |
| NM_017919:XM_024447593:XR_001746332:XM_011518821:XM_011518820 | 10:2:0:0:0 | 3:0:0:0:0 | 45:5:0:4:1 | 1:0:1:0:0 | 17:4:0:1:0 | 1:0:0:0:0 | 117:39:2:6:3 | 13:1:0:1:0 | 11:2:0:0:1 | 3:0:0:0:0 |
| NM_015051 | 181 | 2 | 152 | 0 | 136 | 2 | 916 | 7 | 215 | 1 |

| 3L__WT | 3L__MUT | 3N__WT | 3N__MUT | 3T__WT | 3T__MUT |
|---|---|---|---|---|---|
| 0 | 0 | 4 | 0 | 9 | 0 |
| 0:00 | 0:00 | 0:00 | 0:01 | 0:00 | 1:00 |
| 385:2:24:2 | 0:0:0:0 | 779:27:57:1 | 1:0:0:0 | 1502:50:138:8 | 2:0:0:0 |
| 0:0:0:0 | 1:0:0:0 | 0:0:0:1 | 0:0:0:0 | 0:0:0:0 | 0:0:0:0 |
| 2:62:0:0:3:0:0 | 0:0:0:0:0:0:0 | 2:74:0:0:30:1:1 | 0:0:0:0:0:0:0 | 13:142:5:0:52:1:2 | 0:0:0:0:0:0:1 |
| 2:56:0:0:1:0:0 | 0:0:0:0:0:0:0 | 2:68:0:47:8:0:0 | 0:0:0:0:0:0:0 | 9:136:1:83:14:1:1 | 0:0:0:0:0:0:0 |
| 14:00 | 0:00 | 5:00 | 0:00 | 37:00:00 | 0:00 |
| 14:00 | 0:00 | 3:00 | 0:00 | 34:00:00 | 5:00 |
| 13:00 | 0:00 | 2:00 | 0:00 | 27:00:00 | 0:00 |
| 0 | 0 | 0 | 4 | 0 | 1 |
| 0:0:0:0:0:0 | 3:0:1:2:2:0 | 0:0:0:0:0:0 | 4:1:3:0:7:5 | 0:0:0:0:0:0 | 20:4:10:16:15:14 |
| 2:0:2:1:0:0:0 | 0:0:0:0:0:0:0 | 0:1:2:1:0:1:0 | 0:0:0:0:0:0:0 | 4:0:5:0:1:4:1 | 0:0:0:0:0:0:0 |
| 0:2:0:1:0:0:0 | 0:0:0:0:0:0:0 | 0:1:0:1:0:0:0 | 0:0:0:0:0:0:0 | 0:3:0:1:1:3:1 | 0:0:0:0:0:0:0 |
| 255:2:11:2 | 0:0:0:0 | 961:0:3:7 | 1:0:0:0 | 2094:8:33:21 | 1:0:0:0 |
| 255:2:11:2 | 0:0:0:0 | 954:0:3:7 | 2:0:0:0 | 2088:7:33:21 | 5:0:0:0 |
| 254:2:11:2 | 1:0:0:0 | 950:0:3:7 | 1:0:0:0 | 2092:7:33:21 | 5:0:0:0 |
| 249:2:11:2 | 0:0:0:0 | 839:0:3:7 | 3:0:0:0 | 1969:6:33:20 | 2:0:0:0 |
| 250:2:11:2 | 0:0:0:0 | 818:0:3:7 | 2:0:0:0 | 1943:6:33:20 | 2:0:0:0 |
| 244:2:11:2 | 0:0:0:0 | 736:1:3:7 | 3:0:0:0 | 1776:6:34:20 | 6:0:0:0 |
| 231:11:2:2 | 1:0:0:0 | 478:5:1:7 | 1:0:0:0 | 1280:35:6:21 | 1:0:0:0 |
| 218:2:11:2 | 0:0:0:0 | 371:1:5:7 | 0:0:0:0 | 1067:6:35:21 | 5:0:0:0 |
| 218:2:11:2 | 0:0:0:0 | 348:1:5:7 | 0:0:0:0 | 1023:7:35:21 | 3:0:0:0 |
| 217:2:11:2 | 0:0:0:0 | 321:1:5:7 | 2:0:0:0 | 958:7:35:21 | 1:0:0:0 |
| 193:2:11:2 | 0:0:0:0 | 156:1:7:7 | 0:0:0:0 | 543:8:35:21 | 4:0:0:0 |
| 10:3:10:38 | 0:0:0:0 | 121:64:132:0 | 0:0:0:0 | 76:39:84:52 | 0:0:0:0 |
| 2:7:7:42:17 | 0:0:1:0:1 | 0:39:115:0:58 | 0:0:0:0:0 | 0:25:71:55:68 | 0:0:0:0:0 |
| 2:7:9:42:19 | 0:0:0:0:0 | 0:36:115:0:56 | 0:0:1:0:1 | 0:25:72:55:66 | 0:1:0:0:0 |
| 2:7:9:42:19 | 0:0:0:0:0 | 0:35:116:0:55 | 0:1:0:0:1 | 0:24:71:55:68 | 0:0:1:0:1 |
| 23:9:2:42:8 | 0:0:0:0:0 | 22:91:0:0:22 | 0:0:0:0:0 | 57:56:0:55:20 | 1:0:0:0:1 |
| 2:42:7:33:11 | 0:0:0:0:0 | 0:0:83:35:36 | 0:0:1:2:0 | 0:55:47:86:29 | 0:0:0:0:0 |
| 42:2:6:33:12 | 0:0:1:0:0 | 0:0:73:49:41 | 0:0:1:1:1 | 55:0:44:93:31 | 0:0:0:1:0 |
| 42:2:3:36:13 | 0:0:1:0:0 | 0:0:78:79:59 | 0:0:2:0:0 | 55:0:53:101:37 | 0:0:0:0:0 |
| 41:2:42:3:14 | 0:0:0:0:0 | 168:0:0:79:130 | 0:0:0:3:0 | 149:0:55:58:69 | 1:0:0:1:0 |
| 3:42:2:41:15 | 0:0:0:0:0 | 79:0:0:220:166 | 0:0:0:0:2 | 58:55:0:172:83 | 0:0:0:0:0 |
| 51:2:42:3:17 | 0:0:0:0:0 | 415:0:0:74:347 | 3:0:0:0:1 | 275:0:55:56:172 | 2:0:0:0:0 |
| 1:3:0:3:0 | 1:39:3:48:17 | 0:0:42:216:181 | 0:0:31:220:181 | 0:5:2:19:3 | 0:50:54:268:179 |
| 2:42:57:3:18 | 0:0:0:0:0 | 0:0:548:57:499 | 0:0:3:0:4 | 0:55:351:48:257 | 0:0:3:0:4 |
| 57:2:42:3:18 | 0:0:0:0:0 | 566:0:0:55:523 | 1:0:0:0:3 | 368:0:55:48:270 | 1:0:0:0:0 |
| 67:2:23:42:1 | 0:0:0:0:0 | 754:0:707:0:41 | 2:0:3:0:0 | 467:0:370:55:31 | 1:0:1:0:0 |
| 2:23:42:69:1 | 0:0:0:0:0 | 0:727:0:773:35 | 0:0:0:1:0 | 0:380:55:478:28 | 0:0:0:2:1 |
| 72:2:25:42:2 | 0:0:1:0:0 | 854:0:839:0:21 | 0:0:2:0:0 | 533:0:448:55:19 | 2:0:1:0:0 |
| 2:26:42:72:2 | 0:0:0:0:0 | 0:842:0:852:20 | 0:0:0:2:0 | 0:449:55:533:19 | 0:0:0:2:0 |
| 73:27:2:42:2 | 0:0:0:0:0 | 870:870:0:0:23 | 4:2:0:0:1 | 547:465:0:54:19 | 2:2:0:1:0 |
| 2:29:78:42:2 | 0:0:0:0:0 | 0:913:912:0:20 | 0:1:1:0:1 | 0:485:566:55:21 | 0:1:1:0:0 |
| 42:78:2:30:3 | 0:0:0:0:0 | 0:952:0:958:26 | 0:4:0:4:0 | 55:591:0:515:23 | 0:0:0:1:2 |
| 2:31:7:82:42 | 0:0:0:0:0 | 0:1004:100:1019:0 | 0:0:0:0:0 | 0:536:67:634:55 | 0:0:0:1:0 |
| 2:31:7:81:42 | 0:0:0:1:0 | 0:1003:109:1023:0 | 0:1:1:1:0 | 0:534:68:635:55 | 0:0:0:3:0 |
| 83:2:31:9:42 | 0:0:0:0:0 | 1028:0:1001:126:0 | 3:0:2:1:0 | 640:0:534:73:55 | 1:0:0:0:0 |
| 2:31:9:83:42 | 0:0:0:0:0 | 0:1001:133:1031:0 | 0:1:2:1:0 | 0:533:75:641:55 | 0:0:0:2:0 |
| 2:31:11:88:42 | 0:0:0:0:0 | 0:993:215:1083:0 | 0:1:1:1:0 | 0:524:112:682:55 | 0:0:0:0:0 |
| 2:31:11:89:42 | 0:0:0:0:0 | 0:988:240:1101:0 | 0:1:0:2:0 | 0:521:124:701:55 | 0:1:0:0:0 |
| 13:95:2:29:42 | 0:0:0:0:0 | 334:1206:0:982:0 | 0:0:0:0:0 | 157:754:0:518:55 | 1:0:0:0:0 |
| 0:0:0:0:0:0 | 7:1:0:0:0:0 | 0:0:0:0:0:0 | 6:0:0:1:1:0 | 0:0:0:0:0:0 | 24:7:1:3:0:1 |
| 7 | 0 | 4 | 1 | 6 | 0 |
| 0:3:0:0:0:0:0:0:0 | 0:9:1:0:0:0:0:0:0 | 0:1:4:0:0:0:1:0:0 | 0:0:6:0:1:0:0:0:0 | 0:7:7:0:2:0:0:1:1 | 0:3:1:0:0:0:0:0:0 |
| 0:3:0:1:0:0:0:0:0:0 | 0:7:1:0:0:0:0:0:0:0 | 0:0:5:0:1:0:0:0:1:0 | 0:0:6:3:2:0:0:2:0:0 | 0:6:10:1:3:0:0:6:0:1 | 0:3:1:0:1:0:0:0:0:0 |
| 10:0:1:0:0:0:0:0 | 0:0:0:0:0:0:0:0 | 0:0:2:0:0:3:0:0 | 0:0:0:0:0:0:0:0 | 6:0:6:0:0:3:3:1 | 0:0:0:0:0:0:0:0 |
| 0:00 | 0:02 | 0:00 | 1:05 | 0:00 | 3:05 |
| 1 | 3 | 3 | 8 | 1 | 3 |
| 3 | 1 | 7 | 2 | 3 | 1 |
| 1:00 | 3:00 | 1:01 | 0:00 | 1:00 | 0:00 |

| | | | | | |
|---|---|---|---|---|---|
| 0:0:0:0 | 0:0:0:0 | 0:0:0:0 | 0:0:1:0 | 0:0:0:0 | 0:1:0:1 |
| 1 | 3 | 2 | 1 | 1 | 2 |
| 1:00:00 | 1:00:00 | 0:00:00 | 0:00:00 | 0:00:01 | 2:00:00 |
| 0:00:00 | 2:00:00 | 0:00:00 | 0:00:00 | 0:00:00 | 2:00:01 |
| 2:00 | 0:00 | 0:00 | 0:00 | 2:01 | 0:00 |
| 0:00 | 0:00 | 1:00 | 0:00 | 0:01 | 1:01 |
| 2488:132 | 3:00 | 2903:119 | 7:00 | 16166:1021 | 27:02:00 |
| 2487:132 | 4:00 | 2907:119 | 4:00 | 16179:1023 | 28:00:00 |
| 2492:133 | 0:00 | 2926:120 | 3:00 | 16237:1022 | 10:03 |
| 412:09:00 | 3:00 | 481:39:00 | 1:00 | 3637:263 | 6:02 |
| 415:09:00 | 0:00 | 492:40:00 | 1:00 | 3682:267 | 1:00 |
| 416:08:00 | 0:00 | 492:36:00 | 0:00 | 3678:247 | 3:02 |
| 0 | 1 | 0 | 0 | 3 | 4 |
| 123:2:202:101:64:7:2:1:0:2:0:0 | 0:0:0:0:0:0:0:0:0:0:0:0 | 251:8:348:194:18:10:0:4:1:4:0:1 | 0:0:0:0:0:0:0:0:0:0:0:0 | 876:7:1010:300:111:23:2:2:0:7:0:0 | 1:0:2:0:0:0:0:0:0:0:0:0 |
| 123:2:202:101:64:7:2:1:0:2:0:0 | 0:0:0:0:1:0:0:0:0:0:0:0 | 252:8:347:194:18:10:0:4:1:4:0:1 | 0:0:1:0:0:0:0:0:0:0:0:0 | 881:5:1010:301:112:23:2:2:0:7:0:0 | 0:1:1:1:0:0:0:0:0:0:0:0 |
| 1501:10:3:31:0:9:17:1:0:0:1 | 0:0:0:0:0:0:0:0:0:0:0 | 1724:40:4:7:2:9:3:0:0:0:0 | 3:1:0:0:0:0:0:0:0:0:0 | 4026:108:9:22:1:24:22:2:0:3:0 | 5:2:0:0:0:0:0:0:0:0:0 |
| 1502:10:3:31:0:9:17:1:0:0:1 | 0:0:0:0:0:0:0:0:0:0:0 | 1728:22:4:7:1:9:3:0:0:0:0 | 0:1:0:0:0:0:0:0:0:0:0 | 4032:59:9:22:1:24:22:2:0:3:0 | 3:3:0:0:0:0:0:0:0:0:0 |
| 1481:12:3:31:0:9:17:1:0:0 | 1:0:0:0:0:0:0:0:0:0 | 1629:44:4:7:1:9:3:0:0:0 | 4:0:0:0:0:0:0:0:0:0 | 3813:125:9:22:0:24:22:2:0:1 | 2:0:0:0:0:0:0:0:0:0 |
| 1481:12:3:31:0:9:17:1:0:0 | 1:0:0:0:0:0:0:0:0:0 | 1632:45:4:7:1:9:3:0:0:0 | 2:0:0:0:0:0:0:0:0:0 | 3811:126:9:22:0:24:22:2:0:1 | 2:0:0:0:0:0:0:0:0:0 |
| 1475:13:3:31:0:9:16:1:0:0 | 2:0:0:0:0:0:0:0:0:0 | 1616:47:4:7:1:9:3:0:0:0 | 0:0:0:0:0:0:0:0:0:0 | 3752:127:9:22:0:24:21:2:0:1 | 7:0:0:0:0:0:0:0:0:0 |
| 1469:13:3:31:0:9:16:1:0:0 | 1:0:0:0:0:0:0:0:0:0 | 1595:48:4:6:1:9:3:0:0:0 | 0:0:0:0:0:0:0:0:0:0 | 3718:129:9:22:0:24:19:2:0:1 | 9:0:0:0:0:0:0:0:0:0 |
| 1465:13:3:31:0:9:16:1:0:0 | 1:0:0:0:0:0:0:0:0:0 | 1579:48:4:6:1:9:3:0:0:0 | 0:0:0:0:0:0:0:0:0:0 | 3697:131:9:22:0:24:19:2:0:1 | 2:0:0:0:0:0:0:0:0:0 |
| 1464:13:3:31:0:9:16:1:0:0 | 0:0:0:0:0:0:0:0:0:0 | 1569:48:4:6:1:9:3:0:0:0 | 1:1:0:0:0:0:0:0:0:0 | 3681:131:9:22:0:24:19:2:0:1 | 3:0:0:0:0:0:0:0:0:0 |
| 1461:13:3:31:0:1:16:0:0 | 0:0:0:0:0:0:0:0:0 | 1556:49:4:6:1:5:3:0:0 | 1:0:0:0:0:0:0:0:0 | 3644:128:9:22:0:5:19:2:1 | 1:1:0:0:0:0:0:0:0 |
| 1436:15:11:3:31:0:8:0:1 | 0:0:0:0:0:0:0:0:0 | 1504:2:50:4:5:1:2:0:0 | 4:0:0:0:0:0:1:0:0 | 3576:19:129:8:21:0:19:1:2 | 7:0:0:0:0:0:0:0:0 |
| 1433:11:3:31:0:15:9:0:1 | 0:0:0:0:0:0:0:0:0 | 1506:49:4:5:1:2:6:0:0 | 2:0:0:0:0:0:0:0:0 | 3573:126:8:21:0:19:19:1:2 | 3:0:0:0:0:0:0:0:0 |
| 60 | 0 | 198 | 0 | 356 | 0 |
| 0:0:0:0:0:0 | 0:1:0:1:0:0 | 0:0:0:0:0:0 | 1:0:0:0:0:0 | 0:0:0:0:0:0 | 1:0:0:1:1:2 |
| 0:0:0:0:0:0:0:0 | 1:1:0:0:0:0:0:0 | 0:0:0:0:0:0:0:0 | 0:0:1:1:1:1:0:0 | 1:2:0:0:0:0:1:1 | 1:3:2:0:0:0:2:0 |
| 1:01:00 | 0:00:00 | 0:01:00 | 0:00:00 | 4:03:04 | 0:00:00 |
| 1:00 | 2:00 | 0:00 | 1:01 | 1:00 | 0:01 |
| 0:0:1:0 | 0:0:0:0 | 1:0:0:0 | 0:0:0:0 | 1:6:1:0 | 0:0:0:0 |
| 0:00 | 0:00 | 1:00 | 0:00 | 3:00 | 0:00 |
| 0 | 2 | 1 | 2 | 1 | 7 |
| 0:00 | 3:00 | 0:00 | 2:00 | 0:00 | 6:01 |
| 19:2:0:0 | 0:0:0:0 | 30:4:0:1 | 1:1:0:0 | 76:4:2:1 | 1:0:0:0 |
| 20:0:1:0 | 0:0:0:0 | 34:0:2:1 | 0:0:0:0 | 85:4:2:0 | 0:0:0:0 |
| 20:0:1:0 | 0:0:0:0 | 37:0:3:1 | 0:0:0:0 | 88:4:5:1 | 0:0:0:0 |
| 0:0:0:0:0:0:0:0:0:0 | 0:0:0:0:0:0:0:0:0:0 | 2:0:0:0:0:0:0:0:1:0 | 4:0:1:0:0:0:0:1:0:0 | 2:0:1:0:0:1:0:1:0:0:2 | 0:0:0:0:0:0:0:0:0:0:0 |
| 4 | 0 | 27 | 1 | 21 | 1 |
| 0 | 0 | 8 | 0 | 5 | 0 |
| 0:00 | 8:04 | 0:00 | 0:00 | 0:00 | 17:12 |
| 1:3:0:1:0:1 | 0:0:0:0:0:0 | 3:6:0:0:0:1 | 0:0:0:0:0:0 | 0:0:0:0:0:0 | 0:0:0:0:0:0 |
| 5:1:0:0 | 3:0:0:0 | 9:0:0:0 | 1:0:0:0 | 3:0:0:0 | 0:0:0:0 |
| 0:0:4:0:0:0:0:0:0 | 1:1:4:0:0:0:1:0:1 | 2:0:5:0:0:0:1:0:1 | 1:0:5:0:0:0:0:0:0 | 0:0:1:0:0:0:1:0:0 | 0:0:2:0:0:0:0:0:0 |

| | | | | | |
|---|---|---|---|---|---|
| 0 | 0 | 4718 | 5 | 103 | 0 |
| 0 | 0 | 4662 | 6 | 101 | 0 |
| 0 | 0 | 2 | 4663 | 0 | 101 |
| 0 | 0 | 4642 | 5 | 101 | 0 |
| 0 | 0 | 4660 | 3 | 101 | 0 |
| 2:00:00 | 0:00:00 | 4:01:00 | 1:00:00 | 4:00:02 | 0:00:00 |
| 24:2:0:1 | 17:0:2:0 | 39:1:5:0 | 40:3:4:0 | 107:10:13:0 | 51:2:10:0 |
| 0 | 0 | 0 | 0 | 0 | 2 |
| 0:0:2:0 | 0:0:0:0 | 1:0:1:1 | 0:0:0:0 | 4:0:1:2 | 0:0:0:0 |
| 122 | 0 | 134 | 0 | 482 | 2 |
| 122 | 0 | 131 | 0 | 473 | 3 |
| 1 | 2 | 0 | 0 | 0 | 0 |
| 324 | 0 | 962 | 0 | 1170 | 1 |
| 321 | 0 | 958 | 1 | 1168 | 0 |
| 325 | 0 | 952 | 3 | 1161 | 1 |
| 324 | 0 | 953 | 1 | 1157 | 1 |
| 318 | 1 | 947 | 1 | 1147 | 0 |
| 316 | 2 | 946 | 2 | 1146 | 1 |
| 318 | 0 | 948 | 0 | 1147 | 0 |
| 309 | 2 | 942 | 1 | 1137 | 1 |
| 0:00 | 0:00 | 0:00 | 0:00 | 0:00 | 0:00 |
| 0 | 0 | 1 | 0 | 2 | 3 |
| 0 | 0 | 0 | 0 | 1 | 0 |
| 51:0:7:10:0:0:6:3:0:0:0:5:2:0:1:0:1:0:0 | 0:0:0:0:0:0:0:0:0:0:0:0:0:0:0:0:0:0:0:0 | 67:0:13:13:13:3:52:1:1:0:3:0:1:0:1:0:0:2:0 | 0:0:0:0:0:0:0:0:0:0:0:0:0:0:0:0:0:0:0:0:0 | 113:11:30:75:26:1:127:4:3:0:6:7:2:2:0:0:0:2:1 | 0:0:1:0:0:0:2:0:0:0:0:0:0:0:0:0:0:0:0 |
| 51:0:7:10:0:0:6:3:0:0:0:5:2:0:1:0:1:0:0 | 0:0:0:0:0:0:0:0:0:0:0:0:0:0:0:0:0:0:0:0 | 67:0:13:13:12:3:51:1:1:0:3:0:1:0:1:0:0:2:0 | 0:0:0:0:1:0:1:0:0:0:0:0:0:0:0:0:0:0:0:0:0 | 113:11:31:75:26:1:129:4:3:0:6:7:2:2:0:0:0:3:1 | 0:0:0:0:0:0:0:0:0:0:0:0:0:0:0:0:0:0:0 |
| 51:0:7:10:0:0:6:3:0:0:0:0:5:2:0:1:0:1:0 | 0:0:0:0:0:0:0:0:0:0:0:0:0:0:0:0:0:0:0:0 | 66:0:13:13:13:3:49:1:1:1:0:3:0:2:0:1:0:0:0 | 0:0:0:0:0:0:0:0:0:0:0:0:0:0:0:0:0:0:0:0:0 | 111:11:31:75:26:1:115:4:8:3:0:6:7:2:2:1:0:0:1 | 0:0:0:0:0:0:0:0:0:0:0:0:0:0:0:0:0:0:0 |
| 51:0:7:10:0:0:5:3:0:0:0:0:5:2:0:1:0:1:0 | 0:0:0:0:0:0:0:0:0:0:0:0:0:0:0:0:0:0:0:0 | 64:0:13:13:13:3:46:1:1:1:0:3:0:3:0:1:0:0:0 | 0:0:0:0:0:0:0:0:0:0:0:0:0:0:0:0:0:0:0:0:0 | 109:11:31:75:26:1:112:4:7:3:0:6:7:1:2:1:0:0:1 | 1:0:0:0:0:0:0:0:0:0:0:0:0:0:0:0:0:0:0 |
| 49:0:7:10:0:0:5:3:0:0:0:5:2:0:1:0:0:0:1:0 | 0:0:0:0:0:0:0:0:0:0:0:0:0:0:0:0:0:0:0:0:0 | 61:0:13:13:13:3:33:1:1:0:3:0:4:0:1:0:0:0:0:0 | 0:0:0:0:0:0:0:0:0:0:0:0:0:0:0:0:0:0:0:0:0:0 | 100:11:31:75:26:1:71:4:3:0:6:7:1:2:1:0:1:2:0:1 | 1:0:0:0:0:0:0:0:0:0:0:0:0:0:0:0:0:0:0:0 |
| 49:0:7:10:0:0:3:3:0:0:0:5:2:0:1:0:0:1:0 | 0:0:0:0:0:0:0:0:0:0:0:0:0:0:0:0:0:0:0:0 | 37:0:11:9:8:0:2:1:1:0:2:0:4:0:1:0:0:0:0 | 0:0:0:0:0:0:0:0:0:0:0:0:0:0:0:0:0:0:0:0:0 | 70:11:29:56:21:1:4:1:3:0:4:7:2:2:1:0:1:0:1 | 2:0:0:0:0:0:0:0:0:0:0:0:0:0:0:0:0:0:0 |
| 3:2:0:0:0:0:0:1:0:0 | 0:0:0:0:0:0:0:0:0:0:0 | 4:17:1:0:0:0:1:0:0:0 | 0:0:0:0:0:0:0:0:0:0:0 | 5:22:0:0:0:0:0:1:0:0 | 0:0:0:0:0:0:0:0:0:0 |
| 0:01:00 | 0:00:00 | 0:00:00 | 0:00:00 | 0:02:02 | 0:01:01 |
| 1:00:01 | 0:00:00 | 0:00:00 | 0:00:00 | 5:00:03 | 0:00:00 |
| 0:01:00 | 0:00:00 | 0:00:00 | 0:00:00 | 7:04:00 | 0:00:00 |
| 129 | 0 | 420 | 0 | 783 | 2 |
| 0:00 | 0:00 | 0:00 | 0:00 | 0:00 | 1:01 |
| 0:00 | 0:01 | 0:00 | 2:00 | 0:00 | 2:00 |
| 0:00 | 0:01 | 0:00 | 2:00 | 0:00 | 2:00 |
| 38:00:00 | 0:00 | 62:01:00 | 0:00 | 164:02:00 | 0:00 |
| 36 | 0 | 71 | 0 | 188 | 2 |
| 32 | 3 | 50 | 11 | 153 | 15 |
| 32 | 0 | 44 | 1 | 110 | 2 |
| 437:26:2:0:6:0:1 | 6:1:0:0:0:0:0 | 749:63:3:0:3:0:0 | 8:0:0:0:0:0:0 | 1810:178:9:0:20:1:0 | 14:1:0:0:0:0:0 |
| 443:27:2:0:6:0:1 | 0:0:0:0:0:0:0 | 749:62:3:0:3:0:0 | 9:1:0:0:0:0:0 | 1821:180:9:0:20:1:0 | 5:0:0:0:0:0:0 |
| 448:25:2:0:0:6:0 | 0:0:0:0:0:0:0 | 766:53:3:0:0:3:0 | 1:1:0:0:0:0:0 | 1844:158:9:3:0:20:1 | 2:0:0:0:0:0:0 |
| 447:25:2:0:0:6:0 | 1:0:0:0:0:0:0 | 766:54:3:0:0:3:0 | 2:0:0:0:0:0:0 | 1845:149:9:3:0:20:1 | 1:2:0:0:0:0:0 |
| 449:24:2:0:0:6:0 | 0:0:0:0:0:0:0 | 768:37:3:0:0:3:1 | 2:1:0:0:0:0:0 | 1848:102:9:4:0:20:1 | 0:1:0:0:0:0:0 |
| 450:6:2:0:6:0:6:0:0 | 1:0:0:0:0:0:0:0:0 | 771:7:3:0:28:0:3:1:0 | 4:0:0:0:0:0:0:0:0 | 1854:29:9:5:39:0:20:1:1 | 1:0:0:0:0:0:0:0:0 |
| 0 | 1 | 0 | 0 | 0 | 3 |
| 456:09:07 | 0:00:00 | 738:24:06 | 1:00:00 | 3450:55:25 | 5:00:00 |
| 457:09:07 | 0:00:00 | 741:21:06 | 2:00:00 | 3477:55:25 | 5:00:00 |
| 503:08:07 | 1:00:00 | 932:11:06 | 0:00:00 | 4124:34:25 | 13:00:00 |
| 503:08:07 | 1:00:00 | 934:11:06 | 2:00:00 | 4133:34:25 | 8:00:00 |
| 265:04:00 | 0:00 | 538:02:00 | 0:00 | 2255:08:00 | 7:00 |
| 0 | 0 | 0 | 3 | 0 | 5 |
| 0 | 4 | 0 | 1 | 0 | 7 |
| 0:00 | 0:00 | 0:02 | 0:01 | 0:00 | 0:00 |
| 0 | 0 | 3 | 3 | 1 | 3 |
| 0 | 0 | 3 | 3 | 1 | 3 |
| 87:7:30:6:0:4:0:1:2:0 | 0:0:0:0:0:0:0:0:0:0 | 3:3:9:0:1:2:0:0:0:0 | 0:0:0:0:0:0:0:0:0:0:0 | 90:9:80:9:4:8:2:2:1:1 | 1:0:0:0:0:0:0:0:0:0 |

| | | | | | |
|---|---|---|---|---|---|
| 32:7:96:6:0:2:0:1:2:0 | 0:0:0:0:0:0:0:0:0:0 | 15:3:5:0:1:2:0:0:0:0 | 0:0:0:0:0:0:0:0:0:0 | 93:9:101:10:4:7:2:0:1:1 | 0:0:0:0:0:0:0:0:0:0 |
| 0 | 0 | 1 | 0 | 0 | 0 |
| 0:00:00 | 4:00:00 | 0:00:00 | 2:01:00 | 0:00:00 | 23:00:01 |
| 0:00:00 | 4:00:00 | 0:00:00 | 2:01:00 | 0:00:00 | 33:00:01 |
| 0 | 0 | 0 | 2 | 1 | 1 |
| 0 | 0 | 0 | 0 | 0 | 1 |
| 1 | 0 | 0 | 0 | 0 | 0 |
| 0:00:00 | 0:00:00 | 0:00:00 | 0:00:01 | 0:00:00 | 1:00:01 |
| 0:00:00 | 1:00:00 | 0:00:00 | 0:00:00 | 0:00:00 | 2:00:01 |
| 0:00 | 1:00 | 0:00 | 0:00 | 0:00 | 2:01 |
| 0 | 0 | 0 | 1 | 0 | 0 |
| 5:01 | 0:00 | 50:00:00 | 0:00 | 48:06:00 | 2:01 |
| 5:00 | 0:00 | 50:00:00 | 0:00 | 50:06:00 | 0:00 |
| 5:01 | 0:00 | 50:01:00 | 0:00 | 50:05:00 | 0:00 |
| 5:01 | 0:00 | 49:01:00 | 0:00 | 50:06:00 | 0:00 |
| 0 | 0 | 7 | 0 | 2 | 2 |
| 4:00 | 0:01 | 0:00 | 0:00 | 6:01 | 0:00 |
| 0:2:0:0:0 | 0:0:0:0:0 | 4:1:0:1:1 | 1:1:0:0:0 | 5:1:1:0:2 | 1:0:0:0:0 |
| 0:1:0:0 | 0:1:0:0 | 5:0:0:1 | 0:0:0:0 | 5:0:1:0 | 0:1:0:1 |
| 0:0:1:3:1 | 0:0:0:0:0 | 0:0:0:0:0 | 0:0:0:0:0 | 0:0:4:2:0 | 0:0:0:0:0 |
| 0:00 | 0:00 | 0:00 | 0:00 | 0:01 | 1:01 |
| 12:0:1:0:0:0:0 | 0:0:0:0:0:0:0 | 310:10:28:0:2:4:2 | 0:0:0:0:0:0:0 | 66:0:3:0:0:2:0 | 0:0:0:0:0:0:0 |
| 12:0:2:0:0:0:0 | 0:0:0:0:0:0:0 | 310:10:29:0:2:4:2 | 0:0:0:0:0:0:0 | 66:0:3:0:0:2:0 | 0:0:0:0:0:0:0 |
| 12:2:0:1:0:0:0 | 0:0:0:0:0:0:0 | 308:30:0:10:2:4:2 | 0:0:0:0:0:0:0 | 66:4:0:0:0:2:0 | 0:0:0:0:0:0:0 |
| 0:00 | 0:00 | 0:00 | 0:00 | 0:00 | 0:02 |
| 0 | 5 | 0 | 1 | 0 | 3 |
| 0:00:00 | 0:00:00 | 0:00:00 | 0:01:00 | 1:00:00 | 0:01:02 |
| 0:00:00 | 0:00:00 | 0:00:00 | 0:01:00 | 0:00:00 | 1:01:02 |
| 0:00:00 | 0:00:00 | 0:00:00 | 0:01:00 | 0:00:00 | 1:01:02 |
| 0:00 | 0:00 | 0:00 | 0:00 | 0:00 | 1:02 |
| 0 | 6 | 0 | 10 | 0 | 39 |
| 0 | 5 | 0 | 8 | 0 | 36 |
| 0 | 4 | 0 | 4 | 0 | 25 |
| 0 | 3 | 0 | 2 | 0 | 11 |
| 0:00 | 0:00 | 0:00 | 0:00 | 0:01 | 0:00 |
| 0:00 | 0:00 | 0:00 | 0:00 | 0:01 | 0:00 |
| 0:0:0:0 | 0:0:0:3 | 0:0:0:0 | 1:0:0:0 | 0:0:0:0 | 2:1:0:1 |
| 152:37:1:0:3:1:0:3:1:0:1:0:1:0:0 | 0:0:0:0:0:0:0:0:0:0:0:0:0:0:0 | 167:15:0:4:3:8:1:2:0:2:0:0:2:0:0 | 1:0:0:0:0:0:0:0:0:0:0:0:0:0:0 | 705:99:1:2:30:26:2:2:8:6:2:1:1:1:1 | 3:0:0:0:0:0:0:0:0:0:0:0:0:0:0 |
| 152:37:1:0:3:1:0:3:1:0:1:0:1:0:0 | 0:0:0:0:0:0:0:0:0:0:0:0:0:0:0 | 168:15:0:4:3:7:1:2:0:2:0:0:2:0:0 | 0:0:0:0:0:0:0:0:0:0:0:0:0:0:0 | 709:99:1:2:30:23:2:2:8:6:2:1:1:1:1 | 1:0:0:0:0:0:0:0:0:0:0:0:0:0:0 |
| 39:156:2:0:0:0:0:0:0:0:0 | 0:0:0:0:0:0:0:0:0:0:0 | 21:170:4:0:4:0:0:0:0:0:0 | 0:0:0:0:0:0:0:0:0:0:0 | 113:716:30:4:3:1:0:0:0:1:0 | 1:3:0:0:0:0:0:0:0:0:0 |
| 38:156:2:0:0:0:0:0:0:0:0 | 1:0:0:0:0:0:0:0:0:0:0 | 21:170:4:0:4:0:0:0:0:0:0 | 0:0:0:0:0:0:0:0:0:0:0 | 114:718:27:4:3:1:0:0:0:1:0 | 0:1:0:0:0:0:0:0:0:0:0 |
| 39:156:3:0:0:0:2:0:0:0:0 | 0:0:0:0:0:0:0:0:0:0:0 | 20:170:1:1:4:0:0:0:0:0:0 | 0:0:0:0:0:0:0:0:0:0:0 | 118:720:14:4:3:1:1:0:0:1:0 | 1:0:1:0:0:0:0:0:0:0:0 |
| 40:156:3:0:0:0:2:0:0:0:0 | 0:0:0:0:0:0:0:0:0:0:0 | 20:170:0:1:4:0:0:0:0:0:0 | 0:0:1:0:0:0:0:0:0:0:0 | 119:720:15:4:3:1:1:0:0:1:0 | 0:0:0:0:0:0:0:0:0:0:0 |
| 0:0:0:0:0:0:0:0:0:0:0 | 40:154:2:0:0:0:2:0:0:0:0 | 0:0:0:0:0:0:0:0:0:0:0 | 18:165:1:1:0:1:1:0:0:0:0 | 0:1:0:0:0:0:0:0:0:0:0 | 120:704:6:5:0:1:1:1:1:0:1 |
| 0:00 | 0:08 | 0:00 | 0:00 | 0:00 | 0:50 |
| 0:00 | 0:06 | 0:00 | 0:00 | 0:00 | 0:47 |
| 0:0:0:0:0:0:0 | 0:0:0:0:0:0:0 | 0:0:1:0:1:0:0 | 0:0:1:0:0:0:0 | 0:0:2:0:0:1:1 | 0:0:0:0:0:0:0 |
| 0:1:1:1:1:0 | 0:0:0:0:0:0 | 10:3:7:0:1:0 | 0:0:0:0:0:0 | 12:3:7:0:1:1 | 0:0:0:0:0:0 |
| 0:00 | 0:00 | 0:00 | 0:00 | 0:00 | 0:01 |
| 0 | 1 | 2 | 2 | 0 | 0 |
| 0 | 1 | 2 | 2 | 0 | 0 |
| 15:4:12:1:0 | 0:0:0:0:0 | 18:4:7:1:0 | 0:0:0:0:0 | 30:5:13:1:1 | 0:0:0:0:0 |
| 97 | 0 | 371 | 2 | 538 | 3 |
| 98 | 0 | 374 | 3 | 538 | 5 |
| 0:0:0:0:0 | 1:0:0:1:0 | 1:0:0:0:0 | 2:0:0:0:0 | 0:0:0:0:0 | 6:0:0:2:0 |
| 0:00 | 0:01 | 0:00 | 1:01 | 0:00 | 0:03 |
| 0:00 | 0:01 | 0:00 | 1:01 | 0:00 | 0:03 |
| 0:00 | 1:00 | 0:00 | 1:01 | 0:00 | 3:00 |
| 0 | 2 | 0 | 0 | 0 | 1 |
| 0 | 0 | 0 | 0 | 1 | 2 |
| 21:00 | 0:00 | 175:01:00 | 1:00 | 154:04:00 | 0:00 |
| 25:0:0:0 | 0:0:0:0 | 216:2:0:2 | 1:0:0:0 | 188:6:1:2 | 0:0:0:0 |
| 0 | 2 | 0 | 0 | 0 | 6 |
| 218:02:00 | 1:00 | 671:07:00 | 3:00 | 1176:11:00 | 5:00 |
| 219:02:00 | 0:00 | 673:07:00 | 2:00 | 1183:11:00 | 2:00 |

| | | | | | |
|---|---|---|---|---|---|
| 221:02:00 | 0:00 | 673:07:00 | 4:00 | 1191:11:00 | 1:00 |
| 40:10:220:2:0 | 0:0:1:0:0 | 155:36:675:7:0 | 1:0:0:0:0 | 278:110:1189:11:1 | 0:0:2:0:0 |
| 40:10:222:2:0 | 0:0:0:0:0 | 156:35:671:7:0 | 0:0:0:0:0 | 276:110:1190:11:1 | 1:0:3:0:0 |
| 39:10:218:2:0 | 0:0:0:0:0 | 154:34:658:7:0 | 1:0:5:0:0 | 270:109:1173:11:0 | 0:0:6:0:0 |
| 39:9:218:2:0 | 0:0:0:0:0 | 155:34:659:7:0 | 0:0:3:0:0 | 269:109:1170:11:0 | 0:0:7:0:0 |
| 39:9:214:2:0 | 0:0:1:0:0 | 155:34:677:6:0 | 0:0:5:0:0 | 263:109:1176:11:0 | 0:0:1:0:0 |
| 38:211:9:2:0 | 0:0:0:0:0 | 155:641:32:4:0 | 0:4:0:0:0 | 259:1125:103:11:0 | 0:0:1:0:0 |
| 37:8:204:1:0 | 0:0:0:0:0 | 147:30:544:4:0 | 2:1:2:0:0 | 238:98:953:10:0 | 2:0:3:0:0 |
| 0 | 0 | 0 | 0 | 0 | 3 |
| 159:217 | 0:00 | 866:1046 | 1:01 | 1022:1209 | 0:00 |
| 159:217 | 0:00 | 861:1031 | 0:00 | 1016:1198 | 2:04 |
| 0 | 1 | 0 | 0 | 0 | 0 |
| 0:0:0:0 | 0:0:0:0 | 0:0:0:0 | 0:1:0:0 | 0:0:0:0 | 0:0:0:1 |
| 0:0:0:0 | 0:0:0:0 | 0:0:0:0 | 0:1:0:0 | 0:0:0:0 | 0:0:0:1 |
| 0:0:0:0 | 0:0:0:0 | 0:0:0:0 | 0:1:0:0 | 0:0:0:0 | 0:0:0:1 |
| 0:0:0:0 | 0:0:0:0 | 0:0:0:0 | 0:1:0:0 | 0:0:0:0 | 0:0:0:1 |
| 0:0:0:0 | 0:0:0:0 | 0:0:0:0 | 0:1:0:0 | 0:0:0:0 | 0:0:0:1 |
| 0:0:0:0 | 0:0:0:0 | 0:0:0:0 | 0:1:0:0 | 0:0:0:0 | 0:0:0:1 |
| 30:1:1:2:1:1:1:0:2:0:0:0:0:0:0 | 0:0:0:0:0:1:0:0:0:0:0:0:0:0:0 | 70:2:11:3:3:5:0:3:0:1:0:0:0:0:0 | 0:0:0:0:0:1:0:1:0:0:0:0:0:0:0 | 81:1:13:5:8:6:0:1:1:2:2:0:1:1:1 | 0:0:0:0:0:0:0:0:0:0:0:0:0:0:0 |
| 0 | 0 | 0 | 21 | 0 | 2 |
| 75:6:5:3:2:1:0 | 0:0:0:0:0:0:0 | 116:2:18:60:1:3:1 | 0:0:0:0:0:0:0 | 310:78:32:175:10:8:4 | 0:1:0:1:0:0:0 |
| 75:6:5:4:1:1:0 | 0:0:0:0:0:0:0 | 114:2:18:61:0:3:1 | 1:0:0:0:0:0:0 | 311:79:26:173:12:7:4 | 0:0:0:0:0:0:0 |
| 75:6:5:5:1:1:0 | 0:0:0:0:0:0:0 | 114:2:18:60:0:3:1 | 1:0:0:1:0:0:0 | 311:79:26:173:11:7:4 | 0:0:0:0:1:0:0 |
| 6:0:1:0 | 0:0:0:0 | 1:0:1:1 | 0:0:0:0 | 64:14:7:1 | 2:0:0:0 |
| 3:0:1:0 | 0:0:0:0 | 1:0:1:0 | 0:0:0:0 | 58:14:8:1 | 3:0:1:0 |
| 73:3:5:2:4:1:0 | 0:0:0:0:0:0:0 | 108:1:19:2:53:0:0 | 1:0:0:0:0:0:0 | 297:41:28:22:137:19:1 | 2:1:0:0:2:0:0 |
| 73:3:5:2:3:1:0 | 0:0:0:0:0:0:0 | 108:1:19:2:53:0:0 | 0:0:0:0:0:0:0 | 295:42:31:22:138:19:1 | 1:0:1:0:1:0:0 |
| 72:3:5:2:3:1:0 | 0:0:0:0:0:0:0 | 108:1:20:2:53:0:0 | 0:0:0:0:0:0:0 | 295:40:42:22:138:19:1 | 1:0:0:0:1:0:0 |
| 0 | 0 | 0 | 0 | 0 | 0 |
| 15:00 | 0:00 | 20:01 | 0:00 | 28:02:00 | 0:00 |
| 0 | 0 | 0 | 0 | 0 | 0 |
| 0 | 0 | 0 | 1 | 0 | 1 |
| 0:00 | 1:00 | 2:00 | 2:00 | 0:00 | 6:01 |
| 0:50:3:0:1:1:1:0 | 0:0:0:0:0:0:0:0 | 0:34:0:0:0:0:0:0 | 0:0:0:0:0:0:0:0 | 1:81:7:1:1:2:0:1 | 0:0:0:0:0:0:0:0 |
| 12:10:5:6:5:0:5:0:2:6:3:0:2:0:0:0:1:0:1:3:0:2:1:0 | 0:0:0:0:0:0:0:0:0:0:0:0:0:0:0:0:0:0:0:0:0:0:0:0 | 5:1:7:1:6:0:3:1:14:36:1:0:0:0:0:0:0:2:0:0:0:0:0:0 | 0:0:0:0:0:0:0:0:0:0:0:0:0:0:0:0:0:0:0:0:0:0:0:0 | 26:7:7:12:20:7:12:2:16:72:3:0:0:1:0:0:0:1:0:3:1:0:0:1 | 1:1:0:0:0:0:0:0:0:0:0:0:0:0:0:0:0:0:0:0:0:0:0:0 |
| 12:10:6:5:5:0:5:6:2:0:2:3:0:1:2:0:0:0:1:3:0:2:1:0 | 0:0:0:0:0:0:0:0:0:0:0:0:0:0:0:0:0:0:0:0:0:0:0:0 | 5:1:8:1:5:0:3:59:12:1:14:1:0:5:0:0:0:0:0:0:0:0:0:0 | 0:0:0:0:0:0:0:0:0:0:0:0:0:0:0:0:0:0:0:0:0:0:0:0 | 26:8:8:11:19:7:11:112:43:2:15:3:0:8:0:1:0:0:0:3:1:0:0:1 | 0:0:1:0:0:0:0:1:0:0:0:0:0:0:0:0:0:0:0:0:0:0:0:0 |
| 12:10:6:5:5:0:5:6:2:0:2:3:0:2:0:0:0:1:1:3:0:2:1:0 | 0:0:0:0:0:0:0:0:0:0:0:0:0:0:0:0:0:0:0:0:0:0:0:0 | 5:1:8:1:4:0:3:59:13:1:14:1:0:0:0:0:0:0:4:0:0:0:0:0 | 0:0:0:0:0:0:0:0:0:0:0:0:0:0:0:0:0:1:0:0:0:0:0:0 | 26:8:9:11:19:6:11:112:44:2:15:3:0:0:1:0:0:0:8:3:1:0:0:1 | 0:0:0:0:0:0:0:1:0:0:0:0:0:0:0:0:0:0:0:0:0:0:0:0 |
| 0 | 0 | 0 | 2 | 0 | 3 |
| 0:0:0:2:0:0:2:0:0:0:1:0:0:0:0:0:0:0:0:0:1:0:0:0 | 0:0:0:0:0:0:0:0:0:0:0:0:0:0:0:0:0:0:0:0:0:0:0:0 | 0:2:0:12:0:5:2:0:0:2:0:0:0:0:1:1:0:1:0:0:0:0:0:0 | 0:0:0:0:0:0:0:0:0:0:0:0:0:0:0:0:0:0:0:0:0:0:0:0 | 1:2:0:6:0:1:2:2:0:0:1:0:0:0:2:0:0:0:0:1:0:0:0:1 | 0:0:0:0:0:0:0:0:0:0:0:0:0:0:0:0:0:0:0:0:0:0:0:0 |
| 6505 | 9 | 16842 | 67 | 35799 | 81 |
| 6498 | 19 | 16854 | 67 | 35822 | 94 |
| 6510 | 11 | 16939 | 30 | 35941 | 61 |
| 6647 | 3 | 17993 | 14 | 37319 | 22 |
| 6689 | 2 | 18448 | 13 | 37877 | 22 |
| 6679 | 11 | 18427 | 25 | 37827 | 49 |
| 6667 | 5 | 18415 | 8 | 37796 | 17 |
| 6670 | 4 | 18405 | 10 | 37773 | 16 |
| 1:0:0:0:0:0:0:0 | 0:0:0:0:0:0:0:0 | 0:0:0:0:0:0:0:0 | 0:0:0:0:0:0:0:0 | 9:1:0:0:0:0:0:0 | 0:0:0:0:0:0:0:0 |
| 0:1:0:0:0:0:0:0:1 | 0:0:0:0:0:0:0:0:0 | 0:1:0:0:0:0:0:0:0 | 0:0:0:0:0:0:0:0:0 | 1:10:0:0:0:0:0:0:0 | 0:0:0:0:0:0:0:0:0 |
| 41:21:6:3 | 0:0:0:0 | 19:38:10:3 | 0:0:0:0 | 140:58:13:1 | 0:1:0:0 |
| 41:6:10:3 | 0:0:0:0 | 19:10:24:3 | 0:0:2:0 | 145:13:34:1 | 0:0:1:0 |
| 41:6:10:3 | 0:0:0:0 | 19:10:23:3 | 0:0:3:0 | 145:13:34:1 | 0:0:1:0 |
| 0:00 | 0:00 | 0:00 | 0:00 | 0:00 | 0:01 |
| 0 | 0 | 0 | 13 | 0 | 1 |
| 35:02:01 | 0:00:00 | 77:01:01 | 0:00:00 | 153:05:02 | 1:00:00 |

| | | | | | |
|---|---|---|---|---|---|
| 6:0:1:15:2:0 | 0:0:0:0:0:0 | 24:7:3:8:24:0 | 0:0:0:0:0:0 | 32:24:5:33:87:1 | 0:0:0:0:0:0 |
| 210:2:14:0:0:0:2:0:1:1:0:1:0 | 0:0:0:0:0:0:0:0:0:0:0:0:0 | 838:8:71:1:1:3:8:4:0:1:0:0:0 | 2:0:0:0:0:0:0:0:0:0:0:0:0 | 1174:22:45:2:5:1:10:6:0:0:1:2:1 | 0:1:0:0:0:0:0:0:0:0:0:0:0 |
| 210:2:14:0:0:0:2:0:1:1:0:1:0 | 0:0:0:0:0:0:0:0:0:0:0:0:0 | 837:8:71:1:1:3:8:4:0:1:0:0:0 | 3:0:0:0:0:0:0:0:0:0:0:0:0 | 1173:23:45:2:5:1:10:7:0:0:1:2:1 | 2:0:0:0:0:0:0:0:0:0:0:0:0 |
| 15:7:0:0:2:0:1:0:0 | 0:0:0:0:0:0:0:0:0 | 75:36:6:3:2:0:1:0:0 | 0:0:0:0:0:0:0:0:0 | 48:30:8:6:6:0:0:1:1 | 0:0:0:0:0:0:0:0:0 |
| 29:4:0:0:0 | 0:0:0:0:0 | 192:17:8:1:1 | 0:0:0:0:0 | 107:12:2:2:2 | 0:0:0:0:0 |
| 1:27:4:0:0 | 0:0:0:0:0 | 6:173:16:8:2 | 0:1:0:0:0 | 8:97:12:2:5 | 0:1:0:0:0 |
| 1:25:4:0:0 | 0:0:0:0:0 | 5:163:15:8:7 | 0:0:0:0:0 | 8:95:9:2:9 | 0:0:0:0:0 |
| 1:25:4:0:0 | 0:0:0:0:0 | 5:163:15:8:7 | 0:0:0:0:0 | 8:95:9:2:11 | 0:0:0:0:0 |
| 1:25:4:0:0 | 0:0:0:0:0 | 5:163:15:8:8 | 0:0:0:0:0 | 8:93:9:2:11 | 0:2:0:0:0 |
| 1:25:2:0:0 | 0:0:0:0:0 | 5:163:15:8:9 | 0:0:0:0:0 | 8:95:8:2:13 | 0:0:0:0:0 |
| 1:25:2:0:1 | 0:0:0:0:0 | 5:163:15:8:9 | 0:0:0:0:0 | 7:95:8:2:13 | 0:0:0:0:0 |
| 24:1:3:0:2 | 0:0:0:0:0 | 160:5:15:7:13 | 0:0:0:0:0 | 95:7:8:2:16 | 0:0:0:0:0 |
| 0:0:0:1:2:2:7:117:156:4:3:20:2:0:0:0 | 0:0:0:0:0:0:0:0:0:0:0:0:0:0:0:0 | 0:0:0:0:3:0:9:126:97:1:7:147:2:0:1:0 | 0:0:0:0:0:0:0:0:0:0:0:0:0:0:0:0 | 0:1:0:1:2:1:11:237:251:4:7:254:3:0:0:0 | 0:0:0:0:0:0:0:0:2:0:0:2:0:0:0:0 |
| 0:0:0:2:2:7:117:154:4:3:15:0:1:0:0 | 0:0:0:0:0:0:0:0:0:0:0:0:0:0:0 | 0:1:0:2:0:7:126:97:1:7:142:0:2:1:0 | 0:0:0:0:0:0:0:0:0:0:0:0:0:0:0 | 0:1:1:0:1:8:237:251:4:7:232:0:1:0:0 | 0:0:0:0:0:0:0:0:0:0:0:0:0:0:0 |
| 0:0:0:2:2:7:117:154:4:3:15:0:1:0:0 | 0:0:0:0:0:1:0:0:0:0:0:0:0:0:0 | 0:1:0:2:0:7:125:97:1:7:141:0:2:1:0 | 0:0:0:0:0:0:1:0:0:0:1:0:0:0:0 | 0:1:1:0:1:8:237:250:4:7:231:0:1:0:0 | 0:0:0:0:0:0:0:1:0:0:1:0:0:0:0 |
| 0:0:0:2:2:8:116:154:4:3:15:0:1:0:0 | 0:0:0:0:0:0:1:0:0:0:0:0:0:0:0 | 0:1:0:2:0:7:126:96:1:7:137:0:2:1:0 | 0:0:0:0:0:0:0:1:0:0:3:0:0:0:0 | 0:1:1:0:1:12:237:251:4:7:230:0:1:0:0 | 0:0:0:0:0:0:0:0:0:0:0:0:0:0:0 |
| 0:2:2:2:13:117:149:4:3:8:2:0:0:0 | 0:0:0:0:0:0:0:0:0:0:0:0:0:0 | 3:0:6:0:45:126:97:1:7:99:15:0:0:0 | 0:0:0:0:1:0:0:0:0:0:0:0:0:0 | 2:3:10:1:52:236:242:4:7:150:17:0:0:0 | 0:0:0:0:2:1:0:0:0:0:0:0:0:0 |
| 24:0:2:2:2:117:149:4:3:7:2:0:0:0 | 1:0:0:2:0:0:0:0:0:0:0:0:0:0 | 67:3:3:12:0:126:97:1:7:99:24:0:0:0 | 2:0:0:0:0:0:0:0:0:0:0:0:0:0 | 77:2:5:16:1:237:240:4:7:149:29:0:0:0 | 1:0:0:2:0:0:1:0:0:0:1:0:0:0 |
| 148:0:2:4:2:25:117:4:3:7:2:0:0:0 | 0:0:0:0:0:0:0:0:0:0:0:0:0:0 | 96:3:3:12:0:70:126:1:7:95:24:0:0:0 | 0:0:0:0:0:0:0:0:0:1:0:0:0:0 | 241:2:5:18:1:77:237:4:7:146:30:0:0:0 | 0:0:0:0:0:0:0:0:0:0:0:0:0:0 |
| 0:2:4:2:117:27:150:4:3:4:3:0:0 | 0:0:0:0:0:0:0:0:0:0:0:0:0 | 3:3:13:0:126:72:88:1:7:28:24:0:0 | 0:0:0:0:0:0:0:0:0:0:0:0:0 | 2:5:19:1:237:92:229:4:7:38:33:0:0 | 0:0:1:0:0:0:0:0:0:0:0:0:0 |
| 45:8:2:0:0 | 0:0:0:0:0 | 26:4:2:0:0 | 1:1:0:0:0 | 36:8:2:0:1 | 0:0:0:0:0 |
| 0 | 0 | 0 | 1 | 0 | 2 |
| 0 | 0 | 0 | 1 | 0 | 0 |
| 0 | 0 | 0 | 3 | 0 | 0 |
| 0 | 0 | 0 | 5 | 0 | 0 |
| 4324:01:00 | 3:00 | 3618:04:00 | 1:00 | 22024:9 | 21:00 |
| 4322:01:00 | 6:00 | 3610:04:00 | 8:00 | 22017:9 | 31:00:00 |
| 4316:01:00 | 7:00 | 3600:04:00 | 5:00 | 21913:9 | 45:00:00 |
| 4303:01:00 | 5:00 | 3558:03:00 | 4:00 | 21648:9 | 22:00 |
| 4287:01:00 | 8:00 | 3526:03:00 | 4:00 | 21498:9 | 34:00:00 |
| 4276:01:00 | 2:00 | 3462:03:00 | 2:00 | 21158:9 | 11:00 |
| 4264:01:00 | 2:00 | 3429:03:00 | 2:00 | 20935:9 | 28:00:00 |
| 4261:01:00 | 3:00 | 3420:03:00 | 7:00 | 20909:9 | 39:00:00 |
| 4225:01:00 | 5:00 | 3383:03:00 | 5:00 | 20637:9 | 31:00:00 |
| 4217:01:00 | 7:00 | 3380:03:00 | 4:00 | 20588:9 | 35:00:00 |
| 4198:01:00 | 5:00 | 3331:04:00 | 4:00 | 20355:9 | 28:00:00 |
| 4185:01:00 | 2:00 | 3321:04:00 | 9:00 | 20350:9 | 23:00 |
| 4159:01:00 | 5:00 | 3299:04:00 | 3:00 | 20154:9 | 30:00:00 |
| 4135:01:00 | 6:00 | 3259:04:00 | 9:00 | 19926:9 | 39:00:00 |
| 4131:01:00 | 4:00 | 3255:04:00 | 6:00 | 19908:9 | 21:00 |
| 4130:01:00 | 4:00 | 3249:04:00 | 8:00 | 19866:9 | 30:00:00 |
| 4132:01:00 | 3:00 | 3248:04:00 | 7:00 | 19834:9 | 43:00:00 |
| 4088:01:00 | 5:00 | 3223:04:00 | 5:00 | 19644:9 | 49:00:00 |
| 4083:01:00 | 8:00 | 3216:04:00 | 7:00 | 19671:9 | 37:00:00 |
| 4086:01:00 | 2:00 | 3207:04:00 | 8:00 | 19643:9 | 38:00:00 |
| 4063:01:00 | 3:00 | 3180:04:00 | 11:00 | 19527:9 | 38:00:00 |
| 4049:01:00 | 2:00 | 3132:04:00 | 5:00 | 19342:9 | 33:00:00 |
| 4044:01:00 | 5:00 | 3127:04:00 | 6:00 | 19287:9 | 44:00:00 |
| 4032:01:00 | 5:00 | 3102:04:00 | 3:00 | 19174:9 | 23:00 |
| 4034:01:00 | 5:00 | 3105:04:00 | 8:00 | 19199:9 | 28:00:00 |
| 4045:01:00 | 2:00 | 3114:04:00 | 2:00 | 19238:9 | 24:00:00 |
| 4040:01:00 | 4:00 | 3112:04:00 | 2:00 | 19213:9 | 24:00:00 |

| | | | | | |
|---|---|---|---|---|---|
| 4029:01:00 | 8:00 | 3099:04:00 | 5:00 | 19153:9 | 17:00 |
| 4038:00:01 | 3:00:00 | 3095:00:00 | 4:00:00 | 19141:1:1 | 14:00:00 |
| 4039:00:01 | 2:00:00 | 3100:00:00 | 1:00:00 | 19160:1:1 | 6:00:00 |
| 4030:00:01 | 7:00:00 | 3099:00:00 | 0:00:00 | 19128:1:1 | 20:00:00 |
| 4029:00:01 | 2:00:00 | 3086:00:00 | 4:00:00 | 19089:1:1 | 19:00:00 |
| 4004:00:01 | 3:00:00 | 3076:02:00 | 0:00:00 | 18981:4:1 | 6:00:00 |
| 0 | 0 | 1 | 1 | 0 | 0 |
| 0 | 0 | 1 | 1 | 0 | 0 |
| 0:00:00 | 0:00:00 | 0:00:00 | 0:00:00 | 0:00:00 | 0:00:00 |
| 0:00:00 | 0:00:00 | 0:00:00 | 0:00:00 | 0:00:00 | 0:00:00 |
| 0:00:00 | 0:00:00 | 0:00:00 | 0:00:00 | 0:00:00 | 0:00:00 |
| 3311:07:00 | 8:00 | 6469:21:00 | 15:00 | 13703:24 | 18:00 |
| 3314:07:00 | 6:00 | 6481:21:00 | 5:00 | 13717:24 | 12:00 |
| 3318:07:00 | 2:00 | 6479:21:00 | 9:00 | 13713:24 | 20:00 |
| 3320:07:00 | 1:00 | 6486:21:00 | 5:00 | 13719:24 | 19:00 |
| 3323:07:00 | 2:00 | 6494:21:00 | 2:00 | 13762:24 | 5:00 |
| 3323:07:00 | 3:00 | 6501:21:00 | 0:00 | 13771:24 | 3:00 |
| 3391:07:00 | 5:00 | 6554:21:00 | 7:00 | 13867:22 | 12:01 |
| 3414:07:00 | 2:00 | 6611:20:00 | 6:00 | 13989:21 | 17:00 |
| 3415:07:00 | 2:00 | 6617:19:00 | 6:00 | 13999:21 | 15:00 |
| 3415:07:00 | 3:00 | 6620:19:00 | 3:00 | 14002:20 | 8:00 |
| 3466:05:00 | 5:00 | 6745:15:00 | 5:00 | 14322:12 | 19:00 |
| 3476:05:00 | 1:00 | 6749:14:00 | 9:00 | 14352:9 | 12:00 |
| 3485:03:00 | 4:00 | 6758:10:00 | 11:00 | 14403:9 | 11:00 |
| 3495:01:00 | 1:00 | 6795:00:00 | 3:01 | 14465:2 | 1:00 |
| 3501:01:00 | 3:00 | 6803:00:00 | 2:00 | 14463:0 | 16:00 |
| 3516:00:00 | 0:00 | 6816:01:00 | 9:01 | 14518:0 | 16:04 |
| 3511:03:00 | 3:00 | 6817:13:00 | 3:01 | 14511:24 | 18:03 |
| 3510:09:00 | 3:00 | 6816:23:00 | 6:00 | 14509:54 | 13:03 |
| 3507:17:00 | 0:00 | 6813:33:00 | 1:01 | 14512:135 | 3:03 |
| 3496:28:00 | 4:00 | 6805:59:00 | 4:02 | 14483:193 | 14:03 |
| 3491:43:00 | 3:00 | 6791:122 | 5:01 | 14437:337 | 19:02 |
| 3483:47:00 | 1:00 | 6782:145 | 5:00 | 14409:358 | 13:00 |
| 3468:55:00 | 5:00 | 6771:179 | 8:00 | 14366:412 | 16:02 |
| 3461:62 | 6:00 | 6754:242 | 10:01 | 14371:500 | 8:02 |
| 3458:63 | 1:00 | 6739:266 | 5:00 | 14345:545 | 9:00 |
| 3178:61 | 198:01:00 | 5984:257 | 461:03:00 | 13033:519 | 891:11:00 |
| 3162:57:00 | 0:00 | 3649:170 | 9:00 | 8798:365 | 13:00 |
| 2:2:192:2:2:2:0 | 0:0:0:0:0:0:0 | 7:6:541:5:6:6:1 | 0:0:0:0:0:0:0 | 6:17:607:7:12:5:2 | 0:0:1:0:0:0:0 |
| 2:2:188:2:2:2:0 | 0:0:0:0:0:0:0 | 6:6:535:2:5:6:6:1 | 0:0:2:0:0:0:0 | 6:17:591:7:7:11:5:2 | 0:0:0:0:0:0:0 |
| 0:00:00 | 0:00:00 | 0:00:00 | 0:01:00 | 0:00:00 | 0:01:01 |
| 0 | 0 | 0 | 0 | 0 | 0 |
| 0:0:0:0 | 0:0:0:0 | 0:0:0:0 | 0:1:0:1 | 0:0:0:0 | 0:1:0:0 |
| 0:00 | 1:01 | 0:00 | 1:00 | 0:00 | 4:00 |
| 27:22:7:2 | 0:0:0:0 | 26:24:9:3 | 0:0:0:0 | 101:82:33:6 | 0:0:0:0 |
| 21:24:3:2 | 0:0:0:0 | 19:19:6:2 | 0:0:0:0 | 76:78:33:5 | 0:0:0:0 |
| 25:9:20:2 | 0:0:0:0 | 17:14:15:2 | 0:0:0:0 | 74:46:67:5 | 0:0:0:0 |
| 19:28:11:2 | 0:0:0:0 | 16:16:15:3 | 0:0:0:0 | 64:79:52:5 | 1:0:0:0 |
| 28:18:13:2 | 0:1:0:0 | 17:15:16:2 | 1:0:1:1 | 72:56:54:4 | 5:9:1:1 |
| 0:0:0:0 | 0:0:0:0 | 0:0:0:0 | 0:0:0:0 | 0:0:0:0 | 0:0:0:0 |
| 0 | 1 | 0 | 0 | 0 | 9 |
| 3:3:3:9:2:2:0:0:0:0:1:0:3:0:0:1:0:1:0:1:0:0:0:1:0:1:0:0:0:1:0:0 | 0:0:0:0:0:0:0:0:0:0:0:0:0:0:0:0:0:0:0:0:0:0:0:0:0:0:0:0:0:0:0:0 | 1:2:2:7:0:1:0:0:0:0:0:1:0:0:0:1:0:0:0:1:1:0:0:0:0:0:0:2:0:0:0:0 | 0:0:0:0:0:0:0:0:0:0:0:0:0:0:0:0:0:0:0:0:0:0:0:0:0:0:0:0:0:0:0:0 | 4:4:4:17:4:10:0:1:0:0:1:1:8:0:0:4:1:0:1:0:2:0:0:0:1:3:0:1:0:0:1:1 | 0:0:0:0:0:0:0:0:0:0:0:0:0:0:0:0:0:0:0:0:0:0:0:0:0:0:0:0:0:0:0:0 |
| 8 | 0 | 11 | 0 | 22 | 0 |
| 0 | 9 | 0 | 11 | 0 | 22 |
| 0 | 9 | 0 | 11 | 0 | 22 |
| 0 | 3 | 2 | 1 | 0 | 3 |
| 0:00 | 0:01 | 0:00 | 0:00 | 0:00 | 2:00 |
| 35:02:00 | 1:00 | 45:05:00 | 4:00 | 108:03:00 | 7:00 |
| 0 | 0 | 0 | 0 | 0 | 0 |
| 0 | 0 | 0 | 0 | 0 | 0 |
| 0:00:00 | 0:00:00 | 0:00:00 | 2:01:01 | 0:00:00 | 3:01:03 |
| 0:00 | 0:00 | 0:00 | 0:00 | 1:00 | 0:00 |
| 0:00 | 0:00 | 0:00 | 0:00 | 1:00 | 0:00 |
| 0:0:0:0 | 0:0:0:0 | 0:0:0:1 | 0:0:0:0 | 0:0:0:0 | 0:0:0:0 |
| 0:0:0:0 | 0:0:0:0 | 0:0:0:1 | 0:0:0:0 | 0:0:0:0 | 0:0:0:0 |
| 0 | 0 | 0 | 0 | 0 | 0 |
| 88 | 0 | 62 | 0 | 192 | 0 |
| 3:0:2:1:1:0:0:0:0 | 0:0:0:0:0:0:0:0:0 | 2:8:2:0:0:0:0:0:0 | 3:1:4:0:0:0:0:0:0 | 7:2:7:0:3:0:0:1:2 | 1:1:5:0:0:0:0:0:0 |
| 0:0:0:0:0:0:1:0 | 0:0:0:0:0:0:0:0 | 0:0:0:0:0:0:2:0 | 0:0:0:0:0:0:0:0 | 0:0:0:1:0:1:0:1 | 0:0:0:0:0:0:1:0 |

| | | | | | |
|---|---|---|---|---|---|
| 0:0:0:0:0:0:1:0 | 0:0:0:0:0:0:0:0 | 0:0:0:0:0:0:3:0 | 0:0:0:0:0:0:0:0 | 0:0:0:1:0:1:1:1 | 0:0:0:0:0:0:0:0 |
| 0:0:0:0:2:0:0:0 | 0:0:0:0:0:0:0:0 | 0:0:0:0:3:0:0:1 | 0:0:0:0:0:0:0:0 | 0:0:0:1:0:0:1:1 | 0:0:0:0:1:0:0:0 |
| 0:0:4:0:3:1:0:0:0:0:0:0 | 0:0:0:0:0:0:0:0:0:0:0:0 | 0:0:6:0:25:1:0:0:1:0:1:0 | 0:0:0:0:0:0:0:0:0:0:0:0 | 0:0:8:1:24:0:0:0:1:0:0:1 | 0:0:0:0:0:0:0:0:0:0:0:0 |
| 0:1:0:4:3:0:0:0:0:0:0:0 | 0:0:0:0:0:0:0:0:0:0:0:0 | 0:2:0:6:21:0:0:0:1:1:0:0 | 0:0:0:0:0:0:0:0:0:0:0:0 | 0:0:0:8:22:0:0:1:0:1:0:1 | 0:0:0:0:0:0:0:0:0:0:0:0 |
| 0:00 | 0:00 | 0:00 | 0:01 | 0:00 | 0:03 |
| 189:02:03 | 1:00:00 | 340:00:01 | 1:00:00 | 814:10:09 | 0:00:02 |
| 190:02:03 | 0:00:00 | 340:00:01 | 1:00:00 | 814:10:11 | 2:00:02 |
| 190:08:04 | 0:00:00 | 335:02:07 | 0:00:00 | 804:30:21 | 0:00:00 |
| 188:10:04 | 0:00:00 | 331:03:05 | 2:00:00 | 799:37:19 | 2:01:00 |
| 0:00 | 5:00 | 0:00 | 1:00 | 0:00 | 4:01 |
| 3 | 0 | 3 | 4 | 6 | 0 |
| 3 | 0 | 4 | 3 | 7 | 0 |
| 66:216:20 | 0:00:00 | 96:446:10 | 0:01:00 | 292:848:59 | 0:00:02 |
| 216:57:23 | 0:00:00 | 447:77:8 | 0:00:00 | 853:230:59 | 0:01:00 |
| 216:56:23 | 0:01:00 | 447:77:8 | 0:00:01 | 853:226:61 | 0:00:00 |
| 215:05:18 | 0:00:00 | 447:07:10 | 0:00:00 | 847:11:55 | 1:00:00 |
| 741:32:00 | 1:00 | 672:17:00 | 3:00 | 2338:64 | 7:00 |
| 763:33:00 | 0:00 | 714:21:00 | 1:00 | 2480:71 | 8:00 |
| 762:33:00 | 1:00 | 716:21:00 | 2:00 | 2489:71 | 4:00 |
| 800:33:00 | 1:00 | 770:22:00 | 0:00 | 2680:73 | 4:00 |
| 800:33:00 | 2:00 | 773:22:00 | 1:00 | 2692:74 | 5:00 |
| 806:33:00 | 2:00 | 777:22:00 | 0:00 | 2709:74 | 3:00 |
| 829:35:00 | 0:00 | 816:24:00 | 1:00 | 2882:81 | 4:00 |
| 827:35:00 | 2:00 | 816:24:00 | 1:00 | 2881:81 | 5:00 |
| 831:35:00 | 1:00 | 816:24:00 | 4:00 | 2894:81 | 7:00 |
| 833:35:00 | 0:01 | 822:24:00 | 2:00 | 2914:80 | 5:01 |
| 833:36:00 | 0:00 | 823:24:00 | 1:00 | 2918:80 | 7:00 |
| 841:36:00 | 0:00 | 835:24:00 | 0:00 | 2981:83 | 4:00 |
| 843:36:00 | 0:00 | 836:24:00 | 2:00 | 2998:83 | 4:00 |
| 850:37:00 | 0:00 | 852:24:00 | 0:00 | 3061:85 | 7:00 |
| 858:37:00 | 1:01 | 876:25:00 | 1:00 | 3181:86 | 4:00 |
| 859:38:00 | 0:00 | 875:25:00 | 2:00 | 3180:86 | 6:00 |
| 859:38:00 | 0:00 | 885:25:00 | 1:00 | 3184:86 | 5:00 |
| 858:38:00 | 1:00 | 887:25:00 | 0:00 | 3189:86 | 2:00 |
| 1:00 | 0:01 | 0:00 | 0:00 | 2:00 | 3:01 |
| 0:00 | 1:01 | 0:00 | 0:00 | 0:00 | 3:09 |
| 0:00 | 1:01 | 0:00 | 0:00 | 0:00 | 3:09 |
| 0:00:00 | 0:00:00 | 0:01:00 | 0:00:00 | 0:00:01 | 0:00:00 |
| 0:1:0:0 | 0:0:0:0 | 0:1:1:0 | 0:0:0:0 | 0:1:0:1 | 0:0:0:0 |
| 0:1:0:0 | 0:0:0:0 | 0:1:1:0 | 0:0:0:0 | 0:1:0:1 | 0:0:0:0 |
| 2 | 0 | 3 | 0 | 2 | 0 |
| 0:1:1:1:2:2:1:0:0 | 0:0:0:0:0:0:0:0:0 | 3:11:2:0:2:0:1:0:2 | 0:0:0:0:0:0:0:0:0 | 5:14:1:1:2:3:0:3:8 | 0:0:0:0:0:1:0:0:0 |
| 1:00:00 | 0:00:00 | 0:00:00 | 2:00:00 | 0:00:01 | 0:01:00 |
| 0:0:2:0:0:0:0:0:0 | 0:0:0:0:0:0:0:0:0 | 0:6:3:0:0:1:0:0:0 | 0:0:0:0:0:0:0:0:0 | 0:26:17:0:0:7:0:1:1 | 0:4:0:0:0:1:0:0:0 |
| 0:0:2:0:0:0:0:0:0 | 0:0:0:0:0:0:0:0:0 | 0:6:3:0:0:1:0:0:0 | 0:0:0:0:0:0:0:0:0 | 0:26:17:0:0:6:0:1:1 | 0:4:0:0:0:2:0:0:0 |
| 0:2:0:0:0:0:0:0:0 | 0:0:0:0:0:0:0:0:0 | 5:3:0:0:0:2:0:0:0 | 0:0:0:0:0:0:0:0:0 | 25:13:0:0:0:16:0:1:1 | 0:3:0:0:0:0:0:0:0 |
| 0:0:1:0:0:0:0:0 | 0:0:1:0:0:0:0:0 | 2:0:0:0:0:2:0:0 | 3:0:3:0:0:0:0:0 | 11:0:4:0:0:16:0:0 | 11:0:11:0:0:0:1:1 |
| 0:0:1:0 | 0:0:0:0 | 0:0:3:0 | 0:0:0:0 | 0:1:27:0 | 0:0:2:0 |
| 0:0:0:0:0:0:0:0:0:0:0 | 0:0:1:0:0:0:0:0:0:0:0 | 0:0:0:0:0:0:0:0:0:0:0 | 0:0:0:0:0:0:0:0:0:0:0 | 0:0:0:0:0:0:0:0:0:0:0 | 0:0:0:0:0:0:0:0:0:1:1 |
| 0:0:0:0:0 | 6:2:6:2:1 | 0:0:0:0:0 | 0:0:5:0:0 | 0:0:0:0:0 | 2:2:12:1:2 |
| 1:6:1:1:0:0:0:0:0:0 | 0:1:0:0:0:0:0:0:0:0 | 0:2:1:5:0:0:2:0:2:1 | 0:0:0:2:0:0:0:0:0:0 | 5:11:0:11:0:0:0:0:4:1 | 2:2:1:1:0:0:1:0:0:0 |
| 172:361:0:1:1 | 0:3:0:0:0 | 34:1139:0:10:0 | 0:2:0:0:0 | 178:878:1:7:2 | 0:1:0:0:0 |
| 358:172:0:1:1 | 0:0:0:0:0 | 1043:35:0:10:0 | 4:0:0:0:0 | 841:178:1:7:2 | 3:0:0:0:0 |
| 173:348:0:1:1 | 0:1:0:0:0 | 36:819:0:10:0 | 1:7:0:0:0 | 174:698:1:7:2 | 2:4:0:0:0 |
| 346:173:0:1:1 | 0:0:0:0:0 | 713:37:0:10:0 | 6:1:0:0:0 | 633:176:1:7:2 | 0:0:0:0:0 |
| 345:172:0:1:1 | 1:1:0:0:0 | 704:38:0:10:0 | 8:0:0:0:0 | 628:176:1:7:2 | 0:0:0:0:0 |

| | | | | | |
|---|---|---|---|---|---|
| 178:327:0:1:1 | 0:1:0:0:0 | 61:453:0:10:0 | 34:12:0:0:0 | 190:484:1:6:2 | 14:5:0:0:0 |
| 178:325:0:1:1 | 0:3:0:0:0 | 61:449:0:10:0 | 35:11:0:0:0 | 191:480:1:6:2 | 16:4:0:0:0 |
| 177:322:0:1:1 | 1:0:0:0:0 | 82:429:0:10:0 | 17:8:0:0:0 | 202:467:1:6:2 | 9:5:0:0:0 |
| 178:306:0:1:1 | 0:0:0:0:0 | 103:326:0:9:0 | 1:4:0:0:0 | 215:434:1:6:2 | 0:1:0:0:0 |
| 303:178:0:1:1 | 0:0:0:0:0 | 302:103:0:9:0 | 4:1:0:0:0 | 425:214:1:6:2 | 2:1:0:0:0 |
| 290:177:0:1:1 | 1:1:0:0:0 | 256:103:1:9:0 | 5:1:0:0:0 | 391:215:1:6:2 | 0:0:0:0:0 |
| 281:178:0:1:1 | 0:0:0:0:0 | 198:104:1:8:0 | 4:0:0:0:0 | 347:216:1:6:2 | 2:0:0:0:0 |
| 178:276:0:1:1 | 0:0:0:0:0 | 105:187:1:8:0 | 0:2:0:0:0 | 216:328:1:6:2 | 0:1:0:0:0 |
| 191:182:0:1:1 | 0:0:0:0:0 | 70:121:1:8:0 | 1:0:0:0:0 | 213:226:1:6:2 | 1:1:0:0:0 |
| 189:182:0:1:1 | 0:0:0:0:0 | 69:121:1:8:0 | 1:0:0:0:0 | 213:227:1:6:2 | 1:0:0:0:0 |
| 0 | 2 | 0 | 6 | 0 | 3 |
| 1 | 52 | 0 | 69 | 1 | 176 |
| 0:00:00 | 2:01:01 | 0:00:00 | 1:00:00 | 0:00:00 | 0:05:02 |
| 52:6:3:0 | 0:0:0:0 | 103:3:5:1 | 0:0:0:0 | 301:18:24:1 | 1:1:0:0 |
| 48:4:5:0 | 0:0:0:0 | 78:14:2:0 | 0:0:0:0 | 245:42:12:1 | 0:0:2:0 |
| 46:4:5:0 | 2:0:0:0 | 68:13:1:0 | 4:0:1:0 | 228:41:14:1 | 6:0:0:0 |
| 40:2:2:0 | 0:0:0:0 | 45:9:1:0 | 10:2:1:0 | 134:30:10:1 | 31:2:1:0 |
| 24:0:2:0 | 1:0:0:0 | 29:0:7:0 | 1:0:1:0 | 83:6:18:1 | 8:0:1:0 |
| 0 | 0 | 0 | 0 | 0 | 3 |
| 0:00 | 3:00 | 0:00 | 0:01 | 0:00 | 1:00 |
| 3:0:0:1:1:0 | 0:0:0:0:0:0 | 2:1:0:2:0:1 | 0:0:0:0:0:0 | 5:3:1:1:0:0 | 0:0:0:0:0:0 |
| 0:3:0:0:1:0 | 0:0:0:0:0:0 | 0:3:0:0:2:1 | 0:0:0:0:0:0 | 1:5:2:1:3:0 | 0:0:0:0:0:0 |
| 2:0:0:0:0:0 | 1:0:0:1:1:0 | 2:1:1:1:0:1 | 1:0:0:1:2:0 | 1:0:0:0:0:0 | 4:3:1:1:3:0 |
| 2:0:0:0:0:0 | 1:0:0:1:1:0 | 2:2:2:1:0:1 | 1:0:1:1:2:0 | 1:0:0:0:0:0 | 4:4:3:1:3:0 |
| 0 | 0 | 0 | 1 | 0 | 2 |
| 49:409:7 | 0:00:00 | 42:391:3 | 0:01:00 | 81:797:13 | 1:02:00 |
| 49:409:7 | 0:01:00 | 42:389:3 | 0:02:00 | 82:791:13 | 0:03:00 |
| 406:49:07 | 0:00:00 | 381:42:03 | 2:00:00 | 778:82:13 | 1:00:00 |
| 49:401:7 | 0:01:00 | 42:381:3 | 0:01:00 | 81:779:12 | 0:01:00 |
| 49:399:7 | 0:02:00 | 42:375:3 | 0:02:00 | 81:774:13 | 0:01:00 |
| 49:403:7 | 0:00:00 | 42:359:3 | 0:00:00 | 83:757:15 | 1:01:00 |
| 49:403:7 | 0:01:00 | 42:351:3 | 0:01:00 | 85:752:15 | 0:00:00 |
| 49:404:7 | 0:01:00 | 42:350:3 | 0:01:00 | 85:752:15 | 0:00:00 |
| 49:403:7 | 0:00:00 | 42:334:3 | 0:00:00 | 85:734:16 | 0:00:00 |
| 49:401:7 | 0:03:00 | 42:333:3 | 0:01:00 | 85:735:16 | 0:00:00 |
| 403:49:07 | 0:00:00 | 332:42:03 | 0:00:00 | 723:85:16 | 2:00:00 |
| 404:49:07 | 0:00:00 | 327:41:03 | 0:01:00 | 723:86:16 | 1:00:00 |
| 0.330358796 | 0:00:00 | 0.15787037 | 0:00:00 | 16:86:708 | 0:00:03 |
| 49:387:7 | 0:00:00 | 42:267:3 | 0:03:00 | 86:645:16 | 0:04:00 |
| 49:384:7 | 0:00:00 | 42:259:3 | 0:00:00 | 86:629:16 | 0:02:00 |
| 381:49:07 | 1:00:00 | 249:42:03 | 0:00:00 | 608:86:16 | 0:00:00 |
| 379:49:07 | 1:00:00 | 238:42:03 | 0:00:00 | 583:86:16 | 1:00:00 |
| 46:350:7 | 0:00:00 | 28:159:2 | 0:00:00 | 65:455:12 | 0:04:00 |
| 44:338:7 | 0:00:00 | 0.91806713 | 0:00:00 | 53:417:12 | 2:01:00 |
| 44:333:7 | 0:01:00 | 0.867361111 | 0:02:00 | 51:393:11 | 0:01:00 |
| 44:334:6 | 0:00:00 | 0.825 | 0:00:00 | 52:395:11 | 0:01:00 |
| 338:44:06 | 0:00:00 | 111:20:00 | 0:00:00 | 395:49:11 | 2:00:00 |
| 2:73:1:2:122:6:3:4:0:1:0 | 0:0:1:0:1:0:0:0:0:0:0 | 0:106:1:0:50:1:1:0:0:0:0 | 0:0:0:0:0:0:0:0:0:0 | 2:380:14:8:415:11:2:13:2:2:1 | 0:0:0:0:0:0:0:0:0:0:0 |
| 2:73:2:2:123:6:4:4:0:1:0 | 0:0:0:0:0:0:0:0:0:0:0 | 0:106:1:0:50:1:2:0:0:0:0 | 0:0:0:0:0:0:0:0:0:0 | 2:379:14:8:415:11:5:14:1:2:1 | 0:1:0:0:0:0:0:0:0:0:0 |
| 2:73:2:2:123:6:4:5:0:1:0 | 0:0:0:0:0:0:0:0:0:0:0 | 0:106:1:0:50:1:3:0:0:0:0 | 0:0:0:0:0:0:0:0:0:0 | 2:380:14:8:413:11:5:14:1:2:1 | 0:0:0:0:2:0:0:0:0:0:0 |
| 2:73:2:2:122:6:5:5:0:1:0 | 0:0:0:0:1:0:0:0:0:0:0 | 0:106:1:0:50:1:6:0:0:0:0 | 0:0:0:0:0:0:0:0:0:0 | 2:378:14:8:414:11:17:6:1:2:1 | 0:2:0:0:1:0:0:6:0:0:0 |
| 2:73:2:2:123:6:5:5:1:0 | 0:0:0:0:0:0:0:0:0:0 | 0:106:1:0:50:1:6:0:0:0 | 0:0:0:0:0:0:0:0:0 | 2:379:14:8:413:11:19:5:2:1 | 0:1:0:0:1:0:0:0:0:0 |
| 2:73:2:122:6:6:4:1:0 | 0:0:0:1:0:0:0:0:0 | 0:104:0:49:1:11:0:0:0 | 0:1:0:0:0:0:0:0:0 | 2:379:8:411:11:42:2:2:1 | 0:2:0:1:0:0:0:0:0 |
| 72:2:2:123:3:6:1:0 | 1:0:0:0:0:0:0:0 | 106:1:0:50:1:11:0:0 | 0:0:0:0:0:0:0:0 | 379:14:8:414:10:46:2:1 | 2:0:0:0:0:0:0:0 |
| 0:73:2:2:123:6:1:0:0 | 0:0:0:0:0:0:0:0:0 | 0:106:1:0:49:12:0:0:0 | 0:0:0:0:0:0:0:0:0 | 1:381:14:7:413:50:2:1:2 | 0:0:0:0:0:1:0:0:0 |
| 0:73:2:2:123:6:1:0:0:0 | 0:0:0:0:0:0:0:0:0:0 | 0:106:1:0:49:12:0:0:0:0 | 0:0:0:0:0:0:0:0:0:0 | 1:381:14:5:413:53:2:0:1:2 | 0:0:0:0:0:0:0:0:0:0 |
| 73:1:2:0:123:3:6:1:0:0:0 | 0:0:0:0:0:0:0:0:0:0:0 | 104:0:1:0:48:1:12:0:0:0:0 | 0:0:0:0:0:0:0:0:0:0 | 381:1:14:0:413:9:52:2:0:2:1 | 0:0:0:0:0:0:0:0:0:0:0 |

| | | | | | |
|---|---|---|---|---|---|
| 1:73:2:0:123:3:6:1:0:0:0 | 0:0:0:0:0:0:0:0:0:0:0 | 0:104:1:0:48:1:12:0:0:0:0 | 0:0:0:0:0:0:0:0:0:0 | 1:381:14:0:413:10:51:2:0:2:1 | 0:0:0:0:0:0:0:0:0:0 |
| 1:73:2:0:123:3:6:1:0:0:0 | 0:0:0:0:0:0:0:0:0:0:0 | 0:104:1:0:48:1:12:0:1:0:0 | 0:0:0:0:0:0:0:0:0:0 | 1:379:14:0:411:10:47:2:4:2:1 | 0:0:0:0:0:0:0:0:0:0 |
| 3:04:13 | 0:00:00 | 32:15:05 | 0:00:00 | 36:21:31 | 0:00:03 |
| 2:00 | 0:00 | 1:00 | 0:00 | 1:00 | 0:00 |
| 2:00 | 0:00 | 7:00 | 0:00 | 3:00 | 0:00 |
| 48:4:1:0:0:0 | 0:0:0:0:0:0 | 164:46:1:0:0:0 | 1:1:0:0:0:0 | 191:55:3:1:0:2 | 1:0:0:0:0:0 |
| 0:00 | 77:00:00 | 4:00 | 239:01:00 | 1:00 | 159:02:00 |
| 103:01:00 | 0:00 | 525:02:00 | 1:00 | 317:01:00 | 0:00 |
| 103:01:00 | 0:00 | 525:02:00 | 0:00 | 316:01:00 | 1:00 |
| 103:00:00 | 0:00 | 526:02:00 | 0:00 | 318:01:00 | 0:00 |
| 105:00:00 | 0:00 | 537:03:00 | 0:00 | 321:01:00 | 0:00 |
| 102:01:00 | 0:00 | 540:06:00 | 1:00 | 321:02:00 | 2:00 |
| 103:02:00 | 0:00 | 542:26:00 | 0:00 | 323:10:00 | 1:00 |
| 103:04:00 | 0:00 | 541:30:00 | 0:00 | 323:13:00 | 0:00 |
| 103:04:00 | 0:00 | 535:33:00 | 6:00 | 323:14:00 | 0:00 |
| 102:04:00 | 0:00 | 518:39:00 | 8:00 | 316:16:00 | 2:00 |
| 102:04:00 | 0:00 | 515:40:00 | 11:00 | 314:16:00 | 4:00 |
| 97:04:00 | 0:00 | 480:43:00 | 0:00 | 299:20:00 | 0:00 |
| 97:04:00 | 0:00 | 479:42:00 | 1:00 | 299:20:00 | 0:00 |
| 95:03:00 | 2:00 | 477:39:00 | 1:00 | 291:19:00 | 4:00 |
| 19:0:1:0:1:0:0 | 0:0:0:0:0:0:0 | 44:4:0:2:3:0:0 | 0:0:0:0:0:0:0 | 59:0:5:9:2:1:0 | 0:0:0:0:0:0:0 |
| 0:27:1:0:0:1:1:0:0:0:1 | 0:0:0:0:0:0:0:0:0:0:0 | 0:283:19:1:17:42:3:0:1:0:4 | 0:1:0:0:0:0:0:0:0:0 | 0:43:9:1:0:4:0:1:0:0:1 | 0:0:0:0:0:0:0:0:0:0 |
| 0:27:1:0:0:1:1:0:1:1:4:0:1:0 | 0:0:0:0:0:0:0:0:0:0:0:0:0:0 | 0:283:19:1:4:0:48:4:3:16:9:0:3:0 | 0:1:0:0:10:0:0:0:0:0:0:0:0:0 | 0:43:8:1:0:0:4:2:0:1:3:0:1:1 | 0:0:1:0:0:0:0:0:0:0:0:0:0:0 |
| 0:27:1:0:1:1:0:1:1:5:1:0:0:0:0 | 0:0:0:0:0:0:0:0:0:0:0:0:0:0:0 | 0:282:19:1:0:47:4:3:15:21:5:0:3:1:0 | 0:1:0:0:0:0:0:0:0:1:0:0:0:0:0 | 0:43:8:1:0:4:2:0:1:6:1:0:0:0:1 | 0:0:0:0:0:0:0:0:0:0:0:0:0:0:0 |
| 0:27:1:0:6:1:1:0:1:1:1:0:0:0:0 | 0:0:0:0:0:0:0:0:0:0:0:0:0:0:0 | 0:275:19:1:28:0:47:4:3:15:5:0:3:1:0 | 0:1:0:0:1:0:0:0:0:0:0:0:0:0:0 | 0:43:8:1:7:0:4:2:0:1:1:0:0:0:1 | 0:0:0:0:0:0:0:0:0:0:0:0:0:0:0 |
| 0:27:1:0:1:1:0:1:1:6:1:0:0:0:0 | 0:0:0:0:0:0:0:0:0:0:0:0:0:0:0 | 0:260:18:1:0:47:4:3:15:33:4:0:3:1:0 | 0:0:0:0:0:0:0:0:0:1:0:0:0:0 | 0:43:8:1:0:4:2:0:1:8:1:0:0:0:1 | 0:0:0:0:0:0:0:0:0:0:0:0:0:0:0 |
| 0:00 | 0:00 | 0:00 | 0:00 | 6:00 | 2:01 |
| 0:00 | 1:00 | 4:00 | 5:01 | 3:01 | 23:00 |
| 4:00 | 0:00 | 6:01 | 6:00 | 36:01:00 | 3:01 |
| 4:00 | 0:00 | 6:01 | 6:00 | 41:01:00 | 3:01 |
| 3 | 0 | 3 | 0 | 35 | 0 |
| 0 | 3 | 0 | 3 | 0 | 33 |
| 1 | 2 | 0 | 0 | 5 | 10 |
| 0 | 1 | 0 | 1 | 0 | 5 |
| 0 | 0 | 0 | 0 | 0 | 2 |
| 0:00 | 1:00 | 0:00 | 0:00 | 0:00 | 0:00 |
| 3:00 | 0:00 | 40:01:00 | 0:00 | 53:01:00 | 0:00 |
| 0 | 0 | 0 | 0 | 0 | 0 |
| 0 | 0 | 0 | 0 | 0 | 0 |
| 0 | 0 | 0 | 0 | 0 | 0 |
| 0 | 0 | 10 | 0 | 11 | 0 |
| 0:00:00 | 1:00:00 | 0:00:00 | 0:00:00 | 0:00:00 | 0:01:01 |
| 46 | 0 | 52 | 0 | 138 | 2 |
| 51 | 0 | 71 | 1 | 205 | 2 |
| 42 | 4 | 81 | 1 | 211 | 8 |
| 33 | 1 | 45 | 3 | 137 | 7 |
| 0 | 1 | 0 | 0 | 0 | 1 |
| 0 | 1 | 0 | 4 | 0 | 1 |
| 0:0:0:1 | 0:0:0:0 | 2:4:1:1 | 0:0:0:0 | 0:0:2:0 | 0:0:0:0 |
| 0:0:0:1 | 0:0:0:0 | 3:5:1:1 | 0:0:0:0 | 1:1:3:0 | 0:0:0:0 |
| 1:00 | 0:01 | 0:00 | 0:00 | 0:00 | 2:00 |
| 0:4:2:0:4:0:0:0:0:0:0:0:0:0:0 | 1:6:0:0:0:0:0:0:0:0:0:0:0:0:0 | 0:1:1:0:1:0:0:0:0:1:0:0:0:0:0 | 0:4:2:0:1:0:0:1:0:0:1:0:0:0:0 | 1:2:0:0:0:0:0:0:0:0:0:0:0:0:0 | 4:29:4:0:10:0:0:1:0:0:1:0:1:1:1 |
| 0:0:0:0:0:0:0:0:0:0:0 | 0:0:0:0:0:0:0:0:0:0:0 | 3:0:0:0:1:0:1:1:0:0:0 | 0:0:0:0:0:0:0:0:0:0 | 1:0:0:0:1:0:1:0:0:1:1 | 0:0:0:0:0:0:0:0:1:0:0 |
| 421:18:11:5:0:1:2 | 2:0:0:0:0:0:0 | 684:9:3:5:0:0:6 | 0:0:0:0:0:0:0 | 701:34:28:7:0:1:2 | 0:0:0:0:0:0:0 |
| 426:18:11:5:1:0 | 1:0:0:0:0:0 | 693:9:3:5:0:0 | 2:0:0:0:0:0 | 720:37:29:7:1:0 | 0:0:0:0:0:0 |
| 429:18:11:1 | 0:0:0:0 | 704:9:3:0 | 0:0:0:0 | 726:37:29:1 | 2:0:0:0 |
| 434:18:4:5:0:2 | 1:0:0:0:0:0 | 709:9:2:5:0:6 | 1:0:0:0:0:0 | 743:36:23:7:0:3 | 1:0:0:0:0:0 |
| 436:18:11:5:0:2:2 | 2:0:0:0:0:0:0 | 717:9:3:4:0:0:6 | 0:0:0:0:0:0:0 | 762:37:29:7:0:1:3 | 0:0:0:0:0:0:0 |
| 0:0:0:0 | 0:0:0:0 | 4:0:0:0 | 0:0:0:0 | 2:0:0:0 | 0:0:0:0 |

| | | | | | |
|---|---|---|---|---|---|
| 3 | 0 | 4 | 0 | 12 | 0 |
| 0 | 0 | 0 | 11 | 0 | 5 |
| 0 | 0 | 0 | 11 | 0 | 5 |
| 0:00:00 | 0:00:00 | 0:00:00 | 0:00:01 | 0:00:00 | 0:02:00 |
| 51:01:00 | 2:00 | 18:00 | 0:00 | 107:02:00 | 0:00 |
| 0.222222222 | 0:00 | 0.503472222 | 1:01 | 0.547222222 | 0:00 |
| 0.263888889 | 0:00 | 0.545833333 | 0:00 | 0.547222222 | 0:00 |
| 0:0:0:0:0:0:0:0:0:0:0:0 | 0:0:0:0:0:0:0:0:0:0:0 | 0:0:0:0:0:0:0:0:0:0:0 | 0:0:2:0:1:1:0:0:0:1:2:0 | 0:0:0:0:0:0:0:0:0:0:0 | 1:1:3:0:2:0:0:0:0:0:1:1 |
| 0:00 | 0:00 | 0:00 | 0:00 | 0:00 | 0:00 |
| 0 | 0 | 0 | 0 | 0 | 0 |
| 0 | 0 | 0 | 0 | 0 | 0 |
| 0 | 0 | 0 | 0 | 0 | 0 |
| 0:0:0:0 | 0:1:0:0 | 0:0:0:0 | 3:0:0:0 | 0:0:0:0 | 3:0:0:1 |
| 1:00 | 1:00 | 1:00 | 0:01 | 2:00 | 1:01 |
| 0:00:00 | 0:00:01 | 0:00:00 | 0:01:00 | 0:00:00 | 0:02:00 |
| 89 | 0 | 61 | 1 | 168 | 0 |
| 96 | 0 | 70 | 0 | 154 | 1 |
| 222:48:108:34:27:23:32:3:2:45:7:8:9:2:0:9:1:0:0 | 0:0:0:0:0:0:0:0:0:0:0:0:0:0:0:0:0:0:0 | 53:12:52:42:91:3:1:2:5:73:1:4:9:8:0:41:1:0:0 | 0:0:0:0:0:0:0:0:0:0:0:0:0:0:1:0:0:0 | 216:43:160:111:215:20:13:6:5:159:4:15:55:17:2:89:1:1:1 | 0:0:0:0:0:0:0:0:0:0:0:0:0:0:0:1:0:0:0 |
| 48:107:27:222:27:23:32:2:43:7:11:7:2:0:1:9:1:0:1:0 | 0:0:0:0:0:0:0:0:0:0:0:0:0:0:0:0:0:0:0 | 12:53:37:53:91:5:1:5:68:1:15:8:13:0:1:44:1:0:0:0 | 0:0:0:0:0:0:0:0:1:0:1:0:0:0:0:0:0:0:0 | 43:155:98:216:216:21:13:5:148:3:38:46:26:3:1:90:1:1:0:1 | 0:0:0:0:0:0:0:0:0:0:1:0:0:0:0:0:0:0:0 |
| 48:101:20:222:27:24:32:2:40:7:4:0:4:10:9:0:1:0:1:0 | 0:0:0:0:0:0:0:0:0:0:0:0:0:0:0:0:0:0:0 | 12:49:28:53:91:5:1:5:57:1:9:0:4:14:44:0:1:0:0:0 | 0:0:0:0:0:0:0:0:0:0:0:0:0:0:0:0:0:0:0 | 43:146:74:215:216:21:13:5:129:3:21:3:35:34:90:1:1:1:0:1 | 0:1:0:1:0:0:0:0:0:0:0:0:0:0:0:0:0:0:0 |
| 48:100:19:222:27:23:32:2:38:7:11:7:0:2:9:0:1:0:1:0 | 0:0:0:0:0:0:0:0:0:0:0:0:1:0:0:0:0:0:0 | 12:48:24:53:91:5:1:5:55:1:14:12:0:3:44:0:1:0:0:0 | 0:0:0:0:0:0:0:0:0:0:0:0:0:0:0:0:0:0:0 | 43:145:50:216:216:22:13:5:119:3:38:29:3:27:90:1:1:1:0:1 | 0:0:0:0:0:0:0:0:0:0:0:0:0:0:0:0:0:0:0 |
| 48:100:16:222:27:23:32:2:38:7:12:7:0:2:9:0:1:0:1:0 | 0:0:1:0:0:0:0:0:0:0:0:0:1:0:0:0:0:0:0 | 12:47:23:53:91:5:1:5:55:1:14:14:0:3:44:0:1:0:0:0 | 0:0:1:0:0:0:0:0:0:0:0:0:0:0:0:0:0:0:0 | 43:145:49:216:216:22:13:5:118:3:38:29:3:25:90:1:1:1:0:1 | 0:0:0:0:0:0:0:0:0:0:0:0:0:0:0:0:0:0:0 |
| 48:101:5:222:27:23:32:2:35:7:14:8:2:9:1:1:0:0:1:0 | 0:0:0:0:0:0:0:0:0:0:0:0:0:0:0:0:0:0:0 | 12:44:8:53:91:5:1:5:47:1:16:16:0:44:0:1:0:0:0:0 | 0:0:0:0:0:0:0:0:0:0:0:0:0:0:0:0:0:0:0 | 43:141:18:216:216:23:13:5:104:3:42:36:3:90:9:1:1:0:0:1 | 0:0:0:0:0:0:0:0:0:0:0:0:0:0:0:0:0:0:0 |
| 48:99:6:222:16:23:32:2:28:7:21:5:1:5:1:1:1:1:0 | 0:0:0:0:0:0:0:0:0:0:0:0:0:0:0:0:0:0 | 12:40:2:53:63:5:1:2:30:1:19:14:0:33:0:1:0:0:0 | 0:0:0:0:0:0:0:0:0:0:0:0:0:0:0:0:0:0 | 43:124:7:216:137:27:13:2:78:3:47:25:3:55:4:1:1:0:1 | 0:1:0:0:2:0:0:0:2:0:0:0:0:0:0:0:0:0 |
| 48:99:7:221:25:32:2:25:7:7:22:2:1:1:4:1:0:1:0 | 0:0:0:1:0:0:0:0:0:0:0:0:0:0:0:0:0:0 | 12:35:4:53:5:1:0:16:1:31:19:7:0:1:12:1:0:0:0 | 0:0:0:0:0:0:0:0:0:0:0:0:1:0:0:0:0:0 | 43:112:12:216:27:13:1:45:3:60:48:16:3:4:28:1:0:1:1 | 0:0:0:0:0:0:0:1:0:1:0:0:0:0:0:0:0:0 |
| 98:47:10:221:25:32:1:14:7:7:22:2:1:2:4:1:0:1:0 | 0:1:0:1:0:0:0:1:0:0:0:0:0:0:0:0:0:0 | 25:12:6:53:5:1:0:9:1:10:19:4:0:2:6:1:0:0:0 | 0:0:0:0:0:0:0:0:0:0:0:0:1:0:0:0:0:0 | 106:43:13:215:27:13:1:30:4:22:50:5:3:5:16:1:0:0:1 | 0:0:0:1:0:0:0:0:0:0:0:0:0:0:0:0:0:0 |
| 98:41:11:222:25:32:1:10:7:7:22:1:1:2:0:0:0:0 | 0:0:0:0:0:0:0:0:0:0:0:0:0:0:0:0:0 | 25:7:7:53:5:1:0:5:1:2:19:4:0:2:1:1:1:0 | 0:0:0:0:0:0:0:0:0:0:0:0:0:0:0:0:0 | 103:32:19:216:27:13:0:29:4:10:50:5:3:9:4:0:0:1 | 0:0:1:0:0:0:0:2:0:0:0:0:0:0:0:0:0 |
| 220:22:40:100:11:25:32:1:12:7:9:1:0:2:0:0:0:0 | 0:0:1:0:0:0:0:0:0:0:0:0:0:0:0:0:0 | 49:19:7:24:10:5:1:0:8:1:2:3:0:2:1:1:0:0 | 0:0:0:0:0:0:0:0:0:0:0:0:0:0:0:0:0 | 212:50:32:102:27:27:13:0:35:4:10:4:2:11:0:4:0:1 | 0:0:0:0:0:0:0:0:0:0:0:0:0:0:0:0:0 |
| 549 | 1 | 1204 | 1 | 1353 | 1 |
| 550 | 1 | 1204 | 0 | 1354 | 1 |
| 554 | 0 | 1207 | 2 | 1362 | 0 |
| 555 | 0 | 1213 | 1 | 1361 | 2 |
| 554 | 2 | 1210 | 3 | 1366 | 1 |
| 557 | 0 | 1214 | 2 | 1367 | 4 |
| 557 | 0 | 1216 | 1 | 1369 | 2 |
| 555 | 1 | 1215 | 0 | 1369 | 1 |
| 555 | 1 | 1211 | 2 | 1368 | 3 |
| 0 | 7 | 0 | 9 | 0 | 15 |
| 18:0:17:8:0 | 0:0:0:0:0 | 48:0:5:4:10 | 0:0:0:0:0 | 70:0:11:15:9 | 0:0:0:0:0 |
| 197:48:05 | 0:00:00 | 269:46:09 | 0:00:00 | 764:128:15 | 2:00:00 |
| 197:48:05 | 0:00:00 | 273:47:09 | 0:00:00 | 765:131:15 | 4:00:00 |
| 208:49:08 | 1:00:00 | 352:47:18 | 0:00:00 | 947:132:19 | 2:00:00 |
| 60:46:05 | 149:03:03 | 139:47:10 | 212:00:08 | 169:123:14 | 779:09:05 |
| 149:02:03 | 58:01:05 | 213:00:08 | 136:04:10 | 783:06:05 | 164:01:14 |

| | | | | | |
|---|---|---|---|---|---|
| 206:45:08 | 1:01:00 | 352:46:18 | 0:00:00 | 951:129:18 | 0:00:00 |
| 60:45:05 | 147:01:03 | 138:46:10 | 214:00:08 | 168:121:13 | 782:08:05 |
| 133:4:2:0 | 0:0:0:0 | 118:6:3:1 | 1:0:0:0 | 437:35:25:0 | 1:0:0:0 |
| 137:4:3:0 | 0:0:0:0 | 126:6:6:2 | 0:0:0:0 | 453:34:34:1 | 0:0:1:0 |
| 137:4:3:0 | 0:0:0:0 | 126:6:6:2 | 0:0:0:0 | 450:33:38:1 | 3:1:0:0 |
| 136:4:3:0 | 0:0:0:0 | 126:6:7:2 | 0:0:0:0 | 443:35:45:1 | 1:0:2:0 |
| 73:3:1:0 | 63:1:2:0 | 63:4:4:0 | 63:2:3:2 | 298:27:34:0 | 146:8:11:1 |
| 130:01:00 | 0:00:00 | 114:29:02 | 0:00:00 | 416:81:1 | 0:00:00 |
| 131:02:00 | 0:00:00 | 114:32:02 | 0:00:00 | 413:125:1 | 2:02:00 |
| 131:03:00 | 0:00:00 | 112:41:02 | 2:00:00 | 414:139:1 | 0:01:00 |
| 131:03:00 | 0:00:00 | 114:43:02 | 0:00:00 | 412:144:1 | 3:01:00 |
| 123:4:5:0 | 0:0:0:0 | 113:6:49:2 | 0:0:0:0 | 411:35:171:1 | 0:0:0:0 |
| 123:4:5:0 | 0:0:0:0 | 113:6:52:2 | 0:0:0:0 | 411:35:176:1 | 0:0:3:0 |
| 122:4:5:0 | 1:0:0:0 | 112:6:53:2 | 0:0:2:0 | 409:35:184:1 | 0:0:1:0 |
| 121:3:7:0 | 0:0:0:0 | 108:6:61:2 | 1:0:0:0 | 390:33:207:1 | 1:0:0:0 |
| 122:3:7:0 | 0:0:0:0 | 109:6:64:2 | 0:0:0:0 | 389:33:208:1 | 2:0:0:0 |
| 96:9:32:12:150:88:9:1:2:6:5:16:0:2:4:0:1:0:3:0:1:2:1:4:1 | 0:0:0:0:0:0:0:0:0:0:0:0:0:0:0:0:0:0:0:0:0:0:0 | 75:2:10:7:102:43:4:1:0:8:1:7:0:0:0:3:0:0:1:1:1:1:0:0:0 | 0:0:0:0:0:0:0:0:0:0:0:0:0:0:0:0:0:0:0:0:0:0:0:0:0 | 754:3:121:25:301:385:39:13:3:38:12:38:0:0:10:5:0:0:5:1:3:1:0:3:2 | 0:0:0:0:1:3:1:0:0:0:0:0:0:0:0:0:0:0:0:0:0:0:0:0:0 |
| 97:9:32:11:151:88:9:1:1:6:5:16:0:2:4:0:1:0:0:1:2:1:3:4:1 | 0:0:0:0:1:0:0:0:0:0:0:0:0:0:0:0:0:0:0:0:0:0:0 | 75:2:10:7:105:41:6:1:0:8:1:7:0:0:0:3:0:0:1:1:1:0:1:0:0 | 0:0:0:0:0:0:0:0:0:0:0:0:0:0:0:0:0:0:0:0:0:0:0:0:0 | 770:3:123:25:302:378:40:13:3:39:11:37:0:1:7:6:0:0:1:3:1:0:5:4:2 | 1:0:0:0:1:1:0:0:0:0:0:0:0:0:0:0:0:0:0:0:0:0:0:0:0 |
| 96:9:32:13:152:79:9:1:2:6:5:16:0:2:1:0:1:0:0:1:2:1:4:1:3 | 1:0:0:0:0:1:0:0:0:0:0:0:0:0:0:0:0:0:0:0:0:0:0 | 78:2:10:8:106:40:6:1:0:9:1:7:0:0:0:3:0:0:1:1:1:0:0:0:1 | 0:0:0:0:0:0:0:0:0:0:0:0:0:0:0:0:0:0:0:0:0:0:0:0:0 | 789:3:126:26:306:366:42:13:3:39:13:37:0:1:8:6:0:0:1:4:1:0:4:2:5 | 1:0:0:0:0:0:0:0:0:0:0:0:0:0:0:0:0:0:0:0:0:0:0:0:0 |
| 97:9:32:13:153:80:10:1:2:6:5:14:0:2:1:0:1:0:0:1:2:1:4:1:3 | 0:0:0:0:0:0:0:0:0:0:0:0:0:0:0:0:0:0:0:0:0:0:0 | 79:2:10:8:106:39:6:1:0:9:1:6:0:0:0:3:0:0:1:1:1:0:0:0:0 | 0:0:0:0:0:0:0:0:0:0:0:0:0:0:0:0:0:0:0:0:0:0:0:0:0 | 809:3:128:28:307:360:42:13:2:41:13:29:0:1:8:6:0:0:1:4:0:0:4:2:2 | 0:0:0:0:0:1:0:0:0:0:0:1:0:0:0:0:0:0:0:0:0:0:0:0:0 |
| 32:11:2:5:1:0:1:1:3:1:0:1:4:1:2:0 | 0:0:0:0:0:0:0:0:0:0:0:0:0:0:0:0:0 | 10:2:0:0:3:0:1:2:3:0:0:0:0:0:0:0 | 0:0:0:0:0:0:0:0:0:0:0:0:0:0:0:0 | 127:8:6:14:6:1:4:4:10:0:0:0:2:2:4:1 | 0:0:0:0:0:0:0:0:0:0:0:0:0:0:0:0 |
| 8:4:4:1:0:1:0:1:0:2:1:1:3:0:1:0 | 24:1:7:1:1:0:0:0:0:1:0:0:1:1:1:0 | 3:0:1:0:0:3:0:0:0:1:1:0:0:0:0:0 | 7:0:1:0:1:0:0:0:0:2:1:0:0:0:0:0 | 18:4:0:1:1:6:0:0:0:3:1:0:1:0:2:0 | 109:10:8:5:3:0:1:0:0:7:3:0:1:2:2:1 |
| 31:8:4:2:0:1:5:4:1:0:0:0:1:3:1:1:0:1:0 | 67:18:7:8:1:1:1:1:0:0:0:0:0:1:0:0:1:0:0 | 45:2:1:1:0:0:5:0:3:0:0:4:0:1:1:0:0:0:0 | 37:4:1:4:2:0:4:1:1:0:0:0:0:2:1:0:0:0:0 | 206:13:1:10:6:1:19:4:8:0:2:2:0:2:1:0:1:1:0 | 617:100:7:30:11:5:26:10:7:1:2:5:0:9:3:0:2:4:1 |
| 98:26:11:10:1:2:6:5:1:0:0:0:1:4:1:1:1:1:0 | 0:0:0:0:0:0:0:0:0:0:0:0:0:0:0:0:0:0:0 | 82:6:2:4:2:0:9:1:5:0:0:4:0:3:2:0:0:0:0 | 0:0:0:0:0:0:0:0:0:0:0:0:0:0:0:0:0:0:0 | 822:112:8:39:17:6:45:14:26:1:4:7:0:11:4:0:3:4:1 | 1:0:0:1:0:0:0:0:0:0:0:0:0:0:0:0:0:1:0 |
| 97:8:26:153:11:8:1:2:6:5:18:0:0:5:2:1:0:1:1:4:1:1:1:0:0 | 0:0:0:0:0:0:0:0:0:0:0:0:0:0:0:0:0:0:0:0:0:0:0 | 82:2:7:106:2:2:2:0:9:1:20:0:0:4:0:0:3:0:0:3:2:0:0:0:0 | 0:0:0:0:0:0:0:0:0:0:0:0:0:0:0:0:0:1:0:0:0:0:0:0:0 | 814:3:110:311:8:39:17:6:45:14:104:1:4:31:1:5:7:0:0:11:4:0:3:0:1 | 2:0:0:0:0:0:0:0:0:0:0:0:1:0:0:0:0:0:0:0:0:0:0:0:0 |
| 97:8:26:153:11:8:1:2:6:5:18:0:0:5:2:1:0:1:1:1:1:4:0:1:0 | 0:0:0:0:0:0:0:0:0:0:0:0:0:0:0:0:0:0:0:0:0:0:0 | 82:2:7:106:2:2:2:0:9:1:20:0:0:4:0:0:4:0:0:0:2:3:0:0:0 | 0:0:0:0:0:0:0:0:0:0:0:0:0:0:0:0:0:0:0:0:0:0:0:0:0 | 813:3:110:311:8:39:17:6:45:14:105:1:4:31:1:5:7:0:0:0:4:11:0:3:1 | 2:0:0:0:0:0:0:0:0:0:0:0:0:0:0:0:0:0:0:0:0:0:0:0:0 |
| 96:8:26:152:11:8:1:2:6:5:18:0:0:5:2:1:0:1:1:4:1:1:1:0:0 | 0:0:0:0:0:0:0:0:0:0:0:0:0:0:0:0:0:0:0:0:0:0:0 | 82:2:7:106:2:2:2:0:9:1:20:0:0:4:0:0:4:0:0:3:2:0:0:0:0 | 0:0:0:0:0:0:0:0:0:0:0:0:0:0:0:0:0:0:0:0:0:0:0:0:0 | 813:3:109:310:8:39:17:6:45:14:105:1:4:33:1:2:7:0:0:11:4:0:3:0:1 | 2:0:2:0:0:0:0:0:0:0:0:0:0:0:0:0:0:0:0:0:0:0:0:0:0 |
| 151:96:7:26:11:8:1:2:6:5:18:0:0:6:2:1:0:1:1:4:1:1:1:0:0 | 0:0:0:0:0:0:0:0:0:0:0:0:0:0:0:0:0:0:0:0:0:0:0 | 106:81:2:7:2:2:2:0:9:1:20:0:0:4:0:0:4:0:0:3:2:0:0:0:0 | 0:1:0:0:0:0:0:0:0:0:0:0:0:0:0:0:0:0:0:0:0:0:0:0:0 | 309:814:3:111:9:39:17:6:45:14:104:1:4:35:2:2:7:0:0:11:4:0:3:0:1 | 1:1:0:0:0:0:0:0:0:0:0:0:0:0:0:0:0:0:0:0:0:0:0:0:0 |
| 96:7:26:151:11:8:1:2:6:5:18:0:0:6:2:1:0:1:1:4:1:1:1:0:0 | 0:0:0:0:0:0:0:0:0:0:0:0:0:0:0:0:0:0:0:0:0:0:0 | 82:2:7:106:2:2:2:0:9:1:20:0:0:4:0:0:4:0:0:3:2:0:0:0:0 | 0:0:0:0:0:0:0:0:0:0:0:0:0:0:0:0:0:0:0:0:0:0:0:0:0 | 815:3:112:308:9:39:17:6:45:14:104:1:4:38:2:2:7:0:0:11:4:0:3:0:1 | 0:0:0:0:0:0:0:0:0:0:0:0:0:0:0:0:0:0:0:0:0:0:0:0:0 |
| 0:00:00 | 0:00:01 | 0:00:00 | 0:00:01 | 1:03:01 | 0:02:03 |
| 0:0:1:0 | 0:0:0:0 | 0:0:1:0 | 0:0:0:0 | 1:5:3:1 | 0:0:0:0 |
| 6:48:10 | 0:00:00 | 0.176539352 | 0:00:00 | 0.401111111 | 0:01:03 |
| 0:00:00 | 1:01:00 | 0:00:00 | 0:00:00 | 0:00:00 | 2:00:00 |
| 150:04:00 | 0:00 | 182:62 | 0:00 | 300:76 | 2:00 |
| 141:04:00 | 10:01 | 197:92 | 4:05 | 294:107 | 34:16:00 |
| 13 | 0 | 34 | 0 | 85 | 0 |
| 15 | 1 | 39 | 0 | 93 | 2 |

| | | | | | |
|---|---|---|---|---|---|
| 105:36:1:0:0 | 0:0:0:0:0 | 10:106:1:1:0 | 0:0:0:0:0 | 104:255:3:2:0 | 0:0:0:0:0 |
| 105:58:1:0 | 0:0:0:0 | 27:250:1:0 | 0:0:0:0 | 132:638:3:0 | 0:2:0:0 |
| 105:58:1:0 | 0:0:0:0 | 27:258:1:0 | 0:0:0:0 | 136:654:3:0 | 0:3:0:0 |
| 0:00 | 0:00 | 0:00 | 2:01 | 0:00 | 2:00 |
| 15:2:2:0:1:2:9:1:0:0:0:0 | 0:0:0:0:0:0:0:0:0:0:0:0 | 19:2:1:1:1:2:3:1:1:1:0:0 | 0:0:0:0:0:0:0:0:0:0:0:0 | 40:11:0:4:10:3:13:1:2:0:1:1 | 0:0:0:0:0:0:0:0:0:0:1:0 |
| 10:7:0:0:0:0:0:0 | 0:0:0:0:0:0:0:0 | 12:3:0:0:0:0:0:1 | 0:0:0:0:0:0:0:0 | 22:9:1:0:1:0:1:0 | 0:0:0:0:0:0:0:0 |
| 0:0:0:0 | 1:0:0:0 | 0:0:0:0 | 3:1:0:1 | 0:0:0:0 | 8:1:0:0 |
| 0:00:00 | 1:00:00 | 2:00:00 | 2:01:00 | 1:01:01 | 2:00:00 |
| 0:01 | 0:00 | 0:00 | 1:00 | 1:02 | 1:01 |
| 41:222:28:3:0:0:3:0:2:1:0:0:0 | 0:1:0:0:0:0:0:0:0:0:0:0:0 | 92:497:9:3:0:1:3:0:5:2:0:2:0 | 0:0:0:0:0:0:0:0:0:0:0:0:0 | 211:705:76:7:0:1:7:0:9:4:0:4:1 | 0:4:0:0:0:0:0:0:0:0:0:0:0 |
| 229:28:0:0:1:3:0:1:2:0:0:0 | 0:0:0:0:0:0:0:0:0:0:0:0 | 507:9:0:2:3:3:0:2:3:0:2:0 | 0:0:0:0:0:0:0:0:0:0:0:0 | 725:76:0:2:6:7:2:4:10:0:4:1 | 1:1:0:0:0:0:0:0:0:0:0:0 |
| 42:229:28:0:0:1:3:0:0:2:1:0:0 | 0:0:0:0:0:0:0:0:0:0:0:0:0 | 92:512:9:0:2:3:3:0:0:5:2:0:1 | 0:0:0:0:0:0:0:0:0:0:0:0:0 | 215:734:77:0:3:6:7:1:2:7:4:0:0 | 0:0:0:0:0:0:0:0:0:0:0:0:0 |
| 42:229:28:0:0:1:3:0:0:2:1:0:0:0 | 0:0:0:0:0:0:0:0:0:0:0:0:0:0 | 92:513:9:0:2:3:3:0:0:5:2:0:0:1 | 0:0:0:0:0:0:0:0:0:0:0:0:0:0 | 215:743:77:0:3:5:7:1:2:7:4:0:0:0 | 0:0:0:0:0:0:0:0:0:0:0:0:0:0 |
| 42:229:28:0:0:1:3:0:0:2:1:0:0:0 | 0:0:0:0:0:0:0:0:0:0:0:0:0:0 | 92:513:9:0:2:3:3:0:0:5:2:0:0:1 | 0:0:0:0:0:0:0:0:0:0:0:0:0:0 | 215:743:77:0:3:5:7:1:2:7:4:0:0:0 | 0:0:0:0:0:0:0:0:0:0:0:0:0:0 |
| 0:00:00 | 0:00:00 | 3:09:01 | 0:01:00 | 0:01:00 | 0:00:00 |
| 0:00:00 | 0:00:00 | 0:00:00 | 0:01:00 | 0:00:00 | 2:01:01 |
| 679 | 2 | 992 | 2 | 2036 | 1 |
| 623 | 3 | 866 | 1 | 1742 | 4 |
| 623 | 3 | 864 | 3 | 1745 | 4 |
| 625 | 1 | 866 | 3 | 1760 | 5 |
| 626 | 3 | 877 | 0 | 1767 | 5 |
| 634 | 2 | 888 | 0 | 1801 | 2 |
| 695 | 1 | 1024 | 2 | 2120 | 3 |
| 681 | 0 | 1015 | 1 | 2092 | 4 |
| 103 | 0 | 66 | 0 | 460 | 4 |
| 103 | 0 | 67 | 0 | 462 | 4 |
| 0:00 | 0:00 | 0:00 | 0:00 | 0:00 | 0:00 |
| 0 | 0 | 0 | 1 | 1 | 6 |
| 192:21:3:5 | 1:0:0:0 | 155:41:1:0 | 1:0:0:0 | 399:77:1:3 | 1:0:0:0 |
| 202:20:3:6 | 1:0:0:0 | 162:38:0:0 | 0:1:0:0 | 407:75:0:3 | 3:2:0:0 |
| 204:4:3:6 | 0:0:0:0 | 162:16:0:0 | 1:0:0:0 | 410:26:0:3 | 1:0:0:0 |
| 187:12:03 | 0:00:00 | 110:07:01 | 1:00:00 | 315:28:00 | 0:00:00 |
| 1 | 3 | 0 | 0 | 0 | 5 |
| 0 | 4 | 0 | 0 | 0 | 4 |
| 14:12 | 0:00 | 12:00 | 0:00 | 21:05 | 0:00 |
| 13:12 | 0:00 | 12:00 | 0:00 | 21:05 | 0:00 |
| 1:00 | 0:00 | 0:00 | 0:00 | 2:00 | 0:00 |
| 0:00 | 0:00 | 0:00 | 0:01 | 0:00 | 2:01 |
| 0:00 | 0:00 | 0:00 | 0:01 | 0:00 | 2:01 |
| 0:0:0:0 | 14:0:0:2 | 0:0:0:0 | 28:0:1:0 | 0:0:0:0 | 37:1:1:1 |
| 0 | 0 | 0 | 0 | 0 | 0 |
| 0:0:0:0 | 0:0:0:0 | 0:0:0:0 | 0:0:0:0 | 0:0:0:0 | 0:0:0:1 |
| 0:00 | 0:00 | 0:00 | 0:00 | 0:00 | 0:01 |
| 0:00 | 1:00 | 0:00 | 0:00 | 0:00 | 0:00 |
| 0:00 | 1:01 | 0:00 | 0:00 | 0:00 | 1:01 |
| 278:00:03 | 0:00:00 | 354:02:10 | 0:00:00 | 1117:05:32 | 2:00:00 |
| 277:00:03 | 1:00:00 | 354:02:10 | 0:00:00 | 1118:05:32 | 2:00:00 |
| 276:04:00 | 3:00:00 | 356:05:02 | 1:00:00 | 1125:18:05 | 1:00:00 |
| 280:03:00 | 0:00:00 | 361:02:02 | 0:00:00 | 1134:12:05 | 1:00:00 |
| 280:03:00 | 0:00:00 | 359:04:01 | 0:00:00 | 1131:16:05 | 2:00:00 |
| 0:00 | 0:01 | 0:00 | 0:01 | 0:00 | 1:01 |
| 0 | 31 | 0 | 22 | 2 | 95 |
| 0 | 1 | 1 | 3 | 6 | 0 |
| 1 | 0 | 2 | 2 | 10 | 1 |
| 2:0:0:0 | 0:0:0:0 | 5:1:0:0 | 0:0:0:0 | 8:4:0:0 | 0:0:0:0 |
| 2:0:0:0 | 0:0:0:0 | 7:0:0:0 | 0:0:0:0 | 9:0:3:0 | 0:0:0:0 |
| 6:0:1:0:0 | 0:0:0:0:0 | 14:0:3:0:0 | 1:0:1:0:0 | 12:2:5:0:1 | 0:0:1:0:0 |
| 0:0:0:0 | 0:0:0:1 | 0:0:0:0 | 0:0:0:0 | 0:0:0:0 | 1:0:2:3 |
| 0:0:1:0:0 | 0:0:0:0:0 | 0:0:0:0:0 | 0:0:0:0:0 | 1:1:3:0:1 | 0:0:0:1:0 |
| 0:0:1:0 | 0:0:0:0 | 0:0:0:0 | 0:0:0:0 | 1:0:1:1 | 0:1:2:0 |
| 0:0:1:0 | 0:0:0:0 | 0:0:0:0 | 0:0:0:0 | 1:0:1:1 | 0:1:2:0 |
| 0:1:0:0:0 | 0:0:0:0:0 | 1:0:0:0:0 | 0:0:0:0:0 | 0:2:1:0:0 | 0:0:0:0:0 |
| 1:0:0:0:0:0 | 1:0:1:0:0:0 | 2:0:0:0:0:0 | 2:0:2:0:0:0 | 5:0:3:0:0:0 | 0:1:0:0:0:0 |
| 2:1:0:7 | 0:0:0:0 | 3:3:0:0 | 0:0:0:0 | 5:4:0:17 | 0:0:0:1 |
| 2:1:0:7 | 0:0:0:0 | 3:3:0:0 | 0:0:0:0 | 5:4:0:18 | 0:0:0:0 |

| | | | | | |
|---|---|---|---|---|---|
| 2:1:0:7 | 0:0:0:0 | 2:2:0:0 | 1:1:0:0 | 5:4:0:18 | 0:0:0:0 |
| 2:1:0:7 | 0:0:0:0 | 3:3:0:0 | 0:0:0:0 | 5:4:0:18 | 0:0:0:0 |
| 2:1:0:7 | 0:0:0:0 | 3:3:0:0 | 0:0:0:0 | 5:4:0:18 | 0:0:0:0 |
| 2:1:0:8 | 0:0:0:0 | 3:3:0:0 | 0:0:0:0 | 5:4:0:17 | 0:0:0:0 |
| 125:103:0 | 0:00:00 | 19:18:00 | 0:00:00 | 128:143:1 | 0:00:00 |
| 300:4:14:8:7:1:2:0:0 | 0:0:0:0:0:0:0:0 | 378:5:7:5:9:3:2:2:0 | 0:1:0:0:0:0:0:0:0 | 911:15:29:16:15:5:9:5:2 | 2:0:0:0:0:0:0:0:0 |
| 298:0:15:14:8:11:17:4:0 | 1:0:0:0:0:0:0:0 | 375:0:7:7:5:10:14:2:3 | 0:0:0:0:0:0:0:0:0 | 909:0:36:31:16:49:45:17:7 | 1:0:0:0:0:0:0:0:0 |
| 295:15:13:8:11:18:0:0:0 | 0:0:0:0:0:0:0:0 | 369:8:7:5:10:16:3:0:0 | 1:0:0:0:0:0:0:0:0 | 901:38:27:15:48:55:6:0:1 | 2:0:0:0:0:0:0:0:0 |
| 295:15:13:8:11:17:0:0:0 | 0:0:0:0:0:0:0:0 | 370:8:7:5:10:15:3:0:0 | 0:0:0:0:0:0:0:0:0 | 899:38:27:15:47:57:5:0:1 | 2:0:0:0:0:0:0:0:0 |
| 294:15:14:8:11:5:17:4:0 | 0:0:0:0:0:0:0:0 | 368:8:7:5:10:6:15:2:3 | 0:0:0:0:0:0:0:0:0 | 896:38:28:15:47:16:59:18:4 | 1:0:0:0:0:0:0:0:1 |
| 294:15:14:8:11:5:18:4:0 | 0:0:0:0:0:0:0:0 | 367:8:7:5:10:6:16:2:3 | 0:0:0:0:0:0:0:0:0 | 897:38:28:15:47:16:61:18:4 | 0:0:0:0:0:0:0:0:1 |
| 295:15:14:8:11:5:18:4:0 | 0:0:0:0:0:0:0:0 | 368:8:7:5:10:6:16:2:3 | 1:0:0:0:0:0:0:0:0 | 899:38:28:15:47:16:61:18:4 | 0:0:0:0:0:0:0:0:0 |
| 82:06:02 | 0:00:00 | 38:05:01 | 0:01:00 | 210:12:15 | 0:02:00 |
| 82:5:4:0 | 0:1:0:0 | 38:3:5:0 | 0:0:0:0 | 209:11:17:1 | 2:0:1:0 |
| 0:01 | 0:00 | 0:00 | 0:00 | 0:00 | 0:00 |
| 0:01:00 | 0:01:00 | 0:01:02 | 0:00:00 | 0:00:02 | 1:01:00 |
| 0:03:00 | 0:01:01 | 0:02:02 | 0:08:04 | 1:13:03 | 0:01:01 |
| 6:00:01 | 6:00:00 | 13:00:06 | 24:00:00 | 19:01:01 | 42:00:02 |
| 8:00:01 | 11:00:00 | 17:00:06 | 4:00:00 | 36:01:01 | 12:00:00 |
| 10:00:01 | 10:00:00 | 17:00:06 | 19:00:00 | 32:01:01 | 30:00:01 |
| 0:27:01 | 0:00:00 | 0.062581019 | 0:03:00 | 0.154259259 | 0:00:00 |
| 0:28:01 | 0:03:00 | 0.064664352 | 0:00:00 | 0.155648148 | 0:00:00 |
| 12:00 | 0:00 | 6:01 | 0:00 | 24:01:00 | 0:00 |
| 1:10:2:0:1:2:0:1:0 | 0:0:0:0:0:0:0:0 | 6:9:3:1:1:3:0:0:0 | 0:0:0:0:0:0:0:0:0 | 29:21:6:2:6:4:1:0:1 | 0:0:0:0:0:0:0:0:0 |
| 0:0:0:0:0:0:0:0:0 | 10:1:2:0:1:2:0:1:0 | 0:0:0:0:0:0:0:0:0 | 9:6:4:0:1:3:0:0:0 | 0:0:0:0:0:0:0:0:0 | 21:18:8:1:7:3:1:0:1 |
| 4 | 0 | 3 | 0 | 10 | 0 |
| 47 | 1 | 54 | 2 | 97 | 0 |
| 0:00:01 | 0:00:00 | 1:00:00 | 0:00:00 | 0:00:00 | 0:00:00 |
| 127:78:6:3:4:1:1:4:1:0:1 | 1:0:0:0:0:0:0:0:0:0:0 | 136:95:5:7:8:4:2:4:4:0:0 | 0:0:0:0:0:0:0:0:0:0:0 | 402:203:16:28:12:5:2:17:5:1:3 | 2:0:0:0:0:0:0:0:0:0:0 |
| 130:78:4:3:1:4:1:0:1:1 | 0:0:0:0:0:0:0:0:0:0 | 136:95:13:7:2:4:4:0:0:4 | 0:0:0:0:0:0:0:0:0:0 | 403:203:19:28:2:17:5:1:3:5 | 2:0:0:0:0:0:0:0:0:0 |
| 130:78:4:3:1:4:1:0:1:1 | 0:0:0:0:0:0:0:0:0:0 | 136:95:14:7:2:4:4:0:0:4 | 0:0:0:0:0:0:0:0:0:0 | 405:203:20:28:2:17:5:1:3:5 | 0:0:1:0:0:0:0:0:0:0 |
| 130:78:6:4:3:1:4:1:1:1:0 | 0:0:0:0:0:0:0:0:0:0 | 136:95:5:18:7:2:4:3:0:3:0 | 0:0:0:0:0:0:0:0:0:0 | 404:202:16:25:28:2:17:4:3:2:1 | 1:1:0:0:0:0:0:0:0:0:0 |
| 0:0:0:0:0:0 | 0:0:1:0:0:0 | 0:0:0:0:0:0 | 0:0:0:1:1:0 | 0:0:0:0:0:0 | 0:0:0:0:0:1 |
| 26:6:1:5:3 | 0:0:0:0:0 | 5:0:1:1:0 | 1:0:0:0:0 | 37:13:7:10:1 | 0:0:0:0:0 |
| 9:3:0:0:3 | 17:3:1:5:0 | 2:0:1:0:0 | 3:0:0:1:0 | 5:1:1:2:0 | 31:12:6:10:1 |
| 0 | 2 | 0 | 2 | 0 | 2 |
| 143:11:11 | 0:00:00 | 625:61:17 | 1:00:00 | 432:46:08 | 0:00:00 |
| 145:11:00 | 0:00 | 603:61 | 2:00 | 416:46:00 | 1:00 |
| 0:00 | 0:00 | 0:00 | 0:02 | 0:00 | 0:01 |
| 1:5:1:0:0:0 | 0:2:1:0:0:0 | 0:0:0:0:0:0 | 0:0:0:0:0:0 | 4:8:2:1:0:1 | 0:1:0:1:0:0 |
| 2:01:00 | 0:00:00 | 1:00:01 | 0:00:00 | 1:00:02 | 0:00:00 |
| 0:1:0:0:0 | 0:0:0:0:0 | 1:1:2:1:0 | 0:1:0:0:0 | 1:0:1:0:0 | 8:2:6:1:0 |
| 0:00 | 0:00 | 1:00 | 1:00 | 0:00 | 0:00 |
| 111:02:00 | 0:00 | 119:00:00 | 0:00 | 427:06:00 | 0:00 |
| 111:02:00 | 1:00 | 119:00:00 | 1:00 | 425:06:00 | 2:00 |
| 113:02:00 | 0:00 | 124:00:00 | 0:00 | 431:06:00 | 2:00 |
| 111:02:00 | 2:00 | 125:00:00 | 0:00 | 435:05:00 | 0:00 |
| 112:03:00 | 0:00 | 125:01:00 | 0:00 | 429:05:00 | 2:00 |
| 111:05:00 | 0:00 | 123:02:00 | 0:00 | 414:07:00 | 2:00 |
| 0 | 1 | 0 | 2 | 0 | 2 |
| 0 | 0 | 0 | 2 | 0 | 2 |
| 0 | 1 | 0 | 2 | 0 | 2 |
| 0 | 1 | 0 | 2 | 0 | 1 |
| 0 | 1 | 0 | 2 | 0 | 1 |
| 14:02:04 | 0:00:00 | 5:00:03 | 0:00:00 | 22:03:07 | 0:00:00 |
| 0:00:00 | 0:00:00 | 0:00:00 | 7:00:00 | 0:00:00 | 6:00:00 |
| 0:00:00 | 0:00:00 | 0:00:00 | 7:00:00 | 0:00:00 | 6:00:00 |
| 0:00:00 | 0:00:00 | 0:00:00 | 7:00:00 | 0:00:00 | 6:00:00 |
| 28:26:00 | 0:00 | 29:57:00 | 1:00 | 155:113 | 1:01 |
| 34:26:00 | 0:00 | 41:57:00 | 0:00 | 198:114 | 1:00 |
| 0:00 | 3:00 | 0:00 | 0:00 | 1:00 | 1:00 |
| 0:00 | 3:00 | 0:00 | 0:00 | 1:00 | 1:00 |
| 0:0:0:1:3:1 | 0:0:0:0:0:0 | 0:0:0:5:4:0 | 0:0:0:0:0:0 | 0:0:0:11:1:1 | 0:0:0:0:0:0 |

| | | | | | |
|---|---|---|---|---|---|
| 4:33:0:0 | 0:1:0:0 | 6:49:3:4 | 0:0:0:0 | 11:153:5:4 | 0:3:0:0 |
| 29:0:4:0:0 | 4:0:0:0:0 | 40:5:6:4:1 | 9:0:0:0:0 | 133:6:11:4:0 | 14:0:0:0:0 |
| 30:0:4:0:0 | 2:0:0:0:0 | 41:5:6:4:1 | 8:0:0:0:0 | 138:6:11:4:0 | 10:0:0:0:0 |
| 4:2:0:0:0 | 0:1:0:0:0 | 6:2:4:6:1 | 0:0:0:0:0 | 11:7:4:4:1 | 0:3:0:0:0 |
| 0:0:0:0 | 0:0:0:0 | 0:0:0:0 | 0:0:0:1 | 0:0:0:0 | 0:3:0:0 |
| 0:0:0:0:0 | 0:0:0:0:0 | 0:0:0:0:0 | 0:0:0:0:1 | 0:0:0:0:0 | 0:0:1:0:0 |
| 0:00 | 1:00 | 5:00 | 3:01 | 10:00 | 2:00 |
| 2:0:0:0:1:1 | 5:6:1:1:0:0 | 0:3:0:0:0:0 | 1:4:0:0:0:0 | 11:31:12:1:6:0 | 0:2:0:0:0:0 |
| 5:6:3:1:2:1:0 | 1:0:0:0:0:0:0 | 1:5:7:0:0:0:0 | 0:0:0:0:0:0:0 | 7:28:26:3:4:3:0 | 1:1:0:0:0:0:0 |
| 0:0:0:0 | 0:0:0:0 | 0:0:0:0 | 0:0:1:0 | 0:0:0:0 | 1:0:0:1 |
| 19:00:00 | 82:01:00 | 188:05:04 | 177:01:00 | 504:05:06 | 32:01:00 |
| 103:00:00 | 0:00:00 | 374:04:00 | 0:00:00 | 544:05:01 | 2:00:00 |
| 106:00:00 | 0:00:00 | 389:00:00 | 0:00:00 | 565:00:00 | 1:00:01 |
| 106:01:00 | 0:00:00 | 395:06:07 | 1:00:00 | 571:06:10 | 2:00:00 |
| 105:01:00 | 1:00:00 | 395:06:10 | 0:00:00 | 571:06:16 | 1:00:00 |
| 0:00 | 0:00 | 0:00 | 0:00 | 4:01 | 0:00 |
| 0:0:0:0:0:0 | 0:0:0:1:0:0 | 0:0:0:0:0:0 | 0:0:1:4:0:1 | 0:0:0:0:0:0 | 0:2:2:10:1:0 |
| 1:0:0:0:0:0:0 | 0:0:1:0:0:0:0 | 8:0:0:1:1:0:1 | 8:0:0:0:0:0:0 | 15:0:2:1:2:1:0 | 20:0:3:1:3:0:0 |
| 57:56:1:2:2:2:1:1:3:0:1:1:3:0:0:1:0:1:0 | 0:0:0:0:0:0:0:0:0:0:0:0:0:0:0:0:0:0 | 38:68:5:10:10:10:7:32:55:8:3:4:15:1:9:1:0:0:3 | 0:0:0:0:0:1:0:0:1:0:0:0:0:0:0:0:0:0:0 | 144:141:14:17:11:33:18:103:125:15:4:5:22:5:10:1:8:1:5 | 0:0:0:0:0:0:0:0:1:0:0:0:0:0:0:0:0:0:0:0 |
| 56:56:1:3:2:4:0:6:3:0:1:0:1:0:1:0:1:1:0 | 1:0:0:0:0:0:0:0:0:0:0:0:0:0:0:0:0:0 | 38:68:3:4:10:2:6:33:56:8:3:1:8:9:0:0:0:0:3 | 0:0:0:0:0:0:0:0:0:0:0:0:0:0:0:0:0:0:0 | 144:141:13:9:12:14:18:105:126:17:4:6:8:12:0:8:0:1:6 | 0:0:0:0:0:0:0:0:0:0:0:0:1:0:0:0:0:0:0:0 |
| 57:56:1:3:2:5:6:0:3:0:1:0:1:0:1:1:1:1:0 | 0:0:0:0:0:0:0:0:0:0:0:0:0:0:0:0:0:0 | 38:67:4:4:10:2:32:6:56:7:3:1:4:8:0:0:0:0:3 | 0:1:0:0:0:0:1:0:0:0:0:0:0:0:0:0:0:0:0 | 143:141:11:5:12:11:104:16:126:16:4:4:5:6:0:7:0:1:6 | 1:0:0:0:0:0:1:0:0:0:0:0:0:0:0:0:0:0:0:0 |
| 57:56:1:3:2:5:6:0:3:0:1:0:1:0:1:1:1:1:0 | 0:0:0:0:0:0:0:0:0:0:0:0:0:0:0:0:0:0 | 37:68:4:4:10:2:33:6:56:7:3:1:4:8:0:0:0:0:3 | 1:0:0:0:0:0:0:0:0:0:0:0:0:0:0:0:0:0:0 | 144:141:11:5:12:11:105:16:126:16:4:4:5:6:0:7:0:1:6 | 0:0:0:0:0:0:0:0:0:0:0:0:0:0:0:0:0:0:0:0 |
| 56:55:2:2:1:4:6:0:3:0:1:0:1:0:1:1:1:1:0 | 1:0:0:0:1:0:0:0:0:0:0:0:0:0:0:0:0:0 | 38:67:4:4:10:2:32:6:56:7:3:1:2:7:0:0:0:0:3 | 0:0:0:0:0:0:0:0:0:0:0:0:0:0:0:0:0:0:0 | 143:138:10:6:12:10:104:15:124:15:4:2:4:3:6:0:1:0:6 | 0:1:0:0:0:0:0:0:0:0:0:0:0:0:0:0:0:0:0:0 |
| 79:08:00 | 0:00 | 127:07:00 | 2:00 | 580:25:00 | 2:00 |
| 78:07:00 | 1:00 | 105:05:00 | 0:00 | 496:20:00 | 3:00 |
| 74:07:00 | 0:00 | 73:03:00 | 0:00 | 392:12:00 | 0:00 |
| 70:07:00 | 0:00 | 38:03:00 | 0:00 | 305:10:00 | 1:00 |
| 59:05:00 | 2:02 | 15:01 | 17:02 | 5:00 | 267:10:00 |
| 0:00 | 0:00 | 0:00 | 0:04 | 0:00 | 1:01 |
| 0:00:00 | 0:00:00 | 1:00:00 | 0:00:00 | 0:00:00 | 1:01:01 |
| 8:00:01 | 0:00:00 | 8:02:02 | 0:00:00 | 13:01:05 | 0:00:00 |
| 0:00 | 1:01 | 0:00 | 0:00 | 0:00 | 0:00 |
| 0 | 2 | 0 | 0 | 0 | 0 |
| 0:0:0:0 | 0:0:0:0 | 1:0:0:0 | 0:0:0:0 | 1:0:2:1 | 0:0:0:0 |
| 1 | 0 | 1 | 1 | 0 | 7 |
| 1 | 0 | 2 | 0 | 7 | 0 |
| 0 | 0 | 0 | 2 | 0 | 6 |
| 0 | 0 | 0 | 2 | 0 | 5 |
| 6 | 3 | 0 | 0 | 4 | 3 |
| 0 | 0 | 0 | 0 | 0 | 2 |
| 1846:03:00 | 0:00 | 1299:00:00 | 1:00 | 5990:02:00 | 8:00 |
| 1845:03:00 | 1:00 | 1299:00:00 | 1:00 | 5991:02:00 | 7:00 |
| 1852:03:00 | 3:00 | 1303:00:00 | 2:00 | 6015:02:00 | 4:00 |
| 1853:03:00 | 2:00 | 1303:00:00 | 2:00 | 6009:02:00 | 13:00 |
| 1852:03:00 | 2:00 | 1303:00:00 | 3:00 | 6022:02:00 | 5:00 |
| 1851:02:03 | 2:00:00 | 1294:03:00 | 4:00:00 | 5990:16:02 | 5:00:00 |
| 1850:05:03 | 2:00:00 | 1287:03:00 | 2:00:00 | 5961:25:02 | 2:00:00 |
| 1848:05:03 | 1:00:00 | 1285:03:00 | 1:00:00 | 5952:24:02 | 6:01:00 |
| 1847:06:03 | 2:00:00 | 1285:03:00 | 1:00:00 | 5943:25:02 | 9:00:00 |
| 1840:06:03 | 2:00:00 | 1281:04:00 | 0:00:00 | 5935:28:02 | 7:00:00 |
| 1833:06:03 | 4:00:00 | 1275:04:00 | 1:00:00 | 5932:31:02 | 2:00:00 |
| 1836:07:03 | 1:00:00 | 1273:04:00 | 1:00:00 | 5914:31:02 | 8:00:00 |
| 1835:07:03 | 1:00:00 | 1274:04:00 | 0:00:00 | 5915:31:02 | 5:00:00 |
| 1837:08:03 | 0:00:00 | 1273:05:00 | 0:00:00 | 5909:30:02 | 5:01:00 |
| 1835:08:03 | 2:00:00 | 1273:05:00 | 0:00:00 | 5909:30:02 | 6:01:00 |
| 1827:08:03 | 3:00:00 | 1261:05:00 | 2:00:00 | 5863:29:02 | 8:01:00 |
| 1826:08:03 | 3:00:00 | 1261:04:00 | 0:00:00 | 5849:24:02 | 7:03:00 |
| 1825:04:03 | 2:00:00 | 1248:04:00 | 1:00:00 | 5789:15:02 | 11:00:00 |
| 1817:03:02 | 6:00:00 | 1243:00:02 | 0:00:00 | 5754:02:06 | 10:00:00 |
| 1820:03:02 | 3:00:00 | 1239:00:02 | 4:00:00 | 5756:02:04 | 5:00:00 |
| 1818:03:02 | 3:00:00 | 1242:00:02 | 1:00:00 | 5747:02:04 | 5:00:00 |
| 1815:03:00 | 2:00 | 1239:00:00 | 2:00 | 5719:02:00 | 6:00 |
| 1804:01:03 | 0:00:00 | 1231:01:00 | 0:00:00 | 5683:05:02 | 5:00:00 |
| 1796:01:03 | 3:00:00 | 1228:01:00 | 0:00:00 | 5669:05:02 | 10:00:00 |
| 1794:02:03 | 2:00:00 | 1221:00:00 | 2:01:00 | 5654:06:02 | 5:00:00 |
| 1792:02:03 | 2:00:00 | 1222:01:00 | 0:00:00 | 5646:07:02 | 8:00:00 |
| 1780:02:03 | 3:00:00 | 1215:01:00 | 2:00:00 | 5600:06:02 | 8:01:00 |
| 38:00:00 | 0:00 | 163:02:00 | 0:00 | 122:05:00 | 2:00 |
| 38 | 0 | 170 | 0 | 126 | 0 |
| 38:00:00 | 0:00 | 157:03:00 | 4:00 | 121:03:00 | 1:00 |

| | | | | | |
|---|---|---|---|---|---|
| 2:03 | 0:00 | 15:22 | 0:00 | 35:11:00 | 0:00 |
| 0:06 | 0:00 | 0:17 | 0:02 | 0:07 | 0:02 |
| 0 | 0 | 0 | 4 | 0 | 0 |
| 3 | 0 | 5 | 0 | 8 | 1 |
| 6 | 0 | 5 | 0 | 17 | 5 |
| 6 | 0 | 12 | 0 | 27 | 2 |
| 7 | 0 | 40 | 1 | 48 | 0 |
| 10 | 1 | 56 | 12 | 57 | 25 |
| 9 | 2 | 65 | 3 | 71 | 12 |
| 11 | 3 | 49 | 11 | 52 | 21 |
| 14 | 0 | 78 | 0 | 101 | 2 |
| 4 | 9 | 28 | 24 | 82 | 3 |
| 0:00:00 | 0:00:00 | 6:00:03 | 0:00:00 | 2:00:01 | 0:00:00 |
| 0:00:00 | 0:00:00 | 6:00:03 | 0:00:00 | 2:00:00 | 0:00:00 |
| 3:02:01 | 0:00:00 | 1:01:00 | 0:00:00 | 18:06:03 | 0:00:00 |
| 14 | 0 | 29 | 0 | 36 | 0 |
| 3:16:0:0:2:0:2:1:0 | 0:0:0:0:0:0:0:0:0 | 8:47:1:1:1:1:3:1:1 | 0:0:0:0:0:0:0:0:0 | 10:35:4:2:3:6:2:2:3 | 0:1:0:0:0:0:0:0:0 |
| 1:0:0:0:0 | 0:0:0:0:0 | 1:2:0:0:0 | 0:0:0:0:0 | 0:0:1:0:0 | 0:0:0:0:0 |
| 0 | 2 | 0 | 0 | 0 | 0 |
| 0 | 2 | 0 | 0 | 0 | 0 |
| 0 | 2 | 0 | 0 | 0 | 0 |
| 0 | 2 | 0 | 0 | 0 | 0 |
| 135:150:53:59:16:14:69 | 0:1:0:1:0:0:0 | 188:216:79:93:22:19:115 | 0:0:1:0:0:0:0 | 418:456:175:204:61:43:227 | 3:0:1:0:0:0:0 |
| 153:136:54:63:16:14:70 | 0:0:0:0:0:0:0 | 228:196:82:94:23:21:124 | 0:0:0:1:0:0:0 | 474:429:183:210:60:47:241 | 0:1:0:0:1:0:0 |
| 153:136:55:64:16:14:70 | 0:0:0:0:0:0:0 | 231:198:82:95:23:21:128 | 0:0:0:0:0:0:0 | 475:431:185:209:63:48:242 | 1:0:0:1:0:0:0 |
| 67:137:151:56:18:15:71 | 0:0:0:0:0:0:0 | 98:201:233:84:25:21:132 | 0:0:1:0:0:0:0 | 216:436:484:186:63:48:256 | 1:0:1:0:0:0:0 |
| 57:67:138:153:18:15:71 | 0:0:0:0:0:0:0 | 85:100:206:238:25:21:134 | 0:0:1:1:0:0:0 | 191:222:443:487:67:50:266 | 0:0:1:1:0:0:0 |
| 140:154:57:68:20:15:73 | 0:0:0:0:0:0:0 | 213:261:86:104:29:21:152 | 1:2:0:0:0:0:0 | 441:523:200:231:72:52:291 | 0:0:1:1:0:0:0 |
| 139:154:56:68:20:15:73 | 1:1:0:0:0:0:0 | 212:265:86:102:28:21:152 | 1:0:0:2:0:0:0 | 436:526:195:230:72:52:292 | 0:0:1:0:0:0:0 |
| 140:155:56:68:20:15:72 | 0:0:0:0:0:0:0 | 211:268:86:104:28:21:152 | 1:0:0:0:0:0:0 | 436:522:196:230:72:52:291 | 1:1:0:0:0:0:1 |
| 345:114:9 | 1:03:00 | 620:153:15 | 0:00:00 | 1367:342:29 | 5:00:00 |
| 115:342 | 0:00 | 151:616 | 0:01 | 378:1366 | 0:00 |
| 1:0:1:0 | 0:0:0:0 | 2:0:1:1 | 0:0:0:0 | 4:1:1:0 | 0:0:0:0 |
| 1:2:0:0:0:1 | 0:0:0:0:0:0 | 2:1:2:5:0:0 | 0:0:0:0:0:0 | 10:6:9:9:1:1 | 1:0:0:0:0:0 |
| 6:14 | 0:00 | 10:33 | 0:00 | 26:91 | 0:00 |
| 4 | 0 | 5 | 1 | 15 | 0 |
| 14:24 | 0:01 | 0.386805556 | 0:00 | 43:116 | 0:00 |
| 4 | 0 | 8 | 0 | 19 | 0 |
| 248:9:4:3:0:2:0:0:0:0 | 0:0:0:0:0:0:0:0:0:0 | 436:8:18:0:0:1:0:1:0:0 | 1:0:0:0:0:0:0:0:0:0 | 894:30:42:10:2:1:0:0:0:1 | 2:0:0:0:0:0:0:0:0:0 |
| 248:7:9:3:0:0:2:0:2:1:0 | 0:0:0:0:0:1:0:0:0:0:0 | 437:7:8:16:0:5:0:0:1:1:0 | 2:0:0:0:0:12:0:0:0:0:0 | 893:20:30:37:1:3:7:2:1:3:0 | 2:0:0:0:0:17:0:0:0:0:0 |
| 250:7:9:2:0:3:2:0:2:1 | 0:0:0:0:0:0:0:0:0:0 | 437:7:8:4:0:45:0:0:1:0 | 2:0:0:0:0:0:0:0:0:0 | 890:20:30:12:0:46:5:2:1:2 | 1:0:0:1:0:0:0:0:0:0 |
| 2744:18:152:109:0:0 | 5:2:0:0:0:0 | 4757:27:185:240:0:0 | 7:0:0:0:0:0 | 8923:74:534:426:0:1 | 9:1:1:0:0:0 |
| 2749:154:109:19:0:0 | 2:0:0:1:0:0 | 4766:185:240:41:0:0 | 4:0:1:2:0:0 | 8934:535:427:97:0:1 | 16:1:1:4:0:0 |
| 2746:153:109:19:0:0 | 0:0:0:1:0:0 | 4765:184:240:46:0:0 | 2:1:1:0:0:0 | 8931:534:426:102:0:1 | 4:1:0:4:0:0 |
| 154:2750:21:109:0:0 | 0:1:1:0:0:0 | 185:4765:47:240:0:0 | 0:5:0:1:0:0 | 535:8942:105:428:0:1 | 0:8:4:0:0:0 |
| 2749:154:108:25:0:0 | 2:0:1:0:0:0 | 4756:185:240:52:0:0 | 12:0:1:0:0:0 | 8924:529:428:115:0:1 | 20:3:0:0:0:0 |
| 2751:153:109:35:0:0 | 0:0:0:0:0:0 | 4768:186:240:76:0:0 | 2:0:1:0:0:0 | 8947:535:427:159:0:1 | 3:0:0:0:0:0 |
| 2751:153:109:37:0:0 | 0:0:0:0:0:0 | 4769:186:240:83:0:0 | 1:0:1:1:0:0 | 8946:532:428:179:0:1 | 2:0:0:3:0:0 |
| 2748:152:109:39:0:0 | 3:0:0:0:0:0 | 4765:186:239:86:0:0 | 5:0:2:3:0:0 | 8934:531:426:184:0:1 | 13:0:2:8:0:0 |
| 2751:148:43:109:0:0 | 1:0:0:0:0:0 | 4766:181:103:243:0:0 | 4:0:0:0:0:0 | 8943:527:223:426:0:1 | 7:0:2:2:0:0 |
| 2747:109:46:152:0:0 | 4:0:0:0:0:0 | 4759:242:121:183:0:0 | 12:0:9:0:0:0 | 8934:427:239:530:0:1 | 16:1:6:3:0:0 |
| 153:2749:109:46:0:0 | 0:3:0:0:0:0 | 181:4771:242:134:0:0 | 1:0:0:0:0:0 | 533:8951:430:249:0:1 | 1:3:0:0:0:0 |

| | | | | | |
|---|---|---|---|---|---|
| 2749:46:151:109:0 | 2:0:0:0:0 | 4765:133:174:242:0 | 4:0:0:0:0 | 8943:249:518:430:0 | 10:0:0:0:0 |
| 2751:147:109:46:0 | 0:0:0:0:0 | 4766:164:242:133:0 | 4:1:0:0:0 | 8950:501:429:249:0 | 5:6:1:0:0 |
| 550:62:30:3:7:0 | 2:0:0:0:0:0 | 235:33:26:4:1:0 | 0:0:0:0:0:0 | 840:99:102:23:5:1 | 4:1:1:0:0:0 |
| 549:64:30:3:7:0 | 2:0:0:0:0:0 | 236:33:26:4:1:0 | 0:0:0:0:0:0 | 830:101:104:23:5:1 | 6:1:0:0:0:0 |
| 550:64:30:3:7:0 | 0:0:0:0:0:0 | 235:33:26:4:1:0 | 1:0:0:0:0:0 | 832:100:104:24:5:1 | 1:2:0:0:0:0 |
| 70:4:0:0 | 0:0:0:0 | 220:5:2:0 | 0:0:0:0 | 391:17:0:0 | 2:0:0:0 |
| 0.217361111 | 0:00:00 | 0.364606481 | 0:00:00 | 1.027083333 | 0:02:00 |
| 3:00 | 0:00 | 12:02 | 0:00 | 10:01 | 0:00 |
| 3:0:0:0 | 0:0:0:0 | 12:0:1:2 | 0:0:0:0 | 10:0:1:1 | 0:0:0:0 |
| 0:0:0:0:0:0 | 1:0:0:0:1:0 | 4:1:0:0:0:0 | 2:0:0:0:0:0 | 4:0:0:1:0:1 | 0:0:0:0:0:0 |
| 52:13:3:1 | 0:1:0:0 | 139:17:6:22 | 0:0:0:0 | 138:48:14:9 | 0:0:0:0 |
| 52:14:3:1 | 0:0:0:0 | 139:17:5:20 | 0:0:0:0 | 137:47:14:9 | 1:1:0:0 |
| 10:3:1:0 | 42:12:2:1 | 65:9:2:5 | 74:6:3:10 | 118:43:10:2 | 20:3:3:2 |
| 0:00 | 0:00 | 0:00 | 2:00 | 0:00 | 0:00 |
| 8696:40:52:13:5 | 11:0:1:0:0 | 11105:52:69:11:2 | 11:0:0:0:0 | 39483:213:104:46:39 | 35:0:1:0:0 |
| 8791:48:53:13:2:1 | 1:0:0:0:0:0 | 11235:56:69:11:1:0 | 5:0:0:0:0:0 | 39866:239:105:46:3:2 | 24:0:0:0:0:0 |
| 8852:48:53:13:5:1 | 7:0:0:0:0:0 | 11337:53:69:11:2:0 | 13:0:0:0:0:0 | 40226:231:105:46:38:2 | 71:2:0:0:0:0 |
| 8866:33:53:13:5:1 | 6:0:0:0:0:0 | 11380:34:69:11:2:0 | 20:0:0:0:0:0 | 40377:149:104:46:39:2 | 60:1:0:0:0:0 |
| 8866:32:53:13:5:1 | 6:0:0:0:0:0 | 11381:33:69:11:2:0 | 22:0:0:0:0:0 | 40373:139:104:46:39:2 | 72:1:0:0:0:0 |
| 8890:25:53:13:5:1 | 6:0:0:0:0:0 | 11429:20:69:11:2:0 | 15:0:0:0:0:0 | 40507:95:104:46:38:2 | 70:1:0:0:0:0 |
| 8905:25:53:13:5:1 | 6:0:0:0:0:0 | 11442:20:69:11:2:0 | 21:0:0:0:0:0 | 40577:96:104:46:38:2 | 66:0:0:0:0:0 |
| 8916:20:53:13:5:1 | 11:1:0:0:0:0 | 11483:17:69:11:2:0 | 29:0:0:0:0:0 | 40677:82:104:46:38:2 | 101:0:0:0:0:0 |
| 8958:20:53:13:5:1 | 3:0:0:0:0:0 | 11552:14:69:11:2:0 | 12:0:0:0:0:0 | 40926:75:104:46:38:2 | 55:0:0:0:0:0 |
| 8964:18:53:13:5:1 | 14:0:0:0:0:0 | 11583:14:69:11:2:0 | 19:0:0:0:0:0 | 41037:69:104:46:38:2 | 76:1:0:0:0:0 |
| 9009:53:13:1:5:1 | 4:0:0:1:0:0 | 11636:69:11:4:2:0 | 22:0:0:0:0:0 | 41270:104:46:14:38:2 | 70:0:0:2:0:0 |
| 8999:53:13:0:5:1 | 2:0:0:0:0:0 | 11663:68:10:2:2:0 | 20:0:0:0:0:0 | 41341:104:46:5:35:2 | 50:0:0:2:0:0 |
| 9059:52:13:0:5:1 | 12:0:0:0:0:0 | 11836:68:11:1:2:0 | 36:0:0:0:0:0 | 41952:104:46:3:36:2 | 129:0:0:3:0:0 |
| 9088:51:13:0:4:1 | 12:1:0:0:0:0 | 11910:68:11:0:2:0 | 26:0:0:0:0:0 | 42234:103:46:3:36:2 | 76:0:0:2:0:0 |
| 9153:52:13:0:4:1 | 7:0:0:0:0:0 | 12056:68:11:0:2:0 | 42:0:0:1:0:0 | 42740:103:46:2:38:2 | 121:0:0:1:0:0 |
| 9145:52:13:0:4:1 | 15:0:0:0:0:0 | 12070:68:11:0:2:0 | 31:0:0:1:0:0 | 42758:103:46:2:37:2 | 123:0:0:1:0:0 |
| 9105:51:13:1:5:1 | 20:0:0:0:0:0 | 12077:67:11:1:1:0 | 41:0:0:1:1:0 | 42800:103:46:6:37:2 | 98:0:0:1:0:0 |
| 9115:51:13:1:5:1 | 10:0:0:0:0:0 | 12092:67:11:1:2:0 | 25:0:0:1:0:0 | 42819:103:46:6:37:2 | 81:0:0:1:0:0 |
| 9119:51:13:1:5:1 | 9:0:0:0:0:0 | 12097:67:11:1:2:0 | 30:0:0:1:0:0 | 42838:102:46:6:37:2 | 81:1:0:1:0:0 |
| 0:00 | 1:00 | 2:00 | 3:00 | 5:00 | 0:00 |
| 140:12:03 | 0:00:00 | 200:50:01 | 0:00:00 | 509:156:1 | 0:01:00 |
| 139:17:03 | 0:00:00 | 189:97:1 | 0:00:00 | 493:291:1 | 1:02:00 |
| 131:20:02 | 1:00:00 | 25:43:00 | 0:00:00 | 177:114:1 | 0:00:00 |
| 8:1:1:2:1:0:0:0:0:0 | 0:0:0:0:0:0:0:0:0:0 | 86:1:4:38:0:3:1:1:2:5 | 0:0:0:0:0:0:0:0:0:0 | 15:2:0:3:0:1:0:0:2:0 | 0:0:0:0:0:0:0:0:0:0 |
| 0 | 0 | 2 | 4 | 5 | 1 |
| 0:00 | 0:00 | 2:02 | 0:04 | 1:07 | 0:03 |
| 176:52:28:14 | 0:0:0:0 | 213:86:33:7 | 0:1:0:0 | 618:167:113:39 | 1:1:1:1 |
| 177:53:28:14 | 0:0:0:0 | 240:99:36:10 | 0:1:0:0 | 687:185:132:46 | 4:0:1:0 |
| 175:52:29:14 | 1:0:0:0 | 269:109:41:14 | 1:1:0:0 | 758:203:148:50 | 1:3:0:0 |
| 170:53:29:14 | 1:0:0:0 | 271:110:41:14 | 1:0:0:0 | 745:203:151:50 | 2:0:0:0 |
| 170:53:29:14 | 1:0:0:0 | 271:110:41:14 | 1:0:0:0 | 748:204:150:50 | 2:0:1:0 |
| 170:53:29:14 | 1:0:0:0 | 270:109:41:14 | 2:1:0:0 | 750:202:151:50 | 1:2:0:0 |
| 174:54:29:15 | 0:0:0:0 | 296:121:43:18 | 0:1:0:0 | 793:213:172:56 | 4:2:0:0 |
| 174:54:29:15 | 0:0:0:0 | 294:123:43:18 | 3:1:0:0 | 793:215:172:56 | 6:0:0:0 |
| 194:55:29:14 | 0:0:0:0 | 348:150:43:18 | 1:1:0:0 | 945:256:171:53 | 0:1:1:0 |
| 204:56:29:15 | 0:0:0:0 | 372:159:43:18 | 0:0:0:0 | 1013:272:172:53 | 3:2:0:0 |
| 203:56:29:15 | 1:0:0:0 | 372:157:43:18 | 0:0:0:0 | 1016:273:172:53 | 1:1:0:0 |
| 211:59:29:15 | 1:0:0:0 | 382:164:43:18 | 0:0:0:0 | 1052:299:171:51 | 0:1:1:0 |

| | | | | | |
|---|---|---|---|---|---|
| 211:60:29:15 | 1:0:0:0 | 382:164:43:18 | 0:0:0:0 | 1050:298:172:51 | 2:2:0:0 |
| 212:62:29:15 | 0:0:0:0 | 381:166:43:18 | 0:1:0:0 | 1049:303:172:51 | 2:1:0:0 |
| 212:63:29:15 | 0:0:0:0 | 380:168:43:17 | 1:1:0:1 | 1049:308:172:51 | 2:2:0:0 |
| 212:63:29:15 | 0:0:0:0 | 381:169:43:18 | 0:1:0:0 | 1049:310:172:51 | 2:0:0:0 |
| 211:71:29:15 | 1:0:0:0 | 381:181:43:18 | 0:0:0:0 | 1050:351:171:51 | 1:0:1:0 |
| 212:72:29:15 | 0:0:0:0 | 380:181:43:18 | 1:0:0:0 | 1049:351:172:51 | 1:0:0:0 |
| 5934 | 18 | 326 | 0 | 8429 | 14 |
| 5949 | 18 | 322 | 4 | 8426 | 32 |
| 5970 | 8 | 327 | 1 | 8468 | 17 |
| 6141 | 6 | 333 | 1 | 8744 | 9 |
| 6156 | 5 | 334 | 1 | 8784 | 7 |
| 6172 | 5 | 337 | 0 | 8810 | 7 |
| 6198 | 2 | 336 | 2 | 8854 | 2 |
| 6207 | 7 | 337 | 1 | 8873 | 12 |
| 6254 | 6 | 341 | 1 | 8947 | 19 |
| 90:4:4:0 | 0:0:0:0 | 154:1:6:0 | 0:0:0:0 | 509:16:18:0 | 3:0:0:0 |
| 20:1:13:3:1:1 | 0:0:0:0:0:0 | 48:1:32:1:2:3 | 0:0:0:0:0:0 | 56:3:11:2:3:1 | 0:1:0:0:0:0 |
| 26:1:13:2:2:1 | 1:0:0:0:0:0 | 79:3:32:2:3:2 | 0:0:0:0:0:0 | 87:6:11:7:2:2 | 0:0:0:0:0:0 |
| 0 | 0 | 0 | 0 | 0 | 0 |
| 5:1:0:0:3:0 | 0:0:0:0:0:0 | 2:9:3:0:4:0 | 0:0:0:0:0:0 | 34:38:18:0:10:0 | 0:0:0:0:1:0 |
| 0:0:0:0:0:0 | 2:5:0:0:0:3 | 0:0:0:0:0:0 | 7:2:1:0:0:3 | 0:0:0:0:0:0 | 37:34:17:1:0:11 |
| 2:5:0:0:0:3 | 0:0:0:0:0:0 | 7:2:1:0:0:3 | 0:0:0:0:0:0 | 37:34:17:1:0:11 | 0:0:0:0:0:0 |
| 2:5:0:0:0:3 | 0:0:0:0:0:0 | 7:2:1:0:0:3 | 0:0:0:0:0:0 | 36:34:13:1:0:11 | 1:0:0:0:0:0 |
| 6:01 | 0:00 | 4:00 | 0:00 | 11:00 | 0:00 |
| 6:01 | 0:00 | 4:00 | 0:00 | 11:00 | 0:00 |
| 0 | 2 | 0 | 0 | 2 | 0 |
| 0 | 0 | 0 | 0 | 0 | 2 |
| 43:3:0:0:0:1:0:0:0:0 | 0:0:0:0:0:0:0:0:0:0 | 132:5:1:0:1:0:0:1:0:0 | 0:0:0:0:0:0:0:0:0:0 | 120:4:1:0:1:0:1:3:0:0 | 0:0:0:0:0:0:0:0:0:0 |
| 43:3:0:0:0:0:1:0:0:0:0 | 0:0:0:0:0:0:0:0:0:0:0 | 134:5:1:0:0:1:0:0:1:0:0 | 0:0:0:0:0:0:0:0:0:0:0 | 121:4:1:0:0:1:0:1:3:0:0 | 1:0:0:0:0:0:0:0:0:0:0 |
| 0:00 | 0:00 | 0:00 | 0:00 | 0:00 | 0:00 |
| 9:8:0:1:2:5:1:0:0:0:0:0:0:0:0:1:0:2 | 0:0:0:0:0:0:0:0:0:0:0:0:0:0:0:0:0:0 | 24:30:29:2:2:11:1:0:0:0:0:0:0:3:3:3:1:2 | 0:0:0:0:0:0:0:0:0:0:0:0:0:0:0:0:0:0 | 73:89:68:4:4:18:5:6:2:1:2:1:5:6:6:8:3:2 | 0:1:0:0:0:0:0:1:0:0:0:0:0:0:0:0:0:0 |
| 1:0:10:0:1:0:4:1:0:8:0:1:0:0:4:1:0:1:1:0:0:0:0:1:0:0:1:0:0:0 | 0:0:0:0:0:0:0:0:0:0:0:0:0:0:0:0:0:0:0:0:0:0:0:0:0:0:0:0:0:0 | 4:5:71:2:8:0:15:3:2:47:0:0:0:0:18:4:3:0:0:0:0:0:0:1:0:0:1:1:0:0 | 0:0:0:0:0:0:0:0:0:0:0:0:0:0:0:0:0:0:0:0:0:0:0:0:0:0:0:0:0:0 | 15:12:238:4:14:2:82:9:4:88:20:6:2:0:43:16:3:0:1:0:0:0:2:0:0:0:1:0:2:1 | 0:0:1:0:0:0:0:0:0:0:0:0:0:0:0:0:0:0:0:0:0:0:0:0:0:0:0:0:0:0 |
| 9:2:10:2:1:1:0:3:2:4:1:1:8:10:2:1:0:0:1:1:0:3:1:0:3:4:1:0:1:1:0:0:0:1:0:0:0 | 0:0:0:0:0:0:0:0:0:0:0:0:0:0:0:0:0:0:0:0:0:0:0:0:0:0:0:0:0:0:0:0:0:0:0:0:0 | 65:9:71:4:8:4:0:15:0:14:3:2:47:85:13:2:5:2:6:0:0:1:1:3:1:19:4:4:0:0:1:0:0:1:1:0:0 | 0:0:0:0:0:0:0:0:0:0:1:0:0:0:0:0:0:0:0:0:0:0:0:0:0:0:0:0:0:0:0:0:0:0:0:0:0 | 108:23:239:8:14:15:3:13:3:82:9:5:87:99:17:6:12:21:59:5:2:7:4:10:1:46:18:3:1:2:0:0:2:0:0:1:0 | 0:0:0:0:0:0:0:0:0:0:0:0:0:0:1:0:0:0:0:1:0:0:0:0:0:0:0:0:0:0:0:0:0:0:0:0:0 |
| 2:10:2:1:1:9:0:3:2:4:1:1:8:10:2:1:0:0:1:1:0:3:1:0:3:4:1:0:1:1:0:0:0:1:0:0:0 | 0:0:0:0:0:0:0:0:0:0:0:0:0:0:0:0:0:0:0:0:0:0:0:0:0:0:0:0:0:0:0:0:0:0:0:0:0 | 10:71:4:8:4:65:0:15:0:15:3:2:47:85:13:2:5:2:6:0:0:1:1:3:1:19:4:4:0:0:1:0:0:1:1:0:0 | 0:0:0:0:0:0:0:0:0:0:0:0:0:0:0:0:0:0:0:0:0:0:0:0:0:0:0:0:0:0:0:0:0:0:0:0:0 | 23:239:8:14:15:110:3:13:3:82:9:4:88:99:17:6:12:22:59:5:2:7:4:10:1:46:18:3:1:2:0:0:2:0:0:1:0 | 0:0:0:0:0:0:0:0:0:0:0:0:0:0:0:0:0:0:0:0:0:0:0:0:0:0:0:0:0:0:0:0:0:0:0:0:0 |
| 10:4:8:10:3:7:0:2:1:0:1:1:2:0:0:0:2:4:1:0:1:0:0:0:0 | 0:0:0:0:0:2:0:0:0:0:1:0:0:0:1:0:0:0:0:0:0:0:0:0:0 | 56:13:41:74:10:53:1:9:7:0:0:3:13:5:4:1:1:18:4:3:1:0:0:0:1 | 14:2:6:11:4:10:0:1:0:0:0:0:0:0:1:0:0:1:0:0:0:0:0:0:0 | 208:72:79:84:10:101:19:23:13:2:0:6:17:12:47:3:1:37:14:3:0:0:1:0:0 | 30:6:9:14:3:7:0:0:1:0:3:3:0:0:9:0:1:7:3:0:0:0:0:1:0 |
| 6:2:2:4:1:1:2:4:3:3:0:0:0:2:0:0:0:1:0:2:0:0:0:0:0:1:0:1:0:0:0:1 | 0:1:0:0:0:0:0:1:0:0:0:0:0:0:0:0:0:0:0:0:0:0:0:0:0:0:0:0:0:0:0:0 | 18:12:2:28:1:2:4:15:1:6:1:0:0:13:5:0:0:0:2:8:0:2:3:0:2:0:1:3:0:0:1:1 | 1:1:0:4:0:1:0:3:0:0:0:0:0:0:0:0:0:0:1:0:0:0:0:0:0:0:0:0:0:0:0:0 | 87:23:1:27:2:4:23:33:2:13:7:2:0:17:12:12:1:1:19:21:2:7:1:1:3:2:0:2:0:2:0:0 | 3:1:0:2:0:1:4:3:0:1:0:0:0:0:0:0:0:0:1:0:0:0:1:0:0:0:0:0:0:0:0:0 |
| 3:0:1:0:0:0:1:0:0:0:0:0:0:0:1:1:0:0:0:0:0:0 | 0:0:0:0:0:0:1:0:0:0:0:0:0:0:0:0:0:0:0:0:0:0 | 6:4:12:0:0:0:8:1:0:2:1:1:1:0:0:2:0:0:1:0:0:0 | 0:0:0:0:0:0:0:0:0:0:0:0:0:0:0:0:0:0:0:0:0:0 | 20:7:12:0:3:0:16:3:1:3:1:2:2:2:2:5:0:0:0:1:0:0 | 6:2:0:0:1:0:2:2:0:0:0:3:0:0:0:1:0:1:0:0:0:0 |
| 2:0:5:0:0:0:0:1:1:1:0 | 0:0:0:0:0:0:0:0:0:0:0 | 1:0:2:1:0:0:0:0:0:0:0 | 0:0:0:0:0:0:0:0:0:0:0 | 1:4:9:1:0:0:1:2:0:1:1 | 0:0:0:0:0:0:0:0:0:0:0 |
| 121:1:1:2 | 1:0:0:0 | 278:6:6:36 | 6:0:0:1 | 556:13:7:56 | 1:0:0:1 |
| 125:2:1:4 | 0:0:0:0 | 282:3:6:37 | 2:0:0:0 | 556:7:8:58 | 0:1:0:0 |
| 125:3:4:4 | 0:0:0:0 | 284:7:6:34 | 0:0:0:0 | 555:15:9:50 | 1:0:0:1 |
| 125:3:4:1 | 0:0:0:0 | 283:16:6:27 | 0:3:0:0 | 555:20:9:36 | 0:8:0:0 |
| 125:3:4:0 | 0:0:0:0 | 282:19:6:19 | 0:0:0:0 | 551:27:9:26 | 2:0:0:0 |
| 122:2:2:0 | 0:0:0:0 | 267:19:5:0 | 0:0:0:0 | 516:25:9:7 | 1:0:0:0 |
| 79:00:01 | 0:00:00 | 118:04:00 | 0:00:01 | 426:07:06 | 0:01:00 |
| 78:00:01 | 1:00:00 | 118:04:00 | 0:00:01 | 423:07:05 | 2:01:01 |
| 78:00:01 | 0:00:00 | 118:04:01 | 0:00:00 | 424:08:06 | 2:00:00 |

| | | | | | |
|---|---|---|---|---|---|
| 0:00:00 | 78:00:01 | 0:00:00 | 118:03:00 | 0:00:00 | 422:04:05 |
| 77:00:02 | 0:00:00 | 108:03:00 | 1:00:00 | 387:03:12 | 0:00:00 |
| 0 | 1 | 0 | 0 | 2 | 5 |
| 0:0:0:0:0:0:0 | 0:2:0:0:0:0:0 | 0:0:0:0:0:0:0 | 0:0:0:0:0:2:0 | 0:0:0:0:0:0:0 | 0:3:0:0:0:1:1 |
| 0:1:0:0:0:0:0 | 0:1:0:0:0:0:0 | 0:0:0:0:0:1:0 | 0:0:0:0:0:1:0 | 0:2:0:0:0:0:1 | 1:1:0:0:0:1:0 |
| 0 | 0 | 1 | 0 | 5 | 0 |
| 0:00:00 | 0:00:00 | 0:00:00 | 1:01:00 | 0:00:00 | 1:00:01 |
| 3:3:1:1:0:1:0:0 | 0:0:0:0:0:0:0:0 | 0:1:0:2:0:0:0:2 | 0:0:0:0:0:0:0:0 | 0:9:0:8:0:2:0:1 | 0:0:0:0:0:0:0:0 |
| 7 | 0 | 4 | 0 | 43 | 0 |
| 5 | 1 | 3 | 0 | 35 | 6 |
| 0 | 5 | 0 | 3 | 0 | 34 |
| 2 | 0 | 0 | 1 | 14 | 8 |
| 2 | 0 | 0 | 0 | 9 | 4 |
| 0 | 1 | 0 | 0 | 4 | 0 |
| 1:0:1:0 | 0:0:0:0 | 0:0:1:1 | 0:0:0:0 | 1:0:0:0 | 0:1:1:0 |
| 2:00 | 0:00 | 1:00 | 0:00 | 3:01 | 0:00 |
| 8:1:1:0 | 5:1:0:0 | 4:0:0:0 | 3:0:0:0 | 8:0:1:0 | 17:0:0:0 |
| 8:1:1:0 | 5:1:0:0 | 4:0:0:0 | 3:0:0:0 | 8:0:1:0 | 23:1:1:0 |
| 13:26:1:0 | 0:0:0:0 | 7:14:0:0 | 0:0:0:0 | 30:28:4:0 | 0:0:0:0 |
| 1 | 2 | 0 | 0 | 4 | 3 |
| 7:00 | 7:00 | 29:00:00 | 28:01:00 | 61:01:00 | 42:00:00 |
| 6909:36:17 | 11:00:00 | 10006:64:10 | 16:00:00 | 21882:281:23 | 26:00:00 |
| 6918:36:17 | 4:00:00 | 10019:64:10 | 10:00:00 | 21897:281:23 | 20:00:00 |
| 6914:36:17 | 9:00:00 | 9996:64:10 | 11:00:00 | 21827:280:23 | 19:00:00 |
| 6921:35:17 | 1:00:00 | 9895:65:10 | 2:00:00 | 21691:279:23 | 10:00:00 |
| 6915:35:17 | 6:00:00 | 9888:65:10 | 9:00:00 | 21703:282:23 | 24:00:00 |
| 6913:35:17 | 8:00:00 | 9868:65:10 | 15:00:00 | 21688:282:23 | 27:00:00 |
| 6892:36:17 | 3:00:00 | 9718:66:10 | 14:00:00 | 21579:286:22 | 31:00:00 |
| 6889:36:17 | 5:00:00 | 9696:66:10 | 14:00:00 | 21560:286:22 | 30:00:00 |
| 6880:36:17 | 11:00:00 | 9650:66:9 | 31:00:00 | 21497:286:22 | 58:00:00 |
| 6837:36:17 | 6:00:00 | 9578:65:9 | 13:01:00 | 21231:285:21 | 27:00:00 |
| 6703:35:17 | 7:00:00 | 9384:66:9 | 9:00:00 | 20794:279:19 | 17:00:00 |
| 0:0:0:0 | 0:0:0:0 | 1:0:0:0 | 0:0:0:0 | 5:1:2:0 | 3:0:0:1 |
| 0 | 2 | 0 | 1 | 0 | 5 |
| 227:77:11 | 4:01:00 | 220:119:20 | 173:63:4 | 667:290:66 | 434:197:10 |
| 235:82:9 | 0:00:00 | 404:186:21 | 0:00:00 | 1169:520:66 | 1:00:00 |
| 82:235:9 | 0:00:00 | 185:404:19 | 1:00:01 | 519:1168:64 | 1:01:00 |
| 235:82:8 | 0:00:00 | 404:186:18 | 0:00:00 | 1170:519:55 | 0:01:01 |
| 234:82:8 | 1:00:00 | 404:187:19 | 0:00:00 | 1169:521:57 | 2:00:00 |
| 235:82:5 | 2:01:00 | 406:189:13 | 8:05:00 | 1176:523:43 | 18:06:00 |
| 236:82:5 | 1:01:00 | 412:194:13 | 2:00:00 | 1185:528:42 | 11:02:01 |
| 248:82:5 | 3:01:00 | 455:200:14 | 18:00:01 | 1301:536:44 | 44:00:01 |
| 249:83:5 | 3:00:00 | 474:198:13 | 8:02:02 | 1349:537:43 | 8:00:00 |
| 254:03:00 | 0:00 | 494:06:00 | 2:00 | 1366:10:00 | 2:00 |
| 9:00 | 0:00 | 7:01 | 0:00 | 26:04:00 | 1:00 |
| 6:02 | 1:00 | 5:02 | 1:00 | 20:06 | 1:00 |
| 4:15 | 0:00 | 1:02 | 0:01 | 4:14 | 0:01 |
| 1 | 0 | 6 | 0 | 16 | 9 |
| 0:00 | 0:00 | 0:00 | 0:00 | 0:00 | 0:00 |
| 0 | 0 | 0 | 0 | 0 | 3 |
| 12:2:0:0:1:0:0:0:0:0:0:0:0:0:0 | 0:0:0:0:0:0:0:0:0:0:0:0:0:0:0 | 11:3:0:0:0:0:0:0:0:0:4:0:0:0:0 | 0:0:0:0:0:0:0:0:0:0:0:0:0:0:0 | 17:3:0:2:1:1:0:1:0:0:1:0:1:2:1 | 0:0:0:0:0:0:0:0:0:0:0:0:0:0:0 |
| 12:2:0:0:0:0:1:0:0:1:0:0:0:0:0:0:0 | 0:0:0:0:0:0:0:0:0:0:0:0:0:0:0:0:0 | 12:8:1:0:0:0:0:0:0:1:4:0:0:0:0:0:0 | 0:0:0:0:0:0:0:0:0:0:0:0:0:0:0:0:0 | 17:8:1:0:2:0:2:1:0:3:1:0:0:0:1:2:1 | 0:0:0:0:0:0:0:0:0:0:0:0:0:0:0:0:0 |
| 12:2:0:1:0:0:0:0:0:0:0:0:1:0:0 | 0:0:0:0:0:0:0:0:0:0:0:0:0:0:0 | 12:11:1:3:0:0:0:0:5:0:0:0:0:0:0 | 0:0:0:0:0:0:0:0:0:0:0:0:0:0:0 | 17:11:1:4:0:0:1:1:1:0:0:1:1:1:1 | 0:0:0:0:0:0:0:0:0:0:0:0:0:0:0 |
| 0 | 0 | 0 | 0 | 0 | 1 |
| 1:1:2:1:0:0 | 0:3:0:0:0:0 | 0:0:0:0:0:0 | 0:0:0:0:1:0 | 2:2:2:0:0:1 | 0:2:1:0:0:0 |
| 134:29:00 | 0:00 | 33:260 | 0:02 | 285:1297 | 0:04 |
| 276:39:00 | 0:00:00 | 124:250:1 | 0:02:00 | 841:1315:1 | 0:03:00 |
| 277:37:00 | 0:00:00 | 124:236:1 | 1:01:00 | 841:1224:1 | 1:05:00 |
| 277:37:00 | 0:00:00 | 123:231:1 | 1:01:00 | 843:1163:1 | 2:05:00 |
| 280:36:00 | 2:00:00 | 130:215:1 | 0:00:00 | 864:1055:1 | 0:04:00 |
| 282:36:00 | 0:00:00 | 130:206:1 | 0:01:00 | 861:988:1 | 1:07:00 |
| 282:35:00 | 0:00:00 | 129:197:1 | 1:01:00 | 858:957:1 | 2:01:00 |
| 282:35:00 | 0:00:00 | 131:189:1 | 0:00:00 | 860:913:2 | 4:03:00 |
| 301:26:00 | 1:00:00 | 152:113:1 | 2:02:00 | 996:477:2 | 19:10:01 |
| 303:19:00 | 0:00:00 | 155:71:1 | 0:00:00 | 1020:300:3 | 2:01:00 |
| 303:07:00 | 0:00:00 | 155:10:01 | 0:00:00 | 1023:42:03 | 0:00:00 |
| 0:00 | 0:00 | 0:00 | 0:01 | 0:00 | 0:03 |
| 0 | 0 | 0 | 0 | 2 | 2 |
| 0 | 0 | 0 | 0 | 2 | 2 |
| 0:0:0:0:0:0:0 | 5:0:0:0:2:0:0 | 5:3:1:0:0:0:0 | 2:3:1:0:0:0:0 | 11:5:4:0:0:1:1 | 10:3:2:0:0:0:0 |
| 307:10:00 | 0:00 | 2071:06:00 | 2:00 | 3670:32:00 | 1:00 |
| 4 | 0 | 15 | 0 | 20 | 0 |

| | | | | | |
|---|---|---|---|---|---|
| 0:01:00 | 0:00:00 | 0:00:00 | 0:01:00 | 1:00:00 | 0:05:01 |
| 0:02:01 | 0:00:00 | 0:06:00 | 0:00:00 | 0:08:02 | 2:00:00 |
| 0 | 2 | 0 | 5 | 0 | 5 |
| 0:0:0:0:0:0:0:0 | 1:2:0:0:0:1:1:0 | 0:0:0:0:0:0:0:0 | 0:0:0:0:0:0:0:0 | 1:8:0:0:0:0:0:0 | 1:1:1:0:1:1:0:1 |
| 0:0:0:0:0 | 0:0:0:0:0 | 0:0:0:0:0 | 0:0:0:0:0 | 0:0:0:0:0 | 0:1:0:0:1 |
| 0 | 0 | 2 | 1 | 0 | 0 |
| 0:00 | 2:01 | 0:00 | 9:02 | 0:00 | 16:02 |
| 0 | 1 | 2 | 2 | 3 | 4 |
| 17:02 | 2:01 | 19:00 | 29:01:00 | 41:05:00 | 69:07:00 |
| 18:03 | 0:00 | 45:01:00 | 0:00 | 110:11:00 | 2:00 |
| 1 | 0 | 0 | 0 | 1 | 0 |
| 0 | 0 | 0 | 1 | 0 | 3 |
| 0 | 0 | 0 | 1 | 0 | 3 |
| 0 | 0 | 0 | 1 | 0 | 3 |
| 17:1:0:3 | 1:0:0:0 | 47:3:0:2 | 10:0:2:0 | 87:5:8:6 | 28:6:2:0 |
| 18:1:1:3 | 3:0:0:0 | 66:4:2:2 | 45:1:7:3 | 139:12:11:6 | 99:9:7:1 |
| 1:21:1:3 | 1:15:0:1 | 3:72:4:3 | 3:16:0:0 | 12:150:14:6 | 5:72:5:1 |
| 42:1:2:5 | 2:0:0:0 | 157:6:11:5 | 4:1:1:1 | 356:28:27:9 | 7:0:0:0 |
| 99:71:0 | 0:00:00 | 352:472:0 | 1:00:00 | 356:377:1 | 0:01:00 |
| 98:71:0 | 1:00:00 | 353:473:0 | 0:00:00 | 355:378:1 | 1:00:00 |
| 99:71:0 | 0:00:00 | 352:482:0 | 1:01:00 | 356:382:1 | 0:01:00 |
| 75:99:0 | 0:00:00 | 502:350:0 | 0:01:00 | 395:357:1 | 2:00:00 |
| 0:1:0:0:0:0:0 | 4:0:2:4:0:4:2 | 6:0:8:3:3:28:0 | 5:1:9:2:3:17:0 | 10:4:20:23:11:33:3 | 9:1:10:5:5:18:6 |
| 0:00:00 | 0:00:00 | 0:00:00 | 0:01:01 | 0:00:00 | 0:01:00 |
| 0:00:00 | 0:00:00 | 0:00:00 | 0:01:01 | 0:00:00 | 0:01:00 |
| 0:0:0:0:0:0 | 3:0:1:0:0:0 | 0:0:0:0:0:0 | 1:0:0:0:0:0 | 0:0:0:0:0:0 | 23:5:1:1:2:1 |
| 0:0:0:0 | 0:1:0:0 | 0:0:0:0 | 0:0:0:0 | 0:0:0:0 | 0:0:1:1 |
| 0:00:00 | 1:00:00 | 0:00:00 | 0:00:00 | 0:00:01 | 0:01:01 |
| 27:00:00 | 0:00:00 | 141:01:02 | 0:00:00 | 194:01:07 | 0:00:01 |
| 27:01:00 | 0:00:00 | 141:02:01 | 0:00:00 | 193:08:01 | 1:00:00 |
| 3:01 | 0:00 | 0:02 | 0:01 | 3:08 | 0:01 |
| 88:66:9:5:9:1:4:1:6:0:0:0:0 | 1:1:0:0:0:0:0:0:0:0:0:0 | 204:130:2:12:6:1:3:6:5:2:1:0:2 | 0:2:0:1:0:0:0:0:0:0:0:0 | 243:117:6:9:10:0:11:5:8:0:0:0:1 | 252:146:8:11:7:0:7:7:1:7:0:3:4 |
| 89:67:9:5:9:1:4:1:6:0:0:0:0 | 0:0:0:0:0:0:0:0:0:0:0:0 | 204:132:2:10:6:1:3:6:5:2:1:0:2 | 0:0:0:1:0:0:0:0:0:0:0:0 | 494:263:14:20:17:0:18:12:9:7:0:3:5 | 1:0:0:0:0:0:0:0:0:0:0:0 |
| 89:67:9:5:9:1:4:1:6:9:9:0:0:1:0:0:0:0 | 0:0:0:0:0:0:0:0:0:0:0:0:0:0:0:0:0 | 204:132:2:11:6:1:3:6:5:43:26:2:1:1:0:2:0:0 | 0:0:0:0:0:0:0:0:0:0:0:0:0:0:0:0:0 | 494:262:14:26:17:0:18:18:9:126:65:9:0:4:3:5:2:5 | 2:1:0:0:0:0:0:0:0:0:0:0:0:0:0:0:0 |
| 89:67:9:4:9:1:4:1:6:9:9:0:0:0:0:0:0:0 | 0:0:0:0:0:0:0:0:0:0:0:0:0:0:0:0:0 | 204:132:2:9:6:1:3:7:5:43:26:1:1:0:0:2:0:0 | 0:0:0:0:0:0:0:0:0:0:0:0:0:0:0:0:0 | 494:263:14:21:17:0:18:18:9:126:65:10:0:3:3:5:2:5 | 2:0:0:0:0:0:0:0:0:0:0:0:0:1:0:0:0:0 |
| 4:0:0:0 | 0:0:0:0 | 0:1:1:0 | 0:0:0:0 | 11:10:0:1 | 0:1:0:0 |
| 2:14:11:1:0:1:0:0:0 | 0:0:0:0:0:0:0:0:0 | 1:1:40:7:0:1:0:0:0 | 0:0:0:0:0:0:0:0:0 | 7:28:180:37:0:4:0:8:2 | 0:0:0:0:0:0:0:1:0 |
| 17:2:0:7:3:0:0:0:0 | 0:0:0:0:0:0:0:0:0 | 1:2:6:18:4:0:0:0:0 | 0:0:0:0:0:0:0:0:0 | 27:4:34:71:13:0:3:1:1 | 0:0:1:1:0:0:0:0:0 |
| 17:2:0:3:6:1:0 | 0:0:0:0:0:0:0 | 0:2:3:4:10:1:0 | 0:0:0:0:0:0:0 | 25:6:22:14:26:14:1 | 1:0:0:1:0:0:0 |
| 0:0:0:0:0:0 | 4:1:2:1:1:0 | 0:0:0:0:0:0 | 1:1:12:0:0:0 | 0:0:0:0:0:0 | 4:3:22:2:1:1 |
| 43:12:4:4 | 2:0:0:0 | 159:25:3:6 | 0:1:0:0 | 356:49:9:16 | 6:0:0:0 |
| 12:4:43:4 | 0:0:1:0 | 25:3:154:6 | 1:0:1:0 | 49:8:351:17 | 0:1:5:0 |
| 43:11:4:4 | 0:0:0:0 | 145:22:2:6 | 12:3:1:0 | 332:41:8:16 | 19:8:0:1 |
| 9:35:2:3 | 0:1:0:1 | 11:98:1:6 | 2:7:0:0 | 30:227:6:16 | 0:6:0:1 |
| 6:3:13:1 | 0:0:1:0 | 0:7:13:0 | 0:0:2:0 | 10:15:67:3 | 2:0:3:0 |
| 5:3:4:0 | 7:0:1:1 | 4:7:0:0 | 9:0:0:0 | 11:15:6:3 | 44:0:4:0 |
| 48:00:00 | 0:00 | 74:00:00 | 0:00 | 236:01:00 | 0:00 |
| 5:48:00 | 5:00:00 | 0.340972222 | 19:01:00 | 24:232:0 | 43:01:00 |
| 0:00:00 | 2:01:00 | 0:00:00 | 0:00:00 | 0:00:00 | 6:00:02 |
| 0:0:0:0 | 0:0:0:0 | 0:0:0:0 | 0:0:0:0 | 0:3:0:0 | 0:0:0:0 |
| 0:0:0:0:0:0:0:0:0:0:0:0 | 35:17:0:5:6:0:0:0:0:1:0:0 | 0:0:0:0:0:0:0:0:0:0:0:0 | 152:22:1:20:42:0:5:2:1:3:0:0 | 0:0:0:0:0:0:0:0:0:0:0:0 | 46:34:7:8:16:2:0:1:2:1:0:0 |
| 35:17:0:5:7:0:0:0:0:1:0:0 | 0:0:0:0:0:0:0:0:0:0:0:0 | 151:21:1:20:40:0:5:2:1:3:0:0 | 0:1:0:0:0:0:0:0:0:0:0:0 | 46:35:7:8:17:2:0:1:2:1:0:0 | 0:0:0:0:0:0:0:0:0:0:0:0 |
| 34:17:0:5:7:0:0:0:0:1:0:0 | 1:0:0:0:0:0:0:0:0:0:0:0 | 151:20:1:20:42:0:5:2:1:3:0:0 | 0:0:0:0:0:0:0:0:0:0:0:0 | 46:35:7:8:16:2:0:1:2:1:0:0 | 0:0:0:0:0:0:0:0:0:0:0:0 |

| | | | | | |
|---|---|---|---|---|---|
| 35:17:0:5:7:0:0:0:0:1:0:0 | 0:0:0:0:0:0:0:0:0:0:0:0 | 151:19:1:20:43:0:5:2:1:2:0:0 | 0:0:0:0:0:0:0:0:0:1:0:0 | 46:35:7:8:16:2:0:1:2:1:0:0 | 0:0:0:0:0:0:0:0:0:0:0:0 |
| 17:35:0:5:7:0:0:0:0:1:0:0 | 0:0:0:0:0:0:0:0:0:0:0:0 | 19:151:1:20:43:0:5:2:1:3:0:0 | 0:0:0:0:0:0:0:0:0:0:0:0 | 35:46:7:8:15:2:0:1:2:1:0:0 | 0:0:0:0:1:0:0:0:0:0:0:0 |
| 35:17:0:5:7:0:0:0:0:1:0:0 | 0:0:0:0:0:0:0:0:0:0:0:0 | 151:19:1:20:43:0:5:2:1:3:0:0 | 0:0:0:0:0:0:0:0:0:0:0:0 | 46:35:7:8:15:2:0:1:2:1:0:0 | 0:0:0:0:0:0:0:0:0:0:0:0 |
| 35:17:0:5:7:0:0:0:0:1:0:0 | 0:0:0:0:0:0:0:0:0:0:0:0 | 151:17:1:20:42:0:5:2:1:3:0:0 | 0:0:0:0:0:0:0:0:0:0:0:0 | 46:35:7:8:12:2:0:1:2:1:0:0 | 0:1:0:0:0:0:0:0:0:0:0:0 |
| 35:15:0:5:5:0:0:0:0:1:0:0 | 0:0:0:0:0:0:0:0:0:0:0:0 | 152:14:1:20:23:0:5:2:1:3:0:0 | 0:0:0:0:0:0:0:0:0:0:0:0 | 46:35:7:8:8:2:0:2:2:1:0:0 | 0:0:0:0:0:0:0:0:0:0:0:0 |
| 35:15:0:5:5:0:0:0:0:1:0:0 | 0:0:0:0:0:0:0:0:0:0:0:0 | 151:13:1:20:22:0:5:2:1:3:0:0 | 1:1:0:0:0:0:0:0:0:0:0:0 | 46:36:7:8:6:2:0:2:2:1:0:0 | 0:0:0:0:0:0:0:0:0:0:0:0 |
| 0:0:0:0:0:0:0:0:0:0:0:0 | 35:15:0:5:0:0:5:0:0:1:0:0 | 0:0:0:0:0:0:0:0:0:0:0:0 | 152:14:1:20:0:5:20:2:1:3:0:0 | 0:0:0:0:0:0:0:0:0:0:0:0 | 46:36:7:8:2:0:6:2:2:1:0:0 |
| 35:15:0:5:1:0:0:0:0:1:0:0 | 0:0:0:0:0:0:0:0:0:0:0:0 | 152:12:1:20:3:0:5:2:1:3:0:0 | 0:0:0:0:0:0:0:0:0:0:0:0 | 47:35:7:8:0:2:0:2:2:1:0:0 | 0:0:0:0:0:0:0:0:0:0:0:0 |
| 0:0:2:1:0:0:0 | 0:0:0:0:0:0:0 | 7:2:5:1:2:0:0 | 0:2:2:0:2:0:0 | 8:7:3:4:1:0:2 | 0:0:0:0:0:0:0 |
| 5:9:0:0:0:0 | 0:0:0:0:0:0 | 14:173:0:0:4:0 | 0:9:0:0:0:0 | 66:369:0:2:7:1 | 1:18:0:0:0:0 |
| 4:9:0:0:0 | 0:0:0:0:0 | 9:173:0:4:0 | 0:0:0:0:0 | 55:383:0:6:1 | 1:1:0:0:0 |
| 2:7:0:0 | 0:0:0:0 | 4:141:0:3 | 0:1:0:0 | 42:308:0:6 | 0:2:0:0 |
| 1 | 0 | 0 | 2 | 2 | 3 |
| 84:3:4:4:2:3 | 0:0:0:0:0:0 | 51:20:6:6:2:1 | 1:0:0:0:0:0 | 339:163:29:32:8:2 | 0:1:0:1:0:0 |
| 84:3:4:4:2:3 | 0:0:0:0:0:0 | 50:20:1:6:2:1 | 2:0:3:0:0:0 | 326:164:14:30:11:2 | 13:0:16:0:0:0 |
| 84:3:4:4:3:2 | 0:0:0:0:0:0 | 49:20:6:4:2:1 | 0:0:0:0:0:0 | 325:163:30:19:2:6 | 1:1:0:0:0:0 |
| 0:00 | 0:00 | 0:00 | 1:00 | 0:00 | 1:00 |
| 0 | 20 | 0 | 44 | 0 | 64 |
| 36 | 0 | 11 | 1 | 70 | 1 |
| 36 | 0 | 11 | 0 | 68 | 2 |
| 0 | 42 | 0 | 11 | 1 | 78 |
| 44 | 0 | 6 | 2 | 69 | 7 |
| 2 | 42 | 5 | 2 | 5 | 66 |
| 44 | 0 | 3 | 1 | 65 | 3 |
| 0 | 0 | 122 | 2 | 41 | 0 |
| 0:00 | 0:00 | 0:01 | 0:00 | 0:00 | 0:00 |
| 0:00 | 1:01 | 0:00 | 7:01 | 0:00 | 35:05:00 |
| 0:00:00 | 1:01:00 | 0:00:00 | 7:00:00 | 0:00:00 | 35:01:01 |
| 0:00 | 0:01 | 0:00 | 3:02 | 0:00 | 6:12 |
| 0:00 | 0:01 | 1:02 | 0:00 | 0:01 | 2:14 |
| 0:00 | 0:02 | 0:00 | 0:02 | 0:00 | 0:13 |
| 0:00 | 0:00 | 0:00 | 0:00 | 0:00 | 0:00 |
| 0:00:00 | 0:00:00 | 0:00:02 | 2:00:02 | 0:00:02 | 1:00:06 |
| 0:00:00 | 0:00:00 | 5:01:02 | 0:00:00 | 3:00:01 | 0:00:00 |
| 19048:136:48 | 12:00:00 | 52321:550:40 | 33:01:00 | 49199:405:79 | 24:00:00 |
| 19057:136:48 | 15:00:00 | 52336:553:40 | 24:01:00 | 49219:406:79 | 15:02:00 |
| 19173:135:47 | 32:00:00 | 52445:560:43 | 65:02:00 | 49314:408:80 | 68:00:00 |
| 38518:270:96 | 44:00:00 | 105140:1116:92 | 122:06:00 | 98884:814:158 | 130:00:00 |
| 120:19387:48 | 2:22:00 | 516:52889:47 | 10:52:00 | 380:49729:82 | 4:56:00 |
| 19402:119:49 | 22:02:00 | 52923:503:47 | 48:09:00 | 49762:369:82 | 61:07:00 |
| 39052:54:98 | 18:02:00 | 106382:92:130 | 112:04:00 | 100054:98:196 | 96:02:00 |
| 0 | 0 | 0 | 0 | 0 | 2 |
| 0 | 0 | 0 | 0 | 0 | 2 |
| 0 | 0 | 0 | 0 | 0 | 1 |
| 2 | 0 | 3 | 0 | 0 | 0 |
| 1:23:34:20:0:14:1:0:0:1 | 0:0:0:0:0:0:0:0:0 | 12:120:292:92:13:39:5:0:11:3 | 0:1:1:0:0:0:0:0:0:0 | 20:208:333:147:39:64:18:2:7:1 | 0:0:0:0:0:0:0:0:0:0 |
| 1:23:33:0:20:1:14:0:1:0:1 | 0:0:1:0:0:0:0:0:0:0:0 | 11:121:292:0:91:18:39:0:8:11:3 | 1:0:1:0:0:0:0:0:0:0:0 | 20:206:331:0:146:47:64:2:21:7:1 | 0:0:1:0:0:0:0:0:1:0:0 |
| 1:23:33:0:20:1:14:0:1:0:1 | 0:0:1:0:0:0:0:0:0:0:0 | 11:121:292:0:91:18:39:0:8:11:3 | 1:0:1:0:0:0:0:0:0:0:0 | 20:206:331:0:146:47:64:2:24:7:1 | 0:0:1:0:0:0:0:0:0:0:0 |
| 1:24:30:0:18:1:14:4:0:0:1 | 0:0:0:0:0:0:0:0:0:0:0 | 9:118:270:0:89:18:38:25:0:9:3 | 0:0:0:0:0:0:0:0:0:0:0 | 18:195:309:0:142:41:61:56:2:7:1 | 0:0:0:0:0:0:0:0:0:0:0 |
| 1:25:29:0:18:1:14:5:0:0:1 | 0:0:0:0:0:0:0:0:0:0:0 | 9:116:272:0:89:18:39:41:0:9:3 | 0:0:0:0:0:0:0:0:0:0:0 | 18:199:306:0:141:39:61:89:2:7:1 | 0:0:0:0:0:0:0:0:0:0:0 |

| | | | | | |
|---|---|---|---|---|---|
| 1:25:29:0:18:1:13:5:0:0:1 | 0:0:0:0:0:0:1:0:0:0:0 | 9:117:272:0:89:18:39:46:0:9:3 | 0:0:0:0:0:0:0:0:0:0:0 | 18:197:304:0:141:39:61:98:2:7:1 | 0:0:0:0:0:1:0:0:0:0:0 |
| 1:26:30:0:18:1:14:5:0:0:1 | 0:0:0:0:0:0:0:0:0:0:0 | 9:119:272:0:89:18:39:52:0:9:3 | 0:0:0:0:0:0:0:0:0:0:0 | 18:194:305:0:142:38:60:114:2:7:1 | 0:1:0:0:0:0:0:0:0:0:0 |
| 1:26:30:0:18:1:14:7:0:0:1 | 0:0:0:0:0:0:0:0:0:0:0 | 9:118:270:0:87:17:39:61:0:9:3 | 0:0:0:0:0:0:0:0:0:0:0 | 18:194:301:0:140:37:60:131:2:7:1 | 0:0:0:0:0:0:0:0:0:0:0 |
| 2:52:60:0:36:2:28:14:0:0:2 | 0:0:0:0:0:0:0:0:0:0:0 | 18:236:538:0:174:34:78:118:0:18:4 | 0:0:4:0:0:0:0:0:0:0:0 | 34:388:600:0:278:74:120:258:4:14:2 | 0:2:2:0:2:0:0:0:0:0:0 |
| 27:24:18:1:7:1:0:12:0:1:0 | 3:1:0:0:0:0:0:2:0:0:0 | 263:116:82:17:56:9:0:37:0:3:9 | 8:2:5:0:1:0:0:2:0:0:0 | 285:183:137:34:127:15:0:56:2:1:7 | 17:11:2:3:2:2:0:4:0:0:0 |
| 44:46:2:0:32:22:14:0:0:2:2 | 8:0:0:0:2:2:0:0:0:0:0 | 482:220:18:0:150:68:92:0:16:6:30 | 26:8:0:0:8:2:0:0:0:0:0 | 520:338:24:0:238:96:226:4:12:2:62 | 34:12:0:0:20:10:4:0:2:0:4 |
| 0:0:0:0:5:0:7:0:0 | 0:0:0:0:0:0:0:0:0 | 1:5:0:0:3:0:5:2:0 | 0:1:0:0:1:0:1:0:0 | 2:2:0:2:12:0:11:1:1 | 0:1:0:0:1:0:2:0:0 |
| 0:0:0:0:1:0:0:0:0 | 0:0:0:0:7:7:0:0:0 | 0:0:0:0:9:0:0:0:0 | 0:0:0:0:6:3:0:0:0 | 0:0:0:0:1:0:0:0:0 | 0:1:0:1:14:12:0:1:0 |
| 78:08:04 | 4:00:02 | 422:00:04 | 48:00:02 | 280:10:04 | 36:00:00 |
| 96:08:06 | 10:00:00 | 626:00:16 | 34:00:02 | 450:10:06 | 20:00:02 |
| 54:4:3:0 | 0:0:0:0 | 335:0:10:0 | 0:0:0:0 | 228:5:6:0 | 0:0:0:0 |
| 4:54:3:0 | 0:1:0:0 | 0:331:10:0 | 0:5:2:0 | 5:225:6:0 | 0:5:1:0 |
| 0:00 | 0:00 | 0:00 | 0:00 | 0:00 | 0:00 |
| 0:00:00 | 0:00:00 | 0:00:00 | 0:00:00 | 0:00:00 | 0:00:01 |
| 6:44:43:12:2:0:0:100:1:0:0:2:1:0:0:0:0:0:0:0:0:0 | 0:0:0:0:0:0:0:0:0:0:0:0:1:0:0:0:0:0:0:0:0:0 | 11:10:190:38:5:1:0:342:4:0:3:6:2:5:0:0:1:0:1:0:0:0 | 0:0:0:0:0:0:0:0:0:0:0:0:0:0:0:0:0:0:0:0:0:0 | 61:217:204:132:4:2:1:452:8:0:5:4:4:5:0:2:1:1:0:2:1:4 | 0:0:1:0:0:0:0:2:0:0:0:0:0:0:0:0:0:0:0:0:0:0 |
| 6:44:43:12:2:0:0:100:1:0:0:2:0:0:1:0:0:0:0:0:0:0 | 0:0:0:0:0:0:0:0:0:0:0:0:0:0:0:0:0:0:0:0:0:0 | 11:10:189:38:6:0:0:336:4:0:4:6:5:0:2:0:1:0:1:0:0:0 | 0:0:1:0:0:1:0:0:0:0:0:0:0:0:0:0:0:0:0:0:0:0 | 61:217:205:133:4:2:1:457:8:0:5:4:5:0:3:2:1:1:0:2:1:4 | 0:0:0:0:0:0:0:1:0:0:0:0:0:0:0:0:0:0:0:0:0:0 |
| 6:44:43:12:2:0:0:100:1:0:0:2:0:0:1:0:0:0:0:0:0:0 | 0:0:0:0:0:0:0:0:0:0:0:0:0:0:0:0:0:0:0:0:0:0 | 11:10:190:38:6:0:0:340:4:0:4:6:5:0:2:0:1:0:1:0:0:0 | 0:0:0:0:0:0:0:0:0:0:0:0:0:0:0:0:0:0:0:0:0:0 | 61:217:204:132:4:2:1:464:8:0:5:4:5:0:3:2:1:2:0:2:1:4 | 0:0:1:0:0:0:0:1:0:0:0:0:0:0:0:0:0:0:0:0:0:0 |
| 6:44:43:12:2:0:0:100:1:0:0:1:0:0:1:0:0:0:0:0:0:0 | 0:0:0:0:0:0:0:0:0:0:0:0:0:0:0:0:0:0:0:0:0:0 | 11:10:188:38:6:0:0:343:4:0:4:6:5:0:2:0:1:0:1:0:0:0 | 0:0:2:0:0:0:0:0:0:0:0:0:0:0:0:0:0:0:0:0:0:0 | 61:216:205:134:4:2:1:467:8:0:5:4:5:0:3:2:1:2:0:2:1:4 | 0:1:0:0:0:0:0:0:0:0:0:0:0:0:0:0:0:0:0:0:0:0 |
| 6:43:101:2:0:0:0:1:0:0:0:0:0:0:0 | 0:0:0:0:0:0:0:0:0:0:0:0:0:0:0 | 11:190:359:6:0:0:4:6:4:2:1:1:0:0:0 | 0:0:0:0:0:0:0:0:0:0:0:0:0:0:0 | 61:204:479:4:2:0:6:4:4:2:1:0:3:1:2 | 0:1:1:0:0:0:0:0:0:0:0:0:0:0:0 |
| 44:43:6:12:101:1:0:0:0:0:0 | 0:0:0:0:0:0:0:0:0:0:0 | 10:190:11:38:360:4:12:0:4:0:0 | 0:0:0:0:0:0:0:0:0:0:0 | 217:205:61:135:494:8:14:1:3:3:0 | 0:0:0:0:0:0:0:0:0:0:0 |
| 8 | 0 | 32 | 3 | 63 | 7 |
| 16 | 0 | 68 | 4 | 134 | 10 |
| 7:14:0:2:1:0:0:2:0 | 0:1:0:0:0:0:0:0:0 | 8:37:3:11:2:7:0:4:0 | 0:0:0:0:0:0:0:0:0 | 21:60:0:12:5:7:1:3:0 | 0:0:0:0:2:0:0:0:0 |
| 18 | 0 | 27 | 0 | 64 | 1 |
| 16:2:2:0:0:0 | 0:0:0:0:0:0 | 284:12:62:0:0:0 | 0:0:0:0:0:0 | 648:36:94:0:2:0 | 0:0:0:0:0:0 |
| 8:0:1:0:0:0 | 0:1:0:0:0:0 | 141:6:30:0:0:0 | 1:0:0:0:0:0 | 322:18:43:0:1:0 | 2:0:0:0:0:0 |
| 8:1:1:0:0:0 | 0:0:0:0:0:0 | 142:6:23:0:0:0 | 0:0:0:0:0:0 | 322:18:33:0:1:0 | 2:0:0:0:0:0 |
| 0:0:0:0 | 0:0:0:0 | 0:0:0:0 | 0:0:0:0 | 0:0:0:0 | 4:2:2:2 |
| 36:1:0:0:2:0:0 | 1:0:0:0:0:0:0 | 66:19:0:2:16:0:0 | 0:0:0:0:0:0:0 | 171:43:0:1:39:1:1 | 0:0:0:0:0:0:0 |
| 68:2:4:0:0:0:0 | 4:0:0:0:0:0:0 | 130:38:32:0:4:0:0 | 0:0:0:0:0:0:0 | 330:86:78:0:2:2:2 | 2:0:0:0:0:0:0 |
| 1:00 | 0:00 | 6:02 | 1:00 | 6:04 | 0:00 |
| 3:12:0:2:0:0 | 0:0:0:0:0:0 | 13:0:0:2:0:0 | 1:1:0:2:0:0 | 14:3:0:1:0:1 | 4:0:0:0:0:0 |
| 0:3:26:3:0 | 0:0:0:0:0 | 1:13:21:14:0 | 0:0:0:0:0 | 4:5:48:29:1 | 0:0:0:0:0 |
| 18:0:3:0:1 | 9:0:0:0:2 | 5:1:3:0:4 | 16:0:10:0:9 | 41:3:5:1:24 | 10:2:1:0:3 |
| 0:0:0:0:0 | 18:0:0:2:0 | 0:0:0:0:0 | 28:2:0:0:0 | 0:0:0:0:0 | 40:0:0:0:2 |
| 3:727:76:2:2:3:1:1 | 0:2:0:0:0:0:0:0 | 4:293:128:4:1:1:0:0 | 2:0:0:0:0:0:0:0 | 26:2142:641:7:13:5:0:1 | 4:10:0:0:0:0:0:0 |
| 726:76:2:2:2:3:1:1 | 1:0:0:0:0:0:0:0 | 292:128:4:3:1:1:0:0 | 0:0:0:1:0:0:0:0 | 2127:638:7:20:13:5:0:1 | 14:1:0:5:0:0:0:0 |
| 693:10:59:2:23:0:2:0:1:1 | 2:0:1:0:0:0:0:0:0:0 | 265:11:104:4:4:3:1:0:0:1 | 0:0:2:0:0:0:0:0:0:0 | 2003:86:531:7:34:14:14:0:0:1 | 5:1:2:0:0:0:0:0:0:0 |
| 15:689:54:2:24:1:2:0:1:1 | 0:0:0:0:0:0:0:0:0:0 | 12:257:101:4:5:3:1:0:0:1 | 0:0:0:0:0:0:0:0:0:0 | 113:1985:493:7:34:14:14:0:0:1 | 0:0:1:0:0:0:0:0:0:0 |
| 1360:4:84:44:90:4:2:0:2:2 | 2:0:4:0:0:0:0:0:0:0 | 490:2:70:10:172:8:8:0:0:2 | 4:0:2:0:0:0:0:0:0:0 | 3806:30:430:72:820:14:24:0:0:2 | 10:0:6:0:10:0:2:0:0:0 |
| 673:2:72:22:41:2:1:0:1:1 | 1:0:0:0:0:0:0:0:0:0 | 241:2:56:6:77:4:3:0:0:1 | 0:0:0:0:0:0:0:0:0:0 | 1878:15:346:41:374:7:12:0:0:1 | 1:0:1:0:5:0:0:0:0:0 |

| | | | | | |
|---|---|---|---|---|---|
| 660:122:2:24:34:2:1:0:1:0:1 | 1:3:0:0:1:0:0:0:0:1:0 | 230:88:2:7:53:4:2:0:0:0:1 | 0:1:0:0:1:0:0:0:0:0:0 | 1795:511:15:61:256:7:12:0:1:0:1 | 4:5:0:0:3:0:0:0:0:0:0 |
| 668:128:2:24:34:2:1:0:1:0:1 | 1:0:0:0:0:0:0:0:0:1:0 | 237:88:2:7:52:4:2:0:0:0:1 | 1:1:0:0:2:0:0:0:0:0:0 | 1829:529:15:64:259:7:12:0:1:0:1 | 2:1:0:0:3:0:0:0:0:0:0 |
| 657:155:28:0:29:2:1:0:1:1:1 | 2:0:0:1:0:0:0:0:0:0:0 | 239:100:9:1:49:4:2:0:0:1:0 | 1:0:0:0:0:0:0:0:0:0:0 | 1798:633:92:4:227:7:11:0:1:1:0 | 5:0:0:7:2:0:0:0:0:0:0 |
| 657:155:28:0:29:2:1:0:1:1:1 | 2:0:0:1:0:0:0:0:0:0:0 | 239:100:9:1:49:4:2:0:0:1:0 | 1:0:0:0:0:0:0:0:0:0:0 | 1798:633:92:4:227:7:11:0:1:1:0 | 5:0:0:7:2:0:0:0:0:0:0 |
| 659:155:28:0:28:2:1:0:1:1:1 | 0:0:0:1:1:0:0:0:0:0:0 | 239:100:9:1:49:4:2:0:0:1:0 | 1:0:0:0:0:0:0:0:0:0:0 | 1796:633:92:4:225:7:11:0:1:1:0 | 5:0:0:7:4:0:0:0:0:0:0 |
| 658:179:36:31:2:1:0:1:1:2 | 2:0:0:0:0:0:0:0:0:0 | 231:109:13:36:4:2:0:0:1:0 | 1:0:0:0:0:0:0:0:0:0 | 1766:716:131:182:7:9:0:1:1:0 | 0:2:0:1:0:0:0:0:0:0 |
| 660:177:37:31:2:1:0:1:1:2 | 0:1:0:0:0:0:0:0:0:0 | 234:109:14:35:4:2:0:0:1:0 | 0:0:0:0:0:0:0:0:0:0 | 1755:719:131:183:7:10:0:1:1:0 | 8:4:0:1:0:0:0:0:0:0 |
| 656:178:38:31:2:1:0:1:1:2 | 3:0:0:0:0:0:0:0:0:0 | 232:109:14:34:4:2:0:0:1:0 | 0:0:0:0:0:0:0:0:0:0 | 1757:721:133:176:7:10:0:1:0:0 | 2:3:0:1:0:0:0:0:0:0 |
| 661:39:181:30:2:1:0:1:1:2 | 0:0:0:0:0:0:0:0:0:0 | 231:14:111:24:4:2:0:0:0:0 | 0:0:0:0:0:0:0:0:0:0 | 1753:144:735:167:7:9:0:1:1:0 | 0:0:2:0:0:0:0:0:0:0 |
| 664:45:181:33:2:1:1:0:1:1 | 2:0:0:0:0:0:0:0:0 | 234:18:111:24:4:2:1:0:0:0 | 0:0:0:0:0:0:0:0:0:0 | 1760:175:758:176:7:8:3:0:1:0 | 1:1:0:1:0:0:0:0:0:0 |
| 665:45:179:33:2:1:1:0:1:1 | 1:0:2:0:0:0:0:0:0:0 | 234:18:109:23:4:2:1:0:0:0 | 0:0:1:0:0:0:0:0:0:0 | 1755:180:758:168:7:8:4:0:1:0 | 2:0:1:1:0:0:0:0:0:0 |
| 671:57:188:33:2:1:2:0:1:0:1 | 0:1:0:0:0:0:0:0:0:1:0 | 229:23:117:21:4:2:2:0:0:1:0 | 0:0:0:0:0:0:0:0:0:0:0 | 1763:235:789:164:7:8:13:0:1:1:0 | 2:0:2:0:0:0:0:0:0:0:0 |
| 682:86:2:199:2:26:1:0:1:1:1 | 0:2:0:0:0:0:0:0:0:0:0 | 228:47:2:140:4:19:1:0:0:1:0 | 1:0:0:0:0:0:0:0:0:0:0 | 1795:366:15:889:7:142:5:0:1:1:0 | 2:1:0:0:0:4:1:0:0:0:0 |
| 0:00 | 0:00 | 0:00 | 2:02 | 0:00 | 2:02 |
| 0:0:0:0 | 0:2:0:2 | 0:0:0:0 | 0:0:0:2 | 0:0:0:0 | 4:2:0:2 |
| 0:0:0:0 | 2:0:0:2 | 0:0:0:0 | 0:0:0:2 | 0:0:0:0 | 2:4:0:2 |
| 1:00:00 | 0:00:00 | 4:00:01 | 0:00:00 | 8:01:00 | 0:00:00 |
| 199:03:00 | 0:00:00 | 691:22:01 | 1:00:00 | 886:38:00 | 2:02:00 |
| 200:03:00 | 0:00:00 | 695:10:01 | 0:00:00 | 892:17:00 | 2:00:00 |
| 198:01:00 | 2:00:00 | 697:06:01 | 2:00:00 | 893:04:00 | 3:00:00 |
| 398:00:00 | 2:00:00 | 1392:04:02 | 4:00:00 | 1784:02:00 | 12:00:00 |
| 396:00:00 | 2:00:00 | 1398:04:02 | 0:00:00 | 1776:02:04 | 4:00:00 |
| 0:00 | 3:00 | 0:00 | 1:00 | 0:00 | 6:01 |
| 0:01 | 0:00 | 0:01 | 0:00 | 1:03 | 0:00 |
| 26:02:00 | 24:06:00 | 46:06:00 | 48:04:00 | 180:12:00 | 150:18:00 |
| 26:02:00 | 24:04:00 | 36:06:00 | 44:02:00 | 168:12:00 | 148:14:00 |
| 26:02:00 | 24:04:00 | 36:06:00 | 44:02:00 | 164:12:00 | 148:14:00 |
| 49:00:00 | 0:00:00 | 91:00:02 | 0:00:00 | 159:04:02 | 0:00:00 |
| 0:00 | 0:00 | 0:00 | 1:00 | 0:00 | 1:01 |
| 0:00 | 0:00 | 1:00 | 0:00 | 0:00 | 1:01 |
| 354:1:12:2:5:0:0 | 0:0:0:0:0:0:0 | 667:1:22:1:39:1:3 | 1:0:0:0:1:0:0 | 1186:1:43:14:43:14:4 | 2:0:0:0:0:0:0 |
| 354:1:12:2:5:0:0 | 0:0:0:0:0:0:0 | 668:1:22:1:40:1:3 | 0:0:0:0:0:0:0 | 1187:1:43:14:43:14:4 | 1:0:0:0:0:0:0 |
| 354:1:12:2:5:0:0 | 0:0:0:0:0:0:0 | 667:1:22:1:40:1:2 | 1:0:0:0:0:0:1 | 1188:1:43:13:43:12:4 | 0:0:0:1:0:1:0 |
| 355:1:2:5:2:0 | 0:0:0:0:0:0 | 668:1:1:40:6:3 | 0:0:0:0:0:0 | 1191:1:14:43:4:4 | 0:0:0:0:0:0 |
| 2:347:1:12:5:3:0 | 0:2:0:0:0:0:0 | 1:659:1:23:38:15:3 | 0:0:0:0:0:0:0 | 14:1173:1:43:41:17:4 | 0:1:0:0:0:0:0 |
| 0 | 0 | 9 | 0 | 0 | 2 |
| 0 | 0 | 0 | 4 | 0 | 1 |
| 0 | 0 | 0 | 4 | 0 | 1 |
| 0 | 0 | 0 | 2 | 0 | 0 |
| 0 | 0 | 0 | 2 | 0 | 1 |
| 0:00 | 2:00 | 0:00 | 3:03 | 0:00 | 1:01 |
| 0:00 | 3:00 | 0:00 | 1:02 | 0:00 | 2:01 |
| 0:00:00 | 3:00:00 | 1:00:00 | 7:02:00 | 0:00:00 | 9:01:00 |
| 0:00:00 | 3:00:00 | 1:00:00 | 9:02:00 | 0:00:00 | 10:01:00 |
| 3:00:00 | 0:00:00 | 11:01:00 | 1:00:00 | 10:01:00 | 0:00:00 |
| 0:00:00 | 4:00:00 | 1:00:00 | 28:01:00 | 0:00:00 | 21:00:00 |
| 4:08:00 | 2:04:00 | 76:92:2 | 6:02:00 | 46:80:2 | 10:06:00 |
| 0:00 | 3:00 | 0:00 | 3:00 | 0:00 | 2:00 |
| 3:02 | 0:00 | 10:13 | 0:00 | 2:06 | 0:00 |
| 0:01 | 0:01 | 2:00 | 3:06 | 0:01 | 0:04 |
| 0:0:0:0:0 | 0:2:0:0:0 | 4:0:2:0:2 | 4:2:0:0:0 | 2:0:0:0:0 | 4:2:2:0:0 |
| 8:04 | 0:00 | 4:02 | 0:04 | 8:04 | 0:04 |
| 1:4:10:3:3:1:23:4:2:2:1:0:0 | 0:0:0:0:0:0:0:0:0:0:0:0:0 | 0:1:0:1:0:3:7:3:0:5:0:1:0 | 0:0:0:0:0:0:0:0:0:0:0:0:0 | 3:23:9:4:17:28:35:10:4:30:7:7:1 | 0:0:0:0:0:0:0:0:0:0:0:0:0 |
| 1:3:10:3:3:1:23:2:3:1:1:0:0 | 0:0:0:0:0:0:0:0:0:0:0:0:0 | 0:1:0:1:0:2:7:4:2:0:0:0:1 | 0:0:0:0:0:0:0:0:0:0:0:0:0 | 3:23:9:4:15:19:33:21:10:2:7:1:4 | 0:0:0:0:1:1:0:0:0:0:0:0:0 |
| 0:0:0:0:0:0:0:0:0:0:0 | 2:4:20:6:46:6:4:2:4:2:0 | 0:0:0:0:0:0:0:0:0:0:0:0 | 0:2:0:0:10:0:0:0:2:0:0 | 0:0:0:0:0:0:0:0:0:0:0:0 | 6:34:18:6:94:18:16:4:8:12:2 |

| | | | | | |
|---|---|---|---|---|---|
| 16:36 | 2:00 | 28:14:00 | 6:00 | 84:50:00 | 18:00 |
| 3:18 | 1:00 | 6:07 | 0:00 | 16:25 | 3:00 |
| 5:0:0:0 | 17:2:1:1 | 21:4:1:1 | 25:1:1:0 | 2:0:0:0 | 43:7:2:1 |
| 2:23:1:1:0 | 0:0:0:0:0 | 5:47:2:1:1 | 0:0:0:0:0 | 7:46:2:1:0 | 0:0:0:0:0 |
| 0:00:00 | 0:00:00 | 1:00:00 | 0:00:00 | 1:00:00 | 0:00:00 |
| 4 | 0 | 0 | 0 | 6 | 0 |
| 0:00 | 1:00 | 0:00 | 1:01 | 0:00 | 1:00 |
| 0 | 26 | 0 | 95 | 0 | 1 |
| 0:0:0:0:1 | 0:0:1:0:0 | 0:0:0:0:0 | 1:0:1:0:0 | 0:0:0:0:0 | 0:1:2:0:0 |
| 0:02 | 0:01 | 0:01 | 1:00 | 2:03 | 0:00 |
| 0:00 | 2:00 | 0:00 | 0:00 | 0:00 | 0:01 |
| 0:00 | 2:00 | 0:00 | 0:00 | 0:00 | 0:01 |
| 0:00:00 | 0:00:00 | 1:02:00 | 0:01:00 | 0:01:00 | 0:01:00 |
| 0:00:00 | 0:00:00 | 0:00:00 | 0:00:00 | 0:00:00 | 0:01:01 |
| 0:00:00 | 0:00:00 | 0:00:00 | 0:00:00 | 0:00:00 | 0:01:01 |
| 75:08:00 | 2:00 | 100:07:00 | 0:00 | 179:14:00 | 2:00 |
| 0 | 0 | 0 | 0 | 0 | 1 |
| 0 | 0 | 0 | 0 | 0 | 3 |
| 0 | 0 | 0 | 0 | 0 | 3 |
| 1:12 | 0:00 | 0:13 | 0:00 | 2:14 | 0:00 |
| 0:2:6:0:4:11:0:2:0:3:1 | 0:0:0:0:0:0:0:0:0:0:0 | 6:9:17:7:6:26:5:3:0:3:1 | 0:0:0:0:0:1:0:0:0:0:0 | 10:21:40:5:11:63:10:18:4:24:3 | 0:0:0:0:0:2:0:0:0:0:0 |
| 0:2:6:0:4:11:0:2:0:3:1 | 0:0:0:0:0:0:0:0:0:0:0 | 6:9:17:7:6:26:5:3:0:3:1 | 0:0:0:0:0:2:0:0:0:0:0 | 10:21:40:5:11:62:10:18:4:24:3 | 0:0:0:0:0:3:0:0:0:0:0 |
| 0:2:6:0:4:11:0:2:0:3:1 | 0:0:0:0:0:0:0:0:0:0:0 | 6:9:17:7:6:26:5:3:0:3:1 | 0:0:0:0:0:3:0:0:0:0:0 | 10:21:40:5:11:63:10:18:4:24:3 | 0:0:0:0:0:3:0:0:0:0:0 |
| 0:00 | 1:00 | 0:00 | 0:00 | 0:00 | 0:01 |
| 0 | 0 | 0 | 0 | 0 | 1 |
| 0:00:00 | 0:00:00 | 1:00:01 | 0:00:00 | 0:00:00 | 0:00:00 |
| 0:00:00 | 2:00:00 | 2:00:00 | 0:00:00 | 0:00:00 | 3:00:01 |
| 2:00 | 0:00 | 2:00 | 0:00 | 3:00 | 0:00 |
| 1:00:00 | 0:00:00 | 1:00:00 | 0:00:00 | 0:01:00 | 0:00:00 |
| 0:00:00 | 0:00:00 | 0:00:00 | 1:00:01 | 0:00:00 | 0:00:00 |
| 1:00 | 0:00 | 0:00 | 0:00 | 0:00 | 0:00 |
| 0:00:00 | 1:00:01 | 0:00:00 | 0:00:00 | 0:00:00 | 0:00:00 |
| 0:00 | 0:00 | 0:00 | 79:00:00 | 0:00 | 6:00 |
| 0:00 | 0:00 | 0:00 | 81:00:00 | 0:00 | 5:00 |
| 2953:50:5:10 | 3:0:0:0 | 2910:31:1:7 | 3:0:0:0 | 11255:264:15:11 | 4:2:0:0 |
| 2954:56:5:10 | 1:0:0:0 | 2909:35:1:7 | 3:1:0:0 | 11228:289:15:11 | 13:1:0:0 |
| 2952:58:5:10 | 1:0:0:0 | 2905:37:1:7 | 2:1:0:0 | 11228:289:15:11 | 5:4:0:0 |
| 2948:64:5:10 | 5:0:0:0 | 2896:39:1:7 | 7:2:0:0 | 11199:302:15:10 | 16:2:0:1 |
| 2944:65:5:10 | 2:0:0:0 | 2875:37:1:7 | 5:1:0:0 | 11134:299:15:10 | 17:2:0:0 |
| 2943:65:5:10 | 1:0:0:0 | 2879:39:1:7 | 1:0:0:0 | 11141:303:15:10 | 3:1:0:0 |
| 2938:68:5:10 | 4:0:0:0 | 2873:40:1:7 | 4:0:0:0 | 11127:305:15:10 | 10:1:0:0 |
| 2939:68:5:10 | 2:0:0:0 | 2873:40:1:7 | 3:0:0:0 | 11119:307:15:10 | 13:0:0:0 |
| 2929:76:5:10 | 2:0:0:0 | 2864:41:1:7 | 3:1:0:0 | 11080:335:15:10 | 16:2:0:0 |
| 2930:77:5:10 | 1:0:0:0 | 2866:41:1:7 | 1:1:0:0 | 11076:337:15:10 | 18:2:0:0 |
| 2926:80:5:10 | 4:0:0:0 | 2864:42:1:7 | 2:1:0:0 | 11078:346:15:10 | 14:0:0:0 |
| 2922:78:5:10 | 4:0:0:0 | 2851:49:1:7 | 3:1:0:0 | 11054:362:15:10 | 12:0:0:0 |
| 2922:78:5:10 | 4:0:0:0 | 2851:49:1:7 | 2:1:0:0 | 11059:359:15:10 | 11:3:0:0 |
| 80:2925:5:10 | 0:1:0:0 | 49:2851:1:7 | 2:1:0:0 | 367:11059:15:10 | 1:4:0:0 |
| 2926:82:5:10 | 0:1:0:0 | 2849:50:1:7 | 3:1:0:0 | 11061:369:15:10 | 5:6:0:0 |
| 2923:83:5:10 | 2:0:0:0 | 2845:50:1:7 | 6:1:0:0 | 11053:379:15:10 | 10:1:0:0 |
| 2922:83:5:10 | 1:0:0:0 | 2843:50:1:7 | 3:1:0:0 | 11038:380:15:10 | 15:0:0:0 |
| 2923:85:5:10 | 3:0:0:0 | 2849:50:1:7 | 1:0:0:0 | 11045:390:15:10 | 14:1:0:0 |

| | | | | | |
|---|---|---|---|---|---|
| 2923:93:5:10 | 0:1:0:0 | 2837:52:1:7 | 2:0:0:0 | 11030:420:15:10 | 13:1:0:0 |
| 2916:99:5:10 | 1:1:0:0 | 2826:55:1:7 | 5:0:0:0 | 11008:442:15:10 | 12:3:0:0 |
| 113:2887:5:10 | 1:3:0:0 | 63:2800:1:7 | 1:2:0:0 | 508:10917:15:9 | 3:12:0:0 |
| 2897:123:5:10 | 0:1:0:0 | 2782:73:1:7 | 8:0:0:0 | 10894:556:15:9 | 17:2:0:0 |
| 2895:128:5:10 | 2:1:0:0 | 2778:78:1:6 | 13:0:0:0 | 10875:586:15:9 | 18:0:0:0 |
| 2896:131:5:10 | 3:0:0:0 | 2781:81:1:6 | 8:0:0:0 | 10881:605:15:9 | 20:0:0:0 |
| 2890:132:5:10 | 7:0:0:0 | 2775:85:1:6 | 7:0:0:0 | 10878:623:15:9 | 21:0:0:0 |
| 4:40:20:8:3:0:1:0 | 0:0:0:0:0:0:0:0 | 20:49:22:7:1:0:0:0 | 0:0:0:0:0:0:0:0 | 54:219:101:36:2:2:5:1 | 0:0:0:0:0:0:0:0 |
| 39:20:4:7:2:0:1:0 | 0:0:0:0:0:0:0:0 | 48:22:20:11:1:0:0:0 | 0:0:0:0:0:0:0:0 | 218:102:54:30:2:2:5:1 | 1:1:0:1:0:0:0:0 |
| 39:20:4:7:2:0:1:0 | 0:0:0:0:0:0:0:0 | 48:22:20:11:1:0:0:0 | 0:0:0:0:0:0:0:0 | 219:102:54:31:2:2:5:1 | 1:1:0:0:0:0:0:0 |
| 43:20:4:6:3:0:0:0 | 0:0:0:0:0:0:0:0 | 52:22:20:12:1:0:0:0 | 0:0:0:0:0:0:0:0 | 236:103:54:30:2:2:2:1 | 1:0:0:1:0:0:1:0 |
| 20:43:3:6:0:0:0:0 | 0:0:0:0:0:0:0:0 | 20:52:16:6:1:0:0:0 | 1:0:2:1:0:0:0:0 | 94:240:40:16:1:2:1:1 | 1:0:1:2:0:0:0:0 |
| 19:43:3:1:0:0:0:0 | 1:0:0:0:0:0:0:0 | 19:53:1:4:0:0:0:0 | 0:0:1:3:0:0:1:0 | 84:240:17:19:2:1:0:1 | 2:0:2:1:0:0:0:0 |
| 43:19:14:4:0:0 | 1:0:0:0:0:0 | 26:7:23:3:1:0 | 0:0:0:0:0:0 | 148:45:79:3:8:1 | 2:0:0:0:0:0 |
| 43:19:14:4:0:0:0 | 0:0:0:0:0:0 | 26:8:25:3:2:0:0 | 0:1:0:0:0:0:0 | 149:47:85:3:19:1:0 | 1:1:2:0:0:0:0 |
| 44:19:14:4:2:0:0 | 0:0:0:0:0:0 | 27:9:25:2:10:0:0 | 0:0:0:0:0:0 | 154:52:88:3:33:2:0 | 0:0:0:0:0:0 |
| 0:0:0:0:0:0 | 0:0:0:0:0:0 | 1:0:0:0:0:1 | 0:0:0:0:0:0 | 4:1:1:0:0:0 | 1:0:0:0:0:0 |
| 0:00:00 | 0:01:00 | 0:00:00 | 0:00:00 | 0:00:00 | 0:01:01 |
| 3:0:1:0:0:0:0:0 | 0:0:0:0:0:0:0:0 | 0:6:1:0:0:0:1:0 | 0:0:0:0:0:0:0:0 | 8:0:1:0:0:0:0:1 | 0:0:0:0:0:0:0:0 |
| 0:0:0:0:0:0:0:0:0:0 | 3:0:0:0:0:0:0:0:0:0 | 0:6:1:0:0:0:0:0:1:0 | 0:0:0:0:0:0:0:0:0:0 | 0:0:0:0:0:0:0:0:0:0 | 8:0:0:0:0:0:0:0:0:1 |
| 0:00:00 | 13:00:02 | 10:02:01 | 8:04:00 | 0:01:00 | 28:08:03 |
| 0:00:00 | 8:01:00 | 9:02:00 | 8:03:00 | 0:00:00 | 20:02:01 |
| 0:00:00 | 8:01:00 | 9:01:00 | 8:02:00 | 0:00:00 | 19:02:00 |
| 0:00:00 | 11:00:02 | 10:00:01 | 7:01:00 | 0:01:00 | 29:03:01 |
| 0 | 0 | 0 | 0 | 0 | 0 |
| 0:9:13:3:1:0:0:0:0 | 0:0:0:0:0:0:0:0:0 | 5:6:13:5:1:7:0:1:0 | 0:0:0:0:0:0:0:0:0 | 14:23:52:32:12:4:0:2:1 | 0:0:0:0:0:0:0:0:0 |
| 1 | 0 | 44 | 0 | 10 | 0 |
| 0 | 0 | 2 | 1 | 0 | 0 |
| 2:00 | 0:00 | 0:00 | 0:00 | 1:00 | 0:00 |
| 0 | 0 | 0 | 2 | 0 | 1 |
| 1:02:00 | 0:00:01 | 0:02:00 | 0:00:00 | 0:01:00 | 0:08:00 |
| 2:01 | 1:00 | 0:00 | 0:00 | 1:00 | 6:00 |
| 153:10:2:9:0 | 0:0:0:0:0 | 288:19:14:17:1 | 0:0:0:0:0 | 757:23:16:61:1 | 2:0:0:0:0 |
| 10:153:2:8:0 | 0:0:0:1:0 | 19:287:14:17:1 | 0:0:0:0:0 | 23:758:16:59:1 | 0:0:0:0:0 |
| 10:153:2:9:0 | 0:0:0:0:0 | 19:295:14:13:1 | 0:0:0:0:0 | 24:770:16:58:1 | 0:2:0:0:0 |
| 154:10:2:9:0 | 0:0:0:0:0 | 308:19:14:12:1 | 9:0:0:0:0 | 798:24:19:56:1 | 18:0:0:0:0 |
| 11:155:5:0 | 0:0:0:0 | 20:320:5:1 | 0:0:1:0 | 27:836:21:1 | 0:3:0:0 |
| 11:154:5:0 | 0:0:0:0 | 20:316:6:1 | 0:1:0:0 | 27:832:22:1 | 0:0:0:0 |
| 6:01 | 0:00 | 201:68 | 0:00 | 39:09:00 | 0:00 |
| 0 | 0 | 0 | 0 | 0 | 1 |
| 3:04:03 | 1:00:00 | 13:02:03 | 8:01:00 | 14:09:03 | 8:01:00 |
| 4:02:03 | 0:00:00 | 2:12:03 | 1:09:00 | 9:09:03 | 1:07:00 |
| 2:05:03 | 0:00:00 | 9:02:03 | 0:00:00 | 7:09:02 | 3:00:00 |
| 0:0:0:0 | 0:0:0:0 | 0:0:0:0 | 0:0:0:0 | 0:0:0:0 | 1:0:0:1 |
| 0:00 | 0:00 | 2:03 | 0:00 | 1:00 | 2:02 |
| 7:00 | 0:00 | 39:00:00 | 0:00 | 43:00:00 | 2:00 |
| 7:00 | 0:00 | 32:00:00 | 0:00 | 44:00:00 | 1:00 |
| 0 | 0 | 0 | 2 | 0 | 1 |
| 1:00 | 0:00 | 0:00 | 0:00 | 8:01 | 0:00 |
| 0:1:0:0 | 0:21:1:0 | 0:4:1:0 | 0:9:3:0 | 0:3:1:0 | 0:23:8:1 |
| 23 | 0 | 19 | 2 | 64 | 1 |
| 0:00 | 0:00 | 0:00 | 0:00 | 0:00 | 1:01 |
| 0:01 | 0:00 | 0:00 | 0:00 | 1:00 | 0:00 |
| 0:0:0:0 | 1:0:0:1 | 0:0:0:0 | 0:0:0:0 | 0:0:0:0 | 0:1:0:0 |
| 0:00 | 2:03 | 0:00 | 0:00 | 0:00 | 2:00 |
| 0:00 | 1:00 | 0:00 | 0:00 | 0:00 | 1:00 |
| 0 | 0 | 0 | 3 | 0 | 4 |

| | | | | | |
|---|---|---|---|---|---|
| 2:00 | 0:00 | 3:00 | 0:00 | 1:00 | 0:00 |
| 13:4:0:0:0 | 4:1:0:0:0 | 2:2:0:0:0 | 0:2:0:0:0 | 13:8:0:0:0 | 1:1:0:0:0 |
| 4:1:0:0:0 | 13:4:0:0:0 | 0:2:0:0:0 | 2:2:0:0:0 | 1:0:0:0:0 | 13:9:0:0:0 |
| 4:1:0:0:0 | 13:4:0:0:0 | 0:2:0:0:0 | 2:2:0:0:0 | 1:0:0:0:0 | 13:9:0:0:0 |
| 20:7:12:1:0:0:0 | 2:0:0:0:0:0:0 | 7:10:35:11:0:0:0 | 0:0:0:0:0:0:0 | 22:18:60:6:0:0:0 | 0:0:1:0:0:0:0 |
| 10:0:1:0:0 | 0:0:0:0:0 | 47:0:0:0:0 | 0:0:0:0:0 | 32:1:0:2:1 | 0:0:0:0:0 |
| 0 | 5 | 0 | 9 | 0 | 25 |
| 0 | 4 | 0 | 9 | 0 | 22 |
| 0:0:0:0:0 | 0:0:0:1:0 | 0:0:0:0:0 | 0:0:0:0:0 | 0:0:0:0:0 | 0:0:0:0:1 |
| 0:0:0:0:0:0:0:0:0 | 1:0:0:0:0:0:0:1:0 | 0:0:0:0:0:0:0:0:0 | 0:0:1:0:0:0:0:0:0 | 0:0:0:0:0:0:0:0:0 | 4:3:0:0:0:0:1:0:1 |
| 0:00 | 0:00 | 0:00 | 0:00 | 0:00 | 0:00 |
| 34:04:00 | 6:02 | 32:04:00 | 34:03:00 | 154:19:00 | 6:01 |
| 0:0:0:0:0 | 0:0:0:0:0 | 0:0:0:0:0 | 0:0:0:0:0 | 0:0:0:0:0 | 0:0:0:0:0 |
| 13:34:03 | 0:01:00 | 42:47:05 | 0:01:00 | 89:104:5 | 1:01:00 |
| 20:36:3:0 | 0:0:0:0 | 46:53:5:0 | 0:0:0:0 | 115:106:5:0 | 0:0:0:0 |
| 26:36:03 | 0:00:00 | 56:51:05 | 0:00:00 | 130:103:5 | 0:00:00 |
| 186:3107:11:1 | 0:8:0:0 | 559:4446:51:0 | 3:16:5:0 | 1253:13384:118:6 | 8:44:15:0 |
| 3118:174:6:6 | 3:0:0:0 | 4478:519:8:12 | 4:3:0:0 | 13470:1108:19:22 | 14:7:0:0 |
| 3119:172:6:7 | 1:0:0:0 | 4480:512:11:12 | 1:1:0:0 | 13478:1083:20:22 | 6:2:1:0 |
| 3113:159:12:8 | 8:0:0:0 | 4478:507:15:12 | 3:0:0:0 | 13476:1049:29:30 | 13:3:0:3 |
| 3117:155:12:8 | 4:0:0:0 | 4476:496:15:14 | 6:1:0:0 | 13469:1007:36:35 | 20:3:0:0 |
| 3117:156:12:8 | 4:2:0:0 | 4480:493:15:13 | 3:2:0:1 | 13477:988:37:37 | 14:4:0:0 |
| 3121:158:12:8 | 0:0:0:0 | 4481:494:15:14 | 1:0:0:0 | 13477:982:36:36 | 13:3:1:1 |
| 3123:136:13:22 | 2:0:0:0 | 4486:376:15:29 | 2:1:0:1 | 13492:629:42:82 | 6:7:0:0 |
| 3125:148:22:12 | 0:0:0:0 | 4487:380:29:16 | 1:0:1:0 | 13497:622:83:40 | 1:5:0:0 |
| 3126:152:24:11 | 1:0:0:0 | 4487:349:33:14 | 5:1:10:0 | 13495:568:97:43 | 9:0:12:0 |
| 3126:26:151:12 | 2:0:1:0 | 4486:43:347:13 | 7:5:0:0 | 13492:108:564:44 | 17:8:0:0 |
| 3146:149:56:11 | 4:1:0:0 | 4573:341:296:40 | 1:1:2:0 | 13788:543:713:62 | 10:0:1:2 |
| 3141:11:52:149 | 7:0:3:0 | 4559:39:271:335 | 15:1:32:1 | 13768:61:663:531 | 31:1:39:1 |
| 147:3148:56:10 | 0:3:1:0 | 331:4575:303:39 | 0:1:3:0 | 520:13785:722:59 | 2:14:5:1 |
| 3146:147:55:10 | 5:0:2:0 | 4574:330:305:38 | 2:0:1:1 | 13793:519:723:59 | 8:1:4:1 |
| 3138:141:54:15 | 3:0:0:0 | 4573:300:399:49 | 8:1:0:0 | 13803:451:879:85 | 23:0:0:0 |
| 1:4:0:1:1:0:0:0 | 0:0:0:0:0:0:0:0 | 2:5:0:1:0:0:0:0 | 0:0:0:0:0:0:0:0 | 3:3:2:0:0:0:0:1 | 0:0:0:0:0:0:0:0 |
| 0:0:0:2:0:0:0:0:0:0:0:0:0:0:1:0:0:0:0:0 | 0:0:0:0:0:0:0:0:0:0:0:0:0:0:0:0:0:0:0:0 | 0:0:1:1:1:1:0:3:0:0:0:0:0:0:1:0:0:0:0:0 | 0:0:0:0:1:2:1:1:0:0:0:0:0:0:0:2:1:0:0:0 | 1:2:3:5:0:1:1:1:0:0:0:1:0:0:1:0:0:1:1:1 | 0:0:0:0:3:0:1:0:0:0:0:0:0:0:0:0:0:0:0:0 |
| 2:0:1:0:0:2:0:0:0:0:0:0:2:1:0:0:0:2:0:0:0:0:0:0:0:0:0:0:0:0 | 3:0:0:0:1:2:0:0:0:3:2:2:0:2:0:0:0:0:0:0:0:0:0:1:0:0:0:0:0 | 8:0:2:0:8:9:2:1:0:14:2:3:8:9:1:0:1:1:1:0:0:0:0:0:0:0:0:0:0 | 23:1:9:0:17:17:8:1:0:54:16:8:11:9:1:0:0:2:2:0:0:0:3:0:0:0:1:0:0 | 7:0:8:0:9:7:6:5:1:11:5:17:7:9:0:0:1:1:4:0:1:1:0:1:0:0:0:1:1 | 37:1:19:0:16:31:9:6:0:60:20:32:18:19:0:0:0:5:3:0:0:1:2:1:0:0:1:0:0 |
| 2:17:5:9:11:17:10:5:1:5:4:0:1:1:0:0:4:0:0:3:0:0:0:0:0:0:0:0:1:0:0:0:0 | 5:9:5:4:11:10:6:3:0:2:0:0:0:2:0:0:3:0:0:0:0:0:0:0:0:0:0:0:0:0:0:0:0 | 23:45:3:33:20:99:33:18:6:4:22:0:1:12:2:0:26:1:1:2:0:1:3:0:0:0:0:2:0:2:0:0:0 | 0:3:1:0:1:4:1:2:0:1:0:0:0:3:0:0:2:0:0:0:0:0:1:0:0:0:0:0:0:0:0:0:0 | 40:85:10:52:70:113:59:50:12:22:37:1:3:23:1:0:40:1:2:1:0:1:6:3:0:1:0:0:1:1:1:1:1 | 7:27:3:19:15:29:8:7:2:4:10:0:0:4:0:1:3:0:0:0:1:0:0:0:0:0:0:0:0:0:1:0:0 |
| 0:14:16:8:28:19:1:37:11:4:0:0:29:14:5:1:7:0:0:3:0:0:0:0:0:0:0:0:0:0:0:1:0:0 | 0:0:0:0:0:0:0:0:0:0:0:0:0:0:0:0:0:0:0:0:0:0:0:0:0:0:0:0:0:0:0:0:0:0 | 0:30:69:10:92:67:1:199:35:30:3:0:45:7:48:12:52:2:1:4:0:1:8:2:3:1:0:1:0:2:1:0:0 | 0:1:0:0:0:0:0:0:0:0:0:0:0:0:1:0:1:0:0:0:0:0:0:0:0:0:0:0:0:0:0:0:0 | 1:97:119:47:197:119:4:273:92:49:3:1:154:32:78:22:75:3:3:2:2:2:8:3:3:4:0:2:2:0:2:0:1 | 0:0:0:0:0:0:0:1:0:0:0:0:0:0:1:0:0:1:0:0:0:0:0:0:0:0:0:0:0:0:0:0:1:0 |
| 0:14:16:8:28:19:1:37:11:4:0:0:29:14:5:1:7:0:0:3:0:0:0:0:0:0:0:0:0:0:0:1:0:0 | 0:0:0:0:0:0:0:0:0:0:0:0:0:0:0:0:0:0:0:0:0:0:0:0:0:0:0:0:0:0:0:0:0:0 | 0:31:68:10:92:67:1:199:35:30:3:0:45:7:49:12:53:2:1:4:0:1:8:2:3:1:0:1:0:2:1:0:0 | 0:0:1:0:0:0:0:0:0:0:0:0:0:0:0:0:0:0:0:0:0:0:0:0:0:0:0:0:0:0:0:0:0 | 1:96:119:47:197:119:4:274:92:48:3:1:154:32:80:22:75:4:3:2:2:2:8:3:3:4:0:2:2:0:2:0:1 | 0:1:0:0:0:0:0:0:0:0:1:0:0:0:0:0:0:0:0:0:0:0:0:0:0:0:0:0:0:0:0:0:1:0 |
| 0:14:16:8:28:20:1:37:11:4:0:0:29:14:5:1:8:0:0:3:0:0:0:0:0:0:0:0:0:0:0:1:0 | 0:0:0:0:0:0:0:0:0:0:0:0:0:0:0:0:0:0:0:0:0:0:0:0:0:0:0:0:0:0:0:0:0 | 0:31:69:10:93:67:1:199:35:30:3:0:45:7:50:12:58:2:1:4:0:1:8:2:3:1:0:1:0:2:1:0 | 0:0:0:0:1:0:0:0:0:0:0:0:0:0:0:0:1:0:0:0:0:0:0:0:0:0:0:0:0:0:0:0 | 1:97:120:48:197:119:4:273:92:50:3:1:154:32:81:25:76:4:3:2:2:2:8:3:3:4:0:2:2:0:2:1 | 0:0:1:0:1:1:0:1:0:0:0:0:0:0:0:0:2:0:0:0:0:0:0:0:0:0:0:0:0:0:0:0:0 |

| | | | | | |
|---|---|---|---|---|---|
| 0:15:16:10:29:22:1:37:11:6:0:0:29:15:5:2:9:3:0:0:0:0:0:0:0:0:0:0:0:0:1:0 | 0:0:0:0:0:0:0:0:0:0:0:0:0:0:0:0:0:0:0:0:0:0:0:0:0:0:0:0:0:0:0:0 | 0:36:76:11:100:71:1:199:35:32:3:0:45:7:51:15:65:4:2:1:0:1:9:2:3:1:0:1:0:2:1:0 | 0:0:0:0:0:0:0:0:0:0:0:0:0:0:0:0:0:0:0:0:0:0:0:0:0:0:0:0:0:0:0:0 | 1:100:127:49:204:123:4:273:91:51:3:1:154:32:81:26:87:2:4:3:2:2:9:3:3:4:0:2:2:0:2:1 | 0:0:0:0:0:0:0:1:1:0:0:0:0:0:0:1:1:0:0:0:0:0:0:0:0:0:0:0:0:0:0:0 |
| 0:15:17:10:29:22:1:37:11:6:0:0:29:15:5:3:9:0:0:3:0:0:0:0:0:0:0:0:0:0:1:0 | 0:0:0:0:0:0:0:0:0:0:0:0:0:0:0:0:0:0:0:0:0:0:0:0:0:0:0:0:0:0:0:0 | 0:35:76:11:101:71:1:198:35:32:3:0:45:7:51:15:67:2:1:4:0:1:9:2:3:1:0:1:0:2:1:0 | 0:1:1:0:0:0:0:1:0:0:0:0:0:0:0:0:0:0:0:0:0:0:0:0:0:0:0:0:0:0:0:0 | 1:101:127:48:204:122:4:273:91:51:3:1:154:32:81:28:95:4:3:2:2:2:9:3:3:4:0:2:2:0:2:1 | 0:0:0:0:0:1:0:1:1:0:0:0:0:0:0:0:0:0:0:0:0:0:0:0:0:0:0:0:0:0:0:0 |
| 0:14:17:10:29:22:1:37:11:6:0:0:29:15:5:3:8:0:0:3:0:0:0:0:0:0:0:0:0:0:1:0 | 0:1:0:0:0:0:0:0:0:0:0:0:0:0:0:0:1:0:0:0:0:0:0:0:0:0:0:0:0:0:0:0 | 0:36:76:11:101:71:1:199:35:32:3:0:45:7:51:15:68:2:1:4:0:1:9:2:3:1:0:1:0:2:1:0 | 0:0:1:0:0:0:0:0:0:0:0:0:0:0:0:0:0:0:0:0:0:0:0:0:0:0:0:0:0:0:0:0 | 1:101:126:48:204:123:4:273:92:51:3:1:153:32:81:28:96:4:3:2:2:2:9:3:3:4:0:2:2:0:2:1 | 0:0:1:0:0:0:0:1:0:0:0:0:1:0:0:0:0:0:0:0:0:0:0:0:0:0:0:0:0:0:0:0 |
| 0:14:17:10:29:22:1:37:11:6:0:0:29:15:5:3:10:0:0:3:0:0:0:0:0:0:0:0:0:0:0:1:0 | 0:1:0:0:0:0:0:0:0:0:0:0:0:0:0:0:0:0:0:0:0:0:0:0:0:0:0:0:0:0:0:0 | 0:36:77:11:100:71:1:199:35:32:3:0:45:7:49:15:68:2:1:4:0:1:9:2:3:1:0:1:0:2:1:0 | 0:0:0:0:1:0:0:0:0:0:0:0:0:0:0:0:0:0:0:0:0:0:0:0:0:0:0:0:0:0:0:0 | 1:101:127:47:204:124:4:274:92:51:3:1:154:32:79:27:96:4:3:2:2:2:9:3:3:4:0:2:2:0:2:1 | 0:0:0:1:0:0:0:0:0:0:0:0:0:0:0:0:0:0:0:0:0:0:0:0:0:0:0:0:0:0:0:0 |
| 17:29:15:0:10:22:1:37:11:6:0:0:29:15:5:3:10:0:0:3:0:0:0:0:0:0:0:0:1:0:0:0:0 | 2:2:3:0:1:1:0:0:0:1:0:0:0:0:0:0:0:0:0:0:0:0:0:0:0:0:0:0:0:0:0:0 | 77:102:37:0:12:71:1:199:35:32:3:0:45:7:49:15:68:2:1:4:0:1:9:2:3:1:0:1:1:0:2:0 | 5:19:8:0:1:3:0:0:0:5:0:0:0:0:0:1:3:0:0:0:0:0:1:0:0:0:0:0:0:0:0:0:0 | 128:205:102:1:49:124:4:274:92:51:3:1:154:32:79:27:97:4:3:2:2:2:9:3:3:4:0:2:2:2:0:1 | 11:8:7:0:2:3:0:0:0:3:0:0:0:0:0:1:2:0:0:0:0:0:0:0:0:0:0:0:0:0:0:0:0 |
| 32:21:0:19:12:23:1:37:11:7:0:0:29:15:4:3:9:0:0:0:3:0:0:0:0:0:0:0:0:0:0:1:0 | 1:0:0:0:0:0:0:0:0:0:0:0:0:0:0:0:0:0:0:0:0:0:0:0:0:0:0:0:0:0:0:0 | 135:91:0:52:14:80:1:199:35:38:4:0:45:7:49:16:73:2:2:1:4:0:1:10:2:3:1:0:1:0:1:0 | 0:2:0:0:0:0:0:0:0:0:0:0:0:0:0:0:0:0:0:0:0:0:0:0:0:0:0:0:0:0:0:0 | 224:152:1:120:55:129:4:274:92:55:3:1:154:32:76:30:107:0:4:3:2:2:2:9:3:3:4:0:2:2:2:1 | 2:3:0:0:0:1:0:0:0:0:0:0:0:0:0:1:0:0:0:0:0:0:0:0:0:0:0:0:0:0:0:0 |
| 0:19:21:12:33:23:1:37:11:7:0:0:29:15:4:3:9:0:0:0:3:0:0:0:0:0:0:0:0:0:0:1:0 | 0:0:0:0:0:0:0:0:0:0:0:0:0:0:0:0:0:0:0:0:0:0:0:0:0:0:0:0:0:0:0:0 | 0:52:93:14:134:80:1:199:35:38:4:0:45:7:49:16:73:2:2:1:4:0:1:10:2:3:1:0:1:0:1:0 | 0:0:0:0:1:0:0:0:0:0:0:0:0:0:0:0:0:0:0:0:0:0:0:0:0:0:0:0:0:0:0:0 | 1:120:154:55:226:130:4:274:92:55:3:1:154:32:77:30:103:0:4:3:2:2:2:9:3:3:4:0:2:2:2:1 | 0:0:1:0:0:0:0:0:0:0:0:0:0:0:0:0:0:0:0:0:0:0:0:0:0:0:0:0:0:0:0:0 |
| 19:21:0:12:34:23:1:37:11:8:0:0:29:15:4:3:10:0:0:0:3:0:0:0:0:0:0:0:1:0:0:0 | 2:3:0:1:1:0:0:0:0:0:0:0:0:0:0:0:0:0:0:0:0:0:0:0:0:0:0:0:0:0:0:0 | 53:93:0:14:135:81:1:199:35:38:4:0:45:7:49:16:72:2:10:2:1:4:0:1:2:3:1:0:1:1:0:0 | 10:17:0:0:36:4:0:0:0:1:0:0:0:0:2:1:14:1:2:0:0:0:0:0:0:0:0:0:0:0:0:0:0 | 121:156:1:55:227:130:4:274:92:55:3:1:153:32:75:30:99:0:9:4:3:2:2:2:3:3:4:0:2:2:2:1 | 7:20:0:2:33:6:0:0:0:2:0:0:1:1:1:2:10:0:3:0:0:0:0:0:0:0:0:0:0:1:0:0:0 |
| 0:21:26:14:38:25:1:37:11:10:0:0:29:15:4:4:10:0:0:0:3:0:0:0:0:0:0:0:0:0:0:1:0 | 0:0:0:0:0:0:0:0:0:0:0:0:0:0:0:0:0:0:0:0:0:0:0:0:0:0:0:0:0:0:0:0 | 0:69:115:17:182:85:1:198:35:46:4:0:45:7:48:17:89:4:2:1:4:0:1:13:2:3:1:0:2:0:1:0 | 0:0:0:0:1:1:0:0:0:0:0:0:0:0:0:0:1:0:0:0:0:0:0:0:0:0:0:0:0:0:0:0 | 1:145:194:70:294:145:4:274:92:64:4:1:154:33:71:38:108:0:4:2:2:2:2:11:3:3:4:0:2:2:5:1 | 0:0:0:0:0:0:0:0:0:0:0:0:0:0:0:0:0:0:0:0:0:0:0:0:1:0:0:0:0:0:0:0:0 |
| 0:21:26:14:38:25:1:37:11:9:0:0:29:15:4:4:10:0:0:0:3:0:0:0:0:0:0:0:0:0:1:0 | 0:0:0:0:0:0:0:0:0:1:0:0:0:0:0:0:0:0:0:0:0:0:0:0:0:0:0:0:0:0:0:0 | 0:69:114:17:183:86:1:199:35:46:4:0:45:7:48:17:89:4:2:1:4:0:1:13:2:3:1:0:2:0:1:0 | 0:0:1:0:1:0:0:0:0:0:0:0:0:0:0:0:0:0:0:0:0:0:0:0:0:0:0:0:0:0:0:0 | 1:144:194:70:294:145:4:274:92:64:4:1:154:33:71:37:106:0:4:2:2:2:2:12:3:3:4:0:2:2:5:1 | 0:1:0:0:0:0:0:0:0:0:0:0:0:0:0:0:1:0:0:0:0:0:0:0:0:0:0:0:0:0:0:0 |
| 0:21:26:14:38:26:1:37:11:10:0:0:29:15:3:3:10:0:0:0:3:0:0:0:0:0:0:0:0:0:0:1:0 | 0:0:0:0:0:0:0:0:0:0:0:0:0:0:0:0:0:0:0:0:0:0:0:0:0:0:0:0:0:0:0:0 | 0:69:115:17:184:84:1:199:35:46:4:0:45:7:44:16:82:4:2:1:4:0:1:13:2:3:1:0:2:0:1:0 | 0:0:0:0:0:0:0:0:0:0:0:0:0:0:0:0:0:0:0:0:0:0:0:0:0:0:0:0:0:0:0:0 | 1:145:190:70:294:141:4:273:92:64:4:1:154:33:67:31:100:0:4:2:2:2:2:12:3:3:4:0:2:2:5:1 | 0:1:1:0:0:0:0:1:0:0:0:0:0:0:0:1:0:0:0:0:0:0:0:0:0:0:0:0:0:0:0:0 |
| 0:21:26:14:38:24:1:37:11:10:0:29:15:2:1:6:0:0:0:0:3:0:0:0:0:0:0:0:0:0:1:0 | 0:0:0:0:0:1:0:0:0:0:0:0:0:0:0:0:0:0:0:0:0:0:0:0:0:0:0:0:0:0:0:0 | 0:75:126:17:182:99:1:199:35:46:0:45:7:41:17:74:4:14:2:1:4:0:1:2:3:1:0:5:2:0:1:0 | 0:0:0:0:2:1:0:0:0:0:0:0:0:0:0:0:0:0:0:0:0:0:0:0:0:0:0:0:0:0:0:0 | 1:169:203:70:294:164:4:273:92:64:1:154:33:59:28:93:0:13:4:2:2:2:2:3:3:4:1:4:2:2:5:1 | 0:0:2:0:0:1:0:1:0:0:0:0:0:0:0:1:0:0:0:0:0:0:0:0:0:0:0:1:0:0:0:0 |

| | | | | | |
|---|---|---|---|---|---|
| 0:22:26:14:38:25:1:37:11:10:0:29:15:1:1:7:0:0:0:0:3:0:0:0:0:0:0:0:0:0:1:0 | 0:0:0:0:0:0:0:0:0:0:0:0:0:0:0:0:0:0:0:0:0:0:0:0:0:0:0:0:0:0 | 0:75:126:17:184:100:1:199:35:46:0:45:7:38:15:70:4:14:2:1:4:0:1:2:3:1:0:5:2:0:1:0 | 0:0:0:0:0:0:0:0:0:0:0:0:0:0:0:0:1:0:0:0:0:0:0:0:0:0:0:0:0:0:0:0:0 | 1:170:205:70:292:164:4:273:92:64:1:154:33:53:21:86:0:13:4:2:2:2:2:3:3:4:1:5:2:2:5:1 | 0:0:0:0:2:0:0:1:0:0:0:0:0:0:2:0:0:0:0:0:0:0:0:0:0:0:0:0:0:0:0:0 |
| 38:23:26:14:27:1:37:11:10:0:0:29:15:2:1:7:0:0:0:0:3:0:0:0:0:0:0:0:0:1:0 | 0:0:0:0:0:0:0:0:0:0:0:0:0:0:0:0:0:0:0:0:0:0:0:0:0:0:0:0:0 | 184:75:126:17:102:1:199:35:46:5:0:45:7:10:18:36:4:14:2:1:4:0:1:2:3:1:0:2:0:1:0 | 0:0:0:0:0:0:0:0:0:0:0:0:0:0:0:0:0:0:0:0:0:0:0:0:0:0:0:0:0:0:0:0:0 | 294:171:207:70:158:4:274:92:64:5:1:154:33:12:31:34:0:13:4:2:2:2:2:3:3:4:1:2:2:5:1 | 0:0:0:0:1:0:0:0:0:0:0:0:0:0:1:2:0:0:0:0:0:0:0:0:0:0:0:0:0:0 |
| 23:26:14:38:27:1:37:11:10:0:0:29:15:2:1:7:0:0:0:0:3:0:0:0:0:0:0:0:0:1:0 | 0:0:0:0:0:0:0:0:0:0:0:0:0:0:0:0:0:0:0:0:0:0:0:0:0:0:0:0:0 | 75:126:17:184:101:1:198:35:46:5:0:45:7:10:16:34:4:14:2:1:4:0:1:2:3:1:0:2:0:1:0 | 0:0:0:0:1:0:1:0:0:0:0:0:0:0:1:0:0:0:0:0:0:0:0:0:0:0:0:0:0:0:0:0 | 171:207:70:294:158:4:274:92:64:5:1:154:33:11:32:32:0:13:4:2:2:2:2:3:3:4:1:2:2:5:1 | 0:0:0:0:0:0:0:0:0:0:0:0:0:0:0:0:2:0:0:0:0:0:0:0:0:0:0:0:0:0:0 |
| 6:00 | 0:00 | 8:00 | 1:01 | 30:00:00 | 3:00 |
| 56:1:7:0:1:5:0:1:0 | 0:0:0:0:0:0:0:0:0 | 127:2:19:1:0:5:2:0:2 | 0:0:0:0:0:0:0:0:0 | 231:8:32:4:1:13:4:0:0 | 1:0:1:0:0:0:0:0:0 |
| 0:00 | 0:00 | 0:00 | 0:00 | 0:00 | 3:00 |
| 0:00 | 0:00 | 0:00 | 0:00 | 0:00 | 3:00 |
| 0:01 | 2:00 | 0:00 | 0:00 | 0:00 | 0:00 |
| 0 | 0 | 0 | 0 | 0 | 2 |
| 0:00:00 | 0:00:00 | 0:00:00 | 0:00:00 | 0:00:00 | 0:00:02 |
| 0:00:00 | 0:00:00 | 0:00:00 | 0:00:00 | 0:00:00 | 0:00:02 |
| 0:00:00 | 0:00:00 | 0:00:00 | 0:00:00 | 0:00:02 | 0:00:00 |
| 0:00:00 | 0:00:00 | 0:00:00 | 0:00:00 | 0:00:00 | 0:00:01 |
| 12:0:2:1:4:1:0 | 0:0:0:0:0:0:0 | 17:0:2:1:1:1:0 | 0:0:0:0:0:0:0 | 47:3:5:1:2:1:0 | 1:0:0:0:0:0:0 |
| 0:2:12:0:3:1:0 | 0:0:0:0:0:0:0 | 0:2:18:1:1:1:0 | 0:0:1:0:0:0:0 | 3:5:47:1:1:0:0 | 0:0:2:0:0:1:0 |
| 0 | 2 | 8 | 7 | 10 | 15 |
| 0 | 0 | 0 | 1 | 0 | 7 |
| 25:0:0:0 | 0:0:0:0 | 61:2:2:2 | 1:0:1:0 | 149:8:2:2 | 1:0:0:0 |
| 17:1:1:2:0 | 1:1:0:0:0 | 46:3:2:9:0 | 2:0:0:2:0 | 107:11:7:16:9 | 12:1:0:1:0 |
| 15:1:3:2:0 | 0:0:0:0:0 | 13:1:1:3:0 | 21:2:0:7:0 | 35:6:8:13:8 | 52:2:0:1:1 |
| 288:15:00 | 0:00 | 359:35:00 | 2:00 | 1206:52:00 | 1:00 |
| 288:15:00 | 0:00 | 361:35:00 | 0:00 | 1208:52:00 | 2:00 |
| 290:15:00 | 1:00 | 362:35:00 | 1:00 | 1207:52:00 | 3:00 |
| 291:15:00 | 0:00 | 364:35:00 | 0:00 | 1210:52:00 | 3:00 |
| 291:15:00 | 0:00 | 359:34:00 | 2:00 | 1206:52:00 | 6:00 |
| 292:15:00 | 0:00 | 361:34:00 | 1:00 | 1209:52:00 | 2:00 |
| 293:15:00 | 0:00 | 362:34:00 | 0:00 | 1211:52:00 | 0:00 |
| 291:15:00 | 2:00 | 367:34:00 | 1:00 | 1217:52:00 | 0:00 |
| 293:15:00 | 0:00 | 367:34:00 | 1:00 | 1219:52:00 | 1:00 |
| 0:0:0:2:0:0:0:0:0 | 1:1:1:1:0:1:0:0:0 | 0:0:6:5:0:0:1:0:0 | 0:0:4:1:0:0:0:0:0 | 2:0:2:1:0:0:0:0:0 | 4:1:27:59:3:2:2:0:1 |
| 30:04:03 | 0:00:00 | 110:18:03 | 0:00:00 | 243:46:11 | 2:01:00 |
| 1:21:03 | 0:00:00 | 0.334861111 | 0:00:00 | 27:157:21 | 1:06:00 |
| 2:23:04 | 0:02:00 | 0.438344907 | 0:01:00 | 31:194:25 | 2:02:01 |
| 0:00 | 0:00 | 5:01 | 0:00 | 1:01 | 0:00 |
| 20:1:8:0 | 0:0:0:0 | 25:24:37:0 | 0:0:0:0 | 63:41:63:0 | 0:1:1:0 |
| 20:1:8:0 | 0:0:0:0 | 25:22:37:0 | 0:0:0:0 | 63:37:64:0 | 0:0:0:0 |
| 0:1:1:0 | 20:0:7:0 | 14:6:24:0 | 11:11:13:0 | 3:0:2:0 | 60:25:61:0 |
| 2 | 0 | 0 | 1 | 1 | 2 |
| 0 | 49 | 0 | 271 | 0 | 215 |
| 49 | 0 | 271 | 0 | 216 | 0 |
| 770:01:00 | 0:00:00 | 1616:03:01 | 2:00:00 | 3376:16:00 | 3:00:00 |
| 0:00:00 | 769:01:00 | 0:00:00 | 1624:03:01 | 0:00:00 | 3404:16:00 |
| 779:00:00 | 0:00:00 | 1651:02:01 | 2:00:00 | 3468:03:00 | 3:00:00 |
| 779:00:00 | 0:00:00 | 1654:00:01 | 1:00:00 | 3471:00:00 | 2:01:00 |
| 783:00:00 | 0:00 | 1660:00:00 | 1:00 | 3491:01:00 | 1:00 |
| 778:00:00 | 1:00:00 | 1640:01:00 | 3:00:00 | 3462:00:01 | 6:00:00 |
| 774:00:00 | 0:00:00 | 1629:01:00 | 2:00:00 | 3446:00:01 | 4:00:00 |
| 36:03:10 | 2:00:00 | 150:6:66 | 3:00:00 | 271:27:89 | 7:00:01 |
| 3:10:38 | 0:00:00 | 6:66:152 | 0:00:01 | 27:90:276 | 0:00:02 |
| 17:46:06 | 0:06:02 | 84:208:7 | 2:13:00 | 107:375:30 | 1:18:00 |
| 17:49:06 | 0:03:02 | 86:217:7 | 2:08:00 | 109:387:30 | 0:10:01 |
| 0:00 | 0:00 | 3:00 | 0:00 | 2:00 | 3:00 |
| 8:30:52:4:13 | 0:0:0:0:0 | 90:242:209:40:62 | 0:0:1:0:0 | 62:227:297:38:84 | 1:0:1:1:0 |
| 56:30:12:4:13 | 0:0:0:0:0 | 229:241:110:41:61 | 1:0:1:0:0 | 325:227:87:39:83 | 1:0:0:0:0 |
| 29:56:15:4:13 | 1:0:0:0:0 | 231:232:119:38:57 | 7:7:3:2:2 | 219:331:92:35:78 | 7:5:2:3:2 |
| 29:56:16:4:12 | 0:0:0:0:0 | 229:229:123:37:56 | 0:2:0:0:0 | 216:328:92:34:79 | 1:3:2:0:0 |

| | | | | | |
|---|---|---|---|---|---|
| 28:56:17:4:12 | 0:0:0:0:0 | 229:229:126:37:56 | 0:3:1:0:0 | 215:331:92:34:79 | 0:0:1:0:0 |
| 28:56:17:4:12 | 0:0:0:0:0 | 228:230:130:35:56 | 0:0:0:1:0 | 213:339:97:34:79 | 0:0:1:0:0 |
| 28:56:17:4:12 | 0:0:0:0:0 | 227:230:130:35:56 | 1:0:0:1:0 | 213:339:97:34:79 | 0:0:1:0:0 |
| 28:55:17:4:12 | 0:0:0:0:0 | 228:232:130:35:56 | 0:0:0:1:0 | 213:338:98:34:79 | 0:1:1:0:0 |
| 0:00 | 0:00 | 0:00 | 0:00 | 0:00 | 0:00 |
| 7:2:0:0:0:0:0 | 0:0:0:0:0:0:0 | 7:0:2:2:0:0:1 | 0:0:0:0:0:0:0 | 13:0:1:3:0:1:0 | 0:0:0:0:0:0:0 |
| 26:2:11:1:0 | 0:0:0:0:0 | 92:1:19:1:0 | 0:0:0:0:0 | 217:2:44:3:1 | 1:0:0:0:0 |
| 0:0:0:0:0 | 26:2:11:1:0 | 0:0:0:0:0 | 99:1:19:1:0 | 1:0:0:0:0 | 223:3:45:3:1 |
| 26:2:11:1:0 | 0:0:0:0:0 | 107:1:19:1:0 | 0:0:0:0:0 | 240:3:48:3:1 | 2:0:0:0:0 |
| 0:0:0:0:0 | 3:0:0:3:1 | 0:0:0:0:0 | 5:1:1:0:0 | 0:0:0:0:0 | 19:9:3:6:0 |
| 0:0:0:0 | 0:1:4:0 | 0:0:0:0 | 10:17:49:6 | 0:0:0:0 | 8:4:24:2 |
| 0:0:0:0 | 0:1:4:0 | 0:0:0:0 | 10:17:49:6 | 0:0:0:0 | 8:4:24:2 |
| 20:01 | 0:00 | 8:00 | 0:00 | 88:01:00 | 0:00 |
| 18:00 | 1:00 | 6:01 | 0:00 | 78:04:00 | 2:00 |
| 1:1:10:5:2:0:0 | 0:0:0:0:0:0:0 | 0:0:15:15:0:0:0 | 0:0:0:0:0:0:0 | 10:3:11:11:7:0:1 | 0:0:0:0:0:0:0 |
| 0:0:0:0 | 2:5:1:1 | 0:0:0:0 | 0:0:0:0 | 0:0:0:0 | 0:2:0:0 |
| 0:00:00 | 1:01:01 | 0:00:00 | 0:00:00 | 0:00:00 | 0:02:00 |
| 115 | 0 | 265 | 15 | 782 | 30 |
| 122 | 1 | 285 | 3 | 831 | 6 |
| 123 | 0 | 291 | 4 | 838 | 5 |
| 124 | 0 | 299 | 0 | 840 | 0 |
| 121 | 1 | 267 | 1 | 790 | 1 |
| 121 | 0 | 264 | 0 | 784 | 3 |
| 1048 | 0 | 12718 | 17 | 17017 | 22 |
| 1044 | 3 | 12652 | 12 | 16937 | 26 |
| 1043 | 0 | 12498 | 15 | 16699 | 17 |
| 1041 | 2 | 12454 | 57 | 16662 | 55 |
| 1036 | 0 | 12222 | 13 | 16398 | 15 |
| 1011 | 0 | 11889 | 22 | 15887 | 27 |
| 1004 | 1 | 11833 | 18 | 15839 | 12 |
| 1002 | 3 | 11821 | 13 | 15806 | 24 |
| 976 | 0 | 11406 | 12 | 15330 | 10 |
| 955 | 0 | 11251 | 20 | 15169 | 21 |
| 912 | 1 | 10723 | 12 | 14464 | 28 |
| 909 | 3 | 10570 | 31 | 14280 | 43 |
| 914 | 1 | 10612 | 17 | 14345 | 32 |
| 920 | 3 | 10717 | 32 | 14437 | 55 |
| 914 | 1 | 10630 | 35 | 14382 | 54 |
| 907 | 4 | 10540 | 44 | 14240 | 58 |
| 908 | 3 | 10509 | 46 | 14178 | 65 |
| 0:00:00 | 0:00:00 | 0:00:00 | 0:00:00 | 0:00:02 | 0:00:00 |
| 0:00:00 | 0:00:00 | 0:00:00 | 0:00:00 | 0:00:00 | 0:00:01 |
| 0:00 | 0:00 | 0:00 | 1:00 | 0:00 | 0:00 |
| 0:00 | 0:00 | 0:00 | 1:00 | 0:00 | 0:00 |
| 0:0:0:0 | 1:0:0:0 | 0:0:0:0 | 0:0:0:0 | 0:0:0:0 | 0:2:0:0 |
| 108:1:3:2:0:0 | 1:0:0:0:0:0 | 299:2:12:1:2:2 | 1:0:0:0:0:0 | 507:9:19:6:1:1 | 2:0:0:0:0:0 |
| 3:0:2:1:0 | 0:0:0:0:0 | 0:4:7:0:1 | 0:0:0:0:0 | 8:3:8:0:2 | 0:0:1:0:0 |
| 0:00 | 0:00 | 0:00 | 0:00 | 0:00 | 0:01 |
| 110 | 0 | 206 | 1 | 465 | 0 |
| 109 | 1 | 207 | 0 | 467 | 0 |
| 110 | 0 | 205 | 0 | 467 | 0 |
| 109 | 1 | 205 | 0 | 465 | 2 |
| 0:00 | 1:01 | 0:00 | 0:00 | 0:00 | 1:00 |
| 0:00:01 | 0:00:00 | 0:00:00 | 0:00:00 | 2:01:02 | 0:00:00 |
| 2:2:1:1:1:0 | 0:0:0:0:0:0 | 0:8:1:0:0:0 | 0:0:0:0:0:0 | 2:12:0:0:0:1 | 0:0:0:0:0:0 |
| 0:0:6:2:0:1 | 0:0:0:0:0:0 | 2:0:0:3:0:0 | 0:0:0:0:0:0 | 12:0:11:3:0:1 | 1:0:1:0:0:0 |
| 3:10:1:5:2:0 | 0:0:0:0:0:0 | 1:4:2:1:0:1 | 0:0:0:0:0:0 | 5:56:15:6:0:1 | 0:0:3:0:0:0 |
| 3:10:1:5:2:0 | 0:0:0:0:0:0 | 1:4:2:1:0:1 | 0:0:0:0:0:0 | 5:55:14:6:0:1 | 0:0:0:0:0:0 |
| 3:10:1:5:2:0 | 0:0:0:0:0:0 | 1:4:2:1:0:1 | 0:0:0:0:0:0 | 5:54:14:6:0:1 | 0:1:0:0:0:0 |
| 0 | 0 | 0 | 16 | 0 | 1 |
| 0 | 0 | 0 | 16 | 0 | 1 |
| 3:5:2:0:3:2:1:0:0:2:0:1:0:0 | 0:0:0:0:0:0:0:0:0:0:0:0:0:0 | 0:22:0:0:0:1:1:5:0:0:0:0:1:0 | 0:0:0:0:0:0:0:0:0:0:0:0:0:0 | 2:80:1:1:5:1:1:4:0:0:0:0:0:1 | 0:2:0:0:0:0:0:0:0:0:0:0:0:0 |
| 0 | 0 | 0 | 0 | 1 | 0 |
| 0 | 1 | 0 | 0 | 0 | 2 |

| | | | | | |
|---|---|---|---|---|---|
| 5:31:5:1:0:1:0:0 | 0:0:0:0:0:0:0:0 | 18:46:7:1:3:0:0:0 | 1:1:0:0:0:0:0:0 | 12:65:24:0:4:5:4:4 | 0:0:0:0:0:0:0:0 |
| 27:5:0:3:1:0:0:0 | 0:0:0:0:0:0:0:0 | 40:13:3:13:0:0:0:0 | 0:0:0:0:0:0:0:0 | 61:32:4:14:5:3:3:0 | 0:0:0:0:0:0:0:0 |
| 30:7:4:0:0:0:1:1 | 0:0:1:0:0:0:0:0 | 45:31:12:0:3:0:0:1 | 0:0:0:0:0:0:0:0 | 62:28:30:5:4:3:5:0 | 1:0:0:0:0:0:0:0 |
| 1:1:0:0:0:0:0 | 19:14:1:0:0:0:0 | 4:1:0:1:1:0:0 | 6:4:0:0:0:0:0 | 4:1:0:0:1:0:1 | 64:18:6:2:10:1:0 |
| 0 | 0 | 2 | 0 | 1 | 0 |
| 0 | 0 | 0 | 0 | 1 | 0 |
| 13:15:176:2:21:4:18 | 0:0:0:0:0:0:0 | 167:83:295:14:25:8:79 | 0:0:1:0:0:1:0 | 235:67:648:16:43:20:127 | 2:0:2:0:2:1:2 |
| 12:15:176:2:21:4:17 | 0:0:0:0:0:0:0 | 161:83:296:14:25:10:77 | 0:0:1:0:0:0:0 | 233:67:648:16:45:20:120 | 0:0:1:0:0:1:2 |
| 12:15:176:2:21:4:17 | 0:0:0:0:0:0:0 | 146:84:300:14:25:10:72 | 0:0:0:0:0:0:1 | 203:67:652:17:46:20:118 | 3:0:0:0:0:1:0 |
| 190:02:00 | 0:00 | 456:08:00 | 0:00 | 621:09:00 | 0:00 |
| 193:02:00 | 0:00 | 456:06:00 | 1:00 | 623:10:00 | 2:00 |
| 176:11:15:2:21:4:15 | 1:0:0:0:0:0:0 | 304:135:85:14:26:10:63 | 0:0:0:0:0:0:0 | 657:175:66:17:46:20:100 | 0:2:1:0:0:0:0 |
| 15:178:11:1:21:4:13 | 0:0:0:1:0:0:0 | 85:305:72:14:26:10:36 | 0:1:0:0:0:0:0 | 67:658:112:17:46:20:67 | 1:0:1:0:0:0:0 |
| 15:178:11:2:21:4:13 | 0:0:0:0:0:0:0 | 84:306:60:14:26:10:29 | 1:1:0:0:0:0:0 | 68:658:103:16:46:20:60 | 0:0:0:1:0:0:0 |
| 11:15:178:2:21:4:12 | 0:0:0:0:0:0:0 | 52:85:307:14:26:10:27 | 0:0:0:0:0:0:0 | 91:68:658:17:46:20:55 | 1:0:1:0:0:0:0 |
| 11:15:179:2:21:4:12 | 0:0:0:0:0:0:0 | 46:85:308:14:26:10:24 | 0:0:0:0:0:0:1 | 86:68:658:17:46:20:48 | 2:0:2:0:0:0:0 |
| 21:11:15:178:2:4:11 | 0:0:0:1:0:0:0 | 26:44:86:307:14:10:21 | 0:0:0:1:0:0:0 | 46:72:68:659:17:20:47 | 0:0:0:1:0:0:0 |
| 11:15:179:2:21:4:10 | 0:0:0:0:0:0:1 | 43:86:307:14:26:10:21 | 0:0:1:0:0:0:0 | 72:68:659:17:46:20:46 | 0:0:1:0:0:0:0 |
| 134:11:66:12:17:1 | 1:0:0:0:0:0 | 268:29:103:30:13:2 | 0:0:0:0:0:0 | 451:91:226:44:43:1 | 0:0:0:1:0:0 |
| 135:11:66:12:17:1 | 0:0:0:0:0:0 | 268:29:103:30:13:2 | 0:0:0:0:0:0 | 450:91:225:45:43:1 | 1:0:1:0:0:0 |
| 136:11:66:12:17:1 | 0:0:0:0:0:0 | 266:29:103:30:13:2 | 2:0:0:0:0:0 | 448:91:227:45:43:1 | 3:1:0:0:0:0 |
| 136:11:66:12:17:1 | 0:0:0:0:0:0 | 268:28:103:30:13:2 | 0:0:0:0:0:0 | 449:92:227:45:43:1 | 2:0:0:0:0:0 |
| 136:9:66:12:17:1 | 0:0:0:0:0:0 | 270:24:103:30:13:2 | 1:0:0:0:0:0 | 451:73:229:45:43:1 | 1:0:1:0:0:0 |
| 136:68:4:12:17:1 | 0:0:0:0:0:0 | 272:104:10:31:13:2 | 0:0:0:0:0:0 | 457:235:35:46:43:1 | 0:0:0:0:0:0 |
| 136:7:68:10:1 | 0:0:0:0:0 | 272:13:103:28:2 | 0:0:1:1:0 | 457:45:235:31:1 | 0:0:0:0:0 |
| 0:00 | 0:00 | 0:00 | 0:00 | 0:00 | 0:00 |
| 62 | 1 | 37 | 0 | 127 | 2 |
| 5:1:0:3:0:0 | 0:1:0:0:0:0 | 5:11:0:8:0:0 | 0:6:0:0:0:0 | 6:15:0:11:0:1 | 1:7:0:0:0:0 |
| 5:1:3:0:0:0 | 0:0:0:0:0:0 | 5:11:8:0:0:0 | 0:2:0:0:0:0 | 6:12:11:0:0:1 | 0:4:0:0:0:0 |
| 0:0:0:0:0:0 | 0:0:0:0:1:0 | 1:0:0:0:0:0 | 0:0:0:0:0:1 | 1:0:0:0:0:0 | 0:2:0:0:0:0 |
| 0 | 1 | 0 | 1 | 0 | 0 |
| 6:1:0:0 | 1:0:0:0 | 10:0:0:0 | 16:0:1:1 | 45:1:0:2 | 1:0:1:0 |
| 258 | 0 | 329 | 0 | 772 | 1 |
| 258 | 0 | 329 | 0 | 771 | 2 |
| 258 | 0 | 329 | 0 | 767 | 2 |
| 256 | 0 | 327 | 0 | 760 | 3 |
| 4:0:0:0:0 | 0:0:0:0:0 | 7:1:0:0:2 | 0:0:0:0:0 | 16:0:0:0:3 | 0:0:0:0:0 |
| 4:0:0:0:0 | 0:0:0:0:0 | 8:1:0:0:2 | 0:0:0:0:0 | 16:0:0:0:3 | 0:0:0:0:0 |
| 3:0:0:0 | 0:0:0:0 | 6:1:0:2 | 0:0:0:0 | 11:0:0:2 | 0:0:0:0 |
| 0:0:0:0 | 3:0:0:0 | 0:0:0:0 | 0:1:0:0 | 0:0:0:0 | 1:0:0:0 |
| 0:0:0:0:0 | 0:0:0:0:0 | 0:0:0:0:0 | 0:2:0:0:0 | 0:0:0:0:0 | 2:2:0:0:0 |
| 0:1:0:0:0 | 0:0:0:0:0 | 0:1:0:0:0 | 0:0:0:0:0 | 0:2:1:0:0 | 0:0:0:0:0 |
| 0:0:1:0:0 | 0:0:0:0:0 | 0:0:1:0:0 | 0:0:0:0:0 | 0:0:2:1:0 | 0:0:0:0:0 |
| 0:0:0:0 | 1:1:0:0 | 0:0:0:0 | 1:0:0:0 | 0:0:0:0 | 7:4:1:0 |
| 12:4:2:1:0:0:0:0 | 0:0:0:0:0:0:0:0 | 5:11:2:24:0:0:1:0 | 0:1:0:0:0:0:0:0 | 31:22:2:75:1:0:1:1 | 1:0:0:1:0:0:0:0 |
| 16 | 0 | 15 | 0 | 46 | 1 |
| 0.247268519 | 0:00:00 | 0.878587963 | 0:03:00 | 0.869641204 | 0:01:00 |
| 0:0:1:1:0:0:2:0 | 0:0:0:0:0:0:1:0 | 0:3:0:0:0:0:0:0 | 0:0:1:1:0:0:0:0 | 4:7:7:0:0:1:1:2 | 1:0:1:1:0:0:0:0 |
| 0:0:0:0 | 0:0:0:0 | 0:0:0:0 | 0:2:0:1 | 0:0:0:0 | 1:1:0:0 |

| | | | | | |
|---|---|---|---|---|---|
| 0:0:0:0:0 | 0:0:0:0:0 | 0:0:2:0:0 | 0:0:0:0:0 | 3:0:5:1:1 | 0:0:0:0:0 |
| 0:00 | 1:00 | 0:00 | 0:00 | 0:00 | 1:01 |
| 19:8:0:0:0 | 0:0:0:0:0 | 46:6:5:3:0 | 0:0:0:0:0 | 48:13:1:2:1 | 0:0:0:0:0 |
| 386:0:8:0 | 0:0:0:0 | 1112:4:7:0 | 0:0:0:0 | 1901:4:13:0 | 2:0:0:0 |
| 386:0:8:0 | 0:0:0:0 | 1054:2:7:0 | 1:0:0:0 | 1845:4:13:0 | 5:0:0:0 |
| 8:370:0:0 | 0:0:0:0 | 7:894:0:0 | 0:1:0:0 | 12:1673:2:0 | 0:3:0:0 |
| 366:0:8:0 | 0:0:0:0 | 887:0:7:0 | 0:0:0:0 | 1644:2:12:0 | 5:0:0:0 |
| 365:0:8:0 | 1:0:0:0 | 885:0:7:0 | 0:0:0:0 | 1637:2:12:0 | 2:0:0:0 |
| 365:0:8:0 | 1:0:0:0 | 883:0:7:0 | 2:0:0:0 | 1636:2:12:0 | 2:0:0:0 |
| 359:0:8:0 | 1:0:0:0 | 846:0:7:0 | 1:0:0:0 | 1558:2:12:0 | 3:0:0:0 |
| 342:0:8:0 | 2:0:0:0 | 719:0:7:0 | 3:0:0:0 | 1296:2:13:0 | 3:0:0:0 |
| 343:0:8:0 | 1:0:0:0 | 717:0:7:0 | 1:0:0:0 | 1282:2:13:0 | 5:0:0:0 |
| 0:0:0:0:0 | 0:0:0:0:0 | 0:0:0:0:0 | 0:0:0:0:0 | 0:0:0:0:0 | 0:0:0:1:0 |
| 0:0:0:0 | 0:1:1:0 | 0:1:0:0 | 0:0:0:0 | 0:0:0:0 | 2:0:0:1 |
| 17:3:0:0 | 0:0:0:0 | 25:3:2:0 | 0:0:0:0 | 56:5:0:2 | 1:1:0:0 |
| 38 | 0 | 69 | 2 | 138 | 0 |
| 0:0:0:0 | 0:0:0:0 | 0:0:0:0 | 0:0:0:0 | 0:0:0:0 | 0:0:0:2 |
| 0 | 0 | 0 | 2 | 0 | 0 |
| 3:06:00 | 0:00:00 | 3:10:00 | 7:08:00 | 2:21:01 | 1:00:00 |
| 0:0:0:0:0:0:0:0:0<br>:0:0:0:0:0:0:0:0:<br>0:0:0:0 | 13:3:1:0:1:0:0<br>:0:0:1:0:0:0:1:<br>0:0:1:0:0:0:0 | 0:0:0:0:0:0:0:0:0:0:<br>0:0:0:0:0:0:0:0:0:0 | 7:2:0:0:1:0:0:0:0:<br>2:0:0:1:1:0:0:0:0:<br>0:1:0 | 0:0:0:0:0:0:0:0:0:0<br>:0:0:0:0:0:0:0:0:0:0 | 19:5:0:1:4:0:0:0:3:0:<br>0:0:0:1:0:0:2:0:1:0:1 |
| 1:0:0:0 | 2:4:1:0 | 2:2:1:0 | 2:6:0:0 | 1:0:0:0 | 8:13:0:0 |
| 3:00 | 0:00 | 4:00 | 2:00 | 9:00 | 1:00 |
| 27:0:1:1:0:0:0:2:<br>1:1:0 | 0:0:0:0:0:0:0:<br>0:0:0:0 | 39:1:2:3:1:0:1:0:0:0:<br>0 | 0:0:0:0:0:0:0:0:0:<br>0:0 | 88:7:7:0:3:1:3:0:2:0:<br>1 | 0:0:0:0:0:0:0:0:0:0:0 |
| 1 | 0 | 4 | 3 | 5 | 1 |
| 0:00:00 | 0:00:00 | 0:01:00 | 0:00:00 | 0:00:01 | 0:00:00 |
| 100:02:00 | 0:00 | 306:31:00 | 0:00 | 466:53:00 | 1:00 |
| 0:2:0:0 | 0:0:0:0 | 9:24:0:0 | 0:0:0:0 | 2:9:0:1 | 0:0:0:0 |
| 0:2:0:0 | 0:0:0:0 | 5:25:0:0 | 0:0:0:0 | 2:9:0:1 | 0:0:0:0 |
| 4:02:00 | 0:01:00 | 4:00:00 | 3:00:00 | 13:06:02 | 1:01:00 |
| 0:0:0:0:0 | 4:4:0:0:0 | 0:0:0:0:0 | 7:1:0:0:1 | 0:0:0:0:0 | 14:11:5:0:0 |
| 107:00:00 | 0:00 | 352:11:00 | 0:00 | 507:22:00 | 2:00 |
| 0 | 0 | 0 | 0 | 0 | 6 |
| 0:00 | 0:00 | 0:00 | 0:00 | 0:00 | 0:00 |
| 8573:110:75 | 10:02:00 | 18509:380:118 | 31:03:00 | 30314:700:252 | 43:06:00 |
| 8562:110:75 | 12:03:00 | 18460:387:118 | 34:03:00 | 30230:712:251 | 62:04:00 |
| 8457:111:75 | 15:02:00 | 17979:424:118 | 49:03:00 | 29560:742:250 | 88:02:00 |
| 8389:115:74 | 13:02:01 | 17702:426:118 | 77:01:00 | 29172:759:250 | 109:05:00 |
| 8349:121:75 | 17:03:00 | 17526:446:118 | 83:02:00 | 28870:793:250 | 102:03:00 |
| 8197:128:75 | 17:02:00 | 17031:485:118 | 76:05:00 | 27978:835:251 | 98:03:00 |
| 8059:135:75 | 21:02:00 | 16579:522:119 | 76:04:00 | 27242:867:250 | 124:03:00 |
| 7989:132:75 | 25:01:00 | 16400:525:119 | 60:02:00 | 26965:869:250 | 99:04:00 |
| 7900:136:75 | 19:01:00 | 16011:525:119 | 53:06:00 | 26388:883:252 | 94:09:00 |
| 7856:139:75 | 21:00:00 | 15846:526:119 | 73:05:00 | 26035:883:252 | 114:07:00 |
| 7781:138:75 | 28:00:00 | 15618:525:119 | 60:05:00 | 25554:870:252 | 125:07:01 |
| 7720:137:75 | 29:00:00 | 15340:518:119 | 88:05:00 | 25139:861:253 | 153:08:00 |
| 7679:140:75 | 27:00:00 | 15249:516:119 | 74:02:00 | 24969:851:253 | 148:07:00 |
| 7660:138:75 | 26:00:00 | 15203:514:119 | 84:04:00 | 24882:848:253 | 155:08:00 |
| 7627:136:75 | 27:00:00 | 15108:509:120 | 84:00:00 | 24692:844:253 | 144:07:00 |
| 7638:137:75 | 26:00:00 | 15126:510:119 | 80:04:01 | 24707:846:253 | 134:08:00 |
| 7630:134:75 | 25:00:00 | 15041:506:120 | 109:02:00 | 24601:848:253 | 168:12:00 |
| 7591:131:75 | 20:00:00 | 14872:509:120 | 108:01:00 | 24364:845:252 | 188:11:01 |
| 7486:128:75 | 28:00:00 | 14436:499:120 | 119:01:00 | 23675:826:253 | 178:10:01 |
| 7417:127:75 | 32:00:00 | 14188:490:121 | 123:03:00 | 23298:800:254 | 209:10:00 |
| 7387:127:75 | 32:00:00 | 14115:487:121 | 106:02:00 | 23167:787:254 | 186:09:00 |
| 126:7301:75 | 0:21:00 | 480:13795:121 | 0.125694444 | 770:22716:254 | 0.368055556 |
| 7300:126:75 | 16:00:00 | 13763:476:121 | 60:05:00 | 22647:770:254 | 118:06:00 |
| 7269:125:75 | 23:00:00 | 13653:473:121 | 93:05:00 | 22522:762:254 | 151:07:00 |
| 7260:125:75 | 32:00:00 | 13637:472:120 | 102:05:01 | 22496:761:253 | 166:06:01 |
| 7210:121:75 | 34:00:00 | 13431:471:120 | 110:03:01 | 22232:738:255 | 192:07:00 |
| 7228:119:75 | 34:02:00 | 13434:461:120 | 139:04:01 | 22259:744:255 | 215:11:00 |
| 121:7211:75 | 0:30:00 | 457:13382:121 | 0.197916667 | 748:22188:254 | 0.502094907 |
| 7199:121:75 | 39:00:00 | 13425:460:121 | 110:03:00 | 22221:746:255 | 198:12:00 |
| 7126:118:74 | 30:00:01 | 13314:447:121 | 101:01:00 | 21950:737:255 | 166:08:00 |
| 7073:114:75 | 20:00:00 | 13267:438:121 | 58:00:00 | 21873:711:256 | 121:07:00 |
| 7022:112:75 | 20:01:00 | 13215:433:121 | 79:02:00 | 21757:710:256 | 143:06:00 |
| 6987:109:75 | 24:02:00 | 13188:427:121 | 68:01:00 | 21684:707:256 | 134:04:00 |
| 6988:109:75 | 20:02:00 | 13183:424:121 | 73:04:00 | 21690:707:256 | 120:04:00 |

| | | | | | |
|---|---|---|---|---|---|
| 0 | 0 | 3 | 3 | 0 | 3 |
| 365:14:00 | 0:00:00 | 466:39:04 | 1:00:00 | 972:76:3 | 1:00:00 |
| 363:19:00 | 1:00:00 | 461:54:04 | 1:00:00 | 936:89:2 | 4:01:00 |
| 354:27:00 | 0:00:00 | 416:84:4 | 3:02:00 | 844:147:2 | 4:00:00 |
| 354:29:00 | 0:00:00 | 412:94:4 | 3:00:00 | 827:157:2 | 3:00:00 |
| 334:34:02 | 1:00:00 | 291:104:0 | 2:00:00 | 656:158:9 | 2:00:00 |
| 337:36:02 | 0:00:00 | 292:104:0 | 2:01:00 | 639:158:9 | 0:00:00 |
| 34:3:0:0:1:0:1:0 | 0:0:1:0:0:0:0:0 | 36:5:1:1:0:1:0:0 | 1:2:1:0:0:0:0:0 | 69:16:1:0:0:0:0:0 | 0:5:1:0:0:0:0:1 |
| 37:2:1:1:1:1:1 | 0:0:0:0:0:0:0 | 54:1:1:0:0:1:1 | 4:2:1:0:0:0:0 | 91:8:4:0:0:3:1 | 2:6:0:0:0:0:1 |
| 0:00:00 | 0:00:00 | 0:00:00 | 0:00:00 | 0:00:01 | 0:00:00 |
| 0:00:00 | 0:00:00 | 0:00:00 | 0:01:00 | 0:00:00 | 4:00:02 |
| 0:0:0:0 | 0:0:0:0 | 0:0:0:0 | 0:1:0:0 | 1:0:1:1 | 0:0:0:0 |
| 0:00 | 0:00 | 0:00 | 0:00 | 0:00 | 0:00 |
| 5 | 12 | 7 | 1 | 20 | 22 |
| 25 | 3 | 33 | 4 | 128 | 4 |
| 0:0:0:0:0:0 | 0:0:0:0:0:0 | 2:0:0:0:0:0 | 0:0:0:0:0:0 | 1:2:2:0:0:1 | 0:1:1:0:0:1 |
| 0:0:0:0:0:0 | 0:0:0:0:0:0 | 0:0:0:0:0:0 | 0:0:0:0:0:1 | 0:0:0:0:0:0 | 0:0:0:0:0:0 |
| 4:19:28:9:1:0:1:0:0:0:1:0 | 0:1:1:0:0:0:0:0:0:0:0:0 | 11:23:42:28:0:2:3:0:3:1:0:1 | 0:0:0:0:0:0:0:0:0:0:0 | 12:43:38:24:0:0:7:1:1:0:3:0 | 0:0:0:0:0:0:0:0:0:0:0:0 |
| 3:5:5:4:1:0:0:0:0:0:0:0 | 1:15:24:5:0:0:1:0:0:0:1:0 | 11:23:41:28:0:2:3:0:3:1:0:1 | 0:0:1:0:0:1:0:0:0:0:0:0 | 1:14:10:9:0:0:1:1:0:0:0:0 | 11:29:28:15:0:0:6:0:1:0:3:0 |
| 3:20:29:9:1:0:1:0:0:0:1:0 | 0:0:0:0:0:0:0:0:0:0:0:0 | 11:23:42:28:0:3:3:0:3:1:0:1 | 0:0:0:0:0:0:0:0:0:0:0 | 11:42:38:24:0:0:7:1:1:0:3:0 | 1:1:0:0:0:0:0:0:0:0:0:0 |
| 8:228:2:0 | 0:0:0:0 | 29:653:1:1 | 0:1:0:0 | 22:751:6:0 | 0:1:0:0 |
| 9:228:5:0 | 0:0:0:0 | 28:660:3:1 | 1:0:0:0 | 22:755:10:0 | 0:1:0:0 |
| 6:228:6:0 | 0:0:0:0 | 17:660:21:1 | 0:0:0:0 | 16:756:22:0 | 0:0:0:0 |
| 3:228:5:0 | 0:0:0:0 | 5:658:20:1 | 0:2:0:0 | 7:755:23:0 | 0:1:0:0 |
| 3:228:5:0 | 0:0:0:0 | 3:659:20:1 | 2:1:0:0 | 7:756:23:0 | 0:0:0:0 |
| 0:00 | 0:01 | 3:00 | 1:00 | 2:00 | 2:01 |
| 0:0:0:0:0:0:1:0:0:0 | 0:0:0:0:0:0:0:0:0 | 0:0:0:0:0:0:0:0:0:0 | 0:0:0:0:0:0:0:0:0 | 0:1:1:0:0:1:0:2:1:1 | 0:0:0:0:0:0:0:0:0:0 |
| 0:0:0:0:0:0:0:0 | 0:0:0:1:0:0:0:0 | 0:0:0:0:0:0:0:0 | 0:0:0:0:0:0:0:0 | 0:1:0:0:0:1:0:1 | 0:0:0:0:1:0:1:0 |
| 0 | 1 | 3 | 1 | 4 | 1 |
| 1 | 3 | 3 | 0 | 16 | 6 |
| 0:00 | 0:00 | 0:00 | 0:00 | 5:00 | 0:00 |
| 8:1:1:0 | 0:0:0:0 | 2:0:4:2 | 0:0:0:0 | 22:2:6:2 | 0:0:0:0 |
| 9:31:00 | 6:04:00 | 1:06:04 | 0:05:01 | 2:47:03 | 2:21:02 |
| 15:45:00 | 0:00:00 | 0.138912037 | 0:01:00 | 0.302835648 | 0:02:00 |
| 94:63:10:25:4 | 1:0:0:0:0 | 0:9:0:54:1 | 0:0:0:1:0 | 121:117:14:173:4 | 0:1:0:1:0 |
| 97:66:9:26:5 | 0:0:0:0:0 | 1:13:0:56:1 | 0:0:0:0:0 | 128:126:14:181:5 | 0:0:0:1:0 |
| 102:67:9:27:5 | 0:0:0:0:0 | 1:15:0:59:1 | 0:0:0:0:0 | 134:136:16:183:5 | 0:0:0:0:0 |
| 113:72:10:22:5 | 0:0:0:1:0 | 2:28:1:51:1 | 0:0:0:0:0 | 162:178:21:154:6 | 0:0:0:1:0 |
| 22:113:72:10:5 | 1:0:0:0:0 | 50:2:28:1:1 | 1:0:0:0:0 | 150:162:178:21:6 | 4:0:0:0:0 |
| 133:84:9:11:5 | 0:0:0:0:0 | 12:42:3:41:2 | 0:0:0:0:0 | 233:269:27:82:7 | 0:2:0:0:0 |
| 143:88:11:0:0 | 0:0:0:0:0 | 18:51:38:0:0 | 0:1:1:0:0 | 263:324:73:2:0 | 0:1:1:0:0 |
| 147:93:22:11:4 | 0:0:0:0:0 | 20:71:19:57:4 | 0:0:0:0:0 | 279:405:121:81:6 | 0:3:0:0:0 |
| 147:93:22:12:4 | 0:0:0:1:0 | 20:71:19:57:4 | 0:0:0:0:0 | 279:407:121:83:6 | 0:1:0:0:0 |
| 147:91:22:22:4 | 0:0:0:0:0 | 20:68:19:76:3 | 0:0:0:0:0 | 278:390:119:147:6 | 1:1:0:1:0 |
| 147:79:22:31:4 | 0:0:0:0:0 | 20:72:19:147:3 | 0:0:0:3:0 | 279:357:109:399:6 | 0:3:0:4:0 |
| 0 | 2 | 0 | 1 | 0 | 6 |
| 0:3:0:1 | 0:0:0:0 | 0:3:0:0 | 0:0:0:0 | 1:21:1:0 | 0:0:0:0 |
| 4:2:0:2:1:1:0:0:0:0:0:1 | 0:0:0:0:0:0:0:0:0:0:0 | 16:5:0:2:1:1:0:0:0:0:0:0 | 0:0:0:0:0:0:0:0:0:0:0 | 70:20:3:9:2:7:0:3:0:0:0:0 | 1:1:0:0:0:0:0:0:0:0:0:0 |
| 0:0:0:2:1:0:1 | 0:0:0:0:0:0:0 | 0:0:0:1:0:1:0 | 0:0:1:2:0:0:0 | 1:1:0:2:0:0:0 | 2:1:0:0:2:0:0 |
| 0 | 0 | 0 | 0 | 6 | 0 |
| 7 | 0 | 6 | 0 | 35 | 0 |
| 0 | 0 | 0 | 0 | 1 | 0 |
| 0 | 0 | 0 | 0 | 1 | 0 |

| | | | | | |
|---|---|---|---|---|---|
| 0 | 0 | 0 | 0 | 1 | 0 |
| 0:00 | 0:00 | 0:00 | 0:00 | 0:00 | 1:01 |
| 45 | 0 | 109 | 0 | 227 | 0 |
| 39 | 0 | 72 | 1 | 159 | 2 |
| 40 | 0 | 72 | 3 | 160 | 0 |
| 12 | 1 | 17 | 1 | 48 | 4 |
| 9 | 2 | 10 | 6 | 31 | 13 |
| 0:0:0:0 | 2:1:0:0 | 0:0:0:0 | 1:1:0:0 | 0:0:0:0 | 7:9:0:1 |
| 2 | 0 | 4 | 4 | 12 | 2 |
| 0 | 4 | 0 | 6 | 0 | 6 |
| 0:00 | 0:01 | 0:02 | 1:00 | 2:02 | 2:02 |
| 0:0:0:0 | 0:0:0:0 | 0:0:0:1 | 0:0:0:0 | 0:1:0:0 | 0:1:0:0 |
| 9:3:132:25:19:5:0:0 | 0:0:0:0:0:0:0:0 | 17:8:478:25:8:5:0:0 | 0:0:0:0:0:0:0:0 | 37:28:998:91:63:17:1:1 | 0:0:2:1:0:0:0:0 |
| 9:4:135:10:20:27:20:19:0 | 0:0:0:0:0:0:0:0:0 | 21:10:485:4:57:25:15:7:0 | 0:0:2:0:0:0:0:0:0 | 46:33:994:42:177:92:66:65:1 | 0:0:2:0:0:0:0:0:0 |
| 14:01:02 | 0:00:00 | 28:05:03 | 0:00:00 | 120:16:10 | 0:00:00 |
| 0 | 1 | 0 | 2 | 0 | 1 |
| 0:00 | 0:00 | 1:00 | 0:00 | 1:03 | 1:02 |
| 0 | 0 | 4 | 0 | 0 | 2 |
| 0:00 | 0:00 | 1:00 | 2:00 | 1:01 | 1:00 |
| 7 | 0 | 1 | 0 | 8 | 0 |
| 0:00 | 1:00 | 0:00 | 2:00 | 0:00 | 4:01 |
| 0:00 | 1:00 | 0:00 | 2:00 | 0:00 | 5:01 |
| 8:1:1:0 | 0:0:0:0 | 5:2:1:1 | 0:0:0:0 | 9:2:1:1 | 1:0:0:0 |
| 21:19 | 0:00 | 37:11:00 | 0:00 | 102:45:00 | 0:01 |
| 0:0:0:0 | 0:2:0:0 | 0:0:0:0 | 0:1:2:0 | 0:0:0:0 | 0:1:0:1 |
| 0:00 | 0:00 | 0:00 | 0:00 | 1:01 | 1:01 |
| 10:01:01 | 0:00:00 | 27:06:03 | 0:00:00 | 99:23:05 | 1:02:00 |
| 6:01:01 | 0:00:00 | 23:06:02 | 0:00:01 | 87:23:05 | 2:01:00 |
| 6:01:01 | 0:00:00 | 25:06:02 | 0:00:00 | 93:22:05 | 1:00:00 |
| 6:01:01 | 0:00:00 | 38:06:01 | 0:00:00 | 131:18:03 | 2:00:00 |
| 6:01:01 | 0:00:00 | 37:06:01 | 0:00:00 | 131:17:03 | 3:00:00 |
| 6:01:01 | 0:00:00 | 37:06:01 | 0:00:00 | 129:17:03 | 4:00:00 |
| 6:01:01 | 0:00:00 | 37:06:01 | 0:00:00 | 130:17:03 | 3:00:00 |
| 1:00:00 | 7:00:00 | 5:01:00 | 3:00:00 | 18:03:02 | 61:04:10 |
| 1:08:00 | 0:00:00 | 2:07:00 | 0:00:00 | 0.843171296 | 0:00:00 |
| 0:00:00 | 0:00:00 | 0:00:00 | 0:00:00 | 0:00:01 | 1:02:00 |
| 0:00:00 | 0:00:00 | 9:02:00 | 0:00:00 | 20:00:00 | 1:00:00 |
| 0:00:00 | 0:00:00 | 9:02:00 | 0:00:00 | 20:00:00 | 1:00:00 |
| 0:00 | 0:00 | 13:01 | 0:00 | 27:00:00 | 0:00 |
| 0:00 | 0:00 | 10:00 | 1:00 | 13:00 | 7:00 |
| 1:0:0:0 | 0:0:0:0 | 9:0:0:0 | 2:0:0:0 | 14:0:0:0 | 2:0:0:0 |
| 2:0:0:0:0 | 0:0:0:0:0 | 2:0:0:0:1 | 0:0:0:0:0 | 4:0:0:0:0 | 0:0:0:0:0 |
| 2:0:0:0 | 0:0:0:0 | 4:0:0:0 | 5:0:0:0 | 11:0:1:1 | 5:0:0:0 |
| 2:0:0:0 | 0:0:0:0 | 5:0:0:0 | 4:0:0:0 | 12:0:1:1 | 4:0:0:0 |
| 10:1:0:0:0 | 0:0:0:0:0 | 56:1:1:0:0 | 4:1:0:0:0 | 81:4:3:3:0 | 4:0:0:1:1 |
| 10:1:0:0:0 | 0:0:0:0:0 | 64:2:1:0:0 | 0:0:0:0:0 | 87:4:3:4:1 | 0:0:0:0:0 |
| 16:1:0:0:0:1:0 | 0:0:0:0:0:0:0 | 68:2:1:1:1:0:0 | 0:0:0:0:0:0:0 | 95:5:3:0:0:0:1 | 0:0:0:0:0:0:0 |
| 0:00 | 0:00 | 0:00 | 0:00 | 0:00 | 2:06 |
| 0 | 2 | 0 | 17 | 0 | 11 |
| 19:18:2:2:0:0:0 | 0:0:0:0:0:0:0 | 11:29:17:45:7:0:1 | 0:0:0:0:0:0:0 | 52:95:77:102:9:0:3 | 0:0:1:1:0:0:0 |
| 18:16:2:2:0:0:0 | 1:1:0:0:0:0:0 | 9:28:16:48:7:0:1 | 2:0:1:1:0:0:0 | 48:86:78:121:9:0:2 | 1:5:1:0:0:0:1 |
| 2:00:00 | 8:00:01 | 4:00:00 | 10:01:00 | 19:01:01 | 43:00:02 |
| 2:00:00 | 10:00:01 | 9:00:00 | 12:00:00 | 23:01:01 | 64:00:02 |
| 2:00:00 | 10:00:01 | 9:00:00 | 12:00:00 | 23:01:01 | 64:00:02 |
| 12:00:01 | 0:00:00 | 20:02:00 | 0:00:00 | 81:10:03 | 2:00:00 |
| 12:00:01 | 0:00:00 | 20:02:00 | 0:00:00 | 78:10:03 | 1:00:00 |
| 11:00:01 | 0:00:00 | 4:00:00 | 8:01:00 | 28:07:00 | 5:01:00 |
| 0 | 0 | 0 | 0 | 1 | 2 |
| 0:00 | 3:02 | 0:00 | 1:00 | 0:00 | 8:01 |
| 0:00:00 | 3:00:02 | 0:00:00 | 1:00:00 | 0:00:00 | 8:00:01 |
| 0:0:0:0 | 0:0:0:0 | 0:0:0:0 | 0:0:0:0 | 0:0:0:0 | 1:1:1:1 |
| 0 | 2 | 0 | 3 | 0 | 3 |
| 0:00 | 0:00 | 0:00 | 0:00 | 0:00 | 1:01 |
| 1:0:2:0:0:0 | 0:0:0:0:0:0 | 1:0:3:0:1:0 | 0:0:0:0:0:0 | 1:0:5:0:1:0 | 0:0:0:0:0:0 |
| 0:00 | 0:00 | 0:00 | 0:02 | 0:00 | 0:04 |
| 1333:92:23 | 2:00:00 | 3062:144:33 | 6:02:00 | 5166:281:71 | 5:00:00 |
| 1341:92:23 | 2:00:00 | 3081:146:34 | 4:00:00 | 5195:280:71 | 8:00:00 |
| 1343:92:23 | 3:00:00 | 3086:146:34 | 3:00:00 | 5217:280:71 | 10:00:00 |
| 1347:92 | 0:00 | 3086:145 | 3:01 | 5217:277 | 2:00 |
| 1321:92:23 | 2:00:00 | 3011:146:32 | 3:00:00 | 5107:275:71 | 8:00:00 |
| 2:01:00 | 0:00:00 | 1:03:00 | 0:00:00 | 4:10:00 | 0:02:00 |
| 4:00 | 0:00 | 4:01 | 3:00 | 3:00 | 5:00 |
| 4:0:0:0 | 0:0:0:0 | 7:0:0:0 | 0:0:0:0 | 7:0:0:1 | 0:0:0:0 |
| 39:00:00 | 1:01 | 26:01:00 | 0:00 | 119:02:00 | 0:00 |
| 936 | 1 | 1545 | 11 | 5269 | 47 |
| 917 | 6 | 1507 | 11 | 5148 | 47 |
| 908 | 3 | 1463 | 11 | 5043 | 49 |

| | | | | | |
|---|---|---|---|---|---|
| 650 | 0 | 503 | 5 | 1973 | 14 |
| 661 | 2 | 501 | 1 | 1999 | 11 |
| 699 | 1 | 559 | 3 | 2172 | 15 |
| 735 | 0 | 608 | 3 | 2355 | 11 |
| 740 | 1 | 627 | 2 | 2405 | 7 |
| 743 | 3 | 638 | 7 | 2393 | 14 |
| 757 | 0 | 661 | 5 | 2428 | 13 |
| 756 | 3 | 670 | 6 | 2460 | 14 |
| 748 | 5 | 659 | 5 | 2454 | 10 |
| 769 | 2 | 696 | 3 | 2587 | 18 |
| 823 | 2 | 802 | 5 | 2882 | 14 |
| 826 | 0 | 816 | 3 | 2913 | 9 |
| 837 | 1 | 848 | 2 | 2998 | 6 |
| 852 | 0 | 892 | 5 | 3093 | 10 |
| 844 | 3 | 893 | 1 | 3077 | 6 |
| 0 | 0 | 1 | 0 | 0 | 0 |
| 33:1:0:1 | 1:0:0:0 | 75:0:2:2 | 1:1:0:0 | 210:9:8:8 | 0:0:0:0 |
| 0 | 1 | 1 | 1 | 0 | 3 |
| 0:00 | 0:00 | 3:00 | 0:00 | 1:00 | 0:00 |
| 0:02 | 0:01 | 0:00 | 0:01 | 1:01 | 0:00 |
| 0:01:01 | 0:00:00 | 1:00:01 | 0:00:00 | 5:02:01 | 0:00:00 |
| 0:00 | 0:00 | 0:02 | 0:02 | 0:00 | 1:01 |
| 0:00 | 0:00 | 0:00 | 0:05 | 0:00 | 1:01 |
| 0:00 | 0:00 | 0:00 | 0:05 | 0:00 | 1:01 |
| 0:00 | 0:00 | 0:00 | 0:05 | 0:00 | 1:01 |
| 0 | 0 | 6 | 0 | 1 | 0 |
| 1:1:0:0:0 | 0:0:0:0:0 | 0:0:0:0:0 | 0:0:0:0:0 | 2:1:1:1:1 | 0:0:0:0:0 |
| 0:00 | 0:00 | 0:00 | 0:00 | 0:00 | 0:00 |
| 19:00:01 | 0:00:00 | 3:00:01 | 0:00:00 | 22:00:02 | 0:00:00 |
| 0:0:0:0 | 23:1:3:1 | 0:0:0:0 | 23:1:1:0 | 1:0:0:0 | 111:12:2:4 |
| 23:2:0:1 | 0:0:0:0 | 24:1:0:0 | 0:0:0:0 | 106:3:0:3 | 0:0:2:0 |
| 0:0:0:0:0 | 21:1:2:0:1 | 0:0:0:0:0 | 12:0:1:1:0 | 0:0:0:0:0 | 84:1:3:4:3 |
| 0:00:00 | 0:00:00 | 0:00:00 | 0:00:00 | 0:00:01 | 0:02:00 |
| 1 | 0 | 1 | 2 | 2 | 2 |
| 39:03:01 | 0:00:00 | 73:08:00 | 0:00:00 | 186:10:00 | 1:00:00 |
| 0 | 13 | 0 | 2 | 7 | 13 |
| 0 | 0 | 1 | 0 | 0 | 1 |
| 1 | 0 | 1 | 2 | 5 | 4 |
| 0 | 0 | 0 | 2 | 0 | 2 |
| 0 | 0 | 1 | 0 | 2 | 0 |
| 0:0:0:0 | 0:0:1:1 | 0:0:0:0 | 0:0:1:0 | 1:0:0:0 | 0:0:0:0 |
| 67:0:4:12:2:0:0:0:0 | 0:0:0:0:0:0:0:0:0 | 185:1:26:17:5:2:3:2:0 | 1:0:0:0:0:0:0:0:0 | 403:2:84:30:15:3:9:1:1 | 0:0:0:0:0:0:0:0:0 |
| 67:1:7:11:2:0:0:0:0 | 0:0:0:0:0:0:0:0:0 | 182:0:33:17:5:0:3:2:0 | 2:0:0:0:0:0:0:0:0 | 398:3:103:32:15:1:9:1:1 | 2:0:0:0:0:0:0:0:0 |
| 12:01:01 | 0:00:00 | 10:03:03 | 0:00:00 | 71:11:15 | 0:01:01 |
| 0:00 | 2:00 | 0:00 | 1:01 | 0:00 | 4:02 |
| 0:00 | 2:00 | 0:00 | 1:01 | 0:00 | 4:02 |
| 0:00 | 2:00 | 0:00 | 1:01 | 0:00 | 4:00 |
| 0:00 | 2:00 | 0:00 | 1:01 | 0:00 | 4:00 |
| 4:00:00 | 0:00:00 | 5:00:00 | 0:00:00 | 17:01:01 | 0:00:00 |
| 0:00 | 1:00 | 2:02 | 2:06 | 1:00 | 3:02 |
| 1:01:02 | 0:00:00 | 0:00:00 | 0:00:00 | 0:00:00 | 0:00:01 |
| 1:01:01 | 0:00:00 | 0:00:00 | 0:00:00 | 0:00:00 | 0:00:01 |
| 1:5:1:0 | 0:0:0:0 | 1:4:1:1 | 1:2:0:0 | 0:2:0:1 | 1:0:0:0 |
| 0:00 | 0:00 | 2:02 | 0:00 | 8:02 | 0:00 |
| 0:3:0:0:0:0:0:0:0:0:0:0 | 0:0:0:0:0:0:0:0:0:0:0 | 1:22:0:0:2:4:0:1:0:0:4:0 | 0:0:0:0:0:0:0:0:0:0:0 | 0:7:0:0:0:0:0:0:0:0:1:0 | 0:0:0:0:0:0:0:0:0:0:0:0 |
| 3:0:0:0:0:0:0:0:0:0:0 | 0:0:0:0:0:0:0:0:0:0:0 | 23:1:0:2:0:4:0:1:0:0:4:0 | 1:0:0:0:0:0:0:0:0:0:0 | 7:0:0:0:0:0:0:0:0:1:0 | 3:0:0:0:0:0:0:0:0:0:0 |
| 156:01:00 | 0:00 | 188:01:00 | 1:00 | 562:03:00 | 2:00 |
| 156:01:00 | 0:00 | 183:01:00 | 0:00 | 548:04:00 | 1:00 |
| 156:01:00 | 0:00 | 182:01:00 | 1:00 | 548:04:00 | 1:00 |
| 154:00:00 | 0:00 | 179:02:00 | 0:00 | 539:08:00 | 0:00 |
| 152:01:00 | 0:00 | 174:02:00 | 1:00 | 536:09:00 | 2:00 |
| 152:01:00 | 0:00 | 164:03:00 | 0:00 | 509:09:00 | 0:00 |
| 151:01:00 | 0:00 | 163:03:00 | 0:00 | 504:10:00 | 1:00 |
| 0:00 | 0:00 | 0:01 | 3:02 | 0:00 | 1:01 |
| 0:00 | 0:00 | 0:02 | 3:01 | 0:01 | 1:00 |
| 0:00:00 | 0:00:00 | 0:03:01 | 3:00:00 | 0:02:00 | 1:00:00 |
| 0:00 | 0:00 | 0:01 | 2:02 | 0:01 | 1:01 |
| 1:00 | 0:00 | 1:00 | 0:00 | 0:01 | 0:00 |
| 1 | 0 | 1 | 0 | 0 | 0 |
| 0 | 0 | 0 | 0 | 4 | 3 |
| 0 | 0 | 0 | 0 | 0 | 0 |
| 0:00 | 0:00 | 0:00 | 0:00 | 0:00 | 0:01 |
| 0 | 0 | 0 | 0 | 0 | 0 |
| 0:00 | 3:00 | 2:00 | 1:00 | 2:01 | 0:00 |
| 2:00 | 0:00 | 0:00 | 0:00 | 6:01 | 0:00 |

| | | | | | |
|---|---|---|---|---|---|
| 2:00:00 | 1:00:00 | 0:00:00 | 0:00:00 | 3:00:01 | 6:02:00 |
| 0 | 0 | 0 | 0 | 0 | 3 |
| 2:0:0:0:0:0:0:1:0 | 0:0:0:0:0:0:0:0:0 | 1:0:0:0:2:0:0:0:0 | 0:0:0:0:0:0:0:0:0 | 5:0:0:1:3:0:0:0:0 | 8:3:1:1:2:0:1:4:1 |
| 0 | 0 | 0 | 0 | 0 | 1 |
| 0:00:00 | 0:00:00 | 0:00:00 | 1:00:00 | 0:00:01 | 4:01:00 |
| 0:00:00 | 0:00:00 | 0:00:00 | 1:00:00 | 0:00:00 | 5:00:01 |
| 0:0:0:0:0 | 15:1:1:1:0 | 0:0:0:0:0 | 1:2:0:0:0 | 0:0:0:0:0 | 22:8:3:3:1 |
| 0:01 | 0:02 | 0:00 | 2:03 | 3:01 | 1:09 |
| 0 | 0 | 0 | 0 | 0 | 0 |
| 0 | 0 | 0 | 0 | 0 | 0 |
| 0 | 0 | 0 | 0 | 0 | 0 |
| 0:00 | 2:00 | 2:01 | 1:00 | 1:00 | 5:01 |
| 8:0:0:0:0:0:0 | 4:0:0:0:0:0:0 | 4:0:0:1:0:0:0 | 4:0:0:0:0:0:0 | 30:0:0:0:1:1:2 | 16:0:0:0:0:0:0 |
| 0:00 | 0:00 | 0:00 | 0:01 | 1:00 | 1:00 |
| 0:00 | 0:00 | 1:00 | 0:00 | 2:01 | 2:00 |
| 0:00 | 0:00 | 0:00 | 0:00 | 0:00 | 0:00 |
| 0:00 | 0:00 | 0:01 | 0:00 | 0:00 | 0:00 |
| 0:00 | 0:00 | 0:01 | 0:00 | 0:00 | 0:00 |
| 0:00 | 0:00 | 0:01 | 0:00 | 0:00 | 0:00 |
| 0:0:0:0:0:1 | 0:0:0:0:0:0 | 3:0:1:0:0:0 | 0:0:0:0:0:0 | 1:0:0:0:0:0 | 0:0:0:0:0:0 |
| 10:0:1:0 | 0:0:1:0 | 9:1:13:0 | 0:0:2:0 | 32:9:12:2 | 0:0:1:0 |
| 10:0:1:0 | 0:0:1:0 | 9:1:11:0 | 0:0:3:0 | 32:9:9:2 | 0:0:5:0 |
| 9:0:1:0 | 1:0:1:0 | 9:1:8:0 | 0:0:6:0 | 32:9:8:2 | 0:0:5:0 |
| 0:0:5:0 | 0:1:6:0 | 0:3:7:0 | 1:2:3:0 | 3:2:10:1 | 7:6:25:1 |
| 406:01:00 | 1:00 | 615:03:00 | 2:00 | 243:00:00 | 0:00 |
| 407:01:00 | 0:00 | 613:03:00 | 3:00 | 243:00:00 | 1:00 |
| 404:01:00 | 2:00 | 612:03:00 | 0:00 | 245:00:00 | 1:00 |
| 406:01:00 | 0:00 | 610:03:00 | 0:00 | 245:00:00 | 1:00 |
| 401:01:00 | 0:00 | 575:03:00 | 3:00 | 242:00:00 | 0:00 |
| 398:01:00 | 1:00 | 574:03:00 | 4:00 | 241:00:00 | 0:00 |
| 9:1:4:1 | 33:1:15:0 | 4:0:1:0 | 7:1:1:0 | 26:1:6:1 | 40:11:25:3 |
| 0 | 0 | 0 | 0 | 0 | 2 |
| 5:9:1:1:0:0:1:1 | 0:0:0:0:0:0:0:0 | 13:20:1:0:0:0:0:0 | 0:0:0:0:0:0:0:0 | 59:30:2:0:0:2:3:0 | 0:0:0:0:0:0:0:0 |
| 0 | 0 | 0 | 0 | 0 | 2 |
| 0 | 0 | 0 | 0 | 0 | 2 |
| 0:00 | 1:00 | 0:00 | 1:00 | 0:00 | 2:01 |
| 198:44:0:0:1 | 1:1:0:0:0 | 205:15:0:0:0 | 1:0:0:0:0 | 363:46:1:0:0 | 2:0:0:0:0 |
| 220:49:0:0:1:0:0 | 0:1:0:0:0:0:0 | 578:41:0:0:1:0:0 | 10:2:0:0:0:0:0 | 1057:96:1:0:1:0:1 | 11:1:0:0:0:1:0 |
| 233:52:0:0:1:0:0 | 1:1:0:0:0:0:0 | 696:62:2:0:1:0:0 | 24:6:0:0:0:0:0 | 1210:121:1:1:1:1:1 | 34:4:0:0:2:0:0 |
| 234:52:0:0:1:0:0 | 0:1:0:0:0:0:0 | 717:66:2:0:1:0:0 | 4:2:0:0:0:0:0 | 1241:126:1:1:3:1:1 | 4:1:0:0:0:0:0 |
| 235:53:0:0:1:0:0 | 0:0:0:0:0:0:0 | 725:69:2:0:1:0:0 | 1:0:0:0:0:0:0 | 1249:126:1:1:3:1:1 | 3:1:0:0:0:0:0 |
| 235:53:0:0:1:0:0 | 0:0:0:0:0:0:0 | 734:70:2:0:1:0:0 | 1:0:0:0:0:0:0 | 1248:130:1:1:3:1:1 | 2:0:0:0:0:0:0 |
| 234:53:0:0:1:0:0 | 2:0:0:0:0:0:0 | 699:70:2:0:1:0:0 | 1:0:0:0:0:0:0 | 1171:130:1:1:3:1:1 | 8:1:0:0:0:0:0 |
| 234:53:0:0:1:0:0 | 2:0:0:0:0:0:0 | 697:70:2:0:1:0:0 | 3:0:0:0:0:0:0 | 1168:130:1:1:3:1:1 | 5:1:0:0:0:0:0 |
| 239:53:0:0:1:0:0 | 3:0:0:0:0:0:0 | 677:71:2:0:1:0:0 | 4:0:0:0:0:0:0 | 1106:131:1:1:3:1:1 | 4:3:0:0:0:0:0 |
| 240:52:0:0:1:0:0 | 1:1:0:0:0:0:0 | 655:71:2:0:1:0:0 | 4:0:0:0:0:0:0 | 1077:134:1:1:3:1:1 | 6:0:0:0:0:0:0 |
| 239:53:0:0:1:0:0 | 0:0:0:0:0:0:0 | 633:71:2:0:1:0:0 | 1:0:0:0:0:0:0 | 1036:134:1:1:3:1:1 | 3:0:0:0:0:0:0 |
| 239:53:0:0:1:0:0 | 0:0:0:0:0:0:0 | 602:71:2:0:1:0:0 | 0:0:0:0:0:0:0 | 996:136:1:1:3:1:1 | 1:0:0:0:0:0:0 |

| | | | | | |
|---|---|---|---|---|---|
| 234:53:0:1 | 0:0:0:0 | 469:71:2:1 | 2:0:0:0 | 792:136:1:3 | 4:1:0:0 |
| 231:53:0:0:1:0 | 2:0:0:0:0:0 | 452:70:2:0:1:0 | 2:1:0:0:0:0 | 763:136:1:1:3:1 | 1:1:0:0:0:0 |
| 15:3:0:0:1 | 0:0:0:1:0 | 3:5:0:1:0 | 0:0:0:0:0 | 23:26:5:6:0 | 2:0:0:0:0 |
| 0 | 0 | 0 | 0 | 0 | 0 |
| 116:01:00 | 0:00 | 209:01:00 | 1:00 | 595:11:00 | 1:00 |
| 116:00:00 | 0:00 | 204:02:00 | 0:00 | 571:08:00 | 2:00 |
| 116:00:00 | 0:00 | 203:02:00 | 1:00 | 572:08:00 | 1:00 |
| 0 | 1 | 0 | 1 | 0 | 7 |
| 0 | 0 | 1 | 0 | 0 | 0 |
| 0:0:0:0:0 | 0:3:0:0:1 | 0:0:0:0:0 | 0:4:1:0:0 | 0:0:0:0:0 | 0:4:2:2:0 |
| 0:2:0:0:0 | 0:1:0:0:1 | 0:2:0:0:0 | 0:1:1:0:0 | 0:3:2:2:0 | 0:1:0:0:0 |
| 0 | 0 | 0 | 0 | 0 | 2 |
| 79:03:00 | 0:00 | 134:35:00 | 2:00 | 398:48:00 | 0:01 |
| 79:02:00 | 0:00 | 136:29:00 | 0:00 | 401:37:00 | 0:00 |
| 36:31:1:1:0 | 0:0:0:0:0 | 12:130:0:1:0 | 0:4:0:1:0 | 53:247:2:4:1 | 0:2:0:0:0 |
| 15:31:1:1:0 | 0:0:0:0:0 | 10:133:0:5:0 | 1:0:0:0:0 | 32:246:2:7:1 | 0:3:0:1:0 |
| 31:3:1:3:0 | 0:0:0:0:0 | 153:11:0:6:0 | 1:0:0:0:0 | 310:32:2:14:1 | 2:0:0:0:0 |
| 2:30:1:2:0 | 0:0:0:0:0 | 10:154:0:3:0 | 0:0:0:0:0 | 31:316:2:13:1 | 0:2:0:0:0 |
| 62:00:00 | 2:00 | 59:00:00 | 0:00 | 131:00:00 | 0:00 |
| 1550:161 | 3:00 | 709:615 | 1:02 | 4206:2805 | 2:07 |
| 1011:110 | 541:52:00 | 376:305 | 333:313 | 2835:1854 | 1371:959 |
| 1550:162 | 3:00 | 708:616 | 4:02 | 4203:2813 | 9:01 |
| 1553:161 | 0:00 | 707:618 | 7:00 | 4200:2813 | 11:02 |
| 1555:162 | 1:01 | 717:617 | 1:02 | 4215:2821 | 6:03 |
| 1554:162 | 1:00 | 717:617 | 1:00 | 4216:2811 | 5:04 |
| 1555:162 | 1:00 | 717:615 | 1:00 | 4217:2804 | 6:04 |
| 1563:160 | 0:00 | 753:618 | 0:01 | 4346:2812 | 3:04 |
| 1562:162 | 1:00 | 753:611 | 0:00 | 4348:2788 | 0:01 |
| 1563:162 | 0:00 | 752:610 | 3:01 | 4348:2764 | 8:05 |
| 1633:165 | 0:00 | 800:602 | 0:00 | 4599:2731 | 4:05 |
| 1630:167 | 3:00 | 800:598 | 0:00 | 4598:2699 | 3:04 |
| 1632:166 | 1:01 | 798:595 | 2:01 | 4601:2689 | 1:05 |
| 1633:168 | 0:00 | 798:588 | 2:03 | 4596:2674 | 6:04 |
| 1660:168 | 2:00 | 841:588 | 2:00 | 4733:2658 | 8:01 |
| 1661:167 | 1:00 | 843:584 | 0:01 | 4730:2651 | 10:03 |
| 1832:166 | 3:01 | 1517:628 | 25:00:00 | 7719:2874 | 61:06:00 |
| 1831:164 | 4:03 | 1518:627 | 46:02:00 | 7730:2867 | 100:11:00 |
| 1834:166 | 22:03 | 1528:623 | 53:07:00 | 7733:2849 | 233:32:00 |
| 1852:166 | 5:01 | 1555:625 | 27:03:00 | 7888:2866 | 80:07:00 |
| 1857:162 | 0:01 | 1577:626 | 10:02 | 7962:2858 | 14:06 |
| 1854:161 | 3:00 | 1587:612 | 0:03 | 7979:2830 | 2:03 |
| 1857:159 | 0:00 | 1586:609 | 1:02 | 7976:2820 | 4:02 |
| 1856:160 | 2:00 | 1588:597 | 1:02 | 7980:2732 | 5:03 |
| 142:1859 | 0:00 | 565:1588 | 1:01 | 2559:7984 | 6:01 |
| 1857:141 | 2:00 | 1587:564 | 0:00 | 7983:2553 | 2:06 |
| 1858:138 | 1:01 | 1588:553 | 0:02 | 7977:2492 | 7:13 |
| 1857:106 | 3:00 | 1594:391 | 1:01 | 7997:1658 | 5:07 |
| 101:1858 | 0:01 | 378:1592 | 2:03 | 1623:7994 | 6:06 |
| 1861:96 | 0:00 | 1593:348 | 2:00 | 8009:1490 | 3:01 |
| 1859:86 | 1:00 | 1594:278 | 1:00 | 7995:1196 | 8:03 |
| 1857:84 | 4:00 | 1594:260 | 1:02 | 8004:1090 | 12:04 |
| 1858:83 | 3:01 | 1592:256 | 3:02 | 8008:1055 | 8:06 |
| 1860:81 | 1:00 | 1594:221 | 2:02 | 8012:908 | 9:10 |
| 1858:78 | 3:00 | 1595:200 | 1:01 | 8018:761 | 7:08 |
| 1859:77 | 2:00 | 1598:156 | 0:04 | 8025:628 | 3:12 |
| 1861:74 | 0:01 | 1597:111 | 1:04 | 8023:477 | 8:14 |
| 1861:67 | 1:01 | 1598:105 | 0:00 | 8024:433 | 8:03 |
| 1862:38:00 | 0:04 | 1597:21:00 | 1:03 | 8026:142 | 9:15 |
| 1:11:197:3:13:2:2:3:34:0:64:6:0:3:1:0:4:0:9:0:2:0:0:0:0 | 0:0:2:0:0:0:0:0:0:0:0:0:0:0:0:0:0:0:0:0:0:0:0:0:0 | 1:13:147:1:22:4:0:5:12:0:17:4:0:5:2:1:0:4:6:0:0:0:0:0:0 | 0:0:0:0:0:0:0:0:0:0:0:0:0:0:0:0:0:0:0:0:0:0:0:0:0 | 1:22:338:14:20:9:1:11:16:0:58:5:1:3:10:0:1:5:5:2:0:1:0:1:1 | 0:0:0:0:1:0:0:0:0:0:0:0:0:0:0:0:0:0:0:0:0:0:0:0:0 |
| 11:199:2:3:2:36:0:64:6:0:3:0:2:1:0:0:0:0:0:2:0:0:0 | 0:0:0:0:0:0:0:0:0:0:0:0:0:0:0:0:0:0:0:0:0:0:0 | 13:145:4:1:0:13:0:17:4:0:5:0:2:2:1:0:5:0:0:0:0:0:0 | 0:2:0:0:0:0:0:0:0:0:0:0:0:0:0:0:0:0:0:0:0:0:0 | 22:338:10:10:1:23:0:58:5:0:2:1:4:10:0:0:6:5:2:2:1:0:1 | 0:0:0:0:0:0:0:0:0:0:0:0:0:0:0:0:0:0:0:0:0:0:0 |
| 0:0:0:0 | 0:0:1:0 | 0:0:0:0 | 0:2:0:0 | 0:0:1:0 | 1:1:0:1 |
| 0:1:0:0:0:0:0:0:0 | 0:0:0:0:0:0:0:0:0 | 0:0:2:0:0:0:0:1:1 | 0:0:0:0:0:0:0:0:0 | 2:2:3:0:0:1:0:0:1 | 0:0:0:1:0:0:0:0:0 |
| 4:3:2:2:0:1:0:0:0:2:1:1:0:0:0:0:0:0 | 0:0:0:0:0:0:0:0:0:0:0:0:0:0:0:0:0:0 | 8:2:0:7:3:0:2:1:0:3:2:3:0:0:0:0:0:0 | 0:0:0:0:0:0:0:0:0:0:0:0:0:0:0:0:0:0 | 12:12:5:11:2:4:6:2:3:13:5:4:2:2:1:0:3:1 | 0:0:0:0:0:0:1:0:0:0:0:1:0:0:0:0:0:0 |
| 0:00 | 0:00 | 0:00 | 0:00 | 0:00 | 0:00 |
| 0:00 | 0:00 | 0:00 | 0:00 | 0:00 | 1:01 |
| 73:3:1:0:1:0 | 0:1:0:0:0:0 | 133:10:2:0:0:0 | 0:2:0:0:0:0 | 242:19:3:0:1:1 | 1:1:0:0:0:0 |
| 73:2:0:0:1:0 | 0:2:1:0:0:0 | 133:4:1:0:0:0 | 0:8:1:0:0:0 | 243:16:1:0:1:1 | 0:4:2:0:0:0 |
| 0:0:0:0:1 | 0:0:0:0:0 | 1:0:0:0:0 | 0:3:0:0:0 | 2:0:4:0:1 | 2:2:2:0:0 |
| 0:0:0:0:0 | 0:0:0:0:1 | 0:0:0:0:0 | 2:3:0:0:0 | 0:0:0:0:0 | 4:2:4:0:1 |

| | | | | | |
|---|---|---|---|---|---|
| 0:01 | 0:00 | 1:00 | 0:00 | 2:01 | 2:00 |
| 0:0:0:0:0:0:0:0:0 | 22:5:1:0:11:1:9:1:0 | 0:0:0:0:0:0:0:0:0 | 3:0:0:0:2:0:2:0:0 | 0:0:0:0:0:0:0:0:0 | 19:5:0:0:9:0:5:3:2 |
| 19:5:1:0:11:1:9:1:0 | 2:0:0:0:0:0:0:0:0 | 3:0:0:0:2:0:2:0:0 | 0:0:0:0:0:0:0:0:0 | 18:5:0:0:9:0:5:3:2 | 0:0:0:0:0:0:0:0:0 |
| 2:0:0:0 | 8:0:0:0 | 0:0:0:0 | 1:0:0:0 | 3:0:0:0 | 11:0:0:0 |
| 0:0:0:0 | 0:0:0:12 | 0:0:0:0 | 0:0:0:1 | 0:0:0:0 | 0:0:0:18 |
| 1:4:0:0:0 | 1:4:1:0:0 | 0:0:0:0:0 | 0:0:0:0:0 | 0:2:0:0:0 | 0:1:0:1:1 |
| 0 | 0 | 0 | 0 | 0 | 2 |
| 0 | 0 | 0 | 0 | 0 | 2 |
| 0 | 0 | 0 | 0 | 0 | 2 |
| 0:00 | 2:01 | 0:00 | 0:00 | 0:00 | 3:02 |
| 16:0:1:0:3 | 1:0:0:0:0 | 7:26:8:1:1 | 0:0:0:0:0 | 21:82:11:1:4 | 0:3:0:0:0 |
| 17:0:1:0:3 | 0:0:0:0:0 | 7:23:8:1:1 | 0:0:0:0:0 | 21:77:11:1:4 | 1:0:0:0:0 |
| 17:0:1:0:3 | 0:0:0:0:0 | 11:19:8:1:1 | 1:0:0:0:0 | 33:50:11:1:4 | 0:1:0:0:0 |
| 0 | 0 | 0 | 0 | 0 | 0 |
| 0:00 | 0:00 | 0:00 | 0:00 | 0:00 | 1:00 |
| 0 | 0 | 4 | 6 | 2 | 2 |
| 86 | 1 | 193 | 0 | 723 | 3 |
| 85 | 0 | 174 | 2 | 670 | 1 |
| 85 | 0 | 176 | 1 | 669 | 1 |
| 78 | 0 | 143 | 0 | 508 | 0 |
| 68 | 1 | 70 | 2 | 302 | 4 |
| 68 | 1 | 71 | 1 | 305 | 1 |
| 68 | 1 | 70 | 2 | 303 | 3 |
| 69 | 0 | 70 | 2 | 307 | 0 |
| 0:00 | 0:00 | 0:00 | 0:00 | 0:00 | 0:01 |
| 0:00:00 | 0:00:00 | 0:00:01 | 0:01:00 | 0:00:00 | 0:00:00 |
| 0:00 | 0:00 | 0:00 | 1:00 | 0:01 | 0:00 |
| 0:00:00 | 0:00:00 | 0:00:00 | 0:01:00 | 0:00:00 | 1:00:01 |
| 0:00:00 | 0:00:00 | 0:00:00 | 0:01:00 | 0:00:00 | 1:00:01 |
| 0:00:00 | 0:00:00 | 0:00:00 | 0:01:00 | 0:00:00 | 1:00:02 |
| 266:5:1:2:0:0 | 0:0:0:0:0:0 | 717:15:0:10:3:0 | 0:0:0:0:0:0 | 1282:49:4:19:6:2 | 0:1:0:0:0:0 |
| 266:1:1:2:0:0 | 0:0:0:0:0:0 | 716:7:0:9:3:0 | 1:0:0:0:0:0 | 1280:34:4:19:9:0 | 3:0:0:0:0:0 |
| 262:1:2:0:2:0 | 0:0:0:0:0:0 | 700:0:9:6:5:0 | 1:0:0:0:0:0 | 1236:3:17:15:15:2 | 3:0:0:0:0:0 |
| 262:1:2:0:2:0 | 0:0:0:0:0:0 | 700:0:9:6:5:0 | 0:0:0:0:0:0 | 1232:3:16:15:15:2 | 3:0:0:0:0:0 |
| 261:2:1:0:2:0 | 1:0:0:0:0:0 | 693:5:0:7:6:0 | 4:0:0:0:0:0 | 1227:15:3:13:15:3 | 4:0:0:0:0:0 |
| 262:2:1:0:2:0 | 0:0:0:0:0:0 | 692:5:0:6:6:1 | 0:0:0:0:0:0 | 1225:15:3:14:13:3 | 2:0:0:0:0:0 |
| 262:2:1:2:0:0 | 0:0:0:0:0:0 | 686:5:0:5:2:1 | 1:0:0:0:1:0 | 1211:16:2:15:13:4 | 1:0:0:0:0:0 |
| 257:2:1:2:0:0 | 1:0:0:0:0:0 | 675:5:0:5:3:1 | 2:0:0:0:0:0 | 1192:16:2:14:12:4 | 0:0:0:0:0:0 |
| 257:2:1:2:0:0 | 0:0:0:0:0:0 | 679:5:0:5:3:1 | 1:0:0:0:0:0 | 1195:16:2:14:12:4 | 2:0:0:0:0:0 |
| 1:01 | 0:00 | 1:00 | 0:00 | 2:01 | 0:00 |
| 0:00 | 0:00 | 0:00 | 0:00 | 0:00 | 1:01 |
| 3:08:03 | 0:00:00 | 2:06:03 | 0:00:00 | 6:35:15 | 0:00:00 |
| 5:01:00 | 3:02:03 | 4:01:01 | 2:01:04 | 8:02:09 | 25:04:15 |
| 0:00:00 | 0:00:00 | 0:00:00 | 0:00:00 | 0:00:00 | 0:01:01 |
| 0 | 0 | 0 | 0 | 0 | 2 |
| 126 | 0 | 458 | 2 | 882 | 0 |
| 125 | 0 | 454 | 0 | 872 | 4 |
| 124 | 0 | 449 | 2 | 874 | 1 |
| 123 | 1 | 447 | 0 | 870 | 2 |
| 120 | 3 | 446 | 0 | 858 | 3 |
| 123 | 0 | 440 | 0 | 853 | 1 |
| 122 | 0 | 403 | 1 | 800 | 2 |
| 122 | 0 | 378 | 2 | 756 | 4 |
| 123 | 0 | 373 | 1 | 752 | 4 |
| 122 | 1 | 373 | 1 | 752 | 3 |
| 123 | 0 | 371 | 1 | 749 | 4 |
| 123 | 0 | 367 | 3 | 746 | 2 |
| 122 | 1 | 364 | 1 | 736 | 2 |
| 122 | 0 | 359 | 0 | 722 | 2 |
| 121 | 0 | 346 | 2 | 702 | 1 |
| 133 | 0 | 374 | 4 | 696 | 10 |
| 134 | 0 | 307 | 0 | 578 | 3 |
| 132 | 0 | 292 | 0 | 573 | 3 |
| 0 | 3 | 0 | 3 | 0 | 0 |
| 1 | 2 | 0 | 3 | 0 | 0 |
| 1:01 | 0:00 | 1:00 | 1:00 | 4:00 | 1:00 |

| | | | | | |
|---|---|---|---|---|---|
| 4:9:4:2 | 5:2:2:0 | 7:15:4:0 | 6:0:2:0 | 22:39:13:3 | 15:19:9:0 |
| 6:11:5:2 | 3:0:1:0 | 9:14:7:0 | 5:1:1:0 | 27:48:16:3 | 10:11:7:0 |
| 8:11:6:2 | 1:0:0:0 | 14:14:8:0 | 0:1:0:0 | 35:60:22:3 | 2:0:2:0 |
| 0:00 | 0:00 | 0:00 | 0:01 | 0:00 | 0:02 |
| 60:7:1:0:2 | 0:0:0:0:0 | 70:34:1:0:1 | 0:1:0:0:0 | 208:65:0:0:0 | 0:0:0:0:0 |
| 1:01:01 | 0:02:00 | 1:02:00 | 1:02:00 | 4:11:00 | 1:02:00 |
| 18:2:0:2:0 | 4:1:0:1:0 | 12:16:0:0:1 | 12:15:0:2:1 | 48:42:2:10:3 | 18:19:0:3:0 |
| 126:7:0:1:0:0:2:3:1 | 0:0:0:0:0:0:0:0 | 199:11:3:1:0:0:7:3:2 | 1:0:0:0:0:0:0:0:0 | 349:12:9:0:0:0:15:2:0 | 0:0:0:0:0:0:0:0:0 |
| 126:7:0:1:0:0:3:3:1 | 0:0:0:0:0:0:0:0 | 202:11:3:1:0:0:10:3:2 | 2:0:0:0:0:0:0:0:0 | 352:12:9:0:0:0:17:2:0 | 0:0:0:0:0:0:0:0:0 |
| 126:7:0:1:0:0:3:3:1 | 0:0:0:0:0:0:0:0 | 204:11:3:1:0:0:10:3:2 | 0:0:0:0:0:0:0:0:0 | 352:12:8:0:0:0:17:2:0 | 0:0:1:0:0:0:0:0:0 |
| 0:0:0:0:0:0 | 0:0:0:0:0:0 | 0:2:0:0:0:0 | 0:0:0:0:0:0 | 0:7:1:0:1:0 | 0:1:0:0:1:0 |
| 2:0:0:0:1 | 0:0:0:0:0 | 0:0:0:0:0 | 0:0:0:0:0 | 10:0:0:0:0 | 0:0:0:0:0 |
| 10:0:2:2 | 0:0:0:0 | 43:1:2:0 | 0:0:0:0 | 136:2:0:9 | 0:0:0:0 |
| 0:0:0:0 | 1:0:0:0 | 0:0:0:0 | 2:0:0:0 | 0:0:0:0 | 7:1:0:1 |
| 38:10:0:0:1 | 0:0:0:0:0 | 73:11:1:0:1 | 1:0:0:0:0 | 199:44:15:8:3 | 1:0:1:0:0 |
| 28:10:0:1:0:0 | 0:0:0:0:0:0 | 38:11:1:1:0:0 | 2:0:0:0:0:0 | 111:44:16:4:1:5 | 0:0:0:0:0:0 |
| 0:0:0:0 | 8:0:0:0 | 0:0:0:0 | 5:1:0:0 | 0:0:0:0 | 10:0:5:1 |
| 6:0:0:0 | 3:0:0:0 | 2:2:0:0 | 2:0:0:0 | 2:1:6:2 | 8:0:0:0 |
| 6:0:0:0 | 3:0:0:0 | 2:2:0:0 | 2:0:0:0 | 2:1:6:3 | 8:0:0:0 |
| 1:02 | 0:00 | 3:08 | 0:00 | 1:23 | 0:00 |
| 1:02 | 0:00 | 3:12 | 0:00 | 1:30 | 0:00 |
| 1:02 | 0:00 | 3:14 | 0:00 | 2:37 | 0:00 |
| 2:01 | 0:00 | 19:04 | 0:00 | 38:01:00 | 0:00 |
| 2:01 | 0:00 | 19:04 | 0:00 | 38:01:00 | 0:00 |
| 0:00 | 1:02 | 0:00 | 1:19 | 0:00 | 0:37 |
| 111:05:00 | 0:00 | 93:03:00 | 0:00 | 352:02:00 | 0:00 |
| 2429 | 1 | 6711 | 2 | 13309 | 10 |
| 2447 | 2 | 6775 | 6 | 13442 | 16 |
| 2448 | 1 | 6776 | 6 | 13449 | 13 |
| 1749 | 716 | 3498 | 3358 | 9040 | 4572 |
| 2473 | 1 | 6885 | 4 | 13685 | 7 |
| 2473 | 1 | 6901 | 5 | 13740 | 9 |
| 2473 | 5 | 6928 | 20 | 13769 | 46 |
| 2496 | 4 | 7133 | 2 | 14074 | 14 |
| 2506 | 3 | 7184 | 7 | 14147 | 18 |
| 2508 | 1 | 7182 | 9 | 14148 | 19 |
| 2511 | 3 | 7234 | 5 | 14262 | 12 |
| 2513 | 2 | 7255 | 7 | 14305 | 15 |
| 2516 | 0 | 7260 | 3 | 14316 | 10 |
| 2517 | 1 | 7291 | 12 | 14354 | 24 |
| 2517 | 2 | 7306 | 10 | 14371 | 17 |
| 2525 | 1 | 7414 | 5 | 14511 | 13 |
| 2524 | 2 | 7412 | 6 | 14505 | 17 |
| 2524 | 3 | 7395 | 7 | 14480 | 14 |
| 87:5:32:0:0:0 | 0:0:0:0:0:0 | 5:37:60:0:0:0 | 0:0:0:0:0:0 | 89:132:277:0:1:1 | 0:0:2:0:0:0 |
| 86:4:32:0:0:0 | 0:0:0:0:0:0 | 5:34:68:0:0:0 | 0:1:0:0:0:0 | 87:123:329:0:1:1 | 0:1:0:0:0:0 |
| 85:3:33:0:0:0 | 1:1:0:0:0:0 | 4:34:78:0:0:0 | 1:2:0:0:0:0 | 81:109:430:0:1:2 | 3:6:0:0:0:0 |
| 85:3:34:0:0:0 | 0:0:0:0:0:0 | 3:34:89:0:0:0 | 0:0:0:0:0:0 | 81:108:500:0:1:2 | 0:0:3:0:0:0 |
| 85:3:34:0:0:0 | 0:0:0:0:0:0 | 3:34:89:0:0:0 | 0:0:0:0:0:0 | 81:108:500:0:1:2 | 0:0:4:0:0:0 |
| 85:3:34:0:0:0 | 0:0:0:0:0:0 | 3:5:90:0:0:0 | 0:29:0:0:0:0 | 81:14:526:0:1:2 | 0:97:0:0:0:0 |
| 85:3:35:0:0:0 | 0:0:0:0:0:0 | 3:4:96:0:0:0 | 0:0:0:0:0:0 | 81:12:561:0:1:3 | 0:0:2:0:0:0 |
| 85:3:35:0:0:0 | 0:0:0:0:0:0 | 3:3:97:0:0:0 | 0:0:0:0:0:0 | 81:11:566:0:1:3 | 0:0:2:0:0:0 |
| 85:3:35:0:0:0 | 0:0:0:0:0:0 | 3:3:98:0:0:0 | 0:0:0:0:0:0 | 81:11:572:0:1:3 | 0:0:0:0:0:0 |
| 1:2:2:3 | 0:0:1:0 | 2:7:0:1 | 0:0:0:0 | 5:36:5:4 | 3:0:0:0 |
| 78:01:00 | 0:00:00 | 92:01:00 | 0:00:00 | 283:03:01 | 2:00:00 |
| 12:0:2:0:0:0 | 0:0:0:0:0:0 | 31:0:2:0:0:1 | 0:0:0:0:0:0 | 124:3:6:0:0:0 | 1:0:1:0:0:0 |
| 12:0:2:0:0:0 | 0:0:0:0:0:0 | 31:0:2:0:0:1 | 0:0:0:0:0:0 | 119:3:6:0:0:0 | 0:0:0:0:0:0 |
| 287:01:00 | 0:00 | 214:00:00 | 0:00 | 1049:01:00 | 3:00 |
| 286:01:00 | 1:00 | 214:00:00 | 0:00 | 1052:01:00 | 1:00 |
| 286:01:00 | 1:00 | 214:00:00 | 0:00 | 1048:01:00 | 2:00 |
| 287:01:00 | 0:00 | 213:00:00 | 0:00 | 1046:01:00 | 1:00 |
| 287:01:00 | 0:00 | 213:00:00 | 1:00 | 1044:01:00 | 2:00 |
| 285:01:00 | 0:00 | 211:00:00 | 0:00 | 1039:01:00 | 0:00 |
| 289:01:00 | 0:00 | 204:00:00 | 0:00 | 1031:01:00 | 0:00 |
| 292:01:00 | 1:00 | 204:00:00 | 0:00 | 1026:01:00 | 2:00 |
| 293:01:00 | 0:00 | 203:00:00 | 0:00 | 1025:01:00 | 2:00 |
| 293:01:00 | 1:00 | 202:00:00 | 0:00 | 1010:01:00 | 2:00 |
| 296:01:00 | 0:00 | 199:00:00 | 0:00 | 1001:01:00 | 1:00 |
| 292:01:00 | 0:00 | 183:00:00 | 0:00 | 928:01:00 | 1:00 |
| 0:0:0:0:0:0:0 | 0:0:1:0:0:0:0 | 0:0:1:0:0:0:0 | 0:0:2:0:1:1:0 | 0:1:2:1:0:1:1 | 0:1:2:0:0:2:0 |
| 4:145:47:0 | 0:1:0:0 | 24:27:336:1 | 0:0:2:0 | 25:290:797:2 | 0:1:2:0 |
| 4:147:47:0 | 0:0:0:0 | 24:27:331:1 | 0:0:2:0 | 24:291:796:2 | 0:0:0:0 |
| 3:147:47:0 | 0:0:1:0 | 24:27:333:1 | 0:0:0:0 | 20:293:783:2 | 0:1:3:0 |
| 3:147:48:0 | 0:0:0:0 | 23:27:328:1 | 0:0:1:0 | 20:293:780:2 | 0:1:5:0 |
| 149:47:3:0 | 0:0:0:0 | 29:307:19:1 | 0:0:0:0 | 296:742:17:2 | 1:2:0:0 |

| | | | | | |
|---|---|---|---|---|---|
| 149:46:2:0 | 0:0:0:0 | 29:298:16:1 | 0:3:1:0 | 298:716:16:2 | 0:3:0:0 |
| 149:44:2:0 | 0:0:0:0 | 30:288:14:1 | 0:0:0:0 | 300:703:16:2 | 0:2:0:0 |
| 149:44:2:0 | 0:0:0:0 | 30:286:14:1 | 0:2:0:0 | 300:700:16:2 | 0:1:0:0 |
| 147:44:2:0 | 2:0:0:0 | 30:283:13:1 | 0:1:0:0 | 300:694:16:2 | 0:2:0:0 |
| 149:44:2:0 | 0:0:0:0 | 30:276:11:1 | 0:0:0:0 | 300:683:14:2 | 0:3:0:0 |
| 148:41:2:0 | 0:0:0:0 | 30:269:9:1 | 0:0:0:0 | 298:658:12:2 | 1:3:0:0 |
| 148:37:2:0 | 0:0:0:0 | 30:239:7:1 | 0:2:0:0 | 299:569:5:2 | 0:3:0:0 |
| 148:34:2:0 | 0:0:0:0 | 28:209:5:1 | 2:1:0:0 | 299:493:3:2 | 0:1:0:0 |
| 149:30:0:0 | 0:0:0:0 | 30:144:2:1 | 0:2:0:0 | 299:357:1:2 | 1:1:0:0 |
| 149:28:0:0 | 0:0:0:0 | 30:127:2:1 | 0:1:0:0 | 300:315:0:2 | 0:1:0:0 |
| 149:27:0:0 | 0:0:0:0 | 31:103:1:1 | 0:0:0:0 | 298:260:0:2 | 2:0:0:0 |
| 149:16:00 | 0:00:00 | 30:34:01 | 0:02:00 | 296:101:2 | 0:02:00 |
| 0:00 | 0:00 | 0:02 | 0:00 | 1:02 | 0:00 |
| 13:1507:5:190:19:12:1 | 0:1:0:0:1:0:0 | 29:1513:10:61:27:2:16 | 0:0:1:0:0:0:0 | 33:5258:24:490:94:9:17 | 0:8:0:0:0:0:0 |
| 13:1505:4:189:21:12:1 | 0:3:0:1:0:0:0 | 29:1509:12:61:26:2:16 | 0:3:0:1:1:0:0 | 33:5258:24:486:89:9:17 | 0:8:0:0:1:0:0 |
| 554:5:2:5:5:0:1 | 954:8:2:12:7:0:0 | 787:18:11:9:1:2:9 | 726:11:7:8:1:0:7 | 1901:7:9:26:4:8:6 | 3365:26:15:29:5:10:11 |
| 13:1508:5:192:12:15:1 | 0:0:0:0:0:0:0 | 29:1509:21:61:2:14:16 | 0:3:0:0:0:0:0 | 33:5254:26:474:9:39:17 | 0:8:1:0:0:0:0 |
| 1507:13:5:191:12:15:1 | 1:0:0:1:0:0:0 | 1512:29:22:61:2:12:16 | 1:0:0:0:0:0:0 | 5258:33:29:475:9:39:17 | 4:0:0:2:0:0:0 |
| 1505:13:5:192:12:15:1 | 1:0:0:0:0:0:0 | 1509:29:23:61:2:12:16 | 2:0:0:0:0:1:0 | 5248:33:30:479:9:34:17 | 4:0:0:0:0:2:0 |
| 1505:13:5:192:16:12:1 | 0:0:0:0:0:0:0 | 1509:29:33:63:34:2:16 | 1:0:0:0:1:0:0 | 5245:33:36:484:80:9:16 | 2:0:0:1:3:0:0 |
| 13:1496:6:191:28:12:1 | 0:1:0:0:0:0:0 | 28:1501:41:62:67:2:16 | 0:1:0:0:0:0:0 | 32:5215:42:479:163:9:16 | 1:10:0:0:0:0:0 |
| 13:1496:6:189:30:12:1 | 0:0:0:2:0:0:0 | 27:1502:40:61:79:2:16 | 0:1:2:0:0:0:0 | 32:5198:42:475:195:9:16 | 0:5:0:1:0:0:0 |
| 13:1494:6:191:34:12:1 | 0:2:0:0:0:0:0 | 27:1493:48:61:89:2:16 | 0:1:0:0:0:0:0 | 33:5174:49:475:204:9:16 | 0:11:0:0:1:0:0 |
| 0 | 0 | 1 | 2 | 1 | 3 |
| 0 | 2 | 0 | 0 | 0 | 0 |
| 0:0:0:0 | 0:0:0:0 | 0:2:0:1 | 0:0:0:0 | 0:2:1:1 | 0:0:0:0 |
| 1:03:04 | 0:00:00 | 0:01:01 | 0:00:00 | 2:00:02 | 0:00:00 |
| 1:02:04 | 0:00:00 | 0:01:01 | 0:00:00 | 2:00:02 | 0:00:00 |
| 0 | 2 | 2 | 7 | 5 | 9 |
| 0:00:00 | 0:00:00 | 0:00:00 | 0:00:00 | 0:00:00 | 0:00:00 |
| 1 | 0 | 106 | 0 | 30 | 0 |
| 1 | 0 | 99 | 0 | 30 | 0 |
| 1 | 0 | 105 | 0 | 30 | 0 |
| 0:0:0:0:0:0:0:0 | 0:4:0:0:0:1:0:0 | 0:0:0:0:0:0:0:0 | 0:0:1:0:1:0:0:0 | 0:0:0:0:0:0:0:0 | 0:4:0:0:0:3:0:1 |
| 0:4:0:0:1:0:0 | 0:0:0:0:0:0:0 | 0:2:1:1:0:0:0 | 0:0:0:0:0:0:0 | 0:5:0:0:3:0:1 | 0:0:0:0:0:0:0 |
| 0 | 1 | 0 | 0 | 0 | 2 |
| 0:00 | 0:01 | 0:00 | 0:00 | 0:00 | 0:03 |
| 0 | 0 | 0 | 1 | 0 | 4 |
| 3:26:1:0:0:0:0 | 0:0:0:0:0:0:0 | 10:34:5:0:3:0:2 | 1:0:0:0:0:0:0 | 32:204:5:3:2:0:2 | 1:0:0:0:0:0:0 |
| 29:5:1:1:0:0:0 | 0:0:0:0:0:0:0 | 52:34:5:0:3:0:3 | 1:1:0:0:0:0:0 | 258:110:5:0:5:3:2 | 0:1:0:0:0:0:0 |
| 5:29:0:1:0:0:0 | 0:0:1:0:0:0:0 | 32:51:5:0:3:0:3 | 3:2:0:0:0:0:0 | 110:257:5:0:5:3:2 | 1:1:0:0:0:0:0 |
| 0:5:28:0:1:0:0 | 0:0:1:1:0:0:0 | 3:34:53:5:0:0:3 | 0:0:0:0:0:0:0 | 7:108:255:5:0:3:2 | 0:2:3:0:0:0:0 |
| 0:00 | 3:01 | 0:00 | 0:07 | 0:00 | 14:21 |
| 15:16:7:0:0 | 0:0:0:0:0 | 5:6:46:0:0 | 0:0:0:0:0 | 43:37:72:11:0 | 0:0:1:0:0 |
| 0:0:0:1:0 | 15:16:8:0:0 | 3:5:31:2:0 | 2:1:48:0:0 | 19:22:89:6:0 | 20:15:81:7:0 |
| 15:16:9:1:0 | 0:0:0:0:0 | 5:6:80:2:0 | 0:0:0:0:0 | 37:37:171:14:0 | 0:0:3:0:0 |
| 15:16:9:1:0 | 0:0:0:0:0 | 5:6:85:2:0 | 0:0:0:0:0 | 37:37:186:15:0 | 0:0:0:0:0 |
| 66:04:00 | 0:00:00 | 87:06:00 | 3:01:00 | 605:39:00 | 3:00:00 |
| 8:00 | 4:00 | 18:00 | 22:00 | 27:00:00 | 16:02 |
| 0:0:0:0:0:0:0 | 0:0:0:0:1:0:0 | 0:0:0:0:0:0:0 | 0:0:0:0:0:1:0 | 0:0:0:0:0:0:0 | 0:0:0:0:0:0:1 |
| 3:00 | 0:00 | 2:02 | 0:00 | 10:06 | 0:00 |
| 5:00 | 0:00 | 5:01 | 0:00 | 15:00 | 2:00 |
| 5:00 | 0:00 | 4:00 | 0:00 | 17:00 | 0:00 |
| 53:23:4:3 | 0:0:0:0 | 2:12:2:0 | 0:0:0:0 | 67:149:15:10 | 0:3:0:0 |
| 53:6:18:3 | 2:0:0:0 | 9:8:9:0 | 0:0:0:0 | 134:120:86:7 | 0:1:1:0 |
| 54:8:19:3 | 1:0:0:0 | 9:8:5:0 | 0:0:1:0 | 145:136:79:7 | 0:1:1:0 |
| 51:8:7:2 | 0:0:0:0 | 10:9:1:0 | 0:0:0:0 | 149:156:45:5 | 0:1:1:0 |
| 66 | 0 | 68 | 3 | 348 | 5 |
| 13:2:0:0:0:0:0:0 | 0:0:0:0:0:0:0:0 | 16:4:2:2:0:0:0:0 | 0:0:0:0:0:0:0:0 | 52:18:3:2:2:0:0:1 | 0:0:0:0:0:0:0:0 |
| 1:0:0:0:0:0:0:0:0 | 0:0:0:0:0:0:0:0:0 | 1:0:0:1:0:0:0:0:0 | 1:0:0:0:0:0:0:0:0 | 4:0:0:1:0:0:0:0:1 | 2:0:0:1:0:0:0:1:0 |

| | | | | | |
|---|---|---|---|---|---|
| 1:0:0:0:0:0:0:0:0 | 0:0:0:0:0:0:0:0 | 1:0:1:0:0:0:0:0:0 | 1:0:0:0:0:0:0:0:0 | 2:0:0:0:0:0:0:0:1 | 5:0:2:0:0:0:0:1:0 |
| 0:1:0:0:0:0:0 | 0:0:0:0:0:0:0 | 0:0:0:0:0:0:0 | 0:1:0:0:0:0:0 | 0:1:0:0:0:0:1 | 2:3:0:0:0:1:0 |
| 143:28:00 | 0:00:00 | 311:103:0 | 1:01:00 | 647:240:1 | 0:00:00 |
| 143:28:00 | 0:00:00 | 311:103:0 | 1:01:00 | 647:240:1 | 1:01:00 |
| 143:28:00 | 0:00:00 | 311:103:0 | 1:01:00 | 648:240:1 | 0:02:00 |
| 143:28:00 | 0:00:00 | 311:103:0 | 1:00:00 | 647:242:1 | 1:00:00 |
| 137:26:00 | 0:00:00 | 297:93:0 | 2:00:00 | 625:231:1 | 0:01:00 |
| 142:23:00 | 0:00:00 | 308:86:0 | 0:00:00 | 642:209:1 | 0:04:00 |
| 142:23:00 | 0:00:00 | 308:83:0 | 0:00:00 | 638:206:1 | 3:00:00 |
| 143:21:00 | 0:00:00 | 314:65:0 | 0:01:00 | 647:179:1 | 0:01:00 |
| 144:20:00 | 0:00:00 | 315:63:0 | 0:00:00 | 646:174:1 | 2:00:00 |
| 144:21:00 | 1:00:00 | 315:59:00 | 3:00:00 | 652:169:1 | 4:00:00 |
| 145:22:00 | 0:00:00 | 317:50:00 | 1:00:00 | 658:152:1 | 0:00:00 |
| 145:22:00 | 0:00:00 | 318:50:00 | 0:00:00 | 658:152:1 | 0:00:00 |
| 144:21:00 | 1:00:00 | 318:39:00 | 0:00:00 | 661:120:1 | 1:01:00 |
| 145:22:00 | 0:00:00 | 319:38:00 | 0:00:00 | 664:119:1 | 2:00:00 |
| 145:23:00 | 0:00:00 | 327:44:00 | 0:00:00 | 683:140:1 | 0:01:00 |
| 1.017361111 | 0:00:00 | 46:327:0 | 0:00:00 | 145:681:1 | 0:02:00 |
| 1:0:0:0:0:0:0:0:0 | 13:0:0:0:0:0:0:0:1 | 8:0:0:2:0:0:1:0:1 | 22:3:0:1:0:1:0:0:0 | 4:0:0:0:0:0:0:0:0 | 42:0:0:2:0:0:2:0:2 |
| 0:0:0:0:0 | 0:0:0:0:0 | 0:0:0:0:0 | 0:1:0:0:0 | 0:0:2:0:1 | 0:2:1:3:0 |
| 3:0:0:0 | 0:0:0:0 | 2:0:0:1 | 0:0:0:0 | 8:0:0:0 | 0:0:0:0 |
| 88:108:6:22 | 0:0:0:0 | 119:219:10:18 | 0:0:0:0 | 536:461:41:89 | 0:3:0:0 |
| 0:0:0:0:0 | 0:0:0:0:0 | 0:0:0:0:0 | 0:0:0:0:0 | 0:0:0:0:0 | 0:0:0:1:1 |
| 1 | 0 | 3 | 3 | 1 | 1 |
| 1 | 0 | 3 | 3 | 1 | 1 |
| 0:00:00 | 0:00:00 | 0:00:00 | 6:00:00 | 0:00:00 | 4:01:01 |
| 0 | 0 | 0 | 0 | 0 | 2 |
| 143:6:3:8:0:4:5 | 0:0:0:0:0:0:0 | 134:10:1:1:0:1:1 | 0:0:0:0:0:0:0 | 585:28:22:22:3:8:7 | 1:0:0:0:1:0:0 |
| 143:6:3:8:0:4:5 | 0:0:0:0:0:0:0 | 132:10:1:1:0:1:1 | 1:0:0:0:0:0:0 | 585:28:22:22:2:7:7 | 1:0:0:0:0:0:0 |
| 144:6:3:0:4:5 | 0:0:0:0:0:0 | 136:10:1:0:1:1 | 0:0:0:0:0:0 | 592:29:22:0:6:7 | 0:0:0:0:0:0 |
| 142:6:3:8:1:3:0 | 1:0:0:0:0:0:0 | 136:10:1:1:4:0:0 | 0:0:0:0:0:0:0 | 587:29:22:22:13:6:2 | 1:0:0:0:0:0:0 |
| 142:6:3:8:0:4:5 | 1:0:0:0:0:0:0 | 135:10:1:1:0:5:1 | 0:0:0:0:0:0:0 | 581:28:22:22:3:19:8 | 2:0:0:0:1:1:0 |
| 140:6:3:7:0:4:5 | 1:0:0:0:0:0:0 | 135:10:1:1:2:3:1 | 0:0:0:0:0:0:0 | 570:27:22:22:7:18:8 | 2:1:0:0:0:0:0 |
| 0:00 | 0:00 | 0:00 | 0:00 | 0:00 | 1:01 |
| 180:03:00 | 1:00 | 381:07:00 | 0:00 | 1313:09:00 | 5:00 |
| 181:03:00 | 0:00 | 374:08:00 | 0:00 | 1306:23:00 | 0:00 |
| 181:03:00 | 0:00 | 373:08:00 | 0:00 | 1290:23:00 | 6:00 |
| 0:00 | 1:00 | 0:00 | 3:00 | 0:00 | 9:00 |
| 104:9:6:2:2:0 | 0:0:0:0:0:0 | 108:6:10:2:1:0 | 0:0:0:0:0:0 | 458:47:21:2:19:0 | 1:0:0:0:1:0 |
| 104:9:6:2:4:0 | 0:0:0:0:0:0 | 108:6:10:2:3:0 | 0:0:0:0:0:0 | 459:47:25:2:28:0 | 0:0:0:0:0:0 |
| 104:9:6:2:4:0 | 0:0:0:0:0:0 | 108:6:10:2:3:0 | 0:0:0:0:0:0 | 459:47:25:2:28:0 | 0:0:0:0:0:0 |
| 104:9:6:2:4:0 | 0:0:0:0:0:0 | 108:6:10:2:3:0 | 0:0:0:0:0:0 | 459:47:25:2:28:0 | 0:0:0:0:0:0 |
| 103:9:6:4:2:0 | 1:0:0:0:0:0 | 108:6:8:7:2:0 | 0:0:0:0:0:0 | 457:47:24:41:1:0 | 0:0:1:0:0:0 |
| 104:9:6:4:1:0 | 0:0:0:0:1:0 | 108:6:8:7:2:0 | 0:0:0:0:0:0 | 456:47:24:42:1:0 | 2:0:0:1:0:0 |
| 104:9:3:4:0:0 | 0:0:0:0:0:0 | 108:6:7:9:0:1 | 0:0:0:0:0:0 | 456:47:17:52:0:0 | 2:0:0:0:0:0 |
| 0:00 | 0:00 | 0:00 | 1:01 | 0:00 | 0:00 |
| 13:11:156:0:1:7:9:2:3:0:0:0:0:1:0:0:0:0 | 0:0:0:0:0:0:1:0:0:0:0:0:0:0:0:0:0:0 | 37:33:101:1:1:2:8:5:4:0:2:1:1:1:0:0:1:0 | 0:0:2:0:0:0:0:0:0:0:0:0:0:0:0:0:0:0 | 124:74:381:5:8:13:40:14:24:1:9:1:8:5:0:0:2:2 | 0:0:0:0:0:0:0:0:0:0:0:0:0:0:0:0:0:0 |
| 156:370:182:48:13:11:10:27:0:14:9:14:13:3:1:0:0:2:0:0:0 | 0:1:0:0:0:0:0:0:0:2:0:0:0:0:0:0:0:0:0:0:0 | 103:246:111:58:37:33:12:13:1:20:2:11:8:4:0:2:1:1:0:0:1 | 0:0:0:0:0:0:0:0:0:2:0:0:0:0:0:0:0:0:0:0:0 | 380:873:353:312:124:74:40:99:5:65:15:52:62:24:5:11:8:7:1:2:2 | 0:6:0:0:0:0:0:0:0:0:12:0:0:0:0:0:0:0:0:0:0 |
| 10:21:372:182:48:13:11:26:156:0:9:14:13:3:1:0:0:2:0:0:0 | 0:1:0:0:0:0:0:1:0:0:0:0:0:0:0:0:0:0:0:0:0 | 12:25:247:111:58:37:33:13:101:1:2:11:8:4:0:2:1:1:0:0:1 | 0:1:0:0:0:0:0:0:0:0:0:0:0:0:0:0:0:0:0:0:0 | 40:87:875:353:312:124:74:98:379:5:15:52:62:24:5:11:8:7:1:2:2 | 0:9:5:0:0:0:0:1:0:0:0:0:0:0:0:0:0:0:0:0:0 |
| 22:182:48:13:11:371:10:27:156:0:9:14:13:3:1:0:0:2:0:0:0 | 0:0:0:0:0:1:0:0:0:0:0:0:0:0:0:0:0:0:0:0:0 | 24:111:58:37:33:245:12:13:101:1:2:11:7:4:0:2:1:1:0:0:1 | 2:0:0:0:0:2:0:0:1:0:0:0:1:0:0:0:0:0:0:0:0 | 92:353:312:124:74:876:40:99:379:5:15:52:61:24:5:11:8:7:1:2:2 | 6:0:0:0:0:4:0:0:0:0:0:0:1:0:0:0:0:0:0:0:0 |
| 372:182:48:13:11:10:27:156:21:0:9:14:13:3:1:0:0:2:0:0:0 | 0:0:0:0:0:0:0:0:1:0:0:0:0:0:0:0:0:0:0:0:0 | 246:111:58:37:33:12:13:102:24:1:2:11:8:4:0:2:1:1:0:0:1 | 1:0:0:0:0:0:0:0:3:0:0:0:0:0:0:0:0:0:0:0:0 | 876:353:312:124:74:40:99:378:102:5:15:51:62:24:5:11:8:7:1:2:2 | 4:0:0:0:0:0:0:0:8:0:0:1:0:0:0:0:0:0:0:0:0 |

| | | | | | |
|---|---|---|---|---|---|
| 372:182:48:13:11:10:27:156:30:0:9:14:12:3:1:0:0:2:0:0:0 | 2:0:0:0:0:0:0:0:2:0:0:0:1:0:0:0:0:0:0:0:0 | 248:111:57:37:33:12:13:102:27:1:2:11:8:4:0:2:1:1:0:0:1 | 0:0:1:0:0:0:0:0:1:0:0:0:0:0:0:0:0:0:0:0:0 | 887:353:312:124:73:40:99:378:151:5:15:52:62:24:5:11:8:7:1:2:2 | 6:0:0:0:1:0:0:0:4:0:0:0:0:0:0:0:0:0:0:0:0:0 |
| 374:182:48:13:11:10:27:156:34:0:9:14:13:3:1:0:0:2:0:0:0 | 0:0:0:0:0:0:0:0:0:0:0:0:0:0:0:0:0:0:0:0:0 | 247:111:58:37:33:12:13:101:26:1:2:11:8:4:0:2:1:1:0:0:1 | 1:0:0:0:0:0:0:1:2:0:0:0:0:0:0:0:0:0:0:0:0 | 889:351:312:124:74:40:99:377:175:5:15:52:62:24:5:11:8:7:1:2:2 | 4:2:0:0:0:0:0:0:1:0:0:0:0:0:0:0:0:0:0:0:0:0 |
| 182:48:13:11:10:378:27:156:46:0:9:14:13:3:1:0:0:2:0:0:0 | 0:0:0:0:0:0:0:0:0:0:0:0:0:0:0:0:0:0:0:0:0 | 111:58:37:33:12:247:13:102:29:1:2:11:8:4:0:2:1:1:0:0:1 | 0:0:0:0:0:0:0:1:0:0:0:0:0:0:0:0:0:0:0:0:0 | 352:312:124:73:40:901:99:376:225:5:15:52:62:24:5:11:8:7:1:2:2 | 0:0:0:1:0:2:0:0:0:0:0:0:0:0:0:0:0:0:0:0:0:0 |
| 182:48:13:11:10:378:27:156:58:0:9:14:13:3:1:0:0:2:0:0:0 | 0:0:0:0:0:0:0:0:0:0:0:0:0:0:0:0:0:0:0:0:0 | 111:58:37:33:12:249:12:102:32:1:2:11:8:4:0:2:1:1:0:0:1 | 0:0:0:0:0:1:1:0:0:0:0:0:0:0:0:0:0:0:0:0:0 | 353:312:124:74:40:904:99:375:252:5:15:52:62:24:5:11:8:7:1:2:2 | 0:0:0:0:0:1:0:0:4:0:0:0:0:0:0:0:0:0:0:0:0:0 |
| 58:182:48:13:11:10:378:27:156:0:9:14:13:3:1:0:0:2:0:0:0 | 0:0:0:0:0:0:0:0:0:0:0:0:0:0:0:0:0:0:0:0:0 | 35:111:58:37:32:12:250:13:102:1:2:11:8:4:0:2:1:1:0:0:1 | 0:0:0:0:1:0:1:0:0:0:0:0:0:0:0:0:0:0:0:0:0 | 271:352:311:124:74:40:911:99:375:5:14:52:62:23:5:11:8:7:1:2:2 | 5:0:1:0:0:0:0:0:0:0:1:0:0:1:0:0:0:0:0:0:0:0 |
| 58:182:48:13:11:10:378:27:156:0:9:14:13:3:1:0:0:2:0:0:0 | 0:0:0:0:0:0:0:0:0:0:0:0:0:0:0:0:0:0:0:0:0 | 35:111:58:37:33:12:250:13:102:1:2:11:8:4:0:2:1:1:0:0:1 | 0:0:0:0:0:0:1:0:0:0:0:0:0:0:0:0:0:0:0:0:0 | 272:350:311:124:74:40:910:99:374:5:15:52:62:24:5:11:8:7:1:2:2 | 5:1:1:0:0:0:0:0:1:0:0:0:0:0:0:0:0:0:0:0:0:0 |
| 156:181:48:13:11:10:377:27:58:0:9:14:13:3:1:0:0:2:0:0:0 | 0:1:0:0:0:0:1:0:0:0:0:0:0:0:0:0:0:0:0:0:0 | 102:111:58:37:33:12:251:13:37:1:2:11:8:4:0:2:1:1:0:0:1 | 0:0:0:0:0:0:0:0:0:0:0:0:0:0:0:0:0:0:0:0:0 | 375:350:312:124:74:40:914:99:288:5:15:52:62:24:5:11:8:7:1:2:2 | 0:2:0:0:0:0:1:0:5:0:0:0:0:0:0:0:0:0:0:0:0:0 |
| 182:48:13:11:9:380:27:156:65:0:9:14:13:3:1:0:0:2:0:0:0 | 0:0:0:0:1:0:0:0:0:0:0:0:0:0:0:0:0:0:0:0:0 | 111:58:37:33:12:249:13:102:41:1:2:11:8:4:0:2:1:1:0:0:1 | 0:0:0:0:0:2:0:0:0:0:0:0:0:0:0:0:0:0:0:0:0 | 352:312:124:74:40:920:97:373:314:5:15:52:62:24:5:11:8:7:1:2:2 | 0:0:0:0:0:0:0:1:2:0:0:0:0:0:0:0:0:0:0:0:0:0 |
| 380:182:48:13:11:10:27:156:71:0:9:14:13:3:1:0:0:2:0:0:0 | 0:0:0:0:0:0:0:0:0:0:0:0:0:0:0:0:0:0:0:0:0 | 252:111:58:37:33:12:13:102:45:1:2:11:8:4:0:2:1:1:0:0:1 | 0:0:0:0:0:0:0:0:0:0:0:0:0:0:0:0:0:0:0:0:0 | 920:350:312:124:74:40:97:374:343:5:15:51:62:24:5:11:8:7:1:2:2 | 2:1:0:0:0:0:0:0:3:0:0:0:0:0:0:0:0:0:0:0:0:0 |
| 380:182:48:13:11:10:27:156:72:0:9:14:13:3:1:0:0:2:0:0:0 | 0:0:0:0:0:0:0:0:0:0:0:0:0:0:0:0:0:0:0:0:0 | 252:111:57:37:33:12:13:102:47:1:2:11:8:4:0:2:1:1:0:0:1 | 0:0:1:0:0:0:0:0:0:0:0:0:0:0:0:0:0:0:0:0:0 | 923:350:312:124:74:40:97:374:355:5:15:50:61:24:5:11:8:7:1:2:2 | 1:0:0:0:0:0:0:0:0:0:0:0:0:0:0:0:0:0:0:0:0:0 |
| 182:48:13:11:10:380:27:156:72:0:9:14:13:3:1:0:0:2:0:0:0 | 0:0:0:0:0:0:0:0:0:0:0:0:0:0:0:0:0:0:0:0:0 | 110:58:37:33:12:252:13:102:47:1:2:11:8:4:0:2:1:1:0:0:1 | 1:0:0:0:0:0:0:0:0:0:0:0:0:0:0:0:0:0:0:0:0 | 348:312:124:74:40:924:97:374:355:5:15:50:61:24:5:11:8:7:1:2:2 | 2:0:0:0:0:0:0:0:0:0:0:0:0:0:0:0:0:0:0:0:0:0 |
| 182:48:13:11:381:10:27:156:89:0:9:14:13:3:1:0:0:2:0:0:0 | 0:0:0:0:0:0:0:0:1:0:0:0:0:0:0:0:0:0:0:0:0 | 111:58:37:33:251:12:13:102:55:1:2:11:8:4:0:2:1:1:0:0:1 | 0:0:0:0:1:0:0:0:1:0:0:0:0:0:0:0:0:0:0:0:0 | 347:312:124:74:940:40:94:373:429:5:15:50:60:23:5:11:8:7:1:2:2 | 1:0:0:0:1:0:0:0:4:0:0:0:0:1:0:0:0:0:0:0:0:0 |
| 156:102:182:48:13:11:381:10:27:0:9:14:13:3:1:0:0:2:0:0:0 | 0:2:0:0:0:0:0:0:0:0:0:0:0:0:0:0:0:0:0:0:0 | 102:63:111:58:37:33:254:12:13:1:2:11:8:4:0:2:1:1:0:0:1 | 0:0:0:0:0:0:0:0:0:0:0:0:0:0:0:0:0:0:0:0:0 | 373:495:348:311:124:74:946:40:92:5:15:48:59:24:5:11:8:7:1:2:2 | 0:7:0:1:0:0:4:0:0:0:0:0:0:0:0:0:0:0:0:0:0:0 |
| 107:182:48:13:11:381:10:27:156:0:9:14:13:3:1:0:0:2:0:0:0 | 1:0:0:0:0:0:0:0:0:0:0:0:0:0:0:0:0:0:0:0:0 | 64:110:58:37:33:254:12:13:102:1:2:11:8:4:0:2:1:1:0:0:1 | 1:1:0:0:0:0:0:0:0:0:0:0:0:0:0:0:0:0:0:0:0 | 522:348:312:123:73:950:40:92:373:5:15:48:59:24:5:11:8:7:1:2:2 | 10:0:0:1:0:4:0:0:0:0:0:0:0:0:0:0:0:0:0:0:0:0 |
| 110:182:48:13:11:382:10:27:156:0:9:14:13:3:1:0:0:2:0:0:0 | 1:0:0:0:0:0:0:0:0:0:0:0:0:0:0:0:0:0:0:0:0 | 67:111:58:37:33:255:12:13:102:1:2:11:7:4:0:2:1:1:0:0:1 | 1:0:0:0:0:0:0:0:0:0:0:0:0:0:0:0:0:0:0:0:0 | 537:348:312:124:73:952:40:92:371:5:15:47:59:24:5:11:8:7:1:2:2 | 8:0:0:0:0:3:0:0:0:0:0:0:0:0:0:0:0:0:0:0:0:0 |
| 110:182:48:13:11:381:10:27:156:0:9:14:13:3:1:0:0:2:0:0:0 | 1:0:0:0:0:0:0:0:0:0:0:0:0:0:0:0:0:0:0:0:0 | 68:111:58:37:33:256:12:13:102:1:2:11:7:4:0:2:1:1:0:0:1 | 0:0:0:0:0:0:0:0:0:0:0:0:0:0:0:0:0:0:0:0:0 | 535:348:312:124:73:952:40:92:372:5:15:47:59:24:5:11:8:7:1:2:2 | 11:0:0:0:0:3:0:0:0:0:0:0:0:0:0:0:0:0:0:0:0:0 |
| 115:182:48:13:11:382:10:26:156:0:9:14:13:3:1:0:0:2:0:0:0 | 0:0:0:0:0:0:0:0:0:0:0:0:0:0:0:0:0:0:0:0:0 | 79:111:58:37:33:258:12:13:102:1:2:11:7:4:0:2:1:1:0:0:1 | 2:0:0:0:0:1:0:0:0:0:0:0:0:0:0:0:0:0:0:0:0 | 624:346:312:124:73:976:40:91:372:5:15:47:58:24:5:10:8:7:1:2:2 | 8:0:0:0:0:2:0:0:0:0:0:0:0:0:0:0:0:0:0:0:0:0 |
| 383:121:182:48:13:11:10:26:156:0:9:14:13:3:1:0:0:2:0:0:0 | 0:2:0:0:0:0:0:0:0:0:0:0:0:0:0:0:0:0:0:0:0 | 262:88:111:57:37:33:12:13:102:1:2:11:7:4:0:2:1:1:0:0:1 | 0:1:0:1:0:0:0:0:0:0:0:0:0:0:0:0:0:0:0:0:0 | 982:663:345:310:124:73:40:91:370:5:15:46:57:24:5:10:8:7:1:2:2 | 6:14:0:1:0:0:0:0:1:0:0:0:0:0:0:0:0:0:0:0:0:0 |
| 385:129:181:48:13:11:10:25:156:0:14:13:9:3:1:0:0:2:0:0:0 | 0:0:0:0:0:0:0:1:0:0:0:0:0:0:0:0:0:0:0:0:0 | 264:88:111:58:37:33:12:13:102:1:11:7:2:4:0:2:1:1:0:0:1 | 0:2:0:0:0:0:0:0:0:0:0:0:0:0:0:0:0:0:0:0:0 | 985:708:344:311:124:72:40:90:371:5:46:57:15:24:5:10:8:7:1:2:2 | 4:12:0:0:0:1:0:0:0:0:0:0:0:0:0:0:0:0:0:0:0:0 |

| | | | | | |
|---|---|---|---|---|---|
| 385:129:181:48:13:11:10:26:156:0:9:14:12:3:1:0:0:2:0:0:0 | 0:1:0:0:0:0:0:0:0:0:0:0:0:0:0:0:0:0:0:0:0 | 264:88:111:58:37:32:12:13:102:1:2:11:7:4:0:2:1:1:0:0:1 | 0:2:0:0:0:1:0:0:0:0:0:0:0:0:0:0:0:0:0:0:0 | 985:722:345:311:124:73:40:90:371:5:15:46:57:24:5:10:8:7:1:2:2 | 6:9:0:0:0:0:0:0:0:0:0:0:0:0:0:0:0:0:0:0:0 |
| 181:48:13:11:388:10:26:156:139:0:9:14:12:3:1:0:0:2:0:0:0 | 0:0:0:0:0:0:0:0:0:0:0:0:0:0:0:0:0:0:0:0:0 | 111:58:37:33:266:12:13:102:101:1:2:11:7:4:0:2:1:1:0:0:1 | 0:0:0:0:0:0:0:0:1:0:0:0:0:0:0:0:0:0:0:0:0 | 343:311:124:73:994:40:90:370:787:5:15:46:57:24:5:10:8:7:1:2:2 | 0:0:0:0:6:0:0:0:8:0:0:0:0:0:0:0:0:0:0:0:0 |
| 181:48:13:11:388:10:26:156:139:0:9:14:12:3:1:0:0:2:0:0:0 | 0:0:0:0:0:0:0:0:0:0:0:0:0:0:0:0:0:0:0:0:0 | 111:58:37:33:266:12:13:102:102:1:2:11:7:4:0:2:1:1:0:0:1 | 0:0:0:0:0:0:0:0:2:0:0:0:0:0:0:0:0:0:0:0:0 | 343:310:124:72:999:40:89:370:797:5:15:46:57:24:5:10:8:7:1:2:2 | 0:1:0:1:4:0:0:0:5:0:0:0:0:0:0:0:0:0:0:0:0 |
| 150:180:48:13:11:392:10:26:154:0:9:14:12:3:1:0:0:2:0:0:0 | 2:1:0:0:0:0:0:0:1:0:0:0:0:0:0:0:0:0:0:0:0 | 110:111:58:37:33:271:12:13:102:1:2:11:7:4:0:2:1:1:0:0:1 | 1:0:0:0:0:0:0:0:0:0:0:0:0:0:0:0:0:0:0:0:0 | 838:343:310:124:73:1007:40:89:368:5:15:45:55:24:5:10:8:7:1:2:2 | 1:0:0:0:0:4:0:0:2:0:0:0:0:0:0:0:0:0:0:0:0 |
| 152:181:48:13:11:392:10:25:155:0:9:14:12:3:1:0:0:2:0:0:0 | 1:0:0:0:0:1:0:0:0:0:0:0:0:0:0:0:0:0:0:0:0 | 113:111:58:37:33:271:12:13:102:1:2:11:7:4:0:2:1:1:0:0:1 | 1:0:0:0:0:0:0:0:0:0:0:0:0:0:0:0:0:0:0:0:0 | 855:343:310:124:73:1013:40:88:368:5:15:45:55:24:5:10:8:7:1:2:2 | 13:0:1:0:0:3:0:0:0:0:0:0:0:0:0:0:0:0:0:0:0 |
| 181:48:13:11:398:10:25:155:171:0:9:14:12:3:1:0:0:2:0:0:0 | 0:0:0:0:0:0:0:0:0:0:0:0:0:0:0:0:0:0:0:0:0 | 111:58:37:33:275:12:13:102:123:1:2:11:7:4:0:2:1:1:0:0:1 | 0:0:0:0:0:0:0:0:0:0:0:0:0:0:0:0:0:0:0:0:0 | 341:309:124:73:1035:40:85:367:963:5:15:42:55:24:5:10:8:7:1:2:2 | 0:1:0:0:1:0:1:0:8:0:0:0:0:0:0:0:0:0:0:0:0 |
| 398:170:181:48:13:11:10:25:155:0:9:14:12:3:1:0:0:2:0:0:0 | 0:1:0:0:0:0:0:0:0:0:0:0:0:0:0:0:0:0:0:0:0 | 275:124:111:58:37:33:12:13:102:1:2:11:7:4:0:2:1:1:0:0:1 | 0:0:0:0:0:0:0:0:0:0:0:0:0:0:0:0:0:0:0:0:0 | 1033:966:342:309:123:73:40:86:366:5:15:42:55:24:5:10:8:7:1:2:2 | 2:5:0:1:1:0:0:0:1:0:0:0:0:0:0:0:0:0:0:0:0 |
| 181:48:13:11:398:10:25:155:174:0:9:14:12:3:1:0:0:2:0:0:0 | 0:0:0:0:0:0:0:0:0:0:0:0:0:0:0:0:0:0:0:0:0 | 111:58:37:33:275:12:13:101:125:1:2:11:7:4:0:2:1:1:0:0:1 | 0:0:0:0:0:0:0:1:1:0:0:0:0:0:0:0:0:0:0:0:0 | 342:310:123:73:1041:40:86:367:978:5:15:42:55:24:5:10:8:7:1:2:2 | 0:0:1:0:0:0:0:0:3:0:0:0:0:0:0:0:0:0:0:0:0 |
| 181:48:13:11:397:10:25:154:175:0:9:14:12:3:1:0:0:2:0:0:0 | 0:0:0:0:1:0:0:1:0:0:0:0:0:0:0:0:0:0:0:0:0 | 111:58:37:33:275:12:13:102:126:1:2:11:7:4:0:2:1:1:0:0:1 | 0:0:0:0:0:0:0:0:0:0:0:0:0:0:0:0:0:0:0:0:0 | 341:310:124:72:1041:40:86:367:986:5:15:42:54:24:5:10:8:6:1:2:2 | 1:0:0:0:2:0:0:0:3:0:0:0:1:0:0:0:0:1:0:0:0 |
| 155:181:48:13:11:398:10:25:176:0:9:14:12:3:1:0:0:2:0:0:0 | 0:0:0:0:0:0:0:0:0:0:0:0:0:0:0:0:0:0:0:0:0 | 102:111:58:37:33:275:12:13:126:1:2:11:7:4:0:2:1:1:0:0:1 | 0:0:0:0:0:0:0:0:0:0:0:0:0:0:0:0:0:0:0:0:0 | 364:341:309:123:72:1043:40:86:998:5:15:42:55:24:5:10:8:7:1:2:2 | 2:1:1:1:0:1:0:0:3:0:0:0:0:0:0:0:0:0:0:0:0 |
| 174:181:48:13:11:398:10:25:154:0:9:14:13:3:1:0:0:2:0:0:0 | 0:0:0:0:0:1:0:0:1:0:0:0:0:0:0:0:0:0:0:0:0 | 134:110:58:37:33:276:12:13:102:1:2:11:7:4:0:2:1:1:0:0:1 | 0:1:0:0:0:0:0:0:0:0:0:0:0:0:0:0:0:0:0:0:0 | 1045:342:309:124:72:1050:40:86:367:5:15:41:55:24:5:10:8:7:1:2:2 | 4:0:1:0:0:4:0:0:0:0:0:0:0:0:0:0:0:0:0:0:0 |
| 176:181:48:13:11:401:10:25:155:0:9:14:13:3:1:0:0:2:0:0:0 | 1:0:0:0:0:0:0:0:0:0:0:0:0:0:0:0:0:0:0:0:0 | 135:111:58:37:33:278:12:13:102:1:2:11:7:4:0:2:1:1:0:0:1 | 0:0:0:0:0:0:0:0:0:0:0:0:0:0:0:0:0:0:0:0:0 | 1069:341:310:124:72:1063:40:86:367:5:15:42:55:24:5:10:8:7:1:2:2 | 5:0:0:0:0:3:0:0:0:0:0:0:0:0:0:0:0:0:0:0:0 |
| 181:48:13:11:402:10:25:155:177:0:9:14:13:3:1:0:0:2:0:0:0 | 0:0:0:0:0:0:0:0:2:0:0:0:0:0:0:0:0:0:0:0:0 | 111:58:37:33:278:12:13:102:137:1:2:11:7:4:0:2:1:1:0:0:1 | 0:0:0:0:0:0:0:0:0:0:0:0:0:0:0:0:0:0:0:0:0 | 341:309:124:72:1066:40:86:367:1080:5:15:42:55:24:5:10:8:7:1:2:2 | 0:1:0:0:4:0:0:0:4:0:0:0:0:0:0:0:0:0:0:0:0 |
| 402:181:48:13:11:10:25:155:179:0:9:14:13:3:1:0:0:2:0:0:0 | 0:0:0:0:0:0:0:0:1:0:0:0:0:0:0:0:0:0:0:0:0 | 281:110:58:37:33:12:13:102:140:1:2:11:7:4:0:2:1:1:0:0:1 | 0:1:0:0:0:0:0:0:0:0:0:0:0:0:0:0:0:0:0:0:0 | 1073:340:310:124:72:40:86:367:1095:5:15:42:55:24:5:10:8:7:1:2:2 | 3:1:0:0:0:0:0:0:0:5:0:0:0:0:0:0:0:0:0:0:0 |
| 404:181:48:13:11:10:25:154:181:0:9:14:13:3:1:0:0:2:0:0:0 | 0:0:0:0:0:0:0:1:1:0:0:0:0:0:0:0:0:0:0:0:0 | 282:111:58:37:33:12:13:102:147:1:2:11:7:4:0:2:1:1:0:0:1 | 1:0:0:0:0:0:0:0:0:0:0:0:0:0:0:0:0:0:0:0:0 | 1088:341:310:124:72:40:85:366:1118:5:15:42:55:24:5:10:8:7:1:2:2 | 2:0:0:0:0:0:0:1:6:0:0:0:0:0:0:0:0:0:0:0:0 |
| 155:404:181:48:12:11:10:25:183:0:9:14:13:3:1:0:0:2:0:0:0 | 0:1:0:0:0:0:0:0:1:0:0:0:0:0:0:0:0:0:0:0:0 | 102:284:111:58:37:33:12:13:150:1:2:11:7:4:0:2:1:1:0:0:1 | 0:0:0:0:0:0:0:0:1:0:0:0:0:0:0:0:0:0:0:0:0 | 368:1097:341:310:123:72:40:85:1146:5:15:42:55:24:5:10:8:7:1:2:2 | 0:2:1:0:1:0:0:0:5:0:0:0:0:0:0:0:0:0:0:0:0 |
| 155:181:48:12:11:404:10:25:184:0:9:14:13:3:1:0:0:2:0:0:0 | 0:0:0:0:0:1:0:0:1:0:0:0:0:0:0:0:0:0:0:0:0 | 102:111:58:37:33:284:12:13:152:1:2:11:6:4:0:2:1:1:0:0:1 | 0:0:0:0:0:0:0:0:0:0:0:0:0:0:0:0:0:0:0:0:0 | 368:342:310:124:72:1098:40:85:1157:5:15:42:55:24:5:10:8:7:1:2:2 | 0:0:0:0:0:2:0:0:5:0:0:0:0:0:0:0:0:0:0:0:0 |
| 181:48:12:11:406:10:25:155:181:0:9:14:13:3:1:0:0:2:0:0:0 | 0:0:0:0:0:0:0:0:0:0:0:0:0:0:0:0:0:0:0:0:0 | 111:58:37:33:285:12:13:102:151:1:2:11:6:4:0:2:1:1:0:0:1 | 0:0:0:0:0:0:0:0:2:0:0:0:0:0:0:0:0:0:0:0:0 | 342:310:124:72:1105:40:85:368:1187:5:15:42:55:24:5:10:8:7:1:2:2 | 0:0:0:0:1:0:0:0:2:0:0:0:0:0:0:0:0:0:0:0:0 |
| 181:48:12:11:404:10:25:155:182:0:9:13:13:3:1:0:0:2:0:0:0 | 0:0:0:0:2:0:0:0:0:0:0:0:0:0:0:0:0:0:0:0:0 | 111:58:37:33:285:12:13:102:153:1:2:11:6:4:0:2:1:1:0:0:1 | 0:0:0:0:0:0:0:0:1:0:0:0:0:0:0:0:0:0:0:0:0 | 341:310:124:72:1106:40:85:368:1193:5:15:42:55:24:5:10:8:7:1:2:2 | 1:0:0:0:1:0:0:0:4:0:0:0:0:0:0:0:0:0:0:0:0 |

| | | | | | |
|---|---|---|---|---|---|
| 180:48:12:11:405:10:25:155:180:0:9:13:13:3:1:0:0:2:0:0:0 | 0:0:0:0:0:0:0:0:0:0:0:0:0:0:0:0:0:0:0:0:0:0 | 111:58:37:33:286:12:13:102:154:1:2:11:6:4:0:2:1:1:0:0:1 | 0:0:0:0:0:0:0:0:1:0:0:0:0:0:0:0:0:0:0:0:0:0 | 341:310:124:72:1104:40:85:368:1213:5:15:42:55:24:5:10:8:7:1:2:2 | 1:0:0:0:4:0:0:0:5:0:0:0:0:0:0:0:0:0:0:0:0:0 |
| 180:48:11:11:404:10:25:154:179:0:9:13:13:3:1:0:0:2:0:0:0 | 0:0:0:0:0:0:0:1:0:0:0:0:0:0:0:0:0:0:0:0:0:0 | 111:58:37:33:287:12:13:102:156:1:2:11:6:4:0:2:1:1:0:0:1 | 0:0:0:0:0:0:0:0:0:0:0:0:0:0:0:0:0:0:0:0:0:0 | 342:310:124:72:1110:39:85:369:1223:5:15:42:54:24:5:10:8:7:1:2:2 | 0:0:0:0:2:0:0:0:7:0:0:0:0:0:0:0:0:0:0:0:0:0 |
| 0 | 0 | 49 | 10 | 1 | 0 |
| 0 | 0 | 59 | 12 | 1 | 0 |
| 0 | 0 | 65 | 11 | 1 | 0 |
| 0 | 0 | 71 | 15 | 1 | 0 |
| 0 | 0 | 64 | 15 | 1 | 0 |
| 0:00 | 2:00 | 1:00 | 1:00 | 1:01 | 4:00 |
| 0:00 | 2:00 | 1:00 | 1:00 | 2:00 | 4:01 |
| 0:0:0:0 | 0:0:0:0 | 2:0:0:0 | 0:0:0:0 | 4:2:1:2 | 0:0:0:0 |
| 0 | 0 | 0 | 0 | 0 | 0 |
| 0 | 0 | 0 | 0 | 0 | 0 |
| 0 | 0 | 0 | 0 | 0 | 0 |
| 0 | 1 | 0 | 0 | 0 | 2 |
| 0 | 2 | 0 | 2 | 0 | 1 |
| 1 | 0 | 1 | 0 | 6 | 0 |
| 1:04 | 0:02 | 1:05 | 1:05 | 2:08 | 3:15 |
| 500:1:0:0 | 0:0:0:0 | 2189:0:0:0 | 0:0:0:0 | 3309:0:2:1 | 2:0:0:0 |
| 501:1:0:0 | 0:0:0:0 | 2202:0:0:0 | 2:0:0:0 | 3309:0:2:1 | 5:0:0:0 |
| 501:1:0:0 | 0:0:0:0 | 2204:0:0:0 | 1:0:0:0 | 3313:0:2:1 | 3:0:0:0 |
| 501:1:0:0 | 0:0:0:0 | 2208:0:0:0 | 3:0:0:0 | 3325:0:2:1 | 3:0:0:0 |
| 496:00:00 | 0:00 | 2111:00:00 | 4:00 | 3195:00:00 | 16:00 |
| 375:00:00 | 27:00:00 | 1872:00:00 | 205:00:00 | 2758:00:00 | 259:00:00 |
| 477:00:00 | 2:00 | 1988:00:00 | 5:00 | 2977:00:00 | 21:00 |
| 0:00 | 34:00:00 | 4:00 | 1114:00:00 | 9:00 | 1330:00:00 |
| 0:00 | 10:00 | 3:00 | 1134:00:00 | 9:00 | 1342:00:00 |
| 0 | 1 | 0 | 1 | 0 | 3 |
| 0:00 | 1:00 | 0:00 | 3:00 | 0:00 | 7:00 |
| 0 | 0 | 1 | 0 | 0 | 0 |
| 0 | 0 | 1 | 0 | 0 | 0 |
| 0 | 0 | 0 | 0 | 0 | 0 |
| 94:0:0:0 | 0:0:0:0 | 76:0:2:2:0 | 0:0:0:0 | 221:2:4:2:0 | 0:0:0:0 |
| 0 | 0 | 0 | 0 | 3 | 0 |
| 109:11:00 | 0:00 | 128:08:00 | 0:00 | 201:15:00 | 0:00 |
| 103:12:00 | 0:00 | 129:08:00 | 1:00 | 197:15:00 | 1:00 |
| 4:11 | 0:00 | 6:07 | 0:00 | 2:15 | 0:00 |
| 0:00 | 1:00 | 0:00 | 0:01 | 0:00 | 2:00 |
| 0 | 0 | 0 | 0 | 0 | 2 |
| 0:00 | 2:00 | 0:00 | 1:00 | 0:00 | 3:01 |
| 5:0:0:0 | 0:0:0:0 | 3:1:0:0 | 0:0:0:0 | 13:6:2:1 | 0:0:0:0 |
| 56:00:01 | 0:00:00 | 55:00:00 | 0:00:00 | 258:00:00 | 0:00:00 |
| 53:00:01 | 0:00:00 | 52:00:00 | 0:00:00 | 245:00:00 | 2:00:00 |
| 0:00 | 10:11 | 0:00 | 16:11 | 0:00 | 66:20:00 |
| 19 | 24 | 4 | 9 | 0 | 0 |
| 45 | 4 | 15 | 4 | 0 | 0 |
| 21:00 | 28:01:00 | 8:00 | 11:02 | 0:00 | 1:00 |
| 0:00 | 3:00 | 0:00 | 8:00 | 0:00 | 12:00 |
| 0:00 | 0:00 | 0:00 | 0:00 | 0:00 | 5:01 |
| 59:35:02 | 0:00:00 | 146:113:2 | 1:00:00 | 338:189:9 | 1:00:00 |
| 58:44:01 | 1:01:00 | 149:152:1 | 0:00:00 | 339:223:5 | 0:00:00 |
| 19:1:6:0:0:0:1:0:1:0:0:0 | 0:0:0:0:0:0:0:0:0:0:0:0 | 18:3:2:0:1:0:0:2:0:1:0:0 | 0:0:0:0:0:0:0:0:0:0:0:0 | 117:7:10:2:5:0:1:15:1:0:1:1 | 2:0:0:0:0:0:0:0:0:0:0:0 |
| 68:77:4:3:0:1:3:2:3:4:0:0:3:2:1:1:2:2:0:0:2:3:0:1:0:0:3:0:0:1:0:0:0 | 0:0:0:0:0:0:0:0:0:0:0:0:1:0:0:0:0:0:0:0:0:0:0:0:0:0:0:0:0:0:0:0:0:0 | 67:64:7:3:6:2:6:0:21:9:1:0:11:1:3:8:1:1:0:1:3:1:1:2:0:0:2:0:0:0:0:0:0 | 0:1:0:0:0:0:0:0:0:0:0:0:0:0:0:0:0:0:0:0:0:0:0:0:0:0:0:0:0:0:0:0:0:0 | 223:247:9:12:4:9:11:6:43:12:3:0:27:1:23:9:3:4:0:0:4:1:10:5:2:0:3:2:0:0:1:1:1 | 0:1:0:0:0:0:0:0:0:0:0:0:0:0:0:1:0:0:0:0:0:0:0:0:0:0:0:1:0:0:0:0:0:0 |
| 77:68:4:3:0:1:3:2:3:4:0:0:5:2:1:1:2:2:0:0:2:3:0:1:0:3:0:0:1:0:0:0 | 0:0:0:0:0:0:0:0:0:0:0:0:0:0:0:0:0:0:0:0:0:0:0:0:0:0:0:0:0:0:0:0 | 65:67:7:3:6:2:6:0:19:9:1:0:16:1:5:8:1:1:0:1:3:1:1:2:0:2:0:0:0:0:0:0 | 0:0:0:0:0:0:0:0:0:1:0:0:0:0:0:0:0:0:0:0:0:0:0:0:0:0:0:0:0:0:0:0:0:0 | 247:221:9:12:4:9:11:6:35:12:3:0:31:1:27:9:3:7:0:0:4:1:10:5:2:4:2:0:0:1:1:1 | 1:1:0:0:0:0:0:0:0:1:0:0:0:0:0:0:0:0:0:0:0:0:0:0:0:0:1:0:0:0:0:0:0:0:0 |
| 77:68:2:3:1:0:0:3:1:0:4:0:0:2:5:1:2:0:0:2:1:0:3:0:0:0:1:0:3:0:1:0:0:0:0:0 | 0:0:0:0:0:0:0:0:0:0:0:0:0:0:0:0:0:0:0:0:0:0:0:0:0:0:0:0:0:0:0:0:0:0:0:0:0:0:0:0:0 | 65:64:7:13:3:3:6:6:8:21:9:0:0:14:29:1:1:0:1:8:3:1:1:0:2:1:2:0:2:0:0:0:1:0:0:0 | 0:0:0:0:0:0:0:0:0:0:0:0:0:0:0:0:0:0:0:0:0:0:0:0:0:0:0:0:0:0:0:0:0:0:0:0:0:0:0:0:0:0:0:0 | 246:221:6:37:7:1:5:11:8:37:12:0:0:83:62:3:3:0:1:8:6:2:2:1:1:1:5:1:4:0:0:0:2:1:1:1 | 1:1:0:0:0:0:0:0:0:0:0:0:0:0:0:0:0:0:0:0:0:0:0:0:0:0:0:0:0:0:0:0:0:0:0:0:0:0:0:0 |
| 0:00 | 0:00 | 0:00 | 0:00 | 0:00 | 0:00 |
| 0 | 0 | 0 | 0 | 0 | 0 |
| 0 | 0 | 0 | 0 | 0 | 0 |
| 0 | 0 | 0 | 0 | 0 | 0 |

| | | | | | |
|---|---|---|---|---|---|
| 9:1:2:0:0 | 0:0:0:0:0 | 15:2:3:1:1 | 0:0:0:0:0 | 44:2:6:1:2 | 1:0:1:0:0 |
| 0:1:0:0:0 | 9:0:0:0:0 | 5:1:0:1:0 | 9:0:1:0:0 | 4:0:0:0:0 | 35:0:2:1:4 |
| 0:0:0:0:0 | 9:0:0:0:0 | 5:1:0:1:1 | 9:0:1:1:0 | 4:0:0:0:0 | 35:0:2:8:1 |
| 0:0:0:0:0 | 8:0:0:0:0 | 4:1:1:0:1 | 8:0:1:1:0 | 3:0:0:0:0 | 35:0:8:2:1 |
| 58 | 0 | 41 | 0 | 166 | 2 |
| 45 | 0 | 20 | 0 | 104 | 1 |
| 26 | 1 | 11 | 1 | 56 | 1 |
| 5 | 0 | 6 | 0 | 48 | 0 |
| 2 | 0 | 4 | 0 | 29 | 1 |
| 2 | 0 | 5 | 0 | 16 | 0 |
| 6 | 0 | 18 | 0 | 37 | 0 |
| 35 | 0 | 41 | 0 | 147 | 2 |
| 61 | 1 | 94 | 1 | 266 | 2 |
| 415:22:67:45 | 0:0:0:0 | 1545:109:194:69 | 0:0:1:0 | 1670:100:266:123 | 1:2:0:0 |
| 45:22:415:68 | 0:0:0:0 | 69:107:1545:196 | 0:0:1:0 | 123:93:1671:266 | 0:5:2:0 |
| 415:21:68:46 | 0:0:0:0 | 1553:101:196:69 | 0:0:0:0 | 1677:84:266:123 | 0:2:0:0 |
| 418:67:3:46 | 1:0:0:0 | 1559:180:8:68 | 2:0:0:0 | 1685:236:8:123 | 2:1:0:0 |
| 421:66:3:46 | 0:0:0:0 | 1567:168:7:69 | 1:0:1:0 | 1691:221:8:121 | 2:0:0:1 |
| 0 | 0 | 0 | 0 | 0 | 1 |
| 547 | 1 | 1052 | 2 | 1947 | 4 |
| 547 | 0 | 1055 | 0 | 1949 | 2 |
| 546 | 1 | 1055 | 0 | 1950 | 3 |
| 546 | 1 | 1053 | 2 | 1952 | 1 |
| 542 | 0 | 1036 | 2 | 1927 | 1 |
| 541 | 1 | 1037 | 1 | 1927 | 2 |
| 540 | 0 | 1034 | 1 | 1922 | 1 |
| 532 | 1 | 1018 | 1 | 1880 | 2 |
| 519 | 0 | 978 | 2 | 1813 | 1 |
| 516 | 1 | 972 | 3 | 1797 | 2 |
| 508 | 1 | 909 | 3 | 1695 | 6 |
| 508 | 1 | 905 | 1 | 1678 | 2 |
| 509 | 1 | 905 | 1 | 1683 | 0 |
| 510 | 1 | 900 | 0 | 1679 | 2 |
| 513 | 0 | 874 | 2 | 1646 | 1 |
| 0 | 0 | 0 | 0 | 0 | 0 |
| 0 | 0 | 0 | 0 | 0 | 0 |
| 1:00:00 | 0:00:00 | 1:00:00 | 2:00:00 | 2:00:01 | 0:00:00 |
| 0 | 0 | 1 | 0 | 4 | 0 |
| 17:02 | 0:00 | 141:01:00 | 1:00 | 224:03:00 | 0:00 |
| 2 | 0 | 2 | 0 | 10 | 0 |
| 489 | 2 | 1372 | 1 | 2277 | 1 |
| 489 | 0 | 1368 | 1 | 2275 | 0 |
| 0 | 0 | 0 | 0 | 0 | 1 |
| 10:11:0:0:4 | 0:0:0:0:0 | 21:21:2:0:3 | 0:0:0:0:0 | 37:70:1:0:5 | 0:0:0:0:0 |
| 642:51:00 | 0:00 | 1291:647 | 1:02 | 3406:192 | 1:00 |
| 643:50:00 | 0:00 | 1297:646 | 1:01 | 3419:185 | 6:01 |
| 641:50:00 | 2:00 | 1297:642 | 0:03 | 3425:181 | 0:03 |
| 1 | 1 | 3 | 4 | 0 | 14 |
| 1 | 0 | 0 | 3 | 1 | 2 |
| 0:17:0:0:0:1 | 0:0:0:0:0:0 | 5:11:0:1:0:1 | 0:0:0:0:0:0 | 6:25:0:0:0:0 | 0:0:0:0:0:0 |
| 0:0:0:0 | 3:2:0:0 | 0:0:0:0 | 10:0:1:0 | 0:0:0:0 | 15:1:1:0 |
| 3:2:0:0 | 0:0:0:0 | 10:3:1:0 | 0:0:0:0 | 15:1:1:0 | 0:0:0:0 |
| 2 | 0 | 1 | 0 | 1 | 0 |
| 1 | 0 | 0 | 0 | 0 | 0 |
| 0:00 | 2:00 | 0:00 | 2:01 | 0:00 | 11:02 |
| 0:00 | 0:00 | 0:00 | 1:01 | 0:00 | 1:00 |
| 0:00:00 | 1:00:00 | 0:00:00 | 0:00:00 | 0:00:00 | 4:00:01 |
| 1:00:00 | 0:00:00 | 0:00:00 | 0:00:00 | 4:00:01 | 0:00:00 |
| 4:01 | 0:00 | 7:00 | 0:00 | 59:05:00 | 0:00 |
| 44:585:11:191:16:8:6:8 | 0:1:0:0:0:0:0:0 | 22:662:7:274:26:12:13:4 | 0:2:0:0:0:0:0:0 | 187:1571:76:623:51:30:33:8 | 0:0:0:1:0:0:0:0 |
| 45:589:11:193:16:10:8:8 | 0:0:0:0:0:0:0:0 | 22:669:7:276:26:18:27:4 | 0:0:0:0:0:0:0:0 | 187:1603:76:636:50:46:73:8 | 0:0:0:0:0:0:0:0 |
| 45:588:11:193:16:12:8:8 | 0:0:0:0:0:0:0:0 | 22:670:7:276:26:22:30:4 | 0:1:0:1:0:1:1:0 | 186:1613:76:638:50:52:88:8 | 0:1:0:2:0:0:1:0 |
| 2:01 | 2:00 | 2:00 | 4:00 | 20:04 | 33:02:00 |
| 45:592:11:194:16:16:8:8 | 0:0:0:0:0:1:0:0 | 22:676:6:279:25:30:44:4 | 0:0:0:0:0:0:0:0 | 186:1629:75:644:50:70:119:8 | 0:2:0:0:0:0:0:0 |
| 590:45:11:193:16:17:8:8 | 2:0:0:1:0:0:0:0 | 675:22:6:279:25:30:44:4 | 1:0:0:0:0:0:0:0 | 1627:185:75:643:49:70:119:8 | 4:1:1:1:1:0:0:0 |
| 591:45:11:194:16:16:8:8 | 1:0:0:0:0:0:0:0 | 676:22:6:280:25:31:45:4 | 0:0:0:0:0:0:0:0 | 1633:185:76:645:50:71:124:8 | 1:1:0:0:0:0:0:0 |
| 45:592:11:194:16:16:10:8 | 0:0:0:0:0:0:0:0 | 22:677:6:283:25:32:48:4 | 0:1:0:1:0:0:0:0 | 185:1643:76:649:50:73:134:8 | 1:2:0:3:0:0:0:0 |
| 45:597:11:194:16:16:11:8 | 0:0:0:0:0:0:0:0 | 22:678:6:285:25:35:53:4 | 0:2:0:0:0:0:1:0 | 186:1653:76:653:50:70:143:8 | 0:0:0:1:0:0:0:0 |

| | | | | | |
|---|---|---|---|---|---|
| 596:45:11:193:16:16:13:8 | 0:0:0:0:0:0:0:0 | 678:22:6:284:25:43:65:4 | 1:0:0:0:0:0:0:0 | 1628:186:76:653:50:80:152:8 | 3:0:0:0:0:0:0:0 |
| 600:45:11:190:16:15:12:8 | 0:0:0:2:0:0:0:0 | 678:22:6:283:25:53:64:4 | 1:0:0:0:0:0:0:0 | 1637:183:75:653:50:94:150:8 | 2:1:0:0:0:1:0:0 |
| 45:600:11:192:16:15:12:8 | 0:0:0:0:0:0:0:0 | 22:676:6:283:25:55:64:4 | 0:2:0:0:0:2:0:0 | 184:1642:75:653:50:96:149:8 | 0:1:0:0:0:3:0:0 |
| 600:45:11:194:16:16:11:8 | 0:0:0:0:0:0:0:0 | 678:22:6:284:24:68:56:4 | 0:0:0:0:0:0:1:0 | 1645:185:75:651:50:122:138:8 | 3:0:0:0:0:0:0:0 |
| 45:600:11:194:16:18:11:8 | 0:0:0:0:0:0:0:0 | 22:677:6:284:24:70:55:4 | 0:0:0:0:0:0:1:0 | 185:1646:74:650:50:121:136:8 | 0:2:0:1:0:0:0:0 |
| 328:37:0:5:1:2:1:2:0:0:1 | 0:0:0:0:0:0:0:0:0:0:0 | 610:99:0:3:1:3:3:2:0:0:0 | 2:1:0:0:0:0:0:0:0:0:0 | 1761:287:0:23:5:8:5:2:0:0:0 | 2:0:0:0:0:0:0:0:0:0:0 |
| 328:38:0:5:1:2:1:2:0:0:1 | 0:0:0:0:0:0:0:0:0:0:0 | 613:100:0:3:1:3:3:2:0:0:0 | 0:0:0:0:0:0:0:0:0:0:0 | 1763:288:0:23:5:7:5:2:0:0:0 | 2:2:0:0:0:0:0:0:0:0:0 |
| 327:38:0:5:1:2:1:2:0:0:1 | 1:0:0:0:0:0:0:0:0:0:0 | 610:100:0:3:1:2:3:2:0:0:0 | 2:0:0:0:0:0:0:0:0:0:0 | 1762:290:0:22:5:7:5:2:0:0:0 | 4:0:0:0:0:0:0:0:0:0:0 |
| 328:32:37:0:8:5:1:1:1:2:0:0 | 1:0:0:0:0:0:0:0:0:0:0:0 | 613:145:98:0:19:3:1:2:3:2:0:0 | 0:0:0:0:0:0:0:0:0:0:0:0 | 1764:258:282:0:36:22:5:4:5:2:0:0 | 1:0:2:0:0:0:0:0:0:0:0:0 |
| 329:32:37:0:9:5:1:1:1:2:0:0 | 0:0:0:0:0:0:0:0:0:0:0:0 | 613:145:94:0:19:3:1:2:3:2:0:0 | 0:0:0:0:0:0:0:0:0:0:0:0 | 1762:254:281:0:38:22:5:4:5:2:0:0 | 2:1:0:0:0:0:0:0:0:0:0:0 |
| 328:33:34:0:9:5:1:1:1:2:0:0:0 | 1:0:0:0:0:0:0:0:0:0:0:0:0 | 613:154:71:0:19:3:1:2:3:2:0:0:0 | 1:0:0:0:0:0:0:0:0:0:0:0:0 | 1763:272:209:0:43:23:5:4:5:2:0:0:1 | 1:0:2:0:0:0:0:0:0:0:0:0:0 |
| 328:33:31:0:7:5:1:1:1:2:0:0:0 | 1:0:0:0:0:0:0:0:0:0:0:0:0 | 614:155:67:0:14:3:1:2:3:2:0:0:0 | 0:0:1:0:0:0:0:0:0:0:0:0:0 | 1763:273:185:0:33:23:5:4:5:2:0:0:1 | 2:1:4:0:0:0:0:0:0:0:0:0:0 |
| 328:33:29:0:7:5:1:1:1:2:0:0:0 | 1:0:0:0:0:0:0:0:0:0:0:0:0 | 614:155:64:0:13:3:1:2:3:2:0:0:0 | 0:0:1:0:0:0:0:0:0:0:0:0:0 | 1762:275:183:0:31:23:5:4:5:2:0:0:1 | 3:0:2:0:0:0:0:0:0:0:0:0:0 |
| 6:2:7:0:3:0:0:1:0:2:0:0:0 | 0:0:0:0:0:0:0:0:0:0:0:0:0 | 0:0:21:0:21:6:0:0:0:0:0:0:0 | 0:0:0:0:0:0:0:0:0:0:0:0:0 | 4:1:68:1:19:10:0:2:0:1:0:2:1 | 0:0:1:0:1:0:0:0:0:0:0:0:0 |
| 28:2:0:2:2:2:1:0:0:0:1:0:0:0:2:0:0:0:0 | 0:0:0:0:0:0:0:0:0:0:0:0:0:0:0:0:0:0:0 | 77:0:1:6:0:7:4:0:2:2:0:0:1:2:1:0:1:3:0 | 0:0:0:0:0:0:0:0:0:0:0:0:0:0:0:0:0:0:0 | 270:10:1:9:2:9:13:3:2:6:5:2:10:2:7:2:0:1:1 | 0:0:0:0:0:0:0:0:0:0:0:0:0:0:0:0:0:0:0 |
| 28:2:0:2:2:2:1:0:0:0:1:0:0:0:2:0:0:0:0 | 0:0:0:0:0:0:0:0:0:0:0:0:0:0:0:0:0:0:0 | 77:0:1:6:0:7:4:2:2:2:0:0:3:2:1:0:1:3:0 | 0:0:0:0:0:0:0:0:0:0:0:0:0:0:0:0:0:0:0 | 268:9:1:9:2:9:13:5:2:6:5:2:15:2:7:2:0:1:1 | 1:1:0:0:0:0:0:1:0:0:0:0:0:0:0:0:0:0:0 |
| 28:2:0:2:2:2:1:0:0:0:1:0:0:0:2:0:0:0:0 | 0:0:0:0:0:0:0:0:0:0:0:0:0:0:0:0:0:0:0 | 77:0:1:6:0:7:4:2:2:2:0:0:4:2:1:0:1:3:0 | 0:0:0:0:0:0:0:0:0:0:0:0:0:0:0:0:0:0:0 | 269:10:1:9:2:9:13:5:2:6:5:2:15:2:7:2:0:1:1 | 1:0:0:0:0:0:0:1:0:0:0:0:0:0:0:0:0:0:0 |
| 27:2:0:1:2:2:1:0:0:0:0:3:0:0:0:1:2:0:0:0 | 0:0:0:1:0:0:0:0:0:0:0:0:0:0:0:0:0:0:0:0 | 69:0:1:6:0:6:4:0:2:2:0:16:2:0:1:1:0:3:1:0 | 6:0:0:0:0:1:0:0:0:0:0:1:0:0:0:0:0:0:0:0 | 260:10:1:8:2:7:13:2:2:6:5:41:2:3:0:0:5:1:1:1 | 4:0:0:0:0:0:0:0:0:0:0:1:0:0:0:0:0:0:0:0 |
| 11:1:0:0:2:0:0:0:2:0:0:0:0:0:0:1:2:0:0:0 | 15:1:2:1:0:1:0:0:0:0:0:0:0:0:0:0:0:0:0:0 | 56:5:0:3:0:3:0:0:14:1:2:1:1:1:1:1:0:2:1:0 | 11:1:0:3:0:1:0:0:3:1:0:0:1:0:0:0:0:1:0:0 | 191:4:6:6:2:7:2:2:31:1:5:1:2:3:0:0:4:1:1:1 | 57:3:4:1:0:5:3:0:5:1:1:0:0:0:0:0:1:0:0:0 |
| 0:0:9:2:0:3:1:0:0:0:0:0:0:0:1:2:0:0:0 | 0:0:0:0:0:0:0:0:0:0:0:0:0:0:0:0:0:0:0 | 3:0:32:0:3:8:4:2:1:2:1:0:1:3:1:0:2:0:0 | 0:0:5:0:0:0:0:0:0:0:0:0:0:0:0:0:0:0:0 | 3:3:120:2:5:19:3:4:1:1:5:2:1:1:0:2:1:0:2 | 1:0:11:0:0:3:0:0:0:0:0:0:0:0:0:0:0:1:0 |
| 2:1:0:0:0:0:0:0:0:0:1:1:0:0 | 0:0:0:0:0:0:0:0:0:0:0:0:0:0 | 0:1:2:0:2:2:0:0:0:0:1:1:3:0 | 0:2:0:0:0:0:0:0:0:0:0:0:0:0 | 2:5:0:0:2:7:0:0:0:0:3:0:1:1 | 0:2:0:0:0:0:0:0:0:0:1:0:0:0 |
| 2:1:0:0:0:0:0:0:0:0:0:1:0:0 | 0:0:0:0:0:0:0:0:0:1:0:0:0:0 | 0:1:1:0:2:0:2:0:0:0:0:1:3:0 | 0:2:1:0:0:0:0:0:0:1:0:0:0:0 | 2:2:0:0:2:0:7:0:0:2:0:0:1:1 | 0:5:0:0:0:0:0:0:0:1:0:0:0:0 |
| 4:4:0:0:1:1:1:0:0:0 | 4:3:0:0:0:0:1:0:0:0 | 10:13:6:2:1:1:1:0:0:1 | 16:16:6:2:2:1:4:0:0:0 | 4:6:2:1:0:2:0:0:0:0 | 9:5:0:1:0:2:1:1:1:0 |
| 4:3:1:0:0:0:0:0:0:0 | 4:4:1:1:0:1:0:0:0 | 16:16:23:1:1:1:0:1:0 | 10:13:18:2:1:1:0:0:1 | 9:5:7:2:0:0:1:1:0 | 3:6:3:1:0:0:0:0:0 |
| 5:0:0:1 | 0:0:0:1 | 1:0:0:0 | 0:0:1:2 | 6:7:6:8 | 0:0:1:2 |
| 0 | 0 | 5 | 0 | 1 | 0 |
| 0 | 0 | 0 | 0 | 0 | 0 |
| 0 | 0 | 0 | 0 | 0 | 0 |
| 34:0:2:3:4:1:1:0:0:1:0:2:0:2:0:1:0:0:0:0:0:1:0:0:0 | 0:0:0:0:0:0:0:0:0:0:0:0:0:0:0:0:0:0:0:0:0:0:0:0:0 | 43:0:5:2:13:0:1:2:0:0:0:2:1:0:1:0:0:0:1:0:0:0:0:0:0 | 0:0:0:0:0:0:0:0:0:0:0:0:0:0:0:0:0:0:0:0:0:0:0:0:0 | 167:0:22:6:41:0:3:2:1:1:0:2:10:1:5:2:0:0:5:1:0:0:1:1:1 | 2:0:0:0:0:0:0:0:0:0:0:0:0:0:0:0:0:0:0:0:0:0:0:0:0 |
| 0:0:0:0:0:0 | 18:20:21:3:11:3 | 0:0:0:0:0:0 | 50:26:19:10:4:5 | 0:0:0:0:0:0 | 22:15:26:6:11:2 |
| 0:00:00 | 22:02:00 | 0:00:00 | 4:06:00 | 0:00:00 | 37:00:01 |
| 6 | 0 | 15 | 0 | 17 | 0 |
| 6 | 0 | 0 | 15 | 10 | 7 |
| 0:00 | 0:00 | 1:00 | 6:00 | 3:00 | 15:00 |
| 28:58:1:0:102:9:0:0:1:0 | 0:0:0:0:0:0:0:0:0:0 | 48:177:1:8:172:5:0:0:1:0 | 0:0:0:0:0:0:0:0:0:0 | 170:506:4:21:2329:96:5:13:12:3 | 0:2:0:0:1:0:0:0:0:0 |
| 28:57:1:0:101:9:0:0:1:0 | 0:0:0:0:1:0:0:0:0:0 | 48:178:1:8:170:5:0:0:1:0 | 0:0:0:0:1:0:0:0:0:0 | 171:504:4:21:2325:97:5:13:12:3 | 0:1:0:0:3:0:0:0:0:0 |

| | | | | | |
|---|---|---|---|---|---|
| 28:57:1:0:102:9:0:0:1:0 | 0:0:0:0:0:0:0:0:0:0 | 48:177:1:8:171:5:0:0:1:0 | 0:0:0:0:0:0:0:0:0:0 | 171:507:4:21:2323:97:5:13:12:3 | 0:0:0:0:1:0:0:0:0:0 |
| 28:57:1:0:102:9:0:0:1:0 | 0:0:0:0:0:0:0:0:0:0 | 48:176:1:8:171:5:0:0:1:0 | 0:1:0:0:0:0:0:0:0:0 | 171:505:4:21:2317:97:5:14:12:3 | 0:0:0:0:1:0:0:0:0:0 |
| 28:57:1:0:102:9:0:0:1:0 | 0:0:0:0:0:0:0:0:0:0 | 48:176:1:8:171:5:0:0:1:0 | 0:1:0:0:0:0:0:0:0:0 | 171:505:4:21:2318:97:5:14:12:3 | 0:0:0:0:2:0:0:0:0:0 |
| 57:28:1:0:102:9:0:0:1:0 | 0:0:0:0:0:0:0:0:0:0 | 176:48:1:8:171:5:0:0:1:0 | 0:0:0:0:0:0:0:0:0:0 | 505:171:4:21:2318:97:5:15:12:3 | 0:0:0:0:2:0:0:0:0:0 |
| 28:58:1:0:102:9:0:1:1:0 | 0:0:0:0:0:0:0:0:0:0 | 48:176:1:8:171:5:0:0:1:0 | 0:0:0:0:0:0:0:0:0:0 | 171:507:4:21:2309:95:5:16:12:3 | 0:0:0:0:2:0:0:0:0:0 |
| 57:28:1:0:101:9:0:1:1:0 | 0:0:0:0:1:0:0:0:0:0 | 160:45:1:8:151:4:0:0:1:0 | 15:3:0:0:15:1:0:0:0:0 | 472:168:4:21:2152:89:5:16:12:3 | 34:2:0:0:143:5:0:0:0:0 |
| 28:57:1:0:100:9:0:1:1:0 | 0:1:0:0:2:0:0:0:0:0 | 45:160:1:8:151:4:0:0:1:0 | 3:15:0:0:16:1:0:0:0:0 | 168:473:4:21:2151:89:5:16:12:3 | 3:36:0:0:152:5:0:0:0:0 |
| 28:56:1:0:100:9:0:1:1:0 | 0:0:0:0:0:0:0:0:0:0 | 45:160:1:8:149:4:0:0:1:0 | 0:0:0:0:1:0:0:0:0:0 | 168:472:4:21:2150:88:5:15:12:3 | 0:1:0:0:1:0:0:0:0:0 |
| 28:53:1:0:94:9:0:1:0 | 0:0:0:0:1:0:0:0:0 | 44:159:1:8:149:4:0:1:0 | 0:0:0:0:0:0:0:0:0 | 164:466:4:21:2072:82:5:12:3 | 0:1:0:0:0:0:0:0:0 |
| 28:54:1:0:95:9:0:1:0 | 0:0:0:0:0:0:0:0:0 | 44:159:1:8:149:4:0:1:0 | 0:1:0:0:0:0:0:0:0 | 164:469:4:21:2067:82:5:12:3 | 0:0:0:0:1:0:0:0:0 |
| 28:54:1:0:95:9:0:1:0 | 0:0:0:0:0:0:0:0:0 | 44:159:1:8:149:4:0:1:0 | 0:1:0:0:0:0:0:0:0 | 164:469:4:21:2068:82:5:12:3 | 0:0:0:0:0:0:0:0:0 |
| 28:55:1:0:95:9:0:1:0:0 | 0:0:0:0:0:0:0:0:0:0 | 44:159:1:8:148:4:0:1:0:0 | 0:1:0:0:0:0:0:0:0:0 | 163:466:4:21:2057:78:5:12:3:1 | 0:0:0:0:0:1:0:0:0:0 |
| 28:56:0:95:9:0:1 | 0:0:0:0:0:0:0 | 44:159:8:145:4:0:0 | 0:0:0:1:0:0:0 | 161:470:21:2030:76:5:18 | 0:1:0:1:0:0:0 |
| 28:57:1:0:93:9:0:0 | 0:0:0:0:0:0:0:0 | 45:162:1:8:146:4:0:0 | 0:0:0:0:0:0:0:0 | 165:470:5:21:2076:82:5:17 | 1:6:0:0:5:1:0:0 |
| 28:57:1:0:93:9:0:0 | 0:0:0:0:0:0:0:0 | 45:162:1:8:146:4:0:0 | 0:0:0:0:0:0:0:0 | 166:475:5:21:2084:82:5:17 | 0:0:0:0:0:0:0:0 |
| 0:5:0:0 | 0:0:0:0 | 0:0:0:0 | 0:0:0:0 | 0:5:0:0 | 0:0:0:0 |
| 0 | 0 | 0 | 6 | 0 | 1 |
| 69:8:15:2:7:0:0:0:0:0:0 | 0:0:0:0:0:0:0:0:0:0:0 | 20:0:30:0:133:0:0:0:0:0:0 | 0:0:1:0:1:0:0:0:0:0:0 | 97:16:81:5:270:1:0:1:1:1:1 | 0:0:0:0:0:0:0:0:0:0:0 |
| 69:8:15:3:5:0:0:0:1:0:0:0:0 | 0:0:0:0:0:0:0:0:0:0:0:0:0 | 10:0:32:0:20:0:0:0:1:0:1:0:0 | 0:0:0:0:1:0:0:0:0:0:0:0:0 | 89:17:83:5:58:0:1:0:6:1:2:1:0 | 0:1:0:0:1:1:0:0:0:0:0:0:0 |
| 5:1:0:0:1 | 0:0:0:0:0 | 9:0:0:1:0 | 4:0:0:1:0 | 3:0:0:0:0 | 4:0:0:0:0 |
| 5:1:0:0:1 | 0:0:0:0:0 | 12:0:0:2:0 | 0:0:0:0:0 | 7:0:0:0:0 | 0:0:0:0:0 |
| 0:0:0:0:0:0 | 1:1:0:0:1:1 | 1:0:0:0:0:0 | 2:1:0:0:0:0 | 1:0:0:0:0:0 | 0:0:0:0:0:0 |
| 0:00 | 0:00 | 0:00 | 0:00 | 0:00 | 0:00 |
| 0:00 | 0:00 | 0:00 | 0:00 | 0:00 | 0:00 |
| 57:01:00 | 0:00:00 | 120:01:02 | 0:00:00 | 326:06:00 | 2:00:00 |
| 234:743:700:216:54:5 | 0:1:0:0:1:0 | 338:829:667:206:55:15 | 0:3:1:0:0:0 | 657:2023:1909:599:134:45 | 0:5:1:3:0:0 |
| 234:698:216:742:54:5 | 0:2:0:1:1:0 | 337:667:204:830:55:15 | 1:1:2:2:0:0 | 656:1906:599:2025:134:45 | 0:3:2:4:0:0 |
| 697:216:742:54:234:5 | 2:0:2:1:0:0 | 668:206:832:54:338:15 | 1:0:0:1:0:0 | 1905:602:2026:134:656:45 | 4:1:3:0:1:0 |
| 696:216:745:54:234:5 | 0:0:1:1:0:0 | 663:206:830:55:336:15 | 0:0:3:0:3:0 | 1891:604:2029:134:659:45 | 4:0:2:0:0:0 |
| 686:216:745:54:236:5 | 2:0:1:1:0:0 | 631:206:832:54:339:15 | 1:1:1:1:0:0 | 1833:605:2035:134:663:45 | 2:1:5:0:0:0 |
| 746:216:684:53:236:5 | 0:0:2:2:0:0 | 831:207:626:55:339:15 | 1:0:0:0:1:0 | 2035:606:1823:134:665:45 | 5:0:0:0:0:0 |
| 686:215:745:54:236:5 | 0:1:1:1:0:0 | 626:207:833:55:339:15 | 0:0:0:0:1:0 | 1820:606:2037:134:665:45 | 0:0:3:0:0:0 |
| 683:217:746:54:236:5 | 0:0:0:1:0:0 | 623:207:834:52:338:15 | 1:0:1:0:1:0 | 1812:605:2041:130:663:45 | 3:1:2:0:1:0 |
| 745:676:217:54:231:5 | 0:0:0:1:0:0 | 833:617:207:51:332:15 | 1:0:0:0:0:0 | 2043:1790:604:130:657:45 | 2:3:2:0:0:0 |
| 676:217:745:53:229:5 | 0:0:0:1:1:0 | 615:207:833:50:332:15 | 2:0:1:1:0:0 | 1789:605:2038:130:656:45 | 7:1:5:0:0:0 |
| 0:0:0:0 | 0:2:0:0 | 0:0:0:0 | 0:3:0:0 | 0:0:0:0 | 0:13:0:0 |
| 0:00 | 0:00 | 0:00 | 0:00 | 0:00 | 0:00 |
| 0:0:0:0 | 0:0:1:1 | 0:0:0:1 | 0:0:0:0 | 0:0:0:0 | 0:0:0:0 |
| 0:0:0:0 | 0:7:0:0 | 0:0:0:0 | 0:2:0:0 | 0:0:1:0 | 0:12:1:0 |
| 72:01:03 | 0:00:00 | 70:12:05 | 0:00:00 | 303:40:09 | 1:01:00 |
| 1:8:2:0:0:0 | 0:0:0:0:0:0 | 1:12:2:0:1:0 | 0:2:0:0:0:0 | 6:23:2:0:1:1 | 0:2:0:0:0:0 |
| 0:3:1:2:0:1:0 | 1:7:2:1:4:0:0 | 0:3:2:0:0:2:0 | 0:12:0:0:1:2:1 | 4:11:1:1:0:5:1 | 1:14:1:0:4:1:0 |
| 18:1:3:2:3:3 | 1:0:0:0:0:0 | 29:0:2:4:0:2 | 4:0:0:1:0:0 | 42:5:2:8:1:4 | 6:0:0:0:0:0 |
| 84:14:20 | 0:00:01 | 80:05:20 | 3:00:01 | 146:26:23 | 2:00:00 |
| 83:20:05 | 0:00:00 | 80:19:03 | 0:00:00 | 138:23:20 | 0:00:00 |
| 0:1:0:1:0:4:0:1:0:0:2:0:1:0:0:0 | 0:0:0:0:0:0:0:0:0:0:0:0:0:0:0:0 | 1:18:1:12:0:0:0:0:0:2:0:0:4:0:0:0 | 0:0:0:0:0:0:0:0:0:0:0:0:0:0:0:0 | 0:25:2:17:2:7:0:1:0:5:1:0:9:1:1:1 | 0:0:0:0:0:0:0:0:0:0:0:0:0:0:0:0 |
| 4:2:0:0:0:1:0:1:0:0 | 0:0:0:0:0:0:0:0:0:0 | 2:1:0:0:0:0:1:0:0:1 | 0:0:0:0:0:0:0:0:0:0 | 11:18:1:3:0:0:0:0:2:0 | 0:0:0:0:0:0:0:0:0:0 |

| | | | | | |
|---|---|---|---|---|---|
| 159:15:69 | 0:01:01 | 18:161:198 | 0:01:00 | 90:367:279 | 0:01:00 |
| 159:15:70 | 0:00:00 | 18:160:198 | 0:00:00 | 89:356:279 | 0:00:00 |
| 159:15:69 | 0:00:00 | 18:159:198 | 0:00:00 | 89:353:277 | 0:01:01 |
| 158:11:69 | 0:00:00 | 17:117:189 | 0:00:00 | 89:274:261 | 0:01:01 |
| 159:10:66 | 0:00:01 | 19:89:166 | 0:02:00 | 92:163:230 | 1:02:01 |
| 159:10:67 | 0:00:00 | 19:87:163 | 0:01:00 | 92:154:226 | 1:02:01 |
| 160:10:67 | 5:00:01 | 22:68:154 | 5:01:01 | 96:103:211 | 16:01:03 |
| 170:02:59 | 0:00:00 | 32:2:104 | 0:00:01 | 118:2:139 | 1:01:00 |
| 168:2:51:0 | 2:0:0:0 | 34:5:61:0 | 0:0:0:0 | 121:2:84:0 | 0:0:1:0 |
| 1:1:3:0:0:0:0:0:1:0:0:0 | 0:1:0:0:0:0:0:0:0:0:0:0 | 6:21:17:1:1:0:0:2:6:0:3:0 | 0:4:2:0:0:0:1:0:2:0:0:0 | 7:9:14:0:7:0:0:0:10:0:2:0 | 1:2:4:0:2:0:0:0:1:0:0:0 |
| 1:2:2:0:1:0:0:0:0:0 | 0:0:0:0:0:0:0:0:0:0 | 4:7:6:2:10:1:0:0:0:1 | 0:0:0:0:0:0:0:0:0:0 | 6:5:4:0:13:5:0:0:0:1 | 1:1:0:0:0:0:0:0:0:0 |
| 0:1:1:0:0:0 | 0:0:0:0:0:0 | 0:2:0:1:11:0 | 0:0:0:0:9:1 | 1:0:1:0:15:0 | 0:2:0:0:7:1 |
| 0:00:00 | 6:00:00 | 0:00:00 | 23:09:02 | 0:00:00 | 19:14:01 |
| 5:00 | 0:00 | 6:02 | 0:00 | 12:01 | 0:00 |
| 5:00 | 0:00 | 6:02 | 0:00 | 12:01 | 0:00 |
| 5:00 | 0:00 | 5:02 | 0:00 | 12:01 | 0:00 |
| 1:4:1:0:0 | 2:0:1:0:0 | 37:3:4:0:1 | 7:0:0:0:1 | 59:6:14:2:2 | 13:0:4:0:0 |
| 0:0:1:0:3:0:18:8:1:6:3:0:0:0:0:0:0:0:0:0:2:0 | 2:4:3:1:0:1:0:0:0:1:0:0:0:0:0:0:0:0:1:0:0:0 | 1:2:2:0:5:0:1:13:0:4:2:2:0:0:0:0:0:0:0:0:0:4 | 0:3:3:2:0:1:0:3:0:0:0:0:0:0:0:0:0:0:0:0:0:0 | 6:1:8:1:8:3:19:4:1:13:4:3:0:1:1:3:0:0:0:2:2:1 | 7:10:27:3:0:6:2:1:0:2:2:0:1:0:0:1:0:0:1:1:0:0 |
| 2:4:3:1:18:8:1:7:4:2:3:0:0:0:0:0:0:1:0:2:0:0 | 0:1:0:0:0:0:0:0:0:0:0:0:0:0:0:0:0:0:0:0:0:0 | 1:6:5:1:1:16:0:4:5:2:2:2:0:0:0:0:0:0:0:0:0:4 | 0:0:0:0:0:0:0:0:0:0:0:0:0:0:0:0:0:0:0:0:0:0 | 14:34:8:10:21:6:1:15:11:4:6:3:1:1:4:0:0:1:3:2:1:1 | 1:1:0:0:0:0:0:0:0:0:0:0:0:0:0:0:0:0:0:0:0:0 |
| 0:00 | 0:00 | 0:00 | 0:00 | 3:00 | 0:03 |
| 0:0:0:0 | 2:0:0:0 | 0:0:0:0 | 0:0:0:0 | 0:0:0:0 | 9:0:1:1 |
| 0 | 5 | 0 | 9 | 0 | 1 |
| 53:0:97 | 0:00:00 | 19:02:52 | 2:00:00 | 41:0:148 | 0:00:00 |
| 49:93:0 | 4:02:00 | 10:26:00 | 9:27:02 | 35:144:0 | 4:07:00 |
| 0 | 0 | 0 | 0 | 0 | 0 |
| 0 | 1 | 0 | 2 | 0 | 1 |
| 110:412:30:0 | 0:1:0:0 | 28:171:9:2 | 1:0:0:0 | 240:997:46:0 | 1:1:0:0 |
| 2:00 | 2:00 | 0:00 | 0:00 | 4:00 | 4:00 |
| 7:4:2:6:2:2 | 0:0:0:0:0:0 | 37:14:17:13:2:3 | 2:1:2:1:1:2 | 23:6:10:14:4:5 | 0:1:2:1:0:0 |
| 5:4:2:6:3:2 | 0:0:0:0:0:0 | 40:17:20:15:3:5 | 0:0:0:0:0:0 | 23:7:14:14:4:5 | 0:1:0:1:0:0 |
| 1:1:1:0:0:1 | 3:1:1:5:3:1 | 6:13:13:3:1:3 | 11:7:6:13:2:2 | 4:9:13:4:2:3 | 4:5:4:11:2:2 |
| 0:2:14:1:0:2:2:1:0:0:0 | 0:0:0:0:0:0:0:0:0:0:0 | 0:76:48:22:0:19:0:0:0:0:0 | 0:0:0:0:0:0:0:0:0:0:0 | 1:48:34:9:0:6:0:1:0:0:0 | 0:0:0:0:0:0:0:0:0:0:0 |
| 0:2:13:1:0:0:2:2:1:0:0 | 0:0:0:0:0:0:0:0:0:0:0 | 0:76:42:22:0:0:20:0:0:0:0 | 0:0:0:0:0:0:0:0:0:0:0 | 1:48:30:9:0:0:6:0:1:0:0 | 0:0:0:0:0:0:0:0:0:0:0 |
| 0:0:0:2:0:2:0:0 | 0:0:0:0:0:0:0:0 | 0:0:0:5:23:0:0:0 | 0:0:0:0:0:0:0:0 | 0:0:0:1:13:0:0:0 | 0:0:0:2:0:0:0:0 |
| 2:0:0:0:2:0:0:0:0 | 2:0:0:0:0:0:0:0:0 | 8:0:0:0:0:0:17:0:0 | 19:0:2:0:1:0:1:0:0 | 5:0:1:0:0:0:11:0:0 | 4:0:2:0:0:0:0:0:0 |
| 0:0:0:2:2:0:0:0:0:0 | 0:0:0:2:0:0:0:0:0:0 | 0:0:0:12:0:0:16:0:0:1 | 0:1:0:17:1:0:1:0:0:0 | 0:3:0:6:0:0:10:0:0:0 | 0:0:0:3:0:0:0:0:0:0 |
| 0:0:0:2:2:0:0:0:0 | 0:0:0:2:0:0:0:0:0 | 0:1:0:22:0:16:0:0:0 | 0:1:0:9:1:1:0:0:0 | 0:3:0:7:0:10:0:0:0 | 0:0:0:2:0:0:0:0:0 |
| 1:0:2:0:5:2:0:3:0:0:0 | 0:0:0:0:0:0:0:0:0:0:0 | 18:2:31:0:36:20:0:1:0:0:0 | 1:0:0:0:0:0:1:0:0:0:0 | 6:3:13:0:10:6:1:1:0:0:0 | 1:0:0:0:0:0:0:0:0:0:0 |
| 1:0:0:0:0:0:1:0 | 0:0:2:0:0:0:0:0 | 0:0:3:0:0:0:0:0 | 1:0:2:2:0:0:0:0 | 0:0:3:3:0:0:0:0 | 0:0:4:0:0:0:0:1 |
| 0:01:00 | 1:00:00 | 0:00:00 | 0:00:00 | 8:01:00 | 3:03:01 |
| 0:01 | 1:00 | 0:00 | 0:00 | 8:01 | 3:03 |
| 60:04:00 | 0:00:00 | 379:24:00 | 1:00:00 | 340:22:00 | 0:00:00 |
| 0.210416667 | 0:01:00 | 24:402:0 | 1:16:00 | 1.039583333 | 0:15:00 |
| 68:4:0:0 | 4:0:0:0 | 438:26:0:1 | 9:0:0:0 | 409:19:0:0 | 3:0:0:0 |
| 73:04:00 | 0:00:00 | 454:16:01 | 0:00:00 | 416:14:00 | 0:00:00 |
| 1:00 | 1:01 | 0:00 | 0:00 | 3:00 | 2:00 |
| 22:2:11:33:1:1:0:2:2:0:0:1 | 0:0:0:0:0:0:0:0:0:0:0:0 | 33:6:1:15:3:0:2:0:0:0:2:0 | 0:0:0:0:0:0:0:0:0:0:0:0 | 98:8:32:46:6:1:6:0:1:4:6:2 | 0:0:1:1:0:0:0:0:0:0:0:0 |
| 12:3:17:50:2:1:0:2:2:2:1:0 | 0:0:0:0:0:0:0:0:0:0:0:0 | 5:8:6:28:5:1:2:0:0:1:0:4 | 0:0:0:0:0:0:0:0:0:0:0:0 | 25:13:52:95:14:1:11:2:1:2:5:8 | 0:1:0:1:0:0:0:0:0:0:0:0 |
| 0:00:00 | 4:02:00 | 0:00:00 | 0:00:00 | 0:00:00 | 2:01:00 |
| 1:0:0:1:0 | 0:0:0:0:0 | 1:1:0:0:1 | 0:0:0:0:0 | 0:1:0:0:1 | 0:0:0:0:0 |
| 519:110 | 0:00 | 414:130 | 0:00 | 1202:442 | 0:00 |
| 519:111 | 1:00 | 421:132 | 1:00 | 1208:444 | 5:00 |
| 519:111 | 0:00 | 422:132 | 0:00 | 1215:447 | 6:00 |
| 523:112 | 4:00 | 430:135 | 1:00 | 1257:462 | 0:01 |
| 527:112 | 2:00 | 441:135 | 1:00 | 1290:463 | 2:00 |
| 536:112 | 0:00 | 448:134 | 0:00 | 1323:462 | 1:01 |
| 111:552 | 0:00 | 133:466 | 0:01 | 455:1360 | 0:02 |
| 554:110 | 1:01 | 467:132 | 1:00 | 1366:450 | 1:02 |

| | | | | | |
|---|---|---|---|---|---|
| 557:111 | 2:00 | 468:130 | 2:01 | 1372:449 | 4:01 |
| 556:111 | 1:00 | 472:131 | 0:00 | 1377:449 | 3:02 |
| 558:111 | 0:00 | 472:130 | 0:00 | 1386:451 | 3:00 |
| 558:111 | 0:00 | 473:130 | 0:00 | 1392:451 | 3:00 |
| 556:111 | 0:00 | 471:130 | 1:00 | 1397:447 | 2:03 |
| 559:111 | 0:00 | 478:130 | 0:00 | 1411:447 | 1:01 |
| 558:111 | 0:00 | 480:126 | 0:01 | 1416:446 | 2:02 |
| 0 | 2 | 0 | 0 | 0 | 1 |
| 0:00:00 | 1:01:00 | 0:00:00 | 6:02:02 | 0:00:00 | 10:06:01 |
| 1:01:00 | 0:00:00 | 11:02:03 | 0:00:00 | 23:06:03 | 0:00:00 |
| 2 | 0 | 0 | 0 | 1 | 3 |
| 0 | 1 | 0 | 1 | 0 | 2 |
| 0 | 1 | 0 | 1 | 0 | 2 |
| 0 | 1 | 1 | 0 | 1 | 2 |
| 0 | 1 | 0 | 1 | 0 | 3 |
| 0:00 | 1:00 | 0:00 | 1:00 | 0:00 | 5:00 |
| 0:00 | 1:00 | 0:00 | 1:00 | 0:00 | 5:00 |
| 0 | 1 | 0 | 2 | 0 | 2 |
| 0 | 1 | 1 | 1 | 1 | 0 |
| 0:00 | 0:00 | 0:00 | 1:00 | 0:00 | 1:00 |
| 47:15:00 | 0:00 | 51:12:00 | 0:00 | 131:23:00 | 2:00 |
| 15:34:16:0 | 0:0:0:0 | 20:55:89:2 | 0:0:0:0 | 47:201:224:1 | 0:0:2:0 |
| 15:34:16:0 | 0:0:0:0 | 20:55:89:2 | 0:0:0:0 | 46:200:226:1 | 2:1:0:0 |
| 15:32:16:0 | 0:2:0:0 | 20:55:89:2 | 0:0:0:0 | 47:201:222:1 | 1:3:2:0 |
| 15:34:13:0 | 0:0:0:0 | 20:50:82:2 | 0:0:0:0 | 46:196:207:1 | 0:1:1:0 |
| 15:33:13:0 | 0:0:0:0 | 19:50:81:2 | 0:0:0:0 | 45:197:207:1 | 0:0:0:0 |
| 0:0:0:0 | 15:33:12:0 | 0:0:0:0 | 19:51:81:2 | 0:0:0:0 | 45:197:205:1 |
| 16:34:10:0 | 0:0:0:0 | 22:54:58:2 | 0:0:0:0 | 48:206:140:1 | 1:0:3:0 |
| 17:34:8:0 | 0:1:0:0 | 22:56:30:2 | 0:0:0:0 | 51:208:76:1 | 0:2:0:0 |
| 17:35:9:0 | 0:0:0:0 | 22:56:29:2 | 0:0:0:0 | 51:210:75:1 | 0:0:0:0 |
| 35:08:00 | 0:00:00 | 55:07:00 | 0:00:00 | 205:23:00 | 0:00:00 |
| 35:5:0:1:0:9:2:0:2:0:1:0:1:0:0:1:0:0:0:0:0:0 | 0:0:0:0:0:0:0:0:0:0:0:0:0:0:0:0:0:0:0:0:0:0:0:0:0 | 28:5:13:0:8:3:0:9:1:0:1:5:1:0:0:1:1:0:0:0:0:0 | 0:0:0:0:0:0:0:0:0:0:0:0:0:0:0:0:0:0:0:0:0:0 | 87:24:41:3:7:22:2:31:1:1:2:16:0:5:0:2:3:2:1:1:1:2 | 0:0:0:0:0:0:0:0:0:0:0:0:0:0:0:0:0:0:0:0:0:0:0:0:0 |
| 35:5:0:2:1:0:0:9:0:2:1:0:1:0:0:1:0:0:0:0:0:0 | 0:0:0:0:0:0:0:0:0:0:0:0:0:0:0:0:0:0:0:0:0:0:0:0:0 | 28:5:9:1:0:0:13:3:8:0:1:5:1:0:0:1:1:0:0:0:0:0 | 0:0:0:0:0:0:0:0:0:0:0:0:0:0:0:0:0:0:0:0:0:0 | 86:24:31:1:3:1:41:21:7:2:2:16:0:5:0:2:3:2:1:1:1:2 | 1:0:0:0:0:0:0:1:0:0:0:0:0:0:0:0:0:0:0:0:0:0:0:0:0 |
| 35:5:0:2:1:0:0:9:0:2:0:1:0:0:1:0:0:0:0 | 0:0:0:0:0:0:0:0:0:0:0:0:0:0:0:0:0:0:0:0:0:0 | 28:5:18:1:0:2:21:3:8:0:10:1:0:0:1:1:0:0:0:0 | 0:0:0:0:0:0:0:0:0:0:0:0:0:0:0:0:0:0:0 | 87:26:54:1:3:8:67:22:7:2:17:0:5:0:2:2:2:1:1:1 | 0:0:0:0:0:0:0:0:0:0:0:1:0:0:0:0:0:0:0:0:0:0:0 |
| 35:5:0:2:1:0:0:9:0:2:1:0:1:0:0:1:0:0:0:0:0:0 | 0:0:0:0:0:0:0:0:0:0:0:0:0:0:0:0:0:0:0:0:0:0:0:0:0 | 28:6:18:1:0:9:24:3:8:0:1:7:1:0:0:1:1:1:0:0:0:0 | 0:0:0:0:0:0:0:0:0:0:0:0:0:0:0:0:0:0:0:0:0:0 | 87:27:54:0:3:34:88:22:7:2:2:11:0:4:0:2:3:2:2:1:1:1 | 0:0:0:0:0:0:1:0:0:0:0:0:0:1:0:0:0:0:0:0:0:0:0:0:0 |
| 34:5:0:2:1:0:0:9:0:2:1:0:1:0:0:1:0:0:0:0:0:0 | 1:0:0:0:0:0:0:0:0:0:0:0:0:0:0:0:0:0:0:0:0:0:0:0:0 | 27:7:18:1:0:9:24:3:8:0:1:6:1:0:0:1:1:1:0:0:0:0 | 1:0:0:0:0:0:0:0:0:0:0:0:0:0:0:0:0:0:0:0:0:0 | 87:27:54:0:3:34:91:20:7:2:2:11:0:4:0:2:3:2:2:1:1:1 | 0:0:0:0:0:1:1:0:0:0:0:0:0:0:0:0:0:0:0:0:0:0:0:0:0 |
| 35:5:0:2:1:0:0:9:0:2:1:1:0:0:0:1:0:0:0:0:0:0 | 0:0:0:0:0:0:0:0:0:0:0:0:0:0:0:0:0:0:0:0:0:0:0:0:0 | 28:7:18:0:0:11:25:3:8:0:1:1:0:0:6:1:1:1:0:0:0:0 | 0:0:0:0:0:0:0:0:0:0:0:0:0:0:0:0:0:0:0:0:0:0 | 87:27:54:0:3:45:97:21:7:2:2:0:4:0:10:2:3:2:2:1:1:1 | 0:0:0:0:0:0:0:0:0:0:0:0:0:0:0:0:0:0:0:0:0:0:0:0:0 |
| 35:0:1:0:2:0:8:0:1:1:1:0:0:0:1:0:0:0:0:0 | 0:0:0:0:0:0:0:0:0:0:0:0:0:0:0:0:0:0:0:0:0:0 | 28:18:0:0:16:26:3:8:0:1:1:0:0:5:1:1:1:0:0:0 | 0:0:0:0:0:0:0:0:0:0:0:0:0:0:0:0:0:0:0 | 87:54:0:3:63:103:20:7:2:1:0:4:0:8:2:3:2:1:1:1 | 0:0:0:0:1:1:0:0:0:0:0:0:0:0:0:0:0:0:0:0:0:0:0 |
| 35:2:0:1:0:2:0:8:0:1:1:1:0:0:0:1:0:0:0:0:0 | 0:0:0:0:0:0:0:0:0:0:0:0:0:0:0:0:0:0:0:0:0:0 | 28:5:18:0:0:18:26:3:8:0:1:1:0:0:5:1:1:1:0:0:0 | 0:0:0:0:0:0:0:0:0:0:0:0:0:0:0:0:0:0:0 | 87:12:54:0:3:66:104:20:7:2:1:0:4:0:8:2:3:1:2:1:1 | 0:0:0:0:0:0:0:0:0:0:0:0:0:0:0:0:0:0:0:0:0:0:0:0 |
| 9:1:2:2:0:6:4:0:0:3:0:1:1 | 0:0:0:0:0:0:0:0:0:0:0:0:0:0 | 14:5:4:18:1:3:52:2:0:0:0:0:0 | 0:0:0:0:0:0:0:1:0:0:0:0:0 | 59:20:29:33:8:22:83:8:3:11:0:1:0 | 1:1:0:0:0:0:1:0:0:0:0:0:0 |
| 1:2:2:0:9:6:7:0:0:3:0:1:1 | 0:0:0:0:0:0:0:0:0:0:0:0:0:0 | 5:3:18:1:14:3:63:3:0:0:0:0:0 | 0:0:0:0:0:0:1:0:0:0:0:0:0 | 20:27:31:8:59:22:120:8:2:11:0:1:0 | 0:1:0:0:0:0:1:0:0:0:0:0:0 |
| 8:1:2:0:2:5:8:1:0:0:3:0:1 | 1:0:0:0:0:0:0:1:0:0:0:0:0:0 | 12:2:17:1:1:3:75:0:2:0:0:0:0 | 0:0:0:0:0:0:0:0:0:0:0:0:0 | 41:16:26:5:19:16:166:4:6:2:7:0:0 | 7:2:1:1:0:1:1:0:0:0:1:0:0 |
| 0:2:8:0:2:5:8:1:0:0:3:0:1 | 0:0:0:0:0:0:0:0:0:0:0:0:0:0 | 1:16:6:1:0:1:73:0:1:0:0:0:0 | 1:1:3:0:1:2:2:0:1:0:0:0:0 | 14:23:30:4:11:10:171:4:5:2:6:0:0 | 1:0:9:0:5:3:1:0:0:0:0:0:0 |

| | | | | | |
|---|---|---|---|---|---|
| 0:2:4:0:2:5:7:1:0:0:3:1 | 0:0:0:0:0:0:0:0:0:0:0:0 | 0:13:2:1:0:0:73:0:0:0:0:0 | 0:0:0:0:0:0:0:0:0:0:0 | 7:20:17:3:7:5:172:3:1:3:6:0 | 0:0:0:0:0:0:1:0:0:0:0:0 |
| 0:2:4:0:2:5:7:1:0:0:3:1 | 0:0:0:0:0:0:0:0:0:0:0:0 | 0:13:2:1:0:0:74:0:0:0:0:0 | 0:0:0:0:0:0:0:0:0:0:0 | 7:20:16:3:7:5:173:3:1:3:6:0 | 0:0:1:0:0:0:1:0:0:0:0:0 |
| 1:0:2:4:0:2:5:6:0:0:3:1 | 0:0:0:0:0:0:0:0:0:0:0:0 | 0:0:11:1:1:0:0:72:0:0:0:0 | 0:0:1:1:0:0:0:2:0:0:0:0 | 3:7:18:15:3:6:4:171:1:2:5:0 | 0:0:2:2:0:1:1:4:0:0:1:0 |
| 0:6:0:2:4:2:4:1:0:0:3:1 | 0:0:0:0:0:0:0:0:0:0:0:0 | 1:64:0:10:1:0:0:0:0:0:0:0 | 0:5:0:0:0:0:0:0:0:0:0:0 | 3:146:7:14:15:5:3:2:1:2:5:0 | 0:8:0:2:1:0:1:1:0:0:0:0 |
| 1:0:0:0:0:0:0:0:1:0 | 2:1:0:0:0:0:0:0:0:0 | 19:0:0:1:0:0:0:0:0:0 | 6:0:0:0:0:0:0:0:0:0 | 44:1:1:2:1:0:1:0:1:1 | 6:3:0:0:0:0:0:0:0:0 |
| 1:0:0:0:0:0:0:0:1:0 | 2:1:0:0:0:0:0:0:0:0 | 19:0:0:1:0:0:0:0:0:0 | 7:0:0:0:0:0:0:0:0:0 | 44:1:0:1:1:0:1:0:1:1 | 6:3:0:1:0:0:0:0:0:0 |
| 0:0:0:0:0:0:0:0:0 | 0:0:0:0:0:0:0:0:0 | 0:0:0:0:12:0:0:0:0 | 0:0:0:0:0:0:0:0:0 | 0:0:1:0:34:0:0:1:1 | 1:0:0:0:2:1:0:0:0 |
| 0:0:0:0:0:0:0 | 0:0:0:0:0:0:0 | 0:0:0:10:0:0:0 | 0:0:0:0:0:0:0 | 0:1:0:17:0:1:1 | 0:0:0:3:0:0:0 |
| 0:02 | 0:00 | 0:01 | 0:00 | 0:09 | 0:01 |
| 0:0:0:0 | 0:0:0:0 | 0:0:0:0 | 0:0:0:0 | 1:0:2:0 | 0:0:2:1 |
| 5:34:17:37:1:20:26:3:3:3:1 | 0:0:0:0:0:0:2:0:0:0:0 | 0:8:1:0:0:0:1:2:0:0:0 | 0:0:0:0:0:0:0:0:0:0:0 | 8:130:44:69:5:35:51:2:3:12:3 | 0:3:1:0:0:0:0:1:0:0:0 |
| 37:1:5:20:37:32:2:29:3:3:1 | 0:0:0:0:1:0:0:0:0:0:0 | 0:0:0:0:20:2:1:1:0:0:0 | 0:0:0:0:0:0:0:0:0:0:0 | 69:5:8:35:197:64:2:63:3:12:3 | 0:0:0:0:0:0:0:0:0:0:0 |
| 37:1:5:20:49:36:38:2:3:3:1 | 0:0:0:0:0:0:0:0:0:0:0 | 0:0:0:0:59:12:5:1:0:0:0 | 0:0:0:0:0:0:0:0:0:0:0 | 69:5:8:35:389:128:105:2:3:11:3 | 0:0:0:0:5:2:0:0:0:0:0 |
| 37:37:1:5:20:55:47:1:3:3:1 | 0:0:0:0:0:0:0:0:0:0:0 | 29:0:0:0:0:80:23:1:0:0:0 | 0:0:0:0:0:2:1:0:0:0:0 | 217:70:5:9:35:546:153:1:3:11:4 | 1:0:0:0:0:2:0:0:0:0:0 |
| 37:1:5:20:56:37:49:3:3:1:1 | 0:0:0:0:0:0:0:0:0:0:0 | 0:0:0:0:91:40:31:0:0:0:1 | 0:0:0:0:1:1:1:0:0:0:0 | 69:5:9:35:629:252:173:3:11:4:1 | 1:0:0:0:2:2:1:0:0:0:0 |
| 38:1:5:20:58:41:48:3:3:1:1 | 0:0:0:0:0:0:1:0:0:0:0 | 0:0:0:0:97:49:44:0:0:0:1 | 0:0:0:0:0:2:1:0:0:0:0 | 70:5:10:35:666:269:191:3:11:4:1 | 0:0:0:0:1:1:2:0:0:0:0 |
| 38:1:5:20:62:41:48:2:3:3:1 | 0:0:0:0:0:0:1:0:0:0:0 | 0:0:0:0:105:58:48:0:0:0:0 | 0:0:0:0:1:0:0:0:0:0:0 | 69:5:10:35:712:303:210:0:3:11:4 | 1:0:0:0:1:2:0:0:0:0:0 |
| 62:38:1:5:20:41:49:2:3:3:1 | 0:0:0:0:0:0:0:0:0:0:0 | 106:0:0:0:0:59:49:0:0:0:0 | 0:0:0:0:0:0:0:0:0:0:0 | 719:70:5:10:35:310:215:0:3:11:4 | 5:0:0:0:0:2:2:0:0:0:0 |
| 38:1:5:20:63:42:50:2:3:3:1 | 0:0:0:0:0:0:0:0:0:0:0 | 0:0:0:0:108:63:57:0:0:0:0 | 0:0:0:0:0:0:0:0:0:0:0 | 70:5:10:35:741:325:223:0:3:11:4 | 0:0:0:0:2:0:1:0:0:0:0 |
| 38:1:5:20:64:44:51:2:3:3:1 | 0:0:0:0:0:0:0:0:0:0:0 | 0:0:0:0:120:76:72:0:0:0:0 | 0:0:0:0:0:1:0:0:0:0:0 | 70:5:10:35:798:359:278:0:3:11:4 | 0:0:0:0:1:0:0:0:0:0:0 |
| 38:1:5:20:64:45:2:51:3:3:1 | 0:0:0:0:0:0:0:0:0:0:0 | 0:0:0:0:121:82:0:80:0:0:0 | 0:0:0:0:0:0:0:1:0:0:0 | 70:5:11:35:819:368:0:294:3:11:5 | 0:0:0:0:2:0:0:1:0:0:0 |
| 38:1:5:20:66:47:3:51:3:3:1 | 0:0:0:0:1:0:0:0:0:0:0 | 0:0:0:0:125:91:0:101:0:0:0 | 0:0:0:0:0:0:0:0:0:0:0 | 70:5:12:35:854:414:0:337:3:11:6 | 0:0:0:0:1:3:0:2:0:0:0 |
| 38:1:5:20:74:49:2:53:3:3:2 | 0:0:0:0:0:0:0:0:0:0:0 | 0:0:0:0:128:100:0:120:0:0:0 | 0:0:0:0:1:0:0:0:0:0:0 | 69:5:13:35:892:465:0:404:3:11:6 | 1:0:0:0:2:4:0:0:0:0:0 |
| 20:38:1:5:73:51:2:52:3:3:2 | 0:0:0:0:0:0:0:1:0:0:0 | 0:0:0:0:127:103:0:135:0:0:0 | 0:0:0:0:0:1:0:1:0:0:0 | 35:70:5:13:890:492:0:434:3:11:6 | 0:0:0:0:3:0:0:1:0:0:0 |
| 53:38:1:5:20:76:51:2:3:3:2 | 0:0:0:0:0:1:0:0:0:0:0 | 152:0:0:0:0:124:109:0:0:0:0 | 1:0:0:0:0:0:0:0:0:0:0 | 474:70:5:13:35:914:518:0:3:11:6 | 3:0:0:0:0:1:0:0:0:0:0 |
| 38:1:5:20:78:50:2:53:3:3:2 | 0:0:0:0:1:1:0:0:0:0:0 | 0:0:0:0:126:109:0:160:0:0:0 | 0:0:0:0:0:0:0:0:0:0:0 | 70:5:13:35:919:529:0:493:3:11:7 | 0:0:0:0:2:3:0:0:0:0:0 |
| 38:1:5:20:80:52:2:52:3:3:2 | 0:0:0:0:0:0:0:1:0:0:0 | 0:0:0:0:127:111:0:170:0:0:0 | 0:0:0:0:0:0:0:0:0:0:0 | 70:5:13:35:925:551:0:517:3:11:7 | 0:0:0:0:3:0:0:2:0:0:0 |
| 0:00 | 0:00 | 0:00 | 1:00 | 0:00 | 2:01 |
| 0 | 0 | 0 | 0 | 0 | 0 |
| 0:00 | 0:00 | 0:00 | 0:00 | 0:04 | 0:03 |
| 14:0:1:0:0 | 1:0:0:0:0 | 71:9:2:3:1 | 0:0:0:0:0 | 68:4:4:2:1 | 2:0:0:0:0 |
| 14:0:1:0:0 | 1:0:0:0:0 | 71:10:2:3:1 | 0:0:0:0:0 | 70:4:4:2:1 | 0:0:0:0:0 |
| 2:0:0:0:0 | 13:0:1:0:0 | 34:4:1:2:0 | 37:6:1:1:1 | 38:3:2:0:1 | 32:1:2:2:0 |
| 14:0:1:0:0 | 1:0:0:0:0 | 70:11:2:3:1 | 0:0:0:0:0 | 70:4:4:2:1 | 0:0:0:0:0 |
| 0:00:00 | 0:01:00 | 6:00:00 | 7:00:00 | 0:00:00 | 0:00:00 |
| 0:0:4:0:0:0:0:0:0:0:0 | 0:0:0:0:0:0:0:0:0:0:0 | 1:3:3:2:0:0:0:0:0:0:0 | 3:0:0:0:0:0:0:0:3:0:0 | 3:6:11:2:0:0:0:0:1:4:1 | 12:2:0:0:0:0:1:1:0:0:0 |
| 4:0:0:0:0:0:0:0:0:0:0:0:0 | 0:0:0:0:0:0:0:0:0:0:0:0:0 | 3:4:0:2:2:0:1:2:0:2:1:1:0 | 0:0:0:0:0:0:0:0:0:0:0:0:0 | 9:24:0:5:6:0:2:1:1:1:0:2:4 | 0:0:0:0:0:0:0:1:0:0:0:0:0 |
| 1:1:1:0:0 | 0:0:0:0:0 | 3:0:0:0:0 | 0:0:0:0:0 | 12:0:3:0:0 | 0:0:0:0:0 |
| 0:1:2:1:0:0:0:2:0:5:0:0:0:1:1:1 | 0:0:0:0:0:0:0:0:1:0:0:0:0:0:0 | 0:0:2:0:0:0:0:0:0:3:0:0:0:0:0:0 | 0:0:3:0:0:0:0:0:0:1:1:0:0:1:0:0 | 0:0:8:0:0:1:0:1:1:8:0:1:0:1:0:0 | 0:0:5:0:0:1:0:0:1:10:1:0:0:2:0:0 |
| 2:0:0:2:1:0:0:0:6:0:0:1:0:1:1 | 0:0:0:0:0:0:0:0:0:0:0:0:0:0:0 | 1:0:0:6:0:0:0:0:14:0:1:0:0:1:0 | 0:0:0:0:0:0:0:0:0:0:0:0:0:0:0 | 5:0:0:15:0:0:2:0:67:2:1:2:1:3:0 | 1:0:0:0:0:0:0:0:2:0:0:0:0:0:0 |
| 0:2:0:0:0:0:2:6:0:0:1:0:1:0 | 0:0:0:0:0:0:0:1:0:0:0:0:0:0 | 0:2:0:0:0:0:0:9:0:0:0:0:0:0 | 0:2:0:0:0:0:2:9:0:1:0:0:0:0 | 0:5:0:0:1:0:4:42:0:0:1:0:1:0 | 0:6:0:0:1:0:2:48:0:1:2:1:2:1 |
| 0:2:0:0:0:0:2:8:1:0:0:0:0:0 | 0:0:0:0:0:0:0:0:0:0:0:0:0:0 | 0:3:0:0:0:0:2:20:0:0:0:0:1:0 | 0:1:0:0:0:0:1:29:3:0:0:0:0:0 | 0:10:0:0:2:0:6:90:1:0:0:1:1:1 | 0:2:0:0:0:0:2:66:2:0:0:0:0:0 |

| | | | | | |
|---|---|---|---|---|---|
| 0:3:0:0:0:0:7:43:0:1:0:0:0:0:0:0 | 0:0:0:0:0:0:0:0:0:0:0:0:0:0:0:0 | 0:7:0:0:0:0:5:70:0:3:0:0:0:0:1:0 | 0:0:0:0:0:0:0:0:1:0:1:0:0:0:0:0:0 | 0:25:0:0:2:0:15:314:0:3:0:0:0:5:1:1 | 0:1:0:0:0:0:0:2:0:1:0:0:0:0:0 |
| 0:3:0:0:0:0:7:43:0:1:0:0:0:0:0:0 | 0:0:0:0:0:0:0:0:0:0:0:0:0:0:0:0 | 0:7:0:0:0:0:4:63:0:3:0:0:0:0:1:0 | 0:0:0:0:0:0:0:1:8:0:1:0:0:0:0:0:0 | 0:26:0:0:2:0:13:291:0:4:0:0:0:5:1:1 | 0:0:0:0:0:0:2:30:0:0:0:0:0:0:0:0 |
| 34:0:0:3:0:0:0:0:5:1:0:0:0:0:0:0 | 9:0:0:0:0:0:0:0:2:0:0:0:0:0:0:0 | 69:0:0:7:0:0:0:0:5:4:0:0:0:0:1:0 | 2:0:0:0:0:0:0:0:0:0:0:0:0:0:0:0:0 | 305:0:0:26:0:0:2:0:14:3:0:0:5:0:1:1 | 17:0:0:0:0:0:0:0:1:2:0:0:0:0:0:0 |
| 0:3:0:0:0:0:7:43:0:1:0:0:0:0:0:0 | 0:0:0:0:0:0:0:0:0:0:0:0:0:0:0:0 | 0:6:0:0:0:0:5:70:0:4:0:0:0:0:1:0 | 0:0:0:0:0:0:0:0:2:0:0:0:0:0:0:0:0 | 0:26:0:0:2:0:15:320:0:4:0:0:0:5:1:1 | 0:0:0:0:0:0:0:3:0:1:0:0:0:0:0 |
| 6:0:4:0:1 | 3:0:2:0:0 | 0:0:15:0:0 | 4:0:14:0:0 | 9:1:36:0:0 | 16:1:64:0:0 |
| 3900:16:10:4:6:1:0 | 2:0:0:0:0:0:0 | 7163:19:10:8:7:3:1 | 4:0:0:0:0:0:0 | 23080:83:31:29:61:4:3 | 25:0:0:0:0:0:0 |
| 3900:16:10:4:6:1:0 | 2:0:0:0:0:0:0 | 7159:19:10:8:7:3:1 | 7:0:0:0:0:0:0 | 23075:83:31:29:61:4:3 | 29:0:0:0:0:0:0 |
| 3892:16:10:4:6:1:0 | 7:0:0:0:0:0:0 | 7152:19:10:8:7:3:1 | 15:0:0:0:0:0:0 | 23057:83:31:29:60:4:3 | 45:0:0:0:1:0:0 |
| 3887:16:10:4:6:1:0 | 6:0:0:0:0:0:0 | 7127:19:10:8:7:3:1 | 17:0:0:0:0:0:0 | 22957:84:31:29:61:4:3 | 80:0:0:0:0:0:0 |
| 3876:15:10:4:1:0:0 | 7:0:0:0:0:0:0 | 7107:19:11:8:3:0:1 | 4:0:0:0:0:0:0 | 22927:81:31:29:4:1:3 | 26:0:0:0:0:0:0 |
| 3880:15:10:4:1:0 | 1:0:0:0:0:0 | 7089:18:11:8:3:1 | 3:0:0:0:0:0 | 22860:80:31:29:4:3 | 19:0:0:0:0:0 |
| 3856:15:2:1:0:0 | 8:0:0:0:0:0 | 6993:17:0:3:0:0 | 8:0:0:0:0:0 | 22694:81:7:4:3:0 | 26:0:0:0:0:0 |
| 0:0:0:0 | 0:0:0:0 | 0:0:0:0 | 0:0:0:1 | 0:0:0:0 | 0:0:1:0 |
| 0 | 0 | 0 | 1 | 0 | 1 |
| 59 | 0 | 23 | 0 | 99 | 0 |
| 0:0:0:4:0 | 0:0:0:1:0 | 0:0:0:2:0 | 0:0:0:5:0 | 0:0:0:5:1 | 0:0:1:7:0 |
| 3:0:0:3:1:0:0:0:0:0:0:0 | 0:0:0:0:0:0:0:0:0:0:0:0 | 1:0:0:21:8:0:0:0:0:0:1:0 | 0:0:0:0:1:0:0:0:0:0:0:0 | 22:4:0:100:25:0:1:0:2:0:0:0 | 0:0:0:2:3:0:0:0:0:0:0:1 |
| 11:0:1:3:4:0:3:0:0:0:0:0:1:0:0:1:1:0:0:0 | 0:0:0:0:0:0:0:0:0:0:0:0:0:0:0:0:0:0:0:0 | 13:3:0:0:21:0:3:3:2:2:0:1:0:0:0:0:0:1:0:0 | 0:0:0:0:0:0:0:0:0:0:0:0:1:0:0:0:0:0:0:0 | 69:5:0:4:93:0:15:3:10:2:0:2:0:0:0:1:0:0:1:1 | 2:0:0:0:1:0:0:1:0:0:0:0:0:0:0:0:0:0:0:0 |
| 1.052905093 | 0:00:00 | 1.20974537 | 0:04:00 | 119:2272:28 | 2:07:00 |
| 1.001493056 | 2:14:02 | 0.571550926 | 0.765972222 | 57:1082:11 | 74:1197:17 |
| 1.177905093 | 0:00:00 | 28:546:2 | 0:00:00 | 167:2274:28 | 0:03:00 |
| 28:316:11 | 0:00:00 | 42:545:2 | 1:00:00 | 203:2269:28 | 2:00:00 |
| 29:316:11 | 0:00:00 | 60:544:2 | 0:00:00 | 279:2265:28 | 2:03:00 |
| 312:11:29 | 3:00:00 | 529:2:72 | 10:00:00 | 2240:27:347 | 25:01:01 |
| 33:312:11 | 0:00:00 | 80:528:2 | 1:00:00 | 398:2234:27 | 0:01:00 |
| 34:311:11 | 0:01:00 | 81:525:2 | 0:03:00 | 403:2231:27 | 0:03:00 |
| 32:295:9 | 2:14:02 | 51:273:2 | 56:247:0 | 237:1049:10 | 260:1160:17 |
| 33:308:11 | 1:01:00 | 109:520:2 | 0:01:00 | 497:2205:27 | 3:03:00 |
| 34:309:11 | 0:00:00 | 110:521:2 | 1:00:00 | 503:2201:27 | 2:05:00 |
| 308:35:11 | 0:00:00 | 523:113:2 | 0:00:00 | 2199:516:27 | 2:01:00 |
| 303:36:11 | 6:00:00 | 511:111:2 | 8:02:00 | 2154:516:26 | 34:03:00 |
| 301:35:11 | 2:01:00 | 445:107:2 | 48:05:00 | 2016:507:25 | 92:07:01 |
| 290:35:11 | 14:01:00 | 426:107:2 | 17:02:00 | 1983:504:24 | 36:03:01 |
| 35:287:11 | 0:00:00 | 104:413:2 | 0:01:00 | 500:1942:23 | 2:02:00 |
| 35:286:11 | 0:01:00 | 104:414:2 | 0:00:00 | 502:1940:23 | 1:05:00 |
| 35:288:11 | 0:00:00 | 103:414:2 | 0:00:00 | 494:1940:23 | 2:06:00 |
| 4 | 0 | 6 | 6 | 13 | 17 |
| 2:0:2:0 | 0:0:1:0 | 0:0:0:0 | 0:0:0:0 | 3:1:1:0 | 7:0:3:0 |
| 11:2:0:0:0:2 | 0:0:0:0:0:0 | 46:4:2:0:4:3 | 4:0:1:0:0:0 | 113:15:7:1:4:3 | 0:0:0:0:0:0 |
| 0:00 | 1:00 | 0:00 | 0:00 | 1:01 | 2:02 |
| 0:01 | 0:00 | 0:00 | 0:00 | 6:00 | 0:00 |
| 0:0:1:0:0:0 | 0:0:0:0:0:0 | 0:0:0:0:0:0 | 0:0:0:1:0:0 | 0:0:1:0:0:1 | 0:0:0:1:1:0 |
| 0 | 0 | 3 | 0 | 1 | 1 |
| 0:0:0:0 | 7:2:1:0 | 0:0:0:0 | 13:0:0:0 | 0:0:0:0 | 33:1:0:1 |
| 0:00:00 | 0:02:00 | 0:00:00 | 0:00:00 | 0:00:00 | 0:01:00 |
| 1:7:3:0:2:0:1:0:0:0:0:1:1:0:0:0:0:0:0:0:1:0:0 | 0:0:0:0:0:0:0:0:0:0:0:0:0:0:0:0:0:0:0:0:0:0:0 | 15:42:63:2:10:0:3:18:2:4:1:3:2:1:0:5:0:0:0:1:4:1:0 | 0:0:1:1:0:0:0:0:0:0:0:0:0:0:0:0:0:0:0:0:0:0:0 | 18:137:101:4:30:1:7:29:6:12:0:2:4:3:9:5:0:4:0:1:3:0:1 | 0:1:0:0:0:0:0:0:0:0:0:0:0:0:0:0:0:0:0:0:0:0:0 |
| 1:1:7:1:0:3:1:0:0:0:0:1:0:0:0:0:0:0:1:0:0:0:0 | 0:0:0:0:0:0:0:0:0:0:0:0:0:0:0:0:0:0:0:0:0:0:0 | 2:16:45:61:4:10:4:14:2:4:1:4:1:0:1:6:0:0:4:0:2:1:1 | 0:0:0:2:0:0:1:0:0:0:0:0:0:0:0:0:0:0:0:0:0:0:0 | 4:21:141:89:4:32:7:27:6:9:0:2:3:9:6:5:0:5:3:0:1:1:0 | 0:0:1:0:0:0:0:0:0:0:0:0:0:0:0:0:0:0:0:0:0:0:0 |
| 0 | 0 | 2 | 3 | 4 | 6 |
| 8:9:14:3:0:0:2:1:0:1:0:1:0:0:0:0:0:0:0 | 0:0:0:0:0:0:0:0:0:0:0:0:0:0:0:0:0:0:0 | 16:12:10:5:7:8:4:2:0:2:1:1:2:0:0:0:0:0:0 | 0:0:0:0:0:0:0:0:0:0:0:0:0:0:0:0:0:0:0 | 65:35:57:10:15:19:14:8:1:4:1:4:6:1:0:0:1:0:1 | 0:0:0:0:0:0:0:0:0:0:0:0:0:0:0:0:0:0:0 |

| | | | | | |
|---|---|---|---|---|---|
| 8:15:0:0:2:1:0:0:1:0:1:0:0:0:0:0 | 0:0:0:0:0:0:0:0:0:0:0:0:0:0:0 | 18:12:0:10:10:2:0:0:2:1:1:0:0:0:0:0 | 0:0:0:0:0:0:0:0:0:0:0:0:0:0:0 | 66:61:0:24:21:8:2:0:3:1:3:0:0:1:1:0 | 1:0:0:0:0:0:0:0:1:0:0:0:0:0:0 |
| 7:9:15:4:0:0:2:1:0:1:0:1:0:0:0:0:0:0 | 0:0:0:0:0:0:0:0:0:0:0:0:0:0:0:0:0:0 | 18:12:12:5:7:10:8:2:0:2:1:1:2:0:0:0:0:0 | 0:0:0:0:0:0:0:0:0:0:0:0:0:0:0:0:0:0 | 65:35:59:12:15:25:23:7:2:4:1:3:8:0:0:1:1:0 | 0:1:1:0:0:0:0:0:0:0:0:0:0:0:0:0:0:0 |
| 2:7:0:2:5:4:5:3:1:0:0:0 | 0:0:0:0:0:0:0:0:0:0:0:0 | 0:7:2:0:5:3:16:4:2:1:0:0 | 0:11:0:2:2:2:17:1:1:0:0:0 | 1:14:0:2:8:17:72:9:5:0:0:0 | 3:17:3:1:12:8:67:9:3:1:0:1 |
| 0:00 | 0:00 | 0:00 | 0:00 | 0:00 | 0:01 |
| 412:00:00 | 5:00 | 1023:03:00 | 26:00:00 | 2363:04:00 | 28:00:00 |
| 419:20:00 | 0:00 | 1053:36:00 | 2:00 | 2407:107 | 2:00 |
| 419:20:00 | 0:00 | 1048:39:00 | 4:00 | 2387:118 | 5:00 |
| 419:20:00 | 0:00 | 1048:40:00 | 1:00 | 2381:121 | 1:00 |
| 419:20:00 | 0:00 | 1047:40:00 | 2:00 | 2379:120 | 1:00 |
| 414:22:00 | 0:00 | 1034:43:00 | 3:00 | 2343:131 | 3:00 |
| 410:22:00 | 1:00 | 1028:47:00 | 1:00 | 2318:135 | 3:01 |
| 410:22:00 | 0:00 | 1029:47:00 | 0:00 | 2317:136 | 2:00 |
| 400:23:00 | 0:00 | 998:46:00 | 0:00 | 2260:138 | 3:00 |
| 397:22:00 | 0:00 | 985:44:00 | 0:00 | 2242:133 | 8:00 |
| 395:21:00 | 0:00 | 983:43:00 | 0:00 | 2244:133 | 4:00 |
| 0 | 1 | 0 | 0 | 0 | 0 |
| 0 | 0 | 1 | 1 | 2 | 0 |
| 1:0:0:0 | 2:1:1:0 | 4:14:0:0 | 1:7:0:0 | 2:1:1:0 | 6:14:1:0 |
| 2:1:1:0 | 1:0:0:0 | 1:7:0:0 | 4:14:0:0 | 6:14:1:0 | 2:1:1:0 |
| 0:0:0:0 | 2:1:0:1 | 0:3:0:0 | 4:15:0:0 | 1:1:0:0 | 6:13:0:2 |
| 1 | 1 | 0 | 2 | 0 | 10 |
| 0 | 0 | 0 | 1 | 0 | 7 |
| 30:00:02 | 0:00:00 | 90:00:03 | 0:00:00 | 180:01:02 | 0:00:00 |
| 10:0:2:10:0:2:1:0 | 0:0:0:0:0:0:0:0 | 47:8:0:21:2:24:0:1 | 0:0:0:0:0:0:0:0 | 59:6:1:50:1:22:5:0 | 0:0:0:0:0:0:0:0 |
| 10:0:6:0:0:1:1:0 | 0:0:0:0:0:0:0:0 | 45:1:11:0:5:0:0:1 | 0:0:0:0:0:0:0:0 | 58:1:30:1:4:4:1:0 | 0:0:1:0:1:1:0:0 |
| 6:10:0:0:0:1:1:0 | 0:0:0:0:0:0:0:0 | 11:45:1:0:5:0:0:1 | 0:0:0:0:0:0:0:0 | 30:58:1:1:2:4:1:0 | 1:0:0:0:3:1:0:0 |
| 6:10:0:0:0:1:1:0 | 0:0:0:0:0:0:0:0 | 11:45:1:0:5:0:0:1 | 0:0:0:0:0:0:0:0 | 31:58:1:1:3:4:1:0 | 0:0:0:0:2:1:0:0 |
| 10:0:6:0:0:1:1:0 | 0:0:0:0:0:0:0:0 | 45:1:11:5:0:0:0:1 | 0:0:0:0:0:0:0:0 | 58:1:30:2:1:4:1:0 | 0:0:0:0:0:0:0:0 |
| 9:0:4:0:0:1:1:0 | 1:0:2:0:0:0:0:0 | 42:1:8:0:1:0:0:1 | 1:0:0:0:0:0:0:0 | 56:0:29:1:1:3:0:0 | 0:0:0:0:0:0:0:0 |
| 6:4:0:0:0:1:1:0 | 3:2:0:0:0:0:0:0 | 34:5:0:1:1:0:0:1 | 0:0:0:0:0:0:0:0 | 42:20:1:1:0:2:0:0 | 5:3:0:0:0:0:0:0 |
| 0 | 0 | 2 | 1 | 0 | 1 |
| 21:03:01 | 4:00:00 | 43:02:02 | 19:01:00 | 90:24:05 | 36:04:01 |
| 0 | 45 | 0 | 10 | 0 | 124 |
| 0 | 0 | 0 | 1 | 0 | 2 |
| 0:00:00 | 0:00:00 | 0:00:00 | 1:00:01 | 0:00:00 | 1:00:00 |
| 1:3:2:0:0:2 | 0:0:0:0:0:0 | 1:2:0:0:0:1 | 0:0:0:0:0:0 | 5:6:1:0:1:1 | 0:0:0:0:0:0 |
| 0:0:0:0:0 | 2:0:2:0:0 | 1:0:8:0:0 | 1:0:3:0:0 | 0:0:0:0:0 | 3:0:15:1:1 |
| 0:0:0:0:0 | 2:0:2:0:0 | 1:0:8:0:0 | 1:0:2:0:0 | 0:0:0:0:0 | 3:0:13:2:1 |
| 96:00:00 | 0:00 | 93:00:00 | 1:00 | 208:01:00 | 0:00 |
| 97:00:00 | 0:00 | 94:00:00 | 0:00 | 208:01:00 | 2:00 |
| 0 | 0 | 0 | 0 | 0 | 2 |
| 0 | 0 | 0 | 0 | 0 | 1 |
| 0 | 0 | 0 | 0 | 0 | 1 |
| 0:00 | 0:00 | 0:00 | 0:00 | 0:00 | 0:00 |
| 1 | 0 | 0 | 2 | 0 | 0 |
| 0:0:0:0 | 20:2:0:0 | 0:0:0:0 | 6:4:0:0 | 0:0:0:0 | 37:18:0:3 |
| 0:1:0:0 | 20:3:0:0 | 0:0:0:0 | 7:4:0:0 | 0:0:0:0 | 37:19:0:4 |
| 0:0:0:0:0 | 24:5:1:0:0 | 0:0:0:0:0 | 11:9:0:0:0 | 0:0:0:0:0 | 43:49:2:2:4 |
| 25:8:1:0:0 | 0:1:0:0:0 | 12:14:0:0:0 | 1:0:0:0:0 | 52:72:2:8:3 | 2:2:0:0:1 |
| 48:3:4:4:3:0 | 0:0:0:0:0:0 | 86:5:15:13:2:1 | 0:0:0:0:0:0 | 145:15:30:29:1:1 | 0:0:0:0:0:0 |
| 48:3:4:4:3:0 | 0:0:0:0:0:0 | 85:5:14:13:2:1 | 1:0:1:0:0:0 | 145:15:30:28:1:1 | 0:0:0:1:0:0 |
| 47:3:4:3:3:0 | 1:0:0:1:0:0 | 86:5:14:13:2:1 | 0:0:1:0:0:0 | 145:15:30:29:1:1 | 0:0:0:0:0:0 |
| 4:0:0:2:0:0:0 | 0:0:0:0:0:0:0 | 1:0:0:0:0:0:0 | 0:0:0:0:0:0:0 | 15:2:0:0:0:0:1 | 0:1:0:1:0:0:0 |
| 1:00:00 | 0:00:00 | 5:00:00 | 2:01:00 | 13:03:01 | 0:00:00 |
| 0:00 | 0:00 | 0:00 | 0:00 | 0:00 | 2:00 |
| 0 | 0 | 0 | 1 | 0 | 4 |
| 0 | 0 | 0 | 0 | 0 | 1 |
| 0 | 0 | 1 | 0 | 4 | 0 |
| 0 | 1 | 0 | 0 | 0 | 5 |
| 2 | 0 | 1 | 2 | 8 | 0 |
| 0 | 2 | 1 | 2 | 1 | 6 |
| 0 | 1 | 0 | 2 | 1 | 5 |
| 0 | 0 | 0 | 0 | 0 | 2 |
| 0 | 0 | 0 | 0 | 0 | 2 |
| 0 | 0 | 0 | 0 | 0 | 2 |
| 1 | 1 | 0 | 0 | 2 | 0 |

| | | | | | |
|---|---|---|---|---|---|
| 0:1:1:2:0:0:1 | 0:1:0:0:0:0:0 | 2:0:0:4:1:0:0 | 3:3:0:1:1:0:0 | 0:1:2:1:1:0:0 | 7:0:0:0:4:0:0 |
| 0:00:00 | 4:00:00 | 0:00:00 | 0:00:00 | 0:00:00 | 12:01:01 |
| 83:26:00 | 0:00 | 184:42:00 | 0:00 | 374:126 | 2:00 |
| 82:26:00 | 0:00 | 135:38:00 | 0:00 | 280:121 | 2:01 |
| 0:00 | 0:00 | 0:00 | 1:00 | 0:00 | 0:02 |
| 0:00 | 0:00 | 0:00 | 0:00 | 0:00 | 1:02 |
| 1:0:3:0:0:0:0:0:1:0 | 0:0:1:0:0:1:0:0:0:0 | 1:0:3:0:0:0:0:0:0:0 | 2:0:1:1:0:0:0:0:0:0 | 5:0:30:0:0:3:0:0:0:2 | 2:0:16:0:0:2:0:0:0:0 |
| 0:2:0:0:0:0:1:1:0:0:0 | 0:0:0:0:0:0:0:0:0:0:0 | 0:0:0:0:0:0:0:0:0:0:0 | 0:0:0:0:0:0:0:0:0:0:0 | 0:7:0:0:1:1:0:0:2:1:1 | 0:0:0:0:0:0:0:0:0:0:0 |
| 0:1:0:0:0:0:0:0:0:0 | 0:0:0:0:0:0:1:0:0 | 0:0:0:0:0:0:0:0:0 | 0:0:0:0:0:0:0:0:0 | 0:0:0:0:0:0:0:0:0 | 0:3:4:1:1:1:0:2:1 |
| 0:00:00 | 0:00:00 | 0:00:00 | 0:00:00 | 0:00:00 | 0:00:00 |
| 0:00:00 | 0:00:00 | 0:00:00 | 0:00:00 | 0:00:00 | 0:00:00 |
| 72:4:1:0 | 28:1:0:0 | 76:4:1:0 | 59:3:0:1 | 242:24:1:0 | 271:25:0:1 |
| 100:5:1:0 | 1:0:0:0 | 146:7:1:1 | 0:0:0:0 | 538:49:1:1 | 3:0:0:0 |
| 108:4:1:0 | 3:0:0:0 | 162:8:1:1 | 3:1:0:0 | 601:53:1:1 | 6:0:0:0 |
| 0 | 0 | 2 | 3 | 10 | 3 |
| 0 | 0 | 5 | 0 | 5 | 0 |
| 0 | 0 | 3 | 0 | 4 | 0 |
| 1 | 0 | 4 | 0 | 3 | 2 |
| 1 | 0 | 5 | 0 | 5 | 0 |
| 157:34:1:6:14:0:2:0:0:0 | 0:0:0:0:0:0:0:0:0:0 | 629:81:14:16:88:8:17:3:6:3 | 0:1:0:0:0:0:0:0:0:0 | 744:171:13:25:72:4:9:6:6:8 | 3:0:0:0:0:0:0:0:0:0 |
| 36:157:1:6:14:0:2:0:0:0 | 0:0:0:0:0:0:0:0:0:0 | 95:631:14:16:88:7:17:3:6:3 | 0:0:0:0:0:0:0:0:0:0 | 195:749:13:25:72:3:9:5:6:8 | 0:0:0:0:0:0:0:0:0:0 |
| 38:157:1:6:14:2:0:0:0:0 | 0:0:0:0:0:0:0:0:0:0 | 106:630:14:16:88:17:6:6:2:3 | 0:2:0:0:0:0:0:0:0:0 | 216:748:13:25:72:9:6:3:3:8 | 0:1:0:0:0:0:0:0:0:0 |
| 38:157:1:6:14:2:0:0:0:0 | 0:0:0:0:0:0:0:0:0:0 | 108:629:14:16:88:17:6:6:2:3 | 0:2:0:0:0:0:0:0:0:0 | 216:748:13:25:72:9:6:3:3:8 | 2:1:0:0:0:0:0:0:0:0 |
| 38:157:1:6:14:2:0:0:0:0 | 0:0:0:0:0:0:0:0:0:0 | 121:632:14:16:88:17:6:5:2:3 | 0:0:0:0:0:0:0:0:0:0 | 225:750:13:25:72:9:6:3:1:8 | 0:0:0:0:0:0:0:0:0:0 |
| 21:99:10:0:2:2:0:4:0 | 0:0:0:0:0:0:0:0:0 | 116:327:42:8:8:22:6:5:0 | 0:1:0:0:0:0:0:0:0 | 189:368:41:3:4:29:5:12:0 | 0:1:0:0:0:0:0:0:0 |
| 0:00:00 | 1:00:00 | 0:00:00 | 17:04:01 | 0:00:00 | 4:00:02 |
| 0:0:0:0:0 | 1:0:0:0:0 | 0:0:0:0:0 | 58:5:0:1:2 | 0:0:0:0:0 | 10:0:0:0:0 |
| 4:8:3:9:12:2:8:8:8:3:4:4:3:4:5:0:0:0:0:0 | 0:0:0:0:0:0:0:0:0:0:0:0:0:0:1:0:0:0:0:0 | 9:31:8:31:6:11:10:16:8:13:10:22:6:3:14:2:0:1:0:0 | 0:0:0:0:0:0:0:0:0:0:0:0:0:0:0:0:0:0:0:0 | 29:34:20:47:20:29:55:27:11:28:21:17:20:7:22:8:0:2:0:1 | 0:0:0:0:0:0:0:0:0:0:0:0:1:0:0:0:0:0:0:0 |
| 4:8:3:8:4:8:11:2:8:0:8:3:4:3:0:4:6:0:0:0 | 0:0:0:1:0:0:1:0:0:0:0:0:0:0:0:0:0:0:0:0 | 10:18:7:27:8:28:6:10:10:1:7:12:23:4:0:4:13:1:2:3 | 0:0:0:3:0:4:0:1:0:0:1:0:0:2:0:1:0:0:0:0 | 20:27:20:45:25:32:19:29:51:1:11:29:15:21:0:8:23:5:8:2 | 0:0:0:2:4:1:0:0:3:0:0:0:1:0:0:0:0:0:0:0 |
| 4:8:7:10:1:5:0:3:3:4:7:6:1:0:1:6:0:0:0:4 | 0:0:1:1:0:0:0:0:0:0:0:0:0:0:0:0:0:0:0:0 | 4:14:14:3:8:5:1:6:5:10:6:4:3:0:3:3:2:1:2:0 | 1:1:3:0:0:0:0:1:1:2:1:1:0:0:0:0:0:0:0:0 | 12:29:21:8:14:27:1:11:9:7:9:9:12:0:12:9:3:3:2:3 | 2:2:0:2:0:1:0:2:1:0:0:0:0:0:1:0:0:0:0:1 |
| 1:3:7:5:1:10:5:8:3:4:6:6:0:3:4:1:0:0:0:0 | 1:0:1:2:1:1:3:1:0:0:0:0:0:0:2:0:0:0:0:0 | 6:3:10:4:3:1:5:7:4:0:2:5:1:6:7:1:0:2:1:1 | 2:2:10:2:1:3:2:10:4:1:2:2:0:2:6:2:0:0:1:1 | 13:10:17:7:8:7:25:21:7:3:8:8:1:8:6:11:0:3:3:1 | 2:6:9:2:7:7:10:12:5:1:5:5:0:10:2:3:0:1:2:1 |
| 63:108:6:0 | 0:0:0:0 | 44:152:5:0 | 0:0:0:0 | 311:509:36:1 | 1:0:1:0 |
| 63:108:6:0 | 1:0:0:0 | 47:152:5:0 | 0:0:0:0 | 328:507:38:1 | 1:2:0:0 |
| 67:107:7:0 | 0:0:0:0 | 52:152:5:0 | 0:0:0:0 | 355:508:40:1 | 1:1:0:0 |
| 70:106:7:0 | 0:1:0:0 | 57:152:6:0 | 0:0:0:0 | 373:509:41:1 | 2:0:0:0 |
| 68:101:7:0 | 5:5:0:0 | 30:87:4:0 | 37:65:3:0 | 400:498:41:1 | 8:11:0:0 |
| 73:106:7:0 | 0:0:0:0 | 68:152:7:0 | 0:0:0:0 | 412:509:42:1 | 1:0:1:0 |
| 103:76:6:0 | 2:0:0:0 | 151:72:7:0 | 0:0:0:0 | 508:429:43:1 | 0:1:0:0 |
| 77:104:6:0 | 0:0:0:0 | 73:151:6:0 | 1:0:0:0 | 445:506:41:1 | 3:1:0:0 |
| 0 | 0 | 0 | 1 | 0 | 5 |
| 2:27 | 0:00 | 0.102083333 | 0:00 | 0.195138889 | 0:00 |
| 2:27 | 0:00 | 0.102083333 | 0:00 | 0.19375 | 0:02 |
| 20:00 | 0:00 | 5:00 | 0:00 | 32:00:00 | 0:00 |
| 0 | 0 | 0 | 0 | 0 | 0 |
| 0:00 | 2:00 | 0:00 | 4:01 | 0:00 | 2:00 |
| 0 | 0 | 0 | 0 | 0 | 2 |
| 1 | 0 | 2 | 0 | 8 | 0 |
| 29:01:00 | 0:00 | 123:00:00 | 4:00 | 81:00:00 | 0:00 |
| 14:01 | 15:00 | 54:00:00 | 61:00:00 | 38:00:00 | 37:00:00 |
| 32:01:00 | 0:00 | 138:01:00 | 6:00 | 73:01:00 | 5:00 |
| 32:01:00 | 0:00:00 | 174:02:00 | 10:00:00 | 92:01:00 | 8:00:01 |
| 15:01:00 | 16:00:00 | 93:02:00 | 86:00:00 | 51:01:00 | 45:01:01 |
| 34:01:00 | 0:00:00 | 189:03:00 | 4:00:00 | 100:02:01 | 1:00:00 |
| 16:01:00 | 18:00:00 | 99:03:00 | 94:00:00 | 52:01:00 | 49:01:01 |

| | | | | | |
|---|---|---|---|---|---|
| 0:00 | 0:00 | 0:00 | 2:00 | 0:00 | 2:00 |
| 23:0:0:6:4:6:0:3:0:0:0:1:0 | 0:0:0:0:1:0:0:0:0:0:0:0:0 | 24:3:0:13:7:7:1:5:2:0:0:0:1 | 0:0:0:0:0:0:0:0:0:0:0:0:0 | 87:5:0:30:10:18:2:18:4:0:0:0:0 | 0:1:0:1:0:0:0:0:0:0:0:0:0 |
| 0 | 0 | 0 | 0 | 0 | 0 |
| 0:00 | 0:00 | 9:01 | 4:01 | 0:00 | 0:00 |
| 0 | 0 | 9 | 4 | 0 | 0 |
| 1 | 0 | 2 | 0 | 11 | 4 |
| 1:02:00 | 0:00:00 | 0:06:01 | 0:00:00 | 1:02:00 | 0:00:00 |
| 24 | 30 | 37 | 31 | 82 | 143 |
| 101:1:4:1:0 | 1:0:0:0:0 | 208:12:8:1:2 | 0:0:0:0:0 | 492:14:15:2:1 | 0:0:0:0:0 |
| 102:1:2:3:1:0:0:0 | 0:0:0:0:0:0:0:0 | 212:12:2:0:1:2:1:0 | 0:0:0:0:0:0:0:0 | 491:14:7:4:2:1:0:1 | 2:0:0:0:0:0:0:0 |
| 133:00:00 | 0:00:00 | 256:04:00 | 0:00:00 | 660:01:01 | 0:00:00 |
| 133:00:00 | 0:00:00 | 256:04:00 | 0:00:00 | 658:01:01 | 2:00:00 |
| 133:00:00 | 0:00:00 | 255:04:00 | 1:00:00 | 657:01:01 | 3:00:00 |
| 0 | 0 | 2 | 0 | 2 | 0 |
| 1:16:25:9:2:0:0:1:0:0 | 0:0:0:0:0:0:0:0:0:0 | 3:30:19:8:1:1:0:1:4:0 | 0:0:0:0:0:0:0:0:0:0 | 6:129:66:18:4:1:2:1:14:0 | 1:1:0:0:0:0:0:0:0:0 |
| 16:25:9:2:0:0:0:1:0:1 | 0:1:0:0:0:0:0:0:0:0 | 30:17:8:1:1:0:2:3:0:1 | 0:0:0:0:0:0:0:0:0:0 | 127:60:19:4:1:1:9:10:0:4 | 1:1:1:0:0:0:0:0:0:0 |
| 1:16:26:9:2:0:0:1:0:0 | 0:0:0:0:0:0:0:0:0:0 | 1:30:17:8:1:1:0:2:0:0 | 0:0:0:0:0:0:0:1:0:0 | 5:130:63:20:4:1:1:10:7:0 | 0:0:0:0:0:0:0:0:0:0 |
| 0:00 | 0:00 | 0:00 | 0:00 | 0:00 | 1:01 |
| 0:00:00 | 0:00:00 | 0:00:01 | 0:00:00 | 0:00:00 | 0:00:00 |
| 125:2:11:85:2:4:2:11:1:0:0:0:0:0:0:0:0 | 0:0:0:0:0:0:0:0:0:0:0:0:0:0:0:0:0 | 166:16:30:145:12:3:1:33:1:5:1:0:0:1:0:0:0 | 0:0:0:0:0:0:0:0:0:0:0:0:0:0:0:0:0 | 527:26:93:405:27:7:5:91:1:1:4:0:0:0:2:1:1 | 0:0:0:0:0:0:0:0:1:0:0:0:0:0:0:0:0 |
| 125:11:85:2:4:1:2:11:1:0:0:0:0:0:0:0:0 | 0:0:0:0:0:0:0:0:0:0:0:0:0:0:0:0:0 | 166:30:145:12:3:16:1:34:1:5:1:0:0:1:0:0:0 | 0:0:1:0:0:0:0:0:0:0:0:0:0:0:0:0:0 | 527:93:405:27:7:23:5:92:1:1:4:0:0:0:2:1:1 | 0:0:0:0:0:0:0:0:0:0:0:0:0:0:0:0:0 |
| 7:125:9:84:2:7:2:1:0:0:0:0:0:0:0:0:0:0:0 | 0:0:0:0:0:0:0:0:0:0:0:0:0:0:0:0:0:0:0 | 20:166:23:143:12:6:0:1:0:1:5:1:0:0:1:1:0:0:0 | 1:1:0:1:0:1:0:0:0:0:0:0:0:0:0:0:0:0:0 | 57:528:58:403:27:15:8:1:3:2:1:4:0:0:0:2:1:1:1 | 1:0:0:0:0:0:0:0:0:0:0:0:0:0:0:0:0:0:0 |
| 82:125:5:5:2:0:0:1:5:0:0:0:0:0:0:0:0:0:0 | 0:0:0:0:0:0:0:0:0:0:0:0:0:0:0:0:0:0:0 | 141:167:14:7:1:6:4:1:8:0:5:1:0:0:1:0:0:0:0 | 0:0:0:0:0:0:0:0:0:0:0:0:0:0:0:0:0:0:0 | 397:528:41:13:6:14:10:1:29:2:1:4:0:0:0:4:1:1:1 | 0:0:0:0:0:0:0:0:0:1:0:0:0:0:0:0:0:0:0 |
| 0 | 0 | 0 | 0 | 0 | 1 |
| 0:00 | 0:00 | 1:01 | 0:00 | 0:02 | 0:00 |
| 0 | 0 | 0 | 1 | 0 | 2 |
| 0 | 0 | 0 | 1 | 1 | 0 |
| 2:2:1:1:1:0:1 | 0:0:0:0:0:0:0 | 3:1:1:0:0:0:0 | 0:0:0:0:0:0:0 | 8:9:5:1:0:0:0 | 0:0:0:0:0:0:0 |
| 0:0:0:0:0:0:0 | 5:2:0:1:1:1:0 | 0:0:0:0:0:0:0 | 2:2:0:0:0:0:0 | 0:0:0:0:0:0:0 | 15:7:2:0:1:0:0 |
| 4:2:1:0:1:1:0 | 0:0:0:0:0:0:0 | 2:2:0:2:0:0:0 | 0:0:0:0:0:0:0 | 14:9:0:2:1:0:0 | 0:0:0:0:0:0:0 |
| 0:00 | 0:00 | 1:00 | 0:00 | 1:00 | 0:00 |
| 0 | 0 | 0 | 1 | 0 | 4 |
| 0 | 0 | 0 | 1 | 0 | 3 |
| 0 | 0 | 0 | 0 | 0 | 3 |
| 0 | 0 | 0 | 0 | 0 | 2 |
| 5094:624:4:4:43:6 | 7:0:0:0:0:0 | 10673:925:7:3:65:37 | 3:0:0:0:0:1 | 4425:722:5:4:36:16 | 4:0:0:0:0:1 |
| 5062:618:22:3:43:0 | 2:1:0:0:0:0 | 10527:907:73:7:65:0 | 17:0:3:0:0:0 | 4348:700:26:4:36:0 | 4:0:1:0:0:0 |
| 4983:610:3:41:37:3 | 4:0:0:1:0:0 | 10340:883:7:65:106:8 | 12:0:0:0:0:0 | 4233:680:4:36:41:4 | 2:0:0:0:0:0 |
| 4987:37:611:6:3:42 | 2:2:0:1:0:0 | 10361:109:883:7:7:65 | 6:2:2:2:0:0 | 4238:43:683:4:4:36 | 4:0:0:0:0:0 |
| 4979:607:3:42:40:8 | 9:1:0:0:0:0 | 10338:880:7:65:118:10 | 14:0:0:0:0:0 | 4226:679:4:36:46:4 | 3:1:0:0:0:0 |
| 4863:596:3:9:39:49 | 12:0:0:0:0:0 | 10022:853:7:11:60:121 | 16:1:0:0:0:1 | 4071:650:4:8:32:47 | 12:2:0:0:0:1 |
| 595:4860:3:9:39:49 | 1:3:0:0:0:0 | 848:9999:7:10:60:123 | 1:8:0:0:0:0 | 649:4066:4:8:32:49 | 1:8:0:0:0:1 |
| 4895:51:602:3:9:35 | 7:2:0:0:0:0 | 10069:131:856:7:10:57 | 13:2:1:0:0:0 | 4097:58:651:3:8:31 | 5:2:1:0:0:0 |
| 4988:600:2:9:33:73 | 12:2:0:0:0:0 | 10247:859:7:10:54:230 | 22:1:0:0:0:2 | 4156:655:4:8:30:115 | 14:3:0:0:0:1 |
| 4983:600:2:9:32:70 | 5:0:0:0:0:0 | 10215:850:7:10:55:229 | 9:0:0:0:0:0 | 4149:647:4:8:27:113 | 3:4:0:0:0:0 |
| 4993:599:70:2:9:32 | 5:0:0:0:0:0 | 10293:854:234:7:10:54 | 9:1:1:0:0:1 | 4178:654:116:4:8:27 | 5:1:0:0:0:0 |
| 4975:596:2:9:32:69 | 13:1:0:0:0:0 | 10289:853:7:10:55:239 | 6:0:0:0:0:0 | 4164:654:4:8:27:116 | 9:2:0:0:0:0 |
| 21:0:4:0:2:5:0:0:0:0 | 0:0:0:0:0:0:0:0:0:0 | 14:0:4:1:9:17:0:0:1:0 | 0:0:0:0:0:0:0:0:0:0 | 33:1:1:0:6:16:1:1:0:1 | 0:0:0:0:0:0:0:0:0:0 |

| | | | | | |
|---|---|---|---|---|---|
| 21:0:4:0:5:2:0:0:0:0 | 0:0:0:0:0:0:0:0:0:0 | 15:0:5:1:18:4:0:0:1:0 | 0:0:0:0:0:0:0:0:0:0 | 33:1:1:0:16:3:1:1:0:1 | 1:0:0:0:1:0:0:0:0:0 |
| 21:0:4:0:5:2:0:0:0:0 | 0:0:0:0:0:0:0:0:0:0 | 15:0:5:1:18:4:0:0:1:0 | 0:0:0:0:0:0:0:0:0:0 | 34:1:1:0:16:3:1:1:0:1 | 0:0:0:0:1:0:0:0:0:0 |
| 15:0:3:0:3:2:0:0:0:0 | 6:0:1:0:0:0:0:0:0:0 | 6:0:2:1:4:0:0:0:1:0 | 9:0:3:0:3:1:0:0:0:0 | 31:1:1:0:4:2:1:1:0:1 | 3:0:0:0:2:0:0:0:0:0 |
| 0 | 0 | 0 | 0 | 0 | 2 |
| 8:02:01 | 0:00:00 | 198:24:14 | 0:00:00 | 84:07:00 | 0:00:00 |
| 48:1520:2:6:0:1:0:0 | 1:4:0:0:0:0:0:0 | 58:1824:7:4:0:0:0:0 | 0:3:0:0:0:0:0:0 | 290:6387:14:18:0:0:1:1 | 2:5:0:0:0:0:0:0 |
| 1531:2:2:6:0:0 | 3:0:0:0:0:0 | 1833:7:8:4:0:0 | 1:0:0:0:0:0 | 6415:14:28:18:1:1 | 4:0:0:0:0:0 |
| 1531:43:2:8:6:0:1:0 | 3:0:0:0:0:0:0:0 | 1826:37:7:18:4:0:0:0 | 4:0:0:0:0:0:0:0 | 6401:204:14:55:18:0:0:1 | 1:0:0:0:0:0:0:0 |
| 1531:43:2:9:6:0:1:0 | 2:0:0:0:0:0:0:0 | 1808:36:7:24:4:0:0:0 | 1:0:0:0:0:0:0:0 | 6324:204:13:73:18:0:0:1 | 7:0:0:0:0:0:0:0 |
| 1530:43:2:11:0:0:0 | 1:0:0:0:0:0:0 | 1804:36:7:47:0:0:0 | 2:0:0:0:0:0:0 | 6313:204:13:113:0:2:1 | 4:1:0:0:0:0:0 |
| 1529:43:2:11:0:0:0 | 1:0:0:0:0:0:0 | 1803:36:7:47:0:0:0 | 2:0:0:0:0:0:0 | 6307:205:13:118:0:2:1 | 6:0:0:0:0:0:0 |
| 1527:43:2:11:0:0:0 | 1:0:0:0:0:0:0 | 1802:36:7:47:0:0:0 | 3:0:0:0:0:0:0 | 6311:205:13:118:0:2:1 | 2:0:0:0:0:0:0 |
| 26:2:4:0:2 | 0:0:0:0:0 | 13:3:6:0:0 | 0:0:0:0:0 | 54:2:9:0:1 | 1:0:0:0:0 |
| 0 | 0 | 0 | 0 | 0 | 0 |
| 0 | 2 | 3 | 1 | 0 | 7 |
| 18:35 | 0:00 | 10:28 | 0:00 | 85:90 | 0:00 |
| 194:01:12 | 0:00:00 | 329:04:08 | 0:00:00 | 536:05:36 | 0:00:00 |
| 0.05224537 | 0:00:00 | 0.176030093 | 0:00:00 | 0.239502315 | 0:00:01 |
| 194:01:12 | 0:00:00 | 330:04:08 | 0:00:00 | 534:05:36 | 0:00:00 |
| 0.052222222 | 0:00:00 | 0.175300926 | 0:00:00 | 0.239409722 | 0:00:02 |
| 0.052233796 | 0:00:00 | 0.175277778 | 0:00:01 | 0.239409722 | 0:00:02 |
| 0.052233796 | 0:00:00 | 0.1753125 | 0:00:00 | 0.239398148 | 0:00:01 |
| 0.052233796 | 0:00:00 | 0.1753125 | 0:00:00 | 0.239398148 | 0:00:01 |
| 0.052233796 | 0:00:00 | 0.175289352 | 0:00:02 | 0.239398148 | 0:00:01 |
| 0.052233796 | 0:00:00 | 0.175266204 | 0:00:03 | 0.239398148 | 0:00:01 |
| 0.052233796 | 0:00:00 | 0.175289352 | 0:00:01 | 0.239386574 | 0:00:02 |
| 0.052233796 | 0:00:00 | 0.175300926 | 0:00:00 | 0.239409722 | 0:00:01 |
| 192:01:12 | 0:00:00 | 325:04:07 | 1:00:00 | 524:05:36 | 2:00:00 |
| 0.052222222 | 0:00:00 | 0.175266204 | 0:00:01 | 0.239386574 | 0:00:00 |
| 192:01:12 | 0:00:00 | 321:04:07 | 0:00:00 | 510:05:36 | 2:00:00 |
| 0.052141204 | 0:00:00 | 0.175092593 | 0:00:01 | 0.238958333 | 0:00:00 |
| 0.052222222 | 0:00:00 | 0.175196759 | 0:00:00 | 0.2384375 | 0:00:00 |
| 0.05224537 | 0:00:00 | 0.175219907 | 0:00:02 | 0.239247685 | 0:00:01 |
| 0.05224537 | 0:00:00 | 0.175243056 | 0:00:00 | 0.239282407 | 0:00:00 |
| 0.052233796 | 0:00:01 | 0.175219907 | 0:00:02 | 0.239293981 | 0:00:00 |
| 193:01:11 | 0:00:01 | 322:04:07 | 0:00:00 | 516:05:36 | 0:00:00 |
| 0 | 0 | 0 | 0 | 0 | 0 |
| 0:0:0:0 | 0:0:0:0 | 0:0:0:0 | 0:2:1:0 | 0:0:0:0 | 0:0:0:1 |
| 0:00:00 | 0:00:00 | 0:00:00 | 1:02:00 | 0:00:00 | 0:00:01 |
| 0:00:00 | 0:00:00 | 0:00:00 | 1:02:00 | 0:00:00 | 0:00:01 |
| 9693 | 13 | 16082 | 25 | 45994 | 76 |
| 9697 | 20 | 16111 | 37 | 46082 | 90 |
| 9718 | 9 | 16145 | 19 | 46139 | 49 |
| 9743 | 25 | 16206 | 36 | 46253 | 105 |
| 9731 | 11 | 16129 | 27 | 45911 | 56 |
| 9735 | 9 | 16124 | 18 | 45890 | 49 |
| 9691 | 5 | 15888 | 7 | 45258 | 22 |
| 9585 | 11 | 15528 | 14 | 44288 | 52 |
| 9598 | 10 | 15588 | 20 | 44469 | 61 |
| 9589 | 7 | 15562 | 26 | 44427 | 39 |
| 9553 | 12 | 15664 | 25 | 44520 | 45 |
| 9537 | 9 | 15655 | 24 | 44482 | 49 |
| 40 | 0 | 69 | 0 | 289 | 2 |
| 1:16:61:0 | 0:0:0:0 | 7:28:126:0 | 0:0:0:0 | 18:105:442:6 | 0:0:1:1 |
| 4:01 | 0:00 | 3:02 | 0:02 | 6:05 | 0:01 |
| 2:01 | 1:02 | 0:05 | 1:06 | 3:09 | 0:11 |
| 0:00 | 2:03 | 0:00 | 1:10 | 0:00 | 3:19 |
| 11:01 | 1:00 | 16:00 | 0:00 | 72:01:00 | 4:01 |
| 0:01:00 | 3:00:00 | 7:03:00 | 16:01:00 | 12:08:02 | 30:00:01 |
| 4:01:00 | 0:00:00 | 23:04:00 | 0:00:00 | 43:08:03 | 0:00:00 |
| 0:01:00 | 4:00:00 | 8:03:00 | 15:01:00 | 13:08:02 | 31:00:01 |
| 0:00 | 4:00 | 8:00 | 17:00 | 13:02 | 31:01:00 |
| 0:01:00 | 3:00:00 | 6:03:00 | 11:01:00 | 11:07:01 | 21:01:01 |
| 2:00:00 | 0:00:00 | 11:01:00 | 0:00:00 | 27:16:02 | 0:00:00 |
| 31:1:1:0 | 0:0:0:0 | 41:2:2:2 | 0:0:0:0 | 197:0:9:3 | 2:0:0:0 |
| 31:1:1:0 | 0:0:0:0 | 34:5:1:2 | 0:0:0:0 | 224:11:8:3 | 3:0:1:0 |
| 30:1:1:0 | 0:0:0:0 | 32:6:1:2 | 2:0:0:0 | 223:12:6:3 | 4:0:1:0 |

| | | | | | |
|---|---|---|---|---|---|
| 1:39:4:0 | 0:0:0:0 | 14:194:22:2 | 0:1:0:0 | 8:79:8:4 | 0:0:0:0 |
| 1:39:0:3 | 0:0:0:0 | 14:194:2:18 | 0:0:0:0 | 8:81:4:8 | 0:0:0:0 |
| 39:00:01 | 0:00:00 | 194:02:00 | 0:00:00 | 82:03:00 | 0:00:00 |
| 2:0:0:0:1:0:0:0:0:0:0:0:0:0:0 | 0:0:1:0:0:0:2:0:1:0:0:1:0:0:0 | 0:0:1:0:1:0:0:0:1:0:0:0:0:0:1 | 1:1:0:0:1:0:0:0:0:0:0:2:0:0:0 | 0:1:0:0:0:0:0:0:0:0:0:0:0:0 | 1:4:0:0:3:0:2:0:2:1:0:5:0:0:3 |
| 0:00 | 0:00 | 11:01 | 0:00 | 3:01 | 0:00 |
| 31:1:0:0 | 6:0:1:0 | 78:0:1:1 | 14:1:0:0 | 112:1:3:4 | 24:1:1:0 |
| 40:1:1:0:0 | 0:0:0:0:0 | 108:22:1:0:0 | 1:0:0:0:0 | 146:28:5:1:0 | 1:1:0:0:0 |
| 9:01:02 | 8:03:00 | 1:22:00 | 2:08:00 | 27:65:0 | 19:27:01 |
| 0:0:1:0 | 0:1:2:0 | 0:6:4:2 | 0:7:3:6 | 4:19:6:4 | 0:1:0:0 |
| 1:1:3:18:1:0:4:8:1 | 0:0:0:0:0:0:0:0:0 | 0:0:20:51:0:0:77:1:3 | 0:0:0:0:0:0:0:0:0 | 3:2:27:83:2:1:144:7:1 | 0:0:0:1:0:0:0:0:0 |
| 1:1:3:18:1:0:4:8:1 | 0:0:0:0:0:0:0:0:0 | 0:0:20:51:0:0:84:1:3 | 0:0:0:1:0:0:1:0:0 | 3:2:27:88:2:1:154:7:1 | 0:0:0:0:0:0:2:0:0 |
| 4:02 | 0:00 | 5:01 | 0:00 | 14:00 | 0:00 |
| 4:00 | 0:00 | 9:00 | 0:00 | 27:00:00 | 0:00 |
| 4:00 | 0:00 | 8:00 | 0:00 | 29:00:00 | 0:00 |
| 1 | 0 | 3 | 0 | 5 | 1 |
| 1 | 0 | 1 | 0 | 4 | 0 |
| 0 | 2 | 0 | 2 | 0 | 7 |
| 2 | 0 | 2 | 0 | 5 | 2 |
| 0 | 2 | 0 | 2 | 2 | 5 |
| 9:01:01 | 0:00:00 | 28:00:10 | 0:00:00 | 58:00:04 | 2:00:00 |
| 0:0:0:0 | 7:1:1:0 | 0:0:0:0 | 17:0:7:0 | 0:0:0:0 | 35:0:2:0 |
| 6:0:5:0 | 0:0:0:0 | 9:5:18:1 | 0:0:0:0 | 36:14:83:1 | 0:0:0:0 |
| 0:0:0:0 | 6:0:4:0 | 0:0:0:0 | 8:3:12:1 | 0:0:0:0 | 30:10:57:1 |
| 0:0:0:0 | 1:0:4:0 | 0:0:3:0 | 1:0:3:0 | 0:0:1:0 | 5:4:13:0 |
| 0:0:0:0 | 1:0:4:0 | 0:0:3:0 | 1:0:3:0 | 0:0:1:0 | 5:4:13:0 |
| 59 | 0 | 111 | 0 | 218 | 3 |
| 39:2:0:0 | 0:0:0:0 | 78:1:2:5 | 0:0:0:0 | 206:3:3:9 | 0:0:0:0 |
| 8:0:2:0:0:1:0:0:1:0:0:0:0:0 | 0:0:1:0:0:0:0:0:0:0:0:0:0:0 | 11:1:9:0:1:0:0:0:0:0:0:0:0:0 | 1:0:0:0:0:0:0:0:0:0:0:0:0:0 | 27:0:36:6:6:4:1:1:9:4:0:1:2:1 | 0:0:2:0:0:0:0:0:0:0:0:0:0:0 |
| 2:8:0:0:0:1:0:0:1:0:0:0:0:0 | 1:0:0:0:0:0:0:0:0:0:0:0:0:0 | 10:12:1:0:1:0:0:0:0:0:0:0:0:0 | 0:0:0:0:0:0:0:0:0:0:0:0:0:0 | 41:26:0:6:6:4:1:1:9:4:0:1:2:1 | 4:1:0:0:0:0:0:0:0:0:0:0:0:0 |
| 8:0:3:0:0:1:0:0:1:0:0:0:0:0 | 0:0:1:0:0:0:0:0:0:0:0:0:0:0 | 12:1:10:0:1:0:0:0:0:0:0:0:0:0 | 0:0:0:0:0:0:0:0:0:0:0:0:0:0 | 27:0:44:6:6:4:1:1:9:3:0:1:2:1 | 0:0:1:0:0:0:0:0:0:0:0:0:0:0 |
| 22:01:00 | 26:00:00 | 68:00:00 | 78:00:00 | 224:00:01 | 112:00:01 |
| 48:04:00 | 0:00 | 100:05:00 | 0:00 | 282:11:00 | 0:00 |
| 41:08:00 | 0:00 | 64:16:00 | 0:00 | 179:58:00 | 0:00 |
| 1 | 0 | 5 | 7 | 23 | 15 |
| 1:01 | 5:00 | 3:02 | 2:01 | 7:10 | 2:02 |
| 0 | 1 | 0 | 0 | 0 | 1 |
| 0:00:00 | 0:00:00 | 0:03:01 | 0:00:00 | 0:00:00 | 0:00:00 |
| 0:0:0:0:0 | 0:0:0:0:0 | 0:2:0:0:0 | 0:1:0:0:0 | 0:0:0:0:0 | 0:0:0:1:1 |
| 0:0:0:0 | 0:1:0:0 | 0:0:0:0 | 0:0:0:0 | 0:0:0:0 | 2:0:2:1 |
| 1 | 1 | 7 | 2 | 4 | 6 |
| 1:99:1:4:3:0:2:0:1:0:0:1:0:2:0:4:0:0:1:2:0:0:1:1:0:0:0 | 0:1:0:0:0:0:0:0:0:0:0:0:0:0:0:0:0:0:0:0:0:0:0:0:0:0 | 1:75:0:13:3:1:2:9:9:0:0:0:5:8:1:6:0:1:0:3:0:0:1:0:1:0:0 | 0:2:0:0:0:0:0:0:0:0:0:0:0:0:0:0:0:0:0:0:0:0:0:0:0:0:0 | 7:392:5:41:4:1:9:7:17:0:1:2:4:15:2:51:2:0:0:3:1:1:0:0:0:1:1 | 4:0:0:0:0:0:0:0:0:0:0:0:0:0:0:0:0:1:0:0:0:0:0:0:0:0:0 |
| 5:9:168:130:8:8:15:11:35:5:5:5 | 0:0:0:0:0:0:0:0:0:0:0:0 | 16:4:276:53:12:2:2:24:31:17:0:3 | 0:0:0:0:0:0:0:0:0:0:0:0 | 32:11:653:185:36:30:12:56:64:28:7:8 | 1:0:0:0:0:0:0:0:0:0:0:0 |
| 5:9:168:130:8:8:15:11:35:5:5:5 | 0:0:0:0:0:0:0:0:0:0:0:0 | 16:4:276:53:12:2:2:24:31:17:0:3 | 0:0:0:0:0:0:0:0:0:0:0:0 | 33:11:650:185:36:30:12:56:64:28:7:8 | 0:0:3:0:0:0:0:0:0:0:0:0 |
| 6:9:168:130:8:8:15:11:35:5:4:5 | 0:0:0:0:0:0:0:0:0:0:0:0 | 16:4:276:51:12:2:2:24:30:17:0:3 | 0:0:0:1:0:0:0:0:0:0:0:0 | 33:11:652:183:35:29:12:54:64:28:6:8 | 0:0:1:0:1:0:0:1:0:0:0:0 |
| 5:9:165:8:8:36:6:0:0 | 0:0:0:0:0:0:0:0:0 | 13:4:275:12:3:31:18:0:0 | 0:0:0:0:0:0:0:0:0 | 32:12:657:36:30:64:32:0:1 | 0:0:0:0:0:0:0:0:0 |
| 5:8:168:126:8:8:15:36:6:6:5:5 | 0:0:0:0:0:0:0:0:0:0:0:0 | 11:4:276:46:12:3:2:31:3:18:0:3 | 0:0:0:0:0:0:0:0:0:0:0:0 | 30:11:657:176:36:31:12:63:16:32:7:8 | 0:0:1:0:0:0:0:0:0:0:0:0 |
| 5:8:168:125:8:8:15:36:6:6:5:5 | 0:0:0:1:0:0:0:0:0:0:0:0 | 11:4:276:46:12:3:2:31:3:18:0:3 | 0:0:0:0:0:0:0:0:0:0:0:0 | 30:11:658:175:36:32:12:63:16:32:7:8 | 0:0:0:1:0:0:0:0:0:0:0:0 |
| 5:8:166:126:8:8:15:36:6:6:5:5 | 0:0:2:0:0:0:0:0:0:0:0:0 | 10:4:276:46:12:3:2:31:3:18:0:3 | 0:0:0:0:0:0:0:0:0:0:0:0 | 26:11:658:176:36:32:12:63:12:32:7:8 | 0:0:0:0:0:0:0:0:0:0:0:0 |
| 4:8:168:126:8:8:15:36:6:6:5:5 | 0:0:0:0:0:0:0:0:0:0:0:0 | 8:4:276:46:12:4:2:30:2:18:0:3 | 0:0:0:0:0:0:0:1:0:0:0:0 | 24:10:657:175:36:32:12:62:12:32:7:8 | 0:1:1:0:0:1:0:0:0:0:0:0 |

| | | | | | |
|---|---|---|---|---|---|
| 10:10:0:0:0:0 | 0:5:0:1:0:0 | 8:1:0:11:1:1 | 0:0:0:2:0:0 | 21:22:0:23:0:0 | 0:3:0:2:0:0 |
| 0:38:2:8:1:1:3:0:1:1:1:0:1:0 | 0:20:8:6:1:1:2:1:0:0:0:0:1:0 | 0:51:14:1:15:2:4:0:1:0:3:0:0:0 | 0:1:0:0:0:0:0:0:0:0:0:0:0:0 | 0:63:9:5:9:0:2:0:0:1:0:0:0:0 | 0:101:33:19:10:5:8:0:1:3:2:1:1:1 |
| 3:1:8:11 | 7:0:0:0 | 0:0:1:4 | 1:1:0:0 | 2:0:23:26 | 9:0:0:0 |
| 10:8:11:1 | 0:0:0:0 | 1:1:4:1 | 1:0:0:0 | 11:23:25:0 | 11:0:0:4 |
| 10:8:11:1 | 0:0:0:0 | 1:1:4:1 | 1:0:0:0 | 12:23:25:1 | 11:0:0:3 |
| 10:8:11:1 | 2:0:0:0 | 1:1:4:1 | 1:0:0:0 | 13:23:24:1 | 10:0:0:2 |
| 8:37:9:9 | 0:0:0:0 | 1:19:3:3 | 0:0:0:0 | 23:97:19:11 | 0:0:0:0 |
| 8:41:7:8 | 0:0:0:0 | 1:21:2:3 | 0:0:0:0 | 23:102:14:13 | 0:1:0:0 |
| 0:0:0:0 | 8:40:0:1 | 0:0:0:0 | 1:21:0:1 | 0:0:0:0 | 22:102:1:3 |
| 0:0:0:1:0:1:0:0 | 0:0:0:0:0:1:2:0 | 0:0:0:0:0:0:0:0 | 0:0:0:1:0:0:0:0 | 1:0:0:4:5:1:4:1 | 1:0:1:1:1:3:2:0 |
| 0:24:7:2:4:5:0:0:0:0:1:2:1:0:1:0:0:0:0 | 0:0:0:0:0:0:0:0:0:0:0:0:0:0:0:0:0:0:0 | 1:2:0:1:2:1:0:0:0:0:4:0:0:0:0:1:1:0:0 | 0:0:0:0:0:0:0:0:0:0:0:0:0:0:0:0:0:0:0 | 5:56:11:3:8:9:0:0:2:0:10:0:2:1:0:0:6:1:1 | 0:0:0:0:0:0:0:0:0:0:0:0:0:0:0:0:0:0:0 |
| 0 | 1 | 0 | 0 | 0 | 2 |
| 92 | 1 | 121 | 1 | 290 | 3 |
| 49:0:6:1:0:0:0:0:1:2:0:0:1 | 0:0:0:0:0:0:0:0:0:0:0:0:0 | 29:0:4:1:0:1:0:0:8:23:0:3:0 | 0:0:0:0:0:0:0:0:0:0:0:0:0 | 129:1:32:1:3:9:2:0:26:37:2:4:3 | 1:0:0:0:0:0:0:0:0:1:0:0:0 |
| 0:00 | 0:00 | 0:00 | 0:01 | 0:00 | 0:02 |
| 1:0:0:0:0:0:0 | 1:0:0:0:0:1:1 | 1:0:1:0:0:0:0 | 0:0:1:0:0:0:0 | 4:0:1:0:0:0:0 | 12:1:7:3:1:1:5 |
| 0:0:0:0:0:0:0 | 2:0:0:0:1:0:1 | 0:0:0:0:0:0:0 | 1:2:0:0:0:0:1 | 0:0:0:0:0:0:0 | 16:8:1:1:1:0:5 |
| 0:00 | 2:00 | 0:00 | 2:02 | 0:00 | 24:01:00 |
| 0:00:00 | 0:00:00 | 2:00:00 | 2:00:00 | 1:00:02 | 4:00:00 |
| 0.141666667 | 0:01 | 0.095138889 | 0:00 | 0.504861111 | 0:00 |
| 0.152777778 | 0:00 | 0.100694444 | 0:00 | 0.529861111 | 0:01 |
| 0.151388889 | 0:00 | 0.098611111 | 0:00 | 0.502777778 | 0:02 |
| 1 | 2 | 0 | 0 | 0 | 0 |
| 24:1:1:1:1:0 | 2:0:0:0:0:0 | 22:1:3:0:1:0 | 0:0:0:0:0:0 | 138:1:25:3:12:2 | 0:0:0:0:0:0 |
| 8:9:2:1:0:0:1:0 | 0:2:0:0:0:0:0:0 | 0:0:0:0:0:0:0:0 | 0:0:0:0:0:0:0:0 | 9:8:0:5:0:0:0:1 | 0:0:0:0:0:0:0:0 |
| 0:00:00 | 1:00:00 | 0:00:00 | 0:02:00 | 0:00:00 | 1:01:01 |
| 0:01:00 | 0:00:00 | 2:00:00 | 0:00:00 | 0:01:04 | 0:00:00 |
| 0:01:00 | 0:00:00 | 1:00:00 | 0:00:00 | 0:01:04 | 0:00:00 |
| 0:00 | 0:00 | 0:00 | 0:03 | 0:00 | 0:01 |
| 0:00 | 0:00 | 0:01 | 0:00 | 0:00 | 0:01 |
| 0:01 | 1:00 | 0:00 | 1:02 | 4:02 | 0:01 |
| 69:6:2:1:8 | 13:1:0:0:1 | 114:10:7:1:11 | 2:0:0:0:0 | 263:9:9:9:28 | 15:0:0:1:3 |
| 80:6:2:2:10 | 11:1:1:0:1 | 130:10:8:1:13 | 4:0:0:0:0 | 294:10:10:9:32 | 17:0:1:3:3 |
| 81:6:3:2:11 | 9:1:0:0:0 | 121:10:6:1:13 | 2:0:1:0:0 | 295:9:11:12:34 | 16:1:0:1:1 |
| 92:3:3:11:1 | 0:0:0:0:0 | 135:8:1:13:0 | 0:0:0:0:0 | 312:10:12:35:0 | 2:0:0:0:0 |
| 92:3:3:11:1 | 0:0:0:0:0 | 135:8:1:13:0 | 0:0:0:0:0 | 313:10:12:34:0 | 1:0:0:1:0 |
| 92:7:3:3:5 | 0:0:0:0:0 | 135:11:8:1:7 | 0:0:0:0:0 | 314:14:11:11:21 | 0:0:0:0:0 |
| 92:5:3:2:7 | 0:0:0:0:0 | 135:11:8:1:3 | 0:0:0:0:0 | 314:13:11:12:17 | 0:0:0:0:0 |
| 0:0:2:0:0:0 | 0:0:0:0:0:0 | 0:1:6:0:1:0 | 0:0:0:0:0:0 | 0:2:16:0:1:0 | 0:0:0:0:0:0 |
| 675:279:52 | 2:00:00 | 95:272:316 | 2:01:01 | 585:491:514 | 4:04:04 |
| 702:287:39 | 0:00:01 | 114:245:184 | 0:01:05 | 617:462:354 | 2:00:08 |
| 720:293:40 | 0:00:00 | 128:250:178 | 7:02:06 | 647:479:334 | 8:05:06 |
| 822:343:32 | 2:02:00 | 401:395:157 | 0:01:00 | 1199:663:274 | 1:01:05 |
| 366:871:25 | 1:00:00 | 438:485:48 | 2:01:00 | 727:1421:77 | 1:00:00 |
| 877:367:23 | 0:00:00 | 490:415:27 | 1:00:00 | 1424:702:47 | 1:02:00 |
| 877:364:24 | 2:00:00 | 494:392:25 | 1:04:01 | 1427:675:53 | 2:01:00 |
| 877:364:24 | 2:00:00 | 492:393:26 | 3:03:00 | 1427:673:53 | 2:01:00 |
| 880:356:24 | 0:02:00 | 494:388:24 | 2:02:00 | 1428:664:52 | 2:01:00 |
| 876:357:25 | 4:00:00 | 495:382:22 | 0:02:00 | 1428:653:58 | 3:01:00 |
| 25:879:356 | 0:01:01 | 22:494:380 | 0:01:01 | 60:1431:649 | 0:00:00 |
| 880:355:26 | 0:01:00 | 494:365:22 | 4:04:00 | 1435:621:66 | 0:02:00 |
| 879:329:29 | 1:01:00 | 501:312:23 | 0:00:00 | 1436:534:70 | 1:01:02 |
| 880:314:31 | 0:00:00 | 501:281:24 | 0:00:00 | 1438:498:81 | 1:02:02 |
| 0:00 | 1:00 | 0:00 | 3:00 | 1:00 | 4:00 |
| 0 | 2 | 0 | 0 | 0 | 1 |
| 0:00 | 0:01 | 0:00 | 2:00 | 0:00 | 7:01 |
| 0 | 1 | 0 | 0 | 0 | 1 |
| 97:3:3:11:0:1 | 0:0:1:0:0:0 | 120:5:4:23:1:0 | 0:0:3:0:0:0 | 513:24:6:74:2:0 | 1:0:4:0:0:2 |
| 0:00 | 2:00 | 2:01 | 4:00 | 5:01 | 3:00 |
| 2:0:0:0 | 0:0:0:0 | 2:0:0:0 | 0:0:0:0 | 10:2:2:0 | 2:1:0:0 |
| 1:13:0:1:0:2:0 | 3:5:1:0:1:1:0 | 0:0:0:2:0:0:0 | 0:1:0:0:0:0:0 | 12:17:3:5:0:2:1 | 14:35:11:4:1:6:0 |
| 0:0:0:0:0:0 | 4:20:1:2:1:0 | 0:0:0:0:0:0 | 1:1:0:2:0:0 | 0:0:0:0:0:0 | 31:53:14:10:1:1 |
| 4:20:1:2:1:0 | 0:0:0:0:0:0 | 1:1:0:2:0:0 | 0:0:0:0:0:0 | 31:53:14:10:1:1 | 0:0:0:0:0:0 |
| 3:6:1:1:1:0 | 1:14:0:1:0:0 | 1:1:0:0:0:0 | 0:0:0:2:0:0 | 18:36:12:5:1:0 | 14:17:2:5:0:1 |
| 28:17:14:0:0:0 | 0:0:0:0:0:0 | 34:20:16:0:0:0 | 0:0:1:0:0:0 | 204:66:86:0:1:1 | 6:2:4:0:0:0 |
| 29:15:20:0:0:0:0 | 0:0:0:0:0:0:0 | 49:22:30:18:0:0:0 | 0:0:0:0:0:0:0 | 287:130:95:48:0:1:1 | 1:1:0:0:0:0:0 |
| 29:15:20:0:0:0:0 | 0:0:0:0:0:0:0 | 49:22:30:18:0:0:0 | 0:0:0:0:0:0:0 | 287:131:95:48:0:1:1 | 1:0:0:0:0:0:0 |

| | | | | | |
|---|---|---|---|---|---|
| 0:0:1:2 | 0:0:0:0 | 0:0:1:0 | 0:0:2:1 | 1:0:6:5 | 0:1:3:1 |
| 0:00:00 | 0:00:00 | 0:00:00 | 0:00:01 | 0:00:00 | 0:00:00 |
| 9 | 0 | 11 | 0 | 25 | 0 |
| 135:08:13 | 0:01:00 | 206:08:30 | 0:00:00 | 506:17:69 | 0:00:02 |
| 136:10:11 | 0:00:00 | 206:09:24 | 0:00:00 | 503:19:61 | 3:00:00 |
| 136:11:10 | 0:00:00 | 206:15:17 | 0:00:00 | 506:25:44 | 0:00:00 |
| 136:13:09 | 0:00:00 | 206:16:11 | 0:00:00 | 504:29:34 | 2:00:01 |
| 50:11:149:1:51 | 2:9:2:0:0 | 41:11:25:0:60 | 0:3:1:0:0 | 170:35:199:3:244 | 2:19:3:0:0 |
| 1:150:50:20:51 | 0:1:1:0:0 | 0:26:40:17:60 | 0:0:0:2:0 | 3:201:161:61:244 | 0:1:3:0:0 |
| 151:48:22:51:1 | 0:0:0:0:0 | 27:35:32:60:0 | 0:0:0:0:0 | 203:135:88:244:2 | 0:0:0:0:0 |
| 151:48:22:51:1 | 0:0:0:0:0 | 27:35:32:60:0 | 0:0:0:0:0 | 203:135:88:244:2 | 0:0:0:0:0 |
| 151:42:33:50:0 | 0:0:0:0:0 | 34:16:58:60:0 | 0:0:2:0:0 | 210:101:129:244:0 | 2:0:3:0:0 |
| 150:42:34:51:1 | 0:0:0:0:0 | 34:13:70:60:0 | 0:0:0:0:0 | 216:104:168:243:2 | 1:0:0:1:0 |
| 97:3:5:1:1:5:0 | 0:0:0:0:0:0:0 | 120:5:3:2:5:1:1 | 0:0:0:0:0:0:0 | 514:24:8:7:11:5:3 | 2:0:0:0:0:0:0 |
| 97:2:6:0:1:3:0 | 0:0:0:0:0:0:0 | 119:2:2:5:2:1:1 | 0:0:0:0:0:0:0 | 513:14:6:6:6:7:3 | 2:0:1:0:0:0:0 |
| 0:00:00 | 2:00:00 | 0:00:00 | 0:00:01 | 0:00:00 | 0:00:00 |
| 30 | 0 | 28 | 2 | 73 | 1 |
| 1:0:1:0:0:0:2:0:0 | 1:0:1:0:0:0:0:<br>0:0 | 1:0:0:0:1:0:1:0:0 | 0:0:0:0:0:0:0:1:0 | 3:0:1:0:0:0:1:1:1 | 3:0:0:0:0:0:0:2:0 |
| 1 | 1 | 0 | 0 | 2 | 1 |
| 2 | 0 | 0 | 0 | 3 | 0 |
| 0:00:00 | 0:00:00 | 0:00:00 | 3:00:00 | 0:00:00 | 0:01:01 |
| 0:00 | 0:00 | 0:00 | 0:00 | 0:00 | 0:00 |
| 0:00 | 0:00 | 0:00 | 0:00 | 0:00 | 0:00 |
| 0:00 | 0:00 | 0:00 | 0:00 | 0:00 | 0:00 |
| 0:00 | 0:00 | 0:00 | 0:00 | 0:00 | 0:00 |
| 0:00 | 0:00 | 0:00 | 0:00 | 0:00 | 0:00 |
| 0:00 | 0:00 | 0:00 | 0:00 | 0:00 | 0:00 |
| 0:00 | 0:00 | 0:00 | 0:00 | 0:00 | 0:00 |
| 0:00 | 0:00 | 0:00 | 0:00 | 0:00 | 0:00 |
| 0:00 | 0:00 | 0:00 | 0:00 | 0:00 | 0:00 |
| 0:00 | 0:00 | 0:00 | 0:00 | 0:00 | 0:00 |
| 0:00 | 0:00 | 0:00 | 0:00 | 0:00 | 0:00 |
| 0:00 | 0:00 | 0:00 | 0:00 | 0:00 | 0:00 |
| 0:00 | 0:00 | 0:00 | 0:00 | 0:00 | 0:00 |
| 0:00 | 0:00 | 0:00 | 0:00 | 0:00 | 0:00 |
| 0:00 | 0:00 | 0:00 | 0:00 | 0:00 | 0:00 |
| 0:00 | 0:00 | 0:00 | 0:00 | 0:00 | 0:00 |
| 0:00 | 0:00 | 0:00 | 0:00 | 0:00 | 0:00 |
| 0:00 | 0:00 | 0:00 | 0:00 | 0:00 | 0:00 |
| 0:00 | 0:00 | 0:00 | 0:00 | 0:00 | 0:00 |
| 0:00 | 0:00 | 0:00 | 0:00 | 0:00 | 0:00 |
| 0:00 | 0:00 | 0:00 | 0:00 | 0:00 | 0:00 |
| 0:00 | 0:00 | 0:00 | 0:00 | 0:00 | 0:00 |
| 0:00 | 0:00 | 0:00 | 0:00 | 0:00 | 0:00 |
| 0:00 | 0:00 | 0:00 | 0:00 | 0:00 | 0:00 |
| 0:00 | 0:00 | 0:00 | 0:00 | 0:00 | 0:00 |
| 0:00 | 0:00 | 0:00 | 0:00 | 0:00 | 0:00 |
| 0:00 | 0:00 | 0:00 | 0:00 | 0:00 | 0:00 |
| 0:00 | 0:00 | 0:00 | 0:00 | 0:00 | 0:00 |
| 0:00 | 0:00 | 0:00 | 0:00 | 0:00 | 0:00 |
| 0:00 | 0:00 | 0:00 | 0:00 | 0:00 | 0:00 |
| 0:00 | 0:00 | 0:00 | 0:00 | 0:00 | 0:00 |
| 0:00 | 0:00 | 0:00 | 0:00 | 0:00 | 0:00 |
| 0:00 | 0:00 | 0:00 | 0:00 | 0:00 | 0:00 |
| 0:00 | 0:00 | 0:00 | 0:00 | 0:00 | 0:00 |
| 0:00 | 0:00 | 0:00 | 0:00 | 0:00 | 0:00 |
| 0:00 | 0:00 | 0:00 | 0:00 | 0:00 | 0:00 |
| 0:00 | 0:00 | 0:00 | 0:00 | 0:00 | 0:00 |
| 0:00 | 0:00 | 0:00 | 0:00 | 0:00 | 0:00 |
| 0:00 | 0:00 | 0:00 | 0:00 | 0:00 | 0:00 |
| 0:00 | 0:00 | 0:00 | 0:00 | 0:00 | 0:00 |
| 0:00 | 0:00 | 0:00 | 0:00 | 0:00 | 0:00 |
| 0:00 | 0:00 | 0:00 | 0:00 | 0:00 | 0:00 |
| 0:00 | 0:00 | 0:00 | 0:00 | 0:00 | 0:00 |
| 0:00 | 0:00 | 0:00 | 0:00 | 0:00 | 0:00 |
| 202:132:27:6:4 | 0:0:0:0:0 | 276:131:28:10:7 | 0:0:0:0:0 | 901:334:60:19:13 | 1:1:0:0:0 |
| 201:132:27:6:4 | 0:0:0:0:0 | 276:131:28:10:7 | 0:0:0:0:0 | 899:333:60:18:13 | 3:0:0:1:0 |
| 132:200:27:6:4 | 0:0:0:0:0 | 131:276:28:10:7 | 0:0:0:0:0 | 332:898:60:18:13 | 1:2:0:1:0 |
| 0:00 | 0:00 | 0:00 | 1:00 | 0:00 | 1:01 |
| 0:0:0:0 | 0:0:0:0 | 0:0:1:0 | 0:0:0:0 | 2:0:1:1 | 0:0:0:0 |
| 1:01 | 0:00 | 2:01 | 3:00 | 7:01 | 3:00 |
| 1:00:01 | 0:00:00 | 3:00:00 | 0:00:00 | 4:00:01 | 0:00:00 |
| 0 | 3 | 0 | 3 | 0 | 4 |
| 0:00 | 0:00 | 0:00 | 0:00 | 0:00 | 0:00 |
| 0:00 | 253:03:00 | 0:00 | 636:05:00 | 0:00 | 795:08:00 |
| 1:00 | 253:03:00 | 1:00 | 630:05:00 | 1:00 | 788:08:00 |
| 0:00 | 240:03:00 | 1:00 | 600:04:00 | 2:00 | 735:07:00 |

| | | | | | |
|---|---|---|---|---|---|
| 0:00 | 240:03:00 | 1:00 | 600:04:00 | 2:00 | 735:07:00 |
| 0:00 | 241:03:00 | 0:00 | 601:04:00 | 0:00 | 735:07:00 |
| 119:00:00 | 122:03:00 | 303:00:00 | 298:04:00 | 504:00:00 | 232:07:00 |
| 122:03:00 | 119:00:00 | 298:04:00 | 301:00:00 | 232:07:00 | 504:00:00 |
| 0:00 | 239:03:00 | 0:00 | 596:05:00 | 0:00 | 729:07:00 |
| 0:00 | 238:03:00 | 0:00 | 589:05:00 | 1:00 | 726:07:00 |
| 237:03:00 | 0:00 | 583:05:00 | 2:00 | 718:07:00 | 4:00 |
| 123:03:00 | 119:00:00 | 297:05:00 | 306:00:00 | 230:08:00 | 539:00:00 |
| 0:00 | 242:03:00 | 0:00 | 603:05:00 | 2:00 | 767:08:00 |
| 0:00 | 242:03:00 | 0:00 | 603:05:00 | 0:00 | 769:08:00 |
| 0:00 | 240:03:00 | 0:00 | 602:05:00 | 2:00 | 764:08:00 |
| 0:00 | 238:03:00 | 0:00 | 598:05:00 | 0:00 | 758:08:00 |
| 236:03:00 | 0:00 | 594:05:00 | 1:00 | 750:08:00 | 2:00 |
| 236:03:00 | 1:00 | 582:05:00 | 5:00 | 730:08:00 | 8:00 |
| 121:03:00 | 112:00:00 | 290:05:00 | 292:00:00 | 221:09:00 | 507:00:00 |
| 0:01 | 229:03:00 | 0:00 | 561:05:00 | 6:00 | 683:09:00 |
| 228:05:00 | 1:00 | 549:05:00 | 2:00 | 680:09:00 | 0:00 |
| 228:05:00 | 0:00 | 545:05:00 | 2:00 | 675:09:00 | 1:00 |
| 226:05:00 | 1:00 | 543:05:00 | 3:00 | 671:09:00 | 0:00 |
| 224:05:00 | 0:00 | 537:05:00 | 3:00 | 662:09:00 | 1:00 |
| 222:05:00 | 1:00 | 534:05:00 | 4:00 | 660:09:00 | 3:00 |
| 222:05:00 | 1:00 | 534:05:00 | 4:00 | 662:09:00 | 1:00 |
| 218:03:00 | 0:01 | 536:05:00 | 2:00 | 661:09:00 | 1:00 |
| 1:01 | 217:03:00 | 5:00 | 531:05:00 | 4:00 | 654:09:00 |
| 0:00 | 217:04:00 | 0:00 | 524:05:00 | 0:00 | 653:09:00 |
| 0:00 | 216:04:00 | 1:00 | 516:05:00 | 0:00 | 642:09:00 |
| 212:04:00 | 0:00 | 510:05:00 | 5:00 | 631:09:00 | 1:00 |
| 205:04:00 | 0:00 | 496:05:00 | 3:00 | 619:09:00 | 0:00 |
| 204:03:00 | 0:01 | 494:05:00 | 3:00 | 615:09:00 | 2:00 |
| 103:03:00 | 102:01:00 | 231:05:00 | 264:00:00 | 170:09:00 | 444:00:00 |
| 204:04:00 | 1:00 | 492:05:00 | 3:00 | 614:09:00 | 1:00 |
| 102:03:00 | 102:01:00 | 230:05:00 | 263:00:00 | 172:09:00 | 442:00:00 |
| 204:03:00 | 0:01 | 490:05:00 | 3:00 | 612:09:00 | 2:00 |
| 205:04:00 | 0:00 | 492:05:00 | 4:00 | 609:09:00 | 0:00 |
| 205:03:00 | 0:01 | 488:05:00 | 5:00 | 605:09:00 | 4:00 |
| 0:00 | 205:04:00 | 2:00 | 487:05:00 | 0:00 | 604:09:00 |
| 101:01:00 | 104:03:00 | 255:00:00 | 232:05:00 | 431:00:00 | 170:09:00 |
| 0.309027778 | 0:00 | 0.544444444 | 0:01 | 0.75 | 0:00 |
| 0.309027778 | 0:00 | 0.544444444 | 0:01 | 0.749305556 | 0:01 |
| 0.309027778 | 0:00 | 0.544444444 | 0:01 | 0.749305556 | 0:01 |
| 205:04:00 | 0:00 | 484:05:00 | 0:00 | 597:07:00 | 3:01 |
| 102:00:00 | 108:01:00 | 254:00:00 | 242:03:00 | 429:00:00 | 177:03:00 |
| 215:01:00 | 0:00 | 509:03:00 | 0:00 | 611:03:00 | 2:00 |
| 112:01:00 | 103:00:00 | 249:03:00 | 260:00:00 | 180:03:00 | 433:00:00 |
| 112:01:00 | 103:00:00 | 248:02:00 | 261:00:00 | 179:03:00 | 432:00:00 |
| 215:01:00 | 0:00 | 509:02:00 | 0:00 | 610:03:00 | 2:00 |
| 216:00:00 | 0:00 | 517:01:00 | 0:00 | 613:02:00 | 1:00 |
| 217:00:00 | 0:00 | 516:01:00 | 0:00 | 613:02:00 | 1:00 |
| 218:00:00 | 0:00 | 519:01:00 | 0:00 | 618:01:00 | 2:00 |
| 0.151388889 | 0:00 | 0.402083333 | 0:00 | 0.471527778 | 0:01 |
| 218:00:00 | 1:00 | 521:01:00 | 0:00 | 619:01:00 | 5:00 |
| 221:00:00 | 0:00 | 523:01:00 | 2:00 | 616:01:00 | 4:00 |
| 114:00:00 | 106:00:00 | 254:01:00 | 268:00:00 | 192:01:00 | 428:00:00 |
| 219:00:00 | 0:00 | 520:01:00 | 1:00 | 616:01:00 | 5:01 |
| 218:00:00 | 0:00 | 521:01:00 | 2:00 | 617:02:00 | 3:00 |
| 0:00 | 217:00:00 | 0:00 | 522:01:00 | 1:00 | 615:02:00 |
| 217:00:00 | 0:00 | 521:01:00 | 1:00 | 613:02:00 | 3:00 |
| 113:00:00 | 104:00:00 | 251:01:00 | 271:00:00 | 185:02:00 | 426:00:00 |
| 216:00:00 | 1:00 | 519:01:00 | 1:00 | 605:02:00 | 6:00 |
| 113:00:00 | 104:00:00 | 251:01:00 | 271:00:00 | 187:02:00 | 425:00:00 |
| 112:00:00 | 104:00:00 | 251:01:00 | 270:00:00 | 187:02:00 | 423:00:00 |
| 0:00 | 216:00:00 | 1:00 | 520:01:00 | 2:00 | 608:02:00 |
| 113:00:00 | 103:00:00 | 253:01:00 | 261:00:00 | 186:02:00 | 421:00:00 |
| 0:00 | 217:00:00 | 0:00 | 516:01:00 | 0:00 | 607:02:00 |
| 217:00:00 | 0:00 | 518:01:00 | 0:00 | 605:02:00 | 2:00 |
| 1:00 | 218:00:00 | 0:00 | 522:01:00 | 1:00 | 615:02:00 |
| 1:00 | 218:00:00 | 1:00 | 521:01:00 | 2:00 | 615:02:00 |
| 56:2:2:0:2:0:0 | 0:0:0:0:0:0:0 | 74:0:3:2:1:0:0 | 0:0:0:0:0:0:0 | 237:7:6:8:11:3:0 | 1:0:0:0:0:0:0 |
| 978 | 983 | 2168 | 2503 | 3514 | 1835 |
| 1959 | 5 | 4645 | 16 | 5318 | 11 |
| 1958 | 1 | 4622 | 8 | 5279 | 5 |
| 1957 | 2 | 4623 | 7 | 5285 | 1 |
| 1937 | 2 | 4533 | 12 | 5190 | 6 |
| 1889 | 2 | 4385 | 12 | 4976 | 10 |
| 1848 | 2 | 4275 | 8 | 4777 | 18 |
| 1841 | 4 | 4269 | 12 | 4772 | 19 |
| 1843 | 5 | 4267 | 12 | 4768 | 15 |
| 933 | 921 | 2294 | 1993 | 1632 | 3158 |
| 1830 | 2 | 4221 | 13 | 4697 | 18 |
| 0 | 1808 | 1 | 4163 | 0 | 4618 |

| | | | | | |
|---|---|---|---|---|---|
| 0 | 1808 | 0 | 4163 | 0 | 4620 |
| 0 | 1808 | 1 | 4163 | 0 | 4620 |
| 1807 | 1 | 4170 | 0 | 4622 | 1 |
| 1809 | 0 | 4167 | 3 | 4617 | 4 |
| 904 | 900 | 2207 | 1944 | 1540 | 3049 |
| 1803 | 1 | 4144 | 6 | 4576 | 8 |
| 1801 | 1 | 4140 | 4 | 4575 | 9 |
| 1795 | 5 | 4131 | 8 | 4571 | 9 |
| 2 | 1797 | 6 | 4124 | 4 | 4560 |
| 4 | 1794 | 7 | 4120 | 3 | 4561 |
| 888 | 898 | 1922 | 2190 | 3017 | 1528 |
| 2 | 1784 | 7 | 4109 | 6 | 4541 |
| 0 | 1777 | 4 | 4084 | 1 | 4521 |
| 1774 | 0 | 4068 | 4 | 4500 | 6 |
| 0 | 1758 | 4 | 4012 | 2 | 4442 |
| 1751 | 0 | 3994 | 4 | 4416 | 5 |
| 873 | 878 | 1854 | 2143 | 2936 | 1484 |
| 879 | 873 | 2136 | 1861 | 1477 | 2940 |
| 871 | 883 | 1849 | 2135 | 2926 | 1477 |
| 881 | 873 | 2129 | 1856 | 1475 | 2925 |
| 1744 | 2 | 3967 | 6 | 4371 | 6 |
| 1745 | 2 | 3969 | 5 | 4370 | 7 |
| 1742 | 2 | 3955 | 7 | 4362 | 9 |
| 0 | 1744 | 1 | 3967 | 5 | 4357 |
| 1742 | 3 | 3960 | 7 | 4346 | 13 |
| 1740 | 0 | 3955 | 7 | 4345 | 10 |
| 876 | 864 | 2113 | 1846 | 1458 | 2895 |
| 876 | 864 | 2113 | 1846 | 1458 | 2895 |
| 876 | 864 | 2110 | 1849 | 1454 | 2897 |
| 1744 | 0 | 4012 | 3 | 4364 | 5 |
| 1743 | 2 | 4019 | 1 | 4367 | 5 |
| 2 | 1742 | 11 | 4006 | 9 | 4359 |
| 3 | 1741 | 14 | 4002 | 16 | 4353 |
| 3 | 1741 | 9 | 4001 | 8 | 4354 |
| 3 | 1742 | 7 | 4004 | 15 | 4352 |
| 3 | 1746 | 7 | 4007 | 12 | 4357 |
| 1 | 1749 | 2 | 4028 | 6 | 4381 |
| 1757 | 3 | 4036 | 7 | 4393 | 11 |
| 3 | 1757 | 7 | 4043 | 6 | 4395 |
| 1 | 1758 | 2 | 4042 | 5 | 4402 |
| 0 | 1758 | 3 | 4041 | 4 | 4399 |
| 0 | 1757 | 2 | 4048 | 2 | 4408 |
| 1 | 1760 | 1 | 4053 | 2 | 4415 |
| 895 | 873 | 2195 | 1885 | 1512 | 2915 |
| 895 | 873 | 2197 | 1882 | 1518 | 2910 |
| 886 | 863 | 2157 | 1858 | 1482 | 2867 |
| 884 | 862 | 2151 | 1861 | 1474 | 2870 |
| 880 | 860 | 2141 | 1850 | 1471 | 2851 |
| 878 | 855 | 2136 | 1850 | 1469 | 2840 |
| 1727 | 5 | 3991 | 6 | 4301 | 5 |
| 1729 | 2 | 3992 | 5 | 4294 | 10 |
| 1729 | 2 | 3988 | 6 | 4288 | 9 |
| 879 | 851 | 2141 | 1853 | 1469 | 2827 |
| 880 | 855 | 2144 | 1856 | 1471 | 2834 |
| 1729 | 5 | 3995 | 8 | 4292 | 10 |
| 1725 | 1 | 3987 | 6 | 4294 | 7 |
| 877 | 850 | 2137 | 1859 | 1471 | 2842 |
| 875 | 848 | 2132 | 1859 | 1465 | 2840 |
| 876 | 848 | 2133 | 1858 | 1467 | 2843 |
| 876 | 850 | 2131 | 1853 | 1466 | 2826 |
| 876 | 850 | 2133 | 1854 | 1465 | 2832 |
| 877 | 848 | 2136 | 1862 | 1470 | 2826 |
| 0 | 1 | 0 | 1 | 0 | 0 |
| 0 | 1 | 0 | 4 | 0 | 1 |
| 0 | 1 | 2 | 2 | 1 | 0 |
| 1:0:1:0 | 1:0:0:0 | 0:0:0:0 | 0:0:0:0 | 2:2:0:1 | 1:1:0:0 |
| 15 | 11 | 10 | 12 | 11 | 24 |
| 29 | 0 | 44 | 0 | 47 | 0 |
| 29 | 0 | 52 | 0 | 57 | 0 |
| 27 | 2 | 53 | 0 | 63 | 0 |
| 36 | 0 | 87 | 1 | 113 | 1 |
| 37 | 0 | 86 | 0 | 114 | 1 |
| 0 | 11 | 0 | 19 | 0 | 12 |
| 0:0:0:0 | 0:0:0:0 | 0:0:0:0 | 0:0:0:0 | 0:0:0:0 | 0:0:0:0 |
| 0:0:0:0 | 0:0:0:0 | 0:0:0:0 | 0:0:0:0 | 0:0:0:0 | 0:0:0:0 |
| 0:0:0:0 | 0:0:0:0 | 0:0:0:0 | 0:0:0:0 | 0:0:0:0 | 0:0:0:0 |
| 0:0:0:0 | 0:0:0:0 | 0:0:0:0 | 0:0:0:0 | 0:0:0:0 | 0:0:0:0 |
| 0:00:00 | 0:00:00 | 0:00:00 | 0:00:00 | 0:00:00 | 0:00:00 |
| 0:00:00 | 0:00:00 | 0:00:00 | 0:00:00 | 0:00:00 | 0:00:00 |
| 0:00:00 | 0:00:00 | 0:00:00 | 0:00:00 | 0:00:00 | 0:00:00 |

| | | | | | |
|---|---|---|---|---|---|
| 0:00:00 | 0:00:00 | 0:00:00 | 0:00:00 | 0:00:00 | 0:00:00 |
| 0:00:00 | 0:00:00 | 0:00:00 | 0:00:00 | 0:00:00 | 0:00:00 |
| 0:00:00 | 0:00:00 | 0:00:00 | 0:00:00 | 0:00:00 | 0:00:00 |
| 0:00:00 | 0:00:00 | 0:00:00 | 0:00:00 | 0:00:00 | 0:00:00 |
| 0:00:00 | 0:00:00 | 0:00:00 | 0:00:00 | 0:00:00 | 0:00:00 |
| 0:00:00 | 0:00:00 | 0:00:00 | 0:00:00 | 0:00:00 | 0:00:00 |
| 0:00:00 | 0:00:00 | 0:00:00 | 0:00:00 | 0:00:00 | 0:00:00 |
| 0:00:00 | 0:00:00 | 0:00:00 | 0:00:00 | 0:00:00 | 0:00:00 |
| 0:00:00 | 0:00:00 | 0:00:00 | 0:00:00 | 0:00:00 | 0:00:00 |
| 0:00:00 | 0:00:00 | 0:00:00 | 0:00:00 | 0:00:00 | 0:00:00 |
| 0:00:00 | 0:00:00 | 0:00:00 | 0:00:00 | 0:00:00 | 0:00:00 |
| 0:00:00 | 0:00:00 | 0:00:00 | 0:00:00 | 0:00:00 | 0:00:00 |
| 0:00:00 | 0:00:00 | 0:00:00 | 0:00:00 | 0:00:00 | 0:00:00 |
| 0:00:00 | 0:00:00 | 0:00:00 | 0:00:00 | 0:00:00 | 0:00:00 |
| 0:00:00 | 0:00:00 | 0:00:00 | 0:00:00 | 0:00:00 | 0:00:00 |
| 0:00:00 | 0:00:00 | 0:00:00 | 0:00:00 | 0:00:00 | 0:00:00 |
| 0:00:00 | 0:00:00 | 0:00:00 | 0:00:00 | 0:00:00 | 0:00:00 |
| 0:00:00 | 0:00:00 | 0:00:00 | 0:00:00 | 0:00:00 | 0:00:00 |
| 0:00 | 0:00 | 0:00 | 0:00 | 0:00 | 0:00 |
| 0:00 | 0:00 | 0:00 | 0:00 | 0:00 | 0:00 |
| 0:00 | 0:00 | 0:00 | 0:00 | 0:00 | 0:00 |
| 0:00 | 0:00 | 0:00 | 0:00 | 0:00 | 0:00 |
| 0:00 | 0:00 | 0:00 | 2:00 | 0:00 | 0:00 |
| 0:00 | 0:00 | 0:00 | 2:00 | 0:00 | 0:00 |
| 0:00 | 0:00 | 0:00 | 2:00 | 0:00 | 0:00 |
| 0:00 | 0:00 | 0:00 | 2:00 | 0:00 | 0:00 |
| 0:00 | 0:00 | 0:00 | 2:00 | 0:00 | 0:00 |
| 0:00 | 0:00 | 0:00 | 2:00 | 0:00 | 0:00 |
| 0:00 | 0:00 | 0:00 | 2:00 | 0:00 | 0:00 |
| 0:00 | 0:00 | 0:00 | 2:00 | 0:00 | 0:00 |
| 0:00 | 0:00 | 0:00 | 2:00 | 0:00 | 0:00 |
| 0:00 | 0:00 | 0:00 | 2:00 | 0:00 | 0:00 |
| 0:00 | 0:00 | 0:00 | 2:00 | 0:00 | 0:00 |
| 0:00 | 0:00 | 0:00 | 2:00 | 0:00 | 0:00 |
| 0:00 | 0:00 | 0:00 | 2:00 | 0:00 | 0:00 |
| 0:00 | 0:00 | 0:00 | 2:00 | 0:00 | 0:00 |
| 0:00 | 0:00 | 0:00 | 2:00 | 0:00 | 0:00 |
| 0:00 | 0:00 | 0:00 | 2:00 | 0:00 | 0:00 |
| 0:00 | 0:00 | 0:00 | 2:00 | 0:00 | 0:00 |
| 0 | 0 | 0 | 0 | 0 | 0 |
| 0 | 0 | 0 | 0 | 0 | 0 |
| 0 | 0 | 0 | 2 | 0 | 0 |
| 0 | 0 | 0 | 0 | 0 | 0 |
| 0 | 0 | 0 | 0 | 0 | 0 |
| 0:00 | 0:00 | 0:03 | 0:00 | 0:00 | 0:00 |
| 0:00 | 0:00 | 0:00 | 3:00 | 0:00 | 0:00 |
| 0:00 | 0:00 | 0:00 | 3:00 | 0:00 | 0:00 |
| 0:00 | 0:00 | 0:00 | 3:00 | 0:00 | 0:00 |
| 0:00 | 0:00 | 0:00 | 3:00 | 0:00 | 0:00 |
| 0:00 | 0:00 | 0:03 | 0:00 | 0:00 | 0:00 |
| 0:00 | 0:00 | 0:03 | 0:00 | 0:00 | 0:00 |
| 0:00 | 0:00 | 0:00 | 3:00 | 0:00 | 0:00 |
| 0:00 | 0:00 | 0:00 | 3:00 | 0:00 | 0:00 |
| 0:00 | 0:00 | 0:00 | 3:00 | 0:00 | 0:00 |
| 0:00 | 0:00 | 0:03 | 0:00 | 0:00 | 0:00 |
| 0:00 | 0:00 | 0:00 | 3:00 | 0:00 | 0:00 |
| 0:00 | 0:00 | 0:00 | 3:00 | 0:00 | 0:00 |
| 0:00 | 0:00 | 0:00 | 3:00 | 0:00 | 0:00 |
| 0:00 | 0:00 | 0:00 | 3:00 | 0:00 | 0:00 |
| 0:00 | 0:00 | 0:01 | 0:02 | 0:00 | 0:00 |
| 0:00 | 0:00 | 0:01 | 0:02 | 0:00 | 0:00 |
| 0:00 | 0:00 | 0:01 | 0:02 | 0:00 | 0:00 |
| 0:00 | 0:00 | 0:03 | 0:00 | 0:00 | 0:00 |
| 0:00 | 0:00 | 0:03 | 0:00 | 0:00 | 0:00 |
| 0:00 | 0:00 | 0:00 | 3:00 | 0:00 | 0:00 |
| 0:00 | 0:00 | 0:00 | 0:03 | 0:00 | 0:00 |
| 0:00 | 0:00 | 0:00 | 0:03 | 0:00 | 0:00 |
| 0:00 | 0:00 | 0:00 | 3:00 | 0:00 | 0:00 |
| 0:00 | 0:00 | 0:01 | 0:02 | 0:00 | 0:00 |
| 0:00 | 0:00 | 0:01 | 0:02 | 0:00 | 0:00 |
| 0:00 | 0:00 | 0:01 | 0:02 | 0:00 | 0:00 |
| 0:00 | 0:00 | 0:03 | 0:00 | 0:00 | 0:00 |
| 0:00 | 0:00 | 0:03 | 0:00 | 0:00 | 0:00 |
| 0:00 | 0:00 | 0:03 | 0:00 | 0:00 | 0:00 |
| 0:00 | 0:00 | 0:00 | 0:03 | 0:00 | 0:00 |
| 0:00 | 0:00 | 0:00 | 3:00 | 0:00 | 0:00 |
| 0:00 | 0:00 | 0:00 | 0:03 | 0:00 | 0:00 |
| 0:00 | 0:00 | 0:00 | 3:00 | 0:00 | 0:00 |

| | | | | | |
|---|---|---|---|---|---|
| 0:00 | 0:00 | 0:03 | 0:00 | 0:00 | 0:00 |
| 0:00 | 0:00 | 1:00 | 2:00 | 0:00 | 0:00 |
| 0:00 | 0:00 | 0:00 | 3:00 | 0:00 | 0:00 |
| 0:00 | 0:00 | 0:03 | 0:00 | 0:00 | 0:00 |
| 0 | 0 | 1 | 0 | 2 | 0 |
| 6 | 0 | 1 | 0 | 1 | 0 |
| 8:0:13:1:39:4 | 0:0:0:0:0:0 | 20:0:3:0:127:5 | 0:0:0:0:2:0 | 56:2:29:1:349:20 | 0:0:0:0:5:0 |
| 8:0:13:1:43:4:0 | 0:0:0:0:0:0:0 | 20:0:6:0:141:4:0 | 0:0:0:0:1:1:0 | 55:2:34:1:390:16:0 | 0:0:0:0:2:0:0 |
| 8:0:13:1:43:4:0 | 0:0:0:0:0:0:0 | 20:0:7:0:141:5:0 | 0:0:0:0:1:0:0 | 55:2:36:1:394:16:0 | 0:0:0:0:2:0:0 |
| 8:0:13:1:43:1 | 0:0:0:0:0:0 | 17:0:10:0:142:8 | 1:0:0:0:0:0 | 54:2:42:0:399:20 | 0:0:0:0:2:0 |
| 4 | 2 | 0 | 2 | 0 | 0 |
| 4 | 2 | 0 | 2 | 0 | 0 |
| 4 | 2 | 0 | 2 | 0 | 0 |
| 4 | 2 | 0 | 2 | 0 | 0 |
| 4 | 2 | 0 | 1 | 0 | 0 |
| 4 | 2 | 0 | 1 | 0 | 0 |
| 4 | 2 | 0 | 1 | 0 | 0 |
| 4 | 2 | 0 | 1 | 0 | 0 |
| 1 | 2 | 1 | 2 | 0 | 0 |
| 1 | 2 | 0 | 3 | 0 | 0 |
| 1 | 2 | 0 | 3 | 0 | 0 |
| 1 | 2 | 1 | 2 | 0 | 0 |
| 1 | 2 | 1 | 2 | 0 | 0 |
| 3 | 0 | 3 | 0 | 0 | 0 |
| 1 | 2 | 1 | 2 | 0 | 0 |
| 1 | 2 | 0 | 3 | 0 | 0 |
| 1 | 2 | 0 | 3 | 0 | 0 |
| 1 | 2 | 0 | 3 | 0 | 0 |
| 1 | 2 | 0 | 3 | 0 | 0 |
| 3 | 0 | 3 | 0 | 0 | 0 |
| 1 | 2 | 0 | 3 | 0 | 0 |
| 1 | 2 | 0 | 3 | 0 | 0 |
| 1 | 2 | 0 | 3 | 0 | 0 |
| 1 | 2 | 0 | 3 | 0 | 0 |
| 1 | 2 | 0 | 3 | 0 | 0 |
| 1 | 2 | 0 | 3 | 0 | 0 |
| 1 | 2 | 0 | 2 | 0 | 0 |
| 1 | 2 | 0 | 3 | 0 | 0 |
| 3 | 0 | 3 | 0 | 0 | 0 |
| 0:00 | 0:00 | 0:00 | 0:00 | 0:00 | 0:00 |
| 1:00 | 0:00 | 0:00 | 0:00 | 3:01 | 0:00 |
| 0 | 0 | 1 | 0 | 1 | 0 |
| 5:2:0:0:0:0:0 | 4:1:1:0:0:0:0 | 14:4:0:0:1:2:0 | 9:1:1:0:0:1:0 | 10:1:0:0:2:2:1 | 15:2:1:0:0:0:1 |
| 0 | 1 | 0 | 0 | 0 | 0 |
| 1:0:0:0 | 0:0:0:0 | 1:0:0:0 | 0:0:0:0 | 2:1:0:1 | 0:0:0:0 |
| 0 | 0 | 0 | 0 | 0 | 0 |
| 24:1:2:0 | 0:0:0:0 | 7:0:1:0 | 0:0:0:0 | 129:11:0:9 | 0:0:0:0 |
| 4:14:0:1:4:1:1:2:0:1 | 0:0:0:0:0:0:0:0:0 | 1:6:0:0:0:0:0:3:0:0 | 0:0:0:0:0:0:0:0:0:0 | 23:34:2:0:33:1:16:9:0:0 | 0:0:0:0:0:0:0:0:0:0 |
| 4:13:1:4:1:0:2:0:0:1 | 0:0:0:0:0:0:0:0:0 | 5:5:0:0:0:0:4:0:0:0 | 0:0:0:0:0:0:0:0:0:0 | 42:33:0:31:13:2:20:1:0:0 | 0:0:0:1:0:0:1:0:0:0 |
| 5:10:1:4:1:3:0:0:0:1 | 0:0:0:0:0:0:0:0:0 | 7:4:0:0:0:3:0:0:0:0 | 0:0:0:0:0:0:0:0:0:0 | 55:31:0:30:12:29:2:1:0:0 | 0:1:0:0:0:0:0:0:0:0 |
| 5 | 6 | 0 | 2 | 10 | 1 |
| 0 | 2 | 0 | 7 | 0 | 8 |
| 0:00:00 | 0:00:00 | 1:00:00 | 0:01:00 | 0:00:00 | 0:00:00 |
| 0:0:0:0 | 0:0:0:0 | 0:0:1:0 | 0:0:1:0 | 0:0:0:0 | 0:0:0:0 |
| 0 | 0 | 1 | 0 | 0 | 2 |
| 0:00 | 0:00 | 1:00 | 0:00 | 0:00 | 0:00 |
| 82:07:02 | 0:00:00 | 122:11:07 | 0:00:00 | 373:26:21 | 0:00:00 |
| 82:07:02 | 0:00:00 | 122:11:10 | 0:00:00 | 370:27:21 | 3:00:00 |
| 82:07:03 | 0:00:00 | 123:12:15 | 0:00:00 | 371:27:30 | 0:00:01 |
| 82:07:06 | 0:00:00 | 125:11:30 | 0:00:00 | 376:26:72 | 0:00:01 |
| 82:06:09 | 0:00:00 | 121:11:38 | 2:00:00 | 366:26:120 | 0:00:01 |
| 82:06:09 | 0:00:00 | 123:11:41 | 0:00:00 | 366:26:121 | 0:00:01 |
| 82:06:11 | 0:00:00 | 123:11:52 | 0:00:00 | 366:26:136 | 0:00:00 |
| 0:0:1:0:0:0:0:0:1:0 | 0:0:0:0:0:0:0:0:0 | 0:0:0:0:0:0:0:1:0:0 | 0:0:0:0:0:0:0:0:0:0 | 0:2:0:0:0:1:0:1:0:1 | 0:0:0:0:0:0:0:0:0:0 |

| | | | | | |
|---|---|---|---|---|---|
| 4:00 | 1:00 | 2:00 | 0:00 | 5:01 | 1:01 |
| 0:0:8:0:0:1 | 0:0:0:0:0:0 | 0:0:9:1:0:0 | 0:0:0:0:0:0 | 0:0:18:11:0:0 | 0:0:0:0:0:0 |
| 1 | 1 | 1 | 2 | 7 | 2 |
| 0:00:00 | 0:00:00 | 0:01:00 | 0:01:00 | 1:01:01 | 0:00:00 |
| 0:00:00 | 0:00:00 | 1:00:00 | 0:00:00 | 4:01:01 | 0:00:00 |
| 0:00:00 | 0:00:00 | 1:00:00 | 0:00:00 | 5:00:01 | 0:00:00 |
| 0 | 0 | 0 | 0 | 0 | 2 |
| 348:01:00 | 1:00:00 | 265:08:00 | 1:00:00 | 961:16:01 | 3:00:00 |
| 347:00:00 | 2:00:00 | 266:06:00 | 0:00:00 | 959:14:01 | 1:00:00 |
| 349:00:00 | 0:00:00 | 265:06:00 | 1:00:00 | 959:14:01 | 2:00:00 |
| 0:00:00 | 0:00:00 | 0:00:00 | 1:01:00 | 0:00:00 | 1:00:01 |
| 13:0:32:18:1:1:1:0:0:0:0:3 | 0:0:0:0:0:0:0:0:0:0:0 | 32:0:11:90:9:2:1:1:0:1:2:0 | 0:0:0:0:0:0:0:0:0:0:0 | 54:0:84:140:24:3:11:0:3:2:2:2 | 0:0:0:0:0:0:0:0:0:0:0 |
| 13:0:32:18:1:1:1:0:1:0:0:3 | 0:0:0:0:0:0:0:0:0:0:0 | 32:0:11:90:9:2:1:1:1:1:2:0 | 0:0:0:0:0:0:0:0:0:0:0 | 54:0:84:140:24:4:11:0:6:2:2:2 | 0:0:0:0:0:0:0:0:0:0:0 |
| 13:0:32:18:1:4:1:0:3:0:3:0 | 0:0:0:0:0:0:0:0:0:0:0 | 31:0:11:87:10:5:1:1:4:2:0:0 | 0:0:0:1:0:0:0:0:0:0:0 | 53:0:86:139:24:6:12:0:10:2:2:2 | 0:0:1:0:1:0:0:0:0:0:0 |
| 0:0:0:0:0:0:0:0:0:0:0 | 29:14:6:4:0:0:0:1:0:1 | 0:0:0:0:0:0:0:0:0:0 | 9:13:29:10:0:0:0:0:0:0 | 1:0:0:0:0:0:0:0:0:0 | 86:42:66:24:0:1:0:0:0:1 |
| 0:29:13:5:3:0:1:1:0:0 | 0:0:0:0:0:0:0:0:0:0 | 0:9:19:43:7:0:0:0:0:0 | 0:0:0:0:0:0:0:0:0:0 | 0:86:54:97:22:0:3:0:1:0 | 0:0:1:1:1:0:0:0:0:0 |
| 0:29:13:5:3:0:1:1:0:0 | 0:0:0:0:0:0:0:0:0:0 | 0:9:19:43:7:0:0:0:0:0 | 0:0:0:0:0:0:0:0:0:0 | 0:85:56:99:22:0:3:0:1:0 | 0:1:0:0:0:0:0:0:0:0 |
| 7:12:0:25:3:0:0:0:1:0 | 1:0:0:0:0:0:0:0:0:0 | 65:28:0:5:5:0:0:1:0:0 | 0:1:0:0:0:0:0:0:0:0 | 142:80:0:78:16:0:0:0:0:0 | 1:2:0:0:0:0:0:0:0:0 |
| 0:25:12:8:3:0:0:0:1:0 | 0:0:0:0:0:0:0:0:0:0 | 0:5:28:65:5:0:0:1:0:0 | 0:0:0:0:0:0:0:0:0:0 | 0:78:79:142:16:0:0:0:0:0 | 0:0:3:1:0:0:0:0:0:0 |
| 0 | 1 | 0 | 1 | 0 | 0 |
| 65:3:14:0:0:1 | 0:0:0:0:0:0 | 430:9:203:6:1:4 | 1:0:0:0:0:0 | 694:18:139:7:3:12 | 4:0:0:0:1:0 |
| 3:66:14:0:0:1 | 0:0:0:0:0:0 | 9:437:203:6:1:4 | 0:0:0:0:0:0 | 19:707:139:7:4:12 | 0:1:0:0:0:0 |
| 3:68:14:0:0:1 | 0:0:0:0:0:0 | 9:442:203:6:1:4 | 0:0:0:0:0:0 | 19:715:138:7:4:12 | 0:1:0:0:0:0 |
| 3:68:14:0:0:1 | 0:0:0:0:0:0 | 9:450:200:6:1:4 | 0:1:1:0:0:0 | 18:731:139:7:4:12 | 0:1:0:0:0:0 |
| 3:68:13:0:0:1 | 0:0:1:0:0:0 | 9:454:200:6:1:4 | 0:1:1:0:0:0 | 18:734:139:7:4:12 | 0:1:0:0:0:0 |
| 3:68:14:0:0:1 | 0:1:0:0:0:0 | 13:466:201:6:1:4 | 0:0:0:0:0:0 | 21:749:139:7:4:12 | 0:0:0:0:0:0 |
| 3:68:14:0:0:1 | 0:1:0:0:0:0 | 13:467:201:6:1:4 | 0:1:0:0:0:0 | 22:756:138:7:4:12 | 0:0:0:0:0:0 |
| 2:51:12:0:0:1 | 0:0:0:0:0:0 | 5:300:119:4:1:3 | 0:0:0:0:0:0 | 14:533:90:3:2:7 | 1:1:0:0:0:0 |
| 12:2:49:0:0:1 | 0:0:1:0:0:0 | 117:5:298:4:1:3 | 2:0:1:0:0:0 | 88:15:529:3:2:7 | 0:0:4:0:0:0 |
| 2:47:10:0:0:1 | 0:0:0:0:0:0 | 5:286:113:4:1:3 | 0:2:0:0:0:0 | 14:511:84:2:2:7 | 0:0:0:0:0:0 |
| 1:47:10:0:0:1 | 1:0:0:0:0:0 | 6:272:102:4:1:3 | 0:0:0:0:0:0 | 13:438:73:2:2:6 | 0:3:0:0:0:0 |
| 2:46:10:0:0:1 | 0:0:0:0:0:0 | 6:267:101:4:1:3 | 0:1:0:0:0:0 | 13:438:73:2:2:6 | 0:2:0:0:0:0 |
| 46:2:10:0:0:1 | 0:0:0:0:0:0 | 266:6:100:4:1:3 | 1:0:0:0:0:0 | 424:13:71:2:2:6 | 6:0:1:0:0:0 |
| 44:9:0:2:0:1 | 2:1:0:0:0:0 | 201:78:4:5:1:2 | 32:7:0:0:0:1 | 332:53:2:11:2:5 | 56:9:0:1:0:1 |
| 7:33:0:2:0:1 | 3:13:0:0:0:0 | 93:243:4:5:1:3 | 8:21:0:1:0:0 | 58:364:1:11:2:5 | 14:68:1:2:0:1 |
| 2:22:3:1:0:0 | 0:0:0:0:0:0 | 1:59:18:1:1:1 | 0:3:2:0:0:0 | 5:119:12:1:0:0 | 0:4:1:0:0:0 |
| 0:0:0:0:0:1:0 | 0:0:0:0:0:0:0 | 2:0:0:0:0:0:0 | 1:0:0:0:0:0:0 | 2:0:0:0:0:0:1 | 4:0:0:0:0:0:0 |
| 0:0:0:0:0:1:0 | 0:0:0:0:0:0:0 | 2:0:0:0:0:0:0 | 1:0:0:0:0:0:0 | 3:0:0:0:0:0:1 | 3:0:0:0:0:0:0 |
| 4:1:4:25:4:4:0:0:0:0:0:0 | 0:0:0:0:0:0:0:0:0:0:0 | 4:5:7:56:6:4:0:0:1:0:0:0 | 0:0:0:0:0:0:0:0:0:0:0 | 12:15:7:158:21:14:1:0:4:0:0:1 | 0:1:0:0:1:0:0:0:0:0:0 |
| 48:49:11:8 | 0:0:0:0 | 29:47:4:6 | 0:0:0:0 | 231:212:31:31 | 3:0:0:0 |
| 52:11:47:8 | 0:0:0:0 | 51:4:29:6 | 1:0:0:0 | 216:31:234:31 | 3:0:0:0 |
| 52:11:47:8 | 0:0:0:0 | 54:4:29:6 | 0:0:0:0 | 220:31:234:31 | 1:0:0:0 |
| 2:19:5:0:2:1:10 | 0:0:0:0:0:0:0 | 0:6:9:2:0:0:0 | 0:0:0:0:0:0:0 | 7:35:10:0:0:0:0 | 0:0:0:0:0:0:0 |
| 2:0:2:4:2 | 0:0:0:0:0 | 0:2:0:6:0 | 0:0:0:3:0 | 4:0:3:7:0 | 0:0:0:0:0 |
| 928 | 3 | 1386 | 0 | 2697 | 2 |
| 932 | 0 | 1392 | 2 | 2707 | 3 |
| 940 | 0 | 1494 | 2 | 2860 | 4 |
| 933 | 1 | 1480 | 2 | 2850 | 6 |
| 0:00 | 0:00 | 0:00 | 0:00 | 48:01:00 | 0:02 |
| 0:0:0:0 | 3:1:0:0 | 0:0:0:0 | 0:0:1:0 | 1:1:0:0 | 5:2:0:1 |
| 143:08:00 | 2:01 | 11:09 | 0:00 | 157:36:00 | 0:00 |
| 159:09:00 | 0:01 | 10:10 | 4:00 | 171:37:00 | 6:00 |
| 216:11:00 | 0:00 | 53:11:00 | 0:00 | 299:53:00 | 0:00 |
| 0.572222222 | 0:00 | 0.511805556 | 0:01 | 46:334 | 1:00 |
| 234:09:00 | 1:00 | 107:08:00 | 0:02 | 404:39:00 | 2:02 |
| 0.538888889 | 0:00 | 0.450694444 | 2:01 | 39:412 | 3:02 |
| 242:08:00 | 0:00 | 131:08:00 | 0:01 | 472:39:00 | 0:00 |
| 243:09:00 | 1:00 | 136:08:00 | 1:01 | 490:39:00 | 0:00 |
| 266:00:00 | 2:00 | 287:00:00 | 2:00 | 728:01:00 | 5:00 |
| 268:00:00 | 1:00 | 314:00:00 | 3:00 | 787:01:00 | 10:02 |
| 0.185416667 | 0:02 | 0.221527778 | 0:02 | 0.638194444 | 2:09 |
| 268:00:00 | 1:00 | 320:00:00 | 2:00 | 807:02:00 | 7:02 |

| | | | | | |
|---|---|---|---|---|---|
| 273:00:00 | 0:00 | 340:00:00 | 1:00 | 898:04:00 | 2:00 |
| 272:00:00 | 1:00 | 343:00:00 | 0:00 | 920:04:00 | 0:00 |
| 272:01:00 | 0:00 | 364:00:00 | 1:00 | 973:04:00 | 2:00 |
| 271:01:00 | 0:00 | 368:00:00 | 0:00 | 977:04:00 | 3:00 |
| 272:01:00 | 0:00 | 388:00:00 | 0:01 | 999:06:00 | 1:00 |
| 272:01:00 | 0:00 | 387:00:00 | 1:01 | 999:07:00 | 1:00 |
| 0 | 4 | 4 | 3 | 3 | 10 |
| 1:00:00 | 13:01:00 | 3:01:03 | 9:03:03 | 3:00:00 | 25:01:15 |
| 13:0:0:1:0:0 | 0:0:0:0:0:0 | 33:3:0:6:0:0 | 5:0:0:0:0:0 | 43:3:0:3:0:0 | 0:0:0:0:0:1 |
| 8:0:0:1:0:0 | 5:0:0:0:0:0 | 26:2:0:6:0:0 | 12:1:0:0:0:0 | 33:3:0:3:0:0 | 10:0:0:0:0:1 |
| 0:5:0:1:0 | 0:0:0:0:0 | 1:13:0:6:0 | 1:2:0:0:0 | 1:22:0:3:0 | 0:0:0:0:0 |
| 0:00 | 1:02 | 0:00 | 2:01 | 0:00 | 2:04 |
| 0:00 | 0:00 | 0:00 | 2:00 | 0:00 | 0:00 |
| 1 | 0 | 2 | 0 | 0 | 0 |
| 0 | 0 | 1 | 3 | 0 | 5 |
| 1:0:0:0:1:0:0:0:0 | 0:0:0:0:1:0:0:0:0 | 1:0:0:0:0:1:0:0:0 | 0:0:0:1:0:0:0:0:0 | 5:1:1:0:2:0:0:4:0 | 5:1:4:1:6:1:1:2:1 |
| 1:0:0:0:1:0:0:0:0 | 0:0:0:0:1:0:0:0:0 | 1:0:0:0:0:1:0:0:0 | 0:0:0:1:0:0:0:0:0 | 6:1:0:0:2:0:0:4:0 | 5:4:1:1:6:1:1:2:1 |
| 0:00 | 0:00 | 4:03 | 0:00 | 9:02 | 3:00 |
| 0:00 | 0:00 | 3:04 | 0:00 | 2:11 | 0:01 |
| 0:0:1:1:0 | 9:2:13:1:0 | 5:9:21:3:0 | 10:3:21:2:0 | 3:1:7:0:0 | 37:32:85:12:1 |
| 0:0:3:0:0:0:0 | 1:2:13:2:0:0:0 | 5:2:20:0:0:0:1 | 0:0:0:0:0:0:0 | 0:1:3:0:0:0:0 | 3:5:36:1:0:2:0 |
| 106:03:00 | 0:00 | 800:57:00 | 1:01 | 708:87 | 1:00 |
| 105:03:00 | 0:00 | 798:58:00 | 1:00 | 706:87 | 0:00 |
| 0 | 7 | 0 | 0 | 0 | 2 |
| 0 | 0 | 0 | 0 | 2 | 0 |
| 2 | 0 | 2 | 0 | 16 | 0 |
| 2 | 0 | 2 | 0 | 14 | 2 |
| 12:1:0:0:0:1:0:0:1:1:0:0:2:0:0:0:0:0 | 0:0:0:0:0:0:0:0:0:0:0:0:0:0:0:0:0:0 | 13:1:0:0:0:0:1:1:0:2:0:0:4:0:0:0:0:0 | 0:0:0:0:0:0:0:0:0:0:0:0:0:0:0:0:0:0 | 24:1:9:0:1:9:0:0:2:2:0:0:0:2:2:1:1:1 | 0:0:0:0:0:0:0:0:0:0:0:0:0:0:0:0:0:0 |
| 0:00 | 0:00 | 0:00 | 0:00 | 0:00 | 0:02 |
| 0:00 | 0:00 | 0:00 | 0:00 | 0:00 | 0:02 |
| 0:00 | 0:00 | 0:00 | 0:00 | 0:00 | 0:02 |
| 673:9:56:16:1:24:4 | 0:0:0:0:0:1:0 | 2480:69:126:59:10:143:2 | 6:0:0:0:0:1:0 | 3059:43:295:125:15:172:15 | 4:0:1:0:0:0:0 |
| 675:9:56:16:1:25:4 | 0:0:0:0:0:0:0 | 2493:66:127:59:10:140:2 | 1:0:0:0:0:0:0 | 3070:43:297:125:15:168:15 | 1:0:0:0:0:0:0 |
| 7:678:56:16:1:23:4 | 0:0:0:0:0:0:0 | 65:2512:127:61:10:132:2 | 0:2:0:0:0:0:0 | 42:3091:298:124:15:159:15 | 0:3:0:1:0:2:0 |
| 677:22:7:56:16:1:4 | 1:1:0:0:0:0:0 | 2514:132:65:127:61:10:2 | 1:0:0:0:0:0:0 | 3093:159:42:299:125:15:15 | 1:2:0:0:0:0:0 |
| 22:7:678:56:16:1:4 | 0:0:0:0:0:0:0 | 128:65:2514:127:61:10:2 | 1:0:3:0:0:0:0 | 154:40:3095:299:124:15:15 | 1:0:0:0:1:0:0 |
| 7:678:56:16:1:24:4 | 0:0:0:0:0:0:0 | 65:2515:127:61:10:119:2 | 0:5:0:0:0:0:0 | 41:3095:299:125:15:145:15 | 0:2:0:0:0:0:0 |
| 6:678:56:16:1:24:4 | 0:0:0:0:0:0:0 | 64:2521:128:61:10:112:2 | 0:1:0:0:0:0:0 | 41:3100:299:125:15:142:15 | 0:0:0:0:0:1:0 |
| 83:3:1:0:2:0:0 | 595:53:5:1:14:23:4 | 1196:53:31:8:40:50:1 | 1328:75:29:2:21:51:1 | 202:14:2:0:12:12:0 | 2901:285:39:15:113:122:15 |
| 6:678:56:16:1:22:4 | 0:0:0:0:0:0:0 | 59:2522:128:61:10:102:2 | 0:3:0:0:0:0:0 | 40:3100:299:124:15:127:15 | 0:4:0:0:0:0:0 |
| 6:678:57:15:1:17:4 | 0:0:0:0:0:0:0 | 56:2528:128:59:10:83:2 | 0:2:0:1:0:0:0 | 35:3110:300:118:15:105:15 | 0:1:0:0:0:0:0 |
| 676:5:57:1:14:15:3 | 2:0:0:0:1:0:1 | 2528:56:128:10:51:78:2 | 2:0:0:0:4:0:0 | 3108:36:300:15:108:103:13 | 3:0:0:0:4:0:1 |
| 1:83:3:0:0:0:0 | 4:597:54:1:12:1:0 | 29:1200:53:8:28:0:0 | 26:1338:75:2:26:1:0 | 2:204:14:0:11:1:0 | 30:2913:286:15:69:2:1 |
| 17:6:680:57:1:1:0 | 0:0:1:0:0:0:0 | 58:57:2546:128:10:1:0 | 0:0:2:0:0:0:0 | 83:33:3128:301:15:7:1 | 1:0:2:1:0:0:0 |
| 5:6:685:57:19:1:18 | 0:0:0:0:0:0:1 | 2:58:2561:129:73:10:64 | 0:1:0:0:0:0:4 | 15:38:3150:307:137:15:93 | 1:0:0:0:0:0:0 |
| 685:8:57:20:1:16:5 | 0:0:0:0:0:0:0 | 2560:64:129:81:10:70:3 | 1:0:0:0:0:1:0 | 3148:44:307:146:15:83:17 | 2:0:0:0:0:0:0 |
| 0:0:0:0:0 | 0:0:0:2:1 | 0:1:0:1:0 | 0:1:3:0:0 | 0:0:0:0:0 | 0:3:2:0:0 |
| 0:0:0:0 | 0:0:0:2 | 0:1:0:1 | 0:1:3:0 | 0:0:0:0 | 0:3:2:0 |
| 0:00 | 0:00 | 0:00 | 2:01 | 0:00 | 0:00 |
| 0:00:00 | 0:00:01 | 0:00:00 | 2:01:00 | 0:00:00 | 0:00:00 |
| 0 | 0 | 0 | 0 | 0 | 0 |
| 0 | 2 | 0 | 0 | 0 | 0 |
| 0:0:3:0 | 3:0:3:1 | 1:0:1:0 | 0:0:1:0 | 3:0:1:2 | 0:0:0:0 |

| | | | | | |
|---|---|---|---|---|---|
| 0:0:3:0 | 2:0:3:0 | 1:0:1:0 | 0:0:1:0 | 2:0:1:2 | 0:0:0:0 |
| 61:0:3:7:0 | 0:0:0:0:0 | 97:3:0:29:0 | 0:0:0:0:0 | 267:8:9:60:3 | 0:0:0:1:0 |
| 63:0:3:6:0 | 0:0:0:0:0 | 105:4:0:26:0 | 0:0:0:0:0 | 296:11:9:61:4 | 1:0:0:2:0 |
| 63:0:3:6:0 | 1:0:0:0:0 | 106:4:0:27:0 | 0:0:0:0:0 | 300:11:10:62:4 | 1:0:0:0:0 |
| 5:2:4:2:0:0 | 0:0:0:0:0:0 | 0:0:0:1:0:0 | 0:0:0:0:0:0 | 11:12:61:7:1:1 | 1:1:2:0:0:0 |
| 546:31:29:13:82:4:11:25:3:30:3:4:3:1:1:0:0:0:0:1:0 | 2:0:0:0:0:0:0:0:0:0:0:0:0:0:0:0:0:0:0:0 | 718:51:33:3:101:14:8:48:2:46:1:11:6:2:1:0:0:0:2:2:0 | 0:0:0:0:0:0:0:0:0:0:1:0:0:0:0:0:0:0:0:0 | 2609:180:84:35:854:33:92:154:18:154:12:22:54:1:14:0:0:0:3:1:2 | 6:0:0:0:1:0:0:0:0:0:0:0:0:0:0:0:0:0:0:0:0 |
| 31:29:12:82:4:11:21:3:30:3:546:4:5:1:1:0:0:0:0:1:0 | 0:0:0:0:0:0:0:0:0:0:0:0:0:0:0:0:0:0:0:0 | 50:32:3:101:14:7:46:2:48:1:719:17:8:2:1:0:0:0:2:2:0 | 1:1:0:0:0:0:0:0:0:0:1:0:0:0:0:0:0:0:0:0 | 179:84:36:855:33:92:141:18:154:11:2616:34:65:1:14:0:0:0:4:1:2 | 1:0:0:0:0:0:0:0:0:0:1:4:5:0:0:0:0:0:0:0:0 |
| 31:29:12:82:4:11:21:3:30:3:548:5:5:1:1:0:0:0:0:1:0 | 0:0:0:0:0:0:0:0:0:0:0:0:0:0:0:0:0:0:0:0 | 51:33:3:100:14:7:46:2:48:1:719:17:7:2:1:0:0:0:2:2:0 | 0:0:0:1:0:0:0:0:0:0:0:2:1:0:0:0:0:0:0:0 | 180:84:36:853:33:92:137:18:154:10:2615:34:70:1:14:0:0:0:4:1:2 | 0:0:0:2:0:0:1:0:0:0:2:5:2:0:0:0:0:0:0:0:0 |
| 545:31:28:12:4:10:7:3:30:3:82:6:5:1:0:0:0:0:1:0:0 | 1:0:1:0:0:0:0:0:0:0:0:0:0:0:0:0:0:0:0:0 | 717:51:32:3:14:7:21:2:48:1:101:28:15:2:0:0:0:1:2:1:0 | 0:0:1:0:0:0:0:0:0:0:0:1:0:0:0:0:0:0:0:0 | 2604:180:83:33:33:89:54:18:153:9:855:62:82:1:0:0:0:6:1:0:2 | 1:0:0:0:0:0:1:0:0:0:0:3:2:0:0:0:0:0:0:0:0 |
| 31:29:12:4:10:7:3:30:3:544:82:6:5:1:0:0:0:0:1:0:0 | 0:0:0:0:0:0:0:0:0:1:0:0:0:0:0:0:0:0:0:0 | 51:33:3:14:7:18:2:48:1:717:101:27:15:2:0:0:0:1:2:1:0 | 0:0:0:0:0:0:0:0:0:0:1:0:0:0:0:0:0:0:0:0 | 180:83:33:33:88:44:18:153:9:2598:855:63:85:1:0:0:0:5:1:0:2 | 0:0:0:0:0:0:0:0:0:4:0:5:2:0:0:0:0:1:0:0:0 |
| 31:28:12:4:10:6:3:30:3:544:82:6:5:1:0:0:0:0:1:0:0 | 0:1:0:0:0:0:0:0:0:1:0:0:0:0:0:0:0:0:0:0 | 51:33:3:14:7:16:2:48:1:717:101:27:14:2:0:0:0:1:2:1:0 | 0:0:0:0:0:0:0:0:0:0:2:2:0:0:0:0:0:0:0:0 | 180:83:33:33:88:42:18:151:9:2599:855:59:83:1:0:0:0:5:1:0:2 | 0:0:0:0:0:0:0:0:0:1:0:7:3:0:0:0:0:0:0:0:0 |
| 12:6:31:29:4:10:5:3:3:543:30:82:5:1:0:0:0:1:0:0 | 0:0:0:0:0:0:0:0:0:1:0:0:1:0:0:0:0:0:0 | 3:16:51:32:14:7:13:2:1:714:48:101:30:2:0:0:0:2:1:0 | 0:1:0:0:0:0:0:0:0:0:2:0:0:3:0:0:0:0:0:0:0 | 32:86:180:83:33:88:32:18:9:2588:152:850:64:1:0:0:0:1:0:1 | 0:2:0:0:0:0:0:0:0:5:1:5:6:0:0:0:0:0:0:0 |
| 31:29:12:4:10:2:3:3:542:30:82:7:8:1:0:0:0:1:0 | 0:0:0:0:1:0:0:0:2:0:0:1:0:0:0:0:0:0:0 | 51:32:3:14:7:11:2:1:713:48:101:36:17:2:0:0:0:2:0 | 0:0:0:0:0:0:0:0:1:0:0:0:1:0:0:0:0:0:0 | 180:83:32:33:88:23:18:7:2589:153:854:77:90:1:0:0:0:1:1 | 0:0:0:0:0:0:0:1:1:1:1:1:3:0:0:0:0:0:0 |
| 31:28:12:4:11:1:3:3:543:30:82:10:9:1:0:0:0:0:1:0 | 0:0:0:0:0:0:0:0:0:0:0:0:0:0:0:0:0:0:0 | 51:32:3:14:7:12:2:1:713:48:101:38:17:2:0:0:0:0:2:0 | 0:0:0:0:0:0:0:0:1:0:0:0:0:0:0:0:0:0:0:0 | 178:83:32:33:88:23:18:8:2590:154:855:78:93:1:0:0:0:0:1:1 | 2:0:0:0:0:0:0:0:0:0:0:1:0:0:0:0:0:0:0:0 |
| 542:31:28:12:4:11:1:3:3:30:81:10:9:1:0:0:0:0:1:0 | 1:0:0:0:0:0:0:0:0:1:0:0:0:0:0:0:0:0:0 | 710:51:32:3:14:7:12:2:1:48:101:38:17:2:0:0:0:0:2:0 | 4:0:0:0:0:0:0:0:0:0:0:0:0:0:0:0:0:0:0:0 | 2588:180:83:32:33:88:23:17:8:153:854:77:91:1:0:0:0:0:1:1 | 2:0:0:0:0:0:0:1:0:1:1:2:2:0:0:0:0:0:0:0 |
| 25:31:29:12:4:11:3:30:3:538:82:12:12:1:1:0:0:0:0:1:0 | 0:0:0:0:0:0:0:0:0:1:0:0:0:0:0:0:0:0:0:0 | 39:51:32:3:14:7:2:49:1:704:101:40:20:2:1:0:0:0:2:2:0 | 0:0:0:0:0:0:0:0:0:0:1:0:0:0:0:0:0:0:0:0 | 134:180:81:34:33:88:18:159:9:2578:855:85:117:1:14:0:0:0:4:1:2 | 0:0:0:0:0:0:0:0:0:3:0:0:2:0:0:0:0:0:0:0:0 |
| 26:31:29:11:4:11:3:30:3:538:82:12:12:1:1:0:0:0:0:1:0 | 0:0:0:0:0:0:0:0:0:1:0:0:0:0:0:0:0:0:0:0 | 42:51:32:3:14:7:2:49:1:703:101:40:22:2:1:0:0:0:2:2:0 | 0:0:0:0:0:0:0:0:0:0:0:0:0:0:0:0:0:0:0:0 | 135:180:81:34:33:87:18:159:9:2573:855:85:121:1:14:0:0:0:4:1:2 | 1:0:0:0:0:0:0:0:0:1:0:0:1:0:0:0:0:0:0:0:0 |
| 31:29:11:4:10:32:3:31:3:538:82:12:14:1:1:0:0:0:0:1:0 | 0:0:0:0:0:0:0:0:0:0:0:0:0:0:0:0:0:0:0:0 | 51:32:3:14:6:56:2:49:1:697:101:40:19:2:1:0:0:0:2:2:0 | 0:0:0:0:0:0:0:0:0:2:0:0:0:0:0:0:0:0:0:0 | 180:81:33:33:82:153:18:159:7:2534:855:85:140:1:14:0:0:0:4:1:2 | 0:0:0:0:1:1:0:0:0:2:0:0:0:0:0:0:0:0:0:0:0 |
| 31:29:11:4:10:32:3:31:3:537:82:12:13:1:1:0:0:0:0:1:0 | 0:0:0:0:0:0:0:0:0:1:0:0:1:0:0:0:0:0:0:0 | 51:31:3:14:6:56:2:49:1:698:101:40:19:2:1:0:0:0:2:2:0 | 0:0:0:0:0:0:0:0:0:1:0:0:0:0:0:0:0:0:0:0 | 180:81:33:33:83:153:18:159:7:2522:854:85:137:1:14:0:0:0:4:1:2 | 0:0:0:0:0:1:0:0:0:7:1:0:1:0:0:0:0:0:0:0:0 |
| 536:31:29:11:4:10:32:3:31:3:82:12:14:1:1:0:0:0:0:1:0 | 0:0:0:0:0:0:0:0:0:0:0:0:0:0:0:0:0:0:0:0 | 694:51:31:3:14:6:56:2:49:1:101:40:18:2:1:0:0:0:2:2:0 | 2:0:0:0:0:0:0:0:0:0:0:0:0:0:0:0:0:0:0:0 | 2518:180:81:33:32:81:154:17:159:7:852:85:129:1:14:0:0:0:4:1:2 | 4:0:0:0:1:0:1:1:0:0:2:0:0:0:0:0:0:0:0:0:0 |
| 31:27:11:4:10:32:3:31:3:534:82:12:13:1:1:0:0:0:0:1:0 | 0:1:0:0:0:0:0:0:0:0:0:0:0:0:0:0:0:0:0:0 | 51:31:3:14:5:56:2:49:1:691:101:40:16:2:1:0:0:0:2:2:0 | 0:0:0:0:0:0:0:0:0:1:0:0:0:0:0:0:0:0:0:0 | 180:81:33:33:80:152:18:159:7:2505:852:85:121:1:14:0:0:0:4:1:2 | 0:0:0:0:0:1:0:0:0:1:2:0:0:0:0:0:0:0:0:0:0 |
| 31:27:11:4:10:32:3:31:3:531:82:12:13:1:1:0:0:0:0:1:0 | 0:0:0:0:0:0:0:0:0:1:0:0:0:0:0:0:0:0:0:0 | 51:31:3:14:5:56:2:48:1:687:101:40:16:2:1:0:0:0:2:2:0 | 0:0:0:0:0:0:0:1:0:3:0:0:0:0:0:0:0:0:0:0 | 180:80:32:33:79:152:18:159:7:2497:853:85:121:1:14:0:0:0:4:1:2 | 0:1:0:0:1:1:0:0:0:6:1:0:0:0:0:0:0:0:0:0:0 |

| | | | | | |
|---|---|---|---|---|---|
| 31:525:27:11:4:9:31:3:31:3:82:12:11:1:1:0:0:0:1:0 | 0:0:0:0:0:0:0:0:0:0:0:0:0:0:0:0:0:0:0:0 | 50:668:30:3:14:3:55:2:49:1:101:40:15:2:1:0:0:2:2:0 | 1:5:0:0:0:0:0:0:0:0:0:0:0:0:0:0:0:0:0 | 180:2431:78:32:33:75:146:18:159:7:850:85:102:1:14:0:0:4:1:2 | 0:8:0:0:0:0:1:0:0:0:4:0:1:0:0:0:0:0:0:0 |
| 82:514:31:26:11:4:9:31:3:31:2:12:8:1:1:0:0:0:1:0 | 0:0:0:0:0:0:0:0:0:0:0:0:0:0:0:0:0:0:0:0 | 101:652:51:30:3:14:3:58:2:49:1:39:9:2:1:0:0:2:2:0 | 0:1:0:0:0:0:0:0:0:0:0:1:0:0:0:0:0:0:0:0 | 853:2366:180:75:30:33:68:153:18:158:7:85:92:1:14:0:0:4:1:2 | 1:2:0:0:0:0:0:1:0:1:0:0:0:0:0:0:0:0:0:0 |
| 88:05:00 | 0:00 | 197:24:00 | 1:00 | 590:70 | 5:00 |
| 92:05:00 | 0:00 | 198:22:00 | 0:00 | 606:66 | 0:01 |
| 91:05:00 | 0:00 | 198:11:00 | 0:00 | 605:25:00 | 0:01 |
| 89:03:00 | 1:00 | 193:08:00 | 0:00 | 586:09:00 | 0:00 |
| 80:02:00 | 0:00 | 181:08:00 | 1:00 | 534:07:00 | 4:00 |
| 78:194:6:19 | 0:0:0:0 | 110:92:4:47 | 0:0:0:0 | 269:376:12:113 | 2:1:0:0 |
| 194:78:5:22 | 0:0:0:0 | 91:117:14:82 | 1:1:0:0 | 376:288:35:168 | 1:1:0:1 |
| 193:78:7:22 | 1:0:0:0 | 92:122:16:82 | 0:0:0:0 | 376:291:51:169 | 1:1:1:0 |
| 194:77:7:22 | 0:0:0:0 | 92:121:15:82 | 0:0:0:0 | 375:292:56:168 | 2:0:0:1 |
| 193:80:16:22 | 1:0:0:0 | 92:140:66:82 | 0:0:0:0 | 375:322:183:168 | 2:1:0:1 |
| 194:80:22:22 | 0:0:0:0 | 92:145:95:82 | 0:0:0:0 | 375:326:319:169 | 2:1:3:0 |
| 191:80:23:22 | 3:0:0:0 | 92:145:96:82 | 0:0:0:0 | 376:329:330:169 | 1:0:1:0 |
| 194:79:23:22 | 0:0:0:0 | 92:145:99:82 | 0:0:0:0 | 376:329:339:168 | 1:0:2:1 |
| 194:78:24:22 | 0:1:1:0 | 92:145:103:82 | 0:0:1:0 | 377:326:347:169 | 0:3:0:0 |
| 194:78:25:22 | 0:1:0:0 | 92:138:104:82 | 0:7:0:0 | 377:322:346:169 | 0:7:0:0 |
| 194:76:30:22 | 0:0:0:0 | 92:124:119:81 | 0:2:0:1 | 377:297:378:169 | 0:1:3:0 |
| 194:70:34:22 | 0:0:0:0 | 92:120:154:82 | 0:0:1:0 | 377:278:508:169 | 0:2:2:0 |
| 49:194:37:22 | 9:0:0:0 | 103:92:180:82 | 7:0:0:0 | 226:376:592:169 | 24:1:2:0 |
| 194:43:36:22 | 0:0:0:0 | 92:101:199:81 | 0:0:0:0 | 376:219:638:168 | 0:0:2:0 |
| 194:43:36:22 | 0:0:0:0 | 92:101:199:81 | 0:0:0:0 | 376:219:638:167 | 0:0:1:1 |
| 192:44:36:22 | 1:0:0:0 | 90:102:195:80 | 1:0:3:0 | 372:226:630:166 | 1:2:6:1 |
| 193:44:35:22 | 0:0:0:0 | 89:99:198:80 | 2:0:1:0 | 372:230:631:167 | 1:0:4:0 |
| 193:44:35:22 | 0:0:0:0 | 89:99:198:80 | 2:0:1:0 | 372:230:631:167 | 1:0:4:0 |
| 192:44:34:22 | 1:0:1:0 | 88:100:196:79 | 3:0:0:1 | 371:231:629:166 | 1:0:4:0 |
| 1:00:00 | 0:00:00 | 1:00:01 | 0:00:00 | 2:01:00 | 0:00:00 |
| 0:00:00 | 1:00:00 | 0:00:00 | 1:00:01 | 1:00:00 | 2:01:00 |
| 0:00 | 1:03 | 0:00 | 0:00 | 0:00 | 0:05 |
| 5:1:0:0 | 3:1:0:0 | 6:2:0:0 | 0:2:0:0 | 11:1:0:0 | 8:12:2:0 |
| 1:7:0:0:0:0:0:0 | 0:3:0:0:0:0:1:0 | 3:8:1:1:0:0:0:1 | 1:6:0:0:0:0:0:0 | 1:11:0:0:0:0:0:0 | 1:7:1:0:0:0:0:0 |
| 1:11:0:1:10:5:1:0:2:0:0 | 0:0:0:0:0:0:0:0:0:0:0 | 0:13:0:0:6:7:0:0:1:0:1 | 0:0:0:0:0:0:0:0:0:0:0 | 1:17:3:6:11:10:1:0:1:0:0 | 0:0:0:0:0:0:0:0:0:0:0 |
| 11:1:0:1:9:6:1:0:2:0:0 | 0:0:0:0:0:0:0:0:0:0:0 | 12:0:0:0:6:7:0:0:1:0:1 | 0:0:0:0:0:0:0:0:0:0:0 | 15:1:3:6:9:9:1:0:1:0:0 | 0:0:0:0:0:0:0:0:0:0:0 |
| 5:0:0:0:1:0 | 0:0:0:0:0:0 | 5:0:0:0:0:1 | 0:0:0:0:0:0 | 1:0:0:0:0:0 | 0:0:0:0:0:0 |
| 1:00 | 1:00 | 1:00 | 0:00 | 1:00 | 1:01 |
| 0:00 | 0:00 | 0:00 | 0:00 | 0:00 | 1:00 |
| 0 | 1 | 0 | 1 | 7 | 4 |
| 0:00 | 0:01 | 0:00 | 0:00 | 3:00 | 3:00 |
| 0:02 | 0:00 | 0:00 | 0:00 | 1:03 | 0:00 |
| 338 | 1 | 267 | 1 | 1211 | 10 |
| 332 | 2 | 260 | 4 | 1198 | 7 |
| 330 | 2 | 260 | 2 | 1180 | 10 |
| 327 | 4 | 257 | 2 | 1165 | 9 |
| 303 | 1 | 215 | 0 | 960 | 3 |
| 301 | 2 | 213 | 2 | 954 | 5 |
| 0 | 0 | 0 | 0 | 0 | 0 |
| 0 | 0 | 0 | 0 | 0 | 0 |
| 0 | 0 | 0 | 0 | 0 | 0 |
| 0 | 1 | 0 | 2 | 0 | 2 |
| 0:3:1:3:0:0:0 | 0:0:0:0:0:0:0 | 0:3:1:0:0:1:0 | 0:4:0:0:0:0:0 | 2:3:6:5:0:10:0 | 0:9:0:8:0:0:0 |
| 1 | 1 | 0 | 0 | 3 | 5 |

| | | | | | |
|---|---|---|---|---|---|
| 1 | 1 | 0 | 0 | 3 | 6 |
| 1 | 1 | 0 | 0 | 6 | 3 |
| 1:3:0:0:2:0:0:1:0:0:2:1:1:0 | 0:0:0:0:0:0:0:0:0:0:0:0:0:0 | 0:1:0:5:4:0:0:0:0:0:1:0:0:1 | 0:0:0:0:0:0:0:1:0:0:0:0:0:0 | 2:12:0:12:14:0:2:2:0:0:4:1:1:1 | 0:0:0:0:0:0:0:2:0:0:0:0:0:0 |
| 1:3:0:2:0:0:0:0:1:2:0:0:0:0 | 0:0:0:0:0:0:0:1:0:0:0:0:1:0 | 0:1:2:4:0:0:0:0:0:0:0:0:0:0 | 0:0:3:0:0:0:0:1:0:1:0:0:0:1 | 2:7:9:10:0:1:0:3:1:4:0:0:0:1 | 0:5:3:4:0:1:0:1:0:0:0:0:1:0 |
| 0:0:0:0:0:0:0:0:0:0:0:0:0:0 | 1:3:1:2:0:2:0:0:2:1:0:1:0:0 | 0:0:0:0:0:0:0:0:0:0:0:0:0:0 | 0:1:5:1:0:2:0:0:1:0:0:0:0:1 | 0:0:0:0:0:0:0:0:0:0:0:0:0:0 | 2:12:12:9:0:16:2:0:4:1:0:2:0:1 |
| 0:0:1:0:0:2:0:0:0:0:0:0:1:0 | 1:3:0:0:0:0:0:0:2:1:0:0:0:0 | 0:0:3:0:0:2:0:0:1:0:0:0:0:1 | 0:1:2:1:0:1:0:0:0:0:0:0:0:0 | 0:5:3:2:0:4:1:0:1:0:0:0:1:0 | 2:7:9:2:0:12:1:0:4:1:0:0:1:1 |
| 195:1:0:0:0 | 1:0:0:0:0 | 341:0:1:0:0 | 1:0:0:0:0 | 978:0:2:0:0 | 1:0:0:0:0 |
| 197:17:1:1:0:0 | 0:0:0:0:0:0 | 344:38:0:2:2:0 | 0:0:0:0:0:0 | 985:90:2:4:1:0 | 0:0:0:0:0:0 |
| 197:17:1:1:0 | 0:0:0:0:0 | 341:38:0:2:2 | 1:0:0:0:0 | 980:90:2:4:1 | 2:0:0:0:0 |
| 196:17:1:1:0:0 | 0:0:0:0:0:0 | 342:38:0:2:2:0 | 1:0:0:0:0:0 | 972:85:2:4:1:0 | 0:0:0:0:0:0 |
| 196:17:1:1:0:0 | 0:0:0:0:0:0 | 339:38:0:2:2:0 | 2:0:0:0:0:0 | 969:85:2:4:1:0 | 1:0:0:0:0:0 |
| 196:17:1:1:0:0 | 0:0:0:0:0:0 | 341:38:0:2:2:0 | 0:0:0:0:0:0 | 969:85:2:4:1:0 | 1:0:0:0:0:0 |
| 0 | 0 | 0 | 0 | 0 | 2 |
| 20:01:01 | 0:00:00 | 19:04:02 | 0:00:00 | 74:12:05 | 0:00:02 |
| 0:00 | 0:00 | 0:01 | 0:00 | 0:02 | 0:02 |
| 1 | 0 | 1 | 0 | 2 | 0 |
| 0:00:00 | 2:01:00 | 4:00:00 | 4:00:00 | 0:00:00 | 10:00:01 |
| 0 | 1 | 0 | 1 | 0 | 0 |
| 1:3:0:1:0:0:1:1:0:0:0:0:0:0:1:0:0:1:0:0:0:0:1 | 0:0:0:1:0:0:0:0:0:0:0:0:0:0:0:0:0:1:0:0:0:0:0 | 0:0:2:2:0:0:0:0:1:0:0:0:0:0:2:1:0:0:1:0:1:0:1 | 0:0:1:5:0:0:0:1:2:0:1:0:0:0:0:0:0:0:0:0:0:0:0 | 4:4:15:7:0:0:0:0:2:1:0:0:2:0:3:0:0:1:0:0:0:0:1 | 0:2:1:0:0:0:0:1:0:0:0:0:0:0:0:0:0:0:0:0:0:0:0 |
| 0:0:0:0 | 1:0:0:0 | 0:0:0:0 | 9:0:4:0 | 0:0:0:0 | 13:1:5:2 |
| 1:0:0:1:0:0:0:0 | 0:0:1:0:0:0:0:0 | 4:0:15:23:15:42:3:0 | 2:0:3:4:1:6:0:0 | 4:0:4:8:1:3:1:0 | 0:0:1:0:0:0:0:0 |
| 1:0:0:0:0:0 | 0:0:0:0:0:0 | 2:5:0:1:9:11 | 0:0:0:0:0:0 | 0:2:0:0:2:7 | 0:0:0:0:0:0 |
| 0:00 | 0:00 | 0:00 | 1:00 | 0:00 | 0:00 |
| 0:00:00 | 0:00:00 | 0:01:00 | 0:01:01 | 1:00:01 | 0:01:00 |
| 1:0:0:0:0 | 0:0:0:0:0 | 318:0:200:51:7 | 1:0:3:0:0 | 27:0:13:0:1 | 0:0:0:0:0 |
| 1:0:0:0:0 | 0:0:0:0:0 | 339:0:162:50:7 | 1:0:1:0:0 | 27:0:12:0:1 | 0:0:0:0:0 |
| 1:0:0:0:0 | 0:0:0:0:0 | 341:0:95:48:7 | 0:0:1:0:0 | 27:0:9:0:1 | 0:0:0:0:0 |
| 0 | 0 | 1 | 0 | 3 | 0 |
| 0 | 0 | 0 | 0 | 0 | 0 |
| 1952:64 | 1:00 | 3261:102 | 1:01 | 19415:832 | 13:04 |
| 1946:69 | 2:00 | 3194:131 | 3:01 | 19138:906 | 16:04 |
| 1946:71 | 2:00 | 3185:131 | 5:01 | 19133:907 | 12:06 |
| 1945:70 | 2:01 | 3179:131 | 3:00 | 19093:913 | 25:03:00 |
| 1942:70 | 2:00 | 3173:130 | 3:00 | 19070:907 | 23:05 |
| 1943:67 | 0:00 | 3158:128 | 3:00 | 19007:901 | 20:01 |
| 1943:67 | 0:00 | 3160:128 | 1:00 | 19005:899 | 21:03 |
| 1935:70 | 2:01 | 3130:132 | 0:00 | 18848:913 | 18:02 |
| 1927:64 | 4:02 | 3054:122 | 5:01 | 18584:821 | 30:05:00 |
| 1930:67 | 1:01 | 3044:119 | 5:01 | 18551:807 | 31:10:00 |
| 1925:61 | 3:01 | 3003:112 | 6:00 | 18442:771 | 36:08:00 |
| 1892:62 | 2:03 | 2925:116 | 9:04 | 18052:825 | 34:15:00 |
| 1884:66 | 0:00 | 2906:118 | 9:00 | 17931:832 | 32:03:00 |
| 1872:66 | 3:01 | 2889:115 | 7:00 | 17813:831 | 36:05:00 |
| 1842:77 | 1:00 | 2808:104 | 4:00 | 17376:826 | 21:04 |
| 1812:78 | 4:00 | 2721:100 | 7:01 | 16856:744 | 36:03:00 |
| 1809:79 | 4:00 | 2712:100 | 12:01 | 16834:746 | 34:02:00 |
| 1788:73 | 4:01 | 2637:101 | 15:00 | 16318:729 | 60:10:00 |
| 1787:73 | 6:00 | 2631:100 | 16:01 | 16287:727 | 51:10:00 |
| 1781:72 | 8:00 | 2622:101 | 18:01 | 16228:732 | 45:05:00 |
| 1765:69 | 6:00 | 2582:102 | 17:01 | 15939:770 | 50:03:00 |
| 1752:70 | 4:00 | 2554:103 | 12:00 | 15772:795 | 71:03:00 |
| 1753:71 | 3:00 | 2551:104 | 16:00 | 15778:805 | 73:03:00 |
| 1750:72 | 6:00 | 2551:105 | 11:00 | 15731:820 | 81:01:00 |
| 1738:73 | 8:00 | 2510:106 | 9:00 | 15499:851 | 90:03:00 |
| 1735:73 | 10:00 | 2508:106 | 10:00 | 15490:852 | 79:02:00 |
| 1726:80 | 1:00 | 2466:109 | 2:00 | 15156:872 | 29:01:00 |
| 1717:80 | 7:00 | 2451:108 | 6:00 | 15040:866 | 33:01:00 |
| 1717:80 | 6:00 | 2451:108 | 5:00 | 15009:866 | 40:02:00 |
| 1645:78 | 7:00 | 2239:106 | 5:01 | 13510:866 | 40:04:00 |
| 1640:78 | 8:00 | 2230:107 | 8:00 | 13469:864 | 48:04:00 |
| 1636:77 | 1:00 | 2184:106 | 7:00 | 13279:869 | 22:01 |
| 1632:76 | 1:00 | 2143:106 | 6:01 | 13029:860 | 30:04:00 |
| 1631:76 | 2:00 | 2137:106 | 5:00 | 13031:860 | 19:03 |
| 1585:72 | 2:00 | 2031:104 | 1:00 | 12391:830 | 4:02 |

| | | | | | |
|---|---|---|---|---|---|
| 1617:74 | 3:00 | 2075:106 | 2:00 | 12656:851 | 14:01 |
| 1620:73 | 0:00 | 2068:106 | 7:00 | 12640:851 | 13:01 |
| 1607:73 | 1:00 | 2006:103 | 3:00 | 12247:855 | 31:04:00 |
| 1601:73 | 3:00 | 1996:103 | 1:00 | 12148:850 | 22:03 |
| 1602:73 | 1:00 | 1988:102 | 3:01 | 12094:840 | 19:07 |
| 1591:73 | 1:00 | 1919:99 | 1:00 | 11626:828 | 19:02 |
| 1584:73 | 3:00 | 1874:98 | 5:01 | 11447:827 | 29:01:00 |
| 1585:72 | 1:00 | 1859:99 | 4:00 | 11404:823 | 24:01:00 |
| 1583:72 | 3:01 | 1852:99 | 3:00 | 11364:821 | 19:00 |
| 1571:71 | 5:00 | 1783:95 | 3:00 | 10806:804 | 38:02:00 |
| 1474:63 | 6:00 | 1136:78 | 14:01 | 7773:700 | 48:04:00 |
| 0:0:0:0:0:0:0:0:0:0:0:0:0:1:0:0 | 0:0:0:0:0:0:0:0:0:0:0:0:0:0:0:0 | 0:0:0:0:1:0:1:0:0:0:0:0:1:0:0:0 | 0:0:0:0:0:0:0:0:0:0:0:0:0:0:0:0 | 2:2:0:0:0:0:1:1:0:0:0:1:0:0:0:0 | 0:0:0:0:0:0:0:0:0:0:0:0:0:0:0:0 |
| 0:00:00 | 0:01:00 | 0:00:00 | 0:02:02 | 0:00:00 | 1:08:03 |
| 0:00 | 1:01 | 0:00 | 0:00 | 0:00 | 2:00 |
| 20:03:01 | 0:00:00 | 7:00:00 | 0:00:00 | 31:12:01 | 0:00:00 |
| 0 | 0 | 2 | 0 | 5 | 0 |
| 3:13:0:0:4:2:10:2:0:0:0:1:0 | 0:1:0:0:1:0:0:0:0:0:0:0:0 | 6:0:0:0:3:0:3:2:1:0:0:0:0 | 0:0:0:0:0:0:0:0:0:0:0:0:0 | 10:14:3:0:7:2:19:7:8:0:0:0:1 | 0:0:0:0:0:0:0:0:0:0:0:0:0 |
| 3:16:0:0:5:2:14:2:0:0:0:1:0 | 0:0:0:0:0:0:0:0:0:0:0:0:0 | 6:0:0:0:3:0:5:4:0:0:0:0:0 | 0:0:0:0:0:0:0:0:0:0:0:0:0 | 10:15:2:0:7:2:28:8:0:0:0:0:1 | 0:0:0:0:0:0:0:0:0:0:0:0:0 |
| 106:403:6:25 | 0:1:0:0 | 290:459:4:27 | 0:0:0:0 | 672:2142:24:139 | 0:3:0:0 |
| 106:408:6:26 | 0:0:0:0 | 290:460:4:27 | 0:1:0:0 | 672:2159:24:140 | 0:5:0:0 |
| 106:408:6:26 | 0:0:0:0 | 288:462:4:27 | 2:1:0:0 | 672:2160:28:139 | 0:3:0:1 |
| 106:408:5:26 | 0:0:1:0 | 289:458:5:27 | 1:1:0:0 | 671:2158:32:139 | 1:3:0:0 |
| 106:405:7:26 | 0:2:0:0 | 289:456:5:27 | 1:1:0:0 | 666:2145:39:139 | 2:2:0:0 |
| 1893 | 1 | 1732 | 1 | 7885 | 3 |
| 1892 | 2 | 1732 | 1 | 7885 | 2 |
| 1894 | 1 | 1731 | 2 | 7883 | 5 |
| 1892 | 3 | 1732 | 1 | 7882 | 6 |
| 1904 | 3 | 1740 | 0 | 7921 | 9 |
| 1906 | 0 | 1732 | 3 | 7916 | 10 |
| 1903 | 1 | 1728 | 2 | 7888 | 8 |
| 1903 | 2 | 1729 | 1 | 7887 | 7 |
| 1900 | 2 | 1727 | 1 | 7862 | 8 |
| 1902 | 1 | 1726 | 2 | 7854 | 4 |
| 1905 | 3 | 1737 | 1 | 7953 | 10 |
| 1907 | 0 | 1737 | 0 | 7961 | 0 |
| 1904 | 3 | 1737 | 2 | 7948 | 7 |
| 1907 | 1 | 1729 | 1 | 7914 | 2 |
| 1904 | 1 | 1721 | 0 | 7878 | 3 |
| 1903 | 1 | 1721 | 0 | 7873 | 8 |
| 1900 | 3 | 1720 | 1 | 7871 | 9 |
| 1903 | 1 | 1704 | 2 | 7847 | 5 |
| 1889 | 5 | 1694 | 1 | 7764 | 7 |
| 1887 | 6 | 1694 | 1 | 7746 | 15 |
| 1887 | 4 | 1690 | 3 | 7725 | 25 |
| 1883 | 5 | 1690 | 0 | 7726 | 19 |
| 165 | 0 | 62 | 0 | 373 | 5 |
| 162 | 2 | 51 | 0 | 317 | 0 |
| 164 | 0 | 51 | 0 | 316 | 2 |
| 197 | 1 | 121 | 3 | 693 | 29 |
| 196 | 0 | 115 | 0 | 705 | 1 |
| 192 | 1 | 114 | 0 | 696 | 1 |
| 190 | 0 | 111 | 0 | 666 | 2 |
| 176 | 0 | 104 | 0 | 650 | 0 |
| 169 | 1 | 100 | 0 | 632 | 1 |
| 168 | 1 | 99 | 1 | 632 | 8 |
| 169 | 0 | 99 | 1 | 633 | 8 |
| 161 | 0 | 100 | 0 | 621 | 3 |
| 143 | 0 | 98 | 0 | 609 | 2 |
| 123 | 0 | 95 | 0 | 567 | 4 |
| 91 | 0 | 129 | 0 | 547 | 1 |
| 88 | 1 | 131 | 1 | 549 | 1 |
| 84 | 1 | 119 | 0 | 512 | 3 |
| 84 | 0 | 118 | 0 | 503 | 2 |
| 79 | 0 | 108 | 0 | 431 | 2 |
| 0:893:1:6:55:0 | 0:1:0:0:0:0 | 3:684:1:4:28:0 | 0:0:0:0:0:0 | 4:3470:13:12:218:1 | 0:3:0:0:0:0 |
| 0:893:1:6:55:0:0 | 0:0:0:0:0:0:0 | 3:688:1:4:27:0:0 | 0:2:0:0:0:0:0 | 4:3480:11:12:216:0:1 | 0:2:0:0:0:0:0 |
| 890:0:1:55:1:0:0 | 1:0:0:0:0:0:0 | 690:3:1:28:0:0:0 | 0:0:0:0:0:0:0 | 3471:4:11:217:1:0:0 | 3:0:0:0:0:0:0 |
| 0:891:1:55:1:0:0 | 0:0:0:0:0:0:0 | 3:690:1:28:0:0:0 | 0:0:0:0:0:0:0 | 4:3475:11:217:1:0:0 | 0:1:0:0:0:0:0 |
| 55:894:0:1:1:0:0:0 | 0:1:0:0:0:0:0:0 | 28:695:3:1:0:0:0:0 | 0:2:0:0:0:0:0:0 | 216:3487:4:11:1:0:0:1 | 0:5:0:0:0:0:0:0 |

| | | | | | |
|---|---|---|---|---|---|
| 894:0:1:55:1:0:0:0 | 1:0:0:0:0:0:0:0 | 693:3:1:28:0:0:0:0 | 5:0:0:0:0:0:0:0 | 3483:4:11:216:1:0:0:1 | 9:0:0:0:0:0:0:0 |
| 0:896:1:55:1:0:0:0 | 0:1:0:0:0:0:0:0 | 3:696:1:28:0:0:0:0 | 0:2:0:0:0:0:0:0 | 4:3499:11:216:1:0:0:1 | 0:4:0:0:0:0:0:0 |
| 0:897:0:55:1:0:0:0 | 0:0:0:0:0:0:0:0 | 3:705:1:28:0:0:0:0 | 0:1:0:0:0:0:0:0 | 4:3510:11:216:1:0:0:1 | 0:2:0:0:0:0:0:0 |
| 893:0:0:55:1:0:0:0 | 1:0:0:0:0:0:0:0 | 704:3:1:28:0:0:0:0 | 1:0:0:0:0:0:0:0 | 3513:4:10:216:1:0:0:1 | 4:0:0:0:0:0:0:0 |
| 892:0:0:54:1:0:0:0 | 1:0:0:0:0:0:0:0 | 711:3:1:28:0:0:0:0 | 0:0:0:0:0:0:0:0 | 3518:4:10:216:1:0:0:1 | 7:0:0:0:0:0:0:0 |
| 0:872:0:52:1:0:0:0 | 0:2:0:0:0:0:0:0 | 3:680:1:27:0:0:0:0 | 0:1:0:0:0:0:0:0 | 4:3353:10:210:1:0:0:1 | 0:4:0:0:0:0:0:0 |
| 0:854:1:52:0:1:0:0 | 0:0:0:0:0:0:0:0 | 3:679:1:27:0:0:0:0 | 0:0:0:0:0:0:0:0 | 4:3254:13:205:0:1:0:1 | 0:2:0:0:0:0:0:0 |
| 0:853:1:52:0:1:0:0 | 0:1:0:0:0:0:0:0 | 3:677:1:27:0:0:0:0 | 0:0:0:0:0:0:0:0 | 4:3258:13:205:0:1:0:1 | 0:4:0:0:0:0:0:0 |
| 0:853:1:52:0:1:0:0 | 0:0:0:0:0:0:0:0 | 3:678:1:27:0:0:0:0 | 0:0:0:0:0:0:0:0 | 4:3258:13:205:0:1:0:1 | 0:6:0:0:0:0:0:0 |
| 0:890:1:51:1:0:0:0 | 0:0:0:0:0:0:0:0 | 3:709:1:27:0:0:0:0 | 0:1:0:0:0:0:0:0 | 4:3410:13:207:1:0:0:1 | 0:5:0:1:0:0:0:0 |
| 886:0:1:51:1:0:0:0 | 2:0:0:0:0:0:0:0 | 715:3:1:27:0:0:0:0 | 1:0:0:0:0:0:0:0 | 3444:4:13:207:1:0:0:1 | 5:0:0:0:0:0:0:0 |
| 0:907:1:54:1:0:0:0 | 0:1:0:0:0:0:0:0 | 3:751:1:28:0:0:0:0 | 0:1:0:0:0:0:0:0 | 4:3631:13:211:1:0:0:1 | 0:4:0:0:0:0:0:0 |
| 23:3:3:0:1:0:1:0:0 | 0:0:0:0:0:0:0:0:0 | 29:6:4:0:0:0:0:0:0 | 0:0:0:0:0:0:0:0:0 | 77:10:8:0:0:0:0:0:2 | 0:0:0:0:0:0:0:0:0 |
| 4:2:1:0:0:0:1:0:0 | 19:1:1:0:0:1:0:0:0 | 12:1:5:0:0:0:0:0:0 | 17:3:1:0:0:0:0:0:0 | 8:0:3:0:0:0:0:0:0 | 70:8:5:0:0:0:0:0:2 |
| 3:2:1:0:1:0:1:0:1:1:0 | 3:1:1:0:1:2:0:0:0:0:0 | 0:12:0:0:1:0:6:0:1:2:1 | 0:2:0:0:2:0:5:0:0:1:0 | 1:12:2:0:1:0:6:0:0:0:0 | 1:6:0:0:0:0:1:0:0:2:1 |
| 0:00 | 0:00 | 0:00 | 0:00 | 0:00 | 1:01 |
| 0:0:0:0:0:0 | 1:0:0:0:0:0 | 0:0:0:0:0:0 | 0:0:0:0:0:0 | 0:0:0:0:0:0 | 1:1:3:0:0:0 |
| 0:0:0:0:0:0 | 1:0:0:0:0:0 | 0:0:0:0:0:0 | 0:0:0:0:0:0 | 0:0:0:0:0:0 | 1:1:3:0:0:0 |
| 0:0:0:0:0:0 | 0:1:0:0:0:0 | 0:0:0:0:0:0 | 0:0:0:0:0:0 | 0:0:0:0:0:0 | 1:1:3:0:0:0 |
| 0:0:0:0:0:0 | 1:0:0:0:0:0 | 0:0:0:0:0:0 | 0:0:0:0:0:0 | 0:0:0:0:0:0 | 1:1:3:0:0:0 |
| 0:00 | 0:00 | 0:00 | 1:00 | 0:00 | 1:01 |
| 4166:12:33:1 | 4:0:0:0 | 8397:67:63:3 | 6:0:0:0 | 22554:172:149:11 | 28:0:0:0 |
| 4167:12:33:1 | 6:0:0:0 | 8413:67:63:3 | 10:1:0:0 | 22600:172:150:11 | 26:0:0:0 |
| 4171:12:33:1 | 5:0:0:0 | 8407:68:63:3 | 15:0:0:0 | 22598:172:150:10 | 34:0:0:0 |
| 4169:12:33:1 | 10:0:0:0 | 8429:68:63:3 | 6:0:0:0 | 22649:172:150:9 | 23:0:0:0 |
| 4184:12:18:0 | 2:0:0:1 | 8435:68:25:3 | 7:0:0:0 | 22692:173:42:9 | 12:0:1:0 |
| 4191:34:00 | 6:00:00 | 8454:63:0 | 7:00:00 | 22736:150:2 | 31:00:00 |
| 4193:34:00 | 6:00:00 | 8450:63:0 | 13:00:00 | 22747:150:2 | 36:00:00 |
| 4220:12:34:1 | 4:0:0:0 | 8493:68:63:3 | 8:0:0:0 | 22877:174:150:11 | 18:0:0:0 |
| 4223:12:34:1 | 5:0:0:0 | 8497:68:63:3 | 6:0:0:0 | 22880:174:150:11 | 23:0:0:0 |
| 4226:12:34:1 | 3:0:0:0 | 8505:68:63:3 | 4:0:0:0 | 22905:174:150:11 | 13:0:0:0 |
| 4233:12:34:1 | 3:0:0:0 | 8521:68:63:3 | 16:0:0:0 | 22983:174:150:11 | 19:0:0:0 |
| 4243:12:34:1 | 4:0:0:0 | 8549:68:63:3 | 4:0:0:0 | 23025:174:150:11 | 16:0:0:0 |
| 4246:12:34:1 | 0:0:0:0 | 8542:68:63:3 | 5:0:0:0 | 23017:174:150:11 | 17:0:0:0 |
| 4226:12:34:1 | 7:0:0:0 | 8505:67:63:3 | 7:0:0:0 | 22941:174:150:11 | 27:0:0:0 |
| 4233:12:33:1 | 2:0:1:0 | 8487:67:63:3 | 12:0:0:0 | 22903:174:149:11 | 20:0:1:0 |
| 4222:12:34:1 | 12:0:0:0 | 8455:67:63:3 | 33:0:0:0 | 22820:174:149:11 | 82:0:1:0 |
| 4228:34:12:1 | 2:0:0:0 | 8471:63:67:3 | 12:0:0:0 | 22865:149:174:11 | 22:0:0:0 |
| 4233:12:34:1 | 1:0:0:0 | 8466:67:63:3 | 6:0:0:0 | 22858:174:149:11 | 20:0:0:0 |
| 2638:11:22:1 | 1595:1:12:0 | 4257:33:32:1 | 4212:34:31:2 | 17259:131:105:9 | 5612:43:44:2 |

| | | | | | |
|---|---|---|---|---|---|
| 4231:12:34:1 | 3:0:0:0 | 8453:67:63:2 | 14:0:0:1 | 22832:174:149:11 | 32:0:0:0 |
| 4225:12:34:1 | 4:0:0:0 | 8454:67:63:3 | 5:0:0:0 | 22833:173:147:11 | 22:0:2:0 |
| 4220:12:34:1 | 8:0:0:0 | 8435:66:62:3 | 18:0:0:0 | 22781:173:148:11 | 29:0:0:0 |
| 4222:12:34:1 | 4:0:0:0 | 8447:66:62:3 | 8:0:0:0 | 22785:173:148:11 | 15:0:0:0 |
| 4218:12:34:1 | 5:0:0:0 | 8441:66:62:3 | 10:0:0:0 | 22756:173:147:11 | 27:0:0:0 |
| 4222:12:34:1 | 2:0:0:0 | 8441:66:61:3 | 6:0:1:0 | 22772:173:147:11 | 7:0:0:0 |
| 4210:12:34:1 | 8:0:0:0 | 8416:66:61:3 | 10:0:0:0 | 22678:173:147:11 | 31:0:0:0 |
| 17:02 | 0:00 | 15:02 | 0:00 | 60:08:00 | 0:00 |
| 1:0:0:0:0:0:1:0:1:0:0:0 | 0:0:0:0:0:0:0:0:0:0:0:0 | 1:1:0:0:0:0:0:0:1:0:0:0 | 0:0:0:0:0:0:0:0:0:0:0:0 | 19:3:2:1:8:2:6:0:2:4:1:1 | 0:0:0:0:0:0:0:0:0:0:0:0 |
| 3:2:0:0:4:0:1:1:0:0:0:0:0:0 | 0:0:0:0:0:0:0:0:0:0:0:0:0:0 | 2:2:0:0:1:0:0:0:0:0:0:0:0:0 | 0:0:0:0:0:0:0:0:0:0:0:0:0:0 | 24:82:1:9:9:1:10:3:0:1:4:7:1:1 | 0:0:0:0:0:0:0:0:0:0:0:0:0:0 |
| 98:00:01 | 1:00:00 | 10:00:00 | 0:00:00 | 224:12:05 | 2:00:00 |
| 98:00:02 | 0:00:00 | 10:00:00 | 0:00:00 | 219:09:05 | 0:00:01 |
| 96:00:07 | 0:00:00 | 10:01:00 | 0:00:00 | 214:04:23 | 1:00:00 |
| 85:09:03 | 0:00:00 | 10:02:02 | 0:00:00 | 212:43:23 | 0:00:00 |
| 0 | 0 | 0 | 0 | 0 | 2 |
| 0:00:00 | 8:01:00 | 0:00:00 | 10:00:00 | 0:00:00 | 11:02:00 |
| 8:01:00 | 0:00:00 | 10:00:00 | 0:00:00 | 11:02:00 | 0:00:00 |
| 0 | 0 | 0 | 0 | 0 | 4 |
| 0:00 | 0:00 | 0:00 | 0:00 | 0:00 | 1:02 |
| 0:0:0:0 | 0:0:0:0 | 0:0:0:0 | 0:1:0:0 | 0:0:0:0 | 0:0:1:1 |
| 0 | 0 | 0 | 0 | 0 | 0 |
| 0 | 0 | 0 | 0 | 0 | 0 |
| 0 | 0 | 0 | 0 | 0 | 0 |
| 0 | 0 | 0 | 0 | 0 | 0 |
| 0 | 0 | 0 | 0 | 0 | 0 |
| 0 | 0 | 0 | 0 | 0 | 1 |
| 0:00:00 | 0:00:00 | 0:00:01 | 0:00:00 | 1:00:00 | 0:00:00 |
| 1:0:2:0:0:0:2:0 | 0:0:0:0:0:0:0:0 | 0:1:3:1:0:0:3:1 | 0:0:0:0:0:0:0:0 | 0:1:6:0:0:0:5:0 | 0:0:0:0:0:0:0:0 |
| 0 | 1 | 0 | 0 | 0 | 4 |
| 0 | 2 | 1 | 0 | 2 | 4 |
| 0 | 3 | 1 | 0 | 2 | 4 |
| 0 | 0 | 0 | 0 | 0 | 0 |
| 0 | 0 | 0 | 0 | 0 | 0 |
| 0 | 0 | 0 | 0 | 0 | 0 |
| 0 | 0 | 0 | 0 | 0 | 0 |
| 0:25:0:1:0:0:0:0:0:0:0:0:0 | 0:0:0:0:0:0:0:0:0:0:0:0:0 | 1:28:3:2:8:0:0:0:1:1:0:0:0 | 2:1:0:0:0:0:0:0:0:0:0:0:0 | 5:60:13:2:18:2:4:16:7:4:1:3:1 | 6:2:4:0:0:0:0:0:0:0:0:0:0 |
| 24:0:1:0:0:0:0:0:1:0:0:0 | 1:0:0:0:0:0:0:0:0:0:0:0 | 25:3:1:18:0:0:1:1:1:0:0:0 | 0:0:0:0:0:0:0:0:0:0:0:0 | 59:12:2:49:2:16:6:4:6:1:3:2 | 2:0:0:0:0:0:0:0:0:0:0:0 |
| 2:3:0:1 | 1:0:0:0 | 79:3:3:1 | 0:0:0:0 | 11:7:0:0 | 0:0:0:0 |
| 3:02:00 | 0:00:00 | 95:01:06 | 0:00:00 | 11:03:00 | 0:00:00 |
| 4:00 | 0:00 | 4:00 | 2:00 | 4:00 | 2:00 |
| 0:01:00 | 1:01:00 | 0:00:00 | 2:00:00 | 0:00:00 | 12:02:00 |
| 1:00 | 0:00 | 10:00 | 0:00 | 27:00:00 | 0:00 |
| 0:0:0:0:0:0:1 | 0:0:0:0:0:0:0 | 0:0:0:0:0:0:0 | 0:0:0:0:0:0:0 | 0:0:0:0:0:0:0 | 0:0:0:0:0:0:0 |
| 0:0:0:1:0:0 | 0:0:0:0:0:0 | 0:0:0:0:0:0 | 0:0:0:0:0:0 | 0:0:0:0:1:1 | 0:0:0:0:0:0 |
| 0 | 0 | 9 | 0 | 34 | 2 |
| 0 | 0 | 9 | 2 | 34 | 6 |
| 0 | 2 | 0 | 3 | 0 | 7 |
| 0:00 | 0:00 | 0:00 | 0:00 | 0:00 | 1:01 |
| 52:35:11:3:6:3:2:2:1:5:1:0 | 0:0:0:0:0:0:0:0:0:0:0:0 | 22:15:3:0:1:1:3:1:1:0:0:0 | 0:0:0:0:0:0:0:0:0:0:0:0 | 135:190:22:2:11:10:10:1:5:7:2:2 | 0:2:1:0:0:0:0:0:0:0:0:0 |
| 14 | 0 | 10 | 0 | 61 | 6 |

| | | | | | |
|---|---|---|---|---|---|
| 16:6:2:3:3:2:1:2:1:3:1:1:0:4:0:0:0:0:0:0:0:0:0:3:0:1:0:0:0:0:0:0:1:1:0:1:0:1:0:1:0:0 | 0:0:0:0:0:0:0:0:0:0:0:0:0:0:0:0:0:0:0:0:0:0:0:0:0:0:0:0:0:0:0:0:0:0:0:0:0:0:0:0:0:0 | 0:0:0:0:6:0:0:0:0:2:3:0:0:4:0:0:0:1:0:0:0:0:0:0:0:0:0:0:0:0:0:0:0:0:0:0:1:1:0:0:0:0 | 0:0:0:0:0:0:0:0:0:0:0:0:0:0:0:0:0:0:0:0:0:0:0:0:0:0:0:0:0:0:0:0:0:0:0:0:0:0:0:0:0:0 | 26:19:8:4:28:2:3:1:1:12:2:4:1:13:0:2:5:2:1:0:1:1:1:18:1:1:1:5:0:1:1:0:3:0:0:0:6:5:1:0:2:1 | 1:0:0:0:1:0:0:0:0:0:0:0:0:0:0:0:0:0:0:0:0:0:0:0:0:0:0:0:0:0:0:0:0:0:0:0:0:0:0:0:0:0:0:0 |
| 16:6:2:3:3:2:1:2:1:4:1:3:1:0:4:0:0:0:1:0:0:0:0:0:0:0:1:0:0:0:0:0:0:1:0:1:0:1:0:0:1:0:0 | 0:0:0:0:0:0:0:0:0:0:0:0:0:0:0:0:0:0:0:0:0:0:0:0:0:0:0:0:0:0:0:0:0:0:0:0:0:0:0:0:0:0:0 | 0:2:1:0:6:0:0:0:0:6:3:0:0:0:4:0:0:0:0:1:1:0:0:0:0:0:0:0:0:0:0:0:0:0:0:0:2:1:0:0:0:0:0 | 0:0:0:0:0:0:0:0:0:0:0:0:0:0:0:0:0:0:0:0:0:0:0:0:0:0:0:0:0:0:0:0:0:0:0:0:0:0:0:0:0:0:0 | 27:23:9:6:29:2:3:1:1:45:2:20:4:1:13:0:2:5:5:2:1:0:1:1:1:1:1:1:5:0:1:1:0:0:1:0:9:6:0:1:0:2:1 | 0:0:0:0:0:0:0:0:0:0:0:0:0:0:0:0:0:1:0:0:0:0:0:0:0:0:0:0:0:0:0:0:0:0:0:0:0:0:0:0:0:0:0:0:0 |
| 16:6:2:3:3:2:1:2:5:1:1:3:1:0:4:0:0:0:1:0:0:0:0:0:0:0:1:0:0:0:0:0:0:1:0:1:0:1:0:0:1:0:0 | 0:0:0:0:0:0:0:0:0:0:0:0:0:0:0:0:0:0:0:0:0:0:0:0:0:0:0:0:0:0:0:0:0:0:0:0:0:0:0:0:0:0:0 | 0:3:1:0:6:0:0:0:10:0:3:0:0:0:4:0:0:0:0:1:1:0:0:0:0:0:0:0:0:0:0:0:0:0:0:0:2:1:0:0:0:0:0 | 0:0:0:0:0:0:0:0:0:0:0:0:0:0:0:0:0:0:0:0:0:0:0:0:0:0:0:0:0:0:0:0:0:0:0:0:0:0:0:0:0:0:0 | 27:24:9:6:29:2:3:1:61:1:2:26:4:1:13:0:2:6:6:2:1:0:1:1:1:1:1:1:5:0:1:1:0:0:1:0:11:6:0:1:0:2:1 | 0:0:0:0:0:0:0:0:3:0:0:0:0:0:0:0:0:0:0:0:0:0:0:0:0:0:0:0:0:0:0:0:0:0:0:0:0:0:0:0:0:0:0:0:0 |
| 16:0:10:3:3:2:1:2:6:0:1:3:1:0:0:4:0:0:0:0:0:0:0:0:0:0:1:0:0:0:0:1:0:1:0:0:0:0:0 | 0:0:0:0:0:0:0:0:0:0:0:0:0:0:0:0:0:0:0:0:0:0:0:0:0:0:0:0:0:0:0:0:0:0:0:0:0:0:0 | 0:1:1:0:6:0:0:0:13:1:0:0:18:0:0:4:0:0:1:0:0:0:0:0:0:0:0:0:0:0:1:0:1:0:1:1:0:0:0 | 0:0:0:0:0:0:0:0:0:0:0:0:0:0:0:0:0:0:0:0:0:0:0:0:0:0:0:0:0:0:0:0:0:0:0:0:0:0:0 | 27:1:17:8:29:2:3:1:71:6:1:35:97:2:5:13:0:2:17:13:1:0:1:1:1:1:1:1:5:1:0:0:1:0:5:1:1:1:1 | 0:0:1:0:0:0:0:0:0:0:0:0:0:0:0:0:0:0:0:0:0:0:0:0:0:0:0:0:0:0:0:0:0:0:0:0:0:0:0:0 |
| 16:0:11:3:2:2:1:2:6:0:1:3:2:0:0:4:0:0:0:0:0:0:0:0:0:0:1:0:0:0:0:1:0:1:0:0:0:0:0 | 0:0:0:0:1:0:0:0:0:0:0:0:0:0:0:0:0:0:0:0:0:0:0:0:0:0:0:0:0:0:0:0:0:0:0:0:0:0:0 | 0:1:1:0:6:0:0:0:13:1:0:0:19:0:0:4:0:0:1:0:0:0:0:0:0:0:0:0:0:0:1:0:1:0:0:2:0:0:0 | 0:0:0:0:0:0:0:0:0:0:0:0:0:0:0:0:0:0:0:0:0:0:0:0:0:0:0:0:0:0:0:0:0:0:0:0:0:0:0 | 27:1:19:8:29:2:3:1:71:6:1:37:115:2:6:13:0:2:22:14:2:0:1:1:1:1:1:1:5:1:0:0:1:0:5:2:1:1:1 | 0:0:0:0:0:0:0:0:0:0:0:0:1:0:0:0:0:0:0:0:0:0:0:0:0:0:0:0:0:0:0:0:0:0:0:0:0:0:0:0 |
| 16:0:11:3:3:2:1:2:6:1:0:3:0:2:0:0:4:0:0:0:0:0:0:0:0:0:0:0:0:1:0:1:0:1:0:0:0:0:1 | 0:0:0:0:0:0:0:0:0:0:0:0:0:0:0:0:0:0:0:0:0:0:0:0:0:0:0:0:0:0:0:0:0:0:0:0:0:0:0 | 0:1:5:0:6:0:0:0:13:0:2:0:1:22:0:0:4:0:0:1:0:0:0:0:0:0:0:0:0:0:0:0:1:0:0:2:0:0:0 | 0:0:0:0:0:0:0:0:0:0:0:0:0:0:0:0:0:0:0:0:0:0:0:0:0:0:0:0:0:0:0:0:0:0:0:0:0:0:0 | 27:1:36:8:29:2:3:1:71:1:0:39:6:132:2:8:13:0:1:23:14:2:0:1:1:1:1:1:5:1:1:0:4:0:5:3:1:1:0 | 0:0:0:0:0:0:0:0:0:0:0:0:0:0:0:0:0:0:0:0:0:0:0:0:0:0:0:0:0:0:0:0:0:0:0:0:0:0:0:0 |
| 16:0:2:3:2:2:10:6:1:0:2:0:0:4:0:0:0:0:0:0:1:0:0:0:1:0:0:0:0:1:1:0:0 | 0:0:0:0:0:0:0:0:0:0:0:0:0:0:0:0:0:0:0:0:0:0:0:0:0:0:0:0:0:0:0:0:0 | 0:1:0:6:0:0:3:13:0:2:24:1:0:4:0:1:0:0:0:0:0:0:0:0:0:1:2:0:0:0:0:0:0 | 0:0:0:0:0:0:0:0:0:0:0:0:0:0:0:0:0:0:0:0:0:0:0:0:0:0:0:0:0:0:0:0:0 | 27:1:6:28:2:1:32:69:1:0:137:6:8:13:1:24:1:1:1:1:0:1:5:1:0:4:3:1:1:0:0:1:1 | 0:0:0:1:0:0:0:1:0:0:0:0:0:0:0:0:0:0:0:0:0:0:0:0:0:0:0:0:0:0:0:0:0:0 |
| 3:16:0:11:3:3:2:1:2:6:1:0:3:0:1:0:4:0:0:0:0:0:0:0:0:0:0:1:0:0:0:1:0:1:0:0:0:0:0:1 | 0:0:0:0:0:0:0:0:0:0:0:0:0:0:0:0:0:0:0:0:0:0:0:0:0:0:0:0:0:0:0:0:0:0:0:0:0:0:0:0 | 24:0:1:3:0:6:0:0:0:13:0:2:0:0:1:0:4:0:0:1:0:0:0:0:0:0:0:0:0:0:0:0:1:0:1:2:0:0:0:0 | 0:0:0:0:0:0:0:0:0:0:0:0:0:0:0:0:0:0:0:0:0:0:0:0:0:0:0:0:0:0:0:0:0:0:0:0:0:0:0:0 | 144:26:1:37:7:29:2:3:1:69:1:1:40:2:6:8:13:0:1:24:14:2:0:1:1:1:1:1:1:5:1:0:4:0:4:3:1:0:1:0 | 2:1:0:0:0:0:0:0:0:0:0:0:0:0:0:0:0:0:0:0:0:0:0:0:0:0:0:0:0:0:0:0:0:0:0:0:0:0:0:0:0:0:0 |
| 15:00:01 | 0:00:00 | 32:01:01 | 18:00:00 | 44:04:00 | 3:01:00 |
| 14:00:01 | 1:00:00 | 45:00:00 | 10:01:01 | 49:04:00 | 5:01:00 |
| 16:00:01 | 3:00:00 | 59:01:01 | 2:00:00 | 59:05:01 | 6:01:00 |
| 19:01:00 | 1:00:00 | 65:01:01 | 1:00:00 | 62:03:06 | 4:00:00 |
| 20:00:01 | 0:00:00 | 68:01:01 | 1:00:00 | 66:06:03 | 1:00:00 |
| 0:20:01 | 0:00:00 | 0.089594907 | 0:00:00 | 0.2965625 | 0:00:00 |
| 22:1:3:4:0 | 0:0:0:0:0 | 51:16:8:1:0 | 0:0:0:0:0 | 65:20:20:1:1 | 0:0:0:0:0 |
| 23:3:1:4:0 | 0:0:0:0:0 | 53:8:15:1:0 | 0:0:0:0:0 | 69:19:20:1:1 | 0:0:0:0:0 |
| 22:1:1:4:0 | 0:0:0:0:0 | 53:5:14:1:0 | 0:0:0:0:0 | 69:9:20:1:1 | 0:0:0:0:0 |
| 22:1:1:4:0 | 0:0:0:0:0 | 53:3:14:1:0 | 0:0:0:0:0 | 69:6:20:1:1 | 0:0:0:0:0 |
| 22:1:1:4:0 | 0:0:0:0:0 | 53:3:14:1:0 | 0:0:0:0:0 | 69:5:20:1:1 | 0:0:0:0:0 |
| 23:1:4:0:0 | 0:0:0:0:0 | 53:14:1:1:0 | 0:0:0:0:0 | 69:20:1:2:1 | 0:0:0:0:0 |
| 23:1:4:0:0 | 0:0:0:0:0 | 53:10:1:0:0 | 0:0:0:0:0 | 69:17:1:1:1 | 0:0:0:0:0 |
| 72:17:00 | 0:00:00 | 104:12:01 | 2:00:00 | 481:63:2 | 2:00:01 |
| 80:17:00 | 2:00:00 | 151:12:01 | 1:00:00 | 678:62:2 | 9:00:01 |
| 87:17:00 | 0:00:00 | 194:12:01 | 1:00:00 | 1032:62:3 | 7:00:00 |
| 91:17:00 | 0:00:00 | 204:12:01 | 0:00:00 | 1074:62:3 | 5:00:00 |
| 89:17:00 | 0:00:00 | 199:10:01 | 0:00:00 | 1047:59:03 | 6:00:00 |
| 28:07:00 | 1:00:00 | 100:02:01 | 61:04:00 | 505:18:00 | 264:15:02 |
| 27:07:00 | 0:00:00 | 95:02:01 | 0:00:00 | 479:15:00 | 5:00:00 |
| 24:04:00 | 2:02:00 | 70:02:00 | 24:00:01 | 356:10:00 | 115:03:00 |
| 8:01:00 | 18:05:00 | 84:00:01 | 10:02:00 | 400:13:00 | 70:01:00 |
| 27:07:00 | 0:00:00 | 94:02:01 | 0:00:00 | 476:15:00 | 2:00:00 |
| 5:01 | 2:00 | 20:00 | 4:00 | 84:04:00 | 13:00 |
| 5:01 | 2:00 | 23:00 | 1:00 | 87:04:00 | 9:00 |
| 3:00 | 0:00 | 0:00 | 3:00 | 1:00 | 1:00 |
| 2:00 | 0:00 | 0:00 | 3:00 | 1:00 | 1:00 |

| | | | | | |
|---|---|---|---|---|---|
| 2:00 | 0:00 | 0:00 | 3:00 | 1:00 | 1:00 |
| 4:6:0:6:1:12:1:0:0:0 | 0:0:0:0:0:0:0:0:0:0 | 1:1:0:0:0:1:0:0:0:0 | 0:0:0:0:0:0:0:0:0:0 | 11:13:10:5:5:33:2:0:4:2 | 0:0:0:0:0:0:0:0:0:0 |
| 141 | 1 | 85 | 0 | 421 | 0 |
| 142 | 0 | 86 | 0 | 426 | 0 |
| 143 | 0 | 86 | 0 | 425 | 2 |
| 143 | 0 | 86 | 0 | 425 | 2 |
| 0:00:00 | 2:00:00 | 0:00:00 | 0:01:00 | 0:00:00 | 4:01:01 |
| 12:00 | 0:00 | 58:01:00 | 0:00 | 185:01:00 | 2:00 |
| 11:01 | 1:00 | 55:02:00 | 0:00 | 182:04:00 | 2:00 |
| 0:0:0:0:1 | 0:0:0:0:0 | 3173:221:37:0:0 | 4:0:0:0:0 | 76:2:1:0:0 | 0:0:0:0:0 |
| 0:0:0:0:1 | 0:0:0:0:0 | 3175:220:37:0:0 | 4:2:0:0:0 | 76:2:1:0:0 | 0:0:0:0:0 |
| 0:0:0:0:1 | 0:0:0:0:0 | 3179:222:37:0:0 | 3:1:0:0:0 | 76:2:1:0:0 | 0:0:0:0:0 |
| 0:0:0:0:1 | 0:0:0:0:0 | 3178:223:37:0:0 | 4:0:0:0:0 | 76:2:1:0:0 | 0:0:0:0:0 |
| 0:0:0:0:1 | 0:0:0:0:0 | 3179:219:37:0:0 | 4:1:0:0:0 | 76:2:1:0:0 | 0:0:0:0:0 |
| 0:0:0:0:1 | 0:0:0:0:0 | 3183:220:37:0:0 | 0:0:0:0:0 | 76:2:1:0:0 | 0:0:0:0:0 |
| 0:0:0:0:1 | 0:0:0:0:0 | 3184:202:37:0:0 | 1:0:0:0:0 | 76:2:1:0:0 | 0:0:0:0:0 |
| 0:0:0:0:1 | 0:0:0:0:0 | 3189:204:37:0:0 | 1:0:0:0:0 | 77:2:1:0:0 | 0:0:0:0:0 |
| 0:0:0:1 | 0:0:0:0 | 3190:166:37:0 | 3:0:0:0 | 77:2:1:0 | 0:0:0:0 |
| 0:0:0:1 | 0:0:0:0 | 3193:137:37:0 | 1:4:0:0 | 77:1:1:0 | 0:1:0:0 |
| 0:0:0:1 | 0:0:0:0 | 3195:130:37:0 | 1:3:0:0 | 77:1:1:0 | 0:0:0:0 |
| 0:0:0:1 | 0:0:0:0 | 3201:108:0:0 | 2:3:0:0 | 78:0:0:0 | 0:0:0:0 |
| 0:0:0:1 | 0:0:0:0 | 3199:89:0:0 | 4:2:0:0 | 77:0:0:0 | 1:0:0:0 |
| 0:0:0:1 | 0:0:0:0 | 3197:72:0:0 | 6:0:0:0 | 78:0:0:0 | 0:0:0:0 |
| 0:0:0:1 | 0:0:0:0 | 3202:67:0:0 | 1:0:0:0 | 78:0:0:0 | 0:0:0:0 |
| 0:0:0:1 | 0:0:0:0 | 3201:55:0:0 | 2:2:0:0 | 78:0:0:0 | 0:0:0:0 |
| 0:0:0:1 | 0:0:0:0 | 3197:55:0:0 | 6:0:0:0 | 77:0:0:0 | 1:0:0:0 |
| 0:0:0:1 | 0:0:0:0 | 3203:51:0:0 | 0:2:0:0 | 78:0:0:0 | 0:0:0:0 |
| 0:00:00 | 0:01:00 | 0:00:00 | 0:01:01 | 0:00:00 | 0:02:00 |
| 39:45:04 | 0:01:00 | 39:12:05 | 0:00:00 | 145:248:15 | 1:02:00 |
| 0 | 0 | 0 | 0 | 0 | 2 |
| 9:3:34:5:0:0:0:1 | 1:0:1:0:0:0:0:0 | 23:22:27:2:3:0:0:2 | 0:0:0:0:0:0:0:0 | 94:38:56:12:4:0:0:3 | 0:0:1:0:0:0:0:0 |
| 58:9:3:8:1:2:0:0:0 | 3:0:0:0:0:0:0:0:0 | 42:27:22:7:4:2:0:0:0 | 15:1:1:5:0:0:1:1:0 | 99:95:44:17:4:3:0:0:1 | 34:5:5:8:0:0:0:1:0 |
| 10:4:72:8:1:0:0:2:0 | 0:0:0:0:0:0:0:0:0 | 28:23:58:12:4:1:1:2:0 | 0:0:1:0:0:0:0:0:0 | 102:49:141:25:4:0:1:3:1 | 0:0:4:0:0:0:0:0:0 |
| 9:4:71:8:1:0:0:2:0 | 0:0:1:0:0:0:0:0:0 | 26:23:61:12:5:1:1:2:0 | 0:0:0:0:0:0:0:0:0 | 105:50:152:26:4:0:1:4:1 | 0:0:1:0:0:0:0:0:0 |
| 9:4:72:8:1:0:0:2:0 | 0:0:0:0:0:0:0:0:0 | 26:23:61:12:5:1:1:2:0 | 0:0:0:0:0:0:0:0:0 | 105:50:152:26:4:0:1:4:1 | 0:0:1:0:0:0:0:0:0 |
| 9:4:73:8:0:1:0:2:0 | 0:0:0:0:0:0:0:0:0 | 25:23:61:12:1:5:1:2:0 | 0:0:2:0:0:0:0:0:0 | 104:50:152:26:1:4:0:4:1 | 0:0:1:0:0:0:0:0:0 |
| 10:15:7:4:73:1:0:2:0 | 0:1:0:0:0:0:0:0:0 | 32:22:34:31:140:7:2:2:0 | 0:0:0:0:0:0:0:0:0 | 50:70:113:81:270:10:0:4:1 | 0:1:0:0:0:0:0:0:0 |
| 7:4:73:10:16:1:0:2:0 | 0:0:0:0:0:0:0:0:0 | 34:31:140:32:22:8:2:2:0 | 0:0:0:0:0:0:0:0:0 | 113:84:271:51:72:10:0:4:1 | 0:0:0:0:0:0:0:0:0 |
| 5:4:73:10:16:1:0:2:0 | 0:0:0:0:0:0:0:0:0 | 35:31:140:32:21:8:2:2:0 | 0:0:0:0:0:0:0:0:0 | 108:83:271:51:77:10:0:4:1 | 0:1:0:0:0:0:0:0:0 |
| 4:2:73:10:16:1:0:2:0 | 0:0:0:0:0:0:0:0:0 | 31:28:140:32:22:8:2:2:0 | 0:0:0:0:1:0:0:0:0 | 84:72:270:51:76:10:0:4:1 | 0:0:1:0:1:0:0:0:0 |
| 4:2:73:10:16:1:0:2:0 | 0:0:0:0:0:0:0:0:0 | 31:25:139:32:22:8:2:2:0 | 0:1:1:0:2:0:0:0:0 | 84:70:271:51:76:10:0:4:1 | 0:2:0:0:3:0:0:0:0 |
| 4:2:73:10:17:1:0:2:0 | 0:0:0:0:0:0:0:0:0 | 30:21:140:32:25:8:3:2:0 | 1:0:0:0:0:0:0:0:0 | 84:58:271:53:81:11:0:4:1 | 0:0:0:0:0:0:0:0:0 |
| 4:2:73:10:17:1:0:2:0 | 0:0:0:0:0:0:0:0:0 | 31:16:139:32:25:8:3:2:0 | 0:0:1:0:1:0:0:0:0 | 83:45:271:54:81:11:0:4:1 | 1:0:0:0:1:0:0:0:0 |

| | | | | | |
|---|---|---|---|---|---|
| 4:2:73:10:17:1:0:2:0 | 0:0:0:0:0:0:0:0:0 | 31:14:140:32:26:8:3:2:0 | 0:0:0:0:0:0:0:0:0 | 83:43:271:53:82:11:0:4:1 | 1:0:0:1:0:0:0:0:0 |
| 4:2:73:10:17:1:2:0:0 | 0:0:0:0:0:0:0:0:0 | 31:11:140:32:26:8:2:1:0 | 0:0:0:0:0:0:0:0:0 | 84:28:270:54:82:11:4:0:1 | 0:0:1:0:0:0:0:0:0 |
| 4:73:10:17:1:2:2:0:0 | 0:0:0:0:0:0:0:0:0 | 31:140:32:26:8:4:2:1:0 | 0:0:0:0:0:0:0:0:0 | 84:269:54:82:11:17:4:0:1 | 0:2:0:0:0:0:0:0:0 |
| 549:24:00 | 1:01 | 867:174 | 2:01 | 1494:228 | 2:02 |
| 550:24:00 | 0:01 | 868:166 | 1:01 | 1498:213 | 2:01 |
| 549:19:00 | 0:00 | 874:122 | 3:03 | 1505:169 | 1:01 |
| 551:15:00 | 0:00 | 882:72 | 3:00 | 1515:119 | 1:01 |
| 206:03:00 | 345:12:00 | 429:27:00 | 457:37:00 | 378:19:00 | 1138:92 |
| 0:00 | 551:15:00 | 0:00 | 886:58:00 | 0:00 | 1518:102 |
| 551:02:00 | 0:00 | 867:05:00 | 2:00 | 1483:08:00 | 5:00 |
| 542:16:00 | 0:00 | 700:41:00 | 3:01 | 1217:66 | 2:01 |
| 1.068055556 | 0:01 | 40:602 | 1:03 | 69:1073 | 2:00 |
| 513:18:00 | 1:00 | 563:40:00 | 5:00 | 1020:69 | 3:02 |
| 430:17:00 | 0:00 | 214:24:00 | 1:00 | 494:51:00 | 2:00 |
| 376:15:00 | 0:00 | 115:19:00 | 2:00 | 374:38:00 | 0:00 |
| 0:00 | 254:11:00 | 0:00 | 12:05 | 1:00 | 190:23:00 |
| 197:06:00 | 1:00 | 5:04 | 0:00 | 154:16:00 | 0:00 |
| 15:00 | 0:00 | 0:01 | 0:00 | 38:00:00 | 0:00 |
| 10:00:01 | 0:00:00 | 1:00:00 | 0:00:00 | 12:00:03 | 2:00:00 |
| 16:00:01 | 6:00:00 | 19:00:00 | 23:00:00 | 47:00:08 | 81:00:03 |
| 22:00:01 | 0:00:00 | 39:00:00 | 5:00:00 | 110:00:11 | 21:00:00 |
| 22:00:01 | 0:00:00 | 46:00:01 | 0:00:00 | 130:00:12 | 2:00:00 |
| 998:6:13:2 | 12:0:0:0 | 4667:1:13:0 | 256:0:2:0 | 8480:6:17:5 | 362:1:1:0 |
| 1002:13:6:2 | 26:0:1:0 | 4686:13:1:0 | 3834:13:18:0 | 8525:17:7:5 | 7163:21:22:5 |
| 1026:7:14:2 | 2:0:0:0 | 8801:20:27:0 | 4:0:0:0 | 16065:23:38:10 | 2:0:0:0 |
| 1026:14:14:2 | 3:0:0:0 | 8828:27:138:0 | 6:0:0:0 | 16092:38:186:10 | 13:0:1:0 |
| 1027:14:14:2 | 0:0:0:0 | 8817:27:137:0 | 5:0:1:0 | 16064:38:189:9 | 22:0:0:1 |
| 1024:14:14:2 | 3:0:0:0 | 8817:27:138:0 | 4:0:0:0 | 16080:38:189:10 | 5:0:0:0 |
| 1017:14:14:2 | 0:0:0:0 | 8777:27:136:0 | 7:0:1:0 | 15990:36:185:9 | 14:1:0:0 |
| 1019:14:14:2 | 1:0:0:0 | 8783:27:137:0 | 12:0:0:0 | 16019:37:187:9 | 11:0:1:0 |
| 43 | 0 | 295 | 0 | 69 | 0 |
| 43 | 0 | 304 | 1 | 72 | 0 |
| 62 | 0 | 370 | 0 | 168 | 0 |
| 85 | 0 | 48 | 0 | 291 | 2 |
| 83 | 1 | 52 | 0 | 293 | 0 |
| 183 | 0 | 476 | 1 | 864 | 2 |
| 183 | 0 | 468 | 3 | 861 | 1 |
| 182 | 0 | 473 | 1 | 863 | 0 |
| 180 | 0 | 449 | 0 | 842 | 1 |
| 0:00 | 0:00 | 0:00 | 1:01 | 0:00 | 0:00 |
| 0:00:00 | 0:00:00 | 0:01:00 | 0:00:00 | 0:00:02 | 2:00:00 |
| 0:00:00 | 0:00:00 | 1:00:00 | 0:00:00 | 6:03:04 | 2:00:00 |
| 0:00 | 0:00 | 0:00 | 0:00 | 0:00 | 1:00 |
| 21:1:1:0 | 0:0:0:0 | 64:11:5:0 | 0:0:0:0 | 120:2:8:1 | 1:0:0:0 |
| 1:17:1:0 | 0:0:0:0 | 6:35:12:0 | 5:24:3:0 | 1:50:5:1 | 1:59:24:0 |
| 0:01:00 | 0:00:00 | 0:01:00 | 1:00:01 | 0:00:00 | 1:02:01 |
| 0 | 0 | 0 | 0 | 0 | 3 |
| 0:00 | 3:00 | 0:00 | 0:01 | 0:00 | 9:00 |
| 0:2:0:0:0 | 0:0:0:0:0 | 0:1:0:0:0 | 0:2:0:0:0 | 1:1:1:0:0 | 2:1:0:1:1 |
| 0:2:0:0:0 | 0:0:0:0:0 | 0:1:0:0:0 | 0:2:0:0:0 | 2:1:0:1:0 | 2:1:0:0:1 |
| 0 | 0 | 0 | 0 | 0 | 2 |
| 2 | 0 | 2 | 0 | 8 | 0 |
| 0:00 | 0:00 | 0:00 | 1:00 | 0:00 | 2:01 |
| 1 | 0 | 3 | 0 | 1 | 1 |
| 865 | 1 | 2240 | 8 | 2020 | 2 |
| 869 | 3 | 2259 | 3 | 2039 | 4 |
| 0:00 | 1:01 | 0:00 | 1:00 | 0:00 | 2:00 |
| 0 | 0 | 0 | 1 | 0 | 2 |
| 2:4:0:0:4:0:1 | 0:0:0:0:0:0:0 | 1:0:0:4:2:0:1 | 0:1:0:0:0:0:0 | 5:17:0:8:9:0:1 | 0:0:0:0:0:0:0 |
| 28:1:1:2:0:1:1:2 | 0:0:0:0:0:0:0:0 | 28:3:1:0:4:2:0:0 | 0:0:0:0:0:0:0:0 | 141:7:11:5:10:11:1:2 | 0:0:0:0:0:0:0:0 |
| 0:0:0:0:0:0:0:0 | 2:5:0:0:0:1:0:0 | 5:0:0:0:0:0:0:0 | 3:0:0:1:0:0:0:0 | 9:3:1:2:0:2:1:1 | 8:2:2:2:0:0:1:0 |
| 0:0:1:0:0:0:0:0:0 | 5:2:2:0:0:1:2:0:0 | 0:2:6:0:0:0:0:0:0 | 0:1:1:0:1:0:0:0:0 | 3:4:8:0:1:0:1:1:0 | 2:2:6:0:0:0:1:0:1 |
| 0:0:0:1:0:0:1 | 0:0:0:0:0:0:0 | 0:0:0:2:0:1:0 | 0:1:1:0:1:0:0 | 0:0:0:2:1:2:0 | 0:0:0:1:1:0:0 |
| 0:0:0:0:0:0:0:0:0:0 | 2:1:0:0:2:2:1:0:1:0 | 1:0:6:1:0:1:2:1:0:0 | 0:1:0:1:4:2:1:0:2:0 | 3:0:4:0:3:5:5:2:2:1 | 1:1:4:0:0:4:3:2:0:0 |
| 0 | 0 | 0 | 0 | 0 | 0 |
| 0 | 0 | 0 | 0 | 0 | 0 |
| 0 | 0 | 0 | 0 | 0 | 0 |

| | | | | | |
|---|---|---|---|---|---|
| 0 | 0 | 0 | 0 | 0 | 0 |
| 0 | 1 | 2 | 0 | 2 | 1 |
| 1 | 0 | 0 | 2 | 1 | 2 |
| 0 | 0 | 0 | 0 | 1 | 0 |
| 0:2:0:0 | 0:1:0:0 | 2:7:2:0 | 1:1:0:0 | 4:12:6:0 | 2:6:1:0 |
| 0:3:0:0 | 0:0:0:0 | 3:8:2:0 | 0:0:0:0 | 6:18:8:0 | 0:0:0:0 |
| 0 | 0 | 0 | 0 | 0 | 0 |
| 0:1:0:0 | 0:0:0:0 | 0:0:0:0 | 0:0:0:0 | 3:0:0:0 | 2:0:1:1 |
| 9:11:19:8:8:3:0:1:0 | 4:6:0:0:0:0:0:0:0 | 37:57:10:7:1:0:1:0:1 | 16:12:1:0:0:0:0:0:0 | 104:79:43:11:11:6:0:2:2 | 28:29:0:0:0:0:0:0:0 |
| 19:8:8:14:17:3:0:1:0 | 0:0:0:0:0:0:0:0:0 | 11:7:1:55:72:0:1:0:0 | 0:0:0:3:2:0:0:0:0 | 43:11:11:143:125:6:0:2:2 | 0:0:0:2:4:0:0:0:0 |
| 14:17:19:8:8:3:0:1:0 | 1:0:0:0:0:0:0:0:0 | 55:75:11:7:1:0:1:0:0 | 4:0:0:0:0:0:0:0:0 | 143:126:43:11:11:6:0:2:2 | 6:9:0:0:0:0:0:0:0 |
| 19:8:8:15:17:3:0:1:0 | 0:0:0:0:0:0:0:0:0 | 12:7:1:60:75:0:1:0:0 | 0:0:0:0:3:0:1:0:0 | 42:11:11:152:143:6:0:2:2 | 0:0:0:2:2:0:0:0:0 |
| 4402 | 4 | 7429 | 3 | 17660 | 8 |
| 4401 | 5 | 7429 | 4 | 17655 | 18 |
| 4413 | 1 | 7456 | 5 | 17668 | 8 |
| 4412 | 4 | 7457 | 6 | 17655 | 14 |
| 4398 | 5 | 7413 | 41 | 17536 | 127 |
| 4385 | 7 | 7422 | 14 | 17627 | 8 |
| 4357 | 8 | 7362 | 10 | 17473 | 20 |
| 4344 | 3 | 7361 | 5 | 17464 | 11 |
| 4330 | 2 | 7346 | 6 | 17410 | 17 |
| 0:00 | 0:00 | 0:00 | 3:00 | 0:00 | 9:01 |
| 177 | 0 | 103 | 0 | 493 | 0 |
| 177 | 0 | 106 | 0 | 490 | 1 |
| 175 | 0 | 109 | 0 | 495 | 1 |
| 175 | 0 | 109 | 0 | 494 | 2 |
| 0:0:0:0:0:0:1 | 0:0:0:0:0:0:0 | 0:9:0:0:2:3:1 | 0:0:0:0:0:0:0 | 1:0:0:1:0:0:0 | 0:0:0:0:0:0:0 |
| 0:0:0:0:0:0:0 | 0:0:0:0:0:0:1 | 4:0:1:0:2:0:0 | 5:0:1:0:1:0:1 | 0:1:0:0:0:0:0 | 0:0:0:0:0:0:0 |
| 0 | 2 | 0 | 29 | 0 | 30 |
| 0:00:00 | 2:00:00 | 0:00:00 | 5:00:02 | 0:00:00 | 7:00:00 |
| 0:00:00 | 0:00:00 | 0:00:00 | 0:00:00 | 0:00:04 | 0:00:01 |
| 0 | 0 | 0 | 0 | 3 | 0 |
| 1 | 0 | 2 | 0 | 1 | 0 |
| 29:0:1:0:0 | 0:0:0:0:0 | 49:4:2:1:1 | 0:0:0:0:0 | 100:7:2:4:4 | 2:0:0:0:0 |
| 0:00:00 | 0:00:00 | 0:01:00 | 0:00:00 | 0:00:01 | 0:00:00 |
| 0:00:00 | 0:00:00 | 0:01:00 | 0:00:00 | 0:00:01 | 0:00:00 |
| 0:00:00 | 0:00:01 | 0:00:00 | 0:00:00 | 0:00:00 | 0:00:00 |
| 0:0:0:1:0:0:0:0:0 | 0:0:0:0:0:0:0:0:0 | 0:1:0:1:0:0:1:0:0 | 0:0:0:0:0:0:0:0:0 | 0:1:0:0:1:0:1:0:0 | 0:0:0:0:0:0:0:0:0 |
| 0:0:0:0:0 | 0:0:0:0:0 | 0:0:0:0:0 | 0:0:1:0:0 | 0:0:0:0:0 | 0:0:0:0:1 |
| 0:00 | 1:00 | 3:00 | 3:00 | 2:00 | 3:00 |
| 77:1:0:0 | 0:0:0:0 | 63:0:0:0 | 0:0:0:0 | 277:3:4:1 | 0:0:0:0 |
| 77:1:0:0 | 0:0:0:0 | 63:0:0:0 | 0:0:0:0 | 276:3:4:1 | 1:0:0:0 |
| 76:1:0:0 | 1:0:0:0 | 64:0:0:0 | 0:0:0:0 | 278:3:4:1 | 0:0:0:0 |
| 77:1:0:0 | 0:0:0:0 | 65:0:0:0 | 0:0:0:0 | 276:3:4:2 | 2:0:0:0 |
| 77:1:0:0 | 0:0:0:0 | 65:0:0:0 | 0:0:0:0 | 277:3:4:2 | 1:0:0:0 |
| 77:1:0:0 | 0:0:0:0 | 64:0:0:0 | 0:0:0:0 | 278:3:2:4 | 0:0:0:0 |
| 76:1:0:0 | 1:0:0:0 | 64:0:0:0 | 0:0:0:0 | 276:3:2:4 | 2:0:0:0 |
| 77:1:0:0 | 0:0:0:0 | 64:0:0:0 | 0:0:0:0 | 278:3:2:4 | 0:0:0:0 |
| 77:1:0:0 | 0:0:0:0 | 64:0:0:0 | 0:0:0:0 | 277:3:2:4 | 1:0:0:0 |
| 77:1:0:0 | 0:0:0:0 | 64:0:1:0 | 0:0:0:0 | 278:3:2:4 | 0:0:0:0 |
| 77:1:0:0 | 0:0:0:0 | 64:0:1:0 | 0:0:0:0 | 275:3:2:4 | 2:0:0:0 |
| 77:1:0:0 | 0:0:0:0 | 63:0:1:0 | 1:0:0:0 | 275:3:1:2 | 1:0:0:0 |
| 77:1:0:0 | 0:0:0:0 | 64:0:1:0 | 0:0:0:0 | 272:3:2:2 | 1:0:0:0 |
| 75:1:0:0:0 | 2:0:0:0:0 | 64:0:1:1:0 | 0:0:0:0:0 | 271:3:3:0:2 | 0:0:0:0:0 |
| 77:1:0:0:0 | 0:0:0:0:0 | 63:0:1:1:0 | 0:0:0:0:0 | 270:2:3:0:1 | 0:0:0:0:0 |
| 2:00:00 | 0:00:00 | 1:00:00 | 0:00:00 | 9:00:01 | 0:00:00 |
| 0:0:0:0:0:0:0:0 | 0:0:0:0:1:0:0:0 | 0:2:0:0:0:0:1:0 | 0:0:0:0:0:0:0:0 | 0:0:0:0:0:0:0:0 | 0:0:0:0:0:0:0:0 |
| 6:3:1:9:2:0:0:0 | 0:0:0:0:0:0:0:0 | 15:0:1:15:1:1:0:0 | 0:0:0:0:0:0:0:0 | 44:6:4:43:3:0:4:4 | 0:0:0:0:0:0:0:0 |
| 5:1:0:0:1:0:0:0:0:0:0:0:0 | 0:0:0:0:0:0:0:0:0:0:0:0:0 | 20:15:0:1:0:1:1:0:0:0:0:1:1 | 0:0:0:0:0:0:0:0:0:0:0:0:0 | 96:56:2:1:0:5:1:1:0:0:0:0:0 | 2:0:0:0:0:0:0:0:0:0:0:0:0 |
| 0:0:0:1:0:0:0:0:0 | 2:2:2:6:2:0:0:0:0 | 2:1:0:1:0:0:0:0:0 | 3:1:0:4:0:0:0:0:0 | 10:3:1:14:3:0:2:0:0 | 2:4:3:7:0:1:0:0:1 |
| 0 | 0 | 0 | 0 | 0 | 0 |
| 0:01 | 0:00 | 0:00 | 0:00 | 1:01 | 0:00 |
| 0 | 1 | 0 | 19 | 0 | 11 |
| 10:3:9:1:1 | 0:0:0:0:0 | 68:7:11:2:0 | 0:0:0:0:0 | 138:13:31:7:4 | 0:1:0:1:0 |
| 10:3:9:1:1 | 0:0:0:0:0 | 67:6:9:2:0 | 1:1:1:0:0 | 138:17:29:8:4 | 0:0:2:0:0 |
| 9:2:0:1:0 | 0:0:0:0:1 | 64:11:0:1:0 | 4:0:0:0:0 | 130:38:0:8:3 | 5:0:0:0:0 |
| 181 | 1 | 466 | 8 | 860 | 6 |

Supplementary Table S7: Unevenly distributed SNV isoforms within a gene

| chr | pos | ref | alt | cosmicID | annotation | gene | mutType | AAanno |
|---|---|---|---|---|---|---|---|---|
| chr1 | 112913924 | A | T | COSV63710018 | exonic | SLC16A1 | nonsynonymous SNV | SLC16A1:NM_001166496:exon5:c.T1470A:p.D490E,SLC16A1:NM_003051:exon5:c.T1470A:p.D490E, |
| chr1 | 112914011 | A | T | . | exonic | SLC16A1 | nonsynonymous SNV | SLC16A1:NM_001166496:exon5:c.T1383A:p.S461R,SLC16A1:NM_003051:exon5:c.T1383A:p.S461R, |
| chr1 | 114405659 | A | G | COSV61821233 | exonic | TRIM33 | nonsynonymous SNV | TRIM33:NM_015906:exon15:c.T2519C:p.I840T,TRIM33:NM_033020:exon15:c.T2519C:p.I840T, |
| chr1 | 114510877 | A | G | COSV61824840 | exonic | TRIM33 | nonsynonymous SNV | TRIM33:NM_015906:exon1:c.T200C:p.V67A,TRIM33:NM_033020:exon1:c.T200C:p.V67A, |
| chr1 | 114718665 | C | T | . | exonic | CSDE1 | nonsynonymous SNV | CSDE1:NM_001130523:exon19:c.G2342A:p.S781N,CSDE1:NM_001242892:exon18:c.G2297A:p.S766N,CSDE1:NM_001007553:exon19:c.G2297A:p.S766N,CSDE1:NM_001242893:exon17:c.G2204A:p.S735N,CSDE1:NM_001242891:exon20:c.G2435A:p.S812N,CSDE1:NM_007158:exon18:c.G2204A:p.S735N, |
| chr1 | 114718716 | C | T | COSV54743242 | exonic | CSDE1 | nonsynonymous SNV | CSDE1:NM_001130523:exon19:c.G2291A:p.R764Q,CSDE1:NM_001242892:exon18:c.G2246A:p.R749Q,CSDE1:NM_001007553:exon19:c.G2246A:p.R749Q,CSDE1:NM_001242893:exon17:c.G2153A:p.R718Q,CSDE1:NM_001242891:exon20:c.G2384A:p.R795Q,CSDE1:NM_007158:exon18:c.G2153A:p.R718Q, |
| chr1 | 114723883 | C | T | . | exonic | CSDE1 | nonsynonymous SNV | CSDE1:NM_001130523:exon16:c.G1918A:p.G640S,CSDE1:NM_001242892:exon15:c.G1873A:p.G625S,CSDE1:NM_001007553:exon16:c.G1873A:p.G625S,CSDE1:NM_001242893:exon14:c.G1780A:p.G594S,CSDE1:NM_001242891:exon17:c.G2011A:p.G671S,CSDE1:NM_007158:exon15:c.G1780A:p.G594S, |
| chr1 | 114723979 | C | T | . | exonic | CSDE1 | nonsynonymous SNV | CSDE1:NM_001130523:exon16:c.G1822A:p.D608N,CSDE1:NM_001242892:exon15:c.G1777A:p.D593N,CSDE1:NM_001007553:exon16:c.G1777A:p.D593N,CSDE1:NM_001242893:exon14:c.G1684A:p.D562N,CSDE1:NM_001242891:exon17:c.G1915A:p.D639N,CSDE1:NM_007158:exon15:c.G1684A:p.D562N, |
| chr1 | 114726284 | C | T | . | exonic | CSDE1 | nonsynonymous SNV | CSDE1:NM_001130523:exon14:c.G1612A:p.V538M,CSDE1:NM_001242892:exon13:c.G1567A:p.V523M,CSDE1:NM_001007553:exon14:c.G1567A:p.V523M,CSDE1:NM_001242893:exon12:c.G1474A:p.V492M,CSDE1:NM_001242891:exon15:c.G1705A:p.V569M,CSDE1:NM_007158:exon13:c.G1474A:p.V492M, |
| chr1 | 114730417 | C | T | . | exonic | CSDE1 | nonsynonymous SNV | CSDE1:NM_001130523:exon12:c.G1242A:p.M414I,CSDE1:NM_001242892:exon11:c.G1197A:p.M399I,CSDE1:NM_001007553:exon12:c.G1197A:p.M399I,CSDE1:NM_001242893:exon10:c.G1104A:p.M368I,CSDE1:NM_001242891:exon13:c.G1335A:p.M445I,CSDE1:NM_007158:exon11:c.G1104A:p.M368I, |
| chr1 | 114730647 | C | T | . | exonic | CSDE1 | nonsynonymous SNV | CSDE1:NM_001130523:exon11:c.G1097A:p.G366D,CSDE1:NM_001242892:exon10:c.G1052A:p.G351D,CSDE1:NM_001007553:exon11:c.G1052A:p.G351D,CSDE1:NM_001242893:exon9:c.G959A:p.G320D,CSDE1:NM_001242891:exon12:c.G1190A:p.G397D,CSDE1:NM_007158:exon10:c.G959A:p.G320D, |
| chr1 | 114736763 | A | T | . | exonic | CSDE1 | nonsynonymous SNV | CSDE1:NM_001130523:exon6:c.T540A:p.N180K,CSDE1:NM_001242892:exon5:c.T495A:p.N165K,CSDE1:NM_001007553:exon6:c.T495A:p.N165K,CSDE1:NM_001242893:exon4:c.T402A:p.N134K,CSDE1:NM_001242891:exon7:c.T633A:p.N211K,CSDE1:NM_007158:exon5:c.T402A:p.N134K, |
| chr1 | 116395187 | A | G | COSV55190446 | exonic | ATP1A1 | nonsynonymous SNV | ATP1A1:NM_001160233:exon13:c.A1738G:p.I580V,ATP1A1:NM_001160234:exon13:c.A1645G:p.I549V,ATP1A1:NM_000701:exon13:c.A1738G:p.I580V, |
| chr1 | 116397893 | C | A | . | exonic | ATP1A1 | nonsynonymous SNV | ATP1A1:NM_001160233:exon15:c.C1979A:p.A660D,ATP1A1:NM_001160234:exon15:c.C1886A:p.A629D,ATP1A1:NM_000701:exon15:c.C1979A:p.A660D, |
| chr1 | 13816508 | A | G | COSV52443068 | exonic | PRDM2 | nonsynonymous SNV | PRDM2:NM_001135610:exon7:c.A593G:p.Q198R, |
| chr1 | 147268843 | C | G | COSV63614007 | exonic | CHD1L | nonsynonymous SNV | CHD1L:NM_004284:exon10:c.C1050G:p.H350Q,CHD1L:NM_001256338:exon4:c.C438G:p.H146Q,CHD1L:NM_024568:exon10:c.C711G:p.H237Q,CHD1L:NM_001256336:exon9:c.C750G:p.H250Q,CHD1L:NM_001256337:exon5:c.C207G:p.H69Q, |
| chr1 | 147286485 | A | G | . | exonic | CHD1L | nonsynonymous SNV | CHD1L:NM_004284:exon18:c.A2206G:p.I736V,CHD1L:NM_001256338:exon12:c.A1594G:p.I532V,CHD1L:NM_024568:exon18:c.A1867G:p.I623V,CHD1L:NM_001256336:exon17:c.A1906G:p.I636V,CHD1L:NM_001256337:exon13:c.A1363G:p.I455V, |
| chr1 | 153534727 | G | A | . | exonic | S100A6 | nonsynonymous SNV | S100A6:NM_014624:exon3:c.C242T:p.A81V, |
| chr1 | 153534737 | C | T | . | exonic | S100A6 | nonsynonymous SNV | S100A6:NM_014624:exon3:c.G232A:p.G78R, |
| chr1 | 153534809 | C | T | . | exonic | S100A6 | nonsynonymous SNV | S100A6:NM_014624:exon3:c.G160A:p.A54T, |
| chr1 | 154170403 | C | T | . | exonic | TPM3 | nonsynonymous SNV | TPM3:NM_152263:exon8:c.G772A:p.E258K,TPM3:NM_001278190:exon6:c.G598A:p.E200K,TPM3:NM_001043352:exon7:c.G661A:p.E221K,TPM3:NM_001043351:exon7:c.G661A:p.E221K,TPM3:NM_001043353:exon7:c.G661A:p.E221K,TPM3:NM_001278191:exon7:c.G391A:p.E131K,TPM3:NM_001278188:exon6:c.G463A:p.E155K,TPM3:NM_001278189:exon7:c.G661A:p.E221K,TPM3:NM_153649:exon7:c.G661A:p.E221K, |
| chr1 | 154170448 | C | T | COSV55136361 | exonic | TPM3 | nonsynonymous SNV | TPM3:NM_152263:exon8:c.G727A:p.A243T,TPM3:NM_001278190:exon6:c.G553A:p.A185T,TPM3:NM_001043352:exon7:c.G616A:p.A206T,TPM3:NM_001043351:exon7:c.G616A:p.A206T,TPM3:NM_001043353:exon7:c.G616A:p.A206T,TPM3:NM_001278191:exon7:c.G346A:p.A116T,TPM3:NM_001278188:exon6:c.G418A:p.A140T,TPM3:NM_001278189:exon7:c.G616A:p.A206T,TPM3:NM_153649:exon7:c.G616A:p.A206T, |
| chr1 | 154172915 | C | T | . | exonic | TPM3 | nonsynonymous SNV | TPM3:NM_152263:exon5:c.G559A:p.A187T,TPM3:NM_001278190:exon4:c.G448A:p.A150T,TPM3:NM_001043352:exon4:c.G448A:p.A150T,TPM3:NM_001043351:exon4:c.G448A:p.A150T,TPM3:NM_001043353:exon4:c.G448A:p.A150T,TPM3:NM_001278191:exon4:c.G178A:p.A60T,TPM3:NM_001278188:exon3:c.G250A:p.A84T,TPM3:NM_001278189:exon4:c.G448A:p.A150T,TPM3:NM_153649:exon4:c.G448A:p.A150T, |
| chr1 | 154173103 | G | T | . | exonic | TPM3 | nonsynonymous SNV | TPM3:NM_152263:exon4:c.C476A:p.A159E,TPM3:NM_001278190:exon3:c.C365A:p.A122E,TPM3:NM_001043352:exon3:c.C365A:p.A122E,TPM3:NM_001043351:exon3:c.C365A:p.A122E,TPM3:NM_001043353:exon3:c.C365A:p.A122E,TPM3:NM_001278191:exon3:c.C95A:p.A32E,TPM3:NM_001278188:exon2:c.C167A:p.A56E,TPM3:NM_001278189:exon3:c.C365A:p.A122E,TPM3:NM_153649:exon3:c.C365A:p.A122E, |

| | | | | | | | | |
|---|---|---|---|---|---|---|---|---|
| chr1 | 154173152 | C | T | . | exonic | TPM3 | nonsynonymous SNV | TPM3:NM_152263:exon4:c.G427A:p.E143K,TPM3:NM_001278190:exon3:c.G316A:p.E106K,TPM3:NM_001043352:exon3:c.G316A:p.E106K,TPM3:NM_001043351:exon3:c.G316A:p.E106K,TPM3:NM_001043353:exon3:c.G316A:p.E106K,TPM3:NM_001278191:exon3:c.G46A:p.E16K,TPM3:NM_001278188:exon2:c.G118A:p.E40K,TPM3:NM_001278189:exon3:c.G316A:p.E106K,TPM3:NM_153649:exon3:c.G316A:p.E106K, |
| chr1 | 154183010 | C | T | . | exonic | TPM3 | nonsynonymous SNV | TPM3:NM_001278190:exon1:c.G110A:p.G37E,TPM3:NM_001043352:exon1:c.G110A:p.G37E,TPM3:NM_001043351:exon1:c.G110A:p.G37E,TPM3:NM_001043353:exon1:c.G110A:p.G37E,TPM3:NM_001278188:exon1:c.G46A:p.E16K,TPM3:NM_001278189:exon1:c.G110A:p.G37E,TPM3:NM_153649:exon1:c.G110A:p.G37E, |
| chr1 | 156768177 | C | T | . | exonic | PRCC | nonsynonymous SNV | PRCC:NM_005973:exon1:c.C406T:p.P136S, |
| chr1 | 161781951 | A | G | COSV63406746 | exonic | ATF6 | nonsynonymous SNV | ATF6:NM_007348:exon3:c.A199G:p.M67V, |
| chr1 | 17697195 | A | G | COSV51429640 | exonic | ARHGEF10L | nonsynonymous SNV | ARHGEF10L:NM_001011722:exon27:c.A3538G:p.I1180V,ARHGEF10L:NM_018125:exon29:c.A3655G:p.I1219V, |
| chr1 | 193125191 | C | T | . | exonic | CDC73 | nonsynonymous SNV | CDC73:NM_024529:exon2:c.C211T:p.H71Y, |
| chr1 | 19339555 | G | A | . | exonic | CAPZB | nonsynonymous SNV | CAPZB:NM_004930:exon9:c.C794T:p.A265V,CAPZB:NM_001282162:exon9:c.C881T:p.A294V, |
| chr1 | 19339568 | C | T | . | exonic | CAPZB | nonsynonymous SNV | CAPZB:NM_004930:exon9:c.G781A:p.D261N,CAPZB:NM_001282162:exon9:c.G868A:p.D290N, |
| chr1 | 202011975 | G | A | . | exonic | ELF3 | nonsynonymous SNV | ELF3:NM_001114309:exon3:c.G182A:p.G61E,ELF3:NM_004433:exon3:c.G182A:p.G61E, |
| chr1 | 202012026 | A | G | . | exonic | ELF3 | nonsynonymous SNV | ELF3:NM_001114309:exon3:c.A233G:p.Y78C,ELF3:NM_004433:exon3:c.A233G:p.Y78C, |
| chr1 | 202013945 | T | C | . | exonic | ELF3 | nonsynonymous SNV | ELF3:NM_001114309:exon8:c.T922C:p.S308P,ELF3:NM_004433:exon8:c.T922C:p.S308P, |
| chr1 | 202013948 | G | A | . | exonic | ELF3 | nonsynonymous SNV | ELF3:NM_001114309:exon8:c.G925A:p.E309K,ELF3:NM_004433:exon8:c.G925A:p.E309K, |
| chr1 | 202015319 | A | T | . | exonic | ELF3 | nonsynonymous SNV | ELF3:NM_001114309:exon9:c.A1112T:p.N371I,ELF3:NM_004433:exon9:c.A1112T:p.N371I, |
| chr1 | 212100297 | C | T | . | exonic | DTL | nonsynonymous SNV | DTL:NM_016448:exon14:c.C1307T:p.A436V,DTL:NM_001286230:exon13:c.C1181T:p.A394V,DTL:NM_001286229:exon12:c.C494T:p.A165V, |
| chr1 | 212101071 | A | C | . | exonic | DTL | nonsynonymous SNV | DTL:NM_016448:exon14:c.A2081C:p.K694T,DTL:NM_001286230:exon13:c.A1955C:p.K652T,DTL:NM_001286229:exon12:c.A1268C:p.K423T, |
| chr1 | 225831932 | T | C | COSV55298811 | exonic | EPHX1 | nonsynonymous SNV | EPHX1:NM_001291163:exon3:c.T337C:p.Y113H,EPHX1:NM_000120:exon3:c.T337C:p.Y113H,EPHX1:NM_001136018:exon3:c.T337C:p.Y113H, |
| chr1 | 225838705 | A | G | COSV55299484 | exonic | EPHX1 | nonsynonymous SNV | EPHX1:NM_001291163:exon4:c.A416G:p.H139R,EPHX1:NM_000120:exon4:c.A416G:p.H139R,EPHX1:NM_001136018:exon4:c.A416G:p.H139R, |
| chr1 | 22913770 | G | A | . | exonic | EPHB2 | nonsynonymous SNV | EPHB2:NM_001309193:exon17:c.G3068A:p.G1023E, |
| chr1 | 246507179 | C | G | COSV66438600 | exonic | SMYD3 | nonsynonymous SNV | SMYD3:NM_001167740:exon1:c.G39C:p.K13N, |
| chr1 | 246988255 | A | G | COSV60584996 | exonic | ZNF695 | nonsynonymous SNV | ZNF695:NM_020394:exon4:c.T260C:p.V87A,ZNF695:NM_001204221:exon4:c.T260C:p.V87A, |
| chr1 | 246999356 | C | T | . | exonic | ZNF695 | nonsynonymous SNV | ZNF695:NM_020394:exon3:c.G251A:p.R84K,ZNF695:NM_001204221:exon3:c.G251A:p.R84K, |
| chr1 | 26460136 | G | A | COSV52575895 | exonic | DHDDS | nonsynonymous SNV | DHDDS:NM_001243565:exon7:c.G640A:p.V214M,DHDDS:NM_205861:exon8:c.G757A:p.V253M,DHDDS:NM_001243564:exon7:c.G655A:p.V219M,DHDDS:NM_024887:exon8:c.G757A:p.V253M, |
| chr1 | 28473868 | C | A | COSV65783948 | exonic | PHACTR4 | nonsynonymous SNV | PHACTR4:NM_001048183:exon7:c.C1138A:p.L380I,PHACTR4:NM_023923:exon6:c.C1168A:p.L390I, |
| chr1 | 35184478 | G | T | . | exonic | SFPQ | nonsynonymous SNV | SFPQ:NM_005066:exon10:c.C2102A:p.P701Q, |
| chr1 | 35187987 | C | T | . | exonic | SFPQ | nonsynonymous SNV | SFPQ:NM_005066:exon7:c.G1801A:p.G601S, |
| chr1 | 35188008 | C | T | . | exonic | SFPQ | nonsynonymous SNV | SFPQ:NM_005066:exon7:c.G1780A:p.E594K, |
| chr1 | 46309111 | A | G | COSV61175886 | exonic | UQCRH | nonsynonymous SNV | UQCRH:NM_006004:exon2:c.A65G:p.E22G, |
| chr1 | 51361249 | T | C | COSV65541248 | exonic | EPS15 | nonsynonymous SNV | EPS15:NM_001159969:exon12:c.A1524G:p.I508M,EPS15:NM_001981:exon24:c.A2466G:p.I822M, |
| chr1 | 59321678 | T | G | COSV58028591 | exonic | FGGY | nonsynonymous SNV | FGGY:NM_001244714:exon2:c.T129G:p.N43K,FGGY:NM_001113411:exon2:c.T129G:p.N43K,FGGY:NM_018291:exon2:c.T129G:p.N43K, |
| chr1 | 61990342 | G | A | COSV56249041 | exonic | INADL | nonsynonymous SNV | INADL:NM_176877:exon28:c.G3845A:p.R1282H, |
| chr1 | 62051011 | G | C | COSV56248179 | exonic | INADL | nonsynonymous SNV | INADL:NM_176877:exon31:c.G4078C:p.V1360L, |
| chr1 | 68137903 | C | T | COSV52037496 | exonic | WLS | nonsynonymous SNV | WLS:NM_001002292:exon11:c.G1387A:p.V463I,WLS:NM_024911:exon11:c.G1393A:p.V465I,WLS:NM_001193334:exon10:c.G1120A:p.V374I, |
| chr1 | 6887657 | C | T | COSV57917420 | exonic | CAMTA1 | nonsynonymous SNV | CAMTA1:NM_001242701:exon4:c.C238T:p.L80F, |
| chr1 | 75749507 | A | G | COSV57714736 | exonic | ACADM | nonsynonymous SNV | ACADM:NM_001286043:exon10:c.A896G:p.D299G,ACADM:NM_001286042:exon8:c.A689G:p.D230G,ACADM:NM_001286044:exon6:c.A230G:p.D77G,ACADM:NM_001127328:exon9:c.A809G:p.D270G,ACADM:NM_000016:exon9:c.A797G:p.D266G, |
| chr1 | 77964325 | A | T | . | exonic | FUBP1 | nonsynonymous SNV | FUBP1:NM_003902:exon11:c.T869A:p.V290E,FUBP1:NM_001303433:exon12:c.T932A:p.V311E, |
| chr1 | 8862903 | A | G | . | exonic | ENO1 | nonsynonymous SNV | ENO1:NM_001201483:exon8:c.T940C:p.Y314H,ENO1:NM_001428:exon11:c.T1219C:p.Y407H, |
| chr1 | 8862909 | C | T | . | exonic | ENO1 | nonsynonymous SNV | ENO1:NM_001201483:exon8:c.G934A:p.A312T,ENO1:NM_001428:exon11:c.G1213A:p.A405T, |
| chr1 | 8862933 | C | G | . | exonic | ENO1 | nonsynonymous SNV | ENO1:NM_001201483:exon8:c.G910C:p.A304P,ENO1:NM_001428:exon11:c.G1189C:p.A397P, |
| chr1 | 8863258 | C | T | . | exonic | ENO1 | nonsynonymous SNV | ENO1:NM_001201483:exon7:c.G874A:p.V292I,ENO1:NM_001428:exon10:c.G1153A:p.V385I, |
| chr1 | 8863288 | C | T | . | exonic | ENO1 | nonsynonymous SNV | ENO1:NM_001201483:exon7:c.G844A:p.E282K,ENO1:NM_001428:exon10:c.G1123A:p.E375K, |

| | | | | | | | | |
|---|---|---|---|---|---|---|---|---|
| chr1 | 8863894 | T | C | COSV52304857 | exonic | ENO1 | nonsynonymous SNV | ENO1:NM_001201483:exon6:c.A785G:p.Q262R,ENO1:NM_001428:exon9:c.A1064G:p.Q355R, |
| chr1 | 8863964 | C | T | . | exonic | ENO1 | nonsynonymous SNV | ENO1:NM_001201483:exon6:c.G715A:p.V239M,ENO1:NM_001428:exon9:c.G994A:p.V332M, |
| chr1 | 8863973 | C | T | . | exonic | ENO1 | nonsynonymous SNV | ENO1:NM_001201483:exon6:c.G706A:p.A236T,ENO1:NM_001428:exon9:c.G985A:p.A329T, |
| chr1 | 8864024 | C | T | . | exonic | ENO1 | nonsynonymous SNV | ENO1:NM_001201483:exon6:c.G655A:p.G219R,ENO1:NM_001428:exon9:c.G934A:p.G312R, |
| chr1 | 8864029 | C | T | . | exonic | ENO1 | nonsynonymous SNV | ENO1:NM_001201483:exon6:c.G650A:p.S217N,ENO1:NM_001428:exon9:c.G929A:p.S310N, |
| chr1 | 8864090 | C | T | . | exonic | ENO1 | nonsynonymous SNV | ENO1:NM_001201483:exon6:c.G589A:p.V197M,ENO1:NM_001428:exon9:c.G868A:p.V290M, |
| chr1 | 8865320 | G | T | . | exonic | ENO1 | nonsynonymous SNV | ENO1:NM_001201483:exon5:c.C551A:p.A184D,ENO1:NM_001428:exon8:c.C830A:p.A277D, |
| chr1 | 8865347 | C | T | . | exonic | ENO1 | nonsynonymous SNV | ENO1:NM_001201483:exon5:c.G524A:p.S175N,ENO1:NM_001428:exon8:c.G803A:p.S268N, |
| chr1 | 8865392 | C | T | . | exonic | ENO1 | nonsynonymous SNV | ENO1:NM_001201483:exon5:c.G479A:p.R160K,ENO1:NM_001428:exon8:c.G758A:p.R253K, |
| chr1 | 8865405 | A | G | . | exonic | ENO1 | nonsynonymous SNV | ENO1:NM_001201483:exon5:c.T466C:p.S156P,ENO1:NM_001428:exon8:c.T745C:p.S249P, |
| chr1 | 8865414 | C | T | . | exonic | ENO1 | nonsynonymous SNV | ENO1:NM_001201483:exon5:c.G457A:p.V153I,ENO1:NM_001428:exon8:c.G736A:p.V246I, |
| chr1 | 8865418 | C | T | . | exonic | ENO1 | nonsynonymous SNV | ENO1:NM_001201483:exon5:c.G453A:p.M151I,ENO1:NM_001428:exon8:c.G732A:p.M244I, |
| chr1 | 8866420 | C | T | . | exonic | ENO1 | nonsynonymous SNV | ENO1:NM_001201483:exon4:c.G247A:p.A83T,ENO1:NM_001428:exon7:c.G526A:p.A176T, |
| chr1 | 8866429 | C | T | . | exonic | ENO1 | nonsynonymous SNV | ENO1:NM_001201483:exon4:c.G238A:p.V80I,ENO1:NM_001428:exon7:c.G517A:p.V173I, |
| chr1 | 8866441 | T | C | . | exonic | ENO1 | nonsynonymous SNV | ENO1:NM_001201483:exon4:c.A226G:p.M76V,ENO1:NM_001428:exon7:c.A505G:p.M169V, |
| chr1 | 8866480 | C | T | . | exonic | ENO1 | nonsynonymous SNV | ENO1:NM_001201483:exon4:c.G187A:p.G63S,ENO1:NM_001428:exon7:c.G466A:p.G156S, |
| chr1 | 8867154 | T | C | . | exonic | ENO1 | nonsynonymous SNV | ENO1:NM_001201483:exon3:c.A128G:p.D43G,ENO1:NM_001428:exon6:c.A407G:p.D136G, |
| chr1 | 8867158 | C | T | . | exonic | ENO1 | nonsynonymous SNV | ENO1:NM_001201483:exon3:c.G124A:p.A42T,ENO1:NM_001428:exon6:c.G403A:p.A135T, |
| chr1 | 8868006 | C | T | . | exonic | ENO1 | nonsynonymous SNV | ENO1:NM_001201483:exon2:c.G13A:p.D5N,ENO1:NM_001428:exon5:c.G292A:p.D98N, |
| chr1 | 8868012 | C | T | . | exonic | ENO1 | nonsynonymous SNV | ENO1:NM_001201483:exon2:c.G7A:p.E3K,ENO1:NM_001428:exon5:c.G286A:p.E96K, |
| chr1 | 8868041 | T | C | . | exonic | ENO1 | nonsynonymous SNV | ENO1:NM_001428:exon5:c.A257G:p.E86G, |
| chr1 | 8868048 | C | T | COSV52303828 | exonic | ENO1 | nonsynonymous SNV | ENO1:NM_001428:exon5:c.G250A:p.V84I, |
| chr1 | 8870510 | C | T | . | exonic | ENO1 | nonsynonymous SNV | ENO1:NM_001428:exon4:c.G182A:p.G61D, |
| chr1 | 8871898 | C | T | . | exonic | ENO1 | nonsynonymous SNV | ENO1:NM_001428:exon3:c.G174A:p.M58I, |
| chr1 | 8871903 | A | G | . | exonic | ENO1 | nonsynonymous SNV | ENO1:NM_001428:exon3:c.T169C:p.Y57H, |
| chr1 | 8871921 | C | T | . | exonic | ENO1 | nonsynonymous SNV | ENO1:NM_001428:exon3:c.G151A:p.D51N, |
| chr1 | 8871948 | C | T | . | exonic | ENO1 | nonsynonymous SNV | ENO1:NM_001428:exon3:c.G124A:p.G42S, |
| chr1 | 8874829 | G | T | . | exonic | ENO1 | stopgain | ENO1:NM_001428:exon2:c.C80A:p.S27X, |
| chr1 | 92833389 | G | A | . | exonic | RPL5 | nonsynonymous SNV | RPL5:NM_000969:exon2:c.G4A:p.G2R, |
| chr1 | 92833396 | T | C | . | exonic | RPL5 | nonsynonymous SNV | RPL5:NM_000969:exon2:c.T11C:p.V4A, |
| chr1 | 92833401 | G | A | . | exonic | RPL5 | nonsynonymous SNV | RPL5:NM_000969:exon2:c.G16A:p.V6I, |
| chr1 | 92833404 | G | A | . | exonic | RPL5 | nonsynonymous SNV | RPL5:NM_000969:exon2:c.G19A:p.V7I, |
| chr1 | 92833433 | C | A | COSV59873416 | exonic | RPL5 | stopgain | RPL5:NM_000969:exon2:c.C48A:p.Y16X, |
| chr1 | 92833446 | A | G | . | exonic | RPL5 | nonsynonymous SNV | RPL5:NM_000969:exon2:c.A61G:p.R21G, |
| chr1 | 92833556 | G | A | . | exonic | RPL5 | nonsynonymous SNV | RPL5:NM_000969:exon3:c.G85A:p.D29N, |
| chr1 | 92833632 | G | A | . | exonic | RPL5 | nonsynonymous SNV | RPL5:NM_000969:exon3:c.G161A:p.R54H, |
| chr1 | 92833646 | G | A | . | exonic | RPL5 | nonsynonymous SNV | RPL5:NM_000969:exon3:c.G175A:p.D59N, |
| chr1 | 92833652 | A | T | . | exonic | RPL5 | nonsynonymous SNV | RPL5:NM_000969:exon3:c.A181T:p.I61F, |
| chr1 | 92834818 | G | A | COSV59873410 | exonic | RPL5 | nonsynonymous SNV | RPL5:NM_000969:exon4:c.G229A:p.A77T, |
| chr1 | 92834833 | G | A | COSV59872943 | exonic | RPL5 | nonsynonymous SNV | RPL5:NM_000969:exon4:c.G244A:p.E82K, |
| chr1 | 92834849 | G | A | . | exonic | RPL5 | nonsynonymous SNV | RPL5:NM_000969:exon4:c.G260A:p.G87D, |
| chr1 | 92834879 | C | T | . | exonic | RPL5 | nonsynonymous SNV | RPL5:NM_000969:exon4:c.C290T:p.A97V, |
| chr1 | 92834909 | G | A | . | exonic | RPL5 | nonsynonymous SNV | RPL5:NM_000969:exon4:c.G320A:p.R107H, |
| chr1 | 92836226 | G | A | . | exonic | RPL5 | nonsynonymous SNV | RPL5:NM_000969:exon5:c.G361A:p.G121S, |
| chr1 | 92836244 | G | A | . | exonic | RPL5 | nonsynonymous SNV | RPL5:NM_000969:exon5:c.G379A:p.G127S, |
| chr1 | 92836259 | G | A | . | exonic | RPL5 | nonsynonymous SNV | RPL5:NM_000969:exon5:c.G394A:p.V132M, |
| chr1 | 92836347 | G | A | . | exonic | RPL5 | nonsynonymous SNV | RPL5:NM_000969:exon5:c.G482A:p.G161D, |
| chr1 | 92837488 | G | A | . | exonic | RPL5 | nonsynonymous SNV | RPL5:NM_000969:exon6:c.G560A:p.S187N, |
| chr1 | 92837515 | G | A | COSV59874073 | exonic | RPL5 | nonsynonymous SNV | RPL5:NM_000969:exon6:c.G587A:p.R196Q, |
| chr1 | 92837565 | G | A | . | exonic | RPL5 | nonsynonymous SNV | RPL5:NM_000969:exon6:c.G637A:p.E213K, |
| chr1 | 92837619 | G | A | COSV59872895 | exonic | RPL5 | nonsynonymous SNV | RPL5:NM_000969:exon6:c.G691A:p.V231I, |
| chr1 | 92840600 | T | C | . | exonic | RPL5 | nonsynonymous SNV | RPL5:NM_000969:exon7:c.T755C:p.V252A, |
| chr1 | 92840627 | T | A | . | exonic | RPL5 | nonsynonymous SNV | RPL5:NM_000969:exon7:c.T782A:p.V261D, |
| chr1 | 92841810 | T | C | . | exonic | RPL5 | nonsynonymous SNV | RPL5:NM_000969:exon8:c.T839C:p.V280A, |
| chr10 | 102160718 | T | C | . | exonic | NOLC1 | nonsynonymous SNV | NOLC1:NM_001284388:exon10:c.T1396C:p.S466P,NOLC1:NM_001284389:exon10:c.T1369C:p.S457P,NOLC1:NM_004741:exon10:c.T1366C:p.S456P, |
| chr10 | 102160941 | A | G | . | exonic | NOLC1 | nonsynonymous SNV | NOLC1:NM_001284388:exon10:c.A1619G:p.E540G,NOLC1:NM_001284389:exon10:c.A1592G:p.E531G,NOLC1:NM_004741:exon10:c.A1589G:p.E530G, |
| chr10 | 102161057 | G | A | . | exonic | NOLC1 | nonsynonymous SNV | NOLC1:NM_001284388:exon10:c.G1735A:p.E579K,NOLC1:NM_001284389:exon10:c.G1708A:p.E570K,NOLC1:NM_004741:exon10:c.G1705A:p.E569K, |
| chr10 | 103174952 | T | C | . | exonic | NT5C2 | nonsynonymous SNV | NT5C2:NM_001134373:exon2:c.A7G:p.T3A,NT5C2:NM_012229:exon3:c.A7G:p.T3A, |
| chr10 | 110295351 | A | T | . | exonic | SMNDC1 | stopgain | SMNDC1:NM_005871:exon5:c.T456A:p.Y152X, |
| chr10 | 110881295 | A | G | . | exonic | PDCD4 | nonsynonymous SNV | PDCD4:NM_145341:exon4:c.A73G:p.I25V,PDCD4:NM_014456:exon3:c.A106G:p.I36V,PDCD4:NM_001199492:exon3:c.A64G:p.I22V, |

| | | | | | | | | |
|---|---|---|---|---|---|---|---|---|
| chr10 | 121911030 | A | T | . | exonic | ATE1 | nonsynonymous SNV | ATE1:NM_001288736:exon5:c.T438A:p.S146R,ATE1:NM_001288734:exon4:c.T114A:p.S38R,ATE1:NM_001001976:exon5:c.T459A:p.S153R,ATE1:NM_007041:exon5:c.T459A:p.S153R,ATE1:NM_001288735:exon3:c.T171A:p.S57R, |
| chr10 | 122237461 | G | A | COSV53279154 | exonic | TACC2 | nonsynonymous SNV | TACC2:NM_206861:exon14:c.G2632A:p.A878T,TACC2:NM_206860:exon10:c.G2338A:p.A780T,TACC2:NM_001291876:exon14:c.G7828A:p.A2610T,TACC2:NM_001291879:exon11:c.G1075A:p.A359T,TACC2:NM_001291878:exon11:c.G2374A:p.A792T,TACC2:NM_006997:exon11:c.G2428A:p.A810T,TACC2:NM_001291877:exon14:c.G7975A:p.A2659T,TACC2:NM_206862:exon17:c.G8194A:p.A2732T, |
| chr10 | 122249609 | G | C | COSV53282365 | exonic | TACC2 | nonsynonymous SNV | TACC2:NM_206861:exon19:c.G3164C:p.S1055T,TACC2:NM_206860:exon15:c.G2870C:p.S957T,TACC2:NM_001291876:exon19:c.G8360C:p.S2787T,TACC2:NM_001291879:exon16:c.G1607C:p.S536T,TACC2:NM_001291878:exon16:c.G2906C:p.S969T,TACC2:NM_006997:exon16:c.G2960C:p.S987T,TACC2:NM_001291877:exon19:c.G8507C:p.S2836T,TACC2:NM_206862:exon22:c.G8726C:p.S2909T, |
| chr10 | 127451398 | G | A | COSV54751969 | exonic | DOCK1 | nonsynonymous SNV | DOCK1:NM_001290223:exon52:c.G5632A:p.A1878T,DOCK1:NM_001380:exon52:c.G5569A:p.A1857T, |
| chr10 | 129708019 | C | T | COSV60031127 | exonic | MGMT | nonsynonymous SNV | MGMT:NM_002412:exon3:c.C343T:p.L115F, |
| chr10 | 133352286 | A | G | . | exonic | PRAP1 | nonsynonymous SNV | PRAP1:NM_001145201:exon5:c.A275G:p.H92R,PRAP1:NM_145202:exon5:c.A302G:p.H101R, |
| chr10 | 17229690 | G | A | . | exonic | VIM | nonsynonymous SNV | VIM:NM_003380:exon2:c.G268A:p.D90N, |
| chr10 | 17229886 | G | A | . | exonic | VIM | nonsynonymous SNV | VIM:NM_003380:exon2:c.G464A:p.R155Q, |
| chr10 | 17233635 | G | A | . | exonic | VIM | nonsynonymous SNV | VIM:NM_003380:exon4:c.G673A:p.E225K, |
| chr10 | 17234705 | T | C | . | exonic | VIM | nonsynonymous SNV | VIM:NM_003380:exon6:c.T895C:p.S299P, |
| chr10 | 17234714 | G | A | . | exonic | VIM | nonsynonymous SNV | VIM:NM_003380:exon6:c.G904A:p.A302T, |
| chr10 | 17234739 | G | A | COSV56406122 | exonic | VIM | nonsynonymous SNV | VIM:NM_003380:exon6:c.G929A:p.R310H, |
| chr10 | 17235223 | G | A | . | exonic | VIM | nonsynonymous SNV | VIM:NM_003380:exon7:c.G1063A:p.A355T, |
| chr10 | 17235352 | G | A | . | exonic | VIM | nonsynonymous SNV | VIM:NM_003380:exon7:c.G1192A:p.A398T, |
| chr10 | 17235362 | G | A | . | exonic | VIM | nonsynonymous SNV | VIM:NM_003380:exon7:c.G1202A:p.R401K, |
| chr10 | 20840818 | A | T | . | exonic | NEBL | nonsynonymous SNV | NEBL:NM_006393:exon13:c.T1259A:p.I420K, |
| chr10 | 20840822 | C | T | . | exonic | NEBL | nonsynonymous SNV | NEBL:NM_006393:exon13:c.G1255A:p.E419K, |
| chr10 | 31520330 | T | A | . | exonic | ZEB1 | nonsynonymous SNV | ZEB1:NM_001174093:exon6:c.T935A:p.I312K,ZEB1:NM_001174095:exon6:c.T794A:p.I265K,ZEB1:NM_001128128:exon7:c.T947A:p.I316K,ZEB1:NM_030751:exon7:c.T995A:p.I332K,ZEB1:NM_001174096:exon7:c.T998A:p.I333K,ZEB1:NM_001174094:exon7:c.T944A:p.I315K, |
| chr10 | 31520332 | G | A | . | exonic | ZEB1 | nonsynonymous SNV | ZEB1:NM_001174093:exon6:c.G937A:p.E313K,ZEB1:NM_001174095:exon6:c.G796A:p.E266K,ZEB1:NM_001128128:exon7:c.G949A:p.E317K,ZEB1:NM_030751:exon7:c.G997A:p.E333K,ZEB1:NM_001174096:exon7:c.G1000A:p.E334K,ZEB1:NM_001174094:exon7:c.G946A:p.E316K, |
| chr10 | 31520809 | T | A | . | exonic | ZEB1 | nonsynonymous SNV | ZEB1:NM_001174093:exon6:c.T1414A:p.L472I,ZEB1:NM_001174095:exon6:c.T1273A:p.L425I,ZEB1:NM_001128128:exon7:c.T1426A:p.L476I,ZEB1:NM_030751:exon7:c.T1474A:p.L492I,ZEB1:NM_001174096:exon7:c.T1477A:p.L493I,ZEB1:NM_001174094:exon7:c.T1423A:p.L475I, |
| chr10 | 33263768 | A | G | . | exonic | NRP1 | nonsynonymous SNV | NRP1:NM_001024628:exon4:c.T536C:p.V179A,NRP1:NM_001244973:exon4:c.T536C:p.V179A,NRP1:NM_001024629:exon4:c.T536C:p.V179A,NRP1:NM_003873:exon4:c.T536C:p.V179A,NRP1:NM_001244972:exon4:c.T536C:p.V179A, |
| chr10 | 46010293 | C | T | . | exonic | NCOA4 | nonsynonymous SNV | NCOA4:NM_001145261:exon9:c.G1676A:p.G559E,NCOA4:NM_001145260:exon9:c.G1676A:p.G559E,NCOA4:NM_005437:exon8:c.G1628A:p.G543E,NCOA4:NM_001145263:exon8:c.G1628A:p.G543E,NCOA4:NM_001145262:exon8:c.G1628A:p.G543E, |
| chr10 | 46010365 | C | T | . | exonic | NCOA4 | nonsynonymous SNV | NCOA4:NM_001145261:exon9:c.G1604A:p.G535D,NCOA4:NM_001145260:exon9:c.G1604A:p.G535D,NCOA4:NM_005437:exon8:c.G1556A:p.G519D,NCOA4:NM_001145263:exon8:c.G1556A:p.G519D,NCOA4:NM_001145262:exon8:c.G1556A:p.G519D, |
| chr10 | 46010595 | C | T | . | exonic | NCOA4 | nonsynonymous SNV | NCOA4:NM_001145261:exon9:c.G1374A:p.M458I,NCOA4:NM_001145260:exon9:c.G1374A:p.M458I,NCOA4:NM_005437:exon8:c.G1326A:p.M442I,NCOA4:NM_001145263:exon8:c.G1326A:p.M442I,NCOA4:NM_001145262:exon8:c.G1326A:p.M442I, |
| chr10 | 46010599 | C | T | . | exonic | NCOA4 | nonsynonymous SNV | NCOA4:NM_001145261:exon9:c.G1370A:p.G457E,NCOA4:NM_001145260:exon9:c.G1370A:p.G457E,NCOA4:NM_005437:exon8:c.G1322A:p.G441E,NCOA4:NM_001145263:exon8:c.G1322A:p.G441E,NCOA4:NM_001145262:exon8:c.G1322A:p.G441E, |
| chr10 | 46010618 | C | T | . | exonic | NCOA4 | nonsynonymous SNV | NCOA4:NM_001145261:exon9:c.G1351A:p.E451K,NCOA4:NM_001145260:exon9:c.G1351A:p.E451K,NCOA4:NM_005437:exon8:c.G1303A:p.E435K,NCOA4:NM_001145263:exon8:c.G1303A:p.E435K,NCOA4:NM_001145262:exon8:c.G1303A:p.E435K, |
| chr10 | 46010714 | C | T | . | exonic | NCOA4 | nonsynonymous SNV | NCOA4:NM_001145261:exon9:c.G1255A:p.V419M,NCOA4:NM_001145260:exon9:c.G1255A:p.V419M,NCOA4:NM_005437:exon8:c.G1207A:p.V403M,NCOA4:NM_001145263:exon8:c.G1207A:p.V403M,NCOA4:NM_001145262:exon8:c.G1207A:p.V403M, |
| chr10 | 46010750 | C | T | . | exonic | NCOA4 | nonsynonymous SNV | NCOA4:NM_001145263:exon8:c.G1171A:p.V391I,NCOA4:NM_001145262:exon8:c.G1171A:p.V391I,NCOA4:NM_001145261:exon9:c.G1219A:p.V407I,NCOA4:NM_001145260:exon9:c.G1219A:p.V407I,NCOA4:NM_005437:exon8:c.G1171A:p.V391I, |
| chr10 | 68771301 | C | A | . | exonic | CCAR1 | nonsynonymous SNV | CCAR1:NM_001282959:exon17:c.C2349A:p.D783E,CCAR1:NM_001282960:exon17:c.C2349A:p.D783E,CCAR1:NM_018237:exon18:c.C2394A:p.D798E, |
| chr10 | 68771325 | C | A | COSV56268323 | exonic | CCAR1 | nonsynonymous SNV | CCAR1:NM_001282959:exon17:c.C2373A:p.S791R,CCAR1:NM_001282960:exon17:c.C2373A:p.S791R,CCAR1:NM_018237:exon18:c.C2418A:p.S806R, |
| chr10 | 68771417 | A | G | . | exonic | CCAR1 | nonsynonymous SNV | CCAR1:NM_001282959:exon17:c.A2465G:p.D822G,CCAR1:NM_001282960:exon17:c.A2465G:p.D822G,CCAR1:NM_018237:exon18:c.A2510G:p.D837G, |
| chr10 | 71322806 | A | G | . | exonic | SLC29A3 | nonsynonymous SNV | SLC29A3:NM_001174098:exon2:c.A52G:p.R18G,SLC29A3:NM_018344:exon2:c.A52G:p.R18G, |
| chr10 | 71351651 | C | T | COSV64385485 | exonic | SLC29A3 | nonsynonymous SNV | SLC29A3:NM_001174098:exon4:c.C473T:p.S158F,SLC29A3:NM_018344:exon4:c.C473T:p.S158F, |
| chr10 | 71356185 | G | A | . | exonic | SLC29A3 | nonsynonymous SNV | SLC29A3:NM_001174098:exon5:c.G715A:p.V239I,SLC29A3:NM_018344:exon5:c.G715A:p.V239I, |
| chr10 | 71362156 | A | G | COSV64386549 | exonic | SLC29A3 | nonsynonymous SNV | SLC29A3:NM_018344:exon6:c.A976G:p.I326V, |
| chr10 | 71819553 | C | T | . | exonic | PSAP | nonsynonymous SNV | PSAP:NM_001042465:exon12:c.G1271A:p.R424H,PSAP:NM_002778:exon11:c.G1262A:p.R421H,PSAP:NM_001042466:exon11:c.G1268A:p.R423H, |

| | | | | | | | | |
|---|---|---|---|---|---|---|---|---|
| chr10 | 74398526 | G | A | . | exonic | ADK | nonsynonymous SNV | ADK:NM_001202450:exon6:c.G502A:p.D168N,ADK:NM_006721:exon6:c.G502A:p.D168N,ADK:NM_001202449:exon5:c.G397A:p.D133N,ADK:NM_001123:exon6:c.G451A:p.D151N, |
| chr10 | 86919290 | A | C | . | exonic | BMPR1A | nonsynonymous SNV | BMPR1A:NM_004329:exon10:c.A987C:p.R329S, |
| chr10 | 87864144 | G | C | . | exonic | PTEN | nonsynonymous SNV | PTEN:NM_001304717:exon2:c.G194C:p.C65S, |
| chr10 | 87958004 | C | A | . | exonic | PTEN | nonsynonymous SNV | PTEN:NM_000314:exon7:c.C786A:p.N262K,PTEN:NM_001304717:exon8:c.C1305A:p.N435K,PTEN:NM_001304718:exon7:c.C195A:p.N65K, |
| chr10 | 87960961 | T | A | . | exonic | PTEN | nonsynonymous SNV | PTEN:NM_000314:exon8:c.T869A:p.V290E,PTEN:NM_001304717:exon9:c.T1388A:p.V463E,PTEN:NM_001304718:exon8:c.T278A:p.V93E, |
| chr10 | 87960969 | G | A | . | exonic | PTEN | nonsynonymous SNV | PTEN:NM_000314:exon8:c.G877A:p.G293R,PTEN:NM_001304717:exon9:c.G1396A:p.G466R,PTEN:NM_001304718:exon8:c.G286A:p.G96R, |
| chr10 | 87965468 | T | C | . | exonic | PTEN | nonsynonymous SNV | PTEN:NM_000314:exon9:c.T1208C:p.V403A,PTEN:NM_001304717:exon10:c.T1727C:p.V576A,PTEN:NM_001304718:exon9:c.T617C:p.V206A, |
| chr10 | 88583080 | C | G | COSV59290594 | exonic | RNLS | nonsynonymous SNV | RNLS:NM_018363:exon1:c.G111C:p.E37D,RNLS:NM_001031709:exon1:c.G111C:p.E37D, |
| chr10 | 89719454 | G | T | COSV53306008 | exonic | KIF20B | nonsynonymous SNV | KIF20B:NM_016195:exon13:c.G1470T:p.E490D,KIF20B:NM_001284259:exon13:c.G1470T:p.E490D, |
| chr11 | 117351876 | G | A | COSV54039014 | exonic | CEP164 | nonsynonymous SNV | CEP164:NM_014956:exon5:c.G281A:p.S94N,CEP164:NM_001271933:exon4:c.G281A:p.S94N, |
| chr11 | 119188695 | G | A | COSV52702606 | exonic | PDZD3 | nonsynonymous SNV | PDZD3:NM_001168468:exon9:c.G1115A:p.R372Q,PDZD3:NM_024791:exon9:c.G1073A:p.R358Q, |
| chr11 | 34888850 | C | T | COSV53505953 | exonic | APIP | nonsynonymous SNV | APIP:NM_015957:exon4:c.G227A:p.C76Y, |
| chr11 | 34895110 | C | T | COSV53506741 | exonic | APIP | nonsynonymous SNV | APIP:NM_015957:exon2:c.G58A:p.D20N, |
| chr11 | 34916266 | G | A | COSV53507442 | exonic | APIP | nonsynonymous SNV | APIP:NM_015957:exon1:c.C19T:p.R7W, |
| chr11 | 34916722 | C | T | . | exonic | PDHX | nonsynonymous SNV | PDHX:NM_001166158:exon1:c.C67T:p.R23C,PDHX:NM_003477:exon1:c.C67T:p.R23C, |
| chr11 | 34947565 | A | G | COSV57166122 | exonic | PDHX | nonsynonymous SNV | PDHX:NM_001135024:exon3:c.A256G:p.T86A,PDHX:NM_001166158:exon3:c.A301G:p.T101A,PDHX:NM_003477:exon3:c.A301G:p.T101A, |
| chr11 | 35180332 | G | A | . | exonic | CD44 | nonsynonymous SNV | CD44:NM_001001389:exon3:c.G292A:p.A98T,CD44:NM_001202556:exon3:c.G292A:p.A98T,CD44:NM_001202555:exon3:c.G292A:p.A98T,CD44:NM_001001390:exon3:c.G292A:p.A98T,CD44:NM_000610:exon3:c.G292A:p.A98T,CD44:NM_001202557:exon3:c.G292A:p.A98T,CD44:NM_001001391:exon3:c.G292A:p.A98T, |
| chr11 | 35204608 | A | G | COSV53530507 | exonic | CD44 | nonsynonymous SNV | CD44:NM_001001389:exon9:c.A1121G:p.K374R,CD44:NM_000610:exon10:c.A1250G:p.K417R, |
| chr11 | 35208126 | T | C | . | exonic | CD44 | nonsynonymous SNV | CD44:NM_001001389:exon11:c.T1307C:p.I436T,CD44:NM_000610:exon12:c.T1436C:p.I479T,CD44:NM_001001390:exon6:c.T689C:p.I230T, |
| chr11 | 35208180 | G | A | . | exonic | CD44 | nonsynonymous SNV | CD44:NM_001001389:exon11:c.G1361A:p.R454K,CD44:NM_000610:exon12:c.G1490A:p.R497K,CD44:NM_001001390:exon6:c.G743A:p.R248K, |
| chr11 | 35221726 | G | A | COSV53530853 | exonic | CD44 | nonsynonymous SNV | CD44:NM_001001389:exon16:c.G1889A:p.R630Q,CD44:NM_000610:exon17:c.G2018A:p.R673Q,CD44:NM_001202556:exon7:c.G812A:p.R271Q,CD44:NM_001202555:exon9:c.G1079A:p.R360Q,CD44:NM_001202557:exon8:c.G875A:p.R292Q,CD44:NM_001001390:exon11:c.G1271A:p.R424Q,CD44:NM_001001391:exon8:c.G875A:p.R292Q, |
| chr11 | 43855148 | C | T | COSV53496916 | exonic | HSD17B12 | nonsynonymous SNV | HSD17B12:NM_016142:exon11:c.C839T:p.S280L, |
| chr11 | 46936748 | G | T | COSV53566094 | exonic | C11ORF49 | nonsynonymous SNV | C11orf49:NM_001278222:exon1:c.G19T:p.G7C, |
| chr11 | 47166860 | C | T | COSV53563682 | exonic | ARFGAP2 | nonsynonymous SNV | ARFGAP2:NM_032389:exon13:c.G1232A:p.S411N,ARFGAP2:NM_001242832:exon12:c.G1148A:p.S383N, |
| chr11 | 47618877 | G | C | COSV56766406 | exonic | MTCH2 | nonsynonymous SNV | MTCH2:NM_001317232:exon12:c.C841G:p.P281A,MTCH2:NM_001317233:exon12:c.C424G:p.P142A,MTCH2:NM_014342:exon13:c.C868G:p.P290A, |
| chr11 | 60497529 | C | T | COSV50014135 | exonic | MS4A12 | stopgain | MS4A12:NM_001164470:exon2:c.C211T:p.Q71X,MS4A12:NM_017716:exon2:c.C211T:p.Q71X, |
| chr11 | 64318313 | A | G | COSV50005053 | exonic | PRDX5 | nonsynonymous SNV | PRDX5:NM_012094:exon1:c.A98G:p.Y33C,PRDX5:NM_181652:exon1:c.A98G:p.Y33C,PRDX5:NM_181651:exon1:c.A98G:p.Y33C, |
| chr11 | 68794860 | C | T | . | exonic | CPT1A | nonsynonymous SNV | CPT1A:NM_001031847:exon8:c.G823A:p.A275T,CPT1A:NM_001876:exon8:c.G823A:p.A275T, |
| chr11 | 72195272 | G | A | . | exonic | FOLR1 | nonsynonymous SNV | FOLR1:NM_016729:exon2:c.G170A:p.C57Y,FOLR1:NM_000802:exon3:c.G170A:p.C57Y,FOLR1:NM_016725:exon3:c.G170A:p.C57Y,FOLR1:NM_016724:exon4:c.G170A:p.C57Y, |
| chr11 | 72195331 | G | A | COSV56616313 | exonic | FOLR1 | nonsynonymous SNV | FOLR1:NM_016729:exon2:c.G229A:p.D77N,FOLR1:NM_000802:exon3:c.G229A:p.D77N,FOLR1:NM_016725:exon3:c.G229A:p.D77N,FOLR1:NM_016724:exon4:c.G229A:p.D77N, |
| chr11 | 73808313 | G | T | COSV99063341 | exonic | MRPL48 | nonsynonymous SNV | MRPL48:NM_016055:exon3:c.G75T:p.R25S, |
| chr11 | 77702301 | C | T | COSV57836647 | exonic | RSF1 | nonsynonymous SNV | RSF1:NM_016578:exon6:c.G928A:p.E310K, |

| | | | | | | | | |
|---|---|---|---|---|---|---|---|---|
| chr11 | 77702322 | C | T | . | exonic | RSF1 | nonsynonymous SNV | RSF1:NM_016578:exon6:c.G907A:p.E303K, |
| chr11 | 77702323 | A | T | . | exonic | RSF1 | nonsynonymous SNV | RSF1:NM_016578:exon6:c.T906A:p.N302K, |
| chr11 | 78568744 | T | G | COSV55250015 | exonic | NARS2 | nonsynonymous SNV | NARS2:NM_024678:exon3:c.A260C:p.N87T, |
| chr11 | 87302396 | G | A | COSV59699111 | exonic | TMEM135 | nonsynonymous SNV | TMEM135:NM_022918:exon8:c.G652A:p.G218R,TMEM135:NM_001168724:exon7:c.G586A:p.G196R, |
| chr12 | 100624303 | C | A | . | exonic | GAS2L3 | nonsynonymous SNV | GAS2L3:NM_001303130:exon9:c.C1498A:p.P500T,GAS2L3:NM_001303131:exon10:c.C1186A:p.P396T,GAS2L3:NM_174942:exon10:c.C1498A:p.P500T, |
| chr12 | 108562641 | T | G | COSV57206465 | exonic | ISCU | nonsynonymous SNV | ISCU:NM_001301141:exon1:c.T19G:p.F7V,ISCU:NM_001301140:exon1:c.T19G:p.F7V,ISCU:NM_213595:exon1:c.T19G:p.F7V, |
| chr12 | 108562642 | T | G | COSV57206471 | exonic | ISCU | nonsynonymous SNV | ISCU:NM_001301141:exon1:c.T20G:p.F7C,ISCU:NM_001301140:exon1:c.T20G:p.F7C,ISCU:NM_213595:exon1:c.T20G:p.F7C, |
| chr12 | 108562657 | C | T | COSV57206596 | exonic | ISCU | nonsynonymous SNV | ISCU:NM_001301141:exon1:c.C35T:p.A12V,ISCU:NM_001301140:exon1:c.C35T:p.A12V,ISCU:NM_213595:exon1:c.C35T:p.A12V, |
| chr12 | 112450442 | T | A | . | exonic | PTPN11 | nonsynonymous SNV | PTPN11:NM_002834:exon3:c.T262A:p.L88I,PTPN11:NM_080601:exon3:c.T262A:p.L88I, |
| chr12 | 112453244 | G | A | . | exonic | PTPN11 | nonsynonymous SNV | PTPN11:NM_002834:exon4:c.G382A:p.E128K,PTPN11:NM_080601:exon4:c.G382A:p.E128K, |
| chr12 | 112453250 | G | A | . | exonic | PTPN11 | nonsynonymous SNV | PTPN11:NM_002834:exon4:c.G388A:p.G130R,PTPN11:NM_080601:exon4:c.G388A:p.G130R, |
| chr12 | 11752469 | C | T | . | exonic | ETV6 | nonsynonymous SNV | ETV6:NM_001987:exon2:c.C53T:p.T18I, |
| chr12 | 121888906 | T | C | COSV55838997 | exonic | PSMD9 | nonsynonymous SNV | PSMD9:NM_002813:exon1:c.T50C:p.V17A,PSMD9:NM_001261400:exon1:c.T50C:p.V17A, |
| chr12 | 122327956 | G | C | COSV56798569 | exonic | CLIP1 | nonsynonymous SNV | CLIP1:NM_002956:exon16:c.C3207G:p.D1069E,CLIP1:NM_001247997:exon17:c.C3240G:p.D1080E,CLIP1:NM_198240:exon15:c.C3102G:p.D1034E, |
| chr12 | 122328296 | C | T | . | exonic | CLIP1 | nonsynonymous SNV | CLIP1:NM_002956:exon15:c.G2965A:p.E989K,CLIP1:NM_001247997:exon16:c.G2998A:p.E1000K,CLIP1:NM_198240:exon14:c.G2860A:p.E954K, |
| chr12 | 124863632 | A | C | . | exonic | SCARB1 | nonsynonymous SNV | SCARB1:NM_001082959:exon1:c.T89G:p.V30G,SCARB1:NM_005505:exon1:c.T89G:p.V30G, |
| chr12 | 124863717 | C | T | COSV55547637 | exonic | SCARB1 | nonsynonymous SNV | SCARB1:NM_001082959:exon1:c.G4A:p.G2S,SCARB1:NM_005505:exon1:c.G4A:p.G2S, |
| chr12 | 21635571 | C | T | . | exonic | LDHB | nonsynonymous SNV | LDHB:NM_002300:exon8:c.G976A:p.D326N,LDHB:NM_001174097:exon8:c.G976A:p.D326N, |
| chr12 | 21635572 | C | T | . | exonic | LDHB | stopgain | LDHB:NM_002300:exon8:c.G975A:p.W325X,LDHB:NM_001174097:exon8:c.G975A:p.W325X, |
| chr12 | 21635648 | C | T | . | exonic | LDHB | nonsynonymous SNV | LDHB:NM_002300:exon8:c.G899A:p.G300E,LDHB:NM_001174097:exon8:c.G899A:p.G300E, |
| chr12 | 21635651 | C | T | . | exonic | LDHB | nonsynonymous SNV | LDHB:NM_002300:exon8:c.G896A:p.R299Q,LDHB:NM_001174097:exon8:c.G896A:p.R299Q, |
| chr12 | 21635700 | C | T | . | exonic | LDHB | nonsynonymous SNV | LDHB:NM_002300:exon8:c.G847A:p.G283S,LDHB:NM_001174097:exon8:c.G847A:p.G283S, |
| chr12 | 21637154 | C | T | COSV63364527 | exonic | LDHB | nonsynonymous SNV | LDHB:NM_002300:exon7:c.G754A:p.A252T,LDHB:NM_001174097:exon7:c.G754A:p.A252T, |
| chr12 | 21638389 | T | C | . | exonic | LDHB | nonsynonymous SNV | LDHB:NM_002300:exon6:c.A677G:p.E226G,LDHB:NM_001174097:exon6:c.A677G:p.E226G, |
| chr12 | 21638402 | C | T | . | exonic | LDHB | nonsynonymous SNV | LDHB:NM_002300:exon6:c.G664A:p.D222N,LDHB:NM_001174097:exon6:c.G664A:p.D222N, |
| chr12 | 21638407 | C | T | . | exonic | LDHB | nonsynonymous SNV | LDHB:NM_002300:exon6:c.G659A:p.G220E,LDHB:NM_001174097:exon6:c.G659A:p.G220E, |
| chr12 | 21638426 | C | T | . | exonic | LDHB | nonsynonymous SNV | LDHB:NM_002300:exon6:c.G640A:p.E214K,LDHB:NM_001174097:exon6:c.G640A:p.E214K, |
| chr12 | 21638460 | C | T | . | exonic | LDHB | stopgain | LDHB:NM_002300:exon6:c.G606A:p.W202X,LDHB:NM_001174097:exon6:c.G606A:p.W202X, |
| chr12 | 21641972 | C | T | . | exonic | LDHB | nonsynonymous SNV | LDHB:NM_002300:exon5:c.G575A:p.G192E,LDHB:NM_001174097:exon5:c.G575A:p.G192E, |
| chr12 | 21641973 | C | T | . | exonic | LDHB | nonsynonymous SNV | LDHB:NM_002300:exon5:c.G574A:p.G192R,LDHB:NM_001174097:exon5:c.G574A:p.G192R, |
| chr12 | 21641979 | T | A | . | exonic | LDHB | nonsynonymous SNV | LDHB:NM_002300:exon5:c.A568T:p.I190F,LDHB:NM_001174097:exon5:c.A568T:p.I190F, |
| chr12 | 21641980 | C | T | . | exonic | LDHB | stopgain | LDHB:NM_002300:exon5:c.G567A:p.W189X,LDHB:NM_001174097:exon5:c.G567A:p.W189X, |
| chr12 | 21642032 | C | T | . | exonic | LDHB | nonsynonymous SNV | LDHB:NM_002300:exon5:c.G515A:p.R172H,LDHB:NM_001174097:exon5:c.G515A:p.R172H, |
| chr12 | 21642066 | C | T | COSV63365537 | exonic | LDHB | nonsynonymous SNV | LDHB:NM_002300:exon5:c.G481A:p.G161R,LDHB:NM_001174097:exon5:c.G481A:p.G161R, |
| chr12 | 21643947 | C | T | . | exonic | LDHB | nonsynonymous SNV | LDHB:NM_002300:exon4:c.G409A:p.V137I,LDHB:NM_001174097:exon4:c.G409A:p.V137I, |
| chr12 | 21644036 | C | T | . | exonic | LDHB | nonsynonymous SNV | LDHB:NM_002300:exon4:c.G320A:p.R107Q,LDHB:NM_001174097:exon4:c.G320A:p.R107Q, |
| chr12 | 21644046 | C | T | . | exonic | LDHB | nonsynonymous SNV | LDHB:NM_002300:exon4:c.G310A:p.G104R,LDHB:NM_001174097:exon4:c.G310A:p.G104R, |
| chr12 | 21644057 | C | T | . | exonic | LDHB | nonsynonymous SNV | LDHB:NM_002300:exon4:c.G299A:p.R100H,LDHB:NM_001174097:exon4:c.G299A:p.R100H, |
| chr12 | 21644108 | T | C | . | exonic | LDHB | nonsynonymous SNV | LDHB:NM_002300:exon4:c.A248G:p.D83G,LDHB:NM_001174097:exon4:c.A248G:p.D83G, |
| chr12 | 21646992 | C | T | . | exonic | LDHB | nonsynonymous SNV | LDHB:NM_002300:exon3:c.G154A:p.V52M,LDHB:NM_001174097:exon3:c.G154A:p.V52M, |
| chr12 | 21647010 | C | T | . | exonic | LDHB | nonsynonymous SNV | LDHB:NM_002300:exon3:c.G136A:p.A46T,LDHB:NM_001174097:exon3:c.G136A:p.A46T, |
| chr12 | 21654556 | C | T | . | exonic | LDHB | nonsynonymous SNV | LDHB:NM_002300:exon2:c.G116A:p.S39N,LDHB:NM_001174097:exon2:c.G116A:p.S39N, |
| chr12 | 21654570 | C | T | . | exonic | LDHB | nonsynonymous SNV | LDHB:NM_002300:exon2:c.G102A:p.M34I,LDHB:NM_001174097:exon2:c.G102A:p.M34I, |
| chr12 | 21654669 | C | T | . | exonic | LDHB | nonsynonymous SNV | LDHB:NM_002300:exon2:c.G3A:p.M1I,LDHB:NM_001174097:exon2:c.G3A:p.M1I, |
| chr12 | 25245350 | C | A | COSV55497419 | exonic | KRAS | nonsynonymous SNV | KRAS:NM_004985:exon2:c.G35T:p.G12V,KRAS:NM_033360:exon2:c.G35T:p.G12V, |
| chr12 | 438517 | T | C | COSV53475389 | exonic | CCDC77 | nonsynonymous SNV | CCDC77:NM_032358:exon10:c.T1004C:p.M335T,CCDC77:NM_001130146:exon9:c.T908C:p.M303T,CCDC77:NM_001130147:exon9:c.T908C:p.M303T,CCDC77:NM_001130148:exon8:c.T908C:p.M303T, |
| chr12 | 50819655 | G | A | . | exonic | ATF1 | nonsynonymous SNV | ATF1:NM_005171:exon7:c.G692A:p.R231H, |
| chr12 | 55789103 | C | T | . | exonic | SARNP | nonsynonymous SNV | SARNP:NM_033082:exon9:c.G473A:p.G158D, |
| chr12 | 56671771 | C | T | . | exonic | PTGES3 | nonsynonymous SNV | PTGES3:NM_006601:exon4:c.G263A:p.R88K,PTGES3:NM_001282602:exon4:c.G263A:p.R88K,PTGES3:NM_001282601:exon4:c.G263A:p.R88K,PTGES3:NM_001282605:exon4:c.G263A:p.R88K,PTGES3:NM_001282604:exon5:c.G275A:p.R92K, |
| chr12 | 56712541 | C | T | . | exonic | NACA | nonsynonymous SNV | NACA:NM_001113203:exon11:c.G2775A:p.M925I,NACA:NM_001113202:exon8:c.G645A:p.M215I, |
| chr12 | 56712827 | C | T | . | exonic | NACA | nonsynonymous SNV | NACA:NM_001113203:exon10:c.G2722A:p.A908T,NACA:NM_001113202:exon7:c.G592A:p.A198T, |
| chr12 | 56712838 | T | C | . | exonic | NACA | nonsynonymous SNV | NACA:NM_001113203:exon10:c.A2711G:p.K904R,NACA:NM_001113202:exon7:c.A581G:p.K194R, |
| chr12 | 56712844 | C | T | . | exonic | NACA | nonsynonymous SNV | NACA:NM_001113203:exon10:c.G2705A:p.R902K,NACA:NM_001113202:exon7:c.G575A:p.R192K, |

| | | | | | | | | |
|---|---|---|---|---|---|---|---|---|
| chr12 | 56712902 | C | T | COSV63269367 | exonic | NACA | nonsynonymous SNV | NACA:NM_001113203:exon10:c.G2647A:p.E883K,NACA:NM_001113202:exon7:c.G517A:p.E173K, |
| chr12 | 56713572 | C | T | . | exonic | NACA | nonsynonymous SNV | NACA:NM_001113203:exon8:c.G2476A:p.A826T,NACA:NM_001113202:exon5:c.G346A:p.A116T, |
| chr12 | 56713638 | C | A | . | exonic | NACA | nonsynonymous SNV | NACA:NM_001113203:exon8:c.G2410T:p.V804F,NACA:NM_001113202:exon5:c.G280T:p.V94F, |
| chr12 | 56714379 | C | T | . | exonic | NACA | nonsynonymous SNV | NACA:NM_001113203:exon7:c.G2347A:p.E783K,NACA:NM_001113202:exon4:c.G217A:p.E73K, |
| chr12 | 56714670 | C | T | . | exonic | NACA | nonsynonymous SNV | NACA:NM_001113203:exon6:c.G2218A:p.D740N,NACA:NM_001113202:exon3:c.G88A:p.D30N, |
| chr12 | 56724452 | C | T | . | exonic | NACA | nonsynonymous SNV | NACA:NM_001113203:exon2:c.G70A:p.A24T,NACA:NM_001113202:exon2:c.G70A:p.G24R, |
| chr12 | 56724496 | A | G | . | exonic | NACA | nonsynonymous SNV | NACA:NM_001113203:exon2:c.T26C:p.V9A,NACA:NM_001113202:exon2:c.T26C:p.V9A, |
| chr12 | 57516963 | C | T | . | exonic | DDIT3 | nonsynonymous SNV | DDIT3:NM_001195056:exon3:c.G425A:p.R142H,DDIT3:NM_001195053:exon4:c.G425A:p.R142H,DDIT3:NM_001195057:exon4:c.G356A:p.R119H,DDIT3:NM_001195055:exon4:c.G425A:p.R142H,DDIT3:NM_001195054:exon4:c.G425A:p.R142H,DDIT3:NM_004083:exon4:c.G356A:p.R119H, |
| chr12 | 57517102 | C | T | . | exonic | DDIT3 | nonsynonymous SNV | DDIT3:NM_001195056:exon3:c.G286A:p.V96I,DDIT3:NM_001195053:exon4:c.G286A:p.V96I,DDIT3:NM_001195057:exon4:c.G217A:p.V73I,DDIT3:NM_001195055:exon4:c.G286A:p.V96I,DDIT3:NM_001195054:exon4:c.G286A:p.V96I,DDIT3:NM_004083:exon4:c.G217A:p.V73I, |
| chr12 | 57941843 | A | T | COSV55686289 | exonic | XRCC6BP1 | nonsynonymous SNV | XRCC6BP1:NM_033276:exon1:c.A142T:p.S48C, |
| chr12 | 6333477 | C | T | COSV50610888 | exonic | TNFRSF1A | nonsynonymous SNV | TNFRSF1A:NM_001065:exon4:c.G362A:p.R121Q, |
| chr12 | 6534851 | G | A | . | exonic | GAPDH | nonsynonymous SNV | GAPDH:NM_001289745:exon2:c.G19A:p.G7R,GAPDH:NM_001289746:exon1:c.G19A:p.G7R,GAPDH:NM_002046:exon2:c.G19A:p.G7R, |
| chr12 | 6536591 | A | T | . | exonic | GAPDH | nonsynonymous SNV | GAPDH:NM_001256799:exon2:c.A1T:p.M1L,GAPDH:NM_001289745:exon3:c.A127T:p.M43L,GAPDH:NM_001289746:exon2:c.A127T:p.M43L,GAPDH:NM_002046:exon3:c.A127T:p.M43L, |
| chr12 | 6536756 | G | A | . | exonic | GAPDH | nonsynonymous SNV | GAPDH:NM_001256799:exon3:c.G76A:p.V26I,GAPDH:NM_001289745:exon4:c.G202A:p.V68I,GAPDH:NM_001289746:exon3:c.G202A:p.V68I,GAPDH:NM_002046:exon4:c.G202A:p.V68I, |
| chr12 | 6536957 | G | A | COSV57521174 | exonic | GAPDH | nonsynonymous SNV | GAPDH:NM_001256799:exon4:c.G148A:p.A50T,GAPDH:NM_001289745:exon5:c.G274A:p.A92T,GAPDH:NM_001289746:exon4:c.G274A:p.A92T,GAPDH:NM_002046:exon5:c.G274A:p.A92T, |
| chr12 | 6536969 | G | A | . | exonic | GAPDH | nonsynonymous SNV | GAPDH:NM_001256799:exon4:c.G160A:p.V54M,GAPDH:NM_001289745:exon5:c.G286A:p.V96M,GAPDH:NM_001289746:exon4:c.G286A:p.V96M,GAPDH:NM_002046:exon5:c.G286A:p.V96M, |
| chr12 | 6537101 | G | A | . | exonic | GAPDH | nonsynonymous SNV | GAPDH:NM_001256799:exon5:c.G202A:p.A68T,GAPDH:NM_001289745:exon6:c.G328A:p.A110T,GAPDH:NM_001289746:exon5:c.G328A:p.A110T,GAPDH:NM_002046:exon6:c.G328A:p.A110T, |
| chr12 | 6537116 | G | A | COSV57520634 | exonic | GAPDH | nonsynonymous SNV | GAPDH:NM_001256799:exon5:c.G217A:p.G73R,GAPDH:NM_001289745:exon6:c.G343A:p.G115R,GAPDH:NM_001289746:exon5:c.G343A:p.G115R,GAPDH:NM_002046:exon6:c.G343A:p.G115R, |
| chr12 | 6537140 | G | A | . | exonic | GAPDH | nonsynonymous SNV | GAPDH:NM_001256799:exon5:c.G241A:p.A81T,GAPDH:NM_001289745:exon6:c.G367A:p.A123T,GAPDH:NM_001289746:exon5:c.G367A:p.A123T,GAPDH:NM_002046:exon6:c.G367A:p.A123T, |
| chr12 | 6537158 | C | A | . | exonic | GAPDH | nonsynonymous SNV | GAPDH:NM_001256799:exon5:c.C259A:p.P87T,GAPDH:NM_001289745:exon6:c.C385A:p.P129T,GAPDH:NM_001289746:exon5:c.C385A:p.P129T,GAPDH:NM_002046:exon6:c.C385A:p.P129T, |
| chr12 | 6537167 | G | A | . | exonic | GAPDH | nonsynonymous SNV | GAPDH:NM_001256799:exon5:c.G268A:p.V90I,GAPDH:NM_001289745:exon6:c.G394A:p.V132I,GAPDH:NM_001289746:exon5:c.G394A:p.V132I,GAPDH:NM_002046:exon6:c.G394A:p.V132I, |
| chr12 | 6537216 | G | A | . | exonic | GAPDH | nonsynonymous SNV | GAPDH:NM_001256799:exon5:c.G317A:p.S106N,GAPDH:NM_001289745:exon6:c.G443A:p.S148N,GAPDH:NM_001289746:exon5:c.G443A:p.S148N,GAPDH:NM_002046:exon6:c.G443A:p.S148N, |
| chr12 | 6537361 | G | A | . | exonic | GAPDH | nonsynonymous SNV | GAPDH:NM_001256799:exon6:c.G370A:p.D124N,GAPDH:NM_001289745:exon7:c.G496A:p.D166N,GAPDH:NM_001289746:exon6:c.G496A:p.D166N,GAPDH:NM_002046:exon7:c.G496A:p.D166N, |
| chr12 | 6537660 | G | A | . | exonic | GAPDH | nonsynonymous SNV | GAPDH:NM_001256799:exon7:c.G476A:p.G159E,GAPDH:NM_001289745:exon8:c.G602A:p.G201E,GAPDH:NM_001289746:exon7:c.G602A:p.G201E,GAPDH:NM_002046:exon8:c.G602A:p.G201E, |
| chr12 | 6537734 | G | A | . | exonic | GAPDH | nonsynonymous SNV | GAPDH:NM_001256799:exon7:c.G550A:p.G184R,GAPDH:NM_001289745:exon8:c.G676A:p.G226R,GAPDH:NM_001289746:exon7:c.G676A:p.G226R,GAPDH:NM_002046:exon8:c.G676A:p.G226R, |
| chr12 | 6537820 | A | T | . | exonic | GAPDH | nonsynonymous SNV | GAPDH:NM_001256799:exon7:c.A636T:p.K212N,GAPDH:NM_001289745:exon8:c.A762T:p.K254N,GAPDH:NM_001289746:exon7:c.A762T:p.K254N,GAPDH:NM_002046:exon8:c.A762T:p.K254N, |
| chr12 | 6537821 | T | A | . | exonic | GAPDH | nonsynonymous SNV | GAPDH:NM_001256799:exon7:c.T637A:p.Y213N,GAPDH:NM_001289745:exon8:c.T763A:p.Y255N,GAPDH:NM_001289746:exon7:c.T763A:p.Y255N,GAPDH:NM_002046:exon8:c.T763A:p.Y255N, |
| chr12 | 6537872 | G | A | . | exonic | GAPDH | nonsynonymous SNV | GAPDH:NM_001256799:exon7:c.G688A:p.G230S,GAPDH:NM_001289745:exon8:c.G814A:p.G272S,GAPDH:NM_001289746:exon7:c.G814A:p.G272S,GAPDH:NM_002046:exon8:c.G814A:p.G272S, |
| chr12 | 6537873 | G | A | . | exonic | GAPDH | nonsynonymous SNV | GAPDH:NM_001256799:exon7:c.G689A:p.G230D,GAPDH:NM_001289745:exon8:c.G815A:p.G272D,GAPDH:NM_001289746:exon7:c.G815A:p.G272D,GAPDH:NM_002046:exon8:c.G815A:p.G272D, |
| chr12 | 6537882 | G | A | . | exonic | GAPDH | nonsynonymous SNV | GAPDH:NM_001256799:exon7:c.G698A:p.G233D,GAPDH:NM_001289745:exon8:c.G824A:p.G275D,GAPDH:NM_001289746:exon7:c.G824A:p.G275D,GAPDH:NM_002046:exon8:c.G824A:p.G275D, |
| chr12 | 69253133 | G | A | . | exonic | CPSF6 | nonsynonymous SNV | CPSF6:NM_007007:exon3:c.G353A:p.R118Q,CPSF6:NM_001300947:exon3:c.G353A:p.R118Q, |
| chr12 | 69258629 | G | A | . | exonic | CPSF6 | nonsynonymous SNV | CPSF6:NM_007007:exon6:c.G734A:p.G245D,CPSF6:NM_001300947:exon7:c.G845A:p.G282D, |
| chr12 | 69258638 | G | A | . | exonic | CPSF6 | nonsynonymous SNV | CPSF6:NM_007007:exon6:c.G743A:p.G248D,CPSF6:NM_001300947:exon7:c.G854A:p.G285D, |
| chr12 | 70701175 | C | T | . | exonic | PTPRR | nonsynonymous SNV | PTPRR:NM_001207016:exon4:c.G538A:p.V180I,PTPRR:NM_001207015:exon6:c.G820A:p.V274I,PTPRR:NM_130846:exon3:c.G421A:p.V141I,PTPRR:NM_002849:exon7:c.G1156A:p.V386I, |

| | | | | | | | | |
|---|---|---|---|---|---|---|---|---|
| chr12 | 70745884 | T | C | COSV51736040 | exonic | PTPRR | nonsynonymous SNV | PTPRR:NM_001207016:exon3:c.A323G:p.K108R,PTPRR:NM_001207015:exon5:c.A605G:p.K202R,PTPRR:NM_130846:exon2:c.A206G:p.K69R,PTPRR:NM_002849:exon6:c.A941G:p.K314R, |
| chr12 | 71129354 | A | C | COSV56081147 | exonic | TSPAN8 | nonsynonymous SNV | TSPAN8:NM_004616:exon8:c.T637G:p.S213A, |
| chr12 | 71139754 | C | G | COSV56077446 | exonic | TSPAN8 | nonsynonymous SNV | TSPAN8:NM_004616:exon4:c.G218C:p.G73A, |
| chr13 | 114291534 | T | A | . | exonic | UPF3A | stopgain | UPF3A:NM_023011:exon6:c.T677A:p.L226X,UPF3A:NM_080687:exon5:c.T578A:p.L193X, |
| chr13 | 114291734 | G | A | COSV60894561 | exonic | UPF3A | nonsynonymous SNV | UPF3A:NM_023011:exon7:c.G788A:p.C263Y,UPF3A:NM_080687:exon6:c.G689A:p.C230Y, |
| chr13 | 114298886 | C | A | . | exonic | UPF3A | nonsynonymous SNV | UPF3A:NM_023011:exon8:c.C893A:p.P298Q,UPF3A:NM_080687:exon7:c.C794A:p.P265Q, |
| chr13 | 21510035 | G | T | . | exonic | MICU2 | nonsynonymous SNV | MICU2:NM_152726:exon8:c.C730A:p.Q244K, |
| chr13 | 23837633 | T | C | . | exonic | MIPEP | nonsynonymous SNV | MIPEP:NM_005932:exon13:c.A1462G:p.S488G, |
| chr13 | 31155639 | T | G | COSV99062651 | exonic | HSPH1 | nonsynonymous SNV | HSPH1:NM_001286505:exon3:c.A187C:p.N63H,HSPH1:NM_001286503:exon3:c.A181C:p.N61H,HSPH1:NM_006644:exon3:c.A181C:p.N61H,HSPH1:NM_001286504:exon3:c.A187C:p.N63H, |
| chr13 | 39655754 | G | A | COSV62995992 | exonic | COG6 | nonsynonymous SNV | COG6:NM_020751:exon1:c.G28A:p.A10T,COG6:NM_001145079:exon1:c.G28A:p.A10T, |
| chr13 | 39655820 | T | A | COSV62995068 | exonic | COG6 | nonsynonymous SNV | COG6:NM_020751:exon1:c.T94A:p.C32S,COG6:NM_001145079:exon1:c.T94A:p.C32S, |
| chr13 | 39694708 | T | G | . | exonic | COG6 | nonsynonymous SNV | COG6:NM_020751:exon12:c.T1149G:p.F383L,COG6:NM_001145079:exon12:c.T1149G:p.F383L, |
| chr13 | 40958916 | T | C | COSV53497107 | exonic | ELF1 | nonsynonymous SNV | ELF1:NM_001145353:exon2:c.A173G:p.N58S,ELF1:NM_172373:exon3:c.A173G:p.N58S, |
| chr13 | 45338712 | C | T | . | exonic | TPT1 | nonsynonymous SNV | TPT1:NM_003295:exon5:c.G464A:p.G155D,TPT1:NM_001286272:exon5:c.G464A:p.G155D,TPT1:NM_001286273:exon4:c.G362A:p.G121D, |
| chr13 | 45338721 | C | T | . | exonic | TPT1 | nonsynonymous SNV | TPT1:NM_003295:exon5:c.G455A:p.R152H,TPT1:NM_001286272:exon5:c.G455A:p.R152H,TPT1:NM_001286273:exon4:c.G353A:p.R118H, |
| chr13 | 45338737 | C | T | . | exonic | TPT1 | nonsynonymous SNV | TPT1:NM_003295:exon5:c.G439A:p.A147T,TPT1:NM_001286272:exon5:c.G439A:p.A147T,TPT1:NM_001286273:exon4:c.G337A:p.A113T, |
| chr13 | 45339510 | A | G | . | exonic | TPT1 | nonsynonymous SNV | TPT1:NM_003295:exon4:c.T386C:p.F129S,TPT1:NM_001286272:exon4:c.T386C:p.F129S,TPT1:NM_001286273:exon3:c.T284C:p.F95S, |
| chr13 | 45339541 | C | T | . | exonic | TPT1 | nonsynonymous SNV | TPT1:NM_003295:exon4:c.G355A:p.A119T,TPT1:NM_001286272:exon4:c.G355A:p.A119T,TPT1:NM_001286273:exon3:c.G253A:p.A85T, |
| chr13 | 45339594 | C | T | . | exonic | TPT1 | nonsynonymous SNV | TPT1:NM_003295:exon4:c.G302A:p.G101E,TPT1:NM_001286272:exon4:c.G302A:p.G101E,TPT1:NM_001286273:exon3:c.G200A:p.G67E, |
| chr13 | 45340076 | C | T | . | exonic | TPT1 | nonsynonymous SNV | TPT1:NM_003295:exon3:c.G211A:p.D71N,TPT1:NM_001286272:exon3:c.G211A:p.D71N,TPT1:NM_001286273:exon2:c.G109A:p.D37N, |
| chr13 | 45340082 | C | T | . | exonic | TPT1 | nonsynonymous SNV | TPT1:NM_003295:exon3:c.G205A:p.G69S,TPT1:NM_001286272:exon3:c.G205A:p.G69S,TPT1:NM_001286273:exon2:c.G103A:p.G35S, |
| chr13 | 45340100 | C | T | . | exonic | TPT1 | nonsynonymous SNV | TPT1:NM_003295:exon3:c.G187A:p.E63K,TPT1:NM_001286272:exon3:c.G187A:p.E63K,TPT1:NM_001286273:exon2:c.G85A:p.E29K, |
| chr13 | 45340174 | C | T | . | exonic | TPT1 | nonsynonymous SNV | TPT1:NM_003295:exon3:c.G113A:p.R38K,TPT1:NM_001286272:exon3:c.G113A:p.R38K,TPT1:NM_001286273:exon2:c.G11A:p.R4K, |
| chr13 | 45341067 | C | T | COSV58539175 | exonic | TPT1 | nonsynonymous SNV | TPT1:NM_003295:exon1:c.G3A:p.M1I,TPT1:NM_001286272:exon1:c.G3A:p.M1I, |
| chr13 | 45969538 | G | T | . | exonic | ZC3H13 | nonsynonymous SNV | ZC3H13:NM_015070:exon14:c.C3006A:p.S1002R, |
| chr13 | 48077198 | C | T | . | exonic | MED4 | nonsynonymous SNV | MED4:NM_001270629:exon7:c.G616A:p.E206K,MED4:NM_014166:exon7:c.G754A:p.E252K, |
| chr13 | 48083379 | G | T | . | exonic | MED4 | nonsynonymous SNV | MED4:NM_001270629:exon4:c.C275A:p.A92E,MED4:NM_014166:exon4:c.C413A:p.A138E, |
| chr13 | 48083380 | C | T | . | exonic | MED4 | nonsynonymous SNV | MED4:NM_001270629:exon4:c.G274A:p.A92T,MED4:NM_014166:exon4:c.G412A:p.A138T, |
| chr13 | 48086358 | A | G | . | exonic | MED4 | nonsynonymous SNV | MED4:NM_001270629:exon3:c.T149C:p.L50S,MED4:NM_014166:exon3:c.T287C:p.L96S, |
| chr13 | 48086366 | C | T | . | exonic | MED4 | nonsynonymous SNV | MED4:NM_001270629:exon3:c.G141A:p.M47I,MED4:NM_014166:exon3:c.G279A:p.M93I, |
| chr13 | 72795504 | A | G | COSV58323477 | exonic | PIBF1 | nonsynonymous SNV | PIBF1:NM_006346:exon4:c.A499G:p.I167V, |
| chr14 | 102083681 | C | T | . | exonic | HSP90AA1 | nonsynonymous SNV | HSP90AA1:NM_005348:exon8:c.G1351A:p.E451K,HSP90AA1:NM_001017963:exon9:c.G1717A:p.E573K, |
| chr14 | 102083972 | C | T | . | exonic | HSP90AA1 | nonsynonymous SNV | HSP90AA1:NM_005348:exon7:c.G1159A:p.G387R,HSP90AA1:NM_001017963:exon8:c.G1525A:p.G509R, |
| chr14 | 102084938 | C | T | . | exonic | HSP90AA1 | nonsynonymous SNV | HSP90AA1:NM_005348:exon5:c.G724A:p.E242K,HSP90AA1:NM_001017963:exon6:c.G1090A:p.E364K, |
| chr14 | 102875712 | T | C | COSV61023607 | exonic | TRAF3 | nonsynonymous SNV | TRAF3:NM_001199427:exon4:c.T386C:p.M129T,TRAF3:NM_145725:exon5:c.T386C:p.M129T,TRAF3:NM_145726:exon5:c.T386C:p.M129T,TRAF3:NM_003300:exon4:c.T386C:p.M129T, |
| chr14 | 20456995 | T | G | COSV51657755 | exonic | APEX1 | nonsynonymous SNV | APEX1:NM_080648:exon5:c.T444G:p.D148E,APEX1:NM_080649:exon5:c.T444G:p.D148E,APEX1:NM_001244249:exon5:c.T444G:p.D148E,APEX1:NM_001641:exon5:c.T444G:p.D148E, |
| chr14 | 24145152 | T | G | . | exonic | PSME2 | nonsynonymous SNV | PSME2:NM_002818:exon6:c.A266C:p.H89P, |
| chr14 | 24267755 | T | C | COSV52861986 | exonic | RABGGTA | nonsynonymous SNV | RABGGTA:NM_004581:exon13:c.A1258G:p.T420A,RABGGTA:NM_182836:exon14:c.A1258G:p.T420A, |
| chr14 | 34800273 | C | T | . | exonic | BAZ1A | nonsynonymous SNV | BAZ1A:NM_182648:exon9:c.G1079A:p.R360K,BAZ1A:NM_013448:exon9:c.G1079A:p.R360K, |

| | | | | | | | | |
|---|---|---|---|---|---|---|---|---|
| chr14 | 34800320 | A | T | . | exonic | BAZ1A | nonsynonymous SNV | BAZ1A:NM_182648:exon9:c.T1032A:p.D344E,BAZ1A:NM_013448:exon9:c.T1032A:p.D344E, |
| chr14 | 34800325 | C | T | . | exonic | BAZ1A | nonsynonymous SNV | BAZ1A:NM_182648:exon9:c.G1027A:p.E343K,BAZ1A:NM_013448:exon9:c.G1027A:p.E343K, |
| chr14 | 34801128 | A | T | . | exonic | BAZ1A | nonsynonymous SNV | BAZ1A:NM_182648:exon8:c.T927A:p.D309E,BAZ1A:NM_013448:exon8:c.T927A:p.D309E, |
| chr14 | 35266761 | A | G | COSV51617748 | exonic | KIAA0391 | nonsynonymous SNV | KIAA0391:NM_001256680:exon5:c.A194G:p.N65S,KIAA0391:NM_014672:exon6:c.A1310G:p.N437S,KIAA0391:NM_001256678:exon5:c.A1262G:p.N421S,KIAA0391:NM_001256681:exon4:c.A194G:p.N65S,KIAA0391:NM_001256679:exon6:c.A1025G:p.N342S, |
| chr14 | 50312098 | A | C | COSV55413053 | exonic | L2HGDH | nonsynonymous SNV | L2HGDH:NM_024884:exon1:c.T53G:p.L18R, |
| chr14 | 55138217 | C | A | . | exonic | LGALS3 | nonsynonymous SNV | LGALS3:NM_001177388:exon3:c.C191A:p.P64H,LGALS3:NM_002306:exon3:c.C191A:p.P64H, |
| chr14 | 55138318 | A | C | . | exonic | LGALS3 | nonsynonymous SNV | LGALS3:NM_001177388:exon3:c.A292C:p.T98P,LGALS3:NM_002306:exon3:c.A292C:p.T98P, |
| chr14 | 65078029 | C | T | COSV52414939 | exonic | MAX | nonsynonymous SNV | MAX:NM_145113:exon4:c.G179A:p.R60Q,MAX:NM_145112:exon3:c.G152A:p.R51Q,MAX:NM_002382:exon4:c.G179A:p.R60Q, |
| chr14 | 68874936 | T | A | . | exonic | ACTN1 | nonsynonymous SNV | ACTN1:NM_001130005:exon21:c.A2587T:p.T863S,ACTN1:NM_001130004:exon22:c.A2668T:p.T890S,ACTN1:NM_001102:exon21:c.A2602T:p.T868S, |
| chr14 | 69237836 | G | T | . | exonic | EXD2 | nonsynonymous SNV | EXD2:NM_001193361:exon8:c.G1554T:p.Q518H,EXD2:NM_001193363:exon8:c.G1554T:p.Q518H,EXD2:NM_001193362:exon9:c.G1554T:p.Q518H,EXD2:NM_001193360:exon9:c.G1554T:p.Q518H,EXD2:NM_018199:exon7:c.G1179T:p.Q393H, |
| chr14 | 73105860 | G | A | COSV56201910 | exonic | RBM25 | nonsynonymous SNV | RBM25:NM_021239:exon11:c.G1156A:p.E386K, |
| chr14 | 73105954 | G | A | COSV56202337 | exonic | RBM25 | nonsynonymous SNV | RBM25:NM_021239:exon11:c.G1250A:p.R417Q, |
| chr14 | 73105988 | C | A | . | exonic | RBM25 | nonsynonymous SNV | RBM25:NM_021239:exon11:c.C1284A:p.D428E, |
| chr14 | 73106253 | G | A | . | exonic | RBM25 | nonsynonymous SNV | RBM25:NM_021239:exon12:c.G1435A:p.A479T, |
| chr14 | 73110909 | C | A | . | exonic | RBM25 | nonsynonymous SNV | RBM25:NM_021239:exon15:c.C1771A:p.Q591K, |
| chr14 | 77731069 | C | T | . | exonic | SNW1 | nonsynonymous SNV | SNW1:NM_012245:exon10:c.G952A:p.E318K, |
| chr14 | 91802271 | T | G | . | exonic | TC2N | nonsynonymous SNV | TC2N:NM_001289134:exon4:c.A452C:p.K151T,TC2N:NM_001128595:exon4:c.A452C:p.K151T,TC2N:NM_152332:exon4:c.A452C:p.K151T,TC2N:NM_001128596:exon4:c.A452C:p.K151T, |
| chr14 | 94378506 | T | G | COSV63345434 | exonic | SERPINA1 | nonsynonymous SNV | SERPINA1:NM_001002235:exon5:c.A1200C:p.E400D,SERPINA1:NM_001127704:exon7:c.A1200C:p.E400D,SERPINA1:NM_001127703:exon7:c.A1200C:p.E400D,SERPINA1:NM_001127700:exon5:c.A1200C:p.E400D,SERPINA1:NM_000295:exon5:c.A1200C:p.E400D,SERPINA1:NM_001127706:exon6:c.A1200C:p.E400D,SERPINA1:NM_001127705:exon7:c.A1200C:p.E400D,SERPINA1:NM_001002236:exon7:c.A1200C:p.E400D,SERPINA1:NM_001127702:exon6:c.A1200C:p.E400D,SERPINA1:NM_001127707:exon6:c.A1200C:p.E400D,SERPINA1:NM_001127701:exon7:c.A1200C:p.E400D, |
| chr14 | 94382864 | C | T | COSV63345368 | exonic | SERPINA1 | nonsynonymous SNV | SERPINA1:NM_001002235:exon2:c.G374A:p.R125H,SERPINA1:NM_001127704:exon4:c.G374A:p.R125H,SERPINA1:NM_001127703:exon4:c.G374A:p.R125H,SERPINA1:NM_001127700:exon2:c.G374A:p.R125H,SERPINA1:NM_000295:exon2:c.G374A:p.R125H,SERPINA1:NM_001127706:exon3:c.G374A:p.R125H,SERPINA1:NM_001127705:exon4:c.G374A:p.R125H,SERPINA1:NM_001002236:exon4:c.G374A:p.R125H,SERPINA1:NM_001127702:exon3:c.G374A:p.R125H,SERPINA1:NM_001127707:exon3:c.G374A:p.R125H,SERPINA1:NM_001127701:exon4:c.G374A:p.R125H, |
| chr14 | 95196631 | A | T | . | exonic | CLMN | nonsynonymous SNV | CLMN:NM_024734:exon10:c.T2575A:p.S859T, |
| chr14 | 96847336 | T | A | . | exonic | VRK1 | nonsynonymous SNV | VRK1:NM_003384:exon5:c.T366A:p.N122K, |
| chr15 | 40606445 | G | C | COSV61150583 | exonic | CASC5 | nonsynonymous SNV | CASC5:NM_144508:exon4:c.G128C:p.R43T,CASC5:NM_170589:exon4:c.G128C:p.R43T, |
| chr15 | 40611486 | A | G | COSV61154720 | exonic | CASC5 | nonsynonymous SNV | CASC5:NM_144508:exon7:c.A259G:p.T87A,CASC5:NM_170589:exon8:c.A337G:p.T113A, |
| chr15 | 40621979 | T | C | COSV61151299 | exonic | CASC5 | nonsynonymous SNV | CASC5:NM_144508:exon10:c.T1715C:p.M572T,CASC5:NM_170589:exon11:c.T1793C:p.M598T, |
| chr15 | 40622992 | A | G | COSV61150613 | exonic | CASC5 | nonsynonymous SNV | CASC5:NM_144508:exon10:c.A2728G:p.R910G,CASC5:NM_170589:exon11:c.A2806G:p.R936G, |
| chr15 | 43475576 | G | C | COSV55496034 | exonic | TP53BP1 | nonsynonymous SNV | TP53BP1:NM_005657:exon9:c.C1059G:p.D353E,TP53BP1:NM_001141980:exon9:c.C1074G:p.D358E,TP53BP1:NM_001141979:exon9:c.C1074G:p.D358E, |
| chr15 | 44711577 | G | A | . | exonic | B2M | nonsynonymous SNV | B2M:NM_004048:exon1:c.G31A:p.A11T, |
| chr15 | 44711595 | G | A | . | exonic | B2M | nonsynonymous SNV | B2M:NM_004048:exon1:c.G49A:p.G17S, |
| chr15 | 44715450 | G | A | . | exonic | B2M | nonsynonymous SNV | B2M:NM_004048:exon2:c.G95A:p.R32H, |
| chr15 | 44715489 | G | A | . | exonic | B2M | nonsynonymous SNV | B2M:NM_004048:exon2:c.G134A:p.C45Y, |
| chr15 | 44715585 | G | A | . | exonic | B2M | nonsynonymous SNV | B2M:NM_004048:exon2:c.G230A:p.S77N, |
| chr15 | 44715595 | G | A | . | exonic | B2M | stopgain | B2M:NM_004048:exon2:c.G240A:p.W80X, |
| chr15 | 44716332 | G | A | COSV62564948 | exonic | B2M | nonsynonymous SNV | B2M:NM_004048:exon3:c.G350A:p.R117Q, |
| chr15 | 50459034 | G | A | . | exonic | USP8 | nonsynonymous SNV | USP8:NM_001283049:exon3:c.G139A:p.E47K,USP8:NM_001128610:exon5:c.G370A:p.E124K,USP8:NM_005154:exon5:c.G370A:p.E124K, |
| chr15 | 50481664 | G | A | COSV56164169 | exonic | USP8 | nonsynonymous SNV | USP8:NM_001283049:exon9:c.G1171A:p.E391K,USP8:NM_001128610:exon11:c.G1402A:p.E468K,USP8:NM_005154:exon11:c.G1402A:p.E468K, |
| chr15 | 50481694 | C | A | . | exonic | USP8 | nonsynonymous SNV | USP8:NM_001283049:exon9:c.C1201A:p.Q401K,USP8:NM_001128610:exon11:c.C1432A:p.Q478K,USP8:NM_005154:exon11:c.C1432A:p.Q478K, |

| | | | | | | | | |
|---|---|---|---|---|---|---|---|---|
| chr15 | 50481757 | C | A | . | exonic | USP8 | nonsynonymous SNV | USP8:NM_001283049:exon9:c.C1264A:p.Q422K,USP8:NM_001128610:exon11:c.C1495A:p.Q499K,USP8:NM_005154:exon11:c.C1495A:p.Q499K, |
| chr15 | 50481908 | C | A | . | exonic | USP8 | nonsynonymous SNV | USP8:NM_001128610:exon11:c.C1646A:p.T549K,USP8:NM_005154:exon11:c.C1646A:p.T549K,USP8:NM_001283049:exon9:c.C1415A:p.T472K, |
| chr15 | 50490506 | A | G | . | exonic | USP8 | nonsynonymous SNV | USP8:NM_001283049:exon11:c.A1897G:p.T633A,USP8:NM_001128610:exon14:c.A2215G:p.T739A,USP8:NM_005154:exon14:c.A2215G:p.T739A, |
| chr15 | 55360395 | C | T | . | exonic | CCPG1 | nonsynonymous SNV | CCPG1:NM_004748:exon8:c.G1378A:p.G460R,CCPG1:NM_001204450:exon8:c.G1378A:p.G460R,CCPG1:NM_020739:exon8:c.G1378A:p.G460R, |
| chr15 | 55360686 | G | T | . | exonic | CCPG1 | nonsynonymous SNV | CCPG1:NM_004748:exon8:c.C1087A:p.Q363K,CCPG1:NM_001204450:exon8:c.C1087A:p.Q363K,CCPG1:NM_020739:exon8:c.C1087A:p.Q363K, |
| chr15 | 55377045 | G | T | . | exonic | CCPG1 | nonsynonymous SNV | CCPG1:NM_001204451:exon5:c.C358A:p.Q120K,CCPG1:NM_020739:exon5:c.C358A:p.Q120K,CCPG1:NM_004748:exon5:c.C358A:p.Q120K,CCPG1:NM_001204450:exon5:c.C358A:p.Q120K, |
| chr15 | 67236036 | T | G | COSV56015936 | exonic | AAGAB | nonsynonymous SNV | AAGAB:NM_001271885:exon4:c.A67C:p.I23L,AAGAB:NM_001271886:exon4:c.A67C:p.I23L,AAGAB:NM_024666:exon4:c.A394C:p.I132L, |
| chr15 | 68176535 | C | A | . | exonic | PIAS1 | nonsynonymous SNV | PIAS1:NM_016166:exon11:c.C1362A:p.N454K, |
| chr15 | 70671053 | T | C | . | exonic | UACA | nonsynonymous SNV | UACA:NM_001008224:exon15:c.A1168G:p.M390V,UACA:NM_018003:exon15:c.A1207G:p.M403V, |
| chr15 | 70678124 | T | C | . | exonic | UACA | nonsynonymous SNV | UACA:NM_001008224:exon11:c.A935G:p.N312S,UACA:NM_018003:exon11:c.A974G:p.N325S, |
| chr15 | 72140242 | A | G | . | exonic | SENP8 | nonsynonymous SNV | SENP8:NM_001166340:exon2:c.A619G:p.T207A,SENP8:NM_001172111:exon3:c.A619G:p.T207A,SENP8:NM_001172110:exon3:c.A619G:p.T207A,SENP8:NM_001172109:exon3:c.A619G:p.T207A,SENP8:NM_145204:exon2:c.A619G:p.T207A, |
| chr15 | 72202609 | C | A | . | exonic | PKM | nonsynonymous SNV | PKM:NM_001206797:exon8:c.G930T:p.E310D,PKM:NM_001206798:exon9:c.G1107T:p.E369D,PKM:NM_002654:exon9:c.G1152T:p.E384D, |
| chr15 | 72202620 | T | C | . | exonic | PKM | nonsynonymous SNV | PKM:NM_001206797:exon8:c.A919G:p.I307V,PKM:NM_001206798:exon9:c.A1096G:p.I366V,PKM:NM_002654:exon9:c.A1141G:p.I381V, |
| chr15 | 72206841 | C | T | . | exonic | PKM | nonsynonymous SNV | PKM:NM_001206799:exon8:c.G1042A:p.A348T,PKM:NM_182471:exon8:c.G1027A:p.A343T,PKM:NM_001316318:exon8:c.G1132A:p.A378T,PKM:NM_182470:exon8:c.G1027A:p.A343T,PKM:NM_001206797:exon7:c.G805A:p.A269T,PKM:NM_001206796:exon9:c.G1249A:p.A417T,PKM:NM_001206798:exon8:c.G982A:p.A328T,PKM:NM_002654:exon8:c.G1027A:p.A343T, |
| chr15 | 72208630 | C | T | . | exonic | PKM | nonsynonymous SNV | PKM:NM_001206799:exon6:c.G842A:p.G281E,PKM:NM_182471:exon6:c.G827A:p.G276E,PKM:NM_001316318:exon6:c.G932A:p.G311E,PKM:NM_182470:exon6:c.G827A:p.G276E,PKM:NM_001206797:exon5:c.G605A:p.G202E,PKM:NM_001206796:exon7:c.G1049A:p.G350E,PKM:NM_001206798:exon6:c.G782A:p.G261E,PKM:NM_002654:exon6:c.G827A:p.G276E, |
| chr15 | 72208736 | A | G | . | exonic | PKM | nonsynonymous SNV | PKM:NM_001206799:exon6:c.T736C:p.F246L,PKM:NM_182471:exon6:c.T721C:p.F241L,PKM:NM_001316318:exon6:c.T826C:p.F276L,PKM:NM_182470:exon6:c.T721C:p.F241L,PKM:NM_001206797:exon5:c.T499C:p.F167L,PKM:NM_001206796:exon7:c.T943C:p.F315L,PKM:NM_001206798:exon6:c.T676C:p.F226L,PKM:NM_002654:exon6:c.T721C:p.F241L, |
| chr15 | 72209741 | T | A | . | exonic | PKM | nonsynonymous SNV | PKM:NM_001206799:exon5:c.A512T:p.K171M,PKM:NM_182471:exon5:c.A497T:p.K166M,PKM:NM_001316318:exon5:c.A602T:p.K201M,PKM:NM_182470:exon5:c.A497T:p.K166M,PKM:NM_001206797:exon4:c.A275T:p.K92M,PKM:NM_001206796:exon6:c.A719T:p.K240M,PKM:NM_001206798:exon5:c.A452T:p.K151M,PKM:NM_002654:exon5:c.A497T:p.K166M, |
| chr15 | 72209745 | A | T | . | exonic | PKM | nonsynonymous SNV | PKM:NM_001206799:exon5:c.T508A:p.C170S,PKM:NM_182471:exon5:c.T493A:p.C165S,PKM:NM_001316318:exon5:c.T598A:p.C200S,PKM:NM_182470:exon5:c.T493A:p.C165S,PKM:NM_001206797:exon4:c.T271A:p.C91S,PKM:NM_001206796:exon6:c.T715A:p.C239S,PKM:NM_001206798:exon5:c.T448A:p.C150S,PKM:NM_002654:exon5:c.T493A:p.C165S, |
| chr15 | 72209855 | C | A | COSV99069963 | exonic | PKM | nonsynonymous SNV | PKM:NM_001206799:exon5:c.G398T:p.G133V,PKM:NM_182471:exon5:c.G383T:p.G128V,PKM:NM_001316318:exon5:c.G488T:p.G163V,PKM:NM_182470:exon5:c.G383T:p.G128V,PKM:NM_001206796:exon6:c.G605T:p.G202V,PKM:NM_001206798:exon5:c.G338T:p.G113V,PKM:NM_002654:exon5:c.G383T:p.G128V, |
| chr15 | 72209855 | C | T | . | exonic | PKM | nonsynonymous SNV | PKM:NM_001206799:exon5:c.G398A:p.G133D,PKM:NM_001206798:exon5:c.G338A:p.G113D,PKM:NM_002654:exon5:c.G383A:p.G128D,PKM:NM_182471:exon5:c.G383A:p.G128D,PKM:NM_001316318:exon5:c.G488A:p.G163D,PKM:NM_182470:exon5:c.G383A:p.G128D,PKM:NM_001206796:exon6:c.G605A:p.G202D, |
| chr15 | 72209856 | C | A | . | exonic | PKM | nonsynonymous SNV | PKM:NM_001206799:exon5:c.G397T:p.G133C,PKM:NM_182471:exon5:c.G382T:p.G128C,PKM:NM_001316318:exon5:c.G487T:p.G163C,PKM:NM_182470:exon5:c.G382T:p.G128C,PKM:NM_001206796:exon6:c.G604T:p.G202C,PKM:NM_001206798:exon5:c.G337T:p.G113C,PKM:NM_002654:exon5:c.G382T:p.G128C, |
| chr15 | 72210439 | C | T | . | exonic | PKM | nonsynonymous SNV | PKM:NM_001206799:exon4:c.G301A:p.E101K,PKM:NM_182471:exon4:c.G286A:p.E96K,PKM:NM_001316318:exon4:c.G391A:p.E131K,PKM:NM_182470:exon4:c.G286A:p.E96K,PKM:NM_001206796:exon5:c.G508A:p.E170K,PKM:NM_001206798:exon4:c.G241A:p.E81K,PKM:NM_002654:exon4:c.G286A:p.E96K, |
| chr15 | 72219068 | A | T | . | exonic | PKM | nonsynonymous SNV | PKM:NM_001206798:exon3:c.T77A:p.L26Q, |
| chr15 | 74035775 | A | G | COSV51443885 | exonic | PML | nonsynonymous SNV | PML:NM_033250:exon7:c.A2170G:p.S724G,PML:NM_033239:exon8:c.A2314G:p.S772G, |
| chr15 | 74035800 | G | T | COSV51443888 | exonic | PML | nonsynonymous SNV | PML:NM_033250:exon7:c.G2195T:p.G732V,PML:NM_033239:exon8:c.G2339T:p.G780V, |
| chr15 | 77056128 | A | G | . | exonic | TSPAN3 | nonsynonymous SNV | TSPAN3:NM_005724:exon2:c.T191C:p.L64P,TSPAN3:NM_198902:exon2:c.T191C:p.L64P, |
| chr15 | 79970875 | C | T | COSV51212886 | exonic | BCL2A1 | nonsynonymous SNV | BCL2A1:NM_001114735:exon1:c.G245A:p.G82D,BCL2A1:NM_004049:exon1:c.G245A:p.G82D, |
| chr15 | 79971003 | A | C | COSV51213210 | exonic | BCL2A1 | nonsynonymous SNV | BCL2A1:NM_001114735:exon1:c.T117G:p.N39K,BCL2A1:NM_004049:exon1:c.T117G:p.N39K, |
| chr15 | 79971064 | C | T | COSV51213224 | exonic | BCL2A1 | nonsynonymous SNV | BCL2A1:NM_001114735:exon1:c.G56A:p.C19Y,BCL2A1:NM_004049:exon1:c.G56A:p.C19Y, |

| | | | | | | | | |
|---|---|---|---|---|---|---|---|---|
| chr15 | 85581252 | C | G | COSV63448224 | exonic | AKAP13 | nonsynonymous SNV | AKAP13:NM_006738:exon7:c.C3184G:p.P1062A,AKAP13:NM_007200:exon7:c.C3184G:p.P1062A, |
| chr15 | 85581324 | G | A | COSV63448234 | exonic | AKAP13 | nonsynonymous SNV | AKAP13:NM_006738:exon7:c.G3256A:p.D1086N,AKAP13:NM_007200:exon7:c.G3256A:p.D1086N, |
| chr15 | 85581715 | T | C | COSV63447944 | exonic | AKAP13 | nonsynonymous SNV | AKAP13:NM_006738:exon7:c.T3647C:p.M1216T,AKAP13:NM_007200:exon7:c.T3647C:p.M1216T, |
| chr15 | 85684869 | G | A | . | exonic | AKAP13 | nonsynonymous SNV | AKAP13:NM_006738:exon16:c.G5297A:p.S1766N,AKAP13:NM_007200:exon16:c.G5285A:p.S1762N,AKAP13:NM_001270546:exon8:c.G1151A:p.S384N, |
| chr15 | 89792557 | C | T | COSV55591836 | exonic | ANPEP | nonsynonymous SNV | ANPEP:NM_001150:exon17:c.G2255A:p.S752N, |
| chr15 | 89801121 | T | C | COSV55589989 | exonic | ANPEP | nonsynonymous SNV | ANPEP:NM_001150:exon12:c.A1809G:p.I603M, |
| chr15 | 90360079 | A | G | . | exonic | ZNF774 | nonsynonymous SNV | ZNF774:NM_001004309:exon4:c.A248G:p.N83S, |
| chr15 | 90981611 | G | T | . | exonic | PRC1 | nonsynonymous SNV | PRC1:NM_001267580:exon4:c.C437A:p.A146E,PRC1:NM_199413:exon5:c.C560A:p.A187E,PRC1:NM_003981:exon5:c.C560A:p.A187E, |
| chr15 | 92939625 | G | A | . | exonic | CHD2 | nonsynonymous SNV | CHD2:NM_001271:exon7:c.G599A:p.G200E,CHD2:NM_001042572:exon7:c.G599A:p.G200E, |
| chr15 | 92943036 | C | A | . | exonic | CHD2 | nonsynonymous SNV | CHD2:NM_001271:exon9:c.C1020A:p.N340K,CHD2:NM_001042572:exon9:c.C1020A:p.N340K, |
| chr16 | 10744056 | C | G | COSV51593232 | exonic | NUBP1 | nonsynonymous SNV | NUBP1:NM_002484:exon2:c.C115G:p.P39A,NUBP1:NM_001278506:exon2:c.C115G:p.P39A, |
| chr16 | 25228484 | G | C | COSV54844539 | exonic | AQP8 | nonsynonymous SNV | AQP8:NM_001169:exon6:c.G778C:p.A260P, |
| chr16 | 29779543 | G | A | . | exonic | ZG16 | nonsynonymous SNV | ZG16:NM_152338:exon3:c.G94A:p.G32S, |
| chr16 | 29780400 | G | C | COSV68783707 | exonic | ZG16 | nonsynonymous SNV | ZG16:NM_152338:exon4:c.G485C:p.S162T, |
| chr16 | 30067485 | G | A | . | exonic | ALDOA | nonsynonymous SNV | ALDOA:NM_001127617:exon3:c.G148A:p.E50K,ALDOA:NM_000034:exon8:c.G148A:p.E50K,ALDOA:NM_184043:exon3:c.G148A:p.E50K,ALDOA:NM_184041:exon3:c.G148A:p.E50K,ALDOA:NM_001243177:exon4:c.G310A:p.E104K, |
| chr16 | 30067509 | T | C | . | exonic | ALDOA | nonsynonymous SNV | ALDOA:NM_001127617:exon3:c.T172C:p.F58L,ALDOA:NM_000034:exon8:c.T172C:p.F58L,ALDOA:NM_184043:exon3:c.T172C:p.F58L,ALDOA:NM_184041:exon3:c.T172C:p.F58L,ALDOA:NM_001243177:exon4:c.T334C:p.F112L, |
| chr16 | 30067561 | G | A | . | exonic | ALDOA | nonsynonymous SNV | ALDOA:NM_001127617:exon3:c.G224A:p.G75E,ALDOA:NM_000034:exon8:c.G224A:p.G75E,ALDOA:NM_184043:exon3:c.G224A:p.G75E,ALDOA:NM_184041:exon3:c.G224A:p.G75E,ALDOA:NM_001243177:exon4:c.G386A:p.G129E, |
| chr16 | 30067612 | G | A | . | exonic | ALDOA | nonsynonymous SNV | ALDOA:NM_001127617:exon3:c.G275A:p.R92H,ALDOA:NM_000034:exon8:c.G275A:p.R92H,ALDOA:NM_184043:exon3:c.G275A:p.R92H,ALDOA:NM_184041:exon3:c.G275A:p.R92H,ALDOA:NM_001243177:exon4:c.G437A:p.R146H, |
| chr16 | 30067642 | G | A | . | exonic | ALDOA | nonsynonymous SNV | ALDOA:NM_001127617:exon3:c.G305A:p.G102D,ALDOA:NM_000034:exon8:c.G305A:p.G102D,ALDOA:NM_184043:exon3:c.G305A:p.G102D,ALDOA:NM_184041:exon3:c.G305A:p.G102D,ALDOA:NM_001243177:exon4:c.G467A:p.G156D, |
| chr16 | 30067654 | G | A | . | exonic | ALDOA | nonsynonymous SNV | ALDOA:NM_001127617:exon3:c.G317A:p.G106D,ALDOA:NM_000034:exon8:c.G317A:p.G106D,ALDOA:NM_184043:exon3:c.G317A:p.G106D,ALDOA:NM_184041:exon3:c.G317A:p.G106D,ALDOA:NM_001243177:exon4:c.G479A:p.G160D, |
| chr16 | 30068823 | G | A | COSV57632467 | exonic | ALDOA | nonsynonymous SNV | ALDOA:NM_001127617:exon5:c.G385A:p.D129N,ALDOA:NM_000034:exon10:c.G385A:p.D129N,ALDOA:NM_184043:exon5:c.G385A:p.D129N,ALDOA:NM_184041:exon5:c.G385A:p.D129N,ALDOA:NM_001243177:exon6:c.G547A:p.D183N, |
| chr16 | 30068827 | G | A | . | exonic | ALDOA | nonsynonymous SNV | ALDOA:NM_001127617:exon5:c.G389A:p.G130E,ALDOA:NM_000034:exon10:c.G389A:p.G130E,ALDOA:NM_184043:exon5:c.G389A:p.G130E,ALDOA:NM_184041:exon5:c.G389A:p.G130E,ALDOA:NM_001243177:exon6:c.G551A:p.G184E, |
| chr16 | 30068839 | G | A | . | exonic | ALDOA | nonsynonymous SNV | ALDOA:NM_001127617:exon5:c.G401A:p.R134H,ALDOA:NM_000034:exon10:c.G401A:p.R134H,ALDOA:NM_184043:exon5:c.G401A:p.R134H,ALDOA:NM_184041:exon5:c.G401A:p.R134H,ALDOA:NM_001243177:exon6:c.G563A:p.R188H, |
| chr16 | 30068901 | G | A | . | exonic | ALDOA | nonsynonymous SNV | ALDOA:NM_001127617:exon5:c.G463A:p.G155R,ALDOA:NM_000034:exon10:c.G463A:p.G155R,ALDOA:NM_184043:exon5:c.G463A:p.G155R,ALDOA:NM_184041:exon5:c.G463A:p.G155R,ALDOA:NM_001243177:exon6:c.G625A:p.G209R, |
| chr16 | 30068904 | G | A | . | exonic | ALDOA | nonsynonymous SNV | ALDOA:NM_001127617:exon5:c.G466A:p.E156K,ALDOA:NM_000034:exon10:c.G466A:p.E156K,ALDOA:NM_184043:exon5:c.G466A:p.E156K,ALDOA:NM_184041:exon5:c.G466A:p.E156K,ALDOA:NM_001243177:exon6:c.G628A:p.E210K, |
| chr16 | 30068926 | C | A | . | exonic | ALDOA | nonsynonymous SNV | ALDOA:NM_001127617:exon5:c.C488A:p.A163D,ALDOA:NM_000034:exon10:c.C488A:p.A163D,ALDOA:NM_184043:exon5:c.C488A:p.A163D,ALDOA:NM_184041:exon5:c.C488A:p.A163D,ALDOA:NM_001243177:exon6:c.C650A:p.A217D, |
| chr16 | 30068929 | T | C | . | exonic | ALDOA | nonsynonymous SNV | ALDOA:NM_001127617:exon5:c.T491C:p.I164T,ALDOA:NM_000034:exon10:c.T491C:p.I164T,ALDOA:NM_184043:exon5:c.T491C:p.I164T,ALDOA:NM_184041:exon5:c.T491C:p.I164T,ALDOA:NM_001243177:exon6:c.T653C:p.I218T, |
| chr16 | 30068933 | G | A | . | exonic | ALDOA | nonsynonymous SNV | ALDOA:NM_001127617:exon5:c.G495A:p.M165I,ALDOA:NM_000034:exon10:c.G495A:p.M165I,ALDOA:NM_184043:exon5:c.G495A:p.M165I,ALDOA:NM_184041:exon5:c.G495A:p.M165I,ALDOA:NM_001243177:exon6:c.G657A:p.M219I, |
| chr16 | 30068935 | A | G | . | exonic | ALDOA | nonsynonymous SNV | ALDOA:NM_001127617:exon5:c.A497G:p.E166G,ALDOA:NM_000034:exon10:c.A497G:p.E166G,ALDOA:NM_184043:exon5:c.A497G:p.E166G,ALDOA:NM_184041:exon5:c.A497G:p.E166G,ALDOA:NM_001243177:exon6:c.A659G:p.E220G, |
| chr16 | 30068956 | G | A | . | exonic | ALDOA | nonsynonymous SNV | ALDOA:NM_001127617:exon5:c.G518A:p.R173H,ALDOA:NM_000034:exon10:c.G518A:p.R173H,ALDOA:NM_184043:exon5:c.G518A:p.R173H,ALDOA:NM_184041:exon5:c.G518A:p.R173H,ALDOA:NM_001243177:exon6:c.G680A:p.R227H, |
| chr16 | 30069349 | G | A | COSV57632343 | exonic | ALDOA | nonsynonymous SNV | ALDOA:NM_001127617:exon6:c.G584A:p.G195E,ALDOA:NM_000034:exon11:c.G584A:p.G195E,ALDOA:NM_184043:exon6:c.G584A:p.G195E,ALDOA:NM_184041:exon6:c.G584A:p.G195E,ALDOA:NM_001243177:exon7:c.G746A:p.G249E, |

| | | | | | | | | |
|---|---|---|---|---|---|---|---|---|
| chr16 | 30069505 | G | A | . | exonic | ALDOA | nonsynonymous SNV | ALDOA:NM_001127617:exon7:c.G631A:p.A211T,ALDOA:NM_000034:exon12:c.G631A:p.A211T,ALDOA:NM_184043:exon7:c.G631A:p.A211T,ALDOA:NM_184041:exon7:c.G631A:p.A211T,ALDOA:NM_001243177:exon8:c.G793A:p.A265T, |
| chr16 | 30069664 | G | A | COSV57632104 | exonic | ALDOA | nonsynonymous SNV | ALDOA:NM_001127617:exon7:c.G790A:p.A264T,ALDOA:NM_000034:exon12:c.G790A:p.A264T,ALDOA:NM_184043:exon7:c.G790A:p.A264T,ALDOA:NM_184041:exon7:c.G790A:p.A264T,ALDOA:NM_001243177:exon8:c.G952A:p.A318T, |
| chr16 | 30069881 | T | C | . | exonic | ALDOA | nonsynonymous SNV | ALDOA:NM_001127617:exon8:c.T851C:p.L284P,ALDOA:NM_000034:exon13:c.T851C:p.L284P,ALDOA:NM_184043:exon8:c.T851C:p.L284P,ALDOA:NM_184041:exon8:c.T851C:p.L284P,ALDOA:NM_001243177:exon9:c.T1013C:p.L338P, |
| chr16 | 30069977 | G | A | . | exonic | ALDOA | nonsynonymous SNV | ALDOA:NM_001127617:exon8:c.G947A:p.G316E,ALDOA:NM_000034:exon13:c.G947A:p.G316E,ALDOA:NM_184043:exon8:c.G947A:p.G316E,ALDOA:NM_184041:exon8:c.G947A:p.G316E,ALDOA:NM_001243177:exon9:c.G1109A:p.G370E, |
| chr16 | 30070117 | G | A | . | exonic | ALDOA | nonsynonymous SNV | ALDOA:NM_001127617:exon9:c.G1000A:p.A334T,ALDOA:NM_000034:exon14:c.G1000A:p.A334T,ALDOA:NM_184043:exon9:c.G1000A:p.A334T,ALDOA:NM_184041:exon9:c.G1000A:p.A334T,ALDOA:NM_001243177:exon10:c.G1162A:p.A388T, |
| chr16 | 30070156 | G | A | . | exonic | ALDOA | nonsynonymous SNV | ALDOA:NM_001127617:exon9:c.G1039A:p.G347S,ALDOA:NM_000034:exon14:c.G1039A:p.G347S,ALDOA:NM_184043:exon9:c.G1039A:p.G347S,ALDOA:NM_184041:exon9:c.G1039A:p.G347S,ALDOA:NM_001243177:exon10:c.G1201A:p.G401S, |
| chr16 | 31185078 | T | G | . | exonic | FUS | nonsynonymous SNV | FUS:NM_004960:exon6:c.T663G:p.S221R,FUS:NM_001170634:exon6:c.T660G:p.S220R,FUS:NM_001170937:exon6:c.T651G:p.S217R, |
| chr16 | 3657746 | G | A | COSV55912793 | exonic | DNASE1 | nonsynonymous SNV | DNASE1:NM_005223:exon8:c.G731A:p.R244Q, |
| chr16 | 3658169 | C | T | COSV55914199 | exonic | TRAP1 | nonsynonymous SNV | TRAP1:NM_016292:exon18:c.G2075A:p.R692H,TRAP1:NM_001272049:exon17:c.G1916A:p.R639H, |
| chr16 | 3674464 | G | C | COSV55915372 | exonic | TRAP1 | nonsynonymous SNV | TRAP1:NM_016292:exon9:c.C919G:p.R307G,TRAP1:NM_001272049:exon8:c.C760G:p.R254G, |
| chr16 | 3676074 | G | A | . | exonic | TRAP1 | nonsynonymous SNV | TRAP1:NM_016292:exon7:c.C776T:p.S259F,TRAP1:NM_001272049:exon6:c.C617T:p.S206F, |
| chr16 | 3717472 | G | A | COSV55917281 | exonic | TRAP1 | nonsynonymous SNV | TRAP1:NM_016292:exon1:c.C37T:p.R13C,TRAP1:NM_001272049:exon1:c.C37T:p.R13C, |
| chr16 | 64982184 | A | C | . | exonic | CDH11 | nonsynonymous SNV | CDH11:NM_001308392:exon8:c.T1117G:p.S373A,CDH11:NM_001797:exon8:c.T1117G:p.S373A, |
| chr16 | 64991815 | G | A | COSV51767367 | exonic | CDH11 | nonsynonymous SNV | CDH11:NM_001308392:exon6:c.C764T:p.T255M,CDH11:NM_001797:exon6:c.C764T:p.T255M, |
| chr16 | 66817004 | A | T | . | exonic | NAE1 | nonsynonymous SNV | NAE1:NM_003905:exon10:c.T709A:p.Y237N,NAE1:NM_001286500:exon10:c.T718A:p.Y240N,NAE1:NM_001018159:exon11:c.T691A:p.Y231N,NAE1:NM_001018160:exon9:c.T442A:p.Y148N, |
| chr16 | 67611443 | C | A | COSV50477534 | exonic | CTCF | nonsynonymous SNV | CTCF:NM_006565:exon3:c.C611A:p.T204K, |
| chr16 | 67611452 | C | G | . | exonic | CTCF | nonsynonymous SNV | CTCF:NM_006565:exon3:c.C620G:p.T207S, |
| chr16 | 68738336 | C | A | . | exonic | CDH1 | nonsynonymous SNV | CDH1:NM_001317184:exon2:c.C88A:p.P30T,CDH1:NM_004360:exon2:c.C88A:p.P30T, |
| chr16 | 70514394 | G | A | . | exonic | COG4 | nonsynonymous SNV | COG4:NM_001195139:exon4:c.C485T:p.T162I,COG4:NM_015386:exon4:c.C485T:p.T162I, |
| chr16 | 84745079 | A | G | COSV54746826 | exonic | USP10 | nonsynonymous SNV | USP10:NM_001272075:exon5:c.A610G:p.M204V,USP10:NM_005153:exon4:c.A598G:p.M200V, |
| chr16 | 84745088 | T | C | COSV54751736 | exonic | USP10 | nonsynonymous SNV | USP10:NM_001272075:exon5:c.T619C:p.S207P,USP10:NM_005153:exon4:c.T607C:p.S203P, |
| chr16 | 84745091 | G | C | COSV54751750 | exonic | USP10 | nonsynonymous SNV | USP10:NM_001272075:exon5:c.G622C:p.V208L,USP10:NM_005153:exon4:c.G610C:p.V204L, |
| chr16 | 85801212 | G | A | COSV53665976 | exonic | COX4I1 | nonsynonymous SNV | COX4I1:NM_001861:exon2:c.G7A:p.A3T, |
| chr16 | 89114475 | G | A | COSV58077067 | exonic | ACSF3 | nonsynonymous SNV | ACSF3:NM_001127214:exon5:c.G1114A:p.V372M,ACSF3:NM_001243279:exon6:c.G1114A:p.V372M,ACSF3:NM_174917:exon6:c.G1114A:p.V372M,ACSF3:NM_001284316:exon4:c.G319A:p.V107M, |
| chr16 | 89561665 | G | A | COSV51948976 | exonic | RPL13 | nonsynonymous SNV | RPL13:NM_001243130:exon4:c.G277A:p.A93T,RPL13:NM_033251:exon3:c.G334A:p.A112T,RPL13:NM_000977:exon4:c.G334A:p.A112T, |
| chr17 | 1345458 | C | T | COSV51985336 | exonic | YWHAE | nonsynonymous SNV | YWHAE:NM_006761:exon6:c.G757A:p.E253K, |
| chr17 | 1354316 | C | T | COSV51984710 | exonic | YWHAE | nonsynonymous SNV | YWHAE:NM_006761:exon5:c.G610A:p.A204T, |
| chr17 | 1354342 | G | A | . | exonic | YWHAE | nonsynonymous SNV | YWHAE:NM_006761:exon5:c.C584T:p.A195V, |
| chr17 | 1361101 | C | T | . | exonic | YWHAE | nonsynonymous SNV | YWHAE:NM_006761:exon4:c.G569A:p.R190H, |
| chr17 | 1361143 | T | A | . | exonic | YWHAE | nonsynonymous SNV | YWHAE:NM_006761:exon4:c.A527T:p.N176I, |
| chr17 | 1361156 | C | T | . | exonic | YWHAE | nonsynonymous SNV | YWHAE:NM_006761:exon4:c.G514A:p.G172S, |
| chr17 | 1361161 | C | T | . | exonic | YWHAE | nonsynonymous SNV | YWHAE:NM_006761:exon4:c.G509A:p.R170H, |
| chr17 | 1361927 | C | T | . | exonic | YWHAE | nonsynonymous SNV | YWHAE:NM_006761:exon3:c.G346A:p.E116K, |
| chr17 | 1361947 | A | G | . | exonic | YWHAE | nonsynonymous SNV | YWHAE:NM_006761:exon3:c.T326C:p.I109T, |
| chr17 | 1364864 | G | T | . | exonic | YWHAE | nonsynonymous SNV | YWHAE:NM_006761:exon2:c.C259A:p.Q87K, |
| chr17 | 1364880 | C | T | . | exonic | YWHAE | nonsynonymous SNV | YWHAE:NM_006761:exon2:c.G243A:p.M81I, |
| chr17 | 1364911 | T | C | . | exonic | YWHAE | nonsynonymous SNV | YWHAE:NM_006761:exon2:c.A212G:p.E71G, |
| chr17 | 1364924 | C | T | . | exonic | YWHAE | nonsynonymous SNV | YWHAE:NM_006761:exon2:c.G199A:p.E67K, |
| chr17 | 1364941 | C | T | . | exonic | YWHAE | nonsynonymous SNV | YWHAE:NM_006761:exon2:c.G182A:p.R61K, |

| | | | | | | | | |
|---|---|---|---|---|---|---|---|---|
| chr17 | 1364943 | C | T | . | exonic | YWHAE | stopgain | YWHAE:NM_006761:exon2:c.G180A:p.W60X, |
| chr17 | 1400108 | C | T | . | exonic | YWHAE | nonsynonymous SNV | YWHAE:NM_006761:exon1:c.G3A:p.M1I, |
| chr17 | 16126110 | C | T | . | exonic | NCOR1 | nonsynonymous SNV | NCOR1:NM_001190440:exon14:c.G1606A:p.E536K,NCOR1:NM_001190438:exon12:c.G1279A:p.E427K,NCOR1:NM_006311:exon15:c.G1606A:p.E536K, |
| chr17 | 16126142 | G | T | . | exonic | NCOR1 | nonsynonymous SNV | NCOR1:NM_001190440:exon14:c.C1574A:p.T525K,NCOR1:NM_001190438:exon12:c.C1247A:p.T416K,NCOR1:NM_006311:exon15:c.C1574A:p.T525K, |
| chr17 | 16146392 | G | T | . | exonic | NCOR1 | nonsynonymous SNV | NCOR1:NM_001190440:exon9:c.C1066A:p.Q356K,NCOR1:NM_001190438:exon7:c.C739A:p.Q247K,NCOR1:NM_006311:exon10:c.C1066A:p.Q356K, |
| chr17 | 16146482 | C | T | . | exonic | NCOR1 | nonsynonymous SNV | NCOR1:NM_001190440:exon9:c.G976A:p.E326K,NCOR1:NM_001190438:exon7:c.G649A:p.E217K,NCOR1:NM_006311:exon10:c.G976A:p.E326K, |
| chr17 | 16149456 | G | T | . | exonic | NCOR1 | nonsynonymous SNV | NCOR1:NM_001190440:exon8:c.C904A:p.Q302K,NCOR1:NM_001190438:exon6:c.C577A:p.Q193K,NCOR1:NM_006311:exon9:c.C904A:p.Q302K, |
| chr17 | 16165149 | C | T | COSV51959167 | exonic | NCOR1 | nonsynonymous SNV | NCOR1:NM_001190440:exon4:c.G448A:p.G150R,NCOR1:NM_001190438:exon2:c.G121A:p.G41R,NCOR1:NM_006311:exon5:c.G448A:p.G150R, |
| chr17 | 31993275 | C | A | . | exonic | SUZ12 | nonsynonymous SNV | SUZ12:NM_015355:exon11:c.C1235A:p.T412K, |
| chr17 | 31993308 | G | A | COSV59500263 | exonic | SUZ12 | nonsynonymous SNV | SUZ12:NM_015355:exon11:c.G1268A:p.R423Q, |
| chr17 | 32469044 | T | C | . | exonic | PSMD11 | nonsynonymous SNV | PSMD11:NM_001270482:exon6:c.T494C:p.L165P,PSMD11:NM_002815:exon6:c.T494C:p.L165P, |
| chr17 | 35717225 | A | G | . | exonic | AP2B1 | nonsynonymous SNV | AP2B1:NM_001282:exon20:c.A2615G:p.N872S,AP2B1:NM_001030006:exon21:c.A2657G:p.N886S, |
| chr17 | 35844650 | A | G | . | exonic | TAF15 | nonsynonymous SNV | TAF15:NM_139215:exon15:c.A1351G:p.S451G,TAF15:NM_003487:exon15:c.A1342G:p.S448G, |
| chr17 | 37624364 | C | T | . | exonic | DDX52 | nonsynonymous SNV | DDX52:NM_001291476:exon10:c.G883A:p.V295M,DDX52:NM_007010:exon9:c.G1207A:p.V403M, |
| chr17 | 37626060 | A | G | . | exonic | DDX52 | nonsynonymous SNV | DDX52:NM_001291476:exon9:c.T647C:p.F216S,DDX52:NM_007010:exon8:c.T971C:p.F324S, |
| chr17 | 38753202 | G | A | . | exonic | PSMB3 | nonsynonymous SNV | PSMB3:NM_002795:exon2:c.G56A:p.C19Y, |
| chr17 | 38753246 | A | T | . | exonic | PSMB3 | nonsynonymous SNV | PSMB3:NM_002795:exon2:c.A100T:p.M34L, |
| chr17 | 38755965 | G | A | . | exonic | PSMB3 | nonsynonymous SNV | PSMB3:NM_002795:exon3:c.G271A:p.V91M, |
| chr17 | 38762466 | G | A | . | exonic | PSMB3 | nonsynonymous SNV | PSMB3:NM_002795:exon5:c.G530A:p.R177Q, |
| chr17 | 38764142 | G | A | . | exonic | PSMB3 | nonsynonymous SNV | PSMB3:NM_002795:exon6:c.G593A:p.R198K, |
| chr17 | 38810562 | T | C | . | exonic | CWC25 | nonsynonymous SNV | CWC25:NM_017748:exon5:c.A532G:p.K178E, |
| chr17 | 38812814 | A | T | . | exonic | CWC25 | nonsynonymous SNV | CWC25:NM_017748:exon4:c.T479A:p.M160K, |
| chr17 | 38812820 | A | T | . | exonic | CWC25 | nonsynonymous SNV | CWC25:NM_017748:exon4:c.T473A:p.V158E, |
| chr17 | 39634038 | G | A | . | exonic | PPP1R1B | nonsynonymous SNV | PPP1R1B:NM_181505:exon5:c.G289A:p.E97K,PPP1R1B:NM_001242464:exon5:c.G289A:p.E97K,PPP1R1B:NM_032192:exon5:c.G397A:p.E133K, |
| chr17 | 39634069 | A | T | . | exonic | PPP1R1B | nonsynonymous SNV | PPP1R1B:NM_181505:exon5:c.A320T:p.K107M,PPP1R1B:NM_001242464:exon5:c.A320T:p.K107M,PPP1R1B:NM_032192:exon5:c.A428T:p.K143M, |
| chr17 | 39684607 | G | C | . | exonic | PGAP3 | nonsynonymous SNV | PGAP3:NM_001291728:exon3:c.C422G:p.A141G,PGAP3:NM_001291730:exon3:c.C422G:p.A141G,PGAP3:NM_033419:exon3:c.C422G:p.A141G,PGAP3:NM_001291733:exon3:c.C422G:p.A141G,PGAP3:NM_001291732:exon3:c.C422G:p.A141G, |
| chr17 | 39727784 | C | G | COSV54064104 | exonic | ERBB2 | nonsynonymous SNV | ERBB2:NM_001289936:exon31:c.C3463G:p.P1155A,ERBB2:NM_004448:exon27:c.C3508G:p.P1170A,ERBB2:NM_001005862:exon30:c.C3418G:p.P1140A, |
| chr17 | 39727835 | T | C | . | exonic | ERBB2 | nonsynonymous SNV | ERBB2:NM_001289936:exon31:c.T3514C:p.F1172L,ERBB2:NM_004448:exon27:c.T3559C:p.F1187L,ERBB2:NM_001005862:exon30:c.T3469C:p.F1157L, |

| | | | | | | | | |
|---|---|---|---|---|---|---|---|---|
| chr17 | 39742290 | T | C | . | exonic | GRB7 | nonsynonymous SNV | GRB7:NM_001242442:exon2:c.T58C:p.C20R, |
| chr17 | 39905943 | G | A | COSV58779652 | exonic | GSDMB | nonsynonymous SNV | GSDMB:NM_018530:exon6:c.C865T:p.P289S,GSDMB:NM_001165959:exon8:c.C904T:p.P302S,GSDMB:NM_001042471:exon8:c.C892T:p.P298S,GSDMB:NM_001165958:exon9:c.C931T:p.P311S, |
| chr17 | 39905964 | C | T | COSV58779510 | exonic | GSDMB | nonsynonymous SNV | GSDMB:NM_018530:exon6:c.G844A:p.G282R,GSDMB:NM_001165959:exon8:c.G883A:p.G295R,GSDMB:NM_001042471:exon8:c.G871A:p.G291R,GSDMB:NM_001165958:exon9:c.G910A:p.G304R, |
| chr17 | 40300899 | G | A | COSV52929970 | exonic | CDC6 | nonsynonymous SNV | CDC6:NM_001254:exon10:c.G1321A:p.V441I, |
| chr17 | 47981705 | T | C | COSV58114512 | exonic | CDK5RAP3 | nonsynonymous SNV | CDK5RAP3:NM_001278198:exon13:c.T922C:p.S308P, |
| chr17 | 48184809 | C | T | COSV61146844 | exonic | SKAP1 | nonsynonymous SNV | SKAP1:NM_001075099:exon7:c.G481A:p.G161S,SKAP1:NM_003726:exon7:c.G481A:p.G161S, |
| chr17 | 49706301 | C | T | . | exonic | SLC35B1 | nonsynonymous SNV | SLC35B1:NM_001278784:exon3:c.G41A:p.R14H,SLC35B1:NM_005827:exon3:c.G353A:p.R118H, |
| chr17 | 5335325 | T | A | . | exonic | RABEP1 | stopgain | RABEP1:NM_001083585:exon4:c.T509A:p.L170X,RABEP1:NM_001291581:exon3:c.T380A:p.L127X,RABEP1:NM_001291582:exon4:c.T509A:p.L170X,RABEP1:NM_004703:exon4:c.T509A:p.L170X, |
| chr17 | 5335332 | T | A | COSV52584305 | exonic | RABEP1 | nonsynonymous SNV | RABEP1:NM_001083585:exon4:c.T516A:p.N172K,RABEP1:NM_001291581:exon3:c.T387A:p.N129K,RABEP1:NM_001291582:exon4:c.T516A:p.N172K,RABEP1:NM_004703:exon4:c.T516A:p.N172K, |
| chr17 | 55751132 | A | C | . | exonic | PCTP | nonsynonymous SNV | PCTP:NM_021213:exon1:c.A29C:p.E10A, |
| chr17 | 58357515 | C | G | . | exonic | RNF43 | nonsynonymous SNV | RNF43:NM_017763:exon9:c.G2261C:p.G754A,RNF43:NM_001305545:exon8:c.G1880C:p.G627A,RNF43:NM_001305544:exon9:c.G2261C:p.G754A, |
| chr17 | 58357719 | G | C | . | exonic | RNF43 | nonsynonymous SNV | RNF43:NM_017763:exon9:c.C2057G:p.P686R,RNF43:NM_001305545:exon8:c.C1676G:p.P559R,RNF43:NM_001305544:exon9:c.C2057G:p.P686R, |
| chr17 | 58358524 | G | T | COSV68456749 | exonic | RNF43 | nonsynonymous SNV | RNF43:NM_017763:exon9:c.C1252A:p.L418M,RNF43:NM_001305545:exon8:c.C871A:p.L291M,RNF43:NM_001305544:exon9:c.C1252A:p.L418M, |
| chr17 | 58358748 | C | T | . | exonic | RNF43 | nonsynonymous SNV | RNF43:NM_017763:exon9:c.G1028A:p.R343H,RNF43:NM_001305545:exon8:c.G647A:p.R216H,RNF43:NM_001305544:exon9:c.G1028A:p.R343H, |
| chr17 | 58415439 | T | C | COSV60415571 | exonic | RNF43 | nonsynonymous SNV | RNF43:NM_017763:exon2:c.A139G:p.I47V,RNF43:NM_001305544:exon2:c.A139G:p.I47V, |
| chr17 | 60709264 | A | G | . | exonic | BCAS3 | nonsynonymous SNV | BCAS3:NM_001099432:exon5:c.A260G:p.N87S,BCAS3:NM_017679:exon5:c.A260G:p.N87S, |
| chr17 | 64500188 | C | T | . | exonic | DDX5 | nonsynonymous SNV | DDX5:NM_004396:exon13:c.G1580A:p.G527E, |
| chr17 | 64502200 | C | T | . | exonic | DDX5 | nonsynonymous SNV | DDX5:NM_004396:exon10:c.G1118A:p.G373D, |
| chr17 | 64504015 | C | T | . | exonic | DDX5 | nonsynonymous SNV | DDX5:NM_004396:exon4:c.G409A:p.V137M, |
| chr17 | 64504033 | C | T | . | exonic | DDX5 | nonsynonymous SNV | DDX5:NM_004396:exon4:c.G391A:p.G131R, |
| chr17 | 72947851 | T | C | . | exonic | SLC39A11 | nonsynonymous SNV | SLC39A11:NM_139177:exon5:c.A331G:p.T111A,SLC39A11:NM_001159770:exon5:c.A331G:p.T111A, |
| chr17 | 73209715 | A | G | COSV55428466 | exonic | FAM104A | nonsynonymous SNV | FAM104A:NM_001289411:exon3:c.T293C:p.I98T,FAM104A:NM_001289410:exon2:c.T275C:p.I92T, |
| chr17 | 75225432 | G | A | . | exonic | NUP85 | nonsynonymous SNV | NUP85:NM_001303276:exon8:c.G685A:p.A229T,NUP85:NM_024844:exon9:c.G823A:p.A275T, |
| chr17 | 760232 | C | A | . | exonic | GLOD4 | nonsynonymous SNV | GLOD4:NM_016080:exon9:c.G838T:p.A280S, |
| chr17 | 7676154 | G | C | COSV52666208 | exonic | TP53 | nonsynonymous SNV | TP53:NM_001126118:exon3:c.C98G:p.P33R,TP53:NM_001276696:exon4:c.C98G:p.P33R,TP53:NM_001126113:exon4:c.C215G:p.P72R,TP53:NM_001126112:exon4:c.C215G:p.P72R,TP53:NM_001276695:exon4:c.C98G:p.P33R,TP53:NM_001276761:exon4:c.C98G:p.P33R,TP53:NM_001126114:exon4:c.C215G:p.P72R,TP53:NM_000546:exon4:c.C215G:p.P72R,TP53:NM_001276760:exon4:c.C98G:p.P33R, |
| chr17 | 7703404 | C | G | COSV56832921 | exonic | WRAP53 | nonsynonymous SNV | WRAP53:NM_001143990:exon11:c.C1565G:p.A522G,WRAP53:NM_001143991:exon11:c.C1565G:p.A522G,WRAP53:NM_001143992:exon11:c.C1565G:p.A522G,WRAP53:NM_018081:exon10:c.C1565G:p.A522G, |
| chr17 | 81510694 | A | G | . | exonic | ACTG1 | nonsynonymous SNV | ACTG1:NM_001199954:exon6:c.T1124C:p.F375S,ACTG1:NM_001614:exon6:c.T1124C:p.F375S, |
| chr17 | 81510703 | C | T | . | exonic | ACTG1 | nonsynonymous SNV | ACTG1:NM_001199954:exon6:c.G1115A:p.R372H,ACTG1:NM_001614:exon6:c.G1115A:p.R372H, |
| chr17 | 81510763 | A | G | COSV59510406 | exonic | ACTG1 | nonsynonymous SNV | ACTG1:NM_001199954:exon6:c.T1055C:p.F352S,ACTG1:NM_001614:exon6:c.T1055C:p.F352S, |
| chr17 | 81510790 | C | T | COSV59509690 | exonic | ACTG1 | nonsynonymous SNV | ACTG1:NM_001199954:exon6:c.G1028A:p.G343D,ACTG1:NM_001614:exon6:c.G1028A:p.G343D, |
| chr17 | 81510814 | C | T | . | exonic | ACTG1 | nonsynonymous SNV | ACTG1:NM_001199954:exon6:c.G1004A:p.R335H,ACTG1:NM_001614:exon6:c.G1004A:p.R335H, |
| chr17 | 81510976 | C | T | . | exonic | ACTG1 | nonsynonymous SNV | ACTG1:NM_001199954:exon5:c.G935A:p.R312K,ACTG1:NM_001614:exon5:c.G935A:p.R312K, |
| chr17 | 81511058 | A | G | . | exonic | ACTG1 | nonsynonymous SNV | ACTG1:NM_001199954:exon5:c.T853C:p.C285R,ACTG1:NM_001614:exon5:c.T853C:p.C285R, |
| chr17 | 81511094 | C | T | COSV59510324 | exonic | ACTG1 | nonsynonymous SNV | ACTG1:NM_001199954:exon5:c.G817A:p.G273S,ACTG1:NM_001614:exon5:c.G817A:p.G273S, |
| chr17 | 81511238 | C | T | . | exonic | ACTG1 | nonsynonymous SNV | ACTG1:NM_001199954:exon4:c.G752A:p.G251D,ACTG1:NM_001614:exon4:c.G752A:p.G251D, |
| chr17 | 81511256 | C | T | . | exonic | ACTG1 | nonsynonymous SNV | ACTG1:NM_001199954:exon4:c.G734A:p.G245D,ACTG1:NM_001614:exon4:c.G734A:p.G245D, |
| chr17 | 81511299 | C | T | . | exonic | ACTG1 | nonsynonymous SNV | ACTG1:NM_001199954:exon4:c.G691A:p.A231T,ACTG1:NM_001614:exon4:c.G691A:p.A231T, |
| chr17 | 81511343 | A | G | . | exonic | ACTG1 | nonsynonymous SNV | ACTG1:NM_001199954:exon4:c.T647C:p.L216P,ACTG1:NM_001614:exon4:c.T647C:p.L216P, |

| | | | | | | | | |
|---|---|---|---|---|---|---|---|---|
| chr17 | 81511361 | C | T | . | exonic | ACTG1 | nonsynonymous SNV | ACTG1:NM_001199954:exon4:c.G629A:p.R210H,ACTG1:NM_001614:exon4:c.G629A:p.R210H, |
| chr17 | 81511373 | C | T | . | exonic | ACTG1 | nonsynonymous SNV | ACTG1:NM_001199954:exon4:c.G617A:p.R206Q,ACTG1:NM_001614:exon4:c.G617A:p.R206Q, |
| chr17 | 81511400 | C | T | . | exonic | ACTG1 | nonsynonymous SNV | ACTG1:NM_001199954:exon4:c.G590A:p.G197D,ACTG1:NM_001614:exon4:c.G590A:p.G197D, |
| chr17 | 81511407 | C | T | . | exonic | ACTG1 | nonsynonymous SNV | ACTG1:NM_001199954:exon4:c.G583A:p.E195K,ACTG1:NM_001614:exon4:c.G583A:p.E195K, |
| chr17 | 81511428 | A | G | . | exonic | ACTG1 | nonsynonymous SNV | ACTG1:NM_001199954:exon4:c.T562C:p.Y188H,ACTG1:NM_001614:exon4:c.T562C:p.Y188H, |
| chr17 | 81511455 | C | T | . | exonic | ACTG1 | nonsynonymous SNV | ACTG1:NM_001199954:exon4:c.G535A:p.D179N,ACTG1:NM_001614:exon4:c.G535A:p.D179N, |
| chr17 | 81511523 | C | T | COSV59509822 | exonic | ACTG1 | nonsynonymous SNV | ACTG1:NM_001199954:exon4:c.G467A:p.G156E,ACTG1:NM_001614:exon4:c.G467A:p.G156E, |
| chr17 | 81511571 | A | G | . | exonic | ACTG1 | nonsynonymous SNV | ACTG1:NM_001199954:exon4:c.T419C:p.L140P,ACTG1:NM_001614:exon4:c.T419C:p.L140P, |
| chr17 | 81511594 | C | T | . | exonic | ACTG1 | nonsynonymous SNV | ACTG1:NM_001199954:exon4:c.G396A:p.M132I,ACTG1:NM_001614:exon4:c.G396A:p.M132I, |
| chr17 | 81511926 | C | T | . | exonic | ACTG1 | nonsynonymous SNV | ACTG1:NM_001199954:exon3:c.G340A:p.A114T,ACTG1:NM_001614:exon3:c.G340A:p.A114T, |
| chr17 | 81511934 | T | G | . | exonic | ACTG1 | nonsynonymous SNV | ACTG1:NM_001199954:exon3:c.A332C:p.N111T,ACTG1:NM_001614:exon3:c.A332C:p.N111T, |
| chr17 | 81511943 | G | A | . | exonic | ACTG1 | nonsynonymous SNV | ACTG1:NM_001199954:exon3:c.C323T:p.A108V,ACTG1:NM_001614:exon3:c.C323T:p.A108V, |
| chr17 | 81511944 | C | T | . | exonic | ACTG1 | nonsynonymous SNV | ACTG1:NM_001199954:exon3:c.G322A:p.A108T,ACTG1:NM_001614:exon3:c.G322A:p.A108T, |
| chr17 | 81511980 | C | T | . | exonic | ACTG1 | nonsynonymous SNV | ACTG1:NM_001199954:exon3:c.G286A:p.V96M,ACTG1:NM_001614:exon3:c.G286A:p.V96M, |
| chr17 | 81512025 | C | T | . | exonic | ACTG1 | nonsynonymous SNV | ACTG1:NM_001199954:exon3:c.G241A:p.D81N,ACTG1:NM_001614:exon3:c.G241A:p.D81N, |
| chr17 | 81512045 | C | T | . | exonic | ACTG1 | nonsynonymous SNV | ACTG1:NM_001199954:exon3:c.G221A:p.G74D,ACTG1:NM_001614:exon3:c.G221A:p.G74D, |
| chr17 | 81512088 | T | A | . | exonic | ACTG1 | nonsynonymous SNV | ACTG1:NM_001199954:exon3:c.A178T:p.S60C,ACTG1:NM_001614:exon3:c.A178T:p.S60C, |
| chr17 | 81512130 | C | T | COSV59509545 | exonic | ACTG1 | nonsynonymous SNV | ACTG1:NM_001199954:exon3:c.G136A:p.G46S,ACTG1:NM_001614:exon3:c.G136A:p.G46S, |
| chr17 | 81512285 | C | T | . | exonic | ACTG1 | nonsynonymous SNV | ACTG1:NM_001199954:exon2:c.G70A:p.D24N,ACTG1:NM_001614:exon2:c.G70A:p.D24N, |
| chr17 | 81512315 | A | G | . | exonic | ACTG1 | nonsynonymous SNV | ACTG1:NM_001199954:exon2:c.T40C:p.S14P,ACTG1:NM_001614:exon2:c.T40C:p.S14P, |
| chr17 | 81512336 | C | T | . | exonic | ACTG1 | nonsynonymous SNV | ACTG1:NM_001199954:exon2:c.G19A:p.A7T,ACTG1:NM_001614:exon2:c.G19A:p.A7T, |
| chr17 | 81512339 | C | T | . | exonic | ACTG1 | nonsynonymous SNV | ACTG1:NM_001199954:exon2:c.G16A:p.A6T,ACTG1:NM_001614:exon2:c.G16A:p.A6T, |
| chr17 | 81868822 | A | C | . | exonic | ARHGDIA | nonsynonymous SNV | ARHGDIA:NM_001301242:exon7:c.T557G:p.M186R, |
| chr17 | 81869013 | T | A | . | exonic | ARHGDIA | nonsynonymous SNV | ARHGDIA:NM_001301241:exon6:c.A478T:p.T160S,ARHGDIA:NM_004309:exon6:c.A478T:p.T160S,ARHGDIA:NM_001301243:exon5:c.A613T:p.T205S,ARHGDIA:NM_001301240:exon6:c.A478T:p.T160S,ARHGDIA:NM_001185077:exon6:c.A478T:p.T160S, |
| chr18 | 12274105 | G | T | COSV57599921 | exonic | CIDEA | nonsynonymous SNV | CIDEA:NM_001279:exon4:c.G343T:p.V115F, |
| chr18 | 47896587 | C | T | COSV50994318 | exonic | SMAD2 | nonsynonymous SNV | SMAD2:NM_005901:exon2:c.G170A:p.R57Q,SMAD2:NM_001003652:exon2:c.G170A:p.R57Q,SMAD2:NM_001135937:exon2:c.G170A:p.R57Q, |
| chr19 | 10252575 | A | C | . | exonic | MRPL4 | nonsynonymous SNV | MRPL4:NM_146387:exon4:c.A149C:p.K50T,MRPL4:NM_146388:exon3:c.A149C:p.K50T,MRPL4:NM_015956:exon3:c.A149C:p.K50T, |
| chr19 | 10631494 | A | G | COSV59835474 | exonic | SLC44A2 | nonsynonymous SNV | SLC44A2:NM_020428:exon7:c.A461G:p.Q154R,SLC44A2:NM_001145056:exon7:c.A455G:p.Q152R, |
| chr19 | 10829148 | G | A | COSV53033495 | exonic | DNM2 | nonsynonymous SNV | DNM2:NM_001190716:exon19:c.G2171A:p.R724H,DNM2:NM_001005360:exon19:c.G2171A:p.R724H,DNM2:NM_004945:exon18:c.G2159A:p.R720H,DNM2:NM_001005361:exon19:c.G2171A:p.R724H,DNM2:NM_001005362:exon18:c.G2159A:p.R720H, |
| chr19 | 1360576 | G | A | COSV60900886 | exonic | MUM1 | nonsynonymous SNV | MUM1:NM_032853:exon5:c.G658A:p.G220R, |
| chr19 | 17060075 | C | T | COSV53725996 | exonic | HAUS8 | nonsynonymous SNV | HAUS8:NM_033417:exon5:c.G247A:p.G83R,HAUS8:NM_001011699:exon5:c.G244A:p.G82R, |
| chr19 | 18451628 | C | T | COSV53211465 | exonic | ELL | nonsynonymous SNV | ELL:NM_006532:exon7:c.G890A:p.S297N, |
| chr19 | 2040173 | G | A | . | exonic | MKNK2 | nonsynonymous SNV | MKNK2:NM_199054:exon13:c.C1115T:p.A372V,MKNK2:NM_017572:exon13:c.C1115T:p.A372V, |
| chr19 | 2050824 | G | T | COSV51731205 | exonic | MKNK2 | nonsynonymous SNV | MKNK2:NM_199054:exon2:c.C28A:p.Q10K,MKNK2:NM_017572:exon2:c.C28A:p.Q10K, |
| chr19 | 23359488 | A | G | COSV56084671 | exonic | ZNF91 | nonsynonymous SNV | ZNF91:NM_003430:exon4:c.T3491C:p.L1164P,ZNF91:NM_001300951:exon3:c.T3395C:p.L1132P, |
| chr19 | 23359489 | G | C | COSV56084686 | exonic | ZNF91 | nonsynonymous SNV | ZNF91:NM_003430:exon4:c.C3490G:p.L1164V,ZNF91:NM_001300951:exon3:c.C3394G:p.L1132V, |
| chr19 | 23360293 | T | C | COSV56080094 | exonic | ZNF91 | nonsynonymous SNV | ZNF91:NM_003430:exon4:c.A2686G:p.T896A,ZNF91:NM_001300951:exon3:c.A2590G:p.T864A, |
| chr19 | 23360692 | T | A | . | exonic | ZNF91 | nonsynonymous SNV | ZNF91:NM_003430:exon4:c.A2287T:p.I763F,ZNF91:NM_001300951:exon3:c.A2191T:p.I731F, |
| chr19 | 23362043 | A | T | . | exonic | ZNF91 | nonsynonymous SNV | ZNF91:NM_003430:exon4:c.T936A:p.H312Q,ZNF91:NM_001300951:exon3:c.T840A:p.H280Q, |
| chr19 | 23362045 | G | T | . | exonic | ZNF91 | nonsynonymous SNV | ZNF91:NM_003430:exon4:c.C934A:p.H312N,ZNF91:NM_001300951:exon3:c.C838A:p.H280N, |
| chr19 | 23362645 | A | G | COSV56080539 | exonic | ZNF91 | nonsynonymous SNV | ZNF91:NM_003430:exon4:c.T334C:p.Y112H,ZNF91:NM_001300951:exon3:c.T238C:p.Y80H, |
| chr19 | 23362714 | G | C | . | exonic | ZNF91 | nonsynonymous SNV | ZNF91:NM_003430:exon4:c.C265G:p.H89D,ZNF91:NM_001300951:exon3:c.C169G:p.H57D, |

| | | | | | | | | |
|---|---|---|---|---|---|---|---|---|
| chr19 | 23374700 | G | A | COSV56080556 | exonic | ZNF91 | nonsynonymous SNV | ZNF91:NM_003430:exon2:c.C95T:p.T32I,ZNF91:NM_001300951:exon2:c.C95T:p.T32I, |
| chr19 | 30011135 | T | A | . | exonic | URI1 | nonsynonymous SNV | URI1:NM_003796:exon9:c.T1077A:p.S359R,URI1:NM_001252641:exon9:c.T1023A:p.S341R, |
| chr19 | 33109858 | T | C | COSV50111027 | exonic | GPATCH1 | nonsynonymous SNV | GPATCH1:NM_018025:exon11:c.T1427C:p.L476P, |
| chr19 | 33114394 | A | G | COSV50114971 | exonic | GPATCH1 | nonsynonymous SNV | GPATCH1:NM_018025:exon15:c.A2171G:p.H724R, |
| chr19 | 33114406 | T | C | COSV50114723 | exonic | GPATCH1 | nonsynonymous SNV | GPATCH1:NM_018025:exon15:c.T2183C:p.L728S, |
| chr19 | 33125171 | C | T | . | exonic | GPATCH1 | nonsynonymous SNV | GPATCH1:NM_018025:exon18:c.C2588T:p.A863V, |
| chr19 | 33126632 | T | A | COSV50118646 | exonic | GPATCH1 | nonsynonymous SNV | GPATCH1:NM_018025:exon19:c.T2664A:p.N888K, |
| chr19 | 35157462 | T | G | COSV54343702 | exonic | FXYD5 | nonsynonymous SNV | FXYD5:NM_001164605:exon3:c.T103G:p.S35A,FXYD5:NM_144779:exon3:c.T103G:p.S35A,FXYD5:NM_014164:exon3:c.T103G:p.S35A, |
| chr19 | 35169605 | G | A | . | exonic | FXYD5 | nonsynonymous SNV | FXYD5:NM_001164605:exon9:c.G527A:p.R176H,FXYD5:NM_144779:exon9:c.G527A:p.R176H,FXYD5:NM_014164:exon9:c.G527A:p.R176H, |
| chr19 | 38624111 | G | A | . | exonic | EIF3K | nonsynonymous SNV | EIF3K:NM_001300992:exon3:c.G193A:p.V65I,EIF3K:NM_013234:exon3:c.G193A:p.V65I, |
| chr19 | 38624192 | G | A | . | exonic | EIF3K | nonsynonymous SNV | EIF3K:NM_001300992:exon3:c.G274A:p.A92T,EIF3K:NM_013234:exon3:c.G274A:p.A92T,EIF3K:NM_001308393:exon3:c.G13A:p.A5T, |
| chr19 | 38626047 | G | A | COSV50263701 | exonic | EIF3K | nonsynonymous SNV | EIF3K:NM_001300992:exon4:c.G299A:p.R100Q,EIF3K:NM_013234:exon4:c.G299A:p.R100Q,EIF3K:NM_001308393:exon4:c.G38A:p.R13Q, |
| chr19 | 38636906 | G | A | . | exonic | EIF3K | nonsynonymous SNV | EIF3K:NM_001300992:exon7:c.G565A:p.A189T,EIF3K:NM_013234:exon8:c.G643A:p.A215T,EIF3K:NM_001308393:exon8:c.G382A:p.A128T, |
| chr19 | 38816463 | C | T | COSV55489256 | exonic | ECH1 | nonsynonymous SNV | ECH1:NM_001398:exon7:c.G649A:p.G217R, |
| chr19 | 41686928 | T | A | . | exonic | CEACAM7 | nonsynonymous SNV | CEACAM7:NM_001291485:exon2:c.A358T:p.I120F,CEACAM7:NM_006890:exon2:c.A358T:p.I120F, |
| chr19 | 41761981 | T | G | . | exonic | CEACAM6 | nonsynonymous SNV | CEACAM6:NM_002483:exon4:c.T716G:p.V239G, |
| chr19 | 43356040 | T | G | COSV65120741 | exonic | CD177 | nonsynonymous SNV | CD177:NM_020406:exon5:c.T551G:p.V184G, |
| chr19 | 43356099 | G | A | COSV65120743 | exonic | CD177 | nonsynonymous SNV | CD177:NM_020406:exon5:c.G610A:p.D204N, |
| chr19 | 43356103 | T | G | COSV65120745 | exonic | CD177 | nonsynonymous SNV | CD177:NM_020406:exon5:c.T614G:p.M205R, |
| chr19 | 44908684 | T | C | COSV52985231 | exonic | APOE | nonsynonymous SNV | APOE:NM_000041:exon4:c.T388C:p.C130R,APOE:NM_001302691:exon4:c.T388C:p.C130R,APOE:NM_001302689:exon4:c.T388C:p.C130R,APOE:NM_001302690:exon4:c.T388C:p.C130R,APOE:NM_001302688:exon4:c.T466C:p.C156R, |
| chr19 | 45351661 | T | G | COSV67266431 | exonic | ERCC2 | nonsynonymous SNV | ERCC2:NM_000400:exon23:c.A2251C:p.K751Q, |
| chr19 | 45364001 | C | T | COSV55538614 | exonic | ERCC2 | nonsynonymous SNV | ERCC2:NM_000400:exon10:c.G934A:p.D312N,ERCC2:NM_001130867:exon9:c.G862A:p.D288N, |
| chr19 | 46775602 | C | G | COSV69467128 | exonic | SLC1A5 | nonsynonymous SNV | SLC1A5:NM_001145145:exon7:c.G928C:p.V310L,SLC1A5:NM_005628:exon8:c.G1534C:p.V512L,SLC1A5:NM_001145144:exon8:c.G850C:p.V284L, |
| chr19 | 51024714 | C | T | COSV99031948 | exonic | KLK11 | nonsynonymous SNV | KLK11:NM_144947:exon3:c.G217A:p.E73K,KLK11:NM_001136032:exon3:c.G121A:p.E41K,KLK11:NM_001167605:exon3:c.G121A:p.E41K,KLK11:NM_006853:exon3:c.G121A:p.E41K, |
| chr19 | 8371280 | C | T | COSV56844530 | exonic | ANGPTL4 | nonsynonymous SNV | ANGPTL4:NM_001039667:exon5:c.C683T:p.T228M,ANGPTL4:NM_139314:exon6:c.C797T:p.T266M, |
| chr2 | 100974842 | A | G | COSV59555566 | exonic | NPAS2 | nonsynonymous SNV | NPAS2:NM_002518:exon13:c.A1180G:p.T394A, |
| chr2 | 101308564 | T | C | COSV54863782 | exonic | RNF149 | nonsynonymous SNV | RNF149:NM_173647:exon1:c.A25G:p.S9G, |
| chr2 | 102814945 | T | C | COSV68425595 | exonic | TMEM182 | nonsynonymous SNV | TMEM182:NM_144632:exon5:c.T667C:p.W223R, |
| chr2 | 10790777 | T | C | COSV55349252 | exonic | PDIA6 | nonsynonymous SNV | PDIA6:NM_001282705:exon8:c.A785G:p.K262R,PDIA6:NM_001282707:exon8:c.A632G:p.K211R,PDIA6:NM_001282706:exon8:c.A656G:p.K219R,PDIA6:NM_005742:exon7:c.A641G:p.K214R,PDIA6:NM_001282704:exon9:c.A797G:p.K266R, |
| chr2 | 119372265 | A | G | COSV61055619 | exonic | DBI | nonsynonymous SNV | DBI:NM_020548:exon5:c.A262G:p.M88V,DBI:NM_001282633:exon5:c.A262G:p.M88V,DBI:NM_001282634:exon5:c.A262G:p.M88V,DBI:NM_001178041:exon5:c.A337G:p.M113V,DBI:NM_001079863:exon4:c.A214G:p.M72V,DBI:NM_001282636:exon4:c.A139G:p.M47V,DBI:NM_001178017:exon4:c.A394G:p.M132V,DBI:NM_001282635:exon5:c.A262G:p.M88V,DBI:NM_001178043:exon4:c.A241G:p.M81V,DBI:NM_001079862:exon4:c.A211G:p.M71V,DBI:NM_001178042:exon5:c.A262G:p.M88V, |
| chr2 | 135135700 | C | A | . | exonic | RAB3GAP1 | nonsynonymous SNV | RAB3GAP1:NM_001172435:exon17:c.C1691A:p.T564K,RAB3GAP1:NM_012233:exon17:c.C1691A:p.T564K, |
| chr2 | 135135704 | T | A | . | exonic | RAB3GAP1 | nonsynonymous SNV | RAB3GAP1:NM_001172435:exon17:c.T1695A:p.D565E,RAB3GAP1:NM_012233:exon17:c.T1695A:p.D565E, |
| chr2 | 135135802 | A | G | COSV51482881 | exonic | RAB3GAP1 | nonsynonymous SNV | RAB3GAP1:NM_001172435:exon17:c.A1793G:p.N598S,RAB3GAP1:NM_012233:exon17:c.A1793G:p.N598S, |
| chr2 | 143040620 | A | G | COSV51580314 | exonic | KYNU | nonsynonymous SNV | KYNU:NM_001199241:exon14:c.A1234G:p.K412E,KYNU:NM_003937:exon13:c.A1234G:p.K412E, |
| chr2 | 177230913 | C | T | . | exonic | NFE2L2 | nonsynonymous SNV | NFE2L2:NM_001145413:exon5:c.G1621A:p.V541I,NFE2L2:NM_001313901:exon5:c.G1642A:p.V548I,NFE2L2:NM_001313902:exon4:c.G1600A:p.V534I,NFE2L2:NM_001313900:exon5:c.G1642A:p.V548I,NFE2L2:NM_001313904:exon5:c.G1390A:p.V464I,NFE2L2:NM_001313903:exon5:c.G1471A:p.V491I,NFE2L2:NM_006164:exon5:c.G1690A:p.V564I,NFE2L2:NM_001145412:exon5:c.G1642A:p.V548I, |

| | | | | | | | | |
|---|---|---|---|---|---|---|---|---|
| chr2 | 177231000 | C | T | . | exonic | NFE2L2 | nonsynonymous SNV | NFE2L2:NM_001145413:exon5:c.G1534A:p.E512K,NFE2L2:NM_001313901:exon5:c.G1555A:p.E519K,NFE2L2:NM_001313902:exon4:c.G1513A:p.E505K,NFE2L2:NM_001313900:exon5:c.G1555A:p.E519K,NFE2L2:NM_001313904:exon5:c.G1303A:p.E435K,NFE2L2:NM_001313903:exon5:c.G1384A:p.E462K,NFE2L2:NM_006164:exon5:c.G1603A:p.E535K,NFE2L2:NM_001145412:exon5:c.G1555A:p.E519K, |
| chr2 | 177231062 | C | T | . | exonic | NFE2L2 | nonsynonymous SNV | NFE2L2:NM_001145413:exon5:c.G1472A:p.C491Y,NFE2L2:NM_001313901:exon5:c.G1493A:p.C498Y,NFE2L2:NM_001313902:exon4:c.G1451A:p.C484Y,NFE2L2:NM_001313900:exon5:c.G1493A:p.C498Y,NFE2L2:NM_001313904:exon5:c.G1241A:p.C414Y,NFE2L2:NM_001313903:exon5:c.G1322A:p.C441Y,NFE2L2:NM_006164:exon5:c.G1541A:p.C514Y,NFE2L2:NM_001145412:exon5:c.G1493A:p.C498Y, |
| chr2 | 177231064 | A | T | . | exonic | NFE2L2 | nonsynonymous SNV | NFE2L2:NM_001145413:exon5:c.T1470A:p.N490K,NFE2L2:NM_001313901:exon5:c.T1491A:p.N497K,NFE2L2:NM_001313902:exon4:c.T1449A:p.N483K,NFE2L2:NM_001313900:exon5:c.T1491A:p.N497K,NFE2L2:NM_001313904:exon5:c.T1239A:p.N413K,NFE2L2:NM_001313903:exon5:c.T1320A:p.N440K,NFE2L2:NM_006164:exon5:c.T1539A:p.N513K,NFE2L2:NM_001145412:exon5:c.T1491A:p.N497K, |
| chr2 | 177231095 | C | T | . | exonic | NFE2L2 | nonsynonymous SNV | NFE2L2:NM_001145413:exon5:c.G1439A:p.R480K,NFE2L2:NM_001313901:exon5:c.G1460A:p.R487K,NFE2L2:NM_001313902:exon4:c.G1418A:p.R473K,NFE2L2:NM_001313900:exon5:c.G1460A:p.R487K,NFE2L2:NM_001313904:exon5:c.G1208A:p.R403K,NFE2L2:NM_001313903:exon5:c.G1289A:p.R430K,NFE2L2:NM_006164:exon5:c.G1508A:p.R503K,NFE2L2:NM_001145412:exon5:c.G1460A:p.R487K, |
| chr2 | 177231740 | G | A | . | exonic | NFE2L2 | nonsynonymous SNV | NFE2L2:NM_001145413:exon5:c.C794T:p.A265V,NFE2L2:NM_001313901:exon5:c.C815T:p.A272V,NFE2L2:NM_001313902:exon4:c.C773T:p.A258V,NFE2L2:NM_001313900:exon5:c.C815T:p.A272V,NFE2L2:NM_001313904:exon5:c.C563T:p.A188V,NFE2L2:NM_001313903:exon5:c.C644T:p.A215V,NFE2L2:NM_006164:exon5:c.C863T:p.A288V,NFE2L2:NM_001145412:exon5:c.C815T:p.A272V, |
| chr2 | 177231758 | C | T | . | exonic | NFE2L2 | nonsynonymous SNV | NFE2L2:NM_001145413:exon5:c.G776A:p.G259D,NFE2L2:NM_001313901:exon5:c.G797A:p.G266D,NFE2L2:NM_001313902:exon4:c.G755A:p.G252D,NFE2L2:NM_001313900:exon5:c.G797A:p.G266D,NFE2L2:NM_001313904:exon5:c.G545A:p.G182D,NFE2L2:NM_001313903:exon5:c.G626A:p.G209D,NFE2L2:NM_006164:exon5:c.G845A:p.G282D,NFE2L2:NM_001145412:exon5:c.G797A:p.G266D, |
| chr2 | 177231891 | C | T | . | exonic | NFE2L2 | nonsynonymous SNV | NFE2L2:NM_001313902:exon4:c.G622A:p.E208K,NFE2L2:NM_001313900:exon5:c.G664A:p.E222K,NFE2L2:NM_001313904:exon5:c.G412A:p.E138K,NFE2L2:NM_001145412:exon5:c.G664A:p.E222K,NFE2L2:NM_001145413:exon5:c.G643A:p.E215K,NFE2L2:NM_001313901:exon5:c.G664A:p.E222K,NFE2L2:NM_001313903:exon5:c.G493A:p.E165K,NFE2L2:NM_006164:exon5:c.G712A:p.E238K, |
| chr2 | 177231951 | C | T | . | exonic | NFE2L2 | nonsynonymous SNV | NFE2L2:NM_001145413:exon5:c.G583A:p.A195T,NFE2L2:NM_001313901:exon5:c.G604A:p.A202T,NFE2L2:NM_001313902:exon4:c.G562A:p.A188T,NFE2L2:NM_001313900:exon5:c.G604A:p.A202T,NFE2L2:NM_001313904:exon5:c.G352A:p.A118T,NFE2L2:NM_001313903:exon5:c.G433A:p.A145T,NFE2L2:NM_006164:exon5:c.G652A:p.A218T,NFE2L2:NM_001145412:exon5:c.G604A:p.A202T, |
| chr2 | 177232454 | C | T | . | exonic | NFE2L2 | nonsynonymous SNV | NFE2L2:NM_001145413:exon4:c.G463A:p.D155N,NFE2L2:NM_001313901:exon4:c.G484A:p.D162N,NFE2L2:NM_001313902:exon3:c.G442A:p.D148N,NFE2L2:NM_001313900:exon4:c.G484A:p.D162N,NFE2L2:NM_001313904:exon4:c.G232A:p.D78N,NFE2L2:NM_001313903:exon4:c.G313A:p.D105N,NFE2L2:NM_006164:exon4:c.G532A:p.D178N,NFE2L2:NM_001145412:exon4:c.G484A:p.D162N, |
| chr2 | 177234073 | C | T | . | exonic | NFE2L2 | nonsynonymous SNV | NFE2L2:NM_001313902:exon2:c.G244A:p.E82K,NFE2L2:NM_001313900:exon2:c.G196A:p.E66K,NFE2L2:NM_001145412:exon2:c.G196A:p.E66K,NFE2L2:NM_001145413:exon2:c.G196A:p.E66K,NFE2L2:NM_001313901:exon2:c.G196A:p.E66K,NFE2L2:NM_006164:exon2:c.G244A:p.E82K, |
| chr2 | 177234109 | A | G | . | exonic | NFE2L2 | nonsynonymous SNV | NFE2L2:NM_001145413:exon2:c.T160C:p.F54L,NFE2L2:NM_001313901:exon2:c.T160C:p.F54L,NFE2L2:NM_001313902:exon2:c.T208C:p.F70L,NFE2L2:NM_001313900:exon2:c.T160C:p.F54L,NFE2L2:NM_006164:exon2:c.T208C:p.F70L,NFE2L2:NM_001145412:exon2:c.T160C:p.F54L, |
| chr2 | 190296896 | T | C | COSV62891843 | exonic | HIBCH | nonsynonymous SNV | HIBCH:NM_198047:exon3:c.A136G:p.T46A,HIBCH:NM_014362:exon3:c.A136G:p.T46A, |
| chr2 | 190319749 | A | G | COSV62891757 | exonic | HIBCH | nonsynonymous SNV | HIBCH:NM_198047:exon1:c.T2C:p.M1T,HIBCH:NM_014362:exon1:c.T2C:p.M1T, |
| chr2 | 197487080 | C | G | COSV61445925 | exonic | HSPD1 | nonsynonymous SNV | HSPD1:NM_002156:exon12:c.G1688C:p.G563A,HSPD1:NM_199440:exon12:c.G1688C:p.G563A, |
| chr2 | 197487083 | C | T | . | exonic | HSPD1 | nonsynonymous SNV | HSPD1:NM_002156:exon12:c.G1685A:p.G562D,HSPD1:NM_199440:exon12:c.G1685A:p.G562D, |
| chr2 | 197487094 | C | T | . | exonic | HSPD1 | nonsynonymous SNV | HSPD1:NM_002156:exon12:c.G1674A:p.M558I,HSPD1:NM_199440:exon12:c.G1674A:p.M558I, |
| chr2 | 197488473 | C | T | . | exonic | HSPD1 | nonsynonymous SNV | HSPD1:NM_002156:exon10:c.G1234A:p.V412I,HSPD1:NM_199440:exon10:c.G1234A:p.V412I, |
| chr2 | 197489109 | G | T | COSV61444464 | exonic | HSPD1 | nonsynonymous SNV | HSPD1:NM_002156:exon9:c.C1108A:p.R370S,HSPD1:NM_199440:exon9:c.C1108A:p.R370S, |
| chr2 | 197489115 | C | T | . | exonic | HSPD1 | nonsynonymous SNV | HSPD1:NM_002156:exon9:c.G1102A:p.E368K,HSPD1:NM_199440:exon9:c.G1102A:p.E368K, |
| chr2 | 197489132 | C | T | . | exonic | HSPD1 | nonsynonymous SNV | HSPD1:NM_002156:exon9:c.G1085A:p.G362D,HSPD1:NM_199440:exon9:c.G1085A:p.G362D, |
| chr2 | 197489145 | A | T | . | exonic | HSPD1 | nonsynonymous SNV | HSPD1:NM_002156:exon9:c.T1072A:p.L358I,HSPD1:NM_199440:exon9:c.T1072A:p.L358I, |
| chr2 | 197489178 | C | T | . | exonic | HSPD1 | nonsynonymous SNV | HSPD1:NM_002156:exon9:c.G1039A:p.E347K,HSPD1:NM_199440:exon9:c.G1039A:p.E347K, |
| chr2 | 197493385 | G | T | . | exonic | HSPD1 | nonsynonymous SNV | HSPD1:NM_002156:exon7:c.C808A:p.P270T,HSPD1:NM_199440:exon7:c.C808A:p.P270T, |
| chr2 | 197497334 | A | G | . | exonic | HSPD1 | nonsynonymous SNV | HSPD1:NM_002156:exon3:c.T233C:p.V78A,HSPD1:NM_199440:exon3:c.T233C:p.V78A, |
| chr2 | 197498682 | C | T | COSV52098272 | exonic | HSPD1 | nonsynonymous SNV | HSPD1:NM_002156:exon2:c.G167A:p.G56E,HSPD1:NM_199440:exon2:c.G167A:p.G56E, |
| chr2 | 197498823 | C | T | . | exonic | HSPD1 | nonsynonymous SNV | HSPD1:NM_002156:exon2:c.G26A:p.R9H,HSPD1:NM_199440:exon2:c.G26A:p.R9H, |
| chr2 | 213317301 | G | C | . | exonic | SPAG16 | nonsynonymous SNV | SPAG16:NM_024532:exon5:c.G481C:p.E161Q,SPAG16:NM_001025436:exon5:c.G481C:p.E161Q, |
| chr2 | 213317306 | T | A | . | exonic | SPAG16 | nonsynonymous SNV | SPAG16:NM_024532:exon5:c.T486A:p.N162K,SPAG16:NM_001025436:exon5:c.T486A:p.N162K, |
| chr2 | 215370366 | C | T | . | exonic | FN1 | nonsynonymous SNV | FN1:NM_001306130:exon40:c.G6151A:p.V2051I,FN1:NM_212474:exon39:c.G5878A:p.V1960I,FN1:NM_212476:exon39:c.G6238A:p.V2080I,FN1:NM_001306129:exon42:c.G6688A:p.V2230I,FN1:NM_001306131:exon40:c.G6145A:p.V2049I,FN1:NM_001306132:exon40:c.G6070A:p.V2024I,FN1:NM_002026:exon41:c.G6415A:p.V2139I,FN1:NM_212478:exon41:c.G6340A:p.V2114I,FN1:NM_212482:exon41:c.G6781A:p.V2261I, |

| | | | | | | | | |
|---|---|---|---|---|---|---|---|---|
| chr2 | 215407182 | A | T | . | exonic | FN1 | nonsynonymous SNV | FN1:NM_001306130:exon18:c.T2658A:p.N886K,FN1:NM_212474:exon18:c.T2658A:p.N886K,FN1:NM_212476:exon18:c.T2658A:p.N886K,FN1:NM_001306129:exon18:c.T2658A:p.N886K,FN1:NM_001306131:exon18:c.T2658A:p.N886K,FN1:NM_001306132:exon18:c.T2658A:p.N886K,FN1:NM_002026:exon18:c.T2658A:p.N886K,FN1:NM_212478:exon18:c.T2658A:p.N886K,FN1:NM_212482:exon18:c.T2658A:p.N886K, |
| chr2 | 215408177 | T | G | . | exonic | FN1 | nonsynonymous SNV | FN1:NM_001306130:exon17:c.A2449C:p.T817P,FN1:NM_212474:exon17:c.A2449C:p.T817P,FN1:NM_212476:exon17:c.A2449C:p.T817P,FN1:NM_001306129:exon17:c.A2449C:p.T817P,FN1:NM_001306131:exon17:c.A2449C:p.T817P,FN1:NM_001306132:exon17:c.A2449C:p.T817P,FN1:NM_002026:exon17:c.A2449C:p.T817P,FN1:NM_212478:exon17:c.A2449C:p.T817P,FN1:NM_212482:exon17:c.A2449C:p.T817P, |
| chr2 | 224511367 | A | T | . | exonic | CUL3 | nonsynonymous SNV | CUL3:NM_001257198:exon6:c.T888A:p.N296K,CUL3:NM_001257197:exon5:c.T672A:p.N224K,CUL3:NM_003590:exon6:c.T870A:p.N290K, |
| chr2 | 233204655 | C | T | COSV64037273 | exonic | INPP5D | nonsynonymous SNV | INPP5D:NM_001017915:exon26:c.C3505T:p.H1169Y,INPP5D:NM_005541:exon26:c.C3502T:p.H1168Y, |
| chr2 | 240025297 | G | C | COSV53155269 | exonic | NDUFA10 | nonsynonymous SNV | NDUFA10:NM_004544:exon1:c.C5G:p.A2G, |
| chr2 | 240630832 | A | C | COSV60576322 | exonic | GPR35 | nonsynonymous SNV | GPR35:NM_001195382:exon6:c.A973C:p.S325R,GPR35:NM_005301:exon2:c.A880C:p.S294R,GPR35:NM_001195381:exon6:c.A973C:p.S325R, |
| chr2 | 27342108 | G | C | . | exonic | GTF3C2 | nonsynonymous SNV | GTF3C2:NM_001521:exon5:c.C695G:p.A232G,GTF3C2:NM_001035521:exon4:c.C695G:p.A232G, |
| chr2 | 27342117 | T | A | . | exonic | GTF3C2 | nonsynonymous SNV | GTF3C2:NM_001521:exon5:c.A686T:p.Q229L,GTF3C2:NM_001035521:exon4:c.A686T:p.Q229L, |
| chr2 | 28851255 | C | T | . | exonic | TRMT61B | nonsynonymous SNV | TRMT61B:NM_017910:exon5:c.G1129A:p.E377K, |
| chr2 | 32736791 | A | G | COSV58653535 | exonic | TTC27 | nonsynonymous SNV | TTC27:NM_017735:exon12:c.A1427G:p.Y476C,TTC27:NM_001193509:exon12:c.A1277G:p.Y426C, |
| chr2 | 33360729 | T | C | . | exonic | LTBP1 | nonsynonymous SNV | LTBP1:NM_000627:exon23:c.T3155C:p.V1052A,LTBP1:NM_001166264:exon22:c.T3029C:p.V1010A,LTBP1:NM_001166266:exon22:c.T2870C:p.V957A,LTBP1:NM_001166265:exon23:c.T2996C:p.V999A,LTBP1:NM_206943:exon27:c.T4133C:p.V1378A, |
| chr2 | 42282878 | A | G | COSV59293461 | exonic | EML4 | nonsynonymous SNV | EML4:NM_001145076:exon7:c.A673G:p.K225E,EML4:NM_019063:exon8:c.A847G:p.K283E, |
| chr2 | 42288248 | A | G | COSV59293527 | exonic | EML4 | nonsynonymous SNV | EML4:NM_001145076:exon10:c.A970G:p.I324V,EML4:NM_019063:exon11:c.A1144G:p.I382V, |
| chr2 | 42288297 | A | G | COSV59293537 | exonic | EML4 | nonsynonymous SNV | EML4:NM_001145076:exon10:c.A1019G:p.K340R,EML4:NM_019063:exon11:c.A1193G:p.K398R, |
| chr2 | 44320435 | G | A | COSV53215689 | exonic | SLC3A1 | nonsynonymous SNV | SLC3A1:NM_000341:exon10:c.G1854A:p.M618I, |
| chr2 | 47471001 | T | A | COSV51879859 | exonic | MSH2 | nonsynonymous SNV | MSH2:NM_000251:exon11:c.T1698A:p.N566K,MSH2:NM_001258281:exon12:c.T1500A:p.N500K, |
| chr2 | 47475249 | C | A | . | exonic | MSH2 | nonsynonymous SNV | MSH2:NM_000251:exon12:c.C1984A:p.Q662K,MSH2:NM_001258281:exon13:c.C1786A:p.Q596K, |
| chr2 | 60546080 | C | T | . | exonic | BCL11A | nonsynonymous SNV | BCL11A:NM_138559:exon2:c.G276A:p.M92I,BCL11A:NM_018014:exon2:c.G276A:p.M92I,BCL11A:NM_022893:exon2:c.G276A:p.M92I, |
| chr2 | 69327071 | C | T | COSV61947419 | exonic | GFPT1 | nonsynonymous SNV | GFPT1:NM_002056:exon18:c.G1844A:p.R615Q,GFPT1:NM_001244710:exon19:c.G1898A:p.R633Q, |
| chr2 | 69338463 | A | G | . | exonic | GFPT1 | nonsynonymous SNV | GFPT1:NM_002056:exon13:c.T1252C:p.F418L,GFPT1:NM_001244710:exon14:c.T1306C:p.F436L, |
| chr2 | 69804572 | G | A | . | exonic | ANXA4 | nonsynonymous SNV | ANXA4:NM_001153:exon4:c.G137A:p.R46H, |
| chr2 | 69804622 | G | A | . | exonic | ANXA4 | nonsynonymous SNV | ANXA4:NM_001153:exon4:c.G187A:p.G63S, |
| chr2 | 69806452 | C | T | COSV67848858 | exonic | ANXA4 | nonsynonymous SNV | ANXA4:NM_001153:exon5:c.C260T:p.T87M, |
| chr2 | 69807913 | G | A | . | exonic | ANXA4 | nonsynonymous SNV | ANXA4:NM_001153:exon6:c.G314A:p.G105D, |
| chr2 | 69807924 | G | A | . | exonic | ANXA4 | nonsynonymous SNV | ANXA4:NM_001153:exon6:c.G325A:p.G109S, |
| chr2 | 69807952 | G | A | COSV67848884 | exonic | ANXA4 | nonsynonymous SNV | ANXA4:NM_001153:exon6:c.G353A:p.R118Q, |
| chr2 | 69818664 | G | A | . | exonic | ANXA4 | nonsynonymous SNV | ANXA4:NM_001153:exon10:c.G694A:p.G232S, |
| chr2 | 69820699 | G | A | . | exonic | ANXA4 | nonsynonymous SNV | ANXA4:NM_001153:exon12:c.G784A:p.G262S, |
| chr2 | 69825510 | G | A | COSV57003814 | exonic | ANXA4 | nonsynonymous SNV | ANXA4:NM_001153:exon13:c.G961A:p.D321N, |
| chr2 | 99439044 | A | G | COSV51482947 | exonic | REV1 | nonsynonymous SNV | REV1:NM_016316:exon6:c.T770C:p.F257S,REV1:NM_001037872:exon6:c.T770C:p.F257S, |
| chr20 | 17990067 | G | A | . | exonic | MGME1 | nonsynonymous SNV | MGME1:NM_001310339:exon3:c.G640A:p.E214K, |
| chr20 | 18162473 | T | G | . | exonic | CSRP2BP | nonsynonymous SNV | CSRP2BP:NM_020536:exon6:c.T1199G:p.V400G, |
| chr20 | 18532728 | C | T | COSV52719453 | exonic | SEC23B | nonsynonymous SNV | SEC23B:NM_001172745:exon11:c.C1298T:p.P433L,SEC23B:NM_032985:exon11:c.C1298T:p.P433L,SEC23B:NM_001172746:exon11:c.C1244T:p.P415L,SEC23B:NM_006363:exon11:c.C1298T:p.P433L,SEC23B:NM_032986:exon11:c.C1298T:p.P433L, |
| chr20 | 18542358 | C | G | COSV52719183 | exonic | SEC23B | nonsynonymous SNV | SEC23B:NM_001172745:exon13:c.C1467G:p.H489Q,SEC23B:NM_032985:exon13:c.C1467G:p.H489Q,SEC23B:NM_001172746:exon13:c.C1413G:p.H471Q,SEC23B:NM_006363:exon13:c.C1467G:p.H489Q,SEC23B:NM_032986:exon13:c.C1467G:p.H489Q, |
| chr20 | 23637790 | C | T | COSV65361855 | exonic | CST3 | nonsynonymous SNV | CST3:NM_001288614:exon1:c.G73A:p.A25T,CST3:NM_000099:exon1:c.G73A:p.A25T, |
| chr20 | 25282133 | G | A | COSV53816266 | exonic | PYGB | nonsynonymous SNV | PYGB:NM_002862:exon12:c.G1504A:p.D502N, |
| chr20 | 32432924 | G | A | . | exonic | ASXL1 | nonsynonymous SNV | ASXL1:NM_015338:exon10:c.G1024A:p.E342K, |
| chr20 | 32432949 | G | A | . | exonic | ASXL1 | stopgain | ASXL1:NM_015338:exon10:c.G1049A:p.W350X, |

| | | | | | | | | |
|---|---|---|---|---|---|---|---|---|
| chr20 | 34936982 | A | G | . | exonic | GSS | nonsynonymous SNV | GSS:NM_000178:exon7:c.T650C:p.I217T, |
| chr20 | 35384111 | C | T | . | exonic | UQCC1 | nonsynonymous SNV | UQCC1:NM_199487:exon3:c.G152A:p.R51Q,UQCC1:NM_018244:exon3:c.G152A:p.R51Q, |
| chr20 | 54214409 | G | A | . | exonic | PFDN4 | nonsynonymous SNV | PFDN4:NM_002623:exon2:c.G83A:p.R28Q, |
| chr20 | 54219119 | G | A | . | exonic | PFDN4 | nonsynonymous SNV | PFDN4:NM_002623:exon4:c.G374A:p.S125N, |
| chr20 | 56513348 | A | G | COSV50130089 | exonic | RTFDC1 | nonsynonymous SNV | RTFDC1:NM_001283035:exon7:c.A601G:p.M201V,RTFDC1:NM_016407:exon6:c.A511G:p.M171V,RTFDC1:NM_001283037:exon6:c.A511G:p.M171V,RTFDC1:NM_001283036:exon6:c.A511G:p.M171V, |
| chr20 | 58903571 | A | C | . | exonic | GNAS | nonsynonymous SNV | GNAS:NM_080426:exon3:c.A256C:p.K86Q,GNAS:NM_080425:exon4:c.A2227C:p.K743Q,GNAS:NM_001309840:exon4:c.A121C:p.K41Q,GNAS:NM_001077489:exon3:c.A253C:p.K85Q,GNAS:NM_001309861:exon4:c.A121C:p.K41Q,GNAS:NM_000516:exon4:c.A298C:p.K100Q,GNAS:NM_001077488:exon4:c.A301C:p.K101Q, |
| chr20 | 58903682 | C | A | . | exonic | GNAS | nonsynonymous SNV | GNAS:NM_080426:exon4:c.C281A:p.A94D,GNAS:NM_080425:exon5:c.C2252A:p.A751D,GNAS:NM_001309840:exon5:c.C146A:p.A49D,GNAS:NM_001077489:exon4:c.C278A:p.A93D,GNAS:NM_001309861:exon5:c.C146A:p.A49D,GNAS:NM_000516:exon5:c.C323A:p.A108D,GNAS:NM_001077488:exon5:c.C326A:p.A109D, |
| chr20 | 58903684 | G | A | COSV55672684 | exonic | GNAS | nonsynonymous SNV | GNAS:NM_080426:exon4:c.G283A:p.A95T,GNAS:NM_080425:exon5:c.G2254A:p.A752T,GNAS:NM_001309840:exon5:c.G148A:p.A50T,GNAS:NM_001077489:exon4:c.G280A:p.A94T,GNAS:NM_001309861:exon5:c.G148A:p.A50T,GNAS:NM_000516:exon5:c.G325A:p.A109T,GNAS:NM_001077488:exon5:c.G328A:p.A110T, |
| chr20 | 58903691 | G | A | COSV99039588 | exonic | GNAS | nonsynonymous SNV | GNAS:NM_080426:exon4:c.G290A:p.S97N,GNAS:NM_080425:exon5:c.G2261A:p.S754N,GNAS:NM_001309840:exon5:c.G155A:p.S52N,GNAS:NM_001077489:exon4:c.G287A:p.S96N,GNAS:NM_001309861:exon5:c.G155A:p.S52N,GNAS:NM_000516:exon5:c.G332A:p.S111N,GNAS:NM_001077488:exon5:c.G335A:p.S112N, |
| chr20 | 58903717 | G | A | . | exonic | GNAS | nonsynonymous SNV | GNAS:NM_080426:exon4:c.G316A:p.A106T,GNAS:NM_080425:exon5:c.G2287A:p.A763T,GNAS:NM_001309840:exon5:c.G181A:p.A61T,GNAS:NM_001077489:exon4:c.G313A:p.A105T,GNAS:NM_001309861:exon5:c.G181A:p.A61T,GNAS:NM_000516:exon5:c.G358A:p.A120T,GNAS:NM_001077488:exon5:c.G361A:p.A121T, |
| chr20 | 58903726 | G | A | . | exonic | GNAS | nonsynonymous SNV | GNAS:NM_080426:exon4:c.G325A:p.E109K,GNAS:NM_080425:exon5:c.G2296A:p.E766K,GNAS:NM_001309840:exon5:c.G190A:p.E64K,GNAS:NM_001077489:exon4:c.G322A:p.E108K,GNAS:NM_001309861:exon5:c.G190A:p.E64K,GNAS:NM_000516:exon5:c.G367A:p.E123K,GNAS:NM_001077488:exon5:c.G370A:p.E124K, |
| chr20 | 58903768 | G | A | . | exonic | GNAS | nonsynonymous SNV | GNAS:NM_080426:exon4:c.G367A:p.V123M,GNAS:NM_080425:exon5:c.G2338A:p.V780M,GNAS:NM_001309840:exon5:c.G232A:p.V78M,GNAS:NM_001077489:exon4:c.G364A:p.V122M,GNAS:NM_001309861:exon5:c.G232A:p.V78M,GNAS:NM_000516:exon5:c.G409A:p.V137M,GNAS:NM_001077488:exon5:c.G412A:p.V138M, |
| chr20 | 58905383 | G | A | . | exonic | GNAS | nonsynonymous SNV | GNAS:NM_080426:exon5:c.G391A:p.E131K,GNAS:NM_080425:exon6:c.G2362A:p.E788K,GNAS:NM_001309840:exon6:c.G256A:p.E86K,GNAS:NM_001077489:exon5:c.G388A:p.E130K,GNAS:NM_001309861:exon6:c.G256A:p.E86K,GNAS:NM_000516:exon6:c.G433A:p.E145K,GNAS:NM_001077488:exon6:c.G436A:p.E146K, |
| chr20 | 58905398 | G | A | . | exonic | GNAS | nonsynonymous SNV | GNAS:NM_080426:exon5:c.G406A:p.A136T,GNAS:NM_080425:exon6:c.G2377A:p.A793T,GNAS:NM_001309840:exon6:c.G271A:p.A91T,GNAS:NM_001077489:exon5:c.G403A:p.A135T,GNAS:NM_001309861:exon6:c.G271A:p.A91T,GNAS:NM_000516:exon6:c.G448A:p.A150T,GNAS:NM_001077488:exon6:c.G451A:p.A151T, |
| chr20 | 58905399 | C | T | COSV55688594 | exonic | GNAS | nonsynonymous SNV | GNAS:NM_001309840:exon6:c.C272T:p.A91V,GNAS:NM_000516:exon6:c.C449T:p.A150V,GNAS:NM_080426:exon5:c.C407T:p.A136V,GNAS:NM_080425:exon6:c.C2378T:p.A793V,GNAS:NM_001077489:exon5:c.C404T:p.A135V,GNAS:NM_001309861:exon6:c.C272T:p.A91V,GNAS:NM_001077488:exon6:c.C452T:p.A151V, |
| chr20 | 58905411 | G | A | . | exonic | GNAS | stopgain | GNAS:NM_080426:exon5:c.G419A:p.W140X,GNAS:NM_080425:exon6:c.G2390A:p.W797X,GNAS:NM_001309840:exon6:c.G284A:p.W95X,GNAS:NM_001077489:exon5:c.G416A:p.W139X,GNAS:NM_001309861:exon6:c.G284A:p.W95X,GNAS:NM_000516:exon6:c.G461A:p.W154X,GNAS:NM_001077488:exon6:c.G464A:p.W155X, |
| chr20 | 58905425 | G | A | . | exonic | GNAS | nonsynonymous SNV | GNAS:NM_080426:exon5:c.G433A:p.V145M,GNAS:NM_080425:exon6:c.G2404A:p.V802M,GNAS:NM_001309840:exon6:c.G298A:p.V100M,GNAS:NM_001077489:exon5:c.G430A:p.V144M,GNAS:NM_001309861:exon6:c.G298A:p.V100M,GNAS:NM_000516:exon6:c.G475A:p.V159M,GNAS:NM_001077488:exon6:c.G478A:p.V160M, |

| | | | | | | | | |
|---|---|---|---|---|---|---|---|---|
| chr20 | 58905455 | T | C | . | exonic | GNAS | nonsynonymous SNV | GNAS:NM_080426:exon5:c.T463C:p.Y155H,GNAS:NM_080425:exon6:c.T2434C:p.Y812H,GNAS:NM_001309840:exon6:c.T328C:p.Y110H,GNAS:NM_001077489:exon5:c.T460C:p.Y154H,GNAS:NM_001309861:exon6:c.T328C:p.Y110H,GNAS:NM_000516:exon6:c.T505C:p.Y169H,GNAS:NM_001077488:exon6:c.T508C:p.Y170H, |
| chr20 | 58909211 | G | A | . | exonic | GNAS | nonsynonymous SNV | GNAS:NM_080426:exon6:c.G538A:p.D180N,GNAS:NM_080425:exon7:c.G2509A:p.D837N,GNAS:NM_001309840:exon7:c.G403A:p.D135N,GNAS:NM_001077489:exon6:c.G535A:p.D179N,GNAS:NM_001309861:exon7:c.G403A:p.D135N,GNAS:NM_000516:exon7:c.G580A:p.D194N,GNAS:NM_001077488:exon7:c.G583A:p.D195N, |
| chr20 | 58909380 | G | A | . | exonic | GNAS | nonsynonymous SNV | GNAS:NM_080426:exon7:c.G574A:p.G192R,GNAS:NM_080425:exon8:c.G2545A:p.G849R,GNAS:NM_001309840:exon8:c.G439A:p.G147R,GNAS:NM_001077489:exon7:c.G571A:p.G191R,GNAS:NM_001309861:exon8:c.G439A:p.G147R,GNAS:NM_000516:exon8:c.G616A:p.G206R,GNAS:NM_001077488:exon8:c.G619A:p.G207R, |
| chr20 | 58909398 | T | C | COSV55672955 | exonic | GNAS | nonsynonymous SNV | GNAS:NM_001309840:exon8:c.T457C:p.F153L,GNAS:NM_000516:exon8:c.T634C:p.F212L,GNAS:NM_080426:exon7:c.T592C:p.F198L,GNAS:NM_080425:exon8:c.T2563C:p.F855L,GNAS:NM_001077489:exon7:c.T589C:p.F197L,GNAS:NM_001309861:exon8:c.T457C:p.F153L,GNAS:NM_001077488:exon8:c.T637C:p.F213L, |
| chr20 | 58909410 | A | C | . | exonic | GNAS | nonsynonymous SNV | GNAS:NM_080426:exon7:c.A604C:p.K202Q,GNAS:NM_080425:exon8:c.A2575C:p.K859Q,GNAS:NM_001309840:exon8:c.A469C:p.K157Q,GNAS:NM_001077489:exon7:c.A601C:p.K201Q,GNAS:NM_001309861:exon8:c.A469C:p.K157Q,GNAS:NM_000516:exon8:c.A646C:p.K216Q,GNAS:NM_001077488:exon8:c.A649C:p.K217Q, |
| chr20 | 58909417 | A | G | . | exonic | GNAS | nonsynonymous SNV | GNAS:NM_080426:exon7:c.A611G:p.N204S,GNAS:NM_080425:exon8:c.A2582G:p.N861S,GNAS:NM_001309840:exon8:c.A476G:p.N159S,GNAS:NM_001077489:exon7:c.A608G:p.N203S,GNAS:NM_001309861:exon8:c.A476G:p.N159S,GNAS:NM_000516:exon8:c.A653G:p.N218S,GNAS:NM_001077488:exon8:c.A656G:p.N219S, |
| chr20 | 58909556 | G | A | COSV55674033 | exonic | GNAS | nonsynonymous SNV | GNAS:NM_080426:exon8:c.G653A:p.R218H,GNAS:NM_080425:exon9:c.G2624A:p.R875H,GNAS:NM_001309840:exon9:c.G518A:p.R173H,GNAS:NM_001077489:exon8:c.G650A:p.R217H,GNAS:NM_001309861:exon9:c.G518A:p.R173H,GNAS:NM_000516:exon9:c.G695A:p.R232H,GNAS:NM_001077488:exon9:c.G698A:p.R233H, |
| chr20 | 58909686 | G | A | . | exonic | GNAS | nonsynonymous SNV | GNAS:NM_080426:exon9:c.G679A:p.V227M,GNAS:NM_080425:exon10:c.G2650A:p.V884M,GNAS:NM_001309840:exon10:c.G544A:p.V182M,GNAS:NM_001077489:exon9:c.G676A:p.V226M,GNAS:NM_001309861:exon10:c.G544A:p.V182M,GNAS:NM_000516:exon10:c.G721A:p.V241M,GNAS:NM_001077488:exon10:c.G724A:p.V242M, |
| chr20 | 58909717 | G | A | . | exonic | GNAS | nonsynonymous SNV | GNAS:NM_001309840:exon10:c.G575A:p.S192N,GNAS:NM_000516:exon10:c.G752A:p.S251N,GNAS:NM_080426:exon9:c.G710A:p.S237N,GNAS:NM_080425:exon10:c.G2681A:p.S894N,GNAS:NM_001077489:exon9:c.G707A:p.S236N,GNAS:NM_001309861:exon10:c.G575A:p.S192N,GNAS:NM_001077488:exon10:c.G755A:p.S252N, |
| chr20 | 58909721 | C | A | . | exonic | GNAS | nonsynonymous SNV | GNAS:NM_080426:exon9:c.C714A:p.S238R,GNAS:NM_080425:exon10:c.C2685A:p.S895R,GNAS:NM_001309840:exon10:c.C579A:p.S193R,GNAS:NM_001077489:exon9:c.C711A:p.S237R,GNAS:NM_001309861:exon10:c.C579A:p.S193R,GNAS:NM_000516:exon10:c.C756A:p.S252R,GNAS:NM_001077488:exon10:c.C759A:p.S253R, |
| chr20 | 58909753 | C | A | . | exonic | GNAS | nonsynonymous SNV | GNAS:NM_080426:exon9:c.C746A:p.T249N,GNAS:NM_080425:exon10:c.C2717A:p.T906N,GNAS:NM_001309840:exon10:c.C611A:p.T204N,GNAS:NM_001077489:exon9:c.C743A:p.T248N,GNAS:NM_001309861:exon10:c.C611A:p.T204N,GNAS:NM_000516:exon10:c.C788A:p.T263N,GNAS:NM_001077488:exon10:c.C791A:p.T264N, |
| chr20 | 58909759 | G | A | COSV55672773 | exonic | GNAS | nonsynonymous SNV | GNAS:NM_080426:exon9:c.G752A:p.R251H,GNAS:NM_080425:exon10:c.G2723A:p.R908H,GNAS:NM_001309840:exon10:c.G617A:p.R206H,GNAS:NM_001077489:exon9:c.G749A:p.R250H,GNAS:NM_001309861:exon10:c.G617A:p.R206H,GNAS:NM_000516:exon10:c.G794A:p.R265H,GNAS:NM_001077488:exon10:c.G797A:p.R266H, |
| chr20 | 58909795 | G | A | . | exonic | GNAS | stopgain | GNAS:NM_080426:exon9:c.G788A:p.W263X,GNAS:NM_080425:exon10:c.G2759A:p.W920X,GNAS:NM_001309840:exon10:c.G653A:p.W218X,GNAS:NM_001077489:exon9:c.G785A:p.W262X,GNAS:NM_001309861:exon10:c.G653A:p.W218X,GNAS:NM_000516:exon10:c.G830A:p.W277X,GNAS:NM_001077488:exon10:c.G833A:p.W278X, |

| | | | | | | | | |
|---|---|---|---|---|---|---|---|---|
| chr20 | 58909952 | T | C | . | exonic | GNAS | nonsynonymous SNV | GNAS:NM_001309840:exon11:c.T664C:p.W222R,GNAS:NM_000516:exon11:c.T841C:p.W281R,GNAS:NM_080426:exon10:c.T799C:p.W267R,GNAS:NM_080425:exon11:c.T2770C:p.W924R,GNAS:NM_001077489:exon10:c.T796C:p.W266R,GNAS:NM_001309861:exon11:c.T664C:p.W222R,GNAS:NM_001077488:exon11:c.T844C:p.W282R, |
| chr20 | 58910021 | G | A | . | exonic | GNAS | nonsynonymous SNV | GNAS:NM_080426:exon10:c.G868A:p.G290R,GNAS:NM_080425:exon11:c.G2839A:p.G947R,GNAS:NM_001309840:exon11:c.G733A:p.G245R,GNAS:NM_001077489:exon10:c.G865A:p.G289R,GNAS:NM_001309861:exon11:c.G733A:p.G245R,GNAS:NM_000516:exon11:c.G910A:p.G304R,GNAS:NM_001077488:exon11:c.G913A:p.G305R, |
| chr20 | 58910022 | G | A | . | exonic | GNAS | nonsynonymous SNV | GNAS:NM_080426:exon10:c.G869A:p.G290E,GNAS:NM_080425:exon11:c.G2840A:p.G947E,GNAS:NM_001309840:exon11:c.G734A:p.G245E,GNAS:NM_001077489:exon10:c.G866A:p.G289E,GNAS:NM_001309861:exon11:c.G734A:p.G245E,GNAS:NM_000516:exon11:c.G911A:p.G304E,GNAS:NM_001077488:exon11:c.G914A:p.G305E, |
| chr20 | 58910039 | G | A | . | exonic | GNAS | nonsynonymous SNV | GNAS:NM_080426:exon10:c.G886A:p.D296N,GNAS:NM_080425:exon11:c.G2857A:p.D953N,GNAS:NM_001309840:exon11:c.G751A:p.D251N,GNAS:NM_001077489:exon10:c.G883A:p.D295N,GNAS:NM_001309861:exon11:c.G751A:p.D251N,GNAS:NM_000516:exon11:c.G928A:p.D310N,GNAS:NM_001077488:exon11:c.G931A:p.D311N, |
| chr20 | 58910045 | T | C | . | exonic | GNAS | nonsynonymous SNV | GNAS:NM_080426:exon10:c.T892C:p.F298L,GNAS:NM_080425:exon11:c.T2863C:p.F955L,GNAS:NM_001309840:exon11:c.T757C:p.F253L,GNAS:NM_001077489:exon10:c.T889C:p.F297L,GNAS:NM_001309861:exon11:c.T757C:p.F253L,GNAS:NM_000516:exon11:c.T934C:p.F312L,GNAS:NM_001077488:exon11:c.T937C:p.F313L, |
| chr20 | 58910061 | G | A | COSV55679114 | exonic | GNAS | nonsynonymous SNV | GNAS:NM_080426:exon10:c.G908A:p.R303H,GNAS:NM_080425:exon11:c.G2879A:p.R960H,GNAS:NM_001309840:exon11:c.G773A:p.R258H,GNAS:NM_001077489:exon10:c.G905A:p.R302H,GNAS:NM_001309861:exon11:c.G773A:p.R258H,GNAS:NM_000516:exon11:c.G950A:p.R317H,GNAS:NM_001077488:exon11:c.G953A:p.R318H, |
| chr20 | 58910348 | G | A | COSV55690777 | exonic | GNAS | nonsynonymous SNV | GNAS:NM_080426:exon11:c.G943A:p.G315R,GNAS:NM_080425:exon12:c.G2914A:p.G972R,GNAS:NM_001309840:exon12:c.G808A:p.G270R,GNAS:NM_001077489:exon11:c.G940A:p.G314R,GNAS:NM_001309861:exon12:c.G808A:p.G270R,GNAS:NM_000516:exon12:c.G985A:p.G329R,GNAS:NM_001077488:exon12:c.G988A:p.G330R, |
| chr20 | 58910364 | T | C | . | exonic | GNAS | nonsynonymous SNV | GNAS:NM_080426:exon11:c.T959C:p.V320A,GNAS:NM_080425:exon12:c.T2930C:p.V977A,GNAS:NM_001309840:exon12:c.T824C:p.V275A,GNAS:NM_001077489:exon11:c.T956C:p.V319A,GNAS:NM_001309861:exon12:c.T824C:p.V275A,GNAS:NM_000516:exon12:c.T1001C:p.V334A,GNAS:NM_001077488:exon12:c.T1004C:p.V335A, |
| chr20 | 58910373 | C | A | . | exonic | GNAS | nonsynonymous SNV | GNAS:NM_080426:exon11:c.C968A:p.A323D,GNAS:NM_080425:exon12:c.C2939A:p.A980D,GNAS:NM_001309840:exon12:c.C833A:p.A278D,GNAS:NM_001077489:exon11:c.C965A:p.A322D,GNAS:NM_001309861:exon12:c.C833A:p.A278D,GNAS:NM_000516:exon12:c.C1010A:p.A337D,GNAS:NM_001077488:exon12:c.C1013A:p.A338D, |
| chr20 | 58910385 | T | C | . | exonic | GNAS | nonsynonymous SNV | GNAS:NM_001309840:exon12:c.T845C:p.I282T,GNAS:NM_000516:exon12:c.T1022C:p.I341T,GNAS:NM_080426:exon11:c.T980C:p.I327T,GNAS:NM_080425:exon12:c.T2951C:p.I984T,GNAS:NM_001077489:exon11:c.T977C:p.I326T,GNAS:NM_001309861:exon12:c.T845C:p.I282T,GNAS:NM_001077488:exon12:c.T1025C:p.I342T, |
| chr20 | 58910684 | G | A | . | exonic | GNAS | nonsynonymous SNV | GNAS:NM_080426:exon12:c.G998A:p.R333K,GNAS:NM_080425:exon13:c.G2969A:p.R990K,GNAS:NM_001309840:exon13:c.G863A:p.R288K,GNAS:NM_001077489:exon12:c.G995A:p.R332K,GNAS:NM_001309861:exon13:c.G863A:p.R288K,GNAS:NM_000516:exon13:c.G1040A:p.R347K,GNAS:NM_001077488:exon13:c.G1043A:p.R348K, |
| chr20 | 58910699 | G | A | . | exonic | GNAS | nonsynonymous SNV | GNAS:NM_080426:exon12:c.G1013A:p.S338N,GNAS:NM_080425:exon13:c.G2984A:p.S995N,GNAS:NM_001309840:exon13:c.G878A:p.S293N,GNAS:NM_001077489:exon12:c.G1010A:p.S337N,GNAS:NM_001309861:exon13:c.G878A:p.S293N,GNAS:NM_000516:exon13:c.G1055A:p.S352N,GNAS:NM_001077488:exon13:c.G1058A:p.S353N, |
| chr20 | 58910704 | G | A | COSV55681349 | exonic | GNAS | nonsynonymous SNV | GNAS:NM_080426:exon12:c.G1018A:p.D340N,GNAS:NM_080425:exon13:c.G2989A:p.D997N,GNAS:NM_001309840:exon13:c.G883A:p.D295N,GNAS:NM_001077489:exon12:c.G1015A:p.D339N,GNAS:NM_001309861:exon13:c.G883A:p.D295N,GNAS:NM_000516:exon13:c.G1060A:p.D354N,GNAS:NM_001077488:exon13:c.G1063A:p.D355N, |

| | | | | | | | | |
|---|---|---|---|---|---|---|---|---|
| chr20 | 58910738 | G | A | . | exonic | GNAS | nonsynonymous SNV | GNAS:NM_080426:exon12:c.G1052A:p.C351Y,GNAS:NM_080425:exon13:c.G3023A:p.C1008Y,GNAS:NM_001309840:exon13:c.G917A:p.C306Y,GNAS:NM_001077489:exon12:c.G1049A:p.C350Y,GNAS:NM_001309861:exon13:c.G917A:p.C306Y,GNAS:NM_000516:exon13:c.G1094A:p.C365Y,GNAS:NM_001077488:exon13:c.G1097A:p.C366Y, |
| chr20 | 58910746 | G | A | . | exonic | GNAS | nonsynonymous SNV | GNAS:NM_080426:exon12:c.G1060A:p.D354N,GNAS:NM_080425:exon13:c.G3031A:p.D1011N,GNAS:NM_001309840:exon13:c.G925A:p.D309N,GNAS:NM_001077489:exon12:c.G1057A:p.D353N,GNAS:NM_001309861:exon13:c.G925A:p.D309N,GNAS:NM_000516:exon13:c.G1102A:p.D368N,GNAS:NM_001077488:exon13:c.G1105A:p.D369N, |
| chr20 | 58910762 | G | A | . | exonic | GNAS | nonsynonymous SNV | GNAS:NM_080426:exon12:c.G1076A:p.R359H,GNAS:NM_080425:exon13:c.G3047A:p.R1016H,GNAS:NM_001309840:exon13:c.G941A:p.R314H,GNAS:NM_001077489:exon12:c.G1073A:p.R358H,GNAS:NM_001309861:exon13:c.G941A:p.R314H,GNAS:NM_000516:exon13:c.G1118A:p.R373H,GNAS:NM_001077488:exon13:c.G1121A:p.R374H, |
| chr20 | 58910809 | C | A | . | exonic | GNAS | nonsynonymous SNV | GNAS:NM_001309840:exon13:c.C988A:p.R330S,GNAS:NM_000516:exon13:c.C1165A:p.R389S,GNAS:NM_080426:exon12:c.C1123A:p.R375S,GNAS:NM_080425:exon13:c.C3094A:p.R1032S,GNAS:NM_001077489:exon12:c.C1120A:p.R374S,GNAS:NM_001309861:exon13:c.C988A:p.R330S,GNAS:NM_001077488:exon13:c.C1168A:p.R390S, |
| chr21 | 33343393 | G | C | COSV54251383 | exonic | IFNAR1 | nonsynonymous SNV | IFNAR1:NM_000629:exon4:c.G502C:p.V168L, |
| chr21 | 33543109 | A | T | . | exonic | SON | nonsynonymous SNV | SON:NM_001291411:exon1:c.A17T:p.E6V,SON:NM_001291412:exon1:c.A17T:p.E6V,SON:NM_032195:exon1:c.A17T:p.E6V,SON:NM_138927:exon1:c.A17T:p.E6V, |
| chr21 | 33543145 | G | A | . | exonic | SON | nonsynonymous SNV | SON:NM_001291412:exon1:c.G53A:p.R18Q,SON:NM_138927:exon1:c.G53A:p.R18Q,SON:NM_001291411:exon1:c.G53A:p.R18Q,SON:NM_032195:exon1:c.G53A:p.R18Q, |
| chr21 | 33543159 | G | A | . | exonic | SON | nonsynonymous SNV | SON:NM_001291412:exon1:c.G67A:p.E23K,SON:NM_138927:exon1:c.G67A:p.E23K,SON:NM_001291411:exon1:c.G67A:p.E23K,SON:NM_032195:exon1:c.G67A:p.E23K, |
| chr21 | 33549563 | A | G | . | exonic | SON | nonsynonymous SNV | SON:NM_001291411:exon3:c.A332G:p.H111R,SON:NM_032195:exon3:c.A332G:p.H111R,SON:NM_138927:exon3:c.A332G:p.H111R, |
| chr21 | 33549579 | C | A | . | exonic | SON | nonsynonymous SNV | SON:NM_001291411:exon3:c.C348A:p.N116K,SON:NM_032195:exon3:c.C348A:p.N116K,SON:NM_138927:exon3:c.C348A:p.N116K, |
| chr21 | 33549584 | A | G | . | exonic | SON | nonsynonymous SNV | SON:NM_001291411:exon3:c.A353G:p.K118R,SON:NM_032195:exon3:c.A353G:p.K118R,SON:NM_138927:exon3:c.A353G:p.K118R, |
| chr21 | 33549616 | T | A | COSV51654122 | exonic | SON | nonsynonymous SNV | SON:NM_001291411:exon3:c.T385A:p.Y129N,SON:NM_032195:exon3:c.T385A:p.Y129N,SON:NM_138927:exon3:c.T385A:p.Y129N, |
| chr21 | 33549618 | T | A | . | exonic | SON | stopgain | SON:NM_001291411:exon3:c.T387A:p.Y129X,SON:NM_032195:exon3:c.T387A:p.Y129X,SON:NM_138927:exon3:c.T387A:p.Y129X, |
| chr21 | 33553954 | C | T | COSV51652492 | exonic | SON | nonsynonymous SNV | SON:NM_001291411:exon3:c.C4723T:p.R1575C,SON:NM_032195:exon3:c.C4723T:p.R1575C,SON:NM_138927:exon3:c.C4723T:p.R1575C, |
| chr21 | 41480570 | C | T | COSV59820912 | exonic | TMPRSS2 | nonsynonymous SNV | TMPRSS2:NM_001135099:exon6:c.G589A:p.V197M,TMPRSS2:NM_005656:exon6:c.G478A:p.V160M, |
| chr21 | 41507982 | C | A | COSV59827803 | exonic | TMPRSS2 | nonsynonymous SNV | TMPRSS2:NM_001135099:exon1:c.G23T:p.G8V, |
| chr22 | 19963748 | G | A | COSV52889256 | exonic | COMT | nonsynonymous SNV | COMT:NM_001135162:exon4:c.G472A:p.V158M,COMT:NM_000754:exon4:c.G472A:p.V158M,COMT:NM_007310:exon2:c.G322A:p.V108M,COMT:NM_001135161:exon4:c.G472A:p.V158M, |
| chr22 | 21000681 | C | G | COSV53145905 | exonic | THAP7 | nonsynonymous SNV | THAP7:NM_030573:exon3:c.G343C:p.A115P,THAP7:NM_001008695:exon4:c.G343C:p.A115P, |
| chr22 | 22895680 | A | G | COSV66535329 | exonic | IGLL5 | nonsynonymous SNV | IGLL5:NM_001178126:exon3:c.A631G:p.T211A,IGLL5:NM_001256296:exon2:c.A406G:p.T136A, |
| chr22 | 23285182 | A | G | COSV59932309 | exonic | BCR | nonsynonymous SNV | BCR:NM_021574:exon10:c.A2387G:p.N796S,BCR:NM_004327:exon10:c.A2387G:p.N796S, |
| chr22 | 36191656 | T | C | COSV60654226 | exonic | APOL4 | nonsynonymous SNV | APOL4:NM_030643:exon6:c.A466G:p.M156V,APOL4:NM_145660:exon5:c.A475G:p.M159V, |
| chr22 | 36265284 | G | A | COSV59868177 | exonic | APOL1 | nonsynonymous SNV | APOL1:NM_001136541:exon5:c.G394A:p.E132K,APOL1:NM_145343:exon7:c.G496A:p.E166K,APOL1:NM_003661:exon6:c.G448A:p.E150K,APOL1:NM_001136540:exon6:c.G448A:p.E150K, |
| chr22 | 36265520 | G | A | COSV59868421 | exonic | APOL1 | nonsynonymous SNV | APOL1:NM_001136541:exon5:c.G630A:p.M210I,APOL1:NM_145343:exon7:c.G732A:p.M244I,APOL1:NM_003661:exon6:c.G684A:p.M228I,APOL1:NM_001136540:exon6:c.G684A:p.M228I, |
| chr22 | 36265600 | G | A | COSV59868427 | exonic | APOL1 | nonsynonymous SNV | APOL1:NM_001136541:exon5:c.G710A:p.R237K,APOL1:NM_145343:exon7:c.G812A:p.R271K,APOL1:NM_003661:exon6:c.G764A:p.R255K,APOL1:NM_001136540:exon6:c.G764A:p.R255K, |
| chr22 | 36467860 | C | T | . | exonic | TXN2 | nonsynonymous SNV | TXN2:NM_012473:exon4:c.G445A:p.V149M, |
| chr22 | 36467871 | A | G | . | exonic | TXN2 | nonsynonymous SNV | TXN2:NM_012473:exon4:c.T434C:p.V145A, |
| chr22 | 36467911 | C | T | . | exonic | TXN2 | nonsynonymous SNV | TXN2:NM_012473:exon4:c.G394A:p.A132T, |
| chr22 | 36476789 | C | T | . | exonic | TXN2 | nonsynonymous SNV | TXN2:NM_012473:exon3:c.G331A:p.V111M, |
| chr22 | 36476848 | C | T | COSV53398335 | exonic | TXN2 | nonsynonymous SNV | TXN2:NM_012473:exon3:c.G272A:p.G91E, |
| chr22 | 40156115 | G | A | COSV57292828 | exonic | TNRC6B | nonsynonymous SNV | TNRC6B:NM_001024843:exon4:c.G46A:p.V16M, |
| chr22 | 41532697 | A | G | . | exonic | POLR3H | nonsynonymous SNV | POLR3H:NM_001282884:exon4:c.T257C:p.I86T,POLR3H:NM_001282885:exon4:c.T257C:p.I86T,POLR3H:NM_138338:exon4:c.T257C:p.I86T,POLR3H:NM_001018050:exon3:c.T257C:p.I86T, |
| chr22 | 43162920 | A | G | COSV53355454 | exonic | TSPO | nonsynonymous SNV | TSPO:NM_000714:exon4:c.A439G:p.T147A,TSPO:NM_001256530:exon4:c.A439G:p.T147A,TSPO:NM_001256531:exon4:c.A439G:p.T147A, |

| | | | | | | | | |
|---|---|---|---|---|---|---|---|---|
| chr22 | 43162966 | G | A | COSV53354339 | exonic | TSPO | nonsynonymous SNV | TSPO:NM_000714:exon4:c.G485A:p.R162H,TSPO:NM_001256530:exon4:c.G485A:p.R162H,TSPO:NM_001256531:exon4:c.G485A:p.R162H, |
| chr22 | 45527947 | A | G | . | exonic | FBLN1 | nonsynonymous SNV | FBLN1:NM_001996:exon4:c.A422G:p.Q141R,FBLN1:NM_006485:exon4:c.A422G:p.Q141R,FBLN1:NM_006486:exon4:c.A422G:p.Q141R,FBLN1:NM_006487:exon4:c.A422G:p.Q141R, |
| chr3 | 10064848 | C | T | COSV55033303 | exonic | FANCD2 | nonsynonymous SNV | FANCD2:NM_033084:exon23:c.C2141T:p.P714L,FANCD2:NM_001018115:exon23:c.C2141T:p.P714L, |
| chr3 | 10072973 | G | A | COSV55034331 | exonic | FANCD2 | nonsynonymous SNV | FANCD2:NM_033084:exon27:c.G2597A:p.G866E,FANCD2:NM_001018115:exon27:c.G2597A:p.G866E, |
| chr3 | 100732618 | G | A | . | exonic | TFG | nonsynonymous SNV | TFG:NM_006070:exon5:c.G526A:p.E176K,TFG:NM_001195479:exon5:c.G526A:p.E176K,TFG:NM_001195478:exon5:c.G526A:p.E176K,TFG:NM_001007565:exon5:c.G526A:p.E176K, |
| chr3 | 100744893 | C | T | . | exonic | TFG | nonsynonymous SNV | TFG:NM_006070:exon7:c.C782T:p.P261L,TFG:NM_001195479:exon7:c.C770T:p.P257L,TFG:NM_001195478:exon7:c.C782T:p.P261L,TFG:NM_001007565:exon7:c.C782T:p.P261L, |
| chr3 | 100748190 | T | C | . | exonic | TFG | nonsynonymous SNV | TFG:NM_001195479:exon8:c.T850C:p.F284L,TFG:NM_001195478:exon8:c.T862C:p.F288L,TFG:NM_001007565:exon8:c.T862C:p.F288L,TFG:NM_006070:exon8:c.T862C:p.F288L, |
| chr3 | 10312082 | C | T | . | exonic | SEC13 | stopgain | SEC13:NM_183352:exon5:c.G333A:p.W111X,SEC13:NM_030673:exon6:c.G342A:p.W114X,SEC13:NM_001136232:exon5:c.G291A:p.W97X,SEC13:NM_001278946:exon5:c.G333A:p.W111X,SEC13:NM_001136026:exon6:c.G471A:p.W157X, |
| chr3 | 10312608 | C | T | . | exonic | SEC13 | nonsynonymous SNV | SEC13:NM_030673:exon5:c.G296A:p.S99N,SEC13:NM_001278946:exon4:c.G287A:p.S96N,SEC13:NM_001136026:exon5:c.G425A:p.S142N,SEC13:NM_183352:exon4:c.G287A:p.S96N,SEC13:NM_001136232:exon4:c.G245A:p.S82N, |
| chr3 | 107772704 | T | A | . | exonic | BBX | nonsynonymous SNV | BBX:NM_020235:exon11:c.T983A:p.I328N,BBX:NM_001142568:exon11:c.T983A:p.I328N, |
| chr3 | 107772736 | G | A | . | exonic | BBX | nonsynonymous SNV | BBX:NM_020235:exon11:c.G1015A:p.E339K,BBX:NM_001142568:exon11:c.G1015A:p.E339K, |
| chr3 | 107773061 | C | A | . | exonic | BBX | nonsynonymous SNV | BBX:NM_020235:exon11:c.C1340A:p.P447Q,BBX:NM_001142568:exon11:c.C1340A:p.P447Q, |
| chr3 | 107773304 | A | G | . | exonic | BBX | nonsynonymous SNV | BBX:NM_020235:exon11:c.A1583G:p.K528R,BBX:NM_001142568:exon11:c.A1583G:p.K528R, |
| chr3 | 107773313 | T | A | . | exonic | BBX | nonsynonymous SNV | BBX:NM_020235:exon11:c.T1592A:p.V531E,BBX:NM_001142568:exon11:c.T1592A:p.V531E, |
| chr3 | 108353451 | T | C | COSV63315693 | exonic | HHLA2 | nonsynonymous SNV | HHLA2:NM_001282556:exon4:c.T89C:p.I30T,HHLA2:NM_001282557:exon4:c.T89C:p.I30T,HHLA2:NM_001282558:exon5:c.T89C:p.I30T,HHLA2:NM_007072:exon4:c.T89C:p.I30T, |
| chr3 | 108362430 | C | A | . | exonic | HHLA2 | nonsynonymous SNV | HHLA2:NM_001282556:exon7:c.C1092A:p.S364R,HHLA2:NM_001282559:exon8:c.C900A:p.S300R,HHLA2:NM_001282557:exon7:c.C1092A:p.S364R,HHLA2:NM_001282558:exon8:c.C1092A:p.S364R,HHLA2:NM_007072:exon7:c.C1092A:p.S364R, |
| chr3 | 120670469 | T | A | COSV52199452 | exonic | HGD | nonsynonymous SNV | HGD:NM_000187:exon4:c.A240T:p.Q80H, |
| chr3 | 122106350 | G | A | COSV52608918 | exonic | CD86 | nonsynonymous SNV | CD86:NM_176892:exon4:c.G535A:p.V179I,CD86:NM_175862:exon4:c.G553A:p.V185I,CD86:NM_006889:exon4:c.G535A:p.V179I,CD86:NM_001206924:exon3:c.G217A:p.V73I,CD86:NM_001206925:exon3:c.G307A:p.V103I, |
| chr3 | 12351626 | C | G | COSV55140710 | exonic | PPARG | nonsynonymous SNV | PPARG:NM_015869:exon1:c.C34G:p.P12A, |
| chr3 | 12379892 | G | A | . | exonic | PPARG | nonsynonymous SNV | PPARG:NM_005037:exon2:c.G187A:p.A63T,PPARG:NM_138712:exon3:c.G187A:p.A63T,PPARG:NM_015869:exon2:c.G271A:p.A91T,PPARG:NM_138711:exon3:c.G187A:p.A63T, |
| chr3 | 12392691 | T | A | . | exonic | PPARG | nonsynonymous SNV | PPARG:NM_005037:exon4:c.T474A:p.S158R,PPARG:NM_138712:exon5:c.T474A:p.S158R,PPARG:NM_015869:exon4:c.T558A:p.S186R,PPARG:NM_138711:exon5:c.T474A:p.S158R, |
| chr3 | 12392697 | T | A | . | exonic | PPARG | nonsynonymous SNV | PPARG:NM_005037:exon4:c.T480A:p.N160K,PPARG:NM_138712:exon5:c.T480A:p.N160K,PPARG:NM_015869:exon4:c.T564A:p.N188K,PPARG:NM_138711:exon5:c.T480A:p.N160K, |
| chr3 | 12433907 | G | T | . | exonic | PPARG | nonsynonymous SNV | PPARG:NM_005037:exon7:c.G1196T:p.G399V,PPARG:NM_138712:exon8:c.G1196T:p.G399V,PPARG:NM_015869:exon7:c.G1280T:p.G427V,PPARG:NM_138711:exon8:c.G1196T:p.G399V, |
| chr3 | 126107216 | T | C | COSV56412084 | exonic | ALDH1L1 | nonsynonymous SNV | ALDH1L1:NM_001270365:exon19:c.A2075G:p.D692G,ALDH1L1:NM_012190:exon21:c.A2378G:p.D793G,ALDH1L1:NM_001270364:exon21:c.A2408G:p.D803G, |
| chr3 | 129170475 | C | T | . | exonic | CNBP | nonsynonymous SNV | CNBP:NM_001127195:exon5:c.G494A:p.C165Y,CNBP:NM_001127193:exon5:c.G515A:p.C172Y,CNBP:NM_001127196:exon5:c.G491A:p.C164Y,CNBP:NM_001127192:exon5:c.G518A:p.C173Y,CNBP:NM_003418:exon5:c.G512A:p.C171Y,CNBP:NM_001127194:exon5:c.G497A:p.C166Y, |
| chr3 | 129170479 | C | T | . | exonic | CNBP | nonsynonymous SNV | CNBP:NM_001127195:exon5:c.G490A:p.E164K,CNBP:NM_001127193:exon5:c.G511A:p.E171K,CNBP:NM_001127196:exon5:c.G487A:p.E163K,CNBP:NM_001127192:exon5:c.G514A:p.E172K,CNBP:NM_003418:exon5:c.G508A:p.E170K,CNBP:NM_001127194:exon5:c.G493A:p.E165K, |

| | | | | | | | | |
|---|---|---|---|---|---|---|---|---|
| chr3 | 129170485 | C | T | . | exonic | CNBP | nonsynonymous SNV | CNBP:NM_001127195:exon5:c.G484A:p.A162T,CNBP:NM_001127193:exon5:c.G505A:p.A169T,CNBP:NM_001127196:exon5:c.G481A:p.A161T,CNBP:NM_001127192:exon5:c.G508A:p.A170T,CNBP:NM_003418:exon5:c.G502A:p.A168T,CNBP:NM_001127194:exon5:c.G487A:p.A163T, |
| chr3 | 129170535 | C | T | . | exonic | CNBP | nonsynonymous SNV | CNBP:NM_001127195:exon5:c.G434A:p.S145N,CNBP:NM_001127193:exon5:c.G455A:p.S152N,CNBP:NM_001127196:exon5:c.G431A:p.S144N,CNBP:NM_001127192:exon5:c.G458A:p.S153N,CNBP:NM_003418:exon5:c.G452A:p.S151N,CNBP:NM_001127194:exon5:c.G437A:p.S146N, |
| chr3 | 129171245 | C | T | . | exonic | CNBP | nonsynonymous SNV | CNBP:NM_001127195:exon4:c.G232A:p.A78T,CNBP:NM_001127193:exon4:c.G253A:p.A85T,CNBP:NM_001127196:exon4:c.G229A:p.A77T,CNBP:NM_001127192:exon4:c.G256A:p.A86T,CNBP:NM_003418:exon4:c.G250A:p.A84T,CNBP:NM_001127194:exon4:c.G235A:p.A79T, |
| chr3 | 129171262 | C | T | . | exonic | CNBP | nonsynonymous SNV | CNBP:NM_001127195:exon4:c.G215A:p.G72D,CNBP:NM_001127193:exon4:c.G236A:p.G79D,CNBP:NM_001127196:exon4:c.G212A:p.G71D,CNBP:NM_001127192:exon4:c.G239A:p.G80D,CNBP:NM_003418:exon4:c.G233A:p.G78D,CNBP:NM_001127194:exon4:c.G218A:p.G73D, |
| chr3 | 129171265 | C | T | . | exonic | CNBP | nonsynonymous SNV | CNBP:NM_001127195:exon4:c.G212A:p.C71Y,CNBP:NM_001127193:exon4:c.G233A:p.C78Y,CNBP:NM_001127196:exon4:c.G209A:p.C70Y,CNBP:NM_001127192:exon4:c.G236A:p.C79Y,CNBP:NM_003418:exon4:c.G230A:p.C77Y,CNBP:NM_001127194:exon4:c.G215A:p.C72Y, |
| chr3 | 129171671 | C | T | . | exonic | CNBP | nonsynonymous SNV | CNBP:NM_001127195:exon2:c.G87A:p.M29I,CNBP:NM_001127193:exon2:c.G87A:p.M29I,CNBP:NM_001127196:exon2:c.G87A:p.M29I,CNBP:NM_001127192:exon2:c.G87A:p.M29I,CNBP:NM_003418:exon2:c.G87A:p.M29I,CNBP:NM_001127194:exon2:c.G87A:p.M29I, |
| chr3 | 129171694 | C | T | . | exonic | CNBP | nonsynonymous SNV | CNBP:NM_001127195:exon2:c.G64A:p.G22S,CNBP:NM_001127193:exon2:c.G64A:p.G22S,CNBP:NM_001127196:exon2:c.G64A:p.G22S,CNBP:NM_001127194:exon2:c.G64A:p.G22S,CNBP:NM_001127192:exon2:c.G64A:p.G22S,CNBP:NM_003418:exon2:c.G64A:p.G22S, |
| chr3 | 133623144 | T | C | COSV53439574 | exonic | TOPBP1 | nonsynonymous SNV | TOPBP1:NM_007027:exon19:c.A3125G:p.N1042S, |
| chr3 | 13629036 | A | G | COSV55488788 | exonic | FBLN2 | nonsynonymous SNV | FBLN2:NM_001165035:exon12:c.A2701G:p.T901A,FBLN2:NM_001998:exon11:c.A2560G:p.T854A,FBLN2:NM_001004019:exon12:c.A2701G:p.T901A, |
| chr3 | 136604433 | C | A | . | exonic | STAG1 | nonsynonymous SNV | STAG1:NM_005862:exon4:c.G173T:p.S58I, |
| chr3 | 136623187 | C | T | . | exonic | STAG1 | nonsynonymous SNV | STAG1:NM_005862:exon3:c.G91A:p.V31I, |
| chr3 | 136945895 | C | T | COSV56637558 | exonic | NCK1 | nonsynonymous SNV | NCK1:NM_001190796:exon2:c.C347T:p.A116V,NCK1:NM_006153:exon3:c.C539T:p.A180V,NCK1:NM_001291999:exon3:c.C539T:p.A180V, |
| chr3 | 138295865 | G | C | . | exonic | ARMC8 | nonsynonymous SNV | ARMC8:NM_015396:exon23:c.G1953C:p.K651N,ARMC8:NM_001282342:exon20:c.G1776C:p.K592N,ARMC8:NM_001267041:exon21:c.G1902C:p.K634N,ARMC8:NM_001267042:exon21:c.G1794C:p.K598N, |
| chr3 | 138570379 | C | T | . | exonic | CEP70 | nonsynonymous SNV | CEP70:NM_001288966:exon6:c.G404A:p.S135N,CEP70:NM_001288964:exon6:c.G350A:p.S117N,CEP70:NM_024491:exon6:c.G404A:p.S135N,CEP70:NM_001288965:exon5:c.G344A:p.S115N, |
| chr3 | 139369498 | A | T | . | exonic | COPB2 | nonsynonymous SNV | COPB2:NM_004766:exon11:c.T1252A:p.F418I, |
| chr3 | 141442720 | G | A | . | exonic | ZBTB38 | nonsynonymous SNV | ZBTB38:NM_001080412:exon8:c.G332A:p.G111E, |
| chr3 | 141442835 | T | A | . | exonic | ZBTB38 | nonsynonymous SNV | ZBTB38:NM_001080412:exon8:c.T447A:p.N149K, |
| chr3 | 141444232 | C | T | . | exonic | ZBTB38 | nonsynonymous SNV | ZBTB38:NM_001080412:exon8:c.C1844T:p.T615M, |
| chr3 | 14145949 | G | T | COSV53207797 | exonic | XPC | nonsynonymous SNV | XPC:NM_004628:exon16:c.C2815A:p.Q939K, |
| chr3 | 14158387 | G | A | COSV53203534 | exonic | XPC | nonsynonymous SNV | XPC:NM_004628:exon9:c.C1496T:p.A499V, |
| chr3 | 14158408 | C | T | COSV53203815 | exonic | XPC | nonsynonymous SNV | XPC:NM_004628:exon9:c.G1475A:p.R492H, |
| chr3 | 14158440 | C | A | COSV53206797 | exonic | XPC | nonsynonymous SNV | XPC:NM_004628:exon9:c.G1443T:p.K481N, |
| chr3 | 14170484 | G | T | . | exonic | XPC | nonsynonymous SNV | XPC:NM_004628:exon3:c.C366A:p.S122R, |
| chr3 | 141974094 | C | T | . | exonic | TFDP2 | stopgain | TFDP2:NM_001178140:exon5:c.G533A:p.W178X,TFDP2:NM_001178139:exon8:c.G617A:p.W206X,TFDP2:NM_006286:exon6:c.G437A:p.W146X,TFDP2:NM_001178138:exon9:c.G437A:p.W146X,TFDP2:NM_001178142:exon3:c.G209A:p.W70X,TFDP2:NM_001178141:exon5:c.G326A:p.W109X, |
| chr3 | 142562770 | A | G | COSV63383325 | exonic | ATR | nonsynonymous SNV | ATR:NM_001184:exon4:c.T632C:p.M211T, |
| chr3 | 158602914 | C | A | COSV63035321 | exonic | MLF1 | nonsynonymous SNV | MLF1:NM_001195434:exon8:c.C646A:p.P216T,MLF1:NM_001130156:exon6:c.C601A:p.P201T,MLF1:NM_001195433:exon6:c.C472A:p.P158T,MLF1:NM_001195432:exon8:c.C769A:p.P257T,MLF1:NM_022443:exon6:c.C676A:p.P226T,MLF1:NM_001130157:exon7:c.C601A:p.P201T, |
| chr3 | 168029234 | C | T | COSV58329953 | exonic | GOLIM4 | nonsynonymous SNV | GOLIM4:NM_001308155:exon10:c.G1418A:p.G473E,GOLIM4:NM_014498:exon11:c.G1502A:p.G501E, |
| chr3 | 169116038 | A | T | . | exonic | MECOM | nonsynonymous SNV | MECOM:NM_001105078:exon7:c.T1270A:p.F424I,MECOM:NM_001163999:exon7:c.T1273A:p.F425I,MECOM:NM_001105077:exon8:c.T1465A:p.F489I,MECOM:NM_004991:exon8:c.T1834A:p.F612I,MECOM:NM_001164000:exon7:c.T1270A:p.F424I,MECOM:NM_001205194:exon7:c.T1270A:p.F424I,MECOM:NM_005241:exon7:c.T1270A:p.F424I, |
| chr3 | 169116044 | C | T | . | exonic | MECOM | nonsynonymous SNV | MECOM:NM_001105078:exon7:c.G1264A:p.E422K,MECOM:NM_001163999:exon7:c.G1267A:p.E423K,MECOM:NM_001105077:exon8:c.G1459A:p.E487K,MECOM:NM_004991:exon8:c.G1828A:p.E610K,MECOM:NM_001164000:exon7:c.G1264A:p.E422K,MECOM:NM_001205194:exon7:c.G1264A:p.E422K,MECOM:NM_005241:exon7:c.G1264A:p.E422K, |
| chr3 | 169116050 | C | T | COSV52964849 | exonic | MECOM | nonsynonymous SNV | MECOM:NM_001105078:exon7:c.G1258A:p.D420N,MECOM:NM_004991:exon8:c.G1822A:p.D608N,MECOM:NM_001164000:exon7:c.G1258A:p.D420N,MECOM:NM_005241:exon7:c.G1258A:p.D420N,MECOM:NM_001163999:exon7:c.G1261A:p.D421N,MECOM:NM_001105077:exon8:c.G1453A:p.D485N,MECOM:NM_001205194:exon7:c.G1258A:p.D420N, |
| chr3 | 169381204 | G | A | COSV72109887 | exonic | MECOM | nonsynonymous SNV | MECOM:NM_004991:exon2:c.C358T:p.P120S, |
| chr3 | 172506729 | G | T | . | exonic | TNFSF10 | nonsynonymous SNV | TNFSF10:NM_003810:exon5:c.C609A:p.D203E, |

| | | | | | | | | |
|---|---|---|---|---|---|---|---|---|
| chr3 | 172506761 | G | T | . | exonic | TNFSF10 | nonsynonymous SNV | TNFSF10:NM_003810:exon5:c.C577A:p.Q193K, |
| chr3 | 184178961 | G | A | . | exonic | AP2M1 | nonsynonymous SNV | AP2M1:NM_001025205:exon3:c.G179A:p.R60Q, |
| chr3 | 184179005 | G | A | . | exonic | AP2M1 | nonsynonymous SNV | AP2M1:NM_001025205:exon3:c.G223A:p.A75T, |
| chr3 | 184982638 | C | T | COSV54982221 | exonic | VPS8 | nonsynonymous SNV | VPS8:NM_001009921:exon40:c.C3493T:p.H1165Y,VPS8:NM_015303:exon40:c.C3487T:p.H1163Y, |
| chr3 | 195725928 | C | T | COSV57846901 | exonic | MUC20 | nonsynonymous SNV | MUC20:NM_020790:exon4:c.C869T:p.T290I,MUC20:NM_001291833:exon4:c.C812T:p.T271I,MUC20:NM_001282506:exon2:c.C1325T:p.T442I,MUC20:NM_152673:exon4:c.C812T:p.T271I, |
| chr3 | 195726080 | G | C | COSV57846804 | exonic | MUC20 | nonsynonymous SNV | MUC20:NM_020790:exon4:c.G1021C:p.D341H,MUC20:NM_001291833:exon4:c.G964C:p.D322H,MUC20:NM_001282506:exon2:c.G1477C:p.D493H,MUC20:NM_152673:exon4:c.G964C:p.D322H, |
| chr3 | 32997107 | C | T | . | exonic | GLB1 | nonsynonymous SNV | GLB1:NM_001317040:exon17:c.G2116A:p.E706K,GLB1:NM_000404:exon16:c.G1972A:p.E658K,GLB1:NM_001135602:exon13:c.G1579A:p.E527K,GLB1:NM_001079811:exon16:c.G1882A:p.E628K, |
| chr3 | 33014196 | T | C | . | exonic | GLB1 | nonsynonymous SNV | GLB1:NM_001317040:exon16:c.A1738G:p.S580G,GLB1:NM_000404:exon15:c.A1594G:p.S532G,GLB1:NM_001135602:exon12:c.A1201G:p.S401G,GLB1:NM_001079811:exon15:c.A1504G:p.S502G, |
| chr3 | 33014229 | A | G | . | exonic | GLB1 | nonsynonymous SNV | GLB1:NM_001317040:exon16:c.T1705C:p.C569R,GLB1:NM_000404:exon15:c.T1561C:p.C521R,GLB1:NM_001135602:exon12:c.T1168C:p.C390R,GLB1:NM_001079811:exon15:c.T1471C:p.C491R, |
| chr3 | 33097057 | G | A | COSV56561124 | exonic | GLB1 | nonsynonymous SNV | GLB1:NM_001317040:exon1:c.C29T:p.P10L,GLB1:NM_000404:exon1:c.C29T:p.P10L,GLB1:NM_001135602:exon1:c.C29T:p.P10L, |
| chr3 | 37012077 | A | G | COSV51613800 | exonic | MLH1 | nonsynonymous SNV | MLH1:NM_000249:exon8:c.A655G:p.I219V,MLH1:NM_001258271:exon8:c.A655G:p.I219V,MLH1:NM_001167617:exon8:c.A361G:p.I121V, |
| chr3 | 37323683 | T | A | . | exonic | GOLGA4 | nonsynonymous SNV | GOLGA4:NM_002078:exon14:c.T1797A:p.N599K,GOLGA4:NM_001172713:exon15:c.T1863A:p.N621K, |
| chr3 | 37323837 | C | A | . | exonic | GOLGA4 | nonsynonymous SNV | GOLGA4:NM_002078:exon14:c.C1951A:p.Q651K,GOLGA4:NM_001172713:exon15:c.C2017A:p.Q673K, |
| chr3 | 37324641 | G | A | COSV62710454 | exonic | GOLGA4 | nonsynonymous SNV | GOLGA4:NM_002078:exon14:c.G2755A:p.G919R,GOLGA4:NM_001172713:exon15:c.G2821A:p.G941R, |
| chr3 | 37324792 | C | A | . | exonic | GOLGA4 | nonsynonymous SNV | GOLGA4:NM_002078:exon14:c.C2906A:p.T969K,GOLGA4:NM_001172713:exon15:c.C2972A:p.T991K, |
| chr3 | 37324794 | T | A | . | exonic | GOLGA4 | nonsynonymous SNV | GOLGA4:NM_002078:exon14:c.T2908A:p.L970I,GOLGA4:NM_001172713:exon15:c.T2974A:p.L992I, |
| chr3 | 37324968 | C | A | COSV62709296 | exonic | GOLGA4 | nonsynonymous SNV | GOLGA4:NM_002078:exon14:c.C3082A:p.Q1028K,GOLGA4:NM_001172713:exon15:c.C3148A:p.Q1050K, |
| chr3 | 41225688 | G | A | . | exonic | CTNNB1 | nonsynonymous SNV | CTNNB1:NM_001904:exon6:c.G763A:p.A255T,CTNNB1:NM_001098210:exon6:c.G763A:p.A255T,CTNNB1:NM_001098209:exon6:c.G763A:p.A255T, |
| chr3 | 41233679 | G | A | . | exonic | CTNNB1 | nonsynonymous SNV | CTNNB1:NM_001904:exon9:c.G1336A:p.A446T,CTNNB1:NM_001098210:exon9:c.G1336A:p.A446T,CTNNB1:NM_001098209:exon9:c.G1336A:p.A446T, |
| chr3 | 41235790 | G | A | . | exonic | CTNNB1 | nonsynonymous SNV | CTNNB1:NM_001098210:exon11:c.G1750A:p.V584I,CTNNB1:NM_001098209:exon11:c.G1750A:p.V584I,CTNNB1:NM_001904:exon11:c.G1750A:p.V584I, |
| chr3 | 41235800 | G | A | COSV62694859 | exonic | CTNNB1 | nonsynonymous SNV | CTNNB1:NM_001904:exon11:c.G1760A:p.R587Q,CTNNB1:NM_001098210:exon11:c.G1760A:p.R587Q,CTNNB1:NM_001098209:exon11:c.G1760A:p.R587Q, |
| chr3 | 41236618 | T | G | . | exonic | CTNNB1 | nonsynonymous SNV | CTNNB1:NM_001904:exon13:c.T1985G:p.M662R,CTNNB1:NM_001098210:exon13:c.T1985G:p.M662R,CTNNB1:NM_001098209:exon13:c.T1985G:p.M662R, |
| chr3 | 41238041 | G | A | . | exonic | CTNNB1 | nonsynonymous SNV | CTNNB1:NM_001098210:exon14:c.G2102A:p.G701D,CTNNB1:NM_001098209:exon14:c.G2102A:p.G701D,CTNNB1:NM_001904:exon14:c.G2102A:p.G701D, |
| chr3 | 41239247 | G | A | COSV62706846 | exonic | CTNNB1 | nonsynonymous SNV | CTNNB1:NM_001904:exon15:c.G2251A:p.D751N,CTNNB1:NM_001098210:exon15:c.G2251A:p.D751N,CTNNB1:NM_001098209:exon15:c.G2251A:p.D751N, |
| chr3 | 41239286 | G | A | . | exonic | CTNNB1 | nonsynonymous SNV | CTNNB1:NM_001098210:exon15:c.G2290A:p.D764N,CTNNB1:NM_001098209:exon15:c.G2290A:p.D764N,CTNNB1:NM_001904:exon15:c.G2290A:p.D764N, |
| chr3 | 46408579 | T | A | COSV62193340 | exonic | CCRL2 | nonsynonymous SNV | CCRL2:NM_003965:exon2:c.T500A:p.F167Y,CCRL2:NM_001130910:exon2:c.T536A:p.F179Y, |
| chr3 | 46408581 | G | A | COSV62194015 | exonic | CCRL2 | nonsynonymous SNV | CCRL2:NM_003965:exon2:c.G502A:p.V168M,CCRL2:NM_001130910:exon2:c.G538A:p.V180M, |
| chr3 | 47062226 | C | T | COSV57435899 | exonic | SETD2 | nonsynonymous SNV | SETD2:NM_014159:exon14:c.G6230A:p.R2077Q, |
| chr3 | 47083895 | G | A | COSV57428846 | exonic | SETD2 | nonsynonymous SNV | SETD2:NM_014159:exon12:c.C5885T:p.P1962L, |
| chr3 | 47418189 | C | T | COSV55542934 | exonic | SCAP | nonsynonymous SNV | SCAP:NM_012235:exon16:c.G2392A:p.V798I, |
| chr3 | 47914934 | C | T | COSV53135994 | exonic | MAP4 | nonsynonymous SNV | MAP4:NM_002375:exon8:c.G1882A:p.V628I,MAP4:NM_001134364:exon8:c.G1882A:p.V628I, |

| | | | | | | | | |
|---|---|---|---|---|---|---|---|---|
| chr3 | 47916041 | G | T | . | exonic | MAP4 | nonsynonymous SNV | MAP4:NM_002375:exon7:c.C1786A:p.Q596K,MAP4:NM_001134364:exon7:c.C1786A:p.Q596K, |
| chr3 | 47916547 | G | T | COSV64236753 | exonic | MAP4 | nonsynonymous SNV | MAP4:NM_002375:exon7:c.C1280A:p.S427Y,MAP4:NM_001134364:exon7:c.C1280A:p.S427Y, |
| chr3 | 47916670 | A | T | . | exonic | MAP4 | nonsynonymous SNV | MAP4:NM_002375:exon7:c.T1157A:p.I386K,MAP4:NM_001134364:exon7:c.T1157A:p.I386K, |
| chr3 | 47916700 | T | C | . | exonic | MAP4 | nonsynonymous SNV | MAP4:NM_002375:exon7:c.A1127G:p.N376S,MAP4:NM_001134364:exon7:c.A1127G:p.N376S, |
| chr3 | 47916709 | G | T | . | exonic | MAP4 | nonsynonymous SNV | MAP4:NM_002375:exon7:c.C1118A:p.P373H,MAP4:NM_001134364:exon7:c.C1118A:p.P373H, |
| chr3 | 49030471 | T | C | COSV58708022 | exonic | QRICH1 | nonsynonymous SNV | QRICH1:NM_017730:exon11:c.A2312G:p.N771S,QRICH1:NM_198880:exon10:c.A2312G:p.N771S, |
| chr3 | 49360262 | C | T | . | exonic | RHOA | nonsynonymous SNV | RHOA:NM_001313945:exon5:c.G286A:p.A96T,RHOA:NM_001664:exon5:c.G529A:p.A177T,RHOA:NM_001313946:exon3:c.G277A:p.A93T,RHOA:NM_001313941:exon6:c.G529A:p.A177T,RHOA:NM_001313944:exon5:c.G469A:p.A157T, |
| chr3 | 49360264 | C | T | . | exonic | RHOA | nonsynonymous SNV | RHOA:NM_001313945:exon5:c.G284A:p.R95K,RHOA:NM_001664:exon5:c.G527A:p.R176K,RHOA:NM_001313946:exon3:c.G275A:p.R92K,RHOA:NM_001313941:exon6:c.G527A:p.R176K,RHOA:NM_001313944:exon5:c.G467A:p.R156K, |
| chr3 | 49360277 | C | T | COSV69041753 | exonic | RHOA | nonsynonymous SNV | RHOA:NM_001313945:exon5:c.G271A:p.E91K,RHOA:NM_001664:exon5:c.G514A:p.E172K,RHOA:NM_001313946:exon3:c.G262A:p.E88K,RHOA:NM_001313941:exon6:c.G514A:p.E172K,RHOA:NM_001313944:exon5:c.G454A:p.E152K, |
| chr3 | 49360350 | C | A | . | exonic | RHOA | nonsynonymous SNV | RHOA:NM_001313945:exon5:c.G198T:p.M66I,RHOA:NM_001313946:exon3:c.G189T:p.M63I,RHOA:NM_001313941:exon6:c.G441T:p.M147I,RHOA:NM_001664:exon5:c.G441T:p.M147I,RHOA:NM_001313944:exon5:c.G381T:p.M127I, |
| chr3 | 49362534 | C | T | . | exonic | RHOA | nonsynonymous SNV | RHOA:NM_001313945:exon4:c.G127A:p.D43N,RHOA:NM_001664:exon4:c.G370A:p.D124N,RHOA:NM_001313941:exon5:c.G370A:p.D124N,RHOA:NM_001313947:exon3:c.G249A:p.M83I,RHOA:NM_001313944:exon4:c.G310A:p.D104N,RHOA:NM_001313943:exon4:c.G370A:p.D124N, |
| chr3 | 49362608 | C | T | . | exonic | RHOA | stopgain | RHOA:NM_001313945:exon4:c.G53A:p.W18X,RHOA:NM_001664:exon4:c.G296A:p.W99X,RHOA:NM_001313941:exon5:c.G296A:p.W99X,RHOA:NM_001313944:exon4:c.G236A:p.W79X,RHOA:NM_001313943:exon4:c.G296A:p.W99X, |
| chr3 | 49368531 | C | T | . | exonic | RHOA | stopgain | RHOA:NM_001664:exon3:c.G174A:p.W58X,RHOA:NM_001313941:exon4:c.G174A:p.W58X,RHOA:NM_001313943:exon3:c.G174A:p.W58X, |
| chr3 | 52476576 | G | A | COSV61900229 | exonic | NISCH | nonsynonymous SNV | NISCH:NM_007184:exon8:c.G895A:p.V299I,NISCH:NM_001276293:exon8:c.G895A:p.V299I,NISCH:NM_001276294:exon8:c.G895A:p.V299I, |
| chr3 | 52603593 | C | T | . | exonic | PBRM1 | nonsynonymous SNV | PBRM1:NM_018313:exon18:c.G2707A:p.E903K, |
| chr3 | 52687262 | G | A | . | exonic | GNL3 | nonsynonymous SNV | GNL3:NM_206826:exon3:c.G53A:p.R18Q,GNL3:NM_014366:exon3:c.G89A:p.R30Q,GNL3:NM_206825:exon3:c.G53A:p.R18Q, |
| chr3 | 52687289 | G | A | COSV66920847 | exonic | GNL3 | nonsynonymous SNV | GNL3:NM_206826:exon3:c.G80A:p.R27Q,GNL3:NM_014366:exon3:c.G116A:p.R39Q,GNL3:NM_206825:exon3:c.G80A:p.R27Q, |
| chr3 | 52690686 | T | A | . | exonic | GNL3 | nonsynonymous SNV | GNL3:NM_206826:exon7:c.T600A:p.D200E,GNL3:NM_014366:exon7:c.T636A:p.D212E,GNL3:NM_206825:exon7:c.T600A:p.D200E, |
| chr3 | 52693241 | G | A | COSV56476624 | exonic | GNL3 | nonsynonymous SNV | GNL3:NM_206826:exon11:c.G1063A:p.V355M,GNL3:NM_014366:exon11:c.G1099A:p.V367M,GNL3:NM_206825:exon11:c.G1063A:p.V355M, |
| chr3 | 52693700 | G | A | . | exonic | GNL3 | nonsynonymous SNV | GNL3:NM_206826:exon13:c.G1357A:p.E453K,GNL3:NM_014366:exon13:c.G1393A:p.E465K,GNL3:NM_206825:exon13:c.G1357A:p.E453K, |
| chr3 | 52693728 | C | A | . | exonic | GNL3 | nonsynonymous SNV | GNL3:NM_206826:exon13:c.C1385A:p.P462Q,GNL3:NM_014366:exon13:c.C1421A:p.P474Q,GNL3:NM_206825:exon13:c.C1385A:p.P462Q, |
| chr3 | 52693734 | G | A | COSV56477800 | exonic | GNL3 | nonsynonymous SNV | GNL3:NM_206826:exon13:c.G1391A:p.R464Q,GNL3:NM_014366:exon13:c.G1427A:p.R476Q,GNL3:NM_206825:exon13:c.G1391A:p.R464Q, |
| chr3 | 52694079 | G | A | . | exonic | GNL3 | nonsynonymous SNV | GNL3:NM_206826:exon14:c.G1507A:p.A503T,GNL3:NM_014366:exon14:c.G1543A:p.A515T,GNL3:NM_206825:exon14:c.G1507A:p.A503T, |
| chr3 | 52694211 | C | T | . | exonic | GNL3 | nonsynonymous SNV | GNL3:NM_206826:exon15:c.C1550T:p.S517F,GNL3:NM_014366:exon15:c.C1586T:p.S529F,GNL3:NM_206825:exon15:c.C1550T:p.S517F, |
| chr3 | 52763618 | G | C | COSV51778826 | exonic | NEK4 | nonsynonymous SNV | NEK4:NM_003157:exon5:c.C673G:p.P225A,NEK4:NM_001193533:exon4:c.C406G:p.P136A, |
| chr3 | 69180910 | G | A | COSV68330003 | exonic | FRMD4B | nonsynonymous SNV | FRMD4B:NM_015123:exon21:c.C2840T:p.S947L, |
| chr3 | 9914595 | G | A | . | exonic | IL17RE | nonsynonymous SNV | IL17RE:NM_001193380:exon14:c.G1417A:p.G473R, |
| chr3 | 9964092 | C | T | . | exonic | EMC3 | nonsynonymous SNV | EMC3:NM_018447:exon8:c.G763A:p.E255K, |
| chr3 | 9970612 | T | A | . | exonic | EMC3 | nonsynonymous SNV | EMC3:NM_018447:exon6:c.A544T:p.I182F, |
| chr3 | 9986637 | C | T | COSV55277193 | exonic | EMC3 | nonsynonymous SNV | EMC3:NM_018447:exon1:c.G25A:p.D9N, |
| chr4 | 105235006 | G | A | COSV54398666 | exonic | TET2 | nonsynonymous SNV | TET2:NM_001127208:exon3:c.G1064A:p.G355D,TET2:NM_017628:exon3:c.G1064A:p.G355D, |
| chr4 | 105236187 | C | A | . | exonic | TET2 | nonsynonymous SNV | TET2:NM_001127208:exon3:c.C2245A:p.Q749K,TET2:NM_017628:exon3:c.C2245A:p.Q749K, |
| chr4 | 112647304 | T | A | . | exonic | LARP7 | nonsynonymous SNV | LARP7:NM_016648:exon7:c.T752A:p.M251K,LARP7:NM_001267039:exon9:c.T773A:p.M258K,LARP7:NM_015454:exon7:c.T752A:p.M251K, |

| | | | | | | | | |
|---|---|---|---|---|---|---|---|---|
| chr4 | 112647767 | C | A | . | exonic | LARP7 | nonsynonymous SNV | LARP7:NM_016648:exon8:c.C1075A:p.H359N,LARP7:NM_015454:exon8:c.C1075A:p.H359N,LARP7:NM_001267039:exon10:c.C1096A:p.H366N, |
| chr4 | 121816828 | G | A | . | exonic | EXOSC9 | nonsynonymous SNV | EXOSC9:NM_005033:exon12:c.G1292A:p.R431K,EXOSC9:NM_001034194:exon13:c.G1343A:p.R448K, |
| chr4 | 150277928 | G | A | COSV63949643 | exonic | LRBA | nonsynonymous SNV | LRBA:NM_006726:exon57:c.C8426T:p.S2809L,LRBA:NM_001199282:exon56:c.C8390T:p.S2797L, |
| chr4 | 150285975 | C | T | COSV63966388 | exonic | LRBA | nonsynonymous SNV | LRBA:NM_006726:exon55:c.G8110A:p.A2704T,LRBA:NM_001199282:exon54:c.G8074A:p.A2692T, |
| chr4 | 1728226 | G | A | COSV57603699 | exonic | TACC3 | nonsynonymous SNV | TACC3:NM_006342:exon4:c.G824A:p.C275Y, |
| chr4 | 1728261 | G | A | COSV57603712 | exonic | TACC3 | nonsynonymous SNV | TACC3:NM_006342:exon4:c.G859A:p.G287S, |
| chr4 | 1731251 | G | A | COSV57605388 | exonic | TACC3 | nonsynonymous SNV | TACC3:NM_006342:exon6:c.G1541A:p.G514E, |
| chr4 | 185190485 | T | C | COSV57111528 | exonic | CFAP97 | nonsynonymous SNV | CFAP97:NM_020827:exon2:c.A712G:p.T238A,CFAP97:NM_001292033:exon2:c.A712G:p.T238A, |
| chr4 | 1935170 | C | A | . | exonic | WHSC1 | nonsynonymous SNV | WHSC1:NM_133335:exon7:c.C1582A:p.H528N,WHSC1:NM_007331:exon9:c.C1582A:p.H528N,WHSC1:NM_133330:exon9:c.C1582A:p.H528N,WHSC1:NM_001042424:exon7:c.C1582A:p.H528N,WHSC1:NM_133331:exon8:c.C1582A:p.H528N,WHSC1:NM_133334:exon6:c.C1582A:p.H528N, |
| chr4 | 2042401 | C | T | . | exonic | C4ORF48 | nonsynonymous SNV | C4orf48:NM_001168243:exon2:c.C149T:p.P50L,C4orf48:NM_001141936:exon2:c.C50T:p.P17L, |
| chr4 | 41646183 | G | A | . | exonic | LIMCH1 | nonsynonymous SNV | LIMCH1:NM_001289122:exon10:c.G1123A:p.E375K,LIMCH1:NM_001112719:exon6:c.G697A:p.E233K,LIMCH1:NM_001112720:exon5:c.G661A:p.E221K,LIMCH1:NM_014988:exon11:c.G1159A:p.E387K,LIMCH1:NM_001289124:exon10:c.G646A:p.E216K,LIMCH1:NM_001112717:exon11:c.G1159A:p.E387K,LIMCH1:NM_001112718:exon11:c.G1159A:p.E387K, |
| chr4 | 4274597 | C | T | . | exonic | LYAR | nonsynonymous SNV | LYAR:NM_001145725:exon7:c.G602A:p.R201K,LYAR:NM_017816:exon7:c.G602A:p.R201K, |
| chr4 | 4274649 | C | T | . | exonic | LYAR | nonsynonymous SNV | LYAR:NM_001145725:exon7:c.G550A:p.V184M,LYAR:NM_017816:exon7:c.G550A:p.V184M, |
| chr4 | 48986433 | C | A | COSV56939573 | exonic | CWH43 | nonsynonymous SNV | CWH43:NM_025087:exon1:c.C4A:p.P2T, |
| chr4 | 49061855 | C | A | COSV56936745 | exonic | CWH43 | nonsynonymous SNV | CWH43:NM_001286791:exon16:c.C1984A:p.H662N,CWH43:NM_025087:exon16:c.C2065A:p.H689N, |
| chr4 | 53458720 | G | A | . | exonic | FIP1L1 | nonsynonymous SNV | FIP1L1:NM_030917:exon17:c.G1567A:p.E523K,FIP1L1:NM_001134938:exon14:c.G1345A:p.E449K,FIP1L1:NM_001134937:exon17:c.G1549A:p.E517K, |
| chr4 | 53459377 | C | A | . | exonic | FIP1L1 | nonsynonymous SNV | FIP1L1:NM_030917:exon18:c.C1713A:p.S571R,FIP1L1:NM_001134938:exon15:c.C1491A:p.S497R,FIP1L1:NM_001134937:exon18:c.C1695A:p.S565R, |
| chr4 | 56448739 | A | C | . | exonic | PAICS | nonsynonymous SNV | PAICS:NM_001079525:exon6:c.A624C:p.K208N,PAICS:NM_006452:exon6:c.A603C:p.K201N,PAICS:NM_001079524:exon5:c.A603C:p.K201N, |
| chr4 | 87491654 | T | C | COSV56803313 | exonic | SPARCL1 | nonsynonymous SNV | SPARCL1:NM_001291977:exon3:c.A880G:p.T294A,SPARCL1:NM_001291976:exon6:c.A880G:p.T294A,SPARCL1:NM_001128310:exon6:c.A1255G:p.T419A,SPARCL1:NM_004684:exon5:c.A1255G:p.T419A, |
| chr4 | 87494149 | G | C | . | exonic | SPARCL1 | nonsynonymous SNV | SPARCL1:NM_001291977:exon2:c.C276G:p.N92K,SPARCL1:NM_001291976:exon5:c.C276G:p.N92K,SPARCL1:NM_001128310:exon5:c.C651G:p.N217K,SPARCL1:NM_004684:exon4:c.C651G:p.N217K, |
| chr4 | 87494337 | C | T | . | exonic | SPARCL1 | nonsynonymous SNV | SPARCL1:NM_001291977:exon2:c.G88A:p.E30K,SPARCL1:NM_001291976:exon5:c.G88A:p.E30K,SPARCL1:NM_001128310:exon5:c.G463A:p.E155K,SPARCL1:NM_004684:exon4:c.G463A:p.E155K, |
| chr4 | 87494484 | G | C | COSV56802379 | exonic | SPARCL1 | nonsynonymous SNV | SPARCL1:NM_001128310:exon5:c.C316G:p.H106D,SPARCL1:NM_004684:exon4:c.C316G:p.H106D, |
| chr4 | 87495036 | G | T | COSV56802387 | exonic | SPARCL1 | nonsynonymous SNV | SPARCL1:NM_001128310:exon4:c.C146A:p.A49D,SPARCL1:NM_004684:exon3:c.C146A:p.A49D, |
| chr4 | 99339632 | T | C | COSV72624794 | exonic | ADH1C | nonsynonymous SNV | ADH1C:NM_000669:exon8:c.A1048G:p.I350V, |
| chr5 | 116092684 | T | G | COSV57234444 | exonic | COMMD10 | nonsynonymous SNV | COMMD10:NM_016144:exon4:c.T383G:p.I128S,COMMD10:NM_001308080:exon4:c.T341G:p.I114S, |
| chr5 | 146013801 | C | T | COSV63039708 | exonic | SH3RF2 | nonsynonymous SNV | SH3RF2:NM_152550:exon5:c.C799T:p.R267C, |
| chr5 | 146060084 | T | C | COSV63040514 | exonic | SH3RF2 | nonsynonymous SNV | SH3RF2:NM_152550:exon9:c.T1774C:p.W592R, |
| chr5 | 146062640 | G | C | COSV63040479 | exonic | SH3RF2 | nonsynonymous SNV | SH3RF2:NM_152550:exon10:c.G2129C:p.G710A, |
| chr5 | 150404692 | C | T | . | exonic | CD74 | nonsynonymous SNV | CD74:NM_004355:exon6:c.G613A:p.A205T,CD74:NM_001025159:exon6:c.G613A:p.A205T, |
| chr5 | 150405089 | C | T | . | exonic | CD74 | stopgain | CD74:NM_004355:exon5:c.G533A:p.W178X,CD74:NM_001025159:exon5:c.G533A:p.W178X, |
| chr5 | 150406271 | C | T | . | exonic | CD74 | nonsynonymous SNV | CD74:NM_004355:exon4:c.G429A:p.M143I,CD74:NM_001025158:exon4:c.G429A:p.M143I,CD74:NM_001025159:exon4:c.G429A:p.M143I, |
| chr5 | 150406277 | A | T | . | exonic | CD74 | nonsynonymous SNV | CD74:NM_004355:exon4:c.T423A:p.H141Q,CD74:NM_001025159:exon4:c.T423A:p.H141Q,CD74:NM_001025158:exon4:c.T423A:p.H141Q, |
| chr5 | 150406296 | C | T | . | exonic | CD74 | nonsynonymous SNV | CD74:NM_004355:exon4:c.G404A:p.G135D,CD74:NM_001025158:exon4:c.G404A:p.G135D,CD74:NM_001025159:exon4:c.G404A:p.G135D, |
| chr5 | 150406931 | C | T | . | exonic | CD74 | nonsynonymous SNV | CD74:NM_004355:exon3:c.G328A:p.A110T,CD74:NM_001025158:exon3:c.G328A:p.A110T,CD74:NM_001025159:exon3:c.G328A:p.A110T, |
| chr5 | 150406957 | G | T | . | exonic | CD74 | nonsynonymous SNV | CD74:NM_004355:exon3:c.C302A:p.P101H,CD74:NM_001025158:exon3:c.C302A:p.P101H,CD74:NM_001025159:exon3:c.C302A:p.P101H, |
| chr5 | 150407199 | G | A | . | exonic | CD74 | nonsynonymous SNV | CD74:NM_004355:exon2:c.C251T:p.T84I,CD74:NM_001025158:exon2:c.C251T:p.T84I,CD74:NM_001025159:exon2:c.C251T:p.T84I, |
| chr5 | 150407308 | C | T | . | exonic | CD74 | nonsynonymous SNV | CD74:NM_004355:exon2:c.G142A:p.A48T,CD74:NM_001025158:exon2:c.G142A:p.A48T,CD74:NM_001025159:exon2:c.G142A:p.A48T, |
| chr5 | 150412635 | C | G | . | exonic | CD74 | nonsynonymous SNV | CD74:NM_004355:exon1:c.G115C:p.A39P,CD74:NM_001025158:exon1:c.G115C:p.A39P,CD74:NM_001025159:exon1:c.G115C:p.A39P, |
| chr5 | 150412686 | C | T | COSV50532067 | exonic | CD74 | nonsynonymous SNV | CD74:NM_004355:exon1:c.G64A:p.D22N,CD74:NM_001025158:exon1:c.G64A:p.D22N,CD74:NM_001025159:exon1:c.G64A:p.D22N, |

| | | | | | | | | |
|---|---|---|---|---|---|---|---|---|
| chr5 | 150412727 | C | T | . | exonic | CD74 | nonsynonymous SNV | CD74:NM_004355:exon1:c.G23A:p.S8N,CD74:NM_001025159:exon1:c.G23A:p.S8N,CD74:NM_001025158:exon1:c.G23A:p.S8N, |
| chr5 | 154002288 | C | T | COSV61078046 | exonic | FAM114A2 | nonsynonymous SNV | FAM114A2:NM_018691:exon11:c.G1219A:p.E407K, |
| chr5 | 154002906 | C | T | . | exonic | FAM114A2 | nonsynonymous SNV | FAM114A2:NM_018691:exon10:c.G1057A:p.E353K, |
| chr5 | 154002910 | A | T | . | exonic | FAM114A2 | nonsynonymous SNV | FAM114A2:NM_018691:exon10:c.T1053A:p.N351K, |
| chr5 | 154033830 | C | T | COSV61077353 | exonic | FAM114A2 | nonsynonymous SNV | FAM114A2:NM_018691:exon4:c.G364A:p.G122S, |
| chr5 | 171392799 | G | A | . | exonic | NPM1 | nonsynonymous SNV | NPM1:NM_001037738:exon5:c.G442A:p.G148S,NPM1:NM_199185:exon5:c.G442A:p.G148S,NPM1:NM_002520:exon5:c.G442A:p.G148S, |
| chr5 | 171400846 | G | A | . | exonic | NPM1 | nonsynonymous SNV | NPM1:NM_001037738:exon8:c.G590A:p.R197Q,NPM1:NM_002520:exon8:c.G590A:p.R197Q, |
| chr5 | 171405303 | G | A | . | exonic | NPM1 | nonsynonymous SNV | NPM1:NM_001037738:exon9:c.G671A:p.G224E,NPM1:NM_199185:exon8:c.G584A:p.G195E,NPM1:NM_002520:exon9:c.G671A:p.G224E, |
| chr5 | 171405389 | G | A | . | exonic | NPM1 | nonsynonymous SNV | NPM1:NM_001037738:exon9:c.G757A:p.A253T,NPM1:NM_199185:exon8:c.G670A:p.A224T,NPM1:NM_002520:exon9:c.G757A:p.A253T, |
| chr5 | 171407742 | G | A | . | exonic | NPM1 | nonsynonymous SNV | NPM1:NM_199185:exon9:c.G727A:p.V243M,NPM1:NM_002520:exon10:c.G814A:p.V272M, |
| chr5 | 171407758 | G | A | COSV51564036 | exonic | NPM1 | nonsynonymous SNV | NPM1:NM_199185:exon9:c.G743A:p.R248Q,NPM1:NM_002520:exon10:c.G830A:p.R277Q, |
| chr5 | 171407762 | G | A | . | exonic | NPM1 | nonsynonymous SNV | NPM1:NM_199185:exon9:c.G747A:p.M249I,NPM1:NM_002520:exon10:c.G834A:p.M278I, |
| chr5 | 179824245 | G | A | . | exonic | SQSTM1 | nonsynonymous SNV | SQSTM1:NM_001142299:exon5:c.G343A:p.E115K,SQSTM1:NM_003900:exon4:c.G595A:p.E199K,SQSTM1:NM_001142298:exon5:c.G343A:p.E115K, |
| chr5 | 179833099 | G | C | COSV52975046 | exonic | SQSTM1 | nonsynonymous SNV | SQSTM1:NM_001142299:exon7:c.G570C:p.E190D,SQSTM1:NM_003900:exon6:c.G822C:p.E274D,SQSTM1:NM_001142298:exon7:c.G570C:p.E190D, |
| chr5 | 32400161 | A | G | COSV54053200 | exonic | ZFR | nonsynonymous SNV | ZFR:NM_016107:exon9:c.T1559C:p.I520T, |
| chr5 | 32403241 | C | T | . | exonic | ZFR | nonsynonymous SNV | ZFR:NM_016107:exon8:c.G1381A:p.V461I, |
| chr5 | 32406858 | A | T | . | exonic | ZFR | nonsynonymous SNV | ZFR:NM_016107:exon6:c.T948A:p.N316K, |
| chr5 | 33989413 | C | T | COSV56871911 | exonic | AMACR | nonsynonymous SNV | AMACR:NM_001167595:exon5:c.G829A:p.E277K,AMACR:NM_014324:exon5:c.G829A:p.E277K, |
| chr5 | 33998663 | C | A | COSV59460034 | exonic | AMACR | nonsynonymous SNV | AMACR:NM_001167595:exon4:c.G717T:p.Q239H,AMACR:NM_203382:exon3:c.G556T:p.V186F,AMACR:NM_014324:exon4:c.G717T:p.Q239H, |
| chr5 | 33998778 | A | G | COSV59461114 | exonic | AMACR | nonsynonymous SNV | AMACR:NM_001167595:exon4:c.T602C:p.L201S,AMACR:NM_014324:exon4:c.T602C:p.L201S, |
| chr5 | 34004602 | C | T | COSV59461134 | exonic | AMACR | nonsynonymous SNV | AMACR:NM_001167595:exon3:c.G524A:p.G175D,AMACR:NM_014324:exon3:c.G524A:p.G175D, |
| chr5 | 34007995 | C | T | COSV59461909 | exonic | AMACR | nonsynonymous SNV | AMACR:NM_001167595:exon1:c.G25A:p.V9M,AMACR:NM_203382:exon1:c.G25A:p.V9M,AMACR:NM_014324:exon1:c.G25A:p.V9M, |
| chr5 | 37752562 | C | A | . | exonic | WDR70 | nonsynonymous SNV | WDR70:NM_018034:exon18:c.C1954A:p.R652S, |
| chr5 | 55664845 | C | G | COSV59422666 | exonic | SLC38A9 | nonsynonymous SNV | SLC38A9:NM_001282429:exon7:c.G173C:p.S58T,SLC38A9:NM_001258286:exon6:c.G356C:p.S119T,SLC38A9:NM_173514:exon8:c.G545C:p.S182T,SLC38A9:NM_001258287:exon7:c.G464C:p.S155T, |
| chr5 | 65563377 | G | A | . | exonic | PPWD1 | nonsynonymous SNV | PPWD1:NM_015342:exon1:c.G67A:p.E23K, |
| chr5 | 6599951 | G | T | . | exonic | NSUN2 | nonsynonymous SNV | NSUN2:NM_001193455:exon18:c.C2174A:p.P725Q,NSUN2:NM_017755:exon19:c.C2279A:p.P760Q, |
| chr5 | 69400113 | T | G | COSV51456873 | exonic | RAD17 | nonsynonymous SNV | RAD17:NM_133344:exon15:c.T1637G:p.L546R,RAD17:NM_133342:exon16:c.T1637G:p.L546R,RAD17:NM_133339:exon14:c.T1670G:p.L557R,RAD17:NM_002873:exon15:c.T1637G:p.L546R,RAD17:NM_133340:exon14:c.T1142G:p.L381R,RAD17:NM_133338:exon17:c.T1637G:p.L546R,RAD17:NM_133341:exon14:c.T1379G:p.L460R,RAD17:NM_001278622:exon16:c.T1637G:p.L546R,RAD17:NM_133343:exon16:c.T1637G:p.L546R, |
| chr5 | 71455991 | T | G | COSV62429550 | exonic | BDP1 | nonsynonymous SNV | BDP1:NM_018429:exon1:c.T114G:p.D38E, |
| chr5 | 71504711 | G | A | COSV62429660 | exonic | BDP1 | nonsynonymous SNV | BDP1:NM_018429:exon16:c.G2332A:p.V778M, |
| chr5 | 78129204 | A | T | COSV54884128 | exonic | AP3B1 | nonsynonymous SNV | AP3B1:NM_001271769:exon16:c.T1607A:p.V536E,AP3B1:NM_003664:exon16:c.T1754A:p.V585E, |
| chr5 | 7867917 | T | C | . | exonic | FASTKD3 | nonsynonymous SNV | FASTKD3:NM_024091:exon2:c.A167G:p.K56R, |
| chr5 | 82253397 | C | T | COSV56424284 | exonic | ATG10 | nonsynonymous SNV | ATG10:NM_031482:exon7:c.C635T:p.T212M,ATG10:NM_001131028:exon8:c.C635T:p.T212M, |
| chr5 | 82253421 | C | A | . | exonic | ATG10 | nonsynonymous SNV | ATG10:NM_031482:exon7:c.C659A:p.P220H,ATG10:NM_001131028:exon8:c.C659A:p.P220H, |
| chr5 | 87399457 | A | G | COSV56924372 | exonic | CCNH | nonsynonymous SNV | CCNH:NM_001239:exon7:c.T809C:p.V270A,CCNH:NM_001199189:exon7:c.T650C:p.V217A, |
| chr5 | 891709 | C | T | . | exonic | BRD9 | nonsynonymous SNV | BRD9:NM_001009877:exon2:c.G44A:p.R15K, |
| chr6 | 105278161 | C | T | COSV64870385 | exonic | PREP | nonsynonymous SNV | PREP:NM_002726:exon15:c.G2116A:p.V706I, |
| chr6 | 106640356 | C | T | . | exonic | QRSL1 | nonsynonymous SNV | QRSL1:NM_018292:exon2:c.C32T:p.A11V, |
| chr6 | 107883003 | G | T | . | exonic | SEC63 | nonsynonymous SNV | SEC63:NM_007214:exon17:c.C1818A:p.N606K, |
| chr6 | 122801319 | C | T | COSV63755382 | exonic | SMPDL3A | nonsynonymous SNV | SMPDL3A:NM_006714:exon4:c.C481T:p.P161S,SMPDL3A:NM_001286138:exon3:c.C88T:p.P30S, |
| chr6 | 125999566 | G | A | . | exonic | TRMT11 | nonsynonymous SNV | TRMT11:NM_001031712:exon7:c.G632A:p.G211E, |
| chr6 | 127996951 | G | T | . | exonic | PTPRK | nonsynonymous SNV | PTPRK:NM_002844:exon17:c.C2717A:p.A906D,PTPRK:NM_001135648:exon17:c.C2717A:p.A906D,PTPRK:NM_001291984:exon17:c.C2714A:p.A905D,PTPRK:NM_001291981:exon19:c.C2765A:p.A922D, |

| | | | | | | | | |
|---|---|---|---|---|---|---|---|---|
| chr6 | 128240094 | C | T | COSV63888698 | exonic | PTPRK | nonsynonymous SNV | PTPRK:NM_002844:exon5:c.G634A:p.A212T,PTPRK:NM_001291983:exon4:c.G247A:p.A83T,PTPRK:NM_001135648:exon5:c.G634A:p.A212T,PTPRK:NM_001291984:exon5:c.G634A:p.A212T,PTPRK:NM_001291982:exon5:c.G634A:p.A212T,PTPRK:NM_001291981:exon5:c.G634A:p.A212T, |
| chr6 | 129599245 | A | T | . | exonic | ARHGAP18 | nonsynonymous SNV | ARHGAP18:NM_033515:exon12:c.T1684A:p.Y562N, |
| chr6 | 129599297 | A | T | . | exonic | ARHGAP18 | nonsynonymous SNV | ARHGAP18:NM_033515:exon12:c.T1632A:p.D544E, |
| chr6 | 129599299 | C | T | . | exonic | ARHGAP18 | nonsynonymous SNV | ARHGAP18:NM_033515:exon12:c.G1630A:p.D544N, |
| chr6 | 129599306 | A | T | . | exonic | ARHGAP18 | nonsynonymous SNV | ARHGAP18:NM_033515:exon12:c.T1623A:p.H541Q, |
| chr6 | 129618698 | A | G | . | exonic | ARHGAP18 | nonsynonymous SNV | ARHGAP18:NM_033515:exon6:c.T941C:p.I314T, |
| chr6 | 129710070 | T | C | COSV63763931 | exonic | ARHGAP18 | nonsynonymous SNV | ARHGAP18:NM_033515:exon1:c.A67G:p.T23A, |
| chr6 | 137002076 | G | A | COSV57358376 | exonic | IL20RA | nonsynonymous SNV | IL20RA:NM_014432:exon7:c.C1144T:p.L382F,IL20RA:NM_001278724:exon6:c.C745T:p.L249F,IL20RA:NM_001278723:exon6:c.C811T:p.L271F,IL20RA:NM_001278722:exon7:c.C997T:p.L333F, |
| chr6 | 137004710 | C | T | . | exonic | IL20RA | nonsynonymous SNV | IL20RA:NM_014432:exon6:c.G775A:p.V259I,IL20RA:NM_001278724:exon5:c.G376A:p.V126I,IL20RA:NM_001278723:exon5:c.G442A:p.V148I,IL20RA:NM_001278722:exon6:c.G628A:p.V210I, |
| chr6 | 138092196 | G | C | . | exonic | PERP | nonsynonymous SNV | PERP:NM_022121:exon3:c.C428G:p.P143R, |
| chr6 | 151615542 | G | A | COSV53337542 | exonic | CCDC170 | nonsynonymous SNV | CCDC170:NM_025059:exon10:c.G1810A:p.V604I, |
| chr6 | 151618046 | G | A | COSV53344090 | exonic | CCDC170 | nonsynonymous SNV | CCDC170:NM_025059:exon11:c.G2047A:p.V683I, |
| chr6 | 152975406 | G | C | COSV57670920 | exonic | FBXO5 | nonsynonymous SNV | FBXO5:NM_012177:exon2:c.C319G:p.Q107E,FBXO5:NM_001142522:exon2:c.C181G:p.Q61E, |
| chr6 | 155257877 | C | T | . | exonic | TFB1M | nonsynonymous SNV | TFB1M:NM_016020:exon7:c.G1000A:p.E334K, |
| chr6 | 155257885 | C | T | . | exonic | TFB1M | nonsynonymous SNV | TFB1M:NM_016020:exon7:c.G992A:p.S331N, |
| chr6 | 155257894 | C | T | . | exonic | TFB1M | nonsynonymous SNV | TFB1M:NM_016020:exon7:c.G983A:p.R328Q, |
| chr6 | 158769941 | A | G | . | exonic | EZR | nonsynonymous SNV | EZR:NM_001111077:exon11:c.T1094C:p.L365P,EZR:NM_003379:exon10:c.T1094C:p.L365P, |
| chr6 | 159790604 | G | A | COSV58459001 | exonic | MRPL18 | nonsynonymous SNV | MRPL18:NM_014161:exon1:c.G17A:p.R6Q, |
| chr6 | 167305948 | G | A | COSV57807281 | exonic | UNC93A | nonsynonymous SNV | UNC93A:NM_001143947:exon5:c.G748A:p.V250I,UNC93A:NM_018974:exon6:c.G874A:p.V292I, |
| chr6 | 167315286 | T | C | COSV57806804 | exonic | UNC93A | nonsynonymous SNV | UNC93A:NM_001143947:exon7:c.T1082C:p.M361T,UNC93A:NM_018974:exon8:c.T1208C:p.M403T, |
| chr6 | 167315287 | G | A | COSV57807036 | exonic | UNC93A | nonsynonymous SNV | UNC93A:NM_001143947:exon7:c.G1083A:p.M361I,UNC93A:NM_018974:exon8:c.G1209A:p.M403I, |
| chr6 | 167315303 | G | A | COSV57807274 | exonic | UNC93A | nonsynonymous SNV | UNC93A:NM_001143947:exon7:c.G1099A:p.V367I,UNC93A:NM_018974:exon8:c.G1225A:p.V409I, |
| chr6 | 167827136 | T | G | . | exonic | MLLT4 | nonsynonymous SNV | MLLT4:NM_001040000:exon1:c.T4G:p.S2A,MLLT4:NM_001207008:exon1:c.T4G:p.S2A, |
| chr6 | 18249708 | A | T | . | exonic | DEK | nonsynonymous SNV | DEK:NM_003472:exon7:c.T705A:p.D235E,DEK:NM_001134709:exon6:c.T603A:p.D201E, |
| chr6 | 18256394 | A | G | . | exonic | DEK | nonsynonymous SNV | DEK:NM_003472:exon5:c.T419C:p.V140A,DEK:NM_001134709:exon4:c.T317C:p.V106A, |
| chr6 | 2955568 | T | C | COSV59565275 | exonic | SERPINB6 | nonsynonymous SNV | SERPINB6:NM_001271823:exon3:c.A325G:p.M109V,SERPINB6:NM_001271825:exon3:c.A268G:p.M90V,SERPINB6:NM_001271822:exon3:c.A310G:p.M104V,SERPINB6:NM_001271824:exon3:c.A268G:p.M90V,SERPINB6:NM_001297699:exon3:c.A268G:p.M90V,SERPINB6:NM_001195291:exon4:c.A280G:p.M94V,SERPINB6:NM_001297700:exon3:c.A268G:p.M90V,SERPINB6:NM_004568:exon3:c.A268G:p.M90V, |
| chr6 | 29942581 | C | G | COSV65136471 | exonic | HLA-A | nonsynonymous SNV | HLA-A:NM_001242758:exon1:c.C28G:p.L10V,HLA-A:NM_002116:exon1:c.C28G:p.L10V, |
| chr6 | 29942781 | T | C | COSV65138565 | exonic | HLA-A | nonsynonymous SNV | HLA-A:NM_001242758:exon2:c.T98C:p.F33S,HLA-A:NM_002116:exon2:c.T98C:p.F33S, |
| chr6 | 29942886 | G | A | COSV65139535 | exonic | HLA-A | nonsynonymous SNV | HLA-A:NM_001242758:exon2:c.G203A:p.R68K,HLA-A:NM_002116:exon2:c.G203A:p.R68K, |
| chr6 | 29942939 | C | G | COSV65136733 | exonic | HLA-A | nonsynonymous SNV | HLA-A:NM_001242758:exon2:c.C256G:p.Q86E,HLA-A:NM_002116:exon2:c.C256G:p.Q86E, |
| chr6 | 29942948 | C | G | COSV65136740 | exonic | HLA-A | nonsynonymous SNV | HLA-A:NM_001242758:exon2:c.C265G:p.R89G,HLA-A:NM_002116:exon2:c.C265G:p.R89G, |
| chr6 | 29942953 | T | A | COSV65137266 | exonic | HLA-A | nonsynonymous SNV | HLA-A:NM_001242758:exon2:c.T270A:p.N90K,HLA-A:NM_002116:exon2:c.T270A:p.N90K, |
| chr6 | 29942954 | G | A | . | exonic | HLA-A | nonsynonymous SNV | HLA-A:NM_001242758:exon2:c.G271A:p.V91M,HLA-A:NM_002116:exon2:c.G271A:p.V91M, |
| chr6 | 29942965 | G | C | COSV65140297 | exonic | HLA-A | nonsynonymous SNV | HLA-A:NM_001242758:exon2:c.G282C:p.Q94H,HLA-A:NM_002116:exon2:c.G282C:p.Q94H, |
| chr6 | 29942982 | T | A | COSV65136627 | exonic | HLA-A | nonsynonymous SNV | HLA-A:NM_001242758:exon2:c.T299A:p.V100E,HLA-A:NM_002116:exon2:c.T299A:p.V100E, |
| chr6 | 29942984 | G | A | COSV65136637 | exonic | HLA-A | nonsynonymous SNV | HLA-A:NM_001242758:exon2:c.G301A:p.D101N,HLA-A:NM_002116:exon2:c.G301A:p.D101N, |
| chr6 | 29942990 | G | C | COSV65141826 | exonic | HLA-A | nonsynonymous SNV | HLA-A:NM_001242758:exon2:c.G307C:p.G103R,HLA-A:NM_002116:exon2:c.G307C:p.G103R, |
| chr6 | 29942994 | C | T | COSV65136647 | exonic | HLA-A | nonsynonymous SNV | HLA-A:NM_001242758:exon2:c.C311T:p.T104I,HLA-A:NM_002116:exon2:c.C311T:p.T104I, |
| chr6 | 29942996 | C | G | COSV65136655 | exonic | HLA-A | nonsynonymous SNV | HLA-A:NM_001242758:exon2:c.C313G:p.L105V,HLA-A:NM_002116:exon2:c.C313G:p.L105V, |
| chr6 | 29942997 | T | C | COSV65136680 | exonic | HLA-A | nonsynonymous SNV | HLA-A:NM_001242758:exon2:c.T314C:p.L105P,HLA-A:NM_002116:exon2:c.T314C:p.L105P, |
| chr6 | 29943000 | G | T | COSV65136690 | exonic | HLA-A | nonsynonymous SNV | HLA-A:NM_001242758:exon2:c.G317T:p.R106L,HLA-A:NM_002116:exon2:c.G317T:p.R106L, |
| chr6 | 29943002 | G | C | COSV65141194 | exonic | HLA-A | nonsynonymous SNV | HLA-A:NM_001242758:exon2:c.G319C:p.G107R,HLA-A:NM_002116:exon2:c.G319C:p.G107R, |
| chr6 | 29943024 | C | A | COSV65140734 | exonic | HLA-A | nonsynonymous SNV | HLA-A:NM_001242758:exon2:c.C341A:p.A114D,HLA-A:NM_002116:exon2:c.C341A:p.A114D, |
| chr6 | 29943279 | A | G | COSV65137506 | exonic | HLA-A | nonsynonymous SNV | HLA-A:NM_001242758:exon3:c.A355G:p.I119V,HLA-A:NM_002116:exon3:c.A355G:p.I119V, |
| chr6 | 29943287 | A | G | COSV65136818 | exonic | HLA-A | nonsynonymous SNV | HLA-A:NM_001242758:exon3:c.A363G:p.I121M,HLA-A:NM_002116:exon3:c.A363G:p.I121M, |
| chr6 | 29943292 | A | T | COSV65145084 | exonic | HLA-A | nonsynonymous SNV | HLA-A:NM_001242758:exon3:c.A368T:p.Y123F,HLA-A:NM_002116:exon3:c.A368T:p.Y123F, |
| chr6 | 29943309 | T | C | COSV65136992 | exonic | HLA-A | nonsynonymous SNV | HLA-A:NM_001242758:exon3:c.T385C:p.S129P,HLA-A:NM_002116:exon3:c.T385C:p.S129P, |
| chr6 | 29943337 | G | A | COSV65137272 | exonic | HLA-A | nonsynonymous SNV | HLA-A:NM_001242758:exon3:c.G413A:p.R138Q,HLA-A:NM_002116:exon3:c.G413A:p.R138Q, |
| chr6 | 29943342 | G | T | COSV65143116 | exonic | HLA-A | nonsynonymous SNV | HLA-A:NM_001242758:exon3:c.G418T:p.D140Y,HLA-A:NM_002116:exon3:c.G418T:p.D140Y, |

| | | | | | | | | |
|---|---|---|---|---|---|---|---|---|
| chr6 | 29943377 | C | A | COSV65137229 | exonic | HLA-A | nonsynonymous SNV | HLA-A:NM_001242758:exon3:c.C453A:p.N151K,HLA-A:NM_002116:exon3:c.C453A:p.N151K, |
| chr6 | 29943445 | C | T | COSV65137144 | exonic | HLA-A | nonsynonymous SNV | HLA-A:NM_001242758:exon3:c.C521T:p.A174V,HLA-A:NM_002116:exon3:c.C521T:p.A174V, |
| chr6 | 29943451 | A | T | COSV65136508 | exonic | HLA-A | nonsynonymous SNV | HLA-A:NM_001242758:exon3:c.A527T:p.E176V,HLA-A:NM_002116:exon3:c.A527T:p.E176V, |
| chr6 | 29943463 | T | A | COSV65136709 | exonic | HLA-A | stopgain | HLA-A:NM_001242758:exon3:c.T539A:p.L180X,HLA-A:NM_002116:exon3:c.T539A:p.L180X, |
| chr6 | 29943463 | T | G | COSV65140034 | exonic | HLA-A | nonsynonymous SNV | HLA-A:NM_001242758:exon3:c.T539G:p.L180W,HLA-A:NM_002116:exon3:c.T539G:p.L180W, |
| chr6 | 29943469 | C | T | COSV65137164 | exonic | HLA-A | nonsynonymous SNV | HLA-A:NM_001242758:exon3:c.C545T:p.A182V,HLA-A:NM_002116:exon3:c.C545T:p.A182V, |
| chr6 | 29943479 | T | G | COSV65143681 | exonic | HLA-A | nonsynonymous SNV | HLA-A:NM_001242758:exon3:c.T555G:p.D185E,HLA-A:NM_002116:exon3:c.T555G:p.D185E, |
| chr6 | 29943483 | A | C | COSV65137172 | exonic | HLA-A | nonsynonymous SNV | HLA-A:NM_001242758:exon3:c.A559C:p.T187P,HLA-A:NM_002116:exon3:c.A559C:p.T187P, |
| chr6 | 29943484 | C | G | COSV65137185 | exonic | HLA-A | nonsynonymous SNV | HLA-A:NM_001242758:exon3:c.C560G:p.T187R,HLA-A:NM_002116:exon3:c.C560G:p.T187R, |
| chr6 | 29943494 | G | C | COSV65141379 | exonic | HLA-A | nonsynonymous SNV | HLA-A:NM_001242758:exon3:c.G570C:p.E190D,HLA-A:NM_002116:exon3:c.G570C:p.E190D, |
| chr6 | 29943495 | T | G | COSV65137199 | exonic | HLA-A | nonsynonymous SNV | HLA-A:NM_001242758:exon3:c.T571G:p.W191G,HLA-A:NM_002116:exon3:c.T571G:p.W191G, |
| chr6 | 29944503 | T | C | COSV65136543 | exonic | HLA-A | nonsynonymous SNV | HLA-A:NM_001242758:exon5:c.T899C:p.L300P,HLA-A:NM_002116:exon5:c.T899C:p.L300P, |
| chr6 | 29944520 | A | G | COSV65137836 | exonic | HLA-A | nonsynonymous SNV | HLA-A:NM_001242758:exon5:c.A916G:p.I306V,HLA-A:NM_002116:exon5:c.A916G:p.I306V, |
| chr6 | 29944609 | G | C | COSV65137133 | exonic | HLA-A | nonsynonymous SNV | HLA-A:NM_001242758:exon5:c.G1005C:p.K335N,HLA-A:NM_002116:exon5:c.G1005C:p.K335N, |
| chr6 | 29945079 | A | T | COSV65136576 | exonic | HLA-A | nonsynonymous SNV | HLA-A:NM_001242758:exon6:c.A1033T:p.T345S,HLA-A:NM_002116:exon6:c.A1033T:p.T345S, |
| chr6 | 31269347 | T | C | COSV66117127 | exonic | HLA-C | nonsynonymous SNV | HLA-C:NM_002117:exon7:c.A1087G:p.T363A,HLA-C:NM_001243042:exon7:c.A1087G:p.T363A, |
| chr6 | 31269385 | C | G | COSV66115057 | exonic | HLA-C | nonsynonymous SNV | HLA-C:NM_002117:exon7:c.G1049C:p.C350S,HLA-C:NM_001243042:exon7:c.G1049C:p.C350S, |
| chr6 | 31269990 | T | C | COSV66110395 | exonic | HLA-C | nonsynonymous SNV | HLA-C:NM_002117:exon5:c.A991G:p.M331V,HLA-C:NM_001243042:exon5:c.A991G:p.M331V, |
| chr6 | 31269992 | G | A | COSV66110576 | exonic | HLA-C | nonsynonymous SNV | HLA-C:NM_002117:exon5:c.C989T:p.A330V,HLA-C:NM_001243042:exon5:c.C989T:p.A330V, |
| chr6 | 31269996 | T | C | COSV66110404 | exonic | HLA-C | nonsynonymous SNV | HLA-C:NM_002117:exon5:c.A985G:p.T329A,HLA-C:NM_001243042:exon5:c.A985G:p.T329A, |
| chr6 | 31269999 | C | T | COSV66112417 | exonic | HLA-C | nonsynonymous SNV | HLA-C:NM_002117:exon5:c.G982A:p.V328I,HLA-C:NM_001243042:exon5:c.G982A:p.V328I, |
| chr6 | 31270002 | C | T | COSV66110360 | exonic | HLA-C | nonsynonymous SNV | HLA-C:NM_002117:exon5:c.G979A:p.V327M,HLA-C:NM_001243042:exon5:c.G979A:p.V327M, |
| chr6 | 31270025 | A | G | COSV66111096 | exonic | HLA-C | nonsynonymous SNV | HLA-C:NM_002117:exon5:c.T956C:p.V319A,HLA-C:NM_001243042:exon5:c.T956C:p.V319A, |
| chr6 | 31270056 | T | C | COSV66114266 | exonic | HLA-C | nonsynonymous SNV | HLA-C:NM_002117:exon5:c.A925G:p.M309V,HLA-C:NM_001243042:exon5:c.A925G:p.M309V, |
| chr6 | 31270085 | T | C | COSV66111578 | exonic | HLA-C | nonsynonymous SNV | HLA-C:NM_002117:exon5:c.A896G:p.E299G,HLA-C:NM_001243042:exon5:c.A896G:p.E299G, |
| chr6 | 31270210 | C | T | COSV66110876 | exonic | HLA-C | nonsynonymous SNV | HLA-C:NM_002117:exon4:c.G895A:p.E299K,HLA-C:NM_001243042:exon4:c.G895A:p.E299K, |
| chr6 | 31270214 | G | T | COSV66110582 | exonic | HLA-C | nonsynonymous SNV | HLA-C:NM_002117:exon4:c.C891A:p.S297R,HLA-C:NM_001243042:exon4:c.C891A:p.S297R, |
| chr6 | 31270233 | T | G | COSV66110591 | exonic | HLA-C | nonsynonymous SNV | HLA-C:NM_002117:exon4:c.A872C:p.Q291P,HLA-C:NM_001243042:exon4:c.A872C:p.Q291P, |
| chr6 | 31270252 | T | C | COSV66110598 | exonic | HLA-C | nonsynonymous SNV | HLA-C:NM_002117:exon4:c.A853G:p.M285V,HLA-C:NM_001243042:exon4:c.A853G:p.M285V, |
| chr6 | 31270276 | G | C | COSV66110239 | exonic | HLA-C | nonsynonymous SNV | HLA-C:NM_002117:exon4:c.C829G:p.Q277E,HLA-C:NM_001243042:exon4:c.C829G:p.Q277E, |
| chr6 | 31270359 | G | T | . | exonic | HLA-C | nonsynonymous SNV | HLA-C:NM_002117:exon4:c.C746A:p.T249N,HLA-C:NM_001243042:exon4:c.C746A:p.T249N, |
| chr6 | 31270378 | G | A | COSV66117576 | exonic | HLA-C | nonsynonymous SNV | HLA-C:NM_002117:exon4:c.C727T:p.R243W,HLA-C:NM_001243042:exon4:c.C727T:p.R243W, |
| chr6 | 31270482 | G | T | COSV66109964 | exonic | HLA-C | nonsynonymous SNV | HLA-C:NM_002117:exon4:c.C623A:p.P208H,HLA-C:NM_001243042:exon4:c.C623A:p.P208H, |
| chr6 | 31271091 | C | T | COSV66110610 | exonic | HLA-C | nonsynonymous SNV | HLA-C:NM_002117:exon3:c.G601A:p.E201K,HLA-C:NM_001243042:exon3:c.G601A:p.E201K, |
| chr6 | 31271103 | C | T | COSV66112160 | exonic | HLA-C | nonsynonymous SNV | HLA-C:NM_002117:exon3:c.G589A:p.E197K,HLA-C:NM_001243042:exon3:c.G589A:p.E197K, |
| chr6 | 31271132 | G | A | COSV66110017 | exonic | HLA-C | nonsynonymous SNV | HLA-C:NM_002117:exon3:c.C560T:p.T187M,HLA-C:NM_001243042:exon3:c.C560T:p.T187M, |
| chr6 | 31271133 | T | G | COSV66110024 | exonic | HLA-C | nonsynonymous SNV | HLA-C:NM_002117:exon3:c.A559C:p.T187P,HLA-C:NM_001243042:exon3:c.A559C:p.T187P, |
| chr6 | 31271153 | A | C | COSV66109994 | exonic | HLA-C | nonsynonymous SNV | HLA-C:NM_002117:exon3:c.T539G:p.L180R,HLA-C:NM_001243042:exon3:c.T539G:p.L180R, |
| chr6 | 31271165 | G | T | COSV66110458 | exonic | HLA-C | nonsynonymous SNV | HLA-C:NM_002117:exon3:c.C527A:p.A176E,HLA-C:NM_001243042:exon3:c.C527A:p.A176E, |
| chr6 | 31271180 | A | C | COSV66110087 | exonic | HLA-C | nonsynonymous SNV | HLA-C:NM_002117:exon3:c.T512G:p.L171W,HLA-C:NM_001243042:exon3:c.T512G:p.L171W, |
| chr6 | 31271207 | G | T | COSV66110642 | exonic | HLA-C | nonsynonymous SNV | HLA-C:NM_002117:exon3:c.C485A:p.T162N,HLA-C:NM_001243042:exon3:c.C485A:p.T162N, |
| chr6 | 31271273 | G | A | COSV66110477 | exonic | HLA-C | nonsynonymous SNV | HLA-C:NM_002117:exon3:c.C419T:p.S140F,HLA-C:NM_001243042:exon3:c.C419T:p.S140F, |
| chr6 | 31271280 | C | T | COSV66110483 | exonic | HLA-C | nonsynonymous SNV | HLA-C:NM_002117:exon3:c.G412A:p.D138N,HLA-C:NM_001243042:exon3:c.G412A:p.D138N, |
| chr6 | 31271313 | G | C | COSV66114851 | exonic | HLA-C | nonsynonymous SNV | HLA-C:NM_002117:exon3:c.C379G:p.L127V,HLA-C:NM_001243042:exon3:c.C379G:p.L127V, |
| chr6 | 31271324 | G | T | COSV66110489 | exonic | HLA-C | nonsynonymous SNV | HLA-C:NM_002117:exon3:c.C368A:p.S123Y,HLA-C:NM_001243042:exon3:c.C368A:p.S123Y, |
| chr6 | 31271331 | T | A | COSV66110137 | exonic | HLA-C | nonsynonymous SNV | HLA-C:NM_002117:exon3:c.A361T:p.R121W,HLA-C:NM_001243042:exon3:c.A361T:p.R121W, |
| chr6 | 31271336 | A | C | . | exonic | HLA-C | nonsynonymous SNV | HLA-C:NM_002117:exon3:c.T356G:p.L119R,HLA-C:NM_001243042:exon3:c.T356G:p.L119R, |
| chr6 | 31271337 | G | A | . | exonic | HLA-C | nonsynonymous SNV | HLA-C:NM_002117:exon3:c.C355T:p.L119F,HLA-C:NM_001243042:exon3:c.C355T:p.L119F, |
| chr6 | 31271337 | G | T | COSV66110498 | exonic | HLA-C | nonsynonymous SNV | HLA-C:NM_002117:exon3:c.C355A:p.L119I,HLA-C:NM_001243042:exon3:c.C355A:p.L119I, |
| chr6 | 31271339 | G | A | COSV66110997 | exonic | HLA-C | nonsynonymous SNV | HLA-C:NM_002117:exon3:c.C353T:p.T118I,HLA-C:NM_001243042:exon3:c.C353T:p.T118I, |
| chr6 | 31271601 | T | G | COSV66110096 | exonic | HLA-C | nonsynonymous SNV | HLA-C:NM_002117:exon2:c.A341C:p.D114A,HLA-C:NM_001243042:exon2:c.A341C:p.D114A, |
| chr6 | 31271630 | G | T | COSV66115598 | exonic | HLA-C | nonsynonymous SNV | HLA-C:NM_002117:exon2:c.C312A:p.N104K,HLA-C:NM_001243042:exon2:c.C312A:p.N104K, |
| chr6 | 31271640 | C | T | COSV66115617 | exonic | HLA-C | nonsynonymous SNV | HLA-C:NM_002117:exon2:c.G302A:p.S101N,HLA-C:NM_001243042:exon2:c.G302A:p.S101N, |
| chr6 | 31271653 | C | T | COSV66110668 | exonic | HLA-C | nonsynonymous SNV | HLA-C:NM_002117:exon2:c.G289A:p.A97T,HLA-C:NM_001243042:exon2:c.G289A:p.A97T, |
| chr6 | 31271672 | C | G | COSV66110203 | exonic | HLA-C | nonsynonymous SNV | HLA-C:NM_002117:exon2:c.G270C:p.K90N,HLA-C:NM_001243042:exon2:c.G270C:p.K90N, |
| chr6 | 31271674 | T | G | . | exonic | HLA-C | nonsynonymous SNV | HLA-C:NM_002117:exon2:c.A268C:p.K90Q,HLA-C:NM_001243042:exon2:c.A268C:p.K90Q, |
| chr6 | 31271724 | G | T | COSV66115644 | exonic | HLA-C | nonsynonymous SNV | HLA-C:NM_002117:exon2:c.C218A:p.A73E,HLA-C:NM_001243042:exon2:c.C218A:p.A73E, |
| chr6 | 31271766 | C | T | COSV66117347 | exonic | HLA-C | nonsynonymous SNV | HLA-C:NM_002117:exon2:c.G176A:p.R59Q,HLA-C:NM_001243042:exon2:c.G176A:p.R59Q, |
| chr6 | 31271800 | A | C | COSV66110102 | exonic | HLA-C | nonsynonymous SNV | HLA-C:NM_002117:exon2:c.T142G:p.S48A,HLA-C:NM_001243042:exon2:c.T142G:p.S48A, |
| chr6 | 31271808 | C | T | COSV66110108 | exonic | HLA-C | nonsynonymous SNV | HLA-C:NM_002117:exon2:c.G134A:p.R45H,HLA-C:NM_001243042:exon2:c.G134A:p.R45H, |
| chr6 | 31271830 | G | A | COSV66111027 | exonic | HLA-C | nonsynonymous SNV | HLA-C:NM_002117:exon2:c.C112T:p.R38W,HLA-C:NM_001243042:exon2:c.C112T:p.R38W, |
| chr6 | 31271836 | C | T | COSV66110844 | exonic | HLA-C | nonsynonymous SNV | HLA-C:NM_002117:exon2:c.G106A:p.V36M,HLA-C:NM_001243042:exon2:c.G106A:p.V36M, |
| chr6 | 31271839 | C | A | COSV66109978 | exonic | HLA-C | nonsynonymous SNV | HLA-C:NM_002117:exon2:c.G103T:p.A35S,HLA-C:NM_001243042:exon2:c.G103T:p.A35S, |
| chr6 | 31271844 | T | G | COSV66111042 | exonic | HLA-C | nonsynonymous SNV | HLA-C:NM_002117:exon2:c.A98C:p.D33A,HLA-C:NM_001243042:exon2:c.A98C:p.D33A, |
| chr6 | 31271845 | C | A | COSV66119680 | exonic | HLA-C | nonsynonymous SNV | HLA-C:NM_002117:exon2:c.G97T:p.D33Y,HLA-C:NM_001243042:exon2:c.G97T:p.D33Y, |
| chr6 | 31271999 | A | C | . | exonic | HLA-C | nonsynonymous SNV | HLA-C:NM_002117:exon1:c.T73G:p.C25G,HLA-C:NM_001243042:exon1:c.T73G:p.C25G, |
| chr6 | 31272025 | C | G | COSV66117644 | exonic | HLA-C | nonsynonymous SNV | HLA-C:NM_002117:exon1:c.G47C:p.G16A,HLA-C:NM_001243042:exon1:c.G47C:p.G16A, |
| chr6 | 31272044 | G | T | COSV66117654 | exonic | HLA-C | nonsynonymous SNV | HLA-C:NM_002117:exon1:c.C28A:p.L10I,HLA-C:NM_001243042:exon1:c.C28A:p.L10I, |
| chr6 | 31272050 | C | T | COSV66117662 | exonic | HLA-C | nonsynonymous SNV | HLA-C:NM_002117:exon1:c.G22A:p.A8T,HLA-C:NM_001243042:exon1:c.G22A:p.A8T, |
| chr6 | 31896770 | G | T | . | exonic | EHMT2 | nonsynonymous SNV | EHMT2:NM_001289413:exon2:c.C335A:p.T112N,EHMT2:NM_025256:exon3:c.C164A:p.T55N,EHMT2:NM_006709:exon3:c.C164A:p.T55N, |

| | | | | | | | | |
|---|---|---|---|---|---|---|---|---|
| chr6 | 32976408 | G | A | . | exonic | BRD2 | nonsynonymous SNV | BRD2:NM_001113182:exon6:c.G769A:p.A257T,BRD2:NM_005104:exon6:c.G769A:p.A257T,BRD2:NM_001291986:exon7:c.G409A:p.A137T,BRD2:NM_001199456:exon5:c.G628A:p.A210T,BRD2:NM_001199455:exon5:c.G769A:p.A257T, |
| chr6 | 41936044 | A | C | COSV57827647 | exonic | CCND3 | nonsynonymous SNV | CCND3:NM_001760:exon5:c.T775G:p.S259A,CCND3:NM_001136017:exon5:c.T532G:p.S178A,CCND3:NM_001287434:exon5:c.T187G:p.S63A,CCND3:NM_001136126:exon4:c.T187G:p.S63A,CCND3:NM_001136125:exon4:c.T559G:p.S187A,CCND3:NM_001287427:exon5:c.T625G:p.S209A, |
| chr6 | 75715572 | A | G | . | exonic | SENP6 | nonsynonymous SNV | SENP6:NM_015571:exon24:c.A3317G:p.Y1106C,SENP6:NM_001100409:exon23:c.A3296G:p.Y1099C, |
| chr6 | 99545173 | C | T | . | exonic | CCNC | nonsynonymous SNV | CCNC:NM_005190:exon11:c.G736A:p.E246K,CCNC:NM_001013399:exon11:c.G481A:p.E161K, |
| chr7 | 102274250 | G | A | COSV52922844 | exonic | CUX1 | nonsynonymous SNV | CUX1:NM_001202546:exon15:c.G1273A:p.A425T,CUX1:NM_181500:exon16:c.G1384A:p.A462T,CUX1:NM_001202545:exon15:c.G1252A:p.A418T,CUX1:NM_001202544:exon15:c.G1342A:p.A448T,CUX1:NM_001913:exon16:c.G1390A:p.A464T, |
| chr7 | 102278021 | A | C | . | exonic | CUX1 | nonsynonymous SNV | CUX1:NM_001202546:exon17:c.A1519C:p.K507Q,CUX1:NM_181500:exon18:c.A1630C:p.K544Q,CUX1:NM_001202545:exon17:c.A1498C:p.K500Q,CUX1:NM_001202544:exon17:c.A1588C:p.K530Q,CUX1:NM_001913:exon18:c.A1636C:p.K546Q, |
| chr7 | 102283050 | A | G | . | exonic | CUX1 | nonsynonymous SNV | CUX1:NM_001202546:exon22:c.A1880G:p.E627G,CUX1:NM_181500:exon23:c.A1991G:p.E664G,CUX1:NM_001202545:exon22:c.A1859G:p.E620G,CUX1:NM_001202544:exon22:c.A1949G:p.E650G,CUX1:NM_001913:exon23:c.A1997G:p.E666G, |
| chr7 | 102283059 | A | G | . | exonic | CUX1 | nonsynonymous SNV | CUX1:NM_001202546:exon22:c.A1889G:p.N630S,CUX1:NM_181500:exon23:c.A2000G:p.N667S,CUX1:NM_001202545:exon22:c.A1868G:p.N623S,CUX1:NM_001202544:exon22:c.A1958G:p.N653S,CUX1:NM_001913:exon23:c.A2006G:p.N669S, |
| chr7 | 105062265 | G | T | . | exonic | KMT2E | nonsynonymous SNV | KMT2E:NM_182931:exon4:c.G173T:p.G58V,KMT2E:NM_018682:exon3:c.G173T:p.G58V, |
| chr7 | 105090113 | G | A | . | exonic | KMT2E | nonsynonymous SNV | KMT2E:NM_018682:exon13:c.G1463A:p.R488K,KMT2E:NM_182931:exon14:c.G1463A:p.R488K, |
| chr7 | 105090116 | G | A | COSV57576667 | exonic | KMT2E | nonsynonymous SNV | KMT2E:NM_018682:exon13:c.G1466A:p.R489Q,KMT2E:NM_182931:exon14:c.G1466A:p.R489Q, |
| chr7 | 105090133 | G | A | . | exonic | KMT2E | nonsynonymous SNV | KMT2E:NM_018682:exon13:c.G1483A:p.D495N,KMT2E:NM_182931:exon14:c.G1483A:p.D495N, |
| chr7 | 105091303 | A | T | . | exonic | KMT2E | stopgain | KMT2E:NM_018682:exon14:c.A1711T:p.K571X,KMT2E:NM_182931:exon15:c.A1711T:p.K571X, |
| chr7 | 127686719 | G | A | . | exonic | SND1 | nonsynonymous SNV | SND1:NM_014390:exon2:c.G185A:p.R62Q, |
| chr7 | 128074541 | T | C | . | exonic | SND1 | nonsynonymous SNV | SND1:NM_014390:exon17:c.T1819C:p.F607L, |
| chr7 | 128081366 | G | A | . | exonic | SND1 | nonsynonymous SNV | SND1:NM_014390:exon18:c.G1975A:p.A659T, |
| chr7 | 128081388 | T | C | . | exonic | SND1 | nonsynonymous SNV | SND1:NM_014390:exon18:c.T1997C:p.V666A, |
| chr7 | 128081405 | G | A | . | exonic | SND1 | nonsynonymous SNV | SND1:NM_014390:exon18:c.G2014A:p.V672M, |
| chr7 | 128084805 | G | A | . | exonic | SND1 | nonsynonymous SNV | SND1:NM_014390:exon19:c.G2192A:p.R731H, |
| chr7 | 128086980 | G | A | . | exonic | SND1 | nonsynonymous SNV | SND1:NM_014390:exon21:c.G2347A:p.A783T, |
| chr7 | 128089572 | T | G | . | exonic | SND1 | nonsynonymous SNV | SND1:NM_014390:exon22:c.T2502G:p.S834R, |
| chr7 | 134005051 | G | A | . | exonic | EXOC4 | nonsynonymous SNV | EXOC4:NM_021807:exon16:c.G2488A:p.A830T, |
| chr7 | 134933201 | T | A | . | exonic | CALD1 | nonsynonymous SNV | CALD1:NM_033157:exon5:c.T432A:p.N144K,CALD1:NM_033139:exon3:c.T414A:p.N138K,CALD1:NM_004342:exon5:c.T432A:p.N144K,CALD1:NM_033140:exon3:c.T414A:p.N138K,CALD1:NM_033138:exon5:c.T432A:p.N144K, |
| chr7 | 140098970 | A | T | . | exonic | KDM7A | nonsynonymous SNV | KDM7A:NM_030647:exon14:c.T1827A:p.D609E, |
| chr7 | 140459051 | C | G | COSV55436481 | exonic | MKRN1 | nonsynonymous SNV | MKRN1:NM_001291663:exon6:c.G535C:p.V179L,MKRN1:NM_013446:exon4:c.G727C:p.V243L,MKRN1:NM_001145125:exon4:c.G727C:p.V243L, |
| chr7 | 140753336 | A | T | COSV56056643 | exonic | BRAF | nonsynonymous SNV | BRAF:NM_004333:exon15:c.T1799A:p.V600E, |
| chr7 | 148828812 | C | G | COSV57449162 | exonic | EZH2 | nonsynonymous SNV | EZH2:NM_001203249:exon6:c.G526C:p.D176H,EZH2:NM_004456:exon6:c.G553C:p.D185H,EZH2:NM_001203247:exon6:c.G553C:p.D185H,EZH2:NM_001203248:exon6:c.G526C:p.D176H,EZH2:NM_152998:exon5:c.G436C:p.D146H, |
| chr7 | 150792139 | C | T | . | exonic | TMEM176B | nonsynonymous SNV | TMEM176B:NM_014020:exon6:c.G637A:p.V213I,TMEM176B:NM_001101312:exon6:c.G637A:p.V213I,TMEM176B:NM_001101311:exon6:c.G637A:p.V213I,TMEM176B:NM_001101314:exon5:c.G526A:p.V176I, |
| chr7 | 150793105 | A | G | . | exonic | TMEM176B | nonsynonymous SNV | TMEM176B:NM_014020:exon5:c.T583C:p.Y195H,TMEM176B:NM_001101312:exon5:c.T583C:p.Y195H,TMEM176B:NM_001101311:exon5:c.T583C:p.Y195H,TMEM176B:NM_001101314:exon4:c.T472C:p.Y158H, |
| chr7 | 150793191 | C | T | COSV58441166 | exonic | TMEM176B | nonsynonymous SNV | TMEM176B:NM_014020:exon5:c.G497A:p.R166H,TMEM176B:NM_001101312:exon5:c.G497A:p.R166H,TMEM176B:NM_001101311:exon5:c.G497A:p.R166H,TMEM176B:NM_001101314:exon4:c.G386A:p.R129H, |
| chr7 | 150793200 | A | C | . | exonic | TMEM176B | nonsynonymous SNV | TMEM176B:NM_014020:exon5:c.T488G:p.V163G,TMEM176B:NM_001101312:exon5:c.T488G:p.V163G,TMEM176B:NM_001101311:exon5:c.T488G:p.V163G,TMEM176B:NM_001101314:exon4:c.T377G:p.V126G, |
| chr7 | 150793219 | A | G | . | exonic | TMEM176B | nonsynonymous SNV | TMEM176B:NM_014020:exon5:c.T469C:p.F157L,TMEM176B:NM_001101312:exon5:c.T469C:p.F157L,TMEM176B:NM_001101311:exon5:c.T469C:p.F157L,TMEM176B:NM_001101314:exon4:c.T358C:p.F120L, |
| chr7 | 150793249 | C | T | COSV58439227 | exonic | TMEM176B | nonsynonymous SNV | TMEM176B:NM_014020:exon5:c.G439A:p.V147M,TMEM176B:NM_001101312:exon5:c.G439A:p.V147M,TMEM176B:NM_001101311:exon5:c.G439A:p.V147M,TMEM176B:NM_001101314:exon4:c.G328A:p.V110M, |

| | | | | | | | | |
|---|---|---|---|---|---|---|---|---|
| chr7 | 150793260 | A | G | . | exonic | TMEM176B | nonsynonymous SNV | TMEM176B:NM_014020:exon5:c.T428C:p.V143A,TMEM176B:NM_001101312:exon5:c.T428C:p.V143A,TMEM176B:NM_001101311:exon5:c.T428C:p.V143A,TMEM176B:NM_001101314:exon4:c.T317C:p.V106A, |
| chr7 | 150793288 | C | T | COSV58438904 | exonic | TMEM176B | nonsynonymous SNV | TMEM176B:NM_014020:exon5:c.G400A:p.A134T,TMEM176B:NM_001101312:exon5:c.G400A:p.A134T,TMEM176B:NM_001101311:exon5:c.G400A:p.A134T,TMEM176B:NM_001101314:exon4:c.G289A:p.A97T, |
| chr7 | 150793302 | C | T | . | exonic | TMEM176B | nonsynonymous SNV | TMEM176B:NM_014020:exon5:c.G386A:p.S129N,TMEM176B:NM_001101312:exon5:c.G386A:p.S129N,TMEM176B:NM_001101311:exon5:c.G386A:p.S129N,TMEM176B:NM_001101314:exon4:c.G275A:p.S92N, |
| chr7 | 150793587 | C | T | . | exonic | TMEM176B | nonsynonymous SNV | TMEM176B:NM_014020:exon4:c.G329A:p.G110E,TMEM176B:NM_001101312:exon4:c.G329A:p.G110E,TMEM176B:NM_001101311:exon4:c.G329A:p.G110E,TMEM176B:NM_001101314:exon3:c.G218A:p.G73E, |
| chr7 | 150793996 | T | G | COSV58439821 | exonic | TMEM176B | nonsynonymous SNV | TMEM176B:NM_014020:exon3:c.A280C:p.S94R,TMEM176B:NM_001101312:exon3:c.A280C:p.S94R,TMEM176B:NM_001101311:exon3:c.A280C:p.S94R, |
| chr7 | 150796412 | G | C | . | exonic | TMEM176B | nonsynonymous SNV | TMEM176B:NM_014020:exon2:c.C158G:p.T53S,TMEM176B:NM_001101312:exon2:c.C158G:p.T53S,TMEM176B:NM_001101311:exon2:c.C158G:p.T53S,TMEM176B:NM_001101314:exon2:c.C158G:p.T53S, |
| chr7 | 150796518 | A | G | . | exonic | TMEM176B | nonsynonymous SNV | TMEM176B:NM_014020:exon2:c.T52C:p.S18P,TMEM176B:NM_001101312:exon2:c.T52C:p.S18P,TMEM176B:NM_001101311:exon2:c.T52C:p.S18P,TMEM176B:NM_001101314:exon2:c.T52C:p.S18P, |
| chr7 | 26192560 | C | T | . | exonic | HNRNPA2B1 | nonsynonymous SNV | HNRNPA2B1:NM_002137:exon10:c.G982A:p.G328S,HNRNPA2B1:NM_031243:exon11:c.G1018A:p.G340S, |
| chr7 | 26193286 | C | T | . | exonic | HNRNPA2B1 | nonsynonymous SNV | HNRNPA2B1:NM_002137:exon9:c.G929A:p.G310D,HNRNPA2B1:NM_031243:exon10:c.G965A:p.G322D, |
| chr7 | 26193298 | C | T | . | exonic | HNRNPA2B1 | nonsynonymous SNV | HNRNPA2B1:NM_002137:exon9:c.G917A:p.S306N,HNRNPA2B1:NM_031243:exon10:c.G953A:p.S318N, |
| chr7 | 26193611 | C | T | . | exonic | HNRNPA2B1 | nonsynonymous SNV | HNRNPA2B1:NM_002137:exon8:c.G805A:p.G269R,HNRNPA2B1:NM_031243:exon9:c.G841A:p.G281R, |
| chr7 | 26193623 | C | T | . | exonic | HNRNPA2B1 | nonsynonymous SNV | HNRNPA2B1:NM_002137:exon8:c.G793A:p.G265S,HNRNPA2B1:NM_031243:exon9:c.G829A:p.G277S, |
| chr7 | 26193629 | C | T | . | exonic | HNRNPA2B1 | nonsynonymous SNV | HNRNPA2B1:NM_002137:exon8:c.G787A:p.G263R,HNRNPA2B1:NM_031243:exon9:c.G823A:p.G275R, |
| chr7 | 26193649 | C | A | . | exonic | HNRNPA2B1 | nonsynonymous SNV | HNRNPA2B1:NM_002137:exon8:c.G767T:p.G256V,HNRNPA2B1:NM_031243:exon9:c.G803T:p.G268V, |
| chr7 | 26196437 | C | T | . | exonic | HNRNPA2B1 | nonsynonymous SNV | HNRNPA2B1:NM_002137:exon6:c.G622A:p.G208R,HNRNPA2B1:NM_031243:exon7:c.G658A:p.G220R, |
| chr7 | 26196849 | A | G | . | exonic | HNRNPA2B1 | nonsynonymous SNV | HNRNPA2B1:NM_002137:exon4:c.T433C:p.F145L,HNRNPA2B1:NM_031243:exon5:c.T469C:p.F157L, |
| chr7 | 26196870 | C | T | . | exonic | HNRNPA2B1 | nonsynonymous SNV | HNRNPA2B1:NM_002137:exon4:c.G412A:p.G138R,HNRNPA2B1:NM_031243:exon5:c.G448A:p.G150R, |

| | | | | | | | | |
|---|---|---|---|---|---|---|---|---|
| chr7 | 26196903 | C | T | . | exonic | HNRNPA2B1 | nonsynonymous SNV | HNRNPA2B1:NM_002137:exon4:c.G379A:p.D127N,HNRNPA2B1:NM_031243:exon5:c.G415A:p.D139N, |
| chr7 | 26197017 | C | T | . | exonic | HNRNPA2B1 | nonsynonymous SNV | HNRNPA2B1:NM_002137:exon4:c.G265A:p.E89K,HNRNPA2B1:NM_031243:exon5:c.G301A:p.E101K, |
| chr7 | 27538413 | G | T | . | exonic | HIBADH | nonsynonymous SNV | HIBADH:NM_152740:exon6:c.C623A:p.A208E, |
| chr7 | 27792169 | T | A | . | exonic | TAX1BP1 | nonsynonymous SNV | TAX1BP1:NM_006024:exon9:c.T1202A:p.V401E,TAX1BP1:NM_001206901:exon9:c.T1202A:p.V401E,TAX1BP1:NM_001206902:exon8:c.T731A:p.V244E,TAX1BP1:NM_001079864:exon9:c.T1202A:p.V401E, |
| chr7 | 27800028 | G | A | . | exonic | TAX1BP1 | nonsynonymous SNV | TAX1BP1:NM_006024:exon13:c.G1702A:p.E568K,TAX1BP1:NM_001206901:exon13:c.G1702A:p.E568K,TAX1BP1:NM_001206902:exon12:c.G1231A:p.E411K,TAX1BP1:NM_001079864:exon13:c.G1702A:p.E568K, |
| chr7 | 27828701 | G | A | . | exonic | TAX1BP1 | nonsynonymous SNV | TAX1BP1:NM_006024:exon17:c.G2242A:p.E748K,TAX1BP1:NM_001206901:exon17:c.G2116A:p.E706K,TAX1BP1:NM_001206902:exon16:c.G1645A:p.E549K,TAX1BP1:NM_001079864:exon17:c.G2116A:p.E706K, |
| chr7 | 27828706 | T | G | . | exonic | TAX1BP1 | nonsynonymous SNV | TAX1BP1:NM_006024:exon17:c.T2247G:p.H749Q,TAX1BP1:NM_001079864:exon17:c.T2121G:p.H707Q,TAX1BP1:NM_001206901:exon17:c.T2121G:p.H707Q,TAX1BP1:NM_001206902:exon16:c.T1650G:p.H550Q, |
| chr7 | 27828720 | G | A | . | exonic | TAX1BP1 | stopgain | TAX1BP1:NM_006024:exon17:c.G2261A:p.W754X,TAX1BP1:NM_001206901:exon17:c.G2135A:p.W712X,TAX1BP1:NM_001206902:exon16:c.G1664A:p.W555X,TAX1BP1:NM_001079864:exon17:c.G2135A:p.W712X, |
| chr7 | 30595045 | C | G | COSV66827983 | exonic | GARS | nonsynonymous SNV | GARS:NM_002047:exon1:c.C124G:p.P42A, |
| chr7 | 35670232 | C | T | COSV60943904 | exonic | HERPUD2 | nonsynonymous SNV | HERPUD2:NM_022373:exon4:c.G322A:p.A108T, |
| chr7 | 36513048 | G | A | COSV51735862 | exonic | AOAH | nonsynonymous SNV | AOAH:NM_001177506:exon22:c.C2051T:p.P684L, |
| chr7 | 36616429 | G | C | COSV51741603 | exonic | AOAH | nonsynonymous SNV | AOAH:NM_001637:exon11:c.C797G:p.A266G,AOAH:NM_001177506:exon11:c.C797G:p.A266G,AOAH:NM_001177507:exon10:c.C701G:p.A234G, |
| chr7 | 36724067 | C | T | COSV51752564 | exonic | AOAH | nonsynonymous SNV | AOAH:NM_001637:exon1:c.G82A:p.D28N,AOAH:NM_001177506:exon1:c.G82A:p.D28N,AOAH:NM_001177507:exon1:c.G82A:p.D28N, |
| chr7 | 5725380 | C | T | . | exonic | RNF216 | nonsynonymous SNV | RNF216:NM_207111:exon8:c.G1448A:p.R483K,RNF216:NM_207116:exon8:c.G1277A:p.R426K, |
| chr7 | 5986771 | C | T | . | exonic | PMS2 | nonsynonymous SNV | PMS2:NM_000535:exon11:c.G1994A:p.R665K, |
| chr7 | 5986899 | C | T | COSV56221681 | exonic | PMS2 | nonsynonymous SNV | PMS2:NM_000535:exon11:c.G1866A:p.M622I, |
| chr7 | 5987144 | T | C | COSV56223170 | exonic | PMS2 | nonsynonymous SNV | PMS2:NM_000535:exon11:c.A1621G:p.K541E, |
| chr7 | 5987357 | G | A | COSV56220401 | exonic | PMS2 | nonsynonymous SNV | PMS2:NM_000535:exon11:c.C1408T:p.P470S, |
| chr7 | 76048186 | C | T | COSV50388126 | exonic | MDH2 | nonsynonymous SNV | MDH2:NM_001282403:exon1:c.C26T:p.A9V,MDH2:NM_005918:exon1:c.C26T:p.A9V, |
| chr7 | 900544 | C | T | COSV56214400 | exonic | ADAP1 | nonsynonymous SNV | ADAP1:NM_001284309:exon7:c.G505A:p.G169S,ADAP1:NM_006869:exon7:c.G721A:p.G241S,ADAP1:NM_001284308:exon7:c.G754A:p.G252S,ADAP1:NM_001284311:exon5:c.G436A:p.G146S,ADAP1:NM_001284310:exon6:c.G505A:p.G169S, |
| chr8 | 100704315 | C | T | . | exonic | PABPC1 | nonsynonymous SNV | PABPC1:NM_002568:exon14:c.G1894A:p.G632S, |
| chr8 | 100704992 | C | T | COSV59397438 | exonic | PABPC1 | nonsynonymous SNV | PABPC1:NM_002568:exon13:c.G1752A:p.M584I, |
| chr8 | 100704996 | C | T | COSV59400112 | exonic | PABPC1 | nonsynonymous SNV | PABPC1:NM_002568:exon13:c.G1748A:p.G583D, |
| chr8 | 100705009 | C | T | COSV59397447 | exonic | PABPC1 | nonsynonymous SNV | PABPC1:NM_002568:exon13:c.G1735A:p.G579S, |
| chr8 | 100705025 | C | T | COSV59400560 | exonic | PABPC1 | nonsynonymous SNV | PABPC1:NM_002568:exon13:c.G1719A:p.M573I, |
| chr8 | 100705050 | C | T | . | exonic | PABPC1 | nonsynonymous SNV | PABPC1:NM_002568:exon13:c.G1694A:p.R565Q, |
| chr8 | 100705051 | G | A | COSV59397476 | exonic | PABPC1 | nonsynonymous SNV | PABPC1:NM_002568:exon13:c.C1693T:p.R565W, |
| chr8 | 100705654 | C | T | . | exonic | PABPC1 | nonsynonymous SNV | PABPC1:NM_002568:exon12:c.G1622A:p.G541D, |
| chr8 | 100706700 | C | T | . | exonic | PABPC1 | nonsynonymous SNV | PABPC1:NM_002568:exon11:c.G1553A:p.R518H, |
| chr8 | 100706713 | C | T | . | exonic | PABPC1 | nonsynonymous SNV | PABPC1:NM_002568:exon11:c.G1540A:p.A514T, |
| chr8 | 100706740 | C | T | COSV59397513 | exonic | PABPC1 | nonsynonymous SNV | PABPC1:NM_002568:exon11:c.G1513A:p.V505I, |
| chr8 | 100706910 | C | T | COSV59397547 | exonic | PABPC1 | nonsynonymous SNV | PABPC1:NM_002568:exon10:c.G1424A:p.R475Q, |
| chr8 | 100706929 | C | T | . | exonic | PABPC1 | nonsynonymous SNV | PABPC1:NM_002568:exon10:c.G1405A:p.A469T, |
| chr8 | 100706947 | A | G | . | exonic | PABPC1 | nonsynonymous SNV | PABPC1:NM_002568:exon10:c.T1387C:p.F463L, |
| chr8 | 100709162 | C | T | . | exonic | PABPC1 | nonsynonymous SNV | PABPC1:NM_002568:exon9:c.G1307A:p.R436H, |
| chr8 | 100709490 | G | A | . | exonic | PABPC1 | nonsynonymous SNV | PABPC1:NM_002568:exon8:c.C1214T:p.P405L, |
| chr8 | 100709497 | C | T | . | exonic | PABPC1 | nonsynonymous SNV | PABPC1:NM_002568:exon8:c.G1207A:p.A403T, |
| chr8 | 100709573 | G | T | . | exonic | PABPC1 | nonsynonymous SNV | PABPC1:NM_002568:exon8:c.C1131A:p.H377Q, |
| chr8 | 100709578 | C | T | . | exonic | PABPC1 | nonsynonymous SNV | PABPC1:NM_002568:exon8:c.G1126A:p.A376T, |
| chr8 | 100709587 | C | T | . | exonic | PABPC1 | nonsynonymous SNV | PABPC1:NM_002568:exon8:c.G1117A:p.E373K, |

| | | | | | | | | |
|---|---|---|---|---|---|---|---|---|
| chr8 | 100709641 | C | T | . | exonic | PABPC1 | nonsynonymous SNV | PABPC1:NM_002568:exon8:c.G1063A:p.G355S, |
| chr8 | 100709667 | G | T | . | exonic | PABPC1 | nonsynonymous SNV | PABPC1:NM_002568:exon8:c.C1037A:p.A346D, |
| chr8 | 100709668 | C | T | . | exonic | PABPC1 | nonsynonymous SNV | PABPC1:NM_002568:exon8:c.G1036A:p.A346T, |
| chr8 | 100709674 | C | T | . | exonic | PABPC1 | nonsynonymous SNV | PABPC1:NM_002568:exon8:c.G1030A:p.E344K, |
| chr8 | 100709708 | G | T | . | exonic | PABPC1 | nonsynonymous SNV | PABPC1:NM_002568:exon8:c.C996A:p.S332R, |
| chr8 | 100709709 | C | T | . | exonic | PABPC1 | nonsynonymous SNV | PABPC1:NM_002568:exon8:c.G995A:p.S332N, |
| chr8 | 100712384 | A | G | . | exonic | PABPC1 | nonsynonymous SNV | PABPC1:NM_002568:exon7:c.T950C:p.F317S, |
| chr8 | 100712402 | C | T | COSV59399520 | exonic | PABPC1 | nonsynonymous SNV | PABPC1:NM_002568:exon7:c.G932A:p.R311Q, |
| chr8 | 100712405 | A | G | . | exonic | PABPC1 | nonsynonymous SNV | PABPC1:NM_002568:exon7:c.T929C:p.L310P, |
| chr8 | 100712765 | C | T | COSV59400273 | exonic | PABPC1 | nonsynonymous SNV | PABPC1:NM_002568:exon6:c.G763A:p.E255K, |
| chr8 | 100712771 | C | T | . | exonic | PABPC1 | nonsynonymous SNV | PABPC1:NM_002568:exon6:c.G757A:p.G253R, |
| chr8 | 100713101 | C | T | . | exonic | PABPC1 | nonsynonymous SNV | PABPC1:NM_002568:exon5:c.G724A:p.E242K, |
| chr8 | 100713145 | C | T | . | exonic | PABPC1 | nonsynonymous SNV | PABPC1:NM_002568:exon5:c.G680A:p.S227N, |
| chr8 | 100713148 | T | C | . | exonic | PABPC1 | nonsynonymous SNV | PABPC1:NM_002568:exon5:c.A677G:p.E226G, |
| chr8 | 100715467 | T | C | . | exonic | PABPC1 | nonsynonymous SNV | PABPC1:NM_002568:exon4:c.A638G:p.K213R, |
| chr8 | 100715488 | C | T | COSV59397665 | exonic | PABPC1 | nonsynonymous SNV | PABPC1:NM_002568:exon4:c.G617A:p.R206H, |
| chr8 | 100715492 | C | T | . | exonic | PABPC1 | nonsynonymous SNV | PABPC1:NM_002568:exon4:c.G613A:p.E205K, |
| chr8 | 100715549 | T | C | . | exonic | PABPC1 | nonsynonymous SNV | PABPC1:NM_002568:exon4:c.A556G:p.R186G, |
| chr8 | 100715564 | C | T | COSV59397693 | exonic | PABPC1 | nonsynonymous SNV | PABPC1:NM_002568:exon4:c.G541A:p.A181T, |
| chr8 | 100715573 | C | T | COSV59397705 | exonic | PABPC1 | nonsynonymous SNV | PABPC1:NM_002568:exon4:c.G532A:p.E178K, |
| chr8 | 100717798 | C | T | . | exonic | PABPC1 | nonsynonymous SNV | PABPC1:NM_002568:exon3:c.G478A:p.G160R, |
| chr8 | 100717822 | C | T | COSV59399238 | exonic | PABPC1 | nonsynonymous SNV | PABPC1:NM_002568:exon3:c.G454A:p.E152K, |
| chr8 | 100717827 | G | A | . | exonic | PABPC1 | nonsynonymous SNV | PABPC1:NM_002568:exon3:c.C449T:p.A150V, |
| chr8 | 100717831 | C | T | . | exonic | PABPC1 | nonsynonymous SNV | PABPC1:NM_002568:exon3:c.G445A:p.E149K, |
| chr8 | 100717885 | C | A | . | exonic | PABPC1 | nonsynonymous SNV | PABPC1:NM_002568:exon3:c.G391T:p.V131F, |
| chr8 | 100721459 | A | G | . | exonic | PABPC1 | nonsynonymous SNV | PABPC1:NM_002568:exon1:c.T125C:p.V42A, |
| chr8 | 102305099 | G | T | . | exonic | UBR5 | nonsynonymous SNV | UBR5:NM_001282873:exon21:c.C2813A:p.T938N,UBR5:NM_015902:exon21:c.C2813A:p.T938N, |
| chr8 | 103404817 | C | T | . | exonic | SLC25A32 | nonsynonymous SNV | SLC25A32:NM_030780:exon3:c.G350A:p.R117H, |
| chr8 | 109339385 | G | A | . | exonic | ENY2 | nonsynonymous SNV | ENY2:NM_001193557:exon3:c.G134A:p.C45Y,ENY2:NM_020189:exon3:c.G149A:p.C50Y, |
| chr8 | 109340542 | G | A | . | exonic | ENY2 | nonsynonymous SNV | ENY2:NM_001193557:exon4:c.G193A:p.E65K,ENY2:NM_020189:exon4:c.G208A:p.E70K, |
| chr8 | 11802853 | T | C | COSV55043146 | exonic | FDFT1 | nonsynonymous SNV | FDFT1:NM_001287756:exon1:c.T7C:p.W3R, |
| chr8 | 11808849 | G | A | . | exonic | FDFT1 | nonsynonymous SNV | FDFT1:NM_004462:exon2:c.G155A:p.R52H,FDFT1:NM_001287743:exon3:c.G155A:p.R52H,FDFT1:NM_001287750:exon1:c.G332A:p.R111H,FDFT1:NM_001287742:exon4:c.G155A:p.R52H, |
| chr8 | 11809701 | G | A | . | exonic | FDFT1 | nonsynonymous SNV | FDFT1:NM_004462:exon3:c.G232A:p.A78T,FDFT1:NM_001287743:exon4:c.G232A:p.A78T,FDFT1:NM_001287750:exon2:c.G409A:p.A137T,FDFT1:NM_001287742:exon5:c.G232A:p.A78T,FDFT1:NM_001287747:exon3:c.G40A:p.A14T,FDFT1:NM_001287745:exon4:c.G40A:p.A14T,FDFT1:NM_001287744:exon3:c.G40A:p.A14T,FDFT1:NM_001287749:exon3:c.G40A:p.A14T,FDFT1:NM_001287748:exon3:c.G40A:p.A14T, |
| chr8 | 11809707 | G | A | . | exonic | FDFT1 | nonsynonymous SNV | FDFT1:NM_004462:exon3:c.G238A:p.D80N,FDFT1:NM_001287743:exon4:c.G238A:p.D80N,FDFT1:NM_001287750:exon2:c.G415A:p.D139N,FDFT1:NM_001287742:exon5:c.G238A:p.D80N,FDFT1:NM_001287747:exon3:c.G46A:p.D16N,FDFT1:NM_001287745:exon4:c.G46A:p.D16N,FDFT1:NM_001287744:exon3:c.G46A:p.D16N,FDFT1:NM_001287749:exon3:c.G46A:p.D16N,FDFT1:NM_001287748:exon3:c.G46A:p.D16N, |
| chr8 | 11821811 | G | A | COSV55042913 | exonic | FDFT1 | nonsynonymous SNV | FDFT1:NM_004462:exon4:c.G443A:p.R148Q,FDFT1:NM_001287751:exon3:c.G188A:p.R63Q,FDFT1:NM_001287743:exon5:c.G443A:p.R148Q,FDFT1:NM_001287750:exon3:c.G620A:p.R207Q,FDFT1:NM_001287742:exon6:c.G443A:p.R148Q,FDFT1:NM_001287747:exon4:c.G251A:p.R84Q,FDFT1:NM_001287745:exon5:c.G251A:p.R84Q,FDFT1:NM_001287744:exon4:c.G251A:p.R84Q,FDFT1:NM_001287749:exon4:c.G251A:p.R84Q,FDFT1:NM_001287748:exon4:c.G251A:p.R84Q, |
| chr8 | 11830358 | G | A | . | exonic | FDFT1 | nonsynonymous SNV | FDFT1:NM_004462:exon6:c.G817A:p.V273I,FDFT1:NM_001287751:exon5:c.G562A:p.V188I,FDFT1:NM_001287743:exon7:c.G817A:p.V273I,FDFT1:NM_001287750:exon5:c.G994A:p.V332I,FDFT1:NM_001287742:exon8:c.G817A:p.V273I,FDFT1:NM_001287747:exon6:c.G625A:p.V209I,FDFT1:NM_001287745:exon7:c.G625A:p.V209I,FDFT1:NM_001287744:exon6:c.G625A:p.V209I,FDFT1:NM_001287756:exon3:c.G316A:p.V106I,FDFT1:NM_001287749:exon6:c.G625A:p.V209I,FDFT1:NM_001287748:exon6:c.G625A:p.V209I, |
| chr8 | 11831620 | G | A | . | exonic | FDFT1 | nonsynonymous SNV | FDFT1:NM_004462:exon7:c.G982A:p.A328T,FDFT1:NM_001287750:exon6:c.G1159A:p.A387T,FDFT1:NM_001287742:exon9:c.G982A:p.A328T,FDFT1:NM_001287744:exon7:c.G790A:p.A264T,FDFT1:NM_001287756:exon4:c.G481A:p.A161T,FDFT1:NM_001287749:exon7:c.G790A:p.A264T,FDFT1:NM_001287751:exon6:c.G727A:p.A243T,FDFT1:NM_001287743:exon8:c.G982A:p.A328T,FDFT1:NM_001287747:exon7:c.G790A:p.A264T,FDFT1:NM_001287745:exon8:c.G790A:p.A264T,FDFT1:NM_001287748:exon7:c.G790A:p.A264T, |
| chr8 | 11838530 | T | C | . | exonic | FDFT1 | nonsynonymous SNV | FDFT1:NM_004462:exon8:c.T1175C:p.L392P,FDFT1:NM_001287751:exon7:c.T920C:p.L307P,FDFT1:NM_001287743:exon9:c.T1175C:p.L392P,FDFT1:NM_001287750:exon7:c.T1352C:p.L451P,FDFT1:NM_001287742:exon10:c.T1175C:p.L392P,FDFT1:NM_001287747:exon8:c.T983C:p.L328P,FDFT1:NM_001287745:exon9:c.T983C:p.L328P,FDFT1:NM_001287744:exon8:c.T983C:p.L328P,FDFT1:NM_001287756:exon5:c.T674C:p.L225P,FDFT1:NM_001287749:exon8:c.T983C:p.L328P,FDFT1:NM_001287748:exon8:c.T983C:p.L328P, |

| | | | | | | | | |
|---|---|---|---|---|---|---|---|---|
| chr8 | 11852665 | T | C | . | exonic | CTSB | nonsynonymous SNV | CTSB:NM_001908:exon3:c.A157G:p.S53G,CTSB:NM_147783:exon4:c.A157G:p.S53G,CTSB:NM_147781:exon4:c.A157G:p.S53G,CTSB:NM_147780:exon5:c.A157G:p.S53G,CTSB:NM_147782:exon4:c.A157G:p.S53G, |
| chr8 | 11853379 | G | C | COSV61540513 | exonic | CTSB | nonsynonymous SNV | CTSB:NM_001908:exon2:c.C76G:p.L26V,CTSB:NM_147783:exon3:c.C76G:p.L26V,CTSB:NM_147781:exon3:c.C76G:p.L26V,CTSB:NM_147780:exon4:c.C76G:p.L26V,CTSB:NM_147782:exon3:c.C76G:p.L26V, |
| chr8 | 119583783 | A | G | . | exonic | ENPP2 | nonsynonymous SNV | ENPP2:NM_001040092:exon17:c.T1477C:p.S493P,ENPP2:NM_006209:exon18:c.T1633C:p.S545P,ENPP2:NM_001130863:exon17:c.T1477C:p.S493P, |
| chr8 | 124539205 | G | A | . | exonic | NDUFB9 | nonsynonymous SNV | NDUFB9:NM_001311168:exon1:c.G19A:p.G7R,NDUFB9:NM_005005:exon1:c.G19A:p.G7R, |
| chr8 | 124539232 | G | A | . | exonic | NDUFB9 | nonsynonymous SNV | NDUFB9:NM_001311168:exon1:c.G46A:p.V16M,NDUFB9:NM_005005:exon1:c.G46A:p.V16M, |
| chr8 | 124539242 | T | C | . | exonic | NDUFB9 | nonsynonymous SNV | NDUFB9:NM_001311168:exon1:c.T56C:p.L19P,NDUFB9:NM_005005:exon1:c.T56C:p.L19P, |
| chr8 | 124539253 | G | A | . | exonic | NDUFB9 | nonsynonymous SNV | NDUFB9:NM_001311168:exon1:c.G67A:p.A23T,NDUFB9:NM_005005:exon1:c.G67A:p.A23T, |
| chr8 | 124539280 | G | A | . | exonic | NDUFB9 | nonsynonymous SNV | NDUFB9:NM_001311168:exon1:c.G94A:p.V32I,NDUFB9:NM_005005:exon1:c.G94A:p.V32I, |
| chr8 | 124543098 | G | A | . | exonic | NDUFB9 | nonsynonymous SNV | NDUFB9:NM_005005:exon2:c.G113A:p.R38Q, |
| chr8 | 124543106 | G | A | . | exonic | NDUFB9 | nonsynonymous SNV | NDUFB9:NM_005005:exon2:c.G121A:p.A41T, |
| chr8 | 124543156 | G | A | . | exonic | NDUFB9 | nonsynonymous SNV | NDUFB9:NM_001278646:exon2:c.G42A:p.M14I,NDUFB9:NM_001278645:exon2:c.G3A:p.M1I,NDUFB9:NM_001311168:exon2:c.G138A:p.M46I,NDUFB9:NM_005005:exon2:c.G171A:p.M57I, |
| chr8 | 124543163 | G | A | . | exonic | NDUFB9 | nonsynonymous SNV | NDUFB9:NM_001278646:exon2:c.G49A:p.A17T,NDUFB9:NM_001311168:exon2:c.G145A:p.A49T,NDUFB9:NM_001278645:exon2:c.G10A:p.A4T,NDUFB9:NM_005005:exon2:c.G178A:p.A60T, |
| chr8 | 124543173 | T | C | . | exonic | NDUFB9 | nonsynonymous SNV | NDUFB9:NM_001278646:exon2:c.T59C:p.L20P,NDUFB9:NM_001278645:exon2:c.T20C:p.L7P,NDUFB9:NM_001311168:exon2:c.T155C:p.L52P,NDUFB9:NM_005005:exon2:c.T188C:p.L63P, |
| chr8 | 124543206 | G | A | . | exonic | NDUFB9 | nonsynonymous SNV | NDUFB9:NM_001278646:exon2:c.G92A:p.R31H,NDUFB9:NM_001278645:exon2:c.G53A:p.R18H,NDUFB9:NM_001311168:exon2:c.G188A:p.R63H,NDUFB9:NM_005005:exon2:c.G221A:p.R74H, |
| chr8 | 124543248 | G | A | . | exonic | NDUFB9 | nonsynonymous SNV | NDUFB9:NM_001278646:exon2:c.G134A:p.G45D,NDUFB9:NM_001311168:exon2:c.G230A:p.G77D,NDUFB9:NM_001278645:exon2:c.G95A:p.G32D,NDUFB9:NM_005005:exon2:c.G263A:p.G88D, |
| chr8 | 124543268 | G | A | . | exonic | NDUFB9 | nonsynonymous SNV | NDUFB9:NM_001278646:exon2:c.G154A:p.D52N,NDUFB9:NM_001278645:exon2:c.G115A:p.D39N,NDUFB9:NM_001311168:exon2:c.G250A:p.D84N,NDUFB9:NM_005005:exon2:c.G283A:p.D95N, |
| chr8 | 124547042 | G | A | . | exonic | NDUFB9 | nonsynonymous SNV | NDUFB9:NM_001278646:exon3:c.G208A:p.A70T,NDUFB9:NM_001278645:exon3:c.G169A:p.A57T,NDUFB9:NM_001311168:exon3:c.G304A:p.A102T,NDUFB9:NM_005005:exon3:c.G337A:p.A113T, |
| chr8 | 124547080 | G | A | . | exonic | NDUFB9 | stopgain | NDUFB9:NM_001278646:exon3:c.G246A:p.W82X,NDUFB9:NM_001278645:exon3:c.G207A:p.W69X,NDUFB9:NM_001311168:exon3:c.G342A:p.W114X,NDUFB9:NM_005005:exon3:c.G375A:p.W125X, |
| chr8 | 124547096 | G | A | . | exonic | NDUFB9 | nonsynonymous SNV | NDUFB9:NM_001278646:exon3:c.G262A:p.E88K,NDUFB9:NM_001278645:exon3:c.G223A:p.E75K,NDUFB9:NM_001311168:exon3:c.G358A:p.E120K,NDUFB9:NM_005005:exon3:c.G391A:p.E131K, |
| chr8 | 124547109 | G | A | . | exonic | NDUFB9 | nonsynonymous SNV | NDUFB9:NM_001278646:exon3:c.G275A:p.R92Q,NDUFB9:NM_001278645:exon3:c.G236A:p.R79Q,NDUFB9:NM_001311168:exon3:c.G371A:p.R124Q,NDUFB9:NM_005005:exon3:c.G404A:p.R135Q, |
| chr8 | 124549776 | G | A | . | exonic | NDUFB9 | nonsynonymous SNV | NDUFB9:NM_001278646:exon4:c.G295A:p.E99K,NDUFB9:NM_001278645:exon4:c.G256A:p.E86K,NDUFB9:NM_001311168:exon4:c.G391A:p.E131K,NDUFB9:NM_005005:exon4:c.G424A:p.E142K, |
| chr8 | 124549788 | C | T | . | exonic | NDUFB9 | nonsynonymous SNV | NDUFB9:NM_001278646:exon4:c.C307T:p.P103S,NDUFB9:NM_001278645:exon4:c.C268T:p.P90S,NDUFB9:NM_001311168:exon4:c.C403T:p.P135S,NDUFB9:NM_005005:exon4:c.C436T:p.P146S, |
| chr8 | 124549792 | G | A | . | exonic | NDUFB9 | nonsynonymous SNV | NDUFB9:NM_001278646:exon4:c.G311A:p.G104D,NDUFB9:NM_001278645:exon4:c.G272A:p.G91D,NDUFB9:NM_001311168:exon4:c.G407A:p.G136D,NDUFB9:NM_005005:exon4:c.G440A:p.G147D, |
| chr8 | 124549809 | G | A | . | exonic | NDUFB9 | nonsynonymous SNV | NDUFB9:NM_001278646:exon4:c.G328A:p.A110T,NDUFB9:NM_001278645:exon4:c.G289A:p.A97T,NDUFB9:NM_001311168:exon4:c.G424A:p.A142T,NDUFB9:NM_005005:exon4:c.G457A:p.A153T, |
| chr8 | 124549834 | G | A | . | exonic | NDUFB9 | nonsynonymous SNV | NDUFB9:NM_001278646:exon4:c.G353A:p.G118D,NDUFB9:NM_001278645:exon4:c.G314A:p.G105D,NDUFB9:NM_001311168:exon4:c.G449A:p.G150D,NDUFB9:NM_005005:exon4:c.G482A:p.G161D, |
| chr8 | 124549842 | C | T | . | exonic | NDUFB9 | nonsynonymous SNV | NDUFB9:NM_001278646:exon4:c.C361T:p.P121S,NDUFB9:NM_001278645:exon4:c.C322T:p.P108S,NDUFB9:NM_001311168:exon4:c.C457T:p.P153S,NDUFB9:NM_005005:exon4:c.C490T:p.P164S, |
| chr8 | 124549856 | G | A | . | exonic | NDUFB9 | stopgain | NDUFB9:NM_001278646:exon4:c.G375A:p.W125X,NDUFB9:NM_001278645:exon4:c.G336A:p.W112X,NDUFB9:NM_001311168:exon4:c.G471A:p.W157X,NDUFB9:NM_005005:exon4:c.G504A:p.W168X, |
| chr8 | 124549861 | T | C | . | exonic | NDUFB9 | nonsynonymous SNV | NDUFB9:NM_001278646:exon4:c.T380C:p.I127T,NDUFB9:NM_001278645:exon4:c.T341C:p.I114T,NDUFB9:NM_001311168:exon4:c.T476C:p.I159T,NDUFB9:NM_005005:exon4:c.T509C:p.I170T, |
| chr8 | 124549882 | G | A | COSV52674143 | exonic | NDUFB9 | nonsynonymous SNV | NDUFB9:NM_001278646:exon4:c.G401A:p.R134Q,NDUFB9:NM_001278645:exon4:c.G362A:p.R121Q,NDUFB9:NM_001311168:exon4:c.G497A:p.R166Q,NDUFB9:NM_005005:exon4:c.G530A:p.R177Q, |
| chr8 | 132625413 | G | A | COSV51539603 | exonic | LRRC6 | nonsynonymous SNV | LRRC6:NM_012472:exon6:c.C695T:p.T232I, |
| chr8 | 133284281 | C | T | . | exonic | NDRG1 | nonsynonymous SNV | NDRG1:NM_001135242:exon2:c.G31A:p.A11T,NDRG1:NM_006096:exon2:c.G31A:p.A11T, |
| chr8 | 144095929 | A | G | COSV59644891 | exonic | CYC1 | nonsynonymous SNV | CYC1:NM_001916:exon2:c.A226G:p.M76V, |
| chr8 | 18009676 | A | T | COSV59586229 | exonic | PCM1 | nonsynonymous SNV | PCM1:NM_001315508:exon30:c.A4930T:p.I1644L,PCM1:NM_006197:exon31:c.A5092T:p.I1698L,PCM1:NM_001315507:exon31:c.A5068T:p.I1690L, |
| chr8 | 26354556 | T | C | . | exonic | PPP2R2A | nonsynonymous SNV | PPP2R2A:NM_002717:exon4:c.T269C:p.L90S,PPP2R2A:NM_001177591:exon4:c.T299C:p.L100S, |
| chr8 | 27500988 | A | G | COSV66844518 | exonic | EPHX2 | nonsynonymous SNV | EPHX2:NM_001256482:exon2:c.A5G:p.K2R,EPHX2:NM_001256484:exon2:c.A5G:p.K2R,EPHX2:NM_001979:exon2:c.A164G:p.K55R, |
| chr8 | 27516348 | G | A | COSV66844144 | exonic | EPHX2 | nonsynonymous SNV | EPHX2:NM_001256482:exon8:c.G701A:p.R234Q,EPHX2:NM_001256484:exon8:c.G701A:p.R234Q,EPHX2:NM_001979:exon8:c.G860A:p.R287Q,EPHX2:NM_001256483:exon7:c.G662A:p.R221Q, |

| | | | | | | | | |
|---|---|---|---|---|---|---|---|---|
| chr8 | 31141764 | G | T | COSV53297644 | exonic | WRN | nonsynonymous SNV | WRN:NM_000553:exon26:c.G3222T:p.L1074F, |
| chr8 | 31167138 | T | C | COSV53296936 | exonic | WRN | nonsynonymous SNV | WRN:NM_000553:exon34:c.T4099C:p.C1367R, |
| chr8 | 32595840 | G | A | COSV55177787 | exonic | NRG1 | nonsynonymous SNV | NRG1:NM_001160004:exon2:c.G113A:p.R38Q,NRG1:NM_001160005:exon2:c.G113A:p.R38Q,NRG1:NM_001160002:exon2:c.G113A:p.R38Q,NRG1:NM_013962:exon2:c.G758A:p.R253Q,NRG1:NM_013957:exon2:c.G113A:p.R38Q,NRG1:NM_001160007:exon2:c.G113A:p.R38Q,NRG1:NM_013964:exon2:c.G113A:p.R38Q,NRG1:NM_013958:exon2:c.G113A:p.R38Q,NRG1:NM_004495:exon2:c.G113A:p.R38Q,NRG1:NM_013960:exon2:c.G113A:p.R38Q,NRG1:NM_001160008:exon2:c.G113A:p.R38Q,NRG1:NM_013956:exon2:c.G113A:p.R38Q, |
| chr8 | 51820489 | G | T | . | exonic | PCMTD1 | nonsynonymous SNV | PCMTD1:NM_001286783:exon6:c.C408A:p.N136K,PCMTD1:NM_052937:exon6:c.C936A:p.N312K,PCMTD1:NM_001286782:exon4:c.C708A:p.N236K, |
| chr8 | 51820490 | T | A | COSV62119484 | exonic | PCMTD1 | nonsynonymous SNV | PCMTD1:NM_001286783:exon6:c.A407T:p.N136I,PCMTD1:NM_052937:exon6:c.A935T:p.N312I,PCMTD1:NM_001286782:exon4:c.A707T:p.N236I, |
| chr8 | 51820592 | C | T | . | exonic | PCMTD1 | nonsynonymous SNV | PCMTD1:NM_001286783:exon6:c.G305A:p.R102K,PCMTD1:NM_052937:exon6:c.G833A:p.R278K,PCMTD1:NM_001286782:exon4:c.G605A:p.R202K, |
| chr8 | 51820598 | C | T | COSV62124907 | exonic | PCMTD1 | nonsynonymous SNV | PCMTD1:NM_001286783:exon6:c.G299A:p.R100K,PCMTD1:NM_052937:exon6:c.G827A:p.R276K,PCMTD1:NM_001286782:exon4:c.G599A:p.R200K, |
| chr8 | 51820604 | G | T | . | exonic | PCMTD1 | nonsynonymous SNV | PCMTD1:NM_001286783:exon6:c.C293A:p.P98H,PCMTD1:NM_052937:exon6:c.C821A:p.P274H,PCMTD1:NM_001286782:exon4:c.C593A:p.P198H, |
| chr8 | 60742412 | G | T | . | exonic | CHD7 | nonsynonymous SNV | CHD7:NM_001316690:exon1:c.G980T:p.G327V,CHD7:NM_017780:exon2:c.G980T:p.G327V, |
| chr8 | 60742756 | G | A | . | exonic | CHD7 | nonsynonymous SNV | CHD7:NM_001316690:exon1:c.G1324A:p.A442T,CHD7:NM_017780:exon2:c.G1324A:p.A442T, |
| chr8 | 60742862 | C | A | . | exonic | CHD7 | nonsynonymous SNV | CHD7:NM_001316690:exon1:c.C1430A:p.P477Q,CHD7:NM_017780:exon2:c.C1430A:p.P477Q, |
| chr8 | 60781138 | A | G | . | exonic | CHD7 | nonsynonymous SNV | CHD7:NM_017780:exon3:c.A1804G:p.K602E, |
| chr8 | 60781153 | G | A | . | exonic | CHD7 | nonsynonymous SNV | CHD7:NM_017780:exon3:c.G1819A:p.V607I, |
| chr8 | 60781220 | T | A | . | exonic | CHD7 | nonsynonymous SNV | CHD7:NM_017780:exon3:c.T1886A:p.L629Q, |
| chr8 | 60781249 | G | C | . | exonic | CHD7 | nonsynonymous SNV | CHD7:NM_017780:exon3:c.G1915C:p.E639Q, |
| chr8 | 60781252 | G | A | COSV71113211 | exonic | CHD7 | nonsynonymous SNV | CHD7:NM_017780:exon3:c.G1918A:p.E640K, |
| chr8 | 60781279 | G | A | . | exonic | CHD7 | nonsynonymous SNV | CHD7:NM_017780:exon3:c.G1945A:p.A649T, |
| chr8 | 60781280 | C | T | . | exonic | CHD7 | nonsynonymous SNV | CHD7:NM_017780:exon3:c.C1946T:p.A649V, |
| chr8 | 63038678 | C | T | COSV52649507 | exonic | GGH | nonsynonymous SNV | GGH:NM_003878:exon1:c.G91A:p.A31T, |
| chr8 | 63038753 | A | G | COSV52650077 | exonic | GGH | nonsynonymous SNV | GGH:NM_003878:exon1:c.T16C:p.C6R, |
| chr8 | 85328643 | C | T | . | exonic | CA1 | nonsynonymous SNV | CA1:NM_001164830:exon7:c.G703A:p.E235K,CA1:NM_001291967:exon7:c.G505A:p.E169K,CA1:NM_001128830:exon9:c.G703A:p.E235K,CA1:NM_001128831:exon8:c.G703A:p.E235K,CA1:NM_001291968:exon6:c.G364A:p.E122K,CA1:NM_001738:exon9:c.G703A:p.E235K,CA1:NM_001128829:exon9:c.G703A:p.E235K, |
| chr8 | 85328663 | C | T | . | exonic | CA1 | nonsynonymous SNV | CA1:NM_001164830:exon7:c.G683A:p.R228H,CA1:NM_001291967:exon7:c.G485A:p.R162H,CA1:NM_001128830:exon9:c.G683A:p.R228H,CA1:NM_001128831:exon8:c.G683A:p.R228H,CA1:NM_001291968:exon6:c.G344A:p.R115H,CA1:NM_001738:exon9:c.G683A:p.R228H,CA1:NM_001128829:exon9:c.G683A:p.R228H, |
| chr8 | 85329729 | C | T | . | exonic | CA1 | stopgain | CA1:NM_001164830:exon6:c.G629A:p.W210X,CA1:NM_001291967:exon6:c.G431A:p.W144X,CA1:NM_001128830:exon8:c.G629A:p.W210X,CA1:NM_001128831:exon7:c.G629A:p.W210X,CA1:NM_001291968:exon5:c.G290A:p.W97X,CA1:NM_001738:exon8:c.G629A:p.W210X,CA1:NM_001128829:exon8:c.G629A:p.W210X, |
| chr8 | 85329736 | C | T | . | exonic | CA1 | nonsynonymous SNV | CA1:NM_001164830:exon6:c.G622A:p.V208I,CA1:NM_001291967:exon6:c.G424A:p.V142I,CA1:NM_001128830:exon8:c.G622A:p.V208I,CA1:NM_001128831:exon7:c.G622A:p.V208I,CA1:NM_001291968:exon5:c.G283A:p.V95I,CA1:NM_001738:exon8:c.G622A:p.V208I,CA1:NM_001128829:exon8:c.G622A:p.V208I, |
| chr8 | 85329768 | C | T | . | exonic | CA1 | nonsynonymous SNV | CA1:NM_001164830:exon6:c.G590A:p.G197D,CA1:NM_001291967:exon6:c.G392A:p.G131D,CA1:NM_001128830:exon8:c.G590A:p.G197D,CA1:NM_001128831:exon7:c.G590A:p.G197D,CA1:NM_001291968:exon5:c.G251A:p.G84D,CA1:NM_001738:exon8:c.G590A:p.G197D,CA1:NM_001128829:exon8:c.G590A:p.G197D, |
| chr8 | 85329802 | G | A | COSV56279080 | exonic | CA1 | nonsynonymous SNV | CA1:NM_001164830:exon6:c.C556T:p.L186F,CA1:NM_001291967:exon6:c.C358T:p.L120F,CA1:NM_001128830:exon8:c.C556T:p.L186F,CA1:NM_001128831:exon7:c.C556T:p.L186F,CA1:NM_001291968:exon5:c.C217T:p.L73F,CA1:NM_001738:exon8:c.C556T:p.L186F,CA1:NM_001128829:exon8:c.C556T:p.L186F, |
| chr8 | 85332531 | G | T | . | exonic | CA1 | nonsynonymous SNV | CA1:NM_001164830:exon5:c.C472A:p.L158M,CA1:NM_001291967:exon5:c.C274A:p.L92M,CA1:NM_001128830:exon7:c.C472A:p.L158M,CA1:NM_001128831:exon6:c.C472A:p.L158M,CA1:NM_001291968:exon4:c.C133A:p.L45M,CA1:NM_001738:exon7:c.C472A:p.L158M,CA1:NM_001128829:exon7:c.C472A:p.L158M, |
| chr8 | 85333535 | A | G | . | exonic | CA1 | nonsynonymous SNV | CA1:NM_001164830:exon4:c.T440C:p.V147A,CA1:NM_001291967:exon4:c.T242C:p.V81A,CA1:NM_001128830:exon6:c.T440C:p.V147A,CA1:NM_001128831:exon5:c.T440C:p.V147A,CA1:NM_001291968:exon3:c.T101C:p.V34A,CA1:NM_001738:exon6:c.T440C:p.V147A,CA1:NM_001128829:exon6:c.T440C:p.V147A, |
| chr8 | 85336964 | C | T | . | exonic | CA1 | nonsynonymous SNV | CA1:NM_001164830:exon3:c.G335A:p.G112E,CA1:NM_001291967:exon3:c.G137A:p.G46E,CA1:NM_001128830:exon5:c.G335A:p.G112E,CA1:NM_001128831:exon4:c.G335A:p.G112E,CA1:NM_001738:exon5:c.G335A:p.G112E,CA1:NM_001128829:exon5:c.G335A:p.G112E, |
| chr8 | 85337036 | C | T | . | exonic | CA1 | nonsynonymous SNV | CA1:NM_001164830:exon3:c.G263A:p.S88N,CA1:NM_001291967:exon3:c.G65A:p.S22N,CA1:NM_001128830:exon5:c.G263A:p.S88N,CA1:NM_001128831:exon4:c.G263A:p.S88N,CA1:NM_001738:exon5:c.G263A:p.S88N,CA1:NM_001128829:exon5:c.G263A:p.S88N, |
| chr8 | 85338277 | A | T | . | exonic | CA1 | nonsynonymous SNV | CA1:NM_001164830:exon2:c.T210A:p.N70K,CA1:NM_001128830:exon4:c.T210A:p.N70K,CA1:NM_001128831:exon3:c.T210A:p.N70K,CA1:NM_001738:exon4:c.T210A:p.N70K,CA1:NM_001128829:exon4:c.T210A:p.N70K, |

| | | | | | | | | |
|---|---|---|---|---|---|---|---|---|
| chr8 | 85338318 | C | T | . | exonic | CA1 | nonsynonymous SNV | CA1:NM_001164830:exon2:c.G169A:p.A57T,CA1:NM_001128830:exon4:c.G169A:p.A57T,CA1:NM_001128831:exon3:c.G169A:p.A57T,CA1:NM_001738:exon4:c.G169A:p.A57T,CA1:NM_001128829:exon4:c.G169A:p.A57T, |
| chr8 | 85338375 | C | T | . | exonic | CA1 | nonsynonymous SNV | CA1:NM_001164830:exon2:c.G112A:p.E38K,CA1:NM_001128830:exon4:c.G112A:p.E38K,CA1:NM_001128831:exon3:c.G112A:p.E38K,CA1:NM_001738:exon4:c.G112A:p.E38K,CA1:NM_001128829:exon4:c.G112A:p.E38K, |
| chr8 | 85338380 | G | T | . | exonic | CA1 | nonsynonymous SNV | CA1:NM_001164830:exon2:c.C107A:p.T36N,CA1:NM_001128830:exon4:c.C107A:p.T36N,CA1:NM_001128831:exon3:c.C107A:p.T36N,CA1:NM_001738:exon4:c.C107A:p.T36N,CA1:NM_001128829:exon4:c.C107A:p.T36N, |
| chr8 | 85338411 | C | T | COSV56279142 | exonic | CA1 | nonsynonymous SNV | CA1:NM_001164830:exon2:c.G76A:p.G26R,CA1:NM_001128830:exon4:c.G76A:p.G26R,CA1:NM_001128831:exon3:c.G76A:p.G26R,CA1:NM_001738:exon4:c.G76A:p.G26R,CA1:NM_001128829:exon4:c.G76A:p.G26R, |
| chr8 | 85338417 | C | T | . | exonic | CA1 | nonsynonymous SNV | CA1:NM_001164830:exon2:c.G70A:p.A24T,CA1:NM_001128830:exon4:c.G70A:p.A24T,CA1:NM_001128831:exon3:c.G70A:p.A24T,CA1:NM_001738:exon4:c.G70A:p.A24T,CA1:NM_001128829:exon4:c.G70A:p.A24T, |
| chr8 | 85338423 | G | A | . | exonic | CA1 | nonsynonymous SNV | CA1:NM_001164830:exon2:c.C64T:p.P22S,CA1:NM_001128830:exon4:c.C64T:p.P22S,CA1:NM_001128831:exon3:c.C64T:p.P22S,CA1:NM_001738:exon4:c.C64T:p.P22S,CA1:NM_001128829:exon4:c.C64T:p.P22S, |
| chr8 | 89953445 | A | T | . | exonic | NBN | nonsynonymous SNV | NBN:NM_002485:exon11:c.T1644A:p.N548K,NBN:NM_001024688:exon12:c.T1398A:p.N466K, |
| chr8 | 89970473 | A | G | COSV55378087 | exonic | NBN | nonsynonymous SNV | NBN:NM_002485:exon7:c.T787C:p.F263L,NBN:NM_001024688:exon8:c.T541C:p.F181L, |
| chr8 | 89978251 | C | G | COSV55371261 | exonic | NBN | nonsynonymous SNV | NBN:NM_002485:exon5:c.G553C:p.E185Q,NBN:NM_001024688:exon6:c.G307C:p.E103Q, |
| chr8 | 94128313 | C | T | COSV50327363 | exonic | CDH17 | nonsynonymous SNV | CDH17:NM_001144663:exon18:c.G2426A:p.R809H,CDH17:NM_004063:exon18:c.G2426A:p.R809H, |
| chr8 | 94130926 | C | T | COSV50345009 | exonic | CDH17 | nonsynonymous SNV | CDH17:NM_001144663:exon16:c.G2234A:p.R745H,CDH17:NM_004063:exon16:c.G2234A:p.R745H, |
| chr8 | 94130944 | T | G | COSV50337541 | exonic | CDH17 | nonsynonymous SNV | CDH17:NM_001144663:exon16:c.A2216C:p.E739A,CDH17:NM_004063:exon16:c.A2216C:p.E739A, |
| chr8 | 94130958 | C | G | COSV50336268 | exonic | CDH17 | nonsynonymous SNV | CDH17:NM_001144663:exon16:c.G2202C:p.E734D,CDH17:NM_004063:exon16:c.G2202C:p.E734D, |
| chr8 | 94145972 | C | T | . | exonic | CDH17 | nonsynonymous SNV | CDH17:NM_001144663:exon15:c.G2123A:p.G708D,CDH17:NM_004063:exon15:c.G2123A:p.G708D, |
| chr8 | 94176622 | T | C | COSV50326476 | exonic | CDH17 | nonsynonymous SNV | CDH17:NM_001144663:exon5:c.A343G:p.K115E,CDH17:NM_004063:exon5:c.A343G:p.K115E, |
| chr8 | 94527128 | A | T | . | exonic | KIAA1429 | nonsynonymous SNV | KIAA1429:NM_015496:exon8:c.T1116A:p.D372E,KIAA1429:NM_183009:exon8:c.T1116A:p.D372E, |
| chr8 | 94529205 | C | T | . | exonic | KIAA1429 | nonsynonymous SNV | KIAA1429:NM_015496:exon7:c.G745A:p.E249K,KIAA1429:NM_183009:exon7:c.G745A:p.E249K, |
| chr8 | 94529220 | G | T | . | exonic | KIAA1429 | nonsynonymous SNV | KIAA1429:NM_015496:exon7:c.C730A:p.Q244K,KIAA1429:NM_183009:exon7:c.C730A:p.Q244K, |
| chr8 | 96231068 | C | T | . | exonic | UQCRB | stopgain | UQCRB:NM_001199975:exon5:c.G227A:p.W76X,UQCRB:NM_006294:exon4:c.G323A:p.W108X, |
| chr8 | 96231075 | C | T | . | exonic | UQCRB | nonsynonymous SNV | UQCRB:NM_001199975:exon5:c.G220A:p.E74K,UQCRB:NM_006294:exon4:c.G316A:p.E106K, |
| chr8 | 96231860 | C | T | . | exonic | UQCRB | nonsynonymous SNV | UQCRB:NM_001254752:exon3:c.G172A:p.D58N,UQCRB:NM_001199975:exon4:c.G76A:p.D26N,UQCRB:NM_006294:exon3:c.G172A:p.D58N, |
| chr8 | 96233194 | C | T | . | exonic | UQCRB | nonsynonymous SNV | UQCRB:NM_001254752:exon2:c.G53A:p.R18Q,UQCRB:NM_006294:exon2:c.G53A:p.R18Q, |
| chr8 | 96233201 | C | T | . | exonic | UQCRB | nonsynonymous SNV | UQCRB:NM_001254752:exon2:c.G46A:p.G16S,UQCRB:NM_006294:exon2:c.G46A:p.G16S, |
| chr8 | 96235512 | C | T | . | exonic | UQCRB | nonsynonymous SNV | UQCRB:NM_001254752:exon1:c.G19A:p.V7I,UQCRB:NM_006294:exon1:c.G19A:p.V7I, |
| chr8 | 96235527 | C | T | COSV54629256 | exonic | UQCRB | nonsynonymous SNV | UQCRB:NM_001254752:exon1:c.G4A:p.A2T,UQCRB:NM_006294:exon1:c.G4A:p.A2T, |
| chr9 | 113370054 | C | T | COSV53057170 | exonic | BSPRY | nonsynonymous SNV | BSPRY:NM_017688:exon6:c.C1121T:p.T374I, |
| chr9 | 115030304 | C | G | COSV60785451 | exonic | TNC | nonsynonymous SNV | TNC:NM_002160:exon24:c.G6022C:p.E2008Q, |
| chr9 | 132595027 | T | C | COSV55036973 | exonic | DDX31 | nonsynonymous SNV | DDX31:NM_022779:exon20:c.A2395G:p.I799V, |
| chr9 | 132662629 | C | T | COSV60137948 | exonic | DDX31 | nonsynonymous SNV | DDX31:NM_022779:exon2:c.G457A:p.E153K,DDX31:NM_138620:exon2:c.G457A:p.E153K, |
| chr9 | 134040214 | G | T | . | exonic | BRD3 | nonsynonymous SNV | BRD3:NM_007371:exon9:c.C1463A:p.P488Q, |
| chr9 | 134041792 | G | A | . | exonic | BRD3 | nonsynonymous SNV | BRD3:NM_007371:exon8:c.C1375T:p.R459W, |
| chr9 | 15474113 | C | T | . | exonic | PSIP1 | nonsynonymous SNV | PSIP1:NM_001128217:exon9:c.G754A:p.E252K,PSIP1:NM_021144:exon8:c.G754A:p.E252K,PSIP1:NM_033222:exon9:c.G754A:p.E252K, |
| chr9 | 15474188 | G | T | . | exonic | PSIP1 | nonsynonymous SNV | PSIP1:NM_001128217:exon9:c.C679A:p.P227T,PSIP1:NM_021144:exon8:c.C679A:p.P227T,PSIP1:NM_033222:exon9:c.C679A:p.P227T, |
| chr9 | 15474200 | C | T | . | exonic | PSIP1 | nonsynonymous SNV | PSIP1:NM_001128217:exon9:c.G667A:p.E223K,PSIP1:NM_021144:exon8:c.G667A:p.E223K,PSIP1:NM_033222:exon9:c.G667A:p.E223K, |
| chr9 | 15478498 | G | T | . | exonic | PSIP1 | nonsynonymous SNV | PSIP1:NM_001128217:exon8:c.C608A:p.P203H,PSIP1:NM_021144:exon7:c.C608A:p.P203H,PSIP1:NM_033222:exon8:c.C608A:p.P203H, |
| chr9 | 20988427 | A | G | . | exonic | FOCAD | nonsynonymous SNV | FOCAD:NM_017794:exon43:c.A5002G:p.K1668E, |
| chr9 | 286491 | G | A | COSV66634213 | exonic | DOCK8 | nonsynonymous SNV | DOCK8:NM_203447:exon3:c.G187A:p.D63N, |
| chr9 | 33798019 | A | G | COSV61553087 | exonic | PRSS3 | nonsynonymous SNV | PRSS3:NM_001197098:exon3:c.A370G:p.T124A,PRSS3:NM_007343:exon3:c.A562G:p.T188A,PRSS3:NM_001197097:exon4:c.A433G:p.T145A,PRSS3:NM_002771:exon3:c.A391G:p.T131A, |

| | | | | | | | | |
|---|---|---|---|---|---|---|---|---|
| chr9 | 7799653 | G | T | COSV64065665 | exonic | TMEM261 | nonsynonymous SNV | TMEM261:NM_033428:exon1:c.C82A:p.P28T, |
| chr9 | 92649479 | G | A | . | exonic | IPPK | nonsynonymous SNV | IPPK:NM_022755:exon5:c.C388T:p.R130W, |
| chr9 | 93113099 | G | C | . | exonic | CARD19 | nonsynonymous SNV | CARD19:NM_032310:exon6:c.G544C:p.G182R, |
| chr9 | 95150025 | T | A | . | exonic | FANCC | nonsynonymous SNV | FANCC:NM_001243743:exon7:c.A584T:p.D195V,FANCC:NM_000136:exon7:c.A584T:p.D195V,FANCC:NM_001243744:exon7:c.A584T:p.D195V, |
| chr9 | 99968585 | T | A | . | exonic | STX17 | stopgain | STX17:NM_017919:exon8:c.T821A:p.L274X, |

| Isoform | 1T__WT | 1T__MUT | 1M__WT | 1M__MUT | 2T__WT | 2T__MUT |
|---|---|---|---|---|---|---|
| NM_003051:NM_001166496 | 0:00 | 20:00 | 0:00 | 9:00 | 9:00 | 10:01 |
| NM_003051:NM_001166496 | 20:00 | 0:00 | 9:00 | 1:00 | 18:01 | 1:00 |
| NM_015906:XM_005270936:XM_011541568:XM_024447584:NM_033020:XM_005270937 | 0:0:0:0:0:0 | 7:0:0:3:3:3 | 0:0:0:0:0:0 | 3:1:1:1:1:0 | 0:0:0:0:0:0 | 2:1:0:1:2:2 |
| NM_015906:XM_017001454:XM_011541568:NM_033020:XM_005270936:XM_005270937:XM_024447584 | 1:2:0:1:0:0:0 | 3:2:0:2:0:0:0 | 1:0:1:1:0:0:0 | 1:1:0:0:0:0:0 | 1:1:0:0:0:0:0 | 0:0:0:0:1:0:0 |
| NM_007158:NM_001242893:NM_001007553:NM_001242892 | 242:1:26:6 | 0:0:0:0 | 324:3:32:3 | 0:0:0:0 | 418:0:35:7 | 0:0:0:0 |
| NM_007158:NM_001242893:NM_001007553:NM_001242892 | 241:1:26:6 | 1:0:0:0 | 324:3:32:3 | 0:0:0:0 | 419:0:35:7 | 0:0:0:0 |
| NM_007158:NM_001242893:NM_001007553:NM_001242892 | 244:1:26:6 | 0:0:0:0 | 319:3:32:3 | 1:0:0:0 | 414:0:35:7 | 1:0:0:0 |
| NM_007158:NM_001242893:NM_001007553:NM_001242892 | 242:1:26:6 | 0:0:0:0 | 320:3:32:3 | 1:0:0:0 | 416:0:35:7 | 0:0:0:0 |
| NM_007158:NM_001242893:NM_001007553:NM_001242892 | 241:1:26:6 | 0:0:0:0 | 321:3:32:3 | 0:0:0:0 | 411:0:35:7 | 0:0:0:0 |
| NM_007158:NM_001007553:NM_001242893:NM_001242892 | 237:25:1:6 | 0:0:0:0 | 318:30:3:3 | 1:1:0:0 | 396:35:0:7 | 0:0:0:0 |
| NM_007158:NM_001242893:NM_001007553:NM_001242892 | 232:1:25:6 | 1:0:0:0 | 307:3:31:3 | 0:0:0:0 | 379:0:35:7 | 0:0:0:0 |
| NM_007158:NM_001242893:NM_001007553:NM_001242892 | 211:1:26:6 | 0:0:0:0 | 281:3:32:3 | 1:0:0:0 | 310:0:35:7 | 0:0:0:0 |
| NM_001160233:NM_000701:XR_002956654:NM_001160234:XM_017001361 | 0:128:22:161:52 | 0:0:0:0:0 | 2:124:9:85:18 | 0:0:0:0:0 | 1:152:15:159:58 | 0:0:0:1:1 |
| NM_001160233:NM_000701:NM_001160234:XR_002956654:XM_017001361 | 0:128:184:14:61 | 0:0:1:0:0 | 2:124:106:7:21 | 0:0:0:0:0 | 1:152:190:13:61 | 0:0:0:0:1 |
| XM_017002264:XM_017002263:NM_001135610:XM_017002256:NM_012231:XR_002957562 | 0:0:0:0:0:0 | 3:0:1:0:0:0 | 0:0:0:0:0:0 | 4:3:0:0:0:0 | 0:0:0:0:0:0 | 7:1:2:0:0:0 |
| NM_001348451:NM_004284:NM_024568:NM_001348462:XR_001737550:NR_145682:NR_046070:NM_001348452:NR_145695 | 0:4:0:0:0:0:0:0:0 | 0:0:0:0:0:0:0:0:0 | 1:2:1:0:0:0:0:0:0 | 0:0:0:0:0:0:0:0:0 | 0:4:1:0:0:0:0:0:0 | 0:4:1:0:0:0:0:0:0 |
| NM_001348451:NM_004284:NM_024568:XM_024451051:XR_001737550:NR_145682:NM_001348462:NR_145695:NR_046070:NM_001348452 | 0:4:0:0:1:0:0:0:0:0 | 0:0:0:0:0:0:0:0:0:0 | 1:1:1:0:0:0:0:0:0:0 | 0:0:0:0:0:0:0:0:0:0 | 0:7:2:1:0:0:0:0:0:0 | 0:0:0:0:0:0:0:0:0:0 |
| NM_014624:XM_017002033 | 753:34:00 | 2:00 | 779:40:00 | 0:00 | 1207:64 | 1:01 |
| NM_014624:XM_017002033 | 751:34:00 | 4:00 | 779:40:00 | 0:00 | 1207:65 | 1:00 |
| NM_014624:XM_017002033 | 757:34:00 | 0:00 | 780:40:00 | 1:00 | 1208:64 | 1:01 |
| NM_153649:NR_103461:NM_001043351:NM_001278189:NM_001349679:NM_001278188:NM_001043352:NM_001278190:NM_152263:NM_001278191:NM_001043353 | 968:8:4:4:0:5:7:0:0:0:0 | 0:0:0:0:0:0:0:0:0:1:0 | 590:9:2:7:1:2:3:0:0:0:0 | 0:0:0:0:0:0:0:0:0:0:0 | 1097:7:7:13:0:11:41:0:0:0:0 | 0:0:0:0:0:0:0:0:0:0:0 |
| NM_153649:NR_103461:NM_001043351:NM_001278189:NM_001349679:NM_001278188:NM_001043352:NM_001278190:NM_152263:NM_001278191:NM_001043353 | 968:7:4:4:0:5:7:0:0:1:0 | 0:0:0:0:0:0:0:0:0:0:0 | 591:8:2:7:1:2:3:0:0:0:0 | 0:0:0:0:0:0:0:0:0:0:0 | 1096:7:7:13:0:11:41:0:0:0:0 | 1:0:0:0:0:0:0:0:0:0:0 |
| NM_153649:NR_103461:NM_001043351:NM_001278189:NM_001349679:NM_001278188:NM_001043352:NM_001278190:NM_152263:NM_001278191 | 949:7:4:4:0:5:7:0:0:0 | 0:0:0:0:0:0:0:0:0:0 | 582:7:2:7:1:2:3:0:0:0 | 0:0:0:0:0:0:0:0:0:0 | 1083:8:7:13:0:11:41:0:0:0 | 0:0:0:0:0:0:0:0:0:0 |
| NM_153649:NR_103461:NM_001043351:NM_001278189:NM_001349679:NM_001278188:NM_001043352:NM_001278190:NM_152263:NM_001278191 | 943:9:4:4:0:5:7:0:0:0 | 0:0:0:0:0:0:0:0:0:0 | 575:7:2:7:1:2:3:0:0:0 | 0:0:0:0:0:0:0:0:0:0 | 1073:8:7:13:0:11:40:0:0:0 | 1:0:0:0:0:0:0:0:0:0 |

| | | | | | | |
|---|---|---|---|---|---|---|
| NM_153649:NR_103461:NM_001043351:NM_001278189:NM_001349679:NM_001278188:NM_001043352:NM_001278190:NM_152263:NM_001278191 | 936:10:4:4:0:5:7:0:0:0 | 2:0:0:0:0:0:0:0:0:0 | 575:8:2:7:1:2:3:0:0:0 | 0:0:0:0:0:0:0:0:0:0 | 1068:9:7:13:0:11:40:0:0:0 | 1:0:0:0:0:0:0:0:0:0 |
| NM_153649:NM_001043352:NR_103461:NM_001043351:NM_001278189:NM_001349679:NM_001278188:NM_001278191:NM_001278190 | 920:6:10:4:4:0:3:0:0 | 0:0:0:0:0:0:0:0:0 | 565:2:9:2:7:1:0:0:0 | 1:1:0:0:0:0:0:0:0 | 1041:37:9:7:12:0:10:0:0 | 2:0:0:0:0:0:0:0:0 |
| NM_005973:XR_921868:XM_005245314:XM_005245313 | 15:0:1:0 | 5:3:0:0 | 10:0:1:0 | 1:1:0:0 | 23:0:0:0 | 0:0:0:0 |
| NM_007348:XM_006711224:XM_011509310:XM_011509308 | 29:0:0:1 | 11:1:1:0 | 9:1:1:0 | 10:0:0:0 | 18:0:1:0 | 24:1:2:0 |
| NR_137288:XR_002956989:XR_001737277:XM_017001622:XR_946690:XR_946686:XM_024448062:NM_001328124:NM_001319838:XM_017001620 | 3:4:0:0:0:0:0:1:0:0 | 2:2:0:0:0:0:1:1:1:0 | 2:3:1:1:0:0:0:0:0:0 | 1:2:0:0:0:0:0:0:0:0 | 0:0:0:0:0:0:0:0:0:0 | 3:2:1:0:0:1:2:0:0:1 |
| NM_024529:XM_006711537 | 28:00:00 | 0:00 | 13:00 | 9:00 | 37:01:00 | 0:00 |
| NM_004930:XM_024450097:NM_001206540:XM_011542228:NM_001282162:NM_001206541:XM_017002428 | 871:35:14:0:6:1:0 | 6:0:2:0:0:0:0 | 329:17:2:0:0:0:0 | 3:0:0:0:0:0:0 | 576:41:0:0:1:1:0 | 3:0:0:0:0:0:0 |
| NM_004930:XM_024450097:NM_001206540:XM_011542228:NM_001282162:NM_001206541:XM_017002428 | 877:35:16:0:6:1:0 | 0:0:0:0:0:0:0 | 332:17:2:0:0:0:0 | 1:0:0:0:0:0:0 | 579:41:0:0:1:1:0 | 0:0:0:0:0:0:0 |
| NM_004433:XM_005244942:NM_001114309 | 888:13:12 | 1:00:00 | 314:06:01 | 1:00:00 | 608:06:02 | 0:00:00 |
| NM_004433:XM_005244942:NM_001114309 | 894:12:12 | 1:00:00 | 313:06:01 | 0:00:00 | 609:06:02 | 1:00:00 |
| NM_004433:XM_005244942:NM_001114309 | 1013:04:12 | 1:00:00 | 330:02:01 | 0:00:00 | 674:02:02 | 0:00:00 |
| NM_004433:XM_005244942:NM_001114309 | 1012:04:12 | 2:00:00 | 331:02:01 | 0:00:00 | 673:02:02 | 1:00:00 |
| NM_004433:NM_001114309 | 560:05:00 | 3:00 | 201:01:00 | 0:00 | 411:02:00 | 0:00 |
| NM_016448:NM_001286230:NM_001286229 | 0:00:00 | 2:00:00 | 0:00:00 | 4:00:00 | 0:00:00 | 11:00:00 |
| NM_016448:NM_001286230:NM_001286229 | 0:00:00 | 2:00:00 | 0:00:00 | 5:00:00 | 0:00:00 | 15:00:00 |
| NM_001136018:NM_001291163 | 13:02 | 12:01 | 18:01 | 6:00 | 7:01 | 0:00 |
| NM_001136018:NM_001291163 | 12:01 | 13:02 | 6:00 | 18:01 | 0:00 | 7:00 |
| XM_024453895:NM_017449:XM_006710442:NM_004442 | 0:1:0:0 | 0:3:1:0 | 2:0:0:0 | 1:0:0:0 | 5:0:1:0 | 1:5:0:0 |
| XM_011544253:NM_001167740:XM_024449137:XM_024449166:NM_022743:XM_024449136:XM_011544258:XM_024449149:XM_011544261:XM_011544254:XM_024449162 | 8:43:2:0:0:0:0:0:0:0:0 | 3:54:0:0:0:0:1:0:0:0:0 | 7:68:1:1:0:1:0:0:0:0:0 | 17:108:2:1:1:0:0:0:0:0:0 | 0:0:0:0:0:0:0:0:0:0:0 | 18:131:0:0:0:1:2:0:0:1:0 |
| NM_020394:NR_037892 | 0:01 | 0:00 | 0:00 | 0:00 | 0:01 | 0:02 |
| NM_020394:NR_037892 | 0:00 | 0:01 | 0:00 | 0:00 | 0:00 | 0:02 |
| NM_205861:NM_001243564:NM_024887:NM_001319959:NM_001243565 | 27:4:7:0:0 | 15:3:9:0:0 | 2:0:1:0:0 | 6:1:1:0:0 | 7:3:1:1:0 | 0:0:0:0:0 |
| NM_023923:NM_001350161:NM_001350158:NM_001048183:NM_001350160 | 1:0:0:4:0 | 1:0:2:1:0 | 1:1:0:0:0 | 1:0:0:0:0 | 1:0:0:1:1 | 0:2:0:1:0 |
| XM_017002054:XM_011541950 | 221:02:00 | 1:00 | 228:05:00 | 0:00 | 358:01:00 | 0:00 |
| NR_136702:XM_005271115:XM_017002054:XM_011541950:NR_136703 | 18:7:222:2:0 | 0:0:0:0:0 | 30:19:227:5:0 | 0:0:0:0:0 | 24:15:354:3:0 | 0:0:0:0:0 |
| NR_136702:XM_005271115:XM_017002054:XM_011541950:NR_136703 | 18:7:222:1:0 | 0:0:0:0:0 | 30:19:227:5:0 | 0:0:0:0:0 | 23:15:352:3:0 | 1:0:1:0:0 |
| NM_006004:NM_001297565 | 13:00 | 0:00 | 9:00 | 0:00 | 10:00 | 8:00 |
| XM_017000618:XM_017000620:NM_001981:XM_017000617:XM_017000615:XM_017000619:XM_017000616:XM_005270618 | 2:32:9:0:0:2:0:0 | 2:15:6:0:0:1:0:0 | 1:28:12:0:1:0:0:0 | 1:17:5:1:0:0:0:0 | 0:1:0:0:0:0:0:0 | 3:78:4:0:0:1:0:0 |
| XM_017001646:XM_024448207:XR_001737287:NM_018291:XM_024448229:XM_017001645:XM_017001652:XM_017001679:NM_001350793:NM_001350794:XM_017001656:XM_017001657:XM_011541730:NM_001244714:XR_001737285:NM_001113411:NR_103473:XM_017001649:XM_017001673:XM_024448176:NM_001350799:NM_001350797:XM_024448220:XM_017001655 | 1:2:0:3:0:2:1:0:1:0:0:0:0:0:0:0:0:0:0:0:0:0:0:0 | 1:0:0:5:0:2:0:1:0:0:0:0:0:0:0:0:0:0:0:0:0:0:0:0 | 0:1:0:1:0:0:0:0:0:0:0:0:0:0:0:0:0:0:0:0:0:0:0:0 | 1:1:1:0:0:0:0:0:0:0:0:0:0:0:0:0:0:0:0:0:0:0:0:0 | 0:1:1:2:0:0:0:0:0:0:0:0:0:0:0:0:0:0:0:0:0:0:0:0 | 1:2:0:6:0:3:2:0:0:0:0:0:0:0:0:0:0:1:0:0:0:0:0:0 |
| NM_176877:XM_024448614:XM_011540465:XR_001736900:XM_017000001:XR_002957157:XM_011540469:NM_001350145 | 1:1:1:0:0:0:0:0 | 0:0:0:0:0:0:0:0 | 1:1:0:0:0:0:0:0 | 0:0:0:0:0:0:0:0 | 0:0:0:0:0:0:0:0 | 0:0:2:0:0:0:0:0 |
| XM_024448614:NM_176877:XR_002957157:XM_011540465:NM_001350145:XR_001736900:XM_017000001:XM_011540469:XM_016999999 | 1:1:0:0:0:0:0:0:0 | 0:0:0:1:1:0:0:0:0 | 0:2:1:0:0:0:0:0:0 | 1:0:0:0:0:0:0:0:0 | 0:0:0:0:0:0:0:0:0 | 0:0:0:2:0:0:0:0:0 |
| XM_011542191:XM_011542192:NM_001002292:NM_024911:XM_017002390:NM_001193334 | 6:2:0:24:7:0 | 0:0:0:0:0:0 | 16:7:4:32:6:0 | 0:0:0:0:0:0 | 11:0:1:8:1:1 | 2:2:0:5:6:0 |
| XR_001737065:NR_038934:NM_001242701:XR_001737066:NR_146202:NR_146203:XR_001737061:XR_001737063:XR_001737062 | 66:39:2:2:2:0:1:1:0 | 82:37:3:2:1:0:0:0:0 | 9:5:1:2:1:0:0:0:0 | 13:2:2:0:0:0:0:0:0 | 64:25:8:1:1:1:0:0:0 | 0:0:0:0:0:0:0:0:0 |
| NM_001286043:NM_000016:NM_001286042:NM_001127328:NM_001286044 | 0:32:2:0:3 | 0:31:10:1:0 | 0:5:1:0:0 | 1:17:2:1:1 | 0:87:2:0:2 | 0:0:0:0:0 |
| XR_001737524:NR_146539:XR_001737520:NR_146540:XM_024450628:XM_017002744:NM_003902:NM_001303433:XM_017002742:XR_946791:XR_001737522:XM_011542393:XR_946790:XR_001737517 | 24:2:0:3:0:78:127:1:1:9:9:1:0:0 | 3:0:0:0:0:0:0:0:0:2:0:0:0 | 27:6:1:6:1:69:145:5:3:7:2:0:0:0 | 4:0:0:0:0:0:0:0:0:0:0:0:0 | 31:8:1:5:0:85:124:1:3:7:5:0:0:0 | 3:0:0:1:0:0:0:0:0:0:1:0:0:0 |
| NM_001428:NM_001201483 | 1581:00:00 | 2:00 | 2354:01:00 | 0:00 | 4334:01:00 | 1:00 |
| NM_001428:NM_001201483 | 1583:00:00 | 1:00 | 2350:01:00 | 4:00 | 4333:01:00 | 1:00 |
| NM_001428:NM_001201483 | 1582:00:00 | 0:00 | 2352:01:00 | 1:00 | 4331:01:00 | 1:00 |
| NM_001428:NM_001201483 | 1568:00:00 | 3:00 | 2338:01:00 | 1:00 | 4291:01:00 | 7:00 |
| NM_001428:NM_001201483 | 1564:00:00 | 2:00 | 2330:01:00 | 2:00 | 4276:01:00 | 5:00 |

| | | | | | | |
|---|---|---|---|---|---|---|
| NM_001428:NM_001201483 | 1545:00:00 | 0:00 | 2304:01:00 | 2:00 | 4223:01:00 | 2:00 |
| NM_001428:NM_001201483 | 1520:00:00 | 1:00 | 2271:01:00 | 4:00 | 4191:01:00 | 6:00 |
| NM_001428:NM_001201483 | 1514:00:00 | 3:00 | 2268:01:00 | 4:00 | 4182:01:00 | 12:00 |
| NM_001428:NM_001201483 | 1499:00:00 | 3:00 | 2250:01:00 | 2:00 | 4141:01:00 | 2:00 |
| NM_001428:NM_001201483 | 1495:00:00 | 2:00 | 2248:01:00 | 2:00 | 4134:01:00 | 3:00 |
| NM_001428:NM_001201483 | 1480:00:00 | 1:00 | 2229:01:00 | 4:00 | 4088:01:00 | 4:00 |
| NM_001428:NM_001201483 | 1472:00:00 | 1:00 | 2241:01:00 | 2:00 | 4064:01:00 | 6:00 |
| NM_001428:NM_001201483 | 1460:00:00 | 3:00 | 2224:01:00 | 3:00 | 4050:01:00 | 7:00 |
| NM_001428:NM_001201483 | 1447:00:00 | 0:00 | 2215:01:00 | 3:00 | 4033:01:00 | 4:00 |
| NM_001428:NM_001201483 | 1444:00:00 | 0:00 | 2212:01:00 | 3:00 | 4029:01:00 | 4:00 |
| NM_001428:NM_001201483 | 1442:00:00 | 1:00 | 2208:01:00 | 1:00 | 4023:01:00 | 4:00 |
| NM_001428:NM_001201483 | 1443:00:00 | 0:00 | 2210:01:00 | 0:00 | 4017:01:00 | 8:00 |
| NM_001428:NM_001201483 | 1463:00:00 | 2:00 | 2211:01:00 | 6:00 | 3952:01:00 | 6:00 |
| NM_001428:NM_001201483 | 1464:00:00 | 2:00 | 2226:01:00 | 0:00 | 3949:01:00 | 7:00 |
| NM_001428:NM_001201483 | 1461:00:00 | 0:00 | 2224:01:00 | 1:00 | 3942:01:00 | 6:00 |
| NM_001428:NM_001201483 | 1443:00:00 | 1:00 | 2214:01:00 | 3:00 | 3917:01:00 | 4:00 |
| NM_001428:NM_001201483 | 1434:00:00 | 0:00 | 2203:01:00 | 3:00 | 3895:01:00 | 1:00 |
| NM_001428:NM_001201483 | 1434:00:00 | 1:00 | 2197:01:00 | 4:00 | 3886:01:00 | 8:00 |
| NM_001428:NM_001201483 | 1441:00:00 | 0:00 | 2197:01:00 | 1:00 | 3868:01:00 | 5:00 |
| NM_001428:NM_001201483 | 1443:00:00 | 1:00 | 2199:01:00 | 5:00 | 3869:01:00 | 6:00 |
| NM_001428:NM_001201483 | 1446:00:00 | 1:00 | 2214:01:00 | 2:00 | 3878:01:00 | 5:00 |
| NM_001428:NM_001201483 | 1444:00:00 | 1:00 | 2214:01:00 | 0:00 | 3873:01:00 | 4:00 |
| NM_001428:NM_001201483 | 1455:00:00 | 2:00 | 2220:01:00 | 2:00 | 3861:01:00 | 6:00 |
| NM_001428:NM_001201483:NM_001353346 | 1455:00:00 | 1:00:00 | 2215:01:00 | 4:00:00 | 3859:00:00 | 2:00:00 |
| NM_001428:NM_001201483:NM_001353346 | 1454:00:00 | 3:00:00 | 2220:01:00 | 0:00:00 | 3859:00:00 | 2:00:00 |
| NM_001428:NM_001201483:NM_001353346 | 1457:00:00 | 1:00:00 | 2216:01:00 | 3:00:00 | 3859:00:00 | 3:00:00 |
| NM_001428:NM_001201483:NM_001353346 | 1455:00:00 | 1:00:00 | 2207:01:00 | 6:00:00 | 3855:00:00 | 4:00:00 |
| NM_001428:NM_001201483:NM_001353346 | 1448:00:00 | 0:00:00 | 2198:01:00 | 1:00:00 | 3839:00:00 | 3:00:00 |
| NM_000969:NR_146333 | 1614:04:00 | 10:00 | 4404:17:00 | 9:00 | 4808:18:00 | 8:00 |
| NM_000969:NR_146333 | 1618:04:00 | 8:00 | 4404:17:00 | 9:00 | 4810:18:00 | 8:00 |
| NM_000969:NR_146333 | 1617:04:00 | 9:00 | 4403:17:00 | 13:00 | 4813:18:00 | 7:00 |
| NM_000969:NR_146333 | 1618:04:00 | 11:00 | 4401:17:00 | 15:00 | 4816:18:00 | 7:00 |
| NM_000969:NR_146333 | 1631:04:00 | 6:00 | 4424:17:00 | 7:00 | 4836:18:00 | 3:00 |
| NM_000969:NR_146333 | 1633:04:00 | 5:00 | 4432:17:00 | 4:00 | 4841:18:00 | 2:00 |
| NM_000969:NR_146333 | 1650:04:00 | 3:00 | 4460:17:00 | 9:00 | 4897:18:00 | 2:00 |
| NM_000969:NR_146333 | 1675:04:00 | 2:00 | 4526:18:00 | 8:00 | 4932:18:00 | 9:00 |
| NM_000969:NR_146333 | 1680:04:00 | 0:00 | 4537:18:00 | 4:00 | 4948:18:00 | 6:00 |
| NM_000969:NR_146333 | 1678:04:00 | 2:00 | 4536:18:00 | 4:00 | 4952:18:00 | 1:00 |
| NM_000969:NR_146333 | 1734:01:00 | 2:00 | 4675:14:00 | 4:00 | 5023:09:00 | 2:01 |
| NM_000969:NR_146333 | 1736:01:00 | 2:00 | 4687:12:00 | 2:00 | 5027:08:00 | 3:00 |
| NM_000969:NR_146333 | 1744:01:00 | 2:00 | 4707:09:00 | 3:00 | 5038:06:00 | 7:00 |
| NM_000969:NR_146333 | 1747:00:00 | 1:00 | 4731:01:00 | 1:00 | 5054:00:00 | 4:00 |
| NM_000969:NR_146333 | 1747:00:00 | 1:00 | 4733:01:00 | 2:00 | 5063:00:00 | 3:00 |
| NM_000969:NR_146333 | 1755:00:00 | 1:00 | 4751:00:00 | 2:01 | 5083:00:00 | 3:00 |
| NM_000969:NR_146333 | 1753:02:00 | 1:00 | 4736:07:00 | 11:00 | 5084:05:00 | 2:00 |
| NM_000969:NR_146333 | 1755:06:00 | 0:00 | 4741:12:00 | 1:00 | 5079:07:00 | 4:00 |
| NM_000969:NR_146333 | 1748:11:00 | 0:00 | 4721:46:00 | 5:01 | 5066:26:00 | 0:00 |
| NM_000969:NR_146333 | 1739:27:00 | 2:00 | 4692:82 | 5:01 | 5048:63 | 4:03 |
| NM_000969:NR_146333 | 1732:35:00 | 0:00 | 4670:94 | 3:02 | 5032:77 | 11:01 |
| NM_000969:NR_146333 | 1725:41:00 | 2:00 | 4666:103 | 5:01 | 5036:94 | 5:00 |
| NM_000969:NR_146333 | 1718:51:00 | 3:00 | 4655:120 | 3:00 | 5023:106 | 8:00 |
| NM_000969:NR_146333 | 1715:60 | 1:00 | 4628:127 | 3:01 | 5023:116 | 5:00 |
| NM_000969:NR_146333 | 1559:57:00 | 123:02:00 | 4226:118 | 304:02:00 | 4526:114 | 377:01:00 |
| NM_000969:NR_146333 | 1552:56:00 | 1:00 | 4196:108 | 3:00 | 4501:110 | 2:00 |
| NM_004741:XM_005270273:NM_001284389:NM_001284388 | 1:4:2:0 | 1:1:2:0 | 7:3:6:1 | 12:7:7:4 | 18:9:10:2 | 13:9:9:1 |
| NM_001284389:NM_004741:XM_005270273:NM_001284388 | 4:3:5:0 | 0:0:0:0 | 13:19:12:5 | 0:0:0:0 | 10:19:9:1 | 9:17:9:2 |
| NM_004741:NM_001284389:XM_005270273:NM_001284388 | 3:4:4:0 | 0:0:0:0 | 19:12:13:5 | 2:1:0:0 | 35:17:18:3 | 0:1:0:0 |
| NM_001351171:NM_001351183:NM_001351172:NM_001134373:NM_012229:NM_001351169:XM_024447902:NM_001351181:NM_001351175:NM_001351182:XM_024447904:NM_001351185:NM_001351170:NM_001351190:NM_001351194:XM_024447911:XM_024447905:XM_024447907:XM_024447908:XM_024447910:XM_011539537:XM_017015966:NM_001351189:NM_001351197:NM_001351187:XM_024447901:NM_001351178:XM_024447909:NM_001351180:NM_001351176:NM_001351177:XM_006717723 | 1:1:2:7:3:5:0:0:1:0:1:0:1:0:0:0:0:1:0:1:1:0:2:1:1:0:0:0:0:0:0:0 | 0:1:3:2:2:6:0:0:0:0:0:0:2:0:0:0:1:0:0:0:1:1:0:0:0:0:0:0:0:0:0:0 | 2:1:2:5:3:4:0:2:1:1:1:0:0:0:0:0:0:0:0:0:0:0:0:0:0:0:0:0:0:0:0:0 | 1:2:1:4:1:1:1:0:0:0:0:0:0:0:0:0:0:0:0:0:0:0:0:0:0:0:0:0:0:0:0:0 | 4:1:5:14:3:4:0:0:0:0:1:0:5:0:0:2:0:0:0:1:0:0:0:0:0:0:0:0:0:1:1:0:0 | 0:0:0:0:0:0:0:0:0:0:0:0:0:0:0:0:0:0:0:0:0:0:0:0:0:0:0:0:0:0:0:0 |
| NM_005871:XM_005269382 | 15:01 | 0:00 | 32:01:00 | 0:00 | 25:04:00 | 1:00 |
| NM_014456:NM_145341:NM_001199492 | 0:00:00 | 9:01:01 | 0:00:00 | 0:00:00 | 0:00:00 | 1:00:00 |

| | | | | | | |
|---|---|---|---|---|---|---|
| NM_007041:XM_017015598:XM_017015599:NM_001288736:NM_001001976:XM_017015604:XM_005269459:XM_005269458:NR_110006 | 1:2:1:1:2:0:1:0:0 | 0:0:0:0:0:0:0:0:0 | 4:2:0:0:0:0:0:0:0 | 0:0:0:0:0:0:0:0:0 | 2:2:0:0:0:0:1:0:0 | 0:0:1:0:0:0:0:0:0 |
| NM_001291877:XM_011539134:NM_001291879:XM_006717553:XM_017015495:XM_024447768:XM_011539133:NM_206860:XM_006717554:XM_024447762:XM_006717552:NM_206862 | 1:0:5:0:5:0:1:1:0:0:0:0 | 0:0:0:0:0:0:0:0:0:0:0:0 | 1:1:2:0:0:0:0:0:0:0:0:0 | 0:0:0:0:0:0:0:0:0:0:0:0 | 1:0:0:0:1:0:0:0:0:0:0:0 | 0:0:2:0:7:0:0:0:0:0:0:0 |
| NM_001291877:XM_024447768:XM_011539134:NM_001291879:XM_017015495:XM_011539133:NM_206860:NM_206862:XM_006717552:XM_006717554:XM_017015499:XM_006717553 | 1:0:0:5:5:1:1:1:0:0:0:0 | 0:0:0:0:0:0:0:0:0:0:0:0 | 1:1:1:2:0:0:0:0:0:0:0:0 | 0:0:0:0:0:0:0:0:0:0:0:0 | 0:0:0:2:6:0:0:0:0:0:0:0 | 1:0:0:0:0:0:0:0:0:0:0:0 |
| XM_024447859:NM_001380 | 1:01 | 0:00 | 1:02 | 0:00 | 0:00 | 0:02 |
| NM_002412:XM_017016275:XM_005252682 | 193:07:10 | 204:06:05 | 52:04:00 | 82:02:01 | 18:01:00 | 0:00:00 |
| NM_145202:NM_001145201 | 0:00 | 21:01 | 0:00 | 3:00 | 0:00 | 7:00 |
| XM_006717500:NM_003380 | 191:06:00 | 0:00 | 415:19:00 | 1:00 | 210:11:00 | 0:00 |
| XM_006717500:NM_003380 | 210:07:00 | 1:01 | 431:20:00 | 0:00 | 224:12:00 | 0:00 |
| XM_006717500:NM_003380 | 231:08:00 | 0:00 | 468:22:00 | 4:00 | 239:12:00 | 1:00 |
| XM_006717500:NM_003380 | 249:08:00 | 0:00 | 490:25:00 | 1:00 | 251:12:00 | 0:00 |
| XM_006717500:NM_003380 | 249:08:00 | 0:00 | 494:25:00 | 1:00 | 251:12:00 | 0:00 |
| XM_006717500:NM_003380 | 250:08:00 | 0:00 | 494:25:00 | 2:00 | 252:12:00 | 0:00 |
| XM_006717500:NM_003380 | 261:09:00 | 0:00 | 503:26:00 | 0:00 | 267:12:00 | 0:00 |
| XM_006717500:NM_003380 | 266:09:00 | 0:00 | 511:26:00 | 0:00 | 276:13:00 | 1:00 |
| XM_006717500:NM_003380 | 267:09:00 | 0:00 | 511:26:00 | 1:00 | 277:15:00 | 0:00 |
| XM_005252342:XR_001746996:XR_001746995:XR_242691 | 0:1:0:0 | 0:0:0:0 | 0:0:0:0 | 1:0:0:0 | 0:0:0:0 | 0:0:0:0 |
| XM_005252342:XR_001746996:XR_001746995:XR_242691 | 0:1:0:0 | 0:0:0:0 | 0:0:0:0 | 1:0:0:0 | 0:0:0:0 | 0:0:0:0 |
| NM_001323647:NM_001174096:NM_030751:NM_001174093:NM_001323655:XM_017016597:NM_001323649:NM_001174095:NM_001323653 | 1:9:3:0:0:1:0:0:0 | 1:1:0:0:0:0:1:0:0 | 0:3:2:1:0:0:0:0:0 | 1:0:0:0:0:0:0:0:0 | 0:2:0:0:0:0:0:0:0 | 0:1:0:0:0:0:0:0:0 |
| NM_001323647:NM_001174096:NM_030751:NM_001174093:NM_001323655:XM_017016597:NM_001323649:NM_001174095:NM_001323653 | 1:9:3:0:0:1:0:0:0 | 1:1:0:0:0:0:1:0:0 | 0:3:2:1:0:0:0:0:0 | 1:0:0:0:0:0:0:0:0 | 0:2:0:0:0:0:0:0:0 | 0:1:0:0:0:0:0:0:0 |
| NM_001174096:NM_001174093:NM_030751:NM_001323655:NM_001323647:XM_017016597:NM_001174095:NM_001323653 | 3:0:1:0:0:1:0:0 | 3:0:2:0:1:0:0:0 | 2:1:0:0:0:0:0:0 | 1:0:0:0:0:0:0:0 | 0:0:0:0:0:0:0:0 | 1:0:0:0:0:0:0:0 |
| NM_001024628:NM_001244973:NM_003873:XM_006717526:NM_001024629 | 0:0:0:0:0 | 1:0:1:0:0 | 0:0:0:0:0 | 3:1:1:0:0 | 0:0:0:0:0 | 5:0:4:1:0 |
| NM_001145263:NM_005437:NM_001145261:NM_001145260:NM_001145262 | 801:878:6:3:3 | 9:3:0:0:0 | 348:172:1:1:0 | 3:0:0:0:0 | 270:316:2:0:1 | 2:1:0:0:0 |
| NM_005437:NM_001145263:NM_001145261:NM_001145260:NM_001145262 | 874:808:6:3:3 | 1:2:0:0:0 | 170:353:1:1:0 | 0:0:0:0:0 | 316:273:2:0:1 | 0:0:0:0:0 |
| NM_001145263:NM_005437:NM_001145261:NM_001145260:NM_001145262 | 820:828:6:3:3 | 1:1:0:0:0 | 358:169:1:1:0 | 1:0:0:0:0 | 279:302:2:0:1 | 2:1:0:0:0 |
| NM_001145263:NM_005437:NM_001145261:NM_001145260:NM_001145262 | 818:826:6:3:3 | 11:1:0:0:0 | 359:169:1:1:0 | 1:0:0:0:0 | 279:302:2:0:1 | 5:1:0:0:0 |
| NM_001145263:NM_005437:NM_001145261:NM_001145260:NM_001145262 | 820:816:6:3:3 | 16:6:0:0:0 | 359:167:1:1:0 | 3:0:0:0:0 | 281:301:2:0:1 | 5:1:0:0:0 |
| NM_001145263:NM_005437:NM_001145261:NM_001145260:NM_001145262 | 839:797:6:3:3 | 1:4:0:0:0 | 361:161:1:1:0 | 1:0:0:0:0 | 286:293:2:0:1 | 0:0:0:0:0 |
| NM_005437:NM_001145263:NM_001145261:NM_001145260:NM_001145262 | 789:838:6:3:3 | 2:2:0:0:0 | 160:362:1:1:0 | 0:0:0:0:0 | 290:286:2:0:1 | 0:0:0:0:0 |
| NM_018237:NR_104262:NM_001282960:NM_001282959 | 24:3:0:0 | 0:0:0:0 | 36:1:1:0 | 0:0:0:0 | 25:7:4:1 | 0:0:0:0 |
| NM_018237:NR_104262:NM_001282960:NM_001282959 | 17:3:0:0 | 1:0:0:0 | 27:1:1:0 | 1:0:0:0 | 16:3:2:1 | 1:0:1:0 |
| NM_018237:NM_001282960:NR_104262:NM_001282959 | 14:0:3:0 | 0:0:0:0 | 18:0:0:0 | 1:1:0:0 | 8:2:3:0 | 2:0:0:0 |
| NM_018344:NR_033414:XM_017016378:NR_033413:XM_017016377:NM_001174098 | 3:0:0:0:0:0 | 0:0:1:0:0:0 | 0:0:1:0:0:0 | 2:1:0:0:0:0 | 0:0:0:1:0:0 | 2:0:0:1:0:0 |
| XM_017016378:NM_018344:NR_033414:XM_017016377:NR_033413:NM_001174098 | 1:0:0:1:0:0 | 0:3:0:0:0:0 | 0:2:1:0:0:0 | 1:0:0:0:0:0 | 0:0:0:0:0:0 | 0:3:0:0:2:0 |
| NM_018344:NR_033414:XM_017016378:XM_017016377:NR_033413:NM_001174098 | 0:0:0:0:0:0 | 3:1:2:1:1:0 | 0:0:0:0:0:0 | 2:1:0:0:0:0 | 0:0:0:0:0:0 | 3:0:0:0:2:0 |
| NM_018344:NR_033414:XM_017016378:XM_017016377:NR_033413:NM_001174098 | 0:0:0:0:0:0 | 3:1:1:1:1:0 | 0:0:0:0:0:0 | 2:2:1:0:0:0 | 0:0:0:0:0:0 | 3:0:0:0:2:0 |
| NM_001042465:NM_002778:NM_001042466 | 59:647:5 | 0:00:00 | 93:256:25 | 0:00:00 | 24:254:3 | 0:01:00 |

| | | | | | | |
|---|---|---|---|---|---|---|
| XM_017015704:NM_006721:NM_001202450:XM_017015698:NM_001123:XM_017015702:XM_017015703:XM_017015706:XM_017015705:NM_001202449:XM_017015699 | 0:48:0:0:38:0:0:0:0:0:0 | 0:0:0:0:0:0:0:0:0:0:0 | 1:72:3:2:52:1:5:0:0:0:0 | 0:0:0:0:0:0:0:0:0:0:0 | 1:111:2:0:130:3:9:3:0:1:0 | 0:0:0:0:0:0:0:0:0:0:0 |
| NM_004329:XM_011540104 | 3:00 | 10:00 | 5:01 | 3:00 | 13:01 | 0:00 |
| NM_001304717:NM_001304718 | 0:00 | 147:02:00 | 0:00 | 177:01:00 | 0:00 | 149:03:00 |
| NM_001304717:NM_001304718 | 215:08:00 | 2:00 | 286:09:00 | 0:00 | 240:06:00 | 0:00 |
| NM_001304717:NM_001304718 | 209:08:00 | 4:00 | 284:10:00 | 1:00 | 233:07:00 | 6:00 |
| NM_001304717:NM_001304718 | 209:08:00 | 3:00 | 284:10:00 | 1:00 | 233:07:00 | 6:00 |
| NM_001304717:NM_001304718 | 198:09:00 | 1:00 | 279:08:00 | 0:00 | 228:05:00 | 0:00 |
| NM_001031709:XM_005269946:XM_017016385:XM_005269948:XM_005269947:XM_017016384:XR_001747122 | 16:1:0:1:0:0:0 | 12:0:0:0:0:0:0 | 5:0:0:0:0:0:0 | 6:0:0:0:0:0:0 | 5:0:0:0:0:0:0 | 0:0:0:0:0:0:0 |
| NM_016195:NM_001284259 | 2:00 | 2:00 | 2:00 | 1:00 | 12:00 | 0:00 |
| XM_017017365:XR_428971:XM_017017382:XM_017017384:XM_017017367:XM_017017377:XM_006718794:XM_017017381:XM_017017366:XM_017017380:XM_017017379:XM_005271456:XM_017017385:XR_002957133:XM_017017386 | 1:9:4:0:3:0:1:1:0:0:0:0:0:0:0 | 0:1:0:0:0:0:0:0:0:0:0:0:0:0:0 | 1:8:1:1:0:0:0:0:0:0:0:0:0:0:0 | 0:0:0:0:0:0:0:0:0:0:0:0:0:0:0 | 2:6:1:0:0:0:0:0:0:0:0:0:0:0:0 | 0:0:0:0:0:0:0:0:0:0:0:0:0:0:0 |
| XM_017018334:NM_001168468:XM_017018335:XM_011543005 | 2:0:0:1 | 3:1:1:0 | 0:0:0:0 | 0:0:0:0 | 0:0:0:0 | 1:0:0:0 |
| NM_015957:XM_017017875:XM_011520154 | 100:04:00 | 183:07:05 | 36:03:00 | 102:01:01 | 197:08:03 | 0:00:00 |
| NM_015957:XM_017017875:XM_011520154 | 284:06:05 | 0:01:00 | 138:04:01 | 0:00:00 | 196:06:03 | 1:00:00 |
| NM_015957:XM_017017875:XM_011520154 | 100:04:00 | 183:04:05 | 35:03:00 | 102:01:01 | 197:07:03 | 0:00:00 |
| NM_003477:NM_001166158:XM_011520390:NM_001135024 | 72:3:3:0 | 83:3:2:0 | 30:2:2:0 | 22:1:2:1 | 46:0:2:0 | 52:0:3:0 |
| NM_003477:NM_001166158:XM_011520390:NM_001135024 | 158:7:7:0 | 0:0:0:0 | 50:3:4:1 | 0:0:0:0 | 97:1:5:0 | 0:0:0:0 |
| NM_001001390:NM_001202555:XM_005253238:NM_001001389:NM_001001391:XM_017018585:XM_011520488:XM_011520486:XM_011520483:XM_005253240:XM_005253235:XM_017018583:XM_006718388:XM_011520485:NM_001202557:XM_005253239:NM_000610:XM_011520487:XM_011520484:XM_017018584:XM_005253231:NM_001202556:XM_005253232:XM_011520489:XM_011520482 | 28:1:4:4:121:34:5:0:0:1:0:3:0:0:2:1:0:0:0:0:0:0:0:0:0 | 0:0:0:0:1:0:0:0:0:0:0:0:0:0:0:0:0:0:0:0:0:0:0:0:0 | 85:3:16:3:67:20:2:3:2:4:2:4:1:1:0:0:0:0:0:0:0:0:0:0:0 | 0:0:0:0:0:0:0:0:0:0:0:0:0:0:0:0:0:0:0:0:0:0:0:0:0 | 53:0:46:8:43:32:8:5:0:20:5:12:1:0:1:1:5:1:2:2:3:0:0:0:0 | 0:0:0:0:0:0:0:0:0:0:0:0:0:0:0:0:0:0:0:0:0:0:0:0:0 |
| XM_005253238:XM_005253235:NM_001001389:XM_011520483:XM_011520486:XM_017018583:XM_006718388:NM_000610:XM_011520487:XM_011520484:XM_005253231:XM_005253232:XM_011520489:XM_011520482:XM_011520485:XM_017018585 | 3:0:0:0:0:1:0:0:0:0:0:0:0:0:0:0 | 1:0:0:0:0:0:0:0:0:0:0:0:0:0:0:0 | 7:1:1:1:0:1:1:0:0:0:0:0:0:0:0:0 | 9:1:1:1:1:0:0:0:0:0:0:0:0:0:0:0 | 47:5:4:0:1:5:1:5:1:2:3:0:0:0:0:0 | 0:0:0:0:0:0:0:0:0:0:0:0:0:0:0:0 |
| NM_001001390:XM_005253238:NM_001001389:XM_011520488:XM_011520486:XM_011520483:XM_005253240:XM_005253235:XM_017018583:XM_006718388:XM_011520485:XM_005253239:NM_000610:XM_011520484:XM_005253231:XM_005253232:XM_011520482:XM_017018585:XM_011520489 | 0:0:0:0:0:0:0:0:0:0:0:0:0:0:0:0:0:0:0 | 29:3:0:5:0:0:1:0:2:0:0:1:0:0:0:0:0:0:0 | 0:0:0:0:0:0:0:0:0:0:0:0:0:0:0:0:0:0:0 | 85:16:3:2:3:2:4:2:2:1:1:0:0:0:0:0:0:0:0 | 56:39:4:8:5:0:20:5:5:1:0:1:5:2:3:0:0:0:0 | 0:0:0:0:0:0:0:0:0:0:0:0:0:0:0:0:0:0:0 |
| NM_001001390:XM_005253238:NM_001001389:XM_011520488:XM_011520486:XM_011520483:XM_005253240:XM_005253235:XM_017018583:XM_006718388:XM_011520485:XM_005253239:NM_000610:XM_011520484:XM_005253231:XM_005253232:XM_011520482:XM_017018585:XM_011520489 | 29:3:0:5:0:0:1:0:2:0:0:1:0:0:0:0:0:0:0 | 0:0:0:0:0:0:0:0:0:0:0:0:0:0:0:0:0:0:0 | 85:16:3:2:3:2:4:2:2:1:1:0:0:0:0:0:0:0:0 | 0:0:0:0:0:0:0:0:0:0:0:0:0:0:0:0:0:0:0 | 56:39:4:8:5:0:20:5:5:1:0:1:5:2:3:0:0:0:0 | 0:0:0:0:0:0:0:0:0:0:0:0:0:0:0:0:0:0:0 |
| NM_001001390:NM_001202555:XM_005253238:NM_001001391:NM_001001389:XM_011520488:XM_011520486:XM_011520483:XM_005253240:XM_005253235:XM_017018583:XM_006718388:XM_011520485:XM_017018585:NM_001202556:NM_001202557:XM_005253239:NM_001001392:NM_000610:XM_005253232:XM_005253231:XM_011520484:XM_017018584:XM_011520482:XM_011520489 | 29:2:3:124:0:5:0:0:1:0:4:0:0:5:1:1:1:1:0:0:0:0:0:0:0 | 0:0:0:1:0:0:0:0:0:0:0:0:0:0:0:0:0:0:0:0:0:0:0:0:0 | 85:4:16:71:3:2:3:2:4:2:4:1:1:2:4:0:0:0:0:0:0:0:0:0:0 | 0:0:0:0:0:0:0:0:0:0:0:0:0:0:0:0:0:0:0:0:0:0:0:0:0 | 27:0:16:20:3:4:0:0:8:3:7:1:0:2:0:0:0:1:3:0:2:2:0:0:0 | 28:0:22:24:0:4:5:0:12:2:10:0:0:1:0:0:1:0:2:0:1:0:0:0:0 |
| NM_016142:XM_024448573 | 0:00 | 75:04:00 | 0:00 | 53:02:00 | 239:04:00 | 0:00 |
| NM_024113:XM_006718315:XM_017018282:NM_001003676:NM_001003677:XM_011520364:NM_001278222:NM_001003678:NR_103472:XM_017018281:XM_017018276:XM_017018279 | 26:0:1:1:7:1:3:1:0:0:0:0 | 0:0:0:0:0:0:0:0:0:0:0:0 | 3:5:0:0:0:0:0:0:0:0:0:0 | 0:0:0:0:0:0:0:0:0:0:0:0 | 3:3:0:1:1:1:3:0:1:0:0:0 | 4:1:0:0:1:0:1:0:0:0:0:0 |
| NM_032389:XM_005253168:XM_011520405:XM_005253166:XM_017018415:XM_005253167:XM_017018413:NM_001242832 | 4:1:0:0:0:0:0:0 | 1:0:0:0:0:0:0:0 | 0:1:0:0:0:0:0:0 | 4:2:0:0:0:0:0:0 | 1:4:1:0:0:0:1:0 | 2:3:0:0:0:1:0:0 |
| NM_014342:NM_001317232:XM_017017462:XM_006718172 | 1:1:0:0 | 1:0:0:0 | 0:0:0:0 | 0:0:0:0 | 0:0:0:0 | 5:0:0:1 |
| XM_011545117:NM_017716:NM_001164470 | 6:38:01 | 2:06:00 | 0:00:00 | 0:00:00 | 0:00:00 | 0:00:00 |
| NM_012094:NM_001358511:NM_181651:NM_181652 | 0:0:0:0 | 19:2:1:1 | 0:1:0:0 | 4:0:0:0 | 0:0:0:0 | 57:0:1:0 |
| NM_001876:XM_005273763:NM_001031847:XM_017017220 | 6:0:2:0 | 5:1:0:0 | 0:0:0:0 | 0:0:0:0 | 1:0:0:0 | 0:0:0:0 |
| NM_016725:NM_016724:NM_000802:NM_016729 | 0:1:0:0 | 0:0:0:0 | 14:13:1:37 | 0:0:0:0 | 0:7:1:6 | 0:0:0:0 |
| NM_016725:NM_016724:NM_000802:NM_016729 | 0:1:0:0 | 0:0:0:0 | 14:13:1:37 | 0:0:0:0 | 0:7:1:6 | 0:0:0:0 |
| NM_016055:NM_001318500:NM_001318499:NM_001318498:XM_017017919:NR_134660:NR_134661:NR_134659:NR_134658 | 125:0:4:1:1:1:2:2:0 | 0:0:0:0:0:0:0:0:0 | 200:3:4:4:4:1:1:1:0 | 0:0:0:0:0:0:0:0:0 | 357:8:8:0:6:2:3:1:0 | 0:0:0:0:0:0:0:0:0 |
| XM_005274051:NM_016578:XM_017017923 | 4:02:00 | 5:00:00 | 3:02:00 | 4:00:00 | 6:01:01 | 7:01:00 |

| | | | | | | |
|---|---|---|---|---|---|---|
| XM_005274051:NM_016578:XM_017017923 | 9:02:00 | 3:00:00 | 5:02:00 | 2:00:00 | 12:02:01 | 7:00:00 |
| XM_005274051:NM_016578:XM_017017923 | 5:02:00 | 7:00:00 | 5:02:00 | 4:00:00 | 11:02:01 | 8:00:00 |
| NM_001243251:NM_024678:XM_017018303:XR_001747964:XR_001747966:XM_017018304:XM_011545253:XR_001747965:XR_001747963 | 0:0:0:1:0:0:0:0:0 | 3:1:1:0:0:0:0:0:0 | 0:3:0:0:0:0:0:0:0 | 3:0:0:0:0:0:0:0:0 | 0:0:0:0:0:0:0:0:0 | 7:5:1:0:1:1:1:0:0 |
| NM_022918:NM_001168724:XM_017018141:XM_017018142:NR_033149 | 0:0:0:0:0 | 40:2:2:8:0 | 0:0:0:0:0 | 22:3:2:1:0 | 8:0:0:2:0 | 10:0:2:0:0 |
| XM_017019206:NM_001303130:NM_174942:XM_005268804:XM_011538220 | 0:0:0:3:0 | 0:0:0:0:0 | 0:0:0:0:0 | 0:0:0:0:0 | 2:4:0:0:1 | 0:0:0:0:0 |
| NM_213595:NM_001301141:XM_011538101 | 0:00:00 | 5:00:00 | 0:00:00 | 4:00:00 | 0:00:00 | 8:00:00 |
| NM_213595:NM_001301141:XM_011538101 | 0:00:00 | 5:00:00 | 0:00:00 | 4:00:00 | 0:00:00 | 8:00:00 |
| NM_213595:NM_001301141:XM_011538101 | 0:00:00 | 5:00:00 | 0:00:00 | 4:00:00 | 0:00:00 | 8:00:00 |
| NM_002834:NM_080601:XM_017019722:NM_001330437 | 1:37:4:2 | 0:5:0:0 | 2:19:1:1 | 0:0:0:0 | 1:47:4:0 | 0:1:0:0 |
| NM_080601:XM_017019722:NM_002834:NM_001330437:XM_011538613 | 32:3:1:2:0 | 3:0:0:0:0 | 17:0:2:1:0 | 2:1:0:0:0 | 42:5:1:0:0 | 4:0:0:0:0 |
| NM_080601:XM_017019722:NM_002834:NM_001330437:XM_011538613 | 33:3:1:2:0 | 3:0:0:0:0 | 17:0:2:1:0 | 2:1:0:0:0 | 43:5:1:0:0 | 3:0:0:0:0 |
| XM_011520607:NM_001987:XM_017018990:XM_011520608:XM_017018991:XM_011520611 | 13:14:2:0:0:0 | 0:0:0:0:0:0 | 8:5:0:0:0:0 | 0:0:0:0:0:0 | 3:6:2:0:4:2 | 0:0:0:0:0:0 |
| NM_002813:NM_001261400:NR_048555 | 0:00:00 | 247:02:03 | 0:00:00 | 87:01:01 | 0:00:00 | 161:03:00 |
| XM_006719552:XM_017019788:XM_017019792:XM_017019790:NM_198240:XM_017019786 | 0:0:0:3:0:0 | 0:0:0:2:0:0 | 1:1:0:0:0:0 | 0:0:0:0:0:0 | 0:0:0:0:0:0 | 0:1:0:4:0:0 |
| XM_017019790:XM_006719552:XM_017019788:XM_017019786:XM_017019792:NM_198240:XM_005253593 | 6:1:0:0:0:0:0 | 1:0:3:0:0:0:0 | 1:1:1:0:0:0:0 | 1:0:0:0:0:0:0 | 4:0:1:0:0:0:0 | 0:0:0:0:0:0:0 |
| NM_005505:NM_001082959 | 95:09:00 | 83:03:00 | 46:03:00 | 17:01 | 77:00:00 | 0:00 |
| NM_005505:NM_001082959 | 171:11:00 | 0:00 | 62:04:00 | 0:00 | 72:00:00 | 0:00 |
| NM_002300:NM_001174097 | 378:00:00 | 0:00 | 214:00:00 | 1:00 | 1753:01:00 | 1:00 |
| NM_002300:NM_001174097 | 378:00:00 | 0:00 | 215:00:00 | 0:00 | 1754:01:00 | 1:00 |
| NM_002300:NM_001174097 | 377:00:00 | 0:00 | 215:00:00 | 0:00 | 1761:01:00 | 0:00 |
| NM_002300:NM_001174097 | 377:00:00 | 0:00 | 216:00:00 | 0:00 | 1760:01:00 | 2:00 |
| NM_002300:NM_001174097 | 377:00:00 | 0:00 | 216:00:00 | 0:00 | 1762:01:00 | 0:00 |
| NM_002300:XM_006719074:NM_001174097 | 375:00:00 | 0:00:00 | 217:01:00 | 0:00:00 | 1763:03:01 | 1:00:00 |
| NM_002300:XM_006719074:NM_001174097 | 371:00:00 | 0:00:00 | 217:02:00 | 0:00:00 | 1755:04:01 | 0:00:00 |
| NM_002300:XM_006719074:NM_001174097 | 371:00:00 | 0:00:00 | 216:02:00 | 1:00:00 | 1754:04:01 | 1:00:00 |
| NM_002300:XM_006719074:NM_001174097 | 371:00:00 | 0:00:00 | 217:02:00 | 0:00:00 | 1753:04:01 | 1:00:00 |
| NM_002300:XM_006719074:NM_001174097 | 371:00:00 | 0:00:00 | 216:02:00 | 0:00:00 | 1750:04:01 | 0:00:00 |
| NM_002300:XM_006719074:NM_001174097 | 368:00:00 | 0:00:00 | 216:02:00 | 0:00:00 | 1744:05:01 | 1:00:00 |
| NM_002300:XM_006719074:NM_001174097 | 365:00:00 | 1:00:00 | 214:02:00 | 0:00:00 | 1737:05:01 | 2:00:00 |
| NM_002300:XM_006719074:NM_001174097 | 366:00:00 | 0:00:00 | 213:02:00 | 1:00:00 | 1737:05:01 | 2:00:00 |
| NM_002300:XM_006719074:NM_001174097 | 365:00:00 | 1:00:00 | 214:02:00 | 0:00:00 | 1737:05:01 | 1:00:00 |
| NM_002300:XM_006719074:NM_001174097 | 365:00:00 | 1:00:00 | 214:02:00 | 0:00:00 | 1735:05:01 | 3:00:00 |
| NM_002300:XM_006719074:NM_001174097 | 364:00:00 | 0:00:00 | 214:01:00 | 0:00:00 | 1726:05:01 | 4:00:00 |
| NM_002300:XM_006719074:NM_001174097 | 360:00:00 | 1:00:00 | 213:01:00 | 0:00:00 | 1725:04:01 | 1:00:00 |
| NM_002300:XM_006719074:NM_001174097 | 357:00:00 | 0:00:00 | 211:01:00 | 2:00:00 | 1708:03:01 | 4:00:00 |
| NM_002300:NM_001174097:XM_006719074 | 353:00:00 | 0:00:00 | 212:00:00 | 0:00:00 | 1700:01:03 | 0:00:00 |
| NM_002300:NM_001174097:XM_006719074 | 355:00:00 | 0:00:00 | 212:00:00 | 0:00:00 | 1698:01:02 | 4:00:00 |
| NM_002300:NM_001174097:XM_006719074 | 355:00:00 | 0:00:00 | 212:00:00 | 0:00:00 | 1701:01:01 | 0:00:01 |
| NM_002300:NM_001174097 | 355:00:00 | 0:00 | 212:00:00 | 0:00 | 1694:01:00 | 0:00 |
| NM_002300:XM_006719074:NM_001174097 | 351:01:00 | 0:00:00 | 212:00:00 | 0:00:00 | 1679:00:01 | 3:00:00 |
| NM_002300:XM_006719074:NM_001174097 | 351:01:00 | 0:00:00 | 211:00:00 | 0:00:00 | 1679:00:01 | 1:00:00 |
| NM_002300:XM_006719074:NM_001174097 | 349:01:00 | 0:00:00 | 210:00:00 | 0:00:00 | 1672:00:01 | 2:00:00 |
| NM_002300:XM_006719074:NM_001174097 | 349:01:00 | 0:00:00 | 209:00:00 | 1:00:00 | 1674:00:01 | 0:00:00 |
| NM_002300:XM_006719074:NM_001174097 | 344:01:00 | 1:00:00 | 205:00:00 | 1:00:00 | 1654:00:01 | 1:00:00 |
| NM_004985:NM_033360 | 7:08 | 0:00 | 1:02 | 0:00 | 3:06 | 0:01 |
| NM_032358:NM_001130146:NM_001130147 | 0:01:00 | 0:00:00 | 3:01:00 | 0:00:00 | 0:00:00 | 4:02:01 |
| XM_017019336:NM_005171:XM_011538388:XM_017019333:XM_017019335:XM_017019334:XM_011538387:XM_011538386:XM_017019331 | 5:17:1:3:0:0:2:3:0 | 0:0:0:0:0:0:0:0:0 | 2:11:1:1:1:1:0:0:0 | 0:0:0:0:0:0:0:0:0 | 3:22:0:0:0:2:2:2:0 | 0:0:0:0:0:0:0:0:0 |
| NM_033082:NR_026723:NR_026722 | 594:125:8 | 0:00:00 | 277:95:6 | 2:00:00 | 394:115:3 | 1:00:00 |
| NM_006601:NM_001282601:NM_001282602:XM_005268576:XM_006719199:XM_011537773:NM_001282605:NR_104219:NM_001282604:XM_011537774:XM_017018716 | 761:7:16:14:1:2:4:1:1:0:0 | 0:0:0:0:0:1:0:0:0:0:0 | 254:5:9:7:1:3:0:0:0:0:0 | 0:0:0:0:0:0:0:0:0:0:0 | 519:7:9:12:1:3:3:0:5:0:0 | 1:0:0:1:0:0:0:0:0:0:0 |
| NM_001113202:NM_001320194:NM_005594:NM_001320193:NM_001113201:NM_001113203 | 1448:10:106:73:1:1 | 1:6:1:0:0:0 | 3031:16:193:112:3:0 | 4:1:0:0:0:0 | 3983:16:254:185:2:0 | 2:1:1:0:0:0 |
| NM_001113202:NM_005594:NM_001320193:NM_001320194:NM_001113201:NM_001113203 | 1443:107:73:15:1:1 | 6:0:0:3:0:0 | 3033:193:112:25:3:0 | 4:0:0:0:0:0 | 3984:254:185:20:2:0 | 3:1:0:1:0:0 |
| NM_001113202:NM_005594:NM_001320193:NM_001320194:NM_001113201:NM_001113203 | 1447:106:73:18:1:1 | 0:0:0:1:0:0 | 3031:193:112:27:3:0 | 2:0:0:0:0:0 | 3982:255:185:23:2:0 | 0:0:0:0:0:0 |
| NM_005594:NM_001113202:NM_001320194:NM_001320193:NM_001113201:NM_001113203 | 107:1447:15:73:1:1 | 0:2:4:0:0:0 | 192:3035:26:112:3:0 | 1:2:1:0:0:0 | 255:3980:24:185:2:0 | 0:5:0:0:0:0 |

| | | | | | | |
|---|---|---|---|---|---|---|
| NM_001113202:NM_005594:NM_001320193:NM_001320194:NM_001113201:NM_001113203 | 1445:107:73:21:1:1 | 4:0:0:0:0:0 | 3027:193:112:33:3:0 | 10:0:0:0:0:0 | 3979:257:185:30:2:0 | 6:0:0:0:0:0 |
| NM_001113202:NM_005594:NM_001320193:NM_001320194:NM_001113201:NM_001113203 | 1449:106:73:27:1:1 | 0:0:0:0:0:0 | 3033:189:112:61:3:0 | 4:0:0:0:0:0 | 3983:256:185:58:2:0 | 3:0:0:2:0:0 |
| NM_001113202:NM_005594:NM_001320194:NM_001320193:NM_001113201:NM_001113203 | 1448:106:29:73:1:1 | 1:0:0:0:0:0 | 3035:186:75:113:3:0 | 3:1:1:0:0:0 | 3980:251:74:185:2:0 | 6:2:0:0:0:0 |
| NM_001113202:NM_001320193:NM_001320194:NM_005594:NM_001113201:NM_001113203 | 1447:73:31:105:1:1 | 2:0:0:1:0:0 | 3035:112:86:187:3:0 | 2:1:3:0:0:0 | 3978:186:84:253:2:0 | 8:0:3:0:0:0 |
| NM_005594:NM_001113202:NM_001320193:NM_001320194:NM_001113201:NM_001113203 | 103:1446:73:31:1:1 | 1:3:0:0:0:0 | 185:3033:113:92:3:0 | 1:3:0:0:0:0 | 249:3986:186:88:2:0 | 1:0:0:0:0:0 |
| NM_001113202:NM_001320194:NM_005594:NM_001320193:NM_001113201 | 1448:31:98:73:1 | 1:0:0:0:0 | 3032:91:177:113:3 | 4:1:0:0:0 | 3985:88:237:186:2 | 1:0:0:0:0 |
| NM_001113202:NM_005594:NM_001320193:NM_001320194:NM_001113201 | 1447:94:73:31:1 | 2:0:0:0:0 | 3034:162:113:92:3 | 2:0:0:0:0 | 3985:227:186:88:2 | 1:5:0:0:0 |
| NM_004083:NM_001195057:NM_001195056:NM_001195053:NM_001195055:NM_001195054 | 511:76:117:5:2:0 | 2:0:0:0:0:0 | 159:19:19:2:0:0 | 0:0:0:0:0:0 | 205:39:10:1:1:0 | 0:0:0:0:0:0 |
| NM_004083:NM_001195057:NM_001195056:NM_001195053:NM_001195055:NM_001195054 | 511:76:117:5:2:0 | 4:0:0:0:0:0 | 157:19:19:2:0:0 | 0:0:0:0:0:0 | 203:39:10:1:1:0 | 0:0:0:0:0:0 |
| NM_033276:NM_001320408 | 4:00 | 6:00 | 1:00 | 2:00 | 0:00 | 7:00 |
| NM_001065:NR_144351:NM_001346092:NM_001346091 | 492:11:10:12 | 0:0:0:0 | 51:1:5:2 | 0:0:0:0 | 59:1:4:0 | 0:0:0:0 |
| NM_002046:NR_152150:NM_001289746:NM_001289745:NM_001357943 | 3416:15:23:6:0 | 5:0:0:0:0 | 2571:8:15:0:0 | 2:0:0:0:0 | 10005:48:56:8:1 | 9:0:0:0:0 |
| NM_002046:NR_152150:NM_001289746:NM_001289745:NM_001357943:NM_001256799 | 3476:16:23:6:0:0 | 3:0:0:0:0:0 | 2604:12:15:0:0:0 | 0:0:0:0:0:0 | 10114:56:55:8:0:2 | 5:0:0:0:0:0 |
| NM_002046:NR_152150:NM_001289746:NM_001289745:NM_001357943:NM_001256799 | 3521:17:23:6:0:0 | 5:0:0:0:0:0 | 2626:13:15:0:0:0 | 0:0:0:0:0:0 | 10224:57:55:8:1:2 | 11:1:0:0:0:0 |
| NM_002046:NR_152150:NM_001289746:NM_001289745:NM_001357943:NM_001256799 | 3539:10:23:6:0:0 | 2:0:0:0:0:0 | 2625:8:15:0:0:0 | 4:0:0:0:0:0 | 10267:34:55:8:1:2 | 15:0:0:0:0:0 |
| NM_002046:NR_152150:NM_001289746:NM_001289745:NM_001357943:NM_001256799 | 3532:9:23:6:0:0 | 8:0:0:0:0:0 | 2630:8:15:0:0:0 | 2:0:0:0:0:0 | 10273:34:55:8:1:2 | 14:0:0:0:0:0 |
| NM_002046:NR_152150:NM_001289746:NM_001289745:NM_001357943:NM_001256799 | 3558:7:23:6:0:0 | 2:0:0:0:0:0 | 2638:7:15:0:0:0 | 4:0:0:0:0:0 | 10299:24:55:8:1:2 | 15:0:0:0:0:0 |
| NM_002046:NR_152150:NM_001289746:NM_001289745:NM_001357943:NM_001256799 | 3566:7:23:6:0:0 | 5:0:0:0:0:0 | 2640:7:15:0:0:0 | 3:0:0:0:0:0 | 10323:24:55:8:1:2 | 6:0:0:0:0:0 |
| NM_002046:NR_152150:NM_001289746:NM_001289745:NM_001357943:NM_001256799 | 3572:6:23:6:0:0 | 11:0:0:0:0:0 | 2647:7:15:0:0:0 | 4:0:0:0:0:0 | 10338:18:55:8:1:2 | 21:1:0:0:0:0 |
| NM_002046:NR_152150:NM_001289746:NM_001289745:NM_001357943:NM_001256799 | 3592:6:23:6:0:0 | 2:0:0:0:0:0 | 2663:6:15:0:0:0 | 2:0:0:0:0:0 | 10404:18:55:8:1:2 | 4:0:0:0:0:0 |
| NM_002046:NR_152150:NM_001289746:NM_001289745:NM_001357943:NM_001256799 | 3603:6:23:6:0:0 | 6:0:0:0:0:0 | 2672:6:15:0:0:0 | 2:0:0:0:0:0 | 10409:16:55:8:1:2 | 15:1:0:0:0:0 |
| NM_002046:NM_001289746:NM_001289745:NR_152150:NM_001357943:NM_001256799 | 3629:23:6:0:0:0 | 2:0:0:0:0:0 | 2688:15:0:0:0:0 | 2:0:0:0:0:0 | 10455:55:8:3:1:2 | 8:0:0:1:0:0 |
| NM_002046:NM_001289746:NM_001289745:NR_152150:NM_001357943:NM_001256799 | 3646:23:6:0:0:0 | 4:0:0:0:0:0 | 2684:15:0:0:0:0 | 5:0:0:0:0:0 | 10468:55:8:1:1:2 | 10:0:0:0:0:0 |
| NM_002046:NM_001289746:NM_001289745:NR_152150:NM_001357943:NM_001256799 | 3675:23:6:0:0:0 | 4:0:0:0:0:0 | 2701:14:0:0:0:0 | 6:0:0:0:0:0 | 10569:55:8:0:1:2 | 13:0:0:0:0:0 |
| NM_002046:NM_001289746:NM_001289745:NR_152150:NM_001357943:NM_001256799 | 3678:23:6:0:0:0 | 3:0:0:0:0:0 | 2732:14:0:0:0:0 | 2:0:0:0:0:0 | 10618:55:8:0:1:2 | 10:0:0:0:0:0 |
| NM_002046:NM_001289746:NM_001289745:NR_152150:NM_001357943:NM_001256799 | 3702:23:6:0:0:0 | 3:0:0:0:0:0 | 2737:14:0:0:0:0 | 3:0:0:0:0:0 | 10685:55:8:0:1:2 | 7:0:0:0:0:0 |
| NM_002046:NM_001289746:NM_001289745:NR_152150:NM_001357943:NM_001256799 | 3702:23:6:0:0:0 | 5:0:0:0:0:0 | 2735:14:0:0:0:0 | 5:0:0:0:0:0 | 10684:55:8:0:1:2 | 6:0:0:0:0:0 |
| NM_002046:NM_001289746:NM_001289745:NR_152150:NM_001357943:NM_001256799 | 3679:22:5:0:0:0 | 7:0:1:0:0:0 | 2733:14:0:0:0:0 | 2:0:0:0:0:0 | 10673:55:8:1:1:2 | 15:0:0:0:0:0 |
| NM_002046:NM_001289746:NM_001289745:NR_152150:NM_001357943:NM_001256799 | 3680:22:6:0:0:0 | 6:0:0:0:0:0 | 2734:14:0:0:0:0 | 1:0:0:0:0:0 | 10675:55:8:1:1:2 | 14:0:0:0:0:0 |
| NM_002046:NM_001289746:NM_001289745:NR_152150:NM_001357943:NM_001256799 | 3683:21:6:0:0:0 | 6:1:0:0:0:0 | 2734:14:0:0:0:0 | 2:0:0:0:0:0 | 10682:55:8:1:1:2 | 10:0:0:0:0:0 |
| NM_007007:XM_005268590:NM_001300947:XM_005268588 | 184:50:17:10 | 0:0:0:0 | 111:31:11:1 | 0:0:0:0 | 170:43:36:11 | 0:0:0:0 |
| NM_007007:XM_005268590:NM_001300947:XM_005268588 | 181:48:18:10 | 0:0:0:1 | 114:28:11:1 | 0:0:0:0 | 173:43:39:11 | 0:0:0:0 |
| NM_007007:XM_005268590:NM_001300947:XM_005268588 | 181:48:18:11 | 0:0:0:0 | 113:29:11:1 | 0:0:0:0 | 173:43:39:11 | 1:0:0:0 |
| NM_001207016:NM_130846:XM_011538615:NM_002849:XR_001748831:NR_073474 | 52:145:6:0:14:3 | 38:144:2:0:10:0 | 1:1:1:0:0:0 | 2:2:0:0:0:0 | 0:1:0:0:0:0 | 0:0:0:0:0:0 |

| | | | | | | |
|---|---|---|---|---|---|---|
| NM_130846:NM_001207016:XM_011538615:NR_073474:NM_002849:XR_001748831 | 151:52:7:5:0:13 | 176:38:23:7:0:11 | 2:1:1:1:0:0 | 6:2:0:0:0:0 | 1:0:0:0:0:0 | 0:0:0:0:0:0 |
| NM_004616:XM_006719583 | 4:00 | 6:00 | 1:00 | 2:00 | 12:01 | 0:00 |
| NM_004616:XM_006719583 | 3:00 | 5:00 | 1:00 | 2:00 | 11:01 | 0:00 |
| NM_023011:XR_001749644:NR_148487:NR_148489:XM_017020711:XR_001749647:XM_017020709:NM_080687:XM_006719991:NM_001353644:NR_148485:NM_001353650:NM_001353646:NM_001353647:XM_005266202:NR_148494:XM_024449401:XM_011534848:XM_011534847:XM_024449400:XM_011534845:NM_001353645:XR_941422:NR_148484:NM_001353649 | 27:11:11:24:7:19:2:9:1:0:0:1:7:3:1:0:1:12:1:0:1:1:1:0:0 | 6:0:1:1:0:0:0:0:0:0:0:0:0:0:0:0:0:0:1:0:1:0:0:0:0 | 37:28:56:41:22:35:3:7:5:1:1:7:16:13:14:2:1:6:6:3:0:0:0:0:0 | 6:4:2:2:1:2:1:0:0:0:0:0:0:0:0:0:0:0:0:0:0:0:0:0:0 | 36:13:18:29:27:39:5:10:0:0:2:1:6:4:14:5:2:14:5:0:1:0:1:0:1 | 0:1:0:2:1:0:1:0:0:0:0:0:0:0:0:0:1:2:0:0:0:0:0:0:0 |
| NM_023011:NR_148487:XM_017020711:NR_148489:NM_001353650:XM_006719991:XR_001749644:XR_001749647:XM_024449401:XM_011534848:XM_011534847:NM_001353644:NR_148485:NM_001353646:NM_001353647:XM_017020709:NR_148491:NR_148492:XM_005266202:NM_080687:NR_148490:NR_148494:XM_024449400:NR_148482:NR_148496:NR_148486:NR_148493:XR_941422:NM_001353645:XR_001749649:NM_001353649:XM_011534845 | 12:8:5:10:0:1:5:13:1:7:1:0:0:7:3:0:0:0:0:8:0:2:0:0:0:0:1:1:2:1:0:0 | 2:0:0:2:0:0:0:2:0:1:0:0:0:0:0:1:0:0:0:0:0:0:0:0:0:0:0:0:0:0:0:0 | 19:27:10:21:2:2:10:21:0:1:2:1:1:16:14:2:1:3:7:6:1:2:1:0:0:0:0:0:0:0:0:0 | 5:5:1:2:1:2:3:3:1:1:3:0:0:0:0:0:0:0:0:0:0:0:0:0:0:0:0:0:0:0:0:0 | 21:8:20:12:1:0:4:17:1:7:3:0:0:6:4:1:0:0:8:10:0:5:0:0:1:1:0:3:0:0:1:0 | 1:2:1:3:0:0:0:4:0:0:0:0:0:0:0:0:0:0:2:0:0:0:0:0:0:0:0:0:0:0:0:0 |
| NM_023011:NM_001353647:NM_001353646:NR_148490:XM_017020709:NR_148492:NM_080687:NR_148494:XM_017020711:XM_011534848:XM_024449400:XM_005266202:XM_024449401:XM_006719991:NR_148486:XR_941422:NM_001353645:XR_001749649:NM_001353649:XM_011534847:NR_148484:NR_148488 | 6:2:4:0:0:0:6:0:0:1:0:0:1:1:0:1:1:1:0:0:0:0 | 0:1:0:0:0:0:2:1:0:1:0:0:0:0:0:0:0:0:0:0:0:0 | 9:10:9:0:1:1:5:2:1:1:1:0:0:0:0:0:0:0:0:0:0:0 | 2:2:2:1:0:0:0:0:0:0:0:0:0:0:0:0:0:0:0:0:0:0 | 7:3:4:0:0:0:7:5:2:3:0:2:1:0:1:3:0:0:1:0:0:1 | 1:0:1:0:0:0:1:0:1:0:0:2:0:0:0:0:0:0:0:0:0:0 |
| NM_152726:XR_429214:XR_001749501:XM_017020433 | 142:3:1:3 | 0:0:0:0 | 185:12:7:1 | 0:0:0:0 | 214:13:0:7 | 0:0:0:0 |
| NM_005932:XM_011535097:XM_011535098 | 0:00:00 | 92:01:03 | 0:00:00 | 75:01:00 | 0:00:00 | 123:02:00 |
| NM_001286504:NM_006644:NM_001286503:XM_011534887:XM_017020364:XM_017020363:XM_005266236:XM_017020362 | 1:0:0:1:0:0:0:0 | 0:0:0:0:0:0:0:0 | 3:15:1:2:1:0:0:0 | 0:0:0:0:0:0:0:0 | 8:4:0:2:0:0:0:0 | 1:3:0:0:1:1:0:0 |
| NM_020751:NR_026745:XM_011535168:NM_001145079 | 7:0:0:0 | 0:0:0:0 | 51:2:1:0 | 0:0:0:0 | 3:0:0:0 | 8:2:0:0 |
| NM_020751:NR_026745:XM_011535168:NM_001145079 | 8:0:0:0 | 0:0:0:0 | 51:2:1:0 | 0:0:0:0 | 3:0:0:0 | 10:2:0:0 |
| NM_020751:NR_026745:XM_011535168:NM_001145079 | 6:2:0:0 | 4:4:0:0 | 49:3:1:0 | 3:7:0:0 | 15:15:1:0 | 0:0:0:0 |
| NM_172373:NM_001145353 | 0:00 | 13:00 | 0:00 | 44:00:00 | 0:00 | 32:00:00 |
| NM_003295:NM_001286273:NM_001286272 | 2663:18:08 | 0:00:00 | 9527:66:14 | 14:00:00 | 7409:53:15 | 10:01:00 |
| NM_003295:NM_001286273:NM_001286272 | 2661:18:08 | 3:00:00 | 9543:66:14 | 9:00:00 | 7414:54:15 | 8:00:00 |
| NM_003295:NM_001286273:NM_001286272 | 2663:18:08 | 1:00:00 | 9541:66:14 | 18:00:00 | 7407:54:15 | 6:00:00 |
| NM_003295:NM_001286273:NM_001286272 | 2662:18:08 | 1:00:00 | 9588:66:14 | 5:00:00 | 7393:53:15 | 0:02:00 |
| NM_003295:NM_001286273:NM_001286272 | 2659:19:08 | 2:00:00 | 9593:66:14 | 16:00:00 | 7393:54:15 | 4:01:00 |
| NM_003295:NM_001286273:NM_001286272 | 2657:19:08 | 2:00:00 | 9606:67:14 | 13:00:00 | 7371:54:15 | 13:01:00 |
| NM_003295:NM_001286273:NM_001286272 | 2638:19:08 | 3:00:00 | 9554:67:14 | 13:00:00 | 7347:55:15 | 5:00:00 |
| NM_003295:NM_001286273:NM_001286272 | 2641:19:08 | 2:00:00 | 9548:67:14 | 16:00:00 | 7341:55:15 | 7:00:00 |
| NM_003295:NM_001286273:NM_001286272 | 2635:19:08 | 8:00:00 | 9536:67:14 | 19:00:00 | 7312:54:15 | 26:01:00 |
| NM_003295:NM_001286273:NM_001286272 | 2617:19:08 | 4:00:00 | 9460:67:13 | 8:00:00 | 7288:54:15 | 6:00:00 |
| NM_003295:NM_001286273:NM_001286272 | 2531:19:08 | 6:00:00 | 9150:67:13 | 12:00:00 | 7086:51:15 | 10:00:00 |
| XM_024449333:XM_017020480:NM_015070:NM_001330567 | 1:0:0:0 | 0:0:0:0 | 0:0:0:0 | 1:0:0:0 | 1:0:0:0 | 0:0:1:0 |
| NM_014166:NM_001270629:XM_017020549 | 171:66:8 | 7:02:00 | 569:177:17 | 12:04:00 | 438:157:11 | 8:03:00 |
| NM_014166:NM_001270629:XM_017020549 | 188:73:6 | 6:00:00 | 604:190:15 | 11:00:00 | 476:186:10 | 20:03:00 |
| NM_014166:NM_001270629:XM_017020549 | 192:73:6 | 2:00:00 | 609:190:15 | 8:00:00 | 483:187:10 | 15:02:00 |
| NM_014166:NM_001270629:XM_017020549 | 202:72:6 | 3:00:00 | 636:190:14 | 12:00:00 | 516:190:9 | 7:00:00 |
| NM_014166:NM_001270629:XM_017020549 | 205:73:6 | 0:00:00 | 646:190:14 | 8:00:00 | 523:190:9 | 4:00:00 |
| NM_006346:XM_017020350:XM_011534885:XM_017020354:NR_146206:NM_001349655:XR_001749460:XR_002957449:XM_017020351:XR_001749457:XR_001749459:XM_024449314:NR_146205:XR_941461:XM_017020352 | 2:0:0:0:0:0:0:0:0:0:0:0:0:0:0 | 2:0:0:0:0:0:0:0:0:0:0:0:0:0:0 | 16:1:1:0:0:0:0:0:0:0:0:0:0:0:0 | 1:0:0:0:0:0:0:0:0:0:0:0:0:0:0 | 2:1:0:0:0:0:0:0:0:0:0:0:0:0:0 | 13:1:0:3:0:1:0:1:0:0:0:0:0:0:0 |
| NM_005348:XM_011536718:NM_001017963 | 96:25:00 | 0:00:00 | 134:33:00 | 0:00:00 | 136:41:00 | 0:00:00 |
| NM_005348:XM_011536718:NM_001017963 | 97:24:00 | 0:00:00 | 134:32:00 | 0:00:00 | 137:40:00 | 1:00:00 |
| NM_005348:XM_011536718:NM_001017963 | 105:21:00 | 0:00:00 | 138:23:00 | 0:00:00 | 145:31:00 | 0:00:00 |
| NM_145725:XM_017021618:NM_003300:XM_011537117:NM_145726:XM_017021619:XM_017021620 | 7:0:3:0:0:0:0 | 3:2:1:0:1:0:0 | 1:0:0:0:0:0:0 | 2:1:1:0:0:0:0 | 1:0:0:0:0:0:0 | 3:1:0:0:0:0:0 |
| NM_080648:NM_001641:NM_080649 | 0:00:00 | 0:00:00 | 0:00:00 | 0:00:00 | 0:00:00 | 0:00:00 |
| NM_002818:XM_006720213 | 0:00 | 9:02 | 0:00 | 7:00 | 0:00 | 5:00 |
| NM_182836:NM_004581 | 31:03:00 | 32:03:00 | 2:00 | 16:01 | 20:00 | 5:00 |
| XR_943381:XM_011536374:XM_024449460:NM_182648 | 24:0:1:1 | 2:0:0:0 | 9:1:0:0 | 2:0:0:0 | 21:0:3:0 | 0:0:0:0 |

| | | | | | | |
|---|---|---|---|---|---|---|
| XR_943381:XM_011536374:XM_024449460:NM_182648 | 36:0:2:1 | 5:0:0:1 | 11:1:1:0 | 3:0:0:0 | 25:1:3:1 | 5:0:1:1 |
| XM_024449460:XR_943381:XM_011536374:NM_182648 | 2:40:0:2 | 0:18:3:1 | 1:13:1:0 | 1:8:0:0 | 4:27:1:2 | 2:13:0:1 |
| XR_943381:XM_011536374:XM_024449460:NM_182648 | 66:3:3:3 | 1:0:0:0 | 21:1:2:0 | 0:0:0:0 | 48:1:9:4 | 0:0:0:0 |
| NM_014672:XM_005268237:NM_001256680:NM_001256681:XM_017021836:XM_011537410:NM_001256678 | 2:1:6:4:4:6:1 | 0:0:0:0:0:0:0 | 2:1:7:1:2:0:0 | 0:0:0:0:0:0:0 | 5:5:12:2:1:3:2 | 0:0:0:0:0:1:0 |
| XM_005268075:NM_024884:XM_017021657:XM_011537166:XM_017021658:XM_017021656 | 3:0:1:0:0:0 | 0:0:0:0:0:0 | 2:0:0:0:0:0 | 0:0:0:0:0:0 | 3:0:0:0:0:0 | 0:0:0:0:0:0 |
| NM_002306:NM_001357678:NR_003225 | 125:02:04 | 0:00:00 | 7:00:00 | 0:00:00 | 13:00:00 | 27:01:00 |
| NM_002306:NR_003225:NM_001357678 | 125:08:02 | 0:00:00 | 7:01:00 | 0:00:00 | 13:00:00 | 27:00:01 |
| NM_145112:NM_002382:XM_017021312:NR_073137:NM_001320415:XR_002957553:XR_001750327:XR_943452:NM_145113:XR_943451:XM_017021313:XR_943450:XR_001750326 | 361:233:6:13:12:1:4:4:6:1:3:0:0 | 0:0:0:0:0:0:0:0:0:0:0:0:0 | 93:45:10:3:3:2:3:1:1:0:0:0:0 | 0:0:0:0:0:0:0:0:0:0:0:0:0 | 85:85:3:4:4:1:2:1:1:2:0:3:2 | 0:0:0:0:0:0:0:0:0:0:0:0:0 |
| NM_001130005:NM_001102:XM_011537267:XM_017021728:XM_011537265:XM_017021723:NM_001130004:XM_017021727:XM_017021726 | 3:3:3:2:0:0:0:0:0 | 5:3:9:1:0:0:0:0:0 | 4:10:2:0:0:0:0:0:0 | 1:1:0:0:0:0:0:0:0 | 1:22:8:0:0:1:0:0:0 | 0:0:0:0:0:0:0:0:0 |
| NM_001193360:NM_001193363:XM_017021421:NM_001193361:NM_018199:XM_011536908 | 3:1:0:1:0:0 | 4:0:1:1:0:0 | 0:0:0:0:0:0 | 2:1:1:0:0:0 | 0:0:2:0:0:0 | 2:0:1:0:0:0 |
| XR_943501:NM_021239:XM_011537044:XM_024449680 | 48:11:5:0 | 1:0:0:0 | 30:7:2:2 | 1:0:0:0 | 42:7:1:0 | 0:0:0:0 |
| NM_021239:XM_011537044:XR_943501:XM_024449680 | 10:5:47:0 | 1:0:0:0 | 7:2:31:2 | 0:0:0:0 | 7:1:40:0 | 0:0:1:0 |
| XR_943501:NM_021239:XM_011537044:XM_024449680 | 46:10:4:0 | 2:0:0:0 | 27:7:2:2 | 3:0:0:0 | 38:7:1:0 | 1:0:0:0 |
| NM_021239:XR_943501:XM_011537044:XM_024449680 | 5:32:2:0 | 1:2:0:0 | 5:16:2:3 | 1:3:0:0 | 7:27:1:0 | 0:2:0:0 |
| XR_943501:XM_024449680:NM_021239:XM_011537044 | 4:0:4:1 | 10:0:0:0 | 0:3:2:1 | 5:0:0:0 | 3:0:0:0 | 1:0:1:0 |
| XM_005267413:NM_012245:NM_001318844 | 7:30:01 | 2:00:00 | 7:27:00 | 3:00:00 | 4:42:00 | 3:00:00 |
| NM_001128595:NM_001289134:NM_152332 | 0:00:00 | 0:00:00 | 0:00:00 | 0:00:00 | 0:00:00 | 0:00:00 |
| NM_001002235:NM_001127707:NM_000295:NM_001127706:XM_017021370:NM_001002236:NM_001127702:NM_001127705:NM_001127703:NM_001127700:NM_001127704:NM_001127701 | 659:58:2:45:86:6:2:2:10:15:5:2 | 0:0:0:0:0:0:0:0:0:0:0:0 | 24:46:18:5:4:1:1:0:0:0:0:0 | 0:0:0:0:0:0:0:0:0:0:0:0 | 35:74:25:4:18:0:1:0:1:2:0:0 | 0:0:0:0:0:0:0:0:0:0:0:0 |
| NM_001002235:NM_001127707:NM_000295:NM_001127706:NM_001002236:NM_001127702:XM_017021370:NM_001127705:NM_001127703:NM_001127700:NM_001127704:NM_001127701 | 655:48:2:45:6:2:40:2:10:15:4:2 | 0:0:0:0:0:0:1:0:0:0:0:0 | 24:45:19:5:1:1:1:0:0:0:0:0 | 0:0:0:0:0:0:0:0:0:0:0:0 | 35:73:25:4:0:1:4:0:1:2:0:0 | 0:0:0:0:0:0:0:0:0:0:0:0 |
| NM_024734:XR_429332:XM_017021647:XR_001750558:XM_017021646:XR_429330 | 14:28:1:0:0:0 | 0:3:0:0:0:0 | 1:1:0:0:0:0 | 0:0:0:0:0:0 | 4:10:0:0:0:0 | 0:1:0:0:0:0 |
| NM_003384:XM_006720247:XR_001750539:XM_017021624:XM_017021625:XM_017021626 | 48:0:1:2:0:0 | 0:0:0:0:0:0 | 24:1:0:0:0:0 | 0:0:0:0:0:0 | 82:2:1:4:1:0 | 0:0:0:0:0:0 |
| NM_144508:XM_017022432 | 0:00 | 3:01 | 0:00 | 4:00 | 11:03 | 0:00 |
| NM_144508:XM_017022432:NM_170589 | 0:00:00 | 3:00:00 | 0:00:00 | 4:00:00 | 11:01:00 | 0:00:00 |
| NM_144508:XM_017022432 | 0:01 | 0:00 | 2:00 | 0:00 | 3:00 | 0:00 |
| NM_144508:XM_017022432 | 0:00 | 0:00 | 0:00 | 1:00 | 2:00 | 0:00 |
| NM_001355001:XR_001751382:XR_931899 | 0:00:01 | 0:00:00 | 1:01:00 | 0:00:00 | 0:00:00 | 0:00:00 |
| NM_004048:XR_002957658:XM_005254549 | 136792:1344:127 | 102:06:00 | 13000:137:22 | 7:00:00 | 17778:163:15 | 18:00:00 |
| NM_004048:XR_002957658:XM_005254549 | 136969:1352:127 | 54:04:00 | 13009:138:22 | 6:00:00 | 17799:165:15 | 10:00:00 |
| NM_004048:XR_002957658:XM_005254549 | 137989:1368:129 | 347:15:00 | 13126:139:22 | 23:01:01 | 17913:164:16 | 30:00:00 |
| NM_004048:XR_002957658:XM_005254549 | 278032:2724:258 | 604:28:00 | 26446:282:48 | 30:00:00 | 36048:328:32 | 50:00:00 |
| XR_002957658:NM_004048:XM_005254549 | 1287:140956:132 | 0.756944444 | 132:13357:24 | 5:11:00 | 155:18194:17 | 2:24:00 |
| NM_004048:XR_002957658:XM_005254549 | 141172:1261:132 | 224:17:00 | 13378:130:24 | 10:05:00 | 18212:149:17 | 19:03:00 |
| NM_004048:XR_002957658:XM_005254549 | 284706:378:308 | 412:16:02 | 26998:44:48 | 42:02:00 | 36674:48:38 | 38:00:00 |
| NM_001128610:XM_017022720:XM_017022718:NM_001283049:XM_017022719:XM_017022722:XM_006720762:XM_011522193:XM_024450098:NM_005154:XM_017022721 | 1:22:82:0:34:3:5:0:2:1:2 | 0:0:0:0:0:0:0:0:0:0:0 | 4:12:57:2:20:2:3:1:1:0:0 | 0:0:0:0:0:0:0:0:0:0:0 | 4:31:73:0:41:5:9:0:1:5:4 | 0:0:1:0:1:0:0:0:0:0:0 |
| NM_001128610:XM_017022720:XM_017022718:NM_001283049:XM_017022719:XM_017022722:XM_006720762:XM_024450098:XM_011522193:NM_005154:XM_017022721 | 2:44:154:0:64:8:10:20:0:2:4 | 0:0:4:0:0:0:0:2:0:0:0 | 8:20:98:4:38:4:6:14:2:0:0 | 0:0:0:0:0:0:0:0:0:0:0 | 6:60:136:0:70:8:14:30:0:10:6 | 0:0:0:0:0:0:0:2:0:0:0 |
| XM_017022718:XM_017022720:XM_017022719:XM_017022722:XM_024450098:NM_001128610:NM_001283049:XM_006720762:XM_011522193:XM_017022721:NM_005154 | 72:20:25:3:9:1:0:5:0:2:1 | 7:2:7:1:1:0:0:0:0:0:0 | 43:8:17:0:5:4:2:3:1:0:0 | 6:1:1:1:2:0:0:0:0:0:0 | 64:27:30:4:13:3:0:5:0:3:5 | 3:2:4:0:3:0:0:1:0:0:0 |

| | | | | | | |
|---|---|---|---|---|---|---|
| XM_017022718:XM_017022720:NM_001128610:NM_001283049:XM_017022719:XM_006720762:XM_024450098:XM_011522193:NM_005154:XM_017022721:XM_017022722 | 118:36:2:0:46:10:18:0:2:2:6 | 8:0:0:0:2:0:0:0:0:2:0 | 72:12:8:2:32:6:10:2:0:0:0 | 6:4:0:0:0:0:0:0:0:0:0 | 106:52:4:0:52:10:24:0:6:6:8 | 8:2:2:0:4:0:0:0:2:0:0 |
| XM_017022719:XM_017022718:NM_001128610:XM_011522193:XM_024450098:XM_017022722:XM_017022720:NM_005154:XM_006720762 | 3:1:1:0:2:1:6:0:0 | 2:1:0:0:0:0:0:1:0 | 1:2:1:1:2:0:0:0:0 | 0:0:0:0:0:0:0:0:0 | 0:1:0:0:5:2:5:1:0 | 0:0:0:0:0:0:1:0:0 |
| XM_017022719:XM_017022718:NM_001128610:XM_011522193:XM_024450098:XM_017022720:XM_017022722:NM_005154:XM_006720762 | 2:1:0:0:4:6:1:0:0 | 0:0:0:0:0:0:0:0:0 | 1:1:1:1:3:0:0:0:0 | 0:0:0:0:0:0:0:0:0 | 0:0:0:0:7:5:2:1:0 | 0:0:0:0:0:0:0:0:0 |
| NM_020739:NM_001204450:NM_004748 | 106:26:04 | 6:00:00 | 58:08:00 | 2:00:00 | 54:02:00 | 2:00:00 |
| NM_020739:NM_001204450:NM_004748 | 150:26:04 | 12:00:00 | 72:08:00 | 6:00:00 | 62:02:02 | 4:00:02 |
| NM_001204450:NM_020739:NM_004748:NM_001204451 | 13:91:2:1 | 0:2:0:0 | 4:41:0:0 | 0:0:0:0 | 1:37:2:0 | 0:2:0:0 |
| NM_024666:NM_001271886:XM_011522020:XM_024450052:XM_024450053:NM_001271885 | 158:12:14:0:0:0 | 112:8:14:0:0:0 | 8:0:4:0:0:0 | 80:6:6:2:0:0 | 0:0:0:0:0:0 | 318:14:18:0:0:2 |
| NM_016166:XM_024450094:XM_017022691:XM_017022689:XM_011522127:XM_017022690:XM_011522126 | 84:6:4:0:2:0:0 | 4:4:0:0:0:0:0 | 64:4:0:0:0:0:0 | 0:0:0:0:0:0:0 | 60:2:4:0:0:0:0 | 2:0:0:0:0:0:0 |
| NM_001008224:XM_017022394:NM_018003:XM_017022395:XM_011521754 | 0:3:12:1:0 | 0:2:8:1:0 | 2:1:21:1:1 | 1:2:5:0:0 | 2:3:16:2:0 | 0:0:0:0:0 |
| NM_018003:NM_001008224:XM_017022394:XM_011521754:XM_017022395 | 21:0:7:0:2 | 0:0:0:0:0 | 27:3:3:1:0 | 0:0:0:0:0 | 17:2:3:0:2 | 0:0:0:0:0 |
| NM_145204:XM_005254157:NM_001172111:XM_011521217:XM_011521215 | 0:0:0:0:0 | 26:0:0:0:0 | 0:0:0:0:0 | 4:0:0:0:0 | 0:0:0:0:0 | 18:0:0:0:0 |
| XR_001751314:NM_002654:XM_011521670:XM_005254445:NM_001206797:NM_182471:XM_006720570:NM_001206798 | 2:176:16:0:0:0:0:0 | 0:2:0:0:0:0:0:0 | 4:253:32:1:0:0:0:0 | 1:1:0:0:0:0:0:0 | 4:695:79:3:2:1:1:0 | 1:2:0:0:0:0:0:0 |
| NM_002654:XM_011521670:XM_005254445:XR_001751314:NM_001206797:NM_182471:XM_006720570:NM_001206798 | 177:16:0:1:0:0:0:0 | 1:0:0:0:0:0:0:0 | 251:32:1:5:0:0:0:0 | 2:0:0:0:0:0:0:0 | 695:79:3:4:2:1:1:0 | 0:0:0:1:0:0:0:0 |
| NM_002654:XR_001751314:XM_011521670:XM_005254445:NM_182471:XM_017022313:NM_001206797:NM_182470:XM_006720570:NM_001206798 | 165:3:12:0:7:0:1:0:0:0 | 0:0:0:0:0:0:0:0:0:0 | 236:7:21:1:14:1:0:0:0:0 | 0:0:0:0:0:0:0:0:0:0 | 655:10:61:3:13:4:2:1:1:0 | 0:1:0:0:0:0:0:0:0:0 |
| NM_002654:NM_001206797:XR_001751314:NM_182471:XM_011521670:XM_005254445:XM_017022313:NM_182470:XM_006720570:NM_001206798 | 316:2:18:12:16:0:0:0:0:0 | 0:0:0:0:0:0:0:0:0:0 | 452:2:36:30:34:2:2:0:0:0 | 0:0:0:0:0:0:0:0:0:0 | 1244:4:60:28:106:6:8:2:2:0 | 2:0:2:0:0:0:0:0:0:0 |
| NM_002654:NM_001206797:XR_001751314:NM_182471:XM_011521670:XM_005254445:XM_017022313:NM_182470:XM_006720570:NM_001206798 | 157:1:17:6:6:0:0:0:0:0 | 0:0:0:0:0:0:0:0:0:0 | 222:1:30:16:12:1:1:0:0:0 | 1:0:0:0:0:0:0:0:0:0 | 614:2:51:16:47:3:3:1:1:0 | 0:0:0:0:0:0:0:0:0:0 |
| NM_002654:XR_001751314:NM_001206797:NM_182471:XM_011521670:XM_005254445:XM_017022313:NM_182470:XM_006720570:NM_001206799:NM_001206798 | 149:31:1:7:6:0:0:0:0:0:0 | 0:0:0:0:0:0:0:0:0:0:0 | 213:52:1:16:9:1:0:0:0:0:0 | 0:0:0:0:0:0:0:0:0:0:0 | 599:74:2:23:38:3:3:1:1:0:0 | 1:3:0:0:0:0:0:0:0:0:0 |
| NM_002654:XR_001751314:NM_001206797:NM_182471:XM_011521670:XM_005254445:XM_017022313:NM_182470:XM_006720570:NM_001206799:NM_001206798 | 148:31:1:8:6:0:0:0:0:0:0 | 1:0:0:0:0:0:0:0:0:0:0 | 215:52:1:16:9:1:0:0:0:0:0 | 0:0:0:0:0:0:0:0:0:0:0 | 607:80:2:23:36:3:3:1:1:0:0 | 0:1:0:0:1:0:0:0:0:0:0 |
| NM_002654:XR_001751314:NM_182471:NM_001206797:XM_011521670:XM_005254445:XM_017022313:NM_182470:XM_006720570:NM_001206798:NM_001206799 | 149:34:8:0:5:0:0:0:0:0:0 | 0:0:0:0:0:0:0:0:0:0:0 | 214:60:21:1:6:1:0:0:0:0:0 | 0:0:0:0:0:0:0:0:0:0:0 | 607:103:26:1:30:3:4:1:1:0:0 | 0:0:0:1:0:0:0:0:0:0:0 |
| NM_002654:XR_001751314:NM_182471:NM_001206797:XM_011521670:XM_005254445:XM_017022313:NM_182470:XM_006720570:NM_001206798:NM_001206799 | 149:34:8:0:5:0:0:0:0:0:0 | 0:0:0:0:0:0:0:0:0:0:0 | 214:60:21:1:6:1:0:0:0:0:0 | 0:0:0:0:0:0:0:0:0:0:0 | 607:103:26:1:30:3:4:1:1:0:0 | 0:0:0:1:0:0:0:0:0:0:0 |
| NM_002654:XR_001751314:NM_182471:NM_001206797:XM_011521670:XM_005254445:XM_017022313:NM_182470:XM_006720570:NM_001206798:NM_001206799 | 149:34:8:0:5:0:0:0:0:0:0 | 0:0:0:0:0:0:0:0:0:0:0 | 213:60:21:1:6:1:0:0:0:0:0 | 1:0:0:0:0:0:0:0:0:0:0 | 607:103:26:1:30:3:4:1:1:0:0 | 0:0:0:1:0:0:0:0:0:0:0 |
| NM_002654:XR_001751314:NM_182471:XM_011521670:XM_005254445:XM_017022313:NM_182470:XM_006720570:NM_001206798:NM_001206799 | 150:39:10:5:0:0:0:0:0:0 | 0:1:0:0:0:0:0:0:0:0 | 211:68:21:8:1:1:0:0:0:0 | 2:0:0:0:0:0:0:0:0:0 | 602:117:28:25:3:4:1:1:0:0 | 2:0:0:0:0:0:0:0:0:0 |
| NM_002654:NM_182471:NM_001206797:XR_001751314:XM_005254445:XM_011521670:XM_017022313:NM_182470:XM_006720570:NM_001206798:NM_001206799 | 153:18:1:41:0:4:0:0:0:0:0 | 1:0:0:0:0:0:0:0:0:0:0 | 214:38:1:77:1:11:2:0:0:0:0 | 0:0:0:0:0:0:0:0:0:0:0 | 621:67:2:138:3:25:3:1:1:0:0 | 0:0:0:0:0:0:0:0:0:0:0 |
| NM_033250:NM_033240:NM_033244:NM_033239 | 0:0:0:0 | 2:2:0:0 | 0:0:0:0 | 0:0:0:0 | 0:0:0:0 | 2:0:0:0 |
| NM_033240:NM_033250:NM_033244:NM_033239 | 0:2:0:0 | 2:0:0:0 | 0:0:0:0 | 0:0:0:0 | 0:2:0:0 | 2:0:0:0 |
| NM_005724:XM_017021857:NM_198902 | 1388:02:00 | 4:00:00 | 324:00:00 | 0:00:00 | 402:00:00 | 6:00:00 |
| NM_004049:NM_001114735 | 8:00 | 4:04 | 14:00 | 4:04 | 2:00 | 4:04 |
| NM_004049:NM_001114735 | 8:00 | 4:04 | 14:00 | 4:04 | 2:00 | 4:00 |
| NM_004049:NM_001114735 | 8:00 | 4:04 | 14:00 | 4:04 | 2:00 | 4:00 |

| | | | | | | |
|---|---|---|---|---|---|---|
| NM_007200:NM_001270546:NM_006738 | 0:00:00 | 6:00:00 | 0:00:00 | 2:00:00 | 1:00:00 | 0:00:00 |
| NM_007200:NM_001270546:NM_006738 | 0:00:00 | 6:00:00 | 0:00:00 | 2:00:00 | 1:00:00 | 0:00:00 |
| NM_007200:NM_001270546:NM_006738 | 0:00:00 | 8:00:00 | 0:00:00 | 2:00:00 | 1:00:00 | 0:00:00 |
| NM_001270546:NM_007200:NM_006738 | 10:12:00 | 2:06:00 | 2:08:00 | 4:00:00 | 2:06:02 | 0:00:00 |
| NM_001150:XM_011521473 | 84:51:00 | 101:55:00 | 1:00 | 0:00 | 1:01 | 0:00 |
| NM_001150:XM_011521473 | 75:42:00 | 56:34:00 | 0:00 | 1:00 | 1:00 | 0:00 |
| NM_001004309:XR_001751249:XM_017022127:XM_017022128:XM_017022129 | 2:0:0:0:0 | 0:0:0:0:0 | 0:0:0:0:0 | 0:0:0:0:0 | 2:0:0:0:0 | 0:2:0:0:0 |
| XM_011522187:XM_006720760:XM_005254987:XM_006720759:NM_199413:XM_011522188:NM_001267580:NM_003981:XM_017022717:XM_017022713:XM_011522191 | 0:0:0:0:0:0:0:0:0:0:0 | 0:0:6:0:4:2:0:0:0:0:0 | 0:0:0:0:0:0:0:0:0:0:0 | 2:6:8:2:0:0:0:0:0:0:0 | 0:0:0:0:0:0:0:0:0:0:0 | 0:2:24:4:34:0:6:12:0:6:0 |
| NM_001271:NM_001042572 | 12:20 | 2:00 | 16:20 | 0:00 | 18:18 | 2:00 |
| NM_001271:NM_001042572 | 5:10 | 0:00 | 4:10 | 1:00 | 6:10 | 1:00 |
| NM_002484:NM_001323595:NM_001278506:XM_017023252 | 11:0:2:0 | 0:0:0:0 | 15:2:0:0 | 0:0:0:0 | 10:0:0:0 | 10:0:0:0 |
| XM_011545822:XM_011545823:NM_001169 | 3:00:03 | 8:01:02 | 0:00:00 | 0:00:00 | 0:00:00 | 0:00:00 |
| NM_152338:XM_011545921 | 0:00 | 146:01:00 | 0:00 | 0:00 | 0:00 | 6:00 |
| NM_152338:XM_011545921 | 0:00 | 151:01:00 | 0:00 | 0:00 | 0:00 | 6:00 |
| NM_184041:NM_184043:NM_001127617:NM_001243177 | 3390:44:3:6 | 2:0:0:0 | 1775:32:3:3 | 0:0:0:0 | 2369:38:6:6 | 2:0:0:0 |
| NM_184041:NM_184043:NM_001127617:NM_001243177 | 3387:46:3:6 | 1:0:0:0 | 1770:33:3:3 | 2:1:0:0 | 2368:38:6:6 | 2:0:0:0 |
| NM_184041:NM_184043:NM_001127617:NM_001243177 | 3379:52:3:6 | 5:0:0:0 | 1769:42:3:3 | 1:0:0:0 | 2365:40:6:6 | 3:0:0:0 |
| NM_184041:NM_184043:NM_001127617:NM_001243177 | 3361:48:3:6 | 4:0:0:0 | 1763:48:3:3 | 1:0:0:0 | 2361:46:6:6 | 4:0:0:0 |
| NM_184041:NM_184043:NM_001127617:NM_001243177 | 3350:54:3:6 | 6:0:0:0 | 1758:50:3:3 | 2:0:0:0 | 2362:50:6:6 | 1:0:0:0 |
| NM_184041:NM_184043:NM_001127617:NM_001243177 | 3351:54:3:6 | 2:0:0:0 | 1758:50:3:3 | 2:0:0:0 | 2360:50:6:6 | 2:0:0:0 |
| NM_184041:NM_184043:NM_001127617:NM_001243177 | 3344:72:3:6 | 4:0:0:0 | 1755:57:3:3 | 0:0:0:0 | 2358:62:6:6 | 1:0:0:0 |
| NM_184041:NM_184043:NM_001127617:NM_001243177 | 3344:72:3:6 | 4:0:0:0 | 1755:58:3:3 | 0:0:0:0 | 2359:63:6:6 | 0:0:0:0 |
| NM_184041:NM_184043:NM_001127617:NM_001243177 | 3339:73:3:6 | 9:0:0:0 | 1753:59:3:3 | 2:0:0:0 | 2358:63:6:6 | 1:0:0:0 |
| NM_184041:NM_184043:NM_001127617:NM_001243177 | 3344:84:3:6 | 4:0:0:0 | 1751:64:3:3 | 2:0:0:0 | 2358:66:6:6 | 1:1:0:0 |
| NM_184041:NM_184043:NM_001127617:NM_001243177 | 3344:84:3:6 | 4:0:0:0 | 1750:63:3:3 | 3:1:0:0 | 2358:66:6:6 | 1:1:0:0 |
| NM_184043:NM_184041:NM_001127617:NM_001243177 | 86:3344:3:6 | 1:3:0:0 | 66:1753:3:3 | 1:0:0:0 | 69:2356:6:6 | 1:0:0:0 |
| NM_184041:NM_184043:NM_001127617:NM_001243177 | 3346:87:3:6 | 2:4:0:0 | 1752:68:3:3 | 1:0:0:0 | 2357:70:6:6 | 1:1:0:0 |
| NM_184041:NM_184043:NM_001127617:NM_001243177 | 3342:91:3:6 | 3:0:0:0 | 1749:69:3:3 | 1:0:0:0 | 2357:73:6:6 | 1:1:0:0 |
| NM_184041:NM_184043:NM_001127617:NM_001243177 | 3334:91:3:6 | 5:0:0:0 | 1747:69:3:3 | 3:0:0:0 | 2357:72:6:6 | 1:2:0:0 |
| NM_184041:NM_184043:NM_001127617:NM_001243177 | 3336:97:3:6 | 6:1:0:0 | 1751:69:3:3 | 1:0:0:0 | 2357:75:6:6 | 2:1:0:0 |
| NM_184041:NM_184043:NM_001127617:NM_001243177 | 3333:103:3:5 | 5:1:0:1 | 1750:79:3:3 | 1:0:0:0 | 2354:81:6:6 | 2:1:0:0 |

| | | | | | | |
|---|---|---|---|---|---|---|
| NM_184041:NM_184043:NM_001127617:NM_001243177 | 3331:113:3:6 | 4:0:0:0 | 1745:81:3:3 | 3:2:0:0 | 2350:92:6:6 | 0:0:0:0 |
| NM_184043:NM_184041:NM_001127617:NM_001243177 | 132:3304:3:6 | 1:4:0:0 | 89:1725:3:3 | 2:0:0:0 | 121:2341:6:6 | 0:2:0:0 |
| NM_184041:NM_184043:NM_001127617:NM_001243177 | 3308:153:3:6 | 3:0:0:0 | 1726:96:3:3 | 1:1:0:0 | 2341:137:6:6 | 1:0:0:0 |
| NM_184041:NM_184043:NM_001127617:NM_001243177 | 3303:160:3:6 | 7:1:0:0 | 1729:101:3:3 | 3:0:0:0 | 2342:146:6:6 | 0:0:0:0 |
| NM_184041:NM_184043:NM_001127617:NM_001243177 | 3304:170:3:6 | 9:0:0:0 | 1727:105:3:3 | 4:0:0:0 | 2341:150:6:6 | 2:1:0:0 |
| NM_184041:NM_184043:NM_001127617:NM_001243177 | 3304:172:3:6 | 4:0:0:0 | 1726:108:3:3 | 1:0:0:0 | 2339:156:6:6 | 3:1:0:0 |
| NM_001170634:NM_004960:XM_024450221:NR_028388:XM_005255233:XM_011545781:XM_011545782:NM_001170937 | 25:60:2:1:0:0:0:0 | 0:0:0:1:0:0:0:0 | 12:29:3:7:1:0:0:0 | 2:0:0:0:0:0:0:0 | 24:55:3:1:1:0:0:0 | 2:0:0:0:0:0:0:0 |
| XM_011522393:NM_005223:XM_024450171:XM_024450170:XR_002957783:XM_017022993:XR_002957785:XM_024450172:NM_001351825:XM_017023001 | 0:0:0:0:0:0:0:0:0:0 | 1:0:0:0:0:0:0:0:0:0 | 1:2:1:0:0:0:0:0:0:0 | 3:0:0:0:0:0:0:0:0:0 | 0:0:0:0:1:0:0:1:0:0 | 0:0:0:0:0:0:0:0:0:0 |
| NM_016292:XM_011522345:NM_001272049 | 6:01:01 | 0:00:00 | 3:01:00 | 0:00:00 | 8:00:01 | 7:00:01 |
| NM_016292:XM_011522345:NM_001272049 | 1:00:00 | 3:01:00 | 0:00:00 | 2:00:00 | 6:00:00 | 4:00:00 |
| NM_016292:XM_011522345:NM_001272049 | 4:01:00 | 0:00:00 | 2:00:00 | 0:00:00 | 6:00:00 | 4:00:00 |
| NM_016292:XM_011522345:NM_001272049 | 5:00:01 | 2:00:00 | 3:00:00 | 0:00:00 | 8:00:01 | 7:00:00 |
| NM_001797:XM_024450134:NM_001308392 | 0:00:00 | 0:01:00 | 0:00:00 | 3:01:00 | 0:00:00 | 0:00:00 |
| XM_024450134:NM_001797 | 1:00 | 0:00 | 0:02 | 1:01 | 0:00 | 0:00 |
| NM_003905:NM_001018159:NM_001018160:XM_005256215:NM_001286500 | 145:6:3:3:0 | 0:0:0:0:0 | 118:16:4:4:0 | 1:0:0:0:0 | 225:22:12:3:0 | 4:0:0:0:0 |
| XM_005255775:NM_006565:XM_017022868 | 12:07:01 | 2:01:00 | 2:11:01 | 1:00:00 | 1:08:02 | 2:00:00 |
| NM_006565:XM_005255775:XM_017022868 | 7:10:01 | 0:01:00 | 11:02:01 | 0:00:00 | 8:01:02 | 0:00:00 |
| NM_004360:NM_001317186 | 82:00:00 | 0:00 | 19:00 | 1:00 | 5:00 | 6:00 |
| XM_024450224:NM_015386:XR_933267:NM_001195139 | 0:0:0:0 | 2:15:0:0 | 0:0:0:0 | 3:10:0:0 | 0:0:0:0 | 5:25:4:0 |
| NM_005153:XM_017023869:XM_006721332:NM_001272075:XM_017023864 | 12:0:1:0:0 | 0:0:0:0:0 | 13:1:0:0:0 | 0:0:0:0:0 | 20:3:1:1:0 | 16:0:0:0:0 |
| NM_005153:XM_017023869:XM_006721332:NM_001272075:XM_017023864 | 0:0:0:0:0 | 12:0:1:0:0 | 0:0:0:0:0 | 13:1:0:0:0 | 36:3:1:1:0 | 0:0:0:0:0 |
| NM_005153:XM_017023869:XM_006721332:NM_001272075:XM_017023864 | 0:0:0:0:0 | 12:0:1:0:0 | 0:0:0:0:0 | 13:1:0:0:0 | 36:3:1:1:0 | 0:0:0:0:0 |
| NM_001861:XM_024450156:NM_001318797:NM_001318794:NM_001318786 | 17:1:0:0:0 | 23:0:0:0:0 | 7:0:0:0:0 | 5:1:0:0:0 | 42:1:0:0:0 | 0:0:0:0:0 |
| XM_024450187:NM_174917:XR_933240:NR_147928:XM_024450186:NM_001127214:XM_017023019:NM_001243279:XR_933238 | 0:0:0:0:0:0:0:0:0 | 5:0:0:1:1:0:0:0:0 | 0:0:0:0:0:0:0:0:0 | 2:0:0:0:0:0:0:0:0 | 0:0:0:0:0:0:0:0:0 | 2:0:0:0:0:0:1:1:1 |
| NM_000977:NM_033251 | 10:02 | 14:03 | 25:04:00 | 16:00 | 97:09:00 | 0:00 |
| NR_024058:NM_006761:XM_017025005:XM_005256784 | 573:6002:25:0 | 3:30:2:0 | 162:1720:5:0 | 1:4:0:0 | 202:2375:6:1 | 1:6:1:0 |
| NM_006761:NR_024058:XM_017025005:XM_005256784 | 6035:559:12:16 | 9:2:0:0 | 1728:151:3:1 | 0:0:0:0 | 2386:196:5:3 | 1:0:0:0 |
| NM_006761:NR_024058:XM_017025005:XM_005256784 | 6042:553:12:17 | 3:0:0:0 | 1727:151:3:1 | 0:0:0:0 | 2386:195:5:4 | 1:0:0:0 |
| NM_006761:NR_024058:XM_017025005:XM_005256784 | 6034:544:20:19 | 14:1:0:0 | 1727:151:3:1 | 1:0:0:0 | 2384:196:6:4 | 3:1:0:0 |
| NM_006761:NR_024058:XM_017025005:XM_005256784 | 6038:533:23:27 | 12:0:0:1 | 1727:151:3:1 | 1:0:0:0 | 2383:196:7:5 | 3:0:0:0 |
| NM_006761:NR_024058:XM_017025005:XM_005256784 | 6045:532:23:31 | 6:0:0:0 | 1725:149:3:1 | 3:2:0:0 | 2387:193:7:5 | 0:1:0:0 |
| NM_006761:NR_024058:XM_017025005:XM_005256784 | 6041:532:23:32 | 10:0:0:0 | 1723:150:3:1 | 5:1:0:0 | 2386:193:7:5 | 1:1:0:0 |
| NM_006761:NR_024058:XM_017025005:XM_005256784 | 6054:491:32:65 | 11:1:0:1 | 1729:136:8:8 | 2:0:0:0 | 2386:177:6:7 | 2:0:0:0 |
| NM_006761:NR_024058:XM_005256784:XM_017025005 | 6059:499:66:31 | 6:0:0:0 | 1731:139:8:7 | 0:0:0:0 | 2388:177:7:6 | 0:0:0:0 |
| NM_006761:NR_024058:XM_005256784:XM_017025005 | 6064:499:77:31 | 3:0:3:0 | 1731:143:11:7 | 0:0:0:0 | 2387:174:10:6 | 0:1:3:0 |
| NM_006761:XM_005256784:NR_024058:XM_017025005 | 6056:81:495:31 | 12:3:3:0 | 1730:11:143:7 | 1:3:0:0 | 2385:13:175:5 | 4:1:0:0 |
| NM_006761:NR_024058:XM_005256784:XM_017025005 | 6099:491:255:48 | 2:2:0:0 | 1748:141:56:12 | 1:1:0:0 | 2435:173:84:9 | 0:0:0:0 |
| NM_006761:XM_017025005:XM_005256784:NR_024058 | 6088:46:238:489 | 12:4:14:3 | 1744:9:51:140 | 6:2:3:0 | 2429:8:79:172 | 3:0:2:0 |
| NR_024058:NM_006761:XM_005256784:XM_017025005 | 488:6092:258:51 | 4:10:3:1 | 137:1750:56:11 | 1:0:0:0 | 171:2435:85:9 | 0:0:0:0 |

| | | | | | | |
|---|---|---|---|---|---|---|
| NM_006761:NR_024058:XM_005256784:XM_017025005 | 6092:488:260:52 | 9:3:1:0 | 1748:139:56:11 | 2:0:0:0 | 2435:171:85:9 | 0:0:0:0 |
| NM_006761:NR_024058:XM_005256784:XM_017025005 | 6067:453:253:38 | 17:1:0:0 | 1746:135:58:8 | 2:0:0:0 | 2429:158:86:9 | 1:1:0:0 |
| XM_011524086:XM_017025411:XM_017025409:XM_017025396:NM_006311:XM_017025420:XM_005256866:XM_006721602:XM_017025419:XM_005256871:XM_017025403:XM_005256868:XM_024451043:XM_024451042:XM_017025404:XM_017025405:XM_017025407:XM_006721601:XM_024451041:XM_017025406:NM_001190438:XM_017025413:XM_017025416:XM_017025397:XM_017025400:XM_017025398:XM_017025402:XM_005256872:XM_006721603 | 6:0:1:1:2:2:1:1:0:6:4:1:1:2:0:0:0:0:2:0:0:0:0:0:0:0:0:0:0 | 5:0:3:1:4:0:2:0:0:6:2:0:1:2:0:0:0:0:0:0:0:1:0:0:0:0:0:0:0 | 1:0:2:0:1:1:0:1:1:1:1:0:0:0:0:0:0:0:0:0:0:0:0:0:0:0:0:0:0 | 3:1:3:1:2:1:1:1:0:0:0:0:0:0:0:0:0:0:0:0:0:0:0:0:0:0:0:0:0 | 2:0:1:0:2:0:1:0:0:3:1:0:2:1:0:0:0:0:0:0:0:0:1:0:0:0:1:0:0 | 1:0:1:0:0:0:0:0:0:5:1:4:0:0:0:0:0:1:0:0:0:0:0:0:1:0:0:0:0 |
| XM_017025403:XM_011524086:XM_006721601:NM_006311:XM_005256868:XM_005256871:XM_017025420:XM_017025409:XM_024451041:XM_006721602:XM_024451042:XM_017025419:XM_017025397:XM_005256866:XM_017025411:XM_017025396:XM_024451043:XM_017025406:NM_001190438:XM_017025404:XM_017025405:XM_017025407:XM_017025416:XM_017025413:XM_017025399:XM_017025412:XM_017025398:XM_017025400:XM_017025401:XM_017025402:XM_011524083:XM_005256872:XM_006721603 | 13:26:1:15:6:58:12:12:3:9:10:0:0:4:1:2:7:0:0:0:0:1:0:1:0:0:0:0:0:0:0:0:0 | 1:24:0:19:7:32:8:7:0:0:9:0:0:3:0:0:3:0:1:0:0:0:1:0:0:0:0:0:0:0:0:0:0 | 4:6:1:11:4:6:6:11:0:6:1:1:1:4:1:1:0:0:0:0:0:0:0:0:0:0:0:0:0:0:0:0:0 | 4:10:6:6:4:7:2:2:1:2:1:0:0:0:0:0:0:0:0:0:0:0:0:0:0:0:0:0:0:0:0:0:0 | 6:17:1:15:10:16:8:10:1:3:5:0:1:2:0:0:3:0:0:0:0:0:2:0:0:0:0:2:0:1:0:0:0 | 3:11:4:3:3:10:6:5:0:3:2:0:0:5:0:2:4:0:0:0:0:0:0:0:0:0:0:0:0:1:0:0:0 |
| NM_006311:XM_011524086:XM_017025409:XM_017025419:XM_006721602:XM_017025420:XM_017025397:XM_005256871:XM_017025403:XM_005256866:XM_017025411:XM_017025396:XM_005256868:XM_006721601:XM_024451042:XM_024451041:XM_024451043:XM_017025406:NM_001190438:XM_017025404:XM_017025405:XM_017025407:XM_017025416:XM_011524083:XM_017025402:XM_017025413:XM_017025399:XM_017025401:XM_017025412:XM_017025398:XM_017025400:XM_005256872 | 45:73:22:0:11:26:0:110:25:8:1:2:17:2:21:6:18:1:1:0:0:1:2:0:0:1:0:0:0:0:0:0 | 3:6:2:0:0:0:0:0:0:1:0:0:0:0:0:1:4:0:0:0:0:0:0:0:0:0:0:0:0:0:0:0 | 25:21:14:1:11:8:2:18:10:6:1:1:13:8:3:2:2:0:0:0:0:0:0:0:0:0:0:0:0:0:0:0 | 1:4:2:0:0:0:0:0:0:0:0:0:0:0:0:0:0:0:0:0:0:0:0:0:0:0:0:0:0:0:0:0 | 23:37:25:0:7:19:1:37:13:9:0:2:20:6:8:2:10:0:0:0:0:0:2:0:2:0:0:1:0:0:2:0 | 5:3:4:0:1:4:0:0:0:1:0:0:0:1:0:0:1:0:0:0:0:0:0:0:0:0:0:0:1:0:0:0:0 |
| XM_011524086:NM_006311:XM_017025419:XM_017025409:XM_006721602:XM_017025420:XM_017025397:XM_005256871:XM_017025403:XM_005256866:XM_017025411:XM_017025396:XM_005256868:XM_006721601:XM_024451042:XM_024451041:XM_024451043:XM_017025400:XM_017025406:NM_001190438:XM_017025404:XM_017025405:XM_017025407:XM_017025416:XM_011524083:XM_017025402:XM_017025413:XM_017025399:XM_017025412:XM_017025398:XM_017025401:XM_005256872 | 83:52:0:24:11:27:0:110:25:9:1:2:17:2:21:7:22:0:1:1:0:0:1:2:0:0:1:0:0:0:0:0 | 2:1:0:1:0:2:0:0:0:0:0:0:0:0:0:0:0:0:0:0:0:0:0:0:0:0:0:0:0:0:0:0 | 25:27:1:16:11:8:2:18:10:6:1:1:13:8:3:2:3:1:0:0:0:0:0:0:0:0:0:0:0:0:0:0 | 1:1:0:0:0:0:0:0:0:0:0:0:0:0:0:0:0:0:0:0:0:0:0:0:0:0:0:0:0:0:0:0 | 41:32:0:29:10:24:1:37:13:11:0:2:20:7:8:2:11:2:0:0:0:0:0:3:0:2:0:0:0:0:2:0 | 0:0:0:0:0:0:0:0:0:0:0:0:0:0:0:0:0:1:0:0:0:0:0:0:0:0:0:0:0:0:0:0:0 |
| XM_017025409:NM_006311:XM_017025419:XM_006721602:XM_011524086:XM_017025420:XM_017025397:XM_005256871:XM_017025403:XM_005256866:XM_017025411:XM_017025396:XM_005256868:XM_006721601:XM_024451042:XM_024451041:XM_024451043:XM_017025400:XM_017025416:XM_017025406:NM_001190438:XM_017025404:XM_017025405:XM_017025407:XM_011524083:XM_017025402:XM_017025413:XM_017025399:XM_017025401:XM_017025412:XM_017025398:XM_005256872 | 25:53:0:11:87:30:0:110:25:9:1:2:17:2:20:8:23:0:2:1:1:0:0:1:0:0:1:0:0:0:0:0 | 2:6:0:0:9:2:0:0:0:1:0:0:0:0:0:0:0:0:0:0:0:0:0:0:0:0:0:0:0:0:0:0 | 16:29:1:11:26:8:2:18:10:6:1:1:13:8:3:2:3:1:0:0:0:0:0:0:0:0:0:0:0:0:0:0 | 2:2:0:0:0:0:0:0:0:0:0:0:0:0:0:0:0:0:0:0:0:0:0:0:0:0:0:0:0:0:0:0 | 32:32:0:10:41:24:1:37:13:11:0:2:20:7:8:3:12:2:3:0:0:0:0:0:0:2:0:0:2:0:0:0 | 3:2:0:4:2:0:0:0:0:2:1:0:0:0:0:0:1:1:0:0:0:0:0:0:0:0:0:0:0:0:0:0 |
| XM_017025419:XM_017025409:NM_006311:XM_006721602:XM_011524086:XM_017025420:XM_017025397:XM_005256871:XM_017025403:XM_005256866:XM_017025396:XM_005256868:XM_006721601:XM_024451042:XM_024451041:XM_024451043:XM_017025400:XM_017025416:XM_017025406:NM_001190438:XM_017025404:XM_017025405:XM_017025407:XM_011524083:XM_017025402:XM_017025413:XM_017025399:XM_017025411:XM_017025412:XM_017025398:XM_017025401:XM_005256872 | 0:32:72:11:105:38:0:110:25:10:2:17:2:14:5:19:1:2:1:1:0:0:1:0:0:1:0:1:0:0:0:0 | 0:0:0:0:0:0:0:0:0:0:0:0:0:0:0:0:0:0:0:0:0:0:0:0:0:0:0:0:0:0:0:0 | 1:23:35:11:26:12:2:18:10:6:1:13:8:4:3:2:1:1:0:0:0:0:0:0:0:0:0:0:0:0:0:0 | 0:0:0:0:0:0:0:0:0:0:0:0:0:0:0:0:0:0:0:0:0:0:0:0:0:0:0:0:0:0:0:0 | 0:36:37:14:45:28:1:37:13:16:2:20:7:7:4:11:2:4:0:0:0:0:0:0:2:0:0:1:0:0:2:0 | 0:0:0:0:0:0:0:0:0:0:0:0:0:0:0:0:0:0:0:0:0:0:0:0:0:0:0:0:0:0:0:0 |
| NM_015355:NM_001321207:XR_001752462:XM_006721794:XM_017024409 | 21:2:6:9:2 | 5:0:0:0:0 | 16:2:4:1:0 | 3:1:0:0:0 | 16:2:4:5:1 | 1:0:0:0:0 |
| NM_015355:NM_001321207:XR_001752462:XM_006721794:XM_017024409 | 15:1:6:7:2 | 1:1:0:1:0 | 13:2:4:1:0 | 1:0:0:0:0 | 12:2:4:5:1 | 1:0:0:0:0 |
| NM_002815:NM_001270482 | 384:15:00 | 1:00 | 128:07:00 | 0:00 | 209:11:00 | 0:00 |
| NM_001282:NM_001030006:XM_017024285:XM_017024286:XM_017024284:XM_011524454:XM_011524451:XM_011524449:XM_005257937 | 1:0:0:0:2:2:0:0:0 | 0:0:0:0:0:0:0:0:0 | 2:1:1:1:0:0:0:0:0 | 0:0:0:0:0:0:0:0:0 | 2:3:1:1:3:1:0:1:0 | 0:0:0:0:0:0:0:0:0 |
| NM_003487:NM_139215:XM_024450959 | 7:27:00 | 0:00:00 | 7:40:01 | 0:00:00 | 2:26:01 | 0:01:00 |
| NM_007010:XM_011524233:XM_011524232:NM_001291476 | 14:3:12:0 | 0:0:0:0 | 5:3:12:0 | 0:0:0:0 | 1:0:2:0 | 9:2:11:0 |
| NM_007010:XM_011524233:XM_011524232:NM_001291476 | 14:2:12:0 | 0:0:0:0 | 5:3:12:0 | 0:0:0:0 | 10:2:13:0 | 0:0:0:0 |
| NM_002795:NR_104195:NR_104194 | 1955:05:00 | 7:00:00 | 652:02:00 | 2:00:00 | 1161:01:00 | 1:00:00 |
| NM_002795:NR_104195:NR_104194 | 1976:05:00 | 3:00:00 | 659:02:00 | 0:00:00 | 1172:01:00 | 2:00:00 |
| NM_002795:NR_104195:NR_104194 | 2012:01:00 | 3:00:00 | 671:00:00 | 1:00:00 | 1187:01:00 | 1:00:00 |
| NM_002795:NR_104194:NR_104195 | 1996:00:00 | 5:00:00 | 668:00:00 | 0:00:00 | 1184:00:00 | 3:00:00 |
| NM_002795:NR_104194:NR_104195 | 1985:00:00 | 5:00:00 | 662:00:00 | 0:00:00 | 1178:00:00 | 1:00:00 |
| XR_001752548:NR_073428:NM_017748 | 189:12:108 | 12:01:04 | 33:06:14 | 1:01:01 | 50:03:29 | 3:00:01 |
| NM_017748:XR_001752548:NR_073428 | 130:291:18 | 15:42:00 | 18:40:07 | 4:08:00 | 37:77:5 | 2:16:00 |
| NM_017748:XR_001752548:NR_073428 | 135:308:18 | 10:25:00 | 20:43:07 | 2:05:00 | 38:86:5 | 1:08:00 |
| NM_032192:NM_181505:XM_017025217:XM_017025216:NM_001242464 | 156:238:64:12:28 | 0:0:2:1:0 | 35:41:15:2:8 | 0:0:0:0:0 | 47:94:18:8:21 | 1:2:0:0:0 |
| NM_032192:NM_181505:XM_017025217:XM_017025216:NM_001242464 | 156:239:65:12:28 | 0:2:1:0:0 | 34:41:15:2:8 | 0:0:0:0:0 | 48:95:18:8:21 | 0:0:0:0:0 |
| NM_033419:NM_001291730:XM_011525480:XR_002958086:XM_011525481:NM_001291733:XR_934601 | 43:0:1:1:0:0:0 | 0:0:0:0:0:0:0 | 9:1:0:0:0:0:0 | 0:0:0:0:0:0:0 | 3:0:0:0:0:0:0 | 2:0:0:0:0:0:0 |
| XM_024450643:NM_004448:NR_110535:NM_001289937:NM_001289938 | 68:1:29:0:0 | 50:3:22:0:0 | 21:1:8:0:0 | 0:0:0:0:0 | 0:0:0:0:0 | 32:2:19:0:0 |
| XM_024450643:NM_004448:NR_110535:NM_001289937:NM_001289938 | 119:4:51:0:0 | 0:0:0:0:0 | 21:1:8:0:0 | 0:0:0:0:0 | 33:2:20:0:0 | 0:0:0:0:0 |

| | | | | | | |
|---|---|---|---|---|---|---|
| NM_005310:NM_001030002:NM_001330207:XM_024450729:NM_001242443 | 0:0:0:0:0 | 7:2:0:1:0 | 0:0:0:0:0 | 5:2:0:0:0 | 0:0:0:0:0 | 11:6:1:2:0 |
| NM_001165959:NM_001165958:NM_001042471:NM_018530 | 10:32:13:1 | 3:6:20:1 | 4:2:2:0 | 0:0:0:0 | 2:15:13:0 | 0:1:5:0 |
| NM_001165959:NM_001165958:NM_001042471:NM_018530 | 10:32:13:1 | 3:6:20:1 | 4:2:2:0 | 0:0:0:0 | 2:15:13:0 | 0:1:5:0 |
| NM_001254:XM_011525542 | 3:01 | 0:00 | 10:00 | 0:00 | 11:00 | 5:01 |
| NM_176096:XM_017025166:NM_001278216:NM_001278198:XM_017025165:NM_001278217:XM_011525297:XM_017025164 | 8:0:7:1:1:1:1:0 | 9:1:1:0:0:0:0:2 | 2:1:0:0:0:0:0:0 | 14:3:7:1:0:0:0:0 | 19:10:3:0:0:1:0:0 | 10:2:3:0:0:0:0:1 |
| NM_003726:NM_001075099:XM_024451012:XM_017025260:XM_005257755:XM_017025258:XM_017025259 | 2:2:1:0:0:1:1 | 2:1:0:0:0:0:0 | 0:0:0:0:0:0:0 | 38:22:4:1:1:0:0 | 8:4:0:0:0:0:0 | 14:5:0:0:2:1:0 |
| NM_005827:NM_001278784:XM_011524180:XM_006721632:XM_011524179:XM_005256929 | 118:42:3:5:4:1 | 72:16:2:5:4:0 | 102:35:3:5:2:0 | 12:9:0:0:0:0 | 81:27:6:6:21:1 | 60:30:0:4:7:2 |
| NM_004703:XM_011524060:XM_017025295:NM_001291582:NM_001083585:NM_001291581 | 2:3:1:0:0:0 | 0:0:0:0:0:0 | 2:9:0:0:0:0 | 1:2:0:0:0:0 | 2:3:0:5:0:0 | 0:1:0:0:0:0 |
| NM_004703:XM_011524060:NM_001291582:XM_017025295:NM_001083585:NM_001291581 | 2:3:0:1:0:0 | 0:0:0:0:0:0 | 2:9:0:0:0:0 | 1:1:0:0:0:0 | 2:3:5:0:0:0 | 0:1:0:0:0:0 |
| NM_021213:NM_001330377:NM_001330378:XR_001752577 | 10:0:0:0 | 9:0:0:0 | 11:0:0:0 | 11:1:0:0 | 9:0:0:0 | 9:1:0:0 |
| NM_017763:XM_017024800:NM_001305544:XM_011524955:XM_011524956 | 2:0:0:0:0 | 0:0:0:0:0 | 25:0:0:0:0 | 0:0:0:0:0 | 10:1:0:0:0 | 12:1:0:0:0 |
| NM_017763:XM_017024800:XM_011524955:XM_011524956 | 2:0:0:0 | 0:0:0:0 | 23:0:0:0 | 0:0:0:0 | 10:1:0:0 | 12:1:0:0 |
| NM_017763:XM_017024800:XM_011524956:XM_011524955 | 1:0:0:0 | 0:0:0:0 | 9:0:0:0 | 9:0:0:0 | 5:1:0:0 | 2:1:0:0 |
| NM_017763:XM_017024800:XM_011524956:NM_001305544:XM_011524955 | 1:1:0:0:0 | 0:0:0:0:0 | 8:0:0:0:0 | 9:0:0:0:0 | 7:2:0:0:0 | 0:0:0:0:0 |
| NM_017763:XM_017024800:XM_011524955:NM_001305544 | 1:1:0:0 | 1:0:0:0 | 2:1:2:1 | 9:3:0:0 | 2:0:0:0 | 1:2:0:0 |
| NM_001320470:NM_017679:NM_001330414:NM_001353145 | 0:0:0:0 | 0:5:1:0 | 0:0:0:0 | 1:0:0:0 | 0:0:0:0 | 0:0:0:0 |
| NM_004396:NM_001320596:NM_001320595:NM_001320597 | 298:0:5:1 | 0:0:0:0 | 463:1:6:0 | 0:0:0:0 | 353:3:3:0 | 0:0:0:0 |
| NM_004396:NM_001320596:NM_001320595:NM_001320597 | 287:0:5:1 | 0:0:0:0 | 429:1:6:0 | 0:0:0:0 | 324:2:3:0 | 0:0:0:0 |
| NM_004396:NM_001320596:NM_001320595:NM_001320597 | 267:0:5:1 | 2:0:0:0 | 403:1:6:0 | 1:0:0:0 | 297:2:3:0 | 0:0:0:0 |
| NM_004396:NM_001320596:NM_001320595:NM_001320597 | 269:0:5:1 | 0:0:0:0 | 402:1:6:0 | 1:0:0:0 | 297:1:3:0 | 0:0:0:0 |
| NM_139177:NM_001159770:XM_006721757:NM_001352692:XM_017024342:XM_011524496:XM_017024339:XM_017024330:NM_001352693:XM_017024332:XM_011524494:XM_011524495:XM_006721756:XM_017024337:XM_017024335:XM_006721759:XM_017024340:XM_017024341:XM_011524498:XM_006721758:XM_006721755 | 0:0:0:0:0:0:0:0:0:0:0:0:0:0:0:0:0:0:0:0:0 | 38:7:0:2:4:0:1:1:2:2:1:1:3:1:0:0:0:0:0:0:0 | 0:0:0:0:0:0:0:0:0:0:0:0:0:0:0:0:0:0:0:0:0 | 22:2:1:0:0:0:0:0:0:0:0:0:0:0:0:0:0:0:0:0:0 | 0:0:0:0:0:0:0:0:0:0:0:0:0:0:0:0:0:0:0:0:0 | 12:0:0:2:0:1:0:0:1:0:0:0:1:0:1:0:0:0:0:0:0 |
| NM_001098832:NM_032837:NM_001289410:XM_024451008 | 4:9:0:1 | 4:15:0:0 | 2:2:0:0 | 1:3:1:0 | 1:2:0:0 | 2:0:0:0 |
| NM_024844:XM_017025110:XR_934552:NM_001303276:XM_017025108:XM_011525267:XR_001752628:XR_001752629:XM_024450951:XR_001752627:XR_002958072 | 52:1:1:2:0:0:2:1:0:0:0 | 0:0:0:0:0:0:0:0:0:0:0 | 29:1:2:0:0:0:0:0:0:0:0 | 0:0:0:0:0:0:0:0:0:0:0 | 11:0:0:0:0:0:1:0:0:0:0 | 20:0:1:0:0:0:1:1:0:0:0 |
| NM_016080:XM_024450778 | 229:10:00 | 0:00 | 74:02:00 | 1:00 | 80:05:00 | 15:00 |
| NM_001276761:NM_001276760:NM_001126118:NM_001276696 | 1:13:0:0 | 4:21:1:0 | 1:3:0:0 | 1:2:0:0 | 0:0:0:0 | 4:9:0:0 |
| NM_001143992:NM_018081:XR_001752551:XM_024450824:XM_011523952 | 2:2:0:0:0 | 0:0:0:0:0 | 1:0:0:0:0 | 0:0:0:0:0 | 4:2:0:0:0 | 0:0:0:0:0 |
| NM_001614:NR_037688:NM_001199954 | 10652:168:56 | 22:05:00 | 7765:116:31 | 10:00:00 | 9309:154:65 | 11:01:00 |
| NM_001614:NR_037688:NM_001199954 | 10630:168:56 | 26:03:00 | 7749:116:31 | 12:00:00 | 9289:155:65 | 16:00:00 |
| NM_001614:NR_037688:NM_001199954 | 10310:171:56 | 37:03:00 | 7575:118:30 | 26:00:00 | 9062:156:65 | 30:00:00 |
| NM_001614:NR_037688:NM_001199954 | 10170:168:56 | 47:03:00 | 7468:115:30 | 37:01:00 | 8958:154:65 | 35:00:00 |
| NM_001614:NR_037688:NM_001199954 | 10074:173:56 | 47:04:00 | 7405:121:30 | 32:00:00 | 8883:153:65 | 33:00:00 |
| NM_001614:NR_037688:NM_001199954 | 9675:171:56 | 43:03:00 | 7215:119:30 | 37:02:00 | 8519:158:65 | 25:00:00 |
| NM_001614:NR_037688:NM_001199954 | 9409:171:56 | 49:02:00 | 7073:124:30 | 33:00:00 | 8272:158:65 | 26:00:00 |
| NM_001614:NR_037688:NM_001199954 | 9253:162:56 | 45:01:00 | 6983:125:30 | 28:01:00 | 8182:155:65 | 28:00:00 |
| NM_001614:NR_037688:NM_001199954 | 9045:166:56 | 35:01:00 | 6864:125:31 | 29:00:00 | 8046:156:65 | 19:00:00 |
| NM_001614:NR_037688:NM_001199954 | 8914:166:56 | 41:01:00 | 6810:125:31 | 28:01:00 | 7937:153:65 | 24:01:00 |
| NM_001614:NR_037688:NM_001199954 | 8775:160:56 | 47:05:00 | 6728:123:31 | 28:01:00 | 7818:156:65 | 24:00:00 |
| NM_001614:NR_037688:NM_001199954 | 8635:157:56 | 57:02:00 | 6639:118:31 | 32:02:00 | 7735:153:65 | 19:00:00 |

| | | | | | | |
|---|---|---|---|---|---|---|
| NM_001614:NR_037688:NM_001199954 | 8552:157:56 | 59:01:00 | 6572:112:31 | 40:04:00 | 7690:151:65 | 26:00:00 |
| NM_001614:NR_037688:NM_001199954 | 8547:157:56 | 51:02:00 | 6567:113:31 | 34:03:00 | 7673:151:65 | 27:00:00 |
| NM_001614:NR_037688:NM_001199954 | 8502:156:56 | 38:02:00 | 6518:114:31 | 34:03:00 | 7625:151:65 | 23:00:00 |
| NM_001614:NR_037688:NM_001199954 | 8495:155:56 | 52:02:00 | 6525:116:31 | 28:01:00 | 7629:150:65 | 22:00:00 |
| NM_001614:NR_037688:NM_001199954 | 8433:153:56 | 66:04:00 | 6488:116:31 | 37:02:00 | 7604:150:66 | 23:00:00 |
| NM_001614:NR_037688:NM_001199954 | 8376:152:56 | 64:03:00 | 6428:116:31 | 40:02:00 | 7534:147:66 | 31:01:00 |
| NM_001614:NR_037688:NM_001199954 | 8177:152:56 | 64:01:00 | 6318:116:31 | 45:02:00 | 7362:150:66 | 31:00:00 |
| NM_001614:NR_037688:NM_001199954 | 8060:154:56 | 62:01:00 | 6255:120:31 | 41:03:00 | 7267:146:66 | 30:00:00 |
| NM_001614:NR_037688:NM_001199954 | 8034:153:56 | 57:02:00 | 6232:124:31 | 37:03:00 | 7237:141:66 | 28:01:00 |
| NR_037688:NM_001614:NM_001199954 | 149:7852:56 | 1:35:00 | 125:6094:31 | 2:24:00 | 136:7116:66 | 0:12:00 |
| NM_001614:NR_037688:NM_001199954 | 7821:147:56 | 39:00:00 | 6083:127:31 | 25:00:00 | 7085:135:66 | 15:00:00 |
| NM_001614:NR_037688:NM_001199954 | 7786:144:56 | 52:01:00 | 6056:124:31 | 41:01:00 | 7050:133:66 | 24:00:00 |
| NM_001614:NR_037688:NM_001199954 | 7775:144:56 | 61:01:00 | 6048:125:31 | 47:00:00 | 7043:133:66 | 32:00:00 |
| NM_001614:NR_037688:NM_001199954 | 7715:144:56 | 60:00:00 | 5991:124:31 | 41:00:00 | 6979:135:66 | 37:00:00 |
| NM_001614:NR_037688:NM_001199954 | 7725:140:56 | 70:01:00 | 5979:115:31 | 42:01:00 | 6979:138:66 | 29:00:00 |
| NR_037688:NM_001614:NM_001199954 | 142:7707:56 | 0:46:00 | 113:5948:31 | 1:38:00 | 139:6941:65 | 0:33:01 |
| NM_001614:NR_037688:NM_001199954 | 7711:143:56 | 62:00:00 | 5947:117:31 | 43:00:00 | 6940:136:66 | 33:02:00 |
| NM_001614:NR_037688:NM_001199954 | 7646:139:56 | 42:00:00 | 5890:115:31 | 31:01:00 | 6868:133:66 | 29:00:00 |
| NM_001614:NR_037688:NM_001199954 | 7610:128:56 | 27:00:00 | 5862:102:31 | 30:00:00 | 6836:127:66 | 16:00:00 |
| NM_001614:NR_037688:NM_001199954 | 7574:123:56 | 28:02:00 | 5842:100:31 | 31:00:00 | 6797:125:66 | 19:00:00 |
| NM_001614:NR_037688:NM_001199954 | 7548:124:56 | 29:00:00 | 5812:98:31 | 36:00:00 | 6766:127:66 | 25:01:00 |
| NM_001614:NR_037688:NM_001199954 | 7550:123:56 | 25:01:00 | 5813:96:31 | 33:02:00 | 6770:128:66 | 20:00:00 |
| NM_004309:XM_011523574:NR_125441:NM_001185078:NM_001185077:NM_001301242:NM_001301243:NM_001301241 | 95:4:4:1:1:1:0:0 | 0:2:1:0:0:0:0:0 | 47:2:0:0:0:0:0:0 | 0:0:0:0:0:0:0:0 | 52:1:4:0:0:0:1:0 | 0:2:1:0:0:0:0:0 |
| NM_004309:XM_011523574:NR_125441:NM_001301243:NM_001185077:NM_001301240:NM_001301241 | 104:2:4:0:1:3:0 | 0:1:1:0:0:0:0 | 53:1:0:0:0:0:0 | 0:0:0:0:0:0:0 | 62:0:1:1:0:1:0 | 0:0:0:0:0:0:0 |
| NM_001279:NR_134607 | 2:00 | 5:01 | 6:00 | 10:00 | 0:00 | 0:00 |
| XM_024451173:XM_005258259:NM_005901:NM_001003652:XM_006722451:XM_017025749:XM_017025746:XM_017025750:XM_017025745:XM_011525985:NM_001135937:XM_017025747 | 19:18:43:6:0:2:2:0:2:1:0:0 | 0:0:0:0:0:0:0:0:0:0:0:0 | 6:5:22:5:2:1:0:0:0:0:0:0 | 0:0:0:0:0:0:0:0:0:0:0:0 | 10:14:37:10:1:1:0:0:1:0:0:0 | 0:0:0:0:0:0:0:0:0:0:0:0 |
| NM_146387:NM_015956:XM_011528045:NM_146388 | 7:0:0:2 | 7:0:2:1 | 1:0:0:0 | 3:0:0:0 | 10:0:3:2 | 0:0:0:0 |
| NM_001145056:NM_020428:XM_005259999 | 0:00:00 | 0.163888889 | 0:00:00 | 11:26:01 | 4:13:00 | 2:29:00 |
| NM_001005360:NM_001190716:NM_001005361:NM_004945:NM_001005362 | 218:90:54:541:53 | 0:1:0:2:0 | 62:22:18:13:0 | 0:0:0:0:0 | 84:45:13:27:6 | 0:0:0:0:0 |
| NM_032853:XM_017027401:XM_011528382:XM_011528383 | 0:0:0:0 | 0:2:0:0 | 0:0:0:0 | 2:2:0:0 | 1:0:0:0 | 1:0:0:0 |
| NM_033417:NM_001011699:XM_005260154 | 43:08:03 | 26:06:00 | 1:01:00 | 6:01:01 | 20:04:02 | 0:00:00 |
| NM_006532:XM_017027335:XR_001753769:XM_017027336 | 4:0:0:0 | 5:0:0:0 | 0:0:0:0 | 5:0:0:0 | 4:0:0:0 | 0:0:0:0 |
| NM_017572:NM_199054:XM_024451471 | 24:12:06 | 0:00:00 | 3:03:00 | 0:00:00 | 7:02:01 | 0:00:00 |
| NM_199054:NM_017572:XM_024451471 | 11:10:01 | 8:13:03 | 4:03:00 | 0:00:00 | 3:07:00 | 0:00:00 |
| XR_002958354:NM_003430:XR_001753756 | 2:02:00 | 5:01:00 | 0:00:00 | 0:00:00 | 0:00:00 | 0:01:01 |
| XR_002958354:NM_003430:XR_001753756 | 2:02:00 | 5:01:00 | 0:00:00 | 0:00:00 | 0:00:00 | 0:01:01 |
| XR_002958354:NM_003430 | 8:02 | 0:00 | 0:00 | 0:00 | 0:00 | 1:01 |
| XR_002958354:NM_003430 | 7:01 | 0:00 | 0:00 | 0:00 | 2:00 | 0:00 |
| XR_002958354:XM_011528266:XM_011528265:XM_017027236 | 6:1:0:0 | 2:0:0:0 | 5:1:1:0 | 1:0:0:0 | 3:0:2:0 | 0:0:0:0 |
| XR_002958354:XM_011528266:XM_011528265:XM_017027236 | 7:1:0:0 | 1:0:0:0 | 5:1:1:0 | 1:0:0:0 | 3:0:2:0 | 0:0:0:0 |
| XR_002958354:XM_011528266:XM_011528265:XM_017027236:NM_003430 | 20:1:0:0:0 | 2:0:0:0:0 | 10:1:1:1:0 | 0:0:0:0:0 | 8:2:0:0:0 | 6:0:2:0:0 |
| XR_002958354:XM_011528266:XM_011528265:XM_017027236:NM_003430 | 22:1:0:0:0 | 0:0:0:0:0 | 10:1:1:1:0 | 0:0:0:0:0 | 6:0:2:0:0 | 8:2:0:0:0 |

| | | | | | | |
|---|---|---|---|---|---|---|
| XR_002958354:XM_011528266:XM_011528265:XR_001753754:XM_024451693:XR_001753755:NM_003430 | 25:1:0:1:0:0:0 | 0:0:0:0:0:0:0 | 12:1:1:0:0:0:0 | 0:0:0:0:0:0:0 | 9:3:0:0:1:0:0 | 6:0:2:1:0:0:0 |
| NR_045557:NM_003796:XM_005259363:XM_024451751:XM_005259362:NM_001252641:XM_011527435 | 12:20:2:2:2:0:0 | 0:2:0:0:0:0:0 | 8:19:1:2:0:0:0 | 0:0:0:0:0:0:0 | 16:21:5:2:0:1:0 | 0:1:0:0:0:0:0 |
| NM_018025:NR_135270:XM_006723255 | 11:00:00 | 0:00:00 | 8:01:00 | 0:00:00 | 6:00:01 | 3:01:00 |
| NM_018025:NR_135270:XM_006723255 | 13:00:00 | 0:00:00 | 11:01:00 | 0:00:00 | 7:00:01 | 3:01:00 |
| NM_018025:NR_135270:XM_006723255 | 13:00:00 | 0:00:00 | 11:01:00 | 0:00:00 | 0:00:00 | 10:01:01 |
| NM_018025:NR_135270:XM_006723255 | 3:05:00 | 9:04:00 | 4:00:00 | 6:01:00 | 10:01:01 | 0:00:00 |
| NM_018025:NR_135270:XM_006723255 | 7:08:00 | 1:00:00 | 7:01:00 | 0:00:00 | 7:01:01 | 1:00:00 |
| NM_014164:NM_144779 | 0:00 | 0:00 | 0:00 | 2:00 | 5:00 | 0:00 |
| NM_014164:NR_028406:NM_144779 | 0:00:00 | 0:00:00 | 0:00:00 | 2:01:00 | 4:01:00 | 1:00:00 |
| NM_013234:NM_001308393:NM_001300992 | 1006:45:05 | 2:00:00 | 1033:68:10 | 1:00:00 | 1396:64:18 | 1:00:00 |
| NM_013234:NM_001308393:NM_001300992 | 1009:45:05 | 3:00:00 | 1033:68:10 | 1:00:00 | 1401:64:18 | 2:00:00 |
| NM_013234:NM_001308393:NM_001300992 | 1017:45:05 | 2:00:00 | 1035:68:10 | 2:00:00 | 1406:64:18 | 2:00:00 |
| NM_013234:NM_001308393:NM_001300992 | 990:43:04 | 1:00:00 | 1016:68:9 | 1:00:00 | 1385:62:18 | 1:00:00 |
| NM_001398:XM_024451409 | 2:00 | 4:00 | 1:00 | 1:00 | 0:00 | 4:00 |
| NM_001291485:NM_006890 | 0:00 | 21:05 | 0:00 | 0:00 | 0:00 | 0:00 |
| XM_011526990:NM_002483 | 0:01 | 1:01 | 0:00 | 0:00 | 0:00 | 0:06 |
| XM_017027022:NM_020406 | 0:00 | 2:11 | 0:00 | 0:00 | 0:00 | 0:00 |
| XM_017027022:NM_020406 | 0:00 | 2:11 | 0:00 | 0:00 | 0:00 | 0:00 |
| XM_017027022:NM_020406 | 0:00 | 2:11 | 0:00 | 0:00 | 0:00 | 0:00 |
| NM_000041:NM_001302688:NM_001302691 | 8:00:01 | 0:00:00 | 9:01:01 | 0:00:00 | 1:00:01 | 3:00:00 |
| XR_001753633:XR_001753634:XM_011526611:XM_017026467 | 4:1:0:0 | 5:1:1:0 | 3:0:0:0 | 1:0:0:0 | 9:0:0:0 | 5:0:0:0 |
| XR_001753633:NM_001130867:XM_011526611:XR_001753634:XM_017026467 | 4:0:0:0:0 | 4:0:1:1:0 | 3:0:0:0:0 | 1:1:0:0:0 | 9:2:0:0:0 | 5:0:0:0:0 |
| NM_005628:NM_001145144:NM_001145145 | 26:01:00 | 0:00:00 | 58:01:00 | 0:00:00 | 22:03:00 | 29:04:00 |
| NM_001136032:NM_144947:NM_006853:XM_011526373 | 1:1:0:0 | 0:0:0:0 | 0:0:0:0 | 0:0:0:0 | 1:1:0:0 | 1:4:0:0 |
| NM_139314:NM_001039667:XM_005272484 | 0:00:00 | 0:00:00 | 0:00:00 | 0:00:00 | 0:00:00 | 0:01:00 |
| XM_005263959:XM_011511242:XM_005263960:NM_002518:XM_017004216:XM_005263961:XM_017004215:XM_017004217:XM_017004214 | 1:2:0:1:1:1:0:0:0 | 1:0:0:2:0:0:0:0:0 | 0:0:1:1:0:0:0:0:0 | 1:1:0:0:0:0:0:0:0 | 0:2:0:0:0:0:0:1:0 | 1:0:0:0:0:0:0:0:0 |
| NM_173647:XM_005263920:XR_922908 | 0:00:00 | 0:00:00 | 0:00:00 | 1:00:00 | 2:00:00 | 0:00:00 |
| NM_001321345:NM_144632:NM_001321346:NM_001321344:NM_001321343 | 0:0:0:0:0 | 0:1:1:0:0 | 0:0:0:0:0 | 1:0:0:0:0 | 0:0:0:0:0 | 2:3:1:2:0 |
| XM_011510308:NM_005742 | 0:00 | 0:00 | 0:00 | 0:00 | 0:01 | 0:04 |
| NM_001079862:NM_001178042:NM_001079863:NM_001282634:NM_001178017:NM_001282636:NM_020548 | 12:1:0:0:0:0:0 | 0:0:0:0:0:0:0 | 2:0:0:0:0:0:0 | 0:0:0:0:0:0:0 | 17:0:0:0:0:0:0 | 0:0:0:0:0:0:0 |
| NM_012233:XM_011510825:XR_001738674:XM_011510823 | 7:0:6:0 | 0:0:4:0 | 7:1:3:0 | 0:0:0:0 | 4:0:1:0 | 0:0:2:0 |
| NM_012233:XM_011510825:XR_001738674:XM_011510823 | 7:0:6:0 | 0:0:4:0 | 7:1:3:0 | 0:0:0:0 | 4:0:1:0 | 0:0:2:0 |
| XM_011510825:XR_001738674:NM_012233:XM_011510823 | 0:3:4:0 | 0:0:3:0 | 0:1:3:0 | 1:1:4:0 | 0:1:5:0 | 0:0:0:0 |
| NM_003937:XM_024453208:NM_001199241:XR_923058 | 6:1:0:0 | 0:0:0:0 | 13:1:0:0 | 0:0:0:0 | 0:0:0:0 | 1:0:0:0 |
| NM_006164:NM_001145412:NM_001313901:NM_001313904:NM_001313900 | 332:39:0:0:0 | 2:0:0:0:0 | 178:54:1:2:0 | 0:0:0:0:0 | 247:33:2:1:1 | 0:0:0:0:0 |

| | | | | | | |
|---|---|---|---|---|---|---|
| NM_006164:NM_001145412:NM_001313901:NM_001313904:NM_001313900:NM_001313902:NM_001313903 | 363:42:0:0:0:0:0 | 1:0:0:0:0:0:0 | 192:54:1:2:0:0:0 | 1:0:0:0:0:0:0 | 270:34:2:1:1:0:0 | 1:1:0:0:0:0:0 |
| NM_006164:NM_001145412:NM_001313901:NM_001313904:NM_001313900:NM_001313903:NM_001313902 | 394:44:0:0:0:0:0 | 6:0:0:0:0:0:0 | 213:62:1:2:0:0:0 | 1:0:0:0:0:0:0 | 293:35:2:1:1:0:0 | 0:1:0:0:0:0:0 |
| NM_006164:NM_001145412:NM_001313901:NM_001313904:NM_001313900:NM_001313903:NM_001313902 | 398:44:0:0:0:0:0 | 3:0:0:0:0:0:0 | 213:62:1:2:0:0:0 | 0:0:0:0:0:0:0 | 293:35:2:1:1:0:0 | 1:1:0:0:0:0:0 |
| NM_006164:NM_001145412:NM_001313901:NM_001313904:NM_001313900:NM_001313903:NM_001313902 | 401:44:0:0:0:0:0 | 0:0:0:0:0:0:0 | 215:62:1:2:0:0:0 | 0:0:0:0:0:0:0 | 296:36:2:1:1:0:0 | 0:0:0:0:0:0:0 |
| NM_006164:NM_001145412:NM_001313901:NM_001313904:NM_001313900:NM_001313903:NM_001313902 | 408:45:1:0:0:0:0 | 0:0:0:0:0:0:0 | 211:62:1:2:0:0:0 | 1:0:0:0:0:0:0 | 291:37:2:1:2:0:0 | 0:0:0:0:0:0:0 |
| NM_006164:NM_001145412:NM_001313901:NM_001313904:NM_001313900:NM_001313903:NM_001313902 | 406:45:1:0:0:0:0 | 0:0:0:0:0:0:0 | 211:62:1:2:0:0:0 | 1:0:0:0:0:0:0 | 290:37:2:1:2:0:0 | 0:0:0:0:0:0:0 |
| NM_006164:NM_001145412:NM_001313901:NM_001313904:NM_001313900:NM_001313903:NM_001313902 | 404:46:1:0:0:0:0 | 1:0:0:0:0:0:0 | 210:62:1:2:0:0:0 | 1:0:0:0:0:0:0 | 290:37:2:1:2:0:0 | 1:0:0:0:0:0:0 |
| NM_006164:NM_001145412:NM_001313901:NM_001313904:NM_001313900:NM_001313903:NM_001313902 | 404:46:1:0:0:0:0 | 0:0:0:0:0:0:0 | 209:62:1:2:0:0:0 | 0:0:0:0:0:0:0 | 291:37:2:1:2:0:0 | 0:0:0:0:0:0:0 |
| NM_006164:NM_001145412:NM_001313901:NM_001313904:NM_001313900:NM_001313903:NM_001313902 | 403:46:1:0:0:0:0 | 0:0:0:0:0:0:0 | 209:62:1:2:0:0:0 | 0:0:0:0:0:0:0 | 290:37:2:1:2:0:0 | 0:0:0:0:0:0:0 |
| NM_006164:NM_001145412:NM_001313901:NM_001313900 | 397:46:1:0 | 0:0:0:0 | 200:62:1:0 | 1:0:0:0 | 283:37:2:2 | 1:0:0:0 |
| NM_006164:NM_001145412:NM_001313901:NM_001313904:NM_001313900:NM_001313902 | 394:46:1:0:0:0 | 2:0:0:0:0:0 | 200:62:1:2:0:0 | 1:0:0:0:0:0 | 284:36:2:1:2:0 | 0:1:0:0:0:0 |
| XM_011510954:NM_014362:NM_198047:XR_922903:XM_011510953 | 0:1:0:0:0 | 0:0:1:0:0 | 1:0:0:0:0 | 0:0:0:0:0 | 0:2:2:0:0 | 0:2:0:0:0 |
| XM_011510954:NM_014362:NM_198047:XR_922903:XM_011510953 | 0:1:1:0:0 | 0:0:0:0:0 | 1:0:0:0:0 | 0:0:0:0:0 | 0:2:2:0:0 | 0:2:0:0:0 |
| NM_002156:NM_199440 | 813:110 | 3:00 | 1614:125 | 3:00 | 1924:191 | 1:00 |
| NM_002156:NM_199440 | 815:110 | 1:00 | 1612:125 | 4:00 | 1922:191 | 3:00 |
| NM_002156:NM_199440 | 815:110 | 1:00 | 1617:125 | 1:00 | 1921:190 | 4:00 |
| NM_002156:NM_199440 | 893:109 | 1:01 | 1707:120 | 3:00 | 2085:194 | 2:01 |
| NM_002156:NM_199440 | 986:124 | 0:00 | 1943:121 | 0:00 | 2312:199 | 10:01 |
| NM_002156:NM_199440 | 986:124 | 2:00 | 1940:121 | 4:00 | 2317:198 | 6:02 |
| NM_002156:NM_199440 | 985:122 | 25:01:00 | 1944:120 | 52:03:00 | 2318:198 | 41:03:00 |
| NM_002156:NM_199440 | 1005:124 | 7:00 | 1985:123 | 11:01 | 2347:196 | 12:02 |
| NM_002156:NM_199440 | 1011:125 | 4:00 | 1996:122 | 3:00 | 2364:195 | 2:01 |
| NM_199440:NM_002156 | 109:1017 | 0:01 | 111:2002 | 1:00 | 175:2368 | 0:01 |
| NM_002156:NM_199440 | 1020:51:00 | 0:00 | 2009:58:00 | 1:00 | 2373:79 | 1:00 |
| NM_002156:NM_199440 | 1018:42:00 | 2:00 | 2009:45:00 | 1:00 | 2371:63 | 2:00 |
| NM_002156:NM_199440 | 1019:23:00 | 1:01 | 2009:18:00 | 1:02 | 2373:25:00 | 2:04 |
| NM_001025436:XM_017004899:NM_024532:XM_011511837:XM_017004896:XM_011511814 | 45:3:1:0:0:0 | 0:0:0:0:0:0 | 9:2:0:0:0:0 | 0:0:0:0:0:0 | 58:1:1:0:0:0 | 0:0:0:0:0:0 |
| NM_001025436:XM_017004899:NM_024532:XM_011511837:XM_017004896:XM_011511814 | 45:2:1:0:0:0 | 0:1:0:0:0:0 | 9:2:0:0:0:0 | 0:0:0:0:0:0 | 58:1:1:0:0:0 | 0:0:0:0:0:0 |
| NM_212482:NM_212474:XM_017003692:NM_212476:NM_002026:NM_001306132:NM_212478:NM_001306131:NM_001306130 | 0:0:0:0:0:0:0:0:0 | 1:0:0:0:0:0:0:0:0 | 0:0:0:0:0:0:0:0:0 | 4:9:3:1:4:1:3:1:0 | 0:0:0:0:0:0:0:0:0 | 0:0:0:0:1:0:1:0:0 |

| | | | | | | |
|---|---|---|---|---|---|---|
| NM_212482:NM_212474:NM_212478:NM_212476 | 0:0:0:0 | 1:0:0:0 | 1:2:2:1 | 6:0:0:0 | 0:0:0:0 | 0:0:0:0 |
| NM_212474:NM_212478:NM_212476:NM_212482 | 0:0:0:0 | 0:0:0:1 | 0:0:0:0 | 2:2:1:7 | 0:0:0:0 | 0:0:0:0 |
| NM_003590:XM_011511996:XM_011511995:XM_011511994:NM_001257197 | 18:2:0:1:0 | 1:0:0:0:0 | 3:0:0:0:0 | 1:0:0:0:0 | 11:2:2:0:2 | 1:0:0:0:0 |
| NM_001017915:NM_005541:XM_017004004 | 0:00:00 | 0:00:00 | 0:00:01 | 1:05:00 | 0:01:00 | 0:00:00 |
| NM_004544:NR_136157:NM_001322019:XM_011511228:XR_001738750:NR_136158 | 138:1:0:1:2:0 | 102:0:0:1:1:0 | 51:1:1:0:0:0 | 81:0:0:0:0:0 | 271:2:1:2:0:0 | 0:0:0:0:0:0 |
| NM_001195382:NM_001195381:NM_005301 | 7:06:00 | 0:00:00 | 8:01:00 | 0:00:00 | 7:02:03 | 5:02:00 |
| NM_001521:NM_001035521:XM_005264272:NM_001318909 | 19:18:4:1 | 8:4:5:1 | 5:9:2:0 | 3:2:0:0 | 6:11:2:0 | 3:1:2:0 |
| NM_001521:NM_001035521:XM_005264272:NM_001318909 | 22:20:3:1 | 5:2:5:1 | 6:10:2:0 | 3:1:0:0 | 8:11:2:0 | 2:1:2:0 |
| NM_017910:XM_005264393:XR_244945:XM_017004403:XR_939691 | 17:2:0:3:0 | 0:0:0:0:0 | 19:5:1:1:0 | 0:0:0:0:0 | 22:5:0:5:0 | 1:0:0:0:0 |
| NM_017735:XR_002959314:NM_001193509:XM_011532958 | 11:1:0:0 | 0:0:0:0 | 17:0:0:0 | 0:0:0:0 | 7:0:0:0 | 5:0:0:1 |
| NM_206943:NM_001166265:NM_001166266:XM_011532859 | 0:0:0:0 | 2:1:0:0 | 0:0:0:0 | 0:0:0:0 | 0:0:0:0 | 1:0:0:0 |
| NM_019063:XR_001738710:XR_001738709:XM_006711992 | 0:0:0:0 | 2:0:0:0 | 0:0:0:0 | 6:0:0:0 | 0:0:0:0 | 2:0:0:0 |
| NM_019063:XR_001738710:XR_001738709:XM_006711992 | 0:0:0:0 | 2:0:0:0 | 0:0:0:0 | 6:0:0:0 | 0:0:0:0 | 3:0:0:0 |
| NM_019063:XR_001738710:XR_001738709:XM_006711992 | 0:0:0:0 | 2:0:0:0 | 3:0:0:0 | 3:0:0:0 | 0:0:0:0 | 3:0:0:0 |
| XM_011533047:NM_000341 | 0:00 | 0.484027778 | 0:00 | 0:00 | 0:04 | 1:05 |
| NM_000251:XR_939685:NM_001258281:XM_011532867:XM_005264332:XR_001738747 | 48:5:14:0:0:0 | 1:1:0:0:0:0 | 45:3:6:0:0:0 | 4:0:0:0:0:0 | 91:5:36:0:1:0 | 2:4:0:0:0:0 |
| NM_000251:XR_939685:NM_001258281:XM_011532867:XM_005264332:XR_001738747 | 47:4:15:0:0:0 | 0:0:0:0:0:0 | 45:3:6:0:0:0 | 0:0:0:0:0:0 | 91:5:37:0:1:0 | 0:0:0:0:0:0 |
| NM_138559:XM_011532910:XM_017004335:NM_022893:XM_024452963:XM_017004333 | 8:0:0:0:0:0 | 0:0:0:0:0:0 | 8:1:0:0:0:0 | 0:0:0:0:0:0 | 50:1:2:0:1:0 | 0:0:0:0:0:0 |
| XM_017003801:NM_002056:XM_017003802:NM_001244710 | 7:68:17:0 | 0:0:0:0 | 3:64:8:0 | 0:0:0:0 | 10:225:83:0 | 0:0:0:0 |
| NM_002056:XM_017003802:XM_017003801:NM_001244710 | 68:15:3:0 | 0:0:0:0 | 64:8:1:0 | 0:0:0:0 | 228:64:7:0 | 0:0:0:0 |
| XM_024452835:NM_001153:NM_001320702:NM_001320700:XR_001738734:XM_017003944:NM_001320698 | 28:1638:5:35:14:1:12 | 0:3:1:0:0:0:0 | 7:497:2:20:7:1:0 | 0:0:0:0:0:0:0 | 11:1192:4:64:11:2:2 | 0:1:0:0:0:0:0 |
| XM_024452835:NM_001153:NM_001320702:NM_001320700:XR_001738734:XM_017003944:NM_001320698 | 28:1641:5:32:13:1:12 | 0:0:0:0:1:0:0 | 7:497:2:20:6:1:0 | 0:0:0:0:0:0:0 | 11:1189:4:64:11:2:2 | 0:4:0:0:0:0:0 |
| NM_001153:XM_024452835:NM_001320702:XR_001738734:XM_017003944:NM_001320700:NM_001320698 | 1641:28:5:11:1:8:12 | 0:0:0:0:0:0:0 | 495:7:2:6:1:1:0 | 2:0:0:0:0:0:0 | 1193:11:4:11:2:1:2 | 0:0:0:0:0:0:0 |
| XM_024452835:NM_001153:NM_001320702:NM_001320700:XM_017003944:XR_001738734:NM_001320698 | 28:1640:5:27:1:10:12 | 0:1:0:0:0:0:0 | 7:497:2:19:1:5:0 | 0:0:0:0:0:0:0 | 11:1192:4:64:2:10:2 | 0:1:0:0:0:0:0 |
| NM_001153:XM_024452835:NM_001320702:NM_001320700:XM_017003944:XR_001738734:NM_001320698 | 1638:28:6:31:1:9:12 | 3:0:0:0:0:0:0 | 496:7:2:19:1:5:0 | 1:0:0:0:0:0:0 | 1193:11:4:64:2:10:2 | 0:0:0:0:0:0:0 |
| NM_001153:XM_024452835:NM_001320702:NM_001320700:XM_017003944:XR_001738734:NM_001320698 | 1635:28:6:33:1:8:12 | 2:0:0:0:0:0:0 | 494:7:2:19:1:5:0 | 1:0:0:0:0:0:0 | 1193:11:5:64:2:10:2 | 0:0:0:0:0:0:0 |
| XM_024452835:NM_001153:NM_001320702:NM_001320700:XR_001738734:XM_017003944:NM_001320698 | 28:1634:12:35:21:1:12 | 0:2:0:0:0:0:0 | 7:496:2:19:8:1:0 | 0:0:0:0:0:0:0 | 11:1187:5:64:18:2:2 | 0:2:0:0:0:0:0 |
| XM_024452835:NM_001153:NM_001320702:NM_001320700:XR_001738734:XM_017003944:NM_001320698 | 28:1631:11:35:25:1:12 | 0:3:0:0:0:0:0 | 7:493:2:19:9:1:0 | 0:0:0:0:0:0:0 | 11:1185:6:63:16:2:2 | 0:1:0:1:0:0:0 |
| XM_024452835:NM_001153:NM_001320702:NM_001320700:XR_001738734:XM_017003944:NM_001320698 | 28:1628:12:34:34:1:12 | 0:7:0:0:0:0:0 | 7:495:2:19:12:1:0 | 0:0:0:0:0:0:0 | 11:1183:6:64:18:2:2 | 0:1:0:0:0:0:0 |
| NM_001321459:NR_135653:NR_135650:XM_017004313:NR_135651:XM_024452960:NM_001321454:XM_017004311 | 0:1:0:1:0:0:0:0 | 0:0:0:0:0:0:0:0 | 1:0:0:0:0:0:0:0 | 0:0:0:0:0:0:0:0 | 0:0:0:0:0:0:1:0 | 0:1:2:0:1:1:0:0 |
| XM_024452017:XM_017028128:NM_052865:XM_017028127:NM_001310338:XM_006723663:XM_011529394 | 0:7:33:2:2:0:0 | 0:0:0:0:0:0:0 | 0:8:24:2:2:0:0 | 1:0:0:0:0:0:0 | 1:6:44:0:2:0:0 | 1:2:1:0:0:0:0 |
| NM_020536:NR_028402 | 0:00 | 12:03 | 0:00 | 5:01 | 0:00 | 3:01 |
| NM_006363:NM_032985:XM_017027593:NM_032986:NM_001172745 | 14:9:4:0:0 | 0:0:0:0:0 | 12:14:3:0:0 | 0:0:0:0:0 | 19:42:8:2:2 | 0:0:0:0:0 |
| NM_006363:NM_032985:XM_017027593:NM_032986:NM_001172745 | 6:3:2:0:0 | 8:6:2:0:0 | 1:2:0:0:0 | 11:12:3:0:0 | 19:42:8:2:2 | 0:0:0:0:0 |
| NM_000099:NM_001288614 | 44:01:00 | 0:00 | 6:00 | 0:00 | 15:00 | 0:00 |
| NM_002862:XR_002958496 | 14:00 | 0:00 | 6:00 | 0:00 | 0:00 | 6:00 |
| XM_017027706:XM_006723727:NM_015338:XM_006723728:XM_006723733:XM_011528648:XM_017027705:XM_017027704:XM_006723730 | 0:0:0:0:0:0:0:0:0 | 1:0:0:0:0:0:0:0:0 | 1:0:0:0:0:0:0:0:0 | 0:0:0:0:0:0:0:0:0 | 0:0:0:0:0:0:0:0:0 | 2:0:1:0:0:0:0:0:0 |
| NM_015338:XM_017027706:XM_006723728:XM_011528648:XM_017027705:XM_017027704:XM_006723730 | 0:0:0:0:0:0:0 | 0:0:0:0:0:0:0 | 0:0:0:0:0:0:0 | 0:0:0:0:0:0:0 | 0:0:0:0:0:0:0 | 0:0:0:0:0:0:0 |

| | | | | | | |
|---|---|---|---|---|---|---|
| NM_000178:NM_001322495:NM_001322494 | 243:33:00 | 0:00:00 | 105:17:00 | 1:00:00 | 271:31:01 | 0:00:00 |
| NM_018244:XM_011528881:XM_011528884:XM_011528880:XM_011528877:NM_001184977:XM_011528882:XM_011528879:NM_199487 | 0:0:0:0:0:0:0:0:0 | 25:0:0:0:0:0:0:0:0 | 0:0:0:0:0:0:0:0:0 | 9:1:1:0:0:0:0:0:0 | 17:0:0:2:1:0:0:0:0 | 20:0:0:2:1:0:0:0:1 |
| NM_002623:XM_017027879 | 46:01:00 | 0:00 | 77:02:00 | 0:00 | 243:06:00 | 2:00 |
| NM_002623:XM_017027879 | 45:01:00 | 0:00 | 78:04:00 | 0:00 | 245:05:00 | 0:00 |
| NM_016407:XM_017027872 | 0:00 | 2:00 | 0:00 | 1:00 | 0:00 | 5:00 |
| NM_000516:NM_080426:NM_001077489:XM_017027817:NM_001309840:NM_001077488:XM_017027818:XM_017027813:XM_024451874:XR_002958471:NM_080425:XM_024451875:XM_017027819:XM_024451873:XM_017027815:XM_017027812:XM_024451872:NM_016592:XM_017027820:XM_017027814:NM_001309861 | 111:256:101:42:9:23:3:10:1:5:1:8:7:2:2:2:0:0:0:0:0 | 0:0:0:0:0:0:0:0:0:0:0:0:0:0:0:0:0:0:0:0:0 | 86:266:105:40:12:9:10:17:2:9:2:3:7:0:0:0:0:0:0:0:0 | 1:1:0:0:0:0:0:0:0:0:0:0:0:0:0:0:0:0:0:0:0 | 175:281:156:32:20:24:5:29:1:29:3:11:11:0:0:1:0:1:0:0:1 | 0:1:0:0:0:0:0:0:0:0:0:0:0:0:0:0:0:0:0:0:0 |
| XM_017027818:XR_002958471:NM_080426:NM_001077489:XM_017027817:NM_001309840:NM_001077488:XM_017027813:NM_000516:XM_024451874:NM_080425:XM_024451875:XM_017027819:XM_024451873:XM_017027815:XM_017027812:XM_024451872:NM_016592:XM_017027820:XM_017027814:NM_001309861 | 3:10:257:101:42:9:23:10:110:1:1:8:7:2:2:2:0:0:0:0:0 | 0:0:0:0:0:0:0:0:0:0:0:0:0:0:0:0:0:0:0:0:0 | 9:16:267:105:40:12:9:17:87:2:2:3:7:0:0:0:0:0:0:0:0 | 1:1:1:0:0:0:0:0:0:0:0:0:0:0:0:0:0:0:0:0:0 | 5:31:284:156:32:20:24:29:173:1:3:11:11:0:0:1:0:1:0:0:1 | 0:1:0:0:0:0:0:0:0:0:0:0:0:0:0:0:0:0:0:0:0 |
| XR_002958471:NM_001077489:XM_017027817:NM_001309840:NM_001077488:NM_080426:XM_017027818:XM_017027813:NM_000516:XM_024451874:NM_080425:XM_024451875:XM_017027819:XM_024451873:XM_017027815:XM_017027812:XM_024451872:NM_016592:XM_017027820:XM_017027814:NM_001309861 | 10:101:42:9:23:256:3:10:110:1:1:8:7:2:2:2:0:0:0:0:0 | 0:0:0:0:0:1:0:0:0:0:0:0:0:0:0:0:0:0:0:0:0 | 16:105:40:12:9:268:10:17:87:2:2:3:7:0:0:0:0:0:0:0:0 | 1:0:0:0:0:0:0:0:0:0:0:0:0:0:0:0:0:0:0:0:0 | 31:155:32:20:24:284:5:29:172:1:3:11:11:0:0:1:0:1:0:0:1 | 1:1:0:0:0:0:0:0:1:0:0:0:0:0:0:0:0:0:0:0:0 |
| NM_080426:NM_001077489:XM_017027817:NM_001309840:NM_001077488:XM_017027818:XM_017027813:NM_000516:XR_002958471:XM_024451874:NM_080425:XM_024451875:XM_017027819:XM_024451873:XM_017027815:XM_017027812:XM_024451872:NM_016592:XM_017027820:XM_017027814:NM_001309861 | 257:101:42:9:23:3:10:110:11:1:1:8:7:2:2:2:0:0:0:0:0 | 0:0:0:0:0:0:0:0:0:0:0:0:0:0:0:0:0:0:0:0:0 | 267:105:40:12:9:10:17:87:18:2:2:3:7:0:0:0:0:0:0:0:0 | 1:0:0:0:0:0:0:0:0:0:0:0:0:0:0:0:0:0:0:0:0 | 285:156:32:20:24:5:29:173:34:1:3:11:11:0:0:1:0:1:0:0:1 | 0:0:0:0:0:0:0:0:0:0:0:0:0:0:0:0:0:0:0:0:0 |
| NM_080426:NM_001077489:XM_017027817:NM_001309840:NM_001077488:XM_017027818:XM_017027813:NM_000516:XR_002958471:XM_024451874:NM_080425:XM_024451875:XM_017027819:XM_024451873:XM_017027815:XM_017027812:XM_024451872:NM_016592:XM_017027820:XM_017027814:NM_001309861 | 255:101:42:9:23:3:10:110:14:1:1:8:7:2:2:2:0:0:0:0:0 | 2:0:0:0:0:0:0:0:0:0:0:0:0:0:0:0:0:0:0:0:0 | 266:105:40:12:9:10:17:87:21:2:2:3:7:0:0:0:0:0:0:0:0 | 3:0:0:0:0:0:0:0:0:0:0:0:0:0:0:0:0:0:0:0:0 | 283:156:32:20:24:4:29:173:39:1:3:11:11:0:0:1:0:1:0:0:1 | 1:0:0:0:0:1:0:0:0:0:0:0:0:0:0:0:0:0:0:0:0 |
| NM_080426:NM_001077489:XM_017027817:NM_001309840:NM_001077488:XM_017027818:XM_017027813:NM_000516:XR_002958471:XM_024451874:NM_080425:XM_024451875:XM_017027819:XM_024451873:XM_017027815:XM_017027812:XM_024451872:NM_016592:XM_017027820:XM_017027814:NM_001309861 | 256:101:42:9:23:3:10:110:17:1:1:8:7:2:2:2:0:0:0:0:0 | 1:0:0:0:0:0:0:0:0:0:0:0:0:0:0:0:0:0:0:0:0 | 268:105:40:12:9:10:17:87:23:2:2:3:7:0:0:0:0:0:0:0:0 | 1:0:0:0:0:0:0:0:0:0:0:0:0:0:0:0:0:0:0:0:0 | 285:156:32:20:24:5:29:173:42:1:3:11:11:0:0:1:0:1:0:0:1 | 0:0:0:0:0:0:0:0:0:0:0:0:0:0:0:0:0:0:0:0:0 |
| NM_001077489:XM_017027817:NM_001309840:NM_001077488:XM_017027818:NM_080426:XM_017027813:NM_000516:XR_002958471:XM_024451874:NM_080425:XM_024451875:XM_017027819:XM_024451873:XM_017027815:XM_017027812:XM_024451872:NM_016592:XM_017027820:XM_017027814:NM_001309861 | 100:42:9:23:3:260:10:109:21:1:1:8:7:2:2:2:0:0:0:0:0 | 1:0:0:0:0:0:0:1:0:0:0:0:0:0:0:0:0:0:0:0:0 | 105:40:12:9:10:272:17:87:28:2:2:3:7:0:0:0:0:0:0:0:0 | 0:0:0:0:0:0:0:0:0:0:0:0:0:0:0:0:0:0:0:0:0 | 156:32:20:24:5:288:29:173:53:1:3:11:11:0:0:1:0:1:0:0:1 | 0:0:0:0:0:0:0:0:0:0:0:0:0:0:0:0:0:0:0:0:0 |
| NM_001077489:XM_017027817:NM_001309840:NM_001077488:XM_017027818:NM_080426:XM_017027813:NM_000516:XR_002958471:XM_024451874:NM_080425:XM_024451875:XM_017027819:XM_024451873:XM_017027815:XM_017027812:XM_024451872:NM_016592:XM_017027820:XM_017027814:NM_001309861 | 100:42:9:23:3:260:10:110:24:1:1:8:7:2:2:2:0:0:0:0:0 | 1:0:0:0:0:0:0:0:0:0:0:0:0:0:0:0:0:0:0:0:0 | 105:40:12:9:10:273:17:87:31:2:2:3:7:0:0:0:0:0:0:0:0 | 0:0:0:0:0:0:0:0:0:0:0:0:0:0:0:0:0:0:0:0:0 | 156:32:20:24:5:290:29:172:60:1:3:11:11:0:0:1:0:1:0:0:1 | 0:0:0:0:0:1:0:1:0:0:0:0:0:0:0:0:0:0:0:0:0 |
| XR_002958471:NM_001077489:XM_017027817:NM_001309840:NM_001077488:XM_017027818:NM_080426:XM_017027813:NM_000516:XM_024451874:NM_080425:XM_024451875:XM_017027819:XM_024451873:XM_017027815:XM_017027812:XM_024451872:NM_016592:XM_017027820:XM_017027814:NM_001309861 | 26:101:42:9:23:3:261:10:110:1:1:8:7:2:2:2:0:0:0:0:0 | 0:0:0:0:0:0:0:0:0:0:0:0:0:0:0:0:0:0:0:0:0 | 32:105:40:12:9:10:274:17:87:2:2:3:7:0:0:0:0:0:0:0:0 | 1:0:0:0:0:0:0:0:0:0:0:0:0:0:0:0:0:0:0:0:0 | 63:156:32:20:24:5:292:28:172:1:3:10:11:0:0:1:0:1:0:0:1 | 0:0:0:0:0:0:0:1:1:0:0:1:0:0:0:0:0:0:0:0:0 |
| XR_002958471:NM_001077489:XM_017027817:NM_001309840:NM_001077488:XM_017027818:NM_080426:XM_017027813:NM_000516:XM_024451874:NM_080425:XM_024451875:XM_017027819:XM_024451873:XM_017027815:XM_017027812:XM_024451872:NM_016592:XM_017027820:XM_017027814:NM_001309861 | 26:101:42:9:23:3:260:10:110:1:1:8:7:2:2:2:0:0:0:0:0 | 0:0:0:0:0:0:1:0:0:0:0:0:0:0:0:0:0:0:0:0:0 | 32:105:40:12:9:10:274:17:87:2:2:3:7:0:0:0:0:0:0:0:0 | 1:0:0:0:0:0:0:0:0:0:0:0:0:0:0:0:0:0:0:0:0 | 62:156:32:20:24:5:291:29:173:1:3:11:11:0:0:1:0:1:0:0:1 | 1:0:0:0:0:0:2:0:0:0:0:0:0:0:0:0:0:0:0:0:0 |
| NM_000516:NM_001077489:XM_017027817:NM_001309840:NM_001077488:XM_017027818:NM_080426:XM_017027813:XR_002958471:XM_024451874:NM_080425:XM_024451875:XM_017027819:XM_024451873:XM_017027815:XM_017027812:XM_024451872:NM_016592:XM_017027820:XM_017027814:NM_001309861 | 110:101:42:9:23:3:262:10:28:1:1:8:7:2:2:2:0:0:0:0:0 | 0:0:0:0:0:0:0:0:0:0:0:0:0:0:0:0:0:0:0:0:0 | 86:105:40:12:9:10:274:17:36:2:2:3:7:0:0:0:0:0:0:0:0 | 1:0:0:0:0:0:0:0:0:0:0:0:0:0:0:0:0:0:0:0:0 | 173:156:32:20:24:5:293:29:67:1:3:11:11:0:0:1:0:1:0:0:1 | 0:0:0:0:0:0:0:0:0:0:0:0:0:0:0:0:0:0:0:0:0 |
| NM_001077489:XM_017027817:NM_001309840:NM_001077488:XM_017027818:NM_080426:XM_017027813:NM_000516:XR_002958471:XM_024451874:NM_080425:XM_024451875:XM_017027819:XM_024451873:XM_017027815:XM_017027812:XM_024451872:NM_016592:XM_017027820:XM_017027814:NM_001309861 | 101:42:9:23:3:262:10:110:30:1:1:8:7:2:2:2:0:0:0:0:0 | 0:0:0:0:0:0:0:0:0:0:0:0:0:0:0:0:0:0:0:0:0 | 104:40:12:9:10:276:17:87:38:2:2:3:7:0:0:0:0:0:0:0:0 | 1:0:0:0:0:0:0:0:0:0:0:0:0:0:0:0:0:0:0:0:0 | 156:32:20:24:5:293:29:172:73:1:3:11:11:0:0:1:0:1:0:0:1 | 0:0:0:0:0:0:0:0:0:0:0:0:0:0:0:0:0:0:0:0:0 |

| | | | | | | |
|---|---|---|---|---|---|---|
| NM_080426:NM_001077489:XM_017027817:NM_001309840:NM_001077488:XM_017027818:XM_017027813:NM_000516:XR_002958471:XM_024451874:NM_080425:XM_024451875:XM_017027819:XM_024451873:XM_017027815:XM_017027812:XM_024451872:NM_016592:XM_017027820:XM_017027814:NM_001309861 | 263:101:42:9:23:3:10:109:33:1:1:8:7:2:2:2:0:0:0:0:0 | 0:0:0:0:0:0:0:1:0:0:0:0:0:0:0:0:0:0:0:0:0 | 277:105:40:12:9:10:17:87:44:2:2:3:7:0:0:0:0:0:0:0:0 | 1:0:0:0:0:0:0:0:0:0:0:0:0:0:0:0:0:0:0:0:0 | 294:156:32:20:24:5:29:171:80:1:3:11:11:0:0:1:0:1:0:0:1 | 0:0:0:0:0:0:0:1:1:0:0:0:0:0:0:0:0:0:0:0:0 |
| NM_001077489:XM_017027817:NM_001309840:NM_001077488:NM_080426:XM_017027818:XM_017027813:NM_000516:XR_002958471:XM_024451874:NM_080425:XM_024451875:XM_017027819:XM_024451873:XM_017027815:XM_017027812:XM_024451872:NM_016592:XM_017027820:XM_017027814:NM_001309861 | 101:41:9:23:264:3:10:110:37:1:1:8:7:2:2:2:0:0:0:0:0 | 0:1:0:0:0:0:0:0:0:0:0:0:0:0:0:0:0:0:0:0:0 | 105:40:12:9:277:10:16:87:56:2:2:3:7:0:0:0:0:0:0:0:0 | 0:0:0:0:0:0:0:0:0:0:0:0:0:0:0:0:0:0:0:0:0 | 156:32:20:24:300:5:29:172:92:1:3:11:10:0:0:1:0:1:0:0:1 | 0:0:0:0:0:0:0:0:0:0:0:0:0:0:0:0:0:0:0:0:0 |
| NM_000516:XR_002958471:NM_001077489:XM_017027817:NM_001309840:NM_001077488:NM_080426:XM_017027818:XM_017027813:XM_024451874:NM_080425:XM_024451875:XM_017027819:XM_024451873:XM_017027815:XM_017027812:XM_024451872:NM_016592:XM_017027820:XM_017027814:NM_001309861 | 110:41:101:42:9:23:265:3:10:1:1:8:7:2:2:2:0:0:0:0:0 | 0:0:0:0:0:0:0:0:0:0:0:0:0:0:0:0:0:0:0:0:0 | 86:61:105:40:12:9:277:10:16:2:2:3:7:0:0:0:0:0:0:0:0 | 1:2:0:0:0:0:0:0:0:0:0:0:0:0:0:0:0:0:0:0:0 | 171:103:156:32:20:24:300:5:29:1:3:11:10:0:0:1:0:1:0:0:1 | 1:2:0:0:0:0:1:0:0:0:0:0:0:0:0:0:0:0:0:0:0 |
| XR_002958471:NM_001077489:XM_017027817:NM_001309840:NM_001077488:NM_080426:XM_017027818:XM_017027813:NM_000516:XM_024451874:NM_080425:XM_024451875:XM_017027819:XM_024451873:XM_017027815:XM_017027812:XM_024451872:NM_016592:XM_017027820:XM_017027814:NM_001309861 | 42:101:42:9:23:265:3:10:110:1:1:8:7:2:2:2:0:0:0:0:0 | 0:0:0:0:0:0:0:0:0:0:0:0:0:0:0:0:0:0:0:0:0 | 66:105:40:12:9:278:10:16:87:2:2:3:7:0:0:0:0:0:0:0:0 | 1:0:0:0:0:0:0:0:0:0:0:0:0:0:0:0:0:0:0:0:0 | 107:155:32:20:24:299:5:29:172:1:3:11:10:0:0:1:0:1:0:0:1 | 0:1:0:0:0:1:0:0:0:0:0:0:0:0:0:0:0:0:0:0:0 |
| XR_002958471:NM_001077489:XM_017027817:NM_001309840:NM_001077488:NM_080426:XM_017027818:XM_017027813:NM_000516:XM_024451874:NM_080425:XM_024451875:XM_017027819:XM_024451873:XM_017027815:XM_017027812:XM_024451872:NM_016592:XM_017027820:XM_017027814:NM_001309861 | 42:101:42:9:23:266:3:10:110:1:1:8:7:2:2:2:0:0:0:0:0 | 0:0:0:0:0:0:0:0:0:0:0:0:0:0:0:0:0:0:0:0:0 | 66:105:40:12:9:278:10:15:87:2:2:3:7:0:0:0:0:0:0:0:0 | 1:0:0:0:0:0:0:0:0:0:0:0:0:0:0:0:0:0:0:0:0 | 108:156:32:20:24:300:5:29:172:1:3:10:10:0:0:1:0:1:0:0:1 | 0:0:0:0:0:1:0:0:0:0:0:0:0:0:0:0:0:0:0:0:0 |
| XR_002958471:NM_001077489:XM_017027817:NM_001309840:NM_001077488:NM_080426:XM_017027818:XM_017027813:NM_000516:XM_024451874:NM_080425:XM_024451875:XM_017027819:XM_024451873:XM_017027815:XM_017027812:XM_024451872:NM_016592:XM_017027820:XM_017027814:NM_001309861 | 42:101:42:9:23:266:3:10:110:1:1:8:7:2:2:2:0:0:0:0:0 | 0:0:0:0:0:0:0:0:0:0:0:0:0:0:0:0:0:0:0:0:0 | 66:105:40:12:9:278:10:15:87:2:2:3:7:0:0:0:0:0:0:0:0 | 1:0:0:0:0:0:0:0:0:0:0:0:0:0:0:0:0:0:0:0:0 | 107:156:32:20:24:300:5:29:172:1:3:11:10:0:0:1:0:1:0:0:1 | 1:0:0:0:0:1:0:0:0:0:0:0:0:0:0:0:0:0:0:0:0 |
| XR_002958471:NM_001077489:XM_017027817:NM_001309840:NM_001077488:NM_080426:XM_017027818:XM_017027813:NM_000516:XM_024451874:NM_080425:XM_024451875:XM_017027819:XM_024451873:XM_017027815:XM_017027812:XM_024451872:NM_016592:XM_017027820:XM_017027814:NM_001309861 | 46:101:42:9:22:268:3:10:110:1:1:8:7:2:2:2:0:0:0:0:0 | 0:0:0:0:1:1:0:0:0:0:0:0:0:0:0:0:0:0:0:0:0 | 73:105:40:12:9:281:10:15:87:2:2:3:7:0:0:0:0:0:0:0:0 | 1:0:0:0:0:0:0:0:0:0:0:0:0:0:0:0:0:0:0:0:0 | 119:156:32:20:24:307:5:29:171:1:3:11:10:0:0:1:0:1:0:0:1 | 1:0:0:0:0:3:0:0:1:0:0:0:0:0:0:0:0:0:0:0:0 |
| NM_080426:XR_002958471:NM_001077489:XM_017027817:NM_001309840:NM_001077488:XM_017027818:XM_017027813:NM_000516:XM_024451874:NM_080425:XM_024451875:XM_017027819:XM_024451873:XM_017027815:XM_017027812:XM_024451872:NM_016592:XM_017027820:XM_017027814:NM_001309861 | 268:48:100:42:9:22:3:10:110:1:1:8:7:2:2:2:0:0:0:0:0 | 2:1:0:0:0:0:0:0:0:0:0:0:0:0:0:0:0:0:0:0:0 | 281:78:105:40:12:9:10:14:87:2:2:3:7:0:0:0:0:0:0:0:0 | 2:1:0:0:0:0:0:0:0:0:0:0:0:0:0:0:0:0:0:0:0 | 307:127:156:32:20:24:5:29:171:1:3:11:10:0:0:1:0:1:0:0:1 | 3:1:0:0:0:0:0:0:1:0:0:0:0:0:0:0:0:0:0:0:0 |
| NM_080426:XR_002958471:NM_001077489:XM_017027817:NM_001309840:NM_001077488:XM_017027818:XM_017027813:NM_000516:XM_024451874:XM_024451875:XM_017027819:NM_080425:XM_024451873:XM_017027815:XM_017027812:XM_024451872:NM_016592:XM_017027820:XM_017027814:NM_001309861 | 272:53:100:42:9:22:3:10:110:1:8:7:1:2:2:2:0:0:0:0:0 | 0:1:0:0:0:0:0:0:0:0:0:0:0:0:0:0:0:0:0:0:0 | 282:80:105:40:12:9:10:14:87:2:3:7:1:0:0:0:0:0:0:0:0 | 1:1:0:0:0:0:0:0:0:0:0:0:0:0:0:0:0:0:0:0:0 | 313:135:156:32:20:24:5:29:172:1:11:10:3:0:0:1:0:1:0:0:1 | 2:1:0:0:0:0:0:0:0:0:0:0:0:0:0:0:0:0:0:0:0 |
| NM_080426:XR_002958471:NM_001077489:XM_017027817:NM_001309840:NM_001077488:XM_017027818:XM_017027813:NM_000516:XM_024451874:NM_080425:XM_024451875:XM_017027819:XM_024451873:XM_017027815:XM_017027812:XM_024451872:NM_016592:XM_017027820:XM_017027814:NM_001309861 | 271:54:100:42:9:22:3:10:110:1:1:8:7:2:2:2:0:0:0:0:0 | 1:2:0:0:0:0:0:0:0:0:0:0:0:0:0:0:0:0:0:0:0 | 282:81:105:40:12:9:10:14:87:2:2:3:7:0:0:0:0:0:0:0:0 | 1:1:0:0:0:0:0:0:0:0:0:0:0:0:0:0:0:0:0:0:0 | 314:136:156:32:20:24:5:29:172:1:3:11:10:0:0:1:0:1:0:0:1 | 2:2:0:0:0:0:0:0:0:0:0:0:0:0:0:0:0:0:0:0:0 |
| NM_001077489:XM_017027817:NM_001309840:NM_001077488:NM_080426:XM_017027818:XM_017027813:NM_000516:XR_002958471:XM_024451874:NM_080425:XM_024451875:XM_017027819:XM_024451873:XM_017027815:XM_017027812:XM_024451872:NM_016592:XM_017027820:XM_017027814:NM_001309861 | 100:42:9:22:276:3:10:110:57:1:0:8:7:2:2:2:0:0:0:0:0 | 0:0:0:0:0:0:0:0:2:0:0:0:0:0:0:0:0:0:0:0:0 | 105:40:12:9:285:10:14:87:86:2:2:3:7:0:0:0:0:0:0:0:0 | 0:0:0:0:0:0:0:0:0:0:0:0:0:0:0:0:0:0:0:0:0 | 155:32:20:24:320:5:29:172:145:1:3:11:10:0:0:1:0:1:0:0:1 | 0:0:0:0:2:0:0:0:0:0:0:0:0:0:0:0:0:0:0:0:0 |
| NM_001077489:XM_017027817:NM_001309840:NM_001077488:NM_080426:XM_017027818:XM_017027813:NM_000516:XR_002958471:XM_024451874:NM_080425:XM_024451875:XM_017027819:XM_024451873:XM_017027815:XM_017027812:XM_024451872:NM_016592:XM_017027820:XM_017027814:NM_001309861 | 100:42:9:22:276:3:10:110:58:1:0:7:7:2:2:2:0:0:0:0:0 | 0:0:0:0:0:0:0:0:3:0:0:1:0:0:0:0:0:0:0:0:0 | 105:40:12:9:285:10:14:87:87:2:2:3:7:0:0:0:0:0:0:0:0 | 0:0:0:0:0:0:0:0:0:0:0:0:0:0:0:0:0:0:0:0:0 | 155:32:20:24:321:5:29:172:145:1:3:11:10:0:0:1:0:1:0:0:1 | 0:0:0:0:1:0:0:0:0:0:0:0:0:0:0:0:0:0:0:0:0 |
| XR_002958471:NM_001077489:XM_017027817:NM_001309840:NM_001077488:NM_080426:XM_017027818:XM_017027813:NM_000516:XM_024451874:NM_080425:XM_024451875:XM_017027819:XM_024451873:XM_017027815:XM_017027812:XM_024451872:NM_016592:XM_017027820:XM_017027814:NM_001309861 | 62:100:42:9:22:278:3:10:108:1:0:8:7:2:2:2:0:0:0:0:0 | 0:0:0:0:0:0:0:0:1:0:0:0:0:0:0:0:0:0:0:0:0 | 86:105:40:12:9:286:10:14:87:2:2:2:7:0:0:0:0:0:0:0:0 | 2:0:0:0:0:0:0:0:0:0:0:0:0:0:0:0:0:0:0:0:0 | 144:156:32:20:24:325:5:27:172:1:3:11:10:0:0:1:0:1:0:0:1 | 0:0:0:0:0:0:0:1:0:0:0:0:0:0:0:0:0:0:0:0:0 |

| | | | | | | |
|---|---|---|---|---|---|---|
| XR_002958471:NM_001077489:XM_017027817:NM_001309840:NM_001077488:NM_080426:XM_017027818:XM_017027813:NM_000516:XM_024451874:NM_080425:XM_024451875:XM_017027819:XM_024451873:XM_017027815:XM_017027812:XM_024451872:NM_016592:XM_017027820:XM_017027814:NM_001309861 | 63:100:42:9:22:278:3:10:108:1:0:8:7:2:2:2:0:0:0:0:0 | 1:0:0:0:0:0:0:0:0:0:0:0:0:0:0:0:0:0:0:0 | 91:105:40:12:9:286:10:14:87:2:2:2:7:0:0:0:0:0:0:0:0 | 1:0:0:0:0:0:0:0:0:0:0:0:0:0:0:0:0:0:0:0 | 145:156:32:20:24:326:5:28:172:1:3:11:10:0:0:1:0:1:0:0:1 | 0:0:0:0:0:0:0:0:0:0:0:0:0:0:0:0:0:0:0:0 |
| NM_001077489:XM_017027817:NM_001309840:NM_001077488:NM_080426:XM_017027818:XM_017027813:NM_000516:XR_002958471:XM_024451874:NM_080425:XM_024451875:XM_017027819:XM_024451873:XM_017027815:XM_017027812:XM_024451872:NM_016592:XM_017027820:XM_017027814:NM_001309861 | 98:42:9:22:281:3:10:107:71:1:0:7:7:2:2:2:0:0:0:0:0 | 1:0:0:0:0:0:0:1:0:0:0:0:0:0:0:0:0:0:0:0 | 105:40:12:9:286:10:14:87:94:2:2:2:7:0:0:0:0:0:0:0:0 | 0:0:0:0:0:0:0:0:0:0:0:0:0:0:0:0:0:0:0:0 | 155:32:20:24:327:5:27:170:156:1:3:11:10:0:0:1:0:1:0:0:1 | 0:0:0:0:0:0:0:1:0:0:0:0:0:0:0:0:0:0:0:0 |
| NM_080426:XR_002958471:NM_001077489:XM_017027817:NM_001309840:NM_001077488:XM_017027818:XM_017027813:NM_000516:XM_024451874:NM_080425:XM_024451875:XM_017027819:XM_024451873:XM_017027815:XM_017027812:XM_024451872:NM_016592:XM_017027820:XM_017027814:NM_001309861 | 281:70:100:42:9:22:3:10:107:1:0:7:7:2:2:2:0:0:0:0:0 | 0:0:0:0:0:0:0:0:1:0:0:0:0:0:0:0:0:0:0:0 | 285:93:105:40:12:9:10:14:87:2:2:2:7:0:0:0:0:0:0:0:0 | 1:1:0:0:0:0:0:0:0:0:0:0:0:0:0:0:0:0:0:0 | 327:157:155:32:20:24:5:27:171:1:3:11:10:0:0:1:0:1:0:0:1 | 0:0:0:0:0:0:0:0:0:0:0:0:0:0:0:0:0:0:0:0 |
| NM_001077489:XM_017027817:NM_001309840:NM_001077488:NM_080426:XM_017027818:XM_017027813:NM_000516:XR_002958471:XM_024451874:NM_080425:XM_024451875:XM_017027819:XM_024451873:XM_017027815:XM_017027812:XM_024451872:NM_016592:XM_017027820:XM_017027814:NM_001309861 | 99:42:9:22:281:3:10:107:72:1:0:7:7:2:2:2:0:0:0:0:0 | 1:0:0:0:0:0:0:0:0:0:0:0:0:0:0:0:0:0:0:0 | 105:40:12:9:286:10:14:87:96:2:2:2:7:0:0:0:0:0:0:0:0 | 0:0:0:0:0:0:0:0:0:0:0:0:0:0:0:0:0:0:0:0 | 155:32:20:24:327:5:26:171:157:1:3:11:10:0:0:1:0:1:0:0:1 | 0:0:0:0:0:0:0:0:0:0:0:0:0:0:0:0:0:0:0:0 |
| NM_001077489:XM_017027817:NM_001309840:NM_001077488:NM_080426:XM_017027818:XM_017027813:NM_000516:XR_002958471:XM_024451874:NM_080425:XM_024451875:XM_017027819:XM_024451873:XM_017027815:XM_017027812:XM_024451872:NM_016592:XM_017027820:XM_017027814:NM_001309861 | 100:42:9:22:280:3:10:107:72:1:0:7:7:2:2:2:0:0:0:0:0 | 0:0:0:0:1:0:0:0:0:0:0:0:0:0:0:0:0:0:0:0 | 105:40:12:9:286:10:14:87:96:2:2:2:7:0:0:0:0:0:0:0:0 | 0:0:0:0:0:0:0:0:0:0:0:0:0:0:0:0:0:0:0:0 | 155:32:20:24:326:5:26:171:157:1:3:11:10:0:0:1:0:1:0:0:1 | 0:0:0:0:1:0:0:0:0:0:0:0:0:0:0:0:0:0:0:0 |
| NM_000516:NM_001077489:XM_017027817:NM_001309840:NM_001077488:NM_080426:XM_017027818:XM_017027813:XR_002958471:XM_024451874:NM_080425:XM_024451875:XM_017027819:XM_024451873:XM_017027815:XM_017027812:XM_024451872:NM_016592:XM_017027820:XM_017027814:NM_001309861 | 107:100:42:9:22:282:3:9:72:1:0:7:7:2:2:2:0:0:0:0:0 | 0:0:0:0:0:0:0:1:0:0:0:0:0:0:0:0:0:0:0:0 | 86:104:40:12:9:286:10:14:97:2:2:2:7:0:0:0:0:0:0:0:0 | 1:1:0:0:0:0:0:0:0:0:0:0:0:0:0:0:0:0:0:0 | 171:155:32:20:24:327:5:26:157:1:3:11:10:0:0:1:0:1:0:0:1 | 0:0:0:0:0:0:0:0:0:0:0:0:0:0:0:0:0:0:0:0 |
| XR_002958471:NM_001077489:XM_017027817:NM_001309840:NM_001077488:NM_080426:XM_017027818:XM_017027813:NM_000516:XM_024451874:NM_080425:XM_024451875:XM_017027819:XM_024451873:XM_017027815:XM_017027812:XM_024451872:NM_016592:XM_017027820:XM_017027814:NM_001309861 | 75:100:41:9:22:283:3:10:106:1:0:7:7:2:2:2:0:0:0:0:0 | 1:0:1:0:0:0:0:0:1:0:0:0:0:0:0:0:0:0:0:0 | 96:105:40:12:9:287:10:13:87:2:2:2:7:0:0:0:0:0:0:0:0 | 2:0:0:0:0:0:0:0:0:0:0:0:0:0:0:0:0:0:0:0 | 163:155:32:20:24:329:5:26:170:1:3:11:10:0:0:1:0:1:0:0:1 | 0:0:0:0:0:0:0:0:1:0:0:0:0:0:0:0:0:0:0:0 |
| XR_002958471:NM_001077489:XM_017027817:NM_001309840:NM_001077488:NM_080426:XM_017027818:XM_017027813:NM_000516:XM_024451874:NM_080425:XM_024451875:XM_017027819:XM_024451873:XM_017027815:XM_017027812:XM_024451872:NM_016592:XM_017027820:XM_017027814:NM_001309861 | 76:100:42:9:22:284:3:10:107:1:0:7:7:2:2:2:0:0:0:0:0 | 0:0:0:0:0:0:0:0:0:0:0:0:0:0:0:0:0:0:0:0 | 98:105:40:12:9:287:10:13:87:2:2:2:7:0:0:0:0:0:0:0:0 | 1:0:0:0:0:0:0:0:0:0:0:0:0:0:0:0:0:0:0:0 | 161:155:32:20:24:330:5:25:171:1:3:11:10:0:0:1:0:1:0:0:1 | 0:0:0:0:0:0:0:0:0:0:0:0:0:0:0:0:0:0:0:0 |
| NM_001077489:XM_017027817:NM_001309840:NM_001077488:NM_080426:XM_017027818:XM_017027813:NM_000516:XR_002958471:XM_024451874:NM_080425:XM_024451875:XM_017027819:XM_024451873:XM_017027815:XM_017027812:XM_024451872:NM_016592:XM_017027820:XM_017027814:NM_001309861 | 100:42:9:22:284:3:10:107:76:1:0:7:7:2:2:2:0:0:0:0:0 | 0:0:0:0:0:0:0:0:0:0:0:0:0:0:0:0:0:0:0:0 | 105:40:12:9:286:10:13:87:100:2:2:2:7:0:0:0:0:0:0:0:0 | 0:0:0:0:0:0:0:0:0:0:0:0:0:0:0:0:0:0:0:0 | 155:32:20:24:332:5:25:171:160:1:3:11:10:0:0:1:0:1:0:0:1 | 0:0:0:0:0:0:0:0:0:0:0:0:0:0:0:0:0:0:0:0 |
| NM_080426:NM_001077489:XM_017027817:NM_001309840:NM_001077488:XM_017027818:XM_017027813:NM_000516:XR_002958471:XM_024451874:NM_080425:XM_024451875:XM_017027819:XM_024451873:XM_017027815:XM_017027812:XM_024451872:NM_016592:XM_017027820:XM_017027814:NM_001309861 | 285:100:42:9:22:3:10:107:78:1:0:7:7:2:2:2:0:0:0:0:0 | 0:0:0:0:0:0:0:0:0:0:0:0:0:0:0:0:0:0:0:0 | 287:105:40:12:9:10:13:87:101:2:2:2:7:0:0:0:0:0:0:0:0 | 1:0:0:0:0:0:0:0:0:0:0:0:0:0:0:0:0:0:0:0 | 333:155:32:20:24:5:25:171:160:1:3:11:10:0:0:1:0:1:0:0:1 | 0:0:0:0:0:0:0:0:0:0:0:0:0:0:0:0:0:0:0:0 |
| NM_080426:NM_001077489:XM_017027817:NM_001309840:NM_001077488:XM_017027818:XM_017027813:NM_000516:XR_002958471:XM_024451874:NM_080425:XM_024451875:XM_017027819:XM_024451873:XM_017027815:XM_017027812:XM_024451872:NM_016592:XM_017027820:XM_017027814:NM_001309861 | 283:100:42:9:23:3:10:107:78:1:0:7:7:2:2:2:0:0:0:0:0 | 2:0:0:0:0:0:0:0:0:0:0:0:0:0:0:0:0:0:0:0 | 290:105:40:12:9:10:13:87:104:2:2:2:7:0:0:0:0:0:0:0:0 | 1:0:0:0:0:0:0:0:0:0:0:0:0:0:0:0:0:0:0:0 | 333:155:32:20:24:5:25:171:156:1:3:11:10:0:0:1:0:1:0:0:1 | 0:0:0:0:0:0:0:0:0:0:0:0:0:0:0:0:0:0:0:0 |
| NM_000516:NM_080426:NM_001077489:XM_017027817:NM_001309840:NM_001077488:XM_017027818:XM_017027813:XR_002958471:XM_024451874:NM_080425:XM_024451875:XM_017027819:XM_024451873:XM_017027815:XM_017027812:XM_024451872:NM_016592:XM_017027820:XM_017027814:NM_001309861 | 107:285:100:42:9:23:3:10:78:1:0:7:7:2:2:2:0:0:0:0:0 | 0:0:0:0:0:0:0:0:0:0:0:0:0:0:0:0:0:0:0:0 | 86:291:105:40:12:9:10:13:105:2:2:2:7:0:0:0:0:0:0:0:0 | 1:1:0:0:0:0:0:0:0:0:0:0:0:0:0:0:0:0:0:0 | 171:334:155:32:20:24:5:25:157:1:3:11:10:0:0:1:0:1:0:0:1 | 0:0:0:0:0:0:0:0:0:0:0:0:0:0:0:0:0:0:0:0 |
| NM_000516:NM_001077489:XM_017027817:NM_001309840:NM_001077488:NM_080426:XM_017027818:XM_017027813:XR_002958471:XM_024451874:NM_080425:XM_024451875:XM_017027819:XM_024451873:XM_017027815:XM_017027812:XM_024451872:NM_016592:XM_017027820:XM_017027814:NM_001309861 | 107:100:42:9:23:284:3:10:78:1:0:7:7:2:2:2:0:0:0:0:0 | 0:0:0:0:0:1:0:0:0:0:0:0:0:0:0:0:0:0:0:0 | 86:105:40:12:9:292:10:13:104:2:2:2:7:0:0:0:0:0:0:0:0 | 1:0:0:0:0:0:0:0:0:0:0:0:0:0:0:0:0:0:0:0 | 171:155:32:20:24:334:5:25:157:1:3:11:10:0:0:1:0:1:0:0:1 | 0:0:0:0:0:0:0:0:0:0:0:0:0:0:0:0:0:0:0:0 |

| | | | | | | |
|---|---|---|---|---|---|---|
| NM_001077489:XM_017027817:NM_001309840:NM_001077488:NM_080426:XM_017027818:XM_017027813:NM_000516:XR_002958471:XM_024451874:NM_080425:XM_024451875:XM_017027819:XM_024451873:XM_017027815:XM_017027812:XM_024451872:NM_016592:XM_017027820:XM_017027814:NM_001309861 | 100:42:9:23:285:3:10:107:78:1:0:7:7:2:2:2:0:0:0:0:0 | 0:0:0:0:1:0:0:0:0:0:0:0:0:0:0:0:0:0:0:0:0:0 | 105:40:12:9:295:10:13:87:105:2:2:2:7:0:0:0:0:0:0:0:0 | 0:0:0:0:0:0:0:0:0:0:0:0:0:0:0:0:0:0:0:0:0:0 | 155:32:20:24:337:5:25:170:159:1:3:11:10:0:0:1:0:1:0:0:1 | 0:0:0:0:1:0:0:1:1:0:0:0:0:0:0:0:0:0:0:0:0:0 |
| NM_001077489:XM_017027817:NM_001309840:NM_001077488:NM_080426:XM_017027818:XM_017027813:NM_000516:XR_002958471:XM_024451874:NM_080425:XM_024451875:XM_017027819:XM_024451873:XM_017027815:XM_017027812:XM_024451872:NM_016592:XM_017027820:XM_017027814:NM_001309861 | 100:42:9:22:285:3:10:106:77:1:0:7:7:2:2:2:0:0:0:0:0 | 0:0:0:1:1:0:0:1:1:0:0:0:0:0:0:0:0:0:0:0:0:0 | 105:40:12:9:295:10:13:87:104:2:2:2:7:0:0:0:0:0:0:0:0 | 0:0:0:0:0:0:0:0:0:0:0:0:0:0:0:0:0:0:0:0:0:0 | 155:32:20:24:337:5:25:171:158:1:3:11:10:0:0:1:0:1:0:0:1 | 0:0:0:0:1:0:0:0:1:0:0:0:0:0:0:0:0:0:0:0:0:0 |
| NM_001077489:XM_017027817:NM_001309840:NM_001077488:NM_080426:XM_017027818:XM_017027813:NM_000516:XR_002958471:XM_024451874:NM_080425:XM_024451875:XM_017027819:XM_024451873:XM_017027815:XM_017027812:XM_024451872:NM_016592:XM_017027820:XM_017027814:NM_001309861 | 100:42:9:23:286:3:10:107:78:1:0:7:7:2:2:2:0:0:0:0:0 | 0:0:0:0:0:0:0:0:0:0:0:0:0:0:0:0:0:0:0:0:0:0 | 105:40:12:9:295:10:13:87:104:2:2:2:7:0:0:0:0:0:0:0:0 | 0:0:0:0:0:0:0:0:0:0:0:0:0:0:0:0:0:0:0:0:0:0 | 155:32:20:24:338:5:25:170:164:1:3:11:10:0:0:1:0:1:0:0:1 | 0:0:0:0:0:0:0:1:0:0:0:0:0:0:0:0:0:0:0:0:0:0 |
| NM_001077489:XM_017027817:NM_001309840:NM_001077488:NM_080426:XM_017027818:XM_017027813:NM_000516:XR_002958471:XM_024451874:NM_080425:XM_024451875:XM_017027819:XM_024451873:XM_017027815:XM_017027812:XM_024451872:NM_016592:XM_017027820:XM_017027814:NM_001309861 | 100:42:9:23:285:3:9:107:76:1:0:7:7:2:2:2:0:0:0:0:0 | 0:0:0:0:1:0:0:0:0:0:0:0:0:0:0:0:0:0:0:0:0:0 | 105:40:12:9:296:10:12:87:103:2:2:2:7:0:0:0:0:0:0:0:0 | 0:0:0:0:0:0:0:0:0:0:0:0:0:0:0:0:0:0:0:0:0:0 | 155:32:20:24:338:5:25:171:163:1:3:11:10:0:0:1:0:1:0:0:1 | 0:0:0:0:0:0:0:0:0:0:0:0:0:0:0:0:0:0:0:0:0:0 |
| XM_005260964:NM_000629 | 0:03 | 0:00 | 1:07 | 0:00 | 0:00 | 0:03 |
| NR_103797:NM_001291411:NM_001291412:NM_032195 | 319:0:0:0 | 1:0:0:0 | 340:1:0:0 | 2:0:0:0 | 580:0:0:0 | 0:0:0:0 |
| NR_103797:NM_001291411:NM_001291412:NM_032195 | 319:0:0:0 | 1:0:0:0 | 342:1:0:0 | 0:0:0:0 | 580:0:0:0 | 1:0:0:0 |
| NR_103797:NM_001291411:NM_001291412:NM_032195 | 320:0:0:0 | 0:0:0:0 | 342:1:0:0 | 0:0:0:0 | 581:0:0:0 | 0:0:0:0 |
| NR_103797:NM_001291411 | 310:00:00 | 2:00 | 338:01:00 | 0:00 | 573:00:00 | 0:00 |
| NR_103797:NM_001291411 | 248:00:00 | 18:00 | 258:01:00 | 16:00 | 417:00:00 | 28:00:00 |
| NR_103797:NM_001291411 | 305:00:00 | 1:00 | 320:01:00 | 3:00 | 559:00:00 | 1:00 |
| NR_103797:NM_001291411 | 0:00 | 25:00:00 | 0:01 | 13:00 | 1:00 | 31:00:00 |
| NR_103797:NM_001291411 | 0:00 | 8:00 | 0:01 | 5:00 | 1:00 | 3:00 |
| NR_103797:NM_138927 | 0:00 | 0:00 | 0:00 | 2:01 | 3:00 | 2:00 |
| NM_005656:NM_001135099 | 256:17:00 | 294:19:00 | 21:05 | 28:05:00 | 44:05:00 | 33:08:00 |
| NM_005656:NM_001135099 | 4:18 | 2:18 | 1:06 | 1:05 | 2:13 | 0:00 |
| NM_000754:NM_001135162:NM_001135161 | 0:00:00 | 90:01:00 | 0:00:00 | 99:01:00 | 0:00:00 | 54:01:00 |
| NM_030573:NM_001008695 | 10:08 | 8:05 | 4:03 | 6:01 | 0:00 | 4:04 |
| NM_001178126:NM_001256296 | 11:00 | 11:01 | 1:00 | 4:00 | 3:01 | 0:00 |
| NM_004327:NM_021574 | 0:00 | 2:00 | 0:00 | 3:00 | 0:00 | 5:00 |
| NM_145660:NM_030643 | 0:00 | 0:00 | 2:00 | 0:00 | 0:00 | 0:01 |
| NM_003661:NM_145343:NM_001136540:NM_001136541:XM_011530478 | 6:0:0:0:0 | 5:0:0:0:0 | 4:0:0:0:0 | 2:0:0:0:0 | 0:0:0:0:0 | 3:0:1:1:0 |
| NM_003661:NM_145343:NM_001136540:XM_011530478:NM_001136541 | 6:0:0:0:0 | 5:0:0:0:0 | 4:0:0:0:0 | 2:0:0:0:0 | 0:0:0:0:0 | 3:0:1:0:1 |
| NM_003661:NM_145343:XM_011530478:NM_001136540:NM_001136541 | 5:0:0:0:0 | 5:0:0:0:0 | 4:0:0:0:0 | 2:0:0:0:0 | 0:0:0:0:0 | 3:0:0:1:1 |
| XM_005261508:XM_011530111:XM_006724226:NM_012473 | 1877:123:260:211 | 5:1:0:0 | 534:34:61:66 | 2:0:0:0 | 582:44:85:87 | 1:0:0:0 |
| NM_012473:XM_011530111:XM_005261508:XM_006724226 | 211:121:1884:259 | 0:1:1:1 | 65:32:536:62 | 1:1:1:0 | 87:43:583:86 | 0:0:0:0 |
| XM_005261508:XM_011530111:XM_006724226:NM_012473 | 1886:115:264:211 | 5:1:1:0 | 537:32:62:64 | 1:0:0:0 | 583:39:86:87 | 0:0:0:0 |
| XM_005261508:XM_006724226:XM_011530111:NM_012473 | 1893:247:8:210 | 4:1:0:0 | 536:59:2:62 | 3:0:0:0 | 585:82:3:86 | 1:0:0:0 |
| XM_005261508:XM_006724226:XM_011530111:NM_012473 | 1899:231:12:208 | 1:1:0:0 | 539:52:2:63 | 0:0:0:0 | 586:77:3:86 | 1:0:0:0 |
| NM_001024843:NM_015088 | 40:01:00 | 31:00:00 | 18:00 | 9:00 | 17:00 | 0:00 |
| NM_001018050:NM_138338:NM_001282885:NM_001018052:XR_244356 | 7:4:1:0:1 | 0:0:0:0:0 | 16:9:1:1:3 | 0:0:0:0:0 | 5:3:0:0:1 | 8:2:1:0:5 |
| NM_000714:NR_046308:NM_001256531:NM_001256530 | 6:0:0:0 | 7:0:0:0 | 1:0:0:0 | 2:1:0:0 | 0:0:0:0 | 9:0:0:0 |

| | | | | | | |
|---|---|---|---|---|---|---|
| NM_000714:NR_046308:NM_001256531:NM_001256530 | 6:0:0:0 | 7:0:0:0 | 1:0:0:0 | 2:1:0:0 | 6:0:0:0 | 3:0:0:0 |
| NM_006486:NM_006487 | 0:00 | 17:00 | 0:00 | 4:00 | 0:00 | 1:00 |
| NM_033084:NM_001319984 | 1:01 | 0:00 | 0:00 | 0:00 | 10:01 | 3:00 |
| NM_033084:NM_001319984 | 1:01 | 0:00 | 0:00 | 0:00 | 9:01 | 2:01 |
| NM_001007565:NM_006070:XM_005247066:NM_001195479:NM_001195478:XM_017005527:XM_011512334:XM_006713472 | 19:468:8:174:3:11:9:2 | 0:0:0:0:0:0:0:0 | 14:249:2:99:6:3:4:0 | 1:1:0:0:0:0:0:0 | 17:510:5:147:5:6:10:2 | 0:1:0:1:0:0:0:0 |
| NM_006070:NM_001007565:XM_005247066:NM_001195479:NM_001195478:XM_017005527:XM_011512334:XM_006713472 | 474:19:8:174:3:11:11:2 | 0:0:0:0:0:0:0:0 | 250:15:2:100:6:3:3:0 | 1:0:0:0:0:0:0:0 | 512:17:5:149:5:7:13:2 | 0:0:0:0:0:0:0:0 |
| NM_001007565:NM_006070:XM_005247066:NM_001195479:NM_001195478:XM_017005527:XM_011512334:XM_006713472 | 19:474:8:173:3:11:10:2 | 0:0:0:0:0:0:0:0 | 15:251:2:100:6:4:2:0 | 0:0:0:0:0:0:0:0 | 17:512:5:149:5:7:13:2 | 0:0:0:0:0:0:0:0 |
| NM_183352:XM_017007021:XM_024453704:NM_001136026:XM_024453705:NM_030673:XM_024453703:XM_017007020:NM_001136232:XM_024453702:XM_005265379:XM_024453701:NM_001278946 | 622:120:86:0:15:6:5:0:0:2:1:0:0 | 4:0:0:0:0:0:0:0:0:0:0:0:0 | 336:69:38:1:9:4:2:2:0:0:0:0:0 | 0:0:0:0:0:0:0:0:0:0:0:0:0 | 433:83:35:1:12:12:2:1:0:1:1:0:0 | 0:0:0:0:0:0:0:0:0:0:0:0:0 |
| NM_183352:XM_017007021:XM_024453704:NM_001136026:XM_024453705:NM_030673:XM_024453703:XM_017007020:NM_001136232:XM_024453702:XM_005265379:XM_024453701:NM_001278946 | 626:120:82:0:13:6:5:0:0:2:1:0:0 | 0:0:0:0:0:0:0:0:0:0:0:0:0 | 336:69:37:1:8:4:2:2:0:0:0:0:0 | 0:0:0:0:0:0:0:0:0:0:0:0:0 | 433:82:31:1:12:12:2:1:0:1:1:0:0 | 0:1:0:0:0:0:0:0:0:0:0:0:0 |
| NM_020235:XM_024453659:XM_011513001:XM_024453646:NM_001142568:XM_024453648:XM_024453647:XM_024453653:XM_024453650:XM_005247642:XM_005247643:XM_024453661:XM_017006882:XM_024453655:XM_024453660:XM_024453644:XM_024453666:XM_024453651:XM_024453656:XM_024453667 | 14:1:0:0:2:2:1:0:0:0:0:0:0:0:0:0:0:0:0:0 | 0:0:0:0:0:0:0:0:0:0:0:0:0:0:0:0:0:0:0:0 | 28:1:1:1:1:1:1:0:0:0:0:0:0:0:0:0:0:0:0:0 | 0:0:0:0:0:0:0:0:0:0:0:0:0:0:0:0:0:0:0:0 | 23:1:0:1:1:2:3:0:0:2:0:4:0:0:0:0:0:0:0:0 | 0:0:0:0:0:0:2:0:0:0:0:0:0:0:0:0:0:0:0:0 |
| NM_020235:XM_024453648:XM_024453659:XM_024453646:NM_001142568:XM_024453647:XM_005247643:XM_024453653:XM_024453661:XM_024453650:XM_005247642:XM_011513001:XM_017006882:XM_024453655:XM_024453660:XM_024453644:XM_024453666:XM_024453651:XM_024453656:XM_024453667 | 10:1:1:0:2:0:0:0:0:0:0:0:0:0:0:0:0:0:0:0 | 3:1:0:0:0:1:0:0:0:0:0:0:0:0:0:0:0:0:0:0 | 17:0:1:1:1:1:0:0:0:0:0:0:0:0:0:0:0:0:0:0 | 10:1:0:0:0:0:0:0:0:0:0:0:0:0:0:0:0:0:0:0 | 15:1:0:1:1:2:0:0:0:2:0:0:0:0:0:0:0:0:0:0 | 4:1:1:0:0:1:0:0:4:0:0:0:0:0:0:0:0:0:0:0 |
| XM_024453646:XM_024453659:NM_020235:NM_001142568:XM_024453647:XM_024453661:XM_024453648:XM_005247642:XM_011513001:XM_024453656:XM_024453655:XM_017006882:XM_024453650:XM_024453660:XM_024453644:XM_024453666:XM_024453651:XM_005247643:XM_024453653 | 0:1:6:2:0:0:0:0:0:0:0:0:0:0:0:0:0:0:0 | 0:0:0:0:0:0:0:0:0:0:0:0:0:0:0:0:0:0:0 | 0:1:12:1:1:0:0:0:0:0:0:0:0:0:0:0:0:0:0 | 1:0:0:0:0:0:0:0:0:0:0:0:0:0:0:0:0:0:0 | 1:0:13:1:2:0:1:0:0:0:0:0:0:0:0:0:0:0:0 | 0:0:0:0:0:0:0:0:0:0:0:0:0:0:0:0:0:0:0 |
| NM_001142568:NM_020235:XM_005247642:XM_024453659:XM_024453656:XM_024453655:XM_024453646:XM_017006882:XM_024453648:XM_024453647:XM_024453661:XM_024453644:XM_024453660:XM_024453658 | 2:0:0:0:0:0:0:0:0:0:0:0:0:0 | 0:0:0:0:0:0:0:0:0:0:0:0:0:0 | 1:0:0:0:0:0:0:0:0:0:0:0:0:0 | 0:0:0:0:0:0:0:0:0:0:0:0:0:0 | 1:3:0:0:0:0:0:0:0:0:0:0:0:0 | 0:0:0:0:0:0:0:0:0:0:0:0:0:0 |
| NM_001142568:NM_020235:XM_005247642:XM_024453659:XM_024453656:XM_017006882:XM_024453655:XM_024453648:XM_024453646:XM_024453661:XM_024453647:XM_024453644:XM_024453660:XM_024453658 | 2:0:0:0:0:0:0:0:0:0:0:0:0:0 | 0:0:0:0:0:0:0:0:0:0:0:0:0:0 | 1:0:0:0:0:0:0:0:0:0:0:0:0:0 | 0:0:0:0:0:0:0:0:0:0:0:0:0:0 | 1:1:0:0:0:0:0:0:0:0:0:0:0:0 | 0:2:0:0:0:0:0:0:0:0:0:0:0:0 |
| NM_001282556:XM_005247080:XM_011512367:XM_011512364:XM_011512362:NM_001282558:NM_007072:NM_001282557:XM_024453326:XM_011512363 | 34:52:5:1:2:1:2:0:0:0 | 34:55:4:1:1:0:0:0:0:0 | 0:0:0:0:0:0:0:0:0:0 | 1:1:0:0:0:0:0:0:0:0 | 0:0:0:0:0:0:0:0:0:0 | 6:9:0:0:0:0:0:0:1:0 |
| NM_001282556:XM_005247080:NM_007072:NM_001282558:XM_011512364:XM_011512362:NM_001282557:XM_024453326:XM_011512363 | 68:107:5:1:1:3:0:1:0 | 0:0:0:0:0:0:0:0:0 | 1:1:0:0:0:0:0:0:0 | 0:0:0:0:0:0:0:0:0 | 6:9:0:0:0:0:0:1:0 | 0:0:0:0:0:0:0:0:0 |
| NM_000187:XM_011512746:XM_005247413:XM_005247414:XM_005247412:XM_017006277 | 0:0:0:0:0:0 | 23:4:8:1:2:3 | 0:0:0:0:0:0 | 2:2:1:1:0:0 | 0:0:0:0:0:0 | 10:1:3:1:1:4 |
| NM_175862:NM_006889:NM_176892 | 4:00:00 | 5:00:00 | 4:00:00 | 14:00:00 | 0:00:00 | 0:00:00 |
| NM_015869:NM_138712 | 2:00 | 0:00 | 4:00 | 0:00 | 0:00 | 0:00 |
| NM_005037:NM_138712:NM_001330615:NM_015869:NM_138711:NM_001354667:NM_001354666:NM_001354669:NM_001354670:NM_001354668 | 53:138:1:5:818:50:0:0:0:0 | 0:0:0:0:1:0:0:0:0:0 | 17:30:1:5:18:2:0:0:0:0 | 0:0:0:0:0:0:0:0:0:0 | 31:57:2:1:182:12:0:0:0:0 | 0:0:0:0:0:0:0:0:0:0 |
| NM_138712:NM_005037:NM_001330615:NM_015869:NM_138711:NM_001354667:NM_001354666:NM_001354669:NM_001354670:NM_001354668 | 141:53:1:5:807:50:0:0:0:0 | 1:0:0:0:6:0:0:0:0:0 | 29:16:1:5:18:2:0:0:0:0 | 1:0:0:0:0:0:0:0:0:0 | 57:30:3:1:182:11:0:0:0:0 | 0:0:0:0:0:1:0:0:0:0 |
| NM_005037:NM_138712:NM_001330615:NM_015869:NM_138711:NM_001354667:NM_001354666:NM_001354669:NM_001354670:NM_001354668 | 53:141:1:5:807:50:0:0:0:0 | 0:1:0:0:8:0:0:0:0:0 | 16:29:1:5:18:2:0:0:0:0 | 1:1:0:0:0:0:0:0:0:0 | 30:57:3:1:182:11:0:0:0:0 | 0:0:0:0:0:1:0:0:0:0 |
| NM_005037:NM_138712:NM_001330615:NM_015869:NM_138711:NM_001354667:NM_001354666:NM_001354669 | 52:149:1:5:796:50:0:1 | 0:0:0:0:3:0:0:0 | 16:31:1:5:18:2:0:0 | 0:0:0:0:0:0:0:0 | 30:56:3:1:179:11:0:0 | 0:1:0:0:1:0:0:0 |
| XM_024453325:XM_011512355:XM_017005614:NM_012190:NR_072979 | 1:0:0:0:0 | 2:2:1:1:0 | 0:0:0:0:0 | 0:0:0:0:0 | 1:1:0:0:1 | 4:0:1:2:0 |
| NM_001127195:NM_001127196:NM_003418:NM_001127193:NM_001127194:NM_001127192 | 684:1730:1487:575:111:40 | 1:7:5:2:1:0 | 129:440:460:157:31:5 | 1:1:0:0:0:0 | 192:509:617:178:40:9 | 0:0:1:0:0:0 |
| NM_001127195:NM_003418:NM_001127193:NM_001127196:NM_001127194:NM_001127192 | 683:1485:574:1733:112:40 | 3:6:2:5:0:0 | 128:460:157:441:31:5 | 1:0:0:0:0:0 | 192:616:178:509:40:9 | 0:1:0:1:0:0 |

| | | | | | | |
|---|---|---|---|---|---|---|
| NM_003418:NM_001127193:NM_001127196:NM_001127194:NM_001127195:NM_001127192 | 1484:575:1736:112:685:40 | 5:2:3:0:1:0 | 459:157:441:31:130:5 | 0:0:0:0:0:0 | 616:177:510:39:192:9 | 1:1:0:1:0:0 |
| NM_003418:NM_001127193:NM_001127196:NM_001127194:NM_001127195:NM_001127192 | 1475:575:1737:113:686:41 | 4:4:4:0:2:0 | 456:157:442:31:131:5 | 0:0:0:0:0:0 | 611:179:510:40:192:9 | 4:0:0:0:0:0 |
| NM_003418:NM_001127193:NM_001127196:NM_001127194:NM_001127195:NM_001127192 | 1423:580:1745:114:684:41 | 3:2:4:0:3:0 | 446:157:444:31:131:5 | 0:0:0:0:0:0 | 608:179:511:40:193:9 | 0:0:0:0:0:0 |
| NM_001127196:NM_001127193:NM_003418:NM_001127194:NM_001127195:NM_001127192 | 1745:581:1420:115:688:41 | 4:1:4:0:1:0 | 442:156:445:31:131:5 | 2:1:0:0:0:0 | 510:179:606:40:193:9 | 1:0:1:0:0:0 |
| NM_003418:NM_001127193:NM_001127196:NM_001127194:NM_001127195:NM_001127192 | 1423:581:1743:114:689:41 | 1:1:6:1:0:0 | 445:157:444:31:131:5 | 0:0:0:0:0:0 | 605:179:511:40:193:9 | 1:0:0:0:0:0 |
| NM_001127196:NM_003418:NM_001127193:NM_001127194:NM_001127195:NM_001127192 | 1748:1403:580:113:680:41 | 4:4:0:0:0:0 | 442:446:157:31:130:5 | 2:0:0:0:0:0 | 512:603:179:39:191:9 | 1:0:0:0:1:0 |
| NM_003418:NM_001127193:NM_001127196:NM_001127194:NM_001127195:NM_001127192 | 1401:578:1746:113:679:40 | 5:1:4:0:0:0 | 446:157:444:31:129:5 | 0:0:0:0:0:0 | 603:179:513:39:191:9 | 0:0:0:0:0:0 |
| NM_007027:XM_017005637:XM_017005636:XR_001739988 | 0:0:0:0 | 0:0:1:0 | 1:1:0:0 | 0:0:0:0 | 0:4:0:0 | 1:3:0:0 |
| NM_001004019:NM_001998:XM_006713026:NM_001165035 | 0:3:0:1 | 0:2:0:4 | 0:1:0:1 | 1:1:1:0 | 0:0:0:0 | 0:0:0:0 |
| NM_005862:XR_001739978:XM_017005523:XM_011512329:XM_017005525:XM_011512331:XM_017005524 | 1:26:1:8:0:0:0 | 0:0:0:0:0:0:0 | 3:2:1:0:0:0:0 | 0:0:0:0:0:0:0 | 1:14:1:1:0:0:0 | 0:0:0:0:0:0:0 |
| XR_001739978:NM_005862:XM_017005523:XM_011512331:XM_011512329:XM_017005525 | 38:1:1:1:7:0 | 0:0:0:0:0:0 | 4:3:1:0:0:0 | 1:0:0:0:0:0 | 19:1:1:1:1:0 | 2:0:0:0:0:0 |
| NM_006153:NM_001291999:XM_024453538 | 48:05:03 | 0:00:00 | 33:02:02 | 0:00:00 | 13:03:00 | 13:03:00 |
| XM_017006095:XM_024453441:NM_001267042:XR_001740087:XM_006713566:NM_015396:XM_017006098:NM_001267041:XM_017006099:XR_002959506:XM_006713565:XR_002959505:XM_024453442:XM_017006097:XM_011512632:XM_017006100 | 0:0:0:2:0:5:0:1:0:0:4:0:0:0:0:0 | 0:0:0:0:0:0:0:0:0:0:0:0:0:0:0:0 | 1:1:1:1:0:0:0:0:0:0:0:0:0:0:0:0 | 0:0:0:0:0:0:0:0:0:0:0:0:0:0:0:0 | 1:0:0:2:1:5:0:0:0:0:1:1:0:0:0:0 | 0:0:0:2:0:1:0:0:0:0:0:0:0:0:0:0 |
| NM_001320598:NM_024491:XM_017007271:NM_001288964:XM_017007283:NM_001288965:XM_017007275:NM_001320600:XM_024453781:XM_017007278 | 0:3:0:0:0:1:0:0:0:0 | 0:2:0:0:1:0:0:0:0:0 | 1:0:0:0:0:0:0:0:0:0 | 0:0:0:0:0:0:0:0:0:0 | 1:1:0:0:0:0:0:0:0:0 | 1:2:0:0:0:0:0:0:0:0 |
| NM_004766:XM_024453831:NR_023350 | 94:02:18 | 5:00:01 | 90:09:24 | 12:00:00 | 97:06:33 | 7:00:00 |
| XM_017006058:XM_005247261:XM_024453431:NM_001080412:XM_024453433:XM_005247257:XM_017006050:XM_017006057:XM_024453434:NM_001350099:XM_005247258:NM_001350100 | 0:1:5:0:0:0:0:0:0:0:0:0 | 0:0:0:0:0:0:0:0:0:0:0:0 | 3:1:0:0:0:0:0:0:0:0:0:0 | 0:0:0:0:0:0:0:0:0:0:0:0 | 4:2:12:0:8:0:0:0:3:0:0:0 | 0:0:0:0:0:0:0:0:1:0:0:0 |
| XM_017006058:XM_005247261:XM_024453431:XM_017006057:XM_024453434:XM_024453433:XM_017006050:NM_001350099:NM_001350100:XM_005247258 | 0:1:2:0:0:0:0:0:0:0 | 0:0:2:0:0:0:0:0:0:0 | 2:1:0:0:0:0:0:0:0:0 | 0:0:0:0:0:0:0:0:0:0 | 4:0:4:0:5:4:0:0:0:0 | 0:1:5:0:0:3:0:0:0:0 |
| XM_017006058:XM_005247261:XM_024453431:XM_017006057:XM_024453434:XM_024453433 | 0:1:1:0:0:0 | 0:0:0:0:0:0 | 1:1:0:0:0:0 | 0:0:0:0:0:0 | 1:0:1:0:6:3 | 0:0:0:0:0:0 |
| NM_004628:XR_002959581:XR_002959580 | 0:00:00 | 13:01:00 | 0:00:00 | 10:04:00 | 5:02:00 | 4:02:00 |
| NM_004628:XR_002959581 | 5:00 | 6:00 | 3:00 | 4:00 | 4:01 | 4:00 |
| NM_004628:XR_002959581 | 11:00 | 0:00 | 7:00 | 0:00 | 4:01 | 4:00 |
| NM_004628:XR_002959581 | 6:00 | 5:00 | 4:00 | 3:00 | 8:01 | 0:00 |
| XR_002959581:NM_004628:NR_148951:NR_148950:NM_001354729 | 12:10:1:1:0 | 1:0:0:0:0 | 8:8:1:0:0 | 2:0:0:0:0 | 9:8:0:0:0 | 2:0:1:0:0 |
| XM_017007097:XM_011513107:XM_017007100:XM_011513106:XM_005247731:NM_001178138:NM_001178139:NM_006286:XM_011513111:XM_024453729:XM_017007091:XM_024453730:XM_024453726:XM_011513109:XM_024453728:XM_017007098:XM_024453725:XM_011513102:XM_011513101:NM_001178141:XM_017007102:XM_024453727 | 0:0:0:0:0:0:3:1:0:1:0:1:0:0:0:0:0:0:0:0:0:0 | 1:2:0:0:0:1:0:0:0:0:0:0:0:0:0:0:0:0:0:0:0:0 | 1:0:1:0:1:1:2:6:1:1:0:0:0:0:0:0:0:0:0:0:0:0 | 1:1:1:1:0:0:0:0:0:0:0:0:0:0:0:0:0:0:0:0:0:0 | 2:0:0:1:2:0:9:7:0:2:1:0:0:0:0:0:0:0:0:0:0:0 | 1:0:4:0:0:0:1:0:0:1:1:0:0:0:0:1:0:0:0:0:0:0 |
| XR_924148:NM_001184:XM_011512925:NM_001354579 | 2:0:0:0 | 1:0:0:0 | 1:0:0:0 | 0:0:0:0 | 1:0:0:0 | 2:0:0:0 |
| NM_001130156:NM_022443:NM_001130157 | 4:02:00 | 0:03:00 | 5:07:03 | 9:12:00 | 0:00:00 | 11:10:02 |
| XM_005247364:NM_014498:XM_024453455:XM_017006183:XM_005247365:NM_001308155 | 2:11:7:4:1:1 | 0:0:0:0:0:0 | 1:9:5:5:1:1 | 0:0:0:0:0:0 | 1:9:6:1:1:1 | 0:0:0:0:0:0 |
| XM_011512546:NM_005241:XM_005247214:XM_005247213:NM_001164000:NM_001105078:XM_017005874:XM_005247221:NM_004991 | 12:0:0:1:2:2:5:0:0 | 3:0:0:0:0:0:0:0:0 | 0:1:1:1:0:0:0:0:0 | 1:0:0:0:0:0:0:0:0 | 17:0:0:0:0:1:11:0:1 | 7:0:0:0:0:0:0:0:0 |
| NM_005241:XM_005247214:XM_005247213:XM_011512546:NM_001164000:NM_001105078:XM_017005874:XM_005247221:NM_004991:XM_017005877 | 0:0:1:12:2:2:5:0:0:0 | 0:0:0:3:0:0:0:0:0:0 | 1:1:1:1:0:0:0:0:0:0 | 0:0:0:0:0:0:0:0:0:0 | 0:0:0:17:0:1:11:0:1:0 | 0:0:0:7:0:0:0:0:0:0 |
| NM_005241:XM_005247214:XM_005247213:XM_011512546:NM_001164000:XM_017005874:NM_001105078:XM_005247221:NM_004991 | 0:0:1:13:2:5:2:0:0 | 0:0:0:2:0:0:0:0:0 | 1:1:1:1:0:0:0:0:0 | 0:0:0:0:0:0:0:0:0 | 0:0:0:19:0:11:1:0:1 | 0:0:0:6:0:0:0:0:0 |
| XM_005247225:XM_005247213:XM_017005877:XM_005247214:NM_005241:NM_004991:XM_005247224:XM_017005874 | 1:1:2:0:0:0:0:0 | 0:0:0:0:0:0:0:0 | 1:1:1:1:0:0:0:0 | 0:0:0:0:0:0:0:0 | 1:0:4:0:0:1:0:0 | 0:0:0:0:0:0:0:0 |
| NR_033994:NM_003810:NM_001190942 | 0.849305556 | 0:05:00 | 5:57:00 | 1:00:00 | 0.277083333 | 0:02:00 |

| | | | | | | |
|---|---|---|---|---|---|---|
| NM_003810:NR_033994:NM_001190942:NM_001190943 | 464:12:0:0 | 5:0:0:0 | 59:6:0:0 | 0:0:0:0 | 102:5:0:0 | 1:0:0:0 |
| NM_004068:NM_001025205 | 201:84 | 0:00 | 246:57:00 | 0:00 | 309:74 | 0:00 |
| NM_004068:NM_001025205 | 202:84 | 0:00 | 244:57:00 | 1:00 | 310:73 | 0:00 |
| XM_024453430:NM_015303:NR_146113 | 2:02:00 | 0:02:00 | 1:00:00 | 0:00:00 | 1:00:01 | 0:00:00 |
| NM_152673:NM_001282506 | 0:00 | 2:00 | 0:00 | 0:00 | 0:00 | 0:00 |
| NM_152673:NM_001282506 | 0:00 | 2:00 | 0:00 | 0:00 | 0:00 | 0:00 |
| NM_001079811:NM_000404:NM_001317040:NM_001135602 | 18:68:11:0 | 0:0:0:0 | 10:37:7:0 | 0:0:0:0 | 25:70:16:0 | 0:0:0:0 |
| NM_001079811:NM_000404:NM_001317040:NM_001135602 | 7:26:6:0 | 11:37:2:0 | 2:15:3:0 | 7:19:4:0 | 25:69:17:0 | 0:0:0:0 |
| NM_001079811:NM_000404:NM_001317040:NM_001135602 | 0:0:0:0 | 18:63:7:0 | 0:0:0:0 | 9:34:7:0 | 0:0:0:0 | 25:69:17:0 |
| NM_000404:NM_001317040:NM_001135602 | 0:00:00 | 72:01:00 | 0:00:00 | 40:01:00 | 27:06:00 | 43:03:00 |
| NM_000249:NM_001354628:NM_001258273:NM_001354617:NM_001354619:NM_001354618:NM_001167617:NM_001354629:NM_001354622:XM_017006450:NM_001354627:NM_001354630:NM_001167618:NM_001167619:NM_001354620:NM_001354616:NM_001354624:XM_005265161:NM_001258271:NM_001354621:NM_001258274:NM_001354623 | 19:2:1:0:2:2:0:0:1:0:0:0:0:0:0:0:0:0:0:0:0:0 | 13:0:1:1:0:3:0:0:0:0:1:0:0:1:0:0:0:0:0:0:0:0 | 17:2:1:1:1:7:0:3:1:1:1:0:0:0:0:0:0:0:0:0:0:0 | 9:2:1:1:1:3:1:1:0:0:0:0:0:0:0:0:0:0:0:0:0:0 | 29:3:1:0:1:1:0:1:0:0:0:2:0:1:0:0:1:0:0:0:0:0 | 18:3:0:0:6:3:2:0:0:0:0:1:1:1:0:0:0:0:0:0:0:0 |
| XM_005265074:NM_001172713:XM_024453457:XM_006713110:NM_002078:XM_017006187:XM_024453458:XM_011533598:XM_005265075:XM_024453456:XM_017006188:XM_017006186:XM_017006184 | 11:4:1:0:2:1:1:1:0:1:0:0:0 | 2:0:0:0:1:0:0:0:0:1:0:0:0 | 7:1:3:3:4:4:5:2:0:0:0:0:0 | 2:0:0:0:0:0:0:0:0:0:0:0:0 | 18:2:3:0:2:3:9:0:1:0:0:0:0 | 3:0:0:0:0:0:1:0:0:0:0:0:0 |
| NM_001172713:XM_024453457:XM_005265074:XM_006713110:NM_002078:XM_017006187:XM_024453458:XM_011533598:XM_005265075:XM_024453456:XM_017006188:XM_017006186:XM_017006184 | 3:0:10:0:2:0:1:1:0:0:0:0:0 | 1:0:0:0:0:0:0:0:0:0:0:0:0 | 1:3:5:2:3:4:5:2:0:0:0:0:0 | 0:0:0:0:0:0:0:0:0:0:0:0:0 | 2:3:16:0:1:3:9:0:1:0:0:0:0 | 0:0:0:0:0:0:0:0:0:0:0:0:0 |
| XM_006713110:XM_024453458:NM_001172713:XM_024453457:XM_005265074:NM_002078:XM_017006187:XM_011533598:XM_024453456:XM_005265075:XM_017006188:XM_017006184 | 0:2:1:0:7:0:0:1:0:0:0:0 | 0:0:0:0:0:0:0:0:0:0:0:0 | 1:3:1:2:5:3:4:1:0:0:0:0 | 1:1:0:0:0:0:0:0:0:0:0:0 | 0:7:1:3:9:1:3:0:0:1:0:0 | 0:0:0:0:1:0:0:0:0:0:0:0 |
| XM_024453458:XM_005265074:XM_017006187:XM_024453457:XM_011533598:XM_006713110:NM_001172713:XM_005265075:XM_017006188:NM_002078 | 1:1:0:0:1:0:0:0:0:0 | 0:0:0:0:0:0:0:0:0:0 | 1:3:2:1:1:0:0:0:0:0 | 1:0:0:0:0:0:0:0:0:0 | 2:2:2:1:0:0:0:1:0:0 | 2:0:0:0:0:0:0:0:0:0 |
| XM_024453458:XM_005265074:XM_017006187:XM_024453457:XM_011533598:XM_006713110:NM_001172713:XM_005265075:XM_017006188:NM_002078 | 1:1:0:0:1:0:0:0:0:0 | 0:0:0:0:0:0:0:0:0:0 | 1:3:2:1:1:0:0:0:0:0 | 1:0:0:0:0:0:0:0:0:0 | 2:2:2:1:0:0:0:1:0:0 | 2:0:0:0:0:0:0:0:0:0 |
| XM_005265074:XM_017006187:XM_024453457:XM_011533598:XM_024453458:NM_001172713:XM_005265075:XM_017006188:NM_002078 | 1:0:0:0:1:0:0:0:0 | 0:0:0:1:0:0:0:0:0 | 0:0:1:1:0:0:0:0:0 | 3:2:0:0:0:0:0:0:0 | 0:2:1:0:0:0:0:0:0 | 1:0:0:0:2:0:1:0:0 |
| NM_001904:XM_024453360:XM_024453359:NM_001098210:XM_024453356:NM_001098209:XM_024453357:XM_006712985:XM_017005738:XM_024453358:NM_001330729 | 20:14:36:28:6:20:16:2:0:0:0 | 0:0:0:0:0:0:0:0:0:0:0 | 49:18:37:119:5:25:7:2:1:3:0 | 1:2:1:0:0:0:0:0:0:0:0 | 12:57:30:98:4:30:41:3:4:11:0 | 0:0:0:0:0:0:0:0:0:0:0 |
| NM_001098210:XM_024453356:NM_001904:NM_001098209:XM_024453360:XM_024453357:XM_024453359:XM_006712985:XM_017005738:XM_024453358:NM_001330729 | 28:6:21:20:17:20:55:2:0:0:0 | 0:0:0:0:0:0:0:0:0:0:0 | 119:5:50:25:39:10:49:1:1:3:0 | 0:0:0:0:0:0:0:0:0:0:0 | 98:4:12:30:84:56:49:3:4:11:1 | 0:0:0:0:0:0:1:0:0:0:0 |
| NM_001098210:XM_024453356:NM_001904:NM_001098209:XM_024453360:XM_024453357:XM_024453359:XM_006712985:XM_017005738:XM_024453358:NM_001330729 | 28:6:21:20:22:23:72:2:0:0:1 | 0:0:0:0:0:0:0:0:0:0:0 | 119:5:50:25:53:15:64:1:1:3:0 | 0:0:0:0:0:0:0:0:0:0:0 | 98:4:12:30:117:74:67:2:4:11:1 | 0:0:0:0:0:0:0:0:0:0:0 |
| XM_024453360:NM_001098210:XM_024453356:NM_001904:NM_001098209:XM_024453357:XM_024453359:XM_006712985:XM_017005738:XM_024453358:NM_001330729 | 23:28:6:21:20:23:72:2:0:0:1 | 0:0:0:0:0:0:0:0:0:0:0 | 52:119:5:50:25:15:64:1:1:3:0 | 1:0:0:0:0:0:0:0:0:0:0 | 118:98:4:12:30:74:71:2:4:11:1 | 0:0:0:0:0:0:0:0:0:0:0 |
| NM_001098210:XM_024453356:NM_001904:NM_001098209:XM_024453360:XM_024453357:XM_006712985:XM_024453359:XM_017005738:XM_024453358:NM_001330729 | 28:6:21:20:23:28:2:78:0:0:1 | 0:0:0:0:0:0:0:1:0:0:0 | 119:5:50:25:56:23:2:71:1:3:0 | 0:0:0:0:0:0:0:0:0:0:0 | 98:4:12:30:135:86:2:77:4:11:1 | 0:0:0:0:0:0:0:0:0:0:0 |
| NM_001098210:XM_024453356:NM_001904:NM_001098209:XM_024453360:XM_024453357:XM_006712985:XM_024453359:XM_017005738:XM_024453358:NM_001330729 | 28:6:21:20:27:29:0:81:0:0:2 | 0:0:0:0:0:0:0:0:0:0:0 | 119:5:50:25:59:26:2:79:1:3:0 | 0:0:0:0:0:0:0:0:0:0:0 | 98:4:12:30:149:92:1:86:4:11:2 | 0:0:0:0:0:0:0:0:0:0:0 |
| NM_001098210:XM_024453356:NM_001904:NM_001098209:XM_024453360:XM_024453357:XM_006712985:XM_024453359:XM_017005738:XM_024453358:NM_001330729 | 29:6:21:20:31:31:0:89:0:0:2 | 0:0:0:0:0:0:0:0:0:0:0 | 119:5:50:25:63:30:2:82:1:3:0 | 0:0:0:0:0:0:0:0:0:0:0 | 98:4:14:30:159:100:1:97:4:11:3 | 0:0:0:0:0:0:0:0:0:0:0 |
| NM_001098210:XM_024453356:NM_001904:NM_001098209:XM_024453360:XM_024453357:XM_006712985:XM_024453359:XM_017005738:XM_024453358:NM_001330729 | 29:6:21:20:31:31:1:92:0:0:2 | 0:0:0:0:0:0:0:0:0:0:0 | 119:5:50:25:62:30:2:87:1:3:0 | 0:0:0:0:0:0:0:0:0:0:0 | 98:4:14:30:162:102:1:104:4:11:3 | 0:0:0:0:0:0:0:0:0:0:0 |
| NM_003965:XM_017007436:XM_011534209:XM_011534208:NM_001130910 | 53:5:0:4:5 | 0:0:0:0:0 | 13:1:0:0:0 | 0:0:0:0:0 | 7:5:1:2:1 | 16:1:2:0:0 |
| NM_003965:XM_017007436:XM_011534209:XM_011534208:NM_001130910 | 53:5:0:4:5 | 0:0:0:0:0 | 13:1:0:0:0 | 0:0:0:0:0 | 16:1:2:0:0 | 7:5:1:2:1 |
| XR_002959510:NR_146158:NM_014159:XR_002959515:XR_002959514:XM_011533634:XR_002959512:XR_002959517:XR_002959511:XM_024453488:XM_024453487 | 2:0:9:0:0:0:1:0:0:0:0 | 2:0:0:0:0:0:0:0:0:0:0 | 2:2:3:0:0:0:0:0:0:0:0 | 1:0:0:0:0:0:0:0:0:0:0 | 0:3:6:0:0:0:0:0:0:0:0 | 0:0:0:0:0:0:0:0:0:0:0 |
| NM_014159:XR_002959510:XM_011533634:XR_002959511:XR_002959514:XR_002959515:XR_002959516:NR_146158:XR_002959512:XR_002959517:XR_002959513:XM_024453487:XM_024453488 | 5:3:0:0:1:0:0:0:1:0:0:0:0 | 1:3:0:1:1:0:1:1:0:0:0:0:0 | 2:2:0:0:0:0:0:0:0:0:0:0:0 | 1:2:0:0:0:0:0:0:0:0:0:0:0 | 4:1:0:0:0:0:0:0:0:0:1:0:0 | 3:1:0:0:0:0:1:0:0:0:0:0:0 |
| XM_017005921:XM_005264967:XM_017005918:NM_012235:NM_001320044 | 2:5:0:0:0 | 1:6:0:0:0 | 0:1:0:0:0 | 2:1:0:0:0 | 2:0:0:0:0 | 1:2:2:1:1 |
| NM_002375:XM_024453530:XR_002959534:XM_017006412:XM_017006402:XR_002959526:XR_002959525:XR_002959533:XM_017006410:XR_002959532:XR_002959527:XR_002959524:NM_001134364:XR_002959528:XM_024453528:XM_017006416 | 0:3:0:1:0:0:0:2:0:3:0:0:0:0:0:0 | 0:0:0:0:0:0:0:0:0:0:0:0:0:0:0:0 | 1:3:1:1:1:1:1:1:0:0:0:0:0:0:0:0 | 0:0:0:0:0:0:0:0:0:0:0:0:0:0:0:0 | 0:0:0:0:0:1:0:0:1:3:0:0:0:0:0:0 | 0:1:2:2:0:0:0:0:0:4:0:0:1:0:0:0 |

| | | | | | | |
|---|---|---|---|---|---|---|
| XR_002959533:NM_002375:XM_024453530:XR_002959534:XM_017006412:XM_017006402:XR_002959526:XR_002959525:XR_002959532:XM_017006410:XR_002959527:XR_002959524:XR_002959530:XR_002959528:XM_017006416 | 2:0:3:1:1:0:0:0:8:0:0:0:0:0:0 | 0:0:0:1:0:0:0:0:0:0:0:0:0:0:0 | 2:1:1:1:1:1:1:1:3:0:0:0:0:0:0 | 1:0:0:0:0:0:0:0:0:0:0:0:0:0:0 | 1:0:1:2:1:0:1:0:8:1:0:0:0:0:0 | 0:0:0:0:0:0:0:0:1:0:0:0:0:0:0 |
| NM_002375:XR_002959534:XM_017006412:XM_017006402:XR_002959526:XR_002959525:XR_002959533:XR_002959532:XM_024453530:XR_002959527:XR_002959524:XR_002959530:XR_002959528:XM_017006410 | 0:3:1:0:0:0:2:10:2:0:0:0:0:0 | 0:0:0:0:0:0:0:0:0:0:0:0:0:0 | 1:1:1:1:1:1:3:6:0:0:0:0:0:0 | 0:0:0:0:0:0:0:0:0:0:0:0:0:0 | 0:0:0:0:1:0:1:6:0:0:0:0:0:0 | 0:1:1:0:0:0:1:4:1:0:0:0:0:0 |
| NM_002375:XR_002959534:XM_017006412:XM_017006402:XR_002959526:XR_002959525:XR_002959533:XR_002959532:XR_002959524:XM_017006414:XM_024453530:XR_002959530:XR_002959527:XR_002959528 | 0:3:0:0:0:0:2:12:0:0:2:2:0:0 | 0:0:0:0:0:0:0:0:0:0:0:0:0:0 | 1:1:1:1:1:1:3:6:0:0:0:0:0:0 | 0:0:0:0:0:0:0:0:0:0:0:0:0:0 | 0:1:1:0:1:0:2:12:0:0:0:0:0:0 | 0:0:0:0:0:0:0:0:0:0:0:0:0:0 |
| NM_002375:XR_002959534:XM_017006412:XM_017006402:XR_002959526:XR_002959525:XR_002959533:XR_002959532:XM_017006410:XR_002959524:XM_017006414:XR_002959529:XM_024453530:XR_002959530:XR_002959527:XR_002959528 | 0:6:1:0:0:0:3:79:0:3:0:0:3:5:0:0 | 0:0:0:0:0:0:0:0:0:0:0:0:0:0:0:0 | 1:3:1:1:1:1:6:34:1:1:0:0:0:0:0:0 | 0:0:0:0:0:0:0:0:0:0:0:0:0:0:0:0 | 0:2:1:0:1:0:4:40:0:1:0:0:0:0:0:0 | 0:0:0:0:0:0:0:0:0:0:0:0:0:0:0:0 |
| XR_002959532:XM_017006410:NM_002375:XR_002959534:XM_017006412:XM_017006402:XR_002959526:XR_002959525:XR_002959533:XR_002959524:XM_017006414:XR_002959529:XR_002959530:XM_024453530:XR_002959527:XR_002959528 | 62:0:0:4:1:0:0:0:3:2:0:0:4:3:0:0 | 17:0:0:2:0:0:0:0:0:1:0:0:1:0:0:0 | 28:0:1:3:1:1:1:1:6:1:0:0:0:0:0:0 | 5:1:0:0:0:0:0:0:0:0:0:0:0:0:0:0 | 34:0:0:1:1:0:1:0:3:0:0:0:0:0:0:0 | 6:0:0:1:0:0:0:0:1:1:0:0:0:0:0:0 |
| NM_198880:NM_017730:XM_011533863:NM_001320584:NM_001320581 | 12:3:9:0:0 | 0:0:0:0:0 | 8:1:4:1:0 | 0:0:0:0:0 | 4:0:6:0:0 | 12:0:3:0:0 |
| NM_001664:NM_001313943:NM_001313947:NM_001313944:NM_001313946:NM_001313941:NM_001313945 | 8868:29:9:7:16:2:0 | 18:0:0:0:0:0:0 | 3222:21:8:3:2:1:0 | 0:0:0:0:0:0:0 | 4444:27:4:2:9:5:0 | 2:0:0:0:0:0:0 |
| NM_001664:NM_001313943:NM_001313947:NM_001313944:NM_001313946:NM_001313941:NM_001313945 | 8869:29:9:7:16:2:0 | 17:0:0:0:0:0:0 | 3219:21:8:3:2:1:0 | 3:0:0:0:0:0:0 | 4444:27:4:2:9:5:0 | 4:0:0:0:0:0:0 |
| NM_001664:NM_001313943:NM_001313947:NM_001313944:NM_001313946:NM_001313941:NM_001313945 | 8860:29:9:7:16:2:0 | 22:0:0:0:0:0:0 | 3216:21:8:3:2:1:0 | 4:0:0:0:0:0:0 | 4440:27:4:2:9:5:0 | 4:0:0:0:0:0:0 |
| NM_001664:NM_001313943:NM_001313947:NM_001313944:NM_001313946:NM_001313941:NM_001313945 | 8863:28:9:7:16:1:0 | 13:1:0:0:0:1:0 | 3219:21:8:3:2:1:0 | 2:0:0:0:0:0:0 | 4438:27:4:2:9:5:0 | 10:0:0:0:0:0:0 |
| NM_001664:NM_001313943:NM_001313947:NM_001313944:NM_001313941:NM_001313946:NM_001313945 | 8838:29:9:7:2:0:0 | 12:0:0:0:0:0:0 | 3206:21:8:3:1:0:0 | 3:0:0:0:0:0:0 | 4440:26:4:2:5:1:0 | 4:0:0:0:0:0:0 |
| NM_001664:NM_001313943:NM_001313947:NM_001313944:NM_001313941:NM_001313945 | 8833:29:9:7:2:0 | 5:0:0:0:0:0 | 3208:21:8:3:1:0 | 1:0:0:0:0:0 | 4433:26:4:2:5:0 | 6:0:0:0:0:0 |
| NM_001664:NM_001313943:NM_001313944:NM_001313941:NM_001313945:NM_001313947 | 8787:29:1:2:0:0 | 8:0:0:0:0:0 | 3194:21:1:1:0:0 | 0:0:0:0:0:0 | 4420:26:1:5:0:0 | 3:0:0:0:0:0 |
| XM_005264839:NM_007184:NM_001276293:NM_001276294:XM_006712955 | 0:0:0:0:0 | 0:1:1:6:0 | 0:0:0:0:0 | 1:1:0:0:0 | 0:0:0:0:0 | 0:1:0:1:0 |
| XM_017006750:NM_181042:XM_017006748:XM_017006764:XR_001740194:XM_017006759:XM_017006749:XM_017006745:XM_005265280:XM_024453619:XM_017006732:NM_018313 | 3:2:0:2:1:0:0:1:0:0:0:0 | 0:0:0:0:0:0:0:0:0:0:0:0 | 5:2:1:0:0:0:0:0:0:0:0:0 | 0:0:0:0:0:0:0:0:0:0:0:0 | 9:2:2:4:0:0:1:1:3:0:0:0 | 0:0:0:0:0:0:0:0:0:0:0:0 |
| NM_206826:NM_014366:NM_206825 | 0.208344907 | 0:00:00 | 0.561168981 | 0:00:00 | 35:346:10 | 0:00:00 |
| NM_206826:NM_014366:NM_206825 | 4:28:01 | 1:32:00 | 0.217384259 | 0.343784722 | 36:345:10 | 0:01:00 |
| NM_014366:NM_206825:NM_206826 | 59:01:05 | 1:00:00 | 203:04:15 | 4:01:00 | 338:10:54 | 7:00:00 |
| NM_206826:NM_014366:NM_206825 | 3:27:01 | 2:31:00 | 0.297928241 | 0.925034722 | 61:337:10 | 0:00:00 |
| NM_014366:NM_206826:NM_206825 | 55:08:01 | 3:00:00 | 194:23:04 | 6:00:00 | 332:59:10 | 5:01:00 |
| NM_014366:NM_206826:NM_206825 | 55:08:01 | 0:00:00 | 192:22:04 | 2:01:00 | 327:61:10 | 5:00:00 |
| NM_014366:NM_206826:NM_206825 | 55:08:01 | 0:00:00 | 192:22:04 | 2:01:00 | 318:60:9 | 14:01:01 |
| NM_206826:NM_014366:NM_206825 | 7:55:01 | 0:00:00 | 1.00837963 | 0:00:00 | 57:312:9 | 0:00:00 |
| NM_206826:NM_014366:NM_206825 | 7:55:01 | 0:00:00 | 0.966712963 | 0:00:00 | 57:312:9 | 0:00:00 |
| NM_003157:XM_017007085:NM_001348412:XM_017007086 | 3:0:2:0 | 1:1:0:0 | 0:0:1:0 | 4:1:1:0 | 5:0:2:0 | 0:0:0:0 |
| XM_017005994:XM_005264723:NM_015123:XM_017005997 | 7:1:0:0 | 3:0:0:0 | 2:0:0:0 | 5:0:0:0 | 0:0:0:0 | 1:0:0:0 |
| XM_006712975:NM_153480:XM_011533362:NR_104198:NM_001193380:NM_153481:XM_011533361:XM_006712976:NM_153483:XR_940382:XM_011533363:XR_940380 | 0:0:0:0:0:0:0:0:0:0:0:0 | 4:70:3:6:6:10:16:20:6:2:2:0 | 0:0:0:0:0:0:0:0:0:0:0:0 | 1:12:1:1:1:3:1:1:0:0:0:0 | 0:0:0:0:0:0:0:0:0:0:0:0 | 1:14:0:0:2:1:5:1:2:0:0:0 |
| NM_018447:XM_005265321 | 1106:09:00 | 12:00 | 400:01:00 | 2:00 | 487:03:00 | 4:00 |
| NM_018447:XM_005265321 | 1119:42:00 | 2:00 | 405:09:00 | 0:00 | 490:29:00 | 0:00 |
| NM_018447:XM_005265321 | 1045:44:00 | 0:00 | 376:16:00 | 0:00 | 464:28:00 | 0:00 |
| XM_005263082:XR_938747:NM_001127208:XM_024454102 | 7:6:1:1 | 0:0:0:0 | 1:0:0:0 | 0:0:0:0 | 0:0:0:0 | 0:0:0:0 |
| XM_005263082:XR_938747:XM_024454102:NM_001127208 | 1:1:0:0 | 5:5:1:0 | 0:0:0:0 | 1:0:0:0 | 0:0:0:0 | 0:0:0:0 |
| NM_015454:NM_016648:XM_024454083:XM_024454089:XM_024454086:XM_024454087:NM_001267039:XM_024454084 | 9:18:0:0:1:0:0:0 | 0:0:0:0:0:0:0:0 | 8:6:1:0:0:0:0:0 | 1:0:0:0:0:0:0:0 | 7:8:0:1:0:1:0:0 | 0:0:0:1:0:0:0:0 |

| | | | | | | |
|---|---|---|---|---|---|---|
| NM_016648:NM_015454:XM_024454089:XM_024454086:XM_024454083:XM_024454087:NM_001267039:XM_024454084 | 15:6:0:1:0:0:0:0 | 2:3:0:0:0:0:0:0 | 5:7:0:0:0:0:0:0 | 1:0:0:0:0:0:0:0 | 8:5:1:0:0:1:0:0 | 0:1:0:0:0:0:0:0 |
| NM_005033:NM_001034194:XR_001741241 | 11:02:00 | 1:00:00 | 14:01:01 | 2:00:00 | 28:00:00 | 4:00:00 |
| NM_006726:NM_001199282:XM_017008874:XM_011532434:XM_017008873 | 5:0:3:2:0 | 0:0:0:0:0 | 2:1:0:0:0 | 0:0:0:0:0 | 1:0:4:0:0 | 0:0:0:0:0 |
| NM_006726:NM_001199282:XM_017008874:XM_011532434:XM_017008873 | 5:0:3:2:0 | 0:0:0:0:0 | 2:1:0:0:0 | 0:0:0:0:0 | 1:0:4:0:0 | 0:0:0:0:0 |
| NM_006342:XM_017007653:XM_005247929:XM_011513386 | 5:3:0:2 | 0:0:0:0 | 5:1:0:0 | 0:0:0:0 | 13:2:1:0 | 0:0:0:0 |
| NM_006342:XM_017007653:XM_005247929:XM_011513386 | 5:3:0:2 | 0:0:0:0 | 5:1:0:0 | 0:0:0:0 | 13:3:1:0 | 0:0:0:0 |
| NM_006342:XM_017007653:XM_005247929:XM_011513386:XM_005247930 | 6:7:0:0:0 | 0:0:0:0:0 | 5:1:0:0:0 | 0:0:0:0:0 | 15:8:1:0:1 | 0:0:0:0:0 |
| XM_024454160:XM_017008483:NM_020827 | 0:00:00 | 0:00:00 | 1:01:00 | 0:00:00 | 0:00:00 | 3:00:00 |
| XM_011513557:XM_005248005:NM_007331:XM_006713914:NM_001042424:XM_005248001:XM_005248002 | 0:0:0:0:0:0:0 | 0:0:0:1:0:0:0 | 0:1:1:1:0:0:0 | 1:0:0:0:0:0:0 | 1:0:1:3:0:0:1 | 0:0:0:0:1:1:0 |
| NM_001168243:NM_001141936:XM_011513469 | 1:00:00 | 2:00:00 | 0:00:00 | 2:00:00 | 0:00:00 | 6:00:02 |
| XM_017007900:XM_024453938:XM_024453937:NM_014988:NM_001330790:NM_001330792:NM_001289124:XM_011513646:NM_001112718:NM_001289122 | 0:0:1:0:0:0:0:0:0:0 | 0:0:0:0:0:0:0:0:0:0 | 1:1:1:0:0:0:0:0:0:0 | 1:1:0:0:0:0:0:0:0:0 | 0:0:0:0:0:1:0:0:0:0 | 0:0:0:0:0:1:0:0:0:0 |
| NM_017816:XM_011513506:XM_011513505:NM_001145725 | 28:3:0:0 | 21:0:0:0 | 44:2:0:0 | 23:2:0:0 | 75:3:1:0 | 43:3:0:1 |
| NM_017816:XM_011513506:XM_011513505:NM_001145725 | 57:3:0:0 | 0:0:0:0 | 72:5:0:0 | 2:0:0:0 | 132:7:2:1 | 1:0:0:0 |
| NM_025087:XM_011513758:XM_011513755 | 0:00:00 | 29:02:01 | 0:00:00 | 0:00:00 | 2:00:00 | 0:00:00 |
| NM_025087:XM_011513758:XM_011513755:XM_011513757:XM_011513759 | 0:0:0:0:0 | 31:2:1:0:0 | 0:0:0:0:0 | 0:0:0:0:0 | 2:0:0:0:0 | 0:0:0:0:0 |
| XM_017008662:XM_017008669:XM_005265781:XM_005265774:NM_030917:XM_017008666:XM_017008665:XM_005265769:XM_005265782:XM_005265778:NM_001134938:XM_005265773:XM_017008664:XM_017008663:XM_005265779:XM_017008667:XM_017008668:XM_017008671:NM_001134937:XM_017008670 | 0:6:6:11:6:16:1:3:4:0:5:7:9:1:1:0:3:1:3:0 | 0:0:0:1:1:1:0:0:2:0:0:1:0:0:0:0:0:0:1:0 | 7:7:3:8:2:9:3:3:7:1:6:4:2:2:1:1:3:1:0:0 | 2:1:0:0:0:0:0:0:0:0:0:0:0:0:0:0:0:0:0:0 | 3:3:1:9:4:12:3:2:8:0:5:6:4:2:1:2:8:0:2:0 | 0:0:0:0:0:0:0:0:0:0:0:1:0:0:0:0:0:0:0:0 |
| XM_005265769:NM_030917:XM_017008666:NM_001134938:XM_005265781:XM_017008665:XM_005265782:XM_005265774:XM_017008662:XM_017008667:XM_017008668:XM_017008669:XM_005265778:XM_005265773:XM_017008664:XM_017008663:XM_005265779:XM_017008671:NM_001134937:XM_017008670 | 1:2:12:2:3:0:1:7:0:0:3:4:0:5:5:1:0:1:2:0 | 2:4:2:3:3:1:1:4:0:0:0:0:0:2:4:0:1:0:1:0 | 1:1:4:3:1:2:2:7:5:0:1:4:1:4:2:2:1:1:0:0 | 2:1:4:3:1:1:3:1:1:1:1:1:0:0:0:0:0:0:0:0 | 2:2:7:4:0:3:6:7:2:2:2:2:0:5:2:1:0:0:2:0 | 0:1:5:1:1:0:1:2:0:0:5:1:0:1:1:1:1:0:0:0 |
| NM_006452:NM_001079524:XM_011534366:NM_001079525 | 15:38:5:0 | 0:0:0:0 | 35:88:3:0 | 0:0:0:0 | 76:144:10:0 | 0:0:0:0 |
| NM_004684:NM_001128310 | 50:01:00 | 0:00 | 37:00:00 | 0:00 | 43:00:00 | 0:00 |
| NM_004684:NM_001128310 | 58:02:00 | 2:00 | 44:01:00 | 3:00 | 54:00:00 | 3:01 |
| NM_004684:NM_001128310:NM_001291976 | 65:02:00 | 4:01:00 | 56:01:00 | 0:00:00 | 62:01:00 | 1:01:00 |
| NM_004684:NM_001128310:NM_001291976 | 70:03:00 | 0:00:00 | 54:01:00 | 0:00:00 | 30:02:00 | 33:00:00 |
| NM_004684:NM_001128310:NM_001291976 | 71:03:00 | 0:00:00 | 54:01:00 | 0:00:00 | 30:02:00 | 35:00:00 |
| NM_000669:NR_133005 | 0:00 | 12:01 | 0:00 | 0:00 | 0:00 | 1:00 |
| NM_016144:NR_146220:NM_001308080 | 72:00:00 | 0:00:00 | 110:01:01 | 0:00:00 | 79:01:00 | 72:00:00 |
| XM_011537569:NM_152550:XM_017009136:XM_011537567:XM_006714757:XM_005268380:XM_011537568 | 4:3:0:0:0:0:0 | 3:3:1:1:0:0:0 | 0:0:0:0:0:0:0 | 1:1:0:0:0:0:0 | 0:0:0:0:0:0:0 | 2:5:1:1:0:2:1 |
| NM_152550:XM_011537569:XM_017009136:XM_011537568:XM_011537567:XM_006714757:XM_005268380 | 0:0:0:0:0:0:0 | 7:5:1:1:1:0:0 | 0:0:0:0:0:0:0 | 1:0:0:0:0:0:0 | 0:0:0:0:0:0:0 | 6:2:1:1:1:0:2 |
| NM_152550:XM_011537569:XM_011537568:XM_017009136:XM_011537567:XM_006714757:XM_005268380 | 4:4:1:0:0:0:0 | 3:3:0:1:1:0:0 | 0:0:0:0:0:0:0 | 1:0:0:0:0:0:0 | 0:0:0:0:0:0:0 | 5:2:1:1:1:0:2 |
| NM_004355:NM_001025159:XM_017010089:NM_001025158:XM_017010090:XR_002956198 | 7469:1019:10:3:64:27 | 6:1:0:0:0:1 | 6062:1270:12:3:41:29 | 6:0:0:0:0:0 | 1152:108:0:1:3:0 | 1:0:0:0:0:0 |
| NM_004355:NM_001025159:XR_002956198:XM_017010089:XM_017010090:NM_001025158 | 7398:1010:46:10:64:0 | 9:0:0:0:0:0 | 6013:1257:66:12:40:0 | 10:1:1:0:0:0 | 1140:106:6:0:3:0 | 0:0:0:0:0:0 |
| NM_004355:NM_001025159:XM_017010089:XM_017010090:XR_002956198:NM_001025158 | 7234:981:8:64:62:5 | 7:3:0:0:0:0 | 5932:1233:12:40:92:3 | 4:2:0:0:0:0 | 1122:101:0:3:6:1 | 0:0:0:0:0:0 |
| NM_004355:XR_002956198:NM_001025159:NM_001025158:XM_017010089:XM_017010090 | 7240:65:986:7:8:64 | 8:0:0:0:0:0 | 5939:92:1233:4:12:40 | 4:1:3:1:0:0 | 1123:6:101:1:0:3 | 0:0:0:0:0:0 |
| NM_004355:NM_001025159:XM_017010089:XM_017010090:XR_002956198:NM_001025158 | 7212:979:8:64:70:7 | 20:1:0:0:0:0 | 5931:1232:11:40:99:5 | 6:1:0:0:0:0 | 1121:101:0:3:8:2 | 2:0:0:0:0:0 |
| NM_004355:NM_001025159:XM_017010089:NM_001025158:XM_017010090:XR_002956198 | 7021:949:9:8:61:71 | 11:1:0:0:0:0 | 5816:1211:10:6:35:101 | 7:3:0:0:0:0 | 1092:99:0:2:3:7 | 1:1:0:0:0:0 |
| NM_001025159:NM_004355:XM_017010089:NM_001025158:XM_017010090:XR_002956198 | 947:7007:9:8:58:72 | 0:7:0:0:0:0 | 1208:5807:10:6:34:101 | 4:2:0:0:0:0 | 100:1090:0:2:3:7 | 0:1:0:0:0:0 |
| NM_004355:XR_002956198:NM_001025159:XM_017010089:NM_001025158:XM_017010090 | 7052:78:954:9:8:56 | 6:0:2:0:0:0 | 5876:104:1217:10:6:33 | 2:2:2:0:0:0 | 1096:8:100:0:2:3 | 0:0:0:0:0:0 |
| NM_004355:NM_001025159:XM_017010089:NM_001025158:XM_017010090:XR_002956198 | 7140:960:8:8:50:110 | 29:5:0:0:1:1 | 6000:1223:8:6:28:127 | 12:2:1:0:0:0 | 1115:102:0:2:3:14 | 2:0:0:0:0:0 |
| NM_004355:NM_001025159:XM_017010089:NM_001025158:XM_017010090:XR_002956198 | 7118:958:8:8:47:107 | 8:3:0:0:0:0 | 5991:1219:8:6:28:119 | 6:2:0:0:0:0 | 1113:103:0:2:4:14 | 1:0:0:0:0:0 |
| NM_004355:NM_001025159:XR_002956198:XM_017010089:NM_001025158:XM_017010090 | 7140:956:108:7:8:45 | 15:2:1:0:0:1 | 6024:1221:120:8:6:29 | 9:3:1:0:0:0 | 1115:103:14:0:2:4 | 2:0:0:0:0:0 |

| | | | | | | |
|---|---|---|---|---|---|---|
| NM_004355:NM_001025159:XM_017010089:NM_001025158:XM_017010090:XR_002956198 | 7109:951:7:8:46:104 | 19:3:0:0:0:1 | 6014:1216:7:6:27:116 | 8:0:0:0:0:0 | 1111:101:0:2:4:15 | 2:0:0:0:0:0 |
| NM_018691:NM_001317993:XM_005268360:XM_011537542:XR_001741975:XM_024454339:XM_005268361:NM_001317994:XM_017008967:XM_011537540 | 13:0:1:0:0:1:0:0:0:0 | 12:1:3:0:1:5:0:0:0:0 | 8:1:1:0:0:3:0:0:0:0 | 9:2:2:1:1:1:0:0:0:0 | 19:0:9:2:1:9:2:1:1:0 | 0:0:0:0:0:0:0:0:0:0 |
| NM_018691:NM_001317993:XM_005268360:XM_011537542:XM_024454339:XR_001741975:XM_005268361:NM_001317994:XM_017008967:XM_011537540 | 26:1:4:0:6:1:0:0:0:0 | 1:0:0:0:0:0:0:0:0:0 | 17:3:3:1:5:1:0:0:0:0 | 0:0:0:0:0:0:0:0:0:0 | 20:0:9:2:9:0:2:1:1:0 | 1:0:0:0:0:0:0:0:0:0 |
| NM_018691:NM_001317993:XM_005268360:XM_011537542:XM_024454339:XR_001741975:XM_005268361:NM_001317994:XM_017008967:XM_011537540 | 26:1:4:0:6:1:0:0:0:0 | 1:0:0:0:0:0:0:0:0:0 | 17:3:3:1:5:1:0:0:0:0 | 0:0:0:0:0:0:0:0:0:0 | 19:0:9:2:9:0:2:1:1:0 | 2:0:0:0:0:0:0:0:0:0 |
| NM_018691:NM_001317993:XM_005268360:XM_011537542:XM_024454339:XR_001741975:XM_005268361:NM_001317994:XM_017008967:XM_011537540 | 0:0:0:0:0:0:0:0:0:0 | 27:1:4:0:3:1:0:0:0:0 | 0:0:0:0:0:0:0:0:0:0 | 17:3:3:1:1:1:0:0:0:0 | 12:0:1:0:2:0:2:0:1:0 | 9:0:8:2:2:0:0:1:0:0 |
| NM_199185:NM_002520:NM_001355007:NM_001037738:NM_001355010:NM_001355009:NR_149149:NM_001355006 | 18:477:3:0:0:0:0:0 | 0:1:0:0:0:0:0:0 | 51:1804:5:0:0:0:0:0 | 1:0:0:0:0:0:0:0 | 65:2076:0:2:0:0:0:0 | 0:2:0:0:0:0:0:0 |
| NM_002520:NM_001355007:NR_149149:NM_001037738:NM_199185:NM_001355006 | 478:3:4:0:0:0 | 2:0:0:0:0:0 | 1811:5:7:0:0:0 | 3:0:0:0:0:0 | 2078:0:13:2:0:0 | 4:0:0:0:0:0 |
| NM_002520:NM_199185:NM_001355007:NR_149149:NM_001037738:NM_001355010:NM_001355009:NM_001355006 | 474:12:3:7:0:0:0:0 | 1:0:0:0:0:0:0:0 | 1803:39:5:13:0:0:0:0 | 2:0:0:0:0:0:0:0 | 2077:57:0:24:3:0:0:0 | 3:0:0:0:0:0:0:0 |
| NM_002520:NM_199185:NM_001355007:NR_149149:NM_001037738:NM_001355010:NM_001355009:NM_001355006 | 474:12:3:10:0:0:0:0 | 1:0:0:0:0:0:0:0 | 1794:39:5:16:0:0:0:0 | 2:0:0:0:0:0:0:0 | 2075:56:0:29:3:0:0:0 | 3:0:0:0:0:0:0:0 |
| NM_002520:NM_199185:NM_001355007:NR_149149:NM_001355010:NM_001037738:NM_001355006 | 473:12:3:11:0:0:0 | 1:0:0:0:0:0:0 | 1786:39:5:17:0:0:0 | 3:0:0:0:0:0:0 | 2074:56:0:36:0:0:0 | 0:0:0:0:0:0:0 |
| NM_002520:NM_199185:NM_001355007:NR_149149:NM_001355010:NM_001037738:NM_001355006 | 474:12:3:11:0:0:0 | 0:0:0:0:0:0:0 | 1786:39:5:18:0:0:0 | 2:0:0:0:0:0:0 | 2073:56:0:36:0:0:0 | 0:0:0:0:0:0:0 |
| NM_002520:NM_199185:NM_001355007:NR_149149:NM_001355010:NM_001037738:NM_001355006 | 471:12:3:11:0:0:0 | 2:0:0:0:0:0:0 | 1785:39:5:18:0:0:0 | 3:0:0:0:0:0:0 | 2071:56:0:36:0:0:0 | 2:0:0:0:0:0:0 |
| NM_001142298:NM_001142299:NM_003900 | 0.284027778 | 0:00:00 | 0.050127315 | 0:00:00 | 0.186319444 | 0:01:01 |
| NM_001142298:NM_001142299:NM_003900 | 0.22412037 | 0.059837963 | 0.004895833 | 0.045231481 | 0.145358796 | 0:00:00 |
| NM_016107:NR_144318 | 5:11 | 3:22 | 1:00 | 2:00 | 0:00 | 1:01 |
| NM_016107:NR_144318 | 0:00 | 6:34 | 0:00 | 2:00 | 0:00 | 1:01 |
| NR_144318:NM_016107 | 201:02:00 | 9:00 | 9:00 | 2:00 | 11:00 | 0:00 |
| NM_014324:NM_203382:NM_001167595 | 0:00:00 | 82:04:02 | 0:00:00 | 18:02:00 | 0:00:00 | 33:00:00 |
| NM_014324:NM_203382:NM_001167595 | 84:04:02 | 0:00:00 | 18:02:00 | 0:00:00 | 15:00:00 | 20:00:00 |
| NM_014324:NM_203382:NM_001167595 | 1:00:00 | 83:04:02 | 0:00:00 | 18:02:00 | 0:00:00 | 35:00:00 |
| NM_014324:NM_001167595 | 32:01:00 | 52:01:00 | 11:00 | 7:00 | 20:00 | 15:00 |
| NM_014324:NM_203382:NM_001167595 | 25:04:01 | 25:00:01 | 3:01:00 | 3:00:00 | 7:00:00 | 7:00:00 |
| NM_018034:XM_011514061:NM_001345998:NM_001345999 | 16:0:0:0 | 0:0:0:0 | 12:1:0:0 | 0:0:0:0 | 12:0:1:0 | 0:0:0:0 |
| NM_173514:NM_001349384:XM_017009082:XR_948238:XM_011543174:NM_001349382:NM_001349385:XM_024454369:XM_017009077:NR_146165:XM_011543176:XM_017009080:XR_948237:NM_001258286:NM_001349383 | 0:0:0:0:0:0:0:0:0:0:0:0:0:0:0 | 1:2:0:0:0:2:1:1:2:0:0:0:0:0:0 | 0:0:0:0:0:0:0:0:0:0:0:0:0:0:0 | 1:1:0:0:0:0:0:0:0:0:0:0:0:0:0 | 3:1:0:0:0:0:0:0:0:0:0:0:0:1:0 | 2:1:0:0:0:0:1:0:1:0:1:1:1:0:0 |
| NM_015342:XM_006714583:NM_001278926:XM_017009286 | 20:1:1:0 | 4:0:0:0 | 28:1:1:0 | 3:0:0:0 | 40:2:0:0 | 16:0:0:0 |
| NM_017755:NR_037947:NM_001193455 | 4:01:00 | 0:00:00 | 11:00:00 | 0:00:00 | 13:03:01 | 11:00:00 |
| NM_133343:NM_133338:NM_133344:NM_133342:NM_001278622:NM_002873:XM_017009681:NM_133340:NM_133341 | 2:0:5:13:2:0:2:7:0 | 0:0:0:0:0:0:0:0:0 | 3:1:4:16:5:1:1:0:0 | 0:0:0:0:0:0:0:0:0 | 2:0:2:11:1:0:6:3:3 | 1:1:3:15:0:0:2:3:0 |
| XM_017009634:NM_018429 | 4:00 | 0:00 | 5:00 | 0:00 | 0:00 | 2:00 |
| XM_017009634:NM_018429 | 4:00 | 0:00 | 6:00 | 0:00 | 0:00 | 2:00 |
| NM_003664:XM_005248618:XM_017010001:XM_005248619 | 0:0:0:0 | 11:0:1:0 | 0:0:0:0 | 10:1:0:0 | 0:0:0:0 | 21:1:1:1 |
| NM_024091:XR_427664:XM_006714498:XR_001742253 | 0:0:0:0 | 2:1:8:1 | 0:0:0:0 | 3:1:0:0 | 1:1:0:1 | 3:0:1:0 |
| NM_001131028:XM_017009944:NM_031482:XM_011543660 | 0:0:0:0 | 0:0:0:0 | 0:1:0:0 | 1:0:0:0 | 2:0:1:0 | 0:0:0:0 |
| NM_001131028:XM_017009944:NM_031482:XM_011543660 | 0:0:0:0 | 0:0:0:0 | 0:0:0:0 | 1:1:0:0 | 2:0:1:0 | 0:0:0:0 |
| NM_001239:XR_001742327:XM_005248627:NM_001199189 | 26:1:0:0 | 0:0:0:0 | 25:1:2:0 | 0:0:0:0 | 38:0:0:0 | 37:0:0:0 |
| XR_002956175:NM_023924:NR_134296:XM_024446192:XM_024446199:XM_024446198:XR_002956174:NM_001317951:XM_024446193:XM_024446200:XM_006714494:XM_024446196:NM_001009877:XM_024446197 | 10:9:0:1:5:3:1:0:0:0:0:0:0:0 | 4:0:0:0:0:0:0:0:0:0:0:0:0:0 | 2:3:3:0:0:0:0:0:0:0:0:0:0:0 | 3:0:0:0:0:0:0:0:0:0:0:0:0:0 | 4:5:0:1:0:0:1:0:3:1:1:0:0:0 | 0:0:0:0:0:0:0:0:0:0:0:0:0:0 |
| NM_002726:XM_011535925:XM_005267044 | 17:00:00 | 16:00:00 | 4:00:00 | 16:00:00 | 54:00:00 | 0:00:00 |
| NM_018292:XM_011535924 | 29:02:00 | 0:00 | 36:00:00 | 0:00 | 19:02 | 25:01:00 |
| NM_007214:XM_017010218 | 1:01 | 0:00 | 1:00 | 0:00 | 1:00 | 2:00 |
| XM_005266797:NM_006714:XM_005266799 | 0:00:00 | 0:06:00 | 0:00:00 | 0:00:00 | 0:01:00 | 0:00:00 |
| XR_002956294:NM_001031712:NM_001350582:NR_146799:NM_001350580:NR_146802:NM_001350581:NR_146796:NR_146798:NM_001350583:NR_146800:NM_001350586:NR_146801:NM_001350584:NM_001350588:XM_017011168:NR_146804:NM_001350589:NM_001350592:NM_001350591:NR_146795:NM_001350585:NM_001350596:NM_001350597:NR_146803:NM_001350594:XM_024446516 | 8:76:0:10:1:1:3:4:1:0:1:0:5:2:2:2:1:1:1:1:0:0:0:0:0:0:0 | 0:0:0:0:0:0:0:0:0:0:0:0:0:0:0:0:1:0:0:0:0:0:0:0:0:0:0 | 3:59:1:5:1:1:2:2:1:1:2:1:3:1:1:1:0:0:0:0:0:0:0:0:0:0:0 | 2:0:0:0:0:0:0:0:0:0:0:0:0:0:0:0:0:0:0:0:0:0:0:0:0:0:0 | 2:69:0:7:1:1:7:5:3:0:0:1:0:0:0:3:0:0:2:0:0:1:1:0:0:0:0 | 1:0:0:1:0:0:0:0:0:0:0:0:0:0:0:0:0:0:0:0:0:0:0:0:0:0:0 |
| XM_017011148:NM_002844:XM_006715537:XM_017011146:XM_017011145:XM_017011150 | 7:9:0:0:0:1 | 0:0:0:0:0:0 | 4:4:1:0:0:0 | 0:0:0:0:0:0 | 7:6:0:0:0:0 | 0:0:0:0:0:0 |

| | | | | | | |
|---|---|---|---|---|---|---|
| XM_006715537:XM_011536021:XM_017011148:NM_002844:XM_017011146:XM_017011145:XM_011536016:NM_001291982:XM_011536014:XM_011536020:XM_017011147:XM_017011150:XM_011536015:NM_001291983 | 0:38:3:1:0:0:0:1:0:0:0:0:0:0 | 0:0:0:0:0:0:0:0:0:0:0:0:0:0 | 1:7:1:0:0:0:0:0:0:0:0:0:0:0 | 0:0:0:0:0:0:0:0:0:0:0:0:0:0 | 0:18:1:3:5:0:1:0:0:0:1:0:0:0 | 0:0:0:0:0:0:0:0:0:0:0:0:0:0 |
| XM_005267213:XM_017011494:NM_033515:XM_005267212 | 1:0:8:1 | 8:0:0:0 | 3:0:25:4 | 5:1:0:0 | 1:0:16:4 | 4:0:0:0 |
| XM_005267213:NM_033515:XM_005267212:XM_017011494 | 10:8:1:0 | 0:0:0:0 | 9:25:4:1 | 3:0:0:0 | 5:16:4:0 | 0:0:0:0 |
| XM_005267213:NM_033515:XM_005267212:XM_017011494 | 10:8:1:0 | 0:0:0:0 | 9:25:4:1 | 3:0:0:0 | 5:16:4:0 | 0:0:0:0 |
| XM_005267213:NM_033515:XM_005267212:XM_017011494 | 10:8:1:0 | 0:0:0:0 | 9:25:4:1 | 3:0:0:0 | 5:16:4:0 | 0:0:0:0 |
| NM_033515:XM_005267213:XM_005267212:XM_017011494 | 8:23:1:1 | 0:0:0:0 | 25:32:2:3 | 0:0:0:0 | 16:27:4:1 | 0:1:0:0 |
| NM_033515:XM_005267213:XM_005267212:XM_017011494 | 0:0:0:0 | 7:22:1:0 | 0:0:0:0 | 24:31:0:0 | 0:1:0:0 | 14:27:0:0 |
| NM_014432:XM_017010954:NM_001278723:XM_017010955:NM_001278722:XM_011535904:NM_001278724 | 0:0:1:0:0:0:0 | 1:1:1:0:0:0:0 | 2:0:0:0:0:0:0 | 1:0:0:0:0:0:0 | 6:0:4:1:1:0:0 | 0:0:0:0:0:0:0 |
| NM_014432:NM_001278723:XM_017010954:NM_001278722:XM_011535904:XM_017010955:NM_001278724 | 0:0:0:0:0:0:0 | 1:2:1:0:0:0:0 | 0:0:0:0:0:0:0 | 3:0:0:0:0:0:0 | 0:0:0:0:0:0:0 | 6:4:0:1:0:1:0 |
| NM_022121:XM_024446520 | 0:00 | 7:02 | 0:00 | 4:00 | 0:00 | 11:00 |
| XM_011536147:NM_025059:XM_011536148 | 0:00:04 | 0:00:00 | 1:03:08 | 0:00:00 | 0:00:16 | 0:00:11 |
| XM_011536147:NM_025059:XM_011536148 | 0:00:00 | 0:00:04 | 0:02:06 | 1:01:02 | 0:00:15 | 0:00:09 |
| NM_001142522:NM_012177 | 0:00 | 0:00 | 0:00 | 0:00 | 0:00 | 1:00 |
| NM_016020:XR_001743453:NM_001350502:XR_942460:XM_017010917 | 51:10:2:0:4 | 17:0:1:1:0 | 48:5:2:2:0 | 5:0:0:0:0 | 66:10:2:1:5 | 5:1:0:0:1 |
| NM_016020:XR_001743453:NM_001350502:XR_942460:XM_017010917 | 61:11:2:0:5 | 14:0:1:1:1 | 63:6:2:2:0 | 8:0:0:0:0 | 82:11:2:1:5 | 9:1:0:0:2 |
| NM_016020:XR_001743453:NM_001350502:XR_942460:XM_017010917 | 66:11:3:0:6 | 9:0:0:1:0 | 68:6:2:2:0 | 5:0:0:0:0 | 88:11:2:1:7 | 3:1:0:0:0 |
| NM_001111077:NM_003379:XM_011536110 | 603:702:50 | 1:00:01 | 121:80:10 | 0:00:00 | 273:123:10 | 0:00:00 |
| NM_014161:NM_001318817 | 0:00 | 0:00 | 1:00 | 0:00 | 2:00 | 1:01 |
| XM_017010958:XM_011535905:NM_001143947:NM_018974:XM_011535906:XM_011535908:XM_011535907 | 0:1:0:0:0:0:0 | 0:0:0:0:0:0:0 | 2:6:3:1:0:0:0 | 0:0:0:0:0:0:0 | 0:0:0:0:0:0:0 | 0:3:0:1:0:0:0 |
| XM_017010958:XM_011535905:NM_001143947:NM_018974:XM_011535906:XM_011535907 | 0:1:0:0:0:0 | 0:0:0:0:0:0 | 3:6:3:1:0:0 | 0:0:0:0:0:0 | 0:0:0:0:0:0 | 0:4:0:1:0:0 |
| XM_017010958:XM_011535905:NM_001143947:NM_018974:XM_011535906:XM_011535907 | 0:1:0:0:0:0 | 0:0:0:0:0:0 | 3:6:3:1:0:0 | 0:0:0:0:0:0 | 0:0:0:0:0:0 | 0:4:0:1:0:0 |
| XM_017010958:XM_011535905:NM_001143947:NM_018974:XM_011535906:XM_011535907 | 0:1:0:0:0:0 | 0:0:0:0:0:0 | 3:6:3:1:0:0 | 0:0:0:0:0:0 | 0:4:0:1:0:0 | 0:0:0:0:0:0 |
| XR_001743427:XM_017010885:XM_017010880:XM_017010884:XM_017010887:XM_017010886 | 22:15:8:1:0:0 | 1:0:0:0:0:0 | 21:7:10:0:0:0 | 1:0:0:0:0:0 | 39:10:20:0:0:0 | 0:0:0:0:0:0 |
| XM_024446544:XR_001743635:NM_003472:NM_001134709:XM_017011273 | 11:4:43:0:15 | 1:2:0:0:0 | 13:2:71:1:20 | 1:4:1:0:0 | 54:6:110:1:45 | 4:8:2:0:0 |
| NM_003472:XM_024446544:XR_001743635:XM_017011273:NM_001134709 | 42:9:9:15:0 | 0:0:0:0:0 | 73:12:7:20:0 | 0:0:0:0:0 | 54:32:11:21:0 | 61:26:6:24:0 |
| NM_004568:XM_011514674:NM_001271824:XM_024446465:NM_001297699:NM_001271825:NM_001195291:NM_001271822:XM_024446463 | 0:0:0:0:0:0:0:0:0 | 1:0:1:0:1:0:0:0:0 | 0:0:0:0:0:0:0:0:0 | 3:1:0:0:0:0:0:0:0 | 2:0:0:0:0:1:0:0:0 | 1:0:0:0:0:0:0:0:0 |
| NM_002116:NM_001242758 | 0:00 | 15:00 | 0:00 | 1:00 | 0:04 | 0:00 |
| NM_002116:NM_001242758 | 0:00 | 17:00 | 0:00 | 1:00 | 0:04 | 0:00 |
| NM_002116:NM_001242758 | 16:00 | 0:00 | 1:00 | 0:00 | 0:00 | 0:04 |
| NM_002116:NM_001242758 | 0:00 | 16:00 | 0:00 | 1:00 | 0:05 | 0:00 |
| NM_002116:NM_001242758 | 0:00 | 17:00 | 0:00 | 1:00 | 0:05 | 0:00 |
| NM_002116:NM_001242758 | 0:00 | 17:00 | 0:00 | 1:00 | 0:05 | 0:00 |
| NM_002116:NM_001242758 | 17:00 | 0:00 | 1:00 | 0:00 | 0:00 | 0:05 |
| NM_002116:NM_001242758 | 0:00 | 17:00 | 0:00 | 1:00 | 0:00 | 0:05 |
| NM_002116:NM_001242758 | 0:00 | 17:00 | 0:00 | 1:00 | 0:00 | 0:06 |
| NM_002116:NM_001242758 | 0:00 | 17:00 | 0:00 | 1:00 | 0:00 | 0:06 |
| NM_002116:NM_001242758 | 0:00 | 17:00 | 0:00 | 1:00 | 0:06 | 0:00 |
| NM_002116:NM_001242758 | 0:00 | 17:00 | 0:00 | 1:00 | 0:06 | 0:00 |
| NM_002116:NM_001242758 | 0:00 | 17:00 | 0:00 | 1:00 | 0:06 | 0:00 |
| NM_002116:NM_001242758 | 0:00 | 17:00 | 0:00 | 1:00 | 0:06 | 0:00 |
| NM_002116:NM_001242758 | 0:00 | 17:00 | 0:00 | 1:00 | 0:06 | 0:00 |
| NM_002116:NM_001242758 | 0:00 | 17:00 | 0:00 | 1:00 | 0:06 | 0:00 |
| NM_002116:NM_001242758 | 17:00 | 0:00 | 1:00 | 0:00 | 0:00 | 0:06 |
| NM_002116:NM_001242758 | 0:00 | 18:00 | 0:00 | 1:00 | 0:06 | 0:00 |
| NM_002116:NM_001242758 | 0:00 | 18:00 | 0:00 | 1:00 | 0:06 | 0:00 |
| NM_002116:NM_001242758 | 0:00 | 18:00 | 0:00 | 1:00 | 0:06 | 0:00 |
| NM_002116:NM_001242758 | 18:00 | 0:00 | 1:00 | 0:00 | 0:00 | 0:06 |
| NM_002116:NM_001242758 | 0:00 | 18:00 | 0:00 | 1:00 | 0:06 | 0:00 |
| NM_002116:NM_001242758 | 0:00 | 18:00 | 0:00 | 1:00 | 0:06 | 0:00 |

| | | | | | | |
|---|---|---|---|---|---|---|
| NM_002116:NM_001242758 | 0:00 | 18:00 | 0:00 | 1:00 | 0:06 | 0:00 |
| NM_002116:NM_001242758 | 18:00 | 0:00 | 1:00 | 0:00 | 0:00 | 0:07 |
| NM_002116:NM_001242758 | 0:00 | 18:00 | 0:00 | 1:00 | 0:00 | 0:07 |
| NM_002116:NM_001242758 | 0:00 | 19:00 | 0:00 | 1:00 | 0:00 | 0:07 |
| NM_002116:NM_001242758 | 0:00 | 19:00 | 0:00 | 1:00 | 0:00 | 0:07 |
| NM_002116:NM_001242758 | 19:00 | 0:00 | 1:00 | 0:00 | 0:00 | 0:07 |
| NM_002116:NM_001242758 | 0:00 | 19:00 | 0:00 | 1:00 | 0:00 | 0:08 |
| NM_002116:NM_001242758 | 19:00 | 0:00 | 1:00 | 0:00 | 0:00 | 0:08 |
| NM_002116:NM_001242758 | 19:00 | 0:00 | 1:00 | 0:00 | 0:00 | 0:08 |
| NM_002116:NM_001242758 | 0:00 | 19:00 | 0:00 | 1:00 | 0:00 | 0:08 |
| NM_002116:NM_001242758 | 0:00 | 19:00 | 0:00 | 1:00 | 0:00 | 0:08 |
| NM_002116:NM_001242758 | 0:00 | 19:01 | 0:00 | 1:00 | 0:09 | 0:00 |
| NM_002116:NM_001242758 | 1:00 | 18:01 | 0:00 | 1:00 | 0:09 | 0:00 |
| NM_002116:NM_001242758 | 0:00 | 18:00 | 0:00 | 1:00 | 0:10 | 0:00 |
| NM_002116:NM_001242758 | 0:00 | 17:00 | 0:00 | 1:00 | 0:10 | 0:00 |
| NM_002117:NM_001243042 | 0:26 | 214:04:00 | 0:07 | 61:03:00 | 0:00 | 91:01:00 |
| NM_002117:NM_001243042 | 1:25 | 213:04:00 | 0:07 | 60:03:00 | 0:00 | 91:01:00 |
| NM_002117:NM_001243042 | 0:25 | 206:04:00 | 0:07 | 60:03:00 | 0:00 | 91:01:00 |
| NM_002117:NM_001243042 | 0:25 | 206:04:00 | 0:07 | 60:03:00 | 0:00 | 91:01:00 |
| NM_002117:NM_001243042 | 0:25 | 206:04:00 | 0:07 | 60:03:00 | 0:00 | 91:01:00 |
| NM_002117:NM_001243042 | 0:25 | 206:04:00 | 0:07 | 60:03:00 | 6:01 | 85:00:00 |
| NM_002117:NM_001243042 | 206:29:00 | 0:00 | 60:10:00 | 0:00 | 91:01:00 | 0:00 |
| NM_002117:NM_001243042 | 0:25 | 203:04:00 | 0:07 | 60:03:00 | 0:00 | 90:01:00 |
| NM_002117:NM_001243042 | 0:25 | 202:04:00 | 0:07 | 60:03:00 | 0:00 | 90:01:00 |
| NM_002117:NM_001243042 | 1:24 | 199:04:00 | 0:07 | 60:03:00 | 91:01:00 | 0:00 |
| NM_002117:NM_001243042 | 198:27:00 | 1:00 | 60:10:00 | 0:00 | 93:01:00 | 0:00 |
| NM_002117:NM_001243042 | 0:23 | 199:04:00 | 0:07 | 60:03:00 | 0:00 | 93:01:00 |
| NM_002117:NM_001243042 | 0:23 | 198:04:00 | 0:07 | 60:03:00 | 0:00 | 93:01:00 |
| NM_002117:NM_001243042 | 0:23 | 197:04:00 | 0:07 | 60:03:00 | 0:00 | 91:01:00 |
| NM_002117:NM_001243042 | 0:23 | 195:04:00 | 0:07 | 60:03:00 | 0:00 | 91:01:00 |
| NM_002117:NM_001243042 | 190:28:00 | 3:00 | 58:10:00 | 2:00 | 86:01:00 | 1:00 |
| NM_002117:NM_001243042 | 183:28:00 | 2:00 | 59:10:00 | 1:00 | 78:00:00 | 5:01 |
| NM_002117:NM_001243042 | 3:26 | 178:04:00 | 1:07 | 54:03:00 | 1:01 | 83:01:00 |
| NM_002117:NM_001243042 | 2:27 | 179:04:00 | 1:07 | 55:03:00 | 82:01:00 | 0:00 |
| NM_002117:NM_001243042 | 179:31:00 | 1:00 | 56:10:00 | 0:00 | 76:00:00 | 6:01 |
| NM_002117:NM_001243042 | 178:28:00 | 0:03 | 54:10:00 | 0:00 | 75:00:00 | 6:01 |
| NM_002117:NM_001243042 | 178:28:00 | 0:03 | 54:10:00 | 0:00 | 75:00:00 | 6:01 |
| NM_002117:NM_001243042 | 2:26 | 176:04:00 | 1:07 | 53:03:00 | 6:01 | 73:00:00 |
| NM_002117:NM_001243042 | 0:20 | 177:10:00 | 0:07 | 54:03:00 | 0:00 | 79:01:00 |
| NM_002117:NM_001243042 | 0:20 | 177:10:00 | 0:07 | 54:03:00 | 0:00 | 79:01:00 |
| NM_002117:NM_001243042 | 2:26 | 173:04:00 | 1:08 | 53:03:00 | 79:01:00 | 0:00 |
| NM_002117:NM_001243042 | 0:20 | 171:10:00 | 0:07 | 54:04:00 | 71:00:00 | 6:01 |
| NM_002117:NM_001243042 | 0:20 | 170:10:00 | 0:07 | 53:04:00 | 78:01:00 | 0:00 |
| NM_002117:NM_001243042 | 166:28:00 | 2:02 | 51:11:00 | 1:00 | 72:00:00 | 6:01 |
| NM_002117:NM_001243042 | 0:00 | 167:30:00 | 0:00 | 51:11:00 | 0:00 | 78:01:00 |
| NM_002117:NM_001243042 | 165:27:00 | 2:03 | 51:11:00 | 0:00 | 6:01 | 72:00:00 |
| NM_001243042:NM_002117 | 24:167 | 6:00 | 10:51 | 1:00 | 0.095833333 | 0:00 |
| NM_001243042:NM_002117 | 24:167 | 5:00 | 10:51 | 1:00 | 0.05 | 1:06 |
| NM_001243042:NM_002117 | 24:167 | 5:00 | 10:51 | 1:00 | 0.05 | 1:06 |
| NM_002117:NM_001243042 | 167:28:00 | 0:01 | 51:11:00 | 0:00 | 72:00:00 | 6:01 |
| NM_002117:NM_001243042 | 3:20 | 168:04:00 | 0:06 | 52:03:00 | 0:00 | 80:00:00 |
| NM_002117:NM_001243042 | 173:22:00 | 0:00 | 52:09:00 | 0:00 | 81:00:00 | 0:00 |
| NM_002117:NM_001243042 | 175:21:00 | 1:01 | 51:09:00 | 1:00 | 80:00:00 | 0:00 |
| NM_002117:NM_001243042 | 0:19 | 177:02:00 | 1:06 | 52:03:00 | 75:00:00 | 6:00 |
| NM_001243042:NM_002117 | 0.123611111 | 21:00 | 2:54 | 6:00 | 0.05625 | 0:00 |
| NM_002117:NM_001243042 | 180:21:00 | 0:00 | 54:09:00 | 1:00 | 81:00:00 | 0:00 |
| NM_002117:NM_001243042 | 183:19:00 | 1:00 | 56:09:00 | 0:00 | 79:00:00 | 0:00 |
| NM_002117:NM_001243042 | 0:19 | 183:00:00 | 0:07 | 55:02:00 | 79:00:00 | 0:00 |
| NM_002117:NM_001243042 | 0:19 | 182:00:00 | 0:07 | 55:02:00 | 0:00 | 79:00:00 |
| NM_002117:NM_001243042 | 180:19:00 | 2:00 | 55:09:00 | 0:00 | 74:00:00 | 6:00 |
| NM_002117:NM_001243042 | 180:20:00 | 1:00 | 54:09:00 | 0:00 | 80:00:00 | 0:00 |
| NM_002117:NM_001243042 | 179:20:00 | 2:00 | 54:09:00 | 0:00 | 80:00:00 | 0:00 |
| NM_002117:NM_001243042 | 180:20:00 | 1:00 | 55:09:00 | 0:00 | 80:00:00 | 0:00 |
| NM_002117:NM_001243042 | 180:20:00 | 1:00 | 55:09:00 | 0:00 | 80:00:00 | 0:00 |
| NM_002117:NM_001243042 | 1:20 | 180:00:00 | 0:07 | 55:02:00 | 0:00 | 80:00:00 |
| NM_002117:NM_001243042 | 179:20:00 | 1:00 | 55:09:00 | 0:00 | 73:00:00 | 6:00 |
| NM_002117:NM_001243042 | 0:20 | 181:00:00 | 0:07 | 55:03:00 | 0:00 | 79:00:00 |
| NM_002117:NM_001243042 | 0:20 | 181:00:00 | 0:07 | 55:03:00 | 0:00 | 80:00:00 |
| NM_002117:NM_001243042 | 1:20 | 180:00:00 | 0:07 | 55:03:00 | 0:00 | 80:00:00 |
| NM_025256:XM_017010202:XM_006714976:NM_006709 | 0:0:0:0 | 0:0:0:0 | 1:1:0:0 | 0:0:0:0 | 0:0:0:0 | 1:2:1:0 |

| | | | | | | |
|---|---|---|---|---|---|---|
| NM_001291986:NM_001199456:NM_005104:NM_001113182:XR_001743557:XR_241911 | 9:0:9:0:19:0 | 0:0:0:0:0:0 | 2:2:9:1:16:2 | 0:0:0:0:0:0 | 9:0:9:1:28:2 | 0:0:0:0:1:0 |
| NM_001760:NM_001136017:XM_011514972:NM_001287427:XM_011514971:NM_001136126:NM_001136125 | 17:1:2:0:0:0:1 | 20:1:1:0:0:4:0 | 1:0:0:0:0:0:0 | 3:1:0:0:0:0:0 | 0:0:0:0:0:0:0 | 24:0:0:0:0:0:0 |
| NM_001100409:XM_017010719:XM_024446400:XR_241847:NM_015571:XM_017010724 | 1:0:3:0:1:0 | 0:0:0:0:0:0 | 1:1:2:0:0:0 | 0:0:0:0:0:0 | 0:0:2:1:0:0 | 0:0:4:1:0:0 |
| NM_005190:XR_001743714:XM_017011436 | 266:04:00 | 1:00:00 | 115:02:00 | 0:00:00 | 250:02:00 | 1:00:00 |
| NM_001913:NM_181500:XM_006715855:XM_005250154:NM_001202546:XM_017011760:NM_001202545:NM_001202544:XM_024446668:XM_011515825 | 24:7:1:5:0:0:0:0:0:0 | 34:5:0:3:0:0:1:0:0:0 | 51:16:4:3:2:0:0:0:0:0 | 53:13:8:2:0:0:0:0:0:0 | 37:16:5:1:0:0:0:0:0:0 | 42:10:6:3:0:0:0:0:0:0 |
| NM_001202546:NM_001913:NM_181500:XM_006715855:XM_005250154:NM_001202545:XM_011515825:NM_001202544:XM_017011760:XM_024446668 | 0:57:14:2:7:1:0:0:0:0 | 0:0:0:0:0:0:0:0:0:0 | 2:101:29:14:5:0:0:0:0:0 | 0:0:0:0:0:0:0:0:0:0 | 0:41:13:6:3:0:0:0:0:0 | 0:38:16:5:1:0:0:0:0:0 |
| XM_006715855:NM_181500:NM_001202546:NM_001913:XM_005250154:XM_011515825:NM_001202545:XM_017011760:NM_001202544:XM_024446668 | 7:13:0:55:6:0:1:0:0:0 | 0:1:0:0:0:0:0:0:0:0 | 14:33:2:96:5:0:0:0:0:0 | 3:1:0:0:0:0:0:0:0:0 | 17:33:0:75:4:0:0:1:0:0 | 0:0:0:0:0:0:0:0:0:0 |
| NM_001202546:NM_001913:NM_181500:XM_006715855:XM_005250154:XM_011515825:NM_001202545:XM_017011760:NM_001202544:XM_024446668 | 0:55:13:7:6:0:1:0:0:0 | 0:0:1:0:0:0:0:0:0:0 | 2:96:34:17:5:0:0:0:0:0 | 0:0:0:0:0:0:0:0:0:0 | 0:74:33:17:4:0:0:1:0:0 | 0:0:0:0:0:0:0:0:0:0 |
| XR_001744836:XM_017012435:NM_182931:XM_011516400:NM_018682:XM_024446837 | 196:3:65:7:1:3 | 0:0:2:0:0:0 | 95:3:23:4:1:0 | 1:0:0:0:0:0 | 76:3:15:0:0:1 | 0:0:0:0:0:0 |
| XR_001744836:XM_017012435:NM_182931:XM_011516400:NM_018682:XM_024446837 | 123:1:32:4:0:2 | 0:0:0:0:0:0 | 58:2:18:2:0:0 | 1:0:0:0:0:0 | 52:3:13:0:0:1 | 1:0:0:0:0:0 |
| XR_001744836:NM_182931:XM_011516400:XM_017012435:NM_018682:XM_024446837 | 119:30:4:1:0:2 | 4:2:0:0:0:0 | 53:17:1:2:0:0 | 5:1:1:0:0:0 | 48:11:0:3:0:1 | 4:2:0:0:0:0 |
| NM_182931:XR_001744836:XM_011516400:XM_017012435:NM_018682:XM_024446837 | 21:93:2:1:0:1 | 12:28:2:0:0:1 | 15:41:1:1:0:0 | 3:18:1:1:0:0 | 9:39:0:2:0:1 | 4:13:0:1:0:0 |
| XM_017012435:XR_001744836:NM_182931:XM_024446837:NM_018682:XM_011516400 | 1:52:16:0:0:1 | 0:0:0:0:0:0 | 1:26:10:0:0:0 | 0:0:0:0:0:0 | 1:22:4:1:0:0 | 0:0:0:0:0:0 |
| NM_014390:XM_017011987 | 142:07:00 | 2:00 | 331:12:00 | 2:00 | 251:13:00 | 3:00 |
| NM_014390:XM_017011987 | 248:02:00 | 0:00 | 450:00:00 | 1:00 | 386:02:00 | 1:00 |
| NM_014390:XM_017011987 | 249:02:00 | 1:00 | 449:02:00 | 3:00 | 398:03:00 | 1:00 |
| XM_017011987:NM_014390 | 0.259027778 | 0:00 | 0.397222222 | 1:02 | 0.402083333 | 0:01 |
| NM_014390:XM_017011987 | 253:02:00 | 0:00 | 454:03:00 | 2:00 | 401:03:00 | 0:00 |
| NM_014390:XM_017011987 | 255:01:00 | 0:00 | 454:03:00 | 1:00 | 402:03:00 | 0:00 |
| NM_014390:XM_017011987 | 255:02:00 | 0:00 | 452:02:00 | 0:00 | 406:03:00 | 0:00 |
| NM_014390:XM_017011987 | 258:02:00 | 0:00 | 453:02:00 | 1:00 | 405:03:00 | 0:00 |
| NM_021807:XR_001744845:XM_017012494 | 11:01:01 | 0:00:00 | 18:01:00 | 0:00:00 | 27:02:01 | 0:00:00 |
| XR_927542:NM_033140:XM_017012654:XR_927537:XR_001744879:XR_002956490 | 42:9:0:4:1:0 | 10:1:0:0:0:0 | 13:1:1:0:0:0 | 2:0:0:0:0:0 | 12:2:0:2:0:0 | 2:0:0:0:0:0 |
| NM_030647:XM_011516587 | 2:01 | 2:00 | 1:02 | 1:00 | 3:01 | 0:00 |
| NM_013446:XM_011515998:NM_001145125:NM_001291663:XM_011515997 | 0:0:0:0:0 | 59:20:37:4:3 | 0:1:0:0:0 | 24:6:8:3:0 | 41:8:14:2:0 | 0:0:0:0:0 |
| XM_017012558:XM_017012559:NM_004333:NM_001354609:NR_148928:XR_001744857:XR_001744858 | 0:2:8:4:0:1:0 | 0:0:0:0:0:0:0 | 2:2:8:0:0:0:0 | 0:0:0:0:0:0:0 | 0:0:19:1:0:0:0 | 0:0:0:0:0:0:0 |
| NM_001203247:XM_005249962:XM_024446680:XM_011515884:XR_002956414:NM_004456:XM_017011818:XM_017011821:XM_011515894:XR_002956413:XM_017011819:XM_011515897:NM_152998:XM_011515887:XM_017011820:XM_011515889:XM_011515898:NM_001203248 | 11:0:1:0:0:0:0:1:0:1:0:0:0:0:0:0:0:0 | 0:0:0:0:0:0:0:0:0:0:0:0:0:0:0:0:0:0 | 3:1:1:1:1:1:0:0:0:0:0:0:0:0:0:0:0:0 | 0:0:0:0:0:0:0:0:0:0:0:0:0:0:0:0:0:0 | 9:0:0:0:0:2:0:2:0:0:0:0:0:0:1:0:0:0 | 4:0:0:0:0:4:0:1:0:2:0:0:0:1:0:0:0:0 |
| NM_001101312:NM_014020:NM_001101311:NM_001101314:XM_024446733:XM_024446734:XM_006715933 | 3026:93:136:30:19:141:3 | 3:0:0:0:0:0:0 | 1097:23:88:16:10:52:3 | 1:0:0:0:0:0:0 | 919:23:92:28:15:40:7 | 0:0:1:0:0:0:0 |
| NM_001101312:NM_014020:NM_001101311:NM_001101314:XM_024446733:XM_024446734:XM_006715933 | 3032:88:136:30:19:135:3 | 2:0:0:0:0:0:0 | 1100:22:89:16:10:50:3 | 1:0:0:0:0:0:0 | 922:23:93:28:15:40:7 | 0:0:0:0:0:0:0 |
| NM_014020:NM_001101312:NM_001101311:NM_001101314:XM_024446733:XM_024446734:XM_006715933 | 84:3038:136:30:19:126:3 | 0:7:0:0:0:0:0 | 20:1105:89:16:10:50:3 | 0:0:0:0:0:0:0 | 21:922:94:28:15:39:7 | 0:1:0:0:0:0:0 |
| NM_001101312:XM_024446734:NM_014020:NM_001101311:NM_001101314:XM_024446733:XM_006715933 | 3044:126:85:136:30:18:3 | 1:0:0:0:0:1:0 | 1104:48:20:89:16:10:3 | 1:1:0:0:0:0:0 | 922:38:21:94:28:15:7 | 1:0:0:0:0:0:0 |
| XM_024446734:NM_014020:NM_001101312:NM_001101311:NM_001101314:XM_024446733:XM_006715933 | 122:85:3041:136:31:19:3 | 1:0:5:0:0:0:0 | 44:20:1105:89:16:10:3 | 3:0:0:0:0:0:0 | 38:21:924:94:28:15:7 | 0:0:1:0:0:0:0 |
| NM_014020:NM_001101312:NM_001101311:NM_001101314:XM_024446733:XM_024446734:XM_006715933 | 85:3044:137:31:19:120:3 | 0:2:0:0:0:0:0 | 20:1105:89:16:10:46:3 | 0:0:0:0:0:0:0 | 21:924:94:28:15:34:7 | 0:1:0:0:0:0:0 |

| | | | | | | |
|---|---|---|---|---|---|---|
| NM_014020:NM_001101312:NM_001101311:NM_001101314:XM_024446733:XM_024446734:XM_006715933 | 85:3044:137:31:19:116:3 | 0:3:0:0:0:1:0 | 20:1106:89:16:10:46:3 | 0:0:0:0:0:0:0 | 21:925:94:28:15:33:7 | 0:0:0:0:0:0:0 |
| NM_001101312:NM_001101311:NM_014020:XM_024446733:NM_001101314:XM_024446734:XM_006715933 | 1660:67:39:11:17:69:1 | 1388:70:45:8:14:43:2 | 568:52:9:6:9:26:1 | 538:37:10:4:7:18:2 | 474:57:9:4:17:13:4 | 452:37:10:11:11:19:3 |
| NM_014020:NM_001101312:NM_001101311:NM_001101314:XM_024446733:XM_024446734:XM_006715933 | 84:3043:137:31:19:109:3 | 0:5:0:0:0:0:0 | 19:1106:89:16:10:44:3 | 0:0:0:0:0:0:0 | 19:926:94:28:15:32:7 | 0:0:0:0:0:0:0 |
| NM_001101312:NM_014020:NM_001101311:XM_024446733:NM_001101314:XM_024446734:XM_006715933 | 3051:77:137:19:19:86:3 | 4:0:0:0:2:0:0 | 1107:17:89:10:12:43:2 | 2:0:0:0:0:0:0 | 927:16:93:15:25:26:7 | 1:0:1:0:1:0:0 |
| NM_014020:NM_001101312:NM_001101311:XM_024446733:XM_024446734:NM_001101314:XM_006715933 | 1:1:0:0:1:0:0 | 73:3059:137:19:66:1:1 | 1:0:0:0:1:0:0 | 17:1113:89:10:37:0:0 | 0:0:0:0:0:0:0 | 14:930:95:15:21:2:0 |
| XM_006715933:NM_014020:NM_001101312:NM_001101311:NM_001101314:XM_024446733:XM_024446734 | 2:86:3078:138:37:19:82 | 0:0:1:0:1:0:1 | 3:29:1119:90:19:10:38 | 1:0:0:0:0:0:0 | 7:20:932:96:34:15:23 | 0:1:0:0:0:0:1 |
| NM_001101312:NM_014020:NM_001101311:NM_001101314:XM_024446733:XM_024446734:XM_006715933 | 3078:85:138:40:19:79:2 | 1:5:0:0:0:3:0 | 1115:33:90:21:10:23:4 | 4:0:0:0:0:0:0 | 931:24:96:34:15:21:7 | 1:0:0:0:0:0:0 |
| XR_001744669:XR_001744671:XM_017012110:NM_031243:XR_001744664:XR_001744673:XR_242076:XR_002956425:XM_005249729:NM_002137:XR_242077:XR_001744674:XR_428077:XR_001744672:XR_001744675:XR_001744667:XM_017012109:XR_001744668:XR_002956424:XR_001744670:XR_001744666 | 294:22:14:5:51:3:3:16:2:26:1:3:5:1:1:0:0:0:0:0:0 | 0:0:0:0:0:0:0:0:0:0:1:0:0:0:0:0:0:0:0:0:0 | 411:36:16:16:82:6:6:13:1:19:1:1:3:0:0:0:0:0:0:0 | 1:0:0:0:0:0:0:0:0:0:0:0:0:0:0:0:0:0:0:0:0 | 626:25:2:4:100:2:7:33:1:34:3:5:4:1:1:0:1:0:0:0:0 | 0:0:0:0:0:0:0:0:0:0:0:0:0:0:0:0:0:0:0:0:0 |
| XR_001744671:XM_017012110:NM_031243:XR_001744664:XR_001744673:XR_242076:XR_002956425:XM_005249729:NM_002137:XR_242077:XR_001744669:XR_001744674:XR_428077:XR_001744672:XR_001744675:XR_001744667:XM_017012109:XR_001744668:XR_002956424:XR_001744670:XR_001744666 | 22:14:5:51:3:2:15:2:27:1:294:4:5:1:1:0:0:0:0:0:0 | 0:0:0:0:0:0:0:0:0:0:0:0:0:0:0:0:0:0:0:0:0 | 36:16:16:82:6:6:9:1:20:1:412:2:4:0:0:0:0:0:0:0:0 | 0:0:0:0:0:0:0:0:0:0:0:0:0:0:0:0:0:0:0:0:0 | 25:2:4:99:2:7:28:1:34:3:624:7:6:1:1:0:1:0:0:0:0 | 0:0:0:1:0:0:0:0:0:2:1:0:0:0:0:0:0:0:0:0:0 |
| XR_001744671:XM_017012110:NM_031243:XR_001744664:XR_001744673:XR_242076:XR_002956425:XM_005249729:NM_002137:XR_242077:XR_001744669:XR_001744674:XR_428077:XR_001744672:XR_001744675:XR_001744667:XM_017012109:XR_001744668:XR_002956424:XR_001744670:XR_001744666 | 22:14:5:51:3:2:15:2:27:1:294:5:5:1:1:0:0:0:0:0:0 | 0:0:0:0:0:0:0:0:0:0:0:0:0:0:0:0:0:0:0:0:0 | 36:16:16:82:6:6:8:1:20:1:412:2:4:0:0:0:0:0:0:0:0 | 0:0:0:0:0:0:0:0:0:0:0:0:0:0:0:0:0:0:0:0:0 | 25:2:4:100:2:7:28:1:34:3:626:7:6:1:1:0:1:0:0:0:0 | 0:0:0:0:0:0:0:0:0:1:0:0:0:0:0:0:0:0:0:0:0 |
| XR_001744669:XR_001744671:XM_017012110:NM_031243:XR_001744673:XR_242076:XR_002956425:XM_005249729:NM_002137:XR_242077:XR_001744664:XR_001744674:XR_428077:XR_001744672:XR_001744667:XM_017012109:XR_001744668:XR_001744675:XR_001744670:XR_002956424:XR_001744666 | 291:22:14:5:3:2:8:2:27:1:51:8:5:1:0:0:0:0:0:0:0 | 1:0:0:0:0:0:0:0:0:0:0:0:0:0:0:0:0:0:0:0:0 | 410:36:16:15:6:6:6:1:20:1:81:3:8:0:0:0:0:0:0:0:0 | 1:0:0:0:0:0:0:0:0:0:0:0:0:0:0:0:0:0:0:0:0 | 623:25:2:4:2:7:16:1:34:3:100:12:5:1:0:1:0:1:0:0:0 | 0:0:0:0:0:0:0:0:0:0:0:0:0:0:0:0:0:0:0:0:0 |
| XR_001744671:XM_017012110:NM_031243:XR_001744673:XR_242076:XR_002956425:XM_005249729:NM_002137:XR_242077:XR_001744669:XR_001744664:XR_001744674:XR_428077:XR_001744672:XR_001744667:XM_017012109:XR_001744668:XR_001744675:XR_001744670:XR_002956424:XR_001744666 | 22:14:5:3:2:8:2:27:1:291:51:8:5:1:0:0:0:0:0:0:0 | 0:0:0:0:0:0:0:0:0:0:0:0:0:0:0:0:0:0:0:0:0 | 36:16:15:6:6:5:1:20:1:411:81:3:8:0:0:0:0:0:0:0:0 | 0:0:0:0:0:0:0:0:0:0:0:0:0:0:0:0:0:0:0:0:0 | 25:2:4:2:6:15:1:33:3:622:100:11:6:1:0:1:0:0:0:0:0 | 0:0:0:0:0:0:0:0:0:1:0:0:0:0:0:0:0:0:0:0:0 |
| XR_001744671:XM_017012110:NM_031243:XR_001744673:XR_242076:XR_002956425:XM_005249729:NM_002137:XR_242077:XR_001744669:XR_001744664:XR_001744674:XR_428077:XR_001744672:XR_001744667:XM_017012109:XR_001744668:XR_001744675:XR_001744670:XR_002956424:XR_001744666 | 22:14:5:3:2:7:2:27:1:291:51:8:5:1:0:0:0:0:0:0:0 | 0:0:0:0:0:0:0:0:0:0:0:0:0:0:0:0:0:0:0:0:0 | 36:16:15:6:6:5:1:20:1:411:81:2:8:0:0:0:0:0:0:0:0 | 0:0:0:0:0:0:0:0:0:0:0:0:0:0:0:0:0:0:0:0:0 | 25:2:4:2:6:15:1:33:3:621:100:11:6:1:0:1:0:0:0:0:0 | 0:0:0:0:0:0:0:0:0:1:0:0:0:0:0:0:0:0:0:0:0 |
| NM_031243:XR_428077:XR_001744671:XM_017012110:XR_001744673:XR_242076:XR_002956425:XM_005249729:XR_242077:XR_001744669:NM_002137:XR_001744664:XR_001744674:XR_001744672:XR_001744667:XM_017012109:XR_001744668:XR_001744670:XR_001744675:XR_001744666 | 5:5:22:14:3:2:7:2:1:289:27:51:8:1:0:0:0:0:0:0 | 0:1:0:0:0:0:0:0:0:1:0:0:2:0:0:0:0:0:0:0 | 14:9:36:16:6:6:5:1:1:410:19:81:2:0:0:0:0:0:0:0 | 1:1:0:0:0:0:0:0:0:0:0:0:0:0:0:0:0:0:0:0 | 4:6:25:2:2:6:11:1:3:616:33:99:11:1:0:1:0:0:0:0 | 0:0:0:0:0:0:0:0:3:1:0:4:0:0:0:0:0:0:0:0 |
| XR_002956425:XR_001744671:XM_017012110:NM_031243:XR_001744673:XR_242076:XM_005249729:NM_002137:XR_242077:XR_001744669:XR_001744664:XR_001744674:XR_428077:XR_001744672:XR_001744675:XR_001744667:XM_017012109:XR_001744668:XR_002956424:XR_001744670:XR_001744666 | 13:22:14:5:3:2:2:27:0:284:51:12:7:1:1:0:0:0:0:0:0 | 0:0:0:0:0:0:0:0:0:0:0:0:0:0:0:0:0:0:0:0:0 | 8:36:16:15:6:6:1:20:1:408:81:3:11:0:0:0:0:0:0:0:0 | 1:0:0:0:0:0:0:0:0:0:0:0:0:0:0:0:0:0:0:0:0 | 27:25:2:4:2:6:1:35:3:617:100:20:13:1:1:0:1:0:0:0:0 | 0:0:0:0:0:0:0:0:0:0:0:0:0:0:0:0:0:0:0:0:0 |
| XR_001744671:XM_017012110:NM_031243:XR_001744673:XR_242076:XR_002956425:XM_005249729:NM_002137:XR_242077:XR_001744669:XR_001744664:XR_001744674:XR_428077:XR_001744672:XR_001744675:XR_001744667:XM_017012109:XR_001744668:XR_002956424:XR_001744670:XR_001744666 | 22:13:5:3:2:13:2:28:0:278:51:12:8:1:1:0:0:0:0:0:0 | 0:1:0:0:0:0:0:0:0:0:0:0:0:0:0:0:0:0:0:0:0 | 36:16:15:6:6:11:1:20:1:406:81:3:9:0:0:0:0:0:0:0:0 | 0:0:0:0:0:0:0:0:0:0:0:0:0:0:0:0:0:0:0:0:0 | 25:2:3:2:4:35:1:35:2:605:99:20:12:1:1:0:1:0:0:0:0 | 0:0:0:0:0:0:0:0:1:0:0:0:0:0:0:0:0:0:0:0:0 |
| XR_001744669:XR_001744671:XM_017012110:NM_031243:XR_001744673:XR_242076:XR_002956425:XM_005249729:NM_002137:XR_242077:XR_001744664:XR_001744674:XR_428077:XR_001744672:XR_001744675:XR_001744667:XM_017012109:XR_001744668:XR_002956424:XR_001744670:XR_001744666 | 277:22:13:5:3:2:13:2:28:0:51:12:8:1:1:0:0:0:0:0:0 | 0:0:0:0:0:0:0:0:0:0:0:0:0:0:0:0:0:0:0:0:0 | 403:36:16:15:6:6:11:1:20:1:81:3:9:0:0:0:0:0:0:0:0 | 2:0:0:0:0:0:0:0:0:0:0:0:0:0:0:0:0:0:0:0:0 | 604:25:2:3:2:4:35:1:35:2:100:20:12:1:1:0:1:0:0:0:0 | 1:0:0:0:0:0:0:0:0:0:0:0:0:0:0:0:0:0:0:0:0 |

| | | | | | | |
|---|---|---|---|---|---|---|
| XR_001744671:XM_017012110:NM_031243:XR_001744673:XR_242076:XR_002956425:XM_005249729:NM_002137:XR_242077:XR_001744669:XR_001744664:XR_001744674:XR_428077:XR_001744672:XR_001744675:XR_001744667:XM_017012109:XR_001744668:XR_002956424:XR_001744670:XR_001744666 | 22:13:5:3:2:13:2:28:0:276:51:12:7:1:1:0:0:0:0:0:0 | 0:0:0:0:0:0:0:0:0:0:0:0:0:0:0:0:0:0:0:0:0 | 36:15:15:6:6:11:1:20:1:402:81:3:8:0:0:0:0:0:0:0:0 | 0:0:0:0:0:0:0:0:0:0:0:0:0:0:0:0:0:0:0:0:0 | 25:2:3:2:4:35:1:35:2:600:100:20:13:1:1:0:1:0:0:0:0 | 0:0:0:0:0:1:0:0:0:2:0:0:0:0:0:0:0:0:0:0:0 |
| XR_001744671:XR_001744669:XM_017012110:NM_031243:XR_001744673:XR_242076:XR_002956425:XM_005249729:NM_002137:XR_242077:XR_001744664:XR_001744674:XR_428077:XR_001744672:XR_001744675:XR_001744667:XM_017012109:XR_002956424:XR_001744670:XR_001744666 | 22:273:12:5:3:1:14:2:28:0:51:11:5:1:1:0:0:0:0:0 | 0:0:0:0:0:0:0:0:0:0:1:0:0:0:0:0:0:0:0:0 | 35:397:14:15:6:5:11:1:20:1:81:3:5:0:0:0:0:0:0:0 | 1:1:0:0:0:0:0:0:0:0:0:0:0:0:0:0:0:0:0:0 | 25:586:2:3:2:3:37:1:35:2:99:20:12:1:1:0:1:0:0:0 | 0:2:0:0:0:0:1:0:0:0:1:0:0:0:0:0:0:0:0:0 |
| NM_152740:XM_017011719 | 156:10:00 | 0:00 | 114:08:00 | 0:00 | 90:03:00 | 0:00 |
| NM_001079864:NM_006024:NM_001206902:NM_001206901 | 92:66:7:6 | 0:3:0:0 | 103:46:6:4 | 0:0:0:0 | 65:43:14:8 | 0:2:0:0 |
| NM_006024:NM_001079864:NM_001206902:NM_001206901 | 52:91:14:6 | 7:1:0:0 | 30:103:12:4 | 8:0:0:0 | 27:64:23:8 | 1:1:0:0 |
| NM_001079864:NM_006024:NM_001206902:NM_001206901 | 91:48:15:6 | 0:0:0:0 | 101:30:12:4 | 1:0:0:0 | 64:29:26:8 | 0:0:0:0 |
| NM_001079864:NM_006024:NM_001206902:NM_001206901 | 91:48:15:5 | 0:0:0:1 | 102:30:12:4 | 0:0:0:0 | 64:29:25:8 | 0:0:0:0 |
| NM_001079864:NM_006024:NM_001206902:NM_001206901 | 90:48:15:6 | 1:0:0:0 | 102:30:12:4 | 0:0:0:0 | 64:29:24:8 | 0:0:0:0 |
| NM_002047:NM_001316772 | 0:00 | 0:00 | 0:00 | 0:00 | 0:00 | 1:02 |
| XM_017012506:NM_022373:XR_926944:XR_001744846 | 2:1:0:0 | 3:2:0:0 | 3:1:0:0 | 2:3:0:0 | 2:2:0:0 | 0:0:0:0 |
| XM_017012102:NM_001637:XM_011515336:XM_017012106:XM_011515335:XM_011515340:XM_011515341:NM_001177507 | 0:18:0:0:1:0:0:0 | 0:0:0:0:0:0:0:0 | 2:51:1:0:0:0:0:0 | 0:0:0:0:0:0:0:0 | 0:10:0:0:0:0:0:0 | 0:0:0:0:0:0:0:0 |
| XM_017012102:NM_001637:XM_011515336:XM_011515339:XM_011515340:XM_017012106:XM_011515341:XM_011515335:XM_017012104:XM_011515338:NM_001177507 | 0:19:1:1:7:3:1:1:1:0:0 | 0:0:0:0:0:0:0:0:0:0:0 | 2:52:2:1:16:7:1:0:0:0:0 | 0:0:0:0:0:0:0:0:0:0:0 | 0:4:0:0:6:5:0:0:0:0:0 | 0:6:0:2:23:15:1:0:2:0:0 |
| NM_001637:XM_017012106:XM_011515335:XM_011515340:XM_011515341:NM_001177507 | 0:0:0:0:0:0 | 6:0:1:0:0:0 | 0:0:0:0:0:0 | 6:1:0:0:0:0 | 0:0:0:0:0:0 | 1:0:0:0:0:0 |
| XM_017012363:NM_207116:XM_024446805:NM_207111:XM_017012364:XM_024446807:XM_024446806 | 0:2:1:0:0:0:0 | 0:0:0:0:0:0:0 | 0:1:1:0:0:0:0 | 1:0:0:0:0:0:0 | 0:1:2:0:0:0:0 | 0:0:0:0:0:0:0 |
| NM_001322005:NM_000535:NM_001322006:NM_001322015:XM_024446800:NM_001322004:NM_001322003:NR_136154:NM_001322008:NM_001322007:XM_017012342:NM_001322014:NM_001322012:NM_001322011 | 1:6:0:1:0:0:1:0:0:0:0:0:0:0 | 0:0:0:0:0:1:0:0:0:0:0:0:0:0 | 1:6:0:0:0:0:0:0:0:0:0:0:0:0 | 0:0:0:0:0:0:0:0:0:0:0:0:0:0 | 0:7:0:0:0:0:2:0:0:0:0:0:0:0 | 0:0:0:0:0:0:0:0:0:0:0:0:0:0 |
| NM_001322005:NM_000535:NM_001322015:XM_024446800:NM_001322006:NM_001322003:NM_001322004:NR_136154:NM_001322014:XM_017012342:NM_001322008:NM_001322007:NM_001322012:NM_001322011 | 1:6:1:1:0:1:1:0:0:0:0:0:0:0 | 0:0:0:0:0:0:0:0:0:0:0:0:0:0 | 1:6:0:0:0:0:0:0:0:0:0:0:0:0 | 0:0:0:0:0:0:0:0:0:0:0:0:0:0 | 0:4:0:0:0:2:0:0:0:0:0:0:0:0 | 0:3:0:0:0:0:0:0:0:0:0:0:0:0 |
| NM_001322005:NM_000535:NM_001322015:XM_024446800:NM_001322006:NR_136154:NM_001322003:NM_001322004:XM_017012342:NM_001322014:NM_001322007:NM_001322012:NM_001322008:NM_001322011 | 0:0:0:0:0:0:0:0:0:0:0:0:0:0 | 1:6:1:1:0:0:1:1:0:0:0:0:0:0 | 0:0:0:0:0:0:0:0:0:0:0:0:0:0 | 1:6:0:0:0:0:0:0:0:0:0:0:0:0 | 0:0:0:0:0:0:0:0:0:0:0:0:0:0 | 0:7:0:0:0:1:3:0:1:0:0:0:0:0 |
| NM_001322005:NM_000535:NM_001322015:XM_024446800:NM_001322006:NR_136154:NM_001322003:NM_001322004:XM_017012342:NM_001322014:NM_001322007:NM_001322008:NM_001322012:NM_001322011 | 0:0:0:0:0:0:0:0:0:0:0:0:0:0 | 1:6:1:1:0:0:1:1:0:0:0:0:0:0 | 0:0:0:0:0:0:0:0:0:0:0:0:0:0 | 1:6:0:0:0:0:0:0:0:0:0:0:0:0 | 0:3:0:0:0:0:1:0:0:0:0:0:0:0 | 0:4:0:0:0:1:2:0:1:0:0:0:0:0 |
| NM_005918:NR_104165:NM_001282404 | 2:00:00 | 2:00:00 | 1:00:00 | 0:00:00 | 10:00:00 | 0:00:00 |
| NM_001284309:NM_001284308:NM_006869:NM_001284311 | 0:0:0:0 | 56:0:17:0 | 0:0:0:0 | 16:1:0:0 | 0:0:0:0 | 6:0:4:0 |
| NM_002568:XM_005250861 | 968:34:00 | 0:01 | 4164:129 | 4:02 | 1982:36:00 | 2:00 |
| NM_002568:XM_005250861 | 960:38:00 | 1:00 | 4143:141 | 2:00 | 1963:43:00 | 1:00 |
| NM_002568:XM_005250861 | 959:37:00 | 2:00 | 4135:141 | 8:00 | 1962:43:00 | 2:00 |
| NM_002568:XM_005250861 | 961:36:00 | 0:00 | 4137:144 | 3:00 | 1962:43:00 | 2:00 |
| NM_002568:XM_005250861 | 957:36:00 | 3:00 | 4132:143 | 5:00 | 1957:43:00 | 4:00 |
| NM_002568:XM_005250861 | 958:36:00 | 1:00 | 4128:142 | 5:00 | 1951:42:00 | 4:00 |
| NM_002568:XM_005250861 | 959:36:00 | 0:00 | 4130:142 | 3:00 | 1954:42:00 | 1:00 |
| NM_002568:XM_005250861 | 951:37:00 | 0:00 | 4108:138 | 6:00 | 1937:40:00 | 0:00 |
| NM_002568:XM_005250861 | 943:33:00 | 0:01 | 4075:120 | 4:00 | 1922:37:00 | 2:00 |
| NM_002568:XM_005250861 | 941:33:00 | 2:01 | 4070:116 | 5:03 | 1920:38:00 | 2:00 |
| NM_002568:XM_005250861 | 936:34:00 | 0:01 | 4038:107 | 9:00 | 1912:36:00 | 1:00 |
| NM_002568:XM_005250861 | 909:32:00 | 3:00 | 3967:116 | 10:04 | 1869:32:00 | 2:00 |
| NM_002568:XM_005250861 | 899:31:00 | 1:00 | 3949:119 | 9:00 | 1846:31:00 | 2:00 |
| NM_002568:XM_005250861 | 890:32:00 | 1:00 | 3923:121 | 10:00 | 1823:30:00 | 6:00 |
| NM_002568:XM_005250861 | 869:32:00 | 0:00 | 3857:135 | 4:00 | 1793:29:00 | 3:00 |
| NM_002568:XM_005250861 | 836:27:00 | 2:01 | 3775:129 | 11:00 | 1751:27:00 | 4:00 |
| NM_002568:XM_005250861 | 836:27:00 | 0:00 | 3773:131 | 9:01 | 1751:28:00 | 3:00 |
| NM_002568:XM_005250861 | 822:24:00 | 2:00 | 3712:122 | 17:01 | 1731:32:00 | 2:00 |
| NM_002568:XM_005250861 | 822:24:00 | 1:00 | 3708:122 | 16:01 | 1731:32:00 | 0:00 |
| NM_002568:XM_005250861 | 821:25:00 | 1:00 | 3701:123 | 15:00 | 1728:32:00 | 1:00 |

| | | | | | | |
|---|---|---|---|---|---|---|
| NM_002568:XM_005250861 | 813:25:00 | 1:00 | 3669:124 | 14:01 | 1709:35:00 | 1:00 |
| NM_002568:XM_005250861 | 806:24:00 | 1:00 | 3643:129 | 16:00 | 1693:36:00 | 4:00 |
| NM_002568:XM_005250861 | 805:24:00 | 2:00 | 3640:127 | 19:02 | 1693:37:00 | 4:00 |
| NM_002568:XM_005250861 | 802:25:00 | 4:00 | 3647:129 | 10:01 | 1694:39:00 | 1:00 |
| NM_002568:XM_005250861 | 799:26:00 | 0:00 | 3623:130 | 10:00 | 1684:42:00 | 2:00 |
| NM_002568:XM_005250861 | 799:26:00 | 0:00 | 3615:128 | 18:02 | 1684:42:00 | 2:00 |
| NM_002568:XM_005250861 | 787:31:00 | 1:00 | 3569:138 | 5:02 | 1670:45:00 | 1:00 |
| NM_002568:XM_005250861 | 785:33:00 | 1:01 | 3554:140 | 6:01 | 1668:46:00 | 0:00 |
| NM_002568:XM_005250861 | 784:34:00 | 2:00 | 3551:139 | 8:02 | 1666:46:00 | 2:00 |
| NM_002568:XM_005250861 | 729:35:00 | 2:00 | 3365:138 | 3:00 | 1593:48:00 | 0:00 |
| NM_002568:XM_005250861 | 727:35:00 | 2:00 | 3356:137 | 7:00 | 1589:47:00 | 3:00 |
| NM_002568:XM_005250861 | 721:36:00 | 2:00 | 3340:142 | 5:00 | 1584:50:00 | 3:00 |
| NM_002568:XM_005250861 | 720:36:00 | 0:01 | 3331:136 | 3:01 | 1581:50:00 | 1:00 |
| NM_002568:XM_005250861 | 719:35:00 | 0:01 | 3327:138 | 1:00 | 1579:50:00 | 2:00 |
| NM_002568:XM_005250861 | 684:35:00 | 0:00 | 3221:131 | 1:00 | 1526:48:00 | 0:00 |
| NM_002568:XM_005250861 | 707:35:00 | 2:00 | 3297:134 | 4:00 | 1556:49:00 | 2:00 |
| NM_002568:XM_005250861 | 708:35:00 | 1:00 | 3297:134 | 3:00 | 1556:48:00 | 2:00 |
| NM_002568:XM_005250861 | 698:35:00 | 1:00 | 3258:131 | 7:00 | 1546:51:00 | 2:00 |
| NM_002568:XM_005250861 | 698:34:00 | 0:00 | 3252:131 | 3:00 | 1542:51:00 | 3:00 |
| NM_002568:XM_005250861 | 697:33:00 | 0:00 | 3253:132 | 2:00 | 1544:51:00 | 0:00 |
| NM_002568:XM_005250861 | 692:32:00 | 0:00 | 3215:130 | 2:00 | 1534:50:00 | 2:00 |
| NM_002568:XM_005250861 | 685:32:00 | 1:00 | 3199:133 | 4:00 | 1528:48:00 | 1:01 |
| NM_002568:XM_005250861 | 686:32:00 | 0:00 | 3201:133 | 1:00 | 1528:48:00 | 0:00 |
| NM_002568:XM_005250861 | 684:32:00 | 1:00 | 3197:133 | 1:00 | 1528:48:00 | 0:00 |
| NM_002568:XM_005250861 | 678:32:00 | 1:00 | 3161:129 | 8:01 | 1513:45:00 | 1:01 |
| NM_002568:XM_005250861 | 617:26:00 | 2:00 | 2982:123 | 18:00 | 1449:40:00 | 3:00 |
| NM_015902:XR_001745541:XM_024447179 | 0:00:00 | 0:00:00 | 1:00:00 | 1:00:00 | 0:00:00 | 0:00:00 |
| NM_030780:NR_102338:NR_102337 | 1:00:00 | 3:00:00 | 5:01:00 | 13:00:00 | 12:03:02 | 0:00:00 |
| NM_001193557:NM_020189:NR_036472:NR_036471 | 50:122:2:16 | 0:0:0:0 | 54:346:3:19 | 0:0:0:0 | 82:260:5:28 | 0:0:0:0 |
| NM_001193557:NM_020189:NR_036472:NR_036471 | 51:122:3:15 | 0:0:0:0 | 54:343:5:19 | 0:0:0:0 | 82:258:6:28 | 0:2:0:0 |
| NM_001287743:NM_004462:NM_001287756:NM_001287745:NM_001287742:NM_001287751 | 0:1:0:0:0:0 | 4:2393:10:9:12<br>0:0 | 0:0:0:0:0:0 | 5:461:4:3:22:0 | 5:664:2:2:26:0 | 0:0:0:0:0:0 |
| NM_004462:NM_001287743:NM_001287756:NM_001287742:NM_001287745:NM_001287744:NM_0<br>01287750 | 2406:4:10:120:<br>1:1:0 | 5:0:0:0:0:0:0 | 461:5:4:22:1:0<br>:0 | 1:0:0:0:0:0:0 | 665:5:1:26:0:0<br>:1 | 2:0:0:0:0:0:0 |
| NM_001287742:NM_004462:NM_001287743:NM_001287756:NM_001287745:NM_001287744:NM_0<br>01287750:NM_001287751 | 120:2409:4:10:<br>1:1:0:0 | 0:5:0:0:0:0:0:0 | 21:461:5:4:1:0<br>:0:0 | 1:1:0:0:0:0:0:0 | 26:672:5:1:0:0<br>:1:0 | 0:1:0:0:0:0:0:0 |
| NM_004462:NM_001287743:NM_001287756:NM_001287742:NM_001287745:NM_001287744:NM_0<br>01287750:NM_001287751 | 2413:4:10:120:<br>1:1:0:0 | 3:0:0:0:0:0:0:0 | 461:5:4:22:1:0<br>:0:0 | 1:0:0:0:0:0:0:0 | 672:5:1:26:0:0<br>:1:0 | 1:0:0:0:0:0:0:0 |
| NM_004462:NM_001287743:NM_001287756:NM_001287742:NM_001287745:NM_001287744:NM_0<br>01287750:NM_001287751 | 2395:4:10:118:<br>1:1:0:0 | 13:0:0:0:0:0:0:<br>0 | 460:5:4:22:1:0<br>:0:0 | 1:0:0:0:0:0:0:0 | 677:5:1:26:0:0<br>:1:0 | 0:0:0:0:0:0:0:0 |
| NM_001287743:NM_004462:NM_001287756:NM_001287742:NM_001287744:NM_001287745:NM_0<br>01287750:NM_001287751 | 4:2209:10:111:<br>1:1:0:0 | 0:2:0:0:0:0:0:0 | 3:427:4:22:0:0<br>:0:0 | 0:0:0:0:0:0:0:0 | 5:658:2:26:0:0<br>:1:0 | 0:0:0:0:0:0:0:0 |
| NM_001287743:NM_004462:NM_001287756:NM_001287742:NM_001287745:NM_001287744:NM_0<br>01287750:NM_001287751 | 4:2262:10:111:<br>1:1:0:0 | 0:4:0:0:0:0:0:0 | 3:441:4:22:1:0<br>:0:0 | 0:0:0:0:0:0:0:0 | 5:676:2:26:0:0<br>:1:0 | 0:1:0:0:0:0:0:0 |
| NM_004462:NM_001287743:NM_001287756:NM_001287742:NM_001287745:NM_001287744:NM_0<br>01287750:NM_001287751 | 2269:4:10:111:<br>1:1:0:0 | 7:0:0:0:0:0:0:0 | 436:3:4:22:1:0<br>:0:0 | 1:0:0:0:0:0:0:0 | 671:5:2:26:0:0<br>:1:0 | 1:0:0:0:0:0:0:0 |

| | | | | | | |
|---|---|---|---|---|---|---|
| NM_001908:XM_024447071:NM_147782:NM_147780:NM_147783:XM_017013100:XM_017013097:XM_017013099:NM_001317237 | 11:1:1:0:0:1:1:0:0 | 10:1:2:0:1:0:0:0:0 | 20:2:2:1:0:0:0:0:0 | 8:1:1:0:0:0:0:0:0 | 27:6:3:0:0:0:0:0:0 | 0:0:0:0:0:0:0:0:0 |
| NM_001908:NM_147782:XM_024447071:NM_147780:XM_017013100:NM_147783:XM_017013097:XM_017013099:NM_001317237 | 0:0:0:0:0:0:0:0:0 | 21:3:2:0:1:1:1:0:0 | 0:0:0:0:0:0:0:0:0 | 28:3:5:1:0:0:0:0:0 | 27:3:6:0:0:0:0:0:0 | 0:0:0:0:0:0:0:0:0 |
| NM_001040092:XM_017013574:XM_017013575:NR_045555:NM_001130863:XM_017013572:XM_024447182:XM_006716587:NM_006209:XM_017013573:XM_024447181 | 0:0:3:0:0:0:0:0:0:0:0 | 3:0:1:0:0:0:0:0:1:0:0 | 14:2:10:1:2:2:0:0:0:0:0 | 3:1:1:0:0:0:0:0:0:0:0 | 0:0:0:0:0:0:0:0:0:0:0 | 3:2:1:0:1:0:2:1:0:0:0 |
| NM_005005:NM_001311168:NM_001278646:NM_001278645 | 6666:58:42:4 | 33:0:0:0 | 4752:12:45:1 | 16:0:0:0 | 3507:14:24:1 | 4:0:0:0 |
| NM_005005:NM_001311168:NM_001278646:NM_001278645 | 6682:58:42:4 | 30:0:0:0 | 4772:12:45:1 | 11:0:0:0 | 3522:13:24:1 | 3:1:0:0 |
| NM_005005:NM_001311168:NM_001278646:NM_001278645 | 6705:58:42:4 | 18:0:0:0 | 4778:12:45:1 | 13:0:0:0 | 3524:14:24:1 | 5:0:0:0 |
| NM_005005:NM_001311168:NM_001278646:NM_001278645 | 6717:58:42:4 | 13:0:0:0 | 4795:12:45:1 | 2:0:0:0 | 3533:14:24:1 | 1:0:0:0 |
| NM_005005:NM_001311168:NM_001278646:NM_001278645 | 6744:58:24:4 | 2:0:0:0 | 4803:13:16:1 | 1:0:0:0 | 3540:14:9:1 | 3:0:0:0 |
| NM_005005:NM_001278646:NM_001278645 | 6767:42:00 | 35:00:00 | 4824:45:00 | 7:00:00 | 3551:24:00 | 4:00:00 |
| NM_005005:NM_001278646:NM_001278645 | 6793:41:00 | 21:01:00 | 4832:45:00 | 3:00:00 | 3557:24:00 | 4:00:00 |
| NM_005005:NM_001311168:NM_001278646:NM_001278645 | 6853:58:42:4 | 9:0:0:0 | 4858:13:45:1 | 5:0:0:0 | 3573:14:24:1 | 5:0:0:0 |
| NM_005005:NM_001311168:NM_001278646:NM_001278645 | 6860:58:42:4 | 11:0:0:0 | 4859:13:45:1 | 4:0:0:0 | 3574:14:24:1 | 4:0:0:0 |
| NM_005005:NM_001311168:NM_001278646:NM_001278645 | 6865:58:42:4 | 8:0:0:0 | 4863:13:45:1 | 1:0:0:0 | 3576:14:24:1 | 3:0:0:0 |
| NM_005005:NM_001311168:NM_001278646:NM_001278645 | 6880:58:42:4 | 9:0:0:0 | 4876:13:46:1 | 3:0:0:0 | 3587:14:24:1 | 4:0:0:0 |
| NM_005005:NM_001311168:NM_001278646:NM_001278645 | 6895:58:41:4 | 5:0:1:0 | 4880:13:46:1 | 3:0:0:0 | 3604:14:24:1 | 1:0:0:0 |
| NM_005005:NM_001311168:NM_001278646:NM_001278645 | 6890:58:42:4 | 10:0:0:0 | 4879:13:46:1 | 1:0:0:0 | 3603:14:24:1 | 2:0:0:0 |
| NM_005005:NM_001311168:NM_001278646:NM_001278645 | 6878:58:42:4 | 16:0:0:0 | 4862:13:46:1 | 5:0:0:0 | 3586:14:24:1 | 3:0:0:0 |
| NM_005005:NM_001311168:NM_001278646:NM_001278645 | 6867:58:42:4 | 13:0:0:0 | 4854:13:46:1 | 4:0:0:0 | 3585:14:24:1 | 4:0:0:0 |
| NM_005005:NM_001311168:NM_001278646:NM_001278645 | 6852:58:42:4 | 18:0:0:0 | 4842:13:46:1 | 9:0:0:0 | 3579:14:24:1 | 8:0:0:0 |
| NM_005005:NM_001278646:NM_001311168:NM_001278645 | 6852:42:58:4 | 7:0:0:0 | 4845:45:13:1 | 4:1:0:0 | 3582:24:14:1 | 3:0:0:0 |
| NM_005005:NM_001311168:NM_001278646:NM_001278645 | 6841:57:41:4 | 11:0:1:0 | 4832:13:45:1 | 7:0:0:0 | 3583:14:24:1 | 7:0:0:0 |
| NM_005005:NM_001311168:NM_001278646:NM_001278645 | 6840:57:42:4 | 4:0:0:0 | 4832:13:45:1 | 1:0:0:0 | 3589:14:24:1 | 0:0:0:0 |
| NM_005005:NM_001311168:NM_001278646:NM_001278645 | 6826:57:42:4 | 17:0:0:0 | 4827:13:45:1 | 6:0:0:0 | 3584:14:24:1 | 5:0:0:0 |
| NM_005005:NM_001311168:NM_001278646:NM_001278645 | 6820:56:42:4 | 12:0:0:0 | 4819:13:45:1 | 9:0:0:0 | 3585:14:24:1 | 2:0:0:0 |
| NM_005005:NM_001311168:NM_001278646:NM_001278645 | 6801:56:42:4 | 19:0:0:0 | 4815:13:45:1 | 13:0:0:0 | 3582:14:23:1 | 3:0:1:0 |
| NM_005005:NM_001311168:NM_001278646:NM_001278645 | 6796:56:42:4 | 19:0:0:0 | 4811:13:45:1 | 8:0:0:0 | 3582:14:24:1 | 2:0:0:0 |
| NM_005005:NM_001311168:NM_001278646:NM_001278645 | 6774:56:42:4 | 31:0:0:0 | 4806:13:45:1 | 8:0:0:0 | 3580:14:24:1 | 4:0:0:0 |
| NM_005005:NM_001311168:NM_001278646:NM_001278645 | 6784:56:42:4 | 20:0:0:0 | 4801:13:45:1 | 7:0:0:0 | 3583:14:24:1 | 0:0:0:0 |
| NM_005005:NM_001311168:NM_001278646:NM_001278645 | 6740:56:42:4 | 40:0:0:0 | 4780:13:45:1 | 14:0:0:0 | 3574:14:24:1 | 5:0:0:0 |
| NM_012472:NR_073525:XM_017013296:XM_017013297:NR_135907:NR_135909:NM_001321962:NR_135908:XM_011516950:NR_135913:NR_135906:NM_001321961:XM_017013298:NM_001321963 | 3:8:0:0:0:0:0:0:0:0:0:0:0:0 | 3:8:0:0:0:0:0:0:1:0:0:0:0:0 | 0:2:0:0:0:0:0:0:0:0:0:0:0:0 | 1:1:1:0:0:0:0:0:0:0:0:0:0:0 | 3:2:0:0:2:0:0:0:0:0:0:0:0:0 | 0:2:1:0:0:0:0:0:0:0:0:0:0:0 |
| NM_006096:XM_011516791:XM_011516792 | 89:03:00 | 0:00:00 | 51:01:03 | 0:00:00 | 5:00:00 | 7:00:00 |
| NM_001916:XM_024447072:XM_017013102 | 0:00:00 | 4:01:01 | 0:00:00 | 5:01:00 | 0:00:00 | 6:00:00 |
| XM_024447170:XM_017013496:XM_024447168:XM_024447167:XM_017013493:XM_006716340:XM_024447171:NM_001352641 | 0:3:0:0:1:1:0:0 | 0:0:0:0:0:0:0:0 | 1:0:0:0:0:0:0:0 | 0:0:0:0:0:0:0:0 | 0:0:0:0:0:0:0:0 | 0:0:0:0:0:2:0:0 |
| XR_002956638:NM_002717:XM_011544567:XM_005273559:XM_024447187:XM_011544568:XM_005273560:NM_001177591:XM_011544566:XM_024447186:XM_011544569:XM_011544565:XR_949430 | 0:12:1:0:0:0:0:1:0:0:0:0:0 | 0:0:0:0:0:0:0:0:0:0:0:0:0 | 0:18:2:1:1:1:0:0:0:0:0:0:0 | 2:0:0:0:0:0:0:0:0:0:0:0:0 | 0:18:0:2:0:0:0:0:1:0:0:0:0 | 2:1:0:0:0:0:0:0:0:0:0:0:0 |
| NM_001979:XR_001745491:XM_017013199:XM_017013200 | 52:2:1:2 | 54:3:3:0 | 0:0:0:0 | 23:1:0:0 | 8:0:1:0 | 0:0:0:0 |
| NM_001979:XR_001745491:XM_017013199 | 56:02:02 | 52:00:02 | 22:01:00 | 0:00:00 | 8:00:01 | 0:00:00 |

| | | | | | | |
|---|---|---|---|---|---|---|
| XM_024447265:XR_949472:XM_011544639 | 0:00:01 | 0:01:02 | 1:00:00 | 0:00:00 | 1:00:00 | 0:01:00 |
| XM_024447265:XM_011544639 | 0:02 | 0:00 | 1:00 | 0:00 | 0:00 | 2:00 |
| NM_013958:NM_004495:NM_013956:NM_013964:XM_024447143:NM_001160008:NM_013960 | 0:0:0:1:2:1:0 | 0:0:0:0:0:0:0 | 0:0:0:0:0:0:0 | 1:0:0:0:0:0:0 | 0:2:1:0:0:0:0 | 1:4:0:0:0:0:0 |
| NM_052937:NM_001286783:XR_001745466 | 44:01:02 | 4:00:00 | 55:03:04 | 7:00:00 | 21:00:02 | 0:00:00 |
| NM_052937:NM_001286783:XR_001745466 | 46:01:02 | 2:00:00 | 62:03:04 | 1:00:00 | 10:00:00 | 12:00:02 |
| NM_052937:NM_001286783:XR_001745466 | 65:02:01 | 18:02:01 | 78:04:03 | 11:00:00 | 24:00:01 | 1:00:00 |
| NM_052937:XR_001745466:NM_001286783 | 67:02:03 | 17:00:01 | 78:02:04 | 11:01:00 | 23:01:00 | 2:00:00 |
| NM_052937:NM_001286783:XR_001745466 | 79:04:02 | 6:00:00 | 86:04:02 | 4:00:00 | 24:00:01 | 1:00:00 |
| XM_017013613:XM_011517554:XM_017013612 | 75:09:00 | 2:00:00 | 157:22:00 | 0:00:00 | 110:09:03 | 0:00:00 |
| XM_017013613:XM_011517554:XM_017013612 | 81:09:00 | 1:00:00 | 174:23:00 | 0:00:00 | 114:09:03 | 0:00:00 |
| XM_017013613:XM_011517554:XM_017013612 | 87:09:00 | 0:00:00 | 180:23:00 | 0:00:00 | 119:09:03 | 0:00:00 |
| XM_017013613:XM_011517554:XM_017013612 | 87:09:00 | 0:00:00 | 182:23:00 | 1:00:00 | 116:09:03 | 0:00:00 |
| XM_017013613:XM_011517554:XM_017013612 | 35:03:00 | 1:00:00 | 76:09:00 | 1:01:00 | 30:05:00 | 1:00:00 |
| XM_017013613:XM_011517554:XM_017013612 | 35:03:00 | 0:00:00 | 76:09:00 | 1:00:00 | 30:05:00 | 0:00:00 |
| XM_017013613:XM_011517554:XM_017013612 | 31:03:00 | 2:00:00 | 66:09:00 | 8:00:00 | 23:05:00 | 2:00:00 |
| XM_017013613:XM_011517554:XM_017013612 | 8:01:00 | 25:02:00 | 36:01:00 | 38:08:00 | 11:04:00 | 15:01:00 |
| XM_017013613:XM_011517554 | 8:01 | 0:00 | 31:01:00 | 5:02 | 10:04 | 1:00 |
| XM_017013613:XM_011517554 | 8:01 | 0:00 | 30:01:00 | 5:02 | 9:03 | 2:01 |
| NM_003878:XM_011517623 | 2:00 | 1:00 | 3:00 | 1:00 | 5:00 | 4:00 |
| NM_003878:XM_011517623 | 2:00 | 1:00 | 3:00 | 1:00 | 5:00 | 4:00 |
| NM_001164830:NM_001738:NM_001291967:NM_001291968:NM_001128830 | 14257:667:166:1:1 | 21:2:0:0:0 | 0:0:0:0:0 | 0:0:0:0:0 | 10:0:0:0:0 | 0:0:0:0:0 |
| NM_001164830:NM_001738:NM_001291967:NM_001291968:NM_001128830 | 14253:674:166:1:1 | 26:4:0:0:0 | 0:0:0:0:0 | 0:0:0:0:0 | 10:0:0:0:0 | 0:0:0:0:0 |
| NM_001164830:NM_001738:NM_001291967:NM_001291968:NM_001128830 | 14264:694:167:1:1 | 27:0:0:0:0 | 0:0:0:0:0 | 0:0:0:0:0 | 10:0:0:0:0 | 0:0:0:0:0 |
| NM_001164830:NM_001738:NM_001291967:NM_001291968:NM_001128830 | 14274:701:166:1:1 | 22:0:1:0:0 | 0:0:0:0:0 | 0:0:0:0:0 | 10:0:0:0:0 | 0:0:0:0:0 |
| NM_001164830:NM_001738:NM_001291967:NM_001291968:NM_001128830 | 14267:708:167:1:1 | 33:1:0:0:0 | 0:0:0:0:0 | 0:0:0:0:0 | 10:0:0:0:0 | 0:0:0:0:0 |
| NM_001164830:NM_001738:NM_001291967:NM_001291968:NM_001128830 | 14292:717:167:1:1 | 11:0:0:0:0 | 0:0:0:0:0 | 0:0:0:0:0 | 10:0:0:0:0 | 0:0:0:0:0 |
| NM_001164830:NM_001738:NM_001291967:NM_001291968:NM_001128830 | 14314:712:167:1:1 | 6:1:0:0:0 | 0:0:0:0:0 | 0:0:0:0:0 | 10:0:0:0:0 | 0:0:0:0:0 |
| NM_001164830:NM_001738:NM_001291967:NM_001291968:NM_001128830 | 14322:724:171:1:1 | 9:2:0:0:0 | 0:0:0:0:0 | 0:0:0:0:0 | 10:0:0:0:0 | 0:0:0:0:0 |
| NM_001164830:NM_001738:NM_001291967:NM_001128830 | 14334:666:171:1 | 14:0:1:0 | 0:0:0:0 | 0:0:0:0 | 10:0:0:0 | 0:0:0:0 |
| NM_001164830:NM_001738:NM_001291967:NM_001128830 | 14351:544:171:1 | 17:1:0:0 | 0:0:0:0 | 0:0:0:0 | 10:0:0:0 | 0:0:0:0 |
| NM_001164830:NM_001738:NM_001291967:NM_001128830 | 14382:484:7:1 | 8:3:0:0 | 0:0:0:0 | 0:0:0:0 | 10:0:0:0 | 0:0:0:0 |

| | | | | | | |
|---|---|---|---|---|---|---|
| NM_001164830:NM_001738:NM_001291967:NM_001128830 | 14376:425:7:1 | 16:3:0:0 | 0:0:0:0 | 0:0:0:0 | 10:0:0:0 | 0:0:0:0 |
| NM_001164830:NM_001738:NM_001291967:NM_001128830 | 14366:327:7:1 | 28:12:0:0 | 0:0:0:0 | 0:0:0:0 | 10:0:0:0 | 0:0:0:0 |
| NM_001164830:NM_001738:NM_001291967:NM_001128830 | 14392:314:7:1 | 2:19:0:0 | 0:0:0:0 | 0:0:0:0 | 10:0:0:0 | 0:0:0:0 |
| NM_001164830:NM_001738:NM_001291967:NM_001128830 | 14374:263:7:1 | 20:16:0:0 | 0:0:0:0 | 0:0:0:0 | 10:0:0:0 | 0:0:0:0 |
| NM_001164830:NM_001738:NM_001291967:NM_001128830 | 14383:257:7:1 | 11:18:0:0 | 0:0:0:0 | 0:0:0:0 | 10:0:0:0 | 0:0:0:0 |
| NM_001164830:NM_001738:NM_001291967:NM_001128830 | 14389:249:7:1 | 5:15:0:0 | 0:0:0:0 | 0:0:0:0 | 10:0:0:0 | 0:0:0:0 |
| NM_002485:XM_024447165:XM_024447164:NM_001024688:XM_024447163:XM_011517045:XM_017013462:XM_011517046:XM_017013460 | 8:4:3:0:0:0:0:0:0 | 2:0:0:0:0:0:0:0:0 | 21:16:2:2:0:0:0:0:0 | 6:0:0:0:0:0:0:0:0 | 25:11:0:14:0:0:1:1:0 | 3:0:0:1:1:0:0:0:0 |
| XM_024447164:XM_024447165:NM_002485:NM_001024688:XM_011517046:XM_024447163:XM_017013462:XM_011517045:XM_017013460 | 5:1:11:0:6:0:0:0:0 | 0:0:0:0:0:0:0:0:0 | 2:4:38:6:12:0:0:0:0 | 0:0:0:0:0:0:0:0:0 | 2:3:32:20:17:1:1:0:0 | 0:0:0:0:0:0:0:0:0 |
| XM_024447164:XM_024447165:NM_002485:NM_001024688:XM_011517046:XM_024447163:XM_011517045:XM_017013462:XM_017013460 | 6:1:11:1:6:0:0:0:0 | 0:0:0:0:0:0:0:0:0 | 2:2:38:6:12:1:0:0:0 | 0:0:0:0:0:0:0:0:0 | 0:0:0:0:0:0:0:0:0 | 2:0:32:20:17:1:0:0:0 |
| NM_004063:XM_011516790 | 712:27:00 | 1:00 | 576:17:00 | 2:00 | 379:22:00 | 0:00 |
| NM_004063:XM_011516790 | 715:17:00 | 2:00 | 577:12:00 | 2:00 | 376:17:00 | 2:00 |
| NM_004063:XM_011516790 | 359:06:00 | 358:10:00 | 110:02:00 | 469:10:00 | 145:06:00 | 233:11:00 |
| NM_004063:XM_011516790 | 0:00 | 717:16:00 | 0:00 | 579:12:00 | 144:06:00 | 234:10:00 |
| NM_004063:XM_011516790 | 718:00:00 | 0:00 | 579:02:00 | 0:00 | 381:02:00 | 1:00 |
| NM_004063:XM_011516790 | 461:12:00 | 0:00 | 426:10:00 | 0:00 | 0:00 | 193:09:00 |
| NM_183009:NM_015496:XM_017013308 | 6:00:01 | 0:00:00 | 4:01:03 | 0:00:00 | 8:00:00 | 0:00:00 |
| NM_183009:NM_015496:XM_017013308 | 15:00:01 | 5:00:00 | 14:01:03 | 8:00:00 | 20:00:00 | 8:00:00 |
| NM_183009:NM_015496:XM_017013308 | 19:00:01 | 1:00:00 | 20:01:03 | 2:00:00 | 26:00:00 | 2:00:00 |
| NM_006294:NR_045639:NM_001254752:NM_001199975 | 620:3:2:1 | 4:0:0:0 | 1164:4:10:2 | 14:1:0:0 | 1135:3:10:0 | 9:0:1:0 |
| NM_006294:NM_001254752:NR_045639:NM_001199975 | 621:2:3:1 | 9:0:0:0 | 1170:10:5:2 | 35:0:0:0 | 1140:11:3:0 | 18:0:0:0 |
| NM_006294:NM_001254752:NR_045639:NM_001199975 | 631:2:10:1 | 0:0:0:0 | 1204:10:11:2 | 2:0:0:0 | 1153:11:11:0 | 5:0:0:0 |
| NM_006294:NM_001254752:NR_045639:NM_001199975 | 625:2:10:1 | 2:0:0:0 | 1200:10:11:2 | 2:0:0:0 | 1152:11:10:0 | 0:0:1:0 |
| NM_006294:NM_001254752:NR_045639:NM_001199975 | 625:2:10:1 | 2:0:0:0 | 1201:10:11:2 | 1:0:0:0 | 1151:11:11:0 | 1:0:0:0 |
| NM_006294:NM_001254752:NR_045639:NM_001199975 | 615:2:10:1 | 1:0:0:0 | 1190:10:11:2 | 2:0:0:0 | 1141:10:11:0 | 0:0:0:0 |
| NM_006294:NM_001254752:NR_045639:NM_001199975 | 618:2:10:1 | 0:0:0:0 | 1190:10:11:2 | 3:0:0:0 | 1141:10:11:0 | 1:0:0:0 |
| NM_001317944:NM_017688:XM_006717149:NM_001317943 | 1:22:2:0 | 2:11:1:2 | 0:9:2:0 | 2:12:1:0 | 0:21:5:0 | 0:0:0:0 |
| XM_024447530:NM_002160 | 0:00 | 0:00 | 0:00 | 0:00 | 0:00 | 3:00 |
| XM_005272207:NM_022779:XR_929836:XR_246600:XM_011518921:XM_006717236:NM_001322340:XM_011518922 | 0:2:0:1:0:0:0:0 | 0:2:0:1:0:0:0:0 | 0:1:1:0:0:0:0:0 | 1:0:0:0:0:0:0:0 | 0:0:0:0:0:0:0:0 | 4:6:0:0:0:0:0:0 |
| NM_022779:XR_929836:NM_001322344:XM_011518921:XR_246600:XM_006717236:XM_005272207:NM_001322340:NM_001322341 | 4:0:0:1:0:0:0:0:0 | 0:0:0:0:0:0:0:0:0 | 1:4:1:0:0:0:0:0:0 | 0:0:0:0:0:0:0:0:0 | 6:0:1:0:0:0:0:0:0 | 0:0:0:0:0:0:0:0:0 |
| XM_017015165:NM_007371:XM_017015166:XM_006717291 | 0:3:1:0 | 0:0:0:0 | 1:1:0:0 | 0:0:0:0 | 0:1:0:0 | 0:0:0:0 |
| XM_017015165:NM_007371:XM_017015166:XM_006717291 | 0:3:1:0 | 0:0:0:0 | 1:1:0:0 | 0:0:0:0 | 0:1:0:0 | 0:0:0:0 |
| NM_001317898:XM_024447402:NM_033222:NM_001128217:XM_024447400:XM_024447399:NM_001317900:NM_021144:XM_024447401 | 1:1:9:5:4:0:0:0:0 | 5:4:0:0:0:0:0:0:0 | 7:5:10:8:3:0:0:0:0 | 5:3:1:0:0:0:0:0:0 | 16:15:7:8:14:1:0:1:0 | 7:8:0:0:0:0:0:0:0 |
| NM_033222:NM_001128217:XM_024447400:NM_001317898:XM_024447402:XM_024447399:NM_001317900:NM_021144:XM_024447401 | 9:5:4:7:5:0:0:0:0 | 0:0:0:0:0:0:0:0:0 | 11:8:3:13:8:0:0:0:0 | 0:0:0:0:0:0:0:0:0 | 7:8:14:26:26:1:0:1:0 | 0:0:0:0:0:0:0:0:0 |
| NM_001317898:XM_024447402:NM_033222:NM_001128217:XM_024447400:XM_024447399:NM_001317900:NM_021144:XM_024447401 | 7:5:9:5:4:0:0:0:0 | 1:1:0:0:0:0:0:0:0 | 13:8:11:8:3:0:0:0:0 | 1:1:0:0:0:0:0:0:0 | 26:26:7:8:14:1:0:1:0 | 2:0:0:0:0:0:0:0:0 |
| NM_033222:NM_001128217:XM_024447400:NM_001317898:XM_024447402:XM_024447399:NM_001317900:NM_021144:XM_024447401 | 9:5:4:8:6:0:0:0:0 | 0:0:0:0:2:0:0:0:0 | 11:8:3:14:10:0:0:0:0 | 0:0:0:0:0:0:0:0:0 | 7:8:14:29:26:1:0:1:0 | 0:0:0:0:0:0:0:0:0 |
| XM_024447586:NM_017794 | 0:00 | 0:00 | 0:00 | 0:00 | 0:01 | 0:00 |
| XM_011518045:NM_001193536:XM_011518047:XM_017015174:XM_011518046:NM_203447:XM_011518048 | 0:0:0:0:0:0:0 | 0:1:0:0:0:0:0 | 0:1:0:0:0:0:0 | 1:0:0:0:0:0:0 | 0:0:0:1:0:0:0 | 0:0:0:0:0:0:0 |
| NM_007343:NM_001197098:XM_011517965 | 0:00:00 | 10:01:00 | 0:00:00 | 2:00:00 | 0:00:00 | 0:00:00 |

| | | | | | | |
|---|---|---|---|---|---|---|
| NM_033428:NM_001318058 | 1:00 | 0:00 | 1:00 | 0:00 | 4:00 | 2:01 |
| NM_022755:XR_001746365:XM_017015041 | 10:01:00 | 0:00:00 | 5:01:00 | 0:00:00 | 4:00:00 | 0:00:00 |
| NR_134466:NM_001318011:NR_134468:NM_032310:XR_929848:NR_134465:NR_134467:NM_001318010 | 12:2:5:15:1:0:1:0 | 11:1:0:15:2:1:1:0 | 3:0:1:2:1:0:0:0 | 6:1:1:6:0:0:0:0 | 15:1:0:17:2:0:1:1 | 0:0:0:0:0:0:0:0 |
| XM_024447451:NM_000136:NM_001243744:XM_011518366:XM_006717002:XM_017014453:NM_001243743:XM_017014452:XM_006717004 | 1:2:1:1:3:1:2:0:0 | 0:0:0:0:0:0:0:0:0 | 2:1:2:1:0:0:0:0:0 | 0:0:0:0:0:0:0:0:0 | 1:6:3:4:1:0:0:0:0 | 0:0:0:0:0:0:0:0:0 |
| NM_017919:XM_024447593:XR_001746332:XM_011518821:XM_011518820 | 17:4:0:1:0 | 1:0:0:0:0 | 10:2:0:0:0 | 3:0:0:0:0 | 11:2:0:0:1 | 3:0:0:0:0 |

| 2M__WT | 2M__MUT | 3T__WT | 3T__MUT | 3M__WT | 3M__MUT | 1T__q | 1M__q | 2T__q | 2M__q | 3T__q | 3M__q |
|---|---|---|---|---|---|---|---|---|---|---|---|
| 53:00:00 | 73:00:00 | 37:00:00 | 0:00 | 14:00 | 0:00 | 1.56E-05 | 0.003751524 | 0.010019977 | 5.14E-17 | 1 | 1 |
| 120:00:00 | 26:01:00 | 34:00:00 | 5:00 | 14:00 | 0:00 | 1 | 1 | 1 | 3.28E-06 | 0.031296179 | 1 |
| 0:0:0:0:0:0 | 26:5:8:14:35:1 | 0:0:0:0:0:0 | 20:4:10:16:15: | 0:0:0:0:0:0 | 3:0:1:2:2:0 | 0.033223405 | 0.53837282 | 0.782204594 | 5.95E-07 | 0.048032892 | 0.36927926 |
| 2:0:1:1:0:0:0 | 4:2:0:1:0:0:0 | 0:3:0:1:1:3:1 | 0:0:0:0:0:0:0 | 0:2:0:1:0:0:0 | 0:0:0:0:0:0:0 | 0.1336694 | 1 | 1 | 0.035238333 | 1 | 1 |
| 3544:2:64:46 | 9:0:0:0 | 2088:7:33:21 | 5:0:0:0 | 255:2:11:2 | 0:0:0:0 | 1 | 1 | 1 | 1.18E-05 | 0.002732501 | 1 |
| 3525:2:65:46 | 11:0:0:0 | 2092:7:33:21 | 5:0:0:0 | 254:2:11:2 | 1:0:0:0 | 1 | 1 | 1 | 7.61E-07 | 0.002732501 | 1 |
| 3327:2:67:46 | 6:0:0:0 | 1969:6:33:20 | 2:0:0:0 | 249:2:11:2 | 0:0:0:0 | 1 | 1 | 1 | 0.000699231 | 1 | 1 |
| 3286:2:67:46 | 5:0:0:0 | 1943:6:33:20 | 2:0:0:0 | 250:2:11:2 | 0:0:0:0 | 1 | 1 | 1 | 0.002648305 | 1 | 1 |
| 2959:2:71:47 | 8:0:0:0 | 1776:6:34:20 | 6:0:0:0 | 244:2:11:2 | 0:0:0:0 | 1 | 1 | 1 | 4.60E-05 | 0.000739192 | 1 |
| 2067:72:2:47 | 5:0:0:0 | 1280:35:6:21 | 1:0:0:0 | 231:11:2:2 | 1:0:0:0 | 1 | 1 | 1 | 0.002648305 | 1 | 1 |
| 1688:2:72:48 | 10:0:0:0 | 1067:6:35:21 | 5:0:0:0 | 218:2:11:2 | 0:0:0:0 | 1 | 1 | 1 | 3.04E-06 | 0.002732501 | 1 |
| 812:3:76:48 | 8:0:0:0 | 543:8:35:21 | 4:0:0:0 | 193:2:11:2 | 0:0:0:0 | 1 | 1 | 1 | 4.60E-05 | 1 | 1 |
| 0:47:64:308:251 | 0:0:0:0:2 | 0:5:2:19:3 | 0:50:54:268:179 | 1:3:0:3:0 | 1:39:3:48:17 | 1 | 1 | 1 | 1 | 7.16E-93 | 2.27E-16 |
| 0:47:400:41:341 | 0:0:1:0:1 | 0:55:351:48:257 | 0:0:3:0:4 | 2:42:57:3:18 | 0:0:0:0:0 | 1 | 1 | 1 | 1 | 0.034565431 | 1 |
| 0:0:0:0:0:0 | 12:3:0:1:0:0 | 0:0:0:0:0:0 | 24:7:1:3:0:1 | 0:0:0:0:0:0 | 7:1:0:0:0:0 | 1 | 0.016309875 | 0.000855916 | 1.64E-07 | 4.09E-13 | 3.69E-05 |
| 0:3:3:0:2:1:0:0:0 | 0:7:5:1:1:0:0:0:0 | 0:7:7:0:2:0:0:1:1 | 0:3:1:0:0:0:0:0:0 | 0:3:0:0:0:0:0:0:0 | 0:9:1:0:0:0:0:0:0 | 1 | 1 | 0.002187056 | 5.22E-05 | 1 | 2.96E-10 |
| 0:5:7:5:5:1:1:3:0:0 | 0:0:0:0:0:0:0:0:0:0 | 0:6:10:1:3:0:0:6:0:1 | 0:3:1:0:1:0:0:0:0:0 | 0:3:0:1:0:0:0:0:0:0 | 0:7:1:0:0:0:0:0:0:0 | 1 | 1 | 1 | 1 | 0.057415258 | 4.32E-08 |
| 4343:165 | 2:00 | 16166:1021 | 27:02:00 | 2488:132 | 3:00 | 1 | 1 | 1 | 1 | 7.38E-06 | 1 |
| 4343:165 | 2:00 | 16179:1023 | 28:00:00 | 2487:132 | 4:00 | 1 | 1 | 1 | 1 | 3.00E-07 | 1 |
| 4343:165 | 7:00 | 16237:1022 | 10:03 | 2492:133 | 0:00 | 1 | 1 | 1 | 0.010434472 | 0.061326601 | 1 |
| 2460:56:6:71:3:16:43:2:0:6:0 | 4:0:0:0:0:0:0:0:0:0:0 | 4026:108:9:22:1:24:22:2:0:3:0 | 5:2:0:0:0:0:0:0:0:0:0 | 1501:10:3:31:0:9:17:1:0:0:1 | 0:0:0:0:0:0:0:0:0:0:0 | 1 | 1 | 1 | 1 | 5.85E-05 | 1 |
| 2467:32:6:71:1:16:43:2:0:6:0 | 0:2:0:0:0:0:0:0:0:0:0 | 4032:59:9:22:1:24:22:2:0:3:0 | 3:3:0:0:0:0:0:0:0:0:0 | 1502:10:3:31:0:9:17:1:0:0:1 | 0:0:0:0:0:0:0:0:0:0:0 | 1 | 1 | 1 | 1 | 0.00383779 | 1 |
| 2358:102:6:67:0:16:42:2:0:6 | 6:0:0:0:0:0:0:0:0:0 | 3813:125:9:22:0:24:22:2:0:1 | 2:0:0:0:0:0:0:0:0:0 | 1481:12:3:31:0:9:17:1:0:0 | 1:0:0:0:0:0:0:0:0:0 | 1 | 1 | 1 | 4.92E-08 | 1 | 1 |
| 2328:103:6:67:0:16:42:2:0:5 | 2:0:0:0:0:0:0:0:0:0 | 3752:127:9:22:0:24:21:2:0:1 | 7:0:0:0:0:0:0:0:0:0 | 1475:13:3:31:0:9:16:1:0:0 | 2:0:0:0:0:0:0:0:0:0 | 1 | 1 | 1 | 1 | 1.13E-09 | 1 |

| | | | | | | | | | | | |
|---|---|---|---|---|---|---|---|---|---|---|---|
| 2306:102:6:65:0:16:42:2:0:5 | 4:0:0:1:0:0:0:0:0:0 | 3718:129:9:22:0:24:19:2:0:1 | 9:0:0:0:0:0:0:0:0:0 | 1469:13:3:31:0:9:16:1:0:0 | 1:0:0:0:0:0:0:0:0:0 | 1 | 1 | 1 | 0.001006637 | 4.09E-13 | 1 |
| 2188:36:96:6:60:0:8:4:2 | 4:0:0:0:0:0:0:0:0 | 3576:19:129:8:21:0:19:1:2 | 7:0:0:0:0:0:0:0:0 | 1436:15:11:3:31:0:8:0:1 | 0:0:0:0:0:0:0:0:0 | 1 | 1 | 1 | 1 | 8.30E-09 | 1 |
| 50:6:3:2 | 1:0:0:0 | 76:4:2:1 | 1:0:0:0 | 19:2:0:0 | 0:0:0:0 | 0.034015223 | 1 | 1 | 1 | 1 | 1 |
| 57:5:10:0 | 43:2:8:0 | 107:10:13:0 | 51:2:10:0 | 24:2:0:1 | 17:0:2:0 | 3.17E-05 | 3.23E-06 | 4.19E-12 | 4.16E-19 | 1.13E-22 | 8.77E-09 |
| 0:0:0:0:0:0:0:0:0 | 3:9:0:0:0:0:1:0:0:0 | 5:22:0:0:0:0:1:0:0 | 0:0:0:0:0:0:0:0:0 | 3:2:0:0:0:0:1:0:0 | 0:0:0:0:0:0:0:0:0 | 0.475373601 | 1 | 0.362844892 | 1.39E-08 | 1 | 1 |
| 159:07:00 | 0:00 | 164:02:00 | 0:00 | 38:00:00 | 0:00 | 1 | 0.003751524 | 1 | 1 | 1 | 1 |
| 727:63:1:0:2:0:0 | 1:0:0:0:0:0:0 | 1810:178:9:0:20:1:0 | 14:1:0:0:0:0:0 | 437:26:2:0:6:0:1 | 6:1:0:0:0:0:0 | 0.00023392 | 1 | 1 | 1 | 6.40E-14 | 7.53E-05 |
| 728:64:1:0:2:0:0 | 1:0:0:0:0:0:0 | 1821:180:9:0:20:1:0 | 5:0:0:0:0:0:0 | 443:27:2:0:6:0:1 | 0:0:0:0:0:0:0 | 1 | 1 | 1 | 1 | 7.47E-05 | 1 |
| 1909:46:22 | 6:00:00 | 3450:55:25 | 5:00:00 | 456:09:07 | 0:00:00 | 1 | 1 | 1 | 0.003552529 | 0.009407211 | 1 |
| 1909:46:22 | 8:00:00 | 3477:55:25 | 5:00:00 | 457:09:07 | 0:00:00 | 1 | 1 | 1 | 0.000535794 | 0.009407211 | 1 |
| 2241:24:22 | 4:00:00 | 4124:34:25 | 13:00:00 | 503:08:07 | 1:00:00 | 1 | 1 | 1 | 1 | 4.90E-06 | 1 |
| 2241:24:22 | 7:00:00 | 4133:34:25 | 8:00:00 | 503:08:07 | 1:00:00 | 1 | 1 | 1 | 0.00138895 | 0.000571704 | 1 |
| 1140:09:00 | 3:00 | 2255:08:00 | 7:00 | 265:04:00 | 0:00 | 1 | 1 | 1 | 1 | 0.011137275 | 1 |
| 0:00:00 | 9:00:00 | 0:00:00 | 23:00:01 | 0:00:00 | 4:00:00 | 1 | 1 | 3.88E-05 | 0.000209242 | 2.10E-09 | 1 |
| 0:00:00 | 11:00:00 | 0:00:00 | 33:00:01 | 0:00:00 | 4:00:00 | 1 | 0.008830794 | 8.27E-07 | 3.18E-05 | 1.29E-13 | 1 |
| 124:08:00 | 0:00 | 50:05:00 | 0:00 | 5:01 | 0:00 | 0.003177062 | 0.017552603 | 1 | 1 | 1 | 1 |
| 0:00 | 124:04:00 | 50:06:00 | 0:00 | 5:01 | 0:00 | 0.005948649 | 0.000180834 | 0.011433615 | 1.42E-25 | 1 | 1 |
| 12:1:1:0 | 3:5:1:0 | 5:0:1:0 | 0:1:0:1 | 0:1:0:0 | 0:1:0:0 | 1 | 1 | 0.01363685 | 0.097548727 | 1 | 1 |
| 0:0:0:0:0:0:0:0:0:0 | 62:393:4:6:0:1:3:1:1:0:0 | 0:1:0:0:0:0:0:0:0:0 | 120:704:6:5:0:1:1:1:1:0:1 | 0:0:0:0:0:0:0:0:0:0 | 40:154:2:0:0:0:2:0:0:0:0 | 7.75E-99 | 4.29E-183 | 5.74E-229 | 0 | 0 | 1.14E-251 |
| 0:03 | 0:00 | 0:00 | 0:50 | 0:00 | 0:08 | 1 | 1 | 1 | 1 | 5.62E-12 | 0.006579116 |
| 0:00 | 0:03 | 0:00 | 0:47 | 0:00 | 0:06 | 1 | 1 | 1 | 1 | 2.48E-11 | 0.018161034 |
| 22:4:6:3:0 | 0:0:0:0:0 | 30:5:13:1:1 | 0:0:0:0:0 | 15:4:12:1:0 | 0:0:0:0:0 | 5.55E-06 | 0.004669124 | 1 | 1 | 1 | 1 |
| 0:0:0:0:0 | 7:0:0:0:0 | 0:0:0:0:0 | 6:0:0:2:0 | 0:0:0:0:0 | 1:0:0:1:0 | 1 | 1 | 1 | 2.41E-05 | 0.002903423 | 1 |
| 1115:07:00 | 5:00 | 1176:11:00 | 5:00 | 218:02:00 | 1:00 | 1 | 1 | 1 | 0.030173969 | 0.031296179 | 1 |
| 94:50:1102:7:1 | 0:0:3:0:0 | 270:109:1173:11:0 | 0:0:6:0:0 | 39:10:218:2:0 | 0:0:0:0:0 | 1 | 1 | 1 | 1 | 0.00014768 | 1 |
| 92:49:1094:7:1 | 1:0:8:0:0 | 269:109:1170:11:0 | 0:0:7:0:0 | 39:9:218:2:0 | 0:0:0:0:0 | 1 | 1 | 1 | 3.59E-05 | 2.53E-05 | 1 |
| 12:00 | 8:02 | 28:02:00 | 0:00 | 15:00 | 0:00 | 1 | 1 | 0.007094277 | 0.066678803 | 1 | 1 |
| 0:1:5:1:0:0:0:0 | 0:119:4:0:0:2:0:0 | 1:81:7:1:1:2:0:1 | 0:0:0:0:0:0:0:0 | 0:50:3:0:1:1:1:0 | 0:0:0:0:0:0:0:0 | 5.63E-11 | 5.54E-14 | 4.05E-98 | 5.70E-163 | 1 | 1 |
| 5:0:0:30:2:6:6:0:0:1:0:0:0:0:1:2:0:1:0:1:2:1:1:0 | 7:1:3:35:0:7:11:0:0:7:1:1:1:1:1:2:1:1:1:0:0:0:0:0 | 1:2:0:6:0:1:2:2:0:0:1:0:0:0:2:0:0:0:0:1:0:0:0:1 | 0:0:0:0:0:0:0:0:0:0:0:0:0:0:0:0:0:0:0:0:0:0 | 0:0:0:2:0:0:2:0:0:0:1:0:0:0:0:0:0:0:0:0:1:0:0:0 | 0:0:0:0:0:0:0:0:0:0:0:0:0:0:0:0:0:0:0:0:0:0 | 7.64E-07 | 1 | 1.11E-06 | 1.08E-62 | 1 | 1 |
| 3:0:2:0:0:0:1:1 | 5:0:0:2:1:1:0:0 | 9:1:0:0:0:0:0:0 | 0:0:0:0:0:0:0:0 | 1:0:0:0:0:0:0:0 | 0:0:0:0:0:0:0:0 | 1 | 1 | 1 | 0.012355014 | 1 | 1 |
| 0:4:0:3:2:0:0:1:0 | 0:6:0:0:0:1:1:0:0 | 1:10:0:0:0:0:0:0:0 | 0:0:0:0:0:0:0:0:0 | 0:1:0:0:0:0:0:0:1 | 0:0:0:0:0:0:0:0:0 | 1 | 1 | 1 | 5.44E-05 | 1 | 1 |
| 30:13:3:24:71:1 | 22:10:6:19:58:1 | 32:24:5:33:87:1 | 0:0:0:0:0:0 | 6:0:1:15:2:0 | 0:0:0:0:0:0 | 1 | 1 | 0.032395662 | 3.38E-21 | 1 | 1 |
| 87:39:5:4:1:0:0:0:0 | 0:0:0:0:0:0:0:0 | 48:30:8:6:6:0:0:1:1 | 0:0:0:0:0:0:0:0 | 15:7:0:0:2:0:1:0:0 | 0:0:0:0:0:0:0:0 | 1.69E-92 | 1.44E-12 | 1 | 1 | 1 | 1 |
| 23:506:58:25:45 | 0:0:0:0:0 | 8:95:9:2:11 | 0:0:0:0:0 | 1:25:4:0:0 | 0:0:0:0:0 | 8.07E-17 | 1.02E-08 | 1 | 1 | 1 | 1 |
| 117:22:13:50:3:260:232:2:13:172:26:0:2:1 | 7:0:0:2:0:0:0:0:0:1:4:0:0:0 | 77:2:5:16:1:237:240:4:7:149:29:0:0:0 | 1:0:0:2:0:0:1:0:0:0:1:0:0:0 | 24:0:2:2:2:117:149:4:3:7:2:0:0:0 | 1:0:0:2:0:0:0:0:0:0:0:0:0:0 | 0.003959071 | 1 | 0.022318387 | 6.42E-07 | 0.349682076 | 1 |
| 5157:02:00 | 2:00 | 22024:9 | 21:00 | 4324:01:00 | 3:00 | 1 | 1 | 1 | 1 | 9.75E-06 | 1 |
| 5151:02:00 | 8:00 | 22017:9 | 31:00:00 | 4322:01:00 | 6:00 | 1 | 1 | 1 | 0.006305042 | 6.92E-08 | 0.018161034 |
| 5135:02:00 | 3:00 | 21913:9 | 45:00:00 | 4316:01:00 | 7:00 | 1 | 1 | 1 | 1 | 6.75E-11 | 0.010989758 |
| 5080:02:00 | 3:00 | 21648:9 | 22:00 | 4303:01:00 | 5:00 | 1 | 1 | 0.011433615 | 1 | 5.88E-06 | 0.030032732 |
| 5044:02:00 | 6:00 | 21498:9 | 34:00:00 | 4287:01:00 | 8:00 | 1 | 1 | 0.030109098 | 0.017756017 | 1.57E-08 | 0.006579116 |

| | | | | | | | | | | | |
|---|---|---|---|---|---|---|---|---|---|---|---|
| 4993:02:00 | 0:00 | 21158:9 | 11:00 | 4276:01:00 | 2:00 | 1 | 1 | 1 | 1 | 0.001445472 | 1 |
| 4928:02:00 | 6:00 | 20935:9 | 28:00:00 | 4264:01:00 | 2:00 | 1 | 1 | 0.017871477 | 0.017756017 | 3.00E-07 | 1 |
| 4926:02:00 | 8:00 | 20909:9 | 39:00:00 | 4261:01:00 | 3:00 | 1 | 1 | 0.001007623 | 0.006305042 | 1.35E-09 | 1 |
| 4893:02:00 | 5:00 | 20637:9 | 31:00:00 | 4225:01:00 | 5:00 | 1 | 1 | 1 | 0.030173969 | 6.92E-08 | 0.030032732 |
| 4888:02:00 | 8:00 | 20588:9 | 35:00:00 | 4217:01:00 | 7:00 | 1 | 1 | 1 | 0.006305042 | 9.63E-09 | 0.010989758 |
| 4851:02:00 | 5:00 | 20355:9 | 28:00:00 | 4198:01:00 | 5:00 | 1 | 1 | 1 | 0.030173969 | 3.00E-07 | 0.030032732 |
| 4855:02:00 | 2:00 | 20350:9 | 23:00 | 4185:01:00 | 2:00 | 1 | 1 | 0.017871477 | 1 | 3.55E-06 | 1 |
| 4804:02:00 | 0:00 | 20154:9 | 30:00:00 | 4159:01:00 | 5:00 | 1 | 1 | 0.011433615 | 1 | 1.12E-07 | 0.030032732 |
| 4747:02:00 | 8:00 | 19926:9 | 39:00:00 | 4135:01:00 | 6:00 | 1 | 1 | 1 | 0.006305042 | 1.35E-09 | 0.018161034 |
| 4740:02:00 | 6:00 | 19908:9 | 21:00 | 4131:01:00 | 4:00 | 1 | 1 | 1 | 0.017756017 | 9.75E-06 | 1 |
| 4731:02:00 | 10:00 | 19866:9 | 30:00:00 | 4130:01:00 | 4:00 | 1 | 1 | 1 | 0.002318525 | 1.12E-07 | 1 |
| 4722:02:00 | 8:00 | 19834:9 | 43:00:00 | 4132:01:00 | 3:00 | 1 | 1 | 0.007094277 | 0.006305042 | 1.82E-10 | 1 |
| 4709:02:00 | 12:00 | 19644:9 | 49:00:00 | 4088:01:00 | 5:00 | 1 | 0.017552603 | 0.017871477 | 0.000831463 | 9.18E-12 | 0.030032732 |
| 4710:02:00 | 11:00 | 19671:9 | 37:00:00 | 4083:01:00 | 8:00 | 1 | 1 | 0.011433615 | 0.00138895 | 3.53E-09 | 0.006579116 |
| 4705:02:00 | 10:00 | 19643:9 | 38:00:00 | 4086:01:00 | 2:00 | 1 | 1 | 0.017871477 | 0.002318525 | 2.20E-09 | 1 |
| 4653:02:00 | 12:00 | 19527:9 | 38:00:00 | 4063:01:00 | 3:00 | 1 | 1 | 1 | 0.000831463 | 2.20E-09 | 1 |
| 4630:02:00 | 4:00 | 19342:9 | 33:00:00 | 4049:01:00 | 2:00 | 1 | 1 | 1 | 1 | 2.58E-08 | 1 |
| 4623:02:00 | 8:00 | 19287:9 | 44:00:00 | 4044:01:00 | 5:00 | 1 | 1 | 0.007094277 | 0.006305042 | 1.11E-10 | 0.030032732 |
| 4577:02:00 | 1:00 | 19174:9 | 23:00 | 4032:01:00 | 5:00 | 1 | 1 | 0.030109098 | 1 | 3.55E-06 | 0.030032732 |
| 4578:02:00 | 5:00 | 19199:9 | 28:00:00 | 4034:01:00 | 5:00 | 1 | 0.029932329 | 0.017871477 | 0.030173969 | 3.00E-07 | 0.030032732 |
| 4571:02:00 | 7:00 | 19238:9 | 24:00:00 | 4045:01:00 | 2:00 | 1 | 1 | 0.030109098 | 0.010434472 | 2.16E-06 | 1 |
| 4562:02:00 | 7:00 | 19213:9 | 24:00:00 | 4040:01:00 | 4:00 | 1 | 1 | 1 | 0.010434472 | 2.16E-06 | 1 |
| 4567:02:00 | 4:00 | 19153:9 | 17:00 | 4029:01:00 | 8:00 | 1 | 1 | 0.017871477 | 1 | 7.14E-05 | 0.006579116 |
| 4554:01:00 | 4:00:00 | 19141:1:1 | 14:00:00 | 4038:00:01 | 3:00:00 | 1 | 1 | 1 | 1 | 1.88E-06 | 1 |
| 4558:01:00 | 3:00:00 | 19160:1:1 | 6:00:00 | 4039:00:01 | 2:00:00 | 1 | 1 | 1 | 1 | 0.003657671 | 1 |
| 4547:01:00 | 7:00:00 | 19128:1:1 | 20:00:00 | 4030:00:01 | 7:00:00 | 1 | 1 | 1 | 0.00138895 | 6.12E-09 | 0.001510787 |
| 4536:01:00 | 7:00:00 | 19089:1:1 | 19:00:00 | 4029:00:01 | 2:00:00 | 1 | 0.003558706 | 1 | 0.00138895 | 1.58E-08 | 1 |
| 4500:02:00 | 4:00:00 | 18981:4:1 | 6:00:00 | 4004:00:01 | 3:00:00 | 1 | 1 | 1 | 1 | 0.003657671 | 1 |
| 24487:38 | 24:00:00 | 13703:24 | 18:00 | 3311:07:00 | 8:00 | 0.002247757 | 0.003751524 | 0.007094277 | 2.17E-06 | 4.38E-05 | 0.006579116 |
| 24504:37 | 16:01 | 13717:24 | 12:00 | 3314:07:00 | 6:00 | 0.006139527 | 0.003751524 | 0.007094277 | 0.000445066 | 0.000876481 | 0.018161034 |
| 24502:37 | 27:01:00 | 13713:24 | 20:00 | 3318:07:00 | 2:00 | 0.003657788 | 0.000528313 | 0.011433615 | 2.04E-06 | 1.62E-05 | 1 |
| 24505:38 | 30:00:00 | 13719:24 | 19:00 | 3320:07:00 | 1:00 | 0.00132829 | 0.000197866 | 0.011433615 | 1.08E-07 | 2.64E-05 | 1 |
| 24580:37 | 9:00 | 13762:24 | 5:00 | 3323:07:00 | 2:00 | 0.017290558 | 0.010484149 | 1 | 0.00379461 | 0.031296179 | 1 |
| 24600:37 | 6:00 | 13771:24 | 3:00 | 3323:07:00 | 3:00 | 0.029709415 | 1 | 1 | 0.017756017 | 1 | 1 |
| 25248:37 | 19:00 | 13867:22 | 12:01 | 3391:07:00 | 5:00 | 1 | 0.003751524 | 1 | 2.52E-05 | 0.003418198 | 0.030032732 |
| 25450:37 | 22:00 | 13989:21 | 17:00 | 3414:07:00 | 2:00 | 1 | 0.00624902 | 0.004510635 | 5.73E-06 | 7.14E-05 | 1 |
| 25478:37 | 13:00 | 13999:21 | 15:00 | 3415:07:00 | 2:00 | 1 | 1 | 0.017871477 | 0.000501437 | 0.000196442 | 1 |
| 25479:36 | 10:00 | 14002:20 | 8:00 | 3415:07:00 | 3:00 | 1 | 1 | 1 | 0.002318525 | 0.006667609 | 1 |
| 26089:22 | 23:00 | 14322:12 | 19:00 | 3466:05:00 | 5:00 | 1 | 1 | 1 | 3.50E-06 | 2.64E-05 | 0.030032732 |
| 26124:18 | 24:01:00 | 14352:9 | 12:00 | 3476:05:00 | 1:00 | 1 | 1 | 1 | 8.68E-06 | 0.000876481 | 1 |
| 26191:14 | 24:01:00 | 14403:9 | 11:00 | 3485:03:00 | 4:00 | 1 | 1 | 0.011433615 | 8.68E-06 | 0.001445472 | 1 |
| 26279:6 | 9:01 | 14465:2 | 1:00 | 3495:01:00 | 1:00 | 1 | 1 | 1 | 0.014338921 | 1 | 1 |
| 26293:3 | 20:00 | 14463:0 | 16:00 | 3501:01:00 | 3:00 | 1 | 1 | 1 | 1.52E-05 | 0.000117914 | 1 |
| 26374:4 | 26:04:00 | 14518:0 | 16:04 | 3516:00:00 | 0:00 | 1 | 1 | 1 | 0.00010517 | 0.010139464 | 1 |
| 26364:43 | 30:03:00 | 14511:24 | 18:03 | 3511:03:00 | 3:00 | 1 | 0.001383821 | 1 | 5.49E-06 | 0.001665802 | 1 |
| 26369:92 | 21:01 | 14509:54 | 13:03 | 3510:09:00 | 3:00 | 1 | 1 | 1 | 3.79E-05 | 0.016393516 | 1 |
| 26313:230 | 24:04:00 | 14483:193 | 14:03 | 3496:28:00 | 4:00 | 1 | 0.112198641 | 1 | 0.000264086 | 0.010602306 | 1 |
| 26243:414 | 32:03:00 | 14437:337 | 19:02 | 3491:43:00 | 3:00 | 1 | 0.112198641 | 0.7124885 | 2.14E-06 | 0.000368028 | 1 |
| 26206:466 | 17:01 | 14409:358 | 13:00 | 3483:47:00 | 1:00 | 1 | 0.660447267 | 0.006273661 | 0.000271356 | 0.00053673 | 1 |
| 26175:549 | 33:02:00 | 14366:412 | 16:02 | 3468:55:00 | 5:00 | 1 | 0.112198641 | 0.030109098 | 3.91E-07 | 0.001526854 | 0.030032732 |
| 26108:660 | 32:01:00 | 14371:500 | 8:02 | 3461:62 | 6:00 | 1 | 1 | 0.007094277 | 1.68E-07 | 0.067803829 | 0.018161034 |
| 26019:678 | 23:00 | 14345:545 | 9:00 | 3458:63 | 1:00 | 1 | 1 | 0.030109098 | 3.50E-06 | 0.003951718 | 1 |
| 23298:648 | 1775:09:00 | 13033:519 | 891:11:00 | 3178:61 | 198:01:00 | 1.73E-26 | 2.75E-65 | 1.72E-81 | 0 | 5.02E-187 | 2.93E-43 |
| 15757:404 | 26:02:00 | 8798:365 | 13:00 | 3162:57:00 | 0:00 | 1 | 1 | 1 | 1.17E-05 | 0.00053673 | 1 |
| 33:31:25:4 | 68:29:35:3 | 74:46:67:5 | 0:0:0:0 | 25:9:20:2 | 0:0:0:0 | 1 | 0.232824754 | 0.028624445 | 3.71E-13 | 1 | 1 |
| 35:69:39:3 | 27:33:33:4 | 64:79:52:5 | 1:0:0:0 | 19:28:11:2 | 0:0:0:0 | 1 | 1 | 0.010070087 | 5.70E-05 | 1 | 1 |
| 91:51:75:7 | 11:9:2:0 | 72:56:54:4 | 5:9:1:1 | 28:18:13:2 | 0:1:0:0 | 1 | 1 | 1 | 0.002194616 | 0.015610609 | 1 |
| 1:1:6:4:6:4:2:1:3:0:0:0:6:1:0:2:0:0:0:10:1:0:0:0:0:1:1:6:0:0:0:0 | 0:0:0:0:0:0:0:0:0:0:0:0:0:0:0:0:0:0:0:0:0:0:0:0:0:0:0:0:0:0:0:0 | 4:4:4:17:4:10:0:1:0:0:1:1:8:0:0:4:1:0:1:0:2:0:0:0:1:3:0:1:0:0:1:1 | 0:0:0:0:0:0:0:0:0:0:0:0:0:0:0:0:0:0:0:0:0:0:0:0:0:0:0:0:0:0:0:0 | 3:3:3:9:2:2:0:0:0:0:1:0:3:0:0:1:0:1:0:1:0:0:0:1:0:1:0:0:0:1:0:0 | 0:0:0:0:0:0:0:0:0:0:0:0:0:0:0:0:0:0:0:0:0:0:0:0:0:0:0:0:0:0:0:0 | 2.18E-06 | 0.001310156 | 1 | 1 | 1 | 1 |
| 132:08:00 | 5:00 | 108:03:00 | 7:00 | 35:02:00 | 1:00 | 1 | 1 | 1 | 0.030173969 | 0.011137275 | 1 |
| 0:00:00 | 4:01:01 | 0:00:00 | 3:01:03 | 0:00:00 | 0:00:00 | 0.003985093 | 1 | 1 | 0.238833648 | 0.579075362 | 1 |

| | | | | | | | | | | | |
|---|---|---|---|---|---|---|---|---|---|---|---|
| 4:11:9:1:3:1:0:2:0 | 2:1:4:0:0:0:0:0:0 | 7:2:7:0:3:0:0:1:2 | 1:1:5:0:0:0:0:0:0 | 3:0:2:1:1:0:0:0:0 | 0:0:0:0:0:0:0:0:0 | 1 | 1 | 1 | 0.013067538 | 0.000876481 | 1 |
| 0:1:2:0:15:0:1:0:1:1:1:0 | 1:0:4:0:26:0:1:0:1:0:0:0 | 0:0:8:1:24:0:0:0:1:0:0:1 | 0:0:0:0:0:0:0:0:0:0:0:0 | 0:0:4:0:3:1:0:0:0:0:0:0 | 0:0:0:0:0:0:0:0:0:0:0:0 | 1 | 1 | 1.46E-08 | 4.14E-40 | 1 | 1 |
| 1:2:0:4:23:0:0:0:0:1:1:0 | 0:2:1:2:17:1:0:0:1:1:0:0 | 0:0:0:8:22:0:0:1:0:1:0:1 | 0:0:0:0:0:0:0:0:0:0:0:0 | 0:1:0:4:3:0:0:0:0:0:0:0 | 0:0:0:0:0:0:0:0:0:0:0:0 | 1 | 1 | 1 | 1.02E-19 | 1 | 1 |
| 0:00 | 0:08 | 0:00 | 0:03 | 0:00 | 0:00 | 1 | 1 | 1 | 0.006305042 | 1 | 1 |
| 17:00:00 | 0:00:00 | 804:30:21 | 0:00:00 | 190:08:04 | 0:00:00 | 6.43E-79 | 1.27E-32 | 1 | 1 | 1 | 1 |
| 0:00 | 9:00 | 0:00 | 4:01 | 0:00 | 5:00 | 3.80E-05 | 1 | 0.011433615 | 0.00379461 | 0.194927198 | 0.030032732 |
| 399:05:00 | 3:00 | 2338:64 | 7:00 | 741:32:00 | 1:00 | 1 | 1 | 1 | 1 | 0.011137275 | 1 |
| 437:06:00 | 2:00 | 2480:71 | 8:00 | 763:33:00 | 0:00 | 1 | 1 | 1 | 1 | 0.006667609 | 1 |
| 481:08:00 | 1:00 | 2692:74 | 5:00 | 800:33:00 | 2:00 | 1 | 1 | 1 | 1 | 0.031296179 | 1 |
| 522:10:00 | 2:00 | 2881:81 | 5:00 | 827:35:00 | 2:00 | 1 | 1 | 1 | 1 | 0.031296179 | 1 |
| 525:10:00 | 0:00 | 2894:81 | 7:00 | 831:35:00 | 1:00 | 1 | 1 | 1 | 1 | 0.011137275 | 1 |
| 533:10:00 | 0:00 | 2918:80 | 7:00 | 833:36:00 | 0:00 | 1 | 1 | 1 | 1 | 0.011137275 | 1 |
| 564:12:00 | 0:00 | 3061:85 | 7:00 | 850:37:00 | 0:00 | 1 | 1 | 1 | 1 | 0.011137275 | 1 |
| 593:12:00 | 1:00 | 3180:86 | 6:00 | 859:38:00 | 0:00 | 1 | 1 | 1 | 1 | 0.018436807 | 1 |
| 597:12:00 | 0:00 | 3184:86 | 5:00 | 859:38:00 | 0:00 | 1 | 1 | 1 | 1 | 0.031296179 | 1 |
| 0:7:1:7 | 0:8:0:0 | 0:1:0:1 | 0:0:0:0 | 0:1:0:0 | 0:0:0:0 | 1 | 1 | 1 | 4.60E-05 | 1 | 1 |
| 0:7:1:7 | 0:9:0:0 | 0:1:0:1 | 0:0:0:0 | 0:1:0:0 | 0:0:0:0 | 1 | 1 | 1 | 1.18E-05 | 1 | 1 |
| 0:0:1:0:0:0:1:0:0 | 0:0:0:0:0:0:0:0:0 | 0:26:17:0:0:7:0:1:1 | 0:4:0:0:0:1:0:0:0 | 0:0:2:0:0:0:0:0:0 | 0:0:0:0:0:0:0:0:0 | 1 | 1 | 1 | 1 | 0.001912317 | 1 |
| 0:0:1:0:0:0:1:0:0 | 0:0:0:0:0:0:0:0:0 | 0:26:17:0:0:6:0:1:1 | 0:4:0:0:0:2:0:0:0 | 0:0:2:0:0:0:0:0:0 | 0:0:0:0:0:0:0:0:0 | 1 | 1 | 1 | 1 | 0.00341885 | 1 |
| 0:0:0:0:0:2:0:0 | 0:0:1:0:0:0:0:0 | 11:0:4:0:0:16:0:0 | 11:0:11:0:0:0:1:1 | 0:0:1:0:0:0:0:0 | 0:0:1:0:0:0:0:0 | 0.087611998 | 1 | 1 | 1 | 1.62E-09 | 1 |
| 0:0:0:0:0 | 2:1:1:0:0 | 0:0:0:0:0 | 2:2:12:1:2 | 0:0:0:0:0 | 6:2:6:2:1 | 1 | 0.210077142 | 0.031463429 | 1 | 0.00031109 | 0.157605479 |
| 236:995:4:2:1 | 0:6:0:0:0 | 174:698:1:7:2 | 2:4:0:0:0 | 173:348:0:1:1 | 0:1:0:0:0 | 6.79E-05 | 1 | 1 | 0.000140817 | 0.036957127 | 1 |
| 911:237:4:2:1 | 6:0:0:0:0 | 633:176:1:7:2 | 0:0:0:0:0 | 346:173:0:1:1 | 0:0:0:0:0 | 1 | 1 | 1 | 0.000140817 | 1 | 1 |
| 274:698:3:2:1 | 16:4:1:0:0 | 190:484:1:6:2 | 14:5:0:0:0 | 178:327:0:1:1 | 0:1:0:0:0 | 1 | 1 | 1 | 1.74E-08 | 1.66E-07 | 1 |
| 273:695:4:2:1 | 20:3:0:0:0 | 191:480:1:6:2 | 16:4:0:0:0 | 178:325:0:1:1 | 0:3:0:0:0 | 1.85E-07 | 1 | 0.005703675 | 5.73E-13 | 2.90E-09 | 1 |
| 286:684:4:2:1 | 14:1:0:0:0 | 202:467:1:6:2 | 9:5:0:0:0 | 177:322:0:1:1 | 1:0:0:0:0 | 1.48E-08 | 1 | 0.005703675 | 7.61E-10 | 0.000157209 | 1 |
| 306:623:4:2:1 | 1:1:0:0:0 | 215:434:1:6:2 | 0:1:0:0:0 | 178:306:0:1:1 | 0:0:0:0:0 | 0.020772061 | 1 | 1 | 1 | 1 | 1 |
| 600:306:4:2:1 | 4:1:0:0:0 | 425:214:1:6:2 | 2:1:0:0:0 | 303:178:0:1:1 | 0:0:0:0:0 | 1 | 1 | 1 | 0.021353791 | 1 | 1 |
| 231:43:11:0 | 12:0:1:0 | 228:41:14:1 | 6:0:0:0 | 46:4:5:0 | 2:0:0:0 | 1 | 1 | 1 | 1.47E-06 | 0.000739192 | 1 |
| 138:31:4:0 | 26:6:3:0 | 134:30:10:1 | 31:2:1:0 | 40:2:2:0 | 0:0:0:0 | 1 | 1 | 1 | 8.27E-10 | 1.67E-16 | 1 |
| 86:1:24:0 | 7:1:1:0 | 83:6:18:1 | 8:0:1:0 | 24:0:2:0 | 1:0:0:0 | 1 | 1 | 1 | 0.004727276 | 0.000319008 | 1 |
| 8:0:2:4:0:0 | 5:3:0:4:0:0 | 5:3:1:1:0:0 | 0:0:0:0:0:0 | 3:0:0:1:1:0 | 0:0:0:0:0:0 | 1 | 1 | 1 | 0.028394475 | 1 | 1 |
| 0:0:0:0:1:0 | 0:15:0:3:9:0 | 1:5:2:1:3:0 | 0:0:0:0:0:0 | 0:3:0:0:1:0 | 0:0:0:0:0:0 | 1 | 1 | 0.066914403 | 9.30E-08 | 1 | 1 |
| 0:0:0:0:0:0 | 15:4:3:5:12:0 | 1:0:0:0:0:0 | 4:3:1:1:3:0 | 2:0:0:0:0:0 | 1:0:0:1:1:0 | 0.558277499 | 1 | 0.066914403 | 0.000195629 | 0.322819363 | 1 |
| 0:0:0:0:0:0 | 15:8:9:5:12:0 | 1:0:0:0:0:0 | 4:4:3:1:3:0 | 2:0:0:0:0:0 | 1:0:0:1:1:0 | 0.539746124 | 0.335876219 | 0.066914403 | 0.006163028 | 0.385681441 | 1 |
| 256:1034:65 | 0:07:01 | 83:757:15 | 1:01:00 | 49:403:7 | 0:00:00 | 1 | 1 | 1 | 0.006305042 | 1 | 1 |

| | | | | | | | | | | | |
|---|---|---|---|---|---|---|---|---|---|---|---|
| 3:418:11:19:219:7:19:5:0:2:0 | 0:0:0:0:0:0:1:2:1:0:0 | 2:378:14:8:414:11:17:6:1:2:1 | 0:2:0:0:1:0:0:6:0:0:0 | 2:73:2:2:122:6:5:5:0:1:0 | 0:0:0:0:1:0:0:0:0:0:0 | 1 | 1 | 1 | 1 | 2.22E-05 | 1 |
| 23:07 | 0:00 | 3:00 | 0:00 | 2:00 | 0:00 | 0.002247757 | 1 | 1 | 1 | 1 | 1 |
| 1:00 | 765:08:00 | 1:00 | 159:02:00 | 0:00 | 77:00:00 | 1.10E-31 | 1.78E-38 | 5.40E-31 | 1.49E-161 | 2.53E-34 | 9.69E-18 |
| 1479:69 | 7:00 | 323:14:00 | 0:00 | 103:04:00 | 0:00 | 1 | 1 | 1 | 0.010434472 | 1 | 1 |
| 1463:73 | 6:01 | 316:16:00 | 2:00 | 102:04:00 | 0:00 | 1 | 1 | 0.017871477 | 0.067682005 | 1 | 1 |
| 1458:75 | 11:01 | 314:16:00 | 4:00 | 102:04:00 | 0:00 | 1 | 1 | 0.017871477 | 0.005338322 | 1 | 1 |
| 1368:86 | 10:01 | 291:19:00 | 4:00 | 95:03:00 | 2:00 | 1 | 1 | 1 | 0.008900344 | 1 | 1 |
| 59:2:0:5:3:1:1 | 0:0:0:0:0:0:0 | 59:0:5:9:2:1:0 | 0:0:0:0:0:0:0 | 19:0:1:0:1:0:0 | 0:0:0:0:0:0:0 | 5.52E-13 | 6.19E-06 | 1 | 1 | 1 | 1 |
| 15:00 | 0:00 | 3:01 | 23:00 | 0:00 | 1:00 | 1 | 1 | 1 | 1 | 3.55E-06 | 1 |
| 7:30:6:0:4:1:0:0:2:2:2:1:0:0:0 | 0:0:0:0:0:0:0:0:0:0:0:0:0:0:0 | 1:2:0:0:0:0:0:0:0:0:0:0:0:0:0 | 4:29:4:0:10:0:0:1:0:0:1:0:1:1:1 | 0:4:2:0:4:0:0:0:0:0:0:0:0:0:0 | 1:6:0:0:0:0:0:0:0:0:0:0:0:0:0 | 1 | 1 | 1 | 1 | 2.74E-40 | 2.50E-09 |
| 0:0:0:0 | 11:1:0:1 | 2:0:0:0 | 0:0:0:0 | 0:0:0:0 | 0:0:0:0 | 0.293795914 | 1 | 1 | 3.18E-05 | 1 | 1 |
| 792:21:10 | 0:00:00 | 169:123:14 | 779:09:05 | 60:46:05 | 149:03:03 | 3.63E-69 | 5.45E-42 | 1 | 1 | 0 | 6.13E-59 |
| 791:15:09 | 1:01:00 | 783:06:05 | 164:01:14 | 149:02:03 | 58:01:05 | 1 | 1 | 1 | 1 | 1.89E-59 | 1.48E-20 |
| 790:16:10 | 0:00:00 | 168:121:13 | 782:08:05 | 60:45:05 | 147:01:03 | 1.94E-71 | 5.45E-42 | 1 | 1 | 0 | 1.18E-59 |
| 154:11:8:0 | 158:12:13:0 | 437:35:25:0 | 1:0:0:0 | 133:4:2:0 | 0:0:0:0 | 2.02E-47 | 3.53E-10 | 2.35E-29 | 1.98E-78 | 1 | 1 |
| 324:23:29:2 | 0:0:0:0 | 298:27:34:0 | 146:8:11:1 | 73:3:1:0 | 63:1:2:0 | 1 | 1 | 1 | 1 | 6.56E-75 | 1.12E-36 |
| 316:0:162:47:66:201:16:6:6:124:41:31:5:2:1:112:50:8:9:12:46:1:13:10:4 | 0:0:0:0:0:0:0:0:0:0:0:0:0:0:0:0:0:0:0:0:0:0:0:0:0 | 754:3:121:25:301:385:39:13:3:38:12:38:0:0:10:5:0:0:5:1:3:1:0:3:2 | 0:0:0:0:1:3:1:0:0:0:0:0:0:0:0:0:0:0:0:0:0:0:0:0:0 | 96:9:32:12:150:88:9:1:2:6:5:16:0:2:4:0:1:0:3:0:1:2:1:4:1 | 0:0:0:0:0:0:0:0:0:0:0:0:0:0:0:0:0:0:0:0:0:0:0:0:0 | 1 | 1 | 1 | 1 | 0.002190333 | 1 |
| 171:48:56:18:0:12:5:59:9:45:56:16:9:5:4:2 | 0:0:0:0:0:2:0:0:0:0:0:0:0:0:0:0 | 18:4:0:1:1:6:0:0:0:3:1:0:1:0:2:0 | 109:10:8:5:3:0:1:0:0:7:3:0:1:2:2:1 | 8:4:4:1:0:1:0:1:0:2:1:1:3:0:1:0 | 24:1:7:1:1:0:0:0:0:1:0:0:1:1:1:0 | 1 | 2.81E-12 | 1 | 1 | 5.33E-229 | 4.02E-39 |
| 334:150:55:14:29:16:151:46:20:5:9:132:59:53:56:16:3:1:0 | 0:0:0:0:0:0:0:0:1:0:0:0:0:0:0:0:0:0:0 | 206:13:1:10:6:1:19:4:8:0:2:2:0:2:1:0:1:1:0 | 617:100:7:30:11:5:26:10:7:1:2:5:0:9:3:0:2:4:1 | 31:8:4:2:0:1:5:4:1:0:0:0:1:3:1:1:0:1:0 | 67:18:7:8:1:1:1:1:0:0:0:0:0:1:0:0:1:0:0 | 1.20E-65 | 3.60E-212 | 1 | 1 | 0 | 8.35E-152 |
| 329:150:55:14:29:16:151:46:34:5:9:132:59:52:56:16:3:1:0 | 4:0:0:0:0:0:0:0:1:0:0:0:0:0:0:0:0:0:0 | 822:112:8:39:17:6:45:14:26:1:4:7:0:11:4:0:3:4:1 | 1:0:0:1:0:0:0:0:0:0:0:0:0:0:0:0:0:1:0 | 98:26:11:10:1:2:6:5:1:0:0:0:1:4:1:1:1:1:0 | 0:0:0:0:0:0:0:0:0:0:0:0:0:0:0:0:0:0:0 | 1 | 1 | 1 | 5.05E-06 | 1 | 1 |
| 155:0:75:46:26:8:11:9:73:19:51:3:5:13:1:0:62:0:26:10:34:31:0:1:0 | 173:0:74:20:28:6:17:7:75:27:85:2:4:19:1:0:69:0:31:6:21:19:1:1:0 | 813:3:110:311:8:39:17:6:45:14:105:1:4:31:1:5:7:0:0:0:4:11:0:3:1 | 2:0:0:0:0:0:0:0:0:0:0:0:0:0:0:0:0:0:0:0:0:0:0:0:0 | 97:8:26:153:11:8:1:2:6:5:18:0:0:5:2:1:0:1:1:1:1:4:0:1:0 | 0:0:0:0:0:0:0:0:0:0:0:0:0:0:0:0:0:0:0:0:0:0:0:0:0 | 1 | 1 | 4.10E-61 | 5.56E-281 | 1 | 1 |
| 275:84 | 0:00 | 294:107 | 34:16:00 | 141:04:00 | 10:01 | 5.15E-15 | 2.43E-11 | 1 | 1 | 0.014613092 | 0.009228162 |
| 19:3:1:1:1:0:4:1:1:0:0:0 | 8:1:0:1:1:1:1:0:0:0:0:0 | 40:11:0:4:10:3:13:1:2:0:1:1 | 0:0:0:0:0:0:0:0:0:0:1:0 | 15:2:2:0:1:2:9:1:0:0:0:0 | 0:0:0:0:0:0:0:0:0:0:0:0 | 1 | 1 | 0.011087152 | 1.10E-06 | 1 | 1 |
| 8:6:0:0:1:0:0:0 | 16:4:1:1:2:0:0:0 | 22:9:1:0:1:0:1:0 | 0:0:0:0:0:0:0:0 | 10:7:0:0:0:0:0:0 | 0:0:0:0:0:0:0:0 | 1 | 0.005806493 | 0.088875902 | 8.88E-12 | 1 | 1 |
| 0:0:0:0 | 4:0:1:0 | 0:0:0:0 | 8:1:0:0 | 0:0:0:0 | 1:0:0:0 | 1 | 1 | 0.01363685 | 0.04093502 | 0.000319008 | 1 |
| 0:00:00 | 0:00:00 | 0:01:00 | 0:00:00 | 0:00:00 | 0:00:00 | 0.034668522 | 1 | 1 | 1 | 1 | 1 |
| 0:0:0:0 | 79:3:2:1 | 0:0:0:0 | 37:1:1:1 | 0:0:0:0 | 14:0:0:2 | 1.92E-08 | 1 | 2.38E-34 | 3.17E-44 | 3.27E-20 | 4.87E-07 |
| 5:0:0:1 | 0:0:0:0 | 9:0:3:0 | 0:0:0:0 | 2:0:0:0 | 0:0:0:0 | 0.012449474 | 1 | 1 | 1 | 1 | 1 |
| 140:344:3:1653 | 0:1:0:4 | 5:4:0:18 | 0:0:0:0 | 2:1:0:7 | 0:0:0:0 | 1 | 1 | 1 | 0.04093502 | 1 | 1 |
| 139:345:3:1657 | 1:0:0:5 | 5:4:0:18 | 0:0:0:0 | 2:1:0:7 | 0:0:0:0 | 1 | 1 | 1 | 0.012726037 | 1 | 1 |
| 1254:13:31:6:33:8:11:12:2 | 6:0:1:0:0:0:0:0:0 | 911:15:29:16:15:5:9:5:2 | 2:0:0:0:0:0:0:0:0 | 300:4:14:8:7:1:2:0:0 | 0:0:0:0:0:0:0:0:0 | 1 | 1 | 1 | 5.31E-06 | 1 | 1 |
| 17:02:05 | 39:00:03 | 19:01:01 | 42:00:02 | 6:00:01 | 6:00:00 | 0.0086191 | 1 | 0.007094277 | 9.24E-15 | 1.10E-16 | 0.00371338 |

| | | | | | | | | | | | |
|---|---|---|---|---|---|---|---|---|---|---|---|
| 30:02:07 | 10:00:00 | 36:01:01 | 12:00:00 | 8:00:01 | 11:00:00 | 1 | 1 | 0.001636609 | 8.14E-05 | 1.29E-05 | 3.35E-05 |
| 28:02:07 | 28:00:01 | 32:01:01 | 30:00:01 | 10:00:01 | 10:00:00 | 0.00132829 | 1 | 0.00065969 | 1.47E-11 | 2.36E-12 | 8.62E-05 |
| 0:0:0:0:0:0:0:0:0:0 | 19:16:6:0:8:1:0:0:0 | 0:0:0:0:0:0:0:0:0:0 | 21:18:8:1:7:3:1:0:1 | 0:0:0:0:0:0:0:0:0:0 | 10:1:2:0:1:2:0:1:0 | 0.069647953 | 1 | 0.000954665 | 2.49E-13 | 3.63E-12 | 3.38E-06 |
| 11:1:0:3:0 | 7:2:1:2:0 | 5:1:1:2:0 | 31:12:6:10:1 | 9:3:0:0:3 | 17:3:1:5:0 | 7.07E-22 | 8.43E-12 | 6.60E-06 | 0.019952615 | 2.29E-08 | 6.04E-07 |
| 5:3:6:0:0 | 0:0:0:0:0 | 1:0:1:0:0 | 8:2:6:1:0 | 0:1:0:0:0 | 0:0:0:0:0 | 1 | 1 | 1 | 1 | 0.010654386 | 1 |
| 0:00:00 | 8:01:01 | 0:00:00 | 6:00:00 | 0:00:00 | 0:00:00 | 0.0086191 | 1 | 0.00065969 | 0.009816388 | 0.003657671 | 1 |
| 0:00:00 | 8:01:01 | 0:00:00 | 6:00:00 | 0:00:00 | 0:00:00 | 0.0086191 | 1 | 0.00065969 | 0.009816388 | 0.003657671 | 1 |
| 0:00:00 | 8:01:01 | 0:00:00 | 6:00:00 | 0:00:00 | 0:00:00 | 0.0086191 | 1 | 0.00065969 | 0.009816388 | 0.003657671 | 1 |
| 7:187:10:0 | 0:1:0:0 | 11:153:5:4 | 0:3:0:0 | 4:33:0:0 | 0:1:0:0 | 0.002557517 | 1 | 1 | 1 | 1 | 1 |
| 171:12:7:0:0 | 16:0:0:0:0 | 133:6:11:4:0 | 14:0:0:0:0 | 29:0:4:0:0 | 4:0:0:0:0 | 1 | 1 | 1 | 1.42E-12 | 6.85E-11 | 1 |
| 164:11:7:0:0 | 22:1:0:0:0 | 138:6:11:4:0 | 10:0:0:0:0 | 30:0:4:0:0 | 2:0:0:0:0 | 1 | 1 | 1 | 2.06E-16 | 1.12E-07 | 1 |
| 10:22:3:1:9:0 | 0:0:0:0:0:0 | 11:31:12:1:6:0 | 0:2:0:0:0:0 | 2:0:0:0:1:1 | 5:6:1:1:0:0 | 1 | 1 | 1 | 1 | 1 | 0.009222385 |
| 0:00:00 | 381:03:12 | 504:05:06 | 32:01:00 | 19:00:00 | 82:01:00 | 3.96E-102 | 1.41E-35 | 2.41E-66 | 3.56E-152 | 3.45E-13 | 1.66E-34 |
| 0:0:0:0:0:0 | 0:0:0:6:1:0 | 0:0:0:0:0:0 | 0:2:2:10:1:0 | 0:0:0:0:0:0 | 0:0:0:1:0:0 | 1 | 1 | 0.012154512 | 0.000265566 | 5.42E-05 | 1 |
| 11:0:0:0:1:1:0 | 13:0:0:0:0:0:0 | 15:0:2:1:2:1:0 | 20:0:3:1:3:0:0 | 1:0:0:0:0:0:0 | 0:0:1:0:0:0:0 | 1 | 1 | 1 | 3.35E-14 | 7.17E-15 | 1 |
| 136:04:00 | 0:00 | 305:10:00 | 1:00 | 70:07:00 | 0:00 | 2.63E-17 | 0.000285302 | 1 | 1 | 1 | 1 |
| 122:02:00 | 2:00 | 5:00 | 267:10:00 | 59:05:00 | 2:02 | 1 | 1 | 1 | 1 | 7.86E-53 | 1 |
| 7247:07:00 | 3:00 | 5990:02:00 | 8:00 | 1846:03:00 | 0:00 | 1 | 1 | 1 | 1 | 0.006667609 | 1 |
| 7242:07:00 | 8:00 | 5991:02:00 | 7:00 | 1845:03:00 | 1:00 | 1 | 1 | 1 | 0.006305042 | 0.011137275 | 1 |
| 7258:07:00 | 9:00 | 6015:02:00 | 4:00 | 1852:03:00 | 3:00 | 1 | 1 | 1 | 0.00379461 | 1 | 1 |
| 7257:07:00 | 9:00 | 6009:02:00 | 13:00 | 1853:03:00 | 2:00 | 1 | 1 | 1 | 0.00379461 | 0.00053673 | 1 |
| 7269:07:00 | 0:00 | 6022:02:00 | 5:00 | 1852:03:00 | 2:00 | 1 | 1 | 1 | 1 | 0.031296179 | 1 |
| 7252:13:07 | 8:00:00 | 5990:16:02 | 5:00:00 | 1851:02:03 | 2:00:00 | 1 | 1 | 1 | 0.000535794 | 0.009407211 | 1 |
| 7220:20:07 | 8:00:00 | 5961:25:02 | 2:00:00 | 1850:05:03 | 2:00:00 | 1 | 1 | 1 | 0.000535794 | 1 | 1 |
| 7207:21:07 | 9:00:00 | 5952:24:02 | 6:01:00 | 1848:05:03 | 1:00:00 | 1 | 1 | 1 | 0.000209242 | 0.015787883 | 1 |
| 7205:21:07 | 9:00:00 | 5943:25:02 | 9:00:00 | 1847:06:03 | 2:00:00 | 1 | 1 | 1 | 0.000209242 | 0.000224738 | 1 |
| 7184:22:07 | 15:00:00 | 5935:28:02 | 7:00:00 | 1840:06:03 | 2:00:00 | 1 | 1 | 1 | 7.25E-07 | 0.001445472 | 1 |
| 7173:23:07 | 7:00:00 | 5932:31:02 | 2:00:00 | 1833:06:03 | 4:00:00 | 1 | 1 | 1 | 0.00138895 | 1 | 1 |
| 7158:24:07 | 9:00:00 | 5914:31:02 | 8:00:00 | 1836:07:03 | 1:00:00 | 1 | 1 | 1 | 0.000209242 | 0.000571704 | 1 |
| 7157:24:07 | 10:00:00 | 5915:31:02 | 5:00:00 | 1835:07:03 | 1:00:00 | 1 | 1 | 1 | 8.14E-05 | 0.009407211 | 1 |
| 7153:25:07 | 11:00:00 | 5909:30:02 | 5:01:00 | 1837:08:03 | 0:00:00 | 1 | 1 | 1 | 3.18E-05 | 0.036538834 | 1 |
| 7153:25:07 | 11:00:00 | 5909:30:02 | 6:01:00 | 1835:08:03 | 2:00:00 | 1 | 1 | 1 | 3.18E-05 | 0.015787883 | 1 |
| 7129:26:07 | 9:00:00 | 5863:29:02 | 8:01:00 | 1827:08:03 | 3:00:00 | 1 | 1 | 1 | 0.000209242 | 0.00271654 | 1 |
| 7123:23:07 | 7:00:00 | 5849:24:02 | 7:03:00 | 1826:08:03 | 3:00:00 | 1 | 1 | 1 | 0.00138895 | 0.031234246 | 1 |
| 7055:15:07 | 9:00:00 | 5789:15:02 | 11:00:00 | 1825:04:03 | 2:00:00 | 1 | 1 | 1 | 0.000209242 | 3.36E-05 | 1 |
| 7013:07:06 | 11:00:00 | 5754:02:06 | 10:00:00 | 1817:03:02 | 6:00:00 | 1 | 1 | 1 | 3.18E-05 | 8.55E-05 | 0.00371338 |
| 7007:07:05 | 12:00:00 | 5756:02:04 | 5:00:00 | 1820:03:02 | 3:00:00 | 1 | 1 | 1 | 1.22E-05 | 0.009407211 | 1 |
| 6987:07:04 | 19:00:00 | 5747:02:04 | 5:00:00 | 1818:03:02 | 3:00:00 | 1 | 1 | 1 | 1.52E-08 | 0.009407211 | 1 |
| 6969:07:00 | 3:00 | 5719:02:00 | 6:00 | 1815:03:00 | 2:00 | 1 | 1 | 1 | 1 | 0.018436807 | 1 |
| 6923:06:07 | 7:00:00 | 5683:05:02 | 5:00:00 | 1804:01:03 | 0:00:00 | 1 | 1 | 1 | 0.00138895 | 0.009407211 | 1 |
| 6917:06:07 | 6:00:00 | 5669:05:02 | 10:00:00 | 1796:01:03 | 3:00:00 | 1 | 1 | 1 | 0.003552529 | 8.55E-05 | 1 |
| 6891:06:07 | 9:00:00 | 5654:06:02 | 5:00:00 | 1794:02:03 | 2:00:00 | 1 | 1 | 1 | 0.000209242 | 0.009407211 | 1 |
| 6884:06:07 | 12:00:00 | 5646:07:02 | 8:00:00 | 1792:02:03 | 2:00:00 | 1 | 1 | 1 | 1.22E-05 | 0.000571704 | 1 |
| 6844:06:07 | 13:00:00 | 5600:06:02 | 8:01:00 | 1780:02:03 | 3:00:00 | 1 | 1 | 1 | 4.83E-06 | 0.00271654 | 1 |
| 6:09 | 20:05 | 35:11:00 | 0:00 | 2:03 | 0:00 | 1 | 1 | 1 | 0.00379461 | 1 | 1 |
| 0:00:00 | 16:08:01 | 18:06:03 | 0:00:00 | 3:02:01 | 0:00:00 | 1 | 1 | 0.379895604 | 0.001752603 | 1 | 1 |
| 37:80:2:4:6:22:12:4:1 | 0:5:0:0:0:0:0:0:0 | 10:35:4:2:3:6:2:2:3 | 0:1:0:0:0:0:0:0:0 | 3:16:0:0:2:0:2:1:0 | 0:0:0:0:0:0:0:0:0 | 1 | 1 | 1 | 6.65E-06 | 1 | 1 |
| 901:550:20 | 2:01:00 | 1367:342:29 | 5:00:00 | 345:114:9 | 1:03:00 | 1 | 1 | 1 | 1 | 0.009407211 | 1 |
| 1221:17:51:56:2:12:9:2:9:2:1 | 6:0:0:0:0:29:0:0:0:0:0 | 893:20:30:37:1:3:7:2:1:3:0 | 2:0:0:0:0:17:0:0:0:0:0 | 248:7:9:3:0:0:2:0:2:1:0 | 0:0:0:0:0:1:0:0:0:0:0 | 1 | 1 | 1 | 4.09E-45 | 1.66E-26 | 1 |
| 18048:90:777:931:7:2 | 15:3:2:0:0:0 | 8923:74:534:426:0:1 | 9:1:1:0:0:0 | 2744:18:152:109:0:0 | 5:2:0:0:0:0 | 0.00145773 | 0.011213959 | 1 | 2.04E-09 | 4.59E-06 | 0.004619521 |
| 18055:781:933:134:7:2 | 34:0:0:9:0:0 | 8934:535:427:97:0:1 | 16:1:1:4:0:0 | 2749:154:109:19:0:0 | 2:0:0:1:0:0 | 0.001193772 | 1 | 0.156193138 | 1.47E-25 | 1.22E-09 | 1 |
| 18060:779:933:151:7:2 | 10:0:0:3:0:0 | 8931:534:426:102:0:1 | 4:1:0:4:0:0 | 2746:153:109:19:0:0 | 0:0:0:1:0:0 | 1 | 1 | 1 | 1.21E-06 | 0.029638508 | 1 |
| 781:18076:152:933:7:2 | 0:11:8:0:0:0 | 535:8942:105:428:0:1 | 0:8:4:0:0:0 | 154:2750:21:109:0:0 | 0:1:1:0:0:0 | 0.018731299 | 1 | 0.00028777 | 4.71E-07 | 6.98E-05 | 1 |

| | | | | | | | | | | | |
|---|---|---|---|---|---|---|---|---|---|---|---|
| 18051:777:931:185:7:2 | 31:4:2:0:0:0 | 8924:529:428:115:0:1 | 20:3:0:0:0:0 | 2749:154:108:25:0:0 | 2:0:1:0:0:0 | 1 | 4.32E-09 | 3.48E-05 | 5.68E-24 | 6.41E-16 | 1 |
| 18064:782:932:323:7:2 | 17:2:3:5:0:0 | 8934:531:426:184:0:1 | 13:0:2:8:0:0 | 2748:152:109:39:0:0 | 3:0:0:0:0:0 | 1 | 1 | 0.066914403 | 2.85E-08 | 6.12E-07 | 1 |
| 18065:781:361:934:7:2 | 16:0:0:1:0:0 | 8943:527:223:426:0:1 | 7:0:2:2:0:0 | 2751:148:43:109:0:0 | 1:0:0:0:0:0 | 1 | 0.154481828 | 0.000996291 | 6.24E-14 | 0.001877356 | 1 |
| 18054:935:384:781:7:2 | 26:1:22:3:0:0 | 8934:427:239:530:0:1 | 16:1:6:3:0:0 | 2747:109:46:152:0:0 | 4:0:0:0:0:0 | 1 | 0.165313539 | 5.78E-05 | 7.60E-16 | 7.16E-08 | 1 |
| 773:18077:937:414:7:2 | 3:5:0:0:0:0 | 533:8951:430:249:0:1 | 1:3:0:0:0:0 | 153:2749:109:46:0:0 | 0:3:0:0:0:0 | 1 | 1 | 1 | 0.005016446 | 1 | 1 |
| 18066:412:742:936:7 | 17:2:2:1:0 | 8943:249:518:430:0 | 10:0:0:0:0 | 2749:46:151:109:0 | 2:0:0:0:0 | 1 | 0.021065045 | 1 | 7.81E-09 | 1.12E-07 | 1 |
| 18075:711:937:414:7 | 10:7:0:0:0 | 8950:501:429:249:0 | 5:6:1:0:0 | 2751:147:109:46:0 | 0:0:0:0:0 | 1 | 1 | 0.005703675 | 4.06E-05 | 0.010871201 | 1 |
| 1527:223:80:4:11:1 | 3:1:0:0:0:0 | 840:99:102:23:5:1 | 4:1:1:0:0:0 | 550:62:30:3:7:0 | 2:0:0:0:0:0 | 1 | 1 | 1 | 1 | 0.041747173 | 1 |
| 1517:228:81:4:11:1 | 1:0:0:0:0:0 | 830:101:104:23:5:1 | 6:1:0:0:0:0 | 549:64:30:3:7:0 | 2:0:0:0:0:0 | 1 | 1 | 1 | 1 | 0.000284747 | 1 |
| 0:00 | 23:00 | 10:01 | 0:00 | 3:00 | 0:00 | 0.017290558 | 1 | 0.011433615 | 3.50E-06 | 1 | 1 |
| 241:12:23:23 | 0:0:0:0 | 118:43:10:2 | 20:3:3:2 | 10:3:1:0 | 42:12:2:1 | 1 | 1 | 1 | 1 | 1.06E-06 | 6.24E-16 |
| 24249:95:139:12:3 | 14:1:0:0:0 | 39483:213:104:46:39 | 35:0:1:0:0 | 8696:40:52:13:5 | 11:0:1:0:0 | 0.000755847 | 1 | 7.98E-07 | 7.61E-10 | 2.88E-27 | 1.97E-07 |
| 24509:118:139:12:3:3 | 16:0:0:0:0:0 | 39866:239:105:46:3:2 | 24:0:0:0:0:0 | 8791:48:53:13:2:1 | 1:0:0:0:0:0 | 1 | 1 | 0.00028777 | 3.19E-15 | 1.76E-23 | 1 |
| 24681:116:139:12:3:3 | 41:0:0:0:0:0 | 40226:231:105:46:38:2 | 71:2:0:0:0:0 | 8852:48:53:13:5:1 | 7:0:0:0:0:0 | 0.00022653 | 1 | 7.30E-09 | 1.75E-41 | 1.28E-70 | 3.38E-06 |
| 24694:74:138:12:4:3 | 36:2:0:0:0:0 | 40377:149:104:46:39:2 | 60:1:0:0:0:0 | 8866:33:53:13:5:1 | 6:0:0:0:0:0 | 1 | 1 | 5.17E-14 | 1.72E-33 | 3.04E-60 | 3.01E-05 |
| 24687:72:138:12:4:3 | 36:0:0:0:0:0 | 40373:139:104:46:39:2 | 72:1:0:0:0:0 | 8866:32:53:13:5:1 | 6:0:0:0:0:0 | 3.58E-07 | 1 | 5.14E-13 | 3.47E-36 | 4.51E-73 | 3.01E-05 |
| 24771:47:137:12:4:3 | 43:1:0:0:0:0 | 40507:95:104:46:38:2 | 70:1:0:0:0:0 | 8890:25:53:13:5:1 | 6:0:0:0:0:0 | 1 | 1 | 5.17E-14 | 3.56E-42 | 6.12E-71 | 3.01E-05 |
| 24818:47:136:12:4:3 | 36:1:1:0:0:0 | 40577:96:104:46:38:2 | 66:0:0:0:0:0 | 8905:25:53:13:5:1 | 6:0:0:0:0:0 | 0.00022653 | 1 | 3.48E-05 | 1.99E-33 | 3.96E-68 | 3.01E-05 |
| 24871:43:137:12:4:3 | 55:0:0:0:0:0 | 40677:82:104:46:38:2 | 101:0:0:0:0:0 | 8916:20:53:13:5:1 | 11:1:0:0:0:0 | 3.94E-10 | 1 | 9.48E-19 | 2.59E-56 | 1.27E-105 | 6.89E-09 |
| 25043:38:137:12:4:3 | 27:0:0:0:0:0 | 40926:75:104:46:38:2 | 55:0:0:0:0:0 | 8958:20:53:13:5:1 | 3:0:0:0:0:0 | 1 | 1 | 1 | 1.10E-26 | 2.28E-56 | 1 |
| 25119:34:137:12:4:3 | 39:0:0:0:0:0 | 41037:69:104:46:38:2 | 76:1:0:0:0:0 | 8964:18:53:13:5:1 | 14:0:0:0:0:0 | 2.84E-05 | 1 | 9.12E-13 | 2.31E-39 | 2.40E-77 | 4.55E-13 |
| 25233:137:12:7:4:3 | 34:0:0:1:0:0 | 41270:104:46:14:38:2 | 70:0:0:2:0:0 | 9009:53:13:1:5:1 | 4:0:0:1:0:0 | 1 | 1 | 5.22E-06 | 1.16E-32 | 1.49E-69 | 0.011761049 |
| 25224:137:10:2:4:3 | 34:0:0:2:0:0 | 41341:104:46:5:35:2 | 50:0:0:2:0:0 | 8999:53:13:0:5:1 | 2:0:0:0:0:0 | 1 | 0.000251091 | 4.67E-09 | 2.18E-31 | 3.01E-48 | 1 |
| 25479:135:10:0:4:3 | 49:0:0:1:0:0 | 41952:104:46:3:36:2 | 129:0:0:3:0:0 | 9059:52:13:0:5:1 | 12:0:0:0:0:0 | 1 | 3.04E-05 | 5.01E-12 | 1.58E-48 | 1.99E-131 | 4.53E-11 |
| 25617:135:10:0:4:2 | 36:0:0:1:0:0 | 42234:103:46:3:36:2 | 76:0:0:2:0:0 | 9088:51:13:0:4:1 | 12:1:0:0:0:0 | 1 | 1 | 4.67E-09 | 8.96E-35 | 5.93E-76 | 7.39E-10 |
| 25904:136:11:1:4:2 | 72:0:0:0:0:0 | 42740:103:46:2:38:2 | 121:0:0:1:0:0 | 9153:52:13:0:4:1 | 7:0:0:0:0:0 | 1 | 1 | 3.89E-06 | 2.56E-74 | 1.09E-125 | 3.38E-06 |
| 25922:136:11:1:4:2 | 59:0:0:0:0:0 | 42758:103:46:2:37:2 | 123:0:0:1:0:0 | 9145:52:13:0:4:1 | 15:0:0:0:0:0 | 0.00022653 | 0.000251091 | 3.48E-05 | 1.57E-60 | 7.84E-128 | 4.60E-14 |
| 25862:135:11:2:4:2 | 52:1:0:0:0:0 | 42800:103:46:6:37:2 | 98:0:0:1:0:0 | 9105:51:13:1:5:1 | 20:0:0:0:0:0 | 3.52E-05 | 1 | 5.17E-14 | 1.08E-51 | 6.48E-101 | 3.13E-19 |
| 25881:136:11:2:4:2 | 36:0:0:0:0:0 | 42819:103:46:6:37:2 | 81:0:0:1:0:0 | 9115:51:13:1:5:1 | 10:0:0:0:0:0 | 2.84E-05 | 1 | 5.14E-13 | 3.47E-36 | 1.04E-82 | 4.48E-09 |
| 25883:136:11:2:4:2 | 41:0:0:0:0:0 | 42838:102:46:6:37:2 | 81:1:0:1:0:0 | 9119:51:13:1:5:1 | 9:0:0:0:0:0 | 0.000253109 | 1 | 4.67E-09 | 1.75E-41 | 2.83E-81 | 4.17E-08 |
| 384:85:148:54 | 0:0:0:1 | 687:185:132:46 | 4:0:1:0 | 177:53:28:14 | 0:0:0:0 | 1 | 1 | 1 | 1 | 0.041786793 | 1 |
| 467:96:191:74 | 5:1:0:0 | 793:213:172:56 | 4:2:0:0 | 174:54:29:15 | 0:0:0:0 | 1 | 1 | 1 | 0.012726037 | 0.071750259 | 1 |
| 470:98:191:73 | 4:1:0:1 | 793:215:172:56 | 6:0:0:0 | 174:54:29:15 | 0:0:0:0 | 1 | 1 | 1 | 0.122686968 | 0.000739192 | 1 |
| 0:2:1:0:1:0 | 0:0:0:0:0:0 | 34:38:18:0:10:0 | 0:0:0:0:1:0 | 5:1:0:0:3:0 | 0:0:0:0:0:0 | 3.98E-103 | 1 | 1 | 1 | 1 | 1 |

| | | | | | | | | | | | |
|---|---|---|---|---|---|---|---|---|---|---|---|
| 2:0:2:1:0:1 | 0:0:0:0:0:0 | 0:0:0:0:0:0 | 37:34:17:1:0:11 | 0:0:0:0:0:0 | 2:5:0:0:0:3 | 2.58E-109 | 0.000823477 | 1 | 1 | 2.27E-14 | 0.030032732 |
| 9:00 | 0:00 | 11:00 | 0:00 | 6:01 | 0:00 | 0.017290558 | 1 | 1 | 1 | 1 | 1 |
| 9:00 | 0:00 | 11:00 | 0:00 | 6:01 | 0:00 | 0.029709415 | 1 | 1 | 1 | 1 | 1 |
| 72:34:60:112:129:106:3:30:11:0:0:12:24:9:21:1:6:26:29:2:2:4:1:0:2 | 8:6:7:37:18:14:0:1:4:1:0:6:0:0:4:0:3:14:7:0:0:0:0:1:0 | 208:72:79:84:10:101:19:23:13:2:0:6:17:12:47:3:1:37:14:3:0:0:1:0:0 | 30:6:9:14:3:7:0:0:1:0:3:3:0:0:9:0:1:7:3:0:0:0:0:1:0 | 10:4:8:10:3:7:0:2:1:0:1:1:2:0:0:0:2:4:1:0:1:0:0:0:0 | 0:0:0:0:0:2:0:0:0:0:1:0:0:0:1:0:0:0:0:0:0:0:0:0:0 | 4.06E-09 | 1.45E-06 | 0.21166456 | 1.57E-52 | 5.86E-43 | 1 |
| 36:24:47:41:3:5:12:39:4:14:14:0:0:24:9:2:1:0:11:27:1:4:2:0:4:2:0:9:4:1:2:0 | 2:0:6:2:0:1:2:3:0:2:1:0:0:0:0:0:0:0:1:0:1:1:0:0:0:0:0:0:0:0:0:0 | 87:23:1:27:2:4:23:33:2:13:7:2:0:17:12:12:1:1:19:21:2:7:1:1:3:2:0:2:0:2:0:0 | 3:1:0:2:0:1:4:3:0:1:0:0:0:0:0:0:0:0:1:0:0:0:1:0:0:0:0:0:0:0:0:0 | 6:2:2:4:1:1:2:4:3:3:0:0:0:2:0:0:0:1:0:2:0:0:0:0:0:1:0:1:0:0:0:1 | 0:1:0:0:0:0:0:1:0:0:0:0:0:0:0:0:0:0:0:0:0:0:0:0:0:0:0:0:0:0:0:0 | 0.030773955 | 1.19E-05 | 6.62E-05 | 4.20E-05 | 0.000757719 | 1 |
| 12:8:19:0:0:0:19:2:9:6:2:3:2:1:2:8:3:0:2:2:1:0 | 3:0:2:0:0:0:2:0:2:0:0:1:2:0:0:0:0:0:0:0:0:0 | 20:7:12:0:3:0:16:3:1:3:1:2:2:2:2:5:0:0:0:1:0:0 | 6:2:0:0:1:0:2:2:0:0:0:3:0:0:0:1:0:1:0:0:0:0 | 3:0:1:0:0:0:1:0:0:0:0:0:0:0:1:1:0:0:0:0:0:0 | 0:0:0:0:0:0:1:0:0:0:0:0:0:0:0:0:0:0:0:0:0:0 | 0.224886797 | 0.043435636 | 0.340552607 | 0.028903294 | 0.000117914 | 1 |
| 563:27:5:50 | 0:4:0:0 | 555:20:9:36 | 0:8:0:0 | 125:3:4:1 | 0:0:0:0 | 1 | 1 | 1 | 1 | 4.90E-05 | 1 |
| 2:00:00 | 516:07:07 | 0:00:00 | 422:04:05 | 0:00:00 | 78:00:01 | 1.88E-36 | 2.99E-31 | 8.65E-51 | 3.43E-211 | 1.09E-174 | 8.63E-33 |
| 1:16:0:3:0:2:1:1 | 1:9:0:3:0:1:0:0 | 0:9:0:8:0:2:0:1 | 0:0:0:0:0:0:0:0 | 3:3:1:1:0:1:0:0 | 0:0:0:0:0:0:0:0 | 1 | 1 | 0.199751792 | 5.03E-06 | 1 | 1 |
| 14:0:0:1 | 17:2:2:1 | 8:0:1:0 | 17:0:0:0 | 8:1:1:0 | 5:1:0:0 | 1 | 1 | 0.001211998 | 1.12E-06 | 1.63E-10 | 0.013325074 |
| 15:0:0:1 | 19:2:2:1 | 8:0:1:0 | 23:1:1:0 | 8:1:1:0 | 5:1:0:0 | 1 | 1 | 0.00010415 | 8.47E-08 | 2.27E-12 | 0.013325074 |
| 36:59:5:2 | 0:0:0:0 | 30:28:4:0 | 0:0:0:0 | 13:26:1:0 | 0:0:0:0 | 0.051305088 | 0.00573066 | 1 | 1 | 1 | 1 |
| 0:00 | 187:01:00 | 61:01:00 | 42:00:00 | 7:00 | 7:00 | 0.000492136 | 1.14E-10 | 4.63E-08 | 4.57E-41 | 2.99E-10 | 0.010989758 |
| 49153:459:69 | 41:00:00 | 21882:281:23 | 26:00:00 | 6909:36:17 | 11:00:00 | 1 | 1.97E-06 | 0.000517066 | 6.45E-18 | 1.81E-11 | 3.35E-05 |
| 49168:459:69 | 49:00:00 | 21897:281:23 | 20:00:00 | 6918:36:17 | 4:00:00 | 1 | 0.000225925 | 0.00065969 | 2.45E-21 | 6.12E-09 | 1 |
| 49166:459:69 | 47:00:00 | 21827:280:23 | 19:00:00 | 6914:36:17 | 9:00:00 | 1 | 4.28E-08 | 0.004168205 | 1.74E-20 | 1.58E-08 | 0.000224434 |
| 48876:461:69 | 19:00:00 | 21691:279:23 | 10:00:00 | 6921:35:17 | 1:00:00 | 1 | 0.008830794 | 1 | 1.52E-08 | 8.55E-05 | 1 |
| 48878:460:69 | 62:01:00 | 21703:282:23 | 24:00:00 | 6915:35:17 | 6:00:00 | 1 | 2.91E-07 | 0.082060324 | 4.36E-26 | 1.27E-10 | 0.00371338 |
| 48903:461:67 | 55:00:01 | 21688:282:23 | 27:00:00 | 6913:35:17 | 8:00:00 | 1 | 5.15E-06 | 3.26E-05 | 4.44E-23 | 6.80E-12 | 0.000575289 |
| 48700:461:66 | 55:00:01 | 21579:286:22 | 31:00:00 | 6892:36:17 | 3:00:00 | 1 | 5.15E-06 | 0.010070087 | 4.44E-23 | 1.39E-13 | 1 |
| 48669:461:66 | 41:00:00 | 21560:286:22 | 30:00:00 | 6889:36:17 | 5:00:00 | 1 | 2.91E-07 | 0.001636609 | 6.45E-18 | 3.76E-13 | 0.00927654 |
| 48504:461:65 | 121:00:00 | 21497:286:22 | 58:00:00 | 6880:36:17 | 11:00:00 | 0.000526708 | 1.60E-08 | 1.36E-10 | 2.79E-52 | 3.73E-25 | 3.35E-05 |
| 47991:459:65 | 68:00:00 | 21231:285:21 | 27:00:00 | 6837:36:17 | 6:00:00 | 1 | 0.000555359 | 0.004168205 | 1.66E-29 | 6.80E-12 | 0.00371338 |
| 47226:460:65 | 42:00:00 | 20794:279:19 | 17:00:00 | 6703:35:17 | 7:00:00 | 0.003376462 | 1.30E-05 | 0.000100319 | 2.43E-18 | 1.09E-07 | 0.001510787 |
| 3:0:0:0 | 6:1:0:0 | 5:1:2:0 | 3:0:0:1 | 0:0:0:0 | 0:0:0:0 | 1 | 1 | 1 | 0.003813544 | 1 | 1 |
| 1127:356:67 | 668:223:18 | 667:290:66 | 434:197:10 | 227:77:11 | 4:01:00 | 0.01605427 | 0.001383821 | 0.015717509 | 2.36E-157 | 2.54E-91 | 0.082974197 |
| 1863:607:62 | 37:10:00 | 1176:523:43 | 18:06:00 | 235:82:5 | 2:01:00 | 0.003376462 | 3.40E-05 | 7.19E-07 | 2.09E-10 | 5.38E-05 | 1 |
| 1880:614:62 | 24:04:00 | 1185:528:42 | 11:02:01 | 236:82:5 | 1:01:00 | 1 | 0.000555359 | 2.02E-05 | 5.27E-08 | 0.002319044 | 1 |
| 2106:625:54 | 92:01:00 | 1301:536:44 | 44:00:01 | 248:82:5 | 3:01:00 | 1 | 1.30E-05 | 0.001636609 | 5.46E-39 | 2.66E-18 | 1 |
| 2204:626:54 | 28:01:00 | 1349:537:43 | 8:00:00 | 249:83:5 | 3:00:00 | 1 | 0.000555359 | 1 | 1.47E-11 | 0.000571704 | 1 |
| 8:7:0:0:0:0:0:0:1:1:1:1:0:0:0 | 17:5:1:3:1:1:1:2:0:0:0:0:0:0:0 | 17:3:0:2:1:1:0:1:0:0:1:0:1:2:1 | 0:0:0:0:0:0:0:0:0:0:0:0:0:0:0 | 12:2:0:0:1:0:0:0:0:0:0:0:0:0:0 | 0:0:0:0:0:0:0:0:0:0:0:0:0:0:0 | 1 | 1 | 9.48E-19 | 4.45E-20 | 1 | 1 |
| 306:411:0 | 1:02:00 | 841:1224:1 | 1:05:00 | 277:37:00 | 0:00:00 | 1 | 1 | 1 | 1 | 0.036538834 | 1 |
| 312:312:0 | 1:03:00 | 861:988:1 | 1:07:00 | 282:36:00 | 0:00:00 | 1 | 1 | 1 | 1 | 0.006653127 | 1 |
| 384:159:0 | 5:01:00 | 996:477:2 | 19:10:01 | 301:26:00 | 1:00:00 | 1 | 1 | 1 | 0.035780388 | 0.0005272 | 1 |
| 0:0:0:0:0:0:0 | 8:2:4:0:0:0:0 | 11:5:4:0:0:1:1 | 10:3:2:0:0:0:0 | 0:0:0:0:0:0:0 | 5:0:0:0:2:0:0 | 0.248455623 | 1 | 1 | 0.000164516 | 2.82E-06 | 0.001989292 |
| 0:00:00 | 2:02:01 | 1:00:00 | 0:05:01 | 0:01:00 | 0:00:00 | 1 | 1 | 1 | 0.826026374 | 0.036538834 | 1 |
| 0:00 | 4:00 | 0:00 | 16:02 | 0:00 | 2:01 | 0.039691116 | 0.010484149 | 0.030109098 | 1 | 0.001526854 | 1 |
| 67:03:00 | 0:00 | 41:05:00 | 69:07:00 | 17:02 | 2:01 | 2.15E-06 | 0.000470572 | 0.030109098 | 1 | 4.20E-12 | 1 |
| 41:1:4:5 | 14:0:0:1 | 87:5:8:6 | 28:6:2:0 | 17:1:0:3 | 1:0:0:0 | 1 | 1 | 1 | 8.97E-08 | 1.84E-11 | 1 |

| | | | | | | | | | | | |
|---|---|---|---|---|---|---|---|---|---|---|---|
| 63:1:5:6 | 46:0:9:1 | 139:12:11:6 | 99:9:7:1 | 18:1:1:3 | 3:0:0:0 | 0.012449474 | 1 | 0.044045542 | 4.33E-21 | 6.73E-48 | 1 |
| 5:69:1:6 | 8:39:1:1 | 12:150:14:6 | 5:72:5:1 | 1:21:1:3 | 1:15:0:1 | 5.18E-08 | 0.00031277 | 1.15E-05 | 9.16E-17 | 1.27E-35 | 1.66E-07 |
| 176:3:17:8 | 4:0:0:0 | 356:28:27:9 | 7:0:0:0 | 42:1:2:5 | 2:0:0:0 | 1 | 1 | 1 | 1 | 0.000193006 | 1 |
| 8:0:21:3:4:18:0 | 0:0:0:0:0:0:0 | 10:4:20:23:11:33:3 | 9:1:10:5:5:18:6 | 0:1:0:0:0:0:0 | 4:0:2:4:0:4:2 | 1 | 1 | 1 | 1 | 0.001420221 | 0.215856467 |
| 3:2:0:0:0:0 | 0:0:0:0:0:0 | 0:0:0:0:0:0 | 23:5:1:1:2:1 | 0:0:0:0:0:0 | 3:0:1:0:0:0 | 1 | 1 | 1 | 1 | 6.48E-13 | 1 |
| 0:00:00 | 17:00:00 | 194:01:07 | 0:00:01 | 27:00:00 | 0:00:00 | 1 | 1 | 5.13E-11 | 1.04E-07 | 1 | 1 |
| 0:00:00 | 17:00:00 | 193:08:01 | 1:00:00 | 27:01:00 | 0:00:00 | 1 | 1 | 5.13E-11 | 1.04E-07 | 1 | 1 |
| 164:184:2:16:13:3:6:8:8:5:1:3:6 | 0:0:0:1:0:0:0:0:0:0:0:0:0 | 243:117:6:9:10:0:11:5:8:0:0:0:1 | 252:146:8:11:7:0:7:7:1:7:0:3:4 | 88:66:9:5:9:1:4:1:6:0:0:0:0 | 1:1:0:0:0:0:0:0:0:0:0:0:0 | 1 | 1 | 1 | 1 | 0 | 1 |
| 2:21:61:9:1:1:2:0:0 | 0:0:0:0:0:0:0:0:0 | 7:28:180:37:0:4:0:8:2 | 0:0:0:0:0:0:0:1:0 | 2:14:11:1:0:1:0:0:0 | 0:0:0:0:0:0:0:0:0 | 6.67E-06 | 1 | 1 | 1 | 1 | 1 |
| 0:0:0:1:0:0 | 2:3:19:2:1:0 | 0:0:0:0:0:0 | 4:3:22:2:1:1 | 0:0:0:0:0:0 | 4:1:2:1:1:0 | 0.039691116 | 1 | 1 | 1.37E-10 | 3.12E-11 | 0.288398333 |
| 182:25:10:5 | 3:0:0:0 | 356:49:9:16 | 6:0:0:0 | 43:12:4:4 | 2:0:0:0 | 1 | 1 | 1 | 1 | 0.000739192 | 1 |
| 25:10:183:5 | 0:0:1:0 | 49:8:351:17 | 0:1:5:0 | 12:4:43:4 | 0:0:1:0 | 1 | 1 | 1 | 1 | 0.013499631 | 1 |
| 170:21:8:6 | 11:4:0:0 | 332:41:8:16 | 19:8:0:1 | 43:11:4:4 | 0:0:0:0 | 1 | 1 | 1 | 0.000143304 | 8.08E-07 | 1 |
| 10:104:5:6 | 0:5:1:0 | 30:227:6:16 | 0:6:0:1 | 9:35:2:3 | 0:1:0:1 | 1 | 1 | 1 | 0.012726037 | 0.003971575 | 1 |
| 6:7:1:3 | 19:0:2:0 | 11:15:6:3 | 44:0:4:0 | 5:3:4:0 | 7:0:1:1 | 3.02E-06 | 0.002652154 | 1 | 4.88E-10 | 6.22E-24 | 0.004954386 |
| 0.828518519 | 19:00:00 | 24:232:0 | 43:01:00 | 5:48:00 | 5:00:00 | 1 | 1 | 1 | 1.52E-08 | 7.03E-18 | 0.00927654 |
| 0:00:00 | 1:01:00 | 0:00:00 | 6:00:02 | 0:00:00 | 2:01:00 | 1 | 1 | 1 | 1 | 0.036538834 | 1 |
| 42:73:17:5:27:2:1:0:0:0:1:0 | 0:0:0:0:0:0:0:0:0:0:0:0 | 0:0:0:0:0:0:0:0:0:0:0:0 | 46:34:7:8:16:2:0:1:2:1:0:0 | 0:0:0:0:0:0:0:0:0:0:0:0 | 35:17:0:5:6:0:0:0:0:1:0:0 | 1 | 1 | 1 | 1 | 6.50E-48 | 2.18E-42 |
| 42:72:18:5:2:1:10:0:0:1:1:0 | 0:1:0:0:0:0:0:0:0:0:0:0 | 0:0:0:0:0:0:0:0:0:0:0:0 | 46:36:7:8:2:0:6:2:2:1:0:0 | 0:0:0:0:0:0:0:0:0:0:0:0 | 35:15:0:5:0:0:5:0:0:1:0:0 | 1 | 1 | 1 | 1 | 1.35E-52 | 6.06E-43 |
| 4:41:0:1:0:0 | 0:3:0:0:0:0 | 66:369:0:2:7:1 | 1:18:0:0:0:0 | 5:9:0:0:0:0 | 0:0:0:0:0:0 | 1 | 1 | 1 | 1 | 6.58E-16 | 1 |
| 85:46:2:7:1:4 | 3:0:4:0:0:0 | 326:164:14:30:11:2 | 13:0:16:0:0:0 | 84:3:4:4:2:3 | 0:0:0:0:0:0 | 1 | 1 | 1 | 0.016364524 | 6.88E-11 | 1 |
| 20:05 | 0:00 | 0:00 | 35:05:00 | 0:00 | 1:01 | 1 | 1 | 1 | 1 | 4.59E-06 | 1 |
| 19:01:00 | 0:00:00 | 0:00:00 | 35:01:01 | 0:00:00 | 1:01:00 | 1 | 1 | 1 | 1 | 1.12E-13 | 1 |
| 0:05 | 0:00 | 0:01 | 2:14 | 0:00 | 0:01 | 1 | 1 | 1 | 1 | 0.003951718 | 1 |
| 0:06 | 0:00 | 0:00 | 0:13 | 0:00 | 0:02 | 1 | 1 | 1 | 1 | 0.00053673 | 1 |
| 5:01:09 | 0:00:00 | 0:00:02 | 1:00:06 | 0:00:00 | 0:00:00 | 1 | 1 | 1 | 1 | 0.015787883 | 1 |
| 42988:238:47 | 30:00:00 | 49199:405:79 | 24:00:00 | 19048:136:48 | 12:00:00 | 2.92E-39 | 0.001383821 | 4.60E-08 | 3.30E-13 | 1.27E-10 | 1.31E-05 |
| 43013:240:47 | 15:01:00 | 49219:406:79 | 15:02:00 | 19057:136:48 | 15:00:00 | 2.19E-20 | 0.003558706 | 0.000100319 | 4.03E-06 | 1.72E-05 | 7.25E-07 |
| 43057:239:47 | 69:03:00 | 49314:408:80 | 68:00:00 | 19173:135:47 | 32:00:00 | 2.08E-137 | 1.14E-08 | 4.81E-13 | 1.62E-27 | 1.85E-29 | 5.97E-14 |
| 86286:476:94 | 122:06:00 | 98884:814:158 | 130:00:00 | 38518:270:96 | 44:00:00 | 3.10E-238 | 6.23E-13 | 2.56E-21 | 6.17E-48 | 3.30E-56 | 4.54E-19 |
| 213:43356:47 | 6:52:00 | 380:49729:82 | 4:56:00 | 120:19387:48 | 2:22:00 | 1.50E-96 | 0.004670383 | 4.49E-09 | 2.78E-18 | 3.45E-21 | 2.72E-08 |
| 43383:207:47 | 45:08:00 | 49762:369:82 | 61:07:00 | 19402:119:49 | 22:02:00 | 2.34E-83 | 0.008830794 | 1.75E-06 | 2.50E-14 | 2.43E-21 | 2.72E-08 |
| 87102:66:114 | 98:00:00 | 100054:98:196 | 96:02:00 | 39052:54:98 | 18:02:00 | 5.56E-163 | 2.51E-16 | 2.15E-16 | 2.06E-42 | 7.27E-40 | 1.07E-06 |
| 36:190:667:0:355:33:81:2:33:10:18 | 1:1:1:0:5:0:0:0:1:0:0 | 20:206:331:0:146:47:64:2:24:7:1 | 0:0:1:0:0:0:0:0:0:0:0 | 1:23:33:0:20:1:14:0:1:0:1 | 0:0:1:0:0:0:0:0:0:0:0 | 1 | 1 | 1 | 0.004453753 | 1 | 1 |
| 66:368:1242:0:684:56:152:286:4:18:34 | 0:0:6:0:0:2:0:0:0:0:0 | 34:388:600:0:278:74:120:258:4:14:2 | 0:2:2:0:2:0:0:0:0:0:0 | 2:52:60:0:36:2:28:14:0:0:2 | 0:0:0:0:0:0:0:0:0:0:0 | 0.000989986 | 1 | 1 | 2.14E-06 | 0.112785534 | 1 |
| 602:183:332:29:138:32:0:75:2:14:9 | 20:1:8:0:4:1:0:1:0:3:0 | 285:183:137:34:127:15:0:56:2:1:7 | 17:11:2:3:2:2:0:4:0:0:0 | 27:24:18:1:7:1:0:12:0:1:0 | 3:1:0:0:0:0:0:2:0:0:0 | 3.75E-06 | 0.000656642 | 0.034449441 | 2.83E-17 | 3.03E-12 | 0.038125841 |

| | | | | | | | | | | | |
|---|---|---|---|---|---|---|---|---|---|---|---|
| 1086:324:60:0:610:126:232:4:16:26:54 | 56:18:0:0:24:16:12:0:2:2:2 | 520:338:24:0:238:96:226:4:12:2:62 | 34:12:0:0:20:10:4:0:2:0:4 | 44:46:2:0:32:22:14:0:0:2:2 | 8:0:0:0:2:2:0:0:0:0:0 | 1.21E-07 | 2.04E-06 | 7.84E-05 | 1.04E-44 | 3.87E-26 | 1.33E-07 |
| 0:6:1:2:11:1:11:1:2 | 7:4:0:0:1:0:1:0:0 | 2:2:0:2:12:0:11:1:1 | 0:1:0:0:1:0:2:0:0 | 0:0:0:0:5:0:7:0:0 | 0:0:0:0:0:0:0:0:0 | 1 | 1 | 1 | 9.38E-05 | 1 | 1 |
| 0:1:0:1:31:10:1:1:2 | 0:0:0:0:0:0:0:0:0 | 0:0:0:0:1:0:0:0:0 | 0:1:0:1:14:12:0:1:0 | 0:0:0:0:1:0:0:0:0 | 0:0:0:0:7:7:0:0:0 | 1 | 1 | 1 | 1 | 6.26E-13 | 1.71E-07 |
| 290:06:22 | 60:00:12 | 280:10:04 | 36:00:00 | 78:08:04 | 4:00:02 | 0.003376462 | 1 | 1 | 2.43E-18 | 1.04E-15 | 0.147810323 |
| 552:06:54 | 40:00:00 | 450:10:06 | 20:00:02 | 96:08:06 | 10:00:00 | 1.24E-05 | 0.003558706 | 0.146375793 | 1.71E-17 | 1.68E-07 | 8.62E-05 |
| 3:290:30:0 | 0:4:3:0 | 5:225:6:0 | 0:5:1:0 | 4:54:3:0 | 0:1:0:0 | 1 | 1 | 1 | 0.072263067 | 0.013499631 | 1 |
| 0:0:2:0:0:0 | 352:14:100:4:2:0 | 648:36:94:0:2:0 | 0:0:0:0:0:0 | 16:2:2:0:0:0 | 0:0:0:0:0:0 | 3.12E-91 | 2.38E-65 | 2.96E-294 | 1.85E-262 | 1 | 1 |
| 290:76:82:0:0:0:0 | 10:0:0:0:0:0:0 | 330:86:78:0:2:2:2 | 2:0:0:0:0:0:0 | 68:2:4:0:0:0:0 | 4:0:0:0:0:0:0 | 0.00373202 | 1 | 1 | 1.36E-10 | 1 | 1 |
| 1:1:50:15:0 | 0:0:0:0:0 | 4:5:48:29:1 | 0:0:0:0:0 | 0:3:26:3:0 | 0:0:0:0:0 | 0.000660439 | 0.034743704 | 1 | 1 | 1 | 1 |
| 52:1:2:0:14 | 0:0:0:0:0 | 41:3:5:1:24 | 10:2:1:0:3 | 18:0:3:0:1 | 9:0:0:0:2 | 1 | 1 | 1 | 1 | 0.000969656 | 3.01E-05 |
| 0:0:0:0:0 | 84:4:2:2:0 | 0:0:0:0:0 | 40:0:0:0:2 | 0:0:0:0:0 | 18:0:0:2:0 | 6.77E-21 | 1 | 4.97E-14 | 5.68E-61 | 2.21E-30 | 4.58E-12 |
| 19:1059:350:3:10:3:1:0 | 4:7:0:0:0:0:0:0 | 26:2142:641:7:13:5:0:1 | 4:10:0:0:0:0:0:0 | 3:727:76:2:2:3:1:1 | 0:2:0:0:0:0:0:0 | 1 | 1 | 1 | 1.27E-05 | 1.46E-08 | 1 |
| 1054:349:3:15:10:3:1:0 | 7:0:0:3:0:0:0:0 | 2127:638:7:20:13:5:0:1 | 14:1:0:5:0:0:0:0 | 726:76:2:2:2:3:1:1 | 1:0:0:0:0:0:0:0 | 1 | 1 | 1 | 1.21E-05 | 9.19E-12 | 1 |
| 976:45:279:3:12:2:11:0:1:0 | 0:2:3:0:0:0:0:0:0:0 | 2003:86:531:7:34:14:14:0:0:1 | 5:1:2:0:0:0:0:0:0:0 | 693:10:59:2:23:0:2:0:1:1 | 2:0:1:0:0:0:0:0:0:0 | 1 | 1 | 1 | 0.015816907 | 0.000877508 | 1 |
| 1874:22:246:26:452:6:0:0:2:0 | 0:0:6:0:0:0:0:0:0:0 | 3806:30:430:72:820:14:24:0:0:2 | 10:0:6:0:10:0:2:0:0:0 | 1360:4:84:44:90:4:2:0:2:2 | 2:0:4:0:0:0:0:0:0:0 | 1 | 1 | 1 | 4.92E-08 | 1.07E-08 | 0.002015081 |
| 930:12:175:19:200:3:2:0:1:0 | 4:0:1:0:3:0:0:0:0:0 | 1878:15:346:41:374:7:12:0:0:1 | 1:0:1:0:5:0:0:0:0:0 | 673:2:72:22:41:2:1:0:1:1 | 1:0:0:0:0:0:0:0:0:0 | 1 | 1 | 1 | 0.004939017 | 0.00041677 | 1 |
| 897:265:12:31:153:3:2:0:1:0:0 | 3:4:0:0:3:0:0:0:0:0:0 | 1795:511:15:61:256:7:12:0:1:0:1 | 4:5:0:0:3:0:0:0:0:0:0 | 660:122:2:24:34:2:1:0:1:0:1 | 1:3:0:0:1:0:0:0:0:1:0 | 1 | 1 | 1 | 0.003257345 | 0.000350713 | 0.110671218 |
| 921:282:12:32:157:3:2:0:1:0:0 | 2:2:0:0:0:0:0:0:0:0:0 | 1829:529:15:64:259:7:12:0:1:0:1 | 2:1:0:0:3:0:0:0:0:0:0 | 668:128:2:24:34:2:1:0:1:0:1 | 1:0:0:0:0:0:0:0:0:1:0 | 1 | 1 | 1 | 1 | 0.039232735 | 1 |
| 912:342:49:2:134:3:3:0:1:0:0 | 2:0:0:7:0:0:0:0:0:0:0 | 1798:633:92:4:227:7:11:0:1:1:0 | 5:0:0:7:2:0:0:0:0:0:0 | 657:155:28:0:29:2:1:0:1:1:1 | 2:0:0:1:0:0:0:0:0:0:0 | 1 | 1 | 1 | 5.84E-08 | 1.89E-06 | 1 |
| 912:342:49:2:134:3:3:0:1:0:0 | 2:0:0:7:0:0:0:0:0:0:0 | 1798:633:92:4:227:7:11:0:1:1:0 | 5:0:0:7:2:0:0:0:0:0:0 | 657:155:28:0:29:2:1:0:1:1:1 | 2:0:0:1:0:0:0:0:0:0:0 | 1 | 1 | 1 | 5.84E-08 | 1.89E-06 | 1 |
| 913:342:49:2:134:3:2:0:1:0:0 | 1:0:0:7:1:0:1:0:0:0:0 | 1796:633:92:4:225:7:11:0:1:1:0 | 5:0:0:7:4:0:0:0:0:0:0 | 659:155:28:0:28:2:1:0:1:1:1 | 0:0:0:1:1:0:0:0:0:0:0 | 1 | 1 | 1 | 1.99E-06 | 3.33E-06 | 1 |
| 892:388:61:110:3:3:0:1:0:0 | 3:0:0:0:0:0:0:0:0:0 | 1755:719:131:183:7:10:0:1:1:0 | 8:4:0:1:0:0:0:0:0:0 | 660:177:37:31:2:1:0:1:1:2 | 0:1:0:0:0:0:0:0:0:0 | 1 | 1 | 1 | 1 | 3.56E-07 | 1 |
| 905:152:12:463:3:85:4:0:1:0:0 | 0:1:0:1:0:2:0:0:0:0:0 | 1795:366:15:889:7:142:5:0:1:1:0 | 2:1:0:0:0:4:1:0:0:0:0 | 682:86:2:199:2:26:1:0:1:1:1 | 0:2:0:0:0:0:0:0:0:0:0 | 1 | 1 | 1 | 1 | 0.018102765 | 1 |
| 0:0:0:0 | 16:6:2:0 | 0:0:0:0 | 4:2:0:2 | 0:0:0:0 | 0:2:0:2 | 1 | 1 | 1 | 2.53E-05 | 0.277789695 | 1 |
| 4:8:0:0 | 2:8:2:0 | 0:0:0:0 | 2:4:0:2 | 0:0:0:0 | 2:0:0:2 | 1 | 1 | 1 | 0.009770702 | 0.277789695 | 1 |
| 940:04:02 | 8:02:00 | 1784:02:00 | 12:00:00 | 398:00:00 | 2:00:00 | 1 | 1 | 0.004168205 | 0.007425864 | 1.29E-05 | 1 |
| 20:04 | 22:08 | 180:12:00 | 150:18:00 | 26:02:00 | 24:06:00 | 1 | 1 | 1 | 0.013351848 | 1.32E-23 | 0.001674537 |
| 20:04 | 22:08 | 168:12:00 | 148:14:00 | 26:02:00 | 24:04:00 | 1 | 1 | 1 | 0.013351848 | 3.72E-25 | 0.000277862 |
| 20:04 | 18:08 | 164:12:00 | 148:14:00 | 26:02:00 | 24:04:00 | 1 | 1 | 1 | 0.057801533 | 3.72E-25 | 0.000277862 |

| | | | | | | | | | | | |
|---|---|---|---|---|---|---|---|---|---|---|---|
| 5:00:00 | 0:00:00 | 0:00:00 | 9:01:00 | 0:00:00 | 3:00:00 | 0.003376462 | 1 | 1 | 1 | 0.001091486 | 1 |
| 5:00:00 | 0:00:00 | 0:00:00 | 10:01:00 | 0:00:00 | 3:00:00 | 0.003376462 | 1 | 1 | 1 | 0.000446416 | 1 |
| 9:00:00 | 0:00:00 | 0:00:00 | 21:00:00 | 0:00:00 | 4:00:00 | 0.000526708 | 1 | 1 | 1 | 2.35E-09 | 1 |
| 4:50:04 | 2:06:02 | 46:80:2 | 10:06:00 | 4:08:00 | 2:04:00 | 0.034668522 | 1 | 1 | 0.21701746 | 0.01179186 | 0.147810323 |
| 5:04 | 0:00 | 2:06 | 0:00 | 3:02 | 0:00 | 0.000366557 | 1 | 1 | 1 | 1 | 1 |
| 2:02 | 0:00 | 0:01 | 0:04 | 0:01 | 0:01 | 0.024185881 | 1 | 1 | 1 | 1 | 1 |
| 2:0:0:2:0 | 8:2:2:0:0 | 2:0:0:0:0 | 4:2:2:0:0 | 0:0:0:0:0 | 0:2:0:0:0 | 1 | 1 | 1 | 0.001863031 | 0.150388493 | 1 |
| 0:0:0:0:0:0:0:0:0:0:0 | 0:8:20:2:34:2:10:6:6:2:2 | 0:0:0:0:0:0:0:0:0:0:0 | 6:34:18:6:94:18:16:4:8:12:2 | 0:0:0:0:0:0:0:0:0:0:0 | 2:4:20:6:46:6:4:2:4:2:0 | 4.05E-05 | 1.49E-06 | 1.57E-27 | 3.31E-21 | 1.98E-67 | 4.58E-38 |
| 72:40:00 | 30:00:00 | 84:50:00 | 18:00 | 16:36 | 2:00 | 1 | 1 | 1 | 1.08E-07 | 4.38E-05 | 1 |
| 11:20 | 5:00 | 16:25 | 3:00 | 3:18 | 1:00 | 1 | 1 | 1 | 0.030173969 | 1 | 1 |
| 30:2:2:1 | 26:2:0:0 | 2:0:0:0 | 43:7:2:1 | 5:0:0:0 | 17:2:1:1 | 1 | 1 | 3.59E-06 | 2.40E-14 | 8.15E-19 | 2.88E-07 |
| 0:00:00 | 0:00:00 | 0:00:00 | 0:00:00 | 0:00:00 | 0:00:00 | 0.023899761 | 1 | 1 | 1 | 1 | 1 |
| 0:00 | 0:00 | 0:00 | 6:00 | 0:00 | 0:00 | 4.36E-32 | 1 | 0.017871477 | 1 | 0.018436807 | 1 |
| 0:00 | 0:00 | 0:00 | 5:00 | 0:00 | 0:00 | 3.63E-33 | 1 | 0.017871477 | 1 | 0.031296179 | 1 |
| 4635:84:9:19 | 9:0:0:0 | 11228:289:15:11 | 13:1:0:0 | 2954:56:5:10 | 1:0:0:0 | 1 | 1 | 1 | 1.18E-05 | 3.64E-07 | 1 |
| 4636:82:9:19 | 1:0:0:0 | 11228:289:15:11 | 5:4:0:0 | 2952:58:5:10 | 1:0:0:0 | 1 | 1 | 1 | 1 | 0.032544232 | 1 |
| 4625:83:9:19 | 4:2:0:0 | 11199:302:15:10 | 16:2:0:1 | 2948:64:5:10 | 5:0:0:0 | 0.002557517 | 1 | 1 | 0.070826298 | 1.95E-07 | 0.002841239 |
| 4585:76:9:19 | 5:0:0:0 | 11134:299:15:10 | 17:2:0:0 | 2944:65:5:10 | 2:0:0:0 | 1 | 1 | 1 | 0.002648305 | 8.41E-09 | 1 |
| 4579:76:9:19 | 3:0:0:0 | 11127:305:15:10 | 10:1:0:0 | 2938:68:5:10 | 4:0:0:0 | 0.000677933 | 1 | 1 | 1 | 2.23E-05 | 1 |
| 4575:76:9:19 | 5:0:0:0 | 11119:307:15:10 | 13:0:0:0 | 2939:68:5:10 | 2:0:0:0 | 1 | 1 | 1 | 0.002648305 | 4.74E-08 | 1 |
| 4560:91:9:18 | 5:0:0:0 | 11080:335:15:10 | 16:2:0:0 | 2929:76:5:10 | 2:0:0:0 | 1 | 1 | 1 | 0.002648305 | 3.29E-08 | 1 |
| 4558:91:9:18 | 7:0:0:0 | 11076:337:15:10 | 18:2:0:0 | 2930:77:5:10 | 1:0:0:0 | 1 | 1 | 1 | 0.000182133 | 2.16E-09 | 1 |
| 4558:92:9:18 | 5:0:0:0 | 11078:346:15:10 | 14:0:0:0 | 2926:80:5:10 | 4:0:0:0 | 1.19E-05 | 1 | 1 | 0.002648305 | 1.16E-08 | 1 |
| 4560:100:9:17 | 3:0:0:1 | 11054:362:15:10 | 12:0:0:0 | 2922:78:5:10 | 4:0:0:0 | 1 | 1 | 1 | 1 | 1.91E-07 | 1 |
| 4562:100:9:18 | 1:0:0:0 | 11059:359:15:10 | 11:3:0:0 | 2922:78:5:10 | 4:0:0:0 | 1 | 1 | 1 | 1 | 7.18E-05 | 1 |
| 101:4560:9:18 | 0:1:0:0 | 367:11059:15:10 | 1:4:0:0 | 80:2925:5:10 | 0:1:0:0 | 1 | 1 | 1 | 1 | 0.041786793 | 1 |
| 4559:101:9:18 | 1:4:0:0 | 11061:369:15:10 | 5:6:0:0 | 2926:82:5:10 | 0:1:0:0 | 0.068525294 | 1 | 1 | 0.04093502 | 0.014460454 | 1 |
| 4557:105:9:18 | 3:0:0:0 | 11053:379:15:10 | 10:1:0:0 | 2923:83:5:10 | 2:0:0:0 | 1 | 1 | 1 | 1 | 2.23E-05 | 1 |
| 4551:105:9:18 | 2:0:0:0 | 11038:380:15:10 | 15:0:0:0 | 2922:83:5:10 | 1:0:0:0 | 0.002557517 | 1 | 1 | 1 | 2.85E-09 | 1 |
| 4552:113:9:18 | 4:0:0:0 | 11045:390:15:10 | 14:1:0:0 | 2923:85:5:10 | 3:0:0:0 | 0.003671241 | 1 | 1 | 1 | 9.42E-08 | 1 |
| 4540:125:9:18 | 5:1:0:0 | 11030:420:15:10 | 13:1:0:0 | 2923:93:5:10 | 0:1:0:0 | 0.042679166 | 1 | 1 | 0.012726037 | 3.64E-07 | 1 |

| | | | | | | | | | | | |
|---|---|---|---|---|---|---|---|---|---|---|---|
| 4531:131:9:18 | 5:2:0:0 | 11008:442:15:10 | 12:3:0:0 | 2916:99:5:10 | 1:1:0:0 | 1 | 0.154096241 | 1 | 0.027461316 | 2.17E-05 | 1 |
| 146:4491:9:18 | 1:4:0:0 | 508:10917:15:9 | 3:12:0:0 | 113:2887:5:10 | 1:3:0:0 | 0.039854726 | 1 | 1 | 0.04093502 | 2.17E-05 | 1 |
| 4484:173:9:18 | 1:0:0:0 | 10894:556:15:9 | 17:2:0:0 | 2897:123:5:10 | 0:1:0:0 | 1 | 1 | 1 | 1 | 8.41E-09 | 1 |
| 4475:183:9:18 | 7:2:0:0 | 10875:586:15:9 | 18:0:0:0 | 2895:128:5:10 | 2:1:0:0 | 0.001044569 | 1 | 1 | 0.003244975 | 3.87E-11 | 1 |
| 4478:189:9:18 | 3:0:0:0 | 10881:605:15:9 | 20:0:0:0 | 2896:131:5:10 | 3:0:0:0 | 1.19E-05 | 1 | 1 | 1 | 2.23E-12 | 1 |
| 4476:190:9:18 | 4:0:0:0 | 10878:623:15:9 | 21:0:0:0 | 2890:132:5:10 | 7:0:0:0 | 1 | 1 | 1 | 1 | 5.34E-13 | 0.000193253 |
| 131:223:13:23:3:5:0:0 | 2:0:2:7:0:0:0:0 | 84:240:17:19:2:1:0:1 | 2:0:2:1:0:0:0:0 | 19:43:3:1:0:0:0:0 | 1:0:0:0:0:0:0:0 | 1 | 1 | 1 | 0.000138615 | 0.241187977 | 1 |
| 3:11:1:1:1:0:1:0:0:0 | 2:2:0:1:1:1:0:0:0:0 | 0:0:0:0:0:0:0:0:0:0 | 8:0:0:0:0:0:0:0:0:1 | 0:0:0:0:0:0:0:0:0:0 | 3:0:0:0:0:0:0:0:0:0 | 1 | 1 | 1 | 0.479586441 | 1.03E-09 | 1 |
| 21:06:02 | 23:05:02 | 0:01:00 | 28:08:03 | 0:00:00 | 13:00:02 | 1 | 1 | 0.007094277 | 5.30E-06 | 3.15E-06 | 0.000104742 |
| 15:01:02 | 17:03:01 | 0:00:00 | 20:02:01 | 0:00:00 | 8:01:00 | 1 | 1 | 1 | 3.65E-05 | 7.83E-07 | 0.002822994 |
| 15:01:01 | 17:03:01 | 0:00:00 | 19:02:00 | 0:00:00 | 8:01:00 | 1 | 1 | 1 | 3.65E-05 | 4.16E-07 | 0.002822994 |
| 20:00:01 | 21:00:02 | 0:01:00 | 29:03:01 | 0:00:00 | 11:00:02 | 1 | 1 | 0.001636609 | 6.32E-08 | 7.52E-10 | 0.000618575 |
| 0:00:00 | 0:02:00 | 0:01:00 | 0:08:00 | 1:02:00 | 0:00:01 | 1 | 1 | 1 | 1 | 0.000571704 | 1 |
| 1:00 | 1:00 | 1:00 | 6:00 | 2:01 | 1:00 | 1 | 1 | 1 | 1 | 0.018436807 | 1 |
| 682:113:54:44:1 | 22:1:3:2:0 | 798:24:19:56:1 | 18:0:0:0:0 | 154:10:2:9:0 | 0:0:0:0:0 | 1 | 1 | 1 | 6.08E-12 | 3.59E-14 | 1 |
| 25:09:05 | 16:02:00 | 14:09:03 | 8:01:00 | 3:04:03 | 1:00:00 | 1 | 1 | 1 | 6.56E-06 | 0.00271654 | 1 |
| 9:21:05 | 1:16:00 | 9:09:03 | 1:07:00 | 4:02:03 | 0:00:00 | 1 | 1 | 1 | 1.61E-06 | 0.006653127 | 1 |
| 30:01:00 | 34:00:00 | 43:00:00 | 2:00 | 7:00 | 0:00 | 1 | 1 | 0.017871477 | 1.51E-08 | 1 | 1 |
| 0:0:0:0 | 5:29:7:0 | 0:3:1:0 | 0:23:8:1 | 0:1:0:0 | 0:21:1:0 | 1.21E-07 | 0.000239033 | 4.42E-09 | 5.51E-10 | 1.03E-08 | 5.40E-12 |
| 8:4:0:0:0 | 17:5:0:1:1 | 13:8:0:0:0 | 1:1:0:0:0 | 13:4:0:0:0 | 4:1:0:0:0 | 1 | 1 | 1.95E-12 | 4.74E-08 | 1 | 0.021884983 |
| 25:9:0:1:1 | 0:0:0:0:0 | 1:0:0:0:0 | 13:9:0:0:0 | 4:1:0:0:0 | 13:4:0:0:0 | 3.08E-08 | 5.35E-09 | 1 | 1 | 1.17E-06 | 3.74E-07 |
| 25:10:0:1:1 | 0:0:0:0:0 | 1:0:0:0:0 | 13:9:0:0:0 | 4:1:0:0:0 | 13:4:0:0:0 | 3.08E-08 | 5.35E-09 | 1 | 1 | 1.17E-06 | 3.74E-07 |
| 26:1:0:0:0 | 0:0:0:0:0 | 32:1:0:2:1 | 0:0:0:0:0 | 10:0:1:0:0 | 0:0:0:0:0 | 2.17E-18 | 0.004806476 | 1 | 1 | 1 | 1 |
| 0:0:0:0:0:0:0:0:0 | 3:5:1:0:0:2:1:0:0 | 0:0:0:0:0:0:0:0:0 | 4:3:0:0:0:0:1:0:1 | 0:0:0:0:0:0:0:0:0 | 1:0:0:0:0:0:0:1:0 | 0.000798008 | 1 | 0.486505676 | 0.02590394 | 0.027004257 | 1 |
| 142:18:00 | 0:00 | 154:19:00 | 6:01 | 34:04:00 | 6:02 | 0.009714577 | 0.00012042 | 1 | 1 | 0.068536718 | 0.169900525 |
| 1019:7198:72:1 | 3:30:9:0 | 1253:13384:118:6 | 8:44:15:0 | 186:3107:11:1 | 0:8:0:0 | 2.14E-14 | 0.040765436 | 0.015801098 | 7.97E-11 | 1.36E-13 | 4.84E-05 |
| 7244:964:12:16 | 6:6:0:0 | 13470:1108:19:22 | 14:7:0:0 | 3118:174:6:6 | 3:0:0:0 | 0.00028273 | 1 | 1 | 0.009770702 | 2.28E-05 | 1 |
| 7246:955:12:17 | 4:2:0:0 | 13478:1083:20:22 | 6:2:1:0 | 3119:172:6:7 | 1:0:0:0 | 1 | 1 | 1 | 0.070826298 | 0.032544232 | 1 |
| 7241:945:15:20 | 7:0:0:0 | 13476:1049:29:30 | 13:3:0:3 | 3113:159:12:8 | 8:0:0:0 | 8.94E-08 | 1 | 1 | 0.000182133 | 0.000257537 | 4.84E-05 |
| 7241:923:15:19 | 8:0:1:1 | 13469:1007:36:35 | 20:3:0:0 | 3117:155:12:8 | 4:0:0:0 | 1.44E-06 | 1 | 1 | 0.001427729 | 6.49E-10 | 1 |
| 7240:910:16:20 | 9:2:0:0 | 13477:988:37:37 | 14:4:0:0 | 3117:156:12:8 | 4:2:0:0 | 0.000677933 | 0.154096241 | 1 | 0.000294194 | 4.62E-06 | 0.069659881 |
| 7245:908:17:19 | 4:1:0:1 | 13477:982:36:36 | 13:3:1:1 | 3121:158:12:8 | 0:0:0:0 | 3.02E-06 | 0.012695226 | 1 | 0.122686968 | 0.000121223 | 1 |
| 7244:733:20:25 | 10:0:0:0 | 13492:629:42:82 | 6:7:0:0 | 3123:136:13:22 | 2:0:0:0 | 3.17E-05 | 1 | 1 | 3.04E-06 | 0.006249361 | 1 |
| 7252:723:25:19 | 2:1:0:0 | 13497:622:83:40 | 1:5:0:0 | 3125:148:22:12 | 0:0:0:0 | 0.000677933 | 1 | 1 | 1 | 0.013499631 | 1 |
| 7255:675:28:17 | 1:2:2:0 | 13495:568:97:43 | 9:0:12:0 | 3126:152:24:11 | 1:0:0:0 | 0.119939346 | 1 | 1 | 0.545876526 | 0.000129588 | 1 |
| 7249:29:675:18 | 11:5:1:1 | 13492:108:564:44 | 17:8:0:0 | 3126:26:151:12 | 2:0:1:0 | 0.000677933 | 1 | 0.040878917 | 0.002786391 | 1.54E-06 | 1 |
| 7440:660:357:53 | 4:2:0:0 | 13788:543:713:62 | 10:0:1:2 | 3146:149:56:11 | 4:1:0:0 | 1 | 1 | 1 | 0.070826298 | 0.00041677 | 0.040495738 |
| 7423:50:312:652 | 22:7:40:6 | 13768:61:663:531 | 31:1:39:1 | 3141:11:52:149 | 7:0:3:0 | 0.012951673 | 0.086973632 | 0.156140712 | 2.06E-08 | 1.26E-13 | 0.006116201 |
| 652:7432:356:55 | 3:13:3:1 | 520:13785:722:59 | 2:14:5:1 | 147:3148:56:10 | 0:3:1:0 | 0.022173773 | 1 | 1 | 0.000831463 | 0.000456921 | 1 |

| | | | | | | | | | | | |
|---|---|---|---|---|---|---|---|---|---|---|---|
| 7442:651:356:56 | 3:4:2:0 | 13793:519:723:59 | 8:1:4:1 | 3146:147:55:10 | 5:0:2:0 | 0.002557517 | 1 | 1 | 0.289932809 | 0.030521768 | 0.028211333 |
| 7484:588:471:62 | 7:2:0:1 | 13803:451:879:85 | 23:0:0:0 | 3138:141:54:15 | 3:0:0:0 | 1.32E-09 | 1 | 1 | 0.011333842 | 2.94E-14 | 1 |
| 5:1:5:0:7:8:4:6:0:8:6:9:4:5:0:0:0:4:4:0:0:1:0:2:0:1:0:0:0 | 15:1:16:0:25:17:6:13:0:20:11:24:11:6:2:1:1:9:2:0:0:1:5:1:1:0:0:0:0 | 7:0:8:0:9:7:6:5:1:11:5:17:7:9:0:0:1:1:4:0:1:1:0:1:0:0:0:1:1 | 37:1:19:0:16:31:9:6:0:60:20:32:18:19:0:0:0:5:3:0:0:1:2:1:0:0:1:0:0 | 2:0:1:0:0:2:0:0:0:0:0:0:2:1:0:0:0:2:0:0:0:0:0:0:0:0:0:0:0 | 3:0:0:0:1:2:0:0:0:3:2:2:0:2:0:0:0:0:0:0:0:0:0:0:1:0:0:0:0 | 9.31E-07 | 0.016175488 | 1.14E-06 | 1.04E-40 | 2.94E-113 | 0.0107453 |
| 26:36:22:59:69:49:40:39:11:29:19:0:3:20:2:0:20:1:1:6:1:1:7:2:0:2:1:2:1:0:0:0:0 | 4:9:2:14:10:24:9:5:2:12:6:0:1:8:0:0:3:0:0:0:0:0:1:0:1:0:0:0:0:0:0:0:0 | 40:85:10:52:70:113:59:50:12:22:37:1:3:23:1:0:40:1:2:1:0:1:6:3:0:1:0:0:1:1:1:1:1 | 7:27:3:19:15:29:8:7:2:4:10:0:0:4:0:1:3:0:0:0:1:0:0:0:0:0:0:0:0:0:1:0:0 | 2:17:5:9:11:17:10:5:1:5:4:0:1:1:0:0:4:0:0:3:0:0:0:0:0:0:0:0:1:0:0:0:0 | 5:9:5:4:11:10:6:3:0:2:0:0:0:2:0:0:3:0:0:0:0:0:0:0:0:0:0:0:0:0:0:0 | 9.49E-89 | 2.51E-16 | 4.36E-15 | 6.49E-42 | 8.84E-72 | 4.06E-21 |
| 132:100:91:0:76:83:7:151:60:45:4:0:144:44:47:28:61:1:1:13:1:2:13:1:1:3:3:4:4:1:2:0 | 5:7:9:0:4:3:0:0:0:4:0:0:0:1:0:4:2:0:0:0:0:0:3:0:0:0:0:1:0:0:0:0 | 128:205:102:1:49:124:4:274:92:51:3:1:154:32:79:27:97:4:3:2:2:2:9:3:3:4:0:2:2:2:0:1 | 11:8:7:0:2:3:0:0:0:3:0:0:0:0:0:1:2:0:0:0:0:0:0:0:0:0:0:0:0:0:0:0 | 17:29:15:0:10:22:1:37:11:6:0:0:29:15:5:3:10:0:0:3:0:0:0:0:0:0:0:1:0:0:0:0 | 2:2:3:0:1:1:0:0:0:1:0:0:0:0:0:0:0:0:0:0:0:0:0:0:0:0:0:0:0:0:0:0 | 3.81E-10 | 4.18E-07 | 8.27E-07 | 7.32E-13 | 9.02E-24 | 0.00897458 |
| 115:143:0:107:86:87:7:151:60:52:4:0:144:45:47:33:64:2:1:1:13:1:2:16:1:1:3:3:3:1:5:0 | 1:2:0:1:0:0:0:0:0:1:0:0:0:1:0:0:1:0:0:0:0:0:0:0:0:0:0:0:1:0:0:0 | 224:152:1:120:55:129:4:274:92:55:3:1:154:32:76:30:107:0:4:3:2:2:2:9:3:3:4:0:2:2:2:1 | 2:3:0:0:0:1:0:0:0:0:0:0:0:0:1:0:0:0:0:0:0:0:0:0:0:0:0:0:0:0:0:0 | 32:21:0:19:12:23:1:37:11:7:0:0:29:15:4:3:9:0:0:0:3:0:0:0:0:0:0:0:0:0:1:0 | 1:0:0:0:0:0:0:0:0:0:0:0:0:0:0:0:0:0:0:0:0:0:0:0:0:0:0:0:0:0:0:0 | 0.034884421 | 1 | 1 | 0.432446198 | 0.001405578 | 1 |
| 109:146:0:86:117:87:7:151:60:55:4:0:144:46:46:33:61:2:16:1:1:13:1:2:1:2:3:3:5:4:1:0 | 10:15:0:0:11:5:0:0:0:3:0:0:0:0:0:1:0:0:0:0:0:0:0:0:0:0:0:0:1:0:0:0 | 121:156:1:55:227:130:4:274:92:55:3:1:153:32:75:30:99:0:9:4:3:2:2:2:3:3:4:0:2:2:2:1 | 7:20:0:2:33:6:0:0:0:2:0:0:1:1:1:2:10:0:3:0:0:0:0:0:0:0:0:0:1:0:0:0 | 19:21:0:12:34:23:1:37:11:8:0:0:29:15:4:3:10:0:0:0:0:3:0:0:0:0:0:0:1:0:0:0 | 2:3:0:1:1:0:0:0:0:0:0:0:0:0:0:0:0:0:0:0:0:0:0:0:0:0:0:0:0:0:0:0 | 1.73E-22 | 1 | 0.000860931 | 8.03E-43 | 2.42E-89 | 0.001479193 |
| 0:154:186:96:149:114:7:151:60:64:0:144:46:29:28:45:6:23:1:1:13:1:2:1:2:3:3:4:4:1:6:0 | 0:0:0:0:0:1:0:0:0:0:0:0:0:0:0:1:0:0:0:0:0:0:0:0:0:0:0:0:0:0:0:0 | 1:170:205:70:292:164:4:273:92:64:1:154:33:53:21:86:0:13:4:2:2:2:2:3:3:4:1:5:2:2:5:1 | 0:0:0:0:2:0:0:1:0:0:0:0:0:0:2:0:0:0:0:0:0:0:0:0:0:0:0:0:0:0:0:0 | 0:22:26:14:38:25:1:37:11:10:0:29:15:1:1:7:0:0:0:0:3:0:0:0:0:0:0:0:0:0:1:0 | 0:0:0:0:0:0:0:0:0:0:0:0:0:0:0:0:0:0:0:0:0:0:0:0:0:0:0:0:0:0:0:0 | 1 | 1 | 1 | 1 | 0.012226778 | 1 |
| 58:13:6:11:2 | 8:0:1:1:0 | 107:11:7:16:9 | 12:1:0:1:0 | 17:1:1:2:0 | 1:1:0:0:0 | 0.000755847 | 1 | 1 | 0.000213629 | 2.64E-07 | 1 |
| 17:4:7:10:2 | 22:5:1:1:0 | 35:6:8:13:8 | 52:2:0:1:1 | 15:1:3:2:0 | 0:0:0:0:0 | 1 | 1 | 1 | 1.43E-11 | 2.83E-38 | 1 |
| 455:19:00 | 1:01 | 1206:52:00 | 6:00 | 291:15:00 | 0:00 | 1 | 1 | 1 | 1 | 0.018436807 | 1 |
| 0:1:2:5:1:0:1:0:0 | 0:0:0:0:0:0:0:0:0 | 2:0:2:1:0:0:0:0:0 | 4:1:27:59:3:2:2:0:1 | 0:0:0:2:0:0:0:0:0 | 1:1:1:1:0:1:0:0:0 | 1 | 1 | 1 | 1 | 2.40E-56 | 0.859321213 |
| 0.707789352 | 0:02:00 | 27:157:21 | 1:06:00 | 1:21:03 | 0:00:00 | 1 | 1 | 1 | 1 | 0.015787883 | 1 |
| 0:1:0:0 | 33:20:34:0 | 63:37:64:0 | 0:0:0:0 | 20:1:8:0 | 0:0:0:0 | 1 | 1 | 0.002598276 | 3.59E-07 | 1 | 1 |
| 33:20:35:0 | 0:0:0:0 | 3:0:2:0 | 60:25:61:0 | 0:1:1:0 | 20:0:7:0 | 1 | 1 | 1 | 1 | 8.01E-15 | 3.90E-08 |
| 3821:04:00 | 6:00:00 | 3376:16:00 | 3:00:00 | 770:01:00 | 0:00:00 | 0.00132829 | 1 | 1 | 0.003552529 | 1 | 1 |
| 3847:04:00 | 6:00:00 | 0:00:00 | 3404:16:00 | 0:00:00 | 769:01:00 | 1 | 1 | 1 | 0.003552529 | 0 | 0 |
| 3913:03:00 | 7:00:00 | 3468:03:00 | 3:00:00 | 779:00:00 | 0:00:00 | 1 | 1 | 1 | 0.00138895 | 1 | 1 |
| 3900:00:00 | 3:00:00 | 3462:00:01 | 6:00:00 | 778:00:00 | 1:00:00 | 0.0086191 | 1 | 1 | 1 | 0.003657671 | 1 |
| 3875:00:00 | 3:00:00 | 3446:00:01 | 4:00:00 | 774:00:00 | 0:00:00 | 0.0086191 | 1 | 1 | 1 | 1 | 1 |
| 309:23:102 | 5:00:01 | 271:27:89 | 7:00:01 | 36:03:10 | 2:00:00 | 0.004446886 | 1 | 1 | 0.035780388 | 0.006653127 | 1 |
| 127:415:28 | 8:27:00 | 107:375:30 | 1:18:00 | 17:46:06 | 0:06:02 | 1.39E-10 | 0.022158081 | 7.91E-06 | 1.70E-07 | 2.43E-07 | 0.035456988 |
| 132:431:28 | 3:13:00 | 109:387:30 | 0:10:01 | 17:49:06 | 0:03:02 | 2.85E-06 | 0.074426239 | 0.003124028 | 0.000280402 | 0.000446416 | 0.259890601 |
| 541:477:187:62:111 | 20:10:1:2:1 | 219:331:92:35:78 | 7:5:2:3:2 | 29:56:15:4:13 | 1:0:0:0:0 | 1 | 1 | 1 | 8.97E-08 | 0.308909393 | 1 |
| 538:476:188:61:110 | 0:15:6:0:0 | 216:328:92:34:79 | 1:3:2:0:0 | 29:56:16:4:12 | 0:0:0:0:0 | 1 | 1 | 1 | 6.44E-08 | 0.241187977 | 1 |
| 10:1:1:2:0:1:0 | 9:0:1:2:1:0:0 | 13:0:1:3:0:1:0 | 0:0:0:0:0:0:0 | 7:2:0:0:0:0:0 | 0:0:0:0:0:0:0 | 1 | 1 | 1 | 1.42E-05 | 1 | 1 |
| 0:0:0:0:0 | 300:2:59:1:0 | 1:0:0:0:0 | 223:3:45:3:1 | 0:0:0:0:0 | 26:2:11:1:0 | 3.33E-25 | 1 | 2.96E-15 | 4.89E-198 | 1.85E-141 | 1.02E-11 |
| 309:2:60:1:0 | 4:0:1:0:0 | 240:3:48:3:1 | 2:0:0:0:0 | 26:2:11:1:0 | 0:0:0:0:0 | 1 | 1 | 1 | 0.021353791 | 1 | 1 |

| | | | | | | | | | | | |
|---|---|---|---|---|---|---|---|---|---|---|---|
| 0:0:0:0:0 | 56:31:4:11:4 | 0:0:0:0:0 | 19:9:3:6:0 | 0:0:0:0:0 | 3:0:0:3:1 | 0.002706129 | 0.010591505 | 0.000778496 | 6.37E-19 | 1.77E-05 | 0.172713293 |
| 6:28:11:2 | 6:2:13:1 | 0:0:0:0 | 8:4:24:2 | 0:0:0:0 | 0:1:4:0 | 3.85E-06 | 1 | 0.01363685 | 0.00157572 | 1.54E-06 | 0.040495738 |
| 6:28:11:2 | 6:2:13:1 | 0:0:0:0 | 8:4:24:2 | 0:0:0:0 | 0:1:4:0 | 3.85E-06 | 1 | 0.01363685 | 0.00157572 | 1.54E-06 | 0.040495738 |
| 17:01 | 13:01 | 88:01:00 | 0:00 | 20:01 | 0:00 | 1 | 1 | 0.111416266 | 0.002011949 | 1 | 1 |
| 76:35:10:1:10:2:2:0 | 52:16:9:1:3:0:0:0 | 62:28:30:5:4:3:5:0 | 1:0:0:0:0:0:0:0 | 30:7:4:0:0:0:1:1 | 0:0:1:0:0:0:0:0 | 2.22E-06 | 2.28E-09 | 2.12E-06 | 4.33E-43 | 1 | 1 |
| 66:26:4:0:9:0:0 | 79:23:2:0:10:1:2 | 4:1:0:0:1:0:1 | 64:18:6:2:10:1:0 | 1:1:0:0:0:0:0 | 19:14:1:0:0:0:0 | 1 | 1.19E-26 | 1.56E-08 | 1.54E-59 | 8.87E-43 | 1.18E-14 |
| 301:192:16:30:46:2 | 270:165:14:22:50:3 | 457:235:35:46:43:1 | 0:0:0:0:0:0 | 136:68:4:12:17:1 | 0:0:0:0:0:0 | 3.32E-47 | 8.81E-08 | 5.40E-32 | 1.27E-138 | 1 | 1 |
| 9:20:0:35:1:1 | 2:13:0:0:0:0 | 6:15:0:11:0:1 | 1:7:0:0:0:0 | 5:1:0:3:0:0 | 0:1:0:0:0:0 | 1 | 1 | 1 | 5.61E-10 | 3.70E-05 | 1 |
| 8:19:35:0:1:1 | 1:5:0:0:0:0 | 6:12:11:0:0:1 | 0:4:0:0:0:0 | 5:1:3:0:0:0 | 0:0:0:0:0:0 | 1 | 1 | 1 | 0.001878282 | 1 | 1 |
| 21:1:0:0 | 20:0:0:2 | 45:1:0:2 | 1:0:1:0 | 6:1:0:0 | 1:0:0:0 | 1.19E-05 | 5.95E-06 | 9.93E-05 | 1.19E-10 | 1 | 1 |
| 94:1:0:0:2 | 81:2:1:1:0 | 16:0:0:0:3 | 0:0:0:0:0 | 4:0:0:0:0 | 0:0:0:0:0 | 1 | 1 | 3.54E-08 | 8.93E-63 | 1 | 1 |
| 60:3:0:2 | 51:3:1:0 | 11:0:0:2 | 0:0:0:0 | 3:0:0:0 | 0:0:0:0 | 1 | 1 | 1.67E-06 | 2.63E-28 | 1 | 1 |
| 11:4:1:0 | 17:4:1:1 | 0:0:0:0 | 1:0:0:0 | 0:0:0:0 | 3:0:0:0 | 1 | 1.27E-05 | 1 | 2.56E-06 | 1 | 1 |
| 25:7:1:2:1 | 2:1:0:0:0 | 0:0:0:0:0 | 2:2:0:0:0 | 0:0:0:0:0 | 0:0:0:0:0 | 1 | 7.26E-07 | 1 | 1 | 1 | 1 |
| 32:77:5:17 | 32:80:11:6 | 0:0:0:0 | 7:4:1:0 | 0:0:0:0 | 1:1:0:0 | 1 | 0.000704574 | 1 | 3.66E-22 | 0.023786496 | 1 |
| 0:0:0:0 | 1:5:0:1 | 0:0:0:0 | 1:1:0:0 | 0:0:0:0 | 0:0:0:0 | 0.012449474 | 1 | 1 | 0.044129954 | 1 | 1 |
| 1524:5:27:1 | 1:0:0:0 | 1845:4:13:0 | 5:0:0:0 | 386:0:8:0 | 0:0:0:0 | 1 | 1 | 1 | 1 | 0.002732501 | 1 |
| 1363:3:27:1 | 0:0:0:0 | 1644:2:12:0 | 5:0:0:0 | 366:0:8:0 | 0:0:0:0 | 1 | 1 | 1 | 1 | 0.002732501 | 1 |
| 1111:2:27:1 | 5:0:0:0 | 1296:2:13:0 | 3:0:0:0 | 342:0:8:0 | 2:0:0:0 | 1 | 1 | 1 | 0.002648305 | 1 | 1 |
| 1101:2:27:1 | 8:0:0:0 | 1282:2:13:0 | 5:0:0:0 | 343:0:8:0 | 1:0:0:0 | 1 | 1 | 1 | 4.60E-05 | 0.002732501 | 1 |
| 0:0:0:0:0:0:0:0:0:0:0:0:0:0:0:0:0:0:0:0:0:0 | 21:7:1:2:3:0:0:0:2:5:0:1:5:2:1:3:1:1:2:0:0 | 0:0:0:0:0:0:0:0:0:0:0:0:0:0:0:0:0:0:0:0:0:0 | 19:5:0:1:4:0:0:0:3:0:0:0:0:1:0:0:2:0:1:0:1 | 0:0:0:0:0:0:0:0:0:0:0:0:0:0:0:0:0:0:0:0:0:0 | 13:3:1:0:1:0:0:0:0:1:0:0:0:1:0:0:1:0:0:0:0 | 4.10E-81 | 7.02E-68 | 1.48E-22 | 1.84E-22 | 5.07E-31 | 2.19E-23 |
| 8:9:0:0 | 0:9:0:0 | 1:0:0:0 | 8:13:0:0 | 1:0:0:0 | 2:4:1:0 | 1.36E-06 | 0.294968773 | 1 | 1.18E-05 | 6.47E-05 | 0.184047759 |
| 37:3:1:2:0:1:2:0:0:0:0 | 44:2:5:0:1:1:2:0:1:0:0 | 88:7:7:0:3:1:3:0:2:0:1 | 0:0:0:0:0:0:0:0:0:0:0 | 27:0:1:1:0:0:0:2:1:1:0 | 0:0:0:0:0:0:0:0:0:0:0 | 1 | 1 | 6.46E-30 | 5.98E-64 | 1 | 1 |
| 372:27:00 | 4:00 | 466:53:00 | 1:00 | 100:02:00 | 0:00 | 1 | 1 | 0.000225444 | 1 | 1 | 1 |
| 0:0:0:0 | 2:5:0:0 | 2:9:0:1 | 0:0:0:0 | 0:2:0:0 | 0:0:0:0 | 3.43E-09 | 1 | 0.001426566 | 0.027461316 | 1 | 1 |
| 8:3:1:0:0 | 0:0:0:0:0 | 0:0:0:0:0 | 14:11:5:0:0 | 0:0:0:0:0 | 4:4:0:0:0 | 1 | 1 | 1 | 1 | 3.96E-05 | 0.021884983 |
| 22060:560:143 | 32:11:00 | 30314:700:252 | 43:06:00 | 8573:110:75 | 10:02:00 | 8.91E-07 | 8.82E-05 | 0.000204673 | 2.63E-08 | 1.63E-14 | 0.001510787 |
| 21996:565:143 | 45:10:00 | 30230:712:251 | 62:04:00 | 8562:110:75 | 12:03:00 | 2.33E-09 | 1.30E-05 | 3.26E-07 | 2.13E-13 | 9.68E-24 | 0.000696549 |
| 21526:604:145 | 63:05:00 | 29560:742:250 | 88:02:00 | 8457:111:75 | 15:02:00 | 6.27E-14 | 2.06E-11 | 4.81E-13 | 1.58E-23 | 2.05E-36 | 1.75E-05 |
| 21237:599:145 | 90:06:00 | 29172:759:250 | 109:05:00 | 8389:115:74 | 13:02:01 | 3.89E-18 | 4.92E-15 | 3.93E-15 | 2.67E-34 | 4.10E-43 | 0.000432266 |
| 20992:619:145 | 95:04:00 | 28870:793:250 | 102:03:00 | 8349:121:75 | 17:03:00 | 1.89E-17 | 9.32E-14 | 2.74E-14 | 6.88E-38 | 1.19E-41 | 9.41E-06 |
| 20426:665:145 | 81:05:01 | 27978:835:251 | 98:03:00 | 8197:128:75 | 17:02:00 | 1.87E-16 | 2.76E-14 | 5.65E-11 | 2.16E-30 | 6.33E-40 | 2.78E-06 |
| 19959:709:146 | 117:06:00 | 27242:867:250 | 124:03:00 | 8059:135:75 | 21:02:00 | 1.04E-19 | 3.58E-14 | 2.19E-11 | 8.49E-46 | 4.02E-51 | 6.86E-08 |
| 19741:720:145 | 93:05:01 | 26965:869:250 | 99:04:00 | 7989:132:75 | 25:01:00 | 8.84E-19 | 1.88E-11 | 3.19E-12 | 1.66E-35 | 1.39E-39 | 3.22E-10 |
| 19327:730:146 | 73:06:00 | 26388:883:252 | 94:09:00 | 7900:136:75 | 19:01:00 | 1.58E-14 | 1.57E-12 | 1.74E-08 | 4.43E-27 | 6.34E-34 | 9.88E-08 |
| 19113:726:146 | 88:03:00 | 26035:883:252 | 114:07:00 | 7856:139:75 | 21:00:00 | 4.44E-17 | 1.88E-11 | 8.92E-10 | 1.17E-35 | 8.71E-44 | 2.51E-09 |
| 18837:730:146 | 95:06:00 | 25554:870:252 | 125:07:01 | 7781:138:75 | 28:00:00 | 8.07E-17 | 1.88E-11 | 1.49E-10 | 2.03E-36 | 1.06E-47 | 2.91E-12 |
| 18546:734:146 | 124:05:00 | 25139:861:253 | 153:08:00 | 7720:137:75 | 29:00:00 | 4.29E-23 | 2.80E-12 | 1.74E-08 | 1.56E-49 | 9.99E-60 | 1.12E-12 |

| | | | | | | | | | | | |
|---|---|---|---|---|---|---|---|---|---|---|---|
| 18367:732:146 | 128:03:00 | 24969:851:253 | 148:07:00 | 7679:140:75 | 27:00:00 | 9.47E-25 | 3.35E-14 | 2.19E-11 | 8.69E-53 | 2.49E-58 | 7.50E-12 |
| 18328:732:146 | 132:06:00 | 24882:848:253 | 155:08:00 | 7660:138:75 | 26:00:00 | 1.48E-20 | 1.99E-12 | 8.31E-12 | 3.43E-52 | 1.40E-60 | 1.94E-11 |
| 18251:726:146 | 95:06:00 | 24692:844:253 | 144:07:00 | 7627:136:75 | 27:00:00 | 4.81E-15 | 1.99E-12 | 3.83E-10 | 2.03E-36 | 1.28E-56 | 7.50E-12 |
| 18269:729:146 | 100:04:00 | 24707:846:253 | 134:08:00 | 7638:137:75 | 26:00:00 | 5.64E-21 | 1.88E-11 | 9.97E-10 | 4.88E-40 | 1.31E-51 | 1.94E-11 |
| 18176:726:146 | 133:08:00 | 24601:848:253 | 168:12:00 | 7630:134:75 | 25:00:00 | 1.96E-25 | 2.76E-14 | 3.83E-10 | 3.72E-51 | 2.83E-63 | 5.12E-11 |
| 18013:715:146 | 147:03:00 | 24364:845:252 | 188:11:01 | 7591:131:75 | 20:00:00 | 2.56E-25 | 1.68E-15 | 1.12E-12 | 6.78E-61 | 9.82E-72 | 6.45E-09 |
| 17566:690:146 | 143:05:00 | 23675:826:253 | 178:10:01 | 7486:128:75 | 28:00:00 | 7.50E-27 | 1.44E-17 | 1.84E-13 | 1.18E-57 | 3.46E-68 | 2.91E-12 |
| 17281:671:146 | 168:05:00 | 23298:800:254 | 209:10:00 | 7417:127:75 | 32:00:00 | 5.26E-26 | 3.03E-15 | 4.81E-13 | 2.79E-68 | 2.90E-82 | 5.97E-14 |
| 17178:666:146 | 130:04:00 | 23167:787:254 | 186:09:00 | 7387:127:75 | 32:00:00 | 4.29E-23 | 1.26E-13 | 2.01E-11 | 7.54E-53 | 3.93E-73 | 5.97E-14 |
| 643:16786:146 | 0.222222222 | 770:22716:254 | 0.368055556 | 126:7301:75 | 0:21:00 | 1.58E-14 | 4.09E-09 | 1.51E-05 | 1.82E-31 | 4.43E-42 | 2.51E-09 |
| 16734:641:146 | 93:04:00 | 22647:770:254 | 118:06:00 | 7300:126:75 | 16:00:00 | 4.73E-17 | 5.20E-11 | 8.27E-07 | 4.97E-37 | 3.11E-46 | 2.88E-07 |
| 16600:638:146 | 111:04:00 | 22522:762:254 | 151:07:00 | 7269:125:75 | 23:00:00 | 9.47E-22 | 1.16E-16 | 1.49E-10 | 9.79E-45 | 1.29E-59 | 3.58E-10 |
| 16569:634:145 | 130:08:01 | 22496:761:253 | 166:06:01 | 7260:125:75 | 32:00:00 | 1.39E-25 | 4.96E-20 | 6.94E-14 | 4.15E-49 | 5.07E-66 | 5.97E-14 |
| 16379:627:145 | 142:06:00 | 22232:738:255 | 192:07:00 | 7210:121:75 | 34:00:00 | 5.45E-26 | 1.80E-17 | 5.66E-16 | 1.86E-56 | 3.11E-77 | 8.70E-15 |
| 16372:628:147 | 157:03:00 | 22259:744:255 | 215:11:00 | 7228:119:75 | 34:02:00 | 1.92E-29 | 4.42E-17 | 1.20E-12 | 3.92E-65 | 4.36E-84 | 3.08E-13 |
| 627:16296:147 | 0.222222222 | 748:22188:254 | 0.502094907 | 121:7211:75 | 0:30:00 | 4.87E-20 | 2.01E-15 | 1.70E-13 | 6.18E-58 | 4.72E-71 | 4.25E-13 |
| 16322:620:146 | 129:01:01 | 22221:746:255 | 198:12:00 | 7199:121:75 | 39:00:00 | 7.76E-27 | 2.61E-18 | 8.94E-13 | 5.12E-54 | 4.49E-76 | 6.45E-17 |
| 16131:611:146 | 108:03:01 | 21950:737:255 | 166:08:00 | 7126:118:74 | 30:00:01 | 2.50E-18 | 1.46E-12 | 1.20E-12 | 2.06E-43 | 2.59E-65 | 2.70E-12 |
| 16022:598:146 | 74:00:01 | 21873:711:256 | 121:07:00 | 7073:114:75 | 20:00:00 | 6.36E-12 | 6.23E-13 | 3.26E-07 | 3.06E-31 | 8.74E-47 | 6.45E-09 |
| 15932:586:145 | 82:00:01 | 21757:710:256 | 143:06:00 | 7022:112:75 | 20:01:00 | 8.02E-11 | 2.43E-13 | 1.74E-08 | 1.09E-34 | 6.09E-57 | 3.85E-08 |
| 15877:579:146 | 66:01:01 | 21684:707:256 | 134:04:00 | 6987:109:75 | 24:02:00 | 8.78E-13 | 2.11E-15 | 3.48E-10 | 5.57E-27 | 1.24E-54 | 4.23E-09 |
| 15883:578:145 | 60:02:02 | 21690:707:256 | 120:04:00 | 6988:109:75 | 20:02:00 | 2.81E-10 | 1.18E-12 | 6.83E-09 | 7.18E-23 | 1.34E-48 | 1.75E-07 |
| 124:12:1:1:1:2:0:0 | 2:11:5:0:0:0:0:0 | 69:16:1:0:0:0:0:0 | 0:5:1:0:0:0:0:1 | 34:3:0:0:1:0:1:0 | 0:0:1:0:0:0:0:0 | 1 | 1 | 1 | 6.73E-08 | 0.001883319 | 1 |
| 149:10:4:0:3:2:1 | 2:7:3:0:0:0:0 | 91:8:4:0:0:3:1 | 2:6:0:0:0:0:1 | 37:2:1:1:1:1:1 | 0:0:0:0:0:0:0 | 1 | 1 | 1 | 0.000770016 | 0.001340465 | 1 |
| 0:00 | 0:00 | 0:00 | 0:00 | 0:00 | 0:00 | 0.110352776 | 0.002334609 | 1 | 1 | 1 | 1 |
| 17:41:71:17:0:1:4:1:0:1:0:0 | 0:0:1:0:0:0:0:0:0:0:0:0 | 1:14:10:9:0:0:1:1:0:0:0:0 | 11:29:28:15:0:0:6:0:1:0:3:0 | 3:5:5:4:1:0:0:0:0:0:0:0 | 1:15:24:5:0:0:1:0:0:0:1:0 | 1 | 1 | 1 | 1 | 2.68E-29 | 1.31E-28 |
| 8:1:3:1 | 0:0:0:0 | 22:2:6:2 | 0:0:0:0 | 8:1:1:0 | 0:0:0:0 | 0.011225544 | 1 | 1 | 1 | 1 | 1 |
| 0:00:00 | 2:24:00 | 2:47:03 | 2:21:02 | 9:31:00 | 6:04:00 | 3.72E-48 | 8.32E-06 | 4.00E-11 | 3.68E-09 | 1.23E-06 | 0.068846966 |
| 81:90:20:79:1 | 0:0:0:1:0 | 279:357:109:399:6 | 0:3:0:4:0 | 147:79:22:31:4 | 0:0:0:0:0 | 1 | 1 | 1 | 1 | 0.034565431 | 1 |
| 4:0:1:0 | 0:1:0:0 | 0:0:0:0 | 7:9:0:1 | 0:0:0:0 | 2:1:0:0 | 1 | 1 | 1 | 1 | 0.004606311 | 1 |
| 25:08:00 | 0:00:00 | 120:16:10 | 0:00:00 | 14:01:02 | 0:00:00 | 7.24E-08 | 0.050505578 | 1 | 1 | 1 | 1 |
| 5:1:0:0 | 0:0:0:0 | 9:2:1:1 | 1:0:0:0 | 8:1:1:0 | 0:0:0:0 | 0.002557517 | 0.002652154 | 1 | 1 | 1 | 1 |
| 5:00:03 | 0:00:00 | 18:03:02 | 61:04:10 | 1:00:00 | 7:00:00 | 1 | 1 | 1 | 1 | 4.28E-17 | 0.001510787 |
| 5:05:01 | 0:00:00 | 0.843171296 | 0:00:00 | 1:08:00 | 0:00:00 | 0.049209366 | 1 | 1 | 1 | 1 | 1 |
| 0:00:00 | 3:00:00 | 20:00:00 | 1:00:00 | 0:00:00 | 0:00:00 | 0.034668522 | 1 | 1 | 1 | 1 | 1 |
| 0:00:00 | 3:00:00 | 20:00:00 | 1:00:00 | 0:00:00 | 0:00:00 | 0.034668522 | 1 | 1 | 1 | 1 | 1 |
| 0:00 | 8:00 | 27:00:00 | 0:00 | 0:00 | 0:00 | 1 | 1 | 1 | 0.006305042 | 1 | 1 |
| 8:00 | 0:00 | 13:00 | 7:00 | 0:00 | 0:00 | 1 | 1 | 1 | 1 | 0.011137275 | 1 |
| 23:0:3:2 | 13:0:1:1 | 11:0:1:1 | 5:0:0:0 | 2:0:0:0 | 0:0:0:0 | 1 | 1 | 1 | 2.32E-06 | 0.002732501 | 1 |
| 27:0:4:3 | 9:0:0:0 | 12:0:1:1 | 4:0:0:0 | 2:0:0:0 | 0:0:0:0 | 1 | 1 | 1 | 1.18E-05 | 1 | 1 |
| 1:0:0:0:0 | 61:0:8:4:0 | 81:4:3:3:0 | 4:0:0:1:1 | 10:1:0:0:0 | 0:0:0:0:0 | 1 | 1 | 0.003310176 | 1.35E-38 | 0.070859781 | 1 |
| 61:0:8:4:0 | 1:0:0:0:0 | 87:4:3:4:1 | 0:0:0:0:0 | 10:1:0:0:0 | 0:0:0:0:0 | 1 | 1 | 0.000172782 | 1 | 1 | 1 |

| | | | | | | | | | | | |
|---|---|---|---|---|---|---|---|---|---|---|---|
| 1:0:0:0:1:0:0 | 59:0:8:0:2:0:0 | 95:5:3:0:0:0:1 | 0:0:0:0:0:0:0 | 16:1:0:0:0:1:0 | 0:0:0:0:0:0:0 | 1 | 1 | 0.001559109 | 8.53E-59 | 1 | 1 |
| 26:43:32:52:2:0:0 | 3:4:2:0:0:0:0 | 48:86:78:121:9:0:2 | 1:5:1:0:0:0:1 | 18:16:2:2:0:0:0 | 1:1:0:0:0:0:0 | 1 | 1 | 1 | 0.04093502 | 0.015127731 | 1 |
| 28:01:02 | 0:00:00 | 19:01:01 | 43:00:02 | 2:00:00 | 8:00:01 | 1 | 1 | 1 | 1 | 4.19E-17 | 0.002822994 |
| 39:01:02 | 0:00:00 | 23:01:01 | 64:00:02 | 2:00:00 | 10:00:01 | 1 | 1 | 1 | 1 | 4.32E-26 | 0.000445789 |
| 0:00:00 | 39:01:02 | 23:01:01 | 64:00:02 | 2:00:00 | 10:00:01 | 1 | 1 | 0.002092311 | 1.06E-14 | 4.32E-26 | 0.000445789 |
| 34:02:03 | 1:00:00 | 78:10:03 | 1:00:00 | 12:00:01 | 0:00:00 | 0.011548543 | 0.014957633 | 1 | 1 | 1 | 1 |
| 9:01:00 | 2:00:00 | 28:07:00 | 5:01:00 | 11:00:01 | 0:00:00 | 1 | 1 | 1 | 1 | 0.036538834 | 1 |
| 2:00 | 0:00 | 0:00 | 8:01 | 0:00 | 3:02 | 1 | 1 | 1 | 1 | 0.025061882 | 0.661487987 |
| 2:00:00 | 0:00:00 | 0:00:00 | 8:00:01 | 0:00:00 | 3:00:02 | 1 | 1 | 1 | 1 | 0.00271654 | 0.259890601 |
| 2410:175:50 | 3:01:01 | 5166:281:71 | 5:00:00 | 1333:92:23 | 2:00:00 | 1 | 1 | 1 | 0.464838796 | 0.009407211 | 1 |
| 2419:175:51 | 2:01:00 | 5195:280:71 | 8:00:00 | 1341:92:23 | 2:00:00 | 1 | 1 | 1 | 1 | 0.000571704 | 1 |
| 2431:173:51 | 0:00:00 | 5217:280:71 | 10:00:00 | 1343:92:23 | 3:00:00 | 1 | 1 | 1 | 1 | 8.55E-05 | 1 |
| 2385:168:52 | 5:01:00 | 5107:275:71 | 8:00:00 | 1321:92:23 | 2:00:00 | 1 | 1 | 1 | 0.035780388 | 0.000571704 | 1 |
| 0:00 | 5:00 | 3:00 | 5:00 | 4:00 | 0:00 | 1 | 1 | 1 | 0.030173969 | 0.031296179 | 1 |
| 0:00 | 0:00 | 1:00 | 0:00 | 0:00 | 0:00 | 0.002436771 | 1 | 1 | 1 | 1 | 1 |
| 0:00 | 0:00 | 1:01 | 0:00 | 0:02 | 0:01 | 1 | 1 | 0.017871477 | 1 | 1 | 1 |
| 0:00 | 0:00 | 0:00 | 1:01 | 0:00 | 0:00 | 0.015395813 | 1 | 1 | 1 | 1 | 1 |
| 0:00 | 0:00 | 0:00 | 1:01 | 0:00 | 0:00 | 0.015395813 | 1 | 1 | 1 | 1 | 1 |
| 0:00 | 0:00 | 0:00 | 1:01 | 0:00 | 0:00 | 0.015395813 | 1 | 1 | 1 | 1 | 1 |
| 31:01:02 | 39:01:05 | 22:00:02 | 0:00:00 | 19:00:01 | 0:00:00 | 1 | 1 | 1 | 8.14E-13 | 1 | 1 |
| 39:4:0:3 | 1:0:0:0 | 1:0:0:0 | 111:12:2:4 | 0:0:0:0 | 23:1:3:1 | 0.042679166 | 1 | 0.003124028 | 1 | 1.06E-54 | 4.21E-10 |
| 30:3:0:1:3 | 0:0:0:0:0 | 0:0:0:0:0 | 84:1:3:4:3 | 0:0:0:0:0 | 21:1:2:0:1 | 0.067828499 | 1 | 0.000954665 | 1 | 5.41E-58 | 1.50E-12 |
| 0:00:00 | 255:25:00 | 186:10:00 | 1:00:00 | 39:03:01 | 0:00:00 | 1 | 1 | 6.50E-10 | 3.35E-91 | 1 | 1 |
| 2:3:0:0 | 0:0:0:0 | 0:2:0:1 | 1:0:0:0 | 1:5:1:0 | 0:0:0:0 | 1 | 1 | 0.040878917 | 1 | 1 | 1 |
| 0:00:00 | 0:00:00 | 3:00:01 | 6:02:00 | 2:00:00 | 1:00:00 | 1 | 1 | 1 | 1 | 0.036538834 | 1 |
| 5:0:0:0:0:0:1:3:0 | 7:1:0:0:2:0:1:1:0 | 5:0:0:1:3:0:0:0:0 | 8:3:1:1:2:0:1:4:1 | 2:0:0:0:0:0:0:1:0 | 0:0:0:0:0:0:0:0:0 | 1 | 1 | 1 | 0.000346294 | 0.011432274 | 1 |
| 0:01:00 | 0:00:00 | 0:00:00 | 5:00:01 | 0:00:00 | 0:00:00 | 1 | 1 | 1 | 1 | 0.036538834 | 1 |
| 0:0:0:0:0 | 15:5:7:2:0 | 0:0:0:0:0 | 22:8:3:3:1 | 0:0:0:0:0 | 15:1:1:1:0 | 1 | 1 | 0.528465382 | 0.000195629 | 1.34E-07 | 1.03E-08 |
| 3:01 | 1:01 | 3:01 | 1:09 | 0:01 | 0:02 | 1 | 1 | 1 | 1 | 0.015211392 | 1 |
| 17:0:1:0:0:0:0 | 0:0:0:0:0:0:0 | 30:0:0:0:1:1:2 | 16:0:0:0:0:0:0 | 8:0:0:0:0:0:0 | 4:0:0:0:0:0:0 | 1 | 1 | 1 | 1 | 8.21E-18 | 1 |
| 24:4:20:0 | 0:0:7:0 | 32:9:9:2 | 0:0:5:0 | 10:0:1:0 | 0:0:1:0 | 1 | 1 | 1 | 0.000182133 | 0.002732501 | 1 |
| 24:4:18:0 | 0:0:9:0 | 32:9:8:2 | 0:0:5:0 | 9:0:1:0 | 1:0:1:0 | 1 | 1 | 1 | 1.18E-05 | 0.002732501 | 1 |
| 5:14:26:0 | 0:1:0:0 | 3:2:10:1 | 7:6:25:1 | 0:0:5:0 | 0:1:6:0 | 1 | 0.120678556 | 1 | 1 | 4.94E-07 | 0.004024056 |
| 6:1:1:0 | 5:0:0:0 | 26:1:6:1 | 40:11:25:3 | 9:1:4:1 | 33:1:15:0 | 1 | 1 | 1 | 0.002648305 | 2.65E-08 | 5.40E-12 |
| 764:198:0:1:0 | 5:0:0:0:0 | 363:46:1:0:0 | 2:0:0:0:0 | 198:44:0:0:1 | 1:1:0:0:0 | 1 | 1 | 1 | 0.000787752 | 1 | 1 |

| | | | | | | | | | | | |
|---|---|---|---|---|---|---|---|---|---|---|---|
| 2122:392:1:1:0:0:0 | 10:3:0:0:0:0:0 | 1057:96:1:0:1:0:1 | 11:1:0:0:0:1:0 | 220:49:0:0:1:0:0 | 0:1:0:0:0:0:0 | 1 | 1 | 1 | 8.66E-08 | 3.19E-09 | 1 |
| 2442:497:2:1:1:0:0 | 84:22:0:0:0:0:0 | 1210:121:1:1:1:1:1 | 34:4:0:0:2:0:0 | 233:52:0:0:1:0:0 | 1:1:0:0:0:0:0 | 5.79E-06 | 1 | 1 | 3.52E-80 | 2.32E-32 | 1 |
| 2520:519:2:1:1:0:0 | 11:1:0:0:0:0:0 | 1241:126:1:1:3:1:1 | 4:1:0:0:0:0:0 | 234:52:0:0:1:0:0 | 0:1:0:0:0:0:0 | 1 | 1 | 1 | 2.00E-10 | 0.006512479 | 1 |
| 2550:520:2:1:1:0:0 | 8:1:0:0:0:0:0 | 1249:126:1:1:3:1:1 | 3:1:0:0:0:0:0 | 235:53:0:0:1:0:0 | 0:0:0:0:0:0:0 | 1 | 1 | 1 | 5.40E-07 | 1 | 1 |
| 2428:530:2:1:1:0:0 | 4:2:0:0:0:0:0 | 1171:130:1:1:3:1:1 | 8:1:0:0:0:0:0 | 234:53:0:0:1:0:0 | 2:0:0:0:0:0:0 | 1 | 1 | 1 | 0.010434472 | 5.36E-07 | 1 |
| 2399:531:2:1:1:0:0 | 8:2:0:0:0:0:0 | 1168:130:1:1:3:1:1 | 5:1:0:0:0:0:0 | 234:53:0:0:1:0:0 | 2:0:0:0:0:0:0 | 1 | 1 | 1 | 2.98E-06 | 0.000758491 | 1 |
| 2262:532:2:1:1:0:0 | 12:1:0:0:0:0:0 | 1106:131:1:1:3:1:1 | 4:3:0:0:0:0:0 | 239:53:0:0:1:0:0 | 3:0:0:0:0:0:0 | 1 | 1 | 1 | 1.34E-11 | 0.008848579 | 1 |
| 2189:533:2:1:1:0:0 | 7:1:0:0:0:0:0 | 1077:134:1:1:3:1:1 | 6:0:0:0:0:0:0 | 240:52:0:0:1:0:0 | 1:1:0:0:0:0:0 | 1 | 1 | 1 | 6.42E-06 | 5.92E-06 | 1 |
| 2026:537:2:1:1:0:0 | 7:0:0:0:0:0:0 | 996:136:1:1:3:1:1 | 1:0:0:0:0:0:0 | 239:53:0:0:1:0:0 | 0:0:0:0:0:0:0 | 1 | 1 | 1 | 4.44E-07 | 1 | 1 |
| 1619:537:2:1 | 8:1:0:0 | 792:136:1:3 | 4:1:0:0 | 234:53:0:1 | 0:0:0:0 | 1 | 1 | 1 | 0.000296079 | 0.041786793 | 1 |
| 1550:538:2:1:1:0 | 7:0:0:0:0:0 | 763:136:1:1:3:1 | 1:1:0:0:0:0 | 231:53:0:0:1:0 | 2:0:0:0:0:0 | 1 | 1 | 1 | 3.28E-06 | 1 | 1 |
| 0:9:0:0:0 | 0:5:0:1:0 | 0:0:0:0:0 | 0:4:2:2:0 | 0:0:0:0:0 | 0:3:0:0:1 | 1 | 1 | 1 | 0.00485331 | 0.150388493 | 1 |
| 0:9:0:0:0 | 0:5:0:1:0 | 0:3:2:2:0 | 0:1:0:0:0 | 0:2:0:0:0 | 0:1:0:0:1 | 1 | 1 | 1 | 0.00485331 | 1 | 1 |
| 5429:3492 | 10:10 | 2835:1854 | 1371:959 | 1011:110 | 541:52:00 | 1 | 1 | 1 | 1 | 6.53E-17 | 4.72E-88 |
| 5432:3497 | 17:09 | 4203:2813 | 9:01 | 1550:162 | 3:00 | 1 | 1 | 1 | 0.127966727 | 0.015211392 | 1 |
| 5430:3493 | 24:10:00 | 4200:2813 | 11:02 | 1553:161 | 0:00 | 1 | 1 | 1 | 0.020171785 | 0.016532414 | 1 |
| 6395:3318 | 10:04 | 4733:2658 | 8:01 | 1660:168 | 2:00 | 1 | 1 | 1 | 0.119866821 | 0.025061882 | 1 |
| 10874:3715 | 217:08:00 | 7719:2874 | 61:06:00 | 1832:166 | 3:01 | 1 | 1 | 0.010019977 | 3.09E-43 | 6.27E-11 | 1 |
| 10881:3708 | 460:23:00 | 7730:2867 | 100:11:00 | 1831:164 | 4:03 | 1 | 1 | 0.168359308 | 1.58E-86 | 1.37E-16 | 0.71091155 |
| 10899:3686 | 844:65 | 7733:2849 | 233:32:00 | 1834:166 | 22:03 | 5.37E-06 | 1.32E-10 | 3.13E-08 | 1.31E-145 | 3.47E-34 | 0.000258339 |
| 11353:3724 | 399:20:00 | 7888:2866 | 80:07:00 | 1852:166 | 5:01 | 0.010321833 | 0.005302269 | 0.011087152 | 3.30E-75 | 2.14E-14 | 0.113501247 |
| 11737:3734 | 34:06:00 | 7962:2858 | 14:06 | 1857:162 | 0:01 | 1 | 1 | 1 | 1.86E-05 | 0.084590798 | 1 |
| 3397:11777 | 11:00 | 2559:7984 | 6:01 | 142:1859 | 0:00 | 1 | 1 | 1 | 0.00138895 | 0.068536718 | 1 |
| 11890:929 | 10:19 | 8025:628 | 3:12 | 1859:77 | 2:00 | 1 | 1 | 1 | 0.10520116 | 0.025677895 | 1 |
| 11899:635 | 9:01 | 8024:433 | 8:03 | 1861:67 | 1:01 | 1 | 1 | 1 | 0.014338921 | 0.146312724 | 1 |
| 11901:191 | 8:29 | 8026:142 | 9:15 | 1862:38:00 | 0:04 | 1 | 1 | 0.426673808 | 0.000867113 | 0.237931795 | 1 |
| 267:23:5:2:0:0 | 0:6:2:0:0:0 | 242:19:3:0:1:1 | 1:1:0:0:0:0 | 73:3:1:0:1:0 | 0:1:0:0:0:0 | 1 | 1 | 1 | 0.000822106 | 1 | 1 |
| 267:14:1:2:0:0 | 0:15:6:0:0:0 | 243:16:1:0:1:1 | 0:4:2:0:0:0 | 73:2:0:0:1:0 | 0:2:1:0:0:0 | 1 | 1 | 1 | 7.49E-10 | 0.020069249 | 1 |
| 0:0:0:0:0:0:0:0:0 | 2:1:0:0:0:0:0:0:0 | 0:0:0:0:0:0:0:0:0 | 19:5:0:0:9:0:5:3:2 | 0:0:0:0:0:0:0:0:0 | 22:5:1:0:11:1:9:1:0 | 1 | 0.011416545 | 1 | 1 | 4.48E-10 | 4.36E-13 |

| | | | | | | | | | | | |
|---|---|---|---|---|---|---|---|---|---|---|---|
| 0:0:0:0 | 2:0:0:0 | 3:0:0:0 | 11:0:0:0 | 2:0:0:0 | 8:0:0:0 | 1 | 0.000704574 | 1 | 1 | 7.58E-07 | 4.84E-05 |
| 0:0:0:0 | 0:0:0:2 | 0:0:0:0 | 0:0:0:18 | 0:0:0:0 | 0:0:0:12 | 1 | 0.06987518 | 1 | 1 | 3.87E-11 | 1.97E-07 |
| 27:37:8:0:0 | 5:0:0:0:0 | 21:77:11:1:4 | 1:0:0:0:0 | 17:0:1:0:3 | 0:0:0:0:0 | 1 | 1 | 1 | 0.000787752 | 1 | 1 |
| 0:01:02 | 0:00:00 | 0:00:00 | 0:00:00 | 0:00:00 | 0:00:00 | 1 | 0.035298293 | 1 | 1 | 1 | 1 |
| 904:9:1:10:9:2 | 0:0:0:0:0:0 | 1195:16:2:14:12:4 | 2:0:0:0:0:0 | 257:2:1:2:0:0 | 0:0:0:0:0:0 | 2.47E-103 | 1.07E-83 | 1 | 1 | 1 | 1 |
| 30:08:14 | 29:05:18 | 8:02:09 | 25:04:15 | 5:01:00 | 3:02:03 | 1 | 1 | 0.07339536 | 0.000395051 | 0.000888046 | 0.882496903 |
| 21:32:20:0 | 10:7:15:2 | 22:39:13:3 | 15:19:9:0 | 4:9:4:2 | 5:2:2:0 | 0.144836634 | 0.154096241 | 0.356023712 | 0.01854747 | 0.000465937 | 0.142486432 |
| 27:34:33:2 | 4:5:7:0 | 27:48:16:3 | 10:11:7:0 | 6:11:5:2 | 3:0:1:0 | 0.279887099 | 1 | 0.54250814 | 0.099744641 | 0.018436807 | 1 |
| 77:80:7:13:6 | 0:0:0:1:0 | 48:42:2:10:3 | 18:19:0:3:0 | 18:2:0:2:0 | 4:1:0:1:0 | 1 | 1 | 1 | 1 | 5.13E-09 | 0.068846966 |
| 78:5:1:2 | 67:3:7:3 | 136:2:0:9 | 0:0:0:0 | 10:0:2:2 | 0:0:0:0 | 1 | 1 | 0.01363685 | 4.63E-31 | 1 | 1 |
| 0:0:0:0 | 0:0:1:0 | 0:0:0:0 | 7:1:0:1 | 0:0:0:0 | 1:0:0:0 | 1 | 1 | 1 | 1 | 0.004950569 | 1 |
| 0:0:0:0 | 9:0:0:0 | 0:0:0:0 | 10:0:5:1 | 0:0:0:0 | 8:0:0:0 | 1 | 0.000704574 | 1 | 1.18E-05 | 0.002217522 | 4.84E-05 |
| 0:0:0:0 | 13:0:0:0 | 2:1:6:2 | 8:0:0:0 | 6:0:0:0 | 3:0:0:0 | 1 | 0.000704574 | 1 | 4.58E-08 | 4.90E-05 | 1 |
| 0:0:0:0 | 13:0:0:0 | 2:1:6:3 | 8:0:0:0 | 6:0:0:0 | 3:0:0:0 | 1 | 1 | 1 | 4.58E-08 | 4.90E-05 | 1 |
| 5:50 | 4:37 | 0:00 | 0:37 | 0:00 | 1:02 | 5.27E-43 | 1 | 0.111416266 | 6.09E-07 | 3.53E-09 | 1 |
| 41:49:184:1:2:0 | 7:9:0:0:0:0 | 81:109:430:0:1:2 | 3:6:0:0:0:0 | 85:3:33:0:0:0 | 1:1:0:0:0:0 | 1 | 1 | 0.019323705 | 8.68E-06 | 0.001303035 | 1 |
| 39:10:225:1:3:0 | 2:37:0:0:0:0 | 81:14:526:0:1:2 | 0:97:0:0:0:0 | 85:3:34:0:0:0 | 0:0:0:0:0:0 | 1 | 1 | 1 | 1.52E-34 | 2.59E-101 | 1 |
| 130:4:6:1:1:1 | 6:0:0:0:0:0 | 124:3:6:0:0:0 | 1:0:1:0:0:0 | 12:0:2:0:0:0 | 0:0:0:0:0:0 | 1 | 1 | 1 | 2.84E-05 | 1 | 1 |
| 34:221:660:0 | 0:0:2:0 | 20:293:780:2 | 0:1:5:0 | 3:147:48:0 | 0:0:0:0 | 1 | 1 | 1 | 1 | 0.013499631 | 1 |
| 223:484:20:0 | 1:4:0:0 | 299:569:5:2 | 0:3:0:0 | 148:37:2:0 | 0:0:0:0 | 1 | 1 | 1 | 0.04093502 | 1 | 1 |
| 36:2688:4:101:41:3:7 | 0:1:1:0:0:0:0 | 33:5258:24:490:94:9:17 | 0:8:0:0:0:0:0 | 13:1507:5:190:19:12:1 | 0:1:0:0:1:0:0 | 1 | 1 | 1 | 1 | 3.28E-08 | 1 |
| 36:2688:4:98:39:3:7 | 0:1:0:0:0:0:0 | 33:5258:24:486:89:9:17 | 0:8:0:0:1:0:0 | 13:1505:4:189:21:12:1 | 0:3:0:1:0:0:0 | 1 | 1 | 1 | 1 | 5.36E-07 | 1 |
| 2688:36:6:33:3:8:7 | 1:0:0:1:0:0:0 | 1901:7:9:26:4:8:6 | 3365:26:15:29:5:10:11 | 554:5:2:5:5:0:1 | 954:8:2:12:7:0:0 | 1 | 1 | 1 | 1 | 0 | 0 |
| 36:2687:6:93:3:28:7 | 0:1:0:0:0:0:0 | 33:5254:26:474:9:39:17 | 0:8:1:0:0:0:0 | 13:1508:5:192:12:15:1 | 0:0:0:0:0:0:0 | 1 | 1 | 1 | 1 | 5.36E-07 | 1 |
| 2686:36:6:93:3:25:7 | 3:0:0:1:0:0:0 | 5258:33:29:475:9:39:17 | 4:0:0:2:0:0:0 | 1507:13:5:191:12:15:1 | 1:0:0:1:0:0:0 | 1 | 1 | 1 | 1 | 0.011137275 | 1 |
| 2677:36:6:95:3:22:7 | 1:0:0:0:0:0:0 | 5248:33:30:479:9:34:17 | 4:0:0:0:0:2:0:0 | 1505:13:5:192:12:15:1 | 1:0:0:0:0:0:0 | 1 | 1 | 1 | 1 | 0.011137275 | 1 |
| 36:2668:13:98:76:3:7 | 0:0:0:0:0:0:0 | 32:5215:42:479:163:9:16 | 1:10:0:0:0:0:0 | 13:1496:6:191:28:12:1 | 0:1:0:0:0:0:0 | 1 | 1 | 1 | 1 | 3.15E-09 | 1 |
| 36:2655:14:98:88:3:7 | 0:3:0:0:1:0:0 | 32:5198:42:475:195:9:16 | 0:5:0:1:0:0:0 | 13:1496:6:189:30:12:1 | 0:0:0:2:0:0:0 | 1 | 1 | 1 | 1 | 0.000758491 | 1 |
| 36:2645:19:98:104:3:7 | 0:6:1:0:0:0:0 | 33:5174:49:475:204:9:16 | 0:11:0:0:1:0:0 | 13:1494:6:191:34:12:1 | 0:2:0:0:0:0:0 | 4.37E-07 | 1 | 1 | 7.10E-05 | 2.20E-10 | 1 |
| 0:4:0:0:0:1:0 | 0:2:1:0:1:4:0:0 | 0:0:0:0:0:0:0:0 | 0:4:0:0:0:3:0:1 | 0:0:0:0:0:0:0:0 | 0:4:0:0:0:1:0:0 | 1 | 1 | 0.528465382 | 0.059265681 | 0.015819578 | 0.003640542 |
| 5:47:90:4:1:1:2 | 0:0:2:0:0:0:0 | 7:108:255:5:0:3:2 | 0:2:3:0:0:0:0 | 0:5:28:0:1:0:0 | 0:0:1:1:0:0:0 | 1 | 1 | 1 | 1 | 0.047360969 | 1 |
| 0:00 | 9:18 | 0:00 | 14:21 | 0:00 | 3:01 | 0.023936139 | 0.112198641 | 1 | 0.092970541 | 0.252985249 | 1 |
| 18:27:99:10:0 | 0:0:0:0:0 | 19:22:89:6:0 | 20:15:81:7:0 | 0:0:0:1:0 | 15:16:8:0:0 | 1 | 1 | 1 | 1 | 4.23E-35 | 7.14E-06 |
| 18:25:107:11:0 | 0:0:0:0:0 | 37:37:186:15:0 | 0:0:0:0:0 | 15:16:9:1:0 | 0:0:0:0:0 | 0.003355337 | 4.59E-05 | 1 | 1 | 1 | 1 |
| 11:00 | 0:00 | 27:00:00 | 16:02 | 8:00 | 4:00 | 1 | 1 | 1 | 1 | 0.001526854 | 1 |
| 0:00 | 34:01:00 | 17:00 | 0:00 | 5:00 | 0:00 | 1 | 1 | 0.017871477 | 6.28E-08 | 1 | 1 |
| 7:0:4:0:0:1:1:0:0 | 3:1:3:2:1:0:0:0:0 | 2:0:0:0:0:0:0:0:1 | 5:0:2:0:0:0:0:1:0 | 1:0:0:0:0:0:0:0:0 | 0:0:0:0:0:0:0:0:0 | 1 | 1 | 1 | 0.184247002 | 0.001813262 | 1 |
| 0:3:0:0:1:0:0 | 6:1:1:1:0:0:0 | 0:1:0:0:0:0:1 | 2:3:0:0:0:1:0 | 0:1:0:0:0:0:0 | 0:0:0:0:0:0:0 | 1 | 1 | 1 | 0.002363323 | 0.124057299 | 1 |

| | | | | | | | | | | | |
|---|---|---|---|---|---|---|---|---|---|---|---|
| 580:114:1 | 12:00:00 | 652:169:1 | 4:00:00 | 144:21:00 | 1:00:00 | 1 | 1 | 1 | 1.22E-05 | 1 | 1 |
| 27:0:0:2:1:0:0:0:1 | 131:1:2:12:3:1:3:1:2 | 4:0:0:0:0:0:0:0:0 | 42:0:0:2:0:0:2:0:2 | 1:0:0:0:0:0:0:0:0 | 13:0:0:0:0:0:0:0:1 | 5.86E-38 | 5.69E-09 | 1.02E-22 | 3.96E-175 | 6.00E-56 | 2.15E-16 |
| 3211:38:00 | 5:00 | 1313:09:00 | 5:00 | 180:03:00 | 1:00 | 1 | 1 | 1 | 0.030173969 | 0.031296179 | 1 |
| 3135:51:00 | 5:00 | 1290:23:00 | 6:00 | 181:03:00 | 0:00 | 1 | 1 | 1 | 0.030173969 | 0.018436807 | 1 |
| 0:00 | 1:01 | 0:00 | 9:00 | 0:00 | 1:00 | 1 | 1 | 0.030109098 | 1 | 0.003951718 | 1 |
| 493:1710:922:244:67:58:142:123:3:119:14:46:79:48:10:4:4:3:2:5:1 | 0:7:0:0:0:0:0:0:0:24:0:0:0:0:0:0:0:0:0:0:0 | 380:873:353:312:124:74:40:99:5:65:15:52:62:24:5:11:8:7:1:2:2 | 0:6:0:0:0:0:0:0:0:12:0:0:0:0:0:0:0:0:0:0:0 | 156:370:182:48:13:11:10:27:0:14:9:14:13:3:1:0:0:2:0:0:0 | 0:1:0:0:0:0:0:0:0:2:0:0:0:0:0:0:0:0:0:0:0 | 1 | 1 | 1 | 1.52E-69 | 2.66E-29 | 1 |
| 142:145:1711:922:244:67:57:123:491:3:14:46:79:47:10:4:4:3:2:5:1 | 0:24:6:0:0:0:0:0:0:0:0:0:0:0:0:0:0:0:0:0:0 | 40:87:875:353:312:124:74:98:379:5:15:52:62:24:5:11:8:7:1:2:2 | 0:9:5:0:0:0:0:1:0:0:0:0:0:0:0:0:0:0:0:0:0 | 10:21:372:182:48:13:11:26:156:0:9:14:13:3:1:0:0:2:0:0:0 | 0:1:0:0:0:0:0:1:0:0:0:0:0:0:0:0:0:0:0:0:0 | 1 | 1 | 1 | 8.77E-71 | 2.42E-18 | 1 |
| 152:920:244:67:57:1707:142:123:491:3:14:46:79:47:10:4:4:3:2:5:1 | 19:2:0:0:0:10:0:0:0:0:0:0:0:0:0:0:0:0:0:0:0 | 92:353:312:124:74:876:40:99:379:5:15:52:61:24:5:11:8:7:1:2:2 | 6:0:0:0:0:4:0:0:0:0:0:0:1:0:0:0:0:0:0:0:0 | 22:182:48:13:11:371:10:27:156:0:9:14:13:3:1:0:0:2:0:0:0 | 0:0:0:0:0:1:0:0:0:0:0:0:0:0:0:0:0:0:0:0:0 | 1 | 1 | 1 | 1.24E-47 | 2.27E-10 | 1 |
| 1712:923:243:67:57:142:123:491:158:3:14:46:79:47:10:4:4:3:2:5:1 | 8:0:1:0:0:0:0:0:22:0:0:0:0:0:0:0:0:0:0:0:0 | 876:353:312:124:74:40:99:378:102:5:15:51:62:24:5:11:8:7:1:2:2 | 4:0:0:0:0:0:0:0:8:0:0:1:0:0:0:0:0:0:0:0:0 | 372:182:48:13:11:10:27:156:21:0:9:14:13:3:1:0:0:2:0:0:0 | 0:0:0:0:0:0:0:0:1:0:0:0:0:0:0:0:0:0:0:0:0 | 1 | 1 | 1 | 4.24E-59 | 3.15E-15 | 1 |
| 1721:923:244:67:57:142:123:490:199:3:14:46:79:47:10:4:4:3:2:5:1 | 6:0:0:0:0:0:0:1:15:0:0:0:0:0:0:0:0:0:0:0:0 | 887:353:312:124:73:40:99:378:151:5:15:52:62:24:5:11:8:7:1:2:2 | 6:0:0:0:1:0:0:0:4:0:0:0:0:0:0:0:0:0:0:0:0 | 372:182:48:13:11:10:27:156:30:0:9:14:12:3:1:0:0:2:0:0:0 | 2:0:0:0:0:0:0:0:2:0:0:0:1:0:0:0:0:0:0:0:0 | 1 | 1 | 1 | 1.92E-36 | 2.27E-10 | 0.040807783 |
| 1722:923:244:66:57:140:123:491:217:3:14:46:79:47:10:4:4:3:2:5:1 | 5:0:0:1:0:1:0:0:10:0:0:0:0:0:0:0:0:0:0:0:0 | 889:351:312:124:74:40:99:377:175:5:15:52:62:24:5:11:8:7:1:2:2 | 4:2:0:0:0:0:0:0:1:0:0:0:0:0:0:0:0:0:0:0:0 | 374:182:48:13:11:10:27:156:34:0:9:14:13:3:1:0:0:2:0:0:0 | 0:0:0:0:0:0:0:0:0:0:0:0:0:0:0:0:0:0:0:0:0 | 1 | 1 | 1 | 2.34E-19 | 5.65E-05 | 1 |
| 919:244:67:57:141:1729:122:490:272:3:14:46:79:47:10:4:4:3:2:5:1 | 3:0:0:0:0:1:1:0:1:0:0:0:0:0:0:0:0:0:0:0:0 | 352:312:124:73:40:901:99:376:225:5:15:52:62:24:5:11:8:7:1:2:2 | 0:0:0:1:0:2:0:0:0:0:0:0:0:0:0:0:0:0:0:0:0 | 182:48:13:11:10:378:27:156:46:0:9:14:13:3:1:0:0:2:0:0:0 | 0:0:0:0:0:0:0:0:0:0:0:0:0:0:0:0:0:0:0:0:0 | 1 | 1 | 1 | 0.019044153 | 1 | 1 |
| 921:244:67:57:141:1732:123:490:307:3:14:46:78:47:10:4:4:3:2:5:1 | 1:0:0:0:0:2:0:0:7:0:0:0:1:0:0:0:0:0:0:0:0 | 353:312:124:74:40:904:99:375:252:5:15:52:62:24:5:11:8:7:1:2:2 | 0:0:0:0:0:1:0:0:4:0:0:0:0:0:0:0:0:0:0:0:0 | 182:48:13:11:10:378:27:156:58:0:9:14:13:3:1:0:0:2:0:0:0 | 0:0:0:0:0:0:0:0:0:0:0:0:0:0:0:0:0:0:0:0:0 | 1 | 1 | 1 | 4.56E-11 | 1.58E-06 | 1 |
| 335:922:244:67:57:141:1739:123:490:3:14:46:79:47:10:4:4:3:2:5:1 | 4:0:0:0:0:0:1:0:0:0:0:0:0:0:0:0:0:0:0:0:0 | 271:352:311:124:74:40:911:99:375:5:14:52:62:23:5:11:8:7:1:2:2 | 5:0:1:0:0:0:0:0:0:0:1:0:0:1:0:0:0:0:0:0:0 | 58:182:48:13:11:10:378:27:156:0:9:14:13:3:1:0:0:2:0:0:0 | 0:0:0:0:0:0:0:0:0:0:0:0:0:0:0:0:0:0:0:0:0 | 1 | 1 | 1 | 1.61E-06 | 2.17E-06 | 1 |
| 334:921:244:67:57:141:1738:123:490:3:14:46:79:47:10:4:4:3:2:5:1 | 7:0:0:0:0:0:3:0:0:0:0:0:0:0:0:0:0:0:0:0:0 | 272:350:311:124:74:40:910:99:374:5:15:52:62:24:5:11:8:7:1:2:2 | 5:1:1:0:0:0:0:0:1:0:0:0:0:0:0:0:0:0:0:0:0 | 58:182:48:13:11:10:378:27:156:0:9:14:13:3:1:0:0:2:0:0:0 | 0:0:0:0:0:0:0:0:0:0:0:0:0:0:0:0:0:0:0:0:0 | 1 | 1 | 1 | 3.35E-14 | 2.17E-06 | 1 |
| 490:921:244:67:56:141:1749:123:360:3:14:46:79:47:10:4:4:3:2:5:1 | 0:1:0:0:0:0:0:0:4:0:0:0:0:0:0:0:0:0:0:0:0 | 375:350:312:124:74:40:914:99:288:5:15:52:62:24:5:11:8:7:1:2:2 | 0:2:0:0:0:0:1:0:5:0:0:0:0:0:0:0:0:0:0:0:0 | 156:181:48:13:11:10:377:27:58:0:9:14:13:3:1:0:0:2:0:0:0 | 0:1:0:0:0:0:1:0:0:0:0:0:0:0:0:0:0:0:0:0:0 | 1 | 1 | 1 | 1.61E-06 | 3.37E-07 | 1 |
| 921:244:67:56:140:1752:123:489:387:3:14:46:79:47:10:4:4:3:2:5:1 | 1:0:0:0:1:2:0:0:3:0:0:0:0:0:0:0:0:0:0:0:0 | 352:312:124:74:40:920:97:373:314:5:15:52:62:24:5:11:8:7:1:2:2 | 0:0:0:0:0:0:0:1:2:0:0:0:0:0:0:0:0:0:0:0:0 | 182:48:13:11:9:380:27:156:65:0:9:14:13:3:1:0:0:2:0:0:0 | 0:0:0:0:1:0:0:0:0:0:0:0:0:0:0:0:0:0:0:0:0 | 1 | 1 | 1 | 0.01130807 | 1 | 1 |

| | | | | | | | | | | | |
|---|---|---|---|---|---|---|---|---|---|---|---|
| 1756:921:244:67:56:141:123:489:429:3:14:46:79:47:10:4:4:3:2:5:1 | 4:1:0:0:0:0:0:0:2:0:0:0:0:0:0:0:0:0:0:0 | 920:350:312:124:74:40:97:374:343:5:15:51:62:24:5:11:8:7:1:2:2 | 2:1:0:0:0:0:0:0:3:0:0:0:0:0:0:0:0:0:0:0 | 380:182:48:13:11:10:27:156:71:0:9:14:13:3:1:0:0:2:0:0:0 | 0:0:0:0:0:0:0:0:0:0:0:0:0:0:0:0:0:0:0:0 | 1 | 1 | 1 | 5.34E-05 | 0.003066482 | 1 |
| 919:243:67:56:1777:141:121:488:539:3:14:46:77:47:10:4:4:3:2:5:1 | 1:0:0:0:5:0:0:0:3:0:0:0:0:0:0:0:0:0:0:0 | 347:312:124:74:940:40:94:373:429:5:15:50:60:23:5:11:8:7:1:2:2 | 1:0:0:0:1:0:0:0:4:0:0:0:0:1:0:0:0:0:0:0 | 182:48:13:11:381:10:27:156:89:0:9:14:13:3:1:0:0:2:0:0:0 | 0:0:0:0:0:0:0:0:1:0:0:0:0:0:0:0:0:0:0:0 | 1 | 1 | 1 | 1.64E-07 | 0.000392176 | 1 |
| 487:599:918:243:67:56:1789:141:118:3:14:46:77:47:10:4:4:3:2:5:1 | 0:20:2:0:0:0:8:0:1:0:0:0:0:0:0:0:0:0:0:0 | 373:495:348:311:124:74:946:40:92:5:15:48:59:24:5:11:8:7:1:2:2 | 0:7:0:1:0:0:4:0:0:0:0:0:0:0:0:0:0:0:0:0 | 156:102:182:48:13:11:381:10:27:0:9:14:13:3:1:0:0:2:0:0:0 | 0:2:0:0:0:0:0:0:0:0:0:0:0:0:0:0:0:0:0:0 | 1 | 1 | 1 | 3.55E-48 | 1.14E-12 | 1 |
| 622:919:243:67:56:1794:141:119:487:3:14:46:76:47:10:4:4:3:2:5:1 | 20:0:0:0:0:6:0:0:0:0:0:0:0:0:0:0:0:0:0:0 | 522:348:312:123:73:950:40:92:373:5:15:48:59:24:5:11:8:7:1:2:2 | 10:0:0:0:1:0:4:0:0:0:0:0:0:0:0:0:0:0:0:0 | 107:182:48:13:11:381:10:27:156:0:9:14:13:3:1:0:0:2:0:0:0 | 1:0:0:0:0:0:0:0:0:0:0:0:0:0:0:0:0:0:0:0 | 1 | 1 | 1 | 3.87E-56 | 5.69E-21 | 1 |
| 629:918:243:67:56:1796:139:118:487:3:14:46:76:47:10:4:4:3:2:5:1 | 26:0:0:0:0:6:0:1:0:0:0:0:0:0:0:0:0:0:0:0 | 537:348:312:124:73:952:40:92:371:5:15:47:59:24:5:11:8:7:1:2:2 | 8:0:0:0:0:3:0:0:0:0:0:0:0:0:0:0:0:0:0:0 | 110:182:48:13:11:382:10:27:156:0:9:14:13:3:1:0:0:2:0:0:0 | 1:0:0:0:0:0:0:0:0:0:0:0:0:0:0:0:0:0:0:0 | 1 | 1 | 1 | 2.20E-75 | 3.76E-17 | 1 |
| 634:919:243:67:56:1798:141:118:487:3:14:46:76:47:10:4:4:3:2:5:1 | 21:0:0:0:0:5:0:0:0:0:0:0:0:0:0:0:0:0:0:0 | 535:348:312:124:73:952:40:92:372:5:15:47:59:24:5:11:8:7:1:2:2 | 11:0:0:0:0:3:0:0:0:0:0:0:0:0:0:0:0:0:0:0 | 110:182:48:13:11:381:10:27:156:0:9:14:13:3:1:0:0:2:0:0:0 | 1:0:0:0:0:0:0:0:0:0:0:0:0:0:0:0:0:0:0:0 | 1 | 1 | 1 | 5.48E-61 | 3.74E-27 | 1 |
| 729:919:243:67:56:1812:141:118:487:3:14:45:76:47:10:4:4:3:2:5:1 | 19:0:0:0:0:8:0:0:0:0:0:1:0:0:0:0:0:0:0:0 | 624:346:312:124:73:976:40:91:372:5:15:47:58:24:5:10:8:7:1:2:2 | 8:0:0:0:0:2:0:0:0:0:0:0:0:0:0:0:0:0:0:0 | 115:182:48:13:11:382:10:26:156:0:9:14:13:3:1:0:0:2:0:0:0 | 0:0:0:0:0:0:0:0:0:0:0:0:0:0:0:0:0:0:0:0 | 1 | 1 | 0.005703675 | 3.88E-49 | 5.64E-18 | 1 |
| 1818:782:919:242:65:56:140:117:487:3:14:46:76:47:10:4:4:3:2:5:1 | 5:19:0:0:0:0:0:1:0:0:0:0:0:0:0:0:0:0:0:0 | 982:663:345:310:124:73:40:91:370:5:15:46:57:24:5:10:8:7:1:2:2 | 6:14:0:1:0:0:0:0:1:0:0:0:0:0:0:0:0:0:0:0 | 383:121:182:48:13:11:10:26:156:0:9:14:13:3:1:0:0:2:0:0:0 | 0:2:0:0:0:0:0:0:0:0:0:0:0:0:0:0:0:0:0:0 | 1 | 1 | 0.005703675 | 7.21E-51 | 3.97E-31 | 1 |
| 1821:803:918:242:64:56:141:117:487:3:46:76:13:47:10:4:4:3:2:5:1 | 8:28:0:0:0:0:0:0:0:0:0:0:1:0:0:0:0:0:0:0 | 985:708:344:311:124:72:40:90:371:5:46:57:15:24:5:10:8:7:1:2:2 | 4:12:0:0:0:1:0:0:0:0:0:0:0:0:0:0:0:0:0:0 | 385:129:181:48:13:11:10:25:156:0:14:13:9:3:1:0:0:2:0:0:0 | 0:0:0:0:0:0:0:1:0:0:0:0:0:0:0:0:0:0:0:0 | 1 | 1 | 1 | 2.27E-80 | 2.54E-27 | 1 |
| 1824:816:918:242:64:56:141:116:487:3:14:46:76:47:10:4:4:3:2:5:1 | 6:25:0:0:0:0:0:1:0:0:0:0:0:0:0:0:0:0:0:0 | 985:722:345:311:124:73:40:90:371:5:15:46:57:24:5:10:8:7:1:2:2 | 6:9:0:0:0:0:0:0:0:0:0:0:0:0:0:0:0:0:0:0 | 385:129:181:48:13:11:10:26:156:0:9:14:12:3:1:0:0:2:0:0:0 | 0:1:0:0:0:0:0:0:0:0:0:0:0:0:0:0:0:0:0:0 | 1 | 1 | 1 | 1.30E-71 | 5.69E-21 | 1 |
| 916:242:64:56:1839:141:117:486:891:3:14:46:75:47:10:4:4:3:2:5:1 | 1:0:0:0:6:0:0:0:18:0:0:0:0:0:0:0:0:0:0:0 | 343:311:124:73:994:40:90:370:787:5:15:46:57:24:5:10:8:7:1:2:2 | 0:0:0:0:6:0:0:0:8:0:0:0:0:0:0:0:0:0:0:0 | 181:48:13:11:388:10:26:156:139:0:9:14:12:3:1:0:0:2:0:0:0 | 0:0:0:0:0:0:0:0:0:0:0:0:0:0:0:0:0:0:0:0 | 1 | 1 | 1 | 1.80E-46 | 1.45E-18 | 1 |
| 917:241:64:56:1847:140:117:486:903:3:13:46:75:47:10:4:4:3:2:5:1 | 0:1:0:0:1:1:0:0:16:0:1:0:0:0:0:0:0:0:0:0 | 343:310:124:72:999:40:89:370:797:5:15:46:57:24:5:10:8:7:1:2:2 | 0:1:0:1:4:0:0:0:5:0:0:0:0:0:0:0:0:0:0:0 | 181:48:13:11:388:10:26:156:139:0:9:14:12:3:1:0:0:2:0:0:0 | 0:0:0:0:0:0:0:0:0:0:0:0:0:0:0:0:0:0:0:0 | 1 | 1 | 1 | 2.03E-41 | 3.00E-07 | 1 |
| 977:916:242:64:56:1855:141:117:486:3:14:46:75:47:10:4:4:3:2:5:1 | 2:0:0:0:0:3:0:0:0:0:0:0:0:0:0:0:0:0:0:0 | 838:343:310:124:73:1007:40:89:368:5:15:45:55:24:5:10:8:7:1:2:2 | 1:0:0:0:0:4:0:0:2:0:0:0:0:0:0:0:0:0:0:0 | 150:180:48:13:11:392:10:26:154:0:9:14:12:3:1:0:0:2:0:0:0 | 2:1:0:0:0:0:0:0:1:0:0:0:0:0:0:0:0:0:0:0 | 1 | 1 | 1 | 0.000411219 | 5.65E-05 | 1 |

| | | | | | | | | | | | |
|---|---|---|---|---|---|---|---|---|---|---|---|
| 996:915:242:64:56:1861:141:117:486:3:14:46:75:47:10:4:4:3:2:5:1 | 7:1:0:1:0:2:0:0:0:0:0:0:0:0:0:0:0:0:0:0:0 | 855:343:310:124:73:1013:40:88:368:5:15:45:55:24:5:10:8:7:1:2:2 | 13:0:1:0:0:3:0:0:0:0:0:0:0:0:0:0:0:0:0:0:0 | 152:181:48:13:11:392:10:25:155:0:9:14:12:3:1:0:0:2:0:0:0 | 1:0:0:0:0:1:0:0:0:0:0:0:0:0:0:0:0:0:0:0:0 | 1 | 1 | 1 | 4.56E-11 | 1.14E-31 | 1 |
| 913:242:65:56:1877:141:116:486:1081:3:14:45:75:46:10:4:4:3:2:5:1 | 1:0:0:0:1:0:1:0:7:0:0:0:0:1:0:0:0:0:0:0:0 | 341:309:124:73:1035:40:85:367:963:5:15:42:55:24:5:10:8:7:1:2:2 | 0:1:0:0:1:0:1:0:8:0:0:0:0:0:0:0:0:0:0:0:0 | 181:48:13:11:398:10:25:155:171:0:9:14:12:3:1:0:0:2:0:0:0 | 0:0:0:0:0:0:0:0:0:0:0:0:0:0:0:0:0:0:0:0:0 | 1 | 1 | 1 | 2.06E-10 | 4.65E-15 | 1 |
| 1876:1085:913:242:65:56:141:116:485:3:14:45:75:47:10:4:4:3:2:5:1 | 2:3:2:0:0:0:0:1:1:0:0:0:0:0:0:0:0:0:0:0:0 | 1033:966:342:309:123:73:40:86:366:5:15:42:55:24:5:10:8:7:1:2:2 | 2:5:0:1:1:0:0:0:1:0:0:0:0:0:0:0:0:0:0:0:0 | 398:170:181:48:13:11:10:25:155:0:9:14:12:3:1:0:0:2:0:0:0 | 0:1:0:0:0:0:0:0:0:0:0:0:0:0:0:0:0:0:0:0:0 | 1 | 1 | 1 | 0.022613964 | 3.82E-05 | 1 |
| 912:242:65:56:1879:141:116:485:1105:3:14:45:75:47:10:4:4:3:2:5:1 | 2:0:0:0:2:0:1:0:5:0:0:0:0:0:0:0:0:0:0:0:0 | 342:310:123:73:1041:40:86:367:978:5:15:42:55:24:5:10:8:7:1:2:2 | 0:0:1:0:0:0:0:0:3:0:0:0:0:0:0:0:0:0:0:0:0 | 181:48:13:11:398:10:25:155:174:0:9:14:12:3:1:0:0:2:0:0:0 | 0:0:0:0:0:0:0:0:0:0:0:0:0:0:0:0:0:0:0:0:0 | 1 | 1 | 1 | 8.85E-06 | 1 | 1 |
| 910:241:65:56:1879:140:116:484:1104:3:14:45:74:47:10:4:4:3:2:5:1 | 3:1:0:0:0:0:1:1:8:0:0:0:1:0:0:0:0:0:0:0:0 | 341:310:124:72:1041:40:86:367:986:5:15:42:54:24:5:10:8:6:1:2:2 | 1:0:0:0:2:0:0:0:3:0:0:0:1:0:0:0:0:1:0:0:0 | 181:48:13:11:397:10:25:154:175:0:9:14:12:3:1:0:0:2:0:0:0 | 0:0:0:0:1:0:0:1:0:0:0:0:0:0:0:0:0:0:0:0:0 | 1 | 1 | 1 | 7.36E-11 | 0.032219461 | 1 |
| 485:912:242:65:56:1876:141:116:1123:3:14:44:75:47:10:4:4:3:2:5:1 | 0:1:0:0:0:3:0:1:7:0:0:0:0:0:0:0:0:0:0:0:0 | 364:341:309:123:72:1043:40:86:998:5:15:42:55:24:5:10:8:7:1:2:2 | 2:1:1:1:0:1:0:0:3:0:0:0:0:0:0:0:0:0:0:0:0 | 155:181:48:13:11:398:10:25:176:0:9:14:12:3:1:0:0:2:0:0:0 | 0:0:0:0:0:0:0:0:0:0:0:0:0:0:0:0:0:0:0:0:0 | 1 | 1 | 1 | 6.81E-11 | 0.069588407 | 1 |
| 1142:913:242:65:56:1885:141:116:485:3:14:44:75:47:10:4:4:3:2:5:1 | 8:0:0:0:0:2:0:0:0:0:0:0:0:0:0:0:0:0:0:0:0 | 1045:342:309:124:72:1050:40:86:367:5:15:41:55:24:5:10:8:7:1:2:2 | 4:0:1:0:0:4:0:0:0:0:0:0:0:0:0:0:0:0:0:0:0 | 174:181:48:13:11:398:10:25:154:0:9:14:13:3:1:0:0:2:0:0:0 | 0:0:0:0:0:1:0:0:1:0:0:0:0:0:0:0:0:0:0:0:0 | 1 | 1 | 1 | 4.85E-18 | 8.98E-07 | 1 |
| 1155:913:240:65:55:1890:141:116:485:3:14:44:75:47:10:4:4:3:2:5:1 | 7:0:1:0:0:3:0:0:0:0:0:0:0:0:0:0:0:0:0:0:0 | 1069:341:310:124:72:1063:40:86:367:5:15:42:55:24:5:10:8:7:1:2:2 | 5:0:0:0:0:3:0:0:0:0:0:0:0:0:0:0:0:0:0:0:0 | 176:181:48:13:11:401:10:25:155:0:9:14:13:3:1:0:0:2:0:0:0 | 1:0:0:0:0:0:0:0:0:0:0:0:0:0:0:0:0:0:0:0:0 | 1 | 1 | 1 | 2.16E-12 | 7.10E-09 | 1 |
| 913:240:65:54:1891:141:116:484:1165:3:14:44:75:47:10:4:4:3:2:5:1 | 0:1:0:1:3:0:0:0:5:0:0:0:0:0:0:0:0:0:0:0:0 | 341:309:124:72:1066:40:86:367:1080:5:15:42:55:24:5:10:8:7:1:2:2 | 0:1:0:0:4:0:0:0:4:0:0:0:0:0:0:0:0:0:0:0:0 | 181:48:13:11:402:10:25:155:177:0:9:14:13:3:1:0:0:2:0:0:0 | 0:0:0:0:0:0:0:0:2:0:0:0:0:0:0:0:0:0:0:0:0 | 1 | 1 | 1 | 2.13E-06 | 8.98E-07 | 1 |
| 1898:913:241:65:55:141:116:484:1195:3:14:44:75:47:10:4:4:3:2:5:1 | 1:0:0:0:0:0:0:0:2:0:0:0:0:0:0:0:0:0:0:0:0 | 1073:340:310:124:72:40:86:367:1095:5:15:42:55:24:5:10:8:7:1:2:2 | 3:1:0:0:0:0:0:0:5:0:0:0:0:0:0:0:0:0:0:0:0 | 402:181:48:13:11:10:25:155:179:0:9:14:13:3:1:0:0:2:0:0:0 | 0:0:0:0:0:0:0:0:1:0:0:0:0:0:0:0:0:0:0:0:0 | 1 | 1 | 1 | 1 | 1.70E-07 | 1 |
| 1909:913:241:65:55:141:116:483:1221:3:14:44:75:47:10:4:4:3:2:5:1 | 2:0:0:0:0:0:0:0:4:0:0:0:0:0:0:0:0:0:0:0:0 | 1088:341:310:124:72:40:85:366:1118:5:15:42:55:24:5:10:8:7:1:2:2 | 2:0:0:0:0:0:0:1:6:0:0:0:0:0:0:0:0:0:0:0:0 | 404:181:48:13:11:10:25:154:181:0:9:14:13:3:1:0:0:2:0:0:0 | 0:0:0:0:0:0:0:1:1:0:0:0:0:0:0:0:0:0:0:0:0 | 1 | 1 | 1 | 3.62E-06 | 9.09E-10 | 1 |
| 484:1913:913:241:65:55:141:116:1236:3:14:44:75:47:10:4:4:3:2:5:1 | 0:4:0:0:0:0:0:0:4:0:0:0:0:0:0:0:0:0:0:0:0 | 368:1097:341:310:123:72:40:85:1146:5:15:42:55:24:5:10:8:7:1:2:2 | 0:2:1:0:1:0:0:0:5:0:0:0:0:0:0:0:0:0:0:0:0 | 155:404:181:48:12:11:10:25:183:0:9:14:13:3:1:0:0:2:0:0:0 | 0:1:0:0:0:0:0:0:1:0:0:0:0:0:0:0:0:0:0:0:0 | 1 | 1 | 1 | 4.86E-08 | 4.66E-06 | 1 |
| 484:912:240:65:55:1918:141:116:1249:3:14:44:75:47:10:4:4:3:2:5:1 | 0:1:1:0:0:2:0:0:2:0:0:0:0:0:0:0:0:0:0:0:0 | 368:342:310:124:72:1098:40:85:1157:5:15:42:55:24:5:10:8:7:1:2:2 | 0:0:0:0:0:2:0:0:5:0:0:0:0:0:0:0:0:0:0:0:0 | 155:181:48:12:11:404:10:25:184:0:9:14:13:3:1:0:0:2:0:0:0 | 0:0:0:0:0:1:0:0:1:0:0:0:0:0:0:0:0:0:0:0:0 | 1 | 1 | 1 | 0.097734196 | 1.13E-08 | 1 |

| | | | | | | | | | | | |
|---|---|---|---|---|---|---|---|---|---|---|---|
| 913:241:65:55:1920:141:116:484:1270:3:14:44:75:47:10:4:4:3:2:5:1 | 1:0:0:0:2:0:0:0:5:0:0:0:0:0:0:0:0:0:0:0 | 342:310:124:72:1105:40:85:368:1187:5:15:42:55:24:5:10:8:7:1:2:2 | 0:0:0:0:1:0:0:0:2:0:0:0:0:0:0:0:0:0:0:0 | 181:48:12:11:406:10:25:155:181:0:9:14:13:3:1:0:0:2:0:0:0 | 0:0:0:0:0:0:0:0:0:0:0:0:0:0:0:0:0:0:0:0 | 1 | 1 | 1 | 3.36E-07 | 1 | 1 |
| 914:241:65:55:1920:141:116:484:1271:3:14:44:75:47:10:4:4:3:2:5:1 | 0:0:0:0:3:0:0:0:4:0:0:0:0:0:0:0:0:0:0:0 | 341:310:124:72:1106:40:85:368:1193:5:15:42:55:24:5:10:8:7:1:2:2 | 1:0:0:0:1:0:0:0:4:0:0:0:0:0:0:0:0:0:0:0 | 181:48:12:11:404:10:25:155:182:0:9:13:13:3:1:0:0:2:0:0:0 | 0:0:0:0:2:0:0:0:0:0:0:0:0:0:0:0:0:0:0:0 | 1 | 1 | 1 | 9.06E-07 | 4.09E-05 | 1 |
| 914:240:65:55:1922:141:116:484:1274:3:14:44:74:47:10:4:4:3:2:5:1 | 0:1:0:0:2:0:0:0:5:0:0:0:0:0:0:0:0:0:0:0 | 341:310:124:72:1104:40:85:368:1213:5:15:42:55:24:5:10:8:7:1:2:2 | 1:0:0:0:4:0:0:0:5:0:0:0:0:0:0:0:0:0:0:0 | 180:48:12:11:405:10:25:155:180:0:9:13:13:3:1:0:0:2:0:0:0 | 0:0:0:0:0:0:0:0:0:0:0:0:0:0:0:0:0:0:0:0 | 1 | 1 | 1 | 3.36E-07 | 2.23E-08 | 1 |
| 914:242:65:55:1925:141:116:483:1294:3:14:44:74:47:10:4:4:3:2:5:1 | 0:0:0:0:1:0:0:0:3:0:0:0:0:0:0:0:0:0:0:0 | 342:310:124:72:1110:39:85:369:1223:5:15:42:54:24:5:10:8:7:1:2:2 | 0:0:0:0:2:0:0:0:7:0:0:0:0:0:0:0:0:0:0:0 | 180:48:11:11:404:10:25:154:179:0:9:13:13:3:1:0:0:2:0:0:0 | 0:0:0:0:0:0:0:1:0:0:0:0:0:0:0:0:0:0:0:0 | 1 | 1 | 1 | 1 | 1.18E-14 | 1 |
| 7:11 | 5:13 | 2:08 | 3:15 | 1:04 | 0:02 | 1 | 1 | 1 | 0.068255103 | 0.006667609 | 1 |
| 3876:0:3:1 | 5:0:0:0 | 3309:0:2:1 | 2:0:0:0 | 500:1:0:0 | 0:0:0:0 | 1 | 1 | 1 | 0.002648305 | 1 | 1 |
| 3873:0:3:1 | 8:0:0:0 | 3309:0:2:1 | 5:0:0:0 | 501:1:0:0 | 0:0:0:0 | 1 | 1 | 1 | 4.60E-05 | 0.002732501 | 1 |
| 3874:0:3:1 | 9:0:0:0 | 3313:0:2:1 | 3:0:0:0 | 501:1:0:0 | 0:0:0:0 | 1 | 1 | 1 | 1.18E-05 | 1 | 1 |
| 3672:00:00 | 18:00 | 3195:00:00 | 16:00 | 496:00:00 | 0:00 | 1 | 1 | 1 | 4.15E-05 | 0.000117914 | 1 |
| 3074:00:00 | 362:00:00 | 2758:00:00 | 259:00:00 | 375:00:00 | 27:00:00 | 4.05E-05 | 0.00012042 | 3.46E-07 | 2.34E-79 | 2.81E-57 | 4.97E-07 |
| 3318:00:00 | 21:00 | 2977:00:00 | 21:00 | 477:00:00 | 2:00 | 1 | 1 | 1 | 9.40E-06 | 9.75E-06 | 1 |
| 17:00 | 1320:00:00 | 9:00 | 1330:00:00 | 0:00 | 34:00:00 | 1.32E-06 | 0.000528313 | 7.63E-08 | 5.19E-287 | 2.48E-289 | 1.66E-08 |
| 18:00 | 1324:00:00 | 9:00 | 1342:00:00 | 0:00 | 10:00 | 0.006139527 | 0.029932329 | 1 | 7.79E-288 | 6.56E-292 | 0.002529224 |
| 6:00 | 8:00 | 0:00 | 7:00 | 0:00 | 1:00 | 1 | 1 | 1 | 0.006305042 | 0.011137275 | 1 |
| 70:07:00 | 71:08:00 | 201:15:00 | 0:00 | 109:11:00 | 0:00 | 3.27E-53 | 0.00012042 | 0.000202249 | 4.51E-12 | 1 | 1 |
| 4:17 | 0:00 | 2:15 | 0:00 | 4:11 | 0:00 | 0.000542535 | 0.112198641 | 1 | 1 | 1 | 1 |
| 0:00:00 | 136:01:00 | 258:00:00 | 0:00:00 | 56:00:01 | 0:00:00 | 5.47E-38 | 1.48E-41 | 3.50E-22 | 7.22E-58 | 1 | 1 |
| 0:00 | 37:09:00 | 0:00 | 66:20:00 | 0:00 | 10:11 | 0.416962858 | 0.06690288 | 1 | 6.64E-05 | 1.60E-06 | 0.831512608 |
| 0:00 | 0:00 | 0:00 | 1:00 | 21:00 | 28:01:00 | 0.005176196 | 1 | 1 | 1 | 1 | 1.25E-06 |
| 0:00 | 16:01 | 0:00 | 12:00 | 0:00 | 3:00 | 1 | 1 | 0.030109098 | 0.000445066 | 0.000876481 | 1 |
| 0:00 | 5:00 | 0:00 | 0:00 | 0:00 | 0:00 | 1 | 1 | 1 | 0.030173969 | 1 | 1 |
| 0:0:0:0:0 | 20:0:1:0:0 | 4:0:0:0:0 | 35:0:2:1:4 | 0:1:0:0:0 | 9:0:0:0:0 | 0.000755847 | 1 | 0.209334886 | 9.67E-15 | 2.41E-21 | 6.90E-07 |
| 0:0:0:0:0 | 20:0:1:2:0 | 4:0:0:0:0 | 35:0:2:8:1 | 0:0:0:0:0 | 9:0:0:0:0 | 0.000755847 | 1 | 0.209334886 | 8.59E-13 | 6.88E-19 | 6.90E-07 |
| 0:0:0:0:0 | 18:0:2:1:0 | 3:0:0:0:0 | 35:0:8:2:1 | 0:0:0:0:0 | 8:0:0:0:0 | 0.000755847 | 1 | 0.209334886 | 3.30E-11 | 6.88E-19 | 4.27E-06 |
| 2499:127:420:192 | 4:0:0:0 | 1670:100:266:123 | 1:2:0:0 | 415:22:67:45 | 0:0:0:0 | 0.012449474 | 1 | 1 | 1 | 1 | 1 |
| 192:121:2511:421 | 0:0:1:0 | 123:93:1671:266 | 0:5:2:0 | 45:22:415:68 | 0:0:0:0 | 1 | 1 | 1 | 1 | 0.028663669 | 1 |
| 2516:109:422:193 | 1:2:0:0 | 1677:84:266:123 | 0:2:0:0 | 415:21:68:46 | 0:0:0:0 | 0.042679166 | 1 | 1 | 1 | 1 | 1 |
| 2528:401:13:190 | 5:1:1:0 | 1685:236:8:123 | 2:1:0:0 | 418:67:3:46 | 1:0:0:0 | 0.039854726 | 1 | 1 | 0.044129954 | 1 | 1 |
| 2533:376:16:189 | 7:0:0:0 | 1691:221:8:121 | 2:0:0:1 | 421:66:3:46 | 0:0:0:0 | 1 | 1 | 1 | 0.000182133 | 1 | 1 |
| 236:03:00 | 0:00 | 224:03:00 | 0:00 | 17:02 | 0:00 | 6.60E-08 | 0.003751524 | 1 | 1 | 1 | 1 |
| 7:8:2:0:1 | 29:21:3:0:4 | 37:70:1:0:5 | 0:0:0:0:0 | 10:11:0:0:4 | 0:0:0:0:0 | 1 | 1 | 0.013214975 | 2.85E-11 | 1 | 1 |
| 0:0:0:0 | 7:1:0:1 | 0:0:0:0 | 15:1:1:0 | 0:0:0:0 | 3:2:0:0 | 0.000173965 | 1 | 1.46E-05 | 0.004727276 | 1.58E-07 | 0.15720761 |

| | | | | | | | | | | | |
|---|---|---|---|---|---|---|---|---|---|---|---|
| 3:0:0:1 | 4:1:0:0 | 15:1:1:0 | 0:0:0:0 | 3:2:0:0 | 0:0:0:0 | 0.000173965 | 1 | 1 | 0.04093502 | 1 | 1 |
| 0:00 | 0:00 | 0:00 | 11:02 | 0:00 | 2:00 | 6.43E-05 | 1 | 1 | 1 | 0.016532414 | 1 |
| 15:01 | 7:01 | 59:05:00 | 0:00 | 4:01 | 0:00 | 1 | 1 | 1 | 0.039929313 | 1 | 1 |
| 5:02 | 9:00 | 20:04 | 33:02:00 | 2:01 | 2:00 | 1 | 1 | 1 | 0.00379461 | 3.88E-07 | 1 |
| 46:1896:27:671:24:66:171:10 | 0:5:0:1:0:0:0:0 | 185:1643:76:649:50:73:134:8 | 1:2:0:3:0:0:0:0 | 45:592:11:194:16:16:10:8 | 0:0:0:0:0:0:0:0 | 1 | 1 | 1 | 0.000277059 | 0.092169241 | 1 |
| 1866:46:27:670:24:74:188:10 | 5:0:0:0:0:0:0:0 | 1628:186:76:653:50:80:152:8 | 3:0:0:0:0:0:0:0 | 596:45:11:193:16:16:13:8 | 0:0:0:0:0:0:0:0 | 1 | 1 | 1 | 2.17E-05 | 1 | 1 |
| 46:1887:27:673:24:91:184:9 | 0:4:0:0:1:0:0:0 | 184:1642:75:653:50:96:149:8 | 0:1:0:0:0:3:0:0 | 45:600:11:192:16:15:12:8 | 0:0:0:0:0:0:0:0 | 1 | 1 | 1 | 0.003391368 | 1 | 1 |
| 1734:170:201:2:26:24:9:2:17:3:3:1:0 | 1:0:7:0:0:0:0:0:0:0:0:0:0 | 1763:273:185:0:33:23:5:4:5:2:0:0:1 | 2:1:4:0:0:0:0:0:0:0:0:0:0 | 328:33:31:0:7:5:1:1:1:2:0:0:0 | 1:0:0:0:0:0:0:0:0:0:0:0:0 | 1 | 1 | 1 | 2.34E-10 | 0.002143587 | 1 |
| 1735:169:202:2:23:24:9:2:17:3:3:1:0 | 0:2:4:0:0:0:0:0:0:0:0:0:0 | 1762:275:183:0:31:23:5:4:5:2:0:0:1 | 3:0:2:0:0:0:0:0:0:0:0:0:0 | 328:33:29:0:7:5:1:1:1:2:0:0:0 | 1:0:0:0:0:0:0:0:0:0:0:0:0 | 1 | 1 | 1 | 0.000324681 | 0.006124303 | 1 |
| 226:14:1:8:0:10:19:0:2:8:1:27:10:3:1:1:0:0:0:0 | 10:0:0:0:0:0:0:0:0:0:0:0:0:0:0:0:0:0:0:0 | 260:10:1:8:2:7:13:2:2:6:5:41:2:3:0:0:5:1:1:1 | 4:0:0:0:0:0:0:0:0:0:0:1:0:0:0:0:0:0:0:0 | 27:2:0:1:2:2:1:0:0:0:0:3:0:0:0:1:2:0:0:0 | 0:0:0:1:0:0:0:0:0:0:0:0:0:0:0:0:0:0:0:0 | 1 | 1 | 1 | 1.85E-29 | 2.85E-06 | 1 |
| 163:8:11:8:0:14:1:0:24:2:7:1:5:3:1:1:0:0:0:0 | 54:2:4:0:0:2:0:0:0:0:0:0:3:0:0:0:0:0:0:0 | 191:4:6:6:2:7:2:2:31:1:5:1:2:3:0:0:4:1:1:1 | 57:3:4:1:0:5:3:0:5:1:1:0:0:0:0:0:1:0:0:0 | 11:1:0:0:2:0:0:0:2:0:0:0:0:0:0:1:2:0:0:0 | 15:1:2:1:0:1:0:0:0:0:0:0:0:0:0:0:0:0:0:0 | 0.005858491 | 1.20E-25 | 0.000115477 | 1.18E-164 | 8.26E-144 | 1.28E-33 |
| 6:7:89:0:9:17:4:6:1:0:5:3:1:1:1:0:0:0:0 | 0:0:13:0:1:0:1:0:0:0:1:0:0:0:0:0:0:0:0 | 3:3:120:2:5:19:3:4:1:1:5:2:1:1:0:2:1:0:2 | 1:0:11:0:0:3:0:0:0:0:0:0:0:0:0:0:0:1:0 | 0:0:9:2:0:3:1:0:0:0:0:0:0:0:1:2:0:0:0 | 0:0:0:0:0:0:0:0:0:0:0:0:0:0:0:0:0:0:0 | 1 | 1 | 1 | 1.26E-29 | 2.43E-20 | 1 |
| 0:6:0:1:0:2:2:1:2:1:1:2:0:0 | 0:2:0:0:0:3:0:0:0:0:0:0:0:0 | 2:5:0:0:2:7:0:0:0:0:3:0:1:1 | 0:2:0:0:0:0:0:0:0:0:1:0:0:0 | 2:1:0:0:0:0:0:0:0:0:1:1:0:0 | 0:0:0:0:0:0:0:0:0:0:0:0:0:0 | 1 | 1 | 1 | 0.004123223 | 1 | 1 |
| 0:1:0:0:0:0:2:1:1:0:1:2:0:0 | 0:7:0:0:0:1:3:1:1:1:0:0:0:0 | 2:2:0:0:2:0:7:0:0:2:0:0:1:1 | 0:5:0:0:0:0:0:0:0:1:0:0:0:0 | 2:1:0:0:0:0:0:0:0:0:0:1:0:0 | 0:0:0:0:0:0:0:0:0:1:0:0:0:0 | 1 | 1 | 1 | 1.29E-05 | 1.07E-06 | 1 |
| 0:0:0:0:0:0:0:0:0:0 | 13:19:1:0:0:3:1:1:1:0 | 4:6:2:1:0:2:0:0:0:0 | 9:5:0:1:0:2:1:1:1:0 | 4:4:0:0:1:1:1:0:0:0 | 4:3:0:0:0:0:1:0:0:0 | 5.66E-68 | 1 | 1.17E-08 | 5.58E-17 | 5.15E-05 | 0.005199327 |
| 13:19:7:3:0:0:1:1:0 | 0:0:0:0:0:0:0:0:0 | 9:5:7:2:0:0:1:1:0 | 3:6:3:1:0:0:0:0:0 | 4:3:1:0:0:0:0:0:0 | 4:4:1:1:0:1:0:0:0 | 1 | 1 | 1 | 1 | 0.002323676 | 0.030032732 |
| 0:0:0:0:0:0 | 11:11:17:9:3:2 | 0:0:0:0:0:0 | 22:15:26:6:11:2 | 0:0:0:0:0:0 | 18:20:21:3:11:3 | 3.62E-09 | 0.557471904 | 0.004168205 | 0.004564638 | 1.84E-05 | 0.000109789 |
| 0:00:00 | 9:01:00 | 0:00:00 | 37:00:01 | 0:00:00 | 22:02:00 | 0.0086191 | 1.97E-06 | 1 | 0.001047753 | 2.58E-15 | 2.72E-08 |
| 15:01 | 0:00 | 3:00 | 15:00 | 0:00 | 0:00 | 1 | 1 | 1 | 1 | 0.000196442 | 1 |
| 274:483:2:17:1576:67:1:10:4:1 | 0:0:0:0:5:0:0:0:0:0 | 171:504:4:21:2325:97:5:13:12:3 | 0:1:0:0:3:0:0:0:0:0 | 28:57:1:0:101:9:0:0:1:0 | 0:0:0:0:1:0:0:0:0:0 | 1 | 1 | 1 | 2.10E-06 | 1 | 1 |
| 460:268:2:17:1448:64:1:10:3:1 | 23:6:0:0:110:3:0:0:0:0 | 472:168:4:21:2152:89:5:16:12:3 | 34:2:0:0:143:5:0:0:0:0 | 57:28:1:0:101:9:0:1:1:0 | 0:0:0:0:1:0:0:0:0:0 | 1.45E-06 | 1 | 1 | 3.98E-154 | 5.61E-206 | 1 |
| 268:459:2:17:1448:64:1:10:3:1 | 6:25:0:0:115:3:0:0:0:0 | 168:473:4:21:2151:89:5:16:12:3 | 3:36:0:0:152:5:0:0:0:0 | 28:57:1:0:100:9:0:1:1:0 | 0:1:0:0:2:0:0:0:0:0 | 9.36E-10 | 1 | 1 | 3.42E-161 | 1.19E-218 | 1 |
| 267:462:2:17:1429:61:1:11 | 0:2:0:0:7:0:0:0 | 165:470:5:21:2076:82:5:17 | 1:6:0:0:5:1:0:0 | 28:57:1:0:93:9:0:0 | 0:0:0:0:0:0:0:0 | 1 | 1 | 1 | 6.05E-06 | 0.000908241 | 1 |
| 14:0:3:4:1 | 22:0:0:8:1 | 7:0:0:0:0 | 0:0:0:0:0 | 5:1:0:0:1 | 0:0:0:0:0 | 0.681467576 | 1 | 0.100638069 | 2.99E-11 | 1 | 1 |
| 716:2105:3105:1003:195:65 | 1:3:4:0:0:0 | 657:2023:1909:599:134:45 | 0:5:1:3:0:0 | 234:743:700:216:54:5 | 0:1:0:0:1:0 | 0.018731299 | 1 | 1 | 0.049116069 | 0.017791031 | 1 |
| 716:3105:1001:2103:195:65 | 1:3:2:5:0:0 | 656:1906:599:2025:134:45 | 0:3:2:4:0:0 | 234:698:216:742:54:5 | 0:2:0:1:1:0 | 0.04307984 | 1 | 1 | 0.077182787 | 0.076429239 | 1 |

| | | | | | | | | | | | |
|---|---|---|---|---|---|---|---|---|---|---|---|
| 3101:1003:2105:195:717:65 | 5:0:3:0:0:0 | 1905:602:2026:134:656:45 | 4:1:3:0:1:0 | 697:216:742:54:234:5 | 2:0:2:1:0:0 | 0.074196581 | 1 | 1 | 0.005016446 | 0.121941343 | 0.338281183 |
| 3081:1005:2103:195:717:65 | 8:0:5:0:0:0 | 1891:604:2029:134:659:45 | 4:0:2:0:0:0 | 696:216:745:54:234:5 | 0:0:1:1:0:0 | 0.150747748 | 1 | 1 | 6.40E-05 | 0.020069249 | 1 |
| 3019:1007:2112:195:722:65 | 5:0:1:0:1:0 | 1833:605:2035:134:663:45 | 2:1:5:0:0:0 | 686:216:745:54:236:5 | 2:0:1:1:0:0 | 0.231190676 | 1 | 1 | 0.00866799 | 0.016698024 | 1 |
| 2109:1005:3016:195:723:64 | 3:2:3:0:1:1 | 2035:606:1823:134:665:45 | 5:0:0:0:0:0 | 746:216:684:53:236:5 | 0:0:2:2:0:0 | 0.071157571 | 1 | 1 | 0.507825879 | 0.000252891 | 1 |
| 3010:1004:2110:195:724:65 | 8:3:2:0:0:0 | 1820:606:2037:134:665:45 | 0:0:3:0:0:0 | 686:215:745:54:236:5 | 0:1:1:1:0:0 | 0.005948649 | 1 | 1 | 0.000658376 | 1 | 1 |
| 2118:2971:1001:189:714:65 | 1:3:3:0:1:0 | 2043:1790:604:130:657:45 | 2:3:2:0:0:0 | 745:676:217:54:231:5 | 0:0:0:1:0:0 | 0.008732692 | 1 | 1 | 0.236416902 | 0.196659836 | 1 |
| 2964:1003:2115:189:713:65 | 6:0:3:0:1:0 | 1789:605:2038:130:656:45 | 7:1:5:0:0:0 | 676:217:745:53:229:5 | 0:0:0:1:1:0 | 0.011991799 | 1 | 1 | 0.00485331 | 0.001009902 | 1 |
| 0:1:0:0 | 0:8:0:2 | 0:0:0:0 | 0:13:0:0 | 0:0:0:0 | 0:2:0:0 | 1 | 1 | 1 | 0.001001643 | 4.74E-08 | 1 |
| 0:0:0:0 | 0:8:3:0 | 0:0:1:0 | 0:12:1:0 | 0:0:0:0 | 0:7:0:0 | 0.068525294 | 1 | 1 | 0.002105672 | 1.45E-06 | 0.000193253 |
| 8:39:2:27:0:2:0 | 0:0:0:0:1:0:0 | 4:11:1:1:0:5:1 | 1:14:1:0:4:1:0 | 0:3:1:2:0:1:0 | 1:7:2:1:4:0:0 | 1 | 1 | 1 | 1 | 1.01E-08 | 0.00827895 |
| 68:8:2:2:26:1 | 12:0:0:2:0:1 | 42:5:2:8:1:4 | 6:0:0:0:0:0 | 18:1:3:2:3:3 | 1:0:0:0:0:0 | 1 | 1 | 1 | 4.61E-08 | 2.97E-05 | 1 |
| 82:13:11 | 71:11:10 | 138:23:20 | 0:00:00 | 83:20:05 | 0:00:00 | 1 | 1 | 0.000344915 | 2.10E-17 | 1 | 1 |
| 1:17:0:11:1:3:0:0:0:5:3:0:3:0:0:0 | 0:20:1:12:1:2:0:2:0:4:0:1:6:0:0:0 | 0:25:2:17:2:7:0:1:0:5:1:0:9:1:1:1 | 0:0:0:0:0:0:0:0:0:0:0:0:0:0:0:0 | 0:1:0:1:0:4:0:1:0:0:2:0:1:0:0:0 | 0:0:0:0:0:0:0:0:0:0:0:0:0:0:0:0 | 1 | 1 | 1 | 1.72E-23 | 1 | 1 |
| 2:6:0:0:0:0:0:0:1:0 | 0:6:0:0:0:1:1:1:0:0 | 11:18:1:3:0:0:0:0:2:0 | 0:0:0:0:0:0:0:0:0:0 | 4:2:0:0:0:1:0:1:0:0 | 0:0:0:0:0:0:0:0:0:0 | 1 | 1 | 1 | 0.00013835 | 1 | 1 |
| 107:130:204 | 19:02:01 | 96:103:211 | 16:01:03 | 160:10:67 | 5:00:01 | 0.034668522 | 1.30E-05 | 0.001636609 | 1.99E-06 | 8.96E-05 | 0.035456988 |
| 7:14:38:0:38:0:3:3:7:1:1:1 | 1:0:6:0:1:0:0:0:2:0:0:0 | 7:9:14:0:7:0:0:0:10:0:2:0 | 1:2:4:0:2:0:0:0:1:0:0:0 | 1:1:3:0:0:0:0:0:1:0:0:0 | 0:1:0:0:0:0:0:0:0:0:0:0 | 1 | 1 | 1 | 5.59E-05 | 0.037858102 | 1 |
| 4:5:15:0:11:16:2:1:1:0 | 1:0:0:0:0:1:0:0:0:0 | 6:5:4:0:13:5:0:0:0:1 | 1:1:0:0:0:0:0:0:0:0 | 1:2:2:0:1:0:0:0:0:0 | 0:0:0:0:0:0:0:0:0:0 | 1 | 1 | 0.000884194 | 1 | 1 | 1 |
| 3:2:0:0:41:2 | 0:0:0:0:0:0 | 1:0:1:0:15:0 | 0:2:0:0:7:1 | 0:1:1:0:0:0 | 0:0:0:0:0:0 | 1 | 1 | 1 | 1 | 0.000739192 | 1 |
| 13:11:00 | 18:12:00 | 0:00:00 | 19:14:01 | 0:00:00 | 6:00:00 | 2.62E-05 | 0.005909413 | 0.146375793 | 0.000367801 | 0.000818744 | 0.00371338 |
| 6:01 | 7:01 | 12:01 | 0:00 | 5:00 | 0:00 | 0.017290558 | 1 | 1 | 0.039929313 | 1 | 1 |
| 6:01 | 7:01 | 12:01 | 0:00 | 5:00 | 0:00 | 1 | 1 | 1 | 0.039929313 | 1 | 1 |
| 12:03 | 0:00 | 12:01 | 0:00 | 5:00 | 0:00 | 0.029709415 | 1 | 1 | 1 | 1 | 1 |
| 71:6:12:2:0 | 12:0:1:0:0 | 59:6:14:2:2 | 13:0:4:0:0 | 1:4:1:0:0 | 2:0:1:0:0 | 1 | 1 | 1 | 3.09E-08 | 3.62E-07 | 1 |
| 9:0:4:1:4:3:25:59:0:34:12:4:0:1:1:2:7:2:1:2:1:0 | 7:13:20:2:0:5:1:10:0:2:5:0:1:0:0:0:0:0:0:0:0:0 | 6:1:8:1:8:3:19:4:1:13:4:3:0:1:1:3:0:0:0:2:2:1 | 7:10:27:3:0:6:2:1:0:2:2:0:1:0:0:1:0:0:1:1:0:0 | 0:0:1:0:3:0:18:8:1:6:3:0:0:0:0:0:0:0:0:0:2:0 | 2:4:3:1:0:1:0:0:0:1:0:0:0:0:0:0:0:0:1:0:0:0 | 1 | 1 | 0.006180963 | 4.11E-29 | 4.58E-42 | 0.004834004 |
| 2:0:0:0 | 3:0:0:0 | 0:0:0:0 | 9:0:1:1 | 0:0:0:0 | 2:0:0:0 | 1 | 1 | 1 | 1 | 0.000441095 | 1 |
| 0:00:00 | 63:112:9 | 35:144:0 | 4:07:00 | 49:93:0 | 4:02:00 | 1 | 0.005203732 | 0.0484396 | 6.89E-19 | 0.041599774 | 0.147810323 |
| 17:46:29:13:5:10 | 0:0:0:0:0:0 | 4:9:13:4:2:3 | 4:5:4:11:2:2 | 1:1:1:0:0:1 | 3:1:1:5:3:1 | 1 | 1 | 1 | 1 | 0.043748509 | 0.346541957 |
| 14:0:0:0:2:0:7:0:0 | 48:0:1:0:1:0:1:1:0 | 5:0:1:0:0:0:11:0:0 | 4:0:2:0:0:0:0:0:0 | 2:0:0:0:2:0:0:0:0 | 2:0:0:0:0:0:0:0:0 | 1 | 1 | 9.20E-09 | 5.95E-69 | 0.00341885 | 1 |
| 0:0:0:22:2:0:7:0:0 | 0:1:0:40:1:0:1:1:0:0 | 0:3:0:6:0:0:10:0:0 | 0:0:0:3:0:0:0:0:0 | 0:0:0:2:2:0:0:0:0 | 0:0:0:2:0:0:0:0:0 | 1 | 1 | 1.25E-09 | 1.73E-62 | 1 | 1 |
| 0:1:0:46:2:7:0:1:0 | 0:0:0:23:1:1:0:0:0 | 0:3:0:7:0:10:0:0:0 | 0:0:0:2:0:0:0:0:0 | 0:0:0:2:2:0:0:0:0 | 0:0:0:2:0:0:0:0:0 | 1 | 1 | 2.90E-07 | 4.81E-31 | 1 | 1 |
| 0:0:0:1:0:0:0:0 | 0:0:0:0:0:0:0:0 | 0:0:3:3:0:0:0:0 | 0:0:4:0:0:0:0:1 | 1:0:0:0:0:0:1:0 | 0:0:2:0:0:0:0:0 | 1 | 1 | 1 | 1 | 0.003495428 | 1 |
| 0.959722222 | 0:15:00 | 1.039583333 | 0:15:00 | 0.210416667 | 0:01:00 | 0.0086191 | 1 | 1 | 7.25E-07 | 7.26E-07 | 1 |

| | | | | | | | | | | | |
|---|---|---|---|---|---|---|---|---|---|---|---|
| 347:18:0:0 | 4:0:0:0 | 409:19:0:0 | 3:0:0:0 | 68:4:0:0 | 4:0:0:0 | 0.002557517 | 1 | 1 | 1 | 1 | 1 |
| 1063:213 | 1:00 | 1208:444 | 5:00 | 519:111 | 1:00 | 1 | 1 | 1 | 1 | 0.031296179 | 1 |
| 1070:215 | 0:00 | 1215:447 | 6:00 | 519:111 | 0:00 | 1 | 1 | 1 | 1 | 0.018436807 | 1 |
| 2:02:01 | 0:00:00 | 0:00:00 | 10:06:01 | 0:00:00 | 1:01:00 | 1 | 1 | 1 | 1 | 0.033943254 | 1 |
| 0:01 | 0:00 | 0:00 | 5:00 | 0:00 | 1:00 | 1 | 1 | 1 | 1 | 0.031296179 | 1 |
| 0:01 | 0:00 | 0:00 | 5:00 | 0:00 | 1:00 | 1 | 1 | 1 | 1 | 0.031296179 | 1 |
| 38:157:246:2 | 0:1:4:0 | 47:201:222:1 | 1:3:2:0 | 15:32:16:0 | 0:2:0:0 | 1 | 1 | 1 | 0.04093502 | 0.359192143 | 1 |
| 35:151:235:2 | 0:0:0:0 | 45:197:207:1 | 0:0:0:0 | 15:33:13:0 | 0:0:0:0 | 1.98E-14 | 1.36E-05 | 1 | 1 | 1 | 1 |
| 0:0:0:0 | 36:148:236:2 | 0:0:0:0 | 45:197:205:1 | 0:0:0:0 | 15:33:12:0 | 8.15E-23 | 6.94E-11 | 3.61E-19 | 4.81E-69 | 6.25E-62 | 1.15E-07 |
| 78:18:01 | 93:11:00 | 205:23:00 | 0:00:00 | 35:08:00 | 0:00:00 | 2.63E-30 | 2.88E-16 | 3.20E-16 | 2.69E-32 | 1 | 1 |
| 36:9:7:0:9:23:3:10:1:0:4:7:0:0:1:0:1:0:0:0:0:0 | 20:11:14:1:14:8:3:13:0:0:2:4:0:0:1:1:1:0:0:0:0:0 | 87:24:41:3:7:22:2:31:1:1:2:16:0:5:0:2:3:2:1:1:1:2 | 0:0:0:0:0:0:0:0:0:0:0:0:0:0:0:0:0:0:0:0:0:0 | 35:5:0:1:0:9:2:0:2:0:1:0:1:0:0:1:0:0:0:0:0:0 | 0:0:0:0:0:0:0:0:0:0:0:0:0:0:0:0:0:0:0:0:0:0 | 1.73E-26 | 6.11E-11 | 3.65E-31 | 1.14E-27 | 1 | 1 |
| 41:15:19:4:12:9:108:2:3:1:1:1:0 | 6:0:1:0:2:0:1:0:0:0:0:0:0 | 41:16:26:5:19:16:166:4:6:2:7:0:0 | 7:2:1:1:0:1:1:0:0:0:1:0:0 | 8:1:2:0:2:5:8:1:0:0:3:0:1 | 1:0:0:0:0:0:0:1:0:0:0:0:0 | 1 | 1 | 1 | 2.34E-05 | 0.000140647 | 1 |
| 10:18:31:3:12:6:104:1:1:1:1:1:0 | 2:1:7:1:0:2:2:0:1:0:0:0:0 | 14:23:30:4:11:10:171:4:5:2:6:0:0 | 1:0:9:0:5:3:1:0:0:0:0:0:0 | 0:2:8:0:2:5:8:1:0:0:3:0:1 | 0:0:0:0:0:0:0:0:0:0:0:0:0 | 1 | 1 | 1 | 0.00052074 | 4.00E-08 | 1 |
| 1:83:3:10:9:4:3:1:1:1:0:0 | 0:5:0:1:0:0:0:0:0:0:0:0 | 3:146:7:14:15:5:3:2:1:2:5:0 | 0:8:0:2:1:0:1:1:0:0:0:0 | 0:6:0:2:4:2:4:1:0:0:3:1 | 0:0:0:0:0:0:0:0:0:0:0:0 | 1 | 1 | 1 | 6.70E-06 | 5.22E-07 | 1 |
| 19:1:1:1:1:0:1:0:0:0 | 6:0:0:2:0:1:0:0:0:0 | 44:1:1:2:1:0:1:0:1:1 | 6:3:0:0:0:0:0:0:0:0 | 1:0:0:0:0:0:0:0:1:0 | 2:1:0:0:0:0:0:0:0:0 | 1 | 1 | 1 | 5.76E-05 | 1.06E-05 | 1 |
| 18:1:1:1:1:0:1:0:0:0 | 7:0:0:2:0:1:0:0:0:0 | 44:1:0:1:1:0:1:0:1:1 | 6:3:0:1:0:0:0:0:0:0 | 1:0:0:0:0:0:0:0:1:0 | 2:1:0:0:0:0:0:0:0:0 | 1 | 1 | 1 | 3.10E-06 | 7.52E-05 | 1 |
| 0:0:0:0:5:0:0:0:0 | 1:1:1:0:7:1:0:0:0 | 0:0:1:0:34:0:0:1:1 | 1:0:0:0:2:1:0:0:0 | 0:0:0:0:0:0:0:0:0 | 0:0:0:0:0:0:0:0:0 | 1 | 0.022206229 | 1 | 0.000141071 | 1 | 1 |
| 5:78:32:38:3:13:33:3:3:5:1 | 0:0:0:0:0:0:0:0:0:0:0 | 8:130:44:69:5:35:51:2:3:12:3 | 0:3:1:0:0:0:0:1:0:0:0 | 5:34:17:37:1:20:26:3:3:3:1 | 0:0:0:0:0:0:2:0:0:0:0 | 1 | 1 | 1 | 1 | 0.044883775 | 1 |
| 38:3:5:13:235:102:68:0:3:5:1 | 0:0:0:0:3:0:0:2:0:0:0 | 69:5:8:35:389:128:105:2:3:11:3 | 0:0:0:0:5:2:0:0:0:0:0 | 37:1:5:20:49:36:38:2:3:3:1 | 0:0:0:0:0:0:0:0:0:0:0 | 1 | 1 | 1 | 0.011169271 | 5.85E-05 | 1 |
| 39:3:5:13:416:203:139:0:3:5:1 | 0:0:0:0:4:1:0:0:0:0:0 | 69:5:10:35:712:303:210:0:3:11:4 | 1:0:0:0:1:2:0:0:0:0:0 | 38:1:5:20:62:41:48:2:3:3:1 | 0:0:0:0:0:0:1:0:0:0:0 | 1 | 1 | 1 | 0.000547567 | 1 | 1 |
| 421:39:3:5:13:207:141:0:3:5:1 | 2:0:0:0:0:2:0:0:0:0:0 | 719:70:5:10:35:310:215:0:3:11:4 | 5:0:0:0:0:2:2:0:0:0:0 | 62:38:1:5:20:41:49:2:3:3:1 | 0:0:0:0:0:0:0:0:0:0:0 | 1 | 1 | 1 | 1 | 0.000856572 | 1 |
| 39:3:5:13:482:282:0:202:3:5:2 | 0:0:0:0:3:0:0:0:0:0:0 | 70:5:12:35:854:414:0:337:3:11:6 | 0:0:0:0:1:3:0:2:0:0:0 | 38:1:5:20:66:47:3:51:3:3:1 | 0:0:0:0:1:0:0:0:0:0:0 | 1 | 1 | 1 | 1 | 0.039232735 | 1 |
| 39:3:5:13:487:317:0:240:3:5:3 | 0:0:0:0:0:1:0:0:0:0:0 | 69:5:13:35:892:465:0:404:3:11:6 | 1:0:0:0:2:4:0:0:0:0:0 | 38:1:5:20:74:49:2:53:3:3:2 | 0:0:0:0:0:0:0:0:0:0:0 | 1 | 1 | 1 | 1 | 0.005445271 | 1 |
| 39:3:5:13:490:344:0:281:3:5:6 | 0:0:0:0:1:1:0:0:0:0:0 | 70:5:13:35:919:529:0:493:3:11:7 | 0:0:0:0:2:3:0:0:0:0:0 | 38:1:5:20:78:50:2:53:3:3:2 | 0:0:0:0:1:1:0:0:0:0:0 | 1 | 1 | 1 | 1 | 0.011876933 | 1 |
| 39:3:5:13:490:352:0:294:3:5:6 | 0:0:0:0:0:0:0:2:0:0:0 | 70:5:13:35:925:551:0:517:3:11:7 | 0:0:0:0:3:0:0:2:0:0:0 | 38:1:5:20:80:52:2:52:3:3:2 | 0:0:0:0:0:0:0:1:0:0:0 | 1 | 1 | 1 | 1 | 0.011876933 | 1 |
| 70:15:3:8:1 | 71:14:2:7:1 | 70:4:4:2:1 | 0:0:0:0:0 | 14:0:1:0:0 | 1:0:0:0:0 | 1 | 1 | 1.48E-09 | 9.03E-38 | 1 | 1 |
| 71:14:2:7:1 | 70:15:3:8:1 | 38:3:2:0:1 | 32:1:2:2:0 | 2:0:0:0:0 | 13:0:1:0:0 | 1 | 1 | 0.068053893 | 4.06E-35 | 1.46E-20 | 5.60E-09 |
| 4:11:14:3:0:0:0:0:1:2:0 | 8:1:0:2:0:0:0:1:0:0:0 | 3:6:11:2:0:0:0:0:1:4:1 | 12:2:0:0:0:0:1:1:0:0:0 | 0:0:4:0:0:0:0:0:0:0:0 | 0:0:0:0:0:0:0:0:0:0:0 | 1 | 1 | 1 | 2.61E-07 | 8.26E-14 | 1 |
| 6:8:0:5:3:1:3:2:0:0:0:0:0 | 8:5:0:2:3:1:1:1:0:1:0:0:0 | 9:24:0:5:6:0:2:1:1:1:0:2:4 | 0:0:0:0:0:0:0:1:0:0:0:0:0 | 4:0:0:0:0:0:0:0:0:0:0:0:0 | 0:0:0:0:0:0:0:0:0:0:0:0:0 | 0.265398929 | 1 | 0.028355434 | 0.000100568 | 1 | 1 |
| 11:1:1:0:0 | 9:1:1:0:0 | 12:0:3:0:0 | 0:0:0:0:0 | 1:1:1:0:0 | 0:0:0:0:0 | 0.000965198 | 1 | 0.935177969 | 4.22E-05 | 1 | 1 |
| 0:1:8:0:0:0:0:1:0:5:0:0:0:0:0:0 | 0:2:4:0:1:2:0:0:1:3:2:1:0:0:0:0 | 0:0:8:0:0:1:0:1:1:8:0:1:0:1:0:0 | 0:0:5:0:0:1:0:0:1:10:1:0:0:2:0:0 | 0:1:2:1:0:0:0:2:0:5:0:0:0:1:1:1 | 0:0:0:0:0:0:0:0:0:1:0:0:0:0:0:0 | 1 | 1 | 0.010824061 | 0.074194925 | 2.28E-11 | 1 |

| | | | | | | | | | | | |
|---|---|---|---|---|---|---|---|---|---|---|---|
| 2:0:3:15:0:1:2:0:43:0:2:2:1:0:0 | 0:0:0:1:0:0:0:0:5:0:0:0:0:0:0 | 5:0:0:15:0:0:2:0:67:2:1:2:1:3:0 | 1:0:0:0:0:0:0:0:2:0:0:0:0:0:0 | 2:0:0:2:1:0:0:0:6:0:0:1:0:1:1 | 0:0:0:0:0:0:0:0:0:0:0:0:0:0:0 | 1 | 1 | 1 | 4.26E-07 | 1 | 1 |
| 0:7:0:0:0:0:1:37:0:0:1:1:0:0 | 0:3:0:1:2:0:1:29:0:2:2:0:0:0 | 0:5:0:0:1:0:4:42:0:0:1:0:1:0 | 0:6:0:0:1:0:2:48:0:1:2:1:2:1 | 0:2:0:0:0:0:2:6:0:0:1:0:1:0 | 0:0:0:0:0:0:0:1:0:0:0:0:0:0 | 1 | 1 | 0.016710098 | 1.41E-47 | 4.39E-87 | 1 |
| 0:8:0:1:2:0:3:76:3:0:0:1:2:0 | 0:7:0:0:0:0:3:100:0:0:0:0:0:0 | 0:10:0:0:2:0:6:90:1:0:0:1:1:1 | 0:2:0:0:0:0:2:66:2:0:0:0:0:0 | 0:2:0:0:0:0:2:8:1:0:0:0:0:0 | 0:0:0:0:0:0:0:0:0:0:0:0:0:0 | 1 | 1 | 1 | 3.47E-240 | 3.58E-156 | 1 |
| 0:23:0:1:2:0:11:274:0:10:0:0:0:2:2:0 | 0:1:0:0:0:0:1:42:0:1:0:0:0:0:0:0 | 0:26:0:0:2:0:13:291:0:4:0:0:0:5:1:1 | 0:0:0:0:0:0:2:30:0:0:0:0:0:0:0:0 | 0:3:0:0:0:0:7:43:0:1:0:0:0:0:0:0 | 0:0:0:0:0:0:0:0:0:0:0:0:0:0:0:0 | 1 | 1 | 1 | 4.05E-113 | 5.79E-79 | 1 |
| 300:0:0:24:0:1:2:0:12:11:0:0:2:0:2:0 | 19:0:0:0:0:0:0:0:0:0:0:0:0:0:0:0 | 305:0:0:26:0:0:2:0:14:3:0:0:5:0:1:1 | 17:0:0:0:0:0:0:1:2:0:0:0:0:0:0:0 | 34:0:0:3:0:0:0:0:5:1:0:0:0:0:0:0 | 9:0:0:0:0:0:0:0:2:0:0:0:0:0:0:0 | 2.97E-34 | 1.78E-07 | 6.26E-07 | 6.79E-51 | 1.21E-36 | 2.65E-16 |
| 12:0:24:0:0 | 6:0:22:1:0 | 9:1:36:0:0 | 16:1:64:0:0 | 6:0:4:0:1 | 3:0:2:0:0 | 1 | 1 | 7.98E-07 | 6.35E-12 | 1.21E-38 | 0.102014456 |
| 15023:155:18:16:20:15:2 | 17:0:0:0:0:0:0 | 23080:83:31:29:61:4:3 | 25:0:0:0:0:0:0 | 3900:16:10:4:6:1:0 | 2:0:0:0:0:0:0 | 2.52E-20 | 1 | 1 | 4.16E-19 | 4.76E-29 | 1 |
| 15016:155:18:16:20:15:2 | 23:0:0:0:0:0:0 | 23075:83:31:29:61:4:3 | 29:0:0:0:0:0:0 | 3900:16:10:4:6:1:0 | 2:0:0:0:0:0:0 | 4.32E-19 | 1 | 1 | 1.40E-26 | 4.37E-34 | 1 |
| 15012:155:18:16:20:15:2 | 23:0:0:0:0:0:0 | 23057:83:31:29:60:4:3 | 45:0:0:0:1:0:0 | 3892:16:10:4:6:1:0 | 7:0:0:0:0:0:0 | 2.86E-25 | 1 | 1 | 1.40E-26 | 8.72E-53 | 4.55E-07 |
| 14971:155:18:16:20:15:2 | 22:0:0:0:0:0:0 | 22957:84:31:29:61:4:3 | 80:0:0:0:0:0:0 | 3887:16:10:4:6:1:0 | 6:0:0:0:0:0:0 | 1.56E-11 | 1 | 1.76E-10 | 2.46E-25 | 3.05E-99 | 6.12E-06 |
| 14944:153:18:16:15:1:2 | 14:0:0:0:0:0:0 | 22927:81:31:29:4:1:3 | 26:0:0:0:0:0:0 | 3876:15:10:4:1:0:0 | 7:0:0:0:0:0:0 | 5.52E-13 | 1 | 1 | 2.03E-15 | 2.67E-30 | 4.55E-07 |
| 14906:153:17:16:15:2 | 13:0:0:0:0:0 | 22860:80:31:29:4:3 | 19:0:0:0:0:0 | 3880:15:10:4:1:0 | 1:0:0:0:0:0 | 0.00022653 | 1 | 3.48E-05 | 3.73E-12 | 2.94E-18 | 1 |
| 14754:151:0:15:1:1 | 13:0:0:0:0:0 | 22694:81:7:4:3:0 | 26:0:0:0:0:0 | 3856:15:2:1:0:0 | 8:0:0:0:0:0 | 3.58E-07 | 1 | 1 | 3.73E-12 | 1.40E-25 | 3.74E-07 |
| 0:0:0:0:0 | 2:1:0:14:0 | 0:0:0:5:1 | 0:0:1:7:0 | 0:0:0:4:0 | 0:0:0:1:0 | 0.004498193 | 1 | 1 | 4.21E-08 | 0.000205879 | 1 |
| 2:5:1:56:17:1:1:1:1:1:0:0 | 0:1:0:3:2:0:0:0:0:0:0:0 | 22:4:0:100:25:0:1:0:2:0:0:0 | 0:0:0:2:3:0:0:0:0:0:0:1 | 3:0:0:3:1:0:0:0:0:0:0:0 | 0:0:0:0:0:0:0:0:0:0:0:0 | 1 | 1 | 1 | 0.029531909 | 0.030827483 | 1 |
| 110:1863:29 | 1:03:00 | 119:2272:28 | 2:07:00 | 1.052905093 | 0:00:00 | 1 | 1 | 1 | 1 | 0.017156951 | 1 |
| 120:1865:29 | 1:00:00 | 57:1082:11 | 74:1197:17 | 1.001493056 | 2:14:02 | 3.03E-13 | 1.02E-50 | 1 | 1 | 0 | 0.000575289 |
| 1831:27:288 | 24:02:01 | 2240:27:347 | 25:01:01 | 312:11:29 | 3:00:00 | 1 | 0.083167178 | 0.001636609 | 1.89E-08 | 1.71E-09 | 1 |
| 384:1809:27 | 1:00:00 | 237:1049:10 | 260:1160:17 | 32:295:9 | 2:14:02 | 4.45E-12 | 9.82E-41 | 1 | 1 | 0 | 0.000575289 |
| 1769:391:26 | 29:05:00 | 2154:516:26 | 34:03:00 | 303:36:11 | 6:00:00 | 1 | 0.003558706 | 0.035359329 | 1.77E-09 | 1.27E-12 | 0.00371338 |
| 1659:381:25 | 79:07:01 | 2016:507:25 | 92:07:01 | 301:35:11 | 2:01:00 | 1 | 1 | 0.010070087 | 3.35E-28 | 1.21E-33 | 1 |
| 1624:374:24 | 32:02:01 | 1983:504:24 | 36:03:01 | 290:35:11 | 14:01:00 | 1 | 1 | 5.91E-05 | 9.04E-12 | 1.01E-12 | 1.09E-05 |
| 365:1594:24 | 1:01:00 | 502:1940:23 | 1:05:00 | 35:286:11 | 0:01:00 | 1 | 1 | 1 | 1 | 0.036538834 | 1 |
| 362:1594:24 | 2:01:00 | 494:1940:23 | 2:06:00 | 35:288:11 | 0:00:00 | 1 | 1 | 1 | 1 | 0.036538834 | 1 |
| 5:0:0:3 | 0:0:0:0 | 3:1:1:0 | 7:0:3:0 | 2:0:2:0 | 0:0:1:0 | 1 | 0.120678556 | 1 | 1 | 0.006124303 | 1 |
| 0:0:0:0 | 6:1:1:0 | 0:0:0:0 | 33:1:0:1 | 0:0:0:0 | 7:2:1:0 | 1 | 0.002652154 | 1 | 0.014715107 | 1.28E-18 | 0.011859295 |
| 0:0:0:0:0:0:0:0:0:0:0:0 | 3:40:1:5:11:14:107:14:10:2:0:1 | 1:14:0:2:8:17:72:9:5:0:0:0 | 3:17:3:1:12:8:67:9:3:1:0:1 | 2:7:0:2:5:4:5:3:1:0:0:0 | 0:0:0:0:0:0:0:0:0:0:0:0 | 7.88E-64 | 4.21E-10 | 4.17E-12 | 2.54E-116 | 2.62E-70 | 1 |
| 2041:01:00 | 36:00:00 | 2363:04:00 | 28:00:00 | 412:00:00 | 5:00 | 0.000798008 | 1 | 1 | 5.52E-09 | 3.00E-07 | 0.030032732 |
| 2082:44:00 | 2:00 | 2387:118 | 5:00 | 419:20:00 | 0:00 | 1 | 1 | 1 | 1 | 0.031296179 | 1 |
| 1954:56:00 | 4:00 | 2242:133 | 8:00 | 397:22:00 | 0:00 | 1 | 1 | 1 | 1 | 0.006667609 | 1 |
| 1:21:0:1 | 0:0:0:0 | 2:1:1:0 | 6:14:1:0 | 1:0:0:0 | 2:1:1:0 | 1 | 1 | 1 | 1 | 6.47E-05 | 1 |
| 0:2:0:0 | 0:11:1:0 | 1:1:0:0 | 6:13:0:2 | 0:0:0:0 | 2:1:0:1 | 0.062751851 | 1 | 1 | 5.54E-06 | 0.000520745 | 1 |
| 60:60:2:2:0:0:0:0 | 4:1:1:0:1:0:0:0 | 30:58:1:1:2:4:1:0 | 1:0:0:0:3:1:0:0 | 6:10:0:0:0:1:1:0 | 0:0:0:0:0:0:0:0 | 1 | 1 | 1 | 0.046292247 | 0.094146177 | 1 |

| | | | | | | | | | | | |
|---|---|---|---|---|---|---|---|---|---|---|---|
| 40:42:2:0:1:0:0:0 | 3:3:0:0:0:0:0:0 | 42:20:1:1:0:2:0:0 | 5:3:0:0:0:0:0:0 | 6:4:0:0:0:1:1:0 | 3:2:0:0:0:0:0:0 | 0.031509029 | 1 | 1 | 0.014984561 | 0.000836121 | 0.031903402 |
| 77:03:02 | 19:01:02 | 90:24:05 | 36:04:01 | 21:03:01 | 4:00:00 | 1 | 1 | 1 | 1.99E-06 | 4.04E-12 | 1 |
| 21:0:25:3:0 | 0:0:0:0:0 | 0:0:0:0:0 | 3:0:15:1:1 | 0:0:0:0:0 | 2:0:2:0:0 | 1 | 1 | 1 | 1 | 1.78E-07 | 1 |
| 19:0:21:3:0 | 0:0:0:0:0 | 0:0:0:0:0 | 3:0:13:2:1 | 0:0:0:0:0 | 2:0:2:0:0 | 1 | 1 | 1 | 1 | 1.52E-05 | 1 |
| 13:5:1:1 | 0:0:0:0 | 0:0:0:0 | 37:18:0:3 | 0:0:0:0 | 20:2:0:0 | 1 | 1 | 1 | 1 | 2.97E-12 | 1.40E-10 |
| 13:7:1:1 | 0:0:0:0 | 0:0:0:0 | 37:19:0:4 | 0:1:0:0 | 20:3:0:0 | 1 | 1 | 1 | 1 | 1.22E-11 | 6.85E-10 |
| 15:19:1:1:0 | 0:0:0:0:0 | 0:0:0:0:0 | 43:49:2:2:4 | 0:0:0:0:0 | 24:5:1:0:0 | 1 | 1 | 1 | 1 | 6.60E-23 | 8.98E-14 |
| 0:00:00 | 15:03:01 | 13:03:01 | 0:00:00 | 1:00:00 | 0:00:00 | 1 | 1 | 1 | 0.000201192 | 1 | 1 |
| 2:1:0:9:0:0:0 | 5:4:0:0:2:0:0 | 0:1:2:1:1:0:0 | 7:0:0:0:4:0:0 | 0:1:1:2:0:0:1 | 0:1:0:0:0:0:0 | 1 | 1 | 1 | 0.009576467 | 6.47E-05 | 1 |
| 0:00:00 | 1:02:00 | 0:00:00 | 12:01:01 | 0:00:00 | 4:00:00 | 1 | 1 | 0.035359329 | 1 | 0.000315403 | 1 |
| 2:0:7:0:0:2:1:1:0:0 | 0:0:10:0:0:1:0:0:0:0 | 5:0:30:0:0:3:0:0:0:2 | 2:0:16:0:0:2:0:0:0:0 | 1:0:3:0:0:0:0:0:1:0 | 0:0:1:0:0:1:0:0:0:0 | 1 | 1 | 1 | 3.89E-13 | 2.96E-19 | 1 |
| 226:22:2:0 | 268:9:0:1 | 242:24:1:0 | 271:25:0:1 | 72:4:1:0 | 28:1:0:0 | 4.73E-13 | 2.86E-12 | 9.89E-23 | 5.64E-162 | 4.86E-150 | 2.35E-16 |
| 570:34:2:1 | 8:0:0:0 | 601:53:1:1 | 6:0:0:0 | 108:4:1:0 | 3:0:0:0 | 1 | 1 | 1 | 4.60E-05 | 0.000739192 | 1 |
| 0:01:00 | 0:00:00 | 0:00:00 | 4:00:02 | 0:00:00 | 1:00:00 | 1.66E-10 | 1 | 1 | 1 | 0.150081592 | 1 |
| 2:1:0:1:1 | 0:0:0:0:0 | 0:0:0:0:0 | 10:0:0:0:0 | 0:0:0:0:0 | 1:0:0:0:0 | 9.71E-22 | 1 | 1 | 1 | 1.12E-07 | 1 |
| 39:36:15:39:56:32:34:28:32:2:29:19:28:16:0:12:62:1:16:1 | 2:2:0:0:6:0:3:1:2:0:2:0:0:3:0:1:1:0:0:0 | 20:27:20:45:25:32:19:29:51:1:11:29:15:21:0:8:23:5:8:2 | 0:0:0:2:4:1:0:0:3:0:0:0:1:0:0:0:0:0:0:0 | 4:8:3:8:4:8:11:2:8:0:8:3:4:3:0:4:6:0:0:0 | 0:0:0:1:0:0:1:0:0:0:0:0:0:0:0:0:0:0:0:0 | 0.485902809 | 1 | 1 | 0.003967309 | 0.00099502 | 1 |
| 7:17:13:14:6:8:13:12:13:1:18:10:0:8:6:7:0:1:9:0 | 12:16:9:5:3:10:11:16:10:4:20:7:1:6:8:6:0:0:4:1 | 13:10:17:7:8:7:25:21:7:3:8:8:1:8:6:11:0:3:3:1 | 2:6:9:2:7:7:10:12:5:1:5:5:0:1:0:2:3:0:1:2:1 | 1:3:7:5:1:10:5:8:3:4:6:6:0:3:4:1:0:0:0:0 | 1:0:1:2:1:1:3:1:0:0:0:0:0:0:0:2:0:0:0:0 | 0.050245353 | 0.140328346 | 0.007063949 | 4.02E-09 | 2.25E-05 | 0.184047759 |
| 366:672:49:0 | 0:0:0:0 | 400:498:41:1 | 8:11:0:0 | 68:101:7:0 | 5:5:0:0 | 1 | 1 | 1 | 1 | 0.000311761 | 0.02334199 |
| 25:00:00 | 0:00 | 38:00:00 | 37:00:00 | 14:01 | 15:00 | 1 | 1 | 1 | 1 | 3.53E-09 | 0.000196434 |
| 42:00:00 | 4:00 | 73:01:00 | 5:00 | 32:01:00 | 0:00 | 1 | 1 | 1 | 1 | 0.031296179 | 1 |
| 50:00:00 | 6:00:00 | 92:01:00 | 8:00:01 | 32:01:00 | 0:00:00 | 0.08085072 | 1 | 1 | 0.003552529 | 0.00271654 | 1 |
| 32:00:00 | 26:00:00 | 51:01:00 | 45:01:01 | 15:01:00 | 16:00:00 | 1 | 1 | 2.74E-14 | 1.62E-11 | 6.25E-18 | 2.88E-07 |
| 33:00:00 | 27:00:00 | 52:01:00 | 49:01:01 | 16:01:00 | 18:00:00 | 1 | 1 | 3.93E-15 | 6.16E-12 | 1.24E-19 | 4.32E-08 |
| 0:00 | 0:00 | 0:00 | 0:00 | 0:00 | 0:00 | 0.003177062 | 1 | 1 | 1 | 1 | 1 |
| 151:00:01 | 8:00:00 | 660:01:01 | 0:00:00 | 133:00:00 | 0:00:00 | 1 | 1 | 1.16E-30 | 0.000535794 | 1 | 1 |
| 0:0:0:0:0:0:0 | 12:18:2:0:1:0:0 | 8:9:5:1:0:0:0 | 0:0:0:0:0:0:0 | 2:2:1:1:1:0:1 | 0:0:0:0:0:0:0 | 0.156345239 | 1 | 0.184820174 | 4.76E-12 | 1 | 1 |
| 0:0:0:0:0:0:0 | 27:11:1:0:0:1:0 | 0:0:0:0:0:0:0 | 15:7:2:0:1:0:0 | 0:0:0:0:0:0:0 | 5:2:0:1:1:1:0 | 0.002647618 | 1 | 0.064012353 | 1.42E-20 | 3.35E-09 | 0.060943212 |
| 0:0:0:0:0:0:0 | 28:12:0:2:0:1:0 | 14:9:0:2:1:0:0 | 0:0:0:0:0:0:0 | 4:2:1:0:1:1:0 | 0:0:0:0:0:0:0 | 0.156345239 | 1 | 0.184820174 | 1.57E-20 | 1 | 1 |
| 6548:934:6:2:43:17 | 4:0:0:0:0:2 | 4425:722:5:4:36:16 | 4:0:0:0:0:1 | 5094:624:4:4:43:6 | 7:0:0:0:0:0 | 0.00145773 | 3.04E-05 | 1 | 0.019303392 | 0.011881852 | 3.38E-06 |
| 6396:903:46:5:43:3 | 8:2:0:0:0:0 | 4348:700:26:4:36:0 | 4:0:1:0:0:0 | 5062:618:22:3:43:0 | 2:1:0:0:0:0 | 3.85E-08 | 6.00E-07 | 1 | 1.99E-05 | 0.011881852 | 1 |
| 6233:870:5:43:57:9 | 9:3:0:0:0:0 | 4233:680:4:36:41:4 | 2:0:0:0:0:0 | 4983:610:3:41:37:3 | 4:0:0:1:0:0 | 0.000245212 | 0.019017106 | 1 | 7.78E-06 | 1 | 0.011761049 |
| 6246:58:874:10:5:43 | 1:2:2:1:0:0 | 4238:43:683:4:4:36 | 4:0:0:0:0:0 | 4987:37:611:6:3:42 | 2:2:0:1:0:0 | 3.58E-07 | 0.118667291 | 1 | 0.56323426 | 1 | 0.338281183 |
| 6212:868:5:43:65:11 | 9:2:0:0:0:0 | 4226:679:4:36:46:4 | 3:1:0:0:0:0 | 4979:607:3:42:40:8 | 9:1:0:0:0:0 | 5.05E-18 | 0.000280691 | 1 | 2.83E-06 | 1 | 5.26E-07 |
| 5934:828:5:12:41:68 | 12:0:0:0:0:0 | 4071:650:4:8:32:47 | 12:2:0:0:0:1 | 4863:596:3:9:39:49 | 12:0:0:0:0:0 | 6.18E-09 | 0.000273007 | 1 | 3.80E-11 | 4.76E-08 | 4.53E-11 |
| 823:5899:5:12:41:68 | 1:16:0:0:0:0 | 649:4066:4:8:32:49 | 1:8:0:0:0:1 | 595:4860:3:9:39:49 | 1:3:0:0:0:0 | 3.28E-06 | 0.019017106 | 1 | 6.24E-14 | 3.55E-05 | 1 |
| 5981:72:834:5:12:39 | 11:1:1:0:0:0 | 4097:58:651:3:8:31 | 5:2:1:0:0:0 | 4895:51:602:3:9:35 | 7:2:0:0:0:0 | 0.0007901 | 0.317220796 | 1 | 6.61E-08 | 0.016698024 | 0.00014384 |
| 6117:850:5:11:39:175 | 18:4:0:0:0:0 | 4156:655:4:8:30:115 | 14:3:0:0:0:1 | 4988:600:2:9:33:73 | 12:2:0:0:0:0 | 3.95E-21 | 4.80E-08 | 1 | 2.58E-13 | 3.10E-09 | 5.77E-09 |
| 6083:847:5:11:39:172 | 12:0:0:0:0:1 | 4149:647:4:8:27:113 | 3:4:0:0:0:0 | 4983:600:2:9:32:70 | 5:0:0:0:0:0 | 4.55E-05 | 0.000823477 | 1 | 6.38E-10 | 0.01715236 | 0.000249906 |
| 6156:851:193:5:12:38 | 5:1:0:0:0:0 | 4178:654:116:4:8:27 | 5:1:0:0:0:0 | 4993:599:70:2:9:32 | 5:0:0:0:0:0 | 6.33E-10 | 4.61E-05 | 1 | 0.001878282 | 0.00194145 | 0.000249906 |

| | | | | | | | | | | | |
|---|---|---|---|---|---|---|---|---|---|---|---|
| 6136:847:4:12:37:192 | 9:1:0:0:0:0 | 4164:654:4:8:27:116 | 9:2:0:0:0:0 | 4975:596:2:9:32:69 | 13:1:0:0:0:0 | 6.01E-14 | 3.83E-07 | 1 | 5.20E-07 | 2.83E-06 | 7.65E-11 |
| 80:2:6:0:11:74:1:3:0:0 | 0:0:0:0:0:0:0:0:0:0 | 33:1:1:0:6:16:1:1:0:1 | 0:0:0:0:0:0:0:0:0:0 | 21:0:4:0:2:5:0:0:0:0 | 0:0:0:0:0:0:0:0:0:0 | 4.07E-09 | 8.99E-06 | 1 | 1 | 1 | 1 |
| 81:2:6:0:75:4:1:3:0:0 | 6:0:0:0:1:0:0:0:0:0 | 33:1:1:0:16:3:1:1:0:1 | 1:0:0:0:1:0:0:0:0:0 | 21:0:4:0:5:2:0:0:0:0 | 0:0:0:0:0:0:0:0:0:0 | 1 | 1 | 1 | 1.49E-06 | 1 | 1 |
| 81:2:6:0:75:4:1:3:0:0 | 5:0:0:0:1:0:0:0:0:0 | 34:1:1:0:16:3:1:1:0:1 | 0:0:0:0:1:0:0:0:0:0 | 21:0:4:0:5:2:0:0:0:0 | 0:0:0:0:0:0:0:0:0:0 | 1 | 1 | 1 | 4.28E-05 | 1 | 1 |
| 51:0:3:0:15:3:1:2:0:0 | 36:2:4:0:5:1:0:1:0:0 | 31:1:1:0:4:2:1:1:0:1 | 3:0:0:0:2:0:0:0:0:0 | 15:0:3:0:3:2:0:0:0:0 | 6:0:1:0:0:0:0:0:0:0 | 6.61E-32 | 3.16E-15 | 7.19E-07 | 1.35E-42 | 0.016638054 | 1.49E-06 |
| 306:9109:17:16:2:0:0:0 | 1:5:0:0:0:0:0:0 | 290:6387:14:18:0:0:1:1 | 2:5:0:0:0:0:0:0 | 48:1520:2:6:0:1:0:0 | 1:4:0:0:0:0:0:0 | 1 | 1 | 1 | 0.000277059 | 0.000793207 | 0.003640542 |
| 9149:17:38:16:0:0 | 10:0:0:0:0:0 | 6415:14:28:18:1:1 | 4:0:0:0:0:0 | 1531:2:2:6:0:0 | 3:0:0:0:0:0 | 1 | 1 | 1 | 3.91E-09 | 1 | 1 |
| 9129:195:17:78:16:2:0:0 | 6:0:0:0:0:0:0:0 | 6401:204:14:55:18:0:0:1 | 1:0:0:0:0:0:0:0 | 1531:43:2:8:6:0:1:0 | 3:0:0:0:0:0:0:0 | 1 | 1 | 1 | 1.21E-06 | 1 | 1 |
| 9010:192:16:98:16:2:0:0 | 12:1:0:0:0:0:0:0 | 6324:204:13:73:18:0:0:1 | 7:0:0:0:0:0:0:0 | 1531:43:2:9:6:0:1:0 | 2:0:0:0:0:0:0:0 | 1 | 1 | 1 | 2.86E-13 | 6.12E-08 | 1 |
| 8998:193:16:136:2:0:0 | 9:0:0:0:0:0 | 6313:204:13:113:0:2:1 | 4:1:0:0:0:0 | 1530:43:2:11:0:0:0 | 1:0:0:0:0:0 | 1 | 1 | 1 | 2.09E-09 | 0.006512479 | 1 |
| 8995:192:16:138:2:0:0 | 6:1:0:1:0:0 | 6307:205:13:118:0:2:1 | 6:0:0:0:0:0 | 1529:43:2:11:0:0:0 | 1:0:0:0:0:0 | 1 | 1 | 1 | 0.000495625 | 5.92E-06 | 1 |
| 8992:193:16:139:2:0:0 | 7:0:0:0:0:0 | 6311:205:13:118:0:2:1 | 2:0:0:0:0:0 | 1527:43:2:11:0:0:0 | 1:0:0:0:0:0 | 1 | 1 | 1 | 4.44E-07 | 1 | 1 |
| 18:62:1772 | 0:00:05 | 0.239386574 | 0:00:02 | 0.052233796 | 0:00:00 | 1 | 1 | 1 | 0.008963405 | 1 | 1 |
| 18:61:1763 | 0:00:04 | 0.239386574 | 0:00:00 | 0.052222222 | 0:00:00 | 1.18E-222 | 1.52E-51 | 1 | 1 | 1 | 1 |
| 0:00 | 2:20 | 3:09 | 0:11 | 2:01 | 1:02 | 0.00023392 | 1 | 1 | 0.000209945 | 0.001445472 | 1 |
| 0:00 | 2:19 | 0:00 | 3:19 | 0:00 | 2:03 | 1.91E-05 | 1 | 1 | 0.00034049 | 0.001051209 | 0.661487987 |
| 66:00:00 | 5:00 | 72:01:00 | 4:01 | 11:01 | 1:00 | 0.003657788 | 1 | 1 | 0.030173969 | 0.194927198 | 1 |
| 0:00:00 | 279:10:08 | 12:08:02 | 30:00:01 | 0:01:00 | 3:00:00 | 1.07E-30 | 1.11E-06 | 2.74E-14 | 7.46E-106 | 2.36E-12 | 1 |
| 135:00:03 | 146:10:05 | 43:08:03 | 0:00:00 | 4:01:00 | 0:00:00 | 1 | 1 | 6.83E-09 | 1.54E-51 | 1 | 1 |
| 0:00:00 | 282:10:08 | 13:08:02 | 31:00:01 | 0:01:00 | 4:00:00 | 4.04E-31 | 1.11E-06 | 3.93E-15 | 3.92E-107 | 9.01E-13 | 1 |
| 144:05:00 | 137:03:00 | 13:02 | 31:01:00 | 0:00 | 4:00 | 8.28E-12 | 0.010484149 | 0.000225444 | 5.46E-29 | 2.86E-07 | 1 |
| 84:06:01 | 64:00:02 | 11:07:01 | 21:01:01 | 0:01:00 | 3:00:00 | 2.81E-10 | 1 | 0.001636609 | 3.83E-26 | 7.47E-08 | 1 |
| 63:7:1:1 | 0:0:0:0 | 223:12:6:3 | 4:0:1:0 | 30:1:1:0 | 0:0:0:0 | 1 | 1 | 1 | 1 | 0.041786793 | 1 |
| 5:1:0:1:0:2:0:0:0:2:0:0:0:0:0 | 0:0:0:0:0:0:0:0:0:0:0:0:0:0 | 0:1:0:0:0:0:0:0:0:0:0:0:0:0 | 1:4:0:0:3:0:2:0:2:1:0:5:0:0:3 | 2:0:0:0:1:0:0:0:0:0:0:0:0:0 | 0:0:1:0:0:0:2:0:1:0:0:1:0:0:0 | 0.323648844 | 1 | 0.7124885 | 1 | 0.017082132 | 0.326744913 |
| 58:3:0:0 | 14:1:0:0 | 112:1:3:4 | 24:1:1:0 | 31:1:0:0 | 6:0:1:0 | 1 | 1 | 7.75E-10 | 8.97E-08 | 5.52E-13 | 0.004024056 |
| 13:28:00 | 17:27:00 | 27:65:0 | 19:27:01 | 9:01:02 | 8:03:00 | 1 | 1 | 3.88E-05 | 6.36E-06 | 2.47E-05 | 0.015356793 |
| 0:0:0:2:0:0:0:0:0 | 0:2:16:84:1:2:120:7:2 | 3:2:27:83:2:1:144:7:1 | 0:0:0:1:0:0:0:0:0 | 1:1:3:18:1:0:4:8:1 | 0:0:0:0:0:0:0:0:0 | 1 | 1 | 2.20E-10 | 1.43E-123 | 1 | 1 |
| 0:00 | 6:00 | 14:00 | 0:00 | 4:02 | 0:00 | 1 | 1 | 1 | 0.017756017 | 1 | 1 |
| 0:00 | 9:00 | 29:00:00 | 0:00 | 4:00 | 0:00 | 1 | 1 | 1 | 0.00379461 | 1 | 1 |
| 0:0:0:0 | 12:0:6:0 | 0:0:0:0 | 35:0:2:0 | 0:0:0:0 | 7:1:1:0 | 5.55E-06 | 2.27E-05 | 3.07E-10 | 0.000115802 | 6.27E-20 | 0.004954386 |
| 7:1:5:0 | 6:2:5:0 | 0:0:0:0 | 30:10:57:1 | 0:0:0:0 | 6:0:4:0 | 0.012449474 | 1 | 1 | 0.081585557 | 1.28E-15 | 0.016870733 |
| 3:0:7:1 | 0:0:0:0 | 0:0:1:0 | 5:4:13:0 | 0:0:0:0 | 1:0:4:0 | 1 | 1 | 1 | 1 | 0.001633008 | 0.040495738 |
| 3:0:7:1 | 0:0:0:0 | 0:0:1:0 | 5:4:13:0 | 0:0:0:0 | 1:0:4:0 | 1 | 1 | 1 | 1 | 0.001633008 | 0.040495738 |
| 2:0:0:0 | 142:1:0:5 | 206:3:3:9 | 0:0:0:0 | 39:2:0:0 | 0:0:0:0 | 1 | 1 | 1.02E-22 | 1.84E-84 | 1 | 1 |
| 21:3:1:3:2:2:1:2:1:1:0:0:0:0 | 2:0:0:0:0:0:0:0:0:0:0:0:0:0 | 41:26:0:6:6:4:1:1:9:4:0:1:2:1 | 4:1:0:0:0:0:0:0:0:0:0:0:0:0 | 2:8:0:0:0:1:0:0:1:0:0:0:0:0 | 1:0:0:0:0:0:0:0:0:0:0:0:0:0 | 1 | 1 | 1 | 1 | 9.77E-05 | 1 |
| 123:01:00 | 0:00:00 | 224:00:01 | 112:00:01 | 22:01:00 | 26:00:00 | 2.74E-07 | 2.91E-07 | 1 | 1 | 1.37E-47 | 1.94E-11 |
| 142:16:00 | 92:06:00 | 282:11:00 | 0:00 | 48:04:00 | 0:00 | 1 | 1 | 6.38E-06 | 1.52E-17 | 1 | 1 |
| 8:04 | 3:01 | 7:10 | 2:02 | 1:01 | 5:00 | 1 | 1 | 1 | 1 | 1 | 0.030032732 |
| 0:02:01 | 0:00:00 | 0:00:00 | 0:00:00 | 0:00:00 | 0:00:00 | 0.003376462 | 1 | 1 | 1 | 1 | 1 |
| 4:235:0:43:0:0:10:6:5:0:0:1:0:18:0:42:1:0:0:0:2:1:0:0:0:0:0 | 1:0:0:0:0:0:0:0:0:0:0:0:0:0:0:0:0:0:0:0:0:0:0:0:0:0:0 | 7:392:5:41:4:1:9:7:17:0:1:2:4:15:2:51:2:0:0:3:1:1:0:0:0:1:1 | 4:0:0:0:0:0:0:0:0:0:0:0:0:0:1:0:0:0:0:0:0:0:0:0:0:0:0 | 1:99:1:4:3:0:2:0:1:0:0:1:0:2:0:4:0:0:1:2:0:0:1:1:0:0:0 | 0:1:0:0:0:0:0:0:0:0:0:0:0:0:0:0:0:0:0:0:0:0:0:0:0:0:0 | 1 | 1 | 1 | 1 | 4.91E-08 | 1 |
| 18:7:2:24:0:0 | 0:2:0:6:0:0 | 21:22:0:23:0:0 | 0:3:0:2:0:0 | 10:10:0:0:0:0 | 0:5:0:1:0:0 | 1 | 1 | 1 | 0.000822106 | 0.069632805 | 0.002036019 |

| | | | | | | | | | | | |
|---|---|---|---|---|---|---|---|---|---|---|---|
| 2:126:34:7:22:3:9:0:1:2:2:1:0:0 | 0:1:0:0:0:0:0:0:0:0:0:0:0:0 | 0:63:9:5:9:0:2:0:0:1:0:0:0:0 | 0:101:33:19:10:5:8:0:1:3:2:1:1:1 | 0:38:2:8:1:1:3:0:1:1:1:0:1:0 | 0:20:8:6:1:1:2:1:0:0:0:0:1:0 | 1 | 1 | 1 | 1 | 2.27E-142 | 3.66E-22 |
| 5:1:29:74 | 24:2:0:0 | 2:0:23:26 | 9:0:0:0 | 3:1:8:11 | 7:0:0:0 | 4.53E-05 | 0.012695226 | 1 | 4.17E-13 | 1.24E-05 | 0.000193253 |
| 29:29:73:4 | 24:0:0:2 | 11:23:25:0 | 11:0:0:4 | 10:8:11:1 | 0:0:0:0 | 1 | 1 | 1 | 4.17E-13 | 0.000150553 | 1 |
| 31:29:73:4 | 22:0:0:2 | 12:23:25:1 | 11:0:0:3 | 10:8:11:1 | 0:0:0:0 | 1 | 1 | 1 | 7.10E-12 | 7.18E-05 | 1 |
| 34:29:73:4 | 18:0:0:2 | 13:23:24:1 | 10:0:0:2 | 10:8:11:1 | 2:0:0:0 | 1 | 1 | 1 | 1.97E-09 | 8.91E-05 | 1 |
| 29:261:35:40 | 0:10:0:2 | 23:102:14:13 | 0:1:0:0 | 8:41:7:8 | 0:0:0:0 | 1 | 1 | 1 | 8.49E-05 | 1 | 1 |
| 0:0:0:0 | 29:274:0:7 | 0:0:0:0 | 22:102:1:3 | 0:0:0:0 | 8:40:0:1 | 1.62E-08 | 1.24E-11 | 4.29E-10 | 2.29E-143 | 6.47E-45 | 5.78E-18 |
| 19:0:7:4:2:3:0 | 0:0:0:0:0:0:0 | 4:0:1:0:0:0:0 | 12:1:7:3:1:1:5 | 1:0:0:0:0:0:0 | 1:0:0:0:0:1:1 | 1 | 1 | 1 | 1 | 0.000983287 | 1 |
| 0:0:0:0:0:0:0 | 20:7:0:2:3:0:0 | 0:0:0:0:0:0:0 | 16:8:1:1:1:0:5 | 0:0:0:0:0:0:0 | 2:0:0:0:1:0:1 | 1 | 1 | 0.005630511 | 2.15E-12 | 1.78E-07 | 1 |
| 0:00 | 5:01 | 0:00 | 24:01:00 | 0:00 | 2:00 | 0.103374839 | 1 | 0.001636609 | 0.113250947 | 9.02E-06 | 1 |
| 0.001643519 | 0.043263889 | 0:01:04 | 0:00:00 | 0:01:00 | 0:00:00 | 1 | 1 | 3.88E-05 | 1.05E-58 | 1 | 1 |
| 0.00150463 | 0.043043981 | 0:01:04 | 0:00:00 | 0:01:00 | 0:00:00 | 1 | 1 | 0.000257469 | 1.38E-50 | 1 | 1 |
| 0:00 | 7:01 | 4:02 | 0:01 | 0:01 | 1:00 | 1 | 1 | 1 | 0.039929313 | 1 | 1 |
| 252:12:6:4:33 | 16:0:0:0:1 | 263:9:9:9:28 | 15:0:0:1:3 | 69:6:2:1:8 | 13:1:0:0:1 | 3.06E-11 | 0.000792624 | 0.01911573 | 1.82E-11 | 3.11E-08 | 4.69E-08 |
| 285:13:6:4:33 | 19:1:0:0:1 | 294:10:10:9:32 | 17:0:1:3:3 | 80:6:2:2:10 | 11:1:1:0:1 | 5.42E-08 | 4.44E-06 | 0.000185616 | 7.22E-13 | 1.06E-07 | 9.35E-06 |
| 284:14:6:4:32 | 16:0:0:0:3 | 295:9:11:12:34 | 16:1:0:1:1 | 81:6:3:2:11 | 9:1:0:0:0 | 6.40E-06 | 0.000792624 | 1 | 7.42E-10 | 1.71E-09 | 6.76E-06 |
| 240:217:131 | 0:00:02 | 617:462:354 | 2:00:08 | 702:287:39 | 0:00:01 | 1 | 1 | 1 | 1 | 0.00784274 | 1 |
| 1:00 | 1:00 | 0:00 | 7:01 | 0:00 | 0:01 | 1 | 1 | 1 | 1 | 0.040786183 | 1 |
| 0:0:0:0:0:0:0 | 0:4:0:0:0:0:0 | 12:17:3:5:0:2:1 | 14:35:11:4:1:6:0 | 1:13:0:1:0:2:0 | 3:5:1:0:1:1:0 | 1 | 1 | 1 | 1 | 8.17E-16 | 0.057965639 |
| 0:0:0:0:0:0 | 0:7:0:0:0:0 | 0:0:0:0:0:0 | 31:53:14:10:1:1 | 0:0:0:0:0:0 | 4:20:1:2:1:0 | 1 | 1 | 0.012154512 | 3.28E-06 | 8.98E-22 | 1.49E-11 |
| 0:0:0:0:0:0 | 0:7:0:0:0:0 | 31:53:14:10:1:1 | 0:0:0:0:0:0 | 4:20:1:2:1:0 | 0:0:0:0:0:0 | 1 | 1 | 0.012154512 | 3.28E-06 | 1 | 1 |
| 0:7:0:0:0:0 | 0:0:0:0:0:0 | 18:36:12:5:1:0 | 14:17:2:5:0:1 | 3:6:1:1:1:0 | 1:14:0:1:0:0 | 1 | 1 | 1 | 1 | 3.30E-07 | 1.01E-10 |
| 132:49:62:0:0:0 | 9:4:2:0:0:0 | 204:66:86:0:1:1 | 6:2:4:0:0:0 | 28:17:14:0:0:0 | 0:0:0:0:0:0 | 1 | 1 | 1 | 0.000200727 | 0.009543145 | 1 |
| 107:36:150:2:170 | 3:11:3:0:0 | 170:35:199:3:244 | 2:19:3:0:0 | 50:11:149:1:51 | 2:9:2:0:0 | 1 | 0.069086341 | 0.005023791 | 0.000147906 | 1.82E-10 | 0.00049525 |
| 76:40:30:90:0 | 79:46:32:79:1 | 203:135:88:244:2 | 0:0:0:0:0 | 151:48:22:51:1 | 0:0:0:0:0 | 1 | 1 | 3.80E-19 | 1.57E-18 | 1 | 1 |
| 1:0:0:0:1:0:0:0:0 | 0:0:0:0:0:0:0:0:0 | 3:0:1:0:0:0:1:1:1 | 3:0:0:0:0:0:0:2:0 | 1:0:1:0:0:0:2:0:0 | 1:0:1:0:0:0:0:0:0 | 1 | 1 | 1 | 1 | 0.023626577 | 1 |
| 0:07 | 0:00 | 0:00 | 0:00 | 0:00 | 0:00 | 0.000177077 | 1 | 1 | 1 | 1 | 1 |
| 0:07 | 0:00 | 0:00 | 0:00 | 0:00 | 0:00 | 6.43E-05 | 1 | 1 | 1 | 1 | 1 |
| 0:00 | 0:07 | 0:00 | 0:00 | 0:00 | 0:00 | 1 | 1 | 1 | 0.010434472 | 1 | 1 |
| 0:07 | 0:00 | 0:00 | 0:00 | 0:00 | 0:00 | 0.00010713 | 1 | 1 | 1 | 1 | 1 |
| 0:07 | 0:00 | 0:00 | 0:00 | 0:00 | 0:00 | 6.43E-05 | 1 | 1 | 1 | 1 | 1 |
| 0:07 | 0:00 | 0:00 | 0:00 | 0:00 | 0:00 | 6.43E-05 | 1 | 1 | 1 | 1 | 1 |
| 0:00 | 0:07 | 0:00 | 0:00 | 0:00 | 0:00 | 1 | 1 | 0.030109098 | 0.010434472 | 1 | 1 |
| 0:00 | 0:07 | 0:00 | 0:00 | 0:00 | 0:00 | 6.43E-05 | 1 | 0.030109098 | 0.010434472 | 1 | 1 |
| 0:00 | 0:07 | 0:00 | 0:00 | 0:00 | 0:00 | 6.43E-05 | 1 | 0.017871477 | 0.010434472 | 1 | 1 |
| 0:00 | 0:07 | 0:00 | 0:00 | 0:00 | 0:00 | 6.43E-05 | 1 | 0.017871477 | 0.010434472 | 1 | 1 |
| 0:07 | 0:00 | 0:00 | 0:00 | 0:00 | 0:00 | 6.43E-05 | 1 | 1 | 1 | 1 | 1 |
| 0:07 | 0:00 | 0:00 | 0:00 | 0:00 | 0:00 | 6.43E-05 | 1 | 1 | 1 | 1 | 1 |
| 0:07 | 0:00 | 0:00 | 0:00 | 0:00 | 0:00 | 6.43E-05 | 1 | 1 | 1 | 1 | 1 |
| 0:07 | 0:00 | 0:00 | 0:00 | 0:00 | 0:00 | 6.43E-05 | 1 | 1 | 1 | 1 | 1 |
| 0:07 | 0:00 | 0:00 | 0:00 | 0:00 | 0:00 | 6.43E-05 | 1 | 1 | 1 | 1 | 1 |
| 0:07 | 0:00 | 0:00 | 0:00 | 0:00 | 0:00 | 6.43E-05 | 1 | 1 | 1 | 1 | 1 |
| 0:00 | 0:07 | 0:00 | 0:00 | 0:00 | 0:00 | 1 | 1 | 0.017871477 | 0.010434472 | 1 | 1 |
| 0:07 | 0:00 | 0:00 | 0:00 | 0:00 | 0:00 | 4.05E-05 | 1 | 1 | 1 | 1 | 1 |
| 0:07 | 0:00 | 0:00 | 0:00 | 0:00 | 0:00 | 4.05E-05 | 1 | 1 | 1 | 1 | 1 |
| 0:07 | 0:00 | 0:00 | 0:00 | 0:00 | 0:00 | 4.05E-05 | 1 | 1 | 1 | 1 | 1 |
| 0:00 | 0:07 | 0:00 | 0:00 | 0:00 | 0:00 | 1 | 1 | 0.017871477 | 0.010434472 | 1 | 1 |
| 0:07 | 0:00 | 0:00 | 0:00 | 0:00 | 0:00 | 4.05E-05 | 1 | 1 | 1 | 1 | 1 |
| 0:07 | 0:00 | 0:00 | 0:00 | 0:00 | 0:00 | 4.05E-05 | 1 | 1 | 1 | 1 | 1 |

| | | | | | | | | | | | |
|---|---|---|---|---|---|---|---|---|---|---|---|
| 0:07 | 0:00 | 0:00 | 0:00 | 0:00 | 0:00 | 4.05E-05 | 1 | 1 | 1 | 1 | 1 |
| 0:00 | 0:07 | 0:00 | 0:00 | 0:00 | 0:00 | 1 | 1 | 0.011433615 | 0.010434472 | 1 | 1 |
| 0:00 | 0:07 | 0:00 | 0:00 | 0:00 | 0:00 | 4.05E-05 | 1 | 0.011433615 | 0.010434472 | 1 | 1 |
| 0:00 | 0:07 | 0:00 | 0:00 | 0:00 | 0:00 | 2.55E-05 | 1 | 0.011433615 | 0.010434472 | 1 | 1 |
| 0:00 | 0:07 | 0:00 | 0:00 | 0:00 | 0:00 | 2.55E-05 | 1 | 0.011433615 | 0.010434472 | 1 | 1 |
| 0:00 | 0:07 | 0:00 | 0:00 | 0:00 | 0:00 | 1 | 1 | 0.011433615 | 0.010434472 | 1 | 1 |
| 0:00 | 0:07 | 0:00 | 0:00 | 0:00 | 0:00 | 2.55E-05 | 1 | 0.007094277 | 0.010434472 | 1 | 1 |
| 0:00 | 0:07 | 0:00 | 0:00 | 0:00 | 0:00 | 1 | 1 | 0.007094277 | 0.010434472 | 1 | 1 |
| 0:00 | 0:07 | 0:00 | 0:00 | 0:00 | 0:00 | 1 | 1 | 0.007094277 | 0.010434472 | 1 | 1 |
| 0:00 | 0:07 | 0:00 | 0:00 | 0:00 | 0:00 | 2.55E-05 | 1 | 0.007094277 | 0.010434472 | 1 | 1 |
| 0:00 | 0:07 | 0:00 | 0:00 | 0:00 | 0:00 | 2.55E-05 | 1 | 0.007094277 | 0.010434472 | 1 | 1 |
| 0:07 | 0:00 | 0:00 | 0:00 | 0:00 | 0:00 | 9.67E-05 | 1 | 1 | 1 | 1 | 1 |
| 0:07 | 0:00 | 0:00 | 0:00 | 0:00 | 0:00 | 0.000160489 | 1 | 1 | 1 | 1 | 1 |
| 0:07 | 0:00 | 0:00 | 0:00 | 0:00 | 0:00 | 4.05E-05 | 1 | 1 | 1 | 1 | 1 |
| 0:06 | 0:00 | 0:00 | 0:00 | 0:00 | 0:00 | 6.43E-05 | 1 | 1 | 1 | 1 | 1 |
| 0:00 | 316:07:00 | 0:00 | 795:08:00 | 0:00 | 253:03:00 | 1.07E-44 | 2.44E-12 | 6.58E-20 | 5.16E-65 | 4.22E-168 | 1.37E-53 |
| 0:00 | 312:07:00 | 1:00 | 788:08:00 | 1:00 | 253:03:00 | 1.72E-44 | 3.23E-12 | 6.58E-20 | 3.76E-64 | 1.34E-166 | 1.37E-53 |
| 0:00 | 308:07:00 | 2:00 | 735:07:00 | 0:00 | 240:03:00 | 4.98E-43 | 3.23E-12 | 6.58E-20 | 2.65E-63 | 7.81E-156 | 6.55E-51 |
| 0:00 | 308:07:00 | 2:00 | 735:07:00 | 0:00 | 240:03:00 | 4.98E-43 | 3.23E-12 | 6.58E-20 | 2.65E-63 | 7.81E-156 | 6.55E-51 |
| 0:00 | 307:07:00 | 0:00 | 735:07:00 | 0:00 | 241:03:00 | 4.98E-43 | 3.23E-12 | 6.58E-20 | 4.31E-63 | 7.81E-156 | 4.66E-51 |
| 13:00 | 295:07:00 | 504:00:00 | 232:07:00 | 119:00:00 | 122:03:00 | 4.98E-43 | 3.23E-12 | 2.85E-19 | 1.48E-60 | 4.51E-47 | 1.55E-25 |
| 308:07:00 | 0:00 | 232:07:00 | 504:00:00 | 122:03:00 | 119:00:00 | 1 | 1 | 1 | 1 | 2.49E-110 | 8.90E-27 |
| 1:00 | 303:07:00 | 0:00 | 729:07:00 | 0:00 | 239:03:00 | 2.13E-42 | 3.23E-12 | 1.04E-19 | 3.00E-62 | 1.53E-154 | 1.03E-50 |
| 1:00 | 297:07:00 | 1:00 | 726:07:00 | 0:00 | 238:03:00 | 3.43E-42 | 3.23E-12 | 1.04E-19 | 5.72E-61 | 6.66E-154 | 1.55E-50 |
| 297:07:00 | 0:00 | 718:07:00 | 4:00 | 237:03:00 | 0:00 | 1.47E-41 | 3.23E-12 | 1 | 1 | 1 | 1 |
| 297:07:00 | 0:00 | 230:08:00 | 539:00:00 | 123:03:00 | 119:00:00 | 1 | 1 | 1 | 1 | 6.39E-118 | 8.90E-27 |
| 0:00 | 296:07:00 | 2:00 | 767:08:00 | 0:00 | 242:03:00 | 1.47E-41 | 3.23E-12 | 2.90E-20 | 9.19E-61 | 4.56E-162 | 3.00E-51 |
| 0:00 | 297:07:00 | 0:00 | 769:08:00 | 0:00 | 242:03:00 | 2.37E-41 | 3.23E-12 | 2.90E-20 | 5.72E-61 | 1.74E-162 | 3.00E-51 |
| 0:00 | 295:07:00 | 2:00 | 764:08:00 | 0:00 | 240:03:00 | 3.83E-41 | 3.23E-12 | 6.58E-20 | 1.48E-60 | 1.97E-161 | 6.55E-51 |
| 0:00 | 292:07:00 | 0:00 | 758:08:00 | 0:00 | 238:03:00 | 1.02E-40 | 3.23E-12 | 6.58E-20 | 6.46E-60 | 3.84E-160 | 1.55E-50 |
| 294:06:00 | 1:01 | 730:08:00 | 8:00 | 236:03:00 | 1:00 | 1 | 1 | 1 | 1 | 0.006667609 | 1 |
| 279:07:00 | 10:00 | 221:09:00 | 507:00:00 | 121:03:00 | 112:00:00 | 1 | 1 | 0.111416266 | 0.002318525 | 5.64E-111 | 2.85E-25 |
| 0:00 | 284:08:00 | 6:00 | 683:09:00 | 0:01 | 229:03:00 | 4.10E-37 | 5.20E-11 | 2.96E-18 | 1.24E-57 | 2.51E-143 | 1.31E-48 |
| 281:10:00 | 1:00 | 680:09:00 | 0:00 | 228:05:00 | 1:00 | 2.53E-37 | 3.28E-11 | 1 | 1 | 1 | 1 |
| 272:10:00 | 9:00 | 675:09:00 | 1:00 | 228:05:00 | 0:00 | 1 | 1 | 0.066371669 | 0.00379461 | 1 | 1 |
| 269:10:00 | 9:00 | 660:09:00 | 3:00 | 222:05:00 | 1:00 | 1 | 1 | 0.066371669 | 0.00379461 | 1 | 1 |
| 269:10:00 | 9:00 | 662:09:00 | 1:00 | 222:05:00 | 1:00 | 1 | 1 | 0.066371669 | 0.00379461 | 1 | 1 |
| 11:01 | 261:08:00 | 4:00 | 654:09:00 | 1:01 | 217:03:00 | 1.10E-36 | 8.26E-11 | 9.19E-17 | 1.11E-52 | 4.67E-137 | 4.36E-46 |
| 0:00 | 264:09:00 | 0:00 | 653:09:00 | 0:00 | 217:04:00 | 2.11E-33 | 5.20E-11 | 2.03E-17 | 9.96E-53 | 7.52E-137 | 1.79E-45 |
| 0:00 | 257:09:00 | 0:00 | 642:09:00 | 0:00 | 216:04:00 | 2.11E-33 | 5.20E-11 | 2.03E-17 | 3.08E-51 | 1.81E-134 | 2.87E-45 |
| 249:09:00 | 1:00 | 619:09:00 | 0:00 | 205:04:00 | 0:00 | 4.79E-36 | 8.26E-11 | 1 | 1 | 1 | 1 |
| 237:08:00 | 10:01 | 170:09:00 | 444:00:00 | 103:03:00 | 102:01:00 | 4.04E-32 | 1.73E-10 | 0.066371669 | 0.008900344 | 2.56E-97 | 1.59E-22 |
| 245:08:00 | 2:01 | 172:09:00 | 442:00:00 | 102:03:00 | 102:01:00 | 6.44E-32 | 2.82E-10 | 1 | 1 | 6.87E-97 | 1.59E-22 |
| 241:08:00 | 10:01 | 605:09:00 | 4:00 | 205:03:00 | 0:01 | 1 | 1 | 0.066371669 | 0.008900344 | 1 | 1 |
| 1:00 | 250:09:00 | 0:00 | 604:09:00 | 0:00 | 205:04:00 | 8.70E-22 | 9.33E-07 | 3.31E-17 | 9.73E-50 | 2.76E-126 | 6.55E-43 |
| 9:01 | 242:08:00 | 431:00:00 | 170:09:00 | 101:01:00 | 104:03:00 | 0.662609049 | 1 | 1.50E-16 | 1.29E-48 | 1.58E-32 | 9.97E-22 |
| 0.548611111 | 0:01 | 0.75 | 0:00 | 0.309027778 | 0:00 | 0.017290558 | 1 | 1 | 1 | 1 | 1 |
| 0.542361111 | 0:10 | 0.749305556 | 0:01 | 0.309027778 | 0:00 | 0.029709415 | 1 | 0.066371669 | 0.002318525 | 1 | 1 |
| 0.542361111 | 0:10 | 0.749305556 | 0:01 | 0.309027778 | 0:00 | 0.029709415 | 1 | 0.066371669 | 0.002318525 | 1 | 1 |
| 241:09:00 | 10:00 | 597:07:00 | 3:01 | 205:04:00 | 0:00 | 1 | 1 | 0.066371669 | 0.002318525 | 1 | 1 |
| 1:00 | 254:03:00 | 429:00:00 | 177:03:00 | 102:00:00 | 108:01:00 | 5.80E-35 | 1.32E-10 | 2.96E-18 | 3.19E-54 | 1.31E-37 | 9.19E-24 |
| 261:02:00 | 0:00 | 180:03:00 | 433:00:00 | 112:01:00 | 103:00:00 | 1 | 1 | 1 | 1 | 6.15E-95 | 2.26E-23 |
| 261:02:00 | 0:00 | 179:03:00 | 432:00:00 | 112:01:00 | 103:00:00 | 1 | 1 | 1 | 1 | 9.99E-95 | 2.26E-23 |
| 252:00:00 | 9:00 | 613:02:00 | 1:00 | 216:00:00 | 0:00 | 4.07E-38 | 1.32E-10 | 0.017871477 | 0.00379461 | 1 | 1 |
| 0.181944444 | 0:00 | 0.471527778 | 0:01 | 0.151388889 | 0:00 | 9.53E-06 | 0.017552603 | 1 | 1 | 1 | 1 |
| 264:00:00 | 2:00 | 619:01:00 | 5:00 | 218:00:00 | 1:00 | 1 | 1 | 1 | 1 | 0.031296179 | 1 |
| 266:00:00 | 2:00 | 192:01:00 | 428:00:00 | 114:00:00 | 106:00:00 | 1 | 1 | 1 | 1 | 7.31E-94 | 5.77E-24 |
| 266:00:00 | 0:00 | 617:02:00 | 3:00 | 218:00:00 | 0:00 | 1.18E-40 | 9.72E-12 | 1 | 1 | 1 | 1 |
| 0:00 | 265:00:00 | 1:00 | 615:02:00 | 0:00 | 217:00:00 | 1.90E-40 | 9.72E-12 | 4.73E-18 | 1.69E-58 | 4.31E-133 | 5.50E-48 |
| 255:00:00 | 9:00 | 613:02:00 | 3:00 | 217:00:00 | 0:00 | 1 | 1 | 0.017871477 | 0.00379461 | 1 | 1 |
| 262:00:00 | 1:00 | 185:02:00 | 426:00:00 | 113:00:00 | 104:00:00 | 1 | 1 | 1 | 1 | 1.96E-93 | 1.46E-23 |
| 262:00:00 | 0:00 | 605:02:00 | 6:00 | 216:00:00 | 1:00 | 1 | 1 | 1 | 1 | 0.018436807 | 1 |
| 264:01:00 | 1:00 | 187:02:00 | 425:00:00 | 113:00:00 | 104:00:00 | 1 | 1 | 1 | 1 | 3.19E-93 | 1.46E-23 |
| 263:01:00 | 2:00 | 187:02:00 | 423:00:00 | 112:00:00 | 104:00:00 | 1 | 1 | 1 | 1 | 8.44E-93 | 1.46E-23 |
| 1:00 | 264:01:00 | 2:00 | 608:02:00 | 0:00 | 216:00:00 | 4.79E-40 | 9.72E-12 | 2.96E-18 | 1.18E-57 | 1.41E-131 | 8.79E-48 |
| 252:01:00 | 9:00 | 186:02:00 | 421:00:00 | 113:00:00 | 103:00:00 | 1 | 1 | 0.017871477 | 0.00379461 | 2.27E-92 | 2.26E-23 |
| 0:00 | 266:01:00 | 0:00 | 607:02:00 | 0:00 | 217:00:00 | 3.02E-40 | 3.28E-11 | 4.73E-18 | 4.48E-58 | 2.23E-131 | 5.50E-48 |
| 0:00 | 267:01:00 | 1:00 | 615:02:00 | 1:00 | 218:00:00 | 3.02E-40 | 3.28E-11 | 2.96E-18 | 2.74E-58 | 4.31E-133 | 3.58E-48 |
| 0:00 | 268:01:00 | 2:00 | 615:02:00 | 1:00 | 218:00:00 | 4.79E-40 | 3.28E-11 | 2.96E-18 | 1.69E-58 | 4.31E-133 | 3.58E-48 |
| 0:0:0:0 | 5:2:0:0 | 2:2:0:1 | 1:1:0:0 | 1:0:1:0 | 1:0:0:0 | 1 | 1 | 1 | 0.027461316 | 1 | 1 |

| | | | | | | | | | | | |
|---|---|---|---|---|---|---|---|---|---|---|---|
| 61:3:26:0:381:15 | 0:0:0:0:0:1 | 56:2:29:1:349:20 | 0:0:0:0:5:0 | 8:0:13:1:39:4 | 0:0:0:0:0:0 | 1 | 1 | 1 | 1 | 0.000252891 | 1 |
| 0:0:0:0:0:0:0 | 43:12:4:1:1:9:1 | 10:1:0:0:2:2:1 | 15:2:1:0:0:0:1 | 5:2:0:0:0:0:0 | 4:1:1:0:0:0:0 | 6.09E-16 | 1 | 2.56E-27 | 4.98E-26 | 9.18E-12 | 0.025385819 |
| 0:0:8:3:0:0 | 0:0:5:1:0:0 | 0:0:18:11:0:0 | 0:0:0:0:0:0 | 0:0:8:0:0:1 | 0:0:0:0:0:0 | 1 | 1 | 0.012154512 | 0.001878282 | 1 | 1 |
| 715:14:00 | 6:00:00 | 961:16:01 | 3:00:00 | 348:01:00 | 1:00:00 | 1 | 1 | 1 | 0.003552529 | 1 | 1 |
| 78:59:77:30:0:0:1:0:0:0 | 71:44:85:23:0:0:0:1:1:0 | 1:0:0:0:0:0:0:0:0:0 | 86:42:66:24:0:1:0:0:0:1 | 0:0:0:0:0:0:0:0:0:0 | 29:14:6:4:0:0:0:1:0:1 | 3.65E-44 | 4.11E-63 | 1.59E-47 | 1.64E-85 | 7.62E-84 | 1.70E-25 |
| 0:65:78:149:22:0:1:1:1:1 | 0:74:80:140:25:1:0:0:0:0 | 0:85:56:99:22:0:3:0:1:0 | 0:1:0:0:0:0:0:0:0:0 | 0:29:13:5:3:0:1:1:0:0 | 0:0:0:0:0:0:0:0:0:0 | 1 | 1 | 1.61E-42 | 1.03E-139 | 1 | 1 |
| 427:192:0:119:35:6:1:1:1:1 | 5:5:0:0:0:0:0:0:0:0 | 142:80:0:78:16:0:0:0:0:0 | 1:2:0:0:0:0:0:0:0:0 | 7:12:0:25:3:0:0:0:1:0 | 1:0:0:0:0:0:0:0:0:0 | 1 | 1 | 1 | 1.49E-05 | 1 | 1 |
| 0:119:193:423:35:6:1:1:1:1 | 0:0:5:6:0:0:0:0:0:0 | 0:78:79:142:16:0:0:0:0:0 | 0:0:3:1:0:0:0:0:0:0 | 0:25:12:8:3:0:0:0:1:0 | 0:0:0:0:0:0:0:0:0:0 | 1 | 1 | 1 | 2.60E-06 | 1 | 1 |
| 798:13:248:13:4:9 | 3:0:2:0:0:0 | 694:18:139:7:3:12 | 4:0:0:0:1:0 | 65:3:14:0:0:1 | 0:0:0:0:0:0 | 1 | 1 | 1 | 0.068829746 | 0.011881852 | 1 |
| 472:10:123:9:3:5 | 6:0:3:0:0:0 | 424:13:71:2:2:6 | 6:0:1:0:0:0 | 46:2:10:0:0:1 | 0:0:0:0:0:0 | 1 | 1 | 1 | 0.001254484 | 0.000284747 | 1 |
| 370:91:8:9:2:3 | 59:17:0:1:1:0 | 332:53:2:11:2:5 | 56:9:0:1:0:1 | 44:9:0:2:0:1 | 2:1:0:0:0:0 | 0.018731299 | 0.008580571 | 0.019323705 | 5.45E-43 | 6.36E-45 | 1 |
| 101:401:8:6:3:3 | 22:71:1:4:0:2 | 58:364:1:11:2:5 | 14:68:1:2:0:1 | 7:33:0:2:0:1 | 3:13:0:0:0:0 | 1.04E-16 | 6.91E-12 | 6.90E-08 | 2.27E-47 | 3.45E-51 | 3.20E-09 |
| 1:111:30:0:0:2 | 0:12:1:0:0:1 | 5:119:12:1:0:0 | 0:4:1:0:0:0 | 2:22:3:1:0:0 | 0:0:0:0:0:0 | 1 | 1 | 1 | 7.58E-09 | 0.011881852 | 1 |
| 293:75 | 18:00 | 171:37:00 | 6:00 | 159:09:00 | 0:01 | 1 | 1 | 1 | 4.15E-05 | 0.018436807 | 1 |
| 1430:02:00 | 16:02 | 728:01:00 | 5:00 | 266:00:00 | 2:00 | 1 | 1 | 1 | 0.001469165 | 0.031296179 | 1 |
| 1585:04:00 | 14:00 | 787:01:00 | 10:02 | 268:00:00 | 1:00 | 1 | 1 | 1 | 0.000301919 | 0.026647174 | 1 |
| 1.283333333 | 0:13 | 0.638194444 | 2:09 | 0.185416667 | 0:02 | 1 | 1 | 1 | 0.000501437 | 0.041747173 | 1 |
| 1634:04:00 | 5:00 | 807:02:00 | 7:02 | 268:00:00 | 1:00 | 1 | 1 | 1 | 0.030173969 | 0.108311537 | 1 |
| 1771:04:00 | 10:01 | 898:04:00 | 2:00 | 273:00:00 | 0:00 | 1 | 1 | 1 | 0.008900344 | 1 | 1 |
| 1890:05:00 | 5:00 | 973:04:00 | 2:00 | 272:01:00 | 0:00 | 1 | 1 | 1 | 0.030173969 | 1 | 1 |
| 1952:05:00 | 7:00 | 999:07:00 | 1:00 | 272:01:00 | 0:00 | 1 | 1 | 1 | 0.010434472 | 1 | 1 |
| 40:02:13 | 0:00:00 | 3:00:00 | 25:01:15 | 1:00:00 | 13:01:00 | 1 | 1 | 1 | 1 | 4.76E-05 | 2.82E-05 |
| 3:0:0:0:0:0 | 0:1:0:0:0:0 | 33:3:0:3:0:0 | 10:0:0:0:0:1 | 8:0:0:1:0:0 | 5:0:0:0:0:0 | 5.74E-08 | 1 | 1 | 1 | 6.01E-08 | 0.000249906 |
| 40:05:00 | 9:00 | 9:02 | 3:00 | 0:00 | 0:00 | 1 | 1 | 1 | 0.00379461 | 1 | 1 |
| 76:43:82:10:1 | 0:0:0:0:0 | 3:1:7:0:0 | 37:32:85:12:1 | 0:0:1:1:0 | 9:2:13:1:0 | 2.82E-18 | 3.66E-08 | 1 | 1 | 2.30E-25 | 6.10E-05 |
| 3:9:96:1:1:0:1 | 0:0:0:0:0:0:0 | 0:1:3:0:0:0:0 | 3:5:36:1:0:2:0 | 0:0:3:0:0:0:0 | 1:2:13:2:0:0:0 | 1 | 1 | 1 | 1 | 1.99E-29 | 8.23E-09 |
| 2:0:1:0:1:3:0:2:0:0:1:0:1:0:0:0:0:0 | 4:0:1:0:0:0:0:1:0:1:1:1:0:0:0:0:0:0 | 24:1:9:0:1:9:0:0:2:2:0:0:0:2:2:1:1:1 | 0:0:0:0:0:0:0:0:0:0:0:0:0:0:0:0:0:0 | 12:1:0:0:0:1:0:0:1:1:0:0:2:0:0:0:0:0 | 0:0:0:0:0:0:0:0:0:0:0:0:0:0:0:0:0:0 | 1 | 1 | 0.000495455 | 0.014197495 | 1 | 1 |
| 5632:100:1062:107:174:350:27 | 3:0:3:0:0:1:0 | 3059:43:295:125:15:172:15 | 4:0:1:0:0:0:0 | 673:9:56:16:1:24:4 | 0:0:0:0:0:1:0 | 1 | 1 | 1 | 0.070826298 | 0.006512479 | 1 |
| 5641:99:1066:108:174:336:27 | 7:0:0:0:0:1:0 | 3070:43:297:125:15:168:15 | 1:0:0:0:0:0:0 | 675:9:56:16:1:25:4 | 0:0:0:0:0:0:0 | 1 | 1 | 1 | 6.42E-06 | 1 | 1 |
| 94:5670:1071:108:179:323:27 | 0:10:0:0:0:0:0 | 42:3091:298:124:15:159:15 | 0:3:0:1:0:2:0 | 7:678:56:16:1:23:4 | 0:0:0:0:0:0:0 | 4.37E-07 | 1 | 1 | 1.36E-10 | 0.124057299 | 1 |
| 5675:320:94:1070:108:179:27 | 5:1:0:0:0:0:0 | 3093:159:42:299:125:15:15 | 1:2:0:0:0:0:0 | 677:22:7:56:16:1:4 | 1:1:0:0:0:0:0 | 1 | 1 | 1 | 0.000718566 | 1 | 1 |
| 318:93:5675:1071:108:179:28 | 0:1:5:0:0:0:0 | 154:40:3095:299:124:15:15 | 1:0:0:0:1:0:0 | 22:7:678:56:16:1:4 | 0:0:0:0:0:0:0 | 0.000696723 | 1 | 1 | 0.000718566 | 1 | 1 |
| 94:5675:1073:108:179:304:28 | 0:5:0:0:0:0:0 | 41:3095:299:125:15:145:15 | 0:2:0:0:0:0:0 | 7:678:56:16:1:24:4 | 0:0:0:0:0:0:0 | 1 | 1 | 1 | 7.10E-05 | 1 | 1 |

| | | | | | | | | | | | |
|---|---|---|---|---|---|---|---|---|---|---|---|
| 92:5679:1073:<br>108:179:296:2<br>8 | 0:4:1:0:0:1:0 | 41:3100:299:1<br>25:15:142:15 | 0:0:0:0:0:1:0 | 6:678:56:16:1:<br>24:4 | 0:0:0:0:0:0:0 | 1 | 1 | 1 | 0.024750111 | 1 | 1 |
| 2837:564:41:4<br>1:52:124:12 | 2852:510:49:1<br>39:55:149:16 | 202:14:2:0:12:<br>12:0 | 2901:285:39:1<br>5:113:122:15 | 83:3:1:0:2:0:0 | 595:53:5:1:14:<br>23:4 | 0 | 0 | 0 | 0 | 0 | 0 |
| 90:5687:1074:<br>107:180:262:2<br>8 | 0:2:1:0:0:0:0 | 40:3100:299:1<br>24:15:127:15 | 0:4:0:0:0:0:0 | 6:678:56:16:1:<br>22:4 | 0:0:0:0:0:0:0 | 6.72E-05 | 1 | 1 | 1 | 1 | 1 |
| 5695:81:1074:<br>180:87:196:17 | 6:0:2:0:6:0:0 | 3108:36:300:1<br>5:108:103:13 | 3:0:0:0:4:0:1 | 676:5:57:1:14:<br>15:3 | 2:0:0:0:1:0:1 | 0.010321833 | 1 | 1 | 0.000822106 | 0.028327806 | 1 |
| 2:4:0:0:0:0:0 | 71:5714:1078:<br>180:161:9:1 | 2:204:14:0:11:<br>1:0 | 30:2913:286:1<br>5:69:2:1 | 1:83:3:0:0:0:0 | 4:597:54:1:12:<br>1:0 | 0 | 0 | 0 | 0 | 0 | 0 |
| 31:90:5785:10<br>94:126:183:17<br>7 | 1:3:1:1:0:0:11 | 15:38:3150:30<br>7:137:15:93 | 1:0:0:0:0:0:0 | 5:6:685:57:19:<br>1:18 | 0:0:0:0:0:0:1 | 1 | 1 | 1 | 2.78E-06 | 1 | 1 |
| 5780:106:1095<br>:136:183:179:4<br>0 | 6:0:1:0:0:1:0 | 3148:44:307:1<br>46:15:83:17 | 2:0:0:0:0:0:0 | 685:8:57:20:1:<br>16:5 | 0:0:0:0:0:0:0 | 0.007462076 | 1 | 1 | 0.000495625 | 1 | 1 |
| 1618:143:9:17:<br>191:31:27:66:1<br>:123:5:19:12:4<br>:2:1:1:1:2:0:0 | 0:1:0:0:0:0:0:0<br>:0:0:0:1:0:0:0:<br>0:0:0:0:0:0:0 | 2609:180:84:3<br>5:854:33:92:15<br>4:18:154:12:22<br>:54:1:14:0:0:0:<br>3:1:2 | 6:0:0:0:1:0:0:0<br>:0:0:0:0:0:0:0:<br>0:0:0:0:0:0:0 | 546:31:29:13:8<br>2:4:11:25:3:30<br>:3:4:3:1:1:0:0:<br>0:0:1:0 | 2:0:0:0:0:0:0:0<br>:0:0:0:0:0:0:0:<br>0:0:0:0:0:0:0 | 1 | 1 | 1 | 1 | 9.33E-13 | 1 |
| 144:9:17:191:3<br>1:26:62:1:122:<br>5:1618:24:18:4<br>:2:1:1:1:2:0:0 | 0:0:0:0:0:0:0:0<br>:0:0:0:3:0:0:0:<br>0:0:0:0:0:0:0 | 179:84:36:855:<br>33:92:141:18:1<br>54:11:2616:34:<br>65:1:14:0:0:0:<br>4:1:2 | 1:0:0:0:0:0:0:0<br>:0:0:1:4:5:0:0:<br>0:0:0:0:0:0:0 | 31:29:12:82:4:<br>11:21:3:30:3:5<br>46:4:5:1:1:0:0:<br>0:0:1:0 | 0:0:0:0:0:0:0:0<br>:0:0:0:0:0:0:0:<br>0:0:0:0:0:0:0 | 1 | 1 | 1 | 1 | 3.00E-07 | 1 |
| 144:9:17:191:3<br>1:26:62:1:122:<br>5:1617:24:17:4<br>:2:1:1:1:2:0:0 | 0:0:0:0:0:0:0:0<br>:0:0:1:3:1:0:0:<br>0:0:0:0:0:0:0 | 180:84:36:853:<br>33:92:137:18:1<br>54:10:2615:34:<br>70:1:14:0:0:0:<br>4:1:2 | 0:0:0:2:0:0:1:0<br>:0:0:2:5:2:0:0:<br>0:0:0:0:0:0:0 | 31:29:12:82:4:<br>11:21:3:30:3:5<br>48:5:5:1:1:0:0:<br>0:0:1:0 | 0:0:0:0:0:0:0:0<br>:0:0:0:0:0:0:0:<br>0:0:0:0:0:0:0 | 1 | 1 | 1 | 0.004854451 | 9.14E-05 | 1 |
| 1613:144:9:16:<br>31:26:29:1:123<br>:4:191:35:23:4<br>:1:1:1:1:0:0:0 | 0:0:0:0:0:0:0:0<br>:1:0:0:3:0:0:0:<br>0:0:0:0:0:0:0 | 2604:180:83:3<br>3:33:89:54:18:<br>153:9:855:62:8<br>2:1:0:0:0:6:1:0<br>:2 | 1:0:0:0:0:0:1:0<br>:0:0:0:3:2:0:0:<br>0:0:0:0:0:0:0 | 545:31:28:12:4<br>:10:7:3:30:3:8<br>2:6:5:1:0:0:0:0<br>:1:0:0 | 1:0:1:0:0:0:0:0<br>:0:0:0:0:0:0:0:<br>0:0:0:0:0:0:0 | 1 | 1 | 1 | 1 | 0.011964927 | 1 |
| 144:9:16:31:26<br>:29:1:125:4:16<br>10:191:34:23:4<br>:1:1:1:1:0:0:0 | 0:0:0:0:0:0:0:0<br>:0:2:0:2:0:0:0:<br>0:0:0:0:0:0:0 | 180:83:33:33:8<br>8:44:18:153:9:<br>2598:855:63:8<br>5:1:0:0:0:5:1:0<br>:2 | 0:0:0:0:0:0:0:0<br>:0:4:0:5:2:0:0:<br>0:0:1:0:0:0:0 | 31:29:12:4:10:<br>7:3:30:3:544:8<br>2:6:5:1:0:0:0:0<br>:1:0:0 | 0:0:0:0:0:0:0:0<br>:0:1:0:0:0:0:0:<br>0:0:0:0:0:0:0 | 1 | 1 | 1 | 1 | 7.53E-07 | 1 |
| 144:9:15:31:26<br>:28:1:124:4:16<br>12:191:34:23:4<br>:1:1:1:1:0:0:0 | 0:0:1:0:0:0:0:1<br>:0:0:0:3:0:0:0:<br>0:0:0:0:0:0:0 | 180:83:33:33:8<br>8:42:18:151:9:<br>2599:855:59:8<br>3:1:0:0:0:5:1:0<br>:2 | 0:0:0:0:0:0:0:0<br>:0:1:0:7:3:0:0:<br>0:0:0:0:0:0:0 | 31:28:12:4:10:<br>6:3:30:3:544:8<br>2:6:5:1:0:0:0:0<br>:1:0:0 | 0:1:0:0:0:0:0:0<br>:0:1:0:0:0:0:0:<br>0:0:0:0:0:0:0 | 1 | 1 | 1 | 0.004854451 | 2.40E-12 | 1 |
| 16:26:143:9:31<br>:26:20:1:4:160<br>8:125:191:34:4<br>:1:1:1:0:0:0 | 0:1:1:0:0:0:0:0<br>:0:2:2:0:7:0:0:<br>0:0:0:0:0:0 | 32:86:180:83:3<br>3:88:32:18:9:2<br>588:152:850:6<br>4:1:0:0:0:1:0:1 | 0:2:0:0:0:0:0:0<br>:0:5:1:5:6:0:0:<br>0:0:0:0:0:0 | 12:6:31:29:4:1<br>0:5:3:3:543:30<br>:82:5:1:0:0:0:1<br>:0:0 | 0:0:0:0:0:0:0:0<br>:0:1:0:0:1:0:0:<br>0:0:0:0:0:0 | 1 | 1 | 2.78E-05 | 1.24E-08 | 1.86E-08 | 1 |
| 64:144:9:14:31<br>:25:1:130:4:15<br>98:191:54:27:4<br>:2:1:1:1:2:0:0 | 2:0:0:0:0:0:0:0<br>:0:3:0:0:1:0:0:<br>0:0:0:0:0:0:0 | 134:180:81:34:<br>33:88:18:159:9<br>:2578:855:85:1<br>17:1:14:0:0:0:<br>4:1:2 | 0:0:0:0:0:0:0:0<br>:0:3:0:0:2:0:0:<br>0:0:0:0:0:0:0 | 25:31:29:12:4:<br>11:3:30:3:538:<br>82:12:12:1:1:0<br>:0:0:0:1:0 | 0:0:0:0:0:0:0:0<br>:0:1:0:0:0:0:0:<br>0:0:0:0:0:0:0 | 1 | 1 | 1 | 0.002971703 | 0.000443721 | 1 |
| 143:9:14:31:22<br>:76:1:129:3:15<br>76:191:54:29:4<br>:2:1:1:1:2:0:0 | 1:0:0:0:1:0:0:1<br>:0:0:0:0:0:0:0:<br>0:0:0:0:0:0:0 | 180:81:33:33:8<br>3:153:18:159:7<br>:2522:854:85:1<br>37:1:14:0:0:0:<br>4:1:2 | 0:0:0:0:0:1:0:0<br>:0:7:1:0:1:0:0:<br>0:0:0:0:0:0:0 | 31:29:11:4:10:<br>32:3:31:3:537:<br>82:12:13:1:1:0<br>:0:0:0:1:0 | 0:0:0:0:0:0:0:0<br>:0:1:0:0:1:0:0:<br>0:0:0:0:0:0:0 | 1 | 1 | 1 | 1 | 6.38E-12 | 1 |
| 1566:144:9:14:<br>31:22:76:1:130<br>:3:191:54:26:4<br>:2:1:1:1:2:0:0 | 3:0:0:0:0:0:0:0<br>:0:0:0:0:0:0:0:<br>0:0:0:0:0:0:0 | 2518:180:81:3<br>3:32:81:154:17<br>:159:7:852:85:<br>129:1:14:0:0:0<br>:4:1:2 | 4:0:0:0:1:0:1:1<br>:0:0:2:0:0:0:0:<br>0:0:0:0:0:0:0 | 536:31:29:11:4<br>:10:32:3:31:3:<br>82:12:14:1:1:0<br>:0:0:0:1:0 | 0:0:0:0:0:0:0:0<br>:0:0:0:0:0:0:0:<br>0:0:0:0:0:0:0 | 1 | 1 | 1 | 1 | 0.001907292 | 1 |

| | | | | | | | | | | | |
|---|---|---|---|---|---|---|---|---|---|---|---|
| 144:9:14:31:21:74:1:130:3:1555:191:54:26:4:2:1:1:1:2:0:0 | 0:0:0:0:1:0:0:0:0:1:0:0:0:0:0:0:0:0:0:0:0 | 180:80:32:33:79:152:18:159:7:2497:853:85:121:1:14:0:0:0:4:1:2 | 0:1:0:0:1:1:0:0:0:6:1:0:0:0:0:0:0:0:0:0:0 | 31:27:11:4:10:32:3:31:3:531:82:12:13:1:1:0:0:0:0:1:0 | 0:0:0:0:0:0:0:0:0:1:0:0:0:0:0:0:0:0:0:0:0 | 1 | 1 | 1 | 1 | 1.06E-07 | 1 |
| 144:1520:9:14:31:19:77:1:130:3:191:54:20:4:2:1:1:2:0:0 | 0:3:0:0:0:0:0:0:0:0:0:0:0:0:0:0:0:0:0:0 | 180:2431:78:32:33:75:146:18:159:7:850:85:102:1:14:0:0:4:1:2 | 0:8:0:0:0:0:1:0:0:0:4:0:1:0:0:0:0:0:0:0 | 31:525:27:11:4:9:31:3:31:3:82:12:11:1:1:0:0:0:1:0 | 0:0:0:0:0:0:0:0:0:0:0:0:0:0:0:0:0:0:0:0 | 1 | 1 | 1 | 1 | 5.68E-13 | 1 |
| 448:57:00 | 3:00 | 590:70 | 5:00 | 88:05:00 | 0:00 | 1 | 1 | 1 | 1 | 0.031296179 | 1 |
| 146:158:126:61 | 0:5:1:0 | 377:322:346:169 | 0:7:0:0 | 194:78:25:22 | 0:1:0:0 | 1 | 1 | 1 | 0.012726037 | 0.000193006 | 1 |
| 136:146:219:61 | 6:0:2:0 | 226:376:592:169 | 24:1:2:0 | 49:194:37:22 | 9:0:0:0 | 0.001044569 | 4.88E-05 | 1 | 0.009770702 | 3.43E-12 | 1.26E-05 |
| 143:150:240:60 | 0:1:2:0 | 372:230:631:167 | 1:0:4:0 | 193:44:35:22 | 0:0:0:0 | 1 | 1 | 1 | 1 | 0.041786793 | 1 |
| 143:151:241:60 | 0:0:1:0 | 372:230:631:167 | 1:0:4:0 | 193:44:35:22 | 0:0:0:0 | 1 | 1 | 1 | 1 | 0.041786793 | 1 |
| 143:151:241:59 | 0:0:1:1 | 371:231:629:166 | 1:0:4:0 | 192:44:34:22 | 1:0:1:0 | 1 | 1 | 1 | 1 | 0.041786793 | 1 |
| 0:00 | 1:00 | 0:00 | 0:05 | 0:00 | 1:03 | 1 | 1 | 1 | 1 | 0.031296179 | 1 |
| 13:4:0:0 | 17:7:0:1 | 11:1:0:0 | 8:12:2:0 | 5:1:0:0 | 3:1:0:0 | 0.154229945 | 0.154096241 | 1 | 4.28E-06 | 0.001405523 | 1 |
| 6:24:0:0:1:1:0:0 | 0:0:0:0:0:0:0:0 | 1:11:0:0:0:0:0:0 | 1:7:1:0:0:0:0:0 | 1:7:0:0:0:0:0:0 | 0:3:0:0:0:0:1:0 | 1 | 1 | 1 | 1 | 1.32E-05 | 1 |
| 0:6:0:2:30:27:1:0:3:0:0 | 3:20:1:7:71:67:2:1:15:1:0 | 1:17:3:6:11:10:1:0:1:0:0 | 0:0:0:0:0:0:0:0:0:0:0 | 1:11:0:1:10:5:1:0:2:0:0 | 0:0:0:0:0:0:0:0:0:0:0 | 1 | 1 | 5.39E-22 | 1.72E-80 | 1 | 1 |
| 0:0:0:0:0:0 | 2:0:0:1:0:0 | 1:0:0:0:0:0 | 0:0:0:0:0:0 | 5:0:0:0:1:0 | 0:0:0:0:0:0 | 0.000253109 | 0.000280691 | 1 | 1 | 1 | 1 |
| 1:1:3:3:0:7:0 | 0:4:0:3:0:0:3 | 2:3:6:5:0:10:0 | 0:9:0:8:0:0:0 | 0:3:1:3:0:0:0 | 0:0:0:0:0:0:0 | 1 | 1 | 1 | 0.03781524 | 3.29E-07 | 1 |
| 5:10:0:13:14:0:0:4:0:0:4:2:0:0 | 0:0:0:0:0:0:0:5:1:1:0:0:0:0 | 2:12:0:12:14:0:2:2:0:0:4:1:1:1 | 0:0:0:0:0:0:0:2:0:0:0:0:0:0 | 1:3:0:0:2:0:0:1:0:0:2:1:1:0 | 0:0:0:0:0:0:0:0:0:0:0:0:0:0 | 1 | 1 | 1 | 1.88E-05 | 1 | 1 |
| 3:7:5:6:0:0:0:6:1:3:0:1:0:0 | 2:3:8:7:0:0:0:4:1:1:1:0:0:0 | 2:7:9:10:0:1:0:3:1:4:0:0:0:1 | 0:5:3:4:0:1:0:1:0:0:0:0:1:0 | 1:3:0:2:0:0:0:0:1:2:0:0:0:0 | 0:0:0:0:0:0:0:1:0:0:0:0:1:0 | 1 | 1 | 1 | 1.21E-05 | 0.001610444 | 1 |
| 0:0:0:0:0:0:0:0:0:0:0:0:1:0 | 5:10:13:10:0:22:0:0:4:2:1:1:0:0 | 0:0:0:0:0:0:0:0:0:0:0:0:0:0 | 2:12:12:9:0:16:2:0:4:1:0:2:0:1 | 0:0:0:0:0:0:0:0:0:0:0:0:0:0 | 1:3:1:2:0:2:0:0:2:1:0:1:0:0 | 0.000149562 | 8.43E-09 | 3.43E-07 | 2.89E-18 | 7.33E-13 | 0.390262798 |
| 2:3:8:5:0:9:0:0:1:1:0:1:1:0 | 3:7:5:4:0:13:0:0:3:1:1:0:0:0 | 0:5:3:2:0:4:1:0:1:0:0:0:1:0 | 2:7:9:2:0:12:1:0:4:1:0:0:1:1 | 0:0:1:0:0:2:0:0:0:0:0:0:1:0 | 1:3:0:0:0:0:0:0:2:1:0:0:0:0 | 0.000149562 | 8.43E-09 | 0.006401509 | 4.13E-09 | 1.41E-08 | 0.047647448 |
| 9:00:00 | 0:00:00 | 0:00:00 | 10:00:01 | 0:00:00 | 2:01:00 | 1 | 1 | 1 | 1 | 0.000446416 | 1 |
| 0:0:0:0 | 53:0:19:0 | 0:0:0:0 | 13:1:5:2 | 0:0:0:0 | 1:0:0:0 | 6.07E-24 | 5.88E-09 | 0.017213806 | 9.57E-22 | 0.001192805 | 1 |
| 21216:637 | 12:08 | 19415:832 | 13:04 | 1952:64 | 1:00 | 1 | 0.424020111 | 1 | 0.387840147 | 0.035544253 | 1 |
| 20965:813 | 18:01 | 19138:906 | 16:04 | 1946:69 | 2:00 | 1 | 1 | 1 | 0.000167796 | 0.010139464 | 1 |
| 20958:808 | 15:05 | 19133:907 | 12:06 | 1946:71 | 2:00 | 1 | 0.00624902 | 1 | 0.030173969 | 0.171642121 | 1 |
| 20915:807 | 23:04 | 19093:913 | 25:03:00 | 1945:70 | 2:01 | 1 | 1 | 1 | 0.000415451 | 6.22E-05 | 1 |
| 20892:812 | 28:02:00 | 19070:907 | 23:05 | 1942:70 | 2:00 | 1 | 0.029932329 | 1 | 4.42E-06 | 0.001086938 | 1 |
| 20852:798 | 26:03:00 | 19007:901 | 20:01 | 1943:67 | 0:00 | 1 | 0.029932329 | 1 | 3.68E-05 | 6.49E-05 | 1 |
| 20862:801 | 16:00 | 19005:899 | 21:03 | 1943:67 | 0:00 | 1 | 1 | 1 | 0.000112646 | 0.000420381 | 1 |
| 20697:845 | 26:02:00 | 18848:913 | 18:02 | 1935:70 | 2:01 | 1 | 0.017552603 | 1 | 1.17E-05 | 0.000589794 | 1 |
| 20446:671 | 32:03:00 | 18584:821 | 30:05:00 | 1927:64 | 4:02 | 1 | 1 | 1 | 2.14E-06 | 4.71E-05 | 0.425087994 |
| 20407:643 | 29:07:00 | 18551:807 | 31:10:00 | 1930:67 | 1:01 | 1 | 0.489401305 | 1 | 0.000400643 | 0.001633008 | 1 |
| 20247:583 | 52:07:00 | 18442:771 | 36:08:00 | 1925:61 | 3:01 | 1 | 0.003751524 | 1 | 1.29E-08 | 4.79E-05 | 1 |
| 19856:604 | 61:25:00 | 18052:825 | 34:15:00 | 1892:62 | 2:03 | 1 | 0.118667291 | 1 | 0.000180475 | 0.009407211 | 0.661487987 |
| 19740:630 | 53:01:00 | 17931:832 | 32:03:00 | 1884:66 | 0:00 | 1 | 0.003751524 | 1 | 4.88E-12 | 2.14E-06 | 1 |
| 19616:623 | 49:04:00 | 17813:831 | 36:05:00 | 1872:66 | 3:01 | 1 | 0.002334609 | 0.017871477 | 1.82E-09 | 2.85E-06 | 1 |
| 19183:752 | 19:03 | 17376:826 | 21:04 | 1842:77 | 1:00 | 1 | 1 | 1 | 0.001006164 | 0.001091486 | 1 |
| 18643:740 | 45:06:00 | 16856:744 | 36:03:00 | 1812:78 | 4:00 | 1 | 0.001383821 | 1 | 1.18E-07 | 3.12E-07 | 1 |
| 18623:742 | 35:06:00 | 16834:746 | 34:02:00 | 1809:79 | 4:00 | 1 | 0.014358417 | 1 | 1.18E-05 | 2.44E-07 | 1 |
| 17924:781 | 103:03:00 | 16318:729 | 60:10:00 | 1788:73 | 4:01 | 1 | 0.000285302 | 1 | 1.25E-21 | 6.76E-09 | 0.191464811 |
| 17876:779 | 82:04:00 | 16287:727 | 51:10:00 | 1787:73 | 6:00 | 1 | 0.000470572 | 1 | 1.58E-16 | 3.69E-07 | 0.018161034 |
| 17833:785 | 58:05:00 | 16228:732 | 45:05:00 | 1781:72 | 8:00 | 1 | 0.000197866 | 1 | 7.47E-11 | 4.24E-08 | 0.006579116 |

| | | | | | | | | | | | |
|---|---|---|---|---|---|---|---|---|---|---|---|
| 17450:812 | 64:03:00 | 15939:770 | 50:03:00 | 1765:69 | 6:00 | 1 | 0.001235444 | 1 | 3.24E-13 | 3.52E-10 | 0.018161034 |
| 17242:835 | 73:04:00 | 15772:795 | 71:03:00 | 1752:70 | 4:00 | 1 | 0.00012042 | 1 | 1.39E-14 | 1.16E-14 | 1 |
| 17244:841 | 76:05:00 | 15778:805 | 73:03:00 | 1753:71 | 3:00 | 1 | 0.000358978 | 1 | 1.14E-14 | 4.30E-15 | 1 |
| 17195:841 | 77:05:00 | 15731:820 | 81:01:00 | 1750:72 | 6:00 | 1 | 0.00882314 | 1 | 7.00E-15 | 4.91E-18 | 0.018161034 |
| 16899:842 | 76:02:00 | 15499:851 | 90:03:00 | 1738:73 | 8:00 | 1 | 0.002334609 | 1 | 2.06E-16 | 9.36E-19 | 0.006579116 |
| 16872:838 | 79:02:00 | 15490:852 | 79:02:00 | 1735:73 | 10:00 | 1 | 0.000571097 | 1 | 4.67E-17 | 5.51E-17 | 0.002529224 |
| 16544:890 | 19:02 | 15156:872 | 29:01:00 | 1726:80 | 1:00 | 1 | 0.267669829 | 1 | 0.00034049 | 7.53E-07 | 1 |
| 16412:884 | 33:05:00 | 15040:866 | 33:01:00 | 1717:80 | 7:00 | 1 | 0.06690288 | 1 | 1.14E-05 | 1.07E-07 | 0.010989758 |
| 16396:884 | 32:05:00 | 15009:866 | 40:02:00 | 1717:80 | 6:00 | 1 | 0.066197787 | 1 | 1.77E-05 | 1.30E-08 | 0.018161034 |
| 14557:863 | 51:02:00 | 13510:866 | 40:04:00 | 1645:78 | 7:00 | 1 | 1 | 1 | 5.23E-11 | 1.48E-07 | 0.010989758 |
| 14500:860 | 60:03:00 | 13469:864 | 48:04:00 | 1640:78 | 8:00 | 1 | 0.010484149 | 1 | 2.31E-12 | 3.18E-09 | 0.006579116 |
| 14286:867 | 15:03 | 13279:869 | 22:01 | 1636:77 | 1:00 | 1 | 0.029932329 | 1 | 0.006305042 | 2.43E-05 | 1 |
| 13984:855 | 34:03:00 | 13029:860 | 30:04:00 | 1632:76 | 1:00 | 1 | 1 | 1 | 8.17E-07 | 1.72E-05 | 1 |
| 13965:856 | 17:01 | 13031:860 | 19:03 | 1631:76 | 2:00 | 1 | 1 | 1 | 0.000271356 | 0.001051209 | 1 |
| 13204:814 | 8:01 | 12391:830 | 4:02 | 1585:72 | 2:00 | 1 | 1 | 1 | 0.024013973 | 0.428784618 | 1 |
| 13513:841 | 15:02 | 12656:851 | 14:01 | 1617:74 | 3:00 | 1 | 1 | 1 | 0.002386856 | 0.001271219 | 1 |
| 13485:840 | 20:02 | 12640:851 | 13:01 | 1620:73 | 0:00 | 1 | 1 | 1 | 0.000209945 | 0.002079713 | 1 |
| 12956:831 | 45:02:00 | 12247:855 | 31:04:00 | 1607:73 | 1:00 | 1 | 0.010484149 | 1 | 1.03E-09 | 1.06E-05 | 1 |
| 12817:824 | 40:02:00 | 12148:850 | 22:03 | 1601:73 | 3:00 | 1 | 1 | 1 | 1.25E-08 | 0.000261317 | 1 |
| 12757:821 | 24:00:00 | 12094:840 | 19:07 | 1602:73 | 1:00 | 1 | 1 | 1 | 2.17E-06 | 0.023805629 | 1 |
| 12225:811 | 10:03 | 11626:828 | 19:02 | 1591:73 | 1:00 | 1 | 1 | 1 | 0.060380822 | 0.000368028 | 1 |
| 11992:801 | 30:05:00 | 11447:827 | 29:01:00 | 1584:73 | 3:00 | 1 | 1 | 1 | 4.44E-05 | 7.53E-07 | 1 |
| 11941:800 | 30:03:00 | 11404:823 | 24:01:00 | 1585:72 | 1:00 | 1 | 1 | 1 | 5.49E-06 | 9.02E-06 | 1 |
| 11895:799 | 23:00 | 11364:821 | 19:00 | 1583:72 | 3:01 | 1 | 1 | 1 | 3.50E-06 | 2.64E-05 | 1 |
| 11316:774 | 50:03:00 | 10806:804 | 38:02:00 | 1571:71 | 5:00 | 1 | 0.023584054 | 1 | 3.19E-10 | 3.47E-08 | 0.030032732 |
| 7955:593 | 55:04:00 | 7773:700 | 48:04:00 | 1474:63 | 6:00 | 1 | 4.44E-05 | 1 | 9.59E-11 | 3.18E-09 | 0.018161034 |
| 0:01:01 | 1:07:04 | 0:00:00 | 1:08:03 | 0:00:00 | 0:01:00 | 1 | 1 | 1 | 0.116361657 | 0.045996854 | 1 |
| 36:15:01 | 0:00:00 | 31:12:01 | 0:00:00 | 20:03:01 | 0:00:00 | 1 | 5.15E-06 | 1 | 1 | 1 | 1 |
| 1148:1649:27:137 | 0:0:0:0 | 672:2159:24:140 | 0:5:0:0 | 106:408:6:26 | 0:0:0:0 | 1 | 1 | 1 | 1 | 0.002732501 | 1 |
| 1145:1639:31:137 | 2:7:0:0 | 671:2158:32:139 | 1:3:0:0 | 106:408:5:26 | 0:0:1:0 | 1 | 1 | 1 | 0.003244975 | 1 | 1 |
| 29:964:2:4:49:0 | 0:2:0:0:0:0 | 4:3470:13:12:218:1 | 0:3:0:0:0:0 | 0:893:1:6:55:0 | 0:1:0:0:0:0 | 0 | 0 | 1 | 1 | 1 | 1 |
| 971:29:1:48:2:0:0 | 2:0:0:1:0:0:0 | 3471:4:11:217:1:0:0 | 3:0:0:0:0:0:0 | 890:0:1:55:1:0:0 | 1:0:0:0:0:0:0 | 6.72E-05 | 1 | 1 | 1 | 1 | 1 |
| 48:981:29:1:2:0:0:0 | 0:0:0:0:0:0:0:0 | 216:3487:4:11:1:0:0:1 | 0:5:0:0:0:0:0:0 | 55:894:0:1:1:0:0:0 | 0:1:0:0:0:0:0:0 | 2.23E-05 | 1 | 1 | 1 | 2.28E-05 | 1 |
| 981:29:1:48:2:0:0:0 | 0:0:0:0:0:0:0:0 | 3483:4:11:216:1:0:0:1 | 9:0:0:0:0:0:0:0 | 894:0:1:55:1:0:0:0 | 1:0:0:0:0:0:0:0 | 1 | 1 | 1 | 1 | 1.27E-10 | 1 |
| 981:28:1:48:2:0:0:0 | 2:0:0:0:0:0:0:0 | 3518:4:10:216:1:0:0:1 | 7:0:0:0:0:0:0:0 | 892:0:0:54:1:0:0:0 | 1:0:0:0:0:0:0:0 | 2.99E-16 | 1 | 1 | 1 | 6.12E-08 | 1 |
| 25:928:2:44:0:2:0:0 | 1:2:0:0:0:0:0:0 | 4:3258:13:205:0:1:0:1 | 0:6:0:0:0:0:0:0 | 0:853:1:52:0:1:0:0 | 0:0:0:0:0:0:0:0 | 1 | 1 | 1 | 1 | 1.20E-06 | 1 |
| 27:953:2:43:2:0:0:0 | 0:0:0:0:0:0:0:0 | 4:3410:13:207:1:0:0:1 | 0:5:0:1:0:0:0:0 | 0:890:1:51:1:0:0:0 | 0:0:0:0:0:0:0:0 | 1 | 1 | 1 | 1 | 0.000299023 | 1 |
| 955:27:2:41:2:0:0:0 | 2:0:0:0:0:0:0:0 | 3444:4:13:207:1:0:0:1 | 5:0:0:0:0:0:0:0 | 886:0:1:51:1:0:0:0 | 2:0:0:0:0:0:0:0 | 5.83E-08 | 1 | 1 | 1 | 2.28E-05 | 1 |

| | | | | | | | | | | | |
|---|---|---|---|---|---|---|---|---|---|---|---|
| 111:21:12:0:0:0:0:1:0 | 0:0:0:0:0:0:0:0:0 | 77:10:8:0:0:0:0:0:2 | 0:0:0:0:0:0:0:0:0 | 23:3:3:0:1:0:1:0:0 | 0:0:0:0:0:0:0:0:0 | 1.76E-08 | 1.43E-07 | 1 | 1 | 1 | 1 |
| 111:12:28:0:0:0:0:1:0 | 0:0:0:0:0:0:0:0:0 | 8:0:3:0:0:0:0:0:0 | 70:8:5:0:0:0:0:0:2 | 4:2:1:0:0:0:1:0:0 | 19:1:1:0:0:1:0:0:0 | 4.66E-20 | 8.27E-30 | 1 | 1 | 1.25E-89 | 8.19E-23 |
| 0:0:0:0:0:0:0:0:0:0:0 | 2:2:2:0:0:0:2:1:0:0:0 | 1:12:2:0:1:0:6:0:0:0:0 | 1:6:0:0:0:0:1:0:0:2:1 | 3:2:1:0:1:0:1:0:1:1:0 | 3:1:1:0:1:2:0:0:0:0:0 | 0.042679166 | 0.04358988 | 0.298112523 | 0.317003333 | 0.000679247 | 0.184807975 |
| 32693:119:325:9 | 35:0:0:0 | 22554:172:149:11 | 28:0:0:0 | 4166:12:33:1 | 4:0:0:0 | 1.24E-20 | 6.82E-10 | 1 | 6.19E-22 | 2.03E-17 | 1 |
| 32708:119:323:9 | 48:0:1:0 | 22600:172:150:11 | 26:0:0:0 | 4167:12:33:1 | 6:0:0:0 | 9.68E-19 | 8.02E-07 | 1 | 3.21E-29 | 3.75E-16 | 0.000744507 |
| 32723:119:323:9 | 47:0:0:0 | 22598:172:150:10 | 34:0:0:0 | 4171:12:33:1 | 5:0:0:0 | 3.63E-11 | 4.80E-08 | 0.003124028 | 1.31E-29 | 3.13E-21 | 0.002841239 |
| 32774:119:324:9 | 25:0:0:0 | 22649:172:150:9 | 23:0:0:0 | 4169:12:33:1 | 10:0:0:0 | 4.56E-08 | 1 | 1 | 1.39E-15 | 2.94E-14 | 3.17E-06 |
| 32779:120:130:9 | 25:0:1:0 | 22692:173:42:9 | 12:0:1:0 | 4184:12:18:0 | 2:0:0:1 | 1 | 1 | 1 | 1.36E-14 | 1.45E-06 | 1 |
| 32842:324:0 | 43:00:00 | 22736:150:2 | 31:00:00 | 4191:34:00 | 6:00:00 | 2.41E-15 | 0.001383821 | 1 | 9.06E-19 | 1.39E-13 | 0.00371338 |
| 32870:322:0 | 46:01:00 | 22747:150:2 | 36:00:00 | 4193:34:00 | 6:00:00 | 1.35E-08 | 1 | 1 | 3.19E-19 | 1.04E-15 | 0.00371338 |
| 32985:120:323:9 | 34:0:0:0 | 22877:174:150:11 | 18:0:0:0 | 4220:12:34:1 | 4:0:0:0 | 1.19E-05 | 0.002652154 | 0.003124028 | 2.67E-21 | 3.87E-11 | 1 |
| 32986:120:323:9 | 48:0:0:0 | 22880:174:150:11 | 23:0:0:0 | 4223:12:34:1 | 5:0:0:0 | 7.59E-07 | 1 | 1 | 2.99E-30 | 2.94E-14 | 0.002841239 |
| 33024:120:323:9 | 19:0:0:0 | 22905:174:150:11 | 13:0:0:0 | 4226:12:34:1 | 3:0:0:0 | 4.53E-05 | 1 | 1 | 8.34E-12 | 4.74E-08 | 1 |
| 33096:120:321:9 | 36:0:2:0 | 22983:174:150:11 | 19:0:0:0 | 4233:12:34:1 | 3:0:0:0 | 1.19E-05 | 1 | 1 | 1.25E-20 | 9.18E-12 | 1 |
| 33145:120:323:9 | 28:0:0:0 | 23025:174:150:11 | 16:0:0:0 | 4243:12:34:1 | 4:0:0:0 | 0.012449474 | 1 | 1 | 1.71E-17 | 6.90E-10 | 1 |
| 33134:120:323:9 | 27:0:0:0 | 23017:174:150:11 | 17:0:0:0 | 4246:12:34:1 | 0:0:0:0 | 3.02E-06 | 1 | 1 | 7.37E-17 | 1.63E-10 | 1 |
| 33062:119:322:9 | 30:1:0:0 | 22941:174:150:11 | 27:0:0:0 | 4226:12:34:1 | 7:0:0:0 | 6.31E-10 | 0.002652154 | 1 | 9.39E-18 | 8.68E-17 | 0.000193253 |
| 33004:118:322:9 | 38:1:0:0 | 22903:174:149:11 | 20:0:1:0 | 4233:12:33:1 | 2:0:1:0 | 4.56E-08 | 1 | 1 | 7.82E-23 | 2.00E-11 | 1 |
| 32913:119:322:9 | 108:0:0:0 | 22820:174:149:11 | 82:0:1:0 | 4222:12:34:1 | 12:0:0:0 | 3.63E-11 | 1.27E-05 | 5.77E-05 | 1.18E-68 | 4.80E-51 | 1.97E-07 |
| 32952:322:119:9 | 48:0:0:0 | 22865:149:174:11 | 22:0:0:0 | 4228:34:12:1 | 2:0:0:0 | 0.000173965 | 0.040765436 | 1 | 2.99E-30 | 1.26E-13 | 1 |
| 32949:119:322:9 | 41:0:0:0 | 22858:174:149:11 | 20:0:0:0 | 4233:12:34:1 | 1:0:0:0 | 5.55E-06 | 0.000195836 | 0.000223034 | 8.97E-26 | 2.23E-12 | 1 |
| 32970:118:322:9 | 10:0:0:0 | 17259:131:105:9 | 5612:43:44:2 | 2638:11:22:1 | 1595:1:12:0 | 1 | 1 | 1 | 3.04E-06 | 0 | 0 |
| 32942:118:322:9 | 34:0:0:0 | 22832:174:149:11 | 32:0:0:0 | 4231:12:34:1 | 3:0:0:0 | 1.53E-10 | 0.000704574 | 0.003124028 | 2.67E-21 | 5.89E-20 | 1 |
| 32930:117:322:9 | 32:0:0:0 | 22833:173:147:11 | 22:0:2:0 | 4225:12:34:1 | 4:0:0:0 | 1.84E-07 | 1.27E-05 | 1 | 5.01E-20 | 7.87E-12 | 1 |
| 32887:117:319:9 | 41:0:3:0 | 22781:173:148:11 | 29:0:0:0 | 4220:12:34:1 | 8:0:0:0 | 8.52E-12 | 4.80E-08 | 1 | 6.38E-23 | 4.74E-18 | 4.84E-05 |
| 32887:117:322:9 | 28:0:0:0 | 22785:173:148:11 | 15:0:0:0 | 4222:12:34:1 | 4:0:0:0 | 8.52E-12 | 4.88E-05 | 1 | 1.71E-17 | 2.85E-09 | 1 |
| 32863:116:322:9 | 42:1:0:0 | 22756:173:147:11 | 27:0:0:0 | 4218:12:34:1 | 5:0:0:0 | 2.26E-19 | 4.88E-05 | 1 | 2.17E-25 | 8.68E-17 | 0.002841239 |
| 32879:117:322:9 | 23:0:0:0 | 22772:173:147:11 | 7:0:0:0 | 4222:12:34:1 | 2:0:0:0 | 2.02E-12 | 0.000195836 | 1 | 2.55E-14 | 0.000193006 | 1 |
| 32788:117:322:9 | 41:0:0:0 | 22678:173:147:11 | 31:0:0:0 | 4210:12:34:1 | 8:0:0:0 | 4.42E-25 | 1.16E-08 | 0.003124028 | 8.97E-26 | 2.56E-19 | 4.84E-05 |
| 5:13:0:2:0:0:1:0:0:0:0:0:0:0 | 3:14:16:0:0:0:8:0:0:4:0:0:0:0 | 24:82:1:9:9:1:10:3:0:1:4:7:1:1 | 0:0:0:0:0:0:0:0:0:0:0:0:0:0 | 3:2:0:0:4:0:1:1:0:0:0:0:0:0 | 0:0:0:0:0:0:0:0:0:0:0:0:0:0 | 3.85E-10 | 1 | 1 | 1.97E-19 | 1 | 1 |
| 12:00:05 | 36:08:10 | 212:43:23 | 0:00:00 | 85:09:03 | 0:00:00 | 1 | 1 | 0.001636609 | 2.88E-06 | 1 | 1 |
| 0:00:00 | 12:02:00 | 0:00:00 | 11:02:00 | 0:00:00 | 8:01:00 | 0.233314524 | 0.035298293 | 0.004168205 | 0.000239779 | 0.00061549 | 0.002822994 |
| 0:0:0:0:0:0:1:0 | 0:1:5:1:0:0:8:1 | 0:1:6:0:0:0:5:0 | 0:0:0:0:0:0:0:0 | 1:0:2:0:0:0:2:0 | 0:0:0:0:0:0:0:0 | 1 | 1 | 1 | 0.000165976 | 1 | 1 |
| 4:37:4:1:4:0:0:0:1:3:1:0:0 | 2:2:1:0:0:0:1:0:0:0:0:0:0 | 5:60:13:2:18:2:4:16:7:4:1:3:1 | 6:2:4:0:0:0:0:0:0:0:0:0:0 | 0:25:0:1:0:0:0:0:0:0:0:0:0 | 0:0:0:0:0:0:0:0:0:0:0:0:0 | 1 | 1 | 1 | 0.222252266 | 5.18E-06 | 1 |
| 48:13:3:0 | 0:0:0:0 | 11:7:0:0 | 0:0:0:0 | 2:3:0:1 | 1:0:0:0 | 2.26E-28 | 4.82E-13 | 1 | 1 | 1 | 1 |
| 59:03:03 | 0:00:00 | 11:03:00 | 0:00:00 | 3:02:00 | 0:00:00 | 5.64E-21 | 1 | 1 | 1 | 1 | 1 |

| | | | | | | | | | | | |
|---|---|---|---|---|---|---|---|---|---|---|---|
| 8:01:00 | 0:00:00 | 0:00:00 | 12:02:00 | 0:01:00 | 1:01:00 | 1 | 1 | 1 | 1 | 0.000257537 | 1 |
| 0:00 | 11:00 | 27:00:00 | 0:00 | 1:00 | 0:00 | 1 | 1 | 1 | 0.00138895 | 1 | 1 |
| 0:0:0:0:0:0:0 | 0:0:0:0:0:0:0 | 0:0:0:0:0:0:0 | 0:0:0:0:0:0:0 | 0:0:0:0:0:0:1 | 0:0:0:0:0:0:0 | 1 | 1 | 0.007094277 | 1 | 1 | 1 |
| 276:00:09 | 58:00:00 | 44:04:00 | 3:01:00 | 15:00:01 | 0:00:00 | 1 | 0.001383821 | 1 | 3.25E-25 | 1 | 1 |
| 138:00:02 | 210:00:07 | 49:04:00 | 5:01:00 | 14:00:01 | 1:00:00 | 1 | 1 | 0.000292552 | 1.03E-84 | 0.036538834 | 1 |
| 403:00:08 | 41:00:00 | 59:05:01 | 6:01:00 | 16:00:01 | 3:00:00 | 4.85E-06 | 3.40E-05 | 1 | 6.45E-18 | 0.015787883 | 1 |
| 431:09:00 | 31:01:00 | 62:03:06 | 4:00:00 | 19:01:00 | 1:00:00 | 6.13E-07 | 0.000180834 | 1 | 7.95E-13 | 1 | 1 |
| 478:00:10 | 10:00:00 | 66:06:03 | 1:00:00 | 20:00:01 | 0:00:00 | 0.003376462 | 1 | 1 | 8.14E-05 | 1 | 1 |
| 952:59:08 | 6:00:01 | 678:62:2 | 9:00:01 | 80:17:00 | 2:00:00 | 1 | 1 | 1 | 0.014954762 | 0.001091486 | 1 |
| 1085:58:09 | 3:01:00 | 1032:62:3 | 7:00:00 | 87:17:00 | 0:00:00 | 1 | 1 | 1 | 1 | 0.001445472 | 1 |
| 1112:60:9 | 5:00:00 | 1074:62:3 | 5:00:00 | 91:17:00 | 0:00:00 | 1 | 1 | 1 | 0.008963405 | 0.009407211 | 1 |
| 1045:56:08 | 9:00:00 | 1047:59:03 | 6:00:00 | 89:17:00 | 0:00:00 | 1 | 1 | 1 | 0.000209242 | 0.003657671 | 1 |
| 454:18:01 | 228:16:02 | 505:18:00 | 264:15:02 | 28:07:00 | 1:00:00 | 1 | 1 | 1 | 2.99E-84 | 1.48E-100 | 1 |
| 430:18:00 | 5:00:00 | 479:15:00 | 5:00:00 | 27:07:00 | 0:00:00 | 1 | 1 | 1 | 0.008963405 | 0.009407211 | 1 |
| 321:13:00 | 105:04:00 | 356:10:00 | 115:03:00 | 24:04:00 | 2:02:00 | 1 | 0.000555359 | 1 | 3.56E-42 | 3.01E-47 | 1 |
| 339:14:00 | 87:03:00 | 400:13:00 | 70:01:00 | 8:01:00 | 18:05:00 | 1.40E-09 | 1.83E-11 | 4.81E-06 | 3.13E-35 | 1.78E-29 | 2.70E-05 |
| 82:05:00 | 22:02 | 84:04:00 | 13:00 | 5:01 | 2:00 | 1 | 0.267669829 | 1 | 8.04E-05 | 0.00053673 | 1 |
| 86:06:00 | 17:01 | 87:04:00 | 9:00 | 5:01 | 2:00 | 1 | 0.267669829 | 1 | 0.000271356 | 0.003951718 | 1 |
| 0:00 | 11:01 | 1:00 | 1:00 | 2:00 | 0:00 | 1 | 1 | 1 | 0.005338322 | 1 | 1 |
| 0:00 | 11:01 | 1:00 | 1:00 | 2:00 | 0:00 | 1 | 1 | 1 | 0.005338322 | 1 | 1 |
| 4:1:0:0:0 | 0:0:0:0:0 | 76:2:1:0:0 | 0:0:0:0:0 | 0:0:0:0:1 | 0:0:0:0:0 | 1.37E-14 | 1 | 1 | 1 | 1 | 1 |
| 4:1:0:0:0 | 0:0:0:0:0 | 76:2:1:0:0 | 0:0:0:0:0 | 0:0:0:0:1 | 0:0:0:0:0 | 5.24E-17 | 1 | 1 | 1 | 1 | 1 |
| 4:1:0:0:0 | 0:0:0:0:0 | 76:2:1:0:0 | 0:0:0:0:0 | 0:0:0:0:1 | 0:0:0:0:0 | 9.89E-22 | 1 | 1 | 1 | 1 | 1 |
| 4:1:0:0:0 | 0:0:0:0:0 | 76:2:1:0:0 | 0:0:0:0:0 | 0:0:0:0:1 | 0:0:0:0:0 | 2.07E-16 | 1 | 1 | 1 | 1 | 1 |
| 4:1:0:0:0 | 0:0:0:0:0 | 76:2:1:0:0 | 0:0:0:0:0 | 0:0:0:0:1 | 0:0:0:0:0 | 1.47E-25 | 1 | 1 | 1 | 1 | 1 |
| 4:1:0:0:0 | 0:0:0:0:0 | 76:2:1:0:0 | 0:0:0:0:0 | 0:0:0:0:1 | 0:0:0:0:0 | 1.75E-08 | 1 | 1 | 1 | 1 | 1 |
| 4:1:0:0:0 | 0:0:0:0:0 | 76:2:1:0:0 | 0:0:0:0:0 | 0:0:0:0:1 | 0:0:0:0:0 | 0.000965198 | 1 | 1 | 1 | 1 | 1 |
| 4:1:0:0:0 | 0:0:0:0:0 | 77:2:1:0:0 | 0:0:0:0:0 | 0:0:0:0:1 | 0:0:0:0:0 | 2.84E-05 | 1 | 1 | 1 | 1 | 1 |
| 4:1:0:0 | 0:0:0:0 | 77:2:1:0 | 0:0:0:0 | 0:0:0:1 | 0:0:0:0 | 8.94E-08 | 1 | 1 | 1 | 1 | 1 |
| 4:1:0:0 | 0:0:0:0 | 77:1:1:0 | 0:0:0:0 | 0:0:0:1 | 0:0:0:0 | 1.32E-09 | 1 | 1 | 1 | 1 | 1 |
| 4:1:0:0 | 0:0:0:0 | 78:0:0:0 | 0:0:0:0 | 0:0:0:1 | 0:0:0:0 | 0.002029226 | 1 | 1 | 1 | 1 | 1 |

| | | | | | | | | | | | |
|---|---|---|---|---|---|---|---|---|---|---|---|
| 4:1:0:0 | 0:0:0:0 | 77:0:0:0 | 1:0:0:0 | 0:0:0:1 | 0:0:0:0 | 1.26E-07 | 1 | 1 | 1 | 1 | 1 |
| 4:0:0:0 | 0:0:0:0 | 78:0:0:0 | 0:0:0:0 | 0:0:0:1 | 0:0:0:0 | 6.45E-11 | 1 | 1 | 1 | 1 | 1 |
| 4:0:0:0 | 0:0:0:0 | 78:0:0:0 | 0:0:0:0 | 0:0:0:1 | 0:0:0:0 | 4.91E-10 | 1 | 1 | 1 | 1 | 1 |
| 4:0:0:0 | 0:0:0:0 | 78:0:0:0 | 0:0:0:0 | 0:0:0:1 | 0:0:0:0 | 1.21E-07 | 1 | 1 | 1 | 1 | 1 |
| 4:0:0:0 | 0:0:0:0 | 77:0:0:0 | 1:0:0:0 | 0:0:0:1 | 0:0:0:0 | 1.01E-06 | 1 | 1 | 1 | 1 | 1 |
| 4:0:0:0 | 0:0:0:0 | 78:0:0:0 | 0:0:0:0 | 0:0:0:1 | 0:0:0:0 | 3.02E-06 | 1 | 1 | 1 | 1 | 1 |
| 94:55:30:20:10:3:6:1:0 | 63:10:4:6:0:1:2:0:0 | 99:95:44:17:4:3:0:0:1 | 34:5:5:8:0:0:0:1:0 | 58:9:3:8:1:2:0:0:0 | 3:0:0:0:0:0:0:0:0 | 1 | 2.61E-07 | 0.070151041 | 1.30E-68 | 2.64E-30 | 1 |
| 57:63:348:83:99:17:17:6:0 | 0:1:0:0:2:0:0:0:0 | 84:70:271:51:76:10:0:4:1 | 0:2:0:0:3:0:0:0:0 | 4:2:73:10:16:1:0:2:0 | 0:0:0:0:0:0:0:0:0 | 1 | 1 | 1 | 1 | 0.023626577 | 1 |
| 0:0:0:0:0:0:0:0:0 | 59:18:348:83:104:17:6:0:0 | 84:28:270:54:82:11:4:0:1 | 0:0:1:0:0:0:0:0:0 | 4:2:73:10:17:1:2:0:0 | 0:0:0:0:0:0:0:0:0 | 1 | 1 | 1.09E-25 | 6.67E-293 | 1 | 1 |
| 2649:372 | 8:01 | 1494:228 | 2:02 | 549:24:00 | 1:01 | 1 | 1 | 1 | 0.024013973 | 1 | 1 |
| 2688:140 | 9:02 | 1515:119 | 1:01 | 551:15:00 | 0:00 | 1 | 1 | 1 | 0.04093502 | 1 | 1 |
| 470:11:00 | 2228:105 | 378:19:00 | 1138:92 | 206:03:00 | 345:12:00 | 3.65E-72 | 5.81E-96 | 2.64E-44 | 0 | 9.63E-194 | 6.28E-68 |
| 468:11:00 | 2233:85 | 0:00 | 1518:102 | 0:00 | 551:15:00 | 5.90E-146 | 1.48E-118 | 4.59E-45 | 0 | 2.68E-269 | 1.04E-110 |
| 2643:31:00 | 8:01 | 1483:08:00 | 5:00 | 551:02:00 | 0:00 | 1 | 1 | 1 | 0.024013973 | 0.031296179 | 1 |
| 0:00 | 278:13:00 | 1:00 | 190:23:00 | 0:00 | 254:11:00 | 1 | 1 | 7.25E-37 | 2.13E-53 | 1.63E-29 | 3.56E-49 |
| 18:00:02 | 7:00:00 | 12:00:03 | 2:00:00 | 10:00:01 | 0:00:00 | 1 | 1 | 1 | 0.00138895 | 1 | 1 |
| 116:00:04 | 193:00:02 | 47:00:08 | 81:00:03 | 16:00:01 | 6:00:00 | 0.0086191 | 0.000555359 | 0.00065969 | 1.94E-81 | 1.31E-32 | 0.00371338 |
| 269:00:05 | 40:00:01 | 110:00:11 | 21:00:00 | 22:00:01 | 0:00:00 | 1 | 1 | 1 | 1.14E-16 | 2.35E-09 | 1 |
| 13171:8:104:13 | 398:0:3:0 | 8480:6:17:5 | 362:1:1:0 | 998:6:13:2 | 12:0:0:0 | 1 | 9.59E-08 | 9.93E-05 | 1.89E-253 | 3.63E-231 | 1.97E-07 |
| 13238:104:8:13 | 11600:102:63:1 | 8525:17:7:5 | 7163:21:22:5 | 1002:13:6:2 | 26:0:1:0 | 1.19E-05 | 1.74E-21 | 4.71E-11 | 0 | 0 | 4.32E-15 |
| 25306:211:177:14 | 23:0:0:0 | 16092:38:186:10 | 13:0:1:0 | 1026:14:14:2 | 3:0:0:0 | 1 | 1 | 0.003124028 | 2.55E-14 | 3.64E-07 | 1 |
| 25259:210:180:14 | 28:0:0:0 | 16064:38:189:9 | 22:0:0:1 | 1027:14:14:2 | 0:0:0:0 | 1 | 1 | 1 | 1.71E-17 | 1.16E-12 | 1 |
| 25274:209:180:14 | 13:0:0:0 | 16080:38:189:10 | 5:0:0:0 | 1024:14:14:2 | 3:0:0:0 | 1 | 1 | 1 | 4.58E-08 | 0.002732501 | 1 |
| 25174:207:179:13 | 17:1:0:0 | 15990:36:185:9 | 14:1:0:0 | 1017:14:14:2 | 0:0:0:0 | 1 | 1 | 1 | 1.31E-09 | 9.42E-08 | 1 |
| 25206:208:180:12 | 20:0:0:0 | 16019:37:187:9 | 11:0:1:0 | 1019:14:14:2 | 1:0:0:0 | 1 | 1 | 1 | 1.99E-12 | 5.69E-06 | 1 |
| 3:60:17:0 | 0:0:0:0 | 1:50:5:1 | 1:59:24:0 | 1:17:1:0 | 0:0:0:0 | 0.001315682 | 3.55E-05 | 1 | 1 | 8.89E-23 | 1 |
| 0:00 | 0:00 | 0:00 | 9:00 | 0:00 | 3:00 | 1 | 1 | 1 | 1 | 0.003951718 | 1 |
| 0:0:0:0:0:0:0 | 27:4:0:2:2:1:1:0 | 9:3:1:2:0:2:1:1 | 8:2:2:2:0:0:1:0 | 0:0:0:0:0:0:0:0 | 2:5:0:0:0:1:0:0 | 1 | 1 | 0.000105307 | 1.88E-23 | 0.000817231 | 0.003791228 |
| 4:4:5:1:1:1:4:1:0 | 0:0:0:0:0:0:0:0:0 | 3:4:8:0:1:0:1:1:0 | 2:2:6:0:0:0:1:0:1 | 0:0:1:0:0:0:0:0:0 | 5:2:2:0:0:1:2:0:0 | 1 | 1 | 1 | 1 | 0.005916294 | 0.041000792 |
| 3:12:2:1 | 0:1:1:0 | 4:12:6:0 | 2:6:1:0 | 0:2:0:0 | 0:1:0:0 | 1 | 1 | 1 | 1 | 0.032544232 | 1 |
| 3:5:2:1 | 0:8:1:0 | 6:18:8:0 | 0:0:0:0 | 0:3:0:0 | 0:0:0:0 | 1 | 1 | 1 | 0.000296079 | 1 | 1 |
| 138:95:26:13:11:3:1:0:0 | 42:25:0:0:0:0:0:0:0 | 104:79:43:11:11:6:0:2:2 | 28:29:0:0:0:0:0:0:0 | 9:11:19:8:8:3:0:1:0 | 4:6:0:0:0:0:0:0:0 | 0.000155958 | 0.001586916 | 3.59E-08 | 2.28E-49 | 5.65E-38 | 2.65E-05 |
| 27:13:11:190:132:3:1:0:0 | 0:0:0:8:1:0:0:0:0 | 43:11:11:143:125:6:0:2:2 | 0:0:0:2:4:0:0:0:0 | 19:8:8:14:17:3:0:1:0 | 0:0:0:0:0:0:0:0:0 | 1 | 1 | 1 | 7.83E-09 | 0.00341885 | 1 |
| 193:135:27:13:11:3:1:0:0 | 5:2:0:0:0:0:0:0:0 | 143:126:43:11:11:6:0:2:2 | 6:9:0:0:0:0:0:0:0 | 14:17:19:8:8:3:0:1:0 | 1:0:0:0:0:0:0:0:0 | 1 | 1 | 1 | 0.000310236 | 1.16E-08 | 1 |
| 27:13:11:204:140:3:1:0:0 | 0:0:0:3:2:0:0:0:0 | 42:11:11:152:143:6:0:2:2 | 0:0:0:2:2:0:0:0:0 | 19:8:8:15:17:3:0:1:0 | 0:0:0:0:0:0:0:0:0 | 1 | 1 | 1 | 0.022613964 | 1 | 1 |
| 0:00 | 0:00 | 0:00 | 9:01 | 0:00 | 0:00 | 1 | 1 | 1 | 1 | 0.015211392 | 1 |
| 0:0:0:0:0:1:0 | 0:0:0:0:1:6:0 | 1:0:0:1:0:0:0 | 0:0:0:0:0:0:0 | 0:0:0:0:0:0:1 | 0:0:0:0:0:0:0 | 1 | 1 | 1 | 7.10E-05 | 1 | 1 |
| 0:00:00 | 2:01:00 | 0:00:00 | 7:00:00 | 0:00:00 | 2:00:00 | 0.000404766 | 1 | 1 | 1 | 0.001445472 | 1 |

| | | | | | | | | | | | |
|---|---|---|---|---|---|---|---|---|---|---|---|
| 2:00 | 5:00 | 2:00 | 3:00 | 0:00 | 1:00 | 1 | 1 | 1 | 0.030173969 | 1 | 1 |
| 1:00:00 | 5:00:00 | 9:00:01 | 0:00:00 | 2:00:00 | 0:00:00 | 1 | 1 | 1 | 0.008963405 | 1 | 1 |
| 16:5:1:50:3:0:7:2 | 0:0:0:0:0:0:0:0 | 44:6:4:43:3:0:4:4 | 0:0:0:0:0:0:0:0 | 6:3:1:9:2:0:0:0 | 0:0:0:0:0:0:0:0 | 4.33E-10 | 0.000339323 | 1 | 1 | 1 | 1 |
| 2:2:9:13:2:1:0:1:0 | 0:0:0:0:0:0:0:0:0 | 10:3:1:14:3:0:2:0:0 | 2:4:3:7:0:1:0:0:1 | 0:0:0:1:0:0:0:0:0 | 2:2:2:6:2:0:0:0:0 | 1 | 1 | 1 | 1 | 0.006997882 | 0.016756739 |
| 117:39:2:6:3 | 13:1:0:1:0 | 130:38:0:8:3 | 5:0:0:0:0 | 9:2:0:1:0 | 0:0:0:0:1 | 1 | 1 | 1 | 4.44E-08 | 0.000830299 | 1 |

Supplementary Table S8: Differential expression of SNV isoforms compared with wild type counterparts

| chr | pos | ref | alt | cosmicID | annotation | gene | mutType | AAanno |
|---|---|---|---|---|---|---|---|---|
| chr1 | 51361249 | T | C | COSV65541248 | exonic | EPS15 | nonsynonymous SNV | EPS15:NM_001159969:exon12:c.A1524G:p.I508M,EPS15:NM_001981:exon24:c.A2466G:p.I822M, |
| chr1 | 92836226 | G | A | . | exonic | RPL5 | nonsynonymous SNV | RPL5:NM_000969:exon5:c.G361A:p.G121S, |
| chr1 | 92836244 | G | A | . | exonic | RPL5 | nonsynonymous SNV | RPL5:NM_000969:exon5:c.G379A:p.G127S, |
| chr1 | 92836299 | A | T | . | exonic | RPL5 | nonsynonymous SNV | RPL5:NM_000969:exon5:c.A434T:p.Y145F, |
| chr1 | 92840627 | T | A | . | exonic | RPL5 | nonsynonymous SNV | RPL5:NM_000969:exon7:c.T782A:p.V261D, |
| chr10 | 46010595 | C | T | . | exonic | NCOA4 | nonsynonymous SNV | NCOA4:NM_001145261:exon9:c.G1374A:p.M458I,NCOA4:NM_001145260:exon9:c.G1374A:p.M458I,NCOA4:NM_005437:exon8:c.G1326A:p.M442I,NCOA4:NM_001145263:exon8:c.G1326A:p.M442I,NCOA4:NM_001145262:exon8:c.G1326A:p.M442I, |
| chr10 | 46010599 | C | T | . | exonic | NCOA4 | nonsynonymous SNV | NCOA4:NM_001145261:exon9:c.G1370A:p.G457E,NCOA4:NM_001145260:exon9:c.G1370A:p.G457E,NCOA4:NM_005437:exon8:c.G1322A:p.G441E,NCOA4:NM_001145263:exon8:c.G1322A:p.G441E,NCOA4:NM_001145262:exon8:c.G1322A:p.G441E, |
| chr10 | 46010618 | C | T | . | exonic | NCOA4 | nonsynonymous SNV | NCOA4:NM_001145261:exon9:c.G1351A:p.E451K,NCOA4:NM_001145260:exon9:c.G1351A:p.E451K,NCOA4:NM_005437:exon8:c.G1303A:p.E435K,NCOA4:NM_001145263:exon8:c.G1303A:p.E435K,NCOA4:NM_001145262:exon8:c.G1303A:p.E435K, |
| chr10 | 74398526 | G | A | . | exonic | ADK | nonsynonymous SNV | ADK:NM_001202450:exon6:c.G502A:p.D168N,ADK:NM_006721:exon6:c.G502A:p.D168N,ADK:NM_001202449:exon5:c.G397A:p.D133N,ADK:NM_001123:exon6:c.G451A:p.D151N, |
| chr11 | 34888850 | C | T | COSV53505953 | exonic | APIP | nonsynonymous SNV | APIP:NM_015957:exon4:c.G227A:p.C76Y, |
| chr11 | 34895110 | C | T | COSV53506741 | exonic | APIP | nonsynonymous SNV | APIP:NM_015957:exon2:c.G58A:p.D20N, |
| chr11 | 34916266 | G | A | COSV53507442 | exonic | APIP | nonsynonymous SNV | APIP:NM_015957:exon1:c.C19T:p.R7W, |
| chr12 | 56712541 | C | T | . | exonic | NACA | nonsynonymous SNV | NACA:NM_001113203:exon11:c.G2775A:p.M925I,NACA:NM_001113202:exon8:c.G645A:p.M215I, |
| chr12 | 56712827 | C | T | . | exonic | NACA | nonsynonymous SNV | NACA:NM_001113203:exon10:c.G2722A:p.A908T,NACA:NM_001113202:exon7:c.G592A:p.A198T, |
| chr12 | 56712838 | T | C | . | exonic | NACA | nonsynonymous SNV | NACA:NM_001113203:exon10:c.A2711G:p.K904R,NACA:NM_001113202:exon7:c.A581G:p.K194R, |
| chr12 | 56712844 | C | T | . | exonic | NACA | nonsynonymous SNV | NACA:NM_001113203:exon10:c.G2705A:p.R902K,NACA:NM_001113202:exon7:c.G575A:p.R192K, |
| chr12 | 56713572 | C | T | . | exonic | NACA | nonsynonymous SNV | NACA:NM_001113203:exon8:c.G2476A:p.A826T,NACA:NM_001113202:exon5:c.G346A:p.A116T, |
| chr12 | 56714379 | C | T | . | exonic | NACA | nonsynonymous SNV | NACA:NM_001113203:exon7:c.G2347A:p.E783K,NACA:NM_001113202:exon4:c.G217A:p.E73K, |
| chr13 | 21510035 | G | T | . | exonic | MICU2 | nonsynonymous SNV | MICU2:NM_152726:exon8:c.C730A:p.Q244K, |
| chr13 | 48077198 | C | T | . | exonic | MED4 | nonsynonymous SNV | MED4:NM_001270629:exon7:c.G616A:p.E206K,MED4:NM_014166:exon7:c.G754A:p.E252K, |
| chr13 | 48086358 | A | G | . | exonic | MED4 | nonsynonymous SNV | MED4:NM_001270629:exon3:c.T149C:p.L50S,MED4:NM_014166:exon3:c.T287C:p.L96S, |
| chr14 | 73110909 | C | A | . | exonic | RBM25 | nonsynonymous SNV | RBM25:NM_021239:exon15:c.C1771A:p.Q591K, |
| chr14 | 77731069 | C | T | . | exonic | SNW1 | nonsynonymous SNV | SNW1:NM_012245:exon10:c.G952A:p.E318K, |
| chr14 | 96847336 | T | A | . | exonic | VRK1 | nonsynonymous SNV | VRK1:NM_003384:exon5:c.T366A:p.N122K, |
| chr15 | 44715585 | G | A | . | exonic | B2M | nonsynonymous SNV | B2M:NM_004048:exon2:c.G230A:p.S77N, |
| chr15 | 44715595 | G | A | . | exonic | B2M | stopgain | B2M:NM_004048:exon2:c.G240A:p.W80X, |
| chr15 | 72202620 | T | C | . | exonic | PKM | nonsynonymous SNV | PKM:NM_001206797:exon8:c.A919G:p.I307V,PKM:NM_001206798:exon9:c.A1096G:p.I366V,PKM:NM_002654:exon9:c.A1141G:p.I381V, |
| chr15 | 72209855 | C | A | COSV99069963 | exonic | PKM | nonsynonymous SNV | PKM:NM_001206799:exon5:c.G398T:p.G133V,PKM:NM_182471:exon5:c.G383T:p.G128V,PKM:NM_001316318:exon5:c.G488T:p.G163V,PKM:NM_182470:exon5:c.G383T:p.G128V,PKM:NM_001206796:exon6:c.G605T:p.G202V,PKM:NM_001206798:exon5:c.G338T:p.G113V,PKM:NM_002654:exon5:c.G383T:p.G128V, |
| chr15 | 72209855 | C | T | . | exonic | PKM | nonsynonymous SNV | PKM:NM_001206799:exon5:c.G398A:p.G133D,PKM:NM_001206798:exon5:c.G338A:p.G113D,PKM:NM_002654:exon5:c.G383A:p.G128D,PKM:NM_182471:exon5:c.G383A:p.G128D,PKM:NM_001316318:exon5:c.G488A:p.G163D,PKM:NM_182470:exon5:c.G383A:p.G128D,PKM:NM_001206796:exon6:c.G605A:p.G202D, |
| chr15 | 72209856 | C | A | . | exonic | PKM | nonsynonymous SNV | PKM:NM_001206799:exon5:c.G397T:p.G133C,PKM:NM_182471:exon5:c.G382T:p.G128C,PKM:NM_001316318:exon5:c.G487T:p.G163C,PKM:NM_182470:exon5:c.G382T:p.G128C,PKM:NM_001206796:exon6:c.G604T:p.G202C,PKM:NM_001206798:exon5:c.G337T:p.G113C,PKM:NM_002654:exon5:c.G382T:p.G128C, |

| | | | | | | | | |
|---|---|---|---|---|---|---|---|---|
| chr16 | 30068929 | T | C | . | exonic | ALDOA | nonsynonymous SNV | ALDOA:NM_001127617:exon5:c.T491C:p.I164T,ALDOA:NM_000034:exon10:c.T491C:p.I164T,ALDOA:NM_184043:exon5:c.T491C:p.I164T,ALDOA:NM_184041:exon5:c.T491C:p.I164T,ALDOA:NM_001243177:exon6:c.T653C:p.I218T, |
| chr16 | 31185078 | T | G | . | exonic | FUS | nonsynonymous SNV | FUS:NM_004960:exon6:c.T663G:p.S221R,FUS:NM_001170634:exon6:c.T660G:p.S220R,FUS:NM_001170937:exon6:c.T651G:p.S217R, |
| chr17 | 1345458 | C | T | COSV51985336 | exonic | YWHAE | nonsynonymous SNV | YWHAE:NM_006761:exon6:c.G757A:p.E253K, |
| chr17 | 1361101 | C | T | . | exonic | YWHAE | nonsynonymous SNV | YWHAE:NM_006761:exon4:c.G569A:p.R190H, |
| chr17 | 1361927 | C | T | . | exonic | YWHAE | nonsynonymous SNV | YWHAE:NM_006761:exon3:c.G346A:p.E116K, |
| chr17 | 1361947 | A | G | . | exonic | YWHAE | nonsynonymous SNV | YWHAE:NM_006761:exon3:c.T326C:p.I109T, |
| chr17 | 1364864 | G | T | . | exonic | YWHAE | nonsynonymous SNV | YWHAE:NM_006761:exon2:c.C259A:p.Q87K, |
| chr17 | 1364880 | C | T | . | exonic | YWHAE | nonsynonymous SNV | YWHAE:NM_006761:exon2:c.G243A:p.M81I, |
| chr17 | 1364924 | C | T | . | exonic | YWHAE | nonsynonymous SNV | YWHAE:NM_006761:exon2:c.G199A:p.E67K, |
| chr17 | 31993308 | G | A | COSV59500263 | exonic | SUZ12 | nonsynonymous SNV | SUZ12:NM_015355:exon11:c.G1268A:p.R423Q, |
| chr17 | 81510694 | A | G | . | exonic | ACTG1 | nonsynonymous SNV | ACTG1:NM_001199954:exon6:c.T1124C:p.F375S,ACTG1:NM_001614:exon6:c.T1124C:p.F375S, |
| chr17 | 81510703 | C | T | . | exonic | ACTG1 | nonsynonymous SNV | ACTG1:NM_001199954:exon6:c.G1115A:p.R372H,ACTG1:NM_001614:exon6:c.G1115A:p.R372H, |
| chr17 | 81868822 | A | C | . | exonic | ARHGDIA | nonsynonymous SNV | ARHGDIA:NM_001301242:exon7:c.T557G:p.M186R, |
| chr17 | 81869013 | T | A | . | exonic | ARHGDIA | nonsynonymous SNV | ARHGDIA:NM_001301241:exon6:c.A478T:p.T160S,ARHGDIA:NM_004309:exon6:c.A478T:p.T160S,ARHGDIA:NM_001301243:exon5:c.A613T:p.T205S,ARHGDIA:NM_001301240:exon6:c.A478T:p.T160S,ARHGDIA:NM_001185077:exon6:c.A478T:p.T160S, |
| chr2 | 197489109 | G | T | COSV61444464 | exonic | HSPD1 | nonsynonymous SNV | HSPD1:NM_002156:exon9:c.C1108A:p.R370S,HSPD1:NM_199440:exon9:c.C1108A:p.R370S, |
| chr2 | 197489115 | C | T | . | exonic | HSPD1 | nonsynonymous SNV | HSPD1:NM_002156:exon9:c.G1102A:p.E368K,HSPD1:NM_199440:exon9:c.G1102A:p.E368K, |
| chr2 | 197489132 | C | T | . | exonic | HSPD1 | nonsynonymous SNV | HSPD1:NM_002156:exon9:c.G1085A:p.G362D,HSPD1:NM_199440:exon9:c.G1085A:p.G362D, |
| chr2 | 197489145 | A | T | . | exonic | HSPD1 | nonsynonymous SNV | HSPD1:NM_002156:exon9:c.T1072A:p.L358I,HSPD1:NM_199440:exon9:c.T1072A:p.L358I, |
| chr2 | 197493385 | G | T | . | exonic | HSPD1 | nonsynonymous SNV | HSPD1:NM_002156:exon7:c.C808A:p.P270T,HSPD1:NM_199440:exon7:c.C808A:p.P270T, |
| chr2 | 197497242 | C | T | . | exonic | HSPD1 | nonsynonymous SNV | HSPD1:NM_002156:exon3:c.G325A:p.A109T,HSPD1:NM_199440:exon3:c.G325A:p.A109T, |
| chr2 | 197497287 | C | T | . | exonic | HSPD1 | nonsynonymous SNV | HSPD1:NM_002156:exon3:c.G280A:p.G94R,HSPD1:NM_199440:exon3:c.G280A:p.G94R, |
| chr2 | 197497334 | A | G | . | exonic | HSPD1 | nonsynonymous SNV | HSPD1:NM_002156:exon3:c.T233C:p.V78A,HSPD1:NM_199440:exon3:c.T233C:p.V78A, |
| chr2 | 197497392 | C | T | . | exonic | HSPD1 | nonsynonymous SNV | HSPD1:NM_002156:exon3:c.G175A:p.G59R,HSPD1:NM_199440:exon3:c.G175A:p.G59R, |
| chr2 | 197498823 | C | T | . | exonic | HSPD1 | nonsynonymous SNV | HSPD1:NM_002156:exon2:c.G26A:p.R9H,HSPD1:NM_199440:exon2:c.G26A:p.R9H, |
| chr2 | 213317301 | G | C | . | exonic | SPAG16 | nonsynonymous SNV | SPAG16:NM_024532:exon5:c.G481C:p.E161Q,SPAG16:NM_001025436:exon5:c.G481C:p.E161Q, |
| chr2 | 213317306 | T | A | . | exonic | SPAG16 | nonsynonymous SNV | SPAG16:NM_024532:exon5:c.T486A:p.N162K,SPAG16:NM_001025436:exon5:c.T486A:p.N162K, |
| chr2 | 47471001 | T | A | COSV51879859 | exonic | MSH2 | nonsynonymous SNV | MSH2:NM_000251:exon11:c.T1698A:p.N566K,MSH2:NM_001258281:exon12:c.T1500A:p.N500K, |
| chr2 | 47475249 | C | A | . | exonic | MSH2 | nonsynonymous SNV | MSH2:NM_000251:exon12:c.C1984A:p.Q662K,MSH2:NM_001258281:exon13:c.C1786A:p.Q596K, |
| chr3 | 139369498 | A | T | . | exonic | COPB2 | nonsynonymous SNV | COPB2:NM_004766:exon11:c.T1252A:p.F418I, |
| chr3 | 141974094 | C | T | . | exonic | TFDP2 | stopgain | TFDP2:NM_001178140:exon5:c.G533A:p.W178X,TFDP2:NM_001178139:exon8:c.G617A:p.W206X,TFDP2:NM_006286:exon6:c.G437A:p.W146X,TFDP2:NM_001178138:exon9:c.G437A:p.W146X,TFDP2:NM_001178142:exon3:c.G209A:p.W70X,TFDP2:NM_001178141:exon5:c.G326A:p.W109X, |
| chr5 | 150406277 | A | T | . | exonic | CD74 | nonsynonymous SNV | CD74:NM_004355:exon4:c.T423A:p.H141Q,CD74:NM_001025159:exon4:c.T423A:p.H141Q,CD74:NM_001025158:exon4:c.T423A:p.H141Q, |
| chr6 | 125999566 | G | A | . | exonic | TRMT11 | nonsynonymous SNV | TRMT11:NM_001031712:exon7:c.G632A:p.G211E, |
| chr6 | 129599245 | A | T | . | exonic | ARHGAP18 | nonsynonymous SNV | ARHGAP18:NM_033515:exon12:c.T1684A:p.Y562N, |

| | | | | | | | | |
|---|---|---|---|---|---|---|---|---|
| chr6 | 129599297 | A | T | . | exonic | ARHGAP18 | nonsynonymous SNV | ARHGAP18:NM_033515:exon12:c.T1632A:p.D544E, |
| chr6 | 129599299 | C | T | . | exonic | ARHGAP18 | nonsynonymous SNV | ARHGAP18:NM_033515:exon12:c.G1630A:p.D544N, |
| chr6 | 129599306 | A | T | . | exonic | ARHGAP18 | nonsynonymous SNV | ARHGAP18:NM_033515:exon12:c.T1623A:p.H541Q, |
| chr6 | 1624213 | C | T | . | exonic | GMDS | nonsynonymous SNV | GMDS:NM_001500:exon11:c.G1075A:p.V359M,GMDS:NM_001253846:exon11:c.G985A:p.V329M, |
| chr6 | 18249708 | A | T | . | exonic | DEK | nonsynonymous SNV | DEK:NM_003472:exon7:c.T705A:p.D235E,DEK:NM_001134709:exon6:c.T603A:p.D201E, |
| chr6 | 31269347 | T | C | COSV66117127 | exonic | HLA-C | nonsynonymous SNV | HLA-C:NM_002117:exon7:c.A1087G:p.T363A,HLA-C:NM_001243042:exon7:c.A1087G:p.T363A, |
| chr6 | 31269385 | C | G | COSV66115057 | exonic | HLA-C | nonsynonymous SNV | HLA-C:NM_002117:exon7:c.G1049C:p.C350S,HLA-C:NM_001243042:exon7:c.G1049C:p.C350S, |
| chr6 | 31269990 | T | C | COSV66110395 | exonic | HLA-C | nonsynonymous SNV | HLA-C:NM_002117:exon5:c.A991G:p.M331V,HLA-C:NM_001243042:exon5:c.A991G:p.M331V, |
| chr6 | 31269992 | G | A | COSV66110576 | exonic | HLA-C | nonsynonymous SNV | HLA-C:NM_002117:exon5:c.C989T:p.A330V,HLA-C:NM_001243042:exon5:c.C989T:p.A330V, |
| chr6 | 31269996 | T | C | COSV66110404 | exonic | HLA-C | nonsynonymous SNV | HLA-C:NM_002117:exon5:c.A985G:p.T329A,HLA-C:NM_001243042:exon5:c.A985G:p.T329A, |
| chr6 | 31269999 | C | T | COSV66112417 | exonic | HLA-C | nonsynonymous SNV | HLA-C:NM_002117:exon5:c.G982A:p.V328I,HLA-C:NM_001243042:exon5:c.G982A:p.V328I, |
| chr6 | 31270025 | A | G | COSV66111096 | exonic | HLA-C | nonsynonymous SNV | HLA-C:NM_002117:exon5:c.T956C:p.V319A,HLA-C:NM_001243042:exon5:c.T956C:p.V319A, |
| chr6 | 31270056 | T | C | COSV66114266 | exonic | HLA-C | nonsynonymous SNV | HLA-C:NM_002117:exon5:c.A925G:p.M309V,HLA-C:NM_001243042:exon5:c.A925G:p.M309V, |
| chr6 | 31270085 | T | C | COSV66111578 | exonic | HLA-C | nonsynonymous SNV | HLA-C:NM_002117:exon5:c.A896G:p.E299G,HLA-C:NM_001243042:exon5:c.A896G:p.E299G, |
| chr6 | 31270214 | G | T | COSV66110582 | exonic | HLA-C | nonsynonymous SNV | HLA-C:NM_002117:exon4:c.C891A:p.S297R,HLA-C:NM_001243042:exon4:c.C891A:p.S297R, |
| chr6 | 31270233 | T | G | COSV66110591 | exonic | HLA-C | nonsynonymous SNV | HLA-C:NM_002117:exon4:c.A872C:p.Q291P,HLA-C:NM_001243042:exon4:c.A872C:p.Q291P, |
| chr6 | 31270252 | T | C | COSV66110598 | exonic | HLA-C | nonsynonymous SNV | HLA-C:NM_002117:exon4:c.A853G:p.M285V,HLA-C:NM_001243042:exon4:c.A853G:p.M285V, |
| chr6 | 31270276 | G | C | COSV66110239 | exonic | HLA-C | nonsynonymous SNV | HLA-C:NM_002117:exon4:c.C829G:p.Q277E,HLA-C:NM_001243042:exon4:c.C829G:p.Q277E, |
| chr6 | 31270482 | G | T | COSV66109964 | exonic | HLA-C | nonsynonymous SNV | HLA-C:NM_002117:exon4:c.C623A:p.P208H,HLA-C:NM_001243042:exon4:c.C623A:p.P208H, |
| chr6 | 31271091 | C | T | COSV66110610 | exonic | HLA-C | nonsynonymous SNV | HLA-C:NM_002117:exon3:c.G601A:p.E201K,HLA-C:NM_001243042:exon3:c.G601A:p.E201K, |
| chr6 | 31271153 | A | C | COSV66109994 | exonic | HLA-C | nonsynonymous SNV | HLA-C:NM_002117:exon3:c.T539G:p.L180R,HLA-C:NM_001243042:exon3:c.T539G:p.L180R, |
| chr6 | 31271165 | G | T | COSV66110458 | exonic | HLA-C | nonsynonymous SNV | HLA-C:NM_002117:exon3:c.C527A:p.A176E,HLA-C:NM_001243042:exon3:c.C527A:p.A176E, |
| chr6 | 31271180 | A | C | COSV66110087 | exonic | HLA-C | nonsynonymous SNV | HLA-C:NM_002117:exon3:c.T512G:p.L171W,HLA-C:NM_001243042:exon3:c.T512G:p.L171W, |
| chr6 | 31271207 | G | T | COSV66110642 | exonic | HLA-C | nonsynonymous SNV | HLA-C:NM_002117:exon3:c.C485A:p.T162N,HLA-C:NM_001243042:exon3:c.C485A:p.T162N, |
| chr6 | 31271273 | G | A | COSV66110477 | exonic | HLA-C | nonsynonymous SNV | HLA-C:NM_002117:exon3:c.C419T:p.S140F,HLA-C:NM_001243042:exon3:c.C419T:p.S140F, |
| chr6 | 31271280 | C | T | COSV66110483 | exonic | HLA-C | nonsynonymous SNV | HLA-C:NM_002117:exon3:c.G412A:p.D138N,HLA-C:NM_001243042:exon3:c.G412A:p.D138N, |
| chr6 | 31271336 | A | C | . | exonic | HLA-C | nonsynonymous SNV | HLA-C:NM_002117:exon3:c.T356G:p.L119R,HLA-C:NM_001243042:exon3:c.T356G:p.L119R, |
| chr6 | 31271601 | T | G | COSV66110096 | exonic | HLA-C | nonsynonymous SNV | HLA-C:NM_002117:exon2:c.A341C:p.D114A,HLA-C:NM_001243042:exon2:c.A341C:p.D114A, |
| chr6 | 31271653 | C | T | COSV66110668 | exonic | HLA-C | nonsynonymous SNV | HLA-C:NM_002117:exon2:c.G289A:p.A97T,HLA-C:NM_001243042:exon2:c.G289A:p.A97T, |
| chr6 | 31271672 | C | G | COSV66110203 | exonic | HLA-C | nonsynonymous SNV | HLA-C:NM_002117:exon2:c.G270C:p.K90N,HLA-C:NM_001243042:exon2:c.G270C:p.K90N, |
| chr6 | 31271766 | C | T | COSV66117347 | exonic | HLA-C | nonsynonymous SNV | HLA-C:NM_002117:exon2:c.G176A:p.R59Q,HLA-C:NM_001243042:exon2:c.G176A:p.R59Q, |
| chr6 | 31271800 | A | C | COSV66110102 | exonic | HLA-C | nonsynonymous SNV | HLA-C:NM_002117:exon2:c.T142G:p.S48A,HLA-C:NM_001243042:exon2:c.T142G:p.S48A, |
| chr6 | 31271845 | C | A | COSV66119680 | exonic | HLA-C | nonsynonymous SNV | HLA-C:NM_002117:exon2:c.G97T:p.D33Y,HLA-C:NM_001243042:exon2:c.G97T:p.D33Y, |
| chr6 | 31272025 | C | G | COSV66117644 | exonic | HLA-C | nonsynonymous SNV | HLA-C:NM_002117:exon1:c.G47C:p.G16A,HLA-C:NM_001243042:exon1:c.G47C:p.G16A, |
| chr6 | 31272044 | G | T | COSV66117654 | exonic | HLA-C | nonsynonymous SNV | HLA-C:NM_002117:exon1:c.C28A:p.L10I,HLA-C:NM_001243042:exon1:c.C28A:p.L10I, |
| chr6 | 31272050 | C | T | COSV66117662 | exonic | HLA-C | nonsynonymous SNV | HLA-C:NM_002117:exon1:c.G22A:p.A8T,HLA-C:NM_001243042:exon1:c.G22A:p.A8T, |
| chr7 | 127844417 | G | A | . | exonic | SND1 | nonsynonymous SNV | SND1:NM_014390:exon12:c.G1336A:p.G446R, |
| chr7 | 150793587 | C | T | . | exonic | TMEM176B | nonsynonymous SNV | TMEM176B:NM_014020:exon4:c.G329A:p.G110E,TMEM176B:NM_001101312:exon4:c.G329A:p.G110E,TMEM176B:NM_001101311:exon4:c.G329A:p.G110E,TMEM176B:NM_001101314:exon3:c.G218A:p.G73E, |
| chr7 | 150796412 | G | C | . | exonic | TMEM176B | nonsynonymous SNV | TMEM176B:NM_014020:exon2:c.C158G:p.T53S,TMEM176B:NM_001101312:exon2:c.C158G:p.T53S,TMEM176B:NM_001101311:exon2:c.C158G:p.T53S,TMEM176B:NM_001101314:exon2:c.C158G:p.T53S, |
| chr7 | 150796518 | A | G | . | exonic | TMEM176B | nonsynonymous SNV | TMEM176B:NM_014020:exon2:c.T52C:p.S18P,TMEM176B:NM_001101312:exon2:c.T52C:p.S18P,TMEM176B:NM_001101311:exon2:c.T52C:p.S18P,TMEM176B:NM_001101314:exon2:c.T52C:p.S18P, |

| | | | | | | | | |
|---|---|---|---|---|---|---|---|---|
| chr7 | 27800028 | G | A | . | exonic | TAX1BP1 | nonsynonymous SNV | TAX1BP1:NM_006024:exon13:c.G1702A:p.E568K,TAX1BP1:NM_001206901:exon13:c.G1702A:p.E568K,TAX1BP1:NM_001206902:exon12:c.G1231A:p.E411K,TAX1BP1:NM_001079864:exon13:c.G1702A:p.E568K, |
| chr8 | 100704315 | C | T | . | exonic | PABPC1 | nonsynonymous SNV | PABPC1:NM_002568:exon14:c.G1894A:p.G632S, |
| chr8 | 100706713 | C | T | . | exonic | PABPC1 | nonsynonymous SNV | PABPC1:NM_002568:exon11:c.G1540A:p.A514T, |
| chr8 | 100706910 | C | T | COSV59397547 | exonic | PABPC1 | nonsynonymous SNV | PABPC1:NM_002568:exon10:c.G1424A:p.R475Q, |
| chr8 | 85338375 | C | T | . | exonic | CA1 | nonsynonymous SNV | CA1:NM_001164830:exon2:c.G112A:p.E38K,CA1:NM_001128830:exon4:c.G112A:p.E38K,CA1:NM_001128831:exon3:c.G112A:p.E38K,CA1:NM_001738:exon4:c.G112A:p.E38K,CA1:NM_001128829:exon4:c.G112A:p.E38K, |
| chr8 | 85338380 | G | T | . | exonic | CA1 | nonsynonymous SNV | CA1:NM_001164830:exon2:c.C107A:p.T36N,CA1:NM_001128830:exon4:c.C107A:p.T36N,CA1:NM_001128831:exon3:c.C107A:p.T36N,CA1:NM_001738:exon4:c.C107A:p.T36N,CA1:NM_001128829:exon4:c.C107A:p.T36N, |
| chr8 | 85338411 | C | T | COSV56279142 | exonic | CA1 | nonsynonymous SNV | CA1:NM_001164830:exon2:c.G76A:p.G26R,CA1:NM_001128830:exon4:c.G76A:p.G26R,CA1:NM_001128831:exon3:c.G76A:p.G26R,CA1:NM_001738:exon4:c.G76A:p.G26R,CA1:NM_001128829:exon4:c.G76A:p.G26R, |
| chr8 | 85338417 | C | T | . | exonic | CA1 | nonsynonymous SNV | CA1:NM_001164830:exon2:c.G70A:p.A24T,CA1:NM_001128830:exon4:c.G70A:p.A24T,CA1:NM_001128831:exon3:c.G70A:p.A24T,CA1:NM_001738:exon4:c.G70A:p.A24T,CA1:NM_001128829:exon4:c.G70A:p.A24T, |
| chr8 | 85338423 | G | A | . | exonic | CA1 | nonsynonymous SNV | CA1:NM_001164830:exon2:c.C64T:p.P22S,CA1:NM_001128830:exon4:c.C64T:p.P22S,CA1:NM_001128831:exon3:c.C64T:p.P22S,CA1:NM_001738:exon4:c.C64T:p.P22S,CA1:NM_001128829:exon4:c.C64T:p.P22S, |
| chr8 | 96231075 | C | T | . | exonic | UQCRB | nonsynonymous SNV | UQCRB:NM_001199975:exon5:c.G220A:p.E74K,UQCRB:NM_006294:exon4:c.G316A:p.E106K, |

| Isoform | 1T__WT | 1T__MUT | 1M__WT | 1M__MUT | 2T__WT | 2T__MUT |
|---|---|---|---|---|---|---|
| XM_017000618:XM_017000620:NM_001981:XM_017000617:XM_017000615:XM_017000619:XM_017000616:XM_005270618 | 2:32:9:0:0:2:0:0 | 2:15:6:0:0:1:0:0 | 1:28:12:0:1:0:0:0 | 1:17:5:1:0:0:0:0 | 0:1:0:0:0:0:0:0 | 3:78:4:0:0:1:0:0 |
| NM_000969:NR_146333 | 1755:00:00 | 1:00 | 4751:00:00 | 2:01 | 5083:00:00 | 3:00 |
| NM_000969:NR_146333 | 1753:02:00 | 1:00 | 4736:07:00 | 11:00 | 5084:05:00 | 2:00 |
| NM_000969:NR_146333 | 1753:09:00 | 0:00 | 4737:28:00 | 3:00 | 5075:12:00 | 0:01 |
| NM_000969:NR_146333 | 1559:57:00 | 123:02:00 | 4226:118 | 304:02:00 | 4526:114 | 377:01:00 |
| NM_001145263:NM_005437:NM_001145261:NM_001145260:NM_001145262 | 820:828:6:3:3 | 1:1:0:0:0 | 358:169:1:1:0 | 1:0:0:0:0 | 279:302:2:0:1 | 2:1:0:0:0 |
| NM_001145263:NM_005437:NM_001145261:NM_001145260:NM_001145262 | 818:826:6:3:3 | 11:1:0:0:0 | 359:169:1:1:0 | 1:0:0:0:0 | 279:302:2:0:1 | 5:1:0:0:0 |
| NM_001145263:NM_005437:NM_001145261:NM_001145260:NM_001145262 | 820:816:6:3:3 | 16:6:0:0:0 | 359:167:1:1:0 | 3:0:0:0:0 | 281:301:2:0:1 | 5:1:0:0:0 |
| XM_017015704:NM_006721:NM_001202450:XM_017015698:NM_001123:XM_017015702:XM_017015703:XM_017015706:XM_017015705:NM_001202449:XM_017015699 | 0:48:0:0:38:0:0:0:0:0:0 | 0:0:0:0:0:0:0:0:0:0:0 | 1:72:3:2:52:1:5:0:0:0:0 | 0:0:0:0:0:0:0:0:0:0:0 | 1:111:2:0:130:3:9:3:0:1:0 | 0:0:0:0:0:0:0:0:0:0:0 |
| NM_015957:XM_017017875:XM_011520154 | 100:04:00 | 183:07:05 | 36:03:00 | 102:01:01 | 197:08:03 | 0:00:00 |
| NM_015957:XM_017017875:XM_011520154 | 284:06:05 | 0:01:00 | 138:04:01 | 0:00:00 | 196:06:03 | 1:00:00 |
| NM_015957:XM_017017875:XM_011520154 | 100:04:00 | 183:04:05 | 35:03:00 | 102:01:01 | 197:07:03 | 0:00:00 |
| NM_001113202:NM_001320194:NM_005594:NM_001320193:NM_001113201:NM_001113203 | 1448:10:106:73:1:1 | 1:6:1:0:0:0 | 3031:16:193:112:3:0 | 4:1:0:0:0:0 | 3983:16:254:185:2:0 | 2:1:1:0:0:0 |
| NM_001113202:NM_005594:NM_001320193:NM_001320194:NM_001113201:NM_001113203 | 1443:107:73:15:1:1 | 6:0:0:3:0:0 | 3033:193:112:25:3:0 | 4:0:0:0:0:0 | 3984:254:185:20:2:0 | 3:1:0:1:0:0 |
| NM_001113202:NM_005594:NM_001320193:NM_001320194:NM_001113201:NM_001113203 | 1447:106:73:18:1:1 | 0:0:0:1:0:0 | 3031:193:112:27:3:0 | 2:0:0:0:0:0 | 3982:255:185:23:2:0 | 0:0:0:0:0:0 |
| NM_005594:NM_001113202:NM_001320194:NM_001320193:NM_001113201:NM_001113203 | 107:1447:15:73:1:1 | 0:2:4:0:0:0 | 192:3035:26:112:3:0 | 1:2:1:0:0:0 | 255:3980:24:185:2:0 | 0:5:0:0:0:0 |
| NM_001113202:NM_005594:NM_001320193:NM_001320194:NM_001113201:NM_001113203 | 1449:106:73:27:1:1 | 0:0:0:0:0:0 | 3033:189:112:61:3:0 | 4:0:0:0:0:0 | 3983:256:185:58:2:0 | 3:0:0:2:0:0 |
| NM_001113202:NM_001320193:NM_001320194:NM_005594:NM_001113201:NM_001113203 | 1447:73:31:105:1:1 | 2:0:0:1:0:0 | 3035:112:86:187:3:0 | 2:1:3:0:0:0 | 3978:186:84:253:2:0 | 8:0:3:0:0:0 |
| NM_152726:XR_429214:XR_001749501:XM_017020433 | 142:3:1:3 | 0:0:0:0 | 185:12:7:1 | 0:0:0:0 | 214:13:0:7 | 0:0:0:0 |
| NM_014166:NM_001270629:XM_017020549 | 171:66:8 | 7:02:00 | 569:177:17 | 12:04:00 | 438:157:11 | 8:03:00 |
| NM_014166:NM_001270629:XM_017020549 | 202:72:6 | 3:00:00 | 636:190:14 | 12:00:00 | 516:190:9 | 7:00:00 |
| XR_943501:XM_024449680:NM_021239:XM_011537044 | 4:0:4:1 | 10:0:0:0 | 0:3:2:1 | 5:0:0:0 | 3:0:0:0 | 1:0:1:0 |
| XM_005267413:NM_012245:NM_001318844 | 7:30:01 | 2:00:00 | 7:27:00 | 3:00:00 | 4:42:00 | 3:00:00 |
| NM_003384:XM_006720247:XR_001750539:XM_017021624:XM_017021625:XM_017021626 | 48:0:1:2:0:0 | 0:0:0:0:0:0 | 24:1:0:0:0:0 | 0:0:0:0:0:0 | 82:2:1:4:1:0 | 0:0:0:0:0:0 |
| XR_002957658:NM_004048:XM_005254549 | 1287:140956:132 | 0.756944444 | 132:13357:24 | 5:11:00 | 155:18194:17 | 2:24:00 |
| NM_004048:XR_002957658:XM_005254549 | 141172:1261:132 | 224:17:00 | 13378:130:24 | 10:05:00 | 18212:149:17 | 19:03:00 |
| NM_002654:XM_011521670:XM_005254445:XR_001751314:NM_001206797:NM_182471:XM_006720570:NM_001206798 | 177:16:0:1:0:0:0:0 | 1:0:0:0:0:0:0:0 | 251:32:1:5:0:0:0:0 | 2:0:0:0:0:0:0:0 | 695:79:3:4:2:1:1:0 | 0:0:0:1:0:0:0:0 |
| NM_002654:XR_001751314:NM_182471:NM_001206797:XM_011521670:XM_005254445:XM_017022313:NM_182470:XM_006720570:NM_001206798:NM_001206799 | 149:34:8:0:5:0:0:0:0:0:0 | 0:0:0:0:0:0:0:0:0:0:0 | 214:60:21:1:6:1:0:0:0:0:0 | 0:0:0:0:0:0:0:0:0:0:0 | 607:103:26:1:30:3:4:1:1:0:0 | 0:0:0:1:0:0:0:0:0:0:0 |
| NM_002654:XR_001751314:NM_182471:NM_001206797:XM_011521670:XM_005254445:XM_017022313:NM_182470:XM_006720570:NM_001206798:NM_001206799 | 149:34:8:0:5:0:0:0:0:0:0 | 0:0:0:0:0:0:0:0:0:0:0 | 214:60:21:1:6:1:0:0:0:0:0 | 0:0:0:0:0:0:0:0:0:0:0 | 607:103:26:1:30:3:4:1:1:0:0 | 0:0:0:1:0:0:0:0:0:0:0 |
| NM_002654:XR_001751314:NM_182471:NM_001206797:XM_011521670:XM_005254445:XM_017022313:NM_182470:XM_006720570:NM_001206798:NM_001206799 | 149:34:8:0:5:0:0:0:0:0:0 | 0:0:0:0:0:0:0:0:0:0:0 | 213:60:21:1:6:1:0:0:0:0:0 | 1:0:0:0:0:0:0:0:0:0:0 | 607:103:26:1:30:3:4:1:1:0:0 | 0:0:0:1:0:0:0:0:0:0:0 |

| | | | | | | |
|---|---|---|---|---|---|---|
| NM_184041:NM_184043:NM_001127617:NM_001243177 | 3346:87:3:6 | 2:4:0:0 | 1752:68:3:3 | 1:0:0:0 | 2357:70:6:6 | 1:1:0:0 |
| NM_001170634:NM_004960:XM_024450221:NR_028388:XM_005255233:XM_011545781:XM_011545782:NM_001170937 | 25:60:2:1:0:0:0:0 | 0:0:0:1:0:0:0:0 | 12:29:3:7:1:0:0:0 | 2:0:0:0:0:0:0:0 | 24:55:3:1:1:0:0:0 | 2:0:0:0:0:0:0:0 |
| NR_024058:NM_006761:XM_017025005:XM_005256784 | 573:6002:25:0 | 3:30:2:0 | 162:1720:5:0 | 1:4:0:0 | 202:2375:6:1 | 1:6:1:0 |
| NM_006761:NR_024058:XM_017025005:XM_005256784 | 6034:544:20:19 | 14:1:0:0 | 1727:151:3:1 | 1:0:0:0 | 2384:196:6:4 | 3:1:0:0 |
| NM_006761:NR_024058:XM_017025005:XM_005256784 | 6054:491:32:65 | 11:1:0:1 | 1729:136:8:8 | 2:0:0:0 | 2386:177:6:7 | 2:0:0:0 |
| NM_006761:NR_024058:XM_005256784:XM_017025005 | 6059:499:66:31 | 6:0:0:0 | 1731:139:8:7 | 0:0:0:0 | 2388:177:7:6 | 0:0:0:0 |
| NM_006761:NR_024058:XM_005256784:XM_017025005 | 6064:499:77:31 | 3:0:3:0 | 1731:143:11:7 | 0:0:0:0 | 2387:174:10:6 | 0:1:3:0 |
| NM_006761:XM_005256784:NR_024058:XM_017025005 | 6056:81:495:31 | 12:3:3:0 | 1730:11:143:7 | 1:3:0:0 | 2385:13:175:5 | 4:1:0:0 |
| NM_006761:XM_017025005:XM_005256784:NR_024058 | 6088:46:238:489 | 12:4:14:3 | 1744:9:51:140 | 6:2:3:0 | 2429:8:79:172 | 3:0:2:0 |
| NM_015355:NM_001321207:XR_001752462:XM_006721794:XM_017024409 | 15:1:6:7:2 | 1:1:0:1:0 | 13:2:4:1:0 | 1:0:0:0:0 | 12:2:4:5:1 | 1:0:0:0:0 |
| NM_001614:NR_037688:NM_001199954 | 10652:168:56 | 22:05:00 | 7765:116:31 | 10:00:00 | 9309:154:65 | 11:01:00 |
| NM_001614:NR_037688:NM_001199954 | 10630:168:56 | 26:03:00 | 7749:116:31 | 12:00:00 | 9289:155:65 | 16:00:00 |
| NM_004309:XM_011523574:NR_125441:NM_001185078:NM_001185077:NM_001301242:NM_001301243:NM_001301241 | 95:4:4:1:1:1:0:0 | 0:2:1:0:0:0:0:0 | 47:2:0:0:0:0:0:0 | 0:0:0:0:0:0:0:0 | 52:1:4:0:0:0:1:0 | 0:2:1:0:0:0:0:0 |
| NM_004309:XM_011523574:NR_125441:NM_001301243:NM_001185077:NM_001301240:NM_001301241 | 104:2:4:0:1:3:0 | 0:1:1:0:0:0:0 | 53:1:0:0:0:0:0 | 0:0:0:0:0:0:0 | 62:0:1:1:0:1:0 | 0:0:0:0:0:0:0 |
| NM_002156:NM_199440 | 986:124 | 0:00 | 1943:121 | 0:00 | 2312:199 | 10:01 |
| NM_002156:NM_199440 | 986:124 | 2:00 | 1940:121 | 4:00 | 2317:198 | 6:02 |
| NM_002156:NM_199440 | 985:122 | 25:01:00 | 1944:120 | 52:03:00 | 2318:198 | 41:03:00 |
| NM_002156:NM_199440 | 1005:124 | 7:00 | 1985:123 | 11:01 | 2347:196 | 12:02 |
| NM_199440:NM_002156 | 109:1017 | 0:01 | 111:2002 | 1:00 | 175:2368 | 0:01 |
| NM_002156:NM_199440 | 1018:59:00 | 2:00 | 2002:62 | 3:01 | 2369:90 | 3:00 |
| NM_002156:NM_199440 | 1019:57:00 | 1:00 | 2005:59:00 | 3:01 | 2371:83 | 1:00 |
| NM_002156:NM_199440 | 1020:51:00 | 0:00 | 2009:58:00 | 1:00 | 2373:79 | 1:00 |
| NM_002156:NM_199440 | 1018:46:00 | 2:00 | 2010:48:00 | 0:00 | 2373:68 | 1:00 |
| NM_002156:NM_199440 | 1019:23:00 | 1:01 | 2009:18:00 | 1:02 | 2373:25:00 | 2:04 |
| NM_001025436:XM_017004899:NM_024532:XM_011511837:XM_017004896:XM_011511814 | 45:3:1:0:0:0 | 0:0:0:0:0:0 | 9:2:0:0:0:0 | 0:0:0:0:0:0 | 58:1:1:0:0:0 | 0:0:0:0:0:0 |
| NM_001025436:XM_017004899:NM_024532:XM_011511837:XM_017004896:XM_011511814 | 45:2:1:0:0:0 | 0:1:0:0:0:0 | 9:2:0:0:0:0 | 0:0:0:0:0:0 | 58:1:1:0:0:0 | 0:0:0:0:0:0 |
| NM_000251:XR_939685:NM_001258281:XM_011532867:XM_005264332:XR_001738747 | 48:5:14:0:0:0 | 1:1:0:0:0:0 | 45:3:6:0:0:0 | 4:0:0:0:0:0 | 91:5:36:0:1:0 | 2:4:0:0:0:0 |
| NM_000251:XR_939685:NM_001258281:XM_011532867:XM_005264332:XR_001738747 | 47:4:15:0:0:0 | 0:0:0:0:0:0 | 45:3:6:0:0:0 | 0:0:0:0:0:0 | 91:5:37:0:1:0 | 0:0:0:0:0:0 |
| NM_004766:XM_024453831:NR_023350 | 94:02:18 | 5:00:01 | 90:09:24 | 12:00:00 | 97:06:33 | 7:00:00 |
| XM_017007097:XM_011513107:XM_017007100:XM_011513106:XM_005247731:NM_001178138:NM_001178139:NM_006286:XM_011513111:XM_024453729:XM_017007091:XM_024453730:XM_024453726:XM_011513109:XM_024453728:XM_017007098:XM_024453725:XM_011513102:XM_011513101:NM_001178141:XM_017007102:XM_024453727 | 0:0:0:0:0:0:3:1:0:1:0:1:0:0:0:0:0:0:0:0:0:0 | 1:2:0:0:0:1:0:0:0:0:0:0:0:0:0:0:0:0:0:0:0:0 | 1:0:1:0:1:1:2:6:1:1:0:0:0:0:0:0:0:0:0:0:0:0 | 1:1:1:1:0:0:0:0:0:0:0:0:0:0:0:0:0:0:0:0:0:0 | 2:0:0:1:2:0:9:7:0:2:1:0:0:0:0:0:0:0:0:0:0:0 | 1:0:4:0:0:0:1:0:0:1:1:0:0:0:0:1:0:0:0:0:0:0 |
| NM_004355:XR_002956198:NM_001025159:NM_001025158:XM_017010089:XM_017010090 | 7240:65:986:7:8:64 | 8:0:0:0:0:0 | 5939:92:1233:4:12:40 | 4:1:3:1:0:0 | 1123:6:101:1:0:3 | 0:0:0:0:0:0 |
| XR_002956294:NM_001031712:NM_001350582:NR_146799:NM_001350580:NR_146802:NM_001350581:NR_146796:NR_146798:NM_001350583:NR_146800:NM_001350586:NR_146801:NM_001350584:NM_001350588:XM_017011168:NR_146804:NM_001350589:NM_001350592:NM_001350591:NR_146795:NM_001350585:NM_001350596:NM_001350597:NR_146803:NM_001350594:XM_024446516 | 8:76:0:10:1:1:3:4:1:0:1:0:5:2:2:2:1:1:1:1:0:0:0:0:0:0:0 | 0:0:0:0:0:0:0:0:0:0:0:0:0:0:0:1:0:0:0:0:0:0:0:0:0:0:0 | 3:59:1:5:1:1:2:2:1:1:2:1:3:1:1:1:0:0:0:0:0:0:0:0:0:0:0 | 2:0:0:0:0:0:0:0:0:0:0:0:0:0:0:0:0:0:0:0:0:0:0:0:0:0:0 | 2:69:0:7:1:1:7:5:3:0:0:1:0:0:0:3:0:0:2:0:0:1:1:0:0:0:0 | 1:0:0:1:0:0:0:0:0:0:0:0:0:0:0:0:0:0:0:0:0:0:0:0:0:0:0 |
| XM_005267213:XM_017011494:NM_033515:XM_005267212 | 1:0:8:1 | 8:0:0:0 | 3:0:25:4 | 5:1:0:0 | 1:0:16:4 | 4:0:0:0 |

| | | | | | | |
|---|---|---|---|---|---|---|
| XM_005267213:NM_033515:XM_005267212:XM_017011494 | 10:8:1:0 | 0:0:0:0 | 9:25:4:1 | 3:0:0:0 | 5:16:4:0 | 0:0:0:0 |
| XM_005267213:NM_033515:XM_005267212:XM_017011494 | 10:8:1:0 | 0:0:0:0 | 9:25:4:1 | 3:0:0:0 | 5:16:4:0 | 0:0:0:0 |
| XM_005267213:NM_033515:XM_005267212:XM_017011494 | 10:8:1:0 | 0:0:0:0 | 9:25:4:1 | 3:0:0:0 | 5:16:4:0 | 0:0:0:0 |
| NM_001500:XM_011514500:XR_001743349:XM_017010752:XM_011514507:XR_001743350 | 68:5:4:10:0:0 | 0:0:1:0:0:0 | 36:2:1:2:0:0 | 0:0:0:0:0:0 | 132:2:2:8:0:0 | 1:0:1:1:0:0 |
| XM_024446544:XR_001743635:NM_003472:NM_001134709:XM_017011273 | 11:4:43:0:15 | 1:2:0:0:0 | 13:2:71:1:20 | 1:4:1:0:0 | 54:6:110:1:45 | 4:8:2:0:0 |
| NM_002117:NM_001243042 | 0:26 | 214:04:00 | 0:07 | 61:03:00 | 0:00 | 91:01:00 |
| NM_002117:NM_001243042 | 1:25 | 213:04:00 | 0:07 | 60:03:00 | 0:00 | 91:01:00 |
| NM_002117:NM_001243042 | 0:25 | 206:04:00 | 0:07 | 60:03:00 | 0:00 | 91:01:00 |
| NM_002117:NM_001243042 | 0:25 | 206:04:00 | 0:07 | 60:03:00 | 0:00 | 91:01:00 |
| NM_002117:NM_001243042 | 0:25 | 206:04:00 | 0:07 | 60:03:00 | 0:00 | 91:01:00 |
| NM_002117:NM_001243042 | 0:25 | 206:04:00 | 0:07 | 60:03:00 | 6:01 | 85:00:00 |
| NM_002117:NM_001243042 | 0:25 | 203:04:00 | 0:07 | 60:03:00 | 0:00 | 90:01:00 |
| NM_002117:NM_001243042 | 0:25 | 202:04:00 | 0:07 | 60:03:00 | 0:00 | 90:01:00 |
| NM_002117:NM_001243042 | 1:24 | 199:04:00 | 0:07 | 60:03:00 | 91:01:00 | 0:00 |
| NM_002117:NM_001243042 | 0:23 | 199:04:00 | 0:07 | 60:03:00 | 0:00 | 93:01:00 |
| NM_002117:NM_001243042 | 0:23 | 198:04:00 | 0:07 | 60:03:00 | 0:00 | 93:01:00 |
| NM_002117:NM_001243042 | 0:23 | 197:04:00 | 0:07 | 60:03:00 | 0:00 | 91:01:00 |
| NM_002117:NM_001243042 | 0:23 | 195:04:00 | 0:07 | 60:03:00 | 0:00 | 91:01:00 |
| NM_002117:NM_001243042 | 3:26 | 178:04:00 | 1:07 | 54:03:00 | 1:01 | 83:01:00 |
| NM_002117:NM_001243042 | 2:27 | 179:04:00 | 1:07 | 55:03:00 | 82:01:00 | 0:00 |
| NM_002117:NM_001243042 | 2:26 | 176:04:00 | 1:07 | 53:03:00 | 6:01 | 73:00:00 |
| NM_002117:NM_001243042 | 0:20 | 177:10:00 | 0:07 | 54:03:00 | 0:00 | 79:01:00 |
| NM_002117:NM_001243042 | 0:20 | 177:10:00 | 0:07 | 54:03:00 | 0:00 | 79:01:00 |
| NM_002117:NM_001243042 | 2:26 | 173:04:00 | 1:08 | 53:03:00 | 79:01:00 | 0:00 |
| NM_002117:NM_001243042 | 0:20 | 171:10:00 | 0:07 | 54:04:00 | 71:00:00 | 6:01 |
| NM_002117:NM_001243042 | 0:20 | 170:10:00 | 0:07 | 53:04:00 | 78:01:00 | 0:00 |
| NM_001243042:NM_002117 | 24:167 | 6:00 | 10:51 | 1:00 | 0.095833333 | 0:00 |
| NM_002117:NM_001243042 | 3:20 | 168:04:00 | 0:06 | 52:03:00 | 0:00 | 80:00:00 |
| NM_002117:NM_001243042 | 0:19 | 177:02:00 | 1:06 | 52:03:00 | 75:00:00 | 6:00 |
| NM_001243042:NM_002117 | 0.123611111 | 21:00 | 2:54 | 6:00 | 0.05625 | 0:00 |
| NM_002117:NM_001243042 | 0:19 | 183:00:00 | 0:07 | 55:02:00 | 79:00:00 | 0:00 |
| NM_002117:NM_001243042 | 0:19 | 182:00:00 | 0:07 | 55:02:00 | 0:00 | 79:00:00 |
| NM_002117:NM_001243042 | 1:20 | 180:00:00 | 0:07 | 55:02:00 | 0:00 | 80:00:00 |
| NM_002117:NM_001243042 | 0:20 | 181:00:00 | 0:07 | 55:03:00 | 0:00 | 79:00:00 |
| NM_002117:NM_001243042 | 0:20 | 181:00:00 | 0:07 | 55:03:00 | 0:00 | 80:00:00 |
| NM_002117:NM_001243042 | 1:20 | 180:00:00 | 0:07 | 55:03:00 | 0:00 | 80:00:00 |
| XM_017011987:NM_014390 | 0.521527778 | 1:00 | 0.543055556 | 4:00 | 0.532638889 | 1:01 |
| NM_001101312:NM_014020:NM_001101311:XM_024446733:NM_001101314:XM_024446734:XM_006715933 | 3051:77:137:19:19:86:3 | 4:0:0:0:2:0:0 | 1107:17:89:10:12:43:2 | 2:0:0:0:0:0:0 | 927:16:93:15:25:26:7 | 1:0:1:0:1:0:0 |
| XM_006715933:NM_014020:NM_001101312:NM_001101311:NM_001101314:XM_024446733:XM_024446734 | 2:86:3078:138:37:19:82 | 0:0:1:0:1:0:1 | 3:29:1119:90:19:10:38 | 1:0:0:0:0:0:0 | 7:20:932:96:34:15:23 | 0:1:0:0:0:0:1 |
| NM_001101312:NM_014020:NM_001101311:NM_001101314:XM_024446733:XM_024446734:XM_006715933 | 3078:85:138:40:19:79:2 | 1:5:0:0:0:3:0 | 1115:33:90:21:10:23:4 | 4:0:0:0:0:0:0 | 931:24:96:34:15:21:7 | 1:0:0:0:0:0:0 |

| | | | | | | |
|---|---|---|---|---|---|---|
| NM_006024:NM_001079864:NM_001206902:NM_001206901 | 52:91:14:6 | 7:1:0:0 | 30:103:12:4 | 8:0:0:0 | 27:64:23:8 | 1:1:0:0 |
| NM_002568:XM_005250861 | 968:34:00 | 0:01 | 4164:129 | 4:02 | 1982:36:00 | 2:00 |
| NM_002568:XM_005250861 | 941:33:00 | 2:01 | 4070:116 | 5:03 | 1920:38:00 | 2:00 |
| NM_002568:XM_005250861 | 909:32:00 | 3:00 | 3967:116 | 10:04 | 1869:32:00 | 2:00 |
| NM_001164830:NM_001738:NM_001291967:NM_001128830 | 14366:327:7:1 | 28:12:0:0 | 0:0:0:0 | 0:0:0:0 | 10:0:0:0 | 0:0:0:0 |
| NM_001164830:NM_001738:NM_001291967:NM_001128830 | 14392:314:7:1 | 2:19:0:0 | 0:0:0:0 | 0:0:0:0 | 10:0:0:0 | 0:0:0:0 |
| NM_001164830:NM_001738:NM_001291967:NM_001128830 | 14374:263:7:1 | 20:16:0:0 | 0:0:0:0 | 0:0:0:0 | 10:0:0:0 | 0:0:0:0 |
| NM_001164830:NM_001738:NM_001291967:NM_001128830 | 14383:257:7:1 | 11:18:0:0 | 0:0:0:0 | 0:0:0:0 | 10:0:0:0 | 0:0:0:0 |
| NM_001164830:NM_001738:NM_001291967:NM_001128830 | 14389:249:7:1 | 5:15:0:0 | 0:0:0:0 | 0:0:0:0 | 10:0:0:0 | 0:0:0:0 |
| NM_006294:NM_001254752:NR_045639:NM_001199975 | 621:2:3:1 | 9:0:0:0 | 1170:10:5:2 | 35:0:0:0 | 1140:11:3:0 | 18:0:0:0 |

| 2M__WT | 2M__MUT | 3T__WT | 3T__MUT | 3M__WT | 3M__MUT | 1T__q | 1M__q | 2T__q | 2M__q | 3T__q | 3M__q |
|---|---|---|---|---|---|---|---|---|---|---|---|
| 0:1:5:1:0:0:0:0 | 0:119:4:0:0:2:0:0 | 1:81:7:1:1:2:0:1 | 0:0:0:0:0:0:0:0 | 0:50:3:0:1:1:1:0 | 0:0:0:0:0:0:0:0 | 1 | 1 | 1 | 0.00181137 | 1 | 1 |
| 26374:4 | 26:04:00 | 14518:0 | 16:04 | 3516:00:00 | 0:00 | 1 | 1 | 1 | 3.95E-07 | 1.09E-08 | 1 |
| 26364:43 | 30:03:00 | 14511:24 | 18:03 | 3511:03:00 | 3:00 | 1 | 1 | 1 | 0.10770649 | 0.03085654 | 1 |
| 26351:152 | 12:06 | 14512:135 | 3:03 | 3507:17:00 | 0:00 | 1 | 1 | 1 | 2.98E-06 | 0.0665619 | 1 |
| 23298:648 | 1775:09:00 | 13033:519 | 891:11:00 | 3178:61 | 198:01:00 | 1 | 1 | 1 | 7.25E-08 | 0.03632093 | 1 |
| 274:698:3:2:1 | 16:4:1:0:0 | 190:484:1:6:2 | 14:5:0:0:0 | 178:327:0:1:1 | 0:1:0:0:0 | 1 | 1 | 1 | 0.02381625 | 1 | 1 |
| 273:695:4:2:1 | 20:3:0:0:0 | 191:480:1:6:2 | 16:4:0:0:0 | 178:325:0:1:1 | 0:3:0:0:0 | 1 | 1 | 1 | 0.0010254 | 0.34691846 | 1 |
| 286:684:4:2:1 | 14:1:0:0:0 | 202:467:1:6:2 | 9:5:0:0:0 | 177:322:0:1:1 | 1:0:0:0:0 | 1 | 1 | 1 | 0.02881053 | 1 | 1 |
| 3:418:11:19:219:7:19:5:0:2:0 | 0:0:0:0:0:0:1:2:1:0:0 | 2:378:14:8:414:11:17:6:1:2:1 | 0:2:0:0:1:0:0:6:0:0:0 | 2:73:2:2:122:6:5:5:0:1:0 | 0:0:0:0:1:0:0:0:0:0:0 | 1 | 1 | 1 | 0.00795109 | 0.0003734 | 1 |
| 792:21:10 | 0:00:00 | 169:123:14 | 779:09:05 | 60:46:05 | 149:03:03 | 1 | 1 | 1 | 1 | 3.64E-68 | 1.49E-14 |
| 791:15:09 | 1:01:00 | 783:06:05 | 164:01:14 | 149:02:03 | 58:01:05 | 1 | 1 | 1 | 1 | 0.00132523 | 1 |
| 790:16:10 | 0:00:00 | 168:121:13 | 782:08:05 | 60:45:05 | 147:01:03 | 1 | 1 | 1 | 1 | 6.50E-68 | 1.79E-16 |
| 18048:90:777:931:7:2 | 15:3:2:0:0:0 | 8923:74:534:426:0:1 | 9:1:1:0:0:0 | 2744:18:152:109:0:0 | 5:2:0:0:0:0 | 4.67E-08 | 1 | 1 | 0.909711 | 1 | 1 |
| 18055:781:933:134:7:2 | 34:0:0:9:0:0 | 8934:535:427:97:0:1 | 16:1:1:4:0:0 | 2749:154:109:19:0:0 | 2:0:0:1:0:0 | 1 | 1 | 1 | 1.79E-06 | 1 | 1 |
| 18060:779:933:151:7:2 | 10:0:0:3:0:0 | 8931:534:426:102:0:1 | 4:1:0:4:0:0 | 2746:153:109:19:0:0 | 0:0:0:1:0:0 | 1 | 1 | 1 | 1 | 0.01540288 | 1 |
| 781:18076:152:933:7:2 | 0:11:8:0:0:0 | 535:8942:105:428:0:1 | 0:8:4:0:0:0 | 154:2750:21:109:0:0 | 0:1:1:0:0:0 | 0.00837057 | 1 | 1 | 2.70E-07 | 0.16878358 | 1 |
| 18064:782:932:323:7:2 | 17:2:3:5:0:0 | 8934:531:426:184:0:1 | 13:0:2:8:0:0 | 2748:152:109:39:0:0 | 3:0:0:0:0:0 | 1 | 1 | 1 | 0.26244724 | 7.70E-05 | 1 |
| 18054:935:384:781:7:2 | 26:1:22:3:0:0 | 8934:427:239:530:0:1 | 16:1:6:3:0:0 | 2747:109:46:152:0:0 | 4:0:0:0:0:0 | 1 | 1 | 1 | 1.43E-18 | 0.16421334 | 1 |
| 563:27:5:50 | 0:4:0:0 | 555:20:9:36 | 0:8:0:0 | 125:3:4:1 | 0:0:0:0 | 1 | 1 | 1 | 0.05270185 | 1.03E-07 | 1 |
| 1127:356:67 | 668:223:18 | 667:290:66 | 434:197:10 | 227:77:11 | 4:01:00 | 1 | 1 | 1 | 1 | 0.01849052 | 1 |
| 2106:625:54 | 92:01:00 | 1301:536:44 | 44:00:01 | 248:82:5 | 3:01:00 | 1 | 1 | 1 | 2.06E-05 | 0.00835274 | 1 |
| 6:7:1:3 | 19:0:2:0 | 11:15:6:3 | 44:0:4:0 | 5:3:4:0 | 7:0:1:1 | 1 | 1 | 1 | 0.39269103 | 1.79E-06 | 1 |
| 0.828518519 | 19:00:00 | 24:232:0 | 43:01:00 | 5:48:00 | 5:00:00 | 1 | 1 | 1 | 1.20E-10 | 7.13E-30 | 0.23125997 |
| 85:46:2:7:1:4 | 3:0:4:0:0:0 | 326:164:14:30:11:2 | 13:0:16:0:0:0 | 84:3:4:4:2:3 | 0:0:0:0:0:0 | 1 | 1 | 1 | 0.6657948 | 1.91E-11 | 1 |
| 213:43356:47 | 6:52:00 | 380:49729:82 | 4:56:00 | 120:19387:48 | 2:22:00 | 1 | 0.00505282 | 1 | 0.00575256 | 1 | 1 |
| 43383:207:47 | 45:08:00 | 49762:369:82 | 61:07:00 | 19402:119:49 | 22:02:00 | 1 | 0.00340962 | 1 | 2.05E-06 | 0.00686016 | 1 |
| 1054:349:3:15:10:3:1:0 | 7:0:0:3:0:0:0:0 | 2127:638:7:20:13:5:0:1 | 14:1:0:5:0:0:0:0 | 726:76:2:2:2:3:1:1 | 1:0:0:0:0:0:0:0 | 1 | 1 | 1 | 1 | 0.03556103 | 1 |
| 912:342:49:2:134:3:3:0:1:0:0 | 2:0:0:7:0:0:0:0:0:0:0 | 1798:633:92:4:227:7:11:0:1:1:0 | 5:0:0:7:2:0:0:0:0:0:0 | 657:155:28:0:29:2:1:0:1:1:1 | 2:0:0:1:0:0:0:0:0:0:0 | 1 | 1 | 1 | 7.67E-11 | 9.78E-10 | 1 |
| 912:342:49:2:134:3:3:0:1:0:0 | 2:0:0:7:0:0:0:0:0:0:0 | 1798:633:92:4:227:7:11:0:1:1:0 | 5:0:0:7:2:0:0:0:0:0:0 | 657:155:28:0:29:2:1:0:1:1:1 | 2:0:0:1:0:0:0:0:0:0:0 | 1 | 1 | 1 | 7.67E-11 | 9.78E-10 | 1 |
| 913:342:49:2:134:3:2:0:1:0:0 | 1:0:0:7:1:0:1:0:0:0:0 | 1796:633:92:4:225:7:11:0:1:1:0 | 5:0:0:7:4:0:0:0:0:0:0 | 659:155:28:0:28:2:1:0:1:1:1 | 0:0:0:1:1:0:0:0:0:0:0 | 1 | 1 | 1 | 4.08E-13 | 2.34E-10 | 1 |

| | | | | | | | | | | | |
|---|---|---|---|---|---|---|---|---|---|---|---|
| 4559:101:9:18 | 1:4:0:0 | 11061:369:15:10 | 5:6:0:0 | 2926:82:5:10 | 0:1:0:0 | 0.07769114 | 1 | 1 | 0.01889599 | 0.00942178 | 1 |
| 131:223:13:23:3:5:0:0 | 2:0:2:7:0:0:0:0 | 84:240:17:19:2:1:0:1 | 2:0:2:1:0:0:0:0 | 19:43:3:1:0:0:0:0 | 1:0:0:0:0:0:0:0 | 1 | 1 | 1 | 0.00100057 | 1 | 1 |
| 1019:7198:72:1 | 3:30:9:0 | 1253:13384:118:6 | 8:44:15:0 | 186:3107:11:1 | 0:8:0:0 | 1 | 1 | 1 | 2.07E-06 | 2.22E-13 | 1 |
| 7241:945:15:20 | 7:0:0:0 | 13476:1049:29:30 | 13:3:0:3 | 3113:159:12:8 | 8:0:0:0 | 1 | 1 | 1 | 1 | 0.02824949 | 1 |
| 7244:733:20:25 | 10:0:0:0 | 13492:629:42:82 | 6:7:0:0 | 3123:136:13:22 | 2:0:0:0 | 1 | 1 | 1 | 1 | 0.01748971 | 1 |
| 7252:723:25:19 | 2:1:0:0 | 13497:622:83:40 | 1:5:0:0 | 3125:148:22:12 | 0:0:0:0 | 1 | 1 | 1 | 1 | 0.01752706 | 1 |
| 7255:675:28:17 | 1:2:2:0 | 13495:568:97:43 | 9:0:12:0 | 3126:152:24:11 | 1:0:0:0 | 0.41069488 | 1 | 0.00031796 | 0.02970966 | 4.52E-16 | 1 |
| 7249:29:675:18 | 11:5:1:1 | 13492:108:564:44 | 17:8:0:0 | 3126:26:151:12 | 2:0:1:0 | 1 | 0.01255668 | 1 | 7.65E-06 | 6.10E-07 | 1 |
| 7423:50:312:652 | 22:7:40:6 | 13768:61:663:531 | 31:1:39:1 | 3141:11:52:149 | 7:0:3:0 | 5.00E-13 | 0.04431818 | 1 | 3.50E-40 | 1.48E-28 | 1 |
| 17:4:7:10:2 | 22:5:1:1:0 | 35:6:8:13:8 | 52:2:0:1:1 | 15:1:3:2:0 | 0:0:0:0:0 | 1 | 1 | 1 | 1 | 0.00548791 | 1 |
| 22060:560:143 | 32:11:00 | 30314:700:252 | 43:06:00 | 8573:110:75 | 10:02:00 | 0.37916208 | 1 | 1 | 0.00010099 | 1 | 1 |
| 21996:565:143 | 45:10:00 | 30230:712:251 | 62:04:00 | 8562:110:75 | 12:03:00 | 1 | 1 | 1 | 0.01791562 | 1 | 1 |
| 124:12:1:1:1:2:0:0 | 2:11:5:0:0:0:0:0 | 69:16:1:0:0:0:0:0 | 0:5:1:0:0:0:0:1 | 34:3:0:0:1:0:1:0 | 0:0:1:0:0:0:0:0 | 1 | 1 | 1 | 2.08E-08 | 0.02949837 | 1 |
| 149:10:4:0:3:2:1 | 2:7:3:0:0:0:0 | 91:8:4:0:0:3:1 | 2:6:0:0:0:0:1 | 37:2:1:1:1:1:1 | 0:0:0:0:0:0:0 | 1 | 1 | 1 | 0.00092213 | 0.28834485 | 1 |
| 10874:3715 | 217:08:00 | 7719:2874 | 61:06:00 | 1832:166 | 3:01 | 1 | 1 | 1 | 7.66E-15 | 1 | 1 |
| 10881:3708 | 460:23:00 | 7730:2867 | 100:11:00 | 1831:164 | 4:03 | 1 | 1 | 1 | 3.01E-29 | 0.05369671 | 1 |
| 10899:3686 | 844:65 | 7733:2849 | 233:32:00 | 1834:166 | 22:03 | 1 | 1 | 1 | 7.34E-40 | 3.34E-05 | 1 |
| 11353:3724 | 399:20:00 | 7888:2866 | 80:07:00 | 1852:166 | 5:01 | 1 | 1 | 1 | 8.82E-24 | 0.09811423 | 1 |
| 3397:11777 | 11:00 | 2559:7984 | 6:01 | 142:1859 | 0:00 | 1 | 1 | 1 | 0.00030002 | 1 | 1 |
| 11867:1285 | 17:09 | 8012:908 | 9:10 | 1860:81 | 1:00 | 1 | 1 | 1 | 1 | 0.02015401 | 1 |
| 11873:1118 | 17:16 | 8018:761 | 7:08 | 1858:78 | 3:00 | 1 | 1 | 1 | 1.16E-05 | 0.05064863 | 1 |
| 11890:929 | 10:19 | 8025:628 | 3:12 | 1859:77 | 2:00 | 1 | 1 | 1 | 1.08E-11 | 3.70E-08 | 1 |
| 11896:696 | 12:07 | 8023:477 | 8:14 | 1861:74 | 0:01 | 1 | 1 | 1 | 0.18820854 | 3.25E-09 | 1 |
| 11901:191 | 8:29 | 8026:142 | 9:15 | 1862:38:00 | 0:04 | 1 | 1 | 0.00106254 | 6.81E-41 | 4.17E-17 | 0.00086285 |
| 267:23:5:2:0:0 | 0:6:2:0:0:0 | 242:19:3:0:1:1 | 1:1:0:0:0:0 | 73:3:1:0:1:0 | 0:1:0:0:0:0 | 1 | 1 | 1 | 0.00013638 | 1 | 1 |
| 267:14:1:2:0:0 | 0:15:6:0:0:0 | 243:16:1:0:1:1 | 0:4:2:0:0:0 | 73:2:0:0:1:0 | 0:2:1:0:0:0 | 1 | 1 | 1 | 6.87E-20 | 0.00060886 | 1 |
| 41:49:184:1:2:0 | 7:9:0:0:0:0 | 81:109:430:0:1:2 | 3:6:0:0:0:0 | 85:3:33:0:0:0 | 1:1:0:0:0:0 | 1 | 1 | 1 | 0.00115274 | 0.46908418 | 1 |
| 39:10:225:1:3:0 | 2:37:0:0:0:0 | 81:14:526:0:1:2 | 0:97:0:0:0:0 | 85:3:34:0:0:0 | 0:0:0:0:0:0 | 1 | 1 | 1 | 1.14E-33 | 9.56E-102 | 1 |
| 107:130:204 | 19:02:01 | 96:103:211 | 16:01:03 | 160:10:67 | 5:00:01 | 1 | 1 | 1 | 1.57E-05 | 0.00538893 | 1 |
| 9:0:4:1:4:3:25:59:0:34:12:4:0:1:1:2:7:2:1:2:1:0 | 7:13:20:2:0:5:1:10:0:2:5:0:1:0:0:0:0:0:0:0:0:0 | 6:1:8:1:8:3:19:4:1:13:4:3:0:1:1:3:0:0:0:2:2:1 | 7:10:27:3:0:6:2:1:0:2:2:0:1:0:0:1:0:0:1:1:0:0 | 0:0:1:0:3:0:18:8:1:6:3:0:0:0:0:0:0:0:0:0:2:0 | 2:4:3:1:0:1:0:0:0:1:0:0:0:0:0:0:0:0:1:0:0:0 | 1 | 1 | 1 | 1 | 1 | 0.00030741 |
| 6246:58:874:10:5:43 | 1:2:2:1:0:0 | 4238:43:683:4:4:36 | 4:0:0:0:0:0 | 4987:37:611:6:3:42 | 2:2:0:1:0:0 | 1 | 1 | 1 | 0.01693504 | 1 | 0.06624921 |
| 4:235:0:43:0:0:10:6:5:0:0:1:0:18:0:42:1:0:0:0:2:1:0:0:0:0:0 | 1:0:0:0:0:0:0:0:0:0:0:0:0:0:0:0:0:0:0:0:0:0:0:0:0:0:0 | 7:392:5:41:4:1:9:7:17:0:1:2:4:15:2:51:2:0:0:3:1:1:0:0:0:1:1 | 4:0:0:0:0:0:0:0:0:0:0:0:0:0:0:0:1:0:0:0:0:0:0:0:0:0:0 | 1:99:1:4:3:0:2:0:1:0:0:1:0:2:0:4:0:0:1:2:0:0:1:1:0:0:0 | 0:1:0:0:0:0:0:0:0:0:0:0:0:0:0:0:0:0:0:0:0:0:0:0:0:0:0 | 1 | 1 | 1 | 1 | 0.03633546 | 1 |
| 5:1:29:74 | 24:2:0:0 | 2:0:23:26 | 9:0:0:0 | 3:1:8:11 | 7:0:0:0 | 1 | 0.21923214 | 1 | 7.85E-19 | 1.56E-05 | 0.32214559 |

| | | | | | | | | | | | |
|---|---|---|---|---|---|---|---|---|---|---|---|
| 29:29:73:4 | 24:0:0:2 | 11:23:25:0 | 11:0:0:4 | 10:8:11:1 | 0:0:0:0 | 1 | 1 | 1 | 1.08E-09 | 3.14E-06 | 1 |
| 31:29:73:4 | 22:0:0:2 | 12:23:25:1 | 11:0:0:3 | 10:8:11:1 | 0:0:0:0 | 1 | 1 | 1 | 2.02E-08 | 0.00023607 | 1 |
| 34:29:73:4 | 18:0:0:2 | 13:23:24:1 | 10:0:0:2 | 10:8:11:1 | 2:0:0:0 | 1 | 1 | 1 | 3.44E-06 | 0.00346073 | 1 |
| 125:4:3:19:1:0 | 0:0:3:0:0:0 | 513:24:6:74:2:0 | 1:0:4:0:0:2 | 97:3:3:11:0:1 | 0:0:1:0:0:0 | 1 | 1 | 1 | 0.30651322 | 9.47E-07 | 1 |
| 107:36:150:2:170 | 3:11:3:0:0 | 170:35:199:3:244 | 2:19:3:0:0 | 50:11:149:1:51 | 2:9:2:0:0 | 1 | 0.14506308 | 8.00E-05 | 5.45E-05 | 4.27E-15 | 3.16E-05 |
| 0:00 | 316:07:00 | 0:00 | 795:08:00 | 0:00 | 253:03:00 | 1.57E-27 | 0.00037979 | 1 | 1 | 1 | 1 |
| 0:00 | 312:07:00 | 1:00 | 788:08:00 | 1:00 | 253:03:00 | 3.25E-25 | 0.00042123 | 1 | 1 | 1 | 1 |
| 0:00 | 308:07:00 | 2:00 | 735:07:00 | 0:00 | 240:03:00 | 3.08E-26 | 0.00042123 | 1 | 1 | 1 | 1 |
| 0:00 | 308:07:00 | 2:00 | 735:07:00 | 0:00 | 240:03:00 | 3.08E-26 | 0.00042123 | 1 | 1 | 1 | 1 |
| 0:00 | 307:07:00 | 0:00 | 735:07:00 | 0:00 | 241:03:00 | 3.08E-26 | 0.00042123 | 1 | 1 | 1 | 1 |
| 13:00 | 295:07:00 | 504:00:00 | 232:07:00 | 119:00:00 | 122:03:00 | 3.08E-26 | 0.00042123 | 1 | 1 | 1 | 1 |
| 1:00 | 303:07:00 | 0:00 | 729:07:00 | 0:00 | 239:03:00 | 4.32E-26 | 0.00042123 | 1 | 1 | 1 | 1 |
| 1:00 | 297:07:00 | 1:00 | 726:07:00 | 0:00 | 238:03:00 | 4.84E-26 | 0.00042123 | 1 | 1 | 1 | 1 |
| 297:07:00 | 0:00 | 718:07:00 | 4:00 | 237:03:00 | 0:00 | 1.18E-23 | 0.00042123 | 1 | 1 | 1 | 1 |
| 0:00 | 296:07:00 | 2:00 | 767:08:00 | 0:00 | 242:03:00 | 4.36E-24 | 0.00042123 | 1 | 1 | 1 | 1 |
| 0:00 | 297:07:00 | 0:00 | 769:08:00 | 0:00 | 242:03:00 | 4.85E-24 | 0.00042123 | 1 | 1 | 1 | 1 |
| 0:00 | 295:07:00 | 2:00 | 764:08:00 | 0:00 | 240:03:00 | 5.40E-24 | 0.00042123 | 1 | 1 | 1 | 1 |
| 0:00 | 292:07:00 | 0:00 | 758:08:00 | 0:00 | 238:03:00 | 6.70E-24 | 0.00042123 | 1 | 1 | 1 | 1 |
| 0:00 | 284:08:00 | 6:00 | 683:09:00 | 0:01 | 229:03:00 | 2.93E-22 | 0.00550702 | 1 | 1 | 1 | 1 |
| 281:10:00 | 1:00 | 680:09:00 | 0:00 | 228:05:00 | 1:00 | 4.88E-24 | 0.00492925 | 1 | 1 | 1 | 1 |
| 11:01 | 261:08:00 | 4:00 | 654:09:00 | 1:01 | 217:03:00 | 4.61E-23 | 0.00616505 | 1 | 1 | 1 | 1 |
| 0:00 | 264:09:00 | 0:00 | 653:09:00 | 0:00 | 217:04:00 | 3.79E-17 | 0.00080977 | 1 | 1 | 1 | 1 |
| 0:00 | 257:09:00 | 0:00 | 642:09:00 | 0:00 | 216:04:00 | 3.79E-17 | 0.00080977 | 1 | 1 | 1 | 1 |
| 249:09:00 | 1:00 | 619:09:00 | 0:00 | 205:04:00 | 0:00 | 6.89E-23 | 0.00117508 | 1 | 1 | 1 | 1 |
| 237:08:00 | 10:01 | 170:09:00 | 444:00:00 | 103:03:00 | 102:01:00 | 7.02E-17 | 0.00198515 | 1 | 1 | 0.04800051 | 1 |
| 245:08:00 | 2:01 | 172:09:00 | 442:00:00 | 102:03:00 | 102:01:00 | 7.79E-17 | 0.0022242 | 1 | 1 | 0.05313239 | 1 |
| 0.548611111 | 0:01 | 0.75 | 0:00 | 0.309027778 | 0:00 | 0.03299937 | 1 | 1 | 1 | 1 | 1 |
| 1:00 | 254:03:00 | 429:00:00 | 177:03:00 | 102:00:00 | 108:01:00 | 7.78E-17 | 0.00632511 | 1 | 1 | 1 | 1 |
| 252:00:00 | 9:00 | 613:02:00 | 1:00 | 216:00:00 | 0:00 | 6.03E-21 | 0.03813644 | 1 | 1 | 1 | 1 |
| 0.181944444 | 0:00 | 0.471527778 | 0:01 | 0.151388889 | 0:00 | 1 | 0.00190798 | 1 | 1 | 1 | 1 |
| 266:00:00 | 0:00 | 617:02:00 | 3:00 | 218:00:00 | 0:00 | 1 | 0.00024409 | 1 | 1 | 1 | 1 |
| 0:00 | 265:00:00 | 1:00 | 615:02:00 | 0:00 | 217:00:00 | 1 | 0.00024409 | 1 | 1 | 1 | 1 |
| 1:00 | 264:01:00 | 2:00 | 608:02:00 | 0:00 | 216:00:00 | 4.92E-23 | 0.00024409 | 1 | 1 | 1 | 1 |
| 0:00 | 266:01:00 | 0:00 | 607:02:00 | 0:00 | 217:00:00 | 1 | 0.00072308 | 1 | 1 | 1 | 1 |
| 0:00 | 267:01:00 | 1:00 | 615:02:00 | 1:00 | 218:00:00 | 1 | 0.00072308 | 1 | 1 | 1 | 1 |
| 0:00 | 268:01:00 | 2:00 | 615:02:00 | 1:00 | 218:00:00 | 4.92E-23 | 0.00072308 | 1 | 1 | 1 | 1 |
| 53:827 | 1:02 | 39:412 | 3:02 | 0.538888889 | 0:00 | 1 | 0.00061571 | 1 | 1 | 1 | 1 |
| 5695:81:1074:180:87:196:17 | 6:0:2:0:6:0:0 | 3108:36:300:15:108:103:13 | 3:0:0:0:4:0:1 | 676:5:57:1:14:15:3 | 2:0:0:0:1:0:1 | 1 | 1 | 1 | 0.00403038 | 0.19639467 | 1 |
| 31:90:5785:1094:126:183:177 | 1:3:1:1:0:0:11 | 15:38:3150:307:137:15:93 | 1:0:0:0:0:0:0 | 5:6:685:57:19:1:18 | 0:0:0:0:0:0:1 | 1 | 1 | 1 | 2.55E-14 | 1 | 1 |
| 5780:106:1095:136:183:179:40 | 6:0:1:0:0:1:0 | 3148:44:307:146:15:83:17 | 2:0:0:0:0:0:0 | 685:8:57:20:1:16:5 | 0:0:0:0:0:0:0 | 5.24E-05 | 1 | 1 | 1 | 1 | 1 |

| | | | | | | | | | | | |
|---|---|---|---|---|---|---|---|---|---|---|---|
| 136:146:219:61 | 6:0:2:0 | 226:376:592:169 | 24:1:2:0 | 49:194:37:22 | 9:0:0:0 | 1 | 0.24561009 | 1 | 1 | 2.11E-11 | 0.00374444 |
| 21216:637 | 12:08 | 19415:832 | 13:04 | 1952:64 | 1:00 | 1 | 1 | 1 | 0.00021055 | 1 | 1 |
| 20407:643 | 29:07:00 | 18551:807 | 31:10:00 | 1930:67 | 1:01 | 1 | 1 | 1 | 0.40638236 | 0.02367994 | 1 |
| 19856:604 | 61:25:00 | 18052:825 | 34:15:00 | 1892:62 | 2:03 | 1 | 1 | 1 | 1.97E-14 | 7.25E-06 | 1 |
| 4:0:0:0 | 0:0:0:0 | 78:0:0:0 | 0:0:0:0 | 0:0:0:1 | 0:0:0:0 | 7.84E-07 | 1 | 1 | 1 | 1 | 1 |
| 4:0:0:0 | 0:0:0:0 | 78:0:0:0 | 0:0:0:0 | 0:0:0:1 | 0:0:0:0 | 1.27E-25 | 1 | 1 | 1 | 1 | 1 |
| 4:0:0:0 | 0:0:0:0 | 78:0:0:0 | 0:0:0:0 | 0:0:0:1 | 0:0:0:0 | 2.31E-14 | 1 | 1 | 1 | 1 | 1 |
| 4:0:0:0 | 0:0:0:0 | 77:0:0:0 | 1:0:0:0 | 0:0:0:1 | 0:0:0:0 | 2.90E-20 | 1 | 1 | 1 | 1 | 1 |
| 4:0:0:0 | 0:0:0:0 | 78:0:0:0 | 0:0:0:0 | 0:0:0:1 | 0:0:0:0 | 1.43E-18 | 1 | 1 | 1 | 1 | 1 |
| 13238:104:8:13 | 11600:102:63:1 | 8525:17:7:5 | 7163:21:22:5 | 1002:13:6:2 | 26:0:1:0 | 1 | 1 | 1 | 8.17E-11 | 1 | 1 |

Supplementary Table S9: SNV isoform validation via targeted RNA sequencing
S9A: Gene FAM104A
chr17:73209715 A -> G

| Read count | Loop-seq | | | |
|---|---|---|---|---|
| sample | NM_001098832 | NM_032837 | NM_001289410 | XM_024451008 |
| 1M__WT | 2 | 2 | 0 | 0 |
| 1M__MUT | 1 | 3 | 1 | 0 |
| 1N__WT | 0 | 0 | 0 | 0 |
| 1N__MUT | 0 | 0 | 0 | 0 |
| 1T__WT | 4 | 9 | 0 | 1 |
| 1T__MUT | 4 | 15 | 0 | 0 |
| 2M__WT | 8 | 9 | 0 | 0 |
| 2M__MUT | 0 | 9 | 0 | 0 |
| 2T__WT | 1 | 2 | 0 | 0 |
| 2T__MUT | 2 | 0 | 0 | 0 |
| 2N__WT | | | | |
| 2N__MUT | | | | |
| 3M__WT | 1 | 0 | 0 | 0 |
| 3M__MUT | 2 | 4 | 1 | 0 |
| 3N__WT | 2 | 2 | 1 | 0 |
| 3N__MUT | 2 | 6 | 0 | 0 |
| 3T__WT | 1 | 0 | 0 | 0 |
| 3T__MUT | 8 | 13 | 0 | 0 |

| RNA-seq validation, experiment 1 | | | | |
|---|---|---|---|---|
| sample | NM_001098832 | NM_032837 | NM_001289410 | Undetermine |
| GRM_FAM104A_1M__WT | 5 | 24 | 0 | 11 |
| GRM_FAM10A_1M__MUT | 8 | 13 | 0 | 6 |
| GRM_FAM10A_1T__WT | 3 | 17 | 0 | 3 |
| GRM_FAM104A_1T__MUT | 1 | 19 | 0 | 12 |
| GRM_FAM104A_2M__WT | 5 | 7 | 0 | 0 |
| GRM_FAM104A_2M__MUT | 3 | 17 | 0 | 0 |
| GRM_FAM104A_2T__WT | 9 | 7 | 0 | 1 |
| GRM_FAM104A_2T__MUT | 3 | 9 | 0 | 2 |

| RNA-seq validation, experiment 2 | | | | |
|---|---|---|---|---|
| | NM_001098832 | NM_032837 | NM_001289410 | Undetermine |
| PHU_FAM104A_1M__WT | 7 | 26 | 9 | 5 |
| PHU_FAM104A_1M__MUT | 15 | 28 | 8 | 2 |
| PHU_FAM104A1_1T__WT | 7 | 35 | 7 | 2 |
| PHU_FAM104A_1T__MUT | 7 | 19 | 4 | 4 |
| PHU_FAM104A_2M__WT | 31 | 36 | 18 | 6 |
| PHU_FAM104A_2M__MUT | 21 | 48 | 22 | 3 |
| PHU_FAM104A_2T__WT | 2 | 22 | 20 | 8 |
| PHU_FAM104A_2T__MUT | 4 | 25 | 2 | 14 |

| WES | |
|---|---|
| | Count |
| 1M-T__WT | 39 |
| 1M-T__MUT | 41 |
| 1T-N__WT | 62 |
| 1T-N__MUT | 87 |
| 1T__WT | 68 |
| 1T__MUT | 59 |
| 2M-T__WT | 62 |
| 2M-T__MUT | 71 |
| 2T-T__WT | 62 |
| 2T-T__MUT | 84 |
| 2M-N__WT | 40 |
| 2M-N__MUT | 38 |
| 3M-T__WT | 17 |
| 3M-T__MUT | 102 |
| 3T-N__WT | 38 |
| 3T-N__MUT | 57 |
| 3T-T__WT | 14 |
| 3T-T__MUT | 42 |

Supplementary Table S9: SNV isoform validation via targeted RNA sequencing

S9B: Gene PABPC1

chr8: 100705025 C->T

| Read count | Loop-seq | | RNA-seq validation, experiment 1 | | | | RNA-seq validation, experiment 2 | | | |
|---|---|---|---|---|---|---|---|---|---|---|
| sample | NM_002568 | XM_005250861 | | NM_002568 | XM_005250861 | Undetermine | | NM_002568 | XM_005250861 | Undetermine |
| 1M__WT | 0 | 0 | | | | | | | | |
| 1M__MUT | 0 | 0 | | | | | | | | |
| 1N__WT | 1209 | 38 | | | | | | | | |
| 1N__MUT | 4 | 0 | | | | | | | | |
| 1T__WT | 957 | 36 | | | | | | | | |
| 1T__MUT | 3 | 0 | | | | | | | | |
| 2M__WT | 20892 | 812 | GRM_PABPC1_2M__WT | 30564 | 58 | 1574 | PHU_PABPC1_2M__WT | 4104 | 3 | 366 |
| 2M__MUT | 28 | 2 | GRM_PABPC1_2M__MUT | 26 | 0 | 3 | PHU_PABPC1_2M__MUT | 2 | 0 | 2 |
| 2T__WT | 1957 | 43 | GRM_PABPC1_2T__WT | 6222 | 17 | 354 | PHU_PABPC1_2T__WT | 6165 | 12 | 763 |
| 2T__MUT | 4 | 0 | GRM_PABPC1_2T__MUT | 5 | 0 | 2 | PHU_PABPC1_2T__MUT | 5 | 0 | 0 |
| 2N__WT | | | | | | | | | | |
| 2N__MUT | | | | | | | | | | |
| 3M__WT | 1942 | 70 | GRM_PABPC1_3M__WT | 15305 | 83 | 808 | PHU_PABPC1_3M__WT | 1249 | 10 | 151 |
| 3M__MUT | 2 | 0 | GRM_PABPC1_3M__MUT | 19 | 0 | 2 | PHU_PABPC1_3M__MUT | 0 | 0 | 2 |
| 3N__WT | 0 | 0 | | | | | | | | |
| 3N__MUT | 0 | 0 | | | | | | | | |
| 3T__WT | 19070 | 907 | GRM_PABPC1_3T__WT | 1892 | 12 | 90 | PHU_PABPC1_3T__WT | 1062 | 10 | 124 |
| 3T__MUT | 23 | 5 | GRM_PABPC1_3T__MUT | 2 | 0 | 0 | PHU_PABPC1_3T__MUT | 1 | 0 | 0 |

| WES | |
|---|---|
| | Count |
| 1M-T__WT | 182 |
| 1M-T__MUT | 1 |
| 1T-N__WT | 139 |
| 1T-N__MUT | 3 |
| 1T__WT | 146 |
| 1T__MUT | 1 |
| 2M-T__WT | 354 |
| 2M-T__MUT | 2 |
| 2T-T__WT | 176 |
| 2T-T__MUT | 1 |
| 2M-N__WT | 169 |
| 2M-N__MUT | 0 |
| 3M-T__WT | 223 |
| 3M-T__MUT | 1 |
| 3T-N__WT | 248 |
| 3T-N__MUT | 2 |
| 3T-T__WT | 152 |
| 3T-T__MUT | 0 |

Supplementary Table S10: Switching SNV isoforms between tumor and metastasis

| chr | pos | ref | alt | cosmicID | annotation | gene | mutType |
|---|---|---|---|---|---|---|---|
| chr14 | 22830077 | T | C | . | exonic | MRPL52 | nonsynonymous SNV |
| chr14 | 34800325 | C | T | . | exonic | BAZ1A | nonsynonymous SNV |
| chr17 | 72947851 | T | C | . | exonic | SLC39A11 | nonsynonymous SNV |
| chr3 | 169116038 | A | T | . | exonic | MECOM | nonsynonymous SNV |
| chr6 | 29555899 | C | G | . | exonic | UBD | nonsynonymous SNV |
| chr6 | 33080765 | C | T | COSV69603920 | exonic | HLA-DPB1 | nonsynonymous SNV |
| chr6 | 33080822 | C | A | COSV69602760 | exonic | HLA-DPB1 | nonsynonymous SNV |
| chr1 | 13816508 | A | G | COSV52443068 | exonic | PRDM2 | nonsynonymous SNV |
| chr1 | 42164693 | G | A | COSV63691329 | exonic | GUCA2A | nonsynonymous SNV |
| chr10 | 31520809 | T | A | . | exonic | ZEB1 | nonsynonymous SNV |
| chr11 | 20383067 | T | G | COSV58178278 | exonic | HTATIP2 | nonsynonymous SNV |
| chr13 | 72727590 | T | C | COSV64695018 | exonic | MZT1 | nonsynonymous SNV |
| chr14 | 24440156 | T | C | COSV52162406 | exonic | SDR39U1 | nonsynonymous SNV |
| chr14 | 24440269 | C | G | COSV52161794 | exonic | SDR39U1 | nonsynonymous SNV |
| chr14 | 24441767 | T | G | COSV52162442 | exonic | SDR39U1 | nonsynonymous SNV |
| chr16 | 29779543 | G | A | . | exonic | ZG16 | nonsynonymous SNV |
| chr16 | 29780400 | G | C | COSV68783707 | exonic | ZG16 | nonsynonymous SNV |
| chr16 | 74661181 | G | T | . | exonic | RFWD3 | nonsynonymous SNV |
| chr19 | 990281 | G | A | COSV52120984 | exonic | WDR18 | nonsynonymous SNV |
| chr20 | 32432924 | G | A | . | exonic | ASXL1 | nonsynonymous SNV |
| chr7 | 5987144 | T | C | COSV56223170 | exonic | PMS2 | nonsynonymous SNV |
| chr7 | 5987357 | G | A | COSV56220401 | exonic | PMS2 | nonsynonymous SNV |
| chr8 | 31141764 | G | T | COSV53297644 | exonic | WRN | nonsynonymous SNV |

| AAanno | Isoform | 1T | 2T | 3T |
|---|---|---|---|---|
| MRPL52:NM_181307:exon2:c.T56C:p.V19A,MRPL52:NM_180982:exon2:c.T53C:p.V18A,MRPL52:NM_178336:exon2:c.T56C:p.V19A, | NM_180982 | 0 | 0 | 0.5 |
| BAZ1A:NM_182648:exon9:c.G1027A:p.E343K,BAZ1A:NM_013448:exon9:c.G1027A:p.E343K, | XM_024449460 | 0 | 0.333333333 | 0.294117647 |
| SLC39A11:NM_139177:exon5:c.A331G:p.T111A,SLC39A11:NM_001159770:exon5:c.A331G:p.T111A, | XM_006721757 | 0 | 0 | 0 |
| MECOM:NM_001105078:exon7:c.T1270A:p.F424I,MECOM:NM_001163999:exon7:c.T1273A:p.F425I,MECOM:NM_001105077:exon8:c.T1465A:p.F489I,MECOM:NM_004991:exon8:c.T1834A:p.F612I,MECOM:NM_001164000:exon7:c.T1270A:p.F424I,MECOM:NM_001205194:exon7:c.T1270A:p.F424I,MECOM:NM_005241:exon7:c.T1270A:p.F424I, | XM_011512546 | 0.2 | 0.291666667 | 0.444444444 |
| UBD:NM_006398:exon2:c.G479C:p.C160S, | NM_006398 | 0 | 0 | 0.333333333 |
| HLA-DPB1:NM_002121:exon2:c.C194T:p.A65V, | NM_002121 | 0 | 0 | 0 |
| HLA-DPB1:NM_002121:exon2:c.C251A:p.A84D, | NM_002121 | 0 | 0 | 0 |
| PRDM2:NM_001135610:exon7:c.A593G:p.Q198R, | NM_001135610 | 1 | 1 | 1 |
| GUCA2A:NM_033553:exon1:c.C20T:p.S7F, | NM_033553 | 1 | 0.5 | 1 |
| ZEB1:NM_001174093:exon6:c.T1414A:p.L472I,ZEB1:NM_001174095:exon6:c.T1273A:p.L425I,ZEB1:NM_001128128:exon7:c.T1426A:p.L476I,ZEB1:NM_030751:exon7:c.T1474A:p.L492I,ZEB1:NM_001174096:exon7:c.T1477A:p.L493I,ZEB1:NM_001174094:exon7:c.T1423A:p.L475I, | NM_001174096 | 0.5 | 1 | 0.5 |
| HTATIP2:NM_006410:exon6:c.T591G:p.S197R,HTATIP2:NM_001098522:exon5:c.T591G:p.S197R,HTATIP2:NM_001098520:exon6:c.T693G:p.S231R,HTATIP2:NM_001098521:exon6:c.T591G:p.S197R, | NM_001098522 | 1 | 1 | 1 |
| MZT1:NM_001071775:exon1:c.A13G:p.S5G, | NM_001071775 | 1 | 1 | 1 |
| SDR39U1:NM_001290292:exon5:c.A563G:p.Q188R,SDR39U1:NM_020195:exon6:c.A809G:p.Q270R,SDR39U1:NM_001308075:exon5:c.A563G:p.Q188R,SDR39U1:NM_001290293:exon5:c.A485G:p.Q162R, | NM_020195 | 1 | 1 | 1 |
| SDR39U1:NM_001290292:exon5:c.G450C:p.L150F,SDR39U1:NM_020195:exon6:c.G696C:p.L232F,SDR39U1:NM_001308075:exon5:c.G450C:p.L150F,SDR39U1:NM_001290293:exon5:c.G372C:p.L124F, | NM_020195 | 1 | 1 | 1 |
| SDR39U1:NM_020195:exon4:c.A235C:p.I79L, | NM_020195 | 1 | 1 | 1 |
| ZG16:NM_152338:exon3:c.G94A:p.G32S, | NM_152338 | 1 | 1 | 1 |
| ZG16:NM_152338:exon4:c.G485C:p.S162T, | NM_152338 | 1 | 1 | 1 |
| RFWD3:NM_018124:exon2:c.C269A:p.T90N, | NM_018124 | 1 | 1 | 1 |
| WDR18:NM_024100:exon4:c.G514A:p.A172T, | NM_024100 | 1 | 1 | 1 |
| ASXL1:NM_015338:exon10:c.G1024A:p.E342K, | XM_017027706 | 1 | 1 | 0.714285714 |
| PMS2:NM_000535:exon11:c.A1621G:p.K541E, | NM_001322003 | 1 | 1 | 1 |
| PMS2:NM_000535:exon11:c.C1408T:p.P470S, | NM_001322003 | 1 | 0.666666667 | 0.5 |
| WRN:NM_000553:exon26:c.G3222T:p.L1074F, | XR_949472 | 1 | 1 | 1 |

| 1M | 2M | 3M | aveT | aveL | diffRate |
|---|---|---|---|---|---|
| 1 | 1 | 1 | 0.166666667 | 1 | 0.833333333 |
| 0.5 | 0.615384615 | 0.5 | 0.209150327 | 0.538461538 | 0.329311212 |
| 1 | 1 | 1 | 0 | 1 | 1 |
| 1 | 0.774193548 | 0.5 | 0.312037037 | 0.758064516 | 0.446027479 |
| 1 | 0.666666667 | 0.5 | 0.111111111 | 0.722222222 | 0.611111111 |
| 1 | 1 | 0.666666667 | 0 | 0.888888889 | 0.888888889 |
| 0.75 | 1 | 0.666666667 | 0 | 0.805555556 | 0.805555556 |
| 0 | 0 | 0 | 1 | 0 | -1 |
| 0 | 0 | 0 | 0.833333333 | 0 | -0.833333333 |
| 0.333333333 | 0 | 0 | 0.666666667 | 0.111111111 | -0.555555556 |
| 0 | 0 | 0 | 1 | 0 | -1 |
| 0 | 0 | 0 | 1 | 0 | -1 |
| 0 | 0 | 0 | 1 | 0 | -1 |
| 0 | 0 | 0 | 1 | 0 | -1 |
| 0 | 0 | 0 | 1 | 0 | -1 |
| 0 | 0 | 0 | 1 | 0 | -1 |
| 0 | 0 | 0 | 1 | 0 | -1 |
| 0 | 0 | 0 | 1 | 0 | -1 |
| 0 | 0 | 0 | 1 | 0 | -1 |
| 0 | 0.3 | 0 | 0.904761905 | 0.1 | -0.804761905 |
| 0 | 0 | 0 | 1 | 0 | -1 |
| 0 | 0 | 0 | 0.722222222 | 0 | -0.722222222 |
| 0 | 0 | 0.5 | 1 | 0.166666667 | -0.833333333 |

Supplementary Table S11: Colon cancer related mutations/SNVs

| Mutation | | | | | | | | |
|---|---|---|---|---|---|---|---|---|
| chr | pos | ref | alt | cosmicID | annotation | gene | mutType | AAanno |
| chr1 | 1014228 | G | A | COSV65106391 | exonic | ISG15 | nonsynonymous SNV | ISG15:NM_005101:exon2:c.G248A:p.S83N, |
| chr1 | 114405659 | A | G | COSV61821233 | exonic | TRIM33 | nonsynonymous SNV | TRIM33:NM_015906:exon15:c.T2519C:p.I840T,TRIM33:NM_033020:exon15:c.T2519C:p.I840T, |
| chr1 | 150755063 | G | A | COSV64566034 | exonic | CTSS | nonsynonymous SNV | CTSS:NM_004079:exon4:c.C337T:p.R113W, |
| chr1 | 160610759 | G | T | COSV52502577 | exonic | SLAMF1 | nonsynonymous SNV | SLAMF1:NM_003037:exon7:c.C997A:p.P333T, |
| chr1 | 161781951 | A | G | COSV63406746 | exonic | ATF6 | nonsynonymous SNV | ATF6:NM_007348:exon3:c.A199G:p.M67V, |
| chr1 | 17697195 | A | G | COSV51429640 | exonic | ARHGEF10L | nonsynonymous SNV | ARHGEF10L:NM_001011722:exon27:c.A3538G:p.I1180V,ARHGEF10L:NM_018125:exon29:c.A3655G:p.I1219V, |
| chr1 | 225838705 | A | G | COSV55299484 | exonic | EPHX1 | nonsynonymous SNV | EPHX1:NM_001291163:exon4:c.A416G:p.H139R,EPHX1:NM_000120:exon4:c.A416G:p.H139R,EPHX1:NM_001136018:exon4:c.A416G:p.H139R, |
| chr1 | 230978719 | C | G | COSV59617127 | exonic | TTC13 | nonsynonymous SNV | TTC13:NM_024525:exon1:c.G112C:p.G38R,TTC13:NM_001122835:exon1:c.G112C:p.G38R, |
| chr1 | 236554626 | T | C | COSV63979833 | exonic | HEATR1 | nonsynonymous SNV | HEATR1:NM_018072:exon42:c.A6050G:p.E2017G, |
| chr1 | 236555320 | T | C | COSV63979840 | exonic | HEATR1 | nonsynonymous SNV | HEATR1:NM_018072:exon41:c.A5899G:p.N1967D, |
| chr1 | 236555893 | A | G | COSV63979843 | exonic | HEATR1 | nonsynonymous SNV | HEATR1:NM_018072:exon39:c.T5561C:p.V1854A, |
| chr1 | 246507179 | C | G | COSV66438600 | exonic | SMYD3 | nonsynonymous SNV | SMYD3:NM_001167740:exon1:c.G39C:p.K13N, |
| chr1 | 26320239 | A | G | COSV57485055 | exonic | CD52 | nonsynonymous SNV | CD52:NM_001803:exon2:c.A123G:p.I41M, |
| chr1 | 28473868 | C | A | COSV65783948 | exonic | PHACTR4 | nonsynonymous SNV | PHACTR4:NM_001048183:exon7:c.C1138A:p.L380I,PHACTR4:NM_023923:exon6:c.C1168A:p.L390I, |
| chr1 | 59321678 | T | G | COSV58028591 | exonic | FGGY | nonsynonymous SNV | FGGY:NM_001244714:exon2:c.T129G:p.N43K,FGGY:NM_001113411:exon2:c.T129G:p.N43K,FGGY:NM_018291:exon2:c.T129G:p.N43K, |
| chr1 | 75749507 | A | G | COSV57714736 | exonic | ACADM | nonsynonymous SNV | ACADM:NM_001286043:exon10:c.A896G:p.D299G,ACADM:NM_001286042:exon8:c.A689G:p.D230G,ACADM:NM_001286044:exon6:c.A230G:p.D77G,ACADM:NM_001127328:exon9:c.A809G:p.D270G,ACADM:NM_000016:exon9:c.A797G:p.D266G, |
| chr1 | 92834833 | G | A | COSV59872943 | exonic | RPL5 | nonsynonymous SNV | RPL5:NM_000969:exon4:c.G244A:p.E82K, |
| chr1 | 97515839 | T | C | COSV64593269 | exonic | DPYD | nonsynonymous SNV | DPYD:NM_000110:exon13:c.A1627G:p.I543V, |
| chr10 | 102813206 | T | C | COSV64009277 | exonic | WBP1L | nonsynonymous SNV | WBP1L:NM_017787:exon4:c.T904C:p.S302P,WBP1L:NM_001083913:exon4:c.T967C:p.S323P, |
| chr10 | 104263031 | C | A | COSV63846494 | exonic | GSTO1 | nonsynonymous SNV | GSTO1:NM_001191003:exon4:c.C335A:p.A112D,GSTO1:NM_004832:exon4:c.C419A:p.A140D, |
| chr10 | 133373302 | A | G | COSV63906597 | exonic | ECHS1 | nonsynonymous SNV | ECHS1:NM_004092:exon1:c.T32C:p.V11A, |
| chr10 | 59792934 | G | T | COSV54054295 | exonic | CCDC6 | nonsynonymous SNV | CCDC6:NM_005436:exon9:c.C1408A:p.P470T, |
| chr10 | 71351651 | C | T | COSV64385485 | exonic | SLC29A3 | nonsynonymous SNV | SLC29A3:NM_001174098:exon4:c.C473T:p.S158F,SLC29A3:NM_018344:exon4:c.C473T:p.S158F, |
| chr10 | 86876022 | C | A | COSV64409048 | exonic | BMPR1A | nonsynonymous SNV | BMPR1A:NM_004329:exon3:c.C4A:p.P2T, |
| chr10 | 89719454 | G | T | COSV53306008 | exonic | KIF20B | nonsynonymous SNV | KIF20B:NM_016195:exon13:c.G1470T:p.E490D,KIF20B:NM_001284259:exon13:c.G1470T:p.E490D, |
| chr11 | 20383067 | T | G | COSV58178278 | exonic | HTATIP2 | nonsynonymous SNV | HTATIP2:NM_006410:exon6:c.T591G:p.S197R,HTATIP2:NM_001098522:exon5:c.T591G:p.S197R,HTATIP2:NM_001098520:exon6:c.T693G:p.S231R,HTATIP2:NM_001098521:exon6:c.T591G:p.S197R, |
| chr11 | 308290 | T | C | COSV67714640 | exonic | IFITM2 | nonsynonymous SNV | IFITM2:NM_006435:exon1:c.T98C:p.V33A, |
| chr11 | 314207 | C | G | COSV60248916 | exonic | IFITM1 | nonsynonymous SNV | IFITM1:NM_003641:exon1:c.C37G:p.P13A, |
| chr11 | 35204608 | A | G | COSV53530507 | exonic | CD44 | nonsynonymous SNV | CD44:NM_001001389:exon9:c.A1121G:p.K374R,CD44:NM_000610:exon10:c.A1250G:p.K417R, |
| chr11 | 43855148 | C | T | COSV53496916 | exonic | HSD17B12 | nonsynonymous SNV | HSD17B12:NM_016142:exon11:c.C839T:p.S280L, |
| chr11 | 64318313 | A | G | COSV50005053 | exonic | PRDX5 | nonsynonymous SNV | PRDX5:NM_012094:exon1:c.A98G:p.Y33C,PRDX5:NM_181652:exon1:c.A98G:p.Y33C,PRDX5:NM_181651:exon1:c.A98G:p.Y33C, |
| chr11 | 78079609 | G | C | COSV55247515 | exonic | NDUFC2 | nonsynonymous SNV | NDUFC2-KCTD14:NM_001203261:exon1:c.C136G:p.L46V,NDUFC2:NM_004549:exon1:c.C136G:p.L46V,NDUFC2-KCTD14:NM_001203262:exon1:c.C136G:p.L46V,NDUFC2:NM_001204054:exon1:c.C136G:p.L46V,NDUFC2:NM_001204055:exon1:c.C136G:p.L46V,NDUFC2-KCTD14:NM_001203260:exon1:c.C136G:p.L46V, |
| chr11 | 78568744 | T | G | COSV55250015 | exonic | NARS2 | nonsynonymous SNV | NARS2:NM_024678:exon3:c.A260C:p.N87T, |
| chr11 | 7987468 | C | T | COSV59141570 | exonic | EIF3F | nonsynonymous SNV | EIF3F:NM_003754:exon1:c.C116T:p.P39L, |
| chr11 | 88312415 | A | G | COSV57060752 | exonic | CTSC | nonsynonymous SNV | CTSC:NM_001814:exon3:c.T458C:p.I153T, |
| chr12 | 121888906 | T | C | COSV55838997 | exonic | PSMD9 | nonsynonymous SNV | PSMD9:NM_002813:exon1:c.T50C:p.V17A,PSMD9:NM_001261400:exon1:c.T50C:p.V17A, |
| chr12 | 124950034 | C | T | COSV58132345 | exonic | DHX37 | nonsynonymous SNV | DHX37:NM_032656:exon25:c.G3242A:p.R1081Q, |
| chr12 | 309877 | C | T | COSV66991870 | exonic | KDM5A | nonsynonymous SNV | KDM5A:NM_001042603:exon22:c.G3304A:p.E1102K, |
| chr12 | 318409 | A | G | COSV66991468 | exonic | KDM5A | nonsynonymous SNV | KDM5A:NM_001042603:exon19:c.T2594C:p.M865T, |
| chr12 | 56712902 | C | T | COSV63269367 | exonic | NACA | nonsynonymous SNV | NACA:NM_001113203:exon10:c.G2647A:p.E883K,NACA:NM_001113202:exon7:c.G517A:p.E173K, |
| chr12 | 6347896 | T | C | COSV57434232 | exonic | SCNN1A | nonsynonymous SNV | SCNN1A:NM_001159576:exon12:c.A2164G:p.T722A,SCNN1A:NM_001038:exon13:c.A1987G:p.T663A,SCNN1A:NM_001159575:exon13:c.A2056G:p.T686A, |
| chr12 | 70745884 | T | C | COSV51736040 | exonic | PTPRR | nonsynonymous SNV | PTPRR:NM_001207016:exon3:c.A323G:p.K108R,PTPRR:NM_001207015:exon5:c.A605G:p.K202R,PTPRR:NM_130846:exon2:c.A206G:p.K69R,PTPRR:NM_002849:exon6:c.A941G:p.K314R, |
| chr12 | 71139754 | C | G | COSV56077446 | exonic | TSPAN8 | nonsynonymous SNV | TSPAN8:NM_004616:exon4:c.G218C:p.G73A, |
| chr13 | 27434894 | G | C | COSV63528570 | exonic | GTF3A | nonsynonymous SNV | GTF3A:NM_002097:exon7:c.G733C:p.V245L, |
| chr13 | 40941150 | T | A | COSV53497091 | exonic | ELF1 | nonsynonymous SNV | ELF1:NM_001145353:exon7:c.A955T:p.T319S,ELF1:NM_172373:exon8:c.A1027T:p.T343S, |

| | | | | | | | | |
|---|---|---|---|---|---|---|---|---|
| chr13 | 40958916 | T | C | COSV53497107 | exonic | ELF1 | nonsynonymous SNV | ELF1:NM_001145353:exon2:c.A173G:p.N58S,ELF1:NM_172373:exon3:c.A173G:p.N58S, |
| chr14 | 102875712 | T | C | COSV61023607 | exonic | TRAF3 | nonsynonymous SNV | TRAF3:NM_001199427:exon4:c.T386C:p.M129T,TRAF3:NM_145725:exon5:c.T386C:p.M129T,TRAF3:NM_145726:exon5:c.T386C:p.M129T,TRAF3:NM_003300:exon4:c.T386C:p.M129T, |
| chr14 | 24267755 | T | C | COSV52861986 | exonic | RABGGTA | nonsynonymous SNV | RABGGTA:NM_004581:exon13:c.A1258G:p.T420A,RABGGTA:NM_182836:exon14:c.A1258G:p.T420A, |
| chr14 | 73105860 | G | A | COSV56201910 | exonic | RBM25 | nonsynonymous SNV | RBM25:NM_021239:exon11:c.G1156A:p.E386K, |
| chr15 | 40185630 | G | A | COSV55009718 | exonic | BUB1B | nonsynonymous SNV | BUB1B:NM_001211:exon8:c.G1046A:p.R349Q, |
| chr15 | 40606445 | G | C | COSV61150583 | exonic | CASC5 | nonsynonymous SNV | CASC5:NM_144508:exon4:c.G128C:p.R43T,CASC5:NM_170589:exon4:c.G128C:p.R43T, |
| chr16 | 30069349 | G | A | COSV57632343 | exonic | ALDOA | nonsynonymous SNV | ALDOA:NM_001127617:exon6:c.G584A:p.G195E,ALDOA:NM_000034:exon11:c.G584A:p.G195E,ALDOA:NM_184043:exon6:c.G584A:p.G195E,ALDOA:NM_184041:exon6:c.G584A:p.G195E,ALDOA:NM_001243177:exon7:c.G746A:p.G249E, |
| chr16 | 3657746 | G | A | COSV55912793 | exonic | DNASE1 | nonsynonymous SNV | DNASE1:NM_005223:exon8:c.G731A:p.R244Q, |
| chr16 | 84745091 | G | C | COSV54751750 | exonic | USP10 | nonsynonymous SNV | USP10:NM_001272075:exon5:c.G622C:p.V208L,USP10:NM_005153:exon4:c.G610C:p.V204L, |
| chr16 | 88643420 | A | G | COSV54918013 | exonic | CYBA | nonsynonymous SNV | CYBA:NM_000101:exon6:c.T521C:p.V174A, |
| chr16 | 88646828 | A | G | COSV55368224 | exonic | CYBA | nonsynonymous SNV | CYBA:NM_000101:exon4:c.T214C:p.Y72H, |
| chr16 | 89114475 | G | A | COSV58077067 | exonic | ACSF3 | nonsynonymous SNV | ACSF3:NM_001127214:exon5:c.G1114A:p.V372M,ACSF3:NM_001243279:exon6:c.G1114A:p.V372M,ACSF3:NM_174917:exon6:c.G1114A:p.V372M,ACSF3:NM_001284316:exon4:c.G319A:p.V107M, |
| chr17 | 1345458 | C | T | COSV51985336 | exonic | YWHAE | nonsynonymous SNV | YWHAE:NM_006761:exon6:c.G757A:p.E253K, |
| chr17 | 31993308 | G | A | COSV59500263 | exonic | SUZ12 | nonsynonymous SNV | SUZ12:NM_015355:exon11:c.G1268A:p.R423Q, |
| chr17 | 41528069 | G | C | COSV64237813 | exonic | KRT19 | nonsynonymous SNV | KRT19:NM_002276:exon1:c.C179G:p.A60G, |
| chr17 | 43071077 | T | C | COSV58783381 | exonic | BRCA1 | nonsynonymous SNV | BRCA1:NM_007300:exon16:c.A4900G:p.S1634G,BRCA1:NM_007298:exon14:c.A1525G:p.S509G,BRCA1:NM_007294:exon15:c.A4837G:p.S1613G,BRCA1:NM_007297:exon14:c.A4696G:p.S1566G,BRCA1:NM_007299:exon15:c.A1525G:p.S509G, |
| chr17 | 48184809 | C | T | COSV61146844 | exonic | SKAP1 | nonsynonymous SNV | SKAP1:NM_001075099:exon7:c.G481A:p.G161S,SKAP1:NM_003726:exon7:c.G481A:p.G161S, |
| chr17 | 58415439 | T | C | COSV60415571 | exonic | RNF43 | nonsynonymous SNV | RNF43:NM_017763:exon2:c.A139G:p.I47V,RNF43:NM_001305544:exon2:c.A139G:p.I47V, |
| chr17 | 68452981 | G | A | COSV50929487 | exonic | WIPI1 | nonsynonymous SNV | WIPI1:NM_017983:exon2:c.C92T:p.T31I, |
| chr17 | 73209715 | A | G | COSV55428466 | exonic | FAM104A | nonsynonymous SNV | FAM104A:NM_001289411:exon3:c.T293C:p.I98T,FAM104A:NM_001289410:exon2:c.T275C:p.I92T, |
| chr17 | 75261905 | C | T | COSV55452937 | exonic | MRPS7 | nonsynonymous SNV | MRPS7:NM_015971:exon1:c.C5T:p.A2V, |
| chr17 | 7676154 | G | C | COSV52666208 | exonic | TP53 | nonsynonymous SNV | TP53:NM_001126118:exon3:c.C98G:p.P33R,TP53:NM_001276696:exon4:c.C98G:p.P33R,TP53:NM_001126113:exon4:c.C215G:p.P72R,TP53:NM_001126112:exon4:c.C215G:p.P72R,TP53:NM_001276695:exon4:c.C98G:p.P33R,TP53:NM_001276761:exon4:c.C98G:p.P33R,TP53:NM_001126114:exon4:c.C215G:p.P72R,TP53:NM_000546:exon4:c.C215G:p.P72R,TP53:NM_001276760:exon4:c.C98G:p.P33R, |
| chr18 | 9117869 | T | C | COSV59186794 | exonic | NDUFV2 | nonsynonymous SNV | NDUFV2:NM_021074:exon2:c.T86C:p.V29A, |
| chr19 | 1360576 | G | A | COSV60900886 | exonic | MUM1 | nonsynonymous SNV | MUM1:NM_032853:exon5:c.G658A:p.G220R, |
| chr19 | 17060075 | C | T | COSV53725996 | exonic | HAUS8 | nonsynonymous SNV | HAUS8:NM_033417:exon5:c.G247A:p.G83R,HAUS8:NM_001011699:exon5:c.G244A:p.G82R, |
| chr19 | 18451628 | C | T | COSV53211465 | exonic | ELL | nonsynonymous SNV | ELL:NM_006532:exon7:c.G890A:p.S297N, |
| chr19 | 45364001 | C | T | COSV55538614 | exonic | ERCC2 | nonsynonymous SNV | ERCC2:NM_000400:exon10:c.G934A:p.D312N,ERCC2:NM_001130867:exon9:c.G862A:p.D288N, |
| chr2 | 132783534 | T | A | COSV58444336 | exonic | NCKAP5 | nonsynonymous SNV | NCKAP5:NM_207363:exon14:c.A3277T:p.N1093Y, |
| chr2 | 132783881 | A | G | COSV58434578 | exonic | NCKAP5 | nonsynonymous SNV | NCKAP5:NM_207363:exon14:c.T2930C:p.I977T, |
| chr2 | 135135802 | A | G | COSV51482881 | exonic | RAB3GAP1 | nonsynonymous SNV | RAB3GAP1:NM_001172435:exon17:c.A1793G:p.N598S,RAB3GAP1:NM_012233:exon17:c.A1793G:p.N598S, |
| chr2 | 197487080 | C | G | COSV61445925 | exonic | HSPD1 | nonsynonymous SNV | HSPD1:NM_002156:exon12:c.G1688C:p.G563A,HSPD1:NM_199440:exon12:c.G1688C:p.G563A, |
| chr2 | 240025297 | G | C | COSV53155269 | exonic | NDUFA10 | nonsynonymous SNV | NDUFA10:NM_004544:exon1:c.C5G:p.A2G, |
| chr2 | 240630275 | C | T | COSV60576792 | exonic | GPR35 | nonsynonymous SNV | GPR35:NM_001195382:exon6:c.C416T:p.T139M,GPR35:NM_005301:exon2:c.C323T:p.T108M,GPR35:NM_001195381:exon6:c.C416T:p.T139M, |
| chr2 | 42288248 | A | G | COSV59293527 | exonic | EML4 | nonsynonymous SNV | EML4:NM_001145076:exon10:c.A970G:p.I324V,EML4:NM_019063:exon11:c.A1144G:p.I382V, |
| chr2 | 42288297 | A | G | COSV59293537 | exonic | EML4 | nonsynonymous SNV | EML4:NM_001145076:exon10:c.A1019G:p.K340R,EML4:NM_019063:exon11:c.A1193G:p.K398R, |
| chr20 | 18542358 | C | G | COSV52719183 | exonic | SEC23B | nonsynonymous SNV | SEC23B:NM_001172745:exon13:c.C1467G:p.H489Q,SEC23B:NM_032985:exon13:c.C1467G:p.H489Q,SEC23B:NM_001172746:exon13:c.C1413G:p.H471Q,SEC23B:NM_006363:exon13:c.C1467G:p.H489Q,SEC23B:NM_032986:exon13:c.C1467G:p.H489Q, |
| chr20 | 45409934 | A | G | COSV61919052 | exonic | DBNDD2 | nonsynonymous SNV | DBNDD2:NM_001048221:exon4:c.A280G:p.M94V,DBNDD2:NM_001197140:exon4:c.A280G:p.M94V,DBNDD2:NM_001048223:exon4:c.A280G:p.M94V,DBNDD2:NM_018478:exon4:c.A574G:p.M192V,DBNDD2:NM_001197139:exon4:c.A280G:p.M94V,DBNDD2:NM_001048225:exon3:c.A586G:p.M196V, |
| chr20 | 58903684 | G | A | COSV55672684 | exonic | GNAS | nonsynonymous SNV | GNAS:NM_080426:exon4:c.G283A:p.A95T,GNAS:NM_080425:exon5:c.G2254A:p.A752T,GNAS:NM_001309840:exon5:c.G148A:p.A50T,GNAS:NM_001077489:exon4:c.G280A:p.A94T,GNAS:NM_001309861:exon5:c.G148A:p.A50T,GNAS:NM_000516:exon5:c.G325A:p.A109T,GNAS:NM_001077488:exon5:c.G328A:p.A110T, |
| chr20 | 58905399 | C | T | COSV55688594 | exonic | GNAS | nonsynonymous SNV | GNAS:NM_001309840:exon6:c.C272T:p.A91V,GNAS:NM_000516:exon6:c.C449T:p.A150V,GNAS:NM_080426:exon5:c.C407T:p.A136V,GNAS:NM_080425:exon6:c.C2378T:p.A793V,GNAS:NM_001077489:exon5:c.C404T:p.A135V,GNAS:NM_001309861:exon6:c.C272T:p.A91V,GNAS:NM_001077488:exon6:c.C452T:p.A151V, |

| | | | | | | | | |
|---|---|---|---|---|---|---|---|---|
| chr20 | 58909556 | G | A | COSV55674033 | exonic | GNAS | nonsynonymous SNV | GNAS:NM_080426:exon8:c.G653A:p.R218H,GNAS:NM_080425:exon9:c.G2624A:p.R875H,GNAS:NM_001309840:exon9:c.G518A:p.R173H,GNAS:NM_001077489:exon8:c.G650A:p.R217H,GNAS:NM_001309861:exon9:c.G518A:p.R173H,GNAS:NM_000516:exon9:c.G695A:p.R232H,GNAS:NM_001077488:exon9:c.G698A:p.R233H, |
| chr21 | 33549616 | T | A | COSV51654122 | exonic | SON | nonsynonymous SNV | SON:NM_001291411:exon3:c.T385A:p.Y129N,SON:NM_032195:exon3:c.T385A:p.Y129N,SON:NM_138927:exon3:c.T385A:p.Y129N, |
| chr21 | 41480570 | C | T | COSV59820912 | exonic | TMPRSS2 | nonsynonymous SNV | TMPRSS2:NM_001135099:exon6:c.G589A:p.V197M,TMPRSS2:NM_005656:exon6:c.G478A:p.V160M, |
| chr22 | 19963748 | G | A | COSV52889256 | exonic | COMT | nonsynonymous SNV | COMT:NM_001135162:exon4:c.G472A:p.V158M,COMT:NM_000754:exon4:c.G472A:p.V158M,COMT:NM_007310:exon2:c.G322A:p.V108M,COMT:NM_001135161:exon4:c.G472A:p.V158M, |
| chr22 | 22895680 | A | G | COSV66535329 | exonic | IGLL5 | nonsynonymous SNV | IGLL5:NM_001178126:exon3:c.A631G:p.T211A,IGLL5:NM_001256296:exon2:c.A406G:p.T136A, |
| chr22 | 23285182 | A | G | COSV59932309 | exonic | BCR | nonsynonymous SNV | BCR:NM_021574:exon10:c.A2387G:p.N796S,BCR:NM_004327:exon10:c.A2387G:p.N796S, |
| chr22 | 36518817 | C | T | COSV53403638 | exonic | EIF3D | nonsynonymous SNV | EIF3D:NM_003753:exon9:c.G805A:p.V269I, |
| chr22 | 43162966 | G | A | COSV53354339 | exonic | TSPO | nonsynonymous SNV | TSPO:NM_000714:exon4:c.G485A:p.R162H,TSPO:NM_001256530:exon4:c.G485A:p.R162H,TSPO:NM_001256531:exon4:c.G485A:p.R162H, |
| chr3 | 158602914 | C | A | COSV63035321 | exonic | MLF1 | nonsynonymous SNV | MLF1:NM_001195434:exon8:c.C646A:p.P216T,MLF1:NM_001130156:exon6:c.C601A:p.P201T,MLF1:NM_001195433:exon6:c.C472A:p.P158T,MLF1:NM_001195432:exon8:c.C769A:p.P257T,MLF1:NM_022443:exon6:c.C676A:p.P226T,MLF1:NM_001130157:exon7:c.C601A:p.P201T, |
| chr3 | 37012077 | A | G | COSV51613800 | exonic | MLH1 | nonsynonymous SNV | MLH1:NM_000249:exon8:c.A655G:p.I219V,MLH1:NM_001258271:exon8:c.A655G:p.I219V,MLH1:NM_001167617:exon8:c.A361G:p.I121V, |
| chr3 | 47062226 | C | T | COSV57435899 | exonic | SETD2 | nonsynonymous SNV | SETD2:NM_014159:exon14:c.G6230A:p.R2077Q, |
| chr3 | 47083895 | G | A | COSV57428846 | exonic | SETD2 | nonsynonymous SNV | SETD2:NM_014159:exon12:c.C5885T:p.P1962L, |
| chr3 | 47418189 | C | T | COSV55542934 | exonic | SCAP | nonsynonymous SNV | SCAP:NM_012235:exon16:c.G2392A:p.V798I, |
| chr3 | 52706166 | C | G | COSV51776812 | exonic | SPCS1 | nonsynonymous SNV | SPCS1:NM_014041:exon1:c.C121G:p.P41A, |
| chr4 | 40425993 | T | C | COSV55931179 | exonic | RBM47 | nonsynonymous SNV | RBM47:NM_019027:exon5:c.A1486G:p.M496V,RBM47:NM_001098634:exon7:c.A1693G:p.M565V, |
| chr5 | 32400161 | A | G | COSV54053200 | exonic | ZFR | nonsynonymous SNV | ZFR:NM_016107:exon9:c.T1559C:p.I520T, |
| chr5 | 33989413 | C | T | COSV56871911 | exonic | AMACR | nonsynonymous SNV | AMACR:NM_001167595:exon5:c.G829A:p.E277K,AMACR:NM_014324:exon5:c.G829A:p.E277K, |
| chr5 | 55664845 | C | G | COSV59422666 | exonic | SLC38A9 | nonsynonymous SNV | SLC38A9:NM_001282429:exon7:c.G173C:p.S58T,SLC38A9:NM_001258286:exon6:c.G356C:p.S119T,SLC38A9:NM_173514:exon8:c.G545C:p.S182T,SLC38A9:NM_001258287:exon7:c.G464C:p.S155T, |
| chr5 | 74685445 | T | C | COSV54668597 | exonic | HEXB | nonsynonymous SNV | HEXB:NM_000521:exon1:c.T185C:p.L62S, |
| chr6 | 131950812 | G | C | COSV63466445 | exonic | CTGF | nonsynonymous SNV | CTGF:NM_001901:exon2:c.C247G:p.H83D, |
| chr6 | 137002076 | G | A | COSV57358376 | exonic | IL20RA | nonsynonymous SNV | IL20RA:NM_014432:exon7:c.C1144T:p.L382F,IL20RA:NM_001278724:exon6:c.C745T:p.L249F,IL20RA:NM_001278723:exon6:c.C811T:p.L271F,IL20RA:NM_001278722:exon7:c.C997T:p.L333F, |
| chr6 | 158770806 | G | A | COSV61456427 | exonic | EZR | nonsynonymous SNV | EZR:NM_001111077:exon10:c.C1048T:p.R350W,EZR:NM_003379:exon9:c.C1048T:p.R350W, |
| chr6 | 29942781 | T | C | COSV65138565 | exonic | HLA-A | nonsynonymous SNV | HLA-A:NM_001242758:exon2:c.T98C:p.F33S,HLA-A:NM_002116:exon2:c.T98C:p.F33S, |
| chr6 | 29942939 | C | G | COSV65136733 | exonic | HLA-A | nonsynonymous SNV | HLA-A:NM_001242758:exon2:c.C256G:p.Q86E,HLA-A:NM_002116:exon2:c.C256G:p.Q86E, |
| chr6 | 29942948 | C | G | COSV65136740 | exonic | HLA-A | nonsynonymous SNV | HLA-A:NM_001242758:exon2:c.C265G:p.R89G,HLA-A:NM_002116:exon2:c.C265G:p.R89G, |
| chr6 | 29942953 | T | A | COSV65137266 | exonic | HLA-A | nonsynonymous SNV | HLA-A:NM_001242758:exon2:c.T270A:p.N90K,HLA-A:NM_002116:exon2:c.T270A:p.N90K, |
| chr6 | 29942982 | T | A | COSV65136627 | exonic | HLA-A | nonsynonymous SNV | HLA-A:NM_001242758:exon2:c.T299A:p.V100E,HLA-A:NM_002116:exon2:c.T299A:p.V100E, |
| chr6 | 29942984 | G | A | COSV65136637 | exonic | HLA-A | nonsynonymous SNV | HLA-A:NM_001242758:exon2:c.G301A:p.D101N,HLA-A:NM_002116:exon2:c.G301A:p.D101N, |
| chr6 | 29942994 | C | T | COSV65136647 | exonic | HLA-A | nonsynonymous SNV | HLA-A:NM_001242758:exon2:c.C311T:p.T104I,HLA-A:NM_002116:exon2:c.C311T:p.T104I, |
| chr6 | 29942996 | C | G | COSV65136655 | exonic | HLA-A | nonsynonymous SNV | HLA-A:NM_001242758:exon2:c.C313G:p.L105V,HLA-A:NM_002116:exon2:c.C313G:p.L105V, |
| chr6 | 29942997 | T | C | COSV65136680 | exonic | HLA-A | nonsynonymous SNV | HLA-A:NM_001242758:exon2:c.T314C:p.L105P,HLA-A:NM_002116:exon2:c.T314C:p.L105P, |
| chr6 | 29943000 | G | T | COSV65136690 | exonic | HLA-A | nonsynonymous SNV | HLA-A:NM_001242758:exon2:c.G317T:p.R106L,HLA-A:NM_002116:exon2:c.G317T:p.R106L, |
| chr6 | 29943337 | G | A | COSV65137272 | exonic | HLA-A | nonsynonymous SNV | HLA-A:NM_001242758:exon3:c.G413A:p.R138Q,HLA-A:NM_002116:exon3:c.G413A:p.R138Q, |
| chr6 | 29943377 | C | A | COSV65137229 | exonic | HLA-A | nonsynonymous SNV | HLA-A:NM_001242758:exon3:c.C453A:p.N151K,HLA-A:NM_002116:exon3:c.C453A:p.N151K, |
| chr6 | 29943494 | G | C | COSV65141379 | exonic | HLA-A | nonsynonymous SNV | HLA-A:NM_001242758:exon3:c.G570C:p.E190D,HLA-A:NM_002116:exon3:c.G570C:p.E190D, |
| chr6 | 29943495 | T | G | COSV65137199 | exonic | HLA-A | nonsynonymous SNV | HLA-A:NM_001242758:exon3:c.T571G:p.W191G,HLA-A:NM_002116:exon3:c.T571G:p.W191G, |
| chr6 | 31269990 | T | C | COSV66110395 | exonic | HLA-C | nonsynonymous SNV | HLA-C:NM_002117:exon5:c.A991G:p.M331V,HLA-C:NM_001243042:exon5:c.A991G:p.M331V, |
| chr6 | 31269992 | G | A | COSV66110576 | exonic | HLA-C | nonsynonymous SNV | HLA-C:NM_002117:exon5:c.C989T:p.A330V,HLA-C:NM_001243042:exon5:c.C989T:p.A330V, |
| chr6 | 31269996 | T | C | COSV66110404 | exonic | HLA-C | nonsynonymous SNV | HLA-C:NM_002117:exon5:c.A985G:p.T329A,HLA-C:NM_001243042:exon5:c.A985G:p.T329A, |
| chr6 | 31270025 | A | G | COSV66111096 | exonic | HLA-C | nonsynonymous SNV | HLA-C:NM_002117:exon5:c.T956C:p.V319A,HLA-C:NM_001243042:exon5:c.T956C:p.V319A, |
| chr6 | 31270214 | G | T | COSV66110582 | exonic | HLA-C | nonsynonymous SNV | HLA-C:NM_002117:exon4:c.C891A:p.S297R,HLA-C:NM_001243042:exon4:c.C891A:p.S297R, |
| chr6 | 31270233 | T | G | COSV66110591 | exonic | HLA-C | nonsynonymous SNV | HLA-C:NM_002117:exon4:c.A872C:p.Q291P,HLA-C:NM_001243042:exon4:c.A872C:p.Q291P, |
| chr6 | 31270276 | G | C | COSV66110239 | exonic | HLA-C | nonsynonymous SNV | HLA-C:NM_002117:exon4:c.C829G:p.Q277E,HLA-C:NM_001243042:exon4:c.C829G:p.Q277E, |
| chr6 | 31271109 | A | G | COSV66110011 | exonic | HLA-C | nonsynonymous SNV | HLA-C:NM_002117:exon3:c.T583C:p.Y195H,HLA-C:NM_001243042:exon3:c.T583C:p.Y195H, |
| chr6 | 31271229 | G | T | COSV66110004 | exonic | HLA-C | nonsynonymous SNV | HLA-C:NM_002117:exon3:c.C463A:p.R155S,HLA-C:NM_001243042:exon3:c.C463A:p.R155S, |
| chr6 | 31271337 | G | T | COSV66110498 | exonic | HLA-C | nonsynonymous SNV | HLA-C:NM_002117:exon3:c.C355A:p.L119I,HLA-C:NM_001243042:exon3:c.C355A:p.L119I, |
| chr6 | 31271800 | A | C | COSV66110102 | exonic | HLA-C | nonsynonymous SNV | HLA-C:NM_002117:exon2:c.T142G:p.S48A,HLA-C:NM_001243042:exon2:c.T142G:p.S48A, |
| chr6 | 31271845 | C | A | COSV66119680 | exonic | HLA-C | nonsynonymous SNV | HLA-C:NM_002117:exon2:c.G97T:p.D33Y,HLA-C:NM_001243042:exon2:c.G97T:p.D33Y, |
| chr6 | 31355134 | C | T | COSV69520203 | exonic | HLA-B | nonsynonymous SNV | HLA-B:NM_005514:exon5:c.G985A:p.A329T, |
| chr6 | 31355544 | G | A | COSV69521058 | exonic | HLA-B | nonsynonymous SNV | HLA-B:NM_005514:exon4:c.C668T:p.A223V, |
| chr6 | 31356183 | G | C | COSV69520290 | exonic | HLA-B | nonsynonymous SNV | HLA-B:NM_005514:exon3:c.C603G:p.D201E, |
| chr6 | 31356247 | C | A | COSV69520219 | exonic | HLA-B | nonsynonymous SNV | HLA-B:NM_005514:exon3:c.G539T:p.R180L, |

| | | | | | | | | |
|---|---|---|---|---|---|---|---|---|
| chr6 | 31356259 | T | A | COSV69520258 | exonic | HLA-B | nonsynonymous SNV | HLA-B:NM_005514:exon3:c.A527T:p.E176V, |
| chr6 | 31356377 | G | A | COSV69520464 | exonic | HLA-B | nonsynonymous SNV | HLA-B:NM_005514:exon3:c.C409T:p.H137Y, |
| chr6 | 31356430 | A | C | COSV69520377 | exonic | HLA-B | nonsynonymous SNV | HLA-B:NM_005514:exon3:c.T356G:p.L119R, |
| chr6 | 31356431 | G | A | COSV69520378 | exonic | HLA-B | nonsynonymous SNV | HLA-B:NM_005514:exon3:c.C355T:p.L119F, |
| chr6 | 31356433 | G | A | COSV69520255 | exonic | HLA-B | nonsynonymous SNV | HLA-B:NM_005514:exon3:c.C353T:p.T118I, |
| chr6 | 31356712 | C | G | COSV69520745 | exonic | HLA-B | nonsynonymous SNV | HLA-B:NM_005514:exon2:c.G319C:p.G107R, |
| chr6 | 31356714 | C | A | COSV69520695 | exonic | HLA-B | nonsynonymous SNV | HLA-B:NM_005514:exon2:c.G317T:p.R106L, |
| chr6 | 31356717 | A | G | COSV69520426 | exonic | HLA-B | nonsynonymous SNV | HLA-B:NM_005514:exon2:c.T314C:p.L105P, |
| chr6 | 31356718 | G | C | COSV69520618 | exonic | HLA-B | nonsynonymous SNV | HLA-B:NM_005514:exon2:c.C313G:p.L105V, |
| chr6 | 31356729 | C | T | COSV69520964 | exonic | HLA-B | nonsynonymous SNV | HLA-B:NM_005514:exon2:c.G302A:p.S101N, |
| chr6 | 31356739 | C | A | COSV69520380 | exonic | HLA-B | nonsynonymous SNV | HLA-B:NM_005514:exon2:c.G292T:p.D98Y, |
| chr6 | 31356748 | C | T | COSV69520384 | exonic | HLA-B | nonsynonymous SNV | HLA-B:NM_005514:exon2:c.G283A:p.A95T, |
| chr6 | 31356749 | C | G | COSV69520387 | exonic | HLA-B | nonsynonymous SNV | HLA-B:NM_005514:exon2:c.G282C:p.Q94H, |
| chr6 | 31356751 | G | T | COSV69520389 | exonic | HLA-B | nonsynonymous SNV | HLA-B:NM_005514:exon2:c.C280A:p.Q94K, |
| chr6 | 31356754 | C | T | COSV69520472 | exonic | HLA-B | nonsynonymous SNV | HLA-B:NM_005514:exon2:c.G277A:p.A93T, |
| chr6 | 31356770 | G | C | COSV69520395 | exonic | HLA-B | nonsynonymous SNV | HLA-B:NM_005514:exon2:c.C261G:p.N87K, |
| chr6 | 31356772 | T | C | COSV69520398 | exonic | HLA-B | nonsynonymous SNV | HLA-B:NM_005514:exon2:c.A259G:p.N87D, |
| chr6 | 31356838 | C | T | COSV69520400 | exonic | HLA-B | nonsynonymous SNV | HLA-B:NM_005514:exon2:c.G193A:p.A65T, |
| chr6 | 31356864 | T | A | COSV69520244 | exonic | HLA-B | nonsynonymous SNV | HLA-B:NM_005514:exon2:c.A167T:p.Q56L, |
| chr6 | 31356925 | C | T | COSV69520570 | exonic | HLA-B | nonsynonymous SNV | HLA-B:NM_005514:exon2:c.G106A:p.V36M, |
| chr6 | 31356928 | A | C | COSV69520756 | exonic | HLA-B | nonsynonymous SNV | HLA-B:NM_005514:exon2:c.T103G:p.S35A, |
| chr6 | 31356934 | A | G | COSV69520552 | exonic | HLA-B | nonsynonymous SNV | HLA-B:NM_005514:exon2:c.T97C:p.Y33H, |
| chr6 | 32580249 | G | C | COSV63511877 | exonic | HLA-DRB1 | nonsynonymous SNV | HLA-DRB1:NM_002124:exon5:c.C785G:p.T262R, |
| chr6 | 32581675 | C | G | COSV63511369 | exonic | HLA-DRB1 | nonsynonymous SNV | HLA-DRB1:NM_002124:exon3:c.G534C:p.Q178H, |
| chr6 | 32581704 | C | T | COSV63511807 | exonic | HLA-DRB1 | nonsynonymous SNV | HLA-DRB1:NM_002124:exon3:c.G505A:p.A169T, |
| chr6 | 32581724 | A | C | COSV63513549 | exonic | HLA-DRB1 | nonsynonymous SNV | HLA-DRB1:NM_002124:exon3:c.T485G:p.L162R, |
| chr6 | 32584181 | C | T | COSV63515717 | exonic | HLA-DRB1 | nonsynonymous SNV | HLA-DRB1:NM_002124:exon2:c.G298A:p.A100T, |
| chr6 | 32584252 | A | T | COSV63512106 | exonic | HLA-DRB1 | nonsynonymous SNV | HLA-DRB1:NM_002124:exon2:c.T227A:p.F76Y, |
| chr6 | 32589658 | A | C | COSV63512004 | exonic | HLA-DRB1 | nonsynonymous SNV | HLA-DRB1:NM_002124:exon1:c.T85G:p.S29A, |
| chr6 | 32641349 | T | C | COSV58239937 | exonic | HLA-DQA1 | nonsynonymous SNV | HLA-DQA1:NM_002122:exon2:c.T122C:p.F41S, |
| chr6 | 32641477 | G | T | COSV58237257 | exonic | HLA-DQA1 | nonsynonymous SNV | HLA-DQA1:NM_002122:exon2:c.G250T:p.G84C, |
| chr6 | 32641478 | G | T | COSV58237262 | exonic | HLA-DQA1 | nonsynonymous SNV | HLA-DQA1:NM_002122:exon2:c.G251T:p.G84V, |
| chr6 | 32641487 | G | C | COSV58238124 | exonic | HLA-DQA1 | nonsynonymous SNV | HLA-DQA1:NM_002122:exon2:c.G260C:p.R87T, |
| chr6 | 32641494 | G | C | COSV58238131 | exonic | HLA-DQA1 | nonsynonymous SNV | HLA-DQA1:NM_002122:exon2:c.G267C:p.M89I, |
| chr6 | 32641501 | G | C | COSV58238145 | exonic | HLA-DQA1 | nonsynonymous SNV | HLA-DQA1:NM_002122:exon2:c.G274C:p.A92P, |
| chr6 | 32641502 | C | T | COSV58238153 | exonic | HLA-DQA1 | nonsynonymous SNV | HLA-DQA1:NM_002122:exon2:c.C275T:p.A92V, |
| chr6 | 32641535 | A | C | COSV58237540 | exonic | HLA-DQA1 | nonsynonymous SNV | HLA-DQA1:NM_002122:exon2:c.A308C:p.Y103S, |
| chr6 | 32642096 | G | C | COSV58237000 | exonic | HLA-DQA1 | nonsynonymous SNV | HLA-DQA1:NM_002122:exon3:c.G456C:p.Q152H, |
| chr6 | 32664870 | T | C | COSV66574305 | exonic | HLA-DQB1 | nonsynonymous SNV | HLA-DQB1:NM_001243961:exon2:c.A307G:p.T103A,HLA-DQB1:NM_002123:exon2:c.A307G:p.T103A, |
| chr6 | 32664911 | T | A | COSV66574314 | exonic | HLA-DQB1 | nonsynonymous SNV | HLA-DQB1:NM_001243961:exon2:c.A266T:p.D89V,HLA-DQB1:NM_002123:exon2:c.A266T:p.D89V, |
| chr6 | 32664968 | G | A | COSV66571252 | exonic | HLA-DQB1 | nonsynonymous SNV | HLA-DQB1:NM_001243961:exon2:c.C209T:p.A70V,HLA-DQB1:NM_002123:exon2:c.C209T:p.A70V, |
| chr6 | 32664993 | A | G | COSV66574347 | exonic | HLA-DQB1 | nonsynonymous SNV | HLA-DQB1:NM_001243961:exon2:c.T184C:p.Y62H,HLA-DQB1:NM_002123:exon2:c.T184C:p.Y62H, |
| chr6 | 42938646 | G | T | COSV65709724 | exonic | CNPY3 | nonsynonymous SNV | CNPY3:NM_006586:exon6:c.G692T:p.S231I, |
| chr6 | 69697573 | A | T | COSV65296211 | exonic | LMBRD1 | nonsynonymous SNV | LMBRD1:NM_018368:exon14:c.T1407A:p.D469E, |
| chr7 | 102274250 | G | A | COSV52922844 | exonic | CUX1 | nonsynonymous SNV | CUX1:NM_001202546:exon15:c.G1273A:p.A425T,CUX1:NM_181500:exon16:c.G1384A:p.A462T,CUX1:NM_001202545:exon15:c.G1252A:p.A418T,CUX1:NM_001202544:exon15:c.G1342A:p.A448T,CUX1:NM_001913:exon16:c.G1390A:p.A464T, |
| chr7 | 105090116 | G | A | COSV57576667 | exonic | KMT2E | nonsynonymous SNV | KMT2E:NM_018682:exon13:c.G1466A:p.R489Q,KMT2E:NM_182931:exon14:c.G1466A:p.R489Q, |
| chr7 | 127807503 | G | A | COSV61241417 | exonic | SND1 | nonsynonymous SNV | SND1:NM_014390:exon11:c.G1172A:p.R391H, |
| chr7 | 138627377 | A | C | COSV55991488 | exonic | SVOPL | nonsynonymous SNV | SVOPL:NM_001139456:exon11:c.T1154G:p.F385C,SVOPL:NM_174959:exon8:c.T698G:p.F233C, |
| chr7 | 150793288 | C | T | COSV58438904 | exonic | TMEM176B | nonsynonymous SNV | TMEM176B:NM_014020:exon5:c.G400A:p.A134T,TMEM176B:NM_001101312:exon5:c.G400A:p.A134T,TMEM176B:NM_001101311:exon5:c.G400A:p.A134T,TMEM176B:NM_001101314:exon4:c.G289A:p.A97T, |
| chr7 | 150793996 | T | G | COSV58439821 | exonic | TMEM176B | nonsynonymous SNV | TMEM176B:NM_014020:exon3:c.A280C:p.S94R,TMEM176B:NM_001101312:exon3:c.A280C:p.S94R,TMEM176B:NM_001101311:exon3:c.A280C:p.S94R, |
| chr7 | 35670232 | C | T | COSV60943904 | exonic | HERPUD2 | nonsynonymous SNV | HERPUD2:NM_022373:exon4:c.G322A:p.A108T, |
| chr7 | 5987357 | G | A | COSV56220401 | exonic | PMS2 | nonsynonymous SNV | PMS2:NM_000535:exon11:c.C1408T:p.P470S, |
| chr8 | 100704992 | C | T | COSV59397438 | exonic | PABPC1 | nonsynonymous SNV | PABPC1:NM_002568:exon13:c.G1752A:p.M584I, |
| chr8 | 100705025 | C | T | COSV59400560 | exonic | PABPC1 | nonsynonymous SNV | PABPC1:NM_002568:exon13:c.G1719A:p.M573I, |
| chr8 | 100712402 | C | T | COSV59399520 | exonic | PABPC1 | nonsynonymous SNV | PABPC1:NM_002568:exon7:c.G932A:p.R311Q, |
| chr8 | 100712765 | C | T | COSV59400273 | exonic | PABPC1 | nonsynonymous SNV | PABPC1:NM_002568:exon6:c.G763A:p.E255K, |
| chr8 | 11821811 | G | A | COSV55042913 | exonic | FDFT1 | nonsynonymous SNV | FDFT1:NM_004462:exon4:c.G443A:p.R148Q,FDFT1:NM_001287751:exon3:c.G188A:p.R63Q,FDFT1:NM_001287743:exon5:c.G443A:p.R148Q,FDFT1:NM_001287750:exon3:c.G620A:p.R207Q,FDFT1:NM_001287742:exon6:c.G443A:p.R148Q,FDFT1:NM_001287747:exon4:c.G251A:p.R84Q,FDFT1:NM_001287745:exon5:c.G251A:p.R84Q,FDFT1:NM_001287744:exon4:c.G251A:p.R84Q,FDFT1:NM_001287749:exon4:c.G251A:p.R84Q,FDFT1:NM_001287748:exon4:c.G251A:p.R84Q, |

| | | | | | | | | |
|---|---|---|---|---|---|---|---|---|
| chr8 | 11853379 | G | C | COSV61540513 | exonic | CTSB | nonsynonymous SNV | CTSB:NM_001908:exon2:c.C76G:p.L26V,CTSB:NM_147783:exon3:c.C76G:p.L26V,CTSB:NM_147781:exon3:c.C76G:p.L26V,CTSB:NM_147780:exon4:c.C76G:p.L26V,CTSB:NM_147782:exon3:c.C76G:p.L26V, |
| chr8 | 27500988 | A | G | COSV66844518 | exonic | EPHX2 | nonsynonymous SNV | EPHX2:NM_001256482:exon2:c.A5G:p.K2R,EPHX2:NM_001256484:exon2:c.A5G:p.K2R,EPHX2:NM_001979:exon2:c.A164G:p.K55R, |
| chr8 | 31081188 | G | A | COSV53296756 | exonic | WRN | nonsynonymous SNV | WRN:NM_000553:exon9:c.G1161A:p.M387I, |
| chr8 | 51820490 | T | A | COSV62119484 | exonic | PCMTD1 | nonsynonymous SNV | PCMTD1:NM_001286783:exon6:c.A407T:p.N136I,PCMTD1:NM_052937:exon6:c.A935T:p.N312I,PCMTD1:NM_001286782:exon4:c.A707T:p.N236I, |
| chr8 | 63038678 | C | T | COSV52649507 | exonic | GGH | nonsynonymous SNV | GGH:NM_003878:exon1:c.G91A:p.A31T, |
| chr8 | 63038753 | A | G | COSV52650077 | exonic | GGH | nonsynonymous SNV | GGH:NM_003878:exon1:c.T16C:p.C6R, |
| chr8 | 94130926 | C | T | COSV50345009 | exonic | CDH17 | nonsynonymous SNV | CDH17:NM_001144663:exon16:c.G2234A:p.R745H,CDH17:NM_004063:exon16:c.G2234A:p.R745H, |
| chr8 | 94130958 | C | G | COSV50336268 | exonic | CDH17 | nonsynonymous SNV | CDH17:NM_001144663:exon16:c.G2202C:p.E734D,CDH17:NM_004063:exon16:c.G2202C:p.E734D, |
| chr9 | 113370054 | C | T | COSV53057170 | exonic | BSPRY | nonsynonymous SNV | BSPRY:NM_017688:exon6:c.C1121T:p.T374I, |
| chr9 | 21816759 | G | A | COSV66477132 | exonic | MTAP | nonsynonymous SNV | MTAP:NM_002451:exon3:c.G166A:p.V56I, |
| chr9 | 33264542 | C | G | COSV56303169 | exonic | BAG1 | nonsynonymous SNV | BAG1:NM_004323:exon1:c.G133C:p.G45R, |
| chr1 | 109757819 | G | A | COSV62609036 | exonic | EPS8L3 | nonsynonymous SNV | EPS8L3:NM_139053:exon10:c.C880T:p.H294Y,EPS8L3:NM_133181:exon10:c.C877T:p.H293Y,EPS8L3:NM_024526:exon10:c.C877T:p.H293Y, |
| chr1 | 116401151 | G | A | COSV55193103 | exonic | ATP1A1 | nonsynonymous SNV | ATP1A1:NM_001160233:exon20:c.G2740A:p.V914M,ATP1A1:NM_001160234:exon20:c.G2647A:p.V883M,ATP1A1:NM_000701:exon20:c.G2740A:p.V914M, |
| chr1 | 116403975 | G | A | COSV55190853 | exonic | ATP1A1 | nonsynonymous SNV | ATP1A1:NM_001160233:exon22:c.G3043A:p.G1015S,ATP1A1:NM_001160234:exon22:c.G2950A:p.G984S,ATP1A1:NM_000701:exon22:c.G3043A:p.G1015S, |
| chr1 | 146156641 | G | A | COSV61245135 | exonic | NOTCH2NL | nonsynonymous SNV | NOTCH2NL:NM_203458:exon4:c.C473T:p.T158I, |
| chr1 | 154170448 | C | T | COSV55136361 | exonic | TPM3 | nonsynonymous SNV | TPM3:NM_152263:exon8:c.G727A:p.A243T,TPM3:NM_001278190:exon6:c.G553A:p.A185T,TPM3:NM_001043352:exon7:c.G616A:p.A206T,TPM3:NM_001043351:exon7:c.G616A:p.A206T,TPM3:NM_001043353:exon7:c.G616A:p.A206T,TPM3:NM_001278191:exon7:c.G346A:p.A116T,TPM3:NM_001278188:exon6:c.G418A:p.A140T,TPM3:NM_001278189:exon7:c.G616A:p.A206T,TPM3:NM_153649:exon7:c.G616A:p.A206T, |
| chr1 | 15770439 | C | T | COSV60030004 | exonic | FBLIM1 | nonsynonymous SNV | FBLIM1:NM_001024215:exon5:c.C572T:p.S191F,FBLIM1:NM_001024216:exon2:c.C281T:p.S94F,FBLIM1:NM_017556:exon6:c.C572T:p.S191F, |
| chr1 | 20589208 | A | C | COSV66751560 | exonic | CDA | nonsynonymous SNV | CDA:NM_001785:exon1:c.A79C:p.K27Q, |
| chr1 | 20661380 | G | C | COSV56117446 | exonic | DDOST | nonsynonymous SNV | DDOST:NM_005216:exon1:c.C22G:p.R8G, |
| chr1 | 46668139 | T | C | COSV61304776 | exonic | ATPAF1 | nonsynonymous SNV | ATPAF1:NM_022745:exon1:c.A253G:p.S85G,ATPAF1:NM_001042546:exon1:c.A253G:p.S85G, |
| chr1 | 61990342 | G | A | COSV56249041 | exonic | INADL | nonsynonymous SNV | INADL:NM_176877:exon28:c.G3845A:p.R1282H, |
| chr1 | 68137903 | C | T | COSV52037496 | exonic | WLS | nonsynonymous SNV | WLS:NM_001002292:exon11:c.G1387A:p.V463I,WLS:NM_024911:exon11:c.G1393A:p.V465I,WLS:NM_001193334:exon10:c.G1120A:p.V374I, |
| chr10 | 87933135 | G | A | COSV64288974 | exonic | PTEN | nonsynonymous SNV | PTEN:NM_000314:exon5:c.G376A:p.A126T,PTEN:NM_001304717:exon6:c.G895A:p.A299T, |
| chr10 | 87952143 | G | A | COSV64288961 | exonic | PTEN | nonsynonymous SNV | PTEN:NM_000314:exon6:c.G518A:p.R173H,PTEN:NM_001304717:exon7:c.G1037A:p.R346H, |
| chr11 | 237087 | A | G | COSV61500926 | exonic | PSMD13 | nonsynonymous SNV | PSMD13:NM_175932:exon1:c.A38G:p.N13S,PSMD13:NM_002817:exon1:c.A38G:p.N13S, |
| chr11 | 34895110 | C | T | COSV53506741 | exonic | APIP | nonsynonymous SNV | APIP:NM_015957:exon2:c.G58A:p.D20N, |
| chr11 | 35221726 | G | A | COSV53530853 | exonic | CD44 | nonsynonymous SNV | CD44:NM_001001389:exon16:c.G1889A:p.R630Q,CD44:NM_000610:exon17:c.G2018A:p.R673Q,CD44:NM_001202556:exon7:c.G812A:p.R271Q,CD44:NM_001202555:exon9:c.G1079A:p.R360Q,CD44:NM_001202557:exon8:c.G875A:p.R292Q,CD44:NM_001001390:exon11:c.G1271A:p.R424Q,CD44:NM_001001391:exon8:c.G875A:p.R292Q, |
| chr11 | 62672097 | G | A | COSV53656335 | exonic | UQCC3 | nonsynonymous SNV | UQCC3:NM_001085372:exon2:c.G265A:p.G89S, |
| chr11 | 67585218 | A | G | COSV66992376 | exonic | GSTP1 | nonsynonymous SNV | GSTP1:NM_000852:exon5:c.A313G:p.I105V, |
| chr11 | 95923947 | T | G | COSV60579185 | exonic | MTMR2 | nonsynonymous SNV | MTMR2:NM_016156:exon1:c.A8C:p.K3T, |
| chr12 | 122327956 | G | C | COSV56798569 | exonic | CLIP1 | nonsynonymous SNV | CLIP1:NM_002956:exon16:c.C3207G:p.D1069E,CLIP1:NM_001247997:exon17:c.C3240G:p.D1080E,CLIP1:NM_198240:exon15:c.C3102G:p.D1034E, |
| chr12 | 14503834 | G | C | COSV53694489 | exonic | PLBD1 | nonsynonymous SNV | PLBD1:NM_024829:exon11:c.C1600G:p.P534A, |
| chr12 | 25245350 | C | A | COSV55497419 | exonic | KRAS | nonsynonymous SNV | KRAS:NM_004985:exon2:c.G35T:p.G12V,KRAS:NM_033360:exon2:c.G35T:p.G12V, |
| chr13 | 32332592 | A | C | COSV66448817 | exonic | BRCA2 | nonsynonymous SNV | BRCA2:NM_000059:exon10:c.A1114C:p.N372H, |
| chr13 | 39655754 | G | A | COSV62995992 | exonic | COG6 | nonsynonymous SNV | COG6:NM_020751:exon1:c.G28A:p.A10T,COG6:NM_001145079:exon1:c.G28A:p.A10T, |
| chr13 | 39655820 | T | A | COSV62995068 | exonic | COG6 | nonsynonymous SNV | COG6:NM_020751:exon1:c.T94A:p.C32S,COG6:NM_001145079:exon1:c.T94A:p.C32S, |
| chr14 | 73105954 | G | A | COSV56202337 | exonic | RBM25 | nonsynonymous SNV | RBM25:NM_021239:exon11:c.G1250A:p.R417Q, |
| chr15 | 74035775 | A | G | COSV51443885 | exonic | PML | nonsynonymous SNV | PML:NM_033250:exon7:c.A2170G:p.S724G,PML:NM_033239:exon8:c.A2314G:p.S772G, |
| chr15 | 74035800 | G | T | COSV51443888 | exonic | PML | nonsynonymous SNV | PML:NM_033250:exon7:c.G2195T:p.G732V,PML:NM_033239:exon8:c.G2339T:p.G780V, |
| chr16 | 84745079 | A | G | COSV54746826 | exonic | USP10 | nonsynonymous SNV | USP10:NM_001272075:exon5:c.A610G:p.M204V,USP10:NM_005153:exon4:c.A598G:p.M200V, |
| chr16 | 89100686 | T | C | COSV58076417 | exonic | ACSF3 | nonsynonymous SNV | ACSF3:NM_001127214:exon2:c.T5C:p.L2P,ACSF3:NM_001243279:exon3:c.T5C:p.L2P,ACSF3:NM_174917:exon3:c.T5C:p.L2P, |
| chr17 | 39657827 | G | A | COSV60419443 | exonic | STARD3 | nonsynonymous SNV | STARD3:NM_001165937:exon4:c.G350A:p.R117Q,STARD3:NM_006804:exon4:c.G350A:p.R117Q,STARD3:NM_001165938:exon4:c.G350A:p.R117Q, |
| chr17 | 39727784 | C | G | COSV54064104 | exonic | ERBB2 | nonsynonymous SNV | ERBB2:NM_001289936:exon31:c.C3463G:p.P1155A,ERBB2:NM_004448:exon27:c.C3508G:p.P1170A,ERBB2:NM_001005862:exon30:c.C3418G:p.P1140A, |
| chr17 | 39905943 | G | A | COSV58779652 | exonic | GSDMB | nonsynonymous SNV | GSDMB:NM_018530:exon6:c.C865T:p.P289S,GSDMB:NM_001165959:exon8:c.C904T:p.P302S,GSDMB:NM_001042471:exon8:c.C892T:p.P298S,GSDMB:NM_001165958:exon9:c.C931T:p.P311S, |
| chr17 | 39905964 | C | T | COSV58779510 | exonic | GSDMB | nonsynonymous SNV | GSDMB:NM_018530:exon6:c.G844A:p.G282R,GSDMB:NM_001165959:exon8:c.G883A:p.G295R,GSDMB:NM_001042471:exon8:c.G871A:p.G291R,GSDMB:NM_001165958:exon9:c.G910A:p.G304R, |

| | | | | | | | | |
|---|---|---|---|---|---|---|---|---|
| chr19 | 18386214 | G | C | COSV53250533 | exonic | GDF15 | nonsynonymous SNV | GDF15:NM_004864:exon1:c.G25C:p.V9L, |
| chr19 | 23359488 | A | G | COSV56084671 | exonic | ZNF91 | nonsynonymous SNV | ZNF91:NM_003430:exon4:c.T3491C:p.L1164P,ZNF91:NM_001300951:exon3:c.T3395C:p.L1132P, |
| chr19 | 23359489 | G | C | COSV56084686 | exonic | ZNF91 | nonsynonymous SNV | ZNF91:NM_003430:exon4:c.C3490G:p.L1164V,ZNF91:NM_001300951:exon3:c.C3394G:p.L1132V, |
| chr19 | 23360293 | T | C | COSV56080094 | exonic | ZNF91 | nonsynonymous SNV | ZNF91:NM_003430:exon4:c.A2686G:p.T896A,ZNF91:NM_001300951:exon3:c.A2590G:p.T864A, |
| chr19 | 23362645 | A | G | COSV56080539 | exonic | ZNF91 | nonsynonymous SNV | ZNF91:NM_003430:exon4:c.T334C:p.Y112H,ZNF91:NM_001300951:exon3:c.T238C:p.Y80H, |
| chr19 | 33114406 | T | C | COSV50114723 | exonic | GPATCH1 | nonsynonymous SNV | GPATCH1:NM_018025:exon15:c.T2183C:p.L728S, |
| chr19 | 44908684 | T | C | COSV52985231 | exonic | APOE | nonsynonymous SNV | APOE:NM_000041:exon4:c.T388C:p.C130R,APOE:NM_001302691:exon4:c.T388C:p.C130R,APOE:NM_001302689:exon4:c.T388C:p.C130R,APOE:NM_001302690:exon4:c.T388C:p.C130R,APOE:NM_001302688:exon4:c.T466C:p.C156R, |
| chr19 | 49880379 | T | C | COSV55578602 | exonic | TBC1D17 | nonsynonymous SNV | TBC1D17:NM_001168222:exon3:c.T197C:p.L66P,TBC1D17:NM_024682:exon4:c.T296C:p.L99P, |
| chr19 | 51347036 | G | A | COSV58535703 | exonic | ETFB | nonsynonymous SNV | ETFB:NM_001014763:exon4:c.C734T:p.T245M,ETFB:NM_001985:exon5:c.C461T:p.T154M, |
| chr2 | 143040620 | A | G | COSV51580314 | exonic | KYNU | nonsynonymous SNV | KYNU:NM_001199241:exon14:c.A1234G:p.K412E,KYNU:NM_003937:exon13:c.A1234G:p.K412E, |
| chr2 | 197497182 | C | T | COSV52099976 | exonic | HSPD1 | nonsynonymous SNV | HSPD1:NM_002156:exon3:c.G385A:p.E129K,HSPD1:NM_199440:exon3:c.G385A:p.E129K, |
| chr2 | 32736791 | A | G | COSV58653535 | exonic | TTC27 | nonsynonymous SNV | TTC27:NM_017735:exon12:c.A1427G:p.Y476C,TTC27:NM_001193509:exon12:c.A1277G:p.Y426C, |
| chr2 | 74958729 | G | T | COSV52050939 | exonic | POLE4 | nonsynonymous SNV | POLE4:NM_019896:exon1:c.G50T:p.G17V, |
| chr2 | 99442408 | C | T | COSV51481954 | exonic | REV1 | nonsynonymous SNV | REV1:NM_016316:exon5:c.G412A:p.V138M,REV1:NM_001037872:exon5:c.G412A:p.V138M, |
| chr20 | 1118880 | T | G | COSV55672896 | exonic | PSMF1 | nonsynonymous SNV | PSMF1:NM_178578:exon2:c.T107G:p.F36C,PSMF1:NM_006814:exon1:c.T107G:p.F36C, |
| chr20 | 25278370 | G | T | COSV53816254 | exonic | PYGB | nonsynonymous SNV | PYGB:NM_002862:exon8:c.G907T:p.A303S, |
| chr20 | 58909398 | T | C | COSV55672955 | exonic | GNAS | nonsynonymous SNV | GNAS:NM_001309840:exon8:c.T457C:p.F153L,GNAS:NM_000516:exon8:c.T634C:p.F212L,GNAS:NM_080426:exon7:c.T592C:p.F198L,GNAS:NM_080425:exon8:c.T2563C:p.F855L,GNAS:NM_001077489:exon7:c.T589C:p.F197L,GNAS:NM_001309861:exon8:c.T457C:p.F153L,GNAS:NM_001077488:exon8:c.T637C:p.F213L, |
| chr20 | 58909759 | G | A | COSV55672773 | exonic | GNAS | nonsynonymous SNV | GNAS:NM_080426:exon9:c.G752A:p.R251H,GNAS:NM_080425:exon10:c.G2723A:p.R908H,GNAS:NM_001309840:exon10:c.G617A:p.R206H,GNAS:NM_001077489:exon9:c.G749A:p.R250H,GNAS:NM_001309861:exon10:c.G617A:p.R206H,GNAS:NM_000516:exon10:c.G794A:p.R265H,GNAS:NM_001077488:exon10:c.G797A:p.R266H, |
| chr21 | 33553954 | C | T | COSV51652492 | exonic | SON | nonsynonymous SNV | SON:NM_001291411:exon3:c.C4723T:p.R1575C,SON:NM_032195:exon3:c.C4723T:p.R1575C,SON:NM_138927:exon3:c.C4723T:p.R1575C, |
| chr3 | 124908177 | T | G | COSV60698847 | exonic | MUC13 | nonsynonymous SNV | MUC13:NM_033049:exon11:c.A1509C:p.R503S, |
| chr3 | 126107216 | T | C | COSV56412084 | exonic | ALDH1L1 | nonsynonymous SNV | ALDH1L1:NM_001270365:exon19:c.A2075G:p.D692G,ALDH1L1:NM_012190:exon21:c.A2378G:p.D793G,ALDH1L1:NM_001270364:exon21:c.A2408G:p.D803G, |
| chr3 | 142562770 | A | G | COSV63383325 | exonic | ATR | nonsynonymous SNV | ATR:NM_001184:exon4:c.T632C:p.M211T, |
| chr3 | 185048513 | T | C | COSV54997868 | exonic | VPS8 | nonsynonymous SNV | VPS8:NM_001009921:exon46:c.T4091C:p.I1364T,VPS8:NM_015303:exon46:c.T4085C:p.I1362T, |
| chr3 | 46408579 | T | A | COSV62193340 | exonic | CCRL2 | nonsynonymous SNV | CCRL2:NM_003965:exon2:c.T500A:p.F167Y,CCRL2:NM_001130910:exon2:c.T536A:p.F179Y, |
| chr3 | 47914934 | C | T | COSV53135994 | exonic | MAP4 | nonsynonymous SNV | MAP4:NM_002375:exon8:c.G1882A:p.V628I,MAP4:NM_001134364:exon8:c.G1882A:p.V628I, |
| chr3 | 47916547 | G | T | COSV64236753 | exonic | MAP4 | nonsynonymous SNV | MAP4:NM_002375:exon7:c.C1280A:p.S427Y,MAP4:NM_001134364:exon7:c.C1280A:p.S427Y, |
| chr3 | 52693734 | G | A | COSV56477800 | exonic | GNL3 | nonsynonymous SNV | GNL3:NM_206826:exon13:c.G1391A:p.R464Q,GNL3:NM_014366:exon13:c.G1427A:p.R476Q,GNL3:NM_206825:exon13:c.G1391A:p.R464Q, |
| chr4 | 103092639 | T | C | COSV56477889 | exonic | BDH2 | nonsynonymous SNV | BDH2:NM_020139:exon4:c.A209G:p.N70S, |
| chr4 | 87494484 | G | C | COSV56802379 | exonic | SPARCL1 | nonsynonymous SNV | SPARCL1:NM_001128310:exon5:c.C316G:p.H106D,SPARCL1:NM_004684:exon4:c.C316G:p.H106D, |
| chr4 | 87495036 | G | T | COSV56802387 | exonic | SPARCL1 | nonsynonymous SNV | SPARCL1:NM_001128310:exon4:c.C146A:p.A49D,SPARCL1:NM_004684:exon3:c.C146A:p.A49D, |
| chr5 | 116092684 | T | G | COSV57234444 | exonic | COMMD10 | nonsynonymous SNV | COMMD10:NM_016144:exon4:c.T383G:p.I128S,COMMD10:NM_001308080:exon4:c.T341G:p.I114S, |
| chr5 | 33998663 | C | A | COSV59460034 | exonic | AMACR | nonsynonymous SNV | AMACR:NM_001167595:exon4:c.G717T:p.Q239H,AMACR:NM_203382:exon3:c.G556T:p.V186F,AMACR:NM_014324:exon4:c.G717T:p.Q239H, |
| chr5 | 36985201 | A | G | COSV56947629 | exonic | NIPBL | nonsynonymous SNV | NIPBL:NM_133433:exon10:c.A2021G:p.N674S,NIPBL:NM_015384:exon10:c.A2021G:p.N674S, |
| chr5 | 69400113 | T | G | COSV51456873 | exonic | RAD17 | nonsynonymous SNV | RAD17:NM_133344:exon15:c.T1637G:p.L546R,RAD17:NM_133342:exon16:c.T1637G:p.L546R,RAD17:NM_133339:exon14:c.T1670G:p.L557R,RAD17:NM_002873:exon15:c.T1637G:p.L546R,RAD17:NM_133340:exon14:c.T1142G:p.L381R,RAD17:NM_133338:exon17:c.T1637G:p.L546R,RAD17:NM_133341:exon14:c.T1379G:p.L460R,RAD17:NM_001278622:exon16:c.T1637G:p.L546R,RAD17:NM_133343:exon16:c.T1637G:p.L546R, |
| chr5 | 71504711 | G | A | COSV62429660 | exonic | BDP1 | nonsynonymous SNV | BDP1:NM_018429:exon16:c.G2332A:p.V778M, |
| chr5 | 71510630 | G | A | COSV62429841 | exonic | BDP1 | nonsynonymous SNV | BDP1:NM_018429:exon17:c.G3538A:p.G1180S, |
| chr5 | 71510822 | T | A | COSV62430362 | exonic | BDP1 | nonsynonymous SNV | BDP1:NM_018429:exon17:c.T3730A:p.F1244I, |
| chr6 | 125957084 | G | C | COSV57649843 | exonic | HINT3 | nonsynonymous SNV | HINT3:NM_138571:exon1:c.G107C:p.G36A, |
| chr6 | 143503967 | G | T | COSV50019718 | exonic | FUCA2 | nonsynonymous SNV | FUCA2:NM_032020:exon3:c.C698A:p.A233E, |
| chr6 | 151615542 | G | A | COSV53337542 | exonic | CCDC170 | nonsynonymous SNV | CCDC170:NM_025059:exon10:c.G1810A:p.V604I, |
| chr6 | 152975406 | G | C | COSV57670920 | exonic | FBXO5 | nonsynonymous SNV | FBXO5:NM_012177:exon2:c.C319G:p.Q107E,FBXO5:NM_001142522:exon2:c.C181G:p.Q61E, |
| chr6 | 167305948 | G | A | COSV57807281 | exonic | UNC93A | nonsynonymous SNV | UNC93A:NM_001143947:exon5:c.G748A:p.V250I,UNC93A:NM_018974:exon6:c.G874A:p.V292I, |
| chr6 | 167315286 | T | C | COSV57806804 | exonic | UNC93A | nonsynonymous SNV | UNC93A:NM_001143947:exon7:c.T1082C:p.M361T,UNC93A:NM_018974:exon8:c.T1208C:p.M403T, |
| chr6 | 170553212 | G | C | COSV51529993 | exonic | PSMB1 | nonsynonymous SNV | PSMB1:NM_002793:exon1:c.C31G:p.P11A, |

| | | | | | | | | |
|---|---|---|---|---|---|---|---|---|
| chr6 | 18130729 | C | T | COSV59428760 | exonic | TPMT | nonsynonymous SNV | TPMT:NM_000367:exon9:c.G677A:p.R226Q, |
| chr6 | 31271103 | C | T | COSV66112160 | exonic | HLA-C | nonsynonymous SNV | HLA-C:NM_002117:exon3:c.G589A:p.E197K,HLA-C:NM_001243042:exon3:c.G589A:p.E197K, |
| chr6 | 31271132 | G | A | COSV66110017 | exonic | HLA-C | nonsynonymous SNV | HLA-C:NM_002117:exon3:c.C560T:p.T187M,HLA-C:NM_001243042:exon3:c.C560T:p.T187M, |
| chr6 | 31271133 | T | G | COSV66110024 | exonic | HLA-C | nonsynonymous SNV | HLA-C:NM_002117:exon3:c.A559C:p.T187P,HLA-C:NM_001243042:exon3:c.A559C:p.T187P, |
| chr6 | 31271331 | T | A | COSV66110137 | exonic | HLA-C | nonsynonymous SNV | HLA-C:NM_002117:exon3:c.A361T:p.R121W,HLA-C:NM_001243042:exon3:c.A361T:p.R121W, |
| chr6 | 31271339 | G | A | COSV66110997 | exonic | HLA-C | nonsynonymous SNV | HLA-C:NM_002117:exon3:c.C353T:p.T118I,HLA-C:NM_001243042:exon3:c.C353T:p.T118I, |
| chr6 | 31356274 | C | A | COSV69520985 | exonic | HLA-B | nonsynonymous SNV | HLA-B:NM_005514:exon3:c.G512T:p.W171L, |
| chr6 | 34246545 | C | G | COSV58988465 | exonic | C6ORF1 | nonsynonymous SNV | C6orf1:NM_001287398:exon4:c.G389C:p.G130A,C6orf1:NM_001008703:exon5:c.G449C:p.G150A,C6orf1:NM_001287397:exon4:c.G389C:p.G130A,C6orf1:NM_001008704:exon5:c.G449C:p.G150A,C6orf1:NM_178508:exon5:c.G449C:p.G150A,C6orf1:NM_001287396:exon5:c.G449C:p.G150A, |
| chr6 | 7402828 | G | A | COSV53572602 | exonic | RIOK1 | nonsynonymous SNV | RIOK1:NM_031480:exon8:c.G698A:p.G233D, |
| chr7 | 148828812 | C | G | COSV57449162 | exonic | EZH2 | nonsynonymous SNV | EZH2:NM_001203249:exon6:c.G526C:p.D176H,EZH2:NM_004456:exon6:c.G553C:p.D185H,EZH2:NM_001203247:exon6:c.G553C:p.D185H,EZH2:NM_001203248:exon6:c.G526C:p.D176H,EZH2:NM_152998:exon5:c.G436C:p.D146H, |
| chr7 | 150793191 | C | T | COSV58441166 | exonic | TMEM176B | nonsynonymous SNV | TMEM176B:NM_014020:exon5:c.G497A:p.R166H,TMEM176B:NM_001101312:exon5:c.G497A:p.R166H,TMEM176B:NM_001101311:exon5:c.G497A:p.R166H,TMEM176B:NM_001101314:exon4:c.G386A:p.R129H, |
| chr7 | 150793249 | C | T | COSV58439227 | exonic | TMEM176B | nonsynonymous SNV | TMEM176B:NM_014020:exon5:c.G439A:p.V147M,TMEM176B:NM_001101312:exon5:c.G439A:p.V147M,TMEM176B:NM_001101311:exon5:c.G439A:p.V147M,TMEM176B:NM_001101314:exon4:c.G328A:p.V110M, |
| chr7 | 30595045 | C | G | COSV66827983 | exonic | GARS | nonsynonymous SNV | GARS:NM_002047:exon1:c.C124G:p.P42A, |
| chr7 | 36616429 | G | C | COSV51741603 | exonic | AOAH | nonsynonymous SNV | AOAH:NM_001637:exon11:c.C797G:p.A266G,AOAH:NM_001177506:exon11:c.C797G:p.A266G,AOAH:NM_001177507:exon10:c.C701G:p.A234G, |
| chr7 | 56021132 | T | A | COSV51908933 | exonic | PSPH | nonsynonymous SNV | PSPH:NM_004577:exon4:c.A81T:p.R27S, |
| chr7 | 56021209 | C | T | COSV51908733 | exonic | PSPH | nonsynonymous SNV | PSPH:NM_004577:exon4:c.G4A:p.V2I, |
| chr7 | 5986899 | C | T | COSV56221681 | exonic | PMS2 | nonsynonymous SNV | PMS2:NM_000535:exon11:c.G1866A:p.M622I, |
| chr8 | 18009676 | A | T | COSV59586229 | exonic | PCM1 | nonsynonymous SNV | PCM1:NM_001315508:exon30:c.A4930T:p.I1644L,PCM1:NM_006197:exon31:c.A5092T:p.I1698L,PCM1:NM_001315507:exon31:c.A5068T:p.I1690L, |
| chr8 | 31167138 | T | C | COSV53296936 | exonic | WRN | nonsynonymous SNV | WRN:NM_000553:exon34:c.T4099C:p.C1367R, |
| chr8 | 89978251 | C | G | COSV55371261 | exonic | NBN | nonsynonymous SNV | NBN:NM_002485:exon5:c.G553C:p.E185Q,NBN:NM_001024688:exon6:c.G307C:p.E103Q, |
| chr8 | 94467452 | G | C | COSV52594351 | exonic | RAD54B | nonsynonymous SNV | RAD54B:NM_012415:exon2:c.C88G:p.L30V,RAD54B:NM_001205262:exon2:c.C88G:p.L30V, |
| chr9 | 124414882 | A | G | COSV52330411 | exonic | PSMB7 | nonsynonymous SNV | PSMB7:NM_002799:exon2:c.T116C:p.V39A, |
| chr9 | 7799653 | G | T | COSV64065665 | exonic | TMEM261 | nonsynonymous SNV | TMEM261:NM_033428:exon1:c.C82A:p.P28T, |
| chr1 | 151760851 | T | C | COSV58618941 | exonic | MRPL9 | nonsynonymous SNV | MRPL9:NM_001300733:exon5:c.A535G:p.I179V,MRPL9:NM_031420:exon6:c.A637G:p.I213V, |
| chr1 | 154172926 | C | T | COSV55137972 | exonic | TPM3 | nonsynonymous SNV | TPM3:NM_152263:exon5:c.G548A:p.R183Q,TPM3:NM_001278190:exon4:c.G437A:p.R146Q,TPM3:NM_001043352:exon4:c.G437A:p.R146Q,TPM3:NM_001043351:exon4:c.G437A:p.R146Q,TPM3:NM_001043353:exon4:c.G437A:p.R146Q,TPM3:NM_001278191:exon4:c.G167A:p.R56Q,TPM3:NM_001278188:exon3:c.G239A:p.R80Q,TPM3:NM_001278189:exon4:c.G437A:p.R146Q,TPM3:NM_153649:exon4:c.G437A:p.R146Q, |
| chr1 | 155133751 | A | T | COSV57596245 | exonic | EFNA1 | nonsynonymous SNV | EFNA1:NM_004428:exon4:c.A476T:p.D159V,EFNA1:NM_182685:exon3:c.A410T:p.D137V, |
| chr1 | 156591859 | G | C | COSV63992221 | exonic | APOA1BP | nonsynonymous SNV | APOA1BP:NM_144772:exon1:c.G55C:p.V19L, |
| chr1 | 160646913 | G | T | COSV52498809 | exonic | SLAMF1 | nonsynonymous SNV | SLAMF1:NM_003037:exon1:c.C33A:p.F11L, |
| chr1 | 17044848 | C | T | COSV64965099 | exonic | SDHB | nonsynonymous SNV | SDHB:NM_003000:exon2:c.G113A:p.R38H, |
| chr1 | 201386394 | T | C | COSV66212460 | exonic | LAD1 | nonsynonymous SNV | LAD1:NM_005558:exon3:c.A967G:p.K323E, |
| chr1 | 2193733 | G | A | COSV66031527 | exonic | FAAP20 | nonsynonymous SNV | FAAP20:NM_001282673:exon3:c.C376T:p.P126S,FAAP20:NM_001282670:exon7:c.C685T:p.P229S,FAAP20:NM_001256946:exon3:c.C376T:p.P126S,FAAP20:NM_182533:exon3:c.C376T:p.P126S, |
| chr1 | 35189057 | C | T | COSV61758334 | exonic | SFPQ | nonsynonymous SNV | SFPQ:NM_005066:exon6:c.G1643A:p.R548H, |
| chr1 | 47333967 | G | C | COSV64101982 | exonic | CMPK1 | nonsynonymous SNV | CMPK1:NM_001136140:exon1:c.G22C:p.G8R,CMPK1:NM_016308:exon1:c.G22C:p.G8R, |
| chr1 | 999842 | C | A | COSV59241795 | exonic | HES4 | nonsynonymous SNV | HES4:NM_001142467:exon1:c.G132T:p.R44S, |
| chr10 | 102878966 | T | C | COSV54916951 | exonic | AS3MT | nonsynonymous SNV | AS3MT:NM_020682:exon9:c.T860C:p.M287T, |
| chr10 | 104279427 | A | G | COSV58537765 | exonic | GSTO2 | nonsynonymous SNV | GSTO2:NM_183239:exon5:c.A424G:p.N142D,GSTO2:NM_001191014:exon3:c.A340G:p.N114D, |
| chr10 | 17594310 | G | A | COSV63517260 | exonic | HACD1 | nonsynonymous SNV | HACD1:NM_014241:exon6:c.C679T:p.H227Y, |
| chr10 | 17617132 | C | A | COSV58892476 | exonic | HACD1 | nonsynonymous SNV | HACD1:NM_014241:exon1:c.G208T:p.V70F, |
| chr10 | 17617150 | C | T | COSV58892488 | exonic | HACD1 | nonsynonymous SNV | HACD1:NM_014241:exon1:c.G190A:p.E64K, |
| chr11 | 117351876 | G | A | COSV54039014 | exonic | CEP164 | nonsynonymous SNV | CEP164:NM_014956:exon5:c.G281A:p.S94N,CEP164:NM_001271933:exon4:c.G281A:p.S94N, |
| chr11 | 3038130 | C | T | COSV53457881 | exonic | CARS | nonsynonymous SNV | CARS:NM_001751:exon6:c.G472A:p.A158T,CARS:NM_001194997:exon7:c.G721A:p.A241T,CARS:NM_139273:exon6:c.G472A:p.A158T,CARS:NM_001014437:exon7:c.G721A:p.A241T, |
| chr11 | 34947565 | A | G | COSV57166122 | exonic | PDHX | nonsynonymous SNV | PDHX:NM_001135024:exon3:c.A256G:p.T86A,PDHX:NM_001166158:exon3:c.A301G:p.T101A,PDHX:NM_003477:exon3:c.A301G:p.T101A, |

| | | | | | | | | |
|---|---|---|---|---|---|---|---|---|
| chr11 | 35434513 | T | C | COSV53510283 | exonic | PAMR1 | nonsynonymous SNV | PAMR1:NM_015430:exon11:c.A1676G:p.Q559R,PAMR1:NM_001282675:exon12:c.A1505G:p.Q502R,PAMR1:NM_001001991:exon10:c.A1625G:p.Q542R,PAMR1:NM_001282676:exon8:c.A1292G:p.Q431R, |
| chr11 | 43919052 | C | G | COSV57023880 | exonic | ALKBH3 | nonsynonymous SNV | ALKBH3:NM_139178:exon9:c.C684G:p.D228E, |
| chr11 | 61445981 | G | A | COSV57099212 | exonic | SDHAF2 | nonsynonymous SNV | SDHAF2:NM_017841:exon4:c.G411A:p.M137I, |
| chr12 | 10420495 | C | G | COSV67249876 | exonic | KLRC3 | nonsynonymous SNV | KLRC3:NM_002261:exon1:c.G56C:p.W19S,KLRC3:NM_007333:exon1:c.G56C:p.W19S, |
| chr12 | 109557065 | A | T | COSV57169652 | exonic | MMAB | nonsynonymous SNV | MMAB:NM_052845:exon9:c.T716A:p.M239K, |
| chr12 | 109573425 | C | T | COSV57169511 | exonic | MMAB | nonsynonymous SNV | MMAB:NM_052845:exon1:c.G56A:p.R19H, |
| chr12 | 11884577 | G | A | COSV56765875 | exonic | ETV6 | nonsynonymous SNV | ETV6:NM_001987:exon6:c.G1142A:p.G381E, |
| chr12 | 124863717 | C | T | COSV55547637 | exonic | SCARB1 | nonsynonymous SNV | SCARB1:NM_001082959:exon1:c.G4A:p.G2S,SCARB1:NM_005505:exon1:c.G4A:p.G2S, |
| chr13 | 77613943 | A | G | COSV62993129 | exonic | SCEL | nonsynonymous SNV | SCEL:NM_003843:exon23:c.A1379G:p.K460R,SCEL:NM_001160706:exon22:c.A1313G:p.K438R,SCEL:NM_144777:exon24:c.A1439G:p.K480R, |
| chr14 | 35266761 | A | G | COSV51617748 | exonic | KIAA0391 | nonsynonymous SNV | KIAA0391:NM_001256680:exon5:c.A194G:p.N65S,KIAA0391:NM_014672:exon6:c.A1310G:p.N437S,KIAA0391:NM_001256678:exon5:c.A1262G:p.N421S,KIAA0391:NM_001256681:exon4:c.A194G:p.N65S,KIAA0391:NM_001256679:exon6:c.A1025G:p.N342S, |
| chr14 | 50312098 | A | C | COSV55413053 | exonic | L2HGDH | nonsynonymous SNV | L2HGDH:NM_024884:exon1:c.T53G:p.L18R, |
| chr14 | 53950804 | A | G | COSV55416014 | exonic | BMP4 | nonsynonymous SNV | BMP4:NM_130850:exon4:c.T455C:p.V152A,BMP4:NM_130851:exon4:c.T455C:p.V152A,BMP4:NM_001202:exon4:c.T455C:p.V152A, |
| chr14 | 65078029 | C | T | COSV52414939 | exonic | MAX | nonsynonymous SNV | MAX:NM_145113:exon4:c.G179A:p.R60Q,MAX:NM_145112:exon3:c.G152A:p.R51Q,MAX:NM_002382:exon4:c.G179A:p.R60Q, |
| chr14 | 94378506 | T | G | COSV63345434 | exonic | SERPINA1 | nonsynonymous SNV | SERPINA1:NM_001002235:exon5:c.A1200C:p.E400D,SERPINA1:NM_001127704:exon7:c.A1200C:p.E400D,SERPINA1:NM_001127703:exon7:c.A1200C:p.E400D,SERPINA1:NM_001127700:exon5:c.A1200C:p.E400D,SERPINA1:NM_000295:exon5:c.A1200C:p.E400D,SERPINA1:NM_001127706:exon6:c.A1200C:p.E400D,SERPINA1:NM_001127705:exon7:c.A1200C:p.E400D,SERPINA1:NM_001002236:exon7:c.A1200C:p.E400D,SERPINA1:NM_001127702:exon6:c.A1200C:p.E400D,SERPINA1:NM_001127707:exon6:c.A1200C:p.E400D,SERPINA1:NM_001127701:exon7:c.A1200C:p.E400D, |
| chr14 | 94382864 | C | T | COSV63345368 | exonic | SERPINA1 | nonsynonymous SNV | SERPINA1:NM_001002235:exon2:c.G374A:p.R125H,SERPINA1:NM_001127704:exon4:c.G374A:p.R125H,SERPINA1:NM_001127703:exon4:c.G374A:p.R125H,SERPINA1:NM_001127700:exon2:c.G374A:p.R125H,SERPINA1:NM_000295:exon2:c.G374A:p.R125H,SERPINA1:NM_001127706:exon3:c.G374A:p.R125H,SERPINA1:NM_001127705:exon4:c.G374A:p.R125H,SERPINA1:NM_001002236:exon4:c.G374A:p.R125H,SERPINA1:NM_001127702:exon3:c.G374A:p.R125H,SERPINA1:NM_001127707:exon3:c.G374A:p.R125H,SERPINA1:NM_001127701:exon4:c.G374A:p.R125H, |
| chr15 | 40621979 | T | C | COSV61151299 | exonic | CASC5 | nonsynonymous SNV | CASC5:NM_144508:exon10:c.T1715C:p.M572T,CASC5:NM_170589:exon11:c.T1793C:p.M598T, |
| chr15 | 40622992 | A | G | COSV61150613 | exonic | CASC5 | nonsynonymous SNV | CASC5:NM_144508:exon10:c.A2728G:p.R910G,CASC5:NM_170589:exon11:c.A2806G:p.R936G, |
| chr15 | 50476967 | G | A | COSV56168389 | exonic | USP8 | nonsynonymous SNV | USP8:NM_001283049:exon7:c.G737A:p.R246H,USP8:NM_001128610:exon9:c.G968A:p.R323H,USP8:NM_005154:exon9:c.G968A:p.R323H, |
| chr15 | 85735078 | G | A | COSV63448267 | exonic | AKAP13 | nonsynonymous SNV | AKAP13:NM_006738:exon31:c.G7381A:p.G2461S,AKAP13:NM_007200:exon31:c.G7369A:p.G2457S,AKAP13:NM_001270546:exon23:c.G3232A:p.G1078S, |
| chr16 | 11915422 | A | G | COSV70452514 | exonic | GSPT1 | nonsynonymous SNV | GSPT1:NM_002094:exon1:c.T299C:p.V100A,GSPT1:NM_001130006:exon1:c.T299C:p.V100A, |
| chr16 | 15038105 | A | G | COSV55074188 | exonic | NTAN1 | nonsynonymous SNV | NTAN1:NM_001270767:exon8:c.T544C:p.S182P,NTAN1:NM_001270766:exon9:c.T544C:p.S182P,NTAN1:NM_173474:exon10:c.T859C:p.S287P, |
| chr16 | 15038117 | G | T | COSV55074343 | exonic | NTAN1 | nonsynonymous SNV | NTAN1:NM_001270767:exon8:c.C532A:p.H178N,NTAN1:NM_001270766:exon9:c.C532A:p.H178N,NTAN1:NM_173474:exon10:c.C847A:p.H283N, |
| chr16 | 64991815 | G | A | COSV51767367 | exonic | CDH11 | nonsynonymous SNV | CDH11:NM_001308392:exon6:c.C764T:p.T255M,CDH11:NM_001797:exon6:c.C764T:p.T255M, |
| chr16 | 766977 | G | A | COSV53502132 | exonic | MSLN | nonsynonymous SNV | MSLN:NM_001177355:exon15:c.G1466A:p.G489E,MSLN:NM_005823:exon15:c.G1466A:p.G489E,MSLN:NM_013404:exon14:c.G1490A:p.G497E, |
| chr17 | 40484492 | C | T | COSV54184711 | exonic | TNS4 | nonsynonymous SNV | TNS4:NM_032865:exon6:c.G1493A:p.S498N, |
| chr17 | 47827054 | C | T | COSV51621557 | exonic | MRPL10 | nonsynonymous SNV | MRPL10:NM_145255:exon3:c.G373A:p.V125I,MRPL10:NM_148887:exon4:c.G403A:p.V135I, |
| chr17 | 5422825 | C | G | COSV56705547 | exonic | RPAIN | nonsynonymous SNV | RPAIN:NM_001160266:exon3:c.C309G:p.N103K,RPAIN:NM_001160244:exon3:c.C309G:p.N103K,RPAIN:NM_001033002:exon3:c.C309G:p.N103K,RPAIN:NM_001160246:exon3:c.C309G:p.N103K,RPAIN:NM_001160243:exon3:c.C309G:p.N103K, |
| chr17 | 59886176 | A | G | COSV57872241 | exonic | TUBD1 | nonsynonymous SNV | TUBD1:NM_001193609:exon3:c.T227C:p.M76T,TUBD1:NM_001193611:exon3:c.T227C:p.M76T,TUBD1:NM_016261:exon3:c.T227C:p.M76T,TUBD1:NM_001193610:exon3:c.T227C:p.M76T, |
| chr17 | 623667 | G | T | COSV52126287 | exonic | VPS53 | nonsynonymous SNV | VPS53:NM_018289:exon10:c.C895A:p.L299I,VPS53:NM_001128159:exon11:c.C982A:p.L328I, |
| chr17 | 7703404 | C | G | COSV56832921 | exonic | WRAP53 | nonsynonymous SNV | WRAP53:NM_001143990:exon11:c.C1565G:p.A522G,WRAP53:NM_001143991:exon11:c.C1565G:p.A522G,WRAP53:NM_001143992:exon11:c.C1565G:p.A522G,WRAP53:NM_018081:exon10:c.C1565G:p.A522G, |
| chr19 | 12956978 | G | C | COSV57534227 | exonic | GADD45GIP1 | nonsynonymous SNV | GADD45GIP1:NM_052850:exon1:c.C235G:p.P79A, |
| chr19 | 18568569 | C | G | COSV55889000 | exonic | KXD1 | nonsynonymous SNV | KXD1:NM_001171948:exon6:c.C469G:p.P157A,KXD1:NM_001171949:exon5:c.C469G:p.P157A,KXD1:NM_024069:exon5:c.C469G:p.P157A, |
| chr19 | 8371280 | C | T | COSV56844530 | exonic | ANGPTL4 | nonsynonymous SNV | ANGPTL4:NM_001039667:exon5:c.C683T:p.T228M,ANGPTL4:NM_139314:exon6:c.C797T:p.T266M, |
| chr2 | 201258272 | A | G | COSV51848843 | exonic | CASP8 | nonsynonymous SNV | CASP8:NM_001080125:exon1:c.A41G:p.K14R, |
| chr2 | 214780740 | C | G | COSV53612030 | exonic | BARD1 | nonsynonymous SNV | BARD1:NM_001282543:exon3:c.G1077C:p.R359S,BARD1:NM_000465:exon4:c.G1134C:p.R378S, |

| | | | | | | | | |
|---|---|---|---|---|---|---|---|---|
| chr2 | 215435759 | T | A | COSV60547598 | exonic | FN1 | nonsynonymous SNV | FN1:NM_001306130:exon1:c.A44T:p.Q15L,FN1:NM_212474:exon1:c.A44T:p.Q15L,FN1:NM_212476:exon1:c.A44T:p.Q15L,FN1:NM_001306129:exon1:c.A44T:p.Q15L,FN1:NM_054034:exon1:c.A44T:p.Q15L,FN1:NM_001306131:exon1:c.A44T:p.Q15L,FN1:NM_001306132:exon1:c.A44T:p.Q15L,FN1:NM_002026:exon1:c.A44T:p.Q15L,FN1:NM_212478:exon1:c.A44T:p.Q15L,FN1:NM_212482:exon1:c.A44T:p.Q15L, |
| chr2 | 26191602 | C | T | COSV66106823 | exonic | HADHA | nonsynonymous SNV | HADHA:NM_000182:exon19:c.G2027A:p.R676H, |
| chr2 | 26201217 | C | T | COSV66106966 | exonic | HADHA | nonsynonymous SNV | HADHA:NM_000182:exon13:c.G1324A:p.D442N, |
| chr20 | 23637790 | C | T | COSV65361855 | exonic | CST3 | nonsynonymous SNV | CST3:NM_001288614:exon1:c.G73A:p.A25T,CST3:NM_000099:exon1:c.G73A:p.A25T, |
| chr20 | 58903691 | G | A | COSV99039588 | exonic | GNAS | nonsynonymous SNV | GNAS:NM_080426:exon4:c.G290A:p.S97N,GNAS:NM_080425:exon5:c.G2261A:p.S754N,GNAS:NM_001309840:exon5:c.G155A:p.S52N,GNAS:NM_001077489:exon4:c.G287A:p.S96N,GNAS:NM_001309861:exon5:c.G155A:p.S52N,GNAS:NM_000516:exon5:c.G332A:p.S111N,GNAS:NM_001077488:exon5:c.G335A:p.S112N, |
| chr20 | 63742354 | G | T | COSV55507914 | exonic | SLC2A4RG | nonsynonymous SNV | SLC2A4RG:NM_020062:exon6:c.G699T:p.E233D, |
| chr22 | 17159155 | G | A | COSV50084409 | exonic | CECR5 | nonsynonymous SNV | CECR5:NM_033070:exon1:c.C97T:p.R33C, |
| chr22 | 40967827 | G | A | COSV53430854 | exonic | RBX1 | nonsynonymous SNV | RBX1:NM_014248:exon4:c.G257A:p.R86H, |
| chr3 | 160279273 | T | A | COSV58451332 | exonic | IFT80 | nonsynonymous SNV | IFT80:NM_020800:exon16:c.A1756T:p.T586S,IFT80:NM_001190242:exon15:c.A1345T:p.T449S,IFT80:NM_001190241:exon17:c.A1345T:p.T449S, |
| chr3 | 169796797 | A | T | COSV57186258 | exonic | LRRC34 | nonsynonymous SNV | LRRC34:NM_001172780:exon8:c.T856A:p.L286I,LRRC34:NM_001172779:exon8:c.T856A:p.L286I,LRRC34:NM_153353:exon7:c.T760A:p.L254I, |
| chr4 | 105235006 | G | A | COSV54398666 | exonic | TET2 | nonsynonymous SNV | TET2:NM_001127208:exon3:c.G1064A:p.G355D,TET2:NM_017628:exon3:c.G1064A:p.G355D, |
| chr4 | 105396272 | C | G | COSV58995765 | exonic | PPA2 | nonsynonymous SNV | PPA2:NM_176866:exon5:c.G540C:p.K180N,PPA2:NM_006903:exon8:c.G759C:p.K253N,PPA2:NM_176869:exon9:c.G846C:p.K282N,PPA2:NM_176867:exon3:c.G348C:p.K116N, |
| chr4 | 150277928 | G | A | COSV63949643 | exonic | LRBA | nonsynonymous SNV | LRBA:NM_006726:exon57:c.C8426T:p.S2809L,LRBA:NM_001199282:exon56:c.C8390T:p.S2797L, |
| chr4 | 150285975 | C | T | COSV63966388 | exonic | LRBA | nonsynonymous SNV | LRBA:NM_006726:exon55:c.G8110A:p.A2704T,LRBA:NM_001199282:exon54:c.G8074A:p.A2692T, |
| chr4 | 1728226 | G | A | COSV57603699 | exonic | TACC3 | nonsynonymous SNV | TACC3:NM_006342:exon4:c.G824A:p.C275Y, |
| chr4 | 1731251 | G | A | COSV57605388 | exonic | TACC3 | nonsynonymous SNV | TACC3:NM_006342:exon6:c.G1541A:p.G514E, |
| chr5 | 74697056 | A | G | COSV54667253 | exonic | HEXB | nonsynonymous SNV | HEXB:NM_000521:exon5:c.A619G:p.I207V, |
| chr6 | 110456925 | T | C | COSV57930020 | exonic | SLC22A16 | nonsynonymous SNV | SLC22A16:NM_033125:exon2:c.A146G:p.H49R, |
| chr6 | 128240094 | C | T | COSV63888698 | exonic | PTPRK | nonsynonymous SNV | PTPRK:NM_002844:exon5:c.G634A:p.A212T,PTPRK:NM_001291983:exon4:c.G247A:p.A83T,PTPRK:NM_001135648:exon5:c.G634A:p.A212T,PTPRK:NM_001291984:exon5:c.G634A:p.A212T,PTPRK:NM_001291982:exon5:c.G634A:p.A212T,PTPRK:NM_001291981:exon5:c.G634A:p.A212T, |
| chr6 | 159692840 | A | G | COSV61622991 | exonic | SOD2 | nonsynonymous SNV | SOD2:NM_001024465:exon2:c.T47C:p.V16A,SOD2:NM_001024466:exon2:c.T47C:p.V16A,SOD2:NM_000636:exon2:c.T47C:p.V16A, |
| chr6 | 166326158 | T | C | COSV62277048 | exonic | SFT2D1 | nonsynonymous SNV | SFT2D1:NM_145169:exon5:c.A325G:p.I109V, |
| chr6 | 167315303 | G | A | COSV57807274 | exonic | UNC93A | nonsynonymous SNV | UNC93A:NM_001143947:exon7:c.G1099A:p.V367I,UNC93A:NM_018974:exon8:c.G1225A:p.V409I, |
| chr6 | 31271148 | C | T | COSV66113730 | exonic | HLA-C | nonsynonymous SNV | HLA-C:NM_002117:exon3:c.G544A:p.A182T,HLA-C:NM_001243042:exon3:c.G544A:p.A182T, |
| chr6 | 31271283 | A | G | COSV66110655 | exonic | HLA-C | nonsynonymous SNV | HLA-C:NM_002117:exon3:c.T409C:p.Y137H,HLA-C:NM_001243042:exon3:c.T409C:p.Y137H, |
| chr6 | 31271830 | G | A | COSV66111027 | exonic | HLA-C | nonsynonymous SNV | HLA-C:NM_002117:exon2:c.C112T:p.R38W,HLA-C:NM_001243042:exon2:c.C112T:p.R38W, |
| chr6 | 31871532 | C | T | COSV57666266 | exonic | SLC44A4 | nonsynonymous SNV | SLC44A4:NM_001178045:exon8:c.G331A:p.V111I,SLC44A4:NM_001178044:exon7:c.G433A:p.V145I,SLC44A4:NM_025257:exon8:c.G559A:p.V187I, |
| chr7 | 13939184 | T | C | COSV54136900 | exonic | ETV1 | nonsynonymous SNV | ETV1:NM_001163152:exon3:c.A178G:p.S60G,ETV1:NM_004956:exon7:c.A298G:p.S100G,ETV1:NM_001163151:exon2:c.A124G:p.S42G,ETV1:NM_001163150:exon3:c.A178G:p.S60G,ETV1:NM_001163148:exon6:c.A244G:p.S82G,ETV1:NM_001163147:exon6:c.A298G:p.S100G,ETV1:NM_001163149:exon5:c.A244G:p.S82G, |
| chr7 | 140753336 | A | T | COSV56056643 | exonic | BRAF | nonsynonymous SNV | BRAF:NM_004333:exon15:c.T1799A:p.V600E, |
| chr7 | 27792114 | G | A | COSV55291444 | exonic | TAX1BP1 | nonsynonymous SNV | TAX1BP1:NM_006024:exon9:c.G1147A:p.V383I,TAX1BP1:NM_001206901:exon9:c.G1147A:p.V383I,TAX1BP1:NM_001206902:exon8:c.G676A:p.V226I,TAX1BP1:NM_001079864:exon9:c.G1147A:p.V383I, |
| chr7 | 29566700 | C | A | COSV60501902 | exonic | PRR15 | nonsynonymous SNV | PRR15:NM_175887:exon2:c.C371A:p.P124H, |
| chr7 | 39570578 | A | G | COSV56232709 | exonic | YAE1D1 | nonsynonymous SNV | YAE1D1:NM_001282446:exon2:c.A202G:p.K68E,YAE1D1:NM_020192:exon2:c.A202G:p.K68E, |
| chr7 | 5919854 | G | A | COSV58073320 | exonic | CCZ1 | nonsynonymous SNV | CCZ1:NM_015622:exon12:c.G994A:p.V332I, |
| chr7 | 76048186 | C | T | COSV50388126 | exonic | MDH2 | nonsynonymous SNV | MDH2:NM_001282403:exon1:c.C26T:p.A9V,MDH2:NM_005918:exon1:c.C26T:p.A9V, |
| chr8 | 123427934 | A | G | COSV54873851 | exonic | WDYHV1 | nonsynonymous SNV | WDYHV1:NM_018024:exon2:c.A94G:p.I32V, |
| chr8 | 123436496 | A | G | COSV54874032 | exonic | WDYHV1 | nonsynonymous SNV | WDYHV1:NM_001283024:exon4:c.A98G:p.N33S,WDYHV1:NM_001283027:exon5:c.A74G:p.N25S,WDYHV1:NM_018024:exon4:c.A278G:p.N93S, |
| chr8 | 123436564 | T | A | COSV54873861 | exonic | WDYHV1 | nonsynonymous SNV | WDYHV1:NM_001283024:exon4:c.T166A:p.F56I,WDYHV1:NM_001283027:exon5:c.T142A:p.F48I,WDYHV1:NM_018024:exon4:c.T346A:p.F116I, |
| chr8 | 123437226 | C | T | COSV54874614 | exonic | WDYHV1 | nonsynonymous SNV | WDYHV1:NM_001283024:exon5:c.C220T:p.R74C,WDYHV1:NM_001283027:exon6:c.C196T:p.R66C,WDYHV1:NM_018024:exon5:c.C400T:p.R134C, |
| chr8 | 18069818 | T | C | COSV50501249 | exonic | ASAH1 | nonsynonymous SNV | ASAH1:NM_177924:exon4:c.A277G:p.I93V,ASAH1:NM_004315:exon4:c.A325G:p.I109V, |
| chr8 | 18071302 | C | T | COSV50502085 | exonic | ASAH1 | nonsynonymous SNV | ASAH1:NM_001127505:exon4:c.G283A:p.V95M,ASAH1:NM_177924:exon3:c.G214A:p.V72M,ASAH1:NM_004315:exon3:c.G262A:p.V88M, |
| chr8 | 31141764 | G | T | COSV53297644 | exonic | WRN | nonsynonymous SNV | WRN:NM_000553:exon26:c.G3222T:p.L1074F, |
| chr9 | 115030304 | C | G | COSV60785451 | exonic | TNC | nonsynonymous SNV | TNC:NM_002160:exon24:c.G6022C:p.E2008Q, |
| chr9 | 132911517 | A | G | COSV53763263 | exonic | TSC1 | nonsynonymous SNV | TSC1:NM_001162426:exon10:c.T965C:p.M322T,TSC1:NM_000368:exon10:c.T965C:p.M322T,TSC1:NM_001162427:exon9:c.T812C:p.M271T, |

| | Loop-seq count | | | | |
|---|---|---|---|---|---|
| Isoform | 1T__WT | 1T__MUT | 1M__WT | 1M__MUT | 2T__WT |
| NM_005101 | 3 | 4 | 0 | 2 | 0 |
| NM_015906:XM_005270936:XM_011541568:XM_024447584:NM_033020:XM_005270937 | 0:0:0:0:0:0 | 7:0:0:3:3:3 | 0:0:0:0:0:0 | 3:1:1:1:1:0 | 0:0:0:0:0:0 |
| NM_004079 | 2 | 7 | 0 | 1 | 0 |
| NR_104400:NM_003037:XM_005245456:XM_011509905:NM_001330754:NR_104399 | 0:3:0:0:1:0 | 0:1:0:0:0:0 | 1:3:0:1:0:0 | 2:2:1:0:0:0 | 0:1:0:0:0:0 |
| NM_007348:XM_006711224:XM_011509310:XM_011509308 | 29:0:0:1 | 11:1:1:0 | 9:1:1:0 | 10:0:0:0 | 18:0:1:0 |
| NR_137288:XR_002956989:XR_001737277:XM_017001622:XR_946690:XR_946686:XM_024448062:NM_001328124:NM_001319838:XM_017001620 | 3:4:0:0:0:0:0:1:0:0 | 2:2:0:0:0:0:1:1:1:0 | 2:3:1:1:0:0:0:0:0:0 | 1:2:0:0:0:0:0:0:0:0 | 0:0:0:0:0:0:0:0:0:0 |
| NM_001136018:NM_001291163 | 12:01 | 13:02 | 6:00 | 18:01 | 0:00 |
| XM_006711815:XM_005273262:NM_024525:NM_001122835:XM_005273260 | 0:0:0:0:0 | 0:0:1:0:0 | 0:0:0:0:0 | 1:1:0:0:0 | 0:0:2:0:0 |
| NM_018072 | 0 | 1 | 0 | 1 | 0 |
| NM_018072 | 0 | 1 | 0 | 1 | 0 |
| NM_018072 | 0 | 1 | 0 | 1 | 0 |
| XM_011544253:NM_001167740:XM_024449137:XM_024449166:NM_022743:XM_024449136:XM_011544258:XM_024449149:XM_011544261:XM_011544254:XM_024449162 | 8:43:2:0:0:0:0:0:0:0:0 | 3:54:0:0:0:0:1:0:0:0:0 | 7:68:1:1:0:1:0:0:0:0:0 | 17:108:2:1:1:0:0:0:0:0:0 | 0:0:0:0:0:0:0:0:0:0:0 |
| NM_001803 | 0 | 2 | 0 | 3 | 0 |
| NM_023923:NM_001350161:NM_001350158:NM_001048183:NM_001350160 | 1:0:0:4:0 | 1:0:2:1:0 | 1:1:0:0:0 | 1:0:0:0:0 | 1:0:0:1:1 |
| XM_017001646:XM_024448207:XR_001737287:NM_018291:XM_024448229:XM_017001645:XM_017001652:XM_017001679:NM_001350793:NM_001350794:XM_017001656:XM_017001657:XM_011541730:NM_001244714:XR_001737285:NM_001113411:NR_103473:XM_017001649:XM_017001673:XM_024448176:NM_001350799:NM_001350797:XM_024448220:XM_017001655 | 1:2:0:3:0:2:1:0:1:0:0:0:0:0:0:0:0:0:0:0:0:0:0:0 | 1:0:0:5:0:2:0:1:0:0:0:0:0:0:0:0:0:0:0:0:0:0:0:0 | 0:1:0:1:0:0:0:0:0:0:0:0:0:0:0:0:0:0:0:0:0:0:0:0 | 1:1:1:0:0:0:0:0:0:0:0:0:0:0:0:0:0:0:0:0:0:0:0:0 | 0:1:1:2:0:0:0:0:0:0:0:0:0:0:0:0:0:0:0:0:0:0:0:0 |
| NM_001286043:NM_000016:NM_001286042:NM_001127328:NM_001286044 | 0:32:2:0:3 | 0:31:10:1:0 | 0:5:1:0:0 | 1:17:2:1:1 | 0:87:2:0:2 |
| NM_000969:NR_146333 | 1736:01:00 | 2:00 | 4687:12:00 | 2:00 | 5027:08:00 |
| NM_000110:XM_017000510:XM_017000507:XM_006710397 | 0:0:0:0 | 2:2:0:1 | 0:0:0:0 | 2:2:0:0 | 0:0:0:0 |
| XM_011539913:NM_017787:XM_017016360:NM_001083913 | 0:0:0:1 | 0:1:1:0 | 0:1:0:0 | 1:0:0:0 | 0:0:0:0 |
| NM_004832 | 2 | 2 | 1 | 1 | 3 |
| NM_004092 | 0 | 10 | 0 | 1 | 0 |
| NM_005436 | 29 | 16 | 17 | 11 | 15 |
| XM_017016378:NM_018344:NR_033414:XM_017016377:NR_033413:NM_001174098 | 1:0:0:1:0:0 | 0:3:0:0:0:0 | 0:2:1:0:0:0 | 1:0:0:0:0:0 | 0:0:0:0:0:0 |
| NM_004329:XM_011540104 | 3:00 | 4:00 | 5:00 | 3:00 | 4:00 |
| NM_016195:NM_001284259 | 2:00 | 2:00 | 2:00 | 1:00 | 12:00 |
| NM_001098520:NM_006410:NM_001098521:NM_001098522 | 0:0:0:0 | 3:2:0:1 | 0:0:0:0 | 2:0:0:0 | 0:0:0:0 |
| NM_006435 | 1 | 1044 | 0 | 1384 | 960 |
| NM_003641 | 0 | 10 | 0 | 14 | 0 |
| XM_005253238:XM_005253235:NM_001001389:XM_011520483:XM_011520486:XM_017018583:XM_006718388:NM_000610:XM_011520487:XM_011520484:XM_005253231:XM_005253232:XM_011520489:XM_011520482:XM_011520485:XM_017018585 | 3:0:0:0:0:1:0:0:0:0:0:0:0:0:0:0 | 1:0:0:0:0:0:0:0:0:0:0:0:0:0:0:0 | 7:1:1:1:0:1:1:0:0:0:0:0:0:0:0:0 | 9:1:1:1:1:0:0:0:0:0:0:0:0:0:0:0 | 47:5:4:0:1:5:1:5:1:2:3:0:0:0:0:0 |
| NM_016142:XM_024448573 | 0:00 | 75:04:00 | 0:00 | 53:02:00 | 239:04:00 |
| NM_012094:NM_001358511:NM_181651:NM_181652 | 0:0:0:0 | 19:2:1:1 | 0:1:0:0 | 4:0:0:0 | 0:0:0:0 |
| NM_004549:NM_001204055 | 3:00 | 4:00 | 0:00 | 2:00 | 7:00 |
| NM_001243251:NM_024678:XM_017018303:XR_001747964:XR_001747966:XM_017018304:XM_011545253:XR_001747965:XR_001747963 | 0:0:0:1:0:0:0:0:0 | 3:1:1:0:0:0:0:0:0 | 0:3:0:0:0:0:0:0:0 | 3:0:0:0:0:0:0:0:0 | 0:0:0:0:0:0:0:0:0 |
| NM_003754 | 1 | 3 | 1 | 1 | 10 |
| NM_001814 | 0 | 1 | 0 | 1 | 0 |
| NM_002813:NM_001261400:NR_048555 | 0:00:00 | 247:02:03 | 0:00:00 | 87:01:01 | 0:00:00 |
| NM_032656:XR_001748820 | 0:00 | 1:00 | 0:00 | 1:00 | 1:00 |
| NM_001042603 | 15 | 2 | 8 | 1 | 6 |
| NM_001042603 | 10 | 8 | 5 | 3 | 20 |
| NM_001113202:NM_005594:NM_001320193:NM_001320194:NM_001113201:NM_001113203 | 1445:107:73:21:1:1 | 4:0:0:0:0:0 | 3027:193:112:33:3:0 | 10:0:0:0:0:0 | 3979:257:185:30:2:0 |
| NM_001159576:NM_001038 | 0:00 | 0:01 | 0:00 | 1:00 | 0:00 |
| NM_130846:NM_001207016:XM_011538615:NR_073474:NM_002849:XR_001748831 | 151:52:7:5:0:13 | 176:38:23:7:0:11 | 2:1:1:1:0:0 | 6:2:0:0:0:0 | 1:0:0:0:0:0 |
| NM_004616:XM_006719583 | 3:00 | 5:00 | 1:00 | 2:00 | 11:01 |
| NM_002097 | 1 | 1 | 1 | 1 | 3 |
| NM_172373 | 0 | 4 | 0 | 21 | 0 |

| | | | | | |
|---|---|---|---|---|---|
| NM_172373:NM_001145353 | 0:00 | 13:00 | 0:00 | 44:00:00 | 0:00 |
| NM_145725:XM_017021618:NM_003300:XM_011537117:NM_145726:XM_017021619:XM_017021620 | 7:0:3:0:0:0:0 | 3:2:1:0:1:0:0 | 1:0:0:0:0:0:0 | 2:1:1:0:0:0:0 | 1:0:0:0:0:0:0 |
| NM_182836:NM_004581 | 31:03:00 | 32:03:00 | 2:00 | 16:01 | 20:00 |
| XR_943501:NM_021239:XM_011537044:XM_024449680 | 48:11:5:0 | 1:0:0:0 | 30:7:2:2 | 1:0:0:0 | 42:7:1:0 |
| NM_001211 | 2 | 6 | 0 | 10 | 46 |
| NM_144508:XM_017022432 | 0:00 | 3:01 | 0:00 | 4:00 | 11:03 |
| NM_184041:NM_184043:NM_001127617:NM_001243177 | 3333:103:3:5 | 5:1:0:1 | 1750:79:3:3 | 1:0:0:0 | 2354:81:6:6 |
| XM_011522393:NM_005223:XM_024450171:XM_024450170:XR_002957783:XM_017022993:XR_002957785:XM_024450172:NM_001351825:XM_017023001 | 0:0:0:0:0:0:0:0:0:0 | 1:0:0:0:0:0:0:0:0:0 | 1:2:1:0:0:0:0:0:0:0 | 3:0:0:0:0:0:0:0:0:0 | 0:0:0:0:1:0:0:1:0:0 |
| NM_005153:XM_017023869:XM_006721332:NM_001272075:XM_017023864 | 0:0:0:0:0 | 12:0:1:0:0 | 0:0:0:0:0 | 13:1:0:0:0 | 36:3:1:1:0 |
| NM_000101 | 6 | 4 | 2 | 1 | 5 |
| NM_000101 | 0 | 10 | 0 | 3 | 5 |
| XM_024450187:NM_174917:XR_933240:NR_147928:XM_024450186:NM_001127214:XM_017023019:NM_001243279:XR_933238 | 0:0:0:0:0:0:0:0:0 | 5:0:0:1:1:0:0:0:0 | 0:0:0:0:0:0:0:0:0 | 2:0:0:0:0:0:0:0:0 | 0:0:0:0:0:0:0:0:0 |
| NR_024058:NM_006761:XM_017025005:XM_005256784 | 573:6002:25:0 | 3:30:2:0 | 162:1720:5:0 | 1:4:0:0 | 202:2375:6:1 |
| NM_015355:NM_001321207:XR_001752462:XM_006721794:XM_017024409 | 15:1:6:7:2 | 1:1:0:1:0 | 13:2:4:1:0 | 1:0:0:0:0 | 12:2:4:5:1 |
| NM_002276 | 5 | 4288 | 3 | 1174 | 0 |
| NM_007298:NM_007299:NR_027676 | 0:00:01 | 0:01:01 | 0:00:00 | 1:00:00 | 0:00:02 |
| NM_003726:NM_001075099:XM_024451012:XM_017025260:XM_005257755:XM_017025258:XM_017025259 | 2:2:1:0:0:1:1 | 2:1:0:0:0:0:0 | 0:0:0:0:0:0:0 | 38:22:4:1:1:0:0 | 8:4:0:0:0:0:0 |
| NM_017763:XM_017024800:XM_011524955:NM_001305544 | 1:1:0:0 | 1:0:0:0 | 2:1:2:1 | 9:3:0:0 | 2:0:0:0 |
| NM_017983 | 0 | 1 | 0 | 1 | 0 |
| NM_001098832:NM_032837:NM_001289410:XM_024451008 | 4:9:0:1 | 4:15:0:0 | 2:2:0:0 | 1:3:1:0 | 1:2:0:0 |
| NM_015971 | 0 | 3 | 0 | 1 | 0 |
| NM_001276761:NM_001276760:NM_001126118:NM_001276696 | 1:13:0:0 | 4:21:1:0 | 1:3:0:0 | 1:2:0:0 | 0:0:0:0 |
| NM_021074 | 4 | 4 | 1 | 1 | 0 |
| NM_032853:XM_017027401:XM_011528382:XM_011528383 | 0:0:0:0 | 0:2:0:0 | 0:0:0:0 | 2:2:0:0 | 1:0:0:0 |
| NM_033417:NM_001011699:XM_005260154 | 43:08:03 | 26:06:00 | 1:01:00 | 6:01:01 | 20:04:02 |
| NM_006532:XM_017027335:XR_001753769:XM_017027336 | 4:0:0:0 | 5:0:0:0 | 0:0:0:0 | 5:0:0:0 | 4:0:0:0 |
| XR_001753633:NM_001130867:XM_011526611:XR_001753634:XM_017026467 | 4:0:0:0:0 | 4:0:1:1:0 | 3:0:0:0:0 | 1:1:0:0:0 | 9:2:0:0:0 |
| XM_017003977:XM_017003978 | 1:00 | 1:00 | 0:00 | 1:00 | 0:00 |
| XM_017003977:XM_017003978 | 0:00 | 2:00 | 0:00 | 1:00 | 0:00 |
| XM_011510825:XR_001738674:NM_012233:XM_011510823 | 0:3:4:0 | 0:0:3:0 | 0:1:3:0 | 1:1:4:0 | 0:1:5:0 |
| NM_002156:NM_199440 | 813:110 | 3:00 | 1614:125 | 3:00 | 1924:191 |
| NM_004544:NR_136157:NM_001322019:XM_011511228:XR_001738750:NR_136158 | 138:1:0:1:2:0 | 102:0:0:1:1:0 | 51:1:1:0:0:0 | 81:0:0:0:0:0 | 271:2:1:2:0:0 |
| NM_001195381:NM_001195382:NM_005301 | 1:05:00 | 5:02:00 | 0:06:00 | 1:02:00 | 4:12:03 |
| NM_019063:XR_001738710:XR_001738709:XM_006711992 | 0:0:0:0 | 2:0:0:0 | 0:0:0:0 | 6:0:0:0 | 0:0:0:0 |
| NM_019063:XR_001738710:XR_001738709:XM_006711992 | 0:0:0:0 | 2:0:0:0 | 3:0:0:0 | 3:0:0:0 | 0:0:0:0 |
| NM_006363:NM_032985:XM_017027593:NM_032986:NM_001172745 | 6:3:2:0:0 | 8:6:2:0:0 | 1:2:0:0:0 | 11:12:3:0:0 | 19:42:8:2:2 |
| NM_001048223:NM_001048221:NM_001197140 | 0:00:00 | 1:01:00 | 0:00:00 | 1:00:00 | 1:00:00 |
| XR_002958471:NM_001077489:XM_017027817:NM_001309840:NM_001077488:NM_080426:XM_017027818:XM_017027813:NM_000516:XM_024451874:NM_080425:XM_024451875:XM_017027819:XM_024451873:XM_017027815:XM_017027812:XM_024451872:NM_016592:XM_017027820:XM_017027814:NM_001309861 | 10:101:42:9:23:256:3:10:110:1:1:8:7:2:2:2:0:0:0:0:0 | 0:0:0:0:0:1:0:0:0:0:0:0:0:0:0:0:0:0:0:0:0 | 16:105:40:12:9:268:10:17:87:2:2:3:7:0:0:0:0:0:0:0:0 | 1:0:0:0:0:0:0:0:0:0:0:0:0:0:0:0:0:0:0:0:0 | 31:155:32:20:24:284:5:29:172:1:3:11:11:0:0:1:0:1:0:0:1 |
| XR_002958471:NM_001077489:XM_017027817:NM_001309840:NM_001077488:XM_017027818:NM_080426:XM_017027813:NM_000516:XM_024451874:NM_080425:XM_024451875:XM_017027819:XM_024451873:XM_017027815:XM_017027812:XM_024451872:NM_016592:XM_017027820:XM_017027814:NM_001309861 | 26:101:42:9:23:3:260:10:110:1:1:8:7:2:2:2:0:0:0:0:0 | 0:0:0:0:0:1:0:0:0:0:0:0:0:0:0:0:0:0:0:0:0 | 32:105:40:12:9:10:274:17:87:2:2:3:7:0:0:0:0:0:0:0:0 | 1:0:0:0:0:0:0:0:0:0:0:0:0:0:0:0:0:0:0:0:0 | 62:156:32:20:24:5:291:29:173:1:3:11:11:0:0:1:0:1:0:0:1 |

| | | | | | |
|---|---|---|---|---|---|
| XR_002958471:NM_001077489:XM_017027817:NM_001309840:NM_001077488:NM_080426:XM_017027818:XM_017027813:NM_000516:XM_024451874:NM_080425:XM_024451875:XM_017027819:XM_024451873:XM_017027815:XM_017027812:XM_024451872:NM_016592:XM_017027820:XM_017027814:NM_001309861 | 46:101:42:9:22:268:3:10:110:1:1:8:7:2:2:2:0:0:0:0:0 | 0:0:0:0:1:1:0:0:0:0:0:0:0:0:0:0:0:0:0:0:0 | 73:105:40:12:9:281:10:15:87:2:2:3:7:0:0:0:0:0:0:0:0 | 1:0:0:0:0:0:0:0:0:0:0:0:0:0:0:0:0:0:0:0:0 | 119:156:32:20:24:307:5:29:171:1:3:11:10:0:0:1:0:1:0:0:1 |
| NR_103797:NM_001291411 | 0:00 | 25:00:00 | 0:01 | 13:00 | 1:00 |
| NM_005656:NM_001135099 | 256:17:00 | 294:19:00 | 21:05 | 28:05:00 | 44:05:00 |
| NM_000754:NM_001135162:NM_001135161 | 0:00:00 | 90:01:00 | 0:00:00 | 99:01:00 | 0:00:00 |
| NM_001178126:NM_001256296 | 11:00 | 11:01 | 1:00 | 4:00 | 3:01 |
| NM_004327:NM_021574 | 0:00 | 2:00 | 0:00 | 3:00 | 0:00 |
| NM_003753 | 927 | 2 | 678 | 2 | 544 |
| NM_000714:NR_046308:NM_001256531:NM_001256530 | 6:0:0:0 | 7:0:0:0 | 1:0:0:0 | 2:1:0:0 | 6:0:0:0 |
| NM_001130156:NM_022443:NM_001130157 | 4:02:00 | 0:03:00 | 5:07:03 | 9:12:00 | 0:00:00 |
| NM_000249:NM_001354628:NM_001258273:NM_001354617:NM_001354619:NM_001354618:NM_001167617:NM_001354629:NM_001354622:XM_017006450:NM_001354627:NM_001354630:NM_001167618:NM_001167619:NM_001354620:NM_001354616:NM_001354624:XM_005265161:NM_001258271:NM_001354621:NM_001258274:NM_001354623 | 19:2:1:0:2:2:0:0:1:0:0:0:0:0:0:0:0:0:0:0:0:0 | 13:0:1:1:0:3:0:0:0:0:1:0:0:1:0:0:0:0:0:0:0:0 | 17:2:1:1:1:7:0:3:1:1:1:0:0:0:0:0:0:0:0:0:0:0 | 9:2:1:1:1:3:1:1:0:0:0:0:0:0:0:0:0:0:0:0:0:0 | 29:3:1:0:1:1:0:1:0:0:0:2:0:1:0:0:1:0:0:0:0:0 |
| XR_002959510:NR_146158:NM_014159:XR_002959515:XR_002959514:XM_011533634:XR_002959512:XR_002959517:XR_002959511:XM_024453488:XM_024453487 | 2:0:9:0:0:0:1:0:0:0:0 | 2:0:0:0:0:0:0:0:0:0:0 | 2:2:3:0:0:0:0:0:0:0:0 | 1:0:0:0:0:0:0:0:0:0:0 | 0:3:6:0:0:0:0:0:0:0:0 |
| NM_014159:XR_002959510:XM_011533634:XR_002959511:XR_002959514:XR_002959515:XR_002959516:NR_146158:XR_002959512:XR_002959517:XR_002959513:XM_024453487:XM_024453488 | 5:3:0:0:1:0:0:0:1:0:0:0:0 | 1:3:0:1:1:0:1:1:0:0:0:0:0 | 2:2:0:0:0:0:0:0:0:0:0:0:0 | 1:2:0:0:0:0:0:0:0:0:0:0:0 | 4:1:0:0:0:0:0:0:0:0:1:0:0 |
| XM_017005921:XM_005264967:XM_017005918:NM_012235:NM_001320044 | 2:5:0:0:0 | 1:6:0:0:0 | 0:1:0:0:0 | 2:1:0:0:0 | 2:0:0:0:0 |
| NM_014041 | 3 | 4 | 3 | 4 | 19 |
| NM_001098634:XM_005248103 | 0:00 | 2:03 | 0:02 | 1:00 | 1:01 |
| NM_016107:NR_144318 | 5:11 | 3:22 | 1:00 | 2:00 | 0:00 |
| NM_014324:NM_203382:NM_001167595 | 0:00:00 | 82:04:02 | 0:00:00 | 18:02:00 | 0:00:00 |
| NM_173514:NM_001349384:XM_017009082:XR_948238:XM_011543174:NM_001349382:NM_001349385:XM_024454369:XM_017009077:NR_146165:XM_011543176:XM_017009080:XR_948237:NM_001258286:NM_001349383 | 0:0:0:0:0:0:0:0:0:0:0:0:0:0:0 | 1:2:0:0:0:2:1:1:2:0:0:0:0:0:0 | 0:0:0:0:0:0:0:0:0:0:0:0:0:0:0 | 1:1:0:0:0:0:0:0:0:0:0:0:0:0:0 | 3:1:0:0:0:0:0:0:0:0:0:0:0:1:0 |
| NM_000521 | 0 | 4 | 0 | 2 | 0 |
| NM_001901 | 0 | 1 | 0 | 3 | 0 |
| NM_014432:XM_017010954:NM_001278723:XM_017010955:NM_001278722:XM_011535904:NM_001278724 | 0:0:1:0:0:0:0 | 1:1:1:0:0:0:0 | 2:0:0:0:0:0:0 | 1:0:0:0:0:0:0 | 6:0:4:1:1:0:0 |
| NM_001111077:NM_003379:XM_011536110 | 621:725:51 | 3:01:01 | 122:81:10 | 1:00:00 | 279:126:13 |
| NM_002116:NM_001242758 | 0:00 | 17:00 | 0:00 | 1:00 | 0:04 |
| NM_002116:NM_001242758 | 0:00 | 16:00 | 0:00 | 1:00 | 0:05 |
| NM_002116:NM_001242758 | 0:00 | 17:00 | 0:00 | 1:00 | 0:05 |
| NM_002116:NM_001242758 | 0:00 | 17:00 | 0:00 | 1:00 | 0:05 |
| NM_002116:NM_001242758 | 0:00 | 17:00 | 0:00 | 1:00 | 0:00 |
| NM_002116:NM_001242758 | 0:00 | 17:00 | 0:00 | 1:00 | 0:00 |
| NM_002116:NM_001242758 | 0:00 | 17:00 | 0:00 | 1:00 | 0:06 |
| NM_002116:NM_001242758 | 0:00 | 17:00 | 0:00 | 1:00 | 0:06 |
| NM_002116:NM_001242758 | 0:00 | 17:00 | 0:00 | 1:00 | 0:06 |
| NM_002116:NM_001242758 | 0:00 | 17:00 | 0:00 | 1:00 | 0:06 |
| NM_002116:NM_001242758 | 0:00 | 18:00 | 0:00 | 1:00 | 0:06 |
| NM_002116:NM_001242758 | 0:00 | 18:00 | 0:00 | 1:00 | 0:06 |
| NM_002116:NM_001242758 | 0:00 | 19:00 | 0:00 | 1:00 | 0:00 |
| NM_002116:NM_001242758 | 0:00 | 19:00 | 0:00 | 1:00 | 0:00 |
| NM_002117:NM_001243042 | 0:25 | 206:04:00 | 0:07 | 60:03:00 | 0:00 |
| NM_002117:NM_001243042 | 0:25 | 206:04:00 | 0:07 | 60:03:00 | 0:00 |
| NM_002117:NM_001243042 | 0:25 | 206:04:00 | 0:07 | 60:03:00 | 0:00 |
| NM_002117:NM_001243042 | 0:25 | 203:04:00 | 0:07 | 60:03:00 | 0:00 |
| NM_002117:NM_001243042 | 0:23 | 199:04:00 | 0:07 | 60:03:00 | 0:00 |
| NM_002117:NM_001243042 | 0:23 | 198:04:00 | 0:07 | 60:03:00 | 0:00 |
| NM_002117:NM_001243042 | 0:23 | 195:04:00 | 0:07 | 60:03:00 | 0:00 |
| NM_002117:NM_001243042 | 177:28:00 | 3:03 | 54:10:00 | 1:00 | 82:01:00 |
| NM_002117:NM_001243042 | 171:24:00 | 2:06 | 53:10:00 | 1:01 | 78:01:00 |
| NM_001243042:NM_002117 | 24:167 | 5:00 | 10:51 | 1:00 | 0.05 |
| NM_002117:NM_001243042 | 0:19 | 182:00:00 | 0:07 | 55:02:00 | 0:00 |
| NM_002117:NM_001243042 | 1:20 | 180:00:00 | 0:07 | 55:02:00 | 0:00 |
| NM_005514 | 1751 | 1894 | 687 | 428 | 1024 |
| NM_005514 | 3382 | 7 | 1032 | 3 | 961 |
| NM_005514 | 0 | 3290 | 1 | 990 | 119 |
| NM_005514 | 6 | 3238 | 2 | 971 | 1 |

| | | | | | |
|---|---|---|---|---|---|
| NM_005514 | 3218 | 7 | 963 | 4 | 0 |
| NM_005514 | 3 | 3145 | 0 | 949 | 897 |
| NM_005514 | 1474 | 1649 | 571 | 369 | 887 |
| NM_005514 | 1470 | 1653 | 570 | 370 | 887 |
| NM_005514 | 3115 | 6 | 939 | 1 | 887 |
| NM_005514 | 1503 | 1659 | 583 | 379 | 116 |
| NM_005514 | 1499 | 1658 | 583 | 379 | 117 |
| NM_005514 | 1495 | 1657 | 582 | 379 | 116 |
| NM_005514 | 1498 | 1658 | 584 | 378 | 116 |
| NM_005514 | 1499 | 1678 | 580 | 385 | 117 |
| NM_005514 | 1509 | 1680 | 586 | 380 | 1 |
| NM_005514 | 6 | 3177 | 2 | 966 | 0 |
| NM_005514 | 5 | 3180 | 3 | 967 | 0 |
| NM_005514 | 10 | 3184 | 1 | 969 | 0 |
| NM_005514 | 4 | 3196 | 1 | 971 | 1 |
| NM_005514 | 1534 | 1688 | 590 | 383 | 795 |
| NM_005514 | 1535 | 1688 | 592 | 382 | 794 |
| NM_005514 | 1526 | 1662 | 577 | 376 | 788 |
| NM_005514 | 1522 | 1655 | 576 | 375 | 786 |
| NM_005514 | 1512 | 1631 | 573 | 371 | 781 |
| NM_005514 | 1 | 3145 | 1 | 944 | 778 |
| NM_005514 | 1513 | 1639 | 572 | 375 | 778 |
| NM_001243965:NM_002124:XM_024452554:XM_024452553 | 0:1:0:0 | 0:0:1:1 | 0:1:0:0 | 1:0:0:0 | 0:0:0:0 |
| NM_001243965:NM_002124:XM_024452554:XM_024452553 | 0:1:0:0 | 0:0:1:1 | 0:1:0:0 | 1:0:0:0 | 0:0:0:0 |
| NM_001243965:NM_002124:XM_024452554:XM_024452553 | 0:1:0:0 | 0:0:1:1 | 0:1:0:0 | 1:0:0:0 | 0:0:0:0 |
| NM_002124:XM_024452554:XM_024452553 | 0:00:00 | 1:01:01 | 0:00:00 | 1:00:00 | 0:00:00 |
| NM_002124:XM_024452554:XM_024452553 | 0:00:01 | 1:00:00 | 0:00:00 | 1:00:00 | 0:00:00 |
| NM_002124:XM_024452554:XM_024452553 | 0:00:01 | 1:00:00 | 0:00:00 | 1:00:00 | 0:00:00 |
| NM_002124:XM_024452554 | 0:00 | 1:00 | 0:00 | 1:00 | 0:00 |
| NM_002122:XM_006715079 | 0:00 | 1:00 | 2:00 | 1:00 | 0:00 |
| NM_002122:XM_006715079 | 0:00 | 1:00 | 2:00 | 1:00 | 0:00 |
| NM_002122:XM_006715079 | 0:00 | 1:00 | 2:00 | 1:00 | 0:00 |
| NM_002122:XM_006715079 | 0:00 | 1:00 | 2:00 | 1:00 | 0:00 |
| NM_002122:XM_006715079 | 0:00 | 1:00 | 2:00 | 1:00 | 0:00 |
| NM_002122:XM_006715079 | 0:00 | 1:00 | 2:00 | 1:00 | 0:00 |
| NM_002122:XM_006715079 | 0:00 | 1:00 | 2:00 | 1:00 | 0:00 |
| NM_002122:XM_006715079 | 0:00 | 1:00 | 2:00 | 1:00 | 0:00 |
| NM_002122:XM_006715079 | 0:00 | 1:00 | 2:00 | 1:00 | 0:00 |
| NM_002123:NM_001243962 | 0:00 | 2:00 | 0:01 | 1:00 | 1:00 |
| NM_002123:NM_001243962 | 0:00 | 2:00 | 0:01 | 1:00 | 1:00 |
| NM_002123:NM_001243962 | 0:00 | 2:00 | 0:01 | 1:00 | 1:00 |
| NM_002123:NM_001243962 | 0:00 | 2:00 | 0:01 | 1:00 | 0:00 |
| NM_006586:NM_001318845:NR_134888:NR_134882 | 0:0:0:0 | 0:1:1:0 | 0:0:0:0 | 1:0:0:0 | 0:0:0:0 |
| NM_018368 | 0 | 3 | 0 | 1 | 0 |
| NM_001913:NM_181500:XM_006715855:XM_005250154:NM_001202546:XM_017011760:NM_001202545:NM_001202544:XM_024446668:XM_011515825 | 24:7:1:5:0:0:0:0:0:0 | 34:5:0:3:0:0:1:0:0:0 | 51:16:4:3:2:0:0:0:0:0 | 53:13:8:2:0:0:0:0:0:0 | 37:16:5:1:0:0:0:0:0:0 |
| XR_001744836:NM_182931:XM_011516400:XM_017012435:NM_018682:XM_024446837 | 119:30:4:1:0:2 | 4:2:0:0:0:0 | 53:17:1:2:0:0 | 5:1:1:0:0:0 | 48:11:0:3:0:1 |
| XM_017011987:NM_014390 | 0.514583333 | 0:01 | 0.738194444 | 1:01 | 0.736805556 |
| XM_024446655:NM_001331192 | 0:00 | 1:01 | 0:00 | 1:00 | 0:00 |
| NM_001101312:NM_001101311:NM_014020:XM_024446733:NM_001101314:XM_024446734:XM_006715933 | 1660:67:39:11:17:69:1 | 1388:70:45:8:14:43:2 | 568:52:9:6:9:26:1 | 538:37:10:4:7:18:2 | 474:57:9:4:17:13:4 |
| NM_014020:NM_001101312:NM_001101311:XM_024446733:XM_024446734:NM_001101314:XM_006715933 | 1:1:0:0:1:0:0 | 73:3059:137:19:66:1:1 | 1:0:0:0:1:0:0 | 17:1113:89:10:37:0:0 | 0:0:0:0:0:0:0 |
| XM_017012506:NM_022373:XR_926944:XR_001744846 | 2:1:0:0 | 3:2:0:0 | 3:1:0:0 | 2:3:0:0 | 2:2:0:0 |
| NM_001322005:NM_000535:NM_001322015:XM_024446800:NM_001322006:NR_136154:NM_001322003:NM_001322004:XM_017012342:NM_001322014:NM_001322007:NM_001322008:NM_001322012:NM_001322011 | 0:0:0:0:0:0:0:0:0:0:0:0:0:0 | 1:6:1:1:0:0:1:1:0:0:0:0:0:0 | 0:0:0:0:0:0:0:0:0:0:0:0:0:0 | 1:6:0:0:0:0:0:0:0:0:0:0:0:0 | 0:3:0:0:0:0:1:0:0:0:0:0:0:0 |
| NM_002568:XM_005250861 | 960:38:00 | 1:00 | 4143:141 | 2:00 | 1963:43:00 |
| NM_002568:XM_005250861 | 957:36:00 | 3:00 | 4132:143 | 5:00 | 1957:43:00 |
| NM_002568:XM_005250861 | 785:33:00 | 1:01 | 3554:140 | 6:01 | 1668:46:00 |
| NM_002568:XM_005250861 | 729:35:00 | 2:00 | 3365:138 | 3:00 | 1593:48:00 |
| NM_004462:NM_001287743:NM_001287756:NM_001287742:NM_001287745:NM_001287744:NM_001287750:NM_001287751 | 2395:4:10:118:1:1:0:0 | 13:0:0:0:0:0:0:0 | 460:5:4:22:1:0:0:0 | 1:0:0:0:0:0:0:0 | 677:5:1:26:0:0:1:0 |

| | | | | | |
|---|---|---|---|---|---|
| NM_001908:NM_147782:XM_024447071:NM_147780:XM_017013100:NM_147783:XM_017013097:XM_017013099:NM_001317237 | 0:0:0:0:0:0:0:0:0 | 21:3:2:0:1:1:1:0:0 | 0:0:0:0:0:0:0:0:0 | 28:3:5:1:0:0:0:0:0 | 27:3:6:0:0:0:0:0:0 |
| NM_001979:XR_001745491:XM_017013199:XM_017013200 | 52:2:1:2 | 54:3:3:0 | 0:0:0:0 | 23:1:0:0 | 8:0:1:0 |
| XR_949472:XM_011544639 | 2:02 | 3:01 | 0:00 | 2:00 | 1:00 |
| NM_052937:NM_001286783:XR_001745466 | 46:01:02 | 2:00:00 | 62:03:04 | 1:00:00 | 10:00:00 |
| NM_003878:XM_011517623 | 2:00 | 1:00 | 3:00 | 1:00 | 5:00 |
| NM_003878:XM_011517623 | 2:00 | 1:00 | 3:00 | 1:00 | 5:00 |
| NM_004063:XM_011516790 | 715:17:00 | 2:00 | 577:12:00 | 2:00 | 376:17:00 |
| NM_004063:XM_011516790 | 0:00 | 717:16:00 | 0:00 | 579:12:00 | 144:06:00 |
| NM_001317944:NM_017688:XM_006717149:NM_001317943 | 1:22:2:0 | 2:11:1:2 | 0:9:2:0 | 2:12:1:0 | 0:21:5:0 |
| NM_002451 | 0 | 32 | 0 | 75 | 118 |
| NM_004323 | 1 | 24 | 0 | 11 | 0 |
| NM_133181:XM_011542134:NM_024526:XR_001737407 | 1:0:0:0 | 0:0:0:0 | 0:0:0:0 | 0:0:0:0 | 0:0:0:0 |
| NM_001160233:XM_017001361:XR_002956654:NM_001160234:NM_000701 | 0:110:48:354:128 | 0:0:1:1:0 | 2:39:16:187:124 | 0:0:0:0:0 | 1:105:22:312:151 |
| XR_002956654:NM_001160234:NM_001160233:XM_017001361:NM_000701 | 99:424:0:104:128 | 0:0:0:0:0 | 35:220:2:36:124 | 1:1:0:0:0 | 47:354:1:102:152 |
| NM_203458 | 4 | 0 | 1 | 0 | 0 |
| NM_153649:NR_103461:NM_001043351:NM_001278189:NM_001349679:NM_001278188:NM_001043352:NM_001278190:NM_152263:NM_001278191:NM_001043353 | 968:7:4:4:0:5:7:0:0:1:0 | 0:0:0:0:0:0:0:0:0:0:0 | 591:8:2:7:1:2:3:0:0:0:0 | 0:0:0:0:0:0:0:0:0:0:0 | 1096:7:7:13:0:11:41:0:0:0:0 |
| XM_017001525:XM_006710704:XM_024447751:XM_017001526:XM_017001519:NM_001350151:XM_006710705:XM_011541617:NM_001024216:XM_011541616:NM_017556 | 4:0:1:1:1:1:0:0:0:0:0 | 0:0:0:0:0:0:0:0:0:0:0 | 0:0:0:0:0:0:0:0:0:0:0 | 0:0:0:0:0:0:0:0:0:0:0 | 0:0:0:0:0:0:0:0:0:0:0 |
| NM_001785 | 2 | 0 | 0 | 0 | 1 |
| NM_005216 | 0 | 3 | 0 | 0 | 0 |
| NM_022745 | 0 | 1 | 0 | 0 | 0 |
| NM_176877:XM_024448614:XM_011540465:XR_001736900:XM_017000001:XR_002957157:XM_011540469:NM_001350145 | 1:1:1:0:0:0:0:0 | 0:0:0:0:0:0:0:0 | 1:1:0:0:0:0:0:0 | 0:0:0:0:0:0:0:0 | 0:0:0:0:0:0:0:0 |
| XM_011542191:XM_011542192:NM_001002292:NM_024911:XM_017002390:NM_001193334 | 6:2:0:24:7:0 | 0:0:0:0:0:0 | 16:7:4:32:6:0 | 0:0:0:0:0:0 | 11:0:1:8:1:1 |
| NM_001304717:NM_001304718 | 216:03:00 | 0:00 | 284:02:00 | 0:00 | 243:04:00 |
| NM_001304717:NM_001304718 | 217:02:00 | 0:00 | 285:01:00 | 0:00 | 237:02:00 |
| NM_002817:NM_175932 | 0:00 | 0:00 | 0:00 | 1:00 | 0:00 |
| NM_015957:XM_017017875:XM_011520154 | 284:06:05 | 0:01:00 | 138:04:01 | 0:00:00 | 196:06:03 |
| NM_001001390:NM_001202555:XM_005253238:NM_001001391:NM_001001389:XM_011520488:XM_011520486:XM_011520483:XM_005253240:XM_005253235:XM_017018583:XM_006718388:XM_011520485:XM_017018585:NM_001202556:NM_001202557:XM_005253239:NM_001001392:NM_000610:XM_005253232:XM_005253231:XM_011520484:XM_017018584:XM_011520482:XM_011520489 | 29:2:3:124:0:5:0:0:1:0:4:0:0:5:1:1:1:1:0:0:0:0:0:0:0 | 0:0:0:1:0:0:0:0:0:0:0:0:0:0:0:0:0:0:0:0:0:0:0:0:0 | 85:4:16:71:3:2:3:2:4:2:4:1:1:2:4:0:0:0:0:0:0:0:0:0:0 | 0:0:0:0:0:0:0:0:0:0:0:0:0:0:0:0:0:0:0:0:0:0:0:0:0 | 27:0:16:20:3:4:0:0:8:3:7:1:0:2:0:0:0:1:3:0:2:2:0:0:0 |
| NM_001085372 | 0 | 1 | 2 | 0 | 0 |
| NM_000852 | 14 | 0 | 6 | 1 | 15 |
| NM_016156:NM_201281:NM_201278:XM_005274375:NM_001243571:XM_005274374 | 1:2:0:1:0:0 | 0:0:0:0:0:0 | 5:2:2:1:0:0 | 0:0:0:0:0:0 | 0:1:0:0:0:1 |
| XM_006719552:XM_017019788:XM_017019792:XM_017019790:NM_198240:XM_017019786 | 0:0:0:3:0:0 | 0:0:0:2:0:0 | 1:1:0:0:0:0 | 0:0:0:0:0:0 | 0:0:0:0:0:0 |
| NM_024829 | 0 | 0 | 0 | 0 | 0 |
| NM_004985:NM_033360 | 7:08 | 0:00 | 1:02 | 0:00 | 3:06 |
| NM_000059 | 1 | 0 | 8 | 0 | 13 |
| NM_020751:NR_026745:XM_011535168:NM_001145079 | 7:0:0:0 | 0:0:0:0 | 51:2:1:0 | 0:0:0:0 | 3:0:0:0 |
| NM_020751:NR_026745:XM_011535168:NM_001145079 | 8:0:0:0 | 0:0:0:0 | 51:2:1:0 | 0:0:0:0 | 3:0:0:0 |
| NM_021239:XM_011537044:XR_943501:XM_024449680 | 10:5:47:0 | 1:0:0:0 | 7:2:31:2 | 0:0:0:0 | 7:1:40:0 |
| NM_033250:NM_033240:NM_033244:NM_033239 | 0:0:0:0 | 2:2:0:0 | 0:0:0:0 | 0:0:0:0 | 0:0:0:0 |
| NM_033240:NM_033250:NM_033244:NM_033239 | 0:2:0:0 | 2:0:0:0 | 0:0:0:0 | 0:0:0:0 | 0:2:0:0 |
| NM_005153:XM_017023869:XM_006721332:NM_001272075:XM_017023864 | 12:0:1:0:0 | 0:0:0:0:0 | 13:1:0:0:0 | 0:0:0:0:0 | 20:3:1:1:0 |
| XM_011522943:NM_001127214:NM_174917:NM_001243279:XR_933238 | 0:0:0:0:0 | 0:0:0:0:0 | 0:0:0:0:0 | 0:0:0:0:0 | 0:0:0:0:0 |
| XM_024450549:NM_006804 | 0:00 | 0:00 | 0:00 | 0:00 | 0:00 |
| XM_024450643:NM_004448:NR_110535:NM_001289937:NM_001289938 | 68:1:29:0:0 | 50:3:22:0:0 | 21:1:8:0:0 | 0:0:0:0:0 | 0:0:0:0:0 |
| NM_001165959:NM_001165958:NM_001042471:NM_018530 | 10:32:13:1 | 3:6:20:1 | 4:2:2:0 | 0:0:0:0 | 2:15:13:0 |
| NM_001165959:NM_001165958:NM_001042471:NM_018530 | 10:32:13:1 | 3:6:20:1 | 4:2:2:0 | 0:0:0:0 | 2:15:13:0 |

| | | | | | |
|---|---|---|---|---|---|
| XM_024451789:NM_004864 | 3:00 | 0:00 | 2:00 | 0:00 | 0:00 |
| XR_002958354:NM_003430:XR_001753756 | 2:02:00 | 5:01:00 | 0:00:00 | 0:00:00 | 0:00:00 |
| XR_002958354:NM_003430:XR_001753756 | 2:02:00 | 5:01:00 | 0:00:00 | 0:00:00 | 0:00:00 |
| XR_002958354:NM_003430 | 8:02 | 0:00 | 0:00 | 0:00 | 0:00 |
| XR_002958354:XM_011528266:XM_011528265:XM_017027236:NM_003430 | 20:1:0:0:0 | 2:0:0:0:0 | 10:1:1:1:0 | 0:0:0:0:0 | 8:2:0:0:0 |
| NM_018025:NR_135270:XM_006723255 | 13:00:00 | 0:00:00 | 11:01:00 | 0:00:00 | 0:00:00 |
| NM_000041:NM_001302688:NM_001302691 | 8:00:01 | 0:00:00 | 9:01:01 | 0:00:00 | 1:00:01 |
| NM_024682:XM_011527317:NM_001168222 | 1:00:01 | 0:00:00 | 1:00:00 | 0:00:00 | 0:00:00 |
| NM_001985:XM_024451418 | 1:00 | 0:00 | 1:00 | 0:00 | 0:00 |
| NM_003937:XM_024453208:NM_001199241:XR_923058 | 6:1:0:0 | 0:0:0:0 | 13:1:0:0 | 0:0:0:0 | 0:0:0:0 |
| NM_002156:NM_199440 | 1019:65 | 1:00 | 2005:67 | 0:00 | 2370:97 |
| NM_017735:XR_002959314:NM_001193509:XM_011532958 | 11:1:0:0 | 0:0:0:0 | 17:0:0:0 | 0:0:0:0 | 7:0:0:0 |
| NM_019896 | 0 | 0 | 0 | 0 | 1 |
| NM_001321459:NR_135653:NR_135650:NR_135651:XM_024452960:NM_001321454:XM_017004311 | 0:1:0:0:0:0:0 | 0:0:0:0:0:0:0 | 1:1:0:0:0:0:0 | 0:0:0:0:0:0:0 | 0:0:0:0:0:1:0 |
| NM_006814 | 0 | 0 | 0 | 0 | 0 |
| NM_002862:XR_002958496 | 9:00 | 0:00 | 7:00 | 0:00 | 0:00 |
| XR_002958471:NM_001077489:XM_017027817:NM_001309840:NM_001077488:NM_080426:XM_017027818:XM_017027813:NM_000516:XM_024451874:NM_080425:XM_024451875:XM_017027819:XM_024451873:XM_017027815:XM_017027812:XM_024451872:NM_016592:XM_017027820:XM_017027814:NM_001309861 | 42:101:42:9:23:265:3:10:110:1:1:8:7:2:2:2:0:0:0:0:0 | 0:0:0:0:0:0:0:0:0:0:0:0:0:0:0:0:0:0:0:0:0 | 66:105:40:12:9:278:10:16:87:2:2:3:7:0:0:0:0:0:0:0:0 | 1:0:0:0:0:0:0:0:0:0:0:0:0:0:0:0:0:0:0:0:0 | 107:155:32:20:24:299:5:29:172:1:3:11:10:0:0:1:0:1:0:0:1 |
| NM_001077489:XM_017027817:NM_001309840:NM_001077488:NM_080426:XM_017027818:XM_017027813:NM_000516:XR_002958471:XM_024451874:NM_080425:XM_024451875:XM_017027819:XM_024451873:XM_017027815:XM_017027812:XM_024451872:NM_016592:XM_017027820:XM_017027814:NM_001309861 | 100:42:9:22:276:3:10:110:58:1:0:7:7:2:2:2:0:0:0:0:0 | 0:0:0:0:0:0:0:3:0:0:1:0:0:0:0:0:0:0:0:0:0 | 105:40:12:9:285:10:14:87:87:2:2:3:7:0:0:0:0:0:0:0:0 | 0:0:0:0:0:0:0:0:0:0:0:0:0:0:0:0:0:0:0:0:0 | 155:32:20:24:321:5:29:172:145:1:3:11:10:0:0:1:0:1:0:0:1 |
| NR_103797:NM_138927 | 0:00 | 0:00 | 0:00 | 2:01 | 3:00 |
| NM_033049 | 1 | 1 | 0 | 0 | 1 |
| XM_024453325:XM_011512355:XM_017005614:NM_012190:NR_072979 | 1:0:0:0:0 | 2:2:1:1:0 | 0:0:0:0:0 | 0:0:0:0:0 | 1:1:0:0:1 |
| XR_924148:NM_001184:XM_011512925:NM_001354579 | 2:0:0:0 | 1:0:0:0 | 1:0:0:0 | 0:0:0:0 | 1:0:0:0 |
| XM_024453430:NM_015303:NR_146113 | 6:04:04 | 0:00:00 | 1:01:00 | 0:00:00 | 1:00:00 |
| NM_003965:XM_017007436:XM_011534209:XM_011534208:NM_001130910 | 53:5:0:4:5 | 0:0:0:0:0 | 13:1:0:0:0 | 0:0:0:0:0 | 7:5:1:2:1 |
| NM_002375:XM_024453530:XR_002959534:XM_017006412:XM_017006402:XR_002959526:XR_002959525:XR_002959533:XM_017006410:XR_002959532:XR_002959527:XR_002959524:NM_001134364:XR_002959528:XM_024453528:XM_017006416 | 0:3:0:1:0:0:0:2:0:3:0:0:0:0:0:0 | 0:0:0:0:0:0:0:0:0:0:0:0:0:0:0:0 | 1:3:1:1:1:1:1:1:0:0:0:0:0:0:0:0 | 0:0:0:0:0:0:0:0:0:0:0:0:0:0:0:0 | 0:0:0:0:0:1:0:0:1:3:0:0:0:0:0:0 |
| NM_002375:XR_002959534:XM_017006412:XM_017006402:XR_002959526:XR_002959525:XR_002959533:XR_002959532:XM_024453530:XR_002959527:XR_002959524:XR_002959530:XR_002959528:XM_017006410 | 0:3:1:0:0:0:2:1:0:2:0:0:0:0 | 0:0:0:0:0:0:0:0:0:0:0:0:0:0 | 1:1:1:1:1:1:3:6:0:0:0:0:0:0 | 0:0:0:0:0:0:0:0:0:0:0:0:0:0 | 0:0:0:0:1:0:1:6:0:0:0:0:0:0 |
| NM_014366:NM_206826:NM_206825 | 55:08:01 | 0:00:00 | 192:22:04 | 2:01:00 | 318:60:9 |
| NM_020139 | 0 | 1 | 0 | 0 | 0 |
| NM_004684:NM_001128310:NM_001291976 | 70:03:00 | 0:00:00 | 54:01:00 | 0:00:00 | 30:02:00 |
| NM_004684:NM_001128310:NM_001291976 | 71:03:00 | 0:00:00 | 54:01:00 | 0:00:00 | 30:02:00 |
| NM_016144:NR_146220:NM_001308080 | 72:00:00 | 0:00:00 | 110:01:01 | 0:00:00 | 79:01:00 |
| NM_014324:NM_203382:NM_001167595 | 84:04:02 | 0:00:00 | 18:02:00 | 0:00:00 | 15:00:00 |
| XM_017009329:XM_017009330:XM_006714468 | 1:00:00 | 0:00:00 | 3:01:00 | 0:00:00 | 0:00:00 |
| NM_133343:NM_133338:NM_133344:NM_133342:NM_001278622:NM_002873:XM_017009681:NM_133340:NM_133341 | 2:0:5:13:2:0:2:7:0 | 0:0:0:0:0:0:0:0:0 | 3:1:4:16:5:1:1:0:0 | 0:0:0:0:0:0:0:0:0 | 2:0:2:11:1:0:6:3:3 |
| XM_017009634:NM_018429 | 4:00 | 0:00 | 6:00 | 0:00 | 0:00 |
| XM_017009634 | 1 | 0 | 2 | 0 | 0 |
| XM_017009634 | 1 | 0 | 1 | 0 | 0 |
| NM_138571 | 0 | 0 | 0 | 1 | 0 |
| NM_032020 | 1 | 0 | 0 | 0 | 1 |
| XM_011536147:NM_025059:XM_011536148 | 0:00:04 | 0:00:00 | 1:03:08 | 0:00:00 | 0:00:16 |
| NM_001142522:NM_012177 | 0:00 | 0:00 | 0:00 | 0:00 | 0:00 |
| XM_017010958:XM_011535905:NM_001143947:NM_018974:XM_011535906:XM_011535908:XM_011535907 | 0:1:0:0:0:0:0 | 0:0:0:0:0:0:0 | 2:6:3:1:0:0:0 | 0:0:0:0:0:0:0 | 0:0:0:0:0:0:0 |
| XM_017010958:XM_011535905:NM_001143947:NM_018974:XM_011535906:XM_011535907 | 0:1:0:0:0:0 | 0:0:0:0:0:0 | 3:6:3:1:0:0 | 0:0:0:0:0:0 | 0:0:0:0:0:0 |
| NM_002793 | 2 | 0 | 1 | 1 | 5 |

| | | | | | |
|---|---|---|---|---|---|
| NM_000367:NM_001346818:NM_001346817 | 101:02:05 | 0:00:00 | 34:02:02 | 0:00:00 | 107:02:13 |
| NM_002117:NM_001243042 | 179:31:00 | 1:00 | 56:10:00 | 0:00 | 76:00:00 |
| NM_002117:NM_001243042 | 178:28:00 | 0:03 | 54:10:00 | 0:00 | 75:00:00 |
| NM_002117:NM_001243042 | 178:28:00 | 0:03 | 54:10:00 | 0:00 | 75:00:00 |
| NM_002117:NM_001243042 | 165:27:00 | 2:03 | 51:11:00 | 0:00 | 6:01 |
| NM_002117:NM_001243042 | 167:28:00 | 0:01 | 51:11:00 | 0:00 | 72:00:00 |
| NM_005514 | 3214 | 3 | 961 | 0 | 794 |
| NM_001008703:NM_178508 | 0:00 | 0:00 | 0:00 | 0:00 | 0:00 |
| NM_031480:NM_001348194:XM_011514933 | 45:07:01 | 0:00:00 | 61:03:02 | 0:00:00 | 66:09:08 |
| NM_001203247:XM_005249962:XM_024446680:XM_011515884:XR_002956414:NM_004456:XM_017011818:XM_017011821:XM_011515894:XR_002956413:XM_017011819:XM_011515897:NM_152998:XM_011515887:XM_017011820:XM_011515889:XM_011515898:NM_001203248 | 11:0:1:0:0:0:0:1:0:1:0:0:0:0:0:0:0:0 | 0:0:0:0:0:0:0:0:0:0:0:0:0:0:0:0:0:0 | 3:1:1:1:1:1:0:0:0:0:0:0:0:0:0:0:0:0 | 0:0:0:0:0:0:0:0:0:0:0:0:0:0:0:0:0:0 | 9:0:0:0:0:2:0:2:0:0:0:0:0:0:1:0:0:0 |
| NM_014020:NM_001101312:NM_001101311:NM_001101314:XM_024446733:XM_024446734:XM_006715933 | 84:3038:136:30:19:126:3 | 0:7:0:0:0:0:0 | 20:1105:89:16:10:50:3 | 0:0:0:0:0:0:0 | 21:922:94:28:15:39:7 |
| NM_014020:NM_001101312:NM_001101311:NM_001101314:XM_024446733:XM_024446734:XM_006715933 | 85:3044:137:31:19:120:3 | 0:2:0:0:0:0:0 | 20:1105:89:16:10:46:3 | 0:0:0:0:0:0:0 | 21:924:94:28:15:34:7 |
| NM_002047:NM_001316772 | 0:00 | 0:00 | 0:00 | 0:00 | 0:00 |
| XM_017012102:NM_001637:XM_011515336:XM_011515339:XM_011515340:XM_017012106:XM_011515341:XM_011515335:XM_017012104:XM_011515338:NM_001177507 | 0:19:1:1:7:3:1:1:1:0:0 | 0:0:0:0:0:0:0:0:0:0:0 | 2:52:2:1:16:7:1:0:0:0:0 | 0:0:0:0:0:0:0:0:0:0:0 | 0:4:0:0:6:5:0:0:0:0:0 |
| XR_002956470 | 0 | 0 | 0 | 0 | 0 |
| XR_002956470 | 0 | 0 | 0 | 0 | 0 |
| NM_001322005:NM_000535:NM_001322015:XM_024446800:NM_001322006:NM_001322003:NM_001322004:NR_136154:NM_001322014:XM_017012342:NM_001322008:NM_001322007:NM_001322012:NM_001322011 | 1:6:1:1:0:1:1:0:0:0:0:0:0:0 | 0:0:0:0:0:0:0:0:0:0:0:0:0:0 | 1:6:0:0:0:0:0:0:0:0:0:0:0:0 | 0:0:0:0:0:0:0:0:0:0:0:0:0:0 | 0:4:0:0:0:2:0:0:0:0:0:0:0:0 |
| XM_024447170:XM_017013496:XM_024447168:XM_024447167:XM_017013493:XM_006716340:XM_024447171:NM_001352641 | 0:3:0:0:1:1:0:0 | 0:0:0:0:0:0:0:0 | 1:0:0:0:0:0:0:0 | 0:0:0:0:0:0:0:0 | 0:0:0:0:0:0:0:0 |
| XM_024447265:XM_011544639 | 0:02 | 0:00 | 1:00 | 0:00 | 0:00 |
| XM_024447164:XM_024447165:NM_002485:NM_001024688:XM_011517046:XM_024447163:XM_011517045:XM_017013462:XM_017013460 | 6:1:11:1:6:0:0:0:0 | 0:0:0:0:0:0:0:0:0 | 2:2:38:6:12:1:0:0:0 | 0:0:0:0:0:0:0:0:0 | 0:0:0:0:0:0:0:0:0 |
| NM_012415:NM_001205262 | 2:00 | 0:00 | 3:00 | 0:00 | 7:00 |
| NM_002799 | 4 | 0 | 0 | 0 | 3 |
| NM_033428:NM_001318058 | 1:00 | 0:00 | 1:00 | 0:00 | 4:00 |
| NM_031420:NR_125331:XR_921910 | 0:00:00 | 0:00:00 | 1:00:00 | 0:00:00 | 3:00:00 |
| NM_153649:NR_103461:NM_001043351:NM_001278189:NM_001349679:NM_001278188:NM_001043352:NM_001278190:NM_152263:NM_001278191 | 949:7:4:4:0:5:7:0:0:0 | 0:0:0:0:0:0:0:0:0:0 | 581:7:2:7:1:2:3:0:0:0 | 1:0:0:0:0:0:0:0:0:0 | 1083:8:7:13:0:11:41:0:0:0 |
| NM_004428:NM_182685 | 0:00 | 0:00 | 1:00 | 0:00 | 0:00 |
| NM_144772:XM_017000319 | 0:00 | 2:00 | 0:00 | 0:00 | 0:00 |
| NR_104400:XM_011509905:NM_003037:XM_005245456:NM_001330754:XM_017002131:XM_017002130:NR_104401:NR_104399 | 0:0:5:0:1:0:1:1:0 | 0:0:0:0:0:0:0:0:0 | 3:1:8:1:0:0:0:0:0 | 0:0:0:0:0:0:0:0:0 | 0:0:1:0:0:0:0:0:0 |
| NM_003000 | 2585 | 15 | 266 | 0 | 339 |
| NM_005558 | 1 | 1 | 0 | 0 | 1 |
| NM_182533:NM_001282670:NM_001256947 | 2:00:00 | 0:00:00 | 0:00:00 | 0:00:00 | 4:01:00 |
| NR_136702:XM_005271115:XM_017002054:XM_011541950:NR_136703 | 17:7:221:1:0 | 0:0:0:0:0 | 30:19:225:5:0 | 0:0:0:0:0 | 24:15:349:3:0 |
| NM_016308:NR_046394 | 4:00 | 0:00 | 0:00 | 0:00 | 0:00 |
| XM_005244771:NM_001142467 | 0:00 | 1:00 | 0:00 | 0:00 | 0:01 |
| NM_020682 | 0 | 0 | 0 | 0 | 2 |
| XM_011539270:XM_006717624 | 0:00 | 0:00 | 0:00 | 0:00 | 0:00 |
| NM_014241:XR_428651 | 0:00 | 0:00 | 1:00 | 0:00 | 0:00 |
| NM_014241:XR_428651 | 0:00 | 0:00 | 0:00 | 1:00 | 0:00 |
| NM_014241:XR_428651 | 0:00 | 0:00 | 0:00 | 1:00 | 0:00 |
| XM_017017365:XR_428971:XM_017017382:XM_017017384:XM_017017367:XM_017017377:XM_006718794:XM_017017381:XM_017017366:XM_017017380:XM_017017379:XM_005271456:XM_017017385:XR_002957133:XM_017017386 | 1:9:4:0:3:0:1:1:0:0:0:0:0:0:0 | 0:1:0:0:0:0:0:0:0:0:0:0:0:0:0 | 1:8:1:1:0:0:0:0:0:0:0:0:0:0:0 | 0:0:0:0:0:0:0:0:0:0:0:0:0:0:0 | 2:6:1:0:0:0:0:0:0:0:0:0:0:0:0 |
| XM_006718341:NM_139273:XR_930914:NM_001751:XM_017018389:NM_001014437:XR_001747995:NR_036542:NM_001194997:XM_017018391:XR_001747996:XR_930913:XR_001747997:XR_001747998:XM_017018392:XM_017018390:XR_002957203:XM_024448705:XM_024448703 | 23:60:11:127:24:13:0:7:10:7:23:5:1:1:1:0:0:0:0 | 0:0:0:0:0:0:0:0:0:0:0:0:0:0:0:0:0:0:0 | 28:51:7:100:12:7:3:2:7:4:8:2:1:1:1:0:0:0:0 | 0:0:0:0:0:0:0:0:0:0:0:0:0:0:0:0:0:0:0 | 31:64:3:116:17:17:2:8:3:7:12:2:0:4:3:1:0:0:0 |
| NM_003477:NM_001166158:XM_011520390:NM_001135024 | 158:7:7:0 | 0:0:0:0 | 50:3:4:1 | 0:0:0:0 | 97:1:5:0 |

| | | | | | |
|---|---|---|---|---|---|
| NM_001001991:NM_001282675:NM_015430 | 1:00:00 | 0:00:00 | 0:00:00 | 0:00:00 | 1:00:00 |
| NM_139178 | 27 | 0 | 20 | 0 | 33 |
| NM_017841 | 422 | 1 | 338 | 0 | 425 |
| NM_002261 | 0 | 0 | 0 | 0 | 0 |
| NM_052845:NR_038118 | 0:00 | 0:00 | 0:00 | 2:00 | 0:00 |
| NM_052845:NR_038118 | 0:00 | 0:00 | 0:00 | 2:00 | 0:00 |
| XM_011520607:NM_001987:XM_017018991:XM_011520608:XM_011520611:XM_017018990:XM_011520612 | 10:14:5:0:2:1:1 | 0:0:0:0:0:0:0 | 6:4:1:0:0:0:0 | 0:0:0:0:0:0:0 | 3:5:7:0:1:0:0 |
| NM_005505:NM_001082959 | 171:11:00 | 0:00 | 62:04:00 | 0:00 | 72:00:00 |
| XM_011535289:NM_144777:NM_003843:XM_011535288:XM_011535290:XM_006719885 | 0:0:0:0:0:0 | 0:0:0:0:0:0 | 0:0:0:0:0:0 | 0:0:0:0:0:0 | 0:0:0:0:0:0 |
| NM_014672:XM_005268237:NM_001256680:NM_001256681:XM_017021836:XM_011537410:NM_001256678 | 2:1:6:4:4:6:1 | 0:0:0:0:0:0:0 | 2:1:7:1:2:0:0 | 0:0:0:0:0:0:0 | 5:5:12:2:1:3:2 |
| XM_005268075:NM_024884:XM_017021657:XM_011537166:XM_017021658:XM_017021656 | 3:0:1:0:0:0 | 0:0:0:0:0:0 | 2:0:0:0:0:0 | 0:0:0:0:0:0 | 3:0:0:0:0:0 |
| NM_001202:NM_130850:NM_130851 | 0:00:00 | 0:00:00 | 0:00:00 | 0:00:00 | 0:00:00 |
| NM_145112:NM_002382:XM_017021312:NR_073137:NM_001320415:XR_002957553:XR_001750327:XR_943452:NM_145113:XR_943451:XM_017021313:XR_943450:XR_001750326 | 361:233:6:13:12:1:4:4:6:1:3:0:0 | 0:0:0:0:0:0:0:0:0:0:0:0:0 | 93:45:10:3:3:2:3:1:1:0:0:0:0 | 0:0:0:0:0:0:0:0:0:0:0:0:0 | 85:85:3:4:4:1:2:1:1:2:0:3:2 |
| NM_001002235:NM_001127707:NM_000295:NM_001127706:XM_017021370:NM_001002236:NM_001127702:NM_001127705:NM_001127703:NM_001127700:NM_001127704:NM_001127701 | 659:58:2:45:86:6:2:2:10:15:5:2 | 0:0:0:0:0:0:0:0:0:0:0:0 | 24:46:18:5:4:1:1:0:0:0:0:0 | 0:0:0:0:0:0:0:0:0:0:0:0 | 35:74:25:4:18:0:1:0:1:2:0:0 |
| NM_001002235:NM_001127707:NM_000295:NM_001127706:NM_001002236:NM_001127702:XM_017021370:NM_001127705:NM_001127703:NM_001127700:NM_001127704:NM_001127701 | 655:48:2:45:6:2:40:2:10:15:4:2 | 0:0:0:0:0:0:1:0:0:0:0:0 | 24:45:19:5:1:1:1:0:0:0:0:0 | 0:0:0:0:0:0:0:0:0:0:0:0 | 35:73:25:4:0:1:4:0:1:2:0:0 |
| NM_144508:XM_017022432 | 0:01 | 0:00 | 2:00 | 0:00 | 3:00 |
| NM_144508:XM_017022432 | 0:00 | 0:00 | 0:00 | 1:00 | 2:00 |
| NM_001128610:XM_017022720:XM_017022718:NM_001283049:XM_017022719:XM_017022722:XM_006720762:XM_024450098:XM_011522193:NM_005154:XM_017022721 | 1:23:79:0:32:4:5:9:0:1:2 | 0:0:0:0:0:0:0:0:0:0:0 | 4:10:52:2:19:2:3:8:1:0:0 | 0:0:0:0:0:0:0:0:0:0:0 | 3:30:68:0:34:4:7:11:0:5:3 |
| NM_007200:NM_001270546 | 3:00 | 4:01 | 2:00 | 0:00 | 0:00 |
| NM_002094:NM_001130006 | 0:00 | 0:00 | 0:00 | 0:00 | 0:00 |
| NM_173474:NM_001270766 | 1:00 | 0:00 | 0:00 | 0:00 | 1:01 |
| NM_173474:NM_001270766 | 1:00 | 0:00 | 0:00 | 0:00 | 1:01 |
| XM_024450134:NM_001797 | 1:00 | 0:00 | 0:02 | 1:01 | 0:00 |
| NM_005823:NM_013404 | 0:00 | 0:00 | 0:00 | 0:00 | 0:00 |
| NM_032865:XM_005257744:XM_017025236:XR_001752669 | 0:3:0:0 | 0:0:0:0 | 1:0:0:0 | 0:0:0:0 | 0:4:0:0 |
| NM_145255 | 1 | 0 | 0 | 0 | 2 |
| NM_001033002:NM_001160246:NR_027683:NM_001160266:NM_001160244:XR_001752663 | 1:0:0:0:0:0 | 0:0:0:0:0:0 | 0:0:0:0:0:0 | 0:0:0:0:0:0 | 1:0:2:1:0:0 |
| XM_005257425:NM_001193609:NM_016261:XM_024450784:NM_001193612:XM_024450783:XM_017024720:XM_017024716 | 0:1:1:0:1:0:0:0 | 0:0:0:0:0:0:0:0 | 3:1:0:0:0:0:0:0 | 0:0:0:0:0:0:0:0 | 0:0:4:0:0:0:0:0 |
| NM_018289:XR_001752553 | 0:00 | 0:00 | 0:00 | 0:00 | 0:00 |
| NM_001143992:NM_018081:XR_001752551:XM_024450824:XM_011523952 | 2:2:0:0:0 | 0:0:0:0:0 | 1:0:0:0:0 | 0:0:0:0:0 | 4:2:0:0:0 |
| NM_052850 | 13 | 0 | 9 | 0 | 6 |
| XM_006722883:NM_024069:XM_011528273:NM_001171948 | 0:0:0:0 | 0:0:0:0 | 0:0:0:0 | 0:0:0:0 | 0:0:0:0 |
| NM_139314:NM_001039667:XM_005272484 | 0:00:00 | 0:00:00 | 0:00:00 | 0:00:00 | 0:00:00 |
| NM_001080125:NR_111983:NM_033356:XM_005246885 | 2:0:0:0 | 0:0:0:0 | 1:0:0:0 | 0:0:0:0 | 0:0:0:0 |
| NM_000465:XR_002959322:XM_017004613:XM_017004614:NR_104212 | 0:0:0:0:0 | 0:0:0:0:0 | 0:0:0:0:0 | 0:0:0:0:0 | 0:0:1:1:0 |

| | | | | | |
|---|---|---|---|---|---|
| NM_002026:NM_212482:NM_212478:NM_212474:NM_054034 | 0:1:0:0:0 | 0:0:0:0:0 | 2:1:0:0:0 | 1:2:0:0:0 | 0:0:0:0:0 |
| NM_000182 | 196 | 0 | 126 | 1 | 177 |
| NM_000182 | 188 | 0 | 126 | 0 | 173 |
| NM_000099:NM_001288614 | 44:01:00 | 0:00 | 6:00 | 0:00 | 15:00 |
| NM_080426:NM_001077489:XM_017027817:NM_001309840:NM_001077488:XM_017027818:XM_017027813:NM_000516:XR_002958471:XM_024451874:NM_080425:XM_024451875:XM_017027819:XM_024451873:XM_017027815:XM_017027812:XM_024451872:NM_016592:XM_017027820:XM_017027814:NM_001309861 | 257:101:42:9:23:3:10:110:11:1:1:8:7:2:2:2:0:0:0:0:0 | 0:0:0:0:0:0:0:0:0:0:0:0:0:0:0:0:0:0:0:0:0 | 267:105:40:12:9:10:17:87:18:2:2:3:7:0:0:0:0:0:0:0:0 | 1:0:0:0:0:0:0:0:0:0:0:0:0:0:0:0:0:0:0:0:0 | 285:156:32:20:24:5:29:173:34:1:3:11:11:0:0:1:0:1:0:0:1 |
| NM_020062 | 0 | 1 | 0 | 0 | 0 |
| NM_033070:XR_002958690 | 0:00 | 0:00 | 2:00 | 0:00 | 0:00 |
| NM_014248 | 540 | 1 | 340 | 0 | 434 |
| NM_020800:NM_001190241 | 1:02 | 0:00 | 0:00 | 0:00 | 2:01 |
| NM_001172780:NM_001172779:NM_153353 | 0:00:00 | 0:00:00 | 1:00:00 | 0:00:00 | 0:00:00 |
| XM_005263082:XR_938747:NM_001127208:XM_024454102 | 7:6:1:1 | 0:0:0:0 | 1:0:0:0 | 0:0:0:0 | 0:0:0:0 |
| NM_176869 | 2 | 1 | 2 | 0 | 0 |
| NM_006726:NM_001199282:XM_017008874:XM_011532434:XM_017008873 | 5:0:3:2:0 | 0:0:0:0:0 | 2:1:0:0:0 | 0:0:0:0:0 | 1:0:4:0:0 |
| NM_006726:NM_001199282:XM_017008874:XM_011532434:XM_017008873 | 5:0:3:2:0 | 0:0:0:0:0 | 2:1:0:0:0 | 0:0:0:0:0 | 1:0:4:0:0 |
| NM_006342:XM_017007653:XM_005247929:XM_011513386 | 5:3:0:2 | 0:0:0:0 | 5:1:0:0 | 0:0:0:0 | 13:2:1:0 |
| NM_006342:XM_017007653:XM_005247929:XM_011513386:XM_005247930 | 6:7:0:0:0 | 0:0:0:0:0 | 5:1:0:0:0 | 0:0:0:0:0 | 15:8:1:0:1 |
| NM_000521 | 4 | 0 | 2 | 0 | 2 |
| NM_033125 | 0 | 0 | 0 | 0 | 0 |
| XM_006715537:XM_011536021:XM_017011148:NM_002844:XM_017011146:XM_017011145:XM_011536016:NM_001291982:XM_011536014:XM_011536020:XM_017011147:XM_017011150:XM_011536015:NM_001291983 | 0:38:3:1:0:0:0:1:0:0:0:0:0:0 | 0:0:0:0:0:0:0:0:0:0:0:0:0:0 | 1:7:1:0:0:0:0:0:0:0:0:0:0:0 | 0:0:0:0:0:0:0:0:0:0:0:0:0:0 | 0:18:1:3:5:0:1:0:0:0:1:0:0:0 |
| NM_001024465:NM_000636 | 0:00 | 0:00 | 1:01 | 0:00 | 0:00 |
| NM_145169:NR_130112 | 4:01 | 0:00 | 3:01 | 0:00 | 5:01 |
| XM_017010958:XM_011535905:NM_001143947:NM_018974:XM_011535906:XM_011535907 | 0:1:0:0:0:0 | 0:0:0:0:0:0 | 3:6:3:1:0:0 | 0:0:0:0:0:0 | 0:4:0:1:0:0 |
| NM_002117:NM_001243042 | 178:30:00 | 0:00 | 54:10:00 | 0:00 | 79:01:00 |
| NM_002117:NM_001243042 | 170:30:00 | 0:00 | 53:11:00 | 0:00 | 78:01:00 |
| NM_002117:NM_001243042 | 180:20:00 | 1:00 | 54:09:00 | 0:00 | 80:00:00 |
| NM_025257 | 0 | 1 | 0 | 0 | 0 |
| XM_005249635:XM_011515168:XM_024446679:NM_001163150:NM_004956:NR_120445:XM_011515167:XM_011515170:NM_001163147 | 1:0:1:1:0:0:0:0:0 | 0:0:0:0:0:0:0:0:0 | 1:1:0:0:0:0:0:0:0 | 0:0:0:0:0:0:0:0:0 | 1:0:0:0:1:0:0:0:0 |
| XM_017012558:XM_017012559:NM_004333:NM_001354609:NR_148928:XR_001744857:XR_001744858 | 0:2:8:4:0:1:0 | 0:0:0:0:0:0:0 | 2:2:8:0:0:0:0 | 0:0:0:0:0:0:0 | 0:0:19:1:0:0:0 |
| NM_001079864:NM_006024:NM_001206902:NM_001206901 | 92:69:6:6 | 0:0:0:0 | 103:46:5:4 | 0:0:0:0 | 65:45:13:8 |
| NM_175887:XM_011515199:NM_001329996 | 0:00:00 | 0:00:00 | 0:00:00 | 0:00:00 | 2:01:00 |
| NM_020192:NM_001282446 | 0:00 | 0:00 | 0:00 | 0:00 | 0:00 |
| NM_015622 | 1 | 1 | 1 | 0 | 1 |
| NM_005918:NR_104165:NM_001282404 | 2:00:00 | 2:00:00 | 1:00:00 | 0:00:00 | 10:00:00 |
| NM_001283024:NM_001283027:NM_018024:XM_017013603:XR_002956636:XM_011517151 | 1:5:1:1:0:0 | 0:0:0:0:0:0 | 15:6:19:1:1:0 | 0:0:0:0:0:0 | 5:6:11:0:0:2 |
| NM_001283024:NM_001283027:NM_018024:XM_017013603:XR_002956636:XM_011517151 | 1:5:1:1:0:0 | 0:0:0:0:0:0 | 15:6:19:1:1:0 | 0:0:0:0:0:0 | 5:6:11:0:0:2 |
| NM_001283027:NM_001283024:NM_018024:XM_017013603:XR_002956636:XM_011517151 | 5:1:1:1:0:0 | 0:0:0:0:0:0 | 5:15:19:1:1:0 | 1:0:0:0:0:0 | 6:5:11:0:0:2 |
| NM_001283024:NM_001283027:NM_018024:XM_017013603:XR_002956636:XM_011517151 | 1:5:1:1:0:0 | 0:0:0:0:0:0 | 15:6:19:1:1:0 | 0:0:0:0:0:0 | 5:6:11:0:0:3 |
| NM_177924 | 0 | 0 | 0 | 1 | 0 |
| NM_177924 | 0 | 0 | 0 | 1 | 0 |
| XM_024447265:XR_949472:XM_011544639 | 0:00:01 | 0:01:02 | 1:00:00 | 0:00:00 | 1:00:00 |
| XM_024447530:NM_002160 | 0:00 | 0:00 | 0:00 | 0:00 | 0:00 |
| XM_005272211:XM_006717271:NM_001162426:NM_001162427:XM_017015098:NM_000368:XM_017015100:XM_017015101:XM_017015097:XM_011518979 | 1:0:0:1:1:2:0:0:0:0 | 0:0:0:0:0:0:0:0:0:0 | 2:1:2:0:0:0:0:0:0:0 | 0:0:0:0:0:0:0:0:0:0 | 0:0:0:1:1:5:0:0:0:0 |

| 2T__MUT | 2M__WT | 2M__MUT | 3T__WT | 3T__MUT | 3M__WT | 3M__MUT |
|---|---|---|---|---|---|---|
| 0 | 0 | 0 | 9 | 0 | 0 | 0 |
| 2:1:0:1:2:2 | 0:0:0:0:0:0 | 26:5:8:14:35:13 | 0:0:0:0:0:0 | 20:4:10:16:15:14 | 0:0:0:0:0:0 | 3:0:1:2:2:0 |
| 1 | 1 | 1 | 1 | 3 | 1 | 3 |
| 0:0:0:0:0:0 | 0:0:0:0:0:0 | 0:0:0:0:0:0 | 0:0:0:0:0:0 | 0:0:0:0:0:0 | 1:3:0:1:0:1 | 0:0:0:0:0:0 |
| 24:1:2:0 | 57:5:10:0 | 43:2:8:0 | 107:10:13:0 | 51:2:10:0 | 24:2:0:1 | 17:0:2:0 |
| 3:2:1:0:0:1:2:0:0:1 | 0:0:0:0:0:0:0:0:0:0 | 3:9:0:0:0:0:1:0:0:0 | 5:22:0:0:0:0:0:1:0:0 | 0:0:0:0:0:0:0:0:0:0 | 3:2:0:0:0:0:0:1:0:0 | 0:0:0:0:0:0:0:0:0:0 |
| 7:00 | 0:00 | 124:04:00 | 50:06:00 | 0:00 | 5:01 | 0:00 |
| 0:0:0:0:0 | 0:0:3:0:0 | 0:0:0:0:0 | 0:0:4:2:0 | 0:0:0:0:0 | 0:0:1:3:1 | 0:0:0:0:0 |
| 7 | 0 | 36 | 0 | 39 | 0 | 6 |
| 8 | 0 | 32 | 0 | 36 | 0 | 5 |
| 7 | 0 | 26 | 0 | 25 | 0 | 4 |
| 18:131:0:0:0:1:2:0:0:1:0 | 0:0:0:0:0:0:0:0:0:0:0 | 62:393:4:6:0:1:3:1:1:0:0 | 0:1:0:0:0:0:0:0:0:0:0 | 120:704:6:5:0:1:1:1:1:0:1 | 0:0:0:0:0:0:0:0:0:0:0 | 40:154:2:0:0:0:2:0:0:0:0 |
| 0 | 0 | 1 | 0 | 0 | 0 | 1 |
| 0:2:0:1:0 | 0:0:0:0:0 | 7:0:0:0:0 | 0:0:0:0:0 | 6:0:0:2:0 | 0:0:0:0:0 | 1:0:0:1:0 |
| 1:2:0:6:0:3:2:0:0:0:0:0:0:0:0:0:0:1:0:0:0:0:0:0 | 5:0:0:30:2:6:6:0:0:1:0:0:0:0:1:2:0:1:0:1:2:1:1:0 | 7:1:3:35:0:7:11:0:0:7:1:1:1:1:1:2:1:1:1:0:0:0:0:0 | 1:2:0:6:0:1:2:2:0:0:1:0:0:0:2:0:0:0:0:1:0:0:0:1 | 0:0:0:0:0:0:0:0:0:0:0:0:0:0:0:0:0:0:0:0:0:0:0:0 | 0:0:0:2:0:0:2:0:0:0:1:0:0:0:0:0:0:0:0:0:1:0:0:0 | 0:0:0:0:0:0:0:0:0:0:0:0:0:0:0:0:0:0:0:0:0:0:0:0 |
| 0:0:0:0:0 | 23:506:58:25:45 | 0:0:0:0:0 | 8:95:9:2:11 | 0:0:0:0:0 | 1:25:4:0:0 | 0:0:0:0:0 |
| 3:00 | 26124:18 | 24:01:00 | 14352:9 | 12:00 | 3476:05:00 | 1:00 |
| 0:0:0:0 | 0:0:0:0 | 0:0:0:0 | 0:0:0:0 | 0:1:0:0 | 0:0:0:0 | 0:0:0:0 |
| 0:0:0:0 | 0:0:0:0 | 0:0:1:0 | 0:0:0:0 | 0:0:0:0 | 0:0:0:0 | 0:0:0:0 |
| 0 | 4 | 0 | 0 | 3 | 0 | 3 |
| 6 | 0 | 8 | 7 | 0 | 3 | 0 |
| 32 | 84 | 86 | 1 | 176 | 1 | 52 |
| 0:3:0:0:2:0 | 0:0:0:0:1:0 | 0:15:0:3:9:0 | 1:5:2:1:3:0 | 0:0:0:0:0:0 | 0:3:0:0:1:0 | 0:0:0:0:0:0 |
| 4:01 | 4:03 | 3:04 | 1:00 | 0:00 | 2:00 | 0:00 |
| 0:00 | 15:00 | 0:00 | 3:01 | 23:00 | 0:00 | 1:00 |
| 2:0:1:1 | 0:0:0:0 | 0:1:0:0 | 0:0:0:0 | 3:0:0:1 | 0:0:0:0 | 0:1:0:0 |
| 2 | 3256 | 1 | 1353 | 1 | 549 | 1 |
| 28 | 0 | 66 | 0 | 15 | 0 | 7 |
| 0:0:0:0:0:0:0:0:0:0:0:0:0:0:0:0:0 | 171:48:56:18:0:12:5:59:9:45:56:16:9:5:4:2 | 0:0:0:0:0:2:0:0:0:0:0:0:0:0:0:0:0 | 18:4:0:1:1:6:0:0:0:3:1:0:1:0:2:0 | 109:10:8:5:3:0:1:0:0:7:3:0:1:2:2:1 | 8:4:4:1:0:1:0:1:0:2:1:1:3:0:1:0 | 24:1:7:1:1:0:0:0:0:1:0:0:1:1:1:0 |
| 0:00 | 275:84 | 0:00 | 294:107 | 34:16:00 | 141:04:00 | 10:01 |
| 57:0:1:0 | 0:0:0:0 | 79:3:2:1 | 0:0:0:0 | 37:1:1:1 | 0:0:0:0 | 14:0:0:2 |
| 0:00 | 11:00 | 0:00 | 24:01:00 | 0:00 | 12:00 | 0:00 |
| 7:5:1:0:1:1:1:0:0 | 0:0:0:0:0:0:0:0:0 | 19:16:6:0:8:1:0:0:0 | 0:0:0:0:0:0:0:0:0 | 21:18:8:1:7:3:1:0:1 | 0:0:0:0:0:0:0:0:0 | 10:1:2:0:1:2:0:1:0 |
| 0 | 10 | 0 | 10 | 0 | 4 | 0 |
| 1 | 0 | 0 | 0 | 2 | 0 | 2 |
| 161:03:00 | 0:00:00 | 381:03:12 | 504:05:06 | 32:01:00 | 19:00:00 | 82:01:00 |
| 0:00 | 0:00 | 0:00 | 0:00 | 1:01 | 0:00 | 0:00 |
| 6 | 78 | 39 | 57 | 25 | 10 | 1 |
| 0 | 133 | 0 | 82 | 3 | 4 | 9 |
| 6:0:0:0:0:0 | 18051:777:931:185:7:2 | 31:4:2:0:0:0 | 8924:529:428:115:0:1 | 20:3:0:0:0:0 | 2749:154:108:25:0:0 | 2:0:1:0:0:0 |
| 1:00 | 0:00 | 0:00 | 0:00 | 0:00 | 0:00 | 0:00 |
| 0:0:0:0:0:0 | 2:0:2:1:0:1 | 0:0:0:0:0:0 | 0:0:0:0:0:0 | 37:34:17:1:0:11 | 0:0:0:0:0:0 | 2:5:0:0:0:3 |
| 0:00 | 9:00 | 0:00 | 11:00 | 0:00 | 6:01 | 0:00 |
| 0 | 5 | 0 | 2 | 5 | 0 | 1 |
| 10 | 0 | 13 | 4 | 3 | 1 | 2 |

| | | | | | | |
|---|---|---|---|---|---|---|
| 32:00:00 | 0:00 | 187:01:00 | 61:01:00 | 42:00:00 | 7:00 | 7:00 |
| 3:1:0:0:0:0:0 | 0:0:0:0:0:0:0 | 8:2:4:0:0:0:0 | 11:5:4:0:0:1:1 | 10:3:2:0:0:0:0 | 0:0:0:0:0:0:0 | 5:0:0:0:2:0:0 |
| 5:00 | 67:03:00 | 0:00 | 41:05:00 | 69:07:00 | 17:02 | 2:01 |
| 0:0:0:0 | 182:25:10:5 | 3:0:0:0 | 356:49:9:16 | 6:0:0:0 | 43:12:4:4 | 2:0:0:0 |
| 0 | 36 | 0 | 1 | 78 | 0 | 42 |
| 0:00 | 20:05 | 0:00 | 0:00 | 35:05:00 | 0:00 | 1:01 |
| 2:1:0:0 | 4540:125:9:18 | 5:1:0:0 | 11030:420:15:10 | 13:1:0:0 | 2923:93:5:10 | 0:1:0:0 |
| 0:0:0:0:0:0:0:0:0:0 | 3:11:1:1:1:0:1:0:0:0 | 2:2:0:1:1:1:0:0:0:0 | 0:0:0:0:0:0:0:0:0:0 | 8:0:0:0:0:0:0:0:0:1 | 0:0:0:0:0:0:0:0:0:0 | 3:0:0:0:0:0:0:0:0:0 |
| 0:0:0:0:0 | 25:10:0:1:1 | 0:0:0:0:0 | 1:0:0:0:0 | 13:9:0:0:0 | 4:1:0:0:0 | 13:4:0:0:0 |
| 5 | 5 | 2 | 0 | 25 | 0 | 5 |
| 5 | 2 | 5 | 0 | 22 | 0 | 4 |
| 2:0:0:0:0:0:1:1:1 | 0:0:0:0:0:0:0:0:0 | 3:5:1:0:0:2:1:0:0 | 0:0:0:0:0:0:0:0:0 | 4:3:0:0:0:0:1:0:1 | 0:0:0:0:0:0:0:0:0 | 1:0:0:0:0:0:0:1:0 |
| 1:6:1:0 | 1019:7198:72:1 | 3:30:9:0 | 1253:13384:118:6 | 8:44:15:0 | 186:3107:11:1 | 0:8:0:0 |
| 1:0:0:0:0 | 17:4:7:10:2 | 22:5:1:1:0 | 35:6:8:13:8 | 52:2:0:1:1 | 15:1:3:2:0 | 0:0:0:0:0 |
| 2027 | 10 | 3285 | 14382 | 54 | 914 | 1 |
| 0:00:00 | 0:00:00 | 0:00:01 | 2:01:02 | 0:00:00 | 0:00:01 | 0:00:00 |
| 14:5:0:0:2:1:0 | 66:26:4:0:9:0:0 | 79:23:2:0:10:1:2 | 4:1:0:0:1:0:1 | 64:18:6:2:10:1:0 | 1:1:0:0:0:0:0 | 19:14:1:0:0:0:0 |
| 1:2:0:0 | 32:77:5:17 | 32:80:11:6 | 0:0:0:0 | 7:4:1:0 | 0:0:0:0 | 1:1:0:0 |
| 0 | 0 | 0 | 0 | 0 | 0 | 0 |
| 2:0:0:0 | 8:9:0:0 | 0:9:0:0 | 1:0:0:0 | 8:13:0:0 | 1:0:0:0 | 2:4:1:0 |
| 3 | 0 | 2 | 5 | 1 | 1 | 0 |
| 4:9:0:0 | 0:0:0:0 | 2:5:0:0 | 2:9:0:1 | 0:0:0:0 | 0:2:0:0 | 0:0:0:0 |
| 7 | 0 | 3 | 16 | 6 | 1 | 3 |
| 1:0:0:0 | 4:0:1:0 | 0:1:0:0 | 0:0:0:0 | 7:9:0:1 | 0:0:0:0 | 2:1:0:0 |
| 0:00:00 | 25:08:00 | 0:00:00 | 120:16:10 | 0:00:00 | 14:01:02 | 0:00:00 |
| 0:0:0:0 | 5:1:0:0 | 0:0:0:0 | 9:2:1:1 | 1:0:0:0 | 8:1:1:0 | 0:0:0:0 |
| 5:0:0:0:0 | 30:3:0:1:3 | 0:0:0:0:0 | 0:0:0:0:0 | 84:1:3:4:3 | 0:0:0:0:0 | 21:1:2:0:1 |
| 0:01 | 0:00 | 0:04 | 0:00 | 0:00 | 0:00 | 0:00 |
| 0:01 | 0:00 | 0:02 | 0:00 | 0:00 | 0:00 | 0:00 |
| 0:0:0:0 | 5:14:26:0 | 0:1:0:0 | 3:2:10:1 | 7:6:25:1 | 0:0:5:0 | 0:1:6:0 |
| 1:00 | 5429:3492 | 10:10 | 2835:1854 | 1371:959 | 1011:110 | 541:52:00 |
| 0:0:0:0:0:0 | 904:9:1:10:9:2 | 0:0:0:0:0:0 | 1195:16:2:14:12:4 | 2:0:0:0:0:0 | 257:2:1:2:0:0 | 0:0:0:0:0:0 |
| 0:00:00 | 0.583576389 | 0:00:00 | 6:35:15 | 0:00:00 | 3:08:03 | 0:00:00 |
| 3:0:0:0 | 0:0:0:0 | 13:0:0:0 | 2:1:6:2 | 8:0:0:0 | 6:0:0:0 | 3:0:0:0 |
| 3:0:0:0 | 0:0:0:0 | 13:0:0:0 | 2:1:6:3 | 8:0:0:0 | 6:0:0:0 | 3:0:0:0 |
| 0:0:0:0:0 | 18:25:107:11:0 | 0:0:0:0:0 | 37:37:186:15:0 | 0:0:0:0:0 | 15:16:9:1:0 | 0:0:0:0:0 |
| 0:00:00 | 3:00:00 | 1:00:00 | 0:00:00 | 4:01:01 | 0:00:00 | 0:00:00 |
| 1:1:0:0:0:0:0:0:1:0:0:0:0:0:0:0:0:0:0:0:0 | 152:920:244:67:57:1707:142:123:491:3:14:46:79:47:10:4:4:3:2:5:1 | 19:2:0:0:0:10:0:0:0:0:0:0:0:0:0:0:0:0:0:0:0 | 92:353:312:124:74:876:40:99:379:5:15:52:61:24:5:11:8:7:1:2:2 | 6:0:0:0:0:4:0:0:0:0:0:0:1:0:0:0:0:0:0:0:0 | 22:182:48:13:11:371:10:27:156:0:9:14:13:3:1:0:0:2:0:0:0 | 0:0:0:0:0:1:0:0:0:0:0:0:0:0:0:0:0:0:0:0:0 |
| 1:0:0:0:0:0:2:0:0:0:0:0:0:0:0:0:0:0:0:0:0 | 334:921:244:67:57:141:1738:123:490:3:14:46:79:47:10:4:4:3:2:5:1 | 7:0:0:0:0:0:3:0:0:0:0:0:0:0:0:0:0:0:0:0:0 | 272:350:311:124:74:40:910:99:374:5:15:52:62:24:5:11:8:7:1:2:2 | 5:1:1:0:0:0:0:0:1:0:0:0:0:0:0:0:0:0:0:0:0 | 58:182:48:13:11:10:378:27:156:0:9:14:13:3:1:0:0:2:0:0:0 | 0:0:0:0:0:0:0:0:0:0:0:0:0:0:0:0:0:0:0:0:0 |

| | | | | | | |
|---|---|---|---|---|---|---|
| 1:0:0:0:0:3:0:0:1:0:0:0:0:0:0:0:0:0:0:0:0 | 729:919:243:67:56:1812:141:118:487:3:14:45:76:47:10:4:4:3:2:5:1 | 19:0:0:0:0:8:0:0:0:0:0:1:0:0:0:0:0:0:0:0:0 | 624:346:312:124:73:976:40:91:372:5:15:47:58:24:5:10:8:7:1:2:2 | 8:0:0:0:0:2:0:0:0:0:0:0:0:0:0:0:0:0:0:0:0 | 115:182:48:13:11:382:10:26:156:0:9:14:13:3:1:0:0:2:0:0:0 | 0:0:0:0:0:0:0:0:0:0:0:0:0:0:0:0:0:0:0:0:0 |
| 31:00:00 | 17:00 | 1320:00:00 | 9:00 | 1330:00:00 | 0:00 | 34:00:00 |
| 33:08:00 | 70:07:00 | 71:08:00 | 201:15:00 | 0:00 | 109:11:00 | 0:00 |
| 54:01:00 | 0:00:00 | 136:01:00 | 258:00:00 | 0:00:00 | 56:00:01 | 0:00:00 |
| 0:00 | 0:00 | 0:00 | 0:00 | 1:00 | 21:00 | 28:01:00 |
| 5:00 | 0:00 | 16:01 | 0:00 | 12:00 | 0:00 | 3:00 |
| 1 | 1901 | 2 | 1813 | 1 | 519 | 0 |
| 3:0:0:0 | 3:0:0:1 | 4:1:0:0 | 15:1:1:0 | 0:0:0:0 | 3:2:0:0 | 0:0:0:0 |
| 11:10:02 | 0:00:00 | 63:112:9 | 35:144:0 | 4:07:00 | 49:93:0 | 4:02:00 |
| 18:3:0:0:6:3:2:0:0:0:0:1:1:1:0:0:0:0:0:0:0:0 | 36:9:7:0:9:23:3:10:1:0:4:7:0:0:1:0:1:0:0:0:0:0 | 20:11:14:1:14:8:3:13:0:0:2:4:0:0:1:1:1:0:0:0:0:0 | 87:24:41:3:7:22:2:31:1:1:2:16:0:5:0:2:3:2:1:1:1:2 | 0:0:0:0:0:0:0:0:0:0:0:0:0:0:0:0:0:0:0:0:0:0 | 35:5:0:1:0:9:2:0:2:0:1:0:1:0:0:1:0:0:0:0:0:0 | 0:0:0:0:0:0:0:0:0:0:0:0:0:0:0:0:0:0:0:0:0:0 |
| 0:0:0:0:0:0:0:0:0:0:0 | 4:11:14:3:0:0:0:0:1:2:0 | 8:1:0:2:0:0:0:1:0:0:0 | 3:6:11:2:0:0:0:0:1:4:1 | 12:2:0:0:0:0:1:1:0:0:0 | 0:0:4:0:0:0:0:0:0:0:0 | 0:0:0:0:0:0:0:0:0:0:0 |
| 3:1:0:0:0:0:1:0:0:0:0:0:0:0 | 6:8:0:5:3:1:3:2:0:0:0:0:0:0 | 8:5:0:2:3:1:1:1:0:1:0:0:0:0 | 9:24:0:5:6:0:2:1:1:1:0:2:4 | 0:0:0:0:0:0:0:1:0:0:0:0:0:0 | 4:0:0:0:0:0:0:0:0:0:0:0:0:0 | 0:0:0:0:0:0:0:0:0:0:0:0:0:0 |
| 1:2:2:1:1 | 11:1:1:0:0 | 9:1:1:0:0 | 12:0:3:0:0 | 0:0:0:0:0 | 1:1:1:0:0 | 0:0:0:0:0 |
| 0 | 15 | 0 | 13 | 17 | 4 | 0 |
| 0:00 | 0:00 | 0:00 | 0:00 | 1:02 | 0:00 | 0:00 |
| 1:01 | 0:00 | 2:20 | 3:09 | 0:11 | 2:01 | 1:02 |
| 33:00:00 | 0:00:00 | 279:10:08 | 12:08:02 | 30:00:01 | 0:01:00 | 3:00:00 |
| 2:1:0:0:0:0:1:0:1:0:1:1:1:0:0 | 5:1:0:1:0:2:0:0:0:2:0:0:0:0:0 | 0:0:0:0:0:0:0:0:0:0:0:0:0:0:0 | 0:1:0:0:0:0:0:0:0:0:0:0:0:0:0 | 1:4:0:0:3:0:2:0:2:1:0:5:0:0:3 | 2:0:0:0:1:0:0:0:0:0:0:0:0:0:0 | 0:0:1:0:0:0:2:0:1:0:0:1:0:0:0 |
| 2 | 0 | 1 | 0 | 7 | 0 | 2 |
| 0 | 0 | 1 | 0 | 2 | 0 | 1 |
| 0:0:0:0:0:0:0 | 19:0:7:4:2:3:0 | 0:0:0:0:0:0:0 | 4:0:1:0:0:0:0 | 12:1:7:3:1:1:5 | 1:0:0:0:0:0:0 | 1:0:0:0:0:1:1 |
| 1:00:00 | 261:221:126 | 2:03:03 | 647:479:334 | 8:05:06 | 720:293:40 | 0:00:00 |
| 0:00 | 0:07 | 0:00 | 0:00 | 0:00 | 0:00 | 0:00 |
| 0:00 | 0:07 | 0:00 | 0:00 | 0:00 | 0:00 | 0:00 |
| 0:00 | 0:07 | 0:00 | 0:00 | 0:00 | 0:00 | 0:00 |
| 0:00 | 0:07 | 0:00 | 0:00 | 0:00 | 0:00 | 0:00 |
| 0:06 | 0:00 | 0:07 | 0:00 | 0:00 | 0:00 | 0:00 |
| 0:06 | 0:00 | 0:07 | 0:00 | 0:00 | 0:00 | 0:00 |
| 0:00 | 0:07 | 0:00 | 0:00 | 0:00 | 0:00 | 0:00 |
| 0:00 | 0:07 | 0:00 | 0:00 | 0:00 | 0:00 | 0:00 |
| 0:00 | 0:07 | 0:00 | 0:00 | 0:00 | 0:00 | 0:00 |
| 0:00 | 0:07 | 0:00 | 0:00 | 0:00 | 0:00 | 0:00 |
| 0:00 | 0:07 | 0:00 | 0:00 | 0:00 | 0:00 | 0:00 |
| 0:00 | 0:07 | 0:00 | 0:00 | 0:00 | 0:00 | 0:00 |
| 0:08 | 0:00 | 0:07 | 0:00 | 0:00 | 0:00 | 0:00 |
| 0:08 | 0:00 | 0:07 | 0:00 | 0:00 | 0:00 | 0:00 |
| 91:01:00 | 0:00 | 308:07:00 | 2:00 | 735:07:00 | 0:00 | 240:03:00 |
| 91:01:00 | 0:00 | 308:07:00 | 2:00 | 735:07:00 | 0:00 | 240:03:00 |
| 91:01:00 | 0:00 | 307:07:00 | 0:00 | 735:07:00 | 0:00 | 241:03:00 |
| 90:01:00 | 1:00 | 303:07:00 | 0:00 | 729:07:00 | 0:00 | 239:03:00 |
| 93:01:00 | 0:00 | 296:07:00 | 2:00 | 767:08:00 | 0:00 | 242:03:00 |
| 93:01:00 | 0:00 | 297:07:00 | 0:00 | 769:08:00 | 0:00 | 242:03:00 |
| 91:01:00 | 0:00 | 292:07:00 | 0:00 | 758:08:00 | 0:00 | 238:03:00 |
| 0:00 | 281:10:00 | 0:00 | 671:09:00 | 0:00 | 226:05:00 | 1:00 |
| 0:00 | 249:08:00 | 1:01 | 615:09:00 | 2:00 | 204:03:00 | 0:01 |
| 1:06 | 0.542361111 | 0:10 | 0.749305556 | 0:01 | 0.309027778 | 0:00 |
| 79:00:00 | 0:00 | 265:00:00 | 1:00 | 615:02:00 | 0:00 | 217:00:00 |
| 80:00:00 | 1:00 | 264:01:00 | 2:00 | 608:02:00 | 0:00 | 216:00:00 |
| 0 | 2906 | 3 | 5279 | 5 | 1958 | 1 |
| 1 | 2695 | 9 | 1632 | 3158 | 933 | 921 |
| 807 | 382 | 2213 | 0 | 4620 | 0 | 1808 |
| 917 | 6 | 2517 | 6 | 4541 | 2 | 1784 |

| | | | | | | |
|---|---|---|---|---|---|---|
| 913 | 0 | 2512 | 1 | 4521 | 0 | 1777 |
| 0 | 2463 | 7 | 1475 | 2925 | 881 | 873 |
| 0 | 2444 | 7 | 4345 | 10 | 1740 | 0 |
| 0 | 2444 | 6 | 1458 | 2895 | 876 | 864 |
| 0 | 2445 | 4 | 1454 | 2897 | 876 | 864 |
| 782 | 371 | 2111 | 9 | 4359 | 2 | 1742 |
| 782 | 368 | 2111 | 16 | 4353 | 3 | 1741 |
| 781 | 366 | 2111 | 8 | 4354 | 3 | 1741 |
| 781 | 369 | 2108 | 15 | 4352 | 3 | 1742 |
| 786 | 371 | 2118 | 6 | 4381 | 1 | 1749 |
| 905 | 7 | 2503 | 6 | 4395 | 3 | 1757 |
| 906 | 6 | 2507 | 5 | 4402 | 1 | 1758 |
| 906 | 7 | 2508 | 4 | 4399 | 0 | 1758 |
| 907 | 11 | 2506 | 2 | 4408 | 0 | 1757 |
| 907 | 7 | 2522 | 2 | 4415 | 1 | 1760 |
| 118 | 2172 | 376 | 1512 | 2915 | 895 | 873 |
| 118 | 2171 | 376 | 1518 | 2910 | 895 | 873 |
| 115 | 2159 | 371 | 1474 | 2870 | 884 | 862 |
| 115 | 2158 | 369 | 1471 | 2851 | 880 | 860 |
| 111 | 2138 | 370 | 1469 | 2827 | 879 | 851 |
| 113 | 2135 | 374 | 1471 | 2834 | 880 | 855 |
| 113 | 2137 | 372 | 4292 | 10 | 1729 | 5 |
| 0:0:1:0 | 0:0:0:0 | 0:0:2:0 | 0:0:0:0 | 0:0:0:0 | 0:0:0:0 | 0:0:0:0 |
| 0:0:1:0 | 0:0:0:0 | 0:0:2:0 | 0:0:0:0 | 0:0:0:0 | 0:0:0:0 | 0:0:0:0 |
| 0:0:1:0 | 0:0:0:0 | 0:0:2:0 | 0:0:0:0 | 0:0:0:0 | 0:0:0:0 | 0:0:0:0 |
| 0:01:00 | 0:00:00 | 0:02:00 | 0:00:00 | 0:00:00 | 0:00:00 | 0:00:00 |
| 0:00:00 | 0:01:00 | 0:00:00 | 0:00:00 | 0:00:00 | 0:00:00 | 0:00:00 |
| 0:00:00 | 0:01:00 | 0:00:00 | 0:00:00 | 0:00:00 | 0:00:00 | 0:00:00 |
| 0:00 | 0:00 | 0:01 | 0:00 | 0:00 | 0:00 | 0:00 |
| 0:00 | 0:01 | 0:00 | 0:00 | 0:00 | 0:00 | 0:00 |
| 0:00 | 0:01 | 0:00 | 0:00 | 0:00 | 0:00 | 0:00 |
| 0:00 | 0:01 | 0:00 | 0:00 | 0:00 | 0:00 | 0:00 |
| 0:00 | 0:01 | 0:00 | 0:00 | 0:00 | 0:00 | 0:00 |
| 0:00 | 0:01 | 0:00 | 0:00 | 0:00 | 0:00 | 0:00 |
| 0:00 | 0:01 | 0:00 | 0:00 | 0:00 | 0:00 | 0:00 |
| 0:00 | 0:01 | 0:00 | 0:00 | 0:00 | 0:00 | 0:00 |
| 0:00 | 0:01 | 0:00 | 0:00 | 0:00 | 0:00 | 0:00 |
| 0:00 | 0:00 | 0:01 | 0:00 | 0:00 | 0:00 | 0:00 |
| 0:00 | 3:00 | 0:00 | 0:00 | 0:00 | 0:00 | 0:00 |
| 0:00 | 3:00 | 0:00 | 0:00 | 0:00 | 0:00 | 0:00 |
| 0:00 | 3:00 | 0:00 | 0:00 | 0:00 | 0:00 | 0:00 |
| 1:00 | 0:00 | 3:00 | 0:00 | 0:00 | 0:00 | 0:00 |
| 2:0:0:0 | 0:0:0:0 | 2:0:1:0 | 2:1:0:1 | 0:0:0:0 | 1:0:0:0 | 0:0:0:0 |
| 0 | 0 | 0 | 0 | 2 | 0 | 0 |
| 42:10:6:3:0:0:0:0:0:0 | 78:59:77:30:0:0:1:0:0:0 | 71:44:85:23:0:0:0:1:1:0 | 1:0:0:0:0:0:0:0:0:0 | 86:42:66:24:0:1:0:0:0:1 | 0:0:0:0:0:0:0:0:0 | 29:14:6:4:0:0:0:1:0:1 |
| 4:2:0:0:0:0 | 370:91:8:9:2:3 | 59:17:0:1:1:0 | 332:53:2:11:2:5 | 56:9:0:1:0:1 | 44:9:0:2:0:1 | 2:1:0:0:0:0 |
| 0:00 | 71:689 | 2:02 | 46:334 | 1:00 | 0.572222222 | 0:00 |
| 0:00 | 0:00 | 1:00 | 0:00 | 0:00 | 0:00 | 0:00 |
| 452:37:10:11:11:19:3 | 2837:564:41:41:52:124:12 | 2852:510:49:139:55:149:16 | 202:14:2:0:12:12:0 | 2901:285:39:15:113:122:15 | 83:3:1:0:2:0:0 | 595:53:5:1:14:23:4 |
| 14:930:95:15:21:2:0 | 2:4:0:0:0:0:0 | 71:5714:1078:180:161:9:1 | 2:204:14:0:11:1:0 | 30:2913:286:15:69:2:1 | 1:83:3:0:0:0:0 | 4:597:54:1:12:1:0 |
| 0:0:0:0 | 13:4:0:0 | 17:7:0:1 | 11:1:0:0 | 8:12:2:0 | 5:1:0:0 | 3:1:0:0 |
| 0:4:0:0:0:1:2:0:1:0:0:0:0:0 | 2:3:8:5:0:9:0:0:1:1:0:1:1:0 | 3:7:5:4:0:13:0:0:3:1:1:0:0:0 | 0:5:3:2:0:4:1:0:1:0:0:0:1:0 | 2:7:9:2:0:12:1:0:4:1:0:0:1:1 | 0:0:1:0:0:2:0:0:0:0:0:0:1:0 | 1:3:0:0:0:0:0:0:2:1:0:0:0:0 |
| 1:00 | 20965:813 | 18:01 | 19138:906 | 16:04 | 1946:69 | 2:00 |
| 4:00 | 20892:812 | 28:02:00 | 19070:907 | 23:05 | 1942:70 | 2:00 |
| 0:00 | 16412:884 | 33:05:00 | 15040:866 | 33:01:00 | 1717:80 | 7:00 |
| 0:00 | 14557:863 | 51:02:00 | 13510:866 | 40:04:00 | 1645:78 | 7:00 |
| 0:0:0:0:0:0:0:0 | 981:28:1:48:2:0:0:0 | 2:0:0:0:0:0:0:0 | 3518:4:10:216:1:0:0:1 | 7:0:0:0:0:0:0:0 | 892:0:0:54:1:0:0:0 | 1:0:0:0:0:0:0:0 |

| | | | | | | |
|---|---|---|---|---|---|---|
| 0:0:0:0:0:0:0:0:0 | 111:12:28:0:0:0:0:1:0 | 0:0:0:0:0:0:0:0:0 | 8:0:3:0:0:0:0:0:0 | 70:8:5:0:0:0:0:0:2 | 4:2:1:0:0:0:1:0:0 | 19:1:1:0:0:1:0:0:0 |
| 0:0:0:0 | 48:13:3:0 | 0:0:0:0 | 11:7:0:0 | 0:0:0:0 | 2:3:0:1 | 1:0:0:0 |
| 0:00 | 5:01 | 0:00 | 4:00 | 2:00 | 4:00 | 0:00 |
| 12:00:02 | 138:00:02 | 210:00:07 | 49:04:00 | 5:01:00 | 14:00:01 | 1:00:00 |
| 4:00 | 0:00 | 11:01 | 1:00 | 1:00 | 2:00 | 0:00 |
| 4:00 | 0:00 | 11:01 | 1:00 | 1:00 | 2:00 | 0:00 |
| 2:00 | 2688:140 | 9:02 | 1515:119 | 1:01 | 551:15:00 | 0:00 |
| 234:10:00 | 468:11:00 | 2233:85 | 0:00 | 1518:102 | 0:00 | 551:15:00 |
| 0:0:0:0 | 3:60:17:0 | 0:0:0:0 | 1:50:5:1 | 1:59:24:0 | 1:17:1:0 | 0:0:0:0 |
| 0 | 434 | 1 | 490 | 1 | 177 | 0 |
| 14 | 1 | 36 | 0 | 30 | 0 | 2 |
| 1:0:1:0 | 0:0:0:0 | 1:1:0:0 | 0:0:0:0 | 0:0:0:0 | 0:0:0:0 | 1:0:0:0 |
| 0:1:0:0:1 | 0:689:83:678:47 | 0:2:1:1:0 | 0:536:67:634:55 | 0:0:0:1:0 | 2:31:7:82:42 | 0:0:0:0:0 |
| 0:2:0:0:0 | 201:769:0:678:46 | 0:2:0:1:0 | 157:754:0:518:55 | 1:0:0:0:0 | 13:95:2:29:42 | 0:0:0:0:0 |
| 1 | 1 | 1 | 6 | 0 | 7 | 0 |
| 1:0:0:0:0:0:0:0:0:0 | 2467:32:6:71:1:16:43:2:0:6:0 | 0:2:0:0:0:0:0:0:0:0 | 4032:59:9:22:1:24:22:2:0:3:0 | 3:3:0:0:0:0:0:0:0:0 | 1502:10:3:31:0:9:17:1:0:0:1 | 0:0:0:0:0:0:0:0:0:0 |
| 1:0:0:0:0:1:0:1:1:0:0 | 0:0:0:0:0:0:0:0:0:0 | 2:0:1:1:0:0:1:0:0:0:0 | 2:0:1:0:0:1:0:1:0:0:2 | 0:0:0:0:0:0:0:0:0:0 | 0:0:0:0:0:0:0:0:0:0 | 0:0:0:0:0:0:0:0:0:0 |
| 1 | 0 | 4 | 0 | 5 | 0 | 0 |
| 6 | 0 | 2 | 0 | 7 | 0 | 4 |
| 2 | 0 | 1 | 0 | 1 | 0 | 0 |
| 0:0:2:0:0:0:0:0 | 3:0:2:0:0:0:1:1 | 5:0:0:2:1:1:0:0 | 9:1:0:0:0:0:0:0 | 0:0:0:0:0:0:0:0 | 1:0:0:0:0:0:0:0 | 0:0:0:0:0:0:0:0 |
| 2:2:0:5:6:0 | 30:13:3:24:71:1 | 22:10:6:19:58:1 | 32:24:5:33:87:1 | 0:0:0:0:0:0 | 6:0:1:15:2:0 | 0:0:0:0:0:0 |
| 1:00 | 1459:14:00 | 3:00 | 317:01:00 | 0:00 | 103:01:00 | 0:00 |
| 2:00 | 1470:15:00 | 1:01 | 321:02:00 | 2:00 | 102:01:00 | 0:00 |
| 1:00 | 0:00 | 1:00 | 2:00 | 1:01 | 1:00 | 1:00 |
| 1:00:00 | 791:15:09 | 1:01:00 | 783:06:05 | 164:01:14 | 149:02:03 | 58:01:05 |
| 28:0:22:24:0:4:5:0:12:2:10:0:0:1:0:0:1:0:2:0:1:0:0:0:0 | 155:0:75:46:26:8:11:9:73:19:51:3:5:13:1:0:62:0:26:10:34:31:0:1:0 | 173:0:74:20:28:6:17:7:75:27:85:2:4:19:1:0:69:0:31:6:21:19:1:1:0 | 813:3:110:311:8:39:17:6:45:14:105:1:4:31:1:5:7:0:0:0:4:11:0:3:1 | 2:0:0:0:0:0:0:0:0:0:0:0:0:0:0:0:0:0:0:0:0:0:0:0:0 | 97:8:26:153:11:8:1:2:6:5:18:0:0:5:2:1:0:1:1:1:1:4:0:1:0 | 0:0:0:0:0:0:0:0:0:0:0:0:0:0:0:0:0:0:0:0:0:0:0:0:0 |
| 7 | 0 | 4 | 1 | 6 | 0 | 0 |
| 24 | 25 | 21 | 2 | 95 | 0 | 31 |
| 1:1:0:3:0:0 | 2:2:0:1:1:0 | 1:3:1:0:0:0 | 4:8:2:1:0:1 | 0:1:0:1:0:0 | 1:5:1:0:0:0 | 0:2:1:0:0:0 |
| 0:1:0:4:0:0 | 0:0:0:0:0:0 | 0:0:0:6:1:0 | 0:0:0:0:0:0 | 0:2:2:10:1:0 | 0:0:0:0:0:0 | 0:0:0:1:0:0 |
| 1 | 0 | 3 | 0 | 7 | 1 | 0 |
| 0:01 | 6:09 | 20:05 | 35:11:00 | 0:00 | 2:03 | 0:00 |
| 5 | 23 | 8 | 0 | 34 | 0 | 5 |
| 8:2:0:0 | 14:0:0:1 | 17:2:2:1 | 8:0:1:0 | 17:0:0:0 | 8:1:1:0 | 5:1:0:0 |
| 10:2:0:0 | 15:0:0:1 | 19:2:2:1 | 8:0:1:0 | 23:1:1:0 | 8:1:1:0 | 5:1:0:0 |
| 0:0:1:0 | 25:10:183:5 | 0:0:1:0 | 49:8:351:17 | 0:1:5:0 | 12:4:43:4 | 0:0:1:0 |
| 2:0:0:0 | 0:0:0:0 | 16:6:2:0 | 0:0:0:0 | 4:2:0:2 | 0:0:0:0 | 0:2:0:2 |
| 2:0:0:0 | 4:8:0:0 | 2:8:2:0 | 0:0:0:0 | 2:4:0:2 | 0:0:0:0 | 2:0:0:2 |
| 16:0:0:0:0 | 8:4:0:0:0 | 17:5:0:1:1 | 13:8:0:0:0 | 1:1:0:0:0 | 13:4:0:0:0 | 4:1:0:0:0 |
| 0:0:0:1:1 | 0:0:0:0:0 | 1:2:2:0:0 | 0:0:0:0:0 | 0:0:0:0:1 | 0:0:0:0:0 | 0:0:0:1:0 |
| 0:01 | 0:00 | 1:01 | 0:00 | 0:00 | 0:00 | 0:00 |
| 32:2:19:0:0 | 0:0:0:0:0 | 300:2:59:1:0 | 1:0:0:0:0 | 223:3:45:3:1 | 0:0:0:0:0 | 26:2:11:1:0 |
| 0:1:5:0 | 6:28:11:2 | 6:2:13:1 | 0:0:0:0 | 8:4:24:2 | 0:0:0:0 | 0:1:4:0 |
| 0:1:5:0 | 6:28:11:2 | 6:2:13:1 | 0:0:0:0 | 8:4:24:2 | 0:0:0:0 | 0:1:4:0 |

| | | | | | | |
|---|---|---|---|---|---|---|
| 3:01 | 0:00 | 1:00 | 0:00 | 4:01 | 0:00 | 1:00 |
| 0:01:01 | 0:00:00 | 3:00:00 | 20:00:00 | 1:00:00 | 0:00:00 | 0:00:00 |
| 0:01:01 | 0:00:00 | 3:00:00 | 20:00:00 | 1:00:00 | 0:00:00 | 0:00:00 |
| 1:01 | 0:00 | 8:00 | 27:00:00 | 0:00 | 0:00 | 0:00 |
| 6:0:2:0:0 | 1:0:0:0:0 | 61:0:8:4:0 | 81:4:3:3:0 | 4:0:0:1:1 | 10:1:0:0:0 | 0:0:0:0:0 |
| 10:01:01 | 0:00:00 | 39:01:02 | 23:01:01 | 64:00:02 | 2:00:00 | 10:00:01 |
| 3:00:00 | 31:01:02 | 39:01:05 | 22:00:02 | 0:00:00 | 19:00:01 | 0:00:00 |
| 1:00:00 | 0:00:00 | 3:00:02 | 17:01:01 | 0:00:00 | 4:00:00 | 0:00:00 |
| 4:00 | 0:00 | 1:00 | 8:02 | 0:00 | 0:00 | 0:00 |
| 1:0:0:0 | 6:1:1:0 | 5:0:0:0 | 26:1:6:1 | 40:11:25:3 | 9:1:4:1 | 33:1:15:0 |
| 2:01 | 11866:1484 | 12:10 | 8004:1090 | 12:04 | 1857:84 | 4:00 |
| 5:0:0:1 | 78:5:1:2 | 67:3:7:3 | 136:2:0:9 | 0:0:0:0 | 10:0:2:2 | 0:0:0:0 |
| 4 | 3 | 4 | 1 | 3 | 0 | 0 |
| 0:1:2:1:1:0:0 | 0:4:0:0:0:1:0 | 0:2:1:1:4:0:0 | 0:5:0:0:3:0:1 | 0:0:0:0:0:0:0 | 0:4:0:0:1:0:0 | 0:0:0:0:0:0:0 |
| 1 | 0 | 2 | 0 | 2 | 0 | 1 |
| 1:01 | 0:00 | 23:21 | 10:06 | 0:00 | 3:00 | 0:00 |
| 0:1:0:0:0:1:0:0:0:0:0:0:0:0:0:0:0:0:0:0:0 | 622:919:243:67:56:1794:141:119:487:3:14:46:76:47:10:4:4:3:2:5:1 | 20:0:0:0:0:6:0:0:0:0:0:0:0:0:0:0:0:0:0:0:0 | 522:348:312:123:73:950:40:92:373:5:15:48:59:24:5:11:8:7:1:2:2 | 10:0:0:1:0:4:0:0:0:0:0:0:0:0:0:0:0:0:0:0:0 | 107:182:48:13:11:381:10:27:156:0:9:14:13:3:1:0:0:2:0:0:0 | 1:0:0:0:0:0:0:0:0:0:0:0:0:0:0:0:0:0:0:0:0 |
| 0:0:0:0:1:0:0:0:0:0:0:0:0:0:0:0:0:0:0:0:0 | 917:241:64:56:1847:140:117:486:903:3:13:46:75:47:10:4:4:3:2:5:1 | 0:1:0:0:1:1:0:0:16:0:1:0:0:0:0:0:0:0:0:0:0 | 343:310:124:72:999:40:89:370:797:5:15:46:57:24:5:10:8:7:1:2:2 | 0:1:0:1:4:0:0:0:5:0:0:0:0:0:0:0:0:0:0:0:0 | 181:48:13:11:388:10:26:156:139:0:9:14:12:3:1:0:0:2:0:0:0 | 0:0:0:0:0:0:0:0:0:0:0:0:0:0:0:0:0:0:0:0:0 |
| 2:00 | 6:00 | 8:00 | 0:00 | 7:00 | 0:00 | 1:00 |
| 1 | 0 | 1 | 0 | 1 | 0 | 0 |
| 4:0:1:2:0 | 14:0:3:4:1 | 22:0:0:8:1 | 7:0:0:0:0 | 0:0:0:0:0 | 5:1:0:0:1 | 0:0:0:0:0 |
| 2:0:0:0 | 2:0:0:0 | 3:0:0:0 | 0:0:0:0 | 9:0:1:1 | 0:0:0:0 | 2:0:0:0 |
| 0:00:01 | 1:01:01 | 2:01:00 | 23:06:03 | 0:00:00 | 1:01:00 | 0:00:00 |
| 16:1:2:0:0 | 70:15:3:8:1 | 71:14:2:7:1 | 70:4:4:2:1 | 0:0:0:0:0 | 14:0:1:0:0 | 1:0:0:0:0 |
| 0:1:2:2:0:0:0:0:0:4:0:0:1:0:0:0 | 0:1:8:0:0:0:0:1:0:5:0:0:0:0:0:0 | 0:2:4:0:1:2:0:0:1:3:2:1:0:0:0:0 | 0:0:8:0:0:1:0:1:1:8:0:1:0:1:0:0 | 0:0:5:0:0:1:0:0:1:10:1:0:0:2:0:0 | 0:1:2:1:0:0:0:2:0:5:0:0:0:1:1:1 | 0:0:0:0:0:0:0:0:0:1:0:0:0:0:0:0 |
| 0:1:1:0:0:0:1:4:1:0:0:0:0:0 | 0:7:0:0:0:0:1:37:0:0:1:1:0:0 | 0:3:0:1:2:0:1:29:0:2:2:0:0:0 | 0:5:0:0:1:0:4:42:0:0:1:0:1:0 | 0:6:0:0:1:0:2:48:0:1:2:1:2:1 | 0:2:0:0:0:0:2:6:0:0:1:0:1:0 | 0:0:0:0:0:0:0:1:0:0:0:0:0:0 |
| 14:01:01 | 1624:374:24 | 32:02:01 | 1983:504:24 | 36:03:01 | 290:35:11 | 14:01:00 |
| 1 | 1 | 1 | 2 | 0 | 0 | 0 |
| 33:00:00 | 32:00:00 | 26:00:00 | 51:01:00 | 45:01:01 | 15:01:00 | 16:00:00 |
| 35:00:00 | 33:00:00 | 27:00:00 | 52:01:00 | 49:01:01 | 16:01:00 | 18:00:00 |
| 72:00:00 | 151:00:01 | 8:00:00 | 660:01:01 | 0:00:00 | 133:00:00 | 0:00:00 |
| 20:00:00 | 135:00:03 | 146:10:05 | 43:08:03 | 0:00:00 | 4:01:00 | 0:00:00 |
| 1:00:00 | 11:06:00 | 8:08:01 | 27:16:02 | 0:00:00 | 2:00:00 | 0:00:00 |
| 1:1:3:15:0:0:2:3:0 | 0:0:0:2:0:0:0:0:0 | 0:2:16:84:1:2:120:7:2 | 3:2:27:83:2:1:144:7:1 | 0:0:0:1:0:0:0:0:0 | 1:1:3:18:1:0:4:8:1 | 0:0:0:0:0:0:0:0:0 |
| 2:00 | 0:00 | 9:00 | 29:00:00 | 0:00 | 4:00 | 0:00 |
| 1 | 0 | 2 | 5 | 1 | 1 | 0 |
| 1 | 0 | 2 | 4 | 0 | 1 | 0 |
| 2 | 0 | 19 | 4 | 6 | 1 | 1 |
| 1 | 0 | 2 | 0 | 0 | 1 | 2 |
| 0:00:11 | 0.001643519 | 0.043263889 | 0:01:04 | 0:00:00 | 0:01:00 | 0:00:00 |
| 1:00 | 0:00 | 7:01 | 4:02 | 0:01 | 0:01 | 1:00 |
| 0:3:0:1:0:0:0 | 0:0:0:0:0:0:0 | 0:4:0:0:0:0:0 | 12:17:3:5:0:2:1 | 14:35:11:4:1:6:0 | 1:13:0:1:0:2:0 | 3:5:1:0:1:1:0 |
| 0:4:0:1:0:0 | 0:0:0:0:0:0 | 0:7:0:0:0:0 | 0:0:0:0:0:0 | 31:53:14:10:1:1 | 0:0:0:0:0:0 | 4:20:1:2:1:0 |
| 3 | 2 | 2 | 25 | 0 | 9 | 0 |

| | | | | | | |
|---|---|---|---|---|---|---|
| 1:00:00 | 279:10:31 | 1:00:00 | 506:17:69 | 0:00:02 | 135:08:13 | 0:01:00 |
| 6:01 | 272:10:00 | 9:00 | 675:09:00 | 1:00 | 228:05:00 | 0:00 |
| 6:01 | 269:10:00 | 9:00 | 660:09:00 | 3:00 | 222:05:00 | 1:00 |
| 6:01 | 269:10:00 | 9:00 | 662:09:00 | 1:00 | 222:05:00 | 1:00 |
| 72:00:00 | 9:01 | 242:08:00 | 431:00:00 | 170:09:00 | 101:01:00 | 104:03:00 |
| 6:01 | 241:09:00 | 10:00 | 597:07:00 | 3:01 | 205:04:00 | 0:00 |
| 116 | 2149 | 362 | 4500 | 6 | 1774 | 0 |
| 1:00 | 0:00 | 2:00 | 0:00 | 0:00 | 0:00 | 0:00 |
| 1:00:00 | 310:41:58 | 2:00:01 | 376:26:72 | 0:00:01 | 82:07:06 | 0:00:00 |
| 4:0:0:0:0:4:0:1:0:2:0:0:0:1:0:0:0:0 | 2:0:1:0:1:3:0:2:0:0:1:0:1:0:0:0:0:0 | 4:0:1:0:0:0:0:1:0:1:1:1:0:0:0:0:0:0 | 24:1:9:0:1:9:0:0:2:2:0:0:0:2:2:1:1:1 | 0:0:0:0:0:0:0:0:0:0:0:0:0:0:0:0:0:0 | 12:1:0:0:0:1:0:0:1:1:0:0:2:0:0:0:0:0 | 0:0:0:0:0:0:0:0:0:0:0:0:0:0:0:0:0:0 |
| 0:1:0:0:0:0:0 | 94:5670:1071:108:179:323:27 | 0:10:0:0:0:0:0 | 42:3091:298:124:15:159:15 | 0:3:0:1:0:2:0 | 7:678:56:16:1:23:4 | 0:0:0:0:0:0:0 |
| 0:1:0:0:0:0:0 | 94:5675:1073:108:179:304:28 | 0:5:0:0:0:0:0 | 41:3095:299:125:15:145:15 | 0:2:0:0:0:0:0 | 7:678:56:16:1:24:4 | 0:0:0:0:0:0:0 |
| 1:02 | 0:00 | 1:00 | 0:00 | 0:05 | 0:00 | 1:03 |
| 0:6:0:2:23:15:1:0:2:0:0 | 0:6:0:2:30:27:1:0:3:0:0 | 3:20:1:7:71:67:2:1:15:1:0 | 1:17:3:6:11:10:1:0:1:0:0 | 0:0:0:0:0:0:0:0:0:0:0 | 1:11:0:1:10:5:1:0:2:0:0 | 0:0:0:0:0:0:0:0:0:0:0 |
| 1 | 0 | 1 | 0 | 0 | 0 | 0 |
| 1 | 0 | 1 | 0 | 0 | 0 | 0 |
| 0:3:0:0:0:0:0:0:0:0:0:0:0:0 | 3:7:5:6:0:0:0:6:1:3:0:1:0:0 | 2:3:8:7:0:0:0:4:1:1:1:0:0:0 | 2:7:9:10:0:1:0:3:1:4:0:0:0:1 | 0:5:3:4:0:1:0:1:0:0:0:0:1:0 | 1:3:0:2:0:0:0:0:1:2:0:0:0:0 | 0:0:0:0:0:0:0:1:0:0:0:0:1:0 |
| 0:0:0:0:0:2:0:0 | 0:0:0:0:0:0:1:0 | 0:1:5:1:0:0:8:1 | 0:1:6:0:0:0:5:0 | 0:0:0:0:0:0:0:0 | 1:0:2:0:0:0:2:0 | 0:0:0:0:0:0:0:0 |
| 2:00 | 0:00 | 11:00 | 27:00:00 | 0:00 | 1:00 | 0:00 |
| 2:0:32:20:17:1:0:0:0 | 0:0:0:0:0:0:0:0:0 | 59:18:348:83:104:17:6:0:0 | 84:28:270:54:82:11:4:0:1 | 0:0:1:0:0:0:0:0:0 | 4:2:73:10:17:1:2:0:0 | 0:0:0:0:0:0:0:0:0 |
| 3:00 | 7:00 | 3:00 | 38:00:00 | 0:00 | 15:00 | 0:00 |
| 1 | 4 | 4 | 8 | 0 | 2 | 0 |
| 2:01 | 2:00 | 5:00 | 2:00 | 3:00 | 0:00 | 1:00 |
| 0:00:00 | 4:00:00 | 0:00:00 | 0:00:01 | 2:00:00 | 1:00:00 | 1:00:00 |
| 0:0:0:0:0:0:0:0:0:0 | 2361:102:6:67:0:16:42:2:0:6 | 1:0:0:0:0:0:0:0:0:0 | 3811:126:9:22:0:24:22:2:0:1 | 2:0:0:0:0:0:0:0:0:0 | 1481:12:3:31:0:9:17:1:0:0 | 1:0:0:0:0:0:0:0:0:0 |
| 0:00 | 8:00 | 1:00 | 1:00 | 0:01 | 1:00 | 2:00 |
| 0:00 | 0:00 | 3:00 | 0:00 | 6:01 | 0:00 | 3:00 |
| 0:0:0:0:0:0:0:0:0 | 0:0:1:0:0:0:0:1:0 | 0:0:0:0:0:0:0:0:0 | 0:0:1:0:0:0:1:0:0 | 0:0:2:0:0:0:0:0:0 | 0:0:4:0:0:0:0:0:0 | 1:1:4:0:0:0:1:0:1 |
| 0 | 596 | 0 | 1146 | 1 | 316 | 2 |
| 3 | 0 | 0 | 0 | 3 | 0 | 1 |
| 0:00:00 | 1:00:00 | 0:00:00 | 0:00:00 | 2:00:01 | 0:00:00 | 1:00:00 |
| 0:0:0:0:0 | 92:49:1104:7:1 | 0:0:2:0:0 | 263:109:1176:11:0 | 0:0:1:0:0 | 39:9:214:2:0 | 0:0:1:0:0 |
| 0:00 | 0:00 | 0:00 | 0:00 | 6:01 | 0:00 | 1:00 |
| 0:01 | 0:00 | 0:00 | 0:00 | 4:00 | 0:00 | 1:01 |
| 0 | 7 | 0 | 0 | 9 | 0 | 1 |
| 0:00 | 1:00 | 0:00 | 0:00 | 2:00 | 0:00 | 0:01 |
| 0:00 | 4:00 | 0:00 | 2:00 | 3:01 | 1:00 | 0:01 |
| 0:00 | 0:00 | 3:03 | 0:00 | 3:09 | 0:00 | 1:01 |
| 0:00 | 0:00 | 3:03 | 0:00 | 3:09 | 0:00 | 1:01 |
| 0:0:0:0:0:0:0:0:0:0:0:0:0:0 | 7:30:6:0:4:1:0:0:2:2:2:1:0:0:0 | 0:0:0:0:0:0:0:0:0:0:0:0:0:0 | 1:2:0:0:0:0:0:0:0:0:0:0:0:0 | 4:29:4:0:10:0:0:1:0:0:1:0:1:1:1 | 0:4:2:0:4:0:0:0:0:0:0:0:0:0 | 1:6:0:0:0:0:0:0:0:0:0:0:0:0 |
| 0:0:0:0:0:0:0:0:0:0:0:0:0:0:0:0:0:0:0 | 30:94:11:167:18:13:6:39:6:43:47:20:2:3:25:4:1:0:0 | 0:1:0:1:0:0:0:0:0:0:0:0:0:0:1:0:0:0:0 | 43:112:12:216:27:13:1:45:3:60:48:16:3:4:28:1:0:1:1 | 0:0:0:0:0:0:0:1:0:1:0:0:0:0:0:0:0:0:0 | 48:99:7:221:25:32:2:25:7:7:22:2:1:1:4:1:0:1:0 | 0:0:0:1:0:0:0:0:0:0:0:0:0:0:0:0:0:0:0 |
| 0:0:0:0 | 324:23:29:2 | 0:0:0:0 | 298:27:34:0 | 146:8:11:1 | 73:3:1:0 | 63:1:2:0 |

| | | | | | | |
|---|---|---|---|---|---|---|
| 0:00:00 | 0:00:00 | 0:00:00 | 1:03:01 | 0:02:03 | 0:00:00 | 0:00:01 |
| 0 | 72 | 0 | 93 | 2 | 15 | 1 |
| 0 | 2311 | 4 | 2120 | 3 | 695 | 1 |
| 0 | 1 | 1 | 0 | 1 | 0 | 1 |
| 0:00 | 0:01 | 0:00 | 1:00 | 1:00 | 0:00 | 3:00 |
| 0:00 | 0:01 | 0:00 | 1:00 | 1:00 | 0:00 | 3:00 |
| 0:0:0:0:1:0:0 | 7:21:23:2:1:0:0 | 0:0:0:0:0:0:0 | 7:28:26:3:4:3:0 | 1:1:0:0:0:0:0 | 5:6:3:1:2:1:0 | 1:0:0:0:0:0:0 |
| 0:00 | 122:02:00 | 2:00 | 5:00 | 267:10:00 | 59:05:00 | 2:02 |
| 0:0:0:0:0:0 | 0:0:0:0:0:0 | 0:0:0:0:0:0 | 2:2:2:0:0:1 | 0:2:1:0:0:0 | 1:1:2:1:0:0 | 0:3:0:0:0:0 |
| 0:0:0:0:0:1:0 | 8:0:21:3:4:18:0 | 0:0:0:0:0:0:0 | 10:4:20:23:11:33:3 | 9:1:10:5:5:18:6 | 0:1:0:0:0:0:0 | 4:0:2:4:0:4:2 |
| 0:0:0:0:0:0 | 3:2:0:0:0:0 | 0:0:0:0:0:0 | 0:0:0:0:0:0 | 23:5:1:1:2:1 | 0:0:0:0:0:0 | 3:0:1:0:0:0 |
| 0:00:00 | 0:00:00 | 0:00:00 | 0:00:01 | 0:01:01 | 0:00:00 | 1:00:00 |
| 0:0:0:0:0:0:0:0:0:0:0:0:0 | 164:184:2:16:13:3:6:8:8:5:1:3:6 | 0:0:0:1:0:0:0:0:0:0:0:0:0 | 243:117:6:9:10:0:11:5:8:0:0:0:1 | 252:146:8:11:7:0:7:7:1:7:0:3:4 | 88:66:9:5:9:1:4:1:6:0:0:0:0 | 1:1:0:0:0:0:0:0:0:0:0:0:0 |
| 0:0:0:0:0:0:0:0:0:0:0:0 | 42:73:17:5:27:2:1:0:0:0:1:0 | 0:0:0:0:0:0:0:0:0:0:0:0 | 0:0:0:0:0:0:0:0:0:0:0:0 | 46:34:7:8:16:2:0:1:2:1:0:0 | 0:0:0:0:0:0:0:0:0:0:0:0 | 35:17:0:5:6:0:0:0:0:1:0:0 |
| 0:0:0:0:0:0:0:0:0:0:0:0 | 42:72:18:5:2:1:10:0:0:1:1:0 | 0:1:0:0:0:0:0:0:0:0:0:0 | 0:0:0:0:0:0:0:0:0:0:0:0 | 46:36:7:8:2:0:6:2:2:1:0:0 | 0:0:0:0:0:0:0:0:0:0:0:0 | 35:15:0:5:0:0:5:0:0:1:0:0 |
| 0:00 | 0:05 | 0:00 | 0:01 | 2:14 | 0:00 | 0:01 |
| 0:00 | 0:06 | 0:00 | 0:00 | 0:13 | 0:00 | 0:02 |
| 0:0:0:0:0:0:0:0:0:0:0 | 33:180:634:0:345:30:77:111:2:9:17 | 0:0:1:0:2:0:0:0:0:0:0 | 18:197:304:0:141:39:61:98:2:7:1 | 0:0:0:0:0:1:0:0:0:0:0 | 1:25:29:0:18:1:13:5:0:0:1 | 0:0:0:0:0:0:1:0:0:0:0 |
| 0:00 | 3:00 | 0:00 | 0:00 | 2:00 | 0:00 | 3:00 |
| 1:00 | 0:00 | 0:00 | 0:00 | 1:00 | 0:00 | 1:00 |
| 0:00 | 0:00 | 0:00 | 0:00 | 0:01 | 0:00 | 2:00 |
| 0:00 | 0:00 | 0:00 | 0:00 | 0:01 | 0:00 | 2:00 |
| 0:00 | 1:00 | 1:00 | 1:00 | 6:00 | 2:01 | 1:00 |
| 0:00 | 3:00 | 0:00 | 0:00 | 2:00 | 0:00 | 2:03 |
| 0:0:0:0 | 3:9:2:0 | 0:0:0:0 | 0:0:0:0 | 0:2:0:0 | 0:0:0:0 | 2:5:1:1 |
| 0 | 0 | 0 | 0 | 2 | 0 | 1 |
| 0:0:0:0:0:0 | 1:1:0:0:0:0 | 0:0:0:0:0:0 | 1:0:0:0:0:0 | 0:2:0:0:0:0 | 0:0:0:0:0:0 | 0:0:0:0:1:0 |
| 0:0:0:0:0:0:0 | 5:2:13:1:1:1:2:2 | 0:0:0:0:0:0:0 | 4:7:7:0:0:1:1:2 | 1:0:1:1:0:0:0:0 | 0:0:1:1:0:0:2:0 | 0:0:0:0:0:0:1:0 |
| 0:00 | 2:00 | 0:00 | 0:00 | 1:01 | 0:00 | 1:00 |
| 0:0:0:0:0 | 8:3:1:0:0 | 0:0:0:0:0 | 0:0:0:0:0 | 14:11:5:0:0 | 0:0:0:0:0 | 4:4:0:0:0 |
| 0 | 10 | 0 | 31 | 13 | 9 | 2 |
| 1:0:0:0 | 0:0:0:0 | 0:0:0:0 | 0:0:0:0 | 0:1:0:1 | 0:0:0:0 | 0:2:0:0 |
| 0:01:00 | 0:00:00 | 0:00:00 | 3:00:01 | 6:02:00 | 2:00:00 | 1:00:00 |
| 0:0:0:0 | 0:1:3:0 | 0:2:1:0 | 0:0:1:0 | 1:1:0:1 | 0:0:0:0 | 0:0:1:0 |
| 2:0:0:0:0 | 0:0:0:0:0 | 0:0:0:0:0 | 0:0:0:0:0 | 4:2:4:0:1 | 0:0:0:0:0 | 0:0:0:0:1 |

| | | | | | | |
|---|---|---|---|---|---|---|
| 0:0:0:0:0 | 0:0:0:0:0 | 0:0:0:0:0 | 0:2:0:0:0 | 0:1:0:1:1 | 1:4:0:0:0 | 1:4:1:0:0 |
| 0 | 657 | 2 | 870 | 2 | 123 | 1 |
| 0 | 578 | 3 | 752 | 3 | 122 | 1 |
| 0:00 | 11:00 | 0:00 | 27:00:00 | 16:02 | 8:00 | 4:00 |
| 0:0:0:0:0:0:0:0:0:0:0:0:0:0:0:0:0:0:0:0 | 1712:923:243:67:57:142:123:491:158:3:14:46:79:47:10:4:4:3:2:5:1 | 8:0:1:0:0:0:0:0:22:0:0:0:0:0:0:0:0:0:0:0 | 876:353:312:124:74:40:99:378:102:5:15:51:62:24:5:11:8:7:1:2:2 | 4:0:0:0:0:0:0:0:8:0:0:1:0:0:0:0:0:0:0:0 | 372:182:48:13:11:10:27:156:21:0:9:14:13:3:1:0:0:2:0:0:0 | 0:0:0:0:0:0:0:0:1:0:0:0:0:0:0:0:0:0:0:0 |
| 0 | 0 | 2 | 0 | 2 | 0 | 1 |
| 0:00 | 0:00 | 0:00 | 0:00 | 3:01 | 0:00 | 2:00 |
| 0 | 1368 | 0 | 2277 | 1 | 489 | 2 |
| 0:00 | 0:01 | 0:00 | 4:00 | 4:00 | 2:00 | 2:00 |
| 0:00:00 | 1:00:00 | 0:00:00 | 8:01:00 | 3:03:01 | 0:01:00 | 1:00:00 |
| 0:0:0:0 | 1:21:0:1 | 0:0:0:0 | 2:1:1:0 | 6:14:1:0 | 1:0:0:0 | 2:1:1:0 |
| 3 | 2 | 0 | 0 | 10 | 1 | 1 |
| 0:0:0:0:0 | 21:0:25:3:0 | 0:0:0:0:0 | 0:0:0:0:0 | 3:0:15:1:1 | 0:0:0:0:0 | 2:0:2:0:0 |
| 0:0:0:0:0 | 19:0:21:3:0 | 0:0:0:0:0 | 0:0:0:0:0 | 3:0:13:2:1 | 0:0:0:0:0 | 2:0:2:0:0 |
| 0:0:0:0 | 13:5:1:1 | 0:0:0:0 | 0:0:0:0 | 37:18:0:3 | 0:0:0:0 | 20:2:0:0 |
| 0:0:0:0:0 | 15:19:1:1:0 | 0:0:0:0:0 | 0:0:0:0:0 | 43:49:2:2:4 | 0:0:0:0:0 | 24:5:1:0:0 |
| 0 | 1 | 0 | 2 | 5 | 0 | 2 |
| 0 | 0 | 0 | 0 | 1 | 0 | 1 |
| 0:0:0:0:0:0:0:0:0:0:0:0:0 | 2:126:34:7:22:3:9:0:1:2:2:1:0:0 | 0:1:0:0:0:0:0:0:0:0:0:0:0 | 0:63:9:5:9:0:2:0:0:1:0:0:0:0 | 0:101:33:19:10:5:8:0:1:3:2:1:1:1 | 0:38:2:8:1:1:3:0:1:1:1:0:1:0 | 0:20:8:6:1:1:2:1:0:0:0:0:1:0 |
| 0:00 | 2:01 | 0:00 | 1:00 | 4:00 | 0:00 | 1:00 |
| 0:00 | 2:00 | 0:00 | 5:01 | 3:00 | 0:00 | 2:00 |
| 0:0:0:0:0:0 | 0:7:0:0:0:0 | 0:0:0:0:0:0 | 18:36:12:5:1:0 | 14:17:2:5:0:1 | 3:6:1:1:1:0 | 1:14:0:1:0:0 |
| 0:00 | 273:08:00 | 1:01 | 661:09:00 | 1:00 | 218:03:00 | 0:01 |
| 0:00 | 247:08:00 | 0:01 | 612:09:00 | 2:00 | 204:03:00 | 0:01 |
| 0:00 | 262:00:00 | 1:00 | 185:02:00 | 426:00:00 | 113:00:00 | 104:00:00 |
| 0 | 0 | 0 | 0 | 1 | 0 | 1 |
| 1:0:0:0:0:0:0:0:0 | 0:0:0:0:0:0:0:0:0 | 0:0:0:0:0:0:0:0:0 | 6:1:0:0:2:0:0:4:0 | 5:4:1:1:6:1:1:2:1 | 1:0:0:0:1:0:0:0:0 | 0:0:0:0:1:0:0:0:0 |
| 0:0:0:0:0:0:0 | 3:9:96:1:1:0:1 | 0:0:0:0:0:0:0 | 0:1:3:0:0:0:0 | 3:5:36:1:0:2:0 | 0:0:3:0:0:0:0 | 1:2:13:2:0:0:0 |
| 0:0:0:0 | 145:160:122:61 | 1:0:0:0 | 376:329:330:169 | 1:0:1:0 | 191:80:23:22 | 3:0:0:0 |
| 0:00:00 | 3:01:00 | 0:00:00 | 1:00:00 | 2:01:00 | 0:00:00 | 1:00:00 |
| 0:00 | 3:00 | 0:00 | 1:00 | 1:01 | 1:00 | 1:00 |
| 0 | 0 | 0 | 3 | 6 | 1 | 1 |
| 0:00:00 | 9:00:00 | 0:00:00 | 0:00:00 | 10:00:01 | 0:00:00 | 2:01:00 |
| 0:0:0:0:0:0 | 37:29:46:1:0:15 | 0:0:0:0:0:0 | 0:0:0:0:0:0 | 1:1:3:0:0:0 | 0:0:0:0:0:0 | 1:0:0:0:0:0 |
| 0:0:0:0:0:0 | 37:29:46:1:0:14 | 0:0:0:0:0:0 | 0:0:0:0:0:0 | 1:1:3:0:0:0 | 0:0:0:0:0:0 | 1:0:0:0:0:0 |
| 0:0:0:0:0:0 | 29:37:45:1:0:15 | 0:0:0:0:0:0 | 0:0:0:0:0:0 | 1:1:3:0:0:0 | 0:0:0:0:0:0 | 0:1:0:0:0:0 |
| 0:0:0:0:0:0 | 37:29:45:1:0:18 | 0:0:1:0:0:0 | 0:0:0:0:0:0 | 1:1:3:0:0:0 | 0:0:0:0:0:0 | 1:0:0:0:0:0 |
| 0 | 0 | 0 | 2 | 4 | 0 | 2 |
| 0 | 0 | 0 | 2 | 4 | 0 | 3 |
| 0:01:00 | 8:01:00 | 0:00:00 | 0:00:00 | 12:02:00 | 0:01:00 | 1:01:00 |
| 3:00 | 0:00 | 0:00 | 0:00 | 9:00 | 0:00 | 3:00 |
| 0:0:0:0:0:0:0:0:0:0 | 0:0:9:2:2:2:5:1:0:0 | 0:0:0:0:0:0:0:0:0:0 | 3:0:4:0:3:5:5:2:2:1 | 1:1:4:0:0:4:3:2:0:0 | 0:0:0:0:0:0:0:0:0:0 | 2:1:0:0:2:2:1:0:1:0 |

| WES count | | | | | | | | | | | |
|---|---|---|---|---|---|---|---|---|---|---|---|
| 1M_T__MUT | 1M_T__WT | 1T_T__MUT | 1T_T__WT | 2T_T__MUT | 2T_T__WT | 2M_T__MUT | 2M_T__WT | 3M_T__MUT | 3M_T__WT | 3T_T__MUT | 3T_T__WT |
| 15 | 12 | 33 | 41 | 25 | 21 | 8 | 31 | 1 | 50 | 0 | 0 |
| 48 | 0 | 80 | 0 | 76 | 0 | 81 | 0 | 63 | 0 | 49 | 0 |
| 25 | 35 | 34 | 32 | 19 | 15 | 29 | 26 | 24 | 31 | 21 | 24 |
| 16 | 18 | 26 | 27 | 0 | 0 | 0 | 0 | 0 | 0 | 0 | 0 |
| 32 | 38 | 45 | 46 | 29 | 29 | 46 | 56 | 19 | 35 | 18 | 46 |
| 11 | 24 | 35 | 32 | 60 | 0 | 37 | 0 | 2 | 58 | 0 | 0 |
| 20 | 6 | 26 | 23 | 59 | 1 | 58 | 1 | 0 | 0 | 0 | 0 |
| 29 | 0 | 35 | 0 | 1 | 25 | 0 | 0 | 0 | 0 | 0 | 0 |
| 67 | 0 | 85 | 0 | 61 | 0 | 91 | 0 | 92 | 0 | 0 | 0 |
| 30 | 0 | 36 | 0 | 32 | 0 | 55 | 0 | 36 | 0 | 29 | 0 |
| 48 | 0 | 66 | 0 | 52 | 0 | 73 | 0 | 72 | 0 | 46 | 0 |
| 7 | 12 | 17 | 13 | 25 | 1 | 36 | 0 | 33 | 0 | 19 | 0 |
| 37 | 0 | 57 | 0 | 35 | 41 | 39 | 7 | 20 | 47 | 13 | 36 |
| 37 | 24 | 45 | 47 | 41 | 33 | 46 | 9 | 63 | 28 | 45 | 25 |
| 18 | 27 | 37 | 52 | 44 | 27 | 50 | 43 | 1 | 66 | 0 | 0 |
| 78 | 56 | 81 | 96 | 2 | 111 | 0 | 0 | 0 | 0 | 0 | 0 |
| 1 | 107 | 0 | 0 | 0 | 0 | 0 | 0 | 0 | 0 | 0 | 0 |
| 150 | 1 | 198 | 0 | 64 | 79 | 87 | 110 | 18 | 99 | 31 | 58 |
| 24 | 26 | 31 | 32 | 79 | 0 | 65 | 0 | 1 | 82 | 0 | 0 |
| 35 | 33 | 36 | 31 | 0 | 0 | 1 | 99 | 18 | 5 | 25 | 17 |
| 5 | 0 | 24 | 0 | 13 | 0 | 12 | 0 | 6 | 11 | 2 | 13 |
| 3 | 1 | 7 | 8 | 7 | 7 | 3 | 3 | 8 | 0 | 2 | 0 |
| 25 | 31 | 42 | 56 | 105 | 0 | 89 | 1 | 2 | 77 | 1 | 44 |
| 46 | 46 | 70 | 58 | 35 | 36 | 67 | 66 | 2 | 59 | 2 | 72 |
| 18 | 23 | 30 | 24 | 2 | 31 | 0 | 0 | 19 | 8 | 26 | 9 |
| 64 | 0 | 95 | 0 | 71 | 0 | 104 | 0 | 112 | 0 | 66 | 0 |
| 62 | 0 | 99 | 1 | 1 | 74 | 0 | 0 | 0 | 0 | 0 | 0 |
| 45 | 0 | 77 | 0 | 53 | 0 | 86 | 0 | 94 | 0 | 45 | 0 |
| 29 | 12 | 26 | 38 | 0 | 0 | 0 | 0 | 36 | 21 | 27 | 16 |
| 30 | 0 | 49 | 0 | 1 | 35 | 0 | 0 | 22 | 23 | 17 | 13 |
| 50 | 0 | 63 | 0 | 51 | 0 | 79 | 0 | 82 | 0 | 52 | 0 |
| 14 | 16 | 38 | 26 | 1 | 47 | 0 | 0 | 0 | 0 | 2 | 41 |
| 15 | 22 | 23 | 17 | 31 | 0 | 49 | 1 | 23 | 0 | 34 | 1 |
| 10 | 13 | 17 | 13 | 0 | 0 | 0 | 0 | 0 | 0 | 0 | 0 |
| 95 | 1 | 126 | 1 | 44 | 46 | 66 | 75 | 89 | 1 | 71 | 0 |
| 37 | 0 | 38 | 0 | 30 | 0 | 53 | 0 | 18 | 11 | 11 | 20 |
| 14 | 0 | 44 | 1 | 1 | 41 | 0 | 0 | 35 | 0 | 30 | 0 |
| 0 | 0 | 0 | 0 | 0 | 0 | 1 | 161 | 0 | 0 | 0 | 0 |
| 16 | 24 | 38 | 41 | 0 | 0 | 0 | 0 | 30 | 21 | 9 | 32 |
| 0 | 0 | 0 | 0 | 0 | 0 | 0 | 0 | 0 | 0 | 0 | 0 |
| 39 | 0 | 43 | 0 | 16 | 19 | 29 | 23 | 49 | 0 | 25 | 0 |
| 30 | 37 | 50 | 44 | 54 | 35 | 40 | 48 | 88 | 1 | 41 | 0 |
| 18 | 17 | 19 | 33 | 0 | 0 | 0 | 0 | 0 | 0 | 0 | 0 |
| 82 | 35 | 41 | 55 | 0 | 0 | 0 | 0 | 0 | 0 | 32 | 32 |
| 151 | 0 | 134 | 0 | 240 | 1 | 191 | 0 | 75 | 68 | 38 | 48 |

| | | | | | | | | | | | |
|---|---|---|---|---|---|---|---|---|---|---|---|
| 75 | 0 | 73 | 0 | 93 | 0 | 100 | 0 | 39 | 39 | 21 | 32 |
| 59 | 33 | 76 | 59 | 65 | 20 | 70 | 15 | 84 | 20 | 34 | 54 |
| 15 | 12 | 22 | 34 | 5 | 21 | 2 | 22 | 13 | 36 | 22 | 22 |
| 0 | 0 | 0 | 0 | 0 | 0 | 0 | 0 | 0 | 0 | 1 | 22 |
| 48 | 32 | 74 | 70 | 2 | 126 | 1 | 143 | 106 | 2 | 74 | 0 |
| 39 | 0 | 68 | 0 | 0 | 0 | 0 | 0 | 34 | 1 | 42 | 0 |
| C | 2 | 72 | 0 | 0 | 0 | 0 | 0 | 0 | 0 | 0 | 0 |
| 24 | 41 | 68 | 59 | 69 | 69 | 50 | 48 | 76 | 23 | 44 | 18 |
| 40 | 0 | 71 | 0 | 0 | 0 | 0 | 0 | 43 | 17 | 29 | 14 |
| 11 | 21 | 0 | 0 | 14 | 21 | 0 | 0 | 35 | 1 | 26 | 0 |
| 38 | 0 | 75 | 0 | 37 | 51 | 37 | 42 | 66 | 1 | 43 | 0 |
| 70 | 0 | 106 | 0 | 101 | 0 | 100 | 0 | 106 | 0 | 51 | 0 |
| 0 | 0 | 0 | 0 | 1 | 34 | 0 | 0 | 0 | 0 | 0 | 0 |
| 0 | 0 | 0 | 0 | 0 | 0 | 0 | 0 | 0 | 0 | 0 | 0 |
| 28 | 0 | 58 | 0 | 50 | 0 | 52 | 0 | 0 | 0 | 0 | 0 |
| 16 | 41 | 44 | 51 | 44 | 54 | 49 | 62 | 2 | 72 | 0 | 0 |
| 30 | 22 | 39 | 39 | 29 | 33 | 49 | 42 | 52 | 22 | 27 | 11 |
| 49 | 38 | 45 | 72 | 30 | 36 | 61 | 75 | 83 | 1 | 63 | 1 |
| 42 | 0 | 77 | 0 | 25 | 25 | 18 | 31 | 46 | 0 | 32 | 0 |
| 22 | 22 | 33 | 33 | 44 | 30 | 41 | 33 | 52 | 9 | 23 | 9 |
| 18 | 1 | 34 | 0 | 9 | 0 | 41 | 0 | 3 | 5 | 5 | 19 |
| 12 | 10 | 22 | 28 | 27 | 2 | 32 | 0 | 0 | 0 | 0 | 0 |
| 34 | 21 | 68 | 78 | 35 | 0 | 90 | 1 | 19 | 29 | 45 | 49 |
| 33 | 0 | 70 | 0 | 25 | 26 | 8 | 41 | 53 | 0 | 38 | 0 |
| 11 | 13 | 14 | 17 | 1 | 30 | 0 | 0 | 0 | 0 | 0 | 0 |
| 12 | 14 | 17 | 17 | 0 | 0 | 0 | 0 | 0 | 0 | 0 | 0 |
| 6 | 7 | 8 | 13 | 14 | 19 | 4 | 11 | 32 | 0 | 13 | 0 |
| 6 | 15 | 16 | 14 | 13 | 18 | 11 | 12 | 0 | 0 | 0 | 0 |
| 9 | 0 | 11 | 0 | 14 | 5 | 6 | 7 | 11 | 6 | 4 | 3 |
| 37 | 29 | 66 | 70 | 1 | 85 | 0 | 0 | 46 | 38 | 35 | 43 |
| 0 | 0 | 0 | 0 | 0 | 0 | 0 | 0 | 36 | 99 | 27 | 92 |
| 14 | 11 | 19 | 9 | 1 | 16 | 0 | 0 | 0 | 0 | 0 | 0 |
| 1 | 1 | 7 | 5 | 0 | 0 | 0 | 0 | 0 | 0 | 0 | 0 |
| 42 | 0 | 49 | 0 | 23 | 0 | 57 | 0 | 15 | 13 | 18 | 16 |
| 18 | 31 | 31 | 27 | 23 | 0 | 60 | 0 | 20 | 17 | 16 | 17 |
| 42 | 20 | 60 | 49 | 1 | 69 | 0 | 0 | 0 | 0 | 0 | 0 |
| 23 | 0 | 31 | 0 | 41 | 7 | 51 | 16 | 44 | 0 | 29 | 0 |
| 0 | 0 | 1 | 173 | 0 | 0 | 0 | 0 | 1 | 222 | 1 | 101 |
| 0 | 0 | 0 | 0 | 1 | 174 | 0 | 0 | 0 | 0 | 0 | 0 |

| | | | | | | | | | | | |
|---|---|---|---|---|---|---|---|---|---|---|---|
| 0 | 0 | 0 | 0 | 0 | 0 | 0 | 0 | 0 | 0 | 0 | 0 |
| 0 | 0 | 0 | 0 | 0 | 0 | 0 | 0 | 0 | 0 | 0 | 0 |
| 35 | 31 | 0 | 0 | 46 | 52 | 66 | 53 | 0 | 0 | 2 | 76 |
| 70 | 0 | 79 | 0 | 99 | 0 | 59 | 1 | 0 | 0 | 0 | 0 |
| 25 | 32 | 39 | 47 | 1 | 80 | 1 | 71 | 17 | 66 | 23 | 43 |
| 14 | 0 | 28 | 0 | 25 | 0 | 19 | 0 | 23 | 0 | 14 | 0 |
| 0 | 0 | 0 | 0 | 1 | 52 | 0 | 0 | 0 | 0 | 0 | 0 |
| 15 | 29 | 36 | 31 | 31 | 42 | 28 | 17 | 0 | 0 | 0 | 0 |
| 24 | 17 | 0 | 0 | 43 | 0 | 79 | 0 | 29 | 28 | 30 | 12 |
| 28 | 55 | 73 | 65 | 58 | 60 | 67 | 82 | 1 | 96 | 0 | 0 |
| 0 | 0 | 0 | 0 | 0 | 0 | 0 | 0 | 1 | 69 | 0 | 0 |
| 32 | 56 | 82 | 73 | 58 | 53 | 87 | 75 | 1 | 111 | 0 | 0 |
| 21 | 15 | 32 | 24 | 32 | 36 | 38 | 34 | 0 | 0 | 0 | 0 |
| 23 | 16 | 37 | 24 | 2 | 45 | 0 | 0 | 11 | 31 | 30 | 22 |
| A | 34 | 16 | 0 | 0 | 0 | 0 | 0 | 0 | 0 | 0 | 0 |
| 9 | 14 | 38 | 31 | 42 | 1 | 49 | 0 | 28 | 37 | 27 | 22 |
| 56 | 7 | 96 | 13 | 82 | 2 | 99 | 14 | 31 | 50 | 30 | 35 |
| 56 | 0 | 77 | 0 | 24 | 44 | 11 | 77 | 33 | 11 | 27 | 22 |
| 33 | 0 | 59 | 0 | 37 | 0 | 43 | 0 | 33 | 0 | 34 | 0 |
| 33 | 0 | 0 | 0 | 31 | 0 | 29 | 0 | 29 | 0 | 25 | 0 |
| 33 | 37 | 76 | 68 | 1 | 87 | 1 | 132 | 64 | 64 | 51 | 51 |
| 0 | 0 | 0 | 0 | 1 | 59 | 0 | 0 | 0 | 0 | 0 | 0 |
| A | 20 | 30 | 0 | 0 | 0 | 0 | 0 | 0 | 0 | 0 | 0 |
| 42 | 4 | 79 | 2 | 0 | 0 | 0 | 0 | 24 | 50 | 28 | 27 |
| 27 | 16 | 40 | 27 | 0 | 0 | 0 | 0 | 0 | 0 | 0 | 0 |
| 37 | 0 | 58 | 1 | 0 | 0 | 0 | 0 | 23 | 39 | 20 | 19 |
| 4 | 7 | 12 | 14 | 0 | 0 | 0 | 0 | 0 | 0 | 7 | 13 |
| 3 | 7 | 12 | 17 | 24 | 0 | 38 | 0 | 15 | 9 | 7 | 14 |
| 3 | 5 | 9 | 16 | 9 | 15 | 14 | 21 | 14 | 9 | 7 | 11 |
| 3 | 5 | 10 | 15 | 9 | 15 | 15 | 20 | 13 | 9 | 7 | 9 |
| 3 | 5 | 10 | 14 | 9 | 15 | 15 | 20 | 14 | 9 | 7 | 9 |
| 3 | 5 | 9 | 14 | 9 | 14 | 15 | 20 | 14 | 8 | 7 | 8 |
| 33 | 0 | 46 | 0 | 19 | 16 | 33 | 33 | 33 | 1 | 38 | 0 |
| 50 | 0 | 71 | 0 | 0 | 0 | 0 | 0 | 32 | 32 | 21 | 38 |
| 18 | 38 | 35 | 42 | 18 | 45 | 41 | 58 | 0 | 0 | 1 | 68 |
| 18 | 36 | 34 | 39 | 17 | 46 | 41 | 58 | 0 | 0 | 1 | 64 |
| 11 | 11 | 15 | 20 | 32 | 1 | 41 | 0 | 37 | 0 | 31 | 0 |
| 10 | 11 | 15 | 20 | 32 | 1 | 41 | 0 | 0 | 0 | 30 | 0 |
| 11 | 11 | 15 | 21 | 32 | 1 | 41 | 0 | 37 | 0 | 30 | 0 |
| 11 | 9 | 16 | 20 | 35 | 1 | 42 | 0 | 38 | 0 | 31 | 0 |
| 13 | 7 | 13 | 16 | 14 | 0 | 9 | 0 | 15 | 0 | 9 | 1 |
| 13 | 9 | 18 | 21 | 25 | 0 | 20 | 0 | 31 | 0 | 14 | 1 |
| 20 | 17 | 28 | 42 | 37 | 0 | 40 | 0 | 72 | 0 | 29 | 1 |
| 27 | 86 | 40 | 143 | 0 | 0 | 1 | 113 | 0 | 0 | 0 | 0 |
| 55 | 56 | 91 | 89 | 25 | 35 | 34 | 35 | 46 | 38 | 43 | 28 |
| 0 | 0 | 0 | 0 | 9 | 12 | 9 | 14 | 0 | 0 | 0 | 0 |
| T | 39 | 35 | 65 | 48 | 62 | 0 | 118 | 0 | 102 | 0 | 70 |
| G | 35 | 23 | 51 | 33 | 45 | 0 | 90 | 0 | 0 | 0 | 44 |
| 7 | 15 | 16 | 14 | 1 | 32 | 0 | 0 | 0 | 0 | 0 | 0 |
| 0 | 0 | 0 | 0 | 0 | 0 | 0 | 0 | 28 | 35 | 27 | 24 |
| 26 | 0 | 58 | 0 | 28 | 56 | 29 | 69 | 53 | 0 | 42 | 0 |
| 20 | 24 | 28 | 23 | 100 | 16 | 101 | 31 | 13 | 66 | 11 | 47 |

| | | | | | | | | | | | |
|---|---|---|---|---|---|---|---|---|---|---|---|
| 0 | 0 | 1 | 47 | 66 | 56 | 78 | 68 | 17 | 73 | 17 | 50 |
| 14 | 0 | 22 | 0 | 20 | 50 | 34 | 71 | 70 | 24 | 38 | 18 |
| 0 | 0 | 1 | 3 | 0 | 0 | 0 | 0 | 0 | 0 | 0 | 0 |
| 0 | 0 | 1 | 3 | 0 | 0 | 0 | 0 | 0 | 0 | 0 | 0 |
| 0 | 0 | 0 | 0 | 3 | 43 | 4 | 47 | 26 | 24 | 0 | 0 |
| 8 | 39 | 9 | 46 | 3 | 30 | 7 | 42 | 10 | 0 | 10 | 0 |
| 8 | 39 | 9 | 46 | 3 | 30 | 7 | 43 | 10 | 0 | 10 | 0 |
| 8 | 38 | 10 | 46 | 4 | 30 | 7 | 44 | 10 | 0 | 10 | 0 |
| 8 | 39 | 9 | 48 | 4 | 31 | 7 | 44 | 10 | 0 | 10 | 0 |
| 12 | 38 | 19 | 52 | 13 | 31 | 22 | 44 | 19 | 0 | 24 | 0 |
| 12 | 39 | 21 | 56 | 44 | 0 | 67 | 0 | 18 | 0 | 27 | 0 |
| 44 | 0 | 73 | 0 | 38 | 0 | 57 | 0 | 18 | 0 | 28 | 0 |
| 43 | 0 | 74 | 0 | 38 | 0 | 58 | 0 | 18 | 0 | 28 | 0 |
| 43 | 0 | 73 | 0 | 38 | 0 | 58 | 0 | 18 | 0 | 28 | 0 |
| 43 | 0 | 78 | 0 | 39 | 0 | 59 | 0 | 19 | 0 | 29 | 0 |
| 18 | 40 | 37 | 65 | 26 | 25 | 46 | 40 | 14 | 27 | 21 | 22 |
| 19 | 41 | 37 | 65 | 26 | 25 | 46 | 40 | 14 | 27 | 21 | 23 |
| 24 | 42 | 44 | 62 | 22 | 49 | 44 | 74 | 33 | 62 | 29 | 35 |
| 26 | 37 | 49 | 60 | 15 | 48 | 39 | 76 | 33 | 61 | 25 | 33 |
| 18 | 27 | 28 | 46 | 13 | 39 | 32 | 72 | 30 | 52 | 19 | 26 |
| 42 | 0 | 71 | 0 | 14 | 36 | 34 | 74 | 30 | 51 | 21 | 25 |
| 18 | 25 | 27 | 42 | 12 | 33 | 33 | 72 | 0 | 0 | 0 | 0 |
| 19 | 47 | 42 | 94 | 43 | 54 | 60 | 72 | 0 | 0 | 0 | 0 |
| 33 | 59 | 58 | 82 | 0 | 0 | 0 | 0 | 0 | 0 | 1 | 150 |
| 33 | 69 | 60 | 101 | 133 | 47 | 177 | 47 | 17 | 324 | 7 | 172 |
| 106 | 0 | 171 | 0 | 0 | 0 | 0 | 0 | 0 | 0 | 0 | 0 |
| 56 | 0 | 69 | 0 | 30 | 105 | 33 | 124 | 187 | 19 | 101 | 10 |
| 69 | 17 | 97 | 33 | 84 | 125 | 97 | 133 | 150 | 93 | 65 | 51 |
| 62 | 1 | 95 | 0 | 96 | 0 | 149 | 1 | 168 | 0 | 130 | 0 |
| 21 | 45 | 34 | 68 | 1 | 97 | 0 | 0 | 94 | 1 | 71 | 2 |
| 0 | 0 | 1 | 104 | 0 | 0 | 0 | 0 | 10 | 2 | 8 | 0 |
| 0 | 0 | 1 | 104 | 0 | 0 | 0 | 0 | 10 | 2 | 8 | 0 |
| 3 | 71 | 2 | 108 | 0 | 0 | 0 | 0 | 49 | 3 | 39 | 0 |
| 3 | 70 | 2 | 108 | 0 | 0 | 0 | 0 | 52 | 3 | 44 | 0 |
| 3 | 68 | 2 | 106 | 0 | 0 | 0 | 0 | 53 | 3 | 48 | 0 |
| 3 | 68 | 1 | 107 | 0 | 0 | 0 | 0 | 53 | 3 | 48 | 0 |
| 11 | 61 | 35 | 99 | 0 | 0 | 0 | 0 | 84 | 3 | 69 | 0 |
| 23 | 59 | 42 | 97 | 50 | 70 | 51 | 78 | 39 | 1 | 18 | 0 |
| 38 | 23 | 37 | 42 | 0 | 0 | 0 | 0 | 1 | 38 | 1 | 23 |
| 49 | 23 | 52 | 51 | 45 | 46 | 64 | 59 | 0 | 0 | 0 | 0 |
| 37 | 24 | 49 | 50 | 0 | 0 | 1 | 120 | 29 | 61 | 13 | 20 |
| 40 | 23 | 44 | 42 | 92 | 1 | 119 | 0 | 0 | 0 | 0 | 0 |
| 27 | 1 | 54 | 0 | 47 | 0 | 48 | 0 | 0 | 0 | 2 | 39 |
| 104 | 0 | 116 | 1 | 2 | 63 | 1 | 128 | 38 | 32 | 48 | 37 |
| 20 | 19 | 22 | 15 | 13 | 17 | 20 | 29 | 27 | 1 | 25 | 0 |
| 0 | 0 | 0 | 0 | 0 | 0 | 0 | 0 | 0 | 0 | 1 | 120 |
| 0 | 0 | 0 | 0 | 0 | 0 | 0 | 0 | 0 | 0 | 0 | 0 |
| 49 | 0 | 62 | 0 | 48 | 0 | 74 | 0 | 41 | 0 | 35 | 0 |
| 25 | 25 | 38 | 46 | 32 | 42 | 42 | 48 | 70 | 14 | 36 | 18 |
| 16 | 0 | 30 | 0 | 35 | 0 | 38 | 0 | 26 | 5 | 13 | 9 |
| 15 | 8 | 22 | 19 | 15 | 13 | 15 | 20 | 21 | 6 | 20 | 23 |
| 53 | 1 | 75 | 0 | 43 | 48 | 49 | 54 | 35 | 63 | 33 | 37 |
| 1 | 114 | 2 | 95 | 2 | 117 | 6 | 225 | 9 | 120 | 4 | 89 |
| 0 | 0 | 0 | 0 | 0 | 0 | 0 | 0 | 0 | 0 | 0 | 0 |
| 0 | 0 | 0 | 0 | 0 | 0 | 0 | 0 | 0 | 0 | 0 | 0 |
| 1 | 34 | 0 | 0 | 0 | 0 | 0 | 0 | 0 | 0 | 0 | 0 |
| 1 | 53 | 0 | 0 | 0 | 0 | 0 | 0 | 0 | 0 | 0 | 0 |

| | | | | | | | | | | | |
|---|---|---|---|---|---|---|---|---|---|---|---|
| 18 | 0 | 29 | 0 | 2 | 22 | 0 | 0 | 14 | 10 | 10 | 7 |
| 9 | 6 | 30 | 38 | 0 | 0 | 0 | 0 | 0 | 0 | 0 | 0 |
| 25 | 14 | 37 | 38 | 1 | 31 | 0 | 0 | 0 | 0 | 0 | 0 |
| 0 | 0 | 0 | 0 | 35 | 24 | 118 | 40 | 19 | 25 | 9 | 20 |
| 4 | 15 | 15 | 16 | 16 | 7 | 40 | 8 | 25 | 13 | 14 | 9 |
| 6 | 14 | 10 | 12 | 12 | 7 | 33 | 7 | 20 | 9 | 15 | 6 |
| 0 | 0 | 0 | 0 | 0 | 0 | 0 | 0 | 1 | 92 | 0 | 0 |
| 56 | 0 | 42 | 0 | 29 | 16 | 76 | 29 | 86 | 0 | 44 | 0 |
| 10 | 17 | 22 | 20 | 0 | 0 | 0 | 0 | 10 | 26 | 28 | 13 |
| 44 | 0 | 69 | 1 | 0 | 0 | 1 | 76 | 1 | 33 | 0 | 0 |
| 1 | 0 | 8 | 0 | 3 | 0 | 4 | 0 | 4 | 0 | 6 | 0 |
| 0 | 0 | 0 | 0 | 60 | 3 | 62 | 1 | 54 | 21 | 23 | 13 |
| 0 | 0 | 0 | 0 | 0 | 0 | 1 | 55 | 0 | 0 | 0 | 0 |
| 0 | 0 | 0 | 0 | 0 | 0 | 1 | 40 | 0 | 0 | 0 | 0 |
| 0 | 0 | 0 | 0 | 0 | 0 | 0 | 0 | 0 | 0 | 0 | 0 |
| 0 | 0 | 0 | 0 | 0 | 0 | 0 | 0 | 1 | 171 | 0 | 0 |
| 0 | 0 | 0 | 0 | 30 | 1 | 23 | 1 | 8 | 34 | 7 | 18 |
| 0 | 0 | 0 | 0 | 26 | 41 | 47 | 21 | 57 | 21 | 44 | 12 |
| 15 | 0 | 32 | 0 | 18 | 0 | 19 | 0 | 18 | 0 | 26 | 0 |
| 13 | 0 | 31 | 0 | 28 | 0 | 23 | 0 | 17 | 0 | 18 | 0 |
| 0 | 0 | 1 | 48 | 25 | 28 | 19 | 27 | 7 | 31 | 11 | 16 |
| 0 | 0 | 0 | 0 | 25 | 20 | 38 | 26 | 0 | 0 | 1 | 36 |
| 0 | 0 | 0 | 0 | 0 | 0 | 0 | 0 | 0 | 0 | 0 | 0 |
| 0 | 0 | 0 | 0 | 0 | 0 | 0 | 0 | 0 | 0 | 1 | 57 |
| 45 | 0 | 68 | 0 | 73 | 0 | 71 | 0 | 44 | 38 | 22 | 18 |
| 0 | 0 | 0 | 0 | 1 | 43 | 0 | 0 | 24 | 28 | 11 | 28 |
| 0 | 0 | 0 | 0 | 23 | 21 | 25 | 26 | 0 | 0 | 0 | 0 |
| 11 | 9 | 14 | 20 | 27 | 0 | 42 | 0 | 14 | 9 | 17 | 4 |
| 0 | 0 | 0 | 0 | 0 | 0 | 17 | 23 | 45 | 0 | 20 | 0 |
| 0 | 0 | 0 | 0 | 16 | 15 | 36 | 40 | 19 | 23 | 15 | 34 |
| 14 | 9 | 35 | 23 | 48 | 0 | 51 | 0 | 57 | 0 | 0 | 0 |
| T | 25 | 23 | 0 | 0 | 0 | 0 | 0 | 0 | 0 | 0 | 0 |
| 0 | 0 | 0 | 0 | 32 | 71 | 68 | 116 | 0 | 0 | 0 | 0 |
| 1 | 152 | 0 | 0 | 49 | 119 | 64 | 131 | 84 | 3 | 65 | 2 |
| 0 | 0 | 0 | 0 | 50 | 16 | 53 | 36 | 18 | 23 | 17 | 20 |
| 0 | 0 | 0 | 0 | 65 | 17 | 61 | 40 | 25 | 24 | 20 | 20 |
| 0 | 0 | 0 | 0 | 0 | 0 | 0 | 0 | 0 | 0 | 0 | 0 |
| 31 | 0 | 55 | 0 | 53 | 0 | 67 | 1 | 35 | 14 | 24 | 10 |
| 15 | 13 | 22 | 22 | 19 | 21 | 22 | 26 | 25 | 10 | 20 | 10 |
| 0 | 0 | 0 | 0 | 36 | 34 | 30 | 37 | 17 | 49 | 13 | 34 |
| 29 | 0 | 32 | 0 | 35 | 0 | 44 | 0 | 43 | 0 | 27 | 0 |
| 22 | 20 | 39 | 34 | 66 | 0 | 75 | 0 | 74 | 0 | 31 | 0 |
| 3 | 15 | 8 | 14 | 16 | 0 | 18 | 0 | 28 | 0 | 6 | 1 |
| 13 | 24 | 34 | 37 | 0 | 0 | 22 | 22 | 56 | 0 | 35 | 1 |
| 9 | 22 | 28 | 31 | 19 | 17 | 21 | 21 | 46 | 0 | 29 | 1 |

| | | | | | | | | | | | |
|---|---|---|---|---|---|---|---|---|---|---|---|
| 2 | 7 | 16 | 18 | 16 | 0 | 11 | 0 | 26 | 0 | 13 | 0 |
| 10 | 15 | 20 | 35 | 27 | 25 | 25 | 5 | 1 | 51 | 0 | 0 |
| 10 | 15 | 20 | 37 | 27 | 25 | 25 | 5 | 1 | 52 | 0 | 0 |
| 0 | 0 | 0 | 0 | 20 | 17 | 34 | 9 | 0 | 0 | 0 | 0 |
| 0 | 0 | 0 | 0 | 8 | 6 | 6 | 0 | 0 | 0 | 0 | 0 |
| 0 | 0 | 0 | 0 | 28 | 1 | 25 | 0 | 22 | 18 | 8 | 7 |
| 0 | 0 | 0 | 0 | 53 | 44 | 13 | 53 | 0 | 0 | 1 | 62 |
| 0 | 0 | 0 | 0 | 23 | 29 | 19 | 8 | 0 | 0 | 0 | 0 |
| 17 | 15 | 0 | 0 | 62 | 2 | 49 | 0 | 2 | 85 | 0 | 0 |
| 0 | 0 | 0 | 0 | 18 | 14 | 22 | 18 | 21 | 18 | 11 | 10 |
| 0 | 0 | 0 | 0 | 1 | 67 | 0 | 0 | 0 | 0 | 0 | 0 |
| 0 | 0 | 0 | 0 | 55 | 44 | 68 | 65 | 1 | 85 | 0 | 0 |
| 0 | 0 | 0 | 0 | 2 | 10 | 7 | 8 | 8 | 6 | 11 | 8 |
| 1 | 44 | 0 | 0 | 25 | 25 | 31 | 42 | 0 | 0 | 0 | 0 |
| 24 | 0 | 28 | 0 | 17 | 0 | 24 | 0 | 26 | 0 | 26 | 0 |
| 0 | 0 | 0 | 0 | 31 | 12 | 30 | 8 | 1 | 55 | 0 | 0 |
| A | 0 | 0 | 0 | 0 | 0 | 0 | 0 | 0 | 0 | 0 | 3 |
| 0 | 0 | 0 | 0 | 0 | 0 | 0 | 0 | 0 | 0 | 0 | 0 |
| 71 | 1 | 93 | 0 | 45 | 24 | 49 | 36 | 70 | 1 | 66 | 0 |
| 32 | 36 | 0 | 0 | 37 | 30 | 45 | 47 | 106 | 1 | 49 | 1 |
| 20 | 11 | 27 | 33 | 19 | 18 | 23 | 31 | 0 | 0 | 0 | 0 |
| 9 | 17 | 22 | 20 | 12 | 10 | 24 | 25 | 35 | 0 | 26 | 0 |
| 0 | 0 | 0 | 0 | 39 | 36 | 39 | 41 | 0 | 0 | 0 | 0 |
| 0 | 0 | 0 | 0 | 21 | 27 | 40 | 48 | 1 | 53 | 0 | 0 |
| 0 | 0 | 0 | 0 | 23 | 14 | 25 | 23 | 17 | 15 | 13 | 17 |
| 0 | 0 | 1 | 84 | 0 | 0 | 41 | 52 | 27 | 54 | 31 | 32 |
| 0 | 0 | 0 | 0 | 0 | 0 | 1 | 60 | 0 | 0 | 0 | 0 |
| 57 | 49 | 61 | 65 | 31 | 32 | 66 | 67 | 18 | 38 | 31 | 51 |
| 0 | 0 | 0 | 0 | 50 | 39 | 54 | 58 | 15 | 57 | 22 | 45 |
| 0 | 0 | 1 | 149 | 51 | 55 | 77 | 48 | 22 | 57 | 25 | 57 |
| 0 | 0 | 0 | 0 | 16 | 29 | 9 | 21 | 0 | 0 | 0 | 0 |
| 0 | 0 | 0 | 0 | 6 | 16 | 18 | 32 | 0 | 0 | 0 | 0 |
| 0 | 0 | 0 | 0 | 25 | 26 | 37 | 36 | 0 | 0 | 0 | 0 |
| 0 | 0 | 0 | 0 | 37 | 46 | 53 | 16 | 0 | 0 | 0 | 0 |
| 0 | 0 | 1 | 135 | 0 | 0 | 85 | 2 | 5 | 72 | 1 | 76 |
| 0 | 0 | 0 | 0 | 36 | 1 | 31 | 1 | 0 | 0 | 0 | 0 |
| 0 | 0 | 0 | 0 | 33 | 2 | 52 | 0 | 1 | 28 | 0 | 0 |
| 42 | 0 | 88 | 0 | 43 | 0 | 81 | 0 | 37 | 30 | 24 | 26 |
| 0 | 0 | 0 | 0 | 26 | 21 | 25 | 22 | 22 | 22 | 17 | 29 |
| 0 | 0 | 0 | 0 | 47 | 66 | 77 | 81 | 1 | 127 | 0 | 0 |
| 26 | 30 | 37 | 36 | 37 | 0 | 89 | 0 | 39 | 32 | 24 | 36 |
| 0 | 0 | 0 | 0 | 52 | 2 | 49 | 0 | 32 | 31 | 21 | 11 |
| 1 | 45 | 1 | 80 | 53 | 0 | 70 | 0 | 78 | 0 | 61 | 1 |
| 18 | 29 | 26 | 26 | 20 | 15 | 19 | 21 | 0 | 0 | 0 | 0 |

| | | | | | | | | | | | |
|---|---|---|---|---|---|---|---|---|---|---|---|
| 0 | 0 | 0 | 0 | 0 | 0 | 0 | 0 | 0 | 0 | 0 | 0 |
| 0 | 0 | 0 | 0 | 10 | 57 | 28 | 86 | 0 | 0 | 0 | 0 |
| 28 | 97 | 48 | 144 | 0 | 0 | 0 | 0 | 45 | 110 | 34 | 65 |
| 28 | 97 | 48 | 141 | 9 | 63 | 29 | 83 | 45 | 109 | 33 | 66 |
| 6 | 35 | 9 | 49 | 13 | 10 | 16 | 9 | 18 | 20 | 15 | 12 |
| 0 | 0 | 0 | 0 | 9 | 12 | 9 | 13 | 0 | 0 | 0 | 0 |
| 0 | 0 | 0 | 0 | 57 | 69 | 67 | 86 | 0 | 0 | 0 | 0 |
| 4 | 0 | 4 | 0 | 6 | 0 | 3 | 0 | 10 | 0 | 7 | 0 |
| 1 | 36 | 0 | 0 | 0 | 0 | 0 | 0 | 0 | 0 | 0 | 0 |
| 0 | 0 | 0 | 0 | 43 | 57 | 62 | 67 | 2 | 96 | 0 | 0 |
| 1 | 100 | 0 | 0 | 0 | 0 | 0 | 0 | 0 | 0 | 0 | 0 |
| A | 0 | 0 | 0 | 0 | 0 | 0 | 0 | 0 | 0 | 103 | 0 |
| 60 | 0 | 100 | 0 | 61 | 0 | 64 | 0 | 71 | 0 | 74 | 0 |
| A | 0 | 0 | 0 | 0 | 35 | 0 | 0 | 0 | 0 | 0 | 0 |
| 0 | 0 | 0 | 0 | 42 | 110 | 42 | 172 | 37 | 165 | 43 | 126 |
| 0 | 0 | 0 | 0 | 27 | 109 | 27 | 138 | 22 | 126 | 30 | 110 |
| 0 | 0 | 0 | 0 | 37 | 27 | 45 | 46 | 46 | 30 | 35 | 42 |
| 0 | 0 | 0 | 0 | 10 | 7 | 18 | 3 | 0 | 0 | 0 | 0 |
| 0 | 0 | 0 | 0 | 28 | 12 | 62 | 16 | 1 | 58 | 0 | 0 |
| 0 | 0 | 1 | 164 | 118 | 0 | 320 | 1 | 2 | 134 | 0 | 0 |
| 0 | 0 | 0 | 0 | 22 | 30 | 0 | 0 | 0 | 0 | 0 | 0 |
| 20 | 18 | 25 | 30 | 16 | 14 | 21 | 33 | 0 | 0 | 0 | 0 |
| 0 | 0 | 0 | 0 | 7 | 2 | 6 | 11 | 7 | 1 | 7 | 10 |
| 0 | 0 | 0 | 0 | 0 | 0 | 0 | 0 | 15 | 27 | 10 | 26 |
| 0 | 0 | 0 | 0 | 0 | 0 | 0 | 0 | 1 | 201 | 0 | 0 |
| 0 | 0 | 0 | 0 | 23 | 33 | 36 | 41 | 44 | 31 | 19 | 15 |
| 58 | 0 | 60 | 0 | 57 | 0 | 79 | 0 | 68 | 0 | 59 | 0 |
| 0 | 0 | 0 | 0 | 0 | 0 | 0 | 0 | 42 | 17 | 37 | 30 |
| 0 | 0 | 0 | 0 | 0 | 0 | 1 | 122 | 0 | 0 | 0 | 0 |
| 0 | 0 | 1 | 1 | 0 | 0 | 2 | 1 | 3 | 0 | 1 | 0 |
| 0 | 0 | 0 | 0 | 0 | 0 | 0 | 0 | 18 | 10 | 8 | 8 |
| 1 | 74 | 0 | 0 | 0 | 0 | 0 | 0 | 0 | 0 | 0 | 0 |
| 0 | 0 | 0 | 0 | 11 | 10 | 16 | 17 | 15 | 3 | 14 | 7 |
| 14 | 17 | 29 | 26 | 21 | 32 | 40 | 7 | 46 | 1 | 24 | 1 |
| A | 0 | 0 | 0 | 0 | 0 | 0 | 0 | 0 | 27 | 0 | 0 |
| 22 | 12 | 19 | 27 | 0 | 0 | 0 | 0 | 31 | 15 | 20 | 4 |
| 0 | 0 | 0 | 0 | 0 | 0 | 1 | 80 | 24 | 7 | 27 | 12 |
| 36 | 0 | 58 | 0 | 27 | 0 | 38 | 0 | 27 | 0 | 33 | 0 |
| 35 | 0 | 58 | 0 | 31 | 0 | 38 | 0 | 31 | 0 | 36 | 0 |
| 0 | 0 | 0 | 0 | 0 | 0 | 0 | 0 | 45 | 51 | 61 | 22 |
| 0 | 0 | 0 | 0 | 1 | 75 | 0 | 0 | 0 | 0 | 0 | 0 |
| 0 | 0 | 0 | 0 | 0 | 0 | 1 | 152 | 30 | 52 | 41 | 89 |

| | | | | | | | | | | | |
|---|---|---|---|---|---|---|---|---|---|---|---|
| 0 | 0 | 0 | 0 | 0 | 0 | 0 | 0 | 10 | 11 | 4 | 2 |
| 0 | 0 | 0 | 0 | 24 | 23 | 39 | 32 | 30 | 30 | 30 | 14 |
| 0 | 0 | 0 | 0 | 0 | 0 | 0 | 0 | 0 | 0 | 0 | 0 |
| 39 | 10 | 40 | 13 | 14 | 5 | 43 | 9 | 23 | 5 | 22 | 8 |
| 77 | 0 | 143 | 0 | 2 | 125 | 0 | 0 | 101 | 48 | 37 | 43 |
| 50 | 0 | 65 | 0 | 1 | 46 | 1 | 83 | 36 | 13 | 12 | 33 |
| 0 | 0 | 0 | 0 | 0 | 0 | 1 | 56 | 0 | 0 | 0 | 0 |
| 0 | 0 | 0 | 0 | 0 | 0 | 0 | 0 | 4 | 11 | 7 | 5 |
| 1 | 149 | 0 | 0 | 0 | 0 | 0 | 0 | 47 | 42 | 32 | 38 |
| 0 | 0 | 1 | 44 | 0 | 0 | 0 | 0 | 28 | 14 | 9 | 18 |
| 14 | 40 | 50 | 38 | 14 | 50 | 10 | 48 | 58 | 2 | 64 | 0 |
| 14 | 20 | 28 | 35 | 16 | 5 | 29 | 10 | 32 | 17 | 12 | 34 |
| 0 | 0 | 0 | 0 | 0 | 0 | 0 | 0 | 5 | 87 | 23 | 50 |
| C | 0 | 0 | 0 | 0 | 0 | 0 | 0 | 0 | 66 | 0 | 0 |
| 0 | 0 | 0 | 0 | 0 | 0 | 0 | 0 | 42 | 0 | 27 | 0 |
| 0 | 0 | 0 | 0 | 0 | 0 | 0 | 0 | 26 | 7 | 32 | 10 |
| 41 | 0 | 75 | 0 | 1 | 53 | 1 | 87 | 35 | 0 | 49 | 0 |
| C | T | 0 | 0 | 3 | 69 | 0 | 0 | 0 | 0 | 0 | 0 |
| 15 | 30 | 26 | 27 | 0 | 0 | 0 | 0 | 54 | 0 | 36 | 1 |
| 35 | 0 | 50 | 0 | 46 | 0 | 0 | 0 | 50 | 0 | 42 | 0 |
| 0 | 0 | 0 | 0 | 0 | 0 | 0 | 0 | 26 | 9 | 25 | 13 |
| 0 | 0 | 0 | 0 | 0 | 0 | 0 | 0 | 28 | 11 | 27 | 15 |
| 11 | 13 | 22 | 16 | 0 | 0 | 22 | 20 | 28 | 14 | 9 | 9 |
| 0 | 0 | 0 | 0 | 0 | 0 | 2 | 49 | 40 | 10 | 21 | 10 |
| 0 | 0 | 0 | 0 | 2 | 45 | 0 | 0 | 38 | 11 | 18 | 8 |
| 0 | 0 | 0 | 0 | 0 | 0 | 0 | 0 | 17 | 2 | 8 | 6 |
| 8 | 20 | 27 | 24 | 0 | 0 | 0 | 0 | 0 | 0 | 17 | 13 |
| 0 | 0 | 0 | 0 | 0 | 0 | 0 | 0 | 10 | 22 | 11 | 25 |
| 3 | 17 | 25 | 17 | 0 | 0 | 0 | 0 | 26 | 0 | 24 | 0 |
| 0 | 0 | 0 | 0 | 0 | 0 | 0 | 0 | 73 | 2 | 35 | 0 |
| 1 | 46 | 0 | 0 | 0 | 0 | 0 | 0 | 17 | 40 | 23 | 37 |
| 8 | 21 | 35 | 33 | 77 | 1 | 46 | 0 | 57 | 0 | 51 | 1 |
| 24 | 21 | 37 | 39 | 78 | 0 | 60 | 0 | 30 | 51 | 23 | 31 |
| 0 | 0 | 0 | 0 | 27 | 30 | 66 | 73 | 37 | 1 | 83 | 1 |
| 42 | 58 | 66 | 84 | 48 | 48 | 75 | 84 | 0 | 0 | 0 | 0 |

| | | | | | | | | | | | |
|---|---|---|---|---|---|---|---|---|---|---|---|
| 12 | 14 | 16 | 23 | 1 | 33 | 0 | 0 | 18 | 26 | 12 | 17 |
| 0 | 0 | 0 | 0 | 0 | 0 | 0 | 0 | 0 | 0 | 0 | 0 |
| A | 0 | 0 | 2 | 146 | 0 | 0 | 0 | 0 | 0 | 0 | 0 |
| 0 | 0 | 0 | 0 | 0 | 0 | 0 | 0 | 5 | 12 | 12 | 11 |
| 0 | 0 | 0 | 0 | 0 | 0 | 0 | 0 | 0 | 0 | 0 | 0 |
| 8 | 0 | 21 | 0 | 42 | 0 | 49 | 0 | 38 | 0 | 7 | 0 |
| 0 | 0 | 0 | 0 | 1 | 0 | 0 | 0 | 1 | 0 | 0 | 0 |
| 0 | 0 | 0 | 0 | 0 | 0 | 1 | 40 | 0 | 0 | 0 | 0 |
| 0 | 0 | 0 | 0 | 0 | 0 | 0 | 0 | 47 | 46 | 36 | 48 |
| 0 | 0 | 0 | 0 | 0 | 0 | 0 | 0 | 66 | 52 | 39 | 43 |
| 0 | 0 | 0 | 0 | 0 | 0 | 0 | 0 | 41 | 9 | 30 | 8 |
| 17 | 33 | 32 | 43 | 13 | 14 | 41 | 37 | 20 | 8 | 26 | 18 |
| 0 | 0 | 0 | 0 | 0 | 0 | 0 | 0 | 35 | 16 | 28 | 21 |
| 0 | 0 | 0 | 0 | 0 | 0 | 0 | 0 | 35 | 12 | 37 | 18 |
| 0 | 0 | 0 | 0 | 0 | 0 | 0 | 0 | 49 | 0 | 25 | 0 |
| 0 | 0 | 0 | 0 | 0 | 0 | 0 | 0 | 41 | 0 | 36 | 1 |
| 0 | 0 | 0 | 0 | 0 | 0 | 0 | 0 | 50 | 19 | 59 | 43 |
| 0 | 0 | 0 | 0 | 0 | 0 | 0 | 0 | 14 | 13 | 8 | 7 |
| A | 0 | 0 | 0 | 0 | 0 | 0 | 0 | 0 | 18 | 0 | 0 |
| 0 | 0 | 0 | 0 | 1 | 7 | 7 | 3 | 13 | 0 | 13 | 0 |
| 0 | 0 | 0 | 0 | 0 | 0 | 0 | 0 | 30 | 36 | 15 | 29 |
| 0 | 0 | 0 | 0 | 1 | 55 | 0 | 0 | 35 | 44 | 25 | 35 |
| 0 | 0 | 0 | 0 | 40 | 33 | 51 | 64 | 52 | 100 | 33 | 67 |
| 0 | 0 | 0 | 0 | 3 | 25 | 10 | 31 | 5 | 45 | 6 | 36 |
| 0 | 0 | 0 | 0 | 0 | 0 | 0 | 0 | 26 | 65 | 20 | 36 |
| 23 | 0 | 50 | 0 | 49 | 0 | 53 | 0 | 45 | 0 | 41 | 0 |
| 0 | 0 | 0 | 0 | 40 | 32 | 52 | 66 | 41 | 43 | 38 | 53 |
| 0 | 0 | 1 | 167 | 0 | 0 | 0 | 0 | 39 | 43 | 28 | 60 |
| 1 | 101 | 0 | 0 | 0 | 0 | 0 | 0 | 0 | 0 | 0 | 0 |
| 0 | 0 | 0 | 0 | 0 | 0 | 0 | 0 | 5 | 7 | 4 | 7 |
| 42 | 53 | 0 | 0 | 0 | 0 | 0 | 0 | 0 | 0 | 39 | 41 |
| 25 | 80 | 41 | 152 | 1 | 39 | 0 | 0 | 59 | 55 | 39 | 26 |
| 17 | 20 | 30 | 20 | 0 | 0 | 0 | 0 | 28 | 9 | 18 | 13 |
| 0 | 0 | 0 | 0 | 0 | 0 | 0 | 0 | 56 | 0 | 63 | 1 |
| 0 | 0 | 1 | 50 | 0 | 0 | 1 | 106 | 66 | 0 | 55 | 0 |
| 0 | 0 | 1 | 52 | 0 | 0 | 1 | 126 | 77 | 1 | 52 | 1 |
| 0 | 0 | 0 | 0 | 1 | 54 | 0 | 0 | 91 | 0 | 29 | 0 |
| 34 | 36 | 71 | 70 | 1 | 47 | 0 | 0 | 43 | 18 | 38 | 25 |
| 47 | 24 | 60 | 74 | 1 | 42 | 0 | 0 | 30 | 10 | 59 | 25 |
| 38 | 70 | 92 | 97 | 12 | 43 | 18 | 98 | 46 | 21 | 54 | 30 |
| 30 | 25 | 35 | 29 | 51 | 0 | 58 | 0 | 58 | 0 | 43 | 0 |
| 0 | 0 | 0 | 0 | 0 | 0 | 0 | 0 | 68 | 33 | 39 | 45 |

Supplementary Table S12: LoopSeq targeted sequencing on-target rate

| Sample | On-target rate at gene level | On-target rate at short-read level | On-target rate at long-read level | On-target rate at base level |
|---|---|---|---|---|
| 1N | 31.83% | 91.07% | 91.06% | 93.72% |
| 1T | 28.86% | 91.33% | 91.58% | 95.54% |
| 1M | 36.74% | 84.45% | 83.46% | 95.97% |
| 2T | 27.41% | 85.54% | 84.05% | 93.56% |
| 2M | 26.23% | 85.80% | 86.43% | 95.38% |
| 3N | 26.25% | 89.39% | 89.00% | 93.65% |
| 3T | 20.35% | 87.38% | 87.65% | 94.07% |
| 3M | 31.77% | 88.61% | 87.88% | 94.74% |

**Supplementary Table S13**
**Primers and probes design**

| Gene | Isoform | Primer | Probe |
|---|---|---|---|
| **ATP1A1** | **XR_002956654.1** | **CTCCGGTAACGGTGACGTTTC** /CGCCTCCTCCTCACTTCCTAA | **5'-FAM/CCATGGTGG/ZEN/CGGTGCTCAGT/3IABkFQ** |
| | **XM_017001361.1** | **GTCCTCTCATAACGCATATGCTG** /GCTGACCACCTTACCCTCCT | **5'-FAM/TGTCTCTAA/ZEN/CCAGAGCAATCTGG/3IABkFQ** |
| | **NM_001160233.1** | **GCCCTCCCCCAAAGAAAAAAC /ACACACCCTTTCTGGCACCAG** | **5'-FAM/TTCTAAGTG/ZEN/CGAAGCCGGCTG/3IABkFQ** |
| | **NM_001160234.1** | **TAAAAACATGAAATGAGGTTGGCAC /CCAGCAGTGTATCCTTAGGCAAG** | **5'-FAM/CTTTATCTC/ZEN/CACGCTGTGGAAG/3IABkFQ** |
| | **NM_000701.8/ XR_002956654.1** | **GAGCTGCTCTGTGCTTTTCT /CCATGGTGGCGGTGCTCAGT** | **5'-FAM/TGATTCTCC/ZEN/AGCGACAGGACC/3IABkF** |
| **STAMBPL1 /FAS** | **XM_011539985.2 /XM_011539766.2** | **TTCATCCACACACCAAGGAGC /TGTGCCAGCCTTGTGCACACA** | **5'-FAM/CAGGCTGTT/ZEN/CAGTATGCTCAG/3IABkFQ** |
| **ZNF124 /SMYD3** | **XM_005273257.4 /NM_001375963.1** | **GATGTCGGGACACCCCGGAA /GGCACTGAGAGCATCGCATC** | **5'-FAM/CTGGGAAAT/ZEN/GGAAGGAAAAGCT/3IABkFQ** |
| **PTPRK /ECHDC1** | **XM_011536021.3 /NM_001105544.1** | **GGATCGGCCCAAGGCCAGT /GTGGACGAATCTGATCTTACTC** | **5'-FAM/CTCCGCAGG/ZEN/TTAATGACTCCA/3IABkFQ** |
| **VAPB /GNAS** | **NM_001195677.2 /XM_017027814.2** | **GGATCGGCCCAAGGCCAGT /GTGGACGAATCTGATCTTACTC** | **5'-FAM/CTCCGCAGG/ZEN/TTAATGACTCCA/3IABkFQ** |
